# A small RNA derived from the 5’ end of the IS200 *tnpA* transcript regulates multiple virulence regulons in *Salmonella* Typhimurium

**DOI:** 10.1101/2024.06.26.600842

**Authors:** Ryan S. Trussler, Naomi-Jean Q. Scherba, Michael J. Ellis, Konrad U. Förstner, Matthew Albert, Alexander J. Westermann, David B. Haniford

## Abstract

The insertion sequence IS200 is widely distributed in Eubacteria. Despite its wide-ranging occurrence, IS200 does not appear to be mobile and as such is considered an ancestral component of bacterial genomes. Previous work in *Salmonella enterica* revealed that the IS200 *tnpA* transcript is processed to form a small highly structured RNA (*5’tnpA*) that participates in the post-transcriptional control of *invF* expression, encoding a key transcription factor in this enteropathogen’s invasion regulon. To further examine the scope of *5’tnpA* transcript integration into *Salmonella* gene expression networks, we performed comparative RNA-seq revealing the differential expression of over 200 genes in a *Salmonella* SL1344 *5’tnpA* disruption strain. This includes the genes for the master regulators of both invasion and flagellar regulons (HilD and FlhDC, respectively), plus genes involved in cysteine biosynthesis (cysteine regulon) and an operon (*phsABC*) encoding a thiosulfate reductase complex. These expression changes were accompanied by an 80-fold increase in *Salmonella* invasion of HeLa cells. Likewise, a *phsABC* disruption strain was associated with an increased invasion specifically under anaerobic growth conditions. Based on these findings, we propose that early induction of invasion and motility regulons in the absence of *5’tnpA* causes a metabolic stress resulting in cysteine limitation and activation of CysB, which turns down expression of the *phsABC* operon to increase the flux of thiosulfate in the media towards cysteine production. Taken together, this study provides a powerful new example of bacterial transposon domestication that is based not on the production of a regulatory protein, but of a transposon-derived small RNA.

## Introduction

The Gram-negative bacterium *Salmonella enterica* serovar Typhimurium (hereafter referred to as *Salmonella*) is a major cause of gastrointestinal diseases in humans and a wide range of agriculturally important animals ^1,2^. *Salmonella* typically enters the GI tract through ingestion of contaminated food or water and initially infects epithelial cells of the lower intestine ^3^. Infection is dependent on flagellar-mediated motility of *Salmonella* and the interaction between bacterial fimbria and host cell membranes which promotes adhesion ^4,5^. The subsequent invasion of intestinal epithelial cells is directed by the induction of invasion genes which are mostly encoded on *Salmonella* pathogenicity island 1 (SPI-1), a horizontally acquired genetic element that includes approximately 40 genes arranged in several operons ^6^. The SPI-1 genes include the components of a Type III Secretion System (T3SS), effector proteins, and transcription factors (TF), which control expression of themselves as well as the T3SS and effector genes ^7^. The T3SS transports *Salmonella* effector proteins into the host cell, altering its architecture in a manner that permits *Salmonella* to be taken up^8^. *Salmonella* can replicate in the host epithelial cells and through these cells, gains access to the lymph system where the bacterium can enter macrophage and replicate ^9^.

Because a relatively large number of genes participate in the invasion step, it is expected that efficient invasion involves the precise spatiotemporal regulation of invasion genes. Having the invasion genes turned on only when *Salmonella* is in an optimal position within the GI tract is expected to avoid a metabolic burden that otherwise could reduce the capacity of *Salmonella* to compete for nutrients with other microbes ^10^. Indeed, spatiotemporal regulation of this process is achieved by the invasion genes being organized into a complex positive feed-forward cascade involving several TFs. At a higher level of regulation, this cascade is responsive to several different two-component signal transduction systems and other global regulators ^11,12^. The TF HilD is positioned at the top of this cascade receiving inputs from these signal transduction systems. HilD activates the transcription of two other TFs, HilC and RtsA, as well as of itself. HilC and RtsA also positively self-regulate and control each other, establishing a strong positive feed-forward loop. Upon surpassing a specific threshold of active protein, HilD then activates transcription of *hilA*, encoding a TF that activates transcription of additional TFs InvF and PrgH. The former controls expression of the effector gene operons, while the latter controls the expression of components of the T3SS ^13^. Besides, there is cross-talk between *Salmonella* invasion and flagellar regulons. For example, the FliZ protein, which is a component of the flagellar regulon, binds directly to and activates HilD ^14^. In another linkage, HilD acts as a positive regulator of *flhDC* expression, with FlhDC being the master regulator of the flagellar regulon. Expression of the *flhDC* operon is negatively regulated by a number of TFs including LrhA, YdiV, SlyA, RcsB, RtsB, and RflM, and also positively controlled by Crp-cAMP ^15^.

In addition to receiving regulatory input from proteins, *hilD* expression is subject to control by small non-coding RNAs (sRNA). Bacterial sRNAs constitute a versatile class of regulators that can affect expression of target genes through a variety of different mechanisms (e.g. through modulating the activity of regulatory proteins or binding to complementary sequences in mRNAs, influencing their transcription, translation or stability) ^16^. For example, sRNAs CsrB and CsrC compete with other RNAs for binding the global translational repressor CsrA, effectively titrating CsrA protein, relieving *hilD* repression ^17–19^. Transcription of *csrB* and *csrC* is turned on by the TF SirA and negatively regulated by Crp-cAMP ^20,21^. Spot 42 (encoded by *spf*) is another sRNA which impinges on *hilD* expression. Specifically, Spot 42, whose transcription is negatively regulated by Crp-cAMP, acts as a positive regulator of *hilD* transcription ^22^. While these sRNAs are encoded on *Salmonella’s* core genome, bacterial genomes contain a variety of exogenous DNA elements including plasmids, prophages and transposons ^23^.

IS200 is a small prokaryotic insertion sequence. It is widely conserved in Enterobacteriaceae and found throughout Eubacteria and Archaea. In a few bacteria where IS200 has been studied in detail, including *Escherichia coli* and *Salmonella*, this transposon has been found to be essentially immobile yet conserved in sequence and an active template for transcription producing both a transposase (*tnpA*) mRNA and an antisense transcript (*art200*) that is divergently encoded to *tnpA* (Figure 1) ^24–28^. Given the evidence that multiple independent mechanisms conspire to prevent translation of the *tnpA* transcript ^29^, we previously proposed that one or more IS200-derived transcripts might serve as sRNAs to regulate host gene expression. This led to the discovery that the *tnpA* transcript forms a highly structured and stable sRNA comprised of the first ∼110 nucleotides of the transcript, which we refer to here as *5’tnpA*. Moreover, we demonstrated that the mRNA of the SPI-1 TF InvF, which controls effector gene expression, is a direct target of *5’tnpA*. Over-expression of *5’tnpA* reduced *invF* expression and disrupting all seven copies of IS200 in the pathogenic SL1344 strain of *Salmonella* (referred to here as the Δ*5’tnpA* strain) had the opposite effect ^30^. Once induced, *invF* transcript levels greatly exceed those of *tnpA* and accordingly it is unlikely that *tnpA* regulation of *invF* would have a significant impact on SPI-1 function after the system had been turned on. In contrast, early in growth before SPI-1 induction, *tnpA* transcript levels significantly exceed *invF* mRNA levels and accordingly it is likely that the *5’tnpA* sRNA interferes with SPI-1 induction early in growth ^30^. Interestingly, in a mouse infection model, the Δ*5’tnpA* strain exhibited reduced infectivity in GI tract tissues. As the Δ*5’tnpA* strain displays a slight growth defect in culture, we speculated that the reduced infectivity of this strain in tissues of the mouse GI tract may be due to a reduced ability of this strain to compete with other gut microbes for nutrients ^31^, although another possibility for this observation is presented here.

**Figure 1.**
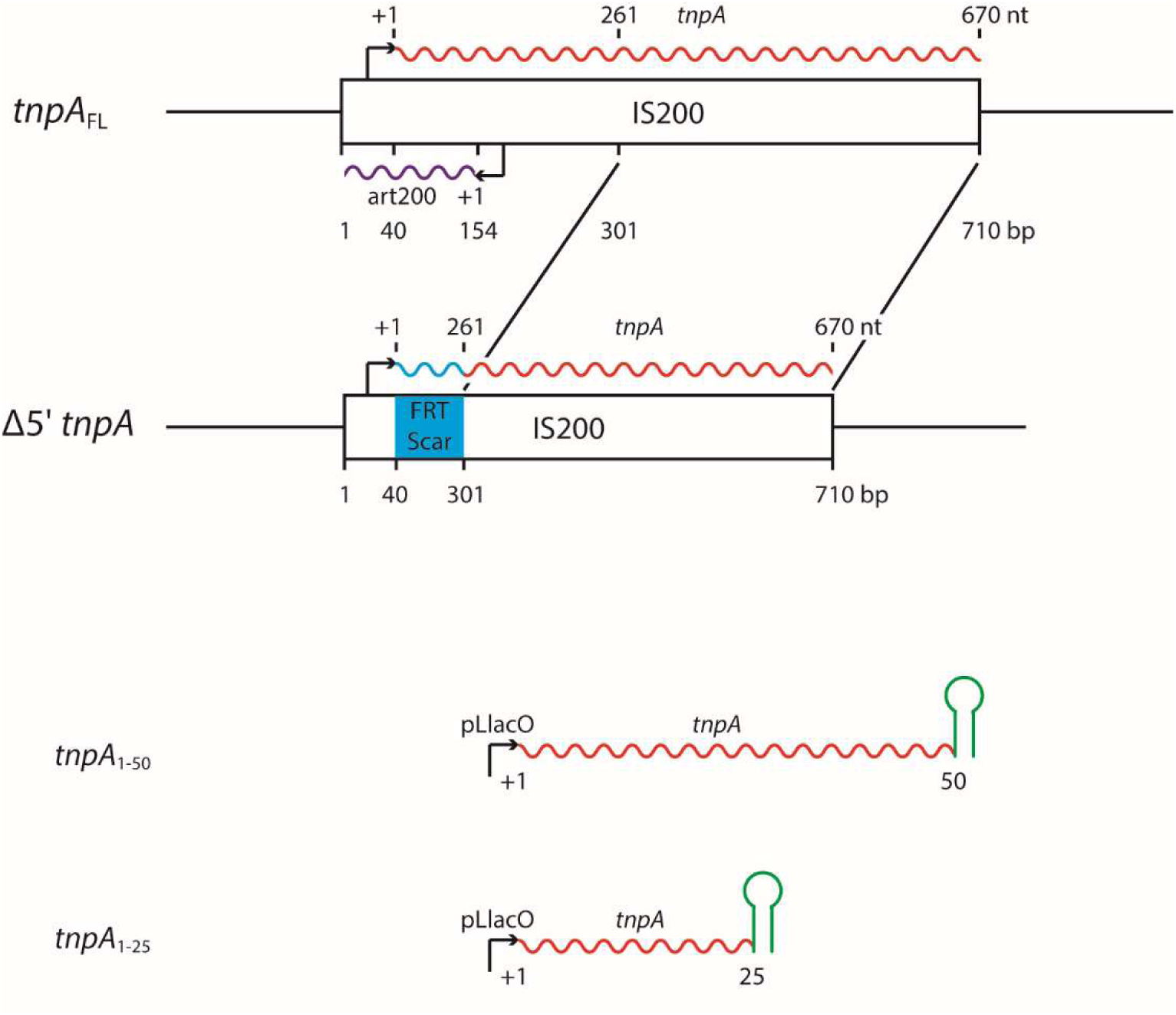
Structure of IS200, position of *5’tnpA* deletion and deletion points for truncated *tnpA* plasmid constructs. The structure of IS200 is shown along with IS200-encoded transcripts including the transposase transcript (red wavy line – *tnpA*) and the antisense transposase transcript (purple wavy line – *art200*). The IS200 insertion sequence is 710 bp in length and the transcription start site for *tnpA* (+1) is located at bp 40. A poorly defined promoter for *tnpA* is located between bp 1 and 40 (arrow). The designation Δ*5’tnpA* defines a 5’ internal deletion in IS200 in which the transcription start site of *tnpA* is fused to bp 301, removing 261 nt from the *tnpA* transcript. *FRT-scar* refers to the Flip-recombinase ‘scar’ sequence left after excision of antibiotic resistance genes used for selection in the construction of the Δ*5’tnpA* strain. IS200 *tnpA* gene derivatives *tnpA_1-50_* and *tnpA_1-25_* encode the first 50 and 25 nt of the *tnpA* transcript, respectively. These truncated *tnpA* genes are present on plasmids pDH1129 and pDH1133 where the IPTG-inducible pLlacO-1 promoter controls expression of the genes and the transcription terminator from the sRNA SgrS (green hairpin structure) controls transcription termination. The native IS200 gene includes a weaker transcription terminator (not shown).

In the current work, we have more extensively characterized the role of the IS200 *5’tnpA* sRNA in *Salmonella* by performing comparative gene expression analysis at multiple growth phases. We show that the deletion of *5’tnpA* from all seven copies of IS200 affects the expression of over 200 genes by 2-fold or more. At least four major pathways are impacted by this absence, including the invasion and flagellar regulons, the cysteine regulon and expression of a thiosulfate reductase. Regarding the SPI-1 regulon, we provide evidence that in addition to its capacity to negatively regulate *invF* expression, *5’tnpA* also regulates the expression of *hilD*, likely by influencing the expression of factors that act upstream of *hilD*, including Crp, CsrC, Spot 42 and LrhA.

## Results

### Differential gene expression analysis by RNA-seq

To better understand the function of the IS200 transposase gene transcript in post-transcriptional regulation of *Salmonella* genes, we removed the 5’ portion (bp 41-300) of the IS200 *tnpA* gene from all seven copies of IS200 in the *Salmonella* SL1344 chromosome (Figure 1 and Figure S1). This was done by successive transduction of independently constructed disruption alleles into a single strain ^30^. The genome of the resulting strain was sequenced, corroborating the successful disruption of all seven IS200 copies and identifying only a single synonymous mutation (in the *ruvB* gene, encoding a Holliday junction DNA helicase) (data not shown). We refer to this strain as Δ*5’tnpA*.

We performed comparative gene expression analysis by RNA-seq using total RNA samples extracted from wild-type (*WT*) and Δ*5’tnpA* strains (three biological replicates of each strain) grown in rich media (LB) at three growth stages; the transition from lag phase to early-exponential phase (EE), mid-exponential phase (ME), and the transition from late-exponential to stationary phase (LE) (see Figure S2 for growth curves and Tables S1-S3 for RNA-seq data). Across all growth stages, a total of 219 genes exhibited differential expression between *WT* and Δ*5’tnpA Salmonella* (FC ≥ |2|; P < 0.05) (Table S3). The data is summarized in the form of MA plots in Figures 2A-C. For all three growth phases, differences in gene expression between the two strains tended towards increases in the Δ*5’tnpA* strain (88%, 94% and 71% of differentially expressed genes for EE, ME and LE, respectively). In EE and ME phases, genes exhibiting the largest increases in expression in the Δ*5’tnpA* strain are involved in invasion (SPI-1 genes), flagellar-dependent cell motility and chemotaxis (blue dots in Figures 2A and B). In LE growth, genes involved in cysteine biosynthesis (cysteine regulon) were the most up-regulated genes in the Δ*5’tnpA* strain (blue dots in Figure 2C). Genes involved in thiosulfate reduction (components of the *phsABC* operon) were the most highly downregulated genes in the entire data set. Results from gene ontology analysis further emphasize the over-representation of genes involved in invasion and flagellar-dependent cell motility in EE and ME growth and genes involved in cysteine biosynthesis in LE growth in the data set (Figure 2D). The magnitude of expression differences between the two strains for each of the three growth phases is also presented in the heat map shown in Figure 3A, and the overlap between differentially expressed genes in the different growth phases is presented in a Venn diagram in Figure 3B. From this analysis it is apparent that EE phase has the highest number of differentially expressed genes that are unique to a specific phase (84 genes). In contrast, ME phase has the fewest number of genes in this category (6 genes). There are 21 differentially expressed genes present in all three growth phases and the majority of these (67%) are constituents of the flagellar regulon. Overlap of differentially expressed genes in EE and ME (35 genes) are mainly genes in the invasion cascade (63%). Gene lists for each of these categories are presented in Table S4. The results support earlier published data implicating *tnpA* mRNA as a negative regulator of SPI-1 (see Introduction). From the current work, further insight into how *invF* expression is affected by *tnpA* expression – i.e., in addition to being directly targeted by the sRNA - can be inferred. The RNA-seq analysis identifies four genes upstream of *invF* in the SPI-1 regulatory cascade as being upregulated in the Δ*5’tnpA* compared to the *WT* strain early in growth, including *hilD* (2.3-fold), *hilC* (2.9-fold), *rtsA* (7.7-fold) and *hilA* (7.7-fold) (Figure 3A). Accordingly, the increased expression of the three genes (*hilD*, *hilC*, and *rtsA*) that make up the positive feed-forward loop that activates the major positive regulator of *invF* transcription (HilA), represents a second mechanism by which *5’tnpA* is linked to *invF* expression. In line with the idea of two additive regulations (direct and indirect) converging at the level InvF, the *invF* mRNA itself is upregulated 16.3-fold in EE growth in the Δ*5’tnpA* strain (Figure 3A).

**Figure 2.**
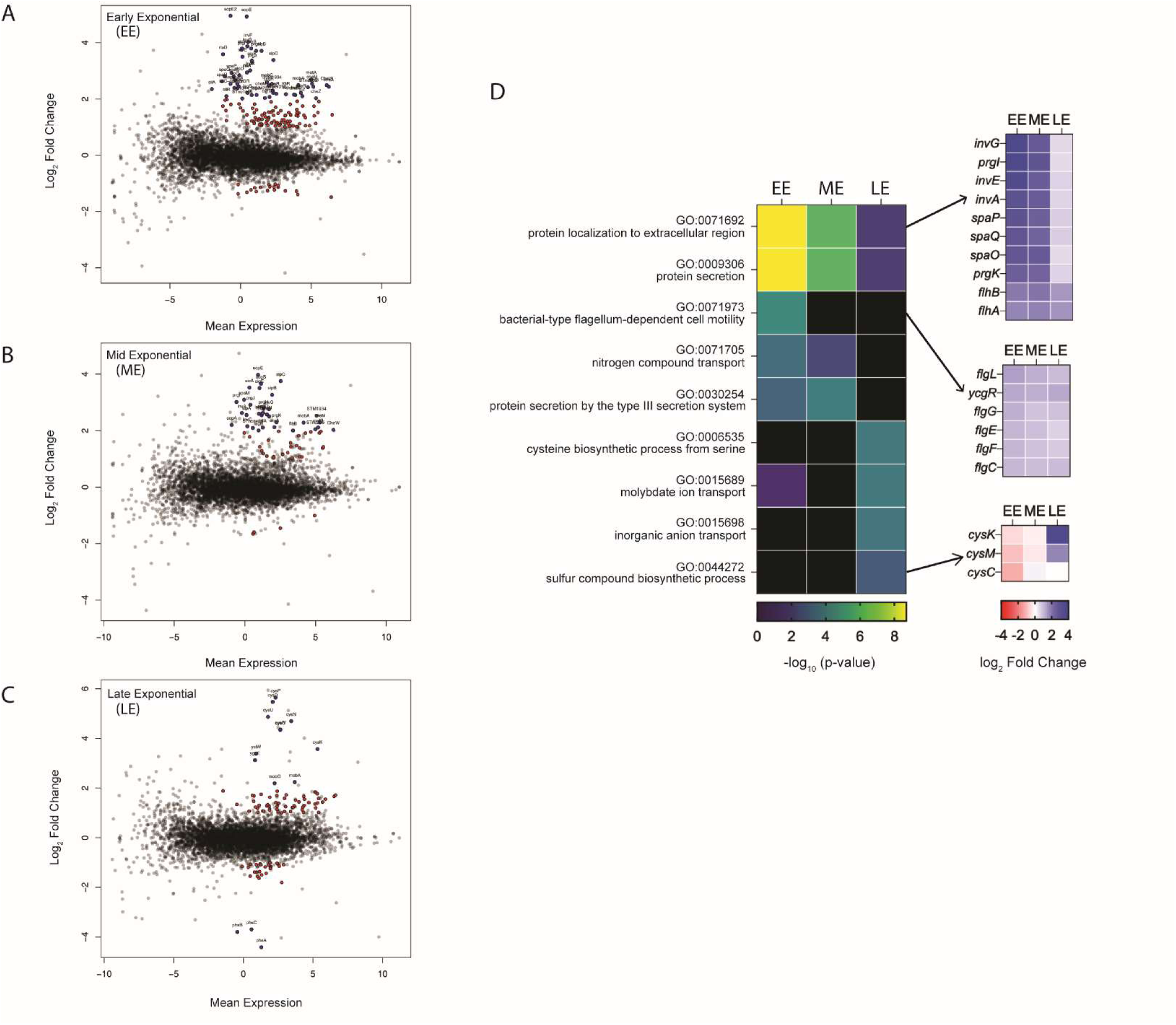
Differential gene expression analysis via RNA-seq. (A-C) MA plots summarizing comparative RNA-seq data (wild-type vs Δ*5’tnpA*) for early (EE), mid (ME) and late (LE) exponential growth phases. For each growth phase, RNA was extracted from 3 clones from each of WT and Δ*5’tnpA* strains. Genes showing differential expression of 2-fold or more are represented by red and blue dots; in panels A and B, blue dots are components of either SPI-1 or flagellar regulons, whereas in panel C, blue dots are components of either the cysteine regulon or the *phsABC* operon. (D) Gene ontology (GO) analysis for the comparative RNA-seq data shows the top GO terms identified along with a heat map showing the extent of differential expression for genes found in 3 selected GO terms.

**Figure 3.**
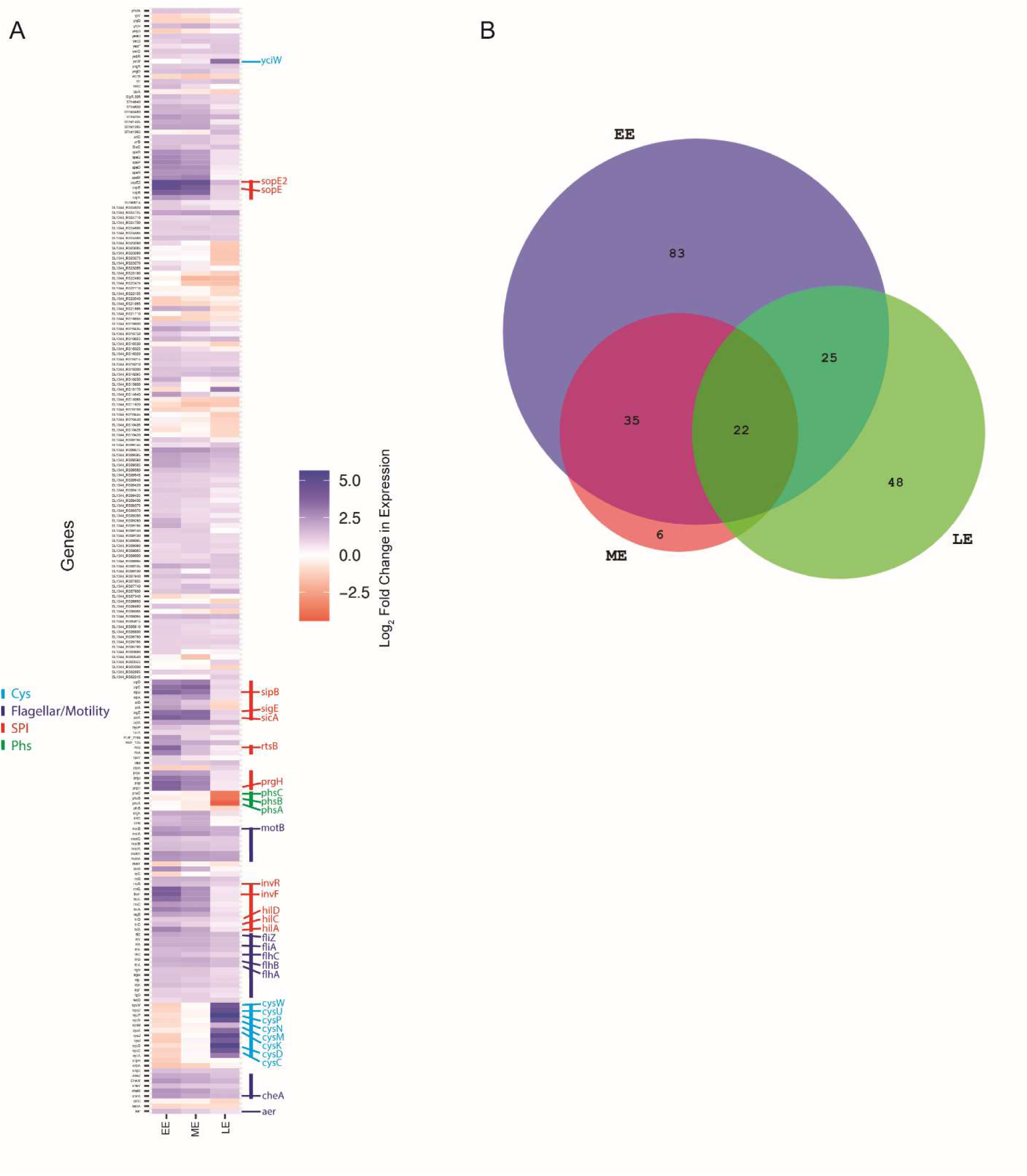
Comparison of differential gene expression over three growth phases and the extent of overlap between phases. (A) Heat map showing expression values of individual differentially expressed genes for EE, ME and LE growth phases. Vertical bars of different color define specific classes of differentially expressed genes with some examples indicated in large font. (B) Venn diagram showing the number of differentially expressed genes that are either growth phase-specific or show overlap between two or more growth phases.

Amongst the most highly upregulated genes in EE growth are those for the SPI-1 effectors (e.g. *sopE2* [31-fold], *sicA* [14-fold] and *sipB* [13-fold]) (Figure 3A). Thus, the increased level of expression of SPI-1 TFs in EE growth is sufficient to induce effector gene expression early in growth. A more detailed example of this is shown in the qRT-PCR experiment shown in Figure S3A where we compare the expression profiles of *sopE* in *WT* and Δ*5’tnpA* strains over 6 growth points. A highlight from this experiment is that during EE growth *sopE* expression is increased 35-fold in the Δ*5’tnpA* strain versus *WT*. Importantly, this increase in effector expression at the transcript level coincides with an increased invasion frequency for the Δ*5’tnpA* strain taken at EE growth. As shown in Figure S3B, Δ*5’tnpA* cells grown to EE are close to 80-fold more competent for invasion of HeLa cells versus *WT* cells taken at the same growth phase. In fact, the invasion frequency of the Δ*5’tnpA* strain taken at EE growth is roughly equivalent to that of the *WT* strain at early stationary phase, where SPI-1 effector gene expression is maximal in LB media ^32^.

For the flagellar regulon, genes for regulators (e.g. *flhDC* [3.1-fold] and *fliA* [3.7-fold]) and structural components of flagella (e.g. *flhB* [3.8-fold], *flhA* [3.4-fold] and *motB* [5.4-fold]), genes involved in chemotaxis (e.g. *cheR* [4.7-fold], *cheA* [5.4-fold] and *aer* [2.9-fold]) and the gene for a post-translational activator of HilD (*fliZ*) [4.3-fold]) are upregulated in the Δ*5’tnpA* strain (Figures 2A, B and D and Figure 3A). In addition, *fliZ* is upregulated 4.3-fold. Notably, both flagellar-dependent motility and chemotaxis are important for invasion ^33,34^.

For LE phase, genes involved in sulfur assimilation/cysteine synthesis show differential expression (increase) of the largest magnitude (Figure 2C). This group comprises 13% of all differentially expressed genes in this growth phase. Components of the cysteine regulon (e.g. *cysP*, *cysD*, *cysU*, *cysN*, *cysW*, *cysC cysK*, and *yciW*) are amongst the most strongly upregulated genes in the entire data set (Figure 3A). In contrast, genes involved in thiosulfate reduction (components of the *phsABC* operon) are amongst the most strongly downregulated genes in the data set (Figure 3A). In LE growth, differential expression of invasion genes is almost completely lost except for relatively small differences in *invR*, *sigE*, and *sopE2* (up to 3.3-fold). This likely reflects the induction of invasion genes in the *WT* strain that is characteristic of this growth phase in rich media ^32^. In contrast to invasion genes, the increased expression of flagellar regulon genes observed in the earlier phases extends into LE growth (Figure 3A).

We used qRT-PCR (Figure 4A-C) to validate the trends in differential gene expression for *WT* and *Δ5’tnpA* strains discussed above. Notably, the magnitude of the decrease in *phsA* expression is smaller in the qRT-PCR (4-fold) versus the RNA-seq analysis (21-fold).

**Figure 4.**
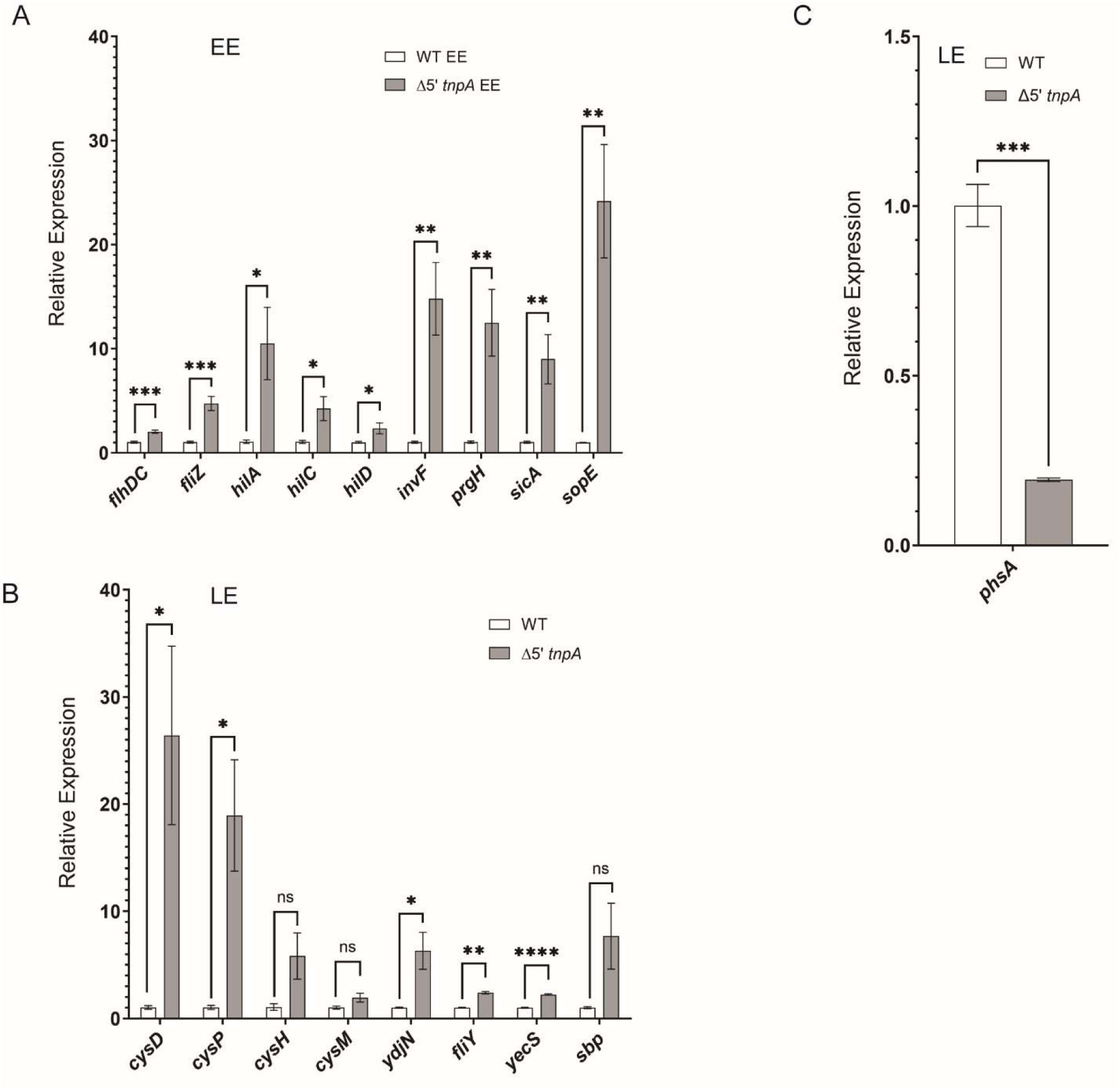
Validation of key hits from RNA-seq analysis. Total RNA was isolated from the indicated strains and subject to qRT-PCR analysis. In (A) strains were grown to EE phase and targets for PCR included select flagellar and SPI-1 regulon genes. In (B) and (C) strains were grown to LE phase and targets included cysteine regulon and *phsABC* operon genes, respectively. For each pairwise comparison, the expression value for the *WT* strain is set to one. Error bars show the standard error of the mean for at least 3 biological replicates. The data shown is representative of at least 3 independent experiments. ns, not statistically significant (P ≥ 0.05); *, **, *** and **** P ≤ 0.05, 0.01, 0.001 and 0.0001, respectively.

### tnpA overexpression suppresses the early induction of hilD and other downstream SPI-1 regulators

To confirm that the *5’tnpA* transcript plays a role in regulating SPI-1 induction via *hilD* expression and to gain insight into what part of the transcript is important for this regulation, we overexpressed various forms of *tnpA* from an IPTG-inducible promoter and assessed the impact of this expression on the transcript levels of *hilD* and other SPI-1-encoded TFs (including *invF* and *hilA*). Constructs included the full-length *tnpA* gene (*tnpA_FL_*) and truncated forms of *tnpA* where only the first 50 (*tnpA_1-50_*) or 25 (*tnpA_1-25_*) nucleotides of the *tnpA* transcript are expressed; notably, we had previously found that constitutive overexpression of the first 50 nucleotides of *tnpA* was sufficient to suppress early induction of *invF* ^30^. The various plasmid constructs were transformed into the Δ*5’tnpA* strain and RNA was prepared for qRT-PCR analysis at EE growth.

Under these conditions, the level of *tnpA_FL_* transcript increased approximately 8-fold relative to uninduced cells (Figure S4). The results in Figure 5A show that induction of *tnpA_FL_* as well as *tnpA_1-50_* suppresses *hilD*, *hilA* and *invF* expression approximately 2-fold, whereas *tnpA_1-25_* expression has no impact on *hilD* or *invF* expression. We also generated a plasmid (pDH1081) where the gene encoding the *FLP-scar-tnpA* chimeric RNA generated from the disrupted form of IS200 (present in the *Δ5’tnpA* strain; Figure 1) was fused to an inducible promoter and the chimeric transcript was expressed in the *WT* strain. Expression of the chimeric transcript did not influence *hilD* or *invF* expression (Figure S5A) even when the expression level of the chimeric transcript was approximately 120-fold higher than in uninduced cells (Figure S5B). Taken together, these results support the contention that the early induction of SPI-1 observed for the Δ*5’tnpA* strain is a consequence of a loss of function (the absence of the 5’ portion of *tnpA*) rather than a gain of function (expression of a chimeric *FLP-scar-tnpA* RNA). Furthermore, the first 50 nt of the *tnpA* transcript are necessary and sufficient for suppression of SPI-1 TF expression.

**Figure 5.**
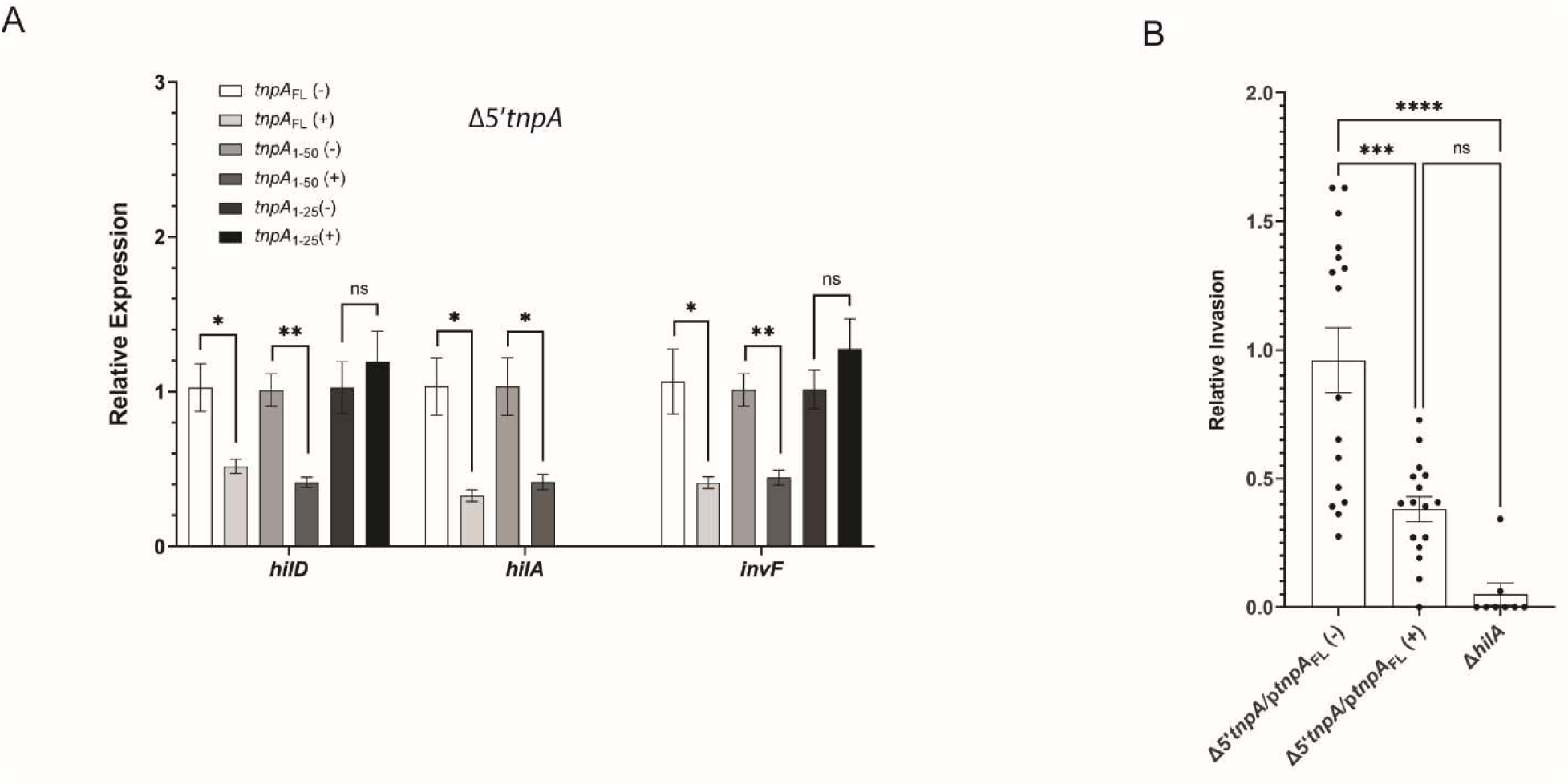
Impact of overexpression of the *tnpA* transcript on select SPI-1 regulators and on *Salmonella* invasion. (A) Plasmids expressing IPTG-inducible full-length *tnpA* (*tnpA_FL_*; pDH968) or truncated forms of *tnpA* (*tnpA_1-50_*; pDH1129 and *tnpA_1-25_*; pDH1133 – Table S6) were transformed into the Δ*5’tnpA* strain. Transformants were grown to EE phase either in the presence (+) or absence (−) of IPTG (0.5 mM) whereupon total RNA was isolated and qRT-PCR performed. For each pair (induced versus uninduced) the relative expression of the uninduced sample was set at 1. Three independent transformants were used for each data set and the results are representative of at least 2 independent experiments. (B) HeLa cells were infected with Δ*5’tnpA* cells transformed with pDH968. Overnight cultures of Δ*5’tnpA*/pDH968 were grown in the presence of ampicillin (Ap). Subcultures were grown in LB-Ap media either in the presence (+) or absence (−) of IPTG (0.5 mM) and grown to ME phase. Bacterial cells were then mixed with HeLa cells as described in Experimental Procedures and the gentamicin protection assay was performed. A non-invasive strain (Δ*hilA*) was included as a negative control. The invasion frequency for Δ*5’tnpA* containing pLacO-1-tnpA_FL_ without IPTG treatment was set to 1. Bars represent the average invasion for two independent experiments each with 8 biological replicates (including technical duplicates). Error bars show the standard error of the mean. ns, not statistically significant (P ≥ 0.05); *, **, *** and **** P ≤ 0.05, 0.01, 0.001 and 0.0001, respectively.

We had previously shown that the 5’ truncation of all seven copies of IS200 increased the capacity of *Salmonella* (SL1344) to invade HeLa cells ^30^. Conversely, we show in Figure 5B that overexpression of *tnpA_FL_* in the Δ*5’tnpA* strain background reduced invasion about 2-fold, supporting the idea that the *tnpA* transcript is a *bona fide* regulator of the invasion machinery presumably due to its ability to downregulate *hilD* expression.

### Profiling differential expression of hilD and its regulators as a function of growth phase in the Δ5’tnpA strain

We carried out a more extensive profiling of *hilD* expression by qRT-PCR to define the point in growth where the largest differential in *hilD* transcript level between *WT* and Δ*5’tnpA* strains exists. As shown in Figure 6A, *hilD* transcript levels gradually increase during growth, peaking in LE phase and decreasing as cells enter early-stationary phase (ES). Consistent with the RNA-seq data where *hilD* was only detected as being differentially expressed in EE, our qRT-PCR analysis shows increased expression of *hilD* in the Δ*5’tnpA* strain from lag phase to LE, with the largest differential occurring in EE and ME phases. As noted above, this early increase in *hilD* expression is biologically relevant because the invasion frequency of the Δ*5’tnpA* strain in EE growth is about 80 times higher than that of the *WT* strain. Thus, we conclude that the early increase in *hilD* expression in the Δ*5’tnpA* strain is sufficient to fully drive SPI-1 induction.

**Figure 6.**
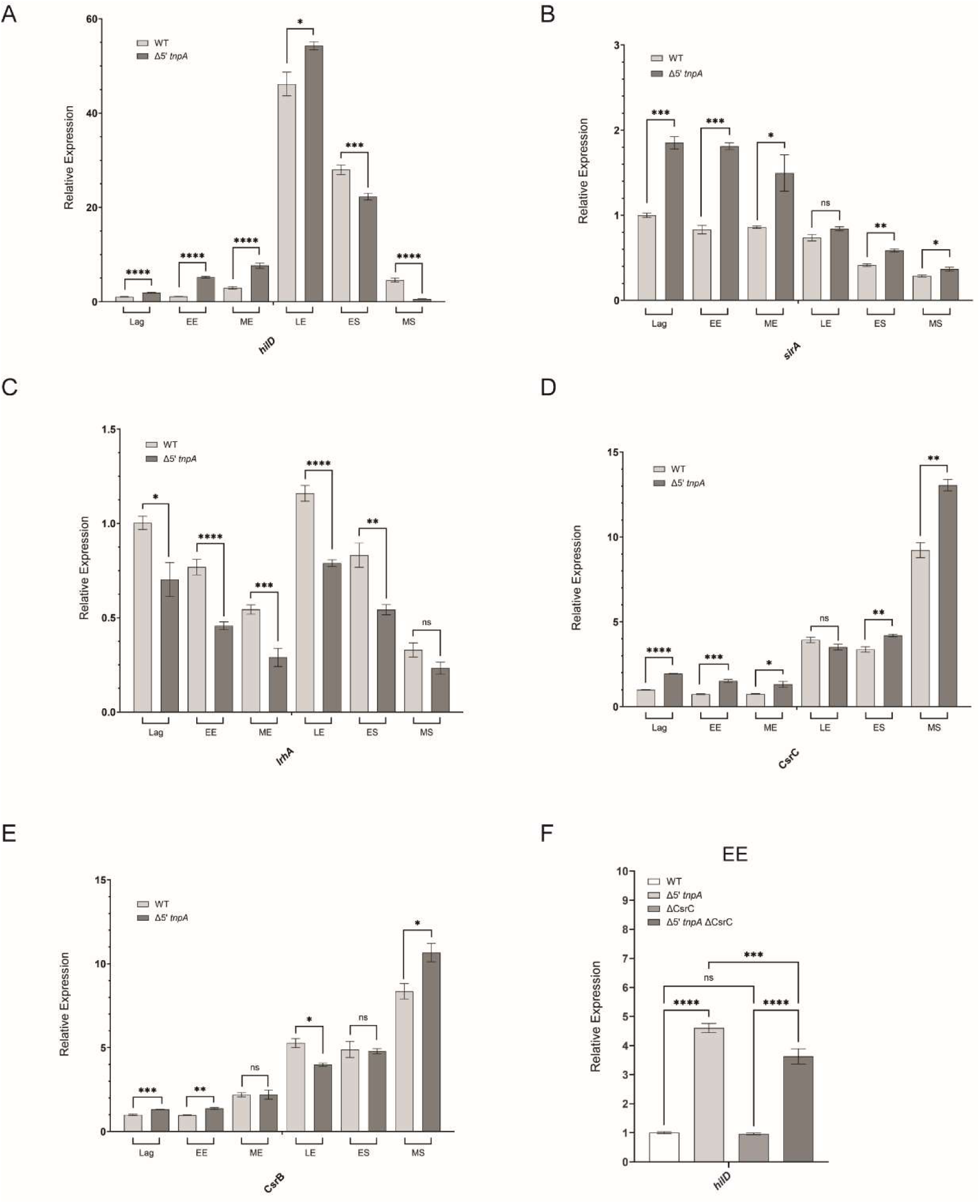
Temporal expression of SPI-1 regulators and the impact of *csrC* disruption on early induction of *hilD* expression. (A-E) Six independent clones of each of the *WT* and Δ*5’tnpA* strains were cultured in LB media and at the indicated growth phases (EE = early-exponential; ME = mid-exponential; LE = late-exponential; ES = early-stationary; MS = mid-stationary), total RNA was isolated and subject to qRT-PCR analysis. For each time course, the relative expression of the *WT* strain in lag phase is set to 1. (F) *hilD* expression levels were assessed by qRT-PCR for the indicated strains. The *hilD* expression level for the *WT* strain was set to 1. Error bars show the standard error of the mean for at least 6 biological replicates. ns, not statistically significant (P ≥ 0.05); *, **, *** and **** P ≤ 0.05, 0.01, 0.001 and 0.0001, respectively.

In considering the possibility that *tnpA* negatively regulates *hilD* expression directly by an RNA-RNA basepairing mechanism, we performed *in silico* pairing analysis using the first 50 nucleotides of *tnpA* as our query subject in RNApredator ^35^. This analysis did not identify *hilD* nor any putative *tnpA* targets that have any obvious connection to SPI-1 or flagellar regulons as top hits (Table S5). To look at the possibility that *tnpA* might act on an upstream regulator of *hilD* (indirect mechanism), we asked if there was evidence of dysregulation of *hilD* regulators early in growth in the Δ*5’tnpA* strain. Note that evidence of this might not have been picked up by the RNA-seq analysis if the extent of altered expression was relatively small (e.g. 2-fold or less). Towards this end, we performed expression profiling (via qRT-PCR) on several genes known to act upstream of *hilD* to regulate its expression, including *hilE* ^36^, *ompR* ^37^, *fur* ^38^, *sirA* ^17^, *rcsB* ^39^, and *lrhA*. The *lrhA* gene was included because it is known to be an early negative regulator of *flhDC* expression which in turn controls the expression of *fliZ* ^15^. In addition, we have already established through RNA-seq analysis that both *flhDC* and *fliZ* are upregulated early in growth (EE and ME) in the Δ*5’tnpA* strain. Of the six genes profiled, only *sirA* and *lrhA* showed differential expression early in growth. The *sirA* gene is upregulated approximately 2-fold starting in lag phase and continuing into ME phase (Figure 6B) and the *lrhA* gene exhibits reduced expression of about 2-fold in EE and ME phases (Figure 6C) in the Δ*5’tnpA* strain.

It was unclear if the small increase in *sirA* transcript early in growth would have a significant biological effect on the SPI-1 system. As part of a two-component phosphorelay system, activation of SirA is dependent on phosphorylation by either acetyl phosphate ^40^ or BarA, which in turn is activated by short chain fatty acids such as formate and acetate ^41^. We looked for evidence of this increase in *sirA* having an impact on SPI-1 by asking if expression levels of SirA-activated genes, *csrB* and *csrC*, also increased in the *Δ5’tnpA* strain. Using the same RNA samples used to investigate the six genes profiled above, we show that from lag to ME phase CsrC is upregulated (up to 1.7-fold) in the Δ*5’tnpA* strain (Figure 6D). By comparison, for CsrB there is only a very small but statistically significant increase in expression (1.1-fold) in the Δ*5’tnpA* strain in EE and ME growth (Figure 6E). While taken together this level of differential expression is relatively small, it should be noted that CsrB and CsrC have 18 and 9 CsrA binding motifs, respectively ^17^. Given that the increase in expression of CsrC is larger than that of CsrB in the *Δ5’tnpA* strain, we anticipated that upregulation of CsrC would be a contributing factor to the early SPI-1 induction phenotype. In support of this we show in Figure 6F that deletion of the *csrC* gene in the *Δ5’tnpA* strain reduces *hilD* expression by roughly 1.2-fold.

### Reduced Crp levels in the Δ5’tnpA strain contribute to early induction of SPI-1

The larger increase in differential expression of CsrC relative to CsrB in the Δ*5’tnpA* strain is suggestive of reduced expression/activity of Crp-cAMP in this strain because Crp-cAMP specifically inhibits CsrC expression early in growth ^21^. We also considered this possibility because of our finding from RNA-seq that the *phsABC* operon is the most downregulated set of genes in the Δ*5’tnpA* strain; earlier published data showed that *phsABC* operon expression is subject to catabolite repression ^42^. Consistent with Crp-cAMP being an important regulator of the *phsABC* operon, we show that *phsABC* expression is strongly reduced in EE growth when the *crp* gene is disrupted (Figure S6A) and that mutations in a putative Crp-cAMP binding site of the *phsABC* operon make expression of the operon insensitive to Crp (Figure S6B-C). To directly test the idea that Crp protein levels are reduced in the Δ*5’tnpA* strain, we compared via western blot analysis, Crp protein levels in *WT* and Δ*5’tnpA* strains at various growth phases. We show in Figure 7A that during lag phase Crp levels are roughly 2-fold lower in the Δ*5’tnpA* strain versus the *WT* strain. Notably, this difference does not manifest at the transcript level at this early growth point (Figure S7A), nor do we see statistically significant differences at later time points for Crp protein levels between the two strains (Figure S7B-C).

**Figure 7.**
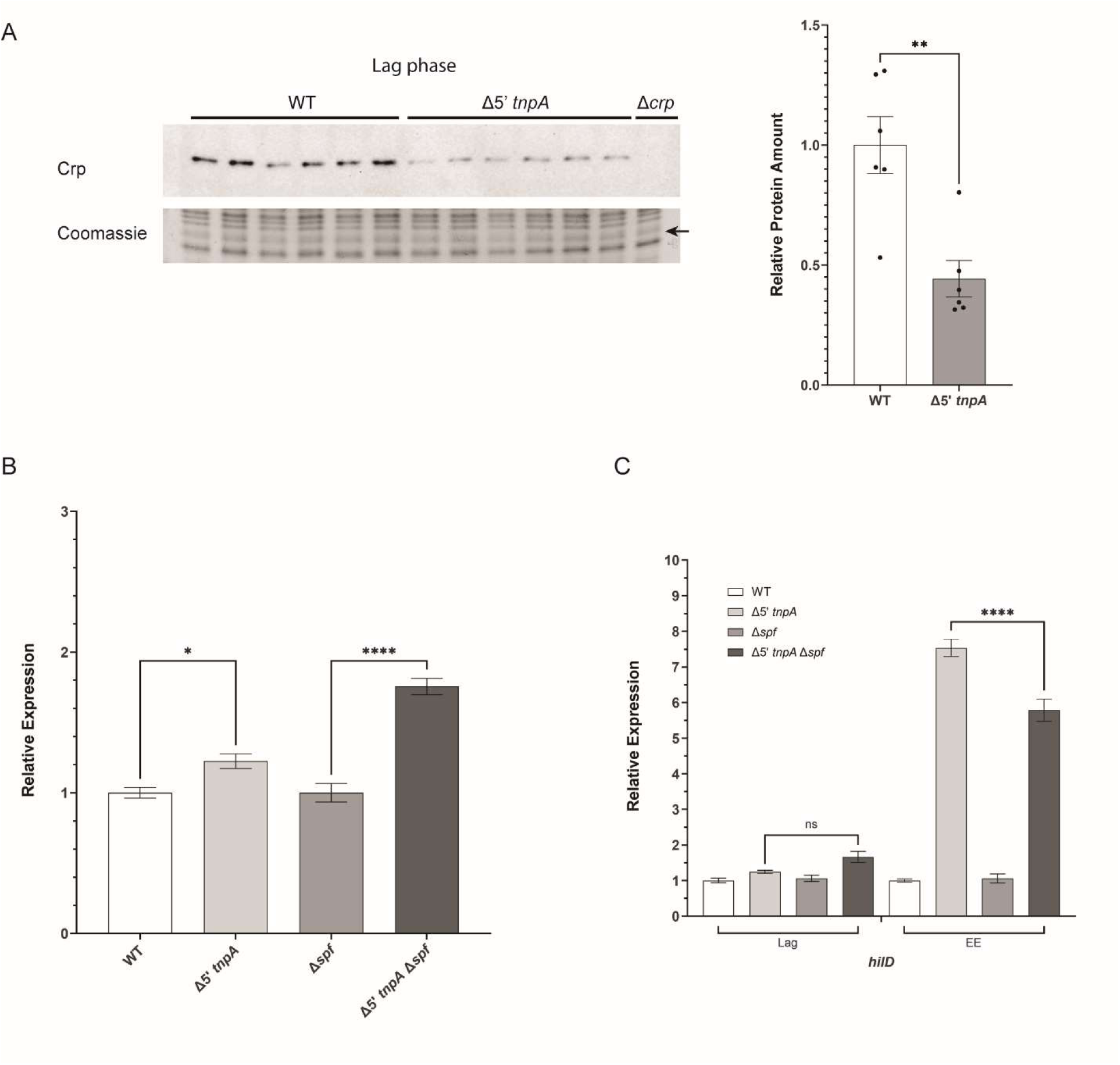
Early induction of SPI-1 in the. Δ*5’tnpA* strain is linked to reduced expression of the catabolite repressor protein (Crp) in lag phase and increased expression of the sRNA Spot 42. (A) Crp western blot from *WT* and *Δ5’tnpA* strains grown to lag phase (OD_600_ = 0.15). DBH569 (Δ*crp*) contains a disruption of the *crp* gene and serves as a negative control for Crp detection. The arrow indicates a protein from a duplicate Coomassie stained gel used as a loading control for quantification purposes; quantification is shown beside the blot with the Crp level in the *WT* strain set to 1. The results for 6 biological replicates are shown. (B) β-galactosidase levels for strains transformed with a Spot 42-lacZ reporter plasmid are shown. Miller units for the *WT* strain were set to 1. (C) *hilD* expression was measured by qRT-PCR for the indicated strains. For (B) and (C) the results for 6 and 3 biological replicates are shown, respectively. ns, not statistically significant (P ≥ 0.05); *, **, *** and **** P ≤ 0.05, 0.01, 0.001 and 0.0001, respectively.

In addition to downregulating CsrC expression early in growth, Crp-cAMP also downregulates expression of the sRNA Spot 42. As indicated in the Introduction, Spot 42 is a positive regulator of *hilD* expression. This led us to ask if Spot 42 levels might be elevated in the Δ*5’tnpA* strain and thereby contribute to the early induction of SPI-1. To investigate this, we generated a Spot 42-lacZ transcriptional fusion and cloned this into a low copy plasmid (pDH1121); one reason for choosing this method of monitoring Spot 42 expression was that we could not reliably perform qRT-PCR on Spot 42 (due to its small size). The pDH1121 plasmid was transformed into WT and Δ*5’tnpA* strains and we used the Miller assay to evaluate Spot 42 expression levels early in growth. The results in Figure 7B show that there is a small increase (1.2-fold) in Spot 42 expression in EE growth in the Δ*5’tnpA* strain. When the same experiment was performed in strain backgrounds where the Spot 42 gene (*spf*) was disrupted (DBH570 and DBH571) the increase in Spot 42 expression from our reporter in the Δ*5’tnpA* background is even higher at approximately 1.8-fold. These results led us to ask if the early *hilD* induction phenotype shows a dependence on Spot 42 expression. As shown in the qRT-PCR experiment in Figure 7C, deleting *spf* specifically in the Δ*5’tnpA* background reduces *hilD* expression approximately 1.25-fold. This is consistent with the early increase in Spot 42 expression contributing to the early induction of *hilD* and SPI-1 in the Δ*5’tnpA* strain.

### Overexpression of LrhA suppresses early SPI-1 induction in the Δ5’tnpA strain

To test the potential significance of the ∼2-fold decrease in *lrhA* mRNA levels in the Δ*5’tnpA* strain on *hilD* expression, we asked if increasing *lrhA* expression would suppress the early increase in *hilD* expression that is characteristic of the Δ*5’tnpA* strain. We were able to increase the *lrhA* transcript level by about 2-fold by integrating a *lrhA-FLAG* allele into the *attTn7* site of the *Salmonella* chromosome in a background where the native *lrhA* gene was disrupted (strains DBH693 and DBH704). We looked at the impact of this overexpression on *hilD* transcript levels for cells grown to EE and ME growth phases. As shown in Figure 8A, increasing *lrhA* expression levels has little effect on *hilD* expression in an otherwise *WT* background (DBH693), but remarkably in the Δ*5’tnpA* strain background (DBH704) full suppression of the ‘*hilD-up*’ phenotype is observed. Notably, this suppression is strongest for cells grown to EE phase. This fits with the idea that LrhA regulation of *flhDC*, and thus *fliZ*, is confined to EE growth ^15^. Invasion studies with the above strains also show that *lrhA* overexpression suppresses invasion to roughly *WT* levels early in growth (Figure 8B). We also show in Figure 8C that while increasing *lrhA* expression does not impact on *sirA* expression, it suppresses CsrC expression specifically in EE growth. This suggests that there is a previously unrecognized interplay between the flagellar pathway and CsrC expression that is independent of SirA. Overall, the results here suggest that the reduced expression of *lrhA* in the *Δ5’tnpA* strain is a key component of the early SPI-1 induction phenotype that is characteristic of this strain.

**Figure 8.**
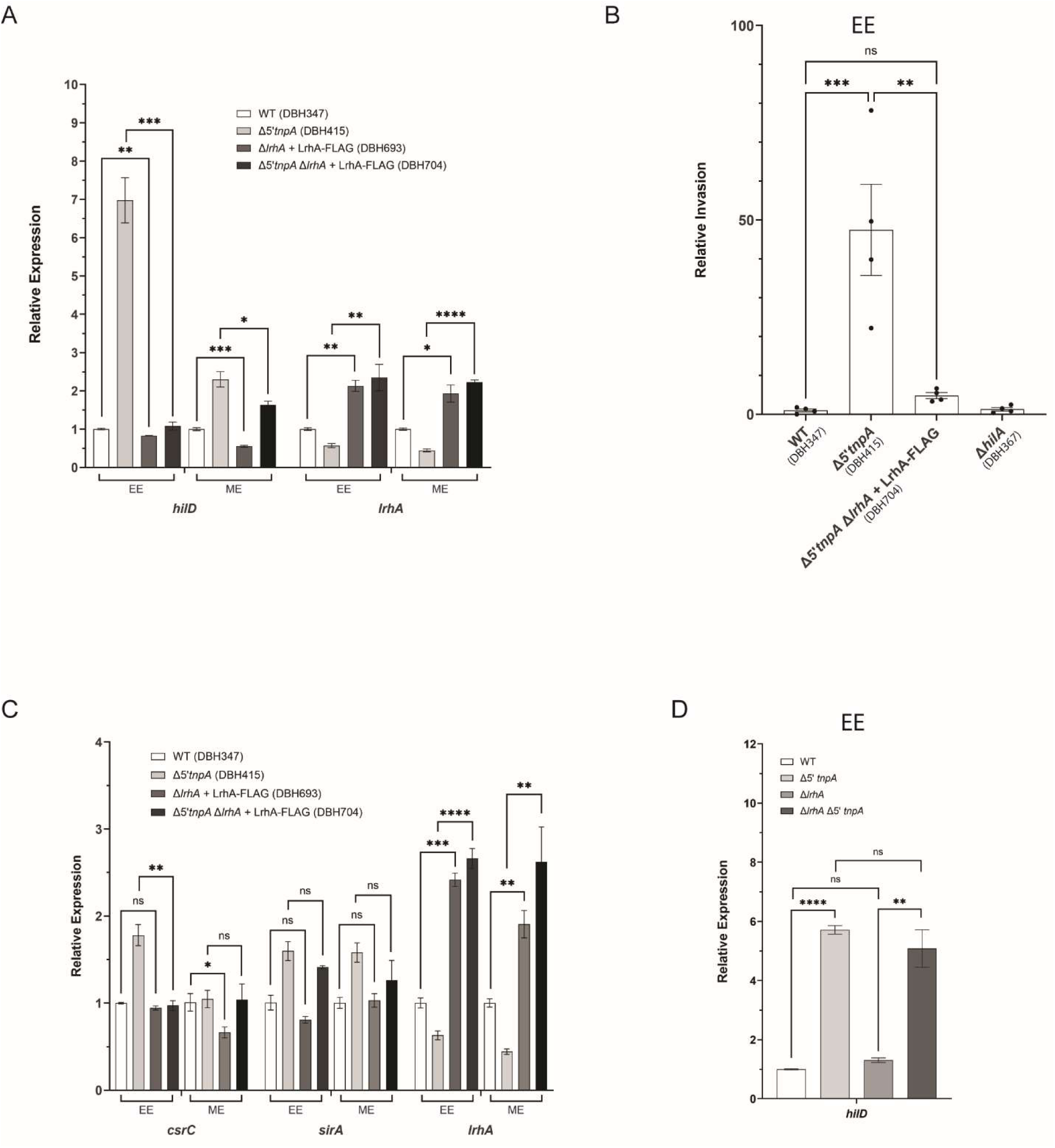
Impact of *lrhA* overexpression and deletion on *hilD* expression and invasion. (A, C and D) The results of qRT-PCR analysis for the indicated strains are shown. For each of the indicated sets the expression of the target gene in the *WT* strain was set to 1. For each of (A), (C) and (D) data is shown for 6 biological replicates. (B) The invasion frequency for the indicated strains is shown with the frequency for the *WT* strain set to 1. Bars represent the average invasion for two independent experiments each with 2 biological replicates (including technical duplicates). Error bars show the standard error of the mean. ns, not statistically significant (P ≥ 0.05); *, **, *** and **** P ≤ 0.05, 0.01, 0.001 and 0.0001, respectively.

### 5’tnpA regulates three parallel pathways that converge on HilD

Based on the above results relating to changes in *lrhA*, CsrC and Spot 42 expression in the Δ*5’tnpA* strain, we suggest that the associated early induction of SPI-1 results from loss of *5’tnpA* regulation of three parallel, additive pathways that converge on *hilD*, including LrhA-FlhDC-FliA-FliZ, Crp-Spot 42 and SirA-CsrC-CsrA (Figure 9). The most straightforward alternative is that *5’tnpA* acts upstream of *lrhA*, *csrC* and *spf* in a single pathway. The main distinguishing feature of these two general models is that in the ‘single pathway’ model, the disruption of any given pathway gene would be fully dominant to the Δ*5’tnpA* disruption. In contrast, in the ‘parallel pathway’ model full dominance would not be seen, because a single gene disruption would never fully abrogate the impact of the Δ*5’tnpA* disruption on the other two pathways. Accordingly, we would expect that the impact on *hilD* expression of a single mutant (either Δ*lrhA*, Δ*csrC* or Δ*spf*) would never equal the impact of the corresponding double mutant. This is borne out by the data shown in Figures 6F, 7C and 8D. For example, for both Δ*csrC* versus Δ*csrC*Δ*5’tnpA* and Δ*spf* versus Δ*spf*Δ*5’tnpA*, the impact of the single mutant on *hilD* expression is much less than that of the double mutant. The results suggest that the relatively large increase in *hilD* expression in the Δ*5’tnpA* strain results from the additive effect of relatively small changes in *lrhA*, CsrC and Spot 42 expression, arising due to the loss of upstream *5’tnpA* function.

**Figure 9.**
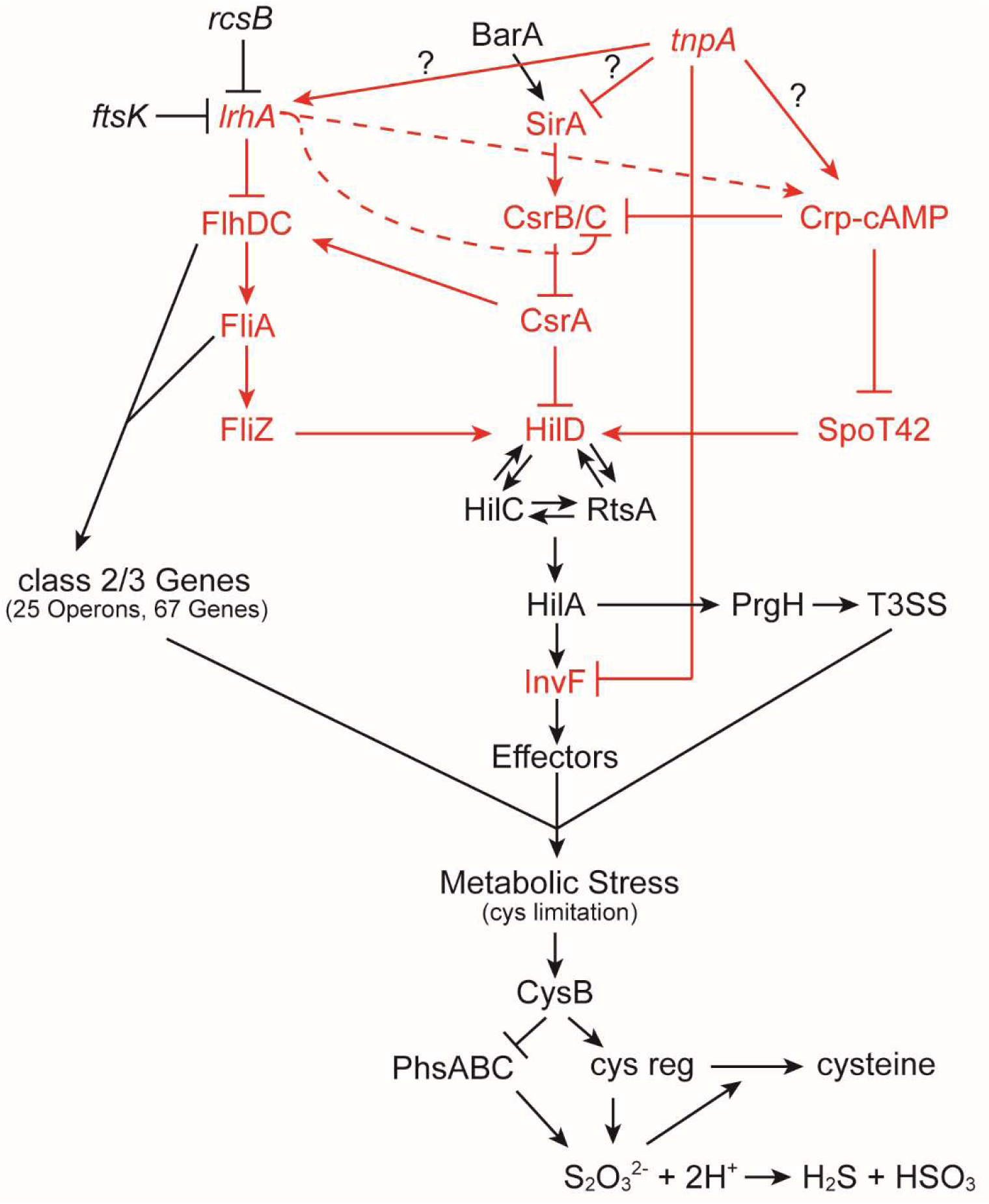
Model for *tnpA* regulation of *hilD* and the impact of *5’tnpA* deletion on the cysteine regulon and the *phsABC* operon. Red lettering indicates pathways predicted to be regulated by *5’tnpA*. Question marks indicate that the regulation of *lrhA*, *crp* and *sirA* by 5’*tnpA* may not be direct. Dotted lines indicate genes that may be regulated by LrhA. Further details are provided in the text.

### Coordinate regulation of the cysteine regulon and phsABC operon through CysB

One feature both the cysteine regulon and the *phsABC* operon have in common is the capacity to influence cysteine biosynthesis. The former because genes in this regulon code for proteins involved in the acquisition of sulfur (in the forms of thiosulfate, sulfate, and cystine) and the assimilation of the sulfur from these compounds into cysteine ^43,44^. The latter because the thiosulfate reductase complex encoded by this operon diverts thiosulfate away from cysteine production ^42^. Given that the expression of these two sets of genes is affected in opposite directions in the Δ*5’tnpA* strain, we considered the possibility that the changes in expression are linked mechanistically. Our working model is that the early induction of invasion and flagellar regulons increases the normal demand for cysteine production as part of the metabolic stress incurred. Accordingly, this would trigger upregulation of the cysteine regulon through activation of CysB, which would also repress expression from the *phsABC* operon; cysteine limitation activates expression of the cysteine regulon through the production of O-acetyl serine, an activator of CysB, the master transcriptional regulator of cysteine regulon genes ^44^. Scanning the *phsABC* operon for a potential CysB binding site revealed a reasonable match to the consensus sequence between the Crp-cAMP binding site and the −35 region of the *phsAB*C promoter (Figure S6B). To test the idea that CysB regulates the *phsABC* operon, we looked at the impact of deleting *cysB* on the expression of a *phsABC-lacZ* reporter. We show in Figure 10A that for cells in a *WT* background grown to EE, LE, and ES phase, disrupting the *cysB* gene consistently results in an increase in *phsABC* expression (up to a maximum of 2-fold), consistent with CysB being a negative regulator of *phsABC* expression.

**Figure 10.**
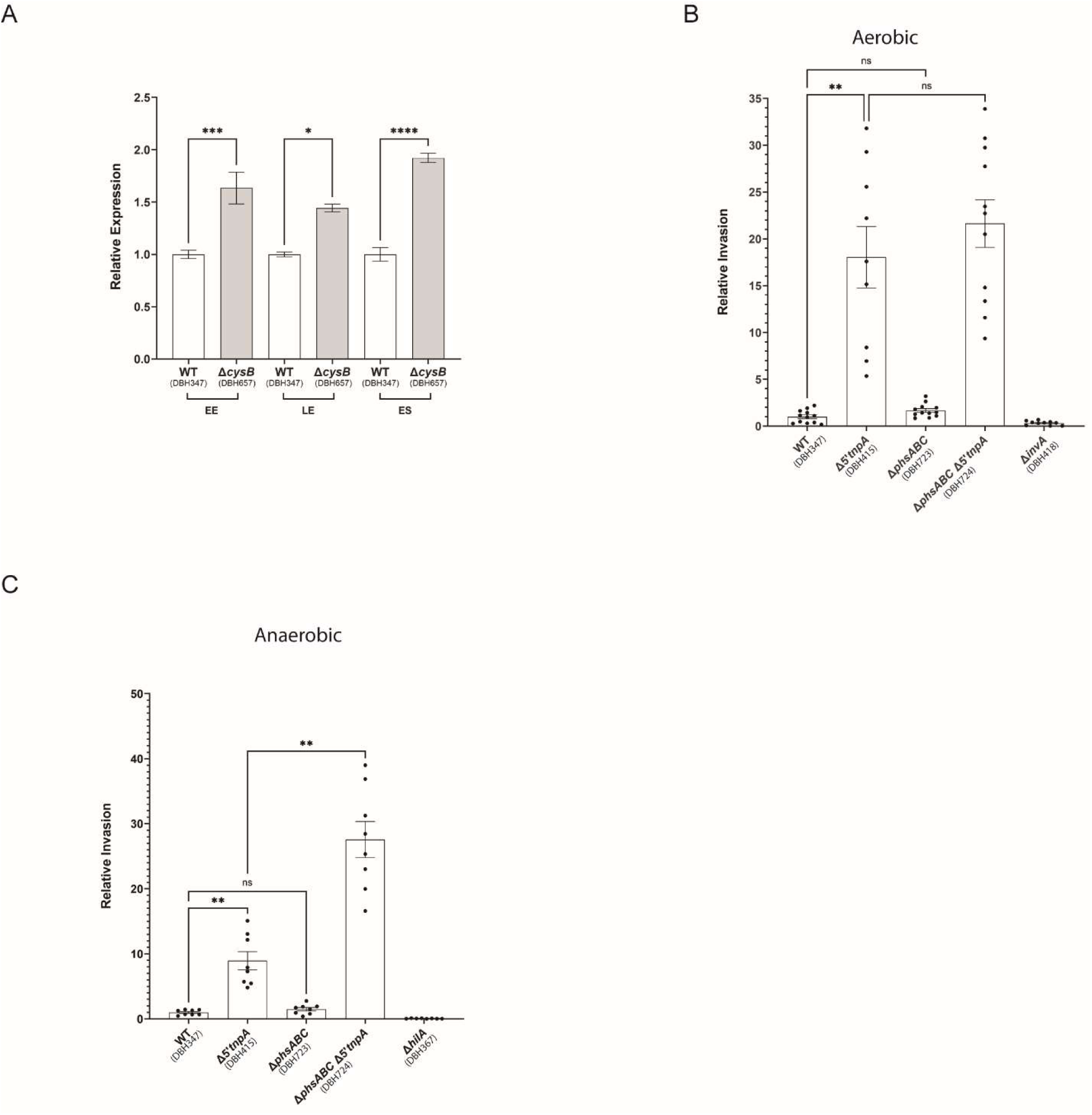
Impact of *cysB* deletion on *phsABC* expression and of *phsABC* deletion on *Salmonella* invasion. (A) qRT-PCR data showing the impact of Δ*cysB* on *phsABC* expression. For each growth phase the relative expression of *phsABC* was set to 1 for the *WT* strain. The results are shown for 6 biological replicates. (B and C) Invasion assays for the indicated strains performed under conditions where the *Salmonella* strains were grown either aerobically or anaerobically. A non-invasive strain (either Δ*invA* or Δ*hilA*) served as a negative control. Bars represent the average invasion for at least 6 biological replicates all measured in technical duplicate. ns, not statistically significant (P ≥ 0.05); *, **, *** and **** P ≤ 0.05, 0.01, 0.001 and 0.0001, respectively.

To further test the general model, we looked at the impact of deleting the *phsABC* operon on *Salmonella* invasion. Our expectation here was that the deletion of the *phsABC* operon would increase the amount of available cysteine for production of proteins necessary for invasion and motility and this would increase the invasion frequency. A corollary of this is that Δ*phsABC* would have to be in a Δ*5’tnpA* strain background to see an increased invasion frequency early in growth because only in this strain background would there be cysteine limitation (due to early induction of SPI-1 and flagellar regulons). Initially, strains were grown under aerobic conditions for testing the efficacy of invading HeLa cells. Under these conditions the *phsABC* disruption in either *WT* or Δ*5’tnpA* strain backgrounds did not have an impact on the invasion frequency (Figure 10B). However, the thiosulfate reductase encoded by the *phsABC* operon is considered an anaerobic enzyme ^42^, so we also monitored invasion frequencies under anaerobic growth conditions where the *phsABC* operon is more highly expressed (Figure S8). Under anaerobic growth conditions with cells in ES phase, Δ*phsABC* increased the invasion frequency approximately 3-fold (Figure 10C). Notably, this increase was only observed when the *phsABC* operon was disrupted in the Δ*5’tnpA* strain background. We also isolated RNA samples from the anaerobically growing strains used for invasion. Their analysis revealed that the increase in invasion observed for the Δ*5’tnpA*Δ*phsABC* strain (DBH724) relative to the Δ*5’tnpA* strain (DBH415) is not due to differences in *hilD* expression levels (Figure S9A).

Based on the above model we reasoned that the increase in invasion frequency in the Δ*5’tnpA*Δ*phsABC* strain background would be dependent on the high level of *hilD* expression that is typical of the Δ*5’tnpA* strain because this would be a major contributor to the metabolic stress these cells are under. To test this idea, we introduced a third mutation, a disruption of the *fur* gene, and repeated the invasion assay. The Fur protein is a TF that is a positive regulator of *hilD* expression ^38^. The results in Figure S10A, where we compare invasion frequencies for Δ*5’tnpA*Δ*phsABC* (DBH724) and Δ*5’tnpA*Δ*phsABC*Δ*fur* (DBH730) strains show that introducing the *fur* gene disruption decreased invasion about 4-fold. This invasion frequency was roughly equivalent to that observed for the Δ*5’tnpA* strain, consistent with the idea that lowering *hilD* expression is sufficient to abrogate the positive impact of the Δ*phsABC* mutation on invasion. We also asked if increasing *hilD* expression would further increase the invasion frequency of a Δ*phsABC* strain. In this case, the third mutation we introduced was a deletion of the *phoPQ* operon as the TF PhoP is a negative regulator of *hilD* expression ^45^. The results in Figure S10B show that the introduction of Δ*phoPQ* into the Δ*5’tnpA*Δ*phsABC* strain (DBH724), creating DBH744, increased the invasion frequency 4-fold over DBH724. This coincides with a 2-fold increase in *hilD* expression (Figure S9B). Finally, we asked if introduction of Δ*phoPQ* into the Δ*phsABC* strain (DBH723), creating DBH749, is sufficient to increase invasion to DBH724 levels. Here we did not see a statistically significant increase in invasion between DBH749 and DBH723 strains (Figure S10C) and this coincided with no significant increase in *hilD* expression (Figure S9B).

## Discussion

In the current work, we investigated the impact of deleting the 5’ portion of the IS200 *tnpA* transcript (*5’tnpA*) on the transcriptome of the pathogenic *Salmonella* strain SL1344 in rich media. Comparative RNA-seq identified over 200 differentially expressed genes throughout exponential growth. Four major groups of differentially expressed genes were identified including genes involved in invasion (SPI-1 regulon), flagellar-directed motility and chemotaxis (flagellar regulon), cysteine biosynthesis (cysteine regulon), and reduction of thiosulfate (*phsABC* operon). Focusing on the SPI-1 and flagellar regulons, additional comparative expression analysis using qRT-PCR identified genes for TFs *crp*, *lrhA*, *sirA*, as well as the sRNAs CsrC and Spot 42 as also being affected by the deletion of *5’tnpA*. An earlier late-stationary phase transcriptomic analysis where *5’tnpA* was overexpressed in the non-pathogenic LT2 strain of *Salmonella* identified 73 genes that were differentially expressed including three SPI-1 effector genes ^30^. Taken together, the results of these transcriptomics data sets are consistent with the IS200 *tnpA* transcript being a regulatory component of multiple *Salmonella* gene expression networks. Importantly, the increased expression of SPI-1 and flagellar regulon genes, which occurred early in growth in the Δ*5’tnpA* strain, resulted in this strain having an increased capacity to invade HeLa cells indicating that the observed gene expression changes are functionally significant. We also identified genes involved in sulfur metabolism (cysteine regulon and the *phsABC* operon) as being coordinately regulated in the Δ*5’tnpA* strain. We suggest that this is an indirect effect of deletion of *5’tnpA* arising due to metabolic stress caused by the early induction of invasion and flagellar regulons. Consistent with a reduced metabolic flux of thiosulfate towards cysteine production, we showed that the disruption of the *phsABC* operon increases invasion by about 3-fold specifically in the Δ*5’tnpA* strain.

### Linking the early invasion phenotype to the absence of *5’tnpA*

The generation of the *Δ5’tnpA* strain involved multiple genetic manipulations (7 independent P22 transductions and 4 rounds of flipping out antibiotic resistance genes used for selection in the transductions). In addition to the removal of the first 261 nt of the *tnpA* transcript from all seven copies of IS200 in the SL1344 strain, our genetic manipulation generated new transcription units in which the *FRT-scar* sequence is fused to the last 409 nt of *tnpA* in 7 locations of the genome. It was therefore important to test the idea that it was the loss of *5’tnpA* sequence rather than the creation of a novel transcript and/or altered transcription at one or more of the modified sites that was responsible for the novel properties of the Δ*5’tnpA* strain. Regarding the latter possibility, one of the disruptions (*tnpA4*) is immediately upstream of the *fliA* gene (Figure S1). The FliA protein, also referred to as σ^28^, drives the expression of *fliZ* ^46^. Accordingly, if the disruption of *tnpA4* were to increase expression of *fliA*, this would increase FliZ levels, which in turn would increase HilD activity and *hilD* expression. We tested the impact of this disruption alone in an otherwise *WT* background on *fliZ* and *hilD* expression via qRT-PCR and found no changes in expression of either of these genes (Figure S11). To directly demonstrate that *5’tnpA* has a regulatory role in the invasion cascade, we performed complementation studies wherein either the full-length or truncated forms of *tnpA* were expressed in the Δ*5’tnpA* strain and we looked at the impact of this on the expression of *hilD* and select downstream genes. We demonstrated that moderate overexpression of either *tnpA_FL_* or *tnpA_1-50_* reduced *hilD*, *hilA*, and *invF* expression by approximately 2-fold and that moderate overexpression of *tnpA_FL_* reduced invasion by about 2-fold. In contrast, increased expression of the chimeric *FLP-scar-tnpA_262-670_* transcript did not influence *hilD* or *invF* expression. One possible reason suppression of *hilD* and downstream factors was limited to 2-fold for *tnpA_FL_* may be that this construct also produces the antisense RNA *art200*, which pairs with the *tnpA* transcript promoting its degradation ^29^. While *tnpA_1-50_* does not produce *art200* it is possible that the 2-fold ceiling in suppression of *hilD* expression observed for this form of *5’tnpA* is due to decreased stability relative to the 5’portion of the full-length transcript, although we have not directly tested this possibility.

### General mechanism of 5’*tnpA* regulatory function

Small regulatory RNAs in bacteria function by multiple mechanisms. The vast majority function through complementary basepairing with target RNAs, influencing translation of the target RNA and/or its stability ^47^. In Gram-negative bacteria, the pairing interaction is typically catalyzed by an RNA-binding chaperone, either Hfq or ProQ ^27,48^, which provides a nucleation center for the RNA-pairing partners. sRNAs can also, through an RNA-RNA pairing mechanism, act as decoys titrating other sRNAs so that they are not available to pair with a partner mRNA ^49^. Finally, sRNAs can titrate RNA-binding proteins, preventing these proteins from binding other transcripts ^50^. We have previously shown that a segment of *tnpA* including nucleotides 1-174 can pair with a segment of the *invF* transcript to reduce *invF* expression. A smaller segment of *tnpA* comprised of nucleotides 1-50 also inhibits *invF* expression ^30^. This shorter form of *tnpA* lacks an Hfq binding site situated from nucleotides 71-87 ^29^ raising the possibility that *5’tnpA* can function in an RNA-RNA pairing capacity independent of Hfq to regulate the expression of target genes. Although the *tnpA_1-50_* transcript has not been directly tested for its ability to pair with the *invF* transcript, it does retain substantial sequence complementarity. For other genes observed in the current work as being regulated by *5’tnpA, in silico* predictions using the RNA-RNA pairing tool RNApredator failed to identify interaction sites with *5’tnpA* (where hits were observed ΔG values were less than 10 kcal/mol). This raises the possibility that the *5’tnpA* transcript might also function through the titration of one or more RNA-binding proteins. At this point, we have only established that the 5’ portion of *tnpA* is capable of binding Hfq ^29^. Other RNA-binding proteins that would be worth investigating for a *5’tnpA* transcript interaction include CsrA and CspC/E, as these proteins have been implicated in SPI-1 and/or flagellar regulon gene expression ^50–53^. With regard to CsrA we note that the minimal binding site, 5’GGA ^54^, is not present in the first 50 nt of *5’tnpA*, nor is the minimal binding site for either CspC or CspE (5’CUG and 5’CAG, respectively) ^55^. On the other hand, ProQ has been shown to bind *art200* and therefore likely plays some role in establishing 5’*tnpA* transcript levels ^56^.

### Misregulation of Crp-Spot 42, SirA-CsrC and LrhA expression drives the early SPI-1 induction phenotype

Previous studies have shown that for *Salmonella* grown in rich media, strong induction of SPI-1 does not occur until the transition from LE to ES phase ^32^. This presumably reflects the build-up of HilD protein above a specific threshold level ^13^. In the current work, we established that the SPI-1 and flagellar regulons are turned on early in growth in the Δ*5’tnpA* strain and this correlates with an ∼80-fold increase in invasive capacity. Strikingly, the invasion frequency for the Δ*5’tnpA* strain in EE growth is roughly equivalent to the *WT* strain at LE growth (i.e. at the transition to stationary phase). While we had previously focused on *invF* as the target of *5’tnpA* in SPI-1 regulation ^30^, the current work establishes *hilD* and *flhDC* gene expression as additional points of *5’tnpA* regulation. Both *hilD* and *flhDC* expression is increased (approximately 5-fold and 2-fold, respectively) in the Δ*5’tnpA* strain early in growth indicating that *5’tnpA* is a negative regulator of these genes.

In our search for how *5’tnpA* regulates SPI-1 and flagellar regulons, we identified several genes upstream of *hilD* as being misregulated in the Δ*5’tnpA* strain. Crp protein levels and *lrhA* transcript levels were reduced and *sirA*, CsrC and Spot 42 levels increased, all early in growth (lag to ME growth). The results of double mutant analysis support the idea that the increase in *hilD* expression in the Δ*5’tnpA* strain early in growth is a consequence of the additive effects of altered regulation of *csrC*, *spf* and *lrhA* genes, with the increased expression of *csrC* and *spf* genes being driven by reduced production of Crp protein in lag phase. The change in *lrhA* gene expression (2-fold decrease) was of a higher magnitude than that observed for either of Spot 42 or CsrC in the Δ*5’tnpA* strain. The downregulation of *lrhA* expression coincided with about a 2-fold increase in the expression of the master regulator of the flagellar regulon *flhDC* and a 4-fold increase in *fliZ*. We suggest that this decrease in *lrhA* expression primarily accounts for the early induction of the flagellar regulon. In addition, because FliZ is a positive post-translational regulator of HilD, there was the potential that a change in *lrhA* expression would also contribute to the early induction of SPI-1. We tested this idea by asking if increasing the expression of *lrhA* in the Δ*5’tnpA* strain would suppress the early induction of *hilD* expression. Remarkably, as little as a 2-fold increase in *lrhA* expression (at the transcript level) fully suppressed the increase in *hilD* expression that is typical of the Δ*5’tnpA* strain early in growth. This coincided with a decrease in invasion of HeLa cells to a level that is typical for the *WT* strain in EE phase. Notably, there is precedent for moderate increases in FliZ expression driving SPI-1 induction. An earlier analysis in which a deletion of *ydiV*, a negative regulator of *flhDC*, resulted in increased expression of *fliZ* also led to the conclusion that increasing FliZ production alone could drive SPI-1 induction ^57^. Overall, the impact of moderately overexpressing *lrhA* on *hilD* expression was significantly greater than the impact of deleting either *csrC* or *spf*. This leads us to predict that while the changes in expression of *lrhA*, *csrC* and *spf* all contribute to the early SPI-1 induction phenotype, the reduced expression of *lrhA* is the major driver of this phenotype.

In further support of the contention that increased expression of CsrC and Spot 42 contribute in an additive manner to the ‘*hilD*-up’ phenotype, we note that the impact of *ΔcsrC* and *Δspf* mutations on *hilD* expression is specific to the Δ*5’tnpA* strain background (Figures 6F and 7C, respectively). These individual changes in gene expression are not sufficient to have an impact on *hilD* expression in an otherwise *WT* background. On the other hand, the *ΔlrhA* mutation did not have a significant impact on *hilD* expression in the Δ*5’tnpA* strain background (Figure 8D). This could be explained by the fact that *lrhA* expression is already suppressed in this strain background (Figure 6C).

Based on the known functions of CsrC, Spot 42 and LrhA, it is straightforward to model the additive impact of altered regulation of these respective genes on *hilD* expression. The increase in Spot 42 levels would increase the amount of *hilD* transcript through Spot 42 binding to the 3’UTR of *hilD* ^21^. The increase in CsrC (and possibly CsrB) levels would increase the amount of *hilD* translation by titrating CsrA. Finally, the decrease in *lrhA* expression would increase *flhDC* expression and ultimately FliZ production increasing the amount of active HilD protein.

One curious observation relating to the *lrhA* overexpression analysis was that it blocked the increase in CsrC expression that is typical of the Δ*5’tnpA* strain early in growth. Furthermore, this occurred in the absence of an impact on *sirA* expression. This leads us to conclude that there is some crosstalk between LrhA and the *csr* pathway that occurs independent of SirA. Potentially, LrhA could be a direct negative regulator of CsrC expression. Alternatively, LrhA could be a positive regulator of Crp expression. Neither of these possibilities have been addressed in the current work (Figure 9).

At this point it remains unclear what the primary target(s) of *5’tnpA* is/are. One attractive possibility is that either *crp* expression or the expression of genes involved in cAMP production are the primary targets. In this scenario, *5’tnpA* would be a positive regulator of *crp* (and/or cAMP production) and Crp-cAMP would be a positive regulator of *lrhA* expression and a negative regulator of both *sirA* and *spf* expression. While the latter is known to be the case, it is unknown if Crp-cAMP regulates *lrhA* expression and there is some evidence that Crp-cAMP acts as a positive not a negative regulator of *sirA* expression ^20^. Currently, known regulators of *lrhA* expression are limited to *rcsB* and *ftsK* ^58^ and neither of these genes were differentially expressed in the Δ*5’tnpA* strain in our RNA-seq analysis (Table S3).

While we have mainly focused on the impact of disrupting *5’tnpA* sRNA expression on invasion, it is worth noting that the increase in flagellar regulon expression observed in the Δ*5’tnpA* strain could provide an explanation for the decreased recovery of this strain from mouse tissues despite the strain having an increased invasive capacity ^31^. Increased expression of the flagellar regulon is expected to increase the number of flagella produced and since the flagellar component FliC is a major target of the immune system ^9^, there may have been selective elimination by immune cells of the Δ*5’tnpA* strain in competitive infection assays.

### An indirect linkage between *5’tnpA* and the coordinate regulation of the cysteine regulon and the *phsABC* operon

We found it curious that the expression of two systems functioning in sulfur metabolism were differentially expressed in opposite directions in the Δ*5’tnpA* strain in LE growth. We suggest that the observed upregulation of the cysteine regulon and the downregulation of the *phsABC* operon are coupled responses to metabolic stress associated with the early induction of invasion and flagellar regulons. Mechanistically, how might changes in cysteine regulon and *phsABC* operon genes be coordinated? The cysteine regulon is induced when cysteine concentrations are low ^59^. Under these conditions, L-serine is converted to O-acetylserine (OAS) by the enzyme CysE, which is inhibited by cysteine. OAS, and its alternate conformer N-acetylserine, bind to the TF CysB, allowing CysB to bind DNA in a sequence-specific manner and activate the transcription of genes containing a CysB binding site ^60^. We have suggested that induction of invasion and flagellar genes would lead to a cysteine shortage and thus induction of the cysteine regulon. One possible way of coordinating cysteine regulon and *phsABC* operon expression might then be through CysB. In support of this possibility, we demonstrated that in a strain where the *cysB* gene was disrupted, *phsABC* expression increased about 2-fold, consistent with CysB acting as a negative regulator of *phsABC* expression.

Based on the above findings, we postulated that deletion of the *phsABC* operon would positively impact on invasion because this deletion would increase sulfur flux towards cysteine (and methionine) production thereby better allowing cells to cope with metabolic stress arising from SPI-1 and flagellar regulon induction. In fact, deletion of the *phsABC* operon did increase invasion about 3-fold. Importantly, this increase is specific to the Δ*5’tnpA* strain under anaerobic growth conditions. The dependence of this response on the absence of the *5’tnpA* fits with the notion that increased invasion is dependent on the metabolic stress linked to induction of invasion and flagellar regulons. Overall, these results are consistent with the idea that downregulating *phsABC* operon expression provides a means for cells to recover from the metabolic stress associated with SPI-1 and flagellar regulon induction.

An alternative model to consider focuses on the hydrogen sulfide produced through reduction of thiosulfate by the products of the *phsABC* operon. Hydrogen sulfide is used in the post-translational modification of cysteine residues in proteins, a reaction called persulfidation ^61^. If persulfidation of one or more proteins involved in invasion plays a negative role in invasion, then depleting hydrogen sulfide production through deletion of the *phsABC* operon would increase invasion. However, at this point there is little known with regards to what proteins in the *Salmonella* proteome are subject to persulfidation, so it is premature to invoke this model.

### Scope of transposon-encoded sRNA regulation of host gene expression

Despite their threat to genome integrity, transposons are ubiquitous in genomes in all three kingdoms of life. It is thought that the successful spread and retention of transposons in genomes reflects their ability to evolve defenses against host mechanisms aimed at suppressing transposon activity, and in some cases providing functions that can contribute to host fitness ^62^. In bacteria, one example of the latter is that transposons have the capacity to acquire and disseminate antibiotic resistance genes to their hosts. Another example is the domestication of transposon-encoded Cas proteins to function in host defense against invading organisms (i.e. the evolution of CRISPR defense systems) ^63–65^. In this and previous work, we have provided evidence for IS200 domestication in *Salmonella*. However, unlike the Cas example, in the case of IS200 it is the function of a transposon-derived sRNA rather than a protein that has been exapted by the host.

Is the integration of *5’tnpA* into *Salmonella* regulatory circuits an isolated example of transposon domestication? IS200 is the founding member of the IS200/IS605 family of insertion sequences. This family is made up of three groups (the IS200, IS605 and IS1341 groups) and has approximately 153 members ^66^. A common feature within the IS200 group, which includes insertion sequences that are widespread in enteric bacteria including *Salmonella*, *E. coli*, *Shigella*, and *Yersinia*, is the presence of an inverted repeat in the 5’UTR of the *tnpA* transcript. In *Salmonella* it has been shown that the 5’ portion of the transcript contains a stable stem-loop structure that sequesters the ribosome binding site, thereby inhibiting translation of the *tnpA* transcript ^24,25^. This is one of several mechanisms that have evolved to reduce *tnpA* expression and to keep IS200 transposition exceedingly low, minimizing negative selection against the transposon ^29^. We propose here that this structural feature of the *tnpA* transcript can benefit the transposon in an additional way, namely by giving rise to a stable sRNA that is available for post-transcriptional regulatory functions in the host. Furthermore, if targets of this sRNA include regulatory molecules that are conserved in enteric bacteria (e.g. Crp), one could anticipate that the example of domestication reported here will be widespread. However, as the copy number of IS200 group insertion sequences is typically quite high, alternate methods of silencing these elements to gene disruption (used here) such as CRISPR interference ^67,68^ would likely need to be applied to uncover such examples.

### Experimental Procedures

#### Growth conditions, strains, and plasmids

For experiments performed under both aerobic and anaerobic conditions, *S.* Typhimurium was grown at 37°C in Lennox Broth (LB; 5 g/L NaCl, 10 g/L tryptone, 5 g/L yeast extract) supplemented with Streptomycin (150 μg/mL). For aerobic growth standard culture tubes (16 x 150 mm) were used and cultures were grown with shaking. For anaerobic growth, cultures were grown without shaking in 2 mL microtubes (Axygen) filled to the top with LB. After approximately 24 hr, anaerobic cultures reached an OD_600_ ≅ 0.5; the culture density did not change significantly with further incubation time indicating that the cultures had reached stationary phase. For all aerobic experiments, overnight cultures were diluted 1:100 and then grown to the desired density (see points on growth curve in Figure S2). For experiments where strains were transformed with plasmids, plasmids were maintained by selection with the appropriate antibiotics; ampicillin, 200 μg/ml; kanamycin, 50 μg/ml; chloramphenicol, 20 μg/ml and tetracycline, 15 μg/ml.

All strains and plasmids used in this study are listed in Table S6 and oligonucleotides are listed in Table S7. *S* Typhimurium str. SL1344 was considered the wild-type strain and derivatives were made in the SL1344 background. *Escherichia coli* DH5α was used for routine cloning and plasmid propagation and *E. coli* MG1655 and SL1344 genomic DNA were used as templates for PCR amplification where indicated.

Mutant strains of SL1344 were constructed by Lambda Red recombineering ^69^, P22 HT105/1 *int-201* (P22)-mediated transduction ^70^ or where indicated mini-Tn7-mediated transposition of genes into the *attTn7* site of the *Salmonella* chromosome ^71^ (see Supplemental Data for details). Colony PCR was used to confirm all new constructs and where indicated antibiotic resistance cassettes were removed using the temperature sensitive plasmid pCP20 (pDH739) carrying the FLP recombinase ^72^.

Each of the disrupted copies of IS200 in the Δ*5’tnpA* strain retains the weak *tnpA* promoter, the *tnpA* transcription start site, an *FRT*-scar sequence and internal *tnpA* sequence extending from bp 301 (nt 261) to bp 710 (nt 670) (Figure 1). We made the disruption in this way to minimize the impact of each IS200 disruption on surrounding genes. The location of each of the seven disrupted copies of IS200, including immediately surrounding transcription units is shown in Figure S1. Based on the raw reads from the RNA-seq analysis, it appeared that a form of *tnpA* for each of the disrupted IS200 copies in the Δ*5’tnpA* strain is expressed (Table S1). We confirmed that the Δ*5’tnpA* strain does produce a truncated *tnpA* transcript using qRT-PCR where primers specific for the 3’ end of *tnpA* were used (Figure S12). In principle, this transcript should be a chimera wherein *FRT*-scar sequence is fused to a portion of the *tnpA* transcript consisting of nt 261 to 670. Cloning of this chimeric form of *tnpA* into a plasmid followed by expression in the wild-type strain did not impact on *hilD* expression (Figure S5). Accordingly, it is unlikely that the production of this transcript is linked to the early induction of SPI-1.

#### RNA isolation

Total RNA was prepared by the hot acid phenol method ^73^ and quantified using a nano-drop spectrophotometer (IMPLEN).

#### RNA-seq and data analysis

Three colonies of each of *WT* and *Δ5’tnpA* strains were grown to EE, ME and LE phases whereupon 600 μL were removed and processed for RNA extraction. Purified RNA was treated with TURBO DNase (Ambion) to remove residual genomic DNA and ∼500 ng of total, DNase-treated RNA was treated with the Ribo-Zero “Bacteria” (Illumina), following the manufacturer’s instructions. rRNA-depleted RNA was precipitated in ethanol for 3 h at −20°C. cDNA libraries for Illumina sequencing were generated by Vertis Biotechnologie AG, Freising-Weihenstephan, Germany. To this end, rRNA-free RNA samples were first sheared via ultrasound sonication (4 pulses of 30 s at 4°C) to generate on average 200-400 nt fragments. Fragments <20 nt were removed using the Agencourt RNAClean XP kit (Beckman Coulter Genomics) and the Illumina TruSeq adapter was ligated to the 3’ end of the remaining fragments. First-strand cDNA synthesis was performed using M-MLV reverse transcriptase (NEB) wherein the 3’ adapter served as a primer. The first-strand cDNA was purified and the 5’ Illumina TruSeq sequencing adapter was ligated to the 3’ end of the antisense cDNA. The resulting cDNA was PCR-amplified to about 10-20 ng/μL using a high fidelity DNA polymerase. The TruSeq barcode sequences were part of the 5’ and 3’ TruSeq sequencing adapters. The cDNA libraries were purified using the Agencourt AMPure XP kit (Beckman Coulter Genomics) and analyzed by capillary electrophoresis (Shimadzu MultiNA microchip). Prior to sequencing, individual cDNA samples were pooled in equimolar amounts. The resulting cDNA pool was size-fractionated in the range of 200-600 bp using a differential clean-up with the Agencourt AMPure kit (Beckman Coulter Genomics). Aliquots of the cDNA pools were analyzed by capillary electrophoresis (Shimadzu MultiNA microchip). Sequencing was performed on a NextSeq 500 platform (Illumina) at Vertis Biotechnologie AG, Freising-Weihenstephan, Germany (single-end mode; 75 cycles). All RNA-seq data discussed in this publication have been deposited in NCBI’s Gene Expression Omnibus and are accessible through GEO Series accession number XXX.

Reads were aligned to the *S*. Typhimurium SL1344 genome (NC_016810.1) with Rockhopper ^74^ (Table S1) and differential expression was analyzed using ALDEx2 ^75^. More detail on data analysis is provided in Supplementary Data.

#### Reverse transcription quantitative polymerase chain reaction (qRT-PCR)

DNase treated RNA (2μg) was converted to cDNA with the High-Capacity cDNA Reverse Transcription Kit (Applied Biosystems); cDNA was diluted to 30 ng/μl in TE [50 mM Tris-HCl, pH 8.0, 1 mM Ethyleneediaminetetraacteticacid (EDTA)] and stored at −20°C. A minimum of three biological replicates were analyzed in technical duplicate in each experiment and the 16S rRNA (*rrsA*) was used as a reference gene for relative quantitation. Reactions (48 μl) contained 360 ng of cDNA, 500 nM of each primer (Table S7) and PowerUP SYBR Green Master Mix (Applied Biosystems). Standard settings on the ViiA 7 Real-Time PCR system were used except for the anneal/extension step, which was performed at 60.5°C. Relative expression of each target was calculated by the efficiency corrected method ^76^.

#### Growth curves

Growth was measured in a Multiskan Go microplate spectrophotometer. Cells from 3 overnight cultures (biological triplicates) of each strain (*WT* and Δ*5’tnpA*) were diluted 100-fold in LB. 200 μL of each dilution was plated in technical replicates in a 96-well microplate. Cultures were grown with continuous shaking at 37°C for 12 hr and absorbance at 600 nm (A_600_) was measured every 15 min. Note that the A_600_ was not adjusted for path length and light scattering from the microplate lid and is therefore not directly comparable to optical density readings measured in a standard cuvette. Absorbance readings from the plate reader were calibrated to a spectrophotometer allowing us to use cuvette readings to identify cultures at specific growth phases for either RNA extraction, western blot analysis or Miller assays.

#### β-galactosidase assays

Overnight cultures were grown in LB supplemented with tetracycline (15 μg/ml) and then diluted (1:40 for assays conducted with Δ*crp* strains, otherwise standard 1:100 dilution were used) and grown to the indicated growth phase. The Miller assay was performed as previously described ^77^.

#### Gentamicin protection (invasion) assay

Tissue culture plates (12-well) were seeded with ∼0.05 X 10^6^ HeLa cells per well in 1 mL of Dulbecco’s Modified Eagle’s Medium (DMEM) supplemented with 10% (v/v) fetal bovine serum (FBS) and kanamycin (300 μg/mL) 20-22 h prior to the invasion assay. At the time of the assay, cells were 70-80% confluent (∼0.1 × 10^6^ cells per well). For invasion assays performed with aerobically and anaerobically grown strains, strains were grown as previously described from freshly streaked colonies. Bacterial cells were washed with 0.85% NaCl saline solution and resuspended in DMEM/10% FBS to a concentration of 1 × 10^7^ cfu/ml with a final volume of 1 mL.

HeLa cells were washed with Dulbecco’s Phosphate Buffered Saline (dPBS) and 1 mL of bacterial suspension (MOI of 40) was added per well. Serial dilutions of the bacterial suspension were plated in technical duplicates on LB agar plates supplemented with 150 μg/ml streptomycin to determine the input number of bacteria.

Bacterial cells were centrifuged onto the HeLa cell monolayer at 100 x g for 3 min at room temperature and then incubated at 37°C for 40 min. Bacterial cells were washed with dPBS, which was then followed by addition of 100 μg/ml gentamicin in DMEM/10% FBS for 1.5 hr incubation at 37°C to kill extracellular bacteria. After washing with dPBS, HeLa cells were resuspended in 250 μL of lysis solution (0.1% Triton X-100 in 1X dPBS) and serial dilutions were plated in technical duplicates on LB with 150 μg/ml of streptomycin to determine the output bacterial cell counts. Invasion was calculated as the ratio of recovered cells to the input and normalized to the indicated strain.

#### Western blot

Strains were grown to the indicated growth phase and 1 ml of culture was pelleted and then resuspended in 100 μL of 1X TE. 15 μL of sample and 15 μL of sodium dodecyl sulphate (SDS) sample buffer (60 mM Tris-HCl, pH 6.8, 2% SDS [w/v], 0.1% bromophenol blue [w/v], 1% β-mercaptoethanol [v/v]) were mixed and boiled for 5 minutes. Samples (10 μL) were then applied to a 15% SDS-polyacrylamide gel and electroblotted to a polyvinylidene difluoride (PVDF) membrane. Membranes were incubated overnight with 5% milk solution in Tris-Buffered Saline Tween 20 (TBST) (1X TBS, 0.5% Tween-20) at 4°C. Membranes were then incubated with primary antibody (1:5000 dilution: mouse α-purified (azide-free) anti-*E. coli* Crp, BioLegend or rabbit α-GroES, Sigma) followed by incubation with the secondary antibody (1:10000 dilution; α-mouse-HRP or α-rabbit-HRP, Promega). Crp and GroES were detected using the Pierce ECL western blotting substrate (Thermoscientific) and imaged using the Bio-Rad ChemiDoc imager; after Crp detection membranes were stripped and re-probed for the loading control (GroES). Protein species (Crp and GroES) were quantified in AlphaView and the Crp level was normalized to the internal standard (GroES or select bands in the Coomassie duplicate gel). It should be noted that for lag phase samples the loading control signal (GroES) was weak making quantification difficult, so randomly selected species from the duplicate Coomassie-stained gel were utilized for loading control normalizations. A total of 5 different species were used in this capacity, the choice of which did not significantly alter (≤ 2%) the level of Crp reported.

#### Statistics

For graphs with one comparison an unpaired t-test (two-tailed) was performed, while for graphs with two or more comparisons a One-way ANOVA by Tukey’s multiple comparisons test was performed using GraphPad Prism version 10.2.3 for Windows, GraphPad Software, Boston, Massachusetts USA, www.graphpad.com

## Acknowledgments

We thank Serena Nath and Lucas Lorimer for the initial characterization of the impact of Δ*lrhA*, Δ*csrC* and Δ*crp* mutations on *hilD* and/or *phsABC* gene expression. We are grateful to J. Slauch for providing Δ*fur* and Δ*phoPQ Salmonella* strains, to S. Gottesman for proving the pLlacO-1 expression plasmid and to A. White for proving plasmids used to introduce genes via Tn7 transposition into the *Salmonella attTn7* site. We thank M. Surette and K. Hoffman for sequencing the Δ*5’tnpA* strain and D. Heinrichs and R. Flanagan for help in performing invasion assays.

## Funding

Natural Sciences and Engineering Research Council of Canada (RGPIN/04753-2016) and Canadian Institutes of Health Research (159655) (to D.H.). Add funding information for A.J.W.

## Author Contributions

D.B.H. conceived of and designed the project. A.J.W. and K.U.F. performed the RNA-seq analysis and associated data analysis. R.S.T. prepared RNA samples for RNA-seq and together with N.-J.Q.S. performed most of the experiments in this study. M.J.E. was involved in the RNA-seq data analysis and prepared figures for the RNA-seq analysis. M.A. performed invasion assays specifically looking at the impact of Δ*phsABC* on invasion in aerobic versus anaerobic growth conditions. D.B.H. discussed the data with all authors and wrote the manuscript primarily with input/editing from A.J.W.

## Notes

### Competing Interest Statement

The authors have declared no competing interest.

