## Supplementary Data for "A small RNA derived from the 5’ end of the IS200 *tnpA* transcript regulates multiple virulence regulons in *Salmonella* Typhimurium"

**Supplementary Experimental Procedures**

**Strain and plasmid construction**

Strains DBH367, 488, 554, 555, 569, 575, 657, 722, 723, 724, 730, 744 and 749 listed in Table S6 were generated by P22 transduction of either *WT* or Δ*5'tnpA* strains. Transducing lysates containing the corresponding disruption alleles used to construct the above strains were generated from strains obtained from other laboratories or generated by the Haniford lab via lambda red recombineering [1]. Strain DBH566 was generated using the modified Tn7 transposition system [2]. Briefly, the *lacI*$^{q}$ gene was cloned into the *EcoRV* site of pDH1004, a derivative of pUC18R6K-mini-Tn7-*PacI* (pDH1001) containing a *kan*$^{R}$ gene cloned into the *HinCII* site. The new plasmid pDH1037 was transformed by electroporation into *WT* containing pDH1002 (a source of Tn7 transposase). After a 2 hr recovery at 30°C cells with Tn7 hops were selected for by plating on LB Kan plates at 37°C. Colony PCR was used to detect the presence of the *lacI*$^{q}$ gene at the *attTn7* site of the Salmonella chromosome. Similarly, DBH689 was constructed by first cloning the *lrhA-3X-FLAG* gene into *SacI-EcoRV* digested pDH1004 producing pDH1100. *WT* containing pDH1002 was then transformed with pDH1100 and selected for Tn7 hops as above generating DBH689. DBH689 was then used to produce a P22 transducing lysate for transducing the *attTn7-lrhA-3X-FLAG-kan*$^{R}$ cassette into DBH554 and DBH555. *Δ5'tnpA* was constructed as previously described [3].

Plasmids expressing *tnpA* under the control of the IPTG-inducible PLlacO-1 promoter, including the full-length gene (pDH968) and truncated forms of *tnpA*$_{1\text{-}50}$ (pDH1129) and *tnpA*$_{1\text{-}25}$ (pDH1133), were constructed as follows. For pDH968 a PCR product encoding *tnpA*$_{full\text{-}length}$ was amplified from DBH347 genomic DNA with oDH747 and oDH753, digested with *AatII* + *MfeI* and cloned into *AatII* + *EcoRI* digested pDH967. This places the *tnpA* gene under the control of the pLlacO-1 promoter. pDH967 also encodes the *lacI*$^q$ gene. pDH967 was derived from a plasmid in which a truncated form of *tnpA* was cloned into pDH763 and subsequently *lacI*$^q$ was cloned into the *HindIII* site of pDH965. Truncated forms of *tnpA* (pDH1129 and pDH1133) containing *tnpA* sequences fused to the pLlacO-1 promoter plus the SgrS transcription terminator immediately following *tnpA* sequence were constructed as follows. For pDH1129 and pDH1133 a PCR product containing either the first 50 or 25 nucleotides of *tnpA* followed by the SgrS terminator was amplified from *WT* genomic DNA using oDH747 and oDH746 or oDH747 and oDH853 and A/T cloned into pGEM-T-easy. For the *tnpA*$_{1\text{-}50}$ construct the new plasmid was digested with *AatII* and *HinCII* and the resulting pLlacO-1-*tnpA*$_{1\text{-}50}$-terminator fragment was cloned into *AatII-EcoRV* cut pDH763. Subsequently, a *lacI*$^q$ PCR product amplified from *E. coli* MG1655 genomic DNA with oDH843 and oDH844 was digested with *StuI* and cloned into the *PhsAI* site generating pDH1129. For the *tnpA*$_{1\text{-}25}$ construct the new plasmid was digested with *AatII* and *EcoRI* and the resulting pLlacO-1-*tnpA*$_{1\text{-}25}$-terminator fragment was cloned into *AatII-EcoRI* cut pDH1129 creating pDH1133. For pDH1081, PCR with oDH856 and oDH857 and *Δ5'tnpA* as template was used to generate a pBAD-*Flp-scar-tnpA*$_{261\text{-}680}$ DNA fragment. A second PCR with this fragment and oDH577 and pBAD24 as template was performed to introduce the pBAD24 *AflII* site into the above fragment. Subsequently, the pBAD-*Flp-scar-tnpA*$_{261\text{-}680}$ fragment was cut with *AflII* and *StuI* and cloned into pBAD24 cut with *AflII* and *SmaI*. The *phsA-lacZ* translational fusion (pDH1120) was constructed by first cloning a PCR product containing a portion of the *phsA* gene (including codons 1-47) into *EcoRI-PstI* digested pBR322. The PCR product was amplified from *WT* with oDH847 and oDH848 and digested with *EcoRI* and *PstI*. Then the *lacZ* gene (starting at codon 11) present on a *PstI* digest product was cloned into the *PstI* site creating pDH1120. Mutations in the putative Crp binding site were added using a modified SOEing technique. A PCR product amplified from pDH1120 with oDH849 (mut2) or oDH850 (mut3) and oDH848 was used with oDH847 in a second PCR reaction with pDH1120 as template. The second PCR product was cloned as in pDH1120. For hopping genes into the *attTn7* site of the SL1344 chromosome we first generated a *mini-Tn7* containing construct (pDH1004) with the $kan^R$ gene from IS903 situated between the Tn7 end sequences. For this the $kan^R$ gene flanked by *PsiI* sites was amplified using oDH841 and oDH842 and the PCR product was cut with *PsiI* and cloned in *HinCII* digested pDH1001. Subsequently, either $lacI^q$ (pDH1037) or *lrhA-3X-FLAG* (pDH1100), was introduced into pDH1004 as follows. For the $lacI^q$ construct a $lacI^q$ PCR product generated from *E. coli* MG1655 genomic DNA and primers oDH843 and oDH844 was cut with *StuI* and cloned into the *EcoRV* site of pDH1004, generating pDH1037. For *lrhA-3X-FLAG*, we first fused the *3X-FLAG* sequence to the C-terminus of the *lrhA* gene by generating a PCR fragment with *WT* genomic DNA as template and primers oDH845 and oDH846. The corresponding PCR product was digested with *EcoRI* and *HindIII* and cloned into *EcoRI-HindIII* cut pDH1093. This *lrhA-3X-FLAG* sequence was then purified after *SacI-NruI* digestion and cloned into *SacI-EcoRV* cut pDH1004 generating pDH1100.

**RNA-seq Data Analysis**

Transcripts predicted by Rockhopper [4] were manually annotated with the genomic interval and categorized into intergenic (predicted RNA, e.g. 4824999-4825049_predicted RNA) or intragenic (antisense, e.g. 1906043-1906092_antisense: fadR) RNAs prior to expression analysis (Table S1). Differential expression was analyzed using ALDEx2 version 1.3.4.0 a tool for analyzing the compositional nature of high-throughput sequencing data [5]. Since ALDEx2 compares two conditions, this analysis was performed on each pairwise comparison (i.e. *WT* EE vs *Δ5'tnpA* EE, *WT* ME vs *Δ5'tnpA* ME, *WT* LE vs *Δ5'tnpA* LE) (Table S2). Briefly, the relative expression (abundance) of each gene within a sample was calculated as the median centered log-ratio (clr) from 1000 Monte-Carlo Dirichlet instances. The ALDEx2 effect size was then calculated for each feature as the median log2 difference between conditions (i.e. strains) divided by the maximum difference within a condition (i.e. difference in relative abundance between biological replicates). Accordingly, the effect size is a measure of the confidence one has in the difference in expression for a gene between two conditions, but does not necessarily inform on the magnitude of the difference. An effect size of 2 means that the difference between conditions (i.e. log2 fold-change between two strains) is twice as large as the greater difference within a condition (variance between biological replicates). We took a more conservative approach for considering genes differentially expressed and used an effect size of 2 as the cutoff for differential expression (Table S3).

**SUPPLEMENTAL TABLES**

**Table S1.** All RNA-seq reads mapped to the *S.* Typhimurium SL1344 genome using Rockhopper. Each pairwise comparison is shown on a separate sheet in Excel (Sheet 1, *WT* EE vs Δ*5'tnpA* EE; Sheet 2, *WT* ME vs Δ*5'tnpA* ME; Sheet 3, *WT* LE vs Δ*5'tnpA* LE).

**Table S2**. Results of ALDEx2 analysis of RNA-seq data. Each pairwise comparison (Sheet 1, *WT* EE vs Δ*5'tnpA* EE; Sheet 2, *WT* ME vs Δ*5'tnpA* ME; Sheet 3, *WT* LE vs Δ*5'tnpA* LE) was made using ALDEx2.

**Table S3**. Genes identified as differentially expressed (|Effect| >2) in our RNA-seq experiment. Each pairwise comparison is shown on a separate sheet in Excel (Sheet 1, *WT* EE vs Δ*5'tnpA* EE; Sheet 2, *WT* ME vs Δ*5'tnpA* ME; Sheet 3, *WT* LE vs Δ*5'tnpA* LE).

**Table S4.** Groupings of differentially expressed genes from Venn diagram. Lists of differentially expressed genes found in 1, 2 or 3 different growth phases depicted in the Venn diagram of Figure 3B. The total number of differentially expressed genes in each category is indicated.

**Table S5**. *In silico* RNA-RNA pairing analysis. The first 50 nt of the *tnpA* transcript was used as the input RNA sequence to search genome-wide for potential pairing partners using the RNApredator search tool [6]. The top 39 hits are shown. Notably, this analysis did not identify the *invF* transcript as a potential pairing partner for *5'tnpA* within the top 39 hits. However, as previously reported, a single extended region of complementarity between the first 63 nucleotides of tnpA and an internal segment of invF (104 to 160 nucleotides upstream of the invF start codon) was identified using the IntaRNA base-pairing interaction tool [7] when the full tnpA sequence was used as a query against the full invF transcript sequence [3].

**Table S6. Bacterial strains and plasmids used in this study.**

| | Relevant genotype/ description | Reference/source |
|---|---|---|
| *S*. Typhimurium | | |
| DBH199 | Wild-type LT2, Str$^{R}$ | Miguel Valvano |
| DBH347 | Wild-type SL1344, Str$^{R}$ | Salmonella Genetic Stock Centre (SGSC438) |
| DBH367 | DBH347 Δ*hilA::kan*$^{R}$, Kan$^{R}$Str$^{R}$ | This study |
| DBH415 | DBH347 Δ*tnpA_2::FRT* Δ*tnpA_7::FRT* Δ*tnpA_6::FRT* Δ*tnpA_4::FRT* Δ*tnpA_1::FRT* Δ*tnpA_5::FRT* Δ*tnpA_3::FRT* | [3] |
| DBH418 | SL1344 Δ*invA* | Brian Coombes |
| DBH457 | JS1068 Δ*phoPQ::cm*$^{R}$, Cm$^{R}$ | James Slauch |
| DBH485 | JS1107 Δ*fur41::cm*$^{R}$ *attλ::pDX1::hilA`-lacZ*, Cm$^{R}$ | James Slauch |
| DBH488 | DBH415 Δ*fur41::cm*$^{R}$, Cm$^{R}$Str$^{R}$ | This study |
| DBH554 | DBH347 Δ*lrhA::kan*$^{R}$, Kan$^{R}$Str$^{R}$ | This study |
| DBH555 | DBH415 Δ*lrhA::kan*$^{R}$, Kan$^{R}$Str$^{R}$ | This study |
| DBH566 | DBH347 *attTn7-lacI*$^{q}$*-kan*$^{R}$, Kan$^{R}$ | This study |
| DBH569 | DBH347 Δ*crp::kan*$^{R}$, Kan$^{R}$Str$^{R}$ | This study |
| DBH575 | DBH415 Δ*crp::kan*$^{R}$, Kan$^{R}$Str$^{R}$ | This study |
| DBH657 | DBH347 Δ*cysB::kan*$^{R}$, Kan$^{R}$Str$^{R}$ | This study |
| DBH693 | DBH347 Δ*lrhA::FRT attTn7-lrhA-3X-FLAG-kan*$^{R}$, Kan$^{R}$Str$^{R}$ | This study |
| DBH704 | DBH415 Δ*lrhA::FRT attTn7-lrhA-3X-FLAG-kan*$^{R}$, Kan$^{R}$Str$^{R}$ | This study |
| DBH722 | DBH199 Δ*phsABC::kan*$^{R}$, Kan$^{R}$Str$^{R}$ | This study |
| DBH723 | DBH347 Δ*phsABC::kan*$^{R}$, Kan$^{R}$Str$^{R}$ | This study |
| DBH724 | DBH415 Δ*phsABC::kan*$^{R}$, Kan$^{R}$Str$^{R}$ | This study |
| DBH730 | DBH415 Δ*phsABC::kan*$^{R}$ *Δfur::cm*$^{R}$*,* Kan$^{R}$Cm$^{R}$Str$^{R}$ | This study |
| DBH744 | DBH415 Δ*phsABC::kan*$^{R}$ *ΔphoPQ::cm*$^{R}$*,* Kan$^{R}$Cm$^{R}$Str$^{R}$ | This study |
| DBH749 | DBH347 Δ*phsABC::kan*$^{R}$ *ΔphoPQ::cm*$^{R}$*,* Kan$^{R}$Cm$^{R}$Str$^{R}$ | This study |
| Plasmids | | |
| pDH337 | pWKS30, low copy plasmid with multiple cloning sites, *pSC101 ori* and Ap$^{R}$ | [8] |
| pDH739 | pCP20, temperature sensitive FLP expression | [1] |

| | | |
|---|---|---|
| | plasmid, $Ap^R Cm^R$ | |
| pDH740 | pKD46, temperature sensitive Lamda Red expression plasmid, $Ap^R$ | [1] |
| pDH742 | pKD4, kanamycin cassette for Lambda Red recombineering, $Ap^R Kn^R$ | [1] |
| pDH763 | pBR322 with *pLlacO-1* promoter and multiple cloning sites, $Ap^R$ | Susan Gottesman |
| pDH859 | pBAD24 | ATTC |
| pDH965 | pDH763 with *tnpA*$_{1\text{-}200}$ fused to the *pLlacO-1* promoter, $Ap^R$ | This study |
| pDH967 | pDH965 with *lacI*$^q$, $Ap^R$ | This study |
| pDH968 | pBR322-derived plasmid expressing *tnpA*$_{Full\text{-}Length}$ from the *pLlacO-1* promoter and $LacI^q$, $Ap^R$ | This study |
| pDH1001 | pUC18*R6K-mini-Tn7-PacI*, $Ap^R$ | Aaron White |
| pDH1002 | pHSG415-t*nsABCD*, $Ap^R$ pSC101*ori*$^{ts}$ | Aaron White |
| pDH1004 | pDH1001 *mini-Tn7-kan*$^R$, $Ap^R$ $Kan^R$ | This study |
| pDH1037 | pDH1004 *mini-Tn7-kan*$^R$*-lacI*$^q$, $Ap^R Kan^R$ | This study |
| pDH1081 | pBAD24-derived plasmid expressing *flp-scar-tnpA*$_{261\text{-}670}$ from the *pAra* promoter, $Ap^R$ | This study |
| pDH1093 | pWKS30 *ΔplacZ*, $Ap^R$ | This study |
| pDH1100 | pDH1004 *mini-Tn7-kan*$^R$*-lrhA-3X-FLAG*, $Ap^R Kan^R$ | This study |
| pDH1120 | pBR322-derived *phsA-lacZ* translational fusion, $Tet^R$ | This study |
| pDH1129 | pBR322-derived plasmid expressing *tnpA*$_{1\text{-}50}$ from the *pLlacO-1* promoter and $LacI^q$, $Ap^R$ | This study |
| pDH1133 | pBR322-derived plasmid expressing *tnpA*$_{1\text{-}25}$ from the *pLlacO-1* promoter and $LacI^q$, $Ap^R$ | This study |
| pDH1138 | pDH1120 with M2 mutation, $Tet^R$ | This study |
| pDH1139 | pDH1120 with M3 mutation, $Tet^R$ | This study |

**Table S7. Oligonucleotides used in this study.**

| Name | Sequence (5' to 3') | Description/use |
|---|---|---|
| Strain and Plasmid Construction | | |
| oDH841 | TATGTCTTATAACAACAAAGCCACGTTGTGTCTC | Amplifying Kan gene with PsiI cut sites |
| oDH842 | TATGTCTTATAAGTTACAACCAATTAACCAATTCTG | |
| oDH843 | ATAAAGGCCTTTGACACCATCGAATGGTGC | Amplifying lacIq gene with StuI cut sites |
| oDH844 | ATAAAGGCCTACATTAATTGCGTTGCGCTC | |
| oDH845 | TTACGAATTCTCATCATTAGTGTCGCGTCG | Amplifying native lrhA with C-term 3xFLAG |
| oDH846 | ACTAAGCTTTTACTATTTATCGTCGTCATCTTTGTAGTCGATATCATGATCTTTATAATCACCGTCATGGTCTTTGTAGTCCTCAAAACCCCCTTCCACC | |
| oDH747 | TACCGAGACGTCTCAGCTTTAAGCCAGTTATTAAA | Amplifying full length tnpA for inducible expression |
| oDH753 | ATCTAACAATTGTAAATATCCTCCGGCATAGCC | |
| oDH847 | TTACGAATTCCTTATCAACATCATTA | Amplifying phsA TLF fragment |
| oDH848 | TTACTGCAGCAAGCTTGGCGTTAGCGTG | |
| oDH849 | CAACTATTTCAAATTATGATTGAAGCCGACATTTC | Used in soeing phsA mut2 |
| oDH850 | CAACTATTTCAAATTATAATTGAAGCCGACATTTC | Used in soeing phsA mut3 |
| oDH857 | AAAGGCCTTAAATATCCTCCGGCATAGCC | Amplifying Δ5' tnpA while adding the arabinose inducible promoter and the repressor araC |
| oDH856 | CGCAACTCTCTACTGTTTCTCCATAAGTGTAGGCTGGAGCTGCTTC | |
| oDH577 | GCCTTTGAGTGAGCTGATACCG | |
| qRT-PCR | | |
| oDH704 | TTCAGTTGGCATACTGCCGC | *sicA* |
| oDH705 | GAAGGCGCCACGCTAAAAGA | |
| oDH815 | GCTGTGTTTCGCTTTCCAGC | *flhDC* |
| oDH816 | TTCGCGCGTCGATGACTTAC | |
| oDH804 | AGGTAATTGCCAAGGCGTCG | *fliZ* |
| oDH805 | CTACTTGACGACGCTGCCTG | |
| oDH661 | TCTGGGAAACTGCCTGATGGAGG | *rrsA* (16S rRNA) |
| oDH662 | CGTAGGAGTCTGGACCGTGTCTC | |
| oDH794 | TGCCGGTGACCATTACGAAG | *hilA* |
| oDH795 | AGAGAGAAGCGGGTTGGTGT | |
| oDH710 | GGTCTCCTGATACTGGTGCG | *invF* |
| oDH711 | AGCCCGGAAGCATGGTTTAT | |
| oDH394 | TATTTGGGCGCGAAAACTATG | *tnpA* |
| oDH428 | GTTGCCAGACCGCAAGATG | |
| oDH764 | GCCAGCTGCGGATAATAGGT | *prgH* |
| oDH765 | TGTCGCTGCGCAAAATGAAA | |
| oDH788 | CGTCCTGCCGCAGATAACTT | *hilD* |
| oDH789 | TGAGCACCAACATCCCAGGT | |
| oDH817 | CCTGTGACCATTTGCGGGTG | *hilC* |
| oDH818 | TGGCATGTCCACGGGTTTG | |

| | | |
|---|---|---|
| oDH819 | AGCCAGACCCGTGAAGCTAT | *sopE* |
| oDH820 | TAAAAGGATTGGCCCCTGCG | |
| oDH798 | GCAACTGGAGGCGGAAAGTA | *cysD* |
| oDH799 | CGCGAAAGGCGTACATCTCA | |
| oDH800 | TGAGCAACAATGGGCGAAGG | *cysP* |
| oDH801 | GTTGTTCGGCAGACGGCTTT | |
| oDH821 | TGTTCCCGGAAACCTACCAGT | *cysH* |
| oDH822 | TTCAGCTCTTTTAGCGCCCG | |
| oDH823 | GCGGCGCTATCGATGATTGT | *cysM* |
| oDH824 | AATCAGTTCCGCGCCATAGG | |
| oDH802 | GCAGGCCCGTCATAAACAGT | *ydjN* |
| oDH803 | CGAATACCAGCGGCATCACA | |
| oDH778 | GAGCTTTGCCGATGAGGGTT | *fliY* |
| oDH779 | CAGCATTCCGTCCCATTTGGT | |
| oDH784 | TCTGCTGCTGGGATTTGTGC | *yecS* |
| oDH785 | GCCGCGGTATTGAGCGATAA | |
| oDH825 | AAAAGTGGGGCGTGGGATTC | *sbp* |
| oDH826 | GTCGCCTGTTTACCCGAACC | |
| oDH827 | AACCTATGAAACGCACCGGG | *phsA* |
| oDH828 | ATTAGGCGAACCAAAGGCCG | |
| oDH829 | GTCGAGCGGGCAATTTTACG | *sirA* |
| oDH830 | TGATGACCACGAACTGGTGC | |
| oDH831 | GATGACGCTTCAGCAGGACA | *csrB* |
| oDH832 | GTTTGCATCCTGCGCACATC | |
| oDH833 | AGCGAGGACGCTAACAGGAT | *csrC* |
| oDH834 | ACGACCCTTAACGGGTTCCA | |
| oDH835 | TCAGTCAGCAAATGCAGCGT | *lrhA* |
| oDH836 | CTGATTCATCAGAAGCGCCGA | |
| oDH837 | CCGTCAAAGAGCACGCTGAT | *crp* |
| oDH838 | GGTTTTCGCACGTACCCAGG | |
| oDH858 | CATGCTTCTGGAGATCCCGC | *3'tnpA* |
| oDH859 | GCTGGTGCTTTATGTAGTCCTGT | |

#### Supplementary Figures

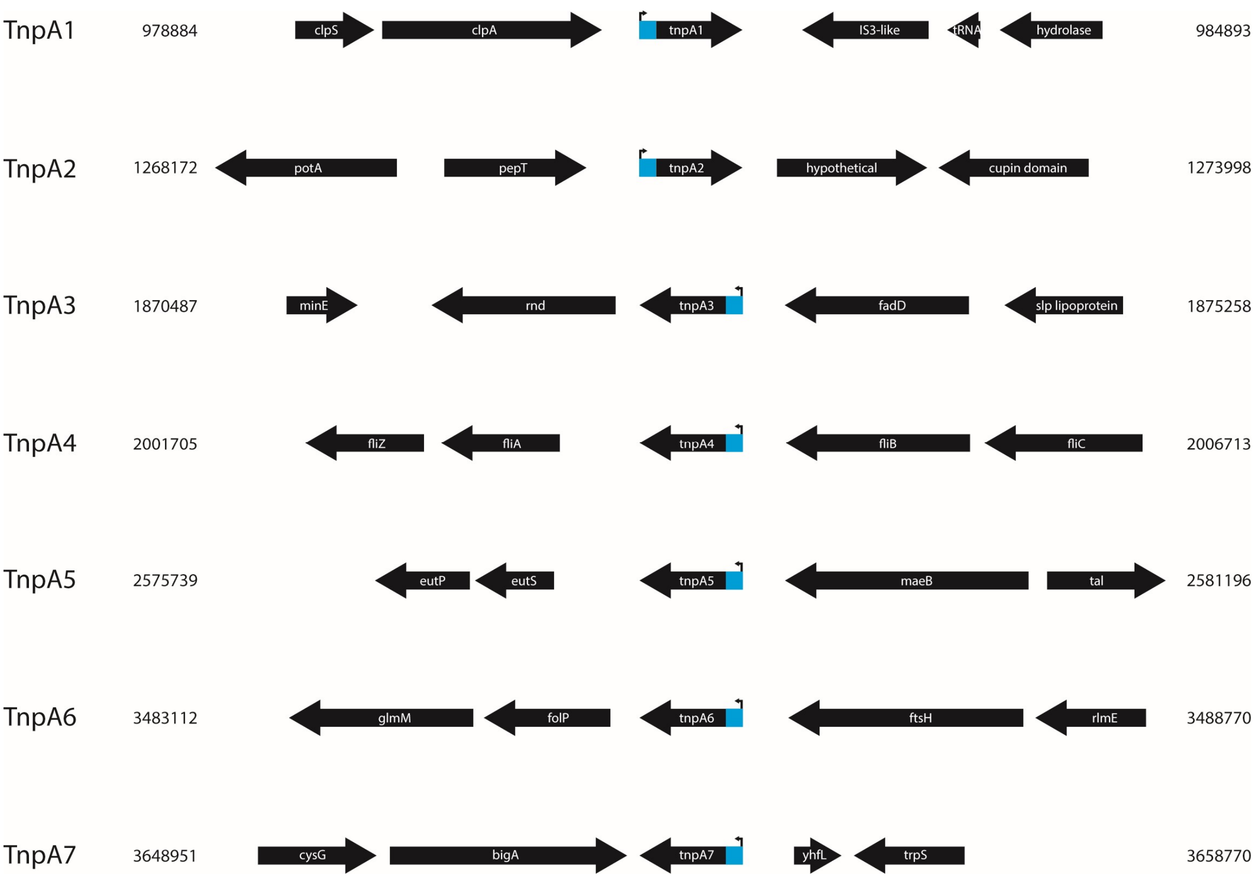

**Figure S1. The position of all 7 disrupted copies of IS200 in the *Salmonella* SL1344 genome is shown.** Arrowheads define the orientation of each disrupted IS200 element along with the direction of transcription of immediately neighboring genes. The small arrow defines the IS200 *tnpA* promoter. The blue rectangle is the *FRT-scar* sequence generated by flipping out antibiotic resistance genes used for selection in the construction of the *Δ5'tnpA* strain. Numbers define the genetic interval where each copy of IS200 is located within the *Salmonella* chromosome.

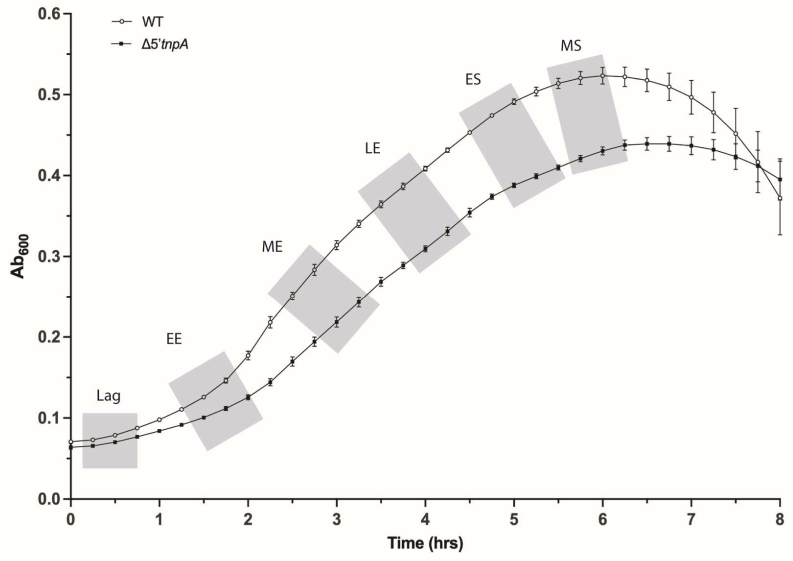

**Figure S2. Growth curves for *WT* and Δ*5'tnpA* strains.** Shaded rectangles define ranges of $A_{600}$ values used for harvesting cells at the specific indicated growth phases for the two strains. The results are shown for 3 biological replicates of each strain. Lag, lag phase; EE, early exponential phase; ME, mid exponential phase; LE, late exponential phase; ES, early stationary phase; MS, mid stationary phase; LS, late stationary phase.

A

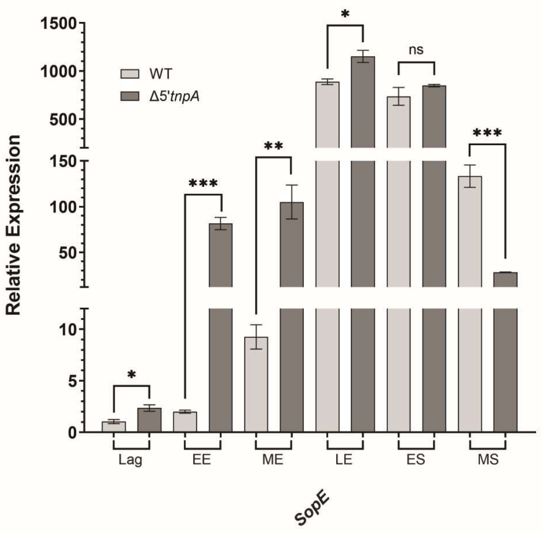

B

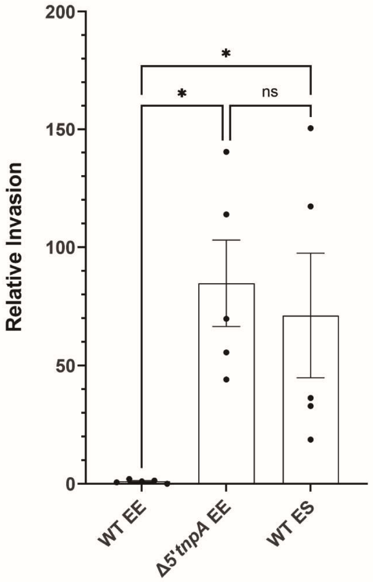

**Figure S3. Profiling *sopE* gene expression and invasion frequencies early in growth in *WT* and Δ*5'tnpA* strains.** (A) Differential expression of the SPI-1 effector gene *sopE* over 6 growth phases in LB media as measured by qRT-PCR is shown. In this experiment, the *sopE* transcript level for the *WT* strain in lag phase was set at 1. n = 6. (B) Relative invasion frequency of *WT* and Δ*5'tnpA* strains measured at the indicated growth phases. The invasion frequency for the *WT* strain at EE growth was set to 1. n = 6. Error bars represent the standard error of the mean. ns, not statistically significant (P ≥ 0.05); *, **, *** and **** P ≤ 0.05, 0.01, 0.001 and 0.0001, respectively.

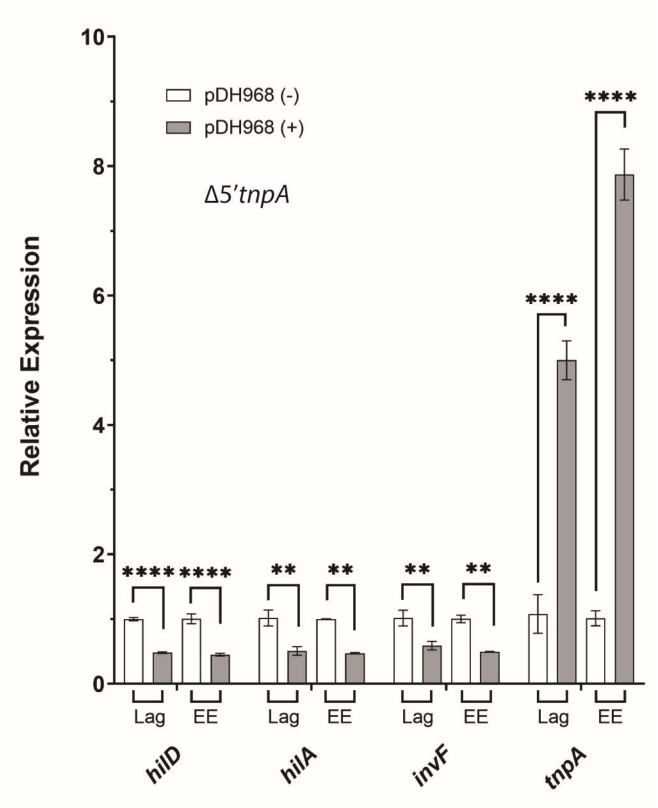

**Figure S4. Impact of *5'tnpA* overexpression on select SPI-1 transcription factors.** The Δ*5'tnpA* strain was transformed with pDH968 (encodes $tnpA_{FL}$ fused to the IPTG-inducible pLlacO-1 promoter). Transformants were subcultured (1:100) from overnight cultures in the presence (+) or absence (-) of IPTG (0.5 mM). Cultures were then grown to the indicated growth phase whereupon total RNA was extracted for qRT-PCR analysis. Note that IPTG treatment increases the *tnpA* transcript level up to 8-fold relative to that seen for uninduced cultures. For each pairwise comparison the expression level for the uninduced sample was set to 1. n = 3. Error bars represent the standard error of the mean. ** and ****, P ≤ 0.01, and 0.0001, respectively.

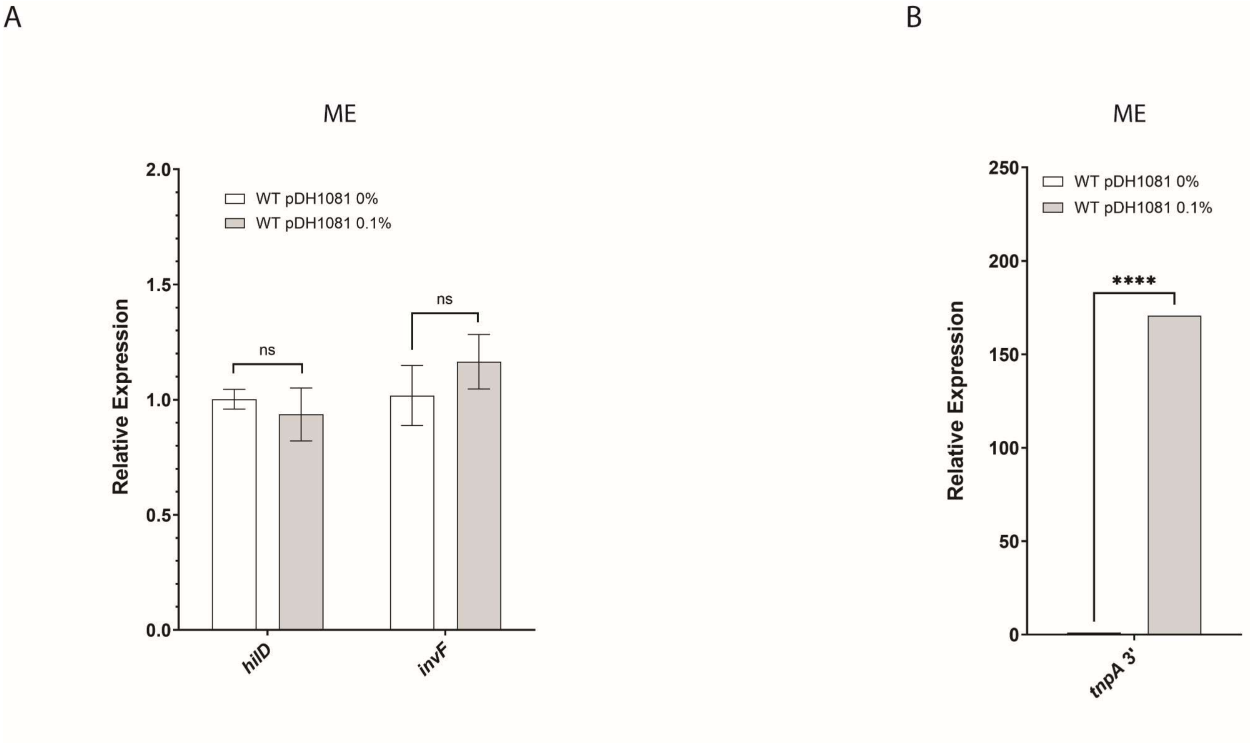

**Figure S5. Impact of overexpressing the chimeric transcript *FLP-scar-tnpA* produced by the Δ*5'tnpA* strain on select SPI-1 TF's.** (A and B) A plasmid (pDH1081) expressing the *FLP-scar-tnpA* transcript from the *pBAD* promoter was transformed into the *WT* strain. Transformants were subcultured (1:100) from overnight cultures in the presence (+) or absence (-) of arabinose (0.1%). Cultures were then grown to the indicated growth phase whereupon total RNA was extracted for qRT-PCR analysis. n = 3. Error bars show the standard error on the mean. ns, not statistically significant ($P \geq 0.05$); *, **, *** and **** $P \leq 0.05$, 0.01, 0.001 and 0.0001, respectively.

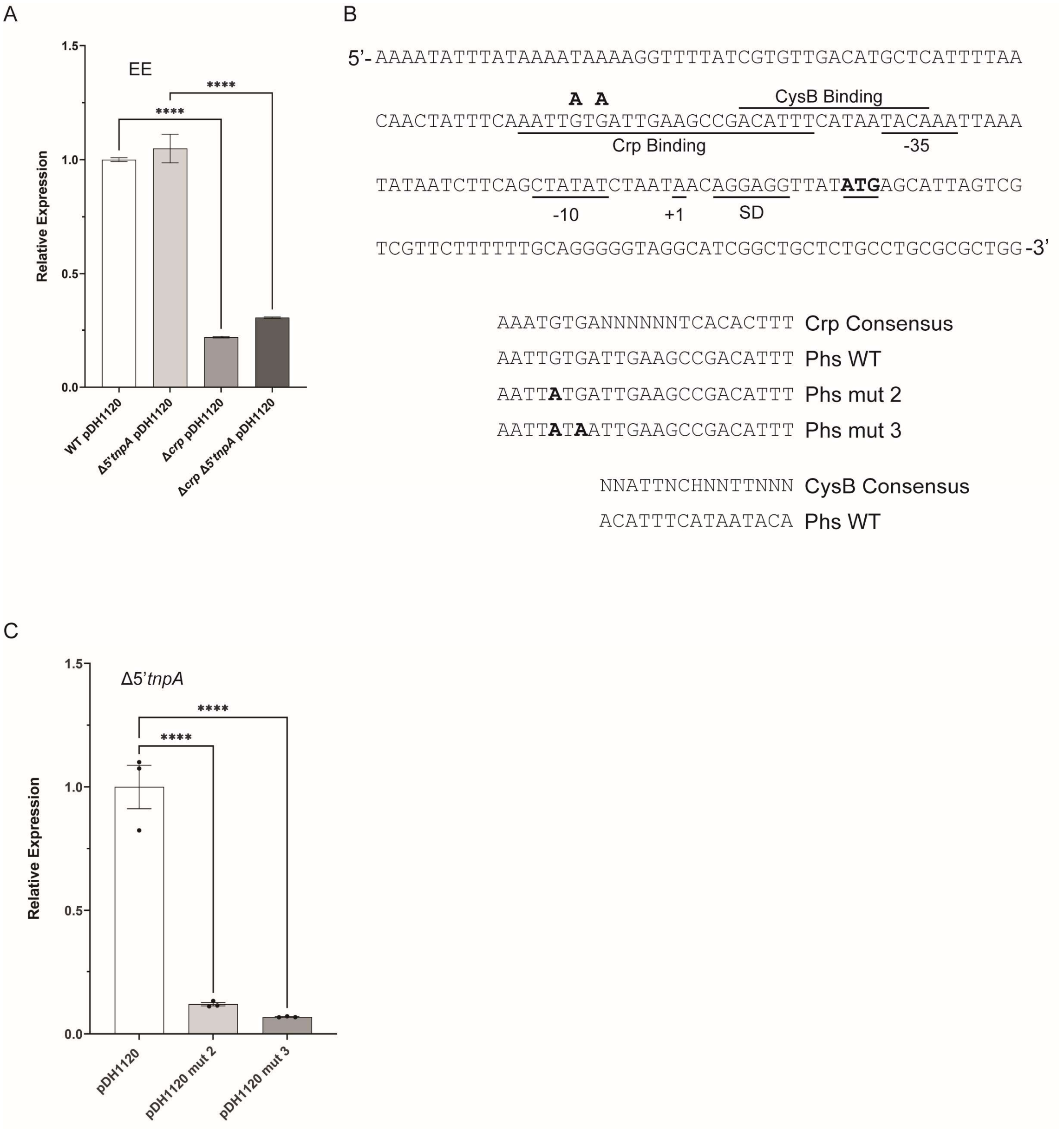

**Figure S6. Impact of Crp on the expression of the *phsABC* operon.** (A) β-galactosidase reporter assay with *phsA-lacZ* translational fusion (present on pDH1120). The indicated strains containing pDH1120 were grown to EE phase prior to processing for the β-galactosidase assay (see Materials and Methods). Note that at this time point we did not see differential expression of the *phsABC* operon in the RNA-seq analysis. Expression of the reporter in the *WT* strain was set to 1. (B) DNA sequence of the *phsABC* operon promoter region with the position of the Crp-cAMP and CysB binding sites indicated as well as the -35, -10, +1, SD (Shine-Delgarno) and start codon. DNA sequence comparisons of putative Crp-cAMP and CysB binding sites and the respective consensus sites are also shown along with the sequence of two mutant forms (mut1 and mut2) of the putative Crp-cAMP site designed to prevent Crp-cAMP binding. (C) β-galactosidase assay comparing *phsA-lacZ* reporter expression for *WT* and mutant forms of the *phsA* gene present in the Δ*5'tnpA* strain. Expression of the *WT* reporter was set to 1. Error bars represent the standard error of the mean. ****, P ≤ 0.0001.

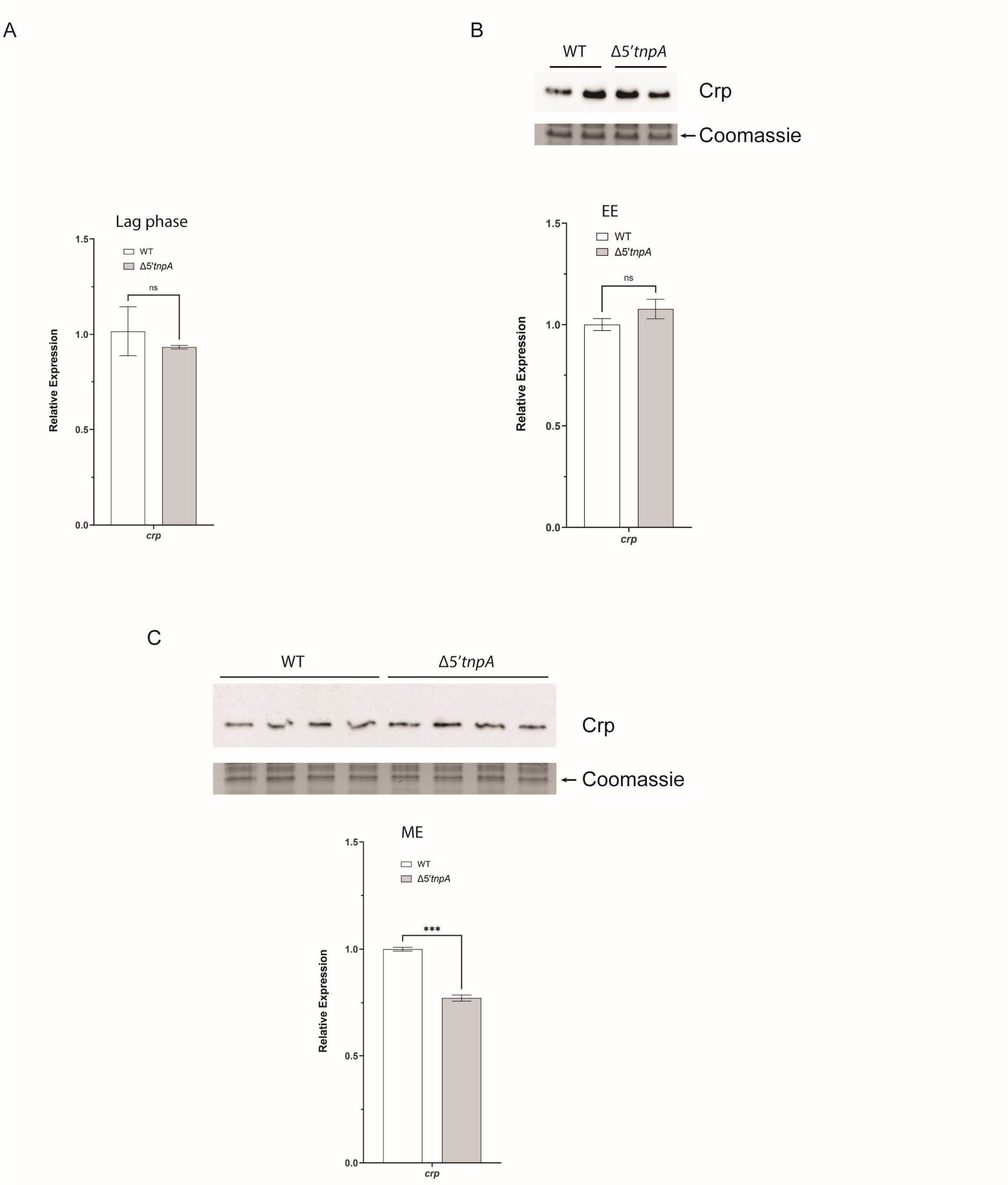

**Figure S7. Crp expression at different growth phases.** (A) Comparison of *crp* transcript levels in lag phase for *WT* and Δ*5'tnpA* strains as measured by qRT-PCR. The level of *crp* expression in the *WT* strain was set to 1. (B and C) Western blot and qRT-PCR analyses comparing Crp protein and transcript levels between *WT* and Δ*5'tnpA* strains at EE and ME growth phases, respectively. Crp protein levels were not significantly different between the two strains at either EE and ME growth phases and there was a small (~20%) decrease in *crp* transcript level in the Δ*5'tnpA* strain at ME growth. In the qRT-PCR the *crp* transcript level for *WT* was set to 1 and n = 3. Error bars represent the standard error of the mean. ns, not statistically significant ($P \geq 0.05$); ***, $P \leq 0.001$.

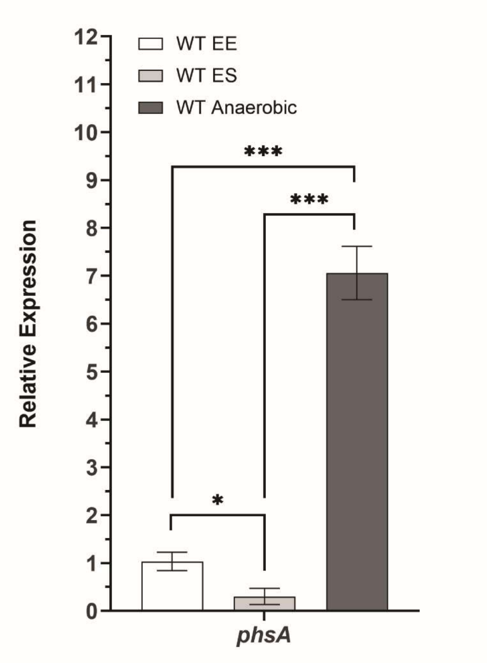

**Figure S8**. **Comparison of *phsA* expression levels in aerobic versus anaerobic growth.** The *WT* strain was grown to EE or ES phase under aerobic conditions or ES phase under anaerobic conditions. Total RNA was extracted and subject to qRT-PCR analysis. The *phsA* expression level for aerobic growth (EE phase) was set to 1. n = 3. Error bars represent the standard error of the mean. * and ***, P ≤ 0.05 and 0.001, respectively.

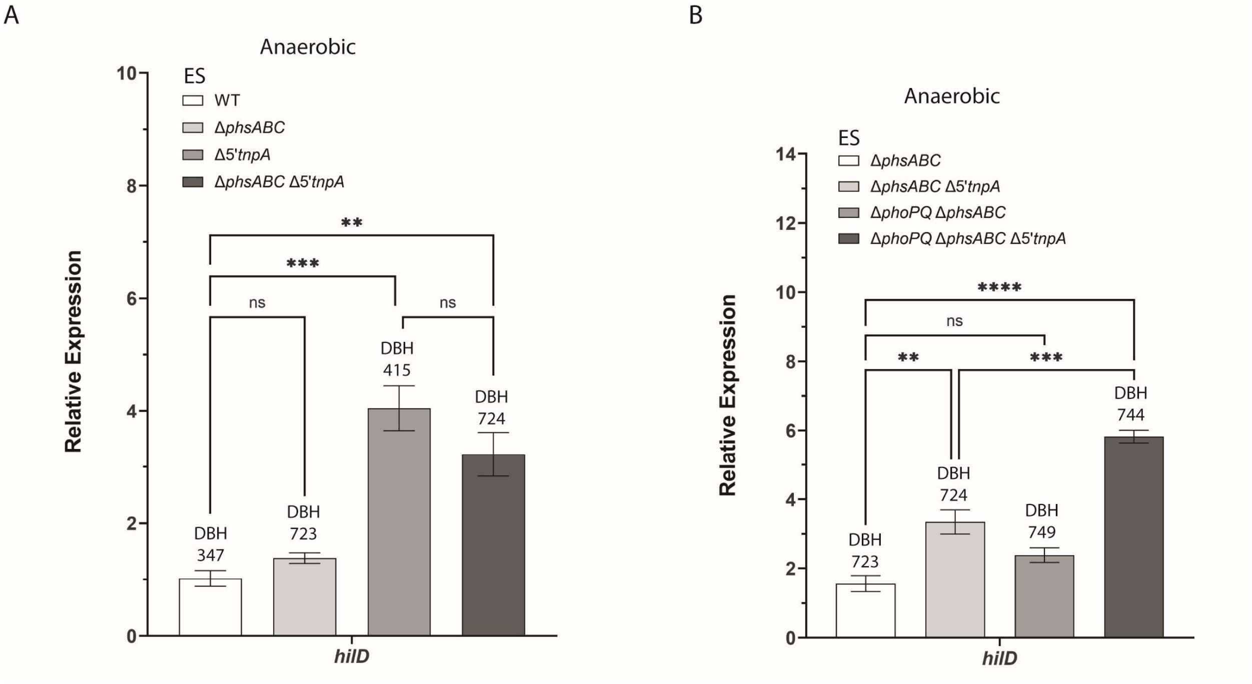

**Figure S9. Expression profiling of *hilD* under anaerobic growth conditions.** (A and B) The indicated strains were grown to ES phase under anaerobic conditions whereupon total RNA was extracted and used for qRT-PCR analysis. In (A) the *hilD* expression level for the *WT* strain was set to 1. In (B) the *hilD* expression level for the Δ*phsABC* strain (DBH723) was set to 1. In each experiment n = 3. Error bars represent the standard error on the mean. ns, not statistically significant ($P \geq 0.05$); **, *** and ****, $P \leq$ 0.01, 0.001 and 0.0001, respectively.

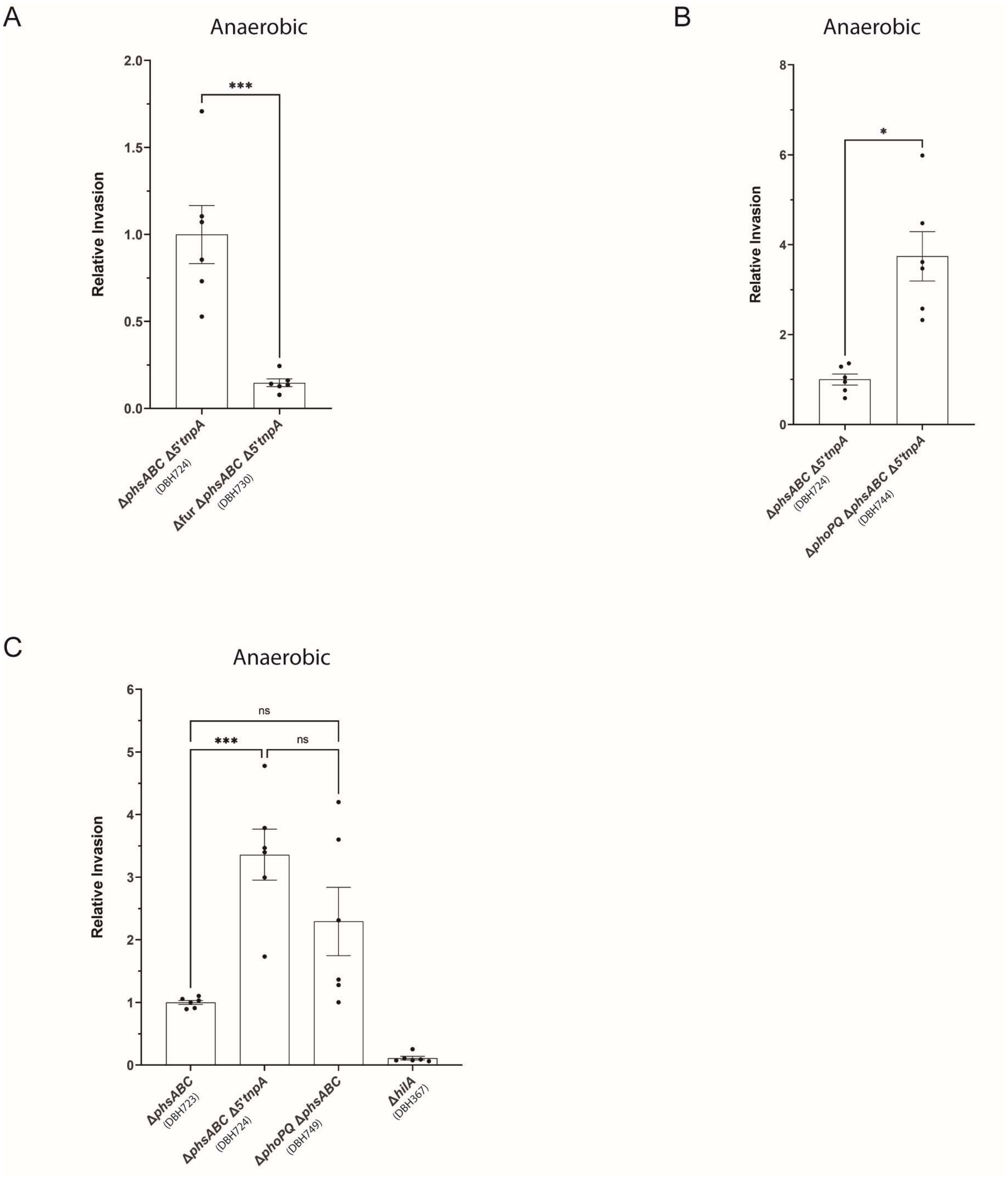

**Figure S10. Invasion assays for Δ*phsABC* strains. (A-C) Relative invasion frequencies for the indicated strains are shown.** All strains were grown under anaerobic conditions to ES phase. In (A) and (B) the invasion frequency for DBH724 was set to 1 and in (C) the invasion frequency of DBH723 was set to 1. n = 6. Error bars represent the standard error of the mean. ns, not statistically significant ($P \geq 0.05$); * and ***, $P \leq 0.05$ and 0.001, respectively.

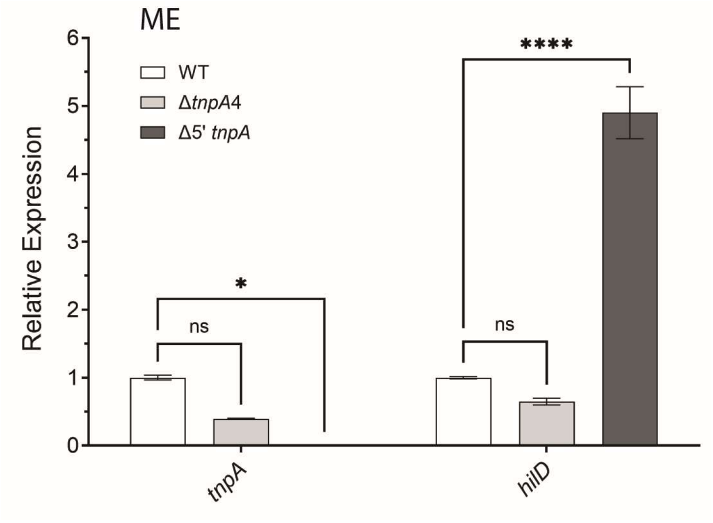

**Figure S11**. **Impact of disruption of *tnpA4* on *hilD* expression.** The indicated strains were grown to ME phase whereupon total RNA was extracted and used for qRT-PCR analysis. The Δ*tnpA4* disruption is in an otherwise *WT* strain background. The Δ*5'tnpA* sample was used as a positive control for *hilD* expression. Note that there is no signal for *tnpA* expression in the Δ*5'tnpA* sample because we used qRT-PCR primers (tnpA – see Table S7) that detect the 5' portion of the *tnpA* transcript. The *tnpA* and *hilD* expression levels for the *WT* strain were set to 1. n = 3. Error bars represent the standard error of the mean. ns, not statistically significant ($P \geq 0.05$); * and ****, $P \leq 0.05$ and 0.0001, respectively.

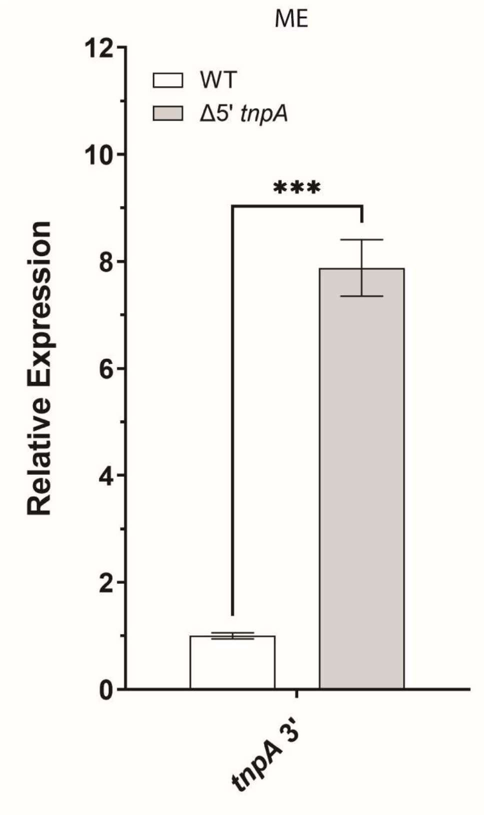

**Figure S12. qRT-PCR analysis showing that the Δ*5'tnpA* strain produces a *tnpA* transcript that includes the 3' portion of the *tnpA* gene.** In other qRT-PCR analyses presented in this work we detected *tnpA* transcript using a primer pair with DNA sequence complementarity to the 5' end of the *tnpA* transcript (nt 32-169). In this experiment the primer pair was complementary to the 3' end of the *tnpA* transcript (nt 287-486) and therefore capable of detecting both the *WT tnpA* transcript and the presumed chimeric *FRT-scar-3'-tnpA* transcript produced specifically by the Δ*5'tnpA* strain. In this experiment the *tnpA* level for the *WT* strain was set at 1. n = 3 and error bars represent the standard error of the mean. *** $P \leq 0.001$ and 0.0001.

### Table S1

## EE

| Gene | WT-1-EE | WT-2-EE | WT-3-EE | 5' tnpA-1-EE | 5' tnpA-2-EE | 5' tnpA-3-EE |
|---|---|---|---|---|---|---|
| aceF_aceF | 16510 | 17611 | 19420 | 17521 | 16793 | 16486 |
| SL1344_RS11990_ackA | 7438 | 7520 | 7591 | 5979 | 5763 | 5896 |
| SL1344_RS08180_acpD | 137 | 134 | 160 | 280 | 262 | 284 |
| acpS_acpS | 231 | 296 | 279 | 298 | 269 | 286 |
| adrA_adrA | 102 | 108 | 79 | 105 | 87 | 78 |
| SL1344_RS16630_aer | 349 | 473 | 406 | 1313 | 1113 | 1329 |
| SL1344_RS13055_alpha/beta hydrolase | 153 | 124 | 155 | 80 | 79 | 96 |
| AmgR_amgR | 8 | 15 | 12 | 12 | 17 | 9 |
| amiC_amiC | 805 | 982 | 905 | 1184 | 1082 | 1134 |
| SL1344_RS04710_amino acid permease | 211 | 228 | 206 | 137 | 120 | 140 |
| SL1344_RS01920_ampH | 193 | 262 | 214 | 120 | 135 | 139 |
| SL1344_RS23585_arcA | 3241 | 3362 | 3186 | 3255 | 3248 | 3243 |
| ArcZ_arcZ | 113 | 134 | 99 | 88 | 91 | 83 |
| STnc490a_art200 | 3 | 0 | 0 | 2 | 0 | 0 |
| STnc490b_art200 | 2 | 0 | 0 | 0 | 0 | 0 |
| STnc490c_art200 | 0 | 7 | 4 | 0 | 0 | 1 |
| STnc490d_art200 | 0 | 0 | 1 | 0 | 0 | 0 |
| STnc490e_art200 | 0 | 0 | 0 | 0 | 0 | 0 |
| STnc490f_art200 | 3 | 1 | 0 | 0 | 0 | 0 |
| STnc490g_art200 | 5 | 0 | 0 | 0 | 0 | 0 |
| artM_artM | 1115 | 1339 | 1310 | 1389 | 1263 | 1386 |
| artP_artP | 95 | 113 | 111 | 133 | 126 | 157 |
| asarfA_asarfA | 360 | 327 | 442 | 357 | 281 | 305 |
| asnB_asnB | 524 | 531 | 568 | 606 | 549 | 583 |
| SL1344_RS13060_asrA | 20 | 12 | 31 | 11 | 5 | 13 |
| SL1344_RS13065_asrB | 24 | 16 | 25 | 32 | 20 | 22 |
| SL1344_RS13070_asrC | 43 | 60 | 44 | 57 | 46 | 43 |
| asyhbQ_asyhbQ | 134 | 147 | 182 | 196 | 152 | 149 |
| atpC_atpC | 10936 | 10961 | 11996 | 8515 | 8483 | 8314 |
| SL1344_RS15305_barA | 693 | 689 | 767 | 658 | 573 | 615 |
| bcsA_bcsA | 849 | 917 | 990 | 482 | 455 | 466 |
| SL1344_RS17950_bigA | 250 | 213 | 222 | 257 | 237 | 238 |
| bssR_bssR | 33 | 38 | 29 | 51 | 49 | 38 |
| bssS_bssS | 736 | 681 | 571 | 660 | 595 | 563 |
| cadB_cadB | 272 | 272 | 294 | 175 | 137 | 161 |
| cbiC_cbiC | 21 | 12 | 10 | 14 | 9 | 11 |
| cbiD_cbiD | 22 | 33 | 25 | 23 | 26 | 17 |
| SL1344_RS11580_ccmA | 4 | 3 | 7 | 9 | 11 | 13 |
| SL1344_RS11575_ccmB | 16 | 8 | 6 | 3 | 0 | 2 |
| cdsA_cdsA | 1112 | 1311 | 1348 | 1265 | 1090 | 1383 |
| SL1344_RS02340_cdsH | 106 | 123 | 116 | 142 | 112 | 112 |
| chaB_chaB | 38 | 29 | 33 | 70 | 67 | 36 |
| SL1344_RS09610_cheA | 4812 | 5708 | 5480 | 31984 | 27014 | 32639 |
| SL1344_RS09600_cheM | 2221 | 2303 | 1915 | 14381 | 12459 | 13827 |
| SL1344_RS09595_chemotaxis protein methyltransferase | 1454 | 1413 | 1305 | 6820 | 6313 | 7450 |
| SL1344_RS09585_chemotaxis regulator | 979 | 1139 | 1141 | 5189 | 4633 | 5437 |
| SL1344_RS09590_chemotaxis response regulator protein-glutamate methylesterase | 953 | 1160 | 1070 | 5159 | 4192 | 5622 |
| SL1344_RS11875_cheV | 1668 | 1708 | 1495 | 5250 | 4586 | 4912 |
| SL1344_RS09605_CheW | 4506 | 4970 | 4761 | 29050 | 25407 | 29529 |
| SL1344_RS09580_cheZ | 2675 | 3293 | 3383 | 13602 | 12502 | 14846 |
| chiP_chiP | 80 | 68 | 74 | 62 | 61 | 57 |
| ChiX_chiX | 359 | 445 | 565 | 353 | 254 | 421 |
| SL1344_RS02290_clpP | 1128 | 1268 | 1209 | 1456 | 1412 | 1379 |
| SL1344_RS02295_clpX | 6335 | 6274 | 6091 | 7305 | 6844 | 6981 |
| SL1344_RS09530_cmoA | 916 | 854 | 871 | 1686 | 1625 | 1694 |
| cobS_cobS | 91 | 101 | 123 | 86 | 81 | 105 |
| copA_copA | 750 | 1069 | 744 | 975 | 865 | 623 |
| SL1344_RS20320_corA | 917 | 1193 | 1359 | 1314 | 1223 | 1147 |
| SL1344_RS20860_cpxA | 1476 | 1793 | 1806 | 1714 | 1598 | 1705 |
| cpxP_cpxP | 572 | 455 | 358 | 450 | 512 | 436 |
| SL1344_RS20865_cpxR | 696 | 729 | 657 | 667 | 599 | 576 |

| | | | | | | |
|---|---|---|---|---|---|---|
| SL1344_RS17890_crp | 3202 | 3292 | 3295 | 3440 | 3188 | 3272 |
| CrpT_crpT | 16 | 15 | 18 | 9 | 16 | 15 |
| csgB_csgB | 0 | 4 | 3 | 5 | 6 | 2 |
| SL1344_RS05640_csgC | 197 | 189 | 144 | 520 | 589 | 504 |
| SL1344_RS18810_cspA | 21479 | 21353 | 25453 | 7946 | 8174 | 9668 |
| SL1344_RS09980_cspB | 307 | 222 | 233 | 91 | 89 | 95 |
| SL1344_RS06145_cspH | 370 | 308 | 305 | 135 | 118 | 170 |
| SL1344_RS14635_csrA | 3122 | 3931 | 4012 | 4414 | 4099 | 4330 |
| CsrB_csrB | 55512 | 65315 | 64096 | 80585 | 76536 | 77058 |
| SL1344_RS15345_csrB | 48940 | 58265 | 54071 | 70154 | 67007 | 67963 |
| CsrC_csrC | 30519 | 35481 | 37984 | 58622 | 62025 | 58815 |
| SL1344_RS17435_csrD | 801 | 832 | 834 | 1369 | 1133 | 1218 |
| CyaR_cyaR | 215 | 213 | 215 | 207 | 191 | 193 |
| SL1344_RS12535_cysA | 1112 | 1223 | 1348 | 650 | 480 | 638 |
| SL1344_RS08565_cysB | 608 | 655 | 560 | 699 | 653 | 691 |
| SL1344_RS15175_cysC | 920 | 893 | 815 | 407 | 296 | 467 |
| SL1344_RS15185_cysD | 1171 | 1243 | 1012 | 692 | 520 | 721 |
| SL1344_RS19060_cysE | 1348 | 1350 | 1416 | 1425 | 1417 | 1524 |
| SL1344_RS17945_cysG | 708 | 768 | 747 | 671 | 609 | 678 |
| SL1344_RS15240_cysH | 908 | 1059 | 1169 | 575 | 359 | 575 |
| SL1344_RS15245_cysI | 3893 | 4241 | 4598 | 1769 | 1284 | 1893 |
| SL1344_RS15250_cysJ | 3544 | 3746 | 4064 | 1703 | 1330 | 1822 |
| SL1344_RS12475_cysK | 4756 | 4785 | 4344 | 3596 | 3098 | 3418 |
| SL1344_RS12530_cysM | 817 | 864 | 887 | 487 | 446 | 508 |
| SL1344_RS15180_cysN | 1592 | 1601 | 1429 | 860 | 640 | 874 |
| SL1344_RS12550_cysP | 1363 | 1552 | 1393 | 916 | 676 | 832 |
| SL1344_RS22605_cysQ | 447 | 516 | 467 | 601 | 577 | 465 |
| SL1344_RS02725_cysS | 2407 | 2415 | 2392 | 2078 | 1959 | 2100 |
| SL1344_RS09720_cysT | 5564 | 5384 | 5143 | 9376 | 8475 | 9594 |
| SL1344_RS12545_cysU | 624 | 855 | 886 | 362 | 315 | 463 |
| SL1344_RS12540_cysW | 1189 | 1294 | 1287 | 586 | 497 | 612 |
| SL1344_RS12470_cysZ | 368 | 291 | 228 | 284 | 250 | 269 |
| SL1344_RS16735_cyuA | 255 | 254 | 260 | 263 | 244 | 255 |
| SL1344_RS16740_cyuP | 127 | 131 | 97 | 132 | 109 | 118 |
| SL1344_RS10610_dacD | 179 | 207 | 234 | 174 | 138 | 188 |
| SL1344_RS17980_dam | 1839 | 1778 | 1986 | 1578 | 1535 | 1576 |
| DapZ_dapZ | 181 | 233 | 256 | 328 | 238 | 322 |
| ddl_ddl | 1315 | 1468 | 1528 | 1434 | 1295 | 1373 |
| SL1344_RS09505_dihydroneopterin triphosphate diphosphatase | 1229 | 1272 | 1391 | 2652 | 2501 | 2688 |
| dinG_dinG | 1012 | 1030 | 1005 | 995 | 957 | 1051 |
| SL1344_RS05725_dinI | 1017 | 989 | 950 | 1191 | 1055 | 1130 |
| SL1344_RS04685_dmsA | 56 | 69 | 66 | 65 | 62 | 71 |
| SL1344_RS04690_dmsB | 23 | 7 | 8 | 16 | 10 | 11 |
| SL1344_RS04695_dmsC | 13 | 21 | 23 | 24 | 24 | 23 |
| SL1344_RS06610_DNA/RNA non-specific endonuclease | 241 | 222 | 192 | 212 | 196 | 200 |
| dnaA_dnaA | 2057 | 2047 | 2005 | 2165 | 2020 | 2190 |
| SL1344_RS00060_dnaK | 27833 | 27701 | 27870 | 36317 | 31763 | 27995 |
| dpiA_dpiA | 60 | 54 | 55 | 60 | 51 | 57 |
| dpiA_dpiA | 61 | 77 | 57 | 66 | 61 | 67 |
| dpiB_dpiB | 53 | 65 | 55 | 69 | 70 | 67 |
| DsrA_dsrA | 11 | 25 | 5 | 7 | 5 | 7 |
| SL1344_RS09330_DUF1441 domain-containing protein | 3 | 3 | 5 | 13 | 13 | 12 |
| SL1344_RS06460_DUF1496 domain-containing protein | 500 | 512 | 406 | 1447 | 1213 | 1430 |
| SL1344_RS16295_DUF2623 domain-containing protein | 343 | 338 | 380 | 1025 | 794 | 1045 |
| emrB_emrB | 836 | 920 | 994 | 707 | 801 | 832 |
| emrD_emrD | 558 | 568 | 454 | 883 | 801 | 749 |
| emtA_emtA | 301 | 356 | 283 | 445 | 413 | 422 |
| SL1344_RS03015_entB | 230 | 317 | 289 | 291 | 313 | 257 |
| entE_entE | 414 | 496 | 431 | 426 | 426 | 398 |
| entF_entF | 598 | 801 | 662 | 689 | 646 | 596 |
| SL1344_RS18065_envZ | 1177 | 1235 | 1206 | 950 | 918 | 1030 |
| eutA_eutA | 20 | 22 | 24 | 31 | 20 | 17 |

| | | | | | | |
|---|---|---|---|---|---|---|
| SL1344_RS12680_eutS | 0 | 2 | 2 | 3 | 2 | 2 |
| fabG_fabG | 3386 | 3913 | 4109 | 4409 | 4092 | 4290 |
| fadA_fadA | 46 | 49 | 42 | 78 | 54 | 63 |
| SL1344_RS09080_fadD | 557 | 691 | 774 | 1249 | 1180 | 1322 |
| fadE_fadE | 228 | 265 | 242 | 326 | 303 | 334 |
| fadI_fadI | 24 | 27 | 32 | 28 | 24 | 27 |
| SL1344_RS13945_Fels-2 prophage protein | 7 | 8 | 12 | 24 | 55 | 6 |
| feoA_feoA | 445 | 539 | 548 | 288 | 268 | 266 |
| ffs_ffs | 1775 | 1727 | 2153 | 1523 | 1172 | 1710 |
| fimA_fimA | 35 | 58 | 48 | 45 | 42 | 44 |
| fimF_fimF | 35 | 42 | 52 | 47 | 43 | 36 |
| SL1344_RS02785_fimZ | 16 | 31 | 15 | 38 | 18 | 63 |
| fis_fis | 3238 | 3891 | 4282 | 3457 | 3014 | 3165 |
| fixB_fixB | 86 | 134 | 113 | 75 | 96 | 95 |
| flgD_flgD | 1703 | 2042 | 2059 | 4422 | 4353 | 4744 |
| flgF_flgF | 2714 | 2826 | 2763 | 5763 | 5979 | 6223 |
| flgI_flgI | 1069 | 1274 | 1443 | 2330 | 2371 | 2646 |
| SL1344_RS05830_flgK | 4037 | 4458 | 4099 | 13126 | 11795 | 13301 |
| flgL_flgL | 3566 | 3835 | 3692 | 10582 | 9599 | 10819 |
| SL1344_RS05775_flgM | 1644 | 1752 | 1412 | 4835 | 4171 | 4699 |
| SL1344_RS05770_flgN | 2134 | 2637 | 2411 | 6666 | 5770 | 6821 |
| SL1344_RS09570_flhA | 1776 | 1867 | 1986 | 6047 | 6888 | 6789 |
| SL1344_RS09575_flhB | 1280 | 1388 | 1392 | 5044 | 5401 | 5580 |
| SL1344_RS09625_flhC | 1465 | 1423 | 1280 | 4924 | 4269 | 4441 |
| SL1344_RS09630_flhD | 640 | 629 | 571 | 1153 | 1094 | 1126 |
| SL1344_RS09565_flhE | 417 | 436 | 509 | 1333 | 1421 | 1516 |
| SL1344_RS09785_fliA | 3259 | 4110 | 4050 | 15564 | 13890 | 16460 |
| SL1344_RS09800_fliC | 10866 | 11389 | 10882 | 143673 | 75208 | 148387 |
| SL1344_RS09845_fliE | 465 | 572 | 549 | 992 | 807 | 1146 |
| fliI_fliI | 2131 | 2061 | 2064 | 2702 | 2923 | 3049 |
| fliJ_fliJ | 737 | 819 | 833 | 1217 | 1333 | 1279 |
| fliL_fliL | 1430 | 1553 | 1447 | 2361 | 2690 | 2583 |
| fliQ_fliQ | 569 | 520 | 522 | 702 | 808 | 769 |
| SL1344_RS09815_fliT | 3209 | 3347 | 2874 | 6134 | 5984 | 6425 |
| SL1344_RS09775_fliY | 902 | 1035 | 985 | 3748 | 3343 | 3782 |
| SL1344_RS09780_fliZ | 1506 | 1674 | 1512 | 7138 | 6672 | 7490 |
| SL1344_RS14340_fljA | 1361 | 1580 | 1483 | 87 | 1492 | 64 |
| flk_flk | 454 | 410 | 395 | 319 | 272 | 302 |
| SL1344_RS08285_fnr | 2517 | 2875 | 2837 | 3511 | 3278 | 3822 |
| FnrS_fnrS | 57 | 61 | 39 | 78 | 58 | 71 |
| SL1344_RS04735_focA | 855 | 821 | 730 | 518 | 458 | 458 |
| folA_folA | 542 | 585 | 666 | 476 | 425 | 484 |
| frsA_frsA | 845 | 880 | 838 | 812 | 751 | 783 |
| SL1344_RS09665_ftnB | 35 | 30 | 21 | 87 | 73 | 86 |
| SL1344_RS06895_fumD | 151 | 168 | 176 | 161 | 148 | 128 |
| SL1344_RS03500_fur | 4195 | 4130 | 3940 | 5160 | 4219 | 4605 |
| gapA_gapA | 1394 | 1705 | 1692 | 1493 | 1229 | 1475 |
| GcvB_gcvB | 31465 | 31984 | 31813 | 25069 | 23273 | 27608 |
| gcvT_gcvT | 622 | 723 | 681 | 1420 | 1225 | 1368 |
| gidB_gidB | 591 | 581 | 622 | 493 | 460 | 474 |
| gldA_gldA | 130 | 125 | 109 | 137 | 146 | 134 |
| gldA_gldA | 342 | 323 | 392 | 380 | 434 | 385 |
| GlmY_glmY | 1698 | 1869 | 1760 | 1131 | 971 | 1253 |
| GlmZ_glmZ | 7476 | 7199 | 6233 | 6460 | 6370 | 6920 |
| glnQ_glnQ | 2188 | 2322 | 2431 | 1842 | 1664 | 1845 |
| SL1344_RS13690_gluconolactonase | 192 | 203 | 169 | 64 | 44 | 32 |
| SL1344_RS09560_glycoside hydrolase family 105 protein | 280 | 309 | 315 | 867 | 917 | 904 |
| gntK_gntK | 88 | 79 | 91 | 91 | 68 | 78 |
| SL1344_RS18265_gntR | 631 | 646 | 633 | 592 | 640 | 642 |
| gpmA_gpmA | 7029 | 8222 | 8362 | 8793 | 8224 | 7925 |
| GraL_graL | 1441 | 1407 | 991 | 1243 | 1176 | 1187 |
| groES_groES | 9764 | 10712 | 10404 | 13206 | 11443 | 11187 |

| | | | | | | |
|---|---|---|---|---|---|---|
| SL1344_RS14595_gshA | 1845 | 1797 | 1892 | 1843 | 1713 | 1860 |
| SL1344_RS16010_gshB | 1183 | 1267 | 1227 | 1228 | 1209 | 1272 |
| SL1344_RS05170_gtgD | 18 | 9 | 14 | 31 | 18 | 40 |
| SL1344_RS05180_gtgE | 147 | 188 | 131 | 225 | 249 | 219 |
| SL1344_RS05185_gtgF | 6 | 7 | 8 | 11 | 5 | 5 |
| SL1344_RS11665_gyrA | 10130 | 10446 | 10908 | 9365 | 9057 | 9950 |
| SL1344_RS13995_head completion/stabilization protein | 13 | 3 | 40 | 1 | 2 | 7 |
| helD_helD | 503 | 597 | 522 | 611 | 590 | 601 |
| hemG_hemG | 309 | 322 | 316 | 379 | 364 | 373 |
| hflB_hflB | 13743 | 14170 | 14778 | 14314 | 13613 | 14078 |
| SL1344_RS22390_hfq | 4616 | 5305 | 5287 | 5590 | 5174 | 5046 |
| SL1344_RS02410_hha | 966 | 995 | 894 | 628 | 645 | 669 |
| SL1344_RS02415_Hha toxicity attenuator | 1128 | 1057 | 858 | 661 | 577 | 620 |
| SL1344_RS14890_hilA | 76 | 81 | 73 | 542 | 746 | 620 |
| SL1344_RS14845_hilC | 382 | 387 | 361 | 1044 | 1321 | 1167 |
| SL1344_RS14885_hilD | 556 | 569 | 481 | 1312 | 1421 | 1223 |
| SL1344_RS23135_hilE | 60 | 65 | 70 | 74 | 68 | 77 |
| SL1344_RS21865_HlyD family type I secretion periplasmic adaptor subunit | 41 | 38 | 37 | 168 | 181 | 135 |
| SL1344_RS08745_hns | 21324 | 20369 | 19610 | 21886 | 20605 | 20344 |
| hscB_hscB | 1062 | 1108 | 1044 | 943 | 951 | 1041 |
| SL1344_RS08210_hslJ | 285 | 306 | 280 | 325 | 290 | 345 |
| SL1344_RS21405_hupA | 22997 | 21521 | 20712 | 23377 | 23069 | 22914 |
| SL1344_RS21410_hupB | 3491 | 3224 | 3629 | 3386 | 3253 | 3254 |
| hycD_hycD | 20 | 10 | 32 | 31 | 27 | 20 |
| hycE_hycE | 50 | 68 | 64 | 53 | 55 | 66 |
| SL1344_RS06065_hypothetical protein | 156 | 173 | 128 | 489 | 523 | 543 |
| SL1344_RS07940_hypothetical protein | 1089 | 1222 | 1152 | 3145 | 3673 | 3326 |
| SL1344_RS09195_hypothetical protein | 36 | 44 | 31 | 126 | 171 | 157 |
| SL1344_RS09675_hypothetical protein | 280 | 308 | 320 | 1737 | 2062 | 1840 |
| SL1344_RS09745_hypothetical protein | 618 | 638 | 512 | 989 | 1097 | 1004 |
| SL1344_RS22135_hypothetical protein | 15 | 15 | 8 | 62 | 79 | 58 |
| iacP_iacP | 185 | 179 | 160 | 220 | 246 | 246 |
| SL1344_RS14895_iagB | 106 | 94 | 77 | 438 | 514 | 417 |
| lamB_iamB | 45 | 54 | 53 | 68 | 72 | 68 |
| idnK_idnK | 35 | 29 | 30 | 48 | 35 | 34 |
| SL1344_RS22990_idnT | 30 | 56 | 28 | 33 | 35 | 42 |
| SL1344_RS06655_ihfA | 4174 | 4475 | 4411 | 5238 | 4909 | 5235 |
| SL1344_RS04775_ihfB | 1852 | 2062 | 2027 | 2334 | 2122 | 2355 |
| lldD_ildD | 129 | 133 | 120 | 177 | 161 | 199 |
| ilvM_ilvM | 158 | 127 | 126 | 196 | 157 | 178 |
| SL1344_RS14990_invA | 85 | 87 | 79 | 675 | 749 | 730 |
| SL1344_RS14980_invC | 55 | 65 | 58 | 318 | 342 | 367 |
| SL1344_RS14995_invE | 37 | 41 | 51 | 490 | 540 | 503 |
| SL1344_RS15005_invF | 50 | 54 | 29 | 785 | 855 | 798 |
| SL1344_RS15000_invG | 49 | 58 | 49 | 834 | 829 | 758 |
| InvR_invR | 511 | 486 | 368 | 1659 | 1642 | 1567 |
| SL1344_RS14355_iroB | 57 | 57 | 50 | 231 | 150 | 224 |
| SL1344_RS24725_IS91 family transposase | 113 | 135 | 114 | 533 | 620 | 552 |
| IsrA_isrA | 7 | 5 | 9 | 3 | 1 | 6 |
| IsrB_isrB | 41 | 43 | 35 | 26 | 24 | 25 |
| IsrC_isrC | 41 | 42 | 33 | 36 | 44 | 40 |
| IsrD_isrD | 0 | 0 | 1 | 0 | 0 | 0 |
| IsrE_isrE | 198 | 193 | 196 | 108 | 95 | 100 |
| IsrF_isrF | 55 | 68 | 53 | 111 | 121 | 106 |
| IsrG_isrG | 40 | 38 | 50 | 53 | 40 | 60 |
| IsrH-1_isrH | 6 | 7 | 5 | 14 | 5 | 17 |
| IsrH-2_isrH2 | 6 | 7 | 5 | 14 | 5 | 16 |
| IsrI_isrI | 25 | 37 | 23 | 26 | 33 | 26 |
| IsrJ_isrJ | 133 | 143 | 116 | 122 | 119 | 134 |
| IsrK_isrK | 350 | 320 | 323 | 300 | 285 | 291 |
| IsrL_isrL | 50 | 37 | 33 | 40 | 41 | 43 |
| IsrM_isrM | 8 | 14 | 8 | 11 | 6 | 5 |

| | | | | | | |
|---|---|---|---|---|---|---|
| IsrN_isrN | 0 | 0 | 2 | 2 | 1 | 1 |
| IsrO_isrO | 108 | 109 | 88 | 126 | 104 | 94 |
| IsrP_isrP | 51 | 60 | 42 | 60 | 56 | 62 |
| IsrQ_isrQ | 6 | 2 | 7 | 6 | 5 | 5 |
| IstR-1_istR1 | 76 | 70 | 53 | 46 | 43 | 40 |
| IstR-2_istR2 | 82 | 76 | 61 | 53 | 51 | 44 |
| ksgA_ksgA | 1422 | 1536 | 1607 | 1378 | 1270 | 1450 |
| SL1344_RS08205_ldhA | 457 | 570 | 556 | 958 | 818 | 842 |
| SL1344_RS00585_leuO | 29 | 15 | 13 | 17 | 16 | 9 |
| SL1344_RS21750_lexA | 2799 | 3097 | 3249 | 3146 | 2949 | 3418 |
| ligA_ligA | 2206 | 2315 | 2368 | 1798 | 1797 | 1729 |
| livF_livF | 58 | 71 | 78 | 92 | 102 | 113 |
| lolA_lolA | 626 | 767 | 778 | 900 | 761 | 857 |
| lolB_lolB | 219 | 253 | 293 | 250 | 278 | 306 |
| lplA_lplA | 511 | 499 | 467 | 531 | 494 | 471 |
| SL1344_RS06600_lpxR | 38 | 49 | 36 | 277 | 282 | 265 |
| SL1344_RS11955_lrhA | 548 | 578 | 645 | 362 | 356 | 401 |
| SL1344_RS04660_lrp | 2049 | 2122 | 2139 | 2975 | 2927 | 2890 |
| SL1344_RS09120_L-serine ammonia-lyase | 2325 | 2419 | 2446 | 4648 | 4558 | 4509 |
| SL1344_RS20930_lsrR | 255 | 255 | 249 | 222 | 184 | 187 |
| SL1344_RS14590_luxS | 2463 | 2556 | 2511 | 2541 | 2229 | 2541 |
| SL1344_RS09795_lysine-N-methylase | 675 | 783 | 860 | 1983 | 1720 | 2125 |
| malF_malF | 34 | 50 | 28 | 39 | 23 | 39 |
| malG_malG | 39 | 39 | 22 | 33 | 36 | 44 |
| SL1344_RS18135_malT | 1435 | 1503 | 1497 | 744 | 628 | 748 |
| SL1344_RS07280_malY* | 119 | 125 | 109 | 123 | 100 | 98 |
| mdoD_mdoD | 1097 | 1181 | 1117 | 1652 | 1509 | 1562 |
| mdoG_mdoG | 3413 | 3386 | 3297 | 3474 | 3366 | 3398 |
| SL1344_RS09375_membrane protein | 606 | 679 | 609 | 1596 | 1398 | 1483 |
| SL1344_RS21870_membrane protein | 1510 | 1546 | 1506 | 2149 | 2112 | 2299 |
| SL1344_RS11835_menC | 446 | 467 | 566 | 481 | 426 | 455 |
| SL1344_RS21470_metA | 96 | 86 | 93 | 105 | 87 | 98 |
| SL1344_RS21065_metB | 485 | 551 | 520 | 523 | 481 | 489 |
| SL1344_RS16345_metC | 154 | 163 | 154 | 247 | 217 | 209 |
| SL1344_RS20385_metE | 330 | 291 | 372 | 416 | 370 | 320 |
| SL1344_RS21095_metF | 189 | 156 | 126 | 223 | 205 | 189 |
| metG_metG | 2452 | 2437 | 2529 | 2075 | 1973 | 2053 |
| SL1344_RS21500_metH | 678 | 732 | 747 | 834 | 738 | 861 |
| SL1344_RS18435_methyl-accepting chemotaxis citrate transducer | 654 | 728 | 661 | 3356 | 3033 | 3319 |
| SL1344_RS08105_methyl-accepting chemotaxis protein | 249 | 267 | 236 | 1152 | 988 | 1135 |
| SL1344_RS16230_methyl-accepting chemotaxis protein | 337 | 391 | 304 | 1401 | 1296 | 1383 |
| SL1344_RS16300_methyl-accepting chemotaxis protein | 2198 | 2180 | 2290 | 6624 | 5652 | 6624 |
| SL1344_RS16625_methyl-accepting chemotaxis protein | 3275 | 3485 | 3304 | 12222 | 10256 | 12408 |
| SL1344_RS21060_metJ | 935 | 1256 | 1298 | 1457 | 1284 | 1565 |
| SL1344_RS21070_metL | 1265 | 1475 | 1481 | 1341 | 1221 | 1274 |
| metQ_metQ | 1576 | 1587 | 1385 | 1888 | 1597 | 1689 |
| SL1344_RS20380_metR | 128 | 110 | 104 | 126 | 111 | 105 |
| MgrR_mgrR | 842 | 896 | 878 | 567 | 437 | 687 |
| MicA_micA | 25 | 35 | 18 | 51 | 43 | 46 |
| MicC_micC | 30 | 26 | 20 | 32 | 48 | 22 |
| MicF_micF | 834 | 732 | 622 | 359 | 350 | 330 |
| SL1344_RS07395_mlc | 426 | 426 | 409 | 361 | 353 | 376 |
| moaA_moaA | 263 | 287 | 300 | 329 | 277 | 324 |
| moaD_moaD | 104 | 125 | 114 | 129 | 104 | 98 |
| moaE_moaE | 172 | 204 | 172 | 186 | 189 | 187 |
| mobA_mobA | 155 | 182 | 180 | 193 | 171 | 193 |
| SL1344_RS24715_mobA | 1050 | 1198 | 1123 | 6555 | 6940 | 6644 |
| SL1344_RS24720_mobC | 182 | 273 | 193 | 1323 | 1197 | 1382 |
| SL1344_RS03955_modA | 655 | 661 | 676 | 2281 | 1917 | 2144 |
| modB_modB | 608 | 646 | 595 | 1931 | 1598 | 1941 |
| SL1344_RS03965_modC | 853 | 940 | 864 | 2552 | 2208 | 2668 |
| SL1344_RS09620_motA | 1787 | 2236 | 1996 | 14622 | 11958 | 14967 |

| | | | | | | |
|---|---|---|---|---|---|---|
| motB_motB | 2364 | 2656 | 2262 | 14590 | 12492 | 14589 |
| SL1344_RS09450_msbB | 649 | 747 | 704 | 1355 | 1307 | 1387 |
| Msr_Msr | 147 | 121 | 113 | 83 | 65 | 72 |
| mtlR_mtlR | 427 | 513 | 426 | 509 | 421 | 532 |
| mukB_mukB | 5551 | 5916 | 6412 | 6149 | 6215 | 6308 |
| murD_murD | 2296 | 2644 | 2727 | 2157 | 2151 | 2432 |
| SL1344_RS00625_murE | 1315 | 1489 | 1773 | 1284 | 1181 | 1489 |
| SL1344_RS09280_N-acetyltransferase | 59 | 89 | 77 | 308 | 244 | 294 |
| napB_napB | 47 | 42 | 34 | 59 | 54 | 64 |
| SL1344_RS08805_narG | 50 | 61 | 59 | 135 | 102 | 116 |
| SL1344_RS08800_narH | 44 | 23 | 35 | 63 | 64 | 65 |
| SL1344_RS08790_narI | 18 | 21 | 19 | 39 | 36 | 32 |
| SL1344_RS08795_narJ | 8 | 9 | 11 | 22 | 18 | 19 |
| SL1344_RS08810_narK | 20 | 20 | 19 | 56 | 40 | 55 |
| SL1344_RS08820_narL | 147 | 153 | 154 | 267 | 283 | 258 |
| nepI_nepI | 313 | 371 | 372 | 394 | 358 | 401 |
| nhaA_nhaA | 641 | 760 | 800 | 835 | 807 | 822 |
| nirD_nirD | 50 | 50 | 77 | 58 | 65 | 56 |
| nlpD_nlpD | 7172 | 7307 | 6763 | 8593 | 8961 | 8491 |
| nrdI_nrdI | 65 | 103 | 89 | 80 | 71 | 63 |
| SL1344_RS21955_nrfB | 85 | 89 | 84 | 338 | 273 | 277 |
| SL1344_RS21960_nrfC | 83 | 107 | 96 | 338 | 265 | 304 |
| SL1344_RS21975_nrfG | 55 | 45 | 25 | 41 | 27 | 30 |
| nudC_nudC | 229 | 209 | 201 | 210 | 189 | 223 |
| nusA_nusA | 8226 | 8545 | 9305 | 7211 | 7250 | 7643 |
| nusB_nusB | 3480 | 3413 | 3158 | 2649 | 2497 | 3058 |
| nusG_nusG | 9790 | 10006 | 10258 | 10643 | 10219 | 10283 |
| obgE_obgE | 1515 | 1880 | 2207 | 1567 | 1446 | 1718 |
| ompL_ompL | 6 | 8 | 5 | 5 | 3 | 7 |
| SL1344_RS18070_ompR | 798 | 824 | 851 | 706 | 732 | 681 |
| ompX_ompX | 32919 | 32624 | 32434 | 26649 | 26880 | 25983 |
| OmrA_omrA | 95 | 86 | 78 | 37 | 27 | 44 |
| OmrB_omrB | 52 | 51 | 50 | 34 | 37 | 25 |
| oppB_oppB | 167 | 136 | 161 | 138 | 113 | 156 |
| SL1344_RS14860_orgA | 319 | 296 | 341 | 1324 | 1558 | 1399 |
| OxyS_oxyS | 81 | 77 | 62 | 80 | 82 | 53 |
| pabB_pabB | 497 | 449 | 443 | 936 | 892 | 872 |
| pagP_pagP | 1494 | 1318 | 1237 | 896 | 838 | 869 |
| SL1344_RS10525_pduA | 2 | 1 | 0 | 3 | 5 | 3 |
| pduC_pduC | 10 | 6 | 12 | 11 | 12 | 12 |
| pduD_pduD | 8 | 6 | 2 | 6 | 5 | 8 |
| pduE_pduE | 2 | 7 | 6 | 7 | 4 | 2 |
| pdxK_pdxK | 115 | 127 | 124 | 173 | 155 | 129 |
| SL1344_RS06590_pfkB | 563 | 539 | 542 | 557 | 574 | 485 |
| SL1344_RS13960_phage virion morphogenesis protein | 2 | 25 | 10 | 7 | 0 | 6 |
| SL1344_RS02035_phoB | 796 | 844 | 754 | 858 | 733 | 795 |
| SL1344_RS11645_phoE | 35271 | 39136 | 38308 | 21916 | 19720 | 21304 |
| SL1344_RS05535_phoH | 111 | 123 | 105 | 222 | 159 | 186 |
| SL1344_RS06080_phoP | 2767 | 2716 | 2621 | 2649 | 2477 | 2584 |
| SL1344_RS06075_phoQ | 2980 | 2903 | 2830 | 2580 | 2447 | 2606 |
| phoR_phoR | 440 | 445 | 456 | 563 | 522 | 500 |
| SL1344_RS13700_phospho-2-dehydro-3-deoxyheptonate aldolase Tyr-sensitive | 416 | 446 | 428 | 275 | 266 | 248 |
| SL1344_RS10625_phsA | 49 | 54 | 44 | 64 | 51 | 54 |
| SL1344_RS10620_phsB | 13 | 17 | 17 | 11 | 5 | 18 |
| SL1344_RS10615_phsC | 30 | 34 | 27 | 36 | 50 | 29 |
| potG_potG | 150 | 201 | 197 | 264 | 205 | 282 |
| SL1344_RS09215_prc | 6151 | 6198 | 6684 | 11643 | 10919 | 11770 |
| SL1344_RS14880_prgH | 76 | 92 | 72 | 1090 | 1054 | 1163 |
| SL1344_RS14875_prgI | 34 | 47 | 25 | 512 | 507 | 479 |
| SL1344_RS14870_prgJ | 86 | 84 | 69 | 826 | 824 | 852 |
| SL1344_RS14865_prgK | 265 | 266 | 249 | 1435 | 1658 | 1551 |
| SL1344_RS09220_proQ | 6069 | 6086 | 6203 | 9890 | 9536 | 10332 |

| | | | | | | |
|---|---|---|---|---|---|---|
| SL1344_RS09255_prpA/pphA | 24 | 28 | 28 | 54 | 57 | 60 |
| psiB_psiB | 34 | 37 | 31 | 23 | 23 | 28 |
| psiB_psiB | 20 | 16 | 17 | 17 | 21 | 18 |
| pssA_pssA | 2528 | 2638 | 2524 | 2356 | 2196 | 2419 |
| SL1344_RS11995_pta | 11922 | 12387 | 14229 | 10384 | 10313 | 10341 |
| SL1344_RS15255_ptpS | 889 | 916 | 988 | 426 | 387 | 414 |
| SL1344_RS02910_PTS mannose/fructose/sorbose transporter subunit IIB | 0 | 8 | 2 | 2 | 3 | 5 |
| SL1344_RS07105_purR | 1240 | 1318 | 1229 | 1015 | 1014 | 1119 |
| putA_putA | 586 | 642 | 634 | 667 | 664 | 692 |
| SL1344_RS23405_putative inner membrane protein | 279 | 273 | 288 | 522 | 479 | 461 |
| SL1344_RS09445_pyruvate kinase II | 814 | 814 | 863 | 1782 | 1671 | 1667 |
| SL1344_RS16430_qseB | 42 | 43 | 45 | 43 | 34 | 44 |
| SL1344_RS16435_qseC | 102 | 123 | 123 | 103 | 97 | 105 |
| SL1344_RS23490_radA | 983 | 1040 | 1028 | 831 | 849 | 899 |
| RaoN_raoN | 42 | 47 | 32 | 31 | 28 | 29 |
| SL1344_RS12895_ratA | 363 | 372 | 379 | 384 | 365 | 421 |
| rcsA_rcsA | 117 | 130 | 99 | 144 | 128 | 150 |
| SL1344_RS11655_rcsB | 1869 | 1825 | 1696 | 1752 | 1679 | 1605 |
| SL1344_RS11660_rcsC | 772 | 834 | 923 | 665 | 610 | 755 |
| SL1344_RS11650_rcsD | 2020 | 2017 | 1951 | 1515 | 1462 | 1541 |
| rcsF_rcsF | 620 | 602 | 630 | 542 | 566 | 518 |
| RdlD_rdlD | 102 | 103 | 119 | 71 | 102 | 85 |
| RdlE_rdlE | 34 | 32 | 49 | 20 | 14 | 23 |
| SL1344_RS14650_recA | 2505 | 3113 | 3042 | 4164 | 3839 | 3663 |
| SL1344_RS13835_recN | 923 | 940 | 892 | 1053 | 1057 | 1058 |
| SL1344_RS06620_rfc | 622 | 581 | 546 | 491 | 442 | 435 |
| SL1344_RS22270_rflM | 721 | 841 | 819 | 1751 | 2040 | 1881 |
| rhtB_rhtB | 39 | 45 | 41 | 47 | 49 | 57 |
| rimM_rimM | 15456 | 15693 | 16536 | 14965 | 13059 | 15499 |
| rimO_rimO | 1693 | 1852 | 1933 | 1755 | 1667 | 1692 |
| SL1344_RS03220_rlpA | 668 | 704 | 727 | 734 | 687 | 753 |
| SL1344_RS05840_rne | 7025 | 6877 | 7093 | 6825 | 6990 | 7097 |
| rnpB_rnpB | 422281 | 420429 | 436257 | 394608 | 373652 | 396378 |
| RnpB_RnpB | 422281 | 420429 | 436257 | 394608 | 373652 | 396378 |
| SL1344_RS15130_rpoS | 3512 | 4034 | 4133 | 5401 | 5247 | 5338 |
| rprA_rprA | 34 | 37 | 33 | 18 | 34 | 27 |
| rpsN_rpsN | 21452 | 21888 | 23621 | 19350 | 19309 | 20041 |
| rpsV_rpsV | 384 | 418 | 363 | 979 | 922 | 800 |
| RseX_rseX | 0 | 0 | 4 | 1 | 2 | 3 |
| rsmD_rsmD | 444 | 380 | 359 | 296 | 294 | 241 |
| SL1344_RS09240_rsmF/yebU | 757 | 767 | 811 | 1307 | 1260 | 1405 |
| SL1344_RS22150_rtsA | 79 | 88 | 51 | 524 | 679 | 598 |
| SL1344_RS22145_rtsB | 18 | 12 | 13 | 187 | 192 | 158 |
| rumA_rumA | 1275 | 1219 | 1110 | 1163 | 1163 | 1169 |
| SL1344_RS09480_ruvA | 432 | 460 | 412 | 906 | 817 | 864 |
| SL1344_RS09475_ruvB | 735 | 770 | 818 | 1440 | 1458 | 1537 |
| RybA_rybA | 821 | 864 | 749 | 1006 | 1063 | 1012 |
| RybB_rybB | 147 | 108 | 144 | 92 | 80 | 99 |
| RybD_rybD | 170 | 196 | 218 | 407 | 323 | 361 |
| RydB_rydB | 20 | 14 | 19 | 19 | 21 | 14 |
| RydC_rydC | 138 | 153 | 172 | 230 | 238 | 238 |
| RyeF_ryeF | 126 | 139 | 125 | 384 | 378 | 357 |
| RyfA_ryfA | 600 | 626 | 655 | 663 | 635 | 788 |
| RyfD_ryfD | 1555 | 1581 | 1459 | 1479 | 1337 | 1345 |
| RyhB_ryhB | 251 | 204 | 122 | 113 | 85 | 85 |
| RyjB_ryjB | 22 | 28 | 32 | 35 | 22 | 29 |
| Sar_sar | 123 | 140 | 137 | 99 | 71 | 93 |
| SL1344_RS10635_sbcB | 638 | 631 | 561 | 482 | 539 | 468 |
| SL1344_RS01925_sbmA | 172 | 206 | 225 | 182 | 168 | 210 |
| SL1344_RS10605_sbmC | 286 | 334 | 372 | 289 | 280 | 298 |
| SL1344_RS20885_sbp | 369 | 392 | 402 | 237 | 181 | 257 |
| sdhC_sdhC | 435 | 420 | 422 | 652 | 622 | 799 |

| | | | | | | |
|---|---|---|---|---|---|---|
| SL1344_RS09755_sdiA | 379 | 366 | 319 | 1888 | 1553 | 1811 |
| SdsR_sdsR | 79 | 76 | 119 | 52 | 52 | 67 |
| sgbH_sgbH | 8 | 1 | 5 | 13 | 11 | 4 |
| SgrS_sgrS | 970 | 862 | 851 | 1257 | 1003 | 1076 |
| SibA_sibA | 265 | 293 | 303 | 186 | 137 | 223 |
| SibC_sibC | 1208 | 1270 | 1201 | 1262 | 1070 | 1406 |
| SibD_sibD | 832 | 956 | 1058 | 819 | 624 | 909 |
| SL1344_RS14940_sicA | 30 | 40 | 42 | 579 | 551 | 536 |
| sifA_sifA | 192 | 182 | 142 | 186 | 148 | 143 |
| sifB_sifB | 32 | 28 | 32 | 28 | 24 | 22 |
| SL1344_RS05355_sigE | 73 | 81 | 84 | 863 | 881 | 848 |
| SL1344_RS21850_siiA | 15 | 16 | 14 | 80 | 93 | 73 |
| SL1344_RS21855_siiB | 31 | 41 | 28 | 153 | 158 | 141 |
| SL1344_RS21875_siiF | 846 | 842 | 644 | 856 | 808 | 790 |
| SL1344_RS01560_sinR | 218 | 264 | 205 | 331 | 326 | 359 |
| SL1344_RS14920_sipA | 147 | 166 | 168 | 750 | 812 | 707 |
| SL1344_RS14935_sipB | 90 | 133 | 108 | 1536 | 1571 | 1441 |
| SL1344_RS14930_sipC | 222 | 250 | 229 | 2520 | 2598 | 2624 |
| SL1344_RS14925_sipD | 44 | 65 | 40 | 322 | 417 | 358 |
| SL1344_RS09740_sirA | 1154 | 1247 | 1203 | 1936 | 1808 | 2090 |
| SL1344_RS10425_sirohydrochlorin cobaltochelatase | 25 | 17 | 25 | 16 | 11 | 14 |
| SL1344_RS09840_SL1896 | 24 | 28 | 13 | 44 | 39 | 37 |
| SL1344_RS07175_slyA | 1118 | 1208 | 1327 | 1332 | 1299 | 1251 |
| SL1344_RS07110_sodB | 1517 | 1927 | 2210 | 2220 | 1935 | 2229 |
| SL1344_RS10630_sopA | 33 | 45 | 29 | 208 | 244 | 201 |
| sopB_sopB | 61 | 72 | 56 | 929 | 1009 | 853 |
| SL1344_RS15235_sopD | 98 | 94 | 74 | 97 | 95 | 96 |
| SL1344_RS13925_sopE | 23 | 38 | 29 | 941 | 1022 | 940 |
| SL1344_RS09265_sopE2 | 15 | 12 | 11 | 427 | 448 | 386 |
| spaK_spaK | 24 | 28 | 15 | 163 | 188 | 153 |
| spaM_spaM | 26 | 30 | 44 | 225 | 209 | 236 |
| spaN_spaN | 51 | 48 | 57 | 292 | 355 | 304 |
| SL1344_RS14965_spaO | 30 | 44 | 35 | 270 | 280 | 248 |
| SL1344_RS14960_spaP | 32 | 34 | 22 | 231 | 244 | 247 |
| SL1344_RS14955_spaQ | 20 | 24 | 26 | 156 | 177 | 192 |
| SL1344_RS14950_spaR | 27 | 28 | 21 | 140 | 184 | 180 |
| SL1344_RS14945_spaS | 30 | 22 | 35 | 163 | 155 | 152 |
| Spf_spf | 46986 | 40347 | 35444 | 40235 | 38908 | 35934 |
| SL1344_RS14840_spi4 transcriptional regulator | 52 | 54 | 59 | 301 | 319 | 292 |
| spiC_spiC | 1 | 4 | 6 | 3 | 4 | 4 |
| spr_spr | 5601 | 5568 | 5446 | 4293 | 4152 | 4447 |
| SraA_sraA | 5 | 4 | 3 | 3 | 4 | 4 |
| SraB_sraB | 312 | 293 | 272 | 296 | 255 | 351 |
| SraC_sraC | 1075 | 940 | 846 | 2769 | 3011 | 2768 |
| SraF_sraF | 963 | 817 | 556 | 630 | 601 | 534 |
| SraG_sraG | 52 | 44 | 40 | 33 | 47 | 48 |
| SraL_sraL | 21 | 25 | 21 | 43 | 32 | 24 |
| SL1344_RS07945_srfB | 1457 | 1478 | 1618 | 3970 | 4500 | 4312 |
| SL1344_RS07950_srfC | 1152 | 1149 | 1157 | 3132 | 3521 | 3340 |
| srlA_srlA | 6 | 8 | 4 | 10 | 11 | 11 |
| srlE_srlE | 20 | 14 | 20 | 20 | 15 | 21 |
| srlR_srlR | 238 | 221 | 203 | 248 | 227 | 229 |
| SroA_sroA | 99 | 102 | 99 | 78 | 77 | 105 |
| SroC_sroC | 3496 | 3027 | 2325 | 3824 | 3330 | 3885 |
| SroD_sroD | 80 | 151 | 163 | 304 | 276 | 284 |
| SroG_sroG | 1020 | 1958 | 2284 | 1706 | 1376 | 1878 |
| SL1344_RS06985_ssaG | 1 | 5 | 9 | 6 | 2 | 3 |
| SL1344_RS07050_ssaR | 49 | 41 | 41 | 36 | 29 | 36 |
| SL1344_RS05160_sseI | 45 | 50 | 41 | 50 | 53 | 60 |
| sspA_sspA | 3477 | 3537 | 3560 | 3470 | 3482 | 3432 |
| SL1344_RS06915_ssrA | 107 | 111 | 91 | 109 | 93 | 85 |
| SsrA_ssrA | 264934 | 283009 | 304498 | 328574 | 281726 | 314845 |

| | | | | | | |
|---|---|---|---|---|---|---|
| SL1344_RS06910_ssrB | 41 | 53 | 47 | 61 | 52 | 27 |
| ssrS_ssrS | 70681 | 77349 | 59124 | 54696 | 45297 | 51849 |
| SsrS_ssrS | 70666 | 77326 | 59104 | 54680 | 45289 | 51836 |
| SL1344_RS16670_sstT | 713 | 825 | 813 | 814 | 768 | 861 |
| SL1344_RS01735_stbB | 0 | 1 | 2 | 5 | 3 | 7 |
| SL1344_RS05145_stfT | 40 | 46 | 35 | 79 | 58 | 68 |
| STnc3140_STnc3140 (sopD) | 301 | 257 | 325 | 79 | 69 | 114 |
| STnc3200_STnc3200 | 5 | 3 | 2 | 3 | 3 | 0 |
| SL1344_RS15350_syd | 719 | 729 | 749 | 591 | 568 | 689 |
| SymR_symR | 40 | 57 | 42 | 51 | 42 | 51 |
| SL1344_RS13990_tail protein | 1 | 3 | 34 | 2 | 1 | 0 |
| tatE_tatE | 251 | 370 | 403 | 479 | 440 | 485 |
| tcuB_tcuB | 23 | 29 | 29 | 26 | 32 | 34 |
| SL1344_RS16770_tdcA | 156 | 69 | 53 | 75 | 45 | 50 |
| SL1344_RS16765_tdcB | 119 | 91 | 104 | 280 | 256 | 310 |
| SL1344_RS16760_tdcC | 194 | 208 | 178 | 674 | 576 | 640 |
| SL1344_RS05150_tfaT | 31 | 39 | 32 | 63 | 39 | 59 |
| Tff_tff | 7714 | 7232 | 6031 | 7631 | 7323 | 7341 |
| thiP_thiP | 555 | 614 | 619 | 550 | 421 | 517 |
| thiQ_thiQ | 287 | 280 | 318 | 193 | 183 | 243 |
| SL1344_RS06625_thrS | 9392 | 9421 | 9657 | 7994 | 8119 | 8179 |
| SL1344_RS04595_tnpA1 | 2 | 4 | 13 | 62 | 66 | 63 |
| SL1344_RS06060_tnpA2 | 12 | 4 | 6 | 41 | 32 | 49 |
| SL1344_RS09075_tnpA3 | 3 | 13 | 8 | 29 | 9 | 34 |
| SL1344_RS09790_tnpA4 | 2 | 2 | 0 | 3 | 1 | 5 |
| SL1344_RS12685_tnpA5 | 3 | 3 | 4 | 5 | 7 | 3 |
| SL1344_RS17020_tnpA6 | 20 | 34 | 13 | 44 | 39 | 51 |
| SL1344_RS17955_tnpA7 | 21 | 17 | 23 | 34 | 44 | 29 |
| tolA_tolA | 544 | 641 | 682 | 612 | 621 | 681 |
| SL1344_RS16475_tolC | 2608 | 2604 | 2547 | 2360 | 2334 | 2378 |
| SL1344_RS21860_TolC family protein | 51 | 47 | 36 | 181 | 115 | 133 |
| tqsA_tqsA | 90 | 105 | 89 | 95 | 79 | 104 |
| SL1344_RS14145_transcriptional regulator | 2 | 1 | 3 | 5 | 41 | 3 |
| trkD_trkD | 1156 | 1170 | 1257 | 919 | 887 | 995 |
| SL1344_RS06175_tRNA arg | 9 | 7 | 5 | 46 | 59 | 37 |
| SL1344_RS13600_tRNA/rRNA methyltransferase | 1856 | 1875 | 1941 | 1174 | 1184 | 1117 |
| truA_truA | 540 | 602 | 607 | 444 | 380 | 452 |
| SL1344_RS23260_tsr | 8411 | 9231 | 8861 | 26369 | 23528 | 26103 |
| SL1344_RS06870_ttrA | 51 | 61 | 80 | 56 | 49 | 73 |
| SL1344_RS06880_ttrB | 6 | 6 | 5 | 2 | 7 | 11 |
| SL1344_RS06875_ttrC | 6 | 4 | 10 | 9 | 8 | 4 |
| SL1344_RS06890_ttrR | 96 | 95 | 70 | 98 | 85 | 77 |
| SL1344_RS06885_ttrS | 206 | 232 | 213 | 197 | 176 | 216 |
| tyrA_tyrA | 442 | 524 | 495 | 391 | 373 | 357 |
| ubiB_ubiB | 2025 | 1995 | 1990 | 1857 | 1818 | 1942 |
| ubiF_ubiF | 346 | 381 | 367 | 447 | 448 | 440 |
| SL1344_RS21190_udhA | 1013 | 1034 | 1018 | 1620 | 1668 | 1596 |
| SL1344_RS03635_UDP-galactopyranose mutase | 7 | 5 | 1 | 27 | 34 | 11 |
| ugpC_ugpC | 76 | 70 | 79 | 87 | 84 | 96 |
| ulaD_ulaD | 14 | 22 | 24 | 22 | 29 | 30 |
| ushA_ushA | 549 | 574 | 574 | 668 | 614 | 638 |
| SL1344_RS09735_uvrC | 1298 | 1460 | 1413 | 2071 | 2031 | 2226 |
| SL1344_RS11530_virulence protein MsgA | 0 | 0 | 1 | 0 | 2 | 3 |
| wecC_wecC | 1547 | 1636 | 1691 | 1339 | 1248 | 1387 |
| SL1344_RS10695_wzzB | 1322 | 1505 | 1640 | 757 | 748 | 718 |
| SL1344_RS02345_ybaO | 311 | 300 | 303 | 306 | 322 | 313 |
| SL1344_RS03185_ybeM | 13 | 34 | 28 | 85 | 56 | 88 |
| SL1344_RS04285_ybiK | 466 | 492 | 492 | 559 | 467 | 544 |
| SL1344_RS08990_ycgO | 835 | 957 | 911 | 2676 | 2263 | 2701 |
| SL1344_RS08975_ycgR | 814 | 914 | 858 | 2157 | 2063 | 2329 |
| SL1344_RS08585_yciK | 349 | 408 | 329 | 464 | 470 | 434 |
| SL1344_RS08495_yciW | 369 | 330 | 296 | 364 | 328 | 344 |

| | | | | | | |
|---|---|---|---|---|---|---|
| SL1344_RS07545_ydeD | 69 | 75 | 72 | 71 | 69 | 90 |
| SL1344_RS06680_ydiV | 113 | 134 | 130 | 91 | 80 | 113 |
| SL1344_RS06595_ydiY | 1466 | 1526 | 1501 | 1168 | 1136 | 1342 |
| SL1344_RS06585_ydiZ | 17 | 26 | 24 | 31 | 20 | 24 |
| SL1344_RS06560_ydjN | 10169 | 11035 | 10896 | 7201 | 6305 | 7437 |
| SL1344_RS09415_yebG | 84 | 110 | 118 | 303 | 285 | 288 |
| SL1344_RS09225_yebR | 1453 | 1490 | 1483 | 3418 | 3081 | 3151 |
| SL1344_RS09230_yebS | 392 | 357 | 293 | 702 | 691 | 669 |
| SL1344_RS09760_yecC | 261 | 319 | 270 | 684 | 749 | 728 |
| SL1344_RS09750_yecF | 1685 | 1789 | 1681 | 2984 | 3173 | 2923 |
| SL1344_RS09765_yecS | 129 | 159 | 170 | 343 | 280 | 395 |
| SL1344_RS09825_yedD | 599 | 633 | 717 | 573 | 509 | 577 |
| SL1344_RS09830_yedE | 302 | 291 | 331 | 189 | 181 | 180 |
| SL1344_RS09835_yedF | 178 | 209 | 233 | 132 | 114 | 98 |
| SL1344_RS09770_yedO | 267 | 312 | 301 | 762 | 653 | 884 |
| SL1344_RS10925_yegD | 561 | 508 | 532 | 242 | 229 | 221 |
| SL1344_RS13585_yfiK | 31 | 28 | 22 | 29 | 32 | 28 |
| SL1344_RS17960_yhfL | 297 | 224 | 216 | 484 | 459 | 455 |
| SL1344_RS18605_yhjH | 1441 | 1698 | 1396 | 5019 | 3991 | 4785 |
| SL1344_RS18650_yhjQ | 606 | 619 | 603 | 300 | 272 | 309 |
| SL1344_RS23255_yjiY | 947 | 931 | 1090 | 440 | 502 | 498 |
| SL1344_RS05645_ymdA | 178 | 237 | 180 | 475 | 548 | 550 |
| SL1344_RS05655_ymdC | 499 | 564 | 590 | 921 | 965 | 932 |
| yopJ_yopJ | 153 | 174 | 157 | 279 | 320 | 324 |
| SL1344_RS15335_yqcB | 539 | 575 | 545 | 446 | 451 | 559 |
| SL1344_RS15340_yqcC (next to csrB) | 266 | 299 | 294 | 221 | 235 | 307 |
| YrlA_yrlA | 12 | 8 | 12 | 10 | 12 | 10 |
| YrlB_yrlB | 9 | 11 | 9 | 14 | 12 | 14 |
| zntA_zntA | 668 | 802 | 668 | 913 | 917 | 949 |
| zntB_zntB | 271 | 232 | 227 | 342 | 351 | 363 |
| zraP_zraP | 29 | 31 | 23 | 25 | 26 | 17 |
| crRNA1_ | 145 | 126 | 162 | 132 | 116 | 154 |
| crRNA2_ | 416 | 413 | 383 | 297 | 330 | 358 |
| ECS021_ | 34 | 33 | 19 | 28 | 30 | 16 |
| ECS173_ | 46 | 50 | 41 | 40 | 45 | 44 |
| Ffs_ | 1784 | 1735 | 2163 | 1535 | 1178 | 1714 |
| fimA-trailer_ | 9 | 13 | 12 | 11 | 11 | 11 |
| GO-10702_ | 82 | 50 | 42 | 65 | 73 | 66 |
| GO-10704_ | 1508 | 1644 | 1344 | 1357 | 1300 | 1448 |
| Ig1085_ | 56 | 46 | 33 | 61 | 35 | 43 |
| Ig1140_ | 26 | 43 | 41 | 60 | 72 | 73 |
| Ig2335_ | 350 | 390 | 355 | 343 | 260 | 293 |
| Ig959_ | 0 | 0 | 0 | 0 | 0 | 0 |
| IS042_ | 16 | 16 | 16 | 7 | 10 | 13 |
| lysC-leader_ | 67 | 42 | 51 | 49 | 58 | 61 |
| mreB-leader_ | 403 | 510 | 484 | 540 | 457 | 501 |
| pRNA_ | 2 | 9 | 3 | 9 | 10 | 2 |
| RprA_ | 34 | 37 | 33 | 18 | 34 | 27 |
| rpsL-leader_ | 3182 | 3227 | 2945 | 3301 | 2739 | 3297 |
| rpsP-leader_ | 3814 | 3812 | 3190 | 3604 | 3457 | 3651 |
| rrf_ | 2 | 2 | 1 | 1 | 3 | 1 |
| rrf_ | 6 | 3 | 3 | 2 | 2 | 4 |
| rrf_ | 11 | 5 | 9 | 5 | 6 | 4 |
| rrf_ | 3 | 5 | 4 | 1 | 1 | 5 |
| rrf_ | 12 | 26 | 19 | 0 | 7 | 5 |
| rrf_ | 1 | 3 | 3 | 0 | 0 | 1 |
| rrf_ | 0 | 6 | 0 | 0 | 0 | 0 |
| rrf_ | 4 | 3 | 0 | 0 | 0 | 0 |
| RUF_107c.1_ | 286 | 313 | 386 | 256 | 244 | 277 |
| RUF_107c.10_ | 214 | 197 | 226 | 235 | 226 | 221 |
| RUF_107c.102_ | 21 | 19 | 15 | 60 | 17 | 24 |
| RUF_107c.105_ | 205 | 270 | 381 | 328 | 288 | 352 |

| | | | | | | |
|---|---|---|---|---|---|---|
| RUF_107c.107_ | 92 | 73 | 69 | 65 | 41 | 49 |
| RUF_107c.108_ | 435 | 519 | 631 | 445 | 348 | 430 |
| RUF_107c.111_ | 770 | 783 | 844 | 719 | 657 | 766 |
| RUF_107c.112_ | 395 | 349 | 407 | 324 | 326 | 329 |
| RUF_107c.18_ | 12 | 10 | 9 | 13 | 19 | 14 |
| RUF_107c.19_ | 226 | 206 | 245 | 192 | 223 | 201 |
| RUF_107c.2_ | 168 | 176 | 204 | 109 | 120 | 129 |
| RUF_107c.23_ | 180 | 173 | 212 | 265 | 231 | 218 |
| RUF_107c.24_ | 80 | 112 | 71 | 92 | 89 | 97 |
| RUF_107c.26_ | 9 | 9 | 11 | 10 | 8 | 7 |
| RUF_107c.30_ | 3 | 1 | 2 | 3 | 4 | 2 |
| RUF_107c.38_ | 81 | 58 | 70 | 76 | 69 | 75 |
| RUF_107c.47_ | 201 | 204 | 301 | 209 | 181 | 189 |
| RUF_107c.50_ | 102 | 86 | 91 | 95 | 82 | 97 |
| RUF_107c.51_ | 107 | 126 | 109 | 110 | 100 | 89 |
| RUF_107c.53_ | 253 | 253 | 291 | 582 | 413 | 577 |
| RUF_107c.56_ | 344 | 347 | 375 | 246 | 235 | 245 |
| RUF_107c.57_ | 75 | 72 | 100 | 75 | 76 | 110 |
| RUF_107c.58_ | 646 | 660 | 788 | 625 | 614 | 642 |
| RUF_107c.59_ | 316 | 332 | 440 | 246 | 251 | 315 |
| RUF_107c.63_ | 128 | 123 | 166 | 86 | 101 | 96 |
| RUF_107c.69_ | 116 | 106 | 88 | 138 | 115 | 120 |
| RUF_107c.70_ | 181 | 199 | 212 | 125 | 143 | 129 |
| RUF_107c.72_ | 105 | 100 | 112 | 71 | 87 | 79 |
| RUF_107c.77_ | 175 | 173 | 246 | 133 | 128 | 139 |
| RUF_107c.8_ | 59 | 71 | 63 | 22 | 23 | 33 |
| RUF_107c.81_ | 13 | 19 | 16 | 11 | 19 | 13 |
| RUF_107c.82_ | 972 | 976 | 953 | 746 | 626 | 803 |
| RUF_107c.83_ | 11 | 5 | 6 | 12 | 14 | 12 |
| RUF_107c.86_ | 66 | 102 | 92 | 106 | 85 | 123 |
| RUF_107c.89_ | 381 | 434 | 517 | 468 | 436 | 419 |
| RUF_107c.90_ | 39 | 26 | 27 | 18 | 25 | 10 |
| RUF_107c.92_ | 81 | 59 | 66 | 62 | 60 | 80 |
| RUF_107c.94_ | 22 | 24 | 17 | 15 | 21 | 17 |
| RUF_107c.95_ | 1363 | 1482 | 1453 | 1404 | 1365 | 1503 |
| RUF_107c.96_ | 124 | 132 | 207 | 120 | 119 | 121 |
| RUF_111_ | 84 | 92 | 90 | 89 | 79 | 100 |
| RUF_121_ | 68 | 88 | 63 | 128 | 119 | 118 |
| RUF_125_ | 343 | 337 | 280 | 1791 | 1592 | 1708 |
| RUF_167_ | 188 | 175 | 177 | 102 | 119 | 95 |
| RUF_178_ | 12 | 7 | 4 | 7 | 3 | 2 |
| RUF_219c_ | 55 | 48 | 42 | 272 | 305 | 231 |
| RUF_220c_ | 91 | 92 | 102 | 149 | 170 | 175 |
| RUF_226_ | 28 | 22 | 16 | 17 | 22 | 20 |
| RUF_247c_ | 441 | 420 | 386 | 384 | 354 | 370 |
| RUF_257c_ | 343 | 320 | 272 | 270 | 240 | 303 |
| RUF_279_ | 1261 | 1243 | 1122 | 705 | 657 | 782 |
| RUF_290_ | 608 | 589 | 587 | 643 | 585 | 670 |
| RUF_321c_ | 0 | 0 | 0 | 0 | 1 | 1 |
| RUF_328c_ | 7 | 4 | 4 | 1 | 4 | 4 |
| RUF_341c.10_ | 141 | 124 | 80 | 76 | 81 | 86 |
| RUF_341c.11_ | 128 | 120 | 138 | 112 | 112 | 114 |
| RUF_341c.2_ | 31 | 37 | 32 | 26 | 30 | 20 |
| SL1344_RS00005_ | 2989 | 2972 | 2447 | 4307 | 4046 | 3808 |
| SL1344_RS00010_ | 855 | 1063 | 958 | 1266 | 1039 | 1152 |
| SL1344_RS00015_ | 625 | 670 | 652 | 828 | 704 | 765 |
| SL1344_RS00020_ | 1083 | 1356 | 1496 | 1569 | 1301 | 1492 |
| SL1344_RS00025_ | 518 | 499 | 482 | 545 | 521 | 512 |
| SL1344_RS00030_ | 524 | 449 | 466 | 465 | 408 | 445 |
| SL1344_RS00035_ | 6819 | 7486 | 7844 | 7839 | 7344 | 7737 |
| SL1344_RS00040_ | 860 | 770 | 828 | 697 | 660 | 664 |
| SL1344_RS00045_ | 2721 | 2912 | 3199 | 1596 | 1536 | 1697 |

| | | | | | | |
|---|---|---|---|---|---|---|
| SL1344_RS00050_ | 16 | 39 | 17 | 22 | 24 | 23 |
| SL1344_RS00055_ | 35 | 25 | 20 | 27 | 30 | 19 |
| SL1344_RS00065_ | 2819 | 3198 | 3020 | 3944 | 3323 | 3370 |
| SL1344_RS00070_ | 116 | 128 | 145 | 114 | 89 | 107 |
| SL1344_RS00075_ | 13 | 14 | 8 | 5 | 5 | 9 |
| SL1344_RS00080_ | 17 | 18 | 8 | 14 | 17 | 18 |
| SL1344_RS00085_ | 0 | 5 | 4 | 8 | 3 | 2 |
| SL1344_RS00090_ | 173 | 169 | 158 | 183 | 186 | 197 |
| SL1344_RS00095_ | 58 | 76 | 69 | 91 | 89 | 100 |
| SL1344_RS00100_ | 18 | 31 | 29 | 33 | 40 | 37 |
| SL1344_RS00105_ | 27 | 29 | 39 | 30 | 29 | 39 |
| SL1344_RS00110_ | 15 | 6 | 14 | 19 | 12 | 15 |
| SL1344_RS00115_ | 53 | 67 | 44 | 59 | 45 | 87 |
| SL1344_RS00120_ | 30 | 46 | 27 | 47 | 37 | 38 |
| SL1344_RS00125_ | 17 | 16 | 16 | 15 | 20 | 15 |
| SL1344_RS00130_ | 12 | 9 | 19 | 14 | 14 | 17 |
| SL1344_RS00135_ | 31 | 21 | 30 | 26 | 20 | 28 |
| SL1344_RS00140_ | 82 | 77 | 93 | 78 | 64 | 76 |
| SL1344_RS00145_ | 78 | 77 | 62 | 53 | 83 | 75 |
| SL1344_RS00150_ | 7 | 5 | 4 | 6 | 5 | 6 |
| SL1344_RS00155_ | 32 | 24 | 36 | 23 | 46 | 31 |
| SL1344_RS00160_ | 9 | 17 | 9 | 8 | 6 | 4 |
| SL1344_RS00165_ | 54 | 83 | 74 | 73 | 56 | 51 |
| SL1344_RS00170_ | 118 | 132 | 113 | 122 | 116 | 126 |
| SL1344_RS00175_ | 1 | 2 | 3 | 2 | 7 | 3 |
| SL1344_RS00180_ | 132 | 110 | 99 | 129 | 115 | 108 |
| SL1344_RS00185_ | 96 | 102 | 74 | 92 | 85 | 77 |
| SL1344_RS00190_ | 43 | 55 | 54 | 42 | 47 | 38 |
| SL1344_RS00195_ | 232 | 202 | 214 | 206 | 209 | 193 |
| SL1344_RS00205_ | 500 | 594 | 540 | 722 | 690 | 797 |
| SL1344_RS00210_ | 137 | 156 | 150 | 162 | 125 | 148 |
| SL1344_RS00215_ | 8 | 15 | 7 | 6 | 16 | 9 |
| SL1344_RS00220_ | 59553 | 57938 | 54758 | 50060 | 52187 | 50330 |
| SL1344_RS00225_ | 112 | 112 | 124 | 146 | 134 | 127 |
| SL1344_RS00230_ | 1185 | 1432 | 1408 | 1391 | 1289 | 1491 |
| SL1344_RS00235_ | 4674 | 5205 | 6017 | 5544 | 5362 | 5486 |
| SL1344_RS00240_ | 1313 | 1304 | 1532 | 1225 | 1176 | 1243 |
| SL1344_RS00245_ | 730 | 795 | 894 | 743 | 782 | 796 |
| SL1344_RS00250_ | 1686 | 1848 | 1820 | 1683 | 1596 | 1784 |
| SL1344_RS00255_ | 519 | 467 | 436 | 500 | 568 | 477 |
| SL1344_RS00260_ | 435 | 468 | 447 | 646 | 700 | 623 |
| SL1344_RS00270_ | 65 | 86 | 94 | 76 | 80 | 81 |
| SL1344_RS00275_ | 11 | 8 | 5 | 9 | 5 | 17 |
| SL1344_RS00280_ | 7 | 6 | 13 | 8 | 1 | 4 |
| SL1344_RS00285_ | 2 | 1 | 0 | 1 | 1 | 0 |
| SL1344_RS00290_ | 27 | 22 | 18 | 31 | 27 | 28 |
| SL1344_RS00295_ | 12 | 9 | 19 | 18 | 13 | 12 |
| SL1344_RS00300_ | 4 | 6 | 5 | 3 | 4 | 9 |
| SL1344_RS00305_ | 21 | 17 | 8 | 17 | 36 | 20 |
| SL1344_RS00310_ | 29 | 27 | 27 | 34 | 48 | 44 |
| SL1344_RS00315_ | 10 | 13 | 14 | 30 | 31 | 41 |
| SL1344_RS00320_ | 27 | 35 | 37 | 59 | 38 | 58 |
| SL1344_RS00325_ | 595 | 763 | 742 | 867 | 720 | 816 |
| SL1344_RS00330_ | 14 | 12 | 9 | 8 | 9 | 10 |
| SL1344_RS00335_ | 277 | 321 | 283 | 261 | 298 | 344 |
| SL1344_RS00340_ | 794 | 778 | 853 | 759 | 754 | 770 |
| SL1344_RS00345_ | 141 | 125 | 139 | 87 | 68 | 83 |
| SL1344_RS00350_ | 15 | 12 | 13 | 20 | 25 | 23 |
| SL1344_RS00355_ | 21 | 23 | 20 | 37 | 22 | 37 |
| SL1344_RS00360_ | 40 | 40 | 37 | 65 | 46 | 56 |
| SL1344_RS00365_ | 35 | 39 | 44 | 53 | 51 | 54 |
| SL1344_RS00370_ | 62 | 67 | 94 | 72 | 77 | 85 |

| | | | | | | |
|---|---|---|---|---|---|---|
| SL1344_RS00375_ | 15 | 10 | 6 | 11 | 18 | 5 |
| SL1344_RS00380_ | 2 | 2 | 2 | 2 | 0 | 7 |
| SL1344_RS00385_ | 22 | 40 | 38 | 38 | 26 | 28 |
| SL1344_RS00390_ | 11 | 14 | 32 | 23 | 23 | 24 |
| SL1344_RS00395_ | 19 | 26 | 23 | 33 | 21 | 26 |
| SL1344_RS00400_ | 16 | 10 | 11 | 14 | 13 | 10 |
| SL1344_RS00405_ | 129 | 107 | 131 | 127 | 97 | 105 |
| SL1344_RS00410_ | 31 | 24 | 31 | 24 | 39 | 33 |
| SL1344_RS00415_ | 152 | 142 | 151 | 122 | 119 | 110 |
| SL1344_RS00420_ | 35 | 35 | 38 | 42 | 49 | 41 |
| SL1344_RS00425_ | 0 | 5 | 5 | 4 | 1 | 3 |
| SL1344_RS00430_ | 199 | 211 | 168 | 175 | 151 | 160 |
| SL1344_RS00435_ | 32 | 30 | 30 | 30 | 20 | 21 |
| SL1344_RS00440_ | 74 | 119 | 115 | 124 | 114 | 117 |
| SL1344_RS00450_ | 862 | 960 | 1057 | 828 | 840 | 869 |
| SL1344_RS00455_ | 855 | 992 | 1088 | 818 | 749 | 906 |
| SL1344_RS00465_ | 1555 | 1678 | 1699 | 1406 | 1276 | 1515 |
| SL1344_RS00470_ | 2968 | 3340 | 3106 | 2780 | 2499 | 2715 |
| SL1344_RS00475_ | 2571 | 2835 | 2835 | 2766 | 2573 | 2747 |
| SL1344_RS00480_ | 428 | 522 | 533 | 433 | 416 | 479 |
| SL1344_RS00485_ | 1227 | 1250 | 1242 | 899 | 936 | 950 |
| SL1344_RS00490_ | 2289 | 2393 | 2514 | 1918 | 1872 | 2021 |
| SL1344_RS00495_ | 519 | 425 | 472 | 479 | 471 | 434 |
| SL1344_RS00500_ | 76 | 67 | 68 | 71 | 48 | 56 |
| SL1344_RS00505_ | 13 | 22 | 7 | 6 | 14 | 4 |
| SL1344_RS00510_ | 17 | 14 | 15 | 19 | 16 | 22 |
| SL1344_RS00515_ | 12 | 30 | 22 | 29 | 22 | 22 |
| SL1344_RS00520_ | 10 | 14 | 11 | 17 | 12 | 20 |
| SL1344_RS00525_ | 166 | 185 | 185 | 179 | 174 | 139 |
| SL1344_RS00530_ | 370 | 473 | 459 | 367 | 390 | 444 |
| SL1344_RS00545_ | 217 | 335 | 298 | 319 | 231 | 306 |
| SL1344_RS00550_ | 182 | 181 | 173 | 155 | 156 | 187 |
| SL1344_RS00555_ | 922 | 816 | 808 | 1184 | 957 | 1026 |
| SL1344_RS00560_ | 56 | 70 | 81 | 91 | 91 | 79 |
| SL1344_RS00565_ | 99 | 118 | 146 | 138 | 161 | 116 |
| SL1344_RS00570_ | 116 | 138 | 102 | 127 | 149 | 101 |
| SL1344_RS00575_ | 99 | 107 | 121 | 96 | 112 | 108 |
| SL1344_RS00580_ | 131 | 122 | 108 | 104 | 78 | 124 |
| SL1344_RS00590_ | 162 | 177 | 184 | 187 | 162 | 202 |
| SL1344_RS00595_ | 151 | 175 | 171 | 233 | 228 | 232 |
| SL1344_RS00600_ | 851 | 901 | 1001 | 823 | 810 | 755 |
| SL1344_RS00605_ | 3155 | 3448 | 3089 | 3063 | 2893 | 3025 |
| SL1344_RS00610_ | 3055 | 3391 | 3419 | 3251 | 3054 | 3214 |
| SL1344_RS00615_ | 1691 | 1831 | 2007 | 1871 | 1662 | 1760 |
| SL1344_RS00620_ | 6989 | 7192 | 6180 | 7259 | 6974 | 7508 |
| SL1344_RS00630_ | 1710 | 1914 | 1976 | 1615 | 1578 | 1753 |
| SL1344_RS00635_ | 1735 | 1936 | 2004 | 1705 | 1550 | 1791 |
| SL1344_RS00645_ | 2018 | 2229 | 2220 | 1967 | 1889 | 2051 |
| SL1344_RS00650_ | 1707 | 1760 | 1871 | 1507 | 1438 | 1611 |
| SL1344_RS00655_ | 2037 | 2256 | 2352 | 2069 | 1979 | 2157 |
| SL1344_RS00665_ | 1049 | 1236 | 1195 | 1130 | 1001 | 1179 |
| SL1344_RS00670_ | 2297 | 2511 | 2531 | 2344 | 2135 | 2548 |
| SL1344_RS00675_ | 2810 | 3085 | 3334 | 3158 | 3011 | 3117 |
| SL1344_RS00680_ | 10181 | 10904 | 11514 | 10874 | 9884 | 11412 |
| SL1344_RS00685_ | 1391 | 1410 | 1245 | 1327 | 1337 | 1268 |
| SL1344_RS00690_ | 6138 | 6270 | 6514 | 5446 | 5455 | 5488 |
| SL1344_RS00695_ | 92 | 146 | 139 | 141 | 105 | 133 |
| SL1344_RS00700_ | 580 | 646 | 576 | 584 | 551 | 614 |
| SL1344_RS00705_ | 447 | 519 | 501 | 541 | 499 | 480 |
| SL1344_RS00710_ | 609 | 631 | 676 | 676 | 599 | 653 |
| SL1344_RS00715_ | 628 | 617 | 595 | 523 | 600 | 622 |
| SL1344_RS00720_ | 1914 | 2093 | 2048 | 2421 | 2109 | 2407 |

| | | | | | | |
|---|---|---|---|---|---|---|
| SL1344_RS00725_ | 22 | 16 | 21 | 29 | 22 | 21 |
| SL1344_RS00730_ | 8 | 6 | 8 | 14 | 12 | 8 |
| SL1344_RS00735_ | 2 | 5 | 3 | 0 | 2 | 6 |
| SL1344_RS00740_ | 453 | 439 | 396 | 410 | 340 | 396 |
| SL1344_RS00745_ | 264 | 285 | 254 | 230 | 218 | 238 |
| SL1344_RS00750_ | 416 | 423 | 448 | 475 | 437 | 505 |
| SL1344_RS00755_ | 90 | 96 | 101 | 98 | 94 | 96 |
| SL1344_RS00760_ | 42 | 45 | 36 | 42 | 44 | 42 |
| SL1344_RS00765_ | 359 | 393 | 430 | 365 | 359 | 393 |
| SL1344_RS00770_ | 6642 | 6759 | 6572 | 5659 | 5636 | 5646 |
| SL1344_RS00775_ | 18742 | 20015 | 21218 | 18524 | 17920 | 18403 |
| SL1344_RS00785_ | 16574 | 17340 | 18028 | 19561 | 18590 | 19103 |
| SL1344_RS00790_ | 94 | 79 | 98 | 92 | 77 | 99 |
| SL1344_RS00795_ | 18 | 25 | 21 | 25 | 28 | 14 |
| SL1344_RS00800_ | 58 | 85 | 81 | 79 | 65 | 79 |
| SL1344_RS00805_ | 1372 | 1777 | 1890 | 1893 | 1878 | 2025 |
| SL1344_RS00810_ | 736 | 558 | 499 | 719 | 659 | 665 |
| SL1344_RS00815_ | 398 | 505 | 467 | 546 | 563 | 495 |
| SL1344_RS00820_ | 41 | 40 | 52 | 62 | 40 | 41 |
| SL1344_RS00825_ | 59 | 73 | 44 | 82 | 68 | 62 |
| SL1344_RS00830_ | 192 | 215 | 216 | 215 | 209 | 221 |
| SL1344_RS00835_ | 866 | 868 | 941 | 868 | 850 | 922 |
| SL1344_RS00840_ | 2621 | 2666 | 2714 | 2737 | 2504 | 2598 |
| SL1344_RS00845_ | 1442 | 1450 | 1373 | 1494 | 1306 | 1447 |
| SL1344_RS00850_ | 226 | 205 | 224 | 212 | 181 | 233 |
| SL1344_RS00855_ | 323 | 587 | 424 | 534 | 518 | 419 |
| SL1344_RS00860_ | 393 | 438 | 446 | 634 | 582 | 614 |
| SL1344_RS00865_ | 1495 | 1676 | 1605 | 1726 | 1731 | 1630 |
| SL1344_RS00870_ | 7309 | 7115 | 6635 | 6174 | 5914 | 5995 |
| SL1344_RS00875_ | 188 | 212 | 208 | 217 | 203 | 234 |
| SL1344_RS00880_ | 257 | 293 | 355 | 287 | 266 | 290 |
| SL1344_RS00885_ | 54 | 72 | 59 | 52 | 49 | 45 |
| SL1344_RS00890_ | 42 | 44 | 48 | 37 | 31 | 27 |
| SL1344_RS00895_ | 6 | 9 | 8 | 8 | 6 | 7 |
| SL1344_RS00900_ | 37 | 17 | 18 | 24 | 24 | 18 |
| SL1344_RS00905_ | 20 | 24 | 23 | 22 | 18 | 13 |
| SL1344_RS00910_ | 401 | 411 | 408 | 414 | 395 | 384 |
| SL1344_RS00915_ | 906 | 911 | 941 | 1007 | 846 | 969 |
| SL1344_RS00920_ | 827 | 784 | 747 | 761 | 666 | 758 |
| SL1344_RS00925_ | 484 | 540 | 632 | 501 | 511 | 538 |
| SL1344_RS00930_ | 1036 | 1062 | 1104 | 884 | 904 | 1046 |
| SL1344_RS00935_ | 4117 | 4088 | 4014 | 3730 | 3607 | 3908 |
| SL1344_RS00940_ | 581 | 578 | 615 | 533 | 508 | 561 |
| SL1344_RS00945_ | 7216 | 8264 | 7819 | 11924 | 11133 | 11384 |
| SL1344_RS00950_ | 315 | 314 | 279 | 293 | 259 | 255 |
| SL1344_RS00955_ | 269 | 238 | 255 | 249 | 242 | 240 |
| SL1344_RS00960_ | 923 | 992 | 941 | 990 | 949 | 976 |
| SL1344_RS00965_ | 2140 | 2319 | 2378 | 2299 | 2227 | 2363 |
| SL1344_RS00970_ | 5186 | 6495 | 6513 | 5095 | 4863 | 5281 |
| SL1344_RS00975_ | 696 | 859 | 1025 | 820 | 782 | 826 |
| SL1344_RS00980_ | 819 | 1033 | 1187 | 813 | 866 | 834 |
| SL1344_RS00985_ | 1886 | 2114 | 2284 | 1594 | 1580 | 1698 |
| SL1344_RS00990_ | 28 | 31 | 32 | 20 | 25 | 25 |
| SL1344_RS00995_ | 82 | 64 | 59 | 70 | 46 | 58 |
| SL1344_RS01000_ | 13 | 14 | 15 | 26 | 15 | 12 |
| SL1344_RS01005_ | 16 | 26 | 11 | 17 | 17 | 16 |
| SL1344_RS01010_ | 14 | 10 | 17 | 30 | 23 | 26 |
| SL1344_RS01015_ | 9 | 17 | 9 | 34 | 20 | 15 |
| SL1344_RS01020_ | 137 | 156 | 154 | 161 | 146 | 169 |
| SL1344_RS01025_ | 1328 | 1495 | 1614 | 1539 | 1583 | 1610 |
| SL1344_RS01030_ | 269 | 292 | 282 | 321 | 290 | 301 |
| SL1344_RS01035_ | 1970 | 2637 | 2668 | 2771 | 2612 | 2746 |

| | | | | | | |
|---|---|---|---|---|---|---|
| SL1344_RS01040_ | 325 | 315 | 325 | 348 | 312 | 316 |
| SL1344_RS01045_ | 435 | 473 | 462 | 502 | 476 | 469 |
| SL1344_RS01050_ | 1042 | 1107 | 1080 | 1289 | 1376 | 1212 |
| SL1344_RS01055_ | 171 | 209 | 198 | 153 | 164 | 189 |
| SL1344_RS01060_ | 1447 | 1470 | 1367 | 1262 | 1228 | 1133 |
| SL1344_RS01065_ | 47 | 68 | 64 | 71 | 68 | 76 |
| SL1344_RS01070_ | 657 | 653 | 653 | 815 | 828 | 783 |
| SL1344_RS01075_ | 328 | 335 | 303 | 334 | 299 | 343 |
| SL1344_RS01080_ | 2220 | 2331 | 2166 | 2421 | 2425 | 2329 |
| SL1344_RS01085_ | 692 | 625 | 709 | 605 | 608 | 592 |
| SL1344_RS01090_ | 2400 | 2225 | 2017 | 2032 | 2094 | 2013 |
| SL1344_RS01095_ | 41180 | 37900 | 38603 | 41757 | 40333 | 41545 |
| SL1344_RS01100_ | 50781 | 53227 | 55287 | 52276 | 50338 | 51976 |
| SL1344_RS01105_ | 2354 | 2582 | 2767 | 2733 | 2597 | 2720 |
| SL1344_RS01110_ | 9525 | 10192 | 9340 | 10590 | 10613 | 10111 |
| SL1344_RS01115_ | 1523 | 1544 | 1696 | 1450 | 1481 | 1572 |
| SL1344_RS01120_ | 1063 | 1346 | 1462 | 1345 | 1242 | 1461 |
| SL1344_RS01130_ | 4429 | 4926 | 5300 | 4847 | 4410 | 4906 |
| SL1344_RS01135_ | 7569 | 8258 | 8434 | 8330 | 7904 | 8473 |
| SL1344_RS01140_ | 8064 | 8251 | 7943 | 9431 | 9124 | 8943 |
| SL1344_RS01145_ | 7908 | 8455 | 9110 | 8917 | 9040 | 9149 |
| SL1344_RS01150_ | 1546 | 1656 | 1748 | 1658 | 1645 | 1704 |
| SL1344_RS01155_ | 3812 | 4287 | 4395 | 4438 | 4543 | 4528 |
| SL1344_RS01160_ | 3709 | 4159 | 4875 | 4433 | 4381 | 4502 |
| SL1344_RS01165_ | 1086 | 1149 | 1192 | 989 | 1040 | 1036 |
| SL1344_RS01170_ | 2973 | 3461 | 3681 | 3164 | 3111 | 3557 |
| SL1344_RS01175_ | 2074 | 1947 | 2018 | 1924 | 1843 | 1952 |
| SL1344_RS01180_ | 263 | 291 | 256 | 326 | 283 | 348 |
| SL1344_RS01185_ | 423 | 435 | 447 | 462 | 376 | 430 |
| SL1344_RS01190_ | 144 | 163 | 167 | 156 | 159 | 162 |
| SL1344_RS01195_ | 506 | 528 | 521 | 415 | 437 | 509 |
| SL1344_RS01200_ | 588 | 589 | 609 | 743 | 684 | 690 |
| SL1344_RS01205_ | 459 | 555 | 430 | 641 | 548 | 610 |
| SL1344_RS01210_ | 255 | 253 | 266 | 265 | 259 | 270 |
| SL1344_RS01215_ | 244 | 228 | 216 | 229 | 218 | 239 |
| SL1344_RS01220_ | 373 | 424 | 431 | 455 | 436 | 442 |
| SL1344_RS01225_ | 4493 | 4887 | 4978 | 4830 | 4655 | 4883 |
| SL1344_RS01230_ | 892 | 892 | 991 | 889 | 818 | 787 |
| SL1344_RS01245_ | 419 | 455 | 439 | 511 | 487 | 499 |
| SL1344_RS01250_ | 831 | 861 | 723 | 842 | 676 | 765 |
| SL1344_RS01255_ | 203 | 214 | 193 | 187 | 194 | 201 |
| SL1344_RS01260_ | 383 | 207 | 147 | 121 | 113 | 166 |
| SL1344_RS01265_ | 4980 | 4666 | 5005 | 4568 | 4766 | 4856 |
| SL1344_RS01270_ | 2361 | 2364 | 2699 | 2602 | 2618 | 3000 |
| SL1344_RS01275_ | 629 | 451 | 463 | 357 | 321 | 453 |
| SL1344_RS01285_ | 127 | 120 | 169 | 154 | 118 | 150 |
| SL1344_RS01290_ | 254 | 317 | 321 | 301 | 285 | 343 |
| SL1344_RS01295_ | 60 | 74 | 94 | 88 | 71 | 77 |
| SL1344_RS01300_ | 59 | 76 | 72 | 57 | 84 | 73 |
| SL1344_RS01305_ | 428 | 441 | 398 | 402 | 360 | 339 |
| SL1344_RS01310_ | 167 | 168 | 171 | 164 | 161 | 178 |
| SL1344_RS01315_ | 4629 | 4608 | 4481 | 6178 | 6063 | 5898 |
| SL1344_RS01320_ | 167 | 156 | 153 | 134 | 149 | 150 |
| SL1344_RS01325_ | 147 | 138 | 121 | 113 | 144 | 167 |
| SL1344_RS01330_ | 357 | 412 | 331 | 303 | 305 | 299 |
| SL1344_RS01335_ | 671 | 656 | 572 | 696 | 588 | 609 |
| SL1344_RS01340_ | 400 | 477 | 446 | 489 | 399 | 430 |
| SL1344_RS01345_ | 32 | 30 | 34 | 40 | 37 | 33 |
| SL1344_RS01350_ | 13 | 27 | 16 | 20 | 14 | 18 |
| SL1344_RS01355_ | 36 | 44 | 28 | 46 | 45 | 51 |
| SL1344_RS01360_ | 9 | 10 | 18 | 17 | 13 | 17 |
| SL1344_RS01365_ | 39 | 35 | 33 | 32 | 32 | 30 |

| | | | | | | |
|---|---|---|---|---|---|---|
| SL1344_RS01370_ | 15 | 15 | 10 | 17 | 12 | 10 |
| SL1344_RS01375_ | 0 | 0 | 1 | 0 | 0 | 0 |
| SL1344_RS01380_ | 167 | 156 | 175 | 166 | 125 | 153 |
| SL1344_RS01385_ | 14 | 36 | 35 | 35 | 28 | 24 |
| SL1344_RS01390_ | 51 | 50 | 57 | 81 | 63 | 61 |
| SL1344_RS01395_ | 29 | 30 | 28 | 32 | 28 | 34 |
| SL1344_RS01400_ | 7 | 16 | 19 | 23 | 9 | 17 |
| SL1344_RS01405_ | 37 | 33 | 22 | 18 | 21 | 14 |
| SL1344_RS01410_ | 8 | 9 | 11 | 13 | 10 | 9 |
| SL1344_RS01415_ | 12 | 22 | 21 | 15 | 14 | 18 |
| SL1344_RS01420_ | 6 | 11 | 8 | 14 | 13 | 9 |
| SL1344_RS01425_ | 32 | 27 | 51 | 40 | 37 | 24 |
| SL1344_RS01430_ | 35 | 36 | 41 | 66 | 53 | 44 |
| SL1344_RS01435_ | 32 | 17 | 12 | 24 | 18 | 10 |
| SL1344_RS01440_ | 4 | 8 | 6 | 33 | 5 | 10 |
| SL1344_RS01445_ | 122 | 140 | 160 | 159 | 133 | 133 |
| SL1344_RS01450_ | 19 | 18 | 14 | 22 | 15 | 29 |
| SL1344_RS01455_ | 6 | 10 | 6 | 8 | 14 | 8 |
| SL1344_RS01460_ | 26 | 48 | 39 | 48 | 53 | 49 |
| SL1344_RS01465_ | 88 | 85 | 86 | 75 | 71 | 84 |
| SL1344_RS01470_ | 9 | 16 | 15 | 12 | 11 | 16 |
| SL1344_RS01475_ | 268 | 274 | 225 | 319 | 271 | 316 |
| SL1344_RS01480_ | 410 | 405 | 383 | 533 | 432 | 476 |
| SL1344_RS01485_ | 221 | 237 | 201 | 253 | 233 | 264 |
| SL1344_RS01490_ | 165 | 160 | 111 | 161 | 141 | 159 |
| SL1344_RS01495_ | 69 | 80 | 62 | 50 | 61 | 56 |
| SL1344_RS01500_ | 2 | 1 | 4 | 2 | 1 | 3 |
| SL1344_RS01505_ | 39 | 30 | 38 | 48 | 35 | 40 |
| SL1344_RS01515_ | 13 | 12 | 21 | 12 | 14 | 17 |
| SL1344_RS01520_ | 5 | 9 | 1 | 6 | 5 | 4 |
| SL1344_RS01525_ | 93 | 118 | 69 | 114 | 162 | 126 |
| SL1344_RS01530_ | 31 | 30 | 31 | 37 | 36 | 33 |
| SL1344_RS01535_ | 139 | 161 | 141 | 160 | 140 | 187 |
| SL1344_RS01540_ | 37 | 50 | 43 | 42 | 40 | 53 |
| SL1344_RS01545_ | 34 | 33 | 25 | 39 | 29 | 37 |
| SL1344_RS01550_ | 17 | 16 | 24 | 17 | 21 | 22 |
| SL1344_RS01565_ | 30 | 28 | 30 | 28 | 35 | 36 |
| SL1344_RS01570_ | 110 | 131 | 139 | 99 | 140 | 128 |
| SL1344_RS01575_ | 103 | 110 | 109 | 88 | 84 | 88 |
| SL1344_RS01580_ | 148 | 152 | 175 | 166 | 168 | 174 |
| SL1344_RS01590_ | 818 | 867 | 813 | 789 | 827 | 724 |
| SL1344_RS01595_ | 583 | 704 | 749 | 645 | 613 | 682 |
| SL1344_RS01600_ | 821 | 970 | 906 | 938 | 925 | 927 |
| SL1344_RS01605_ | 359 | 394 | 381 | 420 | 395 | 417 |
| SL1344_RS01610_ | 113 | 93 | 134 | 102 | 103 | 127 |
| SL1344_RS01615_ | 133 | 122 | 137 | 119 | 128 | 124 |
| SL1344_RS01620_ | 2156 | 2086 | 2191 | 2191 | 2064 | 2140 |
| SL1344_RS01625_ | 2782 | 2631 | 2798 | 2016 | 2030 | 2261 |
| SL1344_RS01635_ | 639 | 755 | 765 | 1120 | 992 | 1012 |
| SL1344_RS01640_ | 77 | 63 | 59 | 64 | 54 | 62 |
| SL1344_RS01645_ | 703 | 747 | 731 | 666 | 659 | 755 |
| SL1344_RS01650_ | 1860 | 1956 | 2125 | 1903 | 1780 | 2009 |
| SL1344_RS01655_ | 302 | 320 | 301 | 225 | 209 | 237 |
| SL1344_RS01660_ | 12 | 12 | 10 | 18 | 8 | 6 |
| SL1344_RS01665_ | 3 | 5 | 7 | 4 | 4 | 3 |
| SL1344_RS01670_ | 16 | 7 | 9 | 12 | 11 | 8 |
| SL1344_RS01675_ | 128 | 127 | 124 | 166 | 124 | 112 |
| SL1344_RS01680_ | 18 | 20 | 19 | 13 | 9 | 13 |
| SL1344_RS01685_ | 18 | 26 | 23 | 27 | 21 | 29 |
| SL1344_RS01690_ | 5 | 12 | 13 | 12 | 6 | 9 |
| SL1344_RS01695_ | 41 | 32 | 33 | 64 | 44 | 48 |
| SL1344_RS01700_ | 88 | 93 | 98 | 141 | 105 | 141 |

| | | | | | | |
|---|---|---|---|---|---|---|
| SL1344_RS01705_ | 148 | 139 | 135 | 158 | 152 | 161 |
| SL1344_RS01710_ | 41 | 31 | 27 | 35 | 42 | 27 |
| SL1344_RS01715_ | 16 | 19 | 16 | 13 | 12 | 19 |
| SL1344_RS01720_ | 80 | 72 | 71 | 74 | 71 | 64 |
| SL1344_RS01725_ | 63 | 58 | 66 | 52 | 65 | 52 |
| SL1344_RS01730_ | 55 | 54 | 56 | 72 | 48 | 64 |
| SL1344_RS01740_ | 3 | 1 | 1 | 1 | 2 | 5 |
| SL1344_RS01745_ | 125 | 137 | 104 | 165 | 171 | 142 |
| SL1344_RS01750_ | 84 | 94 | 74 | 88 | 80 | 93 |
| SL1344_RS01755_ | 113 | 106 | 118 | 124 | 112 | 106 |
| SL1344_RS01760_ | 296 | 377 | 329 | 376 | 310 | 348 |
| SL1344_RS01765_ | 191 | 209 | 198 | 211 | 188 | 210 |
| SL1344_RS01770_ | 15 | 18 | 16 | 9 | 15 | 25 |
| SL1344_RS01775_ | 13 | 17 | 18 | 14 | 19 | 15 |
| SL1344_RS01780_ | 27 | 21 | 22 | 45 | 42 | 15 |
| SL1344_RS01785_ | 80 | 80 | 92 | 64 | 62 | 59 |
| SL1344_RS01790_ | 13 | 28 | 28 | 37 | 27 | 11 |
| SL1344_RS01795_ | 31 | 33 | 47 | 32 | 27 | 37 |
| SL1344_RS01800_ | 5 | 7 | 11 | 7 | 8 | 6 |
| SL1344_RS01805_ | 41 | 43 | 33 | 46 | 48 | 46 |
| SL1344_RS01810_ | 13 | 36 | 17 | 22 | 27 | 17 |
| SL1344_RS01815_ | 27 | 48 | 30 | 48 | 43 | 35 |
| SL1344_RS01820_ | 27 | 16 | 10 | 30 | 16 | 23 |
| SL1344_RS01825_ | 152 | 174 | 135 | 178 | 188 | 195 |
| SL1344_RS01830_ | 614 | 739 | 769 | 719 | 647 | 800 |
| SL1344_RS01835_ | 1433 | 1510 | 1590 | 1422 | 1346 | 1488 |
| SL1344_RS01840_ | 27 | 14 | 15 | 31 | 21 | 16 |
| SL1344_RS01845_ | 47 | 60 | 55 | 64 | 54 | 71 |
| SL1344_RS01850_ | 131 | 144 | 155 | 167 | 162 | 142 |
| SL1344_RS01855_ | 29 | 19 | 27 | 34 | 35 | 26 |
| SL1344_RS01860_ | 532 | 491 | 430 | 436 | 410 | 397 |
| SL1344_RS01865_ | 206 | 201 | 162 | 173 | 164 | 185 |
| SL1344_RS01870_ | 96 | 92 | 87 | 74 | 83 | 72 |
| SL1344_RS01875_ | 134 | 119 | 121 | 204 | 194 | 194 |
| SL1344_RS01880_ | 42 | 63 | 50 | 83 | 65 | 80 |
| SL1344_RS01885_ | 12 | 14 | 16 | 22 | 22 | 29 |
| SL1344_RS01890_ | 26 | 27 | 26 | 40 | 31 | 26 |
| SL1344_RS01895_ | 31 | 33 | 28 | 24 | 31 | 41 |
| SL1344_RS01900_ | 56 | 75 | 50 | 65 | 52 | 69 |
| SL1344_RS01905_ | 1957 | 1982 | 1989 | 1873 | 1794 | 1932 |
| SL1344_RS01910_ | 387 | 403 | 403 | 491 | 483 | 543 |
| SL1344_RS01915_ | 39 | 56 | 44 | 54 | 38 | 42 |
| SL1344_RS01930_ | 174 | 170 | 212 | 172 | 173 | 179 |
| SL1344_RS01935_ | 29 | 26 | 22 | 22 | 18 | 21 |
| SL1344_RS01940_ | 225 | 203 | 167 | 164 | 190 | 146 |
| SL1344_RS01945_ | 879 | 860 | 792 | 1029 | 952 | 904 |
| SL1344_RS01950_ | 18 | 23 | 19 | 25 | 25 | 19 |
| SL1344_RS01955_ | 91 | 72 | 96 | 84 | 107 | 105 |
| SL1344_RS01960_ | 758 | 762 | 705 | 1188 | 1147 | 983 |
| SL1344_RS01965_ | 92 | 56 | 45 | 49 | 51 | 64 |
| SL1344_RS01975_ | 718 | 847 | 798 | 888 | 863 | 912 |
| SL1344_RS01980_ | 217 | 184 | 223 | 225 | 214 | 197 |
| SL1344_RS01985_ | 543 | 633 | 663 | 451 | 438 | 504 |
| SL1344_RS01990_ | 708 | 698 | 719 | 509 | 530 | 492 |
| SL1344_RS01995_ | 390 | 401 | 366 | 297 | 260 | 316 |
| SL1344_RS02000_ | 561 | 596 | 589 | 626 | 613 | 585 |
| SL1344_RS02005_ | 918 | 1071 | 1115 | 992 | 956 | 965 |
| SL1344_RS02010_ | 110 | 133 | 122 | 132 | 142 | 145 |
| SL1344_RS02015_ | 249 | 253 | 212 | 209 | 209 | 241 |
| SL1344_RS02020_ | 818 | 889 | 944 | 818 | 824 | 834 |
| SL1344_RS02025_ | 343 | 382 | 360 | 337 | 326 | 353 |
| SL1344_RS02030_ | 4 | 0 | 1 | 3 | 2 | 3 |

| | | | | | | |
|---|---|---|---|---|---|---|
| SL1344_RS02045_ | 1915 | 2028 | 2239 | 2090 | 2011 | 2103 |
| SL1344_RS02050_ | 1915 | 2102 | 2092 | 1515 | 1483 | 1594 |
| SL1344_RS02055_ | 198 | 256 | 259 | 203 | 132 | 162 |
| SL1344_RS02060_ | 4178 | 4876 | 4611 | 5908 | 5388 | 5690 |
| SL1344_RS02065_ | 910 | 896 | 738 | 760 | 868 | 873 |
| SL1344_RS02070_ | 1368 | 1532 | 1526 | 1149 | 1111 | 1242 |
| SL1344_RS02075_ | 2514 | 2660 | 2650 | 2296 | 2252 | 2303 |
| SL1344_RS02080_ | 1465 | 1866 | 1867 | 1831 | 1679 | 1815 |
| SL1344_RS02085_ | 5281 | 5598 | 5796 | 4694 | 4826 | 5001 |
| SL1344_RS02090_ | 1575 | 2057 | 2411 | 2246 | 2174 | 2193 |
| SL1344_RS02095_ | 94 | 94 | 113 | 102 | 86 | 93 |
| SL1344_RS02100_ | 115 | 146 | 137 | 122 | 87 | 131 |
| SL1344_RS02105_ | 399 | 436 | 454 | 478 | 452 | 442 |
| SL1344_RS02110_ | 1115 | 1067 | 1325 | 1191 | 1147 | 1145 |
| SL1344_RS02115_ | 765 | 1096 | 1243 | 1075 | 922 | 1062 |
| SL1344_RS02120_ | 29 | 33 | 45 | 38 | 46 | 42 |
| SL1344_RS02125_ | 461 | 469 | 462 | 425 | 411 | 442 |
| SL1344_RS02130_ | 756 | 689 | 738 | 626 | 642 | 696 |
| SL1344_RS02135_ | 1272 | 1080 | 979 | 805 | 615 | 1048 |
| SL1344_RS02145_ | 487 | 514 | 553 | 391 | 343 | 455 |
| SL1344_RS02150_ | 465 | 524 | 569 | 367 | 424 | 437 |
| SL1344_RS02155_ | 313 | 309 | 351 | 245 | 274 | 293 |
| SL1344_RS02160_ | 1191 | 1274 | 1460 | 1222 | 1187 | 1292 |
| SL1344_RS02165_ | 1027 | 1035 | 1007 | 921 | 881 | 979 |
| SL1344_RS02170_ | 429 | 420 | 412 | 416 | 395 | 405 |
| SL1344_RS02175_ | 1331 | 1398 | 1539 | 1255 | 1226 | 1341 |
| SL1344_RS02180_ | 36 | 42 | 48 | 64 | 47 | 55 |
| SL1344_RS02185_ | 29 | 27 | 24 | 31 | 31 | 22 |
| SL1344_RS02190_ | 42 | 52 | 58 | 41 | 60 | 53 |
| SL1344_RS02195_ | 71 | 72 | 48 | 77 | 45 | 57 |
| SL1344_RS02200_ | 192 | 212 | 189 | 252 | 221 | 213 |
| SL1344_RS02205_ | 19 | 27 | 16 | 17 | 40 | 14 |
| SL1344_RS02210_ | 19 | 16 | 11 | 16 | 23 | 18 |
| SL1344_RS02215_ | 397 | 410 | 370 | 462 | 442 | 452 |
| SL1344_RS02220_ | 182 | 206 | 199 | 248 | 256 | 224 |
| SL1344_RS02225_ | 1659 | 1992 | 1980 | 2052 | 1895 | 2033 |
| SL1344_RS02230_ | 427 | 474 | 482 | 513 | 443 | 495 |
| SL1344_RS02235_ | 122 | 146 | 111 | 111 | 93 | 117 |
| SL1344_RS02240_ | 256 | 252 | 265 | 222 | 217 | 221 |
| SL1344_RS02245_ | 7683 | 8352 | 8539 | 8199 | 7891 | 7977 |
| SL1344_RS02250_ | 1602 | 1747 | 1801 | 1639 | 1608 | 1687 |
| SL1344_RS02255_ | 2427 | 2723 | 2747 | 2704 | 2504 | 2668 |
| SL1344_RS02260_ | 8999 | 10076 | 10944 | 9867 | 9652 | 9912 |
| SL1344_RS02265_ | 11040 | 11419 | 11876 | 10016 | 9834 | 9923 |
| SL1344_RS02270_ | 1353 | 1490 | 1465 | 1293 | 1245 | 1409 |
| SL1344_RS02275_ | 1037 | 974 | 872 | 936 | 1076 | 908 |
| SL1344_RS02280_ | 840 | 877 | 743 | 1108 | 1071 | 920 |
| SL1344_RS02285_ | 18023 | 19086 | 19528 | 19120 | 17854 | 18401 |
| SL1344_RS02300_ | 7205 | 7882 | 7982 | 10158 | 8521 | 9323 |
| SL1344_RS02305_ | 11144 | 15597 | 15143 | 19327 | 17535 | 19054 |
| SL1344_RS02310_ | 2079 | 2322 | 2396 | 2098 | 1974 | 2359 |
| SL1344_RS02315_ | 4 | 7 | 16 | 18 | 23 | 8 |
| SL1344_RS02320_ | 50 | 44 | 52 | 56 | 55 | 51 |
| SL1344_RS02325_ | 422 | 439 | 514 | 445 | 442 | 446 |
| SL1344_RS02330_ | 42 | 49 | 41 | 50 | 44 | 28 |
| SL1344_RS02335_ | 58 | 46 | 42 | 64 | 55 | 45 |
| SL1344_RS02350_ | 527 | 554 | 575 | 558 | 547 | 527 |
| SL1344_RS02355_ | 563 | 565 | 598 | 563 | 527 | 608 |
| SL1344_RS02360_ | 34 | 32 | 34 | 9 | 10 | 21 |
| SL1344_RS02365_ | 97 | 118 | 149 | 90 | 73 | 97 |
| SL1344_RS02370_ | 460 | 494 | 460 | 466 | 449 | 469 |
| SL1344_RS02375_ | 112 | 107 | 111 | 188 | 210 | 188 |

| | | | | | | |
|---|---|---|---|---|---|---|
| SL1344_RS02380_ | 131 | 109 | 114 | 116 | 141 | 110 |
| SL1344_RS02385_ | 414 | 420 | 360 | 440 | 358 | 383 |
| SL1344_RS02390_ | 88 | 55 | 58 | 51 | 51 | 56 |
| SL1344_RS02395_ | 151 | 140 | 140 | 114 | 138 | 128 |
| SL1344_RS02400_ | 1505 | 1623 | 1654 | 2489 | 2196 | 2467 |
| SL1344_RS02405_ | 1283 | 1155 | 1255 | 960 | 893 | 938 |
| SL1344_RS02420_ | 5966 | 6305 | 6789 | 6008 | 5792 | 5975 |
| SL1344_RS02425_ | 2784 | 2694 | 2651 | 2319 | 2343 | 2342 |
| SL1344_RS02430_ | 559 | 544 | 520 | 571 | 562 | 525 |
| SL1344_RS02435_ | 2006 | 2128 | 2129 | 1844 | 1767 | 1998 |
| SL1344_RS02440_ | 373 | 399 | 407 | 445 | 453 | 487 |
| SL1344_RS02445_ | 134 | 174 | 143 | 163 | 184 | 162 |
| SL1344_RS02450_ | 292 | 267 | 250 | 285 | 300 | 274 |
| SL1344_RS02455_ | 234 | 260 | 235 | 254 | 226 | 229 |
| SL1344_RS02460_ | 1606 | 1870 | 1981 | 1844 | 1773 | 1891 |
| SL1344_RS02465_ | 3243 | 3235 | 3453 | 2717 | 2762 | 2855 |
| SL1344_RS02470_ | 2728 | 2780 | 2883 | 2548 | 2547 | 2487 |
| SL1344_RS02475_ | 4503 | 4385 | 4736 | 3997 | 3978 | 4204 |
| SL1344_RS02480_ | 10674 | 10463 | 10848 | 14207 | 12541 | 11638 |
| SL1344_RS02485_ | 4093 | 4068 | 4327 | 3805 | 3786 | 3713 |
| SL1344_RS02490_ | 405 | 420 | 424 | 510 | 488 | 447 |
| SL1344_RS02495_ | 49 | 20 | 49 | 36 | 37 | 49 |
| SL1344_RS02500_ | 454 | 530 | 584 | 463 | 407 | 531 |
| SL1344_RS02505_ | 972 | 930 | 1027 | 1203 | 1151 | 1163 |
| SL1344_RS02510_ | 429 | 414 | 424 | 431 | 425 | 413 |
| SL1344_RS02520_ | 359 | 374 | 321 | 353 | 316 | 361 |
| SL1344_RS02525_ | 341 | 331 | 315 | 286 | 302 | 256 |
| SL1344_RS02530_ | 112 | 111 | 100 | 87 | 78 | 76 |
| SL1344_RS02540_ | 229 | 244 | 242 | 223 | 211 | 222 |
| SL1344_RS02545_ | 195 | 163 | 158 | 230 | 222 | 232 |
| SL1344_RS02550_ | 143 | 163 | 154 | 227 | 210 | 227 |
| SL1344_RS02555_ | 204 | 181 | 132 | 163 | 129 | 145 |
| SL1344_RS02560_ | 164 | 224 | 188 | 175 | 172 | 140 |
| SL1344_RS02565_ | 1414 | 1428 | 1356 | 1909 | 1765 | 1619 |
| SL1344_RS02570_ | 323 | 292 | 287 | 329 | 286 | 325 |
| SL1344_RS02575_ | 188 | 189 | 167 | 190 | 193 | 162 |
| SL1344_RS02580_ | 45 | 85 | 82 | 86 | 82 | 81 |
| SL1344_RS02585_ | 269 | 300 | 298 | 302 | 304 | 340 |
| SL1344_RS02590_ | 166 | 233 | 202 | 204 | 198 | 230 |
| SL1344_RS02595_ | 97 | 113 | 97 | 119 | 143 | 133 |
| SL1344_RS02600_ | 79 | 114 | 96 | 125 | 91 | 105 |
| SL1344_RS02605_ | 96 | 101 | 101 | 129 | 114 | 143 |
| SL1344_RS02610_ | 996 | 822 | 711 | 806 | 794 | 722 |
| SL1344_RS02615_ | 35 | 34 | 25 | 27 | 26 | 36 |
| SL1344_RS02620_ | 34 | 34 | 33 | 41 | 53 | 46 |
| SL1344_RS02625_ | 335 | 398 | 415 | 489 | 494 | 509 |
| SL1344_RS02630_ | 20 | 23 | 24 | 26 | 30 | 27 |
| SL1344_RS02635_ | 7 | 15 | 15 | 16 | 14 | 18 |
| SL1344_RS02640_ | 23 | 28 | 21 | 14 | 29 | 25 |
| SL1344_RS02645_ | 23 | 32 | 23 | 24 | 27 | 29 |
| SL1344_RS02650_ | 8 | 13 | 10 | 9 | 11 | 10 |
| SL1344_RS02655_ | 24 | 28 | 34 | 59 | 32 | 18 |
| SL1344_RS02660_ | 25 | 15 | 13 | 25 | 37 | 24 |
| SL1344_RS02665_ | 26 | 31 | 35 | 42 | 33 | 28 |
| SL1344_RS02670_ | 161 | 149 | 152 | 176 | 131 | 141 |
| SL1344_RS02675_ | 13 | 21 | 22 | 25 | 24 | 14 |
| SL1344_RS02680_ | 6 | 33 | 5 | 8 | 16 | 26 |
| SL1344_RS02685_ | 34 | 32 | 24 | 30 | 37 | 36 |
| SL1344_RS02690_ | 14 | 15 | 12 | 26 | 16 | 13 |
| SL1344_RS02695_ | 6 | 12 | 18 | 7 | 12 | 13 |
| SL1344_RS02700_ | 32 | 39 | 41 | 43 | 42 | 46 |
| SL1344_RS02705_ | 709 | 702 | 721 | 670 | 631 | 683 |

| | | | | | | |
|---|---|---|---|---|---|---|
| SL1344_RS02710_ | 414 | 465 | 400 | 440 | 411 | 390 |
| SL1344_RS02715_ | 1505 | 1533 | 1686 | 1580 | 1505 | 1537 |
| SL1344_RS02720_ | 5966 | 6042 | 5319 | 6260 | 5757 | 5893 |
| SL1344_RS02730_ | 79 | 73 | 83 | 86 | 70 | 64 |
| SL1344_RS02735_ | 44 | 40 | 43 | 51 | 46 | 42 |
| SL1344_RS02740_ | 61 | 48 | 53 | 63 | 58 | 72 |
| SL1344_RS02745_ | 809 | 927 | 883 | 966 | 929 | 1046 |
| SL1344_RS02750_ | 981 | 1099 | 1105 | 1029 | 972 | 1038 |
| SL1344_RS02760_ | 9 | 12 | 7 | 16 | 20 | 13 |
| SL1344_RS02765_ | 7 | 9 | 6 | 15 | 11 | 7 |
| SL1344_RS02770_ | 22 | 47 | 37 | 68 | 51 | 49 |
| SL1344_RS02775_ | 26 | 26 | 39 | 32 | 37 | 38 |
| SL1344_RS02790_ | 9 | 15 | 29 | 13 | 15 | 12 |
| SL1344_RS02795_ | 1 | 6 | 9 | 4 | 3 | 2 |
| SL1344_RS02800_ | 56 | 34 | 52 | 58 | 41 | 39 |
| SL1344_RS02805_ | 229 | 262 | 285 | 148 | 152 | 209 |
| SL1344_RS02815_ | 177 | 153 | 176 | 312 | 195 | 249 |
| SL1344_RS02820_ | 106 | 111 | 129 | 205 | 122 | 163 |
| SL1344_RS02825_ | 37 | 32 | 41 | 88 | 39 | 58 |
| SL1344_RS02830_ | 8 | 10 | 8 | 13 | 9 | 13 |
| SL1344_RS02835_ | 12 | 8 | 5 | 8 | 4 | 14 |
| SL1344_RS02840_ | 122 | 131 | 115 | 114 | 127 | 95 |
| SL1344_RS02845_ | 51 | 41 | 47 | 48 | 66 | 55 |
| SL1344_RS02850_ | 51 | 56 | 63 | 68 | 70 | 69 |
| SL1344_RS02855_ | 1 | 4 | 7 | 5 | 6 | 5 |
| SL1344_RS02860_ | 19 | 17 | 20 | 22 | 22 | 26 |
| SL1344_RS02865_ | 442 | 541 | 468 | 1268 | 1087 | 1057 |
| SL1344_RS02870_ | 770 | 816 | 894 | 880 | 736 | 871 |
| SL1344_RS02875_ | 545 | 478 | 501 | 455 | 427 | 405 |
| SL1344_RS02880_ | 115 | 131 | 133 | 142 | 115 | 159 |
| SL1344_RS02885_ | 184 | 245 | 194 | 203 | 203 | 229 |
| SL1344_RS02890_ | 34 | 30 | 32 | 25 | 36 | 48 |
| SL1344_RS02895_ | 16 | 19 | 15 | 17 | 10 | 21 |
| SL1344_RS02900_ | 8 | 7 | 7 | 12 | 10 | 8 |
| SL1344_RS02905_ | 4 | 3 | 6 | 1 | 12 | 10 |
| SL1344_RS02915_ | 4 | 2 | 5 | 9 | 5 | 3 |
| SL1344_RS02920_ | 719 | 721 | 728 | 843 | 779 | 775 |
| SL1344_RS02925_ | 105 | 91 | 95 | 123 | 101 | 89 |
| SL1344_RS02930_ | 221 | 210 | 197 | 223 | 210 | 208 |
| SL1344_RS02935_ | 75 | 101 | 67 | 133 | 122 | 102 |
| SL1344_RS02940_ | 85 | 88 | 65 | 84 | 80 | 90 |
| SL1344_RS02945_ | 39 | 64 | 54 | 69 | 48 | 41 |
| SL1344_RS02950_ | 25 | 50 | 44 | 48 | 48 | 32 |
| SL1344_RS02955_ | 536 | 767 | 631 | 641 | 609 | 540 |
| SL1344_RS02960_ | 481 | 619 | 577 | 431 | 331 | 341 |
| SL1344_RS02965_ | 52 | 84 | 71 | 48 | 60 | 64 |
| SL1344_RS02975_ | 943 | 1011 | 1023 | 791 | 766 | 796 |
| SL1344_RS02980_ | 263 | 269 | 292 | 334 | 283 | 283 |
| SL1344_RS02985_ | 435 | 524 | 517 | 505 | 483 | 453 |
| SL1344_RS02990_ | 437 | 481 | 542 | 417 | 410 | 474 |
| SL1344_RS02995_ | 507 | 588 | 735 | 486 | 516 | 517 |
| SL1344_RS03000_ | 324 | 340 | 287 | 274 | 285 | 250 |
| SL1344_RS03005_ | 348 | 569 | 527 | 429 | 438 | 353 |
| SL1344_RS03020_ | 163 | 233 | 214 | 203 | 197 | 236 |
| SL1344_RS03025_ | 117 | 153 | 112 | 139 | 148 | 123 |
| SL1344_RS03030_ | 358 | 390 | 389 | 458 | 420 | 436 |
| SL1344_RS03035_ | 51 | 63 | 48 | 57 | 65 | 69 |
| SL1344_RS03040_ | 182 | 161 | 143 | 213 | 163 | 156 |
| SL1344_RS03045_ | 211 | 215 | 187 | 275 | 188 | 193 |
| SL1344_RS03050_ | 89 | 84 | 89 | 105 | 95 | 114 |
| SL1344_RS03055_ | 122 | 115 | 122 | 120 | 122 | 130 |
| SL1344_RS03060_ | 179 | 139 | 140 | 139 | 114 | 146 |

| | | | | | | |
|---|---|---|---|---|---|---|
| SL1344_RS03065_ | 350 | 331 | 305 | 372 | 350 | 335 |
| SL1344_RS03070_ | 4370 | 4244 | 3899 | 4784 | 4734 | 4263 |
| SL1344_RS03075_ | 1088 | 1227 | 1181 | 1299 | 1186 | 1201 |
| SL1344_RS03080_ | 6 | 2 | 5 | 13 | 6 | 8 |
| SL1344_RS03085_ | 56 | 52 | 50 | 63 | 61 | 69 |
| SL1344_RS03090_ | 35 | 19 | 22 | 35 | 30 | 21 |
| SL1344_RS03095_ | 234 | 208 | 222 | 318 | 325 | 292 |
| SL1344_RS03100_ | 1265 | 1089 | 1049 | 1808 | 1684 | 1623 |
| SL1344_RS03105_ | 43 | 38 | 52 | 35 | 42 | 36 |
| SL1344_RS03110_ | 156 | 214 | 204 | 232 | 229 | 219 |
| SL1344_RS03115_ | 1019 | 951 | 969 | 911 | 880 | 990 |
| SL1344_RS03120_ | 187 | 190 | 183 | 249 | 243 | 292 |
| SL1344_RS03125_ | 5 | 6 | 1 | 8 | 6 | 11 |
| SL1344_RS03130_ | 7 | 10 | 2 | 3 | 9 | 6 |
| SL1344_RS03135_ | 12 | 10 | 14 | 11 | 11 | 8 |
| SL1344_RS03140_ | 4 | 4 | 19 | 0 | 15 | 18 |
| SL1344_RS03145_ | 2 | 1 | 4 | 2 | 2 | 1 |
| SL1344_RS03150_ | 4 | 4 | 0 | 4 | 1 | 2 |
| SL1344_RS03165_ | 115 | 85 | 112 | 52 | 103 | 47 |
| SL1344_RS03175_ | 26305 | 24247 | 24884 | 18480 | 19492 | 18527 |
| SL1344_RS03180_ | 97 | 114 | 90 | 120 | 98 | 108 |
| SL1344_RS03195_ | 2976 | 2883 | 2654 | 2832 | 2881 | 2720 |
| SL1344_RS03200_ | 192 | 187 | 164 | 133 | 99 | 106 |
| SL1344_RS03205_ | 791 | 800 | 819 | 941 | 918 | 950 |
| SL1344_RS03210_ | 2418 | 2594 | 2174 | 3338 | 2940 | 2828 |
| SL1344_RS03215_ | 2510 | 2520 | 2454 | 2377 | 2251 | 2307 |
| SL1344_RS03225_ | 523 | 596 | 671 | 661 | 625 | 661 |
| SL1344_RS03230_ | 1506 | 1607 | 1712 | 1487 | 1402 | 1477 |
| SL1344_RS03235_ | 1568 | 1639 | 1763 | 1422 | 1450 | 1493 |
| SL1344_RS03240_ | 1202 | 1175 | 1144 | 1030 | 1050 | 1125 |
| SL1344_RS03245_ | 544 | 544 | 487 | 415 | 413 | 446 |
| SL1344_RS03250_ | 95 | 126 | 147 | 127 | 129 | 99 |
| SL1344_RS03255_ | 46 | 61 | 48 | 48 | 69 | 58 |
| SL1344_RS03260_ | 380 | 414 | 425 | 389 | 363 | 459 |
| SL1344_RS03265_ | 1345 | 1335 | 1508 | 1276 | 1206 | 1339 |
| SL1344_RS03270_ | 1657 | 1714 | 1650 | 1499 | 1545 | 1496 |
| SL1344_RS03275_ | 5749 | 5655 | 5799 | 5167 | 5170 | 5198 |
| SL1344_RS03280_ | 2 | 2 | 4 | 7 | 1 | 6 |
| SL1344_RS03285_ | 38 | 34 | 25 | 31 | 29 | 31 |
| SL1344_RS03290_ | 80 | 78 | 65 | 62 | 49 | 62 |
| SL1344_RS03295_ | 271 | 248 | 282 | 353 | 316 | 333 |
| SL1344_RS03300_ | 78 | 103 | 93 | 110 | 101 | 93 |
| SL1344_RS03305_ | 60 | 55 | 57 | 45 | 40 | 57 |
| SL1344_RS03310_ | 7 | 11 | 11 | 12 | 12 | 7 |
| SL1344_RS03315_ | 22 | 54 | 24 | 18 | 24 | 31 |
| SL1344_RS03320_ | 17 | 22 | 12 | 20 | 14 | 28 |
| SL1344_RS03325_ | 54 | 79 | 62 | 63 | 74 | 83 |
| SL1344_RS03330_ | 61 | 65 | 71 | 58 | 84 | 84 |
| SL1344_RS03335_ | 105 | 93 | 89 | 82 | 73 | 108 |
| SL1344_RS03340_ | 64 | 78 | 81 | 83 | 63 | 86 |
| SL1344_RS03350_ | 73 | 97 | 109 | 104 | 94 | 104 |
| SL1344_RS03355_ | 148 | 132 | 142 | 132 | 121 | 158 |
| SL1344_RS03360_ | 3180 | 2779 | 2123 | 3519 | 3102 | 3529 |
| SL1344_RS03365_ | 1762 | 1854 | 2012 | 1636 | 1496 | 1688 |
| SL1344_RS03370_ | 1864 | 1846 | 1784 | 1870 | 1632 | 1862 |
| SL1344_RS03375_ | 846 | 861 | 840 | 1093 | 994 | 850 |
| SL1344_RS03380_ | 1311 | 1192 | 1209 | 1528 | 1358 | 1362 |
| SL1344_RS03385_ | 3382 | 3591 | 3566 | 3643 | 3560 | 3556 |
| SL1344_RS03395_ | 56 | 44 | 52 | 53 | 43 | 33 |
| SL1344_RS03400_ | 6722 | 6091 | 6421 | 6304 | 5819 | 6159 |
| SL1344_RS03405_ | 4309 | 3995 | 3997 | 4003 | 3482 | 4292 |
| SL1344_RS03410_ | 4984 | 4693 | 4246 | 4736 | 4422 | 5050 |

| | | | | | | |
|---|---|---|---|---|---|---|
| SL1344_RS03415_ | 8795 | 7768 | 9616 | 7823 | 7338 | 8570 |
| SL1344_RS03420_ | 16069 | 15487 | 17711 | 13255 | 13838 | 15089 |
| SL1344_RS03425_ | 17062 | 16638 | 16765 | 12841 | 13634 | 14381 |
| SL1344_RS03430_ | 13269 | 12789 | 11114 | 8755 | 8467 | 9859 |
| SL1344_RS03440_ | 1055 | 1025 | 1160 | 999 | 985 | 1118 |
| SL1344_RS03445_ | 598 | 655 | 577 | 698 | 664 | 648 |
| SL1344_RS03450_ | 770 | 786 | 698 | 788 | 777 | 809 |
| SL1344_RS03455_ | 215 | 237 | 223 | 236 | 234 | 268 |
| SL1344_RS03460_ | 765 | 882 | 837 | 693 | 651 | 746 |
| SL1344_RS03465_ | 3247 | 3475 | 3463 | 3615 | 3362 | 3413 |
| SL1344_RS03475_ | 35 | 33 | 36 | 35 | 34 | 28 |
| SL1344_RS03480_ | 50 | 52 | 50 | 49 | 45 | 43 |
| SL1344_RS03490_ | 44 | 34 | 45 | 65 | 48 | 55 |
| SL1344_RS03495_ | 228 | 241 | 227 | 223 | 226 | 253 |
| SL1344_RS03505_ | 3661 | 3438 | 3308 | 3104 | 3131 | 3206 |
| SL1344_RS03510_ | 847 | 844 | 777 | 654 | 639 | 699 |
| SL1344_RS03515_ | 629 | 649 | 543 | 649 | 494 | 555 |
| SL1344_RS03520_ | 1100 | 1062 | 1067 | 973 | 1002 | 1062 |
| SL1344_RS03525_ | 4170 | 4204 | 4229 | 4484 | 4102 | 4139 |
| SL1344_RS03530_ | 39 | 32 | 38 | 41 | 26 | 35 |
| SL1344_RS03535_ | 32 | 23 | 18 | 22 | 32 | 41 |
| SL1344_RS03540_ | 40 | 38 | 41 | 45 | 45 | 56 |
| SL1344_RS03545_ | 0 | 1 | 0 | 1 | 0 | 0 |
| SL1344_RS03550_ | 171 | 224 | 188 | 240 | 224 | 227 |
| SL1344_RS03555_ | 255 | 298 | 229 | 281 | 285 | 303 |
| SL1344_RS03560_ | 49 | 50 | 40 | 50 | 48 | 36 |
| SL1344_RS03565_ | 33 | 27 | 18 | 26 | 27 | 26 |
| SL1344_RS03570_ | 12 | 9 | 13 | 16 | 11 | 14 |
| SL1344_RS03575_ | 5 | 2 | 0 | 0 | 1 | 0 |
| SL1344_RS03580_ | 247 | 253 | 331 | 325 | 303 | 319 |
| SL1344_RS03585_ | 111 | 89 | 103 | 88 | 105 | 98 |
| SL1344_RS03590_ | 239 | 294 | 350 | 210 | 197 | 286 |
| SL1344_RS03595_ | 178 | 219 | 219 | 264 | 211 | 278 |
| SL1344_RS03600_ | 145 | 169 | 175 | 196 | 184 | 199 |
| SL1344_RS03605_ | 199 | 177 | 225 | 247 | 201 | 254 |
| SL1344_RS03610_ | 225 | 245 | 254 | 274 | 249 | 266 |
| SL1344_RS03615_ | 12 | 11 | 9 | 5 | 10 | 6 |
| SL1344_RS03620_ | 162 | 165 | 166 | 126 | 143 | 135 |
| SL1344_RS03625_ | 3 | 9 | 4 | 11 | 8 | 13 |
| SL1344_RS03630_ | 17 | 23 | 22 | 27 | 24 | 32 |
| SL1344_RS03640_ | 7 | 5 | 10 | 9 | 8 | 12 |
| SL1344_RS03645_ | 12 | 13 | 11 | 9 | 16 | 9 |
| SL1344_RS03650_ | 6 | 5 | 8 | 31 | 11 | 8 |
| SL1344_RS03655_ | 19 | 10 | 7 | 18 | 13 | 9 |
| SL1344_RS03660_ | 28 | 33 | 26 | 29 | 15 | 28 |
| SL1344_RS03665_ | 10 | 10 | 7 | 11 | 4 | 10 |
| SL1344_RS03670_ | 28 | 24 | 21 | 23 | 39 | 29 |
| SL1344_RS03675_ | 65 | 83 | 88 | 62 | 58 | 62 |
| SL1344_RS03680_ | 399 | 408 | 401 | 355 | 338 | 412 |
| SL1344_RS03685_ | 136 | 118 | 122 | 209 | 176 | 175 |
| SL1344_RS03690_ | 1387 | 1574 | 1350 | 3161 | 3128 | 3076 |
| SL1344_RS03695_ | 17 | 8 | 3 | 14 | 11 | 9 |
| SL1344_RS03705_ | 239 | 268 | 222 | 343 | 330 | 403 |
| SL1344_RS03710_ | 486 | 558 | 575 | 1027 | 986 | 1097 |
| SL1344_RS03715_ | 795 | 935 | 850 | 1389 | 1349 | 1496 |
| SL1344_RS03720_ | 1814 | 2029 | 2027 | 3467 | 3265 | 3522 |
| SL1344_RS03725_ | 1612 | 1817 | 1828 | 3079 | 3089 | 3202 |
| SL1344_RS03730_ | 891 | 1064 | 1127 | 1870 | 1778 | 1958 |
| SL1344_RS03735_ | 893 | 1131 | 1092 | 2192 | 1972 | 2198 |
| SL1344_RS03740_ | 99 | 113 | 138 | 226 | 173 | 201 |
| SL1344_RS03745_ | 2196 | 2038 | 2296 | 1319 | 1267 | 1247 |
| SL1344_RS03750_ | 829 | 962 | 1144 | 666 | 647 | 650 |

| | | | | | | |
|---|---|---|---|---|---|---|
| SL1344_RS03755_ | 703 | 645 | 641 | 411 | 417 | 429 |
| SL1344_RS03760_ | 790 | 738 | 898 | 525 | 489 | 516 |
| SL1344_RS03765_ | 724 | 812 | 975 | 591 | 548 | 584 |
| SL1344_RS03770_ | 526 | 599 | 556 | 578 | 468 | 597 |
| SL1344_RS03775_ | 1079 | 1083 | 1149 | 1002 | 905 | 1022 |
| SL1344_RS03780_ | 624 | 681 | 715 | 622 | 572 | 640 |
| SL1344_RS03790_ | 3786 | 4185 | 3711 | 4738 | 4298 | 4596 |
| SL1344_RS03795_ | 7244 | 8216 | 8129 | 9653 | 8874 | 9928 |
| SL1344_RS03800_ | 4018 | 4617 | 5054 | 5978 | 5333 | 6041 |
| SL1344_RS03805_ | 7015 | 7356 | 6430 | 4911 | 5482 | 5905 |
| SL1344_RS03810_ | 8398 | 8213 | 8470 | 6553 | 6525 | 7474 |
| SL1344_RS03815_ | 4971 | 4575 | 4457 | 3876 | 3852 | 4050 |
| SL1344_RS03820_ | 4846 | 4353 | 4586 | 4448 | 4407 | 4293 |
| SL1344_RS03825_ | 2373 | 2225 | 2322 | 2332 | 2166 | 2201 |
| SL1344_RS03830_ | 25 | 42 | 31 | 30 | 29 | 39 |
| SL1344_RS03835_ | 61 | 62 | 53 | 76 | 67 | 69 |
| SL1344_RS03840_ | 56 | 57 | 65 | 71 | 87 | 90 |
| SL1344_RS03845_ | 22 | 28 | 27 | 36 | 28 | 37 |
| SL1344_RS03850_ | 1172 | 1113 | 1084 | 1230 | 1175 | 1100 |
| SL1344_RS03855_ | 77 | 79 | 83 | 86 | 87 | 82 |
| SL1344_RS03860_ | 39 | 61 | 48 | 45 | 56 | 50 |
| SL1344_RS03865_ | 82 | 105 | 95 | 122 | 95 | 97 |
| SL1344_RS03870_ | 45 | 35 | 29 | 35 | 29 | 37 |
| SL1344_RS03875_ | 8 | 9 | 7 | 4 | 10 | 4 |
| SL1344_RS03880_ | 5 | 3 | 4 | 4 | 4 | 3 |
| SL1344_RS03885_ | 17 | 13 | 22 | 21 | 27 | 18 |
| SL1344_RS03890_ | 17 | 17 | 43 | 23 | 23 | 27 |
| SL1344_RS03895_ | 85 | 97 | 107 | 111 | 109 | 104 |
| SL1344_RS03900_ | 43 | 56 | 49 | 75 | 54 | 62 |
| SL1344_RS03905_ | 21 | 22 | 26 | 27 | 29 | 25 |
| SL1344_RS03915_ | 641 | 578 | 661 | 713 | 755 | 766 |
| SL1344_RS03920_ | 493 | 469 | 434 | 541 | 477 | 451 |
| SL1344_RS03925_ | 216 | 237 | 262 | 215 | 245 | 233 |
| SL1344_RS03930_ | 145 | 173 | 186 | 171 | 166 | 161 |
| SL1344_RS03935_ | 25 | 25 | 25 | 38 | 19 | 29 |
| SL1344_RS03940_ | 676 | 681 | 660 | 693 | 620 | 725 |
| SL1344_RS03945_ | 435 | 419 | 400 | 412 | 406 | 411 |
| SL1344_RS03950_ | 4307 | 4145 | 3382 | 5284 | 5913 | 4662 |
| SL1344_RS03970_ | 297 | 286 | 294 | 294 | 260 | 251 |
| SL1344_RS03975_ | 263 | 286 | 298 | 375 | 310 | 385 |
| SL1344_RS03980_ | 2812 | 3042 | 3011 | 2934 | 2673 | 2912 |
| SL1344_RS03985_ | 237 | 246 | 248 | 239 | 240 | 281 |
| SL1344_RS03990_ | 163 | 148 | 151 | 162 | 165 | 181 |
| SL1344_RS03995_ | 145 | 126 | 138 | 124 | 128 | 138 |
| SL1344_RS04000_ | 80 | 66 | 92 | 87 | 99 | 113 |
| SL1344_RS04005_ | 86 | 98 | 101 | 105 | 127 | 124 |
| SL1344_RS04010_ | 202 | 226 | 227 | 188 | 190 | 188 |
| SL1344_RS04015_ | 45 | 59 | 42 | 43 | 53 | 50 |
| SL1344_RS04020_ | 29 | 23 | 39 | 40 | 25 | 19 |
| SL1344_RS04025_ | 15 | 22 | 22 | 26 | 27 | 25 |
| SL1344_RS04030_ | 12 | 17 | 13 | 10 | 16 | 17 |
| SL1344_RS04035_ | 6 | 10 | 18 | 18 | 13 | 18 |
| SL1344_RS04040_ | 933 | 1169 | 1265 | 1358 | 1225 | 1387 |
| SL1344_RS04045_ | 24 | 26 | 16 | 29 | 25 | 33 |
| SL1344_RS04050_ | 392 | 439 | 434 | 474 | 522 | 503 |
| SL1344_RS04055_ | 278 | 305 | 313 | 276 | 237 | 292 |
| SL1344_RS04065_ | 187 | 213 | 200 | 189 | 207 | 191 |
| SL1344_RS04070_ | 164 | 205 | 169 | 191 | 173 | 171 |
| SL1344_RS04085_ | 1528 | 1506 | 1637 | 1764 | 1621 | 1707 |
| SL1344_RS04090_ | 13 | 14 | 5 | 18 | 20 | 9 |
| SL1344_RS04095_ | 16 | 10 | 11 | 4 | 11 | 16 |
| SL1344_RS04100_ | 17 | 17 | 14 | 14 | 15 | 19 |

| | | | | | | |
|---|---|---|---|---|---|---|
| SL1344_RS04105_ | 51 | 57 | 42 | 48 | 55 | 69 |
| SL1344_RS04110_ | 25 | 16 | 15 | 31 | 24 | 27 |
| SL1344_RS04115_ | 12 | 11 | 15 | 12 | 16 | 21 |
| SL1344_RS04120_ | 512 | 480 | 499 | 491 | 448 | 474 |
| SL1344_RS04125_ | 137 | 171 | 177 | 146 | 144 | 165 |
| SL1344_RS04130_ | 162 | 181 | 196 | 171 | 161 | 158 |
| SL1344_RS04135_ | 263 | 289 | 268 | 259 | 280 | 307 |
| SL1344_RS04140_ | 150 | 157 | 173 | 164 | 157 | 162 |
| SL1344_RS04145_ | 197 | 161 | 145 | 191 | 163 | 143 |
| SL1344_RS04150_ | 795 | 840 | 990 | 631 | 650 | 724 |
| SL1344_RS04160_ | 382 | 433 | 390 | 418 | 363 | 372 |
| SL1344_RS04165_ | 338 | 267 | 196 | 380 | 373 | 261 |
| SL1344_RS04170_ | 179 | 143 | 103 | 191 | 164 | 107 |
| SL1344_RS04175_ | 36 | 53 | 36 | 57 | 31 | 32 |
| SL1344_RS04180_ | 369 | 382 | 358 | 281 | 295 | 283 |
| SL1344_RS04185_ | 313 | 297 | 357 | 173 | 168 | 172 |
| SL1344_RS04195_ | 1014 | 1240 | 1274 | 1001 | 825 | 930 |
| SL1344_RS04200_ | 3680 | 3907 | 3433 | 3302 | 2752 | 3068 |
| SL1344_RS04205_ | 443 | 366 | 303 | 519 | 525 | 403 |
| SL1344_RS04210_ | 225 | 219 | 189 | 188 | 173 | 201 |
| SL1344_RS04220_ | 597 | 601 | 607 | 470 | 428 | 518 |
| SL1344_RS04225_ | 170 | 213 | 217 | 258 | 230 | 246 |
| SL1344_RS04230_ | 235 | 312 | 323 | 363 | 294 | 370 |
| SL1344_RS04235_ | 1857 | 2051 | 1888 | 1692 | 1785 | 1815 |
| SL1344_RS04240_ | 1454 | 1703 | 1798 | 1605 | 1474 | 1753 |
| SL1344_RS04245_ | 32 | 28 | 33 | 31 | 39 | 38 |
| SL1344_RS04250_ | 4 | 10 | 8 | 5 | 5 | 9 |
| SL1344_RS04255_ | 19 | 25 | 34 | 47 | 44 | 48 |
| SL1344_RS04260_ | 1554 | 1487 | 1316 | 1337 | 1240 | 1326 |
| SL1344_RS04265_ | 22 | 23 | 22 | 32 | 32 | 20 |
| SL1344_RS04270_ | 8 | 6 | 7 | 7 | 4 | 11 |
| SL1344_RS04275_ | 457 | 444 | 462 | 506 | 476 | 460 |
| SL1344_RS04280_ | 469 | 420 | 474 | 471 | 422 | 491 |
| SL1344_RS04290_ | 568 | 639 | 663 | 683 | 635 | 654 |
| SL1344_RS04295_ | 307 | 289 | 352 | 310 | 297 | 320 |
| SL1344_RS04300_ | 335 | 273 | 303 | 303 | 278 | 310 |
| SL1344_RS04305_ | 237 | 211 | 226 | 225 | 207 | 226 |
| SL1344_RS04320_ | 15 | 8 | 5 | 5 | 10 | 17 |
| SL1344_RS04325_ | 22 | 13 | 23 | 17 | 29 | 18 |
| SL1344_RS04330_ | 28 | 46 | 25 | 41 | 21 | 38 |
| SL1344_RS04335_ | 4 | 8 | 6 | 5 | 14 | 5 |
| SL1344_RS04340_ | 70 | 102 | 100 | 97 | 74 | 83 |
| SL1344_RS04345_ | 91 | 84 | 104 | 80 | 64 | 67 |
| SL1344_RS04350_ | 112 | 120 | 133 | 110 | 86 | 96 |
| SL1344_RS04355_ | 134 | 145 | 125 | 101 | 116 | 142 |
| SL1344_RS04360_ | 53 | 43 | 54 | 47 | 38 | 37 |
| SL1344_RS04365_ | 238 | 254 | 288 | 339 | 300 | 314 |
| SL1344_RS04370_ | 277 | 296 | 314 | 348 | 350 | 382 |
| SL1344_RS04375_ | 1495 | 1386 | 1402 | 1141 | 1059 | 1206 |
| SL1344_RS04380_ | 473 | 433 | 435 | 246 | 218 | 278 |
| SL1344_RS04385_ | 271 | 353 | 384 | 346 | 262 | 334 |
| SL1344_RS04390_ | 319 | 343 | 342 | 294 | 290 | 292 |
| SL1344_RS04395_ | 364 | 335 | 344 | 297 | 329 | 355 |
| SL1344_RS04400_ | 41 | 42 | 50 | 59 | 51 | 48 |
| SL1344_RS04405_ | 1174 | 1228 | 1200 | 643 | 688 | 718 |
| SL1344_RS04410_ | 17 | 35 | 28 | 25 | 16 | 18 |
| SL1344_RS04415_ | 1665 | 1711 | 1614 | 1918 | 1831 | 2001 |
| SL1344_RS04420_ | 92 | 97 | 85 | 90 | 82 | 99 |
| SL1344_RS04425_ | 128 | 171 | 206 | 166 | 133 | 173 |
| SL1344_RS04430_ | 257 | 361 | 353 | 353 | 340 | 346 |
| SL1344_RS04435_ | 405 | 546 | 562 | 518 | 490 | 532 |
| SL1344_RS04440_ | 226 | 320 | 386 | 425 | 365 | 493 |

| | | | | | | |
|---|---|---|---|---|---|---|
| SL1344_RS04450_ | 56 | 101 | 94 | 109 | 87 | 134 |
| SL1344_RS04455_ | 83 | 110 | 118 | 138 | 134 | 162 |
| SL1344_RS04460_ | 169 | 199 | 168 | 218 | 178 | 224 |
| SL1344_RS04465_ | 677 | 652 | 652 | 565 | 580 | 677 |
| SL1344_RS04470_ | 21 | 28 | 35 | 20 | 19 | 20 |
| SL1344_RS04475_ | 16 | 13 | 11 | 12 | 14 | 4 |
| SL1344_RS04480_ | 63 | 75 | 80 | 90 | 62 | 73 |
| SL1344_RS04485_ | 116 | 151 | 144 | 123 | 101 | 144 |
| SL1344_RS04495_ | 1161 | 1216 | 1392 | 1123 | 993 | 1269 |
| SL1344_RS04500_ | 1666 | 1879 | 2033 | 1825 | 1598 | 2075 |
| SL1344_RS04505_ | 1924 | 2093 | 2234 | 1693 | 1651 | 2197 |
| SL1344_RS04510_ | 541 | 467 | 475 | 533 | 520 | 484 |
| SL1344_RS04515_ | 135 | 177 | 152 | 158 | 168 | 212 |
| SL1344_RS04520_ | 262 | 303 | 289 | 307 | 302 | 318 |
| SL1344_RS04525_ | 379 | 444 | 397 | 422 | 387 | 400 |
| SL1344_RS04530_ | 213 | 215 | 209 | 239 | 257 | 267 |
| SL1344_RS04535_ | 237 | 208 | 200 | 277 | 239 | 249 |
| SL1344_RS04540_ | 58 | 90 | 62 | 104 | 117 | 86 |
| SL1344_RS04545_ | 147 | 171 | 164 | 164 | 178 | 189 |
| SL1344_RS04550_ | 145 | 184 | 172 | 170 | 164 | 198 |
| SL1344_RS04555_ | 351 | 313 | 330 | 386 | 385 | 368 |
| SL1344_RS04560_ | 239 | 226 | 248 | 287 | 275 | 271 |
| SL1344_RS04565_ | 4124 | 4096 | 4074 | 3109 | 3136 | 3463 |
| SL1344_RS04570_ | 362 | 422 | 435 | 435 | 433 | 416 |
| SL1344_RS04575_ | 415 | 438 | 530 | 495 | 531 | 547 |
| SL1344_RS04580_ | 107 | 119 | 90 | 277 | 258 | 262 |
| SL1344_RS04585_ | 399 | 384 | 373 | 464 | 416 | 421 |
| SL1344_RS04590_ | 1341 | 1535 | 1606 | 1966 | 1869 | 1942 |
| SL1344_RS04600_ | 29 | 30 | 29 | 20 | 31 | 20 |
| SL1344_RS04605_ | 105 | 98 | 94 | 86 | 108 | 105 |
| SL1344_RS04610_ | 383 | 396 | 451 | 325 | 300 | 446 |
| SL1344_RS04615_ | 63 | 64 | 81 | 79 | 81 | 89 |
| SL1344_RS04620_ | 16 | 14 | 19 | 30 | 13 | 17 |
| SL1344_RS04625_ | 46 | 43 | 57 | 51 | 61 | 59 |
| SL1344_RS04630_ | 9437 | 9536 | 9718 | 9968 | 9828 | 10239 |
| SL1344_RS04635_ | 6651 | 6574 | 5244 | 6753 | 6712 | 6175 |
| SL1344_RS04640_ | 560 | 524 | 573 | 480 | 412 | 483 |
| SL1344_RS04645_ | 965 | 1014 | 1041 | 857 | 835 | 881 |
| SL1344_RS04650_ | 1303 | 1277 | 1247 | 1124 | 1141 | 1186 |
| SL1344_RS04655_ | 3420 | 3693 | 3346 | 4434 | 4153 | 4544 |
| SL1344_RS04665_ | 3927 | 4064 | 4327 | 4288 | 4189 | 4374 |
| SL1344_RS04675_ | 1185 | 1191 | 1311 | 1214 | 1177 | 1321 |
| SL1344_RS04680_ | 3745 | 3960 | 4296 | 4231 | 3834 | 4141 |
| SL1344_RS04700_ | 4 | 6 | 6 | 5 | 5 | 9 |
| SL1344_RS04705_ | 325 | 398 | 406 | 265 | 187 | 321 |
| SL1344_RS04715_ | 1815 | 2079 | 1928 | 2038 | 1992 | 1997 |
| SL1344_RS04720_ | 88 | 79 | 72 | 55 | 62 | 60 |
| SL1344_RS04725_ | 158 | 137 | 103 | 146 | 126 | 152 |
| SL1344_RS04730_ | 3412 | 3131 | 3049 | 1994 | 1935 | 2077 |
| SL1344_RS04740_ | 1718 | 1832 | 1897 | 1488 | 1412 | 1563 |
| SL1344_RS04745_ | 22 | 28 | 32 | 32 | 26 | 27 |
| SL1344_RS04750_ | 843 | 884 | 799 | 979 | 949 | 952 |
| SL1344_RS04755_ | 713 | 841 | 770 | 817 | 771 | 853 |
| SL1344_RS04760_ | 190 | 220 | 216 | 200 | 200 | 234 |
| SL1344_RS04765_ | 2758 | 2698 | 2558 | 2366 | 2315 | 2662 |
| SL1344_RS04770_ | 50869 | 56136 | 58469 | 57003 | 51979 | 58358 |
| SL1344_RS04780_ | 87 | 103 | 98 | 91 | 84 | 89 |
| SL1344_RS04785_ | 2208 | 2307 | 2218 | 2131 | 2136 | 2288 |
| SL1344_RS04790_ | 1636 | 1755 | 1769 | 1501 | 1408 | 1596 |
| SL1344_RS04795_ | 188 | 201 | 200 | 190 | 173 | 191 |
| SL1344_RS04800_ | 51 | 65 | 61 | 64 | 74 | 82 |
| SL1344_RS04805_ | 443 | 567 | 598 | 577 | 509 | 605 |

| | | | | | | |
|---|---|---|---|---|---|---|
| SL1344_RS04810_ | 96 | 114 | 113 | 110 | 137 | 95 |
| SL1344_RS04815_ | 122 | 120 | 129 | 96 | 96 | 93 |
| SL1344_RS04820_ | 332 | 452 | 540 | 526 | 461 | 505 |
| SL1344_RS04825_ | 792 | 865 | 906 | 956 | 823 | 1002 |
| SL1344_RS04830_ | 888 | 964 | 1009 | 958 | 965 | 956 |
| SL1344_RS04840_ | 891 | 877 | 929 | 1053 | 1042 | 1059 |
| SL1344_RS04845_ | 621 | 645 | 676 | 689 | 624 | 694 |
| SL1344_RS04850_ | 810 | 1039 | 1019 | 1025 | 863 | 1006 |
| SL1344_RS04855_ | 2829 | 2865 | 2856 | 2585 | 2650 | 2683 |
| SL1344_RS04860_ | 730 | 838 | 952 | 1804 | 1695 | 1778 |
| SL1344_RS04865_ | 5317 | 5886 | 5930 | 6065 | 5799 | 6132 |
| SL1344_RS04870_ | 65 | 74 | 72 | 84 | 66 | 81 |
| SL1344_RS04875_ | 7 | 12 | 11 | 12 | 6 | 14 |
| SL1344_RS04880_ | 37 | 40 | 26 | 33 | 20 | 41 |
| SL1344_RS04885_ | 13 | 16 | 19 | 20 | 22 | 21 |
| SL1344_RS04890_ | 1148 | 1194 | 1060 | 1150 | 1122 | 1082 |
| SL1344_RS04895_ | 670 | 641 | 670 | 666 | 639 | 684 |
| SL1344_RS04900_ | 5 | 8 | 11 | 6 | 14 | 12 |
| SL1344_RS04905_ | 1 | 0 | 3 | 2 | 2 | 2 |
| SL1344_RS04910_ | 5 | 2 | 15 | 15 | 10 | 8 |
| SL1344_RS04915_ | 18 | 23 | 16 | 36 | 31 | 28 |
| SL1344_RS04920_ | 2 | 8 | 4 | 8 | 2 | 1 |
| SL1344_RS04925_ | 4 | 8 | 4 | 0 | 5 | 6 |
| SL1344_RS04930_ | 1540 | 1481 | 1254 | 1538 | 1439 | 1509 |
| SL1344_RS04935_ | 6 | 9 | 5 | 18 | 14 | 4 |
| SL1344_RS04940_ | 21 | 15 | 10 | 14 | 13 | 17 |
| SL1344_RS04945_ | 32 | 38 | 34 | 40 | 32 | 32 |
| SL1344_RS04950_ | 8 | 12 | 7 | 10 | 14 | 14 |
| SL1344_RS04955_ | 5 | 6 | 6 | 7 | 7 | 11 |
| SL1344_RS04960_ | 41 | 38 | 44 | 41 | 49 | 46 |
| SL1344_RS04965_ | 741 | 850 | 760 | 934 | 946 | 854 |
| SL1344_RS04970_ | 106 | 114 | 91 | 124 | 86 | 105 |
| SL1344_RS04975_ | 150 | 175 | 136 | 198 | 149 | 174 |
| SL1344_RS04980_ | 102 | 95 | 119 | 95 | 91 | 88 |
| SL1344_RS04985_ | 2 | 2 | 2 | 3 | 4 | 1 |
| SL1344_RS04990_ | 2 | 7 | 9 | 7 | 14 | 14 |
| SL1344_RS04995_ | 1 | 0 | 3 | 0 | 2 | 0 |
| SL1344_RS05000_ | 15 | 15 | 12 | 25 | 13 | 14 |
| SL1344_RS05005_ | 360 | 442 | 432 | 406 | 374 | 438 |
| SL1344_RS05010_ | 3 | 1 | 2 | 2 | 3 | 2 |
| SL1344_RS05015_ | 500 | 447 | 495 | 305 | 337 | 297 |
| SL1344_RS05020_ | 77 | 78 | 41 | 73 | 61 | 64 |
| SL1344_RS05025_ | 2 | 1 | 3 | 2 | 4 | 1 |
| SL1344_RS05030_ | 26 | 23 | 22 | 21 | 27 | 21 |
| SL1344_RS05035_ | 16 | 13 | 17 | 13 | 6 | 9 |
| SL1344_RS05040_ | 27 | 27 | 29 | 20 | 26 | 39 |
| SL1344_RS05045_ | 54 | 52 | 55 | 64 | 56 | 58 |
| SL1344_RS05050_ | 1 | 4 | 4 | 1 | 3 | 4 |
| SL1344_RS05055_ | 41 | 40 | 46 | 43 | 53 | 40 |
| SL1344_RS05060_ | 60 | 78 | 72 | 68 | 69 | 91 |
| SL1344_RS05065_ | 9 | 9 | 16 | 16 | 12 | 14 |
| SL1344_RS05070_ | 3 | 9 | 8 | 8 | 7 | 9 |
| SL1344_RS05075_ | 8 | 17 | 24 | 26 | 27 | 22 |
| SL1344_RS05080_ | 9 | 8 | 14 | 21 | 20 | 14 |
| SL1344_RS05085_ | 39 | 30 | 24 | 34 | 42 | 48 |
| SL1344_RS05090_ | 22 | 26 | 20 | 21 | 31 | 30 |
| SL1344_RS05095_ | 14 | 15 | 13 | 10 | 19 | 17 |
| SL1344_RS05100_ | 105 | 128 | 115 | 138 | 136 | 145 |
| SL1344_RS05105_ | 3 | 13 | 8 | 10 | 11 | 13 |
| SL1344_RS05110_ | 119 | 117 | 81 | 144 | 140 | 159 |
| SL1344_RS05115_ | 432 | 430 | 348 | 555 | 571 | 550 |
| SL1344_RS05120_ | 23 | 24 | 42 | 36 | 41 | 26 |

| | | | | | | |
|---|---|---|---|---|---|---|
| SL1344_RS05125_ | 20 | 24 | 20 | 23 | 21 | 23 |
| SL1344_RS05130_ | 23 | 33 | 32 | 33 | 28 | 41 |
| SL1344_RS05135_ | 125 | 110 | 105 | 140 | 150 | 142 |
| SL1344_RS05140_ | 10 | 14 | 22 | 26 | 23 | 22 |
| SL1344_RS05165_ | 6 | 7 | 10 | 14 | 4 | 11 |
| SL1344_RS05175_ | 52 | 58 | 50 | 88 | 85 | 69 |
| SL1344_RS05190_ | 996 | 1122 | 1082 | 1411 | 1253 | 1428 |
| SL1344_RS05195_ | 452 | 480 | 481 | 504 | 374 | 497 |
| SL1344_RS05200_ | 570 | 587 | 580 | 539 | 502 | 497 |
| SL1344_RS05205_ | 141 | 158 | 168 | 155 | 147 | 181 |
| SL1344_RS05210_ | 1309 | 1626 | 1600 | 1623 | 1470 | 1627 |
| SL1344_RS05215_ | 2233 | 2426 | 2412 | 2360 | 2168 | 2457 |
| SL1344_RS05220_ | 767 | 905 | 888 | 790 | 676 | 801 |
| SL1344_RS05225_ | 837 | 917 | 896 | 898 | 812 | 943 |
| SL1344_RS05230_ | 725 | 793 | 732 | 778 | 746 | 744 |
| SL1344_RS05235_ | 115 | 109 | 121 | 180 | 193 | 173 |
| SL1344_RS05240_ | 2940 | 3823 | 3758 | 5080 | 4696 | 5009 |
| SL1344_RS05245_ | 1449 | 1734 | 1758 | 1675 | 1627 | 1819 |
| SL1344_RS05250_ | 135 | 143 | 153 | 229 | 192 | 201 |
| SL1344_RS05255_ | 60248 | 59572 | 61346 | 50187 | 49717 | 49982 |
| SL1344_RS05260_ | 487 | 385 | 330 | 496 | 444 | 461 |
| SL1344_RS05265_ | 27 | 43 | 27 | 63 | 39 | 49 |
| SL1344_RS05270_ | 430 | 451 | 466 | 493 | 473 | 491 |
| SL1344_RS05275_ | 111 | 110 | 94 | 126 | 108 | 131 |
| SL1344_RS05285_ | 204 | 234 | 193 | 261 | 227 | 235 |
| SL1344_RS05290_ | 10 | 17 | 15 | 23 | 22 | 24 |
| SL1344_RS05295_ | 267 | 386 | 419 | 473 | 384 | 462 |
| SL1344_RS05300_ | 1231 | 1296 | 1202 | 1622 | 1525 | 1435 |
| SL1344_RS05305_ | 766 | 722 | 728 | 659 | 622 | 673 |
| SL1344_RS05310_ | 43 | 42 | 49 | 47 | 46 | 40 |
| SL1344_RS05315_ | 121 | 116 | 100 | 95 | 100 | 104 |
| SL1344_RS05320_ | 24 | 30 | 34 | 34 | 20 | 29 |
| SL1344_RS05325_ | 1551 | 1483 | 1437 | 1612 | 1496 | 1586 |
| SL1344_RS05330_ | 890 | 886 | 824 | 1009 | 1034 | 941 |
| SL1344_RS05335_ | 1760 | 1823 | 1873 | 1834 | 1701 | 1851 |
| SL1344_RS05340_ | 54 | 77 | 65 | 73 | 67 | 52 |
| SL1344_RS05345_ | 16 | 18 | 10 | 34 | 35 | 29 |
| SL1344_RS05350_ | 5 | 1 | 2 | 0 | 1 | 3 |
| SL1344_RS05365_ | 13 | 10 | 13 | 6 | 4 | 9 |
| SL1344_RS05370_ | 13 | 11 | 8 | 6 | 6 | 9 |
| SL1344_RS05375_ | 118 | 109 | 80 | 65 | 70 | 55 |
| SL1344_RS05380_ | 181 | 164 | 150 | 175 | 151 | 169 |
| SL1344_RS05385_ | 134 | 123 | 143 | 126 | 136 | 139 |
| SL1344_RS05390_ | 85 | 92 | 87 | 91 | 100 | 95 |
| SL1344_RS05395_ | 51 | 56 | 33 | 52 | 56 | 56 |
| SL1344_RS05400_ | 23 | 39 | 32 | 27 | 28 | 41 |
| SL1344_RS05405_ | 57 | 60 | 68 | 61 | 62 | 66 |
| SL1344_RS05410_ | 16 | 18 | 19 | 20 | 29 | 25 |
| SL1344_RS05415_ | 27 | 28 | 34 | 23 | 38 | 31 |
| SL1344_RS05420_ | 3 | 10 | 23 | 25 | 19 | 15 |
| SL1344_RS05425_ | 7 | 10 | 10 | 9 | 6 | 13 |
| SL1344_RS05430_ | 15 | 19 | 14 | 15 | 14 | 19 |
| SL1344_RS05435_ | 19 | 19 | 14 | 19 | 18 | 17 |
| SL1344_RS05440_ | 31 | 24 | 20 | 39 | 34 | 46 |
| SL1344_RS05445_ | 24 | 34 | 16 | 28 | 26 | 17 |
| SL1344_RS05450_ | 97 | 108 | 91 | 123 | 111 | 113 |
| SL1344_RS05455_ | 14 | 15 | 15 | 10 | 11 | 9 |
| SL1344_RS05460_ | 141 | 147 | 163 | 239 | 203 | 225 |
| SL1344_RS05465_ | 327 | 278 | 251 | 350 | 309 | 341 |
| SL1344_RS05470_ | 265 | 294 | 303 | 287 | 337 | 315 |
| SL1344_RS05475_ | 130 | 121 | 114 | 109 | 85 | 101 |
| SL1344_RS05480_ | 24 | 32 | 28 | 21 | 23 | 29 |

| | | | | | | |
|---|---|---|---|---|---|---|
| SL1344_RS05485_ | 51 | 57 | 69 | 53 | 47 | 65 |
| SL1344_RS05490_ | 84 | 74 | 79 | 108 | 94 | 101 |
| SL1344_RS05495_ | 140 | 219 | 169 | 261 | 266 | 298 |
| SL1344_RS05500_ | 57 | 93 | 68 | 84 | 77 | 76 |
| SL1344_RS05505_ | 59 | 61 | 54 | 92 | 73 | 99 |
| SL1344_RS05510_ | 218 | 245 | 228 | 244 | 252 | 268 |
| SL1344_RS05515_ | 354 | 373 | 351 | 485 | 429 | 448 |
| SL1344_RS05525_ | 300 | 306 | 304 | 331 | 330 | 353 |
| SL1344_RS05530_ | 105 | 113 | 85 | 269 | 206 | 213 |
| SL1344_RS05540_ | 1207 | 1202 | 1213 | 1060 | 1041 | 1175 |
| SL1344_RS05545_ | 26 | 31 | 26 | 22 | 21 | 27 |
| SL1344_RS05550_ | 12 | 8 | 9 | 12 | 9 | 16 |
| SL1344_RS05555_ | 27 | 29 | 22 | 17 | 17 | 15 |
| SL1344_RS05560_ | 20 | 8 | 6 | 17 | 11 | 16 |
| SL1344_RS05565_ | 23 | 28 | 23 | 31 | 21 | 29 |
| SL1344_RS05570_ | 182 | 162 | 144 | 177 | 133 | 161 |
| SL1344_RS05580_ | 2506 | 2695 | 2490 | 1968 | 1514 | 2210 |
| SL1344_RS05585_ | 5 | 2 | 3 | 6 | 6 | 5 |
| SL1344_RS05590_ | 526 | 632 | 546 | 738 | 737 | 769 |
| SL1344_RS05595_ | 566 | 545 | 500 | 602 | 605 | 574 |
| SL1344_RS05600_ | 342 | 354 | 412 | 361 | 391 | 404 |
| SL1344_RS05605_ | 790 | 880 | 910 | 938 | 856 | 906 |
| SL1344_RS05610_ | 17 | 22 | 20 | 34 | 29 | 51 |
| SL1344_RS05615_ | 3 | 4 | 4 | 6 | 1 | 3 |
| SL1344_RS05620_ | 5 | 5 | 3 | 9 | 7 | 6 |
| SL1344_RS05625_ | 23 | 19 | 16 | 18 | 17 | 16 |
| SL1344_RS05635_ | 13 | 8 | 11 | 18 | 18 | 14 |
| SL1344_RS05650_ | 203 | 293 | 245 | 548 | 476 | 527 |
| SL1344_RS05660_ | 243 | 267 | 250 | 182 | 184 | 176 |
| SL1344_RS05670_ | 4096 | 4451 | 4955 | 4677 | 4378 | 4637 |
| SL1344_RS05675_ | 144 | 151 | 155 | 159 | 133 | 145 |
| SL1344_RS05680_ | 86 | 101 | 121 | 149 | 156 | 106 |
| SL1344_RS05685_ | 173 | 169 | 167 | 129 | 134 | 125 |
| SL1344_RS05690_ | 1107 | 1106 | 1106 | 1109 | 1120 | 1118 |
| SL1344_RS05695_ | 1441 | 1529 | 1508 | 1746 | 1481 | 1728 |
| SL1344_RS05700_ | 45 | 53 | 59 | 86 | 107 | 79 |
| SL1344_RS05705_ | 25 | 31 | 36 | 49 | 34 | 28 |
| SL1344_RS05710_ | 23 | 24 | 17 | 24 | 17 | 18 |
| SL1344_RS05715_ | 602 | 637 | 608 | 591 | 558 | 566 |
| SL1344_RS05730_ | 1493 | 1328 | 1255 | 1562 | 1387 | 1405 |
| SL1344_RS05735_ | 744 | 654 | 685 | 671 | 713 | 691 |
| SL1344_RS05740_ | 263 | 276 | 238 | 393 | 377 | 316 |
| SL1344_RS05745_ | 110 | 122 | 88 | 132 | 106 | 139 |
| SL1344_RS05750_ | 136 | 149 | 169 | 161 | 146 | 161 |
| SL1344_RS05755_ | 234 | 210 | 233 | 214 | 215 | 213 |
| SL1344_RS05760_ | 276 | 339 | 284 | 309 | 320 | 301 |
| SL1344_RS05765_ | 392 | 499 | 515 | 394 | 349 | 422 |
| SL1344_RS05780_ | 1839 | 1974 | 1940 | 3751 | 4210 | 4132 |
| SL1344_RS05785_ | 1692 | 1901 | 1831 | 3644 | 3906 | 3966 |
| SL1344_RS05790_ | 1234 | 1346 | 1284 | 2749 | 2756 | 2886 |
| SL1344_RS05800_ | 3352 | 3498 | 3392 | 7005 | 7367 | 7668 |
| SL1344_RS05810_ | 4378 | 4583 | 4434 | 9822 | 10420 | 10486 |
| SL1344_RS05815_ | 1095 | 1204 | 1304 | 2500 | 2450 | 2776 |
| SL1344_RS05825_ | 953 | 1118 | 1282 | 2179 | 2439 | 2356 |
| SL1344_RS05845_ | 6 | 1 | 3 | 9 | 3 | 1 |
| SL1344_RS05850_ | 1845 | 1864 | 1865 | 1522 | 1606 | 1649 |
| SL1344_RS05855_ | 304 | 324 | 353 | 261 | 250 | 282 |
| SL1344_RS05860_ | 58 | 69 | 79 | 46 | 41 | 60 |
| SL1344_RS05865_ | 238 | 232 | 213 | 312 | 303 | 317 |
| SL1344_RS05870_ | 15915 | 16127 | 15689 | 15262 | 14379 | 15589 |
| SL1344_RS05875_ | 18182 | 19500 | 17841 | 17099 | 16386 | 17190 |
| SL1344_RS05880_ | 7591 | 7849 | 9759 | 8046 | 8029 | 8241 |

| | | | | | | |
|---|---|---|---|---|---|---|
| SL1344_RS05885_ | 2795 | 3097 | 3228 | 2634 | 2395 | 3016 |
| SL1344_RS05890_ | 2891 | 3326 | 3600 | 3108 | 2966 | 3230 |
| SL1344_RS05900_ | 20371 | 26306 | 26675 | 35021 | 30984 | 33591 |
| SL1344_RS05905_ | 15843 | 17839 | 18877 | 18634 | 17653 | 19507 |
| SL1344_RS05910_ | 402 | 453 | 446 | 424 | 396 | 446 |
| SL1344_RS05915_ | 443 | 524 | 491 | 550 | 499 | 582 |
| SL1344_RS05920_ | 477 | 532 | 553 | 585 | 545 | 607 |
| SL1344_RS05925_ | 654 | 628 | 663 | 709 | 643 | 746 |
| SL1344_RS05930_ | 686 | 675 | 778 | 632 | 621 | 720 |
| SL1344_RS05935_ | 4729 | 5129 | 5071 | 4776 | 4520 | 4726 |
| SL1344_RS05940_ | 128 | 165 | 166 | 155 | 164 | 162 |
| SL1344_RS05945_ | 565 | 559 | 501 | 617 | 616 | 598 |
| SL1344_RS05950_ | 547 | 610 | 646 | 668 | 666 | 714 |
| SL1344_RS05955_ | 294 | 343 | 369 | 435 | 485 | 441 |
| SL1344_RS05960_ | 264 | 302 | 334 | 328 | 354 | 341 |
| SL1344_RS05965_ | 226 | 284 | 295 | 382 | 334 | 366 |
| SL1344_RS05970_ | 486 | 528 | 538 | 648 | 633 | 618 |
| SL1344_RS05975_ | 1719 | 1853 | 1835 | 1781 | 1581 | 1880 |
| SL1344_RS05980_ | 50 | 54 | 64 | 56 | 55 | 52 |
| SL1344_RS05985_ | 741 | 797 | 685 | 959 | 890 | 903 |
| SL1344_RS05990_ | 1446 | 1186 | 983 | 1611 | 1635 | 1346 |
| SL1344_RS05995_ | 114 | 108 | 134 | 120 | 127 | 147 |
| SL1344_RS06000_ | 1655 | 1764 | 1769 | 1630 | 1676 | 1636 |
| SL1344_RS06005_ | 957 | 1033 | 1047 | 986 | 929 | 1006 |
| SL1344_RS06010_ | 504 | 513 | 594 | 485 | 447 | 559 |
| SL1344_RS06015_ | 568 | 623 | 699 | 711 | 604 | 703 |
| SL1344_RS06020_ | 647 | 755 | 778 | 762 | 725 | 810 |
| SL1344_RS06025_ | 702 | 796 | 788 | 773 | 738 | 784 |
| SL1344_RS06030_ | 1428 | 1577 | 1568 | 1652 | 1542 | 1489 |
| SL1344_RS06035_ | 687 | 721 | 651 | 617 | 555 | 639 |
| SL1344_RS06045_ | 818 | 922 | 1005 | 934 | 885 | 964 |
| SL1344_RS06050_ | 1004 | 1148 | 1177 | 1240 | 1181 | 1250 |
| SL1344_RS06055_ | 72 | 73 | 89 | 142 | 107 | 98 |
| SL1344_RS06070_ | 1515 | 1388 | 1526 | 977 | 981 | 1092 |
| SL1344_RS06085_ | 1849 | 1898 | 1950 | 1450 | 1402 | 1521 |
| SL1344_RS06090_ | 1017 | 1027 | 1043 | 750 | 759 | 811 |
| SL1344_RS06095_ | 3036 | 3171 | 3247 | 2408 | 2474 | 2554 |
| SL1344_RS06100_ | 214 | 234 | 230 | 179 | 153 | 184 |
| SL1344_RS06105_ | 171 | 189 | 193 | 142 | 155 | 153 |
| SL1344_RS06110_ | 252 | 292 | 217 | 221 | 230 | 244 |
| SL1344_RS06115_ | 3055 | 3008 | 3063 | 4711 | 4475 | 4675 |
| SL1344_RS06120_ | 150 | 148 | 130 | 223 | 206 | 209 |
| SL1344_RS06125_ | 43 | 3 | 1 | 29 | 27 | 2 |
| SL1344_RS06130_ | 25 | 31 | 33 | 26 | 25 | 46 |
| SL1344_RS06135_ | 804 | 959 | 907 | 1073 | 1026 | 936 |
| SL1344_RS06140_ | 146 | 209 | 199 | 131 | 130 | 152 |
| SL1344_RS06150_ | 8 | 2 | 7 | 5 | 2 | 1 |
| SL1344_RS06155_ | 1 | 10 | 1 | 4 | 0 | 1 |
| SL1344_RS06160_ | 8 | 19 | 15 | 13 | 13 | 17 |
| SL1344_RS06165_ | 139 | 193 | 184 | 169 | 141 | 170 |
| SL1344_RS06170_ | 32 | 24 | 24 | 22 | 24 | 17 |
| SL1344_RS06180_ | 217 | 257 | 235 | 180 | 182 | 186 |
| SL1344_RS06185_ | 7 | 6 | 3 | 6 | 4 | 7 |
| SL1344_RS06190_ | 25 | 21 | 13 | 14 | 18 | 11 |
| SL1344_RS06195_ | 94 | 108 | 129 | 125 | 115 | 128 |
| SL1344_RS06200_ | 41 | 35 | 35 | 44 | 39 | 45 |
| SL1344_RS06205_ | 550 | 576 | 468 | 1219 | 1042 | 916 |
| SL1344_RS06210_ | 20 | 27 | 14 | 34 | 23 | 28 |
| SL1344_RS06215_ | 121 | 117 | 101 | 111 | 115 | 114 |
| SL1344_RS06220_ | 202 | 196 | 156 | 147 | 125 | 124 |
| SL1344_RS06225_ | 183 | 225 | 198 | 122 | 126 | 141 |
| SL1344_RS06230_ | 203 | 212 | 186 | 206 | 190 | 197 |

| | | | | | | |
|---|---|---|---|---|---|---|
| SL1344_RS06235_ | 83 | 83 | 61 | 81 | 57 | 96 |
| SL1344_RS06240_ | 34 | 34 | 41 | 39 | 29 | 33 |
| SL1344_RS06245_ | 30 | 33 | 50 | 40 | 32 | 33 |
| SL1344_RS06250_ | 37 | 38 | 45 | 32 | 43 | 35 |
| SL1344_RS06255_ | 88 | 76 | 64 | 82 | 61 | 86 |
| SL1344_RS06260_ | 64 | 68 | 77 | 33 | 41 | 33 |
| SL1344_RS06265_ | 28 | 21 | 24 | 13 | 12 | 16 |
| SL1344_RS06270_ | 140 | 136 | 150 | 106 | 132 | 127 |
| SL1344_RS06275_ | 212 | 191 | 143 | 176 | 157 | 126 |
| SL1344_RS06280_ | 22 | 20 | 15 | 25 | 13 | 19 |
| SL1344_RS06285_ | 381 | 395 | 333 | 222 | 204 | 200 |
| SL1344_RS06290_ | 174 | 171 | 128 | 125 | 139 | 100 |
| SL1344_RS06295_ | 10 | 9 | 2 | 6 | 1 | 6 |
| SL1344_RS06300_ | 217 | 201 | 227 | 90 | 70 | 107 |
| SL1344_RS06305_ | 365 | 489 | 492 | 343 | 270 | 359 |
| SL1344_RS06310_ | 79 | 85 | 66 | 67 | 56 | 67 |
| SL1344_RS06315_ | 212 | 193 | 180 | 212 | 192 | 180 |
| SL1344_RS06320_ | 61 | 46 | 42 | 39 | 40 | 46 |
| SL1344_RS06325_ | 23 | 31 | 20 | 30 | 31 | 35 |
| SL1344_RS06330_ | 134 | 203 | 207 | 264 | 233 | 226 |
| SL1344_RS06335_ | 125 | 123 | 135 | 96 | 101 | 115 |
| SL1344_RS06340_ | 535 | 517 | 444 | 351 | 301 | 328 |
| SL1344_RS06345_ | 300 | 276 | 308 | 229 | 264 | 246 |
| SL1344_RS06350_ | 95 | 120 | 109 | 97 | 105 | 81 |
| SL1344_RS06355_ | 202 | 204 | 181 | 195 | 219 | 204 |
| SL1344_RS06360_ | 73 | 100 | 75 | 48 | 60 | 65 |
| SL1344_RS06365_ | 7 | 11 | 11 | 8 | 6 | 7 |
| SL1344_RS06370_ | 136 | 125 | 138 | 112 | 100 | 94 |
| SL1344_RS06375_ | 111 | 138 | 125 | 108 | 87 | 100 |
| SL1344_RS06380_ | 168 | 196 | 198 | 161 | 158 | 177 |
| SL1344_RS06385_ | 156 | 128 | 124 | 120 | 154 | 141 |
| SL1344_RS06390_ | 2195 | 2478 | 2496 | 2604 | 2262 | 2737 |
| SL1344_RS06395_ | 67 | 68 | 96 | 66 | 67 | 81 |
| SL1344_RS06400_ | 105 | 143 | 107 | 138 | 135 | 121 |
| SL1344_RS06405_ | 479 | 465 | 443 | 406 | 392 | 418 |
| SL1344_RS06410_ | 28846 | 31900 | 29737 | 26423 | 23580 | 25559 |
| SL1344_RS06415_ | 722 | 757 | 596 | 726 | 758 | 713 |
| SL1344_RS06420_ | 705 | 711 | 624 | 905 | 841 | 802 |
| SL1344_RS06425_ | 269 | 307 | 327 | 250 | 221 | 246 |
| SL1344_RS06430_ | 267 | 324 | 359 | 295 | 233 | 256 |
| SL1344_RS06435_ | 998 | 982 | 1005 | 812 | 791 | 754 |
| SL1344_RS06440_ | 317 | 345 | 330 | 358 | 262 | 301 |
| SL1344_RS06445_ | 810 | 892 | 809 | 732 | 676 | 657 |
| SL1344_RS06450_ | 1435 | 1565 | 1608 | 1313 | 1211 | 1452 |
| SL1344_RS06455_ | 363 | 430 | 391 | 310 | 300 | 353 |
| SL1344_RS06465_ | 129 | 133 | 151 | 67 | 84 | 64 |
| SL1344_RS06470_ | 590 | 642 | 686 | 535 | 513 | 513 |
| SL1344_RS06475_ | 24 | 25 | 34 | 28 | 27 | 23 |
| SL1344_RS06480_ | 11 | 16 | 10 | 22 | 13 | 24 |
| SL1344_RS06485_ | 30 | 34 | 31 | 41 | 28 | 39 |
| SL1344_RS06490_ | 35 | 31 | 43 | 40 | 32 | 43 |
| SL1344_RS06495_ | 38 | 29 | 25 | 25 | 30 | 33 |
| SL1344_RS06500_ | 124 | 100 | 108 | 89 | 93 | 90 |
| SL1344_RS06505_ | 278 | 241 | 231 | 255 | 243 | 246 |
| SL1344_RS06510_ | 759 | 698 | 705 | 553 | 583 | 531 |
| SL1344_RS06515_ | 398 | 423 | 402 | 538 | 463 | 419 |
| SL1344_RS06520_ | 176 | 147 | 128 | 153 | 144 | 150 |
| SL1344_RS06525_ | 67 | 78 | 82 | 64 | 74 | 102 |
| SL1344_RS06530_ | 79 | 65 | 71 | 77 | 52 | 60 |
| SL1344_RS06535_ | 132 | 104 | 140 | 126 | 94 | 99 |
| SL1344_RS06540_ | 55 | 66 | 60 | 68 | 60 | 58 |
| SL1344_RS06545_ | 205 | 206 | 162 | 161 | 148 | 181 |

| | | | | | | |
|---|---|---|---|---|---|---|
| SL1344_RS06550_ | 62 | 100 | 73 | 142 | 118 | 113 |
| SL1344_RS06555_ | 175 | 190 | 157 | 132 | 128 | 132 |
| SL1344_RS06565_ | 137 | 138 | 120 | 116 | 91 | 76 |
| SL1344_RS06570_ | 688 | 722 | 618 | 574 | 575 | 589 |
| SL1344_RS06575_ | 202 | 227 | 189 | 181 | 145 | 184 |
| SL1344_RS06580_ | 95 | 91 | 86 | 93 | 85 | 106 |
| SL1344_RS06605_ | 15 | 25 | 19 | 31 | 33 | 21 |
| SL1344_RS06615_ | 22 | 16 | 15 | 13 | 16 | 16 |
| SL1344_RS06630_ | 58412 | 55370 | 54321 | 49917 | 52633 | 49707 |
| SL1344_RS06635_ | 65865 | 65433 | 66157 | 54578 | 62145 | 54728 |
| SL1344_RS06640_ | 63288 | 63763 | 63795 | 59445 | 58533 | 59784 |
| SL1344_RS06645_ | 1496 | 1739 | 1783 | 1502 | 1358 | 1503 |
| SL1344_RS06650_ | 2733 | 3569 | 4046 | 3505 | 3128 | 3531 |
| SL1344_RS06660_ | 453 | 447 | 450 | 360 | 324 | 355 |
| SL1344_RS06665_ | 114 | 125 | 145 | 134 | 130 | 130 |
| SL1344_RS06670_ | 118 | 137 | 158 | 163 | 112 | 150 |
| SL1344_RS06675_ | 211 | 179 | 201 | 121 | 116 | 133 |
| SL1344_RS06685_ | 336 | 315 | 408 | 253 | 284 | 332 |
| SL1344_RS06690_ | 613 | 673 | 592 | 387 | 400 | 391 |
| SL1344_RS06695_ | 702 | 759 | 781 | 580 | 586 | 635 |
| SL1344_RS06700_ | 115 | 123 | 133 | 96 | 98 | 108 |
| SL1344_RS06705_ | 537 | 489 | 531 | 619 | 564 | 587 |
| SL1344_RS06710_ | 65 | 57 | 69 | 55 | 56 | 56 |
| SL1344_RS06715_ | 6 | 8 | 12 | 13 | 1 | 4 |
| SL1344_RS06720_ | 22 | 33 | 22 | 30 | 27 | 23 |
| SL1344_RS06730_ | 9 | 7 | 10 | 11 | 6 | 3 |
| SL1344_RS06735_ | 20 | 20 | 28 | 18 | 9 | 14 |
| SL1344_RS06740_ | 76 | 92 | 88 | 90 | 68 | 78 |
| SL1344_RS06745_ | 39 | 32 | 38 | 29 | 29 | 25 |
| SL1344_RS06750_ | 207 | 238 | 241 | 203 | 205 | 177 |
| SL1344_RS06755_ | 87 | 98 | 76 | 91 | 82 | 79 |
| SL1344_RS06760_ | 77 | 64 | 46 | 78 | 66 | 54 |
| SL1344_RS06765_ | 12 | 11 | 12 | 12 | 21 | 8 |
| SL1344_RS06770_ | 4 | 5 | 7 | 6 | 2 | 5 |
| SL1344_RS06775_ | 395 | 398 | 435 | 370 | 394 | 399 |
| SL1344_RS06780_ | 603 | 655 | 590 | 652 | 569 | 606 |
| SL1344_RS06785_ | 110 | 110 | 113 | 101 | 103 | 96 |
| SL1344_RS06790_ | 320 | 289 | 305 | 337 | 346 | 379 |
| SL1344_RS06795_ | 213 | 200 | 251 | 201 | 156 | 194 |
| SL1344_RS06800_ | 33 | 53 | 63 | 40 | 49 | 38 |
| SL1344_RS06805_ | 194 | 239 | 259 | 265 | 210 | 234 |
| SL1344_RS06810_ | 228 | 284 | 298 | 280 | 235 | 284 |
| SL1344_RS06815_ | 289 | 353 | 312 | 324 | 328 | 312 |
| SL1344_RS06820_ | 215 | 273 | 276 | 239 | 252 | 247 |
| SL1344_RS06825_ | 78 | 115 | 86 | 95 | 99 | 96 |
| SL1344_RS06830_ | 186 | 245 | 233 | 227 | 204 | 245 |
| SL1344_RS06835_ | 361 | 347 | 326 | 363 | 324 | 356 |
| SL1344_RS06840_ | 265022 | 261513 | 246829 | 232284 | 229628 | 242527 |
| SL1344_RS06845_ | 7924 | 8074 | 8220 | 6237 | 6240 | 6093 |
| SL1344_RS06850_ | 66 | 56 | 51 | 62 | 57 | 60 |
| SL1344_RS06855_ | 25 | 22 | 31 | 16 | 22 | 11 |
| SL1344_RS06860_ | 8 | 6 | 10 | 5 | 4 | 7 |
| SL1344_RS06865_ | 27 | 14 | 15 | 5 | 9 | 14 |
| SL1344_RS06900_ | 480 | 485 | 429 | 369 | 364 | 412 |
| SL1344_RS06905_ | 138 | 173 | 173 | 119 | 133 | 116 |
| SL1344_RS06925_ | 13 | 13 | 10 | 17 | 10 | 13 |
| SL1344_RS06930_ | 12 | 16 | 22 | 17 | 12 | 19 |
| SL1344_RS06935_ | 13 | 17 | 15 | 20 | 14 | 13 |
| SL1344_RS06940_ | 29 | 17 | 10 | 10 | 14 | 15 |
| SL1344_RS06945_ | 23 | 26 | 37 | 31 | 21 | 25 |
| SL1344_RS06950_ | 26 | 31 | 28 | 27 | 23 | 26 |
| SL1344_RS06955_ | 58 | 71 | 53 | 28 | 34 | 48 |

| | | | | | | |
|---|---|---|---|---|---|---|
| SL1344_RS06960_ | 18 | 17 | 12 | 15 | 17 | 17 |
| SL1344_RS06965_ | 16 | 16 | 18 | 15 | 8 | 23 |
| SL1344_RS06970_ | 19 | 25 | 31 | 22 | 16 | 7 |
| SL1344_RS06975_ | 27 | 27 | 24 | 21 | 29 | 17 |
| SL1344_RS06980_ | 72 | 71 | 62 | 68 | 58 | 71 |
| SL1344_RS06990_ | 4 | 4 | 11 | 3 | 1 | 4 |
| SL1344_RS06995_ | 4 | 6 | 8 | 4 | 4 | 9 |
| SL1344_RS07000_ | 17 | 15 | 18 | 15 | 16 | 11 |
| SL1344_RS07005_ | 22 | 31 | 21 | 23 | 16 | 25 |
| SL1344_RS07010_ | 18 | 17 | 3 | 7 | 11 | 11 |
| SL1344_RS07015_ | 32 | 46 | 59 | 38 | 38 | 46 |
| SL1344_RS07020_ | 7 | 3 | 5 | 8 | 8 | 10 |
| SL1344_RS07025_ | 42 | 36 | 45 | 29 | 35 | 43 |
| SL1344_RS07030_ | 78 | 96 | 80 | 93 | 76 | 80 |
| SL1344_RS07035_ | 22 | 43 | 38 | 30 | 16 | 22 |
| SL1344_RS07040_ | 19 | 35 | 40 | 25 | 20 | 16 |
| SL1344_RS07045_ | 99 | 94 | 80 | 72 | 70 | 89 |
| SL1344_RS07055_ | 12 | 6 | 10 | 8 | 7 | 9 |
| SL1344_RS07060_ | 21 | 14 | 23 | 16 | 19 | 27 |
| SL1344_RS07065_ | 29 | 31 | 28 | 30 | 24 | 33 |
| SL1344_RS07070_ | 668 | 545 | 615 | 443 | 445 | 471 |
| SL1344_RS07075_ | 1044 | 1114 | 977 | 637 | 576 | 769 |
| SL1344_RS07080_ | 704 | 808 | 679 | 677 | 617 | 672 |
| SL1344_RS07085_ | 303 | 396 | 449 | 348 | 315 | 423 |
| SL1344_RS07090_ | 1034 | 1141 | 1052 | 1558 | 1294 | 1439 |
| SL1344_RS07095_ | 161 | 208 | 188 | 171 | 184 | 181 |
| SL1344_RS07100_ | 60 | 77 | 78 | 77 | 69 | 82 |
| SL1344_RS07115_ | 804 | 832 | 804 | 562 | 552 | 606 |
| SL1344_RS07120_ | 5587 | 5569 | 5502 | 5274 | 4961 | 5229 |
| SL1344_RS07125_ | 776 | 760 | 835 | 569 | 531 | 609 |
| SL1344_RS07130_ | 623 | 722 | 806 | 724 | 675 | 731 |
| SL1344_RS07135_ | 389 | 399 | 380 | 381 | 409 | 381 |
| SL1344_RS07140_ | 176 | 166 | 191 | 156 | 194 | 177 |
| SL1344_RS07145_ | 13 | 15 | 25 | 28 | 18 | 18 |
| SL1344_RS07150_ | 416 | 407 | 394 | 295 | 232 | 264 |
| SL1344_RS07155_ | 245 | 227 | 253 | 149 | 148 | 203 |
| SL1344_RS07160_ | 422 | 374 | 453 | 253 | 217 | 249 |
| SL1344_RS07165_ | 290 | 310 | 326 | 188 | 175 | 186 |
| SL1344_RS07170_ | 104 | 77 | 83 | 60 | 57 | 58 |
| SL1344_RS07180_ | 1623 | 1723 | 1742 | 1389 | 1270 | 1200 |
| SL1344_RS07185_ | 145 | 149 | 159 | 159 | 189 | 198 |
| SL1344_RS07190_ | 169 | 191 | 153 | 197 | 144 | 175 |
| SL1344_RS07195_ | 705 | 663 | 535 | 528 | 521 | 450 |
| SL1344_RS07200_ | 5485 | 5376 | 5386 | 3951 | 3921 | 4040 |
| SL1344_RS07205_ | 788 | 707 | 650 | 620 | 649 | 658 |
| SL1344_RS07210_ | 303 | 330 | 250 | 350 | 336 | 302 |
| SL1344_RS07215_ | 1749 | 1879 | 1781 | 1224 | 1210 | 1307 |
| SL1344_RS07220_ | 946 | 1046 | 1088 | 871 | 823 | 851 |
| SL1344_RS07225_ | 801 | 886 | 786 | 667 | 616 | 684 |
| SL1344_RS07230_ | 659 | 706 | 745 | 552 | 493 | 512 |
| SL1344_RS07235_ | 868 | 927 | 912 | 604 | 687 | 709 |
| SL1344_RS07240_ | 2288 | 2435 | 2710 | 1731 | 1800 | 1963 |
| SL1344_RS07245_ | 620 | 679 | 742 | 532 | 486 | 559 |
| SL1344_RS07250_ | 1488 | 1478 | 1462 | 1236 | 1143 | 1204 |
| SL1344_RS07255_ | 1626 | 1602 | 1569 | 1318 | 1272 | 1417 |
| SL1344_RS07260_ | 728 | 703 | 677 | 584 | 575 | 581 |
| SL1344_RS07265_ | 551 | 490 | 401 | 452 | 431 | 447 |
| SL1344_RS07270_ | 522 | 529 | 535 | 455 | 406 | 421 |
| SL1344_RS07275_ | 884 | 891 | 951 | 697 | 724 | 734 |
| SL1344_RS07285_ | 1229 | 1356 | 1413 | 1191 | 990 | 1154 |
| SL1344_RS07290_ | 717 | 698 | 674 | 586 | 534 | 591 |
| SL1344_RS07295_ | 857 | 996 | 872 | 1010 | 1092 | 1076 |

| | | | | | | |
|---|---|---|---|---|---|---|
| SL1344_RS07300_ | 287 | 309 | 289 | 332 | 285 | 322 |
| SL1344_RS07305_ | 109 | 127 | 102 | 112 | 101 | 99 |
| SL1344_RS07310_ | 135 | 158 | 131 | 109 | 127 | 144 |
| SL1344_RS07315_ | 7 | 11 | 13 | 11 | 26 | 13 |
| SL1344_RS07320_ | 149 | 150 | 152 | 149 | 128 | 139 |
| SL1344_RS07325_ | 4 | 2 | 5 | 13 | 5 | 6 |
| SL1344_RS07330_ | 2144 | 2294 | 2271 | 1617 | 1666 | 1827 |
| SL1344_RS07335_ | 92 | 97 | 94 | 111 | 113 | 122 |
| SL1344_RS07340_ | 1158 | 1209 | 1281 | 611 | 606 | 682 |
| SL1344_RS07345_ | 1694 | 1661 | 1833 | 1324 | 1322 | 1331 |
| SL1344_RS07350_ | 1581 | 1668 | 1648 | 1207 | 1208 | 1183 |
| SL1344_RS07355_ | 1763 | 1933 | 2107 | 1438 | 1402 | 1481 |
| SL1344_RS07365_ | 92 | 80 | 67 | 77 | 72 | 42 |
| SL1344_RS07370_ | 120 | 101 | 83 | 58 | 74 | 44 |
| SL1344_RS07375_ | 84 | 66 | 71 | 94 | 84 | 83 |
| SL1344_RS07380_ | 102 | 108 | 118 | 74 | 76 | 76 |
| SL1344_RS07385_ | 99 | 129 | 107 | 101 | 108 | 105 |
| SL1344_RS07390_ | 59 | 59 | 48 | 61 | 63 | 59 |
| SL1344_RS07400_ | 129 | 111 | 117 | 75 | 78 | 103 |
| SL1344_RS07405_ | 151 | 159 | 134 | 109 | 122 | 118 |
| SL1344_RS07410_ | 51 | 50 | 43 | 75 | 65 | 76 |
| SL1344_RS07415_ | 15 | 16 | 13 | 16 | 9 | 12 |
| SL1344_RS07420_ | 8 | 10 | 6 | 12 | 12 | 14 |
| SL1344_RS07425_ | 8 | 6 | 4 | 7 | 9 | 6 |
| SL1344_RS07430_ | 421 | 422 | 425 | 360 | 471 | 406 |
| SL1344_RS07435_ | 231 | 215 | 226 | 271 | 201 | 252 |
| SL1344_RS07440_ | 19 | 4 | 4 | 7 | 5 | 4 |
| SL1344_RS07445_ | 47 | 32 | 68 | 49 | 47 | 27 |
| SL1344_RS07450_ | 58 | 56 | 50 | 63 | 65 | 81 |
| SL1344_RS07455_ | 195 | 238 | 192 | 200 | 214 | 238 |
| SL1344_RS07460_ | 82 | 85 | 86 | 104 | 73 | 85 |
| SL1344_RS07465_ | 837 | 869 | 811 | 1197 | 1121 | 1072 |
| SL1344_RS07470_ | 776 | 742 | 635 | 941 | 908 | 796 |
| SL1344_RS07475_ | 36 | 52 | 54 | 52 | 54 | 46 |
| SL1344_RS07480_ | 17 | 23 | 22 | 18 | 15 | 27 |
| SL1344_RS07485_ | 18 | 18 | 19 | 29 | 17 | 28 |
| SL1344_RS07490_ | 16 | 40 | 27 | 16 | 23 | 26 |
| SL1344_RS07495_ | 126 | 140 | 148 | 119 | 122 | 118 |
| SL1344_RS07500_ | 176 | 162 | 147 | 153 | 155 | 132 |
| SL1344_RS07505_ | 424 | 455 | 431 | 397 | 370 | 342 |
| SL1344_RS07510_ | 680 | 730 | 667 | 648 | 690 | 640 |
| SL1344_RS07515_ | 330 | 334 | 322 | 360 | 268 | 342 |
| SL1344_RS07520_ | 8 | 19 | 8 | 16 | 14 | 12 |
| SL1344_RS07525_ | 12 | 19 | 15 | 16 | 13 | 16 |
| SL1344_RS07530_ | 23 | 23 | 18 | 22 | 24 | 26 |
| SL1344_RS07535_ | 351 | 300 | 188 | 297 | 312 | 238 |
| SL1344_RS07540_ | 63 | 70 | 72 | 79 | 67 | 69 |
| SL1344_RS07550_ | 146 | 164 | 169 | 150 | 126 | 138 |
| SL1344_RS07555_ | 196 | 189 | 181 | 145 | 148 | 147 |
| SL1344_RS07560_ | 248 | 261 | 214 | 189 | 204 | 188 |
| SL1344_RS07565_ | 202 | 266 | 217 | 198 | 205 | 220 |
| SL1344_RS07570_ | 408 | 393 | 420 | 441 | 460 | 469 |
| SL1344_RS07575_ | 109 | 75 | 75 | 103 | 91 | 83 |
| SL1344_RS07580_ | 94 | 116 | 106 | 92 | 59 | 107 |
| SL1344_RS07585_ | 241 | 278 | 260 | 227 | 213 | 235 |
| SL1344_RS07590_ | 157 | 166 | 159 | 139 | 139 | 146 |
| SL1344_RS07595_ | 1374 | 1395 | 1293 | 804 | 803 | 981 |
| SL1344_RS07600_ | 40 | 41 | 30 | 89 | 101 | 99 |
| SL1344_RS07605_ | 334 | 333 | 293 | 388 | 340 | 381 |
| SL1344_RS07610_ | 33 | 28 | 36 | 47 | 61 | 26 |
| SL1344_RS07615_ | 79 | 68 | 71 | 79 | 96 | 86 |
| SL1344_RS07620_ | 99 | 97 | 90 | 117 | 128 | 130 |

| | | | | | | |
|---|---|---|---|---|---|---|
| SL1344_RS07625_ | 28 | 38 | 33 | 69 | 32 | 52 |
| SL1344_RS07630_ | 32 | 37 | 45 | 48 | 37 | 42 |
| SL1344_RS07635_ | 83 | 108 | 93 | 136 | 111 | 123 |
| SL1344_RS07640_ | 89 | 104 | 171 | 93 | 78 | 139 |
| SL1344_RS07645_ | 131 | 160 | 219 | 142 | 132 | 181 |
| SL1344_RS07650_ | 69 | 89 | 63 | 86 | 90 | 77 |
| SL1344_RS07655_ | 174 | 187 | 157 | 116 | 112 | 114 |
| SL1344_RS07660_ | 283 | 331 | 285 | 363 | 380 | 351 |
| SL1344_RS07665_ | 63 | 43 | 41 | 53 | 45 | 46 |
| SL1344_RS07670_ | 29 | 17 | 17 | 19 | 15 | 29 |
| SL1344_RS07675_ | 16 | 25 | 19 | 23 | 18 | 16 |
| SL1344_RS07680_ | 47 | 34 | 42 | 43 | 37 | 35 |
| SL1344_RS07685_ | 28 | 43 | 39 | 46 | 45 | 29 |
| SL1344_RS07690_ | 193 | 214 | 168 | 294 | 262 | 276 |
| SL1344_RS07695_ | 87 | 78 | 112 | 81 | 113 | 121 |
| SL1344_RS07700_ | 163 | 172 | 158 | 249 | 213 | 231 |
| SL1344_RS07705_ | 258 | 260 | 238 | 304 | 319 | 291 |
| SL1344_RS07710_ | 184 | 177 | 152 | 206 | 184 | 191 |
| SL1344_RS07715_ | 45 | 41 | 29 | 47 | 43 | 33 |
| SL1344_RS07720_ | 46 | 44 | 32 | 33 | 31 | 29 |
| SL1344_RS07725_ | 65 | 69 | 81 | 89 | 105 | 88 |
| SL1344_RS07735_ | 52 | 65 | 52 | 103 | 113 | 99 |
| SL1344_RS07740_ | 314 | 283 | 275 | 657 | 597 | 645 |
| SL1344_RS07745_ | 125 | 116 | 123 | 133 | 139 | 160 |
| SL1344_RS07750_ | 69 | 70 | 75 | 88 | 88 | 105 |
| SL1344_RS07755_ | 118 | 118 | 114 | 145 | 137 | 167 |
| SL1344_RS07760_ | 105 | 101 | 120 | 144 | 190 | 173 |
| SL1344_RS07765_ | 57 | 44 | 66 | 64 | 77 | 73 |
| SL1344_RS07770_ | 40 | 31 | 37 | 57 | 55 | 57 |
| SL1344_RS07775_ | 15 | 13 | 13 | 17 | 27 | 13 |
| SL1344_RS07780_ | 193 | 250 | 172 | 412 | 352 | 363 |
| SL1344_RS07785_ | 36 | 35 | 28 | 51 | 63 | 58 |
| SL1344_RS07790_ | 35 | 32 | 36 | 30 | 41 | 30 |
| SL1344_RS07800_ | 1406 | 1378 | 1447 | 1830 | 1700 | 1773 |
| SL1344_RS07805_ | 265 | 331 | 302 | 685 | 699 | 651 |
| SL1344_RS07810_ | 653 | 708 | 763 | 1186 | 1197 | 1158 |
| SL1344_RS07815_ | 46 | 47 | 56 | 60 | 62 | 73 |
| SL1344_RS07820_ | 196 | 266 | 285 | 265 | 294 | 327 |
| SL1344_RS07825_ | 176 | 214 | 184 | 233 | 261 | 218 |
| SL1344_RS07830_ | 54 | 45 | 49 | 50 | 37 | 59 |
| SL1344_RS07835_ | 26694 | 27295 | 26408 | 32914 | 30219 | 33169 |
| SL1344_RS07840_ | 314 | 354 | 279 | 252 | 276 | 293 |
| SL1344_RS07845_ | 625 | 1884 | 2154 | 1814 | 265 | 1721 |
| SL1344_RS07850_ | 396 | 663 | 560 | 742 | 392 | 691 |
| SL1344_RS07855_ | 34 | 32 | 41 | 44 | 35 | 33 |
| SL1344_RS07860_ | 80 | 85 | 84 | 125 | 124 | 129 |
| SL1344_RS07865_ | 27 | 31 | 38 | 59 | 39 | 47 |
| SL1344_RS07870_ | 19 | 17 | 13 | 30 | 14 | 30 |
| SL1344_RS07875_ | 16 | 30 | 21 | 50 | 42 | 35 |
| SL1344_RS07880_ | 127 | 127 | 110 | 161 | 167 | 171 |
| SL1344_RS07885_ | 238 | 247 | 202 | 218 | 237 | 226 |
| SL1344_RS07890_ | 614 | 648 | 657 | 774 | 724 | 815 |
| SL1344_RS07895_ | 1459 | 1513 | 1735 | 1814 | 1799 | 1730 |
| SL1344_RS07900_ | 49 | 54 | 60 | 39 | 58 | 51 |
| SL1344_RS07905_ | 742 | 788 | 759 | 905 | 827 | 860 |
| SL1344_RS07910_ | 86 | 114 | 98 | 132 | 128 | 126 |
| SL1344_RS07915_ | 136 | 144 | 142 | 129 | 145 | 133 |
| SL1344_RS07920_ | 223 | 274 | 214 | 355 | 359 | 363 |
| SL1344_RS07925_ | 125 | 119 | 101 | 184 | 152 | 186 |
| SL1344_RS07930_ | 133 | 150 | 132 | 211 | 195 | 204 |
| SL1344_RS07935_ | 98 | 121 | 110 | 148 | 167 | 136 |
| SL1344_RS07955_ | 122 | 126 | 131 | 88 | 98 | 93 |

| | | | | | | |
|---|---|---|---|---|---|---|
| SL1344_RS07960_ | 292 | 341 | 296 | 384 | 377 | 379 |
| SL1344_RS07965_ | 317 | 361 | 334 | 456 | 420 | 451 |
| SL1344_RS07970_ | 199 | 192 | 151 | 281 | 258 | 275 |
| SL1344_RS07975_ | 12 | 10 | 14 | 21 | 20 | 14 |
| SL1344_RS07980_ | 23 | 21 | 23 | 36 | 29 | 25 |
| SL1344_RS07990_ | 27 | 27 | 40 | 60 | 59 | 64 |
| SL1344_RS07995_ | 872 | 1081 | 1073 | 1340 | 1274 | 1336 |
| SL1344_RS08000_ | 495 | 441 | 381 | 447 | 448 | 395 |
| SL1344_RS08005_ | 373 | 327 | 310 | 317 | 332 | 326 |
| SL1344_RS08010_ | 806 | 825 | 817 | 1173 | 1076 | 1160 |
| SL1344_RS08015_ | 288 | 322 | 315 | 529 | 440 | 495 |
| SL1344_RS08020_ | 254 | 263 | 236 | 335 | 338 | 344 |
| SL1344_RS08025_ | 25 | 23 | 25 | 39 | 42 | 43 |
| SL1344_RS08030_ | 60 | 57 | 59 | 85 | 100 | 97 |
| SL1344_RS08035_ | 10 | 13 | 16 | 16 | 28 | 19 |
| SL1344_RS08040_ | 1 | 4 | 2 | 4 | 7 | 4 |
| SL1344_RS08045_ | 25 | 15 | 25 | 39 | 47 | 29 |
| SL1344_RS08050_ | 11 | 23 | 14 | 21 | 32 | 32 |
| SL1344_RS08055_ | 13 | 3 | 13 | 13 | 15 | 18 |
| SL1344_RS08060_ | 24 | 13 | 15 | 22 | 30 | 32 |
| SL1344_RS08065_ | 40 | 36 | 34 | 51 | 60 | 36 |
| SL1344_RS08070_ | 30 | 33 | 27 | 59 | 55 | 42 |
| SL1344_RS08075_ | 127 | 124 | 120 | 144 | 144 | 150 |
| SL1344_RS08080_ | 31 | 42 | 36 | 27 | 35 | 42 |
| SL1344_RS08090_ | 200 | 221 | 168 | 261 | 240 | 288 |
| SL1344_RS08095_ | 78 | 75 | 67 | 99 | 86 | 106 |
| SL1344_RS08100_ | 206 | 199 | 183 | 454 | 459 | 480 |
| SL1344_RS08110_ | 236 | 276 | 255 | 248 | 277 | 263 |
| SL1344_RS08115_ | 156 | 138 | 131 | 127 | 106 | 94 |
| SL1344_RS08120_ | 6 | 8 | 4 | 3 | 6 | 2 |
| SL1344_RS08125_ | 16 | 17 | 16 | 15 | 13 | 26 |
| SL1344_RS08130_ | 21 | 21 | 14 | 16 | 19 | 16 |
| SL1344_RS08135_ | 41 | 50 | 40 | 59 | 52 | 64 |
| SL1344_RS08140_ | 34 | 36 | 32 | 63 | 32 | 42 |
| SL1344_RS08145_ | 20 | 26 | 16 | 18 | 23 | 27 |
| SL1344_RS08150_ | 11 | 11 | 6 | 13 | 6 | 16 |
| SL1344_RS08155_ | 88 | 79 | 75 | 89 | 90 | 94 |
| SL1344_RS08160_ | 686 | 745 | 661 | 1379 | 1379 | 1380 |
| SL1344_RS08165_ | 292 | 299 | 258 | 372 | 313 | 422 |
| SL1344_RS08170_ | 180 | 205 | 199 | 252 | 242 | 283 |
| SL1344_RS08175_ | 1192 | 1257 | 1293 | 1688 | 1567 | 1734 |
| SL1344_RS08185_ | 86 | 95 | 110 | 133 | 120 | 132 |
| SL1344_RS08190_ | 75 | 94 | 86 | 111 | 82 | 113 |
| SL1344_RS08195_ | 91 | 113 | 87 | 110 | 112 | 108 |
| SL1344_RS08200_ | 621 | 644 | 665 | 666 | 677 | 745 |
| SL1344_RS08215_ | 247 | 313 | 305 | 393 | 349 | 386 |
| SL1344_RS08220_ | 26 | 36 | 28 | 35 | 43 | 35 |
| SL1344_RS08225_ | 510 | 528 | 529 | 623 | 567 | 644 |
| SL1344_RS08230_ | 33 | 33 | 24 | 35 | 50 | 26 |
| SL1344_RS08235_ | 215 | 201 | 149 | 322 | 285 | 230 |
| SL1344_RS08240_ | 373 | 347 | 267 | 381 | 351 | 383 |
| SL1344_RS08245_ | 101 | 71 | 54 | 77 | 77 | 82 |
| SL1344_RS08250_ | 196 | 149 | 177 | 183 | 170 | 209 |
| SL1344_RS08255_ | 1378 | 1366 | 1339 | 1446 | 1408 | 1507 |
| SL1344_RS08260_ | 468 | 512 | 472 | 481 | 474 | 515 |
| SL1344_RS08270_ | 165 | 163 | 135 | 326 | 277 | 304 |
| SL1344_RS08275_ | 519 | 555 | 493 | 780 | 705 | 705 |
| SL1344_RS08280_ | 188 | 159 | 149 | 263 | 233 | 236 |
| SL1344_RS08290_ | 944 | 994 | 981 | 1214 | 1151 | 1172 |
| SL1344_RS08295_ | 638 | 756 | 635 | 1342 | 1266 | 1388 |
| SL1344_RS08300_ | 151 | 160 | 149 | 182 | 185 | 181 |
| SL1344_RS08305_ | 130 | 168 | 144 | 188 | 178 | 225 |

| | | | | | | |
|---|---|---|---|---|---|---|
| SL1344_RS08310_ | 94 | 109 | 102 | 113 | 103 | 100 |
| SL1344_RS08315_ | 83 | 83 | 79 | 79 | 61 | 83 |
| SL1344_RS08320_ | 9 | 9 | 1 | 14 | 13 | 11 |
| SL1344_RS08325_ | 3 | 8 | 6 | 14 | 11 | 6 |
| SL1344_RS08330_ | 11 | 15 | 16 | 20 | 21 | 20 |
| SL1344_RS08335_ | 13 | 12 | 3 | 16 | 17 | 18 |
| SL1344_RS08340_ | 6 | 7 | 12 | 20 | 8 | 13 |
| SL1344_RS08345_ | 894 | 850 | 724 | 911 | 882 | 975 |
| SL1344_RS08350_ | 108 | 69 | 79 | 119 | 112 | 109 |
| SL1344_RS08355_ | 40 | 52 | 65 | 81 | 66 | 63 |
| SL1344_RS08360_ | 118 | 144 | 134 | 140 | 142 | 134 |
| SL1344_RS08365_ | 151 | 159 | 170 | 158 | 186 | 191 |
| SL1344_RS08370_ | 42 | 49 | 38 | 38 | 42 | 63 |
| SL1344_RS08375_ | 16 | 9 | 9 | 16 | 16 | 14 |
| SL1344_RS08380_ | 589 | 600 | 539 | 840 | 762 | 791 |
| SL1344_RS08385_ | 103 | 101 | 110 | 139 | 127 | 142 |
| SL1344_RS08390_ | 171 | 181 | 167 | 227 | 231 | 212 |
| SL1344_RS08395_ | 4698 | 4562 | 5078 | 5687 | 5593 | 5831 |
| SL1344_RS08400_ | 510 | 575 | 582 | 709 | 558 | 612 |
| SL1344_RS08405_ | 298 | 338 | 301 | 513 | 390 | 415 |
| SL1344_RS08410_ | 246 | 256 | 254 | 437 | 365 | 343 |
| SL1344_RS08415_ | 176 | 144 | 150 | 276 | 271 | 228 |
| SL1344_RS08420_ | 72 | 60 | 58 | 123 | 93 | 88 |
| SL1344_RS08425_ | 85 | 90 | 78 | 146 | 183 | 130 |
| SL1344_RS08430_ | 51 | 37 | 50 | 77 | 77 | 78 |
| SL1344_RS08435_ | 794 | 787 | 707 | 1016 | 1161 | 968 |
| SL1344_RS08440_ | 128 | 154 | 135 | 155 | 166 | 170 |
| SL1344_RS08445_ | 767 | 752 | 671 | 755 | 807 | 858 |
| SL1344_RS08450_ | 390 | 429 | 421 | 437 | 388 | 512 |
| SL1344_RS08455_ | 333 | 332 | 338 | 390 | 356 | 402 |
| SL1344_RS08460_ | 371 | 437 | 421 | 476 | 424 | 460 |
| SL1344_RS08465_ | 254 | 302 | 291 | 299 | 283 | 326 |
| SL1344_RS08470_ | 112 | 127 | 151 | 165 | 136 | 103 |
| SL1344_RS08475_ | 47 | 47 | 37 | 67 | 42 | 54 |
| SL1344_RS08480_ | 1 | 3 | 2 | 2 | 1 | 5 |
| SL1344_RS08485_ | 85 | 60 | 47 | 109 | 99 | 118 |
| SL1344_RS08490_ | 7833 | 7799 | 7704 | 10432 | 10015 | 10341 |
| SL1344_RS08500_ | 2986 | 3156 | 3416 | 3853 | 3818 | 4010 |
| SL1344_RS08505_ | 219 | 210 | 182 | 265 | 254 | 194 |
| SL1344_RS08510_ | 279 | 256 | 252 | 373 | 359 | 339 |
| SL1344_RS08515_ | 368 | 425 | 405 | 599 | 532 | 596 |
| SL1344_RS08520_ | 306 | 326 | 235 | 463 | 443 | 445 |
| SL1344_RS08525_ | 363 | 438 | 360 | 553 | 456 | 546 |
| SL1344_RS08530_ | 435 | 467 | 448 | 520 | 511 | 567 |
| SL1344_RS08535_ | 811 | 817 | 849 | 1197 | 1125 | 1132 |
| SL1344_RS08540_ | 741 | 694 | 618 | 1017 | 967 | 1007 |
| SL1344_RS08545_ | 67 | 87 | 84 | 108 | 92 | 111 |
| SL1344_RS08550_ | 1027 | 1041 | 850 | 1245 | 1184 | 1227 |
| SL1344_RS08555_ | 196 | 246 | 233 | 377 | 380 | 430 |
| SL1344_RS08560_ | 514 | 587 | 553 | 619 | 532 | 650 |
| SL1344_RS08570_ | 3373 | 3546 | 3888 | 4426 | 4264 | 4381 |
| SL1344_RS08575_ | 1148 | 1190 | 991 | 1566 | 1525 | 1432 |
| SL1344_RS08580_ | 289 | 341 | 355 | 462 | 464 | 404 |
| SL1344_RS08590_ | 314 | 380 | 328 | 473 | 474 | 455 |
| SL1344_RS08595_ | 2085 | 2307 | 2332 | 2374 | 2469 | 2455 |
| SL1344_RS08600_ | 409 | 500 | 490 | 788 | 633 | 755 |
| SL1344_RS08605_ | 479 | 456 | 512 | 659 | 642 | 711 |
| SL1344_RS08610_ | 63 | 57 | 64 | 74 | 63 | 54 |
| SL1344_RS08615_ | 99 | 85 | 81 | 120 | 99 | 108 |
| SL1344_RS08620_ | 145 | 174 | 165 | 202 | 199 | 217 |
| SL1344_RS08625_ | 145 | 121 | 132 | 210 | 196 | 206 |
| SL1344_RS08630_ | 197 | 176 | 159 | 231 | 252 | 246 |

| | | | | | | |
|---|---|---|---|---|---|---|
| SL1344_RS08635_ | 6 | 3 | 8 | 4 | 5 | 5 |
| SL1344_RS08640_ | 5 | 2 | 3 | 5 | 2 | 3 |
| SL1344_RS08645_ | 4 | 1 | 7 | 4 | 1 | 3 |
| SL1344_RS08650_ | 24 | 41 | 28 | 51 | 39 | 42 |
| SL1344_RS08655_ | 91 | 75 | 82 | 118 | 115 | 122 |
| SL1344_RS08660_ | 120 | 99 | 99 | 125 | 102 | 138 |
| SL1344_RS08665_ | 941 | 1016 | 1011 | 1328 | 1281 | 1269 |
| SL1344_RS08670_ | 1317 | 1388 | 1317 | 1699 | 1632 | 1660 |
| SL1344_RS08675_ | 745 | 692 | 687 | 916 | 905 | 946 |
| SL1344_RS08680_ | 1643 | 1902 | 1752 | 2325 | 2148 | 2088 |
| SL1344_RS08685_ | 175 | 211 | 197 | 280 | 290 | 290 |
| SL1344_RS08690_ | 250 | 233 | 226 | 389 | 343 | 321 |
| SL1344_RS08695_ | 414 | 551 | 510 | 752 | 715 | 776 |
| SL1344_RS08700_ | 151 | 132 | 133 | 184 | 170 | 151 |
| SL1344_RS08705_ | 260 | 291 | 329 | 473 | 386 | 469 |
| SL1344_RS08710_ | 178 | 183 | 184 | 165 | 148 | 188 |
| SL1344_RS08715_ | 106 | 139 | 146 | 103 | 85 | 118 |
| SL1344_RS08725_ | 460 | 488 | 511 | 554 | 523 | 524 |
| SL1344_RS08730_ | 277 | 316 | 280 | 339 | 257 | 360 |
| SL1344_RS08735_ | 4294 | 4352 | 3984 | 4784 | 4692 | 4636 |
| SL1344_RS08740_ | 361 | 438 | 354 | 540 | 491 | 512 |
| SL1344_RS08750_ | 2361 | 2526 | 2440 | 3607 | 3474 | 3504 |
| SL1344_RS08755_ | 360 | 407 | 465 | 596 | 529 | 560 |
| SL1344_RS08760_ | 380 | 448 | 386 | 613 | 569 | 593 |
| SL1344_RS08765_ | 287 | 302 | 280 | 491 | 421 | 447 |
| SL1344_RS08770_ | 647 | 731 | 698 | 1196 | 1140 | 1218 |
| SL1344_RS08775_ | 2995 | 3321 | 3136 | 3122 | 3085 | 3783 |
| SL1344_RS08780_ | 1341 | 1170 | 1283 | 1505 | 1365 | 1419 |
| SL1344_RS08785_ | 64 | 97 | 77 | 155 | 140 | 136 |
| SL1344_RS08815_ | 380 | 361 | 347 | 447 | 495 | 466 |
| SL1344_RS08825_ | 191 | 224 | 194 | 280 | 305 | 309 |
| SL1344_RS08830_ | 778 | 884 | 836 | 1125 | 1027 | 1121 |
| SL1344_RS08840_ | 218 | 246 | 227 | 231 | 226 | 251 |
| SL1344_RS08845_ | 1874 | 2047 | 2013 | 2403 | 2537 | 2447 |
| SL1344_RS08850_ | 788 | 987 | 989 | 1152 | 993 | 1107 |
| SL1344_RS08855_ | 413 | 415 | 458 | 461 | 431 | 502 |
| SL1344_RS08860_ | 1039 | 1068 | 1119 | 1124 | 1157 | 1201 |
| SL1344_RS08865_ | 1726 | 1659 | 1828 | 1772 | 1855 | 1832 |
| SL1344_RS08870_ | 2112 | 2282 | 2247 | 2569 | 2580 | 2810 |
| SL1344_RS08880_ | 3664 | 3893 | 3778 | 5019 | 5096 | 5236 |
| SL1344_RS08885_ | 10052 | 11285 | 10369 | 13377 | 13486 | 14023 |
| SL1344_RS08890_ | 584 | 652 | 607 | 856 | 879 | 814 |
| SL1344_RS08895_ | 26 | 26 | 14 | 18 | 9 | 9 |
| SL1344_RS08900_ | 358 | 386 | 351 | 442 | 413 | 512 |
| SL1344_RS08905_ | 3618 | 3927 | 3995 | 4515 | 4424 | 4414 |
| SL1344_RS08910_ | 339 | 326 | 331 | 451 | 430 | 473 |
| SL1344_RS08915_ | 6 | 3 | 5 | 14 | 5 | 5 |
| SL1344_RS08920_ | 30 | 24 | 38 | 45 | 38 | 47 |
| SL1344_RS08925_ | 6 | 6 | 6 | 13 | 4 | 6 |
| SL1344_RS08930_ | 1 | 11 | 10 | 6 | 11 | 10 |
| SL1344_RS08935_ | 5 | 5 | 1 | 2 | 7 | 11 |
| SL1344_RS08940_ | 4 | 12 | 13 | 21 | 16 | 24 |
| SL1344_RS08945_ | 25 | 28 | 22 | 40 | 40 | 43 |
| SL1344_RS08950_ | 26 | 27 | 27 | 32 | 37 | 34 |
| SL1344_RS08955_ | 15 | 27 | 14 | 35 | 26 | 26 |
| SL1344_RS08960_ | 1046 | 1016 | 919 | 2028 | 2005 | 1943 |
| SL1344_RS08965_ | 124 | 129 | 152 | 210 | 215 | 218 |
| SL1344_RS08970_ | 21 | 19 | 23 | 31 | 30 | 34 |
| SL1344_RS08985_ | 140 | 131 | 143 | 199 | 182 | 202 |
| SL1344_RS08995_ | 151 | 155 | 162 | 314 | 318 | 316 |
| SL1344_RS09000_ | 143 | 178 | 144 | 304 | 294 | 310 |
| SL1344_RS09005_ | 76 | 72 | 57 | 99 | 98 | 103 |

| | | | | | | |
|---|---|---|---|---|---|---|
| SL1344_RS09010_ | 452 | 514 | 515 | 598 | 604 | 584 |
| SL1344_RS09015_ | 536 | 512 | 481 | 668 | 657 | 607 |
| SL1344_RS09020_ | 1370 | 1441 | 1471 | 1936 | 1867 | 1962 |
| SL1344_RS09025_ | 46 | 45 | 41 | 76 | 69 | 72 |
| SL1344_RS09030_ | 476 | 598 | 571 | 1008 | 964 | 1004 |
| SL1344_RS09035_ | 57 | 53 | 62 | 73 | 75 | 76 |
| SL1344_RS09040_ | 586 | 657 | 656 | 757 | 742 | 774 |
| SL1344_RS09045_ | 728 | 671 | 707 | 777 | 752 | 799 |
| SL1344_RS09050_ | 306 | 302 | 299 | 369 | 330 | 326 |
| SL1344_RS09055_ | 1366 | 1454 | 1357 | 1725 | 1711 | 1748 |
| SL1344_RS09060_ | 1943 | 1897 | 1797 | 2652 | 2336 | 2527 |
| SL1344_RS09065_ | 1444 | 1544 | 1413 | 2092 | 2023 | 2138 |
| SL1344_RS09070_ | 139 | 147 | 148 | 106 | 123 | 132 |
| SL1344_RS09085_ | 587 | 592 | 739 | 1259 | 1133 | 1311 |
| SL1344_RS09090_ | 774 | 908 | 946 | 1673 | 1600 | 1729 |
| SL1344_RS09095_ | 582 | 640 | 619 | 1398 | 1230 | 1382 |
| SL1344_RS09100_ | 1039 | 1017 | 969 | 2210 | 2212 | 1991 |
| SL1344_RS09105_ | 431 | 428 | 348 | 788 | 781 | 623 |
| SL1344_RS09115_ | 255 | 234 | 255 | 520 | 482 | 520 |
| SL1344_RS09125_ | 38 | 20 | 43 | 60 | 69 | 75 |
| SL1344_RS09130_ | 112 | 155 | 188 | 324 | 313 | 329 |
| SL1344_RS09135_ | 59 | 59 | 40 | 106 | 109 | 108 |
| SL1344_RS09140_ | 447 | 528 | 472 | 947 | 738 | 975 |
| SL1344_RS09145_ | 247 | 339 | 321 | 721 | 622 | 736 |
| SL1344_RS09150_ | 672 | 797 | 814 | 1811 | 1539 | 1827 |
| SL1344_RS09155_ | 102 | 82 | 87 | 150 | 144 | 168 |
| SL1344_RS09160_ | 465 | 377 | 356 | 556 | 532 | 478 |
| SL1344_RS09165_ | 363 | 474 | 493 | 923 | 871 | 985 |
| SL1344_RS09170_ | 261 | 281 | 285 | 395 | 368 | 394 |
| SL1344_RS09175_ | 9258 | 9901 | 9986 | 13742 | 13461 | 13779 |
| SL1344_RS09180_ | 4533 | 4741 | 4013 | 7907 | 7456 | 7310 |
| SL1344_RS09185_ | 611 | 751 | 823 | 1348 | 1215 | 1349 |
| SL1344_RS09190_ | 156 | 213 | 209 | 299 | 255 | 348 |
| SL1344_RS09200_ | 1383 | 1440 | 1299 | 2215 | 2256 | 2058 |
| SL1344_RS09205_ | 112 | 98 | 85 | 171 | 175 | 156 |
| SL1344_RS09210_ | 1593 | 1684 | 1590 | 2938 | 2526 | 2665 |
| SL1344_RS09235_ | 871 | 883 | 888 | 1806 | 1672 | 1793 |
| SL1344_RS09245_ | 148 | 192 | 185 | 273 | 328 | 356 |
| SL1344_RS09250_ | 35 | 22 | 24 | 44 | 40 | 65 |
| SL1344_RS09260_ | 87 | 118 | 131 | 259 | 241 | 281 |
| SL1344_RS09270_ | 21 | 19 | 22 | 20 | 22 | 23 |
| SL1344_RS09275_ | 253 | 244 | 218 | 438 | 410 | 453 |
| SL1344_RS09285_ | 40 | 37 | 34 | 70 | 82 | 66 |
| SL1344_RS09290_ | 47 | 50 | 55 | 134 | 128 | 115 |
| SL1344_RS09295_ | 20 | 20 | 20 | 67 | 34 | 44 |
| SL1344_RS09300_ | 8 | 12 | 9 | 11 | 7 | 13 |
| SL1344_RS09305_ | 67 | 55 | 45 | 88 | 81 | 106 |
| SL1344_RS09310_ | 12 | 9 | 12 | 5 | 8 | 8 |
| SL1344_RS09315_ | 38 | 46 | 53 | 61 | 61 | 68 |
| SL1344_RS09320_ | 154 | 133 | 138 | 204 | 267 | 198 |
| SL1344_RS09325_ | 10 | 15 | 13 | 26 | 23 | 23 |
| SL1344_RS09335_ | 17 | 11 | 13 | 24 | 17 | 31 |
| SL1344_RS09340_ | 229 | 293 | 253 | 405 | 436 | 439 |
| SL1344_RS09345_ | 4 | 3 | 7 | 14 | 17 | 7 |
| SL1344_RS09350_ | 2 | 4 | 2 | 5 | 6 | 5 |
| SL1344_RS09355_ | 21 | 15 | 11 | 25 | 29 | 30 |
| SL1344_RS09360_ | 13 | 21 | 29 | 36 | 44 | 39 |
| SL1344_RS09365_ | 131 | 135 | 180 | 99 | 93 | 119 |
| SL1344_RS09370_ | 642 | 724 | 762 | 1545 | 1589 | 1567 |
| SL1344_RS09380_ | 541 | 486 | 435 | 1144 | 1102 | 1130 |
| SL1344_RS09385_ | 276 | 313 | 294 | 592 | 583 | 648 |
| SL1344_RS09390_ | 255 | 236 | 255 | 480 | 431 | 451 |

| | | | | | | |
|---|---|---|---|---|---|---|
| SL1344_RS09395_ | 312 | 329 | 289 | 582 | 588 | 606 |
| SL1344_RS09400_ | 299 | 301 | 286 | 656 | 680 | 711 |
| SL1344_RS09405_ | 177 | 223 | 214 | 465 | 467 | 497 |
| SL1344_RS09410_ | 248 | 241 | 251 | 614 | 638 | 656 |
| SL1344_RS09420_ | 149 | 155 | 166 | 374 | 384 | 316 |
| SL1344_RS09425_ | 694 | 863 | 888 | 1495 | 1458 | 1558 |
| SL1344_RS09430_ | 458 | 464 | 530 | 937 | 838 | 857 |
| SL1344_RS09435_ | 1724 | 2136 | 2013 | 4215 | 3785 | 4114 |
| SL1344_RS09440_ | 196 | 137 | 134 | 354 | 281 | 252 |
| SL1344_RS09455_ | 673 | 775 | 815 | 1292 | 1171 | 1262 |
| SL1344_RS09460_ | 465 | 488 | 491 | 812 | 761 | 756 |
| SL1344_RS09465_ | 297 | 268 | 263 | 462 | 464 | 481 |
| SL1344_RS09470_ | 404 | 380 | 397 | 658 | 722 | 732 |
| SL1344_RS09485_ | 37 | 37 | 35 | 61 | 59 | 56 |
| SL1344_RS09490_ | 35 | 28 | 29 | 62 | 49 | 46 |
| SL1344_RS09495_ | 1677 | 1963 | 1995 | 4051 | 3731 | 4236 |
| SL1344_RS09500_ | 1041 | 1436 | 1473 | 3234 | 2650 | 3193 |
| SL1344_RS09510_ | 3424 | 3598 | 3826 | 7231 | 7146 | 7159 |
| SL1344_RS09515_ | 214 | 220 | 224 | 406 | 444 | 389 |
| SL1344_RS09520_ | 184 | 215 | 193 | 394 | 371 | 352 |
| SL1344_RS09525_ | 533 | 526 | 479 | 1017 | 903 | 983 |
| SL1344_RS09535_ | 1145 | 1175 | 1195 | 2063 | 1975 | 2026 |
| SL1344_RS09540_ | 241 | 254 | 253 | 672 | 610 | 591 |
| SL1344_RS09545_ | 167 | 206 | 193 | 495 | 432 | 464 |
| SL1344_RS09550_ | 2372 | 2637 | 2738 | 4753 | 4632 | 4786 |
| SL1344_RS09555_ | 134 | 143 | 136 | 190 | 198 | 203 |
| SL1344_RS09635_ | 57 | 49 | 34 | 86 | 59 | 66 |
| SL1344_RS09640_ | 147 | 136 | 112 | 220 | 168 | 205 |
| SL1344_RS09645_ | 92 | 98 | 74 | 237 | 205 | 173 |
| SL1344_RS09650_ | 27 | 37 | 26 | 68 | 74 | 65 |
| SL1344_RS09655_ | 3 | 1 | 2 | 0 | 1 | 5 |
| SL1344_RS09660_ | 28 | 29 | 21 | 69 | 66 | 67 |
| SL1344_RS09670_ | 47 | 38 | 53 | 81 | 94 | 72 |
| SL1344_RS09680_ | 1176 | 1140 | 1046 | 2125 | 2058 | 2044 |
| SL1344_RS09685_ | 8 | 4 | 10 | 11 | 8 | 9 |
| SL1344_RS09690_ | 811 | 874 | 843 | 801 | 754 | 772 |
| SL1344_RS09695_ | 384 | 461 | 435 | 708 | 638 | 692 |
| SL1344_RS09700_ | 720 | 780 | 784 | 1226 | 1187 | 1253 |
| SL1344_RS09705_ | 328 | 320 | 296 | 424 | 438 | 433 |
| SL1344_RS09710_ | 162 | 162 | 119 | 203 | 197 | 188 |
| SL1344_RS09715_ | 4997 | 4581 | 4346 | 8872 | 7814 | 8640 |
| SL1344_RS09725_ | 3403 | 3662 | 3125 | 5507 | 5659 | 6470 |
| SL1344_RS09730_ | 492 | 523 | 485 | 850 | 811 | 861 |
| SL1344_RS09805_ | 6971 | 7490 | 6834 | 14096 | 13460 | 14272 |
| SL1344_RS09810_ | 2798 | 2790 | 2434 | 5138 | 5046 | 5280 |
| SL1344_RS09820_ | 526 | 566 | 563 | 835 | 871 | 834 |
| SL1344_RS09850_ | 6304 | 6272 | 6248 | 8276 | 9747 | 9175 |
| SL1344_RS09855_ | 2967 | 2724 | 2832 | 3963 | 4148 | 4138 |
| SL1344_RS09860_ | 2294 | 2318 | 2223 | 3120 | 3565 | 3198 |
| SL1344_RS09875_ | 1599 | 1564 | 1622 | 2484 | 2668 | 2643 |
| SL1344_RS09885_ | 3016 | 3119 | 3054 | 4359 | 5126 | 4959 |
| SL1344_RS09890_ | 1541 | 1555 | 1555 | 2269 | 2590 | 2462 |
| SL1344_RS09895_ | 1253 | 1205 | 1268 | 1693 | 2016 | 1783 |
| SL1344_RS09900_ | 1793 | 1601 | 1673 | 2325 | 2627 | 2660 |
| SL1344_RS09910_ | 706 | 655 | 620 | 732 | 879 | 949 |
| SL1344_RS09920_ | 451 | 457 | 458 | 412 | 361 | 413 |
| SL1344_RS09925_ | 14 | 15 | 21 | 23 | 25 | 10 |
| SL1344_RS09930_ | 15 | 18 | 31 | 19 | 15 | 23 |
| SL1344_RS09935_ | 161 | 178 | 171 | 151 | 129 | 134 |
| SL1344_RS09940_ | 357 | 433 | 399 | 390 | 399 | 406 |
| SL1344_RS09945_ | 183 | 178 | 198 | 207 | 166 | 196 |
| SL1344_RS09950_ | 87 | 108 | 80 | 62 | 58 | 61 |

| | | | | | | |
|---|---|---|---|---|---|---|
| SL1344_RS09955_ | 201 | 235 | 232 | 140 | 145 | 149 |
| SL1344_RS09960_ | 451 | 453 | 449 | 393 | 358 | 428 |
| SL1344_RS09965_ | 188 | 201 | 197 | 135 | 141 | 160 |
| SL1344_RS09970_ | 145 | 144 | 94 | 97 | 107 | 102 |
| SL1344_RS09975_ | 170 | 209 | 157 | 179 | 178 | 212 |
| SL1344_RS09985_ | 161 | 181 | 174 | 128 | 115 | 137 |
| SL1344_RS09990_ | 69 | 77 | 58 | 52 | 26 | 23 |
| SL1344_RS09995_ | 12 | 16 | 9 | 14 | 9 | 14 |
| SL1344_RS10000_ | 23 | 36 | 32 | 17 | 32 | 41 |
| SL1344_RS10005_ | 28 | 24 | 37 | 37 | 30 | 51 |
| SL1344_RS10010_ | 35 | 27 | 31 | 30 | 24 | 45 |
| SL1344_RS10015_ | 4 | 5 | 10 | 10 | 5 | 11 |
| SL1344_RS10020_ | 3 | 9 | 16 | 9 | 11 | 24 |
| SL1344_RS10025_ | 10 | 15 | 13 | 15 | 18 | 16 |
| SL1344_RS10030_ | 7 | 7 | 9 | 8 | 4 | 10 |
| SL1344_RS10035_ | 15 | 27 | 32 | 26 | 18 | 36 |
| SL1344_RS10040_ | 40 | 46 | 57 | 55 | 36 | 48 |
| SL1344_RS10045_ | 26 | 22 | 29 | 38 | 31 | 38 |
| SL1344_RS10050_ | 84 | 87 | 97 | 123 | 125 | 131 |
| SL1344_RS10055_ | 18 | 23 | 17 | 28 | 36 | 29 |
| SL1344_RS10060_ | 23 | 24 | 28 | 23 | 29 | 26 |
| SL1344_RS10065_ | 30 | 27 | 45 | 50 | 42 | 61 |
| SL1344_RS10070_ | 8 | 7 | 10 | 26 | 15 | 15 |
| SL1344_RS10075_ | 13 | 7 | 11 | 22 | 19 | 21 |
| SL1344_RS10080_ | 6 | 7 | 14 | 9 | 5 | 15 |
| SL1344_RS10085_ | 3 | 8 | 20 | 10 | 9 | 14 |
| SL1344_RS10090_ | 6 | 6 | 13 | 21 | 10 | 14 |
| SL1344_RS10095_ | 19 | 21 | 25 | 45 | 42 | 46 |
| SL1344_RS10100_ | 7 | 12 | 13 | 16 | 14 | 19 |
| SL1344_RS10105_ | 14 | 18 | 21 | 30 | 27 | 23 |
| SL1344_RS10110_ | 3 | 1 | 5 | 8 | 6 | 7 |
| SL1344_RS10115_ | 57 | 37 | 59 | 59 | 58 | 58 |
| SL1344_RS10120_ | 16 | 12 | 13 | 26 | 16 | 18 |
| SL1344_RS10130_ | 27 | 35 | 21 | 43 | 33 | 14 |
| SL1344_RS10135_ | 10 | 4 | 7 | 9 | 22 | 25 |
| SL1344_RS10140_ | 20 | 18 | 21 | 25 | 29 | 24 |
| SL1344_RS10145_ | 13 | 16 | 13 | 30 | 19 | 21 |
| SL1344_RS10150_ | 10 | 15 | 21 | 11 | 11 | 8 |
| SL1344_RS10155_ | 89 | 97 | 100 | 77 | 81 | 88 |
| SL1344_RS10160_ | 89 | 92 | 107 | 122 | 109 | 144 |
| SL1344_RS10165_ | 48 | 54 | 47 | 49 | 51 | 64 |
| SL1344_RS10170_ | 29 | 40 | 35 | 48 | 51 | 59 |
| SL1344_RS10175_ | 11 | 19 | 16 | 17 | 24 | 34 |
| SL1344_RS10180_ | 31 | 28 | 31 | 31 | 34 | 36 |
| SL1344_RS10185_ | 16 | 22 | 23 | 30 | 23 | 39 |
| SL1344_RS10190_ | 33 | 46 | 27 | 80 | 64 | 48 |
| SL1344_RS10195_ | 11 | 8 | 16 | 23 | 15 | 24 |
| SL1344_RS10200_ | 992 | 968 | 941 | 754 | 654 | 682 |
| SL1344_RS10205_ | 30 | 34 | 34 | 49 | 52 | 47 |
| SL1344_RS10210_ | 18 | 33 | 19 | 22 | 25 | 16 |
| SL1344_RS10215_ | 1641 | 1581 | 1388 | 1752 | 1719 | 1663 |
| SL1344_RS10225_ | 84 | 83 | 69 | 49 | 68 | 88 |
| SL1344_RS10230_ | 258 | 256 | 226 | 256 | 251 | 242 |
| SL1344_RS10235_ | 9 | 8 | 17 | 15 | 15 | 19 |
| SL1344_RS10240_ | 2 | 5 | 1 | 3 | 5 | 2 |
| SL1344_RS10245_ | 12 | 23 | 14 | 14 | 13 | 7 |
| SL1344_RS10250_ | 8 | 14 | 19 | 13 | 14 | 17 |
| SL1344_RS10255_ | 33 | 57 | 45 | 50 | 37 | 50 |
| SL1344_RS10260_ | 47 | 51 | 40 | 31 | 34 | 21 |
| SL1344_RS10265_ | 22 | 31 | 25 | 32 | 38 | 27 |
| SL1344_RS10270_ | 83 | 74 | 66 | 50 | 63 | 64 |
| SL1344_RS10275_ | 1232 | 1141 | 1109 | 674 | 603 | 746 |

| | | | | | | |
|---|---|---|---|---|---|---|
| SL1344_RS10280_ | 90 | 97 | 110 | 92 | 60 | 84 |
| SL1344_RS10285_ | 545 | 494 | 454 | 352 | 307 | 374 |
| SL1344_RS10290_ | 39 | 43 | 34 | 31 | 40 | 32 |
| SL1344_RS10295_ | 4 | 6 | 3 | 6 | 4 | 1 |
| SL1344_RS10300_ | 325 | 286 | 327 | 176 | 136 | 196 |
| SL1344_RS10305_ | 308 | 342 | 338 | 246 | 226 | 254 |
| SL1344_RS10310_ | 20 | 19 | 27 | 23 | 17 | 22 |
| SL1344_RS10315_ | 43 | 41 | 43 | 44 | 46 | 53 |
| SL1344_RS10320_ | 856 | 793 | 795 | 667 | 575 | 605 |
| SL1344_RS10325_ | 19 | 50 | 38 | 27 | 36 | 28 |
| SL1344_RS10330_ | 503 | 603 | 521 | 519 | 475 | 528 |
| SL1344_RS10335_ | 23 | 27 | 29 | 21 | 24 | 12 |
| SL1344_RS10340_ | 137 | 148 | 151 | 116 | 114 | 113 |
| SL1344_RS10345_ | 205 | 214 | 173 | 149 | 147 | 160 |
| SL1344_RS10350_ | 48 | 42 | 40 | 31 | 33 | 16 |
| SL1344_RS10355_ | 1374 | 1111 | 1307 | 812 | 778 | 949 |
| SL1344_RS10360_ | 84 | 127 | 107 | 80 | 70 | 71 |
| SL1344_RS10365_ | 361 | 354 | 362 | 283 | 264 | 283 |
| SL1344_RS10370_ | 5 | 0 | 2 | 9 | 0 | 1 |
| SL1344_RS10375_ | 95 | 124 | 111 | 129 | 127 | 111 |
| SL1344_RS10380_ | 111 | 135 | 137 | 124 | 129 | 138 |
| SL1344_RS10390_ | 19 | 51 | 45 | 51 | 40 | 32 |
| SL1344_RS10395_ | 90 | 105 | 114 | 109 | 117 | 110 |
| SL1344_RS10400_ | 58 | 70 | 52 | 68 | 63 | 63 |
| SL1344_RS10405_ | 36 | 40 | 29 | 34 | 31 | 36 |
| SL1344_RS10410_ | 8 | 9 | 9 | 5 | 13 | 13 |
| SL1344_RS10415_ | 24 | 39 | 21 | 31 | 18 | 27 |
| SL1344_RS10420_ | 13 | 24 | 26 | 31 | 24 | 17 |
| SL1344_RS10430_ | 25 | 30 | 31 | 26 | 21 | 27 |
| SL1344_RS10435_ | 29 | 35 | 26 | 21 | 34 | 13 |
| SL1344_RS10440_ | 17 | 15 | 20 | 14 | 32 | 25 |
| SL1344_RS10445_ | 11 | 10 | 7 | 8 | 9 | 16 |
| SL1344_RS10450_ | 9 | 13 | 14 | 15 | 12 | 9 |
| SL1344_RS10455_ | 9 | 7 | 15 | 8 | 16 | 7 |
| SL1344_RS10470_ | 30 | 31 | 23 | 19 | 24 | 18 |
| SL1344_RS10475_ | 25 | 26 | 23 | 23 | 22 | 23 |
| SL1344_RS10480_ | 136 | 106 | 112 | 83 | 89 | 143 |
| SL1344_RS10485_ | 63 | 44 | 56 | 40 | 36 | 57 |
| SL1344_RS10490_ | 1 | 2 | 1 | 1 | 0 | 0 |
| SL1344_RS10495_ | 4 | 7 | 5 | 7 | 6 | 1 |
| SL1344_RS10515_ | 12 | 12 | 18 | 26 | 19 | 21 |
| SL1344_RS10520_ | 2 | 2 | 0 | 3 | 0 | 1 |
| SL1344_RS10530_ | 4 | 3 | 5 | 13 | 7 | 6 |
| SL1344_RS10535_ | 10 | 12 | 7 | 13 | 13 | 12 |
| SL1344_RS10540_ | 11 | 14 | 9 | 24 | 18 | 34 |
| SL1344_RS10545_ | 9 | 13 | 9 | 9 | 13 | 15 |
| SL1344_RS10550_ | 8 | 14 | 15 | 22 | 23 | 27 |
| SL1344_RS10555_ | 9 | 24 | 10 | 37 | 16 | 35 |
| SL1344_RS10560_ | 13 | 24 | 15 | 22 | 36 | 22 |
| SL1344_RS10565_ | 17 | 26 | 19 | 30 | 42 | 40 |
| SL1344_RS10570_ | 12 | 6 | 8 | 19 | 22 | 23 |
| SL1344_RS10575_ | 3 | 3 | 5 | 12 | 7 | 7 |
| SL1344_RS10580_ | 4 | 4 | 4 | 8 | 8 | 14 |
| SL1344_RS10585_ | 42 | 60 | 50 | 54 | 65 | 74 |
| SL1344_RS10590_ | 53 | 85 | 70 | 72 | 89 | 77 |
| SL1344_RS10595_ | 4723 | 4694 | 4325 | 3571 | 3507 | 3285 |
| SL1344_RS10600_ | 286 | 239 | 251 | 277 | 220 | 247 |
| SL1344_RS10640_ | 8927 | 9940 | 9898 | 7017 | 6136 | 7067 |
| SL1344_RS10645_ | 162 | 163 | 151 | 153 | 130 | 175 |
| SL1344_RS10650_ | 615 | 592 | 526 | 476 | 470 | 414 |
| SL1344_RS10655_ | 317 | 325 | 329 | 191 | 199 | 217 |
| SL1344_RS10660_ | 295 | 288 | 301 | 202 | 186 | 197 |

| | | | | | | |
|---|---|---|---|---|---|---|
| SL1344_RS10665_ | 287 | 335 | 298 | 186 | 225 | 193 |
| SL1344_RS10670_ | 440 | 420 | 482 | 307 | 301 | 331 |
| SL1344_RS10675_ | 204 | 246 | 235 | 140 | 139 | 156 |
| SL1344_RS10680_ | 279 | 340 | 344 | 266 | 216 | 232 |
| SL1344_RS10685_ | 494 | 536 | 489 | 339 | 348 | 372 |
| SL1344_RS10690_ | 402 | 408 | 376 | 284 | 273 | 285 |
| SL1344_RS10700_ | 452 | 467 | 483 | 220 | 206 | 238 |
| SL1344_RS10705_ | 1726 | 1888 | 1987 | 1543 | 1399 | 1569 |
| SL1344_RS10710_ | 1076 | 1229 | 1230 | 948 | 822 | 1049 |
| SL1344_RS10715_ | 1038 | 1199 | 1198 | 896 | 831 | 1031 |
| SL1344_RS10720_ | 1815 | 1927 | 1934 | 1457 | 1374 | 1549 |
| SL1344_RS10725_ | 1242 | 1278 | 1214 | 928 | 904 | 931 |
| SL1344_RS10730_ | 1140 | 1191 | 1059 | 949 | 847 | 923 |
| SL1344_RS10735_ | 1679 | 1709 | 1771 | 1256 | 1153 | 1358 |
| SL1344_RS10740_ | 1732 | 1854 | 1714 | 1184 | 1025 | 1273 |
| SL1344_RS10745_ | 2956 | 3109 | 3096 | 2157 | 1830 | 2319 |
| SL1344_RS10750_ | 5398 | 6401 | 6007 | 4548 | 4083 | 4645 |
| SL1344_RS10755_ | 1519 | 1902 | 1890 | 1485 | 1175 | 1523 |
| SL1344_RS10760_ | 834 | 1044 | 1095 | 843 | 680 | 921 |
| SL1344_RS10765_ | 1296 | 1518 | 1644 | 1126 | 958 | 1258 |
| SL1344_RS10770_ | 968 | 1051 | 1046 | 779 | 724 | 924 |
| SL1344_RS10775_ | 870 | 1019 | 1073 | 832 | 708 | 879 |
| SL1344_RS10780_ | 1480 | 1707 | 1709 | 1146 | 1131 | 1259 |
| SL1344_RS10785_ | 1876 | 1897 | 1893 | 1466 | 1277 | 1455 |
| SL1344_RS10790_ | 2694 | 2929 | 2825 | 2626 | 2483 | 2658 |
| SL1344_RS10795_ | 32 | 33 | 40 | 31 | 29 | 39 |
| SL1344_RS10800_ | 22 | 17 | 21 | 22 | 18 | 20 |
| SL1344_RS10805_ | 22 | 18 | 21 | 14 | 21 | 16 |
| SL1344_RS10810_ | 23 | 32 | 34 | 33 | 23 | 35 |
| SL1344_RS10815_ | 18 | 11 | 12 | 18 | 21 | 18 |
| SL1344_RS10820_ | 13 | 19 | 13 | 15 | 5 | 15 |
| SL1344_RS10825_ | 25 | 29 | 26 | 32 | 29 | 26 |
| SL1344_RS10830_ | 32 | 30 | 31 | 30 | 27 | 25 |
| SL1344_RS10835_ | 16 | 13 | 11 | 18 | 8 | 9 |
| SL1344_RS10840_ | 12 | 21 | 21 | 23 | 19 | 21 |
| SL1344_RS10845_ | 13 | 26 | 17 | 29 | 25 | 17 |
| SL1344_RS10850_ | 7 | 5 | 2 | 5 | 5 | 4 |
| SL1344_RS10855_ | 7 | 10 | 19 | 15 | 12 | 10 |
| SL1344_RS10860_ | 24 | 25 | 14 | 17 | 16 | 37 |
| SL1344_RS10865_ | 24 | 16 | 32 | 16 | 16 | 16 |
| SL1344_RS10870_ | 20 | 4 | 6 | 7 | 5 | 6 |
| SL1344_RS10875_ | 10 | 6 | 11 | 14 | 18 | 20 |
| SL1344_RS10880_ | 15 | 20 | 19 | 27 | 15 | 28 |
| SL1344_RS10885_ | 1 | 6 | 4 | 7 | 5 | 3 |
| SL1344_RS10890_ | 0 | 3 | 6 | 8 | 7 | 9 |
| SL1344_RS10895_ | 297 | 385 | 385 | 313 | 322 | 313 |
| SL1344_RS10900_ | 2599 | 2768 | 2759 | 1993 | 1912 | 2082 |
| SL1344_RS10905_ | 910 | 970 | 1021 | 718 | 682 | 755 |
| SL1344_RS10910_ | 1500 | 1474 | 1262 | 918 | 980 | 982 |
| SL1344_RS10915_ | 579 | 571 | 572 | 392 | 379 | 384 |
| SL1344_RS10920_ | 9 | 20 | 15 | 14 | 13 | 20 |
| SL1344_RS10930_ | 67 | 68 | 57 | 68 | 68 | 67 |
| SL1344_RS10935_ | 101 | 113 | 116 | 87 | 106 | 102 |
| SL1344_RS10940_ | 110 | 116 | 107 | 102 | 99 | 88 |
| SL1344_RS10945_ | 130 | 127 | 107 | 130 | 129 | 97 |
| SL1344_RS10950_ | 209 | 190 | 147 | 169 | 174 | 177 |
| SL1344_RS10955_ | 317 | 321 | 310 | 284 | 310 | 297 |
| SL1344_RS10960_ | 38 | 34 | 40 | 21 | 35 | 30 |
| SL1344_RS10965_ | 17 | 19 | 26 | 18 | 16 | 8 |
| SL1344_RS10970_ | 80 | 73 | 87 | 71 | 77 | 86 |
| SL1344_RS10975_ | 398 | 565 | 545 | 503 | 530 | 494 |
| SL1344_RS10980_ | 20 | 25 | 30 | 21 | 16 | 23 |

| | | | | | | |
|---|---|---|---|---|---|---|
| SL1344_RS10985_ | 29 | 43 | 32 | 35 | 35 | 23 |
| SL1344_RS10990_ | 38 | 56 | 55 | 38 | 36 | 48 |
| SL1344_RS10995_ | 1 | 4 | 5 | 8 | 5 | 4 |
| SL1344_RS11000_ | 73 | 65 | 59 | 70 | 53 | 53 |
| SL1344_RS11005_ | 88 | 73 | 85 | 103 | 105 | 98 |
| SL1344_RS11010_ | 62 | 79 | 66 | 79 | 62 | 84 |
| SL1344_RS11015_ | 146 | 175 | 115 | 120 | 121 | 153 |
| SL1344_RS11020_ | 96 | 113 | 135 | 92 | 110 | 129 |
| SL1344_RS11025_ | 364 | 345 | 300 | 294 | 236 | 227 |
| SL1344_RS11030_ | 78 | 146 | 156 | 106 | 62 | 98 |
| SL1344_RS11035_ | 104 | 94 | 141 | 97 | 81 | 91 |
| SL1344_RS11040_ | 430 | 460 | 397 | 342 | 338 | 302 |
| SL1344_RS11045_ | 31 | 23 | 24 | 19 | 19 | 18 |
| SL1344_RS11050_ | 131 | 36 | 80 | 112 | 33 | 93 |
| SL1344_RS11055_ | 8 | 4 | 2 | 2 | 1 | 7 |
| SL1344_RS11060_ | 6 | 8 | 11 | 12 | 12 | 5 |
| SL1344_RS11065_ | 287 | 258 | 201 | 227 | 207 | 192 |
| SL1344_RS11070_ | 1785 | 1865 | 1791 | 1652 | 1626 | 1648 |
| SL1344_RS11080_ | 63 | 76 | 65 | 55 | 64 | 69 |
| SL1344_RS11085_ | 44 | 30 | 29 | 22 | 28 | 33 |
| SL1344_RS11090_ | 718 | 706 | 735 | 497 | 515 | 556 |
| SL1344_RS11095_ | 231 | 224 | 284 | 155 | 119 | 146 |
| SL1344_RS11100_ | 383 | 419 | 423 | 233 | 243 | 244 |
| SL1344_RS11105_ | 45 | 43 | 42 | 39 | 38 | 40 |
| SL1344_RS11110_ | 367 | 391 | 312 | 265 | 265 | 305 |
| SL1344_RS11115_ | 58 | 40 | 46 | 30 | 38 | 35 |
| SL1344_RS11120_ | 71 | 61 | 68 | 54 | 56 | 46 |
| SL1344_RS11125_ | 65 | 78 | 83 | 69 | 78 | 66 |
| SL1344_RS11130_ | 60 | 66 | 85 | 76 | 85 | 70 |
| SL1344_RS11135_ | 612 | 676 | 686 | 511 | 492 | 544 |
| SL1344_RS11140_ | 400 | 383 | 413 | 378 | 328 | 357 |
| SL1344_RS11145_ | 1087 | 996 | 987 | 702 | 674 | 655 |
| SL1344_RS11150_ | 16 | 13 | 20 | 14 | 8 | 20 |
| SL1344_RS11155_ | 15 | 22 | 22 | 25 | 19 | 22 |
| SL1344_RS11160_ | 19 | 30 | 53 | 36 | 37 | 40 |
| SL1344_RS11165_ | 75 | 43 | 44 | 34 | 33 | 40 |
| SL1344_RS11170_ | 349 | 359 | 423 | 350 | 311 | 320 |
| SL1344_RS11175_ | 83 | 74 | 83 | 95 | 67 | 71 |
| SL1344_RS11180_ | 19 | 20 | 22 | 25 | 24 | 14 |
| SL1344_RS11185_ | 11 | 4 | 17 | 17 | 20 | 15 |
| SL1344_RS11190_ | 15 | 18 | 11 | 11 | 7 | 9 |
| SL1344_RS11195_ | 26 | 12 | 8 | 13 | 14 | 18 |
| SL1344_RS11200_ | 184 | 217 | 171 | 207 | 214 | 214 |
| SL1344_RS11205_ | 104 | 79 | 61 | 88 | 51 | 64 |
| SL1344_RS11210_ | 159 | 114 | 119 | 139 | 101 | 118 |
| SL1344_RS11215_ | 1234 | 1262 | 1189 | 1070 | 936 | 963 |
| SL1344_RS11220_ | 751 | 797 | 700 | 544 | 501 | 544 |
| SL1344_RS11225_ | 218 | 217 | 205 | 159 | 148 | 158 |
| SL1344_RS11230_ | 43 | 36 | 22 | 14 | 10 | 21 |
| SL1344_RS11235_ | 86 | 72 | 86 | 57 | 71 | 54 |
| SL1344_RS11240_ | 40 | 38 | 35 | 40 | 38 | 42 |
| SL1344_RS11245_ | 65 | 47 | 58 | 56 | 50 | 83 |
| SL1344_RS11250_ | 194 | 218 | 165 | 215 | 156 | 184 |
| SL1344_RS11255_ | 94 | 95 | 95 | 66 | 46 | 68 |
| SL1344_RS11260_ | 617 | 633 | 687 | 551 | 490 | 563 |
| SL1344_RS11265_ | 657 | 735 | 722 | 511 | 530 | 510 |
| SL1344_RS11270_ | 986 | 1045 | 956 | 987 | 897 | 982 |
| SL1344_RS11275_ | 46 | 60 | 36 | 30 | 40 | 27 |
| SL1344_RS11280_ | 91 | 107 | 70 | 62 | 75 | 75 |
| SL1344_RS11285_ | 6 | 7 | 4 | 5 | 7 | 6 |
| SL1344_RS11290_ | 12 | 23 | 20 | 14 | 14 | 20 |
| SL1344_RS11295_ | 278 | 444 | 435 | 305 | 279 | 258 |

| | | | | | | |
|---|---|---|---|---|---|---|
| SL1344_RS11300_ | 854 | 1023 | 1224 | 987 | 932 | 1038 |
| SL1344_RS11305_ | 160 | 228 | 211 | 280 | 265 | 267 |
| SL1344_RS11310_ | 217 | 225 | 189 | 131 | 98 | 126 |
| SL1344_RS11315_ | 571 | 634 | 636 | 509 | 427 | 487 |
| SL1344_RS11320_ | 1711 | 1841 | 1809 | 1107 | 955 | 1145 |
| SL1344_RS11325_ | 976 | 1115 | 1122 | 602 | 570 | 647 |
| SL1344_RS11330_ | 1216 | 1357 | 1401 | 861 | 729 | 858 |
| SL1344_RS11335_ | 136 | 154 | 168 | 88 | 73 | 107 |
| SL1344_RS11340_ | 37 | 54 | 42 | 49 | 42 | 41 |
| SL1344_RS11345_ | 12 | 11 | 20 | 12 | 18 | 12 |
| SL1344_RS11350_ | 59 | 76 | 58 | 56 | 35 | 39 |
| SL1344_RS11355_ | 2345 | 2411 | 2251 | 1700 | 1638 | 1543 |
| SL1344_RS11360_ | 326 | 308 | 314 | 205 | 208 | 245 |
| SL1344_RS11365_ | 2181 | 2110 | 2078 | 1233 | 1183 | 1370 |
| SL1344_RS11375_ | 489 | 466 | 459 | 379 | 357 | 356 |
| SL1344_RS11380_ | 386 | 408 | 417 | 349 | 337 | 380 |
| SL1344_RS11385_ | 304 | 331 | 301 | 239 | 256 | 286 |
| SL1344_RS11390_ | 198 | 214 | 219 | 153 | 164 | 185 |
| SL1344_RS11395_ | 419 | 411 | 440 | 284 | 308 | 316 |
| SL1344_RS11400_ | 1061 | 1206 | 1333 | 1109 | 1078 | 1304 |
| SL1344_RS11405_ | 50 | 50 | 69 | 37 | 33 | 42 |
| SL1344_RS11410_ | 683 | 721 | 789 | 529 | 436 | 536 |
| SL1344_RS11415_ | 684 | 708 | 655 | 485 | 456 | 491 |
| SL1344_RS11420_ | 416 | 443 | 407 | 333 | 339 | 367 |
| SL1344_RS11425_ | 14230 | 14555 | 14294 | 13350 | 12623 | 13298 |
| SL1344_RS11430_ | 360 | 430 | 442 | 375 | 342 | 352 |
| SL1344_RS11435_ | 650 | 753 | 795 | 595 | 607 | 653 |
| SL1344_RS11440_ | 733 | 779 | 736 | 583 | 531 | 631 |
| SL1344_RS11445_ | 2182 | 2086 | 2154 | 1561 | 1526 | 1804 |
| SL1344_RS11450_ | 506 | 498 | 538 | 364 | 319 | 404 |
| SL1344_RS11455_ | 11 | 11 | 30 | 8 | 11 | 6 |
| SL1344_RS11460_ | 2 | 3 | 2 | 5 | 5 | 7 |
| SL1344_RS11465_ | 1364 | 1608 | 1646 | 1138 | 1033 | 1223 |
| SL1344_RS11470_ | 45 | 54 | 58 | 29 | 32 | 17 |
| SL1344_RS11475_ | 10 | 18 | 13 | 10 | 13 | 16 |
| SL1344_RS11480_ | 23 | 18 | 17 | 21 | 11 | 15 |
| SL1344_RS11485_ | 9 | 4 | 9 | 5 | 3 | 4 |
| SL1344_RS11490_ | 2 | 4 | 3 | 4 | 3 | 3 |
| SL1344_RS11495_ | 673 | 640 | 606 | 493 | 419 | 471 |
| SL1344_RS11500_ | 27 | 29 | 23 | 27 | 33 | 22 |
| SL1344_RS11505_ | 34 | 48 | 23 | 19 | 30 | 24 |
| SL1344_RS11510_ | 18 | 23 | 18 | 23 | 9 | 18 |
| SL1344_RS11515_ | 235 | 236 | 221 | 240 | 194 | 222 |
| SL1344_RS11520_ | 0 | 3 | 1 | 2 | 2 | 2 |
| SL1344_RS11525_ | 27 | 29 | 19 | 15 | 28 | 25 |
| SL1344_RS11535_ | 7 | 5 | 3 | 10 | 5 | 4 |
| SL1344_RS11540_ | 231 | 225 | 240 | 267 | 242 | 214 |
| SL1344_RS11545_ | 79 | 97 | 115 | 105 | 78 | 101 |
| SL1344_RS11550_ | 5 | 5 | 8 | 4 | 7 | 8 |
| SL1344_RS11555_ | 13 | 21 | 32 | 25 | 24 | 30 |
| SL1344_RS11560_ | 19 | 23 | 12 | 11 | 16 | 12 |
| SL1344_RS11565_ | 4 | 2 | 1 | 2 | 2 | 2 |
| SL1344_RS11570_ | 4 | 9 | 4 | 9 | 7 | 8 |
| SL1344_RS11585_ | 60 | 58 | 59 | 58 | 72 | 75 |
| SL1344_RS11595_ | 161 | 159 | 188 | 144 | 137 | 172 |
| SL1344_RS11600_ | 70 | 93 | 104 | 74 | 61 | 88 |
| SL1344_RS11605_ | 287 | 351 | 343 | 377 | 303 | 348 |
| SL1344_RS11610_ | 23 | 32 | 29 | 29 | 35 | 29 |
| SL1344_RS11615_ | 51 | 57 | 56 | 73 | 53 | 48 |
| SL1344_RS11620_ | 257 | 200 | 212 | 141 | 125 | 93 |
| SL1344_RS11625_ | 527 | 638 | 670 | 500 | 470 | 554 |
| SL1344_RS11630_ | 120 | 146 | 138 | 159 | 172 | 169 |

| | | | | | | |
|---|---|---|---|---|---|---|
| SL1344_RS11635_ | 298 | 309 | 355 | 302 | 305 | 356 |
| SL1344_RS11640_ | 213 | 253 | 278 | 294 | 245 | 280 |
| SL1344_RS11670_ | 150 | 122 | 127 | 99 | 91 | 110 |
| SL1344_RS11675_ | 10 | 11 | 15 | 19 | 8 | 17 |
| SL1344_RS11680_ | 26 | 24 | 24 | 33 | 66 | 50 |
| SL1344_RS11685_ | 876 | 951 | 872 | 807 | 744 | 839 |
| SL1344_RS11690_ | 4240 | 4604 | 4783 | 4119 | 4056 | 4472 |
| SL1344_RS11695_ | 1782 | 1976 | 2123 | 1774 | 1614 | 1847 |
| SL1344_RS11700_ | 1562 | 1700 | 1821 | 1681 | 1689 | 1697 |
| SL1344_RS11705_ | 182 | 179 | 137 | 107 | 80 | 107 |
| SL1344_RS11710_ | 58 | 74 | 61 | 56 | 62 | 58 |
| SL1344_RS11715_ | 945 | 1063 | 989 | 937 | 851 | 973 |
| SL1344_RS11720_ | 725 | 719 | 694 | 569 | 520 | 584 |
| SL1344_RS11725_ | 96 | 96 | 84 | 78 | 61 | 71 |
| SL1344_RS11730_ | 72 | 80 | 75 | 70 | 53 | 61 |
| SL1344_RS11735_ | 148 | 179 | 188 | 169 | 173 | 129 |
| SL1344_RS11740_ | 1 | 2 | 1 | 0 | 0 | 1 |
| SL1344_RS11745_ | 9 | 15 | 10 | 11 | 13 | 6 |
| SL1344_RS11750_ | 62 | 58 | 53 | 51 | 55 | 51 |
| SL1344_RS11755_ | 15 | 25 | 27 | 18 | 18 | 21 |
| SL1344_RS11760_ | 29 | 27 | 40 | 30 | 32 | 19 |
| SL1344_RS11765_ | 7 | 6 | 6 | 12 | 7 | 5 |
| SL1344_RS11770_ | 454 | 489 | 478 | 435 | 393 | 396 |
| SL1344_RS11775_ | 542 | 680 | 633 | 496 | 479 | 572 |
| SL1344_RS11780_ | 205 | 251 | 218 | 172 | 158 | 148 |
| SL1344_RS11785_ | 601 | 670 | 746 | 279 | 248 | 389 |
| SL1344_RS11790_ | 652 | 645 | 830 | 354 | 369 | 395 |
| SL1344_RS11795_ | 658 | 668 | 758 | 358 | 360 | 388 |
| SL1344_RS11800_ | 1317 | 1241 | 1333 | 755 | 683 | 845 |
| SL1344_RS11805_ | 567 | 560 | 550 | 313 | 300 | 323 |
| SL1344_RS11810_ | 495 | 548 | 591 | 394 | 318 | 373 |
| SL1344_RS11815_ | 170 | 195 | 213 | 146 | 113 | 142 |
| SL1344_RS11820_ | 157 | 176 | 181 | 86 | 89 | 121 |
| SL1344_RS11825_ | 423 | 452 | 435 | 384 | 364 | 454 |
| SL1344_RS11830_ | 709 | 763 | 780 | 614 | 631 | 666 |
| SL1344_RS11840_ | 369 | 434 | 437 | 385 | 296 | 342 |
| SL1344_RS11845_ | 223 | 211 | 218 | 168 | 160 | 187 |
| SL1344_RS11850_ | 455 | 472 | 528 | 333 | 342 | 399 |
| SL1344_RS11855_ | 487 | 452 | 512 | 438 | 443 | 456 |
| SL1344_RS11860_ | 299 | 393 | 392 | 398 | 358 | 353 |
| SL1344_RS11865_ | 41 | 61 | 50 | 49 | 69 | 56 |
| SL1344_RS11870_ | 189 | 183 | 191 | 212 | 195 | 185 |
| SL1344_RS11880_ | 107 | 117 | 95 | 72 | 83 | 79 |
| SL1344_RS11885_ | 1682 | 2181 | 2335 | 2069 | 2067 | 2100 |
| SL1344_RS11890_ | 946 | 1202 | 1388 | 1184 | 1188 | 1155 |
| SL1344_RS11895_ | 3419 | 3770 | 3940 | 3093 | 2963 | 3256 |
| SL1344_RS11900_ | 632 | 738 | 746 | 624 | 577 | 591 |
| SL1344_RS11905_ | 1646 | 1765 | 2077 | 1466 | 1391 | 1501 |
| SL1344_RS11910_ | 1362 | 1416 | 1437 | 1196 | 1177 | 1263 |
| SL1344_RS11915_ | 2176 | 2566 | 2702 | 2155 | 2008 | 2275 |
| SL1344_RS11920_ | 7541 | 7987 | 8227 | 6703 | 6260 | 6913 |
| SL1344_RS11925_ | 3936 | 4160 | 4307 | 3279 | 3093 | 3463 |
| SL1344_RS11930_ | 1688 | 1863 | 1909 | 1553 | 1375 | 1609 |
| SL1344_RS11935_ | 5806 | 5940 | 6033 | 4677 | 4419 | 4920 |
| SL1344_RS11940_ | 2963 | 2728 | 2834 | 2329 | 2159 | 2274 |
| SL1344_RS11945_ | 2595 | 2547 | 2652 | 2112 | 1946 | 2077 |
| SL1344_RS11950_ | 30 | 16 | 37 | 16 | 17 | 14 |
| SL1344_RS11960_ | 2567 | 2644 | 2470 | 1705 | 1665 | 1889 |
| SL1344_RS11965_ | 151 | 142 | 156 | 135 | 143 | 130 |
| SL1344_RS11970_ | 229 | 251 | 293 | 255 | 213 | 289 |
| SL1344_RS11975_ | 1035 | 1108 | 1060 | 881 | 851 | 947 |
| SL1344_RS11980_ | 352 | 404 | 333 | 351 | 316 | 336 |

| | | | | | | |
|---|---|---|---|---|---|---|
| SL1344_RS11985_ | 391 | 364 | 385 | 341 | 367 | 368 |
| SL1344_RS12000_ | 213 | 217 | 254 | 144 | 148 | 184 |
| SL1344_RS12005_ | 61 | 51 | 70 | 63 | 53 | 72 |
| SL1344_RS12010_ | 18 | 17 | 13 | 16 | 14 | 16 |
| SL1344_RS12015_ | 25 | 29 | 15 | 18 | 26 | 31 |
| SL1344_RS12020_ | 6 | 7 | 6 | 1 | 6 | 10 |
| SL1344_RS12025_ | 16 | 14 | 17 | 19 | 17 | 9 |
| SL1344_RS12030_ | 3 | 0 | 2 | 2 | 3 | 3 |
| SL1344_RS12035_ | 154 | 148 | 155 | 147 | 119 | 131 |
| SL1344_RS12040_ | 1231 | 1395 | 1457 | 1486 | 1421 | 1438 |
| SL1344_RS12045_ | 505 | 668 | 554 | 611 | 528 | 569 |
| SL1344_RS12050_ | 91 | 108 | 99 | 103 | 104 | 123 |
| SL1344_RS12055_ | 17 | 20 | 10 | 15 | 12 | 5 |
| SL1344_RS12060_ | 176 | 165 | 160 | 133 | 139 | 135 |
| SL1344_RS12065_ | 85 | 132 | 127 | 127 | 99 | 132 |
| SL1344_RS12070_ | 105 | 142 | 141 | 95 | 91 | 126 |
| SL1344_RS12075_ | 146 | 191 | 206 | 176 | 138 | 163 |
| SL1344_RS12080_ | 1224 | 1417 | 1396 | 1204 | 1050 | 1146 |
| SL1344_RS12085_ | 91 | 92 | 84 | 54 | 57 | 70 |
| SL1344_RS12090_ | 506 | 487 | 492 | 435 | 382 | 432 |
| SL1344_RS12095_ | 538 | 489 | 463 | 412 | 400 | 378 |
| SL1344_RS12100_ | 126 | 131 | 111 | 108 | 101 | 118 |
| SL1344_RS12105_ | 74 | 74 | 53 | 54 | 53 | 55 |
| SL1344_RS12110_ | 61 | 55 | 64 | 47 | 51 | 60 |
| SL1344_RS12115_ | 141 | 123 | 117 | 106 | 105 | 98 |
| SL1344_RS12120_ | 1674 | 1659 | 1532 | 1606 | 1480 | 1467 |
| SL1344_RS12125_ | 781 | 775 | 783 | 724 | 687 | 724 |
| SL1344_RS12130_ | 1037 | 1111 | 1127 | 1082 | 1080 | 1044 |
| SL1344_RS12135_ | 1276 | 1360 | 1350 | 1201 | 1160 | 1111 |
| SL1344_RS12140_ | 5650 | 5676 | 5842 | 4435 | 4205 | 4524 |
| SL1344_RS12145_ | 474 | 428 | 439 | 315 | 293 | 367 |
| SL1344_RS12155_ | 908 | 1104 | 1271 | 852 | 887 | 921 |
| SL1344_RS12160_ | 1728 | 1799 | 1787 | 1533 | 1349 | 1534 |
| SL1344_RS12170_ | 498 | 560 | 574 | 469 | 444 | 493 |
| SL1344_RS12175_ | 9 | 5 | 7 | 5 | 8 | 2 |
| SL1344_RS12180_ | 9 | 10 | 8 | 8 | 4 | 8 |
| SL1344_RS12185_ | 10 | 9 | 2 | 2 | 5 | 5 |
| SL1344_RS12190_ | 13 | 16 | 13 | 4 | 8 | 12 |
| SL1344_RS12195_ | 120 | 146 | 179 | 122 | 102 | 120 |
| SL1344_RS12200_ | 3236 | 3827 | 4015 | 4540 | 4043 | 4442 |
| SL1344_RS12205_ | 500 | 487 | 508 | 454 | 467 | 484 |
| SL1344_RS12210_ | 1051 | 1065 | 1167 | 799 | 724 | 721 |
| SL1344_RS12215_ | 427 | 459 | 488 | 324 | 289 | 305 |
| SL1344_RS12220_ | 472 | 516 | 517 | 346 | 357 | 373 |
| SL1344_RS12225_ | 455 | 491 | 505 | 366 | 342 | 388 |
| SL1344_RS12230_ | 661 | 770 | 809 | 637 | 585 | 638 |
| SL1344_RS12235_ | 928 | 1048 | 952 | 727 | 697 | 747 |
| SL1344_RS12240_ | 155 | 186 | 134 | 146 | 128 | 136 |
| SL1344_RS12245_ | 2349 | 2566 | 2938 | 1944 | 1941 | 2058 |
| SL1344_RS12250_ | 117 | 135 | 130 | 181 | 157 | 133 |
| SL1344_RS12260_ | 769 | 737 | 495 | 698 | 532 | 443 |
| SL1344_RS12265_ | 2906 | 2671 | 2779 | 2234 | 2152 | 2261 |
| SL1344_RS12270_ | 2386 | 2532 | 2372 | 2312 | 2314 | 2307 |
| SL1344_RS12275_ | 21 | 37 | 36 | 19 | 27 | 31 |
| SL1344_RS12280_ | 131 | 165 | 134 | 88 | 88 | 86 |
| SL1344_RS12285_ | 107 | 82 | 61 | 65 | 56 | 68 |
| SL1344_RS12290_ | 159 | 185 | 192 | 157 | 140 | 189 |
| SL1344_RS12295_ | 168 | 143 | 145 | 113 | 107 | 137 |
| SL1344_RS12300_ | 29 | 34 | 33 | 29 | 29 | 31 |
| SL1344_RS12305_ | 6 | 3 | 3 | 5 | 1 | 3 |
| SL1344_RS12310_ | 28 | 30 | 28 | 25 | 21 | 16 |
| SL1344_RS12315_ | 64 | 61 | 43 | 59 | 48 | 39 |

| | | | | | | |
|---|---|---|---|---|---|---|
| SL1344_RS12320_ | 0 | 3 | 1 | 1 | 3 | 0 |
| SL1344_RS12325_ | 180 | 167 | 166 | 97 | 121 | 123 |
| SL1344_RS12330_ | 715 | 817 | 801 | 905 | 850 | 794 |
| SL1344_RS12335_ | 333 | 264 | 319 | 200 | 216 | 216 |
| SL1344_RS12340_ | 96 | 128 | 126 | 121 | 94 | 94 |
| SL1344_RS12345_ | 90 | 109 | 63 | 95 | 99 | 90 |
| SL1344_RS12350_ | 188 | 199 | 190 | 191 | 174 | 197 |
| SL1344_RS12355_ | 143 | 171 | 148 | 142 | 175 | 166 |
| SL1344_RS12360_ | 225 | 225 | 236 | 166 | 135 | 145 |
| SL1344_RS12365_ | 697 | 768 | 747 | 512 | 470 | 464 |
| SL1344_RS12370_ | 623 | 580 | 583 | 518 | 477 | 529 |
| SL1344_RS12375_ | 3414 | 3208 | 3617 | 3512 | 3052 | 4002 |
| SL1344_RS12380_ | 15996 | 14954 | 11375 | 16011 | 14674 | 16443 |
| SL1344_RS12385_ | 135 | 127 | 96 | 105 | 91 | 97 |
| SL1344_RS12390_ | 204 | 221 | 156 | 185 | 196 | 148 |
| SL1344_RS12395_ | 3842 | 3942 | 3907 | 3171 | 2920 | 2999 |
| SL1344_RS12400_ | 8916 | 7967 | 6831 | 5680 | 5473 | 6457 |
| SL1344_RS12405_ | 7102 | 6392 | 7026 | 4277 | 4204 | 5007 |
| SL1344_RS12410_ | 11445 | 10081 | 10912 | 8032 | 7689 | 8679 |
| SL1344_RS12415_ | 5957 | 5062 | 5539 | 3974 | 4043 | 4164 |
| SL1344_RS12420_ | 45 | 43 | 38 | 35 | 44 | 27 |
| SL1344_RS12425_ | 4 | 5 | 3 | 6 | 3 | 2 |
| SL1344_RS12430_ | 31 | 38 | 37 | 62 | 31 | 42 |
| SL1344_RS12435_ | 4 | 3 | 5 | 3 | 3 | 6 |
| SL1344_RS12440_ | 40 | 50 | 20 | 38 | 45 | 24 |
| SL1344_RS12445_ | 109 | 123 | 98 | 92 | 101 | 98 |
| SL1344_RS12450_ | 382 | 401 | 402 | 283 | 297 | 333 |
| SL1344_RS12455_ | 441 | 499 | 483 | 324 | 388 | 363 |
| SL1344_RS12465_ | 3301 | 3128 | 3016 | 2469 | 2454 | 2471 |
| SL1344_RS12480_ | 1 | 4 | 4 | 5 | 4 | 5 |
| SL1344_RS12485_ | 4013 | 3689 | 3556 | 3139 | 2742 | 2966 |
| SL1344_RS12490_ | 25150 | 25164 | 24802 | 20654 | 19045 | 19580 |
| SL1344_RS12495_ | 7109 | 8867 | 9342 | 6878 | 6402 | 6988 |
| SL1344_RS12500_ | 28 | 32 | 42 | 56 | 38 | 32 |
| SL1344_RS12510_ | 128 | 158 | 138 | 129 | 116 | 116 |
| SL1344_RS12515_ | 82 | 115 | 97 | 77 | 80 | 86 |
| SL1344_RS12520_ | 34 | 46 | 46 | 50 | 44 | 38 |
| SL1344_RS12525_ | 77 | 67 | 68 | 65 | 77 | 63 |
| SL1344_RS12555_ | 144 | 151 | 174 | 136 | 125 | 145 |
| SL1344_RS12560_ | 767 | 824 | 824 | 720 | 635 | 715 |
| SL1344_RS12565_ | 381 | 348 | 382 | 340 | 287 | 337 |
| SL1344_RS12570_ | 320 | 367 | 357 | 305 | 291 | 322 |
| SL1344_RS12575_ | 552 | 571 | 569 | 530 | 464 | 453 |
| SL1344_RS12580_ | 410 | 413 | 362 | 309 | 317 | 283 |
| SL1344_RS12585_ | 260 | 291 | 312 | 206 | 209 | 203 |
| SL1344_RS12590_ | 79 | 124 | 109 | 73 | 82 | 85 |
| SL1344_RS12595_ | 199 | 205 | 166 | 172 | 160 | 137 |
| SL1344_RS12600_ | 36 | 47 | 32 | 34 | 25 | 28 |
| SL1344_RS12605_ | 19 | 18 | 18 | 14 | 77 | 29 |
| SL1344_RS12610_ | 7 | 9 | 9 | 2 | 14 | 4 |
| SL1344_RS12615_ | 6 | 11 | 19 | 13 | 10 | 12 |
| SL1344_RS12620_ | 18 | 12 | 17 | 15 | 17 | 16 |
| SL1344_RS12630_ | 13 | 17 | 16 | 14 | 23 | 11 |
| SL1344_RS12635_ | 13 | 8 | 12 | 6 | 13 | 11 |
| SL1344_RS12640_ | 19 | 17 | 24 | 9 | 14 | 15 |
| SL1344_RS12645_ | 28 | 16 | 20 | 19 | 27 | 16 |
| SL1344_RS12650_ | 2 | 1 | 3 | 5 | 3 | 3 |
| SL1344_RS12655_ | 7 | 8 | 8 | 2 | 9 | 19 |
| SL1344_RS12660_ | 10 | 6 | 4 | 4 | 5 | 5 |
| SL1344_RS12665_ | 3 | 5 | 1 | 7 | 4 | 8 |
| SL1344_RS12670_ | 5 | 9 | 4 | 6 | 6 | 5 |
| SL1344_RS12675_ | 2 | 7 | 1 | 5 | 1 | 6 |

| | | | | | | |
|---|---|---|---|---|---|---|
| SL1344_RS12690_ | 699 | 689 | 671 | 768 | 737 | 738 |
| SL1344_RS12695_ | 93 | 72 | 66 | 56 | 75 | 80 |
| SL1344_RS12700_ | 123 | 158 | 136 | 146 | 151 | 179 |
| SL1344_RS12705_ | 145 | 192 | 166 | 160 | 165 | 172 |
| SL1344_RS12710_ | 132 | 159 | 150 | 132 | 109 | 121 |
| SL1344_RS12715_ | 323 | 362 | 342 | 320 | 314 | 289 |
| SL1344_RS12720_ | 130 | 102 | 99 | 80 | 80 | 79 |
| SL1344_RS12725_ | 27 | 61 | 43 | 42 | 38 | 40 |
| SL1344_RS12730_ | 67 | 58 | 59 | 49 | 59 | 68 |
| SL1344_RS12735_ | 328 | 288 | 297 | 293 | 267 | 265 |
| SL1344_RS12740_ | 226 | 252 | 221 | 161 | 187 | 190 |
| SL1344_RS12745_ | 927 | 994 | 965 | 748 | 744 | 741 |
| SL1344_RS12750_ | 480 | 510 | 533 | 420 | 388 | 412 |
| SL1344_RS12755_ | 435 | 426 | 470 | 414 | 413 | 444 |
| SL1344_RS12760_ | 313 | 323 | 322 | 331 | 343 | 361 |
| SL1344_RS12765_ | 347 | 477 | 463 | 523 | 495 | 466 |
| SL1344_RS12770_ | 6031 | 6027 | 6108 | 5866 | 5767 | 5986 |
| SL1344_RS12775_ | 2729 | 2913 | 2880 | 2765 | 2730 | 2765 |
| SL1344_RS12780_ | 706 | 695 | 526 | 644 | 648 | 585 |
| SL1344_RS12785_ | 1828 | 1874 | 1751 | 1835 | 1898 | 1761 |
| SL1344_RS12790_ | 35 | 43 | 51 | 41 | 44 | 53 |
| SL1344_RS12795_ | 280 | 329 | 374 | 397 | 355 | 406 |
| SL1344_RS12800_ | 501 | 507 | 464 | 584 | 518 | 529 |
| SL1344_RS12805_ | 375 | 411 | 381 | 476 | 547 | 458 |
| SL1344_RS12810_ | 1242 | 1360 | 1455 | 1163 | 1292 | 1395 |
| SL1344_RS12815_ | 657 | 676 | 670 | 623 | 564 | 559 |
| SL1344_RS12820_ | 3292 | 3103 | 2970 | 3313 | 3064 | 3212 |
| SL1344_RS12825_ | 704 | 690 | 642 | 837 | 713 | 695 |
| SL1344_RS12830_ | 523 | 569 | 565 | 651 | 573 | 612 |
| SL1344_RS12835_ | 963 | 1038 | 942 | 866 | 871 | 902 |
| SL1344_RS12840_ | 992 | 1017 | 1084 | 831 | 756 | 858 |
| SL1344_RS12845_ | 50 | 30 | 30 | 50 | 23 | 52 |
| SL1344_RS12850_ | 10 | 13 | 22 | 16 | 12 | 9 |
| SL1344_RS12855_ | 397 | 375 | 420 | 423 | 426 | 464 |
| SL1344_RS12860_ | 40 | 42 | 53 | 33 | 27 | 48 |
| SL1344_RS12865_ | 58 | 68 | 69 | 59 | 53 | 65 |
| SL1344_RS12870_ | 3129 | 3927 | 4349 | 3787 | 3427 | 3677 |
| SL1344_RS12875_ | 1935 | 2420 | 2632 | 2261 | 1938 | 2297 |
| SL1344_RS12880_ | 1310 | 1192 | 1369 | 1096 | 1092 | 1160 |
| SL1344_RS12885_ | 230 | 263 | 257 | 290 | 254 | 281 |
| SL1344_RS12890_ | 757 | 882 | 852 | 873 | 822 | 848 |
| SL1344_RS12900_ | 64 | 35 | 40 | 46 | 58 | 44 |
| SL1344_RS12905_ | 30 | 19 | 27 | 40 | 45 | 31 |
| SL1344_RS12910_ | 306 | 281 | 312 | 246 | 233 | 264 |
| SL1344_RS12915_ | 1337 | 1514 | 1559 | 1217 | 1122 | 1334 |
| SL1344_RS12920_ | 1600 | 1821 | 1890 | 1849 | 1728 | 1899 |
| SL1344_RS12925_ | 1161 | 1282 | 1323 | 1258 | 1205 | 1315 |
| SL1344_RS12930_ | 1910 | 2213 | 2271 | 2126 | 2063 | 2143 |
| SL1344_RS12935_ | 1671 | 1742 | 1881 | 1685 | 1717 | 1771 |
| SL1344_RS12940_ | 1289 | 1236 | 1299 | 1037 | 1173 | 1196 |
| SL1344_RS12945_ | 2128 | 2291 | 2322 | 2008 | 2055 | 2060 |
| SL1344_RS12950_ | 2311 | 2260 | 2001 | 2826 | 2636 | 2851 |
| SL1344_RS12955_ | 27 | 31 | 22 | 27 | 24 | 19 |
| SL1344_RS12960_ | 34 | 30 | 37 | 34 | 42 | 44 |
| SL1344_RS12965_ | 23 | 19 | 28 | 34 | 23 | 22 |
| SL1344_RS12970_ | 53 | 79 | 72 | 69 | 87 | 77 |
| SL1344_RS12975_ | 259 | 274 | 308 | 270 | 259 | 272 |
| SL1344_RS12980_ | 1061 | 1097 | 1067 | 1105 | 1081 | 1145 |
| SL1344_RS12985_ | 487 | 475 | 439 | 501 | 432 | 485 |
| SL1344_RS12990_ | 87 | 83 | 76 | 59 | 76 | 79 |
| SL1344_RS12995_ | 524 | 649 | 676 | 578 | 576 | 641 |
| SL1344_RS13000_ | 1001 | 1135 | 1253 | 1176 | 1042 | 1231 |

| | | | | | | |
|---|---|---|---|---|---|---|
| SL1344_RS13005_ | 1189 | 1277 | 1253 | 1170 | 1150 | 1185 |
| SL1344_RS13010_ | 1773 | 1774 | 1784 | 1645 | 1772 | 1637 |
| SL1344_RS13015_ | 2976 | 3137 | 3105 | 2763 | 2737 | 2784 |
| SL1344_RS13025_ | 2770 | 3132 | 3004 | 3174 | 2983 | 3123 |
| SL1344_RS13030_ | 1742 | 1874 | 1701 | 1913 | 1746 | 1895 |
| SL1344_RS13035_ | 4385 | 4681 | 4813 | 4639 | 4677 | 4757 |
| SL1344_RS13040_ | 1945 | 1977 | 1797 | 1802 | 1725 | 1868 |
| SL1344_RS13045_ | 1147 | 1091 | 1269 | 1008 | 944 | 991 |
| SL1344_RS13050_ | 6877 | 6540 | 6775 | 4776 | 4969 | 5205 |
| SL1344_RS13075_ | 82 | 131 | 140 | 132 | 113 | 136 |
| SL1344_RS13080_ | 55 | 74 | 76 | 99 | 69 | 86 |
| SL1344_RS13085_ | 46 | 34 | 28 | 36 | 21 | 24 |
| SL1344_RS13090_ | 649 | 638 | 660 | 652 | 601 | 700 |
| SL1344_RS13095_ | 3398 | 3544 | 3442 | 4195 | 4005 | 3832 |
| SL1344_RS13100_ | 251 | 234 | 226 | 297 | 292 | 307 |
| SL1344_RS13105_ | 852 | 826 | 823 | 693 | 708 | 722 |
| SL1344_RS13115_ | 410 | 494 | 503 | 628 | 617 | 606 |
| SL1344_RS13120_ | 98 | 135 | 150 | 129 | 87 | 135 |
| SL1344_RS13125_ | 889 | 977 | 983 | 886 | 875 | 900 |
| SL1344_RS13130_ | 862 | 955 | 935 | 871 | 825 | 868 |
| SL1344_RS13135_ | 369 | 371 | 392 | 333 | 341 | 364 |
| SL1344_RS13140_ | 362 | 429 | 469 | 428 | 409 | 406 |
| SL1344_RS13145_ | 949 | 1018 | 1080 | 1113 | 1074 | 1140 |
| SL1344_RS13150_ | 139 | 133 | 106 | 110 | 73 | 118 |
| SL1344_RS13155_ | 748 | 750 | 682 | 620 | 578 | 663 |
| SL1344_RS13160_ | 548 | 547 | 508 | 476 | 503 | 461 |
| SL1344_RS13165_ | 276 | 244 | 213 | 265 | 229 | 221 |
| SL1344_RS13170_ | 166 | 176 | 159 | 181 | 131 | 165 |
| SL1344_RS13175_ | 88 | 109 | 121 | 72 | 70 | 79 |
| SL1344_RS13180_ | 248 | 281 | 258 | 263 | 250 | 222 |
| SL1344_RS13185_ | 153 | 176 | 144 | 167 | 169 | 171 |
| SL1344_RS13190_ | 14 | 20 | 19 | 20 | 32 | 22 |
| SL1344_RS13195_ | 23 | 25 | 14 | 22 | 11 | 18 |
| SL1344_RS13200_ | 269 | 283 | 292 | 271 | 266 | 314 |
| SL1344_RS13210_ | 310 | 379 | 411 | 399 | 344 | 389 |
| SL1344_RS13215_ | 1154 | 1265 | 1378 | 916 | 881 | 924 |
| SL1344_RS13220_ | 1982 | 2177 | 2178 | 1690 | 1562 | 1850 |
| SL1344_RS13225_ | 1780 | 1950 | 2076 | 1444 | 1507 | 1517 |
| SL1344_RS13230_ | 3399 | 3764 | 3939 | 3616 | 3339 | 3634 |
| SL1344_RS13235_ | 3584 | 3776 | 3959 | 3417 | 3087 | 3465 |
| SL1344_RS13240_ | 407 | 394 | 378 | 375 | 361 | 405 |
| SL1344_RS13245_ | 155 | 106 | 120 | 113 | 138 | 253 |
| SL1344_RS13250_ | 7 | 8 | 15 | 5 | 4 | 3 |
| SL1344_RS13255_ | 6 | 5 | 3 | 1 | 2 | 4 |
| SL1344_RS13260_ | 16 | 7 | 6 | 18 | 12 | 14 |
| SL1344_RS13265_ | 33 | 32 | 27 | 29 | 50 | 36 |
| SL1344_RS13270_ | 10 | 3 | 6 | 22 | 12 | 17 |
| SL1344_RS13275_ | 62 | 62 | 46 | 50 | 49 | 62 |
| SL1344_RS13280_ | 38 | 37 | 43 | 51 | 32 | 54 |
| SL1344_RS13285_ | 22 | 21 | 36 | 27 | 26 | 24 |
| SL1344_RS13290_ | 36 | 40 | 41 | 30 | 41 | 38 |
| SL1344_RS13295_ | 12 | 6 | 9 | 10 | 9 | 13 |
| SL1344_RS13300_ | 104 | 107 | 118 | 131 | 116 | 95 |
| SL1344_RS13305_ | 2 | 11 | 5 | 8 | 7 | 3 |
| SL1344_RS13310_ | 4 | 2 | 4 | 6 | 9 | 13 |
| SL1344_RS13315_ | 57 | 76 | 69 | 81 | 81 | 64 |
| SL1344_RS13320_ | 18 | 21 | 14 | 29 | 14 | 16 |
| SL1344_RS13325_ | 3421 | 3388 | 2969 | 2586 | 2361 | 2708 |
| SL1344_RS13330_ | 8 | 16 | 25 | 21 | 21 | 25 |
| SL1344_RS13335_ | 16 | 9 | 12 | 12 | 12 | 15 |
| SL1344_RS13340_ | 11 | 9 | 8 | 15 | 11 | 8 |
| SL1344_RS13345_ | 30 | 28 | 32 | 39 | 32 | 32 |

| | | | | | | |
|---|---|---|---|---|---|---|
| SL1344_RS13350_ | 7 | 20 | 16 | 26 | 26 | 25 |
| SL1344_RS13355_ | 38 | 28 | 24 | 39 | 39 | 37 |
| SL1344_RS13360_ | 44 | 53 | 62 | 69 | 65 | 58 |
| SL1344_RS13365_ | 10 | 10 | 16 | 10 | 12 | 9 |
| SL1344_RS13370_ | 42 | 42 | 36 | 51 | 56 | 36 |
| SL1344_RS13375_ | 11 | 4 | 3 | 5 | 5 | 7 |
| SL1344_RS13380_ | 79 | 64 | 57 | 60 | 59 | 65 |
| SL1344_RS13385_ | 40 | 34 | 25 | 29 | 19 | 26 |
| SL1344_RS13390_ | 31 | 23 | 12 | 19 | 10 | 14 |
| SL1344_RS13395_ | 8 | 0 | 0 | 3 | 1 | 2 |
| SL1344_RS13400_ | 144 | 127 | 116 | 165 | 152 | 169 |
| SL1344_RS13405_ | 23 | 34 | 22 | 25 | 32 | 24 |
| SL1344_RS13410_ | 83 | 63 | 67 | 52 | 38 | 37 |
| SL1344_RS13415_ | 198 | 205 | 195 | 261 | 194 | 196 |
| SL1344_RS13420_ | 16 | 21 | 19 | 9 | 10 | 18 |
| SL1344_RS13425_ | 0 | 2 | 3 | 4 | 3 | 2 |
| SL1344_RS13430_ | 1 | 11 | 11 | 16 | 9 | 3 |
| SL1344_RS13435_ | 3 | 4 | 0 | 0 | 1 | 0 |
| SL1344_RS13440_ | 312 | 282 | 280 | 210 | 226 | 249 |
| SL1344_RS13450_ | 129 | 136 | 68 | 109 | 107 | 95 |
| SL1344_RS13455_ | 229 | 207 | 235 | 196 | 194 | 206 |
| SL1344_RS13460_ | 17 | 11 | 10 | 27 | 24 | 21 |
| SL1344_RS13465_ | 38 | 28 | 27 | 43 | 41 | 36 |
| SL1344_RS13470_ | 9 | 13 | 10 | 12 | 15 | 17 |
| SL1344_RS13475_ | 18 | 20 | 16 | 19 | 28 | 17 |
| SL1344_RS13480_ | 4 | 16 | 13 | 17 | 11 | 14 |
| SL1344_RS13485_ | 36 | 17 | 29 | 34 | 26 | 18 |
| SL1344_RS13490_ | 16 | 17 | 8 | 25 | 12 | 16 |
| SL1344_RS13495_ | 29 | 9 | 4 | 7 | 4 | 5 |
| SL1344_RS13500_ | 1610 | 1664 | 1463 | 1568 | 1513 | 1563 |
| SL1344_RS13505_ | 7183 | 6822 | 5988 | 6421 | 5970 | 6524 |
| SL1344_RS13510_ | 5 | 5 | 4 | 10 | 4 | 6 |
| SL1344_RS13515_ | 11 | 10 | 11 | 19 | 15 | 13 |
| SL1344_RS13520_ | 44 | 34 | 44 | 52 | 64 | 49 |
| SL1344_RS13525_ | 6 | 15 | 12 | 58 | 8 | 9 |
| SL1344_RS13530_ | 6 | 4 | 5 | 3 | 4 | 2 |
| SL1344_RS13535_ | 16 | 15 | 11 | 17 | 24 | 18 |
| SL1344_RS13540_ | 249 | 250 | 286 | 207 | 223 | 211 |
| SL1344_RS13545_ | 482 | 470 | 511 | 381 | 438 | 462 |
| SL1344_RS13550_ | 839 | 905 | 891 | 930 | 889 | 940 |
| SL1344_RS13555_ | 732 | 803 | 837 | 927 | 854 | 869 |
| SL1344_RS13560_ | 753 | 788 | 920 | 864 | 813 | 820 |
| SL1344_RS13565_ | 84 | 74 | 70 | 83 | 64 | 70 |
| SL1344_RS13570_ | 538 | 583 | 555 | 498 | 471 | 485 |
| SL1344_RS13575_ | 3524 | 3755 | 3642 | 3260 | 3325 | 3230 |
| SL1344_RS13580_ | 103 | 92 | 107 | 96 | 118 | 124 |
| SL1344_RS13590_ | 504 | 375 | 230 | 556 | 382 | 259 |
| SL1344_RS13595_ | 520 | 569 | 546 | 516 | 492 | 517 |
| SL1344_RS13605_ | 1112 | 1089 | 948 | 1127 | 1007 | 1038 |
| SL1344_RS13610_ | 467 | 416 | 427 | 388 | 393 | 390 |
| SL1344_RS13615_ | 687 | 691 | 740 | 676 | 639 | 643 |
| SL1344_RS13625_ | 265 | 229 | 253 | 178 | 188 | 222 |
| SL1344_RS13630_ | 107 | 103 | 111 | 124 | 144 | 157 |
| SL1344_RS13635_ | 4 | 4 | 6 | 16 | 3 | 21 |
| SL1344_RS13645_ | 6451 | 6313 | 4716 | 5084 | 4827 | 5466 |
| SL1344_RS13650_ | 372 | 327 | 408 | 302 | 257 | 330 |
| SL1344_RS13655_ | 509 | 239 | 199 | 138 | 157 | 227 |
| SL1344_RS13660_ | 3399 | 3754 | 3712 | 5353 | 4522 | 4536 |
| SL1344_RS13665_ | 1403 | 1415 | 1468 | 1243 | 1132 | 1307 |
| SL1344_RS13670_ | 1739 | 1815 | 1691 | 1478 | 1506 | 1598 |
| SL1344_RS13675_ | 1736 | 1716 | 1607 | 1568 | 1491 | 1535 |
| SL1344_RS13680_ | 492 | 387 | 259 | 475 | 456 | 321 |

| | | | | | | |
|---|---|---|---|---|---|---|
| SL1344_RS13685_ | 745 | 721 | 703 | 818 | 765 | 825 |
| SL1344_RS13705_ | 162 | 145 | 190 | 67 | 76 | 85 |
| SL1344_RS13710_ | 312 | 262 | 319 | 171 | 210 | 188 |
| SL1344_RS13715_ | 869 | 938 | 824 | 1061 | 931 | 1039 |
| SL1344_RS13720_ | 712 | 797 | 739 | 948 | 870 | 852 |
| SL1344_RS13725_ | 499 | 564 | 521 | 713 | 632 | 647 |
| SL1344_RS13730_ | 2 | 3 | 3 | 3 | 4 | 3 |
| SL1344_RS13735_ | 4 | 6 | 1 | 1 | 4 | 2 |
| SL1344_RS13745_ | 3 | 2 | 6 | 7 | 1 | 2 |
| SL1344_RS13750_ | 1 | 1 | 1 | 3 | 0 | 1 |
| SL1344_RS13755_ | 0 | 1 | 2 | 0 | 0 | 0 |
| SL1344_RS13760_ | 2 | 1 | 0 | 2 | 2 | 3 |
| SL1344_RS13765_ | 6 | 11 | 7 | 18 | 13 | 13 |
| SL1344_RS13770_ | 17 | 26 | 14 | 19 | 15 | 16 |
| SL1344_RS13775_ | 70 | 58 | 47 | 54 | 64 | 69 |
| SL1344_RS13780_ | 312 | 331 | 281 | 350 | 334 | 332 |
| SL1344_RS13785_ | 22373 | 26370 | 26810 | 24006 | 21532 | 24680 |
| SL1344_RS13790_ | 23460 | 25245 | 26151 | 23426 | 23022 | 24167 |
| SL1344_RS13800_ | 11157 | 11091 | 11206 | 9952 | 9704 | 10458 |
| SL1344_RS13805_ | 1247 | 1573 | 1704 | 1574 | 1513 | 1628 |
| SL1344_RS13810_ | 821 | 821 | 821 | 746 | 738 | 813 |
| SL1344_RS13815_ | 836 | 924 | 920 | 799 | 763 | 917 |
| SL1344_RS13820_ | 120 | 126 | 167 | 103 | 97 | 132 |
| SL1344_RS13825_ | 5763 | 6178 | 6229 | 9063 | 8099 | 7182 |
| SL1344_RS13830_ | 371 | 316 | 278 | 340 | 341 | 302 |
| SL1344_RS13840_ | 1400 | 1464 | 1325 | 1362 | 1395 | 1436 |
| SL1344_RS13845_ | 540 | 572 | 558 | 600 | 593 | 662 |
| SL1344_RS13850_ | 938 | 926 | 903 | 957 | 1043 | 1044 |
| SL1344_RS13855_ | 2249 | 2016 | 1922 | 1904 | 1810 | 1912 |
| SL1344_RS13860_ | 231 | 202 | 181 | 232 | 169 | 208 |
| SL1344_RS13865_ | 15 | 13 | 15 | 21 | 11 | 18 |
| SL1344_RS13870_ | 37 | 29 | 40 | 31 | 41 | 21 |
| SL1344_RS13875_ | 25 | 44 | 200 | 33 | 32 | 28 |
| SL1344_RS13880_ | 30 | 32 | 29 | 29 | 35 | 26 |
| SL1344_RS13885_ | 22 | 17 | 17 | 20 | 20 | 38 |
| SL1344_RS13890_ | 10 | 9 | 16 | 9 | 8 | 15 |
| SL1344_RS13895_ | 90 | 60 | 68 | 61 | 74 | 73 |
| SL1344_RS13900_ | 6 | 4 | 11 | 5 | 7 | 8 |
| SL1344_RS13905_ | 4 | 9 | 5 | 9 | 32 | 6 |
| SL1344_RS13910_ | 17 | 5 | 6 | 4 | 15 | 7 |
| SL1344_RS13915_ | 39 | 21 | 40 | 33 | 46 | 39 |
| SL1344_RS13920_ | 3 | 9 | 10 | 6 | 7 | 8 |
| SL1344_RS13930_ | 36 | 44 | 38 | 92 | 57 | 50 |
| SL1344_RS13935_ | 39 | 41 | 47 | 46 | 37 | 47 |
| SL1344_RS13940_ | 14 | 4 | 8 | 9 | 7 | 30 |
| SL1344_RS13950_ | 1 | 2 | 1 | 2 | 2 | 0 |
| SL1344_RS13955_ | 2 | 1 | 5 | 6 | 6 | 5 |
| SL1344_RS13965_ | 2 | 6 | 5 | 15 | 5 | 10 |
| SL1344_RS13970_ | 16 | 4 | 8 | 11 | 5 | 6 |
| SL1344_RS13975_ | 2 | 4 | 21 | 3 | 9 | 8 |
| SL1344_RS13980_ | 3 | 4 | 5 | 6 | 15 | 1 |
| SL1344_RS13985_ | 3 | 0 | 0 | 0 | 1 | 0 |
| SL1344_RS14000_ | 8 | 10 | 9 | 25 | 7 | 14 |
| SL1344_RS14005_ | 10 | 19 | 33 | 14 | 11 | 18 |
| SL1344_RS14010_ | 41 | 6 | 21 | 22 | 16 | 3 |
| SL1344_RS14015_ | 8 | 19 | 8 | 15 | 5 | 8 |
| SL1344_RS14020_ | 14 | 6 | 8 | 5 | 5 | 20 |
| SL1344_RS14025_ | 440 | 456 | 417 | 382 | 376 | 395 |
| SL1344_RS14030_ | 249 | 310 | 267 | 269 | 249 | 248 |
| SL1344_RS14035_ | 286 | 267 | 303 | 282 | 256 | 299 |
| SL1344_RS14040_ | 1265 | 1274 | 1224 | 1450 | 1308 | 1404 |
| SL1344_RS14045_ | 548 | 526 | 502 | 602 | 516 | 592 |

| | | | | | | |
|---|---|---|---|---|---|---|
| SL1344_RS14050_ | 59 | 81 | 89 | 109 | 83 | 115 |
| SL1344_RS14055_ | 63 | 82 | 86 | 77 | 83 | 102 |
| SL1344_RS14060_ | 60 | 58 | 91 | 57 | 59 | 53 |
| SL1344_RS14065_ | 34 | 23 | 25 | 27 | 24 | 20 |
| SL1344_RS14070_ | 7 | 5 | 3 | 1 | 1 | 3 |
| SL1344_RS14075_ | 9 | 4 | 5 | 4 | 10 | 3 |
| SL1344_RS14080_ | 4 | 9 | 1 | 2 | 6 | 4 |
| SL1344_RS14085_ | 97 | 90 | 65 | 123 | 117 | 114 |
| SL1344_RS14090_ | 5 | 4 | 4 | 18 | 16 | 4 |
| SL1344_RS14095_ | 5 | 9 | 7 | 22 | 12 | 4 |
| SL1344_RS14100_ | 135 | 200 | 163 | 174 | 143 | 224 |
| SL1344_RS14105_ | 265 | 314 | 262 | 299 | 288 | 310 |
| SL1344_RS14110_ | 263 | 273 | 233 | 251 | 246 | 299 |
| SL1344_RS14115_ | 72 | 107 | 88 | 130 | 116 | 93 |
| SL1344_RS14120_ | 437 | 411 | 397 | 456 | 426 | 525 |
| SL1344_RS14125_ | 43 | 44 | 51 | 49 | 36 | 56 |
| SL1344_RS14130_ | 98 | 115 | 100 | 119 | 89 | 104 |
| SL1344_RS14135_ | 173 | 200 | 169 | 174 | 197 | 195 |
| SL1344_RS14140_ | 6 | 7 | 7 | 8 | 13 | 6 |
| SL1344_RS14150_ | 7 | 29 | 29 | 13 | 20 | 15 |
| SL1344_RS14155_ | 13 | 5 | 13 | 19 | 17 | 18 |
| SL1344_RS14165_ | 11 | 15 | 13 | 12 | 11 | 12 |
| SL1344_RS14170_ | 15 | 25 | 18 | 29 | 18 | 19 |
| SL1344_RS14175_ | 278 | 236 | 212 | 277 | 201 | 219 |
| SL1344_RS14180_ | 16 | 11 | 9 | 14 | 16 | 11 |
| SL1344_RS14190_ | 11 | 6 | 0 | 8 | 7 | 7 |
| SL1344_RS14195_ | 561 | 539 | 551 | 703 | 684 | 735 |
| SL1344_RS14200_ | 344 | 306 | 284 | 285 | 260 | 276 |
| SL1344_RS14205_ | 7 | 10 | 2 | 13 | 8 | 7 |
| SL1344_RS14210_ | 32 | 28 | 23 | 23 | 20 | 23 |
| SL1344_RS14215_ | 965 | 844 | 738 | 771 | 727 | 747 |
| SL1344_RS14220_ | 373 | 337 | 339 | 324 | 309 | 328 |
| SL1344_RS14225_ | 488 | 443 | 400 | 490 | 482 | 480 |
| SL1344_RS14230_ | 11 | 7 | 4 | 13 | 11 | 12 |
| SL1344_RS14235_ | 2 | 5 | 2 | 10 | 4 | 9 |
| SL1344_RS14240_ | 2 | 5 | 3 | 0 | 2 | 2 |
| SL1344_RS14245_ | 8 | 3 | 6 | 8 | 28 | 10 |
| SL1344_RS14250_ | 14 | 16 | 39 | 25 | 12 | 11 |
| SL1344_RS14255_ | 410 | 410 | 382 | 323 | 313 | 318 |
| SL1344_RS14260_ | 14 | 17 | 22 | 18 | 23 | 12 |
| SL1344_RS14265_ | 5 | 8 | 12 | 6 | 10 | 8 |
| SL1344_RS14270_ | 11 | 12 | 7 | 22 | 16 | 11 |
| SL1344_RS14275_ | 18 | 24 | 22 | 14 | 14 | 33 |
| SL1344_RS14280_ | 26 | 20 | 34 | 22 | 36 | 17 |
| SL1344_RS14285_ | 5 | 2 | 1 | 4 | 5 | 6 |
| SL1344_RS14290_ | 16 | 12 | 15 | 47 | 17 | 26 |
| SL1344_RS14295_ | 147 | 133 | 135 | 132 | 113 | 114 |
| SL1344_RS14300_ | 162 | 158 | 156 | 154 | 128 | 163 |
| SL1344_RS14305_ | 179 | 182 | 146 | 145 | 162 | 154 |
| SL1344_RS14310_ | 8 | 9 | 12 | 22 | 4 | 7 |
| SL1344_RS14315_ | 7 | 12 | 16 | 16 | 20 | 16 |
| SL1344_RS14320_ | 377 | 357 | 311 | 414 | 281 | 401 |
| SL1344_RS14325_ | 1410 | 1525 | 1381 | 1659 | 1333 | 1595 |
| SL1344_RS14330_ | 50 | 36 | 34 | 41 | 35 | 41 |
| SL1344_RS14335_ | 22 | 30 | 21 | 37 | 27 | 43 |
| SL1344_RS14345_ | 67364 | 74621 | 67456 | 6530 | 96267 | 4747 |
| SL1344_RS14350_ | 30 | 43 | 30 | 21 | 31 | 32 |
| SL1344_RS14360_ | 130 | 161 | 157 | 383 | 290 | 416 |
| SL1344_RS14365_ | 36 | 38 | 37 | 82 | 60 | 109 |
| SL1344_RS14370_ | 51 | 46 | 43 | 114 | 79 | 102 |
| SL1344_RS14375_ | 157 | 177 | 168 | 168 | 186 | 157 |
| SL1344_RS14380_ | 36 | 29 | 42 | 33 | 39 | 27 |

| | | | | | | |
|---|---|---|---|---|---|---|
| SL1344_RS14385_ | 87 | 111 | 124 | 116 | 82 | 133 |
| SL1344_RS14390_ | 2326 | 2026 | 2032 | 1469 | 1457 | 1637 |
| SL1344_RS14395_ | 2439 | 2075 | 2222 | 1944 | 1853 | 1949 |
| SL1344_RS14400_ | 272 | 252 | 281 | 175 | 149 | 166 |
| SL1344_RS14405_ | 154 | 146 | 147 | 159 | 151 | 193 |
| SL1344_RS14410_ | 99 | 80 | 87 | 98 | 87 | 94 |
| SL1344_RS14415_ | 37 | 37 | 31 | 58 | 39 | 44 |
| SL1344_RS14420_ | 6 | 13 | 11 | 19 | 7 | 9 |
| SL1344_RS14425_ | 35 | 28 | 40 | 46 | 32 | 47 |
| SL1344_RS14430_ | 8 | 5 | 9 | 7 | 8 | 9 |
| SL1344_RS14435_ | 89 | 71 | 84 | 79 | 106 | 107 |
| SL1344_RS14440_ | 15 | 12 | 10 | 16 | 8 | 21 |
| SL1344_RS14445_ | 10 | 23 | 22 | 16 | 10 | 19 |
| SL1344_RS14450_ | 18 | 19 | 32 | 12 | 15 | 31 |
| SL1344_RS14455_ | 20 | 24 | 27 | 33 | 30 | 40 |
| SL1344_RS14460_ | 62 | 51 | 56 | 46 | 34 | 38 |
| SL1344_RS14465_ | 48 | 53 | 47 | 43 | 47 | 37 |
| SL1344_RS14470_ | 62 | 47 | 39 | 43 | 41 | 37 |
| SL1344_RS14475_ | 8 | 17 | 15 | 22 | 20 | 19 |
| SL1344_RS14480_ | 42 | 46 | 38 | 35 | 40 | 42 |
| SL1344_RS14485_ | 97 | 102 | 91 | 121 | 113 | 104 |
| SL1344_RS14490_ | 151 | 146 | 130 | 202 | 222 | 193 |
| SL1344_RS14495_ | 4262 | 4733 | 4909 | 4832 | 4129 | 4494 |
| SL1344_RS14500_ | 244 | 211 | 139 | 181 | 161 | 119 |
| SL1344_RS14505_ | 68 | 94 | 78 | 56 | 57 | 58 |
| SL1344_RS14510_ | 34 | 49 | 54 | 81 | 80 | 106 |
| SL1344_RS14515_ | 203 | 192 | 205 | 247 | 234 | 215 |
| SL1344_RS14520_ | 25 | 33 | 27 | 32 | 41 | 40 |
| SL1344_RS14525_ | 4 | 9 | 4 | 9 | 10 | 11 |
| SL1344_RS14530_ | 74 | 93 | 80 | 69 | 75 | 44 |
| SL1344_RS14540_ | 302 | 318 | 286 | 230 | 243 | 229 |
| SL1344_RS14545_ | 266 | 307 | 288 | 233 | 187 | 251 |
| SL1344_RS14550_ | 666 | 615 | 550 | 435 | 422 | 436 |
| SL1344_RS14555_ | 193 | 212 | 226 | 157 | 164 | 173 |
| SL1344_RS14560_ | 284 | 298 | 281 | 268 | 278 | 291 |
| SL1344_RS14565_ | 146 | 159 | 183 | 121 | 121 | 127 |
| SL1344_RS14570_ | 1066 | 1331 | 1299 | 1146 | 1312 | 1166 |
| SL1344_RS14575_ | 842 | 955 | 913 | 803 | 863 | 907 |
| SL1344_RS14585_ | 54 | 45 | 62 | 57 | 29 | 53 |
| SL1344_RS14600_ | 690 | 809 | 822 | 822 | 740 | 828 |
| SL1344_RS14605_ | 833 | 888 | 834 | 954 | 895 | 890 |
| SL1344_RS14610_ | 1410 | 1287 | 1407 | 1239 | 1115 | 1238 |
| SL1344_RS14615_ | 4200 | 4299 | 4657 | 3602 | 3497 | 3983 |
| SL1344_RS14620_ | 6108 | 6160 | 7196 | 5700 | 5278 | 6122 |
| SL1344_RS14625_ | 7667 | 8161 | 9229 | 7860 | 7426 | 9006 |
| SL1344_RS14630_ | 24964 | 28331 | 27582 | 22368 | 23018 | 26601 |
| SL1344_RS14640_ | 3657 | 4377 | 4912 | 4653 | 4221 | 4526 |
| SL1344_RS14645_ | 183 | 148 | 179 | 152 | 157 | 190 |
| SL1344_RS14655_ | 294 | 343 | 349 | 382 | 328 | 355 |
| SL1344_RS14660_ | 585 | 571 | 491 | 514 | 500 | 546 |
| SL1344_RS14675_ | 18 | 8 | 11 | 9 | 4 | 10 |
| SL1344_RS14680_ | 58 | 72 | 51 | 58 | 56 | 63 |
| SL1344_RS14685_ | 79 | 70 | 72 | 85 | 72 | 75 |
| SL1344_RS14690_ | 63 | 62 | 84 | 53 | 69 | 57 |
| SL1344_RS14700_ | 297 | 274 | 265 | 273 | 310 | 272 |
| SL1344_RS14705_ | 155 | 181 | 170 | 210 | 212 | 207 |
| SL1344_RS14710_ | 30 | 31 | 25 | 19 | 25 | 28 |
| SL1344_RS14715_ | 31 | 27 | 26 | 38 | 32 | 44 |
| SL1344_RS14720_ | 558 | 553 | 502 | 535 | 536 | 497 |
| SL1344_RS14725_ | 13 | 20 | 20 | 18 | 25 | 23 |
| SL1344_RS14730_ | 69 | 76 | 63 | 72 | 62 | 70 |
| SL1344_RS14735_ | 30 | 47 | 40 | 50 | 40 | 41 |

| | | | | | | |
|---|---|---|---|---|---|---|
| SL1344_RS14740_ | 33 | 31 | 42 | 47 | 46 | 38 |
| SL1344_RS14745_ | 49 | 38 | 45 | 53 | 51 | 58 |
| SL1344_RS14750_ | 23 | 28 | 32 | 37 | 26 | 35 |
| SL1344_RS14765_ | 45 | 53 | 63 | 55 | 51 | 45 |
| SL1344_RS14770_ | 10 | 12 | 13 | 9 | 6 | 9 |
| SL1344_RS14775_ | 7 | 9 | 11 | 2 | 4 | 8 |
| SL1344_RS14780_ | 122 | 61 | 70 | 107 | 110 | 81 |
| SL1344_RS14785_ | 91 | 122 | 103 | 111 | 109 | 98 |
| SL1344_RS14790_ | 41 | 50 | 51 | 40 | 57 | 60 |
| SL1344_RS14795_ | 116 | 130 | 142 | 174 | 145 | 122 |
| SL1344_RS14800_ | 149 | 163 | 176 | 180 | 154 | 181 |
| SL1344_RS14805_ | 194 | 206 | 224 | 192 | 185 | 190 |
| SL1344_RS14810_ | 41 | 54 | 38 | 46 | 51 | 36 |
| SL1344_RS14815_ | 266 | 262 | 178 | 262 | 207 | 185 |
| SL1344_RS14820_ | 95 | 114 | 122 | 118 | 117 | 122 |
| SL1344_RS14825_ | 53 | 77 | 59 | 66 | 60 | 80 |
| SL1344_RS14830_ | 133 | 159 | 189 | 167 | 165 | 162 |
| SL1344_RS14850_ | 1056 | 964 | 964 | 985 | 1018 | 1116 |
| SL1344_RS14855_ | 974 | 972 | 964 | 1114 | 1080 | 1117 |
| SL1344_RS14900_ | 937 | 1079 | 1079 | 1138 | 1015 | 1128 |
| SL1344_RS14905_ | 301 | 321 | 311 | 314 | 311 | 319 |
| SL1344_RS14910_ | 371 | 376 | 397 | 418 | 391 | 345 |
| SL1344_RS15010_ | 216 | 193 | 155 | 299 | 367 | 340 |
| SL1344_RS15015_ | 491 | 502 | 443 | 498 | 463 | 442 |
| SL1344_RS15020_ | 168 | 179 | 143 | 156 | 146 | 131 |
| SL1344_RS15025_ | 72 | 62 | 42 | 66 | 48 | 42 |
| SL1344_RS15030_ | 303 | 329 | 301 | 326 | 359 | 335 |
| SL1344_RS15035_ | 422 | 451 | 438 | 559 | 482 | 512 |
| SL1344_RS15040_ | 3 | 11 | 6 | 8 | 3 | 9 |
| SL1344_RS15045_ | 126 | 134 | 122 | 111 | 108 | 124 |
| SL1344_RS15050_ | 7 | 7 | 12 | 14 | 12 | 15 |
| SL1344_RS15055_ | 1132 | 1300 | 1386 | 1312 | 1172 | 1289 |
| SL1344_RS15060_ | 29 | 25 | 28 | 29 | 26 | 25 |
| SL1344_RS15065_ | 33 | 28 | 36 | 32 | 36 | 41 |
| SL1344_RS15070_ | 60 | 69 | 64 | 62 | 56 | 60 |
| SL1344_RS15075_ | 119 | 108 | 77 | 139 | 116 | 105 |
| SL1344_RS15080_ | 18 | 10 | 17 | 8 | 11 | 21 |
| SL1344_RS15085_ | 19 | 11 | 18 | 19 | 12 | 20 |
| SL1344_RS15090_ | 16 | 19 | 22 | 20 | 18 | 24 |
| SL1344_RS15095_ | 19 | 29 | 20 | 25 | 28 | 52 |
| SL1344_RS15100_ | 13 | 15 | 20 | 16 | 21 | 24 |
| SL1344_RS15105_ | 246 | 273 | 240 | 220 | 242 | 209 |
| SL1344_RS15110_ | 197 | 221 | 167 | 164 | 160 | 180 |
| SL1344_RS15115_ | 6 | 9 | 6 | 10 | 12 | 8 |
| SL1344_RS15120_ | 32 | 35 | 28 | 34 | 37 | 30 |
| SL1344_RS15125_ | 13 | 17 | 12 | 30 | 20 | 24 |
| SL1344_RS15140_ | 833 | 884 | 945 | 880 | 845 | 854 |
| SL1344_RS15145_ | 717 | 732 | 775 | 680 | 637 | 680 |
| SL1344_RS15150_ | 804 | 940 | 1050 | 880 | 717 | 827 |
| SL1344_RS15155_ | 358 | 414 | 450 | 427 | 340 | 431 |
| SL1344_RS15160_ | 616 | 728 | 749 | 604 | 599 | 670 |
| SL1344_RS15165_ | 820 | 917 | 979 | 809 | 794 | 844 |
| SL1344_RS15170_ | 563 | 516 | 561 | 290 | 283 | 327 |
| SL1344_RS15190_ | 370 | 346 | 328 | 268 | 247 | 265 |
| SL1344_RS15195_ | 31 | 29 | 28 | 38 | 35 | 41 |
| SL1344_RS15200_ | 71 | 69 | 80 | 106 | 84 | 99 |
| SL1344_RS15205_ | 50 | 45 | 45 | 53 | 50 | 32 |
| SL1344_RS15210_ | 39 | 34 | 29 | 47 | 33 | 35 |
| SL1344_RS15215_ | 50 | 53 | 63 | 54 | 53 | 57 |
| SL1344_RS15220_ | 12 | 13 | 10 | 11 | 15 | 23 |
| SL1344_RS15225_ | 33 | 37 | 47 | 40 | 30 | 53 |
| SL1344_RS15230_ | 30 | 36 | 25 | 22 | 30 | 27 |

| | | | | | | |
|---|---|---|---|---|---|---|
| SL1344_RS15260_ | 64 | 82 | 62 | 84 | 78 | 97 |
| SL1344_RS15265_ | 947 | 975 | 876 | 861 | 915 | 927 |
| SL1344_RS15270_ | 13993 | 15597 | 16422 | 14515 | 14092 | 14302 |
| SL1344_RS15275_ | 5963 | 6156 | 6358 | 5719 | 5493 | 5776 |
| SL1344_RS15280_ | 513 | 570 | 634 | 581 | 544 | 576 |
| SL1344_RS15285_ | 575 | 586 | 477 | 636 | 588 | 607 |
| SL1344_RS15290_ | 829 | 889 | 767 | 964 | 864 | 856 |
| SL1344_RS15295_ | 2615 | 2718 | 2776 | 2818 | 2774 | 2803 |
| SL1344_RS15310_ | 81 | 79 | 102 | 107 | 74 | 104 |
| SL1344_RS15315_ | 91 | 136 | 112 | 139 | 119 | 165 |
| SL1344_RS15320_ | 69 | 89 | 89 | 94 | 82 | 71 |
| SL1344_RS15325_ | 26 | 40 | 24 | 25 | 21 | 30 |
| SL1344_RS15330_ | 519 | 655 | 684 | 615 | 519 | 660 |
| SL1344_RS15355_ | 446 | 446 | 423 | 428 | 403 | 378 |
| SL1344_RS15360_ | 1887 | 1709 | 1713 | 1602 | 1605 | 1518 |
| SL1344_RS15365_ | 2862 | 2930 | 2932 | 1991 | 1816 | 1936 |
| SL1344_RS15370_ | 2876 | 2741 | 2796 | 1721 | 1672 | 1670 |
| SL1344_RS15375_ | 374 | 311 | 400 | 274 | 307 | 294 |
| SL1344_RS15380_ | 19 | 19 | 17 | 28 | 26 | 42 |
| SL1344_RS15385_ | 4 | 6 | 5 | 7 | 4 | 9 |
| SL1344_RS15390_ | 12 | 12 | 10 | 5 | 9 | 5 |
| SL1344_RS15395_ | 25 | 15 | 28 | 24 | 20 | 29 |
| SL1344_RS15400_ | 19 | 21 | 16 | 16 | 17 | 14 |
| SL1344_RS15405_ | 13 | 15 | 32 | 22 | 22 | 21 |
| SL1344_RS15410_ | 139 | 119 | 112 | 111 | 106 | 94 |
| SL1344_RS15415_ | 1255 | 1453 | 1462 | 1308 | 1224 | 1340 |
| SL1344_RS15420_ | 318 | 352 | 344 | 349 | 309 | 325 |
| SL1344_RS15425_ | 589 | 563 | 511 | 516 | 501 | 526 |
| SL1344_RS15430_ | 229 | 194 | 170 | 272 | 285 | 242 |
| SL1344_RS15435_ | 552 | 512 | 499 | 429 | 394 | 408 |
| SL1344_RS15440_ | 357 | 329 | 351 | 280 | 278 | 284 |
| SL1344_RS15445_ | 108 | 105 | 91 | 81 | 70 | 56 |
| SL1344_RS15450_ | 568 | 668 | 748 | 619 | 578 | 639 |
| SL1344_RS15455_ | 1224 | 1356 | 1258 | 1252 | 1157 | 1253 |
| SL1344_RS15460_ | 3429 | 3174 | 2688 | 1920 | 1872 | 2386 |
| SL1344_RS15465_ | 10180 | 9155 | 9917 | 5584 | 5362 | 6336 |
| SL1344_RS15475_ | 202 | 219 | 191 | 216 | 186 | 215 |
| SL1344_RS15480_ | 401 | 419 | 490 | 408 | 363 | 418 |
| SL1344_RS15485_ | 891 | 977 | 1098 | 955 | 885 | 1002 |
| SL1344_RS15490_ | 805 | 870 | 824 | 718 | 709 | 789 |
| SL1344_RS15495_ | 811 | 880 | 979 | 828 | 781 | 792 |
| SL1344_RS15500_ | 53 | 57 | 46 | 46 | 43 | 50 |
| SL1344_RS15505_ | 19 | 33 | 29 | 30 | 25 | 39 |
| SL1344_RS15510_ | 35 | 36 | 30 | 26 | 41 | 29 |
| SL1344_RS15515_ | 11 | 21 | 15 | 7 | 10 | 12 |
| SL1344_RS15520_ | 9 | 15 | 7 | 3 | 13 | 2 |
| SL1344_RS15525_ | 1485 | 1526 | 1566 | 1352 | 1297 | 1377 |
| SL1344_RS15530_ | 701 | 769 | 733 | 748 | 662 | 714 |
| SL1344_RS15535_ | 1501 | 1815 | 1821 | 1663 | 1544 | 1786 |
| SL1344_RS15540_ | 957 | 1048 | 914 | 1068 | 910 | 927 |
| SL1344_RS15545_ | 3 | 7 | 5 | 4 | 7 | 10 |
| SL1344_RS15550_ | 907 | 1020 | 895 | 957 | 904 | 844 |
| SL1344_RS15555_ | 1237 | 1181 | 1167 | 1378 | 1260 | 1294 |
| SL1344_RS15560_ | 59 | 78 | 57 | 53 | 56 | 61 |
| SL1344_RS15565_ | 265 | 276 | 266 | 296 | 262 | 322 |
| SL1344_RS15570_ | 282 | 331 | 287 | 287 | 272 | 352 |
| SL1344_RS15575_ | 671 | 794 | 798 | 774 | 667 | 791 |
| SL1344_RS15580_ | 327 | 346 | 283 | 303 | 297 | 335 |
| SL1344_RS15585_ | 295 | 283 | 279 | 272 | 270 | 276 |
| SL1344_RS15590_ | 74 | 45 | 41 | 76 | 90 | 78 |
| SL1344_RS15595_ | 132 | 134 | 147 | 119 | 123 | 126 |
| SL1344_RS15600_ | 198 | 195 | 227 | 243 | 212 | 228 |

| | | | | | | |
|---|---|---|---|---|---|---|
| SL1344_RS15605_ | 77 | 59 | 70 | 65 | 75 | 55 |
| SL1344_RS15610_ | 38 | 36 | 53 | 41 | 40 | 48 |
| SL1344_RS15615_ | 23 | 14 | 30 | 21 | 22 | 17 |
| SL1344_RS15620_ | 118 | 151 | 182 | 196 | 187 | 191 |
| SL1344_RS15625_ | 243 | 218 | 235 | 282 | 316 | 275 |
| SL1344_RS15630_ | 87 | 98 | 112 | 106 | 109 | 83 |
| SL1344_RS15635_ | 762 | 743 | 689 | 469 | 480 | 447 |
| SL1344_RS15640_ | 199 | 210 | 163 | 234 | 229 | 222 |
| SL1344_RS15645_ | 111 | 105 | 85 | 87 | 69 | 78 |
| SL1344_RS15650_ | 24 | 32 | 46 | 28 | 30 | 33 |
| SL1344_RS15655_ | 4 | 8 | 8 | 4 | 13 | 7 |
| SL1344_RS15660_ | 31 | 31 | 34 | 27 | 27 | 26 |
| SL1344_RS15665_ | 15 | 12 | 8 | 11 | 15 | 9 |
| SL1344_RS15670_ | 34 | 23 | 25 | 30 | 24 | 29 |
| SL1344_RS15675_ | 14 | 12 | 17 | 12 | 14 | 20 |
| SL1344_RS15680_ | 174 | 142 | 148 | 184 | 188 | 185 |
| SL1344_RS15685_ | 22 | 29 | 39 | 27 | 14 | 17 |
| SL1344_RS15690_ | 659 | 654 | 662 | 668 | 637 | 695 |
| SL1344_RS15695_ | 562 | 500 | 475 | 523 | 495 | 523 |
| SL1344_RS15700_ | 5 | 6 | 9 | 5 | 3 | 3 |
| SL1344_RS15705_ | 27 | 11 | 21 | 22 | 17 | 15 |
| SL1344_RS15710_ | 1701 | 1420 | 1019 | 1376 | 1272 | 1138 |
| SL1344_RS15715_ | 490 | 567 | 598 | 573 | 595 | 573 |
| SL1344_RS15720_ | 527 | 593 | 505 | 609 | 628 | 556 |
| SL1344_RS15725_ | 3487 | 4443 | 5061 | 5041 | 4760 | 5121 |
| SL1344_RS15730_ | 3245 | 3748 | 4148 | 3695 | 3862 | 3656 |
| SL1344_RS15735_ | 1203 | 1377 | 1459 | 1397 | 1253 | 1415 |
| SL1344_RS15740_ | 682 | 715 | 640 | 748 | 672 | 730 |
| SL1344_RS15745_ | 529 | 531 | 491 | 528 | 458 | 518 |
| SL1344_RS15750_ | 615 | 781 | 809 | 706 | 679 | 734 |
| SL1344_RS15755_ | 607 | 678 | 664 | 697 | 606 | 686 |
| SL1344_RS15760_ | 300 | 369 | 293 | 318 | 291 | 310 |
| SL1344_RS15765_ | 64 | 73 | 62 | 97 | 59 | 71 |
| SL1344_RS15770_ | 831 | 882 | 895 | 871 | 862 | 835 |
| SL1344_RS15775_ | 501 | 525 | 544 | 513 | 495 | 525 |
| SL1344_RS15780_ | 161 | 140 | 156 | 160 | 112 | 192 |
| SL1344_RS15785_ | 302 | 352 | 304 | 304 | 298 | 286 |
| SL1344_RS15790_ | 24 | 27 | 37 | 43 | 46 | 34 |
| SL1344_RS15795_ | 1407 | 1553 | 1545 | 2942 | 2570 | 3021 |
| SL1344_RS15800_ | 377 | 402 | 451 | 943 | 795 | 989 |
| SL1344_RS15810_ | 755 | 872 | 856 | 796 | 791 | 873 |
| SL1344_RS15815_ | 747 | 762 | 925 | 890 | 739 | 853 |
| SL1344_RS15820_ | 1116 | 1210 | 1169 | 1238 | 1190 | 1233 |
| SL1344_RS15825_ | 1252 | 1101 | 1066 | 1097 | 1061 | 1050 |
| SL1344_RS15830_ | 3825 | 3682 | 3288 | 2843 | 2918 | 2993 |
| SL1344_RS15835_ | 625 | 546 | 616 | 446 | 464 | 486 |
| SL1344_RS15840_ | 259 | 273 | 332 | 365 | 364 | 337 |
| SL1344_RS15845_ | 1214 | 1292 | 1373 | 1268 | 1218 | 1330 |
| SL1344_RS15850_ | 129 | 148 | 124 | 126 | 100 | 126 |
| SL1344_RS15855_ | 19 | 18 | 22 | 14 | 13 | 12 |
| SL1344_RS15860_ | 481 | 524 | 497 | 443 | 403 | 427 |
| SL1344_RS15865_ | 97 | 90 | 82 | 77 | 83 | 77 |
| SL1344_RS15870_ | 438 | 487 | 532 | 494 | 429 | 463 |
| SL1344_RS15875_ | 12557 | 13558 | 13302 | 10966 | 10263 | 10811 |
| SL1344_RS15880_ | 2486 | 2918 | 3115 | 2923 | 2546 | 2853 |
| SL1344_RS15890_ | 10 | 6 | 12 | 11 | 9 | 13 |
| SL1344_RS15895_ | 20 | 14 | 23 | 9 | 8 | 14 |
| SL1344_RS15900_ | 21 | 32 | 28 | 21 | 27 | 13 |
| SL1344_RS15905_ | 51 | 47 | 54 | 53 | 54 | 45 |
| SL1344_RS15910_ | 21 | 28 | 24 | 22 | 17 | 16 |
| SL1344_RS15915_ | 7847 | 7429 | 7839 | 7356 | 6988 | 7122 |
| SL1344_RS15920_ | 382 | 411 | 359 | 402 | 382 | 360 |

| | | | | | | |
|---|---|---|---|---|---|---|
| SL1344_RS15925_ | 5154 | 4854 | 5162 | 4061 | 3810 | 3985 |
| SL1344_RS15930_ | 44 | 39 | 35 | 37 | 44 | 38 |
| SL1344_RS15935_ | 28 | 21 | 21 | 27 | 23 | 27 |
| SL1344_RS15940_ | 60 | 84 | 83 | 97 | 81 | 97 |
| SL1344_RS15945_ | 50 | 43 | 52 | 63 | 64 | 64 |
| SL1344_RS15950_ | 48 | 27 | 20 | 30 | 27 | 30 |
| SL1344_RS15955_ | 18 | 20 | 31 | 24 | 25 | 29 |
| SL1344_RS15960_ | 647 | 732 | 619 | 752 | 645 | 706 |
| SL1344_RS15965_ | 131 | 132 | 152 | 84 | 58 | 99 |
| SL1344_RS15970_ | 4321 | 4813 | 4765 | 4305 | 3385 | 4139 |
| SL1344_RS15975_ | 650 | 655 | 630 | 618 | 454 | 579 |
| SL1344_RS15985_ | 9817 | 10109 | 9348 | 9701 | 9031 | 9675 |
| SL1344_RS15990_ | 297 | 252 | 255 | 264 | 269 | 267 |
| SL1344_RS15995_ | 391 | 386 | 375 | 285 | 297 | 308 |
| SL1344_RS16000_ | 461 | 438 | 454 | 447 | 377 | 435 |
| SL1344_RS16005_ | 849 | 830 | 786 | 705 | 617 | 732 |
| SL1344_RS16015_ | 474 | 459 | 428 | 412 | 389 | 448 |
| SL1344_RS16020_ | 437 | 474 | 419 | 396 | 386 | 432 |
| SL1344_RS16025_ | 245 | 239 | 262 | 244 | 198 | 244 |
| SL1344_RS16030_ | 59 | 44 | 34 | 61 | 57 | 56 |
| SL1344_RS16035_ | 551 | 629 | 582 | 547 | 502 | 623 |
| SL1344_RS16040_ | 511 | 566 | 612 | 613 | 569 | 580 |
| SL1344_RS16045_ | 362 | 420 | 434 | 410 | 389 | 416 |
| SL1344_RS16050_ | 609 | 642 | 712 | 581 | 649 | 602 |
| SL1344_RS16055_ | 1414 | 1595 | 1586 | 1294 | 1272 | 1475 |
| SL1344_RS16060_ | 60 | 50 | 46 | 57 | 44 | 61 |
| SL1344_RS16065_ | 219 | 216 | 220 | 201 | 202 | 269 |
| SL1344_RS16070_ | 978 | 1144 | 1237 | 1158 | 1003 | 1069 |
| SL1344_RS16075_ | 1037 | 1348 | 1479 | 1418 | 1189 | 1242 |
| SL1344_RS16080_ | 1783 | 1978 | 2164 | 1674 | 1577 | 1639 |
| SL1344_RS16085_ | 515 | 542 | 561 | 492 | 479 | 488 |
| SL1344_RS16090_ | 407 | 446 | 387 | 393 | 392 | 393 |
| SL1344_RS16095_ | 614 | 609 | 598 | 533 | 530 | 540 |
| SL1344_RS16100_ | 156 | 177 | 190 | 122 | 124 | 136 |
| SL1344_RS16105_ | 329 | 326 | 354 | 342 | 302 | 328 |
| SL1344_RS16110_ | 5 | 9 | 5 | 7 | 3 | 5 |
| SL1344_RS16115_ | 72 | 60 | 75 | 72 | 74 | 40 |
| SL1344_RS16120_ | 389 | 435 | 401 | 351 | 338 | 344 |
| SL1344_RS16125_ | 56 | 59 | 65 | 44 | 52 | 56 |
| SL1344_RS16130_ | 117 | 123 | 114 | 106 | 108 | 110 |
| SL1344_RS16135_ | 48 | 51 | 54 | 52 | 56 | 65 |
| SL1344_RS16140_ | 87 | 103 | 72 | 94 | 85 | 87 |
| SL1344_RS16145_ | 266 | 312 | 255 | 226 | 218 | 244 |
| SL1344_RS16150_ | 162 | 174 | 151 | 148 | 159 | 185 |
| SL1344_RS16155_ | 17 | 15 | 23 | 18 | 18 | 20 |
| SL1344_RS16160_ | 20 | 20 | 19 | 14 | 16 | 11 |
| SL1344_RS16165_ | 100 | 88 | 104 | 83 | 95 | 77 |
| SL1344_RS16170_ | 156 | 152 | 148 | 137 | 151 | 156 |
| SL1344_RS16175_ | 64 | 70 | 61 | 63 | 50 | 71 |
| SL1344_RS16180_ | 127 | 149 | 119 | 147 | 124 | 137 |
| SL1344_RS16185_ | 109 | 124 | 113 | 144 | 108 | 140 |
| SL1344_RS16190_ | 10 | 19 | 20 | 21 | 29 | 25 |
| SL1344_RS16195_ | 22 | 22 | 29 | 42 | 27 | 29 |
| SL1344_RS16200_ | 23 | 33 | 49 | 44 | 27 | 44 |
| SL1344_RS16205_ | 13 | 14 | 13 | 12 | 14 | 18 |
| SL1344_RS16210_ | 59 | 49 | 59 | 69 | 60 | 71 |
| SL1344_RS16215_ | 209 | 195 | 189 | 163 | 184 | 196 |
| SL1344_RS16220_ | 119 | 93 | 104 | 101 | 97 | 88 |
| SL1344_RS16225_ | 222 | 226 | 195 | 211 | 190 | 188 |
| SL1344_RS16235_ | 536 | 590 | 634 | 669 | 655 | 738 |
| SL1344_RS16240_ | 254 | 282 | 201 | 250 | 213 | 212 |
| SL1344_RS16245_ | 36 | 50 | 61 | 47 | 60 | 69 |

| | | | | | | |
|---|---|---|---|---|---|---|
| SL1344_RS16250_ | 36 | 40 | 44 | 47 | 43 | 67 |
| SL1344_RS16255_ | 275 | 295 | 296 | 480 | 453 | 496 |
| SL1344_RS16260_ | 165 | 174 | 134 | 235 | 211 | 253 |
| SL1344_RS16265_ | 147 | 141 | 154 | 241 | 212 | 221 |
| SL1344_RS16270_ | 281 | 290 | 236 | 466 | 410 | 499 |
| SL1344_RS16275_ | 607 | 630 | 682 | 1028 | 982 | 1164 |
| SL1344_RS16280_ | 179 | 203 | 226 | 386 | 355 | 428 |
| SL1344_RS16285_ | 126 | 133 | 156 | 296 | 230 | 342 |
| SL1344_RS16290_ | 369 | 361 | 332 | 584 | 548 | 671 |
| SL1344_RS16305_ | 950 | 1078 | 960 | 1672 | 1402 | 1569 |
| SL1344_RS16310_ | 1054 | 1038 | 892 | 2526 | 2418 | 2464 |
| SL1344_RS16315_ | 355 | 439 | 353 | 1061 | 938 | 1031 |
| SL1344_RS16320_ | 386 | 402 | 401 | 1044 | 989 | 1130 |
| SL1344_RS16325_ | 352 | 401 | 405 | 976 | 853 | 1050 |
| SL1344_RS16330_ | 1652 | 1972 | 2133 | 1697 | 1560 | 1671 |
| SL1344_RS16335_ | 2433 | 2718 | 2378 | 2346 | 2090 | 2072 |
| SL1344_RS16340_ | 1005 | 1055 | 711 | 992 | 784 | 693 |
| SL1344_RS16350_ | 491 | 428 | 418 | 377 | 390 | 411 |
| SL1344_RS16355_ | 280 | 334 | 335 | 371 | 325 | 347 |
| SL1344_RS16360_ | 437 | 421 | 404 | 509 | 445 | 475 |
| SL1344_RS16365_ | 213 | 228 | 259 | 290 | 287 | 296 |
| SL1344_RS16370_ | 82 | 86 | 51 | 81 | 57 | 53 |
| SL1344_RS16375_ | 280 | 231 | 201 | 263 | 238 | 233 |
| SL1344_RS16380_ | 92 | 96 | 96 | 86 | 87 | 66 |
| SL1344_RS16385_ | 2046 | 2061 | 2005 | 1518 | 1556 | 1615 |
| SL1344_RS16390_ | 14 | 11 | 7 | 15 | 13 | 24 |
| SL1344_RS16395_ | 4 | 7 | 7 | 14 | 7 | 5 |
| SL1344_RS16400_ | 52 | 76 | 51 | 54 | 35 | 47 |
| SL1344_RS16405_ | 829 | 1077 | 1260 | 908 | 885 | 878 |
| SL1344_RS16410_ | 646 | 685 | 708 | 530 | 466 | 467 |
| SL1344_RS16415_ | 2887 | 2990 | 3113 | 2644 | 2594 | 2795 |
| SL1344_RS16420_ | 273 | 308 | 332 | 421 | 463 | 476 |
| SL1344_RS16425_ | 371 | 551 | 593 | 677 | 725 | 719 |
| SL1344_RS16440_ | 258 | 296 | 255 | 296 | 277 | 261 |
| SL1344_RS16445_ | 1113 | 1140 | 1269 | 1308 | 1336 | 1293 |
| SL1344_RS16450_ | 3224 | 3358 | 3778 | 2740 | 2712 | 2885 |
| SL1344_RS16455_ | 1019 | 892 | 900 | 831 | 816 | 829 |
| SL1344_RS16460_ | 1390 | 1299 | 1293 | 1316 | 1196 | 1179 |
| SL1344_RS16465_ | 348 | 367 | 305 | 333 | 329 | 334 |
| SL1344_RS16470_ | 193 | 218 | 202 | 259 | 235 | 252 |
| SL1344_RS16480_ | 597 | 631 | 583 | 545 | 497 | 583 |
| SL1344_RS16485_ | 1107 | 1189 | 1221 | 1111 | 1023 | 1181 |
| SL1344_RS16490_ | 440 | 428 | 460 | 498 | 493 | 505 |
| SL1344_RS16495_ | 447 | 412 | 440 | 448 | 406 | 455 |
| SL1344_RS16500_ | 10 | 13 | 5 | 11 | 14 | 10 |
| SL1344_RS16505_ | 32 | 39 | 41 | 38 | 28 | 40 |
| SL1344_RS16510_ | 21 | 27 | 19 | 13 | 27 | 32 |
| SL1344_RS16515_ | 82 | 97 | 87 | 118 | 107 | 103 |
| SL1344_RS16520_ | 1048 | 1412 | 2002 | 1276 | 1327 | 1468 |
| SL1344_RS16525_ | 1621 | 1479 | 1210 | 1358 | 1163 | 1122 |
| SL1344_RS16530_ | 106 | 111 | 85 | 89 | 75 | 72 |
| SL1344_RS16535_ | 29 | 32 | 33 | 28 | 36 | 32 |
| SL1344_RS16540_ | 184 | 160 | 173 | 202 | 189 | 160 |
| SL1344_RS16545_ | 822 | 884 | 958 | 796 | 776 | 833 |
| SL1344_RS16550_ | 693 | 806 | 852 | 808 | 758 | 757 |
| SL1344_RS16555_ | 494 | 518 | 463 | 475 | 437 | 490 |
| SL1344_RS16560_ | 318 | 348 | 322 | 307 | 308 | 307 |
| SL1344_RS16565_ | 768 | 774 | 714 | 655 | 656 | 632 |
| SL1344_RS16570_ | 502 | 599 | 610 | 511 | 506 | 508 |
| SL1344_RS16575_ | 163 | 173 | 172 | 155 | 128 | 172 |
| SL1344_RS16580_ | 492 | 490 | 444 | 434 | 447 | 466 |
| SL1344_RS16585_ | 1027 | 1129 | 1114 | 909 | 897 | 922 |

| | | | | | | |
|---|---|---|---|---|---|---|
| SL1344_RS16590_ | 43382 | 41666 | 41170 | 38673 | 39899 | 39931 |
| SL1344_RS16595_ | 6612 | 6509 | 6530 | 6269 | 6058 | 6549 |
| SL1344_RS16600_ | 5376 | 5923 | 6337 | 6696 | 6557 | 6374 |
| SL1344_RS16605_ | 31 | 67 | 62 | 81 | 79 | 77 |
| SL1344_RS16610_ | 306 | 334 | 283 | 227 | 190 | 256 |
| SL1344_RS16615_ | 364 | 358 | 320 | 351 | 269 | 315 |
| SL1344_RS16620_ | 271 | 290 | 249 | 214 | 191 | 178 |
| SL1344_RS16635_ | 46 | 68 | 86 | 83 | 64 | 73 |
| SL1344_RS16640_ | 54 | 41 | 40 | 56 | 54 | 37 |
| SL1344_RS16645_ | 951 | 1022 | 1171 | 791 | 770 | 774 |
| SL1344_RS16650_ | 324 | 324 | 266 | 265 | 289 | 283 |
| SL1344_RS16655_ | 105 | 112 | 102 | 110 | 81 | 95 |
| SL1344_RS16660_ | 182 | 141 | 132 | 156 | 145 | 165 |
| SL1344_RS16665_ | 353 | 389 | 442 | 307 | 350 | 344 |
| SL1344_RS16675_ | 907 | 831 | 843 | 849 | 791 | 905 |
| SL1344_RS16680_ | 565 | 567 | 566 | 612 | 583 | 630 |
| SL1344_RS16685_ | 1182 | 1098 | 1064 | 1036 | 1213 | 1057 |
| SL1344_RS16690_ | 719 | 703 | 693 | 769 | 753 | 646 |
| SL1344_RS16695_ | 949 | 907 | 1013 | 863 | 963 | 924 |
| SL1344_RS16700_ | 884 | 920 | 836 | 1127 | 1104 | 1080 |
| SL1344_RS16705_ | 27 | 32 | 31 | 28 | 19 | 19 |
| SL1344_RS16710_ | 153 | 112 | 121 | 132 | 117 | 137 |
| SL1344_RS16715_ | 576 | 575 | 564 | 527 | 486 | 525 |
| SL1344_RS16720_ | 151 | 162 | 208 | 177 | 172 | 186 |
| SL1344_RS16725_ | 52 | 50 | 38 | 63 | 48 | 50 |
| SL1344_RS16730_ | 68 | 62 | 60 | 82 | 61 | 65 |
| SL1344_RS16745_ | 123 | 170 | 150 | 323 | 273 | 350 |
| SL1344_RS16750_ | 206 | 221 | 226 | 577 | 511 | 589 |
| SL1344_RS16755_ | 90 | 104 | 133 | 307 | 257 | 301 |
| SL1344_RS16775_ | 72 | 87 | 72 | 70 | 88 | 75 |
| SL1344_RS16780_ | 106 | 85 | 71 | 92 | 90 | 94 |
| SL1344_RS16785_ | 36 | 39 | 32 | 40 | 39 | 33 |
| SL1344_RS16790_ | 38 | 41 | 32 | 52 | 41 | 44 |
| SL1344_RS16795_ | 221 | 237 | 224 | 280 | 245 | 223 |
| SL1344_RS16800_ | 360 | 371 | 277 | 443 | 328 | 292 |
| SL1344_RS16805_ | 20 | 15 | 14 | 22 | 26 | 21 |
| SL1344_RS16810_ | 18 | 13 | 8 | 13 | 16 | 9 |
| SL1344_RS16815_ | 23 | 14 | 24 | 24 | 33 | 18 |
| SL1344_RS16820_ | 9 | 13 | 20 | 14 | 14 | 23 |
| SL1344_RS16825_ | 18 | 27 | 14 | 15 | 27 | 18 |
| SL1344_RS16830_ | 5 | 21 | 3 | 5 | 6 | 5 |
| SL1344_RS16835_ | 12 | 7 | 4 | 13 | 10 | 8 |
| SL1344_RS16840_ | 34 | 41 | 30 | 34 | 43 | 49 |
| SL1344_RS16845_ | 19 | 8 | 20 | 14 | 18 | 16 |
| SL1344_RS16850_ | 356 | 454 | 428 | 426 | 410 | 388 |
| SL1344_RS16855_ | 627 | 610 | 617 | 543 | 531 | 497 |
| SL1344_RS16860_ | 1327 | 1246 | 1365 | 1249 | 1227 | 1308 |
| SL1344_RS16865_ | 563 | 535 | 592 | 579 | 526 | 538 |
| SL1344_RS16870_ | 472 | 508 | 447 | 501 | 463 | 490 |
| SL1344_RS16875_ | 907 | 915 | 853 | 972 | 980 | 1000 |
| SL1344_RS16880_ | 71 | 72 | 72 | 61 | 71 | 74 |
| SL1344_RS16885_ | 5 | 17 | 16 | 20 | 15 | 19 |
| SL1344_RS16890_ | 94 | 82 | 104 | 122 | 107 | 125 |
| SL1344_RS16895_ | 102 | 117 | 113 | 98 | 116 | 104 |
| SL1344_RS16900_ | 304 | 423 | 400 | 447 | 372 | 409 |
| SL1344_RS16905_ | 377 | 410 | 355 | 384 | 357 | 373 |
| SL1344_RS16910_ | 15 | 19 | 20 | 5 | 15 | 10 |
| SL1344_RS16915_ | 18 | 17 | 15 | 14 | 16 | 17 |
| SL1344_RS16920_ | 121 | 132 | 117 | 125 | 118 | 133 |
| SL1344_RS16925_ | 23 | 35 | 24 | 32 | 29 | 30 |
| SL1344_RS16930_ | 8 | 10 | 12 | 9 | 4 | 6 |
| SL1344_RS16935_ | 206 | 204 | 226 | 206 | 144 | 182 |

| | | | | | | |
|---|---|---|---|---|---|---|
| SL1344_RS16940_ | 7096 | 6891 | 8123 | 5590 | 5790 | 6619 |
| SL1344_RS16945_ | 10429 | 11362 | 11390 | 10500 | 10311 | 11167 |
| SL1344_RS16950_ | 6986 | 7724 | 8798 | 7246 | 7110 | 7487 |
| SL1344_RS16955_ | 38366 | 36270 | 35810 | 30173 | 31467 | 29791 |
| SL1344_RS16960_ | 2267 | 2218 | 2276 | 1906 | 2007 | 2003 |
| SL1344_RS16965_ | 1111 | 1150 | 1131 | 979 | 1067 | 1004 |
| SL1344_RS16970_ | 14262 | 15277 | 16982 | 14141 | 13623 | 14148 |
| SL1344_RS16980_ | 2263 | 2432 | 2456 | 2101 | 2207 | 2143 |
| SL1344_RS16985_ | 5042 | 4355 | 3977 | 4477 | 4124 | 4478 |
| SL1344_RS16990_ | 304 | 327 | 362 | 296 | 280 | 370 |
| SL1344_RS16995_ | 208 | 232 | 233 | 327 | 249 | 268 |
| SL1344_RS17000_ | 3988 | 4204 | 5527 | 5347 | 4979 | 5283 |
| SL1344_RS17005_ | 18729 | 21763 | 21609 | 23871 | 22818 | 24245 |
| SL1344_RS17010_ | 1085 | 1281 | 1441 | 1563 | 1464 | 1604 |
| SL1344_RS17015_ | 249 | 313 | 303 | 399 | 334 | 419 |
| SL1344_RS17030_ | 1840 | 1992 | 1886 | 2095 | 1890 | 1902 |
| SL1344_RS17035_ | 4813 | 4674 | 4812 | 3775 | 3995 | 3940 |
| SL1344_RS17040_ | 2038 | 2078 | 2110 | 2269 | 2200 | 2118 |
| SL1344_RS17045_ | 1592 | 1559 | 1131 | 1363 | 1289 | 1324 |
| SL1344_RS17050_ | 1122 | 1086 | 985 | 968 | 980 | 997 |
| SL1344_RS17060_ | 1216 | 1469 | 1641 | 1132 | 1061 | 1347 |
| SL1344_RS17065_ | 7223 | 11420 | 12606 | 11354 | 8458 | 11222 |
| SL1344_RS17070_ | 11581 | 15616 | 16450 | 15449 | 14278 | 16927 |
| SL1344_RS17075_ | 3576 | 3516 | 3497 | 2947 | 2886 | 2953 |
| SL1344_RS17080_ | 211 | 192 | 156 | 157 | 173 | 190 |
| SL1344_RS17085_ | 2128 | 2310 | 2514 | 2397 | 2206 | 2239 |
| SL1344_RS17090_ | 300 | 344 | 353 | 340 | 276 | 337 |
| SL1344_RS17095_ | 857 | 900 | 999 | 963 | 879 | 894 |
| SL1344_RS17100_ | 760 | 768 | 773 | 821 | 715 | 861 |
| SL1344_RS17105_ | 299 | 380 | 373 | 404 | 334 | 367 |
| SL1344_RS17110_ | 391 | 453 | 453 | 480 | 414 | 464 |
| SL1344_RS17115_ | 724 | 686 | 785 | 722 | 631 | 712 |
| SL1344_RS17120_ | 1004 | 1117 | 1096 | 1036 | 983 | 1079 |
| SL1344_RS17125_ | 1059 | 1125 | 1083 | 986 | 1021 | 1058 |
| SL1344_RS17130_ | 1695 | 1567 | 1539 | 1423 | 1412 | 1389 |
| SL1344_RS17135_ | 1377 | 1337 | 1205 | 1147 | 1111 | 1190 |
| SL1344_RS17140_ | 1503 | 1541 | 1320 | 1284 | 1261 | 1308 |
| SL1344_RS17145_ | 1416 | 1345 | 1413 | 1223 | 1200 | 1223 |
| SL1344_RS17150_ | 3165 | 3143 | 3018 | 2658 | 2750 | 2762 |
| SL1344_RS17155_ | 831 | 837 | 743 | 928 | 845 | 885 |
| SL1344_RS17160_ | 899 | 962 | 885 | 860 | 812 | 954 |
| SL1344_RS17165_ | 1636 | 1654 | 1658 | 1523 | 1493 | 1522 |
| SL1344_RS17170_ | 554 | 588 | 572 | 547 | 457 | 531 |
| SL1344_RS17175_ | 84 | 71 | 72 | 70 | 64 | 70 |
| SL1344_RS17180_ | 330 | 403 | 440 | 873 | 728 | 826 |
| SL1344_RS17185_ | 424 | 395 | 397 | 800 | 624 | 780 |
| SL1344_RS17190_ | 841 | 960 | 1010 | 1323 | 1151 | 1253 |
| SL1344_RS17195_ | 64 | 49 | 50 | 58 | 44 | 65 |
| SL1344_RS17200_ | 9 | 14 | 14 | 12 | 3 | 8 |
| SL1344_RS17205_ | 1371 | 1570 | 1680 | 1588 | 1497 | 1663 |
| SL1344_RS17210_ | 474 | 588 | 692 | 714 | 671 | 702 |
| SL1344_RS17215_ | 24 | 22 | 32 | 32 | 30 | 24 |
| SL1344_RS17220_ | 308 | 354 | 397 | 272 | 251 | 290 |
| SL1344_RS17225_ | 773 | 801 | 780 | 717 | 677 | 639 |
| SL1344_RS17230_ | 60 | 83 | 71 | 100 | 87 | 88 |
| SL1344_RS17235_ | 59 | 83 | 52 | 71 | 78 | 63 |
| SL1344_RS17240_ | 58 | 47 | 53 | 49 | 73 | 59 |
| SL1344_RS17245_ | 96 | 108 | 89 | 111 | 96 | 100 |
| SL1344_RS17250_ | 45 | 40 | 50 | 43 | 46 | 46 |
| SL1344_RS17255_ | 1275 | 1302 | 1282 | 1238 | 1200 | 1296 |
| SL1344_RS17260_ | 6090 | 6354 | 6373 | 6556 | 6477 | 6146 |
| SL1344_RS17270_ | 192 | 268 | 277 | 171 | 148 | 181 |

| | | | | | | |
|---|---|---|---|---|---|---|
| SL1344_RS17275_ | 38722 | 46051 | 50282 | 42621 | 38053 | 41552 |
| SL1344_RS17280_ | 19114 | 20734 | 21938 | 19778 | 17441 | 19941 |
| SL1344_RS17285_ | 476 | 528 | 517 | 451 | 457 | 533 |
| SL1344_RS17290_ | 1301 | 1472 | 1356 | 1548 | 1420 | 1409 |
| SL1344_RS17295_ | 890 | 936 | 1072 | 1129 | 994 | 1018 |
| SL1344_RS17300_ | 1076 | 992 | 1066 | 883 | 812 | 814 |
| SL1344_RS17305_ | 38 | 22 | 21 | 22 | 18 | 23 |
| SL1344_RS17310_ | 65 | 45 | 54 | 57 | 65 | 71 |
| SL1344_RS17315_ | 104 | 106 | 109 | 134 | 85 | 91 |
| SL1344_RS17320_ | 8 | 8 | 13 | 2 | 20 | 8 |
| SL1344_RS17325_ | 46 | 42 | 50 | 39 | 38 | 38 |
| SL1344_RS17330_ | 77 | 47 | 57 | 64 | 58 | 54 |
| SL1344_RS17335_ | 27 | 29 | 20 | 34 | 23 | 33 |
| SL1344_RS17340_ | 387 | 414 | 431 | 466 | 398 | 419 |
| SL1344_RS17345_ | 505 | 501 | 479 | 496 | 458 | 457 |
| SL1344_RS17350_ | 1603 | 1936 | 1727 | 3531 | 3401 | 3429 |
| SL1344_RS17355_ | 816 | 896 | 841 | 800 | 848 | 854 |
| SL1344_RS17360_ | 292 | 261 | 205 | 263 | 261 | 247 |
| SL1344_RS17365_ | 415 | 389 | 388 | 432 | 433 | 423 |
| SL1344_RS17370_ | 78 | 104 | 89 | 79 | 51 | 69 |
| SL1344_RS17375_ | 226 | 272 | 312 | 263 | 228 | 263 |
| SL1344_RS17380_ | 121 | 102 | 146 | 112 | 122 | 114 |
| SL1344_RS17385_ | 20 | 21 | 23 | 25 | 24 | 30 |
| SL1344_RS17390_ | 181 | 171 | 141 | 182 | 158 | 155 |
| SL1344_RS17395_ | 702 | 863 | 935 | 804 | 712 | 823 |
| SL1344_RS17400_ | 2106 | 2017 | 2141 | 1983 | 1873 | 2054 |
| SL1344_RS17405_ | 1853 | 2240 | 2260 | 2062 | 2098 | 2115 |
| SL1344_RS17410_ | 1222 | 1366 | 1528 | 1312 | 1229 | 1319 |
| SL1344_RS17415_ | 803 | 1039 | 1054 | 970 | 1053 | 1018 |
| SL1344_RS17420_ | 1926 | 2104 | 2223 | 2100 | 2003 | 2130 |
| SL1344_RS17425_ | 5092 | 5306 | 5435 | 5221 | 5315 | 5227 |
| SL1344_RS17430_ | 9 | 9 | 2 | 4 | 2 | 12 |
| SL1344_RS17440_ | 397 | 334 | 285 | 350 | 338 | 296 |
| SL1344_RS17445_ | 235 | 249 | 215 | 204 | 214 | 182 |
| SL1344_RS17450_ | 163 | 114 | 142 | 151 | 119 | 131 |
| SL1344_RS17455_ | 5334 | 5065 | 5411 | 4176 | 4310 | 4164 |
| SL1344_RS17460_ | 10971 | 10707 | 10927 | 8181 | 8133 | 8407 |
| SL1344_RS17465_ | 65 | 78 | 105 | 83 | 68 | 78 |
| SL1344_RS17470_ | 627 | 590 | 670 | 533 | 523 | 600 |
| SL1344_RS17475_ | 984 | 1009 | 1006 | 854 | 780 | 955 |
| SL1344_RS17480_ | 7264 | 7830 | 8242 | 6002 | 5972 | 6618 |
| SL1344_RS17490_ | 262 | 282 | 288 | 232 | 196 | 239 |
| SL1344_RS17495_ | 327 | 271 | 251 | 261 | 227 | 250 |
| SL1344_RS17500_ | 291 | 309 | 305 | 220 | 244 | 244 |
| SL1344_RS17505_ | 24 | 33 | 24 | 26 | 24 | 27 |
| SL1344_RS17510_ | 21 | 22 | 17 | 18 | 25 | 24 |
| SL1344_RS17515_ | 126 | 161 | 137 | 152 | 142 | 144 |
| SL1344_RS17520_ | 288 | 355 | 290 | 194 | 178 | 174 |
| SL1344_RS17530_ | 124 | 129 | 140 | 91 | 87 | 92 |
| SL1344_RS17540_ | 1367 | 836 | 758 | 557 | 514 | 918 |
| SL1344_RS17545_ | 251 | 210 | 153 | 182 | 127 | 170 |
| SL1344_RS17550_ | 455 | 164 | 150 | 112 | 103 | 196 |
| SL1344_RS17555_ | 256 | 226 | 213 | 325 | 337 | 349 |
| SL1344_RS17560_ | 299 | 366 | 384 | 323 | 321 | 340 |
| SL1344_RS17565_ | 594 | 671 | 677 | 634 | 549 | 677 |
| SL1344_RS17570_ | 470 | 502 | 491 | 460 | 427 | 480 |
| SL1344_RS17575_ | 1338 | 1229 | 1272 | 1240 | 1120 | 1198 |
| SL1344_RS17580_ | 1278 | 1296 | 1290 | 1472 | 1364 | 1383 |
| SL1344_RS17585_ | 306 | 333 | 306 | 343 | 327 | 342 |
| SL1344_RS17590_ | 1601 | 1695 | 1791 | 1552 | 1464 | 1512 |
| SL1344_RS17595_ | 2911 | 3043 | 3169 | 2817 | 2687 | 2904 |
| SL1344_RS17600_ | 294 | 334 | 413 | 324 | 305 | 344 |

| | | | | | | |
|---|---|---|---|---|---|---|
| SL1344_RS17605_ | 517 | 559 | 480 | 353 | 411 | 432 |
| SL1344_RS17610_ | 1485 | 1630 | 1734 | 1541 | 1364 | 1488 |
| SL1344_RS17615_ | 390 | 496 | 502 | 475 | 459 | 414 |
| SL1344_RS17620_ | 283 | 306 | 265 | 284 | 294 | 258 |
| SL1344_RS17625_ | 154 | 146 | 129 | 164 | 126 | 137 |
| SL1344_RS17630_ | 30708 | 31878 | 34411 | 28309 | 27451 | 28399 |
| SL1344_RS17635_ | 58860 | 60249 | 68166 | 57885 | 56251 | 58228 |
| SL1344_RS17640_ | 34026 | 39370 | 47649 | 39535 | 40488 | 38416 |
| SL1344_RS17645_ | 5151 | 7630 | 9430 | 9394 | 8829 | 8714 |
| SL1344_RS17650_ | 13443 | 15415 | 18728 | 15474 | 15633 | 15491 |
| SL1344_RS17655_ | 16478 | 17919 | 17960 | 16483 | 15198 | 16459 |
| SL1344_RS17660_ | 49597 | 54721 | 60476 | 52392 | 48472 | 55395 |
| SL1344_RS17665_ | 13358 | 15976 | 18050 | 15345 | 14030 | 16152 |
| SL1344_RS17670_ | 6370 | 7295 | 7595 | 7319 | 6563 | 7475 |
| SL1344_RS17675_ | 14593 | 16210 | 17484 | 16079 | 14406 | 16210 |
| SL1344_RS17680_ | 14482 | 15438 | 16641 | 14333 | 13433 | 14573 |
| SL1344_RS17685_ | 24042 | 24720 | 26072 | 22610 | 21619 | 23503 |
| SL1344_RS17690_ | 22896 | 23362 | 24869 | 22011 | 21135 | 22433 |
| SL1344_RS17700_ | 37343 | 37458 | 40690 | 34449 | 33264 | 35791 |
| SL1344_RS17705_ | 28850 | 29882 | 30631 | 27743 | 26948 | 28237 |
| SL1344_RS17710_ | 32930 | 33056 | 35970 | 29590 | 29288 | 30396 |
| SL1344_RS17715_ | 49446 | 54243 | 53054 | 44047 | 45074 | 44630 |
| SL1344_RS17720_ | 24732 | 24836 | 23742 | 22348 | 22079 | 21488 |
| SL1344_RS17725_ | 54170 | 51507 | 54076 | 48684 | 51885 | 46868 |
| SL1344_RS17730_ | 43351 | 41207 | 41712 | 38852 | 38024 | 38472 |
| SL1344_RS17735_ | 23938 | 23388 | 23718 | 22640 | 21519 | 22190 |
| SL1344_RS17740_ | 19335 | 20002 | 21397 | 18692 | 18526 | 18630 |
| SL1344_RS17745_ | 27314 | 29233 | 30232 | 29121 | 26613 | 28316 |
| SL1344_RS17750_ | 33731 | 35458 | 41671 | 31141 | 32553 | 32193 |
| SL1344_RS17755_ | 28310 | 29512 | 31867 | 28357 | 27893 | 27798 |
| SL1344_RS17760_ | 33720 | 33714 | 35828 | 32847 | 31578 | 32062 |
| SL1344_RS17765_ | 17023 | 16619 | 17577 | 14817 | 13954 | 15074 |
| SL1344_RS17770_ | 4 | 6 | 2 | 10 | 4 | 6 |
| SL1344_RS17775_ | 1782 | 2264 | 2223 | 2049 | 2060 | 2043 |
| SL1344_RS17780_ | 2386 | 2480 | 2088 | 2225 | 2401 | 2087 |
| SL1344_RS17785_ | 50709 | 58885 | 70138 | 54879 | 52195 | 53301 |
| SL1344_RS17790_ | 93779 | 99082 | 106388 | 91811 | 90274 | 92394 |
| SL1344_RS17795_ | 23440 | 23444 | 23867 | 21560 | 20187 | 22000 |
| SL1344_RS17800_ | 16274 | 15838 | 15718 | 13969 | 13020 | 15113 |
| SL1344_RS17805_ | 644 | 604 | 551 | 805 | 650 | 664 |
| SL1344_RS17810_ | 517 | 486 | 462 | 401 | 393 | 442 |
| SL1344_RS17815_ | 539 | 525 | 479 | 417 | 422 | 438 |
| SL1344_RS17820_ | 975 | 970 | 950 | 834 | 850 | 833 |
| SL1344_RS17825_ | 6494 | 6579 | 6748 | 6434 | 6592 | 6372 |
| SL1344_RS17830_ | 426 | 477 | 395 | 611 | 620 | 590 |
| SL1344_RS17835_ | 4985 | 5090 | 5506 | 4520 | 4408 | 4531 |
| SL1344_RS17840_ | 181 | 227 | 177 | 244 | 202 | 208 |
| SL1344_RS17845_ | 609 | 614 | 587 | 791 | 727 | 727 |
| SL1344_RS17850_ | 176 | 170 | 179 | 242 | 236 | 272 |
| SL1344_RS17855_ | 1752 | 1737 | 1744 | 1423 | 1491 | 1408 |
| SL1344_RS17860_ | 81 | 99 | 105 | 81 | 66 | 63 |
| SL1344_RS17865_ | 129 | 162 | 144 | 195 | 175 | 162 |
| SL1344_RS17870_ | 682 | 659 | 662 | 602 | 530 | 614 |
| SL1344_RS17875_ | 139 | 164 | 163 | 162 | 127 | 170 |
| SL1344_RS17880_ | 461 | 469 | 435 | 449 | 451 | 444 |
| SL1344_RS17885_ | 736 | 705 | 646 | 625 | 592 | 555 |
| SL1344_RS17895_ | 1346 | 1401 | 1397 | 1270 | 1240 | 1384 |
| SL1344_RS17900_ | 412 | 477 | 452 | 452 | 430 | 476 |
| SL1344_RS17905_ | 327 | 343 | 315 | 303 | 264 | 314 |
| SL1344_RS17910_ | 105 | 109 | 102 | 80 | 77 | 82 |
| SL1344_RS17915_ | 70 | 70 | 81 | 65 | 54 | 54 |
| SL1344_RS17920_ | 1298 | 1207 | 1203 | 837 | 856 | 825 |

| | | | | | | |
|---|---|---|---|---|---|---|
| SL1344_RS17925_ | 2888 | 2917 | 2969 | 1882 | 1775 | 1907 |
| SL1344_RS17930_ | 312 | 343 | 348 | 205 | 276 | 233 |
| SL1344_RS17940_ | 137 | 160 | 126 | 94 | 120 | 116 |
| SL1344_RS17965_ | 2820 | 3220 | 3280 | 2968 | 2739 | 3074 |
| SL1344_RS17970_ | 1342 | 1396 | 1429 | 1190 | 1167 | 1263 |
| SL1344_RS17975_ | 1129 | 1116 | 1130 | 924 | 893 | 932 |
| SL1344_RS17985_ | 4245 | 4185 | 4555 | 3809 | 3596 | 3695 |
| SL1344_RS17990_ | 2846 | 3242 | 3550 | 3430 | 3118 | 3418 |
| SL1344_RS17995_ | 2412 | 2896 | 3060 | 2885 | 2647 | 2937 |
| SL1344_RS18000_ | 83 | 99 | 84 | 97 | 74 | 85 |
| SL1344_RS18005_ | 17 | 18 | 24 | 16 | 18 | 21 |
| SL1344_RS18010_ | 18 | 22 | 15 | 18 | 23 | 16 |
| SL1344_RS18015_ | 20 | 16 | 21 | 29 | 24 | 36 |
| SL1344_RS18020_ | 25 | 16 | 22 | 30 | 32 | 26 |
| SL1344_RS18025_ | 1663 | 1696 | 1736 | 1573 | 1495 | 1570 |
| SL1344_RS18030_ | 1030 | 1026 | 975 | 711 | 683 | 743 |
| SL1344_RS18035_ | 2097 | 2022 | 1930 | 2287 | 2165 | 2208 |
| SL1344_RS18040_ | 588 | 586 | 526 | 567 | 530 | 543 |
| SL1344_RS18045_ | 426 | 424 | 412 | 481 | 451 | 427 |
| SL1344_RS18050_ | 1048 | 988 | 1004 | 952 | 881 | 994 |
| SL1344_RS18055_ | 225 | 246 | 215 | 232 | 205 | 221 |
| SL1344_RS18060_ | 1627 | 1584 | 1632 | 1847 | 1742 | 1681 |
| SL1344_RS18075_ | 300 | 366 | 329 | 356 | 363 | 297 |
| SL1344_RS18080_ | 2033 | 2135 | 2291 | 2060 | 1943 | 2009 |
| SL1344_RS18090_ | 2882 | 3104 | 3209 | 1662 | 1616 | 1670 |
| SL1344_RS18095_ | 1188 | 1182 | 1199 | 772 | 738 | 745 |
| SL1344_RS18100_ | 140 | 169 | 143 | 153 | 124 | 138 |
| SL1344_RS18105_ | 152 | 206 | 219 | 187 | 167 | 212 |
| SL1344_RS18110_ | 183 | 177 | 165 | 262 | 214 | 217 |
| SL1344_RS18115_ | 2806 | 2813 | 2700 | 2980 | 2744 | 2758 |
| SL1344_RS18120_ | 253 | 257 | 273 | 167 | 174 | 192 |
| SL1344_RS18125_ | 211 | 221 | 228 | 222 | 222 | 211 |
| SL1344_RS18130_ | 137 | 154 | 142 | 152 | 134 | 144 |
| SL1344_RS18140_ | 417 | 369 | 393 | 458 | 412 | 390 |
| SL1344_RS18145_ | 737 | 688 | 577 | 823 | 727 | 650 |
| SL1344_RS18150_ | 138 | 125 | 150 | 226 | 212 | 209 |
| SL1344_RS18155_ | 189 | 201 | 186 | 263 | 271 | 286 |
| SL1344_RS18160_ | 88 | 96 | 113 | 107 | 111 | 114 |
| SL1344_RS18165_ | 558 | 537 | 435 | 425 | 378 | 416 |
| SL1344_RS18170_ | 479 | 572 | 616 | 602 | 602 | 636 |
| SL1344_RS18175_ | 1184 | 1231 | 1284 | 1188 | 1177 | 1178 |
| SL1344_RS18180_ | 424 | 418 | 439 | 401 | 405 | 378 |
| SL1344_RS18185_ | 20288 | 20805 | 20138 | 22163 | 20369 | 22160 |
| SL1344_RS18190_ | 63 | 50 | 52 | 53 | 56 | 57 |
| SL1344_RS18195_ | 202 | 226 | 208 | 229 | 188 | 225 |
| SL1344_RS18205_ | 15 | 14 | 19 | 12 | 11 | 15 |
| SL1344_RS18210_ | 32 | 21 | 21 | 25 | 23 | 15 |
| SL1344_RS18215_ | 67 | 81 | 45 | 43 | 41 | 38 |
| SL1344_RS18220_ | 542 | 583 | 505 | 663 | 561 | 601 |
| SL1344_RS18225_ | 564 | 592 | 581 | 611 | 621 | 635 |
| SL1344_RS18230_ | 178 | 194 | 194 | 216 | 200 | 249 |
| SL1344_RS18235_ | 162 | 162 | 203 | 195 | 213 | 244 |
| SL1344_RS18240_ | 217 | 244 | 247 | 312 | 269 | 336 |
| SL1344_RS18245_ | 335 | 357 | 391 | 479 | 475 | 455 |
| SL1344_RS18250_ | 482 | 585 | 667 | 742 | 722 | 768 |
| SL1344_RS18255_ | 173 | 192 | 222 | 181 | 203 | 204 |
| SL1344_RS18270_ | 130 | 153 | 127 | 135 | 131 | 129 |
| SL1344_RS18275_ | 179 | 208 | 190 | 211 | 218 | 227 |
| SL1344_RS18280_ | 137 | 137 | 102 | 119 | 120 | 117 |
| SL1344_RS18285_ | 14 | 5 | 7 | 7 | 4 | 3 |
| SL1344_RS18290_ | 5 | 8 | 6 | 5 | 5 | 3 |
| SL1344_RS18295_ | 6 | 8 | 5 | 18 | 3 | 7 |

| | | | | | | |
|---|---|---|---|---|---|---|
| SL1344_RS18300_ | 43 | 61 | 42 | 53 | 39 | 53 |
| SL1344_RS18305_ | 130 | 112 | 117 | 151 | 137 | 146 |
| SL1344_RS18310_ | 105 | 115 | 80 | 142 | 128 | 98 |
| SL1344_RS18315_ | 117 | 132 | 167 | 154 | 173 | 147 |
| SL1344_RS18325_ | 82 | 104 | 96 | 88 | 93 | 106 |
| SL1344_RS18330_ | 91 | 79 | 78 | 97 | 75 | 84 |
| SL1344_RS18335_ | 176 | 191 | 188 | 235 | 248 | 220 |
| SL1344_RS18340_ | 69 | 66 | 63 | 77 | 71 | 75 |
| SL1344_RS18345_ | 41 | 35 | 36 | 64 | 61 | 68 |
| SL1344_RS18355_ | 64 | 67 | 90 | 84 | 72 | 90 |
| SL1344_RS18360_ | 106 | 96 | 89 | 122 | 101 | 103 |
| SL1344_RS18365_ | 56 | 62 | 61 | 82 | 68 | 96 |
| SL1344_RS18370_ | 65 | 94 | 113 | 147 | 116 | 152 |
| SL1344_RS18375_ | 467 | 486 | 404 | 387 | 368 | 394 |
| SL1344_RS18380_ | 612 | 586 | 714 | 529 | 496 | 512 |
| SL1344_RS18385_ | 280 | 284 | 345 | 317 | 360 | 324 |
| SL1344_RS18390_ | 4184 | 4583 | 4760 | 5452 | 5062 | 5158 |
| SL1344_RS18395_ | 1707 | 1833 | 1928 | 1592 | 1539 | 1645 |
| SL1344_RS18400_ | 846 | 841 | 961 | 767 | 758 | 759 |
| SL1344_RS18405_ | 1386 | 1672 | 1721 | 1479 | 1408 | 1355 |
| SL1344_RS18415_ | 489 | 503 | 475 | 349 | 351 | 332 |
| SL1344_RS18420_ | 135 | 184 | 147 | 162 | 163 | 161 |
| SL1344_RS18425_ | 310 | 308 | 277 | 342 | 284 | 295 |
| SL1344_RS18440_ | 558 | 571 | 522 | 394 | 440 | 389 |
| SL1344_RS18445_ | 360 | 428 | 485 | 415 | 310 | 405 |
| SL1344_RS18450_ | 1778 | 1769 | 1698 | 1869 | 1748 | 1831 |
| SL1344_RS18455_ | 214 | 237 | 226 | 273 | 263 | 239 |
| SL1344_RS18460_ | 86 | 92 | 72 | 75 | 63 | 76 |
| SL1344_RS18465_ | 432 | 445 | 396 | 407 | 415 | 429 |
| SL1344_RS18470_ | 370 | 386 | 386 | 302 | 303 | 312 |
| SL1344_RS18475_ | 92 | 103 | 101 | 90 | 105 | 102 |
| SL1344_RS18480_ | 319 | 319 | 304 | 440 | 412 | 405 |
| SL1344_RS18485_ | 144 | 153 | 179 | 170 | 192 | 194 |
| SL1344_RS18490_ | 924 | 992 | 1058 | 902 | 766 | 832 |
| SL1344_RS18495_ | 5147 | 5321 | 5523 | 3769 | 3397 | 3714 |
| SL1344_RS18500_ | 177 | 224 | 227 | 243 | 237 | 245 |
| SL1344_RS18505_ | 1835 | 1865 | 1693 | 3057 | 3053 | 2834 |
| SL1344_RS18510_ | 234 | 278 | 240 | 208 | 202 | 202 |
| SL1344_RS18515_ | 361 | 429 | 427 | 337 | 328 | 371 |
| SL1344_RS18520_ | 680 | 775 | 845 | 1059 | 907 | 890 |
| SL1344_RS18525_ | 418 | 433 | 354 | 427 | 428 | 443 |
| SL1344_RS18530_ | 760 | 668 | 610 | 699 | 623 | 633 |
| SL1344_RS18535_ | 1499 | 1435 | 1473 | 1532 | 1403 | 1491 |
| SL1344_RS18540_ | 107 | 111 | 133 | 144 | 150 | 122 |
| SL1344_RS18545_ | 92 | 109 | 109 | 133 | 98 | 87 |
| SL1344_RS18550_ | 59 | 48 | 56 | 76 | 42 | 55 |
| SL1344_RS18555_ | 113 | 136 | 140 | 131 | 133 | 110 |
| SL1344_RS18560_ | 275 | 260 | 227 | 278 | 220 | 220 |
| SL1344_RS18565_ | 292 | 261 | 283 | 252 | 257 | 277 |
| SL1344_RS18570_ | 397 | 387 | 394 | 767 | 744 | 715 |
| SL1344_RS18575_ | 30 | 21 | 29 | 20 | 23 | 20 |
| SL1344_RS18580_ | 80 | 56 | 75 | 33 | 41 | 31 |
| SL1344_RS18585_ | 152 | 148 | 155 | 152 | 122 | 128 |
| SL1344_RS18590_ | 228 | 188 | 184 | 240 | 228 | 191 |
| SL1344_RS18595_ | 199 | 189 | 138 | 214 | 189 | 216 |
| SL1344_RS18600_ | 612 | 656 | 598 | 1486 | 1290 | 1499 |
| SL1344_RS18610_ | 125 | 141 | 123 | 92 | 78 | 75 |
| SL1344_RS18615_ | 780 | 867 | 839 | 807 | 758 | 718 |
| SL1344_RS18620_ | 171 | 263 | 267 | 245 | 230 | 267 |
| SL1344_RS18625_ | 848 | 879 | 827 | 820 | 782 | 793 |
| SL1344_RS18630_ | 1301 | 1427 | 1333 | 1011 | 945 | 1003 |
| SL1344_RS18635_ | 504 | 546 | 559 | 297 | 308 | 367 |

| | | | | | | |
|---|---|---|---|---|---|---|
| SL1344_RS18640_ | 650 | 674 | 707 | 408 | 406 | 414 |
| SL1344_RS18655_ | 237 | 215 | 203 | 79 | 85 | 102 |
| SL1344_RS18660_ | 546 | 542 | 593 | 269 | 302 | 321 |
| SL1344_RS18665_ | 138 | 121 | 135 | 68 | 76 | 85 |
| SL1344_RS18670_ | 623 | 639 | 589 | 366 | 364 | 384 |
| SL1344_RS18675_ | 794 | 669 | 804 | 874 | 760 | 896 |
| SL1344_RS18680_ | 133 | 130 | 143 | 124 | 141 | 151 |
| SL1344_RS18685_ | 463 | 509 | 478 | 376 | 364 | 409 |
| SL1344_RS18690_ | 123 | 134 | 150 | 230 | 225 | 247 |
| SL1344_RS18695_ | 140 | 150 | 155 | 235 | 248 | 278 |
| SL1344_RS18700_ | 159 | 191 | 177 | 278 | 243 | 318 |
| SL1344_RS18705_ | 86 | 108 | 87 | 140 | 128 | 176 |
| SL1344_RS18710_ | 415 | 422 | 407 | 748 | 667 | 779 |
| SL1344_RS18715_ | 64 | 60 | 47 | 65 | 46 | 47 |
| SL1344_RS18720_ | 133 | 126 | 139 | 152 | 116 | 127 |
| SL1344_RS18725_ | 197 | 181 | 187 | 178 | 154 | 172 |
| SL1344_RS18735_ | 1216 | 1131 | 1141 | 1114 | 881 | 974 |
| SL1344_RS18740_ | 321 | 351 | 305 | 329 | 260 | 277 |
| SL1344_RS18745_ | 59 | 50 | 68 | 60 | 65 | 61 |
| SL1344_RS18750_ | 70 | 62 | 85 | 59 | 62 | 56 |
| SL1344_RS18755_ | 42 | 41 | 36 | 49 | 50 | 46 |
| SL1344_RS18760_ | 6 | 8 | 11 | 6 | 9 | 8 |
| SL1344_RS18765_ | 13 | 9 | 21 | 15 | 13 | 19 |
| SL1344_RS18770_ | 27 | 44 | 38 | 45 | 60 | 42 |
| SL1344_RS18775_ | 232 | 218 | 216 | 242 | 196 | 213 |
| SL1344_RS18780_ | 134 | 127 | 152 | 167 | 156 | 151 |
| SL1344_RS18785_ | 955 | 1044 | 1051 | 910 | 931 | 962 |
| SL1344_RS18790_ | 423 | 432 | 410 | 386 | 373 | 386 |
| SL1344_RS18795_ | 422 | 439 | 400 | 422 | 438 | 461 |
| SL1344_RS18800_ | 1965 | 2423 | 2489 | 2197 | 1948 | 2288 |
| SL1344_RS18805_ | 57 | 61 | 44 | 63 | 92 | 58 |
| SL1344_RS18815_ | 122 | 179 | 101 | 90 | 103 | 86 |
| SL1344_RS18820_ | 348 | 359 | 357 | 400 | 400 | 416 |
| SL1344_RS18825_ | 282 | 293 | 280 | 328 | 307 | 358 |
| SL1344_RS18830_ | 76 | 101 | 87 | 61 | 54 | 62 |
| SL1344_RS18835_ | 4033 | 4317 | 4799 | 3929 | 3716 | 3860 |
| SL1344_RS18840_ | 1236 | 1377 | 1501 | 1316 | 1184 | 1281 |
| SL1344_RS18845_ | 63 | 64 | 49 | 72 | 65 | 64 |
| SL1344_RS18850_ | 128 | 142 | 103 | 136 | 135 | 137 |
| SL1344_RS18855_ | 17 | 28 | 14 | 24 | 23 | 18 |
| SL1344_RS18860_ | 115 | 115 | 130 | 155 | 173 | 177 |
| SL1344_RS18865_ | 146 | 164 | 137 | 235 | 193 | 179 |
| SL1344_RS18870_ | 78 | 94 | 72 | 69 | 56 | 45 |
| SL1344_RS18875_ | 2284 | 2078 | 1906 | 2100 | 2111 | 2078 |
| SL1344_RS18880_ | 18 | 16 | 21 | 26 | 17 | 24 |
| SL1344_RS18885_ | 56 | 55 | 51 | 58 | 41 | 57 |
| SL1344_RS18890_ | 1485 | 1489 | 1569 | 1727 | 1645 | 1832 |
| SL1344_RS18895_ | 129 | 145 | 149 | 214 | 214 | 184 |
| SL1344_RS18900_ | 244 | 299 | 297 | 352 | 331 | 354 |
| SL1344_RS18905_ | 11 | 14 | 13 | 20 | 13 | 23 |
| SL1344_RS18910_ | 7 | 3 | 5 | 12 | 5 | 8 |
| SL1344_RS18915_ | 23 | 16 | 18 | 19 | 17 | 12 |
| SL1344_RS18920_ | 2 | 1 | 1 | 1 | 1 | 2 |
| SL1344_RS18925_ | 8 | 16 | 9 | 14 | 8 | 13 |
| SL1344_RS18930_ | 6 | 12 | 10 | 13 | 8 | 8 |
| SL1344_RS18935_ | 85 | 64 | 77 | 70 | 66 | 71 |
| SL1344_RS18945_ | 7 | 8 | 14 | 23 | 6 | 10 |
| SL1344_RS18950_ | 16 | 19 | 6 | 27 | 26 | 16 |
| SL1344_RS18955_ | 60 | 53 | 37 | 36 | 35 | 36 |
| SL1344_RS18960_ | 62 | 54 | 70 | 81 | 63 | 84 |
| SL1344_RS18965_ | 40 | 45 | 46 | 51 | 57 | 77 |
| SL1344_RS18970_ | 83 | 66 | 78 | 78 | 68 | 79 |

| | | | | | | |
|---|---|---|---|---|---|---|
| SL1344_RS18975_ | 890 | 859 | 846 | 868 | 822 | 837 |
| SL1344_RS18980_ | 1069 | 1094 | 899 | 863 | 834 | 827 |
| SL1344_RS18985_ | 271 | 267 | 269 | 358 | 337 | 301 |
| SL1344_RS18990_ | 532 | 523 | 498 | 656 | 666 | 652 |
| SL1344_RS18995_ | 537 | 633 | 576 | 664 | 605 | 674 |
| SL1344_RS19005_ | 16 | 23 | 21 | 23 | 16 | 10 |
| SL1344_RS19010_ | 1461 | 1544 | 1522 | 1305 | 1249 | 1216 |
| SL1344_RS19015_ | 4 | 13 | 11 | 0 | 5 | 2 |
| SL1344_RS19020_ | 144 | 181 | 157 | 184 | 166 | 167 |
| SL1344_RS19025_ | 91 | 100 | 110 | 170 | 149 | 157 |
| SL1344_RS19030_ | 56 | 41 | 36 | 73 | 60 | 76 |
| SL1344_RS19040_ | 484 | 447 | 452 | 378 | 388 | 377 |
| SL1344_RS19045_ | 323 | 421 | 380 | 392 | 384 | 430 |
| SL1344_RS19050_ | 67 | 62 | 70 | 64 | 66 | 62 |
| SL1344_RS19055_ | 58 | 67 | 57 | 55 | 64 | 53 |
| SL1344_RS19065_ | 2467 | 2843 | 3005 | 3516 | 3262 | 3528 |
| SL1344_RS19070_ | 3060 | 3234 | 3153 | 3482 | 3416 | 3604 |
| SL1344_RS19075_ | 1454 | 1573 | 1483 | 1686 | 1560 | 1644 |
| SL1344_RS19080_ | 2164 | 2392 | 2142 | 2391 | 2231 | 2089 |
| SL1344_RS19085_ | 1206 | 1260 | 1152 | 1436 | 1306 | 1330 |
| SL1344_RS19090_ | 2354 | 2277 | 2151 | 2122 | 1994 | 2036 |
| SL1344_RS19095_ | 1155 | 1225 | 1151 | 965 | 912 | 974 |
| SL1344_RS19100_ | 127 | 122 | 134 | 76 | 61 | 84 |
| SL1344_RS19105_ | 3148 | 3069 | 2857 | 2896 | 2852 | 2756 |
| SL1344_RS19110_ | 1571 | 1628 | 1698 | 1585 | 1446 | 1660 |
| SL1344_RS19115_ | 1414 | 1447 | 1486 | 1440 | 1287 | 1458 |
| SL1344_RS19120_ | 1563 | 1699 | 1614 | 1496 | 1440 | 1493 |
| SL1344_RS19125_ | 1417 | 1439 | 1444 | 1278 | 1200 | 1346 |
| SL1344_RS19130_ | 2389 | 2352 | 2224 | 1995 | 1855 | 1992 |
| SL1344_RS19135_ | 977 | 902 | 926 | 934 | 906 | 954 |
| SL1344_RS19140_ | 965 | 920 | 885 | 541 | 525 | 647 |
| SL1344_RS19145_ | 394 | 467 | 476 | 474 | 464 | 515 |
| SL1344_RS19150_ | 736 | 749 | 674 | 708 | 626 | 755 |
| SL1344_RS19155_ | 959 | 944 | 880 | 1005 | 829 | 999 |
| SL1344_RS19160_ | 921 | 946 | 910 | 905 | 856 | 879 |
| SL1344_RS19165_ | 239 | 293 | 251 | 222 | 186 | 247 |
| SL1344_RS19170_ | 910 | 1006 | 906 | 952 | 819 | 955 |
| SL1344_RS19175_ | 1310 | 1280 | 1202 | 1254 | 1160 | 1309 |
| SL1344_RS19180_ | 1108 | 1126 | 1058 | 1136 | 1060 | 1164 |
| SL1344_RS19185_ | 3097 | 3089 | 2866 | 2252 | 2211 | 2452 |
| SL1344_RS19190_ | 1247 | 1269 | 1251 | 1029 | 958 | 988 |
| SL1344_RS19195_ | 310 | 355 | 341 | 397 | 386 | 361 |
| SL1344_RS19200_ | 36649 | 39321 | 38775 | 39622 | 39417 | 36622 |
| SL1344_RS19205_ | 28314 | 29437 | 25740 | 29685 | 29586 | 26866 |
| SL1344_RS19210_ | 165 | 161 | 146 | 125 | 135 | 150 |
| SL1344_RS19215_ | 1488 | 1452 | 1388 | 1488 | 1567 | 1448 |
| SL1344_RS19220_ | 960 | 1134 | 1126 | 1122 | 1073 | 1165 |
| SL1344_RS19225_ | 1038 | 1147 | 1161 | 1199 | 1071 | 1212 |
| SL1344_RS19230_ | 155 | 180 | 183 | 200 | 211 | 161 |
| SL1344_RS19235_ | 361 | 454 | 513 | 475 | 432 | 507 |
| SL1344_RS19240_ | 1378 | 1516 | 1551 | 1597 | 1584 | 1544 |
| SL1344_RS19245_ | 159 | 155 | 163 | 148 | 138 | 134 |
| SL1344_RS19250_ | 60 | 70 | 58 | 49 | 62 | 53 |
| SL1344_RS19255_ | 395 | 416 | 333 | 359 | 314 | 309 |
| SL1344_RS19260_ | 94 | 99 | 99 | 78 | 90 | 112 |
| SL1344_RS19265_ | 1610 | 1642 | 1480 | 1400 | 1341 | 1446 |
| SL1344_RS19270_ | 2492 | 2686 | 2864 | 2580 | 2265 | 2567 |
| SL1344_RS19275_ | 7019 | 7480 | 7877 | 6716 | 6585 | 7014 |
| SL1344_RS19280_ | 1799 | 1885 | 1961 | 1517 | 1468 | 1683 |
| SL1344_RS19285_ | 1922 | 1966 | 2091 | 1520 | 1510 | 1668 |
| SL1344_RS19290_ | 854 | 907 | 947 | 733 | 707 | 763 |
| SL1344_RS19295_ | 880 | 1030 | 977 | 836 | 751 | 785 |

| | | | | | | |
|---|---|---|---|---|---|---|
| SL1344_RS19300_ | 346 | 382 | 322 | 346 | 299 | 288 |
| SL1344_RS19305_ | 793 | 795 | 770 | 873 | 780 | 794 |
| SL1344_RS19310_ | 86 | 109 | 86 | 108 | 85 | 109 |
| SL1344_RS19315_ | 63 | 58 | 43 | 46 | 43 | 49 |
| SL1344_RS19320_ | 249 | 216 | 260 | 177 | 205 | 206 |
| SL1344_RS19325_ | 35 | 52 | 43 | 26 | 25 | 49 |
| SL1344_RS19330_ | 720 | 716 | 639 | 868 | 890 | 866 |
| SL1344_RS19335_ | 192 | 192 | 193 | 247 | 249 | 243 |
| SL1344_RS19340_ | 661 | 722 | 663 | 710 | 759 | 685 |
| SL1344_RS19345_ | 3 | 4 | 9 | 8 | 11 | 10 |
| SL1344_RS19350_ | 113 | 137 | 125 | 175 | 159 | 144 |
| SL1344_RS19355_ | 92 | 112 | 119 | 77 | 60 | 72 |
| SL1344_RS19360_ | 150 | 115 | 134 | 88 | 77 | 105 |
| SL1344_RS19365_ | 93 | 107 | 74 | 94 | 84 | 80 |
| SL1344_RS19370_ | 155 | 187 | 135 | 195 | 162 | 145 |
| SL1344_RS19375_ | 254 | 242 | 278 | 347 | 356 | 368 |
| SL1344_RS19380_ | 77 | 61 | 82 | 82 | 91 | 114 |
| SL1344_RS19385_ | 377 | 339 | 323 | 295 | 277 | 248 |
| SL1344_RS19390_ | 74 | 88 | 59 | 82 | 88 | 95 |
| SL1344_RS19395_ | 86 | 59 | 63 | 84 | 78 | 84 |
| SL1344_RS19400_ | 44 | 48 | 51 | 64 | 54 | 67 |
| SL1344_RS19405_ | 33 | 25 | 27 | 40 | 31 | 24 |
| SL1344_RS19410_ | 26 | 20 | 15 | 30 | 27 | 19 |
| SL1344_RS19415_ | 7 | 10 | 7 | 13 | 10 | 1 |
| SL1344_RS19420_ | 2 | 6 | 2 | 4 | 6 | 4 |
| SL1344_RS19425_ | 572 | 600 | 585 | 636 | 648 | 631 |
| SL1344_RS19430_ | 6 | 5 | 2 | 10 | 2 | 4 |
| SL1344_RS19435_ | 59 | 69 | 65 | 83 | 73 | 70 |
| SL1344_RS19445_ | 135 | 140 | 105 | 116 | 133 | 100 |
| SL1344_RS19450_ | 173 | 150 | 143 | 181 | 138 | 146 |
| SL1344_RS19455_ | 18 | 27 | 20 | 27 | 19 | 19 |
| SL1344_RS19460_ | 11 | 14 | 9 | 13 | 5 | 21 |
| SL1344_RS19465_ | 168 | 188 | 164 | 123 | 122 | 140 |
| SL1344_RS19470_ | 70 | 75 | 59 | 52 | 51 | 52 |
| SL1344_RS19475_ | 44 | 33 | 36 | 29 | 47 | 46 |
| SL1344_RS19480_ | 75 | 79 | 73 | 53 | 57 | 67 |
| SL1344_RS19485_ | 143 | 131 | 137 | 96 | 100 | 93 |
| SL1344_RS19490_ | 221 | 265 | 275 | 222 | 249 | 219 |
| SL1344_RS19495_ | 63 | 56 | 62 | 53 | 57 | 65 |
| SL1344_RS19500_ | 422 | 449 | 468 | 362 | 329 | 392 |
| SL1344_RS19505_ | 557 | 606 | 626 | 556 | 536 | 555 |
| SL1344_RS19510_ | 311 | 347 | 363 | 352 | 362 | 382 |
| SL1344_RS19515_ | 145 | 179 | 138 | 148 | 135 | 143 |
| SL1344_RS19520_ | 34 | 52 | 58 | 35 | 32 | 49 |
| SL1344_RS19525_ | 42 | 51 | 43 | 52 | 34 | 33 |
| SL1344_RS19530_ | 114 | 116 | 104 | 112 | 104 | 112 |
| SL1344_RS19535_ | 46 | 47 | 41 | 71 | 57 | 66 |
| SL1344_RS19540_ | 225 | 250 | 214 | 259 | 284 | 273 |
| SL1344_RS19545_ | 769 | 880 | 695 | 637 | 584 | 727 |
| SL1344_RS19550_ | 365 | 361 | 345 | 391 | 384 | 410 |
| SL1344_RS19555_ | 144 | 118 | 110 | 115 | 94 | 89 |
| SL1344_RS19565_ | 279 | 248 | 235 | 251 | 228 | 261 |
| SL1344_RS19570_ | 88 | 118 | 88 | 106 | 87 | 107 |
| SL1344_RS19575_ | 47 | 36 | 30 | 49 | 47 | 63 |
| SL1344_RS19580_ | 146 | 170 | 155 | 199 | 145 | 183 |
| SL1344_RS19585_ | 206 | 230 | 228 | 214 | 213 | 241 |
| SL1344_RS19590_ | 175 | 183 | 177 | 206 | 174 | 218 |
| SL1344_RS19595_ | 209 | 229 | 211 | 206 | 207 | 193 |
| SL1344_RS19600_ | 234 | 222 | 193 | 293 | 293 | 273 |
| SL1344_RS19605_ | 772 | 762 | 656 | 1746 | 1512 | 1207 |
| SL1344_RS19610_ | 467 | 469 | 335 | 1271 | 885 | 738 |
| SL1344_RS19615_ | 110 | 172 | 153 | 185 | 169 | 151 |

| | | | | | | |
|---|---|---|---|---|---|---|
| SL1344_RS19620_ | 356 | 335 | 331 | 338 | 305 | 327 |
| SL1344_RS19625_ | 48 | 50 | 48 | 69 | 67 | 53 |
| SL1344_RS19630_ | 2 | 5 | 4 | 5 | 6 | 7 |
| SL1344_RS19635_ | 15 | 23 | 23 | 23 | 10 | 18 |
| SL1344_RS19640_ | 15 | 15 | 10 | 20 | 10 | 9 |
| SL1344_RS19645_ | 5 | 4 | 3 | 8 | 4 | 1 |
| SL1344_RS19650_ | 10 | 11 | 15 | 27 | 12 | 12 |
| SL1344_RS19655_ | 4 | 8 | 1 | 3 | 6 | 5 |
| SL1344_RS19660_ | 3 | 12 | 5 | 10 | 7 | 2 |
| SL1344_RS19665_ | 17 | 22 | 16 | 38 | 17 | 34 |
| SL1344_RS19670_ | 15 | 14 | 16 | 24 | 17 | 27 |
| SL1344_RS19675_ | 81 | 77 | 75 | 81 | 101 | 77 |
| SL1344_RS19680_ | 27 | 26 | 20 | 23 | 26 | 25 |
| SL1344_RS19685_ | 365 | 353 | 362 | 389 | 392 | 374 |
| SL1344_RS19690_ | 62 | 62 | 62 | 71 | 54 | 57 |
| SL1344_RS19695_ | 319 | 343 | 304 | 287 | 298 | 321 |
| SL1344_RS19700_ | 38 | 42 | 37 | 46 | 33 | 42 |
| SL1344_RS19705_ | 54 | 58 | 47 | 38 | 51 | 45 |
| SL1344_RS19710_ | 21 | 14 | 12 | 19 | 18 | 19 |
| SL1344_RS19715_ | 112 | 112 | 128 | 109 | 130 | 133 |
| SL1344_RS19720_ | 73 | 80 | 70 | 78 | 86 | 74 |
| SL1344_RS19725_ | 537 | 562 | 527 | 565 | 542 | 498 |
| SL1344_RS19730_ | 17 | 14 | 18 | 21 | 15 | 21 |
| SL1344_RS19735_ | 27 | 31 | 42 | 61 | 22 | 33 |
| SL1344_RS19740_ | 66 | 62 | 46 | 40 | 57 | 53 |
| SL1344_RS19745_ | 6589 | 7005 | 7166 | 6436 | 6107 | 6543 |
| SL1344_RS19750_ | 1603 | 1612 | 1640 | 1513 | 1517 | 1581 |
| SL1344_RS19755_ | 2503 | 2405 | 2539 | 2398 | 2274 | 2372 |
| SL1344_RS19765_ | 20564 | 19479 | 18685 | 18100 | 18013 | 17995 |
| SL1344_RS19770_ | 6434 | 6075 | 6669 | 5762 | 5521 | 5756 |
| SL1344_RS19775_ | 2676 | 2837 | 2813 | 2426 | 2544 | 2502 |
| SL1344_RS19780_ | 12594 | 12606 | 12918 | 10651 | 10620 | 11190 |
| SL1344_RS19785_ | 835 | 856 | 939 | 711 | 713 | 760 |
| SL1344_RS19790_ | 38 | 36 | 28 | 27 | 34 | 25 |
| SL1344_RS19795_ | 39 | 38 | 37 | 49 | 45 | 32 |
| SL1344_RS19800_ | 1729 | 1560 | 1365 | 1453 | 1398 | 1402 |
| SL1344_RS19805_ | 1043 | 869 | 816 | 730 | 724 | 749 |
| SL1344_RS19810_ | 46 | 69 | 41 | 42 | 46 | 51 |
| SL1344_RS19815_ | 162 | 144 | 126 | 147 | 138 | 147 |
| SL1344_RS19820_ | 106 | 106 | 93 | 85 | 77 | 79 |
| SL1344_RS19825_ | 450 | 480 | 458 | 416 | 489 | 465 |
| SL1344_RS19830_ | 769 | 797 | 812 | 725 | 674 | 760 |
| SL1344_RS19835_ | 487 | 603 | 621 | 650 | 565 | 636 |
| SL1344_RS19840_ | 727 | 840 | 825 | 675 | 649 | 716 |
| SL1344_RS19845_ | 350 | 333 | 345 | 311 | 246 | 297 |
| SL1344_RS19850_ | 185 | 227 | 189 | 188 | 181 | 217 |
| SL1344_RS19855_ | 211 | 221 | 203 | 251 | 190 | 254 |
| SL1344_RS19860_ | 278 | 312 | 274 | 394 | 334 | 348 |
| SL1344_RS19865_ | 39 | 62 | 54 | 80 | 49 | 69 |
| SL1344_RS19870_ | 9 | 10 | 10 | 11 | 7 | 12 |
| SL1344_RS19875_ | 214 | 238 | 216 | 265 | 251 | 240 |
| SL1344_RS19880_ | 7229 | 7211 | 7523 | 7191 | 6915 | 7387 |
| SL1344_RS19885_ | 6862 | 7049 | 6744 | 6644 | 6412 | 6741 |
| SL1344_RS19890_ | 162 | 167 | 205 | 149 | 146 | 126 |
| SL1344_RS19900_ | 13645 | 14390 | 15693 | 11482 | 10715 | 11156 |
| SL1344_RS19905_ | 11035 | 11180 | 11893 | 8401 | 7767 | 8352 |
| SL1344_RS19910_ | 26975 | 28819 | 30404 | 20294 | 20125 | 20675 |
| SL1344_RS19915_ | 9464 | 9874 | 10213 | 8019 | 7489 | 7646 |
| SL1344_RS19920_ | 12440 | 12760 | 13738 | 9831 | 10145 | 9600 |
| SL1344_RS19925_ | 5806 | 6237 | 6319 | 5080 | 4508 | 4758 |
| SL1344_RS19930_ | 5651 | 6232 | 6096 | 4792 | 4268 | 4767 |
| SL1344_RS19935_ | 2586 | 2734 | 2758 | 2120 | 1972 | 2160 |

| | | | | | | |
|---|---|---|---|---|---|---|
| SL1344_RS19945_ | 1611 | 1893 | 2084 | 1499 | 1329 | 1644 |
| SL1344_RS19950_ | 4113 | 4114 | 3917 | 3858 | 3745 | 3935 |
| SL1344_RS19955_ | 265 | 237 | 219 | 248 | 275 | 245 |
| SL1344_RS19960_ | 135 | 165 | 149 | 154 | 143 | 145 |
| SL1344_RS19965_ | 364 | 380 | 394 | 325 | 345 | 353 |
| SL1344_RS19970_ | 582 | 581 | 531 | 534 | 564 | 540 |
| SL1344_RS19980_ | 561 | 594 | 552 | 898 | 923 | 840 |
| SL1344_RS19985_ | 446 | 448 | 387 | 617 | 610 | 539 |
| SL1344_RS19990_ | 338 | 353 | 326 | 476 | 488 | 509 |
| SL1344_RS19995_ | 949 | 1014 | 974 | 1390 | 1318 | 1240 |
| SL1344_RS20000_ | 409 | 456 | 492 | 533 | 540 | 528 |
| SL1344_RS20005_ | 429 | 495 | 472 | 480 | 514 | 474 |
| SL1344_RS20020_ | 534 | 534 | 425 | 429 | 386 | 386 |
| SL1344_RS20025_ | 115 | 52 | 61 | 27 | 45 | 40 |
| SL1344_RS20030_ | 274 | 213 | 139 | 177 | 134 | 157 |
| SL1344_RS20035_ | 939 | 570 | 502 | 450 | 363 | 608 |
| SL1344_RS20045_ | 3555 | 3460 | 3586 | 3011 | 2371 | 3223 |
| SL1344_RS20050_ | 5849 | 5564 | 6536 | 4772 | 3864 | 4995 |
| SL1344_RS20055_ | 323 | 290 | 260 | 292 | 256 | 299 |
| SL1344_RS20060_ | 4690 | 6307 | 6064 | 6303 | 5285 | 5989 |
| SL1344_RS20065_ | 64 | 72 | 82 | 92 | 93 | 113 |
| SL1344_RS20070_ | 585 | 496 | 558 | 539 | 476 | 547 |
| SL1344_RS20075_ | 206 | 250 | 211 | 309 | 293 | 286 |
| SL1344_RS20085_ | 332 | 348 | 349 | 434 | 377 | 449 |
| SL1344_RS20090_ | 584 | 639 | 678 | 710 | 669 | 849 |
| SL1344_RS20095_ | 1042 | 1133 | 1174 | 1204 | 1131 | 1386 |
| SL1344_RS20100_ | 174 | 163 | 158 | 201 | 196 | 186 |
| SL1344_RS20105_ | 153 | 151 | 145 | 186 | 175 | 172 |
| SL1344_RS20110_ | 191 | 198 | 221 | 200 | 217 | 202 |
| SL1344_RS20115_ | 924 | 1045 | 1071 | 2015 | 1731 | 2123 |
| SL1344_RS20120_ | 958 | 1118 | 1281 | 1078 | 1007 | 1057 |
| SL1344_RS20125_ | 88 | 76 | 86 | 75 | 77 | 68 |
| SL1344_RS20130_ | 1332 | 1571 | 1505 | 1486 | 1327 | 1355 |
| SL1344_RS20135_ | 3873 | 3580 | 3828 | 2812 | 2883 | 3110 |
| SL1344_RS20140_ | 3539 | 3191 | 3239 | 2639 | 2514 | 2853 |
| SL1344_RS20145_ | 12812 | 13536 | 12678 | 11884 | 10543 | 12160 |
| SL1344_RS20150_ | 13848 | 14036 | 15090 | 11620 | 11512 | 12625 |
| SL1344_RS20155_ | 1713 | 1741 | 1654 | 1632 | 1434 | 1573 |
| SL1344_RS20160_ | 2002 | 2230 | 2047 | 2067 | 1887 | 2018 |
| SL1344_RS20165_ | 1717 | 1673 | 1667 | 1446 | 1363 | 1520 |
| SL1344_RS20175_ | 1160 | 1255 | 1294 | 1012 | 987 | 1010 |
| SL1344_RS20180_ | 383 | 395 | 424 | 293 | 279 | 278 |
| SL1344_RS20185_ | 582 | 649 | 683 | 557 | 558 | 575 |
| SL1344_RS20190_ | 863 | 968 | 940 | 812 | 763 | 794 |
| SL1344_RS20195_ | 904 | 943 | 979 | 716 | 730 | 778 |
| SL1344_RS20200_ | 630 | 710 | 686 | 582 | 552 | 605 |
| SL1344_RS20205_ | 925 | 998 | 931 | 849 | 822 | 751 |
| SL1344_RS20210_ | 657 | 690 | 661 | 588 | 566 | 592 |
| SL1344_RS20215_ | 1572 | 1751 | 1750 | 1338 | 1188 | 1268 |
| SL1344_RS20220_ | 3965 | 4535 | 4393 | 3969 | 3880 | 4807 |
| SL1344_RS20225_ | 5249 | 5979 | 5973 | 5502 | 5137 | 5986 |
| SL1344_RS20230_ | 11102 | 10720 | 10672 | 10507 | 9693 | 10465 |
| SL1344_RS20235_ | 6433 | 5547 | 7748 | 6091 | 4919 | 5663 |
| SL1344_RS20240_ | 1010 | 1147 | 1170 | 1137 | 1095 | 1109 |
| SL1344_RS20245_ | 1038 | 1021 | 1009 | 984 | 901 | 1003 |
| SL1344_RS20250_ | 970 | 920 | 843 | 764 | 779 | 815 |
| SL1344_RS20255_ | 1111 | 1060 | 1099 | 974 | 969 | 1001 |
| SL1344_RS20260_ | 4646 | 4631 | 4681 | 4270 | 4219 | 4528 |
| SL1344_RS20265_ | 92 | 91 | 77 | 59 | 91 | 98 |
| SL1344_RS20270_ | 58 | 44 | 55 | 40 | 26 | 34 |
| SL1344_RS20275_ | 58 | 46 | 52 | 41 | 30 | 41 |
| SL1344_RS20280_ | 348 | 538 | 576 | 519 | 439 | 549 |

| | | | | | | |
|---|---|---|---|---|---|---|
| SL1344_RS20285_ | 14 | 21 | 12 | 12 | 10 | 11 |
| SL1344_RS20290_ | 369 | 314 | 296 | 301 | 297 | 279 |
| SL1344_RS20295_ | 929 | 985 | 963 | 988 | 955 | 954 |
| SL1344_RS20300_ | 624 | 720 | 816 | 663 | 621 | 672 |
| SL1344_RS20305_ | 684 | 769 | 791 | 675 | 664 | 709 |
| SL1344_RS20310_ | 356 | 404 | 401 | 332 | 313 | 381 |
| SL1344_RS20315_ | 2014 | 2095 | 2273 | 2145 | 2098 | 2141 |
| SL1344_RS20325_ | 39 | 31 | 36 | 35 | 31 | 27 |
| SL1344_RS20330_ | 16 | 14 | 24 | 16 | 8 | 20 |
| SL1344_RS20335_ | 248 | 276 | 290 | 327 | 295 | 325 |
| SL1344_RS20340_ | 45 | 45 | 34 | 57 | 60 | 92 |
| SL1344_RS20345_ | 1399 | 1481 | 1329 | 1715 | 1460 | 1489 |
| SL1344_RS20350_ | 1179 | 1258 | 1309 | 1178 | 1107 | 1205 |
| SL1344_RS20355_ | 317 | 343 | 352 | 325 | 325 | 315 |
| SL1344_RS20365_ | 674 | 689 | 681 | 670 | 636 | 717 |
| SL1344_RS20370_ | 437 | 478 | 507 | 496 | 508 | 481 |
| SL1344_RS20375_ | 152 | 197 | 201 | 177 | 143 | 167 |
| SL1344_RS20390_ | 97 | 88 | 129 | 98 | 84 | 100 |
| SL1344_RS20395_ | 137 | 108 | 106 | 117 | 102 | 134 |
| SL1344_RS20400_ | 3184 | 3595 | 3299 | 3992 | 3809 | 3481 |
| SL1344_RS20405_ | 478 | 513 | 486 | 451 | 454 | 516 |
| SL1344_RS20410_ | 1079 | 1169 | 1056 | 1186 | 1057 | 1210 |
| SL1344_RS20415_ | 768 | 778 | 805 | 841 | 732 | 876 |
| SL1344_RS20425_ | 2362 | 2231 | 2322 | 2482 | 2371 | 2498 |
| SL1344_RS20430_ | 2725 | 2594 | 2810 | 2700 | 2497 | 2530 |
| SL1344_RS20435_ | 2316 | 2394 | 2410 | 2211 | 2093 | 2268 |
| SL1344_RS20440_ | 333 | 322 | 306 | 331 | 310 | 271 |
| SL1344_RS20445_ | 1455 | 1415 | 1244 | 1036 | 1083 | 1074 |
| SL1344_RS20450_ | 3504 | 3545 | 3520 | 3714 | 3485 | 3650 |
| SL1344_RS20455_ | 1694 | 1866 | 1805 | 2092 | 1831 | 1981 |
| SL1344_RS20460_ | 384 | 378 | 349 | 381 | 368 | 371 |
| SL1344_RS20470_ | 94 | 89 | 83 | 132 | 109 | 100 |
| SL1344_RS20475_ | 1322 | 1303 | 1274 | 1430 | 1470 | 1398 |
| SL1344_RS20480_ | 891 | 948 | 887 | 1007 | 877 | 983 |
| SL1344_RS20485_ | 641 | 828 | 828 | 753 | 690 | 852 |
| SL1344_RS20495_ | 99 | 43 | 40 | 33 | 30 | 60 |
| SL1344_RS20500_ | 297 | 273 | 233 | 259 | 201 | 259 |
| SL1344_RS20505_ | 94 | 107 | 111 | 104 | 104 | 130 |
| SL1344_RS20510_ | 222 | 138 | 144 | 127 | 127 | 147 |
| SL1344_RS20520_ | 119 | 151 | 128 | 182 | 148 | 137 |
| SL1344_RS20530_ | 318 | 295 | 276 | 308 | 303 | 253 |
| SL1344_RS20535_ | 1041 | 1002 | 953 | 968 | 879 | 899 |
| SL1344_RS20540_ | 797 | 876 | 933 | 907 | 812 | 869 |
| SL1344_RS20545_ | 1396 | 1300 | 1101 | 1241 | 1168 | 1296 |
| SL1344_RS20550_ | 47 | 44 | 39 | 40 | 45 | 46 |
| SL1344_RS20555_ | 1545 | 1644 | 1715 | 1614 | 1448 | 1645 |
| SL1344_RS20560_ | 1319 | 1331 | 1263 | 1298 | 1802 | 1191 |
| SL1344_RS20565_ | 30472 | 35460 | 37953 | 58588 | 61921 | 58773 |
| SL1344_RS20570_ | 1369 | 1639 | 1632 | 1448 | 1411 | 1458 |
| SL1344_RS20575_ | 1460 | 1577 | 1538 | 1353 | 1292 | 1418 |
| SL1344_RS20580_ | 546 | 559 | 536 | 428 | 416 | 398 |
| SL1344_RS20585_ | 377 | 453 | 407 | 338 | 306 | 352 |
| SL1344_RS20590_ | 4631 | 4573 | 4108 | 3250 | 3016 | 3230 |
| SL1344_RS20595_ | 15880 | 16488 | 16783 | 14931 | 14510 | 15165 |
| SL1344_RS20600_ | 85 | 96 | 77 | 125 | 96 | 107 |
| SL1344_RS20605_ | 31 | 20 | 35 | 28 | 25 | 32 |
| SL1344_RS20610_ | 34 | 41 | 38 | 26 | 29 | 34 |
| SL1344_RS20615_ | 29 | 28 | 33 | 32 | 23 | 37 |
| SL1344_RS20620_ | 24 | 24 | 26 | 29 | 37 | 25 |
| SL1344_RS20625_ | 33 | 39 | 41 | 45 | 58 | 41 |
| SL1344_RS20635_ | 59 | 60 | 55 | 54 | 47 | 50 |
| SL1344_RS20640_ | 122 | 109 | 108 | 103 | 102 | 132 |

| | | | | | | |
|---|---|---|---|---|---|---|
| SL1344_RS20645_ | 61 | 78 | 67 | 68 | 75 | 67 |
| SL1344_RS20650_ | 41 | 27 | 25 | 41 | 41 | 61 |
| SL1344_RS20655_ | 34 | 31 | 41 | 65 | 39 | 48 |
| SL1344_RS20660_ | 19 | 11 | 20 | 25 | 22 | 12 |
| SL1344_RS20665_ | 6 | 2 | 6 | 12 | 8 | 13 |
| SL1344_RS20670_ | 27 | 32 | 17 | 28 | 26 | 30 |
| SL1344_RS20675_ | 445 | 480 | 417 | 466 | 435 | 456 |
| SL1344_RS20680_ | 513 | 579 | 548 | 603 | 575 | 627 |
| SL1344_RS20685_ | 596 | 562 | 557 | 685 | 551 | 623 |
| SL1344_RS20690_ | 454 | 447 | 422 | 478 | 420 | 425 |
| SL1344_RS20695_ | 887 | 924 | 869 | 955 | 950 | 975 |
| SL1344_RS20700_ | 271 | 244 | 201 | 400 | 368 | 320 |
| SL1344_RS20705_ | 226 | 202 | 165 | 341 | 318 | 283 |
| SL1344_RS20710_ | 109 | 102 | 99 | 129 | 114 | 109 |
| SL1344_RS20715_ | 77 | 89 | 85 | 105 | 108 | 108 |
| SL1344_RS20720_ | 87 | 127 | 78 | 119 | 88 | 101 |
| SL1344_RS20725_ | 1536 | 1695 | 1920 | 1662 | 1464 | 1642 |
| SL1344_RS20730_ | 1123 | 1213 | 1278 | 1189 | 1065 | 1159 |
| SL1344_RS20735_ | 1063 | 1076 | 1297 | 993 | 1014 | 1039 |
| SL1344_RS20740_ | 3152 | 3158 | 3371 | 3070 | 2878 | 3103 |
| SL1344_RS20745_ | 2036 | 2087 | 2155 | 1920 | 2019 | 2070 |
| SL1344_RS20750_ | 380 | 367 | 352 | 379 | 374 | 377 |
| SL1344_RS20755_ | 60 | 79 | 58 | 69 | 57 | 67 |
| SL1344_RS20760_ | 41 | 32 | 20 | 43 | 39 | 35 |
| SL1344_RS20765_ | 173 | 198 | 150 | 174 | 200 | 217 |
| SL1344_RS20770_ | 251 | 236 | 256 | 278 | 276 | 281 |
| SL1344_RS20775_ | 543 | 569 | 481 | 516 | 546 | 521 |
| SL1344_RS20780_ | 59 | 69 | 76 | 119 | 94 | 98 |
| SL1344_RS20785_ | 25 | 35 | 27 | 39 | 41 | 35 |
| SL1344_RS20790_ | 14 | 17 | 24 | 20 | 14 | 19 |
| SL1344_RS20795_ | 21 | 28 | 19 | 19 | 31 | 23 |
| SL1344_RS20800_ | 26 | 18 | 20 | 23 | 14 | 20 |
| SL1344_RS20805_ | 39 | 46 | 51 | 91 | 82 | 68 |
| SL1344_RS20810_ | 29 | 30 | 30 | 34 | 26 | 28 |
| SL1344_RS20815_ | 45 | 39 | 39 | 44 | 56 | 43 |
| SL1344_RS20820_ | 1 | 4 | 0 | 5 | 1 | 1 |
| SL1344_RS20825_ | 20 | 16 | 16 | 20 | 15 | 16 |
| SL1344_RS20830_ | 80 | 80 | 77 | 84 | 63 | 66 |
| SL1344_RS20835_ | 5 | 10 | 9 | 10 | 8 | 8 |
| SL1344_RS20840_ | 13 | 11 | 5 | 12 | 16 | 12 |
| SL1344_RS20845_ | 15188 | 15481 | 14945 | 16099 | 15478 | 15819 |
| SL1344_RS20850_ | 218 | 242 | 198 | 240 | 197 | 218 |
| SL1344_RS20855_ | 183 | 211 | 208 | 247 | 227 | 248 |
| SL1344_RS20875_ | 489 | 551 | 522 | 542 | 480 | 542 |
| SL1344_RS20880_ | 788 | 852 | 979 | 769 | 736 | 838 |
| SL1344_RS20890_ | 1947 | 1948 | 1832 | 1991 | 1796 | 1716 |
| SL1344_RS20895_ | 119 | 104 | 99 | 74 | 81 | 59 |
| SL1344_RS20900_ | 69 | 89 | 80 | 89 | 62 | 79 |
| SL1344_RS20905_ | 598 | 645 | 644 | 488 | 499 | 559 |
| SL1344_RS20910_ | 621 | 618 | 582 | 632 | 580 | 633 |
| SL1344_RS20915_ | 21 | 22 | 22 | 25 | 23 | 32 |
| SL1344_RS20920_ | 41 | 58 | 44 | 45 | 43 | 32 |
| SL1344_RS20925_ | 222 | 202 | 213 | 145 | 138 | 164 |
| SL1344_RS20935_ | 16 | 41 | 17 | 37 | 28 | 34 |
| SL1344_RS20940_ | 41 | 54 | 45 | 47 | 47 | 42 |
| SL1344_RS20945_ | 17 | 20 | 28 | 20 | 17 | 17 |
| SL1344_RS20950_ | 69 | 41 | 47 | 55 | 52 | 40 |
| SL1344_RS20955_ | 59 | 53 | 47 | 56 | 48 | 60 |
| SL1344_RS20960_ | 22 | 28 | 38 | 34 | 25 | 28 |
| SL1344_RS20965_ | 118 | 108 | 125 | 101 | 119 | 102 |
| SL1344_RS20970_ | 4069 | 4011 | 4107 | 4019 | 3682 | 3979 |
| SL1344_RS20975_ | 308 | 329 | 260 | 336 | 290 | 262 |

| | | | | | | |
|---|---|---|---|---|---|---|
| SL1344_RS20980_ | 221 | 190 | 186 | 188 | 154 | 168 |
| SL1344_RS20985_ | 622 | 657 | 563 | 764 | 650 | 571 |
| SL1344_RS20990_ | 441 | 573 | 650 | 633 | 480 | 528 |
| SL1344_RS20995_ | 1860 | 2028 | 1932 | 1861 | 1847 | 1799 |
| SL1344_RS21000_ | 569 | 645 | 705 | 538 | 500 | 635 |
| SL1344_RS21005_ | 2542 | 2852 | 2873 | 2690 | 2586 | 2575 |
| SL1344_RS21010_ | 1364 | 1530 | 1500 | 1959 | 1871 | 1922 |
| SL1344_RS21015_ | 506 | 577 | 592 | 559 | 491 | 551 |
| SL1344_RS21020_ | 4893 | 4631 | 4301 | 5261 | 4516 | 4351 |
| SL1344_RS21025_ | 1030 | 1055 | 979 | 1430 | 1076 | 1121 |
| SL1344_RS21030_ | 1854 | 1946 | 1859 | 1608 | 1606 | 1632 |
| SL1344_RS21035_ | 591 | 722 | 701 | 619 | 594 | 621 |
| SL1344_RS21040_ | 1199 | 1219 | 1134 | 948 | 1020 | 1005 |
| SL1344_RS21045_ | 14174 | 14266 | 13970 | 12199 | 12037 | 12651 |
| SL1344_RS21050_ | 94 | 81 | 59 | 74 | 91 | 85 |
| SL1344_RS21055_ | 140 | 145 | 116 | 116 | 105 | 113 |
| SL1344_RS21075_ | 18 | 17 | 14 | 20 | 15 | 14 |
| SL1344_RS21080_ | 17 | 10 | 12 | 14 | 13 | 11 |
| SL1344_RS21085_ | 96 | 81 | 82 | 64 | 59 | 55 |
| SL1344_RS21090_ | 143 | 144 | 150 | 129 | 123 | 138 |
| SL1344_RS21100_ | 324 | 341 | 365 | 437 | 390 | 428 |
| SL1344_RS21105_ | 49 | 46 | 34 | 43 | 40 | 45 |
| SL1344_RS21115_ | 188 | 173 | 157 | 190 | 159 | 178 |
| SL1344_RS21120_ | 192 | 174 | 166 | 202 | 175 | 181 |
| SL1344_RS21125_ | 31 | 40 | 35 | 41 | 38 | 32 |
| SL1344_RS21130_ | 34 | 42 | 37 | 35 | 73 | 54 |
| SL1344_RS21135_ | 267 | 274 | 265 | 301 | 291 | 280 |
| SL1344_RS21140_ | 292 | 334 | 337 | 369 | 363 | 386 |
| SL1344_RS21145_ | 75 | 81 | 99 | 96 | 97 | 85 |
| SL1344_RS21150_ | 173 | 227 | 266 | 165 | 184 | 187 |
| SL1344_RS21155_ | 2073 | 2095 | 2135 | 1518 | 1430 | 1662 |
| SL1344_RS21160_ | 2790 | 2695 | 2655 | 2655 | 2617 | 2593 |
| SL1344_RS21165_ | 229 | 235 | 223 | 270 | 242 | 264 |
| SL1344_RS21170_ | 103 | 114 | 136 | 89 | 115 | 115 |
| SL1344_RS21175_ | 137 | 162 | 195 | 181 | 164 | 188 |
| SL1344_RS21180_ | 351 | 409 | 395 | 434 | 439 | 430 |
| SL1344_RS21185_ | 1167 | 1316 | 1288 | 1328 | 1293 | 1354 |
| SL1344_RS21195_ | 872 | 897 | 762 | 871 | 788 | 758 |
| SL1344_RS21200_ | 334 | 384 | 358 | 396 | 331 | 374 |
| SL1344_RS21205_ | 723 | 806 | 796 | 641 | 598 | 621 |
| SL1344_RS21210_ | 1410 | 1374 | 1418 | 1246 | 1109 | 1240 |
| SL1344_RS21215_ | 1488 | 1397 | 1456 | 1119 | 1091 | 1224 |
| SL1344_RS21220_ | 62 | 24 | 28 | 15 | 17 | 29 |
| SL1344_RS21225_ | 535 | 506 | 438 | 437 | 477 | 430 |
| SL1344_RS21230_ | 80 | 49 | 63 | 80 | 69 | 71 |
| SL1344_RS21235_ | 172 | 66 | 58 | 107 | 75 | 87 |
| SL1344_RS21245_ | 485 | 519 | 466 | 505 | 450 | 489 |
| SL1344_RS21250_ | 587 | 677 | 707 | 625 | 569 | 696 |
| SL1344_RS21255_ | 960 | 1019 | 1080 | 755 | 659 | 783 |
| SL1344_RS21260_ | 1 | 1 | 0 | 0 | 2 | 2 |
| SL1344_RS21265_ | 2248 | 2792 | 2503 | 2034 | 2106 | 2591 |
| SL1344_RS21270_ | 4244 | 5014 | 4974 | 4031 | 4049 | 4897 |
| SL1344_RS21275_ | 11968 | 11506 | 14312 | 11272 | 10364 | 12218 |
| SL1344_RS21280_ | 6164 | 6107 | 6984 | 6363 | 5519 | 6748 |
| SL1344_RS21285_ | 38095 | 42696 | 46969 | 49157 | 47421 | 46721 |
| SL1344_RS21290_ | 4672 | 4967 | 5191 | 5268 | 5330 | 5112 |
| SL1344_RS21300_ | 19659 | 18682 | 19257 | 17284 | 16668 | 17043 |
| SL1344_RS21305_ | 43432 | 41485 | 42621 | 38507 | 36826 | 38651 |
| SL1344_RS21310_ | 62108 | 59772 | 59192 | 54737 | 55352 | 53967 |
| SL1344_RS21315_ | 129347 | 130652 | 138078 | 127595 | 127631 | 120804 |
| SL1344_RS21320_ | 27945 | 31870 | 35781 | 29996 | 29819 | 31234 |
| SL1344_RS21325_ | 32148 | 36783 | 40788 | 36243 | 34222 | 37641 |

| | | | | | | |
|---|---|---|---|---|---|---|
| SL1344_RS21330_ | 143 | 185 | 160 | 141 | 146 | 175 |
| SL1344_RS21335_ | 67 | 45 | 42 | 39 | 21 | 32 |
| SL1344_RS21340_ | 278 | 291 | 233 | 243 | 224 | 252 |
| SL1344_RS21345_ | 74 | 71 | 77 | 60 | 46 | 63 |
| SL1344_RS21350_ | 192 | 199 | 183 | 194 | 127 | 191 |
| SL1344_RS21355_ | 61 | 100 | 99 | 57 | 56 | 68 |
| SL1344_RS21360_ | 28 | 25 | 22 | 21 | 20 | 9 |
| SL1344_RS21365_ | 72 | 87 | 80 | 73 | 52 | 83 |
| SL1344_RS21370_ | 34 | 59 | 64 | 45 | 45 | 49 |
| SL1344_RS21375_ | 140 | 180 | 181 | 192 | 157 | 135 |
| SL1344_RS21380_ | 73 | 82 | 80 | 108 | 91 | 118 |
| SL1344_RS21390_ | 916 | 992 | 958 | 883 | 909 | 828 |
| SL1344_RS21395_ | 597 | 591 | 642 | 588 | 562 | 590 |
| SL1344_RS21400_ | 791 | 819 | 870 | 783 | 777 | 848 |
| SL1344_RS21420_ | 319 | 332 | 349 | 305 | 285 | 350 |
| SL1344_RS21425_ | 386 | 425 | 451 | 438 | 372 | 441 |
| SL1344_RS21430_ | 410 | 364 | 378 | 426 | 353 | 399 |
| SL1344_RS21435_ | 389 | 425 | 385 | 424 | 398 | 423 |
| SL1344_RS21440_ | 108 | 46 | 59 | 52 | 58 | 56 |
| SL1344_RS21445_ | 9 | 11 | 15 | 11 | 12 | 6 |
| SL1344_RS21450_ | 313 | 241 | 189 | 207 | 162 | 263 |
| SL1344_RS21460_ | 52 | 44 | 61 | 38 | 45 | 32 |
| SL1344_RS21465_ | 196 | 288 | 304 | 331 | 309 | 272 |
| SL1344_RS21475_ | 63 | 69 | 64 | 81 | 75 | 72 |
| SL1344_RS21480_ | 119 | 123 | 147 | 121 | 132 | 127 |
| SL1344_RS21485_ | 134 | 125 | 125 | 179 | 131 | 146 |
| SL1344_RS21490_ | 363 | 336 | 300 | 334 | 322 | 331 |
| SL1344_RS21495_ | 614 | 667 | 585 | 707 | 632 | 597 |
| SL1344_RS21505_ | 407 | 452 | 463 | 436 | 439 | 413 |
| SL1344_RS21510_ | 147 | 143 | 141 | 203 | 209 | 191 |
| SL1344_RS21515_ | 16 | 13 | 16 | 11 | 17 | 12 |
| SL1344_RS21520_ | 760 | 913 | 855 | 816 | 772 | 743 |
| SL1344_RS21525_ | 1057 | 1205 | 1174 | 984 | 879 | 901 |
| SL1344_RS21530_ | 281 | 256 | 230 | 206 | 225 | 187 |
| SL1344_RS21535_ | 76 | 96 | 78 | 63 | 72 | 81 |
| SL1344_RS21540_ | 29 | 28 | 17 | 14 | 17 | 15 |
| SL1344_RS21545_ | 13 | 14 | 13 | 9 | 15 | 12 |
| SL1344_RS21550_ | 54 | 55 | 72 | 53 | 56 | 56 |
| SL1344_RS21555_ | 21 | 18 | 26 | 19 | 21 | 24 |
| SL1344_RS21560_ | 58 | 35 | 37 | 36 | 42 | 42 |
| SL1344_RS21565_ | 15 | 10 | 14 | 7 | 9 | 7 |
| SL1344_RS21570_ | 22 | 27 | 28 | 23 | 30 | 13 |
| SL1344_RS21575_ | 10 | 8 | 6 | 3 | 10 | 11 |
| SL1344_RS21580_ | 499 | 522 | 458 | 398 | 394 | 365 |
| SL1344_RS21585_ | 213 | 189 | 202 | 137 | 127 | 150 |
| SL1344_RS21590_ | 106 | 112 | 99 | 67 | 57 | 94 |
| SL1344_RS21595_ | 50 | 64 | 55 | 43 | 76 | 56 |
| SL1344_RS21600_ | 38 | 37 | 25 | 32 | 35 | 25 |
| SL1344_RS21605_ | 6 | 3 | 5 | 14 | 12 | 13 |
| SL1344_RS21610_ | 18 | 25 | 24 | 32 | 28 | 27 |
| SL1344_RS21615_ | 56 | 42 | 55 | 67 | 52 | 43 |
| SL1344_RS21620_ | 25 | 29 | 33 | 44 | 31 | 34 |
| SL1344_RS21625_ | 22 | 23 | 22 | 34 | 25 | 28 |
| SL1344_RS21630_ | 63 | 68 | 54 | 49 | 80 | 59 |
| SL1344_RS21635_ | 5 | 7 | 3 | 3 | 2 | 5 |
| SL1344_RS21640_ | 4 | 4 | 2 | 4 | 3 | 3 |
| SL1344_RS21645_ | 8 | 4 | 19 | 13 | 11 | 11 |
| SL1344_RS21650_ | 18 | 14 | 13 | 18 | 18 | 16 |
| SL1344_RS21655_ | 16 | 18 | 10 | 26 | 21 | 14 |
| SL1344_RS21660_ | 64 | 119 | 111 | 109 | 85 | 99 |
| SL1344_RS21665_ | 450 | 439 | 461 | 431 | 407 | 422 |
| SL1344_RS21670_ | 3840 | 3707 | 3366 | 3013 | 2741 | 2845 |

| | | | | | | |
|---|---|---|---|---|---|---|
| SL1344_RS21675_ | 122 | 112 | 77 | 97 | 133 | 116 |
| SL1344_RS21680_ | 25 | 25 | 29 | 29 | 29 | 37 |
| SL1344_RS21685_ | 20 | 23 | 15 | 15 | 9 | 25 |
| SL1344_RS21690_ | 226 | 256 | 244 | 225 | 194 | 212 |
| SL1344_RS21695_ | 74 | 89 | 82 | 95 | 76 | 86 |
| SL1344_RS21710_ | 59 | 53 | 46 | 61 | 43 | 46 |
| SL1344_RS21715_ | 20 | 26 | 24 | 16 | 22 | 21 |
| SL1344_RS21725_ | 54 | 59 | 74 | 70 | 60 | 58 |
| SL1344_RS21730_ | 468 | 494 | 377 | 404 | 332 | 306 |
| SL1344_RS21735_ | 357 | 417 | 418 | 326 | 313 | 378 |
| SL1344_RS21740_ | 5361 | 5108 | 5187 | 4649 | 4439 | 4750 |
| SL1344_RS21745_ | 415 | 392 | 339 | 310 | 280 | 266 |
| SL1344_RS21755_ | 605 | 733 | 751 | 659 | 585 | 729 |
| SL1344_RS21760_ | 157 | 154 | 168 | 143 | 117 | 161 |
| SL1344_RS21765_ | 1025 | 1139 | 986 | 868 | 850 | 954 |
| SL1344_RS21770_ | 439 | 481 | 494 | 460 | 424 | 466 |
| SL1344_RS21775_ | 129 | 152 | 154 | 212 | 184 | 191 |
| SL1344_RS21780_ | 1772 | 1741 | 1827 | 2006 | 1768 | 1877 |
| SL1344_RS21785_ | 135 | 152 | 120 | 91 | 108 | 107 |
| SL1344_RS21790_ | 208 | 226 | 197 | 257 | 215 | 224 |
| SL1344_RS21795_ | 1639 | 1541 | 1590 | 1483 | 1452 | 1446 |
| SL1344_RS21800_ | 579 | 685 | 790 | 665 | 645 | 666 |
| SL1344_RS21805_ | 442 | 520 | 508 | 389 | 361 | 389 |
| SL1344_RS21810_ | 567 | 571 | 523 | 623 | 634 | 665 |
| SL1344_RS21815_ | 597 | 519 | 505 | 471 | 452 | 438 |
| SL1344_RS21820_ | 185 | 197 | 180 | 201 | 158 | 146 |
| SL1344_RS21825_ | 24 | 15 | 22 | 10 | 11 | 17 |
| SL1344_RS21830_ | 7 | 16 | 17 | 15 | 16 | 16 |
| SL1344_RS21835_ | 1625 | 1729 | 1761 | 1912 | 1882 | 1997 |
| SL1344_RS21840_ | 10 | 13 | 17 | 20 | 15 | 19 |
| SL1344_RS21845_ | 2672 | 2920 | 2962 | 2898 | 2824 | 3009 |
| SL1344_RS21880_ | 414 | 421 | 415 | 433 | 480 | 462 |
| SL1344_RS21885_ | 231 | 200 | 201 | 195 | 135 | 189 |
| SL1344_RS21890_ | 128 | 106 | 97 | 199 | 155 | 126 |
| SL1344_RS21895_ | 136 | 127 | 131 | 161 | 154 | 144 |
| SL1344_RS21900_ | 101 | 83 | 64 | 78 | 84 | 75 |
| SL1344_RS21905_ | 1073 | 1160 | 1224 | 1168 | 1058 | 1139 |
| SL1344_RS21910_ | 333 | 353 | 380 | 401 | 403 | 467 |
| SL1344_RS21915_ | 521 | 541 | 481 | 533 | 503 | 463 |
| SL1344_RS21920_ | 84 | 82 | 84 | 79 | 83 | 78 |
| SL1344_RS21925_ | 110 | 131 | 145 | 109 | 101 | 101 |
| SL1344_RS21930_ | 67 | 70 | 69 | 82 | 71 | 74 |
| SL1344_RS21935_ | 5 | 12 | 18 | 6 | 10 | 16 |
| SL1344_RS21940_ | 41 | 61 | 37 | 111 | 89 | 92 |
| SL1344_RS21945_ | 57 | 52 | 29 | 60 | 76 | 72 |
| SL1344_RS21950_ | 77 | 62 | 57 | 52 | 84 | 53 |
| SL1344_RS21965_ | 35 | 40 | 42 | 78 | 62 | 70 |
| SL1344_RS21970_ | 59 | 52 | 63 | 79 | 66 | 76 |
| SL1344_RS21980_ | 2256 | 2404 | 2497 | 2415 | 2406 | 2416 |
| SL1344_RS21985_ | 364 | 422 | 402 | 195 | 171 | 169 |
| SL1344_RS21990_ | 298 | 280 | 266 | 325 | 397 | 331 |
| SL1344_RS21995_ | 23 | 14 | 26 | 24 | 26 | 19 |
| SL1344_RS22000_ | 540 | 524 | 469 | 569 | 500 | 516 |
| SL1344_RS22005_ | 42 | 45 | 52 | 60 | 66 | 63 |
| SL1344_RS22010_ | 93 | 84 | 95 | 78 | 78 | 76 |
| SL1344_RS22015_ | 1379 | 1522 | 1527 | 1416 | 1369 | 1352 |
| SL1344_RS22020_ | 709 | 875 | 887 | 1097 | 1092 | 1085 |
| SL1344_RS22030_ | 1076 | 1122 | 1148 | 647 | 660 | 646 |
| SL1344_RS22035_ | 799 | 902 | 952 | 475 | 455 | 502 |
| SL1344_RS22040_ | 2033 | 2153 | 2231 | 931 | 873 | 1050 |
| SL1344_RS22045_ | 121 | 142 | 134 | 128 | 119 | 104 |
| SL1344_RS22050_ | 149 | 149 | 109 | 174 | 129 | 145 |

| | | | | | | |
|---|---|---|---|---|---|---|
| SL1344_RS22055_ | 110 | 88 | 64 | 115 | 78 | 95 |
| SL1344_RS22060_ | 61 | 98 | 75 | 79 | 69 | 62 |
| SL1344_RS22065_ | 51 | 45 | 29 | 54 | 56 | 51 |
| SL1344_RS22070_ | 30 | 27 | 41 | 69 | 35 | 31 |
| SL1344_RS22075_ | 45 | 56 | 46 | 55 | 48 | 41 |
| SL1344_RS22080_ | 244 | 228 | 217 | 235 | 250 | 238 |
| SL1344_RS22085_ | 76 | 71 | 81 | 106 | 86 | 87 |
| SL1344_RS22090_ | 13 | 15 | 18 | 30 | 29 | 28 |
| SL1344_RS22095_ | 288 | 292 | 286 | 288 | 265 | 283 |
| SL1344_RS22100_ | 418 | 403 | 397 | 397 | 388 | 380 |
| SL1344_RS22105_ | 56 | 55 | 33 | 57 | 61 | 56 |
| SL1344_RS22110_ | 23 | 25 | 22 | 24 | 20 | 11 |
| SL1344_RS22115_ | 24 | 26 | 26 | 24 | 23 | 29 |
| SL1344_RS22120_ | 61 | 52 | 48 | 71 | 66 | 78 |
| SL1344_RS22125_ | 43 | 22 | 26 | 16 | 26 | 23 |
| SL1344_RS22130_ | 55 | 48 | 40 | 127 | 143 | 125 |
| SL1344_RS22155_ | 2 | 1 | 1 | 2 | 3 | 1 |
| SL1344_RS22160_ | 154 | 193 | 160 | 180 | 194 | 202 |
| SL1344_RS22165_ | 285 | 311 | 266 | 351 | 295 | 317 |
| SL1344_RS22170_ | 537 | 628 | 605 | 532 | 509 | 606 |
| SL1344_RS22175_ | 36 | 28 | 24 | 27 | 29 | 26 |
| SL1344_RS22180_ | 155 | 161 | 105 | 166 | 148 | 121 |
| SL1344_RS22185_ | 136 | 159 | 144 | 131 | 101 | 133 |
| SL1344_RS22190_ | 208 | 250 | 268 | 287 | 230 | 273 |
| SL1344_RS22195_ | 310 | 294 | 295 | 328 | 297 | 314 |
| SL1344_RS22200_ | 203 | 174 | 166 | 178 | 186 | 147 |
| SL1344_RS22205_ | 305 | 267 | 272 | 246 | 250 | 316 |
| SL1344_RS22210_ | 1492 | 1527 | 1515 | 2425 | 2270 | 2322 |
| SL1344_RS22215_ | 211 | 229 | 191 | 291 | 283 | 272 |
| SL1344_RS22220_ | 1109 | 1156 | 1042 | 1692 | 1466 | 1213 |
| SL1344_RS22225_ | 234 | 234 | 246 | 206 | 159 | 202 |
| SL1344_RS22235_ | 39277 | 40814 | 40348 | 45138 | 42778 | 41878 |
| SL1344_RS22240_ | 1543 | 1639 | 1634 | 1819 | 1644 | 1828 |
| SL1344_RS22245_ | 153 | 176 | 172 | 144 | 138 | 118 |
| SL1344_RS22250_ | 717 | 777 | 773 | 570 | 564 | 566 |
| SL1344_RS22255_ | 3501 | 3804 | 4072 | 4103 | 4090 | 4094 |
| SL1344_RS22260_ | 145 | 180 | 172 | 131 | 125 | 134 |
| SL1344_RS22265_ | 320 | 330 | 325 | 329 | 349 | 322 |
| SL1344_RS22275_ | 231 | 219 | 235 | 197 | 210 | 185 |
| SL1344_RS22280_ | 42 | 33 | 21 | 28 | 30 | 25 |
| SL1344_RS22285_ | 99 | 91 | 99 | 118 | 109 | 115 |
| SL1344_RS22290_ | 53 | 42 | 31 | 69 | 43 | 56 |
| SL1344_RS22295_ | 125 | 123 | 127 | 161 | 151 | 158 |
| SL1344_RS22300_ | 297 | 344 | 333 | 383 | 360 | 372 |
| SL1344_RS22305_ | 1011 | 1175 | 1283 | 1042 | 988 | 990 |
| SL1344_RS22310_ | 148 | 165 | 159 | 82 | 90 | 105 |
| SL1344_RS22315_ | 236 | 199 | 200 | 202 | 171 | 184 |
| SL1344_RS22320_ | 1419 | 1351 | 1410 | 1248 | 1107 | 1202 |
| SL1344_RS22325_ | 563 | 602 | 574 | 529 | 449 | 507 |
| SL1344_RS22330_ | 884 | 860 | 863 | 759 | 859 | 768 |
| SL1344_RS22335_ | 358 | 506 | 510 | 490 | 418 | 548 |
| SL1344_RS22340_ | 177 | 151 | 196 | 196 | 213 | 213 |
| SL1344_RS22345_ | 9126 | 8984 | 7762 | 5979 | 6049 | 7208 |
| SL1344_RS22350_ | 3690 | 3462 | 3362 | 3135 | 3176 | 3153 |
| SL1344_RS22355_ | 509 | 429 | 479 | 473 | 458 | 458 |
| SL1344_RS22360_ | 357 | 349 | 371 | 348 | 346 | 363 |
| SL1344_RS22365_ | 292 | 312 | 300 | 330 | 307 | 347 |
| SL1344_RS22370_ | 109 | 133 | 143 | 144 | 134 | 140 |
| SL1344_RS22375_ | 733 | 760 | 841 | 742 | 717 | 745 |
| SL1344_RS22380_ | 1101 | 1133 | 1174 | 1079 | 1019 | 1064 |
| SL1344_RS22385_ | 3366 | 3442 | 3173 | 3598 | 3099 | 3188 |
| SL1344_RS22395_ | 3508 | 3867 | 3922 | 4030 | 3828 | 3887 |

| | | | | | | |
|---|---|---|---|---|---|---|
| SL1344_RS22400_ | 2411 | 2712 | 2808 | 2597 | 2637 | 2636 |
| SL1344_RS22405_ | 4646 | 4803 | 4933 | 4730 | 4497 | 4806 |
| SL1344_RS22410_ | 178 | 202 | 183 | 161 | 147 | 144 |
| SL1344_RS22415_ | 12202 | 12594 | 11735 | 12998 | 11880 | 11660 |
| SL1344_RS22420_ | 485 | 582 | 519 | 495 | 461 | 542 |
| SL1344_RS22425_ | 2630 | 2665 | 2785 | 2777 | 2650 | 2580 |
| SL1344_RS22430_ | 1736 | 1844 | 1745 | 1521 | 1572 | 1661 |
| SL1344_RS22435_ | 8 | 6 | 9 | 9 | 7 | 6 |
| SL1344_RS22440_ | 32 | 39 | 40 | 32 | 28 | 34 |
| SL1344_RS22445_ | 48 | 42 | 33 | 30 | 47 | 46 |
| SL1344_RS22450_ | 19 | 26 | 15 | 18 | 15 | 10 |
| SL1344_RS22455_ | 10 | 12 | 14 | 7 | 14 | 12 |
| SL1344_RS22460_ | 22 | 25 | 27 | 29 | 37 | 25 |
| SL1344_RS22465_ | 98 | 102 | 91 | 86 | 95 | 99 |
| SL1344_RS22470_ | 77 | 103 | 85 | 87 | 70 | 82 |
| SL1344_RS22475_ | 73 | 60 | 87 | 95 | 77 | 58 |
| SL1344_RS22480_ | 134 | 111 | 110 | 123 | 77 | 90 |
| SL1344_RS22485_ | 139 | 136 | 134 | 188 | 174 | 172 |
| SL1344_RS22490_ | 394 | 403 | 322 | 417 | 312 | 333 |
| SL1344_RS22495_ | 87 | 110 | 58 | 104 | 78 | 59 |
| SL1344_RS22500_ | 17 | 18 | 21 | 21 | 26 | 27 |
| SL1344_RS22505_ | 11 | 2 | 9 | 8 | 11 | 15 |
| SL1344_RS22510_ | 12 | 12 | 9 | 19 | 19 | 20 |
| SL1344_RS22520_ | 35 | 24 | 36 | 51 | 53 | 53 |
| SL1344_RS22525_ | 48 | 35 | 54 | 59 | 58 | 66 |
| SL1344_RS22530_ | 79 | 69 | 66 | 78 | 61 | 71 |
| SL1344_RS22535_ | 4201 | 4331 | 3334 | 3336 | 3212 | 3797 |
| SL1344_RS22540_ | 14670 | 14190 | 15863 | 12534 | 12362 | 12894 |
| SL1344_RS22545_ | 17903 | 17400 | 17624 | 16075 | 17054 | 15721 |
| SL1344_RS22550_ | 9993 | 9537 | 9868 | 9048 | 8867 | 9085 |
| SL1344_RS22555_ | 26848 | 28124 | 28667 | 23592 | 22704 | 24354 |
| SL1344_RS22560_ | 123 | 125 | 116 | 86 | 75 | 81 |
| SL1344_RS22565_ | 376 | 379 | 391 | 358 | 344 | 353 |
| SL1344_RS22570_ | 355 | 574 | 955 | 883 | 806 | 820 |
| SL1344_RS22575_ | 1084 | 1244 | 1378 | 1058 | 978 | 1062 |
| SL1344_RS22580_ | 59 | 69 | 65 | 68 | 94 | 56 |
| SL1344_RS22585_ | 163 | 156 | 127 | 220 | 229 | 178 |
| SL1344_RS22590_ | 34 | 23 | 25 | 33 | 39 | 19 |
| SL1344_RS22595_ | 81 | 81 | 86 | 99 | 102 | 83 |
| SL1344_RS22600_ | 215 | 202 | 227 | 241 | 229 | 236 |
| SL1344_RS22610_ | 115 | 133 | 124 | 130 | 117 | 107 |
| SL1344_RS22615_ | 1687 | 1622 | 1407 | 2860 | 2834 | 2405 |
| SL1344_RS22620_ | 1758 | 1643 | 1612 | 1448 | 1508 | 1613 |
| SL1344_RS22625_ | 273 | 268 | 205 | 481 | 372 | 320 |
| SL1344_RS22630_ | 785 | 979 | 996 | 739 | 639 | 690 |
| SL1344_RS22635_ | 2902 | 2963 | 2859 | 2463 | 2289 | 2367 |
| SL1344_RS22640_ | 2436 | 2605 | 2931 | 3176 | 2992 | 2959 |
| SL1344_RS22645_ | 49 | 47 | 55 | 48 | 58 | 62 |
| SL1344_RS22650_ | 15 | 26 | 27 | 57 | 69 | 42 |
| SL1344_RS22655_ | 6024 | 5957 | 5547 | 5697 | 5479 | 5462 |
| SL1344_RS22660_ | 812 | 863 | 753 | 930 | 887 | 920 |
| SL1344_RS22665_ | 1277 | 1532 | 1582 | 1422 | 1418 | 1392 |
| SL1344_RS22670_ | 12 | 19 | 19 | 27 | 26 | 17 |
| SL1344_RS22675_ | 874 | 922 | 930 | 815 | 800 | 897 |
| SL1344_RS22680_ | 108 | 112 | 114 | 105 | 116 | 114 |
| SL1344_RS22685_ | 5 | 3 | 15 | 2 | 20 | 1 |
| SL1344_RS22690_ | 90 | 123 | 103 | 129 | 133 | 136 |
| SL1344_RS22695_ | 93 | 76 | 96 | 147 | 74 | 106 |
| SL1344_RS22700_ | 49 | 68 | 53 | 60 | 56 | 51 |
| SL1344_RS22705_ | 9 | 15 | 8 | 24 | 8 | 13 |
| SL1344_RS22710_ | 10 | 8 | 12 | 5 | 6 | 8 |
| SL1344_RS22715_ | 54 | 56 | 61 | 61 | 65 | 56 |

| | | | | | | |
|---|---|---|---|---|---|---|
| SL1344_RS22720_ | 100 | 84 | 86 | 87 | 77 | 110 |
| SL1344_RS22725_ | 60 | 96 | 96 | 102 | 91 | 105 |
| SL1344_RS22730_ | 30 | 38 | 40 | 42 | 35 | 64 |
| SL1344_RS22735_ | 89 | 60 | 71 | 97 | 78 | 106 |
| SL1344_RS22740_ | 26 | 39 | 41 | 53 | 50 | 50 |
| SL1344_RS22745_ | 104 | 81 | 59 | 60 | 49 | 54 |
| SL1344_RS22750_ | 73 | 72 | 77 | 106 | 91 | 104 |
| SL1344_RS22755_ | 9 | 11 | 5 | 24 | 5 | 17 |
| SL1344_RS22760_ | 21 | 34 | 25 | 50 | 34 | 24 |
| SL1344_RS22765_ | 715 | 717 | 738 | 591 | 602 | 680 |
| SL1344_RS22770_ | 1037 | 1104 | 1046 | 1110 | 1153 | 1027 |
| SL1344_RS22775_ | 1931 | 2068 | 1799 | 2430 | 2230 | 2257 |
| SL1344_RS22780_ | 7 | 3 | 4 | 4 | 8 | 6 |
| SL1344_RS22785_ | 12 | 14 | 5 | 10 | 12 | 19 |
| SL1344_RS22790_ | 3 | 5 | 8 | 8 | 5 | 8 |
| SL1344_RS22795_ | 15 | 15 | 16 | 16 | 15 | 21 |
| SL1344_RS22800_ | 5 | 14 | 17 | 11 | 20 | 22 |
| SL1344_RS22805_ | 23 | 34 | 38 | 36 | 29 | 41 |
| SL1344_RS22810_ | 23 | 30 | 32 | 24 | 30 | 21 |
| SL1344_RS22815_ | 21 | 24 | 37 | 26 | 32 | 43 |
| SL1344_RS22820_ | 261 | 268 | 243 | 262 | 247 | 264 |
| SL1344_RS22825_ | 221 | 196 | 196 | 216 | 221 | 195 |
| SL1344_RS22830_ | 171 | 186 | 161 | 202 | 197 | 162 |
| SL1344_RS22835_ | 21 | 27 | 26 | 23 | 48 | 14 |
| SL1344_RS22840_ | 149 | 102 | 123 | 91 | 123 | 95 |
| SL1344_RS22845_ | 4 | 1 | 2 | 3 | 0 | 3 |
| SL1344_RS22850_ | 56 | 47 | 67 | 51 | 43 | 59 |
| SL1344_RS22855_ | 66 | 85 | 92 | 60 | 73 | 77 |
| SL1344_RS22860_ | 134 | 142 | 121 | 120 | 127 | 127 |
| SL1344_RS22865_ | 464 | 568 | 605 | 426 | 423 | 414 |
| SL1344_RS22870_ | 479 | 434 | 379 | 506 | 431 | 442 |
| SL1344_RS22875_ | 4340 | 4586 | 4285 | 4446 | 4453 | 4384 |
| SL1344_RS22880_ | 73 | 83 | 78 | 86 | 90 | 103 |
| SL1344_RS22885_ | 165 | 138 | 147 | 156 | 198 | 156 |
| SL1344_RS22890_ | 21 | 32 | 22 | 27 | 35 | 37 |
| SL1344_RS22895_ | 14 | 20 | 12 | 28 | 14 | 8 |
| SL1344_RS22900_ | 82 | 94 | 114 | 68 | 77 | 77 |
| SL1344_RS22905_ | 31 | 10 | 21 | 16 | 41 | 9 |
| SL1344_RS22910_ | 8 | 9 | 16 | 10 | 12 | 5 |
| SL1344_RS22915_ | 12 | 33 | 18 | 29 | 9 | 20 |
| SL1344_RS22920_ | 87 | 99 | 96 | 126 | 98 | 89 |
| SL1344_RS22925_ | 92 | 76 | 76 | 87 | 144 | 102 |
| SL1344_RS22930_ | 513 | 626 | 580 | 679 | 612 | 678 |
| SL1344_RS22935_ | 466 | 488 | 477 | 412 | 399 | 448 |
| SL1344_RS22940_ | 128 | 115 | 118 | 88 | 82 | 104 |
| SL1344_RS22945_ | 380 | 355 | 319 | 296 | 249 | 277 |
| SL1344_RS22950_ | 131 | 149 | 111 | 154 | 99 | 127 |
| SL1344_RS22955_ | 7650 | 7620 | 7749 | 7069 | 6756 | 7277 |
| SL1344_RS22960_ | 1081 | 1015 | 872 | 1056 | 966 | 1007 |
| SL1344_RS22965_ | 2581 | 2449 | 2358 | 2873 | 2779 | 2759 |
| SL1344_RS22970_ | 891 | 952 | 953 | 855 | 784 | 942 |
| SL1344_RS22975_ | 897 | 1134 | 1213 | 1147 | 1055 | 1147 |
| SL1344_RS22980_ | 747 | 961 | 1070 | 971 | 907 | 965 |
| SL1344_RS22985_ | 29 | 24 | 32 | 49 | 36 | 34 |
| SL1344_RS22995_ | 25 | 25 | 16 | 15 | 16 | 22 |
| SL1344_RS23000_ | 9 | 9 | 26 | 5 | 11 | 8 |
| SL1344_RS23010_ | 102 | 106 | 95 | 143 | 137 | 130 |
| SL1344_RS23015_ | 504 | 547 | 600 | 408 | 305 | 498 |
| SL1344_RS23020_ | 37 | 37 | 41 | 19 | 21 | 22 |
| SL1344_RS23025_ | 440 | 539 | 445 | 625 | 483 | 550 |
| SL1344_RS23030_ | 300 | 346 | 329 | 526 | 464 | 497 |
| SL1344_RS23035_ | 1405 | 1373 | 1414 | 1360 | 1391 | 1405 |

| | | | | | | |
|---|---|---|---|---|---|---|
| SL1344_RS23040_ | 2000 | 1948 | 1962 | 1889 | 1844 | 2028 |
| SL1344_RS23045_ | 769 | 745 | 719 | 765 | 811 | 788 |
| SL1344_RS23050_ | 1122 | 1121 | 1141 | 1203 | 1024 | 1130 |
| SL1344_RS23055_ | 1622 | 1663 | 1675 | 1599 | 1570 | 1670 |
| SL1344_RS23060_ | 2803 | 2856 | 3016 | 2545 | 2523 | 2580 |
| SL1344_RS23065_ | 517 | 599 | 604 | 535 | 629 | 485 |
| SL1344_RS23070_ | 507 | 624 | 584 | 471 | 554 | 508 |
| SL1344_RS23075_ | 14 | 14 | 18 | 6 | 14 | 14 |
| SL1344_RS23080_ | 826 | 846 | 835 | 960 | 816 | 939 |
| SL1344_RS23085_ | 128 | 145 | 141 | 180 | 149 | 153 |
| SL1344_RS23090_ | 36 | 20 | 18 | 14 | 18 | 24 |
| SL1344_RS23095_ | 20 | 7 | 9 | 17 | 16 | 16 |
| SL1344_RS23100_ | 264 | 292 | 258 | 268 | 290 | 245 |
| SL1344_RS23105_ | 30 | 28 | 28 | 29 | 29 | 42 |
| SL1344_RS23110_ | 42 | 52 | 42 | 33 | 41 | 50 |
| SL1344_RS23115_ | 131 | 132 | 130 | 131 | 124 | 136 |
| SL1344_RS23120_ | 811 | 781 | 639 | 817 | 817 | 705 |
| SL1344_RS23125_ | 51 | 46 | 42 | 56 | 46 | 35 |
| SL1344_RS23130_ | 5 | 6 | 8 | 6 | 4 | 10 |
| SL1344_RS23140_ | 31 | 50 | 62 | 68 | 59 | 66 |
| SL1344_RS23145_ | 160 | 184 | 195 | 221 | 183 | 218 |
| SL1344_RS23150_ | 406 | 356 | 375 | 360 | 350 | 371 |
| SL1344_RS23155_ | 21 | 16 | 32 | 21 | 17 | 34 |
| SL1344_RS23160_ | 8 | 11 | 12 | 9 | 5 | 10 |
| SL1344_RS23165_ | 3 | 4 | 2 | 5 | 12 | 7 |
| SL1344_RS23170_ | 99 | 78 | 73 | 96 | 85 | 155 |
| SL1344_RS23175_ | 140 | 155 | 140 | 103 | 107 | 105 |
| SL1344_RS23180_ | 145 | 139 | 175 | 166 | 141 | 173 |
| SL1344_RS23185_ | 25 | 38 | 45 | 64 | 34 | 46 |
| SL1344_RS23190_ | 80 | 91 | 101 | 107 | 94 | 89 |
| SL1344_RS23195_ | 2 | 5 | 3 | 5 | 6 | 7 |
| SL1344_RS23200_ | 7 | 5 | 4 | 7 | 3 | 8 |
| SL1344_RS23205_ | 77 | 92 | 63 | 129 | 94 | 111 |
| SL1344_RS23210_ | 17 | 17 | 13 | 29 | 22 | 19 |
| SL1344_RS23215_ | 489 | 539 | 569 | 523 | 475 | 562 |
| SL1344_RS23220_ | 609 | 702 | 722 | 723 | 628 | 695 |
| SL1344_RS23225_ | 715 | 659 | 637 | 678 | 729 | 635 |
| SL1344_RS23230_ | 121 | 186 | 207 | 240 | 237 | 209 |
| SL1344_RS23235_ | 115 | 126 | 119 | 190 | 143 | 132 |
| SL1344_RS23240_ | 136 | 131 | 146 | 157 | 189 | 172 |
| SL1344_RS23245_ | 623 | 503 | 616 | 562 | 531 | 519 |
| SL1344_RS23250_ | 170 | 170 | 178 | 171 | 154 | 158 |
| SL1344_RS23265_ | 270 | 282 | 274 | 628 | 558 | 685 |
| SL1344_RS23270_ | 7 | 4 | 14 | 15 | 11 | 15 |
| SL1344_RS23275_ | 2 | 8 | 4 | 4 | 3 | 8 |
| SL1344_RS23280_ | 5 | 7 | 10 | 16 | 7 | 7 |
| SL1344_RS23285_ | 11 | 16 | 11 | 17 | 13 | 13 |
| SL1344_RS23290_ | 4 | 10 | 15 | 17 | 17 | 23 |
| SL1344_RS23295_ | 103 | 98 | 78 | 97 | 118 | 125 |
| SL1344_RS23300_ | 3338 | 3258 | 3162 | 2394 | 2441 | 2582 |
| SL1344_RS23305_ | 327 | 355 | 377 | 294 | 281 | 287 |
| SL1344_RS23310_ | 691 | 772 | 681 | 652 | 625 | 638 |
| SL1344_RS23315_ | 381 | 421 | 419 | 426 | 386 | 368 |
| SL1344_RS23320_ | 61 | 50 | 61 | 52 | 63 | 50 |
| SL1344_RS23325_ | 67 | 81 | 68 | 77 | 80 | 71 |
| SL1344_RS23330_ | 2 | 3 | 3 | 5 | 2 | 9 |
| SL1344_RS23335_ | 7 | 3 | 4 | 5 | 5 | 9 |
| SL1344_RS23340_ | 674 | 647 | 611 | 561 | 510 | 478 |
| SL1344_RS23345_ | 1380 | 1661 | 1675 | 1243 | 1140 | 1220 |
| SL1344_RS23350_ | 431 | 421 | 381 | 324 | 317 | 333 |
| SL1344_RS23355_ | 125 | 59 | 34 | 65 | 26 | 28 |
| SL1344_RS23360_ | 5342 | 5313 | 6886 | 5682 | 5137 | 5292 |

| | | | | | | |
|---|---|---|---|---|---|---|
| SL1344_RS23365_ | 13134 | 14539 | 13663 | 14228 | 14202 | 13655 |
| SL1344_RS23370_ | 5086 | 5461 | 5176 | 5172 | 5113 | 5265 |
| SL1344_RS23375_ | 1259 | 1254 | 1251 | 905 | 883 | 935 |
| SL1344_RS23380_ | 669 | 776 | 758 | 619 | 666 | 750 |
| SL1344_RS23385_ | 398 | 427 | 490 | 430 | 407 | 476 |
| SL1344_RS23390_ | 772 | 831 | 820 | 700 | 713 | 767 |
| SL1344_RS23395_ | 3006 | 3289 | 3463 | 2949 | 2917 | 3000 |
| SL1344_RS23400_ | 244 | 245 | 230 | 316 | 345 | 332 |
| SL1344_RS23410_ | 248 | 248 | 243 | 290 | 270 | 289 |
| SL1344_RS23415_ | 195 | 174 | 219 | 206 | 170 | 196 |
| SL1344_RS23420_ | 22 | 17 | 25 | 26 | 19 | 20 |
| SL1344_RS23425_ | 25 | 30 | 24 | 36 | 25 | 37 |
| SL1344_RS23430_ | 349 | 322 | 358 | 297 | 261 | 273 |
| SL1344_RS23435_ | 340 | 353 | 361 | 316 | 324 | 292 |
| SL1344_RS23440_ | 1462 | 1661 | 1876 | 1966 | 2115 | 2089 |
| SL1344_RS23445_ | 1720 | 2198 | 2289 | 2661 | 2501 | 2603 |
| SL1344_RS23450_ | 129 | 116 | 126 | 93 | 125 | 103 |
| SL1344_RS23455_ | 161 | 176 | 155 | 128 | 136 | 147 |
| SL1344_RS23460_ | 23 | 22 | 20 | 18 | 10 | 13 |
| SL1344_RS23465_ | 52 | 42 | 39 | 40 | 43 | 32 |
| SL1344_RS23470_ | 36 | 49 | 38 | 29 | 34 | 29 |
| SL1344_RS23480_ | 169 | 199 | 174 | 169 | 173 | 163 |
| SL1344_RS23485_ | 695 | 805 | 828 | 732 | 728 | 655 |
| SL1344_RS23495_ | 1069 | 1177 | 1038 | 1187 | 1060 | 1080 |
| SL1344_RS23500_ | 6175 | 6934 | 7530 | 7139 | 7415 | 6949 |
| SL1344_RS23505_ | 461 | 479 | 480 | 455 | 468 | 455 |
| SL1344_RS23510_ | 353 | 288 | 332 | 274 | 275 | 299 |
| SL1344_RS23515_ | 233 | 207 | 237 | 214 | 240 | 206 |
| SL1344_RS23520_ | 626 | 703 | 741 | 734 | 713 | 669 |
| SL1344_RS23525_ | 1918 | 2005 | 1811 | 1832 | 1758 | 1766 |
| SL1344_RS23530_ | 408 | 441 | 415 | 541 | 511 | 483 |
| SL1344_RS23535_ | 596 | 698 | 601 | 702 | 777 | 707 |
| SL1344_RS23540_ | 494 | 505 | 471 | 531 | 503 | 533 |
| SL1344_RS23545_ | 124 | 123 | 132 | 119 | 124 | 111 |
| SL1344_RS23550_ | 60 | 54 | 47 | 45 | 47 | 49 |
| SL1344_RS23555_ | 6 | 16 | 12 | 6 | 14 | 10 |
| SL1344_RS23560_ | 93 | 63 | 71 | 70 | 60 | 54 |
| SL1344_RS23565_ | 25 | 24 | 27 | 10 | 16 | 18 |
| SL1344_RS23570_ | 53 | 63 | 41 | 42 | 33 | 44 |
| SL1344_RS23575_ | 248 | 290 | 275 | 259 | 257 | 242 |
| SL1344_RS23580_ | 294 | 230 | 222 | 150 | 152 | 158 |
| SL1344_RS23590_ | 5 | 6 | 2 | 3 | 1 | 4 |
| SL1344_RS23595_ | 92 | 133 | 125 | 113 | 119 | 109 |
| SL1344_RS23640_ | 163 | 220 | 217 | 280 | 250 | 244 |
| SL1344_RS23645_ | 106 | 117 | 87 | 89 | 101 | 125 |
| SL1344_RS23650_ | 374 | 445 | 442 | 492 | 492 | 506 |
| SL1344_RS23655_ | 250 | 265 | 230 | 290 | 253 | 252 |
| SL1344_RS23660_ | 432 | 466 | 485 | 505 | 486 | 507 |
| SL1344_RS23665_ | 53 | 64 | 66 | 101 | 86 | 118 |
| SL1344_RS23670_ | 1389 | 1785 | 1658 | 2043 | 1854 | 2050 |
| SL1344_RS23675_ | 445 | 475 | 499 | 541 | 465 | 578 |
| SL1344_RS23680_ | 419 | 442 | 423 | 487 | 391 | 443 |
| SL1344_RS23685_ | 42 | 57 | 56 | 47 | 53 | 46 |
| SL1344_RS23690_ | 33 | 26 | 29 | 22 | 33 | 21 |
| SL1344_RS23695_ | 25 | 19 | 15 | 13 | 8 | 19 |
| SL1344_RS23700_ | 26 | 37 | 35 | 39 | 46 | 42 |
| SL1344_RS23705_ | 11 | 12 | 12 | 20 | 12 | 16 |
| SL1344_RS23710_ | 64 | 96 | 80 | 80 | 90 | 79 |
| SL1344_RS23715_ | 14 | 32 | 26 | 36 | 25 | 37 |
| SL1344_RS23720_ | 26 | 29 | 32 | 40 | 33 | 38 |
| SL1344_RS23725_ | 19 | 16 | 13 | 28 | 22 | 19 |
| SL1344_RS23730_ | 34 | 29 | 25 | 20 | 27 | 24 |

| | | | | | | |
|---|---|---|---|---|---|---|
| SL1344_RS23735_ | 25 | 17 | 16 | 22 | 21 | 17 |
| SL1344_RS23740_ | 19 | 14 | 20 | 16 | 20 | 15 |
| SL1344_RS23745_ | 95 | 82 | 87 | 92 | 107 | 94 |
| SL1344_RS23750_ | 5 | 6 | 9 | 9 | 4 | 13 |
| SL1344_RS23755_ | 9 | 8 | 14 | 8 | 13 | 11 |
| SL1344_RS23760_ | 4 | 7 | 5 | 4 | 7 | 6 |
| SL1344_RS23765_ | 21 | 21 | 30 | 34 | 34 | 37 |
| SL1344_RS23770_ | 10 | 13 | 15 | 20 | 17 | 14 |
| SL1344_RS23775_ | 19 | 27 | 17 | 37 | 23 | 40 |
| SL1344_RS23780_ | 16 | 14 | 20 | 40 | 28 | 21 |
| SL1344_RS23785_ | 21 | 27 | 32 | 22 | 23 | 34 |
| SL1344_RS23790_ | 5 | 7 | 11 | 8 | 10 | 7 |
| SL1344_RS23795_ | 6 | 15 | 13 | 16 | 14 | 16 |
| SL1344_RS23800_ | 9 | 12 | 17 | 16 | 10 | 11 |
| SL1344_RS23805_ | 12 | 12 | 13 | 17 | 14 | 20 |
| SL1344_RS23810_ | 69 | 74 | 64 | 115 | 105 | 89 |
| SL1344_RS23815_ | 45 | 53 | 43 | 45 | 36 | 59 |
| SL1344_RS23820_ | 9 | 18 | 14 | 16 | 14 | 12 |
| SL1344_RS23825_ | 32 | 16 | 31 | 33 | 21 | 24 |
| SL1344_RS23830_ | 11 | 16 | 26 | 34 | 26 | 36 |
| SL1344_RS23840_ | 21 | 12 | 20 | 14 | 23 | 22 |
| SL1344_RS23845_ | 1 | 5 | 3 | 13 | 4 | 9 |
| SL1344_RS23850_ | 8 | 5 | 5 | 13 | 5 | 6 |
| SL1344_RS23855_ | 4 | 6 | 8 | 7 | 5 | 6 |
| SL1344_RS23860_ | 90 | 98 | 79 | 133 | 98 | 103 |
| SL1344_RS23865_ | 12 | 12 | 12 | 22 | 17 | 21 |
| SL1344_RS23870_ | 3 | 8 | 9 | 14 | 14 | 11 |
| SL1344_RS23875_ | 11 | 15 | 19 | 26 | 19 | 32 |
| SL1344_RS23880_ | 4 | 4 | 2 | 0 | 0 | 0 |
| SL1344_RS23885_ | 1 | 2 | 0 | 2 | 3 | 2 |
| SL1344_RS23890_ | 2 | 2 | 2 | 3 | 1 | 1 |
| SL1344_RS23895_ | 2 | 8 | 5 | 2 | 3 | 7 |
| SL1344_RS23900_ | 2 | 4 | 4 | 3 | 4 | 6 |
| SL1344_RS23905_ | 0 | 0 | 1 | 2 | 1 | 0 |
| SL1344_RS23910_ | 104 | 116 | 102 | 117 | 115 | 138 |
| SL1344_RS23915_ | 186 | 218 | 207 | 263 | 229 | 243 |
| SL1344_RS23920_ | 1635 | 1738 | 1576 | 1744 | 1630 | 1781 |
| SL1344_RS23925_ | 1166 | 1131 | 1056 | 1089 | 1087 | 1086 |
| SL1344_RS23930_ | 1540 | 1913 | 1949 | 2062 | 2129 | 2114 |
| SL1344_RS23935_ | 431 | 420 | 458 | 389 | 385 | 408 |
| SL1344_RS23940_ | 200 | 234 | 230 | 281 | 253 | 249 |
| SL1344_RS23945_ | 237 | 254 | 209 | 293 | 276 | 294 |
| SL1344_RS23950_ | 844 | 869 | 799 | 1092 | 1034 | 914 |
| SL1344_RS23955_ | 195 | 232 | 208 | 293 | 302 | 238 |
| SL1344_RS23960_ | 1053 | 1232 | 1234 | 1233 | 1279 | 1293 |
| SL1344_RS23965_ | 637 | 709 | 793 | 772 | 827 | 841 |
| SL1344_RS23970_ | 984 | 1096 | 1080 | 1109 | 1133 | 1219 |
| SL1344_RS23975_ | 374 | 474 | 463 | 486 | 398 | 472 |
| SL1344_RS23980_ | 64 | 57 | 60 | 35 | 47 | 45 |
| SL1344_RS23985_ | 8 | 12 | 15 | 7 | 6 | 16 |
| SL1344_RS23990_ | 4 | 8 | 3 | 2 | 4 | 5 |
| SL1344_RS23995_ | 13 | 9 | 15 | 19 | 14 | 10 |
| SL1344_RS24000_ | 43 | 46 | 42 | 68 | 47 | 59 |
| SL1344_RS24005_ | 1 | 4 | 1 | 1 | 3 | 4 |
| SL1344_RS24010_ | 7 | 5 | 4 | 8 | 7 | 2 |
| SL1344_RS24015_ | 85 | 105 | 94 | 96 | 82 | 93 |
| SL1344_RS24020_ | 2836 | 2927 | 2462 | 2703 | 2688 | 2810 |
| SL1344_RS24025_ | 3 | 2 | 7 | 3 | 4 | 9 |
| SL1344_RS24030_ | 14 | 11 | 13 | 13 | 12 | 13 |
| SL1344_RS24035_ | 29 | 46 | 34 | 48 | 43 | 58 |
| SL1344_RS24040_ | 74 | 92 | 96 | 87 | 78 | 107 |
| SL1344_RS24045_ | 284 | 256 | 319 | 238 | 281 | 250 |

| | | | | | | |
|---|---|---|---|---|---|---|
| SL1344_RS24050_ | 448 | 489 | 536 | 458 | 445 | 505 |
| SL1344_RS24055_ | 411 | 435 | 437 | 475 | 451 | 486 |
| SL1344_RS24060_ | 392 | 403 | 433 | 391 | 415 | 422 |
| SL1344_RS24065_ | 426 | 467 | 456 | 407 | 427 | 444 |
| SL1344_RS24070_ | 366 | 408 | 391 | 371 | 418 | 371 |
| SL1344_RS24075_ | 281 | 275 | 221 | 340 | 307 | 312 |
| SL1344_RS24080_ | 71 | 41 | 47 | 29 | 38 | 51 |
| SL1344_RS24085_ | 3212 | 3189 | 3123 | 3140 | 3086 | 3128 |
| SL1344_RS24090_ | 30 | 31 | 41 | 34 | 40 | 36 |
| SL1344_RS24095_ | 58 | 73 | 53 | 54 | 48 | 51 |
| SL1344_RS24100_ | 13 | 17 | 16 | 16 | 6 | 13 |
| SL1344_RS24105_ | 235 | 251 | 222 | 194 | 188 | 225 |
| SL1344_RS24110_ | 41 | 61 | 41 | 47 | 47 | 51 |
| SL1344_RS24115_ | 30 | 29 | 21 | 29 | 24 | 31 |
| SL1344_RS24120_ | 44 | 61 | 40 | 52 | 47 | 41 |
| SL1344_RS24125_ | 2607 | 2472 | 2096 | 2035 | 1843 | 1975 |
| SL1344_RS24130_ | 132 | 152 | 154 | 103 | 109 | 130 |
| SL1344_RS24135_ | 201 | 209 | 202 | 133 | 151 | 155 |
| SL1344_RS24140_ | 67 | 79 | 90 | 80 | 55 | 54 |
| SL1344_RS24145_ | 73 | 64 | 63 | 50 | 41 | 55 |
| SL1344_RS24150_ | 87 | 115 | 140 | 131 | 102 | 121 |
| SL1344_RS24155_ | 149 | 161 | 156 | 171 | 162 | 168 |
| SL1344_RS24160_ | 1 | 4 | 2 | 5 | 0 | 6 |
| SL1344_RS24165_ | 952 | 1011 | 994 | 1140 | 1091 | 982 |
| SL1344_RS24170_ | 740 | 819 | 677 | 881 | 849 | 815 |
| SL1344_RS24175_ | 3675 | 3143 | 3270 | 3398 | 3585 | 3299 |
| SL1344_RS24180_ | 1624 | 1564 | 1386 | 1384 | 1291 | 1342 |
| SL1344_RS24185_ | 105 | 124 | 138 | 120 | 126 | 149 |
| SL1344_RS24190_ | 232 | 276 | 255 | 203 | 213 | 197 |
| SL1344_RS24195_ | 109 | 142 | 117 | 123 | 167 | 132 |
| SL1344_RS24200_ | 1 | 6 | 2 | 4 | 5 | 7 |
| SL1344_RS24205_ | 1838 | 1745 | 1690 | 1676 | 1613 | 1508 |
| SL1344_RS24210_ | 184 | 190 | 148 | 189 | 182 | 177 |
| SL1344_RS24215_ | 186 | 208 | 170 | 257 | 214 | 218 |
| SL1344_RS24220_ | 571 | 612 | 578 | 662 | 643 | 643 |
| SL1344_RS24225_ | 192 | 241 | 219 | 286 | 200 | 224 |
| SL1344_RS24230_ | 103 | 90 | 88 | 95 | 76 | 98 |
| SL1344_RS24235_ | 157 | 191 | 153 | 183 | 151 | 156 |
| SL1344_RS24240_ | 256 | 243 | 253 | 246 | 224 | 247 |
| SL1344_RS24245_ | 262 | 295 | 242 | 329 | 295 | 307 |
| SL1344_RS24250_ | 1122 | 1263 | 1116 | 1658 | 1695 | 1550 |
| SL1344_RS24255_ | 144 | 175 | 148 | 175 | 146 | 187 |
| SL1344_RS24260_ | 134 | 163 | 140 | 147 | 152 | 155 |
| SL1344_RS24265_ | 162 | 209 | 199 | 195 | 198 | 160 |
| SL1344_RS24270_ | 85 | 85 | 88 | 77 | 79 | 77 |
| SL1344_RS24275_ | 39 | 65 | 52 | 59 | 50 | 52 |
| SL1344_RS24280_ | 71 | 83 | 86 | 90 | 60 | 87 |
| SL1344_RS24285_ | 610 | 687 | 629 | 711 | 732 | 669 |
| SL1344_RS24290_ | 522 | 575 | 626 | 751 | 717 | 657 |
| SL1344_RS24295_ | 876 | 911 | 886 | 1032 | 918 | 997 |
| SL1344_RS24300_ | 494 | 554 | 581 | 613 | 568 | 628 |
| SL1344_RS24305_ | 120 | 101 | 105 | 138 | 150 | 138 |
| SL1344_RS24310_ | 32 | 42 | 33 | 55 | 49 | 61 |
| SL1344_RS24315_ | 4 | 6 | 4 | 4 | 5 | 8 |
| SL1344_RS24320_ | 3 | 3 | 1 | 0 | 1 | 0 |
| SL1344_RS24325_ | 3 | 7 | 4 | 8 | 9 | 11 |
| SL1344_RS24330_ | 12 | 17 | 5 | 21 | 15 | 29 |
| SL1344_RS24335_ | 2 | 5 | 5 | 7 | 6 | 11 |
| SL1344_RS24340_ | 4 | 12 | 5 | 11 | 8 | 18 |
| SL1344_RS24345_ | 7 | 4 | 12 | 12 | 11 | 12 |
| SL1344_RS24350_ | 3 | 4 | 4 | 6 | 9 | 7 |
| SL1344_RS24355_ | 4 | 1 | 1 | 4 | 4 | 4 |

| | | | | | | |
|---|---|---|---|---|---|---|
| SL1344_RS24360_ | 4 | 6 | 5 | 5 | 14 | 7 |
| SL1344_RS24365_ | 1 | 4 | 8 | 8 | 2 | 5 |
| SL1344_RS24370_ | 54 | 41 | 48 | 48 | 48 | 56 |
| SL1344_RS24380_ | 50 | 42 | 38 | 49 | 43 | 43 |
| SL1344_RS24385_ | 48 | 54 | 41 | 47 | 54 | 69 |
| SL1344_RS24390_ | 3 | 1 | 7 | 4 | 2 | 2 |
| SL1344_RS24395_ | 24 | 22 | 23 | 25 | 25 | 36 |
| SL1344_RS24400_ | 3 | 9 | 6 | 12 | 7 | 3 |
| SL1344_RS24405_ | 4 | 7 | 7 | 9 | 9 | 9 |
| SL1344_RS24410_ | 189 | 253 | 228 | 254 | 249 | 271 |
| SL1344_RS24415_ | 21 | 8 | 13 | 15 | 14 | 14 |
| SL1344_RS24420_ | 84 | 102 | 97 | 102 | 85 | 105 |
| SL1344_RS24425_ | 9 | 7 | 8 | 21 | 8 | 13 |
| SL1344_RS24430_ | 784 | 882 | 733 | 1122 | 1061 | 1018 |
| SL1344_RS24435_ | 196 | 223 | 220 | 274 | 252 | 255 |
| SL1344_RS24440_ | 1076 | 1187 | 1223 | 1204 | 1192 | 1312 |
| SL1344_RS24445_ | 277 | 201 | 224 | 218 | 221 | 218 |
| SL1344_RS24450_ | 64 | 60 | 61 | 59 | 49 | 60 |
| SL1344_RS24455_ | 28 | 33 | 31 | 32 | 37 | 26 |
| SL1344_RS24460_ | 39074 | 39225 | 42432 | 42143 | 41088 | 44767 |
| SL1344_RS24465_ | 142 | 172 | 204 | 156 | 185 | 167 |
| SL1344_RS24470_ | 9844 | 11690 | 12968 | 11626 | 11449 | 12728 |
| SL1344_RS24475_ | 319 | 402 | 348 | 452 | 359 | 457 |
| SL1344_RS24480_ | 1485 | 1422 | 1326 | 1639 | 1544 | 1582 |
| SL1344_RS24485_ | 144 | 113 | 137 | 129 | 155 | 128 |
| SL1344_RS24490_ | 27 | 46 | 20 | 50 | 34 | 30 |
| SL1344_RS24495_ | 38 | 39 | 35 | 35 | 45 | 31 |
| SL1344_RS24500_ | 165 | 151 | 160 | 164 | 143 | 145 |
| SL1344_RS24505_ | 9 | 21 | 17 | 21 | 18 | 20 |
| SL1344_RS24510_ | 0 | 4 | 1 | 1 | 2 | 3 |
| SL1344_RS24515_ | 8 | 10 | 11 | 11 | 9 | 16 |
| SL1344_RS24520_ | 14 | 16 | 22 | 15 | 17 | 17 |
| SL1344_RS24525_ | 53 | 51 | 56 | 50 | 40 | 53 |
| SL1344_RS24530_ | 31 | 26 | 34 | 32 | 30 | 28 |
| SL1344_RS24535_ | 18 | 21 | 11 | 28 | 26 | 36 |
| SL1344_RS24540_ | 28 | 20 | 18 | 25 | 18 | 25 |
| SL1344_RS24545_ | 6 | 9 | 10 | 9 | 8 | 16 |
| SL1344_RS24550_ | 91 | 91 | 77 | 87 | 78 | 74 |
| SL1344_RS24555_ | 25 | 23 | 30 | 32 | 33 | 32 |
| SL1344_RS24560_ | 42 | 28 | 42 | 40 | 36 | 51 |
| SL1344_RS24565_ | 40 | 32 | 58 | 36 | 39 | 32 |
| SL1344_RS24570_ | 25 | 23 | 30 | 20 | 19 | 20 |
| SL1344_RS24575_ | 30 | 24 | 19 | 51 | 51 | 54 |
| SL1344_RS24580_ | 273 | 266 | 245 | 737 | 720 | 771 |
| SL1344_RS24585_ | 100 | 114 | 93 | 236 | 194 | 233 |
| SL1344_RS24590_ | 277 | 243 | 245 | 240 | 241 | 237 |
| SL1344_RS24595_ | 8 | 4 | 3 | 1 | 2 | 2 |
| SL1344_RS24600_ | 24 | 20 | 12 | 25 | 26 | 26 |
| SL1344_RS24605_ | 29 | 12 | 23 | 28 | 21 | 21 |
| SL1344_RS24610_ | 83 | 94 | 90 | 69 | 56 | 78 |
| SL1344_RS24615_ | 26 | 21 | 18 | 25 | 31 | 30 |
| SL1344_RS24620_ | 18 | 18 | 29 | 22 | 17 | 20 |
| SL1344_RS24625_ | 3 | 5 | 4 | 3 | 3 | 4 |
| SL1344_RS24630_ | 71 | 49 | 56 | 78 | 57 | 73 |
| SL1344_RS24635_ | 29 | 45 | 42 | 46 | 58 | 25 |
| SL1344_RS24640_ | 5 | 8 | 1 | 3 | 1 | 4 |
| SL1344_RS24645_ | 12 | 12 | 7 | 10 | 5 | 9 |
| SL1344_RS24650_ | 20 | 17 | 22 | 17 | 12 | 21 |
| SL1344_RS24655_ | 35 | 39 | 23 | 20 | 22 | 32 |
| SL1344_RS24660_ | 252 | 293 | 257 | 307 | 272 | 282 |
| SL1344_RS24665_ | 210 | 196 | 143 | 213 | 202 | 161 |
| SL1344_RS24670_ | 33 | 31 | 31 | 33 | 30 | 28 |

| | | | | | | |
|---|---|---|---|---|---|---|
| SL1344_RS24675_ | 57 | 61 | 53 | 56 | 69 | 84 |
| SL1344_RS24690_ | 12579 | 12121 | 11965 | 16871 | 15506 | 18666 |
| SL1344_RS24695_ | 933 | 929 | 1004 | 2137 | 2303 | 2269 |
| SL1344_RS24700_ | 1159 | 1148 | 1163 | 2557 | 2724 | 2701 |
| SL1344_RS24705_ | 1095 | 1031 | 920 | 2094 | 2068 | 2127 |
| SL1344_RS24710_ | 865 | 798 | 730 | 1523 | 1533 | 1619 |
| SL1344_RS24730_ | 6823 | 7593 | 7434 | 10569 | 9144 | 11150 |
| SL1344_RS24735_ | 10514 | 11764 | 11962 | 16366 | 14759 | 17363 |
| SL1344_RS24740_ | 9 | 18 | 10 | 18 | 14 | 22 |
| SL1344_RS24745_ | 0 | 1 | 3 | 2 | 1 | 1 |
| SL1344_RS24750_ | 263 | 173 | 192 | 167 | 145 | 201 |
| SL1344_RS24755_ | 360 | 370 | 354 | 342 | 320 | 353 |
| SL1344_RS24760_ | 224 | 201 | 193 | 143 | 153 | 144 |
| SL1344_RS24780_ | 43 | 27 | 18 | 39 | 39 | 35 |
| SL1344_RS24785_ | 13787 | 14155 | 13081 | 10432 | 11584 | 12047 |
| SL1344_RS24790_ | 11171 | 10578 | 10640 | 10511 | 10754 | 9963 |
| SL1344_RS24795_ | 202 | 190 | 186 | 186 | 171 | 158 |
| SL1344_RS24800_ | 12 | 6 | 18 | 16 | 14 | 6 |
| SL1344_RS24805_ | 266 | 246 | 227 | 197 | 219 | 190 |
| SL1344_RS24810_ | 463 | 449 | 471 | 309 | 315 | 394 |
| SL1344_RS24820_ | 1159 | 1209 | 1313 | 2671 | 3118 | 3090 |
| SL1344_RS24825_ | 6629 | 6897 | 7772 | 7518 | 7264 | 8743 |
| SL1344_RS24830_ | 414 | 371 | 417 | 515 | 520 | 515 |
| SL1344_RS24835_ | 66 | 61 | 49 | 59 | 76 | 72 |
| SL1344_RS24840_ | 18 | 18 | 19 | 12 | 23 | 21 |
| SL1344_RS24845_ | 14 | 11 | 10 | 2 | 8 | 19 |
| SL1344_RS24850_ | 4 | 4 | 7 | 3 | 6 | 5 |
| SL1344_RS24860_ | 1197 | 1251 | 1298 | 835 | 799 | 978 |
| SL1344_RS24865_ | 2 | 4 | 2 | 1 | 2 | 4 |
| SL1344_RS24875_ | 2 | 5 | 10 | 4 | 6 | 3 |
| SL1344_RS24885_ | 342 | 350 | 383 | 219 | 223 | 273 |
| SLnc0002_ | 22 | 15 | 13 | 24 | 11 | 16 |
| SLnc0010_ | 14 | 30 | 19 | 23 | 23 | 21 |
| SLnc0010a_ | 110 | 95 | 74 | 65 | 90 | 91 |
| SLnc0014_ | 183 | 178 | 147 | 419 | 419 | 424 |
| SLnc0022_ | 115 | 124 | 115 | 96 | 76 | 105 |
| SLnc0030_ | 1803 | 1805 | 2092 | 1537 | 1350 | 1421 |
| SLnc0030a_ | 70 | 75 | 81 | 73 | 85 | 72 |
| SLnc0031_ | 16 | 14 | 23 | 10 | 16 | 15 |
| SLnc0035_ | 2465 | 2673 | 2392 | 2358 | 2365 | 2291 |
| SLnc0039_ | 258 | 199 | 159 | 136 | 121 | 150 |
| SLnc0048_ | 39 | 57 | 41 | 30 | 57 | 46 |
| SLnc1001_ | 1 | 0 | 0 | 0 | 0 | 0 |
| SLnc1002_ | 62 | 75 | 76 | 70 | 74 | 75 |
| SLnc1006_ | 322 | 344 | 257 | 361 | 321 | 379 |
| SLnc1008_ | 3 | 8 | 8 | 4 | 3 | 2 |
| SLnc1013_ | 43 | 33 | 50 | 41 | 51 | 39 |
| SLnc1014_ | 17 | 17 | 16 | 28 | 17 | 9 |
| SLnc1015_ | 31 | 18 | 32 | 21 | 21 | 22 |
| SLnc1016_ | 33 | 33 | 27 | 24 | 24 | 32 |
| SLnc1017_ | 201 | 240 | 270 | 154 | 134 | 168 |
| SLnc1018_ | 18 | 18 | 14 | 8 | 18 | 15 |
| SLnc1019_ | 25 | 34 | 25 | 34 | 26 | 26 |
| SLnc1021_ | 70 | 73 | 47 | 67 | 52 | 56 |
| SLnc1022_ | 9 | 13 | 9 | 7 | 8 | 5 |
| SLnc1024_ | 59 | 47 | 63 | 32 | 32 | 33 |
| SLnc1025_ | 49 | 53 | 37 | 32 | 20 | 30 |
| SLnc1026_ | 18 | 22 | 11 | 17 | 22 | 24 |
| SLnc1027_ | 25 | 30 | 39 | 28 | 21 | 25 |
| SLnc1028_ | 40 | 44 | 44 | 27 | 42 | 34 |
| SLnc1029_ | 12 | 13 | 7 | 5 | 5 | 9 |
| sRNA1_ | 1855 | 2001 | 2147 | 2112 | 1972 | 1988 |

| | | | | | | |
|---|---|---|---|---|---|---|
| sRNA10_ | 433 | 383 | 448 | 436 | 404 | 415 |
| sRNA11_ | 21 | 41 | 34 | 25 | 11 | 35 |
| sRNA12_ | 1 | 5 | 0 | 1 | 2 | 1 |
| sRNA15_ | 3 | 0 | 0 | 0 | 1 | 2 |
| sRNA16_ | 10 | 12 | 8 | 6 | 16 | 9 |
| sRNA18_ | 5 | 5 | 7 | 2 | 2 | 2 |
| sRNA19_ | 0 | 0 | 0 | 0 | 1 | 0 |
| sRNA2_ | 1628 | 1478 | 1258 | 1073 | 1031 | 1134 |
| sRNA21_ | 45 | 44 | 50 | 87 | 69 | 65 |
| sRNA22_ | 40 | 39 | 22 | 42 | 50 | 38 |
| sRNA23_ | 7 | 21 | 14 | 16 | 6 | 19 |
| sRNA24_ | 5 | 6 | 3 | 6 | 11 | 6 |
| sRNA25_ | 1 | 2 | 3 | 36 | 0 | 3 |
| sRNA26_ | 2 | 4 | 5 | 1 | 4 | 5 |
| sRNA28_ | 2 | 4 | 2 | 5 | 11 | 7 |
| sRNA29_ | 4 | 5 | 4 | 1 | 6 | 3 |
| sRNA31_ | 16 | 5 | 3 | 5 | 4 | 8 |
| sRNA32_ | 11 | 8 | 17 | 5 | 4 | 7 |
| sRNA33_ | 31 | 30 | 20 | 30 | 19 | 27 |
| sRNA5_ | 38 | 18 | 33 | 39 | 24 | 28 |
| sRNA7_ | 121 | 128 | 122 | 105 | 98 | 110 |
| STnc10_ | 1 | 4 | 4 | 2 | 4 | 1 |
| STnc1010_ | 29 | 21 | 14 | 28 | 14 | 18 |
| STnc1030_ | 39 | 27 | 20 | 29 | 47 | 32 |
| STnc1040_ | 8 | 8 | 11 | 9 | 5 | 10 |
| STnc1065_ | 2086 | 2183 | 2628 | 2012 | 1936 | 1957 |
| STnc1070_ | 60 | 56 | 66 | 58 | 75 | 44 |
| STnc1080_ | 14 | 21 | 23 | 18 | 21 | 28 |
| STnc1090_ | 81 | 79 | 85 | 99 | 90 | 88 |
| STnc1100_ | 27 | 21 | 22 | 16 | 15 | 22 |
| STnc1130_ | 81 | 98 | 77 | 156 | 148 | 146 |
| STnc1150_ | 261 | 280 | 245 | 243 | 254 | 272 |
| STnc1160_ | 710 | 649 | 722 | 572 | 546 | 624 |
| STnc1170_ | 18 | 8 | 7 | 9 | 10 | 6 |
| STnc1180_ | 29 | 21 | 19 | 25 | 34 | 29 |
| STnc1190_ | 356 | 384 | 381 | 363 | 384 | 362 |
| STnc1200_ | 43 | 41 | 46 | 37 | 42 | 47 |
| STnc1210_ | 6 | 7 | 5 | 7 | 7 | 2 |
| STnc1220_ | 0 | 1 | 0 | 0 | 0 | 1 |
| STnc1275_ | 85 | 91 | 64 | 105 | 122 | 104 |
| STnc1280_ | 4 | 8 | 5 | 8 | 9 | 8 |
| STnc1285_ | 283 | 441 | 471 | 2186 | 1712 | 2122 |
| STnc1290_ | 608 | 552 | 586 | 422 | 400 | 394 |
| STnc130_ | 10 | 9 | 5 | 9 | 10 | 5 |
| STnc1320_ | 13 | 16 | 13 | 7 | 10 | 14 |
| STnc1340_ | 114 | 133 | 122 | 84 | 66 | 84 |
| STnc1380_ | 19 | 25 | 21 | 16 | 21 | 19 |
| STnc1385_ | 1892 | 1836 | 1467 | 1326 | 1073 | 1326 |
| STnc1390_ | 30 | 35 | 16 | 20 | 13 | 23 |
| STnc140_ | 4 | 1 | 2 | 1 | 4 | 1 |
| STnc1400_ | 298 | 303 | 277 | 245 | 241 | 280 |
| STnc1405_ | 87 | 84 | 84 | 352 | 414 | 344 |
| STnc1410_ | 26 | 43 | 37 | 30 | 28 | 33 |
| STnc1460_ | 20950 | 19272 | 18242 | 14004 | 13868 | 14351 |
| STnc1490_ | 279 | 244 | 209 | 169 | 151 | 201 |
| STnc150_ | 152 | 136 | 110 | 102 | 69 | 105 |
| STnc1500_ | 19 | 24 | 19 | 26 | 17 | 20 |
| STnc1510_ | 89 | 80 | 89 | 96 | 79 | 86 |
| STnc1530_ | 6 | 4 | 4 | 3 | 2 | 2 |
| STnc1540_ | 116 | 160 | 144 | 95 | 77 | 93 |
| STnc1545_ | 414 | 434 | 434 | 349 | 321 | 398 |
| STnc1550_ | 112 | 93 | 85 | 153 | 142 | 125 |

| | | | | | | |
|---|---|---|---|---|---|---|
| STnc1560_ | 3898 | 4077 | 6006 | 3747 | 3490 | 3493 |
| STnc1565_ | 128 | 105 | 107 | 74 | 108 | 115 |
| STnc1580_ | 95 | 113 | 110 | 80 | 101 | 74 |
| STnc1590_ | 1923 | 1583 | 1498 | 1029 | 1123 | 1057 |
| STnc1600_ | 31 | 33 | 16 | 21 | 38 | 22 |
| STnc1640_ | 33 | 27 | 25 | 24 | 19 | 21 |
| STnc1650_ | 11 | 12 | 8 | 9 | 3 | 6 |
| STnc1660_ | 140 | 172 | 135 | 131 | 174 | 137 |
| STnc1670_ | 16 | 17 | 10 | 11 | 9 | 1 |
| STnc1675_ | 30336 | 31330 | 35397 | 29717 | 29427 | 30343 |
| STnc1680_ | 471 | 563 | 653 | 390 | 376 | 416 |
| STnc1685_ | 8214 | 8637 | 7728 | 1869 | 11720 | 1592 |
| STnc1690_ | 14 | 16 | 12 | 34 | 32 | 29 |
| STnc170_ | 7 | 8 | 1 | 22 | 22 | 19 |
| STnc1700_ | 19 | 17 | 28 | 17 | 17 | 26 |
| STnc1710_ | 347 | 329 | 273 | 260 | 268 | 271 |
| STnc1760_ | 27 | 25 | 15 | 19 | 20 | 26 |
| STnc1820_ | 168 | 147 | 157 | 118 | 102 | 99 |
| STnc1830_ | 16 | 28 | 14 | 38 | 26 | 31 |
| STnc1870_ | 6 | 3 | 2 | 8 | 2 | 4 |
| STnc1880_ | 69 | 46 | 48 | 55 | 40 | 54 |
| STnc1890_ | 4 | 0 | 4 | 0 | 5 | 3 |
| STnc190_ | 22 | 26 | 27 | 59 | 65 | 56 |
| STnc1920_ | 111 | 124 | 116 | 88 | 71 | 87 |
| STnc1990_ | 114 | 120 | 114 | 83 | 87 | 88 |
| STnc20_ | 25 | 23 | 33 | 46 | 24 | 27 |
| STnc200_ | 76 | 109 | 103 | 140 | 143 | 172 |
| STnc2000_ | 349 | 323 | 376 | 483 | 380 | 415 |
| STnc2010_ | 88 | 92 | 81 | 70 | 81 | 86 |
| STnc2030_ | 7 | 7 | 9 | 10 | 8 | 4 |
| STnc2040_ | 1 | 0 | 0 | 9 | 3 | 7 |
| STnc205_ | 1777 | 1967 | 1853 | 10806 | 9908 | 10985 |
| STnc2070_ | 162 | 178 | 196 | 154 | 119 | 160 |
| STnc2080_ | 248 | 253 | 239 | 160 | 175 | 170 |
| STnc2090_ | 129 | 108 | 87 | 171 | 215 | 143 |
| STnc210_ | 4 | 0 | 2 | 5 | 0 | 3 |
| STnc2100_ | 66 | 79 | 77 | 72 | 85 | 70 |
| STnc2110_ | 66 | 38 | 56 | 110 | 81 | 75 |
| STnc2130_ | 7 | 6 | 5 | 7 | 8 | 7 |
| STnc2160_ | 1 | 5 | 6 | 35 | 1 | 4 |
| STnc2170_ | 43 | 41 | 50 | 46 | 31 | 37 |
| STnc2175_ | 21 | 27 | 75 | 24 | 17 | 59 |
| STnc2180_ | 1929 | 1782 | 1694 | 3022 | 2705 | 3190 |
| STnc220_ | 32 | 22 | 35 | 10 | 11 | 21 |
| STnc230_ | 2 | 0 | 1 | 1 | 0 | 0 |
| STnc240_ | 219 | 274 | 276 | 125 | 117 | 171 |
| STnc260_ | 3 | 1 | 1 | 2 | 3 | 2 |
| STnc280_ | 4 | 4 | 4 | 4 | 1 | 4 |
| STnc290_ | 213 | 199 | 216 | 254 | 219 | 226 |
| STnc30_ | 350 | 388 | 488 | 320 | 266 | 352 |
| STnc300_ | 0 | 2 | 2 | 3 | 1 | 0 |
| STnc3000_ | 2 | 3 | 4 | 1 | 5 | 4 |
| STnc3020_ | 11 | 19 | 9 | 3 | 7 | 6 |
| STnc3030_ | 18 | 9 | 8 | 9 | 12 | 9 |
| STnc3040_ | 5 | 5 | 9 | 4 | 3 | 8 |
| STnc3060_ | 27 | 23 | 22 | 18 | 21 | 17 |
| STnc3070_ | 39 | 27 | 31 | 33 | 41 | 35 |
| STnc3080_ | 17 | 23 | 23 | 12 | 28 | 10 |
| STnc3090_ | 7 | 4 | 5 | 2 | 4 | 11 |
| STnc310_ | 8 | 8 | 12 | 9 | 8 | 7 |
| STnc3100_ | 271 | 277 | 249 | 250 | 208 | 225 |
| STnc3110_ | 112 | 96 | 105 | 109 | 109 | 131 |

| | | | | | | |
|---|---|---|---|---|---|---|
| STnc3120_ | 98 | 89 | 81 | 91 | 62 | 80 |
| STnc3150_ | 162 | 195 | 128 | 157 | 197 | 154 |
| STnc3160_ | 10 | 15 | 24 | 13 | 16 | 24 |
| STnc3170_ | 21 | 15 | 13 | 22 | 24 | 17 |
| STnc3180_ | 11 | 8 | 8 | 8 | 7 | 8 |
| STnc320_ | 28 | 36 | 30 | 38 | 27 | 31 |
| STnc3210_ | 3 | 4 | 3 | 6 | 9 | 4 |
| STnc3220_ | 149 | 131 | 107 | 116 | 141 | 131 |
| STnc3230_ | 15 | 13 | 10 | 8 | 8 | 11 |
| STnc3270_ | 13 | 20 | 24 | 22 | 16 | 17 |
| STnc3280_ | 23 | 16 | 16 | 11 | 10 | 15 |
| STnc3290_ | 48 | 50 | 48 | 31 | 23 | 31 |
| STnc3300_ | 11 | 17 | 13 | 5 | 9 | 7 |
| STnc3310_ | 1320 | 1484 | 1331 | 1501 | 1435 | 1428 |
| STnc3330_ | 1 | 2 | 2 | 0 | 5 | 1 |
| STnc3340_ | 22 | 24 | 16 | 36 | 26 | 27 |
| STnc3350_ | 34 | 34 | 35 | 27 | 40 | 42 |
| STnc340_ | 60 | 44 | 51 | 69 | 56 | 75 |
| STnc3400_ | 8 | 13 | 5 | 6 | 6 | 13 |
| STnc3410_ | 203 | 176 | 224 | 213 | 157 | 193 |
| STnc3420_ | 30 | 20 | 22 | 21 | 20 | 14 |
| STnc3430_ | 73 | 99 | 89 | 55 | 57 | 61 |
| STnc3440_ | 30 | 21 | 41 | 21 | 26 | 32 |
| STnc3460_ | 697 | 723 | 709 | 585 | 541 | 626 |
| STnc3480_ | 34 | 32 | 38 | 123 | 102 | 150 |
| STnc3490_ | 16 | 17 | 6 | 12 | 19 | 9 |
| STnc350_ | 44 | 37 | 27 | 14 | 9 | 28 |
| STnc3500_ | 9 | 10 | 6 | 7 | 9 | 11 |
| STnc3510_ | 103 | 92 | 92 | 106 | 87 | 88 |
| STnc3540_ | 604 | 666 | 714 | 1127 | 1130 | 1108 |
| STnc3550_ | 21 | 16 | 32 | 9 | 15 | 9 |
| STnc3580_ | 6 | 4 | 7 | 8 | 3 | 11 |
| STnc360_ | 8 | 3 | 4 | 3 | 3 | 3 |
| STnc3600_ | 78 | 69 | 54 | 78 | 88 | 64 |
| STnc3610_ | 130 | 125 | 139 | 178 | 161 | 230 |
| STnc3620_ | 12 | 15 | 10 | 13 | 6 | 14 |
| STnc3630_ | 10 | 15 | 17 | 18 | 16 | 15 |
| STnc3640_ | 36 | 20 | 31 | 31 | 34 | 43 |
| STnc3680_ | 809 | 737 | 758 | 603 | 527 | 539 |
| STnc370_ | 6 | 8 | 5 | 5 | 2 | 5 |
| STnc3700_ | 25 | 30 | 23 | 22 | 17 | 30 |
| STnc3720_ | 19 | 16 | 5 | 8 | 7 | 8 |
| STnc3730_ | 74 | 70 | 86 | 78 | 86 | 87 |
| STnc3750_ | 16 | 22 | 11 | 7 | 14 | 11 |
| STnc3770_ | 23 | 24 | 22 | 21 | 13 | 23 |
| STnc3790_ | 9 | 5 | 5 | 5 | 4 | 22 |
| STnc380_ | 10 | 17 | 12 | 21 | 16 | 14 |
| STnc3800_ | 140 | 120 | 117 | 88 | 84 | 77 |
| STnc3830_ | 19 | 12 | 17 | 12 | 23 | 13 |
| STnc3850_ | 13 | 19 | 15 | 17 | 6 | 13 |
| STnc3860_ | 14 | 21 | 8 | 14 | 8 | 22 |
| STnc3870_ | 37 | 30 | 38 | 33 | 31 | 41 |
| STnc3880_ | 5 | 2 | 2 | 5 | 2 | 2 |
| STnc3890_ | 15 | 12 | 15 | 8 | 14 | 17 |
| STnc390_ | 453 | 447 | 590 | 459 | 454 | 425 |
| STnc3920_ | 0 | 4 | 4 | 2 | 2 | 3 |
| STnc3950_ | 49 | 69 | 42 | 83 | 63 | 63 |
| STnc3960_ | 116 | 104 | 107 | 122 | 124 | 116 |
| STnc3970_ | 12 | 7 | 10 | 4 | 8 | 5 |
| STnc3990_ | 84 | 112 | 87 | 122 | 68 | 108 |
| STnc40_ | 375 | 342 | 342 | 340 | 320 | 312 |
| STnc4000_ | 31 | 29 | 27 | 26 | 25 | 20 |

| | | | | | | |
|---|---|---|---|---|---|---|
| STnc4010_ | 0 | 5 | 1 | 7 | 0 | 3 |
| STnc4020_ | 10 | 1 | 5 | 1 | 7 | 7 |
| STnc4040_ | 66 | 78 | 69 | 64 | 57 | 72 |
| STnc4070_ | 25 | 18 | 11 | 16 | 14 | 18 |
| STnc4080_ | 41 | 37 | 21 | 33 | 24 | 25 |
| STnc410_ | 8 | 10 | 5 | 6 | 6 | 8 |
| STnc4100_ | 10 | 11 | 14 | 28 | 52 | 25 |
| STnc4110_ | 15 | 8 | 12 | 12 | 7 | 14 |
| STnc4120_ | 51 | 48 | 52 | 45 | 37 | 49 |
| STnc4130_ | 27 | 21 | 13 | 23 | 25 | 24 |
| STnc4140_ | 50 | 51 | 67 | 41 | 51 | 59 |
| STnc4150_ | 2 | 0 | 1 | 1 | 1 | 2 |
| STnc4160_ | 24 | 27 | 18 | 24 | 30 | 29 |
| STnc4170_ | 16 | 11 | 21 | 14 | 15 | 11 |
| STnc4190_ | 4 | 10 | 6 | 6 | 5 | 6 |
| STnc420_ | 42 | 37 | 39 | 47 | 35 | 47 |
| STnc4200_ | 33 | 41 | 42 | 32 | 31 | 38 |
| STnc4210_ | 4 | 10 | 6 | 42 | 32 | 30 |
| STnc4220_ | 399 | 392 | 401 | 272 | 267 | 345 |
| STnc4230_ | 99 | 116 | 132 | 104 | 93 | 158 |
| STnc4240_ | 564 | 607 | 525 | 424 | 428 | 429 |
| STnc4250_ | 3888 | 3767 | 4073 | 3405 | 3432 | 3290 |
| STnc4260_ | 11 | 3 | 7 | 14 | 10 | 6 |
| STnc430_ | 4 | 6 | 6 | 6 | 6 | 7 |
| STnc440_ | 24 | 18 | 27 | 20 | 28 | 27 |
| STnc450_ | 187 | 174 | 153 | 149 | 169 | 170 |
| STnc460_ | 1 | 0 | 4 | 1 | 3 | 1 |
| STnc470_ | 28 | 31 | 28 | 33 | 32 | 31 |
| STnc475_ | 5093 | 5249 | 4234 | 5540 | 5295 | 5739 |
| STnc480_ | 12 | 15 | 11 | 18 | 11 | 13 |
| STnc50_ | 8 | 9 | 3 | 5 | 8 | 2 |
| STnc500_ | 11 | 11 | 6 | 9 | 12 | 5 |
| STnc510_ | 23 | 36 | 33 | 32 | 25 | 45 |
| STnc520_ | 62 | 64 | 53 | 57 | 46 | 50 |
| STnc530_ | 2 | 8 | 4 | 4 | 5 | 5 |
| STnc540_ | 2257 | 2333 | 2657 | 2866 | 2653 | 2799 |
| STnc590_ | 32 | 51 | 26 | 74 | 61 | 73 |
| STnc60_ | 0 | 1 | 1 | 2 | 0 | 2 |
| STnc600_ | 180 | 146 | 125 | 530 | 582 | 528 |
| STnc610_ | 1878 | 1665 | 1622 | 1752 | 1631 | 2051 |
| STnc620_ | 694 | 761 | 936 | 869 | 737 | 850 |
| STnc640_ | 46 | 31 | 24 | 45 | 35 | 40 |
| STnc650_ | 5533 | 5337 | 4126 | 4808 | 5012 | 4167 |
| STnc660_ | 456 | 550 | 486 | 631 | 603 | 573 |
| STnc670_ | 148 | 139 | 105 | 326 | 298 | 300 |
| STnc680_ | 874 | 810 | 711 | 815 | 709 | 730 |
| STnc690_ | 476 | 475 | 484 | 263 | 243 | 312 |
| STnc695_ | 51 | 44 | 43 | 30 | 34 | 41 |
| STnc70_ | 1096 | 1166 | 972 | 895 | 858 | 1012 |
| STnc720_ | 28 | 34 | 30 | 27 | 29 | 21 |
| STnc730_ | 865 | 696 | 525 | 765 | 656 | 671 |
| STnc740_ | 1 | 2 | 1 | 1 | 4 | 2 |
| STnc750_ | 355 | 304 | 224 | 246 | 254 | 227 |
| STnc760_ | 69 | 70 | 64 | 86 | 61 | 53 |
| STnc770_ | 834 | 778 | 585 | 914 | 808 | 748 |
| STnc780_ | 492 | 435 | 400 | 464 | 447 | 451 |
| STnc790_ | 78 | 91 | 54 | 88 | 78 | 79 |
| STnc800_ | 2725 | 2534 | 2579 | 1827 | 1793 | 1923 |
| STnc810_ | 7 | 9 | 5 | 5 | 10 | 3 |
| STnc835_ | 1408 | 1506 | 1774 | 1684 | 1489 | 1751 |
| STnc840_ | 358 | 393 | 425 | 1026 | 893 | 1095 |
| STnc850_ | 10 | 19 | 19 | 18 | 19 | 14 |

| | | | | | | |
|---|---|---|---|---|---|---|
| STnc870_ | 90 | 80 | 84 | 95 | 130 | 104 |
| STnc880_ | 89 | 66 | 83 | 58 | 70 | 70 |
| STnc890_ | 120 | 159 | 187 | 143 | 152 | 151 |
| STnc920_ | 396 | 377 | 394 | 295 | 319 | 320 |
| STnc925_ | 115 | 130 | 114 | 151 | 117 | 147 |
| STnc930_ | 5 | 0 | 3 | 5 | 6 | 5 |
| STnc940_ | 7 | 4 | 11 | 12 | 6 | 14 |
| STnc950_ | 9 | 7 | 7 | 7 | 14 | 13 |
| STnc970_ | 208 | 168 | 137 | 249 | 257 | 261 |
| StyR-103a_ | 0 | 0 | 0 | 0 | 1 | 0 |
| StyR-103b_ | 0 | 0 | 2 | 0 | 0 | 0 |
| StyR-103c_ | 0 | 0 | 3 | 1 | 1 | 1 |
| StyR-103d_ | 0 | 0 | 0 | 0 | 0 | 0 |
| StyR-103e_ | 0 | 0 | 2 | 0 | 0 | 0 |
| StyR-103f_ | 0 | 0 | 0 | 0 | 0 | 0 |
| StyR-103g_ | 1 | 0 | 0 | 1 | 0 | 0 |
| StyR-122_ | 11 | 3 | 10 | 6 | 4 | 5 |
| StyR-126_ | 27 | 62 | 70 | 84 | 43 | 72 |
| StyR-128_ | 9 | 4 | 7 | 8 | 12 | 11 |
| StyR-134_ | 277 | 250 | 212 | 258 | 241 | 238 |
| StyR-137_ | 0 | 0 | 0 | 0 | 0 | 0 |
| StyR-150_ | 46 | 29 | 28 | 38 | 34 | 35 |
| StyR-169_ | 134 | 121 | 136 | 99 | 78 | 89 |
| StyR-172_ | 0 | 0 | 0 | 0 | 0 | 1 |
| StyR-178_ | 0 | 1 | 0 | 1 | 0 | 0 |
| StyR-195_ | 27 | 14 | 12 | 20 | 17 | 24 |
| StyR-196_ | 3 | 1 | 1 | 3 | 1 | 1 |
| StyR-199_ | 16 | 18 | 26 | 19 | 15 | 16 |
| StyR-215a_ | 5169 | 4335 | 4130 | 3714 | 2967 | 3886 |
| StyR-215b_ | 3920 | 3332 | 3186 | 2257 | 2231 | 2564 |
| StyR-215c_ | 578 | 472 | 501 | 332 | 357 | 417 |
| StyR-215d_ | 930 | 750 | 719 | 741 | 569 | 674 |
| StyR-215e_ | 589 | 350 | 405 | 341 | 220 | 354 |
| StyR-215f_ | 1629 | 1558 | 1660 | 1169 | 1081 | 1280 |
| StyR-215g_ | 386 | 385 | 344 | 254 | 227 | 352 |
| StyR-219_ | 12 | 7 | 11 | 14 | 8 | 10 |
| StyR-222_ | 1 | 0 | 2 | 2 | 1 | 1 |
| StyR-234_ | 8 | 12 | 9 | 6 | 14 | 13 |
| StyR-243_ | 97 | 113 | 95 | 166 | 167 | 164 |
| StyR-248_ | 20 | 22 | 18 | 25 | 16 | 16 |
| StyR-24a_ | 3 | 3 | 3 | 10 | 6 | 9 |
| StyR-24b_ | 3 | 1 | 5 | 58 | 61 | 79 |
| StyR-24c_ | 17 | 17 | 12 | 24 | 12 | 15 |
| StyR-24d_ | 10 | 12 | 9 | 5 | 2 | 4 |
| StyR-24e_ | 1 | 2 | 2 | 6 | 9 | 5 |
| StyR-24f_ | 13 | 18 | 30 | 35 | 25 | 48 |
| StyR-24g_ | 5 | 5 | 5 | 22 | 28 | 35 |
| StyR-250_ | 18 | 17 | 10 | 10 | 11 | 11 |
| StyR-252_ | 56 | 62 | 72 | 72 | 94 | 89 |
| StyR-254_ | 2 | 1 | 0 | 1 | 3 | 5 |
| StyR-262_ | 80 | 73 | 63 | 76 | 75 | 65 |
| StyR-264_ | 96 | 61 | 76 | 41 | 53 | 52 |
| StyR-27_ | 4 | 10 | 10 | 10 | 7 | 10 |
| StyR-277_ | 137 | 214 | 150 | 151 | 129 | 169 |
| StyR-281_ | 4 | 7 | 1 | 6 | 9 | 7 |
| StyR-285_ | 14 | 17 | 12 | 11 | 7 | 8 |
| StyR-287_ | 51 | 50 | 38 | 57 | 46 | 63 |
| StyR-29_ | 9854 | 10119 | 7675 | 8219 | 8372 | 8739 |
| StyR-291_ | 20 | 29 | 41 | 54 | 57 | 53 |
| StyR-292_ | 4 | 4 | 4 | 4 | 3 | 2 |
| StyR-293_ | 12 | 10 | 12 | 13 | 11 | 12 |
| StyR-295_ | 12 | 13 | 15 | 9 | 11 | 11 |

| | | | | | | |
|---|---|---|---|---|---|---|
| StyR-296_ | 2943 | 2293 | 1564 | 2584 | 2138 | 1824 |
| StyR-3_ | 61 | 74 | 66 | 81 | 99 | 97 |
| StyR-302_ | 0 | 1 | 3 | 1 | 0 | 3 |
| StyR-305_ | 43 | 47 | 40 | 143 | 123 | 151 |
| StyR-306_ | 421 | 356 | 263 | 354 | 277 | 272 |
| StyR-315_ | 1 | 0 | 1 | 2 | 0 | 2 |
| StyR-322_ | 15 | 12 | 16 | 8 | 7 | 9 |
| StyR-327_ | 11 | 4 | 9 | 10 | 5 | 4 |
| StyR-328_ | 3 | 3 | 0 | 1 | 0 | 3 |
| StyR-329a_ | 124 | 150 | 127 | 113 | 108 | 116 |
| StyR-329b_ | 10 | 4 | 16 | 7 | 14 | 10 |
| StyR-336_ | 15 | 6 | 7 | 14 | 14 | 11 |
| StyR-341_ | 8 | 8 | 2 | 8 | 7 | 14 |
| StyR-354_ | 113 | 70 | 60 | 40 | 31 | 19 |
| StyR-358_ | 7 | 20 | 14 | 10 | 18 | 8 |
| StyR-362_ | 9 | 11 | 5 | 10 | 11 | 12 |
| StyR-369_ | 4 | 10 | 1 | 5 | 3 | 11 |
| StyR-377_ | 352 | 476 | 604 | 558 | 529 | 658 |
| StyR-378_ | 0 | 2 | 0 | 31 | 1 | 1 |
| StyR-44a_ | 235 | 266 | 379 | 212 | 175 | 185 |
| StyR-44b_ | 802 | 824 | 883 | 520 | 483 | 661 |
| StyR-44c_ | 181 | 150 | 176 | 88 | 77 | 103 |
| StyR-44d_ | 134 | 132 | 118 | 63 | 91 | 83 |
| StyR-44e_ | 32 | 54 | 38 | 32 | 23 | 41 |
| StyR-44f_ | 26 | 33 | 17 | 20 | 12 | 14 |
| StyR-44g_ | 26 | 22 | 23 | 23 | 14 | 20 |
| StyR-51_ | 1524 | 1530 | 1678 | 1415 | 1293 | 1394 |
| StyR-55_ | 9892 | 9860 | 10723 | 8747 | 8008 | 9300 |
| StyR-8_ | 5730 | 5517 | 5478 | 5070 | 4792 | 5437 |
| StyR-88_ | 533 | 476 | 491 | 490 | 429 | 543 |
| StyR-90_ | 19 | 18 | 15 | 21 | 19 | 17 |
| StyR-99_ | 125 | 119 | 82 | 105 | 88 | 107 |
| tp2_ | 11 | 5 | 4 | 4 | 3 | 1 |
| tp28_ | 330 | 332 | 259 | 384 | 349 | 324 |
| tpk3_ | 776 | 789 | 771 | 446 | 458 | 612 |
| tpke11_ | 3291 | 2973 | 4303 | 4339 | 3803 | 3151 |
| tpke70_ | 33 | 39 | 43 | 38 | 56 | 26 |
| zipA-leader_ | 947 | 850 | 718 | 662 | 685 | 627 |
| _ | | | | | | |
| _ | | | | | | |

# ME

| Gene | WT-1-ME | WT-2-ME | WT-3-ME | 5' tnpA-1-ME | 5' tnpA-2-ME | 5' tnpA-3-ME |
|---|---|---|---|---|---|---|
| aceF_aceF | 7940 | 11939 | 9470 | 5535 | 9297 | 6696 |
| SL1344_RS11990_ackA | 1819 | 2762 | 2461 | 1804 | 1502 | 1637 |
| SL1344_RS08180_acpD | 229 | 296 | 212 | 361 | 383 | 370 |
| acpS_acpS | 332 | 345 | 430 | 339 | 327 | 362 |
| adrA_adrA | 134 | 180 | 140 | 170 | 151 | 133 |
| SL1344_RS16630_aer | 1937 | 2162 | 1658 | 4327 | 3635 | 4818 |
| SL1344_RS13055_alpha/beta hydrolase | 34 | 52 | 61 | 60 | 41 | 48 |
| AmgR_amgR | 8 | 10 | 10 | 6 | 7 | 6 |
| amiC_amiC | 673 | 713 | 661 | 891 | 787 | 819 |
| SL1344_RS04710_amino acid permease | 354 | 299 | 296 | 212 | 198 | 193 |
| SL1344_RS01920_ampH | 248 | 313 | 268 | 192 | 104 | 133 |
| SL1344_RS23585_arcA | 4493 | 5047 | 4987 | 5519 | 4409 | 4491 |
| ArcZ_arcZ | 782 | 969 | 1046 | 825 | 696 | 365 |
| STnc490a_art200 | 2 | 3 | 1 | 4 | 1 | 0 |
| STnc490b_art200 | 6 | 4 | 3 | 0 | 0 | 0 |
| STnc490c_art200 | 3 | 3 | 1 | 0 | 0 | 0 |
| STnc490d_art200 | 0 | 1 | 0 | 0 | 0 | 0 |
| STnc490e_art200 | 1 | 0 | 0 | 0 | 0 | 0 |
| STnc490f_art200 | 1 | 3 | 2 | 0 | 0 | 0 |
| STnc490g_art200 | 0 | 3 | 0 | 0 | 0 | 0 |
| artM_artM | 779 | 849 | 910 | 886 | 692 | 744 |
| artP_artP | 1379 | 1272 | 1103 | 782 | 1958 | 1912 |
| asarfA_asarfA | 362 | 567 | 427 | 263 | 517 | 272 |
| asnB_asnB | 916 | 1142 | 966 | 941 | 931 | 1059 |
| SL1344_RS13060_asrA | 18 | 16 | 20 | 17 | 17 | 16 |
| SL1344_RS13065_asrB | 43 | 27 | 49 | 42 | 32 | 45 |
| SL1344_RS13070_asrC | 48 | 63 | 102 | 92 | 49 | 75 |
| asyhbQ_asyhbQ | 312 | 425 | 270 | 240 | 469 | 295 |
| atpC_atpC | 14364 | 17283 | 16696 | 12517 | 12887 | 11324 |
| SL1344_RS15305_barA | 701 | 760 | 732 | 786 | 681 | 655 |
| bcsA_bcsA | 769 | 1014 | 808 | 487 | 482 | 494 |
| SL1344_RS17950_bigA | 216 | 269 | 271 | 350 | 289 | 296 |
| bssR_bssR | 88 | 107 | 88 | 89 | 65 | 78 |
| bssS_bssS | 709 | 908 | 1009 | 744 | 489 | 662 |
| cadB_cadB | 565 | 1327 | 1297 | 598 | 415 | 676 |
| cbiC_cbiC | 99 | 94 | 94 | 52 | 77 | 89 |
| cbiD_cbiD | 192 | 208 | 157 | 87 | 167 | 152 |
| SL1344_RS11580_ccmA | 43 | 25 | 24 | 13 | 55 | 83 |
| SL1344_RS11575_ccmB | 11 | 16 | 12 | 14 | 9 | 16 |
| cdsA_cdsA | 1189 | 1189 | 1325 | 1182 | 1227 | 1432 |
| SL1344_RS02340_cdsH | 144 | 169 | 150 | 164 | 192 | 155 |
| chaB_chaB | 90 | 74 | 95 | 162 | 120 | 157 |
| SL1344_RS09610_cheA | 7904 | 9380 | 8475 | 38430 | 29512 | 39896 |
| SL1344_RS09600_cheM | 2952 | 4505 | 3232 | 18860 | 15110 | 19105 |
| SL1344_RS09595_chemotaxis protein methyltransferase | 1762 | 2370 | 1973 | 8886 | 6862 | 8633 |
| SL1344_RS09585_chemotaxis regulator | 1595 | 1541 | 1759 | 6118 | 4521 | 7071 |
| SL1344_RS09590_chemotaxis response regulator protein-glutamate methylesterase | 1513 | 1652 | 1627 | 6358 | 5037 | 6401 |
| SL1344_RS11875_cheV | 2849 | 4579 | 3758 | 9439 | 6900 | 10710 |
| SL1344_RS09605_CheW | 8092 | 8303 | 8779 | 39607 | 28354 | 36844 |
| SL1344_RS09580_cheZ | 3850 | 4141 | 4351 | 17430 | 14638 | 16936 |
| chiP_chiP | 1696 | 1391 | 1587 | 1643 | 942 | 770 |
| ChiX_chiX | 248 | 1456 | 480 | 1043 | 1076 | 408 |
| SL1344_RS02290_clpP | 1715 | 1927 | 1800 | 2101 | 1544 | 2042 |
| SL1344_RS02295_clpX | 7470 | 8884 | 9399 | 9599 | 7663 | 9029 |
| SL1344_RS09530_cmoA | 647 | 657 | 672 | 1483 | 1172 | 1161 |
| cobS_cobS | 163 | 170 | 196 | 206 | 170 | 156 |
| copA_copA | 1221 | 2219 | 2402 | 1347 | 1678 | 1616 |
| SL1344_RS20320_corA | 717 | 812 | 604 | 967 | 907 | 711 |
| SL1344_RS20860_cpxA | 1978 | 2365 | 1984 | 1932 | 1944 | 1874 |
| cpxP_cpxP | 693 | 548 | 731 | 478 | 329 | 428 |
| SL1344_RS20865_cpxR | 748 | 905 | 876 | 793 | 600 | 808 |
| SL1344_RS17890_crp | 7560 | 9546 | 8964 | 8867 | 6976 | 8035 |

| | | | | | | |
|---|---|---|---|---|---|---|
| CrpT_crpT | 185 | 196 | 198 | 285 | 104 | 177 |
| csgB_csgB | 7 | 3 | 3 | 5 | 4 | 3 |
| SL1344_RS05640_csgC | 242 | 221 | 349 | 585 | 558 | 994 |
| SL1344_RS18810_cspA | 11183 | 9395 | 12350 | 5125 | 5260 | 4259 |
| SL1344_RS09980_cspB | 171 | 187 | 228 | 113 | 99 | 106 |
| SL1344_RS06145_cspH | 58 | 43 | 38 | 63 | 43 | 26 |
| SL1344_RS14635_csrA | 4658 | 4804 | 4468 | 4943 | 4201 | 5212 |
| CsrB_csrB | 112815 | 110262 | 113428 | 147506 | 124065 | 141167 |
| SL1344_RS15345_csrB | 95851 | 86860 | 94805 | 129678 | 83531 | 122075 |
| CsrC_csrC | 32502 | 30961 | 29003 | 54081 | 46473 | 55231 |
| SL1344_RS17435_csrD | 1087 | 1288 | 1031 | 1794 | 1785 | 1928 |
| CyaR_cyaR | 3763 | 5406 | 4896 | 4906 | 4859 | 3712 |
| SL1344_RS12535_cysA | 377 | 385 | 438 | 552 | 456 | 327 |
| SL1344_RS08565_cysB | 497 | 633 | 703 | 922 | 733 | 826 |
| SL1344_RS15175_cysC | 278 | 278 | 306 | 518 | 329 | 270 |
| SL1344_RS15185_cysD | 546 | 517 | 757 | 1016 | 207 | 522 |
| SL1344_RS19060_cysE | 1952 | 2006 | 2151 | 2075 | 2120 | 2072 |
| SL1344_RS17945_cysG | 909 | 1119 | 1020 | 996 | 985 | 865 |
| SL1344_RS15240_cysH | 196 | 218 | 202 | 306 | 327 | 223 |
| SL1344_RS15245_cysI | 971 | 989 | 1048 | 1421 | 1282 | 734 |
| SL1344_RS15250_cysJ | 960 | 688 | 901 | 1591 | 644 | 633 |
| SL1344_RS12475_cysK | 3000 | 3335 | 3856 | 4133 | 2648 | 2768 |
| SL1344_RS12530_cysM | 531 | 667 | 650 | 497 | 554 | 485 |
| SL1344_RS15180_cysN | 660 | 518 | 673 | 1046 | 471 | 501 |
| SL1344_RS12550_cysP | 629 | 798 | 1163 | 1025 | 239 | 725 |
| SL1344_RS22605_cysQ | 586 | 807 | 694 | 650 | 851 | 698 |
| SL1344_RS02725_cysS | 1781 | 2181 | 2233 | 1945 | 1784 | 1783 |
| SL1344_RS09720_cysT | 2885 | 2812 | 2425 | 8243 | 7743 | 3516 |
| SL1344_RS12545_cysU | 232 | 186 | 264 | 343 | 149 | 189 |
| SL1344_RS12540_cysW | 347 | 299 | 444 | 552 | 289 | 279 |
| SL1344_RS12470_cysZ | 365 | 387 | 450 | 361 | 254 | 281 |
| SL1344_RS16735_cyuA | 259 | 291 | 277 | 240 | 238 | 226 |
| SL1344_RS16740_cyuP | 118 | 120 | 86 | 105 | 90 | 96 |
| SL1344_RS10610_dacD | 55 | 65 | 58 | 53 | 56 | 60 |
| SL1344_RS17980_dam | 1311 | 1455 | 1377 | 1442 | 1161 | 1065 |
| DapZ_dapZ | 330 | 415 | 443 | 403 | 529 | 449 |
| ddl_ddl | 1485 | 1521 | 1497 | 1695 | 1513 | 1558 |
| SL1344_RS09505_dihydroneopterin triphosphate diphosphatase | 847 | 920 | 992 | 1816 | 1698 | 1724 |
| dinG_dinG | 743 | 924 | 983 | 1060 | 891 | 930 |
| SL1344_RS05725_dinI | 671 | 984 | 797 | 1151 | 997 | 833 |
| SL1344_RS04685_dmsA | 91 | 95 | 97 | 114 | 90 | 100 |
| SL1344_RS04690_dmsB | 11 | 15 | 10 | 9 | 3 | 8 |
| SL1344_RS04695_dmsC | 25 | 32 | 47 | 23 | 42 | 43 |
| SL1344_RS06610_DNA/RNA non-specific endonuclease | 158 | 170 | 153 | 248 | 137 | 187 |
| dnaA_dnaA | 1820 | 1951 | 1956 | 2475 | 1721 | 1978 |
| SL1344_RS00060_dnaK | 26742 | 22685 | 27720 | 40964 | 25581 | 25193 |
| dpiA_dpiA | 87 | 78 | 74 | 92 | 64 | 88 |
| dpiA_dpiA | 79 | 116 | 77 | 91 | 108 | 107 |
| dpiB_dpiB | 109 | 88 | 99 | 127 | 119 | 145 |
| DsrA_dsrA | 14 | 42 | 31 | 11 | 7 | 4 |
| SL1344_RS09330_DUF1441 domain-containing protein | 6 | 6 | 6 | 18 | 26 | 22 |
| SL1344_RS06460_DUF1496 domain-containing protein | 606 | 989 | 799 | 2174 | 1629 | 2466 |
| SL1344_RS16295_DUF2623 domain-containing protein | 270 | 355 | 321 | 1084 | 647 | 1014 |
| emrB_emrB | 520 | 659 | 619 | 619 | 530 | 502 |
| emrD_emrD | 619 | 749 | 613 | 1009 | 594 | 696 |
| emtA_emtA | 310 | 307 | 370 | 465 | 347 | 424 |
| SL1344_RS03015_entB | 116 | 114 | 148 | 230 | 148 | 185 |
| entE_entE | 152 | 169 | 218 | 304 | 194 | 238 |
| entF_entF | 239 | 278 | 362 | 556 | 362 | 393 |
| SL1344_RS18065_envZ | 1262 | 1630 | 1384 | 1257 | 1551 | 1259 |
| eutA_eutA | 52 | 58 | 52 | 46 | 25 | 48 |
| SL1344_RS12680_eutS | 2 | 5 | 7 | 3 | 6 | 2 |
| fabG_fabG | 3092 | 3409 | 3399 | 3730 | 3001 | 3618 |

| | | | | | | |
|---|---|---|---|---|---|---|
| fadA_fadA | 917 | 887 | 883 | 962 | 1321 | 1480 |
| SL1344_RS09080_fadD | 666 | 507 | 568 | 1199 | 1325 | 1166 |
| fadE_fadE | 939 | 1072 | 908 | 805 | 1339 | 1051 |
| fadI_fadI | 266 | 237 | 216 | 173 | 224 | 144 |
| SL1344_RS13945_Fels-2 prophage protein | 11 | 29 | 37 | 10 | 19 | 28 |
| feoA_feoA | 482 | 502 | 569 | 624 | 324 | 464 |
| ffs_ffs | 774 | 3632 | 2704 | 1785 | 3785 | 1381 |
| fimA_fimA | 53 | 55 | 36 | 69 | 48 | 50 |
| fimF_fimF | 54 | 50 | 48 | 54 | 58 | 46 |
| SL1344_RS02785_fimZ | 52 | 13 | 4 | 15 | 12 | 25 |
| fis_fis | 870 | 946 | 929 | 942 | 740 | 753 |
| fixB_fixB | 116 | 72 | 100 | 106 | 99 | 91 |
| flgD_flgD | 2294 | 2552 | 2238 | 5469 | 4551 | 5507 |
| flgF_flgF | 3337 | 3922 | 3941 | 7783 | 6882 | 7512 |
| flgI_flgI | 1279 | 1502 | 1454 | 2839 | 2746 | 2881 |
| SL1344_RS05830_flgK | 5198 | 6777 | 5445 | 16440 | 13285 | 18138 |
| flgL_flgL | 4911 | 6627 | 5688 | 14327 | 10507 | 14497 |
| SL1344_RS05775_flgM | 2601 | 3036 | 2728 | 8101 | 4971 | 9188 |
| SL1344_RS05770_flgN | 3139 | 4231 | 3134 | 9531 | 7929 | 10789 |
| SL1344_RS09570_flhA | 1405 | 1636 | 1504 | 5262 | 5879 | 5473 |
| SL1344_RS09575_flhB | 1020 | 1268 | 987 | 4327 | 5318 | 4315 |
| SL1344_RS09625_flhC | 2066 | 2279 | 1983 | 6263 | 5550 | 5590 |
| SL1344_RS09630_flhD | 1071 | 989 | 1114 | 2144 | 1437 | 1665 |
| SL1344_RS09565_flhE | 327 | 413 | 358 | 1232 | 1513 | 1194 |
| SL1344_RS09785_fliA | 4366 | 4807 | 3701 | 17666 | 17063 | 18981 |
| SL1344_RS09800_fliC | 14378 | 17232 | 17478 | 226801 | 78808 | 214223 |
| SL1344_RS09845_fliE | 608 | 616 | 716 | 1243 | 1021 | 1393 |
| fliI_fliI | 1741 | 2057 | 2001 | 3528 | 3558 | 2903 |
| fliJ_fliJ | 877 | 828 | 723 | 1447 | 1284 | 1308 |
| fliL_fliL | 1316 | 1655 | 1592 | 2781 | 2351 | 2792 |
| fliQ_fliQ | 405 | 437 | 464 | 817 | 723 | 746 |
| SL1344_RS09815_fliT | 4574 | 5584 | 5889 | 9870 | 6727 | 9669 |
| SL1344_RS09775_fliY | 1363 | 1618 | 1494 | 4869 | 5069 | 6229 |
| SL1344_RS09780_fliZ | 2659 | 2753 | 2493 | 10941 | 9406 | 11643 |
| SL1344_RS14340_fljA | 1551 | 2092 | 1873 | 116 | 1830 | 78 |
| flk_flk | 287 | 285 | 369 | 284 | 177 | 228 |
| SL1344_RS08285_fnr | 2267 | 2638 | 2906 | 4552 | 3005 | 3893 |
| FnrS_fnrS | 186 | 443 | 492 | 294 | 226 | 318 |
| SL1344_RS04735_focA | 769 | 1256 | 1210 | 1070 | 738 | 1379 |
| folA_folA | 437 | 450 | 491 | 427 | 473 | 362 |
| frsA_frsA | 769 | 876 | 893 | 900 | 682 | 832 |
| SL1344_RS09665_ftnB | 159 | 172 | 140 | 347 | 173 | 268 |
| SL1344_RS06895_fumD | 252 | 263 | 230 | 194 | 222 | 229 |
| SL1344_RS03500_fur | 3519 | 3498 | 3802 | 4665 | 3707 | 3865 |
| gapA_gapA | 1416 | 1478 | 1458 | 1460 | 1195 | 1503 |
| GcvB_gcvB | 23143 | 42325 | 30247 | 48447 | 26989 | 25212 |
| gcvT_gcvT | 5331 | 5739 | 5334 | 6790 | 5985 | 6185 |
| gidB_gidB | 336 | 387 | 455 | 357 | 265 | 335 |
| gldA_gldA | 188 | 186 | 185 | 178 | 212 | 192 |
| gldA_gldA | 862 | 935 | 832 | 814 | 783 | 828 |
| GlmY_glmY | 994 | 3174 | 2010 | 3159 | 1897 | 1134 |
| GlmZ_glmZ | 5415 | 8234 | 8441 | 11045 | 6527 | 7529 |
| glnQ_glnQ | 1449 | 1831 | 1522 | 1212 | 1818 | 1578 |
| SL1344_RS13690_gluconolactonase | 149 | 96 | 79 | 65 | 65 | 79 |
| SL1344_RS09560_glycoside hydrolase family 105 protein | 248 | 245 | 246 | 715 | 729 | 651 |
| gntK_gntK | 64 | 57 | 71 | 46 | 51 | 58 |
| SL1344_RS18265_gntR | 483 | 604 | 580 | 639 | 649 | 539 |
| gpmA_gpmA | 10385 | 10864 | 9375 | 10000 | 11740 | 10396 |
| GraL_graL | 684 | 518 | 776 | 918 | 463 | 690 |
| groES_groES | 12496 | 12264 | 12008 | 19634 | 11498 | 14012 |
| SL1344_RS14595_gshA | 1710 | 1889 | 2010 | 1833 | 1504 | 1809 |
| SL1344_RS16010_gshB | 1130 | 1418 | 1279 | 1248 | 1134 | 1127 |
| SL1344_RS05170_gtgD | 17 | 18 | 13 | 16 | 18 | 16 |

| | | | | | | |
|---|---|---|---|---|---|---|
| SL1344_RS05180_gtgE | 258 | 261 | 322 | 381 | 287 | 363 |
| SL1344_RS05185_gtgF | 14 | 13 | 12 | 9 | 14 | 13 |
| SL1344_RS11665_gyrA | 7674 | 8898 | 9037 | 7994 | 6840 | 6862 |
| SL1344_RS13995_head completion/stabilization protein | 6 | 7 | 5 | 1 | 12 | 8 |
| helD_helD | 723 | 871 | 734 | 773 | 926 | 865 |
| hemG_hemG | 350 | 382 | 393 | 399 | 337 | 440 |
| hflB_hflB | 13772 | 15991 | 15629 | 15885 | 13668 | 13625 |
| SL1344_RS22390_hfq | 5695 | 6914 | 6083 | 6285 | 5984 | 5854 |
| SL1344_RS02410_hha | 915 | 1211 | 1188 | 789 | 553 | 693 |
| SL1344_RS02415_Hha toxicity attenuator | 899 | 1147 | 1374 | 783 | 514 | 704 |
| SL1344_RS14890_hilA | 236 | 246 | 230 | 1212 | 860 | 1143 |
| SL1344_RS14845_hilC | 1027 | 1309 | 1190 | 2336 | 2289 | 2597 |
| SL1344_RS14885_hilD | 808 | 848 | 1014 | 2283 | 1619 | 2384 |
| SL1344_RS23135_hilE | 74 | 88 | 74 | 92 | 61 | 111 |
| SL1344_RS21865_HlyD family type I secretion periplasmic adaptor subunit | 33 | 42 | 56 | 156 | 141 | 171 |
| SL1344_RS08745_hns | 21436 | 24849 | 27930 | 28338 | 28096 | 26676 |
| hscB_hscB | 695 | 629 | 745 | 810 | 564 | 815 |
| SL1344_RS08210_hslJ | 151 | 125 | 135 | 218 | 123 | 223 |
| SL1344_RS21405_hupA | 25260 | 25467 | 29304 | 33495 | 21329 | 31452 |
| SL1344_RS21410_hupB | 3971 | 5387 | 4430 | 4933 | 4987 | 4282 |
| hycD_hycD | 34 | 38 | 20 | 35 | 36 | 24 |
| hycE_hycE | 74 | 86 | 70 | 90 | 65 | 63 |
| SL1344_RS06065_hypothetical protein | 128 | 147 | 133 | 505 | 499 | 467 |
| SL1344_RS07940_hypothetical protein | 1022 | 1304 | 1177 | 3332 | 3561 | 3739 |
| SL1344_RS09195_hypothetical protein | 129 | 108 | 160 | 307 | 190 | 376 |
| SL1344_RS09675_hypothetical protein | 259 | 300 | 260 | 1658 | 1555 | 1637 |
| SL1344_RS09745_hypothetical protein | 243 | 202 | 268 | 538 | 292 | 381 |
| SL1344_RS22135_hypothetical protein | 43 | 42 | 40 | 158 | 99 | 154 |
| iacP_iacP | 171 | 108 | 183 | 302 | 252 | 384 |
| SL1344_RS14895_iagB | 176 | 178 | 220 | 894 | 654 | 868 |
| lamB_iamB | 5670 | 5394 | 4536 | 3310 | 1928 | 2998 |
| idnK_idnK | 86 | 58 | 112 | 80 | 60 | 56 |
| SL1344_RS22990_idnT | 196 | 189 | 182 | 175 | 112 | 158 |
| SL1344_RS06655_ihfA | 4982 | 4587 | 4923 | 5470 | 4072 | 5428 |
| SL1344_RS04775_ihfB | 1413 | 1816 | 1626 | 2163 | 1457 | 1692 |
| lldD_ildD | 3170 | 2873 | 3017 | 3267 | 4519 | 4781 |
| ilvM_ilvM | 260 | 255 | 302 | 246 | 180 | 283 |
| SL1344_RS14990_invA | 169 | 187 | 174 | 1172 | 1019 | 1192 |
| SL1344_RS14980_invC | 95 | 115 | 111 | 496 | 465 | 541 |
| SL1344_RS14995_invE | 134 | 133 | 163 | 997 | 828 | 909 |
| SL1344_RS15005_invF | 197 | 227 | 259 | 1482 | 1302 | 1429 |
| SL1344_RS15000_invG | 172 | 185 | 219 | 1327 | 1053 | 1352 |
| InvR_invR | 684 | 1137 | 1337 | 3217 | 3414 | 3012 |
| SL1344_RS14355_iroB | 43 | 41 | 57 | 225 | 101 | 196 |
| SL1344_RS24725_IS91 family transposase | 120 | 180 | 161 | 643 | 1005 | 792 |
| IsrA_isrA | 2 | 2 | 3 | 1 | 3 | 2 |
| IsrB_isrB | 18 | 61 | 38 | 31 | 38 | 24 |
| IsrC_isrC | 49 | 24 | 41 | 57 | 29 | 37 |
| IsrD_isrD | 0 | 0 | 2 | 0 | 0 | 0 |
| IsrE_isrE | 164 | 345 | 168 | 389 | 271 | 150 |
| IsrF_isrF | 93 | 104 | 119 | 332 | 239 | 326 |
| IsrG_isrG | 26 | 39 | 27 | 55 | 69 | 26 |
| IsrH-1_isrH | 12 | 19 | 14 | 28 | 18 | 16 |
| IsrH-2_isrH2 | 12 | 19 | 10 | 28 | 18 | 16 |
| IsrI_isrI | 28 | 28 | 30 | 30 | 29 | 28 |
| IsrJ_isrJ | 54 | 61 | 54 | 85 | 52 | 42 |
| IsrK_isrK | 264 | 531 | 385 | 533 | 559 | 252 |
| IsrL_isrL | 61 | 45 | 46 | 72 | 54 | 43 |
| IsrM_isrM | 13 | 5 | 10 | 9 | 4 | 8 |
| IsrN_isrN | 0 | 1 | 4 | 2 | 0 | 0 |
| IsrO_isrO | 155 | 115 | 225 | 138 | 100 | 174 |
| IsrP_isrP | 61 | 59 | 59 | 82 | 40 | 65 |
| IsrQ_isrQ | 2 | 1 | 2 | 5 | 2 | 2 |

| | | | | | | |
|---|---|---|---|---|---|---|
| IstR-1_istR1 | 19 | 34 | 49 | 31 | 19 | 24 |
| IstR-2_istR2 | 21 | 39 | 55 | 45 | 23 | 29 |
| ksgA_ksgA | 1412 | 1594 | 1659 | 1430 | 1128 | 1491 |
| SL1344_RS08205_ldhA | 448 | 571 | 463 | 927 | 598 | 729 |
| SL1344_RS00585_leuO | 4 | 13 | 22 | 16 | 11 | 10 |
| SL1344_RS21750_lexA | 2545 | 3040 | 3431 | 3619 | 3738 | 3719 |
| ligA_ligA | 1932 | 2327 | 2149 | 1514 | 1630 | 1559 |
| livF_livF | 66 | 74 | 86 | 76 | 82 | 108 |
| lolA_lolA | 1042 | 1058 | 1200 | 1039 | 760 | 1156 |
| lolB_lolB | 260 | 270 | 301 | 337 | 253 | 370 |
| lplA_lplA | 492 | 590 | 474 | 572 | 575 | 527 |
| SL1344_RS06600_lpxR | 94 | 108 | 132 | 481 | 270 | 433 |
| SL1344_RS11955_lrhA | 564 | 688 | 596 | 433 | 419 | 374 |
| SL1344_RS04660_lrp | 4419 | 5050 | 4781 | 4317 | 4289 | 5309 |
| SL1344_RS09120_L-serine ammonia-lyase | 3266 | 3837 | 3382 | 6563 | 5512 | 5787 |
| SL1344_RS20930_lsrR | 473 | 402 | 499 | 479 | 424 | 460 |
| SL1344_RS14590_luxS | 1899 | 2183 | 2334 | 2224 | 1722 | 1976 |
| SL1344_RS09795_lysine-N-methylase | 977 | 1164 | 1012 | 2342 | 2109 | 2567 |
| malF_malF | 1444 | 1690 | 1490 | 1315 | 947 | 1165 |
| malG_malG | 767 | 986 | 809 | 496 | 463 | 515 |
| SL1344_RS18135_malT | 3953 | 3821 | 4079 | 3564 | 3530 | 2932 |
| SL1344_RS07280_malY* | 90 | 91 | 79 | 104 | 93 | 87 |
| mdoD_mdoD | 1352 | 1582 | 1485 | 1985 | 1759 | 1739 |
| mdoG_mdoG | 2893 | 3215 | 3443 | 3479 | 2426 | 3123 |
| SL1344_RS09375_membrane protein | 989 | 961 | 977 | 1953 | 1770 | 1930 |
| SL1344_RS21870_membrane protein | 1720 | 1733 | 1842 | 2640 | 2212 | 2725 |
| SL1344_RS11835_menC | 615 | 685 | 571 | 485 | 514 | 492 |
| SL1344_RS21470_metA | 109 | 146 | 124 | 149 | 128 | 156 |
| SL1344_RS21065_metB | 401 | 412 | 379 | 450 | 429 | 417 |
| SL1344_RS16345_metC | 298 | 289 | 290 | 404 | 291 | 361 |
| SL1344_RS20385_metE | 305 | 300 | 322 | 414 | 375 | 410 |
| SL1344_RS21095_metF | 118 | 140 | 108 | 235 | 166 | 175 |
| metG_metG | 1927 | 2173 | 2263 | 1981 | 1451 | 1718 |
| SL1344_RS21500_metH | 1217 | 1506 | 1201 | 1271 | 1486 | 1400 |
| SL1344_RS18435_methyl-accepting chemotaxis citrate transducer | 2081 | 2408 | 2118 | 8512 | 5482 | 9373 |
| SL1344_RS08105_methyl-accepting chemotaxis protein | 2563 | 3097 | 2878 | 10182 | 8450 | 11512 |
| SL1344_RS16230_methyl-accepting chemotaxis protein | 1785 | 1822 | 1837 | 2586 | 1878 | 2620 |
| SL1344_RS16300_methyl-accepting chemotaxis protein | 1573 | 2218 | 1906 | 6150 | 4195 | 5602 |
| SL1344_RS16625_methyl-accepting chemotaxis protein | 4820 | 6943 | 5693 | 16267 | 13840 | 18658 |
| SL1344_RS21060_metJ | 1178 | 1180 | 1122 | 1561 | 1533 | 1411 |
| SL1344_RS21070_metL | 1203 | 1373 | 1201 | 1230 | 1268 | 1081 |
| metQ_metQ | 1544 | 1785 | 1708 | 1948 | 1350 | 1775 |
| SL1344_RS20380_metR | 63 | 80 | 81 | 106 | 114 | 82 |
| MgrR_mgrR | 418 | 1200 | 676 | 1181 | 733 | 485 |
| MicA_micA | 38 | 78 | 47 | 58 | 86 | 39 |
| MicC_micC | 24 | 33 | 37 | 41 | 30 | 37 |
| MicF_micF | 395 | 621 | 722 | 198 | 224 | 187 |
| SL1344_RS07395_mlc | 548 | 547 | 507 | 674 | 464 | 544 |
| moaA_moaA | 394 | 446 | 389 | 488 | 438 | 402 |
| moaD_moaD | 122 | 195 | 155 | 158 | 119 | 163 |
| moaE_moaE | 217 | 301 | 262 | 274 | 201 | 262 |
| mobA_mobA | 269 | 365 | 290 | 325 | 302 | 294 |
| SL1344_RS24715_mobA | 1336 | 1990 | 1453 | 7720 | 8196 | 7524 |
| SL1344_RS24720_mobC | 336 | 390 | 391 | 1626 | 1905 | 2244 |
| SL1344_RS03955_modA | 1171 | 1735 | 1259 | 3693 | 3990 | 4093 |
| modB_modB | 1033 | 1610 | 1255 | 3360 | 3373 | 3510 |
| SL1344_RS03965_modC | 1253 | 1988 | 1762 | 3814 | 2981 | 4313 |
| SL1344_RS09620_motA | 2359 | 3521 | 2435 | 15899 | 14535 | 18443 |
| motB_motB | 3092 | 4142 | 3360 | 17578 | 13754 | 17476 |
| SL1344_RS09450_msbB | 628 | 710 | 673 | 1270 | 1142 | 1184 |
| Msr_Msr | 59 | 234 | 146 | 152 | 164 | 77 |
| mtlR_mtlR | 547 | 595 | 567 | 643 | 584 | 595 |
| mukB_mukB | 6514 | 7332 | 7224 | 9059 | 8555 | 9214 |

| | | | | | | |
|---|---|---|---|---|---|---|
| murD_murD | 2646 | 2714 | 3090 | 3029 | 2637 | 2676 |
| SL1344_RS00625_murE | 1712 | 1952 | 1617 | 1671 | 1839 | 1648 |
| SL1344_RS09280_N-acetyltransferase | 231 | 198 | 181 | 339 | 326 | 537 |
| napB_napB | 120 | 139 | 120 | 115 | 117 | 127 |
| SL1344_RS08805_narG | 157 | 329 | 234 | 352 | 328 | 388 |
| SL1344_RS08800_narH | 54 | 73 | 71 | 127 | 103 | 89 |
| SL1344_RS08790_narI | 37 | 41 | 41 | 81 | 49 | 59 |
| SL1344_RS08795_narJ | 27 | 22 | 18 | 62 | 39 | 35 |
| SL1344_RS08810_narK | 89 | 165 | 156 | 197 | 161 | 207 |
| SL1344_RS08820_narL | 170 | 151 | 173 | 301 | 215 | 265 |
| nepI_nepI | 63 | 106 | 58 | 89 | 71 | 89 |
| nhaA_nhaA | 679 | 706 | 686 | 745 | 688 | 760 |
| nirD_nirD | 25 | 23 | 29 | 24 | 16 | 20 |
| nlpD_nlpD | 12344 | 13101 | 15758 | 15804 | 12976 | 15439 |
| nrdI_nrdI | 43 | 64 | 62 | 85 | 55 | 58 |
| SL1344_RS21955_nrfB | 93 | 109 | 94 | 465 | 323 | 385 |
| SL1344_RS21960_nrfC | 98 | 130 | 95 | 464 | 384 | 444 |
| SL1344_RS21975_nrfG | 31 | 34 | 31 | 52 | 31 | 48 |
| nudC_nudC | 350 | 324 | 374 | 319 | 282 | 310 |
| nusA_nusA | 4275 | 5044 | 5201 | 4521 | 4463 | 4369 |
| nusB_nusB | 2490 | 3442 | 4351 | 3503 | 2523 | 2736 |
| nusG_nusG | 6129 | 6569 | 7497 | 7771 | 6121 | 6908 |
| obgE_obgE | 647 | 716 | 696 | 829 | 661 | 762 |
| ompL_ompL | 11 | 5 | 9 | 14 | 4 | 22 |
| SL1344_RS18070_ompR | 1036 | 1340 | 961 | 1022 | 1114 | 1035 |
| ompX_ompX | 10566 | 14833 | 14113 | 9629 | 10915 | 9687 |
| OmrA_omrA | 54 | 127 | 113 | 59 | 70 | 44 |
| OmrB_omrB | 40 | 50 | 36 | 37 | 35 | 26 |
| oppB_oppB | 414 | 512 | 501 | 342 | 545 | 524 |
| SL1344_RS14860_orgA | 357 | 424 | 457 | 1863 | 1942 | 1826 |
| OxyS_oxyS | 66 | 100 | 105 | 95 | 131 | 78 |
| pabB_pabB | 546 | 663 | 642 | 1126 | 999 | 1065 |
| pagP_pagP | 635 | 788 | 846 | 834 | 470 | 632 |
| SL1344_RS10525_pduA | 5 | 8 | 7 | 6 | 5 | 13 |
| pduC_pduC | 10 | 14 | 13 | 24 | 5 | 13 |
| pduD_pduD | 1 | 8 | 6 | 13 | 5 | 4 |
| pduE_pduE | 2 | 7 | 6 | 8 | 2 | 8 |
| pdxK_pdxK | 143 | 165 | 128 | 136 | 146 | 158 |
| SL1344_RS06590_pfkB | 1337 | 1335 | 1481 | 1052 | 862 | 1048 |
| SL1344_RS13960_phage virion morphogenesis protein | 10 | 12 | 10 | 8 | 3 | 7 |
| SL1344_RS02035_phoB | 1377 | 1627 | 1415 | 1253 | 1139 | 1228 |
| SL1344_RS11645_phoE | 40475 | 46916 | 48095 | 32004 | 22656 | 29537 |
| SL1344_RS05535_phoH | 516 | 611 | 459 | 695 | 805 | 779 |
| SL1344_RS06080_phoP | 2970 | 3753 | 3606 | 4131 | 3097 | 3490 |
| SL1344_RS06075_phoQ | 2740 | 3307 | 2995 | 3502 | 2847 | 2595 |
| phoR_phoR | 1060 | 1286 | 1277 | 871 | 671 | 795 |
| SL1344_RS13700_phospho-2-dehydro-3-deoxyheptonate aldolase Tyr-sensitive | 341 | 395 | 348 | 278 | 238 | 270 |
| SL1344_RS10625_phsA | 589 | 585 | 623 | 473 | 352 | 456 |
| SL1344_RS10620_phsB | 148 | 126 | 165 | 130 | 95 | 122 |
| SL1344_RS10615_phsC | 338 | 295 | 349 | 303 | 177 | 248 |
| potG_potG | 128 | 129 | 139 | 89 | 111 | 114 |
| SL1344_RS09215_prc | 5347 | 6138 | 6623 | 10614 | 10344 | 10557 |
| SL1344_RS14880_prgH | 231 | 221 | 286 | 1805 | 1474 | 2002 |
| SL1344_RS14875_prgI | 95 | 80 | 115 | 864 | 581 | 854 |
| SL1344_RS14870_prgJ | 183 | 169 | 212 | 1505 | 1125 | 1426 |
| SL1344_RS14865_prgK | 391 | 448 | 491 | 2467 | 1898 | 2470 |
| SL1344_RS09220_proQ | 3725 | 4364 | 4782 | 7858 | 6301 | 6921 |
| SL1344_RS09255_prpA/pphA | 38 | 29 | 36 | 85 | 76 | 74 |
| psiB_psiB | 35 | 36 | 43 | 36 | 50 | 38 |
| psiB_psiB | 31 | 32 | 30 | 27 | 21 | 26 |
| pssA_pssA | 2233 | 2480 | 2646 | 2849 | 2006 | 2380 |
| SL1344_RS11995_pta | 5633 | 8997 | 6817 | 4208 | 7575 | 3865 |
| SL1344_RS15255_ptpS | 274 | 267 | 342 | 192 | 98 | 137 |

| | | | | | | |
|---|---|---|---|---|---|---|
| SL1344_RS02910_PTS mannose/fructose/sorbose transporter subunit IIB | 3 | 6 | 2 | 3 | 4 | 8 |
| SL1344_RS07105_purR | 633 | 690 | 652 | 735 | 527 | 588 |
| putA_putA | 15021 | 14869 | 14433 | 14983 | 15061 | 15431 |
| SL1344_RS23405_putative inner membrane protein | 773 | 750 | 874 | 1239 | 753 | 971 |
| SL1344_RS09445_pyruvate kinase II | 2603 | 3084 | 2674 | 4561 | 4584 | 4819 |
| SL1344_RS16430_qseB | 70 | 79 | 50 | 67 | 77 | 85 |
| SL1344_RS16435_qseC | 138 | 170 | 153 | 144 | 139 | 125 |
| SL1344_RS23490_radA | 929 | 1068 | 1034 | 782 | 815 | 780 |
| RaoN_raoN | 29 | 30 | 41 | 27 | 24 | 32 |
| SL1344_RS12895_ratA | 420 | 567 | 512 | 542 | 589 | 475 |
| rcsA_rcsA | 67 | 65 | 65 | 75 | 59 | 85 |
| SL1344_RS11655_rcsB | 1993 | 2333 | 2531 | 2025 | 1635 | 1808 |
| SL1344_RS11660_rcsC | 990 | 1201 | 1074 | 812 | 856 | 849 |
| SL1344_RS11650_rcsD | 1894 | 2132 | 2067 | 1710 | 1202 | 1539 |
| rcsF_rcsF | 583 | 772 | 760 | 630 | 450 | 635 |
| RdlD_rdlD | 84 | 195 | 98 | 111 | 257 | 58 |
| RdlE_rdlE | 34 | 79 | 32 | 54 | 233 | 46 |
| SL1344_RS14650_recA | 3693 | 3579 | 3086 | 4088 | 4036 | 4788 |
| SL1344_RS13835_recN | 796 | 1053 | 911 | 1322 | 1093 | 1118 |
| SL1344_RS06620_rfc | 429 | 577 | 661 | 481 | 381 | 412 |
| SL1344_RS22270_rflM | 514 | 690 | 505 | 1408 | 1523 | 1303 |
| rhtB_rhtB | 65 | 76 | 47 | 52 | 54 | 73 |
| rimM_rimM | 6681 | 7648 | 9132 | 8583 | 6353 | 7326 |
| rimO_rimO | 575 | 635 | 702 | 838 | 557 | 633 |
| SL1344_RS03220_rlpA | 626 | 660 | 638 | 804 | 541 | 635 |
| SL1344_RS05840_rne | 8017 | 9052 | 9515 | 8758 | 7053 | 8380 |
| rnpB_rnpB | 372573 | 405259 | 448211 | 479591 | 382987 | 413855 |
| RnpB_RnpB | 372573 | 405259 | 448211 | 479591 | 382987 | 413855 |
| SL1344_RS15130_rpoS | 9556 | 9667 | 8535 | 10260 | 12026 | 10570 |
| rprA_rprA | 30 | 82 | 60 | 55 | 79 | 43 |
| rpsN_rpsN | 9374 | 9443 | 10823 | 12507 | 7376 | 8709 |
| rpsV_rpsV | 991 | 1193 | 1293 | 2067 | 1552 | 1731 |
| RseX_rseX | 2 | 3 | 5 | 4 | 7 | 5 |
| rsmD_rsmD | 246 | 337 | 345 | 298 | 223 | 285 |
| SL1344_RS09240_rsmF/yebU | 374 | 424 | 423 | 920 | 706 | 736 |
| SL1344_RS22150_rtsA | 410 | 387 | 453 | 1494 | 997 | 1298 |
| SL1344_RS22145_rtsB | 121 | 120 | 147 | 455 | 366 | 342 |
| rumA_rumA | 1532 | 1691 | 1842 | 1852 | 1461 | 1519 |
| SL1344_RS09480_ruvA | 330 | 466 | 445 | 914 | 762 | 859 |
| SL1344_RS09475_ruvB | 652 | 877 | 767 | 1500 | 1549 | 1467 |
| RybA_rybA | 1860 | 1737 | 2164 | 1973 | 1750 | 2201 |
| RybB_rybB | 80 | 164 | 140 | 122 | 157 | 58 |
| RybD_rybD | 1638 | 2007 | 1764 | 3665 | 2691 | 2954 |
| RydB_rydB | 24 | 19 | 31 | 21 | 41 | 16 |
| RydC_rydC | 112 | 243 | 146 | 303 | 359 | 229 |
| RyeF_ryeF | 112 | 155 | 151 | 326 | 255 | 304 |
| RyfA_ryfA | 631 | 1095 | 951 | 1174 | 1226 | 980 |
| RyfD_ryfD | 1448 | 2826 | 2418 | 2199 | 1650 | 1419 |
| RyhB_ryhB | 117 | 190 | 86 | 185 | 123 | 77 |
| RyjB_ryjB | 24 | 56 | 27 | 80 | 59 | 65 |
| Sar_sar | 115 | 401 | 180 | 300 | 229 | 112 |
| SL1344_RS10635_sbcB | 615 | 789 | 800 | 681 | 508 | 548 |
| SL1344_RS01925_sbmA | 161 | 180 | 179 | 181 | 162 | 206 |
| SL1344_RS10605_sbmC | 1893 | 1718 | 1958 | 2265 | 2378 | 2005 |
| SL1344_RS20885_sbp | 134 | 116 | 148 | 196 | 74 | 134 |
| sdhC_sdhC | 3568 | 3019 | 3917 | 3925 | 2457 | 3068 |
| SL1344_RS09755_sdiA | 963 | 1310 | 1223 | 4282 | 5875 | 5899 |
| SdsR_sdsR | 72 | 150 | 105 | 109 | 161 | 61 |
| sgbH_sgbH | 3 | 8 | 9 | 13 | 12 | 5 |
| SgrS_sgrS | 2030 | 2282 | 2487 | 1916 | 2588 | 2128 |
| SibA_sibA | 216 | 1228 | 592 | 926 | 600 | 356 |
| SibC_sibC | 892 | 2240 | 1504 | 3329 | 2318 | 1334 |
| SibD_sibD | 634 | 2972 | 1353 | 3861 | 2231 | 1040 |

| | | | | | | |
|---|---|---|---|---|---|---|
| SL1344_RS14940_sicA | 60 | 74 | 80 | 894 | 701 | 1033 |
| sifA_sifA | 147 | 169 | 160 | 163 | 74 | 79 |
| sifB_sifB | 9 | 14 | 7 | 21 | 20 | 18 |
| SL1344_RS05355_sigE | 107 | 111 | 148 | 1698 | 1064 | 1594 |
| SL1344_RS21850_siiA | 32 | 23 | 17 | 65 | 56 | 88 |
| SL1344_RS21855_siiB | 33 | 52 | 64 | 129 | 102 | 139 |
| SL1344_RS21875_siiF | 742 | 671 | 867 | 876 | 655 | 772 |
| SL1344_RS01560_sinR | 305 | 336 | 302 | 416 | 373 | 408 |
| SL1344_RS14920_sipA | 192 | 183 | 187 | 1184 | 956 | 1351 |
| SL1344_RS14935_sipB | 236 | 237 | 255 | 2358 | 2275 | 2795 |
| SL1344_RS14930_sipC | 272 | 358 | 314 | 4313 | 4216 | 4707 |
| SL1344_RS14925_sipD | 57 | 59 | 69 | 545 | 471 | 691 |
| SL1344_RS09740_sirA | 1936 | 2027 | 2079 | 2924 | 2865 | 2746 |
| SL1344_RS10425_sirohydrochlorin cobaltochelatase | 107 | 71 | 132 | 136 | 42 | 115 |
| SL1344_RS09840_SL1896 | 64 | 64 | 86 | 122 | 85 | 89 |
| SL1344_RS07175_slyA | 1864 | 2071 | 2036 | 1927 | 1963 | 1899 |
| SL1344_RS07110_sodB | 3060 | 3194 | 2622 | 2567 | 3273 | 3253 |
| SL1344_RS10630_sopA | 45 | 50 | 49 | 239 | 194 | 269 |
| sopB_sopB | 121 | 103 | 131 | 1732 | 1271 | 1746 |
| SL1344_RS15235_sopD | 71 | 71 | 74 | 161 | 175 | 184 |
| SL1344_RS13925_sopE | 104 | 82 | 90 | 1601 | 1110 | 1916 |
| SL1344_RS09265_sopE2 | 23 | 15 | 28 | 637 | 559 | 688 |
| spaK_spaK | 33 | 31 | 48 | 254 | 204 | 262 |
| spaM_spaM | 49 | 38 | 37 | 316 | 345 | 383 |
| spaN_spaN | 86 | 80 | 77 | 521 | 433 | 540 |
| SL1344_RS14965_spaO | 69 | 59 | 70 | 405 | 432 | 444 |
| SL1344_RS14960_spaP | 59 | 42 | 70 | 363 | 296 | 351 |
| SL1344_RS14955_spaQ | 46 | 45 | 64 | 266 | 301 | 238 |
| SL1344_RS14950_spaR | 53 | 22 | 43 | 213 | 206 | 231 |
| SL1344_RS14945_spaS | 53 | 43 | 53 | 196 | 188 | 215 |
| Spf_spf | 3341 | 8401 | 7952 | 4228 | 5186 | 5161 |
| SL1344_RS14840_spi4 transcriptional regulator | 227 | 310 | 326 | 663 | 622 | 780 |
| spiC_spiC | 2 | 6 | 4 | 0 | 0 | 2 |
| spr_spr | 11842 | 11931 | 11336 | 13402 | 10913 | 9710 |
| SraA_sraA | 1 | 2 | 10 | 9 | 5 | 6 |
| SraB_sraB | 224 | 256 | 298 | 368 | 194 | 245 |
| SraC_sraC | 1755 | 1653 | 1888 | 3763 | 4755 | 3895 |
| SraF_sraF | 526 | 466 | 791 | 585 | 544 | 324 |
| SraG_sraG | 31 | 27 | 21 | 25 | 20 | 25 |
| SraL_sraL | 74 | 106 | 87 | 124 | 89 | 72 |
| SL1344_RS07945_srfB | 1440 | 1802 | 1476 | 4349 | 4792 | 4482 |
| SL1344_RS07950_srfC | 1136 | 1337 | 1224 | 3405 | 3361 | 3523 |
| srlA_srlA | 124 | 118 | 142 | 127 | 106 | 134 |
| srlE_srlE | 157 | 141 | 133 | 147 | 147 | 157 |
| srlR_srlR | 616 | 668 | 696 | 633 | 553 | 518 |
| SroA_sroA | 60 | 84 | 50 | 65 | 56 | 38 |
| SroC_sroC | 27420 | 23158 | 32368 | 41993 | 30709 | 31817 |
| SroD_sroD | 114 | 97 | 90 | 201 | 304 | 231 |
| SroG_sroG | 3398 | 3937 | 2351 | 3835 | 4704 | 3769 |
| SL1344_RS06985_ssaG | 2 | 8 | 8 | 5 | 0 | 3 |
| SL1344_RS07050_ssaR | 43 | 43 | 49 | 32 | 27 | 27 |
| SL1344_RS05160_sseI | 62 | 56 | 74 | 58 | 57 | 47 |
| sspA_sspA | 2633 | 3305 | 3055 | 3137 | 2652 | 2760 |
| SL1344_RS06915_ssrA | 109 | 126 | 90 | 130 | 81 | 121 |
| SsrA_ssrA | 357872 | 351838 | 371130 | 382959 | 319759 | 354848 |
| SL1344_RS06910_ssrB | 62 | 56 | 63 | 52 | 33 | 58 |
| ssrS_ssrS | 54360 | 118559 | 142451 | 103175 | 82605 | 65706 |
| SsrS_ssrS | 54347 | 118528 | 142438 | 103156 | 82590 | 65692 |
| SL1344_RS16670_sstT | 2051 | 1625 | 1560 | 1757 | 2142 | 1514 |
| SL1344_RS01735_stbB | 6 | 2 | 3 | 5 | 5 | 8 |
| SL1344_RS05145_stfT | 42 | 56 | 44 | 78 | 65 | 81 |
| STnc3140_STnc3140 (sopD) | 35 | 40 | 30 | 40 | 64 | 24 |
| STnc3200_STnc3200 | 3 | 2 | 4 | 3 | 10 | 4 |

| | | | | | | |
|---|---|---|---|---|---|---|
| SL1344_RS15350_syd | 465 | 442 | 431 | 719 | 483 | 409 |
| SymR_symR | 28 | 32 | 46 | 30 | 37 | 25 |
| SL1344_RS13990_tail protein | 0 | 0 | 6 | 0 | 1 | 1 |
| tatE_tatE | 388 | 353 | 320 | 349 | 346 | 395 |
| tcuB_tcuB | 42 | 54 | 47 | 55 | 69 | 82 |
| SL1344_RS16770_tdcA | 438 | 305 | 407 | 359 | 239 | 186 |
| SL1344_RS16765_tdcB | 185 | 171 | 139 | 286 | 190 | 216 |
| SL1344_RS16760_tdcC | 302 | 322 | 266 | 658 | 389 | 549 |
| SL1344_RS05150_tfaT | 40 | 67 | 51 | 69 | 99 | 82 |
| Tff_tff | 3708 | 3425 | 4569 | 5356 | 2840 | 4195 |
| thiP_thiP | 399 | 450 | 380 | 457 | 346 | 345 |
| thiQ_thiQ | 150 | 179 | 212 | 206 | 147 | 162 |
| SL1344_RS06625_thrS | 7791 | 8687 | 8601 | 7141 | 5989 | 6561 |
| SL1344_RS04595_tnpA1 | 5 | 3 | 5 | 58 | 76 | 83 |
| SL1344_RS06060_tnpA2 | 24 | 11 | 18 | 38 | 32 | 42 |
| SL1344_RS09075_tnpA3 | 4 | 6 | 4 | 31 | 23 | 34 |
| SL1344_RS09790_tnpA4 | 1 | 3 | 1 | 5 | 8 | 13 |
| SL1344_RS12685_tnpA5 | 9 | 5 | 9 | 13 | 11 | 16 |
| SL1344_RS17020_tnpA6 | 8 | 29 | 15 | 27 | 33 | 43 |
| SL1344_RS17955_tnpA7 | 20 | 12 | 21 | 14 | 19 | 41 |
| tolA_tolA | 830 | 909 | 928 | 684 | 766 | 889 |
| SL1344_RS16475_tolC | 2879 | 3740 | 3453 | 3089 | 2362 | 2860 |
| SL1344_RS21860_TolC family protein | 40 | 51 | 55 | 138 | 90 | 133 |
| tqsA_tqsA | 98 | 99 | 97 | 105 | 80 | 101 |
| SL1344_RS14145_transcriptional regulator | 7 | 8 | 2 | 1 | 4 | 3 |
| trkD_trkD | 806 | 811 | 899 | 798 | 748 | 706 |
| SL1344_RS06175_tRNA arg | 6 | 7 | 4 | 26 | 34 | 22 |
| SL1344_RS13600_tRNA/rRNA methyltransferase | 831 | 843 | 939 | 606 | 564 | 563 |
| truA_truA | 372 | 477 | 414 | 371 | 287 | 296 |
| SL1344_RS23260_tsr | 7588 | 9743 | 8272 | 23477 | 16999 | 22293 |
| SL1344_RS06870_ttrA | 56 | 80 | 67 | 75 | 59 | 55 |
| SL1344_RS06880_ttrB | 2 | 8 | 6 | 9 | 4 | 3 |
| SL1344_RS06875_ttrC | 5 | 11 | 14 | 8 | 4 | 9 |
| SL1344_RS06890_ttrR | 367 | 351 | 342 | 277 | 344 | 326 |
| SL1344_RS06885_ttrS | 1180 | 1257 | 1231 | 1127 | 1257 | 1040 |
| tyrA_tyrA | 507 | 538 | 549 | 425 | 387 | 416 |
| ubiB_ubiB | 2205 | 2522 | 2680 | 2203 | 2342 | 2337 |
| ubiF_ubiF | 495 | 518 | 520 | 530 | 573 | 531 |
| SL1344_RS21190_udhA | 2984 | 3039 | 3166 | 2703 | 3160 | 2767 |
| SL1344_RS03635_UDP-galactopyranose mutase | 10 | 11 | 15 | 13 | 11 | 18 |
| ugpC_ugpC | 173 | 193 | 175 | 131 | 156 | 166 |
| ulaD_ulaD | 29 | 36 | 31 | 46 | 37 | 54 |
| ushA_ushA | 2225 | 2239 | 2347 | 1931 | 2061 | 2077 |
| SL1344_RS09735_uvrC | 1666 | 1836 | 1823 | 2397 | 2858 | 2534 |
| SL1344_RS11530_virulence protein MsgA | 1 | 2 | 7 | 1 | 6 | 2 |
| wecC_wecC | 1279 | 1652 | 1673 | 1344 | 1138 | 1241 |
| SL1344_RS10695_wzzB | 1969 | 2106 | 1856 | 837 | 686 | 770 |
| SL1344_RS02345_ybaO | 295 | 396 | 460 | 340 | 375 | 434 |
| SL1344_RS03185_ybeM | 54 | 45 | 57 | 91 | 76 | 128 |
| SL1344_RS04285_ybiK | 940 | 1131 | 1004 | 953 | 991 | 1013 |
| SL1344_RS08990_ycgO | 1007 | 1529 | 1219 | 3808 | 3190 | 3964 |
| SL1344_RS08975_ycgR | 781 | 1037 | 902 | 2015 | 2115 | 2201 |
| SL1344_RS08585_yciK | 837 | 879 | 712 | 994 | 1045 | 921 |
| SL1344_RS08495_yciW | 211 | 209 | 219 | 422 | 189 | 321 |
| SL1344_RS07545_ydeD | 111 | 144 | 123 | 101 | 93 | 111 |
| SL1344_RS06680_ydiV | 146 | 173 | 152 | 101 | 87 | 87 |
| SL1344_RS06595_ydiY | 797 | 690 | 910 | 762 | 479 | 658 |
| SL1344_RS06585_ydiZ | 43 | 64 | 56 | 49 | 39 | 51 |
| SL1344_RS06560_ydjN | 874 | 1058 | 974 | 1527 | 871 | 919 |
| SL1344_RS09415_yebG | 110 | 97 | 101 | 262 | 221 | 326 |
| SL1344_RS09225_yebR | 1705 | 1906 | 2098 | 3346 | 3217 | 3570 |
| SL1344_RS09230_yebS | 360 | 375 | 417 | 722 | 580 | 644 |
| SL1344_RS09760_yecC | 274 | 330 | 298 | 812 | 685 | 824 |

| | | | | | | |
|---|---|---|---|---|---|---|
| SL1344_RS09750_yecF | 746 | 753 | 878 | 1454 | 1097 | 1036 |
| SL1344_RS09765_yecS | 110 | 146 | 168 | 335 | 401 | 399 |
| SL1344_RS09825_yedD | 474 | 618 | 575 | 513 | 563 | 464 |
| SL1344_RS09830_yedE | 173 | 159 | 219 | 116 | 95 | 99 |
| SL1344_RS09835_yedF | 104 | 117 | 120 | 81 | 68 | 63 |
| SL1344_RS09770_yedO | 321 | 342 | 369 | 729 | 805 | 871 |
| SL1344_RS10925_yegD | 114 | 130 | 170 | 82 | 90 | 98 |
| SL1344_RS13585_yfiK | 43 | 47 | 48 | 37 | 38 | 49 |
| SL1344_RS17960_yhfL | 298 | 385 | 425 | 797 | 558 | 678 |
| SL1344_RS18605_yhjH | 1551 | 2350 | 1471 | 6846 | 5654 | 6413 |
| SL1344_RS18650_yhjQ | 514 | 714 | 619 | 286 | 331 | 266 |
| SL1344_RS23255_yjiY | 10822 | 15623 | 12904 | 4492 | 8816 | 7761 |
| SL1344_RS05645_ymdA | 209 | 278 | 253 | 595 | 610 | 669 |
| SL1344_RS05655_ymdC | 641 | 671 | 593 | 1249 | 1150 | 1088 |
| yopJ_yopJ | 285 | 310 | 292 | 445 | 515 | 604 |
| SL1344_RS15335_yqcB | 376 | 380 | 423 | 460 | 376 | 398 |
| SL1344_RS15340_yqcC (next to csrB) | 154 | 177 | 197 | 206 | 193 | 207 |
| YrlA_yrlA | 6 | 12 | 8 | 18 | 9 | 13 |
| YrlB_yrlB | 12 | 14 | 15 | 25 | 19 | 6 |
| zntA_zntA | 774 | 1131 | 949 | 1341 | 1266 | 1025 |
| zntB_zntB | 276 | 315 | 285 | 399 | 374 | 426 |
| zraP_zraP | 30 | 45 | 27 | 36 | 34 | 35 |
| crRNA1_ | 213 | 236 | 224 | 204 | 223 | 206 |
| crRNA2_ | 438 | 494 | 465 | 443 | 443 | 367 |
| ECS021_ | 18 | 10 | 17 | 19 | 23 | 8 |
| ECS173_ | 42 | 53 | 47 | 69 | 45 | 51 |
| Ffs_ | 776 | 3642 | 2706 | 1789 | 3799 | 1386 |
| fimA-trailer_ | 8 | 15 | 11 | 11 | 4 | 11 |
| GO-10702_ | 91 | 102 | 120 | 141 | 84 | 131 |
| GO-10704_ | 1314 | 1095 | 1014 | 1699 | 1142 | 873 |
| Ig1085_ | 54 | 49 | 77 | 43 | 42 | 51 |
| Ig1140_ | 37 | 38 | 36 | 76 | 69 | 53 |
| Ig2335_ | 312 | 394 | 395 | 304 | 265 | 262 |
| Ig959_ | 0 | 0 | 0 | 0 | 1 | 0 |
| IS042_ | 3 | 12 | 10 | 10 | 21 | 9 |
| lysC-leader_ | 68 | 180 | 109 | 119 | 75 | 97 |
| mreB-leader_ | 412 | 423 | 347 | 622 | 499 | 512 |
| pRNA_ | 7 | 11 | 11 | 14 | 6 | 7 |
| RprA_ | 30 | 82 | 60 | 55 | 79 | 43 |
| rpsL-leader_ | 1490 | 1549 | 1835 | 2917 | 1220 | 1699 |
| rpsP-leader_ | 1541 | 1294 | 1759 | 1997 | 1056 | 1561 |
| rrf_ | 2 | 1 | 4 | 1 | 2 | 2 |
| rrf_ | 0 | 11 | 11 | 3 | 4 | 2 |
| rrf_ | 7 | 13 | 14 | 3 | 5 | 3 |
| rrf_ | 1 | 6 | 1 | 0 | 5 | 2 |
| rrf_ | 30 | 38 | 76 | 14 | 39 | 12 |
| rrf_ | 1 | 9 | 5 | 1 | 4 | 1 |
| rrf_ | 1 | 2 | 2 | 0 | 0 | 0 |
| rrf_ | 0 | 1 | 7 | 0 | 1 | 3 |
| RUF_107c.1_ | 327 | 390 | 378 | 356 | 426 | 329 |
| RUF_107c.10_ | 313 | 388 | 352 | 412 | 505 | 368 |
| RUF_107c.102_ | 32 | 32 | 29 | 27 | 33 | 31 |
| RUF_107c.105_ | 214 | 255 | 205 | 238 | 257 | 217 |
| RUF_107c.107_ | 68 | 75 | 93 | 83 | 99 | 63 |
| RUF_107c.108_ | 582 | 1004 | 752 | 378 | 1149 | 429 |
| RUF_107c.111_ | 653 | 934 | 820 | 727 | 783 | 659 |
| RUF_107c.112_ | 413 | 616 | 491 | 439 | 475 | 316 |
| RUF_107c.18_ | 33 | 31 | 22 | 46 | 64 | 29 |
| RUF_107c.19_ | 198 | 248 | 242 | 215 | 185 | 221 |
| RUF_107c.2_ | 35 | 54 | 45 | 57 | 44 | 34 |
| RUF_107c.23_ | 282 | 371 | 300 | 497 | 348 | 270 |
| RUF_107c.24_ | 365 | 410 | 459 | 342 | 369 | 358 |
| RUF_107c.26_ | 10 | 9 | 5 | 8 | 16 | 8 |

| | | | | | | |
|---|---|---|---|---|---|---|
| RUF_107c.30_ | 3 | 1 | 7 | 2 | 4 | 8 |
| RUF_107c.38_ | 102 | 128 | 108 | 96 | 161 | 94 |
| RUF_107c.47_ | 407 | 511 | 341 | 321 | 436 | 209 |
| RUF_107c.50_ | 113 | 141 | 128 | 118 | 164 | 120 |
| RUF_107c.51_ | 131 | 177 | 165 | 157 | 151 | 156 |
| RUF_107c.53_ | 296 | 381 | 344 | 437 | 633 | 416 |
| RUF_107c.56_ | 291 | 406 | 344 | 262 | 310 | 217 |
| RUF_107c.57_ | 69 | 73 | 75 | 81 | 103 | 100 |
| RUF_107c.58_ | 514 | 1113 | 624 | 713 | 1236 | 600 |
| RUF_107c.59_ | 207 | 376 | 298 | 238 | 573 | 205 |
| RUF_107c.63_ | 97 | 233 | 125 | 94 | 330 | 92 |
| RUF_107c.69_ | 168 | 235 | 162 | 200 | 218 | 204 |
| RUF_107c.70_ | 73 | 78 | 103 | 82 | 59 | 47 |
| RUF_107c.72_ | 79 | 136 | 95 | 89 | 156 | 73 |
| RUF_107c.77_ | 89 | 169 | 142 | 154 | 145 | 84 |
| RUF_107c.8_ | 33 | 57 | 53 | 28 | 27 | 20 |
| RUF_107c.81_ | 11 | 19 | 19 | 13 | 21 | 12 |
| RUF_107c.82_ | 741 | 1126 | 916 | 773 | 1085 | 638 |
| RUF_107c.83_ | 9 | 15 | 10 | 9 | 22 | 9 |
| RUF_107c.86_ | 130 | 227 | 184 | 179 | 254 | 166 |
| RUF_107c.89_ | 334 | 494 | 311 | 375 | 881 | 406 |
| RUF_107c.90_ | 32 | 28 | 32 | 25 | 16 | 19 |
| RUF_107c.92_ | 582 | 665 | 614 | 581 | 883 | 697 |
| RUF_107c.94_ | 10 | 21 | 19 | 21 | 10 | 15 |
| RUF_107c.95_ | 940 | 969 | 1131 | 1042 | 815 | 999 |
| RUF_107c.96_ | 82 | 128 | 80 | 120 | 138 | 99 |
| RUF_111_ | 8497 | 7959 | 11375 | 10365 | 7248 | 8100 |
| RUF_121_ | 40 | 60 | 38 | 81 | 130 | 83 |
| RUF_125_ | 442 | 615 | 544 | 2376 | 1865 | 2332 |
| RUF_167_ | 73 | 97 | 106 | 113 | 63 | 76 |
| RUF_178_ | 461 | 700 | 822 | 517 | 204 | 374 |
| RUF_219c_ | 275 | 278 | 280 | 508 | 542 | 593 |
| RUF_220c_ | 55 | 69 | 52 | 212 | 206 | 173 |
| RUF_226_ | 39 | 42 | 37 | 64 | 32 | 38 |
| RUF_247c_ | 447 | 378 | 474 | 502 | 322 | 390 |
| RUF_257c_ | 362 | 374 | 426 | 442 | 318 | 318 |
| RUF_279_ | 957 | 973 | 1163 | 1264 | 718 | 735 |
| RUF_290_ | 894 | 933 | 970 | 864 | 1027 | 962 |
| RUF_321c_ | 1 | 1 | 0 | 1 | 0 | 1 |
| RUF_328c_ | 4 | 2 | 4 | 5 | 1 | 4 |
| RUF_341c.10_ | 83 | 89 | 85 | 96 | 92 | 70 |
| RUF_341c.11_ | 345 | 394 | 458 | 374 | 311 | 302 |
| RUF_341c.2_ | 24 | 48 | 25 | 36 | 35 | 21 |
| SL1344_RS00005_ | 3577 | 3347 | 3138 | 3389 | 3415 | 3818 |
| SL1344_RS00010_ | 569 | 889 | 759 | 729 | 740 | 805 |
| SL1344_RS00015_ | 563 | 688 | 626 | 620 | 540 | 685 |
| SL1344_RS00020_ | 1090 | 1235 | 1001 | 1162 | 1006 | 1181 |
| SL1344_RS00025_ | 503 | 620 | 577 | 640 | 565 | 588 |
| SL1344_RS00030_ | 367 | 530 | 467 | 492 | 421 | 411 |
| SL1344_RS00035_ | 14959 | 17107 | 15412 | 15653 | 14155 | 15555 |
| SL1344_RS00040_ | 719 | 805 | 913 | 802 | 818 | 682 |
| SL1344_RS00045_ | 302 | 324 | 326 | 290 | 209 | 221 |
| SL1344_RS00050_ | 29 | 20 | 19 | 11 | 13 | 16 |
| SL1344_RS00055_ | 20 | 25 | 22 | 19 | 17 | 25 |
| SL1344_RS00065_ | 3158 | 2194 | 2689 | 4036 | 2513 | 2851 |
| SL1344_RS00070_ | 97 | 84 | 92 | 87 | 71 | 93 |
| SL1344_RS00075_ | 1 | 10 | 0 | 5 | 1 | 4 |
| SL1344_RS00080_ | 17 | 8 | 10 | 15 | 16 | 15 |
| SL1344_RS00085_ | 2 | 5 | 6 | 4 | 4 | 6 |
| SL1344_RS00090_ | 170 | 153 | 146 | 210 | 167 | 153 |
| SL1344_RS00095_ | 74 | 121 | 100 | 129 | 144 | 122 |
| SL1344_RS00100_ | 28 | 30 | 19 | 45 | 23 | 28 |
| SL1344_RS00105_ | 33 | 28 | 29 | 26 | 23 | 23 |

| | | | | | | |
|---|---|---|---|---|---|---|
| SL1344_RS00110_ | 10 | 13 | 19 | 17 | 13 | 19 |
| SL1344_RS00115_ | 55 | 57 | 62 | 85 | 45 | 66 |
| SL1344_RS00120_ | 34 | 37 | 29 | 37 | 35 | 37 |
| SL1344_RS00125_ | 16 | 17 | 17 | 36 | 12 | 14 |
| SL1344_RS00130_ | 17 | 17 | 20 | 12 | 15 | 20 |
| SL1344_RS00135_ | 27 | 29 | 27 | 30 | 25 | 18 |
| SL1344_RS00140_ | 62 | 69 | 60 | 83 | 69 | 64 |
| SL1344_RS00145_ | 51 | 52 | 60 | 73 | 63 | 44 |
| SL1344_RS00150_ | 7 | 3 | 4 | 3 | 1 | 3 |
| SL1344_RS00155_ | 20 | 19 | 25 | 20 | 23 | 32 |
| SL1344_RS00160_ | 17 | 14 | 15 | 19 | 11 | 14 |
| SL1344_RS00165_ | 73 | 54 | 70 | 63 | 51 | 81 |
| SL1344_RS00170_ | 125 | 156 | 129 | 142 | 118 | 124 |
| SL1344_RS00175_ | 11 | 4 | 2 | 7 | 4 | 3 |
| SL1344_RS00180_ | 193 | 199 | 220 | 253 | 180 | 229 |
| SL1344_RS00185_ | 112 | 114 | 102 | 98 | 115 | 105 |
| SL1344_RS00190_ | 44 | 55 | 60 | 56 | 56 | 45 |
| SL1344_RS00195_ | 239 | 234 | 276 | 318 | 212 | 212 |
| SL1344_RS00205_ | 520 | 584 | 615 | 727 | 642 | 726 |
| SL1344_RS00210_ | 165 | 143 | 147 | 151 | 155 | 134 |
| SL1344_RS00215_ | 21 | 19 | 23 | 16 | 10 | 15 |
| SL1344_RS00220_ | 22320 | 22498 | 24906 | 25215 | 21459 | 19166 |
| SL1344_RS00225_ | 86 | 62 | 80 | 133 | 87 | 109 |
| SL1344_RS00230_ | 1248 | 1398 | 1237 | 1378 | 1271 | 1480 |
| SL1344_RS00235_ | 5286 | 6230 | 5190 | 5230 | 5532 | 5563 |
| SL1344_RS00240_ | 1302 | 1718 | 1545 | 1374 | 1667 | 1294 |
| SL1344_RS00245_ | 711 | 856 | 808 | 806 | 722 | 818 |
| SL1344_RS00250_ | 1494 | 1674 | 1557 | 1562 | 1480 | 1494 |
| SL1344_RS00255_ | 643 | 622 | 587 | 717 | 777 | 702 |
| SL1344_RS00260_ | 583 | 690 | 688 | 1127 | 810 | 1027 |
| SL1344_RS00270_ | 137 | 164 | 148 | 124 | 116 | 144 |
| SL1344_RS00275_ | 16 | 14 | 11 | 14 | 7 | 13 |
| SL1344_RS00280_ | 7 | 5 | 12 | 9 | 7 | 6 |
| SL1344_RS00285_ | 1 | 2 | 4 | 4 | 2 | 1 |
| SL1344_RS00290_ | 30 | 37 | 43 | 39 | 44 | 45 |
| SL1344_RS00295_ | 26 | 29 | 28 | 22 | 21 | 19 |
| SL1344_RS00300_ | 7 | 3 | 5 | 5 | 4 | 13 |
| SL1344_RS00305_ | 21 | 34 | 18 | 29 | 13 | 31 |
| SL1344_RS00310_ | 40 | 42 | 57 | 40 | 55 | 39 |
| SL1344_RS00315_ | 28 | 23 | 24 | 43 | 34 | 39 |
| SL1344_RS00320_ | 62 | 59 | 51 | 67 | 51 | 86 |
| SL1344_RS00325_ | 931 | 1065 | 1010 | 924 | 963 | 1074 |
| SL1344_RS00330_ | 32 | 26 | 30 | 20 | 13 | 21 |
| SL1344_RS00335_ | 206 | 288 | 254 | 181 | 170 | 176 |
| SL1344_RS00340_ | 779 | 1049 | 925 | 674 | 651 | 603 |
| SL1344_RS00345_ | 213 | 212 | 222 | 214 | 168 | 164 |
| SL1344_RS00350_ | 33 | 31 | 21 | 49 | 21 | 38 |
| SL1344_RS00355_ | 40 | 57 | 42 | 60 | 44 | 55 |
| SL1344_RS00360_ | 56 | 45 | 57 | 73 | 78 | 83 |
| SL1344_RS00365_ | 61 | 65 | 88 | 81 | 113 | 103 |
| SL1344_RS00370_ | 99 | 138 | 145 | 149 | 172 | 173 |
| SL1344_RS00375_ | 13 | 17 | 16 | 16 | 9 | 14 |
| SL1344_RS00380_ | 0 | 4 | 3 | 1 | 4 | 2 |
| SL1344_RS00385_ | 50 | 47 | 24 | 49 | 25 | 26 |
| SL1344_RS00390_ | 19 | 25 | 13 | 26 | 24 | 31 |
| SL1344_RS00395_ | 28 | 31 | 23 | 27 | 26 | 27 |
| SL1344_RS00400_ | 24 | 26 | 27 | 23 | 21 | 14 |
| SL1344_RS00405_ | 115 | 138 | 107 | 144 | 117 | 92 |
| SL1344_RS00410_ | 56 | 40 | 45 | 61 | 30 | 51 |
| SL1344_RS00415_ | 159 | 178 | 198 | 151 | 146 | 205 |
| SL1344_RS00420_ | 76 | 68 | 89 | 62 | 45 | 65 |
| SL1344_RS00425_ | 2 | 6 | 3 | 6 | 1 | 4 |
| SL1344_RS00430_ | 128 | 143 | 163 | 157 | 115 | 115 |

| | | | | | | |
|---|---|---|---|---|---|---|
| SL1344_RS00435_ | 37 | 41 | 39 | 45 | 36 | 33 |
| SL1344_RS00440_ | 198 | 192 | 196 | 161 | 222 | 168 |
| SL1344_RS00450_ | 962 | 1052 | 1124 | 1074 | 1073 | 972 |
| SL1344_RS00455_ | 737 | 946 | 947 | 808 | 699 | 789 |
| SL1344_RS00465_ | 1562 | 2020 | 1778 | 1763 | 1539 | 1453 |
| SL1344_RS00470_ | 3071 | 3863 | 3751 | 2995 | 2720 | 2990 |
| SL1344_RS00475_ | 2950 | 3458 | 2899 | 3047 | 2868 | 3145 |
| SL1344_RS00480_ | 456 | 513 | 540 | 379 | 402 | 417 |
| SL1344_RS00485_ | 284 | 359 | 380 | 460 | 266 | 262 |
| SL1344_RS00490_ | 687 | 789 | 829 | 971 | 585 | 723 |
| SL1344_RS00495_ | 344 | 396 | 344 | 430 | 340 | 371 |
| SL1344_RS00500_ | 29 | 53 | 60 | 63 | 44 | 46 |
| SL1344_RS00505_ | 15 | 19 | 18 | 10 | 7 | 17 |
| SL1344_RS00510_ | 27 | 29 | 37 | 45 | 30 | 42 |
| SL1344_RS00515_ | 45 | 46 | 62 | 42 | 65 | 51 |
| SL1344_RS00520_ | 32 | 18 | 27 | 34 | 45 | 26 |
| SL1344_RS00525_ | 521 | 519 | 467 | 562 | 587 | 409 |
| SL1344_RS00530_ | 437 | 420 | 388 | 399 | 684 | 394 |
| SL1344_RS00545_ | 231 | 266 | 186 | 234 | 220 | 243 |
| SL1344_RS00550_ | 148 | 205 | 165 | 203 | 186 | 169 |
| SL1344_RS00555_ | 1921 | 2057 | 2357 | 1852 | 2151 | 2040 |
| SL1344_RS00560_ | 117 | 132 | 159 | 110 | 87 | 102 |
| SL1344_RS00565_ | 149 | 178 | 170 | 185 | 109 | 161 |
| SL1344_RS00570_ | 170 | 198 | 177 | 193 | 138 | 162 |
| SL1344_RS00575_ | 116 | 130 | 156 | 93 | 100 | 148 |
| SL1344_RS00580_ | 129 | 325 | 199 | 365 | 287 | 193 |
| SL1344_RS00590_ | 199 | 245 | 191 | 278 | 238 | 275 |
| SL1344_RS00595_ | 342 | 297 | 286 | 277 | 248 | 347 |
| SL1344_RS00600_ | 2575 | 2875 | 2647 | 2344 | 2556 | 2167 |
| SL1344_RS00605_ | 3325 | 3720 | 3905 | 3685 | 3067 | 3747 |
| SL1344_RS00610_ | 3494 | 3813 | 3660 | 3552 | 3705 | 3886 |
| SL1344_RS00615_ | 2064 | 2393 | 2194 | 1927 | 2466 | 2133 |
| SL1344_RS00620_ | 8987 | 10956 | 12805 | 15189 | 9019 | 11508 |
| SL1344_RS00630_ | 2149 | 2541 | 2509 | 2342 | 1838 | 2241 |
| SL1344_RS00635_ | 1948 | 2051 | 2243 | 2087 | 1643 | 2111 |
| SL1344_RS00645_ | 2299 | 2021 | 2559 | 2489 | 1876 | 2149 |
| SL1344_RS00650_ | 1722 | 1593 | 1825 | 2151 | 1627 | 1669 |
| SL1344_RS00655_ | 2216 | 2215 | 2112 | 2765 | 2155 | 2160 |
| SL1344_RS00665_ | 1140 | 1048 | 1033 | 1390 | 1259 | 1172 |
| SL1344_RS00670_ | 2531 | 2620 | 2640 | 2999 | 2463 | 2683 |
| SL1344_RS00675_ | 3465 | 3861 | 3505 | 3986 | 3555 | 3610 |
| SL1344_RS00680_ | 8952 | 11175 | 10177 | 12205 | 12388 | 10392 |
| SL1344_RS00685_ | 1093 | 1270 | 1408 | 1382 | 1091 | 1335 |
| SL1344_RS00690_ | 5791 | 6329 | 6284 | 5183 | 4514 | 5169 |
| SL1344_RS00695_ | 135 | 116 | 94 | 118 | 81 | 163 |
| SL1344_RS00700_ | 431 | 534 | 453 | 472 | 442 | 396 |
| SL1344_RS00705_ | 440 | 487 | 543 | 455 | 398 | 493 |
| SL1344_RS00710_ | 519 | 615 | 610 | 664 | 538 | 525 |
| SL1344_RS00715_ | 550 | 607 | 552 | 702 | 680 | 547 |
| SL1344_RS00720_ | 4316 | 4823 | 4462 | 3808 | 2531 | 3707 |
| SL1344_RS00725_ | 17 | 19 | 15 | 27 | 17 | 23 |
| SL1344_RS00730_ | 9 | 14 | 18 | 11 | 9 | 13 |
| SL1344_RS00735_ | 5 | 2 | 4 | 4 | 3 | 1 |
| SL1344_RS00740_ | 387 | 555 | 496 | 399 | 381 | 449 |
| SL1344_RS00745_ | 258 | 367 | 362 | 333 | 336 | 304 |
| SL1344_RS00750_ | 532 | 587 | 562 | 646 | 763 | 551 |
| SL1344_RS00755_ | 114 | 148 | 131 | 125 | 132 | 110 |
| SL1344_RS00760_ | 41 | 41 | 40 | 39 | 45 | 52 |
| SL1344_RS00765_ | 341 | 453 | 372 | 391 | 441 | 370 |
| SL1344_RS00770_ | 1091 | 1195 | 1154 | 1229 | 1111 | 1511 |
| SL1344_RS00775_ | 6477 | 9229 | 7475 | 5541 | 6453 | 6029 |
| SL1344_RS00785_ | 15990 | 19571 | 17528 | 15923 | 18170 | 17306 |
| SL1344_RS00790_ | 27 | 22 | 33 | 39 | 47 | 40 |

| | | | | | | |
|---|---|---|---|---|---|---|
| SL1344_RS00795_ | 23 | 24 | 25 | 25 | 24 | 24 |
| SL1344_RS00800_ | 83 | 69 | 88 | 102 | 69 | 68 |
| SL1344_RS00805_ | 9232 | 9526 | 7406 | 5658 | 6367 | 7489 |
| SL1344_RS00810_ | 805 | 813 | 1049 | 776 | 598 | 812 |
| SL1344_RS00815_ | 797 | 806 | 905 | 787 | 825 | 850 |
| SL1344_RS00820_ | 1808 | 1930 | 1805 | 3753 | 1611 | 1694 |
| SL1344_RS00825_ | 2479 | 2696 | 2465 | 3882 | 2100 | 2461 |
| SL1344_RS00830_ | 2173 | 2228 | 2158 | 2758 | 1139 | 1859 |
| SL1344_RS00835_ | 2677 | 3134 | 2970 | 3347 | 2393 | 2538 |
| SL1344_RS00840_ | 662 | 808 | 965 | 1026 | 735 | 746 |
| SL1344_RS00845_ | 429 | 522 | 577 | 703 | 446 | 512 |
| SL1344_RS00850_ | 222 | 288 | 242 | 259 | 273 | 297 |
| SL1344_RS00855_ | 590 | 1015 | 1027 | 612 | 722 | 793 |
| SL1344_RS00860_ | 290 | 293 | 269 | 355 | 298 | 303 |
| SL1344_RS00865_ | 893 | 1114 | 1060 | 1133 | 919 | 1054 |
| SL1344_RS00870_ | 4015 | 4866 | 4963 | 5316 | 4162 | 4752 |
| SL1344_RS00875_ | 228 | 215 | 225 | 219 | 203 | 235 |
| SL1344_RS00880_ | 341 | 403 | 307 | 244 | 451 | 320 |
| SL1344_RS00885_ | 61 | 53 | 75 | 70 | 86 | 66 |
| SL1344_RS00890_ | 46 | 57 | 35 | 41 | 62 | 54 |
| SL1344_RS00895_ | 6 | 10 | 6 | 7 | 6 | 2 |
| SL1344_RS00900_ | 24 | 29 | 36 | 32 | 17 | 24 |
| SL1344_RS00905_ | 23 | 31 | 41 | 24 | 23 | 41 |
| SL1344_RS00910_ | 292 | 295 | 315 | 373 | 279 | 308 |
| SL1344_RS00915_ | 997 | 1405 | 1288 | 1168 | 1042 | 1171 |
| SL1344_RS00920_ | 455 | 530 | 560 | 530 | 378 | 452 |
| SL1344_RS00925_ | 326 | 436 | 309 | 311 | 353 | 273 |
| SL1344_RS00930_ | 540 | 773 | 659 | 890 | 593 | 548 |
| SL1344_RS00935_ | 2576 | 3027 | 3211 | 3462 | 2396 | 2756 |
| SL1344_RS00940_ | 609 | 610 | 617 | 768 | 592 | 525 |
| SL1344_RS00945_ | 14353 | 14543 | 13275 | 16175 | 15774 | 16764 |
| SL1344_RS00950_ | 337 | 480 | 470 | 435 | 381 | 411 |
| SL1344_RS00955_ | 276 | 352 | 376 | 389 | 416 | 300 |
| SL1344_RS00960_ | 978 | 1074 | 996 | 1140 | 1154 | 1090 |
| SL1344_RS00965_ | 2328 | 2799 | 2631 | 2660 | 2495 | 2588 |
| SL1344_RS00970_ | 1631 | 2044 | 2319 | 4486 | 2625 | 3486 |
| SL1344_RS00975_ | 261 | 354 | 334 | 633 | 463 | 532 |
| SL1344_RS00980_ | 307 | 420 | 443 | 612 | 679 | 544 |
| SL1344_RS00985_ | 539 | 800 | 893 | 1099 | 1063 | 817 |
| SL1344_RS00990_ | 17 | 42 | 28 | 25 | 11 | 13 |
| SL1344_RS00995_ | 48 | 62 | 82 | 69 | 41 | 56 |
| SL1344_RS01000_ | 5 | 24 | 18 | 15 | 18 | 17 |
| SL1344_RS01005_ | 10 | 17 | 19 | 15 | 22 | 17 |
| SL1344_RS01010_ | 26 | 22 | 20 | 41 | 31 | 37 |
| SL1344_RS01015_ | 13 | 19 | 12 | 26 | 25 | 30 |
| SL1344_RS01020_ | 253 | 312 | 326 | 270 | 454 | 273 |
| SL1344_RS01025_ | 1169 | 1493 | 1319 | 1452 | 1173 | 1296 |
| SL1344_RS01030_ | 327 | 397 | 356 | 396 | 358 | 402 |
| SL1344_RS01035_ | 1635 | 1597 | 1379 | 1845 | 2001 | 1992 |
| SL1344_RS01040_ | 351 | 355 | 337 | 415 | 373 | 254 |
| SL1344_RS01045_ | 322 | 424 | 413 | 513 | 394 | 430 |
| SL1344_RS01050_ | 753 | 951 | 982 | 1186 | 985 | 1058 |
| SL1344_RS01055_ | 275 | 344 | 286 | 243 | 253 | 298 |
| SL1344_RS01060_ | 1774 | 2130 | 2263 | 1615 | 1116 | 1411 |
| SL1344_RS01065_ | 555 | 472 | 485 | 594 | 639 | 502 |
| SL1344_RS01070_ | 5820 | 6143 | 5509 | 6462 | 9189 | 6041 |
| SL1344_RS01075_ | 369 | 400 | 417 | 522 | 518 | 419 |
| SL1344_RS01080_ | 3749 | 4787 | 4784 | 4100 | 3603 | 4260 |
| SL1344_RS01085_ | 600 | 742 | 668 | 680 | 548 | 662 |
| SL1344_RS01090_ | 1654 | 1865 | 1994 | 2001 | 1481 | 1919 |
| SL1344_RS01095_ | 20331 | 23602 | 25878 | 27571 | 23940 | 23018 |
| SL1344_RS01100_ | 30295 | 37725 | 37859 | 30668 | 39349 | 30568 |
| SL1344_RS01105_ | 1713 | 1710 | 1837 | 1824 | 1653 | 1555 |

| | | | | | | |
|---|---|---|---|---|---|---|
| SL1344_RS01110_ | 8552 | 8985 | 9784 | 8418 | 7585 | 8638 |
| SL1344_RS01115_ | 1006 | 1166 | 1219 | 1157 | 1096 | 1102 |
| SL1344_RS01120_ | 1106 | 1013 | 915 | 1186 | 1193 | 1218 |
| SL1344_RS01130_ | 4886 | 5602 | 5235 | 5032 | 5574 | 5116 |
| SL1344_RS01135_ | 9388 | 11038 | 10311 | 9609 | 9396 | 9848 |
| SL1344_RS01140_ | 11804 | 12913 | 14725 | 13963 | 10851 | 14131 |
| SL1344_RS01145_ | 8065 | 9792 | 8683 | 9326 | 9755 | 8839 |
| SL1344_RS01150_ | 1276 | 1398 | 1499 | 1681 | 1372 | 1454 |
| SL1344_RS01155_ | 3080 | 3315 | 3618 | 3740 | 3477 | 3916 |
| SL1344_RS01160_ | 3211 | 3879 | 3548 | 3775 | 5331 | 3693 |
| SL1344_RS01165_ | 928 | 1111 | 1168 | 1248 | 1208 | 1048 |
| SL1344_RS01170_ | 2768 | 2889 | 3026 | 3687 | 2884 | 3111 |
| SL1344_RS01175_ | 2066 | 2254 | 2543 | 2437 | 2078 | 2032 |
| SL1344_RS01180_ | 324 | 355 | 339 | 387 | 345 | 350 |
| SL1344_RS01185_ | 666 | 751 | 751 | 588 | 627 | 668 |
| SL1344_RS01190_ | 223 | 256 | 236 | 269 | 251 | 286 |
| SL1344_RS01195_ | 552 | 750 | 613 | 595 | 617 | 539 |
| SL1344_RS01200_ | 1040 | 1679 | 1104 | 1321 | 2060 | 1289 |
| SL1344_RS01205_ | 856 | 972 | 950 | 1003 | 914 | 1149 |
| SL1344_RS01210_ | 217 | 247 | 240 | 238 | 189 | 211 |
| SL1344_RS01215_ | 245 | 257 | 277 | 238 | 211 | 283 |
| SL1344_RS01220_ | 503 | 578 | 571 | 523 | 444 | 547 |
| SL1344_RS01225_ | 3607 | 4093 | 4111 | 4030 | 3298 | 3773 |
| SL1344_RS01230_ | 891 | 1149 | 974 | 921 | 855 | 862 |
| SL1344_RS01245_ | 290 | 294 | 331 | 381 | 277 | 372 |
| SL1344_RS01250_ | 323 | 451 | 355 | 504 | 338 | 354 |
| SL1344_RS01255_ | 233 | 218 | 237 | 194 | 186 | 263 |
| SL1344_RS01260_ | 190 | 258 | 743 | 182 | 238 | 130 |
| SL1344_RS01265_ | 2719 | 3269 | 3640 | 3395 | 3059 | 2937 |
| SL1344_RS01270_ | 2082 | 2300 | 2991 | 2214 | 2283 | 2355 |
| SL1344_RS01275_ | 437 | 487 | 981 | 467 | 611 | 405 |
| SL1344_RS01285_ | 74 | 168 | 104 | 147 | 233 | 76 |
| SL1344_RS01290_ | 279 | 324 | 265 | 379 | 280 | 344 |
| SL1344_RS01295_ | 84 | 135 | 136 | 124 | 145 | 136 |
| SL1344_RS01300_ | 70 | 78 | 72 | 91 | 75 | 82 |
| SL1344_RS01305_ | 588 | 721 | 628 | 581 | 575 | 580 |
| SL1344_RS01310_ | 216 | 228 | 192 | 197 | 228 | 220 |
| SL1344_RS01315_ | 3014 | 2973 | 2985 | 5321 | 3881 | 3531 |
| SL1344_RS01320_ | 202 | 249 | 262 | 245 | 180 | 263 |
| SL1344_RS01325_ | 117 | 211 | 150 | 187 | 135 | 172 |
| SL1344_RS01330_ | 343 | 327 | 360 | 317 | 336 | 335 |
| SL1344_RS01335_ | 577 | 727 | 746 | 653 | 618 | 648 |
| SL1344_RS01340_ | 276 | 517 | 390 | 537 | 691 | 309 |
| SL1344_RS01345_ | 42 | 51 | 53 | 61 | 53 | 33 |
| SL1344_RS01350_ | 22 | 31 | 18 | 36 | 27 | 25 |
| SL1344_RS01355_ | 60 | 74 | 70 | 59 | 62 | 51 |
| SL1344_RS01360_ | 17 | 24 | 14 | 19 | 19 | 19 |
| SL1344_RS01365_ | 50 | 49 | 45 | 46 | 52 | 47 |
| SL1344_RS01370_ | 22 | 22 | 38 | 25 | 13 | 25 |
| SL1344_RS01375_ | 0 | 1 | 1 | 2 | 0 | 0 |
| SL1344_RS01380_ | 103 | 200 | 175 | 145 | 189 | 158 |
| SL1344_RS01385_ | 37 | 41 | 37 | 40 | 25 | 40 |
| SL1344_RS01390_ | 67 | 58 | 66 | 75 | 66 | 70 |
| SL1344_RS01395_ | 32 | 28 | 33 | 29 | 30 | 31 |
| SL1344_RS01400_ | 21 | 11 | 15 | 17 | 16 | 13 |
| SL1344_RS01405_ | 9 | 15 | 12 | 21 | 15 | 18 |
| SL1344_RS01410_ | 10 | 7 | 7 | 2 | 10 | 5 |
| SL1344_RS01415_ | 15 | 28 | 11 | 21 | 10 | 13 |
| SL1344_RS01420_ | 10 | 10 | 11 | 9 | 15 | 15 |
| SL1344_RS01425_ | 33 | 47 | 36 | 44 | 38 | 26 |
| SL1344_RS01430_ | 42 | 51 | 57 | 49 | 61 | 53 |
| SL1344_RS01435_ | 10 | 24 | 22 | 15 | 15 | 18 |
| SL1344_RS01440_ | 5 | 10 | 5 | 6 | 10 | 4 |

| | | | | | | |
|---|---|---|---|---|---|---|
| SL1344_RS01445_ | 165 | 157 | 177 | 147 | 152 | 187 |
| SL1344_RS01450_ | 20 | 28 | 35 | 26 | 31 | 25 |
| SL1344_RS01455_ | 13 | 13 | 13 | 19 | 7 | 21 |
| SL1344_RS01460_ | 33 | 41 | 34 | 49 | 42 | 42 |
| SL1344_RS01465_ | 101 | 60 | 78 | 90 | 80 | 94 |
| SL1344_RS01470_ | 12 | 7 | 8 | 11 | 6 | 7 |
| SL1344_RS01475_ | 308 | 280 | 348 | 344 | 232 | 334 |
| SL1344_RS01480_ | 352 | 397 | 535 | 376 | 312 | 381 |
| SL1344_RS01485_ | 239 | 212 | 238 | 283 | 263 | 275 |
| SL1344_RS01490_ | 125 | 124 | 118 | 182 | 184 | 127 |
| SL1344_RS01495_ | 45 | 72 | 46 | 39 | 51 | 35 |
| SL1344_RS01500_ | 2 | 1 | 4 | 4 | 6 | 0 |
| SL1344_RS01505_ | 25 | 27 | 29 | 32 | 26 | 26 |
| SL1344_RS01515_ | 17 | 23 | 28 | 29 | 33 | 15 |
| SL1344_RS01520_ | 6 | 4 | 2 | 3 | 8 | 1 |
| SL1344_RS01525_ | 65 | 70 | 77 | 106 | 82 | 157 |
| SL1344_RS01530_ | 26 | 42 | 38 | 32 | 25 | 45 |
| SL1344_RS01535_ | 186 | 209 | 201 | 199 | 186 | 234 |
| SL1344_RS01540_ | 53 | 50 | 48 | 65 | 49 | 52 |
| SL1344_RS01545_ | 34 | 47 | 51 | 56 | 36 | 37 |
| SL1344_RS01550_ | 30 | 28 | 28 | 12 | 20 | 21 |
| SL1344_RS01565_ | 31 | 44 | 34 | 51 | 45 | 38 |
| SL1344_RS01570_ | 209 | 227 | 232 | 250 | 164 | 225 |
| SL1344_RS01575_ | 130 | 143 | 179 | 122 | 54 | 132 |
| SL1344_RS01580_ | 187 | 263 | 261 | 254 | 217 | 298 |
| SL1344_RS01590_ | 1024 | 1314 | 1199 | 1041 | 848 | 1085 |
| SL1344_RS01595_ | 772 | 766 | 704 | 737 | 796 | 725 |
| SL1344_RS01600_ | 1189 | 1198 | 1131 | 1292 | 912 | 1080 |
| SL1344_RS01605_ | 365 | 484 | 545 | 616 | 497 | 511 |
| SL1344_RS01610_ | 99 | 101 | 86 | 111 | 107 | 102 |
| SL1344_RS01615_ | 113 | 119 | 116 | 132 | 152 | 126 |
| SL1344_RS01620_ | 3182 | 4476 | 3778 | 3435 | 2985 | 3716 |
| SL1344_RS01625_ | 1313 | 1470 | 1544 | 1588 | 1084 | 1089 |
| SL1344_RS01635_ | 1333 | 1224 | 1087 | 1501 | 1267 | 1461 |
| SL1344_RS01640_ | 82 | 84 | 88 | 81 | 76 | 75 |
| SL1344_RS01645_ | 897 | 1115 | 948 | 878 | 832 | 969 |
| SL1344_RS01650_ | 2326 | 2991 | 2976 | 2225 | 2736 | 2461 |
| SL1344_RS01655_ | 102 | 163 | 115 | 224 | 162 | 91 |
| SL1344_RS01660_ | 18 | 3 | 8 | 16 | 4 | 12 |
| SL1344_RS01665_ | 6 | 5 | 3 | 6 | 9 | 6 |
| SL1344_RS01670_ | 13 | 8 | 16 | 20 | 11 | 14 |
| SL1344_RS01675_ | 2996 | 3159 | 3405 | 2779 | 2672 | 3192 |
| SL1344_RS01680_ | 46 | 32 | 43 | 58 | 37 | 34 |
| SL1344_RS01685_ | 35 | 40 | 42 | 26 | 24 | 29 |
| SL1344_RS01690_ | 20 | 18 | 8 | 13 | 15 | 13 |
| SL1344_RS01695_ | 33 | 43 | 56 | 66 | 53 | 67 |
| SL1344_RS01700_ | 124 | 139 | 120 | 134 | 108 | 151 |
| SL1344_RS01705_ | 175 | 250 | 233 | 210 | 184 | 219 |
| SL1344_RS01710_ | 35 | 40 | 34 | 41 | 19 | 29 |
| SL1344_RS01715_ | 17 | 15 | 13 | 28 | 14 | 22 |
| SL1344_RS01720_ | 80 | 99 | 73 | 95 | 75 | 78 |
| SL1344_RS01725_ | 58 | 53 | 58 | 60 | 34 | 38 |
| SL1344_RS01730_ | 65 | 63 | 78 | 87 | 77 | 69 |
| SL1344_RS01740_ | 2 | 5 | 4 | 1 | 3 | 2 |
| SL1344_RS01745_ | 121 | 139 | 173 | 171 | 138 | 149 |
| SL1344_RS01750_ | 58 | 60 | 64 | 85 | 65 | 68 |
| SL1344_RS01755_ | 114 | 130 | 116 | 101 | 105 | 124 |
| SL1344_RS01760_ | 256 | 305 | 283 | 279 | 304 | 318 |
| SL1344_RS01765_ | 174 | 165 | 149 | 171 | 152 | 176 |
| SL1344_RS01770_ | 9 | 9 | 9 | 6 | 14 | 7 |
| SL1344_RS01775_ | 13 | 20 | 8 | 7 | 3 | 14 |
| SL1344_RS01780_ | 7 | 11 | 14 | 17 | 14 | 11 |
| SL1344_RS01785_ | 68 | 72 | 78 | 65 | 53 | 62 |

| | | | | | | |
|---|---|---|---|---|---|---|
| SL1344_RS01790_ | 24 | 34 | 36 | 27 | 35 | 29 |
| SL1344_RS01795_ | 50 | 59 | 50 | 68 | 65 | 38 |
| SL1344_RS01800_ | 8 | 14 | 9 | 8 | 13 | 6 |
| SL1344_RS01805_ | 67 | 83 | 84 | 85 | 85 | 68 |
| SL1344_RS01810_ | 32 | 37 | 38 | 48 | 41 | 50 |
| SL1344_RS01815_ | 91 | 103 | 116 | 99 | 83 | 81 |
| SL1344_RS01820_ | 27 | 28 | 33 | 24 | 29 | 10 |
| SL1344_RS01825_ | 494 | 484 | 493 | 316 | 478 | 354 |
| SL1344_RS01830_ | 636 | 683 | 666 | 787 | 586 | 739 |
| SL1344_RS01835_ | 1292 | 1475 | 1552 | 1454 | 1291 | 1278 |
| SL1344_RS01840_ | 25 | 28 | 28 | 15 | 21 | 19 |
| SL1344_RS01845_ | 62 | 81 | 74 | 61 | 56 | 47 |
| SL1344_RS01850_ | 213 | 234 | 209 | 261 | 331 | 186 |
| SL1344_RS01855_ | 39 | 32 | 56 | 41 | 41 | 39 |
| SL1344_RS01860_ | 434 | 571 | 446 | 689 | 594 | 545 |
| SL1344_RS01865_ | 153 | 181 | 196 | 192 | 174 | 163 |
| SL1344_RS01870_ | 560 | 574 | 471 | 547 | 439 | 422 |
| SL1344_RS01875_ | 744 | 704 | 803 | 795 | 492 | 753 |
| SL1344_RS01880_ | 2096 | 1727 | 1938 | 1373 | 2703 | 1477 |
| SL1344_RS01885_ | 19 | 19 | 28 | 38 | 46 | 30 |
| SL1344_RS01890_ | 30 | 30 | 39 | 61 | 33 | 36 |
| SL1344_RS01895_ | 64 | 77 | 48 | 58 | 54 | 63 |
| SL1344_RS01900_ | 96 | 92 | 74 | 90 | 91 | 101 |
| SL1344_RS01905_ | 1687 | 2157 | 2245 | 1996 | 1643 | 1819 |
| SL1344_RS01910_ | 470 | 471 | 478 | 541 | 513 | 596 |
| SL1344_RS01915_ | 61 | 54 | 56 | 47 | 44 | 57 |
| SL1344_RS01930_ | 161 | 197 | 196 | 183 | 173 | 156 |
| SL1344_RS01935_ | 33 | 35 | 51 | 34 | 29 | 26 |
| SL1344_RS01940_ | 946 | 844 | 1307 | 998 | 737 | 749 |
| SL1344_RS01945_ | 1107 | 1395 | 1435 | 1307 | 1214 | 1316 |
| SL1344_RS01950_ | 44 | 55 | 44 | 50 | 28 | 66 |
| SL1344_RS01955_ | 48 | 61 | 50 | 62 | 58 | 58 |
| SL1344_RS01960_ | 3332 | 2893 | 3498 | 3616 | 2284 | 3122 |
| SL1344_RS01965_ | 171 | 185 | 192 | 157 | 128 | 129 |
| SL1344_RS01975_ | 1200 | 1409 | 1364 | 1273 | 1278 | 1356 |
| SL1344_RS01980_ | 181 | 194 | 232 | 225 | 207 | 207 |
| SL1344_RS01985_ | 448 | 483 | 475 | 475 | 421 | 419 |
| SL1344_RS01990_ | 507 | 545 | 532 | 474 | 366 | 448 |
| SL1344_RS01995_ | 451 | 472 | 551 | 486 | 430 | 385 |
| SL1344_RS02000_ | 935 | 1234 | 1158 | 996 | 1272 | 1135 |
| SL1344_RS02005_ | 788 | 1027 | 1020 | 932 | 826 | 752 |
| SL1344_RS02010_ | 224 | 271 | 236 | 261 | 310 | 300 |
| SL1344_RS02015_ | 175 | 195 | 196 | 242 | 202 | 199 |
| SL1344_RS02020_ | 648 | 723 | 813 | 968 | 698 | 770 |
| SL1344_RS02025_ | 254 | 252 | 275 | 353 | 280 | 274 |
| SL1344_RS02030_ | 5 | 0 | 7 | 4 | 5 | 2 |
| SL1344_RS02045_ | 1503 | 1628 | 1376 | 2205 | 2121 | 1683 |
| SL1344_RS02050_ | 908 | 1027 | 1123 | 968 | 852 | 911 |
| SL1344_RS02055_ | 157 | 163 | 126 | 145 | 144 | 147 |
| SL1344_RS02060_ | 4827 | 4938 | 4552 | 5296 | 4245 | 5347 |
| SL1344_RS02065_ | 811 | 950 | 1120 | 1143 | 794 | 994 |
| SL1344_RS02070_ | 552 | 624 | 699 | 743 | 599 | 627 |
| SL1344_RS02075_ | 1571 | 1588 | 1694 | 1691 | 1269 | 1372 |
| SL1344_RS02080_ | 2010 | 2072 | 2093 | 2296 | 1716 | 2200 |
| SL1344_RS02085_ | 4648 | 5504 | 5304 | 4852 | 4414 | 4766 |
| SL1344_RS02090_ | 2026 | 1963 | 1761 | 1927 | 2008 | 2096 |
| SL1344_RS02095_ | 96 | 113 | 105 | 106 | 114 | 96 |
| SL1344_RS02100_ | 98 | 139 | 133 | 141 | 176 | 134 |
| SL1344_RS02105_ | 494 | 557 | 663 | 514 | 557 | 683 |
| SL1344_RS02110_ | 671 | 921 | 870 | 1919 | 1333 | 1510 |
| SL1344_RS02115_ | 3962 | 4515 | 3223 | 4449 | 6947 | 5872 |
| SL1344_RS02120_ | 57 | 45 | 49 | 51 | 50 | 87 |
| SL1344_RS02125_ | 425 | 622 | 711 | 558 | 394 | 588 |

| | | | | | | |
|---|---|---|---|---|---|---|
| SL1344_RS02130_ | 687 | 780 | 880 | 905 | 724 | 730 |
| SL1344_RS02135_ | 683 | 824 | 1345 | 1070 | 708 | 731 |
| SL1344_RS02145_ | 288 | 340 | 308 | 322 | 269 | 271 |
| SL1344_RS02150_ | 283 | 268 | 322 | 275 | 190 | 241 |
| SL1344_RS02155_ | 404 | 425 | 376 | 429 | 378 | 351 |
| SL1344_RS02160_ | 1423 | 1703 | 1508 | 1556 | 1516 | 1517 |
| SL1344_RS02165_ | 1150 | 1371 | 1273 | 1380 | 1162 | 1171 |
| SL1344_RS02170_ | 472 | 594 | 615 | 594 | 444 | 554 |
| SL1344_RS02175_ | 798 | 771 | 889 | 799 | 721 | 804 |
| SL1344_RS02180_ | 53 | 81 | 67 | 62 | 79 | 72 |
| SL1344_RS02185_ | 29 | 43 | 27 | 44 | 41 | 25 |
| SL1344_RS02190_ | 55 | 69 | 51 | 90 | 73 | 57 |
| SL1344_RS02195_ | 76 | 64 | 75 | 68 | 52 | 66 |
| SL1344_RS02200_ | 210 | 218 | 198 | 217 | 187 | 221 |
| SL1344_RS02205_ | 31 | 32 | 45 | 29 | 36 | 46 |
| SL1344_RS02210_ | 28 | 24 | 30 | 25 | 25 | 29 |
| SL1344_RS02215_ | 459 | 554 | 482 | 456 | 620 | 449 |
| SL1344_RS02220_ | 257 | 266 | 259 | 250 | 230 | 298 |
| SL1344_RS02225_ | 1524 | 1631 | 1506 | 1750 | 1583 | 1701 |
| SL1344_RS02230_ | 442 | 485 | 466 | 504 | 482 | 501 |
| SL1344_RS02235_ | 121 | 112 | 117 | 119 | 100 | 100 |
| SL1344_RS02240_ | 280 | 317 | 319 | 255 | 235 | 268 |
| SL1344_RS02245_ | 13962 | 14564 | 14023 | 12638 | 12935 | 13404 |
| SL1344_RS02250_ | 2808 | 3222 | 3177 | 2819 | 2615 | 2973 |
| SL1344_RS02255_ | 4669 | 5320 | 4720 | 4474 | 4408 | 4573 |
| SL1344_RS02260_ | 17992 | 18682 | 16886 | 17318 | 15817 | 16611 |
| SL1344_RS02265_ | 19760 | 20781 | 20150 | 19639 | 17347 | 15690 |
| SL1344_RS02270_ | 1062 | 1274 | 1231 | 1206 | 1061 | 1058 |
| SL1344_RS02275_ | 796 | 866 | 1029 | 995 | 595 | 843 |
| SL1344_RS02280_ | 1570 | 1698 | 1724 | 1718 | 1418 | 1680 |
| SL1344_RS02285_ | 17385 | 19573 | 20337 | 18372 | 17343 | 16879 |
| SL1344_RS02300_ | 8057 | 7748 | 8134 | 12270 | 8494 | 9540 |
| SL1344_RS02305_ | 36311 | 33936 | 31910 | 41060 | 37027 | 44253 |
| SL1344_RS02310_ | 2320 | 2581 | 2473 | 2295 | 2050 | 2536 |
| SL1344_RS02315_ | 36 | 23 | 17 | 22 | 35 | 38 |
| SL1344_RS02320_ | 105 | 112 | 97 | 119 | 109 | 92 |
| SL1344_RS02325_ | 532 | 600 | 509 | 543 | 806 | 610 |
| SL1344_RS02330_ | 513 | 543 | 482 | 453 | 769 | 433 |
| SL1344_RS02335_ | 101 | 147 | 139 | 135 | 118 | 142 |
| SL1344_RS02350_ | 513 | 615 | 558 | 538 | 577 | 521 |
| SL1344_RS02355_ | 454 | 518 | 467 | 578 | 471 | 483 |
| SL1344_RS02360_ | 9 | 11 | 12 | 14 | 10 | 7 |
| SL1344_RS02365_ | 102 | 116 | 103 | 137 | 150 | 107 |
| SL1344_RS02370_ | 433 | 495 | 441 | 440 | 459 | 406 |
| SL1344_RS02375_ | 394 | 428 | 477 | 616 | 349 | 513 |
| SL1344_RS02380_ | 177 | 190 | 196 | 196 | 190 | 196 |
| SL1344_RS02385_ | 442 | 509 | 449 | 537 | 388 | 469 |
| SL1344_RS02390_ | 43 | 48 | 59 | 70 | 37 | 62 |
| SL1344_RS02395_ | 158 | 155 | 172 | 178 | 175 | 167 |
| SL1344_RS02400_ | 1867 | 1795 | 1463 | 2516 | 2129 | 2320 |
| SL1344_RS02405_ | 1085 | 1280 | 1332 | 909 | 814 | 778 |
| SL1344_RS02420_ | 5014 | 5808 | 5391 | 4817 | 4205 | 4488 |
| SL1344_RS02425_ | 2016 | 2328 | 2463 | 1910 | 1559 | 1975 |
| SL1344_RS02430_ | 536 | 704 | 672 | 654 | 590 | 700 |
| SL1344_RS02435_ | 1399 | 1520 | 1676 | 1644 | 1303 | 1544 |
| SL1344_RS02440_ | 535 | 578 | 535 | 651 | 544 | 634 |
| SL1344_RS02445_ | 150 | 173 | 127 | 208 | 159 | 181 |
| SL1344_RS02450_ | 270 | 275 | 262 | 298 | 273 | 305 |
| SL1344_RS02455_ | 247 | 283 | 257 | 345 | 252 | 353 |
| SL1344_RS02460_ | 882 | 924 | 960 | 1058 | 836 | 979 |
| SL1344_RS02465_ | 1916 | 2251 | 2320 | 2761 | 2186 | 2032 |
| SL1344_RS02470_ | 2018 | 2438 | 2656 | 2075 | 2036 | 1864 |
| SL1344_RS02475_ | 2505 | 3491 | 3428 | 3437 | 2873 | 2631 |

| | | | | | | |
|---|---|---|---|---|---|---|
| SL1344_RS02480_ | 12416 | 11000 | 12763 | 18098 | 16225 | 11058 |
| SL1344_RS02485_ | 3844 | 4353 | 4471 | 4028 | 3482 | 3135 |
| SL1344_RS02490_ | 369 | 464 | 439 | 563 | 370 | 475 |
| SL1344_RS02495_ | 85 | 122 | 86 | 116 | 127 | 93 |
| SL1344_RS02500_ | 357 | 398 | 361 | 365 | 371 | 381 |
| SL1344_RS02505_ | 824 | 985 | 1005 | 1427 | 1247 | 1240 |
| SL1344_RS02510_ | 400 | 547 | 516 | 427 | 399 | 413 |
| SL1344_RS02520_ | 358 | 430 | 416 | 412 | 360 | 375 |
| SL1344_RS02525_ | 275 | 314 | 301 | 322 | 325 | 365 |
| SL1344_RS02530_ | 121 | 122 | 183 | 104 | 89 | 85 |
| SL1344_RS02540_ | 246 | 279 | 240 | 251 | 205 | 278 |
| SL1344_RS02545_ | 228 | 287 | 228 | 448 | 353 | 368 |
| SL1344_RS02550_ | 212 | 237 | 224 | 358 | 275 | 381 |
| SL1344_RS02555_ | 111 | 170 | 154 | 153 | 101 | 155 |
| SL1344_RS02560_ | 167 | 235 | 209 | 173 | 148 | 196 |
| SL1344_RS02565_ | 2138 | 2136 | 2396 | 3080 | 1717 | 1951 |
| SL1344_RS02570_ | 366 | 462 | 425 | 428 | 353 | 445 |
| SL1344_RS02575_ | 247 | 324 | 276 | 321 | 220 | 301 |
| SL1344_RS02580_ | 128 | 128 | 123 | 107 | 127 | 160 |
| SL1344_RS02585_ | 443 | 495 | 500 | 538 | 508 | 458 |
| SL1344_RS02590_ | 185 | 239 | 228 | 259 | 203 | 258 |
| SL1344_RS02595_ | 123 | 164 | 153 | 157 | 138 | 259 |
| SL1344_RS02600_ | 112 | 133 | 126 | 165 | 152 | 170 |
| SL1344_RS02605_ | 134 | 201 | 215 | 177 | 191 | 202 |
| SL1344_RS02610_ | 655 | 703 | 672 | 753 | 566 | 703 |
| SL1344_RS02615_ | 50 | 63 | 50 | 79 | 48 | 59 |
| SL1344_RS02620_ | 91 | 119 | 97 | 102 | 102 | 103 |
| SL1344_RS02625_ | 707 | 965 | 754 | 881 | 1034 | 887 |
| SL1344_RS02630_ | 33 | 51 | 33 | 29 | 33 | 35 |
| SL1344_RS02635_ | 20 | 25 | 26 | 21 | 20 | 24 |
| SL1344_RS02640_ | 27 | 34 | 28 | 35 | 26 | 32 |
| SL1344_RS02645_ | 33 | 32 | 38 | 31 | 32 | 27 |
| SL1344_RS02650_ | 29 | 23 | 26 | 16 | 11 | 5 |
| SL1344_RS02655_ | 55 | 46 | 66 | 92 | 48 | 56 |
| SL1344_RS02660_ | 14 | 17 | 17 | 22 | 20 | 21 |
| SL1344_RS02665_ | 41 | 51 | 30 | 31 | 35 | 34 |
| SL1344_RS02670_ | 212 | 291 | 249 | 226 | 182 | 229 |
| SL1344_RS02675_ | 41 | 46 | 33 | 42 | 29 | 34 |
| SL1344_RS02680_ | 32 | 27 | 32 | 33 | 36 | 33 |
| SL1344_RS02685_ | 29 | 42 | 48 | 36 | 43 | 43 |
| SL1344_RS02690_ | 37 | 20 | 22 | 21 | 22 | 22 |
| SL1344_RS02695_ | 23 | 16 | 15 | 16 | 20 | 14 |
| SL1344_RS02700_ | 37 | 57 | 61 | 70 | 74 | 65 |
| SL1344_RS02705_ | 381 | 466 | 476 | 552 | 414 | 420 |
| SL1344_RS02710_ | 196 | 190 | 197 | 301 | 176 | 208 |
| SL1344_RS02715_ | 1230 | 1785 | 1538 | 1434 | 1572 | 1273 |
| SL1344_RS02720_ | 5063 | 5877 | 6500 | 5669 | 4027 | 5128 |
| SL1344_RS02730_ | 97 | 118 | 99 | 142 | 135 | 94 |
| SL1344_RS02735_ | 51 | 71 | 61 | 71 | 67 | 67 |
| SL1344_RS02740_ | 44 | 50 | 38 | 47 | 47 | 43 |
| SL1344_RS02745_ | 636 | 718 | 766 | 900 | 826 | 882 |
| SL1344_RS02750_ | 811 | 1074 | 1012 | 1142 | 896 | 962 |
| SL1344_RS02760_ | 9 | 18 | 8 | 10 | 12 | 19 |
| SL1344_RS02765_ | 9 | 12 | 12 | 9 | 18 | 10 |
| SL1344_RS02770_ | 41 | 59 | 53 | 69 | 36 | 63 |
| SL1344_RS02775_ | 41 | 44 | 45 | 59 | 40 | 47 |
| SL1344_RS02790_ | 10 | 43 | 5 | 6 | 5 | 6 |
| SL1344_RS02795_ | 5 | 2 | 4 | 1 | 2 | 2 |
| SL1344_RS02800_ | 18 | 48 | 38 | 36 | 27 | 39 |
| SL1344_RS02805_ | 79 | 142 | 48 | 243 | 131 | 87 |
| SL1344_RS02815_ | 120 | 81 | 102 | 242 | 92 | 116 |
| SL1344_RS02820_ | 49 | 67 | 89 | 133 | 136 | 75 |
| SL1344_RS02825_ | 26 | 15 | 25 | 83 | 32 | 29 |

| | | | | | | |
|---|---|---|---|---|---|---|
| SL1344_RS02830_ | 11 | 13 | 11 | 9 | 12 | 3 |
| SL1344_RS02835_ | 2 | 4 | 7 | 8 | 6 | 1 |
| SL1344_RS02840_ | 164 | 144 | 105 | 137 | 146 | 109 |
| SL1344_RS02845_ | 63 | 95 | 72 | 93 | 70 | 95 |
| SL1344_RS02850_ | 53 | 68 | 74 | 74 | 62 | 83 |
| SL1344_RS02855_ | 4 | 5 | 1 | 6 | 2 | 4 |
| SL1344_RS02860_ | 16 | 16 | 14 | 23 | 17 | 17 |
| SL1344_RS02865_ | 683 | 928 | 599 | 1532 | 1202 | 1835 |
| SL1344_RS02870_ | 433 | 465 | 545 | 662 | 542 | 563 |
| SL1344_RS02875_ | 337 | 474 | 491 | 365 | 270 | 295 |
| SL1344_RS02880_ | 229 | 189 | 176 | 226 | 168 | 239 |
| SL1344_RS02885_ | 621 | 641 | 557 | 593 | 748 | 570 |
| SL1344_RS02890_ | 23 | 32 | 27 | 29 | 25 | 27 |
| SL1344_RS02895_ | 18 | 23 | 11 | 15 | 26 | 16 |
| SL1344_RS02900_ | 7 | 10 | 17 | 21 | 4 | 16 |
| SL1344_RS02905_ | 9 | 8 | 5 | 16 | 8 | 10 |
| SL1344_RS02915_ | 6 | 0 | 5 | 7 | 6 | 7 |
| SL1344_RS02920_ | 913 | 1133 | 1083 | 1071 | 825 | 1035 |
| SL1344_RS02925_ | 126 | 150 | 146 | 183 | 137 | 138 |
| SL1344_RS02930_ | 246 | 364 | 361 | 332 | 296 | 337 |
| SL1344_RS02935_ | 75 | 70 | 76 | 94 | 76 | 81 |
| SL1344_RS02940_ | 75 | 101 | 106 | 91 | 93 | 72 |
| SL1344_RS02945_ | 64 | 52 | 53 | 71 | 55 | 58 |
| SL1344_RS02950_ | 18 | 16 | 16 | 37 | 19 | 18 |
| SL1344_RS02955_ | 190 | 222 | 267 | 398 | 227 | 310 |
| SL1344_RS02960_ | 208 | 183 | 130 | 416 | 257 | 253 |
| SL1344_RS02965_ | 8 | 12 | 19 | 46 | 16 | 28 |
| SL1344_RS02975_ | 647 | 753 | 838 | 626 | 527 | 584 |
| SL1344_RS02980_ | 76 | 104 | 104 | 237 | 145 | 181 |
| SL1344_RS02985_ | 123 | 207 | 267 | 466 | 410 | 346 |
| SL1344_RS02990_ | 135 | 188 | 185 | 425 | 300 | 310 |
| SL1344_RS02995_ | 155 | 177 | 205 | 399 | 333 | 372 |
| SL1344_RS03000_ | 109 | 164 | 179 | 246 | 189 | 170 |
| SL1344_RS03005_ | 136 | 176 | 179 | 303 | 267 | 217 |
| SL1344_RS03020_ | 74 | 118 | 104 | 165 | 140 | 167 |
| SL1344_RS03025_ | 50 | 73 | 61 | 96 | 67 | 83 |
| SL1344_RS03030_ | 3493 | 3210 | 3058 | 3293 | 4149 | 4164 |
| SL1344_RS03035_ | 343 | 318 | 337 | 306 | 413 | 379 |
| SL1344_RS03040_ | 107 | 120 | 95 | 174 | 127 | 109 |
| SL1344_RS03045_ | 135 | 176 | 172 | 236 | 147 | 188 |
| SL1344_RS03050_ | 104 | 99 | 135 | 104 | 94 | 106 |
| SL1344_RS03055_ | 131 | 136 | 159 | 139 | 88 | 102 |
| SL1344_RS03060_ | 184 | 188 | 227 | 183 | 118 | 200 |
| SL1344_RS03065_ | 548 | 631 | 659 | 631 | 514 | 572 |
| SL1344_RS03070_ | 4911 | 4884 | 5493 | 5629 | 4282 | 5707 |
| SL1344_RS03075_ | 1029 | 1109 | 974 | 1174 | 1098 | 1190 |
| SL1344_RS03080_ | 14 | 6 | 4 | 2 | 7 | 7 |
| SL1344_RS03085_ | 104 | 97 | 109 | 96 | 84 | 82 |
| SL1344_RS03090_ | 29 | 42 | 26 | 41 | 58 | 28 |
| SL1344_RS03095_ | 380 | 414 | 440 | 475 | 447 | 487 |
| SL1344_RS03100_ | 4448 | 4579 | 5292 | 6295 | 4326 | 5104 |
| SL1344_RS03105_ | 56 | 70 | 69 | 54 | 60 | 61 |
| SL1344_RS03110_ | 372 | 422 | 374 | 603 | 388 | 485 |
| SL1344_RS03115_ | 1067 | 1393 | 1448 | 1512 | 987 | 1525 |
| SL1344_RS03120_ | 196 | 243 | 240 | 384 | 326 | 284 |
| SL1344_RS03125_ | 6 | 7 | 4 | 8 | 8 | 2 |
| SL1344_RS03130_ | 5 | 7 | 4 | 15 | 12 | 6 |
| SL1344_RS03135_ | 15 | 26 | 15 | 18 | 8 | 12 |
| SL1344_RS03140_ | 7 | 15 | 15 | 11 | 3 | 6 |
| SL1344_RS03145_ | 1 | 4 | 0 | 3 | 3 | 3 |
| SL1344_RS03150_ | 7 | 5 | 21 | 9 | 13 | 5 |
| SL1344_RS03165_ | 63 | 67 | 66 | 69 | 57 | 59 |
| SL1344_RS03175_ | 36095 | 45188 | 50819 | 41938 | 37718 | 36154 |

| | | | | | | |
|---|---|---|---|---|---|---|
| SL1344_RS03180_ | 93 | 102 | 118 | 137 | 108 | 118 |
| SL1344_RS03195_ | 2103 | 2383 | 2402 | 2211 | 1903 | 1870 |
| SL1344_RS03200_ | 185 | 197 | 230 | 119 | 106 | 121 |
| SL1344_RS03205_ | 459 | 598 | 536 | 713 | 473 | 668 |
| SL1344_RS03210_ | 2188 | 1975 | 2626 | 3165 | 1515 | 2480 |
| SL1344_RS03215_ | 1766 | 2020 | 2096 | 2256 | 1630 | 1728 |
| SL1344_RS03225_ | 483 | 493 | 445 | 525 | 439 | 480 |
| SL1344_RS03230_ | 971 | 1052 | 1068 | 1144 | 896 | 998 |
| SL1344_RS03235_ | 790 | 1040 | 1141 | 991 | 853 | 983 |
| SL1344_RS03240_ | 597 | 728 | 874 | 767 | 578 | 793 |
| SL1344_RS03245_ | 223 | 200 | 277 | 272 | 198 | 186 |
| SL1344_RS03250_ | 165 | 189 | 172 | 200 | 154 | 175 |
| SL1344_RS03255_ | 231 | 248 | 253 | 345 | 242 | 233 |
| SL1344_RS03260_ | 343 | 334 | 332 | 369 | 295 | 340 |
| SL1344_RS03265_ | 1100 | 1310 | 1105 | 1374 | 1100 | 1064 |
| SL1344_RS03270_ | 1340 | 1586 | 1626 | 1648 | 1337 | 1334 |
| SL1344_RS03275_ | 4412 | 4953 | 5100 | 5270 | 4121 | 4322 |
| SL1344_RS03280_ | 6 | 5 | 16 | 5 | 4 | 9 |
| SL1344_RS03285_ | 48 | 45 | 41 | 36 | 34 | 55 |
| SL1344_RS03290_ | 66 | 91 | 61 | 97 | 98 | 82 |
| SL1344_RS03295_ | 399 | 463 | 424 | 527 | 447 | 473 |
| SL1344_RS03300_ | 486 | 492 | 384 | 376 | 431 | 356 |
| SL1344_RS03305_ | 50 | 50 | 53 | 55 | 44 | 61 |
| SL1344_RS03310_ | 17 | 18 | 12 | 13 | 15 | 16 |
| SL1344_RS03315_ | 44 | 56 | 41 | 45 | 32 | 53 |
| SL1344_RS03320_ | 20 | 22 | 22 | 28 | 18 | 20 |
| SL1344_RS03325_ | 97 | 98 | 91 | 86 | 112 | 92 |
| SL1344_RS03330_ | 223 | 310 | 251 | 198 | 284 | 219 |
| SL1344_RS03335_ | 823 | 843 | 823 | 893 | 913 | 869 |
| SL1344_RS03340_ | 520 | 653 | 614 | 710 | 606 | 612 |
| SL1344_RS03350_ | 794 | 837 | 743 | 513 | 1275 | 928 |
| SL1344_RS03355_ | 1329 | 1085 | 975 | 860 | 1867 | 1114 |
| SL1344_RS03360_ | 26379 | 19901 | 29746 | 36932 | 25640 | 29214 |
| SL1344_RS03365_ | 1719 | 1885 | 2115 | 2102 | 1728 | 1805 |
| SL1344_RS03370_ | 1817 | 2132 | 2352 | 2671 | 1750 | 2072 |
| SL1344_RS03375_ | 1050 | 1001 | 1085 | 1303 | 878 | 1036 |
| SL1344_RS03380_ | 1403 | 1446 | 1701 | 1953 | 1154 | 1501 |
| SL1344_RS03385_ | 1860 | 2257 | 2336 | 2343 | 1544 | 1948 |
| SL1344_RS03395_ | 55 | 70 | 72 | 88 | 40 | 73 |
| SL1344_RS03400_ | 4171 | 4156 | 2954 | 4560 | 6690 | 2340 |
| SL1344_RS03405_ | 2286 | 2566 | 2214 | 3417 | 2809 | 1830 |
| SL1344_RS03410_ | 2799 | 2513 | 2728 | 3655 | 2585 | 2479 |
| SL1344_RS03415_ | 5183 | 5847 | 4219 | 6201 | 8052 | 3689 |
| SL1344_RS03420_ | 9525 | 9627 | 9132 | 11560 | 12465 | 7773 |
| SL1344_RS03425_ | 9555 | 9714 | 9534 | 11955 | 10136 | 7654 |
| SL1344_RS03430_ | 5785 | 6356 | 6226 | 9236 | 6511 | 4201 |
| SL1344_RS03440_ | 897 | 1263 | 939 | 1100 | 1209 | 893 |
| SL1344_RS03445_ | 803 | 883 | 820 | 1087 | 750 | 857 |
| SL1344_RS03450_ | 1065 | 1128 | 1280 | 1633 | 930 | 1271 |
| SL1344_RS03455_ | 605 | 635 | 704 | 960 | 502 | 589 |
| SL1344_RS03460_ | 4113 | 4095 | 4439 | 4900 | 3187 | 4023 |
| SL1344_RS03465_ | 2549 | 2920 | 2847 | 3490 | 2706 | 2937 |
| SL1344_RS03475_ | 290 | 267 | 362 | 204 | 161 | 182 |
| SL1344_RS03480_ | 104 | 126 | 124 | 133 | 108 | 131 |
| SL1344_RS03490_ | 82 | 92 | 92 | 88 | 110 | 84 |
| SL1344_RS03495_ | 288 | 347 | 289 | 337 | 290 | 278 |
| SL1344_RS03505_ | 2240 | 2990 | 3011 | 2907 | 2177 | 2707 |
| SL1344_RS03510_ | 426 | 294 | 276 | 496 | 361 | 247 |
| SL1344_RS03515_ | 545 | 692 | 678 | 607 | 513 | 570 |
| SL1344_RS03520_ | 1141 | 1318 | 1374 | 1333 | 970 | 1176 |
| SL1344_RS03525_ | 5453 | 6264 | 5967 | 5718 | 5319 | 5088 |
| SL1344_RS03530_ | 122 | 134 | 169 | 84 | 62 | 72 |
| SL1344_RS03535_ | 235 | 294 | 274 | 124 | 76 | 79 |

| | | | | | | |
|---|---|---|---|---|---|---|
| SL1344_RS03540_ | 463 | 572 | 584 | 203 | 159 | 177 |
| SL1344_RS03545_ | 5 | 0 | 3 | 5 | 3 | 3 |
| SL1344_RS03550_ | 218 | 288 | 172 | 291 | 269 | 254 |
| SL1344_RS03555_ | 341 | 421 | 355 | 425 | 420 | 428 |
| SL1344_RS03560_ | 54 | 50 | 56 | 85 | 62 | 59 |
| SL1344_RS03565_ | 29 | 37 | 34 | 48 | 45 | 36 |
| SL1344_RS03570_ | 15 | 14 | 16 | 26 | 18 | 14 |
| SL1344_RS03575_ | 1 | 3 | 0 | 0 | 0 | 2 |
| SL1344_RS03580_ | 200 | 375 | 268 | 297 | 361 | 302 |
| SL1344_RS03585_ | 177 | 189 | 190 | 171 | 138 | 128 |
| SL1344_RS03590_ | 97 | 122 | 100 | 135 | 127 | 115 |
| SL1344_RS03595_ | 518 | 609 | 483 | 540 | 439 | 601 |
| SL1344_RS03600_ | 402 | 456 | 421 | 392 | 451 | 402 |
| SL1344_RS03605_ | 497 | 604 | 577 | 470 | 492 | 621 |
| SL1344_RS03610_ | 555 | 593 | 609 | 561 | 592 | 627 |
| SL1344_RS03615_ | 12 | 10 | 9 | 10 | 6 | 4 |
| SL1344_RS03620_ | 151 | 176 | 170 | 159 | 129 | 121 |
| SL1344_RS03625_ | 9 | 8 | 9 | 5 | 5 | 6 |
| SL1344_RS03630_ | 19 | 23 | 24 | 32 | 33 | 25 |
| SL1344_RS03640_ | 9 | 11 | 11 | 15 | 9 | 11 |
| SL1344_RS03645_ | 13 | 18 | 11 | 14 | 16 | 20 |
| SL1344_RS03650_ | 8 | 10 | 14 | 16 | 7 | 11 |
| SL1344_RS03655_ | 5 | 11 | 18 | 9 | 9 | 11 |
| SL1344_RS03660_ | 27 | 24 | 35 | 42 | 22 | 26 |
| SL1344_RS03665_ | 10 | 8 | 7 | 8 | 18 | 11 |
| SL1344_RS03670_ | 27 | 29 | 25 | 35 | 36 | 37 |
| SL1344_RS03675_ | 40 | 84 | 72 | 88 | 36 | 46 |
| SL1344_RS03680_ | 368 | 414 | 454 | 479 | 406 | 415 |
| SL1344_RS03685_ | 371 | 400 | 370 | 385 | 503 | 391 |
| SL1344_RS03690_ | 15861 | 15537 | 15642 | 18456 | 15869 | 18804 |
| SL1344_RS03695_ | 20 | 8 | 21 | 24 | 6 | 24 |
| SL1344_RS03705_ | 1687 | 1770 | 1830 | 1868 | 1700 | 1639 |
| SL1344_RS03710_ | 6070 | 6180 | 5026 | 5054 | 7474 | 7614 |
| SL1344_RS03715_ | 9925 | 11331 | 9583 | 7010 | 13378 | 10315 |
| SL1344_RS03720_ | 20094 | 20531 | 20912 | 25041 | 22601 | 27680 |
| SL1344_RS03725_ | 17241 | 16662 | 18085 | 22747 | 21120 | 23386 |
| SL1344_RS03730_ | 11044 | 9499 | 10752 | 16242 | 12349 | 16035 |
| SL1344_RS03735_ | 9724 | 8848 | 10186 | 20032 | 11686 | 17672 |
| SL1344_RS03740_ | 905 | 1249 | 1018 | 1929 | 1847 | 1543 |
| SL1344_RS03745_ | 4210 | 5199 | 4580 | 6654 | 3894 | 4610 |
| SL1344_RS03750_ | 1762 | 2140 | 1496 | 2579 | 1534 | 1864 |
| SL1344_RS03755_ | 1099 | 1165 | 1411 | 1402 | 797 | 1228 |
| SL1344_RS03760_ | 1337 | 1878 | 1893 | 1792 | 1639 | 1535 |
| SL1344_RS03765_ | 1163 | 1563 | 1314 | 2013 | 1891 | 1423 |
| SL1344_RS03770_ | 727 | 605 | 690 | 711 | 554 | 649 |
| SL1344_RS03775_ | 1247 | 1584 | 1505 | 1202 | 1181 | 1280 |
| SL1344_RS03780_ | 778 | 1050 | 892 | 814 | 731 | 845 |
| SL1344_RS03790_ | 5271 | 5498 | 6266 | 6135 | 4319 | 5693 |
| SL1344_RS03795_ | 11577 | 13637 | 14130 | 15147 | 11249 | 13134 |
| SL1344_RS03800_ | 6777 | 8621 | 7906 | 8323 | 9019 | 7940 |
| SL1344_RS03805_ | 2896 | 2202 | 2195 | 4120 | 3087 | 2039 |
| SL1344_RS03810_ | 3864 | 3371 | 3329 | 4539 | 4497 | 2545 |
| SL1344_RS03815_ | 1930 | 1598 | 1809 | 2488 | 2098 | 1439 |
| SL1344_RS03820_ | 2098 | 1618 | 1892 | 2423 | 2300 | 1587 |
| SL1344_RS03825_ | 1231 | 1181 | 905 | 1357 | 1685 | 801 |
| SL1344_RS03830_ | 64 | 79 | 58 | 66 | 73 | 56 |
| SL1344_RS03835_ | 125 | 153 | 112 | 142 | 136 | 130 |
| SL1344_RS03840_ | 150 | 185 | 149 | 176 | 128 | 149 |
| SL1344_RS03845_ | 47 | 66 | 63 | 54 | 45 | 43 |
| SL1344_RS03850_ | 1949 | 2453 | 2630 | 1834 | 1684 | 2134 |
| SL1344_RS03855_ | 105 | 108 | 116 | 94 | 100 | 101 |
| SL1344_RS03860_ | 107 | 96 | 86 | 103 | 70 | 104 |
| SL1344_RS03865_ | 260 | 280 | 283 | 310 | 219 | 272 |

| | | | | | | |
|---|---|---|---|---|---|---|
| SL1344_RS03870_ | 45 | 43 | 44 | 45 | 33 | 34 |
| SL1344_RS03875_ | 18 | 5 | 12 | 15 | 13 | 14 |
| SL1344_RS03880_ | 2 | 2 | 3 | 4 | 5 | 2 |
| SL1344_RS03885_ | 36 | 35 | 37 | 35 | 29 | 25 |
| SL1344_RS03890_ | 41 | 42 | 37 | 31 | 24 | 36 |
| SL1344_RS03895_ | 98 | 151 | 122 | 135 | 137 | 108 |
| SL1344_RS03900_ | 61 | 53 | 42 | 49 | 46 | 56 |
| SL1344_RS03905_ | 37 | 39 | 27 | 30 | 36 | 35 |
| SL1344_RS03915_ | 1606 | 2090 | 1998 | 3144 | 1721 | 2479 |
| SL1344_RS03920_ | 1415 | 1415 | 1449 | 2117 | 1638 | 1713 |
| SL1344_RS03925_ | 1013 | 1050 | 990 | 1680 | 1256 | 1295 |
| SL1344_RS03930_ | 972 | 938 | 963 | 1868 | 770 | 1205 |
| SL1344_RS03935_ | 27 | 41 | 30 | 47 | 38 | 30 |
| SL1344_RS03940_ | 722 | 958 | 789 | 806 | 881 | 854 |
| SL1344_RS03945_ | 390 | 441 | 480 | 471 | 364 | 453 |
| SL1344_RS03950_ | 3347 | 3203 | 5138 | 5014 | 3080 | 5643 |
| SL1344_RS03970_ | 363 | 422 | 451 | 363 | 323 | 368 |
| SL1344_RS03975_ | 535 | 587 | 576 | 648 | 502 | 662 |
| SL1344_RS03980_ | 2867 | 2865 | 3104 | 2723 | 2098 | 2247 |
| SL1344_RS03985_ | 715 | 728 | 681 | 589 | 759 | 588 |
| SL1344_RS03990_ | 515 | 509 | 498 | 411 | 735 | 497 |
| SL1344_RS03995_ | 341 | 387 | 403 | 271 | 487 | 354 |
| SL1344_RS04000_ | 761 | 666 | 652 | 491 | 851 | 660 |
| SL1344_RS04005_ | 775 | 771 | 732 | 516 | 1006 | 800 |
| SL1344_RS04010_ | 302 | 386 | 346 | 324 | 358 | 279 |
| SL1344_RS04015_ | 49 | 44 | 44 | 55 | 49 | 44 |
| SL1344_RS04020_ | 40 | 37 | 36 | 34 | 49 | 51 |
| SL1344_RS04025_ | 41 | 34 | 44 | 33 | 40 | 29 |
| SL1344_RS04030_ | 26 | 44 | 26 | 35 | 92 | 30 |
| SL1344_RS04035_ | 18 | 21 | 20 | 23 | 12 | 24 |
| SL1344_RS04040_ | 1003 | 1110 | 987 | 1293 | 1298 | 1363 |
| SL1344_RS04045_ | 14 | 10 | 23 | 33 | 25 | 28 |
| SL1344_RS04050_ | 524 | 554 | 635 | 560 | 577 | 605 |
| SL1344_RS04055_ | 361 | 439 | 382 | 355 | 353 | 306 |
| SL1344_RS04065_ | 239 | 335 | 305 | 326 | 279 | 277 |
| SL1344_RS04070_ | 222 | 260 | 223 | 268 | 214 | 250 |
| SL1344_RS04085_ | 1881 | 2294 | 2134 | 2001 | 2372 | 1722 |
| SL1344_RS04090_ | 11 | 8 | 8 | 8 | 8 | 8 |
| SL1344_RS04095_ | 9 | 14 | 10 | 20 | 10 | 18 |
| SL1344_RS04100_ | 9 | 15 | 13 | 19 | 15 | 15 |
| SL1344_RS04105_ | 84 | 99 | 88 | 98 | 126 | 64 |
| SL1344_RS04110_ | 52 | 52 | 49 | 32 | 24 | 49 |
| SL1344_RS04115_ | 33 | 23 | 16 | 37 | 21 | 32 |
| SL1344_RS04120_ | 1961 | 1792 | 2430 | 2115 | 1626 | 1827 |
| SL1344_RS04125_ | 211 | 289 | 256 | 238 | 210 | 234 |
| SL1344_RS04130_ | 253 | 305 | 295 | 229 | 277 | 260 |
| SL1344_RS04135_ | 373 | 505 | 448 | 351 | 505 | 403 |
| SL1344_RS04140_ | 218 | 285 | 200 | 215 | 289 | 236 |
| SL1344_RS04145_ | 236 | 237 | 216 | 221 | 209 | 178 |
| SL1344_RS04150_ | 171 | 192 | 200 | 155 | 187 | 225 |
| SL1344_RS04160_ | 388 | 474 | 475 | 425 | 407 | 434 |
| SL1344_RS04165_ | 38 | 67 | 64 | 78 | 33 | 58 |
| SL1344_RS04170_ | 12 | 15 | 13 | 19 | 15 | 18 |
| SL1344_RS04175_ | 79 | 71 | 68 | 94 | 66 | 70 |
| SL1344_RS04180_ | 173 | 213 | 243 | 210 | 174 | 228 |
| SL1344_RS04185_ | 138 | 176 | 163 | 188 | 162 | 156 |
| SL1344_RS04195_ | 802 | 816 | 663 | 755 | 949 | 701 |
| SL1344_RS04200_ | 4441 | 5544 | 4810 | 2984 | 4285 | 3476 |
| SL1344_RS04205_ | 1215 | 1274 | 1478 | 1601 | 844 | 1282 |
| SL1344_RS04210_ | 221 | 254 | 232 | 239 | 201 | 203 |
| SL1344_RS04220_ | 409 | 426 | 437 | 409 | 294 | 305 |
| SL1344_RS04225_ | 188 | 190 | 187 | 215 | 223 | 215 |
| SL1344_RS04230_ | 269 | 306 | 252 | 366 | 362 | 343 |

| | | | | | | |
|---|---|---|---|---|---|---|
| SL1344_RS04235_ | 769 | 968 | 965 | 1046 | 717 | 860 |
| SL1344_RS04240_ | 773 | 890 | 975 | 1158 | 885 | 964 |
| SL1344_RS04245_ | 19 | 26 | 28 | 41 | 34 | 30 |
| SL1344_RS04250_ | 5 | 4 | 6 | 5 | 3 | 3 |
| SL1344_RS04255_ | 74 | 94 | 75 | 100 | 79 | 82 |
| SL1344_RS04260_ | 567 | 769 | 732 | 639 | 624 | 604 |
| SL1344_RS04265_ | 40 | 54 | 29 | 42 | 33 | 35 |
| SL1344_RS04270_ | 19 | 15 | 15 | 12 | 19 | 13 |
| SL1344_RS04275_ | 363 | 454 | 456 | 449 | 387 | 395 |
| SL1344_RS04280_ | 353 | 431 | 409 | 482 | 345 | 388 |
| SL1344_RS04290_ | 1197 | 1457 | 1396 | 1158 | 1403 | 1248 |
| SL1344_RS04295_ | 647 | 762 | 804 | 624 | 531 | 745 |
| SL1344_RS04300_ | 563 | 660 | 714 | 566 | 580 | 631 |
| SL1344_RS04305_ | 396 | 488 | 505 | 433 | 414 | 475 |
| SL1344_RS04320_ | 19 | 12 | 14 | 14 | 24 | 16 |
| SL1344_RS04325_ | 30 | 20 | 38 | 18 | 20 | 27 |
| SL1344_RS04330_ | 28 | 52 | 35 | 30 | 30 | 31 |
| SL1344_RS04335_ | 7 | 9 | 12 | 9 | 2 | 11 |
| SL1344_RS04340_ | 67 | 73 | 67 | 81 | 56 | 74 |
| SL1344_RS04345_ | 66 | 50 | 53 | 57 | 38 | 62 |
| SL1344_RS04350_ | 117 | 100 | 109 | 115 | 64 | 93 |
| SL1344_RS04355_ | 130 | 181 | 191 | 167 | 113 | 173 |
| SL1344_RS04360_ | 70 | 85 | 76 | 62 | 65 | 57 |
| SL1344_RS04365_ | 457 | 560 | 516 | 577 | 562 | 599 |
| SL1344_RS04370_ | 706 | 757 | 683 | 585 | 661 | 691 |
| SL1344_RS04375_ | 1107 | 1138 | 1182 | 1258 | 1079 | 971 |
| SL1344_RS04380_ | 481 | 541 | 567 | 371 | 314 | 345 |
| SL1344_RS04385_ | 236 | 238 | 276 | 228 | 259 | 310 |
| SL1344_RS04390_ | 292 | 362 | 408 | 313 | 295 | 316 |
| SL1344_RS04395_ | 345 | 425 | 441 | 387 | 363 | 388 |
| SL1344_RS04400_ | 42 | 87 | 55 | 61 | 57 | 69 |
| SL1344_RS04405_ | 649 | 884 | 952 | 556 | 576 | 465 |
| SL1344_RS04410_ | 39 | 31 | 25 | 20 | 29 | 31 |
| SL1344_RS04415_ | 1830 | 2770 | 2471 | 2772 | 2722 | 2765 |
| SL1344_RS04420_ | 68 | 91 | 98 | 111 | 89 | 86 |
| SL1344_RS04425_ | 160 | 211 | 179 | 180 | 143 | 183 |
| SL1344_RS04430_ | 345 | 326 | 263 | 360 | 331 | 317 |
| SL1344_RS04435_ | 607 | 570 | 511 | 660 | 474 | 497 |
| SL1344_RS04440_ | 157 | 158 | 166 | 137 | 183 | 138 |
| SL1344_RS04450_ | 80 | 71 | 68 | 61 | 78 | 71 |
| SL1344_RS04455_ | 115 | 122 | 88 | 109 | 138 | 135 |
| SL1344_RS04460_ | 147 | 165 | 143 | 210 | 197 | 196 |
| SL1344_RS04465_ | 314 | 463 | 424 | 433 | 348 | 354 |
| SL1344_RS04470_ | 21 | 24 | 20 | 16 | 12 | 23 |
| SL1344_RS04475_ | 11 | 11 | 14 | 10 | 9 | 2 |
| SL1344_RS04480_ | 95 | 101 | 108 | 117 | 95 | 103 |
| SL1344_RS04485_ | 110 | 130 | 144 | 101 | 84 | 97 |
| SL1344_RS04495_ | 641 | 848 | 770 | 785 | 697 | 609 |
| SL1344_RS04500_ | 1006 | 1438 | 1231 | 1219 | 1040 | 1030 |
| SL1344_RS04505_ | 859 | 949 | 937 | 1005 | 745 | 867 |
| SL1344_RS04510_ | 545 | 735 | 775 | 717 | 440 | 682 |
| SL1344_RS04515_ | 204 | 202 | 219 | 213 | 208 | 270 |
| SL1344_RS04520_ | 396 | 478 | 428 | 489 | 460 | 449 |
| SL1344_RS04525_ | 522 | 552 | 480 | 637 | 593 | 514 |
| SL1344_RS04530_ | 382 | 565 | 462 | 507 | 565 | 480 |
| SL1344_RS04535_ | 403 | 524 | 496 | 470 | 500 | 478 |
| SL1344_RS04540_ | 112 | 154 | 172 | 183 | 109 | 137 |
| SL1344_RS04545_ | 148 | 162 | 167 | 190 | 178 | 157 |
| SL1344_RS04550_ | 161 | 183 | 139 | 188 | 179 | 163 |
| SL1344_RS04555_ | 502 | 707 | 541 | 606 | 563 | 631 |
| SL1344_RS04560_ | 438 | 481 | 410 | 577 | 537 | 476 |
| SL1344_RS04565_ | 2132 | 2490 | 2643 | 2751 | 1833 | 2468 |
| SL1344_RS04570_ | 531 | 642 | 490 | 509 | 566 | 581 |

| | | | | | | |
|---|---|---|---|---|---|---|
| SL1344_RS04575_ | 626 | 777 | 646 | 635 | 644 | 710 |
| SL1344_RS04580_ | 2559 | 2742 | 2518 | 5442 | 3739 | 4326 |
| SL1344_RS04585_ | 632 | 807 | 737 | 690 | 762 | 832 |
| SL1344_RS04590_ | 3074 | 3807 | 3398 | 2935 | 3969 | 4091 |
| SL1344_RS04600_ | 34 | 43 | 55 | 49 | 32 | 37 |
| SL1344_RS04605_ | 48 | 36 | 52 | 68 | 51 | 40 |
| SL1344_RS04610_ | 166 | 175 | 133 | 283 | 439 | 115 |
| SL1344_RS04615_ | 94 | 112 | 126 | 130 | 92 | 148 |
| SL1344_RS04620_ | 26 | 29 | 32 | 33 | 40 | 43 |
| SL1344_RS04625_ | 48 | 57 | 31 | 65 | 58 | 73 |
| SL1344_RS04630_ | 5348 | 5298 | 5776 | 7327 | 5340 | 5396 |
| SL1344_RS04635_ | 3766 | 2817 | 3923 | 5164 | 2687 | 3908 |
| SL1344_RS04640_ | 501 | 534 | 466 | 587 | 429 | 427 |
| SL1344_RS04645_ | 744 | 977 | 860 | 1116 | 728 | 738 |
| SL1344_RS04650_ | 886 | 1151 | 1151 | 1299 | 900 | 1047 |
| SL1344_RS04655_ | 3038 | 3578 | 3660 | 3852 | 2885 | 3762 |
| SL1344_RS04665_ | 4145 | 5474 | 5273 | 4883 | 4673 | 4889 |
| SL1344_RS04675_ | 1410 | 1563 | 1646 | 1529 | 1349 | 1508 |
| SL1344_RS04680_ | 6084 | 7216 | 6578 | 5752 | 5761 | 5990 |
| SL1344_RS04700_ | 11 | 11 | 8 | 8 | 16 | 12 |
| SL1344_RS04705_ | 122 | 239 | 196 | 173 | 156 | 192 |
| SL1344_RS04715_ | 1768 | 1850 | 2060 | 1923 | 1470 | 1897 |
| SL1344_RS04720_ | 44 | 57 | 106 | 85 | 52 | 88 |
| SL1344_RS04725_ | 253 | 272 | 308 | 252 | 226 | 226 |
| SL1344_RS04730_ | 2708 | 3559 | 3739 | 3298 | 1921 | 2468 |
| SL1344_RS04740_ | 502 | 594 | 700 | 682 | 428 | 472 |
| SL1344_RS04745_ | 66 | 58 | 76 | 52 | 50 | 71 |
| SL1344_RS04750_ | 1399 | 1951 | 1816 | 1620 | 1293 | 1729 |
| SL1344_RS04755_ | 971 | 1356 | 1149 | 1033 | 1003 | 1173 |
| SL1344_RS04760_ | 198 | 276 | 260 | 220 | 233 | 233 |
| SL1344_RS04765_ | 1524 | 1554 | 1766 | 2158 | 1260 | 1557 |
| SL1344_RS04770_ | 30407 | 36175 | 38529 | 36456 | 32965 | 33261 |
| SL1344_RS04780_ | 99 | 96 | 112 | 138 | 115 | 152 |
| SL1344_RS04785_ | 1259 | 1474 | 1692 | 1746 | 1348 | 1479 |
| SL1344_RS04790_ | 801 | 916 | 919 | 1068 | 977 | 835 |
| SL1344_RS04795_ | 163 | 151 | 160 | 236 | 147 | 175 |
| SL1344_RS04800_ | 57 | 86 | 51 | 74 | 42 | 88 |
| SL1344_RS04805_ | 490 | 634 | 478 | 489 | 605 | 518 |
| SL1344_RS04810_ | 142 | 193 | 131 | 136 | 175 | 161 |
| SL1344_RS04815_ | 121 | 155 | 146 | 175 | 110 | 120 |
| SL1344_RS04820_ | 525 | 557 | 434 | 648 | 765 | 761 |
| SL1344_RS04825_ | 988 | 1004 | 892 | 1323 | 1448 | 1439 |
| SL1344_RS04830_ | 1065 | 1301 | 1304 | 1428 | 1672 | 1691 |
| SL1344_RS04840_ | 1525 | 2027 | 1802 | 2201 | 2140 | 2212 |
| SL1344_RS04845_ | 1167 | 1114 | 1061 | 1268 | 1127 | 998 |
| SL1344_RS04850_ | 1737 | 1835 | 1762 | 1507 | 1583 | 1630 |
| SL1344_RS04855_ | 4289 | 5265 | 5412 | 4181 | 3929 | 3711 |
| SL1344_RS04860_ | 6061 | 5262 | 5047 | 11505 | 7715 | 11317 |
| SL1344_RS04865_ | 6077 | 7234 | 6987 | 6415 | 5381 | 6081 |
| SL1344_RS04870_ | 168 | 161 | 179 | 184 | 175 | 128 |
| SL1344_RS04875_ | 30 | 27 | 39 | 35 | 20 | 27 |
| SL1344_RS04880_ | 38 | 49 | 57 | 63 | 67 | 43 |
| SL1344_RS04885_ | 21 | 20 | 11 | 26 | 15 | 25 |
| SL1344_RS04890_ | 585 | 800 | 710 | 795 | 700 | 704 |
| SL1344_RS04895_ | 896 | 1076 | 987 | 1073 | 1038 | 996 |
| SL1344_RS04900_ | 18 | 13 | 13 | 11 | 12 | 14 |
| SL1344_RS04905_ | 2 | 8 | 3 | 6 | 2 | 3 |
| SL1344_RS04910_ | 16 | 7 | 7 | 20 | 9 | 21 |
| SL1344_RS04915_ | 28 | 49 | 29 | 29 | 26 | 27 |
| SL1344_RS04920_ | 7 | 14 | 14 | 5 | 4 | 2 |
| SL1344_RS04925_ | 6 | 4 | 4 | 2 | 4 | 3 |
| SL1344_RS04930_ | 1055 | 1218 | 1384 | 1399 | 1235 | 1444 |
| SL1344_RS04935_ | 11 | 18 | 15 | 20 | 6 | 16 |

| | | | | | | |
|---|---|---|---|---|---|---|
| SL1344_RS04940_ | 12 | 22 | 13 | 27 | 15 | 18 |
| SL1344_RS04945_ | 48 | 42 | 45 | 54 | 45 | 46 |
| SL1344_RS04950_ | 10 | 7 | 19 | 9 | 7 | 15 |
| SL1344_RS04955_ | 9 | 5 | 6 | 8 | 5 | 7 |
| SL1344_RS04960_ | 79 | 81 | 81 | 65 | 44 | 64 |
| SL1344_RS04965_ | 1110 | 1111 | 1142 | 1024 | 883 | 1041 |
| SL1344_RS04970_ | 146 | 142 | 152 | 170 | 115 | 127 |
| SL1344_RS04975_ | 212 | 216 | 208 | 245 | 186 | 231 |
| SL1344_RS04980_ | 85 | 127 | 106 | 104 | 110 | 55 |
| SL1344_RS04985_ | 1 | 0 | 6 | 4 | 3 | 4 |
| SL1344_RS04990_ | 7 | 9 | 5 | 6 | 12 | 5 |
| SL1344_RS04995_ | 3 | 1 | 5 | 2 | 1 | 2 |
| SL1344_RS05000_ | 25 | 9 | 18 | 28 | 16 | 28 |
| SL1344_RS05005_ | 240 | 253 | 281 | 313 | 235 | 265 |
| SL1344_RS05010_ | 1 | 2 | 2 | 4 | 2 | 1 |
| SL1344_RS05015_ | 91 | 83 | 102 | 84 | 66 | 90 |
| SL1344_RS05020_ | 98 | 87 | 131 | 97 | 72 | 91 |
| SL1344_RS05025_ | 3 | 1 | 0 | 0 | 0 | 6 |
| SL1344_RS05030_ | 24 | 35 | 34 | 25 | 19 | 21 |
| SL1344_RS05035_ | 12 | 8 | 7 | 8 | 12 | 12 |
| SL1344_RS05040_ | 46 | 38 | 50 | 35 | 43 | 35 |
| SL1344_RS05045_ | 60 | 75 | 65 | 90 | 79 | 97 |
| SL1344_RS05050_ | 3 | 3 | 7 | 5 | 7 | 5 |
| SL1344_RS05055_ | 52 | 74 | 69 | 58 | 69 | 83 |
| SL1344_RS05060_ | 99 | 87 | 75 | 114 | 95 | 109 |
| SL1344_RS05065_ | 7 | 5 | 9 | 12 | 11 | 6 |
| SL1344_RS05070_ | 3 | 3 | 6 | 6 | 7 | 7 |
| SL1344_RS05075_ | 11 | 13 | 14 | 13 | 18 | 19 |
| SL1344_RS05080_ | 11 | 7 | 6 | 20 | 16 | 20 |
| SL1344_RS05085_ | 46 | 40 | 42 | 58 | 38 | 43 |
| SL1344_RS05090_ | 28 | 40 | 39 | 30 | 39 | 29 |
| SL1344_RS05095_ | 19 | 22 | 32 | 17 | 14 | 28 |
| SL1344_RS05100_ | 136 | 148 | 152 | 180 | 158 | 187 |
| SL1344_RS05105_ | 12 | 20 | 16 | 17 | 10 | 18 |
| SL1344_RS05110_ | 309 | 249 | 289 | 374 | 267 | 367 |
| SL1344_RS05115_ | 689 | 694 | 846 | 1089 | 502 | 963 |
| SL1344_RS05120_ | 88 | 92 | 80 | 89 | 97 | 62 |
| SL1344_RS05125_ | 51 | 54 | 61 | 53 | 42 | 49 |
| SL1344_RS05130_ | 76 | 61 | 72 | 66 | 54 | 66 |
| SL1344_RS05135_ | 160 | 167 | 162 | 200 | 183 | 186 |
| SL1344_RS05140_ | 19 | 19 | 22 | 33 | 22 | 40 |
| SL1344_RS05165_ | 10 | 15 | 6 | 9 | 7 | 10 |
| SL1344_RS05175_ | 85 | 101 | 81 | 124 | 154 | 120 |
| SL1344_RS05190_ | 2447 | 2680 | 2446 | 2796 | 2381 | 3147 |
| SL1344_RS05195_ | 245 | 277 | 327 | 330 | 204 | 256 |
| SL1344_RS05200_ | 820 | 821 | 973 | 742 | 708 | 613 |
| SL1344_RS05205_ | 173 | 213 | 226 | 207 | 221 | 239 |
| SL1344_RS05210_ | 997 | 977 | 982 | 1253 | 974 | 1030 |
| SL1344_RS05215_ | 1280 | 1310 | 1547 | 1737 | 1211 | 1316 |
| SL1344_RS05220_ | 422 | 497 | 506 | 597 | 498 | 480 |
| SL1344_RS05225_ | 610 | 667 | 635 | 700 | 601 | 636 |
| SL1344_RS05230_ | 549 | 615 | 638 | 587 | 494 | 584 |
| SL1344_RS05235_ | 835 | 1021 | 1025 | 1232 | 1357 | 1142 |
| SL1344_RS05240_ | 2876 | 2737 | 2675 | 4225 | 2919 | 4064 |
| SL1344_RS05245_ | 1285 | 1326 | 1287 | 1516 | 1155 | 1330 |
| SL1344_RS05250_ | 293 | 333 | 303 | 389 | 267 | 364 |
| SL1344_RS05255_ | 53404 | 66117 | 66250 | 49727 | 49551 | 43132 |
| SL1344_RS05260_ | 398 | 457 | 492 | 705 | 458 | 612 |
| SL1344_RS05265_ | 157 | 141 | 165 | 196 | 184 | 171 |
| SL1344_RS05270_ | 382 | 461 | 433 | 472 | 405 | 419 |
| SL1344_RS05275_ | 106 | 116 | 110 | 124 | 98 | 137 |
| SL1344_RS05285_ | 899 | 1082 | 1144 | 1074 | 793 | 948 |
| SL1344_RS05290_ | 40 | 33 | 39 | 45 | 26 | 37 |

| | | | | | | |
|---|---|---|---|---|---|---|
| SL1344_RS05295_ | 696 | 745 | 569 | 596 | 1219 | 736 |
| SL1344_RS05300_ | 1101 | 1038 | 1116 | 1777 | 961 | 1058 |
| SL1344_RS05305_ | 332 | 371 | 444 | 457 | 334 | 361 |
| SL1344_RS05310_ | 78 | 102 | 98 | 89 | 55 | 61 |
| SL1344_RS05315_ | 186 | 185 | 216 | 157 | 133 | 104 |
| SL1344_RS05320_ | 43 | 46 | 29 | 37 | 34 | 35 |
| SL1344_RS05325_ | 996 | 1041 | 1078 | 1257 | 922 | 1082 |
| SL1344_RS05330_ | 992 | 1251 | 1384 | 1531 | 741 | 1352 |
| SL1344_RS05335_ | 855 | 1855 | 1430 | 2036 | 3198 | 1020 |
| SL1344_RS05340_ | 82 | 65 | 59 | 84 | 52 | 67 |
| SL1344_RS05345_ | 16 | 16 | 31 | 50 | 42 | 60 |
| SL1344_RS05350_ | 3 | 1 | 0 | 2 | 3 | 8 |
| SL1344_RS05365_ | 361 | 488 | 489 | 525 | 337 | 306 |
| SL1344_RS05370_ | 16 | 8 | 15 | 15 | 5 | 11 |
| SL1344_RS05375_ | 360 | 337 | 365 | 334 | 250 | 263 |
| SL1344_RS05380_ | 184 | 197 | 218 | 192 | 180 | 206 |
| SL1344_RS05385_ | 169 | 204 | 214 | 219 | 174 | 204 |
| SL1344_RS05390_ | 126 | 144 | 149 | 138 | 107 | 129 |
| SL1344_RS05395_ | 83 | 52 | 95 | 98 | 104 | 91 |
| SL1344_RS05400_ | 35 | 47 | 32 | 51 | 27 | 46 |
| SL1344_RS05405_ | 71 | 83 | 71 | 123 | 78 | 113 |
| SL1344_RS05410_ | 182 | 167 | 178 | 153 | 224 | 151 |
| SL1344_RS05415_ | 153 | 163 | 132 | 121 | 164 | 176 |
| SL1344_RS05420_ | 79 | 82 | 68 | 99 | 84 | 124 |
| SL1344_RS05425_ | 58 | 71 | 66 | 79 | 76 | 116 |
| SL1344_RS05430_ | 48 | 74 | 81 | 67 | 62 | 81 |
| SL1344_RS05435_ | 31 | 67 | 64 | 61 | 67 | 75 |
| SL1344_RS05440_ | 60 | 56 | 70 | 95 | 63 | 92 |
| SL1344_RS05445_ | 41 | 53 | 50 | 68 | 40 | 53 |
| SL1344_RS05450_ | 132 | 113 | 138 | 172 | 142 | 143 |
| SL1344_RS05455_ | 24 | 27 | 24 | 10 | 12 | 9 |
| SL1344_RS05460_ | 367 | 417 | 323 | 397 | 439 | 486 |
| SL1344_RS05465_ | 543 | 620 | 608 | 572 | 485 | 613 |
| SL1344_RS05470_ | 290 | 333 | 346 | 392 | 363 | 275 |
| SL1344_RS05475_ | 97 | 126 | 96 | 128 | 125 | 89 |
| SL1344_RS05480_ | 24 | 27 | 26 | 19 | 30 | 23 |
| SL1344_RS05485_ | 39 | 72 | 61 | 65 | 85 | 65 |
| SL1344_RS05490_ | 1109 | 1158 | 1175 | 1711 | 1104 | 1457 |
| SL1344_RS05495_ | 706 | 580 | 676 | 674 | 411 | 768 |
| SL1344_RS05500_ | 258 | 248 | 282 | 293 | 171 | 255 |
| SL1344_RS05505_ | 95 | 100 | 107 | 171 | 104 | 153 |
| SL1344_RS05510_ | 128 | 129 | 141 | 184 | 152 | 154 |
| SL1344_RS05515_ | 1809 | 1905 | 1941 | 1979 | 1928 | 2209 |
| SL1344_RS05525_ | 3302 | 2935 | 3165 | 2344 | 3130 | 2970 |
| SL1344_RS05530_ | 456 | 475 | 498 | 535 | 604 | 673 |
| SL1344_RS05540_ | 915 | 1187 | 1186 | 1167 | 854 | 1045 |
| SL1344_RS05545_ | 353 | 185 | 272 | 559 | 153 | 293 |
| SL1344_RS05550_ | 732 | 296 | 535 | 1473 | 230 | 452 |
| SL1344_RS05555_ | 297 | 105 | 184 | 420 | 165 | 152 |
| SL1344_RS05560_ | 172 | 67 | 88 | 183 | 54 | 115 |
| SL1344_RS05565_ | 52 | 33 | 52 | 86 | 34 | 53 |
| SL1344_RS05570_ | 182 | 212 | 181 | 255 | 181 | 207 |
| SL1344_RS05580_ | 624 | 711 | 597 | 1242 | 1258 | 460 |
| SL1344_RS05585_ | 1 | 4 | 9 | 7 | 4 | 4 |
| SL1344_RS05590_ | 678 | 678 | 669 | 807 | 684 | 771 |
| SL1344_RS05595_ | 542 | 646 | 601 | 701 | 484 | 632 |
| SL1344_RS05600_ | 349 | 343 | 374 | 356 | 323 | 400 |
| SL1344_RS05605_ | 488 | 482 | 433 | 562 | 527 | 588 |
| SL1344_RS05610_ | 49 | 59 | 56 | 66 | 53 | 60 |
| SL1344_RS05615_ | 9 | 12 | 15 | 8 | 9 | 11 |
| SL1344_RS05620_ | 9 | 16 | 16 | 9 | 13 | 11 |
| SL1344_RS05625_ | 22 | 38 | 32 | 24 | 21 | 28 |
| SL1344_RS05635_ | 9 | 17 | 18 | 22 | 18 | 23 |

| | | | | | | |
|---|---|---|---|---|---|---|
| SL1344_RS05650_ | 267 | 312 | 268 | 688 | 578 | 587 |
| SL1344_RS05660_ | 221 | 295 | 295 | 254 | 187 | 213 |
| SL1344_RS05670_ | 3541 | 4042 | 3733 | 4236 | 3860 | 3674 |
| SL1344_RS05675_ | 67 | 62 | 82 | 77 | 65 | 94 |
| SL1344_RS05680_ | 312 | 263 | 271 | 296 | 218 | 290 |
| SL1344_RS05685_ | 125 | 142 | 120 | 130 | 113 | 106 |
| SL1344_RS05690_ | 739 | 959 | 799 | 952 | 826 | 843 |
| SL1344_RS05695_ | 667 | 651 | 653 | 909 | 603 | 692 |
| SL1344_RS05700_ | 107 | 120 | 116 | 149 | 113 | 131 |
| SL1344_RS05705_ | 43 | 47 | 45 | 55 | 24 | 45 |
| SL1344_RS05710_ | 25 | 26 | 25 | 16 | 13 | 18 |
| SL1344_RS05715_ | 801 | 1090 | 1043 | 821 | 684 | 781 |
| SL1344_RS05730_ | 1405 | 1601 | 1889 | 1782 | 1561 | 1506 |
| SL1344_RS05735_ | 944 | 1281 | 1257 | 1103 | 1098 | 1033 |
| SL1344_RS05740_ | 556 | 730 | 610 | 861 | 609 | 807 |
| SL1344_RS05745_ | 192 | 225 | 225 | 274 | 244 | 293 |
| SL1344_RS05750_ | 662 | 728 | 789 | 672 | 664 | 749 |
| SL1344_RS05755_ | 931 | 1015 | 1022 | 773 | 1035 | 933 |
| SL1344_RS05760_ | 1112 | 1253 | 1219 | 977 | 1272 | 1278 |
| SL1344_RS05765_ | 415 | 518 | 492 | 458 | 529 | 505 |
| SL1344_RS05780_ | 2142 | 2539 | 2332 | 4970 | 4694 | 4443 |
| SL1344_RS05785_ | 1909 | 2127 | 2196 | 4772 | 4250 | 4770 |
| SL1344_RS05790_ | 1478 | 1628 | 1665 | 3472 | 2839 | 3525 |
| SL1344_RS05800_ | 4226 | 4948 | 4766 | 10053 | 8111 | 9564 |
| SL1344_RS05810_ | 5763 | 6439 | 6937 | 12596 | 12011 | 12316 |
| SL1344_RS05815_ | 1347 | 1575 | 1595 | 3127 | 2988 | 3279 |
| SL1344_RS05825_ | 1206 | 1250 | 1204 | 2595 | 2333 | 2731 |
| SL1344_RS05845_ | 4 | 4 | 8 | 3 | 2 | 2 |
| SL1344_RS05850_ | 873 | 901 | 1013 | 1009 | 856 | 956 |
| SL1344_RS05855_ | 391 | 335 | 346 | 422 | 325 | 313 |
| SL1344_RS05860_ | 63 | 66 | 87 | 55 | 68 | 70 |
| SL1344_RS05865_ | 303 | 362 | 316 | 454 | 332 | 463 |
| SL1344_RS05870_ | 7976 | 8091 | 9032 | 10087 | 7706 | 7566 |
| SL1344_RS05875_ | 10381 | 10982 | 12037 | 10822 | 7855 | 9645 |
| SL1344_RS05880_ | 3854 | 5321 | 5003 | 4517 | 6655 | 3895 |
| SL1344_RS05885_ | 1652 | 1838 | 1799 | 2085 | 1543 | 1745 |
| SL1344_RS05890_ | 1872 | 2058 | 2122 | 2379 | 2155 | 2063 |
| SL1344_RS05900_ | 26986 | 24995 | 25697 | 27025 | 27224 | 28713 |
| SL1344_RS05905_ | 14723 | 16052 | 15152 | 15303 | 15971 | 14899 |
| SL1344_RS05910_ | 333 | 324 | 334 | 396 | 306 | 397 |
| SL1344_RS05915_ | 441 | 492 | 468 | 515 | 531 | 566 |
| SL1344_RS05920_ | 478 | 527 | 492 | 567 | 556 | 572 |
| SL1344_RS05925_ | 498 | 628 | 604 | 669 | 613 | 654 |
| SL1344_RS05930_ | 495 | 540 | 616 | 631 | 431 | 580 |
| SL1344_RS05935_ | 3632 | 4104 | 4142 | 4807 | 4861 | 4640 |
| SL1344_RS05940_ | 64 | 72 | 84 | 165 | 101 | 118 |
| SL1344_RS05945_ | 757 | 807 | 781 | 796 | 671 | 916 |
| SL1344_RS05950_ | 751 | 923 | 920 | 821 | 872 | 777 |
| SL1344_RS05955_ | 591 | 687 | 556 | 614 | 542 | 713 |
| SL1344_RS05960_ | 413 | 500 | 421 | 516 | 531 | 523 |
| SL1344_RS05965_ | 513 | 527 | 477 | 543 | 502 | 640 |
| SL1344_RS05970_ | 1082 | 1274 | 1223 | 1219 | 1362 | 1281 |
| SL1344_RS05975_ | 261 | 437 | 376 | 664 | 437 | 697 |
| SL1344_RS05980_ | 69 | 73 | 45 | 54 | 66 | 52 |
| SL1344_RS05985_ | 663 | 994 | 1038 | 1175 | 820 | 1159 |
| SL1344_RS05990_ | 167 | 206 | 187 | 358 | 303 | 280 |
| SL1344_RS05995_ | 175 | 174 | 192 | 229 | 134 | 173 |
| SL1344_RS06000_ | 1434 | 1714 | 1693 | 1774 | 1579 | 1526 |
| SL1344_RS06005_ | 776 | 850 | 822 | 966 | 804 | 880 |
| SL1344_RS06010_ | 374 | 585 | 492 | 523 | 506 | 526 |
| SL1344_RS06015_ | 591 | 646 | 673 | 769 | 631 | 724 |
| SL1344_RS06020_ | 851 | 935 | 819 | 925 | 1272 | 907 |
| SL1344_RS06025_ | 843 | 940 | 903 | 1036 | 1065 | 962 |

| | | | | | | |
|---|---|---|---|---|---|---|
| SL1344_RS06030_ | 966 | 1014 | 943 | 921 | 703 | 876 |
| SL1344_RS06035_ | 354 | 453 | 450 | 450 | 306 | 325 |
| SL1344_RS06045_ | 349 | 325 | 385 | 479 | 261 | 428 |
| SL1344_RS06050_ | 530 | 548 | 531 | 684 | 528 | 618 |
| SL1344_RS06055_ | 265 | 285 | 287 | 252 | 275 | 349 |
| SL1344_RS06070_ | 1194 | 1440 | 1441 | 1295 | 981 | 941 |
| SL1344_RS06085_ | 980 | 1090 | 1108 | 962 | 597 | 711 |
| SL1344_RS06090_ | 622 | 687 | 750 | 696 | 454 | 530 |
| SL1344_RS06095_ | 2006 | 2087 | 2112 | 1977 | 1697 | 1636 |
| SL1344_RS06100_ | 164 | 175 | 178 | 141 | 131 | 133 |
| SL1344_RS06105_ | 143 | 177 | 169 | 132 | 104 | 124 |
| SL1344_RS06110_ | 193 | 229 | 215 | 216 | 144 | 195 |
| SL1344_RS06115_ | 11613 | 13056 | 14524 | 13430 | 13041 | 14198 |
| SL1344_RS06120_ | 236 | 224 | 301 | 224 | 293 | 327 |
| SL1344_RS06125_ | 2 | 8 | 2 | 43 | 5 | 31 |
| SL1344_RS06130_ | 28 | 21 | 25 | 29 | 14 | 28 |
| SL1344_RS06135_ | 854 | 872 | 686 | 1013 | 961 | 870 |
| SL1344_RS06140_ | 148 | 138 | 149 | 113 | 92 | 103 |
| SL1344_RS06150_ | 8 | 2 | 10 | 10 | 9 | 7 |
| SL1344_RS06155_ | 0 | 0 | 0 | 0 | 1 | 0 |
| SL1344_RS06160_ | 22 | 16 | 26 | 20 | 14 | 28 |
| SL1344_RS06165_ | 253 | 220 | 261 | 258 | 217 | 309 |
| SL1344_RS06170_ | 51 | 45 | 54 | 46 | 56 | 41 |
| SL1344_RS06180_ | 388 | 432 | 469 | 368 | 281 | 351 |
| SL1344_RS06185_ | 12 | 10 | 9 | 8 | 15 | 10 |
| SL1344_RS06190_ | 21 | 14 | 17 | 33 | 14 | 22 |
| SL1344_RS06195_ | 275 | 188 | 237 | 286 | 174 | 236 |
| SL1344_RS06200_ | 62 | 50 | 60 | 31 | 56 | 36 |
| SL1344_RS06205_ | 1114 | 819 | 1022 | 2290 | 983 | 1323 |
| SL1344_RS06210_ | 16 | 16 | 12 | 29 | 17 | 19 |
| SL1344_RS06215_ | 128 | 144 | 157 | 142 | 147 | 120 |
| SL1344_RS06220_ | 380 | 479 | 427 | 353 | 197 | 253 |
| SL1344_RS06225_ | 126 | 177 | 165 | 119 | 84 | 92 |
| SL1344_RS06230_ | 916 | 967 | 979 | 883 | 790 | 860 |
| SL1344_RS06235_ | 239 | 239 | 278 | 216 | 263 | 231 |
| SL1344_RS06240_ | 77 | 75 | 77 | 64 | 85 | 67 |
| SL1344_RS06245_ | 98 | 109 | 87 | 90 | 97 | 102 |
| SL1344_RS06250_ | 89 | 101 | 78 | 80 | 105 | 99 |
| SL1344_RS06255_ | 124 | 128 | 134 | 111 | 81 | 88 |
| SL1344_RS06260_ | 101 | 102 | 124 | 69 | 33 | 65 |
| SL1344_RS06265_ | 36 | 48 | 53 | 25 | 14 | 29 |
| SL1344_RS06270_ | 127 | 142 | 157 | 134 | 100 | 97 |
| SL1344_RS06275_ | 204 | 228 | 224 | 223 | 192 | 175 |
| SL1344_RS06280_ | 32 | 37 | 24 | 23 | 19 | 21 |
| SL1344_RS06285_ | 369 | 396 | 391 | 344 | 233 | 296 |
| SL1344_RS06290_ | 353 | 327 | 456 | 291 | 113 | 216 |
| SL1344_RS06295_ | 3 | 6 | 11 | 4 | 2 | 3 |
| SL1344_RS06300_ | 396 | 381 | 346 | 334 | 171 | 269 |
| SL1344_RS06305_ | 510 | 584 | 528 | 437 | 404 | 443 |
| SL1344_RS06310_ | 57 | 71 | 77 | 78 | 63 | 86 |
| SL1344_RS06315_ | 229 | 287 | 320 | 227 | 218 | 256 |
| SL1344_RS06320_ | 46 | 50 | 62 | 54 | 45 | 44 |
| SL1344_RS06325_ | 41 | 43 | 39 | 48 | 21 | 30 |
| SL1344_RS06330_ | 360 | 314 | 369 | 315 | 217 | 352 |
| SL1344_RS06335_ | 204 | 236 | 235 | 160 | 126 | 197 |
| SL1344_RS06340_ | 522 | 705 | 654 | 535 | 379 | 412 |
| SL1344_RS06345_ | 246 | 322 | 363 | 267 | 213 | 229 |
| SL1344_RS06350_ | 62 | 96 | 95 | 90 | 72 | 75 |
| SL1344_RS06355_ | 130 | 136 | 159 | 181 | 137 | 128 |
| SL1344_RS06360_ | 82 | 91 | 95 | 103 | 91 | 60 |
| SL1344_RS06365_ | 6 | 10 | 10 | 3 | 8 | 5 |
| SL1344_RS06370_ | 118 | 156 | 143 | 85 | 83 | 90 |
| SL1344_RS06375_ | 133 | 146 | 125 | 117 | 70 | 89 |

| | | | | | | |
|---|---|---|---|---|---|---|
| SL1344_RS06380_ | 405 | 439 | 380 | 422 | 360 | 373 |
| SL1344_RS06385_ | 469 | 526 | 533 | 389 | 365 | 362 |
| SL1344_RS06390_ | 2985 | 3539 | 3354 | 3734 | 2935 | 3411 |
| SL1344_RS06395_ | 77 | 54 | 56 | 59 | 49 | 60 |
| SL1344_RS06400_ | 288 | 321 | 302 | 269 | 270 | 327 |
| SL1344_RS06405_ | 399 | 486 | 447 | 523 | 508 | 407 |
| SL1344_RS06410_ | 20209 | 21336 | 22745 | 23416 | 16385 | 20534 |
| SL1344_RS06415_ | 1529 | 1403 | 1737 | 1372 | 1004 | 1313 |
| SL1344_RS06420_ | 1658 | 1664 | 1868 | 1717 | 1460 | 1745 |
| SL1344_RS06425_ | 323 | 295 | 290 | 297 | 259 | 285 |
| SL1344_RS06430_ | 303 | 342 | 322 | 275 | 221 | 276 |
| SL1344_RS06435_ | 859 | 1107 | 985 | 834 | 704 | 803 |
| SL1344_RS06440_ | 451 | 496 | 484 | 398 | 349 | 492 |
| SL1344_RS06445_ | 839 | 996 | 886 | 796 | 761 | 849 |
| SL1344_RS06450_ | 1522 | 1924 | 1801 | 1563 | 1573 | 1506 |
| SL1344_RS06455_ | 517 | 649 | 597 | 526 | 553 | 565 |
| SL1344_RS06465_ | 103 | 105 | 90 | 52 | 54 | 44 |
| SL1344_RS06470_ | 656 | 931 | 689 | 516 | 599 | 558 |
| SL1344_RS06475_ | 210 | 167 | 176 | 101 | 126 | 127 |
| SL1344_RS06480_ | 166 | 135 | 112 | 67 | 91 | 107 |
| SL1344_RS06485_ | 205 | 176 | 194 | 142 | 183 | 190 |
| SL1344_RS06490_ | 145 | 129 | 165 | 96 | 125 | 136 |
| SL1344_RS06495_ | 112 | 118 | 110 | 88 | 101 | 138 |
| SL1344_RS06500_ | 211 | 194 | 199 | 188 | 156 | 166 |
| SL1344_RS06505_ | 226 | 263 | 236 | 314 | 233 | 266 |
| SL1344_RS06510_ | 1019 | 1514 | 1350 | 879 | 859 | 991 |
| SL1344_RS06515_ | 1297 | 1189 | 1426 | 1146 | 694 | 1114 |
| SL1344_RS06520_ | 434 | 375 | 534 | 406 | 226 | 320 |
| SL1344_RS06525_ | 296 | 320 | 330 | 177 | 261 | 149 |
| SL1344_RS06530_ | 83 | 111 | 106 | 88 | 66 | 118 |
| SL1344_RS06535_ | 191 | 193 | 191 | 156 | 122 | 160 |
| SL1344_RS06540_ | 73 | 84 | 68 | 60 | 53 | 66 |
| SL1344_RS06545_ | 181 | 241 | 264 | 226 | 185 | 168 |
| SL1344_RS06550_ | 354 | 286 | 441 | 425 | 196 | 307 |
| SL1344_RS06555_ | 148 | 172 | 163 | 109 | 108 | 99 |
| SL1344_RS06565_ | 99 | 170 | 106 | 106 | 192 | 98 |
| SL1344_RS06570_ | 732 | 813 | 897 | 742 | 545 | 692 |
| SL1344_RS06575_ | 250 | 292 | 248 | 203 | 172 | 207 |
| SL1344_RS06580_ | 534 | 555 | 509 | 513 | 443 | 427 |
| SL1344_RS06605_ | 19 | 31 | 19 | 31 | 16 | 44 |
| SL1344_RS06615_ | 13 | 13 | 9 | 13 | 8 | 7 |
| SL1344_RS06630_ | 37150 | 42652 | 44830 | 43313 | 38362 | 37219 |
| SL1344_RS06635_ | 44824 | 49653 | 49962 | 41953 | 54994 | 40610 |
| SL1344_RS06640_ | 39962 | 49087 | 54265 | 53162 | 53659 | 44396 |
| SL1344_RS06645_ | 1055 | 1338 | 1306 | 1167 | 828 | 1011 |
| SL1344_RS06650_ | 3063 | 3130 | 2840 | 2762 | 2469 | 3033 |
| SL1344_RS06660_ | 402 | 412 | 491 | 424 | 338 | 321 |
| SL1344_RS06665_ | 179 | 160 | 158 | 163 | 145 | 186 |
| SL1344_RS06670_ | 203 | 192 | 147 | 153 | 124 | 185 |
| SL1344_RS06675_ | 283 | 305 | 319 | 203 | 135 | 178 |
| SL1344_RS06685_ | 407 | 468 | 469 | 367 | 422 | 463 |
| SL1344_RS06690_ | 351 | 300 | 341 | 445 | 231 | 417 |
| SL1344_RS06695_ | 517 | 756 | 651 | 591 | 512 | 547 |
| SL1344_RS06700_ | 250 | 324 | 249 | 248 | 225 | 239 |
| SL1344_RS06705_ | 8107 | 7985 | 8559 | 8320 | 6679 | 7139 |
| SL1344_RS06710_ | 54 | 75 | 79 | 82 | 56 | 70 |
| SL1344_RS06715_ | 13 | 8 | 7 | 12 | 7 | 8 |
| SL1344_RS06720_ | 30 | 35 | 33 | 25 | 33 | 28 |
| SL1344_RS06730_ | 7 | 19 | 14 | 7 | 9 | 7 |
| SL1344_RS06735_ | 23 | 33 | 37 | 15 | 19 | 19 |
| SL1344_RS06740_ | 135 | 128 | 129 | 128 | 111 | 96 |
| SL1344_RS06745_ | 29 | 43 | 39 | 44 | 16 | 31 |
| SL1344_RS06750_ | 386 | 515 | 402 | 262 | 317 | 312 |

| | | | | | | |
|---|---|---|---|---|---|---|
| SL1344_RS06755_ | 146 | 139 | 169 | 122 | 76 | 153 |
| SL1344_RS06760_ | 49 | 40 | 65 | 50 | 24 | 68 |
| SL1344_RS06765_ | 13 | 12 | 18 | 17 | 17 | 16 |
| SL1344_RS06770_ | 16 | 11 | 20 | 9 | 7 | 9 |
| SL1344_RS06775_ | 457 | 544 | 493 | 407 | 408 | 411 |
| SL1344_RS06780_ | 934 | 1087 | 1032 | 934 | 842 | 979 |
| SL1344_RS06785_ | 160 | 175 | 196 | 153 | 152 | 188 |
| SL1344_RS06790_ | 377 | 530 | 539 | 550 | 452 | 486 |
| SL1344_RS06795_ | 174 | 255 | 265 | 154 | 125 | 113 |
| SL1344_RS06800_ | 35 | 33 | 31 | 65 | 58 | 71 |
| SL1344_RS06805_ | 172 | 235 | 208 | 249 | 189 | 261 |
| SL1344_RS06810_ | 197 | 281 | 260 | 312 | 265 | 288 |
| SL1344_RS06815_ | 245 | 265 | 293 | 363 | 269 | 347 |
| SL1344_RS06820_ | 173 | 220 | 261 | 292 | 217 | 280 |
| SL1344_RS06825_ | 81 | 92 | 90 | 106 | 92 | 109 |
| SL1344_RS06830_ | 341 | 379 | 465 | 389 | 306 | 321 |
| SL1344_RS06835_ | 331 | 392 | 433 | 421 | 237 | 396 |
| SL1344_RS06840_ | 244756 | 313580 | 321302 | 317834 | 271652 | 281070 |
| SL1344_RS06845_ | 1917 | 2390 | 2307 | 1632 | 1560 | 1546 |
| SL1344_RS06850_ | 83 | 83 | 99 | 76 | 82 | 90 |
| SL1344_RS06855_ | 32 | 19 | 35 | 30 | 15 | 17 |
| SL1344_RS06860_ | 74 | 51 | 81 | 47 | 50 | 60 |
| SL1344_RS06865_ | 80 | 61 | 81 | 68 | 33 | 41 |
| SL1344_RS06900_ | 825 | 1017 | 1068 | 766 | 676 | 809 |
| SL1344_RS06905_ | 141 | 170 | 175 | 171 | 132 | 156 |
| SL1344_RS06925_ | 18 | 49 | 21 | 14 | 8 | 22 |
| SL1344_RS06930_ | 32 | 23 | 17 | 11 | 19 | 26 |
| SL1344_RS06935_ | 18 | 18 | 28 | 11 | 6 | 12 |
| SL1344_RS06940_ | 21 | 15 | 17 | 16 | 16 | 18 |
| SL1344_RS06945_ | 34 | 39 | 39 | 29 | 19 | 24 |
| SL1344_RS06950_ | 29 | 12 | 33 | 34 | 12 | 26 |
| SL1344_RS06955_ | 66 | 65 | 56 | 57 | 37 | 68 |
| SL1344_RS06960_ | 19 | 17 | 12 | 21 | 13 | 21 |
| SL1344_RS06965_ | 22 | 26 | 26 | 16 | 13 | 19 |
| SL1344_RS06970_ | 27 | 33 | 38 | 27 | 26 | 21 |
| SL1344_RS06975_ | 35 | 42 | 30 | 27 | 18 | 11 |
| SL1344_RS06980_ | 63 | 52 | 55 | 69 | 56 | 50 |
| SL1344_RS06990_ | 4 | 4 | 7 | 7 | 3 | 2 |
| SL1344_RS06995_ | 0 | 4 | 7 | 4 | 1 | 8 |
| SL1344_RS07000_ | 21 | 18 | 22 | 16 | 10 | 15 |
| SL1344_RS07005_ | 28 | 20 | 26 | 15 | 31 | 18 |
| SL1344_RS07010_ | 15 | 10 | 10 | 11 | 8 | 11 |
| SL1344_RS07015_ | 56 | 36 | 31 | 36 | 41 | 25 |
| SL1344_RS07020_ | 3 | 6 | 9 | 9 | 3 | 7 |
| SL1344_RS07025_ | 50 | 62 | 60 | 51 | 30 | 58 |
| SL1344_RS07030_ | 90 | 98 | 102 | 86 | 59 | 82 |
| SL1344_RS07035_ | 29 | 53 | 39 | 23 | 26 | 20 |
| SL1344_RS07040_ | 40 | 29 | 34 | 22 | 20 | 30 |
| SL1344_RS07045_ | 106 | 88 | 95 | 75 | 61 | 74 |
| SL1344_RS07055_ | 12 | 9 | 4 | 11 | 8 | 8 |
| SL1344_RS07060_ | 30 | 21 | 28 | 15 | 12 | 17 |
| SL1344_RS07065_ | 29 | 30 | 26 | 27 | 26 | 27 |
| SL1344_RS07070_ | 230 | 279 | 230 | 360 | 444 | 132 |
| SL1344_RS07075_ | 279 | 357 | 290 | 601 | 532 | 222 |
| SL1344_RS07080_ | 549 | 626 | 708 | 621 | 436 | 558 |
| SL1344_RS07085_ | 380 | 465 | 416 | 351 | 338 | 424 |
| SL1344_RS07090_ | 988 | 1143 | 1035 | 1249 | 1002 | 1201 |
| SL1344_RS07095_ | 99 | 137 | 136 | 123 | 87 | 131 |
| SL1344_RS07100_ | 74 | 89 | 88 | 59 | 66 | 113 |
| SL1344_RS07115_ | 1042 | 1073 | 982 | 779 | 675 | 658 |
| SL1344_RS07120_ | 4394 | 4841 | 5111 | 4566 | 4018 | 4287 |
| SL1344_RS07125_ | 477 | 574 | 590 | 466 | 400 | 365 |
| SL1344_RS07130_ | 717 | 853 | 781 | 736 | 614 | 697 |

| | | | | | | |
|---|---|---|---|---|---|---|
| SL1344_RS07135_ | 390 | 454 | 476 | 479 | 327 | 386 |
| SL1344_RS07140_ | 171 | 179 | 168 | 180 | 162 | 162 |
| SL1344_RS07145_ | 44 | 40 | 40 | 41 | 36 | 41 |
| SL1344_RS07150_ | 538 | 670 | 593 | 499 | 404 | 385 |
| SL1344_RS07155_ | 308 | 308 | 337 | 235 | 312 | 219 |
| SL1344_RS07160_ | 232 | 306 | 276 | 197 | 194 | 154 |
| SL1344_RS07165_ | 146 | 192 | 178 | 170 | 123 | 144 |
| SL1344_RS07170_ | 55 | 75 | 62 | 49 | 43 | 49 |
| SL1344_RS07180_ | 2505 | 2341 | 2104 | 1394 | 1330 | 1517 |
| SL1344_RS07185_ | 222 | 280 | 248 | 258 | 276 | 262 |
| SL1344_RS07190_ | 221 | 289 | 268 | 264 | 266 | 262 |
| SL1344_RS07195_ | 579 | 597 | 658 | 606 | 412 | 467 |
| SL1344_RS07200_ | 3546 | 4073 | 4369 | 3539 | 2783 | 2951 |
| SL1344_RS07205_ | 823 | 928 | 1004 | 837 | 615 | 782 |
| SL1344_RS07210_ | 558 | 663 | 670 | 688 | 511 | 681 |
| SL1344_RS07215_ | 2130 | 2419 | 2429 | 1927 | 1559 | 1680 |
| SL1344_RS07220_ | 498 | 382 | 476 | 536 | 345 | 333 |
| SL1344_RS07225_ | 405 | 301 | 436 | 438 | 333 | 298 |
| SL1344_RS07230_ | 316 | 233 | 392 | 332 | 185 | 260 |
| SL1344_RS07235_ | 300 | 357 | 451 | 400 | 230 | 305 |
| SL1344_RS07240_ | 967 | 1074 | 1115 | 1131 | 743 | 856 |
| SL1344_RS07245_ | 370 | 428 | 432 | 414 | 400 | 388 |
| SL1344_RS07250_ | 810 | 801 | 895 | 966 | 845 | 761 |
| SL1344_RS07255_ | 915 | 879 | 1119 | 1075 | 714 | 824 |
| SL1344_RS07260_ | 463 | 398 | 539 | 506 | 228 | 401 |
| SL1344_RS07265_ | 350 | 353 | 379 | 379 | 245 | 290 |
| SL1344_RS07270_ | 799 | 949 | 837 | 555 | 586 | 646 |
| SL1344_RS07275_ | 537 | 694 | 651 | 587 | 562 | 578 |
| SL1344_RS07285_ | 1249 | 1277 | 1423 | 1234 | 939 | 1205 |
| SL1344_RS07290_ | 763 | 929 | 990 | 779 | 580 | 731 |
| SL1344_RS07295_ | 6427 | 5777 | 6286 | 5865 | 4235 | 5933 |
| SL1344_RS07300_ | 1753 | 1774 | 1744 | 1328 | 1642 | 1708 |
| SL1344_RS07305_ | 84 | 121 | 119 | 98 | 87 | 85 |
| SL1344_RS07310_ | 123 | 194 | 168 | 160 | 127 | 114 |
| SL1344_RS07315_ | 11 | 18 | 15 | 16 | 14 | 21 |
| SL1344_RS07320_ | 215 | 211 | 232 | 254 | 184 | 193 |
| SL1344_RS07325_ | 9 | 9 | 6 | 9 | 11 | 4 |
| SL1344_RS07330_ | 1734 | 2038 | 2097 | 2052 | 1517 | 1758 |
| SL1344_RS07335_ | 125 | 119 | 121 | 116 | 91 | 136 |
| SL1344_RS07340_ | 152 | 169 | 227 | 169 | 126 | 107 |
| SL1344_RS07345_ | 1846 | 2608 | 2384 | 1964 | 2057 | 1744 |
| SL1344_RS07350_ | 1567 | 1908 | 1852 | 1295 | 1130 | 1310 |
| SL1344_RS07355_ | 1710 | 2142 | 1986 | 1565 | 1268 | 1520 |
| SL1344_RS07365_ | 181 | 181 | 185 | 65 | 77 | 56 |
| SL1344_RS07370_ | 200 | 238 | 265 | 66 | 93 | 79 |
| SL1344_RS07375_ | 172 | 219 | 205 | 218 | 158 | 186 |
| SL1344_RS07380_ | 133 | 162 | 185 | 81 | 76 | 128 |
| SL1344_RS07385_ | 205 | 270 | 258 | 189 | 164 | 176 |
| SL1344_RS07390_ | 59 | 90 | 81 | 74 | 67 | 79 |
| SL1344_RS07400_ | 171 | 209 | 238 | 126 | 136 | 226 |
| SL1344_RS07405_ | 118 | 168 | 120 | 143 | 150 | 96 |
| SL1344_RS07410_ | 107 | 121 | 108 | 123 | 91 | 113 |
| SL1344_RS07415_ | 27 | 26 | 39 | 35 | 21 | 31 |
| SL1344_RS07420_ | 16 | 12 | 21 | 19 | 20 | 20 |
| SL1344_RS07425_ | 18 | 21 | 13 | 19 | 14 | 5 |
| SL1344_RS07430_ | 327 | 457 | 470 | 423 | 386 | 338 |
| SL1344_RS07435_ | 208 | 236 | 282 | 299 | 208 | 249 |
| SL1344_RS07440_ | 10 | 8 | 6 | 5 | 9 | 15 |
| SL1344_RS07445_ | 46 | 85 | 64 | 54 | 31 | 38 |
| SL1344_RS07450_ | 71 | 107 | 85 | 78 | 96 | 76 |
| SL1344_RS07455_ | 299 | 392 | 375 | 394 | 249 | 334 |
| SL1344_RS07460_ | 124 | 137 | 119 | 149 | 147 | 168 |
| SL1344_RS07465_ | 1064 | 1120 | 1169 | 1601 | 1017 | 1416 |

| | | | | | | |
|---|---|---|---|---|---|---|
| SL1344_RS07470_ | 920 | 1102 | 1202 | 1123 | 787 | 1198 |
| SL1344_RS07475_ | 94 | 86 | 89 | 108 | 114 | 109 |
| SL1344_RS07480_ | 217 | 193 | 187 | 291 | 282 | 274 |
| SL1344_RS07485_ | 130 | 123 | 106 | 197 | 176 | 189 |
| SL1344_RS07490_ | 185 | 169 | 158 | 218 | 264 | 247 |
| SL1344_RS07495_ | 234 | 275 | 280 | 242 | 290 | 290 |
| SL1344_RS07500_ | 117 | 201 | 211 | 124 | 117 | 148 |
| SL1344_RS07505_ | 514 | 674 | 599 | 542 | 466 | 504 |
| SL1344_RS07510_ | 1165 | 1499 | 1392 | 1341 | 1055 | 1196 |
| SL1344_RS07515_ | 437 | 635 | 506 | 430 | 520 | 469 |
| SL1344_RS07520_ | 16 | 14 | 22 | 32 | 11 | 23 |
| SL1344_RS07525_ | 26 | 23 | 27 | 29 | 19 | 32 |
| SL1344_RS07530_ | 43 | 60 | 49 | 52 | 33 | 45 |
| SL1344_RS07535_ | 557 | 501 | 710 | 579 | 450 | 656 |
| SL1344_RS07540_ | 92 | 117 | 93 | 124 | 120 | 114 |
| SL1344_RS07550_ | 90 | 115 | 118 | 92 | 88 | 84 |
| SL1344_RS07555_ | 122 | 138 | 136 | 121 | 96 | 114 |
| SL1344_RS07560_ | 137 | 133 | 148 | 155 | 130 | 124 |
| SL1344_RS07565_ | 266 | 255 | 237 | 266 | 213 | 274 |
| SL1344_RS07570_ | 108 | 133 | 138 | 166 | 141 | 167 |
| SL1344_RS07575_ | 101 | 111 | 92 | 113 | 82 | 97 |
| SL1344_RS07580_ | 110 | 122 | 123 | 142 | 114 | 158 |
| SL1344_RS07585_ | 184 | 159 | 154 | 192 | 185 | 170 |
| SL1344_RS07590_ | 118 | 127 | 101 | 155 | 96 | 96 |
| SL1344_RS07595_ | 180 | 366 | 310 | 177 | 176 | 247 |
| SL1344_RS07600_ | 28 | 41 | 24 | 121 | 76 | 77 |
| SL1344_RS07605_ | 206 | 243 | 232 | 398 | 191 | 277 |
| SL1344_RS07610_ | 51 | 39 | 37 | 45 | 44 | 53 |
| SL1344_RS07615_ | 115 | 128 | 129 | 156 | 123 | 128 |
| SL1344_RS07620_ | 194 | 178 | 195 | 203 | 175 | 189 |
| SL1344_RS07625_ | 57 | 85 | 85 | 59 | 41 | 83 |
| SL1344_RS07630_ | 46 | 66 | 60 | 70 | 47 | 84 |
| SL1344_RS07635_ | 148 | 174 | 197 | 182 | 159 | 192 |
| SL1344_RS07640_ | 173 | 201 | 190 | 178 | 159 | 203 |
| SL1344_RS07645_ | 265 | 309 | 274 | 205 | 203 | 218 |
| SL1344_RS07650_ | 116 | 151 | 114 | 123 | 88 | 124 |
| SL1344_RS07655_ | 180 | 185 | 186 | 125 | 88 | 128 |
| SL1344_RS07660_ | 410 | 481 | 425 | 505 | 452 | 566 |
| SL1344_RS07665_ | 274 | 275 | 305 | 212 | 161 | 258 |
| SL1344_RS07670_ | 170 | 158 | 154 | 114 | 80 | 108 |
| SL1344_RS07675_ | 32 | 35 | 35 | 53 | 27 | 47 |
| SL1344_RS07680_ | 53 | 56 | 48 | 70 | 63 | 42 |
| SL1344_RS07685_ | 36 | 65 | 38 | 60 | 47 | 48 |
| SL1344_RS07690_ | 244 | 256 | 264 | 313 | 273 | 337 |
| SL1344_RS07695_ | 98 | 95 | 113 | 180 | 130 | 161 |
| SL1344_RS07700_ | 165 | 181 | 150 | 314 | 193 | 215 |
| SL1344_RS07705_ | 281 | 323 | 355 | 352 | 234 | 331 |
| SL1344_RS07710_ | 189 | 190 | 230 | 236 | 140 | 239 |
| SL1344_RS07715_ | 64 | 50 | 49 | 61 | 42 | 61 |
| SL1344_RS07720_ | 50 | 55 | 41 | 50 | 52 | 53 |
| SL1344_RS07725_ | 101 | 77 | 85 | 119 | 112 | 120 |
| SL1344_RS07735_ | 88 | 97 | 114 | 317 | 224 | 301 |
| SL1344_RS07740_ | 444 | 562 | 520 | 1789 | 1140 | 1783 |
| SL1344_RS07745_ | 121 | 158 | 150 | 203 | 171 | 183 |
| SL1344_RS07750_ | 88 | 79 | 98 | 179 | 96 | 143 |
| SL1344_RS07755_ | 162 | 152 | 154 | 268 | 230 | 257 |
| SL1344_RS07760_ | 213 | 175 | 192 | 281 | 239 | 269 |
| SL1344_RS07765_ | 117 | 134 | 134 | 161 | 147 | 135 |
| SL1344_RS07770_ | 111 | 101 | 100 | 113 | 96 | 101 |
| SL1344_RS07775_ | 40 | 28 | 37 | 37 | 29 | 32 |
| SL1344_RS07780_ | 251 | 236 | 306 | 421 | 337 | 430 |
| SL1344_RS07785_ | 80 | 100 | 80 | 129 | 91 | 128 |
| SL1344_RS07790_ | 35 | 46 | 35 | 46 | 35 | 46 |

| | | | | | | |
|---|---|---|---|---|---|---|
| SL1344_RS07800_ | 2576 | 2680 | 2902 | 3549 | 2891 | 3349 |
| SL1344_RS07805_ | 548 | 585 | 545 | 999 | 893 | 998 |
| SL1344_RS07810_ | 1184 | 1089 | 1059 | 1849 | 1702 | 1592 |
| SL1344_RS07815_ | 72 | 68 | 67 | 84 | 81 | 85 |
| SL1344_RS07820_ | 274 | 327 | 245 | 357 | 284 | 342 |
| SL1344_RS07825_ | 221 | 255 | 236 | 331 | 258 | 259 |
| SL1344_RS07830_ | 52 | 62 | 60 | 97 | 69 | 85 |
| SL1344_RS07835_ | 65865 | 74172 | 74069 | 95211 | 68949 | 88038 |
| SL1344_RS07840_ | 293 | 357 | 380 | 621 | 456 | 476 |
| SL1344_RS07845_ | 805 | 513 | 217 | 2595 | 327 | 948 |
| SL1344_RS07850_ | 388 | 351 | 222 | 803 | 364 | 496 |
| SL1344_RS07855_ | 48 | 48 | 44 | 69 | 51 | 40 |
| SL1344_RS07860_ | 130 | 122 | 149 | 162 | 146 | 177 |
| SL1344_RS07865_ | 55 | 46 | 64 | 72 | 58 | 84 |
| SL1344_RS07870_ | 19 | 15 | 16 | 39 | 16 | 28 |
| SL1344_RS07875_ | 25 | 40 | 47 | 63 | 36 | 60 |
| SL1344_RS07880_ | 166 | 204 | 201 | 301 | 231 | 310 |
| SL1344_RS07885_ | 180 | 217 | 226 | 289 | 240 | 282 |
| SL1344_RS07890_ | 752 | 756 | 780 | 1191 | 865 | 1124 |
| SL1344_RS07895_ | 323 | 448 | 394 | 589 | 424 | 465 |
| SL1344_RS07900_ | 24 | 21 | 22 | 35 | 32 | 32 |
| SL1344_RS07905_ | 389 | 554 | 555 | 997 | 680 | 772 |
| SL1344_RS07910_ | 192 | 187 | 186 | 244 | 208 | 235 |
| SL1344_RS07915_ | 96 | 95 | 93 | 70 | 54 | 59 |
| SL1344_RS07920_ | 353 | 311 | 344 | 470 | 429 | 563 |
| SL1344_RS07925_ | 191 | 278 | 267 | 320 | 337 | 401 |
| SL1344_RS07930_ | 218 | 287 | 228 | 356 | 471 | 322 |
| SL1344_RS07935_ | 69 | 86 | 73 | 106 | 66 | 123 |
| SL1344_RS07955_ | 96 | 83 | 97 | 85 | 87 | 72 |
| SL1344_RS07960_ | 374 | 405 | 456 | 519 | 434 | 536 |
| SL1344_RS07965_ | 351 | 361 | 343 | 561 | 378 | 451 |
| SL1344_RS07970_ | 193 | 210 | 245 | 397 | 213 | 321 |
| SL1344_RS07975_ | 6 | 13 | 24 | 36 | 20 | 30 |
| SL1344_RS07980_ | 25 | 39 | 30 | 46 | 46 | 42 |
| SL1344_RS07990_ | 25 | 27 | 33 | 60 | 76 | 73 |
| SL1344_RS07995_ | 702 | 729 | 652 | 1052 | 814 | 807 |
| SL1344_RS08000_ | 380 | 433 | 448 | 506 | 383 | 502 |
| SL1344_RS08005_ | 279 | 436 | 333 | 460 | 381 | 328 |
| SL1344_RS08010_ | 1492 | 1619 | 1683 | 2258 | 1613 | 1992 |
| SL1344_RS08015_ | 461 | 519 | 518 | 596 | 591 | 635 |
| SL1344_RS08020_ | 313 | 308 | 376 | 469 | 409 | 439 |
| SL1344_RS08025_ | 50 | 49 | 43 | 58 | 62 | 88 |
| SL1344_RS08030_ | 84 | 78 | 79 | 113 | 95 | 153 |
| SL1344_RS08035_ | 38 | 32 | 34 | 53 | 56 | 54 |
| SL1344_RS08040_ | 6 | 10 | 9 | 11 | 17 | 17 |
| SL1344_RS08045_ | 47 | 46 | 41 | 75 | 111 | 71 |
| SL1344_RS08050_ | 28 | 28 | 41 | 62 | 66 | 80 |
| SL1344_RS08055_ | 22 | 20 | 33 | 41 | 19 | 37 |
| SL1344_RS08060_ | 26 | 37 | 36 | 55 | 40 | 38 |
| SL1344_RS08065_ | 61 | 68 | 55 | 120 | 91 | 79 |
| SL1344_RS08070_ | 58 | 56 | 51 | 96 | 77 | 95 |
| SL1344_RS08075_ | 86 | 102 | 108 | 124 | 67 | 107 |
| SL1344_RS08080_ | 20 | 26 | 18 | 21 | 24 | 26 |
| SL1344_RS08090_ | 366 | 342 | 413 | 413 | 481 | 422 |
| SL1344_RS08095_ | 580 | 501 | 567 | 659 | 503 | 535 |
| SL1344_RS08100_ | 1962 | 1094 | 1747 | 1491 | 2340 | 1236 |
| SL1344_RS08110_ | 180 | 210 | 154 | 219 | 219 | 231 |
| SL1344_RS08115_ | 57 | 69 | 64 | 69 | 51 | 54 |
| SL1344_RS08120_ | 4 | 7 | 2 | 11 | 8 | 10 |
| SL1344_RS08125_ | 14 | 6 | 12 | 20 | 10 | 27 |
| SL1344_RS08130_ | 12 | 15 | 11 | 13 | 10 | 18 |
| SL1344_RS08135_ | 63 | 44 | 54 | 86 | 60 | 97 |
| SL1344_RS08140_ | 28 | 37 | 39 | 56 | 68 | 64 |

| | | | | | | |
|---|---|---|---|---|---|---|
| SL1344_RS08145_ | 10 | 13 | 32 | 31 | 50 | 29 |
| SL1344_RS08150_ | 5 | 11 | 7 | 21 | 9 | 15 |
| SL1344_RS08155_ | 50 | 50 | 43 | 114 | 67 | 63 |
| SL1344_RS08160_ | 653 | 641 | 632 | 1406 | 748 | 1211 |
| SL1344_RS08165_ | 359 | 389 | 369 | 413 | 429 | 482 |
| SL1344_RS08170_ | 142 | 157 | 156 | 279 | 208 | 256 |
| SL1344_RS08175_ | 942 | 1047 | 998 | 1466 | 1132 | 1454 |
| SL1344_RS08185_ | 87 | 106 | 100 | 132 | 108 | 160 |
| SL1344_RS08190_ | 59 | 64 | 70 | 104 | 62 | 87 |
| SL1344_RS08195_ | 74 | 47 | 72 | 102 | 89 | 105 |
| SL1344_RS08200_ | 367 | 527 | 546 | 741 | 705 | 641 |
| SL1344_RS08215_ | 576 | 483 | 678 | 805 | 621 | 795 |
| SL1344_RS08220_ | 12 | 13 | 23 | 20 | 15 | 14 |
| SL1344_RS08225_ | 294 | 387 | 372 | 459 | 387 | 423 |
| SL1344_RS08230_ | 26 | 37 | 39 | 44 | 32 | 38 |
| SL1344_RS08235_ | 2029 | 2018 | 2001 | 4516 | 3108 | 4078 |
| SL1344_RS08240_ | 240 | 406 | 408 | 297 | 276 | 322 |
| SL1344_RS08245_ | 75 | 113 | 118 | 98 | 76 | 96 |
| SL1344_RS08250_ | 89 | 86 | 108 | 183 | 128 | 100 |
| SL1344_RS08255_ | 664 | 834 | 881 | 1041 | 874 | 857 |
| SL1344_RS08260_ | 219 | 253 | 273 | 377 | 294 | 327 |
| SL1344_RS08270_ | 260 | 291 | 271 | 436 | 377 | 417 |
| SL1344_RS08275_ | 530 | 578 | 564 | 801 | 759 | 858 |
| SL1344_RS08280_ | 262 | 304 | 316 | 392 | 330 | 438 |
| SL1344_RS08290_ | 1234 | 1494 | 1504 | 2073 | 1747 | 1887 |
| SL1344_RS08295_ | 911 | 990 | 1131 | 1710 | 1267 | 2154 |
| SL1344_RS08300_ | 100 | 142 | 104 | 154 | 111 | 111 |
| SL1344_RS08305_ | 168 | 204 | 219 | 252 | 198 | 236 |
| SL1344_RS08310_ | 102 | 162 | 137 | 101 | 131 | 127 |
| SL1344_RS08315_ | 70 | 96 | 79 | 69 | 70 | 51 |
| SL1344_RS08320_ | 11 | 9 | 15 | 14 | 15 | 17 |
| SL1344_RS08325_ | 12 | 7 | 3 | 16 | 2 | 12 |
| SL1344_RS08330_ | 23 | 14 | 21 | 37 | 28 | 32 |
| SL1344_RS08335_ | 8 | 7 | 7 | 19 | 13 | 15 |
| SL1344_RS08340_ | 8 | 5 | 9 | 12 | 13 | 5 |
| SL1344_RS08345_ | 791 | 858 | 1050 | 1552 | 758 | 1260 |
| SL1344_RS08350_ | 85 | 101 | 84 | 114 | 110 | 140 |
| SL1344_RS08355_ | 41 | 39 | 49 | 80 | 83 | 69 |
| SL1344_RS08360_ | 181 | 269 | 183 | 164 | 332 | 163 |
| SL1344_RS08365_ | 196 | 217 | 206 | 211 | 175 | 217 |
| SL1344_RS08370_ | 54 | 37 | 42 | 57 | 54 | 73 |
| SL1344_RS08375_ | 15 | 36 | 20 | 27 | 18 | 29 |
| SL1344_RS08380_ | 955 | 989 | 1064 | 1460 | 1099 | 1260 |
| SL1344_RS08385_ | 130 | 187 | 164 | 244 | 252 | 244 |
| SL1344_RS08390_ | 168 | 184 | 204 | 307 | 238 | 321 |
| SL1344_RS08395_ | 4392 | 6836 | 5772 | 7308 | 8995 | 6648 |
| SL1344_RS08400_ | 575 | 627 | 551 | 753 | 694 | 720 |
| SL1344_RS08405_ | 460 | 440 | 455 | 741 | 627 | 588 |
| SL1344_RS08410_ | 430 | 409 | 429 | 717 | 462 | 534 |
| SL1344_RS08415_ | 882 | 760 | 919 | 1447 | 965 | 1212 |
| SL1344_RS08420_ | 74 | 92 | 82 | 125 | 130 | 100 |
| SL1344_RS08425_ | 109 | 90 | 99 | 160 | 162 | 136 |
| SL1344_RS08430_ | 65 | 62 | 55 | 97 | 77 | 60 |
| SL1344_RS08435_ | 726 | 797 | 828 | 963 | 905 | 891 |
| SL1344_RS08440_ | 144 | 180 | 135 | 203 | 150 | 222 |
| SL1344_RS08445_ | 511 | 576 | 617 | 764 | 649 | 690 |
| SL1344_RS08450_ | 254 | 328 | 323 | 343 | 283 | 347 |
| SL1344_RS08455_ | 260 | 267 | 286 | 353 | 329 | 325 |
| SL1344_RS08460_ | 306 | 327 | 348 | 383 | 352 | 358 |
| SL1344_RS08465_ | 197 | 231 | 217 | 335 | 244 | 259 |
| SL1344_RS08470_ | 107 | 112 | 137 | 140 | 130 | 115 |
| SL1344_RS08475_ | 28 | 31 | 50 | 63 | 51 | 48 |
| SL1344_RS08480_ | 3 | 3 | 0 | 2 | 3 | 3 |

| | | | | | | |
|---|---|---|---|---|---|---|
| SL1344_RS08485_ | 68 | 48 | 95 | 117 | 78 | 131 |
| SL1344_RS08490_ | 5250 | 6174 | 6765 | 8665 | 7347 | 7757 |
| SL1344_RS08500_ | 2384 | 2853 | 2858 | 3643 | 2645 | 3073 |
| SL1344_RS08505_ | 329 | 350 | 300 | 398 | 405 | 436 |
| SL1344_RS08510_ | 380 | 508 | 564 | 535 | 565 | 597 |
| SL1344_RS08515_ | 467 | 646 | 630 | 709 | 655 | 890 |
| SL1344_RS08520_ | 381 | 339 | 422 | 601 | 421 | 536 |
| SL1344_RS08525_ | 153 | 168 | 213 | 231 | 179 | 250 |
| SL1344_RS08530_ | 164 | 219 | 249 | 327 | 239 | 260 |
| SL1344_RS08535_ | 803 | 799 | 765 | 1426 | 1205 | 1196 |
| SL1344_RS08540_ | 756 | 778 | 842 | 1372 | 1101 | 1207 |
| SL1344_RS08545_ | 116 | 123 | 122 | 145 | 135 | 186 |
| SL1344_RS08550_ | 658 | 790 | 923 | 1096 | 781 | 1028 |
| SL1344_RS08555_ | 607 | 609 | 678 | 1101 | 1084 | 1170 |
| SL1344_RS08560_ | 648 | 535 | 626 | 657 | 532 | 545 |
| SL1344_RS08570_ | 3048 | 3391 | 3272 | 4207 | 3834 | 3995 |
| SL1344_RS08575_ | 1475 | 1518 | 1768 | 2096 | 1803 | 1964 |
| SL1344_RS08580_ | 395 | 428 | 382 | 483 | 536 | 611 |
| SL1344_RS08590_ | 817 | 821 | 699 | 932 | 1226 | 1006 |
| SL1344_RS08595_ | 616 | 625 | 733 | 1070 | 812 | 879 |
| SL1344_RS08600_ | 508 | 611 | 481 | 737 | 712 | 759 |
| SL1344_RS08605_ | 419 | 530 | 542 | 716 | 577 | 753 |
| SL1344_RS08610_ | 49 | 48 | 36 | 61 | 51 | 55 |
| SL1344_RS08615_ | 68 | 97 | 86 | 127 | 111 | 112 |
| SL1344_RS08620_ | 157 | 198 | 170 | 232 | 182 | 290 |
| SL1344_RS08625_ | 147 | 231 | 165 | 316 | 235 | 277 |
| SL1344_RS08630_ | 177 | 252 | 208 | 377 | 229 | 296 |
| SL1344_RS08635_ | 16 | 14 | 14 | 16 | 13 | 12 |
| SL1344_RS08640_ | 6 | 9 | 9 | 9 | 5 | 9 |
| SL1344_RS08645_ | 2 | 4 | 6 | 5 | 14 | 6 |
| SL1344_RS08650_ | 33 | 40 | 43 | 69 | 48 | 42 |
| SL1344_RS08655_ | 454 | 571 | 458 | 609 | 517 | 632 |
| SL1344_RS08660_ | 208 | 279 | 252 | 272 | 226 | 231 |
| SL1344_RS08665_ | 931 | 1285 | 1211 | 1497 | 1204 | 1319 |
| SL1344_RS08670_ | 1172 | 1180 | 1264 | 1538 | 1306 | 1470 |
| SL1344_RS08675_ | 493 | 594 | 628 | 850 | 613 | 602 |
| SL1344_RS08680_ | 741 | 869 | 913 | 1660 | 1220 | 1545 |
| SL1344_RS08685_ | 353 | 426 | 246 | 493 | 512 | 585 |
| SL1344_RS08690_ | 253 | 298 | 268 | 515 | 434 | 462 |
| SL1344_RS08695_ | 458 | 464 | 401 | 737 | 746 | 791 |
| SL1344_RS08700_ | 179 | 199 | 173 | 217 | 229 | 246 |
| SL1344_RS08705_ | 730 | 742 | 563 | 864 | 965 | 1016 |
| SL1344_RS08710_ | 600 | 566 | 568 | 522 | 689 | 672 |
| SL1344_RS08715_ | 412 | 450 | 370 | 301 | 535 | 411 |
| SL1344_RS08725_ | 2027 | 1803 | 1703 | 1918 | 2060 | 2053 |
| SL1344_RS08730_ | 184 | 175 | 216 | 245 | 222 | 200 |
| SL1344_RS08735_ | 2400 | 3404 | 3321 | 4027 | 3359 | 4171 |
| SL1344_RS08740_ | 277 | 302 | 390 | 444 | 310 | 422 |
| SL1344_RS08750_ | 2722 | 2806 | 3119 | 4031 | 3288 | 4014 |
| SL1344_RS08755_ | 525 | 617 | 595 | 788 | 835 | 834 |
| SL1344_RS08760_ | 616 | 647 | 613 | 914 | 863 | 958 |
| SL1344_RS08765_ | 260 | 305 | 315 | 520 | 364 | 492 |
| SL1344_RS08770_ | 633 | 802 | 791 | 1453 | 1206 | 1434 |
| SL1344_RS08775_ | 1225 | 1139 | 858 | 2548 | 1784 | 1225 |
| SL1344_RS08780_ | 549 | 406 | 340 | 859 | 902 | 445 |
| SL1344_RS08785_ | 134 | 156 | 134 | 179 | 158 | 216 |
| SL1344_RS08815_ | 333 | 350 | 329 | 585 | 476 | 463 |
| SL1344_RS08825_ | 244 | 282 | 215 | 324 | 337 | 410 |
| SL1344_RS08830_ | 762 | 906 | 813 | 1120 | 1064 | 1139 |
| SL1344_RS08840_ | 167 | 186 | 195 | 223 | 121 | 187 |
| SL1344_RS08845_ | 2106 | 2351 | 2451 | 3340 | 2895 | 2912 |
| SL1344_RS08850_ | 623 | 669 | 652 | 940 | 719 | 848 |
| SL1344_RS08855_ | 228 | 242 | 231 | 370 | 260 | 266 |

| | | | | | | |
|---|---|---|---|---|---|---|
| SL1344_RS08860_ | 471 | 590 | 653 | 740 | 557 | 631 |
| SL1344_RS08865_ | 829 | 994 | 1087 | 1204 | 1157 | 1023 |
| SL1344_RS08870_ | 1589 | 1993 | 1958 | 2646 | 2293 | 2387 |
| SL1344_RS08880_ | 3316 | 3117 | 4039 | 5972 | 4218 | 5119 |
| SL1344_RS08885_ | 7780 | 9003 | 9075 | 15053 | 9423 | 10666 |
| SL1344_RS08890_ | 459 | 547 | 492 | 877 | 643 | 651 |
| SL1344_RS08895_ | 287 | 168 | 144 | 367 | 240 | 191 |
| SL1344_RS08900_ | 276 | 269 | 302 | 523 | 319 | 363 |
| SL1344_RS08905_ | 2344 | 2611 | 2687 | 2860 | 2805 | 2961 |
| SL1344_RS08910_ | 272 | 294 | 326 | 437 | 390 | 408 |
| SL1344_RS08915_ | 8 | 6 | 1 | 9 | 6 | 7 |
| SL1344_RS08920_ | 38 | 55 | 44 | 72 | 65 | 73 |
| SL1344_RS08925_ | 3 | 4 | 6 | 7 | 16 | 12 |
| SL1344_RS08930_ | 5 | 4 | 5 | 19 | 17 | 9 |
| SL1344_RS08935_ | 6 | 3 | 2 | 5 | 10 | 12 |
| SL1344_RS08940_ | 12 | 11 | 12 | 19 | 22 | 21 |
| SL1344_RS08945_ | 23 | 32 | 31 | 41 | 45 | 49 |
| SL1344_RS08950_ | 34 | 39 | 27 | 42 | 38 | 36 |
| SL1344_RS08955_ | 28 | 22 | 31 | 39 | 44 | 44 |
| SL1344_RS08960_ | 9921 | 9099 | 10835 | 16477 | 13693 | 15592 |
| SL1344_RS08965_ | 645 | 529 | 632 | 934 | 846 | 966 |
| SL1344_RS08970_ | 57 | 56 | 60 | 100 | 75 | 77 |
| SL1344_RS08985_ | 191 | 204 | 174 | 280 | 254 | 269 |
| SL1344_RS08995_ | 1997 | 1649 | 1696 | 1765 | 4892 | 3441 |
| SL1344_RS09000_ | 1781 | 1534 | 1519 | 2186 | 3266 | 3018 |
| SL1344_RS09005_ | 197 | 241 | 244 | 298 | 280 | 237 |
| SL1344_RS09010_ | 965 | 997 | 1045 | 1450 | 1060 | 1268 |
| SL1344_RS09015_ | 594 | 723 | 696 | 834 | 618 | 740 |
| SL1344_RS09020_ | 1104 | 1347 | 1209 | 2135 | 1714 | 1951 |
| SL1344_RS09025_ | 30 | 45 | 32 | 66 | 54 | 60 |
| SL1344_RS09030_ | 775 | 838 | 863 | 902 | 845 | 1142 |
| SL1344_RS09035_ | 54 | 70 | 92 | 99 | 84 | 117 |
| SL1344_RS09040_ | 477 | 492 | 464 | 646 | 652 | 567 |
| SL1344_RS09045_ | 785 | 1025 | 1053 | 849 | 1001 | 876 |
| SL1344_RS09050_ | 320 | 423 | 490 | 412 | 394 | 449 |
| SL1344_RS09055_ | 2201 | 2208 | 2433 | 3190 | 2681 | 2824 |
| SL1344_RS09060_ | 3865 | 4027 | 4138 | 5204 | 4646 | 5181 |
| SL1344_RS09065_ | 3025 | 3752 | 3427 | 4867 | 4346 | 4500 |
| SL1344_RS09070_ | 74 | 125 | 107 | 215 | 243 | 250 |
| SL1344_RS09085_ | 393 | 447 | 414 | 1029 | 923 | 856 |
| SL1344_RS09090_ | 546 | 585 | 560 | 1333 | 1217 | 1200 |
| SL1344_RS09095_ | 538 | 605 | 593 | 1289 | 1145 | 1295 |
| SL1344_RS09100_ | 1233 | 1614 | 1632 | 3098 | 3144 | 3277 |
| SL1344_RS09105_ | 296 | 258 | 333 | 521 | 445 | 647 |
| SL1344_RS09115_ | 272 | 361 | 333 | 656 | 510 | 648 |
| SL1344_RS09125_ | 45 | 60 | 34 | 124 | 88 | 66 |
| SL1344_RS09130_ | 277 | 274 | 262 | 437 | 507 | 536 |
| SL1344_RS09135_ | 119 | 144 | 138 | 270 | 146 | 194 |
| SL1344_RS09140_ | 2059 | 2159 | 2213 | 4242 | 4184 | 4414 |
| SL1344_RS09145_ | 1331 | 1322 | 1270 | 2851 | 2399 | 3148 |
| SL1344_RS09150_ | 3011 | 3267 | 2934 | 7382 | 6673 | 8381 |
| SL1344_RS09155_ | 168 | 145 | 132 | 235 | 302 | 276 |
| SL1344_RS09160_ | 323 | 394 | 393 | 358 | 413 | 378 |
| SL1344_RS09165_ | 242 | 208 | 210 | 541 | 396 | 448 |
| SL1344_RS09170_ | 241 | 235 | 266 | 517 | 486 | 463 |
| SL1344_RS09175_ | 24266 | 24842 | 27667 | 38558 | 34137 | 36693 |
| SL1344_RS09180_ | 12901 | 10593 | 14949 | 23712 | 14541 | 24022 |
| SL1344_RS09185_ | 587 | 574 | 513 | 913 | 791 | 1042 |
| SL1344_RS09190_ | 232 | 259 | 269 | 303 | 282 | 515 |
| SL1344_RS09200_ | 1660 | 1739 | 1687 | 2427 | 1967 | 2198 |
| SL1344_RS09205_ | 74 | 108 | 104 | 204 | 150 | 179 |
| SL1344_RS09210_ | 1389 | 1517 | 1652 | 2570 | 1721 | 2227 |
| SL1344_RS09235_ | 915 | 973 | 1002 | 1748 | 1616 | 1692 |

| | | | | | | |
|---|---|---|---|---|---|---|
| SL1344_RS09245_ | 220 | 186 | 189 | 513 | 303 | 416 |
| SL1344_RS09250_ | 48 | 45 | 62 | 87 | 76 | 113 |
| SL1344_RS09260_ | 136 | 92 | 97 | 294 | 243 | 291 |
| SL1344_RS09270_ | 12 | 18 | 20 | 19 | 14 | 24 |
| SL1344_RS09275_ | 186 | 265 | 318 | 530 | 313 | 495 |
| SL1344_RS09285_ | 53 | 43 | 62 | 95 | 69 | 95 |
| SL1344_RS09290_ | 96 | 90 | 85 | 191 | 91 | 202 |
| SL1344_RS09295_ | 21 | 31 | 19 | 37 | 52 | 50 |
| SL1344_RS09300_ | 6 | 16 | 6 | 20 | 30 | 18 |
| SL1344_RS09305_ | 60 | 78 | 70 | 142 | 108 | 125 |
| SL1344_RS09310_ | 8 | 3 | 7 | 8 | 11 | 11 |
| SL1344_RS09315_ | 29 | 31 | 25 | 45 | 35 | 41 |
| SL1344_RS09320_ | 172 | 120 | 178 | 220 | 118 | 227 |
| SL1344_RS09325_ | 10 | 16 | 10 | 33 | 31 | 28 |
| SL1344_RS09335_ | 14 | 10 | 18 | 17 | 5 | 14 |
| SL1344_RS09340_ | 163 | 176 | 251 | 814 | 373 | 703 |
| SL1344_RS09345_ | 7 | 2 | 3 | 19 | 14 | 13 |
| SL1344_RS09350_ | 3 | 5 | 3 | 13 | 13 | 14 |
| SL1344_RS09355_ | 15 | 12 | 11 | 17 | 28 | 25 |
| SL1344_RS09360_ | 27 | 25 | 22 | 45 | 44 | 58 |
| SL1344_RS09365_ | 102 | 191 | 152 | 147 | 237 | 90 |
| SL1344_RS09370_ | 1089 | 1061 | 1044 | 1777 | 1907 | 1960 |
| SL1344_RS09380_ | 900 | 895 | 1004 | 1610 | 1159 | 1765 |
| SL1344_RS09385_ | 262 | 199 | 243 | 514 | 338 | 560 |
| SL1344_RS09390_ | 276 | 255 | 261 | 422 | 358 | 409 |
| SL1344_RS09395_ | 375 | 436 | 387 | 781 | 660 | 738 |
| SL1344_RS09400_ | 449 | 512 | 450 | 910 | 834 | 962 |
| SL1344_RS09405_ | 176 | 158 | 207 | 453 | 404 | 453 |
| SL1344_RS09410_ | 313 | 353 | 334 | 901 | 712 | 783 |
| SL1344_RS09420_ | 154 | 142 | 181 | 340 | 358 | 322 |
| SL1344_RS09425_ | 778 | 883 | 910 | 1396 | 1405 | 1581 |
| SL1344_RS09430_ | 382 | 469 | 484 | 725 | 668 | 692 |
| SL1344_RS09435_ | 2342 | 2389 | 2206 | 4548 | 4101 | 5837 |
| SL1344_RS09440_ | 741 | 734 | 792 | 1168 | 912 | 1055 |
| SL1344_RS09455_ | 1051 | 1028 | 1028 | 1904 | 1819 | 1790 |
| SL1344_RS09460_ | 538 | 533 | 498 | 1069 | 844 | 979 |
| SL1344_RS09465_ | 202 | 269 | 294 | 478 | 397 | 414 |
| SL1344_RS09470_ | 260 | 328 | 310 | 600 | 597 | 511 |
| SL1344_RS09485_ | 25 | 21 | 23 | 50 | 41 | 41 |
| SL1344_RS09490_ | 31 | 33 | 32 | 79 | 58 | 105 |
| SL1344_RS09495_ | 1103 | 1089 | 1149 | 2517 | 1859 | 2184 |
| SL1344_RS09500_ | 944 | 856 | 888 | 1967 | 1566 | 1931 |
| SL1344_RS09510_ | 2491 | 2783 | 2877 | 5444 | 4585 | 4801 |
| SL1344_RS09515_ | 264 | 298 | 302 | 438 | 445 | 503 |
| SL1344_RS09520_ | 245 | 275 | 237 | 425 | 439 | 410 |
| SL1344_RS09525_ | 535 | 656 | 732 | 1346 | 998 | 1279 |
| SL1344_RS09535_ | 650 | 833 | 834 | 1468 | 1169 | 1288 |
| SL1344_RS09540_ | 197 | 261 | 251 | 569 | 453 | 490 |
| SL1344_RS09545_ | 148 | 162 | 151 | 356 | 278 | 354 |
| SL1344_RS09550_ | 1525 | 1493 | 1440 | 3517 | 2410 | 2765 |
| SL1344_RS09555_ | 67 | 64 | 79 | 183 | 138 | 116 |
| SL1344_RS09635_ | 68 | 93 | 79 | 166 | 108 | 149 |
| SL1344_RS09640_ | 218 | 351 | 303 | 475 | 412 | 543 |
| SL1344_RS09645_ | 159 | 179 | 182 | 355 | 239 | 328 |
| SL1344_RS09650_ | 57 | 66 | 63 | 131 | 101 | 118 |
| SL1344_RS09655_ | 0 | 3 | 3 | 1 | 1 | 1 |
| SL1344_RS09660_ | 51 | 36 | 36 | 65 | 102 | 83 |
| SL1344_RS09670_ | 93 | 67 | 90 | 76 | 124 | 98 |
| SL1344_RS09680_ | 1066 | 1289 | 1188 | 1556 | 1261 | 1435 |
| SL1344_RS09685_ | 14 | 15 | 21 | 28 | 34 | 30 |
| SL1344_RS09690_ | 257 | 242 | 337 | 453 | 380 | 304 |
| SL1344_RS09695_ | 861 | 875 | 834 | 1428 | 1631 | 1552 |
| SL1344_RS09700_ | 703 | 614 | 583 | 1007 | 892 | 967 |

| | | | | | | |
|---|---|---|---|---|---|---|
| SL1344_RS09705_ | 469 | 474 | 454 | 630 | 582 | 518 |
| SL1344_RS09710_ | 196 | 198 | 228 | 292 | 237 | 294 |
| SL1344_RS09715_ | 2826 | 2429 | 1645 | 7504 | 6475 | 3036 |
| SL1344_RS09725_ | 1729 | 1779 | 2240 | 5152 | 3963 | 3391 |
| SL1344_RS09730_ | 863 | 1019 | 1029 | 1960 | 1497 | 1737 |
| SL1344_RS09805_ | 10289 | 11780 | 11419 | 21852 | 14496 | 21509 |
| SL1344_RS09810_ | 3984 | 4518 | 4669 | 8185 | 6205 | 8016 |
| SL1344_RS09820_ | 511 | 426 | 376 | 847 | 996 | 656 |
| SL1344_RS09850_ | 5145 | 6307 | 5933 | 9309 | 8887 | 8743 |
| SL1344_RS09855_ | 2463 | 2969 | 3061 | 4715 | 4605 | 4129 |
| SL1344_RS09860_ | 1987 | 2285 | 2440 | 3702 | 3605 | 3524 |
| SL1344_RS09875_ | 1837 | 2184 | 1987 | 3315 | 3155 | 2964 |
| SL1344_RS09885_ | 2626 | 3097 | 2985 | 5203 | 4503 | 4673 |
| SL1344_RS09890_ | 1355 | 1553 | 1598 | 2470 | 2608 | 2392 |
| SL1344_RS09895_ | 995 | 1380 | 1251 | 1778 | 2210 | 1694 |
| SL1344_RS09900_ | 1380 | 1618 | 1585 | 2602 | 2428 | 2329 |
| SL1344_RS09910_ | 435 | 453 | 423 | 790 | 733 | 605 |
| SL1344_RS09920_ | 636 | 585 | 701 | 429 | 473 | 616 |
| SL1344_RS09925_ | 54 | 37 | 56 | 44 | 30 | 43 |
| SL1344_RS09930_ | 39 | 63 | 56 | 42 | 47 | 41 |
| SL1344_RS09935_ | 286 | 257 | 248 | 216 | 148 | 223 |
| SL1344_RS09940_ | 1258 | 1052 | 1169 | 929 | 837 | 929 |
| SL1344_RS09945_ | 247 | 269 | 252 | 222 | 209 | 238 |
| SL1344_RS09950_ | 93 | 104 | 108 | 88 | 61 | 85 |
| SL1344_RS09955_ | 172 | 204 | 223 | 190 | 186 | 141 |
| SL1344_RS09960_ | 473 | 446 | 482 | 507 | 364 | 388 |
| SL1344_RS09965_ | 107 | 108 | 92 | 92 | 87 | 91 |
| SL1344_RS09970_ | 14 | 14 | 8 | 10 | 14 | 21 |
| SL1344_RS09975_ | 213 | 210 | 204 | 268 | 222 | 190 |
| SL1344_RS09985_ | 180 | 172 | 184 | 170 | 159 | 139 |
| SL1344_RS09990_ | 60 | 60 | 58 | 56 | 38 | 42 |
| SL1344_RS09995_ | 9 | 10 | 8 | 8 | 8 | 8 |
| SL1344_RS10000_ | 29 | 25 | 39 | 40 | 20 | 30 |
| SL1344_RS10005_ | 48 | 43 | 54 | 72 | 38 | 51 |
| SL1344_RS10010_ | 45 | 34 | 30 | 65 | 44 | 55 |
| SL1344_RS10015_ | 6 | 10 | 18 | 11 | 10 | 23 |
| SL1344_RS10020_ | 19 | 10 | 17 | 19 | 19 | 30 |
| SL1344_RS10025_ | 18 | 11 | 12 | 18 | 18 | 18 |
| SL1344_RS10030_ | 4 | 8 | 6 | 13 | 14 | 18 |
| SL1344_RS10035_ | 30 | 42 | 35 | 36 | 39 | 34 |
| SL1344_RS10040_ | 46 | 62 | 81 | 65 | 52 | 73 |
| SL1344_RS10045_ | 40 | 64 | 46 | 61 | 42 | 58 |
| SL1344_RS10050_ | 135 | 162 | 158 | 193 | 138 | 166 |
| SL1344_RS10055_ | 29 | 32 | 28 | 29 | 37 | 28 |
| SL1344_RS10060_ | 49 | 46 | 49 | 44 | 38 | 44 |
| SL1344_RS10065_ | 49 | 57 | 53 | 80 | 54 | 86 |
| SL1344_RS10070_ | 17 | 17 | 16 | 24 | 22 | 22 |
| SL1344_RS10075_ | 19 | 15 | 10 | 25 | 27 | 26 |
| SL1344_RS10080_ | 12 | 15 | 13 | 18 | 9 | 20 |
| SL1344_RS10085_ | 10 | 18 | 20 | 19 | 10 | 25 |
| SL1344_RS10090_ | 11 | 23 | 13 | 14 | 10 | 21 |
| SL1344_RS10095_ | 43 | 50 | 41 | 75 | 58 | 80 |
| SL1344_RS10100_ | 13 | 14 | 16 | 17 | 24 | 32 |
| SL1344_RS10105_ | 33 | 32 | 22 | 54 | 39 | 50 |
| SL1344_RS10110_ | 4 | 5 | 9 | 6 | 6 | 3 |
| SL1344_RS10115_ | 70 | 73 | 67 | 92 | 56 | 83 |
| SL1344_RS10120_ | 18 | 25 | 34 | 32 | 25 | 29 |
| SL1344_RS10130_ | 65 | 60 | 51 | 63 | 78 | 70 |
| SL1344_RS10135_ | 18 | 13 | 17 | 25 | 20 | 25 |
| SL1344_RS10140_ | 28 | 32 | 25 | 47 | 35 | 37 |
| SL1344_RS10145_ | 20 | 32 | 26 | 29 | 27 | 32 |
| SL1344_RS10150_ | 8 | 11 | 15 | 25 | 12 | 21 |
| SL1344_RS10155_ | 60 | 70 | 97 | 89 | 51 | 78 |

| | | | | | | |
|---|---|---|---|---|---|---|
| SL1344_RS10160_ | 102 | 73 | 102 | 134 | 105 | 128 |
| SL1344_RS10165_ | 48 | 54 | 50 | 80 | 65 | 49 |
| SL1344_RS10170_ | 50 | 39 | 33 | 59 | 42 | 59 |
| SL1344_RS10175_ | 29 | 17 | 20 | 25 | 15 | 25 |
| SL1344_RS10180_ | 34 | 38 | 34 | 43 | 37 | 32 |
| SL1344_RS10185_ | 32 | 34 | 31 | 30 | 28 | 34 |
| SL1344_RS10190_ | 50 | 64 | 49 | 60 | 55 | 62 |
| SL1344_RS10195_ | 16 | 14 | 16 | 20 | 20 | 15 |
| SL1344_RS10200_ | 645 | 984 | 774 | 986 | 981 | 538 |
| SL1344_RS10205_ | 61 | 52 | 55 | 77 | 56 | 66 |
| SL1344_RS10210_ | 29 | 25 | 23 | 51 | 39 | 36 |
| SL1344_RS10215_ | 1895 | 2117 | 2363 | 2273 | 1770 | 2367 |
| SL1344_RS10225_ | 106 | 101 | 114 | 103 | 84 | 101 |
| SL1344_RS10230_ | 311 | 286 | 345 | 434 | 279 | 335 |
| SL1344_RS10235_ | 8 | 26 | 15 | 26 | 17 | 26 |
| SL1344_RS10240_ | 13 | 10 | 12 | 4 | 24 | 10 |
| SL1344_RS10245_ | 25 | 15 | 17 | 14 | 11 | 21 |
| SL1344_RS10250_ | 18 | 21 | 23 | 24 | 16 | 47 |
| SL1344_RS10255_ | 56 | 71 | 64 | 71 | 70 | 67 |
| SL1344_RS10260_ | 66 | 75 | 60 | 64 | 47 | 65 |
| SL1344_RS10265_ | 34 | 35 | 37 | 56 | 31 | 38 |
| SL1344_RS10270_ | 111 | 81 | 108 | 74 | 76 | 83 |
| SL1344_RS10275_ | 554 | 962 | 934 | 616 | 538 | 528 |
| SL1344_RS10280_ | 902 | 872 | 1053 | 1032 | 707 | 653 |
| SL1344_RS10285_ | 164 | 245 | 179 | 261 | 356 | 136 |
| SL1344_RS10290_ | 14 | 11 | 10 | 12 | 7 | 7 |
| SL1344_RS10295_ | 1 | 1 | 3 | 3 | 6 | 5 |
| SL1344_RS10300_ | 131 | 195 | 139 | 127 | 186 | 72 |
| SL1344_RS10305_ | 118 | 79 | 98 | 96 | 78 | 69 |
| SL1344_RS10310_ | 10 | 4 | 7 | 10 | 7 | 12 |
| SL1344_RS10315_ | 50 | 69 | 68 | 41 | 41 | 54 |
| SL1344_RS10320_ | 758 | 988 | 1146 | 798 | 941 | 704 |
| SL1344_RS10325_ | 25 | 24 | 33 | 29 | 23 | 21 |
| SL1344_RS10330_ | 1101 | 1424 | 1232 | 1073 | 1061 | 1283 |
| SL1344_RS10335_ | 37 | 50 | 47 | 34 | 27 | 47 |
| SL1344_RS10340_ | 132 | 157 | 149 | 156 | 96 | 125 |
| SL1344_RS10345_ | 162 | 174 | 194 | 159 | 121 | 178 |
| SL1344_RS10350_ | 149 | 180 | 237 | 151 | 122 | 153 |
| SL1344_RS10355_ | 898 | 998 | 674 | 834 | 1310 | 429 |
| SL1344_RS10360_ | 103 | 143 | 154 | 79 | 81 | 110 |
| SL1344_RS10365_ | 180 | 202 | 200 | 256 | 240 | 124 |
| SL1344_RS10370_ | 9 | 12 | 7 | 11 | 11 | 6 |
| SL1344_RS10375_ | 179 | 176 | 182 | 237 | 163 | 170 |
| SL1344_RS10380_ | 215 | 234 | 218 | 270 | 248 | 201 |
| SL1344_RS10390_ | 80 | 59 | 75 | 105 | 61 | 85 |
| SL1344_RS10395_ | 189 | 176 | 203 | 264 | 199 | 214 |
| SL1344_RS10400_ | 166 | 120 | 126 | 181 | 91 | 128 |
| SL1344_RS10405_ | 111 | 65 | 98 | 132 | 49 | 87 |
| SL1344_RS10410_ | 34 | 21 | 26 | 50 | 18 | 30 |
| SL1344_RS10415_ | 98 | 54 | 88 | 120 | 50 | 103 |
| SL1344_RS10420_ | 136 | 81 | 104 | 125 | 47 | 104 |
| SL1344_RS10430_ | 210 | 105 | 187 | 188 | 109 | 145 |
| SL1344_RS10435_ | 197 | 126 | 209 | 186 | 87 | 185 |
| SL1344_RS10440_ | 147 | 115 | 186 | 114 | 67 | 155 |
| SL1344_RS10445_ | 91 | 81 | 67 | 50 | 66 | 88 |
| SL1344_RS10450_ | 89 | 116 | 79 | 63 | 64 | 93 |
| SL1344_RS10455_ | 99 | 113 | 93 | 55 | 85 | 97 |
| SL1344_RS10470_ | 168 | 184 | 190 | 107 | 179 | 157 |
| SL1344_RS10475_ | 244 | 172 | 204 | 186 | 196 | 157 |
| SL1344_RS10480_ | 198 | 275 | 235 | 242 | 140 | 209 |
| SL1344_RS10485_ | 65 | 74 | 62 | 69 | 62 | 60 |
| SL1344_RS10490_ | 2 | 2 | 2 | 1 | 1 | 3 |
| SL1344_RS10495_ | 6 | 11 | 7 | 16 | 10 | 6 |

| | | | | | | |
|---|---|---|---|---|---|---|
| SL1344_RS10515_ | 20 | 26 | 22 | 35 | 26 | 20 |
| SL1344_RS10520_ | 1 | 5 | 5 | 5 | 5 | 2 |
| SL1344_RS10530_ | 14 | 9 | 9 | 11 | 8 | 13 |
| SL1344_RS10535_ | 8 | 13 | 11 | 13 | 12 | 15 |
| SL1344_RS10540_ | 8 | 13 | 16 | 16 | 12 | 26 |
| SL1344_RS10545_ | 6 | 4 | 8 | 12 | 10 | 10 |
| SL1344_RS10550_ | 10 | 10 | 12 | 27 | 8 | 18 |
| SL1344_RS10555_ | 17 | 14 | 22 | 26 | 10 | 27 |
| SL1344_RS10560_ | 15 | 12 | 31 | 30 | 18 | 25 |
| SL1344_RS10565_ | 21 | 18 | 15 | 39 | 39 | 38 |
| SL1344_RS10570_ | 12 | 9 | 9 | 19 | 11 | 14 |
| SL1344_RS10575_ | 5 | 8 | 5 | 13 | 7 | 3 |
| SL1344_RS10580_ | 5 | 9 | 8 | 7 | 7 | 6 |
| SL1344_RS10585_ | 44 | 37 | 48 | 59 | 47 | 55 |
| SL1344_RS10590_ | 91 | 97 | 115 | 102 | 87 | 66 |
| SL1344_RS10595_ | 2546 | 3587 | 3526 | 2196 | 2266 | 2552 |
| SL1344_RS10600_ | 513 | 509 | 486 | 661 | 545 | 585 |
| SL1344_RS10640_ | 2027 | 2279 | 2516 | 2115 | 1134 | 1427 |
| SL1344_RS10645_ | 262 | 292 | 263 | 229 | 265 | 231 |
| SL1344_RS10650_ | 773 | 749 | 790 | 607 | 630 | 696 |
| SL1344_RS10655_ | 230 | 318 | 304 | 137 | 142 | 172 |
| SL1344_RS10660_ | 231 | 287 | 323 | 188 | 144 | 168 |
| SL1344_RS10665_ | 277 | 280 | 313 | 198 | 141 | 174 |
| SL1344_RS10670_ | 505 | 589 | 567 | 360 | 329 | 324 |
| SL1344_RS10675_ | 235 | 240 | 282 | 162 | 120 | 170 |
| SL1344_RS10680_ | 333 | 415 | 384 | 276 | 223 | 235 |
| SL1344_RS10685_ | 502 | 507 | 569 | 386 | 284 | 351 |
| SL1344_RS10690_ | 435 | 379 | 493 | 283 | 206 | 242 |
| SL1344_RS10700_ | 856 | 877 | 1020 | 516 | 408 | 519 |
| SL1344_RS10705_ | 1804 | 2208 | 1965 | 1266 | 1098 | 1420 |
| SL1344_RS10710_ | 1092 | 1011 | 1315 | 853 | 709 | 958 |
| SL1344_RS10715_ | 1006 | 1068 | 1183 | 785 | 665 | 911 |
| SL1344_RS10720_ | 1627 | 1725 | 1897 | 1312 | 1141 | 1423 |
| SL1344_RS10725_ | 1035 | 1169 | 1312 | 824 | 682 | 919 |
| SL1344_RS10730_ | 1003 | 1078 | 1306 | 746 | 773 | 839 |
| SL1344_RS10735_ | 1277 | 1491 | 1605 | 1039 | 1069 | 1090 |
| SL1344_RS10740_ | 1202 | 1389 | 1538 | 981 | 868 | 937 |
| SL1344_RS10745_ | 2366 | 3022 | 3000 | 1850 | 1863 | 1842 |
| SL1344_RS10750_ | 5412 | 6262 | 6704 | 4120 | 3411 | 4622 |
| SL1344_RS10755_ | 1775 | 1600 | 1929 | 1205 | 1014 | 1480 |
| SL1344_RS10760_ | 995 | 1003 | 1090 | 727 | 649 | 847 |
| SL1344_RS10765_ | 1256 | 1241 | 1490 | 980 | 736 | 1003 |
| SL1344_RS10770_ | 941 | 941 | 1073 | 692 | 551 | 716 |
| SL1344_RS10775_ | 885 | 745 | 822 | 663 | 473 | 681 |
| SL1344_RS10780_ | 1247 | 1231 | 1313 | 959 | 752 | 876 |
| SL1344_RS10785_ | 1533 | 1485 | 1620 | 1251 | 864 | 1074 |
| SL1344_RS10790_ | 2699 | 2843 | 2908 | 2646 | 2168 | 2404 |
| SL1344_RS10795_ | 22 | 37 | 40 | 46 | 38 | 35 |
| SL1344_RS10800_ | 32 | 27 | 38 | 33 | 27 | 14 |
| SL1344_RS10805_ | 22 | 31 | 30 | 26 | 20 | 31 |
| SL1344_RS10810_ | 33 | 31 | 35 | 38 | 28 | 25 |
| SL1344_RS10815_ | 26 | 18 | 23 | 22 | 27 | 33 |
| SL1344_RS10820_ | 12 | 16 | 16 | 26 | 25 | 12 |
| SL1344_RS10825_ | 31 | 33 | 32 | 34 | 31 | 31 |
| SL1344_RS10830_ | 30 | 41 | 34 | 49 | 46 | 45 |
| SL1344_RS10835_ | 14 | 19 | 10 | 13 | 11 | 39 |
| SL1344_RS10840_ | 30 | 33 | 27 | 21 | 24 | 38 |
| SL1344_RS10845_ | 21 | 28 | 25 | 25 | 15 | 21 |
| SL1344_RS10850_ | 6 | 5 | 7 | 9 | 3 | 5 |
| SL1344_RS10855_ | 18 | 10 | 20 | 12 | 7 | 18 |
| SL1344_RS10860_ | 27 | 33 | 42 | 29 | 24 | 46 |
| SL1344_RS10865_ | 21 | 55 | 33 | 32 | 31 | 22 |
| SL1344_RS10870_ | 9 | 13 | 11 | 14 | 16 | 13 |

| | | | | | | |
|---|---|---|---|---|---|---|
| SL1344_RS10875_ | 17 | 16 | 13 | 26 | 19 | 20 |
| SL1344_RS10880_ | 35 | 42 | 50 | 46 | 43 | 44 |
| SL1344_RS10885_ | 8 | 8 | 6 | 7 | 5 | 4 |
| SL1344_RS10890_ | 10 | 8 | 15 | 14 | 10 | 8 |
| SL1344_RS10895_ | 324 | 460 | 367 | 299 | 316 | 316 |
| SL1344_RS10900_ | 2305 | 2584 | 2616 | 1895 | 1962 | 1790 |
| SL1344_RS10905_ | 717 | 938 | 964 | 700 | 725 | 634 |
| SL1344_RS10910_ | 717 | 800 | 901 | 771 | 477 | 596 |
| SL1344_RS10915_ | 540 | 744 | 627 | 513 | 469 | 436 |
| SL1344_RS10920_ | 42 | 27 | 29 | 25 | 31 | 27 |
| SL1344_RS10930_ | 73 | 86 | 73 | 61 | 74 | 73 |
| SL1344_RS10935_ | 122 | 130 | 117 | 136 | 97 | 89 |
| SL1344_RS10940_ | 127 | 127 | 137 | 140 | 102 | 96 |
| SL1344_RS10945_ | 121 | 144 | 144 | 143 | 92 | 115 |
| SL1344_RS10950_ | 142 | 223 | 222 | 226 | 165 | 176 |
| SL1344_RS10955_ | 255 | 397 | 346 | 336 | 318 | 334 |
| SL1344_RS10960_ | 22 | 24 | 27 | 11 | 16 | 7 |
| SL1344_RS10965_ | 16 | 19 | 17 | 19 | 17 | 11 |
| SL1344_RS10970_ | 103 | 104 | 109 | 81 | 66 | 70 |
| SL1344_RS10975_ | 166 | 199 | 153 | 187 | 119 | 165 |
| SL1344_RS10980_ | 14 | 9 | 15 | 22 | 4 | 15 |
| SL1344_RS10985_ | 14 | 34 | 34 | 34 | 24 | 27 |
| SL1344_RS10990_ | 35 | 47 | 38 | 53 | 44 | 37 |
| SL1344_RS10995_ | 6 | 4 | 8 | 12 | 4 | 8 |
| SL1344_RS11000_ | 176 | 188 | 191 | 144 | 108 | 126 |
| SL1344_RS11005_ | 238 | 231 | 259 | 283 | 208 | 237 |
| SL1344_RS11010_ | 173 | 173 | 206 | 233 | 162 | 224 |
| SL1344_RS11015_ | 199 | 210 | 224 | 229 | 207 | 233 |
| SL1344_RS11020_ | 160 | 177 | 203 | 223 | 194 | 189 |
| SL1344_RS11025_ | 310 | 398 | 381 | 331 | 286 | 320 |
| SL1344_RS11030_ | 95 | 150 | 111 | 98 | 85 | 70 |
| SL1344_RS11035_ | 98 | 133 | 95 | 115 | 101 | 67 |
| SL1344_RS11040_ | 446 | 577 | 586 | 386 | 282 | 412 |
| SL1344_RS11045_ | 28 | 26 | 19 | 35 | 13 | 17 |
| SL1344_RS11050_ | 44 | 29 | 30 | 40 | 34 | 28 |
| SL1344_RS11055_ | 13 | 4 | 6 | 5 | 2 | 10 |
| SL1344_RS11060_ | 6 | 20 | 10 | 7 | 8 | 10 |
| SL1344_RS11065_ | 152 | 142 | 145 | 164 | 87 | 129 |
| SL1344_RS11070_ | 1688 | 2068 | 2081 | 1651 | 1438 | 1617 |
| SL1344_RS11080_ | 93 | 67 | 78 | 56 | 83 | 71 |
| SL1344_RS11085_ | 34 | 39 | 39 | 41 | 13 | 21 |
| SL1344_RS11090_ | 569 | 652 | 663 | 600 | 460 | 480 |
| SL1344_RS11095_ | 611 | 590 | 665 | 492 | 407 | 424 |
| SL1344_RS11100_ | 1187 | 1051 | 1401 | 1175 | 889 | 856 |
| SL1344_RS11105_ | 96 | 93 | 109 | 62 | 48 | 50 |
| SL1344_RS11110_ | 271 | 330 | 351 | 299 | 167 | 256 |
| SL1344_RS11115_ | 97 | 124 | 153 | 76 | 65 | 65 |
| SL1344_RS11120_ | 173 | 177 | 167 | 143 | 101 | 131 |
| SL1344_RS11125_ | 264 | 241 | 230 | 166 | 173 | 168 |
| SL1344_RS11130_ | 219 | 234 | 229 | 182 | 144 | 152 |
| SL1344_RS11135_ | 757 | 912 | 880 | 600 | 545 | 605 |
| SL1344_RS11140_ | 689 | 848 | 877 | 680 | 501 | 591 |
| SL1344_RS11145_ | 910 | 1178 | 1246 | 713 | 589 | 692 |
| SL1344_RS11150_ | 28 | 24 | 23 | 26 | 20 | 27 |
| SL1344_RS11155_ | 39 | 53 | 44 | 38 | 35 | 27 |
| SL1344_RS11160_ | 66 | 68 | 69 | 51 | 43 | 54 |
| SL1344_RS11165_ | 42 | 60 | 55 | 54 | 36 | 37 |
| SL1344_RS11170_ | 235 | 233 | 238 | 208 | 227 | 183 |
| SL1344_RS11175_ | 155 | 163 | 170 | 150 | 139 | 159 |
| SL1344_RS11180_ | 53 | 53 | 48 | 35 | 53 | 43 |
| SL1344_RS11185_ | 33 | 27 | 43 | 41 | 26 | 43 |
| SL1344_RS11190_ | 37 | 30 | 27 | 30 | 23 | 41 |
| SL1344_RS11195_ | 45 | 36 | 36 | 28 | 66 | 38 |

| | | | | | | |
|---|---|---|---|---|---|---|
| SL1344_RS11200_ | 218 | 225 | 237 | 210 | 218 | 183 |
| SL1344_RS11205_ | 62 | 89 | 73 | 66 | 66 | 64 |
| SL1344_RS11210_ | 112 | 123 | 104 | 108 | 88 | 103 |
| SL1344_RS11215_ | 379 | 526 | 463 | 566 | 494 | 508 |
| SL1344_RS11220_ | 627 | 765 | 706 | 642 | 493 | 488 |
| SL1344_RS11225_ | 270 | 256 | 278 | 200 | 204 | 156 |
| SL1344_RS11230_ | 158 | 153 | 152 | 100 | 76 | 79 |
| SL1344_RS11235_ | 305 | 311 | 299 | 176 | 191 | 177 |
| SL1344_RS11240_ | 2133 | 2104 | 1855 | 2151 | 1967 | 2556 |
| SL1344_RS11245_ | 7391 | 5847 | 6576 | 6497 | 6919 | 7463 |
| SL1344_RS11250_ | 24149 | 17808 | 22061 | 26318 | 19313 | 22682 |
| SL1344_RS11255_ | 1416 | 1167 | 1380 | 1873 | 1192 | 958 |
| SL1344_RS11260_ | 465 | 638 | 549 | 425 | 434 | 460 |
| SL1344_RS11265_ | 462 | 638 | 601 | 425 | 397 | 475 |
| SL1344_RS11270_ | 1769 | 2078 | 1969 | 1830 | 1360 | 1484 |
| SL1344_RS11275_ | 71 | 65 | 49 | 62 | 63 | 45 |
| SL1344_RS11280_ | 100 | 140 | 136 | 138 | 110 | 110 |
| SL1344_RS11285_ | 10 | 9 | 5 | 13 | 3 | 4 |
| SL1344_RS11290_ | 35 | 33 | 26 | 26 | 43 | 39 |
| SL1344_RS11295_ | 91 | 109 | 98 | 208 | 131 | 140 |
| SL1344_RS11300_ | 934 | 962 | 827 | 957 | 925 | 1001 |
| SL1344_RS11305_ | 340 | 398 | 313 | 489 | 355 | 508 |
| SL1344_RS11310_ | 251 | 268 | 209 | 248 | 153 | 206 |
| SL1344_RS11315_ | 591 | 684 | 704 | 479 | 444 | 516 |
| SL1344_RS11320_ | 118 | 136 | 175 | 129 | 131 | 127 |
| SL1344_RS11325_ | 45 | 73 | 54 | 50 | 56 | 52 |
| SL1344_RS11330_ | 37 | 53 | 60 | 54 | 61 | 48 |
| SL1344_RS11335_ | 45 | 62 | 52 | 61 | 45 | 49 |
| SL1344_RS11340_ | 35 | 38 | 52 | 46 | 35 | 34 |
| SL1344_RS11345_ | 7 | 5 | 15 | 8 | 13 | 7 |
| SL1344_RS11350_ | 61 | 49 | 47 | 53 | 40 | 41 |
| SL1344_RS11355_ | 1849 | 2074 | 2145 | 1604 | 1122 | 1195 |
| SL1344_RS11360_ | 226 | 278 | 300 | 183 | 185 | 171 |
| SL1344_RS11365_ | 802 | 841 | 941 | 819 | 574 | 543 |
| SL1344_RS11375_ | 472 | 630 | 617 | 537 | 447 | 476 |
| SL1344_RS11380_ | 461 | 556 | 464 | 465 | 357 | 433 |
| SL1344_RS11385_ | 367 | 481 | 381 | 389 | 289 | 286 |
| SL1344_RS11390_ | 292 | 332 | 292 | 284 | 220 | 241 |
| SL1344_RS11395_ | 534 | 716 | 675 | 489 | 422 | 442 |
| SL1344_RS11400_ | 708 | 636 | 751 | 575 | 522 | 521 |
| SL1344_RS11405_ | 26 | 37 | 34 | 31 | 29 | 26 |
| SL1344_RS11410_ | 441 | 551 | 608 | 389 | 380 | 393 |
| SL1344_RS11415_ | 434 | 538 | 550 | 409 | 312 | 394 |
| SL1344_RS11420_ | 391 | 492 | 507 | 366 | 316 | 361 |
| SL1344_RS11425_ | 6857 | 7001 | 7973 | 6473 | 4942 | 5507 |
| SL1344_RS11430_ | 358 | 352 | 364 | 292 | 284 | 347 |
| SL1344_RS11435_ | 567 | 554 | 618 | 532 | 400 | 446 |
| SL1344_RS11440_ | 537 | 631 | 655 | 623 | 368 | 479 |
| SL1344_RS11445_ | 1514 | 1857 | 1942 | 1673 | 1197 | 1487 |
| SL1344_RS11450_ | 212 | 421 | 230 | 558 | 697 | 167 |
| SL1344_RS11455_ | 18 | 21 | 13 | 12 | 17 | 12 |
| SL1344_RS11460_ | 5 | 3 | 2 | 5 | 4 | 1 |
| SL1344_RS11465_ | 1133 | 1101 | 1197 | 873 | 681 | 843 |
| SL1344_RS11470_ | 22 | 30 | 26 | 17 | 17 | 17 |
| SL1344_RS11475_ | 24 | 29 | 21 | 33 | 16 | 25 |
| SL1344_RS11480_ | 29 | 39 | 40 | 30 | 17 | 32 |
| SL1344_RS11485_ | 3 | 13 | 15 | 6 | 5 | 6 |
| SL1344_RS11490_ | 7 | 6 | 5 | 10 | 4 | 1 |
| SL1344_RS11495_ | 1230 | 1282 | 1462 | 939 | 545 | 830 |
| SL1344_RS11500_ | 34 | 26 | 28 | 25 | 29 | 44 |
| SL1344_RS11505_ | 39 | 36 | 44 | 30 | 28 | 32 |
| SL1344_RS11510_ | 24 | 18 | 13 | 21 | 18 | 21 |
| SL1344_RS11515_ | 297 | 352 | 328 | 283 | 301 | 321 |

| | | | | | | |
|---|---|---|---|---|---|---|
| SL1344_RS11520_ | 4 | 4 | 3 | 1 | 2 | 4 |
| SL1344_RS11525_ | 30 | 33 | 35 | 19 | 18 | 19 |
| SL1344_RS11535_ | 10 | 10 | 10 | 9 | 6 | 5 |
| SL1344_RS11540_ | 277 | 306 | 292 | 271 | 213 | 213 |
| SL1344_RS11545_ | 187 | 187 | 178 | 155 | 124 | 128 |
| SL1344_RS11550_ | 18 | 13 | 15 | 11 | 24 | 5 |
| SL1344_RS11555_ | 63 | 50 | 56 | 45 | 40 | 45 |
| SL1344_RS11560_ | 47 | 53 | 50 | 29 | 32 | 25 |
| SL1344_RS11565_ | 5 | 4 | 12 | 3 | 2 | 6 |
| SL1344_RS11570_ | 20 | 20 | 31 | 19 | 13 | 23 |
| SL1344_RS11585_ | 183 | 187 | 181 | 147 | 129 | 166 |
| SL1344_RS11595_ | 451 | 540 | 537 | 371 | 411 | 412 |
| SL1344_RS11600_ | 240 | 271 | 268 | 197 | 252 | 210 |
| SL1344_RS11605_ | 1187 | 1475 | 1295 | 1033 | 1010 | 948 |
| SL1344_RS11610_ | 140 | 211 | 207 | 128 | 109 | 122 |
| SL1344_RS11615_ | 485 | 697 | 557 | 357 | 310 | 362 |
| SL1344_RS11620_ | 517 | 537 | 658 | 211 | 171 | 209 |
| SL1344_RS11625_ | 446 | 463 | 444 | 559 | 464 | 506 |
| SL1344_RS11630_ | 140 | 146 | 119 | 155 | 179 | 147 |
| SL1344_RS11635_ | 278 | 286 | 319 | 316 | 397 | 308 |
| SL1344_RS11640_ | 180 | 177 | 190 | 244 | 204 | 208 |
| SL1344_RS11670_ | 114 | 77 | 88 | 76 | 84 | 81 |
| SL1344_RS11675_ | 17 | 27 | 11 | 28 | 56 | 9 |
| SL1344_RS11680_ | 42 | 40 | 35 | 84 | 58 | 66 |
| SL1344_RS11685_ | 801 | 1014 | 900 | 955 | 682 | 903 |
| SL1344_RS11690_ | 3999 | 4946 | 4714 | 4379 | 3403 | 4035 |
| SL1344_RS11695_ | 1761 | 2048 | 2019 | 1730 | 1105 | 1439 |
| SL1344_RS11700_ | 1544 | 1712 | 1622 | 1456 | 1166 | 1227 |
| SL1344_RS11705_ | 124 | 233 | 214 | 175 | 176 | 163 |
| SL1344_RS11710_ | 339 | 368 | 339 | 380 | 436 | 318 |
| SL1344_RS11715_ | 32642 | 31058 | 31460 | 30398 | 31853 | 35008 |
| SL1344_RS11720_ | 24044 | 23118 | 24362 | 21300 | 22076 | 19901 |
| SL1344_RS11725_ | 3364 | 3345 | 2963 | 2423 | 2255 | 2269 |
| SL1344_RS11730_ | 2288 | 2417 | 2347 | 1555 | 1782 | 1727 |
| SL1344_RS11735_ | 2740 | 2971 | 2501 | 1817 | 2101 | 2271 |
| SL1344_RS11740_ | 9 | 2 | 5 | 3 | 8 | 0 |
| SL1344_RS11745_ | 56 | 31 | 46 | 30 | 27 | 29 |
| SL1344_RS11750_ | 110 | 102 | 135 | 96 | 88 | 95 |
| SL1344_RS11755_ | 67 | 54 | 67 | 59 | 35 | 46 |
| SL1344_RS11760_ | 105 | 117 | 117 | 89 | 73 | 70 |
| SL1344_RS11765_ | 100 | 47 | 61 | 69 | 46 | 43 |
| SL1344_RS11770_ | 555 | 727 | 633 | 473 | 480 | 545 |
| SL1344_RS11775_ | 505 | 514 | 493 | 442 | 386 | 433 |
| SL1344_RS11780_ | 391 | 413 | 397 | 329 | 282 | 298 |
| SL1344_RS11785_ | 697 | 704 | 761 | 508 | 427 | 488 |
| SL1344_RS11790_ | 961 | 1174 | 987 | 834 | 642 | 743 |
| SL1344_RS11795_ | 1022 | 1121 | 1074 | 865 | 602 | 754 |
| SL1344_RS11800_ | 2273 | 2625 | 2546 | 1792 | 1167 | 1508 |
| SL1344_RS11805_ | 1035 | 1062 | 1106 | 763 | 462 | 589 |
| SL1344_RS11810_ | 891 | 966 | 956 | 781 | 560 | 653 |
| SL1344_RS11815_ | 292 | 384 | 308 | 293 | 246 | 194 |
| SL1344_RS11820_ | 179 | 197 | 173 | 182 | 147 | 122 |
| SL1344_RS11825_ | 565 | 663 | 620 | 718 | 525 | 694 |
| SL1344_RS11830_ | 758 | 820 | 859 | 637 | 597 | 634 |
| SL1344_RS11840_ | 498 | 495 | 531 | 404 | 386 | 391 |
| SL1344_RS11845_ | 218 | 235 | 264 | 239 | 184 | 168 |
| SL1344_RS11850_ | 556 | 671 | 649 | 549 | 437 | 454 |
| SL1344_RS11855_ | 467 | 538 | 528 | 571 | 414 | 454 |
| SL1344_RS11860_ | 1271 | 1246 | 1519 | 1032 | 845 | 970 |
| SL1344_RS11865_ | 104 | 121 | 74 | 79 | 75 | 110 |
| SL1344_RS11870_ | 283 | 309 | 280 | 320 | 270 | 259 |
| SL1344_RS11880_ | 140 | 132 | 126 | 122 | 124 | 128 |
| SL1344_RS11885_ | 3367 | 3249 | 2897 | 2897 | 2371 | 3133 |

| | | | | | | |
|---|---|---|---|---|---|---|
| SL1344_RS11890_ | 2128 | 2084 | 1732 | 1610 | 1767 | 1790 |
| SL1344_RS11895_ | 5285 | 5933 | 5724 | 4655 | 4137 | 4397 |
| SL1344_RS11900_ | 1049 | 1152 | 1032 | 935 | 728 | 883 |
| SL1344_RS11905_ | 2623 | 3268 | 2974 | 2064 | 2578 | 2070 |
| SL1344_RS11910_ | 2209 | 2209 | 2311 | 1780 | 1516 | 1695 |
| SL1344_RS11915_ | 3489 | 3740 | 3633 | 3063 | 2774 | 2866 |
| SL1344_RS11920_ | 11523 | 12683 | 12969 | 9738 | 9202 | 8784 |
| SL1344_RS11925_ | 6094 | 7049 | 6720 | 4891 | 4736 | 4526 |
| SL1344_RS11930_ | 2703 | 2941 | 2945 | 2154 | 1851 | 2024 |
| SL1344_RS11935_ | 8373 | 10043 | 9472 | 7004 | 6489 | 6363 |
| SL1344_RS11940_ | 3919 | 4774 | 4937 | 3607 | 2748 | 3330 |
| SL1344_RS11945_ | 3800 | 4424 | 4617 | 3431 | 2791 | 3069 |
| SL1344_RS11950_ | 28 | 29 | 37 | 23 | 32 | 14 |
| SL1344_RS11960_ | 2300 | 2491 | 2530 | 1548 | 1576 | 1551 |
| SL1344_RS11965_ | 169 | 228 | 246 | 140 | 134 | 165 |
| SL1344_RS11970_ | 508 | 690 | 562 | 451 | 624 | 545 |
| SL1344_RS11975_ | 1443 | 1668 | 1539 | 1214 | 1246 | 1277 |
| SL1344_RS11980_ | 563 | 719 | 654 | 473 | 472 | 553 |
| SL1344_RS11985_ | 407 | 487 | 493 | 429 | 352 | 397 |
| SL1344_RS12000_ | 65 | 78 | 72 | 84 | 62 | 53 |
| SL1344_RS12005_ | 10269 | 8029 | 8741 | 8980 | 5260 | 9204 |
| SL1344_RS12010_ | 4514 | 3648 | 3735 | 3608 | 2558 | 3507 |
| SL1344_RS12015_ | 8055 | 7066 | 7201 | 6303 | 4908 | 6091 |
| SL1344_RS12020_ | 1810 | 1643 | 1578 | 1537 | 900 | 1355 |
| SL1344_RS12025_ | 2097 | 1844 | 1658 | 2019 | 1061 | 1478 |
| SL1344_RS12030_ | 5 | 4 | 3 | 7 | 1 | 1 |
| SL1344_RS12035_ | 874 | 848 | 717 | 817 | 580 | 554 |
| SL1344_RS12040_ | 1423 | 1811 | 1525 | 1504 | 1336 | 1397 |
| SL1344_RS12045_ | 577 | 635 | 511 | 570 | 507 | 513 |
| SL1344_RS12050_ | 195 | 266 | 231 | 208 | 235 | 214 |
| SL1344_RS12055_ | 30 | 28 | 27 | 22 | 18 | 15 |
| SL1344_RS12060_ | 298 | 302 | 255 | 196 | 278 | 231 |
| SL1344_RS12065_ | 157 | 173 | 139 | 103 | 135 | 151 |
| SL1344_RS12070_ | 124 | 149 | 140 | 113 | 131 | 136 |
| SL1344_RS12075_ | 179 | 177 | 156 | 115 | 222 | 156 |
| SL1344_RS12080_ | 1489 | 1657 | 1690 | 955 | 1272 | 1198 |
| SL1344_RS12085_ | 292 | 203 | 239 | 185 | 202 | 137 |
| SL1344_RS12090_ | 600 | 685 | 554 | 697 | 601 | 517 |
| SL1344_RS12095_ | 501 | 580 | 590 | 574 | 395 | 466 |
| SL1344_RS12100_ | 153 | 173 | 187 | 134 | 125 | 128 |
| SL1344_RS12105_ | 108 | 90 | 93 | 76 | 84 | 70 |
| SL1344_RS12110_ | 74 | 67 | 59 | 89 | 59 | 48 |
| SL1344_RS12115_ | 982 | 966 | 1045 | 869 | 817 | 861 |
| SL1344_RS12120_ | 1044 | 1184 | 1271 | 1326 | 793 | 924 |
| SL1344_RS12125_ | 600 | 539 | 560 | 641 | 460 | 463 |
| SL1344_RS12130_ | 1166 | 1184 | 1202 | 1149 | 967 | 1108 |
| SL1344_RS12135_ | 1328 | 1363 | 1405 | 1249 | 1113 | 1111 |
| SL1344_RS12140_ | 4627 | 5762 | 5668 | 4741 | 4048 | 3704 |
| SL1344_RS12145_ | 321 | 345 | 430 | 295 | 208 | 275 |
| SL1344_RS12155_ | 786 | 878 | 787 | 746 | 648 | 704 |
| SL1344_RS12160_ | 1650 | 2017 | 1816 | 1635 | 1434 | 1325 |
| SL1344_RS12170_ | 503 | 702 | 545 | 530 | 628 | 514 |
| SL1344_RS12175_ | 13 | 13 | 6 | 12 | 5 | 8 |
| SL1344_RS12180_ | 4 | 7 | 4 | 3 | 6 | 4 |
| SL1344_RS12185_ | 5 | 8 | 8 | 6 | 5 | 1 |
| SL1344_RS12190_ | 6 | 12 | 16 | 7 | 7 | 14 |
| SL1344_RS12195_ | 92 | 102 | 104 | 126 | 118 | 124 |
| SL1344_RS12200_ | 6822 | 7697 | 7599 | 6713 | 6520 | 8770 |
| SL1344_RS12205_ | 495 | 582 | 508 | 604 | 482 | 485 |
| SL1344_RS12210_ | 853 | 980 | 1030 | 617 | 622 | 604 |
| SL1344_RS12215_ | 320 | 334 | 437 | 287 | 205 | 263 |
| SL1344_RS12220_ | 354 | 372 | 455 | 343 | 272 | 307 |
| SL1344_RS12225_ | 438 | 602 | 484 | 429 | 393 | 363 |

| | | | | | | |
|---|---|---|---|---|---|---|
| SL1344_RS12230_ | 736 | 880 | 724 | 656 | 602 | 686 |
| SL1344_RS12235_ | 738 | 857 | 812 | 731 | 532 | 627 |
| SL1344_RS12240_ | 161 | 197 | 163 | 147 | 134 | 154 |
| SL1344_RS12245_ | 1838 | 2221 | 1878 | 1802 | 2322 | 1553 |
| SL1344_RS12250_ | 771 | 713 | 665 | 529 | 655 | 541 |
| SL1344_RS12260_ | 717 | 686 | 634 | 673 | 407 | 631 |
| SL1344_RS12265_ | 2310 | 2527 | 2818 | 1923 | 2101 | 1746 |
| SL1344_RS12270_ | 2976 | 3265 | 2878 | 3177 | 2327 | 2500 |
| SL1344_RS12275_ | 61 | 98 | 95 | 49 | 47 | 76 |
| SL1344_RS12280_ | 67 | 85 | 55 | 99 | 64 | 58 |
| SL1344_RS12285_ | 93 | 92 | 101 | 68 | 55 | 86 |
| SL1344_RS12290_ | 396 | 417 | 454 | 313 | 313 | 340 |
| SL1344_RS12295_ | 342 | 377 | 378 | 304 | 323 | 283 |
| SL1344_RS12300_ | 170 | 157 | 148 | 133 | 95 | 77 |
| SL1344_RS12305_ | 32 | 35 | 52 | 31 | 17 | 21 |
| SL1344_RS12310_ | 31 | 31 | 30 | 27 | 28 | 29 |
| SL1344_RS12315_ | 179 | 122 | 202 | 126 | 79 | 160 |
| SL1344_RS12320_ | 2 | 3 | 0 | 2 | 3 | 3 |
| SL1344_RS12325_ | 98 | 110 | 91 | 113 | 101 | 98 |
| SL1344_RS12330_ | 1171 | 1443 | 1191 | 1260 | 1258 | 1273 |
| SL1344_RS12335_ | 1209 | 1451 | 1391 | 933 | 994 | 946 |
| SL1344_RS12340_ | 184 | 242 | 203 | 188 | 176 | 146 |
| SL1344_RS12345_ | 203 | 209 | 286 | 215 | 155 | 228 |
| SL1344_RS12350_ | 353 | 396 | 402 | 371 | 328 | 351 |
| SL1344_RS12355_ | 277 | 312 | 250 | 272 | 229 | 235 |
| SL1344_RS12360_ | 243 | 265 | 256 | 219 | 184 | 180 |
| SL1344_RS12365_ | 1231 | 1458 | 1259 | 1252 | 1202 | 1079 |
| SL1344_RS12370_ | 640 | 679 | 679 | 684 | 539 | 582 |
| SL1344_RS12375_ | 2034 | 1693 | 1164 | 3310 | 4307 | 889 |
| SL1344_RS12380_ | 9490 | 4759 | 5589 | 14149 | 9896 | 4966 |
| SL1344_RS12385_ | 722 | 696 | 934 | 787 | 379 | 549 |
| SL1344_RS12390_ | 1271 | 1127 | 1228 | 1200 | 1040 | 864 |
| SL1344_RS12395_ | 2895 | 3409 | 3360 | 2846 | 2159 | 2449 |
| SL1344_RS12400_ | 4355 | 4151 | 4356 | 5517 | 5838 | 2523 |
| SL1344_RS12405_ | 3722 | 4087 | 3232 | 4124 | 6324 | 1773 |
| SL1344_RS12410_ | 6710 | 5284 | 4989 | 6741 | 8473 | 3331 |
| SL1344_RS12415_ | 3297 | 2365 | 2410 | 3108 | 3612 | 1704 |
| SL1344_RS12420_ | 29 | 52 | 49 | 40 | 41 | 44 |
| SL1344_RS12425_ | 5 | 2 | 2 | 4 | 6 | 2 |
| SL1344_RS12430_ | 47 | 53 | 48 | 45 | 66 | 56 |
| SL1344_RS12435_ | 6 | 6 | 6 | 2 | 7 | 5 |
| SL1344_RS12440_ | 40 | 36 | 27 | 30 | 28 | 20 |
| SL1344_RS12445_ | 88 | 122 | 121 | 115 | 79 | 110 |
| SL1344_RS12450_ | 324 | 371 | 382 | 357 | 261 | 302 |
| SL1344_RS12455_ | 356 | 428 | 404 | 291 | 263 | 307 |
| SL1344_RS12465_ | 3429 | 4215 | 4021 | 3002 | 2888 | 2736 |
| SL1344_RS12480_ | 5 | 7 | 3 | 3 | 1 | 10 |
| SL1344_RS12485_ | 2364 | 2783 | 3019 | 2606 | 1705 | 2457 |
| SL1344_RS12490_ | 18314 | 20045 | 22016 | 17688 | 16907 | 17600 |
| SL1344_RS12495_ | 6085 | 6777 | 6033 | 6072 | 6388 | 5778 |
| SL1344_RS12500_ | 41 | 59 | 39 | 67 | 59 | 45 |
| SL1344_RS12510_ | 121 | 171 | 113 | 163 | 123 | 138 |
| SL1344_RS12515_ | 95 | 105 | 90 | 88 | 83 | 91 |
| SL1344_RS12520_ | 40 | 60 | 48 | 44 | 40 | 59 |
| SL1344_RS12525_ | 72 | 80 | 83 | 79 | 66 | 94 |
| SL1344_RS12555_ | 2691 | 2651 | 2611 | 2093 | 2316 | 2086 |
| SL1344_RS12560_ | 716 | 1007 | 893 | 750 | 585 | 615 |
| SL1344_RS12565_ | 362 | 405 | 427 | 348 | 320 | 337 |
| SL1344_RS12570_ | 345 | 495 | 405 | 332 | 354 | 315 |
| SL1344_RS12575_ | 545 | 761 | 573 | 541 | 573 | 483 |
| SL1344_RS12580_ | 343 | 417 | 399 | 285 | 240 | 266 |
| SL1344_RS12585_ | 230 | 257 | 200 | 203 | 200 | 234 |
| SL1344_RS12590_ | 85 | 82 | 76 | 64 | 59 | 54 |

| | | | | | | |
|---|---|---|---|---|---|---|
| SL1344_RS12595_ | 195 | 247 | 257 | 168 | 130 | 178 |
| SL1344_RS12600_ | 48 | 58 | 58 | 50 | 75 | 45 |
| SL1344_RS12605_ | 73 | 69 | 69 | 74 | 71 | 61 |
| SL1344_RS12610_ | 57 | 64 | 96 | 83 | 84 | 49 |
| SL1344_RS12615_ | 21 | 26 | 18 | 22 | 18 | 10 |
| SL1344_RS12620_ | 38 | 29 | 42 | 30 | 23 | 31 |
| SL1344_RS12630_ | 30 | 66 | 43 | 33 | 41 | 31 |
| SL1344_RS12635_ | 9 | 20 | 26 | 17 | 13 | 13 |
| SL1344_RS12640_ | 26 | 32 | 28 | 37 | 24 | 23 |
| SL1344_RS12645_ | 33 | 34 | 63 | 40 | 44 | 32 |
| SL1344_RS12650_ | 4 | 6 | 12 | 10 | 6 | 12 |
| SL1344_RS12655_ | 16 | 16 | 19 | 20 | 18 | 16 |
| SL1344_RS12660_ | 13 | 14 | 21 | 16 | 10 | 11 |
| SL1344_RS12665_ | 8 | 6 | 7 | 2 | 6 | 2 |
| SL1344_RS12670_ | 14 | 10 | 16 | 10 | 15 | 8 |
| SL1344_RS12675_ | 5 | 10 | 5 | 7 | 7 | 9 |
| SL1344_RS12690_ | 6057 | 6149 | 6111 | 7008 | 5412 | 6176 |
| SL1344_RS12695_ | 192 | 194 | 206 | 173 | 122 | 177 |
| SL1344_RS12700_ | 338 | 345 | 374 | 310 | 315 | 321 |
| SL1344_RS12705_ | 333 | 314 | 295 | 266 | 225 | 333 |
| SL1344_RS12710_ | 114 | 129 | 168 | 118 | 136 | 107 |
| SL1344_RS12715_ | 304 | 299 | 378 | 318 | 269 | 333 |
| SL1344_RS12720_ | 77 | 92 | 94 | 85 | 67 | 60 |
| SL1344_RS12725_ | 53 | 67 | 53 | 73 | 69 | 49 |
| SL1344_RS12730_ | 113 | 141 | 101 | 103 | 72 | 95 |
| SL1344_RS12735_ | 293 | 307 | 310 | 327 | 289 | 300 |
| SL1344_RS12740_ | 319 | 319 | 407 | 277 | 174 | 247 |
| SL1344_RS12745_ | 1169 | 1265 | 1318 | 944 | 790 | 918 |
| SL1344_RS12750_ | 685 | 713 | 759 | 511 | 491 | 518 |
| SL1344_RS12755_ | 524 | 608 | 576 | 586 | 642 | 563 |
| SL1344_RS12760_ | 350 | 392 | 394 | 388 | 373 | 405 |
| SL1344_RS12765_ | 488 | 534 | 448 | 519 | 498 | 495 |
| SL1344_RS12770_ | 6729 | 7727 | 7858 | 6749 | 6047 | 6592 |
| SL1344_RS12775_ | 3067 | 4030 | 3737 | 3577 | 3052 | 3269 |
| SL1344_RS12780_ | 963 | 1252 | 1263 | 1121 | 968 | 1083 |
| SL1344_RS12785_ | 3049 | 4150 | 3867 | 2938 | 3156 | 3008 |
| SL1344_RS12790_ | 34 | 32 | 42 | 43 | 38 | 39 |
| SL1344_RS12795_ | 330 | 350 | 309 | 447 | 446 | 371 |
| SL1344_RS12800_ | 659 | 756 | 743 | 758 | 605 | 777 |
| SL1344_RS12805_ | 588 | 783 | 714 | 661 | 571 | 691 |
| SL1344_RS12810_ | 929 | 1024 | 962 | 1203 | 1134 | 905 |
| SL1344_RS12815_ | 507 | 569 | 576 | 438 | 406 | 311 |
| SL1344_RS12820_ | 1410 | 1458 | 1490 | 1765 | 1290 | 1195 |
| SL1344_RS12825_ | 475 | 472 | 566 | 681 | 560 | 544 |
| SL1344_RS12830_ | 455 | 544 | 488 | 639 | 493 | 539 |
| SL1344_RS12835_ | 1452 | 1711 | 1723 | 1204 | 1021 | 1292 |
| SL1344_RS12840_ | 1392 | 1823 | 1711 | 1051 | 1145 | 1093 |
| SL1344_RS12845_ | 45 | 53 | 49 | 76 | 42 | 47 |
| SL1344_RS12850_ | 25 | 28 | 27 | 30 | 16 | 27 |
| SL1344_RS12855_ | 600 | 762 | 794 | 902 | 536 | 927 |
| SL1344_RS12860_ | 31 | 28 | 38 | 23 | 32 | 35 |
| SL1344_RS12865_ | 16 | 20 | 24 | 17 | 13 | 16 |
| SL1344_RS12870_ | 1890 | 2471 | 1884 | 1576 | 1998 | 1543 |
| SL1344_RS12875_ | 423 | 553 | 491 | 735 | 634 | 542 |
| SL1344_RS12880_ | 468 | 665 | 621 | 544 | 577 | 662 |
| SL1344_RS12885_ | 298 | 319 | 321 | 349 | 292 | 334 |
| SL1344_RS12890_ | 954 | 1131 | 1126 | 1239 | 948 | 1061 |
| SL1344_RS12900_ | 51 | 67 | 49 | 82 | 53 | 63 |
| SL1344_RS12905_ | 33 | 28 | 35 | 42 | 31 | 54 |
| SL1344_RS12910_ | 167 | 190 | 196 | 207 | 141 | 121 |
| SL1344_RS12915_ | 1013 | 1049 | 1110 | 1034 | 783 | 969 |
| SL1344_RS12920_ | 1609 | 1780 | 1852 | 1837 | 1632 | 1956 |
| SL1344_RS12925_ | 1056 | 1179 | 1343 | 1216 | 975 | 1175 |

| | | | | | | |
|---|---|---|---|---|---|---|
| SL1344_RS12930_ | 1843 | 2107 | 1804 | 2012 | 1699 | 1875 |
| SL1344_RS12935_ | 1912 | 2258 | 2099 | 1845 | 2296 | 1955 |
| SL1344_RS12940_ | 1092 | 1311 | 1276 | 1393 | 1291 | 1255 |
| SL1344_RS12945_ | 1168 | 1108 | 1439 | 1484 | 1014 | 1183 |
| SL1344_RS12950_ | 4695 | 3977 | 5128 | 6649 | 4223 | 5115 |
| SL1344_RS12955_ | 28 | 38 | 37 | 36 | 22 | 33 |
| SL1344_RS12960_ | 52 | 45 | 60 | 51 | 36 | 45 |
| SL1344_RS12965_ | 31 | 11 | 22 | 34 | 27 | 28 |
| SL1344_RS12970_ | 118 | 90 | 98 | 135 | 89 | 101 |
| SL1344_RS12975_ | 466 | 592 | 518 | 540 | 664 | 487 |
| SL1344_RS12980_ | 2267 | 2641 | 2343 | 2323 | 2448 | 2108 |
| SL1344_RS12985_ | 922 | 1035 | 1015 | 1077 | 983 | 924 |
| SL1344_RS12990_ | 81 | 89 | 106 | 79 | 64 | 185 |
| SL1344_RS12995_ | 607 | 662 | 489 | 675 | 698 | 755 |
| SL1344_RS13000_ | 967 | 1019 | 976 | 1264 | 1197 | 1218 |
| SL1344_RS13005_ | 1035 | 1002 | 1344 | 994 | 868 | 1048 |
| SL1344_RS13010_ | 1174 | 1296 | 1395 | 1298 | 1033 | 1264 |
| SL1344_RS13015_ | 1991 | 2209 | 2263 | 2396 | 1951 | 2119 |
| SL1344_RS13025_ | 2285 | 2116 | 2531 | 2831 | 2463 | 3057 |
| SL1344_RS13030_ | 1410 | 1370 | 1573 | 1670 | 1204 | 1853 |
| SL1344_RS13035_ | 3418 | 3743 | 3607 | 4261 | 4176 | 4347 |
| SL1344_RS13040_ | 1227 | 1250 | 1284 | 1702 | 1386 | 1613 |
| SL1344_RS13045_ | 961 | 1052 | 918 | 935 | 784 | 815 |
| SL1344_RS13050_ | 1636 | 1242 | 1419 | 2338 | 1401 | 1039 |
| SL1344_RS13075_ | 179 | 193 | 149 | 168 | 189 | 212 |
| SL1344_RS13080_ | 95 | 117 | 89 | 108 | 115 | 107 |
| SL1344_RS13085_ | 324 | 274 | 341 | 185 | 127 | 113 |
| SL1344_RS13090_ | 468 | 512 | 594 | 636 | 565 | 499 |
| SL1344_RS13095_ | 7947 | 9277 | 8402 | 8118 | 7280 | 7618 |
| SL1344_RS13100_ | 321 | 385 | 395 | 469 | 340 | 395 |
| SL1344_RS13105_ | 568 | 636 | 640 | 610 | 539 | 512 |
| SL1344_RS13115_ | 926 | 1102 | 842 | 1154 | 797 | 1011 |
| SL1344_RS13120_ | 160 | 212 | 164 | 167 | 150 | 123 |
| SL1344_RS13125_ | 1166 | 1342 | 1276 | 1220 | 1255 | 989 |
| SL1344_RS13130_ | 940 | 1145 | 1197 | 1060 | 885 | 1039 |
| SL1344_RS13135_ | 394 | 480 | 415 | 478 | 418 | 438 |
| SL1344_RS13140_ | 477 | 560 | 487 | 570 | 585 | 601 |
| SL1344_RS13145_ | 652 | 833 | 730 | 1003 | 801 | 776 |
| SL1344_RS13150_ | 80 | 75 | 47 | 119 | 65 | 56 |
| SL1344_RS13155_ | 446 | 486 | 437 | 506 | 488 | 425 |
| SL1344_RS13160_ | 420 | 406 | 420 | 322 | 428 | 343 |
| SL1344_RS13165_ | 518 | 581 | 636 | 407 | 514 | 499 |
| SL1344_RS13170_ | 538 | 449 | 456 | 279 | 397 | 384 |
| SL1344_RS13175_ | 388 | 377 | 364 | 283 | 345 | 184 |
| SL1344_RS13180_ | 144 | 143 | 245 | 167 | 151 | 136 |
| SL1344_RS13185_ | 146 | 178 | 151 | 192 | 154 | 161 |
| SL1344_RS13190_ | 22 | 15 | 6 | 6 | 17 | 10 |
| SL1344_RS13195_ | 74 | 74 | 105 | 74 | 52 | 52 |
| SL1344_RS13200_ | 94 | 81 | 84 | 113 | 89 | 87 |
| SL1344_RS13210_ | 431 | 499 | 418 | 398 | 389 | 435 |
| SL1344_RS13215_ | 808 | 939 | 846 | 760 | 834 | 612 |
| SL1344_RS13220_ | 1447 | 1604 | 1752 | 1428 | 1131 | 1404 |
| SL1344_RS13225_ | 1224 | 1510 | 1375 | 1190 | 1046 | 1144 |
| SL1344_RS13230_ | 2774 | 3059 | 3067 | 2878 | 2948 | 2494 |
| SL1344_RS13235_ | 2725 | 3272 | 3188 | 2943 | 2383 | 2592 |
| SL1344_RS13240_ | 424 | 496 | 474 | 450 | 437 | 419 |
| SL1344_RS13245_ | 153 | 138 | 176 | 145 | 117 | 156 |
| SL1344_RS13250_ | 4 | 2 | 4 | 5 | 10 | 6 |
| SL1344_RS13255_ | 2 | 13 | 5 | 14 | 11 | 8 |
| SL1344_RS13260_ | 12 | 9 | 9 | 19 | 13 | 15 |
| SL1344_RS13265_ | 53 | 48 | 46 | 55 | 66 | 48 |
| SL1344_RS13270_ | 6 | 11 | 10 | 19 | 13 | 20 |
| SL1344_RS13275_ | 67 | 69 | 64 | 56 | 63 | 56 |

| | | | | | | |
|---|---|---|---|---|---|---|
| SL1344_RS13280_ | 55 | 44 | 38 | 61 | 40 | 48 |
| SL1344_RS13285_ | 25 | 36 | 28 | 36 | 33 | 29 |
| SL1344_RS13290_ | 53 | 52 | 50 | 42 | 41 | 44 |
| SL1344_RS13295_ | 12 | 10 | 10 | 11 | 11 | 19 |
| SL1344_RS13300_ | 166 | 163 | 173 | 168 | 160 | 169 |
| SL1344_RS13305_ | 8 | 15 | 12 | 9 | 10 | 10 |
| SL1344_RS13310_ | 6 | 5 | 8 | 4 | 11 | 8 |
| SL1344_RS13315_ | 113 | 135 | 104 | 112 | 117 | 121 |
| SL1344_RS13320_ | 24 | 29 | 33 | 33 | 22 | 30 |
| SL1344_RS13325_ | 4040 | 5051 | 4926 | 5141 | 5538 | 3725 |
| SL1344_RS13330_ | 23 | 36 | 22 | 30 | 42 | 36 |
| SL1344_RS13335_ | 16 | 18 | 26 | 21 | 20 | 22 |
| SL1344_RS13340_ | 10 | 18 | 14 | 11 | 20 | 25 |
| SL1344_RS13345_ | 30 | 56 | 42 | 47 | 46 | 61 |
| SL1344_RS13350_ | 29 | 28 | 20 | 23 | 24 | 34 |
| SL1344_RS13355_ | 44 | 61 | 69 | 56 | 42 | 61 |
| SL1344_RS13360_ | 66 | 90 | 81 | 76 | 70 | 86 |
| SL1344_RS13365_ | 18 | 8 | 17 | 11 | 17 | 14 |
| SL1344_RS13370_ | 57 | 55 | 61 | 60 | 44 | 62 |
| SL1344_RS13375_ | 13 | 7 | 5 | 12 | 1 | 8 |
| SL1344_RS13380_ | 62 | 78 | 93 | 58 | 46 | 63 |
| SL1344_RS13385_ | 17 | 24 | 22 | 27 | 15 | 18 |
| SL1344_RS13390_ | 16 | 15 | 40 | 17 | 22 | 36 |
| SL1344_RS13395_ | 3 | 5 | 2 | 2 | 1 | 0 |
| SL1344_RS13400_ | 218 | 246 | 251 | 270 | 181 | 278 |
| SL1344_RS13405_ | 26 | 24 | 29 | 27 | 26 | 27 |
| SL1344_RS13410_ | 42 | 50 | 33 | 36 | 38 | 25 |
| SL1344_RS13415_ | 145 | 145 | 120 | 192 | 139 | 150 |
| SL1344_RS13420_ | 13 | 22 | 16 | 21 | 25 | 15 |
| SL1344_RS13425_ | 1 | 1 | 4 | 1 | 0 | 3 |
| SL1344_RS13430_ | 17 | 5 | 13 | 5 | 7 | 7 |
| SL1344_RS13435_ | 1 | 4 | 1 | 1 | 2 | 5 |
| SL1344_RS13440_ | 242 | 394 | 308 | 371 | 350 | 252 |
| SL1344_RS13450_ | 125 | 146 | 154 | 178 | 109 | 163 |
| SL1344_RS13455_ | 389 | 389 | 451 | 281 | 198 | 307 |
| SL1344_RS13460_ | 25 | 28 | 27 | 28 | 33 | 28 |
| SL1344_RS13465_ | 52 | 64 | 43 | 70 | 52 | 60 |
| SL1344_RS13470_ | 19 | 25 | 16 | 24 | 16 | 21 |
| SL1344_RS13475_ | 35 | 32 | 28 | 28 | 28 | 29 |
| SL1344_RS13480_ | 15 | 25 | 16 | 21 | 15 | 15 |
| SL1344_RS13485_ | 38 | 22 | 40 | 40 | 35 | 46 |
| SL1344_RS13490_ | 25 | 25 | 26 | 29 | 21 | 37 |
| SL1344_RS13495_ | 9 | 13 | 12 | 19 | 14 | 14 |
| SL1344_RS13500_ | 1327 | 1397 | 1616 | 1476 | 1140 | 1475 |
| SL1344_RS13505_ | 6291 | 6394 | 7517 | 6826 | 4062 | 5885 |
| SL1344_RS13510_ | 10 | 7 | 7 | 4 | 17 | 15 |
| SL1344_RS13515_ | 15 | 14 | 17 | 22 | 19 | 23 |
| SL1344_RS13520_ | 74 | 87 | 78 | 92 | 60 | 71 |
| SL1344_RS13525_ | 13 | 28 | 16 | 23 | 24 | 19 |
| SL1344_RS13530_ | 5 | 2 | 4 | 9 | 6 | 1 |
| SL1344_RS13535_ | 22 | 32 | 27 | 35 | 32 | 25 |
| SL1344_RS13540_ | 209 | 227 | 215 | 222 | 172 | 178 |
| SL1344_RS13545_ | 451 | 578 | 514 | 430 | 616 | 433 |
| SL1344_RS13550_ | 915 | 1152 | 910 | 963 | 1030 | 1018 |
| SL1344_RS13555_ | 941 | 1142 | 858 | 1043 | 1087 | 1114 |
| SL1344_RS13560_ | 984 | 1318 | 1203 | 1105 | 1201 | 1224 |
| SL1344_RS13565_ | 74 | 98 | 78 | 105 | 112 | 100 |
| SL1344_RS13570_ | 558 | 634 | 601 | 613 | 575 | 518 |
| SL1344_RS13575_ | 2084 | 2048 | 2201 | 1880 | 1419 | 1914 |
| SL1344_RS13580_ | 99 | 160 | 131 | 181 | 150 | 143 |
| SL1344_RS13590_ | 619 | 1253 | 669 | 944 | 874 | 1100 |
| SL1344_RS13595_ | 652 | 838 | 707 | 664 | 768 | 795 |
| SL1344_RS13605_ | 917 | 965 | 983 | 1142 | 829 | 829 |

| | | | | | | |
|---|---|---|---|---|---|---|
| SL1344_RS13610_ | 308 | 356 | 361 | 413 | 277 | 363 |
| SL1344_RS13615_ | 1656 | 1661 | 1674 | 1724 | 1679 | 1529 |
| SL1344_RS13625_ | 210 | 243 | 245 | 225 | 271 | 230 |
| SL1344_RS13630_ | 253 | 288 | 240 | 234 | 254 | 286 |
| SL1344_RS13635_ | 23 | 17 | 16 | 11 | 15 | 16 |
| SL1344_RS13645_ | 2839 | 2043 | 5004 | 3233 | 2620 | 2191 |
| SL1344_RS13650_ | 189 | 272 | 393 | 194 | 161 | 213 |
| SL1344_RS13655_ | 287 | 366 | 994 | 219 | 393 | 167 |
| SL1344_RS13660_ | 5114 | 4298 | 4863 | 8470 | 4566 | 5503 |
| SL1344_RS13665_ | 1100 | 1098 | 1448 | 1235 | 977 | 1096 |
| SL1344_RS13670_ | 1400 | 1592 | 1707 | 1556 | 1318 | 1438 |
| SL1344_RS13675_ | 1729 | 2130 | 2001 | 1884 | 1352 | 1667 |
| SL1344_RS13680_ | 5361 | 7815 | 8954 | 6288 | 5196 | 7682 |
| SL1344_RS13685_ | 967 | 1200 | 1249 | 1015 | 1011 | 1086 |
| SL1344_RS13705_ | 72 | 122 | 93 | 59 | 61 | 59 |
| SL1344_RS13710_ | 198 | 230 | 190 | 159 | 123 | 147 |
| SL1344_RS13715_ | 1281 | 1512 | 1537 | 1425 | 1235 | 1476 |
| SL1344_RS13720_ | 1289 | 1409 | 1601 | 1620 | 1067 | 1600 |
| SL1344_RS13725_ | 891 | 886 | 863 | 992 | 774 | 1034 |
| SL1344_RS13730_ | 3 | 2 | 2 | 4 | 2 | 3 |
| SL1344_RS13735_ | 4 | 2 | 8 | 5 | 9 | 10 |
| SL1344_RS13745_ | 4 | 4 | 8 | 5 | 3 | 7 |
| SL1344_RS13750_ | 1 | 2 | 4 | 3 | 3 | 5 |
| SL1344_RS13755_ | 1 | 6 | 2 | 3 | 1 | 2 |
| SL1344_RS13760_ | 4 | 7 | 1 | 3 | 5 | 3 |
| SL1344_RS13765_ | 17 | 35 | 13 | 16 | 26 | 15 |
| SL1344_RS13770_ | 17 | 25 | 14 | 23 | 28 | 24 |
| SL1344_RS13775_ | 112 | 78 | 82 | 89 | 72 | 77 |
| SL1344_RS13780_ | 1882 | 1805 | 2195 | 1946 | 1386 | 1585 |
| SL1344_RS13785_ | 9964 | 10689 | 10312 | 12600 | 10810 | 9798 |
| SL1344_RS13790_ | 10460 | 11080 | 11630 | 12381 | 10194 | 10356 |
| SL1344_RS13800_ | 4749 | 4549 | 5465 | 5695 | 3784 | 4335 |
| SL1344_RS13805_ | 1408 | 1545 | 1492 | 1397 | 1286 | 1712 |
| SL1344_RS13810_ | 669 | 772 | 800 | 827 | 651 | 755 |
| SL1344_RS13815_ | 642 | 819 | 754 | 711 | 656 | 784 |
| SL1344_RS13820_ | 110 | 117 | 93 | 122 | 112 | 118 |
| SL1344_RS13825_ | 7616 | 6791 | 7455 | 9520 | 8314 | 7614 |
| SL1344_RS13830_ | 373 | 486 | 434 | 487 | 373 | 487 |
| SL1344_RS13840_ | 1244 | 1559 | 1563 | 1836 | 1213 | 1528 |
| SL1344_RS13845_ | 502 | 576 | 504 | 581 | 681 | 588 |
| SL1344_RS13850_ | 865 | 1117 | 1226 | 1113 | 952 | 1251 |
| SL1344_RS13855_ | 1750 | 1901 | 2155 | 1896 | 1383 | 1859 |
| SL1344_RS13860_ | 195 | 235 | 226 | 261 | 198 | 234 |
| SL1344_RS13865_ | 27 | 29 | 34 | 25 | 17 | 30 |
| SL1344_RS13870_ | 41 | 59 | 38 | 52 | 30 | 35 |
| SL1344_RS13875_ | 22 | 29 | 52 | 39 | 27 | 38 |
| SL1344_RS13880_ | 26 | 32 | 33 | 44 | 28 | 27 |
| SL1344_RS13885_ | 41 | 29 | 28 | 69 | 38 | 42 |
| SL1344_RS13890_ | 13 | 16 | 12 | 14 | 16 | 14 |
| SL1344_RS13895_ | 81 | 97 | 103 | 101 | 91 | 83 |
| SL1344_RS13900_ | 7 | 14 | 13 | 12 | 11 | 8 |
| SL1344_RS13905_ | 9 | 13 | 5 | 24 | 7 | 9 |
| SL1344_RS13910_ | 12 | 15 | 6 | 8 | 11 | 10 |
| SL1344_RS13915_ | 20 | 27 | 28 | 42 | 29 | 39 |
| SL1344_RS13920_ | 3 | 6 | 6 | 7 | 7 | 6 |
| SL1344_RS13930_ | 58 | 81 | 73 | 74 | 42 | 69 |
| SL1344_RS13935_ | 49 | 56 | 53 | 53 | 51 | 42 |
| SL1344_RS13940_ | 8 | 7 | 19 | 13 | 7 | 7 |
| SL1344_RS13950_ | 7 | 3 | 1 | 2 | 2 | 3 |
| SL1344_RS13955_ | 11 | 12 | 3 | 4 | 4 | 5 |
| SL1344_RS13965_ | 9 | 17 | 8 | 11 | 13 | 5 |
| SL1344_RS13970_ | 7 | 6 | 13 | 9 | 12 | 9 |
| SL1344_RS13975_ | 8 | 2 | 10 | 5 | 3 | 4 |

| | | | | | | |
|---|---|---|---|---|---|---|
| SL1344_RS13980_ | 11 | 11 | 6 | 3 | 5 | 5 |
| SL1344_RS13985_ | 0 | 7 | 4 | 0 | 0 | 3 |
| SL1344_RS14000_ | 11 | 5 | 7 | 12 | 40 | 8 |
| SL1344_RS14005_ | 21 | 30 | 14 | 14 | 21 | 16 |
| SL1344_RS14010_ | 6 | 8 | 4 | 12 | 1 | 8 |
| SL1344_RS14015_ | 20 | 17 | 18 | 25 | 15 | 13 |
| SL1344_RS14020_ | 6 | 15 | 16 | 9 | 12 | 14 |
| SL1344_RS14025_ | 1530 | 1235 | 1482 | 1107 | 1277 | 1070 |
| SL1344_RS14030_ | 1341 | 1108 | 1340 | 963 | 911 | 1004 |
| SL1344_RS14035_ | 288 | 441 | 328 | 342 | 509 | 336 |
| SL1344_RS14040_ | 1485 | 1563 | 1774 | 1648 | 1401 | 1729 |
| SL1344_RS14045_ | 820 | 1026 | 1038 | 899 | 916 | 996 |
| SL1344_RS14050_ | 131 | 148 | 110 | 109 | 155 | 173 |
| SL1344_RS14055_ | 82 | 96 | 76 | 108 | 104 | 120 |
| SL1344_RS14060_ | 71 | 63 | 61 | 74 | 72 | 100 |
| SL1344_RS14065_ | 34 | 11 | 27 | 25 | 29 | 28 |
| SL1344_RS14070_ | 4 | 3 | 2 | 3 | 6 | 5 |
| SL1344_RS14075_ | 2 | 6 | 2 | 5 | 2 | 7 |
| SL1344_RS14080_ | 4 | 9 | 8 | 5 | 3 | 6 |
| SL1344_RS14085_ | 173 | 186 | 262 | 300 | 148 | 288 |
| SL1344_RS14090_ | 7 | 8 | 14 | 11 | 8 | 10 |
| SL1344_RS14095_ | 10 | 2 | 2 | 11 | 28 | 10 |
| SL1344_RS14100_ | 243 | 291 | 267 | 229 | 237 | 325 |
| SL1344_RS14105_ | 444 | 514 | 484 | 457 | 384 | 528 |
| SL1344_RS14110_ | 496 | 456 | 555 | 448 | 360 | 460 |
| SL1344_RS14115_ | 114 | 134 | 89 | 126 | 132 | 120 |
| SL1344_RS14120_ | 1460 | 1433 | 1784 | 1478 | 1424 | 1780 |
| SL1344_RS14125_ | 148 | 159 | 147 | 134 | 157 | 175 |
| SL1344_RS14130_ | 340 | 287 | 325 | 332 | 253 | 308 |
| SL1344_RS14135_ | 561 | 538 | 768 | 653 | 431 | 566 |
| SL1344_RS14140_ | 11 | 17 | 13 | 5 | 7 | 7 |
| SL1344_RS14150_ | 16 | 15 | 26 | 21 | 22 | 21 |
| SL1344_RS14155_ | 9 | 19 | 14 | 21 | 15 | 16 |
| SL1344_RS14165_ | 13 | 13 | 15 | 15 | 5 | 5 |
| SL1344_RS14170_ | 20 | 21 | 20 | 28 | 19 | 15 |
| SL1344_RS14175_ | 251 | 271 | 313 | 282 | 248 | 255 |
| SL1344_RS14180_ | 101 | 124 | 124 | 157 | 117 | 107 |
| SL1344_RS14190_ | 53 | 36 | 36 | 43 | 44 | 42 |
| SL1344_RS14195_ | 529 | 599 | 639 | 755 | 584 | 696 |
| SL1344_RS14200_ | 370 | 441 | 382 | 336 | 392 | 396 |
| SL1344_RS14205_ | 3 | 9 | 6 | 8 | 2 | 7 |
| SL1344_RS14210_ | 36 | 72 | 27 | 37 | 35 | 27 |
| SL1344_RS14215_ | 981 | 1070 | 1327 | 974 | 782 | 907 |
| SL1344_RS14220_ | 483 | 579 | 669 | 463 | 385 | 424 |
| SL1344_RS14225_ | 506 | 532 | 673 | 698 | 482 | 538 |
| SL1344_RS14230_ | 6 | 13 | 6 | 19 | 9 | 13 |
| SL1344_RS14235_ | 13 | 8 | 5 | 6 | 5 | 4 |
| SL1344_RS14240_ | 2 | 9 | 4 | 4 | 6 | 7 |
| SL1344_RS14245_ | 14 | 12 | 11 | 14 | 14 | 12 |
| SL1344_RS14250_ | 45 | 26 | 15 | 34 | 29 | 26 |
| SL1344_RS14255_ | 558 | 519 | 636 | 658 | 421 | 560 |
| SL1344_RS14260_ | 18 | 23 | 30 | 31 | 34 | 41 |
| SL1344_RS14265_ | 13 | 10 | 9 | 7 | 22 | 10 |
| SL1344_RS14270_ | 43 | 13 | 28 | 15 | 28 | 10 |
| SL1344_RS14275_ | 18 | 20 | 17 | 14 | 21 | 15 |
| SL1344_RS14280_ | 41 | 48 | 41 | 29 | 31 | 33 |
| SL1344_RS14285_ | 2 | 3 | 5 | 1 | 8 | 3 |
| SL1344_RS14290_ | 17 | 21 | 27 | 31 | 28 | 33 |
| SL1344_RS14295_ | 116 | 104 | 131 | 129 | 69 | 103 |
| SL1344_RS14300_ | 153 | 151 | 191 | 177 | 128 | 169 |
| SL1344_RS14305_ | 215 | 348 | 388 | 227 | 237 | 350 |
| SL1344_RS14310_ | 7 | 15 | 6 | 20 | 16 | 8 |
| SL1344_RS14315_ | 19 | 22 | 13 | 20 | 16 | 34 |

| | | | | | | |
|---|---|---|---|---|---|---|
| SL1344_RS14320_ | 629 | 737 | 832 | 821 | 606 | 757 |
| SL1344_RS14325_ | 2267 | 2357 | 2544 | 2464 | 2038 | 2518 |
| SL1344_RS14330_ | 43 | 57 | 64 | 57 | 44 | 68 |
| SL1344_RS14335_ | 38 | 33 | 41 | 44 | 33 | 44 |
| SL1344_RS14345_ | 111238 | 112329 | 115872 | 9976 | 122451 | 7324 |
| SL1344_RS14350_ | 50 | 40 | 60 | 25 | 34 | 31 |
| SL1344_RS14360_ | 138 | 168 | 171 | 429 | 244 | 331 |
| SL1344_RS14365_ | 38 | 64 | 47 | 83 | 53 | 78 |
| SL1344_RS14370_ | 44 | 53 | 51 | 92 | 37 | 73 |
| SL1344_RS14375_ | 167 | 177 | 181 | 256 | 213 | 196 |
| SL1344_RS14380_ | 36 | 36 | 32 | 48 | 55 | 44 |
| SL1344_RS14385_ | 91 | 129 | 85 | 97 | 100 | 90 |
| SL1344_RS14390_ | 1294 | 1531 | 1664 | 1802 | 1040 | 1447 |
| SL1344_RS14395_ | 1832 | 2051 | 2335 | 2285 | 1684 | 2012 |
| SL1344_RS14400_ | 258 | 230 | 222 | 197 | 263 | 202 |
| SL1344_RS14405_ | 335 | 333 | 306 | 302 | 399 | 347 |
| SL1344_RS14410_ | 844 | 695 | 687 | 605 | 1139 | 805 |
| SL1344_RS14415_ | 945 | 612 | 695 | 694 | 1068 | 710 |
| SL1344_RS14420_ | 337 | 261 | 264 | 574 | 198 | 300 |
| SL1344_RS14425_ | 1460 | 986 | 1217 | 2497 | 1279 | 1320 |
| SL1344_RS14430_ | 432 | 303 | 345 | 729 | 397 | 378 |
| SL1344_RS14435_ | 1453 | 1078 | 1214 | 2712 | 1143 | 1869 |
| SL1344_RS14440_ | 102 | 53 | 53 | 74 | 105 | 78 |
| SL1344_RS14445_ | 83 | 49 | 78 | 68 | 126 | 101 |
| SL1344_RS14450_ | 80 | 71 | 96 | 132 | 159 | 179 |
| SL1344_RS14455_ | 121 | 107 | 123 | 245 | 207 | 241 |
| SL1344_RS14460_ | 110 | 100 | 104 | 115 | 124 | 161 |
| SL1344_RS14465_ | 51 | 77 | 53 | 91 | 83 | 66 |
| SL1344_RS14470_ | 79 | 64 | 88 | 79 | 55 | 65 |
| SL1344_RS14475_ | 39 | 46 | 41 | 45 | 41 | 51 |
| SL1344_RS14480_ | 46 | 46 | 47 | 54 | 34 | 42 |
| SL1344_RS14485_ | 79 | 97 | 111 | 130 | 104 | 119 |
| SL1344_RS14490_ | 261 | 318 | 326 | 404 | 330 | 450 |
| SL1344_RS14495_ | 2657 | 2893 | 2804 | 2343 | 2452 | 2409 |
| SL1344_RS14500_ | 671 | 1021 | 1055 | 492 | 220 | 352 |
| SL1344_RS14505_ | 51 | 53 | 63 | 63 | 53 | 61 |
| SL1344_RS14510_ | 200 | 183 | 146 | 240 | 156 | 193 |
| SL1344_RS14515_ | 245 | 268 | 288 | 266 | 252 | 303 |
| SL1344_RS14520_ | 41 | 50 | 49 | 43 | 47 | 31 |
| SL1344_RS14525_ | 11 | 14 | 19 | 10 | 13 | 7 |
| SL1344_RS14530_ | 59 | 45 | 47 | 58 | 50 | 35 |
| SL1344_RS14540_ | 153 | 206 | 206 | 216 | 136 | 186 |
| SL1344_RS14545_ | 133 | 153 | 178 | 182 | 127 | 180 |
| SL1344_RS14550_ | 292 | 226 | 226 | 219 | 241 | 225 |
| SL1344_RS14555_ | 75 | 145 | 158 | 104 | 171 | 107 |
| SL1344_RS14560_ | 200 | 242 | 242 | 242 | 229 | 223 |
| SL1344_RS14565_ | 113 | 138 | 178 | 117 | 129 | 130 |
| SL1344_RS14570_ | 1194 | 1253 | 976 | 1029 | 916 | 1207 |
| SL1344_RS14575_ | 654 | 820 | 805 | 756 | 633 | 776 |
| SL1344_RS14585_ | 69 | 74 | 65 | 49 | 56 | 67 |
| SL1344_RS14600_ | 777 | 849 | 765 | 848 | 814 | 799 |
| SL1344_RS14605_ | 1048 | 1109 | 1053 | 980 | 872 | 1183 |
| SL1344_RS14610_ | 739 | 534 | 356 | 779 | 1008 | 339 |
| SL1344_RS14615_ | 2422 | 1793 | 1490 | 2785 | 3509 | 1296 |
| SL1344_RS14620_ | 5099 | 3981 | 2952 | 5274 | 8855 | 2366 |
| SL1344_RS14625_ | 6429 | 5363 | 5004 | 7320 | 8388 | 4594 |
| SL1344_RS14630_ | 17739 | 16292 | 17081 | 22271 | 21820 | 14661 |
| SL1344_RS14640_ | 5081 | 5416 | 4512 | 4395 | 4331 | 4935 |
| SL1344_RS14645_ | 123 | 118 | 124 | 116 | 137 | 119 |
| SL1344_RS14655_ | 530 | 499 | 529 | 574 | 551 | 469 |
| SL1344_RS14660_ | 640 | 732 | 634 | 679 | 524 | 566 |
| SL1344_RS14675_ | 66 | 61 | 68 | 88 | 59 | 69 |
| SL1344_RS14680_ | 307 | 317 | 348 | 414 | 347 | 367 |

| | | | | | | |
|---|---|---|---|---|---|---|
| SL1344_RS14685_ | 190 | 196 | 264 | 221 | 174 | 188 |
| SL1344_RS14690_ | 158 | 212 | 217 | 164 | 176 | 147 |
| SL1344_RS14700_ | 672 | 793 | 710 | 657 | 889 | 711 |
| SL1344_RS14705_ | 174 | 222 | 188 | 191 | 201 | 230 |
| SL1344_RS14710_ | 19 | 51 | 36 | 48 | 39 | 47 |
| SL1344_RS14715_ | 48 | 46 | 45 | 62 | 48 | 52 |
| SL1344_RS14720_ | 474 | 553 | 484 | 536 | 506 | 445 |
| SL1344_RS14725_ | 18 | 24 | 18 | 24 | 23 | 18 |
| SL1344_RS14730_ | 91 | 91 | 73 | 107 | 101 | 75 |
| SL1344_RS14735_ | 44 | 44 | 52 | 79 | 38 | 56 |
| SL1344_RS14740_ | 28 | 38 | 48 | 53 | 44 | 57 |
| SL1344_RS14745_ | 56 | 51 | 59 | 59 | 52 | 57 |
| SL1344_RS14750_ | 44 | 44 | 35 | 45 | 29 | 31 |
| SL1344_RS14765_ | 75 | 68 | 51 | 64 | 67 | 67 |
| SL1344_RS14770_ | 9 | 11 | 16 | 5 | 9 | 17 |
| SL1344_RS14775_ | 10 | 13 | 11 | 16 | 8 | 11 |
| SL1344_RS14780_ | 233 | 244 | 343 | 223 | 181 | 273 |
| SL1344_RS14785_ | 226 | 242 | 201 | 209 | 167 | 198 |
| SL1344_RS14790_ | 74 | 81 | 89 | 82 | 87 | 93 |
| SL1344_RS14795_ | 239 | 266 | 229 | 261 | 220 | 258 |
| SL1344_RS14800_ | 274 | 323 | 295 | 273 | 351 | 273 |
| SL1344_RS14805_ | 242 | 283 | 281 | 262 | 229 | 234 |
| SL1344_RS14810_ | 39 | 42 | 54 | 42 | 64 | 59 |
| SL1344_RS14815_ | 167 | 160 | 158 | 301 | 161 | 243 |
| SL1344_RS14820_ | 77 | 87 | 90 | 135 | 138 | 145 |
| SL1344_RS14825_ | 44 | 58 | 44 | 83 | 74 | 97 |
| SL1344_RS14830_ | 114 | 192 | 182 | 201 | 195 | 197 |
| SL1344_RS14850_ | 761 | 795 | 1014 | 1339 | 959 | 1352 |
| SL1344_RS14855_ | 735 | 725 | 853 | 1485 | 1053 | 1346 |
| SL1344_RS14900_ | 1006 | 1097 | 1111 | 1127 | 1011 | 1290 |
| SL1344_RS14905_ | 281 | 297 | 365 | 413 | 330 | 411 |
| SL1344_RS14910_ | 362 | 391 | 402 | 453 | 452 | 523 |
| SL1344_RS15010_ | 265 | 197 | 303 | 510 | 286 | 501 |
| SL1344_RS15015_ | 626 | 645 | 726 | 701 | 523 | 791 |
| SL1344_RS15020_ | 216 | 214 | 189 | 242 | 251 | 173 |
| SL1344_RS15025_ | 96 | 65 | 72 | 77 | 70 | 53 |
| SL1344_RS15030_ | 325 | 425 | 462 | 445 | 261 | 434 |
| SL1344_RS15035_ | 487 | 578 | 548 | 607 | 397 | 557 |
| SL1344_RS15040_ | 5 | 12 | 9 | 5 | 4 | 5 |
| SL1344_RS15045_ | 79 | 74 | 77 | 54 | 45 | 53 |
| SL1344_RS15050_ | 10 | 14 | 10 | 9 | 10 | 6 |
| SL1344_RS15055_ | 1497 | 1782 | 1440 | 1430 | 1604 | 1532 |
| SL1344_RS15060_ | 49 | 38 | 37 | 42 | 40 | 31 |
| SL1344_RS15065_ | 37 | 56 | 58 | 54 | 52 | 51 |
| SL1344_RS15070_ | 59 | 80 | 96 | 90 | 82 | 68 |
| SL1344_RS15075_ | 196 | 186 | 190 | 230 | 219 | 261 |
| SL1344_RS15080_ | 57 | 69 | 59 | 81 | 77 | 68 |
| SL1344_RS15085_ | 84 | 85 | 73 | 83 | 68 | 100 |
| SL1344_RS15090_ | 98 | 99 | 91 | 87 | 112 | 93 |
| SL1344_RS15095_ | 154 | 154 | 154 | 160 | 211 | 174 |
| SL1344_RS15100_ | 148 | 153 | 147 | 128 | 174 | 147 |
| SL1344_RS15105_ | 361 | 363 | 411 | 354 | 313 | 364 |
| SL1344_RS15110_ | 203 | 257 | 261 | 252 | 187 | 231 |
| SL1344_RS15115_ | 8 | 7 | 7 | 15 | 11 | 17 |
| SL1344_RS15120_ | 39 | 62 | 60 | 59 | 53 | 59 |
| SL1344_RS15125_ | 22 | 18 | 24 | 26 | 15 | 25 |
| SL1344_RS15140_ | 968 | 1082 | 1058 | 1054 | 999 | 1013 |
| SL1344_RS15145_ | 747 | 916 | 815 | 731 | 760 | 743 |
| SL1344_RS15150_ | 946 | 1022 | 990 | 812 | 821 | 936 |
| SL1344_RS15155_ | 331 | 355 | 336 | 316 | 339 | 344 |
| SL1344_RS15160_ | 530 | 582 | 526 | 563 | 459 | 483 |
| SL1344_RS15165_ | 790 | 776 | 802 | 852 | 722 | 754 |
| SL1344_RS15170_ | 251 | 311 | 326 | 337 | 295 | 290 |

| | | | | | | |
|---|---|---|---|---|---|---|
| SL1344_RS15190_ | 401 | 368 | 466 | 320 | 311 | 372 |
| SL1344_RS15195_ | 52 | 49 | 45 | 58 | 60 | 77 |
| SL1344_RS15200_ | 154 | 175 | 172 | 199 | 179 | 216 |
| SL1344_RS15205_ | 99 | 88 | 95 | 134 | 97 | 104 |
| SL1344_RS15210_ | 78 | 83 | 86 | 112 | 61 | 102 |
| SL1344_RS15215_ | 103 | 101 | 132 | 134 | 85 | 135 |
| SL1344_RS15220_ | 39 | 36 | 44 | 37 | 32 | 48 |
| SL1344_RS15225_ | 85 | 103 | 101 | 115 | 100 | 107 |
| SL1344_RS15230_ | 85 | 79 | 60 | 90 | 98 | 79 |
| SL1344_RS15260_ | 159 | 165 | 146 | 180 | 209 | 179 |
| SL1344_RS15265_ | 755 | 975 | 941 | 989 | 807 | 838 |
| SL1344_RS15270_ | 11263 | 13953 | 12416 | 11156 | 10670 | 10347 |
| SL1344_RS15275_ | 3649 | 4553 | 4201 | 3780 | 3407 | 3616 |
| SL1344_RS15280_ | 496 | 540 | 575 | 519 | 492 | 490 |
| SL1344_RS15285_ | 434 | 455 | 434 | 567 | 417 | 504 |
| SL1344_RS15290_ | 728 | 773 | 712 | 874 | 651 | 839 |
| SL1344_RS15295_ | 3211 | 3682 | 3333 | 3615 | 3265 | 3269 |
| SL1344_RS15310_ | 170 | 172 | 141 | 138 | 143 | 173 |
| SL1344_RS15315_ | 321 | 258 | 273 | 258 | 222 | 239 |
| SL1344_RS15320_ | 250 | 171 | 188 | 171 | 117 | 144 |
| SL1344_RS15325_ | 230 | 134 | 203 | 150 | 64 | 103 |
| SL1344_RS15330_ | 679 | 808 | 663 | 875 | 777 | 862 |
| SL1344_RS15355_ | 391 | 487 | 448 | 428 | 327 | 467 |
| SL1344_RS15360_ | 4740 | 5143 | 5554 | 3726 | 4182 | 4094 |
| SL1344_RS15365_ | 18455 | 18635 | 19975 | 23394 | 16377 | 17244 |
| SL1344_RS15370_ | 14237 | 14842 | 15377 | 15074 | 15339 | 12674 |
| SL1344_RS15375_ | 373 | 407 | 418 | 413 | 446 | 370 |
| SL1344_RS15380_ | 72 | 53 | 55 | 56 | 99 | 83 |
| SL1344_RS15385_ | 27 | 32 | 32 | 25 | 16 | 44 |
| SL1344_RS15390_ | 69 | 85 | 87 | 84 | 44 | 38 |
| SL1344_RS15395_ | 112 | 114 | 95 | 117 | 102 | 94 |
| SL1344_RS15400_ | 77 | 87 | 78 | 58 | 68 | 60 |
| SL1344_RS15405_ | 77 | 69 | 78 | 98 | 72 | 92 |
| SL1344_RS15410_ | 788 | 819 | 824 | 845 | 703 | 458 |
| SL1344_RS15415_ | 801 | 892 | 935 | 916 | 763 | 860 |
| SL1344_RS15420_ | 200 | 210 | 264 | 236 | 194 | 232 |
| SL1344_RS15425_ | 345 | 351 | 415 | 510 | 350 | 386 |
| SL1344_RS15430_ | 528 | 569 | 738 | 695 | 472 | 620 |
| SL1344_RS15435_ | 378 | 455 | 473 | 413 | 377 | 376 |
| SL1344_RS15440_ | 237 | 271 | 325 | 263 | 233 | 231 |
| SL1344_RS15445_ | 53 | 95 | 86 | 57 | 55 | 58 |
| SL1344_RS15450_ | 476 | 575 | 481 | 461 | 557 | 561 |
| SL1344_RS15455_ | 1036 | 1357 | 1237 | 1378 | 1196 | 1356 |
| SL1344_RS15460_ | 980 | 964 | 1098 | 1543 | 1484 | 726 |
| SL1344_RS15465_ | 3435 | 4408 | 3697 | 3868 | 10085 | 2052 |
| SL1344_RS15475_ | 276 | 290 | 302 | 252 | 285 | 242 |
| SL1344_RS15480_ | 529 | 512 | 552 | 486 | 470 | 460 |
| SL1344_RS15485_ | 1244 | 1321 | 1204 | 1249 | 1112 | 1146 |
| SL1344_RS15490_ | 960 | 1128 | 1046 | 951 | 888 | 921 |
| SL1344_RS15495_ | 973 | 1193 | 1000 | 906 | 913 | 956 |
| SL1344_RS15500_ | 51 | 45 | 47 | 55 | 41 | 41 |
| SL1344_RS15505_ | 28 | 27 | 41 | 31 | 30 | 31 |
| SL1344_RS15510_ | 40 | 34 | 37 | 37 | 34 | 39 |
| SL1344_RS15515_ | 11 | 13 | 8 | 24 | 14 | 14 |
| SL1344_RS15520_ | 8 | 4 | 7 | 9 | 6 | 9 |
| SL1344_RS15525_ | 942 | 1223 | 1328 | 1089 | 835 | 1001 |
| SL1344_RS15530_ | 629 | 710 | 708 | 701 | 489 | 714 |
| SL1344_RS15535_ | 2214 | 2356 | 2217 | 2436 | 2545 | 2567 |
| SL1344_RS15540_ | 1298 | 1460 | 1684 | 1534 | 1212 | 1738 |
| SL1344_RS15545_ | 35 | 23 | 37 | 21 | 17 | 24 |
| SL1344_RS15550_ | 802 | 890 | 857 | 860 | 904 | 806 |
| SL1344_RS15555_ | 822 | 919 | 953 | 1207 | 907 | 930 |
| SL1344_RS15560_ | 138 | 130 | 172 | 107 | 125 | 132 |

| | | | | | | |
|---|---|---|---|---|---|---|
| SL1344_RS15565_ | 411 | 526 | 444 | 422 | 394 | 448 |
| SL1344_RS15570_ | 448 | 590 | 574 | 496 | 597 | 530 |
| SL1344_RS15575_ | 945 | 1277 | 1187 | 1117 | 1225 | 1188 |
| SL1344_RS15580_ | 821 | 899 | 919 | 859 | 650 | 762 |
| SL1344_RS15585_ | 680 | 694 | 691 | 570 | 732 | 594 |
| SL1344_RS15590_ | 16 | 46 | 41 | 74 | 88 | 74 |
| SL1344_RS15595_ | 105 | 139 | 145 | 131 | 126 | 160 |
| SL1344_RS15600_ | 337 | 474 | 381 | 367 | 345 | 433 |
| SL1344_RS15605_ | 124 | 119 | 137 | 105 | 115 | 96 |
| SL1344_RS15610_ | 87 | 97 | 75 | 93 | 79 | 67 |
| SL1344_RS15615_ | 52 | 37 | 43 | 41 | 33 | 38 |
| SL1344_RS15620_ | 1790 | 1389 | 1478 | 1307 | 2435 | 1524 |
| SL1344_RS15625_ | 233 | 294 | 240 | 354 | 375 | 271 |
| SL1344_RS15630_ | 68 | 102 | 91 | 104 | 63 | 79 |
| SL1344_RS15635_ | 310 | 368 | 445 | 281 | 216 | 240 |
| SL1344_RS15640_ | 157 | 153 | 199 | 175 | 187 | 214 |
| SL1344_RS15645_ | 78 | 87 | 68 | 95 | 55 | 72 |
| SL1344_RS15650_ | 57 | 51 | 47 | 55 | 52 | 56 |
| SL1344_RS15655_ | 7 | 7 | 15 | 8 | 9 | 8 |
| SL1344_RS15660_ | 28 | 22 | 41 | 37 | 49 | 21 |
| SL1344_RS15665_ | 22 | 23 | 19 | 19 | 23 | 9 |
| SL1344_RS15670_ | 38 | 47 | 34 | 38 | 27 | 41 |
| SL1344_RS15675_ | 9 | 7 | 9 | 14 | 10 | 7 |
| SL1344_RS15680_ | 177 | 185 | 248 | 234 | 170 | 265 |
| SL1344_RS15685_ | 12 | 31 | 18 | 26 | 20 | 17 |
| SL1344_RS15690_ | 598 | 845 | 827 | 698 | 693 | 827 |
| SL1344_RS15695_ | 505 | 699 | 812 | 565 | 459 | 677 |
| SL1344_RS15700_ | 4 | 3 | 3 | 3 | 5 | 7 |
| SL1344_RS15705_ | 37 | 42 | 47 | 36 | 28 | 27 |
| SL1344_RS15710_ | 795 | 431 | 665 | 976 | 838 | 696 |
| SL1344_RS15715_ | 523 | 622 | 535 | 653 | 602 | 636 |
| SL1344_RS15720_ | 799 | 847 | 943 | 837 | 784 | 893 |
| SL1344_RS15725_ | 4325 | 4561 | 3943 | 4355 | 4249 | 4949 |
| SL1344_RS15730_ | 3099 | 3452 | 3125 | 2947 | 3034 | 3487 |
| SL1344_RS15735_ | 1849 | 2192 | 2118 | 1740 | 1712 | 1898 |
| SL1344_RS15740_ | 988 | 1083 | 1130 | 927 | 897 | 1183 |
| SL1344_RS15745_ | 618 | 611 | 686 | 627 | 546 | 699 |
| SL1344_RS15750_ | 494 | 646 | 535 | 589 | 581 | 632 |
| SL1344_RS15755_ | 871 | 833 | 815 | 816 | 921 | 822 |
| SL1344_RS15760_ | 363 | 386 | 449 | 446 | 287 | 448 |
| SL1344_RS15765_ | 80 | 91 | 66 | 109 | 66 | 81 |
| SL1344_RS15770_ | 1404 | 1471 | 1272 | 1316 | 1312 | 1227 |
| SL1344_RS15775_ | 648 | 945 | 980 | 788 | 867 | 897 |
| SL1344_RS15780_ | 235 | 299 | 329 | 260 | 246 | 343 |
| SL1344_RS15785_ | 348 | 405 | 354 | 336 | 272 | 403 |
| SL1344_RS15790_ | 60 | 59 | 71 | 49 | 51 | 57 |
| SL1344_RS15795_ | 11779 | 13982 | 12694 | 13543 | 12785 | 13667 |
| SL1344_RS15800_ | 4949 | 5266 | 5180 | 5746 | 5395 | 5630 |
| SL1344_RS15810_ | 836 | 924 | 800 | 874 | 804 | 884 |
| SL1344_RS15815_ | 870 | 1016 | 925 | 1018 | 1005 | 993 |
| SL1344_RS15820_ | 1259 | 1351 | 1372 | 1388 | 1173 | 1379 |
| SL1344_RS15825_ | 1065 | 1278 | 1382 | 1121 | 1010 | 1191 |
| SL1344_RS15830_ | 2245 | 2581 | 2596 | 2815 | 1969 | 2432 |
| SL1344_RS15835_ | 571 | 596 | 519 | 536 | 562 | 459 |
| SL1344_RS15840_ | 603 | 907 | 660 | 544 | 806 | 829 |
| SL1344_RS15845_ | 1712 | 2100 | 1964 | 1673 | 1485 | 1540 |
| SL1344_RS15850_ | 171 | 253 | 176 | 184 | 191 | 192 |
| SL1344_RS15855_ | 11 | 31 | 30 | 23 | 34 | 12 |
| SL1344_RS15860_ | 626 | 790 | 690 | 581 | 563 | 600 |
| SL1344_RS15865_ | 110 | 98 | 78 | 82 | 76 | 63 |
| SL1344_RS15870_ | 579 | 754 | 682 | 583 | 548 | 582 |
| SL1344_RS15875_ | 12137 | 13625 | 14348 | 12126 | 10266 | 11982 |
| SL1344_RS15880_ | 3123 | 3704 | 2825 | 3266 | 3166 | 3407 |

| | | | | | | |
|---|---|---|---|---|---|---|
| SL1344_RS15890_ | 23 | 53 | 39 | 35 | 20 | 35 |
| SL1344_RS15895_ | 33 | 31 | 29 | 39 | 33 | 20 |
| SL1344_RS15900_ | 53 | 58 | 70 | 54 | 49 | 55 |
| SL1344_RS15905_ | 86 | 88 | 105 | 93 | 101 | 91 |
| SL1344_RS15910_ | 35 | 38 | 26 | 26 | 38 | 30 |
| SL1344_RS15915_ | 11116 | 12997 | 13187 | 12134 | 9154 | 10000 |
| SL1344_RS15920_ | 450 | 522 | 561 | 452 | 458 | 450 |
| SL1344_RS15925_ | 3107 | 4170 | 3868 | 3175 | 4049 | 3026 |
| SL1344_RS15930_ | 42 | 60 | 57 | 68 | 55 | 60 |
| SL1344_RS15935_ | 42 | 33 | 35 | 55 | 37 | 43 |
| SL1344_RS15940_ | 91 | 97 | 89 | 106 | 117 | 89 |
| SL1344_RS15945_ | 90 | 111 | 102 | 145 | 141 | 128 |
| SL1344_RS15950_ | 42 | 37 | 36 | 57 | 29 | 47 |
| SL1344_RS15955_ | 22 | 26 | 23 | 32 | 35 | 30 |
| SL1344_RS15960_ | 808 | 850 | 905 | 822 | 598 | 998 |
| SL1344_RS15965_ | 39 | 26 | 38 | 38 | 25 | 34 |
| SL1344_RS15970_ | 2567 | 2934 | 2894 | 3209 | 2157 | 2536 |
| SL1344_RS15975_ | 450 | 363 | 388 | 527 | 388 | 313 |
| SL1344_RS15985_ | 1948 | 2494 | 2396 | 3482 | 2200 | 2289 |
| SL1344_RS15990_ | 3138 | 3569 | 3462 | 7904 | 5263 | 5562 |
| SL1344_RS15995_ | 215 | 160 | 176 | 240 | 224 | 180 |
| SL1344_RS16000_ | 280 | 300 | 315 | 241 | 180 | 201 |
| SL1344_RS16005_ | 624 | 716 | 753 | 688 | 545 | 637 |
| SL1344_RS16015_ | 391 | 504 | 511 | 592 | 443 | 462 |
| SL1344_RS16020_ | 351 | 489 | 498 | 399 | 431 | 453 |
| SL1344_RS16025_ | 131 | 136 | 109 | 142 | 127 | 134 |
| SL1344_RS16030_ | 39 | 43 | 29 | 68 | 65 | 73 |
| SL1344_RS16035_ | 447 | 468 | 540 | 568 | 373 | 492 |
| SL1344_RS16040_ | 479 | 520 | 516 | 561 | 441 | 570 |
| SL1344_RS16045_ | 349 | 374 | 357 | 412 | 307 | 406 |
| SL1344_RS16050_ | 545 | 618 | 610 | 572 | 418 | 536 |
| SL1344_RS16055_ | 1050 | 1358 | 1270 | 1215 | 1002 | 1090 |
| SL1344_RS16060_ | 366 | 241 | 251 | 300 | 349 | 234 |
| SL1344_RS16065_ | 11436 | 9683 | 9913 | 12501 | 10043 | 10767 |
| SL1344_RS16070_ | 653 | 606 | 614 | 718 | 525 | 649 |
| SL1344_RS16075_ | 853 | 851 | 805 | 793 | 831 | 831 |
| SL1344_RS16080_ | 989 | 850 | 826 | 904 | 891 | 691 |
| SL1344_RS16085_ | 444 | 475 | 434 | 432 | 380 | 371 |
| SL1344_RS16090_ | 492 | 511 | 584 | 424 | 410 | 552 |
| SL1344_RS16095_ | 538 | 732 | 603 | 610 | 413 | 571 |
| SL1344_RS16100_ | 1399 | 1375 | 1398 | 1609 | 1539 | 1359 |
| SL1344_RS16105_ | 394 | 448 | 365 | 419 | 340 | 327 |
| SL1344_RS16110_ | 8 | 10 | 7 | 10 | 2 | 4 |
| SL1344_RS16115_ | 68 | 75 | 75 | 57 | 58 | 67 |
| SL1344_RS16120_ | 152 | 164 | 207 | 285 | 366 | 182 |
| SL1344_RS16125_ | 60 | 50 | 63 | 59 | 61 | 53 |
| SL1344_RS16130_ | 187 | 195 | 201 | 143 | 130 | 175 |
| SL1344_RS16135_ | 93 | 66 | 88 | 56 | 54 | 65 |
| SL1344_RS16140_ | 125 | 163 | 180 | 154 | 98 | 144 |
| SL1344_RS16145_ | 297 | 376 | 298 | 255 | 251 | 299 |
| SL1344_RS16150_ | 184 | 244 | 229 | 229 | 248 | 228 |
| SL1344_RS16155_ | 24 | 23 | 25 | 28 | 21 | 21 |
| SL1344_RS16160_ | 386 | 178 | 265 | 226 | 163 | 103 |
| SL1344_RS16165_ | 98 | 99 | 88 | 91 | 84 | 97 |
| SL1344_RS16170_ | 142 | 173 | 137 | 157 | 161 | 158 |
| SL1344_RS16175_ | 74 | 71 | 81 | 77 | 63 | 60 |
| SL1344_RS16180_ | 152 | 169 | 183 | 169 | 152 | 159 |
| SL1344_RS16185_ | 167 | 158 | 112 | 150 | 163 | 169 |
| SL1344_RS16190_ | 39 | 48 | 44 | 53 | 53 | 54 |
| SL1344_RS16195_ | 45 | 47 | 58 | 57 | 56 | 64 |
| SL1344_RS16200_ | 60 | 55 | 50 | 56 | 55 | 61 |
| SL1344_RS16205_ | 25 | 28 | 25 | 20 | 31 | 13 |
| SL1344_RS16210_ | 1039 | 1105 | 970 | 653 | 1205 | 774 |

| | | | | | | |
|---|---|---|---|---|---|---|
| SL1344_RS16215_ | 2727 | 3181 | 2875 | 1678 | 2655 | 2174 |
| SL1344_RS16220_ | 1308 | 1630 | 1372 | 849 | 1471 | 1236 |
| SL1344_RS16225_ | 1428 | 1836 | 1693 | 1089 | 1646 | 1742 |
| SL1344_RS16235_ | 890 | 1005 | 896 | 747 | 765 | 821 |
| SL1344_RS16240_ | 256 | 362 | 339 | 368 | 284 | 279 |
| SL1344_RS16245_ | 52 | 66 | 49 | 78 | 59 | 85 |
| SL1344_RS16250_ | 48 | 50 | 45 | 44 | 77 | 67 |
| SL1344_RS16255_ | 293 | 310 | 319 | 507 | 357 | 410 |
| SL1344_RS16260_ | 142 | 175 | 165 | 309 | 180 | 217 |
| SL1344_RS16265_ | 146 | 190 | 189 | 307 | 210 | 256 |
| SL1344_RS16270_ | 326 | 417 | 412 | 629 | 415 | 484 |
| SL1344_RS16275_ | 868 | 1011 | 953 | 1441 | 1028 | 1218 |
| SL1344_RS16280_ | 334 | 339 | 314 | 519 | 338 | 520 |
| SL1344_RS16285_ | 211 | 204 | 176 | 420 | 229 | 369 |
| SL1344_RS16290_ | 403 | 494 | 555 | 868 | 475 | 724 |
| SL1344_RS16305_ | 901 | 1029 | 991 | 1491 | 1236 | 1394 |
| SL1344_RS16310_ | 1036 | 1333 | 1537 | 2948 | 2079 | 3009 |
| SL1344_RS16315_ | 452 | 505 | 499 | 1229 | 733 | 1121 |
| SL1344_RS16320_ | 547 | 570 | 622 | 1355 | 1034 | 1400 |
| SL1344_RS16325_ | 569 | 678 | 626 | 1129 | 1062 | 1088 |
| SL1344_RS16330_ | 782 | 1005 | 1057 | 1257 | 1226 | 1035 |
| SL1344_RS16335_ | 1550 | 1415 | 1457 | 2054 | 1438 | 1772 |
| SL1344_RS16340_ | 890 | 539 | 713 | 901 | 608 | 736 |
| SL1344_RS16350_ | 576 | 763 | 896 | 829 | 522 | 622 |
| SL1344_RS16355_ | 303 | 288 | 302 | 336 | 277 | 331 |
| SL1344_RS16360_ | 419 | 499 | 415 | 432 | 367 | 419 |
| SL1344_RS16365_ | 346 | 422 | 332 | 377 | 385 | 322 |
| SL1344_RS16370_ | 79 | 89 | 97 | 75 | 76 | 77 |
| SL1344_RS16375_ | 408 | 492 | 430 | 373 | 444 | 375 |
| SL1344_RS16380_ | 60 | 39 | 49 | 56 | 100 | 46 |
| SL1344_RS16385_ | 1022 | 1067 | 1224 | 1199 | 861 | 887 |
| SL1344_RS16390_ | 59 | 68 | 50 | 82 | 66 | 85 |
| SL1344_RS16395_ | 12 | 20 | 12 | 16 | 16 | 11 |
| SL1344_RS16400_ | 83 | 90 | 72 | 97 | 84 | 76 |
| SL1344_RS16405_ | 901 | 1074 | 889 | 751 | 886 | 810 |
| SL1344_RS16410_ | 477 | 496 | 530 | 424 | 346 | 377 |
| SL1344_RS16415_ | 1888 | 2274 | 2120 | 2310 | 2232 | 2044 |
| SL1344_RS16420_ | 648 | 843 | 599 | 801 | 993 | 806 |
| SL1344_RS16425_ | 1119 | 1069 | 940 | 1220 | 1102 | 1411 |
| SL1344_RS16440_ | 237 | 235 | 180 | 290 | 218 | 224 |
| SL1344_RS16445_ | 1592 | 2227 | 2056 | 1674 | 2119 | 1764 |
| SL1344_RS16450_ | 3488 | 3662 | 4003 | 3036 | 3309 | 3149 |
| SL1344_RS16455_ | 1250 | 1395 | 1546 | 1384 | 1398 | 1223 |
| SL1344_RS16460_ | 2277 | 2437 | 2479 | 2512 | 2125 | 2009 |
| SL1344_RS16465_ | 612 | 724 | 849 | 717 | 445 | 641 |
| SL1344_RS16470_ | 294 | 246 | 272 | 320 | 235 | 273 |
| SL1344_RS16480_ | 980 | 1364 | 1456 | 1049 | 883 | 1183 |
| SL1344_RS16485_ | 2310 | 2666 | 2570 | 2047 | 1955 | 2210 |
| SL1344_RS16490_ | 352 | 368 | 398 | 528 | 469 | 456 |
| SL1344_RS16495_ | 422 | 594 | 480 | 459 | 525 | 484 |
| SL1344_RS16500_ | 20 | 56 | 50 | 25 | 24 | 38 |
| SL1344_RS16505_ | 74 | 78 | 92 | 90 | 62 | 82 |
| SL1344_RS16510_ | 26 | 55 | 37 | 35 | 25 | 26 |
| SL1344_RS16515_ | 108 | 151 | 196 | 156 | 153 | 174 |
| SL1344_RS16520_ | 4922 | 5837 | 4285 | 6018 | 7202 | 5930 |
| SL1344_RS16525_ | 1188 | 1261 | 1207 | 1176 | 773 | 940 |
| SL1344_RS16530_ | 363 | 269 | 202 | 271 | 168 | 178 |
| SL1344_RS16535_ | 28 | 19 | 25 | 42 | 25 | 22 |
| SL1344_RS16540_ | 211 | 245 | 190 | 238 | 235 | 208 |
| SL1344_RS16545_ | 840 | 941 | 1049 | 892 | 779 | 748 |
| SL1344_RS16550_ | 1257 | 1423 | 1346 | 1168 | 1219 | 1315 |
| SL1344_RS16555_ | 627 | 782 | 781 | 649 | 622 | 671 |
| SL1344_RS16560_ | 380 | 410 | 373 | 413 | 328 | 405 |

| | | | | | | |
|---|---|---|---|---|---|---|
| SL1344_RS16565_ | 751 | 937 | 799 | 766 | 1008 | 799 |
| SL1344_RS16570_ | 454 | 593 | 551 | 456 | 463 | 510 |
| SL1344_RS16575_ | 127 | 157 | 140 | 160 | 107 | 132 |
| SL1344_RS16580_ | 642 | 776 | 717 | 769 | 706 | 687 |
| SL1344_RS16585_ | 706 | 900 | 868 | 742 | 742 | 668 |
| SL1344_RS16590_ | 19464 | 20373 | 23711 | 29156 | 22612 | 20933 |
| SL1344_RS16595_ | 3151 | 2872 | 3490 | 4316 | 2516 | 3185 |
| SL1344_RS16600_ | 4200 | 4332 | 4375 | 5927 | 4132 | 4666 |
| SL1344_RS16605_ | 66 | 93 | 75 | 74 | 84 | 129 |
| SL1344_RS16610_ | 104 | 167 | 119 | 235 | 222 | 68 |
| SL1344_RS16615_ | 231 | 300 | 280 | 368 | 320 | 357 |
| SL1344_RS16620_ | 232 | 257 | 219 | 219 | 167 | 186 |
| SL1344_RS16635_ | 106 | 101 | 77 | 115 | 107 | 104 |
| SL1344_RS16640_ | 407 | 371 | 383 | 352 | 533 | 299 |
| SL1344_RS16645_ | 368 | 441 | 455 | 395 | 374 | 425 |
| SL1344_RS16650_ | 240 | 304 | 272 | 286 | 206 | 257 |
| SL1344_RS16655_ | 83 | 98 | 68 | 106 | 84 | 77 |
| SL1344_RS16660_ | 838 | 853 | 981 | 838 | 739 | 679 |
| SL1344_RS16665_ | 236 | 251 | 219 | 283 | 332 | 235 |
| SL1344_RS16675_ | 798 | 1082 | 1022 | 984 | 816 | 912 |
| SL1344_RS16680_ | 507 | 510 | 538 | 672 | 425 | 564 |
| SL1344_RS16685_ | 1584 | 1560 | 1677 | 1454 | 1044 | 1464 |
| SL1344_RS16690_ | 1067 | 1110 | 1049 | 1150 | 728 | 903 |
| SL1344_RS16695_ | 1418 | 1587 | 1554 | 1497 | 1196 | 1240 |
| SL1344_RS16700_ | 1735 | 1632 | 1557 | 1881 | 1294 | 1527 |
| SL1344_RS16705_ | 13 | 22 | 12 | 27 | 13 | 24 |
| SL1344_RS16710_ | 162 | 229 | 181 | 204 | 166 | 148 |
| SL1344_RS16715_ | 843 | 1132 | 1128 | 768 | 763 | 779 |
| SL1344_RS16720_ | 206 | 252 | 178 | 242 | 287 | 256 |
| SL1344_RS16725_ | 150 | 123 | 138 | 182 | 108 | 123 |
| SL1344_RS16730_ | 379 | 422 | 477 | 375 | 367 | 329 |
| SL1344_RS16745_ | 167 | 153 | 138 | 284 | 223 | 241 |
| SL1344_RS16750_ | 201 | 223 | 203 | 433 | 260 | 340 |
| SL1344_RS16755_ | 108 | 124 | 115 | 239 | 145 | 224 |
| SL1344_RS16775_ | 200 | 151 | 160 | 137 | 112 | 145 |
| SL1344_RS16780_ | 344 | 299 | 287 | 234 | 149 | 156 |
| SL1344_RS16785_ | 216 | 148 | 142 | 137 | 44 | 66 |
| SL1344_RS16790_ | 196 | 170 | 187 | 135 | 125 | 113 |
| SL1344_RS16795_ | 459 | 495 | 435 | 498 | 428 | 465 |
| SL1344_RS16800_ | 648 | 633 | 641 | 622 | 468 | 664 |
| SL1344_RS16805_ | 16 | 35 | 28 | 48 | 56 | 34 |
| SL1344_RS16810_ | 9 | 25 | 19 | 16 | 13 | 18 |
| SL1344_RS16815_ | 30 | 22 | 31 | 33 | 23 | 25 |
| SL1344_RS16820_ | 24 | 29 | 25 | 18 | 16 | 17 |
| SL1344_RS16825_ | 19 | 27 | 34 | 26 | 45 | 33 |
| SL1344_RS16830_ | 7 | 11 | 9 | 13 | 10 | 10 |
| SL1344_RS16835_ | 7 | 10 | 14 | 16 | 16 | 11 |
| SL1344_RS16840_ | 61 | 67 | 77 | 101 | 69 | 94 |
| SL1344_RS16845_ | 13 | 22 | 17 | 22 | 17 | 15 |
| SL1344_RS16850_ | 556 | 579 | 620 | 648 | 549 | 491 |
| SL1344_RS16855_ | 384 | 494 | 477 | 448 | 382 | 450 |
| SL1344_RS16860_ | 1564 | 1967 | 2023 | 1948 | 1673 | 1726 |
| SL1344_RS16865_ | 730 | 953 | 901 | 814 | 862 | 807 |
| SL1344_RS16870_ | 550 | 593 | 628 | 753 | 471 | 589 |
| SL1344_RS16875_ | 1109 | 1517 | 1344 | 1392 | 1139 | 1179 |
| SL1344_RS16880_ | 85 | 75 | 87 | 102 | 53 | 69 |
| SL1344_RS16885_ | 68 | 81 | 78 | 81 | 50 | 49 |
| SL1344_RS16890_ | 83 | 103 | 73 | 112 | 116 | 121 |
| SL1344_RS16895_ | 104 | 106 | 119 | 119 | 96 | 131 |
| SL1344_RS16900_ | 744 | 853 | 652 | 619 | 832 | 758 |
| SL1344_RS16905_ | 659 | 872 | 683 | 635 | 636 | 715 |
| SL1344_RS16910_ | 54 | 88 | 56 | 55 | 46 | 67 |
| SL1344_RS16915_ | 36 | 33 | 37 | 26 | 21 | 21 |

| | | | | | | |
|---|---|---|---|---|---|---|
| SL1344_RS16920_ | 223 | 250 | 243 | 165 | 179 | 181 |
| SL1344_RS16925_ | 22 | 20 | 23 | 26 | 14 | 18 |
| SL1344_RS16930_ | 10 | 7 | 7 | 8 | 8 | 10 |
| SL1344_RS16935_ | 146 | 167 | 136 | 152 | 143 | 104 |
| SL1344_RS16940_ | 570 | 807 | 967 | 1091 | 920 | 1025 |
| SL1344_RS16945_ | 8843 | 9131 | 10053 | 11102 | 9758 | 11464 |
| SL1344_RS16950_ | 6198 | 7170 | 6547 | 7164 | 6538 | 6451 |
| SL1344_RS16955_ | 15796 | 17145 | 18543 | 20340 | 14864 | 13708 |
| SL1344_RS16960_ | 1177 | 1434 | 1582 | 1476 | 1200 | 1283 |
| SL1344_RS16965_ | 645 | 741 | 761 | 724 | 634 | 689 |
| SL1344_RS16970_ | 8377 | 8598 | 9029 | 8871 | 7648 | 8566 |
| SL1344_RS16980_ | 1221 | 1278 | 1391 | 1358 | 1154 | 1273 |
| SL1344_RS16985_ | 2528 | 2004 | 1775 | 4018 | 4238 | 1715 |
| SL1344_RS16990_ | 264 | 318 | 313 | 198 | 171 | 211 |
| SL1344_RS16995_ | 224 | 307 | 254 | 402 | 348 | 468 |
| SL1344_RS17000_ | 2825 | 3127 | 2095 | 4389 | 5928 | 2542 |
| SL1344_RS17005_ | 14288 | 9465 | 8801 | 19025 | 16890 | 11204 |
| SL1344_RS17010_ | 1829 | 1867 | 1736 | 1941 | 2004 | 2000 |
| SL1344_RS17015_ | 407 | 366 | 329 | 473 | 426 | 510 |
| SL1344_RS17030_ | 2342 | 2100 | 2366 | 2490 | 1888 | 2221 |
| SL1344_RS17035_ | 2393 | 2914 | 2914 | 2796 | 2432 | 2298 |
| SL1344_RS17040_ | 1221 | 1417 | 1514 | 1674 | 1181 | 1432 |
| SL1344_RS17045_ | 780 | 642 | 851 | 1031 | 525 | 745 |
| SL1344_RS17050_ | 957 | 1171 | 1084 | 1227 | 1069 | 1099 |
| SL1344_RS17060_ | 477 | 541 | 560 | 571 | 424 | 519 |
| SL1344_RS17065_ | 6015 | 5831 | 4943 | 5384 | 5591 | 5607 |
| SL1344_RS17070_ | 8135 | 7206 | 6991 | 9209 | 7627 | 8261 |
| SL1344_RS17075_ | 2742 | 3721 | 3604 | 3141 | 2800 | 2728 |
| SL1344_RS17080_ | 69 | 79 | 101 | 91 | 88 | 82 |
| SL1344_RS17085_ | 2001 | 2330 | 2016 | 2025 | 1858 | 2060 |
| SL1344_RS17090_ | 271 | 275 | 233 | 280 | 237 | 275 |
| SL1344_RS17095_ | 1359 | 1361 | 1427 | 1105 | 1073 | 1034 |
| SL1344_RS17100_ | 892 | 920 | 1028 | 878 | 623 | 823 |
| SL1344_RS17105_ | 329 | 379 | 339 | 382 | 289 | 383 |
| SL1344_RS17110_ | 403 | 467 | 393 | 428 | 413 | 492 |
| SL1344_RS17115_ | 583 | 767 | 706 | 680 | 744 | 613 |
| SL1344_RS17120_ | 865 | 1022 | 1043 | 1144 | 1036 | 1063 |
| SL1344_RS17125_ | 987 | 1129 | 1177 | 1132 | 981 | 1069 |
| SL1344_RS17130_ | 1355 | 1609 | 1630 | 1527 | 1486 | 1464 |
| SL1344_RS17135_ | 1100 | 1245 | 1377 | 1284 | 959 | 1149 |
| SL1344_RS17140_ | 1202 | 1444 | 1577 | 1280 | 898 | 1291 |
| SL1344_RS17145_ | 1263 | 1553 | 1503 | 1381 | 1069 | 1261 |
| SL1344_RS17150_ | 2872 | 3498 | 3242 | 3434 | 2584 | 2769 |
| SL1344_RS17155_ | 1145 | 1219 | 1271 | 1474 | 944 | 1372 |
| SL1344_RS17160_ | 1063 | 1216 | 1238 | 1185 | 937 | 1109 |
| SL1344_RS17165_ | 1902 | 2336 | 2399 | 2185 | 1653 | 2031 |
| SL1344_RS17170_ | 553 | 682 | 746 | 670 | 574 | 606 |
| SL1344_RS17175_ | 45 | 61 | 62 | 51 | 51 | 46 |
| SL1344_RS17180_ | 676 | 831 | 561 | 1184 | 1235 | 1459 |
| SL1344_RS17185_ | 592 | 797 | 747 | 1095 | 1005 | 1334 |
| SL1344_RS17190_ | 1350 | 1465 | 1389 | 1629 | 1668 | 1837 |
| SL1344_RS17195_ | 74 | 82 | 76 | 71 | 72 | 77 |
| SL1344_RS17200_ | 8 | 9 | 9 | 19 | 6 | 7 |
| SL1344_RS17205_ | 1037 | 1575 | 1260 | 827 | 852 | 973 |
| SL1344_RS17210_ | 539 | 549 | 573 | 439 | 333 | 422 |
| SL1344_RS17215_ | 45 | 30 | 19 | 54 | 37 | 35 |
| SL1344_RS17220_ | 126 | 147 | 155 | 187 | 133 | 98 |
| SL1344_RS17225_ | 474 | 621 | 533 | 556 | 481 | 502 |
| SL1344_RS17230_ | 1596 | 1271 | 1513 | 2169 | 1301 | 1732 |
| SL1344_RS17235_ | 1112 | 889 | 982 | 1614 | 918 | 1255 |
| SL1344_RS17240_ | 1152 | 905 | 1071 | 1681 | 904 | 1165 |
| SL1344_RS17245_ | 2811 | 1802 | 1924 | 3670 | 2126 | 2531 |
| SL1344_RS17250_ | 5907 | 3973 | 4452 | 7220 | 4303 | 5243 |

| | | | | | | |
|---|---|---|---|---|---|---|
| SL1344_RS17255_ | 1108 | 1345 | 1171 | 1373 | 1367 | 1286 |
| SL1344_RS17260_ | 4417 | 5229 | 4994 | 4494 | 5268 | 4348 |
| SL1344_RS17270_ | 168 | 176 | 178 | 207 | 137 | 254 |
| SL1344_RS17275_ | 19649 | 20345 | 20331 | 22050 | 19953 | 16617 |
| SL1344_RS17280_ | 8757 | 8765 | 9341 | 11491 | 8216 | 7623 |
| SL1344_RS17285_ | 653 | 860 | 815 | 829 | 781 | 731 |
| SL1344_RS17290_ | 2377 | 2872 | 2443 | 2318 | 2419 | 2840 |
| SL1344_RS17295_ | 1480 | 2134 | 1647 | 1551 | 1641 | 1720 |
| SL1344_RS17300_ | 804 | 1057 | 1004 | 954 | 848 | 857 |
| SL1344_RS17305_ | 23 | 28 | 29 | 26 | 20 | 17 |
| SL1344_RS17310_ | 72 | 74 | 82 | 58 | 53 | 62 |
| SL1344_RS17315_ | 91 | 114 | 119 | 124 | 78 | 99 |
| SL1344_RS17320_ | 6 | 4 | 8 | 6 | 9 | 1 |
| SL1344_RS17325_ | 45 | 63 | 63 | 36 | 31 | 50 |
| SL1344_RS17330_ | 42 | 50 | 61 | 47 | 29 | 54 |
| SL1344_RS17335_ | 34 | 32 | 26 | 37 | 25 | 16 |
| SL1344_RS17340_ | 529 | 528 | 426 | 402 | 463 | 481 |
| SL1344_RS17345_ | 601 | 593 | 630 | 517 | 545 | 568 |
| SL1344_RS17350_ | 11984 | 12067 | 11737 | 11321 | 10490 | 11928 |
| SL1344_RS17355_ | 922 | 1057 | 1011 | 1083 | 770 | 915 |
| SL1344_RS17360_ | 46 | 46 | 47 | 43 | 27 | 39 |
| SL1344_RS17365_ | 109 | 147 | 132 | 151 | 109 | 153 |
| SL1344_RS17370_ | 91 | 132 | 105 | 80 | 78 | 85 |
| SL1344_RS17375_ | 155 | 163 | 159 | 155 | 178 | 189 |
| SL1344_RS17380_ | 77 | 99 | 90 | 79 | 88 | 77 |
| SL1344_RS17385_ | 8 | 14 | 6 | 11 | 10 | 9 |
| SL1344_RS17390_ | 215 | 294 | 246 | 236 | 257 | 255 |
| SL1344_RS17395_ | 990 | 1334 | 1101 | 774 | 1029 | 980 |
| SL1344_RS17400_ | 1945 | 2321 | 2248 | 2214 | 2033 | 1823 |
| SL1344_RS17405_ | 1946 | 1961 | 1958 | 2159 | 1740 | 1992 |
| SL1344_RS17410_ | 1274 | 1169 | 1198 | 1308 | 1129 | 1158 |
| SL1344_RS17415_ | 816 | 867 | 841 | 979 | 748 | 994 |
| SL1344_RS17420_ | 1484 | 1649 | 1636 | 1891 | 1510 | 1654 |
| SL1344_RS17425_ | 3841 | 5099 | 4968 | 5102 | 4285 | 5158 |
| SL1344_RS17430_ | 4 | 2 | 8 | 12 | 8 | 5 |
| SL1344_RS17440_ | 556 | 698 | 604 | 651 | 614 | 648 |
| SL1344_RS17445_ | 335 | 480 | 350 | 310 | 379 | 324 |
| SL1344_RS17450_ | 182 | 242 | 242 | 223 | 185 | 247 |
| SL1344_RS17455_ | 4274 | 4854 | 5261 | 3873 | 3614 | 3869 |
| SL1344_RS17460_ | 8056 | 9731 | 9270 | 7645 | 6596 | 6944 |
| SL1344_RS17465_ | 31 | 42 | 45 | 50 | 29 | 44 |
| SL1344_RS17470_ | 310 | 369 | 386 | 418 | 311 | 323 |
| SL1344_RS17475_ | 543 | 647 | 606 | 700 | 495 | 493 |
| SL1344_RS17480_ | 1977 | 2003 | 2294 | 2592 | 1892 | 2074 |
| SL1344_RS17490_ | 99 | 134 | 111 | 139 | 83 | 104 |
| SL1344_RS17495_ | 308 | 270 | 347 | 357 | 307 | 284 |
| SL1344_RS17500_ | 286 | 305 | 306 | 327 | 267 | 248 |
| SL1344_RS17505_ | 26 | 26 | 20 | 16 | 14 | 12 |
| SL1344_RS17510_ | 19 | 20 | 16 | 16 | 14 | 16 |
| SL1344_RS17515_ | 99 | 128 | 127 | 152 | 138 | 125 |
| SL1344_RS17520_ | 118 | 112 | 132 | 89 | 78 | 110 |
| SL1344_RS17530_ | 58 | 187 | 128 | 110 | 126 | 79 |
| SL1344_RS17540_ | 705 | 1017 | 2304 | 850 | 1162 | 679 |
| SL1344_RS17545_ | 77 | 100 | 210 | 95 | 77 | 106 |
| SL1344_RS17550_ | 178 | 306 | 776 | 177 | 201 | 131 |
| SL1344_RS17555_ | 430 | 522 | 539 | 587 | 454 | 475 |
| SL1344_RS17560_ | 442 | 490 | 488 | 363 | 357 | 471 |
| SL1344_RS17565_ | 802 | 989 | 941 | 683 | 678 | 899 |
| SL1344_RS17570_ | 569 | 675 | 559 | 554 | 569 | 636 |
| SL1344_RS17575_ | 2220 | 2711 | 2362 | 2050 | 2813 | 2333 |
| SL1344_RS17580_ | 2701 | 3008 | 3116 | 2732 | 2829 | 3130 |
| SL1344_RS17585_ | 556 | 707 | 760 | 604 | 553 | 714 |
| SL1344_RS17590_ | 1329 | 1760 | 1759 | 1604 | 1247 | 1646 |

| | | | | | | |
|---|---|---|---|---|---|---|
| SL1344_RS17595_ | 2024 | 2173 | 2376 | 2355 | 1915 | 2220 |
| SL1344_RS17600_ | 293 | 266 | 259 | 301 | 199 | 275 |
| SL1344_RS17605_ | 356 | 396 | 442 | 448 | 251 | 292 |
| SL1344_RS17610_ | 1524 | 1793 | 1796 | 1353 | 1285 | 1477 |
| SL1344_RS17615_ | 429 | 372 | 288 | 478 | 473 | 368 |
| SL1344_RS17620_ | 385 | 385 | 338 | 460 | 419 | 392 |
| SL1344_RS17625_ | 187 | 199 | 181 | 242 | 151 | 201 |
| SL1344_RS17630_ | 14305 | 19330 | 17346 | 14251 | 14913 | 13000 |
| SL1344_RS17635_ | 28426 | 35324 | 33455 | 30820 | 32127 | 29442 |
| SL1344_RS17640_ | 21995 | 27124 | 20576 | 20139 | 28479 | 21669 |
| SL1344_RS17645_ | 4456 | 4633 | 3085 | 4507 | 4260 | 5398 |
| SL1344_RS17650_ | 8935 | 10397 | 8075 | 8802 | 11227 | 8703 |
| SL1344_RS17655_ | 8689 | 11500 | 10665 | 8561 | 10706 | 9207 |
| SL1344_RS17660_ | 23938 | 29921 | 27738 | 28098 | 32223 | 25464 |
| SL1344_RS17665_ | 6572 | 7167 | 6591 | 7572 | 7374 | 6361 |
| SL1344_RS17670_ | 2930 | 3066 | 3249 | 3651 | 2897 | 3059 |
| SL1344_RS17675_ | 6930 | 6978 | 7331 | 8404 | 6668 | 6473 |
| SL1344_RS17680_ | 6748 | 6328 | 7242 | 8477 | 5167 | 5951 |
| SL1344_RS17685_ | 10804 | 9862 | 12289 | 13559 | 8341 | 9623 |
| SL1344_RS17690_ | 10477 | 9519 | 11544 | 12698 | 8462 | 8848 |
| SL1344_RS17700_ | 16189 | 17133 | 18217 | 21482 | 14809 | 15034 |
| SL1344_RS17705_ | 12422 | 12599 | 13783 | 16073 | 12080 | 12046 |
| SL1344_RS17710_ | 14510 | 14962 | 16781 | 18447 | 14618 | 13661 |
| SL1344_RS17715_ | 17884 | 27702 | 24128 | 19164 | 33385 | 20371 |
| SL1344_RS17720_ | 9421 | 11650 | 12094 | 9723 | 15147 | 10415 |
| SL1344_RS17725_ | 21341 | 26006 | 26508 | 20221 | 32306 | 22260 |
| SL1344_RS17730_ | 15984 | 18622 | 19503 | 17846 | 19226 | 16760 |
| SL1344_RS17735_ | 8537 | 9685 | 10465 | 10629 | 9280 | 8847 |
| SL1344_RS17740_ | 7686 | 9116 | 8656 | 9276 | 9789 | 7463 |
| SL1344_RS17745_ | 10798 | 11658 | 11974 | 15136 | 11208 | 11344 |
| SL1344_RS17750_ | 13303 | 16205 | 14234 | 15083 | 19767 | 11504 |
| SL1344_RS17755_ | 11659 | 12115 | 13241 | 16080 | 11649 | 10715 |
| SL1344_RS17760_ | 14776 | 13577 | 16188 | 20514 | 12479 | 12271 |
| SL1344_RS17765_ | 6630 | 6831 | 7296 | 10417 | 5803 | 4988 |
| SL1344_RS17770_ | 15 | 17 | 6 | 17 | 20 | 7 |
| SL1344_RS17775_ | 1635 | 1420 | 1564 | 2238 | 1652 | 1905 |
| SL1344_RS17780_ | 1180 | 1073 | 1818 | 2388 | 1253 | 2541 |
| SL1344_RS17785_ | 35417 | 49314 | 37892 | 34710 | 53632 | 32821 |
| SL1344_RS17790_ | 42005 | 50784 | 48068 | 51001 | 44255 | 40579 |
| SL1344_RS17795_ | 8885 | 9480 | 10840 | 12944 | 7844 | 8640 |
| SL1344_RS17800_ | 6058 | 6923 | 7769 | 9574 | 5798 | 6300 |
| SL1344_RS17805_ | 690 | 720 | 696 | 1246 | 687 | 757 |
| SL1344_RS17810_ | 559 | 788 | 654 | 586 | 605 | 551 |
| SL1344_RS17815_ | 609 | 743 | 738 | 659 | 566 | 591 |
| SL1344_RS17820_ | 1137 | 1365 | 1346 | 1342 | 950 | 1125 |
| SL1344_RS17825_ | 6797 | 8017 | 7803 | 7148 | 7030 | 6842 |
| SL1344_RS17830_ | 378 | 344 | 373 | 524 | 355 | 470 |
| SL1344_RS17835_ | 4589 | 5031 | 5318 | 4439 | 4541 | 4164 |
| SL1344_RS17840_ | 275 | 283 | 308 | 201 | 285 | 301 |
| SL1344_RS17845_ | 868 | 1008 | 854 | 645 | 1059 | 868 |
| SL1344_RS17850_ | 271 | 266 | 226 | 180 | 298 | 195 |
| SL1344_RS17855_ | 1277 | 1438 | 1537 | 1365 | 1169 | 1190 |
| SL1344_RS17860_ | 58 | 44 | 49 | 65 | 42 | 45 |
| SL1344_RS17865_ | 367 | 363 | 318 | 332 | 263 | 424 |
| SL1344_RS17870_ | 557 | 677 | 611 | 662 | 617 | 649 |
| SL1344_RS17875_ | 147 | 157 | 175 | 192 | 119 | 190 |
| SL1344_RS17880_ | 501 | 689 | 609 | 542 | 557 | 636 |
| SL1344_RS17885_ | 1211 | 1250 | 1059 | 1222 | 1276 | 935 |
| SL1344_RS17895_ | 2082 | 2716 | 2472 | 2642 | 2196 | 2197 |
| SL1344_RS17900_ | 331 | 400 | 395 | 396 | 379 | 347 |
| SL1344_RS17905_ | 247 | 264 | 279 | 316 | 229 | 258 |
| SL1344_RS17910_ | 99 | 85 | 95 | 96 | 65 | 88 |
| SL1344_RS17915_ | 70 | 68 | 59 | 62 | 50 | 57 |

| | | | | | | |
|---|---|---|---|---|---|---|
| SL1344_RS17920_ | 1105 | 1190 | 1383 | 920 | 704 | 770 |
| SL1344_RS17925_ | 96 | 136 | 126 | 125 | 123 | 113 |
| SL1344_RS17930_ | 110 | 280 | 318 | 142 | 151 | 200 |
| SL1344_RS17940_ | 174 | 177 | 227 | 180 | 118 | 151 |
| SL1344_RS17965_ | 2494 | 3062 | 2791 | 2601 | 2561 | 2553 |
| SL1344_RS17970_ | 1102 | 1302 | 1277 | 1089 | 1065 | 1030 |
| SL1344_RS17975_ | 789 | 918 | 960 | 809 | 606 | 751 |
| SL1344_RS17985_ | 2829 | 3601 | 3588 | 3223 | 2762 | 2933 |
| SL1344_RS17990_ | 2488 | 2623 | 2511 | 2643 | 2323 | 2937 |
| SL1344_RS17995_ | 1992 | 2046 | 2093 | 2361 | 1967 | 2395 |
| SL1344_RS18000_ | 113 | 99 | 95 | 106 | 120 | 115 |
| SL1344_RS18005_ | 26 | 17 | 27 | 21 | 33 | 36 |
| SL1344_RS18010_ | 22 | 25 | 23 | 25 | 35 | 24 |
| SL1344_RS18015_ | 25 | 18 | 20 | 23 | 34 | 26 |
| SL1344_RS18020_ | 30 | 28 | 29 | 45 | 50 | 38 |
| SL1344_RS18025_ | 990 | 1190 | 969 | 1276 | 1066 | 968 |
| SL1344_RS18030_ | 499 | 600 | 599 | 355 | 493 | 458 |
| SL1344_RS18035_ | 2248 | 2525 | 2299 | 2787 | 2553 | 2566 |
| SL1344_RS18040_ | 422 | 449 | 477 | 607 | 399 | 387 |
| SL1344_RS18045_ | 335 | 304 | 343 | 483 | 269 | 309 |
| SL1344_RS18050_ | 660 | 596 | 764 | 1008 | 391 | 594 |
| SL1344_RS18055_ | 292 | 329 | 236 | 316 | 403 | 298 |
| SL1344_RS18060_ | 8360 | 9186 | 9361 | 9576 | 8655 | 8407 |
| SL1344_RS18075_ | 355 | 378 | 330 | 331 | 313 | 343 |
| SL1344_RS18080_ | 2030 | 2647 | 2349 | 2111 | 2379 | 2016 |
| SL1344_RS18090_ | 2028 | 2382 | 2774 | 2469 | 1711 | 1947 |
| SL1344_RS18095_ | 918 | 1128 | 1274 | 906 | 1052 | 870 |
| SL1344_RS18100_ | 166 | 174 | 147 | 233 | 163 | 182 |
| SL1344_RS18105_ | 209 | 260 | 181 | 183 | 278 | 241 |
| SL1344_RS18110_ | 221 | 152 | 220 | 291 | 130 | 185 |
| SL1344_RS18115_ | 2385 | 2165 | 2407 | 2961 | 2292 | 2399 |
| SL1344_RS18120_ | 343 | 406 | 365 | 345 | 404 | 293 |
| SL1344_RS18125_ | 953 | 1322 | 839 | 623 | 761 | 750 |
| SL1344_RS18130_ | 1502 | 1783 | 1613 | 1064 | 1000 | 1115 |
| SL1344_RS18140_ | 340 | 409 | 451 | 427 | 267 | 420 |
| SL1344_RS18145_ | 631 | 653 | 824 | 728 | 483 | 782 |
| SL1344_RS18150_ | 184 | 216 | 206 | 287 | 326 | 302 |
| SL1344_RS18155_ | 215 | 274 | 208 | 329 | 292 | 306 |
| SL1344_RS18160_ | 104 | 104 | 117 | 152 | 123 | 135 |
| SL1344_RS18165_ | 323 | 339 | 351 | 446 | 320 | 289 |
| SL1344_RS18170_ | 803 | 876 | 702 | 727 | 807 | 908 |
| SL1344_RS18175_ | 1297 | 1607 | 1411 | 1336 | 1374 | 1474 |
| SL1344_RS18180_ | 480 | 545 | 615 | 541 | 483 | 505 |
| SL1344_RS18185_ | 26617 | 27332 | 29301 | 34256 | 27909 | 27628 |
| SL1344_RS18190_ | 81 | 78 | 64 | 85 | 84 | 85 |
| SL1344_RS18195_ | 354 | 283 | 304 | 336 | 318 | 365 |
| SL1344_RS18205_ | 28 | 23 | 24 | 33 | 19 | 36 |
| SL1344_RS18210_ | 41 | 44 | 56 | 53 | 35 | 50 |
| SL1344_RS18215_ | 74 | 66 | 74 | 78 | 62 | 71 |
| SL1344_RS18220_ | 791 | 938 | 916 | 1005 | 834 | 939 |
| SL1344_RS18225_ | 1978 | 2089 | 2040 | 1858 | 2167 | 2096 |
| SL1344_RS18230_ | 636 | 690 | 622 | 649 | 681 | 762 |
| SL1344_RS18235_ | 671 | 640 | 688 | 679 | 673 | 773 |
| SL1344_RS18240_ | 870 | 998 | 1117 | 1118 | 806 | 1018 |
| SL1344_RS18245_ | 1020 | 1150 | 1105 | 1271 | 894 | 1163 |
| SL1344_RS18250_ | 1361 | 1568 | 1307 | 1657 | 1493 | 1676 |
| SL1344_RS18255_ | 191 | 225 | 176 | 209 | 217 | 193 |
| SL1344_RS18270_ | 129 | 183 | 130 | 140 | 181 | 174 |
| SL1344_RS18275_ | 295 | 328 | 272 | 280 | 296 | 352 |
| SL1344_RS18280_ | 180 | 321 | 240 | 251 | 267 | 210 |
| SL1344_RS18285_ | 29 | 21 | 22 | 20 | 28 | 15 |
| SL1344_RS18290_ | 10 | 14 | 8 | 16 | 12 | 3 |
| SL1344_RS18295_ | 10 | 14 | 15 | 7 | 6 | 9 |

| | | | | | | |
|---|---|---|---|---|---|---|
| SL1344_RS18300_ | 54 | 57 | 50 | 71 | 65 | 59 |
| SL1344_RS18305_ | 148 | 185 | 147 | 216 | 165 | 181 |
| SL1344_RS18310_ | 186 | 165 | 140 | 219 | 134 | 230 |
| SL1344_RS18315_ | 250 | 256 | 182 | 232 | 255 | 265 |
| SL1344_RS18325_ | 196 | 180 | 198 | 109 | 115 | 151 |
| SL1344_RS18330_ | 197 | 210 | 203 | 168 | 186 | 148 |
| SL1344_RS18335_ | 1062 | 941 | 976 | 870 | 786 | 860 |
| SL1344_RS18340_ | 64 | 84 | 53 | 69 | 47 | 64 |
| SL1344_RS18345_ | 55 | 52 | 56 | 66 | 33 | 79 |
| SL1344_RS18355_ | 86 | 88 | 71 | 77 | 91 | 76 |
| SL1344_RS18360_ | 91 | 111 | 104 | 73 | 123 | 101 |
| SL1344_RS18365_ | 44 | 68 | 52 | 79 | 76 | 61 |
| SL1344_RS18370_ | 82 | 96 | 102 | 120 | 103 | 110 |
| SL1344_RS18375_ | 305 | 360 | 391 | 358 | 250 | 358 |
| SL1344_RS18380_ | 434 | 654 | 532 | 459 | 814 | 382 |
| SL1344_RS18385_ | 250 | 257 | 225 | 281 | 446 | 365 |
| SL1344_RS18390_ | 6057 | 7087 | 5832 | 7884 | 8252 | 6642 |
| SL1344_RS18395_ | 1458 | 1952 | 1900 | 1574 | 1638 | 1458 |
| SL1344_RS18400_ | 773 | 872 | 852 | 848 | 788 | 784 |
| SL1344_RS18405_ | 1396 | 1470 | 1227 | 1242 | 1234 | 1350 |
| SL1344_RS18415_ | 313 | 433 | 445 | 330 | 297 | 310 |
| SL1344_RS18420_ | 193 | 209 | 228 | 238 | 187 | 241 |
| SL1344_RS18425_ | 400 | 570 | 421 | 556 | 578 | 415 |
| SL1344_RS18440_ | 328 | 288 | 296 | 374 | 295 | 246 |
| SL1344_RS18445_ | 177 | 168 | 135 | 286 | 153 | 218 |
| SL1344_RS18450_ | 1657 | 2010 | 2079 | 2203 | 1457 | 1999 |
| SL1344_RS18455_ | 416 | 412 | 377 | 343 | 425 | 421 |
| SL1344_RS18460_ | 114 | 122 | 110 | 135 | 115 | 100 |
| SL1344_RS18465_ | 336 | 421 | 444 | 479 | 341 | 437 |
| SL1344_RS18470_ | 287 | 418 | 388 | 390 | 399 | 303 |
| SL1344_RS18475_ | 175 | 226 | 190 | 185 | 193 | 234 |
| SL1344_RS18480_ | 543 | 724 | 575 | 727 | 768 | 759 |
| SL1344_RS18485_ | 186 | 250 | 175 | 264 | 316 | 273 |
| SL1344_RS18490_ | 525 | 533 | 530 | 547 | 446 | 467 |
| SL1344_RS18495_ | 2682 | 2782 | 3222 | 3337 | 2405 | 2535 |
| SL1344_RS18500_ | 332 | 345 | 311 | 389 | 329 | 356 |
| SL1344_RS18505_ | 7186 | 8539 | 8525 | 9438 | 6934 | 9819 |
| SL1344_RS18510_ | 1863 | 1861 | 1851 | 1454 | 1336 | 1234 |
| SL1344_RS18515_ | 401 | 412 | 375 | 496 | 455 | 405 |
| SL1344_RS18520_ | 1081 | 1155 | 976 | 1311 | 916 | 1164 |
| SL1344_RS18525_ | 470 | 480 | 417 | 677 | 553 | 479 |
| SL1344_RS18530_ | 447 | 499 | 537 | 555 | 417 | 493 |
| SL1344_RS18535_ | 1290 | 1610 | 1592 | 1698 | 1318 | 1606 |
| SL1344_RS18540_ | 129 | 138 | 111 | 162 | 128 | 132 |
| SL1344_RS18545_ | 103 | 133 | 161 | 137 | 144 | 145 |
| SL1344_RS18550_ | 69 | 60 | 90 | 79 | 55 | 85 |
| SL1344_RS18555_ | 227 | 207 | 201 | 247 | 178 | 242 |
| SL1344_RS18560_ | 228 | 333 | 344 | 276 | 266 | 308 |
| SL1344_RS18565_ | 286 | 384 | 289 | 372 | 310 | 315 |
| SL1344_RS18570_ | 3053 | 3342 | 3219 | 4947 | 5219 | 4406 |
| SL1344_RS18575_ | 18 | 17 | 14 | 31 | 22 | 12 |
| SL1344_RS18580_ | 39 | 40 | 44 | 45 | 34 | 28 |
| SL1344_RS18585_ | 157 | 222 | 170 | 154 | 172 | 182 |
| SL1344_RS18590_ | 295 | 375 | 369 | 305 | 341 | 326 |
| SL1344_RS18595_ | 124 | 190 | 170 | 202 | 163 | 206 |
| SL1344_RS18600_ | 606 | 750 | 523 | 1605 | 1440 | 1469 |
| SL1344_RS18610_ | 377 | 434 | 315 | 335 | 299 | 302 |
| SL1344_RS18615_ | 937 | 1056 | 1028 | 1027 | 1008 | 1004 |
| SL1344_RS18620_ | 6538 | 6032 | 4997 | 5516 | 6659 | 5572 |
| SL1344_RS18625_ | 696 | 829 | 781 | 929 | 701 | 844 |
| SL1344_RS18630_ | 1413 | 1304 | 1501 | 1310 | 1167 | 1112 |
| SL1344_RS18635_ | 535 | 522 | 645 | 435 | 312 | 404 |
| SL1344_RS18640_ | 695 | 769 | 796 | 576 | 393 | 489 |

| | | | | | | |
|---|---|---|---|---|---|---|
| SL1344_RS18655_ | 181 | 188 | 191 | 100 | 92 | 86 |
| SL1344_RS18660_ | 569 | 657 | 654 | 427 | 320 | 336 |
| SL1344_RS18665_ | 122 | 158 | 171 | 107 | 77 | 86 |
| SL1344_RS18670_ | 613 | 788 | 725 | 511 | 484 | 435 |
| SL1344_RS18675_ | 1217 | 1939 | 1745 | 1796 | 2081 | 1243 |
| SL1344_RS18680_ | 41 | 36 | 31 | 72 | 49 | 33 |
| SL1344_RS18685_ | 181 | 183 | 249 | 201 | 141 | 138 |
| SL1344_RS18690_ | 343 | 439 | 376 | 445 | 446 | 473 |
| SL1344_RS18695_ | 363 | 381 | 397 | 439 | 344 | 448 |
| SL1344_RS18700_ | 399 | 459 | 458 | 455 | 426 | 475 |
| SL1344_RS18705_ | 246 | 277 | 291 | 282 | 303 | 313 |
| SL1344_RS18710_ | 1437 | 1474 | 1405 | 1312 | 1413 | 1467 |
| SL1344_RS18715_ | 55 | 51 | 51 | 50 | 38 | 54 |
| SL1344_RS18720_ | 96 | 98 | 69 | 73 | 82 | 83 |
| SL1344_RS18725_ | 97 | 114 | 106 | 116 | 112 | 86 |
| SL1344_RS18735_ | 466 | 449 | 421 | 740 | 1066 | 306 |
| SL1344_RS18740_ | 633 | 609 | 611 | 598 | 547 | 543 |
| SL1344_RS18745_ | 54 | 68 | 68 | 67 | 53 | 46 |
| SL1344_RS18750_ | 70 | 71 | 78 | 90 | 80 | 57 |
| SL1344_RS18755_ | 67 | 68 | 61 | 86 | 72 | 61 |
| SL1344_RS18760_ | 5 | 6 | 10 | 14 | 4 | 12 |
| SL1344_RS18765_ | 21 | 9 | 14 | 26 | 8 | 20 |
| SL1344_RS18770_ | 91 | 127 | 84 | 79 | 74 | 64 |
| SL1344_RS18775_ | 644 | 604 | 738 | 733 | 657 | 621 |
| SL1344_RS18780_ | 392 | 458 | 591 | 487 | 478 | 435 |
| SL1344_RS18785_ | 1003 | 1153 | 1098 | 1103 | 975 | 1094 |
| SL1344_RS18790_ | 387 | 477 | 511 | 410 | 348 | 382 |
| SL1344_RS18795_ | 655 | 863 | 698 | 637 | 747 | 716 |
| SL1344_RS18800_ | 2511 | 3021 | 2560 | 2647 | 2186 | 2349 |
| SL1344_RS18805_ | 321 | 318 | 382 | 320 | 147 | 254 |
| SL1344_RS18815_ | 157 | 278 | 353 | 145 | 137 | 196 |
| SL1344_RS18820_ | 280 | 357 | 399 | 354 | 335 | 374 |
| SL1344_RS18825_ | 268 | 319 | 374 | 359 | 312 | 395 |
| SL1344_RS18830_ | 74 | 54 | 77 | 47 | 39 | 56 |
| SL1344_RS18835_ | 3253 | 3947 | 3792 | 3425 | 3063 | 3090 |
| SL1344_RS18840_ | 1089 | 1325 | 1118 | 1200 | 1103 | 1110 |
| SL1344_RS18845_ | 133 | 117 | 125 | 120 | 81 | 111 |
| SL1344_RS18850_ | 104 | 121 | 95 | 150 | 111 | 133 |
| SL1344_RS18855_ | 35 | 27 | 24 | 24 | 30 | 32 |
| SL1344_RS18860_ | 229 | 279 | 254 | 283 | 259 | 244 |
| SL1344_RS18865_ | 312 | 292 | 299 | 369 | 350 | 351 |
| SL1344_RS18870_ | 307 | 290 | 273 | 166 | 194 | 138 |
| SL1344_RS18875_ | 1680 | 1774 | 1751 | 2299 | 1645 | 1701 |
| SL1344_RS18880_ | 46 | 35 | 47 | 51 | 29 | 28 |
| SL1344_RS18885_ | 125 | 162 | 140 | 151 | 100 | 101 |
| SL1344_RS18890_ | 449 | 510 | 598 | 699 | 743 | 657 |
| SL1344_RS18895_ | 282 | 304 | 268 | 325 | 279 | 311 |
| SL1344_RS18900_ | 317 | 391 | 302 | 394 | 309 | 391 |
| SL1344_RS18905_ | 41 | 56 | 58 | 58 | 62 | 54 |
| SL1344_RS18910_ | 31 | 21 | 19 | 33 | 29 | 15 |
| SL1344_RS18915_ | 46 | 34 | 35 | 33 | 21 | 23 |
| SL1344_RS18920_ | 5 | 1 | 4 | 4 | 1 | 3 |
| SL1344_RS18925_ | 9 | 14 | 7 | 9 | 14 | 19 |
| SL1344_RS18930_ | 15 | 12 | 10 | 19 | 9 | 17 |
| SL1344_RS18935_ | 21 | 55 | 53 | 45 | 40 | 41 |
| SL1344_RS18945_ | 12 | 6 | 15 | 14 | 28 | 23 |
| SL1344_RS18950_ | 12 | 22 | 32 | 28 | 21 | 27 |
| SL1344_RS18955_ | 250 | 250 | 267 | 190 | 178 | 162 |
| SL1344_RS18960_ | 136 | 173 | 126 | 160 | 191 | 173 |
| SL1344_RS18965_ | 194 | 196 | 155 | 172 | 202 | 229 |
| SL1344_RS18970_ | 148 | 136 | 129 | 134 | 128 | 121 |
| SL1344_RS18975_ | 850 | 1033 | 970 | 947 | 889 | 948 |
| SL1344_RS18980_ | 927 | 1229 | 1349 | 1287 | 999 | 996 |

| | | | | | | |
|---|---|---|---|---|---|---|
| SL1344_RS18985_ | 538 | 564 | 549 | 747 | 458 | 665 |
| SL1344_RS18990_ | 1756 | 1790 | 1708 | 1632 | 1530 | 1826 |
| SL1344_RS18995_ | 839 | 902 | 835 | 1079 | 837 | 901 |
| SL1344_RS19005_ | 29 | 35 | 23 | 18 | 13 | 23 |
| SL1344_RS19010_ | 1088 | 1216 | 1297 | 823 | 1327 | 943 |
| SL1344_RS19015_ | 3 | 5 | 8 | 6 | 4 | 3 |
| SL1344_RS19020_ | 142 | 188 | 174 | 191 | 152 | 141 |
| SL1344_RS19025_ | 5562 | 3198 | 3770 | 3093 | 5116 | 3433 |
| SL1344_RS19030_ | 2220 | 1440 | 1531 | 1046 | 2330 | 1999 |
| SL1344_RS19040_ | 501 | 462 | 459 | 479 | 618 | 661 |
| SL1344_RS19045_ | 310 | 331 | 346 | 395 | 269 | 302 |
| SL1344_RS19050_ | 432 | 460 | 467 | 365 | 314 | 400 |
| SL1344_RS19055_ | 180 | 191 | 187 | 130 | 152 | 175 |
| SL1344_RS19065_ | 4837 | 5645 | 4425 | 5237 | 5692 | 5095 |
| SL1344_RS19070_ | 4818 | 5836 | 5756 | 6322 | 4634 | 5714 |
| SL1344_RS19075_ | 1843 | 1817 | 2046 | 1951 | 1395 | 1789 |
| SL1344_RS19080_ | 2168 | 1918 | 2193 | 2185 | 1539 | 2092 |
| SL1344_RS19085_ | 558 | 785 | 674 | 609 | 601 | 707 |
| SL1344_RS19090_ | 1720 | 1739 | 1895 | 1941 | 1631 | 1737 |
| SL1344_RS19095_ | 730 | 844 | 841 | 818 | 656 | 700 |
| SL1344_RS19100_ | 168 | 159 | 194 | 101 | 109 | 120 |
| SL1344_RS19105_ | 7393 | 7244 | 8320 | 6243 | 6026 | 6896 |
| SL1344_RS19110_ | 4534 | 4815 | 4484 | 3731 | 4320 | 3612 |
| SL1344_RS19115_ | 1450 | 1756 | 1695 | 1879 | 1195 | 1562 |
| SL1344_RS19120_ | 1530 | 1700 | 1571 | 1730 | 1538 | 1549 |
| SL1344_RS19125_ | 1143 | 1255 | 1196 | 1267 | 989 | 1167 |
| SL1344_RS19130_ | 1416 | 1741 | 1994 | 1628 | 1203 | 1302 |
| SL1344_RS19135_ | 646 | 784 | 715 | 759 | 1005 | 708 |
| SL1344_RS19140_ | 485 | 659 | 698 | 584 | 492 | 452 |
| SL1344_RS19145_ | 381 | 296 | 343 | 410 | 360 | 467 |
| SL1344_RS19150_ | 551 | 566 | 680 | 741 | 531 | 637 |
| SL1344_RS19155_ | 688 | 703 | 871 | 989 | 622 | 839 |
| SL1344_RS19160_ | 770 | 734 | 921 | 975 | 655 | 836 |
| SL1344_RS19165_ | 198 | 232 | 227 | 238 | 178 | 200 |
| SL1344_RS19170_ | 796 | 676 | 821 | 880 | 519 | 844 |
| SL1344_RS19175_ | 1042 | 972 | 1103 | 1238 | 733 | 1075 |
| SL1344_RS19180_ | 939 | 925 | 1019 | 1190 | 792 | 1150 |
| SL1344_RS19185_ | 1929 | 2189 | 2288 | 2198 | 1691 | 1625 |
| SL1344_RS19190_ | 941 | 1139 | 978 | 985 | 899 | 825 |
| SL1344_RS19195_ | 308 | 219 | 236 | 429 | 189 | 266 |
| SL1344_RS19200_ | 16617 | 19814 | 23304 | 22869 | 18963 | 19898 |
| SL1344_RS19205_ | 13823 | 11664 | 16253 | 16791 | 10471 | 14472 |
| SL1344_RS19210_ | 500 | 420 | 499 | 522 | 438 | 384 |
| SL1344_RS19215_ | 1164 | 1248 | 1172 | 1363 | 1188 | 1212 |
| SL1344_RS19220_ | 923 | 968 | 874 | 1013 | 792 | 1067 |
| SL1344_RS19225_ | 882 | 1113 | 1001 | 1170 | 896 | 998 |
| SL1344_RS19230_ | 377 | 413 | 297 | 245 | 349 | 286 |
| SL1344_RS19235_ | 322 | 282 | 218 | 288 | 281 | 307 |
| SL1344_RS19240_ | 1192 | 1414 | 1397 | 1440 | 1151 | 1386 |
| SL1344_RS19245_ | 161 | 200 | 182 | 179 | 175 | 183 |
| SL1344_RS19250_ | 46 | 52 | 61 | 62 | 43 | 61 |
| SL1344_RS19255_ | 385 | 594 | 543 | 591 | 307 | 502 |
| SL1344_RS19260_ | 152 | 140 | 156 | 158 | 135 | 143 |
| SL1344_RS19265_ | 944 | 1056 | 950 | 1085 | 756 | 1108 |
| SL1344_RS19270_ | 1485 | 1604 | 1517 | 1660 | 1626 | 1367 |
| SL1344_RS19275_ | 4401 | 4773 | 4528 | 4883 | 3913 | 4343 |
| SL1344_RS19280_ | 1054 | 1141 | 1150 | 1256 | 869 | 935 |
| SL1344_RS19285_ | 1190 | 1420 | 1359 | 1500 | 1085 | 1129 |
| SL1344_RS19290_ | 618 | 719 | 705 | 624 | 681 | 613 |
| SL1344_RS19295_ | 667 | 747 | 684 | 686 | 670 | 630 |
| SL1344_RS19300_ | 154 | 180 | 231 | 225 | 134 | 155 |
| SL1344_RS19305_ | 1182 | 1605 | 1278 | 1342 | 1469 | 1259 |
| SL1344_RS19310_ | 136 | 169 | 146 | 145 | 135 | 142 |

| | | | | | | |
|---|---|---|---|---|---|---|
| SL1344_RS19315_ | 67 | 57 | 87 | 64 | 71 | 42 |
| SL1344_RS19320_ | 115 | 264 | 177 | 243 | 336 | 136 |
| SL1344_RS19325_ | 19 | 23 | 31 | 23 | 24 | 24 |
| SL1344_RS19330_ | 722 | 724 | 881 | 847 | 634 | 834 |
| SL1344_RS19335_ | 211 | 184 | 206 | 269 | 223 | 255 |
| SL1344_RS19340_ | 591 | 620 | 751 | 687 | 587 | 709 |
| SL1344_RS19345_ | 3 | 9 | 10 | 5 | 8 | 4 |
| SL1344_RS19350_ | 124 | 147 | 140 | 158 | 151 | 193 |
| SL1344_RS19355_ | 77 | 73 | 74 | 51 | 50 | 46 |
| SL1344_RS19360_ | 89 | 83 | 81 | 69 | 48 | 53 |
| SL1344_RS19365_ | 129 | 125 | 153 | 115 | 94 | 97 |
| SL1344_RS19370_ | 182 | 235 | 197 | 193 | 178 | 198 |
| SL1344_RS19375_ | 217 | 271 | 263 | 350 | 315 | 337 |
| SL1344_RS19380_ | 54 | 45 | 61 | 87 | 53 | 92 |
| SL1344_RS19385_ | 269 | 362 | 363 | 357 | 252 | 328 |
| SL1344_RS19390_ | 118 | 101 | 101 | 106 | 109 | 117 |
| SL1344_RS19395_ | 123 | 164 | 137 | 140 | 215 | 103 |
| SL1344_RS19400_ | 65 | 102 | 100 | 111 | 81 | 93 |
| SL1344_RS19405_ | 31 | 48 | 48 | 54 | 38 | 44 |
| SL1344_RS19410_ | 42 | 48 | 42 | 56 | 33 | 51 |
| SL1344_RS19415_ | 34 | 29 | 38 | 31 | 13 | 20 |
| SL1344_RS19420_ | 27 | 22 | 13 | 27 | 13 | 12 |
| SL1344_RS19425_ | 1123 | 1201 | 1135 | 1220 | 1267 | 1082 |
| SL1344_RS19430_ | 21 | 45 | 28 | 23 | 31 | 14 |
| SL1344_RS19435_ | 511 | 496 | 447 | 610 | 653 | 691 |
| SL1344_RS19445_ | 129 | 141 | 141 | 137 | 116 | 126 |
| SL1344_RS19450_ | 152 | 169 | 174 | 173 | 159 | 145 |
| SL1344_RS19455_ | 41 | 61 | 53 | 51 | 35 | 40 |
| SL1344_RS19460_ | 30 | 22 | 39 | 25 | 16 | 37 |
| SL1344_RS19465_ | 338 | 328 | 308 | 237 | 266 | 262 |
| SL1344_RS19470_ | 247 | 239 | 289 | 129 | 148 | 206 |
| SL1344_RS19475_ | 305 | 266 | 326 | 275 | 334 | 292 |
| SL1344_RS19480_ | 530 | 432 | 480 | 384 | 393 | 367 |
| SL1344_RS19485_ | 1141 | 1042 | 1521 | 852 | 683 | 821 |
| SL1344_RS19490_ | 311 | 327 | 347 | 298 | 326 | 329 |
| SL1344_RS19495_ | 97 | 111 | 99 | 90 | 101 | 95 |
| SL1344_RS19500_ | 191 | 228 | 184 | 244 | 180 | 150 |
| SL1344_RS19505_ | 261 | 337 | 341 | 344 | 292 | 258 |
| SL1344_RS19510_ | 209 | 188 | 212 | 273 | 227 | 180 |
| SL1344_RS19515_ | 270 | 276 | 254 | 279 | 258 | 301 |
| SL1344_RS19520_ | 169 | 187 | 136 | 141 | 152 | 167 |
| SL1344_RS19525_ | 154 | 177 | 166 | 188 | 146 | 170 |
| SL1344_RS19530_ | 135 | 171 | 169 | 155 | 135 | 154 |
| SL1344_RS19535_ | 83 | 83 | 89 | 155 | 91 | 107 |
| SL1344_RS19540_ | 331 | 397 | 415 | 569 | 411 | 502 |
| SL1344_RS19545_ | 2096 | 2848 | 2293 | 4153 | 3972 | 2179 |
| SL1344_RS19550_ | 353 | 418 | 413 | 429 | 345 | 414 |
| SL1344_RS19555_ | 106 | 100 | 127 | 109 | 131 | 99 |
| SL1344_RS19565_ | 617 | 643 | 594 | 621 | 953 | 657 |
| SL1344_RS19570_ | 440 | 379 | 402 | 559 | 481 | 422 |
| SL1344_RS19575_ | 1789 | 1358 | 1349 | 2782 | 2784 | 2656 |
| SL1344_RS19580_ | 3487 | 2621 | 2896 | 7152 | 4747 | 7550 |
| SL1344_RS19585_ | 529 | 612 | 603 | 606 | 582 | 452 |
| SL1344_RS19590_ | 284 | 358 | 308 | 325 | 339 | 314 |
| SL1344_RS19595_ | 260 | 378 | 291 | 245 | 312 | 332 |
| SL1344_RS19600_ | 352 | 396 | 367 | 508 | 467 | 347 |
| SL1344_RS19605_ | 916 | 581 | 692 | 2144 | 746 | 882 |
| SL1344_RS19610_ | 885 | 638 | 707 | 1746 | 545 | 777 |
| SL1344_RS19615_ | 843 | 725 | 728 | 783 | 630 | 627 |
| SL1344_RS19620_ | 255 | 368 | 300 | 357 | 342 | 323 |
| SL1344_RS19625_ | 69 | 78 | 55 | 74 | 97 | 57 |
| SL1344_RS19630_ | 14 | 9 | 9 | 12 | 9 | 12 |
| SL1344_RS19635_ | 30 | 50 | 58 | 48 | 22 | 30 |

| | | | | | | |
|---|---|---|---|---|---|---|
| SL1344_RS19640_ | 13 | 22 | 12 | 12 | 17 | 15 |
| SL1344_RS19645_ | 10 | 11 | 2 | 4 | 0 | 2 |
| SL1344_RS19650_ | 20 | 37 | 23 | 13 | 10 | 20 |
| SL1344_RS19655_ | 12 | 18 | 17 | 9 | 14 | 20 |
| SL1344_RS19660_ | 7 | 19 | 7 | 17 | 11 | 6 |
| SL1344_RS19665_ | 47 | 72 | 43 | 77 | 52 | 51 |
| SL1344_RS19670_ | 26 | 25 | 17 | 28 | 34 | 31 |
| SL1344_RS19675_ | 94 | 125 | 115 | 134 | 110 | 113 |
| SL1344_RS19680_ | 37 | 25 | 35 | 40 | 25 | 31 |
| SL1344_RS19685_ | 727 | 874 | 948 | 875 | 874 | 768 |
| SL1344_RS19690_ | 105 | 122 | 136 | 121 | 92 | 125 |
| SL1344_RS19695_ | 247 | 319 | 339 | 313 | 260 | 251 |
| SL1344_RS19700_ | 54 | 100 | 77 | 89 | 93 | 77 |
| SL1344_RS19705_ | 104 | 99 | 87 | 97 | 92 | 135 |
| SL1344_RS19710_ | 60 | 52 | 38 | 37 | 29 | 59 |
| SL1344_RS19715_ | 165 | 213 | 226 | 193 | 212 | 242 |
| SL1344_RS19720_ | 114 | 128 | 155 | 138 | 153 | 159 |
| SL1344_RS19725_ | 380 | 578 | 497 | 372 | 322 | 473 |
| SL1344_RS19730_ | 31 | 36 | 30 | 20 | 21 | 19 |
| SL1344_RS19735_ | 54 | 78 | 73 | 79 | 66 | 71 |
| SL1344_RS19740_ | 652 | 626 | 786 | 732 | 432 | 454 |
| SL1344_RS19745_ | 4545 | 4830 | 5433 | 5571 | 4275 | 4713 |
| SL1344_RS19750_ | 1315 | 1238 | 1269 | 1472 | 1043 | 1118 |
| SL1344_RS19755_ | 2180 | 2509 | 2684 | 2584 | 2276 | 2237 |
| SL1344_RS19765_ | 8770 | 9713 | 11119 | 12068 | 7770 | 8325 |
| SL1344_RS19770_ | 2580 | 3182 | 2894 | 3766 | 3035 | 2490 |
| SL1344_RS19775_ | 1235 | 1066 | 1187 | 1446 | 935 | 1123 |
| SL1344_RS19780_ | 6742 | 7439 | 8452 | 8413 | 6578 | 6333 |
| SL1344_RS19785_ | 454 | 463 | 525 | 462 | 370 | 374 |
| SL1344_RS19790_ | 20 | 21 | 28 | 39 | 27 | 34 |
| SL1344_RS19795_ | 22 | 30 | 18 | 43 | 42 | 30 |
| SL1344_RS19800_ | 1124 | 913 | 1023 | 1168 | 880 | 861 |
| SL1344_RS19805_ | 678 | 719 | 883 | 773 | 622 | 585 |
| SL1344_RS19810_ | 49 | 52 | 47 | 55 | 52 | 54 |
| SL1344_RS19815_ | 149 | 167 | 136 | 169 | 150 | 143 |
| SL1344_RS19820_ | 134 | 188 | 158 | 122 | 103 | 125 |
| SL1344_RS19825_ | 867 | 1019 | 1068 | 857 | 576 | 762 |
| SL1344_RS19830_ | 513 | 661 | 599 | 660 | 661 | 514 |
| SL1344_RS19835_ | 443 | 604 | 498 | 617 | 896 | 591 |
| SL1344_RS19840_ | 3680 | 2542 | 3761 | 1989 | 1117 | 1207 |
| SL1344_RS19845_ | 1574 | 1294 | 1942 | 990 | 398 | 722 |
| SL1344_RS19850_ | 1057 | 1232 | 1413 | 608 | 378 | 626 |
| SL1344_RS19855_ | 1264 | 1617 | 1568 | 786 | 445 | 820 |
| SL1344_RS19860_ | 2848 | 3805 | 3345 | 1570 | 971 | 1717 |
| SL1344_RS19865_ | 87 | 138 | 85 | 100 | 106 | 92 |
| SL1344_RS19870_ | 18 | 34 | 24 | 5 | 20 | 25 |
| SL1344_RS19875_ | 255 | 406 | 326 | 271 | 283 | 268 |
| SL1344_RS19880_ | 5966 | 7715 | 7096 | 6791 | 6291 | 6541 |
| SL1344_RS19885_ | 5335 | 6617 | 6460 | 5790 | 5282 | 5854 |
| SL1344_RS19890_ | 149 | 153 | 136 | 182 | 155 | 148 |
| SL1344_RS19900_ | 18513 | 20546 | 19576 | 15792 | 14353 | 15345 |
| SL1344_RS19905_ | 13425 | 15889 | 14975 | 11976 | 11772 | 10783 |
| SL1344_RS19910_ | 35291 | 42664 | 39236 | 31722 | 28383 | 28242 |
| SL1344_RS19915_ | 13848 | 14025 | 14827 | 11705 | 10190 | 10748 |
| SL1344_RS19920_ | 18083 | 18949 | 18957 | 15248 | 16404 | 13489 |
| SL1344_RS19925_ | 8666 | 8684 | 8780 | 7353 | 6708 | 6853 |
| SL1344_RS19930_ | 8311 | 8638 | 8890 | 8300 | 6056 | 6910 |
| SL1344_RS19935_ | 3795 | 3228 | 3785 | 3760 | 3075 | 2844 |
| SL1344_RS19945_ | 1076 | 1172 | 1147 | 1114 | 878 | 1095 |
| SL1344_RS19950_ | 2458 | 2421 | 2832 | 3235 | 2295 | 3006 |
| SL1344_RS19955_ | 186 | 192 | 206 | 257 | 121 | 197 |
| SL1344_RS19960_ | 156 | 179 | 137 | 122 | 161 | 155 |
| SL1344_RS19965_ | 464 | 641 | 526 | 512 | 466 | 492 |

| | | | | | | |
|---|---|---|---|---|---|---|
| SL1344_RS19970_ | 747 | 982 | 888 | 833 | 784 | 861 |
| SL1344_RS19980_ | 488 | 663 | 552 | 593 | 1045 | 713 |
| SL1344_RS19985_ | 156 | 171 | 137 | 209 | 220 | 166 |
| SL1344_RS19990_ | 326 | 338 | 381 | 499 | 523 | 508 |
| SL1344_RS19995_ | 4175 | 4147 | 4704 | 6321 | 4956 | 6338 |
| SL1344_RS20000_ | 1096 | 1063 | 1097 | 944 | 1120 | 943 |
| SL1344_RS20005_ | 886 | 946 | 892 | 787 | 931 | 776 |
| SL1344_RS20020_ | 483 | 493 | 500 | 420 | 372 | 414 |
| SL1344_RS20025_ | 63 | 85 | 216 | 51 | 74 | 30 |
| SL1344_RS20030_ | 84 | 123 | 187 | 80 | 110 | 99 |
| SL1344_RS20035_ | 650 | 643 | 1490 | 554 | 800 | 445 |
| SL1344_RS20045_ | 1617 | 1825 | 1676 | 3146 | 2263 | 1226 |
| SL1344_RS20050_ | 2994 | 4341 | 3177 | 5246 | 6259 | 2019 |
| SL1344_RS20055_ | 412 | 680 | 521 | 406 | 621 | 670 |
| SL1344_RS20060_ | 3506 | 3131 | 2958 | 3900 | 2724 | 3275 |
| SL1344_RS20065_ | 78 | 101 | 76 | 133 | 110 | 101 |
| SL1344_RS20070_ | 1081 | 2868 | 1483 | 1800 | 3774 | 1369 |
| SL1344_RS20075_ | 222 | 348 | 259 | 274 | 377 | 388 |
| SL1344_RS20085_ | 553 | 678 | 796 | 625 | 554 | 597 |
| SL1344_RS20090_ | 961 | 1127 | 1352 | 1169 | 783 | 1161 |
| SL1344_RS20095_ | 1354 | 1618 | 1715 | 2004 | 1250 | 1554 |
| SL1344_RS20100_ | 184 | 187 | 194 | 228 | 165 | 176 |
| SL1344_RS20105_ | 224 | 193 | 217 | 213 | 204 | 184 |
| SL1344_RS20110_ | 207 | 280 | 226 | 256 | 263 | 219 |
| SL1344_RS20115_ | 959 | 1546 | 1332 | 1579 | 1700 | 2089 |
| SL1344_RS20120_ | 841 | 1221 | 947 | 1085 | 1431 | 1032 |
| SL1344_RS20125_ | 140 | 143 | 127 | 129 | 127 | 94 |
| SL1344_RS20130_ | 798 | 757 | 789 | 930 | 678 | 781 |
| SL1344_RS20135_ | 2728 | 3409 | 3665 | 2816 | 3028 | 2769 |
| SL1344_RS20140_ | 2536 | 3330 | 3390 | 2915 | 2636 | 2896 |
| SL1344_RS20145_ | 9685 | 11874 | 11225 | 10283 | 9908 | 9763 |
| SL1344_RS20150_ | 10329 | 13422 | 13890 | 11310 | 10998 | 10502 |
| SL1344_RS20155_ | 1377 | 1550 | 1672 | 1570 | 1243 | 1538 |
| SL1344_RS20160_ | 2106 | 2374 | 2291 | 2309 | 2028 | 2006 |
| SL1344_RS20165_ | 1500 | 1779 | 1677 | 1557 | 1335 | 1385 |
| SL1344_RS20175_ | 1007 | 1112 | 1157 | 992 | 670 | 926 |
| SL1344_RS20180_ | 358 | 419 | 428 | 368 | 280 | 311 |
| SL1344_RS20185_ | 614 | 590 | 559 | 553 | 404 | 512 |
| SL1344_RS20190_ | 793 | 856 | 963 | 836 | 544 | 680 |
| SL1344_RS20195_ | 756 | 824 | 881 | 718 | 614 | 645 |
| SL1344_RS20200_ | 541 | 503 | 559 | 568 | 377 | 437 |
| SL1344_RS20205_ | 771 | 787 | 934 | 826 | 636 | 700 |
| SL1344_RS20210_ | 521 | 531 | 481 | 529 | 432 | 449 |
| SL1344_RS20215_ | 1066 | 1204 | 1214 | 947 | 862 | 819 |
| SL1344_RS20220_ | 2692 | 2043 | 1867 | 4296 | 2435 | 1878 |
| SL1344_RS20225_ | 3465 | 2563 | 2221 | 4486 | 4112 | 1964 |
| SL1344_RS20230_ | 6794 | 5038 | 4558 | 8671 | 7291 | 4292 |
| SL1344_RS20235_ | 3476 | 6510 | 4209 | 5869 | 9475 | 2700 |
| SL1344_RS20240_ | 1618 | 2047 | 1856 | 1685 | 1434 | 1724 |
| SL1344_RS20245_ | 1232 | 1607 | 1542 | 1304 | 1247 | 1385 |
| SL1344_RS20250_ | 1233 | 1523 | 1373 | 1219 | 1458 | 1209 |
| SL1344_RS20255_ | 1348 | 1811 | 1668 | 1454 | 1444 | 1465 |
| SL1344_RS20260_ | 2466 | 3856 | 3257 | 3016 | 3010 | 3265 |
| SL1344_RS20265_ | 25 | 32 | 26 | 27 | 37 | 27 |
| SL1344_RS20270_ | 30 | 22 | 27 | 28 | 19 | 18 |
| SL1344_RS20275_ | 32 | 23 | 30 | 34 | 17 | 14 |
| SL1344_RS20280_ | 548 | 661 | 582 | 617 | 623 | 771 |
| SL1344_RS20285_ | 18 | 16 | 21 | 17 | 9 | 6 |
| SL1344_RS20290_ | 473 | 625 | 744 | 463 | 377 | 616 |
| SL1344_RS20295_ | 1266 | 1554 | 1493 | 1333 | 1312 | 1349 |
| SL1344_RS20300_ | 958 | 1362 | 1146 | 897 | 1049 | 1161 |
| SL1344_RS20305_ | 994 | 1328 | 1218 | 1015 | 1063 | 1110 |
| SL1344_RS20310_ | 493 | 675 | 609 | 576 | 482 | 548 |

| | | | | | | |
|---|---|---|---|---|---|---|
| SL1344_RS20315_ | 1620 | 1964 | 1998 | 2121 | 2040 | 2006 |
| SL1344_RS20325_ | 24 | 40 | 43 | 20 | 17 | 27 |
| SL1344_RS20330_ | 14 | 7 | 13 | 10 | 5 | 14 |
| SL1344_RS20335_ | 266 | 255 | 281 | 242 | 373 | 345 |
| SL1344_RS20340_ | 107 | 80 | 77 | 52 | 145 | 99 |
| SL1344_RS20345_ | 1480 | 1655 | 1724 | 1800 | 1371 | 1611 |
| SL1344_RS20350_ | 944 | 1023 | 1066 | 1092 | 933 | 1019 |
| SL1344_RS20355_ | 255 | 295 | 296 | 282 | 244 | 278 |
| SL1344_RS20365_ | 687 | 885 | 780 | 807 | 777 | 829 |
| SL1344_RS20370_ | 510 | 573 | 531 | 526 | 517 | 542 |
| SL1344_RS20375_ | 125 | 159 | 141 | 156 | 165 | 142 |
| SL1344_RS20390_ | 103 | 134 | 128 | 123 | 102 | 119 |
| SL1344_RS20395_ | 351 | 418 | 341 | 415 | 389 | 421 |
| SL1344_RS20400_ | 1821 | 2372 | 1951 | 2572 | 3487 | 2733 |
| SL1344_RS20405_ | 674 | 543 | 660 | 667 | 538 | 645 |
| SL1344_RS20410_ | 1122 | 1229 | 1386 | 1292 | 1163 | 1292 |
| SL1344_RS20415_ | 857 | 925 | 932 | 897 | 923 | 917 |
| SL1344_RS20425_ | 2243 | 2555 | 2960 | 2749 | 2539 | 2470 |
| SL1344_RS20430_ | 2430 | 2777 | 2988 | 2622 | 2594 | 2426 |
| SL1344_RS20435_ | 1631 | 1900 | 1863 | 1996 | 1604 | 1719 |
| SL1344_RS20440_ | 239 | 229 | 212 | 238 | 206 | 247 |
| SL1344_RS20445_ | 691 | 674 | 712 | 698 | 524 | 596 |
| SL1344_RS20450_ | 3551 | 3767 | 3885 | 3886 | 3549 | 3391 |
| SL1344_RS20455_ | 1911 | 1982 | 2050 | 1943 | 1990 | 1820 |
| SL1344_RS20460_ | 277 | 265 | 296 | 273 | 257 | 271 |
| SL1344_RS20470_ | 1704 | 1348 | 1463 | 1438 | 2309 | 2092 |
| SL1344_RS20475_ | 1515 | 1911 | 1940 | 1778 | 1406 | 1829 |
| SL1344_RS20480_ | 1027 | 1346 | 1328 | 1168 | 1197 | 1134 |
| SL1344_RS20485_ | 752 | 791 | 766 | 755 | 799 | 854 |
| SL1344_RS20495_ | 46 | 46 | 118 | 97 | 105 | 41 |
| SL1344_RS20500_ | 183 | 236 | 269 | 192 | 123 | 177 |
| SL1344_RS20505_ | 106 | 174 | 164 | 117 | 76 | 97 |
| SL1344_RS20510_ | 116 | 141 | 406 | 98 | 211 | 87 |
| SL1344_RS20520_ | 190 | 282 | 181 | 208 | 227 | 255 |
| SL1344_RS20530_ | 461 | 655 | 644 | 451 | 395 | 498 |
| SL1344_RS20535_ | 1274 | 1764 | 1484 | 1377 | 1538 | 1341 |
| SL1344_RS20540_ | 1067 | 1210 | 1112 | 925 | 933 | 1020 |
| SL1344_RS20545_ | 976 | 885 | 1127 | 1112 | 836 | 1068 |
| SL1344_RS20550_ | 54 | 41 | 55 | 59 | 32 | 48 |
| SL1344_RS20555_ | 1819 | 2231 | 1957 | 1682 | 1783 | 1916 |
| SL1344_RS20560_ | 928 | 1105 | 1141 | 1060 | 1044 | 1049 |
| SL1344_RS20565_ | 32496 | 30935 | 28960 | 54010 | 46336 | 55168 |
| SL1344_RS20570_ | 1309 | 1296 | 1365 | 1276 | 1016 | 1213 |
| SL1344_RS20575_ | 1358 | 1470 | 1415 | 1441 | 1364 | 1343 |
| SL1344_RS20580_ | 302 | 450 | 370 | 379 | 390 | 395 |
| SL1344_RS20585_ | 190 | 238 | 185 | 325 | 246 | 229 |
| SL1344_RS20590_ | 1982 | 2617 | 2716 | 3219 | 2402 | 2288 |
| SL1344_RS20595_ | 3896 | 4616 | 5007 | 5809 | 3993 | 4310 |
| SL1344_RS20600_ | 91 | 99 | 91 | 137 | 110 | 130 |
| SL1344_RS20605_ | 33 | 28 | 42 | 31 | 26 | 29 |
| SL1344_RS20610_ | 23 | 26 | 33 | 60 | 24 | 24 |
| SL1344_RS20615_ | 43 | 33 | 43 | 38 | 26 | 33 |
| SL1344_RS20620_ | 23 | 31 | 36 | 35 | 48 | 33 |
| SL1344_RS20625_ | 50 | 55 | 49 | 57 | 48 | 64 |
| SL1344_RS20635_ | 64 | 72 | 74 | 122 | 71 | 95 |
| SL1344_RS20640_ | 196 | 190 | 204 | 215 | 176 | 179 |
| SL1344_RS20645_ | 124 | 129 | 144 | 187 | 118 | 178 |
| SL1344_RS20650_ | 71 | 84 | 52 | 116 | 105 | 96 |
| SL1344_RS20655_ | 67 | 77 | 86 | 105 | 91 | 90 |
| SL1344_RS20660_ | 43 | 32 | 49 | 55 | 35 | 38 |
| SL1344_RS20665_ | 24 | 29 | 22 | 48 | 20 | 23 |
| SL1344_RS20670_ | 35 | 44 | 28 | 50 | 51 | 52 |
| SL1344_RS20675_ | 661 | 645 | 636 | 640 | 600 | 577 |

| | | | | | | |
|---|---|---|---|---|---|---|
| SL1344_RS20680_ | 731 | 841 | 745 | 888 | 760 | 897 |
| SL1344_RS20685_ | 782 | 881 | 850 | 898 | 797 | 875 |
| SL1344_RS20690_ | 610 | 697 | 661 | 636 | 601 | 653 |
| SL1344_RS20695_ | 1200 | 1355 | 1320 | 1189 | 1193 | 1276 |
| SL1344_RS20700_ | 326 | 351 | 331 | 368 | 344 | 409 |
| SL1344_RS20705_ | 261 | 290 | 263 | 355 | 313 | 334 |
| SL1344_RS20710_ | 163 | 266 | 226 | 253 | 240 | 279 |
| SL1344_RS20715_ | 125 | 100 | 93 | 134 | 123 | 140 |
| SL1344_RS20720_ | 115 | 108 | 111 | 126 | 108 | 129 |
| SL1344_RS20725_ | 2204 | 2531 | 2203 | 1983 | 2169 | 2052 |
| SL1344_RS20730_ | 2157 | 2169 | 2276 | 1942 | 1722 | 1974 |
| SL1344_RS20735_ | 2122 | 2374 | 2220 | 1802 | 1714 | 1875 |
| SL1344_RS20740_ | 5520 | 5794 | 5827 | 5025 | 4640 | 5074 |
| SL1344_RS20745_ | 3673 | 4160 | 4230 | 3536 | 3646 | 3517 |
| SL1344_RS20750_ | 464 | 431 | 428 | 503 | 494 | 457 |
| SL1344_RS20755_ | 63 | 51 | 60 | 59 | 49 | 41 |
| SL1344_RS20760_ | 24 | 32 | 26 | 27 | 26 | 31 |
| SL1344_RS20765_ | 138 | 165 | 147 | 176 | 154 | 179 |
| SL1344_RS20770_ | 183 | 267 | 216 | 274 | 295 | 282 |
| SL1344_RS20775_ | 375 | 493 | 477 | 442 | 462 | 452 |
| SL1344_RS20780_ | 177 | 198 | 164 | 259 | 254 | 230 |
| SL1344_RS20785_ | 204 | 200 | 196 | 221 | 162 | 202 |
| SL1344_RS20790_ | 64 | 69 | 55 | 65 | 59 | 50 |
| SL1344_RS20795_ | 38 | 40 | 25 | 37 | 39 | 24 |
| SL1344_RS20800_ | 25 | 26 | 16 | 31 | 40 | 28 |
| SL1344_RS20805_ | 101 | 122 | 121 | 212 | 145 | 163 |
| SL1344_RS20810_ | 129 | 142 | 95 | 142 | 123 | 102 |
| SL1344_RS20815_ | 57 | 57 | 70 | 58 | 69 | 50 |
| SL1344_RS20820_ | 3 | 1 | 7 | 2 | 3 | 3 |
| SL1344_RS20825_ | 15 | 12 | 18 | 15 | 18 | 16 |
| SL1344_RS20830_ | 60 | 95 | 90 | 78 | 82 | 68 |
| SL1344_RS20835_ | 16 | 12 | 8 | 10 | 4 | 2 |
| SL1344_RS20840_ | 23 | 19 | 11 | 20 | 14 | 22 |
| SL1344_RS20845_ | 14247 | 16420 | 17370 | 17395 | 13264 | 15449 |
| SL1344_RS20850_ | 297 | 352 | 312 | 312 | 346 | 319 |
| SL1344_RS20855_ | 300 | 392 | 322 | 399 | 574 | 388 |
| SL1344_RS20875_ | 458 | 487 | 486 | 577 | 481 | 624 |
| SL1344_RS20880_ | 599 | 881 | 802 | 563 | 952 | 705 |
| SL1344_RS20890_ | 1503 | 1701 | 1520 | 1894 | 1309 | 1541 |
| SL1344_RS20895_ | 255 | 286 | 268 | 189 | 212 | 173 |
| SL1344_RS20900_ | 210 | 179 | 138 | 120 | 124 | 132 |
| SL1344_RS20905_ | 742 | 940 | 809 | 860 | 763 | 695 |
| SL1344_RS20910_ | 589 | 710 | 622 | 714 | 638 | 580 |
| SL1344_RS20915_ | 18 | 13 | 20 | 40 | 42 | 32 |
| SL1344_RS20920_ | 109 | 85 | 87 | 107 | 133 | 122 |
| SL1344_RS20925_ | 360 | 344 | 334 | 331 | 542 | 353 |
| SL1344_RS20935_ | 247 | 189 | 198 | 252 | 278 | 218 |
| SL1344_RS20940_ | 161 | 142 | 142 | 157 | 261 | 166 |
| SL1344_RS20945_ | 112 | 120 | 109 | 107 | 183 | 175 |
| SL1344_RS20950_ | 204 | 184 | 205 | 252 | 274 | 344 |
| SL1344_RS20955_ | 209 | 173 | 194 | 357 | 290 | 429 |
| SL1344_RS20960_ | 64 | 81 | 83 | 101 | 82 | 118 |
| SL1344_RS20965_ | 240 | 274 | 279 | 352 | 256 | 404 |
| SL1344_RS20970_ | 3642 | 4660 | 4378 | 4401 | 3732 | 3922 |
| SL1344_RS20975_ | 251 | 309 | 285 | 294 | 265 | 261 |
| SL1344_RS20980_ | 179 | 195 | 210 | 188 | 220 | 186 |
| SL1344_RS20985_ | 752 | 841 | 729 | 691 | 828 | 863 |
| SL1344_RS20990_ | 2036 | 1952 | 1887 | 1873 | 2332 | 2628 |
| SL1344_RS20995_ | 52844 | 49705 | 50009 | 57567 | 48966 | 57707 |
| SL1344_RS21000_ | 15056 | 13415 | 16152 | 16813 | 16338 | 14260 |
| SL1344_RS21005_ | 3232 | 3662 | 3500 | 3037 | 2589 | 3585 |
| SL1344_RS21010_ | 1470 | 1641 | 1805 | 2072 | 1966 | 2300 |
| SL1344_RS21015_ | 393 | 332 | 379 | 466 | 316 | 335 |

| | | | | | | |
|---|---|---|---|---|---|---|
| SL1344_RS21020_ | 4168 | 3562 | 4493 | 6402 | 2632 | 4033 |
| SL1344_RS21025_ | 1074 | 837 | 951 | 1651 | 711 | 957 |
| SL1344_RS21030_ | 1905 | 2061 | 1989 | 1718 | 1655 | 1799 |
| SL1344_RS21035_ | 1427 | 1733 | 1546 | 1523 | 1725 | 1733 |
| SL1344_RS21040_ | 743 | 935 | 825 | 941 | 813 | 722 |
| SL1344_RS21045_ | 5552 | 6510 | 8304 | 8810 | 5010 | 5497 |
| SL1344_RS21050_ | 98 | 118 | 96 | 136 | 82 | 92 |
| SL1344_RS21055_ | 132 | 154 | 158 | 126 | 139 | 127 |
| SL1344_RS21075_ | 18 | 20 | 10 | 16 | 13 | 11 |
| SL1344_RS21080_ | 12 | 12 | 17 | 14 | 14 | 15 |
| SL1344_RS21085_ | 443 | 408 | 362 | 464 | 380 | 309 |
| SL1344_RS21090_ | 200 | 201 | 200 | 209 | 203 | 210 |
| SL1344_RS21100_ | 993 | 1202 | 998 | 1274 | 1145 | 1278 |
| SL1344_RS21105_ | 58 | 40 | 51 | 61 | 43 | 54 |
| SL1344_RS21115_ | 388 | 401 | 420 | 356 | 299 | 417 |
| SL1344_RS21120_ | 368 | 412 | 387 | 343 | 402 | 334 |
| SL1344_RS21125_ | 81 | 79 | 81 | 77 | 73 | 64 |
| SL1344_RS21130_ | 102 | 92 | 107 | 89 | 127 | 132 |
| SL1344_RS21135_ | 342 | 410 | 413 | 390 | 420 | 407 |
| SL1344_RS21140_ | 460 | 422 | 398 | 464 | 506 | 433 |
| SL1344_RS21145_ | 94 | 102 | 100 | 110 | 142 | 118 |
| SL1344_RS21150_ | 161 | 201 | 176 | 190 | 176 | 181 |
| SL1344_RS21155_ | 1809 | 2186 | 2193 | 1767 | 1430 | 1617 |
| SL1344_RS21160_ | 2747 | 3814 | 3486 | 2635 | 2927 | 2976 |
| SL1344_RS21165_ | 461 | 601 | 576 | 522 | 428 | 585 |
| SL1344_RS21170_ | 142 | 218 | 154 | 99 | 136 | 127 |
| SL1344_RS21175_ | 185 | 299 | 242 | 172 | 185 | 174 |
| SL1344_RS21180_ | 581 | 808 | 733 | 534 | 477 | 586 |
| SL1344_RS21185_ | 1298 | 1695 | 1429 | 1786 | 1404 | 1601 |
| SL1344_RS21195_ | 1332 | 1291 | 1330 | 1148 | 1298 | 1421 |
| SL1344_RS21200_ | 672 | 899 | 852 | 776 | 813 | 877 |
| SL1344_RS21205_ | 302 | 289 | 352 | 346 | 239 | 291 |
| SL1344_RS21210_ | 2048 | 2457 | 2520 | 2046 | 2021 | 1995 |
| SL1344_RS21215_ | 1395 | 1972 | 1712 | 1514 | 1821 | 1394 |
| SL1344_RS21220_ | 17 | 40 | 103 | 33 | 42 | 20 |
| SL1344_RS21225_ | 387 | 361 | 559 | 328 | 272 | 345 |
| SL1344_RS21230_ | 52 | 77 | 90 | 76 | 51 | 88 |
| SL1344_RS21235_ | 68 | 95 | 206 | 92 | 119 | 57 |
| SL1344_RS21245_ | 467 | 568 | 481 | 491 | 447 | 549 |
| SL1344_RS21250_ | 621 | 661 | 615 | 709 | 729 | 655 |
| SL1344_RS21255_ | 1395 | 1234 | 1399 | 1082 | 1090 | 965 |
| SL1344_RS21260_ | 1 | 0 | 2 | 1 | 1 | 3 |
| SL1344_RS21265_ | 1032 | 1182 | 1065 | 1907 | 1593 | 1046 |
| SL1344_RS21270_ | 2303 | 2314 | 1855 | 3565 | 3073 | 1911 |
| SL1344_RS21275_ | 8943 | 10204 | 8880 | 10803 | 19739 | 6783 |
| SL1344_RS21280_ | 3808 | 4830 | 3952 | 5783 | 7871 | 3893 |
| SL1344_RS21285_ | 35513 | 42391 | 35880 | 44639 | 42626 | 36188 |
| SL1344_RS21290_ | 3466 | 3217 | 3428 | 4004 | 3112 | 3562 |
| SL1344_RS21300_ | 7343 | 7052 | 9061 | 10187 | 6032 | 6460 |
| SL1344_RS21305_ | 15572 | 16482 | 19550 | 20991 | 16469 | 14650 |
| SL1344_RS21310_ | 28830 | 35435 | 36101 | 33013 | 26527 | 29533 |
| SL1344_RS21315_ | 55356 | 77613 | 78773 | 66702 | 71738 | 66597 |
| SL1344_RS21320_ | 16537 | 18849 | 17612 | 20777 | 16184 | 17119 |
| SL1344_RS21325_ | 20292 | 25164 | 22409 | 23424 | 24280 | 21003 |
| SL1344_RS21330_ | 32 | 38 | 35 | 76 | 53 | 41 |
| SL1344_RS21335_ | 12 | 11 | 9 | 10 | 7 | 8 |
| SL1344_RS21340_ | 239 | 254 | 289 | 211 | 196 | 242 |
| SL1344_RS21345_ | 47 | 50 | 39 | 35 | 59 | 29 |
| SL1344_RS21350_ | 154 | 180 | 197 | 228 | 149 | 202 |
| SL1344_RS21355_ | 66 | 71 | 69 | 71 | 67 | 80 |
| SL1344_RS21360_ | 18 | 24 | 22 | 38 | 20 | 29 |
| SL1344_RS21365_ | 69 | 98 | 86 | 66 | 70 | 103 |
| SL1344_RS21370_ | 36 | 57 | 51 | 54 | 58 | 38 |

| | | | | | | |
|---|---|---|---|---|---|---|
| SL1344_RS21375_ | 142 | 148 | 145 | 173 | 158 | 131 |
| SL1344_RS21380_ | 332 | 274 | 279 | 299 | 320 | 263 |
| SL1344_RS21390_ | 933 | 1069 | 1184 | 1039 | 935 | 998 |
| SL1344_RS21395_ | 629 | 717 | 751 | 635 | 674 | 627 |
| SL1344_RS21400_ | 780 | 741 | 739 | 763 | 708 | 765 |
| SL1344_RS21420_ | 466 | 500 | 464 | 457 | 483 | 495 |
| SL1344_RS21425_ | 579 | 587 | 542 | 525 | 710 | 585 |
| SL1344_RS21430_ | 222 | 280 | 265 | 361 | 260 | 230 |
| SL1344_RS21435_ | 270 | 293 | 328 | 385 | 272 | 315 |
| SL1344_RS21440_ | 45 | 82 | 175 | 61 | 46 | 43 |
| SL1344_RS21445_ | 10 | 14 | 14 | 6 | 11 | 12 |
| SL1344_RS21450_ | 181 | 224 | 451 | 231 | 256 | 133 |
| SL1344_RS21460_ | 12 | 12 | 12 | 30 | 21 | 17 |
| SL1344_RS21465_ | 411 | 357 | 333 | 346 | 303 | 431 |
| SL1344_RS21475_ | 156 | 157 | 136 | 123 | 108 | 131 |
| SL1344_RS21480_ | 211 | 235 | 225 | 229 | 201 | 219 |
| SL1344_RS21485_ | 186 | 258 | 211 | 246 | 211 | 225 |
| SL1344_RS21490_ | 450 | 489 | 493 | 404 | 437 | 423 |
| SL1344_RS21495_ | 909 | 955 | 1088 | 803 | 725 | 825 |
| SL1344_RS21505_ | 469 | 516 | 470 | 515 | 604 | 494 |
| SL1344_RS21510_ | 687 | 709 | 558 | 572 | 777 | 738 |
| SL1344_RS21515_ | 35 | 18 | 24 | 26 | 7 | 14 |
| SL1344_RS21520_ | 851 | 759 | 764 | 677 | 544 | 629 |
| SL1344_RS21525_ | 985 | 1035 | 989 | 857 | 790 | 755 |
| SL1344_RS21530_ | 418 | 438 | 490 | 396 | 360 | 369 |
| SL1344_RS21535_ | 104 | 144 | 118 | 124 | 149 | 172 |
| SL1344_RS21540_ | 34 | 29 | 25 | 21 | 12 | 25 |
| SL1344_RS21545_ | 15 | 13 | 14 | 13 | 5 | 12 |
| SL1344_RS21550_ | 74 | 78 | 92 | 81 | 60 | 68 |
| SL1344_RS21555_ | 18 | 37 | 22 | 23 | 13 | 24 |
| SL1344_RS21560_ | 48 | 55 | 34 | 32 | 46 | 53 |
| SL1344_RS21565_ | 7 | 14 | 6 | 10 | 6 | 7 |
| SL1344_RS21570_ | 24 | 23 | 30 | 12 | 22 | 22 |
| SL1344_RS21575_ | 4 | 5 | 9 | 8 | 4 | 6 |
| SL1344_RS21580_ | 255 | 267 | 339 | 291 | 161 | 238 |
| SL1344_RS21585_ | 128 | 111 | 127 | 95 | 75 | 74 |
| SL1344_RS21590_ | 84 | 68 | 88 | 80 | 57 | 57 |
| SL1344_RS21595_ | 74 | 76 | 95 | 86 | 62 | 73 |
| SL1344_RS21600_ | 44 | 51 | 55 | 41 | 41 | 41 |
| SL1344_RS21605_ | 10 | 22 | 15 | 16 | 16 | 12 |
| SL1344_RS21610_ | 37 | 53 | 40 | 32 | 42 | 47 |
| SL1344_RS21615_ | 64 | 64 | 67 | 49 | 54 | 58 |
| SL1344_RS21620_ | 47 | 49 | 67 | 66 | 49 | 56 |
| SL1344_RS21625_ | 24 | 26 | 27 | 34 | 24 | 33 |
| SL1344_RS21630_ | 55 | 58 | 55 | 81 | 64 | 43 |
| SL1344_RS21635_ | 2 | 2 | 4 | 8 | 4 | 2 |
| SL1344_RS21640_ | 3 | 5 | 7 | 12 | 1 | 6 |
| SL1344_RS21645_ | 14 | 8 | 12 | 7 | 13 | 10 |
| SL1344_RS21650_ | 19 | 20 | 11 | 18 | 20 | 9 |
| SL1344_RS21655_ | 14 | 9 | 21 | 14 | 13 | 16 |
| SL1344_RS21660_ | 101 | 98 | 85 | 92 | 73 | 84 |
| SL1344_RS21665_ | 612 | 956 | 747 | 632 | 832 | 666 |
| SL1344_RS21670_ | 2722 | 3320 | 3299 | 2954 | 2459 | 2586 |
| SL1344_RS21675_ | 118 | 132 | 154 | 172 | 162 | 158 |
| SL1344_RS21680_ | 27 | 21 | 27 | 24 | 25 | 29 |
| SL1344_RS21685_ | 20 | 17 | 11 | 28 | 14 | 27 |
| SL1344_RS21690_ | 234 | 328 | 248 | 337 | 296 | 301 |
| SL1344_RS21695_ | 46 | 70 | 65 | 77 | 79 | 81 |
| SL1344_RS21710_ | 7812 | 9746 | 9068 | 5470 | 4120 | 4408 |
| SL1344_RS21715_ | 2993 | 3364 | 3133 | 2246 | 1696 | 1895 |
| SL1344_RS21725_ | 2005 | 2461 | 1939 | 1513 | 962 | 1461 |
| SL1344_RS21730_ | 534 | 486 | 489 | 327 | 348 | 335 |
| SL1344_RS21735_ | 477 | 459 | 417 | 337 | 350 | 331 |

| | | | | | | |
|---|---|---|---|---|---|---|
| SL1344_RS21740_ | 3601 | 4541 | 4675 | 4744 | 4273 | 3936 |
| SL1344_RS21745_ | 276 | 289 | 375 | 292 | 209 | 255 |
| SL1344_RS21755_ | 397 | 472 | 395 | 533 | 492 | 468 |
| SL1344_RS21760_ | 130 | 127 | 146 | 197 | 99 | 182 |
| SL1344_RS21765_ | 854 | 926 | 1122 | 1232 | 790 | 1105 |
| SL1344_RS21770_ | 299 | 397 | 399 | 466 | 403 | 428 |
| SL1344_RS21775_ | 223 | 243 | 229 | 264 | 249 | 278 |
| SL1344_RS21780_ | 1189 | 1238 | 1417 | 1706 | 1246 | 1505 |
| SL1344_RS21785_ | 74 | 56 | 58 | 80 | 65 | 65 |
| SL1344_RS21790_ | 403 | 499 | 379 | 455 | 376 | 470 |
| SL1344_RS21795_ | 1187 | 1403 | 1376 | 1520 | 1279 | 1349 |
| SL1344_RS21800_ | 582 | 781 | 595 | 608 | 750 | 628 |
| SL1344_RS21805_ | 749 | 851 | 763 | 583 | 611 | 673 |
| SL1344_RS21810_ | 1956 | 2098 | 1941 | 2765 | 2659 | 2474 |
| SL1344_RS21815_ | 720 | 775 | 848 | 762 | 668 | 759 |
| SL1344_RS21820_ | 236 | 307 | 344 | 248 | 259 | 296 |
| SL1344_RS21825_ | 12 | 15 | 17 | 9 | 30 | 16 |
| SL1344_RS21830_ | 2 | 6 | 7 | 13 | 9 | 2 |
| SL1344_RS21835_ | 1261 | 1693 | 1513 | 1981 | 1710 | 1609 |
| SL1344_RS21840_ | 18 | 23 | 23 | 21 | 14 | 36 |
| SL1344_RS21845_ | 2114 | 2324 | 2300 | 2552 | 2300 | 2080 |
| SL1344_RS21880_ | 31 | 49 | 43 | 52 | 40 | 59 |
| SL1344_RS21885_ | 175 | 204 | 175 | 186 | 161 | 188 |
| SL1344_RS21890_ | 50 | 72 | 54 | 99 | 55 | 56 |
| SL1344_RS21895_ | 82 | 120 | 89 | 130 | 109 | 123 |
| SL1344_RS21900_ | 136 | 125 | 140 | 136 | 146 | 123 |
| SL1344_RS21905_ | 420 | 520 | 478 | 663 | 430 | 436 |
| SL1344_RS21910_ | 574 | 776 | 640 | 740 | 857 | 763 |
| SL1344_RS21915_ | 496 | 522 | 530 | 538 | 428 | 475 |
| SL1344_RS21920_ | 58 | 70 | 61 | 53 | 46 | 63 |
| SL1344_RS21925_ | 90 | 121 | 96 | 118 | 114 | 100 |
| SL1344_RS21930_ | 266 | 294 | 230 | 270 | 364 | 379 |
| SL1344_RS21935_ | 60 | 55 | 64 | 49 | 97 | 67 |
| SL1344_RS21940_ | 1896 | 1401 | 1457 | 1310 | 2158 | 1810 |
| SL1344_RS21945_ | 167 | 101 | 113 | 122 | 107 | 135 |
| SL1344_RS21950_ | 94 | 81 | 119 | 85 | 122 | 96 |
| SL1344_RS21965_ | 59 | 72 | 64 | 91 | 99 | 87 |
| SL1344_RS21970_ | 63 | 95 | 72 | 111 | 106 | 114 |
| SL1344_RS21980_ | 2002 | 2368 | 2360 | 2376 | 2904 | 2249 |
| SL1344_RS21985_ | 521 | 562 | 650 | 269 | 319 | 283 |
| SL1344_RS21990_ | 327 | 441 | 437 | 506 | 467 | 466 |
| SL1344_RS21995_ | 18 | 38 | 36 | 36 | 36 | 37 |
| SL1344_RS22000_ | 578 | 726 | 581 | 549 | 511 | 463 |
| SL1344_RS22005_ | 59 | 73 | 111 | 77 | 61 | 115 |
| SL1344_RS22010_ | 70 | 76 | 68 | 73 | 76 | 78 |
| SL1344_RS22015_ | 696 | 848 | 832 | 843 | 602 | 659 |
| SL1344_RS22020_ | 621 | 746 | 647 | 538 | 591 | 552 |
| SL1344_RS22030_ | 1067 | 1129 | 1068 | 860 | 580 | 774 |
| SL1344_RS22035_ | 768 | 890 | 825 | 676 | 446 | 631 |
| SL1344_RS22040_ | 1566 | 1793 | 1885 | 1269 | 832 | 1256 |
| SL1344_RS22045_ | 122 | 123 | 146 | 129 | 132 | 112 |
| SL1344_RS22050_ | 192 | 195 | 174 | 216 | 101 | 172 |
| SL1344_RS22055_ | 133 | 112 | 139 | 135 | 89 | 120 |
| SL1344_RS22060_ | 102 | 96 | 75 | 95 | 105 | 91 |
| SL1344_RS22065_ | 485 | 464 | 592 | 641 | 340 | 451 |
| SL1344_RS22070_ | 57 | 54 | 49 | 42 | 41 | 51 |
| SL1344_RS22075_ | 44 | 66 | 55 | 59 | 63 | 48 |
| SL1344_RS22080_ | 515 | 569 | 574 | 815 | 564 | 776 |
| SL1344_RS22085_ | 426 | 398 | 489 | 652 | 517 | 517 |
| SL1344_RS22090_ | 216 | 308 | 328 | 449 | 228 | 360 |
| SL1344_RS22095_ | 432 | 457 | 480 | 550 | 409 | 425 |
| SL1344_RS22100_ | 690 | 691 | 745 | 762 | 588 | 611 |
| SL1344_RS22105_ | 84 | 106 | 101 | 101 | 53 | 93 |

| | | | | | | |
|---|---|---|---|---|---|---|
| SL1344_RS22110_ | 36 | 42 | 31 | 29 | 31 | 35 |
| SL1344_RS22115_ | 44 | 47 | 30 | 31 | 56 | 36 |
| SL1344_RS22120_ | 76 | 105 | 104 | 117 | 139 | 114 |
| SL1344_RS22125_ | 45 | 58 | 39 | 55 | 46 | 24 |
| SL1344_RS22130_ | 82 | 81 | 66 | 178 | 128 | 154 |
| SL1344_RS22155_ | 0 | 3 | 3 | 3 | 3 | 4 |
| SL1344_RS22160_ | 177 | 210 | 203 | 190 | 156 | 237 |
| SL1344_RS22165_ | 275 | 383 | 352 | 287 | 333 | 358 |
| SL1344_RS22170_ | 472 | 540 | 444 | 548 | 390 | 557 |
| SL1344_RS22175_ | 20 | 28 | 18 | 30 | 20 | 31 |
| SL1344_RS22180_ | 143 | 128 | 132 | 183 | 131 | 172 |
| SL1344_RS22185_ | 101 | 118 | 89 | 105 | 169 | 84 |
| SL1344_RS22190_ | 365 | 347 | 393 | 357 | 327 | 435 |
| SL1344_RS22195_ | 345 | 466 | 373 | 425 | 394 | 407 |
| SL1344_RS22200_ | 191 | 243 | 245 | 238 | 204 | 235 |
| SL1344_RS22205_ | 832 | 1165 | 1083 | 1107 | 697 | 915 |
| SL1344_RS22210_ | 27880 | 26904 | 30562 | 31407 | 23768 | 28896 |
| SL1344_RS22215_ | 9182 | 5914 | 8804 | 9665 | 4653 | 8410 |
| SL1344_RS22220_ | 1088 | 1057 | 1139 | 1145 | 768 | 735 |
| SL1344_RS22225_ | 213 | 231 | 223 | 285 | 248 | 272 |
| SL1344_RS22235_ | 46628 | 44791 | 49422 | 74812 | 45044 | 56538 |
| SL1344_RS22240_ | 2921 | 4243 | 3738 | 3672 | 4669 | 4047 |
| SL1344_RS22245_ | 147 | 166 | 123 | 104 | 116 | 130 |
| SL1344_RS22250_ | 543 | 716 | 654 | 535 | 582 | 599 |
| SL1344_RS22255_ | 2628 | 3040 | 2768 | 3298 | 2658 | 2766 |
| SL1344_RS22260_ | 92 | 101 | 49 | 114 | 87 | 56 |
| SL1344_RS22265_ | 927 | 917 | 1196 | 1574 | 840 | 1048 |
| SL1344_RS22275_ | 115 | 197 | 156 | 207 | 218 | 171 |
| SL1344_RS22280_ | 157 | 169 | 176 | 146 | 160 | 136 |
| SL1344_RS22285_ | 1527 | 1785 | 1550 | 1736 | 1887 | 1801 |
| SL1344_RS22290_ | 807 | 724 | 805 | 859 | 724 | 898 |
| SL1344_RS22295_ | 2300 | 2161 | 2154 | 2282 | 2157 | 2670 |
| SL1344_RS22300_ | 5437 | 5016 | 4736 | 5373 | 4704 | 5443 |
| SL1344_RS22305_ | 567 | 611 | 572 | 569 | 493 | 553 |
| SL1344_RS22310_ | 85 | 90 | 95 | 92 | 83 | 78 |
| SL1344_RS22315_ | 238 | 289 | 270 | 296 | 279 | 306 |
| SL1344_RS22320_ | 1154 | 1396 | 1390 | 1329 | 1150 | 1183 |
| SL1344_RS22325_ | 386 | 503 | 504 | 562 | 356 | 504 |
| SL1344_RS22330_ | 878 | 1047 | 1067 | 1091 | 947 | 914 |
| SL1344_RS22335_ | 474 | 549 | 456 | 563 | 476 | 626 |
| SL1344_RS22340_ | 251 | 306 | 281 | 309 | 269 | 385 |
| SL1344_RS22345_ | 3133 | 2570 | 2792 | 4497 | 3417 | 2180 |
| SL1344_RS22350_ | 1834 | 1082 | 1455 | 2126 | 1735 | 1167 |
| SL1344_RS22355_ | 293 | 203 | 215 | 285 | 315 | 188 |
| SL1344_RS22360_ | 195 | 268 | 227 | 276 | 309 | 247 |
| SL1344_RS22365_ | 409 | 478 | 416 | 453 | 453 | 422 |
| SL1344_RS22370_ | 195 | 202 | 175 | 144 | 154 | 206 |
| SL1344_RS22375_ | 852 | 1109 | 1036 | 892 | 929 | 972 |
| SL1344_RS22380_ | 1107 | 1447 | 1277 | 1178 | 1221 | 1170 |
| SL1344_RS22385_ | 4237 | 4208 | 4649 | 4334 | 3085 | 3745 |
| SL1344_RS22395_ | 4820 | 5225 | 5059 | 4713 | 4702 | 4850 |
| SL1344_RS22400_ | 3251 | 3380 | 3274 | 3472 | 2566 | 3127 |
| SL1344_RS22405_ | 5844 | 6432 | 6910 | 6520 | 4616 | 5410 |
| SL1344_RS22410_ | 108 | 77 | 107 | 127 | 84 | 98 |
| SL1344_RS22415_ | 8495 | 9134 | 8937 | 10217 | 8733 | 8864 |
| SL1344_RS22420_ | 461 | 607 | 637 | 595 | 474 | 649 |
| SL1344_RS22425_ | 2967 | 3689 | 3381 | 3118 | 3288 | 3532 |
| SL1344_RS22430_ | 1756 | 2006 | 2348 | 1736 | 1543 | 1801 |
| SL1344_RS22435_ | 6 | 15 | 7 | 9 | 15 | 7 |
| SL1344_RS22440_ | 33 | 52 | 42 | 33 | 31 | 53 |
| SL1344_RS22445_ | 40 | 44 | 37 | 51 | 45 | 42 |
| SL1344_RS22450_ | 13 | 22 | 25 | 23 | 21 | 18 |
| SL1344_RS22455_ | 11 | 20 | 11 | 17 | 14 | 14 |

| | | | | | | |
|---|---|---|---|---|---|---|
| SL1344_RS22460_ | 40 | 42 | 27 | 27 | 43 | 34 |
| SL1344_RS22465_ | 147 | 149 | 137 | 139 | 126 | 137 |
| SL1344_RS22470_ | 268 | 246 | 283 | 236 | 257 | 181 |
| SL1344_RS22475_ | 994 | 639 | 809 | 385 | 285 | 222 |
| SL1344_RS22480_ | 2663 | 2111 | 3325 | 1103 | 513 | 537 |
| SL1344_RS22485_ | 208 | 290 | 191 | 205 | 236 | 232 |
| SL1344_RS22490_ | 363 | 368 | 429 | 379 | 360 | 420 |
| SL1344_RS22495_ | 70 | 93 | 91 | 137 | 86 | 118 |
| SL1344_RS22500_ | 31 | 47 | 31 | 42 | 46 | 46 |
| SL1344_RS22505_ | 17 | 19 | 13 | 15 | 9 | 17 |
| SL1344_RS22510_ | 22 | 28 | 20 | 41 | 31 | 28 |
| SL1344_RS22520_ | 69 | 52 | 49 | 78 | 91 | 80 |
| SL1344_RS22525_ | 81 | 110 | 64 | 114 | 120 | 128 |
| SL1344_RS22530_ | 73 | 71 | 61 | 76 | 56 | 66 |
| SL1344_RS22535_ | 1113 | 1082 | 1246 | 2219 | 864 | 1083 |
| SL1344_RS22540_ | 5141 | 5328 | 5527 | 7234 | 5103 | 4535 |
| SL1344_RS22545_ | 5777 | 5888 | 6602 | 8179 | 5252 | 5598 |
| SL1344_RS22550_ | 3381 | 3368 | 3639 | 4563 | 2897 | 2967 |
| SL1344_RS22555_ | 7835 | 7931 | 9267 | 11932 | 7395 | 6878 |
| SL1344_RS22560_ | 52 | 66 | 71 | 69 | 59 | 83 |
| SL1344_RS22565_ | 516 | 614 | 592 | 582 | 499 | 647 |
| SL1344_RS22570_ | 805 | 965 | 430 | 648 | 1346 | 1032 |
| SL1344_RS22575_ | 1772 | 1452 | 1337 | 1256 | 2191 | 1455 |
| SL1344_RS22580_ | 68 | 58 | 58 | 92 | 101 | 80 |
| SL1344_RS22585_ | 185 | 159 | 198 | 188 | 224 | 139 |
| SL1344_RS22590_ | 37 | 46 | 45 | 56 | 45 | 50 |
| SL1344_RS22595_ | 77 | 101 | 94 | 81 | 99 | 109 |
| SL1344_RS22600_ | 1695 | 1843 | 1765 | 1969 | 2079 | 2044 |
| SL1344_RS22610_ | 344 | 373 | 394 | 457 | 313 | 404 |
| SL1344_RS22615_ | 3337 | 3195 | 3406 | 4941 | 5597 | 4675 |
| SL1344_RS22620_ | 1044 | 1214 | 1255 | 1200 | 1007 | 1156 |
| SL1344_RS22625_ | 536 | 522 | 583 | 828 | 540 | 600 |
| SL1344_RS22630_ | 886 | 1101 | 995 | 647 | 743 | 717 |
| SL1344_RS22635_ | 2929 | 3492 | 3272 | 2470 | 2424 | 2601 |
| SL1344_RS22640_ | 3062 | 3666 | 2992 | 3762 | 5592 | 3316 |
| SL1344_RS22645_ | 44 | 37 | 43 | 54 | 38 | 40 |
| SL1344_RS22650_ | 30 | 20 | 35 | 93 | 64 | 42 |
| SL1344_RS22655_ | 3845 | 4603 | 4893 | 4855 | 3485 | 4149 |
| SL1344_RS22660_ | 3829 | 4923 | 4768 | 4040 | 3386 | 3680 |
| SL1344_RS22665_ | 1354 | 1597 | 1409 | 1330 | 1374 | 1475 |
| SL1344_RS22670_ | 22 | 31 | 21 | 35 | 22 | 36 |
| SL1344_RS22675_ | 619 | 627 | 708 | 631 | 676 | 642 |
| SL1344_RS22680_ | 97 | 112 | 116 | 113 | 107 | 109 |
| SL1344_RS22685_ | 4 | 3 | 2 | 1 | 3 | 16 |
| SL1344_RS22690_ | 164 | 150 | 106 | 181 | 175 | 163 |
| SL1344_RS22695_ | 266 | 213 | 182 | 266 | 314 | 179 |
| SL1344_RS22700_ | 134 | 170 | 141 | 162 | 141 | 163 |
| SL1344_RS22705_ | 48 | 16 | 30 | 40 | 26 | 20 |
| SL1344_RS22710_ | 13 | 7 | 15 | 17 | 6 | 4 |
| SL1344_RS22715_ | 55 | 76 | 60 | 73 | 51 | 59 |
| SL1344_RS22720_ | 93 | 110 | 92 | 123 | 124 | 119 |
| SL1344_RS22725_ | 135 | 162 | 108 | 143 | 215 | 152 |
| SL1344_RS22730_ | 50 | 32 | 55 | 59 | 53 | 64 |
| SL1344_RS22735_ | 146 | 158 | 148 | 172 | 147 | 151 |
| SL1344_RS22740_ | 52 | 89 | 62 | 92 | 104 | 88 |
| SL1344_RS22745_ | 44 | 48 | 41 | 48 | 42 | 55 |
| SL1344_RS22750_ | 62 | 84 | 63 | 111 | 90 | 99 |
| SL1344_RS22755_ | 18 | 28 | 23 | 21 | 20 | 18 |
| SL1344_RS22760_ | 41 | 48 | 59 | 41 | 48 | 47 |
| SL1344_RS22765_ | 435 | 482 | 510 | 426 | 455 | 492 |
| SL1344_RS22770_ | 1273 | 1470 | 1455 | 1372 | 1277 | 1468 |
| SL1344_RS22775_ | 3174 | 3208 | 3232 | 3727 | 2616 | 3435 |
| SL1344_RS22780_ | 44 | 40 | 49 | 28 | 36 | 33 |

| | | | | | | |
|---|---|---|---|---|---|---|
| SL1344_RS22785_ | 93 | 63 | 97 | 48 | 71 | 61 |
| SL1344_RS22790_ | 68 | 42 | 75 | 46 | 46 | 35 |
| SL1344_RS22795_ | 67 | 83 | 98 | 66 | 64 | 68 |
| SL1344_RS22800_ | 53 | 57 | 48 | 43 | 53 | 68 |
| SL1344_RS22805_ | 106 | 83 | 83 | 67 | 90 | 87 |
| SL1344_RS22810_ | 104 | 108 | 82 | 71 | 84 | 101 |
| SL1344_RS22815_ | 88 | 92 | 92 | 82 | 63 | 74 |
| SL1344_RS22820_ | 421 | 531 | 436 | 368 | 424 | 458 |
| SL1344_RS22825_ | 370 | 448 | 390 | 350 | 272 | 338 |
| SL1344_RS22830_ | 342 | 335 | 321 | 304 | 270 | 322 |
| SL1344_RS22835_ | 51 | 38 | 22 | 27 | 17 | 23 |
| SL1344_RS22840_ | 267 | 387 | 375 | 202 | 191 | 250 |
| SL1344_RS22845_ | 2 | 4 | 3 | 5 | 4 | 7 |
| SL1344_RS22850_ | 234 | 217 | 213 | 174 | 177 | 171 |
| SL1344_RS22855_ | 692 | 567 | 601 | 629 | 605 | 477 |
| SL1344_RS22860_ | 103 | 151 | 89 | 133 | 132 | 106 |
| SL1344_RS22865_ | 218 | 248 | 187 | 224 | 225 | 233 |
| SL1344_RS22870_ | 864 | 815 | 828 | 1022 | 800 | 807 |
| SL1344_RS22875_ | 10289 | 11370 | 11300 | 11351 | 9809 | 10044 |
| SL1344_RS22880_ | 109 | 131 | 134 | 124 | 101 | 104 |
| SL1344_RS22885_ | 189 | 244 | 223 | 175 | 252 | 184 |
| SL1344_RS22890_ | 15 | 18 | 25 | 40 | 30 | 14 |
| SL1344_RS22895_ | 49 | 76 | 73 | 48 | 53 | 67 |
| SL1344_RS22900_ | 71 | 78 | 84 | 97 | 103 | 64 |
| SL1344_RS22905_ | 49 | 15 | 12 | 17 | 25 | 9 |
| SL1344_RS22910_ | 9 | 10 | 9 | 44 | 9 | 6 |
| SL1344_RS22915_ | 20 | 18 | 11 | 10 | 15 | 15 |
| SL1344_RS22920_ | 200 | 189 | 158 | 165 | 209 | 185 |
| SL1344_RS22925_ | 104 | 151 | 114 | 131 | 156 | 156 |
| SL1344_RS22930_ | 497 | 506 | 576 | 667 | 574 | 844 |
| SL1344_RS22935_ | 326 | 400 | 375 | 386 | 311 | 336 |
| SL1344_RS22940_ | 130 | 144 | 238 | 126 | 76 | 125 |
| SL1344_RS22945_ | 356 | 465 | 423 | 349 | 345 | 294 |
| SL1344_RS22950_ | 103 | 119 | 94 | 124 | 120 | 108 |
| SL1344_RS22955_ | 5935 | 6917 | 6766 | 6787 | 5412 | 5740 |
| SL1344_RS22960_ | 853 | 936 | 1079 | 1088 | 729 | 911 |
| SL1344_RS22965_ | 3394 | 4214 | 4080 | 4143 | 4235 | 3623 |
| SL1344_RS22970_ | 1045 | 1296 | 1174 | 1110 | 1657 | 1056 |
| SL1344_RS22975_ | 1041 | 1274 | 1004 | 1167 | 1200 | 1195 |
| SL1344_RS22980_ | 851 | 941 | 783 | 935 | 870 | 937 |
| SL1344_RS22985_ | 91 | 105 | 74 | 95 | 97 | 77 |
| SL1344_RS22995_ | 51 | 46 | 32 | 40 | 31 | 37 |
| SL1344_RS23000_ | 23 | 39 | 68 | 30 | 23 | 34 |
| SL1344_RS23010_ | 350 | 327 | 359 | 405 | 217 | 260 |
| SL1344_RS23015_ | 171 | 429 | 136 | 779 | 781 | 171 |
| SL1344_RS23020_ | 10 | 13 | 10 | 15 | 5 | 4 |
| SL1344_RS23025_ | 937 | 958 | 932 | 899 | 897 | 1125 |
| SL1344_RS23030_ | 875 | 992 | 945 | 990 | 919 | 1094 |
| SL1344_RS23035_ | 1509 | 1827 | 1942 | 1953 | 1363 | 1633 |
| SL1344_RS23040_ | 2210 | 2472 | 2530 | 2528 | 1875 | 2341 |
| SL1344_RS23045_ | 841 | 953 | 1055 | 1003 | 802 | 1015 |
| SL1344_RS23050_ | 1231 | 1303 | 1223 | 1428 | 1207 | 1347 |
| SL1344_RS23055_ | 1925 | 1884 | 2070 | 1790 | 1455 | 1865 |
| SL1344_RS23060_ | 2673 | 3106 | 3031 | 2729 | 2230 | 2615 |
| SL1344_RS23065_ | 538 | 580 | 451 | 519 | 523 | 505 |
| SL1344_RS23070_ | 585 | 605 | 564 | 625 | 473 | 529 |
| SL1344_RS23075_ | 5 | 3 | 7 | 11 | 8 | 5 |
| SL1344_RS23080_ | 630 | 668 | 723 | 841 | 620 | 821 |
| SL1344_RS23085_ | 295 | 302 | 315 | 347 | 352 | 347 |
| SL1344_RS23090_ | 154 | 122 | 119 | 146 | 103 | 82 |
| SL1344_RS23095_ | 16 | 10 | 18 | 16 | 17 | 10 |
| SL1344_RS23100_ | 1277 | 1169 | 1562 | 1183 | 1008 | 1149 |
| SL1344_RS23105_ | 25 | 25 | 27 | 30 | 31 | 22 |

| | | | | | | |
|---|---|---|---|---|---|---|
| SL1344_RS23110_ | 57 | 65 | 54 | 51 | 48 | 70 |
| SL1344_RS23115_ | 140 | 180 | 131 | 178 | 97 | 124 |
| SL1344_RS23120_ | 1195 | 1254 | 1431 | 1400 | 1175 | 1231 |
| SL1344_RS23125_ | 36 | 52 | 43 | 45 | 42 | 48 |
| SL1344_RS23130_ | 3 | 3 | 8 | 3 | 4 | 3 |
| SL1344_RS23140_ | 63 | 90 | 60 | 61 | 88 | 73 |
| SL1344_RS23145_ | 161 | 167 | 144 | 220 | 189 | 153 |
| SL1344_RS23150_ | 342 | 434 | 403 | 347 | 337 | 323 |
| SL1344_RS23155_ | 12 | 25 | 28 | 36 | 15 | 27 |
| SL1344_RS23160_ | 7 | 22 | 13 | 19 | 13 | 12 |
| SL1344_RS23165_ | 7 | 8 | 8 | 10 | 21 | 9 |
| SL1344_RS23170_ | 126 | 153 | 126 | 147 | 194 | 162 |
| SL1344_RS23175_ | 151 | 199 | 167 | 157 | 137 | 133 |
| SL1344_RS23180_ | 141 | 203 | 194 | 181 | 198 | 177 |
| SL1344_RS23185_ | 59 | 74 | 76 | 89 | 87 | 108 |
| SL1344_RS23190_ | 294 | 328 | 323 | 225 | 189 | 192 |
| SL1344_RS23195_ | 8 | 6 | 4 | 9 | 11 | 5 |
| SL1344_RS23200_ | 9 | 6 | 5 | 6 | 9 | 4 |
| SL1344_RS23205_ | 103 | 96 | 113 | 191 | 189 | 206 |
| SL1344_RS23210_ | 14 | 28 | 21 | 24 | 22 | 24 |
| SL1344_RS23215_ | 334 | 364 | 388 | 399 | 256 | 284 |
| SL1344_RS23220_ | 461 | 463 | 475 | 466 | 461 | 462 |
| SL1344_RS23225_ | 855 | 1049 | 907 | 969 | 865 | 913 |
| SL1344_RS23230_ | 333 | 324 | 229 | 278 | 274 | 372 |
| SL1344_RS23235_ | 200 | 185 | 200 | 217 | 157 | 184 |
| SL1344_RS23240_ | 161 | 170 | 187 | 218 | 160 | 176 |
| SL1344_RS23245_ | 2872 | 4196 | 3932 | 1656 | 2781 | 2809 |
| SL1344_RS23250_ | 519 | 738 | 675 | 340 | 394 | 568 |
| SL1344_RS23265_ | 653 | 667 | 675 | 954 | 952 | 915 |
| SL1344_RS23270_ | 23 | 12 | 26 | 47 | 21 | 24 |
| SL1344_RS23275_ | 14 | 9 | 10 | 22 | 7 | 9 |
| SL1344_RS23280_ | 17 | 11 | 15 | 19 | 13 | 17 |
| SL1344_RS23285_ | 17 | 19 | 13 | 32 | 14 | 22 |
| SL1344_RS23290_ | 13 | 31 | 23 | 33 | 15 | 18 |
| SL1344_RS23295_ | 83 | 117 | 104 | 128 | 104 | 105 |
| SL1344_RS23300_ | 3296 | 3625 | 3752 | 3542 | 3015 | 2991 |
| SL1344_RS23305_ | 254 | 337 | 280 | 322 | 288 | 275 |
| SL1344_RS23310_ | 575 | 668 | 723 | 645 | 592 | 676 |
| SL1344_RS23315_ | 323 | 418 | 428 | 461 | 328 | 468 |
| SL1344_RS23320_ | 68 | 104 | 105 | 94 | 88 | 93 |
| SL1344_RS23325_ | 96 | 108 | 96 | 92 | 91 | 95 |
| SL1344_RS23330_ | 3 | 2 | 2 | 3 | 3 | 6 |
| SL1344_RS23335_ | 2 | 4 | 9 | 6 | 4 | 4 |
| SL1344_RS23340_ | 847 | 1026 | 874 | 592 | 551 | 567 |
| SL1344_RS23345_ | 678 | 847 | 765 | 1671 | 1221 | 1080 |
| SL1344_RS23350_ | 291 | 330 | 336 | 324 | 250 | 247 |
| SL1344_RS23355_ | 67 | 18 | 16 | 46 | 40 | 29 |
| SL1344_RS23360_ | 4501 | 4448 | 3330 | 4852 | 10062 | 3175 |
| SL1344_RS23365_ | 10445 | 8446 | 8663 | 12595 | 12392 | 9908 |
| SL1344_RS23370_ | 3884 | 2187 | 2544 | 3765 | 3690 | 2295 |
| SL1344_RS23375_ | 613 | 677 | 756 | 625 | 575 | 455 |
| SL1344_RS23380_ | 495 | 467 | 493 | 615 | 562 | 495 |
| SL1344_RS23385_ | 372 | 342 | 356 | 339 | 291 | 325 |
| SL1344_RS23390_ | 533 | 682 | 714 | 663 | 539 | 613 |
| SL1344_RS23395_ | 1784 | 1929 | 2026 | 1936 | 1403 | 1602 |
| SL1344_RS23400_ | 598 | 632 | 679 | 645 | 460 | 656 |
| SL1344_RS23410_ | 304 | 365 | 321 | 367 | 370 | 381 |
| SL1344_RS23415_ | 225 | 304 | 275 | 258 | 318 | 273 |
| SL1344_RS23420_ | 39 | 46 | 30 | 35 | 35 | 31 |
| SL1344_RS23425_ | 52 | 64 | 34 | 73 | 53 | 47 |
| SL1344_RS23430_ | 223 | 292 | 273 | 292 | 344 | 282 |
| SL1344_RS23435_ | 276 | 360 | 307 | 362 | 340 | 343 |
| SL1344_RS23440_ | 936 | 1153 | 1003 | 1354 | 1268 | 1297 |

| | | | | | | |
|---|---|---|---|---|---|---|
| SL1344_RS23445_ | 1453 | 1546 | 1259 | 1965 | 1808 | 1922 |
| SL1344_RS23450_ | 132 | 137 | 148 | 137 | 137 | 128 |
| SL1344_RS23455_ | 158 | 151 | 142 | 126 | 115 | 116 |
| SL1344_RS23460_ | 11 | 10 | 16 | 15 | 10 | 13 |
| SL1344_RS23465_ | 23 | 27 | 47 | 40 | 31 | 36 |
| SL1344_RS23470_ | 21 | 18 | 24 | 26 | 18 | 29 |
| SL1344_RS23480_ | 162 | 185 | 168 | 262 | 189 | 209 |
| SL1344_RS23485_ | 730 | 836 | 743 | 751 | 654 | 715 |
| SL1344_RS23495_ | 1040 | 1154 | 1079 | 1200 | 952 | 1074 |
| SL1344_RS23500_ | 6643 | 7454 | 6946 | 7016 | 7195 | 6895 |
| SL1344_RS23505_ | 948 | 1087 | 995 | 860 | 981 | 808 |
| SL1344_RS23510_ | 275 | 376 | 322 | 310 | 403 | 272 |
| SL1344_RS23515_ | 183 | 224 | 202 | 240 | 191 | 219 |
| SL1344_RS23520_ | 1031 | 1023 | 1092 | 1036 | 929 | 918 |
| SL1344_RS23525_ | 1719 | 2185 | 2091 | 2194 | 1719 | 2128 |
| SL1344_RS23530_ | 561 | 621 | 607 | 690 | 535 | 654 |
| SL1344_RS23535_ | 759 | 981 | 853 | 1061 | 912 | 857 |
| SL1344_RS23540_ | 484 | 603 | 530 | 582 | 659 | 573 |
| SL1344_RS23545_ | 131 | 156 | 153 | 144 | 172 | 143 |
| SL1344_RS23550_ | 42 | 44 | 57 | 42 | 38 | 41 |
| SL1344_RS23555_ | 6 | 16 | 11 | 11 | 4 | 5 |
| SL1344_RS23560_ | 61 | 66 | 58 | 75 | 38 | 45 |
| SL1344_RS23565_ | 18 | 9 | 21 | 7 | 14 | 13 |
| SL1344_RS23570_ | 20 | 32 | 24 | 22 | 14 | 20 |
| SL1344_RS23575_ | 262 | 290 | 276 | 230 | 216 | 246 |
| SL1344_RS23580_ | 165 | 206 | 199 | 166 | 169 | 169 |
| SL1344_RS23590_ | 3 | 2 | 6 | 5 | 3 | 3 |
| SL1344_RS23595_ | 173 | 150 | 160 | 164 | 137 | 124 |
| SL1344_RS23640_ | 261 | 293 | 284 | 282 | 313 | 337 |
| SL1344_RS23645_ | 141 | 151 | 143 | 133 | 168 | 153 |
| SL1344_RS23650_ | 621 | 632 | 686 | 699 | 718 | 770 |
| SL1344_RS23655_ | 215 | 188 | 229 | 254 | 165 | 285 |
| SL1344_RS23660_ | 310 | 447 | 402 | 411 | 453 | 450 |
| SL1344_RS23665_ | 75 | 95 | 89 | 147 | 130 | 151 |
| SL1344_RS23670_ | 4127 | 3941 | 3857 | 4103 | 3287 | 4497 |
| SL1344_RS23675_ | 1331 | 1101 | 1393 | 947 | 849 | 1119 |
| SL1344_RS23680_ | 818 | 752 | 778 | 783 | 628 | 721 |
| SL1344_RS23685_ | 64 | 59 | 67 | 60 | 41 | 72 |
| SL1344_RS23690_ | 23 | 24 | 33 | 22 | 17 | 24 |
| SL1344_RS23695_ | 8 | 25 | 19 | 17 | 24 | 14 |
| SL1344_RS23700_ | 31 | 43 | 48 | 57 | 44 | 50 |
| SL1344_RS23705_ | 18 | 15 | 20 | 19 | 16 | 21 |
| SL1344_RS23710_ | 109 | 129 | 117 | 102 | 112 | 110 |
| SL1344_RS23715_ | 29 | 35 | 40 | 38 | 44 | 31 |
| SL1344_RS23720_ | 39 | 35 | 40 | 44 | 40 | 42 |
| SL1344_RS23725_ | 21 | 13 | 26 | 32 | 29 | 21 |
| SL1344_RS23730_ | 41 | 40 | 39 | 35 | 27 | 44 |
| SL1344_RS23735_ | 18 | 16 | 23 | 29 | 24 | 23 |
| SL1344_RS23740_ | 8 | 17 | 11 | 17 | 15 | 17 |
| SL1344_RS23745_ | 80 | 96 | 76 | 96 | 85 | 73 |
| SL1344_RS23750_ | 2 | 12 | 4 | 12 | 5 | 10 |
| SL1344_RS23755_ | 12 | 13 | 16 | 15 | 8 | 12 |
| SL1344_RS23760_ | 5 | 4 | 8 | 11 | 5 | 3 |
| SL1344_RS23765_ | 39 | 33 | 37 | 44 | 47 | 59 |
| SL1344_RS23770_ | 14 | 11 | 21 | 16 | 26 | 27 |
| SL1344_RS23775_ | 25 | 50 | 32 | 100 | 73 | 64 |
| SL1344_RS23780_ | 28 | 26 | 18 | 17 | 26 | 25 |
| SL1344_RS23785_ | 25 | 39 | 23 | 42 | 32 | 35 |
| SL1344_RS23790_ | 7 | 6 | 10 | 10 | 7 | 8 |
| SL1344_RS23795_ | 11 | 8 | 15 | 10 | 8 | 12 |
| SL1344_RS23800_ | 14 | 11 | 5 | 16 | 4 | 8 |
| SL1344_RS23805_ | 15 | 8 | 19 | 10 | 12 | 20 |
| SL1344_RS23810_ | 57 | 41 | 79 | 117 | 67 | 107 |

| | | | | | | |
|---|---|---|---|---|---|---|
| SL1344_RS23815_ | 59 | 65 | 54 | 108 | 121 | 67 |
| SL1344_RS23820_ | 9 | 10 | 9 | 12 | 11 | 12 |
| SL1344_RS23825_ | 17 | 28 | 33 | 17 | 22 | 18 |
| SL1344_RS23830_ | 28 | 30 | 26 | 39 | 34 | 27 |
| SL1344_RS23840_ | 17 | 31 | 25 | 19 | 32 | 29 |
| SL1344_RS23845_ | 6 | 3 | 3 | 8 | 15 | 10 |
| SL1344_RS23850_ | 7 | 0 | 5 | 9 | 8 | 8 |
| SL1344_RS23855_ | 5 | 1 | 4 | 5 | 1 | 3 |
| SL1344_RS23860_ | 101 | 83 | 75 | 81 | 95 | 91 |
| SL1344_RS23865_ | 10 | 10 | 18 | 20 | 10 | 13 |
| SL1344_RS23870_ | 10 | 6 | 2 | 14 | 16 | 21 |
| SL1344_RS23875_ | 27 | 23 | 16 | 42 | 36 | 29 |
| SL1344_RS23880_ | 3 | 1 | 1 | 9 | 4 | 3 |
| SL1344_RS23885_ | 4 | 2 | 1 | 5 | 2 | 3 |
| SL1344_RS23890_ | 0 | 2 | 1 | 0 | 3 | 2 |
| SL1344_RS23895_ | 4 | 6 | 1 | 3 | 6 | 2 |
| SL1344_RS23900_ | 11 | 6 | 7 | 7 | 7 | 5 |
| SL1344_RS23905_ | 1 | 0 | 2 | 1 | 0 | 0 |
| SL1344_RS23910_ | 99 | 143 | 126 | 159 | 145 | 146 |
| SL1344_RS23915_ | 192 | 245 | 184 | 178 | 219 | 219 |
| SL1344_RS23920_ | 4041 | 4327 | 4597 | 4138 | 4529 | 5083 |
| SL1344_RS23925_ | 2639 | 2616 | 2958 | 2473 | 2553 | 2837 |
| SL1344_RS23930_ | 1751 | 2212 | 1714 | 2251 | 2003 | 2403 |
| SL1344_RS23935_ | 325 | 429 | 457 | 462 | 452 | 381 |
| SL1344_RS23940_ | 219 | 252 | 278 | 314 | 203 | 241 |
| SL1344_RS23945_ | 232 | 278 | 269 | 311 | 255 | 219 |
| SL1344_RS23950_ | 931 | 820 | 961 | 1090 | 702 | 853 |
| SL1344_RS23955_ | 209 | 181 | 142 | 271 | 189 | 224 |
| SL1344_RS23960_ | 1273 | 1269 | 1254 | 1463 | 895 | 1510 |
| SL1344_RS23965_ | 823 | 975 | 823 | 949 | 1102 | 944 |
| SL1344_RS23970_ | 1089 | 1240 | 1191 | 1398 | 1369 | 1323 |
| SL1344_RS23975_ | 541 | 629 | 565 | 545 | 623 | 684 |
| SL1344_RS23980_ | 42 | 41 | 55 | 62 | 48 | 45 |
| SL1344_RS23985_ | 9 | 10 | 7 | 9 | 13 | 6 |
| SL1344_RS23990_ | 4 | 4 | 4 | 2 | 4 | 6 |
| SL1344_RS23995_ | 15 | 12 | 17 | 14 | 6 | 14 |
| SL1344_RS24000_ | 65 | 62 | 90 | 83 | 64 | 93 |
| SL1344_RS24005_ | 4 | 2 | 0 | 1 | 5 | 3 |
| SL1344_RS24010_ | 5 | 8 | 11 | 9 | 9 | 7 |
| SL1344_RS24015_ | 70 | 101 | 89 | 94 | 123 | 72 |
| SL1344_RS24020_ | 2222 | 2820 | 3450 | 2996 | 1830 | 2966 |
| SL1344_RS24025_ | 5 | 3 | 5 | 3 | 4 | 2 |
| SL1344_RS24030_ | 5 | 17 | 8 | 14 | 14 | 10 |
| SL1344_RS24035_ | 47 | 47 | 55 | 64 | 61 | 65 |
| SL1344_RS24040_ | 95 | 112 | 123 | 117 | 116 | 109 |
| SL1344_RS24045_ | 341 | 390 | 383 | 337 | 561 | 302 |
| SL1344_RS24050_ | 582 | 678 | 720 | 564 | 639 | 636 |
| SL1344_RS24055_ | 550 | 650 | 595 | 589 | 561 | 541 |
| SL1344_RS24060_ | 515 | 584 | 593 | 506 | 589 | 472 |
| SL1344_RS24065_ | 513 | 602 | 648 | 546 | 515 | 523 |
| SL1344_RS24070_ | 519 | 547 | 551 | 517 | 404 | 519 |
| SL1344_RS24075_ | 296 | 397 | 373 | 527 | 326 | 434 |
| SL1344_RS24080_ | 45 | 50 | 32 | 27 | 32 | 32 |
| SL1344_RS24085_ | 2947 | 3346 | 3449 | 3217 | 3119 | 3150 |
| SL1344_RS24090_ | 20 | 25 | 38 | 46 | 35 | 40 |
| SL1344_RS24095_ | 78 | 60 | 65 | 59 | 43 | 41 |
| SL1344_RS24100_ | 6 | 9 | 8 | 7 | 1 | 6 |
| SL1344_RS24105_ | 152 | 181 | 202 | 176 | 89 | 138 |
| SL1344_RS24110_ | 48 | 56 | 53 | 62 | 38 | 59 |
| SL1344_RS24115_ | 48 | 45 | 40 | 39 | 45 | 36 |
| SL1344_RS24120_ | 45 | 57 | 44 | 43 | 47 | 76 |
| SL1344_RS24125_ | 1891 | 1796 | 2048 | 1731 | 1377 | 1630 |
| SL1344_RS24130_ | 73 | 91 | 93 | 86 | 71 | 79 |

| | | | | | | |
|---|---|---|---|---|---|---|
| SL1344_RS24135_ | 112 | 153 | 146 | 94 | 114 | 119 |
| SL1344_RS24140_ | 32 | 58 | 59 | 28 | 38 | 40 |
| SL1344_RS24145_ | 28 | 44 | 40 | 30 | 25 | 32 |
| SL1344_RS24150_ | 118 | 116 | 112 | 97 | 85 | 114 |
| SL1344_RS24155_ | 249 | 226 | 211 | 165 | 175 | 205 |
| SL1344_RS24160_ | 3 | 6 | 0 | 1 | 12 | 11 |
| SL1344_RS24165_ | 910 | 863 | 651 | 1196 | 1317 | 839 |
| SL1344_RS24170_ | 771 | 720 | 662 | 1251 | 801 | 794 |
| SL1344_RS24175_ | 4122 | 3817 | 2990 | 4813 | 5206 | 3407 |
| SL1344_RS24180_ | 1293 | 1699 | 2044 | 1619 | 1406 | 1803 |
| SL1344_RS24185_ | 123 | 136 | 135 | 164 | 168 | 163 |
| SL1344_RS24190_ | 256 | 270 | 304 | 196 | 179 | 270 |
| SL1344_RS24195_ | 149 | 130 | 168 | 160 | 137 | 174 |
| SL1344_RS24200_ | 4 | 2 | 2 | 3 | 3 | 5 |
| SL1344_RS24205_ | 1715 | 1453 | 1000 | 2182 | 2114 | 1161 |
| SL1344_RS24210_ | 171 | 228 | 225 | 239 | 173 | 237 |
| SL1344_RS24215_ | 206 | 222 | 253 | 241 | 215 | 234 |
| SL1344_RS24220_ | 636 | 788 | 682 | 833 | 663 | 730 |
| SL1344_RS24225_ | 294 | 362 | 271 | 383 | 329 | 360 |
| SL1344_RS24230_ | 139 | 133 | 130 | 155 | 89 | 150 |
| SL1344_RS24235_ | 210 | 178 | 207 | 239 | 172 | 233 |
| SL1344_RS24240_ | 290 | 359 | 454 | 409 | 252 | 397 |
| SL1344_RS24245_ | 480 | 475 | 490 | 527 | 399 | 499 |
| SL1344_RS24250_ | 2273 | 2359 | 2871 | 4307 | 2831 | 3923 |
| SL1344_RS24255_ | 141 | 163 | 216 | 247 | 166 | 257 |
| SL1344_RS24260_ | 87 | 97 | 124 | 171 | 148 | 162 |
| SL1344_RS24265_ | 165 | 260 | 229 | 246 | 277 | 229 |
| SL1344_RS24270_ | 68 | 93 | 94 | 92 | 68 | 127 |
| SL1344_RS24275_ | 61 | 86 | 68 | 90 | 113 | 92 |
| SL1344_RS24280_ | 118 | 139 | 142 | 196 | 206 | 188 |
| SL1344_RS24285_ | 755 | 864 | 767 | 814 | 707 | 782 |
| SL1344_RS24290_ | 658 | 756 | 691 | 710 | 756 | 901 |
| SL1344_RS24295_ | 835 | 946 | 1025 | 1097 | 824 | 1033 |
| SL1344_RS24300_ | 568 | 637 | 657 | 737 | 566 | 664 |
| SL1344_RS24305_ | 112 | 150 | 139 | 190 | 118 | 171 |
| SL1344_RS24310_ | 48 | 39 | 51 | 66 | 68 | 53 |
| SL1344_RS24315_ | 5 | 2 | 3 | 9 | 6 | 2 |
| SL1344_RS24320_ | 1 | 2 | 1 | 3 | 4 | 2 |
| SL1344_RS24325_ | 15 | 19 | 14 | 24 | 16 | 20 |
| SL1344_RS24330_ | 26 | 20 | 20 | 35 | 45 | 33 |
| SL1344_RS24335_ | 9 | 7 | 6 | 12 | 10 | 10 |
| SL1344_RS24340_ | 11 | 7 | 18 | 12 | 9 | 11 |
| SL1344_RS24345_ | 12 | 12 | 14 | 14 | 13 | 15 |
| SL1344_RS24350_ | 13 | 11 | 6 | 5 | 3 | 9 |
| SL1344_RS24355_ | 14 | 6 | 5 | 11 | 6 | 7 |
| SL1344_RS24360_ | 6 | 3 | 14 | 9 | 3 | 7 |
| SL1344_RS24365_ | 8 | 6 | 6 | 7 | 9 | 9 |
| SL1344_RS24370_ | 76 | 74 | 70 | 78 | 51 | 50 |
| SL1344_RS24380_ | 60 | 71 | 66 | 50 | 47 | 43 |
| SL1344_RS24385_ | 81 | 91 | 92 | 91 | 74 | 76 |
| SL1344_RS24390_ | 6 | 8 | 4 | 6 | 4 | 4 |
| SL1344_RS24395_ | 24 | 35 | 43 | 47 | 36 | 35 |
| SL1344_RS24400_ | 4 | 5 | 5 | 10 | 7 | 5 |
| SL1344_RS24405_ | 5 | 6 | 9 | 12 | 11 | 11 |
| SL1344_RS24410_ | 318 | 448 | 347 | 384 | 323 | 381 |
| SL1344_RS24415_ | 21 | 18 | 31 | 13 | 22 | 23 |
| SL1344_RS24420_ | 102 | 95 | 101 | 121 | 104 | 120 |
| SL1344_RS24425_ | 11 | 15 | 10 | 18 | 18 | 16 |
| SL1344_RS24430_ | 991 | 954 | 1269 | 1410 | 1014 | 1582 |
| SL1344_RS24435_ | 268 | 276 | 229 | 315 | 321 | 377 |
| SL1344_RS24440_ | 1142 | 1258 | 1178 | 1706 | 1344 | 1539 |
| SL1344_RS24445_ | 242 | 267 | 349 | 277 | 213 | 263 |
| SL1344_RS24450_ | 53 | 83 | 87 | 88 | 54 | 65 |

| | | | | | | |
|---|---|---|---|---|---|---|
| SL1344_RS24455_ | 28 | 45 | 32 | 35 | 30 | 20 |
| SL1344_RS24460_ | 54643 | 67855 | 61575 | 63657 | 90963 | 50453 |
| SL1344_RS24465_ | 363 | 291 | 198 | 311 | 227 | 348 |
| SL1344_RS24470_ | 7856 | 13140 | 9443 | 16306 | 13713 | 16108 |
| SL1344_RS24475_ | 390 | 404 | 425 | 610 | 323 | 529 |
| SL1344_RS24480_ | 1553 | 1862 | 2040 | 1921 | 1757 | 1749 |
| SL1344_RS24485_ | 144 | 150 | 152 | 148 | 137 | 138 |
| SL1344_RS24490_ | 40 | 59 | 37 | 50 | 33 | 47 |
| SL1344_RS24495_ | 49 | 51 | 43 | 45 | 41 | 53 |
| SL1344_RS24500_ | 133 | 218 | 216 | 219 | 178 | 157 |
| SL1344_RS24505_ | 22 | 24 | 26 | 16 | 17 | 20 |
| SL1344_RS24510_ | 4 | 4 | 7 | 4 | 2 | 7 |
| SL1344_RS24515_ | 10 | 16 | 12 | 11 | 10 | 7 |
| SL1344_RS24520_ | 19 | 31 | 24 | 35 | 28 | 26 |
| SL1344_RS24525_ | 61 | 79 | 79 | 70 | 64 | 100 |
| SL1344_RS24530_ | 45 | 35 | 33 | 34 | 25 | 32 |
| SL1344_RS24535_ | 10 | 19 | 12 | 32 | 23 | 20 |
| SL1344_RS24540_ | 18 | 24 | 19 | 28 | 16 | 23 |
| SL1344_RS24545_ | 10 | 18 | 11 | 20 | 13 | 13 |
| SL1344_RS24550_ | 80 | 109 | 79 | 101 | 76 | 116 |
| SL1344_RS24555_ | 37 | 35 | 64 | 33 | 31 | 32 |
| SL1344_RS24560_ | 44 | 35 | 35 | 34 | 38 | 37 |
| SL1344_RS24565_ | 20 | 15 | 15 | 27 | 21 | 35 |
| SL1344_RS24570_ | 13 | 16 | 5 | 14 | 11 | 13 |
| SL1344_RS24575_ | 37 | 23 | 36 | 89 | 59 | 90 |
| SL1344_RS24580_ | 422 | 468 | 502 | 1503 | 905 | 1329 |
| SL1344_RS24585_ | 139 | 160 | 166 | 390 | 206 | 346 |
| SL1344_RS24590_ | 198 | 210 | 205 | 205 | 162 | 195 |
| SL1344_RS24595_ | 5 | 4 | 7 | 3 | 5 | 2 |
| SL1344_RS24600_ | 28 | 37 | 30 | 46 | 32 | 28 |
| SL1344_RS24605_ | 19 | 17 | 20 | 29 | 28 | 14 |
| SL1344_RS24610_ | 103 | 66 | 91 | 111 | 79 | 58 |
| SL1344_RS24615_ | 37 | 34 | 34 | 42 | 38 | 30 |
| SL1344_RS24620_ | 35 | 39 | 32 | 42 | 32 | 33 |
| SL1344_RS24625_ | 16 | 8 | 8 | 13 | 9 | 7 |
| SL1344_RS24630_ | 68 | 73 | 62 | 100 | 70 | 75 |
| SL1344_RS24635_ | 40 | 56 | 54 | 61 | 28 | 42 |
| SL1344_RS24640_ | 5 | 8 | 2 | 7 | 4 | 5 |
| SL1344_RS24645_ | 12 | 14 | 6 | 10 | 12 | 10 |
| SL1344_RS24650_ | 17 | 16 | 16 | 22 | 8 | 19 |
| SL1344_RS24655_ | 17 | 26 | 20 | 25 | 13 | 21 |
| SL1344_RS24660_ | 314 | 288 | 306 | 368 | 217 | 364 |
| SL1344_RS24665_ | 203 | 156 | 173 | 186 | 135 | 217 |
| SL1344_RS24670_ | 38 | 47 | 39 | 70 | 37 | 45 |
| SL1344_RS24675_ | 48 | 39 | 42 | 126 | 96 | 79 |
| SL1344_RS24690_ | 11117 | 14878 | 13996 | 24896 | 20173 | 20203 |
| SL1344_RS24695_ | 974 | 1192 | 1057 | 2688 | 2628 | 2529 |
| SL1344_RS24700_ | 1045 | 1551 | 1363 | 3192 | 2854 | 2831 |
| SL1344_RS24705_ | 1027 | 1240 | 1209 | 2266 | 2011 | 2305 |
| SL1344_RS24710_ | 783 | 1010 | 1037 | 1803 | 1712 | 1878 |
| SL1344_RS24730_ | 6062 | 6948 | 6404 | 10875 | 9203 | 10045 |
| SL1344_RS24735_ | 10869 | 12966 | 11910 | 19691 | 17791 | 17568 |
| SL1344_RS24740_ | 12 | 15 | 11 | 14 | 12 | 17 |
| SL1344_RS24745_ | 2 | 3 | 5 | 3 | 2 | 1 |
| SL1344_RS24750_ | 121 | 96 | 136 | 106 | 82 | 92 |
| SL1344_RS24755_ | 315 | 340 | 357 | 348 | 328 | 303 |
| SL1344_RS24760_ | 33 | 56 | 53 | 59 | 53 | 23 |
| SL1344_RS24780_ | 56 | 83 | 69 | 65 | 70 | 64 |
| SL1344_RS24785_ | 6458 | 4516 | 4926 | 7760 | 5820 | 4109 |
| SL1344_RS24790_ | 5472 | 3995 | 4549 | 5645 | 5163 | 3924 |
| SL1344_RS24795_ | 66 | 72 | 53 | 68 | 78 | 44 |
| SL1344_RS24800_ | 6 | 4 | 2 | 2 | 11 | 2 |
| SL1344_RS24805_ | 32 | 28 | 26 | 52 | 45 | 31 |

| | | | | | | |
|---|---|---|---|---|---|---|
| SL1344_RS24810_ | 93 | 69 | 71 | 114 | 112 | 48 |
| SL1344_RS24820_ | 1396 | 1642 | 1756 | 3197 | 4476 | 3095 |
| SL1344_RS24825_ | 2936 | 2950 | 2185 | 5317 | 5620 | 2372 |
| SL1344_RS24830_ | 177 | 119 | 152 | 220 | 263 | 166 |
| SL1344_RS24835_ | 10 | 18 | 15 | 24 | 21 | 17 |
| SL1344_RS24840_ | 5 | 4 | 1 | 10 | 10 | 6 |
| SL1344_RS24845_ | 1 | 2 | 2 | 1 | 5 | 1 |
| SL1344_RS24850_ | 8 | 9 | 10 | 12 | 12 | 10 |
| SL1344_RS24860_ | 738 | 1492 | 959 | 1136 | 1796 | 551 |
| SL1344_RS24865_ | 22 | 15 | 11 | 23 | 24 | 23 |
| SL1344_RS24875_ | 6 | 5 | 8 | 5 | 10 | 6 |
| SL1344_RS24885_ | 104 | 71 | 84 | 105 | 125 | 53 |
| SLnc0002_ | 28 | 13 | 29 | 21 | 14 | 25 |
| SLnc0010_ | 22 | 21 | 28 | 25 | 30 | 16 |
| SLnc0010a_ | 135 | 144 | 196 | 188 | 162 | 132 |
| SLnc0014_ | 321 | 422 | 435 | 869 | 436 | 618 |
| SLnc0022_ | 105 | 142 | 176 | 189 | 174 | 163 |
| SLnc0030_ | 1073 | 1774 | 1407 | 1131 | 2238 | 1026 |
| SLnc0030a_ | 88 | 141 | 105 | 75 | 200 | 78 |
| SLnc0031_ | 18 | 32 | 28 | 22 | 31 | 17 |
| SLnc0035_ | 5900 | 5814 | 6822 | 4666 | 4971 | 5281 |
| SLnc0039_ | 1651 | 2135 | 2473 | 2670 | 1610 | 1339 |
| SLnc0048_ | 76 | 95 | 91 | 88 | 67 | 80 |
| SLnc1001_ | 2 | 3 | 10 | 9 | 7 | 2 |
| SLnc1002_ | 96 | 184 | 161 | 205 | 190 | 167 |
| SLnc1006_ | 177 | 228 | 194 | 360 | 177 | 200 |
| SLnc1008_ | 17 | 37 | 13 | 20 | 23 | 20 |
| SLnc1013_ | 49 | 73 | 68 | 35 | 54 | 41 |
| SLnc1014_ | 38 | 72 | 36 | 35 | 83 | 51 |
| SLnc1015_ | 52 | 63 | 47 | 34 | 105 | 38 |
| SLnc1016_ | 41 | 42 | 39 | 31 | 32 | 34 |
| SLnc1017_ | 107 | 250 | 116 | 159 | 312 | 100 |
| SLnc1018_ | 21 | 36 | 30 | 16 | 19 | 19 |
| SLnc1019_ | 27 | 36 | 25 | 36 | 21 | 40 |
| SLnc1021_ | 20 | 27 | 27 | 40 | 27 | 23 |
| SLnc1022_ | 38 | 68 | 39 | 43 | 42 | 41 |
| SLnc1024_ | 43 | 52 | 51 | 30 | 59 | 27 |
| SLnc1025_ | 51 | 62 | 37 | 48 | 64 | 20 |
| SLnc1026_ | 17 | 19 | 39 | 31 | 27 | 40 |
| SLnc1027_ | 70 | 93 | 73 | 68 | 93 | 50 |
| SLnc1028_ | 15 | 20 | 12 | 28 | 33 | 20 |
| SLnc1029_ | 18 | 19 | 18 | 17 | 22 | 12 |
| sRNA1_ | 1878 | 2519 | 2144 | 1614 | 2658 | 1717 |
| sRNA10_ | 521 | 613 | 608 | 641 | 715 | 599 |
| sRNA11_ | 39 | 47 | 34 | 31 | 36 | 26 |
| sRNA12_ | 2 | 5 | 8 | 4 | 5 | 6 |
| sRNA15_ | 1 | 1 | 2 | 3 | 3 | 5 |
| sRNA16_ | 49 | 49 | 68 | 48 | 23 | 31 |
| sRNA18_ | 3 | 3 | 2 | 6 | 1 | 1 |
| sRNA19_ | 3 | 1 | 0 | 1 | 1 | 2 |
| sRNA2_ | 2904 | 2597 | 2417 | 3269 | 2005 | 1918 |
| sRNA21_ | 60 | 70 | 66 | 101 | 70 | 91 |
| sRNA22_ | 42 | 35 | 34 | 35 | 42 | 54 |
| sRNA23_ | 15 | 8 | 6 | 11 | 6 | 11 |
| sRNA24_ | 3 | 5 | 6 | 5 | 3 | 7 |
| sRNA25_ | 0 | 2 | 0 | 3 | 3 | 0 |
| sRNA26_ | 3 | 6 | 6 | 5 | 3 | 7 |
| sRNA28_ | 5 | 2 | 3 | 14 | 9 | 6 |
| sRNA29_ | 1 | 3 | 7 | 5 | 6 | 7 |
| sRNA31_ | 4 | 8 | 6 | 13 | 4 | 6 |
| sRNA32_ | 5 | 6 | 5 | 10 | 20 | 5 |
| sRNA33_ | 22 | 20 | 24 | 33 | 35 | 19 |
| sRNA5_ | 4 | 7 | 4 | 14 | 25 | 11 |

| | | | | | | |
|---|---|---|---|---|---|---|
| sRNA7_ | 109 | 73 | 87 | 157 | 99 | 72 |
| STnc10_ | 2 | 5 | 1 | 2 | 5 | 8 |
| STnc1010_ | 32 | 38 | 47 | 44 | 32 | 38 |
| STnc1030_ | 47 | 66 | 61 | 55 | 57 | 52 |
| STnc1040_ | 5 | 3 | 6 | 2 | 3 | 3 |
| STnc1065_ | 4139 | 5075 | 4931 | 3832 | 4739 | 3382 |
| STnc1070_ | 84 | 126 | 122 | 136 | 145 | 117 |
| STnc1080_ | 11 | 14 | 13 | 12 | 16 | 18 |
| STnc1090_ | 74 | 99 | 51 | 100 | 103 | 83 |
| STnc1100_ | 9 | 10 | 5 | 16 | 9 | 21 |
| STnc1130_ | 109 | 115 | 144 | 249 | 200 | 238 |
| STnc1150_ | 229 | 284 | 388 | 413 | 245 | 336 |
| STnc1160_ | 602 | 1577 | 1018 | 942 | 2082 | 574 |
| STnc1170_ | 19 | 48 | 38 | 18 | 18 | 17 |
| STnc1180_ | 32 | 35 | 72 | 46 | 53 | 45 |
| STnc1190_ | 999 | 1001 | 1298 | 1714 | 892 | 1113 |
| STnc1200_ | 62 | 54 | 72 | 60 | 65 | 59 |
| STnc1210_ | 8 | 9 | 10 | 17 | 16 | 7 |
| STnc1220_ | 2 | 1 | 2 | 2 | 1 | 2 |
| STnc1275_ | 124 | 105 | 133 | 200 | 145 | 183 |
| STnc1280_ | 81 | 84 | 74 | 48 | 54 | 40 |
| STnc1285_ | 631 | 563 | 386 | 2121 | 2322 | 2881 |
| STnc1290_ | 367 | 619 | 458 | 534 | 675 | 294 |
| STnc130_ | 8 | 10 | 11 | 10 | 2 | 10 |
| STnc1320_ | 8 | 8 | 12 | 12 | 8 | 7 |
| STnc1340_ | 83 | 141 | 228 | 83 | 56 | 85 |
| STnc1380_ | 14 | 14 | 15 | 18 | 23 | 16 |
| STnc1385_ | 2649 | 3259 | 3452 | 3397 | 3690 | 2252 |
| STnc1390_ | 8 | 13 | 5 | 23 | 13 | 6 |
| STnc140_ | 3 | 6 | 5 | 3 | 1 | 3 |
| STnc1400_ | 174 | 307 | 356 | 331 | 348 | 243 |
| STnc1405_ | 115 | 155 | 141 | 627 | 628 | 585 |
| STnc1410_ | 23 | 41 | 38 | 35 | 30 | 30 |
| STnc1460_ | 14795 | 28538 | 31922 | 19725 | 26114 | 15179 |
| STnc1490_ | 194 | 210 | 230 | 198 | 179 | 213 |
| STnc150_ | 192 | 233 | 255 | 132 | 184 | 181 |
| STnc1500_ | 19 | 19 | 22 | 28 | 41 | 31 |
| STnc1510_ | 132 | 201 | 211 | 145 | 343 | 134 |
| STnc1530_ | 3 | 6 | 2 | 3 | 2 | 4 |
| STnc1540_ | 72 | 111 | 123 | 66 | 72 | 78 |
| STnc1545_ | 346 | 451 | 451 | 351 | 264 | 329 |
| STnc1550_ | 344 | 271 | 406 | 506 | 329 | 293 |
| STnc1560_ | 2398 | 5089 | 2900 | 1427 | 7451 | 1366 |
| STnc1565_ | 114 | 165 | 151 | 190 | 114 | 101 |
| STnc1580_ | 95 | 99 | 87 | 79 | 114 | 86 |
| STnc1590_ | 596 | 909 | 691 | 653 | 1396 | 538 |
| STnc1600_ | 63 | 80 | 57 | 97 | 85 | 75 |
| STnc1640_ | 69 | 86 | 69 | 69 | 86 | 62 |
| STnc1650_ | 5 | 9 | 8 | 8 | 4 | 7 |
| STnc1660_ | 124 | 151 | 152 | 183 | 135 | 161 |
| STnc1670_ | 356 | 121 | 151 | 277 | 129 | 132 |
| STnc1675_ | 14240 | 17842 | 16972 | 15851 | 16469 | 14937 |
| STnc1680_ | 343 | 801 | 429 | 482 | 1263 | 248 |
| STnc1685_ | 12871 | 14263 | 13697 | 2870 | 12821 | 2457 |
| STnc1690_ | 14 | 20 | 17 | 29 | 38 | 31 |
| STnc170_ | 4 | 2 | 5 | 16 | 8 | 13 |
| STnc1700_ | 24 | 22 | 17 | 10 | 16 | 8 |
| STnc1710_ | 228 | 251 | 277 | 298 | 244 | 222 |
| STnc1760_ | 27 | 35 | 28 | 19 | 17 | 22 |
| STnc1820_ | 56 | 113 | 88 | 102 | 81 | 56 |
| STnc1830_ | 40 | 35 | 31 | 39 | 32 | 37 |
| STnc1870_ | 5 | 5 | 4 | 5 | 4 | 5 |
| STnc1880_ | 22 | 46 | 55 | 46 | 32 | 43 |

| | | | | | | |
|---|---|---|---|---|---|---|
| STnc1890_ | 4 | 4 | 4 | 7 | 1 | 2 |
| STnc190_ | 40 | 40 | 55 | 132 | 97 | 98 |
| STnc1920_ | 128 | 179 | 206 | 154 | 120 | 141 |
| STnc1990_ | 102 | 99 | 123 | 92 | 80 | 83 |
| STnc20_ | 28 | 22 | 28 | 33 | 18 | 31 |
| STnc200_ | 87 | 134 | 66 | 198 | 190 | 157 |
| STnc2000_ | 240 | 431 | 346 | 356 | 519 | 274 |
| STnc2010_ | 85 | 139 | 164 | 113 | 114 | 90 |
| STnc2030_ | 6 | 7 | 8 | 12 | 7 | 10 |
| STnc2040_ | 0 | 2 | 0 | 6 | 5 | 5 |
| STnc205_ | 2988 | 3153 | 3513 | 15019 | 12301 | 14577 |
| STnc2070_ | 169 | 551 | 276 | 298 | 626 | 180 |
| STnc2080_ | 169 | 281 | 234 | 241 | 226 | 134 |
| STnc2090_ | 2246 | 4079 | 3849 | 2926 | 3569 | 3878 |
| STnc210_ | 1 | 3 | 1 | 3 | 2 | 6 |
| STnc2100_ | 17 | 46 | 22 | 27 | 40 | 20 |
| STnc2110_ | 60 | 69 | 55 | 163 | 95 | 46 |
| STnc2130_ | 180 | 301 | 231 | 192 | 131 | 167 |
| STnc2160_ | 6 | 8 | 6 | 10 | 10 | 15 |
| STnc2170_ | 26 | 38 | 27 | 18 | 36 | 23 |
| STnc2175_ | 34 | 50 | 42 | 38 | 39 | 43 |
| STnc2180_ | 1256 | 1079 | 640 | 3449 | 3060 | 1281 |
| STnc220_ | 8 | 9 | 13 | 9 | 11 | 5 |
| STnc230_ | 4 | 3 | 0 | 1 | 0 | 0 |
| STnc240_ | 65 | 79 | 63 | 98 | 45 | 32 |
| STnc260_ | 4 | 4 | 2 | 1 | 1 | 2 |
| STnc280_ | 8 | 7 | 6 | 4 | 7 | 5 |
| STnc290_ | 243 | 388 | 418 | 312 | 417 | 295 |
| STnc30_ | 310 | 383 | 248 | 298 | 337 | 231 |
| STnc300_ | 3 | 0 | 2 | 4 | 2 | 3 |
| STnc3000_ | 7 | 13 | 7 | 12 | 9 | 14 |
| STnc3020_ | 6 | 5 | 7 | 6 | 1 | 3 |
| STnc3030_ | 35 | 46 | 41 | 28 | 57 | 20 |
| STnc3040_ | 7 | 5 | 3 | 8 | 4 | 13 |
| STnc3060_ | 13 | 14 | 15 | 19 | 28 | 13 |
| STnc3070_ | 40 | 66 | 71 | 65 | 69 | 68 |
| STnc3080_ | 6 | 14 | 12 | 11 | 9 | 5 |
| STnc3090_ | 4 | 6 | 6 | 11 | 6 | 9 |
| STnc310_ | 11 | 8 | 11 | 12 | 15 | 11 |
| STnc3100_ | 340 | 438 | 446 | 315 | 239 | 277 |
| STnc3110_ | 157 | 125 | 162 | 234 | 231 | 130 |
| STnc3120_ | 95 | 112 | 88 | 158 | 69 | 48 |
| STnc3150_ | 132 | 169 | 147 | 281 | 206 | 287 |
| STnc3160_ | 31 | 25 | 32 | 77 | 49 | 17 |
| STnc3170_ | 51 | 55 | 59 | 73 | 80 | 74 |
| STnc3180_ | 10 | 20 | 14 | 10 | 19 | 8 |
| STnc320_ | 38 | 54 | 69 | 43 | 29 | 58 |
| STnc3210_ | 1 | 3 | 9 | 3 | 3 | 5 |
| STnc3220_ | 115 | 148 | 189 | 168 | 108 | 149 |
| STnc3230_ | 34 | 29 | 31 | 21 | 16 | 22 |
| STnc3270_ | 29 | 58 | 24 | 33 | 31 | 28 |
| STnc3280_ | 20 | 17 | 20 | 27 | 16 | 23 |
| STnc3290_ | 30 | 71 | 49 | 59 | 34 | 42 |
| STnc3300_ | 26 | 50 | 28 | 27 | 27 | 16 |
| STnc3310_ | 1276 | 1343 | 1473 | 1372 | 1130 | 1361 |
| STnc3330_ | 15 | 10 | 16 | 17 | 15 | 10 |
| STnc3340_ | 19 | 31 | 22 | 22 | 25 | 24 |
| STnc3350_ | 21 | 37 | 23 | 36 | 47 | 23 |
| STnc340_ | 14 | 16 | 21 | 39 | 22 | 25 |
| STnc3400_ | 8 | 10 | 10 | 8 | 3 | 2 |
| STnc3410_ | 157 | 260 | 202 | 176 | 247 | 171 |
| STnc3420_ | 48 | 44 | 55 | 45 | 53 | 38 |
| STnc3430_ | 65 | 101 | 81 | 53 | 44 | 37 |

| | | | | | | |
|---|---|---|---|---|---|---|
| STnc3440_ | 23 | 32 | 36 | 30 | 29 | 17 |
| STnc3460_ | 414 | 355 | 440 | 389 | 244 | 358 |
| STnc3480_ | 32 | 21 | 29 | 128 | 95 | 96 |
| STnc3490_ | 5 | 9 | 8 | 10 | 5 | 6 |
| STnc350_ | 45 | 50 | 65 | 54 | 38 | 31 |
| STnc3500_ | 9 | 13 | 14 | 17 | 9 | 14 |
| STnc3510_ | 152 | 141 | 157 | 173 | 142 | 117 |
| STnc3540_ | 1146 | 1035 | 984 | 1775 | 1647 | 1511 |
| STnc3550_ | 4 | 15 | 14 | 12 | 14 | 14 |
| STnc3580_ | 19 | 8 | 15 | 9 | 7 | 10 |
| STnc360_ | 4 | 7 | 0 | 1 | 6 | 1 |
| STnc3600_ | 64 | 85 | 93 | 65 | 34 | 54 |
| STnc3610_ | 81 | 139 | 77 | 150 | 153 | 142 |
| STnc3620_ | 14 | 12 | 18 | 21 | 14 | 21 |
| STnc3630_ | 33 | 52 | 41 | 83 | 124 | 44 |
| STnc3640_ | 41 | 28 | 37 | 94 | 67 | 61 |
| STnc3680_ | 693 | 929 | 1082 | 729 | 908 | 635 |
| STnc370_ | 3 | 1 | 6 | 4 | 2 | 2 |
| STnc3700_ | 14 | 22 | 15 | 24 | 16 | 12 |
| STnc3720_ | 19 | 18 | 21 | 24 | 7 | 14 |
| STnc3730_ | 102 | 119 | 100 | 102 | 187 | 88 |
| STnc3750_ | 14 | 24 | 22 | 15 | 13 | 22 |
| STnc3770_ | 21 | 34 | 42 | 32 | 32 | 48 |
| STnc3790_ | 4 | 10 | 6 | 9 | 12 | 9 |
| STnc380_ | 29 | 29 | 25 | 15 | 33 | 28 |
| STnc3800_ | 116 | 149 | 109 | 93 | 72 | 74 |
| STnc3830_ | 30 | 32 | 34 | 33 | 29 | 28 |
| STnc3850_ | 17 | 25 | 23 | 30 | 27 | 21 |
| STnc3860_ | 20 | 17 | 23 | 21 | 22 | 17 |
| STnc3870_ | 47 | 79 | 65 | 69 | 106 | 104 |
| STnc3880_ | 72 | 39 | 48 | 32 | 57 | 19 |
| STnc3890_ | 17 | 19 | 18 | 24 | 23 | 17 |
| STnc390_ | 1307 | 1964 | 1557 | 937 | 2504 | 1007 |
| STnc3920_ | 0 | 2 | 1 | 3 | 1 | 1 |
| STnc3950_ | 60 | 76 | 60 | 96 | 72 | 90 |
| STnc3960_ | 87 | 117 | 123 | 130 | 160 | 108 |
| STnc3970_ | 3 | 8 | 3 | 10 | 7 | 2 |
| STnc3990_ | 103 | 94 | 106 | 82 | 81 | 108 |
| STnc40_ | 309 | 414 | 398 | 323 | 337 | 323 |
| STnc4000_ | 22 | 25 | 27 | 21 | 29 | 23 |
| STnc4010_ | 5 | 1 | 9 | 9 | 4 | 3 |
| STnc4020_ | 8 | 8 | 3 | 6 | 9 | 9 |
| STnc4040_ | 78 | 128 | 99 | 113 | 150 | 110 |
| STnc4070_ | 19 | 24 | 27 | 15 | 18 | 14 |
| STnc4080_ | 19 | 20 | 13 | 28 | 19 | 14 |
| STnc410_ | 8 | 9 | 10 | 13 | 11 | 7 |
| STnc4100_ | 7 | 23 | 14 | 29 | 26 | 19 |
| STnc4110_ | 11 | 25 | 16 | 13 | 17 | 27 |
| STnc4120_ | 44 | 54 | 41 | 61 | 42 | 30 |
| STnc4130_ | 29 | 39 | 28 | 33 | 41 | 51 |
| STnc4140_ | 62 | 56 | 78 | 68 | 81 | 50 |
| STnc4150_ | 5 | 5 | 3 | 9 | 6 | 2 |
| STnc4160_ | 31 | 53 | 58 | 47 | 63 | 47 |
| STnc4170_ | 5 | 5 | 2 | 13 | 10 | 1 |
| STnc4190_ | 0 | 3 | 6 | 3 | 2 | 5 |
| STnc420_ | 9 | 30 | 20 | 29 | 59 | 19 |
| STnc4200_ | 32 | 55 | 18 | 60 | 48 | 26 |
| STnc4210_ | 9 | 16 | 18 | 75 | 52 | 61 |
| STnc4220_ | 143 | 245 | 206 | 235 | 160 | 173 |
| STnc4230_ | 86 | 102 | 105 | 92 | 114 | 102 |
| STnc4240_ | 450 | 446 | 340 | 361 | 387 | 300 |
| STnc4250_ | 2085 | 2430 | 2736 | 2429 | 3043 | 1908 |
| STnc4260_ | 7 | 6 | 6 | 12 | 6 | 9 |

| | | | | | | |
|---|---|---|---|---|---|---|
| STnc430_ | 5 | 3 | 9 | 3 | 4 | 6 |
| STnc440_ | 21 | 47 | 39 | 64 | 44 | 26 |
| STnc450_ | 178 | 214 | 218 | 237 | 141 | 199 |
| STnc460_ | 5 | 2 | 2 | 3 | 6 | 7 |
| STnc470_ | 52 | 52 | 54 | 30 | 46 | 44 |
| STnc475_ | 6854 | 8368 | 10560 | 13019 | 6681 | 9421 |
| STnc480_ | 26 | 34 | 25 | 19 | 48 | 16 |
| STnc50_ | 5 | 9 | 4 | 7 | 5 | 10 |
| STnc500_ | 58 | 47 | 40 | 113 | 36 | 51 |
| STnc510_ | 56 | 40 | 51 | 46 | 38 | 65 |
| STnc520_ | 51 | 104 | 79 | 79 | 116 | 51 |
| STnc530_ | 4 | 6 | 2 | 5 | 9 | 6 |
| STnc540_ | 2760 | 2955 | 2639 | 2774 | 4345 | 2761 |
| STnc590_ | 45 | 47 | 65 | 110 | 104 | 146 |
| STnc60_ | 0 | 0 | 2 | 0 | 1 | 0 |
| STnc600_ | 347 | 368 | 399 | 1490 | 1031 | 1205 |
| STnc610_ | 889 | 812 | 632 | 1589 | 1785 | 697 |
| STnc620_ | 504 | 825 | 657 | 698 | 774 | 530 |
| STnc640_ | 43 | 39 | 44 | 51 | 46 | 63 |
| STnc650_ | 2535 | 2160 | 3141 | 2349 | 1388 | 2355 |
| STnc660_ | 691 | 735 | 570 | 701 | 758 | 599 |
| STnc670_ | 564 | 534 | 620 | 781 | 756 | 938 |
| STnc680_ | 729 | 656 | 759 | 915 | 651 | 930 |
| STnc690_ | 199 | 272 | 206 | 175 | 292 | 111 |
| STnc695_ | 59 | 76 | 62 | 55 | 50 | 60 |
| STnc70_ | 1017 | 1954 | 1901 | 1823 | 1504 | 1232 |
| STnc720_ | 23 | 32 | 35 | 31 | 13 | 23 |
| STnc730_ | 695 | 625 | 811 | 666 | 447 | 801 |
| STnc740_ | 5 | 5 | 5 | 3 | 2 | 4 |
| STnc750_ | 234 | 136 | 239 | 225 | 101 | 205 |
| STnc760_ | 156 | 156 | 176 | 147 | 136 | 120 |
| STnc770_ | 1075 | 1041 | 1113 | 1293 | 904 | 1322 |
| STnc780_ | 366 | 385 | 426 | 401 | 343 | 403 |
| STnc790_ | 596 | 528 | 586 | 867 | 740 | 979 |
| STnc800_ | 1618 | 3126 | 2305 | 2557 | 3406 | 1900 |
| STnc810_ | 276 | 402 | 279 | 180 | 113 | 157 |
| STnc835_ | 2231 | 3767 | 2816 | 2908 | 3959 | 2225 |
| STnc840_ | 525 | 1019 | 682 | 1434 | 1941 | 1431 |
| STnc850_ | 18 | 28 | 11 | 14 | 17 | 12 |
| STnc870_ | 78 | 148 | 123 | 116 | 129 | 77 |
| STnc880_ | 385 | 530 | 474 | 642 | 680 | 292 |
| STnc890_ | 132 | 143 | 139 | 215 | 174 | 139 |
| STnc920_ | 410 | 453 | 451 | 499 | 382 | 357 |
| STnc925_ | 112 | 152 | 112 | 168 | 117 | 107 |
| STnc930_ | 8 | 12 | 9 | 16 | 14 | 11 |
| STnc940_ | 15 | 22 | 19 | 19 | 24 | 10 |
| STnc950_ | 18 | 25 | 13 | 16 | 36 | 14 |
| STnc970_ | 72 | 102 | 108 | 117 | 51 | 85 |
| StyR-103a_ | 0 | 0 | 0 | 0 | 0 | 0 |
| StyR-103b_ | 0 | 1 | 0 | 0 | 1 | 2 |
| StyR-103c_ | 0 | 0 | 1 | 0 | 0 | 0 |
| StyR-103d_ | 0 | 0 | 0 | 0 | 0 | 1 |
| StyR-103e_ | 0 | 0 | 0 | 0 | 0 | 0 |
| StyR-103f_ | 0 | 0 | 0 | 0 | 0 | 3 |
| StyR-103g_ | 0 | 1 | 1 | 1 | 0 | 0 |
| StyR-122_ | 4 | 3 | 14 | 12 | 2 | 8 |
| StyR-126_ | 58 | 51 | 40 | 67 | 42 | 45 |
| StyR-128_ | 7 | 4 | 4 | 9 | 17 | 4 |
| StyR-134_ | 214 | 257 | 306 | 257 | 227 | 276 |
| StyR-137_ | 1 | 0 | 0 | 0 | 4 | 0 |
| StyR-150_ | 15 | 19 | 10 | 23 | 26 | 15 |
| StyR-169_ | 87 | 89 | 126 | 78 | 71 | 87 |
| StyR-172_ | 1 | 0 | 0 | 0 | 0 | 0 |

| | | | | | | |
|---|---|---|---|---|---|---|
| StyR-178_ | 0 | 0 | 0 | 0 | 0 | 0 |
| StyR-195_ | 17 | 14 | 10 | 22 | 23 | 14 |
| StyR-196_ | 3 | 2 | 1 | 3 | 0 | 3 |
| StyR-199_ | 19 | 23 | 10 | 18 | 24 | 19 |
| StyR-215a_ | 1623 | 2078 | 2288 | 2190 | 1706 | 1254 |
| StyR-215b_ | 1089 | 1213 | 1300 | 1250 | 1022 | 722 |
| StyR-215c_ | 109 | 153 | 128 | 143 | 81 | 87 |
| StyR-215d_ | 269 | 341 | 360 | 440 | 226 | 247 |
| StyR-215e_ | 118 | 146 | 196 | 140 | 54 | 85 |
| StyR-215f_ | 527 | 554 | 639 | 518 | 391 | 306 |
| StyR-215g_ | 57 | 72 | 112 | 112 | 54 | 49 |
| StyR-219_ | 5 | 4 | 10 | 15 | 10 | 17 |
| StyR-222_ | 0 | 5 | 2 | 1 | 3 | 1 |
| StyR-234_ | 13 | 4 | 4 | 6 | 6 | 4 |
| StyR-243_ | 286 | 353 | 271 | 239 | 567 | 221 |
| StyR-248_ | 52 | 80 | 51 | 47 | 88 | 73 |
| StyR-24a_ | 11 | 11 | 12 | 10 | 16 | 10 |
| StyR-24b_ | 9 | 5 | 13 | 75 | 46 | 58 |
| StyR-24c_ | 7 | 11 | 9 | 37 | 26 | 49 |
| StyR-24d_ | 5 | 9 | 7 | 5 | 2 | 2 |
| StyR-24e_ | 6 | 3 | 2 | 9 | 13 | 6 |
| StyR-24f_ | 15 | 14 | 8 | 22 | 23 | 32 |
| StyR-24g_ | 5 | 1 | 5 | 27 | 22 | 41 |
| StyR-250_ | 13 | 13 | 13 | 9 | 11 | 10 |
| StyR-252_ | 56 | 54 | 39 | 107 | 72 | 73 |
| StyR-254_ | 2 | 2 | 2 | 4 | 1 | 4 |
| StyR-262_ | 101 | 105 | 100 | 102 | 77 | 93 |
| StyR-264_ | 28 | 19 | 35 | 32 | 17 | 26 |
| StyR-27_ | 14 | 24 | 12 | 11 | 46 | 20 |
| StyR-277_ | 201 | 269 | 287 | 309 | 203 | 244 |
| StyR-281_ | 7 | 3 | 5 | 9 | 16 | 10 |
| StyR-285_ | 18 | 18 | 14 | 12 | 8 | 5 |
| StyR-287_ | 108 | 78 | 87 | 84 | 64 | 84 |
| StyR-29_ | 3479 | 3003 | 3842 | 5262 | 2076 | 3251 |
| StyR-291_ | 52 | 56 | 57 | 58 | 125 | 86 |
| StyR-292_ | 0 | 0 | 0 | 5 | 1 | 0 |
| StyR-293_ | 14 | 28 | 31 | 55 | 19 | 27 |
| StyR-295_ | 12 | 7 | 12 | 13 | 12 | 5 |
| StyR-296_ | 2396 | 2238 | 2665 | 2813 | 2089 | 2505 |
| StyR-3_ | 55 | 72 | 56 | 74 | 103 | 52 |
| StyR-302_ | 0 | 2 | 1 | 0 | 0 | 2 |
| StyR-305_ | 49 | 80 | 58 | 174 | 152 | 135 |
| StyR-306_ | 346 | 412 | 503 | 382 | 226 | 339 |
| StyR-315_ | 0 | 3 | 2 | 0 | 0 | 3 |
| StyR-322_ | 11 | 9 | 3 | 12 | 8 | 12 |
| StyR-327_ | 6 | 9 | 11 | 4 | 0 | 1 |
| StyR-328_ | 3 | 3 | 1 | 1 | 2 | 4 |
| StyR-329a_ | 741 | 501 | 665 | 231 | 246 | 162 |
| StyR-329b_ | 24 | 39 | 30 | 22 | 43 | 17 |
| StyR-336_ | 15 | 16 | 9 | 21 | 19 | 16 |
| StyR-341_ | 4 | 5 | 6 | 5 | 7 | 6 |
| StyR-354_ | 446 | 732 | 572 | 140 | 391 | 446 |
| StyR-358_ | 11 | 13 | 8 | 15 | 12 | 7 |
| StyR-362_ | 4 | 1 | 5 | 6 | 3 | 6 |
| StyR-369_ | 6 | 1 | 6 | 5 | 3 | 4 |
| StyR-377_ | 405 | 405 | 332 | 522 | 773 | 607 |
| StyR-378_ | 4 | 1 | 1 | 2 | 0 | 2 |
| StyR-44a_ | 82 | 246 | 168 | 174 | 251 | 81 |
| StyR-44b_ | 316 | 535 | 694 | 350 | 387 | 370 |
| StyR-44c_ | 40 | 97 | 101 | 70 | 54 | 38 |
| StyR-44d_ | 54 | 115 | 98 | 95 | 40 | 77 |
| StyR-44e_ | 22 | 23 | 38 | 15 | 14 | 11 |
| StyR-44f_ | 5 | 19 | 16 | 8 | 12 | 9 |

| | | | | | | |
|---|---|---|---|---|---|---|
| StyR-44g_ | 11 | 18 | 25 | 8 | 12 | 10 |
| StyR-51_ | 241 | 283 | 271 | 304 | 382 | 326 |
| StyR-55_ | 4423 | 6072 | 5914 | 6107 | 6302 | 4508 |
| StyR-8_ | 2531 | 2371 | 2585 | 3857 | 2242 | 2136 |
| StyR-88_ | 343 | 318 | 334 | 459 | 259 | 403 |
| StyR-90_ | 11 | 6 | 10 | 12 | 6 | 4 |
| StyR-99_ | 86 | 104 | 153 | 118 | 57 | 83 |
| tp2_ | 6 | 1 | 3 | 6 | 2 | 5 |
| tp28_ | 223 | 190 | 245 | 344 | 175 | 282 |
| tpk3_ | 189 | 400 | 278 | 406 | 342 | 236 |
| tpke11_ | 3447 | 4184 | 4014 | 4202 | 7920 | 2967 |
| tpke70_ | 59 | 76 | 45 | 49 | 96 | 53 |
| zipA-leader_ | 1011 | 1159 | 1249 | 898 | 770 | 909 |
| _ | | | | | | |
| _ | | | | | | |

# LE

| Gene | WT-1-LE | WT-2-LE | WT-3-LE | 5' tnpA-1-LE | 5' tnpA-2-LE | 5' tnpA-3-LE |
|---|---|---|---|---|---|---|
| aceF_aceF | 8258 | 7866 | 8504 | 9043 | 8414 | 10345 |
| SL1344_RS11990_ackA | 2531 | 2609 | 2890 | 2954 | 2635 | 3064 |
| SL1344_RS08180_acpD | 193 | 207 | 248 | 264 | 199 | 252 |
| acpS_acpS | 392 | 409 | 372 | 277 | 257 | 328 |
| adrA_adrA | 183 | 178 | 160 | 119 | 133 | 158 |
| SL1344_RS16630_aer | 3208 | 3148 | 2980 | 5592 | 3955 | 5742 |
| SL1344_RS13055_alpha/beta hydrolase | 28 | 16 | 50 | 32 | 27 | 32 |
| AmgR_amgR | 8 | 5 | 2 | 6 | 7 | 2 |
| amiC_amiC | 392 | 405 | 378 | 447 | 506 | 597 |
| SL1344_RS04710_amino acid permease | 179 | 177 | 192 | 148 | 138 | 148 |
| SL1344_RS01920_ampH | 190 | 155 | 180 | 114 | 99 | 90 |
| SL1344_RS23585_arcA | 5562 | 5446 | 6034 | 6441 | 5788 | 6141 |
| ArcZ_arcZ | 561 | 566 | 700 | 546 | 486 | 401 |
| STnc490a_art200 | 2 | 0 | 1 | 0 | 0 | 0 |
| STnc490b_art200 | 0 | 0 | 2 | 0 | 0 | 0 |
| STnc490c_art200 | 0 | 1 | 4 | 0 | 0 | 0 |
| STnc490d_art200 | 0 | 0 | 0 | 0 | 0 | 0 |
| STnc490e_art200 | 1 | 0 | 3 | 0 | 0 | 1 |
| STnc490f_art200 | 2 | 2 | 2 | 0 | 0 | 0 |
| STnc490g_art200 | 0 | 0 | 0 | 0 | 0 | 0 |
| artM_artM | 580 | 582 | 675 | 652 | 619 | 699 |
| artP_artP | 307 | 267 | 346 | 422 | 430 | 457 |
| asarfA_asarfA | 406 | 380 | 472 | 364 | 278 | 330 |
| asnB_asnB | 3229 | 2932 | 1597 | 1492 | 1453 | 1663 |
| SL1344_RS13060_asrA | 148 | 162 | 170 | 191 | 124 | 162 |
| SL1344_RS13065_asrB | 161 | 153 | 172 | 192 | 160 | 221 |
| SL1344_RS13070_asrC | 258 | 294 | 281 | 350 | 274 | 356 |
| asyhbQ_asyhbQ | 759 | 774 | 700 | 514 | 494 | 594 |
| atpC_atpC | 13276 | 13721 | 15278 | 9763 | 10099 | 11060 |
| SL1344_RS15305_barA | 859 | 914 | 938 | 853 | 699 | 892 |
| bcsA_bcsA | 964 | 1024 | 965 | 621 | 511 | 541 |
| SL1344_RS17950_bigA | 176 | 216 | 182 | 204 | 203 | 200 |
| bssR_bssR | 842 | 870 | 768 | 566 | 505 | 630 |
| bssS_bssS | 1113 | 1171 | 1230 | 978 | 911 | 1089 |
| cadB_cadB | 967 | 1465 | 2038 | 411 | 660 | 730 |
| cbiC_cbiC | 567 | 627 | 581 | 287 | 261 | 306 |
| cbiD_cbiD | 747 | 842 | 747 | 479 | 332 | 433 |
| SL1344_RS11580_ccmA | 362 | 373 | 374 | 226 | 178 | 305 |
| SL1344_RS11575_ccmB | 188 | 139 | 202 | 172 | 122 | 145 |
| cdsA_cdsA | 1148 | 1098 | 988 | 1068 | 1061 | 1316 |
| SL1344_RS02340_cdsH | 127 | 129 | 114 | 125 | 124 | 133 |
| chaB_chaB | 238 | 209 | 202 | 299 | 240 | 331 |
| SL1344_RS09610_cheA | 11379 | 11338 | 10612 | 37413 | 30516 | 44372 |
| SL1344_RS09600_cheM | 7493 | 7650 | 7477 | 22929 | 19041 | 26094 |
| SL1344_RS09595_chemotaxis protein methyltransferase | 2407 | 2420 | 2285 | 7921 | 6875 | 8496 |
| SL1344_RS09585_chemotaxis regulator | 2740 | 2782 | 2645 | 6593 | 6088 | 8811 |
| SL1344_RS09590_chemotaxis response regulator protein-glutamate methylesterase | 1972 | 2124 | 1968 | 5537 | 4882 | 6676 |
| SL1344_RS11875_cheV | 4952 | 5332 | 5178 | 12641 | 10298 | 13471 |
| SL1344_RS09605_CheW | 11573 | 11951 | 10838 | 37712 | 33630 | 43602 |
| SL1344_RS09580_cheZ | 5205 | 5464 | 5303 | 14305 | 12048 | 17442 |
| chiP_chiP | 992 | 1029 | 1121 | 757 | 636 | 663 |
| ChiX_chiX | 466 | 667 | 1720 | 1417 | 578 | 661 |
| SL1344_RS02290_clpP | 2402 | 2498 | 2386 | 2011 | 1873 | 2474 |
| SL1344_RS02295_clpX | 6865 | 6741 | 7045 | 6621 | 6494 | 7014 |
| SL1344_RS09530_cmoA | 549 | 492 | 523 | 794 | 739 | 819 |
| cobS_cobS | 529 | 560 | 469 | 293 | 259 | 298 |
| copA_copA | 1911 | 2231 | 1643 | 1205 | 1242 | 1725 |
| SL1344_RS20320_corA | 693 | 626 | 656 | 675 | 532 | 721 |
| SL1344_RS20860_cpxA | 1524 | 1543 | 1524 | 1542 | 1353 | 1495 |
| cpxP_cpxP | 1891 | 1839 | 1572 | 1868 | 1387 | 1522 |
| SL1344_RS20865_cpxR | 788 | 734 | 789 | 776 | 775 | 833 |

| | | | | | | |
|---|---|---|---|---|---|---|
| SL1344_RS17890_crp | 7317 | 7115 | 7892 | 7258 | 6973 | 7987 |
| CrpT_crpT | 90 | 114 | 129 | 140 | 96 | 91 |
| csgB_csgB | 6 | 5 | 9 | 5 | 5 | 15 |
| SL1344_RS05640_csgC | 629 | 648 | 588 | 1329 | 1768 | 1750 |
| SL1344_RS18810_cspA | 1234 | 813 | 873 | 1181 | 657 | 596 |
| SL1344_RS09980_cspB | 107 | 64 | 112 | 87 | 74 | 99 |
| SL1344_RS06145_cspH | 8 | 8 | 15 | 7 | 11 | 10 |
| SL1344_RS14635_csrA | 4757 | 4560 | 4154 | 3876 | 3472 | 4247 |
| CsrB_csrB | 180080 | 185435 | 165941 | 156753 | 134903 | 178784 |
| SL1344_RS15345_csrB | 155086 | 164566 | 143191 | 134224 | 118581 | 156727 |
| CsrC_csrC | 68992 | 76805 | 69782 | 79303 | 61603 | 84261 |
| SL1344_RS17435_csrD | 1197 | 1234 | 1125 | 1497 | 1172 | 1757 |
| CyaR_cyaR | 10269 | 9564 | 12104 | 10911 | 8176 | 8079 |
| SL1344_RS12535_cysA | 468 | 617 | 958 | 4552 | 5202 | 6128 |
| SL1344_RS08565_cysB | 733 | 708 | 784 | 1621 | 2643 | 2508 |
| SL1344_RS15175_cysC | 189 | 213 | 424 | 4069 | 5043 | 5145 |
| SL1344_RS15185_cysD | 94 | 90 | 192 | 3963 | 6274 | 6342 |
| SL1344_RS19060_cysE | 908 | 911 | 898 | 985 | 937 | 1017 |
| SL1344_RS17945_cysG | 1245 | 1160 | 1220 | 1024 | 1006 | 1091 |
| SL1344_RS15240_cysH | 126 | 163 | 397 | 3457 | 3393 | 4208 |
| SL1344_RS15245_cysI | 680 | 818 | 1695 | 11664 | 13686 | 15307 |
| SL1344_RS15250_cysJ | 192 | 241 | 503 | 7441 | 8658 | 10317 |
| SL1344_RS12475_cysK | 1840 | 1926 | 3223 | 22813 | 29737 | 32638 |
| SL1344_RS12530_cysM | 254 | 258 | 311 | 837 | 918 | 1064 |
| SL1344_RS15180_cysN | 280 | 345 | 673 | 7526 | 10324 | 11325 |
| SL1344_RS12550_cysP | 97 | 104 | 213 | 4837 | 8490 | 9796 |
| SL1344_RS22605_cysQ | 818 | 884 | 886 | 734 | 703 | 854 |
| SL1344_RS02725_cysS | 1465 | 1386 | 1534 | 1446 | 1336 | 1594 |
| SL1344_RS09720_cysT | 575 | 695 | 915 | 1219 | 898 | 1032 |
| SL1344_RS12545_cysU | 77 | 92 | 193 | 2360 | 3240 | 4237 |
| SL1344_RS12540_cysW | 183 | 234 | 410 | 4066 | 5193 | 5806 |
| SL1344_RS12470_cysZ | 272 | 249 | 300 | 352 | 301 | 318 |
| SL1344_RS16735_cyuA | 268 | 270 | 244 | 237 | 218 | 248 |
| SL1344_RS16740_cyuP | 105 | 114 | 170 | 128 | 102 | 134 |
| SL1344_RS10610_dacD | 77 | 94 | 67 | 55 | 32 | 44 |
| SL1344_RS17980_dam | 748 | 756 | 871 | 759 | 657 | 731 |
| DapZ_dapZ | 442 | 384 | 455 | 640 | 520 | 519 |
| ddl_ddl | 1818 | 1778 | 1640 | 1689 | 1640 | 1904 |
| SL1344_RS09505_dihydroneopterin triphosphate diphosphatase | 859 | 917 | 1057 | 1491 | 1443 | 1598 |
| dinG_dinG | 580 | 560 | 623 | 698 | 718 | 765 |
| SL1344_RS05725_dinI | 429 | 498 | 496 | 813 | 777 | 722 |
| SL1344_RS04685_dmsA | 3166 | 3418 | 3326 | 2142 | 1759 | 2350 |
| SL1344_RS04690_dmsB | 407 | 470 | 368 | 273 | 172 | 262 |
| SL1344_RS04695_dmsC | 1094 | 1225 | 944 | 571 | 476 | 693 |
| SL1344_RS06610_DNA/RNA non-specific endonuclease | 416 | 397 | 431 | 390 | 317 | 355 |
| dnaA_dnaA | 1289 | 1320 | 1353 | 1534 | 1384 | 1543 |
| SL1344_RS00060_dnaK | 27155 | 26788 | 26504 | 25936 | 26561 | 28958 |
| dpiA_dpiA | 102 | 73 | 84 | 77 | 73 | 109 |
| dpiA_dpiA | 130 | 122 | 111 | 123 | 94 | 139 |
| dpiB_dpiB | 132 | 136 | 130 | 117 | 116 | 131 |
| DsrA_dsrA | 17 | 12 | 19 | 18 | 5 | 7 |
| SL1344_RS09330_DUF1441 domain-containing protein | 11 | 12 | 13 | 20 | 23 | 30 |
| SL1344_RS06460_DUF1496 domain-containing protein | 1431 | 1636 | 1386 | 3105 | 2731 | 3595 |
| SL1344_RS16295_DUF2623 domain-containing protein | 198 | 221 | 202 | 717 | 519 | 785 |
| emrB_emrB | 475 | 474 | 457 | 501 | 481 | 493 |
| emrD_emrD | 446 | 427 | 492 | 538 | 528 | 553 |
| emtA_emtA | 211 | 203 | 210 | 251 | 239 | 277 |
| SL1344_RS03015_entB | 46 | 42 | 44 | 35 | 41 | 43 |
| entE_entE | 46 | 30 | 48 | 63 | 39 | 46 |
| entF_entF | 113 | 129 | 132 | 140 | 128 | 146 |
| SL1344_RS18065_envZ | 1468 | 1563 | 1584 | 1272 | 1052 | 1341 |
| eutA_eutA | 60 | 70 | 78 | 58 | 51 | 71 |

| | | | | | | |
|---|---|---|---|---|---|---|
| SL1344_RS12680_eutS | 2 | 9 | 10 | 12 | 9 | 9 |
| fabG_fabG | 2802 | 2725 | 3012 | 2709 | 2656 | 2923 |
| fadA_fadA | 196 | 223 | 228 | 264 | 307 | 341 |
| SL1344_RS09080_fadD | 358 | 384 | 399 | 774 | 768 | 926 |
| fadE_fadE | 497 | 518 | 466 | 576 | 511 | 584 |
| fadI_fadI | 91 | 84 | 104 | 144 | 110 | 119 |
| SL1344_RS13945_Fels-2 prophage protein | 20 | 20 | 17 | 18 | 19 | 24 |
| feoA_feoA | 120 | 71 | 72 | 79 | 39 | 46 |
| ffs_ffs | 2338 | 2103 | 2593 | 2892 | 2220 | 2631 |
| fimA_fimA | 111 | 136 | 135 | 77 | 70 | 76 |
| fimF_fimF | 36 | 44 | 41 | 38 | 41 | 69 |
| SL1344_RS02785_fimZ | 13 | 26 | 31 | 23 | 25 | 18 |
| fis_fis | 324 | 238 | 351 | 598 | 456 | 536 |
| fixB_fixB | 48 | 41 | 65 | 54 | 51 | 68 |
| flgD_flgD | 4531 | 4664 | 4635 | 6982 | 6789 | 8502 |
| flgF_flgF | 6031 | 6465 | 6487 | 10007 | 10802 | 11852 |
| flgI_flgI | 2540 | 2771 | 2637 | 4050 | 3782 | 4486 |
| SL1344_RS05830_flgK | 8844 | 9105 | 8996 | 19745 | 16192 | 23093 |
| flgL_flgL | 8646 | 8773 | 8688 | 17774 | 14763 | 20277 |
| SL1344_RS05775_flgM | 5505 | 5993 | 5468 | 10302 | 9900 | 12296 |
| SL1344_RS05770_flgN | 6094 | 6476 | 5724 | 11652 | 9615 | 12388 |
| SL1344_RS09570_flhA | 2434 | 2604 | 2568 | 7376 | 7457 | 7961 |
| SL1344_RS09575_flhB | 1592 | 1691 | 1679 | 4835 | 4699 | 4928 |
| SL1344_RS09625_flhC | 3536 | 3458 | 3519 | 8034 | 6316 | 7270 |
| SL1344_RS09630_flhD | 1896 | 1719 | 1848 | 3713 | 3399 | 3728 |
| SL1344_RS09565_flhE | 701 | 765 | 775 | 1907 | 1951 | 2117 |
| SL1344_RS09785_fliA | 6967 | 6895 | 6373 | 19598 | 15454 | 20997 |
| SL1344_RS09800_fliC | 28049 | 27951 | 30684 | 242813 | 127566 | 299967 |
| SL1344_RS09845_fliE | 1280 | 1328 | 1137 | 1856 | 1908 | 2308 |
| fliI_fliI | 2672 | 3019 | 3117 | 4228 | 3938 | 4183 |
| fliJ_fliJ | 1480 | 1697 | 1517 | 1968 | 1667 | 1856 |
| fliL_fliL | 2215 | 2468 | 2364 | 3748 | 3775 | 3914 |
| fliQ_fliQ | 449 | 510 | 555 | 913 | 854 | 927 |
| SL1344_RS09815_fliT | 5987 | 5568 | 5476 | 11106 | 9875 | 11407 |
| SL1344_RS09775_fliY | 3343 | 3432 | 3282 | 8675 | 8339 | 11512 |
| SL1344_RS09780_fliZ | 5236 | 5285 | 5017 | 15637 | 13897 | 18094 |
| SL1344_RS14340_fljA | 1547 | 1435 | 1365 | 84 | 1524 | 87 |
| flk_flk | 169 | 173 | 213 | 191 | 162 | 199 |
| SL1344_RS08285_fnr | 1876 | 2001 | 2299 | 2177 | 2248 | 2653 |
| FnrS_fnrS | 13149 | 10863 | 10827 | 10896 | 11335 | 9658 |
| SL1344_RS04735_focA | 1995 | 2004 | 2114 | 2951 | 2690 | 2809 |
| folA_folA | 163 | 138 | 149 | 223 | 176 | 240 |
| frsA_frsA | 1100 | 1068 | 1174 | 900 | 1023 | 1106 |
| SL1344_RS09665_ftnB | 627 | 620 | 644 | 745 | 677 | 787 |
| SL1344_RS06895_fumD | 212 | 231 | 244 | 161 | 171 | 160 |
| SL1344_RS03500_fur | 1531 | 1534 | 1616 | 2104 | 2070 | 2061 |
| gapA_gapA | 1614 | 1595 | 1542 | 1260 | 1229 | 1475 |
| GcvB_gcvB | 22432 | 27045 | 41572 | 36322 | 20096 | 24882 |
| gcvT_gcvT | 6579 | 6675 | 6819 | 6107 | 5727 | 6811 |
| gidB_gidB | 285 | 237 | 269 | 287 | 251 | 323 |
| gldA_gldA | 141 | 142 | 137 | 153 | 138 | 154 |
| gldA_gldA | 2092 | 2275 | 2081 | 1928 | 1711 | 1859 |
| GlmY_glmY | 1318 | 1640 | 2795 | 2545 | 1609 | 1349 |
| GlmZ_glmZ | 4918 | 5778 | 7102 | 7938 | 5688 | 5867 |
| glnQ_glnQ | 503 | 493 | 688 | 840 | 805 | 938 |
| SL1344_RS13690_gluconolactonase | 126 | 147 | 128 | 83 | 63 | 96 |
| SL1344_RS09560_glycoside hydrolase family 105 protein | 238 | 312 | 282 | 686 | 643 | 675 |
| gntK_gntK | 37 | 50 | 45 | 47 | 46 | 42 |
| SL1344_RS18265_gntR | 471 | 477 | 558 | 523 | 542 | 583 |
| gpmA_gpmA | 5843 | 5758 | 5485 | 5580 | 5191 | 6579 |
| GraL_graL | 292 | 285 | 243 | 420 | 433 | 375 |
| groES_groES | 14976 | 13959 | 13853 | 15160 | 14889 | 17836 |

| | | | | | | |
|---|---|---|---|---|---|---|
| SL1344_RS14595_gshA | 1476 | 1389 | 1431 | 1300 | 1376 | 1536 |
| SL1344_RS16010_gshB | 1101 | 1048 | 1032 | 890 | 902 | 990 |
| SL1344_RS05170_gtgD | 8 | 9 | 12 | 7 | 12 | 25 |
| SL1344_RS05180_gtgE | 486 | 478 | 567 | 672 | 671 | 796 |
| SL1344_RS05185_gtgF | 9 | 7 | 6 | 9 | 11 | 17 |
| SL1344_RS11665_gyrA | 6426 | 6508 | 7368 | 6297 | 6322 | 7472 |
| SL1344_RS13995_head completion/stabilization protein | 8 | 5 | 6 | 11 | 6 | 5 |
| helD_helD | 1078 | 1090 | 1027 | 849 | 814 | 989 |
| hemG_hemG | 344 | 347 | 356 | 351 | 339 | 370 |
| hflB_hflB | 11474 | 11439 | 12054 | 10863 | 10567 | 11839 |
| SL1344_RS22390_hfq | 6629 | 6441 | 6498 | 5215 | 4777 | 5608 |
| SL1344_RS02410_hha | 658 | 649 | 778 | 490 | 450 | 498 |
| SL1344_RS02415_Hha toxicity attenuator | 728 | 883 | 978 | 541 | 561 | 639 |
| SL1344_RS14890_hilA | 3222 | 3193 | 3517 | 6023 | 4215 | 5434 |
| SL1344_RS14845_hilC | 4853 | 4502 | 4142 | 7020 | 4988 | 6292 |
| SL1344_RS14885_hilD | 5838 | 5299 | 5199 | 8888 | 8203 | 9101 |
| SL1344_RS23135_hilE | 213 | 177 | 143 | 186 | 161 | 188 |
| SL1344_RS21865_HlyD family type I secretion periplasmic adaptor subunit | 1201 | 988 | 1118 | 530 | 492 | 679 |
| SL1344_RS08745_hns | 21290 | 20411 | 22225 | 26441 | 26129 | 27777 |
| hscB_hscB | 241 | 285 | 260 | 268 | 274 | 288 |
| SL1344_RS08210_hslJ | 130 | 147 | 120 | 141 | 134 | 183 |
| SL1344_RS21405_hupA | 20444 | 19778 | 20438 | 22076 | 22763 | 23334 |
| SL1344_RS21410_hupB | 2522 | 2131 | 2929 | 3351 | 2732 | 2703 |
| hycD_hycD | 31 | 27 | 33 | 35 | 26 | 37 |
| hycE_hycE | 64 | 63 | 70 | 74 | 66 | 69 |
| SL1344_RS06065_hypothetical protein | 168 | 170 | 181 | 580 | 556 | 594 |
| SL1344_RS07940_hypothetical protein | 1624 | 1684 | 1671 | 4099 | 4159 | 4204 |
| SL1344_RS09195_hypothetical protein | 472 | 475 | 447 | 688 | 497 | 657 |
| SL1344_RS09675_hypothetical protein | 552 | 569 | 623 | 2277 | 2059 | 2192 |
| SL1344_RS09745_hypothetical protein | 84 | 98 | 91 | 247 | 227 | 234 |
| SL1344_RS22135_hypothetical protein | 188 | 158 | 121 | 246 | 141 | 206 |
| iacP_iacP | 658 | 629 | 565 | 990 | 835 | 1409 |
| SL1344_RS14895_iagB | 2630 | 2513 | 2420 | 5081 | 3978 | 4653 |
| lamB_iamB | 2174 | 2092 | 2162 | 1223 | 1065 | 1389 |
| idnK_idnK | 131 | 129 | 134 | 102 | 109 | 103 |
| SL1344_RS22990_idnT | 318 | 328 | 376 | 232 | 240 | 293 |
| SL1344_RS06655_ihfA | 6859 | 6399 | 6108 | 5994 | 5647 | 6382 |
| SL1344_RS04775_ihfB | 2032 | 1964 | 2402 | 2054 | 1714 | 2098 |
| lldD_ildD | 96 | 95 | 121 | 149 | 138 | 150 |
| ilvM_ilvM | 171 | 162 | 178 | 166 | 205 | 226 |
| SL1344_RS14990_invA | 3115 | 2919 | 3177 | 4959 | 3420 | 5288 |
| SL1344_RS14980_invC | 1932 | 1936 | 2099 | 2804 | 2130 | 3295 |
| SL1344_RS14995_invE | 2430 | 2328 | 2583 | 4125 | 3037 | 4218 |
| SL1344_RS15005_invF | 3737 | 3836 | 3968 | 6803 | 5001 | 6660 |
| SL1344_RS15000_invG | 3536 | 3472 | 3638 | 5678 | 3829 | 5697 |
| InvR_invR | 7011 | 5342 | 6085 | 14517 | 12408 | 12004 |
| SL1344_RS14355_iroB | 20 | 29 | 37 | 89 | 46 | 76 |
| SL1344_RS24725_IS91 family transposase | 78 | 135 | 102 | 484 | 467 | 467 |
| IsrA_isrA | 6 | 2 | 4 | 2 | 0 | 3 |
| IsrB_isrB | 34 | 32 | 56 | 48 | 36 | 32 |
| IsrC_isrC | 28 | 31 | 17 | 55 | 34 | 27 |
| IsrD_isrD | 1 | 1 | 0 | 1 | 0 | 0 |
| IsrE_isrE | 37 | 51 | 50 | 110 | 63 | 53 |
| IsrF_isrF | 103 | 85 | 89 | 159 | 140 | 154 |
| IsrG_isrG | 9 | 11 | 5 | 18 | 18 | 23 |
| IsrH-1_isrH | 10 | 11 | 9 | 14 | 10 | 11 |
| IsrH-2_isrH2 | 6 | 10 | 5 | 12 | 7 | 8 |
| IsrI_isrI | 51 | 44 | 50 | 32 | 30 | 33 |
| IsrJ_isrJ | 30 | 22 | 55 | 43 | 26 | 32 |
| IsrK_isrK | 217 | 201 | 299 | 281 | 226 | 217 |
| IsrL_isrL | 45 | 62 | 48 | 66 | 40 | 40 |
| IsrM_isrM | 5 | 7 | 12 | 4 | 5 | 5 |

| | | | | | | |
|---|---|---|---|---|---|---|
| IsrN_isrN | 2 | 0 | 0 | 2 | 2 | 1 |
| IsrO_isrO | 112 | 99 | 78 | 114 | 140 | 114 |
| IsrP_isrP | 34 | 29 | 38 | 50 | 46 | 54 |
| IsrQ_isrQ | 1 | 1 | 0 | 4 | 1 | 1 |
| IstR-1_istR1 | 12 | 11 | 16 | 14 | 16 | 5 |
| IstR-2_istR2 | 14 | 14 | 17 | 19 | 17 | 6 |
| ksgA_ksgA | 1026 | 905 | 981 | 1026 | 873 | 997 |
| SL1344_RS08205_ldhA | 985 | 1098 | 1056 | 1015 | 838 | 1092 |
| SL1344_RS00585_leuO | 15 | 9 | 15 | 21 | 17 | 13 |
| SL1344_RS21750_lexA | 2672 | 2991 | 2892 | 3168 | 3440 | 4113 |
| ligA_ligA | 1472 | 1496 | 1360 | 1160 | 1151 | 1369 |
| livF_livF | 55 | 52 | 51 | 55 | 49 | 70 |
| lolA_lolA | 1274 | 1228 | 1169 | 1087 | 1017 | 1304 |
| lolB_lolB | 274 | 258 | 288 | 273 | 266 | 308 |
| lplA_lplA | 524 | 543 | 488 | 516 | 467 | 545 |
| SL1344_RS06600_lpxR | 1587 | 1473 | 1614 | 1924 | 1484 | 2043 |
| SL1344_RS11955_lrhA | 638 | 592 | 717 | 446 | 396 | 478 |
| SL1344_RS04660_lrp | 4103 | 3609 | 3462 | 3361 | 3246 | 3715 |
| SL1344_RS09120_L-serine ammonia-lyase | 3002 | 3059 | 3498 | 4283 | 3990 | 4601 |
| SL1344_RS20930_lsrR | 1672 | 1671 | 1476 | 1869 | 2063 | 2074 |
| SL1344_RS14590_luxS | 1824 | 1853 | 2044 | 1791 | 1939 | 2041 |
| SL1344_RS09795_lysine-N-methylase | 2483 | 2708 | 2872 | 3270 | 3241 | 3850 |
| malF_malF | 473 | 493 | 516 | 331 | 305 | 323 |
| malG_malG | 310 | 378 | 326 | 189 | 155 | 187 |
| SL1344_RS18135_malT | 4859 | 5320 | 5629 | 3292 | 3169 | 3444 |
| SL1344_RS07280_malY* | 64 | 72 | 84 | 70 | 95 | 78 |
| mdoD_mdoD | 1363 | 1319 | 1456 | 1377 | 1292 | 1460 |
| mdoG_mdoG | 2836 | 2734 | 2810 | 2902 | 2901 | 3365 |
| SL1344_RS09375_membrane protein | 765 | 739 | 766 | 1374 | 1170 | 1419 |
| SL1344_RS21870_membrane protein | 8291 | 7632 | 8010 | 4623 | 3684 | 5743 |
| SL1344_RS11835_menC | 1250 | 1407 | 1356 | 963 | 903 | 1142 |
| SL1344_RS21470_metA | 188 | 167 | 151 | 132 | 145 | 134 |
| SL1344_RS21065_metB | 288 | 277 | 242 | 257 | 251 | 279 |
| SL1344_RS16345_metC | 360 | 319 | 294 | 245 | 278 | 341 |
| SL1344_RS20385_metE | 397 | 364 | 376 | 348 | 398 | 359 |
| SL1344_RS21095_metF | 112 | 105 | 99 | 104 | 122 | 123 |
| metG_metG | 1441 | 1455 | 1546 | 1411 | 1409 | 1602 |
| SL1344_RS21500_metH | 1085 | 1001 | 1066 | 1006 | 839 | 1079 |
| SL1344_RS18435_methyl-accepting chemotaxis citrate transducer | 4935 | 5007 | 4683 | 10465 | 7791 | 11584 |
| SL1344_RS08105_methyl-accepting chemotaxis protein | 4996 | 4716 | 4457 | 12458 | 10531 | 15251 |
| SL1344_RS16230_methyl-accepting chemotaxis protein | 5656 | 5129 | 5632 | 9021 | 7370 | 9790 |
| SL1344_RS16300_methyl-accepting chemotaxis protein | 1598 | 1661 | 1870 | 5145 | 3971 | 6122 |
| SL1344_RS16625_methyl-accepting chemotaxis protein | 6484 | 6183 | 6243 | 20475 | 15323 | 24408 |
| SL1344_RS21060_metJ | 890 | 838 | 767 | 1054 | 951 | 979 |
| SL1344_RS21070_metL | 1026 | 973 | 1091 | 1002 | 908 | 1087 |
| metQ_metQ | 1957 | 1949 | 2075 | 1986 | 1946 | 2147 |
| SL1344_RS20380_metR | 78 | 73 | 72 | 95 | 92 | 110 |
| MgrR_mgrR | 526 | 770 | 1482 | 1100 | 597 | 670 |
| MicA_micA | 244 | 267 | 284 | 224 | 99 | 170 |
| MicC_micC | 31 | 31 | 27 | 46 | 32 | 35 |
| MicF_micF | 457 | 366 | 457 | 379 | 330 | 336 |
| SL1344_RS07395_mlc | 1295 | 1206 | 1084 | 844 | 849 | 939 |
| moaA_moaA | 1027 | 1039 | 1225 | 1564 | 1426 | 1416 |
| moaD_moaD | 527 | 542 | 565 | 565 | 558 | 675 |
| moaE_moaE | 760 | 747 | 890 | 870 | 876 | 966 |
| mobA_mobA | 389 | 318 | 241 | 276 | 253 | 290 |
| SL1344_RS24715_mobA | 1064 | 1232 | 1316 | 5855 | 5795 | 6000 |
| SL1344_RS24720_mobC | 416 | 508 | 346 | 1777 | 1915 | 2147 |
| SL1344_RS03955_modA | 1884 | 1826 | 1953 | 4679 | 3536 | 5042 |
| modB_modB | 1675 | 1627 | 1624 | 4049 | 3239 | 4252 |
| SL1344_RS03965_modC | 1618 | 1666 | 1620 | 3888 | 3383 | 4548 |
| SL1344_RS09620_motA | 4194 | 4174 | 3766 | 15477 | 12079 | 17324 |

| | | | | | | |
|---|---|---|---|---|---|---|
| motB_motB | 4412 | 4208 | 4034 | 15878 | 13149 | 16728 |
| SL1344_RS09450_msbB | 493 | 556 | 564 | 806 | 709 | 787 |
| Msr_Msr | 58 | 88 | 115 | 118 | 69 | 59 |
| mtlR_mtlR | 526 | 570 | 524 | 505 | 431 | 491 |
| mukB_mukB | 6065 | 6334 | 6295 | 8853 | 8091 | 9291 |
| murD_murD | 2120 | 2194 | 2083 | 2153 | 1988 | 2380 |
| SL1344_RS00625_murE | 1677 | 1824 | 1578 | 1425 | 1225 | 1524 |
| SL1344_RS09280_N-acetyltransferase | 90 | 95 | 110 | 198 | 176 | 207 |
| napB_napB | 3119 | 3015 | 2930 | 1922 | 1435 | 2045 |
| SL1344_RS08805_narG | 3027 | 1713 | 3998 | 10632 | 11692 | 12052 |
| SL1344_RS08800_narH | 2568 | 1597 | 3271 | 6426 | 6468 | 6946 |
| SL1344_RS08790_narI | 2267 | 1648 | 2647 | 3916 | 3311 | 3685 |
| SL1344_RS08795_narJ | 1154 | 673 | 1514 | 2376 | 2212 | 2482 |
| SL1344_RS08810_narK | 1331 | 727 | 2094 | 5339 | 5898 | 6358 |
| SL1344_RS08820_narL | 301 | 290 | 334 | 478 | 541 | 576 |
| nepI_nepI | 35 | 50 | 39 | 53 | 54 | 42 |
| nhaA_nhaA | 924 | 871 | 906 | 968 | 813 | 1056 |
| nirD_nirD | 292 | 251 | 305 | 301 | 276 | 340 |
| nlpD_nlpD | 10977 | 9971 | 10913 | 13438 | 13632 | 14091 |
| nrdI_nrdI | 6 | 5 | 5 | 5 | 4 | 11 |
| SL1344_RS21955_nrfB | 789 | 738 | 675 | 800 | 574 | 723 |
| SL1344_RS21960_nrfC | 882 | 859 | 891 | 878 | 644 | 853 |
| SL1344_RS21975_nrfG | 256 | 313 | 267 | 250 | 141 | 200 |
| nudC_nudC | 262 | 255 | 250 | 211 | 232 | 236 |
| nusA_nusA | 3172 | 3141 | 3527 | 3787 | 3685 | 4349 |
| nusB_nusB | 2800 | 2805 | 3034 | 2836 | 2949 | 3037 |
| nusG_nusG | 3076 | 2752 | 3086 | 4562 | 4352 | 4460 |
| obgE_obgE | 610 | 665 | 622 | 699 | 690 | 819 |
| ompL_ompL | 7 | 8 | 6 | 12 | 18 | 8 |
| SL1344_RS18070_ompR | 1375 | 1475 | 1480 | 1107 | 973 | 1202 |
| ompX_ompX | 6771 | 7003 | 7519 | 7210 | 7324 | 8354 |
| OmrA_omrA | 41 | 47 | 79 | 85 | 60 | 47 |
| OmrB_omrB | 15 | 20 | 29 | 28 | 21 | 24 |
| oppB_oppB | 756 | 725 | 776 | 731 | 725 | 783 |
| SL1344_RS14860_orgA | 2993 | 2840 | 3022 | 5432 | 4161 | 5300 |
| OxyS_oxyS | 56 | 31 | 42 | 55 | 75 | 42 |
| pabB_pabB | 499 | 488 | 508 | 787 | 734 | 819 |
| pagP_pagP | 440 | 469 | 483 | 483 | 452 | 444 |
| SL1344_RS10525_pduA | 10 | 20 | 18 | 6 | 6 | 11 |
| pduC_pduC | 83 | 93 | 83 | 29 | 18 | 19 |
| pduD_pduD | 24 | 46 | 42 | 5 | 10 | 10 |
| pduE_pduE | 13 | 27 | 26 | 8 | 8 | 13 |
| pdxK_pdxK | 129 | 130 | 96 | 114 | 92 | 126 |
| SL1344_RS06590_pfkB | 2316 | 2403 | 2095 | 1059 | 979 | 1272 |
| SL1344_RS13960_phage virion morphogenesis protein | 5 | 3 | 5 | 11 | 6 | 12 |
| SL1344_RS02035_phoB | 694 | 699 | 600 | 755 | 664 | 751 |
| SL1344_RS11645_phoE | 45528 | 46403 | 57058 | 40051 | 40947 | 46519 |
| SL1344_RS05535_phoH | 1552 | 1708 | 1519 | 1787 | 1604 | 2073 |
| SL1344_RS06080_phoP | 7973 | 7855 | 8264 | 6766 | 6682 | 8123 |
| SL1344_RS06075_phoQ | 4307 | 4255 | 4736 | 4288 | 3626 | 4136 |
| phoR_phoR | 495 | 500 | 452 | 553 | 503 | 555 |
| SL1344_RS13700_phospho-2-dehydro-3-deoxyheptonate aldolase Tyr-sensitive | 356 | 328 | 321 | 244 | 254 | 261 |
| SL1344_RS10625_phsA | 2749 | 2677 | 1781 | 131 | 116 | 126 |
| SL1344_RS10620_phsB | 658 | 631 | 412 | 39 | 44 | 46 |
| SL1344_RS10615_phsC | 1328 | 1198 | 878 | 101 | 88 | 89 |
| potG_potG | 74 | 69 | 85 | 69 | 71 | 115 |
| SL1344_RS09215_prc | 3585 | 3479 | 3757 | 5687 | 5704 | 6478 |
| SL1344_RS14880_prgH | 4507 | 4248 | 4412 | 6476 | 5508 | 8189 |
| SL1344_RS14875_prgI | 2025 | 1922 | 2104 | 3005 | 2505 | 3698 |
| SL1344_RS14870_prgJ | 3410 | 2988 | 3265 | 5276 | 4106 | 5857 |
| SL1344_RS14865_prgK | 5208 | 5247 | 5709 | 8602 | 6839 | 9562 |
| SL1344_RS09220_proQ | 2056 | 2116 | 2176 | 3624 | 3800 | 4103 |

| | | | | | | |
|---|---|---|---|---|---|---|
| SL1344_RS09255_prpA/pphA | 59 | 66 | 56 | 88 | 66 | 102 |
| psiB_psiB | 50 | 41 | 46 | 38 | 37 | 42 |
| psiB_psiB | 35 | 28 | 24 | 33 | 44 | 39 |
| pssA_pssA | 1328 | 1470 | 1607 | 1581 | 1382 | 1483 |
| SL1344_RS11995_pta | 6185 | 6576 | 7185 | 6152 | 5225 | 6668 |
| SL1344_RS15255_ptpS | 38 | 43 | 49 | 91 | 84 | 100 |
| SL1344_RS02910_PTS mannose/fructose/sorbose transporter subunit IIB | 5 | 1 | 6 | 2 | 5 | 0 |
| SL1344_RS07105_purR | 364 | 377 | 470 | 420 | 434 | 518 |
| putA_putA | 11134 | 10722 | 13492 | 12384 | 11755 | 13613 |
| SL1344_RS23405_putative inner membrane protein | 3462 | 3416 | 3352 | 2679 | 2623 | 3182 |
| SL1344_RS09445_pyruvate kinase II | 2973 | 2988 | 3214 | 4667 | 3795 | 4549 |
| SL1344_RS16430_qseB | 82 | 69 | 62 | 50 | 50 | 58 |
| SL1344_RS16435_qseC | 138 | 122 | 130 | 109 | 102 | 108 |
| SL1344_RS23490_radA | 561 | 597 | 597 | 540 | 563 | 605 |
| RaoN_raoN | 26 | 37 | 40 | 33 | 37 | 38 |
| SL1344_RS12895_ratA | 519 | 510 | 478 | 485 | 458 | 526 |
| rcsA_rcsA | 44 | 43 | 33 | 70 | 59 | 67 |
| SL1344_RS11655_rcsB | 1344 | 1386 | 1455 | 1397 | 1425 | 1553 |
| SL1344_RS11660_rcsC | 697 | 751 | 753 | 670 | 593 | 755 |
| SL1344_RS11650_rcsD | 1314 | 1288 | 1390 | 1204 | 1143 | 1258 |
| rcsF_rcsF | 447 | 495 | 478 | 563 | 467 | 525 |
| RdlD_rdlD | 60 | 63 | 66 | 99 | 69 | 73 |
| RdlE_rdlE | 14 | 19 | 41 | 28 | 30 | 22 |
| SL1344_RS14650_recA | 4378 | 4325 | 4030 | 4026 | 3616 | 4723 |
| SL1344_RS13835_recN | 729 | 804 | 843 | 1329 | 1274 | 1360 |
| SL1344_RS06620_rfc | 522 | 537 | 617 | 444 | 420 | 453 |
| SL1344_RS22270_rflM | 183 | 215 | 229 | 612 | 504 | 659 |
| rhtB_rhtB | 44 | 46 | 61 | 48 | 48 | 54 |
| rimM_rimM | 3818 | 3490 | 4824 | 4961 | 5595 | 6562 |
| rimO_rimO | 327 | 352 | 376 | 400 | 413 | 470 |
| SL1344_RS03220_rlpA | 607 | 734 | 756 | 665 | 554 | 738 |
| SL1344_RS05840_rne | 5145 | 5225 | 5292 | 5496 | 5199 | 5715 |
| rnpB_rnpB | 354304 | 355491 | 380126 | 378564 | 430166 | 443316 |
| RnpB_RnpB | 354304 | 355491 | 380126 | 378564 | 430166 | 443316 |
| SL1344_RS15130_rpoS | 11139 | 11068 | 10056 | 9707 | 8927 | 11130 |
| rprA_rprA | 147 | 157 | 202 | 175 | 95 | 181 |
| rpsN_rpsN | 7148 | 7494 | 9310 | 8721 | 9012 | 10486 |
| rpsV_rpsV | 4416 | 4825 | 4538 | 4854 | 4651 | 5019 |
| RseX_rseX | 5 | 4 | 10 | 9 | 6 | 10 |
| rsmD_rsmD | 210 | 217 | 265 | 185 | 212 | 245 |
| SL1344_RS09240_rsmF/yebU | 181 | 192 | 265 | 361 | 365 | 335 |
| SL1344_RS22150_rtsA | 2997 | 2971 | 2756 | 5297 | 3856 | 4581 |
| SL1344_RS22145_rtsB | 828 | 650 | 606 | 1147 | 835 | 900 |
| rumA_rumA | 1683 | 1849 | 1796 | 1534 | 1536 | 1660 |
| SL1344_RS09480_ruvA | 273 | 281 | 265 | 651 | 634 | 623 |
| SL1344_RS09475_ruvB | 656 | 626 | 632 | 1083 | 1083 | 1160 |
| RybA_rybA | 1697 | 1385 | 1351 | 1796 | 2026 | 2250 |
| RybB_rybB | 232 | 229 | 306 | 248 | 112 | 169 |
| RybD_rybD | 2214 | 2064 | 3337 | 2791 | 2134 | 2354 |
| RydB_rydB | 29 | 17 | 31 | 56 | 38 | 28 |
| RydC_rydC | 136 | 158 | 250 | 261 | 191 | 218 |
| RyeF_ryeF | 231 | 210 | 227 | 406 | 298 | 352 |
| RyfA_ryfA | 1437 | 1479 | 1758 | 1941 | 1737 | 1971 |
| RyfD_ryfD | 2064 | 1876 | 2914 | 2508 | 2064 | 2031 |
| RyhB_ryhB | 29 | 21 | 26 | 28 | 20 | 22 |
| RyjB_ryjB | 38 | 30 | 40 | 45 | 26 | 50 |
| Sar_sar | 133 | 167 | 361 | 228 | 118 | 146 |
| SL1344_RS10635_sbcB | 528 | 648 | 630 | 590 | 560 | 574 |
| SL1344_RS01925_sbmA | 184 | 176 | 202 | 177 | 158 | 175 |
| SL1344_RS10605_sbmC | 4012 | 3815 | 3700 | 3653 | 4289 | 4717 |
| SL1344_RS20885_sbp | 107 | 140 | 156 | 2934 | 10597 | 9263 |
| sdhC_sdhC | 2037 | 1954 | 2415 | 3157 | 3117 | 3688 |

| | | | | | | |
|---|---|---|---|---|---|---|
| SL1344_RS09755_sdiA | 944 | 1011 | 964 | 3264 | 2829 | 4009 |
| SdsR_sdsR | 395 | 502 | 833 | 437 | 316 | 356 |
| sgbH_sgbH | 6 | 8 | 5 | 12 | 7 | 11 |
| SgrS_sgrS | 3623 | 4124 | 4525 | 5473 | 4668 | 6202 |
| SibA_sibA | 473 | 727 | 1427 | 1346 | 612 | 636 |
| SibC_sibC | 854 | 829 | 1179 | 1881 | 1348 | 1137 |
| SibD_sibD | 945 | 1439 | 3314 | 3564 | 1743 | 1726 |
| SL1344_RS14940_sicA | 2165 | 1961 | 1866 | 3695 | 3097 | 4984 |
| sifA_sifA | 117 | 116 | 96 | 121 | 131 | 98 |
| sifB_sifB | 6 | 3 | 6 | 13 | 18 | 13 |
| SL1344_RS05355_sigE | 3159 | 2742 | 2878 | 5904 | 5245 | 7443 |
| SL1344_RS21850_siiA | 565 | 546 | 574 | 255 | 212 | 326 |
| SL1344_RS21855_siiB | 1053 | 934 | 1081 | 538 | 401 | 632 |
| SL1344_RS21875_siiF | 767 | 637 | 679 | 797 | 795 | 811 |
| SL1344_RS01560_sinR | 472 | 365 | 346 | 544 | 430 | 555 |
| SL1344_RS14920_sipA | 2934 | 2740 | 2583 | 4832 | 3598 | 6459 |
| SL1344_RS14935_sipB | 5635 | 5376 | 4894 | 9765 | 7372 | 13326 |
| SL1344_RS14930_sipC | 7764 | 7459 | 7580 | 15256 | 12060 | 20954 |
| SL1344_RS14925_sipD | 1315 | 1204 | 1332 | 2336 | 1710 | 3054 |
| SL1344_RS09740_sirA | 1551 | 1697 | 1664 | 2173 | 2052 | 2482 |
| SL1344_RS10425_sirohydrochlorin cobaltochelatase | 648 | 708 | 726 | 367 | 296 | 364 |
| SL1344_RS09840_SL1896 | 163 | 139 | 148 | 204 | 118 | 147 |
| SL1344_RS07175_slyA | 3043 | 2931 | 2776 | 2861 | 2534 | 3195 |
| SL1344_RS07110_sodB | 11397 | 12322 | 12834 | 7226 | 5623 | 8759 |
| SL1344_RS10630_sopA | 478 | 461 | 511 | 1036 | 708 | 1142 |
| sopB_sopB | 3202 | 2912 | 3133 | 6446 | 4999 | 7873 |
| SL1344_RS15235_sopD | 282 | 249 | 261 | 561 | 415 | 622 |
| SL1344_RS13925_sopE | 2985 | 2789 | 2503 | 6559 | 4803 | 7440 |
| SL1344_RS09265_sopE2 | 681 | 622 | 636 | 2235 | 1634 | 2440 |
| spaK_spaK | 793 | 703 | 789 | 1184 | 817 | 1239 |
| spaM_spaM | 1115 | 1073 | 908 | 1355 | 934 | 1648 |
| spaN_spaN | 1853 | 1981 | 1791 | 2585 | 1742 | 3259 |
| SL1344_RS14965_spaO | 1583 | 1508 | 1500 | 2267 | 1470 | 2666 |
| SL1344_RS14960_spaP | 969 | 905 | 920 | 1600 | 1096 | 1788 |
| SL1344_RS14955_spaQ | 737 | 665 | 708 | 1183 | 912 | 1398 |
| SL1344_RS14950_spaR | 512 | 486 | 468 | 904 | 680 | 889 |
| SL1344_RS14945_spaS | 487 | 468 | 445 | 790 | 514 | 814 |
| Spf_spf | 1785 | 1422 | 1512 | 2718 | 2657 | 2152 |
| SL1344_RS14840_spi4 transcriptional regulator | 1463 | 1412 | 1529 | 2136 | 1528 | 2277 |
| spiC_spiC | 5 | 12 | 4 | 10 | 3 | 3 |
| spr_spr | 5340 | 4820 | 4947 | 8045 | 6915 | 7526 |
| SraA_sraA | 5 | 5 | 5 | 8 | 9 | 2 |
| SraB_sraB | 99 | 111 | 131 | 203 | 135 | 156 |
| SraC_sraC | 685 | 650 | 646 | 1821 | 1869 | 1683 |
| SraF_sraF | 190 | 153 | 180 | 387 | 372 | 297 |
| SraG_sraG | 10 | 11 | 12 | 19 | 16 | 19 |
| SraL_sraL | 331 | 360 | 355 | 288 | 219 | 204 |
| SL1344_RS07945_srfB | 3003 | 3090 | 3296 | 5870 | 5495 | 6318 |
| SL1344_RS07950_srfC | 2054 | 2277 | 2266 | 4215 | 4148 | 4538 |
| srlA_srlA | 166 | 163 | 172 | 163 | 314 | 271 |
| srlE_srlE | 214 | 173 | 195 | 208 | 304 | 260 |
| srlR_srlR | 591 | 558 | 546 | 547 | 602 | 645 |
| SroA_sroA | 23 | 14 | 37 | 36 | 26 | 24 |
| SroC_sroC | 23257 | 23428 | 23334 | 31680 | 42398 | 34311 |
| SroD_sroD | 70 | 69 | 67 | 172 | 114 | 175 |
| SroG_sroG | 1842 | 1809 | 1664 | 1665 | 1008 | 1668 |
| SL1344_RS06985_ssaG | 5 | 9 | 9 | 9 | 15 | 12 |
| SL1344_RS07050_ssaR | 29 | 24 | 30 | 41 | 28 | 33 |
| SL1344_RS05160_sseI | 54 | 48 | 65 | 69 | 63 | 53 |
| sspA_sspA | 1837 | 1621 | 1762 | 2174 | 1954 | 2307 |
| SL1344_RS06915_ssrA | 254 | 253 | 275 | 239 | 235 | 268 |
| SsrA_ssrA | 501265 | 457323 | 530330 | 478744 | 460358 | 570213 |

| | | | | | | |
|---|---|---|---|---|---|---|
| SL1344_RS06910_ssrB | 120 | 132 | 132 | 121 | 110 | 120 |
| ssrS_ssrS | 119189 | 144358 | 151674 | 145083 | 148496 | 135005 |
| SsrS_ssrS | 119179 | 144338 | 151659 | 145064 | 148484 | 134994 |
| SL1344_RS16670_sstT | 1238 | 1250 | 1138 | 1224 | 1145 | 1367 |
| SL1344_RS01735_stbB | 2 | 0 | 4 | 3 | 10 | 2 |
| SL1344_RS05145_stfT | 52 | 67 | 51 | 70 | 80 | 69 |
| STnc3140_STnc3140 (sopD) | 3 | 20 | 22 | 241 | 206 | 180 |
| STnc3200_STnc3200 | 0 | 0 | 2 | 6 | 5 | 6 |
| SL1344_RS15350_syd | 241 | 206 | 223 | 240 | 237 | 248 |
| SymR_symR | 19 | 22 | 27 | 40 | 49 | 32 |
| SL1344_RS13990_tail protein | 0 | 5 | 2 | 4 | 4 | 1 |
| tatE_tatE | 436 | 358 | 362 | 425 | 387 | 516 |
| tcuB_tcuB | 41 | 35 | 35 | 49 | 43 | 47 |
| SL1344_RS16770_tdcA | 393 | 358 | 428 | 178 | 157 | 195 |
| SL1344_RS16765_tdcB | 304 | 334 | 315 | 260 | 218 | 304 |
| SL1344_RS16760_tdcC | 743 | 867 | 889 | 906 | 677 | 1000 |
| SL1344_RS05150_tfaT | 67 | 56 | 45 | 81 | 64 | 72 |
| Tff_tff | 1539 | 1388 | 1752 | 2198 | 2446 | 2411 |
| thiP_thiP | 275 | 297 | 321 | 266 | 263 | 280 |
| thiQ_thiQ | 84 | 102 | 110 | 105 | 78 | 109 |
| SL1344_RS06625_thrS | 8854 | 8836 | 9635 | 7061 | 6404 | 7538 |
| SL1344_RS04595_tnpA1 | 8 | 18 | 4 | 68 | 72 | 103 |
| SL1344_RS06060_tnpA2 | 22 | 26 | 19 | 60 | 48 | 68 |
| SL1344_RS09075_tnpA3 | 7 | 5 | 8 | 37 | 23 | 37 |
| SL1344_RS09790_tnpA4 | 5 | 3 | 3 | 11 | 9 | 19 |
| SL1344_RS12685_tnpA5 | 8 | 9 | 4 | 9 | 21 | 18 |
| SL1344_RS17020_tnpA6 | 17 | 21 | 21 | 44 | 38 | 38 |
| SL1344_RS17955_tnpA7 | 12 | 8 | 7 | 30 | 26 | 31 |
| tolA_tolA | 761 | 753 | 734 | 608 | 617 | 823 |
| SL1344_RS16475_tolC | 3741 | 4089 | 4504 | 3249 | 2910 | 3445 |
| SL1344_RS21860_TolC family protein | 1035 | 879 | 970 | 473 | 339 | 595 |
| tqsA_tqsA | 157 | 128 | 148 | 141 | 106 | 116 |
| SL1344_RS14145_transcriptional regulator | 3 | 1 | 1 | 5 | 2 | 5 |
| trkD_trkD | 653 | 705 | 756 | 684 | 637 | 689 |
| SL1344_RS06175_tRNA arg | 21 | 20 | 26 | 42 | 31 | 44 |
| SL1344_RS13600_tRNA/rRNA methyltransferase | 726 | 667 | 640 | 536 | 594 | 667 |
| truA_truA | 374 | 417 | 406 | 314 | 298 | 330 |
| SL1344_RS23260_tsr | 5906 | 6131 | 6099 | 18467 | 14369 | 21493 |
| SL1344_RS06870_ttrA | 100 | 102 | 102 | 96 | 84 | 81 |
| SL1344_RS06880_ttrB | 7 | 11 | 11 | 4 | 11 | 11 |
| SL1344_RS06875_ttrC | 11 | 11 | 19 | 13 | 11 | 9 |
| SL1344_RS06890_ttrR | 564 | 551 | 421 | 313 | 313 | 391 |
| SL1344_RS06885_ttrS | 1528 | 1653 | 1643 | 1095 | 1022 | 1099 |
| tyrA_tyrA | 419 | 378 | 400 | 307 | 341 | 355 |
| ubiB_ubiB | 1839 | 1881 | 2005 | 1811 | 1630 | 1901 |
| ubiF_ubiF | 904 | 969 | 884 | 684 | 662 | 851 |
| SL1344_RS21190_udhA | 1623 | 1689 | 2089 | 2458 | 2357 | 2582 |
| SL1344_RS03635_UDP-galactopyranose mutase | 11 | 9 | 11 | 10 | 9 | 18 |
| ugpC_ugpC | 205 | 221 | 202 | 156 | 127 | 157 |
| ulaD_ulaD | 63 | 64 | 89 | 60 | 48 | 63 |
| ushA_ushA | 2480 | 2242 | 2231 | 1804 | 1761 | 2083 |
| SL1344_RS09735_uvrC | 1405 | 1449 | 1442 | 1835 | 1747 | 2094 |
| SL1344_RS11530_virulence protein MsgA | 4 | 6 | 7 | 4 | 3 | 4 |
| wecC_wecC | 1097 | 1135 | 1166 | 952 | 1024 | 1110 |
| SL1344_RS10695_wzzB | 1598 | 1597 | 1635 | 839 | 712 | 842 |
| SL1344_RS02345_ybaO | 421 | 455 | 450 | 430 | 390 | 458 |
| SL1344_RS03185_ybeM | 96 | 75 | 98 | 128 | 81 | 116 |
| SL1344_RS04285_ybiK | 850 | 819 | 714 | 1556 | 2148 | 2167 |
| SL1344_RS08990_ycgO | 1476 | 1386 | 1460 | 2875 | 2254 | 3114 |
| SL1344_RS08975_ycgR | 744 | 754 | 637 | 1739 | 1560 | 2044 |
| SL1344_RS08585_yciK | 708 | 699 | 667 | 624 | 562 | 708 |
| SL1344_RS08495_yciW | 101 | 143 | 122 | 980 | 1495 | 1459 |

| | | | | | | |
|---|---|---|---|---|---|---|
| SL1344_RS07545_ydeD | 136 | 149 | 144 | 105 | 80 | 99 |
| SL1344_RS06680_ydiV | 111 | 108 | 108 | 86 | 76 | 76 |
| SL1344_RS06595_ydiY | 285 | 288 | 314 | 233 | 216 | 212 |
| SL1344_RS06585_ydiZ | 150 | 106 | 99 | 61 | 41 | 64 |
| SL1344_RS06560_ydjN | 420 | 491 | 959 | 5048 | 5342 | 6593 |
| SL1344_RS09415_yebG | 197 | 185 | 179 | 302 | 321 | 374 |
| SL1344_RS09225_yebR | 833 | 821 | 816 | 1438 | 1510 | 1526 |
| SL1344_RS09230_yebS | 471 | 488 | 430 | 649 | 577 | 706 |
| SL1344_RS09760_yecC | 444 | 473 | 496 | 1158 | 1071 | 1268 |
| SL1344_RS09750_yecF | 308 | 285 | 313 | 777 | 781 | 728 |
| SL1344_RS09765_yecS | 222 | 265 | 261 | 602 | 518 | 685 |
| SL1344_RS09825_yedD | 315 | 326 | 348 | 346 | 252 | 297 |
| SL1344_RS09830_yedE | 111 | 130 | 179 | 104 | 104 | 108 |
| SL1344_RS09835_yedF | 132 | 102 | 138 | 87 | 75 | 96 |
| SL1344_RS09770_yedO | 499 | 526 | 543 | 1189 | 1046 | 1366 |
| SL1344_RS10925_yegD | 59 | 76 | 101 | 67 | 69 | 77 |
| SL1344_RS13585_yfiK | 29 | 23 | 28 | 36 | 20 | 17 |
| SL1344_RS17960_yhfL | 173 | 157 | 227 | 572 | 481 | 552 |
| SL1344_RS18605_yhjH | 2089 | 1996 | 1800 | 5235 | 3486 | 5352 |
| SL1344_RS18650_yhjQ | 451 | 493 | 463 | 330 | 277 | 341 |
| SL1344_RS23255_yjiY | 109 | 100 | 131 | 102 | 94 | 103 |
| SL1344_RS05645_ymdA | 748 | 745 | 735 | 1627 | 1772 | 1700 |
| SL1344_RS05655_ymdC | 1320 | 1318 | 1318 | 2279 | 2049 | 2106 |
| yopJ_yopJ | 1505 | 1357 | 1196 | 1440 | 1159 | 1626 |
| SL1344_RS15335_yqcB | 242 | 229 | 207 | 284 | 284 | 269 |
| SL1344_RS15340_yqcC (next to csrB) | 77 | 92 | 110 | 118 | 116 | 119 |
| YrlA_yrlA | 11 | 7 | 11 | 12 | 8 | 13 |
| YrlB_yrlB | 13 | 10 | 22 | 17 | 12 | 18 |
| zntA_zntA | 756 | 806 | 920 | 995 | 1011 | 1143 |
| zntB_zntB | 358 | 346 | 322 | 346 | 338 | 401 |
| zraP_zraP | 47 | 41 | 36 | 37 | 37 | 41 |
| crRNA1_ | 141 | 192 | 131 | 173 | 165 | 152 |
| crRNA2_ | 449 | 509 | 543 | 384 | 354 | 390 |
| ECS021_ | 33 | 39 | 14 | 11 | 17 | 21 |
| ECS173_ | 33 | 45 | 49 | 51 | 26 | 42 |
| Ffs_ | 2344 | 2105 | 2596 | 2901 | 2225 | 2635 |
| fimA-trailer_ | 35 | 39 | 36 | 18 | 19 | 23 |
| GO-10702_ | 71 | 58 | 68 | 88 | 67 | 57 |
| GO-10704_ | 515 | 518 | 574 | 626 | 560 | 579 |
| Ig1085_ | 93 | 78 | 65 | 75 | 80 | 81 |
| Ig1140_ | 29 | 31 | 22 | 53 | 40 | 48 |
| Ig2335_ | 440 | 421 | 458 | 349 | 358 | 371 |
| Ig959_ | 1 | 0 | 0 | 0 | 1 | 0 |
| IS042_ | 11 | 11 | 6 | 24 | 12 | 7 |
| lysC-leader_ | 77 | 100 | 106 | 97 | 84 | 72 |
| mreB-leader_ | 240 | 249 | 250 | 386 | 271 | 333 |
| pRNA_ | 10 | 18 | 20 | 23 | 29 | 29 |
| RprA_ | 147 | 157 | 202 | 175 | 95 | 181 |
| rpsL-leader_ | 819 | 822 | 1095 | 1304 | 1224 | 1258 |
| rpsP-leader_ | 721 | 749 | 914 | 1018 | 1204 | 1230 |
| rrf_ | 5 | 1 | 9 | 0 | 0 | 1 |
| rrf_ | 0 | 4 | 28 | 2 | 2 | 8 |
| rrf_ | 6 | 4 | 21 | 2 | 7 | 16 |
| rrf_ | 2 | 2 | 7 | 5 | 2 | 6 |
| rrf_ | 6 | 26 | 114 | 8 | 17 | 61 |
| rrf_ | 2 | 2 | 10 | 1 | 0 | 5 |
| rrf_ | 0 | 1 | 3 | 0 | 2 | 0 |
| rrf_ | 1 | 0 | 8 | 0 | 0 | 0 |
| RUF_107c.1_ | 292 | 346 | 331 | 234 | 245 | 307 |
| RUF_107c.10_ | 238 | 257 | 282 | 435 | 337 | 315 |
| RUF_107c.102_ | 23 | 22 | 24 | 28 | 24 | 20 |
| RUF_107c.105_ | 179 | 166 | 165 | 176 | 150 | 168 |

| | | | | | | |
|---|---|---|---|---|---|---|
| RUF_107c.107_ | 77 | 90 | 83 | 105 | 89 | 64 |
| RUF_107c.108_ | 595 | 573 | 618 | 415 | 398 | 503 |
| RUF_107c.111_ | 760 | 729 | 898 | 765 | 640 | 726 |
| RUF_107c.112_ | 198 | 242 | 316 | 245 | 218 | 250 |
| RUF_107c.18_ | 17 | 28 | 16 | 48 | 37 | 18 |
| RUF_107c.19_ | 154 | 153 | 212 | 178 | 146 | 184 |
| RUF_107c.2_ | 32 | 24 | 48 | 25 | 30 | 28 |
| RUF_107c.23_ | 174 | 187 | 250 | 252 | 199 | 213 |
| RUF_107c.24_ | 490 | 480 | 435 | 337 | 324 | 380 |
| RUF_107c.26_ | 10 | 3 | 6 | 9 | 8 | 10 |
| RUF_107c.30_ | 2 | 3 | 0 | 5 | 2 | 6 |
| RUF_107c.38_ | 90 | 116 | 88 | 97 | 87 | 99 |
| RUF_107c.47_ | 196 | 185 | 276 | 238 | 152 | 185 |
| RUF_107c.50_ | 100 | 112 | 87 | 95 | 86 | 85 |
| RUF_107c.51_ | 138 | 118 | 134 | 87 | 102 | 113 |
| RUF_107c.53_ | 430 | 485 | 454 | 495 | 618 | 638 |
| RUF_107c.56_ | 240 | 267 | 340 | 236 | 243 | 255 |
| RUF_107c.57_ | 98 | 129 | 97 | 56 | 77 | 95 |
| RUF_107c.58_ | 519 | 555 | 798 | 613 | 446 | 541 |
| RUF_107c.59_ | 285 | 383 | 328 | 264 | 239 | 302 |
| RUF_107c.63_ | 102 | 130 | 180 | 128 | 108 | 109 |
| RUF_107c.69_ | 206 | 175 | 171 | 151 | 159 | 181 |
| RUF_107c.70_ | 41 | 51 | 49 | 49 | 53 | 62 |
| RUF_107c.72_ | 60 | 43 | 73 | 57 | 56 | 59 |
| RUF_107c.77_ | 82 | 62 | 89 | 73 | 64 | 72 |
| RUF_107c.8_ | 29 | 29 | 41 | 37 | 21 | 20 |
| RUF_107c.81_ | 9 | 10 | 7 | 7 | 11 | 13 |
| RUF_107c.82_ | 722 | 794 | 894 | 653 | 641 | 716 |
| RUF_107c.83_ | 5 | 10 | 22 | 12 | 11 | 14 |
| RUF_107c.86_ | 188 | 149 | 175 | 157 | 129 | 157 |
| RUF_107c.89_ | 223 | 205 | 220 | 241 | 227 | 200 |
| RUF_107c.90_ | 11 | 13 | 15 | 18 | 13 | 16 |
| RUF_107c.92_ | 762 | 748 | 678 | 705 | 635 | 735 |
| RUF_107c.94_ | 16 | 12 | 16 | 19 | 10 | 21 |
| RUF_107c.95_ | 749 | 786 | 838 | 712 | 792 | 873 |
| RUF_107c.96_ | 76 | 71 | 113 | 58 | 43 | 56 |
| RUF_111_ | 429 | 442 | 447 | 611 | 623 | 537 |
| RUF_121_ | 25 | 36 | 50 | 82 | 76 | 79 |
| RUF_125_ | 936 | 951 | 918 | 2754 | 2620 | 3372 |
| RUF_167_ | 103 | 83 | 118 | 143 | 92 | 91 |
| RUF_178_ | 646 | 691 | 673 | 856 | 792 | 748 |
| RUF_219c_ | 883 | 788 | 788 | 1268 | 860 | 1215 |
| RUF_220c_ | 274 | 233 | 230 | 515 | 368 | 458 |
| RUF_226_ | 47 | 55 | 77 | 74 | 40 | 69 |
| RUF_247c_ | 366 | 344 | 397 | 393 | 333 | 372 |
| RUF_257c_ | 337 | 293 | 305 | 287 | 241 | 264 |
| RUF_279_ | 535 | 527 | 629 | 511 | 496 | 494 |
| RUF_290_ | 485 | 476 | 449 | 428 | 437 | 469 |
| RUF_321c_ | 0 | 3 | 4 | 0 | 1 | 1 |
| RUF_328c_ | 1 | 1 | 0 | 2 | 1 | 0 |
| RUF_341c.10_ | 194 | 241 | 235 | 311 | 234 | 228 |
| RUF_341c.11_ | 301 | 327 | 343 | 312 | 300 | 283 |
| RUF_341c.2_ | 25 | 30 | 48 | 32 | 30 | 29 |
| SL1344_RS00005_ | 3228 | 2767 | 2140 | 2976 | 2795 | 2505 |
| SL1344_RS00010_ | 569 | 538 | 472 | 519 | 557 | 623 |
| SL1344_RS00015_ | 503 | 513 | 511 | 547 | 485 | 539 |
| SL1344_RS00020_ | 978 | 1021 | 1037 | 1027 | 768 | 1032 |
| SL1344_RS00025_ | 673 | 754 | 713 | 636 | 560 | 678 |
| SL1344_RS00030_ | 473 | 523 | 575 | 467 | 450 | 477 |
| SL1344_RS00035_ | 13074 | 12931 | 13927 | 9999 | 8913 | 11060 |
| SL1344_RS00040_ | 721 | 788 | 848 | 674 | 752 | 842 |
| SL1344_RS00045_ | 106 | 127 | 102 | 227 | 173 | 205 |

| | | | | | | |
|---|---|---|---|---|---|---|
| SL1344_RS00050_ | 18 | 13 | 11 | 16 | 15 | 17 |
| SL1344_RS00055_ | 18 | 14 | 25 | 19 | 23 | 16 |
| SL1344_RS00065_ | 2760 | 2701 | 2432 | 2489 | 2545 | 2845 |
| SL1344_RS00070_ | 52 | 39 | 46 | 50 | 40 | 52 |
| SL1344_RS00075_ | 8 | 1 | 5 | 3 | 6 | 5 |
| SL1344_RS00080_ | 6 | 12 | 8 | 11 | 16 | 12 |
| SL1344_RS00085_ | 5 | 5 | 7 | 9 | 5 | 5 |
| SL1344_RS00090_ | 148 | 144 | 128 | 195 | 164 | 149 |
| SL1344_RS00095_ | 109 | 101 | 95 | 120 | 92 | 125 |
| SL1344_RS00100_ | 39 | 44 | 55 | 77 | 73 | 83 |
| SL1344_RS00105_ | 44 | 63 | 43 | 58 | 48 | 55 |
| SL1344_RS00110_ | 9 | 7 | 6 | 23 | 8 | 12 |
| SL1344_RS00115_ | 69 | 48 | 64 | 50 | 53 | 66 |
| SL1344_RS00120_ | 20 | 33 | 34 | 42 | 35 | 29 |
| SL1344_RS00125_ | 14 | 12 | 12 | 21 | 16 | 13 |
| SL1344_RS00130_ | 22 | 24 | 15 | 11 | 20 | 21 |
| SL1344_RS00135_ | 18 | 17 | 20 | 23 | 11 | 23 |
| SL1344_RS00140_ | 46 | 42 | 56 | 45 | 41 | 48 |
| SL1344_RS00145_ | 35 | 26 | 45 | 48 | 42 | 41 |
| SL1344_RS00150_ | 1 | 1 | 4 | 6 | 0 | 1 |
| SL1344_RS00155_ | 18 | 10 | 27 | 24 | 20 | 22 |
| SL1344_RS00160_ | 8 | 9 | 9 | 9 | 9 | 12 |
| SL1344_RS00165_ | 82 | 79 | 86 | 80 | 88 | 83 |
| SL1344_RS00170_ | 171 | 165 | 177 | 146 | 135 | 147 |
| SL1344_RS00175_ | 3 | 5 | 4 | 3 | 6 | 3 |
| SL1344_RS00180_ | 253 | 252 | 237 | 253 | 262 | 264 |
| SL1344_RS00185_ | 100 | 88 | 103 | 91 | 82 | 112 |
| SL1344_RS00190_ | 32 | 41 | 48 | 34 | 45 | 44 |
| SL1344_RS00195_ | 198 | 211 | 198 | 253 | 252 | 211 |
| SL1344_RS00205_ | 863 | 839 | 876 | 893 | 911 | 1035 |
| SL1344_RS00210_ | 169 | 156 | 193 | 135 | 125 | 139 |
| SL1344_RS00215_ | 29 | 23 | 38 | 29 | 13 | 19 |
| SL1344_RS00220_ | 3884 | 3222 | 3714 | 8076 | 7411 | 6611 |
| SL1344_RS00225_ | 36 | 39 | 44 | 50 | 44 | 43 |
| SL1344_RS00230_ | 799 | 801 | 807 | 886 | 748 | 989 |
| SL1344_RS00235_ | 5275 | 5184 | 5567 | 4483 | 3752 | 4976 |
| SL1344_RS00240_ | 1241 | 1321 | 1484 | 1089 | 1039 | 1128 |
| SL1344_RS00245_ | 570 | 632 | 644 | 595 | 481 | 577 |
| SL1344_RS00250_ | 1200 | 1220 | 1265 | 1141 | 967 | 1242 |
| SL1344_RS00255_ | 543 | 570 | 474 | 659 | 676 | 621 |
| SL1344_RS00260_ | 879 | 820 | 857 | 1180 | 1194 | 1168 |
| SL1344_RS00270_ | 158 | 155 | 207 | 138 | 134 | 147 |
| SL1344_RS00275_ | 7 | 12 | 10 | 15 | 8 | 11 |
| SL1344_RS00280_ | 7 | 7 | 8 | 6 | 5 | 7 |
| SL1344_RS00285_ | 2 | 1 | 1 | 1 | 3 | 2 |
| SL1344_RS00290_ | 38 | 38 | 37 | 41 | 45 | 56 |
| SL1344_RS00295_ | 15 | 12 | 25 | 13 | 12 | 18 |
| SL1344_RS00300_ | 12 | 6 | 15 | 8 | 13 | 9 |
| SL1344_RS00305_ | 32 | 22 | 16 | 25 | 20 | 42 |
| SL1344_RS00310_ | 58 | 47 | 49 | 36 | 41 | 45 |
| SL1344_RS00315_ | 36 | 22 | 22 | 33 | 36 | 28 |
| SL1344_RS00320_ | 64 | 50 | 53 | 60 | 47 | 57 |
| SL1344_RS00325_ | 783 | 797 | 709 | 673 | 645 | 794 |
| SL1344_RS00330_ | 16 | 29 | 33 | 31 | 20 | 27 |
| SL1344_RS00335_ | 121 | 106 | 96 | 84 | 84 | 122 |
| SL1344_RS00340_ | 466 | 523 | 560 | 430 | 382 | 422 |
| SL1344_RS00345_ | 1097 | 1452 | 1062 | 569 | 472 | 650 |
| SL1344_RS00350_ | 24 | 26 | 26 | 35 | 26 | 38 |
| SL1344_RS00355_ | 62 | 58 | 88 | 86 | 68 | 98 |
| SL1344_RS00360_ | 55 | 43 | 50 | 59 | 52 | 56 |
| SL1344_RS00365_ | 100 | 119 | 103 | 100 | 110 | 104 |
| SL1344_RS00370_ | 205 | 214 | 214 | 215 | 215 | 204 |

| | | | | | | |
|---|---|---|---|---|---|---|
| SL1344_RS00375_ | 9 | 15 | 13 | 17 | 28 | 13 |
| SL1344_RS00380_ | 2 | 0 | 4 | 3 | 6 | 1 |
| SL1344_RS00385_ | 34 | 31 | 49 | 39 | 22 | 19 |
| SL1344_RS00390_ | 42 | 35 | 41 | 27 | 29 | 32 |
| SL1344_RS00395_ | 48 | 54 | 34 | 30 | 30 | 28 |
| SL1344_RS00400_ | 46 | 43 | 19 | 20 | 19 | 17 |
| SL1344_RS00405_ | 64 | 87 | 75 | 85 | 79 | 64 |
| SL1344_RS00410_ | 84 | 80 | 96 | 82 | 87 | 114 |
| SL1344_RS00415_ | 289 | 277 | 237 | 192 | 225 | 235 |
| SL1344_RS00420_ | 101 | 114 | 110 | 91 | 82 | 102 |
| SL1344_RS00425_ | 3 | 0 | 3 | 1 | 2 | 0 |
| SL1344_RS00430_ | 92 | 79 | 105 | 127 | 100 | 114 |
| SL1344_RS00435_ | 39 | 51 | 44 | 23 | 29 | 40 |
| SL1344_RS00440_ | 292 | 273 | 246 | 178 | 148 | 198 |
| SL1344_RS00450_ | 861 | 905 | 877 | 742 | 694 | 881 |
| SL1344_RS00455_ | 670 | 677 | 676 | 587 | 556 | 629 |
| SL1344_RS00465_ | 1188 | 1257 | 1431 | 1034 | 942 | 1073 |
| SL1344_RS00470_ | 2704 | 2894 | 2835 | 2194 | 2176 | 2437 |
| SL1344_RS00475_ | 2795 | 2835 | 2875 | 2278 | 2061 | 2370 |
| SL1344_RS00480_ | 595 | 634 | 511 | 326 | 415 | 498 |
| SL1344_RS00485_ | 317 | 349 | 377 | 311 | 319 | 346 |
| SL1344_RS00490_ | 583 | 633 | 776 | 687 | 704 | 810 |
| SL1344_RS00495_ | 290 | 358 | 365 | 392 | 377 | 370 |
| SL1344_RS00500_ | 38 | 51 | 44 | 59 | 46 | 57 |
| SL1344_RS00505_ | 11 | 13 | 13 | 15 | 7 | 10 |
| SL1344_RS00510_ | 43 | 48 | 38 | 47 | 49 | 29 |
| SL1344_RS00515_ | 71 | 58 | 42 | 34 | 46 | 51 |
| SL1344_RS00520_ | 34 | 32 | 32 | 23 | 35 | 30 |
| SL1344_RS00525_ | 595 | 532 | 526 | 496 | 455 | 526 |
| SL1344_RS00530_ | 386 | 392 | 392 | 312 | 277 | 276 |
| SL1344_RS00545_ | 173 | 212 | 190 | 186 | 158 | 195 |
| SL1344_RS00550_ | 185 | 196 | 201 | 188 | 178 | 240 |
| SL1344_RS00555_ | 3458 | 3991 | 4341 | 5173 | 4487 | 5960 |
| SL1344_RS00560_ | 91 | 88 | 106 | 98 | 96 | 120 |
| SL1344_RS00565_ | 122 | 122 | 111 | 132 | 116 | 148 |
| SL1344_RS00570_ | 108 | 98 | 90 | 126 | 122 | 147 |
| SL1344_RS00575_ | 81 | 73 | 85 | 71 | 66 | 97 |
| SL1344_RS00580_ | 178 | 207 | 294 | 292 | 223 | 183 |
| SL1344_RS00590_ | 230 | 233 | 239 | 229 | 202 | 246 |
| SL1344_RS00595_ | 292 | 296 | 260 | 265 | 210 | 241 |
| SL1344_RS00600_ | 1690 | 1537 | 1540 | 1487 | 1318 | 1494 |
| SL1344_RS00605_ | 2119 | 1907 | 1679 | 2424 | 2489 | 2404 |
| SL1344_RS00610_ | 2932 | 2940 | 2677 | 2831 | 2786 | 3220 |
| SL1344_RS00615_ | 1964 | 2075 | 1719 | 1737 | 1716 | 1882 |
| SL1344_RS00620_ | 9155 | 8973 | 9536 | 12772 | 11608 | 11331 |
| SL1344_RS00630_ | 1864 | 1922 | 2005 | 1780 | 1719 | 2046 |
| SL1344_RS00635_ | 1770 | 1800 | 1825 | 1650 | 1574 | 1766 |
| SL1344_RS00645_ | 1858 | 1914 | 1806 | 1722 | 1666 | 1974 |
| SL1344_RS00650_ | 1374 | 1550 | 1347 | 1440 | 1410 | 1577 |
| SL1344_RS00655_ | 1747 | 1792 | 1846 | 1914 | 1682 | 1884 |
| SL1344_RS00665_ | 1337 | 1343 | 1283 | 1182 | 1095 | 1244 |
| SL1344_RS00670_ | 2663 | 2579 | 2491 | 2451 | 2283 | 2783 |
| SL1344_RS00675_ | 4479 | 4295 | 4406 | 4056 | 3731 | 4896 |
| SL1344_RS00680_ | 10277 | 10886 | 11878 | 11020 | 10056 | 12265 |
| SL1344_RS00685_ | 693 | 728 | 754 | 889 | 798 | 913 |
| SL1344_RS00690_ | 4578 | 4836 | 4948 | 4614 | 4258 | 5054 |
| SL1344_RS00695_ | 103 | 108 | 104 | 103 | 59 | 104 |
| SL1344_RS00700_ | 260 | 283 | 302 | 299 | 252 | 245 |
| SL1344_RS00705_ | 532 | 555 | 491 | 480 | 478 | 554 |
| SL1344_RS00710_ | 493 | 469 | 530 | 491 | 443 | 522 |
| SL1344_RS00715_ | 366 | 381 | 425 | 519 | 408 | 449 |
| SL1344_RS00720_ | 2567 | 2599 | 2700 | 1910 | 1920 | 2092 |

| | | | | | | |
|---|---|---|---|---|---|---|
| SL1344_RS00725_ | 21 | 21 | 21 | 18 | 21 | 26 |
| SL1344_RS00730_ | 12 | 8 | 8 | 11 | 9 | 5 |
| SL1344_RS00735_ | 2 | 1 | 1 | 0 | 3 | 7 |
| SL1344_RS00740_ | 271 | 277 | 272 | 325 | 273 | 308 |
| SL1344_RS00745_ | 444 | 466 | 489 | 352 | 353 | 414 |
| SL1344_RS00750_ | 716 | 722 | 711 | 554 | 570 | 646 |
| SL1344_RS00755_ | 103 | 103 | 80 | 107 | 92 | 112 |
| SL1344_RS00760_ | 22 | 60 | 34 | 45 | 39 | 37 |
| SL1344_RS00765_ | 274 | 338 | 300 | 302 | 279 | 311 |
| SL1344_RS00770_ | 1052 | 949 | 1013 | 1894 | 1651 | 1835 |
| SL1344_RS00775_ | 8676 | 8087 | 9447 | 9779 | 9256 | 11117 |
| SL1344_RS00785_ | 11538 | 10737 | 12676 | 13393 | 12928 | 15011 |
| SL1344_RS00790_ | 13 | 26 | 10 | 17 | 26 | 35 |
| SL1344_RS00795_ | 21 | 17 | 35 | 13 | 28 | 22 |
| SL1344_RS00800_ | 69 | 86 | 85 | 76 | 66 | 78 |
| SL1344_RS00805_ | 8003 | 7817 | 8591 | 4558 | 3611 | 5301 |
| SL1344_RS00810_ | 612 | 601 | 653 | 742 | 730 | 620 |
| SL1344_RS00815_ | 1375 | 1454 | 1432 | 907 | 1013 | 1241 |
| SL1344_RS00820_ | 105 | 116 | 119 | 125 | 124 | 118 |
| SL1344_RS00825_ | 236 | 264 | 259 | 197 | 208 | 230 |
| SL1344_RS00830_ | 442 | 452 | 428 | 356 | 362 | 398 |
| SL1344_RS00835_ | 1197 | 1355 | 1200 | 1139 | 1205 | 1286 |
| SL1344_RS00840_ | 299 | 293 | 383 | 584 | 558 | 582 |
| SL1344_RS00845_ | 266 | 266 | 393 | 394 | 420 | 488 |
| SL1344_RS00850_ | 110 | 112 | 133 | 141 | 113 | 125 |
| SL1344_RS00855_ | 807 | 834 | 631 | 478 | 519 | 744 |
| SL1344_RS00860_ | 389 | 436 | 493 | 317 | 319 | 368 |
| SL1344_RS00865_ | 495 | 453 | 535 | 620 | 628 | 682 |
| SL1344_RS00870_ | 1447 | 1311 | 1467 | 1947 | 1952 | 2012 |
| SL1344_RS00875_ | 185 | 196 | 198 | 205 | 175 | 200 |
| SL1344_RS00880_ | 395 | 399 | 353 | 272 | 239 | 333 |
| SL1344_RS00885_ | 59 | 65 | 64 | 75 | 63 | 63 |
| SL1344_RS00890_ | 30 | 33 | 36 | 58 | 53 | 45 |
| SL1344_RS00895_ | 8 | 6 | 5 | 6 | 5 | 6 |
| SL1344_RS00900_ | 31 | 39 | 29 | 48 | 46 | 46 |
| SL1344_RS00905_ | 281 | 322 | 261 | 157 | 142 | 231 |
| SL1344_RS00910_ | 244 | 263 | 268 | 279 | 209 | 267 |
| SL1344_RS00915_ | 672 | 700 | 714 | 814 | 693 | 814 |
| SL1344_RS00920_ | 399 | 414 | 399 | 427 | 406 | 446 |
| SL1344_RS00925_ | 357 | 332 | 336 | 372 | 314 | 415 |
| SL1344_RS00930_ | 268 | 308 | 438 | 430 | 385 | 404 |
| SL1344_RS00935_ | 1300 | 1313 | 1437 | 1783 | 1687 | 1929 |
| SL1344_RS00940_ | 207 | 231 | 241 | 312 | 250 | 307 |
| SL1344_RS00945_ | 13135 | 12657 | 12146 | 12771 | 11712 | 14268 |
| SL1344_RS00950_ | 532 | 537 | 529 | 494 | 413 | 470 |
| SL1344_RS00955_ | 251 | 299 | 272 | 298 | 281 | 276 |
| SL1344_RS00960_ | 962 | 1033 | 1017 | 1028 | 972 | 1125 |
| SL1344_RS00965_ | 2268 | 2203 | 2171 | 2221 | 1955 | 2297 |
| SL1344_RS00970_ | 189 | 189 | 197 | 155 | 142 | 166 |
| SL1344_RS00975_ | 34 | 36 | 35 | 43 | 17 | 35 |
| SL1344_RS00980_ | 35 | 41 | 43 | 47 | 46 | 52 |
| SL1344_RS00985_ | 97 | 74 | 81 | 116 | 79 | 90 |
| SL1344_RS00990_ | 27 | 23 | 22 | 25 | 24 | 20 |
| SL1344_RS00995_ | 62 | 51 | 61 | 48 | 55 | 60 |
| SL1344_RS01000_ | 10 | 21 | 19 | 18 | 9 | 20 |
| SL1344_RS01005_ | 21 | 12 | 14 | 15 | 19 | 26 |
| SL1344_RS01010_ | 32 | 15 | 15 | 29 | 36 | 30 |
| SL1344_RS01015_ | 21 | 16 | 15 | 20 | 12 | 22 |
| SL1344_RS01020_ | 296 | 263 | 231 | 282 | 279 | 270 |
| SL1344_RS01025_ | 1120 | 1171 | 1304 | 1135 | 1013 | 1216 |
| SL1344_RS01030_ | 438 | 444 | 456 | 422 | 373 | 434 |
| SL1344_RS01035_ | 484 | 433 | 474 | 549 | 527 | 653 |

| | | | | | | |
|---|---|---|---|---|---|---|
| SL1344_RS01040_ | 149 | 165 | 140 | 228 | 198 | 212 |
| SL1344_RS01045_ | 281 | 283 | 322 | 358 | 315 | 360 |
| SL1344_RS01050_ | 712 | 877 | 882 | 905 | 953 | 1128 |
| SL1344_RS01055_ | 425 | 364 | 377 | 274 | 300 | 337 |
| SL1344_RS01060_ | 2365 | 2335 | 2524 | 1932 | 1840 | 2193 |
| SL1344_RS01065_ | 1118 | 1185 | 1196 | 740 | 807 | 1055 |
| SL1344_RS01070_ | 7327 | 8002 | 6878 | 6874 | 6458 | 7289 |
| SL1344_RS01075_ | 445 | 460 | 443 | 478 | 452 | 451 |
| SL1344_RS01080_ | 2590 | 2496 | 2512 | 2272 | 2446 | 2655 |
| SL1344_RS01085_ | 502 | 545 | 538 | 488 | 430 | 481 |
| SL1344_RS01090_ | 1057 | 942 | 1003 | 1063 | 1039 | 1098 |
| SL1344_RS01095_ | 10146 | 9900 | 11705 | 13940 | 15524 | 16624 |
| SL1344_RS01100_ | 15533 | 14936 | 17029 | 17995 | 20525 | 21914 |
| SL1344_RS01105_ | 1481 | 1340 | 1387 | 1362 | 1334 | 1639 |
| SL1344_RS01110_ | 5802 | 5523 | 5326 | 6067 | 6017 | 6895 |
| SL1344_RS01115_ | 667 | 583 | 708 | 700 | 676 | 777 |
| SL1344_RS01120_ | 548 | 519 | 537 | 687 | 486 | 642 |
| SL1344_RS01130_ | 3682 | 3567 | 3383 | 3697 | 3376 | 4000 |
| SL1344_RS01135_ | 7787 | 7790 | 7850 | 7125 | 6512 | 7943 |
| SL1344_RS01140_ | 8573 | 7655 | 8296 | 7118 | 7931 | 8858 |
| SL1344_RS01145_ | 4111 | 3936 | 4539 | 4886 | 4051 | 4470 |
| SL1344_RS01150_ | 884 | 791 | 857 | 1044 | 999 | 1157 |
| SL1344_RS01155_ | 2341 | 2320 | 2235 | 2727 | 2599 | 3095 |
| SL1344_RS01160_ | 1735 | 1709 | 1736 | 2455 | 2009 | 2291 |
| SL1344_RS01165_ | 684 | 729 | 709 | 852 | 752 | 845 |
| SL1344_RS01170_ | 2107 | 1994 | 2029 | 2286 | 2063 | 2569 |
| SL1344_RS01175_ | 1228 | 1182 | 1190 | 1327 | 1355 | 1461 |
| SL1344_RS01180_ | 245 | 256 | 278 | 303 | 256 | 310 |
| SL1344_RS01185_ | 653 | 559 | 606 | 538 | 434 | 533 |
| SL1344_RS01190_ | 258 | 253 | 239 | 176 | 199 | 266 |
| SL1344_RS01195_ | 393 | 417 | 414 | 351 | 355 | 403 |
| SL1344_RS01200_ | 2232 | 2261 | 2114 | 1824 | 1680 | 1944 |
| SL1344_RS01205_ | 1338 | 1234 | 1078 | 1092 | 1112 | 1323 |
| SL1344_RS01210_ | 130 | 159 | 167 | 192 | 157 | 151 |
| SL1344_RS01215_ | 275 | 319 | 328 | 273 | 217 | 265 |
| SL1344_RS01220_ | 504 | 464 | 474 | 430 | 398 | 490 |
| SL1344_RS01225_ | 2454 | 2379 | 2722 | 2532 | 2639 | 2970 |
| SL1344_RS01230_ | 491 | 455 | 481 | 636 | 477 | 565 |
| SL1344_RS01245_ | 289 | 325 | 336 | 336 | 345 | 335 |
| SL1344_RS01250_ | 331 | 323 | 310 | 406 | 385 | 431 |
| SL1344_RS01255_ | 221 | 198 | 192 | 183 | 153 | 228 |
| SL1344_RS01260_ | 369 | 357 | 636 | 213 | 283 | 580 |
| SL1344_RS01265_ | 1390 | 1350 | 1353 | 1631 | 1730 | 1832 |
| SL1344_RS01270_ | 1226 | 1080 | 1228 | 1426 | 1623 | 1847 |
| SL1344_RS01275_ | 872 | 680 | 1058 | 488 | 685 | 1155 |
| SL1344_RS01285_ | 50 | 50 | 75 | 84 | 70 | 84 |
| SL1344_RS01290_ | 272 | 292 | 284 | 271 | 207 | 300 |
| SL1344_RS01295_ | 143 | 144 | 120 | 114 | 117 | 112 |
| SL1344_RS01300_ | 60 | 65 | 67 | 53 | 51 | 77 |
| SL1344_RS01305_ | 637 | 642 | 659 | 569 | 519 | 534 |
| SL1344_RS01310_ | 190 | 189 | 155 | 148 | 97 | 134 |
| SL1344_RS01315_ | 2817 | 2761 | 2901 | 4466 | 3932 | 4204 |
| SL1344_RS01320_ | 204 | 190 | 187 | 185 | 203 | 201 |
| SL1344_RS01325_ | 210 | 188 | 229 | 179 | 172 | 198 |
| SL1344_RS01330_ | 156 | 126 | 130 | 155 | 150 | 157 |
| SL1344_RS01335_ | 582 | 691 | 547 | 588 | 595 | 672 |
| SL1344_RS01340_ | 210 | 198 | 308 | 273 | 207 | 233 |
| SL1344_RS01345_ | 46 | 64 | 56 | 59 | 47 | 39 |
| SL1344_RS01350_ | 22 | 33 | 35 | 28 | 22 | 33 |
| SL1344_RS01355_ | 69 | 73 | 76 | 69 | 63 | 67 |
| SL1344_RS01360_ | 18 | 25 | 36 | 24 | 34 | 26 |
| SL1344_RS01365_ | 72 | 58 | 74 | 60 | 57 | 68 |

| | | | | | | |
|---|---|---|---|---|---|---|
| SL1344_RS01370_ | 47 | 40 | 53 | 51 | 45 | 51 |
| SL1344_RS01375_ | 3 | 2 | 1 | 2 | 2 | 6 |
| SL1344_RS01380_ | 138 | 130 | 140 | 149 | 157 | 170 |
| SL1344_RS01385_ | 45 | 39 | 37 | 38 | 36 | 33 |
| SL1344_RS01390_ | 60 | 82 | 80 | 77 | 71 | 89 |
| SL1344_RS01395_ | 14 | 23 | 36 | 29 | 23 | 33 |
| SL1344_RS01400_ | 11 | 12 | 17 | 16 | 8 | 17 |
| SL1344_RS01405_ | 4 | 7 | 13 | 6 | 9 | 16 |
| SL1344_RS01410_ | 6 | 4 | 4 | 3 | 9 | 9 |
| SL1344_RS01415_ | 15 | 4 | 13 | 17 | 8 | 9 |
| SL1344_RS01420_ | 3 | 3 | 13 | 11 | 2 | 11 |
| SL1344_RS01425_ | 29 | 29 | 39 | 43 | 29 | 31 |
| SL1344_RS01430_ | 46 | 40 | 37 | 50 | 41 | 49 |
| SL1344_RS01435_ | 11 | 11 | 14 | 16 | 15 | 17 |
| SL1344_RS01440_ | 2 | 1 | 8 | 5 | 5 | 16 |
| SL1344_RS01445_ | 122 | 141 | 118 | 134 | 130 | 147 |
| SL1344_RS01450_ | 31 | 24 | 20 | 22 | 22 | 29 |
| SL1344_RS01455_ | 16 | 18 | 11 | 9 | 17 | 11 |
| SL1344_RS01460_ | 39 | 21 | 31 | 40 | 39 | 37 |
| SL1344_RS01465_ | 67 | 83 | 89 | 90 | 76 | 73 |
| SL1344_RS01470_ | 13 | 6 | 11 | 6 | 10 | 11 |
| SL1344_RS01475_ | 337 | 253 | 264 | 327 | 274 | 340 |
| SL1344_RS01480_ | 402 | 336 | 345 | 423 | 450 | 530 |
| SL1344_RS01485_ | 285 | 268 | 236 | 273 | 246 | 289 |
| SL1344_RS01490_ | 107 | 95 | 94 | 112 | 108 | 122 |
| SL1344_RS01495_ | 21 | 30 | 14 | 17 | 26 | 21 |
| SL1344_RS01500_ | 0 | 1 | 2 | 2 | 1 | 2 |
| SL1344_RS01505_ | 24 | 19 | 15 | 16 | 28 | 16 |
| SL1344_RS01515_ | 10 | 14 | 11 | 25 | 14 | 16 |
| SL1344_RS01520_ | 1 | 2 | 0 | 1 | 5 | 2 |
| SL1344_RS01525_ | 99 | 72 | 68 | 111 | 146 | 137 |
| SL1344_RS01530_ | 32 | 33 | 34 | 36 | 31 | 53 |
| SL1344_RS01535_ | 254 | 243 | 223 | 178 | 199 | 243 |
| SL1344_RS01540_ | 54 | 58 | 36 | 43 | 42 | 48 |
| SL1344_RS01545_ | 45 | 33 | 36 | 48 | 45 | 66 |
| SL1344_RS01550_ | 19 | 14 | 22 | 12 | 18 | 21 |
| SL1344_RS01565_ | 19 | 23 | 23 | 30 | 16 | 20 |
| SL1344_RS01570_ | 324 | 289 | 407 | 306 | 246 | 307 |
| SL1344_RS01575_ | 135 | 141 | 108 | 96 | 69 | 97 |
| SL1344_RS01580_ | 287 | 323 | 350 | 255 | 224 | 304 |
| SL1344_RS01590_ | 980 | 1010 | 1133 | 1008 | 948 | 1058 |
| SL1344_RS01595_ | 677 | 683 | 600 | 560 | 508 | 624 |
| SL1344_RS01600_ | 402 | 406 | 401 | 542 | 421 | 508 |
| SL1344_RS01605_ | 357 | 405 | 407 | 564 | 581 | 616 |
| SL1344_RS01610_ | 85 | 71 | 91 | 104 | 93 | 108 |
| SL1344_RS01615_ | 112 | 105 | 118 | 125 | 95 | 120 |
| SL1344_RS01620_ | 3970 | 3936 | 4312 | 2962 | 2608 | 3331 |
| SL1344_RS01625_ | 420 | 366 | 460 | 625 | 594 | 586 |
| SL1344_RS01635_ | 966 | 862 | 869 | 1024 | 889 | 1061 |
| SL1344_RS01640_ | 80 | 62 | 77 | 91 | 75 | 90 |
| SL1344_RS01645_ | 595 | 532 | 534 | 554 | 586 | 673 |
| SL1344_RS01650_ | 1665 | 1625 | 1565 | 1559 | 1463 | 1780 |
| SL1344_RS01655_ | 77 | 91 | 146 | 112 | 83 | 98 |
| SL1344_RS01660_ | 17 | 13 | 13 | 11 | 14 | 24 |
| SL1344_RS01665_ | 2 | 3 | 8 | 4 | 0 | 9 |
| SL1344_RS01670_ | 13 | 13 | 6 | 7 | 10 | 12 |
| SL1344_RS01675_ | 3379 | 3161 | 3195 | 2306 | 2384 | 2786 |
| SL1344_RS01680_ | 27 | 20 | 26 | 18 | 18 | 33 |
| SL1344_RS01685_ | 39 | 26 | 37 | 42 | 28 | 45 |
| SL1344_RS01690_ | 17 | 15 | 15 | 15 | 11 | 21 |
| SL1344_RS01695_ | 48 | 32 | 37 | 71 | 53 | 49 |
| SL1344_RS01700_ | 122 | 112 | 106 | 127 | 111 | 151 |

| | | | | | | |
|---|---|---|---|---|---|---|
| SL1344_RS01705_ | 192 | 190 | 160 | 163 | 180 | 183 |
| SL1344_RS01710_ | 39 | 32 | 41 | 46 | 41 | 30 |
| SL1344_RS01715_ | 24 | 15 | 16 | 30 | 22 | 30 |
| SL1344_RS01720_ | 55 | 46 | 43 | 53 | 54 | 51 |
| SL1344_RS01725_ | 33 | 49 | 40 | 44 | 30 | 53 |
| SL1344_RS01730_ | 93 | 96 | 99 | 75 | 52 | 64 |
| SL1344_RS01740_ | 4 | 2 | 1 | 9 | 5 | 2 |
| SL1344_RS01745_ | 127 | 114 | 89 | 133 | 112 | 118 |
| SL1344_RS01750_ | 48 | 46 | 51 | 59 | 54 | 48 |
| SL1344_RS01755_ | 173 | 178 | 166 | 163 | 137 | 161 |
| SL1344_RS01760_ | 320 | 348 | 300 | 341 | 282 | 348 |
| SL1344_RS01765_ | 136 | 151 | 135 | 171 | 112 | 174 |
| SL1344_RS01770_ | 12 | 2 | 7 | 13 | 4 | 9 |
| SL1344_RS01775_ | 5 | 5 | 4 | 4 | 11 | 11 |
| SL1344_RS01780_ | 6 | 5 | 2 | 13 | 8 | 14 |
| SL1344_RS01785_ | 60 | 50 | 51 | 50 | 63 | 55 |
| SL1344_RS01790_ | 17 | 43 | 22 | 21 | 28 | 25 |
| SL1344_RS01795_ | 35 | 55 | 35 | 36 | 44 | 46 |
| SL1344_RS01800_ | 7 | 7 | 12 | 5 | 14 | 9 |
| SL1344_RS01805_ | 90 | 88 | 78 | 76 | 53 | 65 |
| SL1344_RS01810_ | 42 | 45 | 42 | 20 | 30 | 36 |
| SL1344_RS01815_ | 109 | 146 | 117 | 129 | 97 | 92 |
| SL1344_RS01820_ | 17 | 19 | 19 | 13 | 24 | 21 |
| SL1344_RS01825_ | 127 | 111 | 144 | 163 | 136 | 177 |
| SL1344_RS01830_ | 457 | 432 | 525 | 502 | 418 | 547 |
| SL1344_RS01835_ | 1184 | 1175 | 1297 | 1148 | 1222 | 1411 |
| SL1344_RS01840_ | 60 | 40 | 50 | 39 | 22 | 32 |
| SL1344_RS01845_ | 120 | 112 | 125 | 81 | 80 | 90 |
| SL1344_RS01850_ | 320 | 357 | 272 | 253 | 230 | 286 |
| SL1344_RS01855_ | 69 | 73 | 62 | 40 | 40 | 44 |
| SL1344_RS01860_ | 125 | 105 | 107 | 137 | 113 | 88 |
| SL1344_RS01865_ | 127 | 90 | 107 | 103 | 91 | 134 |
| SL1344_RS01870_ | 574 | 666 | 568 | 489 | 441 | 437 |
| SL1344_RS01875_ | 2872 | 2842 | 2502 | 1987 | 1780 | 2108 |
| SL1344_RS01880_ | 178 | 154 | 190 | 234 | 193 | 220 |
| SL1344_RS01885_ | 34 | 32 | 29 | 26 | 41 | 33 |
| SL1344_RS01890_ | 44 | 36 | 48 | 38 | 36 | 44 |
| SL1344_RS01895_ | 46 | 61 | 76 | 63 | 51 | 80 |
| SL1344_RS01900_ | 113 | 115 | 135 | 111 | 120 | 125 |
| SL1344_RS01905_ | 1647 | 1691 | 1760 | 1645 | 1741 | 1856 |
| SL1344_RS01910_ | 491 | 444 | 428 | 464 | 432 | 513 |
| SL1344_RS01915_ | 75 | 64 | 49 | 57 | 46 | 65 |
| SL1344_RS01930_ | 223 | 221 | 202 | 168 | 135 | 164 |
| SL1344_RS01935_ | 86 | 84 | 90 | 53 | 38 | 59 |
| SL1344_RS01940_ | 674 | 660 | 760 | 635 | 660 | 626 |
| SL1344_RS01945_ | 781 | 778 | 741 | 763 | 766 | 842 |
| SL1344_RS01950_ | 94 | 53 | 90 | 65 | 45 | 69 |
| SL1344_RS01955_ | 60 | 39 | 70 | 50 | 44 | 49 |
| SL1344_RS01960_ | 3797 | 3459 | 3074 | 3557 | 3703 | 3816 |
| SL1344_RS01965_ | 433 | 524 | 443 | 312 | 282 | 343 |
| SL1344_RS01975_ | 1345 | 1287 | 1253 | 1163 | 1147 | 1361 |
| SL1344_RS01980_ | 153 | 138 | 154 | 190 | 158 | 196 |
| SL1344_RS01985_ | 336 | 352 | 305 | 350 | 336 | 381 |
| SL1344_RS01990_ | 594 | 544 | 534 | 477 | 468 | 491 |
| SL1344_RS01995_ | 345 | 364 | 331 | 322 | 386 | 393 |
| SL1344_RS02000_ | 1004 | 902 | 818 | 908 | 948 | 972 |
| SL1344_RS02005_ | 423 | 456 | 428 | 507 | 494 | 534 |
| SL1344_RS02010_ | 309 | 268 | 282 | 225 | 202 | 257 |
| SL1344_RS02015_ | 142 | 127 | 137 | 176 | 147 | 175 |
| SL1344_RS02020_ | 649 | 670 | 662 | 674 | 640 | 716 |
| SL1344_RS02025_ | 158 | 148 | 137 | 183 | 164 | 197 |
| SL1344_RS02030_ | 1 | 1 | 2 | 3 | 0 | 1 |

| | | | | | | |
|---|---|---|---|---|---|---|
| SL1344_RS02045_ | 970 | 1117 | 1028 | 1385 | 1167 | 1190 |
| SL1344_RS02050_ | 447 | 438 | 518 | 526 | 503 | 588 |
| SL1344_RS02055_ | 118 | 139 | 135 | 112 | 103 | 95 |
| SL1344_RS02060_ | 7477 | 8079 | 7882 | 6891 | 6834 | 8045 |
| SL1344_RS02065_ | 755 | 891 | 995 | 996 | 836 | 946 |
| SL1344_RS02070_ | 180 | 203 | 205 | 291 | 269 | 352 |
| SL1344_RS02075_ | 875 | 875 | 922 | 1077 | 998 | 1091 |
| SL1344_RS02080_ | 1701 | 1461 | 1682 | 1705 | 1592 | 1895 |
| SL1344_RS02085_ | 2960 | 3045 | 3321 | 3163 | 2970 | 3344 |
| SL1344_RS02090_ | 1503 | 1513 | 1690 | 1439 | 1151 | 1461 |
| SL1344_RS02095_ | 112 | 96 | 75 | 102 | 105 | 132 |
| SL1344_RS02100_ | 140 | 128 | 108 | 146 | 115 | 139 |
| SL1344_RS02105_ | 576 | 566 | 541 | 500 | 533 | 562 |
| SL1344_RS02110_ | 3144 | 3341 | 3631 | 3399 | 3187 | 3442 |
| SL1344_RS02115_ | 3005 | 3136 | 3090 | 3493 | 2676 | 4364 |
| SL1344_RS02120_ | 97 | 94 | 114 | 82 | 63 | 72 |
| SL1344_RS02125_ | 564 | 657 | 684 | 515 | 587 | 655 |
| SL1344_RS02130_ | 794 | 859 | 925 | 763 | 756 | 707 |
| SL1344_RS02135_ | 775 | 828 | 753 | 713 | 1075 | 1016 |
| SL1344_RS02145_ | 276 | 374 | 358 | 275 | 309 | 375 |
| SL1344_RS02150_ | 289 | 326 | 340 | 269 | 272 | 334 |
| SL1344_RS02155_ | 455 | 508 | 523 | 548 | 307 | 385 |
| SL1344_RS02160_ | 1344 | 1403 | 1646 | 1400 | 1193 | 1385 |
| SL1344_RS02165_ | 730 | 671 | 797 | 956 | 780 | 812 |
| SL1344_RS02170_ | 439 | 406 | 433 | 453 | 532 | 537 |
| SL1344_RS02175_ | 391 | 368 | 409 | 352 | 359 | 467 |
| SL1344_RS02180_ | 68 | 69 | 75 | 69 | 50 | 59 |
| SL1344_RS02185_ | 28 | 29 | 27 | 39 | 47 | 40 |
| SL1344_RS02190_ | 61 | 68 | 54 | 79 | 56 | 73 |
| SL1344_RS02195_ | 68 | 63 | 56 | 55 | 78 | 83 |
| SL1344_RS02200_ | 210 | 190 | 209 | 215 | 210 | 241 |
| SL1344_RS02205_ | 22 | 11 | 24 | 18 | 21 | 28 |
| SL1344_RS02210_ | 18 | 14 | 18 | 24 | 23 | 22 |
| SL1344_RS02215_ | 598 | 569 | 538 | 431 | 437 | 521 |
| SL1344_RS02220_ | 372 | 385 | 318 | 307 | 262 | 316 |
| SL1344_RS02225_ | 1687 | 1823 | 1523 | 1266 | 1335 | 1785 |
| SL1344_RS02230_ | 431 | 477 | 427 | 403 | 374 | 461 |
| SL1344_RS02235_ | 74 | 63 | 56 | 65 | 74 | 95 |
| SL1344_RS02240_ | 87 | 78 | 68 | 135 | 141 | 129 |
| SL1344_RS02245_ | 2303 | 2350 | 3115 | 5638 | 6148 | 6860 |
| SL1344_RS02250_ | 714 | 679 | 1113 | 1505 | 1767 | 2153 |
| SL1344_RS02255_ | 966 | 846 | 1478 | 2126 | 2426 | 2813 |
| SL1344_RS02260_ | 2672 | 2457 | 4225 | 6933 | 7380 | 8856 |
| SL1344_RS02265_ | 2058 | 1787 | 2895 | 6530 | 7085 | 7463 |
| SL1344_RS02270_ | 523 | 560 | 595 | 682 | 637 | 716 |
| SL1344_RS02275_ | 412 | 392 | 519 | 638 | 635 | 651 |
| SL1344_RS02280_ | 2423 | 2528 | 2083 | 1689 | 1771 | 1738 |
| SL1344_RS02285_ | 7995 | 7453 | 8404 | 9205 | 9838 | 11188 |
| SL1344_RS02300_ | 8223 | 7734 | 7803 | 7519 | 7654 | 9199 |
| SL1344_RS02305_ | 46894 | 46032 | 41931 | 36469 | 32358 | 45146 |
| SL1344_RS02310_ | 1916 | 1920 | 1890 | 1735 | 1581 | 1962 |
| SL1344_RS02315_ | 62 | 48 | 37 | 32 | 24 | 37 |
| SL1344_RS02320_ | 104 | 117 | 117 | 106 | 102 | 101 |
| SL1344_RS02325_ | 294 | 309 | 266 | 324 | 279 | 368 |
| SL1344_RS02330_ | 157 | 144 | 144 | 110 | 101 | 128 |
| SL1344_RS02335_ | 119 | 133 | 122 | 122 | 145 | 169 |
| SL1344_RS02350_ | 468 | 448 | 386 | 459 | 451 | 489 |
| SL1344_RS02355_ | 448 | 527 | 482 | 458 | 453 | 498 |
| SL1344_RS02360_ | 5 | 3 | 9 | 8 | 7 | 6 |
| SL1344_RS02365_ | 101 | 106 | 79 | 102 | 84 | 91 |
| SL1344_RS02370_ | 450 | 398 | 444 | 442 | 397 | 457 |
| SL1344_RS02375_ | 1234 | 1185 | 1337 | 1323 | 1095 | 1422 |

| | | | | | | |
|---|---|---|---|---|---|---|
| SL1344_RS02380_ | 195 | 225 | 227 | 189 | 190 | 217 |
| SL1344_RS02385_ | 409 | 404 | 438 | 420 | 370 | 449 |
| SL1344_RS02390_ | 72 | 86 | 73 | 41 | 65 | 68 |
| SL1344_RS02395_ | 245 | 286 | 246 | 163 | 201 | 227 |
| SL1344_RS02400_ | 667 | 499 | 646 | 1515 | 1171 | 1153 |
| SL1344_RS02405_ | 526 | 545 | 575 | 517 | 483 | 487 |
| SL1344_RS02420_ | 4449 | 4582 | 4788 | 4213 | 3816 | 4555 |
| SL1344_RS02425_ | 1623 | 1538 | 1586 | 1621 | 1542 | 1802 |
| SL1344_RS02430_ | 534 | 582 | 604 | 615 | 605 | 651 |
| SL1344_RS02435_ | 1144 | 1222 | 1331 | 1246 | 1191 | 1432 |
| SL1344_RS02440_ | 597 | 631 | 574 | 642 | 610 | 658 |
| SL1344_RS02445_ | 115 | 117 | 82 | 124 | 76 | 102 |
| SL1344_RS02450_ | 126 | 113 | 104 | 151 | 156 | 138 |
| SL1344_RS02455_ | 73 | 61 | 53 | 75 | 77 | 66 |
| SL1344_RS02460_ | 190 | 172 | 191 | 357 | 264 | 307 |
| SL1344_RS02465_ | 1112 | 1293 | 1381 | 1358 | 1273 | 1391 |
| SL1344_RS02470_ | 1274 | 1320 | 1263 | 1427 | 1376 | 1522 |
| SL1344_RS02475_ | 1357 | 1398 | 1551 | 1742 | 1778 | 1765 |
| SL1344_RS02480_ | 10891 | 10067 | 11029 | 11391 | 11131 | 12888 |
| SL1344_RS02485_ | 2461 | 2380 | 2481 | 2871 | 2828 | 3185 |
| SL1344_RS02490_ | 618 | 533 | 536 | 551 | 499 | 542 |
| SL1344_RS02495_ | 71 | 100 | 82 | 83 | 65 | 71 |
| SL1344_RS02500_ | 159 | 201 | 203 | 183 | 172 | 229 |
| SL1344_RS02505_ | 1348 | 1418 | 1322 | 1855 | 1655 | 1830 |
| SL1344_RS02510_ | 291 | 232 | 272 | 289 | 267 | 288 |
| SL1344_RS02520_ | 184 | 211 | 215 | 243 | 233 | 249 |
| SL1344_RS02525_ | 164 | 194 | 192 | 216 | 172 | 244 |
| SL1344_RS02530_ | 89 | 65 | 69 | 91 | 79 | 86 |
| SL1344_RS02540_ | 353 | 395 | 322 | 286 | 269 | 346 |
| SL1344_RS02545_ | 402 | 491 | 430 | 438 | 412 | 498 |
| SL1344_RS02550_ | 430 | 474 | 461 | 462 | 375 | 576 |
| SL1344_RS02555_ | 146 | 102 | 111 | 104 | 141 | 165 |
| SL1344_RS02560_ | 193 | 214 | 194 | 191 | 169 | 173 |
| SL1344_RS02565_ | 1505 | 1410 | 1473 | 1597 | 1516 | 1670 |
| SL1344_RS02570_ | 263 | 272 | 276 | 326 | 282 | 268 |
| SL1344_RS02575_ | 148 | 167 | 190 | 176 | 155 | 206 |
| SL1344_RS02580_ | 177 | 155 | 124 | 119 | 114 | 156 |
| SL1344_RS02585_ | 666 | 738 | 625 | 604 | 564 | 614 |
| SL1344_RS02590_ | 171 | 218 | 159 | 178 | 160 | 202 |
| SL1344_RS02595_ | 96 | 101 | 112 | 134 | 109 | 154 |
| SL1344_RS02600_ | 83 | 101 | 81 | 113 | 97 | 127 |
| SL1344_RS02605_ | 116 | 145 | 130 | 138 | 142 | 175 |
| SL1344_RS02610_ | 786 | 852 | 783 | 834 | 876 | 833 |
| SL1344_RS02615_ | 62 | 69 | 65 | 58 | 54 | 60 |
| SL1344_RS02620_ | 153 | 156 | 123 | 112 | 107 | 123 |
| SL1344_RS02625_ | 856 | 831 | 865 | 806 | 783 | 947 |
| SL1344_RS02630_ | 64 | 64 | 65 | 47 | 37 | 48 |
| SL1344_RS02635_ | 49 | 33 | 34 | 24 | 19 | 31 |
| SL1344_RS02640_ | 59 | 75 | 54 | 50 | 37 | 55 |
| SL1344_RS02645_ | 31 | 38 | 22 | 32 | 36 | 37 |
| SL1344_RS02650_ | 19 | 18 | 12 | 10 | 15 | 20 |
| SL1344_RS02655_ | 123 | 126 | 120 | 140 | 135 | 132 |
| SL1344_RS02660_ | 27 | 16 | 20 | 21 | 26 | 23 |
| SL1344_RS02665_ | 46 | 43 | 29 | 51 | 34 | 48 |
| SL1344_RS02670_ | 259 | 254 | 318 | 254 | 220 | 256 |
| SL1344_RS02675_ | 75 | 63 | 62 | 45 | 40 | 65 |
| SL1344_RS02680_ | 50 | 36 | 50 | 45 | 32 | 35 |
| SL1344_RS02685_ | 52 | 39 | 42 | 57 | 52 | 49 |
| SL1344_RS02690_ | 23 | 21 | 29 | 20 | 22 | 29 |
| SL1344_RS02695_ | 11 | 14 | 25 | 8 | 13 | 12 |
| SL1344_RS02700_ | 36 | 39 | 42 | 59 | 40 | 38 |
| SL1344_RS02705_ | 295 | 335 | 387 | 451 | 426 | 429 |

| | | | | | | |
|---|---|---|---|---|---|---|
| SL1344_RS02710_ | 88 | 97 | 93 | 149 | 141 | 161 |
| SL1344_RS02715_ | 631 | 491 | 644 | 791 | 660 | 694 |
| SL1344_RS02720_ | 3116 | 2953 | 3175 | 3557 | 3627 | 3664 |
| SL1344_RS02730_ | 107 | 81 | 98 | 116 | 80 | 114 |
| SL1344_RS02735_ | 60 | 37 | 58 | 66 | 70 | 70 |
| SL1344_RS02740_ | 31 | 20 | 28 | 37 | 36 | 45 |
| SL1344_RS02745_ | 538 | 571 | 571 | 600 | 668 | 735 |
| SL1344_RS02750_ | 869 | 798 | 904 | 910 | 878 | 935 |
| SL1344_RS02760_ | 39 | 58 | 45 | 28 | 25 | 27 |
| SL1344_RS02765_ | 22 | 21 | 26 | 15 | 20 | 19 |
| SL1344_RS02770_ | 65 | 66 | 58 | 48 | 32 | 41 |
| SL1344_RS02775_ | 42 | 53 | 37 | 52 | 46 | 48 |
| SL1344_RS02790_ | 5 | 4 | 3 | 3 | 3 | 3 |
| SL1344_RS02795_ | 1 | 4 | 4 | 15 | 6 | 4 |
| SL1344_RS02800_ | 24 | 14 | 22 | 30 | 40 | 15 |
| SL1344_RS02805_ | 15 | 30 | 35 | 43 | 13 | 19 |
| SL1344_RS02815_ | 54 | 44 | 81 | 148 | 107 | 111 |
| SL1344_RS02820_ | 17 | 18 | 41 | 75 | 57 | 59 |
| SL1344_RS02825_ | 5 | 2 | 6 | 20 | 10 | 22 |
| SL1344_RS02830_ | 2 | 1 | 7 | 4 | 9 | 7 |
| SL1344_RS02835_ | 2 | 1 | 8 | 7 | 2 | 5 |
| SL1344_RS02840_ | 140 | 97 | 98 | 120 | 100 | 116 |
| SL1344_RS02845_ | 91 | 102 | 103 | 79 | 76 | 97 |
| SL1344_RS02850_ | 96 | 77 | 82 | 97 | 83 | 94 |
| SL1344_RS02855_ | 6 | 2 | 7 | 2 | 2 | 6 |
| SL1344_RS02860_ | 19 | 11 | 18 | 19 | 26 | 21 |
| SL1344_RS02865_ | 923 | 940 | 949 | 1787 | 1179 | 1892 |
| SL1344_RS02870_ | 336 | 334 | 383 | 432 | 408 | 403 |
| SL1344_RS02875_ | 468 | 428 | 428 | 393 | 399 | 430 |
| SL1344_RS02880_ | 364 | 379 | 400 | 358 | 308 | 333 |
| SL1344_RS02885_ | 582 | 662 | 601 | 502 | 469 | 553 |
| SL1344_RS02890_ | 23 | 19 | 36 | 27 | 34 | 31 |
| SL1344_RS02895_ | 14 | 9 | 14 | 16 | 15 | 14 |
| SL1344_RS02900_ | 8 | 8 | 9 | 6 | 10 | 17 |
| SL1344_RS02905_ | 7 | 5 | 8 | 4 | 7 | 4 |
| SL1344_RS02915_ | 5 | 2 | 7 | 9 | 2 | 5 |
| SL1344_RS02920_ | 729 | 816 | 789 | 756 | 706 | 842 |
| SL1344_RS02925_ | 126 | 116 | 119 | 128 | 112 | 123 |
| SL1344_RS02930_ | 253 | 199 | 211 | 250 | 250 | 279 |
| SL1344_RS02935_ | 119 | 82 | 93 | 85 | 115 | 96 |
| SL1344_RS02940_ | 58 | 49 | 67 | 65 | 61 | 63 |
| SL1344_RS02945_ | 111 | 114 | 90 | 85 | 67 | 81 |
| SL1344_RS02950_ | 9 | 8 | 10 | 10 | 11 | 11 |
| SL1344_RS02955_ | 145 | 142 | 122 | 101 | 107 | 111 |
| SL1344_RS02960_ | 37 | 31 | 26 | 52 | 24 | 30 |
| SL1344_RS02965_ | 1 | 2 | 8 | 5 | 9 | 2 |
| SL1344_RS02975_ | 486 | 504 | 517 | 451 | 391 | 425 |
| SL1344_RS02980_ | 9 | 11 | 14 | 7 | 13 | 14 |
| SL1344_RS02985_ | 31 | 20 | 41 | 45 | 35 | 46 |
| SL1344_RS02990_ | 16 | 25 | 26 | 28 | 24 | 24 |
| SL1344_RS02995_ | 48 | 35 | 39 | 47 | 54 | 51 |
| SL1344_RS03000_ | 44 | 46 | 53 | 48 | 51 | 41 |
| SL1344_RS03005_ | 34 | 28 | 33 | 27 | 25 | 33 |
| SL1344_RS03020_ | 39 | 32 | 34 | 35 | 37 | 34 |
| SL1344_RS03025_ | 18 | 27 | 24 | 26 | 24 | 19 |
| SL1344_RS03030_ | 3637 | 3371 | 3927 | 3411 | 2960 | 3735 |
| SL1344_RS03035_ | 476 | 416 | 421 | 409 | 378 | 419 |
| SL1344_RS03040_ | 114 | 111 | 101 | 131 | 145 | 128 |
| SL1344_RS03045_ | 158 | 165 | 151 | 224 | 231 | 216 |
| SL1344_RS03050_ | 205 | 204 | 190 | 186 | 202 | 220 |
| SL1344_RS03055_ | 120 | 134 | 124 | 175 | 159 | 205 |
| SL1344_RS03060_ | 240 | 244 | 286 | 242 | 272 | 262 |

| | | | | | | |
|---|---|---|---|---|---|---|
| SL1344_RS03065_ | 437 | 442 | 447 | 507 | 452 | 559 |
| SL1344_RS03070_ | 5240 | 5575 | 4985 | 5120 | 5301 | 5943 |
| SL1344_RS03075_ | 818 | 837 | 783 | 867 | 810 | 939 |
| SL1344_RS03080_ | 7 | 2 | 4 | 1 | 5 | 8 |
| SL1344_RS03085_ | 94 | 106 | 87 | 96 | 94 | 77 |
| SL1344_RS03090_ | 40 | 20 | 38 | 25 | 28 | 32 |
| SL1344_RS03095_ | 380 | 314 | 285 | 410 | 431 | 405 |
| SL1344_RS03100_ | 8013 | 8760 | 8604 | 7290 | 8105 | 8818 |
| SL1344_RS03105_ | 103 | 100 | 91 | 78 | 72 | 72 |
| SL1344_RS03110_ | 328 | 334 | 324 | 407 | 324 | 419 |
| SL1344_RS03115_ | 1537 | 1568 | 1861 | 1589 | 1533 | 1794 |
| SL1344_RS03120_ | 82 | 96 | 92 | 231 | 209 | 151 |
| SL1344_RS03125_ | 9 | 7 | 6 | 10 | 8 | 1 |
| SL1344_RS03130_ | 8 | 8 | 2 | 6 | 6 | 3 |
| SL1344_RS03135_ | 13 | 6 | 12 | 12 | 8 | 12 |
| SL1344_RS03140_ | 6 | 2 | 7 | 7 | 10 | 5 |
| SL1344_RS03145_ | 2 | 4 | 0 | 5 | 0 | 1 |
| SL1344_RS03150_ | 4 | 5 | 8 | 7 | 11 | 12 |
| SL1344_RS03165_ | 211 | 262 | 235 | 146 | 128 | 141 |
| SL1344_RS03175_ | 8774 | 8108 | 9446 | 14598 | 12824 | 12277 |
| SL1344_RS03180_ | 200 | 163 | 138 | 117 | 136 | 166 |
| SL1344_RS03195_ | 1142 | 1163 | 1360 | 1511 | 1434 | 1611 |
| SL1344_RS03200_ | 129 | 141 | 155 | 126 | 77 | 85 |
| SL1344_RS03205_ | 186 | 184 | 226 | 331 | 304 | 331 |
| SL1344_RS03210_ | 1758 | 1580 | 1588 | 2216 | 2361 | 2272 |
| SL1344_RS03215_ | 901 | 926 | 1145 | 1111 | 1031 | 1192 |
| SL1344_RS03225_ | 468 | 460 | 436 | 404 | 385 | 495 |
| SL1344_RS03230_ | 625 | 633 | 667 | 678 | 625 | 670 |
| SL1344_RS03235_ | 420 | 353 | 374 | 494 | 487 | 572 |
| SL1344_RS03240_ | 348 | 337 | 347 | 431 | 443 | 486 |
| SL1344_RS03245_ | 56 | 61 | 55 | 94 | 102 | 96 |
| SL1344_RS03250_ | 199 | 155 | 175 | 149 | 161 | 158 |
| SL1344_RS03255_ | 263 | 225 | 263 | 323 | 264 | 285 |
| SL1344_RS03260_ | 225 | 274 | 299 | 214 | 242 | 280 |
| SL1344_RS03265_ | 730 | 815 | 951 | 932 | 814 | 918 |
| SL1344_RS03270_ | 1381 | 1501 | 1596 | 1456 | 1420 | 1657 |
| SL1344_RS03275_ | 3835 | 3855 | 3948 | 3895 | 3752 | 4227 |
| SL1344_RS03280_ | 6 | 2 | 8 | 5 | 7 | 9 |
| SL1344_RS03285_ | 47 | 39 | 51 | 46 | 44 | 38 |
| SL1344_RS03290_ | 102 | 117 | 130 | 98 | 106 | 104 |
| SL1344_RS03295_ | 822 | 823 | 819 | 736 | 653 | 708 |
| SL1344_RS03300_ | 1153 | 1249 | 1138 | 480 | 503 | 644 |
| SL1344_RS03305_ | 58 | 74 | 54 | 60 | 52 | 67 |
| SL1344_RS03310_ | 15 | 20 | 30 | 27 | 10 | 14 |
| SL1344_RS03315_ | 48 | 55 | 46 | 52 | 51 | 39 |
| SL1344_RS03320_ | 22 | 17 | 23 | 15 | 20 | 27 |
| SL1344_RS03325_ | 94 | 69 | 71 | 73 | 61 | 104 |
| SL1344_RS03330_ | 354 | 390 | 384 | 396 | 337 | 457 |
| SL1344_RS03335_ | 485 | 477 | 454 | 530 | 512 | 610 |
| SL1344_RS03340_ | 432 | 476 | 530 | 544 | 527 | 701 |
| SL1344_RS03350_ | 166 | 178 | 218 | 337 | 287 | 335 |
| SL1344_RS03355_ | 175 | 176 | 213 | 414 | 393 | 400 |
| SL1344_RS03360_ | 19803 | 19696 | 18828 | 26565 | 36441 | 29914 |
| SL1344_RS03365_ | 1108 | 1147 | 1232 | 1272 | 1160 | 1331 |
| SL1344_RS03370_ | 1528 | 1614 | 1722 | 1724 | 1676 | 1830 |
| SL1344_RS03375_ | 1014 | 1073 | 985 | 993 | 873 | 1023 |
| SL1344_RS03380_ | 1373 | 1388 | 1378 | 1490 | 1494 | 1567 |
| SL1344_RS03385_ | 1149 | 1217 | 1292 | 1370 | 1389 | 1557 |
| SL1344_RS03395_ | 31 | 34 | 43 | 47 | 35 | 37 |
| SL1344_RS03400_ | 773 | 677 | 812 | 1027 | 796 | 874 |
| SL1344_RS03405_ | 520 | 560 | 815 | 831 | 640 | 727 |
| SL1344_RS03410_ | 631 | 581 | 667 | 741 | 783 | 802 |

| | | | | | | |
|---|---|---|---|---|---|---|
| SL1344_RS03415_ | 775 | 857 | 1180 | 1142 | 967 | 1043 |
| SL1344_RS03420_ | 1639 | 1738 | 2261 | 2308 | 1943 | 2137 |
| SL1344_RS03425_ | 1667 | 1629 | 2134 | 2427 | 1931 | 2045 |
| SL1344_RS03430_ | 871 | 924 | 1135 | 1555 | 1291 | 1179 |
| SL1344_RS03440_ | 506 | 519 | 534 | 659 | 603 | 670 |
| SL1344_RS03445_ | 707 | 786 | 802 | 727 | 640 | 786 |
| SL1344_RS03450_ | 1021 | 988 | 1079 | 1045 | 1055 | 1088 |
| SL1344_RS03455_ | 616 | 610 | 594 | 482 | 484 | 555 |
| SL1344_RS03460_ | 3509 | 3424 | 3271 | 2598 | 2502 | 2975 |
| SL1344_RS03465_ | 2248 | 2202 | 2660 | 2562 | 2431 | 2871 |
| SL1344_RS03475_ | 293 | 274 | 291 | 144 | 201 | 206 |
| SL1344_RS03480_ | 91 | 110 | 91 | 100 | 84 | 97 |
| SL1344_RS03490_ | 64 | 57 | 57 | 50 | 63 | 58 |
| SL1344_RS03495_ | 164 | 175 | 204 | 201 | 152 | 200 |
| SL1344_RS03505_ | 1225 | 1046 | 1116 | 2060 | 1722 | 1647 |
| SL1344_RS03510_ | 30 | 56 | 53 | 58 | 71 | 55 |
| SL1344_RS03515_ | 268 | 316 | 348 | 342 | 325 | 398 |
| SL1344_RS03520_ | 692 | 663 | 695 | 838 | 797 | 863 |
| SL1344_RS03525_ | 4364 | 4473 | 4704 | 4219 | 4079 | 4602 |
| SL1344_RS03530_ | 370 | 423 | 462 | 438 | 523 | 473 |
| SL1344_RS03535_ | 1016 | 1232 | 1594 | 977 | 1094 | 1271 |
| SL1344_RS03540_ | 1404 | 1532 | 2038 | 1368 | 1641 | 1811 |
| SL1344_RS03545_ | 3 | 2 | 4 | 2 | 1 | 6 |
| SL1344_RS03550_ | 192 | 187 | 182 | 192 | 150 | 146 |
| SL1344_RS03555_ | 339 | 396 | 363 | 366 | 332 | 378 |
| SL1344_RS03560_ | 46 | 56 | 52 | 43 | 62 | 57 |
| SL1344_RS03565_ | 38 | 30 | 30 | 30 | 43 | 43 |
| SL1344_RS03570_ | 13 | 15 | 14 | 13 | 16 | 36 |
| SL1344_RS03575_ | 1 | 0 | 0 | 1 | 0 | 0 |
| SL1344_RS03580_ | 454 | 583 | 589 | 464 | 530 | 599 |
| SL1344_RS03585_ | 191 | 202 | 206 | 182 | 142 | 181 |
| SL1344_RS03590_ | 103 | 94 | 96 | 94 | 94 | 101 |
| SL1344_RS03595_ | 573 | 551 | 601 | 468 | 412 | 543 |
| SL1344_RS03600_ | 437 | 464 | 393 | 344 | 372 | 351 |
| SL1344_RS03605_ | 557 | 577 | 580 | 473 | 441 | 508 |
| SL1344_RS03610_ | 424 | 446 | 463 | 415 | 352 | 428 |
| SL1344_RS03615_ | 8 | 7 | 8 | 3 | 7 | 9 |
| SL1344_RS03620_ | 45 | 72 | 49 | 105 | 78 | 83 |
| SL1344_RS03625_ | 7 | 13 | 6 | 12 | 12 | 15 |
| SL1344_RS03630_ | 29 | 19 | 28 | 27 | 22 | 38 |
| SL1344_RS03640_ | 5 | 4 | 7 | 13 | 11 | 6 |
| SL1344_RS03645_ | 12 | 16 | 19 | 18 | 18 | 22 |
| SL1344_RS03650_ | 14 | 6 | 13 | 9 | 19 | 12 |
| SL1344_RS03655_ | 12 | 8 | 14 | 11 | 16 | 9 |
| SL1344_RS03660_ | 22 | 19 | 31 | 29 | 32 | 24 |
| SL1344_RS03665_ | 12 | 6 | 11 | 14 | 12 | 10 |
| SL1344_RS03670_ | 28 | 19 | 31 | 48 | 32 | 24 |
| SL1344_RS03675_ | 67 | 74 | 108 | 92 | 78 | 84 |
| SL1344_RS03680_ | 320 | 321 | 348 | 405 | 409 | 451 |
| SL1344_RS03685_ | 256 | 244 | 238 | 241 | 248 | 267 |
| SL1344_RS03690_ | 8167 | 8248 | 9790 | 10487 | 10722 | 11675 |
| SL1344_RS03695_ | 20 | 8 | 15 | 13 | 24 | 18 |
| SL1344_RS03705_ | 1549 | 1395 | 1960 | 1834 | 1867 | 2255 |
| SL1344_RS03710_ | 5099 | 5379 | 6493 | 5752 | 4465 | 6448 |
| SL1344_RS03715_ | 7078 | 7861 | 8602 | 7467 | 7040 | 8746 |
| SL1344_RS03720_ | 15536 | 15531 | 18846 | 16714 | 16263 | 19764 |
| SL1344_RS03725_ | 15207 | 15096 | 16583 | 15146 | 14941 | 17523 |
| SL1344_RS03730_ | 10321 | 10091 | 10335 | 10245 | 9910 | 12300 |
| SL1344_RS03735_ | 12423 | 12781 | 14850 | 13876 | 12585 | 15354 |
| SL1344_RS03740_ | 1172 | 1138 | 1782 | 1568 | 1173 | 1192 |
| SL1344_RS03745_ | 29053 | 27950 | 30133 | 26449 | 23455 | 28470 |
| SL1344_RS03750_ | 14304 | 13483 | 14627 | 11762 | 8793 | 12859 |

| | | | | | | |
|---|---|---|---|---|---|---|
| SL1344_RS03755_ | 11249 | 10792 | 9972 | 8390 | 8015 | 9911 |
| SL1344_RS03760_ | 11766 | 11492 | 11820 | 10217 | 9108 | 11321 |
| SL1344_RS03765_ | 5446 | 5259 | 5803 | 6958 | 5769 | 6851 |
| SL1344_RS03770_ | 263 | 272 | 252 | 316 | 306 | 356 |
| SL1344_RS03775_ | 706 | 598 | 613 | 784 | 715 | 820 |
| SL1344_RS03780_ | 558 | 504 | 574 | 570 | 481 | 563 |
| SL1344_RS03790_ | 5205 | 4835 | 5372 | 4999 | 5530 | 6177 |
| SL1344_RS03795_ | 12379 | 12765 | 14777 | 13828 | 14851 | 15669 |
| SL1344_RS03800_ | 5052 | 5182 | 6702 | 6307 | 5808 | 6190 |
| SL1344_RS03805_ | 471 | 550 | 618 | 672 | 528 | 570 |
| SL1344_RS03810_ | 705 | 650 | 832 | 842 | 730 | 890 |
| SL1344_RS03815_ | 363 | 373 | 399 | 408 | 389 | 484 |
| SL1344_RS03820_ | 364 | 401 | 433 | 469 | 409 | 523 |
| SL1344_RS03825_ | 241 | 252 | 268 | 306 | 279 | 308 |
| SL1344_RS03830_ | 73 | 66 | 68 | 69 | 42 | 53 |
| SL1344_RS03835_ | 185 | 178 | 187 | 177 | 154 | 168 |
| SL1344_RS03840_ | 249 | 267 | 262 | 223 | 215 | 259 |
| SL1344_RS03845_ | 434 | 502 | 470 | 312 | 231 | 277 |
| SL1344_RS03850_ | 1120 | 1120 | 1031 | 1095 | 1110 | 1196 |
| SL1344_RS03855_ | 121 | 160 | 148 | 149 | 141 | 150 |
| SL1344_RS03860_ | 140 | 164 | 159 | 179 | 175 | 181 |
| SL1344_RS03865_ | 218 | 219 | 208 | 240 | 240 | 225 |
| SL1344_RS03870_ | 47 | 40 | 27 | 41 | 51 | 44 |
| SL1344_RS03875_ | 8 | 5 | 13 | 8 | 15 | 20 |
| SL1344_RS03880_ | 3 | 6 | 6 | 5 | 2 | 9 |
| SL1344_RS03885_ | 48 | 30 | 44 | 42 | 43 | 42 |
| SL1344_RS03890_ | 22 | 36 | 34 | 37 | 29 | 53 |
| SL1344_RS03895_ | 84 | 78 | 84 | 104 | 98 | 116 |
| SL1344_RS03900_ | 27 | 43 | 45 | 38 | 41 | 48 |
| SL1344_RS03905_ | 34 | 38 | 27 | 44 | 26 | 23 |
| SL1344_RS03915_ | 1133 | 1244 | 1355 | 1158 | 1102 | 1230 |
| SL1344_RS03920_ | 778 | 816 | 826 | 669 | 673 | 747 |
| SL1344_RS03925_ | 476 | 481 | 500 | 370 | 355 | 356 |
| SL1344_RS03930_ | 441 | 464 | 588 | 340 | 314 | 427 |
| SL1344_RS03935_ | 28 | 17 | 23 | 19 | 19 | 18 |
| SL1344_RS03940_ | 571 | 601 | 660 | 661 | 582 | 635 |
| SL1344_RS03945_ | 393 | 368 | 368 | 412 | 364 | 445 |
| SL1344_RS03950_ | 2605 | 2401 | 2085 | 3493 | 4412 | 3777 |
| SL1344_RS03970_ | 448 | 430 | 381 | 295 | 358 | 369 |
| SL1344_RS03975_ | 745 | 725 | 726 | 575 | 524 | 623 |
| SL1344_RS03980_ | 1736 | 1530 | 1810 | 1457 | 1653 | 1692 |
| SL1344_RS03985_ | 843 | 859 | 810 | 836 | 785 | 838 |
| SL1344_RS03990_ | 562 | 607 | 622 | 513 | 519 | 539 |
| SL1344_RS03995_ | 360 | 410 | 373 | 340 | 324 | 324 |
| SL1344_RS04000_ | 323 | 281 | 312 | 410 | 408 | 443 |
| SL1344_RS04005_ | 393 | 404 | 468 | 470 | 486 | 557 |
| SL1344_RS04010_ | 351 | 354 | 376 | 348 | 301 | 324 |
| SL1344_RS04015_ | 26 | 30 | 29 | 26 | 30 | 32 |
| SL1344_RS04020_ | 40 | 22 | 20 | 27 | 36 | 36 |
| SL1344_RS04025_ | 20 | 21 | 20 | 32 | 33 | 23 |
| SL1344_RS04030_ | 20 | 12 | 20 | 16 | 18 | 13 |
| SL1344_RS04035_ | 16 | 15 | 16 | 19 | 9 | 11 |
| SL1344_RS04040_ | 1008 | 973 | 1002 | 1141 | 980 | 1259 |
| SL1344_RS04045_ | 41 | 36 | 42 | 47 | 56 | 64 |
| SL1344_RS04050_ | 931 | 873 | 831 | 888 | 811 | 933 |
| SL1344_RS04055_ | 472 | 468 | 458 | 412 | 381 | 391 |
| SL1344_RS04065_ | 1318 | 1496 | 1565 | 1552 | 1502 | 1642 |
| SL1344_RS04070_ | 875 | 855 | 1031 | 1037 | 1004 | 1005 |
| SL1344_RS04085_ | 3725 | 3828 | 4204 | 3796 | 3750 | 4309 |
| SL1344_RS04090_ | 14 | 3 | 10 | 21 | 16 | 16 |
| SL1344_RS04095_ | 19 | 12 | 10 | 16 | 14 | 16 |
| SL1344_RS04100_ | 24 | 19 | 26 | 27 | 22 | 23 |

| | | | | | | |
|---|---|---|---|---|---|---|
| SL1344_RS04105_ | 134 | 116 | 148 | 124 | 87 | 105 |
| SL1344_RS04110_ | 100 | 97 | 105 | 86 | 71 | 83 |
| SL1344_RS04115_ | 40 | 52 | 44 | 47 | 35 | 37 |
| SL1344_RS04120_ | 2181 | 2142 | 1903 | 1719 | 1955 | 2055 |
| SL1344_RS04125_ | 323 | 316 | 283 | 272 | 221 | 258 |
| SL1344_RS04130_ | 322 | 367 | 336 | 254 | 252 | 313 |
| SL1344_RS04135_ | 608 | 671 | 627 | 498 | 434 | 501 |
| SL1344_RS04140_ | 407 | 413 | 393 | 343 | 297 | 376 |
| SL1344_RS04145_ | 315 | 274 | 252 | 264 | 264 | 252 |
| SL1344_RS04150_ | 69 | 57 | 79 | 78 | 68 | 98 |
| SL1344_RS04160_ | 360 | 373 | 363 | 390 | 380 | 441 |
| SL1344_RS04165_ | 55 | 56 | 67 | 75 | 41 | 71 |
| SL1344_RS04170_ | 15 | 13 | 18 | 18 | 10 | 11 |
| SL1344_RS04175_ | 163 | 190 | 187 | 126 | 127 | 123 |
| SL1344_RS04180_ | 148 | 152 | 161 | 169 | 136 | 202 |
| SL1344_RS04185_ | 160 | 174 | 179 | 141 | 139 | 162 |
| SL1344_RS04195_ | 249 | 256 | 316 | 399 | 360 | 456 |
| SL1344_RS04200_ | 1407 | 1382 | 1572 | 1976 | 2361 | 2639 |
| SL1344_RS04205_ | 6632 | 7772 | 7438 | 5617 | 5139 | 6081 |
| SL1344_RS04210_ | 111 | 128 | 132 | 138 | 146 | 134 |
| SL1344_RS04220_ | 199 | 185 | 218 | 225 | 232 | 266 |
| SL1344_RS04225_ | 121 | 102 | 130 | 166 | 147 | 158 |
| SL1344_RS04230_ | 238 | 252 | 290 | 292 | 287 | 327 |
| SL1344_RS04235_ | 365 | 368 | 424 | 556 | 482 | 524 |
| SL1344_RS04240_ | 411 | 436 | 472 | 621 | 516 | 658 |
| SL1344_RS04245_ | 19 | 21 | 26 | 20 | 24 | 27 |
| SL1344_RS04250_ | 5 | 4 | 7 | 5 | 6 | 5 |
| SL1344_RS04255_ | 39 | 52 | 53 | 57 | 59 | 66 |
| SL1344_RS04260_ | 282 | 266 | 300 | 388 | 377 | 352 |
| SL1344_RS04265_ | 89 | 82 | 90 | 72 | 69 | 63 |
| SL1344_RS04270_ | 26 | 30 | 25 | 19 | 39 | 19 |
| SL1344_RS04275_ | 379 | 369 | 391 | 361 | 362 | 446 |
| SL1344_RS04280_ | 332 | 317 | 332 | 340 | 322 | 401 |
| SL1344_RS04290_ | 1052 | 988 | 1072 | 1717 | 1947 | 2192 |
| SL1344_RS04295_ | 845 | 822 | 936 | 1276 | 1400 | 1633 |
| SL1344_RS04300_ | 513 | 544 | 466 | 890 | 1014 | 996 |
| SL1344_RS04305_ | 390 | 438 | 361 | 603 | 569 | 708 |
| SL1344_RS04320_ | 16 | 11 | 9 | 15 | 14 | 21 |
| SL1344_RS04325_ | 21 | 16 | 22 | 27 | 40 | 23 |
| SL1344_RS04330_ | 37 | 24 | 29 | 39 | 61 | 39 |
| SL1344_RS04335_ | 7 | 4 | 13 | 9 | 7 | 10 |
| SL1344_RS04340_ | 93 | 87 | 90 | 98 | 97 | 99 |
| SL1344_RS04345_ | 50 | 46 | 39 | 51 | 50 | 56 |
| SL1344_RS04350_ | 84 | 70 | 63 | 91 | 57 | 69 |
| SL1344_RS04355_ | 140 | 135 | 120 | 156 | 150 | 134 |
| SL1344_RS04360_ | 74 | 78 | 75 | 63 | 57 | 56 |
| SL1344_RS04365_ | 574 | 542 | 501 | 531 | 538 | 688 |
| SL1344_RS04370_ | 1168 | 1148 | 1050 | 828 | 769 | 859 |
| SL1344_RS04375_ | 502 | 545 | 575 | 696 | 637 | 650 |
| SL1344_RS04380_ | 322 | 337 | 391 | 320 | 279 | 351 |
| SL1344_RS04385_ | 232 | 253 | 264 | 212 | 218 | 253 |
| SL1344_RS04390_ | 171 | 190 | 243 | 190 | 184 | 199 |
| SL1344_RS04395_ | 187 | 182 | 185 | 212 | 223 | 223 |
| SL1344_RS04400_ | 50 | 37 | 51 | 63 | 38 | 55 |
| SL1344_RS04405_ | 486 | 479 | 523 | 421 | 354 | 380 |
| SL1344_RS04410_ | 62 | 58 | 60 | 46 | 22 | 35 |
| SL1344_RS04415_ | 1699 | 1590 | 1712 | 2219 | 1933 | 2179 |
| SL1344_RS04420_ | 71 | 56 | 99 | 69 | 62 | 65 |
| SL1344_RS04425_ | 252 | 220 | 265 | 188 | 160 | 201 |
| SL1344_RS04430_ | 251 | 215 | 224 | 236 | 182 | 222 |
| SL1344_RS04435_ | 281 | 315 | 320 | 304 | 275 | 356 |
| SL1344_RS04440_ | 68 | 70 | 85 | 87 | 106 | 122 |

| | | | | | | |
|---|---|---|---|---|---|---|
| SL1344_RS04450_ | 45 | 33 | 39 | 41 | 53 | 59 |
| SL1344_RS04455_ | 121 | 100 | 95 | 105 | 77 | 108 |
| SL1344_RS04460_ | 106 | 101 | 102 | 148 | 99 | 115 |
| SL1344_RS04465_ | 156 | 156 | 174 | 205 | 176 | 225 |
| SL1344_RS04470_ | 24 | 19 | 16 | 30 | 20 | 25 |
| SL1344_RS04475_ | 25 | 16 | 13 | 14 | 11 | 21 |
| SL1344_RS04480_ | 128 | 130 | 111 | 97 | 107 | 115 |
| SL1344_RS04485_ | 91 | 74 | 76 | 60 | 78 | 69 |
| SL1344_RS04495_ | 689 | 668 | 857 | 747 | 690 | 701 |
| SL1344_RS04500_ | 1543 | 1793 | 1997 | 1462 | 1497 | 1695 |
| SL1344_RS04505_ | 613 | 643 | 666 | 772 | 723 | 695 |
| SL1344_RS04510_ | 980 | 937 | 1033 | 913 | 787 | 963 |
| SL1344_RS04515_ | 401 | 348 | 319 | 291 | 295 | 387 |
| SL1344_RS04520_ | 654 | 710 | 643 | 590 | 501 | 622 |
| SL1344_RS04525_ | 287 | 317 | 326 | 362 | 323 | 385 |
| SL1344_RS04530_ | 466 | 511 | 492 | 410 | 400 | 462 |
| SL1344_RS04535_ | 540 | 560 | 639 | 516 | 473 | 586 |
| SL1344_RS04540_ | 336 | 323 | 343 | 290 | 298 | 306 |
| SL1344_RS04545_ | 238 | 248 | 250 | 252 | 215 | 256 |
| SL1344_RS04550_ | 203 | 241 | 224 | 215 | 197 | 255 |
| SL1344_RS04555_ | 444 | 434 | 429 | 510 | 423 | 495 |
| SL1344_RS04560_ | 673 | 712 | 687 | 670 | 605 | 700 |
| SL1344_RS04565_ | 1097 | 1254 | 1392 | 1631 | 1374 | 1724 |
| SL1344_RS04570_ | 533 | 554 | 423 | 497 | 391 | 591 |
| SL1344_RS04575_ | 841 | 910 | 846 | 750 | 623 | 866 |
| SL1344_RS04580_ | 4685 | 4274 | 4420 | 6461 | 5236 | 5151 |
| SL1344_RS04585_ | 790 | 866 | 758 | 765 | 734 | 816 |
| SL1344_RS04590_ | 4211 | 4051 | 4093 | 3877 | 3529 | 4421 |
| SL1344_RS04600_ | 77 | 80 | 62 | 71 | 59 | 64 |
| SL1344_RS04605_ | 60 | 55 | 53 | 56 | 38 | 44 |
| SL1344_RS04610_ | 82 | 56 | 84 | 101 | 73 | 88 |
| SL1344_RS04615_ | 175 | 157 | 189 | 196 | 187 | 179 |
| SL1344_RS04620_ | 47 | 56 | 38 | 52 | 55 | 52 |
| SL1344_RS04625_ | 92 | 87 | 87 | 67 | 57 | 98 |
| SL1344_RS04630_ | 814 | 773 | 930 | 1862 | 1553 | 1704 |
| SL1344_RS04635_ | 967 | 870 | 843 | 1594 | 1434 | 1462 |
| SL1344_RS04640_ | 503 | 462 | 545 | 500 | 426 | 469 |
| SL1344_RS04645_ | 1002 | 1128 | 1267 | 971 | 900 | 982 |
| SL1344_RS04650_ | 988 | 1000 | 1138 | 1138 | 910 | 1030 |
| SL1344_RS04655_ | 2834 | 2718 | 3013 | 2928 | 3023 | 3391 |
| SL1344_RS04665_ | 4496 | 4853 | 4785 | 4483 | 4113 | 4671 |
| SL1344_RS04675_ | 1361 | 1425 | 1477 | 1368 | 1207 | 1433 |
| SL1344_RS04680_ | 4397 | 4402 | 4806 | 4155 | 4039 | 4722 |
| SL1344_RS04700_ | 270 | 277 | 202 | 124 | 114 | 136 |
| SL1344_RS04705_ | 197 | 228 | 251 | 209 | 177 | 237 |
| SL1344_RS04715_ | 1689 | 1551 | 1605 | 1846 | 1896 | 1992 |
| SL1344_RS04720_ | 150 | 164 | 146 | 215 | 201 | 209 |
| SL1344_RS04725_ | 173 | 187 | 174 | 240 | 217 | 260 |
| SL1344_RS04730_ | 16144 | 16944 | 18602 | 14568 | 13610 | 15989 |
| SL1344_RS04740_ | 246 | 247 | 316 | 345 | 318 | 401 |
| SL1344_RS04745_ | 133 | 109 | 122 | 85 | 92 | 116 |
| SL1344_RS04750_ | 1546 | 1471 | 1486 | 1427 | 1430 | 1526 |
| SL1344_RS04755_ | 678 | 599 | 702 | 578 | 520 | 645 |
| SL1344_RS04760_ | 154 | 151 | 159 | 154 | 151 | 155 |
| SL1344_RS04765_ | 569 | 536 | 626 | 792 | 743 | 777 |
| SL1344_RS04770_ | 17946 | 17610 | 21855 | 20863 | 21577 | 25874 |
| SL1344_RS04780_ | 91 | 123 | 91 | 90 | 86 | 120 |
| SL1344_RS04785_ | 882 | 938 | 1120 | 1060 | 1079 | 1282 |
| SL1344_RS04790_ | 436 | 507 | 530 | 599 | 582 | 647 |
| SL1344_RS04795_ | 152 | 142 | 175 | 164 | 149 | 156 |
| SL1344_RS04800_ | 93 | 104 | 121 | 94 | 58 | 89 |
| SL1344_RS04805_ | 523 | 582 | 535 | 513 | 477 | 577 |

| | | | | | | |
|---|---|---|---|---|---|---|
| SL1344_RS04810_ | 786 | 886 | 858 | 718 | 645 | 801 |
| SL1344_RS04815_ | 105 | 130 | 104 | 134 | 104 | 124 |
| SL1344_RS04820_ | 454 | 427 | 400 | 707 | 493 | 820 |
| SL1344_RS04825_ | 960 | 970 | 971 | 1392 | 1232 | 1593 |
| SL1344_RS04830_ | 1016 | 1001 | 1041 | 1599 | 1400 | 1688 |
| SL1344_RS04840_ | 1832 | 1937 | 1843 | 1827 | 1678 | 1858 |
| SL1344_RS04845_ | 947 | 899 | 997 | 895 | 754 | 952 |
| SL1344_RS04850_ | 1634 | 1559 | 1513 | 1348 | 1265 | 1618 |
| SL1344_RS04855_ | 3144 | 3403 | 3691 | 2799 | 2895 | 3166 |
| SL1344_RS04860_ | 8291 | 8113 | 10179 | 9303 | 7796 | 9301 |
| SL1344_RS04865_ | 4500 | 4560 | 5166 | 4569 | 4406 | 5045 |
| SL1344_RS04870_ | 183 | 167 | 150 | 156 | 157 | 167 |
| SL1344_RS04875_ | 31 | 27 | 28 | 28 | 25 | 22 |
| SL1344_RS04880_ | 52 | 60 | 53 | 74 | 54 | 61 |
| SL1344_RS04885_ | 22 | 22 | 20 | 25 | 37 | 32 |
| SL1344_RS04890_ | 486 | 478 | 550 | 526 | 512 | 598 |
| SL1344_RS04895_ | 996 | 934 | 922 | 904 | 837 | 945 |
| SL1344_RS04900_ | 20 | 22 | 19 | 13 | 12 | 16 |
| SL1344_RS04905_ | 5 | 7 | 9 | 11 | 4 | 5 |
| SL1344_RS04910_ | 35 | 25 | 29 | 21 | 16 | 20 |
| SL1344_RS04915_ | 63 | 41 | 57 | 42 | 37 | 35 |
| SL1344_RS04920_ | 52 | 62 | 50 | 11 | 21 | 13 |
| SL1344_RS04925_ | 36 | 29 | 30 | 7 | 7 | 9 |
| SL1344_RS04930_ | 1061 | 928 | 965 | 1191 | 1185 | 1221 |
| SL1344_RS04935_ | 26 | 27 | 27 | 12 | 13 | 14 |
| SL1344_RS04940_ | 34 | 25 | 25 | 24 | 20 | 31 |
| SL1344_RS04945_ | 65 | 43 | 46 | 36 | 48 | 39 |
| SL1344_RS04950_ | 25 | 17 | 29 | 20 | 11 | 14 |
| SL1344_RS04955_ | 18 | 12 | 7 | 12 | 5 | 9 |
| SL1344_RS04960_ | 64 | 50 | 66 | 76 | 53 | 67 |
| SL1344_RS04965_ | 1274 | 1435 | 1345 | 1143 | 1086 | 1216 |
| SL1344_RS04970_ | 90 | 109 | 127 | 158 | 122 | 141 |
| SL1344_RS04975_ | 161 | 132 | 157 | 177 | 163 | 180 |
| SL1344_RS04980_ | 63 | 51 | 91 | 80 | 61 | 68 |
| SL1344_RS04985_ | 0 | 0 | 3 | 2 | 9 | 2 |
| SL1344_RS04990_ | 8 | 7 | 12 | 7 | 9 | 7 |
| SL1344_RS04995_ | 4 | 2 | 3 | 0 | 2 | 0 |
| SL1344_RS05000_ | 44 | 30 | 21 | 22 | 30 | 24 |
| SL1344_RS05005_ | 216 | 234 | 246 | 232 | 220 | 273 |
| SL1344_RS05010_ | 3 | 3 | 1 | 1 | 4 | 0 |
| SL1344_RS05015_ | 145 | 173 | 149 | 158 | 154 | 158 |
| SL1344_RS05020_ | 94 | 76 | 92 | 148 | 110 | 118 |
| SL1344_RS05025_ | 2 | 5 | 0 | 0 | 5 | 3 |
| SL1344_RS05030_ | 34 | 28 | 28 | 23 | 28 | 20 |
| SL1344_RS05035_ | 13 | 13 | 10 | 16 | 13 | 10 |
| SL1344_RS05040_ | 46 | 49 | 43 | 55 | 48 | 46 |
| SL1344_RS05045_ | 102 | 77 | 102 | 80 | 67 | 93 |
| SL1344_RS05050_ | 10 | 7 | 12 | 5 | 3 | 14 |
| SL1344_RS05055_ | 74 | 122 | 76 | 82 | 47 | 75 |
| SL1344_RS05060_ | 183 | 179 | 163 | 122 | 122 | 178 |
| SL1344_RS05065_ | 16 | 16 | 21 | 10 | 14 | 20 |
| SL1344_RS05070_ | 14 | 10 | 9 | 3 | 4 | 8 |
| SL1344_RS05075_ | 22 | 30 | 29 | 20 | 28 | 30 |
| SL1344_RS05080_ | 16 | 34 | 20 | 17 | 12 | 16 |
| SL1344_RS05085_ | 75 | 53 | 52 | 48 | 52 | 57 |
| SL1344_RS05090_ | 49 | 43 | 38 | 29 | 31 | 39 |
| SL1344_RS05095_ | 29 | 24 | 20 | 13 | 13 | 25 |
| SL1344_RS05100_ | 210 | 214 | 223 | 177 | 185 | 226 |
| SL1344_RS05105_ | 27 | 21 | 26 | 24 | 25 | 23 |
| SL1344_RS05110_ | 516 | 455 | 380 | 452 | 519 | 665 |
| SL1344_RS05115_ | 1335 | 1432 | 1471 | 1340 | 1325 | 1504 |
| SL1344_RS05120_ | 78 | 83 | 85 | 66 | 70 | 79 |

| | | | | | | |
|---|---|---|---|---|---|---|
| SL1344_RS05125_ | 54 | 47 | 37 | 44 | 41 | 50 |
| SL1344_RS05130_ | 83 | 93 | 83 | 70 | 69 | 90 |
| SL1344_RS05135_ | 171 | 179 | 156 | 160 | 152 | 193 |
| SL1344_RS05140_ | 20 | 17 | 23 | 43 | 23 | 34 |
| SL1344_RS05165_ | 23 | 22 | 17 | 12 | 12 | 22 |
| SL1344_RS05175_ | 120 | 110 | 166 | 175 | 122 | 147 |
| SL1344_RS05190_ | 3224 | 3278 | 3312 | 2821 | 2569 | 3310 |
| SL1344_RS05195_ | 249 | 249 | 329 | 297 | 287 | 348 |
| SL1344_RS05200_ | 526 | 509 | 547 | 696 | 597 | 714 |
| SL1344_RS05205_ | 178 | 153 | 159 | 175 | 149 | 184 |
| SL1344_RS05210_ | 585 | 625 | 693 | 787 | 680 | 856 |
| SL1344_RS05215_ | 934 | 972 | 1085 | 1169 | 1198 | 1426 |
| SL1344_RS05220_ | 457 | 514 | 593 | 480 | 488 | 551 |
| SL1344_RS05225_ | 759 | 794 | 817 | 672 | 676 | 812 |
| SL1344_RS05230_ | 649 | 587 | 595 | 534 | 585 | 567 |
| SL1344_RS05235_ | 4206 | 4362 | 4201 | 5140 | 4156 | 4724 |
| SL1344_RS05240_ | 950 | 792 | 982 | 1771 | 1413 | 1720 |
| SL1344_RS05245_ | 651 | 671 | 818 | 904 | 802 | 1013 |
| SL1344_RS05250_ | 350 | 377 | 417 | 389 | 370 | 463 |
| SL1344_RS05255_ | 54215 | 59421 | 64210 | 44426 | 45174 | 50887 |
| SL1344_RS05260_ | 477 | 527 | 502 | 766 | 811 | 887 |
| SL1344_RS05265_ | 184 | 213 | 211 | 197 | 176 | 204 |
| SL1344_RS05270_ | 456 | 549 | 577 | 504 | 411 | 449 |
| SL1344_RS05275_ | 90 | 74 | 87 | 106 | 81 | 98 |
| SL1344_RS05285_ | 1285 | 1415 | 1619 | 1079 | 1138 | 1292 |
| SL1344_RS05290_ | 87 | 67 | 73 | 48 | 38 | 46 |
| SL1344_RS05295_ | 1156 | 1162 | 950 | 732 | 669 | 979 |
| SL1344_RS05300_ | 1207 | 1279 | 1285 | 1227 | 1400 | 1356 |
| SL1344_RS05305_ | 329 | 362 | 423 | 377 | 372 | 381 |
| SL1344_RS05310_ | 103 | 112 | 96 | 90 | 89 | 96 |
| SL1344_RS05315_ | 196 | 204 | 205 | 156 | 160 | 168 |
| SL1344_RS05320_ | 87 | 94 | 73 | 45 | 46 | 54 |
| SL1344_RS05325_ | 367 | 376 | 378 | 556 | 486 | 487 |
| SL1344_RS05330_ | 2857 | 2870 | 3431 | 3558 | 3295 | 3768 |
| SL1344_RS05335_ | 556 | 461 | 602 | 865 | 672 | 498 |
| SL1344_RS05340_ | 67 | 72 | 92 | 56 | 42 | 53 |
| SL1344_RS05345_ | 33 | 28 | 32 | 84 | 70 | 116 |
| SL1344_RS05350_ | 2 | 0 | 9 | 9 | 2 | 12 |
| SL1344_RS05365_ | 660 | 501 | 548 | 1123 | 825 | 930 |
| SL1344_RS05370_ | 40 | 25 | 28 | 8 | 23 | 17 |
| SL1344_RS05375_ | 536 | 574 | 567 | 430 | 419 | 394 |
| SL1344_RS05380_ | 133 | 140 | 147 | 140 | 129 | 137 |
| SL1344_RS05385_ | 170 | 153 | 145 | 162 | 132 | 137 |
| SL1344_RS05390_ | 159 | 162 | 153 | 155 | 117 | 138 |
| SL1344_RS05395_ | 60 | 57 | 44 | 59 | 59 | 57 |
| SL1344_RS05400_ | 27 | 37 | 43 | 33 | 34 | 43 |
| SL1344_RS05405_ | 79 | 80 | 105 | 85 | 84 | 105 |
| SL1344_RS05410_ | 85 | 89 | 98 | 87 | 104 | 116 |
| SL1344_RS05415_ | 117 | 105 | 133 | 120 | 114 | 117 |
| SL1344_RS05420_ | 68 | 52 | 63 | 78 | 56 | 69 |
| SL1344_RS05425_ | 68 | 73 | 59 | 64 | 71 | 71 |
| SL1344_RS05430_ | 41 | 47 | 42 | 47 | 49 | 46 |
| SL1344_RS05435_ | 34 | 35 | 34 | 39 | 34 | 50 |
| SL1344_RS05440_ | 61 | 71 | 60 | 60 | 64 | 73 |
| SL1344_RS05445_ | 36 | 52 | 55 | 49 | 45 | 52 |
| SL1344_RS05450_ | 111 | 100 | 130 | 131 | 101 | 138 |
| SL1344_RS05455_ | 13 | 5 | 6 | 18 | 13 | 4 |
| SL1344_RS05460_ | 875 | 997 | 826 | 587 | 553 | 726 |
| SL1344_RS05465_ | 1043 | 1001 | 1065 | 848 | 781 | 895 |
| SL1344_RS05470_ | 234 | 245 | 213 | 291 | 230 | 260 |
| SL1344_RS05475_ | 82 | 73 | 78 | 95 | 81 | 65 |
| SL1344_RS05480_ | 29 | 29 | 39 | 15 | 29 | 34 |

| | | | | | | |
|---|---|---|---|---|---|---|
| SL1344_RS05485_ | 47 | 52 | 58 | 42 | 53 | 46 |
| SL1344_RS05490_ | 1613 | 1740 | 1852 | 1495 | 1398 | 1616 |
| SL1344_RS05495_ | 3794 | 3770 | 2903 | 2016 | 1976 | 2344 |
| SL1344_RS05500_ | 1472 | 1542 | 1461 | 906 | 686 | 1001 |
| SL1344_RS05505_ | 344 | 326 | 313 | 495 | 449 | 491 |
| SL1344_RS05510_ | 96 | 109 | 122 | 127 | 118 | 138 |
| SL1344_RS05515_ | 1701 | 1789 | 1800 | 1836 | 1884 | 2046 |
| SL1344_RS05525_ | 2113 | 2033 | 2500 | 2011 | 2043 | 2184 |
| SL1344_RS05530_ | 689 | 651 | 711 | 1016 | 910 | 1106 |
| SL1344_RS05540_ | 624 | 610 | 631 | 675 | 596 | 705 |
| SL1344_RS05545_ | 138 | 146 | 143 | 125 | 124 | 130 |
| SL1344_RS05550_ | 114 | 106 | 107 | 127 | 114 | 127 |
| SL1344_RS05555_ | 97 | 81 | 110 | 75 | 49 | 67 |
| SL1344_RS05560_ | 103 | 88 | 105 | 64 | 48 | 66 |
| SL1344_RS05565_ | 71 | 58 | 74 | 37 | 32 | 53 |
| SL1344_RS05570_ | 367 | 398 | 372 | 271 | 272 | 340 |
| SL1344_RS05580_ | 158 | 128 | 183 | 210 | 197 | 188 |
| SL1344_RS05585_ | 3 | 3 | 3 | 4 | 5 | 4 |
| SL1344_RS05590_ | 688 | 713 | 740 | 645 | 673 | 784 |
| SL1344_RS05595_ | 605 | 653 | 656 | 630 | 566 | 656 |
| SL1344_RS05600_ | 387 | 391 | 412 | 325 | 335 | 384 |
| SL1344_RS05605_ | 283 | 315 | 314 | 371 | 262 | 319 |
| SL1344_RS05610_ | 48 | 39 | 60 | 53 | 54 | 40 |
| SL1344_RS05615_ | 7 | 5 | 11 | 6 | 8 | 8 |
| SL1344_RS05620_ | 24 | 11 | 17 | 11 | 11 | 10 |
| SL1344_RS05625_ | 41 | 14 | 22 | 32 | 35 | 24 |
| SL1344_RS05635_ | 31 | 30 | 29 | 34 | 33 | 44 |
| SL1344_RS05650_ | 825 | 897 | 865 | 1462 | 1551 | 1559 |
| SL1344_RS05660_ | 118 | 124 | 134 | 112 | 85 | 127 |
| SL1344_RS05670_ | 3746 | 3910 | 4053 | 3685 | 3260 | 4044 |
| SL1344_RS05675_ | 44 | 41 | 38 | 55 | 40 | 54 |
| SL1344_RS05680_ | 747 | 761 | 598 | 556 | 496 | 621 |
| SL1344_RS05685_ | 116 | 149 | 170 | 128 | 107 | 103 |
| SL1344_RS05690_ | 463 | 454 | 527 | 661 | 585 | 679 |
| SL1344_RS05695_ | 216 | 202 | 242 | 259 | 284 | 309 |
| SL1344_RS05700_ | 54 | 50 | 62 | 55 | 58 | 66 |
| SL1344_RS05705_ | 17 | 14 | 14 | 17 | 15 | 25 |
| SL1344_RS05710_ | 8 | 16 | 16 | 16 | 10 | 8 |
| SL1344_RS05715_ | 665 | 657 | 763 | 655 | 609 | 673 |
| SL1344_RS05730_ | 1068 | 1051 | 1151 | 1241 | 1264 | 1349 |
| SL1344_RS05735_ | 1120 | 1173 | 1259 | 1113 | 1063 | 1121 |
| SL1344_RS05740_ | 1579 | 1756 | 1923 | 1449 | 1311 | 1358 |
| SL1344_RS05745_ | 238 | 236 | 213 | 251 | 224 | 258 |
| SL1344_RS05750_ | 814 | 847 | 865 | 563 | 561 | 693 |
| SL1344_RS05755_ | 1195 | 1277 | 1138 | 690 | 721 | 887 |
| SL1344_RS05760_ | 1239 | 1253 | 1232 | 756 | 810 | 930 |
| SL1344_RS05765_ | 409 | 432 | 410 | 399 | 354 | 416 |
| SL1344_RS05780_ | 3640 | 4028 | 3748 | 6544 | 7147 | 7051 |
| SL1344_RS05785_ | 2490 | 2731 | 2769 | 5659 | 5904 | 6250 |
| SL1344_RS05790_ | 2400 | 2470 | 2503 | 4423 | 4665 | 5335 |
| SL1344_RS05800_ | 8800 | 9373 | 10021 | 14611 | 14765 | 17013 |
| SL1344_RS05810_ | 8994 | 9055 | 9081 | 15287 | 16978 | 18338 |
| SL1344_RS05815_ | 2245 | 2522 | 2356 | 3901 | 3918 | 4702 |
| SL1344_RS05825_ | 2072 | 2196 | 2032 | 3358 | 2939 | 3535 |
| SL1344_RS05845_ | 3 | 4 | 3 | 4 | 1 | 3 |
| SL1344_RS05850_ | 276 | 289 | 301 | 366 | 325 | 377 |
| SL1344_RS05855_ | 211 | 207 | 199 | 229 | 199 | 185 |
| SL1344_RS05860_ | 104 | 107 | 109 | 93 | 75 | 78 |
| SL1344_RS05865_ | 436 | 461 | 516 | 482 | 487 | 488 |
| SL1344_RS05870_ | 3645 | 3534 | 4049 | 4400 | 4493 | 5015 |
| SL1344_RS05875_ | 7536 | 7649 | 8838 | 7587 | 7770 | 9271 |
| SL1344_RS05880_ | 2189 | 2003 | 2683 | 2549 | 2452 | 2772 |

| | | | | | | |
|---|---|---|---|---|---|---|
| SL1344_RS05885_ | 934 | 936 | 1049 | 1098 | 979 | 1195 |
| SL1344_RS05890_ | 1694 | 1804 | 1940 | 1742 | 1682 | 2057 |
| SL1344_RS05900_ | 17769 | 16407 | 15625 | 19563 | 17739 | 21643 |
| SL1344_RS05905_ | 10220 | 9819 | 10889 | 10333 | 10548 | 11955 |
| SL1344_RS05910_ | 337 | 332 | 273 | 340 | 314 | 384 |
| SL1344_RS05915_ | 363 | 368 | 375 | 426 | 383 | 432 |
| SL1344_RS05920_ | 428 | 418 | 400 | 476 | 404 | 533 |
| SL1344_RS05925_ | 499 | 485 | 499 | 517 | 516 | 552 |
| SL1344_RS05930_ | 460 | 455 | 550 | 502 | 461 | 543 |
| SL1344_RS05935_ | 9514 | 10120 | 11171 | 6704 | 6202 | 7069 |
| SL1344_RS05940_ | 70 | 47 | 49 | 66 | 57 | 55 |
| SL1344_RS05945_ | 716 | 783 | 681 | 736 | 681 | 811 |
| SL1344_RS05950_ | 725 | 752 | 798 | 746 | 619 | 783 |
| SL1344_RS05955_ | 773 | 736 | 838 | 703 | 555 | 728 |
| SL1344_RS05960_ | 574 | 502 | 558 | 501 | 388 | 473 |
| SL1344_RS05965_ | 712 | 789 | 709 | 657 | 623 | 817 |
| SL1344_RS05970_ | 2601 | 2500 | 2562 | 2460 | 2211 | 2425 |
| SL1344_RS05975_ | 793 | 801 | 830 | 725 | 688 | 869 |
| SL1344_RS05980_ | 112 | 108 | 105 | 92 | 70 | 102 |
| SL1344_RS05985_ | 1239 | 1257 | 1319 | 1554 | 1455 | 1505 |
| SL1344_RS05990_ | 217 | 216 | 141 | 323 | 302 | 255 |
| SL1344_RS05995_ | 286 | 284 | 311 | 324 | 282 | 319 |
| SL1344_RS06000_ | 1682 | 1774 | 1830 | 1534 | 1443 | 1592 |
| SL1344_RS06005_ | 434 | 405 | 467 | 555 | 475 | 570 |
| SL1344_RS06010_ | 491 | 538 | 544 | 477 | 463 | 519 |
| SL1344_RS06015_ | 659 | 685 | 671 | 598 | 536 | 615 |
| SL1344_RS06020_ | 939 | 970 | 957 | 927 | 765 | 949 |
| SL1344_RS06025_ | 1168 | 1275 | 1178 | 1062 | 1029 | 1251 |
| SL1344_RS06030_ | 1199 | 1195 | 1359 | 1106 | 904 | 1206 |
| SL1344_RS06035_ | 203 | 227 | 270 | 267 | 283 | 301 |
| SL1344_RS06045_ | 263 | 217 | 262 | 315 | 303 | 328 |
| SL1344_RS06050_ | 315 | 348 | 341 | 440 | 403 | 464 |
| SL1344_RS06055_ | 2261 | 2147 | 1934 | 1814 | 1419 | 1984 |
| SL1344_RS06070_ | 1135 | 1260 | 1453 | 1019 | 891 | 992 |
| SL1344_RS06085_ | 885 | 948 | 1124 | 809 | 821 | 1005 |
| SL1344_RS06090_ | 536 | 594 | 735 | 539 | 532 | 657 |
| SL1344_RS06095_ | 1013 | 1071 | 1152 | 1152 | 997 | 1242 |
| SL1344_RS06100_ | 216 | 197 | 210 | 179 | 171 | 203 |
| SL1344_RS06105_ | 146 | 182 | 144 | 152 | 130 | 167 |
| SL1344_RS06110_ | 135 | 195 | 205 | 173 | 110 | 172 |
| SL1344_RS06115_ | 13640 | 12905 | 14812 | 13809 | 13822 | 15970 |
| SL1344_RS06120_ | 330 | 298 | 286 | 561 | 401 | 556 |
| SL1344_RS06125_ | 13 | 2 | 27 | 1 | 6 | 4 |
| SL1344_RS06130_ | 16 | 10 | 10 | 33 | 20 | 20 |
| SL1344_RS06135_ | 411 | 378 | 281 | 689 | 544 | 566 |
| SL1344_RS06140_ | 228 | 227 | 217 | 153 | 160 | 216 |
| SL1344_RS06150_ | 16 | 16 | 29 | 14 | 11 | 10 |
| SL1344_RS06155_ | 0 | 2 | 1 | 1 | 1 | 1 |
| SL1344_RS06160_ | 40 | 50 | 45 | 29 | 18 | 26 |
| SL1344_RS06165_ | 1120 | 1128 | 1030 | 551 | 456 | 745 |
| SL1344_RS06170_ | 33 | 41 | 32 | 38 | 25 | 32 |
| SL1344_RS06180_ | 1056 | 1112 | 1198 | 842 | 669 | 940 |
| SL1344_RS06185_ | 23 | 25 | 19 | 21 | 10 | 29 |
| SL1344_RS06190_ | 30 | 42 | 24 | 35 | 29 | 34 |
| SL1344_RS06195_ | 1164 | 1082 | 886 | 720 | 577 | 781 |
| SL1344_RS06200_ | 71 | 107 | 64 | 72 | 58 | 62 |
| SL1344_RS06205_ | 2637 | 2330 | 1988 | 2823 | 2941 | 2750 |
| SL1344_RS06210_ | 9 | 9 | 3 | 11 | 22 | 18 |
| SL1344_RS06215_ | 160 | 168 | 155 | 131 | 134 | 144 |
| SL1344_RS06220_ | 865 | 882 | 989 | 684 | 621 | 696 |
| SL1344_RS06225_ | 592 | 633 | 628 | 146 | 95 | 155 |
| SL1344_RS06230_ | 1053 | 918 | 1077 | 1294 | 1780 | 1399 |

| | | | | | | |
|---|---|---|---|---|---|---|
| SL1344_RS06235_ | 341 | 331 | 369 | 446 | 580 | 468 |
| SL1344_RS06240_ | 91 | 105 | 97 | 114 | 121 | 101 |
| SL1344_RS06245_ | 149 | 152 | 139 | 157 | 134 | 152 |
| SL1344_RS06250_ | 132 | 120 | 140 | 138 | 145 | 134 |
| SL1344_RS06255_ | 123 | 140 | 155 | 115 | 109 | 123 |
| SL1344_RS06260_ | 168 | 194 | 189 | 74 | 80 | 77 |
| SL1344_RS06265_ | 80 | 83 | 118 | 37 | 25 | 23 |
| SL1344_RS06270_ | 107 | 107 | 121 | 83 | 89 | 99 |
| SL1344_RS06275_ | 192 | 232 | 232 | 191 | 166 | 182 |
| SL1344_RS06280_ | 25 | 15 | 26 | 21 | 13 | 20 |
| SL1344_RS06285_ | 201 | 178 | 179 | 204 | 160 | 228 |
| SL1344_RS06290_ | 726 | 752 | 655 | 554 | 523 | 581 |
| SL1344_RS06295_ | 10 | 11 | 12 | 13 | 8 | 10 |
| SL1344_RS06300_ | 351 | 380 | 418 | 371 | 296 | 372 |
| SL1344_RS06305_ | 435 | 453 | 461 | 389 | 308 | 430 |
| SL1344_RS06310_ | 86 | 81 | 73 | 92 | 116 | 120 |
| SL1344_RS06315_ | 437 | 375 | 395 | 372 | 417 | 423 |
| SL1344_RS06320_ | 55 | 49 | 47 | 25 | 42 | 36 |
| SL1344_RS06325_ | 111 | 165 | 127 | 84 | 63 | 87 |
| SL1344_RS06330_ | 1594 | 1575 | 1225 | 716 | 655 | 834 |
| SL1344_RS06335_ | 392 | 415 | 381 | 268 | 278 | 311 |
| SL1344_RS06340_ | 661 | 793 | 849 | 601 | 554 | 550 |
| SL1344_RS06345_ | 258 | 360 | 343 | 289 | 290 | 314 |
| SL1344_RS06350_ | 39 | 23 | 62 | 62 | 52 | 45 |
| SL1344_RS06355_ | 118 | 121 | 121 | 156 | 117 | 135 |
| SL1344_RS06360_ | 74 | 65 | 103 | 72 | 69 | 93 |
| SL1344_RS06365_ | 8 | 6 | 9 | 6 | 5 | 9 |
| SL1344_RS06370_ | 359 | 365 | 346 | 228 | 219 | 259 |
| SL1344_RS06375_ | 392 | 398 | 413 | 281 | 267 | 280 |
| SL1344_RS06380_ | 1097 | 1100 | 1018 | 588 | 505 | 654 |
| SL1344_RS06385_ | 1771 | 1888 | 1980 | 911 | 736 | 953 |
| SL1344_RS06390_ | 5005 | 4658 | 5527 | 4603 | 3896 | 5111 |
| SL1344_RS06395_ | 40 | 55 | 49 | 55 | 52 | 66 |
| SL1344_RS06400_ | 455 | 415 | 342 | 312 | 293 | 348 |
| SL1344_RS06405_ | 627 | 550 | 731 | 628 | 528 | 611 |
| SL1344_RS06410_ | 28213 | 27860 | 28820 | 23130 | 24431 | 28032 |
| SL1344_RS06415_ | 1019 | 953 | 1087 | 1118 | 1120 | 1147 |
| SL1344_RS06420_ | 1452 | 1583 | 1624 | 1579 | 1673 | 1724 |
| SL1344_RS06425_ | 355 | 354 | 294 | 219 | 218 | 260 |
| SL1344_RS06430_ | 273 | 264 | 313 | 224 | 199 | 265 |
| SL1344_RS06435_ | 745 | 785 | 822 | 723 | 592 | 774 |
| SL1344_RS06440_ | 570 | 531 | 546 | 409 | 365 | 576 |
| SL1344_RS06445_ | 913 | 910 | 896 | 698 | 692 | 827 |
| SL1344_RS06450_ | 1567 | 1570 | 1562 | 1380 | 1204 | 1402 |
| SL1344_RS06455_ | 556 | 529 | 539 | 495 | 452 | 549 |
| SL1344_RS06465_ | 76 | 88 | 94 | 60 | 45 | 55 |
| SL1344_RS06470_ | 707 | 743 | 840 | 517 | 527 | 601 |
| SL1344_RS06475_ | 95 | 102 | 116 | 95 | 82 | 85 |
| SL1344_RS06480_ | 69 | 59 | 95 | 53 | 67 | 69 |
| SL1344_RS06485_ | 149 | 143 | 151 | 94 | 123 | 122 |
| SL1344_RS06490_ | 132 | 137 | 127 | 64 | 68 | 115 |
| SL1344_RS06495_ | 121 | 118 | 128 | 68 | 74 | 81 |
| SL1344_RS06500_ | 337 | 371 | 370 | 306 | 318 | 309 |
| SL1344_RS06505_ | 247 | 259 | 245 | 304 | 341 | 325 |
| SL1344_RS06510_ | 1059 | 1072 | 1109 | 826 | 775 | 836 |
| SL1344_RS06515_ | 4157 | 4284 | 4056 | 2163 | 2030 | 2365 |
| SL1344_RS06520_ | 548 | 488 | 493 | 317 | 339 | 368 |
| SL1344_RS06525_ | 456 | 581 | 498 | 283 | 275 | 306 |
| SL1344_RS06530_ | 232 | 238 | 185 | 130 | 131 | 150 |
| SL1344_RS06535_ | 362 | 373 | 341 | 218 | 204 | 219 |
| SL1344_RS06540_ | 144 | 151 | 113 | 83 | 80 | 96 |
| SL1344_RS06545_ | 405 | 510 | 464 | 306 | 285 | 297 |

| | | | | | | |
|---|---|---|---|---|---|---|
| SL1344_RS06550_ | 1804 | 1968 | 1991 | 994 | 915 | 1097 |
| SL1344_RS06555_ | 86 | 80 | 107 | 96 | 85 | 91 |
| SL1344_RS06565_ | 54 | 68 | 74 | 97 | 88 | 71 |
| SL1344_RS06570_ | 859 | 848 | 771 | 724 | 747 | 891 |
| SL1344_RS06575_ | 293 | 309 | 306 | 214 | 237 | 290 |
| SL1344_RS06580_ | 928 | 1063 | 1150 | 583 | 545 | 615 |
| SL1344_RS06605_ | 81 | 77 | 60 | 87 | 92 | 109 |
| SL1344_RS06615_ | 18 | 31 | 26 | 17 | 7 | 21 |
| SL1344_RS06630_ | 26840 | 25137 | 28153 | 29185 | 29913 | 29856 |
| SL1344_RS06635_ | 23903 | 24580 | 21534 | 26022 | 27237 | 26701 |
| SL1344_RS06640_ | 26820 | 26768 | 30899 | 31472 | 33507 | 33616 |
| SL1344_RS06645_ | 873 | 835 | 949 | 894 | 841 | 1078 |
| SL1344_RS06650_ | 3898 | 3744 | 3780 | 3238 | 2796 | 3864 |
| SL1344_RS06660_ | 218 | 256 | 262 | 263 | 242 | 265 |
| SL1344_RS06665_ | 306 | 269 | 269 | 194 | 188 | 207 |
| SL1344_RS06670_ | 275 | 265 | 269 | 196 | 155 | 206 |
| SL1344_RS06675_ | 328 | 335 | 295 | 212 | 203 | 212 |
| SL1344_RS06685_ | 359 | 357 | 404 | 274 | 254 | 341 |
| SL1344_RS06690_ | 69 | 59 | 44 | 81 | 27 | 16 |
| SL1344_RS06695_ | 478 | 461 | 499 | 448 | 394 | 477 |
| SL1344_RS06700_ | 386 | 397 | 419 | 285 | 219 | 323 |
| SL1344_RS06705_ | 12743 | 12670 | 13158 | 10681 | 10402 | 12750 |
| SL1344_RS06710_ | 76 | 75 | 63 | 61 | 64 | 84 |
| SL1344_RS06715_ | 13 | 6 | 14 | 7 | 4 | 8 |
| SL1344_RS06720_ | 30 | 32 | 36 | 26 | 20 | 39 |
| SL1344_RS06730_ | 8 | 8 | 3 | 7 | 8 | 15 |
| SL1344_RS06735_ | 26 | 30 | 22 | 23 | 20 | 25 |
| SL1344_RS06740_ | 115 | 139 | 114 | 138 | 121 | 170 |
| SL1344_RS06745_ | 27 | 40 | 26 | 33 | 30 | 32 |
| SL1344_RS06750_ | 386 | 362 | 345 | 331 | 305 | 300 |
| SL1344_RS06755_ | 140 | 143 | 118 | 120 | 111 | 135 |
| SL1344_RS06760_ | 57 | 35 | 44 | 49 | 34 | 45 |
| SL1344_RS06765_ | 35 | 24 | 23 | 26 | 17 | 21 |
| SL1344_RS06770_ | 9 | 14 | 14 | 10 | 6 | 12 |
| SL1344_RS06775_ | 514 | 484 | 558 | 404 | 355 | 400 |
| SL1344_RS06780_ | 1306 | 1328 | 1351 | 1168 | 1129 | 1205 |
| SL1344_RS06785_ | 354 | 320 | 308 | 228 | 230 | 322 |
| SL1344_RS06790_ | 986 | 1113 | 1085 | 1899 | 1599 | 1543 |
| SL1344_RS06795_ | 270 | 260 | 323 | 170 | 167 | 199 |
| SL1344_RS06800_ | 36 | 39 | 40 | 16 | 17 | 13 |
| SL1344_RS06805_ | 169 | 181 | 176 | 90 | 95 | 102 |
| SL1344_RS06810_ | 173 | 163 | 162 | 147 | 119 | 129 |
| SL1344_RS06815_ | 194 | 210 | 201 | 123 | 135 | 144 |
| SL1344_RS06820_ | 166 | 184 | 173 | 123 | 108 | 141 |
| SL1344_RS06825_ | 82 | 84 | 66 | 58 | 56 | 55 |
| SL1344_RS06830_ | 512 | 450 | 453 | 337 | 332 | 355 |
| SL1344_RS06835_ | 498 | 420 | 534 | 458 | 458 | 474 |
| SL1344_RS06840_ | 322181 | 362201 | 339020 | 293648 | 317587 | 330197 |
| SL1344_RS06845_ | 1816 | 2026 | 2035 | 1373 | 1235 | 1445 |
| SL1344_RS06850_ | 281 | 296 | 304 | 142 | 124 | 137 |
| SL1344_RS06855_ | 65 | 82 | 108 | 44 | 51 | 50 |
| SL1344_RS06860_ | 280 | 256 | 272 | 137 | 96 | 121 |
| SL1344_RS06865_ | 190 | 211 | 252 | 98 | 95 | 120 |
| SL1344_RS06900_ | 1567 | 1661 | 1619 | 1130 | 1040 | 1171 |
| SL1344_RS06905_ | 195 | 178 | 196 | 164 | 149 | 172 |
| SL1344_RS06925_ | 15 | 20 | 24 | 18 | 26 | 29 |
| SL1344_RS06930_ | 29 | 21 | 31 | 25 | 22 | 35 |
| SL1344_RS06935_ | 22 | 28 | 14 | 28 | 16 | 24 |
| SL1344_RS06940_ | 31 | 32 | 31 | 30 | 32 | 62 |
| SL1344_RS06945_ | 61 | 60 | 45 | 57 | 47 | 62 |
| SL1344_RS06950_ | 44 | 34 | 36 | 47 | 34 | 37 |
| SL1344_RS06955_ | 148 | 154 | 123 | 92 | 97 | 107 |

| | | | | | | |
|---|---|---|---|---|---|---|
| SL1344_RS06960_ | 46 | 39 | 35 | 31 | 30 | 25 |
| SL1344_RS06965_ | 42 | 36 | 52 | 28 | 21 | 36 |
| SL1344_RS06970_ | 56 | 46 | 50 | 44 | 50 | 55 |
| SL1344_RS06975_ | 61 | 49 | 48 | 53 | 39 | 46 |
| SL1344_RS06980_ | 80 | 100 | 101 | 63 | 66 | 83 |
| SL1344_RS06990_ | 5 | 5 | 8 | 13 | 10 | 20 |
| SL1344_RS06995_ | 1 | 3 | 2 | 8 | 14 | 22 |
| SL1344_RS07000_ | 11 | 14 | 28 | 37 | 31 | 27 |
| SL1344_RS07005_ | 11 | 17 | 16 | 44 | 37 | 35 |
| SL1344_RS07010_ | 30 | 10 | 17 | 23 | 9 | 20 |
| SL1344_RS07015_ | 72 | 72 | 45 | 63 | 48 | 82 |
| SL1344_RS07020_ | 13 | 9 | 11 | 9 | 6 | 12 |
| SL1344_RS07025_ | 62 | 88 | 77 | 75 | 60 | 75 |
| SL1344_RS07030_ | 141 | 119 | 126 | 81 | 115 | 121 |
| SL1344_RS07035_ | 47 | 65 | 53 | 43 | 35 | 39 |
| SL1344_RS07040_ | 67 | 51 | 55 | 33 | 34 | 44 |
| SL1344_RS07045_ | 179 | 153 | 124 | 115 | 90 | 99 |
| SL1344_RS07055_ | 9 | 9 | 6 | 9 | 5 | 9 |
| SL1344_RS07060_ | 24 | 17 | 19 | 25 | 18 | 27 |
| SL1344_RS07065_ | 50 | 35 | 31 | 34 | 34 | 33 |
| SL1344_RS07070_ | 84 | 100 | 143 | 94 | 90 | 105 |
| SL1344_RS07075_ | 112 | 115 | 201 | 125 | 96 | 102 |
| SL1344_RS07080_ | 801 | 853 | 872 | 756 | 812 | 895 |
| SL1344_RS07085_ | 652 | 731 | 700 | 455 | 446 | 624 |
| SL1344_RS07090_ | 1386 | 1512 | 1586 | 1368 | 1059 | 1275 |
| SL1344_RS07095_ | 154 | 154 | 147 | 149 | 127 | 174 |
| SL1344_RS07100_ | 124 | 121 | 103 | 74 | 80 | 102 |
| SL1344_RS07115_ | 817 | 846 | 766 | 701 | 634 | 637 |
| SL1344_RS07120_ | 3272 | 3226 | 3354 | 3913 | 4321 | 4714 |
| SL1344_RS07125_ | 291 | 297 | 327 | 300 | 283 | 311 |
| SL1344_RS07130_ | 1054 | 1073 | 1190 | 970 | 810 | 996 |
| SL1344_RS07135_ | 596 | 601 | 661 | 522 | 489 | 594 |
| SL1344_RS07140_ | 130 | 125 | 139 | 160 | 150 | 143 |
| SL1344_RS07145_ | 86 | 111 | 85 | 55 | 51 | 56 |
| SL1344_RS07150_ | 551 | 582 | 651 | 387 | 406 | 450 |
| SL1344_RS07155_ | 543 | 608 | 496 | 392 | 340 | 378 |
| SL1344_RS07160_ | 175 | 227 | 230 | 151 | 135 | 156 |
| SL1344_RS07165_ | 132 | 125 | 130 | 121 | 105 | 116 |
| SL1344_RS07170_ | 21 | 36 | 43 | 34 | 25 | 24 |
| SL1344_RS07180_ | 3075 | 2895 | 2444 | 1466 | 1296 | 1506 |
| SL1344_RS07185_ | 286 | 297 | 269 | 273 | 235 | 279 |
| SL1344_RS07190_ | 307 | 313 | 319 | 210 | 195 | 268 |
| SL1344_RS07195_ | 512 | 514 | 495 | 514 | 497 | 508 |
| SL1344_RS07200_ | 2400 | 2250 | 2629 | 2425 | 2300 | 2584 |
| SL1344_RS07205_ | 573 | 569 | 587 | 620 | 607 | 634 |
| SL1344_RS07210_ | 1683 | 1953 | 1653 | 1141 | 1102 | 1329 |
| SL1344_RS07215_ | 1389 | 1445 | 1549 | 1313 | 1294 | 1424 |
| SL1344_RS07220_ | 388 | 426 | 446 | 379 | 363 | 409 |
| SL1344_RS07225_ | 436 | 410 | 409 | 345 | 388 | 421 |
| SL1344_RS07230_ | 283 | 302 | 316 | 290 | 264 | 298 |
| SL1344_RS07235_ | 313 | 307 | 368 | 313 | 293 | 333 |
| SL1344_RS07240_ | 523 | 598 | 751 | 669 | 645 | 678 |
| SL1344_RS07245_ | 215 | 210 | 237 | 275 | 205 | 283 |
| SL1344_RS07250_ | 302 | 277 | 349 | 405 | 371 | 476 |
| SL1344_RS07255_ | 488 | 485 | 477 | 567 | 566 | 560 |
| SL1344_RS07260_ | 261 | 220 | 220 | 290 | 283 | 323 |
| SL1344_RS07265_ | 74 | 66 | 91 | 136 | 116 | 98 |
| SL1344_RS07270_ | 1107 | 1158 | 1101 | 659 | 634 | 839 |
| SL1344_RS07275_ | 708 | 736 | 839 | 656 | 695 | 836 |
| SL1344_RS07285_ | 1726 | 1753 | 1874 | 1303 | 1135 | 1485 |
| SL1344_RS07290_ | 942 | 900 | 1042 | 748 | 658 | 793 |
| SL1344_RS07295_ | 6127 | 6400 | 6465 | 3763 | 3965 | 4404 |

| | | | | | | |
|---|---|---|---|---|---|---|
| SL1344_RS07300_ | 2100 | 2196 | 2194 | 1535 | 1455 | 1732 |
| SL1344_RS07305_ | 134 | 125 | 156 | 134 | 113 | 142 |
| SL1344_RS07310_ | 153 | 155 | 142 | 112 | 136 | 140 |
| SL1344_RS07315_ | 33 | 33 | 36 | 24 | 24 | 25 |
| SL1344_RS07320_ | 236 | 213 | 222 | 256 | 283 | 258 |
| SL1344_RS07325_ | 2 | 6 | 4 | 2 | 9 | 11 |
| SL1344_RS07330_ | 873 | 761 | 898 | 1092 | 927 | 1053 |
| SL1344_RS07335_ | 61 | 58 | 55 | 70 | 58 | 55 |
| SL1344_RS07340_ | 84 | 109 | 122 | 104 | 87 | 113 |
| SL1344_RS07345_ | 1881 | 1943 | 2279 | 1568 | 1507 | 1797 |
| SL1344_RS07350_ | 2431 | 2376 | 2254 | 1490 | 1513 | 1688 |
| SL1344_RS07355_ | 2846 | 2839 | 2906 | 1610 | 1561 | 1923 |
| SL1344_RS07365_ | 186 | 206 | 203 | 156 | 104 | 176 |
| SL1344_RS07370_ | 360 | 428 | 367 | 219 | 156 | 242 |
| SL1344_RS07375_ | 441 | 436 | 456 | 293 | 317 | 369 |
| SL1344_RS07380_ | 122 | 121 | 139 | 83 | 86 | 99 |
| SL1344_RS07385_ | 342 | 392 | 368 | 199 | 163 | 167 |
| SL1344_RS07390_ | 79 | 68 | 76 | 76 | 74 | 99 |
| SL1344_RS07400_ | 1071 | 1194 | 1078 | 613 | 619 | 706 |
| SL1344_RS07405_ | 106 | 100 | 112 | 85 | 103 | 107 |
| SL1344_RS07410_ | 221 | 206 | 201 | 122 | 108 | 147 |
| SL1344_RS07415_ | 66 | 48 | 67 | 40 | 45 | 39 |
| SL1344_RS07420_ | 44 | 38 | 42 | 32 | 18 | 30 |
| SL1344_RS07425_ | 34 | 28 | 36 | 23 | 26 | 31 |
| SL1344_RS07430_ | 303 | 283 | 315 | 300 | 289 | 317 |
| SL1344_RS07435_ | 309 | 269 | 301 | 301 | 264 | 312 |
| SL1344_RS07440_ | 51 | 67 | 60 | 42 | 31 | 47 |
| SL1344_RS07445_ | 250 | 316 | 228 | 162 | 152 | 166 |
| SL1344_RS07450_ | 573 | 774 | 640 | 299 | 257 | 351 |
| SL1344_RS07455_ | 654 | 745 | 764 | 558 | 468 | 573 |
| SL1344_RS07460_ | 251 | 269 | 223 | 197 | 199 | 233 |
| SL1344_RS07465_ | 1431 | 1462 | 1436 | 1321 | 1347 | 1670 |
| SL1344_RS07470_ | 1123 | 1178 | 1210 | 1046 | 1128 | 1371 |
| SL1344_RS07475_ | 44 | 41 | 38 | 39 | 42 | 40 |
| SL1344_RS07480_ | 107 | 105 | 133 | 143 | 181 | 185 |
| SL1344_RS07485_ | 88 | 128 | 97 | 120 | 120 | 156 |
| SL1344_RS07490_ | 144 | 145 | 208 | 193 | 160 | 217 |
| SL1344_RS07495_ | 255 | 280 | 223 | 225 | 244 | 278 |
| SL1344_RS07500_ | 2732 | 2619 | 2664 | 2865 | 2636 | 2921 |
| SL1344_RS07505_ | 361 | 359 | 366 | 347 | 317 | 318 |
| SL1344_RS07510_ | 1646 | 1770 | 1789 | 1283 | 1175 | 1359 |
| SL1344_RS07515_ | 616 | 621 | 608 | 468 | 423 | 550 |
| SL1344_RS07520_ | 54 | 53 | 60 | 45 | 49 | 51 |
| SL1344_RS07525_ | 51 | 58 | 61 | 33 | 35 | 36 |
| SL1344_RS07530_ | 166 | 181 | 182 | 129 | 104 | 123 |
| SL1344_RS07535_ | 954 | 784 | 694 | 845 | 985 | 835 |
| SL1344_RS07540_ | 115 | 144 | 152 | 129 | 113 | 116 |
| SL1344_RS07550_ | 82 | 74 | 81 | 80 | 58 | 51 |
| SL1344_RS07555_ | 81 | 106 | 87 | 81 | 74 | 98 |
| SL1344_RS07560_ | 57 | 74 | 71 | 82 | 86 | 79 |
| SL1344_RS07565_ | 304 | 305 | 319 | 258 | 228 | 280 |
| SL1344_RS07570_ | 79 | 119 | 107 | 142 | 122 | 137 |
| SL1344_RS07575_ | 76 | 83 | 80 | 103 | 103 | 99 |
| SL1344_RS07580_ | 189 | 154 | 165 | 120 | 93 | 145 |
| SL1344_RS07585_ | 101 | 108 | 107 | 105 | 92 | 114 |
| SL1344_RS07590_ | 95 | 98 | 99 | 105 | 87 | 97 |
| SL1344_RS07595_ | 108 | 107 | 133 | 108 | 118 | 119 |
| SL1344_RS07600_ | 35 | 42 | 35 | 166 | 125 | 134 |
| SL1344_RS07605_ | 211 | 200 | 255 | 394 | 301 | 340 |
| SL1344_RS07610_ | 62 | 67 | 41 | 59 | 63 | 71 |
| SL1344_RS07615_ | 137 | 149 | 162 | 194 | 150 | 190 |
| SL1344_RS07620_ | 278 | 251 | 313 | 298 | 263 | 341 |

| | | | | | | |
|---|---|---|---|---|---|---|
| SL1344_RS07625_ | 86 | 88 | 103 | 136 | 99 | 129 |
| SL1344_RS07630_ | 108 | 119 | 123 | 117 | 118 | 138 |
| SL1344_RS07635_ | 251 | 252 | 226 | 286 | 269 | 318 |
| SL1344_RS07640_ | 288 | 251 | 312 | 306 | 307 | 323 |
| SL1344_RS07645_ | 399 | 381 | 493 | 415 | 343 | 450 |
| SL1344_RS07650_ | 200 | 192 | 212 | 199 | 202 | 238 |
| SL1344_RS07655_ | 196 | 163 | 178 | 150 | 132 | 170 |
| SL1344_RS07660_ | 723 | 692 | 679 | 628 | 607 | 762 |
| SL1344_RS07665_ | 245 | 242 | 264 | 166 | 184 | 210 |
| SL1344_RS07670_ | 126 | 129 | 121 | 75 | 72 | 101 |
| SL1344_RS07675_ | 57 | 57 | 62 | 58 | 57 | 50 |
| SL1344_RS07680_ | 32 | 35 | 35 | 37 | 32 | 48 |
| SL1344_RS07685_ | 45 | 38 | 46 | 42 | 41 | 32 |
| SL1344_RS07690_ | 343 | 258 | 245 | 297 | 343 | 347 |
| SL1344_RS07695_ | 203 | 139 | 171 | 183 | 165 | 233 |
| SL1344_RS07700_ | 197 | 185 | 196 | 247 | 183 | 243 |
| SL1344_RS07705_ | 329 | 296 | 255 | 375 | 351 | 367 |
| SL1344_RS07710_ | 189 | 202 | 240 | 236 | 247 | 229 |
| SL1344_RS07715_ | 51 | 41 | 47 | 55 | 57 | 64 |
| SL1344_RS07720_ | 53 | 35 | 43 | 46 | 46 | 48 |
| SL1344_RS07725_ | 99 | 97 | 94 | 102 | 105 | 121 |
| SL1344_RS07735_ | 95 | 79 | 79 | 148 | 136 | 148 |
| SL1344_RS07740_ | 541 | 457 | 521 | 964 | 882 | 1000 |
| SL1344_RS07745_ | 141 | 156 | 141 | 179 | 161 | 168 |
| SL1344_RS07750_ | 104 | 117 | 127 | 99 | 106 | 139 |
| SL1344_RS07755_ | 214 | 200 | 175 | 200 | 214 | 256 |
| SL1344_RS07760_ | 420 | 489 | 432 | 370 | 319 | 400 |
| SL1344_RS07765_ | 379 | 457 | 436 | 295 | 257 | 325 |
| SL1344_RS07770_ | 615 | 555 | 317 | 437 | 185 | 408 |
| SL1344_RS07775_ | 82 | 65 | 79 | 66 | 48 | 70 |
| SL1344_RS07780_ | 1021 | 1000 | 799 | 1069 | 803 | 1054 |
| SL1344_RS07785_ | 459 | 542 | 516 | 353 | 297 | 396 |
| SL1344_RS07790_ | 89 | 67 | 63 | 57 | 51 | 70 |
| SL1344_RS07800_ | 2860 | 2979 | 3063 | 2766 | 3012 | 3471 |
| SL1344_RS07805_ | 640 | 599 | 574 | 846 | 806 | 1042 |
| SL1344_RS07810_ | 3445 | 3087 | 2686 | 3702 | 3365 | 4351 |
| SL1344_RS07815_ | 851 | 913 | 965 | 802 | 630 | 1125 |
| SL1344_RS07820_ | 3176 | 3043 | 3318 | 3515 | 2655 | 4032 |
| SL1344_RS07825_ | 1510 | 1307 | 1523 | 2026 | 1632 | 2206 |
| SL1344_RS07830_ | 79 | 84 | 68 | 88 | 83 | 74 |
| SL1344_RS07835_ | 160433 | 151401 | 158184 | 150459 | 148213 | 193758 |
| SL1344_RS07840_ | 364 | 406 | 369 | 485 | 400 | 602 |
| SL1344_RS07845_ | 866 | 884 | 1991 | 1824 | 2068 | 271 |
| SL1344_RS07850_ | 443 | 514 | 587 | 691 | 623 | 339 |
| SL1344_RS07855_ | 55 | 52 | 61 | 70 | 77 | 66 |
| SL1344_RS07860_ | 226 | 184 | 238 | 292 | 281 | 299 |
| SL1344_RS07865_ | 87 | 94 | 118 | 149 | 139 | 148 |
| SL1344_RS07870_ | 45 | 33 | 46 | 55 | 35 | 57 |
| SL1344_RS07875_ | 63 | 68 | 109 | 101 | 83 | 73 |
| SL1344_RS07880_ | 230 | 241 | 227 | 284 | 257 | 327 |
| SL1344_RS07885_ | 190 | 160 | 172 | 255 | 192 | 245 |
| SL1344_RS07890_ | 1620 | 1719 | 1610 | 1758 | 1681 | 1930 |
| SL1344_RS07895_ | 176 | 206 | 239 | 334 | 324 | 342 |
| SL1344_RS07900_ | 17 | 11 | 9 | 19 | 13 | 24 |
| SL1344_RS07905_ | 143 | 157 | 129 | 161 | 178 | 217 |
| SL1344_RS07910_ | 290 | 280 | 272 | 267 | 208 | 227 |
| SL1344_RS07915_ | 100 | 72 | 97 | 87 | 79 | 81 |
| SL1344_RS07920_ | 857 | 768 | 745 | 691 | 591 | 727 |
| SL1344_RS07925_ | 346 | 375 | 396 | 358 | 385 | 389 |
| SL1344_RS07930_ | 377 | 404 | 349 | 391 | 372 | 390 |
| SL1344_RS07935_ | 118 | 144 | 120 | 122 | 130 | 145 |
| SL1344_RS07955_ | 21 | 26 | 42 | 56 | 41 | 41 |

| | | | | | | |
|---|---|---|---|---|---|---|
| SL1344_RS07960_ | 824 | 809 | 789 | 633 | 629 | 635 |
| SL1344_RS07965_ | 531 | 518 | 566 | 624 | 574 | 648 |
| SL1344_RS07970_ | 408 | 422 | 433 | 518 | 553 | 541 |
| SL1344_RS07975_ | 47 | 48 | 65 | 51 | 54 | 46 |
| SL1344_RS07980_ | 107 | 121 | 129 | 124 | 91 | 123 |
| SL1344_RS07990_ | 41 | 39 | 44 | 61 | 49 | 60 |
| SL1344_RS07995_ | 581 | 598 | 671 | 706 | 619 | 784 |
| SL1344_RS08000_ | 375 | 295 | 338 | 363 | 359 | 416 |
| SL1344_RS08005_ | 198 | 193 | 198 | 238 | 231 | 228 |
| SL1344_RS08010_ | 2166 | 2147 | 2274 | 2346 | 2221 | 2393 |
| SL1344_RS08015_ | 785 | 805 | 731 | 653 | 593 | 887 |
| SL1344_RS08020_ | 390 | 357 | 339 | 430 | 443 | 451 |
| SL1344_RS08025_ | 105 | 100 | 108 | 82 | 89 | 118 |
| SL1344_RS08030_ | 157 | 191 | 153 | 161 | 144 | 180 |
| SL1344_RS08035_ | 138 | 130 | 119 | 121 | 113 | 118 |
| SL1344_RS08040_ | 58 | 54 | 51 | 47 | 34 | 51 |
| SL1344_RS08045_ | 153 | 132 | 134 | 131 | 105 | 152 |
| SL1344_RS08050_ | 88 | 103 | 90 | 82 | 88 | 95 |
| SL1344_RS08055_ | 41 | 39 | 41 | 48 | 37 | 43 |
| SL1344_RS08060_ | 61 | 81 | 85 | 60 | 58 | 72 |
| SL1344_RS08065_ | 112 | 112 | 136 | 130 | 102 | 154 |
| SL1344_RS08070_ | 144 | 130 | 104 | 120 | 107 | 120 |
| SL1344_RS08075_ | 87 | 77 | 67 | 91 | 86 | 94 |
| SL1344_RS08080_ | 11 | 11 | 10 | 17 | 6 | 12 |
| SL1344_RS08090_ | 634 | 699 | 740 | 497 | 399 | 511 |
| SL1344_RS08095_ | 1117 | 1113 | 1125 | 798 | 745 | 748 |
| SL1344_RS08100_ | 228 | 213 | 252 | 466 | 582 | 514 |
| SL1344_RS08110_ | 309 | 293 | 270 | 200 | 161 | 233 |
| SL1344_RS08115_ | 50 | 37 | 43 | 49 | 41 | 41 |
| SL1344_RS08120_ | 13 | 19 | 97 | 12 | 209 | 14 |
| SL1344_RS08125_ | 3 | 9 | 13 | 15 | 7 | 20 |
| SL1344_RS08130_ | 4 | 5 | 5 | 11 | 17 | 10 |
| SL1344_RS08135_ | 67 | 69 | 66 | 123 | 108 | 136 |
| SL1344_RS08140_ | 37 | 34 | 48 | 74 | 65 | 83 |
| SL1344_RS08145_ | 13 | 24 | 18 | 29 | 29 | 22 |
| SL1344_RS08150_ | 8 | 5 | 11 | 14 | 13 | 16 |
| SL1344_RS08155_ | 47 | 48 | 56 | 73 | 54 | 62 |
| SL1344_RS08160_ | 628 | 563 | 646 | 983 | 891 | 1023 |
| SL1344_RS08165_ | 349 | 368 | 397 | 386 | 415 | 434 |
| SL1344_RS08170_ | 391 | 392 | 372 | 309 | 283 | 397 |
| SL1344_RS08175_ | 1403 | 1535 | 1392 | 1402 | 1295 | 1602 |
| SL1344_RS08185_ | 109 | 111 | 114 | 106 | 130 | 116 |
| SL1344_RS08190_ | 60 | 65 | 75 | 124 | 71 | 105 |
| SL1344_RS08195_ | 73 | 74 | 76 | 93 | 76 | 109 |
| SL1344_RS08200_ | 485 | 574 | 525 | 632 | 536 | 646 |
| SL1344_RS08215_ | 657 | 691 | 606 | 671 | 624 | 740 |
| SL1344_RS08220_ | 26 | 25 | 16 | 13 | 14 | 27 |
| SL1344_RS08225_ | 524 | 582 | 585 | 486 | 443 | 525 |
| SL1344_RS08230_ | 31 | 33 | 29 | 52 | 33 | 35 |
| SL1344_RS08235_ | 5863 | 6057 | 6109 | 6993 | 6498 | 7566 |
| SL1344_RS08240_ | 373 | 411 | 438 | 362 | 353 | 383 |
| SL1344_RS08245_ | 109 | 147 | 122 | 118 | 126 | 117 |
| SL1344_RS08250_ | 40 | 40 | 55 | 58 | 55 | 59 |
| SL1344_RS08255_ | 454 | 496 | 529 | 630 | 590 | 682 |
| SL1344_RS08260_ | 319 | 332 | 346 | 374 | 293 | 359 |
| SL1344_RS08270_ | 301 | 254 | 272 | 311 | 339 | 324 |
| SL1344_RS08275_ | 762 | 761 | 636 | 683 | 821 | 848 |
| SL1344_RS08280_ | 517 | 513 | 455 | 511 | 491 | 503 |
| SL1344_RS08290_ | 3314 | 3563 | 3787 | 3077 | 3048 | 3473 |
| SL1344_RS08295_ | 1658 | 1783 | 1486 | 2277 | 2550 | 2815 |
| SL1344_RS08300_ | 233 | 279 | 290 | 276 | 222 | 268 |
| SL1344_RS08305_ | 214 | 230 | 208 | 208 | 247 | 225 |

| | | | | | | |
|---|---|---|---|---|---|---|
| SL1344_RS08310_ | 126 | 139 | 127 | 126 | 105 | 135 |
| SL1344_RS08315_ | 63 | 59 | 55 | 59 | 66 | 59 |
| SL1344_RS08320_ | 16 | 12 | 13 | 18 | 26 | 29 |
| SL1344_RS08325_ | 11 | 7 | 13 | 6 | 8 | 15 |
| SL1344_RS08330_ | 22 | 19 | 24 | 36 | 51 | 33 |
| SL1344_RS08335_ | 19 | 12 | 6 | 14 | 11 | 24 |
| SL1344_RS08340_ | 10 | 6 | 10 | 15 | 12 | 10 |
| SL1344_RS08345_ | 1864 | 1795 | 2037 | 2331 | 2259 | 2445 |
| SL1344_RS08350_ | 162 | 177 | 150 | 139 | 151 | 171 |
| SL1344_RS08355_ | 59 | 68 | 68 | 56 | 62 | 76 |
| SL1344_RS08360_ | 166 | 171 | 149 | 141 | 140 | 156 |
| SL1344_RS08365_ | 268 | 267 | 260 | 243 | 251 | 291 |
| SL1344_RS08370_ | 66 | 87 | 56 | 59 | 64 | 79 |
| SL1344_RS08375_ | 30 | 36 | 32 | 20 | 21 | 35 |
| SL1344_RS08380_ | 1636 | 1732 | 1705 | 1759 | 1719 | 1947 |
| SL1344_RS08385_ | 151 | 161 | 149 | 249 | 170 | 186 |
| SL1344_RS08390_ | 312 | 338 | 304 | 355 | 341 | 411 |
| SL1344_RS08395_ | 3164 | 3386 | 4096 | 4833 | 4542 | 4894 |
| SL1344_RS08400_ | 818 | 893 | 839 | 718 | 684 | 828 |
| SL1344_RS08405_ | 705 | 687 | 659 | 656 | 661 | 738 |
| SL1344_RS08410_ | 482 | 479 | 491 | 543 | 517 | 513 |
| SL1344_RS08415_ | 1017 | 964 | 1058 | 1081 | 954 | 1124 |
| SL1344_RS08420_ | 126 | 119 | 129 | 118 | 117 | 143 |
| SL1344_RS08425_ | 175 | 165 | 153 | 170 | 172 | 213 |
| SL1344_RS08430_ | 92 | 79 | 80 | 96 | 87 | 94 |
| SL1344_RS08435_ | 972 | 1008 | 1013 | 1116 | 1259 | 1253 |
| SL1344_RS08440_ | 215 | 207 | 239 | 218 | 222 | 258 |
| SL1344_RS08445_ | 460 | 495 | 545 | 577 | 542 | 566 |
| SL1344_RS08450_ | 340 | 315 | 392 | 327 | 332 | 356 |
| SL1344_RS08455_ | 241 | 291 | 322 | 294 | 308 | 309 |
| SL1344_RS08460_ | 335 | 391 | 379 | 385 | 349 | 360 |
| SL1344_RS08465_ | 285 | 333 | 347 | 314 | 258 | 359 |
| SL1344_RS08470_ | 115 | 98 | 114 | 96 | 90 | 70 |
| SL1344_RS08475_ | 45 | 36 | 38 | 52 | 56 | 55 |
| SL1344_RS08480_ | 0 | 2 | 2 | 3 | 0 | 2 |
| SL1344_RS08485_ | 70 | 76 | 78 | 94 | 100 | 89 |
| SL1344_RS08490_ | 4840 | 4598 | 5318 | 6246 | 6643 | 7073 |
| SL1344_RS08500_ | 1247 | 1250 | 1493 | 2175 | 2366 | 2705 |
| SL1344_RS08505_ | 448 | 441 | 385 | 391 | 367 | 399 |
| SL1344_RS08510_ | 506 | 521 | 490 | 525 | 614 | 571 |
| SL1344_RS08515_ | 441 | 419 | 392 | 515 | 510 | 570 |
| SL1344_RS08520_ | 1203 | 1088 | 1039 | 1179 | 1041 | 1533 |
| SL1344_RS08525_ | 154 | 136 | 140 | 151 | 199 | 220 |
| SL1344_RS08530_ | 116 | 137 | 175 | 189 | 187 | 198 |
| SL1344_RS08535_ | 780 | 723 | 702 | 1002 | 863 | 950 |
| SL1344_RS08540_ | 849 | 782 | 765 | 1127 | 1072 | 1220 |
| SL1344_RS08545_ | 204 | 224 | 230 | 186 | 175 | 229 |
| SL1344_RS08550_ | 708 | 828 | 842 | 883 | 877 | 946 |
| SL1344_RS08555_ | 839 | 874 | 969 | 1255 | 1167 | 1386 |
| SL1344_RS08560_ | 616 | 688 | 623 | 440 | 445 | 495 |
| SL1344_RS08570_ | 2721 | 2614 | 2602 | 2710 | 2521 | 2902 |
| SL1344_RS08575_ | 1471 | 1487 | 1381 | 2033 | 1993 | 1950 |
| SL1344_RS08580_ | 635 | 687 | 586 | 541 | 505 | 663 |
| SL1344_RS08590_ | 784 | 733 | 753 | 697 | 656 | 735 |
| SL1344_RS08595_ | 465 | 529 | 541 | 597 | 596 | 720 |
| SL1344_RS08600_ | 656 | 683 | 650 | 773 | 751 | 883 |
| SL1344_RS08605_ | 598 | 583 | 621 | 659 | 694 | 707 |
| SL1344_RS08610_ | 49 | 35 | 51 | 43 | 43 | 56 |
| SL1344_RS08615_ | 66 | 76 | 77 | 96 | 104 | 111 |
| SL1344_RS08620_ | 178 | 143 | 156 | 186 | 211 | 192 |
| SL1344_RS08625_ | 168 | 184 | 189 | 220 | 216 | 236 |
| SL1344_RS08630_ | 222 | 212 | 229 | 262 | 246 | 273 |

| | | | | | | |
|---|---|---|---|---|---|---|
| SL1344_RS08635_ | 113 | 139 | 125 | 95 | 86 | 101 |
| SL1344_RS08640_ | 25 | 26 | 26 | 23 | 19 | 16 |
| SL1344_RS08645_ | 27 | 21 | 19 | 16 | 12 | 17 |
| SL1344_RS08650_ | 92 | 116 | 91 | 90 | 72 | 80 |
| SL1344_RS08655_ | 18889 | 20558 | 17848 | 13215 | 10003 | 15878 |
| SL1344_RS08660_ | 333 | 315 | 340 | 344 | 339 | 342 |
| SL1344_RS08665_ | 1027 | 1013 | 1091 | 1427 | 1389 | 1287 |
| SL1344_RS08670_ | 738 | 746 | 770 | 947 | 876 | 893 |
| SL1344_RS08675_ | 181 | 168 | 212 | 339 | 307 | 329 |
| SL1344_RS08680_ | 130 | 143 | 109 | 141 | 119 | 138 |
| SL1344_RS08685_ | 538 | 554 | 494 | 442 | 334 | 458 |
| SL1344_RS08690_ | 339 | 354 | 333 | 430 | 397 | 474 |
| SL1344_RS08695_ | 490 | 447 | 404 | 437 | 370 | 437 |
| SL1344_RS08700_ | 214 | 213 | 191 | 230 | 209 | 221 |
| SL1344_RS08705_ | 1169 | 1109 | 1109 | 1101 | 868 | 1241 |
| SL1344_RS08710_ | 1120 | 1102 | 1108 | 916 | 925 | 1090 |
| SL1344_RS08715_ | 752 | 754 | 711 | 672 | 558 | 638 |
| SL1344_RS08725_ | 3090 | 3009 | 3284 | 3232 | 2914 | 3490 |
| SL1344_RS08730_ | 132 | 127 | 132 | 157 | 149 | 189 |
| SL1344_RS08735_ | 5468 | 5919 | 5714 | 5644 | 5118 | 6168 |
| SL1344_RS08740_ | 396 | 424 | 418 | 408 | 459 | 523 |
| SL1344_RS08750_ | 3082 | 2768 | 2703 | 2883 | 3026 | 3429 |
| SL1344_RS08755_ | 558 | 567 | 587 | 604 | 521 | 654 |
| SL1344_RS08760_ | 690 | 677 | 655 | 723 | 681 | 776 |
| SL1344_RS08765_ | 283 | 245 | 251 | 412 | 360 | 443 |
| SL1344_RS08770_ | 884 | 834 | 896 | 1508 | 1623 | 1676 |
| SL1344_RS08775_ | 167 | 175 | 229 | 292 | 257 | 299 |
| SL1344_RS08780_ | 85 | 81 | 108 | 137 | 127 | 124 |
| SL1344_RS08785_ | 266 | 293 | 265 | 243 | 212 | 251 |
| SL1344_RS08815_ | 342 | 311 | 353 | 656 | 674 | 644 |
| SL1344_RS08825_ | 243 | 249 | 239 | 296 | 239 | 259 |
| SL1344_RS08830_ | 840 | 909 | 867 | 947 | 904 | 979 |
| SL1344_RS08840_ | 169 | 153 | 197 | 195 | 180 | 173 |
| SL1344_RS08845_ | 2561 | 2389 | 2848 | 2815 | 2736 | 3133 |
| SL1344_RS08850_ | 477 | 475 | 504 | 574 | 527 | 642 |
| SL1344_RS08855_ | 125 | 152 | 125 | 180 | 186 | 201 |
| SL1344_RS08860_ | 382 | 450 | 458 | 558 | 512 | 640 |
| SL1344_RS08865_ | 505 | 524 | 634 | 811 | 721 | 857 |
| SL1344_RS08870_ | 1705 | 1707 | 1791 | 2341 | 2092 | 2344 |
| SL1344_RS08880_ | 2505 | 2414 | 2270 | 3278 | 3743 | 3588 |
| SL1344_RS08885_ | 5182 | 5500 | 6389 | 7572 | 7565 | 8006 |
| SL1344_RS08890_ | 429 | 484 | 531 | 744 | 672 | 731 |
| SL1344_RS08895_ | 910 | 1101 | 975 | 664 | 537 | 901 |
| SL1344_RS08900_ | 210 | 190 | 196 | 322 | 255 | 278 |
| SL1344_RS08905_ | 1725 | 1781 | 1876 | 2156 | 2050 | 2278 |
| SL1344_RS08910_ | 286 | 341 | 302 | 404 | 357 | 409 |
| SL1344_RS08915_ | 24 | 16 | 11 | 23 | 16 | 13 |
| SL1344_RS08920_ | 56 | 54 | 63 | 59 | 65 | 60 |
| SL1344_RS08925_ | 8 | 17 | 11 | 15 | 12 | 14 |
| SL1344_RS08930_ | 7 | 12 | 4 | 17 | 17 | 20 |
| SL1344_RS08935_ | 12 | 9 | 5 | 8 | 8 | 12 |
| SL1344_RS08940_ | 15 | 19 | 11 | 20 | 14 | 22 |
| SL1344_RS08945_ | 36 | 35 | 38 | 43 | 31 | 35 |
| SL1344_RS08950_ | 32 | 39 | 29 | 45 | 49 | 51 |
| SL1344_RS08955_ | 26 | 33 | 30 | 47 | 39 | 49 |
| SL1344_RS08960_ | 5017 | 4704 | 6116 | 6892 | 7916 | 8182 |
| SL1344_RS08965_ | 1155 | 1244 | 1324 | 986 | 896 | 1064 |
| SL1344_RS08970_ | 348 | 363 | 279 | 203 | 154 | 242 |
| SL1344_RS08985_ | 177 | 163 | 204 | 194 | 194 | 221 |
| SL1344_RS08995_ | 396 | 370 | 479 | 993 | 982 | 1069 |
| SL1344_RS09000_ | 418 | 381 | 492 | 1033 | 1077 | 1206 |
| SL1344_RS09005_ | 1088 | 1192 | 1083 | 702 | 693 | 706 |

| | | | | | | |
|---|---|---|---|---|---|---|
| SL1344_RS09010_ | 1187 | 1291 | 1521 | 1185 | 1154 | 1350 |
| SL1344_RS09015_ | 634 | 602 | 727 | 704 | 613 | 749 |
| SL1344_RS09020_ | 668 | 656 | 726 | 1064 | 885 | 935 |
| SL1344_RS09025_ | 48 | 50 | 39 | 50 | 73 | 50 |
| SL1344_RS09030_ | 1836 | 1822 | 1626 | 1243 | 1139 | 1551 |
| SL1344_RS09035_ | 207 | 196 | 185 | 172 | 168 | 195 |
| SL1344_RS09040_ | 295 | 305 | 305 | 315 | 304 | 343 |
| SL1344_RS09045_ | 702 | 651 | 693 | 730 | 672 | 758 |
| SL1344_RS09050_ | 321 | 308 | 395 | 341 | 298 | 373 |
| SL1344_RS09055_ | 1291 | 1273 | 1245 | 1886 | 1584 | 1837 |
| SL1344_RS09060_ | 4376 | 4017 | 4151 | 4075 | 4181 | 4861 |
| SL1344_RS09065_ | 3158 | 3189 | 3635 | 3452 | 3167 | 3756 |
| SL1344_RS09070_ | 94 | 81 | 84 | 184 | 174 | 202 |
| SL1344_RS09085_ | 338 | 364 | 344 | 608 | 562 | 667 |
| SL1344_RS09090_ | 287 | 284 | 297 | 577 | 505 | 724 |
| SL1344_RS09095_ | 449 | 442 | 425 | 789 | 769 | 825 |
| SL1344_RS09100_ | 1002 | 1032 | 977 | 1774 | 1886 | 1946 |
| SL1344_RS09105_ | 271 | 310 | 282 | 416 | 539 | 481 |
| SL1344_RS09115_ | 308 | 321 | 329 | 455 | 432 | 501 |
| SL1344_RS09125_ | 71 | 90 | 79 | 99 | 78 | 100 |
| SL1344_RS09130_ | 320 | 329 | 296 | 443 | 365 | 488 |
| SL1344_RS09135_ | 110 | 120 | 134 | 178 | 181 | 192 |
| SL1344_RS09140_ | 4303 | 4628 | 4752 | 4604 | 4478 | 5574 |
| SL1344_RS09145_ | 2954 | 3270 | 3404 | 2972 | 2809 | 3572 |
| SL1344_RS09150_ | 6988 | 7587 | 8059 | 7375 | 6432 | 8091 |
| SL1344_RS09155_ | 119 | 135 | 137 | 205 | 181 | 175 |
| SL1344_RS09160_ | 188 | 193 | 162 | 388 | 332 | 324 |
| SL1344_RS09165_ | 67 | 92 | 119 | 178 | 187 | 210 |
| SL1344_RS09170_ | 152 | 126 | 150 | 281 | 261 | 328 |
| SL1344_RS09175_ | 8988 | 7198 | 8062 | 13060 | 12496 | 12586 |
| SL1344_RS09180_ | 5210 | 4023 | 4528 | 8233 | 8821 | 8633 |
| SL1344_RS09185_ | 601 | 592 | 588 | 722 | 583 | 775 |
| SL1344_RS09190_ | 610 | 661 | 632 | 472 | 483 | 687 |
| SL1344_RS09200_ | 1791 | 1809 | 1755 | 2103 | 2163 | 2488 |
| SL1344_RS09205_ | 75 | 66 | 88 | 157 | 103 | 110 |
| SL1344_RS09210_ | 1803 | 1891 | 2034 | 2844 | 2900 | 3190 |
| SL1344_RS09235_ | 1163 | 1164 | 1222 | 1553 | 1444 | 1623 |
| SL1344_RS09245_ | 374 | 427 | 345 | 646 | 547 | 609 |
| SL1344_RS09250_ | 85 | 75 | 95 | 105 | 98 | 105 |
| SL1344_RS09260_ | 288 | 257 | 196 | 465 | 386 | 542 |
| SL1344_RS09270_ | 21 | 15 | 21 | 31 | 22 | 20 |
| SL1344_RS09275_ | 234 | 220 | 309 | 461 | 479 | 448 |
| SL1344_RS09285_ | 64 | 100 | 67 | 116 | 117 | 130 |
| SL1344_RS09290_ | 159 | 147 | 152 | 188 | 204 | 228 |
| SL1344_RS09295_ | 32 | 24 | 38 | 43 | 30 | 32 |
| SL1344_RS09300_ | 10 | 16 | 16 | 16 | 10 | 13 |
| SL1344_RS09305_ | 71 | 60 | 78 | 119 | 114 | 123 |
| SL1344_RS09310_ | 5 | 9 | 3 | 8 | 10 | 10 |
| SL1344_RS09315_ | 44 | 42 | 29 | 48 | 44 | 49 |
| SL1344_RS09320_ | 709 | 700 | 610 | 759 | 850 | 819 |
| SL1344_RS09325_ | 21 | 15 | 18 | 25 | 22 | 42 |
| SL1344_RS09335_ | 6 | 6 | 4 | 11 | 11 | 14 |
| SL1344_RS09340_ | 51 | 39 | 50 | 122 | 97 | 89 |
| SL1344_RS09345_ | 2 | 5 | 3 | 16 | 7 | 12 |
| SL1344_RS09350_ | 3 | 3 | 5 | 12 | 9 | 9 |
| SL1344_RS09355_ | 15 | 14 | 8 | 21 | 17 | 19 |
| SL1344_RS09360_ | 50 | 58 | 52 | 56 | 69 | 93 |
| SL1344_RS09365_ | 407 | 511 | 832 | 459 | 322 | 371 |
| SL1344_RS09370_ | 496 | 470 | 446 | 954 | 882 | 987 |
| SL1344_RS09380_ | 846 | 816 | 882 | 1376 | 1434 | 1515 |
| SL1344_RS09385_ | 132 | 119 | 90 | 167 | 209 | 230 |
| SL1344_RS09390_ | 199 | 219 | 206 | 286 | 229 | 221 |

| | | | | | | |
|---|---|---|---|---|---|---|
| SL1344_RS09395_ | 374 | 395 | 378 | 572 | 575 | 616 |
| SL1344_RS09400_ | 709 | 810 | 807 | 929 | 865 | 1010 |
| SL1344_RS09405_ | 312 | 309 | 283 | 465 | 395 | 516 |
| SL1344_RS09410_ | 677 | 735 | 689 | 965 | 905 | 1080 |
| SL1344_RS09420_ | 175 | 194 | 155 | 347 | 352 | 367 |
| SL1344_RS09425_ | 1079 | 1170 | 1043 | 1386 | 1269 | 1633 |
| SL1344_RS09430_ | 469 | 497 | 472 | 591 | 586 | 733 |
| SL1344_RS09435_ | 1825 | 1699 | 1678 | 2536 | 2274 | 3195 |
| SL1344_RS09440_ | 695 | 735 | 723 | 765 | 709 | 827 |
| SL1344_RS09455_ | 908 | 875 | 942 | 1493 | 1197 | 1513 |
| SL1344_RS09460_ | 500 | 451 | 457 | 694 | 623 | 731 |
| SL1344_RS09465_ | 202 | 222 | 251 | 371 | 353 | 408 |
| SL1344_RS09470_ | 238 | 280 | 249 | 393 | 412 | 401 |
| SL1344_RS09485_ | 15 | 9 | 11 | 34 | 45 | 15 |
| SL1344_RS09490_ | 52 | 33 | 39 | 70 | 72 | 89 |
| SL1344_RS09495_ | 1000 | 1009 | 1148 | 1572 | 1569 | 1911 |
| SL1344_RS09500_ | 1105 | 1127 | 1119 | 1456 | 1536 | 2050 |
| SL1344_RS09510_ | 2070 | 2231 | 2394 | 3776 | 3661 | 4459 |
| SL1344_RS09515_ | 260 | 267 | 222 | 355 | 375 | 382 |
| SL1344_RS09520_ | 161 | 154 | 178 | 267 | 248 | 227 |
| SL1344_RS09525_ | 690 | 658 | 702 | 1048 | 1063 | 967 |
| SL1344_RS09535_ | 607 | 696 | 734 | 870 | 817 | 935 |
| SL1344_RS09540_ | 320 | 274 | 294 | 542 | 427 | 500 |
| SL1344_RS09545_ | 93 | 90 | 94 | 191 | 161 | 225 |
| SL1344_RS09550_ | 884 | 938 | 1101 | 1986 | 1685 | 2149 |
| SL1344_RS09555_ | 54 | 52 | 53 | 107 | 110 | 70 |
| SL1344_RS09635_ | 123 | 115 | 149 | 149 | 146 | 169 |
| SL1344_RS09640_ | 718 | 801 | 799 | 765 | 943 | 1089 |
| SL1344_RS09645_ | 428 | 460 | 435 | 484 | 415 | 578 |
| SL1344_RS09650_ | 186 | 227 | 198 | 218 | 183 | 281 |
| SL1344_RS09655_ | 1 | 0 | 2 | 1 | 1 | 7 |
| SL1344_RS09660_ | 94 | 98 | 86 | 124 | 137 | 167 |
| SL1344_RS09670_ | 122 | 123 | 127 | 145 | 161 | 169 |
| SL1344_RS09680_ | 5346 | 5265 | 6881 | 7760 | 7740 | 8108 |
| SL1344_RS09685_ | 69 | 80 | 68 | 145 | 109 | 154 |
| SL1344_RS09690_ | 166 | 169 | 197 | 281 | 260 | 297 |
| SL1344_RS09695_ | 1071 | 1105 | 997 | 1270 | 1183 | 1417 |
| SL1344_RS09700_ | 588 | 522 | 484 | 827 | 665 | 818 |
| SL1344_RS09705_ | 399 | 408 | 421 | 508 | 475 | 545 |
| SL1344_RS09710_ | 89 | 104 | 108 | 191 | 178 | 192 |
| SL1344_RS09715_ | 394 | 440 | 644 | 913 | 604 | 714 |
| SL1344_RS09725_ | 659 | 714 | 838 | 989 | 1036 | 1210 |
| SL1344_RS09730_ | 905 | 898 | 982 | 1367 | 1298 | 1575 |
| SL1344_RS09805_ | 17411 | 17940 | 16964 | 31471 | 27405 | 36318 |
| SL1344_RS09810_ | 6495 | 6379 | 6445 | 11358 | 10333 | 12872 |
| SL1344_RS09820_ | 355 | 334 | 329 | 500 | 379 | 412 |
| SL1344_RS09850_ | 6367 | 7216 | 7143 | 11220 | 10999 | 11711 |
| SL1344_RS09855_ | 3930 | 4222 | 4210 | 6047 | 5901 | 6339 |
| SL1344_RS09860_ | 2728 | 3160 | 2978 | 4275 | 4494 | 4659 |
| SL1344_RS09875_ | 3369 | 3849 | 3756 | 5060 | 4684 | 5572 |
| SL1344_RS09885_ | 3531 | 3646 | 4032 | 6557 | 5910 | 6537 |
| SL1344_RS09890_ | 2385 | 2609 | 2545 | 3653 | 3733 | 3879 |
| SL1344_RS09895_ | 1314 | 1394 | 1390 | 2418 | 2069 | 2325 |
| SL1344_RS09900_ | 1452 | 1397 | 1549 | 2871 | 2449 | 2685 |
| SL1344_RS09910_ | 286 | 315 | 391 | 642 | 519 | 550 |
| SL1344_RS09920_ | 732 | 673 | 533 | 471 | 493 | 531 |
| SL1344_RS09925_ | 170 | 142 | 139 | 72 | 68 | 104 |
| SL1344_RS09930_ | 100 | 101 | 88 | 78 | 59 | 72 |
| SL1344_RS09935_ | 316 | 311 | 323 | 221 | 224 | 244 |
| SL1344_RS09940_ | 2836 | 2867 | 2285 | 1487 | 1396 | 1749 |
| SL1344_RS09945_ | 241 | 268 | 210 | 239 | 232 | 245 |
| SL1344_RS09950_ | 92 | 77 | 64 | 69 | 49 | 77 |

| | | | | | | |
|---|---|---|---|---|---|---|
| SL1344_RS09955_ | 226 | 206 | 237 | 149 | 147 | 188 |
| SL1344_RS09960_ | 448 | 479 | 512 | 418 | 385 | 444 |
| SL1344_RS09965_ | 67 | 92 | 83 | 72 | 84 | 76 |
| SL1344_RS09970_ | 17 | 9 | 10 | 19 | 11 | 17 |
| SL1344_RS09975_ | 224 | 178 | 191 | 286 | 270 | 260 |
| SL1344_RS09985_ | 160 | 165 | 185 | 148 | 157 | 189 |
| SL1344_RS09990_ | 51 | 43 | 46 | 40 | 53 | 52 |
| SL1344_RS09995_ | 5 | 8 | 3 | 6 | 6 | 6 |
| SL1344_RS10000_ | 33 | 23 | 23 | 46 | 42 | 38 |
| SL1344_RS10005_ | 78 | 69 | 69 | 90 | 86 | 84 |
| SL1344_RS10010_ | 70 | 68 | 67 | 71 | 66 | 72 |
| SL1344_RS10015_ | 21 | 10 | 18 | 21 | 19 | 23 |
| SL1344_RS10020_ | 23 | 19 | 31 | 22 | 18 | 37 |
| SL1344_RS10025_ | 29 | 32 | 32 | 40 | 31 | 44 |
| SL1344_RS10030_ | 17 | 9 | 18 | 19 | 17 | 26 |
| SL1344_RS10035_ | 30 | 28 | 33 | 41 | 27 | 46 |
| SL1344_RS10040_ | 73 | 66 | 77 | 75 | 75 | 98 |
| SL1344_RS10045_ | 57 | 63 | 66 | 58 | 77 | 80 |
| SL1344_RS10050_ | 232 | 257 | 221 | 230 | 211 | 256 |
| SL1344_RS10055_ | 50 | 62 | 59 | 42 | 48 | 75 |
| SL1344_RS10060_ | 61 | 72 | 83 | 78 | 73 | 97 |
| SL1344_RS10065_ | 131 | 98 | 131 | 150 | 126 | 141 |
| SL1344_RS10070_ | 27 | 26 | 40 | 46 | 43 | 32 |
| SL1344_RS10075_ | 20 | 21 | 29 | 42 | 33 | 36 |
| SL1344_RS10080_ | 16 | 17 | 14 | 20 | 23 | 21 |
| SL1344_RS10085_ | 16 | 24 | 20 | 34 | 27 | 23 |
| SL1344_RS10090_ | 25 | 24 | 9 | 37 | 32 | 24 |
| SL1344_RS10095_ | 99 | 93 | 78 | 125 | 106 | 139 |
| SL1344_RS10100_ | 34 | 31 | 33 | 54 | 32 | 54 |
| SL1344_RS10105_ | 42 | 52 | 49 | 83 | 56 | 85 |
| SL1344_RS10110_ | 10 | 8 | 13 | 18 | 18 | 13 |
| SL1344_RS10115_ | 98 | 89 | 93 | 128 | 139 | 135 |
| SL1344_RS10120_ | 31 | 31 | 24 | 38 | 47 | 39 |
| SL1344_RS10130_ | 82 | 110 | 87 | 119 | 118 | 94 |
| SL1344_RS10135_ | 29 | 33 | 31 | 47 | 43 | 43 |
| SL1344_RS10140_ | 54 | 45 | 32 | 68 | 51 | 57 |
| SL1344_RS10145_ | 37 | 29 | 37 | 69 | 39 | 52 |
| SL1344_RS10150_ | 19 | 18 | 16 | 27 | 40 | 33 |
| SL1344_RS10155_ | 54 | 51 | 67 | 78 | 59 | 73 |
| SL1344_RS10160_ | 89 | 79 | 61 | 116 | 104 | 130 |
| SL1344_RS10165_ | 61 | 60 | 73 | 82 | 70 | 85 |
| SL1344_RS10170_ | 77 | 62 | 81 | 78 | 73 | 102 |
| SL1344_RS10175_ | 32 | 32 | 22 | 26 | 34 | 51 |
| SL1344_RS10180_ | 65 | 56 | 61 | 50 | 43 | 75 |
| SL1344_RS10185_ | 26 | 39 | 44 | 39 | 33 | 30 |
| SL1344_RS10190_ | 68 | 80 | 53 | 75 | 69 | 99 |
| SL1344_RS10195_ | 23 | 18 | 20 | 21 | 19 | 24 |
| SL1344_RS10200_ | 449 | 461 | 549 | 791 | 626 | 584 |
| SL1344_RS10205_ | 125 | 105 | 89 | 120 | 111 | 133 |
| SL1344_RS10210_ | 53 | 57 | 68 | 47 | 67 | 73 |
| SL1344_RS10215_ | 2412 | 2302 | 2381 | 2457 | 2488 | 2614 |
| SL1344_RS10225_ | 187 | 172 | 165 | 169 | 173 | 190 |
| SL1344_RS10230_ | 438 | 498 | 469 | 465 | 467 | 516 |
| SL1344_RS10235_ | 33 | 36 | 41 | 31 | 28 | 31 |
| SL1344_RS10240_ | 16 | 11 | 15 | 13 | 9 | 13 |
| SL1344_RS10245_ | 33 | 36 | 35 | 23 | 32 | 50 |
| SL1344_RS10250_ | 58 | 60 | 65 | 53 | 45 | 50 |
| SL1344_RS10255_ | 96 | 99 | 108 | 95 | 80 | 88 |
| SL1344_RS10260_ | 79 | 72 | 87 | 77 | 60 | 72 |
| SL1344_RS10265_ | 40 | 52 | 36 | 45 | 29 | 58 |
| SL1344_RS10270_ | 106 | 78 | 75 | 82 | 72 | 70 |
| SL1344_RS10275_ | 395 | 492 | 565 | 374 | 342 | 410 |

| | | | | | | |
|---|---|---|---|---|---|---|
| SL1344_RS10280_ | 2072 | 2019 | 2200 | 1578 | 1538 | 1567 |
| SL1344_RS10285_ | 99 | 71 | 150 | 137 | 81 | 99 |
| SL1344_RS10290_ | 6 | 8 | 9 | 9 | 3 | 2 |
| SL1344_RS10295_ | 7 | 1 | 1 | 5 | 4 | 6 |
| SL1344_RS10300_ | 9 | 7 | 25 | 23 | 11 | 15 |
| SL1344_RS10305_ | 26 | 29 | 26 | 34 | 32 | 31 |
| SL1344_RS10310_ | 10 | 1 | 7 | 4 | 9 | 8 |
| SL1344_RS10315_ | 52 | 52 | 50 | 60 | 44 | 48 |
| SL1344_RS10320_ | 812 | 778 | 815 | 798 | 824 | 900 |
| SL1344_RS10325_ | 22 | 21 | 26 | 29 | 24 | 22 |
| SL1344_RS10330_ | 1067 | 1059 | 1020 | 913 | 758 | 985 |
| SL1344_RS10335_ | 34 | 46 | 32 | 37 | 28 | 42 |
| SL1344_RS10340_ | 168 | 172 | 174 | 159 | 109 | 131 |
| SL1344_RS10345_ | 322 | 321 | 334 | 215 | 193 | 242 |
| SL1344_RS10350_ | 551 | 526 | 487 | 372 | 490 | 474 |
| SL1344_RS10355_ | 136 | 89 | 160 | 134 | 86 | 144 |
| SL1344_RS10360_ | 149 | 152 | 139 | 103 | 92 | 101 |
| SL1344_RS10365_ | 74 | 52 | 95 | 84 | 67 | 66 |
| SL1344_RS10370_ | 74 | 71 | 70 | 40 | 38 | 41 |
| SL1344_RS10375_ | 574 | 631 | 579 | 354 | 312 | 371 |
| SL1344_RS10380_ | 761 | 809 | 766 | 488 | 414 | 561 |
| SL1344_RS10390_ | 309 | 385 | 289 | 201 | 140 | 197 |
| SL1344_RS10395_ | 789 | 846 | 763 | 494 | 418 | 491 |
| SL1344_RS10400_ | 545 | 582 | 535 | 343 | 266 | 300 |
| SL1344_RS10405_ | 551 | 630 | 586 | 316 | 223 | 269 |
| SL1344_RS10410_ | 225 | 261 | 240 | 162 | 85 | 120 |
| SL1344_RS10415_ | 648 | 674 | 665 | 340 | 249 | 353 |
| SL1344_RS10420_ | 644 | 770 | 719 | 382 | 288 | 355 |
| SL1344_RS10430_ | 965 | 1062 | 935 | 527 | 424 | 573 |
| SL1344_RS10435_ | 1031 | 1188 | 985 | 496 | 440 | 595 |
| SL1344_RS10440_ | 840 | 957 | 991 | 455 | 340 | 462 |
| SL1344_RS10445_ | 643 | 813 | 659 | 269 | 228 | 355 |
| SL1344_RS10450_ | 467 | 511 | 478 | 261 | 200 | 286 |
| SL1344_RS10455_ | 405 | 486 | 393 | 261 | 177 | 284 |
| SL1344_RS10470_ | 818 | 825 | 734 | 545 | 413 | 450 |
| SL1344_RS10475_ | 857 | 921 | 806 | 541 | 387 | 523 |
| SL1344_RS10480_ | 516 | 685 | 662 | 389 | 288 | 387 |
| SL1344_RS10485_ | 74 | 80 | 83 | 49 | 62 | 48 |
| SL1344_RS10490_ | 37 | 60 | 41 | 6 | 2 | 6 |
| SL1344_RS10495_ | 57 | 84 | 62 | 10 | 12 | 13 |
| SL1344_RS10515_ | 59 | 75 | 90 | 31 | 30 | 24 |
| SL1344_RS10520_ | 12 | 11 | 9 | 2 | 4 | 1 |
| SL1344_RS10530_ | 25 | 25 | 31 | 18 | 18 | 17 |
| SL1344_RS10535_ | 16 | 24 | 36 | 23 | 14 | 26 |
| SL1344_RS10540_ | 14 | 14 | 17 | 23 | 17 | 17 |
| SL1344_RS10545_ | 5 | 5 | 8 | 6 | 7 | 10 |
| SL1344_RS10550_ | 16 | 15 | 24 | 20 | 20 | 23 |
| SL1344_RS10555_ | 26 | 38 | 37 | 27 | 27 | 40 |
| SL1344_RS10560_ | 19 | 32 | 24 | 33 | 26 | 40 |
| SL1344_RS10565_ | 24 | 27 | 38 | 37 | 40 | 48 |
| SL1344_RS10570_ | 13 | 14 | 19 | 13 | 26 | 20 |
| SL1344_RS10575_ | 5 | 5 | 3 | 6 | 17 | 12 |
| SL1344_RS10580_ | 8 | 5 | 9 | 13 | 17 | 19 |
| SL1344_RS10585_ | 62 | 71 | 74 | 70 | 71 | 84 |
| SL1344_RS10590_ | 100 | 136 | 134 | 138 | 123 | 97 |
| SL1344_RS10595_ | 2777 | 2653 | 2798 | 2555 | 2557 | 2744 |
| SL1344_RS10600_ | 581 | 691 | 632 | 745 | 664 | 739 |
| SL1344_RS10640_ | 899 | 993 | 1372 | 1111 | 1193 | 1450 |
| SL1344_RS10645_ | 422 | 387 | 386 | 265 | 215 | 281 |
| SL1344_RS10650_ | 1129 | 1126 | 926 | 723 | 714 | 791 |
| SL1344_RS10655_ | 212 | 218 | 213 | 134 | 131 | 125 |
| SL1344_RS10660_ | 237 | 254 | 280 | 121 | 153 | 157 |

| | | | | | | |
|---|---|---|---|---|---|---|
| SL1344_RS10665_ | 244 | 240 | 288 | 151 | 155 | 160 |
| SL1344_RS10670_ | 459 | 492 | 571 | 365 | 323 | 356 |
| SL1344_RS10675_ | 229 | 255 | 309 | 148 | 136 | 161 |
| SL1344_RS10680_ | 327 | 322 | 329 | 219 | 213 | 261 |
| SL1344_RS10685_ | 436 | 466 | 478 | 292 | 303 | 345 |
| SL1344_RS10690_ | 341 | 326 | 376 | 216 | 227 | 236 |
| SL1344_RS10700_ | 980 | 914 | 1045 | 752 | 683 | 748 |
| SL1344_RS10705_ | 2170 | 2269 | 2591 | 1284 | 1154 | 1465 |
| SL1344_RS10710_ | 1501 | 1362 | 1432 | 934 | 898 | 1118 |
| SL1344_RS10715_ | 1492 | 1340 | 1441 | 839 | 857 | 1062 |
| SL1344_RS10720_ | 1662 | 1610 | 1802 | 1172 | 1157 | 1321 |
| SL1344_RS10725_ | 1137 | 1127 | 1260 | 786 | 908 | 939 |
| SL1344_RS10730_ | 1013 | 1035 | 1103 | 721 | 849 | 948 |
| SL1344_RS10735_ | 1402 | 1329 | 1334 | 983 | 1126 | 1190 |
| SL1344_RS10740_ | 1305 | 1289 | 1364 | 926 | 966 | 1073 |
| SL1344_RS10745_ | 2133 | 2155 | 2464 | 1549 | 1531 | 1725 |
| SL1344_RS10750_ | 5803 | 5238 | 5811 | 3601 | 3750 | 4604 |
| SL1344_RS10755_ | 2142 | 1891 | 2054 | 1118 | 1258 | 1562 |
| SL1344_RS10760_ | 1348 | 1193 | 1390 | 653 | 728 | 981 |
| SL1344_RS10765_ | 1352 | 1295 | 1448 | 916 | 900 | 1168 |
| SL1344_RS10770_ | 1016 | 963 | 980 | 669 | 691 | 885 |
| SL1344_RS10775_ | 777 | 716 | 735 | 534 | 498 | 681 |
| SL1344_RS10780_ | 1059 | 1039 | 1122 | 791 | 690 | 883 |
| SL1344_RS10785_ | 1023 | 974 | 1105 | 874 | 843 | 966 |
| SL1344_RS10790_ | 1975 | 1669 | 1832 | 1836 | 1788 | 2209 |
| SL1344_RS10795_ | 40 | 37 | 39 | 64 | 49 | 45 |
| SL1344_RS10800_ | 37 | 19 | 26 | 33 | 40 | 41 |
| SL1344_RS10805_ | 37 | 32 | 35 | 25 | 21 | 46 |
| SL1344_RS10810_ | 36 | 31 | 30 | 36 | 25 | 31 |
| SL1344_RS10815_ | 27 | 16 | 27 | 24 | 34 | 39 |
| SL1344_RS10820_ | 21 | 20 | 25 | 24 | 20 | 19 |
| SL1344_RS10825_ | 44 | 38 | 44 | 39 | 38 | 41 |
| SL1344_RS10830_ | 31 | 35 | 40 | 41 | 37 | 48 |
| SL1344_RS10835_ | 18 | 16 | 22 | 16 | 13 | 18 |
| SL1344_RS10840_ | 33 | 31 | 32 | 31 | 41 | 48 |
| SL1344_RS10845_ | 42 | 25 | 29 | 40 | 23 | 37 |
| SL1344_RS10850_ | 5 | 6 | 4 | 7 | 5 | 5 |
| SL1344_RS10855_ | 15 | 8 | 10 | 14 | 21 | 15 |
| SL1344_RS10860_ | 31 | 34 | 26 | 24 | 31 | 20 |
| SL1344_RS10865_ | 35 | 30 | 39 | 30 | 36 | 55 |
| SL1344_RS10870_ | 17 | 13 | 11 | 16 | 7 | 19 |
| SL1344_RS10875_ | 33 | 37 | 22 | 26 | 19 | 29 |
| SL1344_RS10880_ | 52 | 53 | 45 | 48 | 42 | 72 |
| SL1344_RS10885_ | 8 | 12 | 6 | 4 | 4 | 7 |
| SL1344_RS10890_ | 29 | 16 | 24 | 13 | 13 | 12 |
| SL1344_RS10895_ | 371 | 370 | 398 | 330 | 222 | 322 |
| SL1344_RS10900_ | 2310 | 2339 | 2220 | 1750 | 1767 | 1965 |
| SL1344_RS10905_ | 682 | 769 | 796 | 611 | 585 | 725 |
| SL1344_RS10910_ | 262 | 275 | 349 | 408 | 323 | 388 |
| SL1344_RS10915_ | 544 | 586 | 657 | 461 | 402 | 447 |
| SL1344_RS10920_ | 41 | 43 | 42 | 35 | 34 | 33 |
| SL1344_RS10930_ | 80 | 102 | 87 | 85 | 82 | 92 |
| SL1344_RS10935_ | 151 | 153 | 197 | 137 | 103 | 114 |
| SL1344_RS10940_ | 136 | 158 | 147 | 127 | 130 | 114 |
| SL1344_RS10945_ | 105 | 134 | 147 | 122 | 135 | 121 |
| SL1344_RS10950_ | 200 | 195 | 221 | 220 | 189 | 230 |
| SL1344_RS10955_ | 275 | 310 | 331 | 285 | 282 | 311 |
| SL1344_RS10960_ | 12 | 15 | 17 | 14 | 13 | 12 |
| SL1344_RS10965_ | 14 | 21 | 38 | 17 | 12 | 20 |
| SL1344_RS10970_ | 125 | 104 | 126 | 94 | 135 | 106 |
| SL1344_RS10975_ | 147 | 138 | 177 | 172 | 131 | 171 |
| SL1344_RS10980_ | 15 | 17 | 16 | 15 | 15 | 9 |

| | | | | | | |
|---|---|---|---|---|---|---|
| SL1344_RS10985_ | 22 | 15 | 20 | 29 | 21 | 32 |
| SL1344_RS10990_ | 42 | 34 | 35 | 49 | 35 | 73 |
| SL1344_RS10995_ | 4 | 3 | 3 | 1 | 3 | 9 |
| SL1344_RS11000_ | 237 | 244 | 242 | 201 | 165 | 178 |
| SL1344_RS11005_ | 1630 | 1674 | 1623 | 780 | 740 | 1109 |
| SL1344_RS11010_ | 168 | 181 | 192 | 277 | 296 | 300 |
| SL1344_RS11015_ | 161 | 203 | 225 | 273 | 257 | 315 |
| SL1344_RS11020_ | 127 | 153 | 173 | 228 | 201 | 251 |
| SL1344_RS11025_ | 348 | 389 | 377 | 334 | 295 | 355 |
| SL1344_RS11030_ | 140 | 123 | 115 | 87 | 65 | 84 |
| SL1344_RS11035_ | 79 | 121 | 126 | 68 | 54 | 70 |
| SL1344_RS11040_ | 214 | 203 | 216 | 219 | 209 | 196 |
| SL1344_RS11045_ | 39 | 38 | 41 | 45 | 27 | 41 |
| SL1344_RS11050_ | 75 | 88 | 73 | 78 | 58 | 89 |
| SL1344_RS11055_ | 30 | 10 | 16 | 20 | 20 | 25 |
| SL1344_RS11060_ | 389 | 326 | 303 | 289 | 281 | 349 |
| SL1344_RS11065_ | 98 | 61 | 90 | 85 | 65 | 86 |
| SL1344_RS11070_ | 1632 | 1603 | 1808 | 1576 | 1471 | 1788 |
| SL1344_RS11080_ | 40 | 55 | 44 | 55 | 65 | 48 |
| SL1344_RS11085_ | 13 | 22 | 20 | 31 | 27 | 24 |
| SL1344_RS11090_ | 253 | 262 | 297 | 343 | 321 | 312 |
| SL1344_RS11095_ | 412 | 404 | 437 | 401 | 396 | 460 |
| SL1344_RS11100_ | 645 | 599 | 603 | 654 | 639 | 725 |
| SL1344_RS11105_ | 176 | 200 | 181 | 109 | 88 | 91 |
| SL1344_RS11110_ | 213 | 251 | 232 | 239 | 235 | 278 |
| SL1344_RS11115_ | 161 | 177 | 162 | 118 | 117 | 108 |
| SL1344_RS11120_ | 283 | 284 | 272 | 186 | 154 | 170 |
| SL1344_RS11125_ | 539 | 514 | 506 | 302 | 251 | 293 |
| SL1344_RS11130_ | 576 | 587 | 515 | 338 | 302 | 395 |
| SL1344_RS11135_ | 763 | 766 | 908 | 578 | 530 | 614 |
| SL1344_RS11140_ | 1290 | 1372 | 1433 | 889 | 893 | 1004 |
| SL1344_RS11145_ | 502 | 515 | 505 | 461 | 445 | 504 |
| SL1344_RS11150_ | 49 | 72 | 59 | 47 | 45 | 51 |
| SL1344_RS11155_ | 55 | 58 | 49 | 51 | 32 | 54 |
| SL1344_RS11160_ | 143 | 159 | 162 | 92 | 84 | 101 |
| SL1344_RS11165_ | 40 | 50 | 49 | 43 | 37 | 42 |
| SL1344_RS11170_ | 101 | 101 | 117 | 95 | 84 | 108 |
| SL1344_RS11175_ | 170 | 163 | 145 | 164 | 124 | 185 |
| SL1344_RS11180_ | 50 | 55 | 51 | 55 | 49 | 71 |
| SL1344_RS11185_ | 38 | 51 | 54 | 49 | 32 | 43 |
| SL1344_RS11190_ | 39 | 31 | 51 | 34 | 21 | 37 |
| SL1344_RS11195_ | 19 | 22 | 25 | 34 | 34 | 60 |
| SL1344_RS11200_ | 82 | 74 | 93 | 156 | 113 | 142 |
| SL1344_RS11205_ | 102 | 108 | 139 | 138 | 122 | 152 |
| SL1344_RS11210_ | 202 | 209 | 195 | 215 | 171 | 186 |
| SL1344_RS11215_ | 448 | 475 | 523 | 570 | 514 | 603 |
| SL1344_RS11220_ | 646 | 693 | 781 | 669 | 636 | 649 |
| SL1344_RS11225_ | 292 | 255 | 295 | 245 | 205 | 223 |
| SL1344_RS11230_ | 384 | 389 | 423 | 262 | 262 | 297 |
| SL1344_RS11235_ | 956 | 967 | 916 | 628 | 587 | 703 |
| SL1344_RS11240_ | 273 | 308 | 324 | 278 | 229 | 259 |
| SL1344_RS11245_ | 446 | 508 | 493 | 415 | 390 | 536 |
| SL1344_RS11250_ | 2138 | 1993 | 1988 | 1834 | 1853 | 2258 |
| SL1344_RS11255_ | 141 | 149 | 110 | 110 | 133 | 131 |
| SL1344_RS11260_ | 363 | 288 | 313 | 320 | 288 | 351 |
| SL1344_RS11265_ | 410 | 402 | 433 | 376 | 340 | 443 |
| SL1344_RS11270_ | 1676 | 1607 | 1910 | 1723 | 1647 | 1766 |
| SL1344_RS11275_ | 56 | 64 | 54 | 52 | 58 | 45 |
| SL1344_RS11280_ | 112 | 122 | 129 | 128 | 107 | 122 |
| SL1344_RS11285_ | 12 | 8 | 25 | 9 | 14 | 12 |
| SL1344_RS11290_ | 49 | 53 | 47 | 21 | 27 | 20 |
| SL1344_RS11295_ | 54 | 58 | 46 | 64 | 61 | 72 |

| | | | | | | |
|---|---|---|---|---|---|---|
| SL1344_RS11300_ | 697 | 746 | 758 | 749 | 614 | 871 |
| SL1344_RS11305_ | 517 | 536 | 463 | 592 | 495 | 656 |
| SL1344_RS11310_ | 591 | 554 | 612 | 459 | 290 | 327 |
| SL1344_RS11315_ | 442 | 497 | 506 | 430 | 381 | 455 |
| SL1344_RS11320_ | 122 | 124 | 96 | 102 | 91 | 128 |
| SL1344_RS11325_ | 50 | 68 | 59 | 43 | 39 | 66 |
| SL1344_RS11330_ | 45 | 36 | 36 | 38 | 42 | 49 |
| SL1344_RS11335_ | 41 | 40 | 55 | 59 | 47 | 50 |
| SL1344_RS11340_ | 14 | 26 | 32 | 26 | 21 | 32 |
| SL1344_RS11345_ | 3 | 2 | 2 | 2 | 6 | 3 |
| SL1344_RS11350_ | 30 | 28 | 21 | 29 | 12 | 17 |
| SL1344_RS11355_ | 800 | 730 | 819 | 913 | 847 | 924 |
| SL1344_RS11360_ | 190 | 227 | 250 | 153 | 154 | 187 |
| SL1344_RS11365_ | 345 | 417 | 464 | 430 | 433 | 417 |
| SL1344_RS11375_ | 610 | 675 | 705 | 579 | 488 | 587 |
| SL1344_RS11380_ | 594 | 617 | 646 | 482 | 422 | 517 |
| SL1344_RS11385_ | 457 | 438 | 537 | 365 | 323 | 367 |
| SL1344_RS11390_ | 351 | 346 | 380 | 256 | 215 | 255 |
| SL1344_RS11395_ | 648 | 704 | 639 | 499 | 418 | 482 |
| SL1344_RS11400_ | 708 | 698 | 767 | 851 | 787 | 998 |
| SL1344_RS11405_ | 30 | 38 | 52 | 29 | 31 | 36 |
| SL1344_RS11410_ | 378 | 359 | 400 | 305 | 343 | 386 |
| SL1344_RS11415_ | 286 | 251 | 297 | 339 | 268 | 337 |
| SL1344_RS11420_ | 282 | 282 | 324 | 265 | 281 | 253 |
| SL1344_RS11425_ | 2186 | 1915 | 2166 | 3001 | 3049 | 2974 |
| SL1344_RS11430_ | 354 | 347 | 330 | 212 | 242 | 255 |
| SL1344_RS11435_ | 354 | 360 | 360 | 328 | 341 | 400 |
| SL1344_RS11440_ | 262 | 295 | 328 | 312 | 280 | 321 |
| SL1344_RS11445_ | 1049 | 1066 | 1279 | 1093 | 969 | 1037 |
| SL1344_RS11450_ | 165 | 180 | 313 | 235 | 152 | 202 |
| SL1344_RS11455_ | 16 | 18 | 12 | 8 | 10 | 17 |
| SL1344_RS11460_ | 1 | 3 | 3 | 6 | 3 | 3 |
| SL1344_RS11465_ | 828 | 789 | 951 | 739 | 655 | 839 |
| SL1344_RS11470_ | 11 | 6 | 8 | 3 | 6 | 15 |
| SL1344_RS11475_ | 25 | 39 | 17 | 28 | 17 | 24 |
| SL1344_RS11480_ | 31 | 33 | 31 | 21 | 25 | 26 |
| SL1344_RS11485_ | 8 | 13 | 9 | 7 | 8 | 11 |
| SL1344_RS11490_ | 8 | 6 | 6 | 2 | 4 | 2 |
| SL1344_RS11495_ | 795 | 731 | 881 | 584 | 557 | 613 |
| SL1344_RS11500_ | 23 | 29 | 30 | 29 | 28 | 33 |
| SL1344_RS11505_ | 36 | 22 | 28 | 23 | 30 | 31 |
| SL1344_RS11510_ | 20 | 15 | 12 | 10 | 14 | 21 |
| SL1344_RS11515_ | 197 | 202 | 235 | 208 | 190 | 212 |
| SL1344_RS11520_ | 5 | 6 | 3 | 2 | 3 | 5 |
| SL1344_RS11525_ | 24 | 30 | 28 | 20 | 19 | 26 |
| SL1344_RS11535_ | 17 | 14 | 15 | 6 | 7 | 18 |
| SL1344_RS11540_ | 566 | 617 | 609 | 619 | 579 | 561 |
| SL1344_RS11545_ | 2246 | 2426 | 2273 | 1667 | 1403 | 1505 |
| SL1344_RS11550_ | 261 | 205 | 186 | 157 | 115 | 148 |
| SL1344_RS11555_ | 791 | 674 | 962 | 739 | 469 | 626 |
| SL1344_RS11560_ | 525 | 447 | 443 | 440 | 323 | 401 |
| SL1344_RS11565_ | 57 | 63 | 78 | 73 | 52 | 48 |
| SL1344_RS11570_ | 219 | 196 | 285 | 254 | 198 | 244 |
| SL1344_RS11585_ | 3442 | 3501 | 3392 | 2279 | 1687 | 2259 |
| SL1344_RS11595_ | 9516 | 9506 | 8769 | 6472 | 5821 | 7054 |
| SL1344_RS11600_ | 4849 | 4885 | 5058 | 3461 | 2908 | 3597 |
| SL1344_RS11605_ | 23557 | 23384 | 25151 | 17344 | 15053 | 18282 |
| SL1344_RS11610_ | 2020 | 1836 | 2057 | 1607 | 1517 | 1809 |
| SL1344_RS11615_ | 4322 | 4162 | 4626 | 3073 | 2823 | 3443 |
| SL1344_RS11620_ | 636 | 633 | 691 | 212 | 233 | 285 |
| SL1344_RS11625_ | 116 | 140 | 123 | 146 | 146 | 143 |
| SL1344_RS11630_ | 171 | 221 | 230 | 223 | 232 | 262 |

| | | | | | | |
|---|---|---|---|---|---|---|
| SL1344_RS11635_ | 380 | 403 | 396 | 441 | 445 | 505 |
| SL1344_RS11640_ | 197 | 179 | 146 | 243 | 249 | 318 |
| SL1344_RS11670_ | 90 | 66 | 81 | 77 | 165 | 85 |
| SL1344_RS11675_ | 13 | 13 | 24 | 34 | 9 | 21 |
| SL1344_RS11680_ | 19 | 26 | 28 | 53 | 31 | 42 |
| SL1344_RS11685_ | 716 | 688 | 690 | 867 | 768 | 927 |
| SL1344_RS11690_ | 3458 | 3555 | 4240 | 3725 | 3348 | 4161 |
| SL1344_RS11695_ | 1618 | 1808 | 1934 | 1594 | 1557 | 1822 |
| SL1344_RS11700_ | 1134 | 1064 | 1264 | 1106 | 1113 | 1202 |
| SL1344_RS11705_ | 139 | 148 | 190 | 174 | 153 | 150 |
| SL1344_RS11710_ | 152 | 174 | 160 | 148 | 158 | 198 |
| SL1344_RS11715_ | 54983 | 53065 | 54262 | 38578 | 38781 | 47628 |
| SL1344_RS11720_ | 31427 | 30410 | 32300 | 23972 | 24258 | 28376 |
| SL1344_RS11725_ | 10500 | 11605 | 12187 | 7482 | 6657 | 8377 |
| SL1344_RS11730_ | 7750 | 8472 | 8685 | 5215 | 4844 | 5441 |
| SL1344_RS11735_ | 8847 | 9737 | 9326 | 6031 | 5215 | 6441 |
| SL1344_RS11740_ | 6 | 10 | 1 | 10 | 6 | 7 |
| SL1344_RS11745_ | 46 | 62 | 55 | 60 | 59 | 55 |
| SL1344_RS11750_ | 187 | 191 | 207 | 115 | 153 | 177 |
| SL1344_RS11755_ | 76 | 83 | 95 | 71 | 40 | 66 |
| SL1344_RS11760_ | 155 | 186 | 193 | 117 | 78 | 137 |
| SL1344_RS11765_ | 83 | 74 | 62 | 74 | 58 | 65 |
| SL1344_RS11770_ | 508 | 502 | 557 | 463 | 385 | 491 |
| SL1344_RS11775_ | 261 | 406 | 288 | 308 | 302 | 335 |
| SL1344_RS11780_ | 293 | 284 | 289 | 231 | 231 | 266 |
| SL1344_RS11785_ | 784 | 849 | 953 | 693 | 613 | 785 |
| SL1344_RS11790_ | 1234 | 1336 | 1453 | 1343 | 1111 | 1381 |
| SL1344_RS11795_ | 1973 | 2227 | 2306 | 1804 | 1514 | 1871 |
| SL1344_RS11800_ | 4098 | 4676 | 5187 | 3393 | 3055 | 3677 |
| SL1344_RS11805_ | 1494 | 1668 | 1745 | 1384 | 1169 | 1378 |
| SL1344_RS11810_ | 1596 | 1781 | 1795 | 1267 | 1174 | 1358 |
| SL1344_RS11815_ | 552 | 614 | 621 | 419 | 403 | 429 |
| SL1344_RS11820_ | 176 | 197 | 204 | 166 | 173 | 199 |
| SL1344_RS11825_ | 730 | 846 | 816 | 770 | 630 | 775 |
| SL1344_RS11830_ | 1209 | 1342 | 1336 | 1168 | 1012 | 1082 |
| SL1344_RS11840_ | 1138 | 1231 | 1247 | 869 | 798 | 945 |
| SL1344_RS11845_ | 452 | 448 | 528 | 410 | 385 | 449 |
| SL1344_RS11850_ | 1199 | 1297 | 1266 | 1172 | 1176 | 1193 |
| SL1344_RS11855_ | 393 | 486 | 523 | 550 | 491 | 510 |
| SL1344_RS11860_ | 3533 | 3654 | 3287 | 2346 | 2042 | 2539 |
| SL1344_RS11865_ | 155 | 127 | 105 | 88 | 72 | 134 |
| SL1344_RS11870_ | 273 | 256 | 242 | 300 | 296 | 280 |
| SL1344_RS11880_ | 94 | 113 | 116 | 106 | 89 | 119 |
| SL1344_RS11885_ | 5117 | 5365 | 5945 | 3651 | 2977 | 4133 |
| SL1344_RS11890_ | 3247 | 3241 | 3704 | 2317 | 1679 | 2627 |
| SL1344_RS11895_ | 6584 | 6717 | 7848 | 5406 | 4962 | 6306 |
| SL1344_RS11900_ | 1171 | 1312 | 1560 | 1062 | 900 | 1175 |
| SL1344_RS11905_ | 2832 | 3091 | 3182 | 2510 | 2215 | 2884 |
| SL1344_RS11910_ | 2415 | 2431 | 2909 | 1986 | 1868 | 2302 |
| SL1344_RS11915_ | 3939 | 4181 | 4725 | 3202 | 2964 | 3735 |
| SL1344_RS11920_ | 11085 | 12026 | 13415 | 10289 | 9917 | 11619 |
| SL1344_RS11925_ | 5603 | 5989 | 7044 | 5120 | 4947 | 5992 |
| SL1344_RS11930_ | 2550 | 2603 | 3105 | 2215 | 2184 | 2655 |
| SL1344_RS11935_ | 6479 | 6746 | 8551 | 7219 | 6363 | 7693 |
| SL1344_RS11940_ | 2781 | 2793 | 3513 | 3440 | 3127 | 3567 |
| SL1344_RS11945_ | 2292 | 2216 | 2633 | 2969 | 2651 | 2958 |
| SL1344_RS11950_ | 21 | 15 | 15 | 18 | 12 | 11 |
| SL1344_RS11960_ | 939 | 890 | 1055 | 926 | 895 | 942 |
| SL1344_RS11965_ | 205 | 181 | 205 | 160 | 151 | 213 |
| SL1344_RS11970_ | 640 | 676 | 704 | 469 | 435 | 550 |
| SL1344_RS11975_ | 1596 | 1619 | 1556 | 1150 | 1093 | 1219 |
| SL1344_RS11980_ | 657 | 632 | 691 | 524 | 535 | 593 |

| | | | | | | |
|---|---|---|---|---|---|---|
| SL1344_RS11985_ | 480 | 568 | 531 | 477 | 491 | 546 |
| SL1344_RS12000_ | 169 | 200 | 212 | 264 | 216 | 283 |
| SL1344_RS12005_ | 797 | 792 | 802 | 695 | 592 | 777 |
| SL1344_RS12010_ | 289 | 293 | 332 | 231 | 211 | 285 |
| SL1344_RS12015_ | 503 | 459 | 564 | 366 | 358 | 433 |
| SL1344_RS12020_ | 108 | 122 | 126 | 85 | 82 | 102 |
| SL1344_RS12025_ | 103 | 108 | 128 | 88 | 90 | 112 |
| SL1344_RS12030_ | 0 | 5 | 2 | 3 | 3 | 2 |
| SL1344_RS12035_ | 384 | 409 | 477 | 333 | 274 | 322 |
| SL1344_RS12040_ | 1134 | 1057 | 1067 | 1162 | 1011 | 1261 |
| SL1344_RS12045_ | 511 | 474 | 488 | 455 | 437 | 523 |
| SL1344_RS12050_ | 372 | 386 | 374 | 308 | 266 | 366 |
| SL1344_RS12055_ | 50 | 63 | 46 | 34 | 20 | 40 |
| SL1344_RS12060_ | 389 | 417 | 401 | 280 | 222 | 290 |
| SL1344_RS12065_ | 128 | 150 | 153 | 105 | 93 | 117 |
| SL1344_RS12070_ | 128 | 105 | 109 | 88 | 86 | 107 |
| SL1344_RS12075_ | 114 | 114 | 137 | 130 | 109 | 105 |
| SL1344_RS12080_ | 1363 | 1380 | 1447 | 888 | 1057 | 1174 |
| SL1344_RS12085_ | 87 | 96 | 75 | 88 | 91 | 94 |
| SL1344_RS12090_ | 432 | 412 | 457 | 468 | 354 | 456 |
| SL1344_RS12095_ | 404 | 380 | 376 | 402 | 441 | 394 |
| SL1344_RS12100_ | 184 | 136 | 191 | 122 | 157 | 168 |
| SL1344_RS12105_ | 65 | 60 | 75 | 71 | 71 | 81 |
| SL1344_RS12110_ | 36 | 51 | 35 | 49 | 41 | 44 |
| SL1344_RS12115_ | 503 | 592 | 617 | 560 | 649 | 668 |
| SL1344_RS12120_ | 901 | 944 | 1157 | 1013 | 1035 | 1129 |
| SL1344_RS12125_ | 382 | 398 | 418 | 425 | 420 | 500 |
| SL1344_RS12130_ | 1199 | 1309 | 1233 | 1230 | 1085 | 1331 |
| SL1344_RS12135_ | 1105 | 1184 | 1081 | 1031 | 1066 | 1176 |
| SL1344_RS12140_ | 2625 | 2656 | 2866 | 2899 | 2735 | 2833 |
| SL1344_RS12145_ | 305 | 309 | 333 | 262 | 278 | 311 |
| SL1344_RS12155_ | 680 | 673 | 681 | 567 | 526 | 704 |
| SL1344_RS12160_ | 1009 | 1029 | 1206 | 1136 | 1038 | 1165 |
| SL1344_RS12170_ | 435 | 449 | 418 | 398 | 380 | 431 |
| SL1344_RS12175_ | 9 | 11 | 10 | 9 | 7 | 7 |
| SL1344_RS12180_ | 3 | 9 | 5 | 8 | 3 | 5 |
| SL1344_RS12185_ | 2 | 3 | 2 | 2 | 7 | 6 |
| SL1344_RS12190_ | 9 | 11 | 9 | 8 | 8 | 5 |
| SL1344_RS12195_ | 72 | 56 | 75 | 82 | 76 | 102 |
| SL1344_RS12200_ | 5774 | 6036 | 5765 | 5281 | 4600 | 6439 |
| SL1344_RS12205_ | 552 | 636 | 602 | 594 | 549 | 640 |
| SL1344_RS12210_ | 414 | 476 | 420 | 408 | 384 | 456 |
| SL1344_RS12215_ | 166 | 209 | 218 | 146 | 157 | 194 |
| SL1344_RS12220_ | 147 | 202 | 206 | 193 | 156 | 242 |
| SL1344_RS12225_ | 343 | 323 | 366 | 307 | 308 | 307 |
| SL1344_RS12230_ | 700 | 723 | 735 | 556 | 514 | 601 |
| SL1344_RS12235_ | 502 | 528 | 554 | 446 | 429 | 549 |
| SL1344_RS12240_ | 141 | 181 | 177 | 134 | 124 | 145 |
| SL1344_RS12245_ | 1251 | 1252 | 1287 | 1248 | 1090 | 1240 |
| SL1344_RS12250_ | 756 | 681 | 692 | 560 | 566 | 660 |
| SL1344_RS12260_ | 2490 | 2440 | 2846 | 2322 | 2178 | 2447 |
| SL1344_RS12265_ | 977 | 1043 | 1530 | 1240 | 1262 | 1376 |
| SL1344_RS12270_ | 1913 | 1789 | 1929 | 2120 | 1907 | 1961 |
| SL1344_RS12275_ | 202 | 213 | 205 | 138 | 127 | 148 |
| SL1344_RS12280_ | 16 | 30 | 48 | 35 | 26 | 19 |
| SL1344_RS12285_ | 126 | 115 | 93 | 101 | 85 | 95 |
| SL1344_RS12290_ | 456 | 447 | 423 | 306 | 292 | 337 |
| SL1344_RS12295_ | 443 | 494 | 477 | 337 | 315 | 335 |
| SL1344_RS12300_ | 179 | 179 | 194 | 144 | 141 | 136 |
| SL1344_RS12305_ | 36 | 39 | 54 | 24 | 35 | 31 |
| SL1344_RS12310_ | 42 | 49 | 64 | 43 | 49 | 42 |
| SL1344_RS12315_ | 356 | 275 | 268 | 196 | 183 | 269 |

| | | | | | | |
|---|---|---|---|---|---|---|
| SL1344_RS12320_ | 3 | 2 | 3 | 3 | 2 | 2 |
| SL1344_RS12325_ | 37 | 50 | 60 | 54 | 51 | 63 |
| SL1344_RS12330_ | 1053 | 1018 | 1062 | 989 | 766 | 978 |
| SL1344_RS12335_ | 1225 | 1129 | 1300 | 1038 | 920 | 1068 |
| SL1344_RS12340_ | 188 | 189 | 218 | 181 | 132 | 151 |
| SL1344_RS12345_ | 413 | 441 | 408 | 338 | 345 | 373 |
| SL1344_RS12350_ | 342 | 380 | 388 | 310 | 281 | 363 |
| SL1344_RS12355_ | 388 | 418 | 381 | 379 | 308 | 409 |
| SL1344_RS12360_ | 219 | 156 | 165 | 145 | 131 | 147 |
| SL1344_RS12365_ | 803 | 783 | 990 | 939 | 787 | 981 |
| SL1344_RS12370_ | 855 | 929 | 954 | 926 | 725 | 878 |
| SL1344_RS12375_ | 245 | 243 | 390 | 354 | 295 | 338 |
| SL1344_RS12380_ | 1078 | 1012 | 1120 | 1306 | 1428 | 1405 |
| SL1344_RS12385_ | 961 | 1024 | 1111 | 628 | 698 | 718 |
| SL1344_RS12390_ | 1209 | 1218 | 980 | 833 | 829 | 793 |
| SL1344_RS12395_ | 2082 | 2280 | 2422 | 2125 | 2104 | 2437 |
| SL1344_RS12400_ | 715 | 762 | 869 | 1028 | 909 | 950 |
| SL1344_RS12405_ | 550 | 661 | 780 | 759 | 640 | 683 |
| SL1344_RS12410_ | 1154 | 1207 | 1482 | 1229 | 1255 | 1433 |
| SL1344_RS12415_ | 588 | 580 | 699 | 594 | 586 | 694 |
| SL1344_RS12420_ | 52 | 59 | 45 | 49 | 63 | 52 |
| SL1344_RS12425_ | 1 | 1 | 2 | 1 | 1 | 4 |
| SL1344_RS12430_ | 59 | 87 | 63 | 72 | 67 | 72 |
| SL1344_RS12435_ | 9 | 2 | 7 | 9 | 7 | 13 |
| SL1344_RS12440_ | 44 | 30 | 26 | 27 | 27 | 27 |
| SL1344_RS12445_ | 87 | 107 | 105 | 132 | 100 | 127 |
| SL1344_RS12450_ | 309 | 347 | 367 | 294 | 295 | 340 |
| SL1344_RS12455_ | 309 | 312 | 344 | 242 | 227 | 317 |
| SL1344_RS12465_ | 2507 | 2587 | 2845 | 2587 | 2376 | 2567 |
| SL1344_RS12480_ | 4 | 0 | 7 | 7 | 10 | 5 |
| SL1344_RS12485_ | 2893 | 2596 | 2872 | 2317 | 2231 | 2520 |
| SL1344_RS12490_ | 21081 | 19553 | 18703 | 14886 | 15104 | 17556 |
| SL1344_RS12495_ | 8180 | 8220 | 8335 | 5742 | 5542 | 6673 |
| SL1344_RS12500_ | 53 | 56 | 61 | 51 | 38 | 35 |
| SL1344_RS12510_ | 99 | 111 | 95 | 103 | 95 | 106 |
| SL1344_RS12515_ | 94 | 88 | 120 | 76 | 79 | 98 |
| SL1344_RS12520_ | 43 | 47 | 48 | 32 | 32 | 68 |
| SL1344_RS12525_ | 62 | 61 | 51 | 43 | 45 | 45 |
| SL1344_RS12555_ | 3318 | 3393 | 3745 | 2313 | 2050 | 2875 |
| SL1344_RS12560_ | 1377 | 1448 | 1534 | 1463 | 1287 | 1402 |
| SL1344_RS12565_ | 357 | 320 | 362 | 376 | 343 | 406 |
| SL1344_RS12570_ | 420 | 380 | 339 | 439 | 384 | 403 |
| SL1344_RS12575_ | 610 | 520 | 509 | 594 | 522 | 596 |
| SL1344_RS12580_ | 130 | 133 | 146 | 145 | 135 | 150 |
| SL1344_RS12585_ | 148 | 133 | 123 | 136 | 100 | 135 |
| SL1344_RS12590_ | 50 | 35 | 50 | 40 | 32 | 24 |
| SL1344_RS12595_ | 248 | 235 | 257 | 242 | 213 | 251 |
| SL1344_RS12600_ | 78 | 94 | 90 | 63 | 65 | 57 |
| SL1344_RS12605_ | 112 | 121 | 88 | 64 | 75 | 84 |
| SL1344_RS12610_ | 100 | 119 | 92 | 69 | 81 | 73 |
| SL1344_RS12615_ | 36 | 36 | 40 | 39 | 24 | 33 |
| SL1344_RS12620_ | 69 | 73 | 58 | 58 | 46 | 67 |
| SL1344_RS12630_ | 68 | 63 | 70 | 61 | 39 | 60 |
| SL1344_RS12635_ | 28 | 14 | 23 | 19 | 14 | 21 |
| SL1344_RS12640_ | 49 | 52 | 48 | 37 | 46 | 41 |
| SL1344_RS12645_ | 59 | 64 | 66 | 52 | 65 | 57 |
| SL1344_RS12650_ | 8 | 16 | 14 | 16 | 11 | 17 |
| SL1344_RS12655_ | 43 | 64 | 47 | 49 | 32 | 38 |
| SL1344_RS12660_ | 32 | 34 | 25 | 39 | 19 | 31 |
| SL1344_RS12665_ | 17 | 12 | 7 | 8 | 10 | 17 |
| SL1344_RS12670_ | 18 | 32 | 20 | 23 | 18 | 27 |
| SL1344_RS12675_ | 16 | 18 | 21 | 6 | 11 | 14 |

| | | | | | | |
|---|---|---|---|---|---|---|
| SL1344_RS12690_ | 7002 | 7423 | 8436 | 6564 | 6172 | 7523 |
| SL1344_RS12695_ | 350 | 362 | 341 | 237 | 214 | 266 |
| SL1344_RS12700_ | 829 | 812 | 761 | 483 | 414 | 515 |
| SL1344_RS12705_ | 752 | 659 | 580 | 558 | 480 | 602 |
| SL1344_RS12710_ | 78 | 74 | 88 | 78 | 78 | 70 |
| SL1344_RS12715_ | 187 | 187 | 153 | 208 | 196 | 216 |
| SL1344_RS12720_ | 142 | 167 | 143 | 113 | 106 | 131 |
| SL1344_RS12725_ | 269 | 296 | 246 | 205 | 150 | 164 |
| SL1344_RS12730_ | 328 | 358 | 348 | 234 | 190 | 260 |
| SL1344_RS12735_ | 217 | 242 | 262 | 267 | 248 | 267 |
| SL1344_RS12740_ | 175 | 185 | 169 | 179 | 187 | 186 |
| SL1344_RS12745_ | 734 | 714 | 764 | 625 | 658 | 758 |
| SL1344_RS12750_ | 570 | 586 | 490 | 391 | 390 | 446 |
| SL1344_RS12755_ | 566 | 620 | 657 | 554 | 506 | 569 |
| SL1344_RS12760_ | 506 | 562 | 496 | 422 | 390 | 521 |
| SL1344_RS12765_ | 723 | 776 | 714 | 622 | 547 | 721 |
| SL1344_RS12770_ | 4512 | 4515 | 4599 | 4575 | 4416 | 4841 |
| SL1344_RS12775_ | 2547 | 2576 | 2579 | 2605 | 2357 | 2812 |
| SL1344_RS12780_ | 729 | 707 | 803 | 758 | 793 | 799 |
| SL1344_RS12785_ | 1936 | 1754 | 1790 | 1940 | 1894 | 1945 |
| SL1344_RS12790_ | 45 | 46 | 55 | 51 | 38 | 45 |
| SL1344_RS12795_ | 339 | 320 | 293 | 317 | 305 | 319 |
| SL1344_RS12800_ | 744 | 805 | 805 | 752 | 677 | 822 |
| SL1344_RS12805_ | 892 | 946 | 854 | 849 | 777 | 880 |
| SL1344_RS12810_ | 424 | 474 | 510 | 590 | 507 | 586 |
| SL1344_RS12815_ | 700 | 698 | 814 | 677 | 731 | 898 |
| SL1344_RS12820_ | 2219 | 2437 | 2516 | 2344 | 2569 | 2479 |
| SL1344_RS12825_ | 501 | 481 | 481 | 567 | 621 | 741 |
| SL1344_RS12830_ | 408 | 419 | 442 | 481 | 451 | 524 |
| SL1344_RS12835_ | 1469 | 1465 | 1497 | 1016 | 1079 | 1153 |
| SL1344_RS12840_ | 895 | 1033 | 1006 | 706 | 664 | 689 |
| SL1344_RS12845_ | 254 | 248 | 260 | 182 | 138 | 144 |
| SL1344_RS12850_ | 70 | 55 | 53 | 35 | 58 | 46 |
| SL1344_RS12855_ | 504 | 427 | 462 | 737 | 571 | 591 |
| SL1344_RS12860_ | 24 | 24 | 19 | 21 | 26 | 26 |
| SL1344_RS12865_ | 8 | 8 | 9 | 6 | 6 | 10 |
| SL1344_RS12870_ | 1531 | 1539 | 1752 | 1627 | 1352 | 1847 |
| SL1344_RS12875_ | 576 | 669 | 801 | 807 | 652 | 958 |
| SL1344_RS12880_ | 235 | 279 | 287 | 351 | 290 | 384 |
| SL1344_RS12885_ | 276 | 296 | 268 | 330 | 332 | 337 |
| SL1344_RS12890_ | 721 | 771 | 774 | 745 | 671 | 751 |
| SL1344_RS12900_ | 64 | 53 | 64 | 72 | 75 | 73 |
| SL1344_RS12905_ | 21 | 23 | 19 | 24 | 31 | 45 |
| SL1344_RS12910_ | 95 | 91 | 103 | 134 | 131 | 142 |
| SL1344_RS12915_ | 708 | 706 | 826 | 773 | 776 | 1007 |
| SL1344_RS12920_ | 1850 | 1682 | 1838 | 1715 | 1540 | 1789 |
| SL1344_RS12925_ | 1362 | 1300 | 1214 | 1148 | 1210 | 1428 |
| SL1344_RS12930_ | 1391 | 1398 | 1406 | 1507 | 1338 | 1677 |
| SL1344_RS12935_ | 2405 | 2630 | 2406 | 1864 | 1870 | 2352 |
| SL1344_RS12940_ | 1116 | 1159 | 1245 | 1162 | 1099 | 1131 |
| SL1344_RS12945_ | 634 | 639 | 740 | 857 | 845 | 932 |
| SL1344_RS12950_ | 1978 | 2092 | 2840 | 3068 | 3318 | 3533 |
| SL1344_RS12955_ | 226 | 195 | 193 | 226 | 179 | 208 |
| SL1344_RS12960_ | 328 | 348 | 373 | 402 | 337 | 398 |
| SL1344_RS12965_ | 218 | 211 | 234 | 238 | 198 | 238 |
| SL1344_RS12970_ | 745 | 769 | 824 | 829 | 723 | 853 |
| SL1344_RS12975_ | 434 | 484 | 439 | 411 | 363 | 399 |
| SL1344_RS12980_ | 2727 | 2891 | 2888 | 2200 | 2075 | 2375 |
| SL1344_RS12985_ | 1553 | 1742 | 1801 | 1128 | 1259 | 1407 |
| SL1344_RS12990_ | 62 | 67 | 82 | 74 | 70 | 62 |
| SL1344_RS12995_ | 332 | 383 | 378 | 462 | 352 | 500 |
| SL1344_RS13000_ | 737 | 717 | 809 | 841 | 773 | 974 |

| | | | | | | |
|---|---|---|---|---|---|---|
| SL1344_RS13005_ | 604 | 520 | 520 | 504 | 524 | 607 |
| SL1344_RS13010_ | 541 | 581 | 587 | 522 | 518 | 608 |
| SL1344_RS13015_ | 856 | 922 | 963 | 923 | 874 | 979 |
| SL1344_RS13025_ | 1058 | 1046 | 947 | 892 | 1018 | 1218 |
| SL1344_RS13030_ | 696 | 700 | 675 | 688 | 735 | 761 |
| SL1344_RS13035_ | 1072 | 1134 | 1268 | 1244 | 1188 | 1391 |
| SL1344_RS13040_ | 281 | 294 | 330 | 373 | 388 | 410 |
| SL1344_RS13045_ | 620 | 688 | 679 | 760 | 758 | 786 |
| SL1344_RS13050_ | 313 | 375 | 431 | 581 | 545 | 639 |
| SL1344_RS13075_ | 184 | 216 | 157 | 170 | 153 | 199 |
| SL1344_RS13080_ | 141 | 136 | 107 | 121 | 126 | 123 |
| SL1344_RS13085_ | 580 | 684 | 681 | 252 | 235 | 280 |
| SL1344_RS13090_ | 279 | 284 | 287 | 351 | 357 | 431 |
| SL1344_RS13095_ | 8975 | 9540 | 9369 | 6556 | 6662 | 7092 |
| SL1344_RS13100_ | 203 | 181 | 201 | 211 | 223 | 230 |
| SL1344_RS13105_ | 413 | 427 | 512 | 442 | 407 | 465 |
| SL1344_RS13115_ | 730 | 967 | 1008 | 796 | 764 | 785 |
| SL1344_RS13120_ | 103 | 132 | 123 | 94 | 112 | 106 |
| SL1344_RS13125_ | 685 | 762 | 690 | 747 | 657 | 706 |
| SL1344_RS13130_ | 965 | 940 | 968 | 914 | 881 | 1019 |
| SL1344_RS13135_ | 362 | 435 | 391 | 390 | 353 | 360 |
| SL1344_RS13140_ | 463 | 452 | 391 | 448 | 395 | 522 |
| SL1344_RS13145_ | 877 | 997 | 1130 | 1016 | 921 | 1121 |
| SL1344_RS13150_ | 35 | 45 | 30 | 43 | 36 | 28 |
| SL1344_RS13155_ | 290 | 307 | 306 | 374 | 290 | 352 |
| SL1344_RS13160_ | 223 | 232 | 208 | 186 | 174 | 198 |
| SL1344_RS13165_ | 570 | 626 | 563 | 420 | 448 | 533 |
| SL1344_RS13170_ | 470 | 481 | 368 | 240 | 302 | 348 |
| SL1344_RS13175_ | 195 | 228 | 199 | 126 | 114 | 165 |
| SL1344_RS13180_ | 92 | 96 | 138 | 116 | 97 | 130 |
| SL1344_RS13185_ | 130 | 116 | 104 | 136 | 156 | 159 |
| SL1344_RS13190_ | 9 | 8 | 9 | 11 | 14 | 15 |
| SL1344_RS13195_ | 58 | 34 | 44 | 40 | 60 | 50 |
| SL1344_RS13200_ | 13 | 12 | 8 | 19 | 10 | 24 |
| SL1344_RS13210_ | 434 | 467 | 485 | 411 | 359 | 424 |
| SL1344_RS13215_ | 491 | 524 | 535 | 453 | 435 | 512 |
| SL1344_RS13220_ | 1126 | 1178 | 1199 | 1015 | 1032 | 1235 |
| SL1344_RS13225_ | 676 | 743 | 806 | 706 | 601 | 815 |
| SL1344_RS13230_ | 1972 | 2017 | 2111 | 1893 | 2012 | 2271 |
| SL1344_RS13235_ | 2081 | 2249 | 2320 | 2158 | 2169 | 2459 |
| SL1344_RS13240_ | 342 | 327 | 346 | 332 | 287 | 309 |
| SL1344_RS13245_ | 443 | 441 | 468 | 355 | 334 | 439 |
| SL1344_RS13250_ | 13 | 7 | 9 | 21 | 9 | 11 |
| SL1344_RS13255_ | 10 | 8 | 5 | 12 | 12 | 16 |
| SL1344_RS13260_ | 18 | 18 | 12 | 13 | 25 | 27 |
| SL1344_RS13265_ | 72 | 57 | 66 | 75 | 52 | 65 |
| SL1344_RS13270_ | 20 | 13 | 8 | 28 | 15 | 19 |
| SL1344_RS13275_ | 103 | 83 | 86 | 63 | 67 | 89 |
| SL1344_RS13280_ | 43 | 45 | 52 | 37 | 35 | 32 |
| SL1344_RS13285_ | 38 | 21 | 28 | 26 | 31 | 32 |
| SL1344_RS13290_ | 82 | 37 | 61 | 30 | 41 | 33 |
| SL1344_RS13295_ | 18 | 12 | 21 | 12 | 16 | 17 |
| SL1344_RS13300_ | 225 | 186 | 180 | 125 | 186 | 157 |
| SL1344_RS13305_ | 13 | 9 | 11 | 7 | 11 | 6 |
| SL1344_RS13310_ | 12 | 15 | 9 | 12 | 13 | 4 |
| SL1344_RS13315_ | 148 | 176 | 148 | 175 | 159 | 191 |
| SL1344_RS13320_ | 46 | 38 | 29 | 36 | 48 | 41 |
| SL1344_RS13325_ | 3050 | 3377 | 3289 | 3541 | 3429 | 3246 |
| SL1344_RS13330_ | 51 | 42 | 40 | 39 | 36 | 46 |
| SL1344_RS13335_ | 45 | 46 | 36 | 35 | 32 | 31 |
| SL1344_RS13340_ | 28 | 32 | 21 | 26 | 18 | 35 |
| SL1344_RS13345_ | 92 | 87 | 92 | 79 | 76 | 73 |

| | | | | | | |
|---|---|---|---|---|---|---|
| SL1344_RS13350_ | 50 | 54 | 59 | 47 | 33 | 44 |
| SL1344_RS13355_ | 100 | 100 | 89 | 62 | 58 | 75 |
| SL1344_RS13360_ | 150 | 128 | 140 | 93 | 95 | 111 |
| SL1344_RS13365_ | 31 | 23 | 25 | 12 | 18 | 30 |
| SL1344_RS13370_ | 89 | 73 | 64 | 60 | 63 | 81 |
| SL1344_RS13375_ | 24 | 20 | 19 | 7 | 13 | 12 |
| SL1344_RS13380_ | 134 | 133 | 129 | 98 | 115 | 121 |
| SL1344_RS13385_ | 21 | 33 | 35 | 14 | 16 | 19 |
| SL1344_RS13390_ | 43 | 39 | 44 | 27 | 43 | 33 |
| SL1344_RS13395_ | 0 | 1 | 3 | 0 | 3 | 2 |
| SL1344_RS13400_ | 352 | 291 | 258 | 418 | 354 | 391 |
| SL1344_RS13405_ | 45 | 42 | 42 | 24 | 29 | 31 |
| SL1344_RS13410_ | 59 | 27 | 52 | 46 | 26 | 24 |
| SL1344_RS13415_ | 59 | 44 | 53 | 67 | 75 | 66 |
| SL1344_RS13420_ | 29 | 32 | 22 | 18 | 27 | 17 |
| SL1344_RS13425_ | 3 | 4 | 4 | 1 | 8 | 4 |
| SL1344_RS13430_ | 13 | 9 | 12 | 4 | 10 | 6 |
| SL1344_RS13435_ | 3 | 6 | 11 | 5 | 3 | 6 |
| SL1344_RS13440_ | 299 | 396 | 470 | 336 | 314 | 343 |
| SL1344_RS13450_ | 140 | 131 | 151 | 202 | 225 | 236 |
| SL1344_RS13455_ | 442 | 447 | 424 | 355 | 403 | 410 |
| SL1344_RS13460_ | 36 | 48 | 34 | 27 | 28 | 47 |
| SL1344_RS13465_ | 68 | 79 | 54 | 49 | 59 | 70 |
| SL1344_RS13470_ | 27 | 32 | 44 | 25 | 21 | 30 |
| SL1344_RS13475_ | 56 | 52 | 53 | 31 | 32 | 54 |
| SL1344_RS13480_ | 38 | 32 | 26 | 17 | 19 | 31 |
| SL1344_RS13485_ | 105 | 97 | 62 | 48 | 57 | 80 |
| SL1344_RS13490_ | 66 | 73 | 70 | 43 | 43 | 63 |
| SL1344_RS13495_ | 39 | 46 | 47 | 25 | 38 | 32 |
| SL1344_RS13500_ | 1079 | 1087 | 1012 | 1190 | 1132 | 1133 |
| SL1344_RS13505_ | 6715 | 6300 | 6562 | 6320 | 6629 | 7098 |
| SL1344_RS13510_ | 16 | 29 | 30 | 13 | 28 | 29 |
| SL1344_RS13515_ | 83 | 66 | 64 | 45 | 45 | 53 |
| SL1344_RS13520_ | 163 | 169 | 173 | 116 | 106 | 150 |
| SL1344_RS13525_ | 48 | 27 | 42 | 32 | 21 | 25 |
| SL1344_RS13530_ | 10 | 9 | 16 | 7 | 6 | 5 |
| SL1344_RS13535_ | 46 | 71 | 63 | 50 | 54 | 35 |
| SL1344_RS13540_ | 113 | 106 | 116 | 103 | 118 | 141 |
| SL1344_RS13545_ | 352 | 389 | 354 | 305 | 273 | 329 |
| SL1344_RS13550_ | 1161 | 1191 | 1213 | 951 | 712 | 946 |
| SL1344_RS13555_ | 1860 | 1790 | 1772 | 1419 | 1018 | 1522 |
| SL1344_RS13560_ | 1694 | 1643 | 1579 | 1509 | 1326 | 1646 |
| SL1344_RS13565_ | 75 | 70 | 90 | 74 | 70 | 82 |
| SL1344_RS13570_ | 207 | 208 | 197 | 241 | 257 | 246 |
| SL1344_RS13575_ | 1393 | 1408 | 2898 | 1286 | 1434 | 1562 |
| SL1344_RS13580_ | 171 | 174 | 151 | 153 | 167 | 153 |
| SL1344_RS13590_ | 17530 | 19409 | 19935 | 22000 | 18304 | 23575 |
| SL1344_RS13595_ | 965 | 1018 | 1007 | 816 | 814 | 894 |
| SL1344_RS13605_ | 376 | 407 | 444 | 571 | 493 | 537 |
| SL1344_RS13610_ | 200 | 199 | 208 | 235 | 204 | 227 |
| SL1344_RS13615_ | 1835 | 1964 | 2092 | 1535 | 1417 | 1508 |
| SL1344_RS13625_ | 169 | 180 | 166 | 160 | 118 | 140 |
| SL1344_RS13630_ | 179 | 179 | 187 | 156 | 164 | 194 |
| SL1344_RS13635_ | 32 | 31 | 29 | 16 | 15 | 9 |
| SL1344_RS13645_ | 3762 | 3117 | 4291 | 2053 | 3215 | 4998 |
| SL1344_RS13650_ | 108 | 80 | 113 | 123 | 124 | 145 |
| SL1344_RS13655_ | 526 | 422 | 918 | 314 | 418 | 813 |
| SL1344_RS13660_ | 8343 | 8356 | 7971 | 7324 | 7155 | 8360 |
| SL1344_RS13665_ | 1136 | 1059 | 1022 | 945 | 1013 | 1045 |
| SL1344_RS13670_ | 1410 | 1439 | 1378 | 1309 | 1400 | 1317 |
| SL1344_RS13675_ | 928 | 863 | 976 | 1175 | 978 | 1033 |
| SL1344_RS13680_ | 21704 | 24572 | 25209 | 17304 | 18080 | 20112 |

| | | | | | | |
|---|---|---|---|---|---|---|
| SL1344_RS13685_ | 715 | 721 | 748 | 723 | 823 | 906 |
| SL1344_RS13705_ | 46 | 56 | 91 | 39 | 34 | 28 |
| SL1344_RS13710_ | 181 | 184 | 169 | 142 | 111 | 111 |
| SL1344_RS13715_ | 1149 | 1038 | 1084 | 1139 | 1116 | 1235 |
| SL1344_RS13720_ | 1222 | 1243 | 1115 | 1207 | 1247 | 1375 |
| SL1344_RS13725_ | 756 | 618 | 573 | 656 | 729 | 768 |
| SL1344_RS13730_ | 1 | 6 | 3 | 4 | 4 | 4 |
| SL1344_RS13735_ | 12 | 5 | 8 | 4 | 5 | 7 |
| SL1344_RS13745_ | 7 | 24 | 23 | 4 | 4 | 3 |
| SL1344_RS13750_ | 1 | 2 | 1 | 7 | 5 | 4 |
| SL1344_RS13755_ | 1 | 2 | 1 | 3 | 2 | 0 |
| SL1344_RS13760_ | 7 | 4 | 4 | 9 | 5 | 4 |
| SL1344_RS13765_ | 28 | 11 | 14 | 16 | 18 | 21 |
| SL1344_RS13770_ | 23 | 24 | 17 | 21 | 22 | 18 |
| SL1344_RS13775_ | 85 | 60 | 98 | 74 | 99 | 84 |
| SL1344_RS13780_ | 1812 | 1748 | 1572 | 1497 | 1457 | 1682 |
| SL1344_RS13785_ | 6228 | 6168 | 7596 | 7435 | 7944 | 9332 |
| SL1344_RS13790_ | 5986 | 5748 | 7192 | 7512 | 7928 | 9270 |
| SL1344_RS13800_ | 2340 | 2250 | 3035 | 3078 | 3479 | 3890 |
| SL1344_RS13805_ | 1860 | 1804 | 1625 | 1420 | 1366 | 1693 |
| SL1344_RS13810_ | 281 | 264 | 283 | 400 | 352 | 367 |
| SL1344_RS13815_ | 357 | 346 | 407 | 393 | 338 | 378 |
| SL1344_RS13820_ | 88 | 89 | 92 | 96 | 82 | 87 |
| SL1344_RS13825_ | 5320 | 4702 | 4233 | 5482 | 5156 | 5692 |
| SL1344_RS13830_ | 521 | 501 | 462 | 465 | 539 | 580 |
| SL1344_RS13840_ | 953 | 1002 | 1085 | 1080 | 1025 | 1054 |
| SL1344_RS13845_ | 677 | 729 | 637 | 736 | 630 | 734 |
| SL1344_RS13850_ | 1184 | 1275 | 1203 | 1299 | 1324 | 1436 |
| SL1344_RS13855_ | 1091 | 930 | 1009 | 1465 | 1382 | 1338 |
| SL1344_RS13860_ | 196 | 220 | 280 | 224 | 220 | 238 |
| SL1344_RS13865_ | 27 | 17 | 25 | 23 | 32 | 25 |
| SL1344_RS13870_ | 37 | 44 | 48 | 54 | 41 | 49 |
| SL1344_RS13875_ | 42 | 42 | 39 | 36 | 26 | 48 |
| SL1344_RS13880_ | 19 | 19 | 27 | 24 | 10 | 31 |
| SL1344_RS13885_ | 25 | 21 | 18 | 35 | 40 | 29 |
| SL1344_RS13890_ | 17 | 12 | 14 | 10 | 10 | 16 |
| SL1344_RS13895_ | 85 | 60 | 68 | 51 | 91 | 73 |
| SL1344_RS13900_ | 9 | 8 | 12 | 8 | 11 | 11 |
| SL1344_RS13905_ | 13 | 18 | 19 | 14 | 23 | 16 |
| SL1344_RS13910_ | 9 | 7 | 12 | 14 | 10 | 5 |
| SL1344_RS13915_ | 14 | 22 | 22 | 18 | 15 | 26 |
| SL1344_RS13920_ | 5 | 3 | 1 | 9 | 10 | 1 |
| SL1344_RS13930_ | 88 | 67 | 84 | 71 | 102 | 68 |
| SL1344_RS13935_ | 35 | 43 | 51 | 59 | 50 | 36 |
| SL1344_RS13940_ | 12 | 9 | 8 | 9 | 13 | 14 |
| SL1344_RS13950_ | 4 | 5 | 7 | 2 | 3 | 2 |
| SL1344_RS13955_ | 8 | 9 | 6 | 3 | 9 | 6 |
| SL1344_RS13965_ | 2 | 4 | 5 | 7 | 5 | 10 |
| SL1344_RS13970_ | 10 | 8 | 10 | 11 | 7 | 6 |
| SL1344_RS13975_ | 5 | 6 | 4 | 7 | 5 | 8 |
| SL1344_RS13980_ | 3 | 13 | 12 | 6 | 6 | 15 |
| SL1344_RS13985_ | 2 | 1 | 7 | 2 | 3 | 4 |
| SL1344_RS14000_ | 14 | 9 | 10 | 22 | 12 | 8 |
| SL1344_RS14005_ | 23 | 14 | 12 | 19 | 17 | 18 |
| SL1344_RS14010_ | 7 | 11 | 16 | 4 | 12 | 10 |
| SL1344_RS14015_ | 22 | 7 | 23 | 19 | 16 | 28 |
| SL1344_RS14020_ | 10 | 11 | 22 | 12 | 8 | 15 |
| SL1344_RS14025_ | 1329 | 1101 | 1150 | 987 | 944 | 942 |
| SL1344_RS14030_ | 1405 | 1285 | 1181 | 945 | 920 | 1099 |
| SL1344_RS14035_ | 366 | 250 | 286 | 464 | 387 | 384 |
| SL1344_RS14040_ | 1825 | 1408 | 1333 | 2370 | 2103 | 2150 |
| SL1344_RS14045_ | 1611 | 1370 | 1252 | 1922 | 1747 | 1809 |

| | | | | | | |
|---|---|---|---|---|---|---|
| SL1344_RS14050_ | 154 | 153 | 115 | 192 | 184 | 236 |
| SL1344_RS14055_ | 106 | 97 | 86 | 166 | 166 | 212 |
| SL1344_RS14060_ | 72 | 74 | 59 | 73 | 77 | 74 |
| SL1344_RS14065_ | 18 | 24 | 23 | 27 | 15 | 26 |
| SL1344_RS14070_ | 1 | 0 | 6 | 5 | 2 | 5 |
| SL1344_RS14075_ | 2 | 2 | 6 | 4 | 4 | 8 |
| SL1344_RS14080_ | 2 | 9 | 6 | 2 | 8 | 13 |
| SL1344_RS14085_ | 561 | 573 | 595 | 604 | 546 | 673 |
| SL1344_RS14090_ | 16 | 8 | 14 | 12 | 8 | 9 |
| SL1344_RS14095_ | 6 | 7 | 3 | 14 | 14 | 12 |
| SL1344_RS14100_ | 242 | 234 | 213 | 209 | 208 | 266 |
| SL1344_RS14105_ | 451 | 435 | 416 | 391 | 365 | 385 |
| SL1344_RS14110_ | 317 | 253 | 272 | 314 | 257 | 312 |
| SL1344_RS14115_ | 127 | 155 | 114 | 135 | 124 | 153 |
| SL1344_RS14120_ | 1675 | 1588 | 1569 | 1293 | 1204 | 1552 |
| SL1344_RS14125_ | 242 | 176 | 195 | 126 | 122 | 186 |
| SL1344_RS14130_ | 439 | 361 | 333 | 287 | 304 | 362 |
| SL1344_RS14135_ | 575 | 555 | 613 | 532 | 585 | 678 |
| SL1344_RS14140_ | 16 | 17 | 10 | 11 | 14 | 15 |
| SL1344_RS14150_ | 20 | 14 | 19 | 15 | 12 | 15 |
| SL1344_RS14155_ | 12 | 13 | 19 | 12 | 14 | 11 |
| SL1344_RS14165_ | 15 | 12 | 11 | 11 | 9 | 17 |
| SL1344_RS14170_ | 31 | 14 | 22 | 15 | 17 | 9 |
| SL1344_RS14175_ | 227 | 244 | 256 | 242 | 231 | 256 |
| SL1344_RS14180_ | 342 | 420 | 431 | 251 | 287 | 266 |
| SL1344_RS14190_ | 129 | 112 | 66 | 62 | 64 | 79 |
| SL1344_RS14195_ | 387 | 344 | 394 | 498 | 515 | 574 |
| SL1344_RS14200_ | 362 | 318 | 287 | 313 | 288 | 309 |
| SL1344_RS14205_ | 6 | 16 | 22 | 13 | 13 | 8 |
| SL1344_RS14210_ | 31 | 33 | 30 | 21 | 30 | 51 |
| SL1344_RS14215_ | 993 | 980 | 1028 | 892 | 933 | 945 |
| SL1344_RS14220_ | 421 | 403 | 377 | 335 | 314 | 343 |
| SL1344_RS14225_ | 439 | 442 | 452 | 468 | 483 | 547 |
| SL1344_RS14230_ | 10 | 14 | 19 | 17 | 7 | 16 |
| SL1344_RS14235_ | 7 | 5 | 11 | 9 | 6 | 10 |
| SL1344_RS14240_ | 5 | 2 | 7 | 3 | 8 | 6 |
| SL1344_RS14245_ | 22 | 12 | 20 | 11 | 10 | 19 |
| SL1344_RS14250_ | 22 | 25 | 17 | 28 | 22 | 30 |
| SL1344_RS14255_ | 332 | 334 | 318 | 394 | 387 | 461 |
| SL1344_RS14260_ | 46 | 41 | 38 | 24 | 23 | 43 |
| SL1344_RS14265_ | 11 | 5 | 10 | 9 | 7 | 11 |
| SL1344_RS14270_ | 13 | 14 | 11 | 13 | 13 | 19 |
| SL1344_RS14275_ | 14 | 17 | 25 | 17 | 39 | 22 |
| SL1344_RS14280_ | 115 | 116 | 114 | 57 | 65 | 97 |
| SL1344_RS14285_ | 14 | 11 | 17 | 10 | 12 | 6 |
| SL1344_RS14290_ | 23 | 21 | 28 | 21 | 22 | 28 |
| SL1344_RS14295_ | 69 | 85 | 96 | 87 | 102 | 94 |
| SL1344_RS14300_ | 165 | 154 | 175 | 178 | 173 | 154 |
| SL1344_RS14305_ | 419 | 403 | 416 | 323 | 333 | 417 |
| SL1344_RS14310_ | 25 | 16 | 13 | 27 | 15 | 10 |
| SL1344_RS14315_ | 27 | 19 | 25 | 32 | 24 | 25 |
| SL1344_RS14320_ | 420 | 385 | 466 | 475 | 453 | 486 |
| SL1344_RS14325_ | 1688 | 1488 | 1545 | 1745 | 1536 | 1841 |
| SL1344_RS14330_ | 75 | 70 | 60 | 68 | 74 | 90 |
| SL1344_RS14335_ | 45 | 47 | 35 | 33 | 41 | 53 |
| SL1344_RS14345_ | 182237 | 185687 | 180066 | 11946 | 191360 | 10800 |
| SL1344_RS14350_ | 70 | 69 | 64 | 45 | 40 | 33 |
| SL1344_RS14360_ | 158 | 121 | 120 | 191 | 187 | 190 |
| SL1344_RS14365_ | 41 | 39 | 64 | 30 | 29 | 41 |
| SL1344_RS14370_ | 27 | 34 | 41 | 34 | 45 | 38 |
| SL1344_RS14375_ | 130 | 122 | 151 | 158 | 178 | 170 |
| SL1344_RS14380_ | 25 | 22 | 35 | 47 | 38 | 43 |

| | | | | | | |
|---|---|---|---|---|---|---|
| SL1344_RS14385_ | 216 | 215 | 200 | 186 | 149 | 248 |
| SL1344_RS14390_ | 1526 | 1554 | 1765 | 1921 | 1804 | 1995 |
| SL1344_RS14395_ | 1899 | 1765 | 2280 | 2233 | 2042 | 2258 |
| SL1344_RS14400_ | 143 | 124 | 131 | 149 | 137 | 123 |
| SL1344_RS14405_ | 155 | 139 | 170 | 190 | 172 | 183 |
| SL1344_RS14410_ | 330 | 322 | 383 | 435 | 400 | 475 |
| SL1344_RS14415_ | 234 | 231 | 234 | 352 | 352 | 381 |
| SL1344_RS14420_ | 26 | 28 | 33 | 33 | 24 | 20 |
| SL1344_RS14425_ | 165 | 155 | 186 | 151 | 170 | 185 |
| SL1344_RS14430_ | 45 | 57 | 57 | 48 | 58 | 54 |
| SL1344_RS14435_ | 234 | 267 | 290 | 278 | 250 | 288 |
| SL1344_RS14440_ | 47 | 52 | 54 | 52 | 62 | 75 |
| SL1344_RS14445_ | 46 | 47 | 73 | 56 | 55 | 52 |
| SL1344_RS14450_ | 62 | 83 | 77 | 92 | 99 | 92 |
| SL1344_RS14455_ | 98 | 90 | 110 | 142 | 122 | 153 |
| SL1344_RS14460_ | 86 | 71 | 85 | 105 | 63 | 103 |
| SL1344_RS14465_ | 43 | 46 | 47 | 51 | 61 | 56 |
| SL1344_RS14470_ | 336 | 352 | 337 | 155 | 155 | 205 |
| SL1344_RS14475_ | 139 | 146 | 142 | 84 | 82 | 93 |
| SL1344_RS14480_ | 43 | 51 | 70 | 57 | 53 | 68 |
| SL1344_RS14485_ | 146 | 146 | 142 | 150 | 133 | 174 |
| SL1344_RS14490_ | 772 | 813 | 834 | 971 | 1094 | 1266 |
| SL1344_RS14495_ | 1667 | 1500 | 1342 | 1461 | 1347 | 1543 |
| SL1344_RS14500_ | 2117 | 2090 | 2305 | 1292 | 1095 | 1155 |
| SL1344_RS14505_ | 48 | 40 | 40 | 44 | 49 | 69 |
| SL1344_RS14510_ | 902 | 999 | 786 | 503 | 391 | 591 |
| SL1344_RS14515_ | 278 | 261 | 249 | 227 | 297 | 302 |
| SL1344_RS14520_ | 36 | 44 | 31 | 39 | 35 | 47 |
| SL1344_RS14525_ | 7 | 2 | 10 | 7 | 2 | 6 |
| SL1344_RS14530_ | 4 | 2 | 5 | 3 | 5 | 7 |
| SL1344_RS14540_ | 39 | 37 | 39 | 42 | 51 | 48 |
| SL1344_RS14545_ | 50 | 55 | 38 | 48 | 46 | 48 |
| SL1344_RS14550_ | 141 | 116 | 129 | 144 | 143 | 179 |
| SL1344_RS14555_ | 125 | 136 | 119 | 90 | 74 | 110 |
| SL1344_RS14560_ | 221 | 216 | 228 | 221 | 225 | 251 |
| SL1344_RS14565_ | 73 | 91 | 99 | 110 | 88 | 136 |
| SL1344_RS14570_ | 699 | 776 | 630 | 821 | 611 | 764 |
| SL1344_RS14575_ | 602 | 616 | 655 | 622 | 581 | 707 |
| SL1344_RS14585_ | 71 | 57 | 45 | 57 | 60 | 42 |
| SL1344_RS14600_ | 734 | 665 | 588 | 642 | 588 | 674 |
| SL1344_RS14605_ | 922 | 844 | 739 | 826 | 811 | 881 |
| SL1344_RS14610_ | 101 | 86 | 133 | 111 | 73 | 95 |
| SL1344_RS14615_ | 185 | 217 | 274 | 312 | 198 | 255 |
| SL1344_RS14620_ | 422 | 448 | 558 | 637 | 399 | 587 |
| SL1344_RS14625_ | 1033 | 846 | 1095 | 1393 | 1015 | 1194 |
| SL1344_RS14630_ | 4447 | 4206 | 4982 | 6051 | 4776 | 5473 |
| SL1344_RS14640_ | 4342 | 4017 | 3985 | 3254 | 2820 | 3896 |
| SL1344_RS14645_ | 70 | 86 | 107 | 96 | 98 | 81 |
| SL1344_RS14655_ | 460 | 385 | 370 | 364 | 352 | 445 |
| SL1344_RS14660_ | 448 | 436 | 459 | 443 | 409 | 439 |
| SL1344_RS14675_ | 105 | 98 | 104 | 136 | 201 | 180 |
| SL1344_RS14680_ | 412 | 457 | 477 | 712 | 922 | 868 |
| SL1344_RS14685_ | 204 | 206 | 224 | 203 | 213 | 192 |
| SL1344_RS14690_ | 154 | 165 | 164 | 152 | 148 | 157 |
| SL1344_RS14700_ | 773 | 826 | 854 | 648 | 648 | 704 |
| SL1344_RS14705_ | 222 | 261 | 208 | 245 | 241 | 279 |
| SL1344_RS14710_ | 41 | 30 | 40 | 38 | 36 | 29 |
| SL1344_RS14715_ | 57 | 60 | 49 | 49 | 65 | 45 |
| SL1344_RS14720_ | 400 | 421 | 431 | 417 | 362 | 466 |
| SL1344_RS14725_ | 14 | 18 | 12 | 20 | 13 | 17 |
| SL1344_RS14730_ | 80 | 69 | 87 | 95 | 88 | 103 |
| SL1344_RS14735_ | 46 | 50 | 58 | 57 | 59 | 52 |

| | | | | | | |
|---|---|---|---|---|---|---|
| SL1344_RS14740_ | 39 | 31 | 52 | 47 | 48 | 53 |
| SL1344_RS14745_ | 46 | 47 | 66 | 55 | 56 | 59 |
| SL1344_RS14750_ | 28 | 25 | 39 | 29 | 19 | 33 |
| SL1344_RS14765_ | 48 | 52 | 50 | 71 | 56 | 78 |
| SL1344_RS14770_ | 10 | 5 | 16 | 8 | 8 | 15 |
| SL1344_RS14775_ | 14 | 15 | 9 | 4 | 7 | 10 |
| SL1344_RS14780_ | 1243 | 1376 | 1067 | 1281 | 1392 | 1370 |
| SL1344_RS14785_ | 972 | 1006 | 1070 | 1005 | 872 | 1070 |
| SL1344_RS14790_ | 417 | 407 | 398 | 429 | 343 | 424 |
| SL1344_RS14795_ | 1095 | 1038 | 1047 | 1028 | 897 | 1107 |
| SL1344_RS14800_ | 1093 | 1177 | 1134 | 1140 | 998 | 1151 |
| SL1344_RS14805_ | 540 | 544 | 563 | 564 | 512 | 596 |
| SL1344_RS14810_ | 38 | 37 | 61 | 47 | 39 | 56 |
| SL1344_RS14815_ | 27 | 32 | 37 | 55 | 44 | 50 |
| SL1344_RS14820_ | 34 | 49 | 34 | 36 | 34 | 39 |
| SL1344_RS14825_ | 23 | 24 | 39 | 38 | 26 | 42 |
| SL1344_RS14830_ | 74 | 72 | 84 | 92 | 81 | 125 |
| SL1344_RS14850_ | 1736 | 1602 | 1585 | 2634 | 2198 | 3095 |
| SL1344_RS14855_ | 1522 | 1498 | 1590 | 2749 | 2058 | 2818 |
| SL1344_RS14900_ | 2544 | 2487 | 2277 | 3096 | 2475 | 4057 |
| SL1344_RS14905_ | 827 | 759 | 778 | 1232 | 1029 | 1516 |
| SL1344_RS14910_ | 983 | 894 | 882 | 1515 | 1232 | 1836 |
| SL1344_RS15010_ | 895 | 882 | 839 | 1387 | 1136 | 1308 |
| SL1344_RS15015_ | 710 | 630 | 531 | 841 | 769 | 822 |
| SL1344_RS15020_ | 198 | 148 | 154 | 207 | 115 | 159 |
| SL1344_RS15025_ | 33 | 23 | 26 | 51 | 42 | 47 |
| SL1344_RS15030_ | 598 | 619 | 653 | 491 | 497 | 636 |
| SL1344_RS15035_ | 785 | 702 | 749 | 707 | 589 | 700 |
| SL1344_RS15040_ | 9 | 1 | 6 | 5 | 5 | 7 |
| SL1344_RS15045_ | 34 | 35 | 50 | 38 | 48 | 48 |
| SL1344_RS15050_ | 7 | 9 | 8 | 8 | 5 | 7 |
| SL1344_RS15055_ | 1387 | 1474 | 1451 | 1263 | 1147 | 1371 |
| SL1344_RS15060_ | 41 | 48 | 33 | 40 | 47 | 39 |
| SL1344_RS15065_ | 46 | 36 | 35 | 59 | 36 | 42 |
| SL1344_RS15070_ | 57 | 63 | 69 | 68 | 71 | 69 |
| SL1344_RS15075_ | 305 | 273 | 317 | 238 | 183 | 274 |
| SL1344_RS15080_ | 120 | 104 | 100 | 77 | 55 | 93 |
| SL1344_RS15085_ | 76 | 106 | 96 | 95 | 53 | 84 |
| SL1344_RS15090_ | 92 | 84 | 129 | 71 | 89 | 73 |
| SL1344_RS15095_ | 208 | 190 | 174 | 173 | 145 | 170 |
| SL1344_RS15100_ | 136 | 166 | 137 | 120 | 138 | 153 |
| SL1344_RS15105_ | 320 | 340 | 277 | 288 | 302 | 309 |
| SL1344_RS15110_ | 269 | 294 | 275 | 191 | 237 | 261 |
| SL1344_RS15115_ | 7 | 11 | 9 | 12 | 4 | 8 |
| SL1344_RS15120_ | 53 | 37 | 53 | 53 | 62 | 47 |
| SL1344_RS15125_ | 20 | 15 | 16 | 28 | 18 | 27 |
| SL1344_RS15140_ | 891 | 777 | 844 | 831 | 756 | 690 |
| SL1344_RS15145_ | 762 | 727 | 794 | 696 | 576 | 711 |
| SL1344_RS15150_ | 927 | 918 | 861 | 799 | 753 | 944 |
| SL1344_RS15155_ | 307 | 287 | 292 | 250 | 247 | 314 |
| SL1344_RS15160_ | 245 | 225 | 276 | 347 | 244 | 314 |
| SL1344_RS15165_ | 395 | 332 | 351 | 410 | 350 | 380 |
| SL1344_RS15170_ | 116 | 91 | 131 | 1007 | 1004 | 1053 |
| SL1344_RS15190_ | 310 | 334 | 300 | 400 | 365 | 440 |
| SL1344_RS15195_ | 53 | 48 | 51 | 48 | 42 | 58 |
| SL1344_RS15200_ | 197 | 182 | 175 | 198 | 166 | 195 |
| SL1344_RS15205_ | 128 | 138 | 123 | 149 | 129 | 145 |
| SL1344_RS15210_ | 119 | 115 | 104 | 125 | 92 | 124 |
| SL1344_RS15215_ | 156 | 209 | 176 | 204 | 196 | 194 |
| SL1344_RS15220_ | 77 | 64 | 71 | 77 | 61 | 83 |
| SL1344_RS15225_ | 125 | 145 | 159 | 175 | 157 | 183 |
| SL1344_RS15230_ | 121 | 128 | 125 | 184 | 143 | 200 |

| | | | | | | |
|---|---|---|---|---|---|---|
| SL1344_RS15260_ | 231 | 218 | 230 | 206 | 153 | 196 |
| SL1344_RS15265_ | 568 | 556 | 542 | 683 | 575 | 664 |
| SL1344_RS15270_ | 11817 | 11838 | 13515 | 11334 | 10224 | 12443 |
| SL1344_RS15275_ | 2950 | 2974 | 3129 | 3212 | 2925 | 3515 |
| SL1344_RS15280_ | 470 | 428 | 452 | 476 | 401 | 520 |
| SL1344_RS15285_ | 412 | 361 | 293 | 362 | 330 | 366 |
| SL1344_RS15290_ | 650 | 634 | 554 | 660 | 616 | 729 |
| SL1344_RS15295_ | 3316 | 3609 | 3457 | 2928 | 2817 | 3224 |
| SL1344_RS15310_ | 767 | 850 | 678 | 487 | 375 | 499 |
| SL1344_RS15315_ | 3710 | 3691 | 3018 | 2064 | 1668 | 2400 |
| SL1344_RS15320_ | 1861 | 2036 | 1863 | 1193 | 1111 | 1234 |
| SL1344_RS15325_ | 3609 | 3630 | 3735 | 2144 | 2378 | 2689 |
| SL1344_RS15330_ | 680 | 656 | 652 | 509 | 504 | 634 |
| SL1344_RS15355_ | 449 | 498 | 487 | 402 | 399 | 468 |
| SL1344_RS15360_ | 3721 | 3631 | 3768 | 2639 | 2748 | 3016 |
| SL1344_RS15365_ | 12743 | 12149 | 14722 | 13490 | 12653 | 13896 |
| SL1344_RS15370_ | 8911 | 9860 | 10733 | 8964 | 8429 | 9388 |
| SL1344_RS15375_ | 216 | 221 | 217 | 229 | 193 | 229 |
| SL1344_RS15380_ | 127 | 135 | 116 | 112 | 72 | 89 |
| SL1344_RS15385_ | 55 | 53 | 53 | 53 | 47 | 47 |
| SL1344_RS15390_ | 54 | 61 | 70 | 34 | 33 | 41 |
| SL1344_RS15395_ | 236 | 226 | 247 | 128 | 104 | 155 |
| SL1344_RS15400_ | 124 | 140 | 150 | 84 | 78 | 84 |
| SL1344_RS15405_ | 123 | 158 | 146 | 81 | 94 | 116 |
| SL1344_RS15410_ | 414 | 408 | 406 | 331 | 317 | 375 |
| SL1344_RS15415_ | 876 | 821 | 840 | 944 | 930 | 998 |
| SL1344_RS15420_ | 139 | 176 | 182 | 220 | 210 | 249 |
| SL1344_RS15425_ | 311 | 305 | 322 | 390 | 337 | 397 |
| SL1344_RS15430_ | 1878 | 2058 | 1780 | 1686 | 1645 | 1864 |
| SL1344_RS15435_ | 315 | 338 | 326 | 321 | 307 | 346 |
| SL1344_RS15440_ | 183 | 208 | 209 | 164 | 150 | 184 |
| SL1344_RS15445_ | 240 | 268 | 284 | 324 | 253 | 290 |
| SL1344_RS15450_ | 339 | 306 | 271 | 310 | 300 | 326 |
| SL1344_RS15455_ | 973 | 880 | 859 | 1048 | 1072 | 1154 |
| SL1344_RS15460_ | 110 | 108 | 136 | 188 | 162 | 177 |
| SL1344_RS15465_ | 300 | 302 | 344 | 424 | 404 | 438 |
| SL1344_RS15475_ | 308 | 310 | 262 | 257 | 252 | 249 |
| SL1344_RS15480_ | 436 | 472 | 489 | 426 | 382 | 417 |
| SL1344_RS15485_ | 1070 | 1125 | 1062 | 957 | 878 | 1071 |
| SL1344_RS15490_ | 761 | 813 | 825 | 795 | 699 | 772 |
| SL1344_RS15495_ | 743 | 744 | 728 | 627 | 591 | 690 |
| SL1344_RS15500_ | 24 | 24 | 27 | 32 | 13 | 27 |
| SL1344_RS15505_ | 21 | 25 | 26 | 21 | 22 | 24 |
| SL1344_RS15510_ | 27 | 33 | 16 | 31 | 30 | 24 |
| SL1344_RS15515_ | 10 | 16 | 11 | 19 | 8 | 18 |
| SL1344_RS15520_ | 7 | 8 | 8 | 3 | 12 | 15 |
| SL1344_RS15525_ | 599 | 668 | 776 | 691 | 682 | 763 |
| SL1344_RS15530_ | 469 | 477 | 506 | 511 | 444 | 616 |
| SL1344_RS15535_ | 2638 | 2742 | 2633 | 2435 | 2150 | 2563 |
| SL1344_RS15540_ | 2174 | 2437 | 2352 | 2026 | 2097 | 2558 |
| SL1344_RS15545_ | 48 | 44 | 43 | 37 | 39 | 55 |
| SL1344_RS15550_ | 600 | 594 | 567 | 813 | 760 | 727 |
| SL1344_RS15555_ | 501 | 453 | 553 | 717 | 648 | 741 |
| SL1344_RS15560_ | 368 | 443 | 359 | 173 | 169 | 239 |
| SL1344_RS15565_ | 435 | 390 | 382 | 335 | 303 | 342 |
| SL1344_RS15570_ | 512 | 500 | 470 | 392 | 361 | 465 |
| SL1344_RS15575_ | 955 | 937 | 936 | 910 | 824 | 892 |
| SL1344_RS15580_ | 380 | 432 | 393 | 343 | 392 | 423 |
| SL1344_RS15585_ | 447 | 436 | 363 | 361 | 318 | 347 |
| SL1344_RS15590_ | 47 | 57 | 45 | 75 | 87 | 114 |
| SL1344_RS15595_ | 149 | 142 | 137 | 112 | 117 | 133 |
| SL1344_RS15600_ | 387 | 397 | 341 | 330 | 285 | 383 |

| | | | | | | |
|---|---|---|---|---|---|---|
| SL1344_RS15605_ | 134 | 115 | 130 | 108 | 92 | 114 |
| SL1344_RS15610_ | 118 | 139 | 107 | 69 | 78 | 84 |
| SL1344_RS15615_ | 55 | 60 | 70 | 46 | 31 | 39 |
| SL1344_RS15620_ | 567 | 557 | 551 | 703 | 652 | 740 |
| SL1344_RS15625_ | 264 | 254 | 236 | 357 | 330 | 355 |
| SL1344_RS15630_ | 58 | 52 | 52 | 87 | 56 | 61 |
| SL1344_RS15635_ | 125 | 119 | 137 | 129 | 137 | 127 |
| SL1344_RS15640_ | 409 | 323 | 288 | 329 | 357 | 424 |
| SL1344_RS15645_ | 71 | 56 | 45 | 49 | 50 | 71 |
| SL1344_RS15650_ | 68 | 66 | 62 | 76 | 32 | 57 |
| SL1344_RS15655_ | 15 | 21 | 12 | 9 | 16 | 12 |
| SL1344_RS15660_ | 23 | 29 | 25 | 30 | 22 | 27 |
| SL1344_RS15665_ | 13 | 23 | 13 | 24 | 17 | 19 |
| SL1344_RS15670_ | 26 | 32 | 41 | 39 | 31 | 35 |
| SL1344_RS15675_ | 6 | 11 | 7 | 15 | 7 | 17 |
| SL1344_RS15680_ | 168 | 160 | 147 | 211 | 175 | 201 |
| SL1344_RS15685_ | 24 | 13 | 19 | 19 | 29 | 15 |
| SL1344_RS15690_ | 1143 | 1120 | 1152 | 925 | 1022 | 1060 |
| SL1344_RS15695_ | 956 | 1011 | 1017 | 798 | 978 | 1094 |
| SL1344_RS15700_ | 1 | 1 | 1 | 3 | 3 | 2 |
| SL1344_RS15705_ | 32 | 39 | 50 | 30 | 23 | 37 |
| SL1344_RS15710_ | 251 | 255 | 315 | 261 | 275 | 368 |
| SL1344_RS15715_ | 522 | 470 | 460 | 530 | 510 | 577 |
| SL1344_RS15720_ | 612 | 591 | 496 | 628 | 672 | 633 |
| SL1344_RS15725_ | 4483 | 4585 | 4861 | 3677 | 3082 | 4391 |
| SL1344_RS15730_ | 1940 | 1875 | 1844 | 2046 | 1773 | 2283 |
| SL1344_RS15735_ | 1674 | 1594 | 1522 | 1457 | 1392 | 1604 |
| SL1344_RS15740_ | 986 | 907 | 782 | 825 | 793 | 943 |
| SL1344_RS15745_ | 462 | 466 | 422 | 493 | 467 | 524 |
| SL1344_RS15750_ | 497 | 460 | 550 | 422 | 389 | 498 |
| SL1344_RS15755_ | 1056 | 1092 | 999 | 887 | 858 | 948 |
| SL1344_RS15760_ | 489 | 534 | 503 | 469 | 452 | 460 |
| SL1344_RS15765_ | 74 | 102 | 97 | 85 | 62 | 102 |
| SL1344_RS15770_ | 1369 | 1501 | 1378 | 1130 | 1073 | 1197 |
| SL1344_RS15775_ | 607 | 617 | 631 | 558 | 525 | 664 |
| SL1344_RS15780_ | 503 | 494 | 482 | 345 | 368 | 451 |
| SL1344_RS15785_ | 392 | 396 | 459 | 330 | 286 | 372 |
| SL1344_RS15790_ | 49 | 50 | 48 | 46 | 43 | 41 |
| SL1344_RS15795_ | 16002 | 16053 | 17549 | 12854 | 11961 | 14332 |
| SL1344_RS15800_ | 7105 | 7510 | 7490 | 5781 | 5268 | 6529 |
| SL1344_RS15810_ | 787 | 762 | 789 | 737 | 615 | 741 |
| SL1344_RS15815_ | 1121 | 1162 | 1181 | 957 | 974 | 1028 |
| SL1344_RS15820_ | 1531 | 1562 | 1553 | 1470 | 1429 | 1560 |
| SL1344_RS15825_ | 964 | 1044 | 1035 | 989 | 942 | 1083 |
| SL1344_RS15830_ | 1996 | 2012 | 1858 | 1897 | 1822 | 1806 |
| SL1344_RS15835_ | 460 | 410 | 562 | 400 | 372 | 430 |
| SL1344_RS15840_ | 808 | 768 | 748 | 636 | 538 | 741 |
| SL1344_RS15845_ | 2276 | 2405 | 2590 | 1599 | 1535 | 1919 |
| SL1344_RS15850_ | 192 | 178 | 175 | 161 | 175 | 163 |
| SL1344_RS15855_ | 14 | 23 | 20 | 12 | 21 | 25 |
| SL1344_RS15860_ | 776 | 756 | 760 | 566 | 570 | 697 |
| SL1344_RS15865_ | 70 | 62 | 76 | 94 | 100 | 99 |
| SL1344_RS15870_ | 766 | 771 | 838 | 623 | 557 | 724 |
| SL1344_RS15875_ | 14734 | 14650 | 14694 | 9602 | 9518 | 10782 |
| SL1344_RS15880_ | 4659 | 4495 | 4920 | 3088 | 2719 | 3775 |
| SL1344_RS15890_ | 124 | 98 | 132 | 155 | 113 | 155 |
| SL1344_RS15895_ | 298 | 321 | 329 | 248 | 217 | 275 |
| SL1344_RS15900_ | 338 | 309 | 261 | 270 | 217 | 230 |
| SL1344_RS15905_ | 379 | 363 | 334 | 341 | 297 | 321 |
| SL1344_RS15910_ | 186 | 167 | 174 | 153 | 118 | 105 |
| SL1344_RS15915_ | 9562 | 9093 | 10776 | 8345 | 8392 | 9434 |
| SL1344_RS15920_ | 542 | 567 | 548 | 492 | 467 | 588 |

| | | | | | | |
|---|---|---|---|---|---|---|
| SL1344_RS15925_ | 2224 | 2182 | 2383 | 1930 | 2233 | 2232 |
| SL1344_RS15930_ | 66 | 48 | 57 | 56 | 52 | 54 |
| SL1344_RS15935_ | 24 | 24 | 36 | 46 | 43 | 39 |
| SL1344_RS15940_ | 85 | 98 | 101 | 92 | 76 | 86 |
| SL1344_RS15945_ | 105 | 99 | 107 | 175 | 110 | 148 |
| SL1344_RS15950_ | 41 | 35 | 53 | 70 | 39 | 38 |
| SL1344_RS15955_ | 43 | 55 | 45 | 47 | 40 | 56 |
| SL1344_RS15960_ | 1119 | 1096 | 1021 | 874 | 868 | 967 |
| SL1344_RS15965_ | 23 | 16 | 31 | 18 | 28 | 21 |
| SL1344_RS15970_ | 1641 | 1615 | 1937 | 1627 | 1625 | 1686 |
| SL1344_RS15975_ | 132 | 144 | 141 | 176 | 183 | 161 |
| SL1344_RS15985_ | 1948 | 2180 | 2315 | 2618 | 3097 | 3218 |
| SL1344_RS15990_ | 601 | 546 | 666 | 448 | 415 | 414 |
| SL1344_RS15995_ | 45 | 61 | 53 | 88 | 53 | 61 |
| SL1344_RS16000_ | 67 | 72 | 91 | 129 | 123 | 112 |
| SL1344_RS16005_ | 412 | 340 | 403 | 400 | 406 | 440 |
| SL1344_RS16015_ | 296 | 270 | 284 | 361 | 284 | 347 |
| SL1344_RS16020_ | 213 | 222 | 211 | 308 | 260 | 228 |
| SL1344_RS16025_ | 58 | 66 | 57 | 76 | 45 | 49 |
| SL1344_RS16030_ | 48 | 47 | 51 | 66 | 49 | 71 |
| SL1344_RS16035_ | 342 | 287 | 299 | 380 | 297 | 361 |
| SL1344_RS16040_ | 477 | 460 | 431 | 419 | 436 | 515 |
| SL1344_RS16045_ | 372 | 333 | 352 | 289 | 300 | 355 |
| SL1344_RS16050_ | 373 | 388 | 391 | 439 | 370 | 438 |
| SL1344_RS16055_ | 525 | 554 | 599 | 665 | 560 | 661 |
| SL1344_RS16060_ | 235 | 226 | 194 | 173 | 184 | 167 |
| SL1344_RS16065_ | 14532 | 14987 | 15210 | 9995 | 9901 | 10478 |
| SL1344_RS16070_ | 878 | 801 | 721 | 690 | 571 | 719 |
| SL1344_RS16075_ | 1401 | 1358 | 1228 | 944 | 787 | 1161 |
| SL1344_RS16080_ | 526 | 509 | 492 | 523 | 428 | 496 |
| SL1344_RS16085_ | 219 | 230 | 205 | 260 | 210 | 247 |
| SL1344_RS16090_ | 585 | 612 | 498 | 442 | 408 | 504 |
| SL1344_RS16095_ | 549 | 507 | 582 | 464 | 416 | 526 |
| SL1344_RS16100_ | 1661 | 1653 | 1826 | 1281 | 1380 | 1481 |
| SL1344_RS16105_ | 325 | 362 | 401 | 298 | 325 | 291 |
| SL1344_RS16110_ | 4 | 7 | 3 | 8 | 5 | 7 |
| SL1344_RS16115_ | 36 | 32 | 59 | 44 | 34 | 37 |
| SL1344_RS16120_ | 70 | 57 | 88 | 121 | 102 | 83 |
| SL1344_RS16125_ | 32 | 29 | 38 | 58 | 45 | 49 |
| SL1344_RS16130_ | 172 | 164 | 176 | 135 | 131 | 185 |
| SL1344_RS16135_ | 84 | 82 | 58 | 50 | 58 | 74 |
| SL1344_RS16140_ | 170 | 135 | 155 | 132 | 152 | 137 |
| SL1344_RS16145_ | 213 | 194 | 188 | 195 | 201 | 215 |
| SL1344_RS16150_ | 254 | 289 | 310 | 231 | 227 | 244 |
| SL1344_RS16155_ | 31 | 24 | 33 | 23 | 33 | 25 |
| SL1344_RS16160_ | 80 | 67 | 68 | 73 | 89 | 72 |
| SL1344_RS16165_ | 83 | 77 | 89 | 87 | 74 | 69 |
| SL1344_RS16170_ | 116 | 121 | 124 | 176 | 124 | 144 |
| SL1344_RS16175_ | 84 | 55 | 70 | 88 | 101 | 84 |
| SL1344_RS16180_ | 129 | 123 | 122 | 139 | 117 | 130 |
| SL1344_RS16185_ | 139 | 130 | 129 | 114 | 92 | 108 |
| SL1344_RS16190_ | 177 | 146 | 125 | 127 | 152 | 152 |
| SL1344_RS16195_ | 187 | 159 | 107 | 170 | 159 | 155 |
| SL1344_RS16200_ | 72 | 80 | 76 | 61 | 62 | 59 |
| SL1344_RS16205_ | 27 | 38 | 36 | 20 | 21 | 29 |
| SL1344_RS16210_ | 721 | 722 | 768 | 453 | 507 | 491 |
| SL1344_RS16215_ | 1136 | 1171 | 1139 | 682 | 825 | 789 |
| SL1344_RS16220_ | 676 | 697 | 734 | 368 | 390 | 414 |
| SL1344_RS16225_ | 906 | 887 | 900 | 515 | 589 | 545 |
| SL1344_RS16235_ | 1045 | 973 | 902 | 660 | 591 | 705 |
| SL1344_RS16240_ | 467 | 458 | 513 | 362 | 327 | 396 |
| SL1344_RS16245_ | 70 | 78 | 83 | 61 | 52 | 57 |

| | | | | | | |
|---|---|---|---|---|---|---|
| SL1344_RS16250_ | 68 | 77 | 53 | 64 | 59 | 81 |
| SL1344_RS16255_ | 829 | 767 | 693 | 801 | 591 | 790 |
| SL1344_RS16260_ | 587 | 540 | 569 | 560 | 467 | 549 |
| SL1344_RS16265_ | 690 | 735 | 694 | 687 | 609 | 739 |
| SL1344_RS16270_ | 1790 | 1855 | 1900 | 1529 | 1327 | 1731 |
| SL1344_RS16275_ | 4493 | 4661 | 4480 | 3851 | 3008 | 4163 |
| SL1344_RS16280_ | 2095 | 2098 | 2002 | 1730 | 1171 | 1877 |
| SL1344_RS16285_ | 1638 | 1751 | 1558 | 1298 | 928 | 1340 |
| SL1344_RS16290_ | 1875 | 2036 | 2079 | 1795 | 1450 | 1756 |
| SL1344_RS16305_ | 964 | 942 | 915 | 1307 | 1213 | 1481 |
| SL1344_RS16310_ | 1563 | 1736 | 1595 | 3054 | 2880 | 3340 |
| SL1344_RS16315_ | 408 | 360 | 375 | 917 | 729 | 836 |
| SL1344_RS16320_ | 526 | 518 | 474 | 1183 | 1012 | 1182 |
| SL1344_RS16325_ | 1601 | 1604 | 1655 | 1344 | 992 | 1402 |
| SL1344_RS16330_ | 115 | 119 | 126 | 158 | 101 | 103 |
| SL1344_RS16335_ | 354 | 268 | 229 | 235 | 139 | 172 |
| SL1344_RS16340_ | 435 | 277 | 189 | 193 | 105 | 117 |
| SL1344_RS16350_ | 647 | 801 | 969 | 642 | 604 | 620 |
| SL1344_RS16355_ | 314 | 315 | 247 | 283 | 262 | 306 |
| SL1344_RS16360_ | 322 | 321 | 360 | 318 | 292 | 341 |
| SL1344_RS16365_ | 521 | 517 | 550 | 349 | 322 | 382 |
| SL1344_RS16370_ | 58 | 65 | 51 | 54 | 63 | 63 |
| SL1344_RS16375_ | 325 | 353 | 327 | 369 | 360 | 372 |
| SL1344_RS16380_ | 20 | 21 | 25 | 17 | 27 | 21 |
| SL1344_RS16385_ | 437 | 482 | 620 | 580 | 583 | 703 |
| SL1344_RS16390_ | 120 | 134 | 111 | 107 | 116 | 142 |
| SL1344_RS16395_ | 27 | 22 | 24 | 20 | 22 | 15 |
| SL1344_RS16400_ | 103 | 98 | 132 | 118 | 103 | 107 |
| SL1344_RS16405_ | 902 | 916 | 946 | 755 | 682 | 815 |
| SL1344_RS16410_ | 264 | 239 | 249 | 300 | 256 | 288 |
| SL1344_RS16415_ | 1679 | 1703 | 1819 | 1671 | 1637 | 1852 |
| SL1344_RS16420_ | 932 | 922 | 971 | 944 | 774 | 1052 |
| SL1344_RS16425_ | 1641 | 1936 | 1542 | 1462 | 1275 | 1871 |
| SL1344_RS16440_ | 225 | 169 | 185 | 204 | 155 | 226 |
| SL1344_RS16445_ | 1415 | 1451 | 1386 | 1422 | 1385 | 1602 |
| SL1344_RS16450_ | 2262 | 2301 | 2252 | 1980 | 2083 | 2344 |
| SL1344_RS16455_ | 890 | 791 | 887 | 897 | 841 | 945 |
| SL1344_RS16460_ | 2023 | 2038 | 2052 | 1908 | 1748 | 1811 |
| SL1344_RS16465_ | 956 | 1087 | 1001 | 682 | 671 | 787 |
| SL1344_RS16470_ | 307 | 241 | 234 | 291 | 245 | 299 |
| SL1344_RS16480_ | 1865 | 2033 | 2038 | 1515 | 1648 | 1944 |
| SL1344_RS16485_ | 2677 | 2688 | 2680 | 1960 | 1866 | 2241 |
| SL1344_RS16490_ | 265 | 232 | 212 | 328 | 292 | 295 |
| SL1344_RS16495_ | 399 | 420 | 454 | 421 | 385 | 414 |
| SL1344_RS16500_ | 353 | 475 | 514 | 186 | 187 | 249 |
| SL1344_RS16505_ | 114 | 125 | 112 | 108 | 79 | 101 |
| SL1344_RS16510_ | 45 | 49 | 53 | 40 | 33 | 40 |
| SL1344_RS16515_ | 198 | 168 | 162 | 186 | 212 | 190 |
| SL1344_RS16520_ | 1369 | 1210 | 1274 | 1338 | 898 | 1350 |
| SL1344_RS16525_ | 1104 | 1142 | 1030 | 1063 | 988 | 1098 |
| SL1344_RS16530_ | 1154 | 1301 | 1158 | 571 | 455 | 607 |
| SL1344_RS16535_ | 39 | 23 | 25 | 37 | 30 | 29 |
| SL1344_RS16540_ | 234 | 201 | 163 | 204 | 202 | 197 |
| SL1344_RS16545_ | 893 | 972 | 897 | 752 | 685 | 772 |
| SL1344_RS16550_ | 1224 | 1254 | 1105 | 1011 | 933 | 1167 |
| SL1344_RS16555_ | 580 | 590 | 586 | 466 | 498 | 534 |
| SL1344_RS16560_ | 300 | 311 | 285 | 347 | 297 | 333 |
| SL1344_RS16565_ | 744 | 666 | 641 | 620 | 558 | 685 |
| SL1344_RS16570_ | 453 | 450 | 432 | 439 | 380 | 522 |
| SL1344_RS16575_ | 99 | 95 | 85 | 74 | 76 | 79 |
| SL1344_RS16580_ | 1032 | 1028 | 1072 | 906 | 884 | 965 |
| SL1344_RS16585_ | 496 | 547 | 495 | 482 | 478 | 501 |

| | | | | | | |
|---|---|---|---|---|---|---|
| SL1344_RS16590_ | 3844 | 3435 | 3865 | 9646 | 7677 | 7632 |
| SL1344_RS16595_ | 1920 | 1996 | 2239 | 2610 | 2492 | 2845 |
| SL1344_RS16600_ | 5151 | 5291 | 5516 | 5510 | 4896 | 6043 |
| SL1344_RS16605_ | 238 | 236 | 185 | 118 | 100 | 160 |
| SL1344_RS16610_ | 23 | 24 | 45 | 38 | 34 | 32 |
| SL1344_RS16615_ | 61 | 57 | 57 | 50 | 48 | 80 |
| SL1344_RS16620_ | 314 | 338 | 279 | 190 | 158 | 190 |
| SL1344_RS16635_ | 128 | 189 | 143 | 124 | 99 | 106 |
| SL1344_RS16640_ | 255 | 256 | 300 | 276 | 259 | 276 |
| SL1344_RS16645_ | 237 | 269 | 301 | 296 | 257 | 312 |
| SL1344_RS16650_ | 295 | 326 | 313 | 274 | 284 | 262 |
| SL1344_RS16655_ | 24 | 32 | 32 | 46 | 38 | 26 |
| SL1344_RS16660_ | 1010 | 1207 | 1242 | 758 | 739 | 847 |
| SL1344_RS16665_ | 215 | 217 | 207 | 212 | 184 | 205 |
| SL1344_RS16675_ | 899 | 898 | 1045 | 915 | 861 | 894 |
| SL1344_RS16680_ | 511 | 483 | 552 | 458 | 458 | 459 |
| SL1344_RS16685_ | 2247 | 2084 | 1750 | 1642 | 1452 | 1445 |
| SL1344_RS16690_ | 1976 | 1867 | 1752 | 1491 | 1421 | 1699 |
| SL1344_RS16695_ | 2818 | 2749 | 2733 | 1960 | 1819 | 2190 |
| SL1344_RS16700_ | 2767 | 2679 | 2586 | 2480 | 2126 | 2299 |
| SL1344_RS16705_ | 15 | 10 | 14 | 11 | 15 | 8 |
| SL1344_RS16710_ | 330 | 365 | 410 | 245 | 209 | 226 |
| SL1344_RS16715_ | 980 | 974 | 967 | 845 | 763 | 810 |
| SL1344_RS16720_ | 230 | 257 | 225 | 188 | 151 | 191 |
| SL1344_RS16725_ | 171 | 134 | 152 | 98 | 108 | 110 |
| SL1344_RS16730_ | 458 | 513 | 432 | 335 | 331 | 330 |
| SL1344_RS16745_ | 279 | 317 | 320 | 387 | 268 | 455 |
| SL1344_RS16750_ | 436 | 459 | 589 | 704 | 499 | 727 |
| SL1344_RS16755_ | 237 | 277 | 247 | 338 | 248 | 452 |
| SL1344_RS16775_ | 1358 | 1497 | 1262 | 957 | 771 | 1061 |
| SL1344_RS16780_ | 5038 | 4964 | 4466 | 2999 | 2674 | 3345 |
| SL1344_RS16785_ | 2405 | 2767 | 3030 | 1848 | 1815 | 2020 |
| SL1344_RS16790_ | 1663 | 1960 | 2037 | 1211 | 1115 | 1352 |
| SL1344_RS16795_ | 710 | 661 | 640 | 494 | 460 | 559 |
| SL1344_RS16800_ | 900 | 811 | 718 | 727 | 735 | 793 |
| SL1344_RS16805_ | 75 | 72 | 55 | 65 | 37 | 65 |
| SL1344_RS16810_ | 48 | 50 | 43 | 35 | 28 | 36 |
| SL1344_RS16815_ | 57 | 65 | 58 | 49 | 32 | 39 |
| SL1344_RS16820_ | 35 | 37 | 30 | 29 | 18 | 23 |
| SL1344_RS16825_ | 21 | 39 | 23 | 32 | 36 | 29 |
| SL1344_RS16830_ | 3 | 6 | 6 | 6 | 9 | 11 |
| SL1344_RS16835_ | 24 | 22 | 19 | 11 | 14 | 13 |
| SL1344_RS16840_ | 75 | 107 | 92 | 87 | 53 | 101 |
| SL1344_RS16845_ | 12 | 17 | 19 | 9 | 23 | 20 |
| SL1344_RS16850_ | 724 | 763 | 745 | 624 | 609 | 688 |
| SL1344_RS16855_ | 304 | 313 | 313 | 325 | 323 | 348 |
| SL1344_RS16860_ | 1621 | 1699 | 1843 | 1713 | 1623 | 1924 |
| SL1344_RS16865_ | 837 | 861 | 860 | 763 | 732 | 860 |
| SL1344_RS16870_ | 537 | 480 | 508 | 546 | 445 | 547 |
| SL1344_RS16875_ | 1301 | 1416 | 1437 | 1394 | 1305 | 1525 |
| SL1344_RS16880_ | 97 | 116 | 97 | 86 | 87 | 100 |
| SL1344_RS16885_ | 214 | 187 | 210 | 136 | 115 | 142 |
| SL1344_RS16890_ | 83 | 91 | 97 | 121 | 78 | 129 |
| SL1344_RS16895_ | 95 | 109 | 84 | 67 | 109 | 119 |
| SL1344_RS16900_ | 1887 | 1987 | 1706 | 1283 | 1219 | 1522 |
| SL1344_RS16905_ | 1718 | 1852 | 1742 | 1261 | 1211 | 1668 |
| SL1344_RS16910_ | 415 | 412 | 472 | 609 | 460 | 543 |
| SL1344_RS16915_ | 703 | 743 | 652 | 695 | 575 | 694 |
| SL1344_RS16920_ | 258 | 305 | 330 | 280 | 254 | 289 |
| SL1344_RS16925_ | 21 | 8 | 21 | 16 | 20 | 20 |
| SL1344_RS16930_ | 14 | 7 | 6 | 10 | 10 | 8 |
| SL1344_RS16935_ | 102 | 97 | 104 | 121 | 94 | 110 |

| | | | | | | |
|---|---|---|---|---|---|---|
| SL1344_RS16940_ | 368 | 436 | 557 | 774 | 675 | 838 |
| SL1344_RS16945_ | 11298 | 11344 | 11517 | 11125 | 11038 | 12983 |
| SL1344_RS16950_ | 5691 | 5922 | 6467 | 6219 | 5760 | 7378 |
| SL1344_RS16955_ | 6431 | 5966 | 7089 | 9681 | 9610 | 9159 |
| SL1344_RS16960_ | 1076 | 1051 | 1147 | 1239 | 1325 | 1445 |
| SL1344_RS16965_ | 607 | 594 | 677 | 629 | 687 | 775 |
| SL1344_RS16970_ | 7991 | 7981 | 8381 | 8175 | 8293 | 9969 |
| SL1344_RS16980_ | 557 | 545 | 576 | 778 | 741 | 765 |
| SL1344_RS16985_ | 602 | 622 | 727 | 1028 | 729 | 751 |
| SL1344_RS16990_ | 178 | 162 | 199 | 192 | 164 | 175 |
| SL1344_RS16995_ | 240 | 226 | 239 | 385 | 334 | 385 |
| SL1344_RS17000_ | 691 | 609 | 893 | 974 | 695 | 819 |
| SL1344_RS17005_ | 3097 | 3122 | 3095 | 3377 | 3282 | 3955 |
| SL1344_RS17010_ | 1688 | 1606 | 1602 | 1291 | 1299 | 1570 |
| SL1344_RS17015_ | 301 | 283 | 246 | 303 | 269 | 352 |
| SL1344_RS17030_ | 2046 | 1823 | 1703 | 1955 | 1685 | 1861 |
| SL1344_RS17035_ | 508 | 484 | 485 | 1230 | 982 | 820 |
| SL1344_RS17040_ | 800 | 792 | 908 | 1206 | 1053 | 1240 |
| SL1344_RS17045_ | 329 | 313 | 291 | 492 | 486 | 410 |
| SL1344_RS17050_ | 577 | 636 | 589 | 748 | 626 | 614 |
| SL1344_RS17060_ | 256 | 242 | 307 | 348 | 309 | 351 |
| SL1344_RS17065_ | 1970 | 1815 | 2064 | 2464 | 2077 | 2636 |
| SL1344_RS17070_ | 2363 | 2351 | 2404 | 3538 | 2890 | 3785 |
| SL1344_RS17075_ | 1826 | 1893 | 2112 | 2070 | 1868 | 2142 |
| SL1344_RS17080_ | 22 | 15 | 25 | 35 | 26 | 23 |
| SL1344_RS17085_ | 1222 | 1177 | 1227 | 1320 | 1181 | 1395 |
| SL1344_RS17090_ | 116 | 91 | 91 | 135 | 105 | 128 |
| SL1344_RS17095_ | 1195 | 1067 | 1189 | 1086 | 979 | 1134 |
| SL1344_RS17100_ | 1074 | 989 | 1013 | 923 | 799 | 978 |
| SL1344_RS17105_ | 364 | 368 | 387 | 330 | 266 | 366 |
| SL1344_RS17110_ | 362 | 356 | 366 | 309 | 287 | 429 |
| SL1344_RS17115_ | 430 | 411 | 518 | 552 | 487 | 501 |
| SL1344_RS17120_ | 683 | 663 | 646 | 813 | 676 | 684 |
| SL1344_RS17125_ | 943 | 983 | 1023 | 867 | 873 | 1057 |
| SL1344_RS17130_ | 1082 | 1103 | 989 | 1027 | 1025 | 1021 |
| SL1344_RS17135_ | 837 | 832 | 887 | 916 | 832 | 884 |
| SL1344_RS17140_ | 1169 | 1153 | 1121 | 1075 | 1117 | 1221 |
| SL1344_RS17145_ | 1034 | 1088 | 1139 | 1023 | 959 | 1088 |
| SL1344_RS17150_ | 2536 | 2645 | 2946 | 2655 | 2421 | 2741 |
| SL1344_RS17155_ | 4609 | 5175 | 4674 | 3752 | 3404 | 4092 |
| SL1344_RS17160_ | 1339 | 1434 | 1409 | 1272 | 1151 | 1324 |
| SL1344_RS17165_ | 2078 | 2034 | 2192 | 1993 | 1733 | 1976 |
| SL1344_RS17170_ | 433 | 401 | 468 | 417 | 383 | 438 |
| SL1344_RS17175_ | 83 | 93 | 90 | 48 | 44 | 56 |
| SL1344_RS17180_ | 898 | 983 | 1008 | 1328 | 903 | 1408 |
| SL1344_RS17185_ | 1032 | 1262 | 1166 | 1424 | 1194 | 1810 |
| SL1344_RS17190_ | 1661 | 1536 | 1364 | 1529 | 1354 | 1733 |
| SL1344_RS17195_ | 113 | 110 | 110 | 169 | 150 | 166 |
| SL1344_RS17200_ | 2 | 7 | 5 | 9 | 7 | 9 |
| SL1344_RS17205_ | 545 | 558 | 557 | 491 | 448 | 516 |
| SL1344_RS17210_ | 282 | 222 | 251 | 246 | 186 | 229 |
| SL1344_RS17215_ | 18 | 33 | 39 | 32 | 20 | 30 |
| SL1344_RS17220_ | 71 | 68 | 112 | 89 | 96 | 92 |
| SL1344_RS17225_ | 360 | 374 | 378 | 403 | 338 | 370 |
| SL1344_RS17230_ | 497 | 506 | 445 | 356 | 389 | 373 |
| SL1344_RS17235_ | 270 | 309 | 330 | 234 | 220 | 258 |
| SL1344_RS17240_ | 260 | 293 | 307 | 221 | 187 | 252 |
| SL1344_RS17245_ | 531 | 574 | 567 | 425 | 470 | 522 |
| SL1344_RS17250_ | 512 | 569 | 509 | 376 | 356 | 467 |
| SL1344_RS17255_ | 1038 | 1158 | 1062 | 1117 | 1014 | 1116 |
| SL1344_RS17260_ | 2999 | 3024 | 2721 | 3450 | 3213 | 3584 |
| SL1344_RS17270_ | 217 | 215 | 233 | 156 | 150 | 184 |

| | | | | | | |
|---|---|---|---|---|---|---|
| SL1344_RS17275_ | 6779 | 6324 | 7396 | 10873 | 9742 | 11300 |
| SL1344_RS17280_ | 2845 | 2881 | 3480 | 4854 | 4714 | 5373 |
| SL1344_RS17285_ | 506 | 479 | 517 | 473 | 470 | 532 |
| SL1344_RS17290_ | 2350 | 2285 | 2188 | 2036 | 1825 | 2219 |
| SL1344_RS17295_ | 2133 | 2212 | 2487 | 1603 | 1416 | 1867 |
| SL1344_RS17300_ | 636 | 755 | 703 | 682 | 609 | 658 |
| SL1344_RS17305_ | 30 | 37 | 33 | 32 | 24 | 48 |
| SL1344_RS17310_ | 67 | 50 | 71 | 56 | 58 | 63 |
| SL1344_RS17315_ | 100 | 71 | 88 | 136 | 113 | 124 |
| SL1344_RS17320_ | 8 | 4 | 12 | 6 | 10 | 9 |
| SL1344_RS17325_ | 60 | 50 | 61 | 38 | 56 | 50 |
| SL1344_RS17330_ | 50 | 43 | 44 | 45 | 51 | 48 |
| SL1344_RS17335_ | 32 | 19 | 32 | 27 | 19 | 28 |
| SL1344_RS17340_ | 323 | 290 | 251 | 253 | 245 | 305 |
| SL1344_RS17345_ | 332 | 283 | 276 | 327 | 320 | 371 |
| SL1344_RS17350_ | 6844 | 6102 | 7874 | 7165 | 6649 | 8220 |
| SL1344_RS17355_ | 821 | 846 | 953 | 894 | 905 | 931 |
| SL1344_RS17360_ | 42 | 27 | 19 | 30 | 27 | 15 |
| SL1344_RS17365_ | 111 | 96 | 97 | 136 | 137 | 139 |
| SL1344_RS17370_ | 165 | 213 | 176 | 102 | 77 | 95 |
| SL1344_RS17375_ | 127 | 133 | 123 | 147 | 118 | 126 |
| SL1344_RS17380_ | 70 | 66 | 73 | 71 | 69 | 74 |
| SL1344_RS17385_ | 8 | 1 | 5 | 5 | 6 | 12 |
| SL1344_RS17390_ | 216 | 239 | 207 | 214 | 180 | 230 |
| SL1344_RS17395_ | 887 | 915 | 911 | 681 | 632 | 764 |
| SL1344_RS17400_ | 1783 | 1733 | 1807 | 1475 | 1464 | 1698 |
| SL1344_RS17405_ | 1500 | 1588 | 1565 | 1516 | 1196 | 1688 |
| SL1344_RS17410_ | 724 | 696 | 638 | 745 | 709 | 779 |
| SL1344_RS17415_ | 531 | 524 | 622 | 565 | 463 | 591 |
| SL1344_RS17420_ | 907 | 909 | 1056 | 1130 | 935 | 1099 |
| SL1344_RS17425_ | 2583 | 2697 | 2763 | 3556 | 3326 | 4081 |
| SL1344_RS17430_ | 2 | 2 | 2 | 3 | 2 | 3 |
| SL1344_RS17440_ | 1332 | 1300 | 1264 | 1198 | 962 | 1171 |
| SL1344_RS17445_ | 359 | 346 | 382 | 346 | 307 | 339 |
| SL1344_RS17450_ | 177 | 166 | 229 | 184 | 154 | 188 |
| SL1344_RS17455_ | 1537 | 1333 | 1330 | 1640 | 1591 | 1719 |
| SL1344_RS17460_ | 3669 | 3575 | 4161 | 3832 | 3536 | 3945 |
| SL1344_RS17465_ | 15 | 9 | 10 | 19 | 22 | 32 |
| SL1344_RS17470_ | 194 | 227 | 226 | 250 | 257 | 276 |
| SL1344_RS17475_ | 332 | 355 | 420 | 392 | 402 | 426 |
| SL1344_RS17480_ | 758 | 781 | 970 | 1564 | 1267 | 1389 |
| SL1344_RS17490_ | 97 | 96 | 136 | 127 | 129 | 125 |
| SL1344_RS17495_ | 183 | 181 | 164 | 242 | 243 | 214 |
| SL1344_RS17500_ | 253 | 235 | 272 | 234 | 263 | 249 |
| SL1344_RS17505_ | 20 | 15 | 17 | 24 | 20 | 25 |
| SL1344_RS17510_ | 10 | 18 | 22 | 11 | 19 | 17 |
| SL1344_RS17515_ | 142 | 158 | 143 | 154 | 157 | 150 |
| SL1344_RS17520_ | 105 | 102 | 125 | 98 | 91 | 88 |
| SL1344_RS17530_ | 118 | 106 | 181 | 147 | 83 | 110 |
| SL1344_RS17540_ | 1883 | 1578 | 2544 | 882 | 1234 | 2491 |
| SL1344_RS17545_ | 21 | 32 | 37 | 48 | 31 | 52 |
| SL1344_RS17550_ | 371 | 298 | 683 | 168 | 281 | 614 |
| SL1344_RS17555_ | 512 | 443 | 526 | 521 | 439 | 569 |
| SL1344_RS17560_ | 373 | 379 | 350 | 347 | 253 | 286 |
| SL1344_RS17565_ | 617 | 653 | 692 | 596 | 519 | 635 |
| SL1344_RS17570_ | 306 | 267 | 309 | 332 | 220 | 263 |
| SL1344_RS17575_ | 2841 | 2590 | 2510 | 2016 | 2005 | 2266 |
| SL1344_RS17580_ | 4315 | 4072 | 3479 | 2974 | 3270 | 3501 |
| SL1344_RS17585_ | 814 | 850 | 866 | 606 | 563 | 726 |
| SL1344_RS17590_ | 1363 | 1476 | 1526 | 1243 | 1193 | 1378 |
| SL1344_RS17595_ | 1292 | 1206 | 1344 | 1312 | 1280 | 1420 |
| SL1344_RS17600_ | 206 | 200 | 236 | 200 | 169 | 201 |

| | | | | | | |
|---|---|---|---|---|---|---|
| SL1344_RS17605_ | 398 | 426 | 467 | 359 | 309 | 409 |
| SL1344_RS17610_ | 1924 | 1984 | 1966 | 1534 | 1365 | 1733 |
| SL1344_RS17615_ | 309 | 286 | 221 | 301 | 215 | 321 |
| SL1344_RS17620_ | 542 | 497 | 453 | 455 | 453 | 483 |
| SL1344_RS17625_ | 235 | 251 | 206 | 233 | 198 | 261 |
| SL1344_RS17630_ | 8275 | 7574 | 10272 | 9854 | 9439 | 10587 |
| SL1344_RS17635_ | 20631 | 19659 | 24494 | 22315 | 22866 | 28548 |
| SL1344_RS17640_ | 13927 | 14217 | 15923 | 16175 | 14113 | 18567 |
| SL1344_RS17645_ | 3222 | 2994 | 3801 | 3337 | 2293 | 3609 |
| SL1344_RS17650_ | 5138 | 5284 | 6148 | 6194 | 5196 | 6842 |
| SL1344_RS17655_ | 7178 | 6563 | 7733 | 7038 | 7641 | 8410 |
| SL1344_RS17660_ | 22088 | 21493 | 26599 | 22673 | 23130 | 28495 |
| SL1344_RS17665_ | 6523 | 6587 | 7671 | 6019 | 6384 | 8444 |
| SL1344_RS17670_ | 2838 | 3035 | 3769 | 2974 | 3118 | 4065 |
| SL1344_RS17675_ | 5981 | 5644 | 7291 | 6218 | 6530 | 8402 |
| SL1344_RS17680_ | 5504 | 5147 | 6683 | 5868 | 6164 | 7418 |
| SL1344_RS17685_ | 8565 | 8058 | 10106 | 9419 | 10473 | 12247 |
| SL1344_RS17690_ | 7071 | 6785 | 8668 | 8347 | 9080 | 10306 |
| SL1344_RS17700_ | 10503 | 10550 | 13926 | 13674 | 13918 | 16526 |
| SL1344_RS17705_ | 7493 | 7696 | 9331 | 9774 | 10057 | 11624 |
| SL1344_RS17710_ | 8132 | 7988 | 10549 | 10759 | 11633 | 13230 |
| SL1344_RS17715_ | 13474 | 11911 | 14901 | 15561 | 17432 | 18438 |
| SL1344_RS17720_ | 6947 | 5954 | 7028 | 7508 | 9415 | 9495 |
| SL1344_RS17725_ | 14233 | 13598 | 14934 | 16728 | 19883 | 20474 |
| SL1344_RS17730_ | 12455 | 11724 | 14901 | 13859 | 16461 | 17820 |
| SL1344_RS17735_ | 6945 | 6164 | 8094 | 7662 | 9410 | 10336 |
| SL1344_RS17740_ | 5924 | 5549 | 7113 | 6709 | 7514 | 8414 |
| SL1344_RS17745_ | 8631 | 8254 | 11403 | 9995 | 11371 | 13311 |
| SL1344_RS17750_ | 8892 | 9289 | 10608 | 10616 | 11384 | 13263 |
| SL1344_RS17755_ | 8272 | 7607 | 9910 | 9497 | 11404 | 12812 |
| SL1344_RS17760_ | 8356 | 7277 | 9190 | 9910 | 12212 | 13156 |
| SL1344_RS17765_ | 3018 | 2898 | 4022 | 4303 | 4629 | 5050 |
| SL1344_RS17770_ | 10 | 8 | 8 | 7 | 6 | 8 |
| SL1344_RS17775_ | 3065 | 3069 | 2689 | 1592 | 1561 | 1873 |
| SL1344_RS17780_ | 79 | 70 | 52 | 70 | 96 | 62 |
| SL1344_RS17785_ | 23755 | 22422 | 27822 | 24788 | 23516 | 28665 |
| SL1344_RS17790_ | 29630 | 28775 | 37341 | 34220 | 35459 | 41853 |
| SL1344_RS17795_ | 5457 | 5505 | 7515 | 7479 | 7960 | 9090 |
| SL1344_RS17800_ | 3523 | 3545 | 5125 | 5271 | 5229 | 6270 |
| SL1344_RS17805_ | 330 | 317 | 330 | 434 | 413 | 358 |
| SL1344_RS17810_ | 332 | 322 | 370 | 329 | 261 | 305 |
| SL1344_RS17815_ | 450 | 534 | 496 | 479 | 367 | 450 |
| SL1344_RS17820_ | 617 | 559 | 609 | 710 | 642 | 665 |
| SL1344_RS17825_ | 4804 | 4429 | 4768 | 4598 | 4556 | 4974 |
| SL1344_RS17830_ | 348 | 282 | 243 | 341 | 363 | 392 |
| SL1344_RS17835_ | 2856 | 2656 | 2817 | 2640 | 2595 | 2811 |
| SL1344_RS17840_ | 148 | 145 | 144 | 174 | 134 | 149 |
| SL1344_RS17845_ | 471 | 430 | 409 | 466 | 473 | 511 |
| SL1344_RS17850_ | 66 | 76 | 78 | 105 | 99 | 114 |
| SL1344_RS17855_ | 758 | 889 | 945 | 875 | 913 | 924 |
| SL1344_RS17860_ | 18 | 19 | 15 | 21 | 25 | 17 |
| SL1344_RS17865_ | 637 | 637 | 556 | 406 | 371 | 513 |
| SL1344_RS17870_ | 601 | 584 | 593 | 554 | 538 | 591 |
| SL1344_RS17875_ | 242 | 233 | 239 | 163 | 161 | 184 |
| SL1344_RS17880_ | 679 | 673 | 669 | 527 | 542 | 612 |
| SL1344_RS17885_ | 765 | 786 | 723 | 860 | 729 | 781 |
| SL1344_RS17895_ | 1559 | 1570 | 1808 | 1726 | 1518 | 1744 |
| SL1344_RS17900_ | 304 | 293 | 318 | 308 | 269 | 317 |
| SL1344_RS17905_ | 208 | 196 | 226 | 197 | 170 | 223 |
| SL1344_RS17910_ | 178 | 154 | 156 | 107 | 86 | 127 |
| SL1344_RS17915_ | 101 | 97 | 99 | 78 | 65 | 60 |
| SL1344_RS17920_ | 733 | 654 | 690 | 890 | 819 | 890 |

| | | | | | | |
|---|---|---|---|---|---|---|
| SL1344_RS17925_ | 62 | 53 | 47 | 60 | 67 | 75 |
| SL1344_RS17930_ | 1037 | 1044 | 1200 | 1468 | 1381 | 1593 |
| SL1344_RS17940_ | 241 | 218 | 226 | 223 | 265 | 244 |
| SL1344_RS17965_ | 2368 | 2415 | 2433 | 1957 | 1864 | 2453 |
| SL1344_RS17970_ | 1087 | 1111 | 1113 | 922 | 847 | 1009 |
| SL1344_RS17975_ | 861 | 790 | 878 | 608 | 647 | 751 |
| SL1344_RS17985_ | 3239 | 3387 | 3644 | 2936 | 3049 | 3532 |
| SL1344_RS17990_ | 2086 | 2004 | 2047 | 1885 | 1864 | 2404 |
| SL1344_RS17995_ | 1256 | 1135 | 1198 | 1423 | 1221 | 1456 |
| SL1344_RS18000_ | 127 | 141 | 113 | 103 | 85 | 121 |
| SL1344_RS18005_ | 30 | 35 | 28 | 20 | 22 | 32 |
| SL1344_RS18010_ | 30 | 20 | 16 | 35 | 26 | 34 |
| SL1344_RS18015_ | 25 | 30 | 26 | 29 | 30 | 30 |
| SL1344_RS18020_ | 28 | 35 | 32 | 33 | 32 | 44 |
| SL1344_RS18025_ | 698 | 769 | 820 | 801 | 769 | 872 |
| SL1344_RS18030_ | 298 | 309 | 289 | 337 | 278 | 345 |
| SL1344_RS18035_ | 1804 | 1809 | 1851 | 2130 | 1820 | 2143 |
| SL1344_RS18040_ | 210 | 260 | 221 | 261 | 278 | 339 |
| SL1344_RS18045_ | 180 | 204 | 201 | 217 | 189 | 247 |
| SL1344_RS18050_ | 525 | 542 | 557 | 516 | 530 | 625 |
| SL1344_RS18055_ | 306 | 310 | 304 | 284 | 280 | 378 |
| SL1344_RS18060_ | 11589 | 11242 | 13705 | 12828 | 11445 | 14176 |
| SL1344_RS18075_ | 229 | 169 | 190 | 265 | 253 | 254 |
| SL1344_RS18080_ | 1037 | 1067 | 1173 | 1189 | 1169 | 1262 |
| SL1344_RS18090_ | 360 | 351 | 385 | 295 | 280 | 299 |
| SL1344_RS18095_ | 178 | 149 | 153 | 129 | 127 | 152 |
| SL1344_RS18100_ | 203 | 194 | 189 | 191 | 197 | 196 |
| SL1344_RS18105_ | 221 | 229 | 205 | 167 | 143 | 196 |
| SL1344_RS18110_ | 289 | 234 | 162 | 223 | 240 | 212 |
| SL1344_RS18115_ | 1203 | 1112 | 991 | 979 | 1088 | 1147 |
| SL1344_RS18120_ | 197 | 184 | 215 | 172 | 169 | 215 |
| SL1344_RS18125_ | 738 | 719 | 640 | 470 | 356 | 482 |
| SL1344_RS18130_ | 916 | 986 | 850 | 572 | 524 | 598 |
| SL1344_RS18140_ | 514 | 492 | 497 | 470 | 498 | 507 |
| SL1344_RS18145_ | 853 | 938 | 779 | 800 | 844 | 995 |
| SL1344_RS18150_ | 210 | 229 | 225 | 279 | 244 | 281 |
| SL1344_RS18155_ | 249 | 254 | 231 | 252 | 221 | 296 |
| SL1344_RS18160_ | 130 | 119 | 138 | 133 | 163 | 167 |
| SL1344_RS18165_ | 267 | 248 | 248 | 242 | 235 | 246 |
| SL1344_RS18170_ | 771 | 726 | 787 | 674 | 507 | 729 |
| SL1344_RS18175_ | 739 | 702 | 667 | 899 | 805 | 933 |
| SL1344_RS18180_ | 313 | 346 | 317 | 379 | 375 | 366 |
| SL1344_RS18185_ | 33153 | 33985 | 37538 | 31812 | 33497 | 37951 |
| SL1344_RS18190_ | 75 | 89 | 74 | 74 | 65 | 70 |
| SL1344_RS18195_ | 378 | 380 | 345 | 313 | 274 | 297 |
| SL1344_RS18205_ | 37 | 28 | 35 | 30 | 28 | 33 |
| SL1344_RS18210_ | 82 | 72 | 60 | 41 | 70 | 58 |
| SL1344_RS18215_ | 75 | 75 | 91 | 67 | 62 | 68 |
| SL1344_RS18220_ | 1321 | 1484 | 1440 | 1254 | 1333 | 1438 |
| SL1344_RS18225_ | 2920 | 3100 | 3021 | 1958 | 1846 | 2110 |
| SL1344_RS18230_ | 1330 | 1363 | 1191 | 773 | 633 | 859 |
| SL1344_RS18235_ | 1339 | 1450 | 1278 | 832 | 786 | 996 |
| SL1344_RS18240_ | 1726 | 1763 | 1806 | 1158 | 1121 | 1339 |
| SL1344_RS18245_ | 2028 | 1941 | 1982 | 1360 | 1256 | 1582 |
| SL1344_RS18250_ | 1678 | 1742 | 1727 | 1437 | 1182 | 1522 |
| SL1344_RS18255_ | 141 | 152 | 175 | 145 | 131 | 175 |
| SL1344_RS18270_ | 151 | 150 | 154 | 136 | 123 | 143 |
| SL1344_RS18275_ | 366 | 315 | 354 | 285 | 250 | 296 |
| SL1344_RS18280_ | 224 | 189 | 197 | 234 | 223 | 230 |
| SL1344_RS18285_ | 52 | 42 | 48 | 30 | 44 | 21 |
| SL1344_RS18290_ | 25 | 23 | 45 | 22 | 27 | 16 |
| SL1344_RS18295_ | 19 | 28 | 19 | 14 | 15 | 22 |

| | | | | | | |
|---|---|---|---|---|---|---|
| SL1344_RS18300_ | 113 | 115 | 97 | 72 | 79 | 89 |
| SL1344_RS18305_ | 354 | 337 | 312 | 286 | 253 | 327 |
| SL1344_RS18310_ | 696 | 791 | 721 | 496 | 462 | 569 |
| SL1344_RS18315_ | 361 | 380 | 324 | 232 | 172 | 237 |
| SL1344_RS18325_ | 173 | 159 | 219 | 138 | 158 | 161 |
| SL1344_RS18330_ | 160 | 169 | 198 | 170 | 179 | 155 |
| SL1344_RS18335_ | 1826 | 1905 | 1784 | 1384 | 1157 | 1380 |
| SL1344_RS18340_ | 125 | 146 | 115 | 101 | 103 | 100 |
| SL1344_RS18345_ | 139 | 149 | 121 | 87 | 99 | 133 |
| SL1344_RS18355_ | 65 | 59 | 59 | 64 | 48 | 68 |
| SL1344_RS18360_ | 72 | 86 | 76 | 78 | 97 | 91 |
| SL1344_RS18365_ | 60 | 41 | 66 | 48 | 52 | 69 |
| SL1344_RS18370_ | 65 | 75 | 69 | 89 | 84 | 91 |
| SL1344_RS18375_ | 276 | 285 | 237 | 259 | 285 | 322 |
| SL1344_RS18380_ | 288 | 288 | 275 | 301 | 276 | 341 |
| SL1344_RS18385_ | 216 | 192 | 186 | 237 | 261 | 259 |
| SL1344_RS18390_ | 6053 | 5603 | 5883 | 6412 | 5893 | 7045 |
| SL1344_RS18395_ | 1821 | 1894 | 1873 | 1432 | 1416 | 1496 |
| SL1344_RS18400_ | 922 | 887 | 840 | 717 | 751 | 807 |
| SL1344_RS18405_ | 1383 | 1378 | 1188 | 1053 | 1032 | 1227 |
| SL1344_RS18415_ | 237 | 272 | 278 | 236 | 283 | 278 |
| SL1344_RS18420_ | 136 | 119 | 109 | 181 | 148 | 158 |
| SL1344_RS18425_ | 489 | 503 | 571 | 516 | 425 | 462 |
| SL1344_RS18440_ | 125 | 140 | 123 | 126 | 155 | 131 |
| SL1344_RS18445_ | 78 | 78 | 104 | 135 | 136 | 103 |
| SL1344_RS18450_ | 1826 | 1868 | 2067 | 2016 | 1942 | 2001 |
| SL1344_RS18455_ | 362 | 340 | 304 | 257 | 227 | 279 |
| SL1344_RS18460_ | 171 | 176 | 179 | 140 | 134 | 128 |
| SL1344_RS18465_ | 269 | 283 | 285 | 278 | 327 | 273 |
| SL1344_RS18470_ | 197 | 194 | 199 | 204 | 223 | 194 |
| SL1344_RS18475_ | 352 | 369 | 370 | 273 | 273 | 299 |
| SL1344_RS18480_ | 1123 | 1183 | 1087 | 979 | 871 | 1007 |
| SL1344_RS18485_ | 718 | 748 | 615 | 541 | 487 | 530 |
| SL1344_RS18490_ | 397 | 369 | 361 | 345 | 315 | 382 |
| SL1344_RS18495_ | 1756 | 1601 | 1959 | 2123 | 2040 | 2265 |
| SL1344_RS18500_ | 759 | 739 | 668 | 454 | 413 | 502 |
| SL1344_RS18505_ | 14140 | 14987 | 14128 | 12127 | 12914 | 14368 |
| SL1344_RS18510_ | 2155 | 2146 | 2419 | 1352 | 1375 | 1633 |
| SL1344_RS18515_ | 327 | 318 | 345 | 252 | 245 | 253 |
| SL1344_RS18520_ | 1569 | 1608 | 1576 | 1111 | 938 | 1154 |
| SL1344_RS18525_ | 502 | 500 | 558 | 572 | 553 | 639 |
| SL1344_RS18530_ | 358 | 334 | 390 | 373 | 393 | 405 |
| SL1344_RS18535_ | 1594 | 1591 | 1635 | 1361 | 1403 | 1555 |
| SL1344_RS18540_ | 386 | 313 | 223 | 278 | 145 | 265 |
| SL1344_RS18545_ | 793 | 574 | 465 | 494 | 304 | 437 |
| SL1344_RS18550_ | 703 | 471 | 355 | 366 | 205 | 309 |
| SL1344_RS18555_ | 1638 | 1263 | 859 | 966 | 510 | 1019 |
| SL1344_RS18560_ | 235 | 248 | 219 | 265 | 245 | 273 |
| SL1344_RS18565_ | 382 | 444 | 395 | 361 | 299 | 351 |
| SL1344_RS18570_ | 3757 | 4097 | 4008 | 3948 | 4162 | 4607 |
| SL1344_RS18575_ | 5 | 3 | 6 | 9 | 7 | 8 |
| SL1344_RS18580_ | 26 | 21 | 29 | 36 | 29 | 27 |
| SL1344_RS18585_ | 128 | 136 | 101 | 128 | 106 | 156 |
| SL1344_RS18590_ | 310 | 281 | 301 | 258 | 250 | 244 |
| SL1344_RS18595_ | 132 | 145 | 157 | 154 | 138 | 140 |
| SL1344_RS18600_ | 946 | 955 | 933 | 1396 | 1037 | 1439 |
| SL1344_RS18610_ | 466 | 527 | 504 | 358 | 293 | 377 |
| SL1344_RS18615_ | 768 | 858 | 800 | 739 | 715 | 904 |
| SL1344_RS18620_ | 9004 | 8010 | 10174 | 7007 | 5794 | 7402 |
| SL1344_RS18625_ | 654 | 652 | 719 | 641 | 653 | 742 |
| SL1344_RS18630_ | 1854 | 1873 | 1749 | 1240 | 1152 | 1417 |
| SL1344_RS18635_ | 649 | 763 | 744 | 431 | 411 | 465 |

| | | | | | | |
|---|---|---|---|---|---|---|
| SL1344_RS18640_ | 906 | 1024 | 1000 | 583 | 502 | 615 |
| SL1344_RS18655_ | 137 | 161 | 152 | 88 | 88 | 92 |
| SL1344_RS18660_ | 565 | 580 | 609 | 390 | 362 | 377 |
| SL1344_RS18665_ | 190 | 189 | 203 | 116 | 104 | 119 |
| SL1344_RS18670_ | 737 | 857 | 875 | 528 | 440 | 515 |
| SL1344_RS18675_ | 2613 | 2557 | 3812 | 3056 | 2569 | 2979 |
| SL1344_RS18680_ | 9 | 3 | 8 | 6 | 9 | 6 |
| SL1344_RS18685_ | 73 | 91 | 109 | 119 | 122 | 130 |
| SL1344_RS18690_ | 351 | 362 | 418 | 321 | 274 | 355 |
| SL1344_RS18695_ | 367 | 350 | 375 | 337 | 314 | 343 |
| SL1344_RS18700_ | 422 | 472 | 451 | 392 | 364 | 374 |
| SL1344_RS18705_ | 270 | 242 | 254 | 204 | 192 | 199 |
| SL1344_RS18710_ | 1727 | 1759 | 1728 | 1327 | 1228 | 1287 |
| SL1344_RS18715_ | 44 | 31 | 41 | 42 | 46 | 46 |
| SL1344_RS18720_ | 60 | 81 | 70 | 67 | 61 | 76 |
| SL1344_RS18725_ | 88 | 92 | 101 | 113 | 116 | 103 |
| SL1344_RS18735_ | 272 | 185 | 314 | 334 | 163 | 234 |
| SL1344_RS18740_ | 566 | 615 | 531 | 515 | 456 | 483 |
| SL1344_RS18745_ | 38 | 45 | 67 | 56 | 48 | 56 |
| SL1344_RS18750_ | 46 | 33 | 53 | 48 | 34 | 44 |
| SL1344_RS18755_ | 61 | 74 | 62 | 84 | 68 | 69 |
| SL1344_RS18760_ | 5 | 5 | 4 | 8 | 12 | 12 |
| SL1344_RS18765_ | 7 | 16 | 20 | 19 | 18 | 13 |
| SL1344_RS18770_ | 121 | 134 | 132 | 104 | 73 | 90 |
| SL1344_RS18775_ | 455 | 420 | 389 | 500 | 480 | 517 |
| SL1344_RS18780_ | 417 | 424 | 383 | 392 | 462 | 455 |
| SL1344_RS18785_ | 815 | 791 | 836 | 859 | 774 | 876 |
| SL1344_RS18790_ | 358 | 397 | 418 | 333 | 277 | 393 |
| SL1344_RS18795_ | 795 | 827 | 694 | 598 | 590 | 621 |
| SL1344_RS18800_ | 1815 | 1757 | 1817 | 1510 | 1444 | 1740 |
| SL1344_RS18805_ | 1154 | 1237 | 1220 | 811 | 700 | 781 |
| SL1344_RS18815_ | 626 | 622 | 470 | 298 | 305 | 417 |
| SL1344_RS18820_ | 231 | 253 | 260 | 283 | 279 | 265 |
| SL1344_RS18825_ | 249 | 302 | 254 | 270 | 341 | 361 |
| SL1344_RS18830_ | 62 | 55 | 52 | 64 | 49 | 61 |
| SL1344_RS18835_ | 2627 | 2578 | 2931 | 2404 | 2365 | 2827 |
| SL1344_RS18840_ | 626 | 627 | 701 | 753 | 597 | 795 |
| SL1344_RS18845_ | 105 | 85 | 80 | 88 | 79 | 70 |
| SL1344_RS18850_ | 69 | 80 | 95 | 109 | 91 | 111 |
| SL1344_RS18855_ | 58 | 39 | 53 | 34 | 30 | 39 |
| SL1344_RS18860_ | 229 | 254 | 216 | 218 | 198 | 251 |
| SL1344_RS18865_ | 232 | 218 | 216 | 203 | 194 | 260 |
| SL1344_RS18870_ | 255 | 281 | 239 | 180 | 170 | 191 |
| SL1344_RS18875_ | 1727 | 1711 | 1695 | 2188 | 2054 | 2084 |
| SL1344_RS18880_ | 28 | 41 | 47 | 28 | 25 | 26 |
| SL1344_RS18885_ | 83 | 74 | 100 | 77 | 63 | 78 |
| SL1344_RS18890_ | 192 | 154 | 181 | 223 | 241 | 261 |
| SL1344_RS18895_ | 566 | 610 | 539 | 531 | 484 | 544 |
| SL1344_RS18900_ | 432 | 375 | 398 | 431 | 374 | 464 |
| SL1344_RS18905_ | 33 | 33 | 43 | 49 | 36 | 36 |
| SL1344_RS18910_ | 10 | 13 | 7 | 20 | 14 | 8 |
| SL1344_RS18915_ | 18 | 25 | 23 | 21 | 21 | 22 |
| SL1344_RS18920_ | 4 | 2 | 4 | 7 | 4 | 3 |
| SL1344_RS18925_ | 4 | 13 | 11 | 6 | 12 | 14 |
| SL1344_RS18930_ | 11 | 13 | 12 | 21 | 15 | 21 |
| SL1344_RS18935_ | 19 | 23 | 26 | 35 | 35 | 28 |
| SL1344_RS18945_ | 12 | 10 | 12 | 17 | 17 | 26 |
| SL1344_RS18950_ | 17 | 20 | 17 | 23 | 22 | 25 |
| SL1344_RS18955_ | 258 | 216 | 234 | 217 | 202 | 189 |
| SL1344_RS18960_ | 153 | 174 | 172 | 194 | 178 | 212 |
| SL1344_RS18965_ | 757 | 785 | 740 | 457 | 351 | 508 |
| SL1344_RS18970_ | 212 | 228 | 195 | 143 | 134 | 173 |

| | | | | | | |
|---|---|---|---|---|---|---|
| SL1344_RS18975_ | 1281 | 1308 | 1357 | 1156 | 1142 | 1179 |
| SL1344_RS18980_ | 1345 | 1393 | 1438 | 1352 | 1546 | 1348 |
| SL1344_RS18985_ | 1055 | 1119 | 1129 | 869 | 849 | 909 |
| SL1344_RS18990_ | 2979 | 2661 | 2817 | 1879 | 1740 | 2252 |
| SL1344_RS18995_ | 986 | 999 | 1016 | 855 | 783 | 962 |
| SL1344_RS19005_ | 114 | 123 | 107 | 21 | 16 | 34 |
| SL1344_RS19010_ | 593 | 533 | 473 | 563 | 505 | 602 |
| SL1344_RS19015_ | 3 | 1 | 3 | 3 | 8 | 4 |
| SL1344_RS19020_ | 182 | 177 | 174 | 169 | 173 | 176 |
| SL1344_RS19025_ | 117 | 120 | 146 | 136 | 174 | 166 |
| SL1344_RS19030_ | 34 | 44 | 40 | 44 | 36 | 57 |
| SL1344_RS19040_ | 91 | 78 | 86 | 94 | 135 | 107 |
| SL1344_RS19045_ | 269 | 240 | 293 | 308 | 248 | 296 |
| SL1344_RS19050_ | 589 | 585 | 596 | 360 | 284 | 391 |
| SL1344_RS19055_ | 267 | 253 | 260 | 139 | 142 | 173 |
| SL1344_RS19065_ | 3480 | 3250 | 3527 | 3202 | 2781 | 3207 |
| SL1344_RS19070_ | 4651 | 4560 | 4969 | 4447 | 4700 | 4528 |
| SL1344_RS19075_ | 1207 | 1166 | 1174 | 1223 | 1156 | 1223 |
| SL1344_RS19080_ | 1045 | 917 | 953 | 1000 | 951 | 1045 |
| SL1344_RS19085_ | 1188 | 1227 | 1207 | 1213 | 1107 | 1415 |
| SL1344_RS19090_ | 1037 | 974 | 1034 | 1231 | 1188 | 1262 |
| SL1344_RS19095_ | 465 | 446 | 494 | 455 | 389 | 408 |
| SL1344_RS19100_ | 106 | 110 | 120 | 122 | 106 | 95 |
| SL1344_RS19105_ | 6614 | 6494 | 6479 | 4269 | 4433 | 4957 |
| SL1344_RS19110_ | 3533 | 3497 | 3814 | 2519 | 2344 | 2789 |
| SL1344_RS19115_ | 1589 | 1667 | 1844 | 1504 | 1505 | 1779 |
| SL1344_RS19120_ | 1404 | 1396 | 1440 | 1329 | 1233 | 1408 |
| SL1344_RS19125_ | 812 | 833 | 806 | 893 | 706 | 809 |
| SL1344_RS19130_ | 1039 | 900 | 1056 | 1079 | 1044 | 1044 |
| SL1344_RS19135_ | 679 | 660 | 674 | 785 | 758 | 764 |
| SL1344_RS19140_ | 464 | 449 | 560 | 401 | 397 | 446 |
| SL1344_RS19145_ | 291 | 282 | 276 | 306 | 284 | 322 |
| SL1344_RS19150_ | 609 | 569 | 669 | 651 | 631 | 687 |
| SL1344_RS19155_ | 724 | 695 | 787 | 762 | 799 | 829 |
| SL1344_RS19160_ | 728 | 731 | 793 | 772 | 827 | 869 |
| SL1344_RS19165_ | 244 | 212 | 275 | 256 | 248 | 256 |
| SL1344_RS19170_ | 806 | 709 | 755 | 720 | 699 | 869 |
| SL1344_RS19175_ | 977 | 937 | 1025 | 975 | 980 | 1110 |
| SL1344_RS19180_ | 717 | 683 | 742 | 876 | 803 | 875 |
| SL1344_RS19185_ | 891 | 928 | 1028 | 1080 | 952 | 1036 |
| SL1344_RS19190_ | 802 | 726 | 719 | 761 | 719 | 762 |
| SL1344_RS19195_ | 249 | 230 | 225 | 244 | 256 | 269 |
| SL1344_RS19200_ | 6999 | 5723 | 7633 | 11324 | 11859 | 12227 |
| SL1344_RS19205_ | 4696 | 3972 | 4468 | 7281 | 8183 | 7930 |
| SL1344_RS19210_ | 180 | 191 | 204 | 196 | 177 | 192 |
| SL1344_RS19215_ | 770 | 714 | 841 | 930 | 811 | 899 |
| SL1344_RS19220_ | 814 | 748 | 808 | 733 | 693 | 866 |
| SL1344_RS19225_ | 542 | 566 | 587 | 735 | 680 | 687 |
| SL1344_RS19230_ | 294 | 339 | 281 | 213 | 181 | 221 |
| SL1344_RS19235_ | 176 | 159 | 196 | 179 | 139 | 187 |
| SL1344_RS19240_ | 1279 | 1175 | 1236 | 1131 | 1208 | 1356 |
| SL1344_RS19245_ | 140 | 154 | 143 | 143 | 129 | 151 |
| SL1344_RS19250_ | 53 | 40 | 48 | 49 | 44 | 31 |
| SL1344_RS19255_ | 179 | 220 | 276 | 249 | 239 | 282 |
| SL1344_RS19260_ | 107 | 123 | 104 | 116 | 113 | 119 |
| SL1344_RS19265_ | 671 | 642 | 664 | 707 | 622 | 682 |
| SL1344_RS19270_ | 890 | 880 | 1029 | 1036 | 876 | 960 |
| SL1344_RS19275_ | 2426 | 2508 | 2814 | 2815 | 2415 | 2940 |
| SL1344_RS19280_ | 589 | 605 | 612 | 689 | 600 | 681 |
| SL1344_RS19285_ | 709 | 781 | 815 | 786 | 780 | 792 |
| SL1344_RS19290_ | 573 | 575 | 622 | 921 | 1105 | 1245 |
| SL1344_RS19295_ | 706 | 677 | 705 | 997 | 1274 | 1519 |

| | | | | | | |
|---|---|---|---|---|---|---|
| SL1344_RS19300_ | 111 | 112 | 118 | 131 | 101 | 137 |
| SL1344_RS19305_ | 1434 | 1452 | 1346 | 1202 | 1186 | 1301 |
| SL1344_RS19310_ | 113 | 102 | 105 | 89 | 76 | 104 |
| SL1344_RS19315_ | 33 | 37 | 34 | 57 | 37 | 32 |
| SL1344_RS19320_ | 105 | 77 | 121 | 127 | 90 | 88 |
| SL1344_RS19325_ | 29 | 33 | 34 | 24 | 39 | 28 |
| SL1344_RS19330_ | 834 | 807 | 868 | 887 | 1088 | 1068 |
| SL1344_RS19335_ | 317 | 344 | 315 | 270 | 251 | 287 |
| SL1344_RS19340_ | 532 | 582 | 562 | 593 | 593 | 649 |
| SL1344_RS19345_ | 7 | 5 | 8 | 5 | 10 | 8 |
| SL1344_RS19350_ | 143 | 104 | 89 | 97 | 111 | 137 |
| SL1344_RS19355_ | 68 | 53 | 60 | 42 | 39 | 51 |
| SL1344_RS19360_ | 52 | 58 | 56 | 51 | 49 | 54 |
| SL1344_RS19365_ | 72 | 64 | 62 | 95 | 85 | 102 |
| SL1344_RS19370_ | 408 | 428 | 335 | 353 | 326 | 393 |
| SL1344_RS19375_ | 212 | 224 | 239 | 270 | 229 | 292 |
| SL1344_RS19380_ | 37 | 27 | 31 | 62 | 66 | 55 |
| SL1344_RS19385_ | 217 | 215 | 207 | 240 | 295 | 263 |
| SL1344_RS19390_ | 103 | 107 | 115 | 92 | 109 | 82 |
| SL1344_RS19395_ | 129 | 139 | 159 | 120 | 151 | 134 |
| SL1344_RS19400_ | 114 | 99 | 112 | 89 | 85 | 107 |
| SL1344_RS19405_ | 68 | 56 | 74 | 46 | 41 | 55 |
| SL1344_RS19410_ | 41 | 63 | 50 | 43 | 23 | 35 |
| SL1344_RS19415_ | 29 | 20 | 32 | 19 | 26 | 23 |
| SL1344_RS19420_ | 8 | 11 | 13 | 16 | 8 | 14 |
| SL1344_RS19425_ | 1067 | 1105 | 1139 | 1060 | 958 | 982 |
| SL1344_RS19430_ | 64 | 100 | 57 | 36 | 23 | 40 |
| SL1344_RS19435_ | 417 | 395 | 407 | 345 | 274 | 377 |
| SL1344_RS19445_ | 159 | 174 | 145 | 123 | 143 | 196 |
| SL1344_RS19450_ | 278 | 261 | 264 | 191 | 240 | 254 |
| SL1344_RS19455_ | 47 | 53 | 63 | 52 | 51 | 45 |
| SL1344_RS19460_ | 40 | 37 | 44 | 41 | 19 | 17 |
| SL1344_RS19465_ | 298 | 260 | 289 | 205 | 209 | 223 |
| SL1344_RS19470_ | 223 | 214 | 235 | 154 | 167 | 189 |
| SL1344_RS19475_ | 169 | 180 | 179 | 172 | 170 | 167 |
| SL1344_RS19480_ | 254 | 239 | 243 | 218 | 241 | 233 |
| SL1344_RS19485_ | 845 | 861 | 835 | 633 | 690 | 846 |
| SL1344_RS19490_ | 192 | 184 | 163 | 160 | 187 | 160 |
| SL1344_RS19495_ | 79 | 69 | 58 | 59 | 49 | 49 |
| SL1344_RS19500_ | 125 | 130 | 115 | 136 | 98 | 135 |
| SL1344_RS19505_ | 120 | 138 | 144 | 171 | 166 | 172 |
| SL1344_RS19510_ | 77 | 71 | 78 | 122 | 101 | 114 |
| SL1344_RS19515_ | 298 | 254 | 323 | 235 | 250 | 277 |
| SL1344_RS19520_ | 224 | 205 | 207 | 153 | 137 | 157 |
| SL1344_RS19525_ | 223 | 227 | 278 | 175 | 179 | 208 |
| SL1344_RS19530_ | 144 | 148 | 157 | 161 | 131 | 170 |
| SL1344_RS19535_ | 86 | 91 | 88 | 100 | 113 | 152 |
| SL1344_RS19540_ | 304 | 281 | 300 | 424 | 422 | 461 |
| SL1344_RS19545_ | 1111 | 1365 | 1373 | 1908 | 1576 | 1071 |
| SL1344_RS19550_ | 472 | 352 | 549 | 713 | 621 | 644 |
| SL1344_RS19555_ | 74 | 67 | 63 | 73 | 76 | 65 |
| SL1344_RS19565_ | 340 | 364 | 335 | 303 | 268 | 296 |
| SL1344_RS19570_ | 99 | 87 | 114 | 89 | 72 | 99 |
| SL1344_RS19575_ | 49 | 51 | 53 | 46 | 38 | 59 |
| SL1344_RS19580_ | 263 | 264 | 265 | 200 | 175 | 225 |
| SL1344_RS19585_ | 805 | 831 | 837 | 785 | 705 | 789 |
| SL1344_RS19590_ | 361 | 347 | 380 | 350 | 314 | 348 |
| SL1344_RS19595_ | 321 | 303 | 271 | 259 | 287 | 285 |
| SL1344_RS19600_ | 848 | 944 | 933 | 626 | 555 | 729 |
| SL1344_RS19605_ | 742 | 622 | 707 | 1646 | 1761 | 1388 |
| SL1344_RS19610_ | 1374 | 1416 | 1353 | 1631 | 1803 | 1623 |
| SL1344_RS19615_ | 516 | 554 | 423 | 411 | 417 | 429 |

| | | | | | | |
|---|---|---|---|---|---|---|
| SL1344_RS19620_ | 375 | 390 | 379 | 398 | 367 | 445 |
| SL1344_RS19625_ | 366 | 382 | 344 | 455 | 363 | 439 |
| SL1344_RS19630_ | 93 | 63 | 74 | 79 | 48 | 74 |
| SL1344_RS19635_ | 404 | 355 | 470 | 351 | 280 | 337 |
| SL1344_RS19640_ | 171 | 146 | 216 | 184 | 148 | 165 |
| SL1344_RS19645_ | 56 | 57 | 58 | 55 | 35 | 48 |
| SL1344_RS19650_ | 302 | 256 | 359 | 344 | 185 | 270 |
| SL1344_RS19655_ | 195 | 160 | 242 | 168 | 112 | 155 |
| SL1344_RS19660_ | 214 | 169 | 228 | 167 | 104 | 142 |
| SL1344_RS19665_ | 1509 | 1618 | 1586 | 2074 | 1808 | 2344 |
| SL1344_RS19670_ | 27 | 27 | 34 | 23 | 25 | 25 |
| SL1344_RS19675_ | 122 | 94 | 121 | 134 | 104 | 123 |
| SL1344_RS19680_ | 47 | 39 | 30 | 35 | 23 | 43 |
| SL1344_RS19685_ | 996 | 1003 | 1064 | 790 | 923 | 958 |
| SL1344_RS19690_ | 234 | 220 | 179 | 208 | 176 | 167 |
| SL1344_RS19695_ | 243 | 248 | 261 | 260 | 227 | 231 |
| SL1344_RS19700_ | 58 | 53 | 58 | 38 | 60 | 54 |
| SL1344_RS19705_ | 94 | 91 | 66 | 67 | 87 | 90 |
| SL1344_RS19710_ | 34 | 47 | 46 | 30 | 29 | 26 |
| SL1344_RS19715_ | 204 | 204 | 189 | 147 | 177 | 199 |
| SL1344_RS19720_ | 165 | 141 | 120 | 111 | 113 | 133 |
| SL1344_RS19725_ | 432 | 443 | 399 | 327 | 336 | 457 |
| SL1344_RS19730_ | 45 | 35 | 54 | 28 | 36 | 48 |
| SL1344_RS19735_ | 128 | 121 | 111 | 112 | 98 | 102 |
| SL1344_RS19740_ | 717 | 828 | 931 | 550 | 517 | 620 |
| SL1344_RS19745_ | 3812 | 3978 | 4119 | 3874 | 3865 | 4342 |
| SL1344_RS19750_ | 631 | 666 | 655 | 685 | 603 | 690 |
| SL1344_RS19755_ | 2051 | 2254 | 2266 | 2120 | 2083 | 2395 |
| SL1344_RS19765_ | 2452 | 2105 | 2892 | 4662 | 4293 | 4098 |
| SL1344_RS19770_ | 569 | 495 | 792 | 1205 | 910 | 866 |
| SL1344_RS19775_ | 167 | 169 | 184 | 328 | 274 | 238 |
| SL1344_RS19780_ | 2775 | 2864 | 3332 | 3843 | 4125 | 4317 |
| SL1344_RS19785_ | 236 | 269 | 269 | 217 | 245 | 299 |
| SL1344_RS19790_ | 70 | 92 | 59 | 127 | 108 | 136 |
| SL1344_RS19795_ | 57 | 66 | 40 | 116 | 72 | 75 |
| SL1344_RS19800_ | 560 | 579 | 584 | 606 | 568 | 528 |
| SL1344_RS19805_ | 617 | 609 | 651 | 571 | 496 | 605 |
| SL1344_RS19810_ | 38 | 24 | 43 | 35 | 40 | 29 |
| SL1344_RS19815_ | 130 | 119 | 118 | 124 | 107 | 128 |
| SL1344_RS19820_ | 166 | 167 | 176 | 94 | 89 | 105 |
| SL1344_RS19825_ | 1100 | 1065 | 1070 | 779 | 780 | 808 |
| SL1344_RS19830_ | 410 | 420 | 470 | 532 | 513 | 497 |
| SL1344_RS19835_ | 332 | 374 | 351 | 442 | 461 | 501 |
| SL1344_RS19840_ | 727 | 736 | 712 | 602 | 580 | 673 |
| SL1344_RS19845_ | 347 | 334 | 350 | 327 | 347 | 315 |
| SL1344_RS19850_ | 220 | 253 | 252 | 211 | 253 | 262 |
| SL1344_RS19855_ | 229 | 213 | 285 | 227 | 249 | 298 |
| SL1344_RS19860_ | 431 | 427 | 449 | 469 | 461 | 549 |
| SL1344_RS19865_ | 84 | 89 | 91 | 86 | 69 | 79 |
| SL1344_RS19870_ | 13 | 25 | 8 | 12 | 10 | 12 |
| SL1344_RS19875_ | 284 | 279 | 309 | 237 | 229 | 262 |
| SL1344_RS19880_ | 5956 | 5944 | 6418 | 5596 | 5595 | 6161 |
| SL1344_RS19885_ | 3752 | 3700 | 3575 | 4111 | 3853 | 4148 |
| SL1344_RS19890_ | 91 | 91 | 112 | 84 | 90 | 120 |
| SL1344_RS19900_ | 18206 | 17171 | 19692 | 13024 | 12623 | 15255 |
| SL1344_RS19905_ | 12990 | 12469 | 15355 | 9835 | 10029 | 11419 |
| SL1344_RS19910_ | 30618 | 31821 | 36628 | 24578 | 24219 | 27760 |
| SL1344_RS19915_ | 9836 | 8387 | 9589 | 8737 | 8781 | 9611 |
| SL1344_RS19920_ | 12250 | 11838 | 12128 | 11282 | 11335 | 11324 |
| SL1344_RS19925_ | 6869 | 5998 | 6456 | 5849 | 6039 | 6200 |
| SL1344_RS19930_ | 5769 | 5084 | 5838 | 5453 | 5380 | 6006 |
| SL1344_RS19935_ | 1765 | 1462 | 1488 | 1686 | 1726 | 1891 |

| | | | | | | |
|---|---|---|---|---|---|---|
| SL1344_RS19945_ | 893 | 933 | 849 | 750 | 743 | 866 |
| SL1344_RS19950_ | 634 | 539 | 497 | 872 | 781 | 803 |
| SL1344_RS19955_ | 183 | 191 | 179 | 177 | 176 | 160 |
| SL1344_RS19960_ | 15919 | 15157 | 4195 | 3971 | 3073 | 3758 |
| SL1344_RS19965_ | 352 | 386 | 424 | 342 | 297 | 372 |
| SL1344_RS19970_ | 804 | 766 | 844 | 764 | 685 | 777 |
| SL1344_RS19980_ | 173 | 187 | 192 | 243 | 291 | 304 |
| SL1344_RS19985_ | 134 | 158 | 161 | 169 | 151 | 164 |
| SL1344_RS19990_ | 303 | 298 | 317 | 299 | 286 | 345 |
| SL1344_RS19995_ | 5766 | 5549 | 5571 | 4824 | 4685 | 4936 |
| SL1344_RS20000_ | 940 | 899 | 908 | 713 | 655 | 797 |
| SL1344_RS20005_ | 704 | 777 | 824 | 601 | 518 | 551 |
| SL1344_RS20020_ | 574 | 509 | 449 | 430 | 458 | 438 |
| SL1344_RS20025_ | 127 | 79 | 171 | 56 | 92 | 142 |
| SL1344_RS20030_ | 40 | 29 | 56 | 67 | 58 | 62 |
| SL1344_RS20035_ | 1567 | 1144 | 1795 | 598 | 839 | 1739 |
| SL1344_RS20045_ | 835 | 1039 | 1378 | 1194 | 1013 | 1204 |
| SL1344_RS20050_ | 1292 | 1339 | 2020 | 2127 | 1691 | 1683 |
| SL1344_RS20055_ | 389 | 425 | 337 | 372 | 400 | 411 |
| SL1344_RS20060_ | 3451 | 3605 | 3735 | 2782 | 2797 | 3651 |
| SL1344_RS20065_ | 62 | 50 | 51 | 78 | 74 | 82 |
| SL1344_RS20070_ | 1484 | 1587 | 1778 | 1863 | 1553 | 1433 |
| SL1344_RS20075_ | 195 | 202 | 211 | 224 | 229 | 279 |
| SL1344_RS20085_ | 562 | 546 | 518 | 468 | 506 | 605 |
| SL1344_RS20090_ | 725 | 698 | 680 | 618 | 696 | 738 |
| SL1344_RS20095_ | 870 | 869 | 955 | 1075 | 960 | 1081 |
| SL1344_RS20100_ | 121 | 126 | 127 | 154 | 147 | 153 |
| SL1344_RS20105_ | 129 | 103 | 139 | 130 | 141 | 125 |
| SL1344_RS20110_ | 279 | 284 | 277 | 242 | 255 | 268 |
| SL1344_RS20115_ | 1440 | 1554 | 1554 | 2030 | 2621 | 2525 |
| SL1344_RS20120_ | 705 | 674 | 684 | 597 | 578 | 694 |
| SL1344_RS20125_ | 95 | 94 | 92 | 90 | 91 | 94 |
| SL1344_RS20130_ | 525 | 545 | 531 | 530 | 494 | 527 |
| SL1344_RS20135_ | 1631 | 1714 | 1779 | 1788 | 1740 | 1825 |
| SL1344_RS20140_ | 1543 | 1559 | 1644 | 1792 | 1737 | 1793 |
| SL1344_RS20145_ | 7107 | 6544 | 6601 | 6861 | 6848 | 7335 |
| SL1344_RS20150_ | 5728 | 5802 | 6382 | 7168 | 6744 | 7351 |
| SL1344_RS20155_ | 1320 | 1249 | 1400 | 1182 | 1200 | 1333 |
| SL1344_RS20160_ | 1790 | 1737 | 1799 | 1716 | 1576 | 1794 |
| SL1344_RS20165_ | 999 | 959 | 987 | 1033 | 896 | 1047 |
| SL1344_RS20175_ | 831 | 930 | 898 | 738 | 728 | 850 |
| SL1344_RS20180_ | 403 | 424 | 437 | 310 | 340 | 383 |
| SL1344_RS20185_ | 431 | 473 | 545 | 428 | 347 | 448 |
| SL1344_RS20190_ | 713 | 733 | 807 | 691 | 581 | 729 |
| SL1344_RS20195_ | 506 | 494 | 537 | 499 | 484 | 487 |
| SL1344_RS20200_ | 393 | 384 | 400 | 328 | 356 | 385 |
| SL1344_RS20205_ | 722 | 675 | 799 | 677 | 645 | 700 |
| SL1344_RS20210_ | 429 | 460 | 442 | 401 | 389 | 435 |
| SL1344_RS20215_ | 824 | 716 | 793 | 656 | 712 | 705 |
| SL1344_RS20220_ | 476 | 407 | 548 | 564 | 450 | 468 |
| SL1344_RS20225_ | 511 | 548 | 712 | 604 | 521 | 691 |
| SL1344_RS20230_ | 1056 | 983 | 1339 | 1362 | 1206 | 1276 |
| SL1344_RS20235_ | 1242 | 1245 | 2022 | 2332 | 1762 | 1586 |
| SL1344_RS20240_ | 1701 | 1799 | 1807 | 1465 | 1421 | 1593 |
| SL1344_RS20245_ | 1282 | 1300 | 1247 | 1148 | 1104 | 1299 |
| SL1344_RS20250_ | 1126 | 1183 | 1210 | 1080 | 966 | 1052 |
| SL1344_RS20255_ | 1527 | 1517 | 1628 | 1466 | 1260 | 1473 |
| SL1344_RS20260_ | 2558 | 2623 | 2693 | 2410 | 2306 | 2686 |
| SL1344_RS20265_ | 7 | 14 | 19 | 11 | 13 | 8 |
| SL1344_RS20270_ | 6 | 5 | 20 | 7 | 7 | 9 |
| SL1344_RS20275_ | 8 | 16 | 13 | 18 | 9 | 13 |
| SL1344_RS20280_ | 547 | 549 | 456 | 436 | 422 | 505 |

| | | | | | | |
|---|---|---|---|---|---|---|
| SL1344_RS20285_ | 24 | 10 | 10 | 8 | 10 | 15 |
| SL1344_RS20290_ | 494 | 558 | 552 | 479 | 618 | 627 |
| SL1344_RS20295_ | 1130 | 1103 | 1034 | 1042 | 981 | 1047 |
| SL1344_RS20300_ | 866 | 845 | 806 | 780 | 700 | 858 |
| SL1344_RS20305_ | 830 | 815 | 842 | 765 | 714 | 786 |
| SL1344_RS20310_ | 325 | 311 | 341 | 322 | 307 | 347 |
| SL1344_RS20315_ | 1472 | 1522 | 1425 | 1646 | 1682 | 1942 |
| SL1344_RS20325_ | 19 | 12 | 15 | 13 | 16 | 16 |
| SL1344_RS20330_ | 8 | 1 | 7 | 3 | 4 | 12 |
| SL1344_RS20335_ | 114 | 99 | 114 | 109 | 112 | 123 |
| SL1344_RS20340_ | 11 | 24 | 17 | 25 | 36 | 31 |
| SL1344_RS20345_ | 1527 | 1402 | 1534 | 1355 | 1400 | 1486 |
| SL1344_RS20350_ | 849 | 793 | 766 | 825 | 767 | 866 |
| SL1344_RS20355_ | 208 | 195 | 207 | 259 | 223 | 266 |
| SL1344_RS20365_ | 561 | 499 | 501 | 560 | 514 | 545 |
| SL1344_RS20370_ | 482 | 425 | 488 | 460 | 377 | 465 |
| SL1344_RS20375_ | 72 | 83 | 79 | 98 | 66 | 72 |
| SL1344_RS20390_ | 105 | 128 | 79 | 125 | 107 | 100 |
| SL1344_RS20395_ | 568 | 577 | 579 | 528 | 486 | 623 |
| SL1344_RS20400_ | 2345 | 2299 | 2282 | 2482 | 2247 | 2718 |
| SL1344_RS20405_ | 423 | 464 | 401 | 432 | 474 | 459 |
| SL1344_RS20410_ | 1039 | 972 | 1007 | 962 | 1017 | 1106 |
| SL1344_RS20415_ | 843 | 878 | 847 | 663 | 658 | 785 |
| SL1344_RS20425_ | 2691 | 2630 | 2670 | 2660 | 2962 | 3135 |
| SL1344_RS20430_ | 2704 | 2511 | 2436 | 2485 | 2522 | 2646 |
| SL1344_RS20435_ | 1052 | 965 | 1035 | 1329 | 1069 | 1142 |
| SL1344_RS20440_ | 205 | 230 | 219 | 216 | 179 | 235 |
| SL1344_RS20445_ | 196 | 192 | 191 | 260 | 221 | 199 |
| SL1344_RS20450_ | 2281 | 2241 | 2451 | 2787 | 2683 | 2911 |
| SL1344_RS20455_ | 1330 | 1255 | 1416 | 1347 | 1336 | 1397 |
| SL1344_RS20460_ | 216 | 219 | 211 | 234 | 238 | 222 |
| SL1344_RS20470_ | 322 | 318 | 377 | 479 | 554 | 527 |
| SL1344_RS20475_ | 1784 | 1960 | 1887 | 1727 | 1668 | 1871 |
| SL1344_RS20480_ | 1336 | 1425 | 1327 | 1126 | 1236 | 1324 |
| SL1344_RS20485_ | 541 | 575 | 558 | 515 | 458 | 578 |
| SL1344_RS20495_ | 57 | 57 | 105 | 55 | 59 | 84 |
| SL1344_RS20500_ | 101 | 42 | 81 | 103 | 145 | 128 |
| SL1344_RS20505_ | 69 | 46 | 96 | 96 | 66 | 107 |
| SL1344_RS20510_ | 200 | 216 | 267 | 116 | 169 | 310 |
| SL1344_RS20520_ | 299 | 287 | 234 | 240 | 197 | 262 |
| SL1344_RS20530_ | 807 | 744 | 763 | 659 | 743 | 810 |
| SL1344_RS20535_ | 1457 | 1302 | 1425 | 1321 | 1140 | 1291 |
| SL1344_RS20540_ | 1376 | 1271 | 1370 | 1155 | 1039 | 1367 |
| SL1344_RS20545_ | 505 | 517 | 418 | 597 | 663 | 586 |
| SL1344_RS20550_ | 44 | 34 | 41 | 51 | 56 | 59 |
| SL1344_RS20555_ | 2074 | 2087 | 2113 | 1423 | 1500 | 1788 |
| SL1344_RS20560_ | 867 | 825 | 767 | 910 | 877 | 919 |
| SL1344_RS20565_ | 68898 | 76732 | 69603 | 79152 | 61515 | 84095 |
| SL1344_RS20570_ | 1066 | 1064 | 937 | 936 | 903 | 1047 |
| SL1344_RS20575_ | 1418 | 1381 | 1480 | 1454 | 1384 | 1618 |
| SL1344_RS20580_ | 286 | 325 | 359 | 328 | 301 | 347 |
| SL1344_RS20585_ | 155 | 168 | 187 | 173 | 145 | 201 |
| SL1344_RS20590_ | 1409 | 1623 | 1659 | 1616 | 1758 | 1913 |
| SL1344_RS20595_ | 2395 | 2438 | 2761 | 3332 | 3411 | 3840 |
| SL1344_RS20600_ | 78 | 69 | 66 | 86 | 87 | 106 |
| SL1344_RS20605_ | 21 | 16 | 26 | 27 | 16 | 35 |
| SL1344_RS20610_ | 25 | 15 | 31 | 18 | 26 | 15 |
| SL1344_RS20615_ | 31 | 39 | 41 | 36 | 36 | 42 |
| SL1344_RS20620_ | 31 | 26 | 16 | 28 | 28 | 30 |
| SL1344_RS20625_ | 59 | 58 | 68 | 51 | 37 | 56 |
| SL1344_RS20635_ | 57 | 76 | 75 | 92 | 80 | 94 |
| SL1344_RS20640_ | 205 | 196 | 223 | 196 | 171 | 163 |

| | | | | | | |
|---|---|---|---|---|---|---|
| SL1344_RS20645_ | 149 | 133 | 174 | 147 | 152 | 151 |
| SL1344_RS20650_ | 68 | 56 | 59 | 73 | 58 | 72 |
| SL1344_RS20655_ | 61 | 64 | 84 | 85 | 94 | 97 |
| SL1344_RS20660_ | 36 | 37 | 50 | 36 | 41 | 32 |
| SL1344_RS20665_ | 21 | 24 | 16 | 32 | 28 | 32 |
| SL1344_RS20670_ | 60 | 67 | 62 | 64 | 53 | 61 |
| SL1344_RS20675_ | 702 | 733 | 703 | 585 | 638 | 682 |
| SL1344_RS20680_ | 652 | 698 | 675 | 756 | 760 | 833 |
| SL1344_RS20685_ | 810 | 794 | 807 | 791 | 774 | 834 |
| SL1344_RS20690_ | 500 | 435 | 387 | 429 | 413 | 460 |
| SL1344_RS20695_ | 962 | 966 | 834 | 927 | 920 | 1119 |
| SL1344_RS20700_ | 415 | 374 | 341 | 421 | 462 | 403 |
| SL1344_RS20705_ | 260 | 216 | 191 | 278 | 300 | 270 |
| SL1344_RS20710_ | 170 | 189 | 187 | 209 | 201 | 257 |
| SL1344_RS20715_ | 146 | 139 | 132 | 109 | 114 | 140 |
| SL1344_RS20720_ | 197 | 172 | 143 | 163 | 130 | 184 |
| SL1344_RS20725_ | 1933 | 1827 | 2094 | 2241 | 2247 | 2642 |
| SL1344_RS20730_ | 2591 | 2748 | 2703 | 2906 | 3129 | 3854 |
| SL1344_RS20735_ | 2308 | 2367 | 2506 | 2600 | 2832 | 3305 |
| SL1344_RS20740_ | 5231 | 5101 | 5976 | 6393 | 6392 | 7688 |
| SL1344_RS20745_ | 2907 | 3049 | 3333 | 3996 | 4080 | 4459 |
| SL1344_RS20750_ | 375 | 350 | 317 | 377 | 331 | 366 |
| SL1344_RS20755_ | 54 | 53 | 49 | 61 | 49 | 64 |
| SL1344_RS20760_ | 30 | 17 | 27 | 28 | 34 | 20 |
| SL1344_RS20765_ | 66 | 63 | 66 | 77 | 85 | 77 |
| SL1344_RS20770_ | 104 | 106 | 96 | 151 | 143 | 156 |
| SL1344_RS20775_ | 334 | 362 | 336 | 335 | 409 | 387 |
| SL1344_RS20780_ | 235 | 267 | 220 | 234 | 208 | 227 |
| SL1344_RS20785_ | 233 | 274 | 273 | 189 | 194 | 231 |
| SL1344_RS20790_ | 83 | 70 | 77 | 69 | 51 | 63 |
| SL1344_RS20795_ | 33 | 21 | 33 | 28 | 28 | 35 |
| SL1344_RS20800_ | 21 | 22 | 17 | 22 | 22 | 32 |
| SL1344_RS20805_ | 148 | 157 | 156 | 150 | 119 | 141 |
| SL1344_RS20810_ | 153 | 161 | 150 | 110 | 94 | 104 |
| SL1344_RS20815_ | 36 | 31 | 34 | 28 | 34 | 29 |
| SL1344_RS20820_ | 1 | 1 | 2 | 2 | 2 | 2 |
| SL1344_RS20825_ | 15 | 11 | 16 | 11 | 24 | 13 |
| SL1344_RS20830_ | 40 | 34 | 38 | 49 | 52 | 38 |
| SL1344_RS20835_ | 10 | 9 | 14 | 6 | 11 | 9 |
| SL1344_RS20840_ | 25 | 17 | 20 | 16 | 17 | 8 |
| SL1344_RS20845_ | 575 | 598 | 685 | 486 | 694 | 651 |
| SL1344_RS20850_ | 289 | 246 | 282 | 280 | 286 | 305 |
| SL1344_RS20855_ | 474 | 449 | 443 | 511 | 457 | 510 |
| SL1344_RS20875_ | 536 | 531 | 570 | 505 | 479 | 551 |
| SL1344_RS20880_ | 1017 | 1129 | 1009 | 808 | 726 | 1039 |
| SL1344_RS20890_ | 1458 | 1334 | 1610 | 1359 | 1400 | 1457 |
| SL1344_RS20895_ | 152 | 143 | 130 | 124 | 148 | 147 |
| SL1344_RS20900_ | 130 | 123 | 131 | 107 | 81 | 103 |
| SL1344_RS20905_ | 585 | 610 | 741 | 596 | 566 | 592 |
| SL1344_RS20910_ | 494 | 500 | 556 | 553 | 507 | 518 |
| SL1344_RS20915_ | 17 | 12 | 15 | 24 | 15 | 22 |
| SL1344_RS20920_ | 749 | 833 | 789 | 1102 | 1050 | 1060 |
| SL1344_RS20925_ | 1793 | 1825 | 1662 | 2300 | 2360 | 2336 |
| SL1344_RS20935_ | 1906 | 1880 | 1917 | 2198 | 2249 | 2474 |
| SL1344_RS20940_ | 1391 | 1342 | 1231 | 1468 | 1523 | 1688 |
| SL1344_RS20945_ | 1203 | 1132 | 1175 | 1620 | 1576 | 1703 |
| SL1344_RS20950_ | 2410 | 2546 | 2480 | 3823 | 3585 | 4117 |
| SL1344_RS20955_ | 3208 | 3394 | 2969 | 4994 | 4110 | 5484 |
| SL1344_RS20960_ | 618 | 588 | 621 | 846 | 786 | 843 |
| SL1344_RS20965_ | 1755 | 1874 | 1708 | 3081 | 2648 | 2947 |
| SL1344_RS20970_ | 4352 | 4520 | 5005 | 4002 | 3644 | 4244 |
| SL1344_RS20975_ | 168 | 129 | 134 | 159 | 132 | 147 |

| | | | | | | |
|---|---|---|---|---|---|---|
| SL1344_RS20980_ | 206 | 245 | 239 | 195 | 180 | 229 |
| SL1344_RS20985_ | 792 | 857 | 763 | 786 | 687 | 796 |
| SL1344_RS20990_ | 1443 | 1569 | 1665 | 1701 | 1568 | 1806 |
| SL1344_RS20995_ | 76601 | 79051 | 88973 | 67755 | 63178 | 72945 |
| SL1344_RS21000_ | 18652 | 16718 | 21009 | 17667 | 17091 | 20734 |
| SL1344_RS21005_ | 3688 | 3881 | 3595 | 2880 | 2514 | 3432 |
| SL1344_RS21010_ | 2501 | 2600 | 2641 | 2618 | 2784 | 3207 |
| SL1344_RS21015_ | 164 | 178 | 230 | 206 | 188 | 232 |
| SL1344_RS21020_ | 3369 | 3464 | 3586 | 3492 | 3587 | 3721 |
| SL1344_RS21025_ | 675 | 551 | 643 | 711 | 686 | 773 |
| SL1344_RS21030_ | 1119 | 1056 | 1020 | 969 | 914 | 984 |
| SL1344_RS21035_ | 906 | 900 | 857 | 893 | 850 | 992 |
| SL1344_RS21040_ | 645 | 639 | 696 | 735 | 666 | 628 |
| SL1344_RS21045_ | 2836 | 2871 | 3951 | 3937 | 4250 | 4018 |
| SL1344_RS21050_ | 52 | 61 | 60 | 64 | 82 | 57 |
| SL1344_RS21055_ | 103 | 98 | 108 | 116 | 102 | 112 |
| SL1344_RS21075_ | 12 | 18 | 12 | 14 | 17 | 13 |
| SL1344_RS21080_ | 17 | 9 | 17 | 22 | 22 | 18 |
| SL1344_RS21085_ | 267 | 289 | 271 | 292 | 274 | 245 |
| SL1344_RS21090_ | 168 | 143 | 154 | 186 | 184 | 182 |
| SL1344_RS21100_ | 2683 | 2780 | 2725 | 2109 | 1892 | 2326 |
| SL1344_RS21105_ | 43 | 41 | 41 | 44 | 48 | 64 |
| SL1344_RS21115_ | 626 | 610 | 560 | 563 | 509 | 534 |
| SL1344_RS21120_ | 457 | 437 | 479 | 340 | 324 | 305 |
| SL1344_RS21125_ | 95 | 95 | 66 | 62 | 46 | 84 |
| SL1344_RS21130_ | 204 | 194 | 136 | 109 | 98 | 133 |
| SL1344_RS21135_ | 508 | 541 | 498 | 433 | 358 | 421 |
| SL1344_RS21140_ | 482 | 460 | 414 | 426 | 370 | 412 |
| SL1344_RS21145_ | 156 | 161 | 146 | 109 | 88 | 138 |
| SL1344_RS21150_ | 171 | 166 | 155 | 141 | 159 | 153 |
| SL1344_RS21155_ | 1213 | 1266 | 1462 | 1222 | 1206 | 1278 |
| SL1344_RS21160_ | 2064 | 2047 | 2013 | 1881 | 1706 | 1970 |
| SL1344_RS21165_ | 568 | 601 | 604 | 413 | 394 | 464 |
| SL1344_RS21170_ | 72 | 62 | 59 | 62 | 59 | 70 |
| SL1344_RS21175_ | 100 | 139 | 108 | 141 | 110 | 144 |
| SL1344_RS21180_ | 367 | 380 | 353 | 434 | 321 | 399 |
| SL1344_RS21185_ | 2050 | 2277 | 2118 | 1840 | 1756 | 2058 |
| SL1344_RS21195_ | 1265 | 1201 | 1145 | 1049 | 1165 | 1300 |
| SL1344_RS21200_ | 897 | 829 | 837 | 725 | 701 | 895 |
| SL1344_RS21205_ | 153 | 148 | 182 | 184 | 184 | 186 |
| SL1344_RS21210_ | 1679 | 1575 | 1904 | 1290 | 1444 | 1493 |
| SL1344_RS21215_ | 988 | 922 | 1149 | 934 | 957 | 960 |
| SL1344_RS21220_ | 68 | 55 | 98 | 42 | 42 | 97 |
| SL1344_RS21225_ | 118 | 98 | 146 | 172 | 200 | 165 |
| SL1344_RS21230_ | 22 | 25 | 42 | 48 | 34 | 86 |
| SL1344_RS21235_ | 190 | 170 | 257 | 93 | 136 | 300 |
| SL1344_RS21245_ | 412 | 429 | 394 | 442 | 359 | 463 |
| SL1344_RS21250_ | 639 | 604 | 646 | 556 | 475 | 609 |
| SL1344_RS21255_ | 460 | 394 | 408 | 414 | 369 | 412 |
| SL1344_RS21260_ | 3 | 2 | 5 | 1 | 4 | 5 |
| SL1344_RS21265_ | 417 | 374 | 582 | 517 | 420 | 484 |
| SL1344_RS21270_ | 500 | 531 | 685 | 693 | 513 | 602 |
| SL1344_RS21275_ | 2785 | 2319 | 2910 | 3713 | 3177 | 3331 |
| SL1344_RS21280_ | 1526 | 1256 | 1732 | 2098 | 1704 | 1749 |
| SL1344_RS21285_ | 35763 | 34889 | 41135 | 36999 | 33289 | 41164 |
| SL1344_RS21290_ | 1756 | 1629 | 1581 | 2207 | 1991 | 2262 |
| SL1344_RS21300_ | 3974 | 3781 | 5123 | 4646 | 5853 | 6284 |
| SL1344_RS21305_ | 10409 | 10400 | 13193 | 11884 | 14579 | 16502 |
| SL1344_RS21310_ | 14162 | 13602 | 16772 | 18442 | 19905 | 20298 |
| SL1344_RS21315_ | 34448 | 31203 | 35760 | 42843 | 47898 | 48729 |
| SL1344_RS21320_ | 15906 | 16364 | 19276 | 17828 | 16800 | 20786 |
| SL1344_RS21325_ | 22279 | 22317 | 26443 | 22519 | 22110 | 27832 |

| | | | | | | |
|---|---|---|---|---|---|---|
| SL1344_RS21330_ | 27 | 46 | 72 | 58 | 32 | 65 |
| SL1344_RS21335_ | 9 | 3 | 5 | 6 | 11 | 4 |
| SL1344_RS21340_ | 197 | 176 | 168 | 163 | 168 | 196 |
| SL1344_RS21345_ | 20 | 21 | 24 | 30 | 23 | 28 |
| SL1344_RS21350_ | 152 | 184 | 196 | 198 | 181 | 157 |
| SL1344_RS21355_ | 67 | 65 | 79 | 70 | 67 | 55 |
| SL1344_RS21360_ | 21 | 24 | 21 | 25 | 24 | 20 |
| SL1344_RS21365_ | 53 | 53 | 61 | 58 | 62 | 43 |
| SL1344_RS21370_ | 32 | 47 | 69 | 46 | 34 | 35 |
| SL1344_RS21375_ | 96 | 116 | 117 | 85 | 102 | 96 |
| SL1344_RS21380_ | 265 | 286 | 288 | 256 | 234 | 243 |
| SL1344_RS21390_ | 998 | 978 | 924 | 880 | 944 | 925 |
| SL1344_RS21395_ | 589 | 620 | 566 | 556 | 502 | 578 |
| SL1344_RS21400_ | 653 | 651 | 604 | 630 | 516 | 663 |
| SL1344_RS21420_ | 500 | 506 | 482 | 354 | 350 | 461 |
| SL1344_RS21425_ | 702 | 654 | 593 | 473 | 493 | 587 |
| SL1344_RS21430_ | 294 | 370 | 397 | 345 | 340 | 364 |
| SL1344_RS21435_ | 257 | 279 | 328 | 354 | 353 | 374 |
| SL1344_RS21440_ | 102 | 72 | 161 | 55 | 62 | 149 |
| SL1344_RS21445_ | 5 | 5 | 9 | 2 | 3 | 8 |
| SL1344_RS21450_ | 379 | 342 | 491 | 210 | 252 | 518 |
| SL1344_RS21460_ | 7 | 9 | 7 | 12 | 10 | 9 |
| SL1344_RS21465_ | 510 | 454 | 410 | 310 | 304 | 411 |
| SL1344_RS21475_ | 150 | 131 | 148 | 107 | 85 | 138 |
| SL1344_RS21480_ | 181 | 143 | 165 | 162 | 151 | 164 |
| SL1344_RS21485_ | 153 | 178 | 168 | 184 | 185 | 173 |
| SL1344_RS21490_ | 303 | 329 | 270 | 232 | 261 | 248 |
| SL1344_RS21495_ | 493 | 468 | 382 | 403 | 487 | 493 |
| SL1344_RS21505_ | 523 | 550 | 538 | 465 | 456 | 482 |
| SL1344_RS21510_ | 1624 | 1599 | 1227 | 1356 | 1259 | 1579 |
| SL1344_RS21515_ | 10 | 10 | 14 | 8 | 13 | 9 |
| SL1344_RS21520_ | 331 | 306 | 329 | 372 | 351 | 353 |
| SL1344_RS21525_ | 798 | 755 | 838 | 687 | 681 | 747 |
| SL1344_RS21530_ | 394 | 509 | 476 | 330 | 412 | 403 |
| SL1344_RS21535_ | 123 | 112 | 98 | 148 | 116 | 145 |
| SL1344_RS21540_ | 56 | 62 | 45 | 56 | 50 | 37 |
| SL1344_RS21545_ | 5 | 8 | 12 | 11 | 4 | 4 |
| SL1344_RS21550_ | 101 | 91 | 73 | 80 | 85 | 88 |
| SL1344_RS21555_ | 38 | 51 | 37 | 27 | 38 | 37 |
| SL1344_RS21560_ | 69 | 49 | 58 | 40 | 47 | 45 |
| SL1344_RS21565_ | 13 | 10 | 9 | 14 | 9 | 13 |
| SL1344_RS21570_ | 24 | 16 | 15 | 25 | 14 | 31 |
| SL1344_RS21575_ | 12 | 5 | 8 | 3 | 5 | 6 |
| SL1344_RS21580_ | 167 | 167 | 202 | 221 | 208 | 245 |
| SL1344_RS21585_ | 41 | 51 | 53 | 81 | 68 | 80 |
| SL1344_RS21590_ | 29 | 48 | 41 | 63 | 39 | 49 |
| SL1344_RS21595_ | 96 | 86 | 91 | 63 | 82 | 96 |
| SL1344_RS21600_ | 56 | 39 | 44 | 57 | 43 | 41 |
| SL1344_RS21605_ | 7 | 8 | 19 | 19 | 15 | 16 |
| SL1344_RS21610_ | 41 | 40 | 56 | 41 | 40 | 58 |
| SL1344_RS21615_ | 71 | 46 | 55 | 67 | 45 | 67 |
| SL1344_RS21620_ | 64 | 64 | 66 | 72 | 73 | 93 |
| SL1344_RS21625_ | 21 | 26 | 27 | 21 | 41 | 25 |
| SL1344_RS21630_ | 53 | 56 | 76 | 81 | 71 | 63 |
| SL1344_RS21635_ | 7 | 7 | 20 | 7 | 3 | 6 |
| SL1344_RS21640_ | 8 | 3 | 7 | 5 | 9 | 5 |
| SL1344_RS21645_ | 12 | 10 | 7 | 6 | 22 | 7 |
| SL1344_RS21650_ | 7 | 13 | 10 | 7 | 16 | 7 |
| SL1344_RS21655_ | 11 | 9 | 13 | 11 | 10 | 25 |
| SL1344_RS21660_ | 150 | 174 | 140 | 142 | 130 | 142 |
| SL1344_RS21665_ | 597 | 626 | 609 | 503 | 468 | 453 |
| SL1344_RS21670_ | 2993 | 3023 | 3136 | 2506 | 2445 | 2794 |

| | | | | | | |
|---|---|---|---|---|---|---|
| SL1344_RS21675_ | 125 | 167 | 184 | 243 | 237 | 261 |
| SL1344_RS21680_ | 23 | 16 | 28 | 24 | 17 | 28 |
| SL1344_RS21685_ | 15 | 13 | 20 | 19 | 20 | 21 |
| SL1344_RS21690_ | 224 | 254 | 211 | 216 | 214 | 235 |
| SL1344_RS21695_ | 61 | 71 | 69 | 88 | 71 | 71 |
| SL1344_RS21710_ | 2921 | 2989 | 3059 | 1920 | 1891 | 2186 |
| SL1344_RS21715_ | 585 | 595 | 585 | 356 | 341 | 400 |
| SL1344_RS21725_ | 727 | 719 | 772 | 471 | 370 | 505 |
| SL1344_RS21730_ | 361 | 341 | 305 | 267 | 268 | 335 |
| SL1344_RS21735_ | 331 | 315 | 303 | 261 | 210 | 251 |
| SL1344_RS21740_ | 3371 | 3612 | 4016 | 3437 | 3462 | 3731 |
| SL1344_RS21745_ | 190 | 196 | 201 | 177 | 202 | 209 |
| SL1344_RS21755_ | 307 | 254 | 322 | 370 | 308 | 370 |
| SL1344_RS21760_ | 445 | 585 | 546 | 483 | 358 | 487 |
| SL1344_RS21765_ | 3980 | 4186 | 3447 | 3099 | 2744 | 3292 |
| SL1344_RS21770_ | 252 | 264 | 266 | 293 | 268 | 333 |
| SL1344_RS21775_ | 347 | 305 | 282 | 303 | 280 | 340 |
| SL1344_RS21780_ | 1176 | 1157 | 1216 | 1335 | 1321 | 1531 |
| SL1344_RS21785_ | 79 | 47 | 67 | 63 | 67 | 70 |
| SL1344_RS21790_ | 840 | 783 | 753 | 577 | 500 | 567 |
| SL1344_RS21795_ | 795 | 695 | 764 | 773 | 811 | 862 |
| SL1344_RS21800_ | 477 | 434 | 479 | 460 | 344 | 469 |
| SL1344_RS21805_ | 614 | 533 | 530 | 460 | 435 | 485 |
| SL1344_RS21810_ | 2501 | 2459 | 2383 | 2554 | 2606 | 2871 |
| SL1344_RS21815_ | 880 | 873 | 812 | 689 | 706 | 719 |
| SL1344_RS21820_ | 528 | 547 | 493 | 392 | 364 | 437 |
| SL1344_RS21825_ | 13 | 11 | 8 | 10 | 16 | 10 |
| SL1344_RS21830_ | 3 | 5 | 1 | 7 | 2 | 6 |
| SL1344_RS21835_ | 1258 | 1318 | 1499 | 1597 | 1507 | 1748 |
| SL1344_RS21840_ | 18 | 11 | 20 | 13 | 18 | 19 |
| SL1344_RS21845_ | 1959 | 1877 | 2014 | 2022 | 2017 | 2320 |
| SL1344_RS21880_ | 33 | 39 | 26 | 38 | 42 | 30 |
| SL1344_RS21885_ | 125 | 105 | 109 | 131 | 119 | 159 |
| SL1344_RS21890_ | 92 | 83 | 86 | 108 | 114 | 106 |
| SL1344_RS21895_ | 79 | 85 | 94 | 99 | 101 | 100 |
| SL1344_RS21900_ | 257 | 219 | 233 | 174 | 176 | 185 |
| SL1344_RS21905_ | 265 | 312 | 306 | 378 | 360 | 432 |
| SL1344_RS21910_ | 863 | 786 | 838 | 743 | 664 | 797 |
| SL1344_RS21915_ | 533 | 505 | 497 | 453 | 457 | 550 |
| SL1344_RS21920_ | 10 | 14 | 25 | 22 | 16 | 12 |
| SL1344_RS21925_ | 40 | 39 | 40 | 58 | 46 | 53 |
| SL1344_RS21930_ | 237 | 229 | 292 | 228 | 221 | 242 |
| SL1344_RS21935_ | 28 | 33 | 27 | 21 | 29 | 57 |
| SL1344_RS21940_ | 636 | 557 | 747 | 783 | 791 | 858 |
| SL1344_RS21945_ | 118 | 131 | 126 | 85 | 93 | 85 |
| SL1344_RS21950_ | 4120 | 4056 | 4020 | 3302 | 2578 | 3239 |
| SL1344_RS21965_ | 1281 | 1345 | 1284 | 937 | 726 | 886 |
| SL1344_RS21970_ | 936 | 994 | 877 | 678 | 473 | 618 |
| SL1344_RS21980_ | 1572 | 1503 | 1377 | 1650 | 1639 | 1659 |
| SL1344_RS21985_ | 246 | 256 | 278 | 154 | 188 | 177 |
| SL1344_RS21990_ | 982 | 1012 | 924 | 1133 | 981 | 1100 |
| SL1344_RS21995_ | 212 | 222 | 198 | 191 | 188 | 216 |
| SL1344_RS22000_ | 263 | 249 | 331 | 363 | 305 | 352 |
| SL1344_RS22005_ | 135 | 101 | 109 | 97 | 115 | 100 |
| SL1344_RS22010_ | 117 | 122 | 113 | 102 | 62 | 93 |
| SL1344_RS22015_ | 896 | 871 | 958 | 897 | 953 | 1145 |
| SL1344_RS22020_ | 968 | 972 | 902 | 611 | 473 | 605 |
| SL1344_RS22030_ | 1182 | 1338 | 1271 | 850 | 811 | 966 |
| SL1344_RS22035_ | 1081 | 1115 | 1239 | 770 | 788 | 954 |
| SL1344_RS22040_ | 1232 | 1325 | 1546 | 1048 | 1032 | 1200 |
| SL1344_RS22045_ | 343 | 323 | 359 | 265 | 225 | 228 |
| SL1344_RS22050_ | 1212 | 1165 | 1101 | 867 | 718 | 924 |

| | | | | | | |
|---|---|---|---|---|---|---|
| SL1344_RS22055_ | 485 | 421 | 415 | 356 | 322 | 374 |
| SL1344_RS22060_ | 145 | 124 | 134 | 106 | 90 | 111 |
| SL1344_RS22065_ | 497 | 594 | 633 | 394 | 380 | 501 |
| SL1344_RS22070_ | 330 | 316 | 341 | 148 | 99 | 123 |
| SL1344_RS22075_ | 107 | 134 | 115 | 72 | 51 | 57 |
| SL1344_RS22080_ | 2928 | 3279 | 3067 | 3012 | 2422 | 2898 |
| SL1344_RS22085_ | 2178 | 2301 | 2198 | 2095 | 1754 | 1824 |
| SL1344_RS22090_ | 1171 | 1327 | 1601 | 984 | 916 | 1100 |
| SL1344_RS22095_ | 518 | 489 | 444 | 476 | 436 | 435 |
| SL1344_RS22100_ | 631 | 659 | 616 | 649 | 646 | 580 |
| SL1344_RS22105_ | 1532 | 1691 | 1569 | 850 | 719 | 835 |
| SL1344_RS22110_ | 375 | 446 | 436 | 232 | 177 | 208 |
| SL1344_RS22115_ | 443 | 500 | 432 | 217 | 171 | 255 |
| SL1344_RS22120_ | 443 | 500 | 492 | 252 | 274 | 275 |
| SL1344_RS22125_ | 32 | 26 | 36 | 42 | 38 | 43 |
| SL1344_RS22130_ | 217 | 190 | 194 | 374 | 282 | 301 |
| SL1344_RS22155_ | 1 | 3 | 4 | 2 | 2 | 3 |
| SL1344_RS22160_ | 235 | 224 | 204 | 230 | 255 | 268 |
| SL1344_RS22165_ | 386 | 402 | 339 | 422 | 399 | 515 |
| SL1344_RS22170_ | 746 | 747 | 768 | 733 | 625 | 848 |
| SL1344_RS22175_ | 11 | 15 | 9 | 24 | 18 | 21 |
| SL1344_RS22180_ | 85 | 61 | 62 | 76 | 82 | 88 |
| SL1344_RS22185_ | 39 | 36 | 56 | 54 | 33 | 38 |
| SL1344_RS22190_ | 451 | 385 | 384 | 355 | 326 | 405 |
| SL1344_RS22195_ | 350 | 332 | 381 | 347 | 306 | 362 |
| SL1344_RS22200_ | 209 | 182 | 193 | 196 | 170 | 197 |
| SL1344_RS22205_ | 4789 | 5215 | 5624 | 3011 | 2324 | 2894 |
| SL1344_RS22210_ | 48993 | 48424 | 51183 | 37982 | 37049 | 42964 |
| SL1344_RS22215_ | 10852 | 9574 | 8542 | 9148 | 10095 | 10158 |
| SL1344_RS22220_ | 917 | 811 | 847 | 1156 | 1163 | 1190 |
| SL1344_RS22225_ | 315 | 307 | 325 | 240 | 232 | 281 |
| SL1344_RS22235_ | 63991 | 60347 | 60633 | 60421 | 61514 | 70597 |
| SL1344_RS22240_ | 4213 | 3871 | 3904 | 3769 | 3954 | 4428 |
| SL1344_RS22245_ | 104 | 103 | 76 | 87 | 106 | 74 |
| SL1344_RS22250_ | 263 | 253 | 254 | 273 | 237 | 304 |
| SL1344_RS22255_ | 1389 | 1279 | 1648 | 1563 | 1546 | 1842 |
| SL1344_RS22260_ | 26 | 33 | 43 | 50 | 27 | 30 |
| SL1344_RS22265_ | 3625 | 4341 | 4074 | 4184 | 4217 | 4102 |
| SL1344_RS22275_ | 82 | 65 | 77 | 98 | 100 | 84 |
| SL1344_RS22280_ | 307 | 341 | 273 | 216 | 210 | 226 |
| SL1344_RS22285_ | 6809 | 6783 | 7617 | 5323 | 4081 | 4990 |
| SL1344_RS22290_ | 3566 | 3469 | 3745 | 2604 | 2275 | 2719 |
| SL1344_RS22295_ | 9943 | 9466 | 9779 | 6882 | 6318 | 7883 |
| SL1344_RS22300_ | 20686 | 20047 | 20144 | 15444 | 13144 | 16672 |
| SL1344_RS22305_ | 359 | 411 | 388 | 384 | 339 | 435 |
| SL1344_RS22310_ | 77 | 68 | 74 | 83 | 86 | 93 |
| SL1344_RS22315_ | 266 | 270 | 257 | 233 | 213 | 264 |
| SL1344_RS22320_ | 1517 | 1595 | 1553 | 1269 | 1224 | 1308 |
| SL1344_RS22325_ | 770 | 782 | 815 | 619 | 607 | 676 |
| SL1344_RS22330_ | 731 | 782 | 830 | 699 | 650 | 804 |
| SL1344_RS22335_ | 716 | 698 | 652 | 471 | 474 | 576 |
| SL1344_RS22340_ | 493 | 489 | 394 | 314 | 498 | 349 |
| SL1344_RS22345_ | 779 | 876 | 921 | 809 | 774 | 959 |
| SL1344_RS22350_ | 337 | 367 | 377 | 356 | 348 | 467 |
| SL1344_RS22355_ | 67 | 73 | 88 | 65 | 55 | 64 |
| SL1344_RS22360_ | 157 | 176 | 177 | 202 | 174 | 201 |
| SL1344_RS22365_ | 543 | 495 | 490 | 412 | 399 | 442 |
| SL1344_RS22370_ | 190 | 205 | 218 | 150 | 133 | 158 |
| SL1344_RS22375_ | 894 | 929 | 891 | 820 | 747 | 905 |
| SL1344_RS22380_ | 1155 | 1175 | 1136 | 1014 | 931 | 1090 |
| SL1344_RS22385_ | 4603 | 4280 | 4072 | 3399 | 3434 | 3586 |
| SL1344_RS22395_ | 4242 | 4298 | 3918 | 3633 | 3162 | 3871 |

| | | | | | | |
|---|---|---|---|---|---|---|
| SL1344_RS22400_ | 3283 | 3275 | 3217 | 2661 | 2343 | 2888 |
| SL1344_RS22405_ | 5340 | 5651 | 5607 | 4615 | 4574 | 5153 |
| SL1344_RS22410_ | 41 | 39 | 38 | 40 | 42 | 49 |
| SL1344_RS22415_ | 6965 | 6910 | 7088 | 7529 | 7360 | 7977 |
| SL1344_RS22420_ | 623 | 667 | 685 | 573 | 517 | 588 |
| SL1344_RS22425_ | 3966 | 4141 | 3695 | 3051 | 3074 | 3585 |
| SL1344_RS22430_ | 1165 | 1148 | 960 | 1049 | 934 | 1188 |
| SL1344_RS22435_ | 12 | 13 | 13 | 10 | 12 | 10 |
| SL1344_RS22440_ | 60 | 44 | 40 | 43 | 37 | 50 |
| SL1344_RS22445_ | 38 | 32 | 42 | 46 | 37 | 52 |
| SL1344_RS22450_ | 26 | 21 | 25 | 21 | 15 | 23 |
| SL1344_RS22455_ | 23 | 16 | 15 | 15 | 13 | 11 |
| SL1344_RS22460_ | 50 | 20 | 24 | 40 | 31 | 30 |
| SL1344_RS22465_ | 174 | 123 | 143 | 150 | 133 | 117 |
| SL1344_RS22470_ | 491 | 500 | 413 | 300 | 276 | 302 |
| SL1344_RS22475_ | 708 | 730 | 878 | 279 | 221 | 263 |
| SL1344_RS22480_ | 2249 | 2712 | 2897 | 635 | 831 | 843 |
| SL1344_RS22485_ | 309 | 306 | 262 | 203 | 171 | 243 |
| SL1344_RS22490_ | 392 | 443 | 414 | 411 | 430 | 423 |
| SL1344_RS22495_ | 99 | 76 | 90 | 87 | 86 | 87 |
| SL1344_RS22500_ | 69 | 85 | 77 | 71 | 53 | 63 |
| SL1344_RS22505_ | 32 | 34 | 30 | 21 | 16 | 21 |
| SL1344_RS22510_ | 63 | 49 | 56 | 24 | 40 | 39 |
| SL1344_RS22520_ | 80 | 95 | 98 | 72 | 63 | 88 |
| SL1344_RS22525_ | 132 | 112 | 146 | 128 | 95 | 125 |
| SL1344_RS22530_ | 66 | 72 | 87 | 59 | 59 | 57 |
| SL1344_RS22535_ | 611 | 668 | 921 | 914 | 901 | 1083 |
| SL1344_RS22540_ | 2212 | 2375 | 3344 | 3199 | 3275 | 3996 |
| SL1344_RS22545_ | 2797 | 3047 | 4017 | 4159 | 4210 | 4825 |
| SL1344_RS22550_ | 1600 | 1686 | 2349 | 2265 | 2246 | 2570 |
| SL1344_RS22555_ | 4349 | 4355 | 5961 | 6017 | 6402 | 7081 |
| SL1344_RS22560_ | 94 | 92 | 83 | 91 | 95 | 113 |
| SL1344_RS22565_ | 884 | 929 | 936 | 643 | 600 | 695 |
| SL1344_RS22570_ | 585 | 501 | 551 | 431 | 246 | 547 |
| SL1344_RS22575_ | 825 | 776 | 721 | 832 | 850 | 925 |
| SL1344_RS22580_ | 106 | 81 | 89 | 89 | 93 | 90 |
| SL1344_RS22585_ | 92 | 66 | 80 | 110 | 121 | 123 |
| SL1344_RS22590_ | 33 | 39 | 40 | 46 | 46 | 50 |
| SL1344_RS22595_ | 104 | 106 | 92 | 90 | 103 | 111 |
| SL1344_RS22600_ | 2055 | 2157 | 2125 | 2094 | 1872 | 2218 |
| SL1344_RS22610_ | 300 | 280 | 239 | 277 | 307 | 278 |
| SL1344_RS22615_ | 4972 | 4222 | 4618 | 8419 | 6796 | 8305 |
| SL1344_RS22620_ | 895 | 944 | 995 | 1014 | 878 | 1070 |
| SL1344_RS22625_ | 649 | 707 | 766 | 853 | 857 | 891 |
| SL1344_RS22630_ | 910 | 1039 | 965 | 551 | 551 | 637 |
| SL1344_RS22635_ | 2457 | 2673 | 2401 | 1799 | 1710 | 1856 |
| SL1344_RS22640_ | 1238 | 1073 | 1040 | 1473 | 1195 | 1214 |
| SL1344_RS22645_ | 39 | 30 | 44 | 48 | 33 | 42 |
| SL1344_RS22650_ | 23 | 21 | 29 | 43 | 29 | 30 |
| SL1344_RS22655_ | 2701 | 2455 | 2945 | 3148 | 2991 | 3320 |
| SL1344_RS22660_ | 4066 | 4219 | 5142 | 3305 | 3292 | 3944 |
| SL1344_RS22665_ | 1218 | 1064 | 1167 | 965 | 893 | 1178 |
| SL1344_RS22670_ | 31 | 29 | 20 | 31 | 23 | 15 |
| SL1344_RS22675_ | 799 | 850 | 746 | 845 | 850 | 1007 |
| SL1344_RS22680_ | 110 | 115 | 138 | 110 | 90 | 88 |
| SL1344_RS22685_ | 2 | 0 | 2 | 0 | 3 | 2 |
| SL1344_RS22690_ | 198 | 198 | 212 | 177 | 140 | 161 |
| SL1344_RS22695_ | 161 | 162 | 138 | 159 | 158 | 147 |
| SL1344_RS22700_ | 118 | 113 | 147 | 120 | 102 | 84 |
| SL1344_RS22705_ | 59 | 44 | 34 | 26 | 22 | 31 |
| SL1344_RS22710_ | 13 | 9 | 8 | 10 | 28 | 5 |
| SL1344_RS22715_ | 88 | 62 | 71 | 78 | 61 | 62 |

| | | | | | | |
|---|---|---|---|---|---|---|
| SL1344_RS22720_ | 66 | 67 | 71 | 71 | 78 | 101 |
| SL1344_RS22725_ | 146 | 182 | 149 | 106 | 95 | 141 |
| SL1344_RS22730_ | 48 | 58 | 44 | 42 | 60 | 46 |
| SL1344_RS22735_ | 191 | 208 | 188 | 147 | 173 | 190 |
| SL1344_RS22740_ | 80 | 77 | 68 | 76 | 72 | 69 |
| SL1344_RS22745_ | 41 | 45 | 36 | 50 | 45 | 44 |
| SL1344_RS22750_ | 72 | 70 | 80 | 112 | 95 | 100 |
| SL1344_RS22755_ | 49 | 32 | 40 | 21 | 32 | 41 |
| SL1344_RS22760_ | 58 | 60 | 57 | 55 | 37 | 58 |
| SL1344_RS22765_ | 339 | 333 | 341 | 300 | 259 | 321 |
| SL1344_RS22770_ | 1420 | 1469 | 1451 | 1300 | 1263 | 1465 |
| SL1344_RS22775_ | 1539 | 1462 | 1411 | 1640 | 1714 | 1763 |
| SL1344_RS22780_ | 12 | 18 | 16 | 13 | 10 | 16 |
| SL1344_RS22785_ | 56 | 45 | 32 | 32 | 35 | 29 |
| SL1344_RS22790_ | 29 | 22 | 32 | 28 | 23 | 20 |
| SL1344_RS22795_ | 41 | 42 | 69 | 36 | 38 | 45 |
| SL1344_RS22800_ | 42 | 38 | 49 | 35 | 34 | 41 |
| SL1344_RS22805_ | 52 | 55 | 63 | 63 | 54 | 58 |
| SL1344_RS22810_ | 78 | 64 | 74 | 47 | 39 | 51 |
| SL1344_RS22815_ | 71 | 72 | 73 | 38 | 61 | 59 |
| SL1344_RS22820_ | 370 | 380 | 356 | 317 | 270 | 367 |
| SL1344_RS22825_ | 223 | 223 | 208 | 245 | 262 | 282 |
| SL1344_RS22830_ | 240 | 224 | 234 | 236 | 238 | 258 |
| SL1344_RS22835_ | 178 | 227 | 241 | 175 | 164 | 184 |
| SL1344_RS22840_ | 2084 | 2011 | 2000 | 1672 | 1416 | 1543 |
| SL1344_RS22845_ | 9 | 5 | 4 | 2 | 4 | 8 |
| SL1344_RS22850_ | 141 | 145 | 211 | 155 | 147 | 140 |
| SL1344_RS22855_ | 372 | 380 | 376 | 373 | 356 | 360 |
| SL1344_RS22860_ | 70 | 69 | 80 | 67 | 71 | 92 |
| SL1344_RS22865_ | 201 | 190 | 207 | 184 | 179 | 181 |
| SL1344_RS22870_ | 692 | 700 | 678 | 650 | 630 | 597 |
| SL1344_RS22875_ | 11523 | 12535 | 12034 | 9316 | 9118 | 10121 |
| SL1344_RS22880_ | 141 | 162 | 146 | 107 | 112 | 114 |
| SL1344_RS22885_ | 193 | 218 | 216 | 139 | 144 | 190 |
| SL1344_RS22890_ | 13 | 7 | 19 | 20 | 9 | 15 |
| SL1344_RS22895_ | 60 | 76 | 51 | 36 | 57 | 47 |
| SL1344_RS22900_ | 50 | 43 | 66 | 50 | 35 | 50 |
| SL1344_RS22905_ | 23 | 33 | 44 | 10 | 21 | 16 |
| SL1344_RS22910_ | 8 | 18 | 13 | 10 | 4 | 6 |
| SL1344_RS22915_ | 15 | 18 | 35 | 7 | 13 | 9 |
| SL1344_RS22920_ | 265 | 291 | 251 | 202 | 180 | 280 |
| SL1344_RS22925_ | 71 | 76 | 69 | 116 | 87 | 110 |
| SL1344_RS22930_ | 955 | 944 | 885 | 638 | 644 | 921 |
| SL1344_RS22935_ | 273 | 280 | 281 | 224 | 200 | 250 |
| SL1344_RS22940_ | 114 | 84 | 85 | 69 | 78 | 107 |
| SL1344_RS22945_ | 253 | 207 | 195 | 297 | 254 | 274 |
| SL1344_RS22950_ | 71 | 78 | 64 | 84 | 97 | 76 |
| SL1344_RS22955_ | 4341 | 4453 | 4864 | 4635 | 4573 | 5103 |
| SL1344_RS22960_ | 715 | 718 | 712 | 681 | 703 | 767 |
| SL1344_RS22965_ | 3109 | 3214 | 3568 | 3319 | 3338 | 3641 |
| SL1344_RS22970_ | 776 | 817 | 802 | 845 | 871 | 850 |
| SL1344_RS22975_ | 1152 | 1151 | 1112 | 999 | 791 | 1080 |
| SL1344_RS22980_ | 856 | 889 | 939 | 755 | 599 | 801 |
| SL1344_RS22985_ | 121 | 116 | 115 | 100 | 90 | 95 |
| SL1344_RS22995_ | 62 | 62 | 59 | 50 | 32 | 50 |
| SL1344_RS23000_ | 46 | 41 | 60 | 47 | 45 | 58 |
| SL1344_RS23010_ | 483 | 515 | 556 | 415 | 374 | 450 |
| SL1344_RS23015_ | 62 | 61 | 164 | 128 | 67 | 89 |
| SL1344_RS23020_ | 3 | 6 | 3 | 0 | 6 | 6 |
| SL1344_RS23025_ | 1231 | 1180 | 1104 | 886 | 918 | 1154 |
| SL1344_RS23030_ | 1246 | 1209 | 1114 | 968 | 921 | 1151 |
| SL1344_RS23035_ | 1538 | 1645 | 1634 | 1535 | 1466 | 1734 |

| | | | | | | |
|---|---|---|---|---|---|---|
| SL1344_RS23040_ | 1798 | 1767 | 1845 | 1994 | 1697 | 2054 |
| SL1344_RS23045_ | 695 | 727 | 668 | 852 | 723 | 835 |
| SL1344_RS23050_ | 922 | 865 | 887 | 936 | 873 | 1051 |
| SL1344_RS23055_ | 1748 | 1764 | 1757 | 1600 | 1406 | 1726 |
| SL1344_RS23060_ | 2834 | 2878 | 2951 | 2512 | 2484 | 2675 |
| SL1344_RS23065_ | 494 | 446 | 420 | 421 | 347 | 433 |
| SL1344_RS23070_ | 351 | 288 | 308 | 369 | 316 | 377 |
| SL1344_RS23075_ | 7 | 2 | 7 | 4 | 4 | 1 |
| SL1344_RS23080_ | 636 | 556 | 661 | 757 | 768 | 809 |
| SL1344_RS23085_ | 519 | 586 | 546 | 406 | 430 | 486 |
| SL1344_RS23090_ | 130 | 138 | 179 | 129 | 98 | 107 |
| SL1344_RS23095_ | 33 | 35 | 27 | 47 | 33 | 37 |
| SL1344_RS23100_ | 1704 | 1630 | 1560 | 1104 | 1231 | 1288 |
| SL1344_RS23105_ | 44 | 31 | 50 | 43 | 30 | 38 |
| SL1344_RS23110_ | 108 | 94 | 62 | 83 | 76 | 94 |
| SL1344_RS23115_ | 95 | 96 | 113 | 103 | 94 | 123 |
| SL1344_RS23120_ | 1242 | 1263 | 1255 | 1304 | 1302 | 1512 |
| SL1344_RS23125_ | 43 | 49 | 38 | 50 | 43 | 65 |
| SL1344_RS23130_ | 9 | 14 | 13 | 9 | 11 | 9 |
| SL1344_RS23140_ | 144 | 140 | 93 | 121 | 63 | 111 |
| SL1344_RS23145_ | 109 | 87 | 119 | 98 | 92 | 120 |
| SL1344_RS23150_ | 347 | 362 | 402 | 290 | 259 | 280 |
| SL1344_RS23155_ | 34 | 43 | 27 | 28 | 13 | 22 |
| SL1344_RS23160_ | 34 | 29 | 13 | 10 | 23 | 15 |
| SL1344_RS23165_ | 12 | 8 | 6 | 7 | 8 | 9 |
| SL1344_RS23170_ | 49 | 46 | 57 | 78 | 66 | 51 |
| SL1344_RS23175_ | 246 | 267 | 234 | 239 | 226 | 211 |
| SL1344_RS23180_ | 416 | 402 | 349 | 473 | 420 | 472 |
| SL1344_RS23185_ | 98 | 119 | 92 | 73 | 70 | 87 |
| SL1344_RS23190_ | 1109 | 1301 | 1278 | 548 | 466 | 598 |
| SL1344_RS23195_ | 12 | 13 | 13 | 7 | 11 | 9 |
| SL1344_RS23200_ | 10 | 12 | 2 | 7 | 4 | 1 |
| SL1344_RS23205_ | 118 | 131 | 120 | 212 | 280 | 192 |
| SL1344_RS23210_ | 42 | 28 | 32 | 31 | 42 | 39 |
| SL1344_RS23215_ | 364 | 374 | 389 | 404 | 338 | 400 |
| SL1344_RS23220_ | 491 | 535 | 483 | 484 | 430 | 538 |
| SL1344_RS23225_ | 2107 | 2259 | 2281 | 1989 | 1909 | 2062 |
| SL1344_RS23230_ | 446 | 424 | 405 | 316 | 252 | 354 |
| SL1344_RS23235_ | 291 | 283 | 279 | 213 | 250 | 261 |
| SL1344_RS23240_ | 295 | 280 | 259 | 220 | 260 | 254 |
| SL1344_RS23245_ | 723 | 806 | 722 | 819 | 763 | 898 |
| SL1344_RS23250_ | 112 | 87 | 101 | 104 | 135 | 85 |
| SL1344_RS23265_ | 828 | 907 | 977 | 895 | 771 | 1006 |
| SL1344_RS23270_ | 575 | 555 | 528 | 195 | 230 | 226 |
| SL1344_RS23275_ | 516 | 451 | 487 | 146 | 171 | 189 |
| SL1344_RS23280_ | 670 | 644 | 698 | 236 | 240 | 222 |
| SL1344_RS23285_ | 877 | 801 | 961 | 312 | 298 | 345 |
| SL1344_RS23290_ | 974 | 950 | 1139 | 363 | 396 | 400 |
| SL1344_RS23295_ | 3311 | 3269 | 3356 | 1476 | 1837 | 1588 |
| SL1344_RS23300_ | 3313 | 3386 | 3407 | 2853 | 2621 | 2933 |
| SL1344_RS23305_ | 350 | 299 | 365 | 305 | 279 | 314 |
| SL1344_RS23310_ | 771 | 753 | 717 | 725 | 763 | 797 |
| SL1344_RS23315_ | 399 | 393 | 417 | 483 | 370 | 418 |
| SL1344_RS23320_ | 318 | 324 | 287 | 290 | 274 | 254 |
| SL1344_RS23325_ | 220 | 243 | 228 | 251 | 224 | 232 |
| SL1344_RS23330_ | 1 | 0 | 1 | 2 | 2 | 2 |
| SL1344_RS23335_ | 2 | 7 | 4 | 4 | 6 | 4 |
| SL1344_RS23340_ | 575 | 541 | 512 | 425 | 376 | 395 |
| SL1344_RS23345_ | 43 | 41 | 33 | 46 | 13 | 44 |
| SL1344_RS23350_ | 136 | 129 | 136 | 141 | 160 | 181 |
| SL1344_RS23355_ | 13 | 9 | 6 | 12 | 16 | 9 |
| SL1344_RS23360_ | 1384 | 1231 | 1324 | 1627 | 1407 | 1664 |

| | | | | | | |
|---|---|---|---|---|---|---|
| SL1344_RS23365_ | 5229 | 4639 | 4181 | 5630 | 5317 | 5939 |
| SL1344_RS23370_ | 432 | 426 | 453 | 582 | 538 | 526 |
| SL1344_RS23375_ | 203 | 222 | 240 | 242 | 233 | 236 |
| SL1344_RS23380_ | 166 | 171 | 166 | 215 | 174 | 207 |
| SL1344_RS23385_ | 176 | 152 | 155 | 204 | 159 | 238 |
| SL1344_RS23390_ | 465 | 443 | 532 | 519 | 466 | 566 |
| SL1344_RS23395_ | 897 | 972 | 1002 | 1024 | 1083 | 1138 |
| SL1344_RS23400_ | 2844 | 2833 | 2673 | 2160 | 1996 | 2523 |
| SL1344_RS23410_ | 404 | 445 | 371 | 343 | 318 | 354 |
| SL1344_RS23415_ | 212 | 222 | 264 | 222 | 189 | 224 |
| SL1344_RS23420_ | 474 | 466 | 458 | 433 | 398 | 434 |
| SL1344_RS23425_ | 803 | 832 | 922 | 808 | 698 | 830 |
| SL1344_RS23430_ | 405 | 449 | 483 | 401 | 406 | 472 |
| SL1344_RS23435_ | 466 | 463 | 539 | 451 | 460 | 475 |
| SL1344_RS23440_ | 1488 | 1572 | 1597 | 1488 | 1317 | 1833 |
| SL1344_RS23445_ | 2133 | 2454 | 2401 | 2239 | 1825 | 2451 |
| SL1344_RS23450_ | 79 | 91 | 84 | 99 | 86 | 78 |
| SL1344_RS23455_ | 89 | 96 | 94 | 86 | 94 | 86 |
| SL1344_RS23460_ | 14 | 10 | 7 | 3 | 7 | 4 |
| SL1344_RS23465_ | 37 | 24 | 26 | 31 | 22 | 35 |
| SL1344_RS23470_ | 21 | 20 | 24 | 20 | 24 | 17 |
| SL1344_RS23480_ | 97 | 85 | 95 | 127 | 105 | 120 |
| SL1344_RS23485_ | 380 | 350 | 396 | 397 | 348 | 402 |
| SL1344_RS23495_ | 1160 | 1110 | 1128 | 910 | 935 | 1073 |
| SL1344_RS23500_ | 5217 | 5206 | 5355 | 4800 | 4784 | 5682 |
| SL1344_RS23505_ | 1047 | 1008 | 1024 | 796 | 802 | 864 |
| SL1344_RS23510_ | 327 | 358 | 379 | 311 | 275 | 347 |
| SL1344_RS23515_ | 123 | 152 | 109 | 182 | 174 | 162 |
| SL1344_RS23520_ | 573 | 596 | 559 | 707 | 592 | 736 |
| SL1344_RS23525_ | 2326 | 2516 | 2489 | 2045 | 1995 | 2349 |
| SL1344_RS23530_ | 655 | 694 | 594 | 558 | 583 | 724 |
| SL1344_RS23535_ | 964 | 985 | 950 | 941 | 869 | 1099 |
| SL1344_RS23540_ | 367 | 409 | 396 | 418 | 366 | 433 |
| SL1344_RS23545_ | 126 | 140 | 130 | 131 | 128 | 157 |
| SL1344_RS23550_ | 39 | 50 | 36 | 32 | 33 | 42 |
| SL1344_RS23555_ | 12 | 12 | 8 | 11 | 8 | 10 |
| SL1344_RS23560_ | 50 | 46 | 59 | 38 | 43 | 52 |
| SL1344_RS23565_ | 9 | 23 | 13 | 13 | 27 | 8 |
| SL1344_RS23570_ | 28 | 17 | 27 | 23 | 29 | 30 |
| SL1344_RS23575_ | 260 | 217 | 238 | 332 | 256 | 346 |
| SL1344_RS23580_ | 69 | 79 | 99 | 88 | 72 | 59 |
| SL1344_RS23590_ | 0 | 4 | 4 | 6 | 1 | 6 |
| SL1344_RS23595_ | 156 | 130 | 140 | 140 | 108 | 120 |
| SL1344_RS23640_ | 403 | 427 | 335 | 313 | 324 | 448 |
| SL1344_RS23645_ | 112 | 97 | 90 | 117 | 124 | 126 |
| SL1344_RS23650_ | 637 | 649 | 611 | 632 | 552 | 753 |
| SL1344_RS23655_ | 308 | 351 | 329 | 304 | 309 | 357 |
| SL1344_RS23660_ | 595 | 641 | 702 | 593 | 592 | 650 |
| SL1344_RS23665_ | 91 | 113 | 90 | 171 | 133 | 158 |
| SL1344_RS23670_ | 8228 | 8302 | 7835 | 5473 | 5193 | 6674 |
| SL1344_RS23675_ | 2099 | 1995 | 1703 | 1272 | 1344 | 1535 |
| SL1344_RS23680_ | 1008 | 1097 | 977 | 735 | 716 | 923 |
| SL1344_RS23685_ | 118 | 111 | 99 | 107 | 67 | 111 |
| SL1344_RS23690_ | 46 | 39 | 26 | 41 | 34 | 41 |
| SL1344_RS23695_ | 31 | 25 | 36 | 43 | 27 | 39 |
| SL1344_RS23700_ | 44 | 58 | 64 | 62 | 45 | 60 |
| SL1344_RS23705_ | 21 | 24 | 32 | 30 | 11 | 21 |
| SL1344_RS23710_ | 165 | 147 | 147 | 117 | 92 | 140 |
| SL1344_RS23715_ | 40 | 32 | 40 | 36 | 30 | 39 |
| SL1344_RS23720_ | 37 | 38 | 50 | 57 | 33 | 59 |
| SL1344_RS23725_ | 19 | 18 | 24 | 34 | 28 | 37 |
| SL1344_RS23730_ | 41 | 48 | 42 | 44 | 46 | 52 |

| | | | | | | |
|---|---|---|---|---|---|---|
| SL1344_RS23735_ | 15 | 19 | 26 | 21 | 26 | 32 |
| SL1344_RS23740_ | 9 | 19 | 19 | 8 | 15 | 21 |
| SL1344_RS23745_ | 96 | 87 | 83 | 93 | 92 | 115 |
| SL1344_RS23750_ | 6 | 11 | 19 | 12 | 18 | 12 |
| SL1344_RS23755_ | 16 | 12 | 13 | 19 | 14 | 33 |
| SL1344_RS23760_ | 11 | 11 | 5 | 12 | 6 | 14 |
| SL1344_RS23765_ | 67 | 81 | 65 | 72 | 83 | 91 |
| SL1344_RS23770_ | 34 | 34 | 25 | 40 | 35 | 34 |
| SL1344_RS23775_ | 36 | 26 | 52 | 107 | 82 | 98 |
| SL1344_RS23780_ | 28 | 24 | 30 | 23 | 18 | 28 |
| SL1344_RS23785_ | 54 | 60 | 34 | 34 | 26 | 47 |
| SL1344_RS23790_ | 11 | 11 | 16 | 9 | 9 | 25 |
| SL1344_RS23795_ | 19 | 25 | 30 | 15 | 15 | 16 |
| SL1344_RS23800_ | 17 | 34 | 23 | 15 | 12 | 19 |
| SL1344_RS23805_ | 14 | 18 | 7 | 12 | 22 | 17 |
| SL1344_RS23810_ | 47 | 37 | 31 | 99 | 96 | 89 |
| SL1344_RS23815_ | 73 | 70 | 79 | 142 | 93 | 114 |
| SL1344_RS23820_ | 7 | 8 | 8 | 16 | 7 | 6 |
| SL1344_RS23825_ | 18 | 13 | 21 | 13 | 17 | 22 |
| SL1344_RS23830_ | 30 | 30 | 32 | 27 | 31 | 31 |
| SL1344_RS23840_ | 26 | 27 | 22 | 25 | 26 | 31 |
| SL1344_RS23845_ | 7 | 7 | 4 | 14 | 18 | 9 |
| SL1344_RS23850_ | 8 | 6 | 7 | 7 | 15 | 8 |
| SL1344_RS23855_ | 3 | 5 | 1 | 1 | 2 | 0 |
| SL1344_RS23860_ | 66 | 52 | 41 | 65 | 65 | 66 |
| SL1344_RS23865_ | 10 | 15 | 12 | 13 | 18 | 30 |
| SL1344_RS23870_ | 9 | 10 | 11 | 11 | 9 | 8 |
| SL1344_RS23875_ | 34 | 41 | 33 | 38 | 32 | 49 |
| SL1344_RS23880_ | 3 | 4 | 3 | 2 | 3 | 5 |
| SL1344_RS23885_ | 0 | 3 | 3 | 2 | 3 | 2 |
| SL1344_RS23890_ | 1 | 4 | 2 | 1 | 3 | 0 |
| SL1344_RS23895_ | 3 | 1 | 1 | 3 | 5 | 2 |
| SL1344_RS23900_ | 2 | 8 | 8 | 3 | 2 | 2 |
| SL1344_RS23905_ | 0 | 2 | 0 | 1 | 3 | 0 |
| SL1344_RS23910_ | 121 | 144 | 142 | 131 | 148 | 109 |
| SL1344_RS23915_ | 180 | 141 | 142 | 139 | 135 | 158 |
| SL1344_RS23920_ | 2010 | 2027 | 2074 | 1743 | 1928 | 2124 |
| SL1344_RS23925_ | 1026 | 882 | 1030 | 936 | 1029 | 1081 |
| SL1344_RS23930_ | 2523 | 2766 | 2733 | 2190 | 1888 | 2427 |
| SL1344_RS23935_ | 410 | 438 | 437 | 411 | 440 | 464 |
| SL1344_RS23940_ | 358 | 307 | 334 | 241 | 315 | 381 |
| SL1344_RS23945_ | 344 | 389 | 325 | 299 | 293 | 310 |
| SL1344_RS23950_ | 1288 | 1251 | 1124 | 1111 | 1149 | 1376 |
| SL1344_RS23955_ | 206 | 205 | 202 | 185 | 184 | 247 |
| SL1344_RS23960_ | 1645 | 1652 | 1723 | 1516 | 1412 | 1659 |
| SL1344_RS23965_ | 1204 | 1277 | 1250 | 1227 | 1102 | 1317 |
| SL1344_RS23970_ | 1843 | 2104 | 2037 | 1758 | 1660 | 1973 |
| SL1344_RS23975_ | 1158 | 1179 | 1063 | 892 | 808 | 1072 |
| SL1344_RS23980_ | 39 | 38 | 38 | 46 | 28 | 41 |
| SL1344_RS23985_ | 7 | 9 | 5 | 7 | 8 | 9 |
| SL1344_RS23990_ | 0 | 4 | 4 | 4 | 5 | 0 |
| SL1344_RS23995_ | 36 | 40 | 46 | 22 | 21 | 21 |
| SL1344_RS24000_ | 112 | 91 | 119 | 84 | 81 | 87 |
| SL1344_RS24005_ | 7 | 1 | 4 | 8 | 4 | 3 |
| SL1344_RS24010_ | 8 | 2 | 9 | 7 | 12 | 5 |
| SL1344_RS24015_ | 52 | 62 | 53 | 63 | 37 | 48 |
| SL1344_RS24020_ | 3437 | 3620 | 3995 | 3220 | 3322 | 3903 |
| SL1344_RS24025_ | 4 | 13 | 6 | 4 | 2 | 3 |
| SL1344_RS24030_ | 8 | 7 | 3 | 13 | 12 | 6 |
| SL1344_RS24035_ | 72 | 69 | 88 | 64 | 51 | 65 |
| SL1344_RS24040_ | 103 | 112 | 163 | 115 | 97 | 122 |
| SL1344_RS24045_ | 383 | 396 | 431 | 370 | 296 | 323 |

| | | | | | | |
|---|---|---|---|---|---|---|
| SL1344_RS24050_ | 610 | 652 | 616 | 580 | 532 | 623 |
| SL1344_RS24055_ | 572 | 576 | 608 | 526 | 497 | 608 |
| SL1344_RS24060_ | 547 | 533 | 539 | 529 | 487 | 561 |
| SL1344_RS24065_ | 687 | 793 | 666 | 573 | 600 | 682 |
| SL1344_RS24070_ | 535 | 606 | 479 | 528 | 549 | 667 |
| SL1344_RS24075_ | 470 | 481 | 487 | 564 | 613 | 640 |
| SL1344_RS24080_ | 44 | 45 | 49 | 41 | 25 | 35 |
| SL1344_RS24085_ | 3013 | 3014 | 3123 | 3065 | 2806 | 3080 |
| SL1344_RS24090_ | 12 | 10 | 15 | 14 | 21 | 19 |
| SL1344_RS24095_ | 59 | 50 | 34 | 28 | 39 | 52 |
| SL1344_RS24100_ | 4 | 3 | 2 | 8 | 7 | 7 |
| SL1344_RS24105_ | 160 | 157 | 171 | 141 | 135 | 166 |
| SL1344_RS24110_ | 54 | 57 | 55 | 40 | 47 | 55 |
| SL1344_RS24115_ | 59 | 38 | 39 | 49 | 27 | 40 |
| SL1344_RS24120_ | 69 | 62 | 44 | 45 | 68 | 90 |
| SL1344_RS24125_ | 1237 | 1184 | 1127 | 1202 | 1220 | 1190 |
| SL1344_RS24130_ | 46 | 49 | 51 | 51 | 37 | 67 |
| SL1344_RS24135_ | 61 | 61 | 75 | 61 | 39 | 81 |
| SL1344_RS24140_ | 20 | 29 | 24 | 26 | 29 | 43 |
| SL1344_RS24145_ | 17 | 22 | 39 | 21 | 13 | 32 |
| SL1344_RS24150_ | 162 | 189 | 177 | 113 | 84 | 148 |
| SL1344_RS24155_ | 253 | 219 | 196 | 162 | 174 | 195 |
| SL1344_RS24160_ | 7 | 5 | 3 | 5 | 4 | 6 |
| SL1344_RS24165_ | 717 | 667 | 689 | 1062 | 729 | 884 |
| SL1344_RS24170_ | 691 | 698 | 619 | 1209 | 849 | 856 |
| SL1344_RS24175_ | 3003 | 2956 | 2586 | 3876 | 3590 | 3301 |
| SL1344_RS24180_ | 1344 | 1382 | 1352 | 1195 | 1435 | 1554 |
| SL1344_RS24185_ | 234 | 218 | 218 | 201 | 156 | 189 |
| SL1344_RS24190_ | 256 | 251 | 194 | 195 | 215 | 222 |
| SL1344_RS24195_ | 170 | 204 | 179 | 139 | 138 | 149 |
| SL1344_RS24200_ | 6 | 6 | 3 | 11 | 3 | 7 |
| SL1344_RS24205_ | 789 | 749 | 705 | 1334 | 927 | 982 |
| SL1344_RS24210_ | 330 | 397 | 363 | 371 | 300 | 430 |
| SL1344_RS24215_ | 428 | 422 | 383 | 358 | 349 | 418 |
| SL1344_RS24220_ | 1027 | 1113 | 1008 | 932 | 892 | 1037 |
| SL1344_RS24225_ | 485 | 456 | 473 | 447 | 407 | 478 |
| SL1344_RS24230_ | 234 | 274 | 211 | 181 | 209 | 216 |
| SL1344_RS24235_ | 389 | 462 | 379 | 318 | 350 | 376 |
| SL1344_RS24240_ | 690 | 803 | 783 | 673 | 623 | 791 |
| SL1344_RS24245_ | 847 | 787 | 849 | 726 | 669 | 782 |
| SL1344_RS24250_ | 1134 | 1129 | 1076 | 1354 | 1433 | 1721 |
| SL1344_RS24255_ | 54 | 63 | 57 | 85 | 71 | 75 |
| SL1344_RS24260_ | 56 | 57 | 45 | 45 | 38 | 40 |
| SL1344_RS24265_ | 239 | 309 | 271 | 311 | 232 | 268 |
| SL1344_RS24270_ | 125 | 108 | 132 | 117 | 101 | 126 |
| SL1344_RS24275_ | 117 | 123 | 132 | 135 | 122 | 152 |
| SL1344_RS24280_ | 190 | 214 | 186 | 230 | 220 | 231 |
| SL1344_RS24285_ | 845 | 970 | 843 | 900 | 850 | 1037 |
| SL1344_RS24290_ | 817 | 738 | 749 | 836 | 666 | 869 |
| SL1344_RS24295_ | 1014 | 1039 | 1024 | 1187 | 1103 | 1377 |
| SL1344_RS24300_ | 628 | 669 | 694 | 646 | 637 | 838 |
| SL1344_RS24305_ | 138 | 142 | 111 | 288 | 247 | 243 |
| SL1344_RS24310_ | 38 | 38 | 34 | 57 | 40 | 63 |
| SL1344_RS24315_ | 2 | 1 | 2 | 4 | 3 | 3 |
| SL1344_RS24320_ | 4 | 2 | 1 | 2 | 1 | 3 |
| SL1344_RS24325_ | 12 | 17 | 9 | 19 | 15 | 15 |
| SL1344_RS24330_ | 25 | 28 | 25 | 32 | 30 | 44 |
| SL1344_RS24335_ | 16 | 12 | 12 | 6 | 7 | 11 |
| SL1344_RS24340_ | 4 | 6 | 12 | 17 | 7 | 13 |
| SL1344_RS24345_ | 11 | 18 | 16 | 12 | 15 | 23 |
| SL1344_RS24350_ | 12 | 11 | 8 | 6 | 5 | 8 |
| SL1344_RS24355_ | 10 | 4 | 4 | 5 | 3 | 2 |

| | | | | | | |
|---|---|---|---|---|---|---|
| SL1344_RS24360_ | 8 | 11 | 11 | 5 | 9 | 19 |
| SL1344_RS24365_ | 8 | 4 | 7 | 10 | 7 | 11 |
| SL1344_RS24370_ | 69 | 88 | 83 | 83 | 66 | 91 |
| SL1344_RS24380_ | 80 | 74 | 72 | 66 | 68 | 71 |
| SL1344_RS24385_ | 104 | 122 | 119 | 122 | 97 | 133 |
| SL1344_RS24390_ | 2 | 4 | 7 | 4 | 3 | 3 |
| SL1344_RS24395_ | 47 | 28 | 20 | 44 | 40 | 54 |
| SL1344_RS24400_ | 8 | 8 | 3 | 14 | 6 | 11 |
| SL1344_RS24405_ | 4 | 3 | 11 | 2 | 2 | 5 |
| SL1344_RS24410_ | 548 | 636 | 558 | 523 | 520 | 623 |
| SL1344_RS24415_ | 74 | 89 | 69 | 51 | 36 | 55 |
| SL1344_RS24420_ | 147 | 160 | 114 | 127 | 147 | 148 |
| SL1344_RS24425_ | 12 | 14 | 6 | 20 | 11 | 14 |
| SL1344_RS24430_ | 1160 | 1073 | 1051 | 1446 | 1672 | 1776 |
| SL1344_RS24435_ | 353 | 300 | 305 | 377 | 315 | 390 |
| SL1344_RS24440_ | 1513 | 1403 | 1479 | 1578 | 1543 | 1817 |
| SL1344_RS24445_ | 352 | 327 | 373 | 288 | 306 | 312 |
| SL1344_RS24450_ | 132 | 114 | 136 | 73 | 86 | 116 |
| SL1344_RS24455_ | 57 | 49 | 61 | 36 | 36 | 42 |
| SL1344_RS24460_ | 55018 | 55925 | 64121 | 78587 | 76941 | 85798 |
| SL1344_RS24465_ | 240 | 218 | 175 | 280 | 156 | 212 |
| SL1344_RS24470_ | 25071 | 26785 | 29579 | 26730 | 20776 | 26766 |
| SL1344_RS24475_ | 591 | 492 | 535 | 711 | 654 | 762 |
| SL1344_RS24480_ | 2692 | 2654 | 2880 | 2510 | 2492 | 2840 |
| SL1344_RS24485_ | 227 | 300 | 300 | 204 | 194 | 232 |
| SL1344_RS24490_ | 79 | 75 | 79 | 54 | 52 | 84 |
| SL1344_RS24495_ | 114 | 96 | 125 | 97 | 69 | 109 |
| SL1344_RS24500_ | 334 | 303 | 372 | 265 | 195 | 250 |
| SL1344_RS24505_ | 124 | 109 | 94 | 85 | 69 | 103 |
| SL1344_RS24510_ | 17 | 17 | 20 | 11 | 7 | 11 |
| SL1344_RS24515_ | 139 | 127 | 173 | 79 | 54 | 104 |
| SL1344_RS24520_ | 61 | 63 | 81 | 56 | 46 | 49 |
| SL1344_RS24525_ | 187 | 169 | 162 | 159 | 98 | 165 |
| SL1344_RS24530_ | 188 | 187 | 203 | 107 | 74 | 123 |
| SL1344_RS24535_ | 102 | 100 | 118 | 65 | 60 | 70 |
| SL1344_RS24540_ | 93 | 105 | 104 | 61 | 69 | 73 |
| SL1344_RS24545_ | 53 | 68 | 71 | 33 | 47 | 45 |
| SL1344_RS24550_ | 479 | 480 | 463 | 338 | 268 | 351 |
| SL1344_RS24555_ | 116 | 110 | 109 | 85 | 68 | 81 |
| SL1344_RS24560_ | 126 | 119 | 129 | 119 | 81 | 123 |
| SL1344_RS24565_ | 94 | 86 | 100 | 75 | 56 | 87 |
| SL1344_RS24570_ | 46 | 28 | 40 | 36 | 19 | 39 |
| SL1344_RS24575_ | 51 | 46 | 36 | 127 | 99 | 96 |
| SL1344_RS24580_ | 1083 | 1096 | 1214 | 2568 | 2593 | 3118 |
| SL1344_RS24585_ | 272 | 312 | 336 | 587 | 624 | 709 |
| SL1344_RS24590_ | 124 | 127 | 114 | 155 | 146 | 146 |
| SL1344_RS24595_ | 10 | 7 | 13 | 5 | 8 | 10 |
| SL1344_RS24600_ | 86 | 80 | 72 | 53 | 50 | 75 |
| SL1344_RS24605_ | 239 | 236 | 243 | 166 | 137 | 189 |
| SL1344_RS24610_ | 215 | 223 | 208 | 189 | 165 | 202 |
| SL1344_RS24615_ | 209 | 186 | 169 | 174 | 131 | 192 |
| SL1344_RS24620_ | 246 | 222 | 242 | 169 | 141 | 197 |
| SL1344_RS24625_ | 81 | 95 | 86 | 62 | 43 | 84 |
| SL1344_RS24630_ | 207 | 199 | 223 | 181 | 155 | 192 |
| SL1344_RS24635_ | 260 | 282 | 264 | 196 | 153 | 230 |
| SL1344_RS24640_ | 60 | 50 | 51 | 36 | 23 | 40 |
| SL1344_RS24645_ | 126 | 125 | 121 | 102 | 73 | 119 |
| SL1344_RS24650_ | 34 | 22 | 31 | 22 | 23 | 27 |
| SL1344_RS24655_ | 29 | 32 | 26 | 25 | 19 | 20 |
| SL1344_RS24660_ | 251 | 257 | 224 | 312 | 302 | 323 |
| SL1344_RS24665_ | 232 | 210 | 188 | 227 | 184 | 253 |
| SL1344_RS24670_ | 227 | 231 | 247 | 308 | 281 | 335 |

| | | | | | | |
|---|---|---|---|---|---|---|
| SL1344_RS24675_ | 49 | 56 | 41 | 77 | 73 | 64 |
| SL1344_RS24690_ | 8304 | 8645 | 9509 | 16312 | 15791 | 15228 |
| SL1344_RS24695_ | 886 | 1007 | 1026 | 1946 | 1944 | 2116 |
| SL1344_RS24700_ | 1036 | 1113 | 1201 | 2317 | 2332 | 2492 |
| SL1344_RS24705_ | 1113 | 1140 | 1133 | 1807 | 2094 | 2161 |
| SL1344_RS24710_ | 774 | 857 | 886 | 1561 | 1764 | 1941 |
| SL1344_RS24730_ | 4195 | 4082 | 4087 | 6518 | 6061 | 6361 |
| SL1344_RS24735_ | 11187 | 11330 | 11567 | 15026 | 14794 | 15802 |
| SL1344_RS24740_ | 7 | 20 | 13 | 6 | 7 | 19 |
| SL1344_RS24745_ | 3 | 0 | 0 | 7 | 3 | 4 |
| SL1344_RS24750_ | 68 | 52 | 44 | 63 | 62 | 58 |
| SL1344_RS24755_ | 267 | 234 | 228 | 274 | 242 | 264 |
| SL1344_RS24760_ | 7 | 6 | 19 | 25 | 32 | 20 |
| SL1344_RS24780_ | 47 | 36 | 43 | 76 | 53 | 56 |
| SL1344_RS24785_ | 957 | 1096 | 1222 | 1311 | 1167 | 1245 |
| SL1344_RS24790_ | 1044 | 1042 | 1003 | 1160 | 1167 | 1380 |
| SL1344_RS24795_ | 11 | 9 | 13 | 18 | 8 | 21 |
| SL1344_RS24800_ | 0 | 2 | 2 | 1 | 0 | 1 |
| SL1344_RS24805_ | 10 | 6 | 11 | 15 | 13 | 8 |
| SL1344_RS24810_ | 5 | 6 | 19 | 15 | 16 | 17 |
| SL1344_RS24820_ | 3181 | 2964 | 2566 | 4404 | 4438 | 4426 |
| SL1344_RS24825_ | 314 | 366 | 419 | 562 | 485 | 549 |
| SL1344_RS24830_ | 18 | 35 | 24 | 45 | 35 | 37 |
| SL1344_RS24835_ | 3 | 4 | 3 | 8 | 6 | 8 |
| SL1344_RS24840_ | 2 | 0 | 2 | 5 | 3 | 5 |
| SL1344_RS24845_ | 1 | 2 | 2 | 1 | 4 | 3 |
| SL1344_RS24850_ | 15 | 10 | 18 | 15 | 18 | 14 |
| SL1344_RS24860_ | 319 | 362 | 508 | 403 | 317 | 357 |
| SL1344_RS24865_ | 61 | 77 | 53 | 50 | 34 | 52 |
| SL1344_RS24875_ | 8 | 2 | 2 | 4 | 3 | 1 |
| SL1344_RS24885_ | 9 | 14 | 7 | 11 | 6 | 11 |
| SLnc0002_ | 33 | 36 | 29 | 24 | 21 | 25 |
| SLnc0010_ | 27 | 19 | 23 | 41 | 25 | 31 |
| SLnc0010a_ | 131 | 159 | 119 | 174 | 172 | 162 |
| SLnc0014_ | 1421 | 1507 | 1675 | 2448 | 1953 | 2295 |
| SLnc0022_ | 165 | 185 | 162 | 244 | 226 | 190 |
| SLnc0030_ | 806 | 722 | 847 | 723 | 811 | 840 |
| SLnc0030a_ | 69 | 83 | 103 | 84 | 84 | 71 |
| SLnc0031_ | 40 | 22 | 23 | 21 | 14 | 24 |
| SLnc0035_ | 4219 | 4142 | 4359 | 2671 | 2955 | 3159 |
| SLnc0039_ | 2678 | 2969 | 3586 | 2984 | 2767 | 2572 |
| SLnc0048_ | 152 | 162 | 136 | 111 | 110 | 118 |
| SLnc1001_ | 9 | 7 | 9 | 12 | 6 | 12 |
| SLnc1002_ | 21 | 43 | 41 | 75 | 71 | 44 |
| SLnc1006_ | 197 | 173 | 242 | 337 | 301 | 285 |
| SLnc1008_ | 14 | 18 | 15 | 25 | 6 | 21 |
| SLnc1013_ | 48 | 36 | 59 | 30 | 36 | 50 |
| SLnc1014_ | 53 | 66 | 63 | 43 | 39 | 55 |
| SLnc1015_ | 57 | 61 | 59 | 44 | 25 | 41 |
| SLnc1016_ | 38 | 38 | 42 | 41 | 33 | 39 |
| SLnc1017_ | 70 | 69 | 81 | 88 | 55 | 61 |
| SLnc1018_ | 31 | 26 | 19 | 11 | 15 | 16 |
| SLnc1019_ | 10 | 22 | 26 | 13 | 19 | 18 |
| SLnc1021_ | 74 | 104 | 59 | 57 | 53 | 47 |
| SLnc1022_ | 43 | 35 | 22 | 33 | 33 | 31 |
| SLnc1024_ | 34 | 38 | 55 | 32 | 32 | 39 |
| SLnc1025_ | 36 | 34 | 56 | 45 | 27 | 24 |
| SLnc1026_ | 32 | 34 | 42 | 28 | 40 | 30 |
| SLnc1027_ | 26 | 14 | 26 | 52 | 32 | 31 |
| SLnc1028_ | 56 | 86 | 79 | 56 | 45 | 51 |
| SLnc1029_ | 4 | 2 | 9 | 8 | 15 | 13 |
| sRNA1_ | 1091 | 924 | 1135 | 1241 | 1137 | 1227 |

| | | | | | | |
|---|---|---|---|---|---|---|
| sRNA10_ | 547 | 484 | 519 | 542 | 530 | 560 |
| sRNA11_ | 14 | 31 | 28 | 22 | 21 | 18 |
| sRNA12_ | 10 | 4 | 5 | 7 | 5 | 7 |
| sRNA15_ | 2 | 2 | 4 | 2 | 0 | 2 |
| sRNA16_ | 80 | 112 | 160 | 66 | 107 | 114 |
| sRNA18_ | 3 | 4 | 6 | 3 | 3 | 0 |
| sRNA19_ | 0 | 0 | 1 | 0 | 0 | 0 |
| sRNA2_ | 965 | 783 | 799 | 1459 | 1203 | 1141 |
| sRNA21_ | 206 | 187 | 164 | 237 | 181 | 225 |
| sRNA22_ | 38 | 52 | 29 | 39 | 43 | 41 |
| sRNA23_ | 1 | 3 | 3 | 2 | 6 | 8 |
| sRNA24_ | 4 | 6 | 5 | 2 | 6 | 2 |
| sRNA25_ | 0 | 6 | 1 | 4 | 1 | 0 |
| sRNA26_ | 3 | 1 | 1 | 6 | 2 | 3 |
| sRNA28_ | 3 | 1 | 2 | 8 | 2 | 8 |
| sRNA29_ | 8 | 4 | 5 | 4 | 8 | 2 |
| sRNA31_ | 12 | 9 | 5 | 11 | 11 | 8 |
| sRNA32_ | 3 | 7 | 6 | 5 | 8 | 5 |
| sRNA33_ | 23 | 29 | 20 | 26 | 21 | 21 |
| sRNA5_ | 3 | 10 | 6 | 5 | 5 | 3 |
| sRNA7_ | 37 | 48 | 23 | 55 | 27 | 31 |
| STnc10_ | 0 | 0 | 0 | 3 | 2 | 0 |
| STnc1010_ | 34 | 38 | 61 | 40 | 50 | 38 |
| STnc1030_ | 33 | 19 | 21 | 32 | 22 | 36 |
| STnc1040_ | 5 | 6 | 2 | 4 | 2 | 3 |
| STnc1065_ | 447 | 357 | 512 | 1244 | 1433 | 1511 |
| STnc1070_ | 225 | 256 | 248 | 237 | 259 | 254 |
| STnc1080_ | 8 | 8 | 7 | 9 | 9 | 1 |
| STnc1090_ | 95 | 107 | 120 | 105 | 93 | 92 |
| STnc1100_ | 15 | 25 | 11 | 14 | 30 | 14 |
| STnc1130_ | 140 | 107 | 127 | 209 | 216 | 205 |
| STnc1150_ | 511 | 461 | 557 | 526 | 619 | 545 |
| STnc1160_ | 649 | 498 | 689 | 846 | 668 | 643 |
| STnc1170_ | 87 | 75 | 87 | 73 | 55 | 55 |
| STnc1180_ | 53 | 54 | 63 | 44 | 41 | 44 |
| STnc1190_ | 3848 | 4588 | 4354 | 4383 | 4391 | 4301 |
| STnc1200_ | 37 | 54 | 55 | 48 | 51 | 50 |
| STnc1210_ | 30 | 22 | 33 | 28 | 16 | 25 |
| STnc1220_ | 1 | 1 | 7 | 2 | 0 | 3 |
| STnc1275_ | 64 | 75 | 79 | 132 | 129 | 139 |
| STnc1280_ | 56 | 58 | 60 | 50 | 48 | 43 |
| STnc1285_ | 1151 | 1025 | 868 | 2672 | 1983 | 3229 |
| STnc1290_ | 196 | 202 | 277 | 377 | 304 | 264 |
| STnc130_ | 11 | 8 | 5 | 11 | 14 | 5 |
| STnc1320_ | 14 | 16 | 11 | 10 | 7 | 12 |
| STnc1340_ | 270 | 263 | 266 | 324 | 318 | 386 |
| STnc1380_ | 16 | 12 | 12 | 8 | 12 | 19 |
| STnc1385_ | 2002 | 2317 | 2234 | 2397 | 2361 | 2079 |
| STnc1390_ | 6 | 5 | 8 | 11 | 7 | 9 |
| STnc140_ | 11 | 14 | 16 | 4 | 8 | 9 |
| STnc1400_ | 158 | 124 | 172 | 248 | 237 | 219 |
| STnc1405_ | 1435 | 1533 | 1706 | 2333 | 1802 | 2644 |
| STnc1410_ | 33 | 35 | 37 | 21 | 22 | 17 |
| STnc1460_ | 11199 | 10906 | 11905 | 14281 | 15457 | 13073 |
| STnc1490_ | 136 | 128 | 119 | 169 | 138 | 146 |
| STnc150_ | 204 | 176 | 136 | 157 | 138 | 138 |
| STnc1500_ | 26 | 29 | 32 | 30 | 32 | 29 |
| STnc1510_ | 122 | 111 | 92 | 137 | 157 | 123 |
| STnc1530_ | 5 | 4 | 4 | 6 | 2 | 4 |
| STnc1540_ | 59 | 51 | 57 | 51 | 66 | 65 |
| STnc1545_ | 341 | 394 | 467 | 397 | 391 | 473 |
| STnc1550_ | 151 | 112 | 123 | 378 | 347 | 238 |

| | | | | | | |
|---|---|---|---|---|---|---|
| STnc1560_ | 2250 | 2080 | 2422 | 2236 | 1631 | 2187 |
| STnc1565_ | 169 | 234 | 195 | 254 | 244 | 243 |
| STnc1580_ | 50 | 45 | 44 | 48 | 36 | 46 |
| STnc1590_ | 1085 | 1157 | 972 | 930 | 1005 | 941 |
| STnc1600_ | 83 | 110 | 100 | 102 | 102 | 98 |
| STnc1640_ | 101 | 102 | 105 | 92 | 88 | 71 |
| STnc1650_ | 3 | 3 | 8 | 2 | 4 | 2 |
| STnc1660_ | 203 | 247 | 196 | 204 | 188 | 198 |
| STnc1670_ | 206 | 199 | 193 | 152 | 171 | 180 |
| STnc1675_ | 9926 | 9509 | 12168 | 10917 | 11245 | 13836 |
| STnc1680_ | 155 | 171 | 264 | 232 | 159 | 184 |
| STnc1685_ | 21345 | 21079 | 22302 | 3591 | 23310 | 3752 |
| STnc1690_ | 10 | 21 | 24 | 26 | 22 | 20 |
| STnc170_ | 1 | 1 | 4 | 18 | 21 | 16 |
| STnc1700_ | 10 | 8 | 9 | 13 | 10 | 8 |
| STnc1710_ | 218 | 202 | 193 | 252 | 253 | 184 |
| STnc1760_ | 25 | 23 | 24 | 24 | 21 | 34 |
| STnc1820_ | 38 | 54 | 61 | 63 | 57 | 49 |
| STnc1830_ | 38 | 26 | 31 | 31 | 28 | 35 |
| STnc1870_ | 1 | 1 | 4 | 6 | 0 | 0 |
| STnc1880_ | 18 | 19 | 25 | 21 | 31 | 25 |
| STnc1890_ | 3 | 8 | 8 | 8 | 5 | 3 |
| STnc190_ | 108 | 83 | 100 | 223 | 117 | 130 |
| STnc1920_ | 215 | 182 | 192 | 228 | 217 | 193 |
| STnc1990_ | 72 | 84 | 89 | 101 | 83 | 84 |
| STnc20_ | 49 | 49 | 35 | 29 | 32 | 30 |
| STnc200_ | 44 | 59 | 78 | 101 | 67 | 87 |
| STnc2000_ | 173 | 152 | 166 | 294 | 313 | 292 |
| STnc2010_ | 231 | 297 | 257 | 166 | 182 | 155 |
| STnc2030_ | 55 | 62 | 37 | 52 | 28 | 40 |
| STnc2040_ | 82 | 77 | 276 | 452 | 247 | 284 |
| STnc205_ | 4573 | 4665 | 3834 | 14815 | 14138 | 17749 |
| STnc2070_ | 154 | 147 | 257 | 359 | 249 | 185 |
| STnc2080_ | 284 | 261 | 388 | 323 | 251 | 225 |
| STnc2090_ | 10572 | 12933 | 14255 | 9127 | 8448 | 10058 |
| STnc210_ | 0 | 1 | 0 | 1 | 4 | 2 |
| STnc2100_ | 16 | 24 | 35 | 31 | 16 | 26 |
| STnc2110_ | 51 | 43 | 48 | 128 | 65 | 75 |
| STnc2130_ | 96 | 76 | 112 | 96 | 79 | 96 |
| STnc2160_ | 7 | 5 | 11 | 11 | 11 | 5 |
| STnc2170_ | 18 | 15 | 20 | 20 | 16 | 8 |
| STnc2175_ | 53 | 51 | 61 | 55 | 65 | 63 |
| STnc2180_ | 135 | 158 | 228 | 343 | 183 | 224 |
| STnc220_ | 7 | 4 | 6 | 12 | 8 | 7 |
| STnc230_ | 2 | 0 | 1 | 1 | 2 | 6 |
| STnc240_ | 41 | 37 | 53 | 42 | 43 | 51 |
| STnc260_ | 3 | 1 | 0 | 2 | 1 | 3 |
| STnc280_ | 8 | 4 | 5 | 4 | 4 | 2 |
| STnc290_ | 159 | 136 | 183 | 201 | 166 | 186 |
| STnc30_ | 151 | 153 | 194 | 165 | 149 | 148 |
| STnc300_ | 1 | 0 | 2 | 0 | 2 | 1 |
| STnc3000_ | 8 | 8 | 13 | 7 | 7 | 9 |
| STnc3020_ | 2 | 2 | 6 | 4 | 5 | 8 |
| STnc3030_ | 14 | 12 | 17 | 35 | 24 | 19 |
| STnc3040_ | 5 | 3 | 7 | 6 | 2 | 8 |
| STnc3060_ | 12 | 17 | 9 | 14 | 11 | 11 |
| STnc3070_ | 94 | 99 | 73 | 51 | 67 | 67 |
| STnc3080_ | 2 | 1 | 2 | 0 | 1 | 4 |
| STnc3090_ | 0 | 5 | 6 | 8 | 13 | 7 |
| STnc310_ | 4 | 5 | 4 | 9 | 13 | 8 |
| STnc3100_ | 429 | 396 | 546 | 395 | 341 | 414 |
| STnc3110_ | 174 | 172 | 166 | 221 | 234 | 180 |

| | | | | | | |
|---|---|---|---|---|---|---|
| STnc3120_ | 73 | 82 | 107 | 73 | 65 | 51 |
| STnc3150_ | 36 | 33 | 28 | 52 | 73 | 49 |
| STnc3160_ | 20 | 13 | 10 | 15 | 13 | 10 |
| STnc3170_ | 138 | 114 | 84 | 104 | 140 | 106 |
| STnc3180_ | 19 | 11 | 10 | 17 | 9 | 14 |
| STnc320_ | 48 | 29 | 35 | 41 | 51 | 44 |
| STnc3210_ | 3 | 1 | 3 | 3 | 1 | 2 |
| STnc3220_ | 122 | 124 | 92 | 104 | 119 | 140 |
| STnc3230_ | 17 | 30 | 34 | 30 | 21 | 27 |
| STnc3270_ | 16 | 20 | 25 | 25 | 19 | 24 |
| STnc3280_ | 18 | 30 | 16 | 21 | 25 | 17 |
| STnc3290_ | 25 | 52 | 53 | 55 | 48 | 42 |
| STnc3300_ | 19 | 16 | 30 | 25 | 20 | 22 |
| STnc3310_ | 1086 | 973 | 964 | 1188 | 1300 | 1319 |
| STnc3330_ | 4 | 10 | 7 | 14 | 6 | 3 |
| STnc3340_ | 7 | 13 | 14 | 11 | 13 | 7 |
| STnc3350_ | 26 | 25 | 27 | 37 | 25 | 25 |
| STnc340_ | 5 | 3 | 2 | 21 | 14 | 7 |
| STnc3400_ | 5 | 5 | 9 | 3 | 5 | 7 |
| STnc3410_ | 158 | 183 | 198 | 161 | 134 | 129 |
| STnc3420_ | 33 | 41 | 28 | 38 | 22 | 32 |
| STnc3430_ | 264 | 270 | 272 | 62 | 35 | 72 |
| STnc3440_ | 27 | 22 | 27 | 20 | 8 | 18 |
| STnc3460_ | 207 | 199 | 189 | 178 | 186 | 165 |
| STnc3480_ | 14 | 11 | 12 | 27 | 12 | 35 |
| STnc3490_ | 1 | 4 | 8 | 5 | 7 | 10 |
| STnc350_ | 50 | 59 | 65 | 44 | 46 | 64 |
| STnc3500_ | 12 | 21 | 24 | 20 | 17 | 23 |
| STnc3510_ | 133 | 119 | 109 | 149 | 176 | 129 |
| STnc3540_ | 2848 | 2473 | 2118 | 3089 | 2842 | 3528 |
| STnc3550_ | 58 | 80 | 83 | 38 | 36 | 39 |
| STnc3580_ | 29 | 27 | 23 | 24 | 11 | 18 |
| STnc360_ | 2 | 5 | 1 | 3 | 2 | 3 |
| STnc3600_ | 24 | 34 | 20 | 123 | 173 | 146 |
| STnc3610_ | 109 | 108 | 113 | 118 | 119 | 123 |
| STnc3620_ | 21 | 20 | 25 | 20 | 26 | 20 |
| STnc3630_ | 103 | 129 | 102 | 111 | 94 | 89 |
| STnc3640_ | 52 | 27 | 39 | 87 | 72 | 69 |
| STnc3680_ | 748 | 692 | 739 | 739 | 771 | 826 |
| STnc370_ | 0 | 4 | 0 | 2 | 4 | 4 |
| STnc3700_ | 18 | 12 | 17 | 18 | 19 | 13 |
| STnc3720_ | 30 | 27 | 30 | 22 | 17 | 18 |
| STnc3730_ | 86 | 89 | 90 | 102 | 85 | 98 |
| STnc3750_ | 25 | 22 | 23 | 22 | 14 | 24 |
| STnc3770_ | 28 | 49 | 42 | 38 | 28 | 32 |
| STnc3790_ | 5 | 6 | 4 | 11 | 9 | 9 |
| STnc380_ | 11 | 5 | 6 | 6 | 4 | 7 |
| STnc3800_ | 88 | 97 | 93 | 90 | 84 | 91 |
| STnc3830_ | 48 | 47 | 51 | 33 | 24 | 32 |
| STnc3850_ | 25 | 21 | 18 | 16 | 15 | 15 |
| STnc3860_ | 30 | 43 | 27 | 29 | 26 | 42 |
| STnc3870_ | 117 | 146 | 102 | 134 | 182 | 181 |
| STnc3880_ | 16 | 13 | 14 | 16 | 13 | 11 |
| STnc3890_ | 16 | 13 | 19 | 34 | 18 | 18 |
| STnc390_ | 854 | 720 | 989 | 620 | 523 | 552 |
| STnc3920_ | 4 | 0 | 0 | 2 | 3 | 2 |
| STnc3950_ | 55 | 48 | 55 | 70 | 86 | 82 |
| STnc3960_ | 57 | 61 | 65 | 86 | 46 | 51 |
| STnc3970_ | 8 | 6 | 10 | 8 | 9 | 4 |
| STnc3990_ | 90 | 103 | 70 | 89 | 80 | 72 |
| STnc40_ | 303 | 259 | 350 | 351 | 307 | 295 |
| STnc4000_ | 15 | 12 | 15 | 12 | 11 | 8 |

| | | | | | | |
|---|---|---|---|---|---|---|
| STnc4010_ | 14 | 15 | 16 | 5 | 9 | 6 |
| STnc4020_ | 1 | 1 | 0 | 1 | 1 | 1 |
| STnc4040_ | 96 | 91 | 131 | 85 | 67 | 75 |
| STnc4070_ | 13 | 14 | 9 | 21 | 13 | 8 |
| STnc4080_ | 13 | 6 | 13 | 11 | 14 | 8 |
| STnc410_ | 4 | 3 | 4 | 2 | 4 | 2 |
| STnc4100_ | 10 | 7 | 14 | 29 | 32 | 24 |
| STnc4110_ | 9 | 18 | 10 | 16 | 21 | 11 |
| STnc4120_ | 38 | 49 | 43 | 41 | 32 | 38 |
| STnc4130_ | 28 | 30 | 37 | 67 | 46 | 56 |
| STnc4140_ | 34 | 35 | 36 | 46 | 41 | 29 |
| STnc4150_ | 1 | 4 | 2 | 3 | 3 | 1 |
| STnc4160_ | 59 | 57 | 59 | 65 | 63 | 77 |
| STnc4170_ | 1 | 5 | 2 | 5 | 2 | 5 |
| STnc4190_ | 0 | 0 | 2 | 2 | 1 | 3 |
| STnc420_ | 1 | 2 | 2 | 12 | 4 | 5 |
| STnc4200_ | 14 | 18 | 41 | 29 | 17 | 15 |
| STnc4210_ | 278 | 267 | 311 | 316 | 240 | 325 |
| STnc4220_ | 107 | 145 | 194 | 142 | 128 | 154 |
| STnc4230_ | 51 | 44 | 71 | 64 | 37 | 51 |
| STnc4240_ | 80 | 108 | 78 | 104 | 91 | 83 |
| STnc4250_ | 1023 | 1109 | 1162 | 1355 | 1476 | 1438 |
| STnc4260_ | 0 | 3 | 4 | 6 | 5 | 4 |
| STnc430_ | 0 | 4 | 6 | 6 | 5 | 3 |
| STnc440_ | 85 | 69 | 120 | 133 | 85 | 92 |
| STnc450_ | 142 | 124 | 182 | 214 | 141 | 167 |
| STnc460_ | 0 | 0 | 0 | 1 | 1 | 3 |
| STnc470_ | 69 | 60 | 80 | 63 | 54 | 69 |
| STnc475_ | 7216 | 6822 | 7616 | 10927 | 9939 | 9313 |
| STnc480_ | 26 | 27 | 36 | 21 | 15 | 31 |
| STnc50_ | 1 | 2 | 3 | 4 | 1 | 4 |
| STnc500_ | 53 | 59 | 69 | 61 | 50 | 65 |
| STnc510_ | 120 | 104 | 124 | 83 | 63 | 75 |
| STnc520_ | 104 | 100 | 169 | 134 | 105 | 95 |
| STnc530_ | 21 | 13 | 8 | 11 | 7 | 8 |
| STnc540_ | 3313 | 2867 | 2767 | 3140 | 2896 | 3169 |
| STnc590_ | 277 | 260 | 255 | 325 | 217 | 385 |
| STnc60_ | 0 | 0 | 0 | 0 | 0 | 1 |
| STnc600_ | 2459 | 2168 | 2496 | 4476 | 3268 | 3474 |
| STnc610_ | 235 | 252 | 306 | 469 | 309 | 330 |
| STnc620_ | 497 | 458 | 662 | 600 | 549 | 606 |
| STnc640_ | 35 | 35 | 31 | 88 | 48 | 81 |
| STnc650_ | 1894 | 2014 | 1870 | 1975 | 2075 | 2432 |
| STnc660_ | 876 | 927 | 876 | 793 | 597 | 689 |
| STnc670_ | 1065 | 1033 | 1099 | 1706 | 1616 | 1809 |
| STnc680_ | 1847 | 2057 | 2077 | 1392 | 1424 | 1512 |
| STnc690_ | 21 | 15 | 30 | 31 | 17 | 25 |
| STnc695_ | 125 | 140 | 99 | 75 | 60 | 74 |
| STnc70_ | 997 | 1215 | 1509 | 1473 | 1383 | 1371 |
| STnc720_ | 133 | 93 | 149 | 38 | 38 | 58 |
| STnc730_ | 512 | 404 | 393 | 539 | 544 | 430 |
| STnc740_ | 2 | 9 | 3 | 2 | 3 | 4 |
| STnc750_ | 93 | 75 | 110 | 124 | 127 | 93 |
| STnc760_ | 276 | 341 | 261 | 261 | 298 | 259 |
| STnc770_ | 2068 | 1598 | 1627 | 1719 | 1718 | 1683 |
| STnc780_ | 199 | 166 | 198 | 248 | 194 | 204 |
| STnc790_ | 202 | 201 | 233 | 274 | 336 | 330 |
| STnc800_ | 1097 | 1277 | 1525 | 1597 | 1497 | 1607 |
| STnc810_ | 103 | 116 | 87 | 78 | 60 | 60 |
| STnc835_ | 2159 | 2278 | 3362 | 3016 | 2588 | 2638 |
| STnc840_ | 754 | 758 | 972 | 1938 | 1338 | 1734 |
| STnc850_ | 25 | 20 | 29 | 29 | 19 | 26 |

| | | | | | | |
|---|---|---|---|---|---|---|
| STnc870_ | 470 | 380 | 533 | 497 | 334 | 352 |
| STnc880_ | 74 | 83 | 122 | 119 | 94 | 76 |
| STnc890_ | 171 | 121 | 163 | 181 | 186 | 149 |
| STnc920_ | 205 | 213 | 206 | 226 | 217 | 190 |
| STnc925_ | 131 | 160 | 191 | 138 | 148 | 176 |
| STnc930_ | 12 | 5 | 3 | 14 | 5 | 9 |
| STnc940_ | 20 | 31 | 32 | 33 | 24 | 19 |
| STnc950_ | 5 | 8 | 16 | 6 | 7 | 23 |
| STnc970_ | 104 | 97 | 95 | 103 | 96 | 101 |
| StyR-103a_ | 0 | 2 | 0 | 0 | 0 | 1 |
| StyR-103b_ | 0 | 0 | 0 | 1 | 0 | 1 |
| StyR-103c_ | 0 | 0 | 0 | 0 | 3 | 2 |
| StyR-103d_ | 0 | 0 | 0 | 0 | 0 | 0 |
| StyR-103e_ | 0 | 0 | 0 | 0 | 0 | 0 |
| StyR-103f_ | 0 | 0 | 0 | 0 | 0 | 0 |
| StyR-103g_ | 0 | 0 | 1 | 0 | 1 | 0 |
| StyR-122_ | 6 | 11 | 13 | 5 | 9 | 7 |
| StyR-126_ | 63 | 69 | 61 | 66 | 58 | 72 |
| StyR-128_ | 5 | 7 | 5 | 7 | 8 | 6 |
| StyR-134_ | 143 | 162 | 136 | 141 | 149 | 155 |
| StyR-137_ | 1 | 0 | 0 | 0 | 0 | 1 |
| StyR-150_ | 8 | 7 | 11 | 10 | 5 | 4 |
| StyR-169_ | 109 | 103 | 113 | 82 | 73 | 98 |
| StyR-172_ | 2 | 0 | 1 | 0 | 0 | 1 |
| StyR-178_ | 0 | 0 | 1 | 0 | 1 | 0 |
| StyR-195_ | 20 | 22 | 24 | 17 | 13 | 18 |
| StyR-196_ | 5 | 2 | 2 | 1 | 0 | 4 |
| StyR-199_ | 28 | 14 | 17 | 22 | 15 | 15 |
| StyR-215a_ | 386 | 333 | 583 | 518 | 541 | 519 |
| StyR-215b_ | 291 | 327 | 368 | 417 | 352 | 345 |
| StyR-215c_ | 14 | 19 | 25 | 30 | 19 | 19 |
| StyR-215d_ | 58 | 53 | 91 | 101 | 104 | 92 |
| StyR-215e_ | 18 | 18 | 35 | 32 | 45 | 21 |
| StyR-215f_ | 112 | 128 | 173 | 121 | 124 | 104 |
| StyR-215g_ | 14 | 13 | 17 | 12 | 23 | 23 |
| StyR-219_ | 5 | 5 | 5 | 9 | 8 | 7 |
| StyR-222_ | 4 | 4 | 3 | 4 | 0 | 1 |
| StyR-234_ | 4 | 3 | 3 | 4 | 6 | 1 |
| StyR-243_ | 91 | 117 | 166 | 184 | 168 | 170 |
| StyR-248_ | 75 | 70 | 99 | 58 | 53 | 55 |
| StyR-24a_ | 7 | 17 | 12 | 17 | 6 | 14 |
| StyR-24b_ | 23 | 14 | 23 | 102 | 88 | 94 |
| StyR-24c_ | 9 | 5 | 11 | 33 | 33 | 42 |
| StyR-24d_ | 10 | 9 | 11 | 6 | 8 | 6 |
| StyR-24e_ | 4 | 7 | 4 | 21 | 19 | 9 |
| StyR-24f_ | 27 | 19 | 13 | 28 | 36 | 35 |
| StyR-24g_ | 3 | 3 | 6 | 19 | 19 | 22 |
| StyR-250_ | 8 | 6 | 5 | 12 | 8 | 11 |
| StyR-252_ | 65 | 83 | 57 | 66 | 56 | 72 |
| StyR-254_ | 10 | 6 | 8 | 7 | 1 | 6 |
| StyR-262_ | 118 | 134 | 154 | 125 | 118 | 135 |
| StyR-264_ | 26 | 17 | 23 | 17 | 16 | 19 |
| StyR-27_ | 45 | 59 | 56 | 40 | 20 | 23 |
| StyR-277_ | 337 | 370 | 326 | 262 | 310 | 314 |
| StyR-281_ | 13 | 4 | 11 | 11 | 10 | 9 |
| StyR-285_ | 12 | 18 | 20 | 18 | 12 | 14 |
| StyR-287_ | 242 | 202 | 124 | 133 | 154 | 165 |
| StyR-29_ | 586 | 616 | 717 | 1379 | 1223 | 1092 |
| StyR-291_ | 64 | 55 | 65 | 77 | 53 | 57 |
| StyR-292_ | 2 | 2 | 0 | 2 | 2 | 2 |
| StyR-293_ | 25 | 34 | 44 | 29 | 26 | 34 |
| StyR-295_ | 12 | 9 | 9 | 8 | 10 | 13 |

| | | | | | | |
|---|---|---|---|---|---|---|
| StyR-296_ | 2223 | 1848 | 1862 | 2200 | 2444 | 2135 |
| StyR-3_ | 51 | 56 | 71 | 73 | 66 | 51 |
| StyR-302_ | 0 | 1 | 1 | 0 | 1 | 0 |
| StyR-305_ | 54 | 53 | 84 | 153 | 89 | 126 |
| StyR-306_ | 496 | 509 | 453 | 404 | 397 | 459 |
| StyR-315_ | 3 | 0 | 3 | 0 | 2 | 4 |
| StyR-322_ | 3 | 5 | 2 | 4 | 4 | 7 |
| StyR-327_ | 7 | 18 | 8 | 13 | 5 | 12 |
| StyR-328_ | 3 | 4 | 0 | 4 | 2 | 5 |
| StyR-329a_ | 122 | 91 | 125 | 99 | 70 | 77 |
| StyR-329b_ | 20 | 21 | 10 | 17 | 11 | 20 |
| StyR-336_ | 29 | 21 | 19 | 15 | 10 | 13 |
| StyR-341_ | 1 | 0 | 2 | 6 | 1 | 0 |
| StyR-354_ | 0 | 2 | 1 | 1 | 2 | 1 |
| StyR-358_ | 8 | 9 | 5 | 8 | 7 | 9 |
| StyR-362_ | 8 | 4 | 5 | 4 | 6 | 1 |
| StyR-369_ | 1 | 1 | 0 | 1 | 6 | 5 |
| StyR-377_ | 623 | 704 | 560 | 567 | 560 | 727 |
| StyR-378_ | 1 | 0 | 1 | 1 | 0 | 1 |
| StyR-44a_ | 53 | 41 | 85 | 85 | 71 | 56 |
| StyR-44b_ | 180 | 197 | 268 | 247 | 235 | 239 |
| StyR-44c_ | 16 | 38 | 32 | 24 | 27 | 39 |
| StyR-44d_ | 20 | 28 | 37 | 38 | 44 | 34 |
| StyR-44e_ | 5 | 10 | 9 | 6 | 9 | 11 |
| StyR-44f_ | 7 | 3 | 20 | 2 | 13 | 9 |
| StyR-44g_ | 7 | 5 | 12 | 17 | 5 | 16 |
| StyR-51_ | 376 | 263 | 369 | 546 | 408 | 464 |
| StyR-55_ | 2274 | 2400 | 3362 | 3246 | 3431 | 3816 |
| StyR-8_ | 653 | 672 | 908 | 1103 | 965 | 1043 |
| StyR-88_ | 146 | 101 | 130 | 171 | 180 | 172 |
| StyR-90_ | 4 | 4 | 2 | 4 | 4 | 9 |
| StyR-99_ | 55 | 83 | 109 | 82 | 83 | 96 |
| tp2_ | 2 | 2 | 4 | 7 | 1 | 3 |
| tp28_ | 94 | 83 | 103 | 169 | 145 | 140 |
| tpk3_ | 105 | 120 | 147 | 216 | 145 | 163 |
| tpke11_ | 2666 | 2477 | 3110 | 3116 | 2643 | 2875 |
| tpke70_ | 65 | 68 | 70 | 70 | 32 | 48 |
| zipA-leader_ | 596 | 637 | 626 | 739 | 759 | 705 |
| _ | | | | | | |
| _ | | | | | | |

### Table S2

## EE

| | rab.all | rab.win.tnpA | rab.win.WT | diff.btw | diff.win | effect | overlap | we.ep | we.eBH | wi.ep | wi.eBH |
|---|---|---|---|---|---|---|---|---|---|---|---|
| aceF_aceF | 6.699346691 | 6.653324906 | 6.739925654 | 0.134113493 | 0.100390016 | 1.420443132 | 0.002668394 | 0.124725206 | 0.299410825 | 0.101 | 0.24442054 |
| SL1344_RS11990_ackA | 5.336787792 | 5.110420972 | 5.553882304 | 0.436818029 | 0.045185654 | 9.692985971 | 4.68E-05 | 0.000627291 | 0.016374181 | 0.1 | 0.243021319 |
| SL1344_RS08180_acpD | 0.324121973 | 0.696697028 | -0.166385433 | -0.857124057 | 0.191503065 | -4.440007828 | 4.68E-05 | 0.010537768 | 0.064180614 | 0.1 | 0.243021319 |
| acpS_acpS | 0.749461467 | 0.740820936 | 0.764279065 | 0.008512767 | 0.175763173 | 0.042204138 | 0.483677549 | 0.638008752 | 0.764374145 | 0.7402 | 0.831125874 |
| adrA_adrA | -0.817449159 | -0.912700812 | -0.698801082 | 0.184547515 | 0.355347069 | 0.494904249 | 0.290000009 | 0.473864142 | 0.632827189 | 0.5719 | 0.702701883 |
| SL1344_RS16630_aer | 2.165888621 | 2.900965209 | 1.338086331 | -1.538151745 | 0.200264204 | -7.644209677 | 4.68E-05 | 0.001856391 | 0.027769122 | 0.1 | 0.243021319 |
| SL1344_RS13055_alpha.beta.hydrolase | -0.625659622 | -1.006633953 | -0.123184411 | 0.853644144 | 0.309950664 | 2.650119655 | 0.002666616 | 0.021950368 | 0.101082527 | 0.1012 | 0.244693004 |
| AmgR_amgR | -3.75450313 | -3.7618372 | -3.748182612 | -0.001324353 | 0.830478476 | -0.002423522 | 0.499000666 | 0.632999381 | 0.760608835 | 0.7482 | 0.834470031 |
| amiC_amiC | 2.641567651 | 2.735800427 | 2.490312041 | -0.252357917 | 0.112416025 | -2.171598327 | 0.001335878 | 0.053947457 | 0.172934772 | 0.102 | 0.245825856 |
| SL1344_RS04710_amino.acid.permease | 0.050821637 | -0.361171454 | 0.418901976 | 0.778932112 | 0.184335056 | 4.172545139 | 4.68E-05 | 0.007073435 | 0.051739726 | 0.1 | 0.243021319 |
| SL1344_RS01920_ampH | 0.046019814 | -0.357002567 | 0.438688661 | 0.828056329 | 0.27331662 | 2.994555734 | 4.68E-05 | 0.014814008 | 0.081140251 | 0.1001 | 0.243167487 |
| SL1344_RS23585_arcA | 4.308777357 | 4.245992526 | 4.345317514 | 0.089449981 | 0.061490015 | 1.393977373 | 0.081333385 | 0.120960482 | 0.275522598 | 0.1962 | 0.360656956 |
| ArcZ_arcZ | -0.759677557 | -0.954884441 | -0.466179436 | 0.482589127 | 0.287893689 | 1.582722669 | 0.050632998 | 0.089537907 | 0.225030175 | 0.1557 | 0.310511071 |
| STnc490a_art200 | -8.055667142 | -8.094746637 | -7.974703134 | 0.295102779 | 4.400510704 | 0.053114741 | 0.463690874 | 0.612894598 | 0.74956645 | 0.7584 | 0.843707167 |
| STnc490b_art200 | -8.936593949 | -9.683595655 | -8.093368096 | 1.483145896 | 4.267625335 | 0.315665187 | 0.344437048 | 0.489023929 | 0.637158452 | 0.6039 | 0.717741729 |
| STnc490c_art200 | -7.304401268 | -8.548643145 | -5.381619117 | 2.679191628 | 3.953792189 | 0.628141312 | 0.203197885 | 0.402930811 | 0.562622313 | 0.4368 | 0.593781575 |
| STnc490d_art200 | -9.086225705 | -9.611060749 | -8.540019544 | 0.960683114 | 3.992211739 | 0.198444132 | 0.394666671 | 0.50199085 | 0.649784638 | 0.6122 | 0.724216549 |
| STnc490f_art200 | -8.297128233 | -9.604728126 | -6.987699788 | 2.373312892 | 3.831114645 | 0.560435182 | 0.220666682 | 0.379504974 | 0.536644487 | 0.4601 | 0.59839236 |
| STnc490g_art200 | -8.952917894 | -9.679456539 | -7.952838825 | 1.956528013 | 4.657147695 | 0.38119941 | 0.321119261 | 0.504077194 | 0.647422838 | 0.607 | 0.719470971 |
| artM_artM | 2.989829913 | 2.985760835 | 2.995585585 | -0.003151913 | 0.112329826 | -0.028092271 | 0.488 | 0.679040945 | 0.799118249 | 0.799 | 0.874762387 |
| artP_artP | -0.454788767 | -0.308845932 | -0.594423777 | -0.289626838 | 0.25695993 | -1.098996351 | 0.100666708 | 0.170113131 | 0.326595555 | 0.2369 | 0.392658382 |
| asarfA_asarfA | 1.012191688 | 0.859714872 | 1.19739355 | 0.33510275 | 0.272306199 | 1.173611701 | 0.094603641 | 0.147717867 | 0.322353118 | 0.2035 | 0.374818766 |
| asnB_asnB | 1.757815822 | 1.767150697 | 1.745248723 | -0.016089109 | 0.111994913 | -0.131939319 | 0.438000002 | 0.579460435 | 0.71456952 | 0.6835 | 0.78392828 |
| SL1344_RS13060_asrA | -3.625831349 | -4.091493127 | -2.980468078 | 1.168501081 | 1.062374165 | 1.015128181 | 0.11792142 | 0.179071363 | 0.352494305 | 0.2532 | 0.422145054 |
| SL1344_RS13065_asrB | -2.835880926 | -2.820333038 | -2.850676047 | -0.071482636 | 0.62375611 | -0.115532538 | 0.449700202 | 0.611644049 | 0.744645349 | 0.7364 | 0.825645407 |
| SL1344_RS13070_asrC | -1.775183295 | -1.802645882 | -1.744004967 | 0.06485328 | 0.419085023 | 0.150545851 | 0.426666669 | 0.591886796 | 0.724665098 | 0.7055 | 0.800229242 |
| asyhbQ_asyhbQ | -0.088486137 | -0.069445946 | -0.111352502 | -0.032515263 | 0.319263263 | -0.102488073 | 0.460666668 | 0.715622477 | 0.824594094 | 0.793 | 0.869223113 |
| atpC_atpC | 5.877444835 | 5.61547653 | 6.123416114 | 0.501269157 | 0.096518514 | 5.161734003 | 4.68E-05 | 0.002442019 | 0.034432785 | 0.1 | 0.243021319 |
| SL1344_RS15305_barA | 1.990257411 | 1.850899377 | 2.148406887 | 0.300400784 | 0.132691642 | 2.199139612 | 0.003998486 | 0.036138321 | 0.134654688 | 0.1044 | 0.249099031 |
| bcsA_bcsA | 2.004876475 | 1.462831503 | 2.491151912 | 1.042287125 | 0.122861587 | 8.608564075 | 4.68E-05 | 0.001328365 | 0.023263476 | 0.1 | 0.243021319 |
| SL1344_RS17950_bigA | 0.508092278 | 0.5225828 | 0.483059791 | -0.026040474 | 0.194800096 | -0.115582754 | 0.454363759 | 0.643072362 | 0.767437165 | 0.7424 | 0.834427179 |
| bssR_bssR | -2.089713635 | -1.868080408 | -2.257775773 | -0.376225147 | 0.446835941 | -0.782192992 | 0.186542324 | 0.290445178 | 0.458645416 | 0.3852 | 0.539302758 |
| bssS_bssS | 1.901326552 | 1.848117426 | 2.044468726 | 0.208219281 | 0.21961861 | 0.888248323 | 0.172551653 | 0.245217804 | 0.438355878 | 0.3485 | 0.528436328 |
| cadB_cadB | 0.369519748 | -0.108927712 | 0.797489411 | 0.908172587 | 0.198048352 | 4.579589453 | 4.68E-05 | 0.006955841 | 0.052096764 | 0.1 | 0.243021319 |
| cbiC_cbiC | -3.774438469 | -3.900336004 | -3.605556357 | 0.339008513 | 0.817293515 | 0.395290881 | 0.334000007 | 0.490129711 | 0.643564068 | 0.6129 | 0.728937452 |
| cbiD_cbiD | -2.75435414 | -2.934178606 | -2.610837732 | 0.331749439 | 0.572095202 | 0.551764775 | 0.268487686 | 0.397909929 | 0.560499537 | 0.5087 | 0.640742828 |
| SL1344_RS11580_ccmA | -4.419957581 | -3.938639086 | -5.113035212 | -1.182158645 | 1.008353689 | -1.099799568 | 0.098000042 | 0.177056296 | 0.338990218 | 0.2457 | 0.402551615 |
| SL1344_RS11575_ccmB | -5.165740818 | -6.576592922 | -4.158131725 | 2.646670852 | 1.878793205 | 1.252553278 | 0.027981506 | 0.136188946 | 0.298440847 | 0.1319 | 0.283210226 |
| cdsA_cdsA | 2.925595779 | 2.864679762 | 2.988978881 | 0.097139319 | 0.180898458 | 0.553231462 | 0.278666677 | 0.448766916 | 0.630995385 | 0.567 | 0.71334971 |
| SL1344_RS02340_cdsH | -0.486907305 | -0.500569814 | -0.47756503 | 0.012556492 | 0.258875599 | 0.042827865 | 0.483333334 | 0.608269291 | 0.737807024 | 0.7097 | 0.805510258 |
| chaB_chaB | -2.057662321 | -1.415201744 | -2.260274812 | -0.727706241 | 0.639151066 | -1.035309371 | 0.161892095 | 0.213833279 | 0.395528667 | 0.3374 | 0.507277466 |
| SL1344_RS09610_cheA | 6.251374771 | 7.521963039 | 5.087450782 | -2.436357858 | 0.15604642 | -15.71489294 | 4.68E-05 | 1.12E-05 | 0.004262471 | 0.1 | 0.243021319 |
| SL1344_RS09600_cheM | 5.064598122 | 6.324974701 | 3.7916604 | -2.547998652 | 0.118317071 | -21.58098578 | 4.68E-05 | 0.000231692 | 0.012004357 | 0.1 | 0.243021319 |
| SL1344_RS09595_chemotaxis.protein.methyltransferase | 4.276969012 | 5.29559874 | 3.102008224 | -2.218814439 | 0.141643529 | -15.70235333 | 4.68E-05 | 3.55E-05 | 0.005571355 | 0.1 | 0.243021319 |
| SL1344_RS09585_chemotaxis.regulator | 3.875979251 | 4.897820896 | 2.777772463 | -2.142883283 | 0.123922316 | -17.20640767 | 4.68E-05 | 6.78E-05 | 0.006634798 | 0.1 | 0.243021319 |
| SL1344_RS09590_chemotaxis.response.regulator.protein.glutamate.methylesterase | 3.785640167 | 4.890560207 | 2.734251455 | -2.155477294 | 0.194043159 | -11.13886326 | 4.68E-05 | 0.000323221 | 0.013810541 | 0.1 | 0.243021319 |
| SL1344_RS11875_cheV | 4.125388988 | 4.837653773 | 3.362715758 | -1.506334705 | 0.109813924 | -13.90262856 | 4.68E-05 | 0.000351264 | 0.013115267 | 0.1 | 0.243021319 |
| SL1344_RS09605_CheW | 6.108829296 | 7.383245277 | 4.886034766 | -2.49046073 | 0.080004277 | -31.17887468 | 4.68E-05 | 4.49E-05 | 0.005890127 | 0.1 | 0.243021319 |
| SL1344_RS09580_cheZ | 5.344095707 | 6.288628513 | 4.319599453 | -2.027674712 | 0.173074143 | -11.90077242 | 4.68E-05 | 0.000257995 | 0.012860661 | 0.1 | 0.243021319 |
| chiP_chiP | -1.31563092 | -1.490164992 | -1.123621743 | 0.370105533 | 0.305677308 | 1.177903257 | 0.099267197 | 0.159334385 | 0.311953944 | 0.2375 | 0.39432276 |
| ChiX_chiX | 1.243972948 | 1.020168717 | 1.437563481 | 0.505273785 | 0.433187296 | 1.111562829 | 0.103930752 | 0.142030609 | 0.320605419 | 0.206 | 0.383630524 |
| SL1344_RS02290_clpP | 2.97654408 | 3.06137346 | 2.903497984 | -0.158144083 | 0.08938313 | -1.755507765 | 0.027981506 | 0.0667775 | 0.191633387 | 0.1285 | 0.280114402 |

| | | | | | | | | | | | |
|---|---|---|---|---|---|---|---|---|---|---|---|
| SL1344_RS02295_clpX | 5.338217471 | 5.377167422 | 5.260315393 | -0.101580679 | 0.057885353 | -1.597596496 | 0.050632998 | 0.087215167 | 0.232726968 | 0.1537 | 0.31495274 |
| SL1344_RS09530_cmoA | 2.931536335 | 3.295253213 | 2.446781119 | -0.844024095 | 0.095909724 | -8.768938823 | 4.68E-05 | 0.002576465 | 0.031661555 | 0.1 | 0.243021319 |
| cobS_cobS | -0.776051844 | -0.927540236 | -0.649926298 | 0.287851527 | 0.32157762 | 0.846813597 | 0.17721521 | 0.267317249 | 0.442069583 | 0.3579 | 0.516778737 |
| copA_copA | 2.336889075 | 2.397545798 | 2.274663715 | 0.197306113 | 0.489592882 | 0.40403371 | 0.441333336 | 0.613546564 | 0.764743975 | 0.9907 | 0.994549324 |
| SL1344_RS20320_corA | 2.871116054 | 2.881456651 | 2.856395423 | -0.014339158 | 0.244656789 | -0.054152981 | 0.483333334 | 0.860552919 | 0.922730428 | 0.9688 | 0.982242082 |
| SL1344_RS20860_cpxA | 3.31357113 | 3.296361469 | 3.437086599 | 0.133095148 | 0.106009296 | 0.757144143 | 0.290666676 | 0.383153329 | 0.574709638 | 0.6075 | 0.751608704 |
| cpxP_cpxP | 1.429097855 | 1.401979392 | 1.468015347 | 0.046855801 | 0.359677816 | 0.113054333 | 0.465689542 | 0.821108264 | 0.898212115 | 0.9613 | 0.977563323 |
| SL1344_RS20865_cpxR | 1.987939705 | 1.864438216 | 2.116366443 | 0.253969642 | 0.134486944 | 1.853079865 | 0.01666694 | 0.05899454 | 0.17958926 | 0.1168 | 0.26542874 |
| SL1344_RS17890_crp | 4.312989963 | 4.281257678 | 4.343223867 | 0.063181325 | 0.050398778 | 1.213603594 | 0.086000049 | 0.145664669 | 0.299834644 | 0.2141 | 0.372249821 |
| CrpT_crpT | -3.43347787 | -3.61906783 | -3.285001158 | 0.36140422 | 0.703590875 | 0.468162017 | 0.300000009 | 0.437193432 | 0.595999852 | 0.5459 | 0.672244928 |
| csgB_csgB | -5.573893878 | -5.227131509 | -5.985642026 | -0.862899039 | 2.231632979 | -0.306715495 | 0.315333341 | 0.466598061 | 0.636495783 | 0.5906 | 0.714046248 |
| SL1344_RS05640_csgC | 0.976996543 | 1.608132002 | 0.178561915 | -1.511012943 | 0.300181865 | -5.149707983 | 4.68E-05 | 0.003875939 | 0.039620996 | 0.1 | 0.243021319 |
| SL1344_RS18810_cspA | 6.419062741 | 5.642287591 | 7.097225009 | 1.488111047 | 0.208209484 | 7.06767878 | 4.68E-05 | 0.000281842 | 0.014046512 | 0.1 | 0.243021319 |
| SL1344_RS09980_cspB | -0.109168214 | -0.89066433 | 0.554017648 | 1.488708228 | 0.30374354 | 5.005721512 | 4.68E-05 | 0.006889056 | 0.055138485 | 0.1 | 0.243021319 |
| SL1344_RS06145_cspH | 0.453623017 | -0.326425646 | 0.974281324 | 1.31680604 | 0.31948141 | 4.138720827 | 4.68E-05 | 0.005416229 | 0.048052754 | 0.1 | 0.243021319 |
| SL1344_RS14635_csrA | 4.634736053 | 4.653539814 | 4.576555142 | -0.079677998 | 0.0894491 | -0.877387749 | 0.133910755 | 0.297405684 | 0.502714527 | 0.2841 | 0.448412565 |
| CsrB_csrB | 8.722164343 | 8.854940881 | 8.628602698 | -0.231744645 | 0.066713452 | -3.28673971 | 4.68E-05 | 0.031747426 | 0.133584268 | 0.1 | 0.243021319 |
| SL1344_RS15345_csrB | 8.544452069 | 8.655586599 | 8.395580068 | -0.259365276 | 0.079051202 | -3.249237511 | 4.68E-05 | 0.026945005 | 0.121704318 | 0.1 | 0.243021319 |
| CsrC_csrC | 8.14419993 | 8.414608123 | 7.749186883 | -0.681751551 | 0.154135474 | -4.57142706 | 4.68E-05 | 0.003282148 | 0.040548079 | 0.1 | 0.243021319 |
| SL1344_RS17435_csrD | 2.620934542 | 2.836444736 | 2.356553923 | -0.49597265 | 0.109257166 | -4.604438896 | 4.68E-05 | 0.008658774 | 0.061191649 | 0.1 | 0.243021319 |
| CyaR_cyaR | 0.317568022 | 0.211979399 | 0.417639208 | 0.20400607 | 0.157075388 | 1.260457388 | 0.091938753 | 0.15045815 | 0.294796585 | 0.2182 | 0.371500426 |
| SL1344_RS12535_cysA | 2.374848878 | 1.854442429 | 2.896759145 | 1.130127519 | 0.233502511 | 4.926656611 | 4.68E-05 | 0.003413582 | 0.038562095 | 0.1 | 0.243021319 |
| SL1344_RS08565_cysB | 1.977878246 | 2.001796741 | 1.935883946 | -0.075833602 | 0.118342464 | -0.591470261 | 0.248501012 | 0.364720949 | 0.536860506 | 0.4877 | 0.620905325 |
| SL1344_RS15175_cysC | 1.904217328 | 1.232306554 | 2.439197815 | 1.227346364 | 0.268985273 | 4.591229131 | 4.68E-05 | 0.011999921 | 0.075781897 | 0.1 | 0.243021319 |
| SL1344_RS15185_cysD | 2.363870289 | 1.988431613 | 2.879528939 | 0.894293065 | 0.26149826 | 3.580556534 | 4.68E-05 | 0.007180176 | 0.05758161 | 0.1 | 0.243021319 |
| SL1344_RS19060_cysE | 3.099494727 | 3.104951346 | 3.096006951 | -0.005677811 | 0.091552658 | -0.058067408 | 0.473017989 | 0.6680492 | 0.788669836 | 0.7704 | 0.852839817 |
| SL1344_RS17945_cysG | 2.072548686 | 1.941842239 | 2.203467499 | 0.261832971 | 0.09026446 | 2.852393165 | 0.001335878 | 0.019183315 | 0.089409639 | 0.1003 | 0.243437012 |
| SL1344_RS15240_cysH | 2.155599367 | 1.690897048 | 2.685380409 | 1.085405039 | 0.337883717 | 3.365677907 | 4.68E-05 | 0.016404544 | 0.090712329 | 0.1 | 0.243021319 |
| SL1344_RS15245_cysI | 4.056164126 | 3.347055128 | 4.686254003 | 1.395832738 | 0.204092842 | 6.906043779 | 4.68E-05 | 0.004497541 | 0.046492757 | 0.1 | 0.243021319 |
| SL1344_RS15250_cysJ | 3.961420778 | 3.292039642 | 4.518189381 | 1.276365694 | 0.1891843 | 6.836986975 | 4.68E-05 | 0.002575783 | 0.034873732 | 0.1 | 0.243021319 |
| SL1344_RS12475_cysK | 4.575137911 | 4.310434746 | 4.858385316 | 0.541618046 | 0.102763001 | 5.21872179 | 4.68E-05 | 0.002135894 | 0.031642425 | 0.1 | 0.243021319 |
| SL1344_RS12530_cysM | 1.990876789 | 1.498943585 | 2.410669098 | 0.915782863 | 0.10979517 | 8.441351807 | 4.68E-05 | 0.001230735 | 0.021409605 | 0.1 | 0.243021319 |
| SL1344_RS15180_cysN | 2.758271062 | 2.288870927 | 3.276711903 | 1.010740893 | 0.205968424 | 5.00317438 | 4.68E-05 | 0.006208818 | 0.052857387 | 0.1 | 0.243021319 |
| SL1344_RS12550_cysP | 2.768317888 | 2.270845043 | 3.145568046 | 0.896287022 | 0.168448007 | 5.369776557 | 4.68E-05 | 0.007295123 | 0.057399822 | 0.1 | 0.243021319 |
| SL1344_RS22605_cysQ | 1.606340792 | 1.760942927 | 1.561612067 | -0.164176824 | 0.209599918 | -0.626872617 | 0.306666675 | 0.49730488 | 0.665255898 | 0.6254 | 0.764568548 |
| SL1344_RS02725_cysS | 3.739413314 | 3.58932446 | 3.901736316 | 0.312483744 | 0.055374581 | 5.564684343 | 4.68E-05 | 0.003363622 | 0.036034422 | 0.1 | 0.243021319 |
| SL1344_RS09720_cysT | 5.424373154 | 5.752143842 | 5.03149464 | -0.694917981 | 0.093865116 | -7.428264967 | 4.68E-05 | 0.000892049 | 0.02083602 | 0.1 | 0.243021319 |
| SL1344_RS12545_cysU | 1.717787977 | 1.06724671 | 2.372531399 | 1.115910755 | 0.383183808 | 3.169447553 | 4.68E-05 | 0.00719197 | 0.058865818 | 0.1 | 0.243021319 |
| SL1344_RS12540_cysW | 2.395311822 | 1.74293836 | 2.967392289 | 1.231334168 | 0.124414141 | 9.766974013 | 4.68E-05 | 0.001730374 | 0.02611489 | 0.1 | 0.243021319 |
| SL1344_RS12470_cysZ | 0.692217857 | 0.655493607 | 0.817450326 | 0.16231155 | 0.333441581 | 0.455958579 | 0.314000008 | 0.473184486 | 0.650892847 | 0.6715 | 0.790161142 |
| SL1344_RS16735_cyuA | 0.626135706 | 0.579021663 | 0.675414218 | 0.09254963 | 0.143853023 | 0.608715776 | 0.246502344 | 0.358213327 | 0.515013557 | 0.4629 | 0.599452065 |
| SL1344_RS16740_cyuP | -0.46723147 | -0.511753336 | -0.396678206 | 0.07974062 | 0.288634508 | 0.2589893 | 0.390666671 | 0.635476245 | 0.762591871 | 0.7317 | 0.825437265 |
| SL1344_RS10610_dacD | 0.156671821 | -0.017527382 | 0.337809712 | 0.385896167 | 0.277314091 | 1.348004272 | 0.064623651 | 0.113908269 | 0.26759572 | 0.1747 | 0.334084627 |
| SL1344_RS17980_dam | 3.336009843 | 3.200270304 | 3.550408116 | 0.343456855 | 0.099725632 | 3.268492468 | 4.68E-05 | 0.021886538 | 0.103567822 | 0.1 | 0.243021319 |
| DapZ_dapZ | 0.62217728 | 0.855743701 | 0.496407065 | -0.324876221 | 0.336050318 | -0.992630494 | 0.139240534 | 0.192551743 | 0.376562997 | 0.2747 | 0.45549747 |
| ddl_ddl | 3.058769724 | 3.006853223 | 3.153880596 | 0.149249061 | 0.101270557 | 1.400805094 | 0.060000072 | 0.120721022 | 0.27997827 | 0.1682 | 0.326327442 |
| SL1344_RS09505_dihydroneopterin.triphosphate.diphosphatase | 3.535895544 | 3.941912922 | 2.99058105 | -0.946125408 | 0.085666409 | -10.80911478 | 4.68E-05 | 0.002213078 | 0.030709849 | 0.1 | 0.243021319 |
| dinG_dinG | 2.613601996 | 2.554191245 | 2.659037716 | 0.103160452 | 0.083886925 | 1.173805303 | 0.099267197 | 0.158712024 | 0.313983844 | 0.2357 | 0.394108201 |
| SL1344_RS05725_dinI | 2.681604247 | 2.725296022 | 2.605190046 | -0.114823632 | 0.096809496 | -1.106821827 | 0.13200003 | 0.188162525 | 0.357211133 | 0.2796 | 0.444983521 |
| SL1344_RS04685_dmsA | -1.3472268 | -1.358403952 | -1.336356286 | 0.024006964 | 0.288798756 | 0.078826909 | 0.471019322 | 0.557624691 | 0.696972702 | 0.6701 | 0.77244591 |
| SL1344_RS04690_dmsB | -3.888190765 | -3.791997726 | -4.044947268 | -0.135368121 | 1.062339776 | -0.117809338 | 0.459027317 | 0.705658697 | 0.818106354 | 0.803 | 0.881350058 |
| SL1344_RS04695_dmsC | -2.92063189 | -2.839085004 | -3.052584516 | -0.246718811 | 0.584324616 | -0.383081524 | 0.334666674 | 0.498395993 | 0.648877717 | 0.6092 | 0.725552206 |
| SL1344_RS06610_DNA.RNA.non.specific.endonuclease | 0.319024425 | 0.257064738 | 0.432391359 | 0.180708081 | 0.214358556 | 0.800863477 | 0.196666684 | 0.312390668 | 0.486168369 | 0.42 | 0.568109472 |

| | | | | | | | | | | | |
|---|---|---|---|---|---|---|---|---|---|---|---|
| dnaA_dnaA | 3.648888472 | 3.641643793 | 3.658035009 | 0.019084922 | 0.063611066 | 0.283397616 | 0.382666671 | 0.556643826 | 0.696620885 | 0.675 | 0.77544766 |
| SL1344_RS00060_dnaK | 7.45607739 | 7.600068974 | 7.438320837 | -0.16028949 | 0.114682686 | -0.734192968 | 0.33333334 | 0.397542445 | 0.598169944 | 0.7 | 0.829598552 |
| dpiA_dpiA | -1.551764031 | -1.596138913 | -1.506244586 | 0.08477088 | 0.311304625 | 0.239640991 | 0.380666671 | 0.496047781 | 0.642024721 | 0.6063 | 0.719191507 |
| dpiA_dpiA.1 | -1.364911569 | -1.396604276 | -1.317990301 | 0.080243345 | 0.314489216 | 0.24231787 | 0.393737512 | 0.526568408 | 0.670812277 | 0.6344 | 0.744683849 |
| dpiB_dpiB | -1.392171592 | -1.310387522 | -1.487506002 | -0.18241095 | 0.297538504 | -0.574517102 | 0.259333345 | 0.388217845 | 0.544342365 | 0.4909 | 0.624533527 |
| DsrA_dsrA | -4.426122268 | -4.711751866 | -3.794061522 | 0.968660896 | 1.296410234 | 0.695727499 | 0.238507675 | 0.368111791 | 0.542215283 | 0.4917 | 0.63527515 |
| SL1344_RS09330_DUF1441.domain.containing.protein | -4.381640087 | -3.733663915 | -5.410315802 | -1.680073429 | 0.921547088 | -1.70741769 | 0.022651765 | 0.071131977 | 0.192098261 | 0.1247 | 0.273579664 |
| SL1344_RS06460_DUF1496.domain.containing.protein | 2.338065514 | 3.029339879 | 1.607537276 | -1.422534346 | 0.192821339 | -7.493069104 | 4.68E-05 | 0.002111022 | 0.029723294 | 0.1 | 0.243021319 |
| SL1344_RS16295_DUF2623.domain.containing.protein | 1.760898319 | 2.544147984 | 1.132006326 | -1.362099174 | 0.215723239 | -6.205765485 | 4.68E-05 | 0.00179716 | 0.028346021 | 0.1 | 0.243021319 |
| emrB_emrB | 2.36085831 | 2.245237981 | 2.491461758 | 0.306070678 | 0.189160828 | 1.64197069 | 0.004664544 | 0.064039152 | 0.194736029 | 0.1069 | 0.25218677 |
| emrD_emrD | 1.979738539 | 2.278284975 | 1.761625496 | -0.530493076 | 0.218288178 | -2.518438804 | 4.68E-05 | 0.023387798 | 0.107754648 | 0.1 | 0.243021319 |
| emtA_emtA | 1.197378126 | 1.32795096 | 0.949606302 | -0.371294236 | 0.17130213 | -2.112656155 | 0.009327606 | 0.053863396 | 0.168640463 | 0.1084 | 0.253989452 |
| SL1344_RS03015_entB | 0.787346074 | 0.744213468 | 0.831035007 | 0.040336248 | 0.278409616 | 0.14011597 | 0.443037977 | 0.730112052 | 0.836122945 | 0.8332 | 0.896845594 |
| entE_entE | 1.380111903 | 1.295105056 | 1.46462423 | 0.17462245 | 0.160088475 | 1.064261403 | 0.11133337 | 0.183114536 | 0.349292482 | 0.2658 | 0.426710986 |
| entF_entF | 1.987373964 | 1.945462944 | 2.048216043 | 0.154346012 | 0.219490137 | 0.69807415 | 0.218000015 | 0.296906358 | 0.49075159 | 0.4353 | 0.596162361 |
| SL1344_RS18065_envZ | 2.742381415 | 2.499590067 | 2.907893242 | 0.406455676 | 0.084500494 | 4.665293549 | 4.68E-05 | 0.008092854 | 0.056368445 | 0.1 | 0.243021319 |
| eutA_eutA | -2.891072726 | -2.942049642 | -2.863910886 | 0.068635282 | 0.585643678 | 0.09371855 | 0.456362427 | 0.604148199 | 0.736168832 | 0.7166 | 0.811173504 |
| SL1344_RS12680_eutS | -6.349635673 | -6.074158484 | -6.766916917 | -0.759608726 | 2.029220338 | -0.302812195 | 0.328000007 | 0.45086177 | 0.617584014 | 0.5845 | 0.702293754 |
| fabG_fabG | 4.635559009 | 4.647994935 | 4.5701235 | -0.079395909 | 0.108775766 | -0.709009249 | 0.271152576 | 0.346737138 | 0.546790256 | 0.5496 | 0.700826191 |
| fadA_fadA | -1.627103653 | -1.410618485 | -1.807300826 | -0.413437452 | 0.358822193 | -1.100377511 | 0.121918754 | 0.186991629 | 0.346437404 | 0.2661 | 0.422452846 |
| SL1344_RS09080_fadD | 2.560984218 | 2.871594338 | 2.065660179 | -0.808582246 | 0.196260775 | -4.190220812 | 4.68E-05 | 0.018412901 | 0.096134061 | 0.1 | 0.243021319 |
| fadE_fadE | 0.779466531 | 0.915846922 | 0.607985609 | -0.310045844 | 0.147156192 | -2.041822254 | 0.013990999 | 0.047772916 | 0.151316826 | 0.1154 | 0.262168102 |
| fadI_fadI | -2.606412577 | -2.680709757 | -2.518866695 | 0.155157587 | 0.451808556 | 0.327267034 | 0.352431718 | 0.494962322 | 0.640097239 | 0.5963 | 0.713395409 |
| SL1344_RS13945_Fels.2.prophage.protein | -3.891311935 | -2.838848431 | -4.151806461 | -1.31435387 | 1.473440154 | -0.727941335 | 0.262000011 | 0.390508332 | 0.577872018 | 0.5407 | 0.694231646 |
| feoA_feoA | 1.125978025 | 0.691506157 | 1.693554211 | 0.985219589 | 0.174875986 | 5.597389103 | 4.68E-05 | 0.003802179 | 0.039520722 | 0.1 | 0.243021319 |
| ffs_ffs | 3.350876745 | 3.129558585 | 3.501221104 | 0.443474931 | 0.287293784 | 1.544253821 | 0.007329074 | 0.067944984 | 0.20710276 | 0.1068 | 0.252557941 |
| fimA_fimA | -1.888289796 | -1.951555786 | -1.760704782 | 0.175403124 | 0.421813716 | 0.372769285 | 0.34733334 | 0.536666542 | 0.68063239 | 0.6457 | 0.757815119 |
| fimF_fimF | -1.970317851 | -2.009043752 | -1.921158422 | 0.109721786 | 0.447024476 | 0.23040352 | 0.396000004 | 0.58840225 | 0.722096771 | 0.706 | 0.799080659 |
| SL1344_RS02785_fimZ | -2.713987814 | -2.20607604 | -3.101793879 | -0.815343153 | 1.010860772 | -0.747175077 | 0.216666682 | 0.328460283 | 0.518543066 | 0.4302 | 0.594904851 |
| fis_fis | 4.339702889 | 4.218573608 | 4.560408604 | 0.337009711 | 0.176161797 | 1.704766497 | 0.006667356 | 0.085712568 | 0.238220048 | 0.1053 | 0.25045222 |
| fixB_fixB | -0.766072791 | -0.910719971 | -0.526078016 | 0.385307419 | 0.393329918 | 0.920474008 | 0.160000023 | 0.230440857 | 0.410767699 | 0.3191 | 0.490311209 |
| flgD_flgD | 4.188347548 | 4.731515601 | 3.627937578 | -1.119249887 | 0.121873544 | -9.311773336 | 4.68E-05 | 0.001496583 | 0.026210129 | 0.1 | 0.243021319 |
| flgF_flgF | 4.59508316 | 5.169695436 | 4.105600206 | -1.053106889 | 0.067903461 | -14.99793395 | 4.68E-05 | 0.000993561 | 0.022081965 | 0.1 | 0.243021319 |
| flgI_flgI | 3.458947489 | 3.858084948 | 2.951592712 | -0.890748704 | 0.200503187 | -4.380729816 | 4.68E-05 | 0.007732166 | 0.06050503 | 0.1 | 0.243021319 |
| SL1344_RS05830_flgK | 5.48249124 | 6.237171908 | 4.696100979 | -1.520414367 | 0.069572651 | -21.95421887 | 4.68E-05 | 2.28E-05 | 0.004533891 | 0.1 | 0.243021319 |
| flgL_flgL | 5.222227736 | 5.926007174 | 4.523908198 | -1.400794137 | 0.053833249 | -26.19795106 | 4.68E-05 | 0.000109265 | 0.007909151 | 0.1 | 0.243021319 |
| SL1344_RS05775_flgM | 4.062044313 | 4.764763807 | 3.37721498 | -1.396879636 | 0.140937201 | -10.02597513 | 4.68E-05 | 0.001471838 | 0.026335032 | 0.1 | 0.243021319 |
| SL1344_RS05770_flgN | 4.588268723 | 5.260829108 | 3.910110795 | -1.341469121 | 0.138971468 | -9.61821181 | 4.68E-05 | 0.000271905 | 0.012752666 | 0.1 | 0.243021319 |
| SL1344_RS09570_flhA | 4.396241638 | 5.298935349 | 3.523909141 | -1.750764731 | 0.148069545 | -11.65176313 | 4.68E-05 | 0.000428853 | 0.015431941 | 0.1 | 0.243021319 |
| SL1344_RS09575_flhB | 4.011615192 | 5.013690734 | 3.074764659 | -1.917285917 | 0.115611269 | -16.39679935 | 4.68E-05 | 0.000226597 | 0.010977844 | 0.1 | 0.243021319 |
| SL1344_RS09625_flhC | 3.984253677 | 4.711947848 | 3.110802381 | -1.627266893 | 0.132592009 | -12.20765094 | 4.68E-05 | 0.000266321 | 0.011773641 | 0.1 | 0.243021319 |
| SL1344_RS09630_flhD | 2.380496477 | 2.724765493 | 1.929330577 | -0.794384852 | 0.108692898 | -7.307482247 | 4.68E-05 | 0.004029365 | 0.039495802 | 0.1 | 0.243021319 |
| SL1344_RS09565_flhE | 2.301557338 | 3.103185093 | 1.453344377 | -1.58235673 | 0.196109509 | -8.013426434 | 4.68E-05 | 0.000743449 | 0.018292334 | 0.1 | 0.243021319 |
| SL1344_RS09785_fliA | 5.545836614 | 6.482516976 | 4.631671583 | -1.910807628 | 0.170556176 | -11.34425348 | 4.68E-05 | 0.0003298 | 0.014360643 | 0.1 | 0.243021319 |
| SL1344_RS09800_fliC | 7.490694266 | 9.689576672 | 6.104445851 | -3.583802955 | 0.066313402 | -47.55187159 | 4.68E-05 | 0.007555559 | 0.0617673 | 0.1 | 0.243021319 |
| SL1344_RS09845_fliE | 2.080991186 | 2.511346325 | 1.744827998 | -0.799126926 | 0.239055849 | -3.321751411 | 4.68E-05 | 0.012228363 | 0.076683883 | 0.1 | 0.243021319 |
| fliI_fliI | 3.861285553 | 4.132682645 | 3.690467026 | -0.404450467 | 0.125678729 | -3.094077594 | 4.68E-05 | 0.015822922 | 0.087855004 | 0.1 | 0.243021319 |
| fliJ_fliJ | 2.609177995 | 2.891998106 | 2.310813565 | -0.596899928 | 0.135832418 | -4.370007871 | 4.68E-05 | 0.005179351 | 0.047422276 | 0.1 | 0.243021319 |
| fliL_fliL | 3.496150376 | 3.906478883 | 3.198315502 | -0.707594316 | 0.131425194 | -5.434630496 | 4.68E-05 | 0.008598774 | 0.06387548 | 0.1 | 0.243021319 |
| fliQ_fliQ | 1.94545653 | 2.157179983 | 1.724972786 | -0.414199311 | 0.185880562 | -2.23820499 | 0.004664544 | 0.031110967 | 0.127055449 | 0.1023 | 0.246246308 |
| SL1344_RS09815_fliT | 4.75548426 | 5.189612209 | 4.330260667 | -0.870651018 | 0.091897305 | -9.566668058 | 4.68E-05 | 0.002174887 | 0.03178968 | 0.1 | 0.243021319 |
| SL1344_RS09775_fliY | 3.519804505 | 4.423625937 | 2.605492461 | -1.8120605 | 0.091714775 | -19.84384309 | 4.68E-05 | 0.000100675 | 0.007299907 | 0.1 | 0.243021319 |
| SL1344_RS09780_fliZ | 4.374024066 | 5.365909006 | 3.275929827 | -2.10218249 | 0.082471561 | -25.50883399 | 4.68E-05 | 2.82E-05 | 0.004634667 | 0.1 | 0.243021319 |

| | | | | | | | | | | | |
|---|---|---|---|---|---|---|---|---|---|---|---|
| SL1344_RS14340_fljA | 3.147666376 | -0.990673958 | 3.205012293 | 4.182871554 | 0.585608155 | 1.024932835 | 0.154563649 | 0.184051199 | 0.379810022 | 0.2846 | 0.465746146 |
| flk_flk | 1.084715624 | 0.809162428 | 1.35373462 | 0.567096299 | 0.174819949 | 3.244039655 | 4.68E-05 | 0.014663847 | 0.080244225 | 0.1 | 0.243021319 |
| SL1344_RS08285_fnr | 4.223920212 | 4.345544631 | 4.114424909 | -0.288474706 | 0.120556159 | -2.518089422 | 4.68E-05 | 0.01762485 | 0.093950775 | 0.1 | 0.243021319 |
| FnrS_fnrS | -1.417457161 | -1.301961309 | -1.558471075 | -0.279044016 | 0.422009296 | -0.634418697 | 0.226000015 | 0.339916372 | 0.517688203 | 0.4416 | 0.588511489 |
| SL1344_RS04735_focA | 1.905226926 | 1.488224926 | 2.315865098 | 0.828295815 | 0.162920238 | 5.02361683 | 4.68E-05 | 0.003823458 | 0.040300441 | 0.1 | 0.243021319 |
| folA_folA | 1.64133848 | 1.442738651 | 1.849198284 | 0.429557416 | 0.159962011 | 2.730412819 | 4.68E-05 | 0.026594996 | 0.114356098 | 0.1 | 0.243021319 |
| frsA_frsA | 2.302483137 | 2.200461033 | 2.408967021 | 0.208521495 | 0.080480833 | 2.505390488 | 0.005330642 | 0.024840249 | 0.103128247 | 0.103 | 0.247020634 |
| SL1344_RS09665_ftnB | -1.605445912 | -1.045997887 | -2.488545934 | -1.443731839 | 0.449817361 | -3.146626455 | 4.68E-05 | 0.025839166 | 0.10750159 | 0.1 | 0.243021319 |
| SL1344_RS06895_fumD | -0.075055436 | -0.201237556 | 0.037225053 | 0.250205876 | 0.242403759 | 0.989615735 | 0.126000031 | 0.204224718 | 0.372719387 | 0.2868 | 0.443654221 |
| SL1344_RS03500_fur | 4.710030419 | 4.740073362 | 4.65141692 | -0.095486591 | 0.124173353 | -0.810075109 | 0.172551653 | 0.253237447 | 0.452347119 | 0.338 | 0.523156463 |
| gapA_gapA | 3.135405979 | 3.073331763 | 3.359677017 | 0.279656657 | 0.19264059 | 1.510805695 | 0.039307241 | 0.071817369 | 0.210682177 | 0.1399 | 0.294239524 |
| GcvB_gcvB | 7.462615304 | 7.171518261 | 7.629663212 | 0.445384415 | 0.051020993 | 7.450867497 | 4.68E-05 | 0.012474594 | 0.07962287 | 0.1 | 0.243021319 |
| gcvT_gcvT | 2.540364004 | 2.981206738 | 2.07701553 | -0.903675882 | 0.113497875 | -7.940305465 | 4.68E-05 | 0.001166367 | 0.021407831 | 0.1 | 0.243021319 |
| gidB_gidB | 1.665818231 | 1.485314696 | 1.900434713 | 0.414727673 | 0.114450305 | 3.591275958 | 4.68E-05 | 0.011038117 | 0.066554591 | 0.1001 | 0.243157234 |
| gldA_gldA | -0.346518274 | -0.294299227 | -0.408128039 | -0.125404941 | 0.251442893 | -0.454941025 | 0.308000008 | 0.46819397 | 0.621625499 | 0.5824 | 0.702623653 |
| gldA_gldA.1 | 1.171575127 | 1.195605905 | 1.130606484 | -0.101123372 | 0.228123553 | -0.40627876 | 0.326666674 | 0.518772221 | 0.677983397 | 0.6593 | 0.772435697 |
| GlmY_glmY | 3.151337348 | 2.6999528 | 3.463654215 | 0.763861198 | 0.134616176 | 5.534388322 | 4.68E-05 | 0.009587502 | 0.06716527 | 0.1 | 0.243021319 |
| GlmZ_glmZ | 5.308449701 | 5.282722384 | 5.448488466 | 0.165718888 | 0.140440539 | 1.038098755 | 0.166666689 | 0.200220705 | 0.397329826 | 0.3096 | 0.499933722 |
| glnQ_glnQ | 3.619675013 | 3.392823245 | 3.830339714 | 0.449680219 | 0.080023865 | 5.711843326 | 4.68E-05 | 0.003029456 | 0.035531519 | 0.1 | 0.243021319 |
| SL1344_RS13690_gluconolactonase | -0.594518061 | -1.904568727 | 0.240026745 | 2.123237157 | 0.474360342 | 4.465993536 | 4.68E-05 | 0.015195225 | 0.083755543 | 0.1 | 0.243021319 |
| SL1344_RS09560_glycoside.hydrolase.family.105.protein | 1.706991799 | 2.397269187 | 0.908972887 | -1.491281532 | 0.143101581 | -10.47049142 | 4.68E-05 | 0.000477279 | 0.014373563 | 0.1 | 0.243021319 |
| gntK_gntK | -0.999353424 | -1.113585966 | -0.896142463 | 0.218713075 | 0.301969009 | 0.675088092 | 0.23466668 | 0.358956433 | 0.52160335 | 0.4584 | 0.600520175 |
| SL1344_RS18265_gntR | 1.949042238 | 1.887384408 | 1.986336451 | 0.10491758 | 0.126233341 | 0.786023248 | 0.199333351 | 0.296689173 | 0.471089913 | 0.3989 | 0.545654032 |
| gpmA_gpmA | 5.642131771 | 5.644006769 | 5.639122926 | -0.001159005 | 0.138188653 | -0.010214192 | 0.495333334 | 0.896463347 | 0.943962908 | 0.9514 | 0.971789517 |
| GraL_graL | 2.842430244 | 2.82102837 | 3.093488157 | 0.266368573 | 0.162305201 | 0.646544079 | 0.332666674 | 0.486555279 | 0.668867983 | 0.697 | 0.82727603 |
| groES_groES | 6.025948585 | 6.127856586 | 6.009726569 | -0.12081589 | 0.1051626 | -1.137010492 | 0.11133337 | 0.179197291 | 0.371750094 | 0.2366 | 0.411407804 |
| SL1344_RS14595_gshA | 3.44494442 | 3.408314707 | 3.53488908 | 0.117460505 | 0.075917943 | 1.41253958 | 0.074000058 | 0.124693837 | 0.281762235 | 0.2036 | 0.36515667 |
| SL1344_RS16010_gshB | 2.900526881 | 2.864372044 | 2.930114981 | 0.067454175 | 0.071851735 | 0.896559848 | 0.159893428 | 0.23960233 | 0.403327941 | 0.3299 | 0.481182172 |
| SL1344_RS05170_gtgD | -3.090346874 | -2.51050779 | -3.51668178 | -0.99148724 | 0.846257121 | -1.11664737 | 0.105333372 | 0.164790157 | 0.333945938 | 0.2321 | 0.399769989 |
| SL1344_RS05180_gtgE | 0.251721386 | 0.419450101 | -0.087386481 | -0.512043253 | 0.290649575 | -1.685363487 | 0.028000161 | 0.073245988 | 0.208014962 | 0.1295 | 0.281089504 |
| SL1344_RS05185_gtgF | -4.573772526 | -4.67133256 | -4.503753286 | 0.157002144 | 1.00344907 | 0.144342927 | 0.442000002 | 0.571809209 | 0.710257088 | 0.6871 | 0.786216678 |
| SL1344_RS11665_gyrA | 5.920760982 | 5.790961144 | 6.014924637 | 0.231546235 | 0.073303564 | 3.204576163 | 4.68E-05 | 0.008005341 | 0.061822444 | 0.1 | 0.243021319 |
| SL1344_RS13995_head.completion.stabilization.protein | -4.927840793 | -5.961878472 | -3.587088934 | 2.366015824 | 2.375786142 | 0.943612365 | 0.159333357 | 0.222805676 | 0.410714784 | 0.3031 | 0.481529961 |
| helD_helD | 1.793275209 | 1.822399829 | 1.733322978 | -0.080391187 | 0.122284924 | -0.595245451 | 0.273333344 | 0.424207083 | 0.585548215 | 0.531 | 0.668739265 |
| hemG_hemG | 1.053940299 | 1.132329045 | 0.976551275 | -0.155381576 | 0.116653986 | -1.283612782 | 0.079280533 | 0.142217409 | 0.282529052 | 0.2165 | 0.368471076 |
| hflB_hflB | 6.402006087 | 6.363629663 | 6.454803406 | 0.093597823 | 0.046498279 | 2.018226342 | 0.00067292 | 0.065918828 | 0.200464392 | 0.1002 | 0.243304955 |
| SL1344_RS22390_hfq | 4.987915366 | 4.977564388 | 5.004772222 | 0.025526089 | 0.121363075 | 0.238118463 | 0.354430386 | 0.739058328 | 0.845908185 | 0.7159 | 0.822916833 |
| SL1344_RS02410_hha | 2.228535687 | 1.932907898 | 2.579350378 | 0.640498108 | 0.120629573 | 5.309720561 | 4.68E-05 | 0.002982697 | 0.035021198 | 0.1 | 0.243021319 |
| SL1344_RS02415_Hha.toxicity.attenuator | 2.16294156 | 1.860618058 | 2.682776365 | 0.815178571 | 0.194026539 | 3.897867277 | 4.68E-05 | 0.018157876 | 0.095343769 | 0.1 | 0.243021319 |
| SL1344_RS14890_hilA | 0.46216544 | 1.844773165 | -1.068878221 | -2.94633598 | 0.320253329 | -9.255629251 | 4.68E-05 | 0.000927276 | 0.020626093 | 0.1 | 0.243021319 |
| SL1344_RS14845_hilC | 2.001293974 | 2.758533573 | 1.22961975 | -1.534440364 | 0.213840994 | -7.266082718 | 4.68E-05 | 0.003249642 | 0.038548633 | 0.1 | 0.243021319 |
| SL1344_RS14885_hilD | 2.377187541 | 2.917304772 | 1.765622162 | -1.221617672 | 0.210001924 | -5.902644703 | 4.68E-05 | 0.001153433 | 0.023782471 | 0.1 | 0.243021319 |
| SL1344_RS23135_hilE | -1.26335931 | -1.218387259 | -1.31030363 | -0.092153903 | 0.274091303 | -0.311298187 | 0.362000006 | 0.475029446 | 0.624207123 | 0.5775 | 0.697152386 |
| SL1344_RS21865_HlyD.family.type.I.secretion.periplasmic.adaptor.subunit | -0.983469416 | -0.056688377 | -2.054224364 | -1.9778188 | 0.358244068 | -5.521630883 | 4.68E-05 | 0.002793396 | 0.033906772 | 0.1 | 0.243021319 |
| SL1344_RS08745_hns | 6.959490777 | 6.968185268 | 6.949507425 | 0.043001775 | 0.101526875 | 0.377630873 | 0.438666669 | 0.500449466 | 0.680762075 | 0.9685 | 0.981068718 |
| hscB_hscB | 2.647570669 | 2.530673937 | 2.737342295 | 0.211028937 | 0.098437291 | 2.113212608 | 0.005996769 | 0.048877323 | 0.159818857 | 0.1085 | 0.253986835 |
| SL1344_RS08210_hslJ | 0.880074109 | 0.910487585 | 0.855930331 | -0.053342057 | 0.15118374 | -0.337163827 | 0.356429053 | 0.497517714 | 0.646442971 | 0.6096 | 0.725580842 |
| SL1344_RS21405_hupA | 7.062685431 | 7.07149906 | 7.028943964 | -0.035735358 | 0.087914649 | -0.305499961 | 0.336442379 | 0.882073261 | 0.936151033 | 0.712 | 0.8364456 |
| SL1344_RS21410_hupB | 4.306937313 | 4.279613937 | 4.469483431 | 0.177237242 | 0.093973094 | 1.200460564 | 0.127914754 | 0.159371953 | 0.343048072 | 0.2678 | 0.44319325 |
| hycD_hycD | -2.764111108 | -2.678993192 | -2.974285561 | -0.300493317 | 0.917706059 | -0.312286811 | 0.365089945 | 0.567448205 | 0.718904682 | 0.731 | 0.823890131 |
| hycE_hycE | -1.474055622 | -1.551930832 | -1.387112826 | 0.139736779 | 0.367981478 | 0.370459004 | 0.338441046 | 0.509898409 | 0.656600672 | 0.6145 | 0.730376926 |
| SL1344_RS06065_hypothetical.protein | 0.844358877 | 1.625823591 | -0.048497354 | -1.672955444 | 0.238843919 | -7.081164764 | 4.68E-05 | 0.002938375 | 0.033251317 | 0.1 | 0.243021319 |
| SL1344_RS07940_hypothetical.protein | 3.551981359 | 4.268887735 | 2.844849036 | -1.432641682 | 0.146668364 | -9.811859936 | 4.68E-05 | 0.001896968 | 0.030125021 | 0.1 | 0.243021319 |

| | | | | | | | | | | | |
|---|---|---|---|---|---|---|---|---|---|---|---|
| SL1344_RS09195_hypothetical.protein | -1.094272672 | -0.143684507 | -2.119651457 | -1.959326192 | 0.428207162 | -4.589435772 | 4.68E-05 | 0.004349267 | 0.042394955 | 0.1 | 0.243021319 |
| SL1344_RS09675_hypothetical.protein | 2.22939568 | 3.414896258 | 0.914725207 | -2.535839001 | 0.205338606 | -12.48046964 | 4.68E-05 | 0.000361019 | 0.013495271 | 0.1 | 0.243021319 |
| SL1344_RS09745_hypothetical.protein | 2.268070837 | 2.558971168 | 1.923815754 | -0.730340641 | 0.228230339 | -3.340146053 | 4.68E-05 | 0.008260527 | 0.061433139 | 0.1 | 0.243021319 |
| SL1344_RS22135_hypothetical.protein | -2.278463207 | -1.399696978 | -3.609969862 | -2.277543122 | 0.635364332 | -3.576322579 | 4.68E-05 | 0.017072513 | 0.083500441 | 0.1 | 0.243021319 |
| iacP_iacP | 0.30301757 | 0.490664581 | 0.126704095 | -0.360412978 | 0.219632955 | -1.60940732 | 0.043304566 | 0.081527489 | 0.215080137 | 0.146 | 0.301391375 |
| SL1344_RS14895_iagB | 0.490593634 | 1.358890311 | -0.795690406 | -2.218400274 | 0.332700585 | -6.737920779 | 4.68E-05 | 0.001435808 | 0.024330525 | 0.1 | 0.243021319 |
| lamB_iamB | -1.456848225 | -1.286931236 | -1.664325144 | -0.377692014 | 0.295821026 | -1.190872317 | 0.09726853 | 0.155608477 | 0.303633849 | 0.228 | 0.38190972 |
| idnK_idnK | -2.249588037 | -2.134341149 | -2.36050403 | -0.213708284 | 0.46057775 | -0.447478278 | 0.315123259 | 0.459529115 | 0.613555658 | 0.5786 | 0.69803771 |
| SL1344_RS22990_idnT | -2.234247079 | -2.215172593 | -2.268403629 | 0.008151532 | 0.586635429 | 0.011944941 | 0.495669554 | 0.68308566 | 0.799817114 | 0.7907 | 0.868891148 |
| SL1344_RS06655_ihfA | 4.844557468 | 4.914060905 | 4.759995301 | -0.154833441 | 0.038653881 | -3.999862429 | 4.68E-05 | 0.009076171 | 0.059906001 | 0.1 | 0.243021319 |
| SL1344_RS04775_ihfB | 3.682993304 | 3.740485188 | 3.629052826 | -0.113925815 | 0.074376748 | -1.511638153 | 0.036666788 | 0.088695453 | 0.228213292 | 0.1428 | 0.297327544 |
| lldD_ildD | -0.137678638 | 0.06726181 | -0.326642231 | -0.39887106 | 0.218966549 | -1.768115699 | 0.01866691 | 0.062971066 | 0.180379182 | 0.1247 | 0.274512213 |
| ilvM_ilvM | -0.049992384 | 0.057352121 | -0.274627348 | -0.297920508 | 0.271991312 | -1.042665927 | 0.141905424 | 0.210746497 | 0.386252218 | 0.3034 | 0.470171346 |
| SL1344_RS14990_invA | 0.682471625 | 2.080826505 | -0.936556786 | -3.018579431 | 0.204182441 | -14.85897979 | 4.68E-05 | 0.000234203 | 0.010021995 | 0.1 | 0.243021319 |
| SL1344_RS14980_invC | -0.126215919 | 1.025649803 | -1.436247212 | -2.442809027 | 0.249646491 | -9.791595798 | 4.68E-05 | 0.000815466 | 0.017076322 | 0.1 | 0.243021319 |
| SL1344_RS14995_invE | 0.132476992 | 1.56917593 | -1.92046863 | -3.502716262 | 0.304574765 | -11.45221763 | 4.68E-05 | 0.001305495 | 0.021052468 | 0.1 | 0.243021319 |
| SL1344_RS15005_invF | 0.526054695 | 2.230655994 | -1.758653107 | -4.028525175 | 0.394680239 | -10.26070057 | 4.68E-05 | 0.004065763 | 0.041715541 | 0.1 | 0.243021319 |
| SL1344_RS15000_invG | 0.481355901 | 2.254505709 | -1.625204761 | -3.874918355 | 0.234489853 | -16.4774541 | 4.68E-05 | 0.000416905 | 0.012041437 | 0.1 | 0.243021319 |
| InvR_invR | 2.491244036 | 3.254172862 | 1.557807708 | -1.707186946 | 0.220992542 | -7.970731659 | 4.68E-05 | 0.004999603 | 0.048147822 | 0.1 | 0.243021319 |
| SL1344_RS14355_iroB | -0.683541091 | 0.325388138 | -1.548743051 | -1.807573259 | 0.389868399 | -4.567343227 | 4.68E-05 | 0.004825612 | 0.045446087 | 0.1 | 0.243021319 |
| SL1344_RS24725_IS91.family.transposase | 0.791772113 | 1.694577913 | -0.418879083 | -2.142373223 | 0.239066419 | -9.026182196 | 4.68E-05 | 0.000603919 | 0.016423523 | 0.1 | 0.243021319 |
| IsrA_isrA | -4.953580661 | -5.732263456 | -4.501868808 | 1.209370376 | 1.434731123 | 0.775237563 | 0.172000021 | 0.26124276 | 0.439790025 | 0.3454 | 0.499424656 |
| IsrB_isrB | -2.388008966 | -2.771530479 | -2.019899283 | 0.746345398 | 0.415156579 | 1.725526587 | 0.031333476 | 0.071651794 | 0.188842286 | 0.1396 | 0.289586912 |
| IsrC_isrC | -2.056862534 | -2.076860994 | -2.031991097 | 0.03333814 | 0.413482432 | 0.080744341 | 0.464666668 | 0.588955251 | 0.723044318 | 0.6921 | 0.790936947 |
| IsrD_isrD | -9.086921354 | -9.633147224 | -8.521899827 | 0.979153087 | 4.013947396 | 0.223638074 | 0.390000004 | 0.501827828 | 0.64934662 | 0.6189 | 0.730247305 |
| IsrE_isrE | -0.173774023 | -0.751933646 | 0.285604687 | 1.039982266 | 0.196107162 | 5.201415676 | 4.68E-05 | 0.004079637 | 0.038049167 | 0.1 | 0.243021319 |
| IsrF_isrF | -0.976304385 | -0.601572987 | -1.452732697 | -0.860661871 | 0.304637141 | -2.752679657 | 0.00067292 | 0.021339286 | 0.096797503 | 0.1009 | 0.244281757 |
| IsrG_isrG | -1.83043039 | -1.730877265 | -1.924353092 | -0.182150162 | 0.438557266 | -0.386279825 | 0.339107268 | 0.509759065 | 0.659753235 | 0.6224 | 0.738593652 |
| IsrH.1_isrH | -4.394213021 | -3.744679169 | -4.717608948 | -0.884665786 | 1.088927693 | -0.720780148 | 0.22918056 | 0.373068716 | 0.536298782 | 0.465 | 0.613615133 |
| IsrH.2_isrH2 | -4.405144797 | -3.744249111 | -4.714444512 | -0.862546798 | 1.071192406 | -0.712171557 | 0.239173897 | 0.392535838 | 0.556192584 | 0.4872 | 0.635019659 |
| IsrI_isrI | -2.574121212 | -2.60351219 | -2.542692404 | 0.05233741 | 0.550090968 | 0.099908884 | 0.453031314 | 0.616981017 | 0.746420952 | 0.7177 | 0.812143847 |
| IsrJ_isrJ | -0.379050518 | -0.443179521 | -0.287536605 | 0.143075171 | 0.234688906 | 0.575060237 | 0.264000011 | 0.404316606 | 0.563874338 | 0.5091 | 0.644334104 |
| IsrK_isrK | 0.89480888 | 0.783756739 | 1.037346269 | 0.261286622 | 0.155302934 | 1.615325442 | 0.031333476 | 0.083079015 | 0.213537964 | 0.1381 | 0.288088038 |
| IsrL_isrL | -2.040740341 | -2.034676318 | -2.052156096 | 0.002148506 | 0.43483937 | 0.003117863 | 0.498334444 | 0.620235205 | 0.751206913 | 0.742 | 0.83192029 |
| IsrM_isrM | -4.279429233 | -4.576903412 | -4.020308754 | 0.566922604 | 0.972366373 | 0.55309847 | 0.272485021 | 0.413499548 | 0.574880223 | 0.5216 | 0.655920941 |
| IsrN_isrN | -7.275331547 | -6.896878563 | -8.000164462 | -1.126462272 | 2.965621206 | -0.318505685 | 0.344437048 | 0.471612497 | 0.63373002 | 0.6089 | 0.729125482 |
| IsrO_isrO | -0.64555733 | -0.649293671 | -0.64357883 | -0.001590136 | 0.324948178 | -0.003780137 | 0.497668221 | 0.66839104 | 0.786777829 | 0.7712 | 0.851703429 |
| IsrP_isrP | -1.57394056 | -1.517352931 | -1.655145992 | -0.135902808 | 0.357225827 | -0.364883178 | 0.348666673 | 0.489391227 | 0.639101371 | 0.6035 | 0.720519959 |
| IsrQ_isrQ | -4.939757476 | -4.947990011 | -4.927847318 | -0.010870438 | 1.200013063 | -0.009918897 | 0.494 | 0.60845074 | 0.741698474 | 0.7222 | 0.815963449 |
| IstR.1_istR1 | -1.696591602 | -1.980170249 | -1.257188533 | 0.706860132 | 0.38937272 | 1.717979479 | 0.03530992 | 0.076084305 | 0.204283233 | 0.1426 | 0.2950307 |
| IstR.2_istR2 | -1.494495001 | -1.785799812 | -1.133611692 | 0.643861811 | 0.36217218 | 1.665792549 | 0.029980163 | 0.075251678 | 0.202725244 | 0.1385 | 0.291179224 |
| ksgA_ksgA | 3.125013605 | 2.999470321 | 3.228902911 | 0.233170912 | 0.102895605 | 2.248773992 | 0.002668394 | 0.031231851 | 0.125780525 | 0.1019 | 0.245554858 |
| SL1344_RS08205_ldhA | 2.05522387 | 2.337243125 | 1.750187787 | -0.63280124 | 0.173588597 | -3.745868929 | 4.68E-05 | 0.008666814 | 0.061107289 | 0.1 | 0.243021319 |
| SL1344_RS00585_leuO | -3.435367624 | -3.575206156 | -3.276713621 | 0.474118352 | 0.963720171 | 0.485874301 | 0.306000008 | 0.466911825 | 0.627999076 | 0.6188 | 0.730572986 |
| SL1344_RS21750_lexA | 4.210338266 | 4.191835338 | 4.23005645 | 0.022058671 | 0.12403255 | 0.190497305 | 0.398000004 | 0.734246394 | 0.840793358 | 0.8212 | 0.888978148 |
| ligA_ligA | 3.62289373 | 3.374591988 | 3.830860957 | 0.453369396 | 0.082195504 | 5.473118047 | 4.68E-05 | 0.003735567 | 0.038916736 | 0.1 | 0.243021319 |
| livF_livF | -0.970161396 | -0.721054838 | -1.221668523 | -0.482707027 | 0.326111033 | -1.443799616 | 0.049966777 | 0.098892886 | 0.239839567 | 0.1591 | 0.316139601 |
| lolA_lolA | 2.255258418 | 2.310460799 | 2.208624256 | -0.119273922 | 0.164500713 | -0.721195087 | 0.209193887 | 0.30257313 | 0.492928085 | 0.405 | 0.565658397 |
| lolB_lolB | 0.690919312 | 0.73962598 | 0.632960294 | -0.061619116 | 0.265165153 | -0.229671702 | 0.40866667 | 0.689237413 | 0.805866384 | 0.8074 | 0.876940138 |
| lplA_lplA | 1.586028306 | 1.563012292 | 1.611064786 | 0.060377303 | 0.149110868 | 0.377101385 | 0.335109934 | 0.497465292 | 0.651646877 | 0.637 | 0.747503774 |
| SL1344_RS06600_lpxR | -0.384177069 | 0.685957349 | -1.984439309 | -2.680526135 | 0.300368482 | -8.940706809 | 4.68E-05 | 0.00267493 | 0.030479233 | 0.1 | 0.243021319 |
| SL1344_RS11955_lrhA | 1.512034271 | 1.131662089 | 1.850469302 | 0.738810491 | 0.162198148 | 4.497525777 | 4.68E-05 | 0.004608406 | 0.043904807 | 0.1 | 0.243021319 |
| SL1344_RS04660_lrp | 3.917735211 | 4.102127263 | 3.710195807 | -0.397984998 | 0.063211513 | -6.285331528 | 4.68E-05 | 0.002529007 | 0.031685306 | 0.1 | 0.243021319 |

| | | | | | | | | | | | |
|---|---|---|---|---|---|---|---|---|---|---|---|
| SL1344_RS09120_L.serine.ammonia.lyase | 4.322489391 | 4.741895176 | 3.896183427 | -0.849087602 | 0.062984947 | -13.53608585 | 4.68E-05 | 0.00025976 | 0.011063581 | 0.1 | 0.243021319 |
| SL1344_RS20930_lsrR | 0.47371537 | 0.206500043 | 0.654822143 | 0.439076306 | 0.181001742 | 2.373688417 | 0.004001152 | 0.036531917 | 0.132052649 | 0.1049 | 0.249692349 |
| SL1344_RS14590_luxS | 3.915643615 | 3.855127633 | 3.963984095 | 0.115509877 | 0.066147895 | 1.744718686 | 0.011326186 | 0.081195225 | 0.218445583 | 0.1122 | 0.258701254 |
| SL1344_RS09795_lysine.N.methylase | 2.939225739 | 3.509861644 | 2.245828739 | -1.255046319 | 0.184938424 | -6.792064962 | 4.68E-05 | 0.001202548 | 0.023670409 | 0.1 | 0.243021319 |
| malF_malF | -2.232659695 | -2.297376743 | -2.159289228 | 0.213095124 | 0.624749543 | 0.321444315 | 0.367754835 | 0.555859148 | 0.703265079 | 0.6987 | 0.79640764 |
| malG_malG | -2.174220995 | -2.155967132 | -2.201762511 | -0.084742111 | 0.550402618 | -0.144104462 | 0.434666669 | 0.620377584 | 0.753131819 | 0.752 | 0.836652781 |
| SL1344_RS18135_malT | 2.670235292 | 2.074507441 | 3.201441739 | 1.135073597 | 0.101569693 | 11.26541326 | 4.68E-05 | 0.001708485 | 0.026045236 | 0.1 | 0.243021319 |
| SL1344_RS07280_malY. | -0.55492506 | -0.682846943 | -0.450861491 | 0.223034717 | 0.256522368 | 0.817840462 | 0.185876102 | 0.288510538 | 0.457332971 | 0.3871 | 0.539483653 |
| mdoD_mdoD | 3.000327218 | 3.209997221 | 2.814250411 | -0.396049529 | 0.069437847 | -5.654978387 | 4.68E-05 | 0.003179894 | 0.034405981 | 0.1 | 0.243021319 |
| mdoG_mdoG | 4.351350635 | 4.324610231 | 4.378090146 | 0.057859826 | 0.064425213 | 0.889924228 | 0.159333357 | 0.246400644 | 0.426428985 | 0.3391 | 0.499905915 |
| SL1344_RS09375_membrane.protein | 2.59139072 | 3.123232828 | 1.968837202 | -1.161655302 | 0.104941061 | -10.95922902 | 4.68E-05 | 0.000487246 | 0.0140794 | 0.1 | 0.243021319 |
| SL1344_RS21870_membrane.protein | 3.454233325 | 3.687609089 | 3.242217386 | -0.443514942 | 0.072100657 | -6.140412394 | 4.68E-05 | 0.002810418 | 0.033401303 | 0.1 | 0.243021319 |
| SL1344_RS11835_menC | 1.478127794 | 1.416561139 | 1.550712881 | 0.154604333 | 0.178034613 | 0.889596801 | 0.128666697 | 0.240569805 | 0.428165581 | 0.3004 | 0.45380288 |
| SL1344_RS21470_metA | -0.810965236 | -0.815779605 | -0.805042532 | 0.008530227 | 0.261604423 | 0.0273906 | 0.488341106 | 0.577561581 | 0.713227833 | 0.6852 | 0.785449052 |
| SL1344_RS21065_metB | 1.615367853 | 1.548487932 | 1.687902145 | 0.136402244 | 0.117314711 | 1.128163085 | 0.104000039 | 0.164897708 | 0.320219274 | 0.2436 | 0.400966758 |
| SL1344_RS16345_metC | 0.164635292 | 0.399403441 | -0.032286338 | -0.42955419 | 0.185474971 | -2.268913926 | 0.009327606 | 0.035280901 | 0.128237009 | 0.1066 | 0.251742757 |
| SL1344_RS20385_metE | 1.114027734 | 1.162999675 | 1.070395697 | -0.069896438 | 0.289872878 | -0.245891197 | 0.372418393 | 0.674809095 | 0.797619159 | 0.7605 | 0.848278813 |
| SL1344_RS21095_metF | 0.187063384 | 0.28795048 | -0.073121558 | -0.327930646 | 0.311088006 | -0.964493305 | 0.166000022 | 0.234227214 | 0.423479436 | 0.3319 | 0.501970165 |
| metG_metG | 3.741432694 | 3.579067433 | 3.952952197 | 0.368167621 | 0.06367557 | 5.661373461 | 4.68E-05 | 0.004252582 | 0.040648978 | 0.1 | 0.243021319 |
| SL1344_RS21500_metH | 2.203271737 | 2.255820301 | 2.158988993 | -0.09083985 | 0.111648844 | -0.77311544 | 0.200532995 | 0.307544289 | 0.478611192 | 0.4051 | 0.557496316 |
| SL1344_RS18435_methyl.accepting.chemotaxis.citrate.transducer | 3.215709554 | 4.252169129 | 2.080109521 | -2.171417527 | 0.082611949 | -26.50373454 | 4.68E-05 | 0.000136908 | 0.007834257 | 0.1 | 0.243021319 |
| SL1344_RS08105_methyl.accepting.chemotaxis.protein | 1.724570599 | 2.694411608 | 0.646934955 | -2.035191477 | 0.142868669 | -14.25616253 | 4.68E-05 | 0.00029721 | 0.010960007 | 0.1 | 0.243021319 |
| SL1344_RS16230_methyl.accepting.chemotaxis.protein | 2.16548682 | 3.000576917 | 1.100448581 | -1.901004377 | 0.154751023 | -12.36421511 | 4.68E-05 | 0.002233728 | 0.031494751 | 0.1 | 0.243021319 |
| SL1344_RS16300_methyl.accepting.chemotaxis.protein | 4.491344177 | 5.243814121 | 3.800061179 | -1.425787671 | 0.113734546 | -12.49423159 | 4.68E-05 | 0.000142518 | 0.009288144 | 0.1 | 0.243021319 |
| SL1344_RS16625_methyl.accepting.chemotaxis.protein | 5.202906586 | 6.133739964 | 4.383048865 | -1.745146533 | 0.072053801 | -23.23335138 | 4.68E-05 | 0.000735113 | 0.019931416 | 0.1 | 0.243021319 |
| SL1344_RS21060_metJ | 2.993766704 | 3.064132613 | 2.928099348 | -0.180567541 | 0.218173037 | -0.860409362 | 0.13000003 | 0.231944456 | 0.431516293 | 0.2719 | 0.4482914 |
| SL1344_RS21070_metL | 2.982510621 | 2.909481966 | 3.150871624 | 0.228825444 | 0.10354743 | 1.985060791 | 0.017322048 | 0.065458394 | 0.193549869 | 0.1221 | 0.271391918 |
| metQ_metQ | 3.291671186 | 3.313318546 | 3.262398125 | -0.097248275 | 0.146322343 | -0.673510164 | 0.248501012 | 0.318306577 | 0.512805125 | 0.5084 | 0.654253708 |
| SL1344_RS20380_metR | -0.548187626 | -0.578777153 | -0.514159111 | 0.085760655 | 0.278942803 | 0.268508703 | 0.390666671 | 0.560601397 | 0.701281737 | 0.6831 | 0.782017366 |
| MgrR_mgrR | 2.203929442 | 1.706014366 | 2.440267947 | 0.740699783 | 0.258135595 | 2.922009338 | 4.68E-05 | 0.046188446 | 0.164481683 | 0.1 | 0.243021319 |
| MicA_micA | -2.13285521 | -1.857079211 | -2.651845266 | -0.794986868 | 0.525171118 | -1.408502722 | 0.050000088 | 0.119078308 | 0.274277634 | 0.1611 | 0.315881751 |
| MicC_micC | -2.567799605 | -2.400479712 | -2.666811146 | -0.291139967 | 0.721652925 | -0.386675887 | 0.341105936 | 0.549051574 | 0.698637924 | 0.6829 | 0.788369817 |
| MicF_micF | 1.542441927 | 1.032401039 | 2.150052462 | 1.125827451 | 0.235044572 | 4.813429211 | 4.68E-05 | 0.007677837 | 0.059459554 | 0.1 | 0.243021319 |
| SL1344_RS07395_mlc | 1.241535941 | 1.093409242 | 1.385069017 | 0.290706828 | 0.124261094 | 2.304024147 | 0.005330642 | 0.032247194 | 0.120298648 | 0.1055 | 0.250400663 |
| moaA_moaA | 0.840424092 | 0.867394895 | 0.811787212 | -0.05286633 | 0.16382396 | -0.293446158 | 0.378666671 | 0.559950516 | 0.700307694 | 0.666 | 0.771415961 |
| moaD_moaD | -0.55022863 | -0.633766069 | -0.489396306 | 0.144353852 | 0.267030478 | 0.493058852 | 0.307128589 | 0.47090785 | 0.625413079 | 0.5884 | 0.709947684 |
| moaE_moaE | 0.155400802 | 0.141582971 | 0.174690034 | 0.039742906 | 0.19457969 | 0.188266791 | 0.409726852 | 0.55894659 | 0.696179577 | 0.6743 | 0.77577109 |
| mobA_mobA | 0.113591003 | 0.12248687 | 0.102871148 | -0.020213804 | 0.187583307 | -0.106784647 | 0.449333335 | 0.556504569 | 0.694664021 | 0.6552 | 0.761977567 |
| SL1344_RS24715_mobA | 4.095642886 | 5.270092658 | 2.803286786 | -2.492733262 | 0.119804114 | -20.84744921 | 4.68E-05 | 4.28E-05 | 0.005594591 | 0.1 | 0.243021319 |
| SL1344_RS24720_mobC | 1.866079893 | 2.930481108 | 0.318497624 | -2.602745094 | 0.237277291 | -10.60012456 | 4.68E-05 | 0.003041214 | 0.037093481 | 0.1 | 0.243021319 |
| SL1344_RS03955_modA | 2.852457744 | 3.636485974 | 2.048243761 | -1.587469869 | 0.104869295 | -15.04353678 | 4.68E-05 | 0.000204978 | 0.010146909 | 0.1 | 0.243021319 |
| modB_modB | 2.661557259 | 3.458710901 | 1.939182771 | -1.49350576 | 0.135045782 | -10.9038722 | 4.68E-05 | 0.000669588 | 0.017374703 | 0.1 | 0.243021319 |
| SL1344_RS03965_modC | 3.162052666 | 3.87228866 | 2.45883454 | -1.404521791 | 0.110874414 | -12.68339556 | 4.68E-05 | 0.00043757 | 0.014375108 | 0.1 | 0.243021319 |
| SL1344_RS09620_motA | 5.007755774 | 6.393933904 | 3.634214683 | -2.700483576 | 0.19828057 | -13.68946538 | 4.68E-05 | 2.26E-05 | 0.005279752 | 0.1 | 0.243021319 |
| motB_motB | 5.1606932 | 6.386341578 | 3.915279527 | -2.434202409 | 0.133152282 | -18.32463247 | 4.68E-05 | 1.61E-05 | 0.004453934 | 0.1 | 0.243021319 |
| SL1344_RS09450_msbB | 2.592627692 | 2.989444918 | 2.123878505 | -0.867938165 | 0.089051127 | -9.736842299 | 4.68E-05 | 0.001332566 | 0.021909314 | 0.1 | 0.243021319 |
| Msr_Msr | -0.786431835 | -1.219246538 | -0.38402759 | 0.861816507 | 0.325496187 | 2.634149137 | 4.68E-05 | 0.021513112 | 0.100085283 | 0.1001 | 0.243161305 |
| mtlR_mtlR | 1.50783261 | 1.538258353 | 1.477520383 | -0.020629334 | 0.200316375 | -0.098619031 | 0.458361094 | 0.718768649 | 0.826931882 | 0.8447 | 0.901688031 |
| mukB_mukB | 5.181592097 | 5.19398542 | 5.169432728 | -0.001013486 | 0.104278711 | -0.008142543 | 0.495669554 | 0.794591041 | 0.881371504 | 0.9157 | 0.950946681 |
| murD_murD | 3.846063515 | 3.716274526 | 4.003973603 | 0.269030316 | 0.141979259 | 1.893255514 | 0.012000381 | 0.043983971 | 0.157710567 | 0.1115 | 0.259201311 |
| SL1344_RS00625_murE | 3.089532061 | 2.897260205 | 3.178993734 | 0.280413397 | 0.254795368 | 1.167536212 | 0.097333376 | 0.134162973 | 0.310210194 | 0.1991 | 0.374480822 |
| SL1344_RS09280_N.acetyltransferase | -0.103957665 | 0.744164262 | -1.091435017 | -1.825437165 | 0.305126502 | -6.010539767 | 4.68E-05 | 0.004758531 | 0.042203885 | 0.1 | 0.243021319 |
| napB_napB | -1.708834525 | -1.525358464 | -1.969106163 | -0.444464372 | 0.371892396 | -1.12314375 | 0.110666703 | 0.17465583 | 0.332811715 | 0.2546 | 0.41106957 |

| | | | | | | | | | | | |
|---|---|---|---|---|---|---|---|---|---|---|---|
| SL1344_RS08805_narG | -0.996702404 | -0.544658001 | -1.502054373 | -0.966320688 | 0.301912617 | -3.185919755 | 4.68E-05 | 0.01428869 | 0.07746253 | 0.1002 | 0.24330454 |
| SL1344_RS08800_narH | -1.670000872 | -1.410283835 | -2.22683854 | -0.81930534 | 0.509074715 | -1.547032477 | 0.026666836 | 0.109326797 | 0.262964592 | 0.1326 | 0.281655629 |
| SL1344_RS08790_narI | -2.624773208 | -2.244432912 | -3.036344594 | -0.786475519 | 0.456652918 | -1.625499871 | 0.03264504 | 0.072933052 | 0.194201575 | 0.1365 | 0.287828933 |
| SL1344_RS08795_narJ | -3.520518113 | -3.110608173 | -4.089318056 | -0.989123617 | 0.623723459 | -1.486559453 | 0.039307241 | 0.094976904 | 0.225384223 | 0.1581 | 0.310107048 |
| SL1344_RS08810_narK | -2.398951964 | -1.744786073 | -3.00660269 | -1.247689784 | 0.452873064 | -2.671097611 | 0.000673369 | 0.021511398 | 0.096638211 | 0.102 | 0.245711257 |
| SL1344_RS08820_narL | 0.298539221 | 0.653270491 | -0.085115557 | -0.744747049 | 0.180973709 | -4.103233621 | 4.68E-05 | 0.007257305 | 0.053180999 | 0.1 | 0.243021319 |
| nepI_nepI | 1.162882022 | 1.174836148 | 1.14232866 | -0.042349334 | 0.153074715 | -0.245722929 | 0.392000004 | 0.56311194 | 0.703110657 | 0.677 | 0.779298096 |
| nhaA_nhaA | 2.250948617 | 2.271942759 | 2.202004398 | -0.072177279 | 0.140894281 | -0.491728042 | 0.299800142 | 0.441875271 | 0.616665433 | 0.5979 | 0.723119583 |
| nirD_nirD | -1.5282415 | -1.513608938 | -1.549322387 | 0.004418828 | 0.437246782 | 0.010931698 | 0.495333334 | 0.682596668 | 0.799494985 | 0.7778 | 0.858494453 |
| nlpD_nlpD | 5.575370738 | 5.63418692 | 5.46895191 | -0.208649708 | 0.112024932 | -2.036316059 | 4.68E-05 | 0.035627052 | 0.141073701 | 0.1 | 0.243021319 |
| nrdI_nrdI | -1.119887593 | -1.2439935 | -0.880653446 | 0.344971118 | 0.395156177 | 0.817862991 | 0.195333351 | 0.298277824 | 0.476107608 | 0.4019 | 0.561360719 |
| SL1344_RS21955_nrfB | -0.006362656 | 0.770519689 | -0.897808298 | -1.689144703 | 0.229714663 | -7.299945164 | 4.68E-05 | 0.00124447 | 0.022467006 | 0.1 | 0.243021319 |
| SL1344_RS21960_nrfC | 0.174866838 | 0.824665373 | -0.753517624 | -1.576726583 | 0.248267655 | -6.269139336 | 4.68E-05 | 0.002367804 | 0.029574253 | 0.1 | 0.243021319 |
| SL1344_RS21975_nrfG | -2.235535419 | -2.405134637 | -1.884486279 | 0.456049914 | 0.666331911 | 0.595383264 | 0.271333344 | 0.44499746 | 0.61343206 | 0.5508 | 0.690493779 |
| nudC_nudC | 0.335921767 | 0.283717661 | 0.39918374 | 0.11432035 | 0.193721142 | 0.568370674 | 0.268666678 | 0.405385689 | 0.566660631 | 0.5155 | 0.647516733 |
| nusA_nusA | 5.583967316 | 5.457880398 | 5.716364934 | 0.311106184 | 0.109109132 | 3.092897002 | 4.68E-05 | 0.011915856 | 0.076641341 | 0.1 | 0.243021319 |
| nusB_nusB | 4.221903115 | 3.94989243 | 4.371509049 | 0.384892396 | 0.158745425 | 2.490817464 | 4.68E-05 | 0.019914675 | 0.101261563 | 0.1 | 0.243021319 |
| nusG_nusG | 5.947153284 | 5.934983465 | 5.964489654 | 0.029917762 | 0.050768358 | 0.569961609 | 0.28000001 | 0.432804937 | 0.611706048 | 0.5557 | 0.70121652 |
| obgE_obgE | 3.295348819 | 3.185307452 | 3.511751297 | 0.318174029 | 0.240484415 | 1.231989445 | 0.095333377 | 0.163325353 | 0.351145705 | 0.1886 | 0.364383651 |
| ompL_ompL | -4.841064149 | -5.084693862 | -4.640982542 | 0.416032427 | 1.067024751 | 0.372208862 | 0.327115264 | 0.471947246 | 0.621448387 | 0.5821 | 0.699009443 |
| SL1344_RS18070_ompR | 2.237256969 | 2.038489107 | 2.358649782 | 0.313558209 | 0.115446035 | 2.594122186 | 0.001336769 | 0.02995657 | 0.120714778 | 0.1022 | 0.24603291 |
| ompX_ompX | 7.49578982 | 7.258794465 | 7.658363904 | 0.389416937 | 0.082553859 | 4.791791787 | 4.68E-05 | 0.00244139 | 0.034527245 | 0.1 | 0.243021319 |
| OmrA_omrA | -1.458028411 | -2.23862361 | -0.899842798 | 1.337292861 | 0.409714221 | 3.188255517 | 4.68E-05 | 0.018528375 | 0.089158401 | 0.1 | 0.243021319 |
| OmrB_omrB | -1.929119484 | -2.392473348 | -1.644227874 | 0.746777074 | 0.423798442 | 1.692587318 | 0.026000173 | 0.083032021 | 0.215516843 | 0.1323 | 0.282610587 |
| oppB_oppB | -0.192872169 | -0.328607767 | -0.029606417 | 0.276745024 | 0.307156305 | 0.893638066 | 0.171219209 | 0.247714888 | 0.426377205 | 0.3366 | 0.504345271 |
| SL1344_RS14860_orgA | 2.052963109 | 3.020835543 | 1.004755978 | -2.073637942 | 0.221278451 | -9.457268339 | 4.68E-05 | 0.000328059 | 0.013502444 | 0.1 | 0.243021319 |
| OxyS_oxyS | -1.145601018 | -1.172120361 | -1.117758048 | 0.108035266 | 0.444028292 | 0.223610809 | 0.401732182 | 0.623197637 | 0.755746082 | 0.7515 | 0.83632538 |
| pabB_pabB | 2.016884096 | 2.410189233 | 1.501570743 | -0.894290693 | 0.143685866 | -6.117339159 | 4.68E-05 | 0.003952025 | 0.039559292 | 0.1 | 0.243021319 |
| pagP_pagP | 2.668592147 | 2.351317599 | 3.006611402 | 0.6712693 | 0.137909529 | 4.921185831 | 4.68E-05 | 0.013423683 | 0.080852273 | 0.1 | 0.243021319 |
| SL1344_RS10525_pduA | -6.138558005 | -5.518160215 | -7.214527247 | -1.727523899 | 2.073472507 | -0.722673403 | 0.157228538 | 0.270800896 | 0.446943406 | 0.3434 | 0.493689378 |
| pduC_pduC | -3.930402524 | -3.847064083 | -4.055463983 | -0.230994027 | 0.795783863 | -0.265190043 | 0.37441706 | 0.511867217 | 0.65931157 | 0.6309 | 0.740552208 |
| pduD_pduD | -4.770823815 | -4.697620432 | -4.867404395 | -0.184426815 | 1.231957002 | -0.131144604 | 0.435709529 | 0.571748884 | 0.716483904 | 0.7116 | 0.808179182 |
| pduE_pduE | -5.084100213 | -5.315549463 | -4.894013801 | 0.332738059 | 1.562597136 | 0.201022282 | 0.40373085 | 0.620406664 | 0.75032903 | 0.7196 | 0.813197717 |
| pdxK_pdxK | -0.316225534 | -0.116834778 | -0.392449729 | -0.247840574 | 0.264084237 | -0.856347002 | 0.196000018 | 0.28996978 | 0.46412275 | 0.4045 | 0.561914438 |
| SL1344_RS06590_pfkB | 1.732130808 | 1.67225307 | 1.763460002 | 0.100485843 | 0.171779855 | 0.533306427 | 0.299333342 | 0.433491036 | 0.608665653 | 0.5804 | 0.712049269 |
| SL1344_RS13960_phage.virion.morphogenesis.protein | -4.720324242 | -5.078093283 | -3.947725489 | 1.618339124 | 3.180857389 | 0.476914427 | 0.248501012 | 0.408162646 | 0.595790615 | 0.481 | 0.647823457 |
| SL1344_RS02035_phoB | 2.266895091 | 2.219160849 | 2.318698589 | 0.089477382 | 0.119450933 | 0.712651516 | 0.219187224 | 0.342102907 | 0.514698089 | 0.4334 | 0.587125789 |
| SL1344_RS11645_phoE | 7.398916002 | 6.949009149 | 7.887977465 | 0.925074494 | 0.061275098 | 15.09835297 | 4.68E-05 | 6.32E-05 | 0.007056709 | 0.1 | 0.243021319 |
| SL1344_RS05535_phoH | -0.209757835 | 0.126034042 | -0.505609975 | -0.652607511 | 0.270507975 | -2.388223791 | 0.001336769 | 0.031665003 | 0.124559037 | 0.1029 | 0.247027129 |
| SL1344_RS06080_phoP | 3.979847632 | 3.916764822 | 4.056081022 | 0.143826973 | 0.066481567 | 2.206668664 | 0.001336769 | 0.049965567 | 0.163916264 | 0.1017 | 0.245273689 |
| SL1344_RS06075_phoQ | 4.021757955 | 3.903985687 | 4.157019721 | 0.261463855 | 0.066828481 | 3.919962815 | 4.68E-05 | 0.013606748 | 0.078186339 | 0.1 | 0.243021319 |
| phoR_phoR | 1.541965226 | 1.646419241 | 1.478027018 | -0.163726255 | 0.126916236 | -1.217273472 | 0.10326453 | 0.154775332 | 0.312711792 | 0.2378 | 0.398078907 |
| SL1344_RS13700_phospho.2.dehydro.3.deoxyheptonate.aldolase.Tyr.sensitive | 1.080504995 | 0.633641369 | 1.421750441 | 0.788294186 | 0.14288148 | 5.479442568 | 4.68E-05 | 0.00501528 | 0.042421746 | 0.1 | 0.243021319 |
| SL1344_RS10625_phsA | -1.646765311 | -1.587468313 | -1.707358351 | -0.125739779 | 0.335488076 | -0.338059813 | 0.348666673 | 0.492948277 | 0.637824103 | 0.6037 | 0.714796732 |
| SL1344_RS10620_phsB | -3.53992726 | -3.953204354 | -3.350875154 | 0.603864423 | 0.923512453 | 0.576780278 | 0.24266668 | 0.363291355 | 0.54446924 | 0.4867 | 0.629251172 |
| SL1344_RS10615_phsC | -2.331539106 | -2.221400594 | -2.404994336 | -0.219441125 | 0.550572918 | -0.360431166 | 0.352666673 | 0.529148669 | 0.676199052 | 0.6582 | 0.763656914 |
| potG_potG | 0.346415952 | 0.585550605 | 0.223831472 | -0.367376615 | 0.302788841 | -1.16315629 | 0.09466671 | 0.136618044 | 0.302016713 | 0.2148 | 0.384205158 |
| SL1344_RS09215_prc | 5.729604743 | 6.070939844 | 5.293451612 | -0.778109666 | 0.067249864 | -11.39949854 | 4.68E-05 | 0.002032728 | 0.031157062 | 0.1 | 0.243021319 |
| SL1344_RS14880_prgH | 1.097127461 | 2.692927343 | -1.013791909 | -3.712893743 | 0.208255375 | -17.90356227 | 4.68E-05 | 0.00076078 | 0.017272341 | 0.1 | 0.243021319 |
| SL1344_RS14875_prgI | 0.06611795 | 1.556241612 | -2.181916865 | -3.75119875 | 0.432206482 | -8.557957982 | 4.68E-05 | 0.004496431 | 0.043340865 | 0.1 | 0.243021319 |
| SL1344_RS14870_prgJ | 0.822571265 | 2.295626704 | -1.005174376 | -3.299913064 | 0.213358023 | -15.38494333 | 4.68E-05 | 0.001174153 | 0.02107073 | 0.1 | 0.243021319 |
| SL1344_RS14865_prgK | 2.009988149 | 3.171092611 | 0.695641671 | -2.484969702 | 0.179259219 | -13.95448227 | 4.68E-05 | 0.000282095 | 0.011865632 | 0.1 | 0.243021319 |
| SL1344_RS09220_proQ | 5.56585493 | 5.865015666 | 5.260272937 | -0.614606632 | 0.056574697 | -10.78646336 | 4.68E-05 | 0.000228959 | 0.011395112 | 0.1 | 0.243021319 |

| | | | | | | | | | | | |
|---|---|---|---|---|---|---|---|---|---|---|---|
| SL1344_RS09255_prpA.pphA | -2.032147298 | -1.571395532 | -2.58000419 | -1.013906377 | 0.366247259 | -2.731026727 | 0.002668394 | 0.023755354 | 0.101284612 | 0.1015 | 0.245050878 |
| psiB_psiB | -2.49909085 | -2.788728887 | -2.235201992 | 0.5412055 | 0.433375069 | 1.186094074 | 0.093271197 | 0.155692133 | 0.301588833 | 0.2325 | 0.387379078 |
| psiB_psiB.1 | -3.176861497 | -3.168681023 | -3.183940856 | -0.003447853 | 0.562835447 | -0.007067078 | 0.497001999 | 0.557427873 | 0.695512606 | 0.6665 | 0.770484724 |
| pssA_pssA | 3.889443607 | 3.770335325 | 3.995049317 | 0.223765543 | 0.055065832 | 4.077539892 | 4.68E-05 | 0.00838194 | 0.057898662 | 0.1 | 0.243021319 |
| SL1344_RS11995_pta | 6.105894785 | 5.913706599 | 6.250933435 | 0.344759499 | 0.094220243 | 3.586640354 | 4.68E-05 | 0.027841185 | 0.123703193 | 0.1 | 0.243021319 |
| SL1344_RS15255_ptpS | 1.914132541 | 1.267214467 | 2.522176035 | 1.26413147 | 0.116482635 | 10.82576655 | 4.68E-05 | 0.000503012 | 0.014528149 | 0.1 | 0.243021319 |
| SL1344_RS02910_PTS.mannose.fructose.sorbose.transporter.subunit.IIB | -5.762169465 | -5.611110012 | -6.128104386 | -0.483946587 | 2.493231774 | -0.130727215 | 0.423717525 | 0.596605608 | 0.743783282 | 0.8062 | 0.8809008 |
| SL1344_RS07105_purR | 2.818457198 | 2.627488582 | 2.976209336 | 0.345637776 | 0.094923311 | 3.631305414 | 4.68E-05 | 0.013264071 | 0.07528128 | 0.1 | 0.243021319 |
| putA_putA | 1.969803875 | 1.988663352 | 1.951463064 | -0.038966341 | 0.100470731 | -0.364623235 | 0.347768161 | 0.49227888 | 0.639790135 | 0.5955 | 0.714351186 |
| SL1344_RS23405_putative.inner.membrane.protein | 1.169551582 | 1.525629702 | 0.801500124 | -0.718394736 | 0.150438647 | -4.722536393 | 4.68E-05 | 0.004387577 | 0.041365963 | 0.1 | 0.243021319 |
| SL1344_RS09445_pyruvate.kinase.II | 2.879179313 | 3.332251413 | 2.369856112 | -0.959211434 | 0.091419964 | -10.39769574 | 4.68E-05 | 0.000653497 | 0.016340312 | 0.1 | 0.243021319 |
| SL1344_RS16430_qseB | -1.971392313 | -2.075799039 | -1.886463139 | 0.185652226 | 0.361095296 | 0.485266354 | 0.286666676 | 0.41798378 | 0.570676722 | 0.5241 | 0.649832545 |
| SL1344_RS16435_qseC | -0.620681777 | -0.741885637 | -0.457277855 | 0.276418114 | 0.237329869 | 1.127960721 | 0.114590308 | 0.184733287 | 0.339542056 | 0.2701 | 0.426225198 |
| SL1344_RS23490_radA | 2.516972186 | 2.347615012 | 2.661933002 | 0.318189838 | 0.094356148 | 3.361796318 | 4.68E-05 | 0.01502849 | 0.079765513 | 0.1 | 0.243021319 |
| RaoN_raoN | -2.304640836 | -2.521281671 | -1.981021607 | 0.521180909 | 0.43968718 | 1.115228645 | 0.114666702 | 0.182637138 | 0.340347426 | 0.2674 | 0.425726647 |
| SL1344_RS12895_ratA | 1.196795193 | 1.184079702 | 1.20390164 | 0.013611993 | 0.133979951 | 0.106125829 | 0.459333335 | 0.594174261 | 0.727072716 | 0.6966 | 0.795062683 |
| rcsA_rcsA | -0.354687651 | -0.275715355 | -0.459846772 | -0.191107547 | 0.257397274 | -0.673756542 | 0.197201883 | 0.312094896 | 0.484822061 | 0.422 | 0.566920136 |
| SL1344_RS11655_rcsB | 3.384875032 | 3.318167734 | 3.469872237 | 0.176546599 | 0.11934661 | 1.453297317 | 0.036000124 | 0.085189494 | 0.22975402 | 0.1374 | 0.290800004 |
| SL1344_RS11660_rcsC | 2.215374778 | 1.958147863 | 2.358179928 | 0.39739656 | 0.182259105 | 2.216166032 | 0.0020023 | 0.029039335 | 0.121940072 | 0.1011 | 0.244565918 |
| SL1344_RS11650_rcsD | 3.374690145 | 3.146288109 | 3.628927379 | 0.486917462 | 0.069133582 | 7.113154273 | 4.68E-05 | 0.001857992 | 0.026537083 | 0.1 | 0.243021319 |
| rcsF_rcsF | 1.827129086 | 1.657238129 | 1.945636688 | 0.27483735 | 0.135713603 | 1.941397701 | 0.021985549 | 0.055637159 | 0.172722499 | 0.1214 | 0.27167093 |
| RdlD_rdlD | -0.724601181 | -0.997341399 | -0.578593713 | 0.409424067 | 0.361658501 | 1.075232118 | 0.116588975 | 0.189488171 | 0.364236377 | 0.2721 | 0.4363557 |
| RdlE_rdlE | -2.600479699 | -3.155745143 | -2.112773456 | 1.088899664 | 0.603726719 | 1.787540932 | 0.020653117 | 0.062005973 | 0.1817165 | 0.1186 | 0.266232521 |
| SL1344_RS14650_recA | 4.329896148 | 4.547583189 | 4.223761953 | -0.335274735 | 0.181042726 | -2.122396951 | 4.68E-05 | 0.032444061 | 0.133485259 | 0.1 | 0.243021319 |
| SL1344_RS13835_recN | 2.573542558 | 2.632183214 | 2.515248816 | -0.11919832 | 0.087065385 | -1.306596573 | 0.067333397 | 0.128563774 | 0.277280552 | 0.1972 | 0.353310283 |
| SL1344_RS06620_rfc | 1.629274181 | 1.427974041 | 1.836808206 | 0.429246048 | 0.160615476 | 2.695998052 | 4.68E-05 | 0.020655214 | 0.09823538 | 0.1 | 0.243021319 |
| SL1344_RS22270_rflM | 2.869984061 | 3.445287596 | 2.319415566 | -1.164674856 | 0.179126237 | -6.557579275 | 4.68E-05 | 0.001698805 | 0.027991014 | 0.1 | 0.243021319 |
| rhtB_rhtB | -1.843672703 | -1.738307849 | -1.945916108 | -0.214604746 | 0.346225135 | -0.563324771 | 0.262666678 | 0.368555037 | 0.525142552 | 0.4757 | 0.60890333 |
| rimM_rimM | 6.534473713 | 6.426005813 | 6.623201622 | 0.210127085 | 0.098393424 | 2.202700664 | 4.68E-05 | 0.031893438 | 0.132907718 | 0.1 | 0.243021319 |
| rimO_rimO | 3.393993542 | 3.325232271 | 3.494137054 | 0.175018585 | 0.087743275 | 1.919698626 | 0.006000767 | 0.061142227 | 0.185355616 | 0.1081 | 0.25399026 |
| SL1344_RS03220_rlpA | 2.102952827 | 2.090049957 | 2.119405569 | 0.032779586 | 0.09532073 | 0.316243462 | 0.357333339 | 0.496957074 | 0.645151907 | 0.6095 | 0.7228225 |
| SL1344_RS05840_rne | 5.400589118 | 5.362662672 | 5.462367778 | 0.086893087 | 0.083594565 | 1.088211441 | 0.124583643 | 0.167915119 | 0.350686448 | 0.2454 | 0.427708072 |
| rnpB_rnpB | 11.24254911 | 11.15660436 | 11.39498064 | 0.235242639 | 0.02505745 | 7.953807709 | 4.68E-05 | 0.014196107 | 0.08548482 | 0.1 | 0.243021319 |
| RnpB_RnpB | 11.24154794 | 11.1566316 | 11.39473664 | 0.235383401 | 0.025681783 | 7.852931967 | 4.68E-05 | 0.014184669 | 0.085453515 | 0.1 | 0.243021319 |
| SL1344_RS15130_rpoS | 4.824818126 | 4.966888237 | 4.612579245 | -0.360169204 | 0.090176273 | -4.123837799 | 4.68E-05 | 0.017521592 | 0.093809837 | 0.1 | 0.243021319 |
| rprA_rprA | -2.355438434 | -2.675249833 | -2.196808401 | 0.493502279 | 0.565419066 | 0.811353381 | 0.177333354 | 0.281987288 | 0.460906099 | 0.3816 | 0.530843978 |
| rpsN_rpsN | 6.970366588 | 6.860453667 | 7.096943254 | 0.25749716 | 0.085977266 | 3.159609648 | 4.68E-05 | 0.012381204 | 0.078712232 | 0.1 | 0.243021319 |
| rpsV_rpsV | 1.805865304 | 2.458976333 | 1.278198915 | -1.145222536 | 0.197587593 | -5.724977832 | 4.68E-05 | 0.002683597 | 0.034093618 | 0.1 | 0.243021319 |
| RseX_rseX | -6.671970802 | -6.309209768 | -7.899633625 | -1.385622185 | 3.228250513 | -0.358251144 | 0.330446376 | 0.467966918 | 0.638597768 | 0.6509 | 0.767784153 |
| rsmD_rsmD | 1.006994104 | 0.739611794 | 1.239621813 | 0.587241327 | 0.273156954 | 2.143436815 | 0.001336769 | 0.031412469 | 0.127401023 | 0.1007 | 0.244003204 |
| SL1344_RS09240_rsmF.yebU | 2.622026954 | 2.955055137 | 2.277007983 | -0.682291945 | 0.104435087 | -6.63534974 | 4.68E-05 | 0.001813364 | 0.026859503 | 0.1 | 0.243021319 |
| SL1344_RS22150_rtsA | 0.452902666 | 1.792610775 | -1.062321105 | -2.941557787 | 0.437518983 | -6.899931821 | 4.68E-05 | 0.002582569 | 0.031997126 | 0.1 | 0.243021319 |
| SL1344_RS22145_rtsB | -1.2442888 | 0.089942354 | -3.49905731 | -3.589880254 | 0.489967752 | -7.179369027 | 4.68E-05 | 0.004147625 | 0.03777489 | 0.1 | 0.243021319 |
| rumA_rumA | 2.816983048 | 2.775071615 | 2.886550901 | 0.115055812 | 0.125822271 | 0.882053255 | 0.16322454 | 0.249810858 | 0.438115893 | 0.3418 | 0.507937586 |
| SL1344_RS09480_ruvA | 1.938169207 | 2.343555669 | 1.440215335 | -0.904044842 | 0.107669568 | -8.493239187 | 4.68E-05 | 0.001518273 | 0.023084096 | 0.1 | 0.243021319 |
| SL1344_RS09475_ruvB | 2.720612275 | 3.132343298 | 2.259719655 | -0.857758325 | 0.106956425 | -7.938240881 | 4.68E-05 | 0.001021099 | 0.020600061 | 0.1 | 0.243021319 |
| RybA_rybA | 2.469230304 | 2.572208771 | 2.354729137 | -0.25863299 | 0.140388889 | -1.833702664 | 0.006000767 | 0.050572941 | 0.167056657 | 0.1036 | 0.247909693 |
| RybB_rybB | -0.683901244 | -0.910664652 | -0.211473169 | 0.657305531 | 0.334290559 | 1.814342149 | 0.031312601 | 0.068150744 | 0.195400424 | 0.1335 | 0.285753668 |
| RybD_rybD | 0.740039424 | 1.077118382 | 0.262969954 | -0.81850875 | 0.220867063 | -3.664833143 | 4.68E-05 | 0.009777973 | 0.064284058 | 0.1 | 0.243021319 |
| RydB_rydB | -3.196358547 | -3.210279108 | -3.181885277 | 0.03295844 | 0.651744014 | 0.044532557 | 0.481678881 | 0.606240176 | 0.738886581 | 0.7164 | 0.811067973 |
| RydC_rydC | 0.248645556 | 0.47063186 | -0.073035795 | -0.532900732 | 0.20687663 | -2.50747397 | 0.003332494 | 0.030985642 | 0.120623766 | 0.1027 | 0.246665876 |
| RyeF_ryeF | 0.453566074 | 1.133138641 | -0.307320228 | -1.439201608 | 0.171474545 | -8.381320097 | 4.68E-05 | 0.001184678 | 0.02085438 | 0.1 | 0.243021319 |
| RyfA_ryfA | 1.968878143 | 1.981772481 | 1.961513312 | -0.038285432 | 0.157903636 | -0.228999364 | 0.395333337 | 0.587254325 | 0.730425677 | 0.7236 | 0.818508633 |

| | | | | | | | | | | | |
|---|---|---|---|---|---|---|---|---|---|---|---|
| RyfD_ryfD | 3.135426928 | 3.028940196 | 3.254968368 | 0.223584484 | 0.098796783 | 2.246804752 | 0.005330642 | 0.032809252 | 0.128656802 | 0.1022 | 0.246112609 |
| RyhB_ryhB | -0.510614971 | -0.878871058 | 0.311752426 | 1.135123649 | 0.520742135 | 1.929434053 | 0.009333825 | 0.067941931 | 0.204708531 | 0.112 | 0.259476077 |
| RyjB_ryjB | -2.569896578 | -2.575287656 | -2.563338191 | 0.030527448 | 0.527350111 | 0.053706764 | 0.475682879 | 0.597880051 | 0.729972263 | 0.7074 | 0.802329087 |
| Sar_sar | -0.567565653 | -0.942826522 | -0.2690599 | 0.680237975 | 0.26922451 | 2.46959017 | 0.001335878 | 0.034442606 | 0.128210183 | 0.1015 | 0.245048551 |
| SL1344_RS10635_sbcB | 1.762366129 | 1.503212831 | 1.930972053 | 0.386952725 | 0.196643083 | 1.892426694 | 0.026666836 | 0.051333554 | 0.169062911 | 0.119 | 0.269352381 |
| SL1344_RS01925_sbmA | 0.21334256 | 0.12249792 | 0.324269686 | 0.187096901 | 0.246256967 | 0.731079331 | 0.221185891 | 0.33525192 | 0.511930916 | 0.4409 | 0.593378342 |
| SL1344_RS10605_sbmC | 0.854326667 | 0.764342457 | 1.020268244 | 0.263169159 | 0.187087602 | 1.329286004 | 0.070000061 | 0.140758168 | 0.304124769 | 0.1949 | 0.352071598 |
| SL1344_RS20885_sbp | 0.942163511 | 0.435868121 | 1.269416241 | 0.842837069 | 0.217242129 | 3.873013773 | 4.68E-05 | 0.017928394 | 0.091064694 | 0.1 | 0.243021319 |
| sdhC_sdhC | 1.718983787 | 1.954409462 | 1.405902426 | -0.57905747 | 0.181384904 | -3.258192353 | 4.68E-05 | 0.018522185 | 0.094648023 | 0.1 | 0.243021319 |
| SL1344_RS09755_sdiA | 2.316461944 | 3.388031482 | 1.145221925 | -2.217953869 | 0.17763552 | -12.44066321 | 4.68E-05 | 0.000295005 | 0.011619053 | 0.1 | 0.243021319 |
| SdsR_sdsR | -1.251637054 | -1.587223238 | -0.937377605 | 0.711306438 | 0.451337276 | 1.522234185 | 0.025316634 | 0.085771414 | 0.228114168 | 0.1273 | 0.278467506 |
| sgbH_sgbH | -4.54649964 | -4.073845099 | -5.044450261 | -0.890434866 | 1.743941128 | -0.478001224 | 0.255333345 | 0.416435022 | 0.593471612 | 0.4985 | 0.64655802 |
| SgrS_sgrS | 2.60872081 | 2.660967153 | 2.429635593 | -0.229893681 | 0.201538678 | -1.214921338 | 0.118000034 | 0.130791699 | 0.300280865 | 0.24 | 0.416347905 |
| SibA_sibA | 0.569696499 | 0.098698117 | 0.834707357 | 0.739780226 | 0.295026131 | 2.446572602 | 0.000673369 | 0.048559607 | 0.165498635 | 0.1001 | 0.243161531 |
| SibC_sibC | 2.914955607 | 2.857416339 | 2.932773376 | 0.075895571 | 0.144153455 | 0.441179778 | 0.348666673 | 0.561762769 | 0.719771086 | 0.7321 | 0.840295169 |
| SibD_sibD | 2.401337341 | 2.235316534 | 2.537898941 | 0.356086436 | 0.268830711 | 1.314290622 | 0.054666746 | 0.11103711 | 0.276055282 | 0.1502 | 0.312052792 |
| SL1344_RS14940_sicA | 0.104307709 | 1.710310908 | -2.088841248 | -3.800894935 | 0.299777447 | -12.76025132 | 4.68E-05 | 0.001764519 | 0.025408169 | 0.1 | 0.243021319 |
| sifA_sifA | -0.012460345 | -0.12024883 | 0.134585355 | 0.20397986 | 0.313317982 | 0.610097227 | 0.261825461 | 0.418542779 | 0.591374233 | 0.5215 | 0.666893714 |
| sifB_sifB | -2.580427801 | -2.777144165 | -2.390500595 | 0.391283328 | 0.470507063 | 0.785588597 | 0.186666686 | 0.276055721 | 0.43621223 | 0.3715 | 0.516409101 |
| SL1344_RS05355_sigE | 0.800489393 | 2.337659121 | -1.01830937 | -3.366385307 | 0.186194022 | -18.14525321 | 4.68E-05 | 0.000388788 | 0.011740449 | 0.1 | 0.243021319 |
| SL1344_RS21850_siiA | -2.030717933 | -1.072829764 | -3.417798385 | -2.352665201 | 0.461687872 | -5.099693169 | 4.68E-05 | 0.004310731 | 0.039649776 | 0.1 | 0.243021319 |
| SL1344_RS21855_siiB | -1.006959861 | -0.17372108 | -2.273669372 | -2.099873268 | 0.352077018 | -5.987093527 | 4.68E-05 | 0.005442632 | 0.044391213 | 0.1 | 0.243021319 |
| SL1344_RS21875_siiF | 2.292282255 | 2.270947785 | 2.345469199 | 0.057561165 | 0.184130622 | 0.220930605 | 0.398666671 | 0.828115522 | 0.902339445 | 0.7987 | 0.885041349 |
| SL1344_RS01560_sinR | 0.809038995 | 0.994833844 | 0.494024913 | -0.492579851 | 0.198530554 | -2.407434388 | 0.003332494 | 0.037686547 | 0.137916289 | 0.1025 | 0.246429524 |
| SL1344_RS14920_sipA | 1.1345437 | 2.117925767 | -0.002838095 | -2.146530203 | 0.201971667 | -10.84802867 | 4.68E-05 | 0.000392117 | 0.013636878 | 0.1 | 0.243021319 |
| SL1344_RS14935_sipB | 1.488832336 | 3.145242582 | -0.557600678 | -3.712649722 | 0.252036627 | -14.72291981 | 4.68E-05 | 0.000865723 | 0.018904594 | 0.1 | 0.243021319 |
| SL1344_RS14930_sipC | 2.327392381 | 3.927122235 | 0.538687681 | -3.384909767 | 0.123010798 | -27.62603278 | 4.68E-05 | 6.29E-05 | 0.005578486 | 0.1 | 0.243021319 |
| SL1344_RS14925_sipD | -0.085266264 | 1.066234888 | -1.769775083 | -2.842701841 | 0.430816665 | -6.442256975 | 4.68E-05 | 0.002337971 | 0.030177049 | 0.1 | 0.243021319 |
| SL1344_RS09740_sirA | 3.188132179 | 3.494721225 | 2.902412834 | -0.606158324 | 0.086080915 | -6.992386501 | 4.68E-05 | 0.002602032 | 0.032243747 | 0.1 | 0.243021319 |
| SL1344_RS10425_sirohydrochlorin.cobaltochelatase | -3.269586203 | -3.618498887 | -2.816003342 | 0.784404289 | 0.639067819 | 1.141476795 | 0.105929419 | 0.170837131 | 0.325522605 | 0.2535 | 0.410809338 |
| SL1344_RS09840_SL1896 | -2.374650784 | -2.075063933 | -2.808937992 | -0.767373675 | 0.604605505 | -1.277993622 | 0.052631662 | 0.144213268 | 0.308776774 | 0.1614 | 0.315209059 |
| SL1344_RS07175_slyA | 2.913276313 | 2.932671199 | 2.884463616 | -0.023892808 | 0.130302724 | -0.173346389 | 0.437708197 | 0.763271746 | 0.858325764 | 0.8468 | 0.908261912 |
| SL1344_RS07110_sodB | 3.620996994 | 3.660737483 | 3.547203421 | -0.108450495 | 0.238142801 | -0.418842327 | 0.372418393 | 0.548522285 | 0.717561088 | 0.7969 | 0.883552275 |
| SL1344_RS10630_sopA | -0.71800026 | 0.308877974 | -2.191397152 | -2.537772168 | 0.405457535 | -6.278963215 | 4.68E-05 | 0.003472446 | 0.035709128 | 0.1 | 0.243021319 |
| sopB_sopB | 0.751940032 | 2.419716275 | -1.34041714 | -3.797656393 | 0.280755375 | -13.55305934 | 4.68E-05 | 0.000261169 | 0.010570109 | 0.1 | 0.243021319 |
| SL1344_RS15235_sopD | -0.827915815 | -0.818069905 | -0.84068217 | -0.030038586 | 0.297230742 | -0.085170291 | 0.461333335 | 0.607189478 | 0.739179625 | 0.707 | 0.805289769 |
| SL1344_RS13925_sopE | 0.434435894 | 2.474587091 | -2.428513668 | -4.933570915 | 0.361997062 | -13.68190033 | 4.68E-05 | 0.001543996 | 0.023873407 | 0.1 | 0.243021319 |
| SL1344_RS09265_sopE2 | -0.715698333 | 1.299861847 | -3.649321815 | -4.959373015 | 0.432247897 | -11.43210645 | 4.68E-05 | 0.00189165 | 0.025318374 | 0.1 | 0.243021319 |
| spaK_spaK | -1.096604811 | -0.055809521 | -2.804056057 | -2.787252316 | 0.497202884 | -5.584551035 | 4.68E-05 | 0.007683772 | 0.053647497 | 0.1 | 0.243021319 |
| spaM_spaM | -0.562425059 | 0.397357341 | -2.339012856 | -2.723728827 | 0.418581325 | -6.384504248 | 4.68E-05 | 0.007915656 | 0.057804479 | 0.1 | 0.243021319 |
| spaN_spaN | -0.220006965 | 0.844193922 | -1.618936189 | -2.525143153 | 0.318283055 | -7.911793137 | 4.68E-05 | 0.00098792 | 0.019945195 | 0.1 | 0.243021319 |
| SL1344_RS14965_spaO | -0.47875027 | 0.642431312 | -2.157408938 | -2.799956436 | 0.33273923 | -8.456398132 | 4.68E-05 | 0.002530327 | 0.029330153 | 0.1 | 0.243021319 |
| SL1344_RS14960_spaP | -0.63790011 | 0.507624045 | -2.419606163 | -2.928857998 | 0.386126113 | -7.61693687 | 4.68E-05 | 0.004982833 | 0.04304549 | 0.1 | 0.243021319 |
| SL1344_RS14955_spaQ | -1.133876649 | 0.068431665 | -2.771170661 | -2.810845869 | 0.375017287 | -7.508039557 | 4.68E-05 | 0.002205075 | 0.027067618 | 0.1 | 0.243021319 |
| SL1344_RS14950_spaR | -1.318185988 | 0.027909043 | -2.652955992 | -2.626061566 | 0.42807728 | -6.115346478 | 4.68E-05 | 0.002477192 | 0.030399477 | 0.1 | 0.243021319 |
| SL1344_RS14945_spaS | -1.004024046 | -0.115831626 | -2.435931272 | -2.339541751 | 0.38985271 | -5.887681688 | 4.68E-05 | 0.009277777 | 0.06071528 | 0.1 | 0.243021319 |
| Spf_spf | 7.873000849 | 7.852927228 | 7.934588459 | 0.090794196 | 0.188650307 | 0.596607955 | 0.225849448 | 0.332576066 | 0.539371552 | 0.4 | 0.598503476 |
| SL1344_RS14840_spi4.transcriptional.regulator | -0.212758664 | 0.828219433 | -1.546407687 | -2.387181168 | 0.244206649 | -9.693109221 | 4.68E-05 | 0.001081187 | 0.019394548 | 0.1 | 0.243021319 |
| spiC_spiC | -5.451188066 | -5.478809411 | -5.405698301 | -0.003607282 | 1.55539174 | -0.002762451 | 0.498 | 0.617743598 | 0.74983303 | 0.7298 | 0.821423309 |
| spr_spr | 4.885315312 | 4.662393352 | 5.094453594 | 0.443810552 | 0.06094391 | 7.275749064 | 4.68E-05 | 0.001070208 | 0.02159536 | 0.1 | 0.243021319 |
| SraA_sraA | -5.373023198 | -5.464367237 | -5.272864142 | 0.201500447 | 1.133595809 | 0.156995616 | 0.422000003 | 0.527970685 | 0.671700628 | 0.6443 | 0.753162555 |
| SraB_sraB | 0.824389828 | 0.779357865 | 0.853854261 | 0.067290358 | 0.234836413 | 0.256122479 | 0.40266667 | 0.678649616 | 0.797984497 | 0.7692 | 0.855456377 |
| SraC_sraC | 3.412784354 | 4.017645085 | 2.511625464 | -1.517567166 | 0.219345377 | -6.832444608 | 4.68E-05 | 0.001363065 | 0.025623652 | 0.1 | 0.243021319 |

| | | | | | | | | | | |
|---|---|---|---|---|---|---|---|---|---|---|
| SraF_sraF | 1.874294056 | 1.826888241 | 2.30989864 | 0.490277694 | 0.361651924 | 1.093395443 | 0.184543657 | 0.19868413 | 0.39487119 | 0.3406 | 0.529605194 |
| SraG_sraG | -1.886913161 | -1.949058299 | -1.830642807 | 0.154069794 | 0.466804107 | 0.293272042 | 0.364000005 | 0.53215812 | 0.681158922 | 0.678 | 0.778107485 |
| SraL_sraL | -2.651745817 | -2.367192137 | -2.830708397 | -0.44065933 | 0.55135606 | -0.749559327 | 0.20733335 | 0.334670364 | 0.501909397 | 0.4421 | 0.586992644 |
| SL1344_RS07945_srfB | 3.941243697 | 4.645787207 | 3.222339444 | -1.412616277 | 0.149043599 | -9.380226472 | 4.68E-05 | 0.000480284 | 0.016223079 | 0.1 | 0.243021319 |
| SL1344_RS07950_srfC | 3.547016733 | 4.27518789 | 2.842379854 | -1.43997097 | 0.12664869 | -11.39233849 | 4.68E-05 | 0.001117817 | 0.023149596 | 0.1 | 0.243021319 |
| srlA_srlA | -4.286092987 | -3.957186217 | -4.74256528 | -0.75838823 | 0.84729071 | -0.799678726 | 0.16322454 | 0.264376737 | 0.428202656 | 0.3515 | 0.500147679 |
| srlE_srlE | -3.154272508 | -3.187319005 | -3.126717632 | 0.046325309 | 0.594513028 | 0.072628928 | 0.465333335 | 0.590365059 | 0.724139626 | 0.6944 | 0.793542283 |
| srlR_srlR | 0.457024536 | 0.464264868 | 0.445169811 | -0.012206655 | 0.191264244 | -0.061399465 | 0.473333334 | 0.620140405 | 0.749630268 | 0.7281 | 0.820782922 |
| SroA_sroA | -0.785236179 | -1.006224987 | -0.68962483 | 0.30863946 | 0.288462291 | 0.96235234 | 0.164000023 | 0.239411153 | 0.411456527 | 0.3416 | 0.500879323 |
| SroC_sroC | 4.400811251 | 4.455895706 | 4.198859418 | -0.250742101 | 0.291183233 | -0.803439707 | 0.226515671 | 0.291044784 | 0.497800556 | 0.4327 | 0.606802069 |
| SroD_sroD | 0.416431628 | 0.758594047 | -0.141821682 | -0.922625794 | 0.346029762 | -2.74601483 | 4.68E-05 | 0.06721691 | 0.204594765 | 0.1 | 0.243021319 |
| SroG_sroG | 3.369588338 | 3.295806206 | 3.566913536 | 0.275701762 | 0.378924105 | 0.430569074 | 0.334000007 | 0.808699067 | 0.894536999 | 0.7012 | 0.830290551 |
| SL1344_RS06985_ssaG | -5.340662727 | -5.57546937 | -4.998096671 | 0.449944548 | 1.807696028 | 0.224785106 | 0.392000004 | 0.65493223 | 0.77576741 | 0.7417 | 0.830683021 |
| SL1344_RS07050_ssaR | -2.116954406 | -2.334830207 | -1.886453921 | 0.452727388 | 0.399385595 | 1.054356467 | 0.107261863 | 0.185052296 | 0.336225453 | 0.2715 | 0.419813841 |
| SL1344_RS05160_sseI | -1.727052401 | -1.640711677 | -1.822036323 | -0.176546139 | 0.341981672 | -0.48582564 | 0.294666676 | 0.4167671 | 0.569757301 | 0.5247 | 0.651424172 |
| sspA_sspA | 4.416523251 | 4.33401693 | 4.458121056 | 0.114115284 | 0.06350925 | 1.626754961 | 0.053297883 | 0.086924697 | 0.228933479 | 0.1503 | 0.308409759 |
| SL1344_RS06915_ssrA | -0.729089864 | -0.823871483 | -0.637772835 | 0.187658542 | 0.287013695 | 0.602420735 | 0.251333346 | 0.396274644 | 0.558911056 | 0.5058 | 0.640311348 |
| SsrA_ssrA | 10.79125487 | 10.83476761 | 10.74487127 | -0.027297466 | 0.131899309 | -0.337000688 | 0.343770826 | 0.607599016 | 0.763020892 | 0.7231 | 0.833925104 |
| SL1344_RS06910_ssrB | -1.751158509 | -1.719380148 | -1.769884149 | 0.026505186 | 0.558897217 | 0.038665631 | 0.484000001 | 0.678175621 | 0.80049006 | 0.8021 | 0.876711276 |
| ssrS_ssrS | 8.404742517 | 8.232744725 | 8.815736139 | 0.572853126 | 0.187466993 | 2.581703129 | 4.68E-05 | 0.023629647 | 0.113390193 | 0.1 | 0.243021319 |
| SsrS_ssrS | 8.411211544 | 8.231331319 | 8.815722183 | 0.573153395 | 0.1881292 | 2.585352975 | 4.68E-05 | 0.023742838 | 0.113687019 | 0.1 | 0.243021319 |
| SL1344_RS16670_sstT | 2.273339 | 2.255263825 | 2.298200749 | 0.028082626 | 0.115280463 | 0.239914122 | 0.389073955 | 0.625900372 | 0.755053969 | 0.7224 | 0.81660833 |
| SL1344_RS01735_stbB | -5.920636846 | -5.046364587 | -7.262718831 | -2.19143208 | 2.020961964 | -0.996154962 | 0.080666719 | 0.183705683 | 0.354880055 | 0.2072 | 0.364963105 |
| SL1344_RS05145_stfT | -1.646743229 | -1.311117952 | -1.988077243 | -0.693677578 | 0.365539923 | -1.783377086 | 0.023333527 | 0.06312414 | 0.181428995 | 0.1257 | 0.275580363 |
| STnc3140_STnc3140..sopD. | 0.088839041 | -1.069519508 | 0.916063519 | 1.895326195 | 0.403641889 | 4.59355628 | 4.68E-05 | 0.005932426 | 0.050040424 | 0.1 | 0.243021319 |
| STnc3200_STnc3200 | -5.866588037 | -6.288095108 | -5.551686495 | 0.869717007 | 2.11837602 | 0.298751328 | 0.312666675 | 0.453017227 | 0.623334711 | 0.5905 | 0.709057724 |
| SL1344_RS15350_syd | 2.069106267 | 1.820478142 | 2.189315567 | 0.352806398 | 0.137006745 | 2.296840734 | 0.005996769 | 0.041771041 | 0.148451206 | 0.1028 | 0.246878313 |
| SymR_symR | -1.822552284 | -1.822945366 | -1.822499898 | 0.004686265 | 0.398021794 | 0.010960049 | 0.497001999 | 0.605727233 | 0.737484298 | 0.7191 | 0.81137253 |
| SL1344_RS13990_tail.protein | -6.454850342 | -7.305410628 | -5.560112183 | 2.622431223 | 3.394502661 | 0.696590402 | 0.204000017 | 0.304448344 | 0.488860201 | 0.427 | 0.576910755 |
| tatE_tatE | 1.364931841 | 1.458607259 | 1.160670184 | -0.306842938 | 0.255661119 | -1.243936343 | 0.03331126 | 0.177350157 | 0.366287184 | 0.1366 | 0.289334499 |
| tcuB_tcuB | -2.516692705 | -2.459102206 | -2.569624456 | -0.095219711 | 0.478036275 | -0.17562878 | 0.40866667 | 0.557729336 | 0.695265092 | 0.6593 | 0.764149445 |
| SL1344_RS16770_tdcA | -1.43001076 | -1.677813144 | -1.243931863 | 0.504058546 | 0.811141002 | 0.665528883 | 0.211192554 | 0.321851593 | 0.521256048 | 0.4325 | 0.598156364 |
| SL1344_RS16765_tdcB | 0.212770665 | 0.723829525 | -0.62407887 | -1.348977114 | 0.271456564 | -4.97165038 | 4.68E-05 | 0.00745054 | 0.053995151 | 0.1 | 0.243021319 |
| SL1344_RS16760_tdcC | 1.130860337 | 1.886877934 | 0.273382915 | -1.616080292 | 0.161161287 | -10.01936348 | 4.68E-05 | 0.001021304 | 0.019257218 | 0.1 | 0.243021319 |
| SL1344_RS05150_tfaT | -2.009158716 | -1.618295059 | -2.250608033 | -0.582302722 | 0.453636783 | -1.223638397 | 0.101333374 | 0.159040677 | 0.318527501 | 0.2433 | 0.405632401 |
| Tff_tff | 5.453222988 | 5.452390792 | 5.454710588 | 0.00481523 | 0.176168994 | 0.025849767 | 0.488 | 0.895706374 | 0.944109895 | 0.9544 | 0.974069181 |
| thiP_thiP | 1.740827806 | 1.575617976 | 1.894642084 | 0.33618971 | 0.161700116 | 2.057438454 | 0.003998486 | 0.055323495 | 0.175922454 | 0.103 | 0.247219807 |
| thiQ_thiQ | 0.630043202 | 0.225573254 | 0.875467142 | 0.622940215 | 0.249070468 | 2.377407151 | 0.000673369 | 0.030569258 | 0.1231092 | 0.102 | 0.24580822 |
| SL1344_RS06625_thrS | 5.736350628 | 5.568594015 | 5.897047379 | 0.310334051 | 0.067841902 | 4.591695914 | 4.68E-05 | 0.003530533 | 0.039842814 | 0.1 | 0.243021319 |
| SL1344_RS04595_tnpA1 | -2.336457835 | -1.412404038 | -5.071438826 | -3.675838987 | 1.361754802 | -2.571498728 | 4.68E-05 | 0.049686795 | 0.167974388 | 0.1 | 0.243021319 |
| SL1344_RS06060_tnpA2 | -2.921673677 | -2.062337646 | -4.554530252 | -2.476258748 | 0.930507729 | -2.605344595 | 0.00067292 | 0.038507355 | 0.139057948 | 0.1001 | 0.243166243 |
| SL1344_RS09075_tnpA3 | -3.809621416 | -2.658071577 | -4.319590207 | -1.504554179 | 1.489116321 | -0.991987153 | 0.137908089 | 0.192436047 | 0.374938815 | 0.2734 | 0.45240566 |
| SL1344_RS09790_tnpA4 | -6.29367199 | -5.812294472 | -6.754452329 | -0.988500693 | 2.280751076 | -0.347049131 | 0.291805472 | 0.462010243 | 0.62565581 | 0.5801 | 0.702995692 |
| SL1344_RS12685_tnpA5 | -5.317137662 | -5.078646992 | -5.514062205 | -0.437492609 | 1.221819098 | -0.323746558 | 0.360666672 | 0.501766684 | 0.647563542 | 0.6151 | 0.726407494 |
| SL1344_RS17020_tnpA6 | -2.221208205 | -1.936811535 | -2.941630784 | -1.009881338 | 0.690402479 | -1.3068228 | 0.051333419 | 0.13279263 | 0.299515157 | 0.1667 | 0.322729738 |
| SL1344_RS17955_tnpA7 | -2.634487378 | -2.285914394 | -2.956030771 | -0.694757838 | 0.570292851 | -1.208861859 | 0.088000048 | 0.149015452 | 0.30570396 | 0.2214 | 0.379762333 |
| tolA_tolA | 1.932772252 | 1.91523569 | 1.957879824 | 0.04528356 | 0.175906339 | 0.245760228 | 0.406000004 | 0.686101484 | 0.803388398 | 0.7907 | 0.868255885 |
| SL1344_RS16475_tolC | 3.9125572 | 3.790906161 | 4.003891412 | 0.212922957 | 0.066706022 | 3.223999291 | 4.68E-05 | 0.013306532 | 0.076002455 | 0.1 | 0.243021319 |
| SL1344_RS21860_TolC.family.protein | -1.019082678 | -0.342011265 | -1.841148001 | -1.577772295 | 0.447384351 | -3.563251754 | 4.68E-05 | 0.00830247 | 0.060148511 | 0.1 | 0.243021319 |
| tqsA_tqsA | -0.818137916 | -0.873855327 | -0.767303512 | 0.119205242 | 0.27863135 | 0.403795919 | 0.320666674 | 0.493643111 | 0.641267438 | 0.6008 | 0.71556238 |
| SL1344_RS14145_transcriptional.regulator | -5.614060631 | -4.885832452 | -6.249949909 | -1.842885664 | 2.433392909 | -0.808440164 | 0.170000022 | 0.267291773 | 0.453498379 | 0.3661 | 0.517548504 |
| trkD_trkD | 2.666182234 | 2.450152255 | 2.88712139 | 0.435965995 | 0.108419211 | 3.976168503 | 4.68E-05 | 0.006860586 | 0.053856726 | 0.1 | 0.243021319 |
| SL1344_RS06175_tRNA.arg | -2.975682369 | -1.87766518 | -4.50135315 | -2.637057895 | 0.756434578 | -3.486217831 | 4.68E-05 | 0.013730271 | 0.07402146 | 0.1 | 0.243021319 |

| | | | | | | | | | | | |
|---|---|---|---|---|---|---|---|---|---|---|---|
| SL1344_RS13600_tRNA.rRNA.methyltransferase | 3.18514808 | 2.759351651 | 3.559159119 | 0.791302691 | 0.097916287 | 8.007779639 | 4.68E-05 | 0.001562361 | 0.024886813 | 0.1 | 0.243021319 |
| truA_truA | 1.588339022 | 1.326091191 | 1.863542522 | 0.537415087 | 0.138920894 | 3.862882217 | 4.68E-05 | 0.008834042 | 0.060304335 | 0.1 | 0.243021319 |
| SL1344_RS23260_tsr | 6.495358272 | 7.236024179 | 5.784158144 | -1.44105851 | 0.058443698 | -24.56445079 | 4.68E-05 | 2.09E-05 | 0.004366308 | 0.1 | 0.243021319 |
| SL1344_RS06870_ttrA | -1.462331206 | -1.560723845 | -1.375301127 | 0.184040679 | 0.465943491 | 0.366513465 | 0.341772158 | 0.536572621 | 0.688553998 | 0.6585 | 0.770216112 |
| SL1344_RS06880_ttrB | -4.726775867 | -4.591587931 | -4.790417926 | -0.136518281 | 1.356143924 | -0.087254641 | 0.458361094 | 0.670880034 | 0.791234553 | 0.7701 | 0.855599733 |
| SL1344_RS06875_ttrC | -4.573452139 | -4.533360219 | -4.620355549 | -0.026838881 | 1.15370869 | -0.023061444 | 0.491005996 | 0.61798664 | 0.750296064 | 0.7378 | 0.826649172 |
| SL1344_RS06890_ttrR | -0.915194917 | -0.965779575 | -0.842581808 | 0.103561272 | 0.35203507 | 0.272429338 | 0.37708195 | 0.625188495 | 0.753365193 | 0.7162 | 0.813687917 |
| SL1344_RS06885_ttrS | 0.331790397 | 0.198992445 | 0.429414027 | 0.229154286 | 0.180520295 | 1.211529727 | 0.096666709 | 0.15818341 | 0.313548181 | 0.2343 | 0.392693951 |
| tyrA_tyrA | 1.350254619 | 1.139825459 | 1.607747397 | 0.464193498 | 0.149778162 | 3.025492112 | 0.00067292 | 0.015247966 | 0.0811413 | 0.1002 | 0.243297329 |
| ubiB_ubiB | 3.548678928 | 3.46441818 | 3.635764773 | 0.178650019 | 0.075322756 | 2.340234169 | 0.004664544 | 0.030372069 | 0.121529056 | 0.1022 | 0.246092658 |
| ubiF_ubiF | 1.285120027 | 1.385137041 | 1.183423305 | -0.202144399 | 0.121620928 | -1.633006576 | 0.037333453 | 0.080516203 | 0.206766756 | 0.1426 | 0.294115622 |
| SL1344_RS21190_udhA | 2.968867811 | 3.239574702 | 2.667258779 | -0.585093113 | 0.092052805 | -6.388594091 | 4.68E-05 | 0.003204055 | 0.036022656 | 0.1 | 0.243021319 |
| SL1344_RS03635_UDP.galactopyranose.mutase | -4.133144165 | -2.699188445 | -5.145681728 | -2.383768793 | 1.632555413 | -1.412029333 | 0.038641021 | 0.100055927 | 0.250860341 | 0.1375 | 0.292672541 |
| ugpC_ugpC | -1.004303178 | -0.934278105 | -1.095588423 | -0.166221186 | 0.275137907 | -0.575526795 | 0.257161904 | 0.378588855 | 0.536554018 | 0.4858 | 0.618511485 |
| ulaD_ulaD | -2.805567142 | -2.639118471 | -2.991584933 | -0.353591858 | 0.601743051 | -0.578822282 | 0.258494349 | 0.399617103 | 0.565686542 | 0.5078 | 0.642767957 |
| ushA_ushA | 1.864416124 | 1.912079691 | 1.814644214 | -0.098093254 | 0.09265714 | -0.985756161 | 0.1352432 | 0.21904045 | 0.373345653 | 0.3142 | 0.463964741 |
| SL1344_RS09735_uvrC | 3.361471403 | 3.633279826 | 3.116291089 | -0.523188376 | 0.088255787 | -5.863317806 | 4.68E-05 | 0.002103819 | 0.029521281 | 0.1 | 0.243021319 |
| SL1344_RS11530_virulence.protein.MsgA | -7.52889106 | -6.569315056 | -8.556902716 | -1.818168259 | 3.793663726 | -0.423814761 | 0.28000001 | 0.46951917 | 0.62043645 | 0.5465 | 0.675560472 |
| wecC_wecC | 3.161038422 | 2.959782746 | 3.32975385 | 0.373795733 | 0.082263533 | 4.473419156 | 4.68E-05 | 0.005227051 | 0.046339405 | 0.1 | 0.243021319 |
| SL1344_RS10695_wzzB | 2.656564197 | 2.123876797 | 3.189828041 | 1.075360559 | 0.156566045 | 6.930816896 | 4.68E-05 | 0.001879762 | 0.028953611 | 0.1 | 0.243021319 |
| SL1344_RS02345_ybaO | 0.902392155 | 0.879108876 | 0.919983115 | 0.043179273 | 0.153235438 | 0.251509879 | 0.388407733 | 0.554057237 | 0.693887365 | 0.6611 | 0.765923718 |
| SL1344_RS03185_ybeM | -1.887247457 | -1.102001843 | -2.607254619 | -1.514277192 | 0.71371507 | -2.108678307 | 0.000673369 | 0.050172967 | 0.165856267 | 0.101 | 0.24440773 |
| SL1344_RS04285_ybiK | 1.606464561 | 1.632296426 | 1.591856387 | -0.032933113 | 0.12303337 | -0.235921989 | 0.39573618 | 0.581026713 | 0.718198033 | 0.6907 | 0.794395477 |
| SL1344_RS08990_ycgO | 3.185836132 | 3.935023308 | 2.491779411 | -1.424764616 | 0.124239944 | -11.36350759 | 4.68E-05 | 0.000290642 | 0.012360327 | 0.1 | 0.243021319 |
| SL1344_RS08975_ycgR | 3.077798169 | 3.666727476 | 2.419497872 | -1.256569859 | 0.095413197 | -13.16561674 | 4.68E-05 | 0.000238665 | 0.01071182 | 0.1 | 0.243021319 |
| SL1344_RS08585_yciK | 1.313476995 | 1.420710421 | 1.161670652 | -0.255220221 | 0.180711205 | -1.357535138 | 0.065289873 | 0.1122879 | 0.265015854 | 0.1777 | 0.34004615 |
| SL1344_RS08495_yciW | 1.017857694 | 1.020426355 | 1.011737794 | 0.003529325 | 0.186466025 | 0.015075008 | 0.493670886 | 0.702493599 | 0.813419021 | 0.8018 | 0.87631496 |
| SL1344_RS07545_ydeD | -1.161240013 | -1.167427637 | -1.157296003 | 0.00642917 | 0.303585023 | 0.020634378 | 0.491672219 | 0.587933101 | 0.722280779 | 0.697 | 0.79476312 |
| SL1344_RS06680_ydiV | -0.545039009 | -0.874311945 | -0.353613433 | 0.511133587 | 0.297510664 | 1.610245482 | 0.04266677 | 0.085838076 | 0.223389298 | 0.1445 | 0.299163453 |
| SL1344_RS06595_ydiY | 3.073073808 | 2.805737784 | 3.220567522 | 0.40733851 | 0.105448533 | 3.4744444 | 4.68E-05 | 0.024512372 | 0.110815771 | 0.1 | 0.243021319 |
| SL1344_RS06585_ydiZ | -2.800518492 | -2.779647922 | -2.826011141 | -0.078015565 | 0.563217498 | -0.120266838 | 0.439706864 | 0.583849072 | 0.719823547 | 0.7024 | 0.797801298 |
| SL1344_RS06560_ydjN | 5.729003017 | 5.370988817 | 6.061999662 | 0.689197385 | 0.076561088 | 9.036771989 | 4.68E-05 | 0.001856761 | 0.029485877 | 0.1 | 0.243021319 |
| SL1344_RS09415_yebG | 0.216907789 | 0.783913677 | -0.609827271 | -1.401990134 | 0.243525202 | -5.771971741 | 4.68E-05 | 0.008505593 | 0.058776868 | 0.1 | 0.243021319 |
| SL1344_RS09225_yebR | 3.718558062 | 4.234652987 | 3.199604481 | -1.04112789 | 0.069281914 | -15.02926838 | 4.68E-05 | 0.00026248 | 0.010918994 | 0.1 | 0.243021319 |
| SL1344_RS09230_yebS | 1.647379006 | 2.015673258 | 1.11496096 | -0.906059481 | 0.211713066 | -4.241561031 | 4.68E-05 | 0.015228001 | 0.084821437 | 0.1 | 0.243021319 |
| SL1344_RS09760_yecC | 1.481362513 | 2.078202304 | 0.798515317 | -1.274841766 | 0.180394433 | -7.082569435 | 4.68E-05 | 0.001320243 | 0.023568646 | 0.1 | 0.243021319 |
| SL1344_RS09750_yecF | 3.78655754 | 4.111512795 | 3.418497672 | -0.70686113 | 0.094817808 | -7.543837724 | 4.68E-05 | 0.004196892 | 0.043724461 | 0.1 | 0.243021319 |
| SL1344_RS09765_yecS | 0.439080206 | 0.981402571 | -0.069315648 | -1.07047457 | 0.296412966 | -3.613829995 | 4.68E-05 | 0.00787162 | 0.059433634 | 0.1 | 0.243021319 |
| SL1344_RS09825_yedD | 1.835190538 | 1.702391764 | 1.97472871 | 0.29665231 | 0.152354127 | 1.976380753 | 0.003332494 | 0.053284898 | 0.171483102 | 0.1037 | 0.247874373 |
| SL1344_RS09830_yedE | 0.523488141 | 0.105985583 | 0.940747231 | 0.831544859 | 0.173651501 | 4.707648344 | 4.68E-05 | 0.0050299 | 0.043624116 | 0.1 | 0.243021319 |
| SL1344_RS09835_yedF | -0.111833943 | -0.546350574 | 0.354530253 | 0.917862438 | 0.303792537 | 3.068579684 | 4.68E-05 | 0.014506919 | 0.080603672 | 0.1 | 0.243021319 |
| SL1344_RS09770_yedO | 1.528714396 | 2.132566892 | 0.873812395 | -1.279905843 | 0.211983096 | -6.145200452 | 4.68E-05 | 0.002698803 | 0.033918072 | 0.1 | 0.243021319 |
| SL1344_RS10925_yegD | 1.105670054 | 0.448425334 | 1.732708717 | 1.286800292 | 0.160963891 | 7.967381483 | 4.68E-05 | 0.001097312 | 0.021527603 | 0.1 | 0.243021319 |
| SL1344_RS13585_yfiK | -2.536914442 | -2.509714361 | -2.566560426 | -0.053444848 | 0.481720847 | -0.108296608 | 0.449700202 | 0.56352012 | 0.700851394 | 0.6709 | 0.773113429 |
| SL1344_RS17960_yhfL | 1.187263171 | 1.457500222 | 0.504355929 | -0.933515925 | 0.231316694 | -3.377929899 | 4.68E-05 | 0.030880083 | 0.127955344 | 0.1 | 0.243021319 |
| SL1344_RS18605_yhjH | 4.006014591 | 4.795318718 | 3.200422236 | -1.517188192 | 0.189557008 | -8.08753487 | 4.68E-05 | 0.000210227 | 0.01215761 | 0.1 | 0.243021319 |
| SL1344_RS18650_yhjQ | 1.410054478 | 0.787398733 | 1.923803022 | 1.137689652 | 0.119280818 | 9.489672369 | 4.68E-05 | 0.001069255 | 0.019448682 | 0.1 | 0.243021319 |
| SL1344_RS23255_yjiY | 2.067857536 | 1.522075098 | 2.595633757 | 1.118925448 | 0.198855483 | 5.605948099 | 4.68E-05 | 0.00166455 | 0.027689682 | 0.1 | 0.243021319 |
| SL1344_RS05645_ymdA | 1.040439866 | 1.66012414 | 0.249280718 | -1.340417466 | 0.267435362 | -4.971683887 | 4.68E-05 | 0.003178714 | 0.036502328 | 0.1 | 0.243021319 |
| SL1344_RS05655_ymdC | 2.152516379 | 2.451716326 | 1.776058336 | -0.689659866 | 0.144953581 | -4.759563375 | 4.68E-05 | 0.003999481 | 0.040590722 | 0.1 | 0.243021319 |
| yopJ_yopJ | 0.396320993 | 0.884194712 | 0.005289087 | -0.860521148 | 0.210180776 | -3.99597295 | 4.68E-05 | 0.007951172 | 0.057595449 | 0.1 | 0.243021319 |
| SL1344_RS15335_yqcB | 1.712908696 | 1.473212104 | 1.786034636 | 0.301329472 | 0.169248072 | 1.490552033 | 0.055296548 | 0.115798878 | 0.276727546 | 0.1646 | 0.320430888 |
| SL1344_RS15340_yqcC..next.to.csrB. | 0.771136177 | 0.528658311 | 0.835854045 | 0.287201217 | 0.251808523 | 0.989409788 | 0.177881433 | 0.23569637 | 0.424710989 | 0.3538 | 0.519226869 |

| | | | | | | | | | | | |
|---|---|---|---|---|---|---|---|---|---|---|---|
| YrlA_yrlA | -3.936647917 | -3.978035203 | -3.880133158 | 0.082839239 | 0.750592024 | 0.101156641 | 0.456000002 | 0.564908063 | 0.701885748 | 0.6744 | 0.77606904 |
| YrlB_yrlB | -3.824360724 | -3.645539216 | -4.018178559 | -0.369894388 | 0.648492982 | -0.530209974 | 0.265333344 | 0.398354504 | 0.55212955 | 0.5045 | 0.633156219 |
| zntA_zntA | 2.338744455 | 2.448860742 | 2.111697314 | -0.31986372 | 0.136049603 | -2.127111729 | 0.00733396 | 0.049579465 | 0.164229985 | 0.1083 | 0.254073575 |
| zntB_zntB | 0.881808952 | 1.056353076 | 0.560932824 | -0.474140517 | 0.195387176 | -2.235754654 | 0.009333825 | 0.044829854 | 0.152799156 | 0.1087 | 0.254620548 |
| zraP_zraP | -2.689176596 | -2.874103679 | -2.527512239 | 0.352438876 | 0.558032394 | 0.576648893 | 0.240506342 | 0.372257163 | 0.538174056 | 0.4838 | 0.621158762 |
| crRNA1_ | -0.256643545 | -0.367182119 | -0.131409196 | 0.201554541 | 0.303460021 | 0.627202454 | 0.246502344 | 0.396532166 | 0.567398786 | 0.513 | 0.655518244 |
| crRNA2_ | 1.156714493 | 0.977528218 | 1.332783439 | 0.377470722 | 0.180712103 | 2.060931378 | 0.003332494 | 0.049664633 | 0.16255365 | 0.1056 | 0.250536002 |
| ECS021_ | -2.579663009 | -2.712560873 | -2.428195388 | 0.270809343 | 0.758062555 | 0.34621152 | 0.342666673 | 0.557186301 | 0.703744593 | 0.6456 | 0.761302786 |
| ECS173_ | -1.891841173 | -1.968762406 | -1.808898818 | 0.159045464 | 0.363035963 | 0.415132672 | 0.309333342 | 0.450293689 | 0.602839296 | 0.5586 | 0.682049803 |
| Ffs_ | 3.354922381 | 3.140785394 | 3.506990827 | 0.436780482 | 0.286235434 | 1.590772553 | 0.004001152 | 0.066660642 | 0.204836557 | 0.1044 | 0.249213916 |
| fimA.trailer_ | -3.888911698 | -3.94369004 | -3.818509608 | 0.104856107 | 0.667909368 | 0.139242379 | 0.431712194 | 0.510448666 | 0.655926508 | 0.6221 | 0.733342683 |
| GO.10702_ | -1.403540298 | -1.323481413 | -1.662807591 | -0.29513406 | 0.517473103 | -0.468671632 | 0.342438381 | 0.604076856 | 0.746703271 | 0.7112 | 0.824409362 |
| GO.10704_ | 3.068739513 | 3.000949692 | 3.259553203 | 0.232949746 | 0.134205524 | 1.45705394 | 0.071285869 | 0.111770439 | 0.273598784 | 0.1742 | 0.33942923 |
| Ig1085_ | -1.885185222 | -1.930120319 | -1.836787407 | 0.046786014 | 0.587373006 | 0.077640339 | 0.464357097 | 0.706620457 | 0.818832961 | 0.8273 | 0.890983865 |
| Ig1140_ | -1.688010064 | -1.302548949 | -2.08737041 | -0.813408039 | 0.46163274 | -1.741200159 | 0.01666694 | 0.071352006 | 0.200663838 | 0.1165 | 0.264878279 |
| Ig2335_ | 1.048776422 | 0.787390721 | 1.183459315 | 0.384569648 | 0.182737981 | 2.010726562 | 0.013333676 | 0.058924989 | 0.179652121 | 0.1128 | 0.259611394 |
| IS042_ | -3.603122147 | -4.041803371 | -3.309943125 | 0.743711894 | 0.700664402 | 1.027398797 | 0.114666702 | 0.209517833 | 0.371342828 | 0.2916 | 0.440257595 |
| lysC.leader_ | -1.606210094 | -1.590811372 | -1.641875998 | -0.012247292 | 0.461208484 | -0.024929297 | 0.491005996 | 0.680495189 | 0.798640632 | 0.7978 | 0.872809234 |
| mreB.leader_ | 1.555643046 | 1.549241768 | 1.564674583 | -0.005535736 | 0.168983484 | -0.032381373 | 0.487674884 | 0.688630944 | 0.804993748 | 0.8153 | 0.883888644 |
| pRNA_ | -4.900091749 | -4.456707697 | -5.36090626 | -0.568494283 | 1.809435847 | -0.302342839 | 0.362000006 | 0.622970569 | 0.755417176 | 0.7236 | 0.818097871 |
| RprA_ | -2.372098401 | -2.690611666 | -2.208357433 | 0.479213222 | 0.56021251 | 0.794270558 | 0.18254499 | 0.280202179 | 0.459323674 | 0.3836 | 0.534823679 |
| rpsL.leader_ | 4.250659144 | 4.233744498 | 4.289364107 | 0.078345628 | 0.134597358 | 0.615925139 | 0.240506342 | 0.346227302 | 0.54271127 | 0.4638 | 0.635062739 |
| rpsP.leader_ | 4.404785056 | 4.392899119 | 4.530173097 | 0.136917463 | 0.102629514 | 0.710181296 | 0.326666674 | 0.416595378 | 0.609940113 | 0.6815 | 0.814997383 |
| rrf_ | -6.538672303 | -6.6209936 | -6.46375635 | 0.143096309 | 1.935851317 | 0.066155268 | 0.469020654 | 0.558917393 | 0.700162496 | 0.6704 | 0.774884306 |
| rrf_.1 | -5.657757855 | -5.952165529 | -5.339715647 | 0.638325524 | 1.407063366 | 0.419358013 | 0.313790814 | 0.46237539 | 0.612647547 | 0.5666 | 0.688032164 |
| rrf_.2 | -4.698392521 | -5.053300232 | -4.246189682 | 0.796592401 | 1.0681367 | 0.696221185 | 0.222000015 | 0.35885056 | 0.517312442 | 0.4552 | 0.596814293 |
| rrf_.3 | -5.621115021 | -6.440971441 | -5.275984124 | 1.105310205 | 1.747794541 | 0.555469492 | 0.255333345 | 0.382785223 | 0.555968264 | 0.5091 | 0.645959192 |
| rrf_.4 | -3.999171857 | -5.219654416 | -3.084515034 | 2.233091818 | 1.924317347 | 1.188845912 | 0.016000285 | 0.187114462 | 0.37213155 | 0.1171 | 0.265702125 |
| rrf_.5 | -7.036465159 | -8.554943962 | -6.007787863 | 2.46486847 | 2.759420358 | 0.80673725 | 0.138666695 | 0.237529489 | 0.408474602 | 0.3018 | 0.455011599 |
| rrf_.6 | -8.916885004 | -9.581721582 | -8.016197201 | 2.040390936 | 4.584728469 | 0.385254301 | 0.335109934 | 0.500202394 | 0.645514098 | 0.6146 | 0.726203783 |
| rrf_.7 | -7.961759533 | -9.486558288 | -5.991024132 | 3.105200306 | 4.02132213 | 0.701786159 | 0.17800002 | 0.329669513 | 0.489750545 | 0.3754 | 0.53262821 |
| RUF_107c.1_ | 0.774438808 | 0.60634114 | 0.955852195 | 0.379449447 | 0.22462847 | 1.678972621 | 0.011326186 | 0.081807508 | 0.223205411 | 0.111 | 0.257192692 |
| RUF_107c.10_ | 0.417201614 | 0.423798015 | 0.408583478 | -0.019126662 | 0.184802505 | -0.099765955 | 0.459027317 | 0.595044997 | 0.729465294 | 0.7143 | 0.807822808 |
| RUF_107c.102_ | -2.978492087 | -2.766868375 | -3.115831682 | -0.479455577 | 0.927368983 | -0.492221362 | 0.294470362 | 0.419100755 | 0.59795308 | 0.5757 | 0.699863267 |
| RUF_107c.105_ | 0.868939187 | 0.921797268 | 0.710279986 | -0.207281417 | 0.384706657 | -0.439554875 | 0.357095276 | 0.652491048 | 0.788411394 | 0.754 | 0.858912769 |
| RUF_107c.107_ | -1.346250581 | -1.743300368 | -1.086894232 | 0.674194779 | 0.435667684 | 1.490125227 | 0.049966777 | 0.087746135 | 0.228552183 | 0.1502 | 0.305789187 |
| RUF_107c.108_ | 1.422837794 | 1.289334428 | 1.654424676 | 0.431818621 | 0.284049952 | 1.541129319 | 0.01866691 | 0.086152409 | 0.23499428 | 0.1204 | 0.270295413 |
| RUF_107c.111_ | 2.198934784 | 2.061529858 | 2.307034958 | 0.243496909 | 0.12284441 | 1.914740657 | 0.016000285 | 0.051222692 | 0.164436197 | 0.1168 | 0.265172023 |
| RUF_107c.112_ | 1.049840187 | 0.942632216 | 1.290572834 | 0.328808397 | 0.172698967 | 1.721988733 | 0.037333453 | 0.078237034 | 0.212333652 | 0.1431 | 0.296461547 |
| RUF_107c.18_ | -3.715917651 | -3.486115089 | -3.944461565 | -0.476765919 | 0.732863377 | -0.609408341 | 0.247333346 | 0.373782905 | 0.530994849 | 0.4684 | 0.605950834 |
| RUF_107c.19_ | 0.385308453 | 0.253399743 | 0.507875088 | 0.228409695 | 0.243052203 | 0.890167826 | 0.17800002 | 0.260292417 | 0.440309381 | 0.3476 | 0.513616971 |
| RUF_107c.2_ | -0.150911162 | -0.495438186 | 0.165645612 | 0.682000998 | 0.256287369 | 2.586668412 | 0.000673369 | 0.022134179 | 0.100616716 | 0.1006 | 0.243873778 |
| RUF_107c.23_ | 0.368046632 | 0.493372961 | 0.220588621 | -0.266581066 | 0.241783924 | -1.061427262 | 0.133910755 | 0.192755897 | 0.367128181 | 0.2729 | 0.442191283 |
| RUF_107c.24_ | -0.89137854 | -0.873527441 | -0.943898258 | -0.05285809 | 0.342606215 | -0.128366958 | 0.445702867 | 0.675283436 | 0.793927347 | 0.7668 | 0.853622827 |
| RUF_107c.26_ | -4.174040148 | -4.330125838 | -4.036228959 | 0.304382441 | 0.759628035 | 0.388810226 | 0.33533334 | 0.452657252 | 0.603854251 | 0.5634 | 0.68309248 |
| RUF_107c.30_ | -5.956793026 | -5.73373004 | -6.214306457 | -0.493126162 | 1.545166494 | -0.271700635 | 0.364666672 | 0.490879321 | 0.642560761 | 0.6066 | 0.721806132 |
| RUF_107c.38_ | -1.207907335 | -1.212613083 | -1.198770368 | 0.0095659 | 0.338113588 | 0.026367494 | 0.489333334 | 0.624671576 | 0.753933458 | 0.7275 | 0.822547517 |
| RUF_107c.47_ | 0.283056807 | 0.180823739 | 0.407429079 | 0.260860488 | 0.303074147 | 0.927867643 | 0.110000037 | 0.235418954 | 0.427016778 | 0.2661 | 0.423029503 |
| RUF_107c.50_ | -0.848937681 | -0.891638085 | -0.792012241 | 0.102158461 | 0.275309302 | 0.344895871 | 0.353097941 | 0.512459614 | 0.656874461 | 0.6133 | 0.727760934 |
| RUF_107c.51_ | -0.616411036 | -0.757270338 | -0.498326053 | 0.267456251 | 0.263733424 | 0.961307458 | 0.129247199 | 0.214977262 | 0.376301422 | 0.2944 | 0.446680081 |
| RUF_107c.53_ | 1.117557755 | 1.700702576 | 0.711130174 | -0.916572475 | 0.276970579 | -3.202406588 | 4.68E-05 | 0.013503996 | 0.079985273 | 0.1 | 0.243021319 |
| RUF_107c.56_ | 0.835578135 | 0.511979262 | 1.14439197 | 0.635136766 | 0.144779007 | 4.323344578 | 4.68E-05 | 0.006023999 | 0.047709598 | 0.1 | 0.243021319 |
| RUF_107c.57_ | -1.024479121 | -1.039716855 | -1.009413184 | 0.010416801 | 0.437930728 | 0.019837462 | 0.487674884 | 0.720904185 | 0.828243376 | 0.8029 | 0.876155561 |

| | | | | | | | | | | | |
|---|---|---|---|---|---|---|---|---|---|---|---|
| RUF_107c.58_ | 1.961944132 | 1.88356638 | 2.061737621 | 0.195169778 | 0.158309353 | 1.270583818 | 0.041972124 | 0.143559062 | 0.31504372 | 0.1455 | 0.298542888 |
| RUF_107c.59_ | 0.922534463 | 0.631479459 | 1.066658502 | 0.487231603 | 0.333245032 | 1.537194055 | 0.019333568 | 0.076308512 | 0.217597387 | 0.1223 | 0.27237909 |
| RUF_107c.63_ | -0.54772614 | -0.839594643 | -0.259939098 | 0.618371457 | 0.333104144 | 1.815016545 | 0.010659989 | 0.055224733 | 0.174193866 | 0.1113 | 0.258099393 |
| RUF_107c.69_ | -0.525013144 | -0.452296858 | -0.637836072 | -0.185008938 | 0.292972771 | -0.626010186 | 0.252498347 | 0.392536875 | 0.557653004 | 0.506 | 0.639914129 |
| RUF_107c.70_ | 0.012754962 | -0.374994418 | 0.291908891 | 0.661204988 | 0.231725981 | 2.821443183 | 0.002000966 | 0.021910653 | 0.098641098 | 0.1011 | 0.244554453 |
| RUF_107c.72_ | -0.814709587 | -1.11648042 | -0.59923885 | 0.5066723 | 0.290789266 | 1.652077758 | 0.029980163 | 0.077032879 | 0.207100612 | 0.1403 | 0.293625003 |
| RUF_107c.77_ | -0.095811182 | -0.35525049 | 0.196639762 | 0.58143803 | 0.299960159 | 2.04712886 | 0.0020023 | 0.062824584 | 0.190534027 | 0.1025 | 0.24645647 |
| RUF_107c.8_ | -1.851641843 | -2.735873939 | -1.329228552 | 1.397744343 | 0.432668706 | 3.166439023 | 4.68E-05 | 0.020792898 | 0.095418672 | 0.1001 | 0.243159876 |
| RUF_107c.81_ | -3.438791765 | -3.599875293 | -3.32117349 | 0.24867771 | 0.712028481 | 0.32300376 | 0.362666672 | 0.543581798 | 0.687133566 | 0.6575 | 0.765440032 |
| RUF_107c.82_ | 2.396588463 | 2.102448662 | 2.588201587 | 0.491611942 | 0.14440893 | 3.381339562 | 4.68E-05 | 0.021597144 | 0.102540208 | 0.1 | 0.243021319 |
| RUF_107c.83_ | -4.008000603 | -3.737754426 | -4.509673476 | -0.789631058 | 0.875536453 | -0.804673261 | 0.187208547 | 0.286894867 | 0.456542475 | 0.4001 | 0.546548336 |
| RUF_107c.86_ | -0.778417885 | -0.7080058 | -0.844753058 | -0.183642128 | 0.409267189 | -0.451885428 | 0.313124592 | 0.493153197 | 0.651925622 | 0.6276 | 0.74260402 |
| RUF_107c.89_ | 1.392887617 | 1.384902423 | 1.404037952 | 0.051746387 | 0.213470584 | 0.249250246 | 0.405063295 | 0.608834394 | 0.749362189 | 0.7903 | 0.8654648 |
| RUF_107c.90_ | -2.742391141 | -3.295004889 | -2.423051156 | 0.910671783 | 0.794422155 | 1.094604048 | 0.112591641 | 0.174625385 | 0.350570039 | 0.2409 | 0.405618067 |
| RUF_107c.92_ | -1.310237943 | -1.358950712 | -1.249980164 | 0.101584767 | 0.399545515 | 0.23376877 | 0.39840107 | 0.60955007 | 0.73926478 | 0.7132 | 0.806125083 |
| RUF_107c.94_ | -3.080020301 | -3.262077551 | -2.924750209 | 0.332414944 | 0.603743751 | 0.515883241 | 0.280666677 | 0.427782393 | 0.580276167 | 0.5328 | 0.659060559 |
| RUF_107c.95_ | 3.117511072 | 3.062432051 | 3.156547706 | 0.090828018 | 0.080292693 | 1.040131436 | 0.13000003 | 0.191419905 | 0.360030766 | 0.2683 | 0.432759611 |
| RUF_107c.96_ | -0.392326041 | -0.500978388 | -0.239817647 | 0.302308851 | 0.348478273 | 0.886501808 | 0.114666702 | 0.244657525 | 0.436525398 | 0.2758 | 0.428088726 |
| RUF_111_ | -0.890771891 | -0.936952323 | -0.852519129 | 0.080486222 | 0.261950535 | 0.285621589 | 0.375333338 | 0.512191987 | 0.652873016 | 0.6143 | 0.726354084 |
| RUF_121_ | -0.729722711 | -0.482350892 | -1.153806584 | -0.667538337 | 0.29262648 | -2.210925113 | 0.006000767 | 0.044834892 | 0.149914346 | 0.1065 | 0.251488113 |
| RUF_125_ | 2.27289672 | 3.314760255 | 1.014857074 | -2.303885846 | 0.161421238 | -14.35085656 | 4.68E-05 | 0.00130998 | 0.023903588 | 0.1 | 0.243021319 |
| RUF_167_ | -0.159771849 | -0.724965848 | 0.161230642 | 0.873977902 | 0.26512481 | 3.187649841 | 4.68E-05 | 0.019105239 | 0.092229702 | 0.1001 | 0.243159523 |
| RUF_178_ | -4.927453849 | -5.453769525 | -4.43987603 | 1.036212191 | 1.467203167 | 0.678586237 | 0.230666681 | 0.350902645 | 0.524397481 | 0.4548 | 0.601459461 |
| RUF_219c_ | -0.437915969 | 0.645632625 | -1.737375986 | -2.395092641 | 0.373030012 | -6.346402666 | 4.68E-05 | 0.001669561 | 0.026116961 | 0.1 | 0.243021319 |
| RUF_220c_ | -0.418580384 | -0.026306144 | -0.754734664 | -0.714499738 | 0.251246686 | -2.731649712 | 0.002000966 | 0.020524271 | 0.094915896 | 0.1018 | 0.245333087 |
| RUF_226_ | -3.01065542 | -3.108570944 | -2.864379392 | 0.234163649 | 0.639798827 | 0.337400135 | 0.347768161 | 0.538791207 | 0.681119399 | 0.6474 | 0.756101906 |
| RUF_247c_ | 1.216215001 | 1.120097743 | 1.363223359 | 0.240112146 | 0.150700616 | 1.561810921 | 0.030666813 | 0.092381585 | 0.23194359 | 0.1439 | 0.296819463 |
| RUF_257c_ | 0.787572205 | 0.660772533 | 0.951109158 | 0.286535902 | 0.242334327 | 1.128165166 | 0.104000039 | 0.163501011 | 0.337181071 | 0.2365 | 0.408342729 |
| RUF_279_ | 2.479955851 | 2.048648102 | 2.911798785 | 0.845240743 | 0.149687114 | 5.625546393 | 4.68E-05 | 0.001817634 | 0.028522642 | 0.1 | 0.243021319 |
| RUF_290_ | 1.88974005 | 1.89345049 | 1.88518176 | -0.006066246 | 0.114731316 | -0.04835646 | 0.481333334 | 0.622852625 | 0.751033084 | 0.7363 | 0.824042627 |
| RUF_321c_ | -8.42572738 | -7.659578586 | -9.399215251 | -1.617132937 | 3.691032591 | -0.369760942 | 0.297135252 | 0.442544755 | 0.594921209 | 0.5362 | 0.661215491 |
| RUF_328c_ | -5.322414752 | -5.705424578 | -5.012957187 | 0.727554321 | 1.465172051 | 0.454872969 | 0.271818798 | 0.412308907 | 0.578596789 | 0.5396 | 0.665947044 |
| RUF_341c.10_ | -0.926514128 | -1.062891765 | -0.417949982 | 0.611237158 | 0.419423122 | 1.206123277 | 0.124000032 | 0.176990587 | 0.356502736 | 0.2791 | 0.446768944 |
| RUF_341c.11_ | -0.468461157 | -0.590748739 | -0.319580271 | 0.268961405 | 0.220362813 | 1.146894726 | 0.105929419 | 0.176383748 | 0.333247958 | 0.2557 | 0.412497983 |
| RUF_341c.2_ | -2.458745807 | -2.746544574 | -2.262819028 | 0.468915457 | 0.488902076 | 0.916263417 | 0.165333356 | 0.243468024 | 0.41058288 | 0.3367 | 0.488904199 |
| SL1344_RS00005_ | 4.363603505 | 4.613396395 | 4.172407938 | -0.436089784 | 0.187971192 | -2.444846653 | 4.68E-05 | 0.026659143 | 0.119814126 | 0.1 | 0.243021319 |
| SL1344_RS00010_ | 2.675461171 | 2.748559106 | 2.577978865 | -0.18433307 | 0.152688057 | -1.193108969 | 0.11133337 | 0.150767229 | 0.324814554 | 0.2221 | 0.39428575 |
| SL1344_RS00015_ | 2.076889887 | 2.163805249 | 2.012818583 | -0.152880717 | 0.105061055 | -1.411981697 | 0.059293877 | 0.117358544 | 0.263514314 | 0.1729 | 0.326842894 |
| SL1344_RS00020_ | 3.077539067 | 3.111634098 | 3.039995155 | -0.06703757 | 0.210518002 | -0.273328347 | 0.405729517 | 0.631798698 | 0.775057813 | 0.8882 | 0.933517271 |
| SL1344_RS00025_ | 1.630155547 | 1.631486559 | 1.627695325 | 0.003518917 | 0.125700828 | 0.024059429 | 0.484666667 | 0.626042433 | 0.752925584 | 0.7342 | 0.823126532 |
| SL1344_RS00030_ | 1.441126933 | 1.368269082 | 1.550571993 | 0.195488536 | 0.168111886 | 1.134867611 | 0.094603641 | 0.170937154 | 0.344387463 | 0.2232 | 0.385037954 |
| SL1344_RS00035_ | 5.493241189 | 5.489150441 | 5.50519008 | 0.016360747 | 0.075401836 | 0.19919858 | 0.40906063 | 0.619799899 | 0.764471273 | 0.807 | 0.886841201 |
| SL1344_RS00040_ | 2.130670391 | 1.987780238 | 2.365788372 | 0.370193037 | 0.12871736 | 2.749245316 | 0.000673369 | 0.028725353 | 0.118984065 | 0.1001 | 0.243159399 |
| SL1344_RS00045_ | 3.731343523 | 3.238141872 | 4.151980811 | 0.934461095 | 0.122322146 | 7.76007301 | 4.68E-05 | 0.00146045 | 0.025540288 | 0.1 | 0.243021319 |
| SL1344_RS00050_ | -2.921624063 | -2.882236107 | -3.00209558 | -0.037165361 | 0.724249452 | -0.039352579 | 0.482666667 | 0.701630726 | 0.812913389 | 0.7906 | 0.869618511 |
| SL1344_RS00055_ | -2.682823231 | -2.720258099 | -2.636576346 | 0.134575205 | 0.666035876 | 0.209717038 | 0.408000004 | 0.61255948 | 0.744736455 | 0.735 | 0.8237099 |
| SL1344_RS00065_ | 4.284699042 | 4.344171657 | 4.230789301 | -0.135569677 | 0.114715634 | -1.198118474 | 0.051965441 | 0.138563311 | 0.314336419 | 0.1539 | 0.313372002 |
| SL1344_RS00070_ | -0.522126749 | -0.708258498 | -0.322578772 | 0.399655222 | 0.275881506 | 1.388362642 | 0.054630326 | 0.100317442 | 0.241688337 | 0.1605 | 0.31531355 |
| SL1344_RS00075_ | -4.232061993 | -4.732875893 | -3.752612655 | 0.963552878 | 0.924634706 | 0.99564547 | 0.147333359 | 0.220627398 | 0.385918012 | 0.3068 | 0.463788016 |
| SL1344_RS00080_ | -3.374243652 | -3.364943488 | -3.398936076 | -0.101623264 | 0.781874858 | -0.117351226 | 0.448000002 | 0.621942884 | 0.754385165 | 0.7382 | 0.828546735 |
| SL1344_RS00085_ | -5.53116664 | -5.439517479 | -5.651270434 | -0.44181781 | 2.37659161 | -0.148498863 | 0.401333337 | 0.575307451 | 0.726622707 | 0.7713 | 0.850124045 |
| SL1344_RS00090_ | 0.107350618 | 0.154194758 | 0.054751771 | -0.100157223 | 0.187783712 | -0.506265738 | 0.282666677 | 0.442156141 | 0.592860044 | 0.5507 | 0.672983867 |
| SL1344_RS00095_ | -1.032284528 | -0.858013344 | -1.246542761 | -0.384796732 | 0.281561497 | -1.320225368 | 0.065956094 | 0.125325434 | 0.271074629 | 0.1897 | 0.34461426 |

| | | | | | | | | | | | |
|---|---|---|---|---|---|---|---|---|---|---|---|
| SL1344_RS00100_ | -2.378643558 | -2.204856782 | -2.606811303 | -0.420444106 | 0.514601787 | -0.760987223 | 0.195203216 | 0.298520098 | 0.471679156 | 0.3974 | 0.544234318 |
| SL1344_RS00105_ | -2.38036053 | -2.394313527 | -2.37012291 | 0.012944902 | 0.491953821 | 0.027368007 | 0.484666667 | 0.607110658 | 0.737476882 | 0.7195 | 0.810943923 |
| SL1344_RS00110_ | -3.573501069 | -3.462270161 | -3.711923473 | -0.28479613 | 0.860656399 | -0.312586237 | 0.349100606 | 0.518584651 | 0.669964693 | 0.651 | 0.75584543 |
| SL1344_RS00115_ | -1.538833027 | -1.521142085 | -1.554669255 | -0.08269222 | 0.552921556 | -0.144975023 | 0.441705532 | 0.691141381 | 0.806939125 | 0.831 | 0.893446083 |
| SL1344_RS00120_ | -2.141219423 | -2.055159079 | -2.287099218 | -0.215758896 | 0.492181973 | -0.392888997 | 0.340439713 | 0.54008892 | 0.687456627 | 0.6463 | 0.761447176 |
| SL1344_RS00125_ | -3.302263519 | -3.335532492 | -3.275901382 | 0.062930339 | 0.6001792 | 0.106122743 | 0.449700202 | 0.555179621 | 0.693921377 | 0.6642 | 0.767369529 |
| SL1344_RS00130_ | -3.535378953 | -3.486698361 | -3.63161886 | -0.138079078 | 0.742616927 | -0.169291502 | 0.425333336 | 0.587388368 | 0.722778739 | 0.6981 | 0.798059877 |
| SL1344_RS00135_ | -2.664206847 | -2.777519877 | -2.514367174 | 0.245750919 | 0.562065391 | 0.398509472 | 0.327115264 | 0.505703009 | 0.652896626 | 0.5998 | 0.719963467 |
| SL1344_RS00140_ | -1.095973526 | -1.228124404 | -0.94155926 | 0.291522191 | 0.305933117 | 0.910850578 | 0.152564982 | 0.238776809 | 0.401642729 | 0.3261 | 0.479298351 |
| SL1344_RS00145_ | -1.173748908 | -1.213895276 | -1.143994305 | 0.091768298 | 0.432494249 | 0.206139417 | 0.413724187 | 0.620072789 | 0.753221893 | 0.757 | 0.84057351 |
| SL1344_RS00150_ | -4.875403012 | -4.875938772 | -4.872820142 | 0.005184958 | 0.970728362 | 0.004520991 | 0.496666667 | 0.540944898 | 0.683098193 | 0.6493 | 0.758860292 |
| SL1344_RS00155_ | -2.400070926 | -2.440917002 | -2.369187831 | -0.002330007 | 0.714451526 | -0.003505333 | 0.498666667 | 0.720593419 | 0.828223225 | 0.8339 | 0.896313214 |
| SL1344_RS00160_ | -4.324828809 | -4.787790994 | -3.847798182 | 1.003232922 | 1.011390098 | 0.937736594 | 0.139240534 | 0.222726088 | 0.38895601 | 0.3053 | 0.460099964 |
| SL1344_RS00165_ | -1.360468876 | -1.523626583 | -1.168481303 | 0.321631751 | 0.428517985 | 0.694759018 | 0.233844118 | 0.359898809 | 0.532937771 | 0.4564 | 0.607007523 |
| SL1344_RS00170_ | -0.44735115 | -0.484038268 | -0.407827214 | 0.077717965 | 0.209192132 | 0.337987816 | 0.349333339 | 0.485257416 | 0.63195194 | 0.5894 | 0.705790968 |
| SL1344_RS00175_ | -5.904571454 | -5.544830797 | -6.235892718 | -0.746536015 | 1.714636113 | -0.412099113 | 0.311792147 | 0.480446884 | 0.633320209 | 0.5962 | 0.71445775 |
| SL1344_RS00180_ | -0.532026846 | -0.523454788 | -0.5441823 | 0.008493108 | 0.312697633 | 0.023966658 | 0.492338441 | 0.677438097 | 0.795592074 | 0.7913 | 0.866814102 |
| SL1344_RS00185_ | -0.918514581 | -0.998118759 | -0.794046288 | 0.186252171 | 0.324268699 | 0.509111327 | 0.28800001 | 0.47052144 | 0.625780458 | 0.5734 | 0.703290749 |
| SL1344_RS00190_ | -1.821272371 | -2.001351474 | -1.657467336 | 0.345847228 | 0.38400132 | 0.830370616 | 0.1798801 | 0.279342071 | 0.443035978 | 0.3703 | 0.522161579 |
| SL1344_RS00195_ | 0.330478637 | 0.254482332 | 0.420699558 | 0.171876035 | 0.205107194 | 0.797390719 | 0.194666685 | 0.294115106 | 0.464714307 | 0.3969 | 0.549135146 |
| SL1344_RS00205_ | 1.953685226 | 2.102870892 | 1.755772712 | -0.35334166 | 0.142205602 | -2.445239248 | 0.002000966 | 0.024484865 | 0.107915981 | 0.1005 | 0.243714178 |
| SL1344_RS00210_ | -0.166918342 | -0.232522041 | -0.120257464 | 0.116854923 | 0.215179917 | 0.50709704 | 0.28000001 | 0.437336792 | 0.594061863 | 0.5426 | 0.670177252 |
| SL1344_RS00215_ | -4.094310188 | -4.147109949 | -4.056064711 | 0.059928022 | 1.087817916 | 0.053844917 | 0.479013991 | 0.676664931 | 0.796921227 | 0.7967 | 0.871389146 |
| SL1344_RS00220_ | 8.366048083 | 8.189279664 | 8.456807025 | 0.255186432 | 0.127315254 | 2.238930921 | 4.68E-05 | 0.018373695 | 0.098498816 | 0.1 | 0.243021319 |
| SL1344_RS00225_ | -0.396458474 | -0.329452504 | -0.469553921 | -0.139524128 | 0.22477518 | -0.578567791 | 0.253164569 | 0.39067064 | 0.550009733 | 0.488 | 0.624568431 |
| SL1344_RS00230_ | 3.052152575 | 3.016068901 | 3.097498098 | 0.038716172 | 0.138205529 | 0.259312118 | 0.404397072 | 0.721753779 | 0.830041104 | 0.8186 | 0.888799069 |
| SL1344_RS00235_ | 4.997431632 | 5.004522583 | 4.980250698 | -0.025168325 | 0.101189205 | -0.159346822 | 0.409333337 | 0.800665318 | 0.887696289 | 0.8383 | 0.908156656 |
| SL1344_RS00240_ | 2.923093272 | 2.83916994 | 3.06768061 | 0.236785459 | 0.127162031 | 1.882628417 | 0.002668394 | 0.080450385 | 0.226042615 | 0.1018 | 0.245480486 |
| SL1344_RS00245_ | 2.236304934 | 2.195132988 | 2.281141602 | 0.118559075 | 0.164605313 | 0.729549964 | 0.192666685 | 0.288531878 | 0.47869219 | 0.4099 | 0.564504313 |
| SL1344_RS00250_ | 3.398987566 | 3.302818744 | 3.474603738 | 0.165428092 | 0.080240989 | 1.986539518 | 0.021319333 | 0.0481902 | 0.158350877 | 0.1174 | 0.266588827 |
| SL1344_RS00255_ | 1.534426502 | 1.544736087 | 1.524826701 | -0.032992858 | 0.24260286 | -0.140296279 | 0.421333336 | 0.741188789 | 0.843853706 | 0.7949 | 0.872046122 |
| SL1344_RS00260_ | 1.717074528 | 1.91322311 | 1.486495516 | -0.445603214 | 0.146705243 | -3.044389072 | 4.68E-05 | 0.021303629 | 0.100119501 | 0.1 | 0.243021319 |
| SL1344_RS00270_ | -1.054033921 | -1.10321135 | -0.966505017 | 0.12912763 | 0.330576174 | 0.368751257 | 0.351333339 | 0.556892107 | 0.696678987 | 0.6599 | 0.768944542 |
| SL1344_RS00275_ | -4.266246002 | -4.170835491 | -4.34108484 | -0.216602019 | 1.145165346 | -0.165168816 | 0.427048637 | 0.642065254 | 0.767892307 | 0.7667 | 0.848897563 |
| SL1344_RS00280_ | -4.70165551 | -5.344642626 | -4.288161888 | 1.168087579 | 1.540502241 | 0.689925625 | 0.198534328 | 0.299459204 | 0.485581474 | 0.3984 | 0.552607098 |
| SL1344_RS00285_ | -7.475357692 | -7.720024074 | -7.196511909 | 0.502877472 | 3.040186248 | 0.123221114 | 0.42266667 | 0.571929347 | 0.714045191 | 0.6991 | 0.795363348 |
| SL1344_RS00290_ | -2.682468982 | -2.559619668 | -2.854856632 | -0.291374484 | 0.507999492 | -0.534555114 | 0.275816133 | 0.407316714 | 0.564580889 | 0.5176 | 0.645770926 |
| SL1344_RS00295_ | -3.593005595 | -3.569668691 | -3.621629134 | -0.032373489 | 0.799651401 | -0.037963256 | 0.483677549 | 0.635929688 | 0.763359127 | 0.7454 | 0.833248956 |
| SL1344_RS00300_ | -5.006731514 | -5.095261393 | -4.952906777 | 0.094155447 | 1.243688594 | 0.067085293 | 0.468666668 | 0.59521194 | 0.728737164 | 0.7086 | 0.80380008 |
| SL1344_RS00305_ | -3.14751033 | -2.994559558 | -3.329078298 | -0.531707395 | 0.998057936 | -0.495999248 | 0.29113925 | 0.419939014 | 0.597039969 | 0.5848 | 0.712081561 |
| SL1344_RS00310_ | -2.305519023 | -2.000459201 | -2.542872541 | -0.525890666 | 0.458444079 | -1.043899603 | 0.126666698 | 0.197482421 | 0.357706046 | 0.2816 | 0.440740884 |
| SL1344_RS00315_ | -2.946166309 | -2.331525426 | -3.669873088 | -1.358446053 | 0.575869694 | -2.311935705 | 0.003998486 | 0.03126524 | 0.119570199 | 0.1024 | 0.246375902 |
| SL1344_RS00320_ | -2.03283962 | -1.684772071 | -2.278713278 | -0.563232381 | 0.477303675 | -1.151436354 | 0.108594308 | 0.173391299 | 0.335799948 | 0.2588 | 0.422328864 |
| SL1344_RS00325_ | 2.202709037 | 2.239060851 | 2.175050761 | -0.095608211 | 0.185338311 | -0.493636234 | 0.274483688 | 0.397796952 | 0.583161472 | 0.5475 | 0.684611689 |
| SL1344_RS00330_ | -3.994400753 | -4.200195085 | -3.75734362 | 0.421614027 | 0.734412761 | 0.534617712 | 0.279813468 | 0.413844335 | 0.564570023 | 0.5203 | 0.647068659 |
| SL1344_RS00335_ | 0.863801583 | 0.858601197 | 0.866882703 | 0.030939951 | 0.220028511 | 0.136429561 | 0.445333335 | 0.674589259 | 0.795766129 | 0.8097 | 0.880374809 |
| SL1344_RS00340_ | 2.227753607 | 2.161994718 | 2.337177689 | 0.169108349 | 0.114901887 | 1.413270666 | 0.059293877 | 0.114490924 | 0.263535768 | 0.185 | 0.342536963 |
| SL1344_RS00345_ | -0.671442305 | -1.090245513 | -0.244969708 | 0.852891537 | 0.269517662 | 3.095202543 | 0.00067292 | 0.015661517 | 0.081364965 | 0.1002 | 0.243304006 |
| SL1344_RS00350_ | -3.218094601 | -2.903636279 | -3.564479895 | -0.684789063 | 0.581171403 | -1.106115186 | 0.114000035 | 0.177686938 | 0.330466343 | 0.2537 | 0.408692034 |
| SL1344_RS00355_ | -2.710480223 | -2.360035209 | -2.909867633 | -0.512886078 | 0.540483498 | -0.861351202 | 0.185209879 | 0.287160609 | 0.453471459 | 0.384 | 0.539238141 |
| SL1344_RS00360_ | -1.840100741 | -1.610090027 | -2.033136773 | -0.430015153 | 0.37223255 | -1.061504983 | 0.111259197 | 0.180474273 | 0.337822577 | 0.2587 | 0.415616109 |
| SL1344_RS00365_ | -1.837320485 | -1.676503917 | -2.027989842 | -0.346022001 | 0.344371063 | -0.959422149 | 0.147235203 | 0.223042562 | 0.380895997 | 0.3118 | 0.462664479 |
| SL1344_RS00370_ | -1.148065115 | -1.126943419 | -1.191482415 | -0.011363873 | 0.396279916 | -0.03010481 | 0.487674884 | 0.673311447 | 0.792741645 | 0.7778 | 0.859642665 |

| | | | | | | | | | | | |
|---|---|---|---|---|---|---|---|---|---|---|---|
| SL1344_RS00375_ | -3.974687202 | -3.951946695 | -3.987816491 | -0.012900137 | 1.231711361 | -0.011179327 | 0.495333334 | 0.7162428 | 0.825514958 | 0.8333 | 0.896778068 |
| SL1344_RS00380_ | -6.225999689 | -6.240066357 | -6.214710962 | 0.022173267 | 2.478659259 | 0.007053549 | 0.497001999 | 0.633717119 | 0.768363748 | 0.7906 | 0.87104738 |
| SL1344_RS00385_ | -2.370231902 | -2.486904124 | -2.203168119 | 0.227537343 | 0.572124645 | 0.368834054 | 0.347101938 | 0.589118677 | 0.724838547 | 0.6894 | 0.793028005 |
| SL1344_RS00390_ | -2.963739589 | -2.85013591 | -3.401977921 | -0.480525177 | 0.826369484 | -0.48562468 | 0.339773491 | 0.552120055 | 0.704888134 | 0.6895 | 0.807324608 |
| SL1344_RS00395_ | -2.747454788 | -2.678584009 | -2.805114148 | -0.145974352 | 0.528591505 | -0.262084114 | 0.381745508 | 0.555444723 | 0.694082777 | 0.6591 | 0.763905605 |
| SL1344_RS00400_ | -3.736223779 | -3.770905472 | -3.702808174 | 0.051646521 | 0.734137269 | 0.06993311 | 0.467688209 | 0.566693909 | 0.703557321 | 0.6783 | 0.777522241 |
| SL1344_RS00405_ | -0.52277892 | -0.64935979 | -0.367726523 | 0.255182442 | 0.305592017 | 0.79152939 | 0.195333351 | 0.311469523 | 0.484938626 | 0.4076 | 0.564797661 |
| SL1344_RS00410_ | -2.434347956 | -2.398381827 | -2.461641968 | -0.06997658 | 0.576770529 | -0.106932369 | 0.453333335 | 0.64581671 | 0.771285901 | 0.7408 | 0.831358633 |
| SL1344_RS00415_ | -0.326047926 | -0.53656605 | -0.11165172 | 0.427665423 | 0.2193188 | 1.90272573 | 0.01798826 | 0.054308377 | 0.1650598 | 0.1204 | 0.268681824 |
| SL1344_RS00420_ | -2.05811619 | -1.953316683 | -2.149565515 | -0.197998802 | 0.385230227 | -0.491739742 | 0.282666677 | 0.429387558 | 0.583048362 | 0.5369 | 0.662469787 |
| SL1344_RS00425_ | -5.713642496 | -5.917322379 | -5.416764279 | 0.251912798 | 2.6095707 | 0.07648774 | 0.444666669 | 0.677433641 | 0.800929202 | 0.784 | 0.866795813 |
| SL1344_RS00430_ | 0.057731449 | -0.074234921 | 0.288727467 | 0.342890416 | 0.221955428 | 1.407813163 | 0.070000061 | 0.11286174 | 0.258120336 | 0.1899 | 0.348779969 |
| SL1344_RS00435_ | -2.566426812 | -2.871714474 | -2.368601148 | 0.494532 | 0.494875814 | 0.92889567 | 0.148567647 | 0.241870159 | 0.402861119 | 0.3295 | 0.480898101 |
| SL1344_RS00440_ | -0.535634906 | -0.516706536 | -0.560959431 | -0.077718831 | 0.332667487 | -0.204355098 | 0.404397072 | 0.558911593 | 0.709130964 | 0.7054 | 0.800110553 |
| SL1344_RS00450_ | 2.415422331 | 2.319177533 | 2.544179402 | 0.244394193 | 0.147965998 | 1.629705122 | 0.013324793 | 0.079561999 | 0.220437091 | 0.1169 | 0.264885001 |
| SL1344_RS00455_ | 2.420434937 | 2.251395232 | 2.590441123 | 0.321145076 | 0.187686061 | 1.69226962 | 0.025333511 | 0.066794896 | 0.200448246 | 0.125 | 0.277794021 |
| SL1344_RS00465_ | 3.221272583 | 3.020798151 | 3.35126263 | 0.32370249 | 0.105113294 | 3.031622988 | 4.68E-05 | 0.019340184 | 0.095425189 | 0.1 | 0.243021319 |
| SL1344_RS00470_ | 4.114777786 | 3.971638216 | 4.279412566 | 0.314810041 | 0.065299414 | 4.774604768 | 4.68E-05 | 0.005052064 | 0.045650751 | 0.1 | 0.243021319 |
| SL1344_RS00475_ | 4.021857879 | 3.986955051 | 4.101304918 | 0.11141858 | 0.062175732 | 1.729889722 | 0.042000105 | 0.088996995 | 0.228553128 | 0.1491 | 0.30317612 |
| SL1344_RS00480_ | 1.468464956 | 1.372698568 | 1.65217098 | 0.249141433 | 0.179804718 | 1.255778452 | 0.106595641 | 0.14314386 | 0.307957675 | 0.2185 | 0.387064539 |
| SL1344_RS00485_ | 2.738341491 | 2.45686777 | 2.947743747 | 0.492239092 | 0.094314002 | 5.258235782 | 4.68E-05 | 0.005947974 | 0.047949051 | 0.1 | 0.243021319 |
| SL1344_RS00490_ | 3.71692736 | 3.511638851 | 3.883729159 | 0.384385304 | 0.083792586 | 4.674237626 | 4.68E-05 | 0.004234865 | 0.042031792 | 0.1 | 0.243021319 |
| SL1344_RS00495_ | 1.482514174 | 1.447079365 | 1.555723135 | 0.118253801 | 0.20297178 | 0.525226825 | 0.307794812 | 0.474895236 | 0.643637686 | 0.609 | 0.738966076 |
| SL1344_RS00500_ | -1.344592889 | -1.566792591 | -1.204443892 | 0.351820198 | 0.362186986 | 0.933213413 | 0.153231204 | 0.23421322 | 0.404063384 | 0.3256 | 0.484092648 |
| SL1344_RS00505_ | -4.003941801 | -4.635610304 | -3.559419526 | 0.956470989 | 1.354006909 | 0.674653089 | 0.233844118 | 0.353957311 | 0.537272846 | 0.4598 | 0.617459682 |
| SL1344_RS00510_ | -3.26023585 | -3.15394626 | -3.366850486 | -0.211167366 | 0.566898771 | -0.361644556 | 0.342438381 | 0.477494642 | 0.623944349 | 0.5867 | 0.703747783 |
| SL1344_RS00515_ | -2.826704516 | -2.802424125 | -2.87640323 | -0.127820332 | 0.695015332 | -0.156044482 | 0.435043307 | 0.617653797 | 0.753615665 | 0.7459 | 0.834604816 |
| SL1344_RS00520_ | -3.586890272 | -3.35897806 | -3.784427111 | -0.40523 | 0.686789449 | -0.549741356 | 0.266666678 | 0.410955033 | 0.564693155 | 0.5151 | 0.642857121 |
| SL1344_RS00525_ | 0.083793445 | -0.010160377 | 0.147939276 | 0.176953295 | 0.252597603 | 0.672686931 | 0.199333351 | 0.313383263 | 0.491681162 | 0.4186 | 0.562491861 |
| SL1344_RS00530_ | 1.322427427 | 1.251400032 | 1.476285381 | 0.198128829 | 0.227729506 | 0.862238412 | 0.179333354 | 0.260541027 | 0.449345087 | 0.3457 | 0.52468368 |
| SL1344_RS00545_ | 0.828025615 | 0.794620061 | 0.883949634 | 0.075512183 | 0.359433943 | 0.219412866 | 0.383077953 | 0.743803117 | 0.846737665 | 0.7608 | 0.852068537 |
| SL1344_RS00550_ | 0.078718483 | -0.039053008 | 0.158641923 | 0.19573422 | 0.210011337 | 0.873412208 | 0.166666689 | 0.258808711 | 0.427797085 | 0.3457 | 0.501075668 |
| SL1344_RS00555_ | 2.539371196 | 2.592930408 | 2.357216581 | -0.229111242 | 0.194106219 | -1.237771272 | 0.101265863 | 0.125149926 | 0.291496846 | 0.224 | 0.397007913 |
| SL1344_RS00560_ | -1.069979953 | -0.961838813 | -1.230773508 | -0.267859247 | 0.347836823 | -0.700943717 | 0.215856111 | 0.328631667 | 0.500894559 | 0.4341 | 0.580595068 |
| SL1344_RS00565_ | -0.386483224 | -0.326781023 | -0.458385191 | -0.116975686 | 0.395654563 | -0.301070432 | 0.364666672 | 0.631455323 | 0.764620078 | 0.7446 | 0.83690174 |
| SL1344_RS00570_ | -0.442614354 | -0.450679188 | -0.437828164 | 0.001488832 | 0.373039884 | 0.003105226 | 0.498666667 | 0.746876551 | 0.8474297 | 0.8628 | 0.916736487 |
| SL1344_RS00575_ | -0.62228474 | -0.682563059 | -0.566692766 | 0.12029189 | 0.274568145 | 0.417480915 | 0.320453039 | 0.489081655 | 0.638404323 | 0.5979 | 0.714347027 |
| SL1344_RS00580_ | -0.53687953 | -0.740397085 | -0.418539539 | 0.323056286 | 0.340253127 | 0.886048237 | 0.163333356 | 0.246365674 | 0.431027309 | 0.3378 | 0.501824761 |
| SL1344_RS00590_ | 0.112531864 | 0.105051702 | 0.118826825 | 0.006286208 | 0.205248459 | 0.032558462 | 0.484666667 | 0.590500963 | 0.725131106 | 0.6881 | 0.790941469 |
| SL1344_RS00595_ | 0.265279495 | 0.442479579 | 0.046503614 | -0.397382111 | 0.169321597 | -2.281795468 | 0.005330642 | 0.035753118 | 0.127695942 | 0.1049 | 0.249259525 |
| SL1344_RS00600_ | 2.369854162 | 2.235440459 | 2.476869135 | 0.27160169 | 0.149209183 | 1.827178855 | 0.008000574 | 0.052505316 | 0.170561367 | 0.1062 | 0.251453756 |
| SL1344_RS00605_ | 4.207256098 | 4.138746634 | 4.329450704 | 0.189394266 | 0.062946031 | 2.875731436 | 4.68E-05 | 0.0298208 | 0.121609055 | 0.1003 | 0.243442874 |
| SL1344_RS00610_ | 4.262457042 | 4.222727878 | 4.361616937 | 0.136333367 | 0.063916035 | 1.872189524 | 0.018654474 | 0.075034834 | 0.211738503 | 0.1167 | 0.264141392 |
| SL1344_RS00615_ | 3.424660354 | 3.368243285 | 3.479261795 | 0.121943782 | 0.11377947 | 1.109605899 | 0.083277865 | 0.172818472 | 0.355250344 | 0.1988 | 0.363484701 |
| SL1344_RS00620_ | 5.425624512 | 5.413928066 | 5.444629028 | 0.022417208 | 0.077174597 | 0.184346028 | 0.404397072 | 0.811246627 | 0.89317645 | 0.8437 | 0.910952937 |
| SL1344_RS00630_ | 3.393625367 | 3.273215401 | 3.536659094 | 0.257589528 | 0.109671229 | 2.324611552 | 0.003334715 | 0.030652481 | 0.125327611 | 0.1022 | 0.246108658 |
| SL1344_RS00635_ | 3.42207186 | 3.298857231 | 3.554368417 | 0.249766257 | 0.111667829 | 2.199339594 | 0.002666616 | 0.032367149 | 0.129429308 | 0.1024 | 0.246388978 |
| SL1344_RS00645_ | 3.626878209 | 3.534212977 | 3.751041742 | 0.212739117 | 0.074769465 | 2.813876785 | 0.000673369 | 0.020419654 | 0.096477144 | 0.1003 | 0.243434945 |
| SL1344_RS00650_ | 3.318365439 | 3.149316219 | 3.456113949 | 0.310159054 | 0.097060018 | 3.179139199 | 4.68E-05 | 0.011316419 | 0.071534527 | 0.1 | 0.243021319 |
| SL1344_RS00655_ | 3.672312934 | 3.602122205 | 3.774730277 | 0.174141925 | 0.093506034 | 1.840615715 | 0.011326186 | 0.067276227 | 0.199618206 | 0.1106 | 0.257461681 |
| SL1344_RS00665_ | 2.763343305 | 2.699655701 | 2.874201759 | 0.158273487 | 0.127688104 | 1.186694889 | 0.111259197 | 0.153349293 | 0.324541426 | 0.2306 | 0.404526467 |
| SL1344_RS00670_ | 3.878288407 | 3.757111307 | 3.927889675 | 0.154954267 | 0.107525723 | 1.347618556 | 0.077333388 | 0.118137169 | 0.282232248 | 0.1922 | 0.362192176 |
| SL1344_RS00675_ | 4.197931201 | 4.186190124 | 4.225096411 | 0.041634414 | 0.090659463 | 0.403345626 | 0.299333342 | 0.419095884 | 0.607345822 | 0.5932 | 0.720033135 |

| | | | | | | | | | | | |
|---|---|---|---|---|---|---|---|---|---|---|---|
| SL1344_RS00680_ | 6.030437326 | 5.966127201 | 6.04871734 | 0.098955166 | 0.108269524 | 0.966010559 | 0.165223207 | 0.178211515 | 0.368412366 | 0.2948 | 0.482927347 |
| SL1344_RS00685_ | 2.997067355 | 2.937919066 | 3.088223379 | 0.123388554 | 0.124708508 | 0.928320792 | 0.166666689 | 0.235706448 | 0.423098546 | 0.3148 | 0.493563595 |
| SL1344_RS00690_ | 5.153585105 | 4.996365521 | 5.2914751 | 0.287526325 | 0.067814783 | 4.230661394 | 4.68E-05 | 0.003962119 | 0.042145135 | 0.1 | 0.243021319 |
| SL1344_RS00695_ | -0.36141847 | -0.413246741 | -0.272956623 | 0.095915925 | 0.382062467 | 0.233665819 | 0.397068625 | 0.72383071 | 0.830391655 | 0.7656 | 0.856143464 |
| SL1344_RS00700_ | 1.835472563 | 1.777043777 | 1.897712023 | 0.120476986 | 0.115114318 | 1.016681368 | 0.13000003 | 0.205846799 | 0.368285796 | 0.2905 | 0.447158938 |
| SL1344_RS00705_ | 1.598433358 | 1.584040365 | 1.610888684 | 0.027721552 | 0.139189285 | 0.195136852 | 0.42066667 | 0.61899717 | 0.74990701 | 0.7182 | 0.813531781 |
| SL1344_RS00710_ | 1.946317386 | 1.916009846 | 1.984513238 | 0.072898314 | 0.11121602 | 0.604221672 | 0.250666679 | 0.384336463 | 0.546530895 | 0.5008 | 0.631492956 |
| SL1344_RS00715_ | 1.878153673 | 1.818778779 | 1.928509346 | 0.137576922 | 0.161023082 | 0.848917867 | 0.151333358 | 0.248231227 | 0.436267274 | 0.3274 | 0.480220814 |
| SL1344_RS00720_ | 3.689991731 | 3.782287361 | 3.654215641 | -0.120336476 | 0.080167791 | -1.34436164 | 0.091272531 | 0.122681328 | 0.281247693 | 0.2132 | 0.379720413 |
| SL1344_RS00725_ | -2.916243025 | -2.823760157 | -3.021159627 | -0.213386888 | 0.556892642 | -0.350474338 | 0.348434383 | 0.501449747 | 0.650708183 | 0.6192 | 0.731569588 |
| SL1344_RS00730_ | -4.131208713 | -3.859170757 | -4.416293628 | -0.546835003 | 0.83708342 | -0.611073866 | 0.246000013 | 0.37866695 | 0.536764364 | 0.4723 | 0.611076004 |
| SL1344_RS00735_ | -5.796091657 | -6.189301595 | -5.583711933 | 0.613523285 | 2.415628768 | 0.202958912 | 0.387075288 | 0.520731303 | 0.684112399 | 0.7072 | 0.80648001 |
| SL1344_RS00740_ | 1.279914763 | 1.175717684 | 1.416146791 | 0.257748222 | 0.168523343 | 1.428892902 | 0.055296548 | 0.100471653 | 0.246407228 | 0.1628 | 0.319995242 |
| SL1344_RS00745_ | 0.581006935 | 0.435622451 | 0.738918913 | 0.303376853 | 0.150537303 | 1.933338221 | 0.014000326 | 0.05211979 | 0.159048981 | 0.1205 | 0.268330588 |
| SL1344_RS00750_ | 1.440103138 | 1.466938143 | 1.412905609 | -0.055600897 | 0.129368798 | -0.421364396 | 0.332445044 | 0.489866145 | 0.639475168 | 0.6002 | 0.718271945 |
| SL1344_RS00755_ | -0.787404418 | -0.823884067 | -0.746732077 | 0.082303604 | 0.226361881 | 0.332793921 | 0.355762831 | 0.475893145 | 0.62158126 | 0.5835 | 0.701389306 |
| SL1344_RS00760_ | -1.968894225 | -1.978271492 | -1.95561447 | 0.011770337 | 0.355845854 | 0.028707889 | 0.488341106 | 0.568345001 | 0.705881593 | 0.6728 | 0.776453194 |
| SL1344_RS00765_ | 1.195388327 | 1.12991549 | 1.272263008 | 0.150683025 | 0.156450873 | 0.913307708 | 0.155896093 | 0.230708903 | 0.40417689 | 0.3183 | 0.475201201 |
| SL1344_RS00770_ | 5.229793116 | 5.043324134 | 5.369423773 | 0.321617598 | 0.05467599 | 5.850817256 | 4.68E-05 | 0.002786026 | 0.033627172 | 0.1 | 0.243021319 |
| SL1344_RS00775_ | 6.832298423 | 6.743919686 | 6.924631203 | 0.183199903 | 0.051935312 | 3.323592047 | 4.68E-05 | 0.035295675 | 0.14116235 | 0.1 | 0.243021319 |
| SL1344_RS00785_ | 6.798992061 | 6.811823276 | 6.72857929 | -0.07548958 | 0.044530726 | -1.124311406 | 0.159893428 | 0.187068179 | 0.376823516 | 0.3211 | 0.498366225 |
| SL1344_RS00790_ | -0.875846099 | -0.924731568 | -0.816498123 | 0.09924638 | 0.315246913 | 0.285726065 | 0.373084615 | 0.571289114 | 0.709627192 | 0.6783 | 0.782659236 |
| SL1344_RS00795_ | -2.888719117 | -2.874970809 | -2.894278193 | 0.017908923 | 0.692436477 | 0.024312339 | 0.487008661 | 0.651379912 | 0.776273645 | 0.7671 | 0.850147129 |
| SL1344_RS00800_ | -1.144122753 | -1.187852552 | -1.073877335 | 0.099994154 | 0.349480572 | 0.26027573 | 0.391738845 | 0.609048465 | 0.741474994 | 0.7056 | 0.807229267 |
| SL1344_RS00805_ | 3.483215736 | 3.513602157 | 3.430999577 | -0.083395622 | 0.172268264 | -0.441565316 | 0.299333342 | 0.413777146 | 0.608218166 | 0.6217 | 0.750407227 |
| SL1344_RS00810_ | 1.969744187 | 2.000944725 | 1.763103525 | -0.220422498 | 0.216038117 | -0.571651377 | 0.334000007 | 0.585238203 | 0.741617816 | 0.7006 | 0.829353277 |
| SL1344_RS00815_ | 1.581474989 | 1.649021732 | 1.525011134 | -0.145701587 | 0.20204255 | -0.700376422 | 0.213333349 | 0.318589419 | 0.505295745 | 0.4378 | 0.598139136 |
| SL1344_RS00820_ | -1.883766231 | -1.900331965 | -1.871527308 | 0.007348873 | 0.484906507 | 0.013412234 | 0.492666667 | 0.666295691 | 0.786064749 | 0.7674 | 0.850174251 |
| SL1344_RS00825_ | -1.326313811 | -1.258415234 | -1.436083657 | -0.197520485 | 0.439970375 | -0.420679217 | 0.322000008 | 0.494788057 | 0.655014375 | 0.6243 | 0.743230329 |
| SL1344_RS00830_ | 0.352628361 | 0.337597475 | 0.370419038 | 0.030877923 | 0.158218392 | 0.180852491 | 0.419333336 | 0.537392176 | 0.678720287 | 0.6523 | 0.758779601 |
| SL1344_RS00835_ | 2.41407632 | 2.372759088 | 2.46742251 | 0.098388913 | 0.10814084 | 0.846405499 | 0.18254499 | 0.264000423 | 0.437534411 | 0.3685 | 0.523386357 |
| SL1344_RS00840_ | 3.995633891 | 3.940793688 | 4.054666182 | 0.111345313 | 0.056920286 | 1.835427031 | 0.026649069 | 0.059347266 | 0.177285818 | 0.1268 | 0.27762221 |
| SL1344_RS00845_ | 3.100360771 | 3.061127386 | 3.141354346 | 0.0859678 | 0.087104044 | 0.973616304 | 0.145902758 | 0.217991268 | 0.390973374 | 0.3106 | 0.470089046 |
| SL1344_RS00850_ | 0.376855483 | 0.284011381 | 0.450257695 | 0.150274818 | 0.215857028 | 0.642990895 | 0.225849448 | 0.35806791 | 0.528244496 | 0.4635 | 0.608873238 |
| SL1344_RS00855_ | 1.510719534 | 1.593965746 | 1.40733992 | -0.138984271 | 0.408890184 | -0.284776093 | 0.430379749 | 0.739047087 | 0.847366119 | 0.961 | 0.977490221 |
| SL1344_RS00860_ | 1.658001599 | 1.84239048 | 1.405268289 | -0.435817332 | 0.11448948 | -3.722677137 | 4.68E-05 | 0.011545925 | 0.067537588 | 0.1 | 0.243021319 |
| SL1344_RS00865_ | 3.310975381 | 3.310804591 | 3.311129255 | -0.00874727 | 0.098654599 | -0.087566144 | 0.470000001 | 0.700158333 | 0.814860433 | 0.8231 | 0.889744272 |
| SL1344_RS00870_ | 5.275762183 | 5.148507312 | 5.430925542 | 0.284378024 | 0.083300425 | 3.542788103 | 4.68E-05 | 0.019966668 | 0.101001089 | 0.1 | 0.243021319 |
| SL1344_RS00875_ | 0.348259732 | 0.356748455 | 0.34039462 | -0.022017697 | 0.17102024 | -0.130998806 | 0.444000002 | 0.574491932 | 0.710531395 | 0.6852 | 0.783581165 |
| SL1344_RS00880_ | 0.768745535 | 0.72424595 | 0.848788234 | 0.143591936 | 0.223365926 | 0.631111827 | 0.234000014 | 0.350098632 | 0.534148127 | 0.4651 | 0.606639172 |
| SL1344_RS00885_ | -1.605858214 | -1.801666967 | -1.394201511 | 0.415781929 | 0.349995006 | 1.134605818 | 0.102666707 | 0.163360518 | 0.317441241 | 0.2375 | 0.394186021 |
| SL1344_RS00890_ | -2.102529597 | -2.41764228 | -1.850584349 | 0.573940683 | 0.39821892 | 1.330906015 | 0.063291207 | 0.117433768 | 0.259711456 | 0.1789 | 0.331512382 |
| SL1344_RS00895_ | -4.452029977 | -4.572551961 | -4.337301541 | 0.218802652 | 0.832731636 | 0.246563566 | 0.389333338 | 0.492148761 | 0.63603688 | 0.5992 | 0.712783713 |
| SL1344_RS00900_ | -2.938011655 | -2.921763682 | -2.974811564 | 0.041242743 | 0.761187045 | 0.053400995 | 0.481333334 | 0.672953271 | 0.791790663 | 0.8 | 0.872682733 |
| SL1344_RS00905_ | -3.011688339 | -3.237880545 | -2.848365671 | 0.406582395 | 0.580405726 | 0.667423876 | 0.225183226 | 0.341625512 | 0.503254303 | 0.4423 | 0.57875572 |
| SL1344_RS00910_ | 1.288103876 | 1.226775461 | 1.341379904 | 0.112350628 | 0.118779484 | 0.919695589 | 0.156562316 | 0.251853807 | 0.409428779 | 0.344 | 0.491057574 |
| SL1344_RS00915_ | 2.501450002 | 2.476503245 | 2.520492954 | 0.049749483 | 0.108330645 | 0.434990546 | 0.316000008 | 0.490071847 | 0.646320756 | 0.6139 | 0.73015156 |
| SL1344_RS00920_ | 2.181330355 | 2.102748873 | 2.268496013 | 0.186612059 | 0.127284054 | 1.432938814 | 0.040666776 | 0.102037394 | 0.249276362 | 0.1459 | 0.299114234 |
| SL1344_RS00925_ | 1.655197262 | 1.6093908 | 1.724360585 | 0.136847551 | 0.188625161 | 0.750259819 | 0.190000019 | 0.289759137 | 0.480988532 | 0.4003 | 0.553143264 |
| SL1344_RS00930_ | 2.642383589 | 2.469169646 | 2.730030122 | 0.261155577 | 0.129937759 | 1.840821229 | 0.011326186 | 0.067838006 | 0.200724555 | 0.1114 | 0.258276238 |
| SL1344_RS00935_ | 4.56669497 | 4.461194382 | 4.653547018 | 0.198893593 | 0.065959542 | 2.99865663 | 4.68E-05 | 0.013017223 | 0.076876068 | 0.1 | 0.243021319 |
| SL1344_RS00940_ | 1.755581173 | 1.650607615 | 1.881014753 | 0.22733555 | 0.112979912 | 1.950612112 | 0.018654474 | 0.053028567 | 0.162801651 | 0.1213 | 0.269280704 |
| SL1344_RS00945_ | 5.853839188 | 6.083427413 | 5.60538654 | -0.478185612 | 0.062829233 | -7.658838397 | 4.68E-05 | 0.002190781 | 0.03106436 | 0.1 | 0.243021319 |

| | | | | | | | | | | |
|---|---|---|---|---|---|---|---|---|---|---|
| SL1344_RS00950_ | 0.783627686 | 0.661302052 | 0.918934641 | 0.252953592 | 0.177464336 | 1.40270555 | 0.071952091 | 0.115590888 | 0.262904728 | 0.1844 | 0.344309279 |
| SL1344_RS00955_ | 0.577862136 | 0.519743009 | 0.664734294 | 0.144335293 | 0.171196381 | 0.77390726 | 0.197333351 | 0.318170823 | 0.484856668 | 0.4147 | 0.563773555 |
| SL1344_RS00960_ | 2.53945692 | 2.515560916 | 2.56642643 | 0.049770913 | 0.073718077 | 0.640205318 | 0.238000013 | 0.353180181 | 0.509595133 | 0.4494 | 0.58610773 |
| SL1344_RS00965_ | 3.780947164 | 3.755828397 | 3.818597052 | 0.06741154 | 0.075152184 | 0.856344039 | 0.164666689 | 0.253410366 | 0.433418718 | 0.3574 | 0.51450488 |
| SL1344_RS00970_ | 4.991726263 | 4.897061792 | 5.298235843 | 0.389766174 | 0.087855173 | 2.245515699 | 4.68E-05 | 0.064025895 | 0.200589634 | 0.1 | 0.243021319 |
| SL1344_RS00975_ | 2.27303093 | 2.249451425 | 2.378622953 | 0.126819558 | 0.234627349 | 0.557852836 | 0.309333342 | 0.442276056 | 0.629791459 | 0.6495 | 0.780855259 |
| SL1344_RS00980_ | 2.395348374 | 2.286913478 | 2.647601176 | 0.350481268 | 0.236314294 | 1.364031546 | 0.071952091 | 0.138634347 | 0.315834061 | 0.172 | 0.339442947 |
| SL1344_RS00985_ | 3.436891531 | 3.261941835 | 3.680596969 | 0.428654755 | 0.128001974 | 3.418716439 | 4.68E-05 | 0.015472961 | 0.086540704 | 0.1 | 0.243021319 |
| SL1344_RS00990_ | -2.601715849 | -2.84906249 | -2.393159349 | 0.461988399 | 0.465090236 | 0.93065047 | 0.148567647 | 0.227341883 | 0.385815224 | 0.3079 | 0.459851986 |
| SL1344_RS00995_ | -1.412798391 | -1.562655349 | -1.281874881 | 0.312797096 | 0.433585682 | 0.679818913 | 0.227181893 | 0.334115361 | 0.51003879 | 0.4527 | 0.60064708 |
| SL1344_RS01000_ | -3.448821496 | -3.367409497 | -3.496605404 | -0.169033626 | 0.750002813 | -0.222985542 | 0.410000003 | 0.578744627 | 0.7160386 | 0.696 | 0.794097822 |
| SL1344_RS01005_ | -3.307508857 | -3.343314981 | -3.240470521 | 0.114438606 | 0.738926589 | 0.136982077 | 0.440000002 | 0.635431172 | 0.761812804 | 0.7455 | 0.834320566 |
| SL1344_RS01010_ | -3.034034071 | -2.683774237 | -3.537075206 | -0.842499053 | 0.613526361 | -1.319467723 | 0.057961434 | 0.131956836 | 0.282056574 | 0.1926 | 0.346896656 |
| SL1344_RS01015_ | -3.391500154 | -2.971375685 | -3.852275158 | -0.897650182 | 0.902968361 | -0.951655324 | 0.150000025 | 0.215226452 | 0.388644407 | 0.3044 | 0.468997461 |
| SL1344_RS01020_ | -0.102891817 | -0.098473947 | -0.107702619 | -0.013028437 | 0.18807042 | -0.068247144 | 0.471685544 | 0.54993646 | 0.689419904 | 0.6561 | 0.763225279 |
| SL1344_RS01025_ | 3.204647274 | 3.219497292 | 3.181301466 | -0.023416229 | 0.135910264 | -0.167936122 | 0.433044639 | 0.773016871 | 0.866112825 | 0.8795 | 0.928357333 |
| SL1344_RS01030_ | 0.822039477 | 0.839055752 | 0.807009629 | -0.032297618 | 0.133130619 | -0.221722654 | 0.407061962 | 0.497588546 | 0.642885182 | 0.611 | 0.722516491 |
| SL1344_RS01035_ | 3.994426164 | 3.993960825 | 3.996239844 | -0.000956013 | 0.094157539 | -0.004720065 | 0.495669554 | 0.549297285 | 0.717726622 | 0.8209 | 0.894437972 |
| SL1344_RS01040_ | 0.969069911 | 0.934716933 | 1.002480902 | 0.068872591 | 0.145114939 | 0.436914087 | 0.320666674 | 0.470825383 | 0.620447452 | 0.573 | 0.694292607 |
| SL1344_RS01045_ | 1.506208243 | 1.50431025 | 1.50759286 | 0.00247056 | 0.115454866 | 0.02163502 | 0.486666667 | 0.554800904 | 0.692563647 | 0.6618 | 0.765583554 |
| SL1344_RS01050_ | 2.796627857 | 2.891730912 | 2.742120135 | -0.158370012 | 0.117118379 | -1.348292412 | 0.047968113 | 0.148231473 | 0.323361639 | 0.1477 | 0.301087027 |
| SL1344_RS01055_ | 0.133941653 | -0.005063165 | 0.263334804 | 0.266359749 | 0.235953322 | 1.076204125 | 0.116666701 | 0.181028452 | 0.348098989 | 0.2501 | 0.413484829 |
| SL1344_RS01060_ | 2.999997804 | 2.848812226 | 3.152607215 | 0.315493521 | 0.114256209 | 2.691109515 | 4.68E-05 | 0.023059834 | 0.105943124 | 0.1 | 0.243021319 |
| SL1344_RS01065_ | -1.307494286 | -1.243056936 | -1.401991891 | -0.173235558 | 0.342003533 | -0.485175903 | 0.290666676 | 0.433321248 | 0.592912019 | 0.5411 | 0.666147156 |
| SL1344_RS01070_ | 2.129779416 | 2.244653685 | 2.026609062 | -0.221454766 | 0.113776085 | -1.955528462 | 0.009993796 | 0.052863822 | 0.165442779 | 0.1136 | 0.260669035 |
| SL1344_RS01075_ | 0.968227476 | 0.933998626 | 1.004014879 | 0.065759602 | 0.146430228 | 0.442269666 | 0.321333341 | 0.465895244 | 0.618091171 | 0.5746 | 0.695674552 |
| SL1344_RS01080_ | 3.804358386 | 3.803921436 | 3.804900023 | -0.010874521 | 0.08349084 | -0.123737286 | 0.447035312 | 0.686530155 | 0.802857573 | 0.8141 | 0.881282744 |
| SL1344_RS01085_ | 1.910825312 | 1.821906084 | 2.105110186 | 0.262315838 | 0.148262425 | 1.607875202 | 0.059293877 | 0.092339473 | 0.238165854 | 0.1581 | 0.317736172 |
| SL1344_RS01090_ | 3.662955306 | 3.568075818 | 3.75542322 | 0.178875534 | 0.152388416 | 1.18839695 | 0.080612977 | 0.146592522 | 0.325853176 | 0.1827 | 0.355151765 |
| SL1344_RS01095_ | 7.913594164 | 7.916296505 | 7.909718636 | -0.00890103 | 0.073636799 | -0.099291882 | 0.447035312 | 0.882589934 | 0.936455635 | 0.9052 | 0.946204991 |
| SL1344_RS01100_ | 8.301890763 | 8.238805012 | 8.342123799 | 0.106625487 | 0.03869561 | 2.62843294 | 4.68E-05 | 0.044839178 | 0.161931267 | 0.1 | 0.243021319 |
| SL1344_RS01105_ | 3.977580349 | 3.980344101 | 3.970326741 | -0.007219059 | 0.088351499 | -0.064107856 | 0.468354432 | 0.76500281 | 0.860346038 | 0.8338 | 0.903321999 |
| SL1344_RS01110_ | 5.925024186 | 5.926750403 | 5.923254753 | -0.021426502 | 0.0828564 | -0.275797863 | 0.398000004 | 0.627891606 | 0.77129161 | 0.8465 | 0.907662609 |
| SL1344_RS01115_ | 3.21548448 | 3.157264463 | 3.285411822 | 0.159071703 | 0.121086326 | 1.262141374 | 0.057295212 | 0.111758788 | 0.269890509 | 0.1612 | 0.317972616 |
| SL1344_RS01120_ | 2.992903248 | 2.963866664 | 3.032213272 | 0.048353353 | 0.204874416 | 0.194655014 | 0.437333336 | 0.864026924 | 0.924729947 | 0.9424 | 0.967141375 |
| SL1344_RS01130_ | 4.828713935 | 4.79899155 | 4.899456204 | 0.11139491 | 0.100970693 | 1.083071935 | 0.109926752 | 0.193559472 | 0.386633137 | 0.2169 | 0.389123923 |
| SL1344_RS01135_ | 5.610837983 | 5.597328208 | 5.644569544 | 0.048950535 | 0.058158413 | 0.763939734 | 0.212000016 | 0.294803086 | 0.490594616 | 0.4018 | 0.565468771 |
| SL1344_RS01140_ | 5.691776027 | 5.759983831 | 5.648008466 | -0.104406864 | 0.058740425 | -1.68360192 | 0.040000111 | 0.074371784 | 0.213746933 | 0.1309 | 0.285827427 |
| SL1344_RS01145_ | 5.708779579 | 5.729277978 | 5.682340023 | -0.023938151 | 0.109998185 | -0.242786574 | 0.402398405 | 0.835186845 | 0.90766165 | 0.8542 | 0.916079801 |
| SL1344_RS01150_ | 3.317506597 | 3.297524145 | 3.341087251 | 0.056215388 | 0.089968562 | 0.567034644 | 0.250666679 | 0.377188059 | 0.55514239 | 0.5228 | 0.654366592 |
| SL1344_RS01155_ | 4.710216759 | 4.717780739 | 4.700660207 | -0.033661267 | 0.10329058 | -0.314090726 | 0.37974684 | 0.62222088 | 0.764861662 | 0.7867 | 0.869296914 |
| SL1344_RS01160_ | 4.690375219 | 4.707604598 | 4.654558728 | -0.051165888 | 0.115594 | -0.261542728 | 0.367088613 | 0.899909313 | 0.946007951 | 0.7906 | 0.88107036 |
| SL1344_RS01165_ | 2.725071214 | 2.58675579 | 2.823837355 | 0.240468003 | 0.107561871 | 2.219441379 | 0.003332494 | 0.038074002 | 0.138789477 | 0.1054 | 0.25025769 |
| SL1344_RS01170_ | 4.315847166 | 4.247861904 | 4.389842426 | 0.123561725 | 0.153420034 | 0.749237871 | 0.213857444 | 0.305548256 | 0.507568559 | 0.3851 | 0.564412548 |
| SL1344_RS01175_ | 3.5368104 | 3.487437498 | 3.649819438 | 0.162530101 | 0.084959811 | 1.75424025 | 0.023317982 | 0.079643928 | 0.218972633 | 0.1211 | 0.269630378 |
| SL1344_RS01180_ | 0.818820727 | 0.906911403 | 0.745475237 | -0.158722712 | 0.171721662 | -0.867902572 | 0.176666687 | 0.26208812 | 0.432483916 | 0.3617 | 0.514961999 |
| SL1344_RS01185_ | 1.386944833 | 1.313647459 | 1.436329823 | 0.126209499 | 0.138529889 | 0.864832909 | 0.169886764 | 0.266362403 | 0.438226303 | 0.3617 | 0.51326248 |
| SL1344_RS01190_ | -0.062500497 | -0.094374702 | -0.024290299 | 0.067949464 | 0.192078015 | 0.335073402 | 0.358000006 | 0.493230882 | 0.639250923 | 0.5997 | 0.714368718 |
| SL1344_RS01195_ | 1.603600826 | 1.418937868 | 1.689983659 | 0.272560993 | 0.157190073 | 1.615187429 | 0.026649069 | 0.096578109 | 0.244179016 | 0.1325 | 0.283145005 |
| SL1344_RS01200_ | 1.973086671 | 2.054693743 | 1.889552745 | -0.165689845 | 0.105521461 | -1.526714362 | 0.047968113 | 0.089009081 | 0.220458867 | 0.1549 | 0.308408576 |
| SL1344_RS01205_ | 1.739368581 | 1.818774776 | 1.557612952 | -0.252631923 | 0.175941329 | -1.269738267 | 0.086000049 | 0.140077523 | 0.309543104 | 0.2045 | 0.370993798 |
| SL1344_RS01210_ | 0.660723741 | 0.639726971 | 0.685855557 | 0.044794289 | 0.145207385 | 0.282975722 | 0.366666672 | 0.498553298 | 0.643230924 | 0.6045 | 0.717807944 |
| SL1344_RS01215_ | 0.462538884 | 0.428166923 | 0.505063978 | 0.082968732 | 0.175737125 | 0.439674736 | 0.310666675 | 0.461357084 | 0.613293949 | 0.5774 | 0.69563331 |

| | | | | | | | | | | | |
|---|---|---|---|---|---|---|---|---|---|---|---|
| SL1344_RS01220_ | 1.373063658 | 1.387150795 | 1.353536459 | -0.03516668 | 0.12976983 | -0.245161548 | 0.38774151 | 0.548447617 | 0.689445199 | 0.6558 | 0.762816137 |
| SL1344_RS01225_ | 4.84084855 | 4.817464591 | 4.889224643 | 0.076842221 | 0.058480236 | 1.202387078 | 0.081333385 | 0.153301982 | 0.327678626 | 0.1987 | 0.361081306 |
| SL1344_RS01230_ | 2.399020581 | 2.305325216 | 2.511265741 | 0.231325001 | 0.140242749 | 1.644890336 | 0.022651765 | 0.071335772 | 0.20410273 | 0.1236 | 0.273848943 |
| SL1344_RS01245_ | 1.500329923 | 1.552627844 | 1.443994877 | -0.107966368 | 0.104483025 | -1.0124885 | 0.135909422 | 0.208361689 | 0.359543129 | 0.2899 | 0.439108909 |
| SL1344_RS01250_ | 2.23272748 | 2.152705089 | 2.362296181 | 0.169807288 | 0.180355385 | 0.918597415 | 0.166000022 | 0.231477445 | 0.420537912 | 0.3033 | 0.484214485 |
| SL1344_RS01255_ | 0.267762155 | 0.19360833 | 0.33786848 | 0.142470352 | 0.172847607 | 0.79962898 | 0.192000018 | 0.292604766 | 0.453976113 | 0.389 | 0.534304547 |
| SL1344_RS01260_ | -0.087768166 | -0.437894658 | 0.331157675 | 0.755193861 | 0.668778708 | 1.114780645 | 0.071952091 | 0.169727908 | 0.358434158 | 0.1756 | 0.346192109 |
| SL1344_RS01265_ | 4.845446687 | 4.81546467 | 4.957706691 | 0.12772246 | 0.127055632 | 1.085922447 | 0.148666692 | 0.155986077 | 0.338050801 | 0.2718 | 0.457965308 |
| SL1344_RS01270_ | 3.958899919 | 4.003000522 | 3.914291188 | -0.066293684 | 0.153811423 | -0.434292448 | 0.313333341 | 0.530150537 | 0.69887539 | 0.6181 | 0.757934887 |
| SL1344_RS01275_ | 1.417442794 | 1.058198968 | 1.542328818 | 0.517506195 | 0.414307032 | 1.386067124 | 0.039333446 | 0.089247783 | 0.241598181 | 0.138 | 0.293202459 |
| SL1344_RS01285_ | -0.266398676 | -0.259504852 | -0.279504334 | 0.033098427 | 0.338634888 | 0.094776498 | 0.456666668 | 0.696408686 | 0.810754618 | 0.8373 | 0.896792428 |
| SL1344_RS01290_ | 0.87817082 | 0.849933375 | 0.912500353 | 0.031666923 | 0.216058436 | 0.135993978 | 0.449333335 | 0.687789248 | 0.804665528 | 0.7999 | 0.872831629 |
| SL1344_RS01295_ | -1.120600825 | -1.11655585 | -1.130729862 | 0.009211127 | 0.375450903 | 0.020755912 | 0.491333334 | 0.682751977 | 0.799433929 | 0.7901 | 0.866180792 |
| SL1344_RS01300_ | -1.220282308 | -1.241236832 | -1.207932324 | 0.030466589 | 0.410901882 | 0.062836929 | 0.472666668 | 0.664891078 | 0.785696509 | 0.7764 | 0.856025983 |
| SL1344_RS01305_ | 1.264388494 | 1.122009923 | 1.400394218 | 0.280943391 | 0.164312834 | 1.716171845 | 0.026666836 | 0.072174138 | 0.199811472 | 0.1288 | 0.279462085 |
| SL1344_RS01310_ | 0.02899759 | -0.019585395 | 0.074822177 | 0.090246238 | 0.180349646 | 0.473974816 | 0.297135252 | 0.417519908 | 0.572480013 | 0.5275 | 0.654558875 |
| SL1344_RS01315_ | 5.005446541 | 5.149832741 | 4.819137001 | -0.322101432 | 0.07684763 | -4.269094497 | 4.68E-05 | 0.004151417 | 0.043059795 | 0.1 | 0.243021319 |
| SL1344_RS01320_ | -0.12204947 | -0.22990211 | -0.024171257 | 0.210689519 | 0.217860939 | 0.928269147 | 0.144000027 | 0.231922135 | 0.397436017 | 0.3245 | 0.476948956 |
| SL1344_RS01325_ | -0.232371288 | -0.208299629 | -0.252538823 | -0.003310102 | 0.341239974 | -0.009829501 | 0.496 | 0.715605135 | 0.82477064 | 0.8207 | 0.888405313 |
| SL1344_RS01330_ | 0.974453947 | 0.82797676 | 1.189853062 | 0.358634964 | 0.173780689 | 1.963876593 | 0.013333676 | 0.053869835 | 0.169070982 | 0.1151 | 0.263029587 |
| SL1344_RS01335_ | 1.924727197 | 1.874848744 | 1.991266335 | 0.08901962 | 0.173474041 | 0.492406338 | 0.28600001 | 0.486980055 | 0.651410014 | 0.5964 | 0.724382627 |
| SL1344_RS01340_ | 1.410658078 | 1.349912771 | 1.461097758 | 0.094764438 | 0.160842376 | 0.544993221 | 0.28047969 | 0.451623105 | 0.616249646 | 0.5621 | 0.695833781 |
| SL1344_RS01345_ | -2.260629536 | -2.205073233 | -2.315892481 | -0.109091194 | 0.405812081 | -0.239506484 | 0.382000005 | 0.51011188 | 0.653570552 | 0.6166 | 0.728128623 |
| SL1344_RS01350_ | -3.256259797 | -3.284230079 | -3.209601415 | 0.123205172 | 0.702210613 | 0.171818034 | 0.430000003 | 0.598611004 | 0.733436076 | 0.7218 | 0.815164913 |
| SL1344_RS01355_ | -1.955147933 | -1.832092607 | -2.14125157 | -0.313758814 | 0.423814335 | -0.695419065 | 0.219333349 | 0.331322836 | 0.499795162 | 0.4401 | 0.58148485 |
| SL1344_RS01360_ | -3.560526634 | -3.424189421 | -3.785699692 | -0.31532213 | 0.775259043 | -0.382693282 | 0.339773491 | 0.509984536 | 0.661665726 | 0.627 | 0.744598318 |
| SL1344_RS01365_ | -2.298984866 | -2.425503596 | -2.162286664 | 0.267714487 | 0.393995384 | 0.647367278 | 0.238507675 | 0.353955089 | 0.513403223 | 0.4532 | 0.592359792 |
| SL1344_RS01370_ | -3.63178556 | -3.689912254 | -3.570778895 | 0.124598107 | 0.738974577 | 0.151432853 | 0.432378417 | 0.574053986 | 0.712355538 | 0.6834 | 0.784969223 |
| SL1344_RS01375_ | -9.019843638 | -9.527388243 | -8.557659054 | 0.875683042 | 3.993200118 | 0.191118794 | 0.39840107 | 0.503222392 | 0.6500066 | 0.6168 | 0.726904396 |
| SL1344_RS01380_ | -0.063436998 | -0.191567932 | 0.052682207 | 0.246305437 | 0.238541707 | 0.965448012 | 0.124666699 | 0.209057542 | 0.376694939 | 0.2881 | 0.446388726 |
| SL1344_RS01385_ | -2.482542493 | -2.554048271 | -2.357596421 | 0.093968756 | 0.726317778 | 0.111983652 | 0.447035312 | 0.732235226 | 0.836977952 | 0.8132 | 0.887061331 |
| SL1344_RS01390_ | -1.478279365 | -1.330380492 | -1.607776397 | -0.298478073 | 0.35178703 | -0.774673723 | 0.196535661 | 0.298641797 | 0.462825067 | 0.4026 | 0.547147036 |
| SL1344_RS01395_ | -2.454442295 | -2.436115276 | -2.469583871 | -0.022653763 | 0.405548164 | -0.057758303 | 0.474666668 | 0.510366867 | 0.655459625 | 0.6248 | 0.735444395 |
| SL1344_RS01400_ | -3.411803618 | -3.370366527 | -3.44634044 | -0.153701361 | 1.037587058 | -0.135035935 | 0.436375752 | 0.683123918 | 0.800921776 | 0.786 | 0.863838784 |
| SL1344_RS01405_ | -2.868574408 | -3.260702164 | -2.374089989 | 0.870603218 | 0.640395942 | 1.275695811 | 0.082000051 | 0.130641916 | 0.282623995 | 0.2034 | 0.365550545 |
| SL1344_RS01410_ | -4.026079663 | -3.982368708 | -4.081309129 | -0.08841973 | 0.735983508 | -0.111225695 | 0.451032647 | 0.551455771 | 0.689976315 | 0.6588 | 0.763928768 |
| SL1344_RS01415_ | -3.296608085 | -3.424833968 | -3.094630261 | 0.294381377 | 0.651076827 | 0.416664133 | 0.332445044 | 0.51782127 | 0.661113963 | 0.6154 | 0.733099638 |
| SL1344_RS01420_ | -4.023489195 | -3.792294907 | -4.248373182 | -0.454746631 | 0.830973903 | -0.502620457 | 0.282478358 | 0.427763289 | 0.585930348 | 0.5383 | 0.665808084 |
| SL1344_RS01425_ | -2.256435998 | -2.275518368 | -2.229925409 | 0.151280647 | 0.672263333 | 0.221136416 | 0.41266667 | 0.634659375 | 0.764175119 | 0.7816 | 0.857844694 |
| SL1344_RS01430_ | -1.9093081 | -1.636108919 | -2.098296252 | -0.458477841 | 0.423546645 | -0.983739899 | 0.146000026 | 0.221056364 | 0.385966961 | 0.3073 | 0.465216272 |
| SL1344_RS01435_ | -3.218134132 | -3.242402332 | -3.181299596 | 0.255703381 | 1.030566531 | 0.238617906 | 0.403333337 | 0.639963602 | 0.76927514 | 0.8038 | 0.873522763 |
| SL1344_RS01440_ | -4.460053853 | -4.018594518 | -4.729909181 | -0.908850567 | 1.514257243 | -0.557408221 | 0.271333344 | 0.41099044 | 0.586585048 | 0.5505 | 0.684617988 |
| SL1344_RS01445_ | -0.243944286 | -0.265633176 | -0.212052301 | 0.068449303 | 0.262806165 | 0.234417976 | 0.403333337 | 0.606443579 | 0.738444145 | 0.7222 | 0.814422431 |
| SL1344_RS01450_ | -3.121425368 | -2.965725024 | -3.227698101 | -0.257023872 | 0.665715578 | -0.369462741 | 0.33733334 | 0.530972626 | 0.675214608 | 0.6377 | 0.748946666 |
| SL1344_RS01455_ | -4.294696245 | -4.120991328 | -4.449819533 | -0.360771775 | 0.946062097 | -0.3546905 | 0.345769493 | 0.522566818 | 0.66810357 | 0.6344 | 0.743369358 |
| SL1344_RS01460_ | -1.875306348 | -1.760676156 | -2.071991861 | -0.340725842 | 0.478598595 | -0.624718063 | 0.244503677 | 0.356577126 | 0.533278559 | 0.4797 | 0.620508198 |
| SL1344_RS01465_ | -1.016105388 | -1.149270564 | -0.898273587 | 0.255677669 | 0.270951794 | 0.926647789 | 0.153897426 | 0.241994896 | 0.39766847 | 0.336 | 0.484992461 |
| SL1344_RS01470_ | -3.625439586 | -3.697172425 | -3.549588136 | 0.107719662 | 0.719266301 | 0.120706491 | 0.439040642 | 0.585658764 | 0.720067683 | 0.6886 | 0.789582316 |
| SL1344_RS01475_ | 0.766032814 | 0.83548531 | 0.696085644 | -0.152556929 | 0.201960113 | -0.760523221 | 0.197201883 | 0.307506659 | 0.483870059 | 0.4106 | 0.560945719 |
| SL1344_RS01480_ | 1.391358874 | 1.484835925 | 1.312942663 | -0.179585889 | 0.155212811 | -1.12969434 | 0.113333369 | 0.178546433 | 0.34524688 | 0.2503 | 0.411523424 |
| SL1344_RS01485_ | 0.513310187 | 0.556408159 | 0.460613205 | -0.102755619 | 0.177372187 | -0.551712559 | 0.258666678 | 0.408783057 | 0.567804254 | 0.5302 | 0.65525465 |
| SL1344_RS01490_ | -0.12073583 | -0.138093473 | -0.072636054 | 0.048870037 | 0.307059329 | 0.14375051 | 0.442666669 | 0.748610322 | 0.848546098 | 0.8064 | 0.88499454 |
| SL1344_RS01495_ | -1.39198102 | -1.600857161 | -1.189521161 | 0.407485678 | 0.342905809 | 1.140240037 | 0.106000038 | 0.164868833 | 0.324935606 | 0.2426 | 0.404266728 |

| | | | | | | | | | | | |
|---|---|---|---|---|---|---|---|---|---|---|---|
| SL1344_RS01500_ | -6.190576253 | -6.31800008 | -6.039104819 | 0.314413099 | 1.805603667 | 0.155562065 | 0.428381082 | 0.572699527 | 0.710525186 | 0.6894 | 0.787592386 |
| SL1344_RS01505_ | -2.098956078 | -2.056386258 | -2.142728872 | -0.092183474 | 0.441554749 | -0.197733234 | 0.40639574 | 0.562092389 | 0.701756655 | 0.6734 | 0.775051366 |
| SL1344_RS01515_ | -3.502069809 | -3.549794851 | -3.439750398 | 0.137749899 | 0.725613216 | 0.18084601 | 0.418000003 | 0.578709758 | 0.714236657 | 0.69 | 0.788946141 |
| SL1344_RS01520_ | -5.032816031 | -5.063311196 | -4.963911687 | 0.009999281 | 1.535604532 | 0.005686928 | 0.498 | 0.643248359 | 0.773930079 | 0.7713 | 0.856028833 |
| SL1344_RS01525_ | -0.565193045 | -0.425959017 | -0.75462512 | -0.440784982 | 0.489505425 | -0.871413227 | 0.169220541 | 0.22811219 | 0.418599487 | 0.3352 | 0.505677736 |
| SL1344_RS01530_ | -2.319597852 | -2.258184946 | -2.380540099 | -0.117054852 | 0.392775104 | -0.276120484 | 0.383077953 | 0.508507256 | 0.649377863 | 0.6151 | 0.72470811 |
| SL1344_RS01535_ | -0.117908982 | -0.094281882 | -0.132063375 | -0.047246566 | 0.248614243 | -0.165449106 | 0.420386412 | 0.605393583 | 0.738588391 | 0.726 | 0.817080515 |
| SL1344_RS01540_ | -1.910164829 | -1.927687202 | -1.888693736 | 0.026967061 | 0.402329194 | 0.064773689 | 0.471019322 | 0.592587625 | 0.728295838 | 0.707 | 0.804129508 |
| SL1344_RS01545_ | -2.324522043 | -2.27367427 | -2.376865371 | -0.094423242 | 0.450269479 | -0.195072389 | 0.40866667 | 0.547218967 | 0.688821633 | 0.663 | 0.767232885 |
| SL1344_RS01550_ | -3.071194708 | -3.064945591 | -3.078024595 | 0.003062415 | 0.602333107 | 0.004795063 | 0.496666667 | 0.586331143 | 0.720565376 | 0.6871 | 0.788190861 |
| SL1344_RS01565_ | -2.398927795 | -2.350323905 | -2.441402748 | -0.09484892 | 0.429938973 | -0.195544012 | 0.411333337 | 0.540427063 | 0.681870739 | 0.6506 | 0.756728496 |
| SL1344_RS01570_ | -0.378666462 | -0.434252063 | -0.33943607 | 0.119756294 | 0.344991026 | 0.317653551 | 0.366000005 | 0.564833734 | 0.713181845 | 0.7158 | 0.812634941 |
| SL1344_RS01575_ | -0.774335021 | -0.972164917 | -0.578515848 | 0.391625967 | 0.213947865 | 1.781680125 | 0.030646382 | 0.065129441 | 0.178289247 | 0.1305 | 0.280576105 |
| SL1344_RS01580_ | -0.014692765 | -0.001961595 | -0.032288541 | -0.027806723 | 0.205192842 | -0.118555306 | 0.448666669 | 0.599416359 | 0.73202177 | 0.7053 | 0.802012222 |
| SL1344_RS01590_ | 2.325851941 | 2.181786459 | 2.372706364 | 0.184903675 | 0.136606495 | 1.173121935 | 0.105333372 | 0.168575514 | 0.345288745 | 0.2469 | 0.410796454 |
| SL1344_RS01595_ | 1.964943514 | 1.925424123 | 2.095348631 | 0.15902842 | 0.1626498 | 0.825693989 | 0.221185891 | 0.308278186 | 0.498620093 | 0.4535 | 0.612842701 |
| SL1344_RS01600_ | 2.462862865 | 2.450513965 | 2.486911515 | 0.029752923 | 0.106911096 | 0.275849329 | 0.392000004 | 0.616722085 | 0.748996447 | 0.7345 | 0.826237394 |
| SL1344_RS01605_ | 1.252182898 | 1.269296344 | 1.234517729 | -0.0350689 | 0.112953894 | -0.294211317 | 0.371085948 | 0.484438342 | 0.629625227 | 0.591 | 0.706681981 |
| SL1344_RS01610_ | -0.57073276 | -0.624041016 | -0.475931812 | 0.123971315 | 0.36017186 | 0.318348035 | 0.36908728 | 0.62070635 | 0.75368995 | 0.7289 | 0.823225643 |
| SL1344_RS01615_ | -0.3795145 | -0.459482067 | -0.289932615 | 0.163159979 | 0.231031717 | 0.680319724 | 0.235842785 | 0.370579541 | 0.5285108 | 0.465 | 0.602745289 |
| SL1344_RS01620_ | 3.673660762 | 3.649095697 | 3.753321408 | 0.098613972 | 0.072696762 | 1.184595853 | 0.135333362 | 0.181259057 | 0.354798894 | 0.2883 | 0.453483603 |
| SL1344_RS01625_ | 3.85223798 | 3.631135977 | 4.114717958 | 0.463613188 | 0.123534573 | 3.757213166 | 4.68E-05 | 0.005230891 | 0.049098043 | 0.1 | 0.243021319 |
| SL1344_RS01635_ | 2.423127456 | 2.609114272 | 2.185084306 | -0.440519096 | 0.136641359 | -3.282688641 | 4.68E-05 | 0.014818436 | 0.081415096 | 0.1 | 0.243021319 |
| SL1344_RS01640_ | -1.414267859 | -1.499473057 | -1.304826195 | 0.213030958 | 0.340222586 | 0.578032441 | 0.260666678 | 0.390095671 | 0.552672033 | 0.5097 | 0.639974034 |
| SL1344_RS01645_ | 2.126432385 | 2.015981737 | 2.178919613 | 0.158205827 | 0.114890652 | 1.262508738 | 0.082611643 | 0.149994086 | 0.311862959 | 0.2232 | 0.382416109 |
| SL1344_RS01650_ | 3.545968292 | 3.468338573 | 3.589551685 | 0.131541993 | 0.108784581 | 1.276372442 | 0.065289873 | 0.127763369 | 0.293437379 | 0.1809 | 0.343631185 |
| SL1344_RS01655_ | 0.683105271 | 0.390663096 | 0.938377879 | 0.547823042 | 0.146194017 | 3.701043984 | 4.68E-05 | 0.009945986 | 0.062261697 | 0.1 | 0.243021319 |
| SL1344_RS01660_ | -3.917457855 | -4.177700523 | -3.810957726 | 0.316556398 | 0.973670564 | 0.290553769 | 0.379080618 | 0.58471539 | 0.725540491 | 0.7124 | 0.813344338 |
| SL1344_RS01665_ | -5.240904504 | -5.473503373 | -4.966487334 | 0.509021419 | 1.153890865 | 0.395819197 | 0.328000007 | 0.471028614 | 0.620005533 | 0.5724 | 0.693423484 |
| SL1344_RS01670_ | -4.01339369 | -4.007630166 | -4.022355021 | 0.032178621 | 0.919593041 | 0.027822412 | 0.488341106 | 0.628202487 | 0.758136129 | 0.7486 | 0.83654426 |
| SL1344_RS01675_ | -0.354160078 | -0.3808072 | -0.343126402 | 0.022901198 | 0.307984442 | 0.067188632 | 0.471333334 | 0.701698324 | 0.813864642 | 0.798 | 0.874730563 |
| SL1344_RS01680_ | -3.421289537 | -3.840024369 | -3.087303075 | 0.769321327 | 0.607301478 | 1.225461648 | 0.075949423 | 0.145322696 | 0.29137367 | 0.2119 | 0.365247348 |
| SL1344_RS01685_ | -2.780480826 | -2.722377891 | -2.839076706 | -0.123459988 | 0.506045844 | -0.217482184 | 0.400000004 | 0.527155469 | 0.671618019 | 0.6376 | 0.746377005 |
| SL1344_RS01690_ | -4.11629579 | -4.253036206 | -3.940072926 | 0.275370105 | 0.995346006 | 0.245762001 | 0.386666671 | 0.612540759 | 0.74172363 | 0.7036 | 0.80085467 |
| SL1344_RS01695_ | -1.951358921 | -1.741928212 | -2.198251107 | -0.467581344 | 0.443578329 | -0.995489878 | 0.141905424 | 0.212176806 | 0.378120901 | 0.2958 | 0.453715672 |
| SL1344_RS01700_ | -0.632114884 | -0.373726081 | -0.784521562 | -0.387856656 | 0.272458355 | -1.34910247 | 0.078614311 | 0.131851517 | 0.282245552 | 0.2098 | 0.369596533 |
| SL1344_RS01705_ | -0.146802961 | -0.112365575 | -0.188419133 | -0.069282722 | 0.202044565 | -0.334567939 | 0.349333339 | 0.493357904 | 0.639226443 | 0.6012 | 0.716462499 |
| SL1344_RS01710_ | -2.313703251 | -2.299008891 | -2.323286377 | 0.007683709 | 0.569043025 | 0.01332448 | 0.496 | 0.669664632 | 0.789594194 | 0.7751 | 0.855576528 |
| SL1344_RS01715_ | -3.365156772 | -3.542547113 | -3.22968312 | 0.310804259 | 0.611971859 | 0.457025011 | 0.312000008 | 0.441818614 | 0.596045891 | 0.559 | 0.682643101 |
| SL1344_RS01720_ | -1.194651961 | -1.279810204 | -1.116406867 | 0.165457652 | 0.297099206 | 0.532499797 | 0.261159239 | 0.398065153 | 0.556465681 | 0.5104 | 0.639133426 |
| SL1344_RS01725_ | -1.463745813 | -1.617798306 | -1.358443135 | 0.233386338 | 0.361403508 | 0.601520755 | 0.255163236 | 0.394653594 | 0.557142355 | 0.4928 | 0.629195454 |
| SL1344_RS01730_ | -1.514879861 | -1.464380689 | -1.546145229 | -0.069276698 | 0.35296327 | -0.174866005 | 0.423051302 | 0.598459561 | 0.729145732 | 0.6979 | 0.796341764 |
| SL1344_RS01740_ | -6.309516044 | -6.042560001 | -6.548696545 | -0.456257792 | 2.035312487 | -0.190367787 | 0.392000004 | 0.56697819 | 0.708051212 | 0.6822 | 0.783448038 |
| SL1344_RS01745_ | -0.243311716 | -0.091445931 | -0.375333644 | -0.296679953 | 0.284191148 | -0.981259233 | 0.131245866 | 0.200853168 | 0.37445017 | 0.2846 | 0.448865788 |
| SL1344_RS01750_ | -0.95203099 | -0.9725828 | -0.924838276 | 0.045399381 | 0.27590886 | 0.154681968 | 0.431045972 | 0.579234658 | 0.714781007 | 0.6833 | 0.785005937 |
| SL1344_RS01755_ | -0.540735018 | -0.568549347 | -0.510764496 | 0.05989591 | 0.244668474 | 0.233821234 | 0.40266667 | 0.546362041 | 0.685788317 | 0.6515 | 0.758593835 |
| SL1344_RS01760_ | 1.02789418 | 1.016732085 | 1.045229221 | 0.035577977 | 0.191954671 | 0.171529577 | 0.423717525 | 0.658954954 | 0.782434838 | 0.7792 | 0.858191801 |
| SL1344_RS01765_ | 0.285595872 | 0.255040381 | 0.315121947 | 0.061867375 | 0.160738772 | 0.361757478 | 0.344000006 | 0.466928899 | 0.61540361 | 0.5744 | 0.692810862 |
| SL1344_RS01770_ | -3.323206253 | -3.411339926 | -3.283364654 | 0.128154548 | 0.835372754 | 0.134301803 | 0.439333336 | 0.635838933 | 0.764792243 | 0.7537 | 0.841854768 |
| SL1344_RS01775_ | -3.360037146 | -3.417156226 | -3.312943142 | 0.083320678 | 0.599106631 | 0.135732075 | 0.443037977 | 0.561759731 | 0.702256819 | 0.676 | 0.77907602 |
| SL1344_RS01780_ | -2.631282344 | -2.096641036 | -2.781854057 | -0.555557697 | 0.792083546 | -0.515338359 | 0.323784151 | 0.616299055 | 0.752271414 | 0.6694 | 0.798476473 |
| SL1344_RS01785_ | -1.196589476 | -1.464353302 | -0.937697979 | 0.523611594 | 0.287458961 | 1.764168253 | 0.026666836 | 0.06467624 | 0.180202904 | 0.1312 | 0.281480643 |
| SL1344_RS01790_ | -2.667993602 | -2.616959734 | -2.707370276 | -0.034359283 | 1.077657306 | -0.031084912 | 0.485009994 | 0.77209531 | 0.865036001 | 0.8343 | 0.899347083 |

| | | | | | | | | | | | |
|---|---|---|---|---|---|---|---|---|---|---|---|
| SL1344_RS01795_ | -2.303065311 | -2.403544783 | -2.172590299 | 0.250213487 | 0.493233489 | 0.495801601 | 0.29380414 | 0.444851387 | 0.601361995 | 0.5622 | 0.685099296 |
| SL1344_RS01800_ | -4.507841828 | -4.578729363 | -4.42375206 | 0.187307971 | 0.948293429 | 0.17673812 | 0.419333336 | 0.550641072 | 0.689949843 | 0.6577 | 0.763776324 |
| SL1344_RS01805_ | -1.937587183 | -1.856876171 | -2.034858226 | -0.187436613 | 0.369470819 | -0.472949479 | 0.305333342 | 0.441201092 | 0.592534951 | 0.548 | 0.670132359 |
| SL1344_RS01810_ | -3.012966771 | -2.95071826 | -3.113235822 | -0.052255253 | 0.852184638 | -0.0454578 | 0.477333334 | 0.734232246 | 0.838048698 | 0.8291 | 0.895665596 |
| SL1344_RS01815_ | -2.104521123 | -2.000035263 | -2.289427905 | -0.222676778 | 0.552435005 | -0.403077705 | 0.342666673 | 0.546531589 | 0.694146417 | 0.6608 | 0.773330688 |
| SL1344_RS01820_ | -3.049958073 | -2.891142573 | -3.317415154 | -0.351478927 | 0.885831469 | -0.359737258 | 0.348000006 | 0.552098143 | 0.702904874 | 0.6883 | 0.795635495 |
| SL1344_RS01825_ | 0.052114306 | 0.142394655 | -0.058690691 | -0.203651833 | 0.228368449 | -0.820215551 | 0.167333355 | 0.269570106 | 0.443615468 | 0.359 | 0.514712892 |
| SL1344_RS01830_ | 2.106221457 | 2.059821872 | 2.159663431 | 0.058079894 | 0.186689721 | 0.30791198 | 0.381745508 | 0.669447515 | 0.793411364 | 0.7836 | 0.863025289 |
| SL1344_RS01835_ | 3.144277412 | 3.055373196 | 3.218023547 | 0.167480342 | 0.091335482 | 1.799232518 | 0.015323415 | 0.059039547 | 0.179704419 | 0.1138 | 0.261073887 |
| SL1344_RS01840_ | -3.066958711 | -2.93660534 | -3.24520038 | -0.236199528 | 0.826163725 | -0.283820984 | 0.389740178 | 0.64144353 | 0.768772517 | 0.7515 | 0.83965384 |
| SL1344_RS01845_ | -1.50032576 | -1.434435379 | -1.568055763 | -0.130126119 | 0.327701871 | -0.358320165 | 0.33933334 | 0.495542001 | 0.640851507 | 0.6016 | 0.717265194 |
| SL1344_RS01850_ | -0.136240186 | -0.098335833 | -0.167755571 | -0.060154087 | 0.237399631 | -0.242011011 | 0.404397072 | 0.582492526 | 0.718605147 | 0.6924 | 0.792652328 |
| SL1344_RS01855_ | -2.520959131 | -2.398299402 | -2.658130876 | -0.261528437 | 0.554676648 | -0.4315875 | 0.304463699 | 0.472839703 | 0.627854638 | 0.5777 | 0.700391974 |
| SL1344_RS01860_ | 1.392568424 | 1.286181664 | 1.576910219 | 0.295423372 | 0.188205555 | 1.47055163 | 0.03597614 | 0.099204212 | 0.250741034 | 0.1442 | 0.298854508 |
| SL1344_RS01865_ | 0.105451644 | 0.034532689 | 0.265814856 | 0.21827942 | 0.237513213 | 0.841170878 | 0.20186544 | 0.302774076 | 0.477825599 | 0.4232 | 0.578586363 |
| SL1344_RS01870_ | -0.956833657 | -1.160411815 | -0.811592552 | 0.349371514 | 0.279819623 | 1.154951763 | 0.106000038 | 0.164724863 | 0.320817724 | 0.2415 | 0.399346311 |
| SL1344_RS01875_ | -0.011704524 | 0.217383191 | -0.372537423 | -0.585649716 | 0.205616333 | -2.809594034 | 0.002668394 | 0.024194341 | 0.101381195 | 0.101 | 0.244363689 |
| SL1344_RS01880_ | -1.363485993 | -1.150248391 | -1.663401808 | -0.503200569 | 0.370408523 | -1.27860991 | 0.07533339 | 0.136325238 | 0.289647334 | 0.2022 | 0.361259896 |
| SL1344_RS01885_ | -3.134661938 | -2.788925155 | -3.51053277 | -0.717006333 | 0.564987023 | -1.201021997 | 0.100599641 | 0.159274602 | 0.307664111 | 0.2314 | 0.385267369 |
| SL1344_RS01890_ | -2.52082709 | -2.396001879 | -2.606294018 | -0.204191412 | 0.48178177 | -0.387919765 | 0.332000007 | 0.514274204 | 0.65727269 | 0.6185 | 0.730355026 |
| SL1344_RS01895_ | -2.3869209 | -2.400304242 | -2.380902904 | 0.025138298 | 0.523685331 | 0.044280512 | 0.483677549 | 0.62959851 | 0.757119887 | 0.7496 | 0.836401787 |
| SL1344_RS01900_ | -1.449348207 | -1.4487357 | -1.450162453 | 0.017567882 | 0.389472864 | 0.043374872 | 0.487008661 | 0.651182872 | 0.776057072 | 0.777 | 0.856494638 |
| SL1344_RS01905_ | 3.536536524 | 3.456020437 | 3.622014969 | 0.165143747 | 0.061284401 | 2.657586643 | 0.004664544 | 0.023625946 | 0.102552166 | 0.1021 | 0.245890261 |
| SL1344_RS01910_ | 1.423545157 | 1.569816432 | 1.306764695 | -0.264194153 | 0.126488255 | -2.013493073 | 0.010659989 | 0.048033018 | 0.154386612 | 0.1117 | 0.257911593 |
| SL1344_RS01915_ | -1.872151327 | -1.946822785 | -1.799466818 | 0.124738672 | 0.433873832 | 0.264701222 | 0.374000005 | 0.551019531 | 0.693113373 | 0.6635 | 0.769142612 |
| SL1344_RS01930_ | 0.094066019 | 0.040001551 | 0.175877592 | 0.137735123 | 0.227327167 | 0.580301446 | 0.255829459 | 0.378978968 | 0.547725979 | 0.5001 | 0.634221384 |
| SL1344_RS01935_ | -2.845594708 | -3.048493152 | -2.630339924 | 0.406919624 | 0.49359999 | 0.787903401 | 0.189207214 | 0.292530985 | 0.451991591 | 0.3875 | 0.533213249 |
| SL1344_RS01940_ | 0.136057255 | -0.06501404 | 0.296384021 | 0.336883789 | 0.343189305 | 0.939381028 | 0.15866669 | 0.220540604 | 0.405269429 | 0.2999 | 0.477299205 |
| SL1344_RS01945_ | 2.445299665 | 2.519684869 | 2.385479119 | -0.110067533 | 0.140063271 | -0.784575402 | 0.200532995 | 0.310815695 | 0.497238735 | 0.3964 | 0.565037234 |
| SL1344_RS01950_ | -2.940080649 | -2.86623289 | -3.003441182 | -0.127049319 | 0.534892454 | -0.231216072 | 0.397333337 | 0.531352561 | 0.673513416 | 0.642 | 0.750266916 |
| SL1344_RS01955_ | -0.809590475 | -0.751673142 | -0.861489047 | -0.113764158 | 0.370005414 | -0.298851343 | 0.358427721 | 0.589161517 | 0.72812552 | 0.6896 | 0.792508559 |
| SL1344_RS01960_ | 2.401767888 | 2.760320689 | 2.20605343 | -0.52401049 | 0.172593751 | -2.821337789 | 4.68E-05 | 0.024775137 | 0.112455779 | 0.1 | 0.243021319 |
| SL1344_RS01965_ | -1.591680923 | -1.63569445 | -1.515483359 | 0.184773461 | 0.578374876 | 0.313166673 | 0.373750838 | 0.552973429 | 0.705888939 | 0.7164 | 0.813361844 |
| SL1344_RS01975_ | 2.350130313 | 2.384245423 | 2.301528108 | -0.089933129 | 0.106879596 | -0.800654099 | 0.186666686 | 0.285236572 | 0.458473749 | 0.3844 | 0.532960394 |
| SL1344_RS01980_ | 0.358063516 | 0.323318514 | 0.411638971 | 0.066630182 | 0.239161641 | 0.260270601 | 0.379080618 | 0.635382605 | 0.763704077 | 0.718 | 0.819545693 |
| SL1344_RS01985_ | 1.665696582 | 1.440738491 | 1.935283003 | 0.481190727 | 0.166401013 | 2.802737248 | 0.000673369 | 0.019682486 | 0.095637652 | 0.1007 | 0.24400578 |
| SL1344_RS01990_ | 1.887915107 | 1.574791339 | 2.1459086 | 0.560507686 | 0.129230138 | 4.242540014 | 4.68E-05 | 0.008225304 | 0.057532812 | 0.1 | 0.243021319 |
| SL1344_RS01995_ | 1.033198528 | 0.777162262 | 1.26089734 | 0.484872537 | 0.161308339 | 2.985006669 | 4.68E-05 | 0.01709331 | 0.085936527 | 0.1 | 0.243021319 |
| SL1344_RS02000_ | 1.850361442 | 1.841018913 | 1.857288199 | 0.014758293 | 0.106824335 | 0.132550909 | 0.440000002 | 0.578963356 | 0.714786803 | 0.6938 | 0.792751773 |
| SL1344_RS02005_ | 2.561076816 | 2.514277854 | 2.69851075 | 0.180095141 | 0.123990331 | 1.261115422 | 0.108666704 | 0.160303565 | 0.334531396 | 0.2381 | 0.402018821 |
| SL1344_RS02010_ | -0.33996526 | -0.277139082 | -0.400607699 | -0.127192038 | 0.226074046 | -0.532265896 | 0.282666677 | 0.429455217 | 0.585570626 | 0.5367 | 0.664835113 |
| SL1344_RS02015_ | 0.457265866 | 0.366080219 | 0.586459474 | 0.200374135 | 0.210579788 | 0.896737399 | 0.165889429 | 0.258739846 | 0.435762803 | 0.3546 | 0.519227035 |
| SL1344_RS02020_ | 2.3568116 | 2.28006507 | 2.443685848 | 0.168658713 | 0.115942321 | 1.415950648 | 0.054000081 | 0.107204598 | 0.255532936 | 0.1592 | 0.315237341 |
| SL1344_RS02025_ | 1.082631156 | 0.99573671 | 1.16733443 | 0.173613075 | 0.1227763 | 1.358945496 | 0.070619647 | 0.125580259 | 0.26664754 | 0.1953 | 0.350027218 |
| SL1344_RS02030_ | -6.197287327 | -5.926340791 | -6.900209737 | -0.905811587 | 2.278493533 | -0.309841912 | 0.345103271 | 0.471475462 | 0.639747685 | 0.6379 | 0.75368239 |
| SL1344_RS02045_ | 3.617520547 | 3.604721711 | 3.63661298 | 0.041475419 | 0.09067852 | 0.453802 | 0.281333343 | 0.403574353 | 0.588023665 | 0.5391 | 0.664087361 |
| SL1344_RS02050_ | 3.420542321 | 3.172068754 | 3.667766162 | 0.493453262 | 0.076695259 | 6.405707953 | 4.68E-05 | 0.001968558 | 0.028276171 | 0.1 | 0.243021319 |
| SL1344_RS02055_ | 0.278724878 | -0.08723705 | 0.596067057 | 0.607049682 | 0.335183469 | 1.789523723 | 0.023317982 | 0.057210692 | 0.181040336 | 0.1219 | 0.273472717 |
| SL1344_RS02060_ | 4.96831257 | 5.054561711 | 4.842415456 | -0.220908564 | 0.072920516 | -3.055509094 | 4.68E-05 | 0.027227396 | 0.118493349 | 0.1 | 0.243021319 |
| SL1344_RS02065_ | 2.369620629 | 2.330355541 | 2.440463125 | 0.103936668 | 0.235078134 | 0.462863538 | 0.28580947 | 0.513252985 | 0.682388625 | 0.5586 | 0.713198319 |
| SL1344_RS02070_ | 2.966136562 | 2.772508347 | 3.209391876 | 0.421605505 | 0.102016102 | 4.078616118 | 4.68E-05 | 0.005780341 | 0.04950809 | 0.1 | 0.243021319 |
| SL1344_RS02075_ | 3.889669312 | 3.74593465 | 4.019094896 | 0.272003355 | 0.054825727 | 4.996029121 | 4.68E-05 | 0.004388281 | 0.04095637 | 0.1 | 0.243021319 |
| SL1344_RS02080_ | 3.400933072 | 3.385697711 | 3.492898032 | 0.10011353 | 0.107978003 | 0.47503324 | 0.337774823 | 0.736877173 | 0.843469914 | 0.7069 | 0.83313208 |

| | | | | | | | | | | | |
|---|---|---|---|---|---|---|---|---|---|---|---|
| SL1344_RS02085_ | 4.982884264 | 4.852401428 | 5.093262091 | 0.279889422 | 0.093211657 | 3.116430003 | 4.68E-05 | 0.009468086 | 0.066822358 | 0.1 | 0.243021319 |
| SL1344_RS02090_ | 3.682250672 | 3.695970841 | 3.641458804 | -0.056017594 | 0.271498463 | -0.203055862 | 0.376415728 | 0.717982643 | 0.83618857 | 0.7936 | 0.882659587 |
| SL1344_RS02095_ | -0.788077043 | -0.865752687 | -0.693695015 | 0.172173556 | 0.267788769 | 0.611730864 | 0.244503677 | 0.375965965 | 0.535407847 | 0.4789 | 0.612229495 |
| SL1344_RS02100_ | -0.402192983 | -0.551962321 | -0.268732057 | 0.296899413 | 0.326564283 | 0.874682658 | 0.155229871 | 0.242462773 | 0.424540034 | 0.3237 | 0.484598151 |
| SL1344_RS02105_ | 1.424854871 | 1.433584322 | 1.41604893 | -0.011550833 | 0.133042754 | -0.080132891 | 0.458361094 | 0.595446193 | 0.730423615 | 0.7037 | 0.803251505 |
| SL1344_RS02110_ | 2.789097778 | 2.771657674 | 2.833642992 | 0.064562439 | 0.161048439 | 0.364887423 | 0.357761498 | 0.524567662 | 0.691896325 | 0.7252 | 0.826595448 |
| SL1344_RS02115_ | 2.626046826 | 2.597893789 | 2.732836718 | 0.140734989 | 0.261489243 | 0.426015061 | 0.34333334 | 0.739961375 | 0.848754522 | 0.724 | 0.843190516 |
| SL1344_RS02120_ | -2.091786607 | -2.013187883 | -2.221275834 | -0.175947695 | 0.464315796 | -0.348592803 | 0.346000006 | 0.536888241 | 0.684435559 | 0.6469 | 0.759927127 |
| SL1344_RS02125_ | 1.433066529 | 1.325867563 | 1.529582423 | 0.204253453 | 0.106541493 | 1.851044065 | 0.020666886 | 0.058229892 | 0.168913227 | 0.124 | 0.273497937 |
| SL1344_RS02130_ | 2.048256894 | 1.956735475 | 2.193521479 | 0.236470546 | 0.148842939 | 1.527930997 | 0.050632998 | 0.085071516 | 0.227112164 | 0.1498 | 0.307375146 |
| SL1344_RS02135_ | 2.604407708 | 2.212544866 | 2.714908502 | 0.515927993 | 0.380692214 | 1.388753858 | 0.065956094 | 0.103836534 | 0.267506736 | 0.1562 | 0.319789172 |
| SL1344_RS02145_ | 1.508922466 | 1.180268869 | 1.675983263 | 0.486228234 | 0.192743748 | 2.421209519 | 0.002666616 | 0.036001326 | 0.136380767 | 0.1014 | 0.244978706 |
| SL1344_RS02150_ | 1.481725235 | 1.306945459 | 1.677329457 | 0.415465826 | 0.215332831 | 1.923463197 | 0.005334196 | 0.044934266 | 0.156036969 | 0.1036 | 0.247785776 |
| SL1344_RS02155_ | 0.849621108 | 0.697592554 | 1.009306791 | 0.341642281 | 0.207874114 | 1.558158102 | 0.029980163 | 0.079397017 | 0.215458085 | 0.1257 | 0.275387513 |
| SL1344_RS02160_ | 2.912158649 | 2.857819434 | 2.96653155 | 0.128243759 | 0.129802514 | 1.005141605 | 0.081945421 | 0.202689132 | 0.391559213 | 0.2187 | 0.378180165 |
| SL1344_RS02165_ | 2.56911661 | 2.441613858 | 2.667107522 | 0.225461428 | 0.086876726 | 2.530225487 | 0.004664544 | 0.028922801 | 0.115001223 | 0.1034 | 0.247601558 |
| SL1344_RS02170_ | 1.313796205 | 1.255236292 | 1.384759515 | 0.133793072 | 0.117043619 | 1.07906492 | 0.117333368 | 0.191601615 | 0.343733214 | 0.2797 | 0.428366056 |
| SL1344_RS02175_ | 3.013006523 | 2.904398004 | 3.110228797 | 0.227424395 | 0.116200689 | 1.991522061 | 0.002000966 | 0.052217142 | 0.171640838 | 0.1031 | 0.247324909 |
| SL1344_RS02180_ | -1.776902935 | -1.619018643 | -1.940044558 | -0.309789664 | 0.383588109 | -0.76399316 | 0.197868106 | 0.301499412 | 0.467831913 | 0.4067 | 0.553350614 |
| SL1344_RS02185_ | -2.58929142 | -2.586154635 | -2.591054053 | 0.009710002 | 0.505952626 | 0.015171551 | 0.493333334 | 0.586958625 | 0.722027713 | 0.7029 | 0.799073602 |
| SL1344_RS02190_ | -1.689861289 | -1.716189031 | -1.66441701 | 0.065650488 | 0.457775385 | 0.13976102 | 0.447035312 | 0.640238679 | 0.766834987 | 0.747 | 0.834354087 |
| SL1344_RS02195_ | -1.408156913 | -1.560706009 | -1.280208082 | 0.195786185 | 0.518978643 | 0.358934742 | 0.33933334 | 0.572192882 | 0.718526988 | 0.6895 | 0.792988251 |
| SL1344_RS02200_ | 0.355260598 | 0.421974149 | 0.303562116 | -0.119422869 | 0.185918862 | -0.611506125 | 0.245169899 | 0.382337482 | 0.545226572 | 0.4891 | 0.625885901 |
| SL1344_RS02205_ | -3.057027096 | -3.208457945 | -2.976251254 | 0.103861142 | 0.928317112 | 0.097349845 | 0.459333335 | 0.752693602 | 0.852245617 | 0.8271 | 0.897111072 |
| SL1344_RS02210_ | -3.264856528 | -3.159627727 | -3.378429353 | -0.236614148 | 0.685408722 | -0.330425021 | 0.358427721 | 0.524751979 | 0.672099262 | 0.6445 | 0.752962394 |
| SL1344_RS02215_ | 1.35916082 | 1.411054755 | 1.289951035 | -0.12355641 | 0.123782628 | -0.929437297 | 0.14533336 | 0.228624805 | 0.388375158 | 0.311 | 0.461185689 |
| SL1344_RS02220_ | 0.383546912 | 0.515932991 | 0.289419987 | -0.232943134 | 0.196257035 | -1.116747564 | 0.104596974 | 0.17379678 | 0.336200329 | 0.2484 | 0.406101215 |
| SL1344_RS02225_ | 3.562160938 | 3.55233857 | 3.587526246 | 0.026326051 | 0.098624818 | 0.184767052 | 0.411333337 | 0.788619862 | 0.876774016 | 0.7987 | 0.884784474 |
| SL1344_RS02230_ | 1.513681904 | 1.508337906 | 1.519913309 | 0.00877424 | 0.125340546 | 0.064154276 | 0.472351767 | 0.614209861 | 0.745258065 | 0.7262 | 0.817914255 |
| SL1344_RS02235_ | -0.527592446 | -0.6705182 | -0.352175617 | 0.312041331 | 0.280943497 | 1.077003232 | 0.105333372 | 0.181663451 | 0.345823582 | 0.2591 | 0.418799255 |
| SL1344_RS02240_ | 0.523285648 | 0.37420984 | 0.683073693 | 0.308394079 | 0.151084143 | 1.935420446 | 0.017322048 | 0.051914812 | 0.155858878 | 0.1232 | 0.270293287 |
| SL1344_RS02245_ | 5.602472113 | 5.559553578 | 5.661730959 | 0.107028796 | 0.062487552 | 1.656081203 | 0.020653117 | 0.080014008 | 0.223547482 | 0.1175 | 0.266910504 |
| SL1344_RS02250_ | 3.32971062 | 3.276198836 | 3.408201822 | 0.137373138 | 0.084593086 | 1.601377998 | 0.029313944 | 0.087491119 | 0.226972332 | 0.1381 | 0.290268154 |
| SL1344_RS02255_ | 3.96906259 | 3.948294875 | 4.041559467 | 0.088929701 | 0.074078024 | 1.050725094 | 0.161892095 | 0.221471226 | 0.403925027 | 0.3385 | 0.501178607 |
| SL1344_RS02260_ | 5.866744655 | 5.847666576 | 5.932889367 | 0.083063098 | 0.102465096 | 0.790112913 | 0.185333353 | 0.287706684 | 0.493113922 | 0.3658 | 0.54336699 |
| SL1344_RS02265_ | 6.014362226 | 5.85750797 | 6.139420932 | 0.283311825 | 0.063290892 | 4.620021985 | 4.68E-05 | 0.003137257 | 0.037907782 | 0.1 | 0.243021319 |
| SL1344_RS02270_ | 3.070356513 | 2.942023797 | 3.160502341 | 0.209781898 | 0.092874428 | 2.170568162 | 0.007995244 | 0.039982882 | 0.143976542 | 0.108 | 0.254008322 |
| SL1344_RS02275_ | 2.508673555 | 2.444849081 | 2.56223029 | 0.070738222 | 0.232901496 | 0.322537082 | 0.34733334 | 0.666296527 | 0.796009786 | 0.6991 | 0.809961551 |
| SL1344_RS02280_ | 2.438582782 | 2.656595031 | 2.382275824 | -0.263340482 | 0.20914706 | -1.24295698 | 0.112000036 | 0.125020242 | 0.292632818 | 0.233 | 0.407659579 |
| SL1344_RS02285_ | 6.814818687 | 6.766222161 | 6.858768404 | 0.103985335 | 0.047726751 | 2.394813869 | 4.68E-05 | 0.036184966 | 0.140396213 | 0.1 | 0.243021319 |
| SL1344_RS02300_ | 5.668068995 | 5.75635597 | 5.57931961 | -0.192640752 | 0.100964366 | -1.956099323 | 0.000673369 | 0.040276248 | 0.151368425 | 0.1002 | 0.243295929 |
| SL1344_RS02305_ | 6.657728872 | 6.783788965 | 6.552322332 | -0.236925472 | 0.061072176 | -3.698984144 | 4.68E-05 | 0.122424331 | 0.297261424 | 0.1 | 0.243021319 |
| SL1344_RS02310_ | 3.750479825 | 3.613042055 | 3.813985129 | 0.167640782 | 0.138992679 | 1.242057098 | 0.113257864 | 0.135568803 | 0.308976891 | 0.2188 | 0.39911018 |
| SL1344_RS02315_ | -3.778066783 | -3.305499623 | -4.373621452 | -0.853492627 | 1.356032731 | -0.572337818 | 0.248501012 | 0.397870681 | 0.579276949 | 0.4961 | 0.647899488 |
| SL1344_RS02320_ | -1.67789119 | -1.646983703 | -1.708138028 | -0.068207941 | 0.342781451 | -0.1878621 | 0.41972019 | 0.532001669 | 0.672581803 | 0.6329 | 0.740986602 |
| SL1344_RS02325_ | 1.418720794 | 1.385046048 | 1.46800476 | 0.101892245 | 0.160000072 | 0.621625698 | 0.233844118 | 0.359647409 | 0.537416071 | 0.4822 | 0.618023954 |
| SL1344_RS02330_ | -1.912439526 | -1.989219941 | -1.865356408 | 0.167888769 | 0.493154656 | 0.317987571 | 0.358000006 | 0.520318352 | 0.673153037 | 0.6525 | 0.759274186 |
| SL1344_RS02335_ | -1.685677469 | -1.627737394 | -1.75226273 | -0.084097537 | 0.450267177 | -0.184280377 | 0.417721522 | 0.619593606 | 0.748839963 | 0.7273 | 0.820812859 |
| SL1344_RS02350_ | 1.728285935 | 1.679114102 | 1.776719761 | 0.101485903 | 0.118089498 | 0.832915718 | 0.185333353 | 0.285494359 | 0.449684728 | 0.3788 | 0.528690053 |
| SL1344_RS02355_ | 1.790458944 | 1.72677542 | 1.840273553 | 0.107765114 | 0.123887826 | 0.81358501 | 0.191333352 | 0.290587704 | 0.458945376 | 0.3855 | 0.539884629 |
| SL1344_RS02360_ | -2.711228776 | -3.814557431 | -2.261084368 | 1.531957656 | 0.719545596 | 1.944213644 | 0.005996769 | 0.066859183 | 0.194925054 | 0.1078 | 0.253274383 |
| SL1344_RS02365_ | -0.746015141 | -0.971948634 | -0.461548162 | 0.546344192 | 0.36135084 | 1.442297255 | 0.049966777 | 0.103306179 | 0.252854358 | 0.1601 | 0.316925798 |
| SL1344_RS02370_ | 1.49528688 | 1.442789401 | 1.551646545 | 0.108609652 | 0.107111364 | 0.981244856 | 0.138574312 | 0.217665613 | 0.374059586 | 0.302 | 0.453743401 |

| | | | | | | | | | | | |
|---|---|---|---|---|---|---|---|---|---|---|---|
| SL1344_RS02375_ | -0.16726352 | 0.183673841 | -0.541094958 | -0.734649659 | 0.22743881 | -3.167549573 | 0.00067292 | 0.014003258 | 0.076876245 | 0.1001 | 0.243155301 |
| SL1344_RS02380_ | -0.487731171 | -0.521608429 | -0.457064666 | 0.04127611 | 0.336386159 | 0.111844951 | 0.443333335 | 0.680139762 | 0.79832858 | 0.7727 | 0.855281224 |
| SL1344_RS02385_ | 1.259466532 | 1.190565592 | 1.325759166 | 0.103845521 | 0.187199183 | 0.548220732 | 0.291805472 | 0.460140524 | 0.623194136 | 0.5714 | 0.702127569 |
| SL1344_RS02390_ | -1.565101319 | -1.690359274 | -1.383519531 | 0.359668494 | 0.450989402 | 0.77494615 | 0.187874769 | 0.292527792 | 0.472017883 | 0.3916 | 0.538345206 |
| SL1344_RS02395_ | -0.274562384 | -0.423412539 | -0.166476866 | 0.264672822 | 0.252718046 | 0.964760457 | 0.14533336 | 0.22010957 | 0.388710514 | 0.3082 | 0.464898298 |
| SL1344_RS02400_ | 3.556955383 | 3.81941615 | 3.305787975 | -0.50085008 | 0.083873709 | -5.874181447 | 4.68E-05 | 0.001934315 | 0.02842366 | 0.1 | 0.243021319 |
| SL1344_RS02405_ | 2.638192563 | 2.454753618 | 2.962057232 | 0.498277179 | 0.114233659 | 4.126570407 | 4.68E-05 | 0.01468012 | 0.082724139 | 0.1 | 0.243021319 |
| SL1344_RS02420_ | 5.196150964 | 5.121297116 | 5.265895832 | 0.153701827 | 0.060372418 | 2.462286882 | 4.68E-05 | 0.061747247 | 0.194318347 | 0.1 | 0.243021319 |
| SL1344_RS02425_ | 3.936957269 | 3.771994817 | 4.058099277 | 0.292966152 | 0.089211418 | 3.286593146 | 4.68E-05 | 0.009648069 | 0.065892504 | 0.1 | 0.243021319 |
| SL1344_RS02430_ | 1.725786434 | 1.706645554 | 1.744010723 | 0.045809836 | 0.135974278 | 0.319545608 | 0.359760166 | 0.544331846 | 0.690736455 | 0.6611 | 0.766444312 |
| SL1344_RS02435_ | 3.605113855 | 3.444006424 | 3.698048617 | 0.246967691 | 0.079828677 | 2.954234198 | 4.68E-05 | 0.022182541 | 0.1020953 | 0.1004 | 0.243575287 |
| SL1344_RS02440_ | 1.359961625 | 1.446880313 | 1.288387543 | -0.15366535 | 0.132379305 | -1.119648153 | 0.107928086 | 0.172066362 | 0.330608521 | 0.2524 | 0.413086657 |
| SL1344_RS02445_ | -0.060915412 | -0.022754365 | -0.115574712 | -0.095395014 | 0.265561402 | -0.342287634 | 0.362425055 | 0.547442231 | 0.694837961 | 0.6626 | 0.772153257 |
| SL1344_RS02450_ | 0.734522712 | 0.741548202 | 0.728380964 | -0.006546933 | 0.204010281 | -0.026553546 | 0.486342439 | 0.661553297 | 0.783547473 | 0.7594 | 0.844817449 |
| SL1344_RS02455_ | 0.537313594 | 0.475866379 | 0.594082555 | 0.120878428 | 0.160154667 | 0.686029996 | 0.220666682 | 0.343670632 | 0.499616054 | 0.4407 | 0.577945967 |
| SL1344_RS02460_ | 3.448940133 | 3.433328033 | 3.503793023 | 0.067809923 | 0.121896867 | 0.48236485 | 0.339107268 | 0.53856351 | 0.700266457 | 0.6904 | 0.809182319 |
| SL1344_RS02465_ | 4.189010779 | 4.041611407 | 4.369328714 | 0.332992286 | 0.090497429 | 3.66424645 | 4.68E-05 | 0.006734242 | 0.055093593 | 0.1 | 0.243021319 |
| SL1344_RS02470_ | 4.018432182 | 3.8826523 | 4.121010147 | 0.231288951 | 0.078494308 | 2.900636463 | 0.000673369 | 0.016325364 | 0.086957769 | 0.1 | 0.243021319 |
| SL1344_RS02475_ | 4.672218484 | 4.59084098 | 4.843627406 | 0.249306022 | 0.095184339 | 2.553418733 | 4.68E-05 | 0.020718893 | 0.10192971 | 0.1 | 0.243021319 |
| SL1344_RS02480_ | 6.088414406 | 6.259218036 | 6.073015482 | -0.188282728 | 0.121957259 | -1.243538331 | 0.144666693 | 0.148455902 | 0.332610938 | 0.2708 | 0.453009821 |
| SL1344_RS02485_ | 4.577653723 | 4.458211637 | 4.706126863 | 0.226837358 | 0.085594422 | 2.62788471 | 4.68E-05 | 0.018375336 | 0.094656213 | 0.1003 | 0.243434916 |
| SL1344_RS02490_ | 1.420445366 | 1.520202263 | 1.371038219 | -0.140085726 | 0.141519036 | -0.908773777 | 0.177881433 | 0.279261188 | 0.450402978 | 0.3881 | 0.544351038 |
| SL1344_RS02495_ | -1.983614124 | -2.081315991 | -1.830825669 | 0.134743707 | 0.709590492 | 0.155673333 | 0.433044639 | 0.771890619 | 0.8656883 | 0.8264 | 0.898735856 |
| SL1344_RS02500_ | 1.577015563 | 1.423415215 | 1.683628392 | 0.246417154 | 0.2331668 | 1.053518787 | 0.127914754 | 0.177238157 | 0.359231291 | 0.2547 | 0.434440295 |
| SL1344_RS02505_ | 2.71447715 | 2.786862424 | 2.623541928 | -0.17356112 | 0.103579497 | -1.590996797 | 0.025982851 | 0.092142773 | 0.236758131 | 0.1229 | 0.27203209 |
| SL1344_RS02510_ | 1.354273044 | 1.317617799 | 1.392616408 | 0.076020049 | 0.12954479 | 0.552225891 | 0.2778148 | 0.415991107 | 0.572869598 | 0.5251 | 0.653647035 |
| SL1344_RS02520_ | 1.062856961 | 1.01619609 | 1.14168453 | 0.119337007 | 0.151934397 | 0.72153526 | 0.221333348 | 0.341450065 | 0.51021595 | 0.4461 | 0.594486038 |
| SL1344_RS02525_ | 0.913262639 | 0.724810741 | 1.031017838 | 0.307197306 | 0.191865084 | 1.53169954 | 0.04266677 | 0.093647746 | 0.236999133 | 0.1439 | 0.298752367 |
| SL1344_RS02530_ | -0.825241673 | -1.076415121 | -0.572803259 | 0.501835518 | 0.247797497 | 1.970475157 | 0.017333596 | 0.051331126 | 0.156594609 | 0.1196 | 0.266903819 |
| SL1344_RS02540_ | 0.468767947 | 0.362686497 | 0.5703744 | 0.20579177 | 0.143908966 | 1.363440268 | 0.056000078 | 0.114334378 | 0.245993478 | 0.1913 | 0.342182139 |
| SL1344_RS02545_ | 0.32763408 | 0.426912183 | 0.054725147 | -0.347931298 | 0.218423392 | -1.41539835 | 0.077948089 | 0.124552308 | 0.279844346 | 0.2011 | 0.358836028 |
| SL1344_RS02550_ | 0.175042892 | 0.381702076 | -0.069839847 | -0.453933664 | 0.164327058 | -2.674052648 | 0.002666616 | 0.022237587 | 0.097229659 | 0.1013 | 0.244845254 |
| SL1344_RS02555_ | -0.140258319 | -0.234459017 | 0.134111803 | 0.343711586 | 0.341779534 | 0.877031017 | 0.208527665 | 0.28999357 | 0.479844674 | 0.418 | 0.583563752 |
| SL1344_RS02560_ | 0.084075146 | -0.030387645 | 0.244451225 | 0.315297028 | 0.304129916 | 1.012056829 | 0.120586309 | 0.182339853 | 0.36006655 | 0.2671 | 0.434201717 |
| SL1344_RS02565_ | 3.203785075 | 3.417392943 | 3.121289154 | -0.274554237 | 0.12904842 | -1.867161179 | 0.015989625 | 0.065234892 | 0.197447779 | 0.1159 | 0.264600254 |
| SL1344_RS02570_ | 0.886772094 | 0.886772094 | 0.886780847 | 0.014865428 | 0.175702941 | 0.081651568 | 0.463024652 | 0.631499221 | 0.759069351 | 0.7483 | 0.833591682 |
| SL1344_RS02575_ | 0.144548728 | 0.110803059 | 0.17415119 | 0.070568521 | 0.230756848 | 0.276692402 | 0.370419725 | 0.582979265 | 0.71982949 | 0.7034 | 0.799133732 |
| SL1344_RS02580_ | -1.043971988 | -1.03377867 | -1.061511175 | -0.066065959 | 0.422504218 | -0.131652202 | 0.428000003 | 0.570205564 | 0.721142923 | 0.7307 | 0.821729551 |
| SL1344_RS02585_ | 0.870626933 | 0.893780883 | 0.84946292 | -0.049814092 | 0.15633076 | -0.298207955 | 0.365756168 | 0.537354456 | 0.678488982 | 0.6376 | 0.748678258 |
| SL1344_RS02590_ | 0.315368573 | 0.306838397 | 0.331413113 | 0.007211981 | 0.252672227 | 0.024930463 | 0.486666667 | 0.69363154 | 0.807079607 | 0.8037 | 0.876602355 |
| SL1344_RS02595_ | -0.521368935 | -0.364974497 | -0.648683468 | -0.280995007 | 0.268190849 | -1.016923658 | 0.135333362 | 0.214427083 | 0.381534246 | 0.2954 | 0.454397408 |
| SL1344_RS02600_ | -0.715203007 | -0.685543461 | -0.745066227 | -0.071391229 | 0.347304159 | -0.184236935 | 0.41266667 | 0.642773292 | 0.769569462 | 0.7644 | 0.84623359 |
| SL1344_RS02605_ | -0.561067943 | -0.405630575 | -0.691616141 | -0.282495277 | 0.231638175 | -1.136009206 | 0.094603641 | 0.16719891 | 0.318655023 | 0.2449 | 0.39907167 |
| SL1344_RS02610_ | 2.243606854 | 2.204183111 | 2.319878447 | 0.144141053 | 0.256067448 | 0.608273297 | 0.242000013 | 0.353066954 | 0.551863993 | 0.4856 | 0.648142558 |
| SL1344_RS02615_ | -2.429168201 | -2.518719548 | -2.335070848 | 0.169020032 | 0.502286608 | 0.316598757 | 0.360666672 | 0.534329959 | 0.679463966 | 0.6408 | 0.753120839 |
| SL1344_RS02620_ | -2.077480155 | -1.867367738 | -2.246916632 | -0.379286256 | 0.395406635 | -0.900283604 | 0.147235203 | 0.246352742 | 0.404573145 | 0.3382 | 0.487761655 |
| SL1344_RS02625_ | 1.431635465 | 1.548416693 | 1.261588312 | -0.290034774 | 0.161924249 | -1.759547876 | 0.015323415 | 0.074385842 | 0.205765152 | 0.1178 | 0.265767992 |
| SL1344_RS02630_ | -2.722949722 | -2.616208767 | -2.828537152 | -0.224926293 | 0.44987431 | -0.439446146 | 0.305796144 | 0.429024049 | 0.581080779 | 0.529 | 0.653293391 |
| SL1344_RS02635_ | -3.492271221 | -3.395875451 | -3.654353654 | -0.308054431 | 0.75603036 | -0.373796019 | 0.327333341 | 0.475649539 | 0.631622875 | 0.5956 | 0.712717644 |
| SL1344_RS02640_ | -2.775487962 | -2.827113452 | -2.745808401 | 0.141950234 | 0.668802269 | 0.185564648 | 0.41466667 | 0.597564962 | 0.735800511 | 0.7296 | 0.821650929 |
| SL1344_RS02645_ | -2.651658071 | -2.654750594 | -2.648846603 | 0.018942211 | 0.491308073 | 0.03412191 | 0.484000001 | 0.584064324 | 0.72005113 | 0.6975 | 0.795218967 |
| SL1344_RS02650_ | -4.0214395 | -4.099805386 | -3.933827178 | 0.140384231 | 0.74817051 | 0.166987902 | 0.425716192 | 0.538735454 | 0.678193251 | 0.6449 | 0.751712729 |
| SL1344_RS02655_ | -2.443760927 | -2.347090356 | -2.48338651 | -0.135719025 | 0.823882009 | -0.140978437 | 0.444370422 | 0.749872632 | 0.849436994 | 0.8301 | 0.899050862 |

| | | | | | | | | | | | |
|---|---|---|---|---|---|---|---|---|---|---|---|
| SL1344_RS02660_ | -2.872538091 | -2.635291254 | -3.269967097 | -0.647668202 | 0.743800088 | -0.822371992 | 0.189333352 | 0.276241861 | 0.455632292 | 0.3803 | 0.543208831 |
| SL1344_RS02665_ | -2.351829957 | -2.300297416 | -2.387747172 | -0.071552538 | 0.490923099 | -0.144746574 | 0.435043307 | 0.59399898 | 0.7261159 | 0.6973 | 0.794574806 |
| SL1344_RS02670_ | -0.115879871 | -0.22139357 | -0.057801953 | 0.138440981 | 0.25591149 | 0.506126787 | 0.29380414 | 0.459566989 | 0.620568168 | 0.5776 | 0.706865001 |
| SL1344_RS02675_ | -3.027235775 | -2.964534588 | -3.08483521 | -0.08706337 | 0.726149227 | -0.108453538 | 0.445702867 | 0.652336128 | 0.774875444 | 0.7496 | 0.837317911 |
| SL1344_RS02680_ | -3.766380434 | -3.340613047 | -4.425368816 | -0.594356193 | 1.778765855 | -0.331602461 | 0.39573618 | 0.670649247 | 0.798209088 | 0.8081 | 0.886152701 |
| SL1344_RS02685_ | -2.346232784 | -2.292538818 | -2.411187274 | -0.114042658 | 0.48348614 | -0.21013745 | 0.40866667 | 0.54182156 | 0.684982292 | 0.6548 | 0.761703745 |
| SL1344_RS02690_ | -3.430987116 | -3.264905405 | -3.549052436 | -0.297979205 | 0.687714577 | -0.392648347 | 0.331112599 | 0.506074205 | 0.650729914 | 0.616 | 0.728070896 |
| SL1344_RS02695_ | -3.886237084 | -3.949114229 | -3.777755871 | 0.199177338 | 1.022768927 | 0.169061481 | 0.424000003 | 0.647234218 | 0.771547955 | 0.7539 | 0.839655191 |
| SL1344_RS02700_ | -2.022960286 | -1.956669392 | -2.102090111 | -0.141851154 | 0.372823683 | -0.364163557 | 0.344437048 | 0.468517095 | 0.616077738 | 0.5802 | 0.696457 |
| SL1344_RS02705_ | 2.046145904 | 1.959355733 | 2.147572552 | 0.186049018 | 0.092208728 | 1.948556403 | 0.015989625 | 0.053344081 | 0.161714127 | 0.1229 | 0.271428386 |
| SL1344_RS02710_ | 1.346100291 | 1.293793247 | 1.402198031 | 0.11793741 | 0.152831903 | 0.751621164 | 0.205862775 | 0.315288353 | 0.487858666 | 0.4191 | 0.568147207 |
| SL1344_RS02715_ | 3.215730275 | 3.182153899 | 3.271333717 | 0.095650884 | 0.095653566 | 0.995919656 | 0.112591641 | 0.204167722 | 0.386168285 | 0.2496 | 0.410416345 |
| SL1344_RS02720_ | 5.153160007 | 5.136463758 | 5.19598697 | 0.0519961 | 0.089683101 | 0.448310498 | 0.351333339 | 0.675958691 | 0.804094687 | 0.7333 | 0.845407601 |
| SL1344_RS02730_ | -1.109892296 | -1.213732755 | -1.026654302 | 0.178181307 | 0.327174354 | 0.510578954 | 0.272666677 | 0.436095066 | 0.592097166 | 0.5449 | 0.671562098 |
| SL1344_RS02735_ | -1.896331111 | -1.863373736 | -1.929170485 | -0.06438551 | 0.354204353 | -0.174685119 | 0.425716192 | 0.526137451 | 0.66947132 | 0.6341 | 0.743502732 |
| SL1344_RS02740_ | -1.476616381 | -1.402193959 | -1.57864766 | -0.175874662 | 0.351222008 | -0.464376744 | 0.304000009 | 0.451684793 | 0.606451317 | 0.5632 | 0.687245674 |
| SL1344_RS02745_ | 2.482306867 | 2.520795172 | 2.445334476 | -0.086444579 | 0.10931095 | -0.768254619 | 0.194666685 | 0.297023227 | 0.472986992 | 0.4193 | 0.568090051 |
| SL1344_RS02750_ | 2.62897641 | 2.572958804 | 2.73038529 | 0.152072863 | 0.090672399 | 1.586737048 | 0.045303228 | 0.094725444 | 0.233293711 | 0.1621 | 0.315748158 |
| SL1344_RS02760_ | -3.713872559 | -3.36415363 | -4.097372375 | -0.731906527 | 0.777029032 | -0.901569538 | 0.154000025 | 0.237236469 | 0.401726168 | 0.3225 | 0.476724637 |
| SL1344_RS02765_ | -4.211880508 | -3.939524135 | -4.417853479 | -0.456641604 | 0.912096283 | -0.452295582 | 0.300666675 | 0.454644827 | 0.608491613 | 0.5581 | 0.684873972 |
| SL1344_RS02770_ | -1.840821408 | -1.635576097 | -2.137634116 | -0.567390016 | 0.567852231 | -0.985634912 | 0.119920087 | 0.207625353 | 0.390655745 | 0.2764 | 0.439677691 |
| SL1344_RS02775_ | -2.327010945 | -2.246467954 | -2.447072132 | -0.179117476 | 0.495696318 | -0.347112237 | 0.358427721 | 0.540328471 | 0.685232015 | 0.6574 | 0.765674471 |
| SL1344_RS02790_ | -3.557096177 | -3.644201122 | -3.372136056 | 0.299174127 | 0.913892496 | 0.307202169 | 0.373750838 | 0.584244235 | 0.723111816 | 0.7042 | 0.801271919 |
| SL1344_RS02795_ | -5.457071102 | -5.769732947 | -4.848463463 | 0.824101517 | 1.741973975 | 0.393109552 | 0.33133334 | 0.582239961 | 0.716140437 | 0.6525 | 0.767515643 |
| SL1344_RS02800_ | -1.820266224 | -1.918110942 | -1.693227733 | 0.147828801 | 0.565222791 | 0.234029828 | 0.398666671 | 0.653211906 | 0.777093111 | 0.7519 | 0.839993511 |
| SL1344_RS02805_ | 0.413803654 | -0.074117456 | 0.677790603 | 0.720390015 | 0.305831282 | 2.265346126 | 0.002666616 | 0.037776882 | 0.14090982 | 0.1015 | 0.245131823 |
| SL1344_RS02815_ | 0.241459416 | 0.534029286 | 0.083241527 | -0.484275449 | 0.330575704 | -1.353830507 | 0.05266675 | 0.111105888 | 0.270626674 | 0.1626 | 0.32186589 |
| SL1344_RS02820_ | -0.347386512 | -0.088106602 | -0.48826887 | -0.400586258 | 0.364228417 | -1.035114487 | 0.145236535 | 0.206331223 | 0.38906954 | 0.2868 | 0.455352029 |
| SL1344_RS02825_ | -1.933075969 | -1.562193163 | -2.129481234 | -0.595117924 | 0.612035459 | -0.930719492 | 0.151333358 | 0.232269236 | 0.417229962 | 0.3194 | 0.485779879 |
| SL1344_RS02830_ | -4.017321674 | -3.848347126 | -4.196229676 | -0.349453224 | 0.723296828 | -0.444530271 | 0.310459702 | 0.436724291 | 0.589079312 | 0.5421 | 0.666700319 |
| SL1344_RS02835_ | -4.322804586 | -4.3651876 | -4.280867867 | 0.114631101 | 1.210024546 | 0.076061273 | 0.465689542 | 0.661702422 | 0.783646428 | 0.7814 | 0.859135079 |
| SL1344_RS02840_ | -0.461668156 | -0.608022575 | -0.390745637 | 0.21338001 | 0.293515485 | 0.654573895 | 0.227333348 | 0.351685605 | 0.526871451 | 0.4641 | 0.609695746 |
| SL1344_RS02845_ | -1.707675729 | -1.605778553 | -1.786962651 | -0.185156061 | 0.43344534 | -0.410799337 | 0.323784151 | 0.498977882 | 0.649909519 | 0.6206 | 0.733227926 |
| SL1344_RS02850_ | -1.391673489 | -1.294776498 | -1.508820491 | -0.202749522 | 0.304408622 | -0.648577229 | 0.23866668 | 0.370328968 | 0.52606556 | 0.4736 | 0.607178789 |
| SL1344_RS02855_ | -5.092339238 | -4.966856027 | -5.339949873 | -0.468474274 | 1.422099984 | -0.255588059 | 0.372000005 | 0.521855821 | 0.676718822 | 0.6586 | 0.767854296 |
| SL1344_RS02860_ | -2.971089804 | -2.859572483 | -3.091360162 | -0.240518441 | 0.490505116 | -0.461427388 | 0.301798809 | 0.425842405 | 0.576880367 | 0.5258 | 0.651190627 |
| SL1344_RS02865_ | 2.191308179 | 2.728549631 | 1.56526536 | -1.154670559 | 0.175387203 | -6.533385724 | 4.68E-05 | 0.001159871 | 0.023184017 | 0.1 | 0.243021319 |
| SL1344_RS02870_ | 2.320723489 | 2.305629482 | 2.335940969 | 0.069789284 | 0.149730451 | 0.441532687 | 0.324666674 | 0.466221806 | 0.634120239 | 0.6771 | 0.776263831 |
| SL1344_RS02875_ | 1.473907622 | 1.340998649 | 1.647346499 | 0.324069016 | 0.174420004 | 1.828749864 | 0.015333631 | 0.057038731 | 0.177724915 | 0.1139 | 0.261251404 |
| SL1344_RS02880_ | -0.323498732 | -0.291330092 | -0.343998374 | -0.052952364 | 0.260488592 | -0.189614783 | 0.412391742 | 0.615265862 | 0.745501247 | 0.7292 | 0.821195285 |
| SL1344_RS02885_ | 0.31880992 | 0.31656017 | 0.323492661 | 0.027279187 | 0.234621037 | 0.106684766 | 0.456000002 | 0.648957627 | 0.773534922 | 0.7593 | 0.843677114 |
| SL1344_RS02890_ | -2.274495875 | -2.198815329 | -2.316223205 | -0.099911446 | 0.545819326 | -0.16399628 | 0.434000002 | 0.660843862 | 0.781486472 | 0.7648 | 0.848540388 |
| SL1344_RS02895_ | -3.307573318 | -3.379225039 | -3.262446813 | 0.118488006 | 0.645795237 | 0.175621337 | 0.41266667 | 0.557532567 | 0.699536427 | 0.678 | 0.780398409 |
| SL1344_RS02900_ | -4.232314224 | -4.063986894 | -4.408182824 | -0.349416489 | 0.768061821 | -0.413192122 | 0.330000007 | 0.456941796 | 0.605221478 | 0.562 | 0.683587682 |
| SL1344_RS02905_ | -4.88433965 | -4.243148618 | -5.19610599 | -0.689009007 | 1.839959476 | -0.298843705 | 0.350000006 | 0.703316747 | 0.813792131 | 0.7257 | 0.832982619 |
| SL1344_RS02915_ | -5.175671836 | -4.90679598 | -5.384211889 | -0.494277233 | 1.396910289 | -0.339785513 | 0.353097941 | 0.539280181 | 0.681020712 | 0.6506 | 0.756139464 |
| SL1344_RS02920_ | 2.19888412 | 2.235788177 | 2.169444165 | -0.067433763 | 0.096532855 | -0.645592473 | 0.23451034 | 0.375621374 | 0.538889919 | 0.4725 | 0.615678148 |
| SL1344_RS02925_ | -0.71908535 | -0.695767535 | -0.732323369 | -0.017983957 | 0.320119029 | -0.055685795 | 0.474350434 | 0.644480105 | 0.771342131 | 0.758 | 0.843717413 |
| SL1344_RS02930_ | 0.350089993 | 0.329829985 | 0.375822429 | 0.047207082 | 0.178247909 | 0.252922828 | 0.372000005 | 0.527454356 | 0.672309656 | 0.639 | 0.747944609 |
| SL1344_RS02935_ | -0.728985569 | -0.493919343 | -1.035734367 | -0.498167037 | 0.3909854 | -1.19103192 | 0.093937419 | 0.138273135 | 0.301981617 | 0.2084 | 0.378027554 |
| SL1344_RS02940_ | -0.992827149 | -0.997050167 | -0.986807892 | -0.002894462 | 0.306535708 | -0.007903409 | 0.497001999 | 0.634614607 | 0.759997321 | 0.7438 | 0.831795597 |
| SL1344_RS02945_ | -1.665755395 | -1.744690543 | -1.60667116 | 0.083645014 | 0.52799858 | 0.153247172 | 0.432378417 | 0.693120677 | 0.807972502 | 0.803 | 0.875324766 |
| SL1344_RS02950_ | -1.945415109 | -1.948459872 | -1.942688674 | -0.037429129 | 0.64546259 | -0.047361546 | 0.476349101 | 0.717155006 | 0.826213454 | 0.8023 | 0.876246173 |

| | | | | | | | | | | |
|---|---|---|---|---|---|---|---|---|---|---|
| SL1344_RS02955_ | 1.887613827 | 1.851624303 | 1.976020403 | 0.165583308 | 0.256290851 | 0.663379056 | 0.239333347 | 0.343221358 | 0.538977117 | 0.4814 | 0.648728954 |
| SL1344_RS02960_ | 1.463908791 | 1.056686911 | 1.832239618 | 0.695345694 | 0.252975107 | 2.787738151 | 0.000673369 | 0.014630078 | 0.083298893 | 0.1001 | 0.243164097 |
| SL1344_RS02965_ | -1.404168399 | -1.544228739 | -1.193795644 | 0.3501174 | 0.443952283 | 0.73936396 | 0.215856111 | 0.327918029 | 0.505211204 | 0.4323 | 0.587889659 |
| SL1344_RS02975_ | 2.409456721 | 2.206960508 | 2.624203397 | 0.41693028 | 0.081459645 | 5.030839976 | 4.68E-05 | 0.004323475 | 0.040213711 | 0.1 | 0.243021319 |
| SL1344_RS02980_ | 0.789938311 | 0.810979527 | 0.768651597 | -0.046845817 | 0.177004056 | -0.259049416 | 0.386000004 | 0.583252309 | 0.718409985 | 0.6929 | 0.790074998 |
| SL1344_RS02985_ | 1.557164043 | 1.510796313 | 1.637666253 | 0.117202469 | 0.1650574 | 0.671162798 | 0.231179228 | 0.376800581 | 0.549221412 | 0.4686 | 0.62295402 |
| SL1344_RS02990_ | 1.461721682 | 1.340387465 | 1.562711885 | 0.242948693 | 0.189958873 | 1.234259409 | 0.079946755 | 0.134221418 | 0.298128503 | 0.1936 | 0.358057069 |
| SL1344_RS02995_ | 1.679784431 | 1.576699027 | 1.837442111 | 0.286229427 | 0.229842748 | 1.265835246 | 0.058666741 | 0.157480186 | 0.339268273 | 0.158 | 0.318011769 |
| SL1344_RS03000_ | 0.823263712 | 0.661973527 | 0.996657505 | 0.316867613 | 0.199901827 | 1.533671789 | 0.057333409 | 0.09343049 | 0.236018228 | 0.1612 | 0.32148187 |
| SL1344_RS03005_ | 1.358636966 | 1.294158624 | 1.702880214 | 0.350749772 | 0.415526235 | 0.803185366 | 0.235333347 | 0.3007337 | 0.501180793 | 0.4305 | 0.617576782 |
| SL1344_RS03020_ | 0.341154096 | 0.311229475 | 0.386483542 | 0.032674708 | 0.277511893 | 0.10723002 | 0.453697537 | 0.73242588 | 0.83792986 | 0.8428 | 0.904339429 |
| SL1344_RS03025_ | -0.343473651 | -0.317383733 | -0.375089854 | -0.034726753 | 0.31055162 | -0.113805205 | 0.454000002 | 0.665343933 | 0.787384809 | 0.7787 | 0.857842589 |
| SL1344_RS03030_ | 1.301732369 | 1.361614041 | 1.23833522 | -0.123160325 | 0.118387254 | -1.009166835 | 0.127914754 | 0.214444658 | 0.369654591 | 0.3003 | 0.450249312 |
| SL1344_RS03035_ | -1.493763074 | -1.409236466 | -1.575867496 | -0.159955302 | 0.347275438 | -0.422515978 | 0.316455704 | 0.481003496 | 0.629997944 | 0.5842 | 0.704572425 |
| SL1344_RS03040_ | 0.006308204 | 0.017474424 | -0.007633019 | -0.034051833 | 0.323449856 | -0.102790517 | 0.457694872 | 0.703736042 | 0.816450597 | 0.8057 | 0.878514465 |
| SL1344_RS03045_ | 0.315206866 | 0.252468403 | 0.351574097 | 0.043227645 | 0.30552301 | 0.13181581 | 0.446000002 | 0.755114589 | 0.85389247 | 0.8448 | 0.908896071 |
| SL1344_RS03050_ | -0.783205657 | -0.70024086 | -0.869954371 | -0.178961959 | 0.245714244 | -0.675755094 | 0.226000015 | 0.346432793 | 0.50574667 | 0.4488 | 0.586983062 |
| SL1344_RS03055_ | -0.435556248 | -0.449280096 | -0.421887367 | 0.028738116 | 0.21291669 | 0.122450376 | 0.444000002 | 0.548288239 | 0.6864606 | 0.6606 | 0.764598805 |
| SL1344_RS03060_ | -0.244953633 | -0.354970926 | -0.13555498 | 0.259333882 | 0.303375974 | 0.833026451 | 0.175216543 | 0.255763004 | 0.436311287 | 0.3551 | 0.509987201 |
| SL1344_RS03065_ | 1.043351487 | 1.058253508 | 1.022458443 | -0.024525375 | 0.169836163 | -0.129198929 | 0.445702867 | 0.641118236 | 0.768212846 | 0.7544 | 0.839848505 |
| SL1344_RS03070_ | 4.733289658 | 4.781121854 | 4.683914063 | -0.056856444 | 0.150243443 | -0.411864822 | 0.326666674 | 0.552733143 | 0.718478189 | 0.6806 | 0.802346987 |
| SL1344_RS03075_ | 2.857682335 | 2.853614161 | 2.861190236 | 0.005993472 | 0.089738938 | 0.064383257 | 0.473333334 | 0.641145917 | 0.767880448 | 0.7539 | 0.84017955 |
| SL1344_RS03080_ | -4.672176329 | -4.268381734 | -5.147830233 | -0.927771435 | 1.197796495 | -0.697223069 | 0.19920055 | 0.302921092 | 0.475769346 | 0.3973 | 0.546002173 |
| SL1344_RS03085_ | -1.494576055 | -1.399903808 | -1.60217216 | -0.202707532 | 0.302964493 | -0.646619774 | 0.242000013 | 0.353556234 | 0.508705077 | 0.4508 | 0.585297213 |
| SL1344_RS03090_ | -2.619358117 | -2.522353293 | -2.756009079 | -0.143342977 | 0.711596655 | -0.193574016 | 0.425716192 | 0.665235915 | 0.788271941 | 0.7871 | 0.865549599 |
| SL1344_RS03095_ | 0.676526543 | 0.872554952 | 0.462855727 | -0.409418202 | 0.198881934 | -2.005117242 | 0.010667096 | 0.04577322 | 0.152093208 | 0.1112 | 0.257770764 |
| SL1344_RS03100_ | 3.131346363 | 3.346687746 | 2.745789579 | -0.551010132 | 0.171124215 | -2.830824375 | 4.68E-05 | 0.02353053 | 0.110169735 | 0.1 | 0.243021319 |
| SL1344_RS03105_ | -2.02104754 | -2.181494683 | -1.864110401 | 0.308107869 | 0.435837919 | 0.670634407 | 0.224666681 | 0.354732761 | 0.517909666 | 0.4605 | 0.602014083 |
| SL1344_RS03110_ | 0.366444576 | 0.417464832 | 0.294876754 | -0.149109368 | 0.240546482 | -0.590325445 | 0.23451034 | 0.36197804 | 0.54054691 | 0.4815 | 0.618994654 |
| SL1344_RS03115_ | 2.523149754 | 2.440467055 | 2.600532814 | 0.160964572 | 0.119855252 | 1.283377004 | 0.074616979 | 0.125896055 | 0.284406239 | 0.1884 | 0.353274307 |
| SL1344_RS03120_ | 0.402597512 | 0.603761962 | 0.21170175 | -0.403523228 | 0.189328387 | -2.095959744 | 0.008000574 | 0.044973594 | 0.149642441 | 0.1078 | 0.253204966 |
| SL1344_RS03125_ | -4.722885304 | -4.344930104 | -5.236945269 | -0.941618736 | 1.350754791 | -0.647031084 | 0.195333351 | 0.309673033 | 0.492862213 | 0.4018 | 0.548319503 |
| SL1344_RS03130_ | -4.727863891 | -4.815324991 | -4.593102451 | 0.080081129 | 1.502493713 | 0.053180004 | 0.473017989 | 0.679015204 | 0.797411251 | 0.7773 | 0.85767602 |
| SL1344_RS03135_ | -3.889320911 | -4.057662705 | -3.723601467 | 0.33270231 | 0.720304642 | 0.411642776 | 0.317121927 | 0.454704198 | 0.604821052 | 0.5619 | 0.682709023 |
| SL1344_RS03140_ | -4.122973773 | -3.604857862 | -4.816353215 | -0.064753321 | 3.158415928 | -0.015002681 | 0.492666667 | 0.714561333 | 0.830302157 | 0.9535 | 0.973224257 |
| SL1344_RS03145_ | -6.313709072 | -6.503569078 | -6.073507256 | 0.406655461 | 1.795850291 | 0.188220646 | 0.404397072 | 0.539091104 | 0.682646857 | 0.6525 | 0.758154209 |
| SL1344_RS03150_ | -6.026711614 | -6.198831482 | -5.783634365 | 0.183224098 | 2.500021449 | 0.05154728 | 0.461333335 | 0.655336454 | 0.783125451 | 0.786 | 0.865467771 |
| SL1344_RS03165_ | -0.810508783 | -1.655686252 | -0.574272184 | 0.94068568 | 0.64841806 | 1.150191231 | 0.145902758 | 0.182428222 | 0.372064577 | 0.2731 | 0.453499542 |
| SL1344_RS03175_ | 7.048664488 | 6.749813261 | 7.275974068 | 0.494188795 | 0.1420281 | 3.64915083 | 4.68E-05 | 0.003129793 | 0.039646696 | 0.1 | 0.243021319 |
| SL1344_RS03180_ | -0.665471606 | -0.649451994 | -0.682440178 | -0.03340753 | 0.265192635 | -0.122238549 | 0.445333335 | 0.578669885 | 0.714719342 | 0.6855 | 0.786106782 |
| SL1344_RS03195_ | 4.086210385 | 4.027621721 | 4.127857556 | 0.092015529 | 0.128546821 | 0.710638044 | 0.213857444 | 0.312291869 | 0.510051689 | 0.4104 | 0.587866193 |
| SL1344_RS03200_ | -0.168128623 | -0.616886616 | 0.177625634 | 0.775272322 | 0.270836752 | 2.826285345 | 0.001335878 | 0.020364596 | 0.095914445 | 0.1007 | 0.244008578 |
| SL1344_RS03205_ | 2.397256258 | 2.462630774 | 2.322850187 | -0.141307501 | 0.081146387 | -1.626089722 | 0.039333446 | 0.083228724 | 0.209074335 | 0.1468 | 0.29749863 |
| SL1344_RS03210_ | 4.007797964 | 4.167807661 | 3.93723535 | -0.260452328 | 0.170628938 | -1.6187155 | 0.010659989 | 0.060195284 | 0.191141329 | 0.1084 | 0.254551828 |
| SL1344_RS03215_ | 3.853538221 | 3.766885628 | 3.952147721 | 0.188641939 | 0.058163584 | 3.19072314 | 4.68E-05 | 0.014059608 | 0.077450259 | 0.1 | 0.243021319 |
| SL1344_RS03225_ | 1.915932322 | 1.933417402 | 1.858342929 | -0.070116974 | 0.157072637 | -0.356635302 | 0.376666672 | 0.660638612 | 0.786812506 | 0.7543 | 0.852671689 |
| SL1344_RS03230_ | 3.187793004 | 3.097572684 | 3.299528644 | 0.210888941 | 0.084359272 | 2.504440155 | 4.68E-05 | 0.040363343 | 0.144304025 | 0.1003 | 0.243441169 |
| SL1344_RS03235_ | 3.227578186 | 3.10530549 | 3.343378287 | 0.258970586 | 0.109071568 | 2.411227051 | 0.00067292 | 0.025544801 | 0.111935157 | 0.1003 | 0.243448682 |
| SL1344_RS03240_ | 2.763996298 | 2.664999749 | 2.857367062 | 0.212696615 | 0.109318709 | 1.904855042 | 0.008661422 | 0.047894369 | 0.15939626 | 0.1072 | 0.252273812 |
| SL1344_RS03245_ | 1.498484924 | 1.32513184 | 1.715020761 | 0.384132971 | 0.139424134 | 2.729272794 | 0.001335878 | 0.022916867 | 0.102391207 | 0.1012 | 0.244632215 |
| SL1344_RS03250_ | -0.448020946 | -0.483121681 | -0.38834413 | 0.118828658 | 0.398529135 | 0.294934485 | 0.377333338 | 0.631538497 | 0.764661613 | 0.742 | 0.834161356 |
| SL1344_RS03255_ | -1.605709307 | -1.550328959 | -1.644696119 | -0.090852756 | 0.429958413 | -0.202941863 | 0.413057965 | 0.608887056 | 0.742084696 | 0.7106 | 0.807419078 |
| SL1344_RS03260_ | 1.298181743 | 1.206046719 | 1.334753815 | 0.10472073 | 0.182702524 | 0.536518393 | 0.292471695 | 0.471422083 | 0.637250232 | 0.5831 | 0.721678859 |

| | | | | | | | | | | | |
|---|---|---|---|---|---|---|---|---|---|---|---|
| SL1344_RS03265_ | 2.988327677 | 2.900488549 | 3.102936843 | 0.203011426 | 0.114401984 | 1.723158739 | 0.016655836 | 0.075800956 | 0.213884804 | 0.1147 | 0.263191575 |
| SL1344_RS03270_ | 3.300005949 | 3.136765998 | 3.379599376 | 0.237523714 | 0.083883442 | 2.664408124 | 4.68E-05 | 0.032663829 | 0.126752829 | 0.1018 | 0.245416171 |
| SL1344_RS03275_ | 5.047334684 | 4.919401521 | 5.169705816 | 0.230401924 | 0.072014987 | 3.180720103 | 4.68E-05 | 0.009583176 | 0.066672484 | 0.1 | 0.243021319 |
| SL1344_RS03280_ | -5.545734402 | -5.100915288 | -5.83158431 | -0.594806818 | 1.799794127 | -0.279901274 | 0.36109261 | 0.607693461 | 0.738446652 | 0.6897 | 0.793563601 |
| SL1344_RS03285_ | -2.40409941 | -2.468696287 | -2.3031124 | 0.164815665 | 0.474958958 | 0.314589992 | 0.355762831 | 0.544203159 | 0.684154843 | 0.6546 | 0.762324202 |
| SL1344_RS03290_ | -1.342892264 | -1.544796646 | -1.103310312 | 0.444374028 | 0.335789253 | 1.306434289 | 0.074616979 | 0.127471737 | 0.275708286 | 0.199 | 0.356276051 |
| SL1344_RS03295_ | 0.877409738 | 0.974569647 | 0.750317088 | -0.231793319 | 0.16472049 | -1.342769974 | 0.050666753 | 0.119352984 | 0.269106126 | 0.1699 | 0.323088948 |
| SL1344_RS03300_ | -0.770081504 | -0.736368383 | -0.808006612 | -0.074496853 | 0.284511734 | -0.246250858 | 0.390000004 | 0.547019452 | 0.689055169 | 0.6608 | 0.765212922 |
| SL1344_RS03305_ | -1.638173832 | -1.863609822 | -1.488165051 | 0.357591352 | 0.374880276 | 0.895364032 | 0.157894761 | 0.247094703 | 0.414415718 | 0.3448 | 0.498150847 |
| SL1344_RS03310_ | -4.017932128 | -4.005589707 | -4.02623694 | -0.010086301 | 0.834906947 | -0.012475467 | 0.493670886 | 0.604033539 | 0.734480157 | 0.7049 | 0.801765081 |
| SL1344_RS03315_ | -2.694629855 | -2.788683619 | -2.592730196 | 0.38607521 | 0.881128844 | 0.431574704 | 0.326449041 | 0.470516074 | 0.637264711 | 0.6519 | 0.758196966 |
| SL1344_RS03320_ | -3.167313205 | -3.0809723 | -3.253026228 | -0.190801761 | 0.757106425 | -0.225208684 | 0.389740178 | 0.603474065 | 0.738873566 | 0.7232 | 0.817692613 |
| SL1344_RS03325_ | -1.261447088 | -1.195900801 | -1.335829677 | -0.108695702 | 0.380749271 | -0.272441941 | 0.38774151 | 0.59428994 | 0.729600388 | 0.7103 | 0.805727034 |
| SL1344_RS03330_ | -1.23277491 | -1.10265941 | -1.292452509 | -0.148732607 | 0.377714146 | -0.322863495 | 0.360000006 | 0.626958727 | 0.756177979 | 0.7045 | 0.811189995 |
| SL1344_RS03335_ | -0.847662946 | -1.009057492 | -0.758917659 | 0.230557118 | 0.352628263 | 0.60096255 | 0.257333345 | 0.409486579 | 0.578502246 | 0.5245 | 0.665618539 |
| SL1344_RS03340_ | -1.1157957 | -1.120691979 | -1.112782552 | 0.014981294 | 0.332310875 | 0.03929534 | 0.481012659 | 0.630073089 | 0.756480165 | 0.7393 | 0.827827196 |
| SL1344_RS03350_ | -0.762796303 | -0.754076971 | -0.785119663 | -0.032203472 | 0.306835907 | -0.093177032 | 0.460359762 | 0.634001219 | 0.762058237 | 0.7526 | 0.837937693 |
| SL1344_RS03355_ | -0.249276855 | -0.338591869 | -0.188632348 | 0.133207718 | 0.26383632 | 0.485951827 | 0.303333342 | 0.469378714 | 0.628849576 | 0.5798 | 0.708087495 |
| SL1344_RS03360_ | 4.289818105 | 4.329834475 | 4.07543641 | -0.250603331 | 0.286165834 | -0.841013374 | 0.210526332 | 0.275453708 | 0.481923976 | 0.3983 | 0.573652001 |
| SL1344_RS03365_ | 3.379523309 | 3.237546945 | 3.51350687 | 0.294094265 | 0.105302237 | 2.83891638 | 4.68E-05 | 0.020002927 | 0.098168824 | 0.1 | 0.243021319 |
| SL1344_RS03370_ | 3.453627398 | 3.406516739 | 3.502422953 | 0.11415253 | 0.090372132 | 1.213896188 | 0.070666727 | 0.131775853 | 0.294133964 | 0.1912 | 0.348589888 |
| SL1344_RS03375_ | 2.424976864 | 2.594337708 | 2.402158859 | -0.173241732 | 0.139385655 | -0.749122334 | 0.307128589 | 0.419660827 | 0.605981283 | 0.6327 | 0.773736237 |
| SL1344_RS03380_ | 3.012755191 | 3.054412502 | 2.917384421 | -0.121142772 | 0.134475809 | -0.856472529 | 0.200000017 | 0.274401816 | 0.465747672 | 0.3679 | 0.546250686 |
| SL1344_RS03385_ | 4.429145019 | 4.394382977 | 4.449515845 | 0.052884097 | 0.05423185 | 0.921829476 | 0.16322454 | 0.250934754 | 0.425086612 | 0.3525 | 0.512461806 |
| SL1344_RS03395_ | -1.800133435 | -1.966627611 | -1.660450264 | 0.315562853 | 0.477447621 | 0.623843191 | 0.238507675 | 0.357846232 | 0.531505235 | 0.4748 | 0.616578588 |
| SL1344_RS03400_ | 5.193640011 | 5.162380226 | 5.321008039 | 0.156334852 | 0.097872505 | 1.467175246 | 0.024000188 | 0.131412702 | 0.307297725 | 0.1245 | 0.274999403 |
| SL1344_RS03405_ | 4.611403058 | 4.524304982 | 4.642293165 | 0.143918303 | 0.146661653 | 1.043925256 | 0.15866669 | 0.173460556 | 0.362206739 | 0.2794 | 0.4657132 |
| SL1344_RS03410_ | 4.800697924 | 4.776644876 | 4.830514533 | 0.048734931 | 0.132945151 | 0.31478073 | 0.40373085 | 0.604525626 | 0.756482196 | 0.8935 | 0.934829563 |
| SL1344_RS03415_ | 5.598642331 | 5.49998897 | 5.809314991 | 0.26710217 | 0.167796261 | 1.181142459 | 0.106595641 | 0.158907679 | 0.346659165 | 0.2016 | 0.385110679 |
| SL1344_RS03420_ | 6.502822219 | 6.401131279 | 6.679382736 | 0.302645616 | 0.147863787 | 2.024001891 | 4.68E-05 | 0.029693541 | 0.128486127 | 0.1 | 0.243021319 |
| SL1344_RS03425_ | 6.52042451 | 6.372575076 | 6.7068054 | 0.372839309 | 0.108971212 | 3.629027625 | 4.68E-05 | 0.010306779 | 0.07147594 | 0.1 | 0.243021319 |
| SL1344_RS03430_ | 5.971998186 | 5.692839855 | 6.277430278 | 0.561004736 | 0.161578892 | 3.379151285 | 4.68E-05 | 0.009181666 | 0.067812038 | 0.1 | 0.243021319 |
| SL1344_RS03440_ | 2.666816643 | 2.595723605 | 2.751110761 | 0.142580338 | 0.142978952 | 0.942878867 | 0.149333359 | 0.208731409 | 0.394224041 | 0.2882 | 0.46553064 |
| SL1344_RS03445_ | 1.954487169 | 1.980474663 | 1.92799511 | -0.056132462 | 0.120028357 | -0.454977779 | 0.315789482 | 0.499452114 | 0.650423379 | 0.6031 | 0.722436041 |
| SL1344_RS03450_ | 2.226306079 | 2.219428811 | 2.238888484 | 0.013324651 | 0.113966822 | 0.09017449 | 0.459333335 | 0.66319716 | 0.784453239 | 0.7619 | 0.847492174 |
| SL1344_RS03455_ | 0.503600825 | 0.531491833 | 0.481908578 | -0.048818807 | 0.163763423 | -0.271025356 | 0.37974684 | 0.515983848 | 0.662105551 | 0.6295 | 0.742310446 |
| SL1344_RS03460_ | 2.202612916 | 2.024775716 | 2.371939685 | 0.334797976 | 0.117901168 | 2.729710625 | 0.002000966 | 0.020053181 | 0.09551869 | 0.1009 | 0.244282848 |
| SL1344_RS03465_ | 4.376370803 | 4.351786124 | 4.398378887 | 0.05105222 | 0.054255536 | 0.886627433 | 0.165333356 | 0.240621861 | 0.414440874 | 0.3326 | 0.490882863 |
| SL1344_RS03475_ | -2.291945064 | -2.38344623 | -2.220490105 | 0.1782627 | 0.414654082 | 0.400658405 | 0.326449041 | 0.472976418 | 0.623125416 | 0.5857 | 0.703017935 |
| SL1344_RS03480_ | -1.772893885 | -1.888536692 | -1.665669853 | 0.241625956 | 0.3241457 | 0.679173392 | 0.226666681 | 0.328547547 | 0.484253113 | 0.4223 | 0.559780601 |
| SL1344_RS03490_ | -1.76426017 | -1.605283279 | -1.942751204 | -0.34892628 | 0.407070394 | -0.82551066 | 0.174000021 | 0.274347253 | 0.442912917 | 0.3716 | 0.518784138 |
| SL1344_RS03495_ | 0.500821916 | 0.459068693 | 0.529938667 | 0.07324504 | 0.164581798 | 0.395648746 | 0.327781486 | 0.487921701 | 0.635026241 | 0.5983 | 0.713076634 |
| SL1344_RS03505_ | 4.312368075 | 4.213997095 | 4.391490185 | 0.202263484 | 0.113542776 | 1.951748797 | 4.68E-05 | 0.050411557 | 0.170804729 | 0.1 | 0.243021319 |
| SL1344_RS03510_ | 2.146730694 | 1.965188013 | 2.357505863 | 0.390795287 | 0.119910931 | 3.24503317 | 4.68E-05 | 0.013044046 | 0.074916591 | 0.1 | 0.243021319 |
| SL1344_RS03515_ | 1.83671024 | 1.697064557 | 1.951069718 | 0.187642363 | 0.215119059 | 0.903306316 | 0.15866669 | 0.249145329 | 0.440369185 | 0.3134 | 0.49138921 |
| SL1344_RS03520_ | 2.661967225 | 2.588168125 | 2.739742413 | 0.166874893 | 0.111566329 | 1.474080594 | 0.045303228 | 0.095034261 | 0.239679795 | 0.1481 | 0.302327265 |
| SL1344_RS03525_ | 4.680147153 | 4.646599038 | 4.711507795 | 0.067651346 | 0.063941346 | 1.006921409 | 0.136000029 | 0.194097284 | 0.374806639 | 0.2688 | 0.438548022 |
| SL1344_RS03530_ | -2.221742584 | -2.316896131 | -2.141884629 | 0.204243296 | 0.477801882 | 0.39315245 | 0.33333334 | 0.502561117 | 0.651815021 | 0.6104 | 0.724776601 |
| SL1344_RS03535_ | -2.591651233 | -2.395281853 | -2.762521027 | -0.309751843 | 0.718469911 | -0.40945621 | 0.323333341 | 0.545414214 | 0.693706804 | 0.6507 | 0.764384245 |
| SL1344_RS03540_ | -1.914370932 | -1.804410715 | -2.016383804 | -0.20844651 | 0.364239458 | -0.511737878 | 0.273151243 | 0.411098323 | 0.563358267 | 0.5191 | 0.643957246 |
| SL1344_RS03545_ | -8.468044837 | -8.532396384 | -8.398602851 | 0.096781951 | 3.861177592 | 0.024105731 | 0.485333334 | 0.543703402 | 0.688494884 | 0.6693 | 0.771627084 |
| SL1344_RS03550_ | 0.380504374 | 0.436992669 | 0.253761676 | -0.181051966 | 0.20110167 | -0.824912064 | 0.200666684 | 0.305548696 | 0.477987819 | 0.417 | 0.566569035 |
| SL1344_RS03555_ | 0.746786124 | 0.772141287 | 0.696903609 | -0.071586311 | 0.203373133 | -0.346754567 | 0.359760166 | 0.559287168 | 0.703793646 | 0.6859 | 0.789483306 |

| | | | | | | | | | | | |
|---|---|---|---|---|---|---|---|---|---|---|---|
| SL1344_RS03560_ | -1.834273189 | -1.896400877 | -1.78182124 | 0.134082848 | 0.412303099 | 0.297325993 | 0.378414395 | 0.550892713 | 0.693991931 | 0.6662 | 0.770515916 |
| SL1344_RS03565_ | -2.661970829 | -2.684078465 | -2.619192196 | 0.0425273 | 0.571985311 | 0.060411861 | 0.473333334 | 0.632645008 | 0.761496843 | 0.7501 | 0.836565971 |
| SL1344_RS03570_ | -3.697222256 | -3.607134002 | -3.798158174 | -0.183286078 | 0.677901002 | -0.254871329 | 0.388666671 | 0.521253794 | 0.664265534 | 0.6293 | 0.740128903 |
| SL1344_RS03575_ | -7.525582139 | -8.516499128 | -6.277669871 | 2.080676115 | 3.82838656 | 0.482232414 | 0.260666678 | 0.447599267 | 0.602792479 | 0.5184 | 0.655768203 |
| SL1344_RS03580_ | 0.859714294 | 0.896125137 | 0.701599886 | -0.17100521 | 0.222629241 | -0.603962767 | 0.315333341 | 0.506914754 | 0.669162627 | 0.647 | 0.778392143 |
| SL1344_RS03585_ | -0.736229257 | -0.812951734 | -0.655769052 | 0.143290721 | 0.31715007 | 0.436469705 | 0.315789482 | 0.510574577 | 0.660846215 | 0.6147 | 0.733268178 |
| SL1344_RS03590_ | 0.665998383 | 0.338621662 | 0.837629913 | 0.433889001 | 0.376399418 | 1.190314637 | 0.113333369 | 0.142328256 | 0.316811009 | 0.2217 | 0.400966166 |
| SL1344_RS03595_ | 0.447586994 | 0.583215607 | 0.366594579 | -0.211636055 | 0.244301486 | -0.812592944 | 0.191872104 | 0.276831255 | 0.456946004 | 0.3651 | 0.532173664 |
| SL1344_RS03600_ | 0.118532014 | 0.185469949 | 0.02827826 | -0.157367913 | 0.186472804 | -0.804666448 | 0.183333353 | 0.286182575 | 0.450753026 | 0.382 | 0.527782321 |
| SL1344_RS03605_ | 0.411603845 | 0.478963297 | 0.335976561 | -0.133833382 | 0.261723557 | -0.496238031 | 0.277333344 | 0.450371094 | 0.617324107 | 0.5655 | 0.697970849 |
| SL1344_RS03610_ | 0.607598987 | 0.628817119 | 0.580641382 | -0.04283585 | 0.148628969 | -0.268560535 | 0.379333338 | 0.513701043 | 0.656972317 | 0.6302 | 0.738772475 |
| SL1344_RS03615_ | -4.186009645 | -4.615815826 | -3.895620217 | 0.714312391 | 0.889941792 | 0.726592637 | 0.201199218 | 0.316845363 | 0.483386508 | 0.4226 | 0.567223786 |
| SL1344_RS03620_ | -0.123457946 | -0.341505693 | 0.034055039 | 0.374154876 | 0.206205696 | 1.700438164 | 0.036666788 | 0.075585974 | 0.200494275 | 0.1363 | 0.287063349 |
| SL1344_RS03625_ | -4.335932681 | -3.994029055 | -5.011629038 | -0.999707179 | 1.079150269 | -0.848889775 | 0.167888097 | 0.250241135 | 0.424976897 | 0.3479 | 0.505560852 |
| SL1344_RS03630_ | -2.774425854 | -2.618837199 | -2.943047468 | -0.338734571 | 0.501293938 | -0.597333738 | 0.233333347 | 0.362630658 | 0.522894943 | 0.4704 | 0.604360342 |
| SL1344_RS03640_ | -4.254858778 | -4.099000694 | -4.420513686 | -0.337971745 | 0.893366831 | -0.34443585 | 0.334000007 | 0.498199281 | 0.647635516 | 0.6164 | 0.729539029 |
| SL1344_RS03645_ | -3.82866956 | -3.97077721 | -3.736751252 | 0.207904409 | 0.790069367 | 0.235633598 | 0.394666671 | 0.563527844 | 0.702449314 | 0.6786 | 0.780690147 |
| SL1344_RS03650_ | -4.258196611 | -3.755951677 | -4.630942228 | -1.05238067 | 1.20153973 | -0.829954804 | 0.183333353 | 0.26942167 | 0.449056113 | 0.369 | 0.524067269 |
| SL1344_RS03655_ | -3.762828474 | -3.666206627 | -3.886869493 | -0.120346022 | 1.033431904 | -0.124233738 | 0.453031314 | 0.666404632 | 0.788179661 | 0.7755 | 0.856243452 |
| SL1344_RS03660_ | -2.594316036 | -2.784833976 | -2.456555739 | 0.35486181 | 0.577084342 | 0.55838211 | 0.249167234 | 0.375426429 | 0.54734041 | 0.4834 | 0.618700583 |
| SL1344_RS03665_ | -4.202833178 | -4.275097149 | -4.138354429 | 0.182967307 | 0.977830376 | 0.15972205 | 0.420386412 | 0.57087146 | 0.711642004 | 0.685 | 0.786110955 |
| SL1344_RS03670_ | -2.636823567 | -2.526332781 | -2.715549355 | -0.205356263 | 0.597530871 | -0.324822366 | 0.356000006 | 0.556578239 | 0.69739831 | 0.6744 | 0.777117394 |
| SL1344_RS03675_ | -1.296040846 | -1.486199664 | -1.024482661 | 0.456801662 | 0.309403566 | 1.366958656 | 0.069333395 | 0.125360992 | 0.27064832 | 0.1981 | 0.354441288 |
| SL1344_RS03680_ | 1.255325086 | 1.090231386 | 1.325288565 | 0.225157347 | 0.152181409 | 1.322492434 | 0.086608976 | 0.135949817 | 0.294648481 | 0.2196 | 0.380277574 |
| SL1344_RS03685_ | -0.088807337 | 0.127214608 | -0.363626257 | -0.488000069 | 0.244713563 | -1.987590469 | 0.014666978 | 0.04977532 | 0.158860521 | 0.1168 | 0.264925214 |
| SL1344_RS03690_ | 3.728976152 | 4.190727357 | 3.147925467 | -1.045723005 | 0.103720696 | -10.01729337 | 4.68E-05 | 0.000858938 | 0.019069151 | 0.1 | 0.243021319 |
| SL1344_RS03695_ | -4.010788089 | -3.899897726 | -4.341566158 | -0.410876135 | 1.253987762 | -0.285532854 | 0.389073955 | 0.586361543 | 0.732662517 | 0.7451 | 0.839028075 |
| SL1344_RS03705_ | 0.838371163 | 1.047200117 | 0.601981012 | -0.470825172 | 0.203912504 | -2.250925607 | 0.002666616 | 0.032026908 | 0.125737455 | 0.1023 | 0.246037291 |
| SL1344_RS03710_ | 2.23552372 | 2.604742573 | 1.748809036 | -0.858901343 | 0.131307306 | -6.563301485 | 4.68E-05 | 0.002846519 | 0.033176105 | 0.1 | 0.243021319 |
| SL1344_RS03715_ | 2.751954625 | 3.047079696 | 2.411506801 | -0.636924265 | 0.111360713 | -5.722948469 | 4.68E-05 | 0.002679454 | 0.032993499 | 0.1 | 0.243021319 |
| SL1344_RS03720_ | 3.994723168 | 4.328405965 | 3.61507307 | -0.719060849 | 0.067385348 | -10.70863188 | 4.68E-05 | 0.001532123 | 0.024556555 | 0.1 | 0.243021319 |
| SL1344_RS03725_ | 3.835626936 | 4.205879679 | 3.454470638 | -0.752098689 | 0.087162537 | -8.5576977 | 4.68E-05 | 0.001191836 | 0.022089566 | 0.1 | 0.243021319 |
| SL1344_RS03730_ | 3.13147998 | 3.455204187 | 2.688924868 | -0.773456281 | 0.140232258 | -5.591703482 | 4.68E-05 | 0.008850727 | 0.063765437 | 0.1 | 0.243021319 |
| SL1344_RS03735_ | 3.198225572 | 3.64525896 | 2.743932665 | -0.911436519 | 0.124285077 | -7.605013734 | 4.68E-05 | 0.005729979 | 0.050789893 | 0.1 | 0.243021319 |
| SL1344_RS03740_ | -0.06715373 | 0.230644831 | -0.502970284 | -0.713572425 | 0.293447927 | -2.35214302 | 0.004664544 | 0.034010119 | 0.131040804 | 0.1037 | 0.248213628 |
| SL1344_RS03745_ | 3.292520164 | 2.913322246 | 3.80126867 | 0.866988597 | 0.125810142 | 6.631217569 | 4.68E-05 | 0.002508658 | 0.033004201 | 0.1 | 0.243021319 |
| SL1344_RS03750_ | 2.198349807 | 1.944755773 | 2.545600741 | 0.606873257 | 0.191921553 | 3.269806394 | 4.68E-05 | 0.030955987 | 0.129514999 | 0.1 | 0.243021319 |
| SL1344_RS03755_ | 1.67422377 | 1.302902986 | 2.020292095 | 0.734293309 | 0.142035955 | 5.118957457 | 4.68E-05 | 0.00392936 | 0.040162383 | 0.1 | 0.243021319 |
| SL1344_RS03760_ | 1.889894588 | 1.586539389 | 2.334541442 | 0.747531352 | 0.15873918 | 4.695974701 | 4.68E-05 | 0.011174968 | 0.071748223 | 0.1 | 0.243021319 |
| SL1344_RS03765_ | 2.015285582 | 1.756152231 | 2.303842351 | 0.560144241 | 0.165422591 | 3.406165128 | 4.68E-05 | 0.031861265 | 0.131528824 | 0.1 | 0.243021319 |
| SL1344_RS03770_ | 1.764580627 | 1.718209601 | 1.800072113 | 0.097032359 | 0.156666281 | 0.592376197 | 0.235842785 | 0.355328084 | 0.538115796 | 0.4837 | 0.623472948 |
| SL1344_RS03775_ | 2.64622145 | 2.523415671 | 2.781140334 | 0.258161466 | 0.097102836 | 2.642839761 | 0.0020023 | 0.022644128 | 0.102144193 | 0.1007 | 0.243989838 |
| SL1344_RS03780_ | 1.942167332 | 1.845097771 | 2.056809942 | 0.218868784 | 0.119600413 | 1.793691664 | 0.024000188 | 0.065683762 | 0.18826577 | 0.1224 | 0.271844698 |
| SL1344_RS03790_ | 4.68940722 | 4.738458134 | 4.596353864 | -0.143424868 | 0.070053684 | -1.932907039 | 0.009993796 | 0.063537141 | 0.193105207 | 0.1095 | 0.255865936 |
| SL1344_RS03795_ | 5.709631931 | 5.794660726 | 5.635800453 | -0.185631775 | 0.083595777 | -2.346420001 | 4.68E-05 | 0.028026666 | 0.121844973 | 0.1 | 0.243021319 |
| SL1344_RS03800_ | 5.003112295 | 5.09996399 | 4.808214897 | -0.28366207 | 0.140596241 | -1.834744609 | 0.004001152 | 0.08056875 | 0.229452841 | 0.1049 | 0.249906759 |
| SL1344_RS03805_ | 5.214377114 | 5.063938508 | 5.468735578 | 0.410347482 | 0.176380594 | 2.502999192 | 4.68E-05 | 0.021890821 | 0.107719883 | 0.1 | 0.243021319 |
| SL1344_RS03810_ | 5.536940024 | 5.316571292 | 5.719169574 | 0.383675356 | 0.106736356 | 3.45484841 | 4.68E-05 | 0.012253222 | 0.077675857 | 0.1 | 0.243021319 |
| SL1344_RS03815_ | 4.678666674 | 4.540537703 | 4.810165687 | 0.301320387 | 0.09786867 | 3.148301131 | 4.68E-05 | 0.024466486 | 0.113214211 | 0.1 | 0.243021319 |
| SL1344_RS03820_ | 4.735233572 | 4.678315072 | 4.838138314 | 0.1515766 | 0.116387795 | 1.205554647 | 0.104596974 | 0.152634984 | 0.335340065 | 0.1975 | 0.374802241 |
| SL1344_RS03825_ | 3.755701749 | 3.717290784 | 3.852913369 | 0.129976346 | 0.091377185 | 1.321837217 | 0.081279199 | 0.132977464 | 0.299295927 | 0.2 | 0.36468211 |
| SL1344_RS03830_ | -2.357494691 | -2.381034832 | -2.324094028 | 0.057820983 | 0.508227208 | 0.10910718 | 0.455333335 | 0.623614406 | 0.754028246 | 0.731 | 0.82417803 |
| SL1344_RS03835_ | -1.341613812 | -1.257785209 | -1.445014922 | -0.189025903 | 0.290814719 | -0.599827594 | 0.245333346 | 0.363005532 | 0.520858398 | 0.4641 | 0.599966469 |

| | | | | | | | | | | | |
|---|---|---|---|---|---|---|---|---|---|---|---|
| SL1344_RS03840_ | -1.256539737 | -1.012842556 | -1.430601796 | -0.389459686 | 0.331410796 | -1.139368072 | 0.113333369 | 0.184721618 | 0.345309804 | 0.2664 | 0.426246211 |
| SL1344_RS03845_ | -2.48247065 | -2.320996851 | -2.644182779 | -0.315011477 | 0.44684001 | -0.664113601 | 0.222000015 | 0.343842398 | 0.502141997 | 0.4375 | 0.577162137 |
| SL1344_RS03850_ | 2.793040031 | 2.803864843 | 2.783090217 | 0.018089997 | 0.134488509 | 0.1340546 | 0.452666668 | 0.696754032 | 0.812679235 | 0.872 | 0.919880315 |
| SL1344_RS03855_ | -1.003688856 | -0.997335913 | -1.010125469 | -0.012664466 | 0.250378534 | -0.044791374 | 0.482345104 | 0.538281346 | 0.678667386 | 0.6474 | 0.752968683 |
| SL1344_RS03860_ | -1.744044757 | -1.764855629 | -1.716441105 | 0.038084132 | 0.430625595 | 0.080470733 | 0.460359762 | 0.650303707 | 0.773628613 | 0.7565 | 0.841212974 |
| SL1344_RS03865_ | -0.745608156 | -0.723612243 | -0.769462272 | -0.066624533 | 0.288629941 | -0.217446369 | 0.40106596 | 0.588280048 | 0.723191516 | 0.7051 | 0.800903503 |
| SL1344_RS03870_ | -2.264615293 | -2.331159599 | -2.169107625 | 0.174466793 | 0.500396723 | 0.33249582 | 0.361333339 | 0.521609341 | 0.667751783 | 0.636 | 0.744917277 |
| SL1344_RS03875_ | -4.493669836 | -4.941622405 | -4.274305282 | 0.626591107 | 1.088758327 | 0.518192798 | 0.281333343 | 0.432343029 | 0.595582885 | 0.567 | 0.693518359 |
| SL1344_RS03880_ | -5.369775887 | -5.461950834 | -5.271383348 | 0.150932355 | 1.134794923 | 0.134561627 | 0.439706864 | 0.525059789 | 0.670368615 | 0.6307 | 0.742761264 |
| SL1344_RS03885_ | -3.08066188 | -2.969295758 | -3.199423657 | -0.247840082 | 0.670269738 | -0.353129732 | 0.339107268 | 0.522294623 | 0.670422119 | 0.64 | 0.750061124 |
| SL1344_RS03890_ | -2.853755289 | -2.795412199 | -2.987221727 | -0.093750961 | 0.79836958 | -0.10399578 | 0.459693539 | 0.735725706 | 0.839237398 | 0.8087 | 0.886398405 |
| SL1344_RS03895_ | -0.69081084 | -0.654073049 | -0.743327573 | -0.095109227 | 0.260720528 | -0.345856755 | 0.355096608 | 0.512374354 | 0.659191896 | 0.6212 | 0.737437287 |
| SL1344_RS03900_ | -1.568097947 | -1.421256888 | -1.703131609 | -0.280240343 | 0.368419801 | -0.725555983 | 0.20333335 | 0.316906625 | 0.482135374 | 0.4139 | 0.560199731 |
| SL1344_RS03905_ | -2.719418307 | -2.640910083 | -2.803981333 | -0.157755212 | 0.473596776 | -0.310832346 | 0.353097941 | 0.486008085 | 0.634956745 | 0.5888 | 0.708590319 |
| SL1344_RS03915_ | 2.057365759 | 2.138958711 | 1.999155409 | -0.15829443 | 0.163586754 | -0.957248659 | 0.131333363 | 0.201425897 | 0.383162346 | 0.2782 | 0.448694929 |
| SL1344_RS03920_ | 1.52883417 | 1.536564878 | 1.521465915 | 0.006569008 | 0.180874362 | 0.034945323 | 0.487008661 | 0.711792883 | 0.822221003 | 0.8292 | 0.892476494 |
| SL1344_RS03925_ | 0.50318582 | 0.440290268 | 0.555923657 | 0.122955513 | 0.216187735 | 0.544502017 | 0.282478358 | 0.438032803 | 0.602406689 | 0.5619 | 0.691093955 |
| SL1344_RS03930_ | 0.004902471 | -0.031119543 | 0.064221197 | 0.095848829 | 0.228287687 | 0.391445074 | 0.32933334 | 0.524914103 | 0.670413391 | 0.6247 | 0.740497688 |
| SL1344_RS03935_ | -2.636189586 | -2.56137924 | -2.676769415 | -0.088016713 | 0.569092465 | -0.152266344 | 0.437041974 | 0.632756517 | 0.759027115 | 0.7314 | 0.824280119 |
| SL1344_RS03940_ | 2.036547703 | 1.996596275 | 2.064311818 | 0.066504113 | 0.112013562 | 0.552278168 | 0.261333345 | 0.395513248 | 0.560513209 | 0.5111 | 0.645447696 |
| SL1344_RS03945_ | 1.315232851 | 1.268183734 | 1.371680418 | 0.106965952 | 0.130143097 | 0.767706468 | 0.201199218 | 0.290897632 | 0.458695794 | 0.392 | 0.538177295 |
| SL1344_RS03950_ | 4.767639447 | 4.924247296 | 4.652574505 | -0.354489893 | 0.267803498 | -1.259393125 | 0.08400005 | 0.110893991 | 0.279274059 | 0.1779 | 0.352469678 |
| SL1344_RS03970_ | 0.770447545 | 0.662949945 | 0.865447033 | 0.201654016 | 0.167936453 | 1.160931255 | 0.091938753 | 0.158550958 | 0.316278556 | 0.2264 | 0.385014194 |
| SL1344_RS03975_ | 0.918463619 | 1.083008616 | 0.81532244 | -0.259483166 | 0.17877769 | -1.392224043 | 0.079946755 | 0.120105931 | 0.272379696 | 0.1943 | 0.355089374 |
| SL1344_RS03980_ | 4.124214586 | 4.063142482 | 4.201931953 | 0.140579161 | 0.054781192 | 2.48130295 | 0.004664544 | 0.028061621 | 0.113627434 | 0.1032 | 0.247374927 |
| SL1344_RS03985_ | 0.589551491 | 0.56792257 | 0.602751076 | 0.030064408 | 0.1727882 | 0.15046124 | 0.433710862 | 0.596670854 | 0.730805765 | 0.7001 | 0.80041333 |
| SL1344_RS03990_ | -0.030884618 | 9.16E-05 | -0.068112019 | -0.060763668 | 0.211123972 | -0.265717572 | 0.38507662 | 0.538879674 | 0.681937401 | 0.6505 | 0.758337157 |
| SL1344_RS03995_ | -0.319358355 | -0.388351231 | -0.231545706 | 0.154926585 | 0.238740759 | 0.616456313 | 0.252498347 | 0.387224458 | 0.548115828 | 0.4983 | 0.633690542 |
| SL1344_RS04000_ | -0.873219109 | -0.755264152 | -1.005494864 | -0.251114592 | 0.381029222 | -0.621132769 | 0.23184545 | 0.367123888 | 0.54291087 | 0.4719 | 0.620182692 |
| SL1344_RS04005_ | -0.650999435 | -0.504619157 | -0.756300402 | -0.248036498 | 0.274134054 | -0.82693881 | 0.191205881 | 0.286332961 | 0.453270708 | 0.3787 | 0.532156008 |
| SL1344_RS04010_ | 0.297578934 | 0.150430928 | 0.446238114 | 0.294936007 | 0.171398627 | 1.655709499 | 0.041333441 | 0.079123702 | 0.200481854 | 0.1523 | 0.303924254 |
| SL1344_RS04015_ | -1.773643096 | -1.804629777 | -1.740763706 | 0.069620614 | 0.40528691 | 0.177230042 | 0.41972019 | 0.586277267 | 0.721343835 | 0.6925 | 0.791257682 |
| SL1344_RS04020_ | -2.526938307 | -2.673835536 | -2.418552128 | 0.210873411 | 0.714510984 | 0.286919679 | 0.376666672 | 0.599544719 | 0.736985627 | 0.7203 | 0.815535818 |
| SL1344_RS04025_ | -2.848287438 | -2.707669193 | -3.020169746 | -0.322189681 | 0.509826397 | -0.598880508 | 0.258494349 | 0.38058511 | 0.538697165 | 0.4878 | 0.619373897 |
| SL1344_RS04030_ | -3.517222071 | -3.537055335 | -3.499840125 | 0.0389567 | 0.714658004 | 0.053032518 | 0.473017989 | 0.590859758 | 0.726679213 | 0.7029 | 0.799335322 |
| SL1344_RS04035_ | -3.536224453 | -3.359992955 | -3.935786629 | -0.552989225 | 0.89340124 | -0.548886823 | 0.28047969 | 0.41195685 | 0.583136385 | 0.5424 | 0.677493908 |
| SL1344_RS04040_ | 2.926342023 | 2.960916331 | 2.826559795 | -0.133812702 | 0.17975406 | -0.716437759 | 0.205196552 | 0.305627799 | 0.506764163 | 0.4206 | 0.587079513 |
| SL1344_RS04045_ | -2.677228111 | -2.548953922 | -2.838207087 | -0.315800819 | 0.540031159 | -0.515642174 | 0.262666678 | 0.397824027 | 0.560025258 | 0.5176 | 0.644735054 |
| SL1344_RS04050_ | 1.454741988 | 1.552703124 | 1.390764301 | -0.167302183 | 0.152287095 | -1.030483857 | 0.128000031 | 0.199085033 | 0.370331326 | 0.2768 | 0.438243128 |
| SL1344_RS04055_ | 0.787986889 | 0.656805836 | 0.891104166 | 0.236575251 | 0.169076566 | 1.318226573 | 0.063957429 | 0.121978717 | 0.271902346 | 0.1835 | 0.341020479 |
| SL1344_RS04065_ | 0.263099096 | 0.192912294 | 0.314558409 | 0.118240535 | 0.191751043 | 0.560752838 | 0.27600001 | 0.405282619 | 0.565217481 | 0.5177 | 0.651906819 |
| SL1344_RS04070_ | 0.097609476 | 0.068527253 | 0.135826046 | 0.08316692 | 0.214814117 | 0.352857722 | 0.351099273 | 0.518402106 | 0.663528089 | 0.6251 | 0.736296287 |
| SL1344_RS04085_ | 3.308262532 | 3.319753312 | 3.278605411 | -0.04117166 | 0.087944047 | -0.4347077 | 0.321333341 | 0.500601094 | 0.658363967 | 0.6387 | 0.751718581 |
| SL1344_RS04090_ | -3.626644548 | -3.361472496 | -3.842401996 | -0.507414769 | 1.105533329 | -0.437922376 | 0.295333342 | 0.493540756 | 0.653578059 | 0.586 | 0.716022866 |
| SL1344_RS04095_ | -3.799510469 | -3.96497516 | -3.694870845 | 0.3546825 | 1.096348745 | 0.288364538 | 0.362666672 | 0.523321326 | 0.681231318 | 0.6785 | 0.781410996 |
| SL1344_RS04100_ | -3.361198852 | -3.399796163 | -3.322939179 | 0.092626555 | 0.572570912 | 0.145164658 | 0.438000002 | 0.547819857 | 0.688016107 | 0.6535 | 0.761036694 |
| SL1344_RS04105_ | -1.63335198 | -1.573390928 | -1.68365528 | -0.116927214 | 0.417133475 | -0.255057292 | 0.37974684 | 0.56743196 | 0.707326287 | 0.6788 | 0.781721961 |
| SL1344_RS04110_ | -2.816855596 | -2.627853289 | -3.159606212 | -0.507935103 | 0.589883906 | -0.780198227 | 0.20333335 | 0.302004694 | 0.473142089 | 0.4065 | 0.558518523 |
| SL1344_RS04115_ | -3.531521043 | -3.381812271 | -3.668508057 | -0.289428367 | 0.690435339 | -0.393506484 | 0.33533334 | 0.510421616 | 0.658734019 | 0.6293 | 0.73975108 |
| SL1344_RS04120_ | 1.536025733 | 1.473084715 | 1.631890364 | 0.154851367 | 0.123360612 | 1.1929379 | 0.095333377 | 0.157142832 | 0.311183389 | 0.2311 | 0.386595455 |
| SL1344_RS04125_ | -0.092238358 | -0.164601724 | 0.018729686 | 0.176199599 | 0.240054678 | 0.693383534 | 0.248666679 | 0.380457765 | 0.546930004 | 0.4856 | 0.62960167 |
| SL1344_RS04130_ | 0.038740162 | -0.053342346 | 0.154602356 | 0.212449628 | 0.198338266 | 1.027910703 | 0.134000029 | 0.2058861 | 0.368463015 | 0.2895 | 0.443498082 |
| SL1344_RS04135_ | 0.757961642 | 0.745720522 | 0.767829441 | 0.028544674 | 0.179441815 | 0.144290388 | 0.436375752 | 0.602373936 | 0.735575713 | 0.7199 | 0.812321325 |

| | | | | | | | | | | | |
|---|---|---|---|---|---|---|---|---|---|---|---|
| SL1344_RS04140_ | -0.048123079 | -0.075198684 | -0.012479399 | 0.067908179 | 0.19202701 | 0.331390342 | 0.3637575 | 0.494359342 | 0.64131107 | 0.6077 | 0.721022119 |
| SL1344_RS04145_ | -0.006987723 | -0.013948557 | -0.000714336 | 0.083174727 | 0.34139352 | 0.244722363 | 0.402000004 | 0.640815452 | 0.769128412 | 0.8018 | 0.871860599 |
| SL1344_RS04150_ | 2.206256333 | 1.982065943 | 2.380341416 | 0.451978587 | 0.202123478 | 2.405259026 | 4.68E-05 | 0.027053382 | 0.118287053 | 0.1 | 0.243021319 |
| SL1344_RS04160_ | 1.2515306 | 1.171433125 | 1.315595903 | 0.142007346 | 0.141450864 | 0.932817205 | 0.143237868 | 0.222216877 | 0.389881482 | 0.306 | 0.465209275 |
| SL1344_RS04165_ | 0.902778835 | 1.105200399 | 0.698605053 | -0.27821651 | 0.527290532 | -0.51109103 | 0.272485021 | 0.437651517 | 0.626297023 | 0.5422 | 0.692305363 |
| SL1344_RS04170_ | -0.09536138 | -0.016575068 | -0.203843035 | -0.032263574 | 0.633985648 | -0.053437295 | 0.475016656 | 0.829738203 | 0.903181607 | 0.9211 | 0.952735544 |
| SL1344_RS04175_ | -2.082058895 | -2.232462968 | -1.986571486 | 0.192575501 | 0.6046493 | 0.295440364 | 0.375333338 | 0.613712357 | 0.749288743 | 0.7057 | 0.810657484 |
| SL1344_RS04180_ | 1.00720322 | 0.745490597 | 1.202775624 | 0.452995978 | 0.137518384 | 3.210178963 | 0.00067292 | 0.016344623 | 0.082729306 | 0.1004 | 0.243586648 |
| SL1344_RS04185_ | 0.482125585 | 0.017551849 | 1.003992334 | 0.984950975 | 0.197110356 | 4.979382619 | 4.68E-05 | 0.004855848 | 0.044254493 | 0.1 | 0.243021319 |
| SL1344_RS04195_ | 2.609403915 | 2.432620119 | 2.908784803 | 0.452717727 | 0.175508 | 2.424001324 | 0.000673369 | 0.02749835 | 0.119378785 | 0.1002 | 0.243306123 |
| SL1344_RS04200_ | 4.334521814 | 4.154465946 | 4.541899663 | 0.355638056 | 0.12410193 | 2.92571138 | 4.68E-05 | 0.010240609 | 0.070109227 | 0.1 | 0.243021319 |
| SL1344_RS04205_ | 1.360808805 | 1.57104152 | 1.149393689 | -0.305489278 | 0.383442898 | -0.861452601 | 0.158560983 | 0.259635519 | 0.459399018 | 0.2971 | 0.481700628 |
| SL1344_RS04210_ | 0.242395284 | 0.13830122 | 0.401587545 | 0.254754394 | 0.207591565 | 1.157624913 | 0.110666703 | 0.171598449 | 0.334950077 | 0.2556 | 0.417888438 |
| SL1344_RS04220_ | 1.7260619 | 1.46086777 | 1.906546936 | 0.441568978 | 0.132475713 | 3.214004148 | 4.68E-05 | 0.018763869 | 0.090716422 | 0.1001 | 0.243156847 |
| SL1344_RS04225_ | 0.450150816 | 0.526412435 | 0.33668093 | -0.201239594 | 0.198185707 | -0.96756249 | 0.135333362 | 0.222825003 | 0.394161522 | 0.3 | 0.451646726 |
| SL1344_RS04230_ | 0.955528127 | 1.029957797 | 0.89751805 | -0.146986843 | 0.268413279 | -0.559121811 | 0.259160571 | 0.399128649 | 0.582126673 | 0.5033 | 0.654661247 |
| SL1344_RS04235_ | 3.498615172 | 3.391748428 | 3.581967221 | 0.203281137 | 0.096069859 | 2.084774914 | 0.002000966 | 0.045992336 | 0.157896541 | 0.1022 | 0.245976996 |
| SL1344_RS04240_ | 3.285244701 | 3.216065014 | 3.36608743 | 0.118658628 | 0.163342129 | 0.700432306 | 0.221852113 | 0.338089342 | 0.534133027 | 0.442 | 0.607035379 |
| SL1344_RS04245_ | -2.29920272 | -2.219247408 | -2.366486411 | -0.137845918 | 0.442161089 | -0.283310994 | 0.383333338 | 0.510198647 | 0.655963745 | 0.6244 | 0.736172081 |
| SL1344_RS04250_ | -4.574364509 | -4.710927545 | -4.418258779 | 0.26591968 | 1.081133971 | 0.228898893 | 0.394000004 | 0.568826215 | 0.707585706 | 0.684 | 0.785092107 |
| SL1344_RS04255_ | -2.123945107 | -1.873102486 | -2.654110867 | -0.773149074 | 0.485123029 | -1.481265386 | 0.048634334 | 0.107741815 | 0.255248991 | 0.1617 | 0.315407643 |
| SL1344_RS04260_ | 2.99842291 | 2.935010072 | 3.171580189 | 0.239741167 | 0.129156357 | 1.679005213 | 0.013324793 | 0.09588644 | 0.250136772 | 0.1149 | 0.262262163 |
| SL1344_RS04265_ | -2.730732632 | -2.550221329 | -2.838876215 | -0.250591945 | 0.54372073 | -0.424515963 | 0.324666674 | 0.506423602 | 0.654927494 | 0.6131 | 0.732549094 |
| SL1344_RS04270_ | -4.518020204 | -4.574650901 | -4.491047203 | 0.077654704 | 1.008558245 | 0.075136966 | 0.470353099 | 0.602281455 | 0.734203285 | 0.712 | 0.807213922 |
| SL1344_RS04275_ | 1.505734501 | 1.504491695 | 1.506657749 | 2.78E-06 | 0.128884212 | -0.000231121 | 0.5 | 0.607483363 | 0.737579727 | 0.7098 | 0.805451404 |
| SL1344_RS04280_ | 1.472873568 | 1.439433792 | 1.525622876 | 0.066679958 | 0.163153504 | 0.384691807 | 0.34133334 | 0.576392031 | 0.719081441 | 0.6732 | 0.78488722 |
| SL1344_RS04290_ | 1.951857297 | 1.950525901 | 1.955126557 | 0.002196792 | 0.11811786 | 0.014281469 | 0.493004664 | 0.659653146 | 0.780579105 | 0.7587 | 0.844886383 |
| SL1344_RS04295_ | 0.896534925 | 0.863684909 | 0.969620909 | 0.10640568 | 0.186196073 | 0.525999113 | 0.285333343 | 0.454210542 | 0.615350262 | 0.5858 | 0.709591194 |
| SL1344_RS04300_ | 0.834543872 | 0.801002923 | 0.921029764 | 0.118518487 | 0.208800944 | 0.518776508 | 0.306000008 | 0.476692038 | 0.640004239 | 0.6161 | 0.737458145 |
| SL1344_RS04305_ | 0.418889596 | 0.372104195 | 0.486801644 | 0.116848833 | 0.176919032 | 0.592097501 | 0.255163236 | 0.398308453 | 0.557640975 | 0.5048 | 0.640205016 |
| SL1344_RS04320_ | -4.130277476 | -4.015539961 | -4.255285503 | -0.114702915 | 1.318655942 | -0.080965783 | 0.461692207 | 0.704847783 | 0.817782059 | 0.8238 | 0.890147712 |
| SL1344_RS04325_ | -3.04316048 | -3.085177899 | -2.999098478 | -0.038079385 | 0.778036073 | -0.055115892 | 0.477333334 | 0.664506811 | 0.786734381 | 0.8044 | 0.87440805 |
| SL1344_RS04330_ | -2.328006246 | -2.291553383 | -2.378974577 | 0.04622886 | 0.694167818 | 0.075029421 | 0.474000001 | 0.713040215 | 0.821676751 | 0.846 | 0.90277908 |
| SL1344_RS04335_ | -4.702628329 | -4.672288999 | -4.721340381 | -0.198933522 | 1.289555666 | -0.1371098 | 0.438666669 | 0.631401383 | 0.758671937 | 0.7417 | 0.829668297 |
| SL1344_RS04340_ | -0.930882384 | -1.013975821 | -0.7830919 | 0.190441852 | 0.359281435 | 0.494489006 | 0.305129922 | 0.507869531 | 0.660345917 | 0.6058 | 0.730831543 |
| SL1344_RS04345_ | -1.036253869 | -1.275216051 | -0.780818579 | 0.497515951 | 0.321643141 | 1.457065722 | 0.051299219 | 0.094587913 | 0.230369888 | 0.1675 | 0.322697212 |
| SL1344_RS04350_ | -0.593123072 | -0.809569338 | -0.410567735 | 0.407291945 | 0.260901464 | 1.488328209 | 0.048634334 | 0.093019533 | 0.227447346 | 0.1558 | 0.308808153 |
| SL1344_RS04355_ | -0.337209138 | -0.499155869 | -0.249403881 | 0.256671779 | 0.296264298 | 0.814685978 | 0.192538326 | 0.281852166 | 0.463772153 | 0.3823 | 0.539472673 |
| SL1344_RS04360_ | -1.873400495 | -2.060061221 | -1.665161705 | 0.379026108 | 0.405012913 | 0.899375854 | 0.155896093 | 0.247367584 | 0.411902972 | 0.332 | 0.490256914 |
| SL1344_RS04365_ | 0.823142776 | 0.899828016 | 0.672834524 | -0.218393051 | 0.181310446 | -1.118552255 | 0.120000033 | 0.178951658 | 0.345671086 | 0.2618 | 0.425827396 |
| SL1344_RS04370_ | 0.982268951 | 1.081084435 | 0.875511832 | -0.200918198 | 0.153372043 | -1.265161055 | 0.077281867 | 0.134316159 | 0.286558677 | 0.2096 | 0.370089765 |
| SL1344_RS04375_ | 2.94761388 | 2.7302936 | 3.131693285 | 0.407992541 | 0.119000222 | 3.436808164 | 4.68E-05 | 0.01037029 | 0.067912167 | 0.1 | 0.243021319 |
| SL1344_RS04380_ | 1.037025602 | 0.519118471 | 1.460372886 | 0.941317217 | 0.198544487 | 4.753114415 | 4.68E-05 | 0.004487027 | 0.042714326 | 0.1 | 0.243021319 |
| SL1344_RS04385_ | 0.970320959 | 0.917307389 | 1.095465877 | 0.17676632 | 0.299913142 | 0.599953669 | 0.268487686 | 0.427898587 | 0.608525974 | 0.5228 | 0.681422716 |
| SL1344_RS04390_ | 0.92363654 | 0.777985258 | 1.059841677 | 0.280571026 | 0.125795762 | 2.170020398 | 0.011326186 | 0.041464054 | 0.1386707 | 0.1115 | 0.257663456 |
| SL1344_RS04395_ | 1.041498662 | 0.970925885 | 1.108267382 | 0.159189608 | 0.192048236 | 0.811464504 | 0.182666686 | 0.272890162 | 0.452157887 | 0.3763 | 0.529740023 |
| SL1344_RS04400_ | -1.76905609 | -1.687181886 | -1.854522061 | -0.164392049 | 0.358474469 | -0.426060064 | 0.321785484 | 0.48404413 | 0.631026977 | 0.6044 | 0.71858809 |
| SL1344_RS04405_ | 2.509360943 | 2.029266597 | 2.89991579 | 0.878481648 | 0.105592025 | 8.424080595 | 4.68E-05 | 0.003613315 | 0.038040287 | 0.1 | 0.243021319 |
| SL1344_RS04410_ | -2.910585791 | -3.1351505 | -2.565892684 | 0.516747377 | 0.688865755 | 0.679559242 | 0.23451034 | 0.362028416 | 0.53083945 | 0.4595 | 0.608879565 |
| SL1344_RS04415_ | 3.435081873 | 3.492002687 | 3.372082513 | -0.124070166 | 0.072930019 | -1.655424772 | 0.028000161 | 0.067566071 | 0.191730428 | 0.1287 | 0.279426343 |
| SL1344_RS04420_ | -0.863712067 | -0.914929478 | -0.809849153 | 0.10555461 | 0.254880127 | 0.391497858 | 0.331112599 | 0.475852726 | 0.624280104 | 0.579 | 0.699752931 |
| SL1344_RS04425_ | -0.049933782 | -0.104503549 | 0.057052611 | 0.163877937 | 0.350663667 | 0.446498756 | 0.330666674 | 0.532467806 | 0.690161104 | 0.671 | 0.784938184 |
| SL1344_RS04430_ | 1.046140357 | 1.028378572 | 1.083614597 | 0.030595879 | 0.203250731 | 0.119326844 | 0.443704199 | 0.736843135 | 0.840680088 | 0.7857 | 0.872698158 |

| | | | | | | | | | | | |
|---|---|---|---|---|---|---|---|---|---|---|---|
| SL1344_RS04435_ | 1.61935558 | 1.591618876 | 1.717646425 | 0.10740503 | 0.200620535 | 0.364031519 | 0.361333339 | 0.74178399 | 0.844534607 | 0.742 | 0.851253821 |
| SL1344_RS04440_ | 1.197787618 | 1.294176717 | 0.959846338 | -0.361968436 | 0.374565682 | -0.952642199 | 0.140666694 | 0.206305462 | 0.401156452 | 0.2744 | 0.456787391 |
| SL1344_RS04450_ | -0.754301414 | -0.654387139 | -0.838759561 | -0.288966956 | 0.503473295 | -0.539491079 | 0.269333344 | 0.378801886 | 0.565923333 | 0.546 | 0.683297722 |
| SL1344_RS04455_ | -0.399185946 | -0.240702523 | -0.609392713 | -0.400970701 | 0.304939486 | -1.258188634 | 0.069333395 | 0.132768967 | 0.292910421 | 0.1851 | 0.343698032 |
| SL1344_RS04460_ | 0.21547635 | 0.290938524 | 0.140671 | -0.14321621 | 0.218912318 | -0.613698623 | 0.260000012 | 0.416502123 | 0.578047827 | 0.5264 | 0.659847533 |
| SL1344_RS04465_ | 1.96862721 | 1.826255969 | 2.038086251 | 0.208021205 | 0.15446438 | 1.23946534 | 0.090606308 | 0.140655062 | 0.308864665 | 0.2036 | 0.37000968 |
| SL1344_RS04470_ | -2.860732541 | -3.110360186 | -2.530021323 | 0.564350947 | 0.532126381 | 0.984640669 | 0.137333362 | 0.225466521 | 0.383828174 | 0.3128 | 0.464615307 |
| SL1344_RS04475_ | -3.741871474 | -3.978688244 | -3.577846109 | 0.480433368 | 1.01210666 | 0.418269616 | 0.292471695 | 0.428304914 | 0.603892535 | 0.5819 | 0.704202246 |
| SL1344_RS04480_ | -1.166585397 | -1.200355288 | -1.143688627 | 0.045401882 | 0.350647442 | 0.11690586 | 0.450000002 | 0.62411406 | 0.752364219 | 0.7391 | 0.826953958 |
| SL1344_RS04485_ | -0.340751234 | -0.484363758 | -0.215418783 | 0.244687631 | 0.306033176 | 0.763481163 | 0.211192554 | 0.318775999 | 0.496316382 | 0.4079 | 0.567330095 |
| SL1344_RS04495_ | 2.866572716 | 2.694966233 | 2.915567564 | 0.240007538 | 0.207315039 | 1.21135833 | 0.094000044 | 0.120372163 | 0.287437697 | 0.2135 | 0.384551113 |
| SL1344_RS04500_ | 3.463173717 | 3.389778895 | 3.51044301 | 0.10449092 | 0.194216581 | 0.549579289 | 0.269153908 | 0.437505339 | 0.625668221 | 0.521 | 0.688082116 |
| SL1344_RS04505_ | 3.635485192 | 3.340318627 | 3.670751025 | 0.315742594 | 0.194409811 | 1.185746046 | 0.156000024 | 0.157703011 | 0.343299316 | 0.2958 | 0.483025372 |
| SL1344_RS04510_ | 1.592258861 | 1.598047966 | 1.584075194 | 0.014261825 | 0.184565526 | 0.063424646 | 0.473684212 | 0.702923995 | 0.816124235 | 0.8373 | 0.898387977 |
| SL1344_RS04515_ | -0.012632521 | 0.048951194 | -0.055288263 | -0.118159906 | 0.295795191 | -0.387983578 | 0.33333334 | 0.523298346 | 0.677324297 | 0.657 | 0.767654415 |
| SL1344_RS04520_ | 0.844033121 | 0.859580583 | 0.826838015 | -0.038705929 | 0.142798304 | -0.235106188 | 0.388407733 | 0.527618737 | 0.668243988 | 0.6385 | 0.744521716 |
| SL1344_RS04525_ | 1.285078234 | 1.249382173 | 1.333629276 | 0.088586406 | 0.131747259 | 0.650403369 | 0.247168567 | 0.369371898 | 0.52811049 | 0.4716 | 0.604682019 |
| SL1344_RS04530_ | 0.479286812 | 0.588359626 | 0.399849288 | -0.183881963 | 0.172620661 | -1.017796413 | 0.139906757 | 0.220774144 | 0.382912316 | 0.3128 | 0.47031267 |
| SL1344_RS04535_ | 0.515129744 | 0.581829763 | 0.389105532 | -0.182141782 | 0.207333346 | -0.801491035 | 0.208000016 | 0.312477192 | 0.486112954 | 0.4206 | 0.575879273 |
| SL1344_RS04540_ | -0.955300637 | -0.735694691 | -1.279654414 | -0.496424147 | 0.431353252 | -1.119958621 | 0.11792142 | 0.168376757 | 0.343131326 | 0.2469 | 0.421533871 |
| SL1344_RS04545_ | 0.03373625 | 0.071005682 | 0.003208655 | -0.060977442 | 0.210394323 | -0.285093178 | 0.378414395 | 0.55390787 | 0.695937201 | 0.6594 | 0.767591217 |
| SL1344_RS04550_ | 0.057302028 | 0.049312878 | 0.068179259 | 0.000980925 | 0.237873767 | 0.006045719 | 0.497668221 | 0.641304181 | 0.76766298 | 0.7683 | 0.848881543 |
| SL1344_RS04555_ | 1.109837314 | 1.158256915 | 1.042497991 | -0.116808044 | 0.163870747 | -0.689307244 | 0.230513005 | 0.359337653 | 0.529281845 | 0.4653 | 0.608720101 |
| SL1344_RS04560_ | 0.647302236 | 0.709278318 | 0.569484747 | -0.145894302 | 0.16509244 | -0.83190313 | 0.172000021 | 0.27370761 | 0.438873899 | 0.3695 | 0.518019481 |
| SL1344_RS04565_ | 4.476363206 | 4.261269985 | 4.668815784 | 0.415891674 | 0.088127978 | 4.72515511 | 4.68E-05 | 0.006718794 | 0.054649215 | 0.1 | 0.243021319 |
| SL1344_RS04570_ | 1.334771698 | 1.326098905 | 1.346294055 | 0.009851339 | 0.156388706 | 0.050412672 | 0.476349101 | 0.670099263 | 0.788302762 | 0.78 | 0.857053735 |
| SL1344_RS04575_ | 1.584791227 | 1.641723778 | 1.452730815 | -0.135689748 | 0.215620928 | -0.578074315 | 0.287808137 | 0.467631603 | 0.640376361 | 0.5929 | 0.73068677 |
| SL1344_RS04580_ | 0.12578395 | 0.648500673 | -0.589433157 | -1.239891067 | 0.228718789 | -5.439024414 | 4.68E-05 | 0.008067757 | 0.055576185 | 0.1 | 0.243021319 |
| SL1344_RS04585_ | 1.307895672 | 1.35262884 | 1.25778947 | -0.091065364 | 0.136413605 | -0.628074304 | 0.243837455 | 0.379702761 | 0.541770665 | 0.4914 | 0.627698254 |
| SL1344_RS04590_ | 3.409582995 | 3.502013514 | 3.219844642 | -0.28223851 | 0.102048497 | -2.65499443 | 4.68E-05 | 0.045754859 | 0.159846392 | 0.1 | 0.243021319 |
| SL1344_RS04600_ | -2.59091274 | -2.911282038 | -2.44032241 | 0.43277289 | 0.531518645 | 0.731920507 | 0.20933335 | 0.324211571 | 0.493329072 | 0.4372 | 0.58449803 |
| SL1344_RS04605_ | -0.719002732 | -0.745521994 | -0.698354525 | 0.065936826 | 0.29487418 | 0.187128398 | 0.404397072 | 0.586677349 | 0.723580076 | 0.7135 | 0.804933057 |
| SL1344_RS04610_ | 1.269460701 | 0.946335751 | 1.322454596 | 0.3492109 | 0.293943095 | 1.000436768 | 0.200000017 | 0.221303092 | 0.416009816 | 0.3753 | 0.559844442 |
| SL1344_RS04615_ | -1.11322531 | -1.02877811 | -1.237806473 | -0.19794182 | 0.307061539 | -0.60561441 | 0.253830791 | 0.394066457 | 0.555992008 | 0.4963 | 0.636062106 |
| SL1344_RS04620_ | -3.247405958 | -3.214821178 | -3.265098112 | -0.113360731 | 0.766715898 | -0.144329562 | 0.440373087 | 0.633593685 | 0.758660277 | 0.7425 | 0.828515848 |
| SL1344_RS04625_ | -1.639632056 | -1.572252381 | -1.729009737 | -0.139469349 | 0.394389272 | -0.319538966 | 0.365089945 | 0.520053365 | 0.666708197 | 0.6307 | 0.742715854 |
| SL1344_RS04630_ | 5.89576214 | 5.888703172 | 5.90469281 | 0.014179194 | 0.052804593 | 0.269083391 | 0.359760166 | 0.652172773 | 0.780945049 | 0.7063 | 0.810327451 |
| SL1344_RS04635_ | 5.297855092 | 5.277822838 | 5.317099496 | 0.027220909 | 0.196099252 | 0.12132199 | 0.448000002 | 0.897113371 | 0.94487012 | 0.9802 | 0.988792033 |
| SL1344_RS04640_ | 1.587530886 | 1.438556602 | 1.797482088 | 0.352404263 | 0.148217473 | 2.316829215 | 0.004664544 | 0.032331031 | 0.126332528 | 0.1045 | 0.249078621 |
| SL1344_RS04645_ | 2.494129832 | 2.335167603 | 2.643200278 | 0.313143391 | 0.088699006 | 3.478905916 | 4.68E-05 | 0.010756861 | 0.065679566 | 0.1 | 0.243021319 |
| SL1344_RS04650_ | 2.873316791 | 2.764266605 | 2.982981312 | 0.231327302 | 0.09578301 | 2.364652892 | 0.001336769 | 0.028664021 | 0.117080464 | 0.1014 | 0.244977598 |
| SL1344_RS04655_ | 4.558317081 | 4.684148066 | 4.444144409 | -0.246578676 | 0.063489174 | -3.848695977 | 4.68E-05 | 0.009399432 | 0.063591091 | 0.1 | 0.243021319 |
| SL1344_RS04665_ | 4.655429707 | 4.657519598 | 4.652199213 | 0.006937024 | 0.071246163 | 0.08699952 | 0.470353099 | 0.688643694 | 0.807095058 | 0.8358 | 0.898728019 |
| SL1344_RS04675_ | 2.888758908 | 2.854413703 | 2.923265632 | 0.067259645 | 0.117297912 | 0.582267562 | 0.272000011 | 0.419273173 | 0.592970414 | 0.5334 | 0.675723476 |
| SL1344_RS04680_ | 4.58989907 | 4.582893894 | 4.598551701 | 0.030079531 | 0.081086944 | 0.331810055 | 0.333777489 | 0.443766284 | 0.626661132 | 0.6581 | 0.764205831 |
| SL1344_RS04700_ | -4.825414465 | -4.784921801 | -4.871741549 | -0.12063756 | 1.005645866 | -0.103417265 | 0.449700202 | 0.553751198 | 0.692284248 | 0.6662 | 0.768291326 |
| SL1344_RS04705_ | 0.974094632 | 0.608784991 | 1.248554519 | 0.628501503 | 0.337985186 | 1.783789542 | 0.009333825 | 0.075428712 | 0.21781627 | 0.1083 | 0.254671526 |
| SL1344_RS04715_ | 3.57180098 | 3.559432227 | 3.587975815 | 0.030372104 | 0.078616021 | 0.340188845 | 0.357761498 | 0.563096858 | 0.706710798 | 0.6795 | 0.783383919 |
| SL1344_RS04720_ | -1.264792452 | -1.520483549 | -1.022975428 | 0.507978966 | 0.319893685 | 1.518457483 | 0.042638345 | 0.08566138 | 0.217163935 | 0.1521 | 0.304789336 |
| SL1344_RS04725_ | -0.262960557 | -0.261203299 | -0.265595512 | -0.007353799 | 0.333342843 | -0.022248833 | 0.489673551 | 0.736693534 | 0.839898225 | 0.8414 | 0.904333696 |
| SL1344_RS04730_ | 3.91873596 | 3.560346758 | 4.265628006 | 0.727102724 | 0.104686614 | 7.058569876 | 4.68E-05 | 0.003991394 | 0.042502425 | 0.1 | 0.243021319 |
| SL1344_RS04740_ | 3.313916314 | 3.124199986 | 3.486971392 | 0.367122905 | 0.088208979 | 4.222624073 | 4.68E-05 | 0.00576914 | 0.049288935 | 0.1 | 0.243021319 |
| SL1344_RS04745_ | -2.563904733 | -2.574490055 | -2.547958021 | 0.035314675 | 0.484000337 | 0.075354444 | 0.471333334 | 0.571866835 | 0.708523299 | 0.6873 | 0.785839269 |

| | | | | | | | | | | | |
|---|---|---|---|---|---|---|---|---|---|---|---|
| SL1344_RS04750_ | 2.451372701 | 2.495650289 | 2.397687183 | -0.102744455 | 0.096285089 | -1.019729108 | 0.115922753 | 0.193046175 | 0.35902206 | 0.2703 | 0.427435736 |
| SL1344_RS04755_ | 2.258791526 | 2.25572346 | 2.26324521 | 0.009022169 | 0.11238575 | 0.070422116 | 0.470000001 | 0.661436103 | 0.782497156 | 0.7671 | 0.849288813 |
| SL1344_RS04760_ | 0.346523547 | 0.30092868 | 0.379734294 | 0.065696152 | 0.194698255 | 0.323612846 | 0.365756168 | 0.542468697 | 0.686133678 | 0.6544 | 0.762784216 |
| SL1344_RS04765_ | 3.97022112 | 3.82301601 | 4.03666401 | 0.210286543 | 0.121035924 | 1.768492171 | 0.01866691 | 0.056439806 | 0.182082379 | 0.1161 | 0.264738326 |
| SL1344_RS04770_ | 8.38048061 | 8.355932647 | 8.41154649 | 0.057404793 | 0.087930486 | 0.670830173 | 0.232666681 | 0.332606508 | 0.5379652 | 0.4453 | 0.628048381 |
| SL1344_RS04780_ | -0.845793071 | -0.945280949 | -0.739498478 | 0.204065983 | 0.23929911 | 0.794158087 | 0.185333353 | 0.28915408 | 0.445911834 | 0.3819 | 0.525358827 |
| SL1344_RS04785_ | 3.756764076 | 3.696455174 | 3.803919038 | 0.113910435 | 0.068398258 | 1.638113805 | 0.026649069 | 0.091956264 | 0.234025531 | 0.129 | 0.278431432 |
| SL1344_RS04790_ | 3.296284705 | 3.131430384 | 3.417494521 | 0.278570338 | 0.084193785 | 3.274532487 | 4.68E-05 | 0.013645579 | 0.077151794 | 0.1001 | 0.243160919 |
| SL1344_RS04795_ | 0.203945913 | 0.117744623 | 0.288079569 | 0.171612508 | 0.153053285 | 1.051783184 | 0.124583643 | 0.204500531 | 0.355875672 | 0.2872 | 0.438116909 |
| SL1344_RS04800_ | -1.332854403 | -1.195331611 | -1.442147846 | -0.236932722 | 0.335852286 | -0.672484216 | 0.228000014 | 0.365041691 | 0.529926248 | 0.4617 | 0.605603629 |
| SL1344_RS04805_ | 1.745638486 | 1.73015271 | 1.775617733 | 0.022300924 | 0.201532979 | 0.099809132 | 0.455029982 | 0.790361799 | 0.877583894 | 0.8717 | 0.924586551 |
| SL1344_RS04810_ | -0.595160378 | -0.639517969 | -0.57342993 | 0.026793026 | 0.348629425 | 0.066651627 | 0.475016656 | 0.723710167 | 0.830305953 | 0.8217 | 0.891112713 |
| SL1344_RS04815_ | -0.600661858 | -0.835965716 | -0.374108838 | 0.46447432 | 0.221492519 | 2.034494258 | 0.015333631 | 0.046576118 | 0.146252048 | 0.1187 | 0.266210199 |
| SL1344_RS04820_ | 1.52251966 | 1.546549767 | 1.454910832 | -0.090603231 | 0.301765646 | -0.261721294 | 0.392405067 | 0.625000252 | 0.768886016 | 0.8012 | 0.885145616 |
| SL1344_RS04825_ | 2.423423429 | 2.455691911 | 2.401186717 | -0.037408826 | 0.138902041 | -0.256002792 | 0.394403735 | 0.65857448 | 0.783066193 | 0.7735 | 0.855575853 |
| SL1344_RS04830_ | 2.523762494 | 2.493413106 | 2.55657255 | 0.062745349 | 0.106886366 | 0.561305583 | 0.256495681 | 0.414075339 | 0.580695359 | 0.5265 | 0.661037369 |
| SL1344_RS04840_ | 2.565630101 | 2.629400991 | 2.489399018 | -0.143371992 | 0.093281707 | -1.444129326 | 0.040000111 | 0.098560099 | 0.240879006 | 0.1511 | 0.304638578 |
| SL1344_RS04845_ | 1.990400589 | 1.976068714 | 2.007718203 | 0.032815061 | 0.103541724 | 0.313357399 | 0.368666672 | 0.534191174 | 0.676606935 | 0.649 | 0.754417212 |
| SL1344_RS04850_ | 2.551811602 | 2.52049767 | 2.629381274 | 0.087312689 | 0.182094673 | 0.431327466 | 0.321333341 | 0.622294194 | 0.762698425 | 0.6446 | 0.780450018 |
| SL1344_RS04855_ | 4.070753996 | 3.959986212 | 4.148501652 | 0.188654387 | 0.071223827 | 2.612754238 | 4.68E-05 | 0.02941265 | 0.119979316 | 0.1006 | 0.24386533 |
| SL1344_RS04860_ | 3.00971956 | 3.371265478 | 2.347414532 | -1.02364616 | 0.15384341 | -6.394766598 | 4.68E-05 | 0.009591344 | 0.068088502 | 0.1 | 0.243021319 |
| SL1344_RS04865_ | 5.146255313 | 5.141602551 | 5.157573575 | 0.013177434 | 0.05904785 | 0.183233325 | 0.432000003 | 0.743590859 | 0.845929195 | 0.8249 | 0.896841176 |
| SL1344_RS04870_ | -1.16559015 | -1.143510057 | -1.186906427 | -0.04277728 | 0.283898848 | -0.144566208 | 0.439333336 | 0.544790637 | 0.683386802 | 0.6416 | 0.748962845 |
| SL1344_RS04875_ | -3.975021911 | -3.959541636 | -3.987989733 | 0.020686477 | 0.896756307 | 0.018431377 | 0.492666667 | 0.643182516 | 0.767028463 | 0.7481 | 0.83373461 |
| SL1344_RS04880_ | -2.298971465 | -2.419058718 | -2.199734462 | 0.224296205 | 0.661887696 | 0.332465105 | 0.345103271 | 0.553885016 | 0.701903417 | 0.6842 | 0.785739164 |
| SL1344_RS04885_ | -3.147535251 | -2.998277867 | -3.325043542 | -0.32256141 | 0.558888828 | -0.547458657 | 0.273333344 | 0.400392586 | 0.555971441 | 0.5044 | 0.635043816 |
| SL1344_RS04890_ | 2.763640737 | 2.720981153 | 2.836233736 | 0.104363856 | 0.112242056 | 0.87307052 | 0.175333354 | 0.261835272 | 0.443920231 | 0.3583 | 0.527228504 |
| SL1344_RS04895_ | 1.994435735 | 1.96324835 | 2.043996452 | 0.076386212 | 0.105339592 | 0.66497956 | 0.236000014 | 0.365443488 | 0.529468738 | 0.4712 | 0.613301818 |
| SL1344_RS04900_ | -4.13597835 | -3.933403731 | -4.312274396 | -0.32991069 | 1.052472 | -0.30268 | 0.373333338 | 0.57882856 | 0.71615276 | 0.684 | 0.785588506 |
| SL1344_RS04905_ | -6.508753637 | -6.289226884 | -6.985215241 | -0.783893339 | 2.292981768 | -0.262243993 | 0.365756168 | 0.496983429 | 0.658488034 | 0.642 | 0.756696038 |
| SL1344_RS04910_ | -4.188017633 | -3.953463758 | -4.838332581 | -0.821981519 | 1.480110934 | -0.465002539 | 0.307794812 | 0.446444695 | 0.624964078 | 0.6241 | 0.753300873 |
| SL1344_RS04915_ | -2.728172182 | -2.417044835 | -3.093709804 | -0.679884715 | 0.50320363 | -1.282331907 | 0.077281867 | 0.130196079 | 0.275865851 | 0.1986 | 0.354751057 |
| SL1344_RS04920_ | -5.468157396 | -5.942032928 | -5.17526255 | 0.666577124 | 1.990973506 | 0.323290182 | 0.362666672 | 0.578858926 | 0.722367969 | 0.7175 | 0.814519358 |
| SL1344_RS04925_ | -5.10036529 | -5.317040309 | -4.94292362 | 0.571835323 | 2.127172555 | 0.216723786 | 0.34333334 | 0.479511233 | 0.654788129 | 0.6753 | 0.774284948 |
| SL1344_RS04930_ | 3.143448984 | 3.136063568 | 3.164850868 | 0.028662028 | 0.141867778 | 0.182467609 | 0.431045972 | 0.821170625 | 0.89806393 | 0.8315 | 0.903738111 |
| SL1344_RS04935_ | -4.328834445 | -3.658242854 | -4.591539777 | -0.769828799 | 1.315924141 | -0.519491274 | 0.303131254 | 0.539180726 | 0.685947679 | 0.5974 | 0.731827523 |
| SL1344_RS04940_ | -3.492084911 | -3.532131153 | -3.428637591 | 0.08608124 | 0.744284384 | 0.103518021 | 0.451032647 | 0.61625777 | 0.74554735 | 0.7237 | 0.816615865 |
| SL1344_RS04945_ | -2.253351758 | -2.294811957 | -2.216633919 | 0.088795642 | 0.391549621 | 0.199860551 | 0.41266667 | 0.52652708 | 0.669267985 | 0.6275 | 0.739915667 |
| SL1344_RS04950_ | -3.920540319 | -3.719286625 | -4.162312931 | -0.437946325 | 0.796620562 | -0.530416141 | 0.285143248 | 0.424001793 | 0.579667909 | 0.5288 | 0.656786692 |
| SL1344_RS04955_ | -4.570759491 | -4.351360516 | -4.801453038 | -0.430497481 | 0.901063021 | -0.437463516 | 0.301132587 | 0.426332308 | 0.578965344 | 0.5377 | 0.662709011 |
| SL1344_RS04960_ | -1.934676081 | -1.902289891 | -1.958503823 | -0.060591943 | 0.37911224 | -0.145158113 | 0.437333336 | 0.557732414 | 0.69581776 | 0.6618 | 0.765347263 |
| SL1344_RS04965_ | 2.340923224 | 2.427743666 | 2.278253007 | -0.136213914 | 0.138888407 | -0.96642268 | 0.14656898 | 0.220987665 | 0.40388454 | 0.3045 | 0.47569557 |
| SL1344_RS04970_ | -0.656752229 | -0.708567796 | -0.618936547 | 0.075308393 | 0.323600698 | 0.218067164 | 0.401732182 | 0.611027733 | 0.743944387 | 0.7125 | 0.810047525 |
| SL1344_RS04975_ | -0.021331304 | 0.022483268 | -0.065329854 | -0.092144741 | 0.26284833 | -0.329597348 | 0.359760166 | 0.563175386 | 0.707838114 | 0.6915 | 0.792949138 |
| SL1344_RS04980_ | -0.775990505 | -0.897893033 | -0.611088884 | 0.285966545 | 0.274433256 | 0.994103713 | 0.134666696 | 0.212258128 | 0.374984474 | 0.2928 | 0.448364485 |
| SL1344_RS04985_ | -6.081944211 | -5.893782529 | -6.224980053 | -0.277382992 | 1.628591751 | -0.146322695 | 0.424383747 | 0.575643514 | 0.711711852 | 0.6842 | 0.784406784 |
| SL1344_RS04990_ | -4.232924096 | -3.802443357 | -4.66890062 | -0.85740947 | 1.290221989 | -0.61934007 | 0.210000016 | 0.328597918 | 0.512790947 | 0.4116 | 0.568621103 |
| SL1344_RS04995_ | -7.517573249 | -8.126112748 | -6.961648064 | 0.951365774 | 3.889081888 | 0.213939334 | 0.36642239 | 0.573532876 | 0.715305198 | 0.698 | 0.797863873 |
| SL1344_RS05000_ | -3.458759313 | -3.387870961 | -3.496218205 | -0.158467337 | 0.723745215 | -0.201483477 | 0.413724187 | 0.591461505 | 0.724818599 | 0.6996 | 0.796313206 |
| SL1344_RS05005_ | 1.29829565 | 1.251259224 | 1.378249964 | 0.104841258 | 0.171163859 | 0.571985169 | 0.27514991 | 0.455087477 | 0.61770893 | 0.5578 | 0.693967026 |
| SL1344_RS05010_ | -6.127563627 | -6.08979534 | -6.181803432 | -0.146054099 | 1.576663437 | -0.089788467 | 0.464000001 | 0.543083159 | 0.685838954 | 0.6608 | 0.765012633 |
| SL1344_RS05015_ | 1.244752182 | 0.85125714 | 1.598163722 | 0.708319018 | 0.201725069 | 3.443657993 | 4.68E-05 | 0.00963576 | 0.064720785 | 0.1 | 0.243021319 |
| SL1344_RS05020_ | -1.309954772 | -1.368784686 | -1.167171718 | 0.133630173 | 0.482835051 | 0.218478714 | 0.398666671 | 0.747661131 | 0.846678424 | 0.7823 | 0.870642382 |

| | | | | | | | | | | |
|---|---|---|---|---|---|---|---|---|---|---|
| SL1344_RS05025_ | -6.205252423 | -6.187140329 | -6.212876271 | -0.066260015 | 1.826742967 | -0.034855035 | 0.480346436 | 0.595487686 | 0.729761263 | 0.7047 | 0.801417823 |
| SL1344_RS05030_ | -2.817533417 | -2.892830832 | -2.756999707 | 0.133564674 | 0.515956592 | 0.248061714 | 0.396000004 | 0.533713414 | 0.676360796 | 0.6493 | 0.756181858 |
| SL1344_RS05035_ | -3.716259282 | -4.173141874 | -3.361120412 | 0.813141018 | 0.764114129 | 1.04675126 | 0.124000032 | 0.201564082 | 0.367632682 | 0.2795 | 0.435616784 |
| SL1344_RS05040_ | -2.55631258 | -2.630070724 | -2.527666421 | 0.074278799 | 0.575931128 | 0.113840748 | 0.447035312 | 0.634162621 | 0.7629592 | 0.7459 | 0.837094086 |
| SL1344_RS05045_ | -1.545477685 | -1.516808416 | -1.578700945 | -0.06894896 | 0.285925351 | -0.21381132 | 0.405333337 | 0.506570211 | 0.649006059 | 0.6066 | 0.721663543 |
| SL1344_RS05050_ | -5.77106244 | -5.917261453 | -5.616800483 | 0.244069749 | 1.745053088 | 0.124587968 | 0.432378417 | 0.589439318 | 0.724824131 | 0.6832 | 0.784414917 |
| SL1344_RS05055_ | -1.92688976 | -1.929092728 | -1.924454212 | -0.011849514 | 0.406376708 | -0.025717681 | 0.487674884 | 0.604870837 | 0.735502778 | 0.7143 | 0.807392763 |
| SL1344_RS05060_ | -1.185667747 | -1.185525592 | -1.185912284 | -0.015573597 | 0.348183376 | -0.045492598 | 0.482666667 | 0.630926687 | 0.756461593 | 0.7385 | 0.826090409 |
| SL1344_RS05065_ | -3.696595116 | -3.584893007 | -3.884144151 | -0.268987651 | 0.760931046 | -0.321053645 | 0.351765496 | 0.5155129 | 0.663725338 | 0.6312 | 0.744326963 |
| SL1344_RS05070_ | -4.449107549 | -4.386109298 | -4.548437866 | -0.200077506 | 1.035874324 | -0.175558322 | 0.41066667 | 0.573156326 | 0.714455517 | 0.692 | 0.791334156 |
| SL1344_RS05075_ | -2.919211749 | -2.760486657 | -3.285080192 | -0.549243617 | 0.795513978 | -0.607875328 | 0.222518336 | 0.33649905 | 0.522959948 | 0.4589 | 0.606668434 |
| SL1344_RS05080_ | -3.549582605 | -3.177679386 | -3.980991806 | -0.766501664 | 0.744967655 | -0.96424576 | 0.140666694 | 0.224609157 | 0.389301659 | 0.3114 | 0.468129088 |
| SL1344_RS05085_ | -2.181151576 | -2.015427594 | -2.399490876 | -0.367833687 | 0.546288465 | -0.650685391 | 0.249333346 | 0.389170726 | 0.559813959 | 0.5031 | 0.646765262 |
| SL1344_RS05090_ | -2.729750679 | -2.614801583 | -2.826242633 | -0.18855669 | 0.54807821 | -0.312958664 | 0.359093943 | 0.542423724 | 0.682458369 | 0.6424 | 0.75212926 |
| SL1344_RS05095_ | -3.47293181 | -3.423401348 | -3.505007934 | -0.062711226 | 0.710415875 | -0.081707012 | 0.463024652 | 0.61066008 | 0.741482852 | 0.7165 | 0.81153676 |
| SL1344_RS05100_ | -0.366674329 | -0.282019211 | -0.471238698 | -0.185466176 | 0.217983957 | -0.796361819 | 0.172551653 | 0.273066674 | 0.434665373 | 0.3706 | 0.517796101 |
| SL1344_RS05105_ | -4.0458924 | -3.899596594 | -4.326159728 | -0.466614454 | 1.088788208 | -0.383846753 | 0.320000008 | 0.46108376 | 0.627256079 | 0.5959 | 0.716223475 |
| SL1344_RS05110_ | -0.346887589 | -0.206113708 | -0.538657625 | -0.360738189 | 0.31269831 | -1.132870115 | 0.070000061 | 0.168450572 | 0.343216627 | 0.1955 | 0.35201273 |
| SL1344_RS05115_ | 1.56745783 | 1.706532586 | 1.367581574 | -0.366709485 | 0.186535398 | -1.974144803 | 0.002666616 | 0.060345101 | 0.185509407 | 0.103 | 0.247171109 |
| SL1344_RS05120_ | -2.406768084 | -2.279503292 | -2.577316321 | -0.191169106 | 0.685134172 | -0.257611898 | 0.381745508 | 0.624523039 | 0.756684781 | 0.7413 | 0.831546884 |
| SL1344_RS05125_ | -2.918344971 | -2.929959728 | -2.904183275 | 0.039513689 | 0.473664151 | 0.074194362 | 0.466355764 | 0.511002896 | 0.6550302 | 0.6135 | 0.726498839 |
| SL1344_RS05130_ | -2.386271135 | -2.326325737 | -2.437432844 | -0.124213565 | 0.492603839 | -0.242494247 | 0.397068625 | 0.55504406 | 0.695266805 | 0.6662 | 0.770624262 |
| SL1344_RS05135_ | -0.354929708 | -0.245097518 | -0.517215456 | -0.266473214 | 0.247691596 | -1.023851223 | 0.140666694 | 0.211264748 | 0.376648805 | 0.3021 | 0.461378583 |
| SL1344_RS05140_ | -3.047568349 | -2.850501839 | -3.446190328 | -0.615321541 | 0.687094763 | -0.803987257 | 0.20186544 | 0.291847991 | 0.467751271 | 0.4033 | 0.556242137 |
| SL1344_RS05165_ | -4.25263239 | -4.055197435 | -4.382528905 | -0.25235469 | 1.114924699 | -0.203283886 | 0.41466667 | 0.63887579 | 0.764723749 | 0.736 | 0.82817318 |
| SL1344_RS05175_ | -1.363645444 | -1.053002543 | -1.590299543 | -0.520312922 | 0.329943574 | -1.497625945 | 0.05266675 | 0.099460201 | 0.236971601 | 0.1701 | 0.326075098 |
| SL1344_RS05190_ | 2.857749288 | 3.010555691 | 2.73524174 | -0.274454146 | 0.095373683 | -2.788479741 | 4.68E-05 | 0.01796977 | 0.09020814 | 0.1002 | 0.243294587 |
| SL1344_RS05195_ | 1.523910014 | 1.486502638 | 1.549452267 | 0.084784963 | 0.174323702 | 0.461222946 | 0.283333343 | 0.410997261 | 0.591955888 | 0.5539 | 0.675493055 |
| SL1344_RS05200_ | 1.730713426 | 1.593840284 | 1.849088443 | 0.256728588 | 0.104399732 | 2.375550822 | 0.004001152 | 0.03102096 | 0.118134622 | 0.1034 | 0.24746661 |
| SL1344_RS05205_ | -0.07002558 | -0.098791943 | -0.04472397 | 0.042977088 | 0.229471426 | 0.183348438 | 0.423333336 | 0.590493388 | 0.724300213 | 0.6883 | 0.788076039 |
| SL1344_RS05210_ | 3.22958306 | 3.210420803 | 3.284633704 | 0.06195251 | 0.117502318 | 0.374149657 | 0.361758833 | 0.771535513 | 0.865465027 | 0.7496 | 0.856171186 |
| SL1344_RS05215_ | 3.82926959 | 3.767365737 | 3.875300766 | 0.101949629 | 0.075005199 | 1.303779661 | 0.080612977 | 0.124931046 | 0.28219364 | 0.1934 | 0.358702337 |
| SL1344_RS05220_ | 2.257525529 | 2.161894585 | 2.432964188 | 0.262033265 | 0.14290376 | 1.737367532 | 0.027981506 | 0.05915706 | 0.181801072 | 0.1257 | 0.276733805 |
| SL1344_RS05225_ | 2.42626246 | 2.375232372 | 2.459213818 | 0.080867734 | 0.10223268 | 0.743846067 | 0.204666683 | 0.311841658 | 0.483838019 | 0.4189 | 0.570674048 |
| SL1344_RS05230_ | 2.185980204 | 2.152529344 | 2.221068776 | 0.070336977 | 0.09299104 | 0.704872856 | 0.222518336 | 0.327421203 | 0.487812692 | 0.4266 | 0.567001777 |
| SL1344_RS05235_ | -0.193198942 | 0.085932509 | -0.478784658 | -0.578640594 | 0.220815237 | -2.548785866 | 0.000673369 | 0.026183445 | 0.108844402 | 0.1015 | 0.245036603 |
| SL1344_RS05240_ | 4.700490933 | 4.857677451 | 4.525312056 | -0.339064357 | 0.068995288 | -5.018620076 | 4.68E-05 | 0.055175104 | 0.18393579 | 0.1 | 0.243021319 |
| SL1344_RS05245_ | 3.347879494 | 3.316454218 | 3.391474146 | 0.042243978 | 0.134695728 | 0.274373604 | 0.396666671 | 0.741527148 | 0.844884114 | 0.8201 | 0.892302378 |
| SL1344_RS05250_ | 0.073512156 | 0.27836451 | -0.161842463 | -0.440774005 | 0.203147729 | -2.130796312 | 0.011326186 | 0.038554895 | 0.135995019 | 0.1093 | 0.25511716 |
| SL1344_RS05255_ | 8.370289525 | 8.181899428 | 8.575813256 | 0.344655759 | 0.07266483 | 5.05286737 | 4.68E-05 | 0.000982451 | 0.023126482 | 0.1 | 0.243021319 |
| SL1344_RS05260_ | 1.422331779 | 1.456499646 | 1.22664873 | -0.21842415 | 0.256433862 | -0.606163319 | 0.324000007 | 0.49221369 | 0.668430136 | 0.6765 | 0.808489936 |
| SL1344_RS05265_ | -2.021534372 | -1.771151919 | -2.368985727 | -0.570643794 | 0.549883063 | -0.991204677 | 0.134666696 | 0.201455252 | 0.375601855 | 0.2863 | 0.455852692 |
| SL1344_RS05270_ | 1.498382158 | 1.514629844 | 1.477133014 | -0.034555909 | 0.107001215 | -0.285621534 | 0.366000005 | 0.49485632 | 0.642410483 | 0.6056 | 0.721742802 |
| SL1344_RS05275_ | -0.544380692 | -0.484165327 | -0.613946117 | -0.130636034 | 0.251284457 | -0.473972176 | 0.298666676 | 0.44596152 | 0.59819775 | 0.5591 | 0.679642854 |
| SL1344_RS05285_ | 0.449125745 | 0.503438551 | 0.384848449 | -0.116831559 | 0.189328781 | -0.585730924 | 0.257828126 | 0.397657912 | 0.558878798 | 0.508 | 0.642018807 |
| SL1344_RS05290_ | -3.12684126 | -2.875347174 | -3.495024975 | -0.648418084 | 0.592858973 | -1.03767757 | 0.118587642 | 0.193299066 | 0.352892388 | 0.2684 | 0.421658149 |
| SL1344_RS05295_ | 1.312591919 | 1.387961286 | 1.216951554 | -0.176744454 | 0.302301168 | -0.567198872 | 0.228514338 | 0.343502856 | 0.541259745 | 0.4591 | 0.616740632 |
| SL1344_RS05300_ | 3.024309242 | 3.196530075 | 2.953233481 | -0.230866014 | 0.108306855 | -1.902571241 | 0.013333676 | 0.056963479 | 0.17942736 | 0.114 | 0.261749018 |
| SL1344_RS05305_ | 2.04998644 | 1.938584757 | 2.192372588 | 0.258440952 | 0.109847245 | 2.31691566 | 0.001336769 | 0.036022356 | 0.131297515 | 0.1036 | 0.247601121 |
| SL1344_RS05310_ | -1.885938928 | -1.93114095 | -1.845248046 | 0.090483865 | 0.357641659 | 0.231408678 | 0.390666671 | 0.515368915 | 0.661350151 | 0.625 | 0.735994718 |
| SL1344_RS05315_ | -0.656976188 | -0.766606273 | -0.512251878 | 0.250039004 | 0.258616667 | 0.876855882 | 0.150000025 | 0.251508053 | 0.413994229 | 0.34 | 0.490716348 |
| SL1344_RS05320_ | -2.513721081 | -2.606633492 | -2.439110209 | 0.166342556 | 0.539721249 | 0.288028743 | 0.36908728 | 0.545471274 | 0.68593631 | 0.6531 | 0.758692651 |
| SL1344_RS05325_ | 3.198305063 | 3.202083689 | 3.190341844 | -9.50E-05 | 0.086148394 | -0.001411438 | 0.499000666 | 0.678590156 | 0.797538352 | 0.7753 | 0.859083412 |

| | | | | | | | | | | | |
|---|---|---|---|---|---|---|---|---|---|---|---|
| SL1344_RS05330_ | 2.479843286 | 2.538890729 | 2.428988034 | -0.115806375 | 0.134652231 | -0.838814155 | 0.189873436 | 0.268541509 | 0.452367786 | 0.371 | 0.535497747 |
| SL1344_RS05335_ | 3.446329024 | 3.400765847 | 3.497631838 | 0.098940724 | 0.064197156 | 1.48475363 | 0.053297883 | 0.103224874 | 0.242542156 | 0.165 | 0.320046463 |
| SL1344_RS05340_ | -1.337923625 | -1.377281938 | -1.306120025 | 0.105211634 | 0.403961389 | 0.246923093 | 0.399733515 | 0.596866876 | 0.732718327 | 0.7178 | 0.810382256 |
| SL1344_RS05345_ | -2.820588403 | -2.371306955 | -3.450714052 | -1.082944082 | 0.60411771 | -1.726580526 | 0.021333546 | 0.075428881 | 0.202080451 | 0.1219 | 0.270590339 |
| SL1344_RS05350_ | -6.46565753 | -7.099936275 | -6.001724179 | 1.188199726 | 2.63932618 | 0.394281619 | 0.299333342 | 0.455973188 | 0.621848731 | 0.5861 | 0.708715553 |
| SL1344_RS05365_ | -4.156078596 | -4.763923985 | -3.71512739 | 1.01352206 | 0.87721263 | 1.105057796 | 0.098666708 | 0.170968027 | 0.330972204 | 0.2432 | 0.40126475 |
| SL1344_RS05370_ | -4.230402892 | -4.58261844 | -3.901467056 | 0.686324241 | 0.85823008 | 0.749051373 | 0.197201883 | 0.300804311 | 0.464345879 | 0.3978 | 0.54440951 |
| SL1344_RS05375_ | -1.099097094 | -1.408481512 | -0.61360976 | 0.772723898 | 0.419466047 | 1.759627524 | 0.021985549 | 0.060099183 | 0.182972021 | 0.1266 | 0.278041627 |
| SL1344_RS05380_ | -0.016707274 | -0.040870358 | 0.01969857 | 0.068575357 | 0.223457734 | 0.272461553 | 0.385742843 | 0.54612673 | 0.689012533 | 0.6671 | 0.771821669 |
| SL1344_RS05385_ | -0.302766637 | -0.337222778 | -0.260043962 | 0.079253417 | 0.235682415 | 0.303629955 | 0.370419725 | 0.547114281 | 0.687388421 | 0.6549 | 0.759917528 |
| SL1344_RS05390_ | -0.849920664 | -0.828151947 | -0.864034784 | -0.026050752 | 0.248648563 | -0.100913077 | 0.456000002 | 0.537445996 | 0.677885079 | 0.6444 | 0.750280551 |
| SL1344_RS05395_ | -1.661333633 | -1.627223737 | -1.722853949 | -0.121178857 | 0.452167111 | -0.247580273 | 0.389333338 | 0.549816106 | 0.697308092 | 0.6919 | 0.78920915 |
| SL1344_RS05400_ | -2.399957554 | -2.43118335 | -2.365174496 | 0.050303203 | 0.561113136 | 0.083752583 | 0.463690874 | 0.657992877 | 0.780881897 | 0.7747 | 0.854570147 |
| SL1344_RS05405_ | -1.409405708 | -1.426757671 | -1.388321714 | 0.044100795 | 0.299950265 | 0.143462182 | 0.437333336 | 0.540478906 | 0.680778695 | 0.6523 | 0.757574308 |
| SL1344_RS05410_ | -2.995199384 | -2.776256287 | -3.177187791 | -0.394827987 | 0.551119819 | -0.679605396 | 0.23184545 | 0.344058736 | 0.503917817 | 0.4418 | 0.581924286 |
| SL1344_RS05415_ | -2.45887636 | -2.473728616 | -2.450045595 | 0.031845835 | 0.563363302 | 0.061978076 | 0.472351767 | 0.654483385 | 0.776182463 | 0.7587 | 0.842879427 |
| SL1344_RS05420_ | -3.32076555 | -3.107024351 | -4.028710438 | -0.874519398 | 1.332246678 | -0.578274435 | 0.272485021 | 0.368728097 | 0.562921749 | 0.5345 | 0.683061696 |
| SL1344_RS05425_ | -4.171522599 | -4.216790971 | -4.134940462 | 0.068620792 | 0.861411438 | 0.079722307 | 0.468000001 | 0.579061098 | 0.71394836 | 0.6905 | 0.787986662 |
| SL1344_RS05430_ | -3.351057734 | -3.404771576 | -3.301868567 | 0.109337958 | 0.57545503 | 0.16563036 | 0.430666669 | 0.527946927 | 0.670924851 | 0.6453 | 0.752128572 |
| SL1344_RS05435_ | -3.212922918 | -3.221570769 | -3.202045968 | 0.012269278 | 0.559854153 | 0.024327264 | 0.492666667 | 0.551399983 | 0.692269184 | 0.6656 | 0.769526476 |
| SL1344_RS05440_ | -2.348197655 | -2.096328507 | -2.708685579 | -0.60524124 | 0.52536472 | -1.088790931 | 0.111259197 | 0.177670719 | 0.341278897 | 0.2588 | 0.419715662 |
| SL1344_RS05445_ | -2.771490979 | -2.811855612 | -2.709702611 | 0.105970754 | 0.719487612 | 0.135481817 | 0.439706864 | 0.661369275 | 0.783443494 | 0.7663 | 0.848277332 |
| SL1344_RS05450_ | -0.622685156 | -0.555139977 | -0.705173971 | -0.15229856 | 0.23278315 | -0.614613177 | 0.254666679 | 0.376125873 | 0.532600852 | 0.4796 | 0.611666504 |
| SL1344_RS05455_ | -3.715845278 | -4.050476596 | -3.438877727 | 0.608624361 | 0.621698803 | 0.899416003 | 0.152564982 | 0.236016192 | 0.391577331 | 0.3199 | 0.470184604 |
| SL1344_RS05460_ | 0.169699804 | 0.386660665 | -0.101438682 | -0.485149543 | 0.192399826 | -2.417258863 | 0.004667653 | 0.031071371 | 0.118737326 | 0.1038 | 0.248236918 |
| SL1344_RS05465_ | 0.93013722 | 0.971934348 | 0.772339759 | -0.184276081 | 0.218453421 | -0.656043968 | 0.279147245 | 0.409219787 | 0.588883042 | 0.5698 | 0.712024807 |
| SL1344_RS05470_ | 0.854005667 | 0.876713008 | 0.841082985 | -0.042602849 | 0.196304633 | -0.188490542 | 0.424000003 | 0.644486186 | 0.771552866 | 0.7667 | 0.849303977 |
| SL1344_RS05475_ | -0.589944632 | -0.790273046 | -0.40742911 | 0.389491687 | 0.26247991 | 1.4379221 | 0.053964105 | 0.100535096 | 0.239829814 | 0.1617 | 0.31648112 |
| SL1344_RS05480_ | -2.653240499 | -2.800283254 | -2.522077578 | 0.298258584 | 0.500843137 | 0.541780836 | 0.266489018 | 0.39907656 | 0.556365381 | 0.5028 | 0.634518042 |
| SL1344_RS05485_ | -1.556868274 | -1.652924913 | -1.46751962 | 0.192471226 | 0.387703547 | 0.480098463 | 0.297135252 | 0.463256135 | 0.620414979 | 0.5755 | 0.702034776 |
| SL1344_RS05490_ | -0.8744648 | -0.752084455 | -1.020989219 | -0.26679456 | 0.264035973 | -0.965394585 | 0.135333362 | 0.220949186 | 0.377902687 | 0.3073 | 0.458676941 |
| SL1344_RS05495_ | 0.501673843 | 0.69470672 | 0.07648663 | -0.597949173 | 0.28905117 | -1.876511115 | 0.012658588 | 0.070289495 | 0.204783007 | 0.1124 | 0.25983287 |
| SL1344_RS05500_ | -1.132105522 | -1.107289521 | -1.193068504 | -0.076259839 | 0.371422732 | -0.186597393 | 0.428666669 | 0.650990193 | 0.774912687 | 0.7516 | 0.842609614 |
| SL1344_RS05505_ | -1.233474929 | -0.935467258 | -1.470122685 | -0.534175917 | 0.314875536 | -1.6258781 | 0.045333431 | 0.082917658 | 0.212186928 | 0.1538 | 0.308322169 |
| SL1344_RS05510_ | 0.550058919 | 0.588820744 | 0.518036602 | -0.066538854 | 0.157903883 | -0.397721117 | 0.329780154 | 0.473591067 | 0.624123298 | 0.5796 | 0.700449184 |
| SL1344_RS05515_ | 1.289115751 | 1.413960559 | 1.163738899 | -0.25233417 | 0.122550107 | -2.039717895 | 0.012658588 | 0.047324655 | 0.149639085 | 0.1148 | 0.261613923 |
| SL1344_RS05525_ | 0.956033836 | 0.995720043 | 0.919010192 | -0.072330317 | 0.131513826 | -0.514680566 | 0.28314458 | 0.408038539 | 0.560244886 | 0.5151 | 0.642721861 |
| SL1344_RS05530_ | -0.045807819 | 0.378368962 | -0.640275235 | -1.072326212 | 0.315320067 | -3.46202112 | 4.68E-05 | 0.009720904 | 0.063704403 | 0.1 | 0.243021319 |
| SL1344_RS05540_ | 2.81851838 | 2.673501727 | 2.912521066 | 0.232736825 | 0.100293463 | 2.192155799 | 0.00733396 | 0.041434709 | 0.145890197 | 0.1072 | 0.252968086 |
| SL1344_RS05545_ | -2.68181787 | -2.863874077 | -2.526335189 | 0.321445668 | 0.452860221 | 0.657068809 | 0.223333348 | 0.333584965 | 0.491937784 | 0.4335 | 0.572114343 |
| SL1344_RS05550_ | -3.917190188 | -3.785842595 | -4.048278251 | -0.25151189 | 0.803447073 | -0.299373436 | 0.365756168 | 0.520194547 | 0.667514239 | 0.6385 | 0.74958051 |
| SL1344_RS05555_ | -2.998663406 | -3.370579391 | -2.603800208 | 0.749446935 | 0.505427589 | 1.421159493 | 0.069953426 | 0.115029659 | 0.253786773 | 0.1825 | 0.337142106 |
| SL1344_RS05560_ | -3.679036912 | -3.515354093 | -4.152617615 | -0.548737489 | 1.031556602 | -0.471578343 | 0.342000007 | 0.533268214 | 0.688681615 | 0.6761 | 0.791180541 |
| SL1344_RS05565_ | -2.664987625 | -2.634396554 | -2.694560326 | -0.060596702 | 0.486525541 | -0.116765224 | 0.450666669 | 0.566392107 | 0.7057647 | 0.6848 | 0.787121499 |
| SL1344_RS05570_ | -0.055077454 | -0.109677316 | -0.000157885 | 0.132260102 | 0.279666499 | 0.432047595 | 0.314000008 | 0.482764978 | 0.643269428 | 0.6145 | 0.732592067 |
| SL1344_RS05580_ | 3.816347059 | 3.497314132 | 3.996782212 | 0.498763995 | 0.207126378 | 2.52869756 | 4.68E-05 | 0.056441592 | 0.186079711 | 0.1 | 0.243021319 |
| SL1344_RS05585_ | -5.156486157 | -4.869597786 | -5.566668105 | -0.673498084 | 1.195081547 | -0.543944593 | 0.267821463 | 0.386588599 | 0.548003171 | 0.4922 | 0.626195753 |
| SL1344_RS05590_ | 2.013800448 | 2.142401611 | 1.79929368 | -0.326820554 | 0.131552694 | -2.357088177 | 0.006667356 | 0.036366997 | 0.134533266 | 0.1036 | 0.247949541 |
| SL1344_RS05595_ | 1.771252803 | 1.800235213 | 1.728204804 | -0.069436742 | 0.146642206 | -0.452307078 | 0.317121927 | 0.515008089 | 0.667424432 | 0.6367 | 0.7522286 |
| SL1344_RS05600_ | 1.180445142 | 1.193326565 | 1.163200063 | 0.008442603 | 0.202157409 | 0.036858266 | 0.486666667 | 0.705322594 | 0.817719027 | 0.8495 | 0.905808926 |
| SL1344_RS05605_ | 2.40922643 | 2.403334874 | 2.418101937 | 0.013177333 | 0.104953896 | 0.113608611 | 0.452666668 | 0.630310084 | 0.758357654 | 0.7462 | 0.834682066 |
| SL1344_RS05610_ | -2.655361205 | -2.249353714 | -3.020064901 | -0.811973749 | 0.582839259 | -1.370704621 | 0.054000081 | 0.116020391 | 0.261658556 | 0.1742 | 0.328053782 |
| SL1344_RS05615_ | -5.500503646 | -5.703494075 | -5.379115081 | 0.360985357 | 1.522528536 | 0.206494219 | 0.399067292 | 0.547271406 | 0.694419838 | 0.672 | 0.778771947 |

| | | | | | | | | | | | |
|---|---|---|---|---|---|---|---|---|---|---|---|
| SL1344_RS05620_ | -4.808433592 | -4.500727902 | -5.162999594 | -0.63398498 | 1.018239393 | -0.592987108 | 0.249167234 | 0.36076272 | 0.518826472 | 0.4614 | 0.596675101 |
| SL1344_RS05625_ | -3.183567662 | -3.290375887 | -3.050571644 | 0.252825047 | 0.578376975 | 0.411071339 | 0.323333341 | 0.45104221 | 0.601147989 | 0.5602 | 0.67882816 |
| SL1344_RS05635_ | -3.574768771 | -3.335482035 | -3.87137889 | -0.550551928 | 0.708038957 | -0.719407546 | 0.20333335 | 0.312087934 | 0.474629005 | 0.4132 | 0.556173625 |
| SL1344_RS05650_ | 1.211190853 | 1.604775331 | 0.61968781 | -0.986025875 | 0.218896044 | -4.419840208 | 4.68E-05 | 0.014168972 | 0.080750917 | 0.1 | 0.243021319 |
| SL1344_RS05660_ | 0.399291889 | 0.087287051 | 0.657467797 | 0.572613131 | 0.166790492 | 3.400413035 | 0.00067292 | 0.012604907 | 0.070516668 | 0.1001 | 0.243160671 |
| SL1344_RS05670_ | 4.748621812 | 4.745977302 | 4.75539618 | 0.013137794 | 0.07392316 | 0.127487753 | 0.421718857 | 0.533160822 | 0.702916895 | 0.7721 | 0.857085054 |
| SL1344_RS05675_ | -0.157654266 | -0.224490512 | -0.100281821 | 0.127565449 | 0.194869308 | 0.61414726 | 0.252666679 | 0.379949816 | 0.535845791 | 0.4797 | 0.613044317 |
| SL1344_RS05680_ | -0.513730235 | -0.239347387 | -0.670321049 | -0.348536723 | 0.428418512 | -0.789623879 | 0.203197885 | 0.31051129 | 0.498522394 | 0.3966 | 0.567027893 |
| SL1344_RS05685_ | -0.134937902 | -0.396653501 | 0.081187999 | 0.474776103 | 0.199528499 | 2.350224476 | 0.004667653 | 0.032056706 | 0.120138304 | 0.1055 | 0.25015447 |
| SL1344_RS05690_ | 2.751047776 | 2.708024409 | 2.782782291 | 0.070840504 | 0.085730532 | 0.781707358 | 0.206000017 | 0.313377028 | 0.482853683 | 0.4078 | 0.560947027 |
| SL1344_RS05695_ | 3.235059996 | 3.302169856 | 3.215399392 | -0.077193605 | 0.088715563 | -0.714400484 | 0.265156573 | 0.401859675 | 0.576906384 | 0.5415 | 0.689199304 |
| SL1344_RS05700_ | -1.282088825 | -0.960793442 | -1.621543668 | -0.695585971 | 0.383556141 | -1.769781235 | 0.012658588 | 0.06155729 | 0.183392118 | 0.1174 | 0.26564813 |
| SL1344_RS05705_ | -2.329642875 | -2.239325 | -2.386535206 | -0.152769608 | 0.552698285 | -0.261903277 | 0.387075288 | 0.595603708 | 0.730433391 | 0.7108 | 0.805722274 |
| SL1344_RS05710_ | -3.010095739 | -3.127280688 | -2.903257792 | 0.206356407 | 0.56737755 | 0.352733751 | 0.34533334 | 0.50733581 | 0.653566764 | 0.6114 | 0.728127648 |
| SL1344_RS05715_ | 1.846757326 | 1.749704278 | 1.937536858 | 0.188528953 | 0.094603571 | 1.899360136 | 0.019333568 | 0.052231832 | 0.158027011 | 0.1225 | 0.2707227 |
| SL1344_RS05730_ | 3.061094462 | 3.086303688 | 3.019642704 | -0.044453802 | 0.150974559 | -0.235008584 | 0.407061962 | 0.804113459 | 0.886670298 | 0.8221 | 0.897300974 |
| SL1344_RS05735_ | 2.053468256 | 2.015989967 | 2.097818786 | 0.087450416 | 0.164379514 | 0.510929323 | 0.29646903 | 0.47416515 | 0.641472726 | 0.6103 | 0.73490524 |
| SL1344_RS05740_ | 0.834007064 | 1.140538591 | 0.699681612 | -0.417176136 | 0.228590378 | -1.708756091 | 0.031978821 | 0.068530175 | 0.199181166 | 0.1334 | 0.285809738 |
| SL1344_RS05745_ | -0.493428934 | -0.422354358 | -0.56482852 | -0.143201373 | 0.318654051 | -0.447516205 | 0.29380414 | 0.477842202 | 0.637890669 | 0.5947 | 0.718688657 |
| SL1344_RS05750_ | -0.114626073 | -0.12047762 | -0.105308015 | 0.024801949 | 0.2178848 | 0.11871007 | 0.453697537 | 0.592152425 | 0.727313898 | 0.7065 | 0.802525158 |
| SL1344_RS05755_ | 0.401161299 | 0.332131235 | 0.501135113 | 0.161921142 | 0.184076077 | 0.801352624 | 0.188000019 | 0.29820425 | 0.464481753 | 0.3938 | 0.544532792 |
| SL1344_RS05760_ | 0.866502605 | 0.861402581 | 0.873567786 | 0.022516356 | 0.185273925 | 0.125201327 | 0.455029982 | 0.652383067 | 0.775746283 | 0.7579 | 0.842033397 |
| SL1344_RS05765_ | 1.310171723 | 1.186194114 | 1.585482201 | 0.369108576 | 0.213609295 | 1.576595718 | 0.058627656 | 0.089317007 | 0.235630609 | 0.1605 | 0.321595865 |
| SL1344_RS05780_ | 4.029801777 | 4.583639094 | 3.576962075 | -1.008607651 | 0.11579271 | -8.494163025 | 4.68E-05 | 0.003575647 | 0.041110794 | 0.1 | 0.243021319 |
| SL1344_RS05785_ | 3.973523968 | 4.523586055 | 3.496875889 | -1.016581265 | 0.109591592 | -9.207515444 | 4.68E-05 | 0.000799649 | 0.019511564 | 0.1 | 0.243021319 |
| SL1344_RS05790_ | 3.537710028 | 4.05199206 | 3.001742434 | -1.041910424 | 0.074261154 | -13.98464132 | 4.68E-05 | 0.000250907 | 0.010865385 | 0.1 | 0.243021319 |
| SL1344_RS05800_ | 4.885416065 | 5.470073318 | 4.408407516 | -1.053790504 | 0.064668177 | -15.79503915 | 4.68E-05 | 0.001541289 | 0.027562432 | 0.1 | 0.243021319 |
| SL1344_RS05810_ | 5.322672406 | 5.926962459 | 4.795727729 | -1.130926504 | 0.077553843 | -14.29196198 | 4.68E-05 | 0.001334214 | 0.026137317 | 0.1 | 0.243021319 |
| SL1344_RS05815_ | 3.445989793 | 3.906542298 | 2.874819218 | -1.023076772 | 0.136631169 | -7.460242717 | 4.68E-05 | 0.000889432 | 0.020330782 | 0.1 | 0.243021319 |
| SL1344_RS05825_ | 3.324514754 | 3.774338263 | 2.757897954 | -0.993910481 | 0.218904143 | -4.518065053 | 4.68E-05 | 0.004493682 | 0.045767159 | 0.1 | 0.243021319 |
| SL1344_RS05845_ | -5.589013582 | -5.54247057 | -5.629167811 | -0.156265252 | 2.171701135 | -0.063094075 | 0.467021987 | 0.682321664 | 0.801365233 | 0.8123 | 0.881337066 |
| SL1344_RS05850_ | 3.40120245 | 3.248029019 | 3.53190096 | 0.291768865 | 0.090730621 | 3.243702295 | 4.68E-05 | 0.02120553 | 0.099946435 | 0.1 | 0.243021319 |
| SL1344_RS05855_ | 0.823530599 | 0.634419709 | 1.0149171 | 0.388348955 | 0.163018452 | 2.296195213 | 0.003334715 | 0.032693106 | 0.123529416 | 0.1036 | 0.24775754 |
| SL1344_RS05860_ | -1.489114258 | -1.821898375 | -1.236100834 | 0.567656955 | 0.404910834 | 1.343414222 | 0.063333401 | 0.109861474 | 0.258390277 | 0.174 | 0.333005623 |
| SL1344_RS05865_ | 0.714302097 | 0.871501016 | 0.501598382 | -0.368137677 | 0.157047889 | -2.276257049 | 0.006000767 | 0.038313816 | 0.134364533 | 0.1066 | 0.251828462 |
| SL1344_RS05870_ | 6.5633892 | 6.464311042 | 6.619659876 | 0.157795355 | 0.042308563 | 3.743489909 | 4.68E-05 | 0.006173412 | 0.05239475 | 0.1 | 0.243021319 |
| SL1344_RS05875_ | 6.723579622 | 6.635436212 | 6.857244979 | 0.220648969 | 0.043571913 | 4.763804205 | 4.68E-05 | 0.010262475 | 0.069114766 | 0.1 | 0.243021319 |
| SL1344_RS05880_ | 5.590019004 | 5.578082669 | 5.598849221 | 0.044604365 | 0.09514035 | 0.305118794 | 0.294666676 | 0.396506705 | 0.596180133 | 0.5958 | 0.730571708 |
| SL1344_RS05885_ | 4.140808748 | 3.921851124 | 4.230580323 | 0.278511622 | 0.163004999 | 1.714421617 | 0.027333498 | 0.063597834 | 0.197730611 | 0.125 | 0.277816062 |
| SL1344_RS05890_ | 4.218905631 | 4.185333185 | 4.333502339 | 0.144554727 | 0.126204232 | 1.095843239 | 0.122000033 | 0.204054545 | 0.398960724 | 0.2527 | 0.425799728 |
| SL1344_RS05900_ | 7.474322435 | 7.605849858 | 7.31820046 | -0.291612433 | 0.093132039 | -2.83311551 | 4.68E-05 | 0.071768852 | 0.215600303 | 0.1 | 0.243021319 |
| SL1344_RS05905_ | 6.756949771 | 6.756650287 | 6.757374772 | -0.004257898 | 0.103887256 | -0.051239122 | 0.480346436 | 0.890612072 | 0.940840107 | 0.9574 | 0.975758212 |
| SL1344_RS05910_ | 1.369519968 | 1.311140919 | 1.435524898 | 0.121448902 | 0.125843397 | 0.920840059 | 0.166666689 | 0.259636698 | 0.421528765 | 0.3508 | 0.502533202 |
| SL1344_RS05915_ | 1.638045762 | 1.671911084 | 1.601445667 | -0.07664337 | 0.137057413 | -0.533017993 | 0.266666678 | 0.425592532 | 0.589197195 | 0.5486 | 0.677094781 |
| SL1344_RS05920_ | 1.735823559 | 1.765478154 | 1.69121975 | -0.077584135 | 0.126657015 | -0.552191443 | 0.275816133 | 0.416696648 | 0.578911257 | 0.5318 | 0.663327095 |
| SL1344_RS05925_ | 2.027157722 | 2.035554034 | 2.018561087 | -0.022032652 | 0.124199462 | -0.175031947 | 0.421052635 | 0.598860833 | 0.733512274 | 0.7269 | 0.816491994 |
| SL1344_RS05930_ | 2.044523201 | 1.935023987 | 2.136444957 | 0.195915749 | 0.168433469 | 1.142616738 | 0.110592975 | 0.153547305 | 0.326097825 | 0.2251 | 0.397730822 |
| SL1344_RS05935_ | 4.851892741 | 4.780541009 | 4.954322706 | 0.172485802 | 0.041399255 | 4.060196418 | 4.68E-05 | 0.011518487 | 0.068757839 | 0.1 | 0.243021319 |
| SL1344_RS05940_ | -0.065410704 | -0.079808115 | -0.044408714 | 0.017198151 | 0.243764334 | 0.063423481 | 0.473017989 | 0.649716959 | 0.77376356 | 0.7606 | 0.844958425 |
| SL1344_RS05945_ | 1.806534834 | 1.84205634 | 1.754768932 | -0.08984941 | 0.137907118 | -0.637074811 | 0.241172565 | 0.374038049 | 0.545287285 | 0.4955 | 0.634985856 |
| SL1344_RS05950_ | 1.966940976 | 2.009850169 | 1.890344701 | -0.111945709 | 0.132841043 | -0.790310013 | 0.19920055 | 0.303814259 | 0.481185045 | 0.4061 | 0.562484942 |
| SL1344_RS05955_ | 1.253527538 | 1.386489759 | 1.05833838 | -0.358554223 | 0.209843206 | -1.654056218 | 0.021333546 | 0.068396237 | 0.199107195 | 0.125 | 0.275214264 |
| SL1344_RS05960_ | 0.956928533 | 1.001924198 | 0.886080156 | -0.111788044 | 0.203520807 | -0.532421187 | 0.287808137 | 0.444616358 | 0.613307695 | 0.5739 | 0.70561202 |

| | | | | | | | | | | | |
|---|---|---|---|---|---|---|---|---|---|---|---|
| SL1344_RS05965_ | 0.952232395 | 1.084518745 | 0.767977552 | -0.328748271 | 0.186186728 | -1.726787109 | 0.016655836 | 0.084290176 | 0.224805338 | 0.1181 | 0.266582459 |
| SL1344_RS05970_ | 1.794238703 | 1.899338894 | 1.68706441 | -0.212178377 | 0.112352499 | -1.785888065 | 0.021333546 | 0.061251524 | 0.178089236 | 0.1275 | 0.277577125 |
| SL1344_RS05975_ | 3.446270425 | 3.355833452 | 3.487792691 | 0.130111847 | 0.097412867 | 1.236414151 | 0.081333385 | 0.14643012 | 0.317305814 | 0.2082 | 0.372892374 |
| SL1344_RS05980_ | -1.588488969 | -1.638950491 | -1.524288233 | 0.106591084 | 0.33511507 | 0.318033817 | 0.363091278 | 0.51601389 | 0.65975491 | 0.6321 | 0.740240262 |
| SL1344_RS05985_ | 2.333905657 | 2.432410434 | 2.217228762 | -0.223597888 | 0.115199887 | -1.928946229 | 0.012658588 | 0.059609367 | 0.18064842 | 0.1093 | 0.255369433 |
| SL1344_RS05990_ | 3.081646656 | 3.214459038 | 2.844259382 | -0.325133007 | 0.329967309 | -0.874575647 | 0.168000022 | 0.262100073 | 0.466291183 | 0.2969 | 0.485573634 |
| SL1344_RS05995_ | -0.403890327 | -0.365045031 | -0.446170454 | -0.073737885 | 0.286181534 | -0.233790157 | 0.392000004 | 0.586808092 | 0.722648981 | 0.693 | 0.793108776 |
| SL1344_RS06000_ | 3.372742289 | 3.261846612 | 3.426108545 | 0.157701403 | 0.084013901 | 1.693820446 | 0.03730858 | 0.077446313 | 0.210705256 | 0.1443 | 0.298243856 |
| SL1344_RS06005_ | 2.579303134 | 2.51770656 | 2.653199833 | 0.136925879 | 0.08372443 | 1.5743743 | 0.041333441 | 0.08985535 | 0.220560233 | 0.1513 | 0.302970634 |
| SL1344_RS06010_ | 1.647132693 | 1.512579522 | 1.709145762 | 0.197870601 | 0.20092524 | 0.978341745 | 0.150000025 | 0.208224366 | 0.391653932 | 0.3011 | 0.480000236 |
| SL1344_RS06015_ | 1.968053682 | 1.992645875 | 1.931221683 | -0.025737879 | 0.167451449 | -0.142309112 | 0.448367757 | 0.743887089 | 0.84538411 | 0.8608 | 0.915766222 |
| SL1344_RS06020_ | 2.173372304 | 2.166905888 | 2.188130474 | 0.009270452 | 0.133148505 | 0.062725 | 0.466666668 | 0.719978955 | 0.827641738 | 0.8262 | 0.892488905 |
| SL1344_RS06025_ | 2.199235873 | 2.168908117 | 2.249470358 | 0.07908221 | 0.100871103 | 0.705511436 | 0.235333347 | 0.359615873 | 0.524221673 | 0.4689 | 0.611228356 |
| SL1344_RS06030_ | 3.233782165 | 3.218526033 | 3.247984112 | 0.03990818 | 0.098744298 | 0.389929756 | 0.338666673 | 0.50936392 | 0.666239023 | 0.6313 | 0.746826867 |
| SL1344_RS06035_ | 1.954584254 | 1.828187629 | 2.09924641 | 0.265679577 | 0.11464244 | 2.27833319 | 0.006667356 | 0.033862982 | 0.127388749 | 0.1045 | 0.249338341 |
| SL1344_RS06045_ | 2.457943898 | 2.446013849 | 2.485188094 | 0.046335558 | 0.13003741 | 0.32123529 | 0.378666671 | 0.571408811 | 0.719825364 | 0.7251 | 0.822109565 |
| SL1344_RS06050_ | 2.831335568 | 2.8472184 | 2.796819213 | -0.057947945 | 0.10305234 | -0.503815993 | 0.286666676 | 0.445442294 | 0.612938665 | 0.5846 | 0.707042672 |
| SL1344_RS06055_ | -0.826837664 | -0.586435829 | -1.051879309 | -0.468410122 | 0.357736804 | -1.257507183 | 0.086666715 | 0.131422811 | 0.29033916 | 0.2007 | 0.364321022 |
| SL1344_RS06070_ | 2.861326668 | 2.584695081 | 3.234793718 | 0.622779881 | 0.14819443 | 4.172537857 | 4.68E-05 | 0.004021914 | 0.042361453 | 0.1 | 0.243021319 |
| SL1344_RS06085_ | 3.340792371 | 3.099274762 | 3.562178479 | 0.461685518 | 0.073009477 | 6.368866439 | 4.68E-05 | 0.002067523 | 0.028661356 | 0.1 | 0.243021319 |
| SL1344_RS06090_ | 2.43589244 | 2.191812761 | 2.680367879 | 0.489823543 | 0.098406374 | 5.05390472 | 4.68E-05 | 0.005267231 | 0.044855508 | 0.1 | 0.243021319 |
| SL1344_RS06095_ | 4.094690944 | 3.881012428 | 4.286247954 | 0.419426517 | 0.079144409 | 5.359040283 | 4.68E-05 | 0.003952082 | 0.040342255 | 0.1 | 0.243021319 |
| SL1344_RS06100_ | 0.274304641 | 0.018334242 | 0.494223203 | 0.47973381 | 0.172885924 | 2.737518509 | 0.002000966 | 0.024947583 | 0.103808498 | 0.1009 | 0.244287376 |
| SL1344_RS06105_ | 0.025630264 | -0.176328801 | 0.203604212 | 0.376323931 | 0.198435131 | 1.811260186 | 0.020000227 | 0.060289314 | 0.176942206 | 0.1263 | 0.275853209 |
| SL1344_RS06110_ | 0.520618972 | 0.453420289 | 0.676433185 | 0.217269624 | 0.233551711 | 0.82915335 | 0.200666684 | 0.28453004 | 0.467768269 | 0.4055 | 0.56692862 |
| SL1344_RS06115_ | 4.522265734 | 4.763969225 | 4.249544598 | -0.51582505 | 0.05276388 | -9.858793537 | 4.68E-05 | 0.002055235 | 0.028299392 | 0.1 | 0.243021319 |
| SL1344_RS06120_ | 0.110487906 | 0.326634817 | -0.167405909 | -0.492154632 | 0.194947939 | -2.486226667 | 0.001335878 | 0.031430388 | 0.119277369 | 0.1024 | 0.246405611 |
| SL1344_RS06125_ | -3.739881254 | -2.773496839 | -5.534383341 | -1.059773244 | 3.72707229 | -0.29378852 | 0.441705532 | 0.647661557 | 0.783962935 | 0.9121 | 0.949227961 |
| SL1344_RS06130_ | -2.466421165 | -2.529719383 | -2.436348309 | 0.049430995 | 0.587462022 | 0.073971536 | 0.470666668 | 0.655305182 | 0.778134273 | 0.7629 | 0.848033642 |
| SL1344_RS06135_ | 2.521472273 | 2.606273603 | 2.485729665 | -0.110244245 | 0.151463423 | -0.712437633 | 0.225849448 | 0.344506386 | 0.532407977 | 0.4303 | 0.605871735 |
| SL1344_RS06140_ | -0.132571618 | -0.308062268 | 0.260040914 | 0.519231628 | 0.295470587 | 1.621885352 | 0.048634334 | 0.08859871 | 0.230853656 | 0.1601 | 0.318850574 |
| SL1344_RS06150_ | -5.416414261 | -6.048717476 | -4.726210309 | 1.192617377 | 1.847745888 | 0.591640591 | 0.230000014 | 0.390064575 | 0.558022973 | 0.4581 | 0.608787738 |
| SL1344_RS06155_ | -6.571652244 | -6.936394061 | -6.264682051 | 1.140739915 | 3.222270296 | 0.327184675 | 0.351333339 | 0.532567404 | 0.686038675 | 0.6792 | 0.785776812 |
| SL1344_RS06160_ | -3.55400914 | -3.578123732 | -3.501120127 | 0.036422413 | 0.796317515 | 0.04660978 | 0.480346436 | 0.638600662 | 0.764930123 | 0.748 | 0.836711915 |
| SL1344_RS06165_ | -0.014767358 | -0.079010337 | 0.139583697 | 0.189446097 | 0.269204669 | 0.629860696 | 0.252000012 | 0.422181592 | 0.590849188 | 0.5288 | 0.674117357 |
| SL1344_RS06170_ | -2.794811114 | -3.005371524 | -2.594856205 | 0.438795804 | 0.568920112 | 0.734831803 | 0.216000016 | 0.325229183 | 0.492915019 | 0.4381 | 0.580332019 |
| SL1344_RS06180_ | 0.314930908 | 0.104021125 | 0.555845516 | 0.449400863 | 0.173070151 | 2.541559046 | 0.001335878 | 0.026540576 | 0.107609351 | 0.1025 | 0.246457709 |
| SL1344_RS06185_ | -4.857082601 | -4.856417659 | -4.857572392 | -0.029852215 | 1.105182054 | -0.022792812 | 0.489007329 | 0.582528382 | 0.720415481 | 0.6931 | 0.792801697 |
| SL1344_RS06190_ | -3.32402273 | -3.586632152 | -3.010607819 | 0.528644537 | 0.761316185 | 0.660438996 | 0.23866668 | 0.378402199 | 0.546270757 | 0.4749 | 0.620296528 |
| SL1344_RS06195_ | -0.502032961 | -0.469848296 | -0.568388253 | -0.080738661 | 0.268290921 | -0.273440767 | 0.387075288 | 0.571487869 | 0.713466329 | 0.688 | 0.79220189 |
| SL1344_RS06200_ | -2.044869655 | -1.982412766 | -2.125283622 | -0.139378838 | 0.37670489 | -0.333716389 | 0.355096608 | 0.483571204 | 0.630913174 | 0.6056 | 0.717755097 |
| SL1344_RS06205_ | 2.13097504 | 2.667890725 | 1.764787216 | -0.917064458 | 0.252741579 | -3.60048746 | 4.68E-05 | 0.006343937 | 0.054643203 | 0.1 | 0.243021319 |
| SL1344_RS06210_ | -2.754051255 | -2.587817325 | -2.98496425 | -0.419423662 | 0.616692484 | -0.617950776 | 0.246666679 | 0.372477172 | 0.541559092 | 0.4869 | 0.624735443 |
| SL1344_RS06215_ | -0.553453516 | -0.583520893 | -0.508567687 | 0.075396853 | 0.24631044 | 0.282655189 | 0.373750838 | 0.540743077 | 0.683012232 | 0.6516 | 0.758515393 |
| SL1344_RS06220_ | -0.136279878 | -0.36941085 | 0.227433062 | 0.570050019 | 0.270172278 | 2.013232197 | 0.012000381 | 0.048576063 | 0.160122903 | 0.1138 | 0.261581563 |
| SL1344_RS06225_ | -0.013336038 | -0.390860301 | 0.325896192 | 0.715123488 | 0.223567815 | 3.177621731 | 0.00067292 | 0.014117484 | 0.076838 | 0.1 | 0.243021319 |
| SL1344_RS06230_ | 0.265121656 | 0.216571269 | 0.320371162 | 0.102462985 | 0.171566963 | 0.571830514 | 0.270000011 | 0.401058202 | 0.552414094 | 0.5008 | 0.632903977 |
| SL1344_RS06235_ | -1.082386865 | -1.116862064 | -1.055008134 | 0.048381315 | 0.439539129 | 0.111620973 | 0.449700202 | 0.695470406 | 0.810680125 | 0.8076 | 0.879321508 |
| SL1344_RS06240_ | -2.238642624 | -2.342664676 | -2.144968493 | 0.200377907 | 0.409378112 | 0.471227141 | 0.301132587 | 0.438936653 | 0.591987505 | 0.5492 | 0.673772422 |
| SL1344_RS06245_ | -2.236757968 | -2.273664552 | -2.177649579 | 0.136760057 | 0.505542985 | 0.244579886 | 0.393333337 | 0.570076833 | 0.710568463 | 0.6906 | 0.79074108 |
| SL1344_RS06250_ | -2.097119922 | -2.226738099 | -2.004431628 | 0.215027715 | 0.440749092 | 0.466126614 | 0.301333342 | 0.466021517 | 0.620071642 | 0.573 | 0.697005887 |
| SL1344_RS06255_ | -1.122880277 | -1.141859007 | -1.092694948 | 0.06872526 | 0.384342871 | 0.184951823 | 0.428381082 | 0.644179189 | 0.770186297 | 0.7601 | 0.844143635 |
| SL1344_RS06260_ | -1.63565243 | -2.250546934 | -1.213342325 | 1.040651475 | 0.372572164 | 2.702790226 | 0.001335878 | 0.024642346 | 0.104240854 | 0.1017 | 0.245346965 |

| | | | | | | | | | | | |
|---|---|---|---|---|---|---|---|---|---|---|---|
| SL1344_RS06265_ | -3.157927996 | -3.625775929 | -2.718163623 | 0.914850393 | 0.583001288 | 1.499997693 | 0.048634334 | 0.098444027 | 0.231115465 | 0.166 | 0.318415943 |
| SL1344_RS06270_ | -0.301906348 | -0.471163474 | -0.178364273 | 0.297109067 | 0.272019729 | 1.053293329 | 0.116666701 | 0.188865984 | 0.357553634 | 0.265 | 0.423199781 |
| SL1344_RS06275_ | -0.010001864 | -0.100254047 | 0.209440908 | 0.333980219 | 0.394770805 | 0.830422431 | 0.197868106 | 0.272683853 | 0.465121776 | 0.3729 | 0.552397792 |
| SL1344_RS06280_ | -3.11073832 | -3.177683219 | -3.056653327 | 0.111613395 | 0.67338113 | 0.152016845 | 0.435333336 | 0.61349473 | 0.745081796 | 0.722 | 0.816226908 |
| SL1344_RS06285_ | 0.730934817 | 0.298966091 | 1.222938271 | 0.90995067 | 0.188944766 | 4.769350668 | 4.68E-05 | 0.004500856 | 0.041923422 | 0.1 | 0.243021319 |
| SL1344_RS06290_ | -0.269118991 | -0.479529577 | 0.022426173 | 0.473354459 | 0.394566773 | 1.173522968 | 0.114666702 | 0.149679324 | 0.321478158 | 0.2214 | 0.395822737 |
| SL1344_RS06295_ | -4.845530006 | -5.179023368 | -4.365513293 | 0.767121333 | 1.959352574 | 0.370832787 | 0.29646903 | 0.532953202 | 0.687088019 | 0.5896 | 0.722521274 |
| SL1344_RS06300_ | -0.17499263 | -0.950063216 | 0.430774655 | 1.36817524 | 0.307732214 | 4.395619214 | 4.68E-05 | 0.010078837 | 0.064723926 | 0.1 | 0.243021319 |
| SL1344_RS06305_ | 1.142264227 | 0.963266673 | 1.548899791 | 0.555707306 | 0.294877673 | 1.903357327 | 0.008000574 | 0.044268517 | 0.157032761 | 0.1077 | 0.253650613 |
| SL1344_RS06310_ | -1.264466702 | -1.424571696 | -1.056294507 | 0.369931542 | 0.31510664 | 1.097146907 | 0.124000032 | 0.183264731 | 0.342007762 | 0.2676 | 0.427784488 |
| SL1344_RS06315_ | 0.232822525 | 0.203386963 | 0.263025706 | 0.075958092 | 0.219955638 | 0.346974348 | 0.355096608 | 0.538797946 | 0.685226872 | 0.6736 | 0.775428231 |
| SL1344_RS06320_ | -1.902267233 | -2.022484456 | -1.741200597 | 0.313207915 | 0.430441978 | 0.699064467 | 0.217854779 | 0.334997612 | 0.503886206 | 0.4495 | 0.589396649 |
| SL1344_RS06325_ | -2.521205139 | -2.390493505 | -2.700504708 | -0.311918063 | 0.490282107 | -0.612319323 | 0.254666679 | 0.374677742 | 0.537832533 | 0.4866 | 0.623229723 |
| SL1344_RS06330_ | 0.388006802 | 0.502692219 | 0.263939101 | -0.274152798 | 0.302115189 | -0.942245207 | 0.112000036 | 0.224201316 | 0.414926416 | 0.2507 | 0.413427127 |
| SL1344_RS06335_ | -0.503554063 | -0.711996634 | -0.329530939 | 0.374553051 | 0.24236858 | 1.494790532 | 0.045969449 | 0.09458681 | 0.228998066 | 0.1563 | 0.309535503 |
| SL1344_RS06340_ | 1.259563801 | 0.93987263 | 1.644930393 | 0.69658374 | 0.180678054 | 3.772024064 | 4.68E-05 | 0.010843003 | 0.069050483 | 0.1 | 0.243021319 |
| SL1344_RS06345_ | 0.718039249 | 0.520111081 | 0.885847332 | 0.351645544 | 0.203602988 | 1.639191661 | 0.048634334 | 0.081248598 | 0.216781678 | 0.1437 | 0.300394102 |
| SL1344_RS06350_ | -0.692137418 | -0.838821979 | -0.571135166 | 0.269947127 | 0.306776712 | 0.84305785 | 0.17800002 | 0.27184269 | 0.446192935 | 0.3636 | 0.520528609 |
| SL1344_RS06355_ | 0.279427072 | 0.268113396 | 0.288720581 | 0.012053645 | 0.212280386 | 0.054552845 | 0.473684212 | 0.641609894 | 0.765878542 | 0.7442 | 0.832949726 |
| SL1344_RS06360_ | -1.260474986 | -1.533074306 | -0.995646339 | 0.568585252 | 0.397230601 | 1.41841166 | 0.041305903 | 0.09817155 | 0.241305902 | 0.1522 | 0.305772038 |
| SL1344_RS06365_ | -4.306746368 | -4.58096403 | -4.023182725 | 0.543214993 | 0.827525491 | 0.588880505 | 0.24266668 | 0.37738965 | 0.533191663 | 0.4753 | 0.611323522 |
| SL1344_RS06370_ | -0.501477989 | -0.730892918 | -0.265998044 | 0.463998444 | 0.24634671 | 1.825539318 | 0.021333546 | 0.058656138 | 0.172529011 | 0.1236 | 0.27316986 |
| SL1344_RS06375_ | -0.589642752 | -0.792340441 | -0.364131579 | 0.423971257 | 0.252389921 | 1.648232033 | 0.03664236 | 0.078161779 | 0.202641626 | 0.1441 | 0.296702957 |
| SL1344_RS06380_ | 0.083711977 | -0.045508994 | 0.228325218 | 0.268972468 | 0.199419702 | 1.289643624 | 0.086666715 | 0.134448618 | 0.283857194 | 0.2092 | 0.367754171 |
| SL1344_RS06385_ | -0.285834653 | -0.289200657 | -0.283072576 | 0.046474569 | 0.326352232 | 0.14474185 | 0.444000002 | 0.677987512 | 0.797424796 | 0.795 | 0.868796733 |
| SL1344_RS06390_ | 3.903993018 | 3.903773527 | 3.904332936 | -0.002668628 | 0.122123359 | -0.018439298 | 0.49 | 0.810345621 | 0.890987828 | 0.9044 | 0.943096909 |
| SL1344_RS06395_ | -1.201490174 | -1.257003954 | -1.128210247 | 0.159701117 | 0.385745006 | 0.377235624 | 0.334666674 | 0.508655568 | 0.658667135 | 0.6366 | 0.744784992 |
| SL1344_RS06400_ | -0.409898556 | -0.363834678 | -0.487265876 | -0.097351804 | 0.297198007 | -0.285742226 | 0.376415728 | 0.602445781 | 0.73883042 | 0.7116 | 0.811867176 |
| SL1344_RS06405_ | 1.377194852 | 1.254795128 | 1.518772828 | 0.268745806 | 0.129665252 | 2.040694527 | 0.008000574 | 0.046846228 | 0.151083112 | 0.112 | 0.258137899 |
| SL1344_RS06410_ | 7.383843088 | 7.211974654 | 7.536049854 | 0.342027415 | 0.057554755 | 6.022806984 | 4.68E-05 | 0.000695966 | 0.019631777 | 0.1 | 0.243021319 |
| SL1344_RS06415_ | 2.123300412 | 2.090000066 | 2.168468798 | 0.026515099 | 0.19783223 | 0.127632123 | 0.456666668 | 0.779953492 | 0.87108966 | 0.8873 | 0.934125026 |
| SL1344_RS06420_ | 2.198738733 | 2.336390765 | 2.098907504 | -0.234827994 | 0.14850161 | -1.576469795 | 0.038000117 | 0.077930163 | 0.215282452 | 0.1369 | 0.291327204 |
| SL1344_RS06425_ | 0.667050053 | 0.489380396 | 0.904792145 | 0.413391855 | 0.171205802 | 2.376312146 | 0.005330642 | 0.035677236 | 0.131285015 | 0.1035 | 0.247812696 |
| SL1344_RS06430_ | 0.764103687 | 0.606835515 | 0.97486426 | 0.356375114 | 0.247682309 | 1.361478169 | 0.066000065 | 0.111651022 | 0.267958631 | 0.1733 | 0.336848037 |
| SL1344_RS06435_ | 2.419947956 | 2.214891817 | 2.634377025 | 0.421134412 | 0.105205203 | 4.023281259 | 4.68E-05 | 0.008528024 | 0.058921287 | 0.1 | 0.243021319 |
| SL1344_RS06440_ | 0.985569403 | 0.817843271 | 1.038781774 | 0.210769404 | 0.190822484 | 0.962565315 | 0.175216543 | 0.249088003 | 0.432815619 | 0.3599 | 0.522557468 |
| SL1344_RS06445_ | 2.206732635 | 2.026260925 | 2.376885051 | 0.357354593 | 0.111587527 | 3.183934689 | 4.68E-05 | 0.01476944 | 0.079836975 | 0.1 | 0.243021319 |
| SL1344_RS06450_ | 3.13251069 | 2.930792572 | 3.250751822 | 0.302072966 | 0.122158021 | 2.358198353 | 0.001336769 | 0.029940977 | 0.123754964 | 0.1022 | 0.246110526 |
| SL1344_RS06455_ | 1.120016391 | 0.903674041 | 1.293004609 | 0.383453529 | 0.168016916 | 2.270862866 | 0.008000574 | 0.0343158 | 0.129480106 | 0.1053 | 0.250354372 |
| SL1344_RS06465_ | -0.623270824 | -1.281739433 | -0.225893602 | 1.045108224 | 0.321681015 | 3.077746513 | 4.68E-05 | 0.019980688 | 0.094527374 | 0.1002 | 0.243296074 |
| SL1344_RS06470_ | 1.781958417 | 1.614639511 | 1.977768195 | 0.366939091 | 0.126763705 | 2.891655906 | 4.68E-05 | 0.020345612 | 0.095222454 | 0.1006 | 0.243852856 |
| SL1344_RS06475_ | -2.633753292 | -2.697201324 | -2.558224273 | 0.171775233 | 0.499713283 | 0.316081541 | 0.362000006 | 0.519223193 | 0.6651141 | 0.6279 | 0.74023766 |
| SL1344_RS06480_ | -3.410770018 | -3.069410991 | -3.709327532 | -0.591416125 | 0.732161827 | -0.756432899 | 0.199866773 | 0.304759131 | 0.474916716 | 0.4021 | 0.556215246 |
| SL1344_RS06485_ | -2.297515902 | -2.232198764 | -2.34550976 | -0.098860201 | 0.437429746 | -0.200335091 | 0.41066667 | 0.551754244 | 0.691252298 | 0.6644 | 0.769122785 |
| SL1344_RS06490_ | -2.145552071 | -2.150047269 | -2.141377238 | 0.002600593 | 0.442119843 | 0.004889764 | 0.495333334 | 0.60817022 | 0.739020298 | 0.7173 | 0.811006508 |
| SL1344_RS06495_ | -2.477172027 | -2.524441454 | -2.425223524 | 0.122155645 | 0.534372811 | 0.208818384 | 0.40906063 | 0.584779823 | 0.719524897 | 0.7004 | 0.794353824 |
| SL1344_RS06500_ | -0.747961961 | -0.905396135 | -0.555138203 | 0.364577616 | 0.280611544 | 1.226694126 | 0.079333387 | 0.140771619 | 0.29703368 | 0.2062 | 0.364969184 |
| SL1344_RS06505_ | 0.566019241 | 0.543832863 | 0.600187219 | 0.070831079 | 0.197981932 | 0.318278992 | 0.356666672 | 0.524429961 | 0.673960455 | 0.6594 | 0.76331685 |
| SL1344_RS06510_ | 1.968907294 | 1.687214737 | 2.147791493 | 0.457335405 | 0.161682893 | 2.743934318 | 0.000673369 | 0.01715956 | 0.089209102 | 0.1002 | 0.243303507 |
| SL1344_RS06515_ | 1.378814278 | 1.502394073 | 1.347377635 | -0.143404824 | 0.172156362 | -0.698585196 | 0.260000012 | 0.398231606 | 0.575167177 | 0.5439 | 0.68572803 |
| SL1344_RS06520_ | -0.174762168 | -0.189066305 | -0.144085075 | 0.060631965 | 0.280317295 | 0.193204086 | 0.42066667 | 0.628365264 | 0.75677466 | 0.7347 | 0.826053588 |
| SL1344_RS06525_ | -1.102258061 | -1.126516441 | -1.086658112 | 0.029617602 | 0.406201992 | 0.063718372 | 0.475016656 | 0.695770757 | 0.810061642 | 0.8031 | 0.8776529 |
| SL1344_RS06530_ | -1.281564465 | -1.456667222 | -1.163016496 | 0.282289975 | 0.377242235 | 0.739508712 | 0.215333349 | 0.33003581 | 0.50400736 | 0.4434 | 0.594541599 |

| | | | | | | | | | | | |
|---|---|---|---|---|---|---|---|---|---|---|---|
| SL1344_RS06535_ | -0.540358647 | -0.713506975 | -0.306764423 | 0.335681096 | 0.346027974 | 0.928668948 | 0.175333354 | 0.252515459 | 0.432417128 | 0.3397 | 0.50708182 |
| SL1344_RS06540_ | -1.436673066 | -1.461842868 | -1.409556228 | 0.051948981 | 0.301482972 | 0.166405802 | 0.428666669 | 0.53579309 | 0.677097286 | 0.6577 | 0.759560502 |
| SL1344_RS06545_ | 0.06156413 | -0.063256155 | 0.282658676 | 0.321956086 | 0.25972263 | 1.135250495 | 0.121918754 | 0.173406659 | 0.343030466 | 0.2581 | 0.426998288 |
| SL1344_RS06550_ | -0.674905196 | -0.447044522 | -1.099593163 | -0.630934491 | 0.37062924 | -1.567409399 | 0.043970787 | 0.089766137 | 0.233598306 | 0.1499 | 0.307240639 |
| SL1344_RS06555_ | -0.162848775 | -0.378360884 | 0.123961312 | 0.500866418 | 0.205699667 | 2.298034237 | 0.005996769 | 0.033303428 | 0.12339955 | 0.1072 | 0.252258407 |
| SL1344_RS06565_ | -0.502623933 | -0.84808471 | -0.281387901 | 0.573967508 | 0.338464191 | 1.620240799 | 0.026666836 | 0.088864166 | 0.231586138 | 0.1338 | 0.285537008 |
| SL1344_RS06570_ | 1.882719952 | 1.770246197 | 2.096729469 | 0.311625761 | 0.133461653 | 2.192524779 | 0.006662914 | 0.042264179 | 0.146424317 | 0.109 | 0.254765734 |
| SL1344_RS06575_ | 0.181944388 | 0.010399707 | 0.362944901 | 0.359413315 | 0.22425502 | 1.552813339 | 0.043970787 | 0.089284579 | 0.225373351 | 0.1476 | 0.300763647 |
| SL1344_RS06580_ | -0.83713908 | -0.849296218 | -0.823207721 | 0.028803224 | 0.261642271 | 0.099760178 | 0.456000002 | 0.56062459 | 0.70009311 | 0.6788 | 0.781507997 |
| SL1344_RS06605_ | -2.814099571 | -2.561121318 | -3.022866493 | -0.460179917 | 0.624306549 | -0.717613098 | 0.230513005 | 0.361080366 | 0.530604704 | 0.4562 | 0.60512766 |
| SL1344_RS06615_ | -3.351446524 | -3.486346047 | -3.186893252 | 0.302885935 | 0.627816757 | 0.46671902 | 0.309333342 | 0.43467503 | 0.589048412 | 0.5425 | 0.667213677 |
| SL1344_RS06630_ | 8.358683843 | 8.174330078 | 8.403690983 | 0.225518468 | 0.147081775 | 1.690535807 | 4.68E-05 | 0.037749116 | 0.147643548 | 0.1 | 0.243021319 |
| SL1344_RS06635_ | 8.603194107 | 8.310738555 | 8.686672994 | 0.342260723 | 0.083112401 | 1.824090604 | 4.68E-05 | 0.075435876 | 0.222082075 | 0.1 | 0.243021319 |
| SL1344_RS06640_ | 8.538086351 | 8.438154705 | 8.634592138 | 0.183471185 | 0.045764161 | 4.055169387 | 4.68E-05 | 0.00326388 | 0.039781293 | 0.1 | 0.243021319 |
| SL1344_RS06645_ | 3.193451813 | 3.098723365 | 3.3975363 | 0.292403613 | 0.112941645 | 2.35424953 | 0.002666616 | 0.044544846 | 0.156712953 | 0.1014 | 0.244929331 |
| SL1344_RS06650_ | 4.342975796 | 4.325203714 | 4.435121674 | 0.112781539 | 0.238710921 | 0.492018264 | 0.334666674 | 0.619853959 | 0.770250669 | 0.7024 | 0.830957576 |
| SL1344_RS06660_ | 1.251304146 | 1.027291404 | 1.486486464 | 0.457867257 | 0.117444914 | 3.849209333 | 4.68E-05 | 0.008773473 | 0.058228898 | 0.1 | 0.243021319 |
| SL1344_RS06665_ | -0.361829608 | -0.372913787 | -0.345660601 | 0.03329469 | 0.236982481 | 0.130784242 | 0.449700202 | 0.593682285 | 0.725962429 | 0.6972 | 0.792991919 |
| SL1344_RS06670_ | -0.233530974 | -0.222825997 | -0.242847542 | 0.029957825 | 0.322376334 | 0.088876446 | 0.463333335 | 0.691057965 | 0.806367468 | 0.822 | 0.886966121 |
| SL1344_RS06675_ | -0.10710813 | -0.465283616 | 0.308073112 | 0.75826137 | 0.236496385 | 3.149313691 | 0.000673369 | 0.015716998 | 0.082856731 | 0.1002 | 0.243304792 |
| SL1344_RS06685_ | 0.942052453 | 0.784689801 | 1.103266236 | 0.363250809 | 0.299353901 | 1.220088452 | 0.082000051 | 0.131975038 | 0.300223963 | 0.1966 | 0.36453451 |
| SL1344_RS06690_ | 1.605872596 | 1.205568128 | 1.963015562 | 0.753064164 | 0.13334317 | 5.642622342 | 4.68E-05 | 0.002982105 | 0.033853103 | 0.1 | 0.243021319 |
| SL1344_RS06695_ | 2.022769265 | 1.824391481 | 2.213777223 | 0.395393673 | 0.121124301 | 3.257988148 | 4.68E-05 | 0.012307896 | 0.072727703 | 0.1001 | 0.243159376 |
| SL1344_RS06700_ | -0.564758325 | -0.758502616 | -0.377695877 | 0.380914936 | 0.237434286 | 1.56896333 | 0.041305903 | 0.085351929 | 0.212938943 | 0.1504 | 0.301874044 |
| SL1344_RS06705_ | 1.759565172 | 1.793009293 | 1.709308872 | -0.096226827 | 0.13627017 | -0.675904925 | 0.214000016 | 0.336689926 | 0.510492344 | 0.4596 | 0.598064277 |
| SL1344_RS06710_ | -1.477026957 | -1.602322212 | -1.329760385 | 0.274952812 | 0.320306405 | 0.773814219 | 0.187874769 | 0.291138759 | 0.451180993 | 0.3861 | 0.533033805 |
| SL1344_RS06715_ | -4.481970802 | -5.271389456 | -4.213220712 | 1.00339626 | 1.750228394 | 0.510338024 | 0.289140582 | 0.411039424 | 0.595090888 | 0.5857 | 0.721351133 |
| SL1344_RS06720_ | -2.668698982 | -2.669463575 | -2.666683472 | 0.007656052 | 0.524210439 | 0.013422624 | 0.494666667 | 0.602235618 | 0.732252021 | 0.7122 | 0.805428479 |
| SL1344_RS06730_ | -4.377889711 | -4.744913682 | -4.181986574 | 0.547703327 | 1.121937804 | 0.452652414 | 0.303797477 | 0.452050573 | 0.614787612 | 0.5885 | 0.710909851 |
| SL1344_RS06735_ | -3.194729782 | -3.621940232 | -2.84266403 | 0.799694893 | 0.692747555 | 1.144693155 | 0.100666708 | 0.163769544 | 0.327746631 | 0.2234 | 0.384521725 |
| SL1344_RS06740_ | -1.005808643 | -1.117010938 | -0.913267677 | 0.204939569 | 0.293614291 | 0.649629947 | 0.232000014 | 0.355103467 | 0.51688372 | 0.4548 | 0.59806491 |
| SL1344_RS06745_ | -2.375967911 | -2.601415561 | -2.125495597 | 0.465204611 | 0.435299832 | 1.005641781 | 0.123333366 | 0.200775729 | 0.355063931 | 0.2879 | 0.440730188 |
| SL1344_RS06750_ | 0.360349768 | 0.20125816 | 0.513232071 | 0.310133669 | 0.215422447 | 1.391248894 | 0.06200007 | 0.110374638 | 0.256549472 | 0.1734 | 0.33099331 |
| SL1344_RS06755_ | -0.953235715 | -1.009103381 | -0.876314883 | 0.129507393 | 0.285250668 | 0.415950893 | 0.311125924 | 0.488349182 | 0.634501137 | 0.5975 | 0.71375445 |
| SL1344_RS06760_ | -1.35201324 | -1.343742497 | -1.361609051 | -0.009394562 | 0.490577735 | -0.01598249 | 0.493004664 | 0.717341897 | 0.826372367 | 0.8297 | 0.894222696 |
| SL1344_RS06765_ | -3.774127969 | -3.803654649 | -3.762249127 | -0.020217276 | 0.914420251 | -0.019699154 | 0.491333334 | 0.670098145 | 0.78886018 | 0.7815 | 0.860335976 |
| SL1344_RS06770_ | -5.071081644 | -5.229314719 | -4.90829474 | 0.368982475 | 1.236743838 | 0.267720337 | 0.371333338 | 0.522767704 | 0.670493964 | 0.6389 | 0.746929346 |
| SL1344_RS06775_ | 1.26931416 | 1.196224999 | 1.342652916 | 0.152398684 | 0.153119575 | 0.967958116 | 0.148567647 | 0.221906368 | 0.391508515 | 0.3072 | 0.465984864 |
| SL1344_RS06780_ | 1.887850716 | 1.836222251 | 1.939887152 | 0.100544391 | 0.112785084 | 0.851558249 | 0.183877435 | 0.285208144 | 0.4502295 | 0.3818 | 0.532912619 |
| SL1344_RS06785_ | -0.638996681 | -0.771391551 | -0.528435112 | 0.234925727 | 0.23400091 | 0.939209211 | 0.139240534 | 0.224715405 | 0.382252462 | 0.3121 | 0.464214652 |
| SL1344_RS06790_ | 0.995227892 | 1.062264988 | 0.923151879 | -0.131006715 | 0.178613843 | -0.71078406 | 0.208527665 | 0.332612348 | 0.5030401 | 0.4271 | 0.577014449 |
| SL1344_RS06795_ | 0.268228978 | 0.125133445 | 0.442691996 | 0.349374183 | 0.255576998 | 1.300268857 | 0.065956094 | 0.117872178 | 0.27345533 | 0.1792 | 0.337442503 |
| SL1344_RS06800_ | -1.893938553 | -2.029330754 | -1.652585472 | 0.319048763 | 0.546943769 | 0.530904134 | 0.290666676 | 0.485786402 | 0.643618238 | 0.5861 | 0.713926503 |
| SL1344_RS06805_ | 0.490134452 | 0.461919591 | 0.532094373 | 0.042833577 | 0.242386 | 0.164063321 | 0.427048637 | 0.676971403 | 0.794978656 | 0.778 | 0.859222996 |
| SL1344_RS06810_ | 0.692322371 | 0.652664126 | 0.76376332 | 0.10524899 | 0.224226078 | 0.426338681 | 0.332000007 | 0.541668039 | 0.692226138 | 0.6456 | 0.765373767 |
| SL1344_RS06815_ | 0.941003922 | 0.915942217 | 0.97415784 | 0.062300652 | 0.167331073 | 0.335998595 | 0.357095276 | 0.542325811 | 0.687629621 | 0.6623 | 0.767351428 |
| SL1344_RS06820_ | 0.581059477 | 0.531683224 | 0.691792227 | 0.13478773 | 0.214353801 | 0.579577484 | 0.269333344 | 0.455675274 | 0.618392494 | 0.5698 | 0.703587267 |
| SL1344_RS06825_ | -0.822739703 | -0.812681466 | -0.842909488 | -0.014131635 | 0.318173241 | -0.041137993 | 0.484343771 | 0.660236248 | 0.78333122 | 0.7608 | 0.847655823 |
| SL1344_RS06830_ | 0.435598473 | 0.397983782 | 0.494979064 | 0.069837953 | 0.222231564 | 0.275754094 | 0.379080618 | 0.620070163 | 0.750016083 | 0.7109 | 0.811222908 |
| SL1344_RS06835_ | 1.060536717 | 1.03229207 | 1.097377768 | 0.067949291 | 0.145239889 | 0.448331279 | 0.317333341 | 0.466555373 | 0.61798892 | 0.5867 | 0.703712092 |
| SL1344_RS06840_ | 10.52038526 | 10.45185955 | 10.63123951 | 0.205464849 | 0.079686643 | 2.709754246 | 4.68E-05 | 0.01637949 | 0.092374592 | 0.1 | 0.243021319 |
| SL1344_RS06845_ | 5.435554916 | 5.167453953 | 5.65608414 | 0.471702235 | 0.066484679 | 7.102240159 | 4.68E-05 | 0.001735799 | 0.027412287 | 0.1 | 0.243021319 |
| SL1344_RS06850_ | -1.499612049 | -1.50433163 | -1.493350809 | 0.01744481 | 0.337745616 | 0.039878016 | 0.480666667 | 0.572042385 | 0.709214978 | 0.6911 | 0.790083448 |

| | | | | | | | | | | | |
|---|---|---|---|---|---|---|---|---|---|---|---|
| SL1344_RS06855_ | -2.930215339 | -3.405977712 | -2.623305755 | 0.776133399 | 0.714373182 | 1.039712651 | 0.124666699 | 0.191655435 | 0.362590843 | 0.2683 | 0.430563035 |
| SL1344_RS06860_ | -4.622252533 | -4.961563393 | -4.307003696 | 0.627606497 | 0.96029508 | 0.612597855 | 0.234000014 | 0.356195702 | 0.517582124 | 0.4555 | 0.594933889 |
| SL1344_RS06865_ | -3.641245593 | -4.182282848 | -3.213552177 | 1.040009662 | 1.007104975 | 1.005674308 | 0.114666702 | 0.186161374 | 0.362015404 | 0.2618 | 0.424325615 |
| SL1344_RS06900_ | 1.339026983 | 1.164809555 | 1.541319274 | 0.365484491 | 0.156750176 | 2.266296309 | 0.006662914 | 0.03514442 | 0.131938055 | 0.1066 | 0.252050652 |
| SL1344_RS06905_ | -0.239055799 | -0.479351601 | 0.024779584 | 0.486709303 | 0.269964414 | 1.740058892 | 0.030666813 | 0.066690549 | 0.190950299 | 0.1337 | 0.287519984 |
| SL1344_RS06925_ | -3.701224774 | -3.663562384 | -3.727960523 | -0.067873452 | 0.697085048 | -0.078280101 | 0.461692207 | 0.554433473 | 0.693503796 | 0.6636 | 0.768019008 |
| SL1344_RS06930_ | -3.362157719 | -3.419184948 | -3.294873086 | 0.129772881 | 0.703497059 | 0.163712852 | 0.42238508 | 0.582555648 | 0.718658767 | 0.6892 | 0.789440567 |
| SL1344_RS06935_ | -3.439647373 | -3.450450267 | -3.43422914 | 0.028149342 | 0.618127407 | 0.036304448 | 0.484666667 | 0.554161988 | 0.69429775 | 0.6625 | 0.766982973 |
| SL1344_RS06940_ | -3.518484795 | -3.686215186 | -3.23545854 | 0.504323263 | 0.945726852 | 0.49414843 | 0.297333342 | 0.478363146 | 0.63702764 | 0.6029 | 0.72324906 |
| SL1344_RS06945_ | -2.633235757 | -2.723480818 | -2.513082364 | 0.216440078 | 0.569839405 | 0.35546589 | 0.34133334 | 0.500819599 | 0.650808135 | 0.6254 | 0.736540146 |
| SL1344_RS06950_ | -2.616386126 | -2.72669991 | -2.507071383 | 0.225063432 | 0.428526628 | 0.474100238 | 0.297135252 | 0.412636892 | 0.56253347 | 0.5046 | 0.633787786 |
| SL1344_RS06955_ | -1.735913714 | -2.245376911 | -1.414569876 | 0.836257946 | 0.493059818 | 1.575203881 | 0.041305903 | 0.090114218 | 0.230770355 | 0.1455 | 0.299757596 |
| SL1344_RS06960_ | -3.353347568 | -3.358458816 | -3.346208001 | 0.002442528 | 0.601591596 | 0.00525701 | 0.497001999 | 0.575302061 | 0.712106289 | 0.6935 | 0.792374539 |
| SL1344_RS06965_ | -3.328135966 | -3.488164622 | -3.256541256 | 0.253351659 | 0.830607154 | 0.303119842 | 0.376666672 | 0.555298984 | 0.703186516 | 0.6863 | 0.7912808 |
| SL1344_RS06970_ | -2.995772184 | -3.410503466 | -2.694638074 | 0.776848391 | 0.888253028 | 0.8198784 | 0.137333362 | 0.236306928 | 0.422613504 | 0.2995 | 0.461574191 |
| SL1344_RS06975_ | -2.738837981 | -2.97414983 | -2.617580133 | 0.329315471 | 0.571605374 | 0.533818294 | 0.27600001 | 0.424458921 | 0.586440056 | 0.547 | 0.67765373 |
| SL1344_RS06980_ | -1.306516751 | -1.373899384 | -1.231463908 | 0.146270518 | 0.295502813 | 0.475663064 | 0.298467697 | 0.445071103 | 0.599432674 | 0.5542 | 0.678440882 |
| SL1344_RS06990_ | -5.382401482 | -5.937269538 | -4.826162787 | 1.217042099 | 1.589760145 | 0.708887156 | 0.194000018 | 0.29539495 | 0.470283929 | 0.4046 | 0.553350179 |
| SL1344_RS06995_ | -4.83403562 | -4.971011782 | -4.717389644 | 0.226070463 | 1.165357081 | 0.176105774 | 0.41972019 | 0.579548311 | 0.717678115 | 0.7022 | 0.799243582 |
| SL1344_RS07000_ | -3.411305054 | -3.598604684 | -3.260118678 | 0.348788221 | 0.612464286 | 0.511916707 | 0.284477025 | 0.417785958 | 0.57157152 | 0.5309 | 0.655194745 |
| SL1344_RS07005_ | -2.849528252 | -2.97936586 | -2.72586118 | 0.268950586 | 0.585158844 | 0.453108922 | 0.299333342 | 0.460190152 | 0.614892413 | 0.5685 | 0.691063983 |
| SL1344_RS07010_ | -3.889681996 | -4.096995873 | -3.431890684 | 0.493197944 | 1.343244781 | 0.296660684 | 0.359760166 | 0.744893381 | 0.844245664 | 0.7325 | 0.839896125 |
| SL1344_RS07015_ | -1.982903294 | -2.060714867 | -1.829343199 | 0.210896009 | 0.503735488 | 0.401122399 | 0.335776156 | 0.542847504 | 0.686693348 | 0.6471 | 0.759199818 |
| SL1344_RS07020_ | -4.546962482 | -4.2712582 | -4.970610038 | -0.725417484 | 0.979647704 | -0.669704069 | 0.216666682 | 0.332400092 | 0.498694495 | 0.4404 | 0.579125194 |
| SL1344_RS07025_ | -2.086861572 | -2.237653657 | -1.961400851 | 0.280489561 | 0.457642962 | 0.575059604 | 0.256666678 | 0.392297609 | 0.555903667 | 0.509 | 0.643046155 |
| SL1344_RS07030_ | -0.98549371 | -1.032386785 | -0.931177509 | 0.107861865 | 0.279571857 | 0.368849645 | 0.352431718 | 0.498973834 | 0.645935098 | 0.6143 | 0.726451683 |
| SL1344_RS07035_ | -2.629380564 | -2.92968948 | -2.15714594 | 0.707955527 | 0.699428476 | 0.968961502 | 0.159227205 | 0.237667724 | 0.413753796 | 0.3206 | 0.487771347 |
| SL1344_RS07040_ | -2.800373231 | -3.060721733 | -2.294955111 | 0.700585219 | 0.719499037 | 0.913325437 | 0.173333354 | 0.252354769 | 0.43007404 | 0.352 | 0.51823026 |
| SL1344_RS07045_ | -0.989038787 | -1.151680868 | -0.818383431 | 0.325895481 | 0.316371176 | 0.95877217 | 0.144570313 | 0.227047037 | 0.396639094 | 0.3141 | 0.475090986 |
| SL1344_RS07055_ | -4.265883707 | -4.400204096 | -4.084410911 | 0.300392351 | 0.874168314 | 0.310742763 | 0.362000006 | 0.522910805 | 0.66763783 | 0.6296 | 0.741711924 |
| SL1344_RS07060_ | -3.035610585 | -3.051280868 | -3.023122666 | -0.001219531 | 0.674277564 | -0.002134715 | 0.499000666 | 0.641405204 | 0.76612656 | 0.7516 | 0.835532243 |
| SL1344_RS07065_ | -2.486981549 | -2.53367378 | -2.44448006 | 0.098578973 | 0.435216776 | 0.22549695 | 0.399733515 | 0.503085094 | 0.648786207 | 0.6178 | 0.729274244 |
| SL1344_RS07070_ | 1.597590838 | 1.416378799 | 1.941890606 | 0.522215832 | 0.19111509 | 2.56081084 | 4.68E-05 | 0.035741767 | 0.137682161 | 0.1002 | 0.243308475 |
| SL1344_RS07075_ | 2.396762794 | 1.890460031 | 2.715288212 | 0.786132546 | 0.180295504 | 4.048925459 | 4.68E-05 | 0.011776914 | 0.073774509 | 0.1 | 0.243021319 |
| SL1344_RS07080_ | 2.034122021 | 1.947632304 | 2.170287192 | 0.227358573 | 0.123846564 | 1.771487107 | 0.017333596 | 0.073055934 | 0.203658887 | 0.1198 | 0.2676495 |
| SL1344_RS07085_ | 1.138564051 | 1.030979455 | 1.264172249 | 0.166928527 | 0.323855765 | 0.494708479 | 0.308000008 | 0.497333127 | 0.669214613 | 0.6433 | 0.765955016 |
| SL1344_RS07090_ | 2.88980148 | 3.06215052 | 2.740429421 | -0.325926138 | 0.107484899 | -2.938817851 | 4.68E-05 | 0.019146558 | 0.095037378 | 0.1 | 0.243021319 |
| SL1344_RS07095_ | 0.126187113 | 0.070985861 | 0.209298303 | 0.131747197 | 0.225700981 | 0.554549567 | 0.268000011 | 0.422861168 | 0.585707839 | 0.5354 | 0.668663594 |
| SL1344_RS07100_ | -1.157716193 | -1.164080354 | -1.149779138 | 0.004198542 | 0.300831305 | 0.01105237 | 0.493004664 | 0.60090777 | 0.733248377 | 0.7152 | 0.80906849 |
| SL1344_RS07115_ | 2.03320522 | 1.752513927 | 2.33861406 | 0.587995498 | 0.097079724 | 6.04589412 | 4.68E-05 | 0.003628373 | 0.036291759 | 0.1 | 0.243021319 |
| SL1344_RS07120_ | 4.998567059 | 4.92224556 | 5.103648529 | 0.184023735 | 0.044841053 | 4.133882355 | 4.68E-05 | 0.011578716 | 0.070502751 | 0.1 | 0.243021319 |
| SL1344_RS07125_ | 2.034942656 | 1.736571238 | 2.300370931 | 0.558855827 | 0.131882715 | 4.177000344 | 4.68E-05 | 0.005885442 | 0.049281021 | 0.1 | 0.243021319 |
| SL1344_RS07130_ | 2.080811883 | 2.061979147 | 2.129431576 | 0.074969788 | 0.16017903 | 0.415663506 | 0.334000007 | 0.505736448 | 0.668891409 | 0.6637 | 0.776327721 |
| SL1344_RS07135_ | 1.240966793 | 1.183322277 | 1.275153693 | 0.08235946 | 0.144658728 | 0.536143903 | 0.28847436 | 0.460224089 | 0.616265533 | 0.5727 | 0.700550187 |
| SL1344_RS07140_ | 0.104031839 | 0.043475385 | 0.146144607 | 0.088745164 | 0.261730376 | 0.328276698 | 0.357333339 | 0.567379116 | 0.710800656 | 0.6886 | 0.790695168 |
| SL1344_RS07145_ | -3.126674584 | -3.024535085 | -3.254767563 | -0.204861436 | 0.728556866 | -0.270500483 | 0.376415728 | 0.571052645 | 0.713490439 | 0.6828 | 0.788530706 |
| SL1344_RS07150_ | 1.04427275 | 0.622560176 | 1.330220169 | 0.711635951 | 0.172194056 | 4.019373871 | 4.68E-05 | 0.009371803 | 0.061345307 | 0.1 | 0.243021319 |
| SL1344_RS07155_ | 0.350120983 | -0.095669567 | 0.59543049 | 0.659478769 | 0.280888009 | 2.208194342 | 0.005996769 | 0.044844142 | 0.155458105 | 0.1056 | 0.250848126 |
| SL1344_RS07160_ | 0.85903484 | 0.499859448 | 1.407552256 | 0.888903178 | 0.200217286 | 4.31305096 | 4.68E-05 | 0.00711584 | 0.054968588 | 0.1 | 0.243021319 |
| SL1344_RS07165_ | 0.529823947 | 0.10847007 | 0.938134819 | 0.837118254 | 0.152956392 | 5.378023748 | 4.68E-05 | 0.003449003 | 0.035724413 | 0.1 | 0.243021319 |
| SL1344_RS07170_ | -1.260229741 | -1.545545472 | -0.901295536 | 0.6495027 | 0.336783803 | 1.881734825 | 0.016000285 | 0.060212212 | 0.178586507 | 0.1146 | 0.261608673 |
| SL1344_RS07180_ | 3.171736887 | 2.945757616 | 3.396525608 | 0.463144812 | 0.105332324 | 4.434632883 | 4.68E-05 | 0.010805317 | 0.069460586 | 0.1 | 0.243021319 |
| SL1344_RS07185_ | -0.017178525 | 0.138096331 | -0.094678565 | -0.206123004 | 0.242881421 | -0.767346707 | 0.213333349 | 0.331221594 | 0.504104301 | 0.4384 | 0.592651138 |

| | | | | | | | | | | | |
|---|---|---|---|---|---|---|---|---|---|---|---|
| SL1344_RS07190_ | 0.059442373 | 0.017552053 | 0.096385305 | 0.078170559 | 0.26530277 | 0.2724613 | 0.382000005 | 0.587961061 | 0.726531433 | 0.6975 | 0.79770912 |
| SL1344_RS07195_ | 1.710334755 | 1.585543289 | 2.008663286 | 0.429381275 | 0.267710688 | 1.596519895 | 0.036000124 | 0.074745693 | 0.215603544 | 0.1331 | 0.287560077 |
| SL1344_RS07200_ | 4.808545605 | 4.546622942 | 5.069139054 | 0.528048419 | 0.066624251 | 7.990427687 | 4.68E-05 | 0.000807273 | 0.01919361 | 0.1 | 0.243021319 |
| SL1344_RS07205_ | 2.006019742 | 1.920649967 | 2.106056863 | 0.208766459 | 0.182618362 | 1.151706113 | 0.067288538 | 0.145865999 | 0.319900588 | 0.1795 | 0.340164169 |
| SL1344_RS07210_ | 0.94774445 | 0.973842411 | 0.920277643 | -0.066678948 | 0.243959035 | -0.282350064 | 0.369753503 | 0.585049099 | 0.730191298 | 0.6976 | 0.802026793 |
| SL1344_RS07215_ | 3.208214586 | 2.880013785 | 3.487616371 | 0.611332103 | 0.075379569 | 8.109409257 | 4.68E-05 | 0.001812932 | 0.025815613 | 0.1 | 0.243021319 |
| SL1344_RS07220_ | 2.45746022 | 2.318957645 | 2.670703415 | 0.353308692 | 0.102971006 | 3.382143185 | 4.68E-05 | 0.016001807 | 0.083509873 | 0.1 | 0.243021319 |
| SL1344_RS07225_ | 2.155930127 | 1.945326081 | 2.355816832 | 0.409046561 | 0.103876416 | 3.901933417 | 4.68E-05 | 0.00810823 | 0.057109932 | 0.1 | 0.243021319 |
| SL1344_RS07230_ | 1.897344427 | 1.609480389 | 2.11960193 | 0.517364405 | 0.119316429 | 4.345762271 | 4.68E-05 | 0.005884844 | 0.048678706 | 0.1 | 0.243021319 |
| SL1344_RS07235_ | 2.283267922 | 2.015699215 | 2.488502246 | 0.485372964 | 0.131806624 | 3.751947021 | 4.68E-05 | 0.023702783 | 0.108417622 | 0.1 | 0.243021319 |
| SL1344_RS07240_ | 3.680506512 | 3.454824132 | 3.897506971 | 0.520805247 | 0.16575035 | 3.224129655 | 4.68E-05 | 0.006823452 | 0.056871643 | 0.1 | 0.243021319 |
| SL1344_RS07245_ | 1.838492092 | 1.626183027 | 2.056266523 | 0.443555761 | 0.149488922 | 2.995700481 | 4.68E-05 | 0.016863154 | 0.088117231 | 0.1 | 0.243021319 |
| SL1344_RS07250_ | 3.014282208 | 2.812119926 | 3.195512469 | 0.385531479 | 0.070765538 | 5.471623489 | 4.68E-05 | 0.003587348 | 0.036755406 | 0.1 | 0.243021319 |
| SL1344_RS07255_ | 3.172116448 | 2.966279466 | 3.306172058 | 0.34180443 | 0.091141386 | 3.773269667 | 4.68E-05 | 0.008290835 | 0.059331011 | 0.1 | 0.243021319 |
| SL1344_RS07260_ | 1.95321489 | 1.769198078 | 2.11771274 | 0.353566448 | 0.112596652 | 3.130616192 | 4.68E-05 | 0.015578738 | 0.082200344 | 0.1002 | 0.243292142 |
| SL1344_RS07265_ | 1.417547679 | 1.384000458 | 1.567562523 | 0.184252694 | 0.222143976 | 0.738935113 | 0.255163236 | 0.352144231 | 0.545130618 | 0.534 | 0.681536652 |
| SL1344_RS07270_ | 1.535128756 | 1.327097473 | 1.717719217 | 0.390291116 | 0.110965019 | 3.477097183 | 4.68E-05 | 0.011670222 | 0.068212006 | 0.1002 | 0.243296832 |
| SL1344_RS07275_ | 2.286420912 | 2.08321553 | 2.499893647 | 0.420468654 | 0.115222237 | 3.636686902 | 4.68E-05 | 0.009578036 | 0.063789344 | 0.1 | 0.243021319 |
| SL1344_RS07285_ | 2.878716557 | 2.729299137 | 3.044298655 | 0.340277094 | 0.130077032 | 2.577517055 | 4.68E-05 | 0.02029502 | 0.099377044 | 0.1 | 0.243021319 |
| SL1344_RS07290_ | 1.92361669 | 1.747129719 | 2.108183048 | 0.367764569 | 0.111580418 | 3.310091198 | 4.68E-05 | 0.013191375 | 0.074038455 | 0.1 | 0.243021319 |
| SL1344_RS07295_ | 2.565578379 | 2.63922176 | 2.476313487 | -0.14395437 | 0.137736746 | -1.019253294 | 0.130666697 | 0.191664904 | 0.372321218 | 0.267 | 0.44284809 |
| SL1344_RS07300_ | 0.881569013 | 0.883754385 | 0.879339075 | -0.00074575 | 0.138315127 | -0.007376837 | 0.497333333 | 0.574740179 | 0.713460395 | 0.6862 | 0.787162528 |
| SL1344_RS07305_ | -0.619328001 | -0.706877425 | -0.520177679 | 0.185814144 | 0.252173385 | 0.709300703 | 0.206666683 | 0.32654541 | 0.488387062 | 0.4307 | 0.572239296 |
| SL1344_RS07310_ | -0.28699771 | -0.39574082 | -0.191702874 | 0.229600628 | 0.272724704 | 0.795115715 | 0.180666687 | 0.287587542 | 0.464033828 | 0.3891 | 0.538660033 |
| SL1344_RS07315_ | -3.783788658 | -3.558670639 | -3.955544789 | -0.545993652 | 0.974498047 | -0.545961447 | 0.29380414 | 0.425119347 | 0.591330425 | 0.5682 | 0.691020801 |
| SL1344_RS07320_ | -0.189782072 | -0.2961902 | -0.096000273 | 0.202345216 | 0.184934391 | 1.022100053 | 0.129247199 | 0.212306402 | 0.366477574 | 0.3003 | 0.452059866 |
| SL1344_RS07325_ | -4.979053513 | -4.559989492 | -5.401695319 | -0.90598816 | 1.278048026 | -0.67192767 | 0.211858777 | 0.329077931 | 0.498038395 | 0.4354 | 0.58033785 |
| SL1344_RS07330_ | 3.590793898 | 3.343863229 | 3.798850402 | 0.46133395 | 0.09236961 | 5.022470156 | 4.68E-05 | 0.010126217 | 0.067514845 | 0.1 | 0.243021319 |
| SL1344_RS07335_ | -0.664764255 | -0.556954326 | -0.763643775 | -0.208380726 | 0.217570764 | -0.887518138 | 0.161225873 | 0.249981105 | 0.404655301 | 0.3347 | 0.482763466 |
| SL1344_RS07340_ | 2.452740525 | 1.894330745 | 2.907362981 | 1.024775617 | 0.121449548 | 8.453049834 | 4.68E-05 | 0.001039873 | 0.020437285 | 0.1 | 0.243021319 |
| SL1344_RS07345_ | 3.166723044 | 2.963646585 | 3.431980333 | 0.463356305 | 0.107616921 | 4.225021288 | 4.68E-05 | 0.007332422 | 0.05640975 | 0.1 | 0.243021319 |
| SL1344_RS07350_ | 3.115494409 | 2.810770653 | 3.344039444 | 0.530504016 | 0.075381629 | 6.930157347 | 4.68E-05 | 0.003064994 | 0.033559515 | 0.1 | 0.243021319 |
| SL1344_RS07355_ | 3.306948035 | 3.084160295 | 3.555787575 | 0.483504476 | 0.109776511 | 4.447135387 | 4.68E-05 | 0.012069266 | 0.07514676 | 0.1 | 0.243021319 |
| SL1344_RS07365_ | -1.165742875 | -1.29631277 | -1.022856145 | 0.364736337 | 0.531328484 | 0.664847748 | 0.20933335 | 0.311242253 | 0.502003648 | 0.4393 | 0.588513299 |
| SL1344_RS07370_ | -1.047437783 | -1.568665029 | -0.691157646 | 0.879116662 | 0.504667509 | 1.698520702 | 0.025316634 | 0.067072008 | 0.198023601 | 0.1226 | 0.273557989 |
| SL1344_RS07375_ | -1.035277823 | -0.962799093 | -1.136275506 | -0.173687219 | 0.313251194 | -0.528831627 | 0.303797477 | 0.456175632 | 0.612184124 | 0.5621 | 0.689938845 |
| SL1344_RS07380_ | -0.855001225 | -1.174824078 | -0.556191331 | 0.612327944 | 0.252643277 | 2.376447451 | 0.003998486 | 0.030442768 | 0.115865588 | 0.1051 | 0.249354135 |
| SL1344_RS07385_ | -0.624476405 | -0.69224857 | -0.541540479 | 0.15374117 | 0.26641222 | 0.550594923 | 0.265156573 | 0.405010245 | 0.563510589 | 0.5165 | 0.646810526 |
| SL1344_RS07390_ | -1.500014648 | -1.474359225 | -1.532520939 | -0.053715417 | 0.320134357 | -0.162540658 | 0.424383747 | 0.555727767 | 0.69512591 | 0.6763 | 0.776635922 |
| SL1344_RS07400_ | -0.662145544 | -1.024853838 | -0.432532979 | 0.58439048 | 0.327048049 | 1.70915025 | 0.029313944 | 0.072827252 | 0.20289162 | 0.1332 | 0.286144997 |
| SL1344_RS07405_ | -0.328308445 | -0.547529003 | -0.111190081 | 0.431610897 | 0.243838948 | 1.71971133 | 0.03664236 | 0.071506071 | 0.193174296 | 0.1386 | 0.29025554 |
| SL1344_RS07410_ | -1.461712877 | -1.235888578 | -1.7406373 | -0.504406779 | 0.314198459 | -1.530591743 | 0.043970787 | 0.090461756 | 0.220475591 | 0.1535 | 0.305390388 |
| SL1344_RS07415_ | -3.586720799 | -3.774143406 | -3.431830958 | 0.350474616 | 0.677206761 | 0.484287071 | 0.292000009 | 0.42520575 | 0.582753752 | 0.5322 | 0.661972689 |
| SL1344_RS07420_ | -3.961854424 | -3.718619689 | -4.285220513 | -0.559201909 | 0.740939443 | -0.708163685 | 0.201199218 | 0.313873591 | 0.475183106 | 0.4076 | 0.551455861 |
| SL1344_RS07425_ | -4.62139644 | -4.514407058 | -4.741860077 | -0.227660089 | 0.975472434 | -0.215477084 | 0.402398405 | 0.53894481 | 0.683072653 | 0.6509 | 0.759090824 |
| SL1344_RS07430_ | 1.361613695 | 1.235087526 | 1.391579885 | 0.146400533 | 0.220840682 | 0.570984243 | 0.321333341 | 0.497721943 | 0.667855796 | 0.6603 | 0.787670677 |
| SL1344_RS07435_ | 0.5004721 | 0.529127544 | 0.484138191 | -0.030490229 | 0.221182774 | -0.129975467 | 0.443037977 | 0.669875766 | 0.790144411 | 0.7638 | 0.850308645 |
| SL1344_RS07440_ | -4.911401793 | -4.958331073 | -4.843689931 | 0.31419048 | 1.528819346 | 0.196494765 | 0.426666669 | 0.610997057 | 0.744106813 | 0.7322 | 0.821428174 |
| SL1344_RS07445_ | -1.866616279 | -1.938038709 | -1.731491776 | 0.320308231 | 0.777151304 | 0.392310295 | 0.32933334 | 0.546825959 | 0.703600751 | 0.6716 | 0.786445883 |
| SL1344_RS07450_ | -1.433070775 | -1.299450208 | -1.548571494 | -0.255160535 | 0.337175085 | -0.714346859 | 0.212000016 | 0.33432984 | 0.497626762 | 0.4335 | 0.576814982 |
| SL1344_RS07455_ | 0.36080707 | 0.364969004 | 0.357536886 | 0.013530938 | 0.229535024 | 0.053994844 | 0.478347769 | 0.670840221 | 0.790642468 | 0.7927 | 0.865790083 |
| SL1344_RS07460_ | -0.944384725 | -0.978071521 | -0.927229471 | 0.043727954 | 0.298929246 | 0.14293908 | 0.440000002 | 0.592575725 | 0.727672743 | 0.7084 | 0.805745815 |
| SL1344_RS07465_ | 2.5289325 | 2.749930989 | 2.385902222 | -0.356475637 | 0.103191892 | -3.332165309 | 4.68E-05 | 0.014329011 | 0.079596888 | 0.1 | 0.243021319 |

| | | | | | | | | | | | |
|---|---|---|---|---|---|---|---|---|---|---|---|
| SL1344_RS07470_ | 2.259718103 | 2.419104335 | 2.167991947 | -0.217017474 | 0.224638928 | -0.972591496 | 0.137333362 | 0.190594587 | 0.377641184 | 0.265 | 0.44901904 |
| SL1344_RS07475_ | -1.744283832 | -1.744967483 | -1.743660636 | -0.016003231 | 0.42117288 | -0.033249936 | 0.484343771 | 0.622282894 | 0.75124191 | 0.735 | 0.824790909 |
| SL1344_RS07480_ | -3.015904567 | -3.132596844 | -2.938066558 | 0.140810677 | 0.602036637 | 0.230330036 | 0.404000004 | 0.574396862 | 0.71086015 | 0.686 | 0.786318267 |
| SL1344_RS07485_ | -2.976335388 | -2.769967052 | -3.115385859 | -0.3196577 | 0.575443855 | -0.512799534 | 0.289140582 | 0.449574337 | 0.602808651 | 0.5513 | 0.681587238 |
| SL1344_RS07490_ | -2.791377105 | -2.944333976 | -2.559628581 | 0.408660051 | 0.793224126 | 0.478914523 | 0.290473027 | 0.496963447 | 0.654823829 | 0.6054 | 0.732201381 |
| SL1344_RS07495_ | -0.363822802 | -0.50435533 | -0.225595341 | 0.277581986 | 0.221310122 | 1.230601036 | 0.088607642 | 0.149746556 | 0.297138165 | 0.226 | 0.379937059 |
| SL1344_RS07500_ | -0.097138091 | -0.196185899 | -0.001956546 | 0.213163179 | 0.257726976 | 0.793850727 | 0.194000018 | 0.284663059 | 0.460937879 | 0.3944 | 0.547396499 |
| SL1344_RS07505_ | 1.309059499 | 1.135348801 | 1.44418984 | 0.317539853 | 0.142868872 | 2.245733943 | 0.004001152 | 0.046525997 | 0.153527558 | 0.1041 | 0.248523237 |
| SL1344_RS07510_ | 2.05245534 | 1.93092804 | 2.107890239 | 0.163099077 | 0.129660379 | 1.124537529 | 0.124000032 | 0.183852574 | 0.356590677 | 0.2806 | 0.447273142 |
| SL1344_RS07515_ | 1.003056368 | 0.958952971 | 1.03066542 | 0.086830556 | 0.182640381 | 0.447551191 | 0.306666675 | 0.439181046 | 0.60925499 | 0.5857 | 0.706730055 |
| SL1344_RS07520_ | -3.702844002 | -3.571329055 | -3.955457173 | -0.326100923 | 0.866008592 | -0.334108147 | 0.37175217 | 0.565241609 | 0.709961241 | 0.687 | 0.795227688 |
| SL1344_RS07525_ | -3.446523434 | -3.494416178 | -3.367577714 | 0.115518248 | 0.601618251 | 0.176789395 | 0.426382415 | 0.547895369 | 0.686982123 | 0.6601 | 0.763807933 |
| SL1344_RS07530_ | -2.84698451 | -2.816119915 | -2.878747393 | -0.076505178 | 0.493002727 | -0.134081326 | 0.435709529 | 0.545823032 | 0.684067013 | 0.6511 | 0.756943923 |
| SL1344_RS07535_ | 0.810440289 | 0.767399028 | 0.867702863 | 0.091469098 | 0.461723163 | 0.167665677 | 0.431333336 | 0.855280727 | 0.919399057 | 0.9043 | 0.944594519 |
| SL1344_RS07540_ | -1.232841773 | -1.241516858 | -1.222283261 | 0.014054314 | 0.276880534 | 0.051573445 | 0.476666668 | 0.551856699 | 0.690113635 | 0.6553 | 0.761394664 |
| SL1344_RS07550_ | -0.155368339 | -0.301548263 | -0.007105389 | 0.294175585 | 0.205747184 | 1.368330582 | 0.068000063 | 0.116516586 | 0.259738643 | 0.185 | 0.341938541 |
| SL1344_RS07555_ | 0.006675949 | -0.215203118 | 0.228221008 | 0.442554861 | 0.183104917 | 2.392785775 | 0.005330642 | 0.031023134 | 0.118493013 | 0.1056 | 0.250182605 |
| SL1344_RS07560_ | 0.368494645 | 0.181410975 | 0.604143487 | 0.398176854 | 0.222592234 | 1.745068619 | 0.03397748 | 0.06510371 | 0.188679718 | 0.1311 | 0.284383684 |
| SL1344_RS07565_ | 0.374395935 | 0.296287138 | 0.472046117 | 0.196576939 | 0.222888 | 0.85460033 | 0.158560983 | 0.247495153 | 0.42417413 | 0.3493 | 0.500496045 |
| SL1344_RS07570_ | 1.388638925 | 1.430361765 | 1.348656609 | -0.082201348 | 0.139020292 | -0.545800584 | 0.265822796 | 0.410682035 | 0.574457521 | 0.5209 | 0.656574228 |
| SL1344_RS07575_ | -0.914162279 | -0.874468258 | -0.998640236 | -0.065881502 | 0.39471756 | -0.165817043 | 0.437708197 | 0.703525763 | 0.816304566 | 0.8057 | 0.88124166 |
| SL1344_RS07580_ | -0.731982136 | -0.928950988 | -0.611429184 | 0.34413902 | 0.390901707 | 0.854604797 | 0.151333358 | 0.256522279 | 0.445517783 | 0.3333 | 0.494304836 |
| SL1344_RS07585_ | 0.545671991 | 0.40956859 | 0.690667397 | 0.284193434 | 0.155857706 | 1.738928176 | 0.026000173 | 0.064671739 | 0.181168411 | 0.1325 | 0.283351104 |
| SL1344_RS07590_ | -0.126313865 | -0.263002641 | -0.00080024 | 0.266339999 | 0.181109789 | 1.376622051 | 0.063291207 | 0.113491539 | 0.248342822 | 0.1808 | 0.333235879 |
| SL1344_RS07595_ | 2.766378603 | 2.298560165 | 3.076892544 | 0.759475425 | 0.151355957 | 4.797758948 | 4.68E-05 | 0.009230037 | 0.0647266 | 0.1 | 0.243021319 |
| SL1344_RS07600_ | -1.394194636 | -0.809806051 | -2.099209964 | -1.292230056 | 0.372917663 | -3.438042427 | 4.68E-05 | 0.014946869 | 0.078118184 | 0.1001 | 0.243161826 |
| SL1344_RS07605_ | 1.067978868 | 1.119218359 | 0.999277746 | -0.126126804 | 0.156347121 | -0.730732025 | 0.200532995 | 0.307812966 | 0.477847513 | 0.4142 | 0.560763558 |
| SL1344_RS07610_ | -2.185811276 | -1.909513397 | -2.296006642 | -0.361653444 | 0.676708597 | -0.463752945 | 0.328666674 | 0.554439133 | 0.70329643 | 0.661 | 0.780655791 |
| SL1344_RS07615_ | -1.060682181 | -0.971491776 | -1.150499995 | -0.181772267 | 0.312119399 | -0.526778576 | 0.267333344 | 0.41417994 | 0.573911789 | 0.5242 | 0.654093316 |
| SL1344_RS07620_ | -0.598186575 | -0.438506157 | -0.75256529 | -0.315347224 | 0.238823092 | -1.246954109 | 0.082666718 | 0.144562667 | 0.291663437 | 0.2219 | 0.377359139 |
| SL1344_RS07625_ | -2.122007592 | -1.734752511 | -2.283034363 | -0.529561926 | 0.564463823 | -0.822130455 | 0.206528997 | 0.298516534 | 0.479341028 | 0.4154 | 0.574163287 |
| SL1344_RS07630_ | -2.041269103 | -1.99886138 | -2.081942819 | -0.079273859 | 0.415614762 | -0.183749506 | 0.419333336 | 0.56599656 | 0.705350507 | 0.6833 | 0.78466612 |
| SL1344_RS07635_ | -0.60600728 | -0.466334767 | -0.765227827 | -0.301672003 | 0.261817802 | -1.092674178 | 0.121333366 | 0.184524507 | 0.344701099 | 0.2622 | 0.41945441 |
| SL1344_RS07640_ | -0.718785107 | -0.866158385 | -0.623522759 | 0.292758057 | 0.665589157 | 0.466470401 | 0.287141915 | 0.486813188 | 0.662273157 | 0.5685 | 0.717841737 |
| SL1344_RS07645_ | -0.120942588 | -0.213220766 | -0.035305638 | 0.199635189 | 0.417358262 | 0.481394818 | 0.298467697 | 0.457807622 | 0.633483326 | 0.5951 | 0.727285986 |
| SL1344_RS07650_ | -1.068598604 | -1.008364509 | -1.140959189 | -0.120722763 | 0.336756614 | -0.336836461 | 0.352666673 | 0.549778147 | 0.692423337 | 0.6609 | 0.768687682 |
| SL1344_RS07655_ | -0.259948507 | -0.568145156 | 0.11080238 | 0.678789624 | 0.212288615 | 3.155719654 | 0.00067292 | 0.014446165 | 0.077541247 | 0.1002 | 0.243297893 |
| SL1344_RS07660_ | 1.001689015 | 1.090906413 | 0.888332956 | -0.203226649 | 0.171240983 | -1.180410517 | 0.10326453 | 0.159050002 | 0.323747066 | 0.2288 | 0.392785853 |
| SL1344_RS07665_ | -1.811033079 | -1.819147431 | -1.794632143 | 0.056773007 | 0.438321194 | 0.130806424 | 0.450666669 | 0.62894 | 0.757325629 | 0.7458 | 0.833076596 |
| SL1344_RS07670_ | -3.043663495 | -3.058767055 | -3.025355016 | 0.064869691 | 0.754949554 | 0.08390796 | 0.460359762 | 0.669531316 | 0.789756027 | 0.7784 | 0.857615699 |
| SL1344_RS07675_ | -3.08436978 | -3.14994429 | -3.012692439 | 0.161484651 | 0.58109447 | 0.244445996 | 0.394666671 | 0.553407148 | 0.693716214 | 0.6607 | 0.76667614 |
| SL1344_RS07680_ | -2.067305254 | -2.140572899 | -1.965636019 | 0.175320123 | 0.430582575 | 0.375685526 | 0.33333334 | 0.52017086 | 0.664135793 | 0.627 | 0.738885839 |
| SL1344_RS07685_ | -2.07267973 | -2.02439835 | -2.117597535 | -0.060126529 | 0.552469624 | -0.092559051 | 0.459027317 | 0.661933048 | 0.78503327 | 0.7551 | 0.843494048 |
| SL1344_RS07690_ | 0.518412977 | 0.706135396 | 0.272494945 | -0.443952909 | 0.192869841 | -2.250009592 | 0.004664544 | 0.046272072 | 0.151678889 | 0.103 | 0.246775529 |
| SL1344_RS07695_ | -0.722809868 | -0.608839266 | -0.835319809 | -0.110699146 | 0.470308711 | -0.254586571 | 0.39573618 | 0.680338768 | 0.800797026 | 0.7744 | 0.857523109 |
| SL1344_RS07700_ | 0.241722861 | 0.442873864 | 0.030435084 | -0.412551661 | 0.170555517 | -2.418049969 | 0.006000767 | 0.030660471 | 0.116971509 | 0.1044 | 0.248728994 |
| SL1344_RS07705_ | 0.739297662 | 0.835501674 | 0.650270596 | -0.18959569 | 0.173478112 | -1.018417691 | 0.115922753 | 0.192318448 | 0.35539687 | 0.2713 | 0.426336649 |
| SL1344_RS07710_ | 0.155766949 | 0.188202373 | 0.100695447 | -0.095261458 | 0.217599895 | -0.418850698 | 0.331112599 | 0.48145938 | 0.635798449 | 0.6082 | 0.722878798 |
| SL1344_RS07715_ | -2.039247605 | -2.029514833 | -2.047305941 | -0.022658153 | 0.517993228 | -0.041774582 | 0.482345104 | 0.667079721 | 0.786764557 | 0.7659 | 0.84789319 |
| SL1344_RS07720_ | -2.252642972 | -2.431660933 | -1.976708082 | 0.464285852 | 0.433881322 | 0.972919669 | 0.155896093 | 0.240724725 | 0.402139503 | 0.3305 | 0.486492375 |
| SL1344_RS07725_ | -1.016699699 | -0.879285094 | -1.171587419 | -0.305262036 | 0.329290191 | -0.885114162 | 0.159333357 | 0.246009237 | 0.414834969 | 0.3342 | 0.489840697 |
| SL1344_RS07735_ | -1.058179992 | -0.70807557 | -1.521662789 | -0.813322191 | 0.31299085 | -2.561700934 | 0.002000966 | 0.025147255 | 0.106227456 | 0.1025 | 0.246335998 |
| SL1344_RS07740_ | 1.475745489 | 1.897774495 | 0.832106621 | -1.056958012 | 0.149692772 | -6.850728617 | 4.68E-05 | 0.004451093 | 0.041195799 | 0.1 | 0.243021319 |

| | | | | | | | | | | | |
|---|---|---|---|---|---|---|---|---|---|---|---|
| SL1344_RS07745_ | -0.32609125 | -0.240239005 | -0.395798254 | -0.158032898 | 0.238356877 | -0.633621634 | 0.23984012 | 0.376075131 | 0.53812198 | 0.4853 | 0.622670824 |
| SL1344_RS07750_ | -1.022178676 | -0.858762854 | -1.17176481 | -0.323880932 | 0.270745806 | -1.143064631 | 0.116666701 | 0.182226825 | 0.334819206 | 0.2601 | 0.41425784 |
| SL1344_RS07755_ | -0.337491072 | -0.195043895 | -0.458432033 | -0.27281753 | 0.218139761 | -1.191302604 | 0.087333381 | 0.156861707 | 0.30726831 | 0.232 | 0.386097862 |
| SL1344_RS07760_ | -0.333219409 | 2.54E-05 | -0.563399469 | -0.557824993 | 0.308402321 | -1.743469129 | 0.024650416 | 0.070560046 | 0.200964254 | 0.1314 | 0.283680476 |
| SL1344_RS07765_ | -1.358250961 | -1.251872131 | -1.499171665 | -0.263581228 | 0.412723362 | -0.608081451 | 0.247168567 | 0.377500049 | 0.549749418 | 0.498 | 0.636045524 |
| SL1344_RS07770_ | -1.815020213 | -1.587886051 | -2.148083737 | -0.558093784 | 0.357098029 | -1.489730182 | 0.050666753 | 0.102707828 | 0.240848491 | 0.1623 | 0.315339934 |
| SL1344_RS07775_ | -3.427271481 | -3.262431855 | -3.535338428 | -0.310444672 | 0.75907573 | -0.395128887 | 0.335109934 | 0.515185854 | 0.6639874 | 0.6218 | 0.736681758 |
| SL1344_RS07780_ | 0.854775327 | 1.135479743 | 0.31056829 | -0.821289108 | 0.257275744 | -3.028106398 | 4.68E-05 | 0.025354004 | 0.111719585 | 0.1 | 0.243021319 |
| SL1344_RS07785_ | -1.911711766 | -1.57559211 | -2.265712114 | -0.696731618 | 0.410735702 | -1.650485665 | 0.031312601 | 0.073677571 | 0.198885631 | 0.1371 | 0.289044295 |
| SL1344_RS07790_ | -2.275820302 | -2.374350341 | -2.213768877 | 0.132336271 | 0.46001522 | 0.265083414 | 0.389740178 | 0.555830987 | 0.697179419 | 0.6682 | 0.773753896 |
| SL1344_RS07800_ | 3.275942241 | 3.377454493 | 3.144271615 | -0.237287056 | 0.072090789 | -3.317521744 | 4.68E-05 | 0.018957491 | 0.091520422 | 0.1 | 0.243021319 |
| SL1344_RS07805_ | 1.503038193 | 1.987865449 | 0.90073481 | -1.101896684 | 0.16912114 | -6.46362015 | 4.68E-05 | 0.002183076 | 0.029102611 | 0.1 | 0.243021319 |
| SL1344_RS07810_ | 2.50612927 | 2.784748808 | 2.120358824 | -0.662977896 | 0.126569027 | -5.193070573 | 4.68E-05 | 0.004010128 | 0.040440076 | 0.1 | 0.243021319 |
| SL1344_RS07815_ | -1.536693932 | -1.383608222 | -1.704168432 | -0.310455116 | 0.338345447 | -0.83708192 | 0.173884098 | 0.267097587 | 0.431522624 | 0.3588 | 0.508771436 |
| SL1344_RS07820_ | 0.750461975 | 0.821662878 | 0.680496942 | -0.155221265 | 0.301715244 | -0.497180074 | 0.285143248 | 0.441539771 | 0.621141848 | 0.5675 | 0.706033154 |
| SL1344_RS07825_ | 0.343768799 | 0.449321837 | 0.241523082 | -0.221066341 | 0.240817202 | -0.906915932 | 0.149333359 | 0.229700395 | 0.407338826 | 0.3184 | 0.482401017 |
| SL1344_RS07830_ | -1.751344149 | -1.812162535 | -1.703082367 | 0.102076847 | 0.433915164 | 0.229559489 | 0.396666671 | 0.583390957 | 0.720635635 | 0.6957 | 0.794534722 |
| SL1344_RS07835_ | 7.472800887 | 7.563152686 | 7.373934114 | -0.17836003 | 0.040197915 | -4.467341757 | 4.68E-05 | 0.003094358 | 0.03780411 | 0.1 | 0.243021319 |
| SL1344_RS07840_ | 0.811870706 | 0.703256765 | 0.988312109 | 0.28737065 | 0.221691922 | 1.264660445 | 0.089273864 | 0.135398844 | 0.29730211 | 0.214 | 0.381402057 |
| SL1344_RS07845_ | 3.352397508 | 3.318528308 | 3.51396858 | 0.357635821 | 1.758415796 | 0.248061065 | 0.223333348 | 0.590543779 | 0.751248321 | 0.4021 | 0.600129325 |
| SL1344_RS07850_ | 1.899333842 | 1.998197985 | 1.806205805 | -0.093810837 | 0.637818917 | -0.178213119 | 0.380666671 | 0.836681839 | 0.909148024 | 0.7906 | 0.873930489 |
| SL1344_RS07855_ | -2.188592267 | -2.196190669 | -2.179994955 | 0.008191705 | 0.432087717 | 0.01513912 | 0.492 | 0.585933883 | 0.7221764 | 0.6966 | 0.796799053 |
| SL1344_RS07860_ | -0.673402438 | -0.432444683 | -0.951579645 | -0.524722097 | 0.219699001 | -2.287188002 | 0.005996769 | 0.031135126 | 0.116113252 | 0.106 | 0.250928843 |
| SL1344_RS07865_ | -2.070906539 | -1.83329634 | -2.33695442 | -0.501851372 | 0.476008475 | -0.994857662 | 0.133910755 | 0.20927871 | 0.375597037 | 0.2946 | 0.454740281 |
| SL1344_RS07870_ | -3.088277634 | -2.687968437 | -3.279499763 | -0.510666494 | 0.745976651 | -0.640838079 | 0.254497014 | 0.42741666 | 0.591675281 | 0.5139 | 0.661077419 |
| SL1344_RS07875_ | -2.373537568 | -1.999752432 | -2.891727084 | -0.858954092 | 0.570762814 | -1.42633969 | 0.054000081 | 0.104216172 | 0.251809529 | 0.1581 | 0.315429019 |
| SL1344_RS07880_ | -0.194400908 | -0.031449616 | -0.398921559 | -0.367132534 | 0.216799839 | -1.639416876 | 0.036000124 | 0.077493829 | 0.205171655 | 0.1366 | 0.287234102 |
| SL1344_RS07885_ | 0.459159786 | 0.407589793 | 0.52971688 | 0.101664604 | 0.214805348 | 0.44476497 | 0.330666674 | 0.521427889 | 0.670630534 | 0.6232 | 0.743169071 |
| SL1344_RS07890_ | 2.083122924 | 2.177827388 | 1.991508548 | -0.187671554 | 0.099217184 | -1.789712496 | 0.022651765 | 0.062139412 | 0.178535554 | 0.1248 | 0.27413252 |
| SL1344_RS07895_ | 3.353500133 | 3.383766603 | 3.232469479 | -0.128882144 | 0.141295375 | -0.800599678 | 0.245836122 | 0.335321625 | 0.531107995 | 0.466 | 0.640563845 |
| SL1344_RS07900_ | -1.642327493 | -1.767313553 | -1.564069121 | 0.204373262 | 0.421890703 | 0.469070108 | 0.304463699 | 0.454765598 | 0.614900665 | 0.5797 | 0.702495546 |
| SL1344_RS07905_ | 2.295748555 | 2.346085935 | 2.246940109 | -0.101505243 | 0.080757134 | -1.169433244 | 0.097333376 | 0.161914596 | 0.30979811 | 0.2461 | 0.397625423 |
| SL1344_RS07910_ | -0.517910942 | -0.392803271 | -0.700815785 | -0.314632639 | 0.250775617 | -1.154377676 | 0.098000042 | 0.164484813 | 0.322209911 | 0.2429 | 0.399712368 |
| SL1344_RS07915_ | -0.248691167 | -0.330134554 | -0.191053455 | 0.13488717 | 0.214677454 | 0.586368734 | 0.262666678 | 0.390598257 | 0.547093398 | 0.4953 | 0.630095093 |
| SL1344_RS07920_ | 0.88495841 | 1.078462113 | 0.529448137 | -0.535411642 | 0.183146888 | -2.802943098 | 0.000673369 | 0.03031792 | 0.12065215 | 0.1008 | 0.244143072 |
| SL1344_RS07925_ | -0.209220027 | 0.038622749 | -0.481629407 | -0.515536357 | 0.251386261 | -2.018064097 | 0.013324793 | 0.050099276 | 0.160282814 | 0.1127 | 0.259819375 |
| SL1344_RS07930_ | 0.058664411 | 0.260442686 | -0.21407227 | -0.476304475 | 0.178899489 | -2.601370825 | 0.0020023 | 0.026625157 | 0.108026988 | 0.1024 | 0.24633893 |
| SL1344_RS07935_ | -0.385693 | -0.194852174 | -0.550275932 | -0.367383221 | 0.272700952 | -1.307188787 | 0.066666731 | 0.124833891 | 0.276920508 | 0.1877 | 0.345617474 |
| SL1344_RS07955_ | -0.574572269 | -0.864455476 | -0.344784049 | 0.525032765 | 0.234329308 | 2.141864342 | 0.009993796 | 0.043485913 | 0.141798761 | 0.1139 | 0.260132778 |
| SL1344_RS07960_ | 1.075425919 | 1.158445033 | 0.938402846 | -0.219830507 | 0.143221529 | -1.470398602 | 0.055333412 | 0.108872765 | 0.249361005 | 0.1848 | 0.339781365 |
| SL1344_RS07965_ | 1.24186956 | 1.382668426 | 1.071539833 | -0.308395913 | 0.122764809 | -2.428493714 | 0.003998486 | 0.027964974 | 0.111253642 | 0.1034 | 0.247457681 |
| SL1344_RS07970_ | 0.497136351 | 0.677976117 | 0.203102294 | -0.484329496 | 0.226464743 | -2.12734882 | 0.002668394 | 0.060800539 | 0.184307335 | 0.1022 | 0.245969832 |
| SL1344_RS07975_ | -3.461897072 | -3.188066988 | -3.711630781 | -0.50046829 | 0.683129765 | -0.687525439 | 0.210666683 | 0.333855185 | 0.494940014 | 0.4296 | 0.572035562 |
| SL1344_RS07980_ | -2.678702518 | -2.488934939 | -2.83176665 | -0.336695095 | 0.480579981 | -0.667912439 | 0.224666681 | 0.358172677 | 0.517225177 | 0.4577 | 0.597123883 |
| SL1344_RS07990_ | -1.783007227 | -1.473178304 | -2.404547936 | -0.925986401 | 0.412701934 | -2.074554136 | 0.011992386 | 0.054833922 | 0.168365681 | 0.1151 | 0.261431655 |
| SL1344_RS07995_ | 2.839192934 | 2.952869911 | 2.69593244 | -0.266451195 | 0.114111199 | -2.350225629 | 4.68E-05 | 0.065317459 | 0.198221945 | 0.1 | 0.243021319 |
| SL1344_RS08000_ | 1.379391115 | 1.349594712 | 1.416954053 | 0.103968811 | 0.240077552 | 0.404551512 | 0.33733334 | 0.544984388 | 0.702719029 | 0.6987 | 0.804134966 |
| SL1344_RS08005_ | 0.976477236 | 0.934342993 | 1.027364517 | 0.114279988 | 0.19408001 | 0.574699848 | 0.257828126 | 0.393379318 | 0.566317684 | 0.5149 | 0.647985033 |
| SL1344_RS08010_ | 2.556188153 | 2.740957578 | 2.343163789 | -0.397776048 | 0.073982087 | -5.336691084 | 4.68E-05 | 0.003366574 | 0.035134648 | 0.1 | 0.243021319 |
| SL1344_RS08015_ | 1.194834046 | 1.52393186 | 0.940757097 | -0.583000147 | 0.145024429 | -3.991801157 | 4.68E-05 | 0.007522241 | 0.054870243 | 0.1 | 0.243021319 |
| SL1344_RS08020_ | 0.83491095 | 0.99672486 | 0.648916951 | -0.354023497 | 0.149058222 | -2.327231314 | 0.004667653 | 0.034358367 | 0.125305306 | 0.1044 | 0.248999876 |
| SL1344_RS08025_ | -2.343827573 | -2.033610534 | -2.709560384 | -0.673391335 | 0.408558631 | -1.591913478 | 0.036666788 | 0.08545842 | 0.210373172 | 0.1478 | 0.299862037 |
| SL1344_RS08030_ | -1.169742223 | -0.844438425 | -1.45408324 | -0.600618109 | 0.30080232 | -1.958409231 | 0.019333568 | 0.051406443 | 0.158006125 | 0.1197 | 0.268053721 |

| | | | | | | | | | | | |
|---|---|---|---|---|---|---|---|---|---|---|---|
| SL1344_RS08035_ | -3.353880998 | -3.061862552 | -3.608110239 | -0.571910187 | 0.726064935 | -0.734161675 | 0.195869438 | 0.30177964 | 0.472168683 | 0.4086 | 0.55599245 |
| SL1344_RS08040_ | -5.525490118 | -5.08468385 | -6.112724726 | -1.081822619 | 1.484599684 | -0.647691777 | 0.223850781 | 0.329104961 | 0.499765525 | 0.4224 | 0.569075926 |
| SL1344_RS08045_ | -2.513271763 | -2.153455319 | -2.845191779 | -0.725202874 | 0.665485899 | -1.06845959 | 0.10326453 | 0.173309683 | 0.342958866 | 0.2436 | 0.407931815 |
| SL1344_RS08050_ | -2.931954258 | -2.538117593 | -3.397838446 | -0.792381659 | 0.75244472 | -1.02940591 | 0.131912088 | 0.203294549 | 0.376655774 | 0.2833 | 0.450553978 |
| SL1344_RS08055_ | -3.630383968 | -3.449010587 | -3.885338625 | -0.550150233 | 1.052327131 | -0.44926781 | 0.258666678 | 0.394275789 | 0.578105486 | 0.5278 | 0.657554944 |
| SL1344_RS08060_ | -2.852875595 | -2.571018451 | -3.304006542 | -0.677587304 | 0.680054507 | -0.909317535 | 0.170552986 | 0.25031256 | 0.427703029 | 0.3439 | 0.507149039 |
| SL1344_RS08065_ | -2.01317058 | -1.780346136 | -2.12919355 | -0.334295416 | 0.505394898 | -0.588305126 | 0.266489018 | 0.426432703 | 0.591133287 | 0.54 | 0.67537298 |
| SL1344_RS08070_ | -2.098088136 | -1.680582626 | -2.401128971 | -0.69804514 | 0.4432351 | -1.516147896 | 0.053333415 | 0.093521353 | 0.229620444 | 0.1656 | 0.322043159 |
| SL1344_RS08075_ | -0.295234777 | -0.220245285 | -0.378284856 | -0.152846296 | 0.203802471 | -0.722617396 | 0.202531663 | 0.306750727 | 0.462837299 | 0.4045 | 0.546065994 |
| SL1344_RS08080_ | -2.194373368 | -2.271002222 | -2.135575097 | 0.138297085 | 0.48884615 | 0.266545012 | 0.375083283 | 0.545935557 | 0.688125637 | 0.6632 | 0.766978927 |
| SL1344_RS08090_ | 0.482004982 | 0.617165196 | 0.313335494 | -0.335542425 | 0.23582717 | -1.370363844 | 0.042638345 | 0.109165868 | 0.260706491 | 0.1556 | 0.309969591 |
| SL1344_RS08095_ | -0.96415073 | -0.807907586 | -1.127787125 | -0.322027166 | 0.2814555 | -1.079924761 | 0.119253864 | 0.188372599 | 0.342844507 | 0.2655 | 0.421725991 |
| SL1344_RS08100_ | 0.966232427 | 1.452500709 | 0.28584774 | -1.165156381 | 0.157404971 | -7.326805296 | 4.68E-05 | 0.00251976 | 0.030017308 | 0.1 | 0.243021319 |
| SL1344_RS08110_ | 0.649976968 | 0.625298979 | 0.669431245 | 0.038893781 | 0.186197852 | 0.20869213 | 0.407333337 | 0.602078794 | 0.734747706 | 0.715 | 0.808894408 |
| SL1344_RS08115_ | -0.392762967 | -0.6279424 | -0.197530972 | 0.453920405 | 0.307812732 | 1.466943735 | 0.043970787 | 0.097334864 | 0.241316476 | 0.1573 | 0.313050436 |
| SL1344_RS08120_ | -5.074819072 | -5.55060521 | -4.722625348 | 0.797793456 | 1.310466987 | 0.585884492 | 0.252000012 | 0.387603924 | 0.552366048 | 0.4941 | 0.63137904 |
| SL1344_RS08125_ | -3.301168976 | -3.319042967 | -3.287730924 | -0.001613871 | 0.672969984 | -0.002711155 | 0.499333333 | 0.621303344 | 0.75080874 | 0.7247 | 0.819786488 |
| SL1344_RS08130_ | -3.200077307 | -3.299156875 | -3.062734205 | 0.219646747 | 0.602374316 | 0.349222939 | 0.348666673 | 0.53476319 | 0.673790059 | 0.6421 | 0.749404076 |
| SL1344_RS08135_ | -1.692856873 | -1.545839609 | -1.883341259 | -0.346245376 | 0.344361825 | -0.927348941 | 0.153333358 | 0.237258251 | 0.396956232 | 0.319 | 0.47246334 |
| SL1344_RS08140_ | -2.144771382 | -1.990290339 | -2.226675887 | -0.258469147 | 0.527349694 | -0.468235247 | 0.305333342 | 0.470643275 | 0.63088768 | 0.6058 | 0.724700426 |
| SL1344_RS08145_ | -2.932823574 | -2.893196643 | -2.971439356 | -0.061188477 | 0.611727521 | -0.091432644 | 0.462000001 | 0.600451156 | 0.734202617 | 0.7135 | 0.808370057 |
| SL1344_RS08150_ | -3.962225519 | -3.812971438 | -4.086445199 | -0.241193189 | 0.998424739 | -0.218082187 | 0.40266667 | 0.617999035 | 0.74764763 | 0.7165 | 0.812104664 |
| SL1344_RS08155_ | -0.942417031 | -0.895897382 | -1.01257712 | -0.10657059 | 0.270160903 | -0.374404936 | 0.337774823 | 0.495196889 | 0.64174206 | 0.596 | 0.715940835 |
| SL1344_RS08160_ | 2.600596624 | 3.01851301 | 2.12007831 | -0.903116739 | 0.09881674 | -9.097000283 | 4.68E-05 | 0.000936733 | 0.019360192 | 0.1 | 0.243021319 |
| SL1344_RS08165_ | 0.935392789 | 1.101072139 | 0.823317762 | -0.29071554 | 0.226873795 | -1.228021524 | 0.082666718 | 0.134301116 | 0.298770023 | 0.1982 | 0.360602244 |
| SL1344_RS08170_ | 0.437141543 | 0.601258637 | 0.279781857 | -0.32233138 | 0.178086411 | -1.787011146 | 0.020653117 | 0.06231646 | 0.179688297 | 0.1261 | 0.275359981 |
| SL1344_RS08175_ | 3.136586261 | 3.289297757 | 2.95190701 | -0.335038732 | 0.076231082 | -4.30500177 | 4.68E-05 | 0.006404437 | 0.050291884 | 0.1 | 0.243021319 |
| SL1344_RS08185_ | -0.529564232 | -0.406029008 | -0.748507115 | -0.337147212 | 0.251983897 | -1.252529302 | 0.092666712 | 0.149924543 | 0.30352619 | 0.2325 | 0.390132373 |
| SL1344_RS08190_ | -0.841632744 | -0.71638663 | -0.920958677 | -0.184474315 | 0.308368152 | -0.566498568 | 0.268000011 | 0.436002645 | 0.596263908 | 0.5419 | 0.676043521 |
| SL1344_RS08195_ | -0.672715435 | -0.628707725 | -0.744630899 | -0.11038978 | 0.266428722 | -0.390978693 | 0.337108601 | 0.502497157 | 0.653957456 | 0.6123 | 0.730718647 |
| SL1344_RS08200_ | 2.016119717 | 2.039714032 | 2.001144468 | -0.036196248 | 0.122465487 | -0.27173043 | 0.3904064 | 0.600372105 | 0.734700914 | 0.6996 | 0.800518176 |
| SL1344_RS08215_ | 1.020825657 | 1.145403719 | 0.872707585 | -0.286393508 | 0.176423939 | -1.564099853 | 0.023333527 | 0.093943449 | 0.237629643 | 0.1259 | 0.275056355 |
| SL1344_RS08220_ | -2.298032186 | -2.178397276 | -2.425987666 | -0.259972914 | 0.461335143 | -0.523141594 | 0.272666677 | 0.426476241 | 0.582473637 | 0.5438 | 0.668690595 |
| SL1344_RS08225_ | 1.767642877 | 1.843351523 | 1.704497878 | -0.141026076 | 0.104608466 | -1.253366326 | 0.081945421 | 0.137524736 | 0.285863425 | 0.2089 | 0.365262272 |
| SL1344_RS08230_ | -2.360436099 | -2.281487222 | -2.415953424 | -0.193821854 | 0.630603057 | -0.269140057 | 0.395069957 | 0.604654665 | 0.740638716 | 0.7363 | 0.826012897 |
| SL1344_RS08235_ | 0.441844652 | 0.782109421 | 0.278093272 | -0.494526337 | 0.401872184 | -1.23742364 | 0.096666709 | 0.120490676 | 0.286175731 | 0.1968 | 0.368337875 |
| SL1344_RS08240_ | 1.114056048 | 1.129063642 | 1.072693846 | -0.061639241 | 0.240344384 | -0.240360619 | 0.40066667 | 0.580703197 | 0.730592605 | 0.7511 | 0.8418676 |
| SL1344_RS08245_ | -1.135060704 | -1.111627016 | -1.201689756 | -0.067941066 | 0.451153165 | -0.12472574 | 0.453697537 | 0.750187512 | 0.850612911 | 0.8185 | 0.894238913 |
| SL1344_RS08250_ | 0.128390712 | 0.126340816 | 0.131940695 | -0.016475083 | 0.286113636 | -0.05349338 | 0.475682879 | 0.698863716 | 0.811685018 | 0.8102 | 0.879772043 |
| SL1344_RS08255_ | 3.088541146 | 3.096815538 | 3.079795423 | -0.013114603 | 0.079479401 | -0.158349318 | 0.432000003 | 0.624418477 | 0.753167403 | 0.7343 | 0.825229408 |
| SL1344_RS08260_ | 1.561690166 | 1.529117334 | 1.590908217 | 0.060139704 | 0.117088641 | 0.474656229 | 0.290473027 | 0.434703886 | 0.586607524 | 0.5363 | 0.661161426 |
| SL1344_RS08270_ | 0.4866068 | 0.825872907 | -0.043132346 | -0.880416423 | 0.208660333 | -4.224060959 | 4.68E-05 | 0.009878303 | 0.063031713 | 0.1 | 0.243021319 |
| SL1344_RS08275_ | 1.899587292 | 2.102423991 | 1.704640209 | -0.400483499 | 0.123489829 | -3.233952282 | 4.68E-05 | 0.012582836 | 0.0733251 | 0.1 | 0.243021319 |
| SL1344_RS08280_ | 0.354083425 | 0.522599129 | 0.003383976 | -0.504114872 | 0.223020096 | -2.05822714 | 0.017333596 | 0.058803687 | 0.179470652 | 0.1175 | 0.265669463 |
| SL1344_RS08290_ | 2.697617577 | 2.794620778 | 2.598126097 | -0.194857052 | 0.070376537 | -2.784566687 | 0.002000966 | 0.020912283 | 0.09317221 | 0.1017 | 0.2452759 |
| SL1344_RS08295_ | 2.58956415 | 2.968070983 | 2.046924536 | -0.914198095 | 0.115449761 | -7.718632504 | 4.68E-05 | 0.003492548 | 0.036947791 | 0.1 | 0.243021319 |
| SL1344_RS08300_ | 0.014379328 | 0.107076961 | -0.068775304 | -0.177226895 | 0.177337474 | -0.943815982 | 0.147235203 | 0.239032661 | 0.392384723 | 0.3284 | 0.47434605 |
| SL1344_RS08305_ | 0.04756037 | 0.18761718 | -0.133860865 | -0.33732725 | 0.263744698 | -1.26032239 | 0.07866672 | 0.132223442 | 0.289275117 | 0.2074 | 0.368261589 |
| SL1344_RS08310_ | -0.671203937 | -0.687420901 | -0.657962066 | 0.028435933 | 0.235999659 | 0.121150917 | 0.444370422 | 0.54136636 | 0.681340809 | 0.6438 | 0.749931482 |
| SL1344_RS08315_ | -1.067024741 | -1.181470901 | -0.972099587 | 0.207220621 | 0.296464898 | 0.688425414 | 0.205862775 | 0.320986858 | 0.488415188 | 0.4205 | 0.56389224 |
| SL1344_RS08320_ | -4.022367884 | -3.743634337 | -4.456488481 | -0.785632249 | 1.276500635 | -0.569221789 | 0.186000019 | 0.319827545 | 0.514281214 | 0.3863 | 0.535509715 |
| SL1344_RS08325_ | -4.400738568 | -3.998356841 | -4.797102989 | -0.768765139 | 1.141612484 | -0.643128532 | 0.23184545 | 0.369226616 | 0.53801486 | 0.4669 | 0.611629106 |
| SL1344_RS08330_ | -3.261590808 | -3.060147887 | -3.511656468 | -0.470318193 | 0.574305965 | -0.769473276 | 0.195869438 | 0.290955599 | 0.451411134 | 0.3854 | 0.529099698 |

| | | | | | | | | | | | |
|---|---|---|---|---|---|---|---|---|---|---|---|
| SL1344_RS08335_ | -3.556074449 | -3.314369212 | -3.973400105 | -0.739412619 | 1.029420722 | -0.661505005 | 0.172551653 | 0.300879955 | 0.487914232 | 0.3728 | 0.518563739 |
| SL1344_RS08340_ | -3.996324711 | -3.673405764 | -4.317227434 | -0.595218202 | 0.980477523 | -0.581205256 | 0.253333345 | 0.406377517 | 0.570865754 | 0.5178 | 0.654994406 |
| SL1344_RS08345_ | 2.415834846 | 2.437105763 | 2.365159638 | -0.07822873 | 0.164980489 | -0.436313738 | 0.329780154 | 0.495577479 | 0.666014565 | 0.6717 | 0.785575873 |
| SL1344_RS08350_ | -0.671202434 | -0.588462304 | -1.003759991 | -0.377679138 | 0.367316157 | -0.903655642 | 0.210000016 | 0.285260703 | 0.472639234 | 0.4251 | 0.583983298 |
| SL1344_RS08355_ | -1.411552213 | -1.282610359 | -1.645409814 | -0.349503196 | 0.429616598 | -0.777865316 | 0.205196552 | 0.308711671 | 0.485592719 | 0.4131 | 0.566716999 |
| SL1344_RS08360_ | -0.286620724 | -0.292378793 | -0.27975349 | 0.008167571 | 0.222420078 | 0.038630453 | 0.483011326 | 0.581929146 | 0.716336314 | 0.6888 | 0.788415304 |
| SL1344_RS08365_ | 0.030312846 | 0.100118546 | -0.012207966 | -0.078434854 | 0.240010978 | -0.320351896 | 0.36109261 | 0.564466841 | 0.707232912 | 0.6702 | 0.781080213 |
| SL1344_RS08370_ | -1.902378669 | -1.907317737 | -1.899703699 | -0.026298788 | 0.489552984 | -0.050645818 | 0.480346436 | 0.657334486 | 0.780408971 | 0.7747 | 0.855339139 |
| SL1344_RS08375_ | -3.609135102 | -3.445552146 | -3.89689564 | -0.427006691 | 0.748155298 | -0.528340692 | 0.289140582 | 0.440771096 | 0.599326711 | 0.5562 | 0.685064771 |
| SL1344_RS08380_ | 2.064972035 | 2.229171218 | 1.851303793 | -0.382538868 | 0.113956496 | -3.32394705 | 4.68E-05 | 0.015126454 | 0.080091931 | 0.1 | 0.243021319 |
| SL1344_RS08385_ | -0.456476235 | -0.321083306 | -0.615134341 | -0.296909923 | 0.214088144 | -1.332935231 | 0.077333388 | 0.130725218 | 0.271420919 | 0.2009 | 0.353916863 |
| SL1344_RS08390_ | 0.238421548 | 0.39410931 | 0.108756988 | -0.292264167 | 0.180201556 | -1.575220754 | 0.040666776 | 0.089595842 | 0.221906362 | 0.1567 | 0.310503799 |
| SL1344_RS08395_ | 5.007136488 | 5.073675931 | 4.905638048 | -0.164288295 | 0.093973479 | -1.696516555 | 0.006000767 | 0.08155762 | 0.229136289 | 0.1064 | 0.251902049 |
| SL1344_RS08400_ | 1.816041717 | 1.844867822 | 1.792425634 | -0.077230117 | 0.169526738 | -0.455585272 | 0.316455704 | 0.472017125 | 0.636968427 | 0.6336 | 0.745809021 |
| SL1344_RS08405_ | 1.141176347 | 1.310659093 | 0.956043142 | -0.382363262 | 0.197590136 | -1.928138808 | 0.003998486 | 0.055767734 | 0.177623406 | 0.1045 | 0.249083092 |
| SL1344_RS08410_ | 0.847348664 | 1.158223026 | 0.648987465 | -0.509726918 | 0.19163219 | -2.581921828 | 4.68E-05 | 0.032773064 | 0.127883233 | 0.1007 | 0.244001247 |
| SL1344_RS08415_ | 0.295641004 | 0.632535176 | -0.061265647 | -0.644944971 | 0.280764912 | -2.249891848 | 0.005330642 | 0.031301463 | 0.125429641 | 0.1046 | 0.249440256 |
| SL1344_RS08420_ | -1.051118278 | -0.780494006 | -1.356363138 | -0.576829037 | 0.362761547 | -1.541850013 | 0.0379748 | 0.081973257 | 0.217923657 | 0.1459 | 0.301127577 |
| SL1344_RS08425_ | -0.602294266 | -0.232541367 | -0.924106012 | -0.740462868 | 0.343662612 | -2.131626536 | 0.002666616 | 0.045071484 | 0.15495722 | 0.1031 | 0.247222376 |
| SL1344_RS08430_ | -1.394951154 | -1.132803652 | -1.764610691 | -0.646591246 | 0.364485535 | -1.729488051 | 0.01866691 | 0.077142282 | 0.206555584 | 0.1219 | 0.268957193 |
| SL1344_RS08435_ | 2.418261396 | 2.553335385 | 2.253260274 | -0.351191514 | 0.205432995 | -1.80973187 | 0.002666616 | 0.059594055 | 0.188313347 | 0.1018 | 0.245489565 |
| SL1344_RS08440_ | -0.127196932 | -0.047849719 | -0.211800523 | -0.158698649 | 0.208665106 | -0.709464501 | 0.219333349 | 0.337394773 | 0.499923976 | 0.44 | 0.581494101 |
| SL1344_RS08445_ | 2.228666513 | 2.276511933 | 2.186954701 | -0.067613301 | 0.159188796 | -0.400492521 | 0.320666674 | 0.559855528 | 0.710790972 | 0.6514 | 0.769810082 |
| SL1344_RS08450_ | 1.357907949 | 1.34794102 | 1.361981641 | -0.00467589 | 0.17508929 | -0.025048506 | 0.488341106 | 0.70870645 | 0.817893582 | 0.7953 | 0.872369162 |
| SL1344_RS08455_ | 1.112054026 | 1.168943411 | 1.055678577 | -0.110123159 | 0.126245407 | -0.833568188 | 0.177333354 | 0.276834244 | 0.433766876 | 0.3634 | 0.51148239 |
| SL1344_RS08460_ | 1.391203311 | 1.415986055 | 1.358539863 | -0.061469185 | 0.133215516 | -0.438340093 | 0.312000008 | 0.462400041 | 0.61767819 | 0.5814 | 0.700922213 |
| SL1344_RS08465_ | 0.821060139 | 0.826901846 | 0.815007896 | -0.017127089 | 0.16552853 | -0.096756647 | 0.458361094 | 0.603649579 | 0.735025851 | 0.7199 | 0.810903295 |
| SL1344_RS08470_ | -0.312717669 | -0.281402355 | -0.336710061 | 0.011567119 | 0.408723082 | 0.027038685 | 0.485009994 | 0.746570768 | 0.847070216 | 0.8722 | 0.922131246 |
| SL1344_RS08475_ | -1.779497782 | -1.654102611 | -1.87061836 | -0.22076604 | 0.436846174 | -0.469469537 | 0.302666675 | 0.472743964 | 0.63022083 | 0.5906 | 0.712312552 |
| SL1344_RS08480_ | -6.15231732 | -6.061114394 | -6.246332223 | -0.226608744 | 1.848082863 | -0.101234073 | 0.444370422 | 0.573697012 | 0.713260304 | 0.6901 | 0.789785048 |
| SL1344_RS08485_ | -0.839452494 | -0.655142899 | -1.431395666 | -0.745867784 | 0.453112586 | -1.428534566 | 0.046000096 | 0.117123235 | 0.279057003 | 0.1546 | 0.310985466 |
| SL1344_RS08490_ | 5.782670098 | 5.914153648 | 5.58704086 | -0.322793033 | 0.043990772 | -7.253628137 | 4.68E-05 | 0.002751885 | 0.033538081 | 0.1 | 0.243021319 |
| SL1344_RS08500_ | 4.440966618 | 4.52637841 | 4.272839008 | -0.22564156 | 0.0914715 | -2.165701334 | 0.004667653 | 0.0458148 | 0.160615147 | 0.1052 | 0.250221134 |
| SL1344_RS08505_ | 0.41864231 | 0.557992 | 0.344504142 | -0.16014998 | 0.299751194 | -0.495352251 | 0.308000008 | 0.546562573 | 0.698558688 | 0.6224 | 0.75318493 |
| SL1344_RS08510_ | 0.903708978 | 1.074526823 | 0.696102804 | -0.37291654 | 0.17358481 | -2.067748805 | 0.014657206 | 0.044412571 | 0.150195769 | 0.1136 | 0.261105854 |
| SL1344_RS08515_ | 1.545965147 | 1.760810983 | 1.319112705 | -0.447246656 | 0.123027043 | -3.510763062 | 4.68E-05 | 0.011268838 | 0.067745065 | 0.1 | 0.243021319 |
| SL1344_RS08520_ | 1.199057482 | 1.407962447 | 0.922273548 | -0.50364782 | 0.210542056 | -2.474279518 | 4.68E-05 | 0.05658178 | 0.181176771 | 0.1 | 0.243021319 |
| SL1344_RS08525_ | 1.442429351 | 1.625631268 | 1.241956963 | -0.345717911 | 0.183722801 | -1.898603262 | 0.023984199 | 0.056088482 | 0.175870222 | 0.1267 | 0.279653379 |
| SL1344_RS08530_ | 1.553897687 | 1.647635509 | 1.482898364 | -0.164488731 | 0.115795487 | -1.35910544 | 0.069333395 | 0.125438792 | 0.269032354 | 0.1953 | 0.348703444 |
| SL1344_RS08535_ | 2.55539142 | 2.762972846 | 2.362390408 | -0.399578136 | 0.088544863 | -4.43615086 | 4.68E-05 | 0.006075757 | 0.048700189 | 0.1 | 0.243021319 |
| SL1344_RS08540_ | 2.406676584 | 2.552185641 | 2.075613355 | -0.472378442 | 0.139644898 | -3.202657239 | 4.68E-05 | 0.030442208 | 0.124732684 | 0.1 | 0.243021319 |
| SL1344_RS08545_ | -0.84615862 | -0.710424136 | -1.008567507 | -0.301907526 | 0.284390443 | -1.028976698 | 0.129333364 | 0.201417865 | 0.366769365 | 0.2754 | 0.434620158 |
| SL1344_RS08550_ | 2.769435361 | 2.842124315 | 2.646876139 | -0.203075372 | 0.126696432 | -1.62209793 | 0.006667356 | 0.121567944 | 0.289447318 | 0.1085 | 0.254151598 |
| SL1344_RS08555_ | 0.876308176 | 1.216603239 | 0.497943985 | -0.727822674 | 0.191953938 | -3.727791157 | 4.68E-05 | 0.009306833 | 0.062030134 | 0.1 | 0.243021319 |
| SL1344_RS08560_ | 1.798582478 | 1.826053141 | 1.780249654 | -0.042911486 | 0.142249242 | -0.27866186 | 0.373750838 | 0.610961643 | 0.743635767 | 0.7165 | 0.812702327 |
| SL1344_RS08570_ | 4.629456326 | 4.678432289 | 4.442729399 | -0.227638763 | 0.072214966 | -2.038027503 | 0.0020023 | 0.070515219 | 0.209433097 | 0.1032 | 0.247319216 |
| SL1344_RS08575_ | 2.965312494 | 3.164260894 | 2.832023215 | -0.34904986 | 0.156958765 | -2.303762445 | 4.68E-05 | 0.032471918 | 0.130763088 | 0.1002 | 0.243300986 |
| SL1344_RS08580_ | 1.191499805 | 1.398195444 | 1.03638133 | -0.356856052 | 0.208528759 | -1.65992919 | 0.031333476 | 0.066988048 | 0.196680887 | 0.1305 | 0.283514038 |
| SL1344_RS08590_ | 1.303753112 | 1.456287593 | 1.069265494 | -0.387231266 | 0.155955408 | -2.356747474 | 0.002668394 | 0.030537619 | 0.121024859 | 0.1024 | 0.246386704 |
| SL1344_RS08595_ | 3.818341083 | 3.831442958 | 3.805208108 | -0.035351398 | 0.099443628 | -0.361705312 | 0.353097941 | 0.569993088 | 0.720007114 | 0.7136 | 0.814537726 |
| SL1344_RS08600_ | 1.799782669 | 2.118170493 | 1.560368492 | -0.556664847 | 0.183915488 | -2.976069918 | 4.68E-05 | 0.013366649 | 0.078291274 | 0.1 | 0.243021319 |
| SL1344_RS08605_ | 1.803258513 | 1.982013342 | 1.594809169 | -0.392111822 | 0.137777814 | -2.812984743 | 0.00067292 | 0.021089434 | 0.098243539 | 0.1001 | 0.243158387 |
| SL1344_RS08610_ | -1.383798246 | -1.399603448 | -1.370441799 | 0.02828293 | 0.349758618 | 0.069677385 | 0.471685544 | 0.605819837 | 0.735656344 | 0.7159 | 0.807907453 |

| | | | | | | | | | | | |
|---|---|---|---|---|---|---|---|---|---|---|---|
| SL1344_RS08615_ | -0.734484977 | -0.64303685 | -0.878370666 | -0.22623828 | 0.283582285 | -0.737797438 | 0.20533335 | 0.316614563 | 0.482544076 | 0.4196 | 0.567720406 |
| SL1344_RS08620_ | 0.154052987 | 0.279925388 | 0.004584281 | -0.273253979 | 0.187316725 | -1.405934037 | 0.050000088 | 0.10948536 | 0.247831652 | 0.1729 | 0.325088335 |
| SL1344_RS08625_ | 0.065193505 | 0.263081551 | -0.285057402 | -0.543282937 | 0.215887613 | -2.473050937 | 0.002000966 | 0.037620491 | 0.134254459 | 0.1023 | 0.246098182 |
| SL1344_RS08630_ | 0.362952412 | 0.51738849 | 0.124645162 | -0.384343587 | 0.235053165 | -1.526725713 | 0.048666757 | 0.089577255 | 0.229778263 | 0.1521 | 0.308428708 |
| SL1344_RS08635_ | -5.00758235 | -5.160602244 | -4.803010247 | 0.352502928 | 1.132695304 | 0.271510466 | 0.382666671 | 0.540309809 | 0.682564108 | 0.6396 | 0.751845324 |
| SL1344_RS08640_ | -5.62145318 | -5.654467709 | -5.591718049 | 0.079599551 | 1.416011609 | 0.045240301 | 0.475016656 | 0.59502381 | 0.729144251 | 0.7109 | 0.805577002 |
| SL1344_RS08645_ | -5.656767562 | -5.910400058 | -5.27753877 | 0.615201135 | 1.909802514 | 0.286028339 | 0.364423723 | 0.596279555 | 0.731329686 | 0.6911 | 0.793615839 |
| SL1344_RS08650_ | -2.125948618 | -1.954092326 | -2.426334952 | -0.462456183 | 0.485066353 | -0.878650783 | 0.167333355 | 0.261232992 | 0.434677086 | 0.3608 | 0.516865778 |
| SL1344_RS08655_ | -0.689569528 | -0.518002024 | -0.958302175 | -0.439331449 | 0.265536841 | -1.571115787 | 0.040000111 | 0.093821761 | 0.229348183 | 0.1511 | 0.303508331 |
| SL1344_RS08660_ | -0.547972263 | -0.477048939 | -0.627488683 | -0.126143543 | 0.316929031 | -0.378436564 | 0.342666673 | 0.549869878 | 0.696062324 | 0.6611 | 0.770514889 |
| SL1344_RS08665_ | 2.765760088 | 2.92769418 | 2.620505591 | -0.308263152 | 0.081733622 | -3.693118368 | 4.68E-05 | 0.009185902 | 0.060871779 | 0.1 | 0.243021319 |
| SL1344_RS08670_ | 3.172889908 | 3.290037532 | 3.062538783 | -0.228064931 | 0.063316544 | -3.511492809 | 4.68E-05 | 0.010108478 | 0.062865872 | 0.1 | 0.243021319 |
| SL1344_RS08675_ | 2.321708187 | 2.442853901 | 2.121555567 | -0.308793629 | 0.11478881 | -2.540438738 | 0.001335878 | 0.032194302 | 0.126130784 | 0.1016 | 0.245264602 |
| SL1344_RS08680_ | 3.564015209 | 3.704274331 | 3.452097233 | -0.228767864 | 0.107031848 | -2.144561283 | 0.007995244 | 0.034410314 | 0.133946045 | 0.107 | 0.252820378 |
| SL1344_RS08685_ | 0.527734455 | 0.753462609 | 0.277961671 | -0.479155803 | 0.177609269 | -2.64297989 | 0.002000966 | 0.023514499 | 0.101545523 | 0.1008 | 0.244128891 |
| SL1344_RS08690_ | 0.791936466 | 1.057037293 | 0.547121457 | -0.49082074 | 0.198280958 | -2.400481136 | 0.002666616 | 0.026975404 | 0.113486798 | 0.1025 | 0.246516707 |
| SL1344_RS08695_ | 1.931087024 | 2.133958859 | 1.653997054 | -0.498955837 | 0.159760209 | -3.162910939 | 4.68E-05 | 0.03163779 | 0.127646084 | 0.1 | 0.243021319 |
| SL1344_RS08700_ | -0.111475073 | 0.005617432 | -0.223699672 | -0.208978253 | 0.248506367 | -0.791460716 | 0.199333351 | 0.311406488 | 0.483827307 | 0.4051 | 0.56117119 |
| SL1344_RS08705_ | 1.135410669 | 1.396519548 | 0.841980974 | -0.520401658 | 0.206141148 | -2.441580884 | 0.003334715 | 0.028086432 | 0.117428788 | 0.1032 | 0.247515067 |
| SL1344_RS08710_ | 0.098475965 | -0.043608975 | 0.182752916 | 0.213016766 | 0.205902139 | 1.002232529 | 0.143904091 | 0.217503128 | 0.383633043 | 0.3062 | 0.464658072 |
| SL1344_RS08715_ | -0.547612578 | -0.746084427 | -0.275122536 | 0.451958815 | 0.334219878 | 1.284425759 | 0.088607642 | 0.132590588 | 0.292780132 | 0.2085 | 0.375848396 |
| SL1344_RS08725_ | 1.625476561 | 1.652063753 | 1.591317506 | -0.05758571 | 0.115633127 | -0.470247664 | 0.311125924 | 0.461345499 | 0.614844425 | 0.5665 | 0.691148281 |
| SL1344_RS08730_ | 0.882037211 | 0.949103175 | 0.853346272 | -0.065683863 | 0.216320631 | -0.265865604 | 0.398666671 | 0.701574956 | 0.814298841 | 0.7838 | 0.867220334 |
| SL1344_RS08735_ | 4.758954343 | 4.784750944 | 4.722690969 | -0.075755055 | 0.083927437 | -0.892824406 | 0.155896093 | 0.226726895 | 0.417695412 | 0.3232 | 0.497416918 |
| SL1344_RS08740_ | 1.478634941 | 1.598032993 | 1.228964665 | -0.356984368 | 0.156530685 | -2.113343238 | 0.010659989 | 0.053494861 | 0.16975208 | 0.1093 | 0.255223417 |
| SL1344_RS08750_ | 4.147613265 | 4.374802789 | 3.925712926 | -0.450065714 | 0.049658297 | -9.065194934 | 4.68E-05 | 0.001009089 | 0.019584886 | 0.1 | 0.243021319 |
| SL1344_RS08755_ | 1.606332306 | 1.721529185 | 1.315077256 | -0.399157909 | 0.167873878 | -2.058712722 | 0.011992386 | 0.061052587 | 0.186703247 | 0.1087 | 0.254287913 |
| SL1344_RS08760_ | 1.612024373 | 1.797555376 | 1.320730813 | -0.472534023 | 0.124196647 | -3.660792731 | 4.68E-05 | 0.014862546 | 0.078518821 | 0.1 | 0.243021319 |
| SL1344_RS08765_ | 1.149104003 | 1.406892189 | 0.853020353 | -0.560340712 | 0.134066138 | -4.135981467 | 4.68E-05 | 0.006936121 | 0.051604969 | 0.1 | 0.243021319 |
| SL1344_RS08770_ | 2.485850174 | 2.800872564 | 2.107157273 | -0.696065804 | 0.085153683 | -8.161217386 | 4.68E-05 | 0.001881003 | 0.025587866 | 0.1 | 0.243021319 |
| SL1344_RS08775_ | 4.274484713 | 4.235708914 | 4.29051094 | 0.047276156 | 0.121972425 | 0.290120701 | 0.378414395 | 0.861195124 | 0.923210782 | 0.8071 | 0.890322561 |
| SL1344_RS08780_ | 3.048180944 | 3.068980412 | 2.999705954 | -0.077273039 | 0.127849105 | -0.549650069 | 0.260493016 | 0.383766211 | 0.572225911 | 0.5315 | 0.673636253 |
| SL1344_RS08785_ | -0.537653377 | -0.236735091 | -1.044190817 | -0.795204914 | 0.307910057 | -2.47150003 | 0.001335878 | 0.037130115 | 0.1360149 | 0.1013 | 0.244794092 |
| SL1344_RS08815_ | 1.316030024 | 1.449625178 | 1.164902938 | -0.289118175 | 0.171984329 | -1.643872653 | 0.032666804 | 0.074668123 | 0.207511477 | 0.1328 | 0.285614365 |
| SL1344_RS08825_ | 0.57213566 | 0.819748997 | 0.333026591 | -0.476884443 | 0.18901385 | -2.490648534 | 0.002668394 | 0.026543862 | 0.110841687 | 0.1019 | 0.245690487 |
| SL1344_RS08830_ | 2.53023263 | 2.682987337 | 2.373218829 | -0.310176002 | 0.087500802 | -3.451796889 | 4.68E-05 | 0.012608297 | 0.071513668 | 0.1 | 0.243021319 |
| SL1344_RS08840_ | 0.494505562 | 0.470669953 | 0.517800707 | 0.047385308 | 0.157386312 | 0.271938563 | 0.378666671 | 0.505116239 | 0.64798333 | 0.6045 | 0.719249465 |
| SL1344_RS08845_ | 3.719417259 | 3.832566811 | 3.623624842 | -0.222226295 | 0.094470689 | -2.457058876 | 0.00067292 | 0.035594305 | 0.136901448 | 0.1003 | 0.243445865 |
| SL1344_RS08850_ | 2.61933853 | 2.675458986 | 2.567127386 | -0.126170441 | 0.148415942 | -0.882596539 | 0.148567647 | 0.246243001 | 0.439877523 | 0.3031 | 0.468215906 |
| SL1344_RS08855_ | 1.427818071 | 1.441599336 | 1.411786266 | -0.032162511 | 0.150156261 | -0.214430724 | 0.415333337 | 0.616105297 | 0.747115544 | 0.7356 | 0.82528319 |
| SL1344_RS08860_ | 2.756076339 | 2.781303096 | 2.737322111 | -0.031275945 | 0.100625928 | -0.30328186 | 0.377748173 | 0.598872962 | 0.733932673 | 0.7053 | 0.805268566 |
| SL1344_RS08865_ | 3.435564946 | 3.413280387 | 3.45710199 | 0.016638803 | 0.121801614 | 0.149889337 | 0.444000002 | 0.751927472 | 0.851713682 | 0.8563 | 0.912643474 |
| SL1344_RS08870_ | 3.849742401 | 3.974267901 | 3.785019069 | -0.184543546 | 0.080877599 | -2.183798929 | 0.008661422 | 0.048174916 | 0.16182919 | 0.1052 | 0.250192576 |
| SL1344_RS08880_ | 4.709506699 | 4.923340035 | 4.554948432 | -0.363714479 | 0.056964719 | -6.185453324 | 4.68E-05 | 0.005509424 | 0.048686579 | 0.1 | 0.243021319 |
| SL1344_RS08885_ | 6.187607842 | 6.343690904 | 6.018716955 | -0.280916559 | 0.080568629 | -3.744789223 | 4.68E-05 | 0.003707266 | 0.041928473 | 0.1 | 0.243021319 |
| SL1344_RS08890_ | 2.109843563 | 2.310217702 | 1.933818041 | -0.38079724 | 0.120695047 | -3.189378476 | 4.68E-05 | 0.01570943 | 0.083707961 | 0.1 | 0.243021319 |
| SL1344_RS08895_ | -3.364560316 | -3.914560276 | -2.823205939 | 0.9832072 | 0.883885009 | 1.065157348 | 0.124583643 | 0.185870308 | 0.357660379 | 0.2595 | 0.429995873 |
| SL1344_RS08900_ | 1.282728409 | 1.386845152 | 1.182002394 | -0.224037086 | 0.161953887 | -1.376121501 | 0.055296548 | 0.12038684 | 0.274375425 | 0.1643 | 0.318246575 |
| SL1344_RS08905_ | 4.656919614 | 4.703736258 | 4.57687272 | -0.132492379 | 0.071700177 | -1.778301694 | 0.019986902 | 0.056954848 | 0.179598846 | 0.1171 | 0.265606589 |
| SL1344_RS08910_ | 1.245939094 | 1.408050224 | 1.047292566 | -0.362556002 | 0.127577409 | -2.762009118 | 0.001336769 | 0.02090913 | 0.094862239 | 0.1011 | 0.244486479 |
| SL1344_RS08915_ | -4.877371144 | -4.63211973 | -5.045466975 | -0.547991171 | 1.285724385 | -0.386254677 | 0.333777489 | 0.503700302 | 0.653059294 | 0.6381 | 0.744435308 |
| SL1344_RS08920_ | -2.133916114 | -1.978191614 | -2.377712462 | -0.398845855 | 0.464808286 | -0.825362207 | 0.174550321 | 0.283935776 | 0.454306372 | 0.3849 | 0.534296864 |
| SL1344_RS08925_ | -4.68675605 | -4.641674756 | -4.717622953 | -0.148602456 | 1.101879193 | -0.117898909 | 0.451032647 | 0.612406893 | 0.74319729 | 0.7294 | 0.8222336 |

| | | | | | | | | | | | |
|---|---|---|---|---|---|---|---|---|---|---|---|
| SL1344_RS08930_ | -4.199799965 | -4.202454233 | -4.196898353 | -0.150991357 | 1.552583864 | -0.071879504 | 0.450366424 | 0.598475627 | 0.74594746 | 0.7798 | 0.859117982 |
| SL1344_RS08935_ | -5.033573126 | -4.640841001 | -5.331978209 | -0.694720375 | 1.877847632 | -0.337945109 | 0.316455704 | 0.544673055 | 0.69505475 | 0.6299 | 0.750111561 |
| SL1344_RS08940_ | -3.42563713 | -3.04993118 | -3.976605075 | -0.974204285 | 0.885683183 | -1.054006417 | 0.088000048 | 0.188042309 | 0.362453422 | 0.2212 | 0.377991037 |
| SL1344_RS08945_ | -2.323509464 | -2.040332186 | -2.6758265 | -0.635926399 | 0.404023219 | -1.482989223 | 0.047968113 | 0.093643924 | 0.225646018 | 0.1524 | 0.304552545 |
| SL1344_RS08950_ | -2.444929137 | -2.303792584 | -2.583214103 | -0.280823505 | 0.409356581 | -0.649357742 | 0.240000013 | 0.360017908 | 0.513542013 | 0.4589 | 0.592919322 |
| SL1344_RS08955_ | -2.768995931 | -2.551836494 | -3.192690995 | -0.615975237 | 0.636597004 | -0.880782525 | 0.164666689 | 0.251873534 | 0.428518779 | 0.3541 | 0.513781466 |
| SL1344_RS08960_ | 3.133663021 | 3.545117608 | 2.626335816 | -0.924796048 | 0.122288414 | -7.576835669 | 4.68E-05 | 0.001777088 | 0.027095825 | 0.1 | 0.243021319 |
| SL1344_RS08965_ | 0.108541217 | 0.340844758 | -0.269103842 | -0.600355542 | 0.214291672 | -2.723140271 | 0.001335878 | 0.027874266 | 0.112250623 | 0.102 | 0.245690051 |
| SL1344_RS08970_ | -2.646559406 | -2.410660068 | -2.911662571 | -0.521780954 | 0.446366071 | -1.08470318 | 0.119253864 | 0.182843498 | 0.33404766 | 0.2608 | 0.413053123 |
| SL1344_RS08985_ | 0.005587117 | 0.193734005 | -0.21244511 | -0.407790333 | 0.186519077 | -2.148118156 | 0.007995244 | 0.043039458 | 0.141411448 | 0.1103 | 0.255953142 |
| SL1344_RS08995_ | 0.482534916 | 0.890832805 | -0.041901002 | -0.93658282 | 0.160215215 | -5.785785575 | 4.68E-05 | 0.003174511 | 0.033615475 | 0.1 | 0.243021319 |
| SL1344_RS09000_ | 0.483176976 | 0.831240234 | -0.077041551 | -0.899646518 | 0.188845971 | -4.67933933 | 4.68E-05 | 0.009170126 | 0.05976714 | 0.1 | 0.243021319 |
| SL1344_RS09005_ | -0.9464828 | -0.764071725 | -1.21555579 | -0.465502645 | 0.304916612 | -1.454751503 | 0.048666757 | 0.109545756 | 0.25419319 | 0.1562 | 0.307931117 |
| SL1344_RS09010_ | 1.718520998 | 1.807886016 | 1.627357276 | -0.186954679 | 0.128091026 | -1.395862471 | 0.050666753 | 0.10547643 | 0.247566837 | 0.1664 | 0.321498705 |
| SL1344_RS09015_ | 1.797811964 | 1.931494521 | 1.654174264 | -0.263281484 | 0.153524093 | -1.642071571 | 0.0440001 | 0.072780229 | 0.204265979 | 0.1377 | 0.292710305 |
| SL1344_RS09020_ | 3.362873319 | 3.499919413 | 3.14895777 | -0.350063186 | 0.066522691 | -5.285905265 | 4.68E-05 | 0.004371134 | 0.04047163 | 0.1 | 0.243021319 |
| SL1344_RS09025_ | -1.511628863 | -1.223028837 | -1.864862296 | -0.633636094 | 0.301329547 | -2.01612839 | 0.009993796 | 0.047491262 | 0.148447402 | 0.1154 | 0.261787476 |
| SL1344_RS09030_ | 2.20312251 | 2.543980802 | 1.800009048 | -0.752913856 | 0.131575748 | -5.864604239 | 4.68E-05 | 0.008857995 | 0.061945042 | 0.1 | 0.243021319 |
| SL1344_RS09035_ | -1.31896573 | -1.177706306 | -1.481050079 | -0.305783281 | 0.289658345 | -1.014649367 | 0.129333364 | 0.213289352 | 0.369668095 | 0.2983 | 0.450042002 |
| SL1344_RS09040_ | 2.078732682 | 2.158393523 | 1.979181462 | -0.179630754 | 0.102833081 | -1.686562238 | 0.030646382 | 0.074556201 | 0.200185475 | 0.1321 | 0.282633648 |
| SL1344_RS09045_ | 2.171856927 | 2.192243054 | 2.133355157 | -0.063806811 | 0.114308751 | -0.491092058 | 0.287333343 | 0.447057507 | 0.610554192 | 0.571 | 0.695552378 |
| SL1344_RS09050_ | 0.954587507 | 1.007892846 | 0.910604405 | -0.099359104 | 0.148086269 | -0.625412995 | 0.251832124 | 0.3839177 | 0.542510654 | 0.4965 | 0.62902783 |
| SL1344_RS09055_ | 3.227215503 | 3.346050205 | 3.115018962 | -0.232091558 | 0.07124256 | -3.234300752 | 4.68E-05 | 0.013119113 | 0.074290317 | 0.1 | 0.243021319 |
| SL1344_RS09060_ | 3.740339944 | 3.877934103 | 3.532832157 | -0.336404808 | 0.092423748 | -3.556911665 | 4.68E-05 | 0.010454469 | 0.067866203 | 0.1 | 0.243021319 |
| SL1344_RS09065_ | 3.415022073 | 3.616660978 | 3.192374551 | -0.427991345 | 0.067987423 | -6.257063489 | 4.68E-05 | 0.003009437 | 0.033432027 | 0.1 | 0.243021319 |
| SL1344_RS09070_ | -0.28482864 | -0.473446843 | -0.148403446 | 0.336195446 | 0.236152613 | 1.354363165 | 0.046000096 | 0.120885675 | 0.270546874 | 0.1661 | 0.319487772 |
| SL1344_RS09085_ | 2.51288704 | 2.857597904 | 1.918866911 | -0.91726807 | 0.172812962 | -4.873720198 | 4.68E-05 | 0.010146183 | 0.06851198 | 0.1 | 0.243021319 |
| SL1344_RS09090_ | 2.917299206 | 3.292425705 | 2.458579654 | -0.840312177 | 0.125393972 | -6.793371308 | 4.68E-05 | 0.005407128 | 0.048207438 | 0.1 | 0.243021319 |
| SL1344_RS09095_ | 2.462091382 | 2.978931712 | 1.933616787 | -1.041181092 | 0.094314296 | -11.05197723 | 4.68E-05 | 0.000519536 | 0.014451685 | 0.1 | 0.243021319 |
| SL1344_RS09100_ | 3.144017303 | 3.66649123 | 2.639373618 | -1.006263192 | 0.132748776 | -7.564152963 | 4.68E-05 | 0.001206352 | 0.023325851 | 0.1 | 0.243021319 |
| SL1344_RS09105_ | 1.675415859 | 2.170637143 | 1.358538654 | -0.788441755 | 0.298089645 | -2.761337769 | 4.68E-05 | 0.013839821 | 0.082445809 | 0.1 | 0.243021319 |
| SL1344_RS09115_ | 1.161198876 | 1.576689421 | 0.635446308 | -0.946184053 | 0.143017777 | -6.684148535 | 4.68E-05 | 0.004163884 | 0.038717086 | 0.1 | 0.243021319 |
| SL1344_RS09125_ | -1.675330397 | -1.309162516 | -2.131758717 | -0.864301139 | 0.595684055 | -1.442479417 | 0.018000252 | 0.116303257 | 0.277716007 | 0.1217 | 0.270956013 |
| SL1344_RS09130_ | 0.592516155 | 0.920787522 | -0.085224536 | -1.009064323 | 0.311774907 | -3.165286632 | 4.68E-05 | 0.036174709 | 0.140393132 | 0.1 | 0.243021319 |
| SL1344_RS09135_ | -1.024397558 | -0.655358378 | -1.561176753 | -0.924657451 | 0.358033684 | -2.536875684 | 4.68E-05 | 0.03919606 | 0.139382275 | 0.1002 | 0.243303449 |
| SL1344_RS09140_ | 1.94030072 | 2.434570533 | 1.574446914 | -0.814355596 | 0.193895377 | -4.043010607 | 4.68E-05 | 0.008121063 | 0.060381075 | 0.1 | 0.243021319 |
| SL1344_RS09145_ | 1.535908499 | 2.037534247 | 0.969574386 | -1.080238103 | 0.21359365 | -5.200206154 | 4.68E-05 | 0.007582553 | 0.056890755 | 0.1 | 0.243021319 |
| SL1344_RS09150_ | 2.807353952 | 3.368920386 | 2.265611805 | -1.096496043 | 0.16147347 | -6.822095241 | 4.68E-05 | 0.001163497 | 0.022676963 | 0.1 | 0.243021319 |
| SL1344_RS09155_ | -0.413934542 | -0.146292812 | -0.848622876 | -0.692114323 | 0.271942113 | -2.448556779 | 0.0020023 | 0.031921274 | 0.122575912 | 0.1021 | 0.245900554 |
| SL1344_RS09160_ | 1.51712622 | 1.647365756 | 1.224417859 | -0.347843034 | 0.258154498 | -1.184370078 | 0.113333369 | 0.153472792 | 0.331241361 | 0.2293 | 0.407249891 |
| SL1344_RS09165_ | 2.047203059 | 2.439006289 | 1.516556856 | -0.947925297 | 0.195492181 | -4.95777024 | 4.68E-05 | 0.011346715 | 0.072096915 | 0.1 | 0.243021319 |
| SL1344_RS09170_ | 1.004583908 | 1.181601922 | 0.776801769 | -0.403380693 | 0.125341988 | -3.172701495 | 0.000673369 | 0.015581134 | 0.077997979 | 0.1006 | 0.243858189 |
| SL1344_RS09175_ | 6.132930754 | 6.322983467 | 5.910355509 | -0.412616115 | 0.049457863 | -8.2845554 | 4.68E-05 | 0.00060345 | 0.016995151 | 0.1 | 0.243021319 |
| SL1344_RS09180_ | 5.129292468 | 5.494505052 | 4.83394547 | -0.667370862 | 0.11528253 | -5.855240316 | 4.68E-05 | 0.003852667 | 0.042423933 | 0.1 | 0.243021319 |
| SL1344_RS09185_ | 2.633990886 | 2.941477197 | 2.187731624 | -0.755536105 | 0.178982485 | -4.274180575 | 4.68E-05 | 0.017936969 | 0.094607154 | 0.1 | 0.243021319 |
| SL1344_RS09190_ | 0.530587647 | 0.792078263 | 0.305913782 | -0.55788519 | 0.301458082 | -1.8507772 | 0.00733396 | 0.051193636 | 0.168340487 | 0.107 | 0.252142983 |
| SL1344_RS09200_ | 3.365292196 | 3.669607772 | 3.109138199 | -0.578509584 | 0.121302371 | -4.73832088 | 4.68E-05 | 0.004192421 | 0.043044201 | 0.1 | 0.243021319 |
| SL1344_RS09205_ | -0.302397667 | -0.024758489 | -0.720438475 | -0.692969173 | 0.291757116 | -2.31876265 | 0.005330642 | 0.035461604 | 0.131896452 | 0.1043 | 0.248993242 |
| SL1344_RS09210_ | 3.664071595 | 3.967377715 | 3.3357661 | -0.646697321 | 0.086165921 | -7.66437199 | 4.68E-05 | 0.001891701 | 0.028024496 | 0.1 | 0.243021319 |
| SL1344_RS09235_ | 2.939390739 | 3.3682906 | 2.455528195 | -0.913135257 | 0.06721055 | -13.61110977 | 4.68E-05 | 0.000459347 | 0.012936933 | 0.1 | 0.243021319 |
| SL1344_RS09245_ | 0.457128198 | 0.970070313 | 0.145976514 | -0.796278529 | 0.299818433 | -2.653162251 | 4.68E-05 | 0.019720079 | 0.09744169 | 0.1001 | 0.24316205 |
| SL1344_RS09250_ | -2.149892803 | -1.8528523 | -2.613310462 | -0.782762361 | 0.619713571 | -1.265292969 | 0.081945421 | 0.124708583 | 0.282927806 | 0.1952 | 0.359025573 |
| SL1344_RS09260_ | 0.174611189 | 0.610712655 | -0.495695319 | -1.126901211 | 0.284139568 | -3.955723495 | 4.68E-05 | 0.016737287 | 0.086559691 | 0.1 | 0.243021319 |

| | | | | | | | | | | | |
|---|---|---|---|---|---|---|---|---|---|---|---|
| SL1344_RS09270_ | -2.964028606 | -2.971489194 | -2.958302438 | 0.00410835 | 0.492441519 | 0.008765184 | 0.494337109 | 0.531629642 | 0.675092789 | 0.654 | 0.757315338 |
| SL1344_RS09275_ | 1.033523524 | 1.348828013 | 0.564854417 | -0.785132461 | 0.160308914 | -4.775029634 | 4.68E-05 | 0.007907732 | 0.055056801 | 0.1 | 0.243021319 |
| SL1344_RS09285_ | -1.656503437 | -1.243944674 | -2.120469812 | -0.881034557 | 0.379572622 | -2.26196245 | 0.003334715 | 0.034288018 | 0.126451983 | 0.1054 | 0.250083844 |
| SL1344_RS09290_ | -1.004119291 | -0.426129966 | -1.665272554 | -1.23456574 | 0.280478715 | -4.304154955 | 4.68E-05 | 0.006455287 | 0.049350378 | 0.1 | 0.243021319 |
| SL1344_RS09295_ | -2.51671499 | -1.927488051 | -2.985984469 | -1.114851467 | 0.592057017 | -1.850646925 | 0.014657206 | 0.060430175 | 0.181391743 | 0.1175 | 0.264963111 |
| SL1344_RS09300_ | -4.031724237 | -4.031065036 | -4.032735881 | 0.004245007 | 0.817037401 | 0.005407385 | 0.497668221 | 0.579433229 | 0.71633127 | 0.69 | 0.789654488 |
| SL1344_RS09305_ | -1.150310765 | -0.904847337 | -1.548922593 | -0.645826899 | 0.38643845 | -1.578098233 | 0.034000131 | 0.084739525 | 0.222430561 | 0.1443 | 0.298338942 |
| SL1344_RS09310_ | -4.172301774 | -4.564994987 | -3.841346122 | 0.724272769 | 0.866394604 | 0.79939648 | 0.171885431 | 0.277138364 | 0.439432873 | 0.3703 | 0.516266624 |
| SL1344_RS09315_ | -1.589889427 | -1.416408137 | -1.809504327 | -0.386600074 | 0.348408625 | -1.063336057 | 0.119253864 | 0.195901739 | 0.357894921 | 0.2813 | 0.436543754 |
| SL1344_RS09320_ | 0.076537413 | 0.288888237 | -0.190657831 | -0.539123474 | 0.334675203 | -1.636304296 | 0.013324793 | 0.075601932 | 0.212989718 | 0.1159 | 0.263814494 |
| SL1344_RS09325_ | -3.177605597 | -2.824232244 | -3.653844097 | -0.83505152 | 0.556175812 | -1.412496354 | 0.056628991 | 0.111967592 | 0.249983377 | 0.1774 | 0.329796413 |
| SL1344_RS09335_ | -3.19781221 | -2.842411029 | -3.543925751 | -0.691062247 | 0.685125749 | -0.973533958 | 0.145902758 | 0.223116312 | 0.390346562 | 0.3036 | 0.461165427 |
| SL1344_RS09340_ | 1.034032169 | 1.335507714 | 0.671960512 | -0.655361571 | 0.193251504 | -3.309662721 | 4.68E-05 | 0.012939061 | 0.074713086 | 0.1 | 0.243021319 |
| SL1344_RS09345_ | -4.440012151 | -3.68191699 | -5.125933467 | -1.345468562 | 1.174923899 | -1.112809402 | 0.111259197 | 0.171491063 | 0.333212624 | 0.2524 | 0.414548858 |
| SL1344_RS09350_ | -5.3382586 | -4.948960491 | -5.874439002 | -0.896327996 | 1.209560089 | -0.711040322 | 0.205196552 | 0.317517081 | 0.480183833 | 0.4085 | 0.552663132 |
| SL1344_RS09355_ | -2.888972183 | -2.595791544 | -3.379205426 | -0.758569593 | 0.642101578 | -1.126877946 | 0.100666708 | 0.168414385 | 0.334966962 | 0.2341 | 0.394212144 |
| SL1344_RS09360_ | -2.381924251 | -2.093392043 | -2.934013026 | -0.873953527 | 0.626295761 | -1.267385179 | 0.06795476 | 0.134662336 | 0.296701843 | 0.1819 | 0.340175028 |
| SL1344_RS09365_ | -0.437091859 | -0.732341639 | -0.193557751 | 0.575232919 | 0.340902976 | 1.695646223 | 0.014666978 | 0.067950434 | 0.196328044 | 0.121 | 0.269776561 |
| SL1344_RS09370_ | 2.746360326 | 3.193945875 | 2.136551719 | -1.064875792 | 0.130324294 | -8.211022743 | 4.68E-05 | 0.001346583 | 0.022955514 | 0.1 | 0.243021319 |
| SL1344_RS09380_ | 2.272947878 | 2.725497074 | 1.564489995 | -1.155667367 | 0.158136407 | -6.995611041 | 4.68E-05 | 0.006607337 | 0.054483304 | 0.1 | 0.243021319 |
| SL1344_RS09385_ | 1.354627895 | 1.836316388 | 0.871483353 | -0.965617746 | 0.134944288 | -7.137973362 | 4.68E-05 | 0.001431748 | 0.023553869 | 0.1 | 0.243021319 |
| SL1344_RS09390_ | 1.056330201 | 1.418613176 | 0.637808486 | -0.78335935 | 0.146160926 | -5.32448807 | 4.68E-05 | 0.005306809 | 0.044573569 | 0.1 | 0.243021319 |
| SL1344_RS09395_ | 1.373410967 | 1.804084815 | 0.949449252 | -0.852633509 | 0.134732443 | -6.338496668 | 4.68E-05 | 0.002871873 | 0.032475907 | 0.1 | 0.243021319 |
| SL1344_RS09400_ | 1.428854248 | 2.016499122 | 0.877220891 | -1.131281496 | 0.131544266 | -8.591019714 | 4.68E-05 | 0.000925601 | 0.019198097 | 0.1 | 0.243021319 |
| SL1344_RS09405_ | 0.969512581 | 1.491535214 | 0.366512837 | -1.131027984 | 0.176294543 | -6.541322189 | 4.68E-05 | 0.004183122 | 0.039388961 | 0.1 | 0.243021319 |
| SL1344_RS09410_ | 1.312456198 | 1.91404377 | 0.619513186 | -1.29144454 | 0.145487336 | -8.950364423 | 4.68E-05 | 0.000949139 | 0.019047414 | 0.1 | 0.243021319 |
| SL1344_RS09420_ | 0.483272819 | 1.102424262 | -0.036455482 | -1.122801263 | 0.222942407 | -4.9093638 | 4.68E-05 | 0.004883652 | 0.045013561 | 0.1 | 0.243021319 |
| SL1344_RS09425_ | 2.812817003 | 3.148533192 | 2.380397793 | -0.773895681 | 0.143218657 | -5.60215665 | 4.68E-05 | 0.010873958 | 0.071358449 | 0.1 | 0.243021319 |
| SL1344_RS09430_ | 2.067372187 | 2.361864186 | 1.56831759 | -0.787470761 | 0.144673592 | -5.348477812 | 4.68E-05 | 0.004887871 | 0.044035475 | 0.1 | 0.243021319 |
| SL1344_RS09435_ | 4.118965747 | 4.573755214 | 3.647024218 | -0.932621091 | 0.10335076 | -9.114534933 | 4.68E-05 | 0.003658313 | 0.041094424 | 0.1 | 0.243021319 |
| SL1344_RS09440_ | 0.443107343 | 0.774236745 | -0.182899155 | -0.86955294 | 0.405390549 | -2.076154537 | 0.005330642 | 0.03839136 | 0.144959402 | 0.1045 | 0.2492693 |
| SL1344_RS09455_ | 2.599490014 | 2.868834619 | 2.230572036 | -0.637277286 | 0.119726721 | -5.256023644 | 4.68E-05 | 0.009188998 | 0.063342514 | 0.1 | 0.243021319 |
| SL1344_RS09460_ | 1.880701856 | 2.195523296 | 1.579682683 | -0.612330691 | 0.102232463 | -5.936669334 | 4.68E-05 | 0.002657927 | 0.031531619 | 0.1 | 0.243021319 |
| SL1344_RS09465_ | 1.178652233 | 1.46733494 | 0.760727265 | -0.693105984 | 0.156082118 | -4.335265097 | 4.68E-05 | 0.009644246 | 0.062451839 | 0.1 | 0.243021319 |
| SL1344_RS09470_ | 1.681161102 | 2.072466556 | 1.296648205 | -0.761107139 | 0.159173412 | -4.69556547 | 4.68E-05 | 0.004206417 | 0.042077148 | 0.1 | 0.243021319 |
| SL1344_RS09485_ | -1.818904935 | -1.536446051 | -2.148686953 | -0.605678814 | 0.327726222 | -1.790863006 | 0.023333527 | 0.062648965 | 0.175090599 | 0.1315 | 0.280921862 |
| SL1344_RS09490_ | -2.020831154 | -1.70866495 | -2.392280285 | -0.671938366 | 0.43622647 | -1.499341021 | 0.046000096 | 0.092234127 | 0.228336343 | 0.1556 | 0.310476952 |
| SL1344_RS09495_ | 4.091007811 | 4.544778112 | 3.56961674 | -0.99903274 | 0.115360182 | -8.789931087 | 4.68E-05 | 0.001364273 | 0.024548487 | 0.1 | 0.243021319 |
| SL1344_RS09500_ | 3.62162753 | 4.195881457 | 3.118952117 | -1.077466466 | 0.227845906 | -4.572219172 | 4.68E-05 | 0.007773963 | 0.0611981 | 0.1 | 0.243021319 |
| SL1344_RS09510_ | 4.986757924 | 5.388601984 | 4.46529123 | -0.918980635 | 0.082188409 | -11.16547688 | 4.68E-05 | 0.000516457 | 0.015746681 | 0.1 | 0.243021319 |
| SL1344_RS09515_ | 0.855891889 | 1.248840604 | 0.452219964 | -0.81679726 | 0.180338523 | -4.592297756 | 4.68E-05 | 0.006341372 | 0.05092827 | 0.1 | 0.243021319 |
| SL1344_RS09520_ | 0.742517997 | 1.138869595 | 0.291468884 | -0.841372072 | 0.16905059 | -4.945979876 | 4.68E-05 | 0.00424353 | 0.040292309 | 0.1 | 0.243021319 |
| SL1344_RS09525_ | 2.149353138 | 2.509514245 | 1.67325198 | -0.836664433 | 0.116818096 | -7.119431725 | 4.68E-05 | 0.002768526 | 0.032094238 | 0.1 | 0.243021319 |
| SL1344_RS09535_ | 3.230997253 | 3.570463016 | 2.865726429 | -0.705165315 | 0.063287182 | -11.09750871 | 4.68E-05 | 0.000761037 | 0.01663654 | 0.1 | 0.243021319 |
| SL1344_RS09540_ | 1.242112218 | 1.886644314 | 0.631870796 | -1.243544764 | 0.139455819 | -8.841092661 | 4.68E-05 | 0.000870317 | 0.018737903 | 0.1 | 0.243021319 |
| SL1344_RS09545_ | 0.905197678 | 1.444045942 | 0.240927699 | -1.207413411 | 0.169032331 | -7.150695522 | 4.68E-05 | 0.002899771 | 0.032247258 | 0.1 | 0.243021319 |
| SL1344_RS09550_ | 4.445557312 | 4.795290178 | 3.999911543 | -0.795758405 | 0.084393284 | -9.276606097 | 4.68E-05 | 0.002375559 | 0.032306732 | 0.1 | 0.243021319 |
| SL1344_RS09555_ | -0.010831399 | 0.214614013 | -0.221995595 | -0.433769404 | 0.181594346 | -2.344865279 | 0.004667653 | 0.030752806 | 0.116543274 | 0.1059 | 0.250787593 |
| SL1344_RS09635_ | -1.495169343 | -1.309387725 | -1.7555458 | -0.500236837 | 0.504532811 | -0.988609033 | 0.131333363 | 0.208043265 | 0.386329936 | 0.2943 | 0.458263837 |
| SL1344_RS09640_ | -0.019363914 | 0.226366748 | -0.277420297 | -0.50531083 | 0.290760034 | -1.715638283 | 0.02466685 | 0.069817622 | 0.199459297 | 0.1267 | 0.278216115 |
| SL1344_RS09645_ | -0.312520904 | 0.31235652 | -0.843764996 | -1.127941347 | 0.347657345 | -3.270010542 | 4.68E-05 | 0.011079601 | 0.070047697 | 0.1 | 0.243021319 |
| SL1344_RS09650_ | -1.769479288 | -1.305392248 | -2.443222865 | -1.139653337 | 0.407469844 | -2.744992822 | 0.002668394 | 0.024889926 | 0.105270623 | 0.1009 | 0.244200748 |
| SL1344_RS09655_ | -6.418680292 | -6.810300069 | -6.221618015 | 0.480625034 | 2.747562615 | 0.146182005 | 0.41172552 | 0.583410319 | 0.728657825 | 0.7444 | 0.83445694 |

| | | | | | | | | | | | |
|---|---|---|---|---|---|---|---|---|---|---|---|
| SL1344_RS09660_ | -1.822338072 | -1.337251639 | -2.609590063 | -1.272453261 | 0.381970555 | -3.284931289 | 0.00067292 | 0.020580214 | 0.091819415 | 0.1 | 0.243021319 |
| SL1344_RS09670_ | -1.403592094 | -1.070478742 | -1.78798808 | -0.748835161 | 0.411030943 | -1.766879023 | 0.018000252 | 0.061783207 | 0.182929829 | 0.1202 | 0.268703844 |
| SL1344_RS09680_ | 3.261540946 | 3.608837832 | 2.792100547 | -0.81320824 | 0.107068909 | -7.439694053 | 4.68E-05 | 0.002745213 | 0.033439895 | 0.1 | 0.243021319 |
| SL1344_RS09685_ | -4.269158997 | -4.191922583 | -4.38145754 | -0.232968427 | 0.930845463 | -0.227452213 | 0.395069957 | 0.532821038 | 0.680074338 | 0.6521 | 0.759883803 |
| SL1344_RS09690_ | 2.294395905 | 2.192259406 | 2.389815055 | 0.198153543 | 0.080315714 | 2.43531541 | 0.004667653 | 0.029096688 | 0.112985679 | 0.1045 | 0.248934686 |
| SL1344_RS09695_ | 1.718805549 | 2.000215141 | 1.417622035 | -0.584226199 | 0.123907292 | -4.613397609 | 4.68E-05 | 0.007749999 | 0.054809744 | 0.1 | 0.243021319 |
| SL1344_RS09700_ | 2.574989394 | 2.843658206 | 2.245615501 | -0.599147752 | 0.08251576 | -7.25789761 | 4.68E-05 | 0.00207296 | 0.026920768 | 0.1 | 0.243021319 |
| SL1344_RS09705_ | 1.174925428 | 1.344584107 | 0.967433681 | -0.377975528 | 0.150499016 | -2.477018628 | 0.003332494 | 0.02776696 | 0.112937295 | 0.1028 | 0.246755598 |
| SL1344_RS09710_ | 0.089196271 | 0.20598161 | -0.06442854 | -0.29145875 | 0.272172988 | -1.045964913 | 0.083333384 | 0.178909198 | 0.354478029 | 0.2118 | 0.370245335 |
| SL1344_RS09715_ | 5.297970824 | 5.645030126 | 4.798329923 | -0.81870755 | 0.103625135 | -7.353038884 | 4.68E-05 | 0.004190671 | 0.044598177 | 0.1 | 0.243021319 |
| SL1344_RS09725_ | 4.737435962 | 5.111239089 | 4.435593407 | -0.709539514 | 0.151777549 | -4.630057479 | 4.68E-05 | 0.002229703 | 0.033477864 | 0.1 | 0.243021319 |
| SL1344_RS09730_ | 2.016622601 | 2.304739961 | 1.640627511 | -0.665351184 | 0.091356286 | -7.331104421 | 4.68E-05 | 0.001971603 | 0.025781234 | 0.1 | 0.243021319 |
| SL1344_RS09805_ | 5.930168386 | 6.358827518 | 5.473790267 | -0.887348839 | 0.046555013 | -19.20664293 | 4.68E-05 | 0.000442533 | 0.014643682 | 0.1 | 0.243021319 |
| SL1344_RS09810_ | 4.527793966 | 4.927293166 | 4.081545962 | -0.844864116 | 0.104873773 | -8.204212388 | 4.68E-05 | 0.004199594 | 0.044131285 | 0.1 | 0.243021319 |
| SL1344_RS09820_ | 2.043700478 | 2.301446782 | 1.775864492 | -0.531950684 | 0.113581982 | -4.702259169 | 4.68E-05 | 0.005698158 | 0.047471428 | 0.1 | 0.243021319 |
| SL1344_RS09850_ | 5.447346291 | 5.733008949 | 5.284217692 | -0.44883498 | 0.156294841 | -3.014466278 | 4.68E-05 | 0.034704773 | 0.140420595 | 0.1 | 0.243021319 |
| SL1344_RS09855_ | 4.384799557 | 4.58606054 | 4.139853585 | -0.44591134 | 0.120312509 | -3.683673081 | 4.68E-05 | 0.006150707 | 0.053651611 | 0.1 | 0.243021319 |
| SL1344_RS09860_ | 4.026899908 | 4.216047159 | 3.821853701 | -0.402716847 | 0.114150866 | -3.495293044 | 4.68E-05 | 0.027840779 | 0.122749877 | 0.1 | 0.243021319 |
| SL1344_RS09875_ | 3.605256114 | 3.939716606 | 3.319323905 | -0.623345596 | 0.109615118 | -5.61016549 | 4.68E-05 | 0.00282137 | 0.034960059 | 0.1 | 0.243021319 |
| SL1344_RS09885_ | 4.474329707 | 4.846376046 | 4.252455365 | -0.597383862 | 0.140799121 | -3.971723346 | 4.68E-05 | 0.024033142 | 0.113876283 | 0.1 | 0.243021319 |
| SL1344_RS09890_ | 3.512191465 | 3.835169597 | 3.272110252 | -0.565634194 | 0.129955629 | -4.332496894 | 4.68E-05 | 0.014376299 | 0.084324684 | 0.1 | 0.243021319 |
| SL1344_RS09895_ | 3.141644438 | 3.369775503 | 2.961031568 | -0.439953567 | 0.166242609 | -2.720630445 | 4.68E-05 | 0.031910654 | 0.13182336 | 0.1 | 0.243021319 |
| SL1344_RS09900_ | 3.638520194 | 3.945924015 | 3.382103691 | -0.505307056 | 0.190319162 | -2.764209839 | 4.68E-05 | 0.012130553 | 0.077354544 | 0.1 | 0.243021319 |
| SL1344_RS09910_ | 2.121180779 | 2.406416274 | 2.014399149 | -0.313763051 | 0.244840276 | -1.203354638 | 0.138574312 | 0.160666496 | 0.342891956 | 0.241 | 0.421599592 |
| SL1344_RS09920_ | 1.368574751 | 1.221146667 | 1.501779706 | 0.282971535 | 0.11997862 | 2.292838861 | 0.003998486 | 0.034258151 | 0.126264839 | 0.1044 | 0.248755028 |
| SL1344_RS09925_ | -3.179481998 | -3.016022326 | -3.287486108 | -0.157078512 | 0.835111165 | -0.155610503 | 0.433333336 | 0.701849209 | 0.812693516 | 0.7943 | 0.871110911 |
| SL1344_RS09930_ | -3.105671891 | -3.161418194 | -3.027335562 | 0.201718295 | 0.722273896 | 0.252484682 | 0.393333337 | 0.590157255 | 0.72600668 | 0.7123 | 0.805764103 |
| SL1344_RS09935_ | -0.092792665 | -0.305652907 | 0.083722953 | 0.39006919 | 0.192084295 | 1.965635283 | 0.019320688 | 0.05127033 | 0.156457077 | 0.1201 | 0.268061571 |
| SL1344_RS09940_ | 1.262730195 | 1.233205927 | 1.30474581 | 0.072898071 | 0.147800755 | 0.453676083 | 0.313124592 | 0.508201545 | 0.657975626 | 0.6253 | 0.738969203 |
| SL1344_RS09945_ | 0.186700999 | 0.160194444 | 0.216365522 | 0.061762345 | 0.206479774 | 0.277935213 | 0.376666672 | 0.540201392 | 0.684920098 | 0.6541 | 0.76062667 |
| SL1344_RS09950_ | -1.185898005 | -1.492861331 | -0.820963945 | 0.678957591 | 0.306493151 | 2.166496867 | 0.00733396 | 0.041681086 | 0.141258971 | 0.1088 | 0.254395909 |
| SL1344_RS09955_ | 0.114803869 | -0.225235733 | 0.475633859 | 0.69469954 | 0.192699119 | 3.574418136 | 4.68E-05 | 0.010028678 | 0.063238246 | 0.1001 | 0.243166684 |
| SL1344_RS09960_ | 1.373346933 | 1.202228979 | 1.48986969 | 0.284582147 | 0.133646074 | 2.010032596 | 0.012000381 | 0.048662734 | 0.156849848 | 0.1146 | 0.262000058 |
| SL1344_RS09965_ | 0.069871568 | -0.222356063 | 0.282844292 | 0.505366958 | 0.19755488 | 2.504994481 | 0.000673369 | 0.031676056 | 0.120460375 | 0.1029 | 0.246881641 |
| SL1344_RS09970_ | -0.633331791 | -0.741835291 | -0.238629709 | 0.463945422 | 0.368247468 | 1.018283119 | 0.191333352 | 0.234690501 | 0.418900827 | 0.3725 | 0.539832614 |
| SL1344_RS09975_ | 0.143536914 | 0.14665694 | 0.138278204 | -0.011155477 | 0.257289813 | -0.038762955 | 0.487333334 | 0.676828114 | 0.795251528 | 0.7822 | 0.861185076 |
| SL1344_RS09985_ | -0.152207882 | -0.423230707 | 0.106134135 | 0.522548587 | 0.197925292 | 2.560273567 | 0.003332494 | 0.025977827 | 0.106063045 | 0.1033 | 0.247208821 |
| SL1344_RS09990_ | -1.601803041 | -2.569418298 | -1.229415027 | 1.26753781 | 0.618355842 | 1.784701971 | 0.022651765 | 0.080445714 | 0.223766287 | 0.1206 | 0.270753536 |
| SL1344_RS09995_ | -3.740529771 | -3.760665843 | -3.717517502 | 0.057748669 | 0.745110738 | 0.069318586 | 0.465689542 | 0.582849294 | 0.718846281 | 0.6878 | 0.787605906 |
| SL1344_RS10000_ | -2.400545701 | -2.413885394 | -2.392708632 | 0.077570952 | 0.722926353 | 0.097958551 | 0.455696204 | 0.670523654 | 0.793252816 | 0.791 | 0.867439536 |
| SL1344_RS10005_ | -2.294326541 | -2.159112219 | -2.442389356 | -0.311666554 | 0.571725299 | -0.514127081 | 0.276482355 | 0.439800255 | 0.599979042 | 0.5567 | 0.684101974 |
| SL1344_RS10010_ | -2.403281803 | -2.450586106 | -2.365402532 | 0.045704687 | 0.57626708 | 0.079402706 | 0.468666668 | 0.664904748 | 0.786572979 | 0.7734 | 0.855853868 |
| SL1344_RS10015_ | -4.475060068 | -4.26471634 | -4.744313785 | -0.395356794 | 1.132362505 | -0.323608215 | 0.351333339 | 0.541459668 | 0.687129179 | 0.6524 | 0.762231663 |
| SL1344_RS10020_ | -3.911711755 | -3.733835049 | -4.183019117 | -0.616987082 | 1.406655276 | -0.426649234 | 0.320453039 | 0.480047222 | 0.650140181 | 0.6466 | 0.764275202 |
| SL1344_RS10025_ | -3.507502605 | -3.365843208 | -3.658156138 | -0.311578346 | 0.647279581 | -0.434824898 | 0.307128589 | 0.434004615 | 0.58968124 | 0.5468 | 0.671922204 |
| SL1344_RS10030_ | -4.42701802 | -4.526062666 | -4.359450819 | 0.212191323 | 0.960601219 | 0.188147161 | 0.40906063 | 0.541711998 | 0.684483876 | 0.6531 | 0.759398028 |
| SL1344_RS10035_ | -2.678273949 | -2.700726206 | -2.663374028 | -0.035227415 | 0.753412628 | -0.046518585 | 0.478347769 | 0.696518669 | 0.810036827 | 0.8143 | 0.881892941 |
| SL1344_RS10040_ | -1.814794852 | -1.876442723 | -1.769082399 | 0.117652678 | 0.439583567 | 0.257333173 | 0.380666671 | 0.575146391 | 0.712512708 | 0.6934 | 0.789883921 |
| SL1344_RS10045_ | -2.41834139 | -2.243676366 | -2.636428941 | -0.38656379 | 0.447754782 | -0.832739697 | 0.177881433 | 0.271402398 | 0.431371883 | 0.3662 | 0.512973307 |
| SL1344_RS10050_ | -0.609404509 | -0.420389452 | -0.850534387 | -0.422620181 | 0.237013782 | -1.716334831 | 0.031333476 | 0.069249703 | 0.187623706 | 0.1341 | 0.285282676 |
| SL1344_RS10055_ | -2.749240762 | -2.460466451 | -3.044714942 | -0.579372128 | 0.521611283 | -1.069573313 | 0.115922753 | 0.186377673 | 0.343322476 | 0.2659 | 0.420444735 |
| SL1344_RS10060_ | -2.695084217 | -2.70556383 | -2.686364498 | 0.030191523 | 0.472214494 | 0.056838099 | 0.474666668 | 0.566654566 | 0.703367526 | 0.6661 | 0.7702558 |
| SL1344_RS10065_ | -1.952926412 | -1.746759376 | -2.307435932 | -0.534164613 | 0.532610765 | -0.954946644 | 0.168000022 | 0.233901852 | 0.411192489 | 0.3246 | 0.489975348 |

| | | | | | | | | | | | |
|---|---|---|---|---|---|---|---|---|---|---|---|
| SL1344_RS10070_ | -3.728432176 | -3.245850568 | -4.260702595 | -1.037639437 | 0.770257697 | -1.32205723 | 0.064000067 | 0.124086992 | 0.270463062 | 0.192 | 0.346944693 |
| SL1344_RS10075_ | -3.384814964 | -3.030342848 | -3.931928442 | -0.921293537 | 0.696863316 | -1.23788715 | 0.077333388 | 0.14381481 | 0.299098098 | 0.1972 | 0.351936672 |
| SL1344_RS10080_ | -4.234983906 | -4.212913773 | -4.261603485 | 0.004141871 | 1.180397709 | 0.003072723 | 0.499333333 | 0.673706138 | 0.792985 | 0.7904 | 0.866354478 |
| SL1344_RS10085_ | -4.044972742 | -3.9516995 | -4.342538997 | -0.353253429 | 1.359719357 | -0.231285185 | 0.422000003 | 0.645659305 | 0.77770976 | 0.7972 | 0.876698411 |
| SL1344_RS10090_ | -3.873105406 | -3.538511519 | -4.366373188 | -0.791764842 | 0.979015047 | -0.754207589 | 0.203864107 | 0.309357165 | 0.483665208 | 0.4041 | 0.56031889 |
| SL1344_RS10095_ | -2.330100296 | -1.935975015 | -2.894502692 | -0.957455116 | 0.403649082 | -2.341094912 | 0.007329074 | 0.038426832 | 0.13164087 | 0.108 | 0.253256085 |
| SL1344_RS10100_ | -3.591679149 | -3.363496967 | -3.86982423 | -0.517442169 | 0.708956472 | -0.675465797 | 0.209193887 | 0.318078827 | 0.484847303 | 0.4232 | 0.564497171 |
| SL1344_RS10105_ | -2.902949789 | -2.659132075 | -3.184640849 | -0.518497533 | 0.546940421 | -0.888879913 | 0.157333357 | 0.252524087 | 0.417082264 | 0.3443 | 0.497147649 |
| SL1344_RS10110_ | -4.978880974 | -4.584604459 | -5.689595523 | -1.163627288 | 1.312496433 | -0.774344165 | 0.162558317 | 0.260152006 | 0.437405962 | 0.342 | 0.493583745 |
| SL1344_RS10115_ | -1.548833051 | -1.536885127 | -1.570742966 | -0.073200038 | 0.432830947 | -0.150814204 | 0.430666669 | 0.598435038 | 0.734836742 | 0.7276 | 0.817884981 |
| SL1344_RS10120_ | -3.330383828 | -3.110793298 | -3.532117209 | -0.422143264 | 0.639720594 | -0.635989144 | 0.239333347 | 0.368399401 | 0.527920265 | 0.4653 | 0.601342298 |
| SL1344_RS10130_ | -2.465417933 | -2.334120922 | -2.548250214 | -0.091252805 | 0.845150421 | -0.094315887 | 0.456000002 | 0.758927447 | 0.856249263 | 0.8391 | 0.904124916 |
| SL1344_RS10135_ | -3.994520239 | -3.030830369 | -4.500318013 | -1.322870522 | 1.187488553 | -1.06633538 | 0.136000029 | 0.189653048 | 0.363384493 | 0.2743 | 0.445480742 |
| SL1344_RS10140_ | -2.863234734 | -2.715501869 | -3.01646317 | -0.298874689 | 0.488478369 | -0.572527361 | 0.262000011 | 0.379603283 | 0.536033001 | 0.4862 | 0.61811931 |
| SL1344_RS10145_ | -3.203697939 | -2.893385315 | -3.504890811 | -0.609249037 | 0.584027197 | -0.977239413 | 0.141905424 | 0.214094711 | 0.374201004 | 0.2866 | 0.441764868 |
| SL1344_RS10150_ | -3.789054323 | -4.059165189 | -3.402987133 | 0.637743136 | 0.802063197 | 0.763142394 | 0.199866773 | 0.317851267 | 0.482241668 | 0.4145 | 0.560841223 |
| SL1344_RS10155_ | -0.892678019 | -1.045751336 | -0.75444466 | 0.287970744 | 0.246732209 | 1.124671702 | 0.103333373 | 0.176308791 | 0.330789758 | 0.2523 | 0.408289871 |
| SL1344_RS10160_ | -0.604335974 | -0.460650237 | -0.750298128 | -0.296128218 | 0.28615335 | -0.966836082 | 0.144570313 | 0.222068368 | 0.389252642 | 0.3145 | 0.471247632 |
| SL1344_RS10165_ | -1.670199943 | -1.640898365 | -1.693748106 | -0.053672373 | 0.35029932 | -0.143339587 | 0.437708197 | 0.568402054 | 0.705696964 | 0.6835 | 0.782852995 |
| SL1344_RS10170_ | -1.930994034 | -1.689074222 | -2.199210575 | -0.51479841 | 0.396851651 | -1.278413788 | 0.072000059 | 0.130755859 | 0.27829486 | 0.1985 | 0.354915597 |
| SL1344_RS10175_ | -3.118207906 | -2.771121771 | -3.374344213 | -0.605527904 | 0.753691557 | -0.76701427 | 0.200000017 | 0.302693319 | 0.477790056 | 0.4035 | 0.559150796 |
| SL1344_RS10180_ | -2.361559008 | -2.325897641 | -2.408853724 | -0.076744721 | 0.399104917 | -0.180135466 | 0.419053967 | 0.512901114 | 0.658067318 | 0.6185 | 0.731918464 |
| SL1344_RS10185_ | -2.739986257 | -2.495242979 | -2.966057494 | -0.475711093 | 0.573290309 | -0.78765886 | 0.188540992 | 0.289629323 | 0.458791826 | 0.3897 | 0.53820037 |
| SL1344_RS10190_ | -1.821338143 | -1.373137368 | -2.202491611 | -0.764541589 | 0.564603105 | -1.285356503 | 0.069287204 | 0.117473571 | 0.273559883 | 0.1833 | 0.34924087 |
| SL1344_RS10195_ | -3.346781355 | -3.013301207 | -3.784386613 | -0.743964791 | 0.765684523 | -0.927185733 | 0.150666692 | 0.228651939 | 0.399619245 | 0.3079 | 0.469328745 |
| SL1344_RS10200_ | 2.31962214 | 2.024665672 | 2.582300319 | 0.55940876 | 0.113581422 | 4.902509897 | 4.68E-05 | 0.004165376 | 0.041016196 | 0.1 | 0.243021319 |
| SL1344_RS10205_ | -2.030148177 | -1.773210099 | -2.284285943 | -0.51473084 | 0.362961625 | -1.379235943 | 0.070000061 | 0.120964135 | 0.260621765 | 0.1907 | 0.344464052 |
| SL1344_RS10210_ | -2.944872657 | -3.002874189 | -2.869992154 | 0.203685844 | 0.700260606 | 0.262468202 | 0.382000005 | 0.578162445 | 0.718790345 | 0.7061 | 0.801808184 |
| SL1344_RS10215_ | 3.307280632 | 3.331626767 | 3.259560451 | -0.073866134 | 0.142201978 | -0.483032824 | 0.313333341 | 0.47228149 | 0.649350546 | 0.637 | 0.762988632 |
| SL1344_RS10225_ | -1.115278695 | -1.270087556 | -1.020793272 | 0.26934365 | 0.452712357 | 0.56169469 | 0.257333345 | 0.390574767 | 0.571157616 | 0.5208 | 0.659766646 |
| SL1344_RS10230_ | 0.583507355 | 0.552193457 | 0.622826837 | 0.064925932 | 0.172300099 | 0.358820898 | 0.344666673 | 0.519799336 | 0.665807241 | 0.6296 | 0.742714332 |
| SL1344_RS10235_ | -3.54438638 | -3.348913706 | -3.900266806 | -0.495812541 | 0.789233378 | -0.580821915 | 0.262000011 | 0.402202833 | 0.568064183 | 0.5262 | 0.661764436 |
| SL1344_RS10240_ | -5.79791755 | -5.662359079 | -5.964288225 | -0.321490629 | 1.741352906 | -0.173217248 | 0.419053967 | 0.592174518 | 0.729648247 | 0.7197 | 0.811363973 |
| SL1344_RS10245_ | -3.610843215 | -3.865047719 | -3.363477991 | 0.58877659 | 0.879210632 | 0.617631859 | 0.224000015 | 0.360245075 | 0.531195618 | 0.4855 | 0.620546831 |
| SL1344_RS10250_ | -3.539503439 | -3.521298262 | -3.567636462 | -0.057226822 | 0.788745064 | -0.06197305 | 0.465689542 | 0.631764005 | 0.760460391 | 0.7454 | 0.833750232 |
| SL1344_RS10255_ | -1.869426253 | -1.886359343 | -1.839633868 | 0.046167596 | 0.477046871 | 0.081446191 | 0.457028649 | 0.654901966 | 0.778697341 | 0.7713 | 0.8530621 |
| SL1344_RS10260_ | -2.121999533 | -2.545739506 | -1.786002843 | 0.766885538 | 0.489893574 | 1.468668784 | 0.041333441 | 0.103618148 | 0.248883934 | 0.1533 | 0.307617361 |
| SL1344_RS10265_ | -2.52363656 | -2.406440459 | -2.628063047 | -0.231853971 | 0.506823734 | -0.418837491 | 0.308666675 | 0.471605133 | 0.623198328 | 0.5739 | 0.696287729 |
| SL1344_RS10270_ | -1.308829008 | -1.501064895 | -1.114472694 | 0.406979954 | 0.361317366 | 1.082596813 | 0.114666702 | 0.176446038 | 0.340498536 | 0.2541 | 0.411713364 |
| SL1344_RS10275_ | 2.4678773 | 1.965315643 | 2.817071736 | 0.86196155 | 0.167375694 | 5.210184818 | 4.68E-05 | 0.002276066 | 0.031655431 | 0.1 | 0.243021319 |
| SL1344_RS10280_ | -0.873196068 | -1.085488242 | -0.709005921 | 0.407873296 | 0.351500983 | 1.116936308 | 0.097333376 | 0.168072371 | 0.336643912 | 0.2298 | 0.389255932 |
| SL1344_RS10285_ | 1.312158565 | 1.018663803 | 1.598277971 | 0.602922602 | 0.197278296 | 3.101697583 | 4.68E-05 | 0.014339072 | 0.080953537 | 0.1 | 0.243021319 |
| SL1344_RS10290_ | -2.171436542 | -2.31940323 | -2.050414075 | 0.245418625 | 0.455438507 | 0.521815137 | 0.28000001 | 0.425817932 | 0.583740922 | 0.5403 | 0.669954609 |
| SL1344_RS10295_ | -5.302046874 | -5.4862969 | -5.17231476 | 0.378044632 | 1.558135061 | 0.20471861 | 0.387333338 | 0.555925353 | 0.702741243 | 0.686 | 0.786222218 |
| SL1344_RS10300_ | 0.499420417 | 0.011698361 | 0.98708002 | 0.963526712 | 0.280029394 | 3.48420717 | 4.68E-05 | 0.011583793 | 0.071052948 | 0.1 | 0.243021319 |
| SL1344_RS10305_ | 0.781020006 | 0.508681698 | 1.034717254 | 0.52352308 | 0.142285492 | 3.630728278 | 4.68E-05 | 0.010076776 | 0.062769959 | 0.1 | 0.243021319 |
| SL1344_RS10310_ | -2.957469002 | -3.037553823 | -2.870272235 | 0.183230045 | 0.554910398 | 0.309013865 | 0.365756168 | 0.520429567 | 0.664946298 | 0.6323 | 0.743816435 |
| SL1344_RS10315_ | -1.875759678 | -1.82986189 | -1.917898414 | -0.091719138 | 0.343976511 | -0.255730655 | 0.390666671 | 0.523731488 | 0.664561737 | 0.6328 | 0.740194551 |
| SL1344_RS10320_ | 2.107127947 | 1.84781393 | 2.324586112 | 0.481823693 | 0.13526551 | 3.524096364 | 4.68E-05 | 0.009331379 | 0.063544165 | 0.1 | 0.243021319 |
| SL1344_RS10325_ | -2.373764898 | -2.501285112 | -2.095384476 | 0.302309139 | 0.742867964 | 0.361033828 | 0.361758833 | 0.636172647 | 0.766114226 | 0.7262 | 0.82730322 |
| SL1344_RS10330_ | 1.646707917 | 1.581158833 | 1.732142189 | 0.169027482 | 0.131639095 | 1.234452946 | 0.085942754 | 0.154003595 | 0.315513798 | 0.2099 | 0.367866876 |
| SL1344_RS10335_ | -2.805398289 | -3.122555767 | -2.600772062 | 0.557441435 | 0.657736303 | 0.801878991 | 0.18254499 | 0.276571078 | 0.453843544 | 0.3677 | 0.516827301 |
| SL1344_RS10340_ | -0.362347406 | -0.573648703 | -0.145298458 | 0.428471353 | 0.193046626 | 2.149765485 | 0.012667027 | 0.043050846 | 0.140882593 | 0.1129 | 0.259418738 |

| | | | | | | | | | | | |
|---|---|---|---|---|---|---|---|---|---|---|---|
| SL1344_RS10345_ | 0.020104391 | -0.154034421 | 0.315826618 | 0.454249189 | 0.224384453 | 2.014672641 | 0.01798826 | 0.053958193 | 0.166279569 | 0.1217 | 0.270696703 |
| SL1344_RS10350_ | -2.167796192 | -2.566300149 | -1.888349933 | 0.733891451 | 0.591969859 | 1.203956097 | 0.049333422 | 0.155532428 | 0.32647274 | 0.1726 | 0.327942166 |
| SL1344_RS10355_ | 2.60867805 | 2.271029504 | 3.022667797 | 0.664674065 | 0.241981001 | 2.759541845 | 4.68E-05 | 0.014822021 | 0.085612926 | 0.1 | 0.243021319 |
| SL1344_RS10360_ | -0.983113085 | -1.199401742 | -0.59242299 | 0.591085354 | 0.321348574 | 1.726255161 | 0.032666804 | 0.076163314 | 0.20707666 | 0.1349 | 0.287474839 |
| SL1344_RS10365_ | 0.929603345 | 0.704517951 | 1.161885046 | 0.454393357 | 0.131956169 | 3.387499958 | 0.00067292 | 0.010974474 | 0.065322112 | 0.1 | 0.243021319 |
| SL1344_RS10370_ | -6.502985655 | -6.890980557 | -6.1797549 | -0.007396226 | 3.943088874 | -0.001615002 | 0.499000666 | 0.697324184 | 0.814645298 | 0.8743 | 0.924004251 |
| SL1344_RS10375_ | -0.50317839 | -0.463670501 | -0.541591909 | -0.074440847 | 0.269893306 | -0.273295655 | 0.375333338 | 0.551863602 | 0.694609053 | 0.6643 | 0.770978078 |
| SL1344_RS10380_ | -0.355389658 | -0.380457766 | -0.324524256 | 0.048697123 | 0.24065588 | 0.173738084 | 0.419333336 | 0.577912866 | 0.714181156 | 0.6937 | 0.792001091 |
| SL1344_RS10390_ | -1.982708821 | -2.036348734 | -1.909818632 | 0.014467541 | 0.768639328 | 0.018771921 | 0.492338441 | 0.741525758 | 0.845322175 | 0.8335 | 0.899718005 |
| SL1344_RS10395_ | -0.623705671 | -0.605676986 | -0.639756352 | -0.034848925 | 0.264795814 | -0.120292809 | 0.441039309 | 0.585351188 | 0.724006656 | 0.698 | 0.799062239 |
| SL1344_RS10400_ | -1.40148225 | -1.386567979 | -1.424225964 | -0.032877951 | 0.323124172 | -0.092819953 | 0.461692207 | 0.569667175 | 0.707598803 | 0.6788 | 0.780601811 |
| SL1344_RS10405_ | -2.267470533 | -2.325545681 | -2.178750279 | 0.144863447 | 0.410432329 | 0.315566664 | 0.358427721 | 0.517747385 | 0.659819859 | 0.6221 | 0.732886207 |
| SL1344_RS10410_ | -4.102719295 | -3.931153553 | -4.200979591 | -0.157911397 | 0.964357244 | -0.140442236 | 0.432666669 | 0.635753796 | 0.761238392 | 0.7182 | 0.816453479 |
| SL1344_RS10415_ | -2.676190198 | -2.724813769 | -2.611320635 | 0.175329779 | 0.661701561 | 0.257752358 | 0.399333337 | 0.594868453 | 0.730950842 | 0.7278 | 0.818759581 |
| SL1344_RS10420_ | -2.840301182 | -2.808260609 | -2.869690267 | -0.104787475 | 0.756602843 | -0.134752085 | 0.434666669 | 0.656510235 | 0.780022396 | 0.7618 | 0.844312867 |
| SL1344_RS10430_ | -2.623467055 | -2.784471166 | -2.472852156 | 0.309879214 | 0.448220382 | 0.620355263 | 0.24250501 | 0.354658553 | 0.510821285 | 0.4484 | 0.585143781 |
| SL1344_RS10435_ | -2.575981092 | -3.027248781 | -2.408692013 | 0.55675357 | 0.775910672 | 0.646841362 | 0.252666679 | 0.357736691 | 0.543648225 | 0.5 | 0.649546725 |
| SL1344_RS10440_ | -3.077154967 | -2.792199799 | -3.214826333 | -0.363630794 | 0.773651351 | -0.446621444 | 0.328000007 | 0.559380738 | 0.701488605 | 0.6552 | 0.768708713 |
| SL1344_RS10445_ | -4.04568526 | -4.0035985 | -4.079407486 | -0.120143851 | 0.866372588 | -0.124878941 | 0.441039309 | 0.593942237 | 0.729460996 | 0.7177 | 0.812341018 |
| SL1344_RS10450_ | -3.766065439 | -3.807016075 | -3.731186826 | 0.08668999 | 0.763636147 | 0.111098819 | 0.447701534 | 0.574084427 | 0.71055363 | 0.6947 | 0.791579064 |
| SL1344_RS10455_ | -4.075127915 | -4.168251839 | -4.001466392 | 0.108839658 | 1.053493847 | 0.091319092 | 0.454363759 | 0.663315083 | 0.784655189 | 0.7725 | 0.853532919 |
| SL1344_RS10470_ | -2.783428425 | -3.071826058 | -2.510761802 | 0.545291572 | 0.534087344 | 0.984550015 | 0.145236535 | 0.227940097 | 0.389822381 | 0.3193 | 0.474500424 |
| SL1344_RS10475_ | -2.790907759 | -2.882323032 | -2.686968326 | 0.21310809 | 0.440191432 | 0.445494304 | 0.319333341 | 0.431568318 | 0.580854483 | 0.5305 | 0.654693671 |
| SL1344_RS10480_ | -0.568495302 | -0.846569927 | -0.474101783 | 0.336576754 | 0.463178654 | 0.625691585 | 0.266666678 | 0.41208015 | 0.595382425 | 0.5328 | 0.689281395 |
| SL1344_RS10485_ | -1.760728757 | -1.998653615 | -1.552509893 | 0.404080847 | 0.497709653 | 0.756783697 | 0.210000016 | 0.317783274 | 0.497085251 | 0.41 | 0.569990543 |
| SL1344_RS10490_ | -7.423873728 | -8.436866824 | -6.714391916 | 1.71526666 | 2.831015399 | 0.546304791 | 0.225333348 | 0.342121365 | 0.513808988 | 0.4389 | 0.579747196 |
| SL1344_RS10495_ | -4.9501405 | -5.055660427 | -4.880699008 | 0.271499788 | 1.465305393 | 0.146976846 | 0.415333337 | 0.566297657 | 0.714608916 | 0.7217 | 0.814280508 |
| SL1344_RS10515_ | -3.201562026 | -2.969021887 | -3.531928809 | -0.577701048 | 0.603236133 | -0.87464276 | 0.16322454 | 0.259681987 | 0.422037167 | 0.3508 | 0.500430943 |
| SL1344_RS10520_ | -6.933452443 | -7.122547988 | -6.78730096 | 0.186063456 | 3.340351785 | 0.043042198 | 0.467688209 | 0.621663452 | 0.755225806 | 0.7474 | 0.834284983 |
| SL1344_RS10530_ | -4.832982801 | -4.343825828 | -5.274120058 | -0.954361131 | 1.090187966 | -0.818209231 | 0.175882766 | 0.271775631 | 0.434511164 | 0.3672 | 0.514366675 |
| SL1344_RS10535_ | -3.850671241 | -3.720303215 | -4.023714391 | -0.318032878 | 0.712802102 | -0.387650999 | 0.319333341 | 0.443330272 | 0.598851816 | 0.5502 | 0.675389833 |
| SL1344_RS10540_ | -3.303221654 | -2.782161157 | -3.806121415 | -1.035761596 | 0.716380871 | -1.398858749 | 0.055296548 | 0.107597291 | 0.249665802 | 0.1745 | 0.330522711 |
| SL1344_RS10545_ | -3.843437227 | -3.729902612 | -3.941137296 | -0.174696447 | 0.775554125 | -0.20638932 | 0.407333337 | 0.553889044 | 0.691984017 | 0.664 | 0.766696302 |
| SL1344_RS10550_ | -3.182370927 | -2.821558775 | -3.675141464 | -0.86056612 | 0.641656574 | -1.296101632 | 0.063291207 | 0.131888555 | 0.282173196 | 0.1919 | 0.345951351 |
| SL1344_RS10555_ | -3.03015777 | -2.424272116 | -3.745697335 | -1.054523584 | 1.058550659 | -1.004455605 | 0.14533336 | 0.214136653 | 0.398459264 | 0.2753 | 0.452894654 |
| SL1344_RS10560_ | -2.973832444 | -2.768249031 | -3.251257709 | -0.529059991 | 0.751922303 | -0.67595053 | 0.224666681 | 0.336945911 | 0.513221113 | 0.4479 | 0.598583209 |
| SL1344_RS10565_ | -2.563395086 | -2.154290938 | -2.954874502 | -0.782259928 | 0.555755505 | -1.32219781 | 0.068620982 | 0.115314381 | 0.262375218 | 0.1856 | 0.345090131 |
| SL1344_RS10570_ | -3.446327143 | -2.979980078 | -4.220705331 | -1.228651551 | 0.760125424 | -1.578848077 | 0.03597614 | 0.093492876 | 0.231355434 | 0.1432 | 0.293947264 |
| SL1344_RS10575_ | -4.818354595 | -4.317702914 | -5.420000016 | -1.119148682 | 1.086310574 | -0.949047534 | 0.129913421 | 0.216853008 | 0.378578996 | 0.3009 | 0.455068235 |
| SL1344_RS10580_ | -4.67764184 | -4.125445109 | -5.254602077 | -1.155061718 | 0.971773271 | -1.111854467 | 0.099267197 | 0.175438955 | 0.327446728 | 0.2532 | 0.406766854 |
| SL1344_RS10585_ | -1.531895983 | -1.373231359 | -1.667588006 | -0.267696763 | 0.409203822 | -0.626405497 | 0.243837455 | 0.392968963 | 0.55979993 | 0.4952 | 0.636727529 |
| SL1344_RS10590_ | -1.149260009 | -1.10664543 | -1.209689552 | -0.132291031 | 0.402030731 | -0.310991664 | 0.375083283 | 0.575695214 | 0.719809892 | 0.7154 | 0.810339466 |
| SL1344_RS10595_ | 4.597575551 | 4.358916541 | 4.830189544 | 0.48736106 | 0.11540844 | 4.217757541 | 4.68E-05 | 0.00318468 | 0.03901832 | 0.1 | 0.243021319 |
| SL1344_RS10600_ | 0.596895264 | 0.538278391 | 0.665174908 | 0.140672248 | 0.234844317 | 0.561177648 | 0.261333345 | 0.407564166 | 0.580101544 | 0.531 | 0.669291425 |
| SL1344_RS10640_ | 5.598135439 | 5.330398749 | 5.912185074 | 0.587042835 | 0.09287346 | 6.483625264 | 4.68E-05 | 0.000704793 | 0.019449969 | 0.1 | 0.243021319 |
| SL1344_RS10645_ | -0.07135843 | -0.167326345 | -0.018459326 | 0.144853135 | 0.237337772 | 0.555306686 | 0.261825461 | 0.4084827 | 0.573354007 | 0.5124 | 0.649837273 |
| SL1344_RS10650_ | 1.619857665 | 1.438297809 | 1.84665217 | 0.426038891 | 0.189611197 | 2.221185786 | 0.001335878 | 0.033233801 | 0.130857551 | 0.1019 | 0.245623577 |
| SL1344_RS10655_ | 0.700215749 | 0.256575141 | 1.010601776 | 0.754247957 | 0.163371834 | 4.560929381 | 4.68E-05 | 0.007718295 | 0.053718169 | 0.1 | 0.243021319 |
| SL1344_RS10660_ | 0.519172809 | 0.195650067 | 0.87837928 | 0.681819027 | 0.148280904 | 4.519648028 | 4.68E-05 | 0.005570491 | 0.045689779 | 0.1 | 0.243021319 |
| SL1344_RS10665_ | 0.668216835 | 0.19975459 | 0.927098024 | 0.711416658 | 0.214670081 | 3.130420216 | 4.68E-05 | 0.021930972 | 0.1009933 | 0.1002 | 0.243300468 |
| SL1344_RS10670_ | 1.153365581 | 0.879485682 | 1.483456451 | 0.599198399 | 0.161561228 | 3.608021579 | 4.68E-05 | 0.010787368 | 0.068197245 | 0.1 | 0.243021319 |
| SL1344_RS10675_ | 0.129689532 | -0.228330509 | 0.514817844 | 0.736732439 | 0.205342418 | 3.570392837 | 4.68E-05 | 0.009923152 | 0.063443673 | 0.1 | 0.243021319 |
| SL1344_RS10680_ | 0.729147972 | 0.466901762 | 1.019567873 | 0.521312909 | 0.208595723 | 2.437497388 | 0.003334715 | 0.026926174 | 0.11397479 | 0.1037 | 0.247962433 |

| | | | | | | | | | | | |
|---|---|---|---|---|---|---|---|---|---|---|---|
| SL1344_RS10685_ | 1.343441971 | 1.057032365 | 1.657975637 | 0.597373404 | 0.128615915 | 4.614867475 | 4.68E-05 | 0.006816925 | 0.051091662 | 0.1 | 0.243021319 |
| SL1344_RS10690_ | 0.999586937 | 0.723293542 | 1.299536658 | 0.57561927 | 0.134022429 | 4.22902194 | 4.68E-05 | 0.006624041 | 0.04988847 | 0.1 | 0.243021319 |
| SL1344_RS10700_ | 1.00308898 | 0.380413972 | 1.537990928 | 1.15610963 | 0.14200896 | 8.162686998 | 4.68E-05 | 0.001738009 | 0.024494651 | 0.1 | 0.243021319 |
| SL1344_RS10705_ | 3.319415523 | 3.144599698 | 3.52140695 | 0.388814084 | 0.102074766 | 3.760294678 | 4.68E-05 | 0.009403034 | 0.06412959 | 0.1 | 0.243021319 |
| SL1344_RS10710_ | 2.694371216 | 2.443953419 | 2.885323936 | 0.418957708 | 0.156838087 | 2.53539943 | 0.00067292 | 0.025860068 | 0.114489039 | 0.1004 | 0.243589453 |
| SL1344_RS10715_ | 2.653594169 | 2.389154399 | 2.84827693 | 0.407775265 | 0.15867208 | 2.500930386 | 0.000673369 | 0.02454225 | 0.111607967 | 0.1012 | 0.244721541 |
| SL1344_RS10720_ | 3.352551054 | 3.091565207 | 3.555084267 | 0.458082232 | 0.076351748 | 5.940350877 | 4.68E-05 | 0.003249151 | 0.035485495 | 0.1 | 0.243021319 |
| SL1344_RS10725_ | 2.699746638 | 2.438164587 | 2.953789912 | 0.515632454 | 0.078345958 | 6.528502946 | 4.68E-05 | 0.002043253 | 0.027488419 | 0.1 | 0.243021319 |
| SL1344_RS10730_ | 2.581249669 | 2.416466709 | 2.829267914 | 0.405549351 | 0.103042753 | 3.77921371 | 4.68E-05 | 0.010072528 | 0.065214124 | 0.1 | 0.243021319 |
| SL1344_RS10735_ | 3.186236219 | 2.865912459 | 3.420116975 | 0.544957866 | 0.100965124 | 5.331278037 | 4.68E-05 | 0.004690596 | 0.043467634 | 0.1 | 0.243021319 |
| SL1344_RS10740_ | 3.155227033 | 2.767565657 | 3.459107192 | 0.687179657 | 0.121813647 | 5.669982852 | 4.68E-05 | 0.006503598 | 0.05334519 | 0.1 | 0.243021319 |
| SL1344_RS10745_ | 3.988425168 | 3.63155481 | 4.247361748 | 0.616726546 | 0.117810759 | 5.251312198 | 4.68E-05 | 0.012843768 | 0.079702846 | 0.1 | 0.243021319 |
| SL1344_RS10750_ | 4.932730014 | 4.707410456 | 5.224895168 | 0.507205675 | 0.101993305 | 4.89453874 | 4.68E-05 | 0.003221317 | 0.038505617 | 0.1 | 0.243021319 |
| SL1344_RS10755_ | 3.211424995 | 3.085357836 | 3.515687218 | 0.424589683 | 0.242435454 | 1.943639884 | 0.00067292 | 0.034833343 | 0.138929747 | 0.1006 | 0.243866822 |
| SL1344_RS10760_ | 2.413430745 | 2.277477173 | 2.662109733 | 0.366768615 | 0.267549008 | 1.414090804 | 0.063291207 | 0.085274578 | 0.234752948 | 0.1585 | 0.323708533 |
| SL1344_RS10765_ | 2.961007477 | 2.692961184 | 3.204499159 | 0.502025167 | 0.213663097 | 2.312749771 | 4.68E-05 | 0.022919858 | 0.109316401 | 0.1002 | 0.243305132 |
| SL1344_RS10770_ | 2.526668956 | 2.188227822 | 2.669497262 | 0.464877477 | 0.14221581 | 2.708944978 | 4.68E-05 | 0.036363447 | 0.139591368 | 0.1004 | 0.243514337 |
| SL1344_RS10775_ | 2.412608039 | 2.255550487 | 2.626026797 | 0.37518079 | 0.178964508 | 2.040506491 | 0.004001152 | 0.036225616 | 0.139579464 | 0.1037 | 0.248196582 |
| SL1344_RS10780_ | 3.071600374 | 2.791783812 | 3.361836294 | 0.551330408 | 0.124191149 | 4.398068004 | 4.68E-05 | 0.003771396 | 0.040682142 | 0.1 | 0.243021319 |
| SL1344_RS10785_ | 3.31270946 | 3.051211007 | 3.55487941 | 0.509235594 | 0.079467543 | 6.406985993 | 4.68E-05 | 0.003354331 | 0.035870193 | 0.1 | 0.243021319 |
| SL1344_RS10790_ | 4.026209228 | 3.927692047 | 4.13233438 | 0.202494482 | 0.049690599 | 4.003671636 | 4.68E-05 | 0.007864876 | 0.054849107 | 0.1 | 0.243021319 |
| SL1344_RS10795_ | -2.279774981 | -2.365779047 | -2.205367702 | 0.158684588 | 0.430325568 | 0.334291947 | 0.346666673 | 0.511895118 | 0.658275803 | 0.6211 | 0.733733402 |
| SL1344_RS10800_ | -3.044462046 | -3.087264667 | -2.996488075 | 0.091610441 | 0.515196289 | 0.170510718 | 0.425716192 | 0.532772374 | 0.674204336 | 0.6338 | 0.743690524 |
| SL1344_RS10805_ | -3.120652623 | -3.312983626 | -2.972803178 | 0.336371477 | 0.601955166 | 0.525697846 | 0.276482355 | 0.419451285 | 0.578055191 | 0.5317 | 0.661769915 |
| SL1344_RS10810_ | -2.444802655 | -2.468939407 | -2.422857472 | 0.038533325 | 0.525497835 | 0.067759082 | 0.470666668 | 0.611875125 | 0.742790444 | 0.7245 | 0.815413746 |
| SL1344_RS10815_ | -3.31718091 | -3.154258694 | -3.570959355 | -0.417602012 | 0.654127763 | -0.558413488 | 0.262666678 | 0.385544969 | 0.548297139 | 0.4947 | 0.632369093 |
| SL1344_RS10820_ | -3.565175381 | -3.755471094 | -3.43400293 | 0.408136126 | 0.931857682 | 0.412627923 | 0.300666675 | 0.447528202 | 0.616048079 | 0.591 | 0.707446935 |
| SL1344_RS10825_ | -2.560204945 | -2.547729132 | -2.572881854 | -0.023590676 | 0.425497576 | -0.053827201 | 0.473684212 | 0.542402903 | 0.684031251 | 0.6566 | 0.762477779 |
| SL1344_RS10830_ | -2.498704413 | -2.624029241 | -2.367615242 | 0.25044527 | 0.417020425 | 0.547770211 | 0.256666678 | 0.369867756 | 0.526998901 | 0.4784 | 0.61210485 |
| SL1344_RS10835_ | -3.728486223 | -3.986532942 | -3.584984164 | 0.344964751 | 0.854659457 | 0.388140416 | 0.339107268 | 0.519557141 | 0.667852808 | 0.6303 | 0.745200879 |
| SL1344_RS10840_ | -3.05843765 | -3.001850641 | -3.132353669 | -0.137508184 | 0.607746081 | -0.213372128 | 0.403064627 | 0.550906055 | 0.690965275 | 0.6607 | 0.763344133 |
| SL1344_RS10845_ | -2.974414839 | -2.810252847 | -3.147580679 | -0.316484695 | 0.747055278 | -0.370857291 | 0.342438381 | 0.532020406 | 0.681027975 | 0.6464 | 0.758830178 |
| SL1344_RS10850_ | -5.104078118 | -5.13866668 | -5.06203094 | 0.035972912 | 1.270703097 | 0.022700898 | 0.486000001 | 0.608420754 | 0.739651466 | 0.7182 | 0.809802656 |
| SL1344_RS10855_ | -3.802564409 | -3.762133072 | -3.872424749 | -0.046896907 | 0.922295273 | -0.043590525 | 0.479333334 | 0.649738151 | 0.77530519 | 0.7642 | 0.850126036 |
| SL1344_RS10860_ | -2.978510464 | -3.118796593 | -2.878059498 | 0.068441117 | 0.872849302 | 0.073174275 | 0.471019322 | 0.719188749 | 0.827914681 | 0.8505 | 0.908169916 |
| SL1344_RS10865_ | -3.157870577 | -3.385068175 | -2.709451958 | 0.64194935 | 0.667678058 | 0.905973539 | 0.182666686 | 0.275324641 | 0.445422514 | 0.3844 | 0.53986617 |
| SL1344_RS10870_ | -4.704905136 | -4.784267235 | -4.543647692 | 0.427293281 | 1.375173279 | 0.287211854 | 0.39307129 | 0.580774237 | 0.71833788 | 0.6896 | 0.790319592 |
| SL1344_RS10875_ | -3.649573987 | -3.27081134 | -4.101957784 | -0.831598158 | 0.755199191 | -1.040462791 | 0.114590308 | 0.198389349 | 0.359727267 | 0.2693 | 0.42564276 |
| SL1344_RS10880_ | -3.01204664 | -2.822425443 | -3.152014315 | -0.294245063 | 0.632216971 | -0.425625573 | 0.326449041 | 0.52623238 | 0.670134652 | 0.6086 | 0.729518332 |
| SL1344_RS10885_ | -5.220724271 | -5.067379397 | -5.428121368 | -0.38255147 | 1.546857562 | -0.224012191 | 0.378414395 | 0.558719147 | 0.703799615 | 0.6862 | 0.784955633 |
| SL1344_RS10890_ | -4.793636817 | -4.3823714 | -5.742091962 | -1.417839386 | 1.872049053 | -0.668392938 | 0.140666694 | 0.276604043 | 0.470130673 | 0.3183 | 0.474712457 |
| SL1344_RS10895_ | 0.967239582 | 0.892221716 | 1.191427991 | 0.267539448 | 0.207175544 | 1.145117675 | 0.144666693 | 0.19249032 | 0.369225958 | 0.3043 | 0.471748131 |
| SL1344_RS10900_ | 3.824166743 | 3.550984845 | 4.072480072 | 0.52339997 | 0.06004495 | 8.602345484 | 4.68E-05 | 0.001165591 | 0.021279472 | 0.1 | 0.243021319 |
| SL1344_RS10905_ | 2.341024417 | 2.080068889 | 2.57720473 | 0.504708 | 0.107111821 | 4.729479606 | 4.68E-05 | 0.00462757 | 0.042801277 | 0.1 | 0.243021319 |
| SL1344_RS10910_ | 2.788199835 | 2.50453003 | 3.15929737 | 0.640940965 | 0.166187545 | 3.767607116 | 4.68E-05 | 0.007951515 | 0.060485533 | 0.1 | 0.243021319 |
| SL1344_RS10915_ | 1.518270128 | 1.181904198 | 1.83653487 | 0.656992481 | 0.108445134 | 6.039938777 | 4.68E-05 | 0.002521897 | 0.030565007 | 0.1 | 0.243021319 |
| SL1344_RS10920_ | -3.45288781 | -3.451300872 | -3.454761774 | -0.019499963 | 0.798069575 | -0.023348985 | 0.489333334 | 0.639047396 | 0.76596679 | 0.7578 | 0.8421831 |
| SL1344_RS10930_ | -1.321078827 | -1.321829996 | -1.320333434 | 0.005452327 | 0.295821033 | 0.017372373 | 0.491672219 | 0.555146746 | 0.691472448 | 0.6598 | 0.764686039 |
| SL1344_RS10935_ | -0.644171912 | -0.775160923 | -0.544433774 | 0.229117085 | 0.277244298 | 0.794734533 | 0.185209879 | 0.278612022 | 0.448966544 | 0.3695 | 0.519861327 |
| SL1344_RS10940_ | -0.661631772 | -0.813917346 | -0.526532787 | 0.292257356 | 0.235428689 | 1.1571382 | 0.09466671 | 0.163045601 | 0.314095702 | 0.2332 | 0.385501023 |
| SL1344_RS10945_ | -0.430310734 | -0.465470981 | -0.396928849 | 0.100549982 | 0.328086657 | 0.28529784 | 0.375083283 | 0.57814457 | 0.719822579 | 0.7003 | 0.798346029 |
| SL1344_RS10950_ | 0.078647169 | 0.034633527 | 0.208028355 | 0.155793275 | 0.287186568 | 0.465664075 | 0.319786816 | 0.528978117 | 0.682087007 | 0.6507 | 0.770374566 |
| SL1344_RS10955_ | 0.905182211 | 0.79836007 | 0.973392446 | 0.175927916 | 0.156651298 | 1.039177562 | 0.127333364 | 0.208196012 | 0.372390643 | 0.3016 | 0.457075691 |

| | | | | | | | | | | | |
|---|---|---|---|---|---|---|---|---|---|---|---|
| SL1344_RS10960_ | -2.277202449 | -2.542264247 | -2.100754779 | 0.457099863 | 0.533594439 | 0.802024295 | 0.171333355 | 0.2690433 | 0.445407266 | 0.3609 | 0.514116549 |
| SL1344_RS10965_ | -3.225694325 | -3.523578032 | -2.979108828 | 0.602299011 | 0.825683995 | 0.718345633 | 0.185209879 | 0.292104212 | 0.470148998 | 0.3893 | 0.538769311 |
| SL1344_RS10970_ | -1.061416252 | -1.113617802 | -1.00095815 | 0.10529523 | 0.303890139 | 0.324714116 | 0.352666673 | 0.514865785 | 0.66027806 | 0.6178 | 0.732481871 |
| SL1344_RS10975_ | 1.62153841 | 1.566459462 | 1.725747718 | 0.103502611 | 0.235268574 | 0.371290048 | 0.372000005 | 0.752413726 | 0.852180824 | 0.7645 | 0.862712734 |
| SL1344_RS10980_ | -2.893671924 | -3.069495034 | -2.680884465 | 0.404692503 | 0.551710213 | 0.684092626 | 0.217854779 | 0.342044845 | 0.502557949 | 0.4401 | 0.579804711 |
| SL1344_RS10985_ | -2.321563336 | -2.414601503 | -2.237143324 | 0.223007918 | 0.542253931 | 0.397371734 | 0.32933334 | 0.498917083 | 0.650966248 | 0.6273 | 0.736905804 |
| SL1344_RS10990_ | -1.891887477 | -2.064829393 | -1.66928011 | 0.376012806 | 0.432350984 | 0.786732111 | 0.194666685 | 0.308451852 | 0.47709508 | 0.3994 | 0.553753194 |
| SL1344_RS10995_ | -5.181744352 | -4.882965582 | -5.528542155 | -0.719199133 | 1.42680161 | -0.427619676 | 0.285143248 | 0.431301197 | 0.596632375 | 0.5578 | 0.680805465 |
| SL1344_RS11000_ | -1.409519085 | -1.549141256 | -1.298964351 | 0.248474428 | 0.36234934 | 0.621722588 | 0.239173897 | 0.364442355 | 0.531914631 | 0.4789 | 0.618759676 |
| SL1344_RS11005_ | -0.828075905 | -0.735184012 | -0.950983268 | -0.220413144 | 0.285120884 | -0.717987984 | 0.198534328 | 0.301494195 | 0.468794014 | 0.4016 | 0.545558443 |
| SL1344_RS11010_ | -1.201526653 | -1.173106245 | -1.225448519 | -0.047583798 | 0.32527886 | -0.129690148 | 0.445333335 | 0.606684189 | 0.736923183 | 0.7172 | 0.810877764 |
| SL1344_RS11015_ | -0.287866964 | -0.39195774 | -0.113692653 | 0.229118783 | 0.357414652 | 0.607420511 | 0.260000012 | 0.420105875 | 0.595087878 | 0.5369 | 0.680057381 |
| SL1344_RS11020_ | -0.555260852 | -0.597727162 | -0.50716355 | 0.127102368 | 0.375453771 | 0.332961788 | 0.347768161 | 0.563816721 | 0.710232003 | 0.7097 | 0.80557127 |
| SL1344_RS11025_ | 0.828609049 | 0.541250127 | 1.06406674 | 0.504082001 | 0.250212442 | 1.934286161 | 0.012667027 | 0.045459449 | 0.15737471 | 0.1116 | 0.259167752 |
| SL1344_RS11030_ | -0.752438996 | -0.867631208 | -0.199269636 | 0.606790375 | 0.688217312 | 0.875595315 | 0.195203216 | 0.240148672 | 0.434409364 | 0.373 | 0.562608505 |
| SL1344_RS11035_ | -0.791319724 | -0.916818779 | -0.569799872 | 0.377360369 | 0.357714758 | 1.006330798 | 0.122584976 | 0.203415397 | 0.380557256 | 0.2772 | 0.43578982 |
| SL1344_RS11040_ | 1.17715922 | 0.958848261 | 1.425075222 | 0.467046274 | 0.167431728 | 2.748951219 | 0.000673369 | 0.021655094 | 0.099940071 | 0.1008 | 0.244134396 |
| SL1344_RS11045_ | -2.919556829 | -3.172341732 | -2.633789947 | 0.545813725 | 0.520776765 | 1.011821904 | 0.129913421 | 0.207172533 | 0.36316885 | 0.297 | 0.44746884 |
| SL1344_RS11050_ | -0.941582342 | -0.901128789 | -0.992149731 | 0.135096004 | 1.368926604 | 0.106452749 | 0.454000002 | 0.843843594 | 0.912830355 | 0.925 | 0.955966804 |
| SL1344_RS11055_ | -5.527191587 | -5.954278779 | -5.202058825 | 0.675647652 | 1.980031944 | 0.320428943 | 0.352431718 | 0.547327679 | 0.697831762 | 0.6785 | 0.786415449 |
| SL1344_RS11060_ | -4.154222424 | -4.055734824 | -4.245215387 | -0.116708515 | 1.063983616 | -0.097471909 | 0.455333335 | 0.638068843 | 0.76347588 | 0.7489 | 0.834844459 |
| SL1344_RS11065_ | 0.392954592 | 0.305788989 | 0.640447893 | 0.330562213 | 0.295928103 | 1.028536743 | 0.153897426 | 0.213206669 | 0.399446283 | 0.3138 | 0.487480546 |
| SL1344_RS11070_ | 3.396474299 | 3.270301652 | 3.496110348 | 0.223786727 | 0.060776132 | 3.615635647 | 4.68E-05 | 0.011184589 | 0.066180405 | 0.1002 | 0.243300933 |
| SL1344_RS11080_ | -1.324580179 | -1.423038948 | -1.243579019 | 0.18550795 | 0.324919444 | 0.559113863 | 0.270000011 | 0.400847222 | 0.561076017 | 0.5044 | 0.635217158 |
| SL1344_RS11085_ | -2.453705196 | -2.594885274 | -2.2937191 | 0.339098559 | 0.565786157 | 0.565170703 | 0.270666677 | 0.390724217 | 0.558136066 | 0.5135 | 0.645789897 |
| SL1344_RS11090_ | 1.899477214 | 1.632504517 | 2.167892942 | 0.536829442 | 0.127147615 | 4.21554589 | 4.68E-05 | 0.008061768 | 0.057059922 | 0.1 | 0.243021319 |
| SL1344_RS11095_ | 0.157801163 | -0.26890329 | 0.578565552 | 0.893302069 | 0.279964726 | 3.163996493 | 4.68E-05 | 0.012094431 | 0.073768781 | 0.1 | 0.243021319 |
| SL1344_RS11100_ | 0.953134156 | 0.496649248 | 1.343628392 | 0.848771077 | 0.144084018 | 5.788751361 | 4.68E-05 | 0.003153326 | 0.033746601 | 0.1 | 0.243021319 |
| SL1344_RS11105_ | -2.00720828 | -2.117886832 | -1.893946468 | 0.225821498 | 0.342818518 | 0.620410512 | 0.239173897 | 0.358302598 | 0.511278148 | 0.4531 | 0.586431513 |
| SL1344_RS11110_ | 0.898422714 | 0.709084368 | 1.190225141 | 0.445430528 | 0.209236668 | 2.055729613 | 0.015989625 | 0.044518625 | 0.152919896 | 0.1144 | 0.262468467 |
| SL1344_RS11115_ | -2.048462544 | -2.29440589 | -1.764002817 | 0.556109567 | 0.470942071 | 1.124338294 | 0.101333374 | 0.170235324 | 0.332368333 | 0.2448 | 0.404335355 |
| SL1344_RS11120_ | -1.473622604 | -1.712602731 | -1.261537025 | 0.455393223 | 0.33704845 | 1.269468459 | 0.085333383 | 0.137130062 | 0.2882452 | 0.2047 | 0.362181255 |
| SL1344_RS11125_ | -1.177522595 | -1.270263733 | -1.088115679 | 0.175284385 | 0.321814332 | 0.505048614 | 0.287141915 | 0.453872158 | 0.610177326 | 0.5688 | 0.692013792 |
| SL1344_RS11130_ | -1.185148418 | -1.151594928 | -1.237147328 | -0.065128465 | 0.369906645 | -0.168117817 | 0.423333336 | 0.624251279 | 0.752783528 | 0.7246 | 0.818981166 |
| SL1344_RS11135_ | 1.816353849 | 1.597875273 | 2.035822824 | 0.435308516 | 0.117090137 | 3.685892281 | 4.68E-05 | 0.009336768 | 0.062222959 | 0.1 | 0.243021319 |
| SL1344_RS11140_ | 1.171356552 | 1.053636092 | 1.320391003 | 0.258574442 | 0.144650493 | 1.778150701 | 0.029333486 | 0.066068396 | 0.185322944 | 0.134 | 0.285595774 |
| SL1344_RS11145_ | 2.338848136 | 1.996780865 | 2.640835684 | 0.665158918 | 0.135464705 | 5.007638812 | 4.68E-05 | 0.004291201 | 0.042501896 | 0.1 | 0.243021319 |
| SL1344_RS11150_ | -3.442053321 | -3.651583822 | -3.29447097 | 0.362327544 | 0.822828714 | 0.407598191 | 0.323333341 | 0.491128542 | 0.648432759 | 0.6218 | 0.738559541 |
| SL1344_RS11155_ | -2.96955153 | -2.931846178 | -3.016453033 | -0.075699423 | 0.551424388 | -0.139499185 | 0.443333335 | 0.558936198 | 0.699551914 | 0.6848 | 0.784865325 |
| SL1344_RS11160_ | -2.234467682 | -2.164701286 | -2.429555987 | -0.222535684 | 0.687955524 | -0.30647193 | 0.399067292 | 0.669728448 | 0.794679527 | 0.7849 | 0.8711778 |
| SL1344_RS11165_ | -2.026720788 | -2.253065752 | -1.758334625 | 0.573643322 | 0.566379059 | 1.01569193 | 0.103333373 | 0.192017629 | 0.369350948 | 0.2502 | 0.408675499 |
| SL1344_RS11170_ | 1.061836472 | 0.943253987 | 1.186267984 | 0.264872806 | 0.180659898 | 1.39965933 | 0.03730858 | 0.105173905 | 0.254488877 | 0.1456 | 0.298531642 |
| SL1344_RS11175_ | -1.066132504 | -1.174957048 | -1.003408154 | 0.153190903 | 0.340760389 | 0.428293581 | 0.330446376 | 0.522539317 | 0.670210584 | 0.6372 | 0.750777425 |
| SL1344_RS11180_ | -2.973574986 | -2.971741334 | -2.975216473 | 0.033124585 | 0.63595116 | 0.049229243 | 0.481012659 | 0.612788746 | 0.744270032 | 0.7221 | 0.816125521 |
| SL1344_RS11185_ | -3.482284416 | -3.279456458 | -3.863568966 | -0.626582342 | 1.030248588 | -0.564164769 | 0.245333346 | 0.36410247 | 0.550923423 | 0.4944 | 0.633341061 |
| SL1344_RS11190_ | -3.837427027 | -4.227273061 | -3.43974031 | 0.78127043 | 0.738194997 | 0.98411799 | 0.138574312 | 0.216177728 | 0.373613804 | 0.3029 | 0.455073686 |
| SL1344_RS11195_ | -3.550289906 | -3.486457685 | -3.706418731 | -0.119187896 | 0.968871195 | -0.113511952 | 0.459333335 | 0.706524866 | 0.818897253 | 0.8011 | 0.879729138 |
| SL1344_RS11200_ | 0.287232098 | 0.31705781 | 0.235776714 | -0.072032638 | 0.211904827 | -0.336522201 | 0.36642239 | 0.54669943 | 0.690147111 | 0.6607 | 0.767410249 |
| SL1344_RS11205_ | -1.198937328 | -1.383629591 | -1.054718484 | 0.333289596 | 0.545034516 | 0.594483752 | 0.260000012 | 0.393274682 | 0.576477499 | 0.509 | 0.663074448 |
| SL1344_RS11210_ | -0.452710219 | -0.526093408 | -0.378982544 | 0.192057893 | 0.374450617 | 0.506757582 | 0.297135252 | 0.442655185 | 0.613360539 | 0.579 | 0.707507976 |
| SL1344_RS11215_ | 2.763482877 | 2.530230201 | 2.934675061 | 0.397172668 | 0.099313179 | 4.013589744 | 4.68E-05 | 0.008246395 | 0.058134816 | 0.1 | 0.243021319 |
| SL1344_RS11220_ | 1.908295183 | 1.638618644 | 2.231308677 | 0.586485341 | 0.113141015 | 5.106074333 | 4.68E-05 | 0.004699681 | 0.042723264 | 0.1 | 0.243021319 |
| SL1344_RS11225_ | 0.139384034 | -0.134044758 | 0.409348936 | 0.546638671 | 0.166179116 | 3.241616349 | 0.000673369 | 0.013494222 | 0.073058737 | 0.1001 | 0.243159503 |

| | | | | | | | | | | | |
|---|---|---|---|---|---|---|---|---|---|---|---|
| SL1344_RS11230_ | -2.923735538 | -3.54236346 | -2.219607483 | 1.260064364 | 0.816933635 | 1.500733392 | 0.051965441 | 0.088826595 | 0.229842026 | 0.1565 | 0.315726569 |
| SL1344_RS11235_ | -1.202784423 | -1.518506354 | -0.963453174 | 0.522339321 | 0.38314904 | 1.324075447 | 0.075283201 | 0.127218569 | 0.281490804 | 0.2008 | 0.362861648 |
| SL1344_RS11240_ | -2.081923563 | -2.082462314 | -2.081623704 | -0.005454172 | 0.36040617 | -0.013879694 | 0.494 | 0.521233938 | 0.665009293 | 0.6358 | 0.745570941 |
| SL1344_RS11245_ | -1.518130653 | -1.54617482 | -1.496553855 | -0.030978642 | 0.511600484 | -0.061416487 | 0.479013991 | 0.702122132 | 0.81423217 | 0.8207 | 0.886796263 |
| SL1344_RS11250_ | 0.189765933 | 0.100852531 | 0.284799197 | 0.150675416 | 0.284541096 | 0.491462135 | 0.298000009 | 0.482369216 | 0.643243154 | 0.5843 | 0.715050853 |
| SL1344_RS11255_ | -1.065982602 | -1.473161709 | -0.761629288 | 0.727285854 | 0.327377418 | 2.208394142 | 0.0020023 | 0.049517124 | 0.158968576 | 0.1025 | 0.246394241 |
| SL1344_RS11260_ | 1.826033023 | 1.656864978 | 1.999064402 | 0.353747541 | 0.126709411 | 2.751599002 | 4.68E-05 | 0.019774905 | 0.094663568 | 0.1003 | 0.243434455 |
| SL1344_RS11265_ | 1.872887998 | 1.59711475 | 2.134182713 | 0.530714207 | 0.119121506 | 4.306700175 | 4.68E-05 | 0.007037745 | 0.053341405 | 0.1 | 0.243021319 |
| SL1344_RS11270_ | 2.550953534 | 2.490369068 | 2.635206592 | 0.141490592 | 0.083934132 | 1.574325039 | 0.041305903 | 0.081731331 | 0.210413446 | 0.1503 | 0.304623452 |
| SL1344_RS11275_ | -2.089094613 | -2.418640081 | -1.758660683 | 0.627042457 | 0.561410415 | 1.09304906 | 0.129913421 | 0.178818843 | 0.347635594 | 0.2565 | 0.423468337 |
| SL1344_RS11280_ | -1.099672631 | -1.254046313 | -0.825424858 | 0.417413023 | 0.393204799 | 0.970949152 | 0.144666693 | 0.219184083 | 0.39608702 | 0.3115 | 0.480622509 |
| SL1344_RS11285_ | -4.781036444 | -4.779736996 | -4.784075069 | -0.027457345 | 0.943792106 | -0.02636804 | 0.489673551 | 0.548350898 | 0.688513237 | 0.6659 | 0.769140039 |
| SL1344_RS11290_ | -3.272303987 | -3.405418667 | -3.106812153 | 0.269977024 | 0.712109698 | 0.34155022 | 0.353097941 | 0.536222683 | 0.67781009 | 0.6362 | 0.748920549 |
| SL1344_RS11295_ | 0.843012601 | 0.734666307 | 1.380788916 | 0.604259907 | 0.318339022 | 1.359574483 | 0.080666719 | 0.121924847 | 0.290646244 | 0.1997 | 0.366172017 |
| SL1344_RS11300_ | 2.556670709 | 2.531774982 | 2.632303255 | 0.099880389 | 0.208273463 | 0.470498724 | 0.328000007 | 0.456851674 | 0.641924246 | 0.6687 | 0.789636949 |
| SL1344_RS11305_ | 0.545813664 | 0.671099427 | 0.361289008 | -0.326322395 | 0.232389357 | -1.430471234 | 0.025316634 | 0.120815905 | 0.280422591 | 0.1317 | 0.28220811 |
| SL1344_RS11310_ | -0.069622187 | -0.507936108 | 0.40073898 | 0.906495689 | 0.250921141 | 3.650617152 | 4.68E-05 | 0.01101441 | 0.06791527 | 0.1 | 0.243021319 |
| SL1344_RS11315_ | 1.70278518 | 1.486313825 | 1.931452965 | 0.448075465 | 0.12995254 | 3.379665669 | 4.68E-05 | 0.012183784 | 0.072061324 | 0.1 | 0.243021319 |
| SL1344_RS11320_ | 3.088856199 | 2.664424067 | 3.474823143 | 0.814286185 | 0.100913753 | 8.116621912 | 4.68E-05 | 0.003014204 | 0.034802361 | 0.1 | 0.243021319 |
| SL1344_RS11325_ | 2.282425787 | 1.831058453 | 2.750555279 | 0.904594969 | 0.123161668 | 7.320412754 | 4.68E-05 | 0.001345235 | 0.023770429 | 0.1 | 0.243021319 |
| SL1344_RS11330_ | 2.656920456 | 2.277788248 | 3.042306663 | 0.778199697 | 0.130187271 | 5.898786977 | 4.68E-05 | 0.001786635 | 0.027870351 | 0.1 | 0.243021319 |
| SL1344_RS11335_ | -0.43090074 | -0.956818316 | -0.081763029 | 0.870075616 | 0.307173329 | 2.721607113 | 0.001335878 | 0.025258537 | 0.108854103 | 0.1005 | 0.243714213 |
| SL1344_RS11340_ | -1.915384338 | -1.942106912 | -1.875955581 | 0.064625465 | 0.394790091 | 0.14917313 | 0.427714859 | 0.569524647 | 0.706974118 | 0.6801 | 0.781207793 |
| SL1344_RS11345_ | -3.582444856 | -3.623627608 | -3.530568119 | 0.111656855 | 0.797734445 | 0.134353081 | 0.437708197 | 0.633333339 | 0.760383703 | 0.7322 | 0.824064388 |
| SL1344_RS11350_ | -1.591007927 | -2.020908709 | -1.332352934 | 0.663041517 | 0.439123734 | 1.425003117 | 0.057295212 | 0.10539365 | 0.253559053 | 0.1688 | 0.327900838 |
| SL1344_RS11355_ | 3.57707718 | 3.278504656 | 3.863878927 | 0.594049525 | 0.09669041 | 6.244668275 | 4.68E-05 | 0.002859926 | 0.03409659 | 0.1 | 0.243021319 |
| SL1344_RS11360_ | 0.711314586 | 0.362382654 | 0.974186853 | 0.609300858 | 0.189532159 | 3.177286436 | 4.68E-05 | 0.016361922 | 0.08419186 | 0.1001 | 0.24316068 |
| SL1344_RS11365_ | 3.331588727 | 2.869480187 | 3.708742721 | 0.836230188 | 0.113310189 | 7.243024383 | 4.68E-05 | 0.001103896 | 0.021988381 | 0.1 | 0.243021319 |
| SL1344_RS11375_ | 1.31501647 | 1.097930566 | 1.543483954 | 0.453563145 | 0.132314626 | 3.435393073 | 0.00067292 | 0.011997072 | 0.069787778 | 0.1002 | 0.243302229 |
| SL1344_RS11380_ | 1.206665529 | 1.058614495 | 1.32523654 | 0.265384392 | 0.12744234 | 1.993012468 | 0.017322048 | 0.050582726 | 0.157693301 | 0.1167 | 0.26420219 |
| SL1344_RS11385_ | 0.813576054 | 0.629720688 | 0.951902642 | 0.33189518 | 0.174880876 | 1.844045946 | 0.017333596 | 0.064535016 | 0.187271336 | 0.1165 | 0.263771819 |
| SL1344_RS11390_ | 0.219521938 | -0.016125841 | 0.389167706 | 0.41284849 | 0.202416643 | 1.982743888 | 0.010659989 | 0.054905848 | 0.168646226 | 0.1122 | 0.259184355 |
| SL1344_RS11395_ | 1.129048533 | 0.841935968 | 1.400837656 | 0.561512757 | 0.157244025 | 3.504443449 | 4.68E-05 | 0.01196876 | 0.070903055 | 0.1 | 0.243021319 |
| SL1344_RS11400_ | 2.81677075 | 2.732618653 | 2.872113685 | 0.125403142 | 0.202183245 | 0.639570571 | 0.240506342 | 0.372930253 | 0.566390417 | 0.463 | 0.635512036 |
| SL1344_RS11405_ | -1.878649128 | -2.191407105 | -1.543369335 | 0.661397624 | 0.426191456 | 1.551607462 | 0.041333441 | 0.091461251 | 0.227294555 | 0.1492 | 0.301781598 |
| SL1344_RS11410_ | 1.892895573 | 1.572190147 | 2.164269709 | 0.623889602 | 0.165882255 | 3.833742779 | 4.68E-05 | 0.007267464 | 0.055739147 | 0.1 | 0.243021319 |
| SL1344_RS11415_ | 1.7744063 | 1.489670033 | 2.087929274 | 0.596862465 | 0.102059684 | 5.801845988 | 4.68E-05 | 0.002801416 | 0.032390587 | 0.1 | 0.243021319 |
| SL1344_RS11420_ | 1.216142428 | 1.031551446 | 1.393446288 | 0.362180582 | 0.135956904 | 2.64599605 | 0.002666616 | 0.026169129 | 0.108008158 | 0.1016 | 0.245194685 |
| SL1344_RS11425_ | 6.37041855 | 6.266516754 | 6.479503963 | 0.214156626 | 0.026842359 | 7.875411908 | 4.68E-05 | 0.001790473 | 0.026027245 | 0.1 | 0.243021319 |
| SL1344_RS11430_ | 1.168353686 | 1.067983532 | 1.372700097 | 0.290352941 | 0.16156803 | 1.674707829 | 0.038000117 | 0.083417339 | 0.218745208 | 0.1481 | 0.301652136 |
| SL1344_RS11435_ | 1.993489286 | 1.868283035 | 2.191486616 | 0.330579448 | 0.153817931 | 2.061264027 | 0.011333737 | 0.047022155 | 0.15848901 | 0.109 | 0.255197131 |
| SL1344_RS11440_ | 2.036895702 | 1.762640401 | 2.221300041 | 0.449075618 | 0.112305868 | 3.893858771 | 4.68E-05 | 0.012229443 | 0.070769775 | 0.1 | 0.243021319 |
| SL1344_RS11445_ | 3.52927014 | 3.223128297 | 3.74371683 | 0.492681862 | 0.131603176 | 3.540867444 | 4.68E-05 | 0.009586804 | 0.066448602 | 0.1 | 0.243021319 |
| SL1344_RS11450_ | 1.414771243 | 1.077555461 | 1.680079848 | 0.58898969 | 0.171074058 | 3.41449453 | 4.68E-05 | 0.01287784 | 0.074572942 | 0.1 | 0.243021319 |
| SL1344_RS11455_ | -3.976466164 | -4.351792493 | -3.553840238 | 0.938818472 | 1.100138894 | 0.830549428 | 0.16266669 | 0.258293409 | 0.436903984 | 0.3493 | 0.506008667 |
| SL1344_RS11460_ | -5.383698757 | -4.870811551 | -6.02435916 | -1.139420752 | 1.176173793 | -0.914096375 | 0.144666693 | 0.237988703 | 0.394122486 | 0.3194 | 0.465769181 |
| SL1344_RS11465_ | 2.990571849 | 2.717468713 | 3.283198606 | 0.52589643 | 0.152186987 | 3.478969171 | 4.68E-05 | 0.00864887 | 0.062903941 | 0.1 | 0.243021319 |
| SL1344_RS11470_ | -2.043370687 | -2.651067467 | -1.61240336 | 1.059942854 | 0.571761508 | 1.81603179 | 0.006667356 | 0.07285682 | 0.204021192 | 0.108 | 0.253585381 |
| SL1344_RS11475_ | -3.62449693 | -3.688877533 | -3.557385198 | 0.137398227 | 0.748088069 | 0.169483198 | 0.421718857 | 0.592828819 | 0.726317448 | 0.7024 | 0.799181109 |
| SL1344_RS11480_ | -3.223555363 | -3.443640091 | -3.05809133 | 0.411511239 | 0.688419639 | 0.570886037 | 0.263157906 | 0.402419353 | 0.564016633 | 0.5152 | 0.647073274 |
| SL1344_RS11485_ | -4.948720806 | -5.361312019 | -4.388323237 | 0.913079651 | 1.164201515 | 0.742388798 | 0.206666683 | 0.329831675 | 0.493688382 | 0.4237 | 0.572921814 |
| SL1344_RS11490_ | -5.650049781 | -5.630110091 | -5.674346527 | -0.06731952 | 1.298592539 | -0.043202466 | 0.479013991 | 0.529119981 | 0.671655955 | 0.6434 | 0.749885478 |
| SL1344_RS11495_ | 1.710728465 | 1.441328257 | 1.980495082 | 0.555103595 | 0.143419785 | 3.855948859 | 4.68E-05 | 0.007998659 | 0.057890382 | 0.1 | 0.243021319 |

| | | | | | | | | | | | |
|---|---|---|---|---|---|---|---|---|---|---|---|
| SL1344_RS11500_ | -2.614680538 | -2.6358232 | -2.599996187 | 0.013799049 | 0.521404879 | 0.020455899 | 0.489007329 | 0.595576218 | 0.73017903 | 0.7018 | 0.800438161 |
| SL1344_RS11505_ | -2.562224974 | -2.809668799 | -2.193725988 | 0.578833183 | 0.717920227 | 0.780647457 | 0.207861442 | 0.302805712 | 0.481320546 | 0.4064 | 0.566788999 |
| SL1344_RS11510_ | -3.135598788 | -3.317186078 | -3.024138962 | 0.327271805 | 0.740737478 | 0.403132136 | 0.305796144 | 0.462550534 | 0.62620403 | 0.597 | 0.716655703 |
| SL1344_RS11515_ | 0.450094243 | 0.364518805 | 0.519702291 | 0.153850492 | 0.183836139 | 0.793410564 | 0.184543657 | 0.287132728 | 0.45356545 | 0.3889 | 0.539725468 |
| SL1344_RS11520_ | -6.518961918 | -6.281054039 | -6.952526964 | -0.725545045 | 2.27246242 | -0.271827783 | 0.357095276 | 0.488125804 | 0.650332552 | 0.6411 | 0.751763619 |
| SL1344_RS11525_ | -2.743764851 | -2.871150527 | -2.643929731 | 0.218920298 | 0.689201117 | 0.288011709 | 0.360426388 | 0.550976501 | 0.697505945 | 0.6626 | 0.771007891 |
| SL1344_RS11535_ | -4.884807048 | -4.795888269 | -4.980495992 | -0.242605325 | 1.204708053 | -0.192994719 | 0.411333337 | 0.586121088 | 0.721647113 | 0.6967 | 0.794128363 |
| SL1344_RS11540_ | 0.53368483 | 0.529681991 | 0.53608676 | 0.01930179 | 0.20993683 | 0.093484039 | 0.463690874 | 0.646409879 | 0.771781748 | 0.7551 | 0.84020441 |
| SL1344_RS11545_ | -0.798859784 | -0.83059816 | -0.754234189 | 0.103530926 | 0.353610831 | 0.271874926 | 0.37974684 | 0.604153303 | 0.738442832 | 0.7273 | 0.817501354 |
| SL1344_RS11550_ | -4.711186267 | -4.701178984 | -4.722010931 | 0.037449139 | 1.041858761 | 0.027724485 | 0.487333334 | 0.595554969 | 0.727587605 | 0.7117 | 0.806099688 |
| SL1344_RS11555_ | -2.760530738 | -2.680777698 | -2.945310103 | -0.261852261 | 0.716629068 | -0.323816589 | 0.377333338 | 0.564190136 | 0.712157521 | 0.7006 | 0.805977266 |
| SL1344_RS11560_ | -3.448194807 | -3.711939064 | -3.104281437 | 0.560653367 | 0.758849488 | 0.702809355 | 0.230000014 | 0.361032741 | 0.526673435 | 0.4553 | 0.601636114 |
| SL1344_RS11565_ | -6.206902633 | -6.313939265 | -6.075766475 | 0.213794673 | 1.691890004 | 0.116653443 | 0.444370422 | 0.561238493 | 0.699505276 | 0.6719 | 0.775040666 |
| SL1344_RS11570_ | -4.577090362 | -4.386295446 | -4.887394379 | -0.484798847 | 0.999661229 | -0.428479904 | 0.306462367 | 0.450978348 | 0.607180436 | 0.5635 | 0.687522339 |
| SL1344_RS11585_ | -1.384722765 | -1.295982803 | -1.445975351 | -0.144542835 | 0.335431687 | -0.399334608 | 0.333777489 | 0.506161286 | 0.655057243 | 0.6145 | 0.732587315 |
| SL1344_RS11595_ | -0.052036918 | -0.196475182 | 0.065801685 | 0.259806482 | 0.258154062 | 0.958139513 | 0.146666693 | 0.218716867 | 0.389561106 | 0.3054 | 0.469424159 |
| SL1344_RS11600_ | -1.036392665 | -1.200914029 | -0.842365429 | 0.335683781 | 0.394691485 | 0.803590766 | 0.190539659 | 0.28572873 | 0.467213249 | 0.3774 | 0.541852917 |
| SL1344_RS11605_ | 1.027889151 | 1.010698536 | 1.046041067 | 0.022089818 | 0.191835018 | 0.112129207 | 0.448666669 | 0.670927752 | 0.789255139 | 0.7741 | 0.853925399 |
| SL1344_RS11610_ | -2.48929958 | -2.463241963 | -2.517261735 | -0.052200902 | 0.489309233 | -0.099422027 | 0.455333335 | 0.576808355 | 0.712553775 | 0.6876 | 0.785954403 |
| SL1344_RS11615_ | -1.567689982 | -1.583855704 | -1.56101466 | 0.009269603 | 0.387222437 | 0.018654652 | 0.492338441 | 0.636472001 | 0.762580591 | 0.7526 | 0.838485542 |
| SL1344_RS11620_ | 0.001969852 | -0.430693363 | 0.42283531 | 0.95445671 | 0.381393645 | 2.665662448 | 4.68E-05 | 0.024688418 | 0.11055866 | 0.1 | 0.243021319 |
| SL1344_RS11625_ | 1.718875035 | 1.568398312 | 1.950190325 | 0.352970797 | 0.188120745 | 1.771793489 | 0.030646382 | 0.064726121 | 0.194052158 | 0.1258 | 0.278672045 |
| SL1344_RS11630_ | -0.139273066 | -0.023573522 | -0.256110439 | -0.238627944 | 0.220988297 | -1.018258725 | 0.131912088 | 0.200876273 | 0.362189432 | 0.2865 | 0.441988472 |
| SL1344_RS11635_ | 0.943104727 | 0.914073781 | 0.97063225 | 0.072508185 | 0.204790747 | 0.351886305 | 0.361333339 | 0.5597445 | 0.705526897 | 0.6846 | 0.790958838 |
| SL1344_RS11640_ | 0.659241565 | 0.686833462 | 0.615723397 | -0.069572108 | 0.219214684 | -0.294425543 | 0.376000005 | 0.602350495 | 0.738792631 | 0.7246 | 0.81911802 |
| SL1344_RS11670_ | -0.533932199 | -0.768380561 | -0.299285558 | 0.482234502 | 0.276371624 | 1.703417123 | 0.020000227 | 0.070918251 | 0.196845998 | 0.1275 | 0.277368385 |
| SL1344_RS11675_ | -3.623151511 | -3.483657852 | -3.736517914 | -0.213909251 | 0.841198042 | -0.220818974 | 0.40066667 | 0.610971774 | 0.742528426 | 0.717 | 0.811707582 |
| SL1344_RS11680_ | -2.41688844 | -1.780900495 | -2.706625407 | -0.90175565 | 0.609692279 | -1.361734405 | 0.071333393 | 0.12847403 | 0.289546175 | 0.1946 | 0.354314875 |
| SL1344_RS11685_ | 2.359897564 | 2.226191734 | 2.484275393 | 0.257164528 | 0.093340203 | 2.735306882 | 0.001336769 | 0.021691062 | 0.098155781 | 0.101 | 0.244434617 |
| SL1344_RS11690_ | 4.727428577 | 4.630810387 | 4.804277479 | 0.189522826 | 0.094139488 | 2.013911235 | 0.003334715 | 0.039563178 | 0.147657339 | 0.1028 | 0.246877276 |
| SL1344_RS11695_ | 3.463829827 | 3.356564213 | 3.584011312 | 0.237050979 | 0.122532536 | 1.933778042 | 0.004667653 | 0.053247369 | 0.174584364 | 0.1053 | 0.250261576 |
| SL1344_RS11700_ | 3.338769434 | 3.309337335 | 3.373133801 | 0.071490834 | 0.10989171 | 0.642894194 | 0.223850781 | 0.339116305 | 0.526684892 | 0.4451 | 0.600807467 |
| SL1344_RS11705_ | -0.422888604 | -0.772401762 | 0.082838455 | 0.847747033 | 0.337074843 | 2.454812398 | 0.001335878 | 0.024996723 | 0.110219166 | 0.1008 | 0.244091018 |
| SL1344_RS11710_ | -1.434896006 | -1.528599452 | -1.329380332 | 0.199310493 | 0.326643213 | 0.594577344 | 0.258494349 | 0.39175198 | 0.549271972 | 0.501 | 0.632633172 |
| SL1344_RS11715_ | 2.542039177 | 2.436001576 | 2.634134762 | 0.198398428 | 0.099625967 | 1.958777803 | 0.011333737 | 0.048040045 | 0.157017491 | 0.1121 | 0.259293163 |
| SL1344_RS11720_ | 1.960082856 | 1.714434572 | 2.142415221 | 0.434158222 | 0.103710537 | 4.098102793 | 4.68E-05 | 0.007296505 | 0.054259173 | 0.1 | 0.243021319 |
| SL1344_RS11725_ | -1.026644275 | -1.278631815 | -0.802494807 | 0.476999268 | 0.288586414 | 1.611586617 | 0.03597614 | 0.078767321 | 0.205950205 | 0.1425 | 0.295268844 |
| SL1344_RS11730_ | -1.253888963 | -1.471634492 | -1.087219437 | 0.387126794 | 0.2960998 | 1.244214485 | 0.085333383 | 0.14873861 | 0.299206077 | 0.2237 | 0.380620827 |
| SL1344_RS11735_ | 0.018187693 | -0.064272881 | 0.100120437 | 0.19075058 | 0.314906904 | 0.592327572 | 0.23984012 | 0.376759045 | 0.55699596 | 0.4916 | 0.637241141 |
| SL1344_RS11740_ | -7.411576763 | -8.517498511 | -6.76297888 | 1.786681739 | 2.721326009 | 0.561125077 | 0.230000014 | 0.349738735 | 0.517823723 | 0.4491 | 0.58696427 |
| SL1344_RS11745_ | -3.927828369 | -4.051159482 | -3.830900315 | 0.242426312 | 0.904614654 | 0.248970227 | 0.383333338 | 0.560342117 | 0.702734664 | 0.6828 | 0.784690707 |
| SL1344_RS11750_ | -1.595399039 | -1.704349748 | -1.48427112 | 0.227835297 | 0.328954563 | 0.624949012 | 0.240506342 | 0.357462973 | 0.516241169 | 0.4575 | 0.59684106 |
| SL1344_RS11755_ | -3.02934949 | -3.165487158 | -2.818655992 | 0.310095763 | 0.610514552 | 0.458650964 | 0.303131254 | 0.495743689 | 0.641729072 | 0.5858 | 0.708207019 |
| SL1344_RS11760_ | -2.477623885 | -2.61644237 | -2.355559244 | 0.304454316 | 0.616096817 | 0.454824176 | 0.290473027 | 0.462335205 | 0.62185273 | 0.6017 | 0.716872981 |
| SL1344_RS11765_ | -4.549278242 | -4.443888468 | -4.638646225 | -0.204679184 | 0.99361912 | -0.178902417 | 0.41266667 | 0.569151503 | 0.704727785 | 0.6778 | 0.780248774 |
| SL1344_RS11770_ | 1.424738158 | 1.262264583 | 1.558206828 | 0.295293043 | 0.117405984 | 2.505754317 | 0.002668394 | 0.028222951 | 0.111596483 | 0.1033 | 0.247521264 |
| SL1344_RS11775_ | 1.760376668 | 1.57474308 | 1.964871904 | 0.351705839 | 0.188199344 | 1.839484824 | 0.032666804 | 0.053510787 | 0.173091913 | 0.1251 | 0.278079568 |
| SL1344_RS11780_ | 0.189852745 | -0.092451649 | 0.473139608 | 0.573225023 | 0.209359622 | 2.678674813 | 0.001335878 | 0.02304525 | 0.101661217 | 0.1008 | 0.244143429 |
| SL1344_RS11785_ | 1.57364507 | 0.708606772 | 2.031796316 | 1.301844523 | 0.314623241 | 3.798551176 | 4.68E-05 | 0.011469944 | 0.073547491 | 0.1 | 0.243021319 |
| SL1344_RS11790_ | 1.611567833 | 1.142724627 | 2.061987769 | 0.971327297 | 0.233291994 | 4.316805725 | 4.68E-05 | 0.007787592 | 0.059680597 | 0.1 | 0.243021319 |
| SL1344_RS11795_ | 1.659652618 | 1.120386926 | 2.085516344 | 0.985293543 | 0.160109787 | 6.220089941 | 4.68E-05 | 0.002016223 | 0.028739167 | 0.1 | 0.243021319 |
| SL1344_RS11800_ | 2.610243255 | 2.129647601 | 3.034343808 | 0.86665417 | 0.156016126 | 5.470614518 | 4.68E-05 | 0.002334088 | 0.031647301 | 0.1 | 0.243021319 |
| SL1344_RS11805_ | 1.354175588 | 0.876419694 | 1.794344818 | 0.921191915 | 0.116589588 | 7.886584376 | 4.68E-05 | 0.001402203 | 0.022729385 | 0.1 | 0.243021319 |

| | | | | | | | | | | | |
|---|---|---|---|---|---|---|---|---|---|---|---|
| SL1344_RS11810_ | 1.434024106 | 1.10094704 | 1.740370377 | 0.66096193 | 0.176298609 | 3.726240826 | 4.68E-05 | 0.007526293 | 0.056640198 | 0.1 | 0.243021319 |
| SL1344_RS11815_ | -0.046241068 | -0.337785684 | 0.249241823 | 0.608769687 | 0.257991109 | 2.332531222 | 0.002666616 | 0.030731594 | 0.122099633 | 0.1022 | 0.245910839 |
| SL1344_RS11820_ | -0.254999395 | -0.828645066 | 0.091313099 | 0.906402006 | 0.296297682 | 2.876341971 | 0.00067292 | 0.026960763 | 0.113550649 | 0.1003 | 0.243445977 |
| SL1344_RS11825_ | 1.381906093 | 1.197052559 | 1.441742172 | 0.230729897 | 0.155741578 | 1.254748068 | 0.10926053 | 0.162320675 | 0.330447734 | 0.2538 | 0.415363758 |
| SL1344_RS11830_ | 2.071445693 | 1.913604671 | 2.219851146 | 0.313304428 | 0.113877203 | 2.684341526 | 4.68E-05 | 0.023351065 | 0.102520919 | 0.1008 | 0.244080673 |
| SL1344_RS11840_ | 1.190399728 | 0.995798194 | 1.377805949 | 0.361856527 | 0.193819139 | 1.810168725 | 0.018000252 | 0.059224089 | 0.181272344 | 0.1236 | 0.274910851 |
| SL1344_RS11845_ | 0.232331849 | 0.010665529 | 0.436991456 | 0.421312535 | 0.190696273 | 2.226836634 | 0.008000574 | 0.038533919 | 0.13515599 | 0.1086 | 0.254163921 |
| SL1344_RS11850_ | 1.359234584 | 1.067972032 | 1.570676355 | 0.518277674 | 0.194418641 | 2.655817393 | 4.68E-05 | 0.018877932 | 0.093998217 | 0.1001 | 0.243157981 |
| SL1344_RS11855_ | 1.465338201 | 1.393304637 | 1.605683883 | 0.207144454 | 0.14956489 | 1.26186373 | 0.096000043 | 0.141775406 | 0.301761633 | 0.2272 | 0.391655637 |
| SL1344_RS11860_ | 1.156052108 | 1.124748214 | 1.21862275 | 0.067780774 | 0.210463722 | 0.278092875 | 0.389740178 | 0.707519409 | 0.817878356 | 0.7618 | 0.854397238 |
| SL1344_RS11865_ | -1.623073858 | -1.574206613 | -1.665485405 | -0.111020539 | 0.462074043 | -0.239448232 | 0.404397072 | 0.61451852 | 0.743814926 | 0.7313 | 0.818968172 |
| SL1344_RS11870_ | 0.226276536 | 0.221941525 | 0.230015037 | 0.008772439 | 0.18923474 | 0.049063553 | 0.478666667 | 0.570165967 | 0.707832529 | 0.6869 | 0.786016419 |
| SL1344_RS11880_ | -0.861675026 | -1.123319288 | -0.594140176 | 0.525602535 | 0.288824395 | 1.754857023 | 0.026666836 | 0.06458767 | 0.183531398 | 0.1321 | 0.282690962 |
| SL1344_RS11885_ | 3.63306184 | 3.610798625 | 3.725301116 | 0.109314856 | 0.160987507 | 0.481705684 | 0.34133334 | 0.707475773 | 0.827684672 | 0.7204 | 0.84122138 |
| SL1344_RS11890_ | 2.809770525 | 2.785522181 | 2.86103286 | 0.076212898 | 0.240049803 | 0.301242491 | 0.388666671 | 0.71257444 | 0.829983909 | 0.8374 | 0.907484224 |
| SL1344_RS11895_ | 4.339121666 | 4.184945348 | 4.514050935 | 0.331280116 | 0.093917904 | 3.546995711 | 4.68E-05 | 0.011784699 | 0.074317437 | 0.1 | 0.243021319 |
| SL1344_RS11900_ | 1.931624516 | 1.81253095 | 2.145700547 | 0.322595827 | 0.124899519 | 2.47593502 | 0.003998486 | 0.036419721 | 0.134058064 | 0.1049 | 0.249407196 |
| SL1344_RS11905_ | 3.263684243 | 3.095535113 | 3.432599943 | 0.349730597 | 0.110929182 | 3.085670841 | 4.68E-05 | 0.04614771 | 0.164170658 | 0.1 | 0.243021319 |
| SL1344_RS11910_ | 2.998447525 | 2.836460338 | 3.127255907 | 0.291843037 | 0.075762965 | 3.821767513 | 4.68E-05 | 0.009555407 | 0.062552945 | 0.1001 | 0.243160514 |
| SL1344_RS11915_ | 3.759961817 | 3.647080279 | 3.958167963 | 0.303147315 | 0.134483183 | 2.06516139 | 0.005996769 | 0.058933655 | 0.18822226 | 0.1042 | 0.248921523 |
| SL1344_RS11920_ | 5.462903294 | 5.275962746 | 5.604963499 | 0.334238616 | 0.066282528 | 5.166291996 | 4.68E-05 | 0.00239561 | 0.032764275 | 0.1 | 0.243021319 |
| SL1344_RS11925_ | 4.486658643 | 4.255457676 | 4.668644069 | 0.41510892 | 0.080977373 | 5.141183869 | 4.68E-05 | 0.001929069 | 0.029788035 | 0.1 | 0.243021319 |
| SL1344_RS11930_ | 3.321425203 | 3.157038159 | 3.499338026 | 0.34632399 | 0.108850884 | 3.183840658 | 4.68E-05 | 0.011315668 | 0.071543813 | 0.1 | 0.243021319 |
| SL1344_RS11935_ | 5.006402599 | 4.766716217 | 5.206392959 | 0.428339214 | 0.059425331 | 7.107943886 | 4.68E-05 | 0.001548622 | 0.025345773 | 0.1 | 0.243021319 |
| SL1344_RS11940_ | 3.885164527 | 3.728695283 | 4.140763014 | 0.412868103 | 0.089765888 | 4.564632713 | 4.68E-05 | 0.01375812 | 0.081226449 | 0.1 | 0.243021319 |
| SL1344_RS11945_ | 3.777765527 | 3.58844568 | 4.02809386 | 0.43211123 | 0.06762789 | 6.374505974 | 4.68E-05 | 0.003547646 | 0.037406716 | 0.1 | 0.243021319 |
| SL1344_RS11950_ | -3.143776129 | -3.412583689 | -2.447290289 | 0.903596682 | 0.733604837 | 1.078198912 | 0.161225873 | 0.217114032 | 0.389592443 | 0.3234 | 0.486320194 |
| SL1344_RS11960_ | 3.703546278 | 3.351987747 | 3.998841991 | 0.634789748 | 0.093763347 | 6.67359556 | 4.68E-05 | 0.002488058 | 0.031752807 | 0.1 | 0.243021319 |
| SL1344_RS11965_ | -0.208101753 | -0.325936833 | -0.097393976 | 0.2187597 | 0.224599562 | 0.930360068 | 0.15866669 | 0.241691626 | 0.406394636 | 0.3352 | 0.491319034 |
| SL1344_RS11970_ | 0.605523736 | 0.559768844 | 0.640404279 | 0.112728218 | 0.267339641 | 0.410017907 | 0.328447709 | 0.531554934 | 0.687762782 | 0.6725 | 0.783655542 |
| SL1344_RS11975_ | 2.586114224 | 2.390512754 | 2.73201554 | 0.340065349 | 0.086254932 | 3.790483263 | 4.68E-05 | 0.010982039 | 0.066212906 | 0.1 | 0.243021319 |
| SL1344_RS11980_ | 1.04768265 | 0.977192886 | 1.168463893 | 0.190318617 | 0.158485068 | 1.120763394 | 0.101265863 | 0.16761782 | 0.333080754 | 0.2375 | 0.397602473 |
| SL1344_RS11985_ | 1.16326687 | 1.080461536 | 1.246352148 | 0.167393928 | 0.164400388 | 0.967964003 | 0.143904091 | 0.214239524 | 0.384268381 | 0.3023 | 0.462251212 |
| SL1344_RS12000_ | 0.242352945 | -0.124445877 | 0.487429009 | 0.615594879 | 0.264439197 | 2.272270327 | 0.003998486 | 0.033140225 | 0.128961335 | 0.1024 | 0.24624559 |
| SL1344_RS12005_ | -1.415860713 | -1.436120139 | -1.388431986 | 0.047458587 | 0.392842634 | 0.105102808 | 0.453697537 | 0.64530614 | 0.770233641 | 0.7482 | 0.836036678 |
| SL1344_RS12010_ | -3.377349658 | -3.445450801 | -3.298902338 | 0.130730842 | 0.585973953 | 0.196813636 | 0.408394407 | 0.525759894 | 0.666966674 | 0.6294 | 0.739834233 |
| SL1344_RS12015_ | -2.736526717 | -2.731342984 | -2.744899224 | -0.025217019 | 0.723152112 | -0.028628799 | 0.483677549 | 0.687326062 | 0.802762424 | 0.7841 | 0.8616003 |
| SL1344_RS12020_ | -4.681110545 | -4.794442802 | -4.623292953 | 0.25680898 | 1.475478439 | 0.160498184 | 0.407333337 | 0.560451685 | 0.713452845 | 0.7182 | 0.815553632 |
| SL1344_RS12025_ | -3.388595941 | -3.436384352 | -3.357219183 | 0.120445055 | 0.734979165 | 0.143823411 | 0.431712194 | 0.585660536 | 0.725341736 | 0.7061 | 0.802561376 |
| SL1344_RS12030_ | -6.140168716 | -5.92988654 | -6.504751903 | -0.669055364 | 2.091105299 | -0.260698425 | 0.34733334 | 0.49007201 | 0.65393325 | 0.63 | 0.74153378 |
| SL1344_RS12035_ | -0.208850095 | -0.369922994 | -0.070634424 | 0.292809146 | 0.216784561 | 1.303289779 | 0.073333391 | 0.132492689 | 0.281029298 | 0.2075 | 0.3645118 |
| SL1344_RS12040_ | 3.087939145 | 3.091725554 | 3.07926902 | -0.010435914 | 0.106236863 | -0.091523432 | 0.462358429 | 0.723584006 | 0.831045728 | 0.8326 | 0.899961958 |
| SL1344_RS12045_ | 1.7607758 | 1.740945158 | 1.793819196 | 0.076456958 | 0.179481883 | 0.38936203 | 0.338666673 | 0.508085607 | 0.671273336 | 0.6768 | 0.783223946 |
| SL1344_RS12050_ | -0.664346868 | -0.632925124 | -0.692758398 | -0.060092686 | 0.260725616 | -0.220752627 | 0.398666671 | 0.545544754 | 0.686821996 | 0.6602 | 0.764955051 |
| SL1344_RS12055_ | -3.625823482 | -3.911527547 | -3.336660667 | 0.614431519 | 1.026970046 | 0.554486088 | 0.238000013 | 0.38685413 | 0.561954977 | 0.4867 | 0.630451718 |
| SL1344_RS12060_ | -0.134576406 | -0.325551329 | 0.055438645 | 0.384266709 | 0.212652384 | 1.722962094 | 0.021333546 | 0.063762312 | 0.180946151 | 0.1282 | 0.277326215 |
| SL1344_RS12065_ | -0.463306387 | -0.493195256 | -0.414950556 | 0.046377935 | 0.378263987 | 0.125288793 | 0.449033979 | 0.739702726 | 0.841837041 | 0.8037 | 0.879725987 |
| SL1344_RS12070_ | -0.506546182 | -0.749925182 | -0.284859 | 0.407889444 | 0.359238301 | 1.089493487 | 0.123333366 | 0.177231174 | 0.3507267 | 0.2514 | 0.423647699 |
| SL1344_RS12075_ | 0.000767266 | -0.097939489 | 0.194554663 | 0.269459807 | 0.288390079 | 0.860201118 | 0.185209879 | 0.269994896 | 0.450278531 | 0.3703 | 0.537091346 |
| SL1344_RS12080_ | 2.880548305 | 2.735461902 | 3.082911558 | 0.334877434 | 0.104879854 | 3.041729805 | 0.00067292 | 0.016856054 | 0.088036727 | 0.1001 | 0.243158976 |
| SL1344_RS12085_ | -1.121413407 | -1.501647853 | -0.84788504 | 0.655591817 | 0.303794636 | 2.081160854 | 0.010667096 | 0.050150155 | 0.157050011 | 0.1153 | 0.261536343 |
| SL1344_RS12090_ | 1.451445266 | 1.292500583 | 1.622422054 | 0.332647178 | 0.128190203 | 2.585104197 | 0.002668394 | 0.025042721 | 0.106080392 | 0.1023 | 0.246042076 |
| SL1344_RS12095_ | 1.416322153 | 1.2291478 | 1.595637607 | 0.390880968 | 0.176457198 | 2.237646291 | 0.001336769 | 0.034468201 | 0.131773962 | 0.1012 | 0.244725383 |

| | | | | | | | | | | |
|---|---|---|---|---|---|---|---|---|---|---|
| SL1344_RS12100_ | -0.521017877 | -0.641499454 | -0.384227842 | 0.252085471 | 0.231908728 | 1.039548482 | 0.1352432 | 0.211009625 | 0.370789386 | 0.299 | 0.453690491 |
| SL1344_RS12105_ | -1.506812685 | -1.645442169 | -1.232725857 | 0.396416571 | 0.357949932 | 0.99609599 | 0.150666692 | 0.237022698 | 0.40192541 | 0.3383 | 0.494636221 |
| SL1344_RS12110_ | -1.542914484 | -1.686571195 | -1.413590253 | 0.275564129 | 0.355614912 | 0.729401941 | 0.198534328 | 0.319377491 | 0.484438559 | 0.4123 | 0.557151524 |
| SL1344_RS12115_ | -0.547443942 | -0.724061732 | -0.356052535 | 0.378055702 | 0.259407419 | 1.382907155 | 0.050666753 | 0.102410983 | 0.243155167 | 0.1604 | 0.315659602 |
| SL1344_RS12120_ | 3.232410522 | 3.164355451 | 3.330739474 | 0.175634012 | 0.105880228 | 1.620416243 | 0.029980163 | 0.076353167 | 0.212282554 | 0.1306 | 0.282418149 |
| SL1344_RS12125_ | 2.163436399 | 2.06472333 | 2.276829848 | 0.212884466 | 0.087630504 | 2.38276322 | 0.004664544 | 0.031052664 | 0.118163858 | 0.1048 | 0.249417136 |
| SL1344_RS12130_ | 2.716197576 | 2.645577384 | 2.762133649 | 0.11401186 | 0.092958983 | 1.132456953 | 0.115333368 | 0.168673399 | 0.331541525 | 0.2544 | 0.4161868 |
| SL1344_RS12135_ | 2.924699324 | 2.775258311 | 3.046653051 | 0.275311205 | 0.088625137 | 3.148792555 | 4.68E-05 | 0.018629388 | 0.090541708 | 0.1001 | 0.243166291 |
| SL1344_RS12140_ | 4.916562749 | 4.689167899 | 5.164640325 | 0.469409224 | 0.053607317 | 8.660956966 | 4.68E-05 | 0.001132137 | 0.021689409 | 0.1 | 0.243021319 |
| SL1344_RS12145_ | 1.239443965 | 0.903099913 | 1.469444146 | 0.554258191 | 0.194220752 | 2.790741234 | 4.68E-05 | 0.018633409 | 0.093334819 | 0.1006 | 0.243870611 |
| SL1344_RS12155_ | 2.491465515 | 2.397231223 | 2.742874494 | 0.349541711 | 0.215099185 | 1.632042855 | 0.016000285 | 0.095677605 | 0.252529436 | 0.1109 | 0.257716817 |
| SL1344_RS12160_ | 3.307299094 | 3.123693569 | 3.462049992 | 0.342882196 | 0.074324595 | 4.702951038 | 4.68E-05 | 0.008036498 | 0.056677206 | 0.1 | 0.243021319 |
| SL1344_RS12170_ | 1.594004959 | 1.458259677 | 1.763682156 | 0.300349509 | 0.128545061 | 2.237774955 | 0.007995244 | 0.038783756 | 0.137329103 | 0.108 | 0.25329884 |
| SL1344_RS12175_ | -4.74686653 | -5.078359432 | -4.485865893 | 0.630264382 | 1.322048319 | 0.429410283 | 0.289333343 | 0.442949049 | 0.60952042 | 0.5685 | 0.694987275 |
| SL1344_RS12180_ | -4.364473323 | -4.637549022 | -4.130162364 | 0.520960784 | 0.907729683 | 0.535650213 | 0.276666677 | 0.390271336 | 0.549016398 | 0.4969 | 0.626953735 |
| SL1344_RS12185_ | -4.986062432 | -5.329731884 | -4.382709178 | 0.818619016 | 1.649917395 | 0.457623334 | 0.29380414 | 0.527042536 | 0.674075452 | 0.5862 | 0.719827416 |
| SL1344_RS12190_ | -3.826570693 | -4.39429315 | -3.499821334 | 0.915605466 | 0.919567863 | 0.902978106 | 0.137908089 | 0.2286949 | 0.40172352 | 0.3033 | 0.455540805 |
| SL1344_RS12195_ | -0.413822795 | -0.565124837 | -0.171550897 | 0.430927552 | 0.31383324 | 1.292849256 | 0.069287204 | 0.135314715 | 0.296956666 | 0.1936 | 0.35051004 |
| SL1344_RS12200_ | 4.635824176 | 4.681334901 | 4.537332689 | -0.140808768 | 0.124556788 | -1.08959627 | 0.090666713 | 0.191299788 | 0.385529511 | 0.1936 | 0.365707036 |
| SL1344_RS12205_ | 1.548070638 | 1.46773141 | 1.638329824 | 0.170858043 | 0.126858323 | 1.298612685 | 0.084666716 | 0.135077762 | 0.28682835 | 0.208 | 0.36731151 |
| SL1344_RS12210_ | 2.44877863 | 2.139014361 | 2.753611927 | 0.629002066 | 0.12203532 | 5.092221374 | 4.68E-05 | 0.003347031 | 0.037419613 | 0.1 | 0.243021319 |
| SL1344_RS12215_ | 1.169312022 | 0.848391356 | 1.503227478 | 0.662959356 | 0.139307384 | 4.766479325 | 4.68E-05 | 0.004833214 | 0.043212658 | 0.1 | 0.243021319 |
| SL1344_RS12220_ | 1.364271902 | 1.083291884 | 1.642021184 | 0.561282354 | 0.127676884 | 4.296928054 | 4.68E-05 | 0.007013439 | 0.052066042 | 0.1 | 0.243021319 |
| SL1344_RS12225_ | 1.364012154 | 1.099609907 | 1.589304799 | 0.492077357 | 0.13001063 | 3.742694821 | 4.68E-05 | 0.008994431 | 0.060518982 | 0.1 | 0.243021319 |
| SL1344_RS12230_ | 1.986364306 | 1.86666061 | 2.222379256 | 0.353381043 | 0.135975097 | 2.387138716 | 0.003334715 | 0.042172224 | 0.149034315 | 0.1022 | 0.245976624 |
| SL1344_RS12235_ | 2.341193916 | 2.090570799 | 2.595350354 | 0.504741946 | 0.093576528 | 5.390808042 | 4.68E-05 | 0.003585384 | 0.036962175 | 0.1 | 0.243021319 |
| SL1344_RS12240_ | -0.218402579 | -0.320950908 | -0.017036043 | 0.296700274 | 0.260340083 | 1.062939067 | 0.13200003 | 0.2063387 | 0.376828443 | 0.2977 | 0.457528934 |
| SL1344_RS12245_ | 3.751033568 | 3.553342882 | 3.963225055 | 0.436621024 | 0.134254834 | 3.390914866 | 4.68E-05 | 0.023518998 | 0.11096013 | 0.1 | 0.243021319 |
| SL1344_RS12250_ | -0.256416306 | -0.083197425 | -0.334784159 | -0.233749919 | 0.268064415 | -0.800862116 | 0.212000016 | 0.322429985 | 0.498434004 | 0.4412 | 0.5943902 |
| SL1344_RS12260_ | 1.864367147 | 1.698132888 | 2.156939655 | 0.327064213 | 0.504455165 | 0.749646661 | 0.206000017 | 0.301267356 | 0.504906199 | 0.3663 | 0.559869953 |
| SL1344_RS12265_ | 3.875573646 | 3.704637248 | 4.115443107 | 0.411390761 | 0.090613691 | 4.489870626 | 4.68E-05 | 0.013805899 | 0.081409411 | 0.1 | 0.243021319 |
| SL1344_RS12270_ | 3.853287588 | 3.757601219 | 3.919822358 | 0.155395552 | 0.066668045 | 2.178651749 | 0.005996769 | 0.038850947 | 0.139747534 | 0.1092 | 0.25558621 |
| SL1344_RS12275_ | -2.522192202 | -2.691406433 | -2.306969924 | 0.350078728 | 0.64727226 | 0.514317509 | 0.274483688 | 0.453039359 | 0.615018472 | 0.533 | 0.673040759 |
| SL1344_RS12280_ | -0.567124567 | -0.956227957 | -0.188321046 | 0.776475087 | 0.249085125 | 3.126794871 | 4.68E-05 | 0.014380458 | 0.078113893 | 0.1001 | 0.243167968 |
| SL1344_RS12285_ | -1.298530181 | -1.429110532 | -1.006082248 | 0.443300851 | 0.459412366 | 0.92432492 | 0.167888097 | 0.252928342 | 0.436681423 | 0.3425 | 0.502484462 |
| SL1344_RS12290_ | 0.057865642 | -0.10170664 | 0.154983613 | 0.237216996 | 0.25667625 | 0.87158887 | 0.183333353 | 0.269375008 | 0.447066419 | 0.3637 | 0.526410063 |
| SL1344_RS12295_ | -0.280000455 | -0.53302587 | -0.092683999 | 0.432003138 | 0.280177179 | 1.51906596 | 0.040000111 | 0.089374765 | 0.228184135 | 0.1541 | 0.310634014 |
| SL1344_RS12300_ | -2.422209817 | -2.507837928 | -2.323072255 | 0.1823881 | 0.405023143 | 0.423490542 | 0.313124592 | 0.439595433 | 0.589161131 | 0.5414 | 0.664823611 |
| SL1344_RS12305_ | -5.534497633 | -5.837789889 | -5.279032075 | 0.615634502 | 1.542758254 | 0.336720427 | 0.343104603 | 0.495316731 | 0.649461793 | 0.6362 | 0.743948987 |
| SL1344_RS12310_ | -2.692149855 | -3.018755291 | -2.467429803 | 0.560047391 | 0.50505043 | 1.058208389 | 0.111259197 | 0.195013943 | 0.353493966 | 0.2644 | 0.418583912 |
| SL1344_RS12315_ | -1.660152838 | -1.791354798 | -1.495516183 | 0.284861236 | 0.48532695 | 0.556671262 | 0.275333344 | 0.435904557 | 0.600339492 | 0.5363 | 0.673635826 |
| SL1344_RS12320_ | -7.037393793 | -7.104442583 | -6.982545875 | 0.101654872 | 3.388935512 | 0.027632407 | 0.482000001 | 0.628576691 | 0.76278989 | 0.7852 | 0.861943061 |
| SL1344_RS12325_ | -0.193713406 | -0.541831705 | 0.087470926 | 0.655118896 | 0.270024902 | 2.391854747 | 0.001336769 | 0.037122056 | 0.135539926 | 0.1007 | 0.243937133 |
| SL1344_RS12330_ | 2.306591171 | 2.344104562 | 2.283214519 | -0.053118768 | 0.133314178 | -0.387832641 | 0.342000007 | 0.590719652 | 0.731633384 | 0.671 | 0.7858917 |
| SL1344_RS12335_ | 0.549608989 | 0.310481874 | 0.971285478 | 0.629780778 | 0.247771057 | 2.437208376 | 0.002000966 | 0.032136357 | 0.127647464 | 0.1021 | 0.245877253 |
| SL1344_RS12340_ | -0.597659485 | -0.749762293 | -0.433890559 | 0.279765637 | 0.319169623 | 0.824667992 | 0.188666685 | 0.294105909 | 0.469208877 | 0.3842 | 0.544953709 |
| SL1344_RS12345_ | -0.837965575 | -0.843386775 | -0.827324703 | -0.02007685 | 0.401559756 | -0.045660262 | 0.481678881 | 0.702336121 | 0.81711981 | 0.8219 | 0.892044858 |
| SL1344_RS12350_ | 0.202828137 | 0.140665433 | 0.262352326 | 0.122841543 | 0.163018965 | 0.717635223 | 0.210000016 | 0.323862427 | 0.479081342 | 0.4257 | 0.564395679 |
| SL1344_RS12355_ | -0.06735043 | -0.066914124 | -0.06744226 | 0.005575635 | 0.252739527 | 0.022210202 | 0.489007329 | 0.665320407 | 0.786169094 | 0.7679 | 0.84940417 |
| SL1344_RS12360_ | 0.203418943 | -0.198138386 | 0.509954295 | 0.710012725 | 0.193314459 | 3.574379685 | 4.68E-05 | 0.011519394 | 0.068108241 | 0.1 | 0.243021319 |
| SL1344_RS12365_ | 1.844937629 | 1.504099826 | 2.196764312 | 0.696318215 | 0.110985605 | 6.268906487 | 4.68E-05 | 0.002836798 | 0.032238857 | 0.1 | 0.243021319 |
| SL1344_RS12370_ | 1.717045605 | 1.582125868 | 1.87733913 | 0.304525904 | 0.123466482 | 2.429901744 | 0.001335878 | 0.032263673 | 0.123617618 | 0.1017 | 0.245337994 |
| SL1344_RS12375_ | 4.388493821 | 4.335654895 | 4.443786301 | 0.0563375 | 0.192191896 | 0.282910161 | 0.436375752 | 0.737208702 | 0.84727296 | 0.9532 | 0.972906608 |

| | | | | | | | | | | | |
|---|---|---|---|---|---|---|---|---|---|---|---|
| SL1344_RS12380_ | 6.514225559 | 6.524622189 | 6.503236618 | -0.024372303 | 0.180729426 | -0.105657771 | 0.429713527 | 0.629302812 | 0.778607398 | 0.9424 | 0.967311066 |
| SL1344_RS12385_ | -0.681449477 | -0.799926531 | -0.392922804 | 0.382649791 | 0.306070827 | 1.132599759 | 0.138574312 | 0.197909538 | 0.370011937 | 0.302 | 0.464449213 |
| SL1344_RS12390_ | 0.172099125 | 0.075392015 | 0.331543554 | 0.215962681 | 0.364290143 | 0.579327516 | 0.252000012 | 0.415828917 | 0.59435446 | 0.4917 | 0.651304854 |
| SL1344_RS12395_ | 4.396687684 | 4.155386189 | 4.598987712 | 0.444015903 | 0.049845202 | 8.839252207 | 4.68E-05 | 0.001135032 | 0.020662024 | 0.1 | 0.243021319 |
| SL1344_RS12400_ | 5.31610449 | 5.064404117 | 5.595785095 | 0.530158636 | 0.207609206 | 2.333605204 | 4.68E-05 | 0.035014714 | 0.141206946 | 0.1 | 0.243021319 |
| SL1344_RS12405_ | 5.072867244 | 4.682617177 | 5.450410353 | 0.669795199 | 0.186782995 | 3.874804383 | 4.68E-05 | 0.002725067 | 0.036911418 | 0.1 | 0.243021319 |
| SL1344_RS12410_ | 5.799547294 | 5.555155497 | 6.08579925 | 0.519037428 | 0.127695447 | 3.816071655 | 4.68E-05 | 0.010436162 | 0.072057127 | 0.1 | 0.243021319 |
| SL1344_RS12415_ | 4.782331061 | 4.591549706 | 5.110138283 | 0.523127533 | 0.145418076 | 3.598718703 | 4.68E-05 | 0.01826637 | 0.097224661 | 0.1 | 0.243021319 |
| SL1344_RS12420_ | -2.061144205 | -2.283025703 | -1.931644828 | 0.346513181 | 0.491044063 | 0.651149253 | 0.23717523 | 0.363100279 | 0.533101214 | 0.4725 | 0.614692674 |
| SL1344_RS12425_ | -5.384885707 | -5.560846069 | -5.248570511 | 0.29732633 | 1.320775247 | 0.210040691 | 0.40373085 | 0.537851178 | 0.682520517 | 0.6506 | 0.75879631 |
| SL1344_RS12430_ | -2.11402106 | -2.011998355 | -2.177913401 | -0.204654096 | 0.530715056 | -0.341729123 | 0.356429053 | 0.544819796 | 0.690525583 | 0.6647 | 0.771063523 |
| SL1344_RS12435_ | -5.345405339 | -5.394879029 | -5.28959962 | 0.114000043 | 1.200774536 | 0.084973963 | 0.452000002 | 0.558031873 | 0.698643897 | 0.6647 | 0.771395921 |
| SL1344_RS12440_ | -2.121790216 | -2.212114066 | -2.003066424 | 0.130783485 | 0.865057934 | 0.145326745 | 0.425333336 | 0.762472074 | 0.857753625 | 0.8182 | 0.888297563 |
| SL1344_RS12445_ | -0.68360896 | -0.8098403 | -0.544232747 | 0.259677289 | 0.270797344 | 0.945922199 | 0.154000025 | 0.234373774 | 0.398911007 | 0.3197 | 0.476304629 |
| SL1344_RS12450_ | 1.105882589 | 0.844581304 | 1.297306011 | 0.455080126 | 0.156251963 | 2.820852995 | 0.001335878 | 0.025796566 | 0.108491807 | 0.1002 | 0.243299502 |
| SL1344_RS12455_ | 1.373662489 | 1.074961219 | 1.560684045 | 0.485421619 | 0.187701074 | 2.492778356 | 0.001336769 | 0.03730312 | 0.138488807 | 0.1001 | 0.243164005 |
| SL1344_RS12465_ | 4.072607197 | 3.852913882 | 4.256415346 | 0.415650982 | 0.098376868 | 4.373066836 | 4.68E-05 | 0.008021794 | 0.05996621 | 0.1 | 0.243021319 |
| SL1344_RS12480_ | -5.34028876 | -5.117119788 | -5.672052606 | -0.637594959 | 1.361360751 | -0.400766329 | 0.304463699 | 0.449207664 | 0.609140159 | 0.5775 | 0.694118745 |
| SL1344_RS12485_ | 4.323672006 | 4.109347136 | 4.494166053 | 0.404639779 | 0.125835881 | 3.411323736 | 4.68E-05 | 0.013845377 | 0.082135629 | 0.1 | 0.243021319 |
| SL1344_RS12490_ | 7.070854777 | 6.863379312 | 7.273037735 | 0.421795083 | 0.050627817 | 8.364355505 | 4.68E-05 | 0.000325242 | 0.013912454 | 0.1 | 0.243021319 |
| SL1344_RS12495_ | 5.416904139 | 5.30984894 | 5.748326144 | 0.437598791 | 0.128429522 | 2.392121277 | 4.68E-05 | 0.063144163 | 0.199555377 | 0.1 | 0.243021319 |
| SL1344_RS12500_ | -2.169086973 | -2.055631945 | -2.263363688 | -0.203785208 | 0.581814182 | -0.347909307 | 0.356429053 | 0.554648723 | 0.697418859 | 0.6685 | 0.773539319 |
| SL1344_RS12510_ | -0.353372503 | -0.497579562 | -0.186400583 | 0.315294691 | 0.230894618 | 1.264854404 | 0.080666719 | 0.137199993 | 0.283634308 | 0.2085 | 0.363896858 |
| SL1344_RS12515_ | -0.922498424 | -1.065687874 | -0.71255295 | 0.354575362 | 0.299895045 | 1.12395405 | 0.112591641 | 0.181794193 | 0.343573888 | 0.2663 | 0.425702679 |
| SL1344_RS12520_ | -1.925425252 | -1.925327565 | -1.925727688 | 0.001638325 | 0.413057447 | 0.004151762 | 0.499000666 | 0.610514468 | 0.740504567 | 0.7187 | 0.810685059 |
| SL1344_RS12525_ | -1.249967054 | -1.32848637 | -1.191106166 | 0.130870369 | 0.338776088 | 0.369148338 | 0.346666673 | 0.518615325 | 0.663622661 | 0.6251 | 0.737528937 |
| SL1344_RS12555_ | -0.200817429 | -0.334746537 | -0.052551206 | 0.290627773 | 0.220571435 | 1.233536277 | 0.076666722 | 0.143615192 | 0.29584934 | 0.2134 | 0.369508105 |
| SL1344_RS12560_ | 2.174028239 | 2.023637353 | 2.324002159 | 0.302575271 | 0.096426287 | 3.078936533 | 4.68E-05 | 0.016648861 | 0.083810863 | 0.1003 | 0.243439343 |
| SL1344_RS12565_ | 1.04511419 | 0.924620567 | 1.220710077 | 0.289932594 | 0.171491911 | 1.634149236 | 0.0379748 | 0.076506088 | 0.207254229 | 0.1422 | 0.295472979 |
| SL1344_RS12570_ | 0.96971538 | 0.847266297 | 1.118431622 | 0.267438668 | 0.140214557 | 1.810565907 | 0.026666836 | 0.061631819 | 0.176231901 | 0.131 | 0.28109818 |
| SL1344_RS12575_ | 1.688512059 | 1.505400743 | 1.811801897 | 0.305865382 | 0.13060247 | 2.270474063 | 0.004001152 | 0.041407472 | 0.144747432 | 0.1034 | 0.247411102 |
| SL1344_RS12580_ | 1.066867254 | 0.834244383 | 1.307793115 | 0.460198311 | 0.186794295 | 2.400670554 | 0.002668394 | 0.027113303 | 0.113805751 | 0.1035 | 0.247941624 |
| SL1344_RS12585_ | 0.559373132 | 0.277103633 | 0.826850261 | 0.556984755 | 0.182228251 | 3.032356642 | 0.00067292 | 0.016646578 | 0.084955351 | 0.1001 | 0.243159559 |
| SL1344_RS12590_ | -0.925102373 | -1.069500077 | -0.577502896 | 0.463226989 | 0.36824771 | 1.163547589 | 0.125333365 | 0.174102102 | 0.343868236 | 0.2684 | 0.435525777 |
| SL1344_RS12595_ | 0.053031316 | -0.089753703 | 0.263373485 | 0.35683125 | 0.26416013 | 1.30604392 | 0.079333387 | 0.125980211 | 0.283611767 | 0.1979 | 0.362298092 |
| SL1344_RS12600_ | -2.329929224 | -2.561749025 | -2.075403058 | 0.477799572 | 0.459310178 | 0.981975846 | 0.133333363 | 0.21657808 | 0.380214393 | 0.3031 | 0.458594427 |
| SL1344_RS12605_ | -2.988624408 | -2.552876816 | -3.125272785 | -0.589365091 | 1.163450006 | -0.516557547 | 0.305129922 | 0.459983913 | 0.639338489 | 0.6251 | 0.754035862 |
| SL1344_RS12610_ | -4.448874954 | -5.201098708 | -4.251018541 | 0.843677984 | 1.506728286 | 0.484556724 | 0.340666673 | 0.493008446 | 0.660494236 | 0.6646 | 0.786222986 |
| SL1344_RS12615_ | -3.855890841 | -3.853368406 | -3.860682133 | 0.020326431 | 0.940626916 | 0.021206829 | 0.489673551 | 0.65026812 | 0.775582009 | 0.7647 | 0.850385613 |
| SL1344_RS12620_ | -3.370560786 | -3.389386805 | -3.34757601 | 0.036772439 | 0.618572312 | 0.049271416 | 0.478666667 | 0.567902745 | 0.705338279 | 0.6722 | 0.77626326 |
| SL1344_RS12630_ | -3.423255258 | -3.515805676 | -3.37401278 | 0.105917272 | 0.762154327 | 0.123190473 | 0.447333335 | 0.620548893 | 0.750889899 | 0.7205 | 0.817656299 |
| SL1344_RS12635_ | -3.941089778 | -4.043634177 | -3.849252221 | 0.201772553 | 0.904806664 | 0.199215695 | 0.408394407 | 0.579064013 | 0.716116966 | 0.6851 | 0.785948913 |
| SL1344_RS12640_ | -3.340597288 | -3.722662271 | -2.996559054 | 0.747028441 | 0.682096891 | 1.028632983 | 0.114000035 | 0.191406595 | 0.352916696 | 0.2696 | 0.423070124 |
| SL1344_RS12645_ | -3.015797129 | -3.093148298 | -2.925940786 | 0.153340827 | 0.751191235 | 0.188906156 | 0.42466667 | 0.638451212 | 0.765057904 | 0.7406 | 0.829417496 |
| SL1344_RS12650_ | -5.80357613 | -5.456330244 | -6.278051231 | -0.768851349 | 1.475212765 | -0.466049967 | 0.277333344 | 0.413336707 | 0.571133759 | 0.5195 | 0.645244952 |
| SL1344_RS12655_ | -4.309809068 | -4.156292755 | -4.365217067 | -0.155577903 | 1.437888965 | -0.085500969 | 0.461025984 | 0.727085438 | 0.835406234 | 0.8258 | 0.897161164 |
| SL1344_RS12660_ | -4.90997103 | -5.132181177 | -4.605396033 | 0.534440892 | 1.144993614 | 0.419853576 | 0.317121927 | 0.468897192 | 0.618257671 | 0.5828 | 0.700670125 |
| SL1344_RS12665_ | -5.112787706 | -4.716566561 | -5.685396782 | -0.969236533 | 1.455835333 | -0.612783781 | 0.212524999 | 0.329502935 | 0.50602685 | 0.4299 | 0.575988472 |
| SL1344_RS12670_ | -4.831515409 | -4.883109929 | -4.762441289 | 0.1353295 | 1.033977274 | 0.124425046 | 0.452000002 | 0.557028913 | 0.695885147 | 0.6705 | 0.773497327 |
| SL1344_RS12675_ | -5.539541019 | -5.305237577 | -5.841422675 | -0.253703794 | 2.075588274 | -0.117383727 | 0.437041974 | 0.648670649 | 0.776341159 | 0.7794 | 0.857888373 |
| SL1344_RS12690_ | 2.116249475 | 2.138202996 | 2.088882276 | -0.045369659 | 0.097012906 | -0.452785268 | 0.325333341 | 0.485714086 | 0.634400234 | 0.5905 | 0.709521539 |
| SL1344_RS12695_ | -1.175950924 | -1.223023345 | -1.127511672 | 0.213589019 | 0.455885701 | 0.430779527 | 0.34133334 | 0.504473906 | 0.663721756 | 0.6846 | 0.784567181 |
| SL1344_RS12700_ | -0.16202075 | -0.104459219 | -0.21377434 | -0.112484565 | 0.25651056 | -0.420821084 | 0.327333341 | 0.516334165 | 0.665217619 | 0.6289 | 0.743229898 |

| | | | | | | | | | | | |
|---|---|---|---|---|---|---|---|---|---|---|---|
| SL1344_RS12705_ | -0.00082478 | -0.036090762 | 0.052011391 | 0.089104399 | 0.228320481 | 0.354442126 | 0.355333339 | 0.524716179 | 0.670146087 | 0.6274 | 0.741478202 |
| SL1344_RS12710_ | -0.311569879 | -0.497138363 | -0.125827178 | 0.373267581 | 0.21685555 | 1.617963671 | 0.038000117 | 0.07882033 | 0.204546883 | 0.1449 | 0.2970118 |
| SL1344_RS12715_ | 0.985370089 | 0.863133611 | 1.09122393 | 0.227837769 | 0.154126737 | 1.408231125 | 0.050000088 | 0.106052343 | 0.248095909 | 0.1646 | 0.318579619 |
| SL1344_RS12720_ | -0.864194027 | -1.095602288 | -0.598621564 | 0.528850973 | 0.313589904 | 1.68055367 | 0.019333568 | 0.077855135 | 0.210220197 | 0.1227 | 0.270848222 |
| SL1344_RS12725_ | -2.02828902 | -2.083689507 | -1.899045202 | 0.164469418 | 0.56980261 | 0.250801656 | 0.389740178 | 0.6575793 | 0.780636273 | 0.7506 | 0.843463173 |
| SL1344_RS12730_ | -1.448196742 | -1.508478052 | -1.400418978 | 0.129409204 | 0.375520771 | 0.316600445 | 0.360666672 | 0.533330686 | 0.677357942 | 0.6543 | 0.758624392 |
| SL1344_RS12735_ | 0.785195411 | 0.690639387 | 0.910124563 | 0.231763643 | 0.180746139 | 1.207824676 | 0.096602308 | 0.152452405 | 0.312533551 | 0.2232 | 0.38050328 |
| SL1344_RS12740_ | 0.339938528 | 0.099023876 | 0.531770732 | 0.442545007 | 0.210760667 | 2.008926569 | 0.005330642 | 0.05033523 | 0.161481178 | 0.1079 | 0.253765801 |
| SL1344_RS12745_ | 2.356834816 | 2.127786595 | 2.581496066 | 0.45241227 | 0.084508857 | 5.285286007 | 4.68E-05 | 0.004453218 | 0.040340551 | 0.1 | 0.243021319 |
| SL1344_RS12750_ | 1.443813881 | 1.255354032 | 1.654154077 | 0.401705388 | 0.114389656 | 3.46086872 | 4.68E-05 | 0.012165958 | 0.069356084 | 0.1 | 0.243021319 |
| SL1344_RS12755_ | 1.382255785 | 1.317266693 | 1.467143986 | 0.144885607 | 0.140250782 | 0.978959806 | 0.140000028 | 0.220141496 | 0.386460049 | 0.3082 | 0.466778967 |
| SL1344_RS12760_ | 1.007539376 | 1.028713175 | 0.990388167 | -0.03677784 | 0.139358504 | -0.259337083 | 0.399733515 | 0.545308651 | 0.68712073 | 0.6572 | 0.764801701 |
| SL1344_RS12765_ | 1.519041585 | 1.552555146 | 1.486457638 | -0.091976218 | 0.215165598 | -0.403753821 | 0.308666675 | 0.444183159 | 0.624050038 | 0.5985 | 0.722684353 |
| SL1344_RS12770_ | 5.166153111 | 5.113831609 | 5.245105972 | 0.126912394 | 0.054460704 | 2.271348128 | 0.004667653 | 0.030494332 | 0.124127732 | 0.1024 | 0.246303169 |
| SL1344_RS12775_ | 4.089828558 | 4.013941412 | 4.143186169 | 0.127050674 | 0.05651562 | 2.195731462 | 0.011992386 | 0.040156832 | 0.14147401 | 0.1085 | 0.254529568 |
| SL1344_RS12780_ | 1.935704782 | 1.885708512 | 2.073898684 | 0.158383552 | 0.244341718 | 0.529493038 | 0.311792147 | 0.554002808 | 0.714509897 | 0.65 | 0.784157374 |
| SL1344_RS12785_ | 3.474708583 | 3.404498618 | 3.501339089 | 0.081463833 | 0.104706781 | 0.706024545 | 0.246666679 | 0.3811738 | 0.566581323 | 0.5108 | 0.666806973 |
| SL1344_RS12790_ | -1.899222096 | -1.883190641 | -1.919843795 | -0.024564282 | 0.425106908 | -0.054755052 | 0.474350434 | 0.62569927 | 0.754049357 | 0.7328 | 0.823035001 |
| SL1344_RS12795_ | 1.137289841 | 1.185779621 | 1.001290302 | -0.171794005 | 0.202161488 | -0.738924365 | 0.230666681 | 0.334083699 | 0.521945663 | 0.4755 | 0.63092454 |
| SL1344_RS12800_ | 1.643839781 | 1.676974297 | 1.61173501 | -0.06781083 | 0.125657203 | -0.516144794 | 0.280666677 | 0.437316742 | 0.597340027 | 0.557 | 0.680559451 |
| SL1344_RS12805_ | 1.366171335 | 1.479810307 | 1.274551552 | -0.224511445 | 0.183668734 | -1.284063775 | 0.048634334 | 0.143218884 | 0.313345914 | 0.1577 | 0.31154312 |
| SL1344_RS12810_ | 3.005339235 | 2.969755722 | 3.048668423 | 0.14534932 | 0.187887314 | 0.788595142 | 0.18254499 | 0.245454891 | 0.441914344 | 0.3779 | 0.542279577 |
| SL1344_RS12815_ | 1.933968192 | 1.77237894 | 2.05567095 | 0.282579369 | 0.104157063 | 2.646353681 | 0.002668394 | 0.027873492 | 0.111972862 | 0.1009 | 0.244290894 |
| SL1344_RS12820_ | 4.235497888 | 4.23189066 | 4.242041716 | 0.017101449 | 0.077240252 | 0.177799387 | 0.402398405 | 0.532103083 | 0.696028129 | 0.7233 | 0.816178968 |
| SL1344_RS12825_ | 2.09655117 | 2.12617445 | 2.071887166 | -0.061560226 | 0.158675263 | -0.346738613 | 0.356666672 | 0.581679652 | 0.726312113 | 0.7145 | 0.813500023 |
| SL1344_RS12830_ | 1.811519651 | 1.845212579 | 1.781900074 | -0.065303664 | 0.107413761 | -0.592026979 | 0.256666678 | 0.403660615 | 0.558836108 | 0.5082 | 0.637613647 |
| SL1344_RS12835_ | 2.486461907 | 2.37439728 | 2.612563615 | 0.234243137 | 0.095953248 | 2.356834823 | 0.003334715 | 0.029552314 | 0.117478065 | 0.1041 | 0.24862726 |
| SL1344_RS12840_ | 2.484955922 | 2.261430621 | 2.676294135 | 0.422173176 | 0.105080328 | 3.98516102 | 4.68E-05 | 0.007585925 | 0.055803865 | 0.1 | 0.243021319 |
| SL1344_RS12845_ | -2.0868627 | -1.903217954 | -2.270283365 | -0.093107444 | 0.787045502 | -0.12772299 | 0.452000002 | 0.745338279 | 0.846499161 | 0.8623 | 0.914534957 |
| SL1344_RS12850_ | -3.663611967 | -3.778938133 | -3.516561056 | 0.34981388 | 0.852806461 | 0.388341966 | 0.350000006 | 0.539227367 | 0.682770389 | 0.6597 | 0.763932081 |
| SL1344_RS12855_ | 1.343527747 | 1.367875217 | 1.317978557 | -0.055151838 | 0.15938961 | -0.346419635 | 0.340666673 | 0.529460662 | 0.678379192 | 0.6444 | 0.755113255 |
| SL1344_RS12860_ | -2.0131594 | -2.318518094 | -1.853129954 | 0.440050008 | 0.512960374 | 0.813500579 | 0.192538326 | 0.29359963 | 0.471655072 | 0.4097 | 0.568111682 |
| SL1344_RS12865_ | -1.413141953 | -1.516574411 | -1.312443226 | 0.211938637 | 0.309144681 | 0.642989724 | 0.246666679 | 0.358152175 | 0.518774081 | 0.4607 | 0.599795808 |
| SL1344_RS12870_ | 4.430974072 | 4.415687125 | 4.573431139 | 0.15950047 | 0.197286733 | 0.72262101 | 0.328000007 | 0.4066502 | 0.604279833 | 0.6922 | 0.823583693 |
| SL1344_RS12875_ | 3.71847909 | 3.694502941 | 3.875488038 | 0.18639963 | 0.196881649 | 0.861873407 | 0.235333347 | 0.282679354 | 0.486258575 | 0.4171 | 0.607723961 |
| SL1344_RS12880_ | 2.811119042 | 2.720339161 | 3.050227049 | 0.311035837 | 0.137356924 | 2.015882013 | 0.006000767 | 0.053750728 | 0.175425337 | 0.1064 | 0.251817708 |
| SL1344_RS12885_ | 0.666258925 | 0.69060546 | 0.637379765 | -0.053667794 | 0.151721326 | -0.343224849 | 0.358666672 | 0.484890578 | 0.632174445 | 0.5988 | 0.712770864 |
| SL1344_RS12890_ | 2.338575205 | 2.318057316 | 2.38129361 | 0.055998899 | 0.099386709 | 0.496409906 | 0.309127257 | 0.49328501 | 0.643904438 | 0.5984 | 0.722774468 |
| SL1344_RS12900_ | -1.856289449 | -1.805723265 | -1.931287172 | -0.067024039 | 0.584643331 | -0.106821362 | 0.455029982 | 0.722662034 | 0.829722904 | 0.8083 | 0.882194436 |
| SL1344_RS12905_ | -2.375806281 | -2.135906174 | -2.636093742 | -0.526926492 | 0.566868963 | -0.856458226 | 0.170552986 | 0.253931922 | 0.427230648 | 0.3461 | 0.503128864 |
| SL1344_RS12910_ | 0.698857305 | 0.539271446 | 0.912176172 | 0.368000959 | 0.171146454 | 2.043477158 | 0.017333596 | 0.048623191 | 0.15739708 | 0.1156 | 0.263242539 |
| SL1344_RS12915_ | 3.023266337 | 2.82236842 | 3.197690367 | 0.348553455 | 0.135455203 | 2.498169494 | 0.000673369 | 0.020135342 | 0.099610661 | 0.1006 | 0.243863562 |
| SL1344_RS12920_ | 3.434166598 | 3.421333769 | 3.464226311 | 0.037510293 | 0.106618865 | 0.316412738 | 0.37441706 | 0.638219882 | 0.771064774 | 0.7672 | 0.854795281 |
| SL1344_RS12925_ | 2.915980829 | 2.887684836 | 2.959628004 | 0.072885566 | 0.097824376 | 0.693926666 | 0.227848116 | 0.339004791 | 0.51401537 | 0.4498 | 0.597028708 |
| SL1344_RS12930_ | 3.65254262 | 3.634699665 | 3.746124167 | 0.108824101 | 0.100311361 | 0.84004845 | 0.247333346 | 0.313578295 | 0.507216217 | 0.482 | 0.639921362 |
| SL1344_RS12935_ | 3.39199855 | 3.351767597 | 3.432599948 | 0.105433509 | 0.106504246 | 0.979260691 | 0.125249865 | 0.191697328 | 0.371535462 | 0.2765 | 0.438059667 |
| SL1344_RS12940_ | 2.874494358 | 2.786007078 | 3.000565092 | 0.229750281 | 0.150289399 | 1.548350986 | 0.022666866 | 0.088396456 | 0.237496622 | 0.1187 | 0.267948317 |
| SL1344_RS12945_ | 3.705528553 | 3.580112787 | 3.801902171 | 0.220465343 | 0.083940907 | 2.609523015 | 0.000673369 | 0.022986698 | 0.10459804 | 0.1003 | 0.243436937 |
| SL1344_RS12950_ | 3.946576756 | 4.026128706 | 3.776474309 | -0.252130063 | 0.108453716 | -2.259178928 | 0.00067292 | 0.060617265 | 0.191193186 | 0.1001 | 0.243163658 |
| SL1344_RS12955_ | -2.711597528 | -2.843323828 | -2.581276215 | 0.277917403 | 0.541958673 | 0.479306064 | 0.286475693 | 0.44681224 | 0.599058451 | 0.5564 | 0.679094434 |
| SL1344_RS12960_ | -2.154057271 | -2.069207947 | -2.24582342 | -0.170401987 | 0.424758036 | -0.363323908 | 0.335776156 | 0.495634477 | 0.642615827 | 0.6036 | 0.718236736 |
| SL1344_RS12965_ | -2.754953261 | -2.716516691 | -2.77863002 | -0.071745356 | 0.575157275 | -0.116280604 | 0.448666669 | 0.601119616 | 0.732496568 | 0.7103 | 0.804005902 |
| SL1344_RS12970_ | -1.173598613 | -1.138373116 | -1.212367653 | -0.112700662 | 0.397227537 | -0.251621431 | 0.396000004 | 0.58134785 | 0.721779332 | 0.7205 | 0.811534926 |

| | | | | | | | | | | | |
|---|---|---|---|---|---|---|---|---|---|---|---|
| SL1344_RS12975_ | 0.706291179 | 0.652173093 | 0.786252013 | 0.13954553 | 0.168354481 | 0.791665061 | 0.18254499 | 0.282320208 | 0.450044894 | 0.3823 | 0.525999463 |
| SL1344_RS12980_ | 2.725345063 | 2.708074715 | 2.740107403 | 0.032382643 | 0.072967872 | 0.421430596 | 0.304000009 | 0.447953984 | 0.600328459 | 0.5626 | 0.682809238 |
| SL1344_RS12985_ | 1.501047267 | 1.478152795 | 1.530667154 | 0.061511678 | 0.142101925 | 0.420559069 | 0.325116596 | 0.490016692 | 0.644135627 | 0.6084 | 0.726122623 |
| SL1344_RS12990_ | -1.083304962 | -1.221060269 | -0.970973449 | 0.275299325 | 0.344059455 | 0.748404125 | 0.193870771 | 0.29759382 | 0.4700419 | 0.403 | 0.549213835 |
| SL1344_RS12995_ | 1.859104143 | 1.815590478 | 1.97445848 | 0.138210607 | 0.187853751 | 0.654143042 | 0.266000011 | 0.413185945 | 0.594021816 | 0.5332 | 0.685551193 |
| SL1344_RS13000_ | 2.768976507 | 2.756704895 | 2.785754979 | 0.043292265 | 0.161250251 | 0.267936983 | 0.401732182 | 0.638364736 | 0.772836874 | 0.806 | 0.877930459 |
| SL1344_RS13005_ | 2.866373214 | 2.781594752 | 2.947809297 | 0.166685507 | 0.069460833 | 2.279801396 | 0.006000767 | 0.033698498 | 0.124178195 | 0.1072 | 0.252520554 |
| SL1344_RS13010_ | 3.426578722 | 3.27024596 | 3.46659284 | 0.181574852 | 0.111542718 | 1.309031985 | 0.096602308 | 0.135137038 | 0.305648197 | 0.2229 | 0.388948579 |
| SL1344_RS13015_ | 4.14699435 | 4.018459405 | 4.256458537 | 0.23659387 | 0.054581515 | 4.260473927 | 4.68E-05 | 0.00773985 | 0.055161431 | 0.1 | 0.243021319 |
| SL1344_RS13025_ | 4.19485876 | 4.185267033 | 4.21721578 | 0.027153556 | 0.059030619 | 0.392085811 | 0.342438381 | 0.592401723 | 0.728478002 | 0.6866 | 0.795253692 |
| SL1344_RS13030_ | 3.453164022 | 3.444884778 | 3.467087382 | 0.020367457 | 0.071858715 | 0.270485338 | 0.388407733 | 0.596859357 | 0.729607387 | 0.6982 | 0.798083062 |
| SL1344_RS13035_ | 4.814518355 | 4.785677985 | 4.832596262 | 0.059244584 | 0.074525897 | 0.752401936 | 0.206000017 | 0.281013986 | 0.471799086 | 0.3965 | 0.564978565 |
| SL1344_RS13040_ | 3.46261784 | 3.402779968 | 3.579015228 | 0.169000676 | 0.084968773 | 1.869776725 | 0.01866691 | 0.065997092 | 0.193933464 | 0.1238 | 0.273954553 |
| SL1344_RS13045_ | 2.637597137 | 2.527484665 | 2.871520908 | 0.342391473 | 0.120384532 | 2.669083062 | 0.00067292 | 0.042753364 | 0.152659239 | 0.1001 | 0.243160715 |
| SL1344_RS13050_ | 5.126440234 | 4.906365346 | 5.398807567 | 0.515956772 | 0.11771765 | 4.533494259 | 4.68E-05 | 0.001987474 | 0.031335994 | 0.1 | 0.243021319 |
| SL1344_RS13075_ | -0.4022123 | -0.418659623 | -0.36769111 | 0.017880505 | 0.35147379 | 0.040736643 | 0.482000001 | 0.727784545 | 0.834538813 | 0.8255 | 0.895419811 |
| SL1344_RS13080_ | -1.114822268 | -1.00403978 | -1.215405101 | -0.218231629 | 0.371572121 | -0.565809281 | 0.260666678 | 0.411254996 | 0.575373626 | 0.5265 | 0.658104049 |
| SL1344_RS13085_ | -2.435305729 | -2.707837988 | -2.212504167 | 0.493456142 | 0.604055327 | 0.772253745 | 0.196535661 | 0.294322679 | 0.473197858 | 0.3925 | 0.553285335 |
| SL1344_RS13090_ | 1.980292841 | 1.929235184 | 2.018263629 | 0.083197523 | 0.118752942 | 0.669474511 | 0.23466668 | 0.364872205 | 0.534416715 | 0.4655 | 0.610840242 |
| SL1344_RS13095_ | 4.460704165 | 4.586050063 | 4.428324472 | -0.149710057 | 0.06812079 | -1.675671042 | 0.03664236 | 0.095435112 | 0.24786406 | 0.1391 | 0.291982871 |
| SL1344_RS13100_ | 0.711369419 | 0.814163357 | 0.550134371 | -0.25841281 | 0.161207673 | -1.517652508 | 0.052631662 | 0.094095336 | 0.232132069 | 0.1609 | 0.317078935 |
| SL1344_RS13105_ | 2.218723612 | 2.061447131 | 2.372000049 | 0.314512715 | 0.107878573 | 2.877990821 | 0.000673369 | 0.018449128 | 0.090370545 | 0.1008 | 0.244148643 |
| SL1344_RS13115_ | 1.733769358 | 1.86044293 | 1.564682935 | -0.308223553 | 0.150463184 | -1.977907939 | 0.005334196 | 0.056996491 | 0.176987312 | 0.1051 | 0.249668282 |
| SL1344_RS13120_ | -0.40883319 | -0.476698434 | -0.294426846 | 0.211111888 | 0.44837964 | 0.472100137 | 0.292000009 | 0.489831448 | 0.656640992 | 0.5893 | 0.726074604 |
| SL1344_RS13125_ | 2.46447309 | 2.38506291 | 2.563339632 | 0.177800917 | 0.092518628 | 1.84939495 | 0.019320688 | 0.060147213 | 0.175881108 | 0.1244 | 0.274135474 |
| SL1344_RS13130_ | 2.408490232 | 2.331384835 | 2.515846607 | 0.182622811 | 0.082040305 | 2.160621792 | 0.014000326 | 0.04596808 | 0.147939938 | 0.1162 | 0.263089751 |
| SL1344_RS13135_ | 1.132924439 | 1.028907706 | 1.229746876 | 0.202252052 | 0.139422753 | 1.409606507 | 0.059333406 | 0.112358704 | 0.254251245 | 0.1771 | 0.331459759 |
| SL1344_RS13140_ | 1.313391204 | 1.287645018 | 1.378456255 | 0.092257169 | 0.186732059 | 0.473901115 | 0.321785484 | 0.509344285 | 0.666265025 | 0.6381 | 0.756917768 |
| SL1344_RS13145_ | 2.686931035 | 2.706784407 | 2.642855468 | -0.053468225 | 0.098893161 | -0.497561246 | 0.304666675 | 0.498575051 | 0.651681291 | 0.6148 | 0.737088693 |
| SL1344_RS13150_ | -0.543462537 | -0.703354926 | -0.335442037 | 0.402709626 | 0.400087856 | 0.962920686 | 0.133244533 | 0.199168066 | 0.380831212 | 0.2662 | 0.437997991 |
| SL1344_RS13155_ | 2.017267122 | 1.859259247 | 2.180644876 | 0.314448334 | 0.129280147 | 2.411368307 | 0.003998486 | 0.027874613 | 0.115340684 | 0.1038 | 0.248229959 |
| SL1344_RS13160_ | 1.637006578 | 1.481506145 | 1.731331514 | 0.237972067 | 0.146837011 | 1.545711643 | 0.061958764 | 0.092827654 | 0.232954758 | 0.1632 | 0.320906645 |
| SL1344_RS13165_ | 0.522380084 | 0.484762872 | 0.574746322 | 0.111300242 | 0.255693542 | 0.398563292 | 0.330666674 | 0.528837033 | 0.682946003 | 0.671 | 0.778300758 |
| SL1344_RS13170_ | 0.001844182 | -0.081538931 | 0.055295939 | 0.14723591 | 0.240080177 | 0.57333204 | 0.252666679 | 0.389871318 | 0.5582673 | 0.5092 | 0.641590146 |
| SL1344_RS13175_ | -0.952231694 | -1.202176739 | -0.596617295 | 0.603358094 | 0.30474321 | 1.896490119 | 0.019986902 | 0.057930289 | 0.172505861 | 0.1234 | 0.272481298 |
| SL1344_RS13180_ | 0.633082349 | 0.545827413 | 0.703363587 | 0.170153466 | 0.183489514 | 0.89027152 | 0.153333358 | 0.245603441 | 0.416746326 | 0.338 | 0.488475705 |
| SL1344_RS13185_ | -0.016393876 | -0.006140658 | -0.02955854 | -0.020109134 | 0.204951267 | -0.099088595 | 0.453697537 | 0.594519366 | 0.727040128 | 0.7044 | 0.800742804 |
| SL1344_RS13190_ | -3.018132596 | -2.827128518 | -3.175079424 | -0.379392092 | 0.628776835 | -0.582683109 | 0.251832124 | 0.401624075 | 0.560999041 | 0.5211 | 0.647297877 |
| SL1344_RS13195_ | -3.113294147 | -3.297173353 | -2.924362395 | 0.355435383 | 0.774887371 | 0.437562469 | 0.305333342 | 0.482222472 | 0.640842554 | 0.5877 | 0.715216216 |
| SL1344_RS13200_ | 0.775975367 | 0.727895512 | 0.808429416 | 0.070030912 | 0.166105829 | 0.39614102 | 0.335109934 | 0.507714817 | 0.658409224 | 0.6252 | 0.742825817 |
| SL1344_RS13210_ | 1.167969286 | 1.154384094 | 1.200963618 | 0.04326297 | 0.196744103 | 0.198600564 | 0.42504997 | 0.695947464 | 0.809886963 | 0.7903 | 0.869671485 |
| SL1344_RS13215_ | 2.67734059 | 2.415084169 | 2.943428118 | 0.542024255 | 0.11568326 | 4.771016062 | 4.68E-05 | 0.009910161 | 0.066555164 | 0.1 | 0.243021319 |
| SL1344_RS13220_ | 3.539007675 | 3.294276172 | 3.719749141 | 0.405295443 | 0.103105373 | 3.782087651 | 4.68E-05 | 0.010265354 | 0.067596196 | 0.1 | 0.243021319 |
| SL1344_RS13225_ | 3.35717513 | 3.136316923 | 3.568194988 | 0.454494417 | 0.126987924 | 3.630282537 | 4.68E-05 | 0.006926113 | 0.05585181 | 0.1 | 0.243021319 |
| SL1344_RS13230_ | 4.419967492 | 4.376348079 | 4.511068277 | 0.133703757 | 0.085523834 | 1.534447204 | 0.020000227 | 0.105672973 | 0.265612048 | 0.1216 | 0.271339574 |
| SL1344_RS13235_ | 4.418929803 | 4.292491064 | 4.530955582 | 0.260763053 | 0.082713304 | 3.191021458 | 4.68E-05 | 0.010802232 | 0.070445547 | 0.1 | 0.243021319 |
| SL1344_RS13240_ | 1.222275014 | 1.161522503 | 1.286164212 | 0.125070332 | 0.138648429 | 0.863181247 | 0.171219209 | 0.265719082 | 0.432682818 | 0.3599 | 0.512434863 |
| SL1344_RS13245_ | -0.327735214 | -0.250284301 | -0.401894634 | -0.181784338 | 0.6451541 | -0.283992736 | 0.389740178 | 0.590681827 | 0.743820273 | 0.7998 | 0.875951395 |
| SL1344_RS13250_ | -4.74798783 | -5.354344904 | -4.093935723 | 1.325224996 | 1.106943974 | 1.155774107 | 0.095269864 | 0.161150846 | 0.31518147 | 0.2417 | 0.396310841 |
| SL1344_RS13255_ | -5.540963133 | -6.143989067 | -5.077783018 | 1.083548042 | 1.531628382 | 0.627553019 | 0.213191222 | 0.329516327 | 0.499363339 | 0.4304 | 0.573425141 |
| SL1344_RS13260_ | -3.725592659 | -3.517371723 | -4.263314184 | -0.677630426 | 0.981048438 | -0.645424771 | 0.263824128 | 0.393446836 | 0.567383143 | 0.5232 | 0.665424988 |
| SL1344_RS13265_ | -2.325518492 | -2.221506709 | -2.389838208 | -0.188478236 | 0.552875109 | -0.304186442 | 0.367754835 | 0.554392987 | 0.695929664 | 0.675 | 0.776987543 |
| SL1344_RS13270_ | -3.854230193 | -3.329375377 | -4.692137838 | -1.344503023 | 1.0562952 | -1.195243091 | 0.070666727 | 0.1406298 | 0.30337223 | 0.1901 | 0.350014208 |

| | | | | | | | | | | | |
|---|---|---|---|---|---|---|---|---|---|---|---|
| SL1344_RS13275_ | -1.585026931 | -1.671253218 | -1.478966663 | 0.167297422 | 0.390380762 | 0.40028492 | 0.325782819 | 0.51003193 | 0.657179736 | 0.6186 | 0.732929822 |
| SL1344_RS13280_ | -1.959259058 | -1.849120622 | -2.024120575 | -0.13334895 | 0.450666666 | -0.267102286 | 0.380000005 | 0.620952664 | 0.749779505 | 0.7098 | 0.810246113 |
| SL1344_RS13285_ | -2.708885508 | -2.716553052 | -2.694070055 | 0.053141513 | 0.581747051 | 0.079767238 | 0.468666668 | 0.625230755 | 0.753415077 | 0.7273 | 0.820844988 |
| SL1344_RS13290_ | -2.11496569 | -2.217378541 | -2.042874385 | 0.177311491 | 0.427219461 | 0.385077673 | 0.33133334 | 0.477688428 | 0.628607955 | 0.6006 | 0.716385057 |
| SL1344_RS13295_ | -4.041951759 | -3.977691227 | -4.129512447 | -0.143030208 | 0.840339658 | -0.151668038 | 0.42504997 | 0.562690756 | 0.703784674 | 0.6725 | 0.776442662 |
| SL1344_RS13300_ | -0.549135424 | -0.541651306 | -0.553133106 | 0.009241639 | 0.295799461 | 0.026672237 | 0.490666667 | 0.65079171 | 0.77525479 | 0.7655 | 0.849053349 |
| SL1344_RS13305_ | -4.801728698 | -4.755092553 | -4.879698064 | -0.024029471 | 1.563841003 | -0.017448517 | 0.490339774 | 0.676634019 | 0.796785293 | 0.8033 | 0.874964544 |
| SL1344_RS13310_ | -4.842268058 | -4.185323773 | -5.529570528 | -1.327317204 | 1.151551066 | -1.063221636 | 0.106595641 | 0.176271287 | 0.335579186 | 0.2435 | 0.401738557 |
| SL1344_RS13315_ | -1.202775687 | -1.153153679 | -1.247455626 | -0.082860635 | 0.358428649 | -0.223613594 | 0.407333337 | 0.598119384 | 0.731318933 | 0.7042 | 0.800812804 |
| SL1344_RS13320_ | -3.198944998 | -3.239336383 | -3.172639679 | -0.002883788 | 0.726724627 | -0.004231247 | 0.498 | 0.649996404 | 0.773837134 | 0.7763 | 0.855326924 |
| SL1344_RS13325_ | 4.089935372 | 3.898248246 | 4.359348876 | 0.445499035 | 0.124465293 | 3.447654949 | 4.68E-05 | 0.014169045 | 0.083079173 | 0.1 | 0.243021319 |
| SL1344_RS13330_ | -3.041585846 | -2.924295128 | -3.348014375 | -0.460615177 | 0.800083857 | -0.50192558 | 0.304463699 | 0.444175353 | 0.617786468 | 0.6012 | 0.726003276 |
| SL1344_RS13335_ | -3.697332792 | -3.69709606 | -3.698425818 | -0.004302447 | 0.722410909 | -0.007202334 | 0.497001999 | 0.592318241 | 0.727039859 | 0.7099 | 0.80539883 |
| SL1344_RS13340_ | -3.995476968 | -3.88812505 | -4.086915532 | -0.186917967 | 0.811411486 | -0.222130515 | 0.395069957 | 0.540029815 | 0.682383759 | 0.6559 | 0.762023045 |
| SL1344_RS13345_ | -2.356984867 | -2.300306712 | -2.414969466 | -0.121609661 | 0.415093151 | -0.258318612 | 0.383744175 | 0.514998539 | 0.658670705 | 0.6141 | 0.728216763 |
| SL1344_RS13350_ | -2.978243228 | -2.719302136 | -3.416552667 | -0.740134416 | 0.741762645 | -0.946864676 | 0.119333367 | 0.221142176 | 0.402123678 | 0.2662 | 0.420630001 |
| SL1344_RS13355_ | -2.253790213 | -2.143104128 | -2.461499881 | -0.304179986 | 0.491952307 | -0.572895083 | 0.264490351 | 0.413473204 | 0.577287071 | 0.5322 | 0.665275196 |
| SL1344_RS13360_ | -1.488728932 | -1.395639405 | -1.611251825 | -0.209567755 | 0.358959761 | -0.560515013 | 0.271152576 | 0.421619915 | 0.581149633 | 0.5297 | 0.662122096 |
| SL1344_RS13365_ | -3.896536676 | -4.016977192 | -3.757284088 | 0.262985151 | 0.775899815 | 0.328912823 | 0.353764163 | 0.503665743 | 0.648939089 | 0.6088 | 0.722157328 |
| SL1344_RS13370_ | -1.929382955 | -1.802531864 | -1.995075575 | -0.17425456 | 0.462261645 | -0.328698195 | 0.359093943 | 0.573983993 | 0.711162577 | 0.6691 | 0.777335691 |
| SL1344_RS13375_ | -4.92739262 | -4.879820009 | -5.019986934 | -0.015047481 | 1.289352608 | -0.009556471 | 0.495333334 | 0.646998641 | 0.772944585 | 0.7573 | 0.844336589 |
| SL1344_RS13380_ | -1.401341122 | -1.463947006 | -1.302658907 | 0.180748385 | 0.343920487 | 0.495499669 | 0.303797477 | 0.452208361 | 0.606906191 | 0.5689 | 0.687157319 |
| SL1344_RS13385_ | -2.557615949 | -2.767971449 | -2.293935319 | 0.480030098 | 0.574956075 | 0.759598711 | 0.191333352 | 0.286000891 | 0.45760346 | 0.3841 | 0.537311815 |
| SL1344_RS13390_ | -3.305730526 | -3.581197253 | -2.85380557 | 0.679685549 | 0.900454627 | 0.709064561 | 0.238000013 | 0.372998024 | 0.544817336 | 0.4718 | 0.621238756 |
| SL1344_RS13395_ | -6.724171518 | -6.363916397 | -7.972404823 | -1.404260588 | 3.782290635 | -0.291811131 | 0.388407733 | 0.577796138 | 0.728766281 | 0.7757 | 0.868376264 |
| SL1344_RS13400_ | -0.158562651 | -0.063899051 | -0.333125598 | -0.258814451 | 0.236899567 | -1.02247915 | 0.140572979 | 0.216448831 | 0.383062933 | 0.3114 | 0.469778327 |
| SL1344_RS13405_ | -2.657088972 | -2.670473881 | -2.639604744 | 0.026740646 | 0.544594942 | 0.059210932 | 0.471685544 | 0.617078131 | 0.74734379 | 0.731 | 0.822863779 |
| SL1344_RS13410_ | -1.566570847 | -2.02541223 | -1.204111656 | 0.83144303 | 0.438207146 | 1.895826187 | 0.018000252 | 0.053253775 | 0.166684698 | 0.1186 | 0.26689111 |
| SL1344_RS13415_ | 0.303102582 | 0.288206976 | 0.31098557 | -0.005313893 | 0.230436936 | -0.019843386 | 0.492666667 | 0.668807449 | 0.789246143 | 0.7716 | 0.853958762 |
| SL1344_RS13420_ | -3.350609396 | -3.845716611 | -3.085392619 | 0.728979125 | 0.740206079 | 0.913754086 | 0.153333358 | 0.238678623 | 0.409742498 | 0.3306 | 0.487326133 |
| SL1344_RS13425_ | -6.011331473 | -5.719809853 | -6.504122363 | -0.848346717 | 2.161540155 | -0.320001206 | 0.311792147 | 0.448667899 | 0.617141267 | 0.5741 | 0.693926496 |
| SL1344_RS13430_ | -4.174764452 | -4.167024062 | -4.183767275 | -0.218277026 | 2.143087037 | -0.092408186 | 0.437708197 | 0.706323608 | 0.821705957 | 0.8272 | 0.892180832 |
| SL1344_RS13435_ | -7.396662542 | -8.595749316 | -5.979663638 | 2.198079309 | 3.789289241 | 0.502463683 | 0.23717523 | 0.431824094 | 0.588679277 | 0.4779 | 0.624740973 |
| SL1344_RS13440_ | 0.651743817 | 0.437871539 | 0.841258692 | 0.42740302 | 0.208649083 | 2.025743684 | 0.00733396 | 0.042976305 | 0.148696188 | 0.1056 | 0.250564043 |
| SL1344_RS13450_ | -0.63404671 | -0.706026151 | -0.358145304 | 0.28609183 | 0.461286929 | 0.418926686 | 0.336000007 | 0.723115639 | 0.831996984 | 0.7084 | 0.830505734 |
| SL1344_RS13455_ | 0.327479082 | 0.22287477 | 0.492850322 | 0.267490651 | 0.187792461 | 1.295940986 | 0.087333381 | 0.136941752 | 0.2871545 | 0.2143 | 0.372657965 |
| SL1344_RS13460_ | -3.155419493 | -2.831182446 | -3.696011919 | -0.833349062 | 0.644256079 | -1.229595789 | 0.079280533 | 0.144728537 | 0.299806249 | 0.2189 | 0.377808057 |
| SL1344_RS13465_ | -2.200332153 | -2.072413942 | -2.401013383 | -0.321899793 | 0.459470801 | -0.630898736 | 0.243171232 | 0.372648092 | 0.536843751 | 0.4831 | 0.621778606 |
| SL1344_RS13470_ | -3.704142494 | -3.509505009 | -3.885656831 | -0.382259177 | 0.667705283 | -0.508782413 | 0.272000011 | 0.41486056 | 0.568311909 | 0.5111 | 0.641847257 |
| SL1344_RS13475_ | -3.102122645 | -3.046095075 | -3.139746539 | -0.133791018 | 0.626504291 | -0.193376388 | 0.415722855 | 0.572466669 | 0.709971184 | 0.6896 | 0.788628305 |
| SL1344_RS13480_ | -3.657281084 | -3.605636159 | -3.74163608 | -0.253790904 | 1.009386805 | -0.208252263 | 0.395333337 | 0.552224622 | 0.705235369 | 0.7095 | 0.803812545 |
| SL1344_RS13485_ | -2.597092694 | -2.677406363 | -2.495012799 | 0.161784484 | 0.801188982 | 0.186487123 | 0.418000003 | 0.680530215 | 0.798776821 | 0.7735 | 0.855864475 |
| SL1344_RS13490_ | -3.404972491 | -3.317472975 | -3.495633625 | -0.248968933 | 0.862005312 | -0.274063128 | 0.375083283 | 0.559378445 | 0.704746713 | 0.7049 | 0.800416014 |
| SL1344_RS13495_ | -4.652876852 | -4.964523486 | -4.119446825 | 1.004912139 | 1.575226404 | 0.621598679 | 0.257828126 | 0.393694355 | 0.571618867 | 0.5291 | 0.666567105 |
| SL1344_RS13500_ | 3.213838983 | 3.186078384 | 3.322524888 | 0.122733066 | 0.092353572 | 1.104780953 | 0.151333358 | 0.201594501 | 0.382961778 | 0.3217 | 0.487150133 |
| SL1344_RS13505_ | 5.233429329 | 5.210239503 | 5.371050859 | 0.159288672 | 0.140478416 | 1.062199005 | 0.124583643 | 0.206827674 | 0.405300525 | 0.2676 | 0.441292719 |
| SL1344_RS13510_ | -4.892615699 | -4.698453218 | -5.06092436 | -0.35870229 | 1.077390341 | -0.313122606 | 0.373333338 | 0.515372379 | 0.657475169 | 0.6198 | 0.72909887 |
| SL1344_RS13515_ | -3.665892182 | -3.43369266 | -3.89051253 | -0.44815914 | 0.650368475 | -0.643379232 | 0.230513005 | 0.353807914 | 0.511583509 | 0.4486 | 0.586669633 |
| SL1344_RS13520_ | -1.814087296 | -1.661716161 | -1.964476517 | -0.341974404 | 0.440654187 | -0.734397184 | 0.193204549 | 0.299151556 | 0.471492667 | 0.423 | 0.565600709 |
| SL1344_RS13525_ | -3.883942485 | -3.969656071 | -3.824044455 | -0.201380832 | 1.71432655 | -0.117583506 | 0.450000002 | 0.62868121 | 0.765887751 | 0.8369 | 0.896667131 |
| SL1344_RS13530_ | -5.305570355 | -5.761540308 | -4.964682631 | 0.813805258 | 1.172002621 | 0.621603199 | 0.231179228 | 0.351474115 | 0.510572838 | 0.4488 | 0.587244852 |
| SL1344_RS13535_ | -3.313655583 | -3.14103574 | -3.508871236 | -0.382084265 | 0.665826201 | -0.531368454 | 0.258494349 | 0.402954606 | 0.560840878 | 0.5128 | 0.641697528 |
| SL1344_RS13540_ | 0.518198539 | 0.322827113 | 0.691637852 | 0.368365757 | 0.199571913 | 1.760826644 | 0.019320688 | 0.058885798 | 0.177482895 | 0.1229 | 0.272835025 |

| | | | | | | | | | | | |
|---|---|---|---|---|---|---|---|---|---|---|---|
| SL1344_RS13545_ | 1.484852376 | 1.375137947 | 1.607759999 | 0.258836625 | 0.186595997 | 1.357894543 | 0.036000124 | 0.114723689 | 0.272433131 | 0.1468 | 0.301801281 |
| SL1344_RS13550_ | 2.443834506 | 2.43647331 | 2.452759367 | 0.013651807 | 0.076315427 | 0.166014856 | 0.42066667 | 0.523387528 | 0.666196058 | 0.6223 | 0.734695969 |
| SL1344_RS13555_ | 2.345452071 | 2.376937795 | 2.291155413 | -0.082359266 | 0.106654237 | -0.715021206 | 0.226666681 | 0.348294725 | 0.519280386 | 0.4556 | 0.60295637 |
| SL1344_RS13560_ | 2.29459734 | 2.294115057 | 2.295308507 | 0.024094672 | 0.148496735 | 0.140106579 | 0.433710862 | 0.610046261 | 0.749508448 | 0.7602 | 0.844145201 |
| SL1344_RS13565_ | -1.159424426 | -1.235459168 | -1.080416022 | 0.157098254 | 0.318345922 | 0.483148242 | 0.301333342 | 0.453700242 | 0.60793509 | 0.5671 | 0.689591001 |
| SL1344_RS13570_ | 1.651877062 | 1.515747121 | 1.796854119 | 0.285287453 | 0.09635601 | 2.865384157 | 0.001336769 | 0.016982518 | 0.082994327 | 0.1009 | 0.244227313 |
| SL1344_RS13575_ | 4.411792983 | 4.244968042 | 4.500758193 | 0.248460685 | 0.064032777 | 3.416717804 | 4.68E-05 | 0.023585892 | 0.107714461 | 0.1 | 0.243021319 |
| SL1344_RS13580_ | -0.622795157 | -0.56448744 | -0.6644225 | -0.082825584 | 0.302998288 | -0.258092582 | 0.38507662 | 0.599062077 | 0.733093148 | 0.6956 | 0.799092523 |
| SL1344_RS13590_ | 1.205716811 | 1.225406511 | 1.189455845 | -0.020629654 | 0.724810122 | -0.026557846 | 0.475682879 | 0.907648837 | 0.950499699 | 0.922 | 0.955606076 |
| SL1344_RS13595_ | 1.672217319 | 1.580126275 | 1.762369792 | 0.181615036 | 0.097675082 | 1.797961266 | 0.025333511 | 0.059710761 | 0.168585695 | 0.1294 | 0.278121683 |
| SL1344_RS13605_ | 2.658763661 | 2.632571623 | 2.720962559 | 0.076756552 | 0.136224513 | 0.500319685 | 0.324450374 | 0.52917061 | 0.688334914 | 0.6603 | 0.779030139 |
| SL1344_RS13610_ | 1.298394933 | 1.197674809 | 1.427205649 | 0.238218499 | 0.163562965 | 1.39610387 | 0.045333431 | 0.105649156 | 0.253591654 | 0.1595 | 0.315950989 |
| SL1344_RS13615_ | 2.028075717 | 1.939858177 | 2.132648013 | 0.19452665 | 0.104660194 | 1.789158818 | 0.02466685 | 0.066560022 | 0.187129411 | 0.1276 | 0.27718808 |
| SL1344_RS13625_ | 0.417217542 | 0.201338606 | 0.641385378 | 0.431941502 | 0.242446203 | 1.714669936 | 0.027981506 | 0.063777514 | 0.18997759 | 0.1246 | 0.275828481 |
| SL1344_RS13630_ | -0.454850141 | -0.236137479 | -0.584426238 | -0.338186904 | 0.270268472 | -1.148094356 | 0.102598308 | 0.165772119 | 0.325165735 | 0.25 | 0.411327083 |
| SL1344_RS13635_ | -4.700725194 | -3.482792172 | -5.054511553 | -1.34684587 | 1.54947947 | -0.742262837 | 0.236000014 | 0.390922605 | 0.561112814 | 0.4952 | 0.650142951 |
| SL1344_RS13645_ | 4.934829497 | 4.891215566 | 5.258795392 | 0.351550496 | 0.141643593 | 0.989911902 | 0.210666683 | 0.226056146 | 0.428470248 | 0.4242 | 0.59633021 |
| SL1344_RS13650_ | 0.970417814 | 0.797530688 | 1.234814601 | 0.406066507 | 0.246581658 | 1.59339889 | 0.048634334 | 0.076031696 | 0.214040291 | 0.1438 | 0.301781177 |
| SL1344_RS13655_ | 0.352897843 | -0.052811839 | 0.539730834 | 0.688194748 | 0.786088026 | 1.006837593 | 0.097934752 | 0.186099418 | 0.379973785 | 0.2093 | 0.385301657 |
| SL1344_RS13660_ | 4.620685682 | 4.787931539 | 4.502621639 | -0.303364913 | 0.110032944 | -2.774242068 | 4.68E-05 | 0.028139241 | 0.122869211 | 0.1 | 0.243021319 |
| SL1344_RS13665_ | 3.017209822 | 2.844560993 | 3.153542767 | 0.307149136 | 0.093862591 | 3.105808541 | 0.00067292 | 0.013919744 | 0.078109507 | 0.1 | 0.243021319 |
| SL1344_RS13670_ | 3.308522585 | 3.181608431 | 3.448193963 | 0.271185476 | 0.087992476 | 3.079070158 | 4.68E-05 | 0.016746505 | 0.087118427 | 0.1 | 0.243021319 |
| SL1344_RS13675_ | 3.254272295 | 3.170794243 | 3.382399422 | 0.215319211 | 0.08190168 | 2.560779404 | 0.000673369 | 0.034681734 | 0.131592276 | 0.1006 | 0.243860803 |
| SL1344_RS13680_ | 1.339976991 | 1.417829392 | 1.234242798 | -0.120738497 | 0.552916916 | -0.192988443 | 0.444000002 | 0.805700847 | 0.890150945 | 0.9841 | 0.990562498 |
| SL1344_RS13685_ | 2.20998862 | 2.239677471 | 2.16245582 | -0.070759592 | 0.09739212 | -0.672658151 | 0.245169899 | 0.368223507 | 0.532323299 | 0.4764 | 0.61811501 |
| SL1344_RS13705_ | -0.571771737 | -1.146732198 | 0.045027101 | 1.196991512 | 0.332867957 | 3.642524076 | 4.68E-05 | 0.008905554 | 0.061854524 | 0.1 | 0.243021319 |
| SL1344_RS13710_ | 0.519712758 | 0.136955673 | 0.935617093 | 0.7392554 | 0.289039473 | 2.561352211 | 0.000673369 | 0.02040866 | 0.0992993 | 0.1005 | 0.243721165 |
| SL1344_RS13715_ | 2.51503635 | 2.573389006 | 2.455447663 | -0.119403262 | 0.105646831 | -1.098716895 | 0.11133337 | 0.17568249 | 0.341969805 | 0.2503 | 0.413218409 |
| SL1344_RS13720_ | 2.293508592 | 2.394112466 | 2.216253797 | -0.171997347 | 0.106946841 | -1.518687821 | 0.051333419 | 0.090081005 | 0.226557604 | 0.1565 | 0.313214699 |
| SL1344_RS13725_ | 1.840484929 | 1.95852618 | 1.714155578 | -0.24985643 | 0.116248909 | -2.111736696 | 0.010667096 | 0.041843908 | 0.143978433 | 0.1093 | 0.255194095 |
| SL1344_RS13730_ | -5.716460446 | -5.616929015 | -5.816002624 | -0.182116571 | 1.255448352 | -0.123770655 | 0.436666669 | 0.494478781 | 0.643260604 | 0.6059 | 0.720695753 |
| SL1344_RS13735_ | -5.796876353 | -6.168819661 | -5.39101419 | 0.672962991 | 1.999201985 | 0.305811093 | 0.351333339 | 0.571003316 | 0.70929127 | 0.6713 | 0.775897566 |
| SL1344_RS13745_ | -5.663439436 | -5.977468807 | -5.483405515 | 0.32971084 | 1.853585845 | 0.167399116 | 0.423051302 | 0.588100044 | 0.727784733 | 0.7323 | 0.822777665 |
| SL1344_RS13750_ | -7.057087961 | -7.002402295 | -7.102258825 | 0.004444632 | 2.613935879 | 0.001664366 | 0.498666667 | 0.576430798 | 0.719552679 | 0.7231 | 0.816454722 |
| SL1344_RS13755_ | -8.348699981 | -9.530343666 | -7.289858663 | 2.254772373 | 3.646527674 | 0.554475401 | 0.248666679 | 0.394682864 | 0.55068756 | 0.4718 | 0.608490092 |
| SL1344_RS13760_ | -6.507655941 | -6.111148359 | -7.191030563 | -1.198014155 | 2.2011478 | -0.44596213 | 0.264490351 | 0.395153306 | 0.565057828 | 0.4975 | 0.631258547 |
| SL1344_RS13765_ | -3.870638069 | -3.527002254 | -4.32957359 | -0.818645287 | 0.785693962 | -0.989164163 | 0.129913421 | 0.211070205 | 0.372935441 | 0.2973 | 0.450951089 |
| SL1344_RS13770_ | -3.253952308 | -3.340622114 | -3.131221275 | 0.237587961 | 0.645708785 | 0.343168293 | 0.366000005 | 0.526273916 | 0.671303591 | 0.6399 | 0.750034593 |
| SL1344_RS13775_ | -1.450769328 | -1.427774223 | -1.480162346 | -0.03457858 | 0.413373662 | -0.075747666 | 0.467021987 | 0.658300256 | 0.781439555 | 0.7803 | 0.858946133 |
| SL1344_RS13780_ | 0.977854016 | 0.993712202 | 0.954131513 | -0.047794207 | 0.155524157 | -0.283853507 | 0.368421058 | 0.537530761 | 0.68236695 | 0.6572 | 0.763438978 |
| SL1344_RS13785_ | 7.158109986 | 7.108852638 | 7.321670344 | 0.207150723 | 0.122001257 | 1.470475427 | 0.072618313 | 0.097847667 | 0.258927182 | 0.1639 | 0.332723566 |
| SL1344_RS13790_ | 7.179728581 | 7.126497156 | 7.257838287 | 0.152785112 | 0.070697081 | 2.302512017 | 4.68E-05 | 0.030849824 | 0.130003866 | 0.1 | 0.243021319 |
| SL1344_RS13800_ | 5.999451988 | 5.888903202 | 6.124180827 | 0.23157921 | 0.060450182 | 3.890853455 | 4.68E-05 | 0.004721696 | 0.046530063 | 0.1 | 0.243021319 |
| SL1344_RS13805_ | 3.220602799 | 3.209154101 | 3.253685162 | 0.047042983 | 0.174546936 | 0.222573788 | 0.407333337 | 0.860473937 | 0.922466932 | 0.8314 | 0.904284604 |
| SL1344_RS13810_ | 2.275512299 | 2.170918323 | 2.353260269 | 0.184628399 | 0.104426214 | 1.686464402 | 0.03264504 | 0.072558261 | 0.198983458 | 0.1316 | 0.282976536 |
| SL1344_RS13815_ | 2.409437892 | 2.243781729 | 2.474320435 | 0.20902577 | 0.13291937 | 1.41589588 | 0.07533339 | 0.117221906 | 0.276674555 | 0.1858 | 0.350056441 |
| SL1344_RS13820_ | -0.437113948 | -0.663622202 | -0.297096394 | 0.391365233 | 0.366668491 | 1.061759286 | 0.12391742 | 0.178667791 | 0.35484183 | 0.2625 | 0.432362251 |
| SL1344_RS13825_ | 5.331025357 | 5.628786634 | 5.230678572 | -0.394681963 | 0.097732472 | -2.150185432 | 4.68E-05 | 0.069268876 | 0.210466153 | 0.1 | 0.243021319 |
| SL1344_RS13830_ | 0.953991328 | 0.956631434 | 0.946659244 | 0.032843969 | 0.255572445 | 0.112122672 | 0.453031314 | 0.728130261 | 0.835037842 | 0.8643 | 0.916609761 |
| SL1344_RS13840_ | 3.080910578 | 3.048906081 | 3.128496811 | 0.079590124 | 0.100666204 | 0.728422349 | 0.201333351 | 0.309967332 | 0.491271301 | 0.4044 | 0.569429068 |
| SL1344_RS13845_ | 1.818693243 | 1.85965179 | 1.791082331 | -0.069543137 | 0.111751297 | -0.581774388 | 0.254000012 | 0.388201307 | 0.548329631 | 0.5089 | 0.640235721 |
| SL1344_RS13850_ | 2.545243786 | 2.595930905 | 2.517764725 | -0.065293631 | 0.128065566 | -0.446358843 | 0.315333341 | 0.532088542 | 0.686562915 | 0.6478 | 0.766985384 |
| SL1344_RS13855_ | 3.530316465 | 3.463463981 | 3.622312156 | 0.16728819 | 0.093990548 | 1.757567256 | 0.003334715 | 0.122039649 | 0.292056613 | 0.1026 | 0.246549036 |

| | | | | | | | | | | | |
|---|---|---|---|---|---|---|---|---|---|---|---|
| SL1344_RS13860_ | 0.289607814 | 0.265083676 | 0.317659523 | 0.100895647 | 0.29243463 | 0.323625898 | 0.36908728 | 0.588326413 | 0.729945262 | 0.7309 | 0.821944491 |
| SL1344_RS13865_ | -3.412909913 | -3.333872053 | -3.474584831 | -0.141222084 | 0.686000222 | -0.187656542 | 0.420386412 | 0.575766085 | 0.711944045 | 0.6703 | 0.775404639 |
| SL1344_RS13870_ | -2.281524991 | -2.470015521 | -2.16846941 | 0.291143307 | 0.623673924 | 0.441386835 | 0.315333341 | 0.501159713 | 0.660587853 | 0.63 | 0.748478471 |
| SL1344_RS13875_ | -2.310553471 | -2.441089972 | -1.916200788 | 0.555467846 | 0.997055965 | 0.524557282 | 0.254666679 | 0.37773349 | 0.576498733 | 0.5267 | 0.673798785 |
| SL1344_RS13880_ | -2.454724448 | -2.521329688 | -2.398404863 | 0.116379821 | 0.461251808 | 0.230056472 | 0.395333337 | 0.545295683 | 0.685763147 | 0.6508 | 0.75868489 |
| SL1344_RS13885_ | -2.987465178 | -2.843697991 | -3.107364225 | -0.307213561 | 0.687320799 | -0.431202405 | 0.319786816 | 0.488773761 | 0.641718437 | 0.6104 | 0.726168032 |
| SL1344_RS13890_ | -3.922068509 | -4.03183833 | -3.817567377 | 0.224826511 | 0.870056293 | 0.243144426 | 0.400000004 | 0.565502725 | 0.706523328 | 0.6728 | 0.777849598 |
| SL1344_RS13895_ | -1.248737843 | -1.282022949 | -1.201985117 | 0.117536244 | 0.427607442 | 0.284124783 | 0.391738845 | 0.597982857 | 0.735125942 | 0.7401 | 0.827585858 |
| SL1344_RS13900_ | -4.608339595 | -4.63426882 | -4.563282938 | 0.065621447 | 1.108463153 | 0.054418511 | 0.476666668 | 0.608221223 | 0.740266698 | 0.7308 | 0.820294375 |
| SL1344_RS13905_ | -4.474596725 | -4.111074416 | -4.772821612 | -0.947642186 | 1.547142095 | -0.600163311 | 0.253830791 | 0.376559042 | 0.557138418 | 0.5191 | 0.658361868 |
| SL1344_RS13910_ | -4.465012678 | -4.49092758 | -4.432481926 | 0.188363833 | 1.517652056 | 0.111779614 | 0.454666668 | 0.720080541 | 0.828942097 | 0.8221 | 0.889411502 |
| SL1344_RS13915_ | -2.141177516 | -2.124742287 | -2.162972391 | -0.134920385 | 0.631391295 | -0.193815324 | 0.41466667 | 0.618945055 | 0.754092839 | 0.7685 | 0.846798039 |
| SL1344_RS13920_ | -4.503177537 | -4.582353227 | -4.387664401 | 0.105292504 | 1.107380482 | 0.076053946 | 0.462358429 | 0.639612791 | 0.766397021 | 0.743 | 0.832998777 |
| SL1344_RS13930_ | -1.799524857 | -1.475887543 | -2.030417798 | -0.595422545 | 0.502960233 | -1.170239587 | 0.08400005 | 0.163828878 | 0.333738687 | 0.2153 | 0.374445367 |
| SL1344_RS13935_ | -1.941023445 | -1.959963243 | -1.920675152 | 0.058453259 | 0.37596949 | 0.141652201 | 0.436375752 | 0.546143082 | 0.686374122 | 0.6594 | 0.763770086 |
| SL1344_RS13940_ | -4.161897711 | -4.066177539 | -4.272519013 | -0.549178948 | 1.539136941 | -0.333026329 | 0.376000005 | 0.562457441 | 0.712402861 | 0.732 | 0.824282731 |
| SL1344_RS13950_ | -6.800492567 | -6.849076344 | -6.766892236 | 0.205714287 | 2.474582332 | 0.064221939 | 0.459693539 | 0.576300895 | 0.717630195 | 0.7057 | 0.800785828 |
| SL1344_RS13955_ | -5.251324079 | -4.843761978 | -5.993583251 | -1.137348883 | 1.421487517 | -0.699623601 | 0.196535661 | 0.298709615 | 0.474296069 | 0.4048 | 0.55218495 |
| SL1344_RS13965_ | -4.68576113 | -4.117442209 | -5.163263951 | -1.026045971 | 1.318783797 | -0.747045634 | 0.194000018 | 0.306900763 | 0.478856289 | 0.3982 | 0.551491375 |
| SL1344_RS13970_ | -4.44651674 | -4.575817781 | -4.269723314 | 0.343276041 | 1.272526874 | 0.246929403 | 0.405333337 | 0.63725549 | 0.764072608 | 0.7423 | 0.83151105 |
| SL1344_RS13975_ | -4.764326334 | -4.566678672 | -5.167073868 | -0.153760481 | 2.091601995 | -0.073575366 | 0.478666667 | 0.75692893 | 0.85578033 | 0.8791 | 0.929734007 |
| SL1344_RS13980_ | -5.125045111 | -4.842750687 | -5.265208396 | -0.444465615 | 1.849755971 | -0.198606502 | 0.413724187 | 0.71300817 | 0.821028307 | 0.7914 | 0.87125581 |
| SL1344_RS13985_ | -8.352724746 | -8.616266228 | -8.033573212 | 0.713891137 | 4.343310258 | 0.144511339 | 0.421333336 | 0.583404543 | 0.722699211 | 0.7137 | 0.808685725 |
| SL1344_RS14000_ | -3.950654791 | -3.582185141 | -4.127508149 | -0.557037445 | 1.002903755 | -0.491553882 | 0.297801475 | 0.49171526 | 0.64429836 | 0.6014 | 0.722687712 |
| SL1344_RS14005_ | -3.390846077 | -3.555560846 | -3.091083956 | 0.490985209 | 0.945863938 | 0.468368247 | 0.309333342 | 0.500480569 | 0.656781661 | 0.6251 | 0.743561834 |
| SL1344_RS14010_ | -3.232257424 | -3.41639448 | -2.92812013 | 0.798029549 | 2.086236717 | 0.383096879 | 0.306666675 | 0.55725293 | 0.717263554 | 0.6243 | 0.759160994 |
| SL1344_RS14015_ | -4.112389546 | -4.297537542 | -3.956266314 | 0.425046439 | 1.196987038 | 0.332537026 | 0.34533334 | 0.569575164 | 0.714184534 | 0.6915 | 0.793745595 |
| SL1344_RS14020_ | -4.335104632 | -4.640976278 | -4.165406918 | 0.229057824 | 1.385681978 | 0.16280387 | 0.437708197 | 0.704742159 | 0.818472321 | 0.7928 | 0.873055783 |
| SL1344_RS14025_ | 1.298502459 | 1.175939261 | 1.447862496 | 0.269833715 | 0.117765736 | 2.191817809 | 0.007329074 | 0.040540294 | 0.137941681 | 0.1108 | 0.257513869 |
| SL1344_RS14030_ | 0.661924768 | 0.587471654 | 0.761084202 | 0.185407499 | 0.178359297 | 0.993764428 | 0.142666694 | 0.217829966 | 0.387988236 | 0.2997 | 0.456243923 |
| SL1344_RS14035_ | 0.764395899 | 0.709036868 | 0.837517942 | 0.120591083 | 0.174952711 | 0.65094948 | 0.241172565 | 0.388289011 | 0.552496608 | 0.492 | 0.63250401 |
| SL1344_RS14040_ | 2.998629632 | 3.027505307 | 2.962009191 | -0.067114384 | 0.076411563 | -0.823543869 | 0.183333353 | 0.285497804 | 0.4500906 | 0.3767 | 0.527296587 |
| SL1344_RS14045_ | 1.727820986 | 1.751827341 | 1.699422478 | -0.039644049 | 0.137325319 | -0.271219901 | 0.381745508 | 0.590101717 | 0.727267534 | 0.6991 | 0.799596311 |
| SL1344_RS14050_ | -0.887270302 | -0.721684992 | -1.042551598 | -0.339760347 | 0.380797281 | -0.833403075 | 0.173217876 | 0.265975144 | 0.445519207 | 0.358 | 0.519312678 |
| SL1344_RS14055_ | -1.010677311 | -0.968179154 | -1.052891396 | -0.092665862 | 0.354471202 | -0.247696469 | 0.38507662 | 0.577870021 | 0.71742054 | 0.7114 | 0.806549097 |
| SL1344_RS14060_ | -1.4785383 | -1.589598305 | -1.316828714 | 0.325022071 | 0.437551815 | 0.714907521 | 0.213333349 | 0.311583766 | 0.492439926 | 0.4265 | 0.572553067 |
| SL1344_RS14065_ | -2.719282306 | -2.835786381 | -2.580685256 | 0.280058136 | 0.542706086 | 0.483744181 | 0.29913392 | 0.450204803 | 0.603972173 | 0.5704 | 0.689342951 |
| SL1344_RS14070_ | -5.69438548 | -6.630101004 | -4.993064022 | 1.652381727 | 1.591387901 | 0.953246943 | 0.132578311 | 0.210826841 | 0.376764 | 0.2838 | 0.442607273 |
| SL1344_RS14075_ | -4.916699771 | -5.113398874 | -4.780028528 | 0.287117568 | 1.402791178 | 0.187297409 | 0.419053967 | 0.618496765 | 0.751720422 | 0.7333 | 0.826576081 |
| SL1344_RS14080_ | -5.312694173 | -5.377271081 | -5.217384871 | 0.178282757 | 1.857780729 | 0.084019384 | 0.459027317 | 0.665213158 | 0.787524076 | 0.786 | 0.861947563 |
| SL1344_RS14085_ | -0.661352764 | -0.52109664 | -0.894349605 | -0.40211533 | 0.340526922 | -1.130878692 | 0.084610309 | 0.17265911 | 0.344340433 | 0.2214 | 0.377494085 |
| SL1344_RS14090_ | -4.75612261 | -3.539155527 | -5.155529791 | -1.420030577 | 1.319450924 | -0.904058848 | 0.188000019 | 0.285923929 | 0.458890282 | 0.3926 | 0.554608236 |
| SL1344_RS14095_ | -4.281251357 | -3.775743233 | -4.489105408 | -0.667953582 | 1.3283626 | -0.435993164 | 0.329780154 | 0.592923136 | 0.728149114 | 0.6701 | 0.784318913 |
| SL1344_RS14100_ | 0.019743586 | 0.013625847 | 0.026826157 | -0.027293608 | 0.385591241 | -0.07669208 | 0.468354432 | 0.764942331 | 0.860089399 | 0.8681 | 0.919530735 |
| SL1344_RS14105_ | 0.807566113 | 0.819885714 | 0.785594681 | -0.026263929 | 0.158019999 | -0.137695427 | 0.437041974 | 0.604856588 | 0.737291319 | 0.7001 | 0.800200874 |
| SL1344_RS14110_ | 0.653072227 | 0.618242746 | 0.684988773 | 0.037659918 | 0.209589676 | 0.192290736 | 0.423717525 | 0.647660593 | 0.773706014 | 0.7617 | 0.845480861 |
| SL1344_RS14115_ | -0.716859648 | -0.545131124 | -0.861389764 | -0.28347108 | 0.388479915 | -0.709062665 | 0.226000015 | 0.346166119 | 0.525367313 | 0.4408 | 0.599447996 |
| SL1344_RS14120_ | 1.397958908 | 1.430757063 | 1.362998756 | -0.080603691 | 0.182440164 | -0.407338312 | 0.302666675 | 0.489913866 | 0.648493404 | 0.6093 | 0.731138481 |
| SL1344_RS14125_ | -1.818037565 | -1.834075466 | -1.804588425 | 0.046339472 | 0.410388701 | 0.111944663 | 0.451333335 | 0.615861251 | 0.745468225 | 0.728 | 0.818760007 |
| SL1344_RS14130_ | -0.670952175 | -0.714489773 | -0.633131613 | 0.084130739 | 0.267815739 | 0.275329799 | 0.372000005 | 0.53604305 | 0.678017599 | 0.6485 | 0.75694178 |
| SL1344_RS14135_ | 0.161640472 | 0.16031088 | 0.162602944 | 0.014460343 | 0.217393921 | 0.072182152 | 0.467333335 | 0.636879924 | 0.762872453 | 0.7505 | 0.836279167 |
| SL1344_RS14140_ | -4.439145155 | -4.289186484 | -4.55916946 | -0.287543821 | 0.969027513 | -0.285789077 | 0.371333338 | 0.514867543 | 0.658816651 | 0.6239 | 0.735872156 |
| SL1344_RS14150_ | -3.214724266 | -3.415900556 | -2.66044512 | 0.582638786 | 1.071661325 | 0.423693656 | 0.326000007 | 0.668659923 | 0.789790712 | 0.6821 | 0.80561654 |

| | | | | | | | | | | | |
|---|---|---|---|---|---|---|---|---|---|---|---|
| SL1344_RS14155_ | -3.474643667 | -3.230612687 | -3.852089057 | -0.658775952 | 0.804112791 | -0.733161912 | 0.167221874 | 0.276726463 | 0.458197745 | 0.3589 | 0.505059254 |
| SL1344_RS14165_ | -3.736740307 | -3.860848237 | -3.605971612 | 0.252931663 | 0.642324615 | 0.358790919 | 0.334666674 | 0.462777242 | 0.609105892 | 0.5692 | 0.687779033 |
| SL1344_RS14170_ | -3.03444828 | -3.006982077 | -3.058272367 | -0.08402639 | 0.624624444 | -0.136365418 | 0.440373087 | 0.616303935 | 0.746365798 | 0.7245 | 0.816767795 |
| SL1344_RS14175_ | 0.478017237 | 0.392834032 | 0.53806729 | 0.137943556 | 0.313520234 | 0.435751911 | 0.308461034 | 0.513239143 | 0.676067089 | 0.602 | 0.736981064 |
| SL1344_RS14180_ | -3.691140691 | -3.632167846 | -3.763395823 | -0.127373378 | 0.774442705 | -0.151393107 | 0.426666669 | 0.585308488 | 0.72046165 | 0.6905 | 0.791068561 |
| SL1344_RS14190_ | -4.574009245 | -4.507577204 | -4.748086841 | -0.3546595 | 2.008137084 | -0.111593892 | 0.408394407 | 0.51224634 | 0.684959252 | 0.7525 | 0.843426963 |
| SL1344_RS14195_ | 1.928165314 | 2.055264299 | 1.774637267 | -0.282312016 | 0.109145357 | -2.492925671 | 0.0020023 | 0.027434073 | 0.112394105 | 0.1026 | 0.246600807 |
| SL1344_RS14200_ | 0.784404289 | 0.685773268 | 0.917111423 | 0.24756648 | 0.188011626 | 1.294560824 | 0.059333406 | 0.136507833 | 0.295419022 | 0.1773 | 0.332264639 |
| SL1344_RS14205_ | -4.351457891 | -4.199098818 | -4.583643343 | -0.47302346 | 1.248129875 | -0.33069726 | 0.333777489 | 0.494982471 | 0.655007478 | 0.6437 | 0.75119513 |
| SL1344_RS14210_ | -2.753798643 | -2.940648639 | -2.532668428 | 0.405534463 | 0.485791902 | 0.803659525 | 0.189333352 | 0.295827008 | 0.457533153 | 0.3848 | 0.530682609 |
| SL1344_RS14215_ | 2.19467141 | 2.13893524 | 2.357277489 | 0.222386673 | 0.186357082 | 1.137546373 | 0.066000065 | 0.180564377 | 0.368438572 | 0.1885 | 0.346799196 |
| SL1344_RS14220_ | 0.998065657 | 0.918044167 | 1.108640339 | 0.199881022 | 0.155502578 | 1.217924541 | 0.076000056 | 0.146098723 | 0.300050973 | 0.2092 | 0.361813942 |
| SL1344_RS14225_ | 1.491364938 | 1.51103969 | 1.434114088 | -0.068970857 | 0.17031766 | -0.345851739 | 0.373084615 | 0.63773205 | 0.767998384 | 0.7417 | 0.839047704 |
| SL1344_RS14230_ | -4.037260686 | -3.823083967 | -4.471492841 | -0.655584555 | 0.93714935 | -0.644110059 | 0.232511673 | 0.360973563 | 0.530190126 | 0.4748 | 0.615778826 |
| SL1344_RS14235_ | -5.044776986 | -4.447465535 | -5.708789027 | -1.222415015 | 1.330201951 | -0.850116007 | 0.166666689 | 0.258127939 | 0.42490686 | 0.3353 | 0.491196129 |
| SL1344_RS14240_ | -6.128561263 | -6.871769156 | -5.565544217 | 1.335211103 | 2.041838127 | 0.562294118 | 0.208000016 | 0.338091917 | 0.516378009 | 0.4273 | 0.570458942 |
| SL1344_RS14245_ | -4.347818995 | -3.902649034 | -4.77955986 | -1.145513047 | 1.402140363 | -0.808130297 | 0.180666687 | 0.266590603 | 0.448913349 | 0.3779 | 0.532073787 |
| SL1344_RS14250_ | -3.324001165 | -3.586765781 | -3.152822553 | 0.534031672 | 1.103403034 | 0.463563524 | 0.303333342 | 0.465533051 | 0.635282124 | 0.5938 | 0.725887565 |
| SL1344_RS14255_ | 1.097416685 | 0.903516988 | 1.314810702 | 0.411884173 | 0.129100917 | 3.178679631 | 4.68E-05 | 0.015036947 | 0.079006415 | 0.1002 | 0.24329913 |
| SL1344_RS14260_ | -3.216957178 | -3.257833578 | -3.191105137 | 0.070805738 | 0.729788389 | 0.094458904 | 0.458000002 | 0.632328919 | 0.760450817 | 0.7513 | 0.836739108 |
| SL1344_RS14265_ | -4.352291767 | -4.404579714 | -4.299703393 | 0.078596861 | 0.987922962 | 0.074547044 | 0.468666668 | 0.596803964 | 0.730685736 | 0.7029 | 0.801588891 |
| SL1344_RS14270_ | -3.704249806 | -3.34251309 | -3.991046843 | -0.616941038 | 0.836815005 | -0.716055634 | 0.217333349 | 0.34077348 | 0.506968167 | 0.4392 | 0.585182435 |
| SL1344_RS14275_ | -3.019159687 | -3.334999781 | -2.899203899 | 0.360896873 | 0.759998599 | 0.395896125 | 0.364423723 | 0.591482715 | 0.734374699 | 0.7151 | 0.8223932 |
| SL1344_RS14280_ | -2.714642644 | -2.895144426 | -2.599219806 | 0.19972512 | 0.803259955 | 0.228204688 | 0.399333337 | 0.63830207 | 0.768178226 | 0.7521 | 0.84054184 |
| SL1344_RS14285_ | -5.386316724 | -5.042626319 | -5.977866707 | -0.930341577 | 1.479694719 | -0.585898188 | 0.242000013 | 0.372820822 | 0.543972813 | 0.4916 | 0.630125682 |
| SL1344_RS14290_ | -3.15822657 | -2.690820426 | -3.466270796 | -0.852349571 | 0.826629445 | -1.008655558 | 0.14333336 | 0.217007571 | 0.396333371 | 0.3011 | 0.462576578 |
| SL1344_RS14295_ | -0.360506801 | -0.512601786 | -0.219807346 | 0.29743738 | 0.233612738 | 1.198693045 | 0.096602308 | 0.148583876 | 0.302170433 | 0.2249 | 0.383356116 |
| SL1344_RS14300_ | -0.09624881 | -0.192089443 | -0.017320842 | 0.182450467 | 0.225167919 | 0.756828715 | 0.184666686 | 0.284265331 | 0.452996003 | 0.3786 | 0.527586995 |
| SL1344_RS14305_ | -0.052194437 | -0.147800669 | 0.098214245 | 0.22468707 | 0.254684579 | 0.838768837 | 0.196000018 | 0.296791499 | 0.471474393 | 0.4011 | 0.560734365 |
| SL1344_RS14310_ | -4.167549896 | -4.471592794 | -4.03905708 | 0.370513002 | 1.345048402 | 0.252625382 | 0.412391742 | 0.677313092 | 0.799998754 | 0.7942 | 0.877681294 |
| SL1344_RS14315_ | -3.486347821 | -3.280124738 | -3.770452582 | -0.50051283 | 0.78724267 | -0.597699599 | 0.24266668 | 0.372279966 | 0.543392468 | 0.4885 | 0.626654837 |
| SL1344_RS14320_ | 1.147178577 | 1.183513739 | 1.115065525 | -0.006417454 | 0.305419069 | -0.023855902 | 0.489007329 | 0.786447844 | 0.87473014 | 0.881 | 0.929084104 |
| SL1344_RS14325_ | 3.178445976 | 3.203761136 | 3.163043705 | -0.023093472 | 0.122996565 | -0.147609657 | 0.432378417 | 0.783789227 | 0.872618571 | 0.8303 | 0.899812327 |
| SL1344_RS14330_ | -2.090428277 | -2.111097149 | -2.054482555 | 0.072474099 | 0.449052028 | 0.178266969 | 0.423717525 | 0.591153982 | 0.725217698 | 0.7092 | 0.804179113 |
| SL1344_RS14335_ | -2.499575041 | -2.244210933 | -2.723529519 | -0.4596642 | 0.517794365 | -0.846679202 | 0.174550321 | 0.278596298 | 0.446778534 | 0.3731 | 0.526566654 |
| SL1344_RS14345_ | 8.730707084 | 5.2300533 | 8.746709675 | 3.51870913 | 0.468627303 | 0.900617294 | 0.33333334 | 0.235159373 | 0.439697595 | 0.7 | 0.829598552 |
| SL1344_RS14350_ | -2.404339286 | -2.578292874 | -2.240248899 | 0.362004194 | 0.546188407 | 0.652007798 | 0.223333348 | 0.353791871 | 0.521667639 | 0.4643 | 0.601698721 |
| SL1344_RS14360_ | 0.387939469 | 1.133152856 | -0.09345076 | -1.202023667 | 0.292267724 | -4.077321986 | 4.68E-05 | 0.006276635 | 0.05243307 | 0.1 | 0.243021319 |
| SL1344_RS14365_ | -1.67728103 | -1.07440458 | -2.115699921 | -1.056316458 | 0.455361151 | -2.287901963 | 0.005996769 | 0.041677549 | 0.14706213 | 0.1045 | 0.249015851 |
| SL1344_RS14370_ | -1.317661438 | -0.778412201 | -1.793229813 | -0.999772986 | 0.353066851 | -2.736815669 | 0.0020023 | 0.02209367 | 0.099720646 | 0.1019 | 0.245617891 |
| SL1344_RS14375_ | 0.031713327 | -0.016207711 | 0.06316568 | 0.067790068 | 0.220201171 | 0.282217052 | 0.380000005 | 0.565461956 | 0.704558199 | 0.6733 | 0.778588974 |
| SL1344_RS14380_ | -2.268624969 | -2.379415843 | -2.164432129 | 0.189978866 | 0.522357794 | 0.338079618 | 0.354666673 | 0.53110819 | 0.677684587 | 0.6413 | 0.752337717 |
| SL1344_RS14385_ | -0.588127942 | -0.598452526 | -0.578318935 | 0.039274714 | 0.411807227 | 0.088500288 | 0.462358429 | 0.71241487 | 0.823669973 | 0.8335 | 0.896041648 |
| SL1344_RS14390_ | 3.434211995 | 3.156750741 | 3.667108622 | 0.542618244 | 0.167482064 | 3.429343595 | 4.68E-05 | 0.010683507 | 0.071352666 | 0.1 | 0.243021319 |
| SL1344_RS14395_ | 3.582727238 | 3.493583549 | 3.791122629 | 0.295284357 | 0.141658509 | 2.117731117 | 4.68E-05 | 0.071264303 | 0.212304096 | 0.1001 | 0.24316291 |
| SL1344_RS14400_ | 0.327214349 | -0.05673378 | 0.755642789 | 0.803558515 | 0.191786707 | 4.168797949 | 4.68E-05 | 0.006622982 | 0.051331222 | 0.1 | 0.243021319 |
| SL1344_RS14405_ | -0.082953213 | -0.042494506 | -0.111037522 | -0.079288358 | 0.236337915 | -0.303150037 | 0.366000005 | 0.536339976 | 0.683048662 | 0.6545 | 0.761621254 |
| SL1344_RS14410_ | -0.868478753 | -0.872127935 | -0.86128216 | 0.005134066 | 0.287081063 | 0.0186309 | 0.490339774 | 0.605826873 | 0.737012183 | 0.7134 | 0.807595324 |
| SL1344_RS14415_ | -2.055097758 | -1.892892476 | -2.187290393 | -0.305463672 | 0.431136573 | -0.693644243 | 0.222518336 | 0.348271293 | 0.511364573 | 0.4504 | 0.588396903 |
| SL1344_RS14420_ | -4.011602789 | -4.02449127 | -4.000140795 | -0.078453218 | 1.055444765 | -0.079147027 | 0.469686877 | 0.66412948 | 0.784866987 | 0.7793 | 0.857136665 |
| SL1344_RS14425_ | -2.107793335 | -2.003799134 | -2.210782358 | -0.196788512 | 0.485831274 | -0.375880686 | 0.334666674 | 0.525234565 | 0.670634492 | 0.6147 | 0.732533121 |
| SL1344_RS14430_ | -4.39786965 | -4.38020661 | -4.415714203 | -0.070893587 | 0.893495922 | -0.083151633 | 0.467333335 | 0.559097686 | 0.697173495 | 0.6718 | 0.77246571 |
| SL1344_RS14435_ | -0.881910665 | -0.760772283 | -0.972081838 | -0.180365473 | 0.380677315 | -0.450711076 | 0.312458369 | 0.519189731 | 0.672119944 | 0.6143 | 0.739647076 |

| | | | | | | | | | | | |
|---|---|---|---|---|---|---|---|---|---|---|---|
| SL1344_RS14440_ | -3.612390566 | -3.467731753 | -3.693968334 | -0.191470067 | 0.864655251 | -0.201756668 | 0.410393075 | 0.650749759 | 0.771492855 | 0.7467 | 0.833697671 |
| SL1344_RS14445_ | -3.312799593 | -3.495028474 | -3.054063915 | 0.349329437 | 0.866756064 | 0.385012186 | 0.331112599 | 0.553648114 | 0.697562 | 0.6216 | 0.745755179 |
| SL1344_RS14450_ | -3.059821919 | -3.330774553 | -2.907093258 | 0.400377749 | 0.907781528 | 0.427587895 | 0.325116596 | 0.526624112 | 0.684476801 | 0.6411 | 0.763228118 |
| SL1344_RS14455_ | -2.520583843 | -2.311272525 | -2.759274605 | -0.442734094 | 0.475180337 | -0.876591812 | 0.152666692 | 0.243088737 | 0.406006426 | 0.3323 | 0.483777168 |
| SL1344_RS14460_ | -1.810904461 | -2.114954997 | -1.518171413 | 0.609123382 | 0.38717897 | 1.539933117 | 0.048666757 | 0.090688939 | 0.223571737 | 0.1561 | 0.309996748 |
| SL1344_RS14465_ | -1.835373507 | -2.005159425 | -1.702144484 | 0.306988658 | 0.376658179 | 0.765908997 | 0.191205881 | 0.294317789 | 0.45903613 | 0.4052 | 0.550515441 |
| SL1344_RS14470_ | -1.945946783 | -2.073079853 | -1.755486184 | 0.345921194 | 0.463439256 | 0.707968114 | 0.217854779 | 0.33269132 | 0.502463908 | 0.4445 | 0.587147473 |
| SL1344_RS14475_ | -3.251812782 | -3.051080824 | -3.558952394 | -0.53162804 | 0.684325331 | -0.719573739 | 0.192000018 | 0.30535069 | 0.47672781 | 0.3962 | 0.540224361 |
| SL1344_RS14480_ | -2.016161651 | -2.108831744 | -1.931120877 | 0.183026503 | 0.374800612 | 0.473266974 | 0.306000008 | 0.44772308 | 0.599538509 | 0.5434 | 0.669306587 |
| SL1344_RS14485_ | -0.66616791 | -0.596093708 | -0.72686477 | -0.138183022 | 0.235409318 | -0.515581246 | 0.278666677 | 0.421889985 | 0.577174452 | 0.5237 | 0.651753894 |
| SL1344_RS14490_ | 0.056642871 | 0.258609541 | -0.171966595 | -0.444377829 | 0.244208217 | -1.807348849 | 0.016655836 | 0.05703888 | 0.174283908 | 0.1179 | 0.265991405 |
| SL1344_RS14495_ | 4.77999127 | 4.703988966 | 4.842404372 | 0.129937599 | 0.108927673 | 1.179945101 | 0.104000039 | 0.133406029 | 0.306892865 | 0.1994 | 0.376745182 |
| SL1344_RS14500_ | 0.017734283 | -0.067285281 | 0.352889085 | 0.442418634 | 0.530523555 | 0.831157613 | 0.210000016 | 0.274443736 | 0.472472997 | 0.386 | 0.576555278 |
| SL1344_RS14505_ | -1.320198805 | -1.57569408 | -1.012230153 | 0.560403299 | 0.325798151 | 1.678665843 | 0.03264504 | 0.073348377 | 0.197572088 | 0.1356 | 0.286766972 |
| SL1344_RS14510_ | -1.323382728 | -0.96594508 | -1.79490202 | -0.880548863 | 0.440788516 | -1.991084273 | 0.006662914 | 0.050607796 | 0.163149121 | 0.1062 | 0.251101309 |
| SL1344_RS14515_ | 0.381898617 | 0.455872455 | 0.317991837 | -0.134513228 | 0.190655151 | -0.681105367 | 0.230666681 | 0.357445926 | 0.521954667 | 0.4654 | 0.6060786 |
| SL1344_RS14520_ | -2.332572991 | -2.151286319 | -2.50798247 | -0.353259012 | 0.446565097 | -0.719208698 | 0.216000016 | 0.325599448 | 0.490632353 | 0.4282 | 0.573874031 |
| SL1344_RS14525_ | -4.371556042 | -4.066001379 | -4.89555764 | -0.79636772 | 0.980667475 | -0.744552062 | 0.188000019 | 0.300887191 | 0.469881121 | 0.401 | 0.549263183 |
| SL1344_RS14530_ | -1.128641386 | -1.364645558 | -0.96833121 | 0.439283579 | 0.441230036 | 0.949978081 | 0.127248532 | 0.217391109 | 0.402037767 | 0.2712 | 0.432717377 |
| SL1344_RS14540_ | 0.707438893 | 0.457228011 | 0.910441232 | 0.448293185 | 0.164215721 | 2.659544993 | 0.003334715 | 0.023353924 | 0.10173555 | 0.1014 | 0.244989191 |
| SL1344_RS14545_ | 0.652828533 | 0.411012485 | 0.836065185 | 0.432388451 | 0.203523928 | 2.070786194 | 0.005334196 | 0.052683155 | 0.167940067 | 0.1061 | 0.251151328 |
| SL1344_RS14550_ | 1.579177747 | 1.340847729 | 1.902566269 | 0.571270309 | 0.170265366 | 3.34409885 | 4.68E-05 | 0.01909011 | 0.094767705 | 0.1 | 0.243021319 |
| SL1344_RS14555_ | 0.167446188 | -0.046333954 | 0.383671364 | 0.433860906 | 0.198476576 | 2.162426536 | 0.006662914 | 0.038545291 | 0.136782509 | 0.1065 | 0.251750865 |
| SL1344_RS14560_ | 0.785950691 | 0.717805671 | 0.839823204 | 0.124623809 | 0.147439722 | 0.792074213 | 0.189333352 | 0.282571132 | 0.444471527 | 0.3837 | 0.528405404 |
| SL1344_RS14565_ | -0.244931223 | -0.461183622 | -0.001916895 | 0.474946132 | 0.238873779 | 1.983911395 | 0.011326186 | 0.052247921 | 0.162698746 | 0.1122 | 0.257574781 |
| SL1344_RS14570_ | 2.884018531 | 2.770770566 | 2.987994722 | 0.087350166 | 0.229785927 | 0.522185278 | 0.271818798 | 0.467760174 | 0.648836154 | 0.5672 | 0.704623668 |
| SL1344_RS14575_ | 2.426088155 | 2.363817782 | 2.495012213 | 0.143215563 | 0.127687832 | 1.08679404 | 0.091333379 | 0.167988708 | 0.34104889 | 0.2259 | 0.388657234 |
| SL1344_RS14585_ | -1.678518258 | -1.790240677 | -1.564051191 | 0.283859879 | 0.554750889 | 0.477748469 | 0.284666676 | 0.421408113 | 0.596347111 | 0.5533 | 0.686252407 |
| SL1344_RS14600_ | 2.247337814 | 2.230465884 | 2.287661385 | 0.050502373 | 0.126993621 | 0.354509714 | 0.368421058 | 0.630434058 | 0.761348284 | 0.7187 | 0.821702006 |
| SL1344_RS14605_ | 2.415394703 | 2.429036978 | 2.405249124 | -0.019594458 | 0.088793127 | -0.218101175 | 0.402398405 | 0.554973574 | 0.695378637 | 0.6673 | 0.771575069 |
| SL1344_RS14610_ | 2.909665698 | 2.818628667 | 3.120081143 | 0.291661253 | 0.113705059 | 2.309846113 | 0.001336769 | 0.043432373 | 0.153875475 | 0.1016 | 0.245263478 |
| SL1344_RS14615_ | 4.618685003 | 4.417840008 | 4.745438851 | 0.330512101 | 0.11911946 | 2.83258371 | 4.68E-05 | 0.010993259 | 0.073069623 | 0.1 | 0.243021319 |
| SL1344_RS14620_ | 5.185871602 | 5.038609276 | 5.283745318 | 0.245813127 | 0.147880934 | 1.792224125 | 0.00067292 | 0.062574077 | 0.197261642 | 0.1006 | 0.243869209 |
| SL1344_RS14625_ | 5.620093539 | 5.512394687 | 5.633870873 | 0.121687112 | 0.203803572 | 0.640296451 | 0.224517003 | 0.291126535 | 0.496765326 | 0.3998 | 0.598285418 |
| SL1344_RS14630_ | 7.291746357 | 7.135819841 | 7.417542942 | 0.279520272 | 0.129481556 | 1.912525781 | 4.68E-05 | 0.063145472 | 0.19943728 | 0.1 | 0.243021319 |
| SL1344_RS14640_ | 4.71978715 | 4.715227132 | 4.729129198 | 0.016347804 | 0.183201031 | 0.080744976 | 0.449333335 | 0.858984535 | 0.922322829 | 0.8908 | 0.937993581 |
| SL1344_RS14645_ | 0.029673955 | -0.052529731 | 0.116111616 | 0.124852733 | 0.301225426 | 0.4103058 | 0.320000008 | 0.542303764 | 0.690841051 | 0.6442 | 0.760676638 |
| SL1344_RS14655_ | 1.050543409 | 1.06121175 | 1.036182898 | -0.029317847 | 0.157679052 | -0.179176673 | 0.426000003 | 0.59237919 | 0.727883424 | 0.7036 | 0.799713953 |
| SL1344_RS14660_ | 1.660385143 | 1.613278473 | 1.787538653 | 0.163167595 | 0.157437976 | 0.929891193 | 0.170552986 | 0.24786871 | 0.432190137 | 0.3584 | 0.524975719 |
| SL1344_RS14675_ | -4.12687416 | -4.428106799 | -3.787180015 | 0.708087202 | 1.056265991 | 0.62877746 | 0.230666681 | 0.348844511 | 0.522772222 | 0.4718 | 0.612794353 |
| SL1344_RS14680_ | -1.478872916 | -1.516143929 | -1.421081326 | 0.109148547 | 0.340237263 | 0.282734479 | 0.375333338 | 0.539515595 | 0.682088042 | 0.6528 | 0.759643741 |
| SL1344_RS14685_ | -1.134484079 | -1.140873575 | -1.129297889 | 0.015035266 | 0.290356878 | 0.04701752 | 0.480346436 | 0.549088008 | 0.68825125 | 0.6501 | 0.757605602 |
| SL1344_RS14690_ | -1.374799575 | -1.535636293 | -1.236913815 | 0.306319323 | 0.416988297 | 0.69665419 | 0.218666682 | 0.341315439 | 0.515198531 | 0.4479 | 0.599558581 |
| SL1344_RS14700_ | 0.753925225 | 0.713353567 | 0.783130708 | 0.05471306 | 0.212100471 | 0.249216975 | 0.388666671 | 0.634244828 | 0.761773163 | 0.7264 | 0.821511907 |
| SL1344_RS14705_ | 0.192618679 | 0.299421915 | 0.070379404 | -0.235782735 | 0.179388117 | -1.222371958 | 0.08400005 | 0.142223313 | 0.288722798 | 0.2154 | 0.369436659 |
| SL1344_RS14710_ | -2.626005479 | -2.803955654 | -2.475047316 | 0.338262865 | 0.503763911 | 0.619653688 | 0.236000014 | 0.379273282 | 0.538159286 | 0.4788 | 0.612767174 |
| SL1344_RS14715_ | -2.337877776 | -2.157906731 | -2.513222171 | -0.341676165 | 0.441425863 | -0.747384725 | 0.202000017 | 0.309617435 | 0.469998019 | 0.4042 | 0.550282732 |
| SL1344_RS14720_ | 1.678816961 | 1.617936305 | 1.744681115 | 0.128328229 | 0.151535562 | 0.796596004 | 0.201199218 | 0.298042638 | 0.473479025 | 0.3993 | 0.55643218 |
| SL1344_RS14725_ | -3.04292616 | -2.924243351 | -3.168666062 | -0.235456812 | 0.609624789 | -0.361378434 | 0.336666673 | 0.500578769 | 0.650119448 | 0.6158 | 0.730277909 |
| SL1344_RS14730_ | -1.265679312 | -1.320789213 | -1.200629785 | 0.114845848 | 0.28626717 | 0.373661018 | 0.338441046 | 0.466356883 | 0.615225888 | 0.567 | 0.687763439 |
| SL1344_RS14735_ | -1.992524475 | -1.965215659 | -2.035056719 | -0.083716325 | 0.431854333 | -0.185144483 | 0.420000003 | 0.590005797 | 0.724709999 | 0.7026 | 0.798321647 |
| SL1344_RS14740_ | -2.065417206 | -1.952342015 | -2.208916924 | -0.246425869 | 0.429584703 | -0.535318397 | 0.28047969 | 0.43053281 | 0.587284392 | 0.5345 | 0.663954573 |
| SL1344_RS14745_ | -1.739591752 | -1.656565573 | -1.855998001 | -0.207591225 | 0.360454887 | -0.525432613 | 0.28200001 | 0.416721124 | 0.57347306 | 0.5156 | 0.64463604 |

| | | | | | | | | | | | |
|---|---|---|---|---|---|---|---|---|---|---|---|
| SL1344_RS14750_ | -2.453036948 | -2.369506943 | -2.53651552 | -0.155233739 | 0.480252666 | -0.301021743 | 0.36908728 | 0.521615826 | 0.666965222 | 0.6335 | 0.74332761 |
| SL1344_RS14765_ | -1.679229622 | -1.747265984 | -1.593980275 | 0.176141377 | 0.377159879 | 0.428186911 | 0.315123259 | 0.4720421 | 0.624074165 | 0.591 | 0.707605371 |
| SL1344_RS14770_ | -4.051718903 | -4.378732236 | -3.757428731 | 0.613451099 | 0.739316613 | 0.759438705 | 0.189333352 | 0.289277812 | 0.446505621 | 0.3787 | 0.522306403 |
| SL1344_RS14775_ | -4.538814604 | -5.250063936 | -4.145679251 | 1.103588716 | 1.201790641 | 0.845352832 | 0.161225873 | 0.24487056 | 0.421936879 | 0.3236 | 0.480252504 |
| SL1344_RS14780_ | -0.873793232 | -0.732987388 | -1.168370202 | -0.291032589 | 0.57604076 | -0.445054885 | 0.359093943 | 0.628472207 | 0.76776453 | 0.7588 | 0.855977291 |
| SL1344_RS14785_ | -0.653070778 | -0.679295853 | -0.616543475 | 0.068966973 | 0.276562588 | 0.234823157 | 0.401333337 | 0.569404945 | 0.709128609 | 0.6851 | 0.786489054 |
| SL1344_RS14790_ | -1.715203577 | -1.632845799 | -1.750516975 | -0.098991549 | 0.428201459 | -0.200287665 | 0.426382415 | 0.648564245 | 0.771452277 | 0.7354 | 0.828155484 |
| SL1344_RS14795_ | -0.265469081 | -0.186826082 | -0.315991571 | -0.104347885 | 0.302746229 | -0.327824799 | 0.375333338 | 0.613387519 | 0.747243722 | 0.7276 | 0.822830886 |
| SL1344_RS14800_ | 0.012232864 | 0.014274507 | 0.010061868 | 0.000864099 | 0.206920298 | 0.003683426 | 0.498 | 0.600421121 | 0.731963238 | 0.7117 | 0.805778468 |
| SL1344_RS14805_ | 0.251929899 | 0.15413732 | 0.361923238 | 0.212194682 | 0.174954969 | 1.170098088 | 0.092666712 | 0.166477193 | 0.316181532 | 0.2408 | 0.393665555 |
| SL1344_RS14810_ | -1.897216394 | -1.923414763 | -1.877977875 | 0.062158373 | 0.468344626 | 0.117353762 | 0.456666668 | 0.640705574 | 0.766578164 | 0.7621 | 0.844616457 |
| SL1344_RS14815_ | 0.468985864 | 0.345447231 | 0.646041861 | 0.204884622 | 0.410430899 | 0.515280609 | 0.294666676 | 0.520071065 | 0.686221702 | 0.5968 | 0.737490298 |
| SL1344_RS14820_ | -0.523371102 | -0.513620095 | -0.538964096 | -0.028088505 | 0.242432058 | -0.114872889 | 0.445036644 | 0.58301495 | 0.719648978 | 0.6939 | 0.793559311 |
| SL1344_RS14825_ | -1.352427207 | -1.330704371 | -1.379436244 | -0.059319687 | 0.376866676 | -0.140794134 | 0.436000002 | 0.637055144 | 0.762300593 | 0.7389 | 0.828099769 |
| SL1344_RS14830_ | -0.040366882 | -0.040122812 | -0.041173607 | 0.012333664 | 0.264468436 | 0.046797072 | 0.483333334 | 0.688563986 | 0.803467019 | 0.7915 | 0.86857883 |
| SL1344_RS14850_ | 2.614047776 | 2.6281581 | 2.59907421 | 0.017071828 | 0.154185202 | 0.102909203 | 0.463024652 | 0.738641278 | 0.841788737 | 0.8861 | 0.929862872 |
| SL1344_RS14855_ | 2.650602936 | 2.696873419 | 2.592938568 | -0.105077119 | 0.077681229 | -1.307356865 | 0.079333387 | 0.130431894 | 0.277445207 | 0.2036 | 0.359364971 |
| SL1344_RS14900_ | 2.692436375 | 2.687995823 | 2.699962689 | 0.003839561 | 0.10976651 | 0.032947675 | 0.485676216 | 0.700956489 | 0.813177024 | 0.7876 | 0.867283349 |
| SL1344_RS14905_ | 0.922952434 | 0.888068459 | 0.954396388 | 0.069979554 | 0.123185753 | 0.537950189 | 0.279147245 | 0.389786699 | 0.54062005 | 0.489 | 0.619272664 |
| SL1344_RS14910_ | 1.231830746 | 1.208843177 | 1.246829815 | 0.054625217 | 0.174747759 | 0.308722518 | 0.366666672 | 0.542159687 | 0.692104011 | 0.6785 | 0.779626662 |
| SL1344_RS15010_ | 0.635471403 | 0.979444536 | 0.225047073 | -0.753773422 | 0.318716298 | -2.325711578 | 4.68E-05 | 0.028486867 | 0.120099817 | 0.101 | 0.244419225 |
| SL1344_RS15015_ | 1.519282942 | 1.470471427 | 1.582062989 | 0.111534024 | 0.147014025 | 0.705689486 | 0.218521001 | 0.333373999 | 0.506665874 | 0.4314 | 0.584892638 |
| SL1344_RS15020_ | -0.117614116 | -0.224987417 | 0.038539894 | 0.254932424 | 0.25277472 | 0.96064557 | 0.148000026 | 0.226337773 | 0.399212185 | 0.3191 | 0.48341315 |
| SL1344_RS15025_ | -1.602152145 | -1.735655318 | -1.42577897 | 0.25160957 | 0.543246984 | 0.439677898 | 0.317121927 | 0.517263218 | 0.673288271 | 0.6259 | 0.748005705 |
| SL1344_RS15030_ | 0.968703636 | 0.988506917 | 0.955705002 | -0.039168289 | 0.161038987 | -0.226888315 | 0.399067292 | 0.57644268 | 0.713417375 | 0.6913 | 0.789949242 |
| SL1344_RS15035_ | 1.517873619 | 1.600874187 | 1.444452021 | -0.160275716 | 0.117158112 | -1.326251368 | 0.074666724 | 0.132466785 | 0.277295017 | 0.1996 | 0.35341194 |
| SL1344_RS15040_ | -4.637357152 | -4.638677144 | -4.635297749 | 0.061881387 | 1.423320564 | 0.042687776 | 0.481012659 | 0.661147617 | 0.784248811 | 0.7831 | 0.859824803 |
| SL1344_RS15045_ | -0.444074605 | -0.570493782 | -0.335802135 | 0.238006587 | 0.215822039 | 1.047843637 | 0.123251198 | 0.19511501 | 0.348527359 | 0.2788 | 0.431216209 |
| SL1344_RS15050_ | -3.870328725 | -3.638089981 | -4.253528652 | -0.600893801 | 0.75484153 | -0.733202131 | 0.210000016 | 0.318941059 | 0.478336401 | 0.4242 | 0.564242067 |
| SL1344_RS15055_ | 2.91370517 | 2.888627438 | 2.981833994 | 0.097464811 | 0.129116086 | 0.695398526 | 0.249333346 | 0.361565001 | 0.550594786 | 0.5054 | 0.658239216 |
| SL1344_RS15060_ | -2.607477734 | -2.670770608 | -2.540604226 | 0.13376631 | 0.440711656 | 0.275497618 | 0.38241173 | 0.508423421 | 0.651628241 | 0.6199 | 0.731029878 |
| SL1344_RS15065_ | -2.260528522 | -2.218403714 | -2.304764009 | -0.088678664 | 0.436058789 | -0.195151549 | 0.415333337 | 0.558963064 | 0.696666712 | 0.6681 | 0.771278175 |
| SL1344_RS15070_ | -1.4198456 | -1.511478527 | -1.320878575 | 0.189538823 | 0.284542216 | 0.650244052 | 0.237841452 | 0.342029628 | 0.497351264 | 0.4361 | 0.575689926 |
| SL1344_RS15075_ | -0.555110681 | -0.501223512 | -0.616787375 | -0.147326584 | 0.393154582 | -0.34321478 | 0.356666672 | 0.547102955 | 0.699778938 | 0.7013 | 0.802558239 |
| SL1344_RS15080_ | -3.539410247 | -3.773399439 | -3.380956261 | 0.311091414 | 0.967855457 | 0.299904283 | 0.382666671 | 0.588760593 | 0.73107395 | 0.7188 | 0.816053532 |
| SL1344_RS15085_ | -3.299257492 | -3.299257492 | -3.299432946 | 0.005730785 | 0.740206854 | 0.007382491 | 0.498334444 | 0.632463293 | 0.758953873 | 0.7445 | 0.830792067 |
| SL1344_RS15090_ | -3.058893042 | -3.034664547 | -3.087628551 | -0.042385094 | 0.541487695 | -0.074001605 | 0.465023319 | 0.550949618 | 0.689516547 | 0.6611 | 0.764866481 |
| SL1344_RS15095_ | -2.670283025 | -2.466447513 | -2.838119428 | -0.448632712 | 0.717292996 | -0.607267046 | 0.248501012 | 0.361412974 | 0.539904523 | 0.481 | 0.625685546 |
| SL1344_RS15100_ | -3.180089329 | -3.028541968 | -3.332203605 | -0.27057891 | 0.632454424 | -0.389952535 | 0.327115264 | 0.481469213 | 0.632101081 | 0.5933 | 0.712268648 |
| SL1344_RS15105_ | 0.556114162 | 0.373703507 | 0.653888918 | 0.264943355 | 0.204064657 | 1.220628765 | 0.103333373 | 0.151973405 | 0.313936297 | 0.2338 | 0.39711175 |
| SL1344_RS15110_ | 0.08925828 | -0.018102618 | 0.302205597 | 0.303831968 | 0.234297195 | 1.182036492 | 0.113257864 | 0.164413238 | 0.330061115 | 0.2518 | 0.416553017 |
| SL1344_RS15115_ | -4.280119218 | -4.082644302 | -4.507805339 | -0.422864086 | 0.855113639 | -0.455982547 | 0.302666675 | 0.445876789 | 0.598140927 | 0.5579 | 0.680410466 |
| SL1344_RS15120_ | -2.333992039 | -2.337484713 | -2.330651525 | 0.00615732 | 0.43647742 | 0.013758187 | 0.494 | 0.55913282 | 0.698870607 | 0.6647 | 0.770348118 |
| SL1344_RS15125_ | -3.131605713 | -2.784376165 | -3.49636039 | -0.703631116 | 0.567962289 | -1.170266401 | 0.08933338 | 0.156503107 | 0.307593844 | 0.2307 | 0.386172901 |
| SL1344_RS15140_ | 2.386990052 | 2.337397225 | 2.448092176 | 0.117646919 | 0.105127338 | 1.076658225 | 0.098600974 | 0.178068355 | 0.344466259 | 0.2408 | 0.396513337 |
| SL1344_RS15145_ | 2.073201997 | 1.966885258 | 2.205286006 | 0.239315295 | 0.096108413 | 2.415982638 | 0.003332494 | 0.032510246 | 0.120898076 | 0.1049 | 0.249498197 |
| SL1344_RS15150_ | 2.352244747 | 2.26031818 | 2.511189531 | 0.278517405 | 0.201030243 | 1.331104042 | 0.056628991 | 0.108801023 | 0.269612788 | 0.1559 | 0.31697945 |
| SL1344_RS15155_ | 1.289149532 | 1.257811696 | 1.332171623 | 0.107150523 | 0.215913082 | 0.470355733 | 0.311792147 | 0.467733935 | 0.636834904 | 0.6142 | 0.7387787 |
| SL1344_RS15160_ | 1.970621438 | 1.877392824 | 2.134613233 | 0.245670697 | 0.160294743 | 1.444798705 | 0.066000065 | 0.100587145 | 0.251744307 | 0.1669 | 0.329357624 |
| SL1344_RS15165_ | 2.348079877 | 2.262676058 | 2.480903455 | 0.219997514 | 0.124788538 | 1.716415269 | 0.016655836 | 0.076288358 | 0.21184696 | 0.1191 | 0.267142058 |
| SL1344_RS15170_ | 1.303878065 | 0.815341679 | 1.782778641 | 0.953650992 | 0.168225851 | 5.569602928 | 4.68E-05 | 0.002626405 | 0.032562351 | 0.1 | 0.243021319 |
| SL1344_RS15190_ | 0.828265239 | 0.615256167 | 1.10834728 | 0.499555776 | 0.158319972 | 3.172467139 | 4.68E-05 | 0.016325769 | 0.083529824 | 0.1001 | 0.243160901 |
| SL1344_RS15195_ | -2.290282213 | -2.152131602 | -2.444664848 | -0.293040538 | 0.388415863 | -0.704207106 | 0.217333349 | 0.320417878 | 0.477851294 | 0.417 | 0.558288419 |

| | | | | | | | | | | | |
|---|---|---|---|---|---|---|---|---|---|---|---|
| SL1344_RS15200_ | -0.968085642 | -0.816053601 | -1.126218462 | -0.304745618 | 0.278300007 | -1.038341266 | 0.13200003 | 0.20588054 | 0.364507028 | 0.2896 | 0.443983059 |
| SL1344_RS15205_ | -1.804775736 | -1.845648423 | -1.781934006 | 0.112791511 | 0.476456066 | 0.217277679 | 0.405063295 | 0.579554983 | 0.720179765 | 0.715 | 0.807455852 |
| SL1344_RS15210_ | -2.216628881 | -2.172981681 | -2.256937068 | -0.080819315 | 0.468150157 | -0.168661552 | 0.427333336 | 0.583362272 | 0.720863528 | 0.6958 | 0.795327899 |
| SL1344_RS15215_ | -1.593955759 | -1.632868308 | -1.546853607 | 0.085210961 | 0.330804302 | 0.237486536 | 0.392405067 | 0.527441748 | 0.668409516 | 0.6399 | 0.746810465 |
| SL1344_RS15220_ | -3.619486086 | -3.407998057 | -3.764410529 | -0.34677456 | 0.771293072 | -0.416483793 | 0.316666675 | 0.496880177 | 0.6453513 | 0.6017 | 0.720633577 |
| SL1344_RS15225_ | -2.07760404 | -2.089806401 | -2.068382477 | 0.023054669 | 0.534813476 | 0.038040323 | 0.481333334 | 0.664304688 | 0.786289379 | 0.7836 | 0.860762701 |
| SL1344_RS15230_ | -2.541086153 | -2.68064844 | -2.404458434 | 0.263587032 | 0.51227037 | 0.491025932 | 0.291333343 | 0.440999647 | 0.597458035 | 0.5502 | 0.677797847 |
| SL1344_RS15260_ | -1.086267526 | -0.980323919 | -1.230801276 | -0.244724574 | 0.326644423 | -0.700627544 | 0.227181893 | 0.339184105 | 0.507311073 | 0.4419 | 0.589036662 |
| SL1344_RS15265_ | 2.47591215 | 2.418588737 | 2.547663024 | 0.129418446 | 0.128450699 | 0.972880975 | 0.139240534 | 0.208097554 | 0.388605025 | 0.2854 | 0.457873649 |
| SL1344_RS15270_ | 6.453816562 | 6.387708238 | 6.562877256 | 0.174333062 | 0.089763931 | 1.862475446 | 0.00067292 | 0.079997835 | 0.228828743 | 0.1003 | 0.243446024 |
| SL1344_RS15275_ | 5.14942754 | 5.058740046 | 5.251925581 | 0.197455644 | 0.048697109 | 4.083893452 | 4.68E-05 | 0.010227071 | 0.066617774 | 0.1 | 0.243021319 |
| SL1344_RS15280_ | 1.759945102 | 1.739037123 | 1.803554942 | 0.074286339 | 0.145700981 | 0.483011579 | 0.305333342 | 0.44802667 | 0.61577084 | 0.5862 | 0.705011843 |
| SL1344_RS15285_ | 1.831102879 | 1.844932382 | 1.808459941 | -0.049304507 | 0.160382544 | -0.296230317 | 0.357095276 | 0.505251925 | 0.668908472 | 0.6668 | 0.771191105 |
| SL1344_RS15290_ | 2.391876237 | 2.399871043 | 2.38431333 | -0.0289142 | 0.135147091 | -0.20261338 | 0.404397072 | 0.66253502 | 0.784675228 | 0.7704 | 0.852019097 |
| SL1344_RS15295_ | 4.053636682 | 4.035823779 | 4.069137356 | 0.030212376 | 0.062102275 | 0.466091581 | 0.294470362 | 0.476665836 | 0.634228147 | 0.5914 | 0.716814362 |
| SL1344_RS15310_ | -0.854910669 | -0.798011417 | -0.905578286 | -0.054803381 | 0.367925763 | -0.141606941 | 0.434377084 | 0.661521172 | 0.784459134 | 0.7742 | 0.855328006 |
| SL1344_RS15315_ | -0.388389745 | -0.296668801 | -0.514706085 | -0.231975057 | 0.358362556 | -0.612516033 | 0.247168567 | 0.364517028 | 0.544390606 | 0.4944 | 0.640706981 |
| SL1344_RS15320_ | -0.987866361 | -1.024097939 | -0.951006082 | 0.072865172 | 0.338901781 | 0.199797065 | 0.40466667 | 0.594716786 | 0.730208546 | 0.7003 | 0.799317895 |
| SL1344_RS15325_ | -2.632076044 | -2.752111965 | -2.473851862 | 0.321653911 | 0.5539856 | 0.541665168 | 0.27514991 | 0.433494171 | 0.591210157 | 0.5468 | 0.673813238 |
| SL1344_RS15330_ | 1.877239147 | 1.817885506 | 1.98722815 | 0.139968924 | 0.231341299 | 0.584678445 | 0.267821463 | 0.441186408 | 0.622626177 | 0.5327 | 0.689679546 |
| SL1344_RS15355_ | 1.358094617 | 1.255446706 | 1.447136445 | 0.201453347 | 0.145018418 | 1.380524975 | 0.057295212 | 0.118276126 | 0.267049224 | 0.1741 | 0.330619458 |
| SL1344_RS15360_ | 3.334381921 | 3.205859969 | 3.421371423 | 0.240494686 | 0.140727342 | 1.798735492 | 0.006000767 | 0.05711927 | 0.183046989 | 0.1073 | 0.25302155 |
| SL1344_RS15365_ | 3.839056528 | 3.493557944 | 4.178839317 | 0.684798078 | 0.052239427 | 13.04479741 | 4.68E-05 | 0.000364606 | 0.012300044 | 0.1 | 0.243021319 |
| SL1344_RS15370_ | 3.71200539 | 3.310111182 | 4.123645282 | 0.81282908 | 0.084060801 | 9.617973299 | 4.68E-05 | 0.000848723 | 0.019184168 | 0.1 | 0.243021319 |
| SL1344_RS15375_ | 0.917551124 | 0.775075114 | 1.231733337 | 0.406631031 | 0.25458991 | 1.534796987 | 0.059293877 | 0.09502007 | 0.244627162 | 0.1679 | 0.332008258 |
| SL1344_RS15380_ | -2.815154626 | -2.466652211 | -3.130301405 | -0.689193079 | 0.550305638 | -1.243615466 | 0.086666715 | 0.148443757 | 0.299673345 | 0.2227 | 0.375780084 |
| SL1344_RS15385_ | -4.817938005 | -4.633991419 | -4.958715974 | -0.298077363 | 1.010125091 | -0.26935374 | 0.385742843 | 0.531406709 | 0.671831749 | 0.6425 | 0.747735409 |
| SL1344_RS15390_ | -4.175477241 | -4.745414943 | -3.795667818 | 0.946948499 | 0.863152466 | 0.985655747 | 0.124583643 | 0.208190211 | 0.36896116 | 0.2911 | 0.444316433 |
| SL1344_RS15395_ | -2.789642252 | -2.807487024 | -2.758926923 | 0.007471594 | 0.640465604 | 0.010844837 | 0.493333334 | 0.653479855 | 0.777482478 | 0.7562 | 0.843624692 |
| SL1344_RS15400_ | -3.254167543 | -3.429122535 | -3.083158947 | 0.348778972 | 0.554409855 | 0.581885337 | 0.266489018 | 0.390020536 | 0.545638574 | 0.4876 | 0.619526666 |
| SL1344_RS15405_ | -3.050349778 | -2.970894309 | -3.236771744 | -0.216895124 | 0.761685907 | -0.253010632 | 0.405333337 | 0.655405358 | 0.781758103 | 0.7538 | 0.848686557 |
| SL1344_RS15410_ | -0.562448493 | -0.705436691 | -0.411292162 | 0.315453205 | 0.296239042 | 1.030002658 | 0.111925419 | 0.186439775 | 0.351366224 | 0.2696 | 0.425744548 |
| SL1344_RS15415_ | 2.980936752 | 2.923500523 | 3.130643166 | 0.194745684 | 0.100966469 | 1.748553473 | 0.0346437 | 0.08645309 | 0.22926947 | 0.1411 | 0.294707877 |
| SL1344_RS15420_ | 1.011368585 | 0.945913242 | 1.074394458 | 0.126975248 | 0.127600891 | 0.926649384 | 0.157228538 | 0.247798985 | 0.404816101 | 0.3436 | 0.490194545 |
| SL1344_RS15425_ | 1.662746626 | 1.597224505 | 1.776406749 | 0.18398722 | 0.141163151 | 1.261314101 | 0.081333385 | 0.144576014 | 0.305796057 | 0.2003 | 0.358779623 |
| SL1344_RS15430_ | 0.51270641 | 0.647231007 | 0.25264034 | -0.36490839 | 0.286556989 | -1.194126285 | 0.106000038 | 0.148707439 | 0.320644528 | 0.2243 | 0.396836123 |
| SL1344_RS15435_ | 1.47582792 | 1.27238065 | 1.677472316 | 0.41563587 | 0.137606955 | 3.03805158 | 4.68E-05 | 0.018411405 | 0.090521626 | 0.1 | 0.243021319 |
| SL1344_RS15440_ | 0.896252079 | 0.726712511 | 1.107313658 | 0.380548383 | 0.153927139 | 2.406655612 | 0.005996769 | 0.030157388 | 0.117346865 | 0.1046 | 0.249212152 |
| SL1344_RS15445_ | -0.946361475 | -1.278071246 | -0.658319247 | 0.633372882 | 0.356779323 | 1.727144295 | 0.016655836 | 0.069347459 | 0.197549684 | 0.1187 | 0.267039658 |
| SL1344_RS15450_ | 1.893857479 | 1.847144876 | 2.017733828 | 0.17695812 | 0.174290176 | 0.918299343 | 0.169886764 | 0.247189787 | 0.438789504 | 0.3432 | 0.505881725 |
| SL1344_RS15455_ | 2.911285393 | 2.844129905 | 2.988927326 | 0.146439 | 0.071524245 | 1.975517987 | 0.011326186 | 0.050538321 | 0.157194657 | 0.1152 | 0.262059811 |
| SL1344_RS15460_ | 3.926584854 | 3.519301427 | 4.266274168 | 0.663585921 | 0.267719787 | 2.614356105 | 4.68E-05 | 0.012879067 | 0.080865747 | 0.1 | 0.243021319 |
| SL1344_RS15465_ | 5.501973483 | 5.036883894 | 5.949155245 | 0.823177891 | 0.167539183 | 5.150224961 | 4.68E-05 | 0.000928563 | 0.022777491 | 0.1 | 0.243021319 |
| SL1344_RS15475_ | 0.309546757 | 0.276442653 | 0.346205924 | 0.073967211 | 0.181094288 | 0.382622453 | 0.336442379 | 0.488471801 | 0.636578248 | 0.5892 | 0.708186092 |
| SL1344_RS15480_ | 1.303964167 | 1.221907931 | 1.40084509 | 0.20745926 | 0.177068928 | 1.153492718 | 0.088666714 | 0.163978327 | 0.333476732 | 0.2148 | 0.372357293 |
| SL1344_RS15485_ | 2.522785704 | 2.474012287 | 2.575851921 | 0.116276631 | 0.143744561 | 0.832257854 | 0.173884098 | 0.26475405 | 0.456402727 | 0.3446 | 0.507481735 |
| SL1344_RS15490_ | 2.269105809 | 2.117285863 | 2.372854327 | 0.257307145 | 0.101072851 | 2.429555929 | 0.003334715 | 0.033754744 | 0.126489577 | 0.1027 | 0.246620959 |
| SL1344_RS15495_ | 2.322411872 | 2.237968862 | 2.431265872 | 0.211726849 | 0.130198138 | 1.594716154 | 0.015989625 | 0.091170879 | 0.237606508 | 0.1161 | 0.262715999 |
| SL1344_RS15500_ | -1.757300968 | -1.870007919 | -1.625385443 | 0.2357609 | 0.336169489 | 0.649344663 | 0.225849448 | 0.356749425 | 0.510823135 | 0.4533 | 0.588365273 |
| SL1344_RS15505_ | -2.504213486 | -2.459382481 | -2.546461216 | -0.127578662 | 0.574619055 | -0.199525565 | 0.409333337 | 0.593263551 | 0.728604775 | 0.7123 | 0.806249859 |
| SL1344_RS15510_ | -2.323191745 | -2.472348977 | -2.248619009 | 0.18892707 | 0.526593012 | 0.327654668 | 0.365333339 | 0.564900193 | 0.705164757 | 0.6731 | 0.779601773 |
| SL1344_RS15515_ | -3.758821129 | -4.099116773 | -3.388526049 | 0.733841283 | 0.830733542 | 0.811740452 | 0.168554319 | 0.264739911 | 0.434331378 | 0.3578 | 0.510554868 |
| SL1344_RS15520_ | -4.313644899 | -5.423833097 | -4.004360695 | 1.321498304 | 1.603420026 | 0.706731351 | 0.23666668 | 0.335493109 | 0.524839418 | 0.486 | 0.645196613 |

| | | | | | | | | | | | |
|---|---|---|---|---|---|---|---|---|---|---|---|
| SL1344_RS15525_ | 3.109179345 | 2.981773524 | 3.245192256 | 0.265967167 | 0.06792981 | 3.88451816 | 4.68E-05 | 0.008765367 | 0.058745459 | 0.1 | 0.243021319 |
| SL1344_RS15530_ | 2.128839979 | 2.057821804 | 2.192811101 | 0.133374957 | 0.092960725 | 1.378363793 | 0.057333409 | 0.117605558 | 0.255284243 | 0.1884 | 0.342763531 |
| SL1344_RS15535_ | 3.321967242 | 3.271834565 | 3.453813241 | 0.135999025 | 0.149629891 | 0.854154406 | 0.210526332 | 0.290852309 | 0.488048778 | 0.3978 | 0.570576702 |
| SL1344_RS15540_ | 2.555662432 | 2.486785775 | 2.602247501 | 0.09332482 | 0.136378307 | 0.635044083 | 0.243837455 | 0.383177871 | 0.563662079 | 0.4816 | 0.637031635 |
| SL1344_RS15545_ | -4.786675023 | -4.573512609 | -4.981294972 | -0.389877118 | 1.179933361 | -0.311069308 | 0.359333339 | 0.538557508 | 0.682720118 | 0.6505 | 0.757365597 |
| SL1344_RS15550_ | 2.482457198 | 2.429804255 | 2.538508939 | 0.132219719 | 0.129776342 | 1.001007944 | 0.121918754 | 0.195563275 | 0.374985498 | 0.2851 | 0.447555987 |
| SL1344_RS15555_ | 2.92244772 | 2.94430843 | 2.880579209 | -0.059892875 | 0.092209079 | -0.58530203 | 0.28800001 | 0.457897205 | 0.619251785 | 0.569 | 0.70351321 |
| SL1344_RS15560_ | -1.46906205 | -1.577944077 | -1.328492532 | 0.259614188 | 0.35257114 | 0.687944396 | 0.217333349 | 0.331262877 | 0.496757421 | 0.4397 | 0.581185096 |
| SL1344_RS15565_ | 0.759355198 | 0.782620559 | 0.743608451 | -0.042404797 | 0.160640967 | -0.233161627 | 0.395333337 | 0.555232419 | 0.695069148 | 0.6642 | 0.7681407 |
| SL1344_RS15570_ | 0.856127337 | 0.783903245 | 0.891047073 | 0.08193032 | 0.215814841 | 0.365833056 | 0.362425055 | 0.598861879 | 0.737742537 | 0.6836 | 0.796577178 |
| SL1344_RS15575_ | 2.178920404 | 2.140451744 | 2.253662721 | 0.104592229 | 0.154847761 | 0.66203076 | 0.237841452 | 0.389925841 | 0.567216191 | 0.4862 | 0.645087307 |
| SL1344_RS15580_ | 0.922292707 | 0.87355316 | 1.016580925 | 0.120085533 | 0.188531495 | 0.600423576 | 0.263157906 | 0.427620175 | 0.592284708 | 0.533 | 0.673877013 |
| SL1344_RS15585_ | 0.74929668 | 0.680061743 | 0.827493997 | 0.152168793 | 0.147072839 | 0.959847929 | 0.151333358 | 0.22003033 | 0.379562808 | 0.3044 | 0.457869925 |
| SL1344_RS15590_ | -1.217175158 | -1.080117779 | -1.762817803 | -0.638652697 | 0.504882672 | -1.074571194 | 0.16066669 | 0.203470363 | 0.388934206 | 0.3261 | 0.49844832 |
| SL1344_RS15595_ | -0.345580554 | -0.468747037 | -0.217788843 | 0.248581189 | 0.215372231 | 1.10138131 | 0.10933337 | 0.187282488 | 0.337751012 | 0.2663 | 0.417987654 |
| SL1344_RS15600_ | 0.396330898 | 0.423739813 | 0.355332783 | -0.064155132 | 0.190606156 | -0.313958882 | 0.367333339 | 0.54220121 | 0.686328485 | 0.6567 | 0.763878478 |
| SL1344_RS15605_ | -1.305299734 | -1.40263427 | -1.217359876 | 0.174900786 | 0.404604216 | 0.401434056 | 0.336000007 | 0.538130422 | 0.686472008 | 0.6482 | 0.762315495 |
| SL1344_RS15610_ | -1.97608614 | -1.986661424 | -1.958230791 | 0.033758219 | 0.444181033 | 0.071788579 | 0.472000001 | 0.615796915 | 0.746914454 | 0.7277 | 0.820376467 |
| SL1344_RS15615_ | -2.982362406 | -3.080421221 | -2.810055012 | 0.225957712 | 0.707584008 | 0.284220162 | 0.37708195 | 0.596186014 | 0.730012633 | 0.6947 | 0.797164804 |
| SL1344_RS15620_ | 0.101363374 | 0.169253721 | -0.125617992 | -0.287850451 | 0.298525409 | -0.856536446 | 0.196535661 | 0.27333027 | 0.462585811 | 0.3991 | 0.557902695 |
| SL1344_RS15625_ | 0.645119853 | 0.746835949 | 0.536124866 | -0.234832168 | 0.219064638 | -1.059081205 | 0.110666703 | 0.181405633 | 0.351013179 | 0.2539 | 0.414176932 |
| SL1344_RS15630_ | -0.726176039 | -0.74948225 | -0.706282449 | 0.07405046 | 0.336494599 | 0.192798006 | 0.416000003 | 0.613804199 | 0.745968607 | 0.7444 | 0.830518883 |
| SL1344_RS15635_ | 1.845651873 | 1.444085356 | 2.179040992 | 0.731099189 | 0.144934544 | 5.001268891 | 4.68E-05 | 0.003556945 | 0.038204052 | 0.1 | 0.243021319 |
| SL1344_RS15640_ | 0.365282573 | 0.424028523 | 0.277638534 | -0.161863491 | 0.216852434 | -0.701903225 | 0.202531663 | 0.307850276 | 0.484666731 | 0.4047 | 0.549823552 |
| SL1344_RS15645_ | -0.936457019 | -1.128380048 | -0.667004133 | 0.455883832 | 0.322398181 | 1.342730466 | 0.072666725 | 0.122003312 | 0.272097857 | 0.1943 | 0.354474431 |
| SL1344_RS15650_ | -2.417712219 | -2.476126607 | -2.306898497 | 0.203731097 | 0.565457377 | 0.333178857 | 0.358000006 | 0.568690971 | 0.707375476 | 0.6883 | 0.78804522 |
| SL1344_RS15655_ | -4.551639923 | -4.517566337 | -4.575648434 | -0.11693269 | 1.222144976 | -0.081491458 | 0.465333335 | 0.654808599 | 0.777423887 | 0.765 | 0.848648228 |
| SL1344_RS15660_ | -2.488384187 | -2.651070066 | -2.326786935 | 0.331885629 | 0.411214202 | 0.760711383 | 0.192000018 | 0.286567247 | 0.444321379 | 0.3819 | 0.527596459 |
| SL1344_RS15665_ | -3.835235598 | -3.883621278 | -3.783286628 | 0.073302672 | 0.838902732 | 0.076896019 | 0.462000001 | 0.622071268 | 0.749688963 | 0.7355 | 0.823476316 |
| SL1344_RS15670_ | -2.590093711 | -2.603089434 | -2.568317053 | 0.054244833 | 0.508233887 | 0.094017975 | 0.462666668 | 0.595871568 | 0.728608729 | 0.7007 | 0.797779365 |
| SL1344_RS15675_ | -3.482490918 | -3.489990363 | -3.478644401 | -0.015859356 | 0.684860023 | -0.022260397 | 0.491005996 | 0.596201618 | 0.729671722 | 0.6978 | 0.796455318 |
| SL1344_RS15680_ | 0.062756819 | 0.126432876 | -0.082861631 | -0.200082755 | 0.237820609 | -0.778491153 | 0.215189889 | 0.323247169 | 0.495594401 | 0.4478 | 0.594991632 |
| SL1344_RS15685_ | -2.785403166 | -3.178545786 | -2.453882407 | 0.708381992 | 0.701649319 | 0.971466718 | 0.14656898 | 0.214050193 | 0.389158194 | 0.3014 | 0.467501973 |
| SL1344_RS15690_ | 2.00236545 | 1.972013577 | 2.035764837 | 0.061787388 | 0.096139205 | 0.630510227 | 0.248501012 | 0.379134653 | 0.537937716 | 0.4865 | 0.621620529 |
| SL1344_RS15695_ | 1.607652565 | 1.594374586 | 1.630223022 | 0.051357253 | 0.151943743 | 0.304813884 | 0.360426388 | 0.521504557 | 0.675933563 | 0.6633 | 0.765203124 |
| SL1344_RS15700_ | -5.03473682 | -5.48891941 | -4.598936278 | 0.870762766 | 1.100077866 | 0.718959739 | 0.196666684 | 0.295372134 | 0.455851395 | 0.3915 | 0.53555125 |
| SL1344_RS15705_ | -3.134528029 | -3.223412022 | -2.962619513 | 0.220640058 | 0.81597775 | 0.23997612 | 0.403064627 | 0.659448808 | 0.781839205 | 0.7472 | 0.838498731 |
| SL1344_RS15710_ | 2.97026317 | 2.943530621 | 3.108641103 | 0.173142998 | 0.354527915 | 0.535310176 | 0.312458369 | 0.5137035 | 0.691645168 | 0.6484 | 0.789037202 |
| SL1344_RS15715_ | 1.769086038 | 1.762684423 | 1.779112699 | 0.00511274 | 0.159957166 | 0.031983748 | 0.486342439 | 0.719033747 | 0.826864808 | 0.8271 | 0.891342424 |
| SL1344_RS15720_ | 1.780268659 | 1.810165991 | 1.753481901 | -0.062013566 | 0.168576795 | -0.342752803 | 0.350000006 | 0.577788199 | 0.722459692 | 0.702 | 0.80378038 |
| SL1344_RS15725_ | 4.85782761 | 4.869182397 | 4.750825401 | -0.116561794 | 0.23624185 | -0.545193589 | 0.33333334 | 0.454009657 | 0.645187507 | 0.7003 | 0.829776116 |
| SL1344_RS15730_ | 4.460256043 | 4.422977857 | 4.507693747 | 0.075720879 | 0.169242529 | 0.356745353 | 0.400399737 | 0.585342542 | 0.743406849 | 0.8746 | 0.922675881 |
| SL1344_RS15735_ | 3.017144909 | 2.996315726 | 3.062656678 | 0.068765764 | 0.127748665 | 0.48013789 | 0.312458369 | 0.474463957 | 0.645776735 | 0.6302 | 0.755238636 |
| SL1344_RS15740_ | 2.078781553 | 2.073985001 | 2.087333256 | 0.004821669 | 0.104195898 | 0.034635725 | 0.485009994 | 0.616483193 | 0.747709061 | 0.7245 | 0.819489712 |
| SL1344_RS15745_ | 1.621089096 | 1.562982869 | 1.68667041 | 0.124965697 | 0.127292163 | 0.93558705 | 0.149900092 | 0.233494922 | 0.401068384 | 0.3193 | 0.476491922 |
| SL1344_RS15750_ | 2.082146312 | 2.054857403 | 2.237777842 | 0.173855996 | 0.167196428 | 0.731658708 | 0.28314458 | 0.389294918 | 0.578257879 | 0.5839 | 0.732018556 |
| SL1344_RS15755_ | 1.993803406 | 1.967898232 | 2.019637573 | 0.052422479 | 0.112525429 | 0.453262374 | 0.316666675 | 0.47393269 | 0.626116903 | 0.5774 | 0.699006053 |
| SL1344_RS15760_ | 0.894420471 | 0.849858259 | 0.961041725 | 0.122916168 | 0.178387615 | 0.647028657 | 0.220519669 | 0.3379888 | 0.514229882 | 0.4514 | 0.592985358 |
| SL1344_RS15765_ | -1.260714371 | -1.229994702 | -1.279602307 | -0.08043888 | 0.391150759 | -0.178272519 | 0.426382415 | 0.62746302 | 0.756320777 | 0.7495 | 0.835718739 |
| SL1344_RS15770_ | 2.389566945 | 2.331889907 | 2.434811973 | 0.10320606 | 0.097807864 | 0.977497174 | 0.134000029 | 0.208251077 | 0.373752197 | 0.2993 | 0.456631265 |
| SL1344_RS15775_ | 1.642061113 | 1.588342873 | 1.701273931 | 0.114610809 | 0.107515073 | 1.020782518 | 0.136575645 | 0.207025352 | 0.362325217 | 0.2942 | 0.444533317 |
| SL1344_RS15780_ | -0.086694699 | -0.123725279 | -0.068658242 | 0.053004107 | 0.360338191 | 0.12802736 | 0.439040642 | 0.705137979 | 0.819072277 | 0.8401 | 0.903486812 |
| SL1344_RS15785_ | 0.887548883 | 0.803620042 | 0.979230747 | 0.184037291 | 0.16137929 | 1.102696242 | 0.101333374 | 0.171173674 | 0.331450019 | 0.2515 | 0.407971539 |

| | | | | | | | | | | | |
|---|---|---|---|---|---|---|---|---|---|---|---|
| SL1344_RS15790_ | -2.242610653 | -2.046617387 | -2.487053893 | -0.427049808 | 0.500107857 | -0.799909457 | 0.193204549 | 0.298672182 | 0.472473061 | 0.4028 | 0.557914068 |
| SL1344_RS15795_ | 3.634003125 | 4.076873507 | 3.227464627 | -0.842117188 | 0.100192758 | -8.40075486 | 4.68E-05 | 0.000813916 | 0.019104589 | 0.1 | 0.243021319 |
| SL1344_RS15800_ | 1.941878303 | 2.43282326 | 1.327121114 | -1.069604813 | 0.180625722 | -5.903380874 | 4.68E-05 | 0.001796684 | 0.028144936 | 0.1 | 0.243021319 |
| SL1344_RS15810_ | 2.318461256 | 2.271763637 | 2.379674401 | 0.098144028 | 0.124565246 | 0.756202904 | 0.211192554 | 0.333210783 | 0.51000121 | 0.4321 | 0.591518729 |
| SL1344_RS15815_ | 2.284041618 | 2.295928461 | 2.270298482 | 0.027691812 | 0.19218315 | 0.150249217 | 0.444000002 | 0.674833596 | 0.799633557 | 0.873 | 0.921604524 |
| SL1344_RS15820_ | 2.849511162 | 2.843492122 | 2.85711619 | 0.014019202 | 0.068559377 | 0.190122204 | 0.409726852 | 0.532202189 | 0.673497712 | 0.6472 | 0.752971714 |
| SL1344_RS15825_ | 2.707905587 | 2.65066843 | 2.765287193 | 0.132846589 | 0.134931808 | 1.023081348 | 0.082611643 | 0.202955612 | 0.391763054 | 0.2104 | 0.369624301 |
| SL1344_RS15830_ | 4.248567205 | 4.113032961 | 4.481956674 | 0.377131307 | 0.130308622 | 2.893889383 | 4.68E-05 | 0.021422761 | 0.10509043 | 0.1 | 0.243021319 |
| SL1344_RS15835_ | 1.62677253 | 1.460570296 | 1.920375735 | 0.445508493 | 0.174736415 | 2.473471431 | 0.0020023 | 0.028091938 | 0.117931509 | 0.101 | 0.244415958 |
| SL1344_RS15840_ | 0.986288959 | 1.065306815 | 0.787654475 | -0.247183384 | 0.222628094 | -1.00136445 | 0.160000023 | 0.212862218 | 0.396301315 | 0.3188 | 0.491307418 |
| SL1344_RS15845_ | 2.943866707 | 2.902275739 | 2.988459288 | 0.096296633 | 0.099025225 | 0.947253228 | 0.128000031 | 0.209605311 | 0.388840589 | 0.2825 | 0.444748488 |
| SL1344_RS15850_ | -0.396319789 | -0.522666825 | -0.269066627 | 0.259993386 | 0.247434129 | 1.003278116 | 0.123333366 | 0.20464235 | 0.369032653 | 0.2905 | 0.445468048 |
| SL1344_RS15855_ | -3.343600644 | -3.688442778 | -3.030550257 | 0.660354574 | 0.576099178 | 1.116449073 | 0.112591641 | 0.179607706 | 0.329161725 | 0.2557 | 0.408002762 |
| SL1344_RS15860_ | 1.47882772 | 1.318581967 | 1.638348085 | 0.322099141 | 0.103639735 | 3.043252129 | 0.00067292 | 0.015183425 | 0.078835279 | 0.1003 | 0.243436432 |
| SL1344_RS15865_ | -0.979229294 | -1.108827173 | -0.851104819 | 0.261847542 | 0.279458811 | 0.874152085 | 0.156000024 | 0.257625558 | 0.419703419 | 0.3528 | 0.503052717 |
| SL1344_RS15870_ | 1.496217464 | 1.439526658 | 1.574379379 | 0.146882778 | 0.154962599 | 0.877767566 | 0.164556985 | 0.24500659 | 0.422943037 | 0.3352 | 0.494688463 |
| SL1344_RS15875_ | 6.152703566 | 5.973089924 | 6.356764046 | 0.381865251 | 0.030011943 | 12.51241549 | 4.68E-05 | 0.000733318 | 0.017352249 | 0.1 | 0.243021319 |
| SL1344_RS15880_ | 4.059011707 | 4.039226627 | 4.144144346 | 0.11117019 | 0.141754121 | 0.73411944 | 0.24250501 | 0.336510134 | 0.537095165 | 0.4665 | 0.644163127 |
| SL1344_RS15890_ | -3.987578202 | -3.930422738 | -4.061704079 | -0.134085536 | 0.843737088 | -0.150346522 | 0.426382415 | 0.550848368 | 0.695242192 | 0.6647 | 0.771358114 |
| SL1344_RS15895_ | -3.543852615 | -4.060250863 | -3.047310961 | 0.981500874 | 0.750935584 | 1.253180725 | 0.085942754 | 0.143417834 | 0.296066284 | 0.2187 | 0.378174988 |
| SL1344_RS15900_ | -2.761329189 | -3.063501119 | -2.573633142 | 0.474825014 | 0.68549105 | 0.650348441 | 0.222000015 | 0.340211792 | 0.516917834 | 0.4481 | 0.595686437 |
| SL1344_RS15905_ | -1.69588904 | -1.735590331 | -1.657506995 | 0.082762942 | 0.349876213 | 0.213041584 | 0.402398405 | 0.544651258 | 0.684825897 | 0.6459 | 0.753186413 |
| SL1344_RS15910_ | -2.929712172 | -3.206249698 | -2.718411741 | 0.487617494 | 0.553168332 | 0.847195917 | 0.173884098 | 0.271075784 | 0.431135424 | 0.3658 | 0.513093415 |
| SL1344_RS15915_ | 5.455282117 | 5.396276593 | 5.608337474 | 0.206169118 | 0.068765167 | 2.342708847 | 0.00067292 | 0.046797224 | 0.164728813 | 0.1002 | 0.243307454 |
| SL1344_RS15920_ | 1.215646557 | 1.172614975 | 1.262688842 | 0.094032514 | 0.154419185 | 0.579097633 | 0.256666678 | 0.411642038 | 0.574659309 | 0.5145 | 0.650820233 |
| SL1344_RS15925_ | 4.717620582 | 4.538587754 | 5.005874305 | 0.459330221 | 0.062648276 | 6.597623848 | 4.68E-05 | 0.009918878 | 0.068904706 | 0.1 | 0.243021319 |
| SL1344_RS15930_ | -2.068432502 | -2.099648125 | -2.032085392 | 0.066556222 | 0.412229488 | 0.14162459 | 0.428666669 | 0.565561826 | 0.70293033 | 0.6663 | 0.77027138 |
| SL1344_RS15935_ | -2.749138808 | -2.718138972 | -2.788612857 | -0.063838243 | 0.487318901 | -0.122065434 | 0.442000002 | 0.562382 | 0.697784661 | 0.6694 | 0.771319452 |
| SL1344_RS15940_ | -0.965551287 | -0.896499621 | -1.063739612 | -0.188460165 | 0.317350563 | -0.528460954 | 0.258494349 | 0.405439837 | 0.570399641 | 0.5191 | 0.649539767 |
| SL1344_RS15945_ | -1.546154784 | -1.414401979 | -1.719381834 | -0.30437402 | 0.336365755 | -0.862648318 | 0.162000023 | 0.255197049 | 0.41457295 | 0.3442 | 0.493142871 |
| SL1344_RS15950_ | -2.551657604 | -2.548600469 | -2.558036458 | 0.053579704 | 0.664082742 | 0.070781721 | 0.474350434 | 0.691522774 | 0.805936766 | 0.7803 | 0.86204151 |
| SL1344_RS15955_ | -2.771156707 | -2.720114688 | -2.872792802 | -0.134302626 | 0.56510922 | -0.219378391 | 0.40466667 | 0.593905908 | 0.730490156 | 0.7055 | 0.805642351 |
| SL1344_RS15960_ | 2.044117599 | 2.041240378 | 2.046570281 | 0.005444394 | 0.138507344 | 0.037721107 | 0.487333334 | 0.687891092 | 0.804283727 | 0.7984 | 0.871873255 |
| SL1344_RS15965_ | -0.554873852 | -1.063185858 | -0.218609208 | 0.85908618 | 0.359613698 | 2.356205329 | 4.68E-05 | 0.046026763 | 0.157364799 | 0.1005 | 0.243730926 |
| SL1344_RS15970_ | 4.697603421 | 4.583586553 | 4.86265947 | 0.283805024 | 0.123210619 | 2.408729793 | 4.68E-05 | 0.044185883 | 0.160076024 | 0.1 | 0.243021319 |
| SL1344_RS15975_ | 1.899512335 | 1.737119224 | 2.003358903 | 0.281553938 | 0.169788046 | 1.659692195 | 0.010000458 | 0.098113344 | 0.251349924 | 0.11 | 0.256195084 |
| SL1344_RS15985_ | 5.837103655 | 5.799229908 | 5.937411899 | 0.13378309 | 0.050069325 | 2.304283369 | 0.001336769 | 0.052895434 | 0.174791412 | 0.1013 | 0.244783188 |
| SL1344_RS15990_ | 0.671106845 | 0.648053121 | 0.70081124 | 0.074885062 | 0.198106501 | 0.353574343 | 0.365333339 | 0.537488018 | 0.685716142 | 0.6644 | 0.7702621 |
| SL1344_RS15995_ | 1.052216026 | 0.810175176 | 1.252672246 | 0.448934644 | 0.144110829 | 3.057350311 | 4.68E-05 | 0.01590625 | 0.081761366 | 0.1001 | 0.243160558 |
| SL1344_RS16000_ | 1.397527933 | 1.311547207 | 1.493166921 | 0.183899814 | 0.146165598 | 1.21300832 | 0.077333388 | 0.137374829 | 0.293007787 | 0.2058 | 0.365328558 |
| SL1344_RS16005_ | 2.181820889 | 2.010181808 | 2.347947709 | 0.345034854 | 0.132332023 | 2.60793196 | 0.000673369 | 0.022279601 | 0.102749474 | 0.1003 | 0.243454608 |
| SL1344_RS16015_ | 1.393831583 | 1.28497641 | 1.49104594 | 0.207127493 | 0.147815725 | 1.334440051 | 0.077333388 | 0.119071773 | 0.268916354 | 0.1887 | 0.347863168 |
| SL1344_RS16020_ | 1.356233508 | 1.251595165 | 1.465897675 | 0.211485298 | 0.140300894 | 1.45590315 | 0.060000072 | 0.102365107 | 0.242229921 | 0.1736 | 0.330334442 |
| SL1344_RS16025_ | 0.540168264 | 0.440823969 | 0.632451135 | 0.200119462 | 0.20016798 | 0.963751802 | 0.128666697 | 0.203430018 | 0.37296897 | 0.2799 | 0.438588335 |
| SL1344_RS16030_ | -1.634669179 | -1.543767693 | -1.882632475 | -0.316107008 | 0.452234412 | -0.604974491 | 0.27514991 | 0.409754294 | 0.581966921 | 0.5493 | 0.684861218 |
| SL1344_RS16035_ | 1.813205406 | 1.679069987 | 1.868221394 | 0.173462053 | 0.150455257 | 1.054296714 | 0.145902758 | 0.208905657 | 0.388563192 | 0.305 | 0.475854972 |
| SL1344_RS16040_ | 1.790766671 | 1.790397962 | 1.792144973 | 0.006205103 | 0.137613556 | 0.044898995 | 0.484343771 | 0.66395223 | 0.784638557 | 0.7738 | 0.854596068 |
| SL1344_RS16045_ | 1.28153567 | 1.251414518 | 1.336500065 | 0.080275441 | 0.145994499 | 0.51438504 | 0.302465032 | 0.477354031 | 0.631863666 | 0.6022 | 0.722158266 |
| SL1344_RS16050_ | 1.944963319 | 1.822007383 | 1.99774035 | 0.181133114 | 0.174581007 | 0.998493977 | 0.148666692 | 0.201331501 | 0.383283226 | 0.295 | 0.472760973 |
| SL1344_RS16055_ | 3.132458718 | 2.959928642 | 3.263051889 | 0.270777225 | 0.129962656 | 2.032350613 | 0.009333825 | 0.041110883 | 0.149057325 | 0.1098 | 0.256642092 |
| SL1344_RS16060_ | -1.641629734 | -1.641423132 | -1.642187443 | 0.018052569 | 0.403332869 | 0.044774198 | 0.481333334 | 0.609458062 | 0.740077537 | 0.7206 | 0.812126085 |
| SL1344_RS16065_ | 0.411101183 | 0.336691788 | 0.440636324 | 0.078355221 | 0.243872406 | 0.283190842 | 0.398000004 | 0.67166059 | 0.793366821 | 0.7602 | 0.852816565 |
| SL1344_RS16070_ | 2.697274037 | 2.654318025 | 2.795943944 | 0.137766602 | 0.156058071 | 0.778521036 | 0.217188556 | 0.304681546 | 0.499199048 | 0.4287 | 0.593415999 |

| | | | | | | | | | | | |
|---|---|---|---|---|---|---|---|---|---|---|---|
| SL1344_RS16075_ | 2.94231586 | 2.884260781 | 3.030147823 | 0.126715225 | 0.228553092 | 0.421395181 | 0.377748173 | 0.68097391 | 0.808124803 | 0.8383 | 0.907194462 |
| SL1344_RS16080_ | 3.38978213 | 3.262093668 | 3.584776682 | 0.330077196 | 0.111281743 | 2.967539296 | 4.68E-05 | 0.036214496 | 0.140907837 | 0.1 | 0.243021319 |
| SL1344_RS16085_ | 1.626291614 | 1.517736949 | 1.742658929 | 0.226450751 | 0.104477284 | 2.089952954 | 0.009333825 | 0.046023641 | 0.147857656 | 0.1129 | 0.259239812 |
| SL1344_RS16090_ | 1.274068419 | 1.205450728 | 1.363738747 | 0.155667202 | 0.134763892 | 1.076149612 | 0.129913421 | 0.196846664 | 0.358913787 | 0.2826 | 0.441083123 |
| SL1344_RS16095_ | 1.787786726 | 1.651534265 | 1.916595627 | 0.26533049 | 0.102423915 | 2.523170752 | 0.005330642 | 0.027253201 | 0.110008516 | 0.1031 | 0.246971489 |
| SL1344_RS16100_ | -0.158232583 | -0.411940235 | 0.113102707 | 0.529304265 | 0.219942184 | 2.337291528 | 0.005996769 | 0.032305207 | 0.122739227 | 0.1044 | 0.248756821 |
| SL1344_RS16105_ | 0.987540296 | 0.928504843 | 1.063255424 | 0.132964894 | 0.142364823 | 0.90341658 | 0.161333357 | 0.254983431 | 0.418506992 | 0.349 | 0.499636512 |
| SL1344_RS16110_ | -4.867137986 | -5.067983622 | -4.674643293 | 0.432240653 | 1.070702654 | 0.351871743 | 0.344000006 | 0.487760381 | 0.636055487 | 0.6026 | 0.716700458 |
| SL1344_RS16115_ | -1.263394565 | -1.327721024 | -1.209329025 | 0.176486389 | 0.527403874 | 0.297434313 | 0.345769493 | 0.500215503 | 0.666547003 | 0.6564 | 0.766615006 |
| SL1344_RS16120_ | 1.177733434 | 1.017887541 | 1.346340761 | 0.327349347 | 0.119079042 | 2.733765839 | 0.002000966 | 0.021954168 | 0.097239909 | 0.1021 | 0.245757807 |
| SL1344_RS16125_ | -1.560606151 | -1.72274821 | -1.413113616 | 0.318085452 | 0.342260655 | 0.894977103 | 0.14923387 | 0.245108108 | 0.408226997 | 0.3376 | 0.486464008 |
| SL1344_RS16130_ | -0.545301046 | -0.649799422 | -0.439280866 | 0.215315011 | 0.214905412 | 0.942688538 | 0.142571646 | 0.228205668 | 0.379996608 | 0.3142 | 0.463538449 |
| SL1344_RS16135_ | -1.6064552 | -1.553553721 | -1.653672803 | -0.093526926 | 0.331567487 | -0.269733109 | 0.374000005 | 0.535583712 | 0.676190539 | 0.6421 | 0.750732225 |
| SL1344_RS16140_ | -0.912548386 | -0.936068493 | -0.864991661 | 0.06160888 | 0.311375409 | 0.179213423 | 0.42504997 | 0.618138832 | 0.747618393 | 0.716 | 0.812347062 |
| SL1344_RS16145_ | 0.591433097 | 0.431292235 | 0.780643257 | 0.356873295 | 0.184899903 | 1.852167188 | 0.018000252 | 0.057707068 | 0.173307443 | 0.1227 | 0.271535479 |
| SL1344_RS16150_ | -0.009119502 | -0.046862728 | 0.016378228 | 0.060070666 | 0.240949366 | 0.236725032 | 0.393333337 | 0.570468477 | 0.709784638 | 0.6818 | 0.78441728 |
| SL1344_RS16155_ | -3.149869935 | -3.166048966 | -3.134944427 | 0.028184394 | 0.578145367 | 0.044485477 | 0.478347769 | 0.560088194 | 0.698549208 | 0.6714 | 0.774449624 |
| SL1344_RS16160_ | -3.281584241 | -3.609567424 | -3.022372669 | 0.604021602 | 0.582574945 | 0.971236619 | 0.138000028 | 0.218314416 | 0.378424744 | 0.2997 | 0.451609767 |
| SL1344_RS16165_ | -0.850502862 | -1.017956943 | -0.708725835 | 0.286341328 | 0.31163424 | 0.826842952 | 0.186000019 | 0.276495558 | 0.446886179 | 0.3664 | 0.52259432 |
| SL1344_RS16170_ | -0.130618582 | -0.19043565 | -0.078514256 | 0.120152441 | 0.212002967 | 0.523891241 | 0.28314458 | 0.431983704 | 0.587043887 | 0.5415 | 0.664354472 |
| SL1344_RS16175_ | -1.377421919 | -1.468555108 | -1.311397622 | 0.156509871 | 0.335188258 | 0.443692343 | 0.304463699 | 0.456372357 | 0.611206534 | 0.5636 | 0.687286895 |
| SL1344_RS16180_ | -0.309421334 | -0.320431605 | -0.294499786 | 0.031104831 | 0.235414252 | 0.120066025 | 0.448367757 | 0.588330761 | 0.724500177 | 0.6998 | 0.797655477 |
| SL1344_RS16185_ | -0.435457517 | -0.364351199 | -0.485803663 | -0.10812634 | 0.246289762 | -0.392799901 | 0.319333341 | 0.515788688 | 0.663155561 | 0.6166 | 0.735308489 |
| SL1344_RS16190_ | -2.99272069 | -2.766259953 | -3.254884267 | -0.516853661 | 0.68840874 | -0.735699492 | 0.193204549 | 0.302430636 | 0.478116178 | 0.4018 | 0.551022676 |
| SL1344_RS16195_ | -2.580114665 | -2.417301445 | -2.724025409 | -0.325825101 | 0.543004069 | -0.581142737 | 0.260666678 | 0.399014431 | 0.560123328 | 0.5087 | 0.642144107 |
| SL1344_RS16200_ | -2.172456415 | -2.101552408 | -2.278027383 | -0.07035289 | 0.681217155 | -0.101367846 | 0.457333335 | 0.704361633 | 0.81632681 | 0.8373 | 0.897350116 |
| SL1344_RS16205_ | -3.549475267 | -3.532974112 | -3.563075358 | -0.014848648 | 0.621449723 | -0.026646336 | 0.488 | 0.552650895 | 0.690732727 | 0.6627 | 0.766167416 |
| SL1344_RS16210_ | -1.427469832 | -1.343022687 | -1.522287139 | -0.180850397 | 0.33596389 | -0.494931108 | 0.271818798 | 0.419232021 | 0.576115842 | 0.5268 | 0.65340065 |
| SL1344_RS16215_ | 0.205653096 | 0.118153113 | 0.289030956 | 0.19549155 | 0.226172377 | 0.825685256 | 0.173884098 | 0.259754879 | 0.434702148 | 0.369 | 0.519602152 |
| SL1344_RS16220_ | -0.737812995 | -0.828353262 | -0.608500102 | 0.227882803 | 0.297281272 | 0.709465203 | 0.228000014 | 0.35343711 | 0.521663658 | 0.4649 | 0.607620992 |
| SL1344_RS16225_ | 0.301274693 | 0.209860266 | 0.420268025 | 0.20649638 | 0.193502786 | 1.022408683 | 0.133910755 | 0.203909739 | 0.367129594 | 0.2875 | 0.446623935 |
| SL1344_RS16235_ | 1.951338151 | 2.010734882 | 1.852424338 | -0.157385875 | 0.143518726 | -1.029852797 | 0.145902758 | 0.202548592 | 0.379423468 | 0.2869 | 0.458264402 |
| SL1344_RS16240_ | 0.468138305 | 0.392994191 | 0.664812937 | 0.228816314 | 0.26680649 | 0.749287371 | 0.227333348 | 0.350022506 | 0.533897278 | 0.4726 | 0.630699551 |
| SL1344_RS16245_ | -1.606521524 | -1.499860512 | -1.716169375 | -0.182980595 | 0.514010551 | -0.335362629 | 0.353097941 | 0.580328236 | 0.721780024 | 0.6834 | 0.788699196 |
| SL1344_RS16250_ | -1.900459669 | -1.758730712 | -1.998917784 | -0.261209819 | 0.427579808 | -0.601225093 | 0.260000012 | 0.398784191 | 0.56078155 | 0.5145 | 0.646342889 |
| SL1344_RS16255_ | 1.23220061 | 1.485994337 | 0.846905724 | -0.641933912 | 0.121041768 | -5.255736386 | 4.68E-05 | 0.003759285 | 0.036713471 | 0.1 | 0.243021319 |
| SL1344_RS16260_ | 0.241471639 | 0.448572635 | -0.004604545 | -0.469280662 | 0.228744768 | -2.014037337 | 0.007329074 | 0.055684302 | 0.173270659 | 0.1067 | 0.252010855 |
| SL1344_RS16265_ | 0.158871753 | 0.399281277 | -0.124104882 | -0.521888166 | 0.186038081 | -2.750738347 | 0.0020023 | 0.020615551 | 0.093644543 | 0.1012 | 0.244640002 |
| SL1344_RS16270_ | 1.084360499 | 1.423770034 | 0.774995863 | -0.665515503 | 0.196699915 | -3.35842767 | 4.68E-05 | 0.013388118 | 0.076546694 | 0.1 | 0.243021319 |
| SL1344_RS16275_ | 2.313231517 | 2.606392305 | 1.98090462 | -0.639013398 | 0.147822589 | -4.404087479 | 4.68E-05 | 0.004327098 | 0.043455438 | 0.1 | 0.243021319 |
| SL1344_RS16280_ | 0.785177295 | 1.181053766 | 0.3202536 | -0.868012701 | 0.207308574 | -4.132170458 | 4.68E-05 | 0.008175492 | 0.057811174 | 0.1 | 0.243021319 |
| SL1344_RS16285_ | 0.256397123 | 0.766557316 | -0.238301076 | -0.986907404 | 0.305007054 | -3.15670722 | 4.68E-05 | 0.013218107 | 0.078471776 | 0.1 | 0.243021319 |
| SL1344_RS16290_ | 1.532999375 | 1.786924466 | 1.136585859 | -0.675774358 | 0.177193434 | -3.848883523 | 4.68E-05 | 0.00691616 | 0.054644317 | 0.1 | 0.243021319 |
| SL1344_RS16305_ | 2.883690354 | 3.185350825 | 2.618634494 | -0.554524384 | 0.115531802 | -4.766040296 | 4.68E-05 | 0.003557734 | 0.039084788 | 0.1 | 0.243021319 |
| SL1344_RS16310_ | 3.323443559 | 3.861409786 | 2.654324924 | -1.209591146 | 0.122264025 | -10.05082747 | 4.68E-05 | 0.003278256 | 0.038820858 | 0.1 | 0.243021319 |
| SL1344_RS16315_ | 1.991665067 | 2.572824279 | 1.214730231 | -1.346353041 | 0.149194096 | -8.877880664 | 4.68E-05 | 0.00295462 | 0.034752948 | 0.1 | 0.243021319 |
| SL1344_RS16320_ | 1.99112977 | 2.622152166 | 1.304017801 | -1.326039822 | 0.113655994 | -11.71801523 | 4.68E-05 | 0.000369208 | 0.012745591 | 0.1 | 0.243021319 |
| SL1344_RS16325_ | 1.882296643 | 2.48915854 | 1.263606555 | -1.230675199 | 0.160152667 | -7.705366021 | 4.68E-05 | 0.000888664 | 0.01994067 | 0.1 | 0.243021319 |
| SL1344_RS16330_ | 3.344030766 | 3.272099251 | 3.581079348 | 0.308095033 | 0.153158839 | 1.897046584 | 0.003332494 | 0.088657765 | 0.241618759 | 0.1018 | 0.245543488 |
| SL1344_RS16335_ | 3.824255788 | 3.677286839 | 3.95754554 | 0.290267698 | 0.114551615 | 2.482456607 | 4.68E-05 | 0.019600243 | 0.098842688 | 0.1002 | 0.243296937 |
| SL1344_RS16340_ | 2.392498933 | 2.258892083 | 2.647871561 | 0.226129642 | 0.439006283 | 0.703399504 | 0.220666682 | 0.364309265 | 0.565026843 | 0.4052 | 0.599412829 |
| SL1344_RS16350_ | 1.316636392 | 1.213817588 | 1.425609373 | 0.239884404 | 0.185866847 | 1.302391201 | 0.046000096 | 0.123712772 | 0.282585959 | 0.1693 | 0.323788829 |
| SL1344_RS16355_ | 1.016161559 | 1.030710698 | 0.993380894 | -0.051203083 | 0.158596968 | -0.281845672 | 0.374666672 | 0.540993721 | 0.68632661 | 0.6646 | 0.767801863 |

| | | | | | | | | | | |
|---|---|---|---|---|---|---|---|---|---|---|
| SL1344_RS16360_ | 1.439533879 | 1.484278434 | 1.382719357 | -0.09805929 | 0.132419984 | -0.695823904 | 0.222666682 | 0.350403771 | 0.513925632 | 0.4538 | 0.596582534 |
| SL1344_RS16365_ | 0.689846275 | 0.776459478 | 0.518792256 | -0.249427785 | 0.177508498 | -1.31170426 | 0.087275198 | 0.135869037 | 0.291464012 | 0.2142 | 0.374069617 |
| SL1344_RS16370_ | -1.27796643 | -1.465259994 | -1.044337794 | 0.277700732 | 0.549946443 | 0.523789855 | 0.29113925 | 0.491624599 | 0.654255301 | 0.5793 | 0.714184779 |
| SL1344_RS16375_ | 0.516993767 | 0.527643625 | 0.494346068 | -0.011959615 | 0.264001308 | -0.038001471 | 0.481678881 | 0.748659099 | 0.848393869 | 0.8486 | 0.909536512 |
| SL1344_RS16380_ | -0.888361706 | -1.063732807 | -0.765013849 | 0.303804279 | 0.304469805 | 0.964534377 | 0.128580977 | 0.216537185 | 0.386640002 | 0.2994 | 0.451997065 |
| SL1344_RS16385_ | 3.436513593 | 3.212205588 | 3.660680254 | 0.45581387 | 0.080730502 | 5.65417951 | 4.68E-05 | 0.004060917 | 0.040230301 | 0.1 | 0.243021319 |
| SL1344_RS16390_ | -3.604583533 | -3.347935194 | -3.897758346 | -0.592558489 | 0.847068477 | -0.654468152 | 0.223850781 | 0.345371477 | 0.514900514 | 0.4569 | 0.600689734 |
| SL1344_RS16395_ | -4.58055684 | -4.371412693 | -4.735283315 | -0.394295878 | 1.118164434 | -0.317648717 | 0.355096608 | 0.542670125 | 0.685477109 | 0.6614 | 0.766114214 |
| SL1344_RS16400_ | -1.687120035 | -1.898965031 | -1.481329215 | 0.461507446 | 0.469275569 | 0.951181006 | 0.135909422 | 0.211366639 | 0.385287782 | 0.3038 | 0.46333812 |
| SL1344_RS16405_ | 2.43241069 | 2.387806193 | 2.708841488 | 0.318351184 | 0.26601745 | 1.042313664 | 0.151232537 | 0.20863281 | 0.405614845 | 0.3016 | 0.470607886 |
| SL1344_RS16410_ | 1.844203176 | 1.521404592 | 2.076906357 | 0.558078601 | 0.121414412 | 4.553676061 | 4.68E-05 | 0.006479984 | 0.050876507 | 0.1 | 0.243021319 |
| SL1344_RS16415_ | 4.103209163 | 3.979801954 | 4.211586157 | 0.242431929 | 0.074688651 | 3.239003018 | 4.68E-05 | 0.011661474 | 0.072116529 | 0.1 | 0.243021319 |
| SL1344_RS16420_ | 1.168100125 | 1.438358383 | 0.914839797 | -0.501666056 | 0.19467003 | -2.510742529 | 0.002666616 | 0.023993641 | 0.106857581 | 0.1019 | 0.245685923 |
| SL1344_RS16425_ | 1.9253856 | 2.058474853 | 1.735879687 | -0.338857249 | 0.248550282 | -1.352107572 | 0.024650416 | 0.155322273 | 0.339424171 | 0.1216 | 0.272040011 |
| SL1344_RS16440_ | 0.728639412 | 0.718616798 | 0.740319884 | 0.033751922 | 0.175818306 | 0.183577392 | 0.426382415 | 0.594293471 | 0.728753452 | 0.7193 | 0.810935875 |
| SL1344_RS16445_ | 2.909034161 | 2.937809339 | 2.837222128 | -0.091453019 | 0.130368521 | -0.653821542 | 0.242000013 | 0.382966426 | 0.565904268 | 0.4806 | 0.646349965 |
| SL1344_RS16450_ | 4.210506511 | 4.038290199 | 4.3741613 | 0.362323438 | 0.109490042 | 3.533953789 | 4.68E-05 | 0.019181894 | 0.098333715 | 0.1 | 0.243021319 |
| SL1344_RS16455_ | 2.377558991 | 2.278817841 | 2.49843741 | 0.234961534 | 0.138479689 | 1.726515808 | 0.007329074 | 0.084163492 | 0.228373259 | 0.1072 | 0.252899003 |
| SL1344_RS16460_ | 2.944702963 | 2.861317356 | 3.021609208 | 0.186690901 | 0.123727688 | 1.493467687 | 0.026000173 | 0.081934595 | 0.223196376 | 0.1302 | 0.281774666 |
| SL1344_RS16465_ | 1.010844916 | 0.969073238 | 1.105849345 | 0.122700757 | 0.165267529 | 0.663446583 | 0.245333346 | 0.388225775 | 0.554559889 | 0.4945 | 0.639616888 |
| SL1344_RS16470_ | 0.454200072 | 0.549941988 | 0.342250971 | -0.201461196 | 0.152199924 | -1.249089732 | 0.078614311 | 0.150459258 | 0.290488099 | 0.224 | 0.375980444 |
| SL1344_RS16480_ | 1.807280738 | 1.662333755 | 1.90914217 | 0.24297489 | 0.118888843 | 1.94259581 | 0.018000252 | 0.055141122 | 0.170798985 | 0.1182 | 0.266808913 |
| SL1344_RS16485_ | 2.795194521 | 2.689294326 | 2.861244733 | 0.171850444 | 0.103962374 | 1.589097591 | 0.04266677 | 0.079244758 | 0.215123368 | 0.1429 | 0.298090702 |
| SL1344_RS16490_ | 1.516645891 | 1.5548603 | 1.465688029 | -0.092367993 | 0.12151003 | -0.751811423 | 0.205196552 | 0.315397431 | 0.482911096 | 0.4208 | 0.565425917 |
| SL1344_RS16495_ | 1.387872576 | 1.358809655 | 1.438796546 | 0.078871376 | 0.137147515 | 0.530176669 | 0.294470362 | 0.460408076 | 0.615665286 | 0.5663 | 0.693083604 |
| SL1344_RS16500_ | -3.962287221 | -3.894754402 | -4.06365745 | -0.219963476 | 0.903669019 | -0.228899622 | 0.395069957 | 0.556194304 | 0.698468664 | 0.6833 | 0.782168009 |
| SL1344_RS16505_ | -2.171176016 | -2.253873951 | -2.095075725 | 0.17342654 | 0.424338735 | 0.377375159 | 0.33333334 | 0.491385329 | 0.641294085 | 0.6068 | 0.720982013 |
| SL1344_RS16510_ | -2.780321795 | -2.690390482 | -2.839588378 | -0.070036893 | 0.74180047 | -0.077642161 | 0.466666668 | 0.704985198 | 0.818606765 | 0.7957 | 0.875057602 |
| SL1344_RS16515_ | -0.739929866 | -0.633703526 | -0.849162254 | -0.209883889 | 0.244781517 | -0.806743765 | 0.184543657 | 0.276944032 | 0.437204152 | 0.3717 | 0.521426675 |
| SL1344_RS16520_ | 3.051324988 | 3.020852206 | 3.103162485 | 0.081296363 | 0.402582072 | 0.197715264 | 0.392000004 | 0.596336416 | 0.753928637 | 0.8275 | 0.901351612 |
| SL1344_RS16525_ | 2.940653676 | 2.828189942 | 3.166551704 | 0.334246936 | 0.238710621 | 1.290975966 | 0.095269864 | 0.130438946 | 0.305042777 | 0.1936 | 0.369700644 |
| SL1344_RS16530_ | -0.918668428 | -1.120480084 | -0.649986165 | 0.453284032 | 0.312290019 | 1.358804561 | 0.078614311 | 0.12544772 | 0.275733541 | 0.1964 | 0.356753648 |
| SL1344_RS16535_ | -2.374627944 | -2.399902735 | -2.349189057 | 0.057773817 | 0.434764813 | 0.124257486 | 0.448367757 | 0.570023131 | 0.707009679 | 0.6736 | 0.776138949 |
| SL1344_RS16540_ | 0.124176417 | 0.144237908 | 0.105045323 | -0.023412584 | 0.258771437 | -0.090333877 | 0.463333335 | 0.681498924 | 0.79956755 | 0.782 | 0.859945379 |
| SL1344_RS16545_ | 2.330720825 | 2.239518168 | 2.444299549 | 0.217586712 | 0.116375469 | 1.81692392 | 0.008661422 | 0.06707331 | 0.194228843 | 0.1126 | 0.259425594 |
| SL1344_RS16550_ | 2.216723424 | 2.1893813 | 2.28939932 | 0.096078413 | 0.140479448 | 0.611465947 | 0.265822796 | 0.406473005 | 0.584190748 | 0.5464 | 0.687761716 |
| SL1344_RS16555_ | 1.528416462 | 1.455940716 | 1.616295518 | 0.156929891 | 0.122322897 | 1.209802703 | 0.102598308 | 0.155458823 | 0.309505896 | 0.2377 | 0.396827853 |
| SL1344_RS16560_ | 0.948438058 | 0.854838766 | 1.03542084 | 0.17720661 | 0.13530182 | 1.266818697 | 0.074616979 | 0.133375603 | 0.27561072 | 0.2066 | 0.3613498 |
| SL1344_RS16565_ | 2.074892949 | 1.925417725 | 2.227732174 | 0.296827914 | 0.113636246 | 2.520779757 | 0.003334715 | 0.025573841 | 0.109453021 | 0.1022 | 0.245977624 |
| SL1344_RS16570_ | 1.665216083 | 1.580603595 | 1.847204595 | 0.255192977 | 0.147712837 | 1.564812618 | 0.050000088 | 0.095868856 | 0.239607399 | 0.1589 | 0.315231741 |
| SL1344_RS16575_ | -0.014147054 | -0.16728889 | 0.079161874 | 0.251299485 | 0.221945826 | 1.032583544 | 0.125916087 | 0.194511551 | 0.362215677 | 0.2753 | 0.433551876 |
| SL1344_RS16580_ | 1.4765619 | 1.40604091 | 1.56785201 | 0.162037619 | 0.142808798 | 1.073776173 | 0.113257864 | 0.18463795 | 0.353777295 | 0.2694 | 0.432349695 |
| SL1344_RS16585_ | 2.58881556 | 2.419347989 | 2.761653851 | 0.342390135 | 0.084368755 | 3.984967698 | 4.68E-05 | 0.007959533 | 0.056169987 | 0.1 | 0.243021319 |
| SL1344_RS16590_ | 7.95656745 | 7.855695201 | 8.002313462 | 0.176551366 | 0.106582587 | 1.818578573 | 4.68E-05 | 0.040093874 | 0.152516426 | 0.1 | 0.243021319 |
| SL1344_RS16595_ | 5.27474531 | 5.210138071 | 5.347691323 | 0.138446319 | 0.06216208 | 2.216357705 | 0.002000966 | 0.031457362 | 0.128176639 | 0.1017 | 0.245388334 |
| SL1344_RS16600_ | 5.239980091 | 5.265911311 | 5.167678032 | -0.086501683 | 0.112573695 | -0.73326577 | 0.245333346 | 0.35436226 | 0.554024577 | 0.4626 | 0.640335838 |
| SL1344_RS16605_ | -1.244547843 | -1.105045002 | -1.452522191 | -0.393704535 | 0.469959641 | -0.835934818 | 0.116588975 | 0.255020917 | 0.448523001 | 0.2678 | 0.424412374 |
| SL1344_RS16610_ | 0.691445645 | 0.390576628 | 0.945417442 | 0.545733012 | 0.225672912 | 2.319642603 | 0.002000966 | 0.033523137 | 0.129778392 | 0.1027 | 0.246801977 |
| SL1344_RS16615_ | 1.004316397 | 0.872989515 | 1.114969595 | 0.24297303 | 0.20519644 | 1.117113944 | 0.114590308 | 0.16334071 | 0.332846434 | 0.2348 | 0.40279975 |
| SL1344_RS16620_ | 0.471226605 | 0.197046012 | 0.756997481 | 0.555732555 | 0.20489916 | 2.699199572 | 0.001335878 | 0.022318185 | 0.101093885 | 0.1006 | 0.24384956 |
| SL1344_RS16635_ | -1.241520653 | -1.22138814 | -1.284280009 | -0.065873147 | 0.469513483 | -0.127194894 | 0.450000002 | 0.689722655 | 0.807431392 | 0.8361 | 0.899145197 |
| SL1344_RS16640_ | -1.805492095 | -1.744808039 | -1.86524585 | -0.049278067 | 0.510625924 | -0.095091921 | 0.458361094 | 0.687198764 | 0.80382061 | 0.793 | 0.868860435 |
| SL1344_RS16645_ | 2.402126387 | 2.195968081 | 2.648821732 | 0.471756575 | 0.133978526 | 3.583035975 | 4.68E-05 | 0.020918938 | 0.101520985 | 0.1 | 0.243021319 |

| | | | | | | | | | | | |
|---|---|---|---|---|---|---|---|---|---|---|---|
| SL1344_RS16650_ | 0.798038931 | 0.717476328 | 0.951780936 | 0.214130367 | 0.21750409 | 0.926773862 | 0.175333354 | 0.253023332 | 0.435033598 | 0.3515 | 0.520699199 |
| SL1344_RS16655_ | -0.69199122 | -0.837351702 | -0.594210082 | 0.243492598 | 0.267459905 | 0.856382445 | 0.16322454 | 0.262580846 | 0.431981169 | 0.3532 | 0.5064267 |
| SL1344_RS16660_ | -0.144513063 | -0.133556133 | -0.167127686 | -7.53E-05 | 0.277081289 | -0.000385884 | 0.5 | 0.69381182 | 0.80969571 | 0.7948 | 0.87296919 |
| SL1344_RS16665_ | 1.133400274 | 0.989923602 | 1.262837234 | 0.314327847 | 0.222194985 | 1.369948744 | 0.04266677 | 0.096951167 | 0.245057325 | 0.1506 | 0.305839485 |
| SL1344_RS16675_ | 2.357100574 | 2.310834718 | 2.401721146 | 0.096604656 | 0.129476091 | 0.728329714 | 0.215856111 | 0.317389524 | 0.50015704 | 0.4266 | 0.583118334 |
| SL1344_RS16680_ | 1.829840047 | 1.841006044 | 1.817550259 | -0.022594205 | 0.096437541 | -0.21472886 | 0.397734847 | 0.502862088 | 0.648455589 | 0.6094 | 0.723887414 |
| SL1344_RS16685_ | 2.732184977 | 2.630138954 | 2.760446043 | 0.116225187 | 0.212799419 | 0.548349953 | 0.266489018 | 0.480333229 | 0.657889954 | 0.5252 | 0.692002281 |
| SL1344_RS16690_ | 2.131108859 | 2.131355215 | 2.130925175 | 0.023313754 | 0.160912816 | 0.122771668 | 0.448000002 | 0.650047853 | 0.778821776 | 0.8084 | 0.878519663 |
| SL1344_RS16695_ | 2.506178934 | 2.425690103 | 2.588223853 | 0.139691177 | 0.165375662 | 0.826108975 | 0.17800002 | 0.258667493 | 0.449777397 | 0.34 | 0.517445823 |
| SL1344_RS16700_ | 2.579383148 | 2.697819385 | 2.45854126 | -0.245605285 | 0.099270246 | -2.410024132 | 0.0020023 | 0.028249033 | 0.114925724 | 0.1018 | 0.24549496 |
| SL1344_RS16705_ | -2.645239129 | -2.976694014 | -2.419194282 | 0.545960565 | 0.487216449 | 1.055086019 | 0.124583643 | 0.201517844 | 0.361384215 | 0.2843 | 0.439195524 |
| SL1344_RS16710_ | -0.392786516 | -0.398865001 | -0.376281861 | 0.051668281 | 0.300345417 | 0.17145892 | 0.430000003 | 0.643390346 | 0.769565525 | 0.765 | 0.847482434 |
| SL1344_RS16715_ | 1.707798147 | 1.59162074 | 1.832405146 | 0.242850483 | 0.100210404 | 2.364235572 | 0.005996769 | 0.030713184 | 0.116198822 | 0.1052 | 0.249831364 |
| SL1344_RS16720_ | 0.061751844 | 0.071473787 | 0.040843707 | 0.001050992 | 0.25864942 | 0.003979897 | 0.498 | 0.684194487 | 0.800356893 | 0.7751 | 0.859230788 |
| SL1344_RS16725_ | -1.715762547 | -1.669050308 | -1.774607434 | -0.110146654 | 0.411734128 | -0.241595232 | 0.395333337 | 0.555847653 | 0.697701459 | 0.677 | 0.778086611 |
| SL1344_RS16730_ | -1.329476851 | -1.308818585 | -1.342816555 | -0.032739993 | 0.337169154 | -0.082962829 | 0.462666668 | 0.609216199 | 0.739418193 | 0.7246 | 0.815621384 |
| SL1344_RS16745_ | 0.400673225 | 0.884927566 | -0.116904895 | -1.012141227 | 0.262314893 | -3.883563421 | 4.68E-05 | 0.00862398 | 0.060035002 | 0.1 | 0.243021319 |
| SL1344_RS16750_ | 1.128497179 | 1.716721458 | 0.43723571 | -1.274162927 | 0.145058153 | -8.757947792 | 4.68E-05 | 0.001133298 | 0.020263742 | 0.1 | 0.243021319 |
| SL1344_RS16755_ | 0.220631352 | 0.767915728 | -0.619981776 | -1.357960782 | 0.296989896 | -4.453036555 | 4.68E-05 | 0.011388286 | 0.07039113 | 0.1 | 0.243021319 |
| SL1344_RS16775_ | -1.102544789 | -1.145812548 | -1.072709162 | 0.067966436 | 0.327156643 | 0.19513975 | 0.415056632 | 0.582799047 | 0.719441604 | 0.691 | 0.792455622 |
| SL1344_RS16780_ | -0.899301523 | -0.88551957 | -0.928032497 | -0.027634748 | 0.344656751 | -0.072463192 | 0.469020654 | 0.670334314 | 0.792283053 | 0.7721 | 0.858943594 |
| SL1344_RS16785_ | -2.170559185 | -2.174758588 | -2.16672547 | 0.009158514 | 0.401649153 | 0.016670202 | 0.492338441 | 0.563110594 | 0.69971503 | 0.6639 | 0.769252462 |
| SL1344_RS16790_ | -1.995599323 | -1.897619551 | -2.116940285 | -0.226428593 | 0.385306129 | -0.529404237 | 0.269153908 | 0.408372859 | 0.56388657 | 0.5128 | 0.641213677 |
| SL1344_RS16795_ | 0.52217223 | 0.566562069 | 0.50080239 | -0.056179562 | 0.187945899 | -0.276540835 | 0.393333337 | 0.609132637 | 0.740101912 | 0.7019 | 0.803982102 |
| SL1344_RS16800_ | 1.07865494 | 1.003683278 | 1.133108283 | 0.020101938 | 0.375458326 | 0.053007441 | 0.483677549 | 0.824692326 | 0.900472853 | 0.9598 | 0.976094139 |
| SL1344_RS16805_ | -3.063365496 | -2.878371433 | -3.302393834 | -0.42069826 | 0.591049072 | -0.678140238 | 0.225183226 | 0.348738 | 0.509868378 | 0.4511 | 0.589676741 |
| SL1344_RS16810_ | -3.705330636 | -3.751754128 | -3.64640494 | 0.103984178 | 0.873105186 | 0.10503591 | 0.453333335 | 0.651780397 | 0.775682202 | 0.7603 | 0.844478183 |
| SL1344_RS16815_ | -2.879970093 | -2.811109784 | -2.938046186 | -0.212999438 | 0.728925808 | -0.260204256 | 0.388666671 | 0.593830743 | 0.732848845 | 0.7287 | 0.820625932 |
| SL1344_RS16820_ | -3.460473702 | -3.378211439 | -3.582700863 | -0.209798546 | 0.834760149 | -0.243086707 | 0.394666671 | 0.594527548 | 0.731005551 | 0.7043 | 0.801636792 |
| SL1344_RS16825_ | -3.094546733 | -3.144123107 | -3.046240309 | 0.063814214 | 0.730937672 | 0.075740376 | 0.467688209 | 0.66611068 | 0.786741964 | 0.7719 | 0.852654952 |
| SL1344_RS16830_ | -4.891036108 | -4.972503886 | -4.732808769 | 0.409536233 | 1.561892716 | 0.236548325 | 0.402000004 | 0.613349038 | 0.744983294 | 0.7347 | 0.824612946 |
| SL1344_RS16835_ | -4.169587795 | -4.025713937 | -4.456854265 | -0.409687022 | 1.067719723 | -0.367370236 | 0.340000007 | 0.522887347 | 0.673273194 | 0.6515 | 0.760694344 |
| SL1344_RS16840_ | -2.111920301 | -1.993523498 | -2.209321228 | -0.197138441 | 0.459528233 | -0.40089767 | 0.327781486 | 0.507643889 | 0.655997239 | 0.6167 | 0.734342847 |
| SL1344_RS16845_ | -3.352072278 | -3.412589979 | -3.239344476 | 0.072798586 | 0.846312091 | 0.075109159 | 0.462000001 | 0.684912562 | 0.800725696 | 0.7738 | 0.856775928 |
| SL1344_RS16850_ | 1.309225254 | 1.270864651 | 1.388603624 | 0.101924449 | 0.183922861 | 0.523359139 | 0.297801475 | 0.491689951 | 0.652783499 | 0.5871 | 0.722504663 |
| SL1344_RS16855_ | 1.794258901 | 1.630392979 | 1.944575554 | 0.317123062 | 0.12450368 | 2.517517642 | 0.002000966 | 0.028487119 | 0.115732763 | 0.1012 | 0.244707848 |
| SL1344_RS16860_ | 2.93440008 | 2.894040288 | 3.056968719 | 0.148713482 | 0.106398498 | 1.237806471 | 0.113924086 | 0.153313919 | 0.323181914 | 0.2526 | 0.421559612 |
| SL1344_RS16865_ | 1.735742348 | 1.688301424 | 1.822147878 | 0.128102044 | 0.129356803 | 0.882951161 | 0.179213878 | 0.25898875 | 0.432091025 | 0.3643 | 0.524017203 |
| SL1344_RS16870_ | 1.53577386 | 1.512844552 | 1.571747044 | 0.056674032 | 0.11962233 | 0.437347825 | 0.317333341 | 0.477174645 | 0.624209864 | 0.5754 | 0.696183725 |
| SL1344_RS16875_ | 2.502787701 | 2.53207516 | 2.471725055 | -0.060866952 | 0.09976074 | -0.589877468 | 0.263157906 | 0.405457004 | 0.569678374 | 0.5147 | 0.649538743 |
| SL1344_RS16880_ | -1.22275695 | -1.301042758 | -1.164299031 | 0.142909626 | 0.29976507 | 0.450991141 | 0.307128589 | 0.442196872 | 0.59725935 | 0.5493 | 0.675403468 |
| SL1344_RS16885_ | -3.343621351 | -3.224513633 | -3.541785088 | -0.392193115 | 0.882575805 | -0.407388948 | 0.304463699 | 0.430474803 | 0.605422936 | 0.5717 | 0.690969502 |
| SL1344_RS16890_ | -0.627603108 | -0.527293002 | -0.774877957 | -0.253806744 | 0.274261701 | -0.885274077 | 0.16055965 | 0.255689008 | 0.425258196 | 0.3486 | 0.498338528 |
| SL1344_RS16895_ | -0.596317683 | -0.686171539 | -0.53583613 | 0.137342263 | 0.259570822 | 0.517797106 | 0.291805472 | 0.443344066 | 0.598651533 | 0.5432 | 0.672673129 |
| SL1344_RS16900_ | 1.273128639 | 1.261382579 | 1.288545156 | -0.012888305 | 0.217976739 | -0.048048494 | 0.476349101 | 0.696439196 | 0.812769423 | 0.8428 | 0.903221585 |
| SL1344_RS16905_ | 1.176070795 | 1.126626397 | 1.248679969 | 0.117985283 | 0.133769719 | 0.850182967 | 0.183877435 | 0.289174211 | 0.454278278 | 0.3913 | 0.541175969 |
| SL1344_RS16910_ | -3.470550927 | -4.091596961 | -3.1475604 | 0.943033176 | 0.8877456 | 0.986013289 | 0.123251198 | 0.205564797 | 0.383937292 | 0.2726 | 0.432761911 |
| SL1344_RS16915_ | -3.348083504 | -3.435815843 | -3.251618419 | 0.18225687 | 0.566711521 | 0.315319721 | 0.365333339 | 0.487033179 | 0.63300626 | 0.592 | 0.707754832 |
| SL1344_RS16920_ | -0.410063445 | -0.439045702 | -0.38501856 | 0.053029499 | 0.206005088 | 0.23676523 | 0.39573618 | 0.501361741 | 0.64568754 | 0.6154 | 0.727467384 |
| SL1344_RS16925_ | -2.516331816 | -2.468577442 | -2.591322377 | -0.108150297 | 0.480981241 | -0.214546394 | 0.41266667 | 0.5624425 | 0.701429378 | 0.6847 | 0.784226421 |
| SL1344_RS16930_ | -4.336060448 | -4.758767703 | -4.00206438 | 0.74117599 | 0.923005436 | 0.755810255 | 0.189873436 | 0.293064002 | 0.459210542 | 0.3842 | 0.532900983 |
| SL1344_RS16935_ | 0.271277632 | 0.067210242 | 0.403371037 | 0.344656569 | 0.25499883 | 1.299003876 | 0.068666729 | 0.130905913 | 0.291238993 | 0.186 | 0.345917212 |
| SL1344_RS16940_ | 5.324814609 | 5.143482689 | 5.498478907 | 0.37791427 | 0.169930636 | 2.06664495 | 4.68E-05 | 0.028114826 | 0.12443594 | 0.1 | 0.243021319 |

| | | | | | | | | | | | |
|---|---|---|---|---|---|---|---|---|---|---|---|
| SL1344_RS16945_ | 6.036233123 | 5.976197878 | 6.106726366 | 0.133399128 | 0.067832927 | 1.927405323 | 0.004667653 | 0.039136091 | 0.147450164 | 0.104 | 0.24861331 |
| SL1344_RS16950_ | 5.463041609 | 5.430175681 | 5.550111829 | 0.126622927 | 0.098612412 | 1.272584046 | 0.015323415 | 0.180075033 | 0.373671395 | 0.118 | 0.266449439 |
| SL1344_RS16955_ | 7.683874077 | 7.442598046 | 7.801314826 | 0.353609849 | 0.141967804 | 2.718390603 | 4.68E-05 | 0.007644491 | 0.061850405 | 0.1 | 0.243021319 |
| SL1344_RS16960_ | 3.685787917 | 3.536519693 | 3.818643894 | 0.276687041 | 0.10185435 | 2.629625629 | 4.68E-05 | 0.021085328 | 0.101396029 | 0.1002 | 0.243297692 |
| SL1344_RS16965_ | 2.744579315 | 2.554462315 | 2.81450922 | 0.253786178 | 0.110942849 | 1.971999165 | 0.012658588 | 0.067901561 | 0.199323363 | 0.1128 | 0.259869637 |
| SL1344_RS16970_ | 6.442709685 | 6.359994693 | 6.534315145 | 0.177153782 | 0.050982127 | 3.298234715 | 4.68E-05 | 0.076825259 | 0.223946988 | 0.1 | 0.243021319 |
| SL1344_RS16980_ | 3.802466846 | 3.641836129 | 3.888693521 | 0.236908941 | 0.091723862 | 2.428340406 | 0.001335878 | 0.033507226 | 0.131952914 | 0.1015 | 0.245145479 |
| SL1344_RS16985_ | 4.691875068 | 4.681061974 | 4.722815695 | 0.042942228 | 0.109904089 | 0.295118204 | 0.34333334 | 0.45155078 | 0.641743865 | 0.7057 | 0.817479564 |
| SL1344_RS16990_ | 0.969312575 | 0.829891131 | 1.024307225 | 0.173582854 | 0.234953897 | 0.684541959 | 0.247834789 | 0.37934277 | 0.561163073 | 0.4786 | 0.641690478 |
| SL1344_RS16995_ | 0.574252196 | 0.681343712 | 0.482356158 | -0.219580458 | 0.207774137 | -1.064661796 | 0.10926053 | 0.187424499 | 0.358627618 | 0.2507 | 0.408507754 |
| SL1344_RS17000_ | 4.926122406 | 4.935358027 | 4.68587045 | -0.245235548 | 0.059039615 | -0.626808556 | 0.33333334 | 0.497415974 | 0.67971703 | 0.7 | 0.829598552 |
| SL1344_RS17005_ | 7.086131993 | 7.121588073 | 7.043186459 | -0.080535673 | 0.05118309 | -1.472614876 | 0.005330642 | 0.161502768 | 0.349705583 | 0.1043 | 0.248986093 |
| SL1344_RS17010_ | 3.14713786 | 3.181508777 | 2.956880519 | -0.223774644 | 0.170157643 | -1.119957285 | 0.12658231 | 0.191975359 | 0.38496798 | 0.2659 | 0.43562186 |
| SL1344_RS17015_ | 0.997160432 | 1.18620586 | 0.863472706 | -0.328890904 | 0.209893627 | -1.517913837 | 0.039307241 | 0.084792085 | 0.224267092 | 0.1418 | 0.296678241 |
| SL1344_RS17030_ | 3.552815618 | 3.529961449 | 3.566698493 | 0.036961627 | 0.076186155 | 0.46477253 | 0.313333341 | 0.487783255 | 0.64653462 | 0.6125 | 0.733259832 |
| SL1344_RS17035_ | 4.720363751 | 4.515407666 | 4.903847577 | 0.372410333 | 0.101325761 | 3.601299444 | 4.68E-05 | 0.007529837 | 0.058756005 | 0.1 | 0.243021319 |
| SL1344_RS17040_ | 3.696480866 | 3.701880816 | 3.692727902 | -0.003494523 | 0.077639315 | -0.039019325 | 0.483011326 | 0.711269376 | 0.820799827 | 0.8096 | 0.881995631 |
| SL1344_RS17045_ | 2.98062005 | 2.962918228 | 3.240714592 | 0.273446939 | 0.149193697 | 0.717446316 | 0.33133334 | 0.409093771 | 0.604562901 | 0.6912 | 0.822692581 |
| SL1344_RS17050_ | 2.601854278 | 2.529857127 | 2.722022805 | 0.194687373 | 0.125720607 | 1.486375344 | 0.045969449 | 0.104804599 | 0.257825708 | 0.153 | 0.310663416 |
| SL1344_RS17060_ | 2.959864789 | 2.729543126 | 3.157156901 | 0.37201443 | 0.246525472 | 1.5283142 | 0.069287204 | 0.078005265 | 0.223660378 | 0.1576 | 0.323783791 |
| SL1344_RS17065_ | 6.025973688 | 6.015884194 | 6.113748013 | 0.103208356 | 0.349689068 | 0.343302753 | 0.33333334 | 0.820797099 | 0.902308818 | 0.7 | 0.829598552 |
| SL1344_RS17070_ | 6.520356023 | 6.47339693 | 6.565658775 | 0.060297174 | 0.174673416 | 0.186619392 | 0.441705532 | 0.866378989 | 0.928384289 | 0.9997 | 0.999830616 |
| SL1344_RS17075_ | 4.290369735 | 4.105119239 | 4.44881519 | 0.345815686 | 0.064634072 | 5.398150697 | 4.68E-05 | 0.003357903 | 0.037218702 | 0.1 | 0.243021319 |
| SL1344_RS17080_ | 0.111969519 | 0.044798315 | 0.220164883 | 0.180746239 | 0.293856135 | 0.568390561 | 0.273333344 | 0.416402263 | 0.591881457 | 0.5424 | 0.683635649 |
| SL1344_RS17085_ | 3.776633098 | 3.748370864 | 3.812006741 | 0.079650248 | 0.106745496 | 0.75776331 | 0.17721521 | 0.277134543 | 0.472916848 | 0.3595 | 0.523440556 |
| SL1344_RS17090_ | 0.980266683 | 0.916970917 | 1.051649028 | 0.141999089 | 0.187920111 | 0.719901206 | 0.20333335 | 0.309226997 | 0.48506103 | 0.4149 | 0.56695279 |
| SL1344_RS17095_ | 2.45016796 | 2.422446633 | 2.483370204 | 0.078817137 | 0.123272458 | 0.628255221 | 0.241333346 | 0.35921894 | 0.540830644 | 0.4977 | 0.633347833 |
| SL1344_RS17100_ | 2.248407897 | 2.235204869 | 2.256160156 | 0.019558048 | 0.115905179 | 0.154072581 | 0.431045972 | 0.63680264 | 0.76366209 | 0.751 | 0.837005526 |
| SL1344_RS17105_ | 1.12954071 | 1.105500164 | 1.161066812 | 0.023969488 | 0.198069647 | 0.113353905 | 0.454666668 | 0.702756135 | 0.815508494 | 0.8036 | 0.876996036 |
| SL1344_RS17110_ | 1.422598603 | 1.414526214 | 1.435432969 | 0.014661612 | 0.141750333 | 0.103215934 | 0.458000002 | 0.611311541 | 0.743153835 | 0.72 | 0.813264519 |
| SL1344_RS17115_ | 2.075827785 | 2.020867163 | 2.202875652 | 0.174776306 | 0.136559723 | 1.180718048 | 0.112000036 | 0.169954689 | 0.341758035 | 0.2539 | 0.422346324 |
| SL1344_RS17120_ | 2.66413726 | 2.600553678 | 2.741407684 | 0.137255071 | 0.085572572 | 1.48300234 | 0.057295212 | 0.095111501 | 0.233770335 | 0.1672 | 0.324362476 |
| SL1344_RS17125_ | 2.689930844 | 2.599057773 | 2.759277288 | 0.165114281 | 0.091820245 | 1.774783659 | 0.012658588 | 0.070974202 | 0.196628681 | 0.117 | 0.264108856 |
| SL1344_RS17130_ | 3.173096242 | 3.044413479 | 3.281039244 | 0.25197534 | 0.11591637 | 2.258427513 | 0.000673369 | 0.039971778 | 0.146326936 | 0.101 | 0.244353037 |
| SL1344_RS17135_ | 2.847257449 | 2.755695668 | 3.01697729 | 0.26684175 | 0.114898786 | 2.182466897 | 0.003998486 | 0.051531778 | 0.170319825 | 0.1016 | 0.245275619 |
| SL1344_RS17140_ | 2.998330063 | 2.91655481 | 3.214021921 | 0.283173172 | 0.110981084 | 2.13590368 | 0.011992386 | 0.060087492 | 0.187246704 | 0.1089 | 0.255039485 |
| SL1344_RS17145_ | 2.952181819 | 2.837805654 | 3.126631462 | 0.282039943 | 0.09404063 | 2.872236779 | 0.00067292 | 0.021709997 | 0.100394242 | 0.1003 | 0.243440998 |
| SL1344_RS17150_ | 4.152501871 | 4.00078956 | 4.261057878 | 0.270739924 | 0.089202896 | 3.060889002 | 4.68E-05 | 0.013577536 | 0.079488574 | 0.1 | 0.243021319 |
| SL1344_RS17155_ | 2.363618319 | 2.378272424 | 2.334765685 | -0.049954094 | 0.113212794 | -0.42253048 | 0.33133334 | 0.491073667 | 0.650960377 | 0.6363 | 0.748926327 |
| SL1344_RS17160_ | 2.458476306 | 2.340793674 | 2.51334372 | 0.154448642 | 0.111764322 | 1.287607523 | 0.100599641 | 0.143614032 | 0.308784149 | 0.2264 | 0.393472569 |
| SL1344_RS17165_ | 3.259850595 | 3.152744649 | 3.359698179 | 0.206686369 | 0.06644453 | 3.065429599 | 0.000673369 | 0.016178729 | 0.08225392 | 0.1007 | 0.244004454 |
| SL1344_RS17170_ | 1.72584438 | 1.597113037 | 1.832888996 | 0.238756367 | 0.118329781 | 2.008827343 | 0.008661422 | 0.054583525 | 0.167411829 | 0.1108 | 0.256800614 |
| SL1344_RS17175_ | -1.221525047 | -1.318212357 | -1.10016194 | 0.220948695 | 0.299075152 | 0.709590626 | 0.214523667 | 0.315474029 | 0.477576619 | 0.4157 | 0.558131081 |
| SL1344_RS17180_ | 1.839058828 | 2.25168069 | 1.286409797 | -0.956516008 | 0.199903386 | -4.84473855 | 4.68E-05 | 0.007735369 | 0.057311005 | 0.1 | 0.243021319 |
| SL1344_RS17185_ | 1.697483153 | 2.150396493 | 1.326190287 | -0.785741462 | 0.185195264 | -4.17593021 | 4.68E-05 | 0.006319334 | 0.052342375 | 0.1 | 0.243021319 |
| SL1344_RS17190_ | 2.734139881 | 2.866693333 | 2.540870536 | -0.325207304 | 0.131127562 | -2.46279483 | 0.000673369 | 0.03310612 | 0.129997913 | 0.1011 | 0.244565873 |
| SL1344_RS17195_ | -1.60033116 | -1.610664684 | -1.590560696 | 0.048373526 | 0.429059374 | 0.110723357 | 0.454000002 | 0.638570837 | 0.764468527 | 0.759 | 0.840895129 |
| SL1344_RS17200_ | -3.983880311 | -4.450438291 | -3.680507704 | 0.769996047 | 1.048032777 | 0.678763344 | 0.193333352 | 0.312565863 | 0.495000074 | 0.4028 | 0.554388496 |
| SL1344_RS17205_ | 3.22583938 | 3.21361199 | 3.251436056 | 0.034988762 | 0.126069954 | 0.257252995 | 0.413724187 | 0.675742813 | 0.799252488 | 0.8268 | 0.894362631 |
| SL1344_RS17210_ | 2.004475347 | 2.031171449 | 1.836603442 | -0.192741737 | 0.236403534 | -0.725806354 | 0.286666676 | 0.358810954 | 0.556290053 | 0.5823 | 0.728209324 |
| SL1344_RS17215_ | -2.594840723 | -2.546053884 | -2.651705374 | -0.068437534 | 0.530530938 | -0.124244482 | 0.442000002 | 0.598355313 | 0.73013305 | 0.7061 | 0.799523178 |
| SL1344_RS17220_ | 0.862352688 | 0.668580394 | 1.111753262 | 0.454034209 | 0.20266974 | 2.218341643 | 0.004664544 | 0.039672083 | 0.142383396 | 0.1033 | 0.247538956 |
| SL1344_RS17225_ | 2.164740284 | 2.007889457 | 2.286288704 | 0.283332121 | 0.105050773 | 2.663438577 | 0.00067292 | 0.032269572 | 0.123943105 | 0.1003 | 0.243439983 |

| | | | | | | | | | | | |
|---|---|---|---|---|---|---|---|---|---|---|---|
| SL1344_RS17230_ | -1.005268803 | -0.888030646 | -1.17156571 | -0.293435353 | 0.307049908 | -0.896231315 | 0.170000022 | 0.253561346 | 0.42082629 | 0.3383 | 0.492364822 |
| SL1344_RS17235_ | -1.305852839 | -1.266417281 | -1.363133288 | -0.07056073 | 0.425050468 | -0.162781426 | 0.429047304 | 0.658088766 | 0.781767168 | 0.7649 | 0.84970329 |
| SL1344_RS17240_ | -1.574022478 | -1.531033903 | -1.602116762 | -0.095401781 | 0.447149607 | -0.200104521 | 0.408394407 | 0.634016947 | 0.7619069 | 0.7498 | 0.835772762 |
| SL1344_RS17245_ | -0.721979397 | -0.734126873 | -0.710412953 | 0.018638102 | 0.248329952 | 0.073285615 | 0.466666668 | 0.568080181 | 0.70372693 | 0.6808 | 0.779883753 |
| SL1344_RS17250_ | -1.86819653 | -1.90452916 | -1.820471262 | 0.082952734 | 0.37012981 | 0.202949539 | 0.403064627 | 0.544836349 | 0.684770259 | 0.6496 | 0.755725488 |
| SL1344_RS17255_ | 2.941998214 | 2.872951019 | 3.001244636 | 0.128723179 | 0.072488933 | 1.727986014 | 0.026666836 | 0.072703575 | 0.19372383 | 0.1372 | 0.287605247 |
| SL1344_RS17260_ | 5.272382198 | 5.23537649 | 5.284117652 | 0.050727533 | 0.073814035 | 0.632550532 | 0.269153908 | 0.373770519 | 0.567512798 | 0.5305 | 0.684872478 |
| SL1344_RS17270_ | 0.190972693 | -0.028809789 | 0.680943711 | 0.67532085 | 0.280943209 | 2.108673871 | 0.010659989 | 0.05043909 | 0.166767108 | 0.11 | 0.256246246 |
| SL1344_RS17275_ | 7.941898619 | 7.912648388 | 8.125048893 | 0.209349802 | 0.166598509 | 1.290610472 | 0.018000252 | 0.152730564 | 0.339524415 | 0.1164 | 0.265925571 |
| SL1344_RS17280_ | 6.890526647 | 6.828352448 | 6.974374901 | 0.192703147 | 0.11083919 | 1.901477439 | 4.68E-05 | 0.041078328 | 0.154258336 | 0.1 | 0.243021319 |
| SL1344_RS17285_ | 1.599693273 | 1.482490796 | 1.65937534 | 0.169705782 | 0.150378631 | 1.042389162 | 0.137241867 | 0.210249607 | 0.385643268 | 0.3053 | 0.467085997 |
| SL1344_RS17290_ | 3.099207051 | 3.108523909 | 3.092524945 | -0.007613641 | 0.094141626 | -0.080923242 | 0.469020654 | 0.674011162 | 0.79327564 | 0.7805 | 0.859308709 |
| SL1344_RS17295_ | 2.588083643 | 2.612784309 | 2.539551854 | -0.054657374 | 0.150114034 | -0.330363733 | 0.356666672 | 0.688755069 | 0.807389854 | 0.7404 | 0.842680838 |
| SL1344_RS17300_ | 2.470922827 | 2.299071326 | 2.720862064 | 0.412699039 | 0.123083366 | 3.234325115 | 4.68E-05 | 0.01463092 | 0.080816927 | 0.1 | 0.243021319 |
| SL1344_RS17305_ | -2.885272942 | -3.005120009 | -2.710580048 | 0.347872239 | 0.644496783 | 0.526666841 | 0.277333344 | 0.415922859 | 0.580028139 | 0.5373 | 0.664155153 |
| SL1344_RS17310_ | -1.458328425 | -1.394440594 | -1.55604052 | -0.152802152 | 0.406560995 | -0.349852308 | 0.355096608 | 0.55318887 | 0.69672836 | 0.673 | 0.777625396 |
| SL1344_RS17315_ | -0.650318403 | -0.828630097 | -0.594296163 | 0.205863408 | 0.345506972 | 0.501966723 | 0.334000007 | 0.547704978 | 0.699054271 | 0.6619 | 0.785940421 |
| SL1344_RS17320_ | -4.167549762 | -4.397026242 | -4.076231427 | 0.330398384 | 1.588575863 | 0.177725315 | 0.430000003 | 0.658751171 | 0.788987393 | 0.8349 | 0.901768322 |
| SL1344_RS17325_ | -1.97582167 | -2.147072857 | -1.794934633 | 0.352813395 | 0.352677462 | 0.957368463 | 0.152666692 | 0.234972355 | 0.387472176 | 0.3177 | 0.467035858 |
| SL1344_RS17330_ | -1.506376842 | -1.53214991 | -1.462296996 | 0.078174228 | 0.430872727 | 0.162941527 | 0.432666669 | 0.647326486 | 0.772233298 | 0.7602 | 0.844769994 |
| SL1344_RS17335_ | -2.565409452 | -2.489428947 | -2.643208005 | -0.141815739 | 0.54944453 | -0.238625888 | 0.394000004 | 0.567401764 | 0.705231828 | 0.6698 | 0.774472013 |
| SL1344_RS17340_ | 1.335741633 | 1.319888564 | 1.350048602 | 0.029381513 | 0.140062792 | 0.197188911 | 0.417721522 | 0.584277174 | 0.720062838 | 0.6875 | 0.788900664 |
| SL1344_RS17345_ | 1.544623149 | 1.473430826 | 1.620373664 | 0.151278832 | 0.122267415 | 1.146222704 | 0.094603641 | 0.152559818 | 0.307546769 | 0.2283 | 0.385856584 |
| SL1344_RS17350_ | 3.954038116 | 4.343140998 | 3.429939106 | -0.905783009 | 0.097776725 | -9.148048648 | 4.68E-05 | 0.002450983 | 0.032811768 | 0.1 | 0.243021319 |
| SL1344_RS17355_ | 2.362382402 | 2.301558415 | 2.403327802 | 0.107962445 | 0.108198831 | 0.947492034 | 0.14923387 | 0.2393886 | 0.412246076 | 0.329 | 0.482096793 |
| SL1344_RS17360_ | 0.614928306 | 0.59682361 | 0.658334523 | 0.057119675 | 0.268254799 | 0.184582772 | 0.431712194 | 0.739865286 | 0.842548759 | 0.826 | 0.896399072 |
| SL1344_RS17365_ | 1.318460593 | 1.334250001 | 1.297744415 | -0.032385195 | 0.142950208 | -0.225397637 | 0.409333337 | 0.58478418 | 0.719780185 | 0.6916 | 0.791059017 |
| SL1344_RS17370_ | -1.07438073 | -1.349707373 | -0.8357572 | 0.524597167 | 0.36922525 | 1.360905559 | 0.052631662 | 0.112739395 | 0.263194299 | 0.1697 | 0.327229602 |
| SL1344_RS17375_ | 0.618407922 | 0.565770286 | 0.727876591 | 0.174968237 | 0.238106689 | 0.660130169 | 0.246502344 | 0.367007978 | 0.547117458 | 0.5068 | 0.650088124 |
| SL1344_RS17380_ | -0.496114207 | -0.556120705 | -0.375388544 | 0.167012652 | 0.321563785 | 0.46916744 | 0.319120594 | 0.515268105 | 0.667283447 | 0.64 | 0.754091857 |
| SL1344_RS17385_ | -2.793666965 | -2.685969816 | -2.908672759 | -0.229665625 | 0.477752142 | -0.449425723 | 0.308461034 | 0.43696994 | 0.588336751 | 0.5454 | 0.66873469 |
| SL1344_RS17390_ | -0.011019048 | -0.044672748 | 0.045174477 | 0.076598747 | 0.257173422 | 0.28217097 | 0.38241173 | 0.594021475 | 0.730908586 | 0.6979 | 0.799296946 |
| SL1344_RS17395_ | 2.238629826 | 2.199401094 | 2.389059609 | 0.186711344 | 0.185599407 | 0.86887434 | 0.216666682 | 0.289827973 | 0.486313873 | 0.4317 | 0.60252399 |
| SL1344_RS17400_ | 3.593492266 | 3.532301013 | 3.720472416 | 0.175460391 | 0.085547652 | 1.815078556 | 0.026000173 | 0.065073775 | 0.192607164 | 0.1277 | 0.280026071 |
| SL1344_RS17405_ | 3.649426178 | 3.619156238 | 3.761391897 | 0.116053688 | 0.133004075 | 0.725382581 | 0.271152576 | 0.387323464 | 0.579267249 | 0.547 | 0.704309501 |
| SL1344_RS17410_ | 2.960405967 | 2.919312063 | 3.051643598 | 0.136778695 | 0.128381155 | 1.084796757 | 0.084666716 | 0.19899709 | 0.388894962 | 0.2116 | 0.369493352 |
| SL1344_RS17415_ | 2.604723528 | 2.565742873 | 2.648239025 | 0.028495929 | 0.210135584 | 0.137456543 | 0.443333335 | 0.830199877 | 0.903874494 | 0.9004 | 0.942493195 |
| SL1344_RS17420_ | 3.632680204 | 3.610052572 | 3.677758064 | 0.072650459 | 0.085538373 | 0.808623587 | 0.169886764 | 0.27187998 | 0.462435579 | 0.3575 | 0.514318435 |
| SL1344_RS17425_ | 5.006471261 | 4.930410045 | 5.028687233 | 0.090522638 | 0.075162164 | 1.156568687 | 0.137908089 | 0.158578629 | 0.336526545 | 0.2658 | 0.445036086 |
| SL1344_RS17430_ | -4.776369272 | -5.209872689 | -4.446484118 | 0.357978862 | 1.842023967 | 0.188184877 | 0.413333337 | 0.674794213 | 0.795958254 | 0.8072 | 0.878261427 |
| SL1344_RS17440_ | 0.989560523 | 0.970465726 | 1.026009246 | 0.112300126 | 0.292621529 | 0.351599385 | 0.360666672 | 0.565302913 | 0.718869446 | 0.7488 | 0.838528587 |
| SL1344_RS17445_ | 0.399150282 | 0.234952905 | 0.54378901 | 0.297444086 | 0.208430291 | 1.370941421 | 0.067288538 | 0.119705371 | 0.271337866 | 0.1811 | 0.341799628 |
| SL1344_RS17450_ | -0.302503307 | -0.361207044 | -0.183382804 | 0.158009826 | 0.333008741 | 0.431261659 | 0.332000007 | 0.561238514 | 0.70910449 | 0.6872 | 0.794822889 |
| SL1344_RS17455_ | 4.840169435 | 4.604310736 | 5.067129871 | 0.394914085 | 0.123290533 | 3.50521191 | 4.68E-05 | 0.004903098 | 0.0482805 | 0.1 | 0.243021319 |
| SL1344_RS17460_ | 5.831824503 | 5.608108692 | 6.087954395 | 0.480687989 | 0.066722373 | 7.199167152 | 4.68E-05 | 0.000865375 | 0.020629366 | 0.1 | 0.243021319 |
| SL1344_RS17465_ | -1.111723681 | -1.154709478 | -1.043210035 | 0.135537637 | 0.383329408 | 0.335850745 | 0.363333339 | 0.536278889 | 0.686134952 | 0.6774 | 0.777959139 |
| SL1344_RS17470_ | 1.820955416 | 1.691649691 | 1.990417701 | 0.27866248 | 0.159551692 | 1.673645304 | 0.040639683 | 0.070706727 | 0.201423072 | 0.1392 | 0.295772802 |
| SL1344_RS17475_ | 2.536971152 | 2.31224642 | 2.63577907 | 0.314399396 | 0.118519981 | 2.3651204 | 0.004667653 | 0.045383749 | 0.156706278 | 0.102 | 0.245819794 |
| SL1344_RS17480_ | 5.396413887 | 5.188054863 | 5.569921356 | 0.407498953 | 0.111118946 | 3.750625832 | 4.68E-05 | 0.003986842 | 0.043961483 | 0.1 | 0.243021319 |
| SL1344_RS17490_ | 0.604362622 | 0.393794985 | 0.787464134 | 0.399832357 | 0.168832616 | 2.30425163 | 0.004667653 | 0.035411892 | 0.128013607 | 0.1036 | 0.248009252 |
| SL1344_RS17495_ | 0.622349412 | 0.530928596 | 0.748436431 | 0.250147871 | 0.226845237 | 1.054129184 | 0.093937419 | 0.18821093 | 0.367930647 | 0.2327 | 0.389820445 |
| SL1344_RS17500_ | 0.724306035 | 0.480603209 | 0.907670078 | 0.429458917 | 0.166198285 | 2.558133402 | 0.001335878 | 0.031886674 | 0.12175048 | 0.1017 | 0.245343993 |
| SL1344_RS17505_ | -2.664350215 | -2.72330613 | -2.596008284 | 0.137671888 | 0.454603454 | 0.287773493 | 0.370666672 | 0.496776474 | 0.640295074 | 0.5926 | 0.708502166 |

| | | | | | | | | | | | |
|---|---|---|---|---|---|---|---|---|---|---|---|
| SL1344_RS17510_ | -2.949638255 | -2.901509452 | -2.996234099 | -0.074872701 | 0.565475576 | -0.132424155 | 0.444000002 | 0.571133321 | 0.70895474 | 0.6853 | 0.786213629 |
| SL1344_RS17515_ | -0.209247323 | -0.222840737 | -0.190981296 | 0.032040315 | 0.217973345 | 0.132885544 | 0.444370422 | 0.56948403 | 0.705956737 | 0.6768 | 0.780919068 |
| SL1344_RS17520_ | 0.490562947 | 0.106350125 | 0.921441803 | 0.835063869 | 0.20134128 | 4.12225317 | 4.68E-05 | 0.006510912 | 0.051556571 | 0.1 | 0.243021319 |
| SL1344_RS17530_ | -0.604009545 | -0.918870167 | -0.297729386 | 0.622853738 | 0.222623009 | 2.750227304 | 0.001335878 | 0.020681104 | 0.093003532 | 0.1011 | 0.244566881 |
| SL1344_RS17540_ | 2.29033824 | 1.707120518 | 2.345388245 | 0.65125305 | 0.753999659 | 0.902180186 | 0.20533335 | 0.160708467 | 0.349163252 | 0.3698 | 0.565910239 |
| SL1344_RS17545_ | 0.04793555 | -0.052395482 | 0.347081463 | 0.432186453 | 0.42270265 | 0.961409919 | 0.15866669 | 0.214453699 | 0.405140904 | 0.3177 | 0.499838551 |
| SL1344_RS17550_ | -0.06088896 | -0.556168652 | 0.021793944 | 0.645755009 | 0.967910964 | 0.75538514 | 0.20719522 | 0.260553339 | 0.464693154 | 0.3772 | 0.573421677 |
| SL1344_RS17555_ | 0.80710821 | 0.993427623 | 0.500750459 | -0.472883463 | 0.200206273 | -2.301117207 | 0.005334196 | 0.039216561 | 0.141172425 | 0.107 | 0.252546089 |
| SL1344_RS17560_ | 1.002311493 | 0.952035895 | 1.140176606 | 0.178692046 | 0.183388536 | 0.853934096 | 0.195203216 | 0.288219857 | 0.466435843 | 0.4102 | 0.567351707 |
| SL1344_RS17565_ | 1.948420378 | 1.864658457 | 2.017374759 | 0.146030307 | 0.150138319 | 0.915557563 | 0.157333357 | 0.23509385 | 0.414656567 | 0.3128 | 0.480916944 |
| SL1344_RS17570_ | 1.518124541 | 1.420190723 | 1.599792742 | 0.180103119 | 0.107978281 | 1.599502096 | 0.046635671 | 0.087362258 | 0.215588252 | 0.1547 | 0.307728525 |
| SL1344_RS17575_ | 2.869286651 | 2.803125762 | 2.987754799 | 0.189098749 | 0.107003428 | 1.657955176 | 0.03000015 | 0.085227772 | 0.226658813 | 0.1261 | 0.276451856 |
| SL1344_RS17580_ | 3.025323726 | 3.049646951 | 3.002803317 | -0.045799817 | 0.071573006 | -0.609033657 | 0.248666679 | 0.392789951 | 0.549471054 | 0.4962 | 0.629668945 |
| SL1344_RS17585_ | 0.977552561 | 0.985020237 | 0.969687528 | -0.019379632 | 0.123031076 | -0.14308959 | 0.440373087 | 0.531053404 | 0.672075816 | 0.6361 | 0.744823969 |
| SL1344_RS17590_ | 3.258554658 | 3.152026607 | 3.381044258 | 0.237023062 | 0.082588941 | 2.884103591 | 4.68E-05 | 0.023856518 | 0.105443054 | 0.1 | 0.243021319 |
| SL1344_RS17595_ | 4.14072793 | 4.04255019 | 4.228560689 | 0.193962462 | 0.06716141 | 2.916193863 | 4.68E-05 | 0.019442977 | 0.095009984 | 0.1 | 0.243021319 |
| SL1344_RS17600_ | 0.967584005 | 0.929297744 | 1.030820097 | 0.123781184 | 0.232269182 | 0.519491615 | 0.262666678 | 0.396683883 | 0.579474076 | 0.5456 | 0.676020498 |
| SL1344_RS17605_ | 1.467609942 | 1.279623005 | 1.697195955 | 0.439671025 | 0.200271549 | 2.213197476 | 4.68E-05 | 0.039515521 | 0.144978898 | 0.1007 | 0.244003918 |
| SL1344_RS17610_ | 3.195778431 | 3.106839121 | 3.308762809 | 0.215299202 | 0.105875864 | 2.030387079 | 0.004667653 | 0.054603426 | 0.175090999 | 0.1056 | 0.250340278 |
| SL1344_RS17615_ | 1.465247531 | 1.419557071 | 1.568380526 | 0.13136046 | 0.216771632 | 0.559437112 | 0.269153908 | 0.465010281 | 0.635567824 | 0.5385 | 0.692389602 |
| SL1344_RS17620_ | 0.778797609 | 0.70710372 | 0.833343032 | 0.120227744 | 0.188598183 | 0.581223242 | 0.267155241 | 0.416720241 | 0.580138804 | 0.5289 | 0.664178155 |
| SL1344_RS17625_ | -0.223313863 | -0.271613387 | -0.172481473 | 0.087993766 | 0.268419497 | 0.315277629 | 0.367754835 | 0.570941252 | 0.711535833 | 0.6714 | 0.778211744 |
| SL1344_RS17630_ | 7.493402737 | 7.364753982 | 7.614432926 | 0.257132878 | 0.050756585 | 5.108274592 | 4.68E-05 | 0.019773423 | 0.102102878 | 0.1 | 0.243021319 |
| SL1344_RS17635_ | 8.469012762 | 8.400013467 | 8.552064733 | 0.152259164 | 0.054532038 | 2.365632051 | 4.68E-05 | 0.092581994 | 0.250869374 | 0.1 | 0.243021319 |
| SL1344_RS17640_ | 7.863604604 | 7.828270421 | 7.899393794 | 0.069257385 | 0.158050713 | 0.234839047 | 0.449333335 | 0.545181874 | 0.717585054 | 0.9991 | 0.999476865 |
| SL1344_RS17645_ | 5.709014476 | 5.743429559 | 5.532869376 | -0.206975673 | 0.358789508 | -0.497400271 | 0.33333334 | 0.428182255 | 0.624399292 | 0.7 | 0.829598552 |
| SL1344_RS17650_ | 6.518822176 | 6.492628478 | 6.546927793 | 0.050959885 | 0.136052638 | 0.179234579 | 0.442371754 | 0.54416711 | 0.716472418 | 0.9874 | 0.992830027 |
| SL1344_RS17655_ | 6.647480395 | 6.562112744 | 6.763947168 | 0.202330189 | 0.046121839 | 4.372094822 | 4.68E-05 | 0.009141899 | 0.065046625 | 0.1 | 0.243021319 |
| SL1344_RS17660_ | 8.316768456 | 8.23429957 | 8.374096461 | 0.139279493 | 0.118319162 | 1.188323946 | 0.104596974 | 0.165722458 | 0.356302705 | 0.1975 | 0.379912744 |
| SL1344_RS17665_ | 6.510077338 | 6.462654447 | 6.598509675 | 0.135094294 | 0.193768597 | 0.614093404 | 0.298000009 | 0.381129742 | 0.583341442 | 0.6003 | 0.752582856 |
| SL1344_RS17670_ | 5.415340624 | 5.39473632 | 5.466341901 | 0.072727332 | 0.110975256 | 0.636518105 | 0.266489018 | 0.400925855 | 0.596454563 | 0.5106 | 0.68031311 |
| SL1344_RS17675_ | 6.549019527 | 6.529118994 | 6.618536816 | 0.098565979 | 0.104618409 | 1.025864519 | 0.114666702 | 0.189416836 | 0.384618244 | 0.2332 | 0.412406536 |
| SL1344_RS17680_ | 6.462040532 | 6.37057194 | 6.55081753 | 0.184415033 | 0.055538052 | 3.069622444 | 4.68E-05 | 0.046998514 | 0.16697051 | 0.1 | 0.243021319 |
| SL1344_RS17685_ | 7.160952313 | 7.046000115 | 7.260695099 | 0.219127786 | 0.06774256 | 3.37463823 | 4.68E-05 | 0.010463556 | 0.071400656 | 0.1 | 0.243021319 |
| SL1344_RS17690_ | 7.085349319 | 7.010276034 | 7.190286404 | 0.183564645 | 0.058299956 | 3.210664046 | 4.68E-05 | 0.026209539 | 0.118713131 | 0.1 | 0.243021319 |
| SL1344_RS17700_ | 7.77022489 | 7.666591714 | 7.89574598 | 0.231423263 | 0.077050152 | 3.068017749 | 4.68E-05 | 0.021976051 | 0.108145469 | 0.1 | 0.243021319 |
| SL1344_RS17705_ | 7.431772041 | 7.352115977 | 7.52391986 | 0.185206651 | 0.045030032 | 4.254636601 | 4.68E-05 | 0.005327282 | 0.049405278 | 0.1 | 0.243021319 |
| SL1344_RS17710_ | 7.564814689 | 7.460977769 | 7.71415641 | 0.257045027 | 0.079566899 | 3.34540163 | 4.68E-05 | 0.015808658 | 0.090242453 | 0.1 | 0.243021319 |
| SL1344_RS17715_ | 8.202879182 | 8.016436158 | 8.359298312 | 0.317641145 | 0.070206223 | 4.419304461 | 4.68E-05 | 0.004334725 | 0.045781874 | 0.1 | 0.243021319 |
| SL1344_RS17720_ | 7.141263525 | 7.004821097 | 7.23435309 | 0.233888502 | 0.076154853 | 3.137709116 | 4.68E-05 | 0.007610562 | 0.060907075 | 0.1 | 0.243021319 |
| SL1344_RS17725_ | 8.297306087 | 8.128516056 | 8.395011155 | 0.199236456 | 0.148461699 | 1.535534851 | 0.108000038 | 0.082939377 | 0.234808615 | 0.197 | 0.37920865 |
| SL1344_RS17730_ | 7.915217096 | 7.808476352 | 8.021264076 | 0.209421875 | 0.069735841 | 2.852408251 | 4.68E-05 | 0.024760186 | 0.115487544 | 0.1 | 0.243021319 |
| SL1344_RS17735_ | 7.093103802 | 7.023473236 | 7.20691229 | 0.183585721 | 0.051014812 | 3.544417793 | 4.68E-05 | 0.02134065 | 0.10489442 | 0.1 | 0.243021319 |
| SL1344_RS17740_ | 6.868652428 | 6.759910346 | 6.946836507 | 0.190799493 | 0.082813913 | 2.470767477 | 4.68E-05 | 0.025362622 | 0.116542652 | 0.1 | 0.243021319 |
| SL1344_RS17745_ | 7.415526053 | 7.361184166 | 7.469783503 | 0.110959097 | 0.050284294 | 2.281632214 | 4.68E-05 | 0.054633826 | 0.18146223 | 0.1 | 0.243021319 |
| SL1344_RS17750_ | 7.688803618 | 7.545253089 | 7.754159924 | 0.255608189 | 0.152391668 | 1.828921745 | 4.68E-05 | 0.069983762 | 0.212343321 | 0.1 | 0.243021319 |
| SL1344_RS17755_ | 7.452553766 | 7.349983437 | 7.49798833 | 0.153923624 | 0.082032184 | 2.037731886 | 4.68E-05 | 0.052043919 | 0.177241596 | 0.1 | 0.243021319 |
| SL1344_RS17760_ | 7.633151189 | 7.559954243 | 7.748695623 | 0.184453045 | 0.061442609 | 2.842584126 | 4.68E-05 | 0.027420872 | 0.122005612 | 0.1 | 0.243021319 |
| SL1344_RS17765_ | 6.55360714 | 6.421152944 | 6.759113644 | 0.326720917 | 0.053172674 | 5.373541369 | 4.68E-05 | 0.009579369 | 0.068060985 | 0.1 | 0.243021319 |
| SL1344_RS17770_ | -4.989153671 | -4.681580903 | -5.289571524 | -0.6160566 | 1.27361152 | -0.461153109 | 0.298467697 | 0.453327979 | 0.61032708 | 0.5731 | 0.694171214 |
| SL1344_RS17775_ | 3.615753493 | 3.584989466 | 3.76395735 | 0.150617351 | 0.120964798 | 0.722935927 | 0.312000008 | 0.423905317 | 0.614106412 | 0.6432 | 0.783707827 |
| SL1344_RS17780_ | 3.794631501 | 3.677946428 | 3.895895474 | 0.11753444 | 0.205441386 | 0.781488285 | 0.177881433 | 0.325220409 | 0.526407598 | 0.3391 | 0.518532289 |
| SL1344_RS17785_ | 8.326704173 | 8.300536795 | 8.480179814 | 0.180533124 | 0.148276306 | 1.2432799 | 0.005330642 | 0.209061694 | 0.409728859 | 0.1043 | 0.249060579 |

| | | | | | | | | | | |
|---|---|---|---|---|---|---|---|---|---|---|
| SL1344_RS17790_ | 9.168041902 | 9.065984535 | 9.232605891 | 0.178184612 | 0.065994095 | 2.586527622 | 4.68E-05 | 0.038734417 | 0.149775187 | 0.1 | 0.243021319 |
| SL1344_RS17795_ | 7.078776104 | 6.957185265 | 7.212237747 | 0.233488117 | 0.04952441 | 4.734412374 | 4.68E-05 | 0.003106305 | 0.037661211 | 0.1 | 0.243021319 |
| SL1344_RS17800_ | 6.522230074 | 6.330608309 | 6.613573421 | 0.272568221 | 0.105745176 | 2.730084386 | 4.68E-05 | 0.012128101 | 0.077937838 | 0.1 | 0.243021319 |
| SL1344_RS17805_ | 1.965831792 | 2.007923348 | 1.880622211 | -0.146487437 | 0.198845466 | -0.744927791 | 0.20933335 | 0.305168464 | 0.495651393 | 0.4177 | 0.588772235 |
| SL1344_RS17810_ | 1.43644241 | 1.273024245 | 1.585849967 | 0.322409309 | 0.155067172 | 2.049263707 | 0.007329074 | 0.043230958 | 0.148941462 | 0.1072 | 0.252350329 |
| SL1344_RS17815_ | 1.478862665 | 1.327898778 | 1.675149422 | 0.350172529 | 0.144349608 | 2.362793505 | 0.0020023 | 0.032019986 | 0.12366222 | 0.1034 | 0.247321416 |
| SL1344_RS17820_ | 2.460370496 | 2.297040418 | 2.582911099 | 0.284417458 | 0.100378798 | 2.814736722 | 0.00067292 | 0.020292085 | 0.095452 | 0.1006 | 0.243866673 |
| SL1344_RS17825_ | 5.325433223 | 5.220500186 | 5.367929407 | 0.124341079 | 0.081213753 | 1.420009037 | 0.090000046 | 0.108577089 | 0.270036092 | 0.1843 | 0.35563197 |
| SL1344_RS17830_ | 1.654639068 | 1.831148975 | 1.43599248 | -0.406373876 | 0.151446988 | -2.6084466 | 4.68E-05 | 0.024125187 | 0.106595809 | 0.1002 | 0.243308097 |
| SL1344_RS17835_ | 4.855475312 | 4.718926478 | 4.990766674 | 0.277087527 | 0.07240482 | 3.890091503 | 4.68E-05 | 0.016391757 | 0.089441407 | 0.1 | 0.243021319 |
| SL1344_RS17840_ | 0.306387482 | 0.348134083 | 0.251261834 | -0.086699005 | 0.23854266 | -0.342010575 | 0.343104603 | 0.567672888 | 0.710765386 | 0.6769 | 0.784706721 |
| SL1344_RS17845_ | 2.022867872 | 2.14002811 | 1.907122163 | -0.230350637 | 0.107499016 | -2.072642909 | 0.011326186 | 0.042778577 | 0.143898284 | 0.1137 | 0.260849677 |
| SL1344_RS17850_ | 0.341104649 | 0.551237346 | 0.125506141 | -0.432044568 | 0.179107331 | -2.338679284 | 0.006667356 | 0.031701303 | 0.120277446 | 0.1043 | 0.248628698 |
| SL1344_RS17855_ | 3.293156795 | 3.057910803 | 3.439609269 | 0.367039923 | 0.099539216 | 3.494418917 | 4.68E-05 | 0.01411075 | 0.079856766 | 0.1 | 0.243021319 |
| SL1344_RS17860_ | -1.021978229 | -1.281118403 | -0.745368307 | 0.5245581 | 0.326629731 | 1.542822294 | 0.044666765 | 0.087910074 | 0.221620578 | 0.153 | 0.308116283 |
| SL1344_RS17865_ | -0.042745139 | 0.070969585 | -0.153536189 | -0.213088448 | 0.235371139 | -0.863311986 | 0.18000002 | 0.270332729 | 0.440412505 | 0.3655 | 0.521147218 |
| SL1344_RS17870_ | 1.918937313 | 1.778773001 | 2.051864212 | 0.280278373 | 0.117728135 | 2.287959781 | 0.003998486 | 0.033723679 | 0.128146304 | 0.1036 | 0.247818579 |
| SL1344_RS17875_ | -0.084084226 | -0.133878414 | -0.045129049 | 0.099589682 | 0.250032434 | 0.366072105 | 0.344000006 | 0.517581891 | 0.666967589 | 0.6328 | 0.743581548 |
| SL1344_RS17880_ | 1.447286631 | 1.395356831 | 1.500978724 | 0.104583348 | 0.122997141 | 0.798335122 | 0.1798801 | 0.295173836 | 0.457555175 | 0.3939 | 0.539210554 |
| SL1344_RS17885_ | 1.943239861 | 1.812480733 | 2.105692899 | 0.312791919 | 0.157042275 | 1.973295539 | 0.008667196 | 0.044828252 | 0.154477629 | 0.1067 | 0.251956539 |
| SL1344_RS17895_ | 3.040751224 | 2.927158484 | 3.104042528 | 0.174136918 | 0.083237426 | 2.001803489 | 0.013990999 | 0.058145379 | 0.177380703 | 0.1161 | 0.263999664 |
| SL1344_RS17900_ | 1.441550066 | 1.410304199 | 1.480527851 | 0.066500138 | 0.126080961 | 0.498609491 | 0.29380414 | 0.455426221 | 0.609987925 | 0.5615 | 0.687001016 |
| SL1344_RS17905_ | 0.919233886 | 0.788525521 | 1.033880683 | 0.244067249 | 0.149494767 | 1.583622134 | 0.040639683 | 0.087668699 | 0.221016926 | 0.1476 | 0.300702533 |
| SL1344_RS17910_ | -0.840766554 | -1.090734178 | -0.600707311 | 0.4883324 | 0.225222842 | 2.121858992 | 0.009993796 | 0.041270386 | 0.136027719 | 0.114 | 0.259401317 |
| SL1344_RS17915_ | -1.341951263 | -1.567094664 | -1.132376747 | 0.437576397 | 0.313971805 | 1.395304231 | 0.070000061 | 0.122207729 | 0.264221828 | 0.1912 | 0.345398522 |
| SL1344_RS17920_ | 2.646862322 | 2.291666041 | 2.919159625 | 0.63472564 | 0.126747281 | 5.088711087 | 4.68E-05 | 0.003101527 | 0.036619422 | 0.1 | 0.243021319 |
| SL1344_RS17925_ | 3.810468996 | 3.447699249 | 4.189672419 | 0.738882127 | 0.056431438 | 13.07983067 | 4.68E-05 | 0.000326144 | 0.011787503 | 0.1 | 0.243021319 |
| SL1344_RS17930_ | 0.8712338 | 0.443440218 | 1.055596314 | 0.606212401 | 0.25304048 | 2.164491311 | 0.003998486 | 0.058751481 | 0.184702819 | 0.1038 | 0.248272234 |
| SL1344_RS17940_ | -0.390684284 | -0.6130704 | -0.19118912 | 0.43064282 | 0.3050693 | 1.381431206 | 0.057333409 | 0.113944101 | 0.265615081 | 0.1755 | 0.334101095 |
| SL1344_RS17965_ | 4.164135904 | 4.099462762 | 4.2861648 | 0.174525178 | 0.097155965 | 1.610454623 | 0.042000105 | 0.086612145 | 0.23446672 | 0.1448 | 0.303631356 |
| SL1344_RS17970_ | 2.97844432 | 2.829157967 | 3.10985226 | 0.280532014 | 0.082610934 | 3.416472871 | 4.68E-05 | 0.012017381 | 0.071385859 | 0.1 | 0.243021319 |
| SL1344_RS17975_ | 2.601606553 | 2.430662132 | 2.810626953 | 0.378460211 | 0.080876352 | 4.604472176 | 4.68E-05 | 0.00576952 | 0.046765585 | 0.1 | 0.243021319 |
| SL1344_RS17985_ | 4.567335957 | 4.445407502 | 4.758297634 | 0.311881211 | 0.075831887 | 3.996930959 | 4.68E-05 | 0.016872291 | 0.090975866 | 0.1 | 0.243021319 |
| SL1344_RS17990_ | 4.2929791 | 4.290747127 | 4.297662769 | 0.009038805 | 0.130425616 | 0.060214184 | 0.467688209 | 0.758898467 | 0.858798915 | 0.8959 | 0.940128572 |
| SL1344_RS17995_ | 4.07300093 | 4.053182062 | 4.135234029 | 0.079426674 | 0.138713915 | 0.491247791 | 0.343104603 | 0.605669737 | 0.756451651 | 0.7291 | 0.844277928 |
| SL1344_RS18000_ | -0.924937306 | -1.002462834 | -0.856735546 | 0.145824168 | 0.285994613 | 0.467615737 | 0.302000009 | 0.449815841 | 0.605917319 | 0.5472 | 0.674565248 |
| SL1344_RS18005_ | -3.112210793 | -3.19589523 | -3.02435576 | 0.181015641 | 0.572863423 | 0.284030031 | 0.366666672 | 0.518451827 | 0.661651422 | 0.6313 | 0.740296203 |
| SL1344_RS18010_ | -3.146029612 | -3.163646776 | -3.122688468 | 0.048963605 | 0.602679623 | 0.068150695 | 0.470353099 | 0.581856937 | 0.718612622 | 0.6934 | 0.792525006 |
| SL1344_RS18015_ | -2.799202681 | -2.531769433 | -3.07133861 | -0.549669292 | 0.543698236 | -0.924969312 | 0.141905424 | 0.218498637 | 0.379062345 | 0.302 | 0.455977202 |
| SL1344_RS18020_ | -2.696249965 | -2.523484891 | -2.915721791 | -0.408933389 | 0.544987691 | -0.696196363 | 0.219187224 | 0.33200478 | 0.500200176 | 0.4389 | 0.57989479 |
| SL1344_RS18025_ | 3.281255748 | 3.184137609 | 3.401109281 | 0.216406998 | 0.065332036 | 3.251566572 | 4.68E-05 | 0.013246435 | 0.074088845 | 0.1001 | 0.243156879 |
| SL1344_RS18030_ | 2.36671036 | 2.06660219 | 2.650360938 | 0.584364566 | 0.097710426 | 5.9233616 | 4.68E-05 | 0.002434767 | 0.030768561 | 0.1 | 0.243021319 |
| SL1344_RS18035_ | 3.690215215 | 3.707889102 | 3.62552639 | -0.071916798 | 0.084291802 | -0.709205998 | 0.269153908 | 0.399627089 | 0.578142669 | 0.5415 | 0.689885471 |
| SL1344_RS18040_ | 1.731894213 | 1.681701972 | 1.823633245 | 0.137710936 | 0.128077118 | 0.978813468 | 0.144570313 | 0.217150304 | 0.387952105 | 0.3115 | 0.470351844 |
| SL1344_RS18045_ | 1.402989104 | 1.427006494 | 1.386422757 | -0.034032446 | 0.13356711 | -0.244590695 | 0.40266667 | 0.588429391 | 0.722529943 | 0.6969 | 0.796521696 |
| SL1344_RS18050_ | 2.553172835 | 2.467456964 | 2.649671138 | 0.185933166 | 0.107268053 | 1.67035431 | 0.031333476 | 0.075791487 | 0.208070869 | 0.1347 | 0.288165205 |
| SL1344_RS18055_ | 0.434671947 | 0.3657871 | 0.511378798 | 0.149382554 | 0.164155998 | 0.853871373 | 0.175333354 | 0.276316569 | 0.4376492 | 0.3721 | 0.518300296 |
| SL1344_RS18060_ | 3.355932631 | 3.385952228 | 3.337271578 | -0.04214477 | 0.100949765 | -0.413216545 | 0.324450374 | 0.543346971 | 0.694757925 | 0.6213 | 0.749900009 |
| SL1344_RS18075_ | 1.036234978 | 1.028222765 | 1.041211498 | 0.035467768 | 0.226100666 | 0.134919056 | 0.440373087 | 0.685158652 | 0.802072056 | 0.8271 | 0.890414225 |
| SL1344_RS18080_ | 3.631622647 | 3.558050594 | 3.71790098 | 0.167832775 | 0.083221336 | 2.086105272 | 0.0020023 | 0.058093002 | 0.181538874 | 0.102 | 0.245319422 |
| SL1344_RS18090_ | 3.766820705 | 3.278034218 | 4.239346699 | 0.97033948 | 0.075714497 | 12.93098745 | 4.68E-05 | 0.000462831 | 0.01438357 | 0.1 | 0.243021319 |
| SL1344_RS18095_ | 2.531813245 | 2.145729705 | 2.889386689 | 0.743501158 | 0.085098796 | 8.726940084 | 4.68E-05 | 0.000947607 | 0.018830364 | 0.1 | 0.243021319 |
| SL1344_RS18100_ | -0.199468102 | -0.296140296 | -0.10462787 | 0.193103619 | 0.225747004 | 0.779721812 | 0.188540992 | 0.279902676 | 0.445786058 | 0.38 | 0.530807671 |

| | | | | | | | | | | | |
|---|---|---|---|---|---|---|---|---|---|---|---|
| SL1344_RS18105_ | 0.193571729 | 0.138153694 | 0.297675449 | 0.117526234 | 0.28826916 | 0.393975832 | 0.354430386 | 0.611203867 | 0.747850505 | 0.7095 | 0.813680491 |
| SL1344_RS18110_ | 0.274279984 | 0.424270677 | 0.120823537 | -0.313005812 | 0.210370494 | -1.462556255 | 0.054630326 | 0.100876774 | 0.244750123 | 0.1645 | 0.320667467 |
| SL1344_RS18115_ | 4.082505317 | 4.061216192 | 4.102486867 | 0.053185733 | 0.075901515 | 0.662384267 | 0.23184545 | 0.331437058 | 0.51327104 | 0.4706 | 0.615364266 |
| SL1344_RS18120_ | 0.401250112 | 0.069407254 | 0.69885209 | 0.633597381 | 0.186027572 | 3.375524584 | 4.68E-05 | 0.013783924 | 0.075404509 | 0.1001 | 0.243158543 |
| SL1344_RS18125_ | 0.404452343 | 0.356230411 | 0.452009593 | 0.092724371 | 0.165297511 | 0.512467902 | 0.282666677 | 0.414995523 | 0.566466477 | 0.5174 | 0.644489126 |
| SL1344_RS18130_ | -0.20365145 | -0.253793933 | -0.154348703 | 0.097593379 | 0.186958753 | 0.477067603 | 0.299800142 | 0.426385707 | 0.574586336 | 0.5334 | 0.657997577 |
| SL1344_RS18140_ | 1.303255584 | 1.315258831 | 1.289116008 | -0.01914873 | 0.184493279 | -0.095171017 | 0.460000001 | 0.703946451 | 0.814703876 | 0.8089 | 0.878605513 |
| SL1344_RS18145_ | 2.10321796 | 2.145067732 | 2.059980332 | -0.055538164 | 0.25914239 | -0.211606772 | 0.40266667 | 0.718793062 | 0.831581152 | 0.83 | 0.896718079 |
| SL1344_RS18150_ | 0.105278763 | 0.344293542 | -0.218711259 | -0.565518154 | 0.216398076 | -2.518116435 | 0.001336769 | 0.030487998 | 0.118175982 | 0.1015 | 0.245063477 |
| SL1344_RS18155_ | 0.468311953 | 0.690277776 | 0.255619838 | -0.430026686 | 0.158065119 | -2.714006554 | 0.003334715 | 0.023538377 | 0.101327974 | 0.1024 | 0.246101009 |
| SL1344_RS18160_ | -0.65558081 | -0.61536493 | -0.712945462 | -0.091563828 | 0.267133717 | -0.308008719 | 0.359093943 | 0.535519765 | 0.678940707 | 0.6433 | 0.753079667 |
| SL1344_RS18165_ | 1.375043628 | 1.256113365 | 1.69922011 | 0.432044783 | 0.196152032 | 1.920210717 | 0.015989625 | 0.075209605 | 0.213625157 | 0.1163 | 0.264380835 |
| SL1344_RS18170_ | 1.834041079 | 1.852662158 | 1.793630371 | -0.062576701 | 0.165691201 | -0.330894931 | 0.365756168 | 0.573825124 | 0.72230284 | 0.7366 | 0.829046201 |
| SL1344_RS18175_ | 2.866157451 | 2.793656124 | 2.933572059 | 0.141824919 | 0.087542698 | 1.56195132 | 0.039307241 | 0.085923241 | 0.222405808 | 0.1451 | 0.299634728 |
| SL1344_RS18180_ | 1.321261632 | 1.213598069 | 1.41389116 | 0.193492226 | 0.137798327 | 1.346469753 | 0.069953426 | 0.122552427 | 0.270358902 | 0.196 | 0.356305962 |
| SL1344_RS18185_ | 6.986980276 | 6.990848614 | 6.983204397 | 0.001669016 | 0.035830171 | 0.052409197 | 0.481678881 | 0.747390865 | 0.84776703 | 0.8828 | 0.927524143 |
| SL1344_RS18190_ | -1.588163456 | -1.616131561 | -1.556560049 | 0.080879617 | 0.347770637 | 0.19253979 | 0.407333337 | 0.55930607 | 0.698659423 | 0.6741 | 0.774862191 |
| SL1344_RS18195_ | 0.373055197 | 0.339661827 | 0.399493807 | 0.065665703 | 0.177236004 | 0.33341883 | 0.350666673 | 0.49528942 | 0.64453464 | 0.61 | 0.723748009 |
| SL1344_RS18205_ | -3.532734172 | -3.737632396 | -3.327006973 | 0.40354339 | 0.638422003 | 0.613471283 | 0.247834789 | 0.370077347 | 0.529144154 | 0.4709 | 0.606200361 |
| SL1344_RS18210_ | -2.877840399 | -2.990133694 | -2.768091405 | 0.280083006 | 0.670384167 | 0.400143518 | 0.328666674 | 0.48047947 | 0.637164022 | 0.6182 | 0.729390035 |
| SL1344_RS18215_ | -1.82852982 | -2.05520093 | -1.244602842 | 0.761582661 | 0.489393084 | 1.393507897 | 0.082611643 | 0.122789696 | 0.279722043 | 0.2007 | 0.361704058 |
| SL1344_RS18220_ | 1.800091218 | 1.827689956 | 1.768711408 | -0.081053437 | 0.139219096 | -0.548448017 | 0.29113925 | 0.42476747 | 0.592048663 | 0.5755 | 0.698576164 |
| SL1344_RS18225_ | 1.860969134 | 1.872261081 | 1.849969964 | -0.022598897 | 0.09901642 | -0.202051989 | 0.405333337 | 0.544935229 | 0.685010931 | 0.6495 | 0.756287253 |
| SL1344_RS18230_ | 0.288844518 | 0.363417851 | 0.231086381 | -0.143328923 | 0.199836127 | -0.677943322 | 0.225183226 | 0.344555589 | 0.508110786 | 0.4493 | 0.586462027 |
| SL1344_RS18235_ | 0.243619107 | 0.363445654 | 0.095780896 | -0.230669069 | 0.278141255 | -0.807407724 | 0.200000017 | 0.295522991 | 0.478642143 | 0.3867 | 0.55244041 |
| SL1344_RS18240_ | 0.681098451 | 0.846381019 | 0.554536834 | -0.290258733 | 0.186579779 | -1.495584101 | 0.049966777 | 0.095199088 | 0.234065 | 0.1594 | 0.315050784 |
| SL1344_RS18245_ | 1.344361901 | 1.467440173 | 1.153822342 | -0.305604311 | 0.149096378 | -1.914969597 | 0.021985549 | 0.054460488 | 0.169019239 | 0.12 | 0.269243579 |
| SL1344_RS18250_ | 2.064638227 | 2.131806134 | 1.830488596 | -0.303142896 | 0.199156263 | -1.362713762 | 0.061333404 | 0.146219721 | 0.324019321 | 0.1696 | 0.328113441 |
| SL1344_RS18255_ | 0.235378262 | 0.220126412 | 0.253400544 | 0.058011464 | 0.248393674 | 0.218894208 | 0.40466667 | 0.60209216 | 0.737009261 | 0.7376 | 0.824546645 |
| SL1344_RS18270_ | -0.311416284 | -0.372676288 | -0.241517132 | 0.131965369 | 0.217146041 | 0.582756269 | 0.264490351 | 0.39542846 | 0.55011083 | 0.4912 | 0.625721698 |
| SL1344_RS18275_ | 0.313626538 | 0.371915682 | 0.256587382 | -0.107044817 | 0.174466208 | -0.589684405 | 0.255829459 | 0.393399498 | 0.551004503 | 0.4989 | 0.633552388 |
| SL1344_RS18280_ | -0.449225189 | -0.512343965 | -0.315226603 | 0.170015371 | 0.284853836 | 0.54161306 | 0.295136585 | 0.488263272 | 0.643589622 | 0.5863 | 0.717839478 |
| SL1344_RS18285_ | -4.780053278 | -5.19181043 | -4.335473204 | 0.906279475 | 1.26836448 | 0.69693913 | 0.223850781 | 0.342108101 | 0.512890986 | 0.4475 | 0.594246621 |
| SL1344_RS18290_ | -4.918245615 | -5.244503258 | -4.624365124 | 0.632130255 | 1.051853617 | 0.555140716 | 0.253164569 | 0.383220826 | 0.53957172 | 0.4816 | 0.616581116 |
| SL1344_RS18295_ | -4.601386559 | -4.54383342 | -4.622195851 | -0.169202582 | 1.382080129 | -0.114542355 | 0.452365092 | 0.690292376 | 0.803765243 | 0.7952 | 0.869993119 |
| SL1344_RS18300_ | -1.779709819 | -1.799887629 | -1.758703653 | 0.077405662 | 0.43188368 | 0.166491662 | 0.431333336 | 0.611437852 | 0.742768058 | 0.7233 | 0.813730381 |
| SL1344_RS18305_ | -0.307477608 | -0.230787802 | -0.432656852 | -0.197358914 | 0.228098344 | -0.820989124 | 0.184666686 | 0.285537809 | 0.449290632 | 0.3861 | 0.533903042 |
| SL1344_RS18310_ | -0.556831678 | -0.405960519 | -0.649553078 | -0.217848982 | 0.410278529 | -0.523107908 | 0.273333344 | 0.463811656 | 0.631632762 | 0.5421 | 0.689885375 |
| SL1344_RS18315_ | -0.180564557 | -0.126584159 | -0.274167547 | -0.135804148 | 0.32161483 | -0.38738904 | 0.340666673 | 0.56812189 | 0.71400021 | 0.6676 | 0.782294471 |
| SL1344_RS18325_ | -0.795616923 | -0.831597871 | -0.766415725 | 0.057652774 | 0.282379895 | 0.190661334 | 0.416000003 | 0.587723886 | 0.721969212 | 0.6925 | 0.790899347 |
| SL1344_RS18330_ | -0.987976052 | -1.004939464 | -0.969834379 | 0.03378259 | 0.301527983 | 0.108619802 | 0.448367757 | 0.603598177 | 0.735705399 | 0.7148 | 0.808731146 |
| SL1344_RS18335_ | 0.316359878 | 0.454791577 | 0.208077235 | -0.256473345 | 0.181839458 | -1.345189516 | 0.069287204 | 0.125468233 | 0.272087114 | 0.1918 | 0.345181836 |
| SL1344_RS18340_ | -1.238804743 | -1.194584975 | -1.28479871 | -0.085186749 | 0.269414771 | -0.312358506 | 0.363333339 | 0.488779861 | 0.635739909 | 0.592 | 0.709593164 |
| SL1344_RS18345_ | -1.69577446 | -1.393451759 | -2.10802139 | -0.712861556 | 0.342825166 | -1.997341151 | 0.014666978 | 0.049411138 | 0.155453555 | 0.1134 | 0.259693519 |
| SL1344_RS18355_ | -1.104148195 | -1.057187759 | -1.179443381 | -0.107098704 | 0.357143308 | -0.284940805 | 0.389333338 | 0.60116053 | 0.736364549 | 0.7194 | 0.815849503 |
| SL1344_RS18360_ | -0.693295193 | -0.653598422 | -0.739099705 | -0.085814174 | 0.271861281 | -0.29995783 | 0.377748173 | 0.54551722 | 0.689737151 | 0.65 | 0.761702034 |
| SL1344_RS18365_ | -1.276304503 | -1.061946541 | -1.426488893 | -0.357870358 | 0.318377243 | -1.05881221 | 0.108666704 | 0.188151615 | 0.347980771 | 0.276 | 0.431722242 |
| SL1344_RS18370_ | -0.499959166 | -0.287070621 | -0.807231298 | -0.513265535 | 0.400284628 | -1.242846724 | 0.070666727 | 0.144095531 | 0.315822592 | 0.1825 | 0.346976826 |
| SL1344_RS18375_ | 1.27747501 | 1.173588232 | 1.523760574 | 0.337811361 | 0.15060619 | 1.984855949 | 0.016655836 | 0.060949067 | 0.183003165 | 0.1224 | 0.270805912 |
| SL1344_RS18380_ | 1.723495387 | 1.590196446 | 1.965435861 | 0.380926817 | 0.166790679 | 2.247648173 | 0.002000966 | 0.050876995 | 0.168737036 | 0.1009 | 0.244215984 |
| SL1344_RS18385_ | 0.909949996 | 0.940924938 | 0.871683523 | -0.068786289 | 0.23938901 | -0.26581477 | 0.376415728 | 0.65739889 | 0.783320943 | 0.7224 | 0.82293024 |
| SL1344_RS18390_ | 4.89663046 | 4.946053177 | 4.796543211 | -0.137971594 | 0.077461891 | -1.616883059 | 0.039333446 | 0.089824297 | 0.240494785 | 0.137 | 0.293509612 |
| SL1344_RS18395_ | 3.349973528 | 3.228042412 | 3.485561196 | 0.267648659 | 0.08680094 | 3.093370136 | 4.68E-05 | 0.01900102 | 0.09330065 | 0.1 | 0.243021319 |

| | | | | | | | | | | |
|---|---|---|---|---|---|---|---|---|---|---|
| SL1344_RS18400_ | 2.287178575 | 2.16242643 | 2.437575888 | 0.283827624 | 0.131218034 | 2.109948994 | 0.002668394 | 0.047920479 | 0.16109272 | 0.1035 | 0.2478275 |
| SL1344_RS18405_ | 3.13167521 | 3.070523997 | 3.340041081 | 0.266891606 | 0.149771687 | 1.571164783 | 0.047968113 | 0.092769348 | 0.245429805 | 0.1558 | 0.314671165 |
| SL1344_RS18415_ | 1.349114563 | 1.017560515 | 1.604286679 | 0.589296765 | 0.131244725 | 4.415293754 | 4.68E-05 | 0.00672292 | 0.050381381 | 0.1 | 0.243021319 |
| SL1344_RS18420_ | -0.074044777 | -0.067040015 | -0.086322483 | -0.004544819 | 0.242960268 | -0.015299881 | 0.492 | 0.655347079 | 0.778850122 | 0.7719 | 0.853587087 |
| SL1344_RS18425_ | 0.866727616 | 0.83705453 | 0.895963118 | 0.05017712 | 0.182066854 | 0.269867669 | 0.390666671 | 0.601896055 | 0.733696823 | 0.6977 | 0.795384796 |
| SL1344_RS18440_ | 1.561733975 | 1.226809627 | 1.781465127 | 0.530771996 | 0.167570938 | 3.039201984 | 4.68E-05 | 0.020261201 | 0.097253108 | 0.1 | 0.243021319 |
| SL1344_RS18445_ | 1.255949916 | 1.194565588 | 1.377745595 | 0.253155293 | 0.281356954 | 0.894525063 | 0.167221874 | 0.224031756 | 0.414672595 | 0.3361 | 0.511474025 |
| SL1344_RS18450_ | 3.424622635 | 3.416866101 | 3.437035855 | 0.023513149 | 0.068880753 | 0.297413328 | 0.366000005 | 0.52829223 | 0.675947083 | 0.6458 | 0.756187514 |
| SL1344_RS18455_ | 0.536205837 | 0.617873029 | 0.490269568 | -0.122130267 | 0.169397716 | -0.682878494 | 0.23866668 | 0.371168525 | 0.533055236 | 0.4799 | 0.617728235 |
| SL1344_RS18460_ | -1.115707841 | -1.253902361 | -0.93696934 | 0.320792035 | 0.298370467 | 1.004234365 | 0.146000026 | 0.228569045 | 0.390676899 | 0.3191 | 0.475835673 |
| SL1344_RS18465_ | 1.345689789 | 1.298642075 | 1.408464097 | 0.104258695 | 0.138096061 | 0.705758556 | 0.216000016 | 0.337498695 | 0.505294617 | 0.4401 | 0.586231484 |
| SL1344_RS18470_ | 1.064666989 | 0.848555334 | 1.246530832 | 0.3938117 | 0.122440117 | 3.138204521 | 4.68E-05 | 0.014563786 | 0.075337473 | 0.1002 | 0.243301146 |
| SL1344_RS18475_ | -0.728719075 | -0.769388794 | -0.697065711 | 0.074773364 | 0.263865264 | 0.256329076 | 0.381333338 | 0.532285697 | 0.676553268 | 0.644 | 0.754023695 |
| SL1344_RS18480_ | 1.140559851 | 1.306968941 | 0.963790981 | -0.338531177 | 0.131857157 | -2.497436311 | 0.003998486 | 0.027541598 | 0.111081343 | 0.1043 | 0.248859268 |
| SL1344_RS18485_ | 0.05405281 | 0.143468655 | -0.043291236 | -0.156482305 | 0.245844275 | -0.594361792 | 0.263333345 | 0.410391849 | 0.576434316 | 0.5087 | 0.649846556 |
| SL1344_RS18490_ | 2.46745108 | 2.285498315 | 2.608863383 | 0.332767377 | 0.123822935 | 2.637000124 | 0.00067292 | 0.020966356 | 0.099495659 | 0.1008 | 0.244144828 |
| SL1344_RS18495_ | 4.733102706 | 4.419332204 | 5.040637184 | 0.634602515 | 0.064691324 | 9.836746027 | 4.68E-05 | 0.000456399 | 0.014725572 | 0.1 | 0.243021319 |
| SL1344_RS18500_ | 0.468113807 | 0.503909045 | 0.406910209 | -0.108895599 | 0.198838179 | -0.485393719 | 0.273151243 | 0.415637795 | 0.58026031 | 0.5458 | 0.668378877 |
| SL1344_RS18505_ | 3.804825742 | 4.13559655 | 3.495279589 | -0.649788713 | 0.117126767 | -5.511153544 | 4.68E-05 | 0.001755816 | 0.028831759 | 0.1 | 0.243021319 |
| SL1344_RS18510_ | 0.441030169 | 0.262025835 | 0.629367969 | 0.369916549 | 0.177428715 | 2.025435901 | 0.011333737 | 0.045100652 | 0.147839582 | 0.1101 | 0.256206735 |
| SL1344_RS18515_ | 1.156826818 | 1.017382826 | 1.351258469 | 0.320672883 | 0.161162632 | 1.883378074 | 0.023984199 | 0.05724941 | 0.174164466 | 0.1288 | 0.280450898 |
| SL1344_RS18520_ | 2.378056705 | 2.473635049 | 2.237196764 | -0.23226861 | 0.181235242 | -1.222187434 | 0.088607642 | 0.127564007 | 0.295149029 | 0.2029 | 0.376130136 |
| SL1344_RS18525_ | 1.351204229 | 1.350681826 | 1.351675762 | -0.01272074 | 0.16587912 | -0.068559898 | 0.471019322 | 0.650855154 | 0.777717075 | 0.7679 | 0.851621469 |
| SL1344_RS18530_ | 1.972323348 | 1.935311069 | 2.026229774 | 0.109912392 | 0.183147554 | 0.561762576 | 0.253164569 | 0.363127845 | 0.552913005 | 0.4963 | 0.640754016 |
| SL1344_RS18535_ | 3.14440792 | 3.117526957 | 3.197500947 | 0.077835176 | 0.081621349 | 0.865247308 | 0.198534328 | 0.275484524 | 0.451827894 | 0.3861 | 0.543383635 |
| SL1344_RS18540_ | -0.385793559 | -0.282870149 | -0.476766308 | -0.174008452 | 0.293419563 | -0.587733183 | 0.278666677 | 0.439854075 | 0.602975251 | 0.5481 | 0.681555668 |
| SL1344_RS18545_ | -0.658585616 | -0.724138854 | -0.627497781 | 0.071969766 | 0.342719621 | 0.19477707 | 0.427048637 | 0.664000659 | 0.788014777 | 0.7805 | 0.862942955 |
| SL1344_RS18550_ | -1.580769059 | -1.623619943 | -1.556330485 | 0.038712079 | 0.469466466 | 0.078539433 | 0.464000001 | 0.673165467 | 0.79420551 | 0.7766 | 0.858768995 |
| SL1344_RS18555_ | -0.365495546 | -0.432274572 | -0.304677437 | 0.128552216 | 0.278336355 | 0.432301014 | 0.306462367 | 0.486939282 | 0.642087743 | 0.5985 | 0.718835276 |
| SL1344_RS18560_ | 0.573874098 | 0.454462228 | 0.659867478 | 0.174775807 | 0.242623374 | 0.691354623 | 0.233333347 | 0.366170593 | 0.545215098 | 0.4692 | 0.622941812 |
| SL1344_RS18565_ | 0.701487505 | 0.630729901 | 0.800402442 | 0.168702093 | 0.180810731 | 0.872093541 | 0.16055965 | 0.260718468 | 0.432030208 | 0.3635 | 0.521326166 |
| SL1344_RS18570_ | 1.738124865 | 2.129891371 | 1.287847978 | -0.83860059 | 0.123051116 | -6.874804029 | 4.68E-05 | 0.001847384 | 0.026720464 | 0.1 | 0.243021319 |
| SL1344_RS18575_ | -2.81447896 | -3.003291606 | -2.557933527 | 0.415358478 | 0.529876012 | 0.736843899 | 0.216666682 | 0.341177918 | 0.503874946 | 0.4422 | 0.584840673 |
| SL1344_RS18580_ | -1.755293024 | -2.293206989 | -1.148440722 | 1.097347687 | 0.484993924 | 2.200370615 | 0.008661422 | 0.036576613 | 0.134945677 | 0.1101 | 0.25676252 |
| SL1344_RS18585_ | -0.196572516 | -0.357629118 | -0.077165166 | 0.276149812 | 0.216654536 | 1.181394517 | 0.100666708 | 0.158840892 | 0.313365029 | 0.2381 | 0.39642295 |
| SL1344_RS18590_ | 0.339945997 | 0.405541547 | 0.264799501 | -0.070410998 | 0.298403787 | -0.247645885 | 0.413333337 | 0.691167452 | 0.806815097 | 0.8125 | 0.880407679 |
| SL1344_RS18595_ | 0.246702791 | 0.278780078 | 0.182260882 | -0.123235494 | 0.273671441 | -0.396098196 | 0.32933334 | 0.467316085 | 0.637684792 | 0.6296 | 0.742638564 |
| SL1344_RS18600_ | 2.515971863 | 3.080778592 | 1.953137607 | -1.117755114 | 0.109643032 | -10.22339074 | 4.68E-05 | 0.000526713 | 0.015306698 | 0.1 | 0.243021319 |
| SL1344_RS18610_ | -0.637037428 | -1.051295994 | -0.312710163 | 0.744704125 | 0.258028607 | 2.781701642 | 0.001336769 | 0.020520293 | 0.093326692 | 0.1008 | 0.244139961 |
| SL1344_RS18615_ | 2.274724112 | 2.180354883 | 2.368114444 | 0.194223131 | 0.114656883 | 1.672668358 | 0.022000206 | 0.076104747 | 0.208749164 | 0.1232 | 0.273155618 |
| SL1344_RS18620_ | 0.568842612 | 0.535018201 | 0.644943297 | 0.071502037 | 0.274596932 | 0.185895141 | 0.414000003 | 0.779880819 | 0.870879507 | 0.8123 | 0.89189059 |
| SL1344_RS18625_ | 2.315551814 | 2.233088874 | 2.408373893 | 0.174179921 | 0.085741964 | 1.973046168 | 0.01866691 | 0.051479547 | 0.159050364 | 0.1191 | 0.267341307 |
| SL1344_RS18630_ | 2.797142865 | 2.535625 | 3.071767864 | 0.537107435 | 0.069812172 | 7.678669612 | 4.68E-05 | 0.001403254 | 0.022801433 | 0.1 | 0.243021319 |
| SL1344_RS18635_ | 1.392186053 | 0.918744049 | 1.737907833 | 0.820025196 | 0.180328973 | 4.450222701 | 4.68E-05 | 0.009783339 | 0.064183334 | 0.1 | 0.243021319 |
| SL1344_RS18640_ | 1.689361591 | 1.271789442 | 2.072111321 | 0.800476127 | 0.110688267 | 7.152782822 | 4.68E-05 | 0.001628739 | 0.024705402 | 0.1 | 0.243021319 |
| SL1344_RS18655_ | -0.175635645 | -0.936960747 | 0.420255151 | 1.371876014 | 0.284105542 | 4.805558158 | 4.68E-05 | 0.005185521 | 0.044874286 | 0.1 | 0.243021319 |
| SL1344_RS18660_ | 1.339727386 | 0.833049685 | 1.804873608 | 0.986744391 | 0.180411962 | 5.430780999 | 4.68E-05 | 0.004220738 | 0.040768397 | 0.1 | 0.243021319 |
| SL1344_RS18665_ | -0.686007452 | -1.14480771 | -0.282399469 | 0.860315887 | 0.282183681 | 2.945965032 | 4.68E-05 | 0.018258031 | 0.088346505 | 0.1001 | 0.243160479 |
| SL1344_RS18670_ | 1.54317145 | 1.130480877 | 1.946326254 | 0.812313463 | 0.121500699 | 6.735784643 | 4.68E-05 | 0.001880314 | 0.026472274 | 0.1 | 0.243021319 |
| SL1344_RS18675_ | 2.307688499 | 2.320039695 | 2.296166643 | -0.04619991 | 0.183572452 | -0.226302227 | 0.387333338 | 0.574545923 | 0.727364065 | 0.7404 | 0.83170092 |
| SL1344_RS18680_ | -0.256632422 | -0.270116864 | -0.244768307 | 0.041464454 | 0.241618516 | 0.16124317 | 0.429713527 | 0.58444247 | 0.721324484 | 0.6983 | 0.79731279 |
| SL1344_RS18685_ | 1.406168918 | 1.169040502 | 1.589933885 | 0.418706105 | 0.122608015 | 3.357302361 | 4.68E-05 | 0.01317594 | 0.072396701 | 0.1001 | 0.243162177 |
| SL1344_RS18690_ | 0.157288387 | 0.460397682 | -0.254410295 | -0.715349802 | 0.20154754 | -3.47899647 | 4.68E-05 | 0.014116816 | 0.076014734 | 0.1 | 0.243021319 |

| | | | | | | | | | | | |
|---|---|---|---|---|---|---|---|---|---|---|---|
| SL1344_RS18695_ | 0.211837104 | 0.585129695 | -0.114718361 | -0.695390459 | 0.201722001 | -3.307199739 | 4.68E-05 | 0.011917125 | 0.070157423 | 0.1001 | 0.243164384 |
| SL1344_RS18700_ | 0.400843293 | 0.693779871 | 0.13326622 | -0.581812689 | 0.225777776 | -2.558538189 | 0.000673369 | 0.024768001 | 0.108571049 | 0.1005 | 0.243729018 |
| SL1344_RS18705_ | -0.493691751 | -0.25147571 | -0.791036217 | -0.564602827 | 0.309292012 | -1.779375624 | 0.016655836 | 0.060292935 | 0.180767695 | 0.1207 | 0.269787419 |
| SL1344_RS18710_ | 1.753550349 | 2.100110205 | 1.365924635 | -0.73432234 | 0.119253298 | -6.090954807 | 4.68E-05 | 0.002413175 | 0.030793109 | 0.1 | 0.243021319 |
| SL1344_RS18715_ | -1.597829687 | -1.723659467 | -1.486156702 | 0.209272606 | 0.432687343 | 0.461222078 | 0.295802807 | 0.489839983 | 0.642664066 | 0.6064 | 0.725504704 |
| SL1344_RS18720_ | -0.32047165 | -0.39268043 | -0.267328454 | 0.101926582 | 0.24873762 | 0.396695728 | 0.326449041 | 0.51102646 | 0.661331437 | 0.6112 | 0.729889087 |
| SL1344_RS18725_ | 0.094959749 | -0.019610183 | 0.227526667 | 0.248402974 | 0.192169613 | 1.242579507 | 0.081945421 | 0.144478584 | 0.292365705 | 0.2171 | 0.37142081 |
| SL1344_RS18735_ | 2.725597014 | 2.502866681 | 2.840281349 | 0.328832334 | 0.162650784 | 1.948826874 | 0.007995244 | 0.042559491 | 0.153123079 | 0.1057 | 0.250853111 |
| SL1344_RS18740_ | 0.915600024 | 0.732945704 | 1.018545466 | 0.265552378 | 0.19306304 | 1.285962009 | 0.074616979 | 0.127251787 | 0.284570115 | 0.201 | 0.365611367 |
| SL1344_RS18745_ | -1.448255455 | -1.453678778 | -1.440619734 | 0.007509785 | 0.360595777 | 0.018482639 | 0.490666667 | 0.625210191 | 0.753105432 | 0.7328 | 0.822891375 |
| SL1344_RS18750_ | -1.375810768 | -1.524040079 | -1.161002001 | 0.360397627 | 0.367388657 | 0.94287279 | 0.152666692 | 0.236180308 | 0.403518208 | 0.3231 | 0.480262659 |
| SL1344_RS18755_ | -1.898327311 | -1.808643805 | -2.005618619 | -0.196084827 | 0.355027859 | -0.515505067 | 0.283333343 | 0.390140002 | 0.548058883 | 0.4981 | 0.630796939 |
| SL1344_RS18760_ | -4.35024291 | -4.448244375 | -4.245499212 | 0.175401682 | 0.88880689 | 0.18875615 | 0.41266667 | 0.559582327 | 0.699007617 | 0.6677 | 0.771940446 |
| SL1344_RS18765_ | -3.473921222 | -3.434511517 | -3.537117738 | -0.100932493 | 0.787949435 | -0.111404403 | 0.450666669 | 0.64158665 | 0.767582667 | 0.756 | 0.841262912 |
| SL1344_RS18770_ | -1.983987426 | -1.854900499 | -2.127838059 | -0.335853362 | 0.513197477 | -0.604130253 | 0.250000012 | 0.371460261 | 0.544531156 | 0.5003 | 0.63757532 |
| SL1344_RS18775_ | 0.407744057 | 0.342713768 | 0.459849511 | 0.11054222 | 0.19144053 | 0.561042274 | 0.260493016 | 0.416681172 | 0.576345044 | 0.5181 | 0.651459825 |
| SL1344_RS18780_ | -0.154639209 | -0.104034954 | -0.226696617 | -0.120100228 | 0.226264052 | -0.484359622 | 0.295333342 | 0.447538872 | 0.604202232 | 0.548 | 0.677341228 |
| SL1344_RS18785_ | 2.561603893 | 2.464565941 | 2.663524931 | 0.198525597 | 0.100534379 | 1.939935393 | 0.010000458 | 0.049546871 | 0.159131817 | 0.1131 | 0.259233652 |
| SL1344_RS18790_ | 1.27805353 | 1.167160023 | 1.393713181 | 0.224234423 | 0.113618659 | 1.904129585 | 0.019320688 | 0.052220328 | 0.157267765 | 0.1229 | 0.271410708 |
| SL1344_RS18795_ | 1.385075593 | 1.380682084 | 1.389185858 | 0.010174242 | 0.141120943 | 0.068616143 | 0.468000001 | 0.61045426 | 0.741055654 | 0.7245 | 0.815666722 |
| SL1344_RS18800_ | 3.698573337 | 3.659925642 | 3.878813361 | 0.200705313 | 0.159119039 | 1.021186023 | 0.191872104 | 0.212128252 | 0.408108692 | 0.3416 | 0.530370439 |
| SL1344_RS18805_ | -1.471867142 | -1.385680355 | -1.547182761 | -0.275218575 | 0.515934095 | -0.505090783 | 0.290473027 | 0.423434124 | 0.597989871 | 0.6023 | 0.718963668 |
| SL1344_RS18815_ | -0.661961367 | -0.887875062 | -0.360894923 | 0.553081091 | 0.440977954 | 1.212496157 | 0.082000051 | 0.143908084 | 0.314679002 | 0.1922 | 0.358230405 |
| SL1344_RS18820_ | 1.1961072 | 1.256213118 | 1.141923327 | -0.114635067 | 0.116570343 | -0.93657587 | 0.159227205 | 0.246579212 | 0.399784274 | 0.3373 | 0.485787565 |
| SL1344_RS18825_ | 0.884675132 | 0.954338161 | 0.825484039 | -0.131019205 | 0.145673376 | -0.844869983 | 0.15866669 | 0.255787412 | 0.417921226 | 0.3476 | 0.497029419 |
| SL1344_RS18830_ | -1.227736463 | -1.522440514 | -0.872407768 | 0.64718982 | 0.304958975 | 2.090053797 | 0.010000458 | 0.040562595 | 0.139775141 | 0.1088 | 0.254760843 |
| SL1344_RS18835_ | 4.589745577 | 4.496303525 | 4.714477432 | 0.225275451 | 0.072942413 | 2.930987692 | 4.68E-05 | 0.056800887 | 0.185762214 | 0.1 | 0.243021319 |
| SL1344_RS18840_ | 2.954373181 | 2.889073147 | 3.063769603 | 0.18308492 | 0.12353731 | 1.480279164 | 0.029980163 | 0.111706824 | 0.272361316 | 0.1316 | 0.28305086 |
| SL1344_RS18845_ | -1.37915612 | -1.341910355 | -1.424344336 | -0.098103279 | 0.325362366 | -0.279519668 | 0.383077953 | 0.518835647 | 0.6644242 | 0.6372 | 0.745883336 |
| SL1344_RS18850_ | -0.328457825 | -0.323122489 | -0.340103678 | -0.035432525 | 0.266532526 | -0.117024025 | 0.442666669 | 0.617316256 | 0.748403174 | 0.7361 | 0.82748538 |
| SL1344_RS18855_ | -3.017366113 | -2.961609302 | -3.106479773 | -0.105099945 | 0.679918586 | -0.144099737 | 0.439333336 | 0.626007514 | 0.756388834 | 0.7412 | 0.83044542 |
| SL1344_RS18860_ | -0.218710981 | -0.005657115 | -0.426332065 | -0.409160501 | 0.238892924 | -1.697491633 | 0.033333467 | 0.076102174 | 0.201785432 | 0.1423 | 0.295531334 |
| SL1344_RS18865_ | 0.036794169 | 0.241461507 | -0.105740081 | -0.350462488 | 0.25158644 | -1.375201614 | 0.072000059 | 0.118086886 | 0.269673426 | 0.1848 | 0.34479157 |
| SL1344_RS18870_ | -1.248901294 | -1.568270947 | -0.989856681 | 0.58496985 | 0.399217698 | 1.454283982 | 0.045969449 | 0.104909848 | 0.250844496 | 0.1595 | 0.314798219 |
| SL1344_RS18875_ | 3.631334347 | 3.612236412 | 3.657378209 | 0.049504511 | 0.136187273 | 0.312150956 | 0.362000006 | 0.524265499 | 0.694033571 | 0.7531 | 0.840796589 |
| SL1344_RS18880_ | -3.025672747 | -2.911047007 | -3.12747012 | -0.196281422 | 0.569107598 | -0.317419974 | 0.359760166 | 0.506053619 | 0.652638679 | 0.6162 | 0.730146279 |
| SL1344_RS18885_ | -1.630888246 | -1.699029 | -1.577636689 | 0.139277957 | 0.360532588 | 0.372721703 | 0.337108601 | 0.494873432 | 0.645129419 | 0.6087 | 0.723654605 |
| SL1344_RS18890_ | 3.294693994 | 3.342500308 | 3.239896286 | -0.10906129 | 0.090649924 | -1.177856985 | 0.082611643 | 0.136855845 | 0.298393016 | 0.2017 | 0.362821219 |
| SL1344_RS18895_ | 0.012796917 | 0.279405914 | -0.191253654 | -0.463554215 | 0.232173631 | -1.939802571 | 0.019320688 | 0.052384938 | 0.164089416 | 0.1183 | 0.266655022 |
| SL1344_RS18900_ | 0.943098074 | 1.024733288 | 0.824908708 | -0.213189937 | 0.164579992 | -1.19698031 | 0.064666733 | 0.144637007 | 0.303780753 | 0.1885 | 0.341923622 |
| SL1344_RS18905_ | -3.434658685 | -3.166484985 | -3.645322534 | -0.463414203 | 0.654134547 | -0.659672141 | 0.228000014 | 0.374209557 | 0.531578265 | 0.4726 | 0.609312538 |
| SL1344_RS18910_ | -4.660175945 | -4.36628554 | -4.97152276 | -0.627067202 | 1.146106956 | -0.498156778 | 0.281812135 | 0.45088018 | 0.607265243 | 0.5579 | 0.683437642 |
| SL1344_RS18915_ | -3.22902945 | -3.380166107 | -3.087758 | 0.319179263 | 0.656659589 | 0.458191052 | 0.300000009 | 0.453886944 | 0.608924296 | 0.565 | 0.687379605 |
| SL1344_RS18920_ | -6.778673138 | -6.833700645 | -6.71869921 | 0.054312386 | 1.881889509 | 0.029446722 | 0.487008661 | 0.535190939 | 0.675207184 | 0.6404 | 0.745990286 |
| SL1344_RS18925_ | -3.87281234 | -3.829585913 | -3.924668529 | -0.031130368 | 0.843122618 | -0.033417046 | 0.484000001 | 0.616683514 | 0.748157029 | 0.7418 | 0.82948126 |
| SL1344_RS18930_ | -4.097697398 | -4.125352595 | -4.073690687 | 0.049599535 | 0.853468125 | 0.052269741 | 0.474666668 | 0.601277054 | 0.734616548 | 0.7041 | 0.802306488 |
| SL1344_RS18935_ | -1.214600792 | -1.294068574 | -1.086082097 | 0.203811395 | 0.321151959 | 0.607171144 | 0.271152576 | 0.424559482 | 0.581604703 | 0.5357 | 0.665204935 |
| SL1344_RS18945_ | -4.047198479 | -3.977461591 | -4.090538267 | -0.192610139 | 1.195003001 | -0.159761174 | 0.435043307 | 0.676306329 | 0.794478264 | 0.7817 | 0.860668279 |
| SL1344_RS18950_ | -3.144649512 | -2.836664069 | -3.442219544 | -0.662251084 | 0.945446738 | -0.684415216 | 0.194536993 | 0.31799155 | 0.504295012 | 0.4022 | 0.560625663 |
| SL1344_RS18955_ | -2.04859903 | -2.243164459 | -1.661740951 | 0.553537025 | 0.456852594 | 1.124028139 | 0.125249865 | 0.192820556 | 0.358035606 | 0.2827 | 0.441718034 |
| SL1344_RS18960_ | -1.248454816 | -1.152909096 | -1.353490771 | -0.200427996 | 0.353682361 | -0.558753298 | 0.264000011 | 0.429898715 | 0.588675942 | 0.5406 | 0.668425742 |
| SL1344_RS18965_ | -1.72100768 | -1.489538911 | -1.876341551 | -0.396365892 | 0.417308001 | -0.918537378 | 0.159333357 | 0.247838091 | 0.41505471 | 0.3346 | 0.489029246 |
| SL1344_RS18970_ | -1.133904553 | -1.18726522 | -1.06502296 | 0.11194539 | 0.304675689 | 0.335112894 | 0.363333339 | 0.534373699 | 0.678939729 | 0.6386 | 0.750996363 |

| | | | | | | | | | | | |
|---|---|---|---|---|---|---|---|---|---|---|---|
| SL1344_RS18975_ | 2.356726745 | 2.308056765 | 2.421919658 | 0.117002436 | 0.092988961 | 1.185443564 | 0.088666714 | 0.158571784 | 0.317374607 | 0.2186 | 0.375489213 |
| SL1344_RS18980_ | 2.421056855 | 2.30720228 | 2.714781976 | 0.392247993 | 0.148168686 | 2.190810154 | 0.005330642 | 0.050860489 | 0.170117385 | 0.1037 | 0.248010171 |
| SL1344_RS18985_ | 0.823680972 | 0.990454369 | 0.743204329 | -0.231942814 | 0.170718997 | -1.262154001 | 0.102666707 | 0.153550052 | 0.315237734 | 0.2425 | 0.405305061 |
| SL1344_RS18990_ | 1.831713051 | 1.948831841 | 1.683944822 | -0.266032461 | 0.118659719 | -2.161333634 | 0.008000574 | 0.038668873 | 0.137751752 | 0.1087 | 0.254643101 |
| SL1344_RS18995_ | 1.898828183 | 1.929138743 | 1.852691434 | -0.073941183 | 0.116940618 | -0.597944356 | 0.262491684 | 0.406303832 | 0.571216654 | 0.5157 | 0.653074998 |
| SL1344_RS19005_ | -3.137068104 | -3.366683394 | -2.987845481 | 0.392235542 | 0.738838077 | 0.488102438 | 0.287141915 | 0.42855503 | 0.596094998 | 0.5464 | 0.676074754 |
| SL1344_RS19010_ | 3.069155479 | 2.89348727 | 3.230256545 | 0.340719726 | 0.076543077 | 4.446148284 | 4.68E-05 | 0.008594398 | 0.058500067 | 0.1 | 0.243021319 |
| SL1344_RS19015_ | -5.06429579 | -6.326418761 | -3.999611993 | 2.210808217 | 2.244305757 | 0.898422127 | 0.089940086 | 0.20261293 | 0.388078912 | 0.2099 | 0.378577925 |
| SL1344_RS19020_ | 0.009351026 | 0.021683683 | -0.009552992 | -0.026191996 | 0.205525325 | -0.127766437 | 0.451032647 | 0.598950453 | 0.730658088 | 0.7116 | 0.805327475 |
| SL1344_RS19025_ | -0.339775695 | -0.098406615 | -0.682954979 | -0.5870764 | 0.226002284 | -2.536170468 | 0.002668394 | 0.028793506 | 0.113049502 | 0.1023 | 0.246039822 |
| SL1344_RS19030_ | -1.485109994 | -1.288291038 | -1.928008406 | -0.618679923 | 0.434162217 | -1.297958015 | 0.089940086 | 0.137755339 | 0.298919007 | 0.211 | 0.373918286 |
| SL1344_RS19040_ | 1.341304122 | 1.160032254 | 1.513158857 | 0.355499067 | 0.148162583 | 2.363095135 | 0.003334715 | 0.03206047 | 0.12362206 | 0.1038 | 0.248198716 |
| SL1344_RS19045_ | 1.238738436 | 1.238843977 | 1.238161449 | -0.017125697 | 0.183101948 | -0.083696938 | 0.461025984 | 0.68526731 | 0.803583736 | 0.81 | 0.881868859 |
| SL1344_RS19050_ | -1.344958749 | -1.404236469 | -1.276360092 | 0.123730484 | 0.300272454 | 0.373963484 | 0.328000007 | 0.456700379 | 0.607091202 | 0.5625 | 0.684199363 |
| SL1344_RS19055_ | -1.482308047 | -1.576970722 | -1.408633912 | 0.162917001 | 0.335407405 | 0.437844726 | 0.306000008 | 0.461585711 | 0.615085433 | 0.5766 | 0.698927702 |
| SL1344_RS19065_ | 4.270556916 | 4.33574243 | 4.108675769 | -0.226907981 | 0.116857629 | -1.954393263 | 0.00067292 | 0.083631511 | 0.233312674 | 0.1015 | 0.245059336 |
| SL1344_RS19070_ | 4.318842782 | 4.369542535 | 4.292092815 | -0.073876257 | 0.048197795 | -1.441449421 | 0.071285869 | 0.116840869 | 0.263933806 | 0.1869 | 0.345054728 |
| SL1344_RS19075_ | 3.244401671 | 3.261449765 | 3.224507965 | -0.034215756 | 0.060738076 | -0.525327753 | 0.262491684 | 0.395822208 | 0.548611771 | 0.4918 | 0.623590491 |
| SL1344_RS19080_ | 3.769992583 | 3.754706535 | 3.789958927 | 0.06641666 | 0.119224437 | 0.556722368 | 0.274483688 | 0.370196117 | 0.560908708 | 0.555 | 0.691107439 |
| SL1344_RS19085_ | 2.954220952 | 2.997683885 | 2.913385747 | -0.088162954 | 0.088683495 | -0.967185184 | 0.148567647 | 0.220508865 | 0.392067419 | 0.305 | 0.463854192 |
| SL1344_RS19090_ | 3.679561435 | 3.593650076 | 3.791039711 | 0.207530456 | 0.086161902 | 2.384360622 | 0.000673369 | 0.041785443 | 0.1500931 | 0.1003 | 0.243446814 |
| SL1344_RS19095_ | 2.688802023 | 2.483735608 | 2.872168497 | 0.388100997 | 0.070174777 | 5.518301536 | 4.68E-05 | 0.003465522 | 0.035574882 | 0.1 | 0.243021319 |
| SL1344_RS19100_ | -0.711765486 | -1.203854309 | -0.330217206 | 0.881363924 | 0.292468915 | 2.984033272 | 4.68E-05 | 0.0206952 | 0.094383449 | 0.1001 | 0.243155678 |
| SL1344_RS19105_ | 4.138633596 | 4.056583504 | 4.218225723 | 0.171605577 | 0.104870841 | 1.602690726 | 0.024000188 | 0.073650888 | 0.212670662 | 0.1238 | 0.276029768 |
| SL1344_RS19110_ | 3.279071965 | 3.195324922 | 3.338158008 | 0.143728845 | 0.092483393 | 1.517932937 | 0.041972124 | 0.085785958 | 0.225190091 | 0.1495 | 0.305938178 |
| SL1344_RS19115_ | 3.105871463 | 3.040707413 | 3.171513509 | 0.13583128 | 0.082528067 | 1.598035951 | 0.029980163 | 0.076168355 | 0.206439918 | 0.1346 | 0.286375049 |
| SL1344_RS19120_ | 3.231269056 | 3.11906605 | 3.336903008 | 0.218301681 | 0.061997859 | 3.506134119 | 4.68E-05 | 0.011080409 | 0.065702721 | 0.1 | 0.243021319 |
| SL1344_RS19125_ | 3.04314825 | 2.900542963 | 3.157829961 | 0.252432017 | 0.076655799 | 3.212520815 | 0.000673369 | 0.014978468 | 0.080530679 | 0.1 | 0.243021319 |
| SL1344_RS19130_ | 3.668265051 | 3.516363496 | 3.840411846 | 0.3287269 | 0.076162254 | 4.3002097 | 4.68E-05 | 0.010533367 | 0.067646279 | 0.1 | 0.243021319 |
| SL1344_RS19135_ | 2.481243574 | 2.454040262 | 2.533078023 | 0.082971769 | 0.10718077 | 0.696339211 | 0.219853446 | 0.328402804 | 0.505205891 | 0.4386 | 0.583593986 |
| SL1344_RS19140_ | 2.18890894 | 1.707685429 | 2.504606941 | 0.795182255 | 0.169716039 | 4.599340975 | 4.68E-05 | 0.005602301 | 0.048576858 | 0.1 | 0.243021319 |
| SL1344_RS19145_ | 1.499535923 | 1.511196901 | 1.486479235 | -0.037627813 | 0.15510909 | -0.23245949 | 0.413057965 | 0.60908835 | 0.742521541 | 0.7332 | 0.8231396 |
| SL1344_RS19150_ | 2.105099426 | 2.029689969 | 2.169612825 | 0.133340963 | 0.139871199 | 0.930296503 | 0.155229871 | 0.24159109 | 0.421004454 | 0.3268 | 0.494134243 |
| SL1344_RS19155_ | 2.510594435 | 2.501002813 | 2.522868929 | 0.052544544 | 0.140655042 | 0.342587529 | 0.352666673 | 0.548941054 | 0.699532194 | 0.7125 | 0.807070646 |
| SL1344_RS19160_ | 2.44808316 | 2.371726746 | 2.526180962 | 0.154502405 | 0.076164224 | 1.96213841 | 0.016655836 | 0.052014012 | 0.156728032 | 0.1211 | 0.269112163 |
| SL1344_RS19165_ | 0.546633218 | 0.353971269 | 0.680869274 | 0.338547712 | 0.233405952 | 1.399411826 | 0.059293877 | 0.104426541 | 0.253773155 | 0.1638 | 0.321302771 |
| SL1344_RS19170_ | 2.487304203 | 2.430529749 | 2.547795727 | 0.125273964 | 0.110771696 | 1.090224694 | 0.101265863 | 0.165902297 | 0.333377881 | 0.2347 | 0.396151097 |
| SL1344_RS19175_ | 2.912458818 | 2.86267281 | 2.963883125 | 0.105211438 | 0.106278885 | 0.973683296 | 0.144570313 | 0.208614647 | 0.38587755 | 0.2935 | 0.458469511 |
| SL1344_RS19180_ | 2.741718253 | 2.719784274 | 2.771866126 | 0.052758547 | 0.084823923 | 0.585090579 | 0.256666678 | 0.385886842 | 0.547336234 | 0.4911 | 0.627243471 |
| SL1344_RS19185_ | 3.998287823 | 3.756879256 | 4.228248787 | 0.470771841 | 0.099714571 | 4.667853572 | 4.68E-05 | 0.003012323 | 0.037140628 | 0.1 | 0.243021319 |
| SL1344_RS19190_ | 2.750679778 | 2.544743943 | 2.96641553 | 0.421566823 | 0.07004455 | 6.05254957 | 4.68E-05 | 0.002603356 | 0.030727851 | 0.1 | 0.243021319 |
| SL1344_RS19195_ | 1.110158846 | 1.174644911 | 1.059274739 | -0.111656403 | 0.147292726 | -0.682401167 | 0.221852113 | 0.345567281 | 0.510889937 | 0.4426 | 0.586409479 |
| SL1344_RS19200_ | 7.883695768 | 7.831292853 | 7.897033995 | 0.065743823 | 0.084370737 | 0.74856362 | 0.246000013 | 0.316961597 | 0.523014005 | 0.481 | 0.654338472 |
| SL1344_RS19205_ | 7.445872979 | 7.414505507 | 7.478390041 | 0.041178595 | 0.149635725 | 0.279722551 | 0.376000005 | 0.699745473 | 0.825472819 | 0.8314 | 0.898804748 |
| SL1344_RS19210_ | -0.159796612 | -0.300659426 | -0.029492697 | 0.284235126 | 0.236726351 | 1.165629685 | 0.090666713 | 0.163139129 | 0.321777121 | 0.2358 | 0.393277984 |
| SL1344_RS19215_ | 3.134755704 | 3.107524217 | 3.15151557 | 0.030985304 | 0.131742851 | 0.239350106 | 0.386666671 | 0.719005934 | 0.828314046 | 0.7601 | 0.851316002 |
| SL1344_RS19220_ | 2.733572666 | 2.718948544 | 2.765023105 | 0.0314716 | 0.112719592 | 0.2408809 | 0.405729517 | 0.700801765 | 0.812116144 | 0.7747 | 0.861182922 |
| SL1344_RS19225_ | 2.783610345 | 2.774893627 | 2.795043943 | 0.021472232 | 0.101534587 | 0.204139807 | 0.405729517 | 0.622373177 | 0.753389471 | 0.7316 | 0.823105841 |
| SL1344_RS19230_ | 0.135793196 | 0.194798624 | 0.103182143 | -0.081085452 | 0.271071129 | -0.268024272 | 0.389740178 | 0.65505712 | 0.780691258 | 0.7542 | 0.844937735 |
| SL1344_RS19235_ | 1.463800792 | 1.464395227 | 1.461719998 | -0.014504618 | 0.228624551 | -0.055706953 | 0.482000001 | 0.782181628 | 0.872044506 | 0.8922 | 0.937098793 |
| SL1344_RS19240_ | 3.204988622 | 3.205886463 | 3.203762523 | -0.009091587 | 0.097794263 | -0.090077488 | 0.458000002 | 0.699147887 | 0.813078842 | 0.8127 | 0.882289653 |
| SL1344_RS19245_ | -0.14403874 | -0.275410019 | -0.015500193 | 0.258191934 | 0.191998612 | 1.305776734 | 0.079946755 | 0.138831177 | 0.283415141 | 0.2091 | 0.363876125 |
| SL1344_RS19250_ | -1.479598591 | -1.650126458 | -1.365050771 | 0.278283708 | 0.35551584 | 0.747826511 | 0.205862775 | 0.312249457 | 0.479449995 | 0.411 | 0.560464951 |

| | | | | | | | | | | | |
|---|---|---|---|---|---|---|---|---|---|---|---|
| SL1344_RS19255_ | 1.058622379 | 0.942147187 | 1.283327258 | 0.310626388 | 0.21209668 | 1.37058216 | 0.070000061 | 0.110382629 | 0.263782986 | 0.1866 | 0.351690956 |
| SL1344_RS19260_ | -0.765722589 | -0.860269605 | -0.720588878 | 0.142146711 | 0.315978619 | 0.406676095 | 0.331778821 | 0.498270857 | 0.654846058 | 0.6293 | 0.744112645 |
| SL1344_RS19265_ | 3.137370938 | 3.037402932 | 3.307781922 | 0.261231427 | 0.088801485 | 2.720492605 | 4.68E-05 | 0.028220138 | 0.117289329 | 0.1004 | 0.243588578 |
| SL1344_RS19270_ | 3.946952009 | 3.873242333 | 4.033285214 | 0.192412358 | 0.109977176 | 1.818143355 | 0.003332494 | 0.05252274 | 0.173410259 | 0.102 | 0.245768612 |
| SL1344_RS19275_ | 5.409280092 | 5.32546081 | 5.507155336 | 0.20805638 | 0.080107309 | 2.774118502 | 4.68E-05 | 0.020996831 | 0.103193818 | 0.1 | 0.243021319 |
| SL1344_RS19280_ | 3.413162817 | 3.174883802 | 3.538281326 | 0.355964866 | 0.10568911 | 3.319415867 | 4.68E-05 | 0.012062162 | 0.073766272 | 0.1 | 0.243021319 |
| SL1344_RS19285_ | 3.426033737 | 3.207655507 | 3.622177924 | 0.427824182 | 0.105802854 | 4.067667461 | 4.68E-05 | 0.005074067 | 0.046947202 | 0.1 | 0.243021319 |
| SL1344_RS19290_ | 2.295728455 | 2.109621717 | 2.481710456 | 0.375761796 | 0.098801548 | 3.794661096 | 4.68E-05 | 0.008773997 | 0.059217966 | 0.1 | 0.243021319 |
| SL1344_RS19295_ | 2.378852448 | 2.21523889 | 2.591532399 | 0.368171687 | 0.110042465 | 3.257794689 | 0.000673369 | 0.014333153 | 0.078547008 | 0.1001 | 0.243158083 |
| SL1344_RS19300_ | 0.998507882 | 0.868889336 | 1.127901262 | 0.256971427 | 0.194846823 | 1.259613457 | 0.071285869 | 0.123985909 | 0.281080368 | 0.1904 | 0.352849955 |
| SL1344_RS19305_ | 2.275694699 | 2.259601387 | 2.288850767 | 0.031616632 | 0.100853328 | 0.294128853 | 0.37441706 | 0.569710977 | 0.707845359 | 0.6851 | 0.784308839 |
| SL1344_RS19310_ | -0.772448401 | -0.748782435 | -0.802216558 | -0.036636332 | 0.304150385 | -0.126986919 | 0.443704199 | 0.628151343 | 0.753363261 | 0.7293 | 0.819525441 |
| SL1344_RS19315_ | -1.759361322 | -1.882660575 | -1.535920374 | 0.332786027 | 0.400184251 | 0.763884795 | 0.20186544 | 0.316433289 | 0.483646063 | 0.4287 | 0.576077466 |
| SL1344_RS19320_ | 0.372802381 | 0.226215028 | 0.623438379 | 0.386063086 | 0.260564047 | 1.411816638 | 0.058627656 | 0.098806203 | 0.245442555 | 0.1694 | 0.330699541 |
| SL1344_RS19325_ | -2.073680759 | -2.527441715 | -1.887206073 | 0.55906773 | 0.617189061 | 0.805926604 | 0.214666682 | 0.308448857 | 0.493085292 | 0.4258 | 0.588234395 |
| SL1344_RS19330_ | 2.255134654 | 2.354330267 | 2.115509563 | -0.254790481 | 0.13565364 | -1.809765209 | 0.010000458 | 0.056816712 | 0.176463386 | 0.1119 | 0.25850326 |
| SL1344_RS19335_ | 0.397595509 | 0.535873174 | 0.259280132 | -0.27122597 | 0.158350663 | -1.689454164 | 0.027315288 | 0.074067245 | 0.192542574 | 0.1384 | 0.288617224 |
| SL1344_RS19340_ | 2.070466945 | 2.046310852 | 2.085092999 | 0.022569067 | 0.133064862 | 0.1458759 | 0.434666669 | 0.702915753 | 0.815080551 | 0.7849 | 0.867577092 |
| SL1344_RS19345_ | -4.417377157 | -4.110978311 | -5.041085601 | -0.888269103 | 1.104778317 | -0.758312707 | 0.20933335 | 0.308424378 | 0.483556728 | 0.4174 | 0.568239168 |
| SL1344_RS19350_ | -0.235451302 | -0.079298704 | -0.367752472 | -0.279144979 | 0.240448145 | -1.074299243 | 0.116588975 | 0.188203624 | 0.348764295 | 0.2726 | 0.431897466 |
| SL1344_RS19355_ | -0.95032085 | -1.278477173 | -0.56763665 | 0.704283201 | 0.309924859 | 2.203887699 | 0.006662914 | 0.036213988 | 0.13226109 | 0.1076 | 0.253279443 |
| SL1344_RS19360_ | -0.610726476 | -0.937758809 | -0.268476655 | 0.648706099 | 0.343742941 | 1.834337819 | 0.024650416 | 0.055841069 | 0.173635368 | 0.1229 | 0.274105062 |
| SL1344_RS19365_ | -0.912856505 | -0.977586052 | -0.788247926 | 0.170518124 | 0.334197996 | 0.470962783 | 0.305796144 | 0.498344934 | 0.652054319 | 0.6 | 0.726058344 |
| SL1344_RS19370_ | -0.015162206 | -0.012996403 | -0.017157127 | 0.004139213 | 0.31709 | 0.011829595 | 0.496 | 0.722467764 | 0.830111429 | 0.8285 | 0.893363565 |
| SL1344_RS19375_ | 0.89809128 | 1.071255861 | 0.682873666 | -0.3873344 | 0.169039043 | -2.228309711 | 0.003334715 | 0.0393605 | 0.138346249 | 0.1048 | 0.249598661 |
| SL1344_RS19380_ | -0.976797428 | -0.835330557 | -1.09342116 | -0.292871395 | 0.394390259 | -0.703304421 | 0.225849448 | 0.325696264 | 0.503242695 | 0.4395 | 0.588318142 |
| SL1344_RS19385_ | 0.897895573 | 0.703237298 | 1.080613732 | 0.416234228 | 0.223447442 | 1.899931105 | 0.006000767 | 0.04718777 | 0.157638495 | 0.1069 | 0.251787818 |
| SL1344_RS19390_ | -1.005251688 | -0.939983191 | -1.102467821 | -0.183299748 | 0.343109314 | -0.508889216 | 0.29113925 | 0.444953102 | 0.607797544 | 0.566 | 0.691308721 |
| SL1344_RS19395_ | -1.112385935 | -1.048414473 | -1.282349109 | -0.210081055 | 0.347105288 | -0.531712868 | 0.320666674 | 0.506207832 | 0.661306391 | 0.6214 | 0.746663548 |
| SL1344_RS19400_ | -1.606185804 | -1.464892959 | -1.750959902 | -0.293593847 | 0.325988075 | -0.848775865 | 0.175216543 | 0.269626443 | 0.425566275 | 0.3577 | 0.503002869 |
| SL1344_RS19405_ | -2.465439106 | -2.414384808 | -2.503020486 | -0.083576327 | 0.552472264 | -0.130845344 | 0.442000002 | 0.636116665 | 0.762515898 | 0.7437 | 0.832316586 |
| SL1344_RS19410_ | -2.846813351 | -2.719851588 | -3.00401653 | -0.240306171 | 0.670025573 | -0.345454248 | 0.34133334 | 0.532235178 | 0.678462628 | 0.6478 | 0.758511303 |
| SL1344_RS19415_ | -4.264961868 | -4.164091393 | -4.3181501 | -0.002282975 | 1.600417587 | -0.000797095 | 0.498666667 | 0.628086944 | 0.766938807 | 0.7934 | 0.872173854 |
| SL1344_RS19420_ | -5.344686009 | -5.153095291 | -5.674445022 | -0.501240486 | 1.365422961 | -0.329221433 | 0.346666673 | 0.500033961 | 0.653034338 | 0.6235 | 0.738450069 |
| SL1344_RS19425_ | 1.88205544 | 1.905443325 | 1.864785126 | -0.04371978 | 0.101949433 | -0.404853267 | 0.331112599 | 0.477751486 | 0.625549188 | 0.5822 | 0.700187827 |
| SL1344_RS19430_ | -5.18347101 | -5.205822152 | -5.168999953 | -0.058464118 | 1.565828629 | -0.037297721 | 0.487333334 | 0.669518409 | 0.789976721 | 0.7893 | 0.865235706 |
| SL1344_RS19435_ | -1.242750037 | -1.164334534 | -1.31789579 | -0.153738572 | 0.284854084 | -0.485976787 | 0.300466365 | 0.432857754 | 0.583323921 | 0.5365 | 0.662241579 |
| SL1344_RS19445_ | -0.431869233 | -0.570424851 | -0.310836972 | 0.210734745 | 0.358434073 | 0.564743034 | 0.272485021 | 0.440451648 | 0.61062909 | 0.5473 | 0.686239102 |
| SL1344_RS19450_ | -0.11666515 | -0.167758352 | -0.074941562 | 0.08805513 | 0.275019068 | 0.305902123 | 0.368000005 | 0.576083805 | 0.716907511 | 0.6659 | 0.775779142 |
| SL1344_RS19455_ | -2.940382441 | -2.975624567 | -2.904617607 | 0.066834966 | 0.567480834 | 0.117465833 | 0.447035312 | 0.577589426 | 0.713266105 | 0.6948 | 0.794181946 |
| SL1344_RS19460_ | -3.775899362 | -3.744268545 | -3.794524818 | -0.011522408 | 1.067368104 | -0.014531315 | 0.493333334 | 0.70136619 | 0.815431941 | 0.8238 | 0.89138558 |
| SL1344_RS19465_ | -0.132292624 | -0.403215845 | 0.110119199 | 0.510608305 | 0.209326516 | 2.428968344 | 0.004664544 | 0.029164033 | 0.114626251 | 0.104 | 0.248305803 |
| SL1344_RS19470_ | -1.505040953 | -1.721357002 | -1.228561515 | 0.482152594 | 0.309159097 | 1.48896544 | 0.063333401 | 0.106522388 | 0.242351548 | 0.1813 | 0.336757879 |
| SL1344_RS19475_ | -2.061829154 | -1.993622354 | -2.103811025 | -0.05090457 | 0.543026973 | -0.086911691 | 0.466355764 | 0.698099818 | 0.810541588 | 0.7986 | 0.872872129 |
| SL1344_RS19480_ | -1.277317096 | -1.523574074 | -1.086855068 | 0.440020926 | 0.297012299 | 1.422432866 | 0.051333419 | 0.115661968 | 0.255260228 | 0.1788 | 0.332471828 |
| SL1344_RS19485_ | -0.516583986 | -0.810958288 | -0.227210251 | 0.584801844 | 0.231019518 | 2.472692639 | 0.004667653 | 0.027437084 | 0.111054873 | 0.1037 | 0.247988916 |
| SL1344_RS19490_ | 0.549379753 | 0.415586619 | 0.667668631 | 0.224733415 | 0.226662517 | 0.950942711 | 0.161225873 | 0.244174626 | 0.422755931 | 0.3323 | 0.500147913 |
| SL1344_RS19495_ | -1.46834058 | -1.534482407 | -1.407949948 | 0.118719049 | 0.330131623 | 0.337828148 | 0.353764163 | 0.504261467 | 0.649731865 | 0.6156 | 0.726718536 |
| SL1344_RS19500_ | 1.300645281 | 1.075622784 | 1.467240848 | 0.395299519 | 0.145736375 | 2.599331001 | 0.002666616 | 0.025025876 | 0.106954777 | 0.1018 | 0.24547649 |
| SL1344_RS19505_ | 1.778402918 | 1.693821543 | 1.885366318 | 0.192343526 | 0.107292197 | 1.747924981 | 0.027333498 | 0.071918806 | 0.194398754 | 0.1346 | 0.284636159 |
| SL1344_RS19510_ | 1.095429672 | 1.1055369 | 1.080124895 | -0.024362951 | 0.157652903 | -0.139849423 | 0.440666669 | 0.623794397 | 0.751584123 | 0.7347 | 0.824940134 |
| SL1344_RS19515_ | -0.185725325 | -0.253611608 | -0.080042272 | 0.18804161 | 0.226635396 | 0.756539763 | 0.203864107 | 0.312567048 | 0.48184273 | 0.409 | 0.55615996 |
| SL1344_RS19520_ | -1.975402941 | -2.180390783 | -1.705835467 | 0.403655208 | 0.553660226 | 0.690076628 | 0.224517003 | 0.353439575 | 0.529618696 | 0.4528 | 0.606213249 |

| | | | | | | | | | | | |
|---|---|---|---|---|---|---|---|---|---|---|---|
| SL1344_RS19525_ | -1.94195908 | -2.157375396 | -1.8317495 | 0.309553216 | 0.461077171 | 0.591318376 | 0.266489018 | 0.410772148 | 0.574994815 | 0.5246 | 0.662576568 |
| SL1344_RS19530_ | -0.584818976 | -0.632630116 | -0.529253019 | 0.105464913 | 0.213880176 | 0.458588548 | 0.301798809 | 0.421544612 | 0.575487153 | 0.5218 | 0.649265397 |
| SL1344_RS19535_ | -1.611930168 | -1.394852382 | -1.837039218 | -0.450214487 | 0.325410342 | -1.301486359 | 0.075283201 | 0.12841805 | 0.269849081 | 0.1966 | 0.35172208 |
| SL1344_RS19540_ | 0.591074575 | 0.671321094 | 0.518447381 | -0.159542588 | 0.18796749 | -0.821821883 | 0.174666688 | 0.2798765 | 0.448937397 | 0.3775 | 0.530869301 |
| SL1344_RS19545_ | 2.092506174 | 1.896228203 | 2.292053183 | 0.349253829 | 0.204322605 | 1.697329173 | 0.03331126 | 0.061144497 | 0.189768254 | 0.1319 | 0.287507557 |
| SL1344_RS19550_ | 1.185367841 | 1.21629561 | 1.150811572 | -0.06392273 | 0.128083009 | -0.45330358 | 0.29113925 | 0.442882424 | 0.598393929 | 0.5501 | 0.678277814 |
| SL1344_RS19555_ | -0.593092064 | -0.787408316 | -0.416116132 | 0.392112849 | 0.33076397 | 1.197548653 | 0.083277865 | 0.152678102 | 0.315710654 | 0.2235 | 0.385925828 |
| SL1344_RS19565_ | 0.574040707 | 0.534804529 | 0.625913845 | 0.110031392 | 0.200351897 | 0.507470601 | 0.279813468 | 0.422597422 | 0.5870282 | 0.5506 | 0.675169908 |
| SL1344_RS19570_ | -0.754196079 | -0.754859362 | -0.753688004 | 0.028648492 | 0.29975837 | 0.094119301 | 0.460000001 | 0.62214693 | 0.751975193 | 0.7464 | 0.832663642 |
| SL1344_RS19575_ | -1.863501019 | -1.695315903 | -2.124337557 | -0.422339762 | 0.473788852 | -0.849987911 | 0.186000019 | 0.275763464 | 0.451015665 | 0.3745 | 0.533754346 |
| SL1344_RS19580_ | 0.000996793 | 0.05950356 | -0.033898897 | -0.079100188 | 0.239850099 | -0.296677906 | 0.377748173 | 0.583921692 | 0.722871669 | 0.6922 | 0.796742331 |
| SL1344_RS19585_ | 0.431469259 | 0.389391546 | 0.463532426 | 0.072673913 | 0.170376539 | 0.397167499 | 0.333111266 | 0.478749583 | 0.627173575 | 0.5838 | 0.704250454 |
| SL1344_RS19590_ | 0.187072601 | 0.234204017 | 0.155939308 | -0.080869892 | 0.193753377 | -0.39100362 | 0.347768161 | 0.497052078 | 0.643658569 | 0.6009 | 0.717663367 |
| SL1344_RS19595_ | 0.348655951 | 0.248136815 | 0.431618915 | 0.181262214 | 0.167612844 | 1.020219016 | 0.13000003 | 0.212452961 | 0.370874341 | 0.2869 | 0.443179926 |
| SL1344_RS19600_ | 0.620299691 | 0.755534901 | 0.429314219 | -0.325396038 | 0.205958081 | -1.531479619 | 0.041333441 | 0.091277031 | 0.231607168 | 0.1463 | 0.300812569 |
| SL1344_RS19605_ | 2.56107338 | 3.205585279 | 2.20604272 | -0.977307042 | 0.291884669 | -3.153543537 | 4.68E-05 | 0.014962508 | 0.086701417 | 0.1 | 0.243021319 |
| SL1344_RS19610_ | 1.83297602 | 2.43401771 | 1.49029376 | -1.049553401 | 0.459001086 | -2.339738397 | 4.68E-05 | 0.022645184 | 0.109847932 | 0.1 | 0.243021319 |
| SL1344_RS19615_ | -0.037939571 | 0.006512667 | -0.094536012 | -0.128345862 | 0.341144043 | -0.344402619 | 0.348666673 | 0.533942689 | 0.689479479 | 0.6698 | 0.777115698 |
| SL1344_RS19620_ | 0.996952203 | 0.929596069 | 1.078245637 | 0.148063105 | 0.142245585 | 0.980980812 | 0.125916087 | 0.213709547 | 0.371759833 | 0.293 | 0.443941895 |
| SL1344_RS19625_ | -1.592807096 | -1.41095857 | -1.715439292 | -0.294662047 | 0.347766412 | -0.785549817 | 0.209860109 | 0.310670819 | 0.473195599 | 0.4083 | 0.556515471 |
| SL1344_RS19630_ | -5.056503712 | -4.789883446 | -5.402816015 | -0.630117074 | 1.142306364 | -0.484374248 | 0.281333343 | 0.40342469 | 0.561823846 | 0.5097 | 0.639714111 |
| SL1344_RS19635_ | -3.107302988 | -3.307554928 | -2.945624243 | 0.343468896 | 0.757243791 | 0.452321217 | 0.303131254 | 0.470994721 | 0.630164773 | 0.5779 | 0.70220092 |
| SL1344_RS19640_ | -3.655954377 | -3.806012427 | -3.560391966 | 0.206934163 | 0.850703651 | 0.223821237 | 0.402000004 | 0.605702922 | 0.738267435 | 0.722 | 0.81571348 |
| SL1344_RS19645_ | -5.282801455 | -5.332707349 | -5.260861056 | 0.117969616 | 1.626835451 | 0.074455502 | 0.460000001 | 0.617958641 | 0.753683787 | 0.748 | 0.836512473 |
| SL1344_RS19650_ | -3.650351864 | -3.522446652 | -3.74312626 | -0.282884964 | 0.87264752 | -0.309474232 | 0.362000006 | 0.54912105 | 0.690915374 | 0.6686 | 0.770138852 |
| SL1344_RS19655_ | -5.176996264 | -5.161576773 | -5.210760245 | -0.064430317 | 1.631066629 | -0.037839881 | 0.480346436 | 0.640137632 | 0.770950748 | 0.7739 | 0.855516089 |
| SL1344_RS19660_ | -4.718331861 | -4.657645917 | -4.796041195 | 0.102153156 | 1.638572898 | 0.060313023 | 0.466000001 | 0.69165982 | 0.807390434 | 0.8373 | 0.89665222 |
| SL1344_RS19665_ | -2.915146468 | -2.429611285 | -3.134122418 | -0.629580983 | 0.679426179 | -0.822072083 | 0.20933335 | 0.317853828 | 0.491835404 | 0.4232 | 0.581301343 |
| SL1344_RS19670_ | -3.1512517 | -2.876938551 | -3.391968844 | -0.516129594 | 0.578084896 | -0.848144626 | 0.178547655 | 0.274241207 | 0.433293454 | 0.3657 | 0.515263973 |
| SL1344_RS19675_ | -1.038391095 | -1.024916425 | -1.046111565 | -0.049903557 | 0.327214513 | -0.148840069 | 0.441039309 | 0.607546974 | 0.740564991 | 0.7239 | 0.816332636 |
| SL1344_RS19680_ | -2.743077955 | -2.776038158 | -2.706354202 | 0.059928774 | 0.491218147 | 0.118698281 | 0.447333335 | 0.534532295 | 0.677930772 | 0.637 | 0.748380285 |
| SL1344_RS19685_ | 1.17210532 | 1.17797929 | 1.166136153 | -0.016263959 | 0.138500998 | -0.115711841 | 0.453697537 | 0.585838086 | 0.721849123 | 0.7014 | 0.797303011 |
| SL1344_RS19690_ | -1.423991603 | -1.500977137 | -1.366380796 | 0.123684794 | 0.313667595 | 0.377086321 | 0.344000006 | 0.476918129 | 0.62722406 | 0.583 | 0.704513994 |
| SL1344_RS19695_ | 0.921959579 | 0.836505244 | 1.008835054 | 0.175453491 | 0.158627131 | 1.067705121 | 0.119920087 | 0.190611988 | 0.352909317 | 0.2712 | 0.429757118 |
| SL1344_RS19700_ | -2.06010107 | -2.074241662 | -2.04763724 | 0.044529352 | 0.389399106 | 0.110769489 | 0.455029982 | 0.556048422 | 0.69563715 | 0.6662 | 0.769956198 |
| SL1344_RS19705_ | -1.740748922 | -1.934911005 | -1.589795196 | 0.339222747 | 0.394685182 | 0.79037664 | 0.186000019 | 0.288673274 | 0.455470752 | 0.3812 | 0.531186104 |
| SL1344_RS19710_ | -3.249769222 | -3.175106828 | -3.378899573 | -0.197957824 | 0.634302403 | -0.287248983 | 0.376000005 | 0.527030673 | 0.672462325 | 0.6354 | 0.749212148 |
| SL1344_RS19715_ | -0.441591129 | -0.428112548 | -0.454433891 | -0.005185292 | 0.277092548 | -0.013851178 | 0.492338441 | 0.653147721 | 0.776670025 | 0.7493 | 0.837414328 |
| SL1344_RS19720_ | -1.105199182 | -1.102414838 | -1.106450473 | -0.006986375 | 0.286761176 | -0.01795801 | 0.493333334 | 0.57870659 | 0.714558126 | 0.683 | 0.785993772 |
| SL1344_RS19725_ | 1.720290334 | 1.670729027 | 1.75721383 | 0.09302635 | 0.12983932 | 0.664531884 | 0.20333335 | 0.333501701 | 0.506745915 | 0.4467 | 0.588050848 |
| SL1344_RS19730_ | -3.213634929 | -3.143025331 | -3.282901658 | -0.13643738 | 0.573919166 | -0.22130521 | 0.40639574 | 0.530747099 | 0.672646409 | 0.6434 | 0.750436119 |
| SL1344_RS19735_ | -2.319333931 | -2.36292572 | -2.296509362 | 0.010840037 | 0.764056508 | 0.013808941 | 0.495003331 | 0.760250041 | 0.857052686 | 0.8604 | 0.916172353 |
| SL1344_RS19740_ | -1.605998731 | -1.732590453 | -1.445167143 | 0.282801682 | 0.481127413 | 0.564363304 | 0.250499679 | 0.410958823 | 0.581325513 | 0.522 | 0.661841683 |
| SL1344_RS19745_ | 5.318379376 | 5.224607993 | 5.414275004 | 0.195480419 | 0.05159816 | 3.850513064 | 4.68E-05 | 0.011051274 | 0.070118373 | 0.1 | 0.243021319 |
| SL1344_RS19750_ | 3.255377395 | 3.183657469 | 3.333674124 | 0.151629776 | 0.078297756 | 1.948094892 | 0.013990999 | 0.051063436 | 0.162738103 | 0.1147 | 0.262076513 |
| SL1344_RS19755_ | 3.836349347 | 3.788236752 | 3.968998754 | 0.175550759 | 0.070861291 | 2.059134264 | 0.011326186 | 0.062151137 | 0.188822104 | 0.1133 | 0.260177877 |
| SL1344_RS19765_ | 6.822299096 | 6.711503021 | 6.885522894 | 0.176577229 | 0.08906766 | 2.128507745 | 4.68E-05 | 0.050603497 | 0.174386034 | 0.1 | 0.243021319 |
| SL1344_RS19770_ | 5.139867646 | 5.06267269 | 5.353092092 | 0.284800118 | 0.059836219 | 3.25011264 | 4.68E-05 | 0.040992365 | 0.153008123 | 0.1 | 0.243021319 |
| SL1344_RS19775_ | 4.028292941 | 3.860419022 | 4.109251475 | 0.242222143 | 0.078051157 | 2.893276889 | 4.68E-05 | 0.02980918 | 0.123161351 | 0.1002 | 0.243306834 |
| SL1344_RS19780_ | 6.137863455 | 6.011231779 | 6.319939495 | 0.311551812 | 0.071037333 | 4.533136484 | 4.68E-05 | 0.002168476 | 0.032236664 | 0.1 | 0.243021319 |
| SL1344_RS19785_ | 2.254474708 | 2.101876844 | 2.432160283 | 0.344873663 | 0.123121873 | 2.769388542 | 4.68E-05 | 0.019292488 | 0.094722755 | 0.1 | 0.243021319 |
| SL1344_RS19790_ | -2.393522285 | -2.582152699 | -2.234113339 | 0.324699345 | 0.493298271 | 0.608804046 | 0.231333347 | 0.387209195 | 0.548400969 | 0.4984 | 0.632399684 |
| SL1344_RS19795_ | -2.040406002 | -1.981536335 | -2.077053547 | -0.06013159 | 0.443778411 | -0.120103404 | 0.440666669 | 0.61750466 | 0.74634763 | 0.7251 | 0.817638596 |

| | | | | | | | | | | |
|---|---|---|---|---|---|---|---|---|---|---|
| SL1344_RS19800_ | 3.099862403 | 3.059699667 | 3.242150129 | 0.183753105 | 0.172214282 | 1.035181578 | 0.10933337 | 0.206355536 | 0.401506558 | 0.2472 | 0.413689621 |
| SL1344_RS19805_ | 2.251430627 | 2.111418549 | 2.409151662 | 0.318286276 | 0.159547372 | 1.981670648 | 0.001336769 | 0.085685709 | 0.236079461 | 0.1007 | 0.243998444 |
| SL1344_RS19810_ | -1.800955748 | -1.865311092 | -1.708016122 | 0.189017647 | 0.477979552 | 0.401018243 | 0.349100606 | 0.511871112 | 0.664392632 | 0.641 | 0.752332885 |
| SL1344_RS19815_ | -0.218652695 | -0.243441082 | -0.175227012 | 0.078776507 | 0.246443953 | 0.286248585 | 0.376000005 | 0.56757159 | 0.707095003 | 0.6782 | 0.780952843 |
| SL1344_RS19820_ | -0.880094009 | -1.080003446 | -0.65614925 | 0.420659059 | 0.245635843 | 1.677856983 | 0.033333467 | 0.08166397 | 0.206580659 | 0.1473 | 0.299701554 |
| SL1344_RS19825_ | 1.498682588 | 1.430615682 | 1.526955456 | 0.09510312 | 0.164901585 | 0.52007409 | 0.297333342 | 0.439686683 | 0.612193679 | 0.591 | 0.718583821 |
| SL1344_RS19830_ | 2.204529595 | 2.077115755 | 2.301665173 | 0.222755264 | 0.097130985 | 2.239526093 | 0.00733396 | 0.037954243 | 0.133731133 | 0.109 | 0.254832421 |
| SL1344_RS19835_ | 1.861762432 | 1.861601504 | 1.862138072 | -0.01702809 | 0.172374092 | -0.097668777 | 0.461333335 | 0.699401455 | 0.814334771 | 0.8186 | 0.887130091 |
| SL1344_RS19840_ | 2.133521049 | 1.99794785 | 2.322499965 | 0.314623355 | 0.112220748 | 2.700598294 | 0.002666616 | 0.024631897 | 0.106204797 | 0.1026 | 0.246608253 |
| SL1344_RS19845_ | 0.93123451 | 0.758058052 | 1.096827212 | 0.351413812 | 0.180978051 | 1.908995994 | 0.012667027 | 0.054707705 | 0.169967406 | 0.1121 | 0.258653626 |
| SL1344_RS19850_ | 0.248372122 | 0.190519158 | 0.303919436 | 0.114467132 | 0.215398959 | 0.516769701 | 0.28847436 | 0.453863965 | 0.614168677 | 0.554 | 0.68581383 |
| SL1344_RS19855_ | 0.427321302 | 0.478172039 | 0.399286135 | -0.059089107 | 0.218242758 | -0.241843148 | 0.401333337 | 0.646125843 | 0.770509046 | 0.7309 | 0.828675879 |
| SL1344_RS19860_ | 0.959230807 | 1.069172371 | 0.841095119 | -0.233588543 | 0.160806875 | -1.367340958 | 0.059293877 | 0.10740908 | 0.249057662 | 0.1709 | 0.326672494 |
| SL1344_RS19865_ | -1.495032515 | -1.346854355 | -1.615264843 | -0.26789759 | 0.489803938 | -0.521596443 | 0.276666677 | 0.443281671 | 0.610454388 | 0.5509 | 0.688964874 |
| SL1344_RS19870_ | -4.038130308 | -4.058722792 | -4.018211947 | 0.06927948 | 0.735061667 | 0.078064432 | 0.465023319 | 0.532070204 | 0.67459047 | 0.6452 | 0.752876716 |
| SL1344_RS19875_ | 0.516736233 | 0.5689393 | 0.470957737 | -0.093645541 | 0.15994979 | -0.543873832 | 0.270486353 | 0.408860066 | 0.566273941 | 0.5192 | 0.648226134 |
| SL1344_RS19880_ | 5.43655708 | 5.399278624 | 5.524161409 | 0.116821748 | 0.057696522 | 1.93268256 | 0.015323415 | 0.062330519 | 0.190990021 | 0.1189 | 0.26847146 |
| SL1344_RS19885_ | 5.344053394 | 5.280601316 | 5.419529469 | 0.141344375 | 0.04161904 | 3.368979394 | 4.68E-05 | 0.01209941 | 0.072321351 | 0.1 | 0.243021319 |
| SL1344_RS19890_ | -0.073285062 | -0.264608221 | 0.106422064 | 0.410566896 | 0.274758579 | 1.455389903 | 0.038641021 | 0.095144206 | 0.238282325 | 0.1489 | 0.301536695 |
| SL1344_RS19900_ | 6.244292191 | 6.031485663 | 6.454799699 | 0.431180893 | 0.046921348 | 9.553744309 | 4.68E-05 | 0.011307643 | 0.075464424 | 0.1 | 0.243021319 |
| SL1344_RS19905_ | 5.84425542 | 5.588262702 | 6.137293059 | 0.551179396 | 0.060920302 | 9.131967261 | 4.68E-05 | 0.002527369 | 0.034176384 | 0.1 | 0.243021319 |
| SL1344_RS19910_ | 7.184357358 | 6.906391835 | 7.449755826 | 0.563600601 | 0.07825705 | 7.407162146 | 4.68E-05 | 0.001523218 | 0.027606569 | 0.1 | 0.243021319 |
| SL1344_RS19915_ | 5.712652948 | 5.508702181 | 5.92129915 | 0.428425156 | 0.057636434 | 7.526555645 | 4.68E-05 | 0.001164532 | 0.022578123 | 0.1 | 0.243021319 |
| SL1344_RS19920_ | 6.113237314 | 5.823190419 | 6.309833585 | 0.476821437 | 0.125825678 | 4.001129754 | 4.68E-05 | 0.002382406 | 0.034575374 | 0.1 | 0.243021319 |
| SL1344_RS19925_ | 5.039745902 | 4.798638243 | 5.243560885 | 0.439915358 | 0.066906097 | 6.488626734 | 4.68E-05 | 0.00103222 | 0.022179661 | 0.1 | 0.243021319 |
| SL1344_RS19930_ | 4.985472775 | 4.772040636 | 5.2304443 | 0.458199806 | 0.06593548 | 6.908290827 | 4.68E-05 | 0.000997485 | 0.021344727 | 0.1 | 0.243021319 |
| SL1344_RS19935_ | 3.842447978 | 3.612950514 | 4.062930598 | 0.450806888 | 0.059282523 | 7.620402499 | 4.68E-05 | 0.001314439 | 0.022801341 | 0.1 | 0.243021319 |
| SL1344_RS19945_ | 3.308578198 | 3.110015925 | 3.522855134 | 0.397831135 | 0.191957895 | 2.026684341 | 0.000673369 | 0.039146822 | 0.148493305 | 0.1014 | 0.244987352 |
| SL1344_RS19950_ | 4.565748817 | 4.501610327 | 4.643959278 | 0.149014276 | 0.060829185 | 2.436764737 | 0.00067292 | 0.034170443 | 0.131422975 | 0.1006 | 0.24386232 |
| SL1344_RS19955_ | 0.562984636 | 0.568390528 | 0.558575695 | -0.011918573 | 0.240398004 | -0.050861135 | 0.483333334 | 0.699809354 | 0.812902542 | 0.7913 | 0.867602076 |
| SL1344_RS19960_ | -0.162001132 | -0.20402539 | -0.106160212 | 0.092636176 | 0.199657705 | 0.41945094 | 0.312666675 | 0.450635046 | 0.599463262 | 0.5501 | 0.672189172 |
| SL1344_RS19965_ | 1.131398239 | 1.01425318 | 1.235900195 | 0.22756006 | 0.141462472 | 1.554061657 | 0.040639683 | 0.090713176 | 0.224989919 | 0.1476 | 0.299144558 |
| SL1344_RS19970_ | 1.750686876 | 1.674239537 | 1.81424412 | 0.135491811 | 0.140712518 | 0.888912624 | 0.178666687 | 0.256199421 | 0.433023531 | 0.3507 | 0.513909561 |
| SL1344_RS19980_ | 2.09378709 | 2.370073328 | 1.824687706 | -0.556772182 | 0.128295906 | -4.316726477 | 4.68E-05 | 0.007621366 | 0.056711632 | 0.1 | 0.243021319 |
| SL1344_RS19985_ | 1.584014571 | 1.811559567 | 1.425443917 | -0.380899272 | 0.192259706 | -1.944324535 | 0.012000381 | 0.044643871 | 0.154820457 | 0.1102 | 0.256957179 |
| SL1344_RS19990_ | 1.298844629 | 1.537740269 | 1.078520238 | -0.45640003 | 0.127272124 | -3.565547432 | 0.00067292 | 0.010449439 | 0.065192963 | 0.1 | 0.243021319 |
| SL1344_RS19995_ | 2.747322132 | 2.978039328 | 2.606776402 | -0.358298903 | 0.100841643 | -3.320059543 | 0.00067292 | 0.019047851 | 0.09397061 | 0.1 | 0.243021319 |
| SL1344_RS20000_ | 1.589430677 | 1.646564896 | 1.481485266 | -0.164535397 | 0.150155891 | -1.025988745 | 0.140666694 | 0.199830327 | 0.375435247 | 0.2887 | 0.45416688 |
| SL1344_RS20005_ | 1.522759888 | 1.505723582 | 1.537104864 | 0.012707553 | 0.152828774 | 0.073258015 | 0.467688209 | 0.650675108 | 0.777044232 | 0.7651 | 0.848968562 |
| SL1344_RS20020_ | 1.35796228 | 1.235110856 | 1.681479081 | 0.421678109 | 0.204884096 | 1.798934225 | 0.023984199 | 0.070463791 | 0.204440362 | 0.124 | 0.274246344 |
| SL1344_RS20025_ | -1.745844262 | -2.151580935 | -1.364832578 | 0.999218309 | 0.805075139 | 1.304492105 | 0.040000111 | 0.112107694 | 0.274111913 | 0.147 | 0.303649034 |
| SL1344_RS20030_ | -0.027799057 | -0.126456095 | 0.371233587 | 0.492549 | 0.491885377 | 0.862090663 | 0.208527665 | 0.271900995 | 0.470985702 | 0.4145 | 0.586939342 |
| SL1344_RS20035_ | 1.711694244 | 1.369367431 | 1.795323373 | 0.510521051 | 0.649413689 | 0.891918661 | 0.178547655 | 0.202891882 | 0.400440898 | 0.322 | 0.513818147 |
| SL1344_RS20045_ | 4.315111832 | 4.115065513 | 4.474047524 | 0.353675802 | 0.146180123 | 2.310256083 | 4.68E-05 | 0.059314954 | 0.191391676 | 0.1 | 0.243021319 |
| SL1344_RS20050_ | 4.968484094 | 4.778158251 | 5.221573937 | 0.481802626 | 0.209862644 | 2.385964349 | 4.68E-05 | 0.016805817 | 0.093423888 | 0.1 | 0.243021319 |
| SL1344_RS20055_ | 0.770253819 | 0.7326477 | 0.831744206 | 0.1209642 | 0.206918081 | 0.534202722 | 0.284477025 | 0.435710319 | 0.602353625 | 0.5713 | 0.694580106 |
| SL1344_RS20060_ | 5.148692012 | 5.117459225 | 5.234973232 | 0.084889382 | 0.172261082 | 0.371236576 | 0.335776156 | 0.825983001 | 0.90363318 | 0.7042 | 0.831970939 |
| SL1344_RS20065_ | -0.951866037 | -0.784523149 | -1.150999425 | -0.370749049 | 0.307624287 | -1.12041406 | 0.102666707 | 0.164046724 | 0.324704009 | 0.2373 | 0.399280812 |
| SL1344_RS20070_ | 1.662896621 | 1.616667291 | 1.787922891 | 0.161097465 | 0.173181866 | 0.818264397 | 0.211858777 | 0.306573404 | 0.492503201 | 0.4264 | 0.591481301 |
| SL1344_RS20075_ | 0.662288454 | 0.80083247 | 0.45955662 | -0.334461469 | 0.17540701 | -1.834180116 | 0.023984199 | 0.062591264 | 0.182199673 | 0.1279 | 0.278290304 |
| SL1344_RS20085_ | 1.181418575 | 1.309849362 | 1.093914449 | -0.210851606 | 0.13650084 | -1.475086371 | 0.057333409 | 0.107135389 | 0.247095059 | 0.1765 | 0.332054478 |
| SL1344_RS20090_ | 2.027324928 | 2.067842096 | 1.966248188 | -0.124646991 | 0.187889218 | -0.706663981 | 0.212666683 | 0.308266705 | 0.499866987 | 0.405 | 0.569222513 |
| SL1344_RS20095_ | 2.804735738 | 2.817382146 | 2.787785388 | -0.051151816 | 0.14572155 | -0.354946829 | 0.33133334 | 0.51642963 | 0.680677334 | 0.6802 | 0.787606774 |

| | | | | | | | | | | | |
|---|---|---|---|---|---|---|---|---|---|---|---|
| SL1344_RS20100_ | 0.120140956 | 0.19620369 | 0.031035445 | -0.157356803 | 0.19433457 | -0.758326407 | 0.197333351 | 0.304621834 | 0.469149608 | 0.3973 | 0.544456159 |
| SL1344_RS20105_ | -0.013559231 | 0.062499268 | -0.098762649 | -0.168585894 | 0.185891926 | -0.840411208 | 0.1798801 | 0.27459154 | 0.43477289 | 0.3647 | 0.51122938 |
| SL1344_RS20110_ | 0.300240138 | 0.266077835 | 0.332718725 | 0.063288066 | 0.2088337 | 0.298407432 | 0.366666672 | 0.566265551 | 0.703914157 | 0.6682 | 0.772397922 |
| SL1344_RS20115_ | 3.085407045 | 3.532054938 | 2.663878868 | -0.867405413 | 0.14330121 | -6.044705199 | 4.68E-05 | 0.001442786 | 0.02599311 | 0.1 | 0.243021319 |
| SL1344_RS20120_ | 2.650946206 | 2.623312966 | 2.762330921 | 0.138984793 | 0.174036539 | 0.81400929 | 0.193870771 | 0.29031526 | 0.489600379 | 0.4068 | 0.566574744 |
| SL1344_RS20125_ | -1.079145477 | -1.205375128 | -0.937628117 | 0.266182917 | 0.300960856 | 0.848459635 | 0.17721521 | 0.276169279 | 0.440516155 | 0.3731 | 0.525213111 |
| SL1344_RS20130_ | 3.092166947 | 3.026144077 | 3.213485474 | 0.165948414 | 0.11180866 | 1.425517015 | 0.07533339 | 0.108922571 | 0.26420619 | 0.1823 | 0.348905864 |
| SL1344_RS20135_ | 4.302345212 | 4.134159093 | 4.573954122 | 0.439661513 | 0.136117928 | 3.310701205 | 4.68E-05 | 0.006416139 | 0.055221239 | 0.1 | 0.243021319 |
| SL1344_RS20140_ | 4.164669426 | 3.952994813 | 4.334425449 | 0.386639036 | 0.130329688 | 3.08276708 | 4.68E-05 | 0.014509851 | 0.084367851 | 0.1 | 0.243021319 |
| SL1344_RS20145_ | 6.220479943 | 6.092940645 | 6.346909253 | 0.251701684 | 0.065655908 | 3.85093769 | 4.68E-05 | 0.012428243 | 0.077576673 | 0.1 | 0.243021319 |
| SL1344_RS20150_ | 6.30102635 | 6.136538185 | 6.46518782 | 0.346970609 | 0.096370504 | 3.587369229 | 4.68E-05 | 0.004570252 | 0.047196271 | 0.1 | 0.243021319 |
| SL1344_RS20155_ | 3.299443561 | 3.185863392 | 3.403724308 | 0.222090445 | 0.080858683 | 2.699762528 | 0.00067292 | 0.020139943 | 0.095109477 | 0.1004 | 0.243587716 |
| SL1344_RS20160_ | 3.617404588 | 3.549983836 | 3.694523533 | 0.148461436 | 0.068535771 | 2.151207921 | 0.004667653 | 0.0422113 | 0.14473815 | 0.105 | 0.249646349 |
| SL1344_RS20165_ | 3.245298931 | 3.079167437 | 3.385681602 | 0.303815593 | 0.086254098 | 3.487087687 | 4.68E-05 | 0.009867445 | 0.065355761 | 0.1 | 0.243021319 |
| SL1344_RS20175_ | 2.734723437 | 2.560038294 | 2.937353283 | 0.380648732 | 0.090366422 | 4.229051968 | 4.68E-05 | 0.007258491 | 0.054122121 | 0.1 | 0.243021319 |
| SL1344_RS20180_ | 1.036377227 | 0.740828964 | 1.310326864 | 0.574655563 | 0.140212972 | 4.082523223 | 4.68E-05 | 0.007509942 | 0.054196953 | 0.1 | 0.243021319 |
| SL1344_RS20185_ | 1.837252082 | 1.730652199 | 1.982773398 | 0.255825743 | 0.129285211 | 1.872106483 | 0.014666978 | 0.057367375 | 0.175134165 | 0.1157 | 0.2637231 |
| SL1344_RS20190_ | 2.344853993 | 2.215523383 | 2.525436957 | 0.307277676 | 0.090028886 | 3.406225283 | 0.00067292 | 0.012660566 | 0.072017532 | 0.1001 | 0.243160103 |
| SL1344_RS20195_ | 2.358079644 | 2.132788266 | 2.549707274 | 0.426981318 | 0.107112403 | 3.918213746 | 4.68E-05 | 0.00800073 | 0.057442735 | 0.1 | 0.243021319 |
| SL1344_RS20200_ | 1.911790088 | 1.769305305 | 2.072553418 | 0.297669643 | 0.101453275 | 2.858361926 | 0.0020023 | 0.018975854 | 0.090067601 | 0.1008 | 0.244134193 |
| SL1344_RS20205_ | 2.42872277 | 2.268717367 | 2.563774692 | 0.30465107 | 0.116387896 | 2.621640123 | 4.68E-05 | 0.033917076 | 0.130575739 | 0.1001 | 0.243165697 |
| SL1344_RS20210_ | 1.919688494 | 1.777862117 | 2.058398618 | 0.27970571 | 0.087928782 | 3.141735423 | 0.000673369 | 0.014118864 | 0.074271916 | 0.1002 | 0.243301466 |
| SL1344_RS20215_ | 3.139873722 | 2.890514536 | 3.401379005 | 0.504398466 | 0.087865684 | 5.730590671 | 4.68E-05 | 0.00268867 | 0.032869889 | 0.1 | 0.243021319 |
| SL1344_RS20220_ | 4.717968192 | 4.567162169 | 4.762698233 | 0.147784694 | 0.13773275 | 0.799668416 | 0.28400001 | 0.348407136 | 0.550726355 | 0.5756 | 0.727809282 |
| SL1344_RS20225_ | 5.091563379 | 4.991853996 | 5.178658699 | 0.120293123 | 0.124960821 | 1.260224858 | 0.114666702 | 0.135738555 | 0.311410609 | 0.2012 | 0.383703715 |
| SL1344_RS20230_ | 5.969434354 | 5.911155735 | 6.055140896 | 0.150092157 | 0.054733237 | 2.849815407 | 4.68E-05 | 0.038757238 | 0.147333677 | 0.1 | 0.243021319 |
| SL1344_RS20235_ | 5.1014384 | 5.037813471 | 5.357375883 | 0.317380469 | 0.245605175 | 1.159980054 | 0.118000034 | 0.160450958 | 0.349205723 | 0.2163 | 0.400009862 |
| SL1344_RS20240_ | 2.735066111 | 2.710198873 | 2.794665194 | 0.076232306 | 0.103326493 | 0.676290321 | 0.247333346 | 0.382644124 | 0.556612358 | 0.5112 | 0.653460672 |
| SL1344_RS20245_ | 2.582446698 | 2.500642141 | 2.663733942 | 0.163862387 | 0.088172741 | 1.82889802 | 0.019320688 | 0.058136537 | 0.173882071 | 0.1211 | 0.270040425 |
| SL1344_RS20250_ | 2.332824706 | 2.217351603 | 2.483213663 | 0.286428476 | 0.140544174 | 2.026175205 | 0.005330642 | 0.049748766 | 0.164618598 | 0.1046 | 0.249159153 |
| SL1344_RS20255_ | 2.627111263 | 2.52762521 | 2.766943137 | 0.234289306 | 0.098732727 | 2.25583584 | 0.005334196 | 0.034443163 | 0.13057067 | 0.1039 | 0.248086046 |
| SL1344_RS20260_ | 4.763310876 | 4.682916562 | 4.860623337 | 0.181082533 | 0.066852599 | 2.668942315 | 0.00067292 | 0.019256299 | 0.095911306 | 0.1002 | 0.243305234 |
| SL1344_RS20265_ | -0.902933829 | -0.922227769 | -0.889510079 | 0.098693205 | 0.434182492 | 0.206082115 | 0.401732182 | 0.575527756 | 0.723473933 | 0.7234 | 0.816328419 |
| SL1344_RS20270_ | -1.989613552 | -2.358971953 | -1.594857718 | 0.75100315 | 0.463323193 | 1.543917255 | 0.049333422 | 0.087116389 | 0.223578529 | 0.1491 | 0.305013091 |
| SL1344_RS20275_ | -1.905535069 | -2.176915842 | -1.621074072 | 0.567583061 | 0.411824594 | 1.302069307 | 0.076666722 | 0.130598696 | 0.278695653 | 0.2017 | 0.357852729 |
| SL1344_RS20280_ | 1.615841142 | 1.568761085 | 1.706369008 | 0.11850861 | 0.28515587 | 0.296727387 | 0.376000005 | 0.861011029 | 0.922707766 | 0.7993 | 0.885643639 |
| SL1344_RS20285_ | -3.677902201 | -3.936141697 | -3.372542296 | 0.564444475 | 0.679565471 | 0.784298929 | 0.200000017 | 0.301349868 | 0.462442696 | 0.3995 | 0.544208252 |
| SL1344_RS20290_ | 0.871038161 | 0.78360552 | 0.965819597 | 0.208682316 | 0.21559926 | 0.9456041 | 0.119920087 | 0.207716998 | 0.389736398 | 0.2721 | 0.43314435 |
| SL1344_RS20295_ | 2.546654519 | 2.506297498 | 2.578097298 | 0.072124111 | 0.076725102 | 0.899289795 | 0.164666689 | 0.261136627 | 0.419869723 | 0.3506 | 0.499786197 |
| SL1344_RS20300_ | 1.995131467 | 1.941461339 | 2.127504826 | 0.193587141 | 0.166579572 | 1.10461764 | 0.08933338 | 0.18567871 | 0.36975995 | 0.2227 | 0.382830894 |
| SL1344_RS20305_ | 2.090622354 | 2.007047727 | 2.218942122 | 0.213645118 | 0.117358321 | 1.768282196 | 0.028647725 | 0.073642007 | 0.203398467 | 0.1311 | 0.282150932 |
| SL1344_RS20310_ | 1.162563406 | 0.98725429 | 1.270495185 | 0.272197252 | 0.168389815 | 1.515033624 | 0.047333426 | 0.093239782 | 0.23451218 | 0.1577 | 0.315660109 |
| SL1344_RS20315_ | 3.668854013 | 3.646073681 | 3.696774682 | 0.059776845 | 0.090325673 | 0.679521243 | 0.19920055 | 0.308121004 | 0.497276952 | 0.4107 | 0.563518045 |
| SL1344_RS20325_ | -2.314389643 | -2.441793388 | -2.174488035 | 0.257754735 | 0.444757908 | 0.540283843 | 0.270486353 | 0.398175634 | 0.555888902 | 0.4992 | 0.631777856 |
| SL1344_RS20330_ | -3.334728434 | -3.489792561 | -3.182622359 | 0.373402071 | 0.844686767 | 0.411916187 | 0.323117929 | 0.489601918 | 0.647941666 | 0.6324 | 0.74411384 |
| SL1344_RS20335_ | 0.834785891 | 0.895174122 | 0.753412041 | -0.143543947 | 0.153276989 | -0.893993647 | 0.168554319 | 0.259005665 | 0.422090205 | 0.3483 | 0.497484778 |
| SL1344_RS20340_ | -1.679248023 | -1.396440554 | -1.942672447 | -0.628012931 | 0.503735199 | -1.210331789 | 0.073950757 | 0.133907037 | 0.296455669 | 0.1918 | 0.350808058 |
| SL1344_RS20345_ | 3.149805772 | 3.167175206 | 3.137687532 | -0.061428299 | 0.121677483 | -0.488554517 | 0.32933334 | 0.460226127 | 0.631681247 | 0.6552 | 0.762413953 |
| SL1344_RS20350_ | 2.855687377 | 2.77432493 | 2.948210847 | 0.176604731 | 0.085240264 | 2.056790627 | 0.008000574 | 0.048461901 | 0.156119829 | 0.1108 | 0.256597659 |
| SL1344_RS20355_ | 0.99601845 | 0.918147908 | 1.068507288 | 0.144648851 | 0.140802469 | 0.978147916 | 0.123251198 | 0.212846179 | 0.372813883 | 0.2934 | 0.448542842 |
| SL1344_RS20365_ | 2.042419679 | 1.983427667 | 2.083178198 | 0.096894495 | 0.098236898 | 0.937719055 | 0.161225873 | 0.240488131 | 0.404364234 | 0.338 | 0.492471673 |
| SL1344_RS20370_ | 1.542645628 | 1.534350242 | 1.551297639 | 0.018450553 | 0.147674847 | 0.114944878 | 0.455029982 | 0.648578908 | 0.773470179 | 0.7541 | 0.84006434 |
| SL1344_RS20375_ | 0.032804515 | -0.062592632 | 0.224750765 | 0.268803208 | 0.262924953 | 0.952381368 | 0.170000022 | 0.232979028 | 0.409722575 | 0.3349 | 0.501408554 |

| | | | | | | | | | | | |
|---|---|---|---|---|---|---|---|---|---|---|---|
| SL1344_RS20390_ | -0.791685739 | -0.859356738 | -0.675040397 | 0.212410566 | 0.336713202 | 0.593700733 | 0.261825461 | 0.392340278 | 0.565416365 | 0.5236 | 0.657504816 |
| SL1344_RS20395_ | -0.526613901 | -0.541447258 | -0.508811233 | 0.059459266 | 0.323775686 | 0.1670467 | 0.4170553 | 0.626837729 | 0.757449771 | 0.7272 | 0.821372626 |
| SL1344_RS20400_ | 4.403596997 | 4.51271428 | 4.371075053 | -0.101026844 | 0.120321432 | -0.786815018 | 0.267333344 | 0.361817382 | 0.55901802 | 0.526 | 0.685304352 |
| SL1344_RS20405_ | 1.563608855 | 1.475369964 | 1.614251508 | 0.13440195 | 0.133445423 | 0.99371455 | 0.142666694 | 0.227085942 | 0.396444966 | 0.3294 | 0.483472556 |
| SL1344_RS20410_ | 2.767611567 | 2.759894661 | 2.777028117 | 0.016342016 | 0.097802367 | 0.152408503 | 0.426000003 | 0.62619789 | 0.756085066 | 0.7436 | 0.831236009 |
| SL1344_RS20415_ | 2.28134036 | 2.270396622 | 2.286948913 | 0.01845647 | 0.11719963 | 0.143389951 | 0.438000002 | 0.645536781 | 0.770163906 | 0.7596 | 0.842708051 |
| SL1344_RS20425_ | 3.851460608 | 3.851011632 | 3.85234254 | -0.001429528 | 0.079828666 | -0.011035656 | 0.494666667 | 0.716633991 | 0.826429349 | 0.8146 | 0.888514749 |
| SL1344_RS20430_ | 3.970044804 | 3.924377928 | 4.10511607 | 0.168127601 | 0.095014031 | 1.571018631 | 0.051299219 | 0.088897646 | 0.237248184 | 0.1552 | 0.313487569 |
| SL1344_RS20435_ | 3.789738034 | 3.687356222 | 3.884529277 | 0.196792999 | 0.055075073 | 3.450160307 | 4.68E-05 | 0.010670539 | 0.065598783 | 0.1001 | 0.243165414 |
| SL1344_RS20440_ | 0.93670137 | 0.872936557 | 0.99186364 | 0.141715571 | 0.195788044 | 0.686797017 | 0.202531663 | 0.305160306 | 0.484952728 | 0.4177 | 0.563857675 |
| SL1344_RS20445_ | 2.838192798 | 2.643255878 | 3.101716762 | 0.44780053 | 0.144745382 | 2.9906459 | 4.68E-05 | 0.01719847 | 0.091188844 | 0.1 | 0.243021319 |
| SL1344_RS20450_ | 4.428547129 | 4.410179224 | 4.453806215 | 0.042779049 | 0.045798385 | 0.881336251 | 0.147901425 | 0.245629181 | 0.405710096 | 0.3403 | 0.489207683 |
| SL1344_RS20455_ | 3.499959942 | 3.526936027 | 3.478534049 | -0.054101716 | 0.076052806 | -0.699442857 | 0.224000015 | 0.340588632 | 0.51255846 | 0.4591 | 0.599081215 |
| SL1344_RS20460_ | 1.161866794 | 1.134783454 | 1.200205338 | 0.069892568 | 0.139315627 | 0.480531244 | 0.306666675 | 0.453903901 | 0.607766774 | 0.5715 | 0.691818908 |
| SL1344_RS20470_ | -0.741126169 | -0.582074123 | -0.858890571 | -0.264113582 | 0.290873992 | -0.880423706 | 0.169333355 | 0.260215467 | 0.429977366 | 0.3552 | 0.50970663 |
| SL1344_RS20475_ | 3.036255171 | 3.058479126 | 3.011240777 | -0.054011401 | 0.103170112 | -0.519387097 | 0.274666677 | 0.433822194 | 0.603067716 | 0.553 | 0.686235347 |
| SL1344_RS20480_ | 2.497430544 | 2.494866504 | 2.500016803 | 0.006253868 | 0.090356293 | 0.053639701 | 0.472666668 | 0.604749869 | 0.736964915 | 0.7168 | 0.810647914 |
| SL1344_RS20485_ | 2.210543623 | 2.131490305 | 2.312424883 | 0.091939434 | 0.227707271 | 0.451493856 | 0.349766828 | 0.600889704 | 0.746809355 | 0.7196 | 0.821517349 |
| SL1344_RS20495_ | -1.951704821 | -2.247470216 | -1.829476126 | 0.528603881 | 0.981655249 | 0.573454252 | 0.256666678 | 0.375521864 | 0.567965126 | 0.4831 | 0.652374361 |
| SL1344_RS20500_ | 0.599838804 | 0.522333939 | 0.731894737 | 0.240041014 | 0.267119287 | 0.856852486 | 0.162558317 | 0.241130478 | 0.425231277 | 0.3469 | 0.514872912 |
| SL1344_RS20505_ | -0.619453587 | -0.611623665 | -0.625162678 | -0.02954588 | 0.27583193 | -0.104315272 | 0.453697537 | 0.596771596 | 0.730115537 | 0.7049 | 0.801652836 |
| SL1344_RS20510_ | -0.24277555 | -0.347497728 | -0.116301869 | 0.278702635 | 0.386448749 | 0.782300235 | 0.15866669 | 0.280099318 | 0.474930911 | 0.3438 | 0.500887699 |
| SL1344_RS20520_ | -0.217790089 | -0.133376371 | -0.287570004 | -0.152179519 | 0.282030188 | -0.518805111 | 0.295136585 | 0.470630967 | 0.628102966 | 0.5791 | 0.704100321 |
| SL1344_RS20530_ | 0.833599231 | 0.795580365 | 0.87036259 | 0.116454418 | 0.228517374 | 0.470952374 | 0.303131254 | 0.459987597 | 0.626640404 | 0.6262 | 0.739010055 |
| SL1344_RS20535_ | 2.521981328 | 2.428643198 | 2.619261205 | 0.201491067 | 0.111285391 | 1.776705969 | 0.014657206 | 0.06178043 | 0.185700744 | 0.1139 | 0.261135563 |
| SL1344_RS20540_ | 2.372453911 | 2.341369019 | 2.421259667 | 0.083233069 | 0.112408017 | 0.672323318 | 0.217188556 | 0.342624538 | 0.517326073 | 0.457 | 0.600227482 |
| SL1344_RS20545_ | 2.88249447 | 2.857266968 | 2.976640428 | 0.12260693 | 0.17211732 | 0.602215559 | 0.309793479 | 0.439109007 | 0.625426176 | 0.6415 | 0.774241737 |
| SL1344_RS20550_ | -1.920631968 | -1.94975289 | -1.887717211 | 0.069878901 | 0.371937128 | 0.180319876 | 0.426382415 | 0.553188187 | 0.692178106 | 0.6596 | 0.765610404 |
| SL1344_RS20555_ | 3.272088681 | 3.207751147 | 3.3355696 | 0.139470855 | 0.094036769 | 1.467614413 | 0.042638345 | 0.089353119 | 0.230868 | 0.1413 | 0.294794033 |
| SL1344_RS20560_ | 2.993303531 | 2.899098319 | 3.018359095 | 0.113574982 | 0.170184729 | 0.28070677 | 0.349333339 | 0.880693223 | 0.935188611 | 0.7306 | 0.846958979 |
| SL1344_RS20565_ | 8.144469399 | 8.41354117 | 7.74793765 | -0.681957543 | 0.15383945 | -4.570755881 | 4.68E-05 | 0.003372469 | 0.041062676 | 0.1 | 0.243021319 |
| SL1344_RS20570_ | 3.124636023 | 3.08208992 | 3.303096086 | 0.207731535 | 0.110214662 | 1.433170524 | 0.076000056 | 0.124708809 | 0.290769386 | 0.1923 | 0.353449362 |
| SL1344_RS20575_ | 3.131573263 | 2.991113413 | 3.247210475 | 0.255614876 | 0.070576713 | 3.532040303 | 0.00067292 | 0.011654182 | 0.06901336 | 0.1001 | 0.243161305 |
| SL1344_RS20580_ | 1.544750384 | 1.288382945 | 1.767188493 | 0.479701402 | 0.117877321 | 4.061527222 | 4.68E-05 | 0.008718287 | 0.057635115 | 0.1001 | 0.243163773 |
| SL1344_RS20585_ | 1.149103781 | 0.965682306 | 1.354058331 | 0.390840223 | 0.152374535 | 2.500396908 | 0.002666616 | 0.026737153 | 0.110686737 | 0.1021 | 0.24589176 |
| SL1344_RS20590_ | 4.4488739 | 4.219478561 | 4.794222808 | 0.574482756 | 0.096006859 | 5.929431905 | 4.68E-05 | 0.008373596 | 0.063277527 | 0.1 | 0.243021319 |
| SL1344_RS20595_ | 6.558985319 | 6.454478784 | 6.663865497 | 0.219436808 | 0.043047424 | 5.110423753 | 4.68E-05 | 0.002571301 | 0.033726016 | 0.1 | 0.243021319 |
| SL1344_RS20600_ | -0.771103752 | -0.648399064 | -0.89479714 | -0.261481906 | 0.288622389 | -0.86592039 | 0.174000021 | 0.260893748 | 0.428756595 | 0.3547 | 0.506079811 |
| SL1344_RS20605_ | -2.529841201 | -2.584794399 | -2.430070669 | 0.112234211 | 0.574299369 | 0.182288825 | 0.422000003 | 0.630348334 | 0.758510996 | 0.7287 | 0.82173271 |
| SL1344_RS20610_ | -2.274619611 | -2.498091664 | -2.088405203 | 0.412377082 | 0.414399309 | 0.950333042 | 0.145902758 | 0.230151364 | 0.389503168 | 0.3227 | 0.472567329 |
| SL1344_RS20615_ | -2.44178354 | -2.469846696 | -2.416712536 | 0.053084846 | 0.47869416 | 0.103537713 | 0.447333335 | 0.583451807 | 0.719102999 | 0.6985 | 0.796002289 |
| SL1344_RS20620_ | -2.619189553 | -2.502108748 | -2.70290665 | -0.202400326 | 0.507940426 | -0.378011179 | 0.340439713 | 0.50004637 | 0.645675474 | 0.6086 | 0.72250483 |
| SL1344_RS20625_ | -1.995794916 | -1.857906201 | -2.092851359 | -0.248980977 | 0.44826123 | -0.499034841 | 0.283810803 | 0.423323532 | 0.58275025 | 0.5454 | 0.669657263 |
| SL1344_RS20635_ | -1.608219157 | -1.757904461 | -1.461104255 | 0.295996779 | 0.307971693 | 0.876324121 | 0.155896093 | 0.242114842 | 0.398803383 | 0.3301 | 0.480581078 |
| SL1344_RS20640_ | -0.561536024 | -0.629860017 | -0.516329292 | 0.103680127 | 0.293731644 | 0.331312696 | 0.359760166 | 0.55571679 | 0.700049207 | 0.6762 | 0.782766384 |
| SL1344_RS20645_ | -1.257333503 | -1.284268786 | -1.227032883 | 0.048834126 | 0.309506941 | 0.155567491 | 0.425716192 | 0.552853925 | 0.691756885 | 0.658 | 0.762955957 |
| SL1344_RS20650_ | -2.090705701 | -1.893508619 | -2.45763528 | -0.564024139 | 0.582983111 | -0.907821826 | 0.174000021 | 0.243263604 | 0.423913155 | 0.3426 | 0.511919208 |
| SL1344_RS20655_ | -2.005874913 | -1.785414681 | -2.182611685 | -0.409394752 | 0.473889267 | -0.834746357 | 0.185209879 | 0.285518746 | 0.459148932 | 0.393 | 0.545192842 |
| SL1344_RS20660_ | -3.129686786 | -3.0339368 | -3.218049151 | -0.144712024 | 0.831966275 | -0.163550877 | 0.421718857 | 0.659528813 | 0.780725591 | 0.7418 | 0.833233933 |
| SL1344_RS20665_ | -4.389920021 | -3.922104241 | -4.980810982 | -1.120042065 | 1.117618751 | -0.90455525 | 0.115333368 | 0.211389108 | 0.383344212 | 0.2646 | 0.421420824 |
| SL1344_RS20670_ | -2.599085359 | -2.591827455 | -2.615912756 | -0.05218229 | 0.562681337 | -0.090535637 | 0.460000001 | 0.605567076 | 0.738774346 | 0.7162 | 0.810124892 |
| SL1344_RS20675_ | 1.440347935 | 1.412669354 | 1.484378123 | 0.067586462 | 0.123090314 | 0.506306858 | 0.293333343 | 0.454039194 | 0.607487902 | 0.5584 | 0.686870799 |
| SL1344_RS20680_ | 1.797340212 | 1.822841243 | 1.766845316 | -0.056121625 | 0.103542307 | -0.509811839 | 0.292666676 | 0.422302778 | 0.580688404 | 0.5365 | 0.665201994 |

| | | | | | | | | | | | |
|---|---|---|---|---|---|---|---|---|---|---|---|
| SL1344_RS20685_ | 1.832214277 | 1.853524368 | 1.818188169 | -0.033038363 | 0.156513005 | -0.197117359 | 0.41866667 | 0.663571174 | 0.785078865 | 0.7687 | 0.84981961 |
| SL1344_RS20690_ | 1.415439234 | 1.371864497 | 1.451916326 | 0.082878215 | 0.142500756 | 0.582027769 | 0.268487686 | 0.418095943 | 0.580708799 | 0.5375 | 0.668995568 |
| SL1344_RS20695_ | 2.487446571 | 2.498235126 | 2.477369037 | -0.022943891 | 0.083069892 | -0.259271542 | 0.398666671 | 0.523358327 | 0.668273117 | 0.6289 | 0.740890967 |
| SL1344_RS20700_ | 0.845767756 | 1.133837274 | 0.566592406 | -0.530396701 | 0.287022001 | -1.872427469 | 0.018654474 | 0.056874665 | 0.180843987 | 0.1197 | 0.270587556 |
| SL1344_RS20705_ | 0.635205314 | 0.91517676 | 0.29472758 | -0.584244785 | 0.279259316 | -2.075001716 | 0.006667356 | 0.047048386 | 0.160209187 | 0.1073 | 0.253116822 |
| SL1344_RS20710_ | -0.587055987 | -0.531068928 | -0.637608849 | -0.104614106 | 0.238784273 | -0.405822254 | 0.32933334 | 0.474618474 | 0.624324351 | 0.5876 | 0.707360914 |
| SL1344_RS20715_ | -0.802279224 | -0.665237187 | -0.938742614 | -0.276922223 | 0.239492001 | -1.099988622 | 0.105929419 | 0.175074965 | 0.323961921 | 0.2584 | 0.410003437 |
| SL1344_RS20720_ | -0.769936235 | -0.745668978 | -0.811337254 | -0.032378617 | 0.409084534 | -0.075128899 | 0.470000001 | 0.726040924 | 0.832654064 | 0.8264 | 0.894325681 |
| SL1344_RS20725_ | 3.283545885 | 3.233369383 | 3.364803521 | 0.156618774 | 0.141016627 | 1.156980663 | 0.05266675 | 0.163229493 | 0.346894718 | 0.1638 | 0.322303478 |
| SL1344_RS20730_ | 2.808958825 | 2.743025772 | 2.890910721 | 0.157097882 | 0.097734886 | 1.56153208 | 0.029313944 | 0.087382017 | 0.227767602 | 0.1263 | 0.276368316 |
| SL1344_RS20735_ | 2.671654592 | 2.582411858 | 2.769368866 | 0.215327728 | 0.163295488 | 1.387027188 | 0.021319333 | 0.119927022 | 0.286202767 | 0.1228 | 0.272599766 |
| SL1344_RS20740_ | 4.215468151 | 4.148797682 | 4.328628973 | 0.179268331 | 0.069663507 | 2.437226658 | 0.000673369 | 0.037976943 | 0.140705456 | 0.1006 | 0.243864096 |
| SL1344_RS20745_ | 3.643004912 | 3.579197179 | 3.699531165 | 0.131956166 | 0.093160408 | 1.421251432 | 0.03664236 | 0.106876209 | 0.262104785 | 0.1395 | 0.292614117 |
| SL1344_RS20750_ | 1.164159718 | 1.149152737 | 1.183721464 | 0.038508398 | 0.137259226 | 0.26163404 | 0.388000004 | 0.538684301 | 0.679690726 | 0.6533 | 0.758700881 |
| SL1344_RS20755_ | -1.361615553 | -1.402837953 | -1.315567041 | 0.101164067 | 0.332287379 | 0.284587215 | 0.380666671 | 0.535848903 | 0.679696309 | 0.6523 | 0.758517593 |
| SL1344_RS20760_ | -2.200999343 | -2.124332846 | -2.373179591 | -0.255410931 | 0.581284923 | -0.412074117 | 0.34333334 | 0.510853837 | 0.668531477 | 0.6518 | 0.764913514 |
| SL1344_RS20765_ | 0.180765381 | 0.242001499 | 0.123832602 | -0.103249802 | 0.287563092 | -0.34882109 | 0.34133334 | 0.572363965 | 0.716436291 | 0.6622 | 0.775135757 |
| SL1344_RS20770_ | 0.675147779 | 0.70943443 | 0.630675386 | -0.081527781 | 0.158027131 | -0.488208538 | 0.295802807 | 0.438370041 | 0.589544816 | 0.55 | 0.671690057 |
| SL1344_RS20775_ | 1.675964865 | 1.621123615 | 1.751704906 | 0.095937866 | 0.162307619 | 0.54975896 | 0.287808137 | 0.460545977 | 0.624531258 | 0.5651 | 0.698834636 |
| SL1344_RS20780_ | -0.963109586 | -0.726254635 | -1.242663473 | -0.52773389 | 0.307676547 | -1.715724206 | 0.03000015 | 0.074335967 | 0.199906377 | 0.1394 | 0.290973621 |
| SL1344_RS20785_ | -2.301269871 | -2.157059379 | -2.477761954 | -0.32303016 | 0.437304669 | -0.72502872 | 0.215189889 | 0.335900371 | 0.498019035 | 0.4338 | 0.575401797 |
| SL1344_RS20790_ | -3.210617559 | -3.246642659 | -3.174945573 | 0.084005955 | 0.667022973 | 0.11336086 | 0.445036644 | 0.585235498 | 0.720330086 | 0.6931 | 0.791072688 |
| SL1344_RS20795_ | -2.830519747 | -2.844751288 | -2.820473429 | 0.000775607 | 0.608196071 | 0.001455229 | 0.5 | 0.640692884 | 0.766052901 | 0.7463 | 0.834833219 |
| SL1344_RS20800_ | -3.040533483 | -3.149956111 | -2.926536785 | 0.254948597 | 0.624808568 | 0.366049952 | 0.332445044 | 0.497660097 | 0.648471745 | 0.6097 | 0.724754673 |
| SL1344_RS20805_ | -1.465506596 | -1.059813072 | -1.826685824 | -0.744406874 | 0.386208592 | -1.863753175 | 0.019333568 | 0.053366498 | 0.164922446 | 0.1238 | 0.273727831 |
| SL1344_RS20810_ | -2.478239334 | -2.529301198 | -2.430559527 | 0.08773461 | 0.422224791 | 0.202928224 | 0.406000004 | 0.484733341 | 0.632718308 | 0.5957 | 0.711310384 |
| SL1344_RS20815_ | -1.923102583 | -1.868124895 | -1.967042413 | -0.117381203 | 0.407572493 | -0.281540267 | 0.380413063 | 0.551740858 | 0.693680266 | 0.6773 | 0.777767189 |
| SL1344_RS20820_ | -6.64679529 | -6.447978145 | -6.872260114 | -0.563329102 | 2.865882419 | -0.159235861 | 0.41666667 | 0.604798241 | 0.74433521 | 0.7527 | 0.838015484 |
| SL1344_RS20825_ | -3.269261834 | -3.330945244 | -3.209635562 | 0.112106265 | 0.578086067 | 0.191897878 | 0.416000003 | 0.540606368 | 0.679570862 | 0.658 | 0.760538568 |
| SL1344_RS20830_ | -1.117230174 | -1.281228182 | -1.023011465 | 0.257387702 | 0.299488171 | 0.783140496 | 0.199333351 | 0.304187734 | 0.469167983 | 0.4064 | 0.556184451 |
| SL1344_RS20835_ | -4.277335431 | -4.267868215 | -4.286864731 | -0.051881708 | 0.852865378 | -0.049385964 | 0.476000001 | 0.570432053 | 0.705226035 | 0.6785 | 0.778390704 |
| SL1344_RS20840_ | -3.797750727 | -3.659644815 | -3.999890893 | -0.378204833 | 0.897479968 | -0.397957226 | 0.321333341 | 0.473377903 | 0.63346765 | 0.6109 | 0.726733271 |
| SL1344_RS20845_ | 6.54709906 | 6.534682285 | 6.556278229 | 0.024583326 | 0.038621735 | 0.608890326 | 0.23866668 | 0.373956393 | 0.557445777 | 0.4858 | 0.640443172 |
| SL1344_RS20850_ | 0.403344491 | 0.35744754 | 0.459098482 | 0.095454299 | 0.20207163 | 0.433576148 | 0.317788149 | 0.497838865 | 0.649877969 | 0.606 | 0.72528661 |
| SL1344_RS20855_ | 0.419644115 | 0.502972947 | 0.320771532 | -0.184546399 | 0.158609682 | -1.093564856 | 0.112591641 | 0.188046341 | 0.342348855 | 0.2646 | 0.418520489 |
| SL1344_RS20875_ | 1.656710349 | 1.618979537 | 1.695099479 | 0.080753357 | 0.115951478 | 0.656256304 | 0.234000014 | 0.361484533 | 0.521672807 | 0.4725 | 0.60869249 |
| SL1344_RS20880_ | 2.299385699 | 2.188593387 | 2.387291973 | 0.221251028 | 0.164609645 | 1.351138103 | 0.041972124 | 0.118149292 | 0.281204963 | 0.149 | 0.305687898 |
| SL1344_RS20890_ | 3.51358683 | 3.45177986 | 3.566929894 | 0.134326313 | 0.119154944 | 1.114492827 | 0.098600974 | 0.157465858 | 0.334831602 | 0.2137 | 0.381301758 |
| SL1344_RS20895_ | -0.874453646 | -1.241469529 | -0.594070418 | 0.676068552 | 0.355647637 | 1.826565666 | 0.012658588 | 0.059754674 | 0.180148227 | 0.114 | 0.261650053 |
| SL1344_RS20900_ | -1.079820372 | -1.151124251 | -1.023193725 | 0.12857628 | 0.337626034 | 0.371476085 | 0.347768161 | 0.505707548 | 0.655888242 | 0.623 | 0.735485422 |
| SL1344_RS20905_ | 1.820721712 | 1.601120299 | 1.967416416 | 0.368974105 | 0.132739233 | 2.726594139 | 0.000673369 | 0.026809778 | 0.111586576 | 0.1008 | 0.244142222 |
| SL1344_RS20910_ | 1.880834871 | 1.856534326 | 1.915632916 | 0.062229321 | 0.10602287 | 0.544626726 | 0.28314458 | 0.416952454 | 0.570872826 | 0.5215 | 0.647048692 |
| SL1344_RS20915_ | -2.784154351 | -2.682393192 | -2.876789896 | -0.192633108 | 0.456440612 | -0.393100944 | 0.335776156 | 0.460090684 | 0.609576111 | 0.5669 | 0.68826369 |
| SL1344_RS20920_ | -1.917330837 | -2.063757432 | -1.767023133 | 0.311117394 | 0.466580782 | 0.656156772 | 0.224000015 | 0.341640197 | 0.512049494 | 0.4626 | 0.602738315 |
| SL1344_RS20925_ | 0.125827703 | -0.195810787 | 0.40318407 | 0.594334907 | 0.214152611 | 2.765062435 | 0.000673369 | 0.019723728 | 0.093413337 | 0.1006 | 0.243874204 |
| SL1344_RS20935_ | -2.488767083 | -2.351617024 | -2.999193629 | -0.574110986 | 0.740740856 | -0.632851285 | 0.290473027 | 0.434096623 | 0.610753422 | 0.5927 | 0.732625084 |
| SL1344_RS20940_ | -1.849748507 | -1.902810356 | -1.786640956 | 0.123773679 | 0.372929815 | 0.305015008 | 0.368421058 | 0.505499446 | 0.649231091 | 0.6097 | 0.722549472 |
| SL1344_RS20945_ | -3.088218389 | -3.215146381 | -2.924017722 | 0.311329559 | 0.573604885 | 0.509910684 | 0.29380414 | 0.43168665 | 0.588472795 | 0.5506 | 0.674385526 |
| SL1344_RS20950_ | -1.750452237 | -1.77193308 | -1.725863425 | 0.130075234 | 0.539000442 | 0.226895676 | 0.415333337 | 0.619850715 | 0.753019083 | 0.7698 | 0.850074971 |
| SL1344_RS20955_ | -1.61504908 | -1.628501984 | -1.597080509 | 0.032143722 | 0.351472368 | 0.09047391 | 0.457694872 | 0.585897245 | 0.718827707 | 0.6915 | 0.788362756 |
| SL1344_RS20960_ | -2.533104363 | -2.556841509 | -2.500451113 | 0.072019064 | 0.547195424 | 0.122810394 | 0.445702867 | 0.626547559 | 0.753621317 | 0.7319 | 0.822612023 |
| SL1344_RS20965_ | -0.551588539 | -0.687225002 | -0.445407684 | 0.213671741 | 0.278209842 | 0.705122052 | 0.221333348 | 0.347763431 | 0.516787416 | 0.444 | 0.59349821 |
| SL1344_RS20970_ | 4.56982799 | 4.518919853 | 4.669824276 | 0.147010642 | 0.058035432 | 2.355391571 | 0.003334715 | 0.035670712 | 0.133902071 | 0.1041 | 0.248592092 |

| | | | | | | | | | | | |
|---|---|---|---|---|---|---|---|---|---|---|---|
| SL1344_RS20975_ | 0.871978483 | 0.818396812 | 0.929471749 | 0.096009468 | 0.257189775 | 0.366089956 | 0.316666675 | 0.577499505 | 0.723054181 | 0.6666 | 0.780168586 |
| SL1344_RS20980_ | 0.140347211 | -0.002604013 | 0.278904425 | 0.29665014 | 0.233024551 | 1.208366565 | 0.069953426 | 0.134885492 | 0.291937827 | 0.1996 | 0.359624457 |
| SL1344_RS20985_ | 1.967646091 | 1.992333314 | 1.954265639 | -0.037657985 | 0.21647736 | -0.161772804 | 0.433333336 | 0.781291817 | 0.871715422 | 0.8518 | 0.9117147 |
| SL1344_RS20990_ | 1.704981555 | 1.629954003 | 1.800477672 | 0.113439362 | 0.3008052 | 0.350641177 | 0.373333338 | 0.648708309 | 0.783285172 | 0.83 | 0.894434353 |
| SL1344_RS20995_ | 3.529900066 | 3.426358829 | 3.592006688 | 0.160982965 | 0.074175794 | 2.112165623 | 0.009333825 | 0.047371749 | 0.156427412 | 0.1114 | 0.257944522 |
| SL1344_RS21000_ | 1.867677452 | 1.666394965 | 1.970486635 | 0.27954022 | 0.210683997 | 1.33518674 | 0.085333383 | 0.110227372 | 0.269636991 | 0.1921 | 0.362828049 |
| SL1344_RS21005_ | 4.003349144 | 3.947159746 | 4.110108014 | 0.157177641 | 0.08490173 | 1.787114488 | 0.01798826 | 0.064903517 | 0.194583808 | 0.1227 | 0.27334291 |
| SL1344_RS21010_ | 3.359311412 | 3.496018866 | 3.194757186 | -0.303594217 | 0.072459542 | -4.224706032 | 4.68E-05 | 0.009622993 | 0.061938109 | 0.1 | 0.243021319 |
| SL1344_RS21015_ | 1.710050319 | 1.652829475 | 1.802381269 | 0.146492042 | 0.137798045 | 1.043936243 | 0.134576978 | 0.199510457 | 0.368050584 | 0.2862 | 0.446619533 |
| SL1344_RS21020_ | 4.799734285 | 4.786750521 | 4.812512348 | 0.052997457 | 0.164543311 | 0.312587265 | 0.355762831 | 0.634803731 | 0.779160768 | 0.7462 | 0.850889221 |
| SL1344_RS21025_ | 2.704500395 | 2.740189598 | 2.672722728 | -0.095280679 | 0.152745598 | -0.619409465 | 0.202000017 | 0.321793949 | 0.520826232 | 0.4179 | 0.566564272 |
| SL1344_RS21030_ | 3.417274782 | 3.246873542 | 3.55331258 | 0.305435986 | 0.064295856 | 4.638920248 | 4.68E-05 | 0.006290183 | 0.049091657 | 0.1 | 0.243021319 |
| SL1344_RS21035_ | 1.90857068 | 1.84777487 | 2.093169402 | 0.231694909 | 0.128201447 | 1.547680539 | 0.077948089 | 0.115264591 | 0.270230712 | 0.1939 | 0.352445753 |
| SL1344_RS21040_ | 2.731064182 | 2.546702402 | 2.881276751 | 0.336050149 | 0.120247897 | 2.723144949 | 0.00067292 | 0.022448864 | 0.10293135 | 0.1001 | 0.243163644 |
| SL1344_RS21045_ | 6.320638826 | 6.189624152 | 6.448438025 | 0.282188657 | 0.054974235 | 5.22336633 | 4.68E-05 | 0.001796732 | 0.028968845 | 0.1 | 0.243021319 |
| SL1344_RS21050_ | -1.020826636 | -1.016248414 | -1.026183195 | -0.022279932 | 0.418301261 | -0.04403281 | 0.480000001 | 0.703078657 | 0.815610771 | 0.821 | 0.888549464 |
| SL1344_RS21055_ | -0.473139806 | -0.606049359 | -0.250568505 | 0.343465481 | 0.248930085 | 1.313343058 | 0.078614311 | 0.136568792 | 0.287569945 | 0.2199 | 0.378127077 |
| SL1344_RS21075_ | -3.32690253 | -3.376772145 | -3.285563512 | 0.097287414 | 0.606024757 | 0.154777667 | 0.438666669 | 0.552439701 | 0.691010992 | 0.6678 | 0.770158928 |
| SL1344_RS21080_ | -3.686512645 | -3.734610322 | -3.628042498 | 0.103501171 | 0.741107467 | 0.139856672 | 0.435043307 | 0.573970242 | 0.709723188 | 0.6775 | 0.780076736 |
| SL1344_RS21085_ | -1.211726564 | -1.515366957 | -0.915357168 | 0.613929778 | 0.311888778 | 1.916020971 | 0.012667027 | 0.052751147 | 0.162937883 | 0.1151 | 0.261999024 |
| SL1344_RS21090_ | -0.259926402 | -0.385718838 | -0.140876718 | 0.245366459 | 0.196538527 | 1.221481545 | 0.091333379 | 0.153724812 | 0.300773051 | 0.2312 | 0.38618354 |
| SL1344_RS21100_ | 1.212475373 | 1.300589301 | 1.089395145 | -0.207221412 | 0.136956377 | -1.475955913 | 0.060626321 | 0.107587041 | 0.246895471 | 0.1791 | 0.335800433 |
| SL1344_RS21105_ | -1.954195415 | -1.989852454 | -1.891932061 | 0.088274127 | 0.416835123 | 0.203095073 | 0.41066667 | 0.592104806 | 0.723186412 | 0.6963 | 0.794026456 |
| SL1344_RS21115_ | 0.063762858 | 0.044322248 | 0.095992695 | 0.06038862 | 0.221768768 | 0.240633981 | 0.391072622 | 0.57181101 | 0.70852781 | 0.6869 | 0.786233718 |
| SL1344_RS21120_ | 0.127914036 | 0.126582835 | 0.129672717 | 0.012474261 | 0.209597162 | 0.051251666 | 0.477015324 | 0.600753606 | 0.735302526 | 0.7141 | 0.809626774 |
| SL1344_RS21125_ | -2.189010923 | -2.192168811 | -2.185431939 | 0.011063942 | 0.408915826 | 0.024809611 | 0.487674884 | 0.572542401 | 0.709583935 | 0.6792 | 0.78115145 |
| SL1344_RS21130_ | -1.971161954 | -1.67734319 | -2.08839166 | -0.400516466 | 0.608022153 | -0.576557177 | 0.28314458 | 0.433824157 | 0.60604866 | 0.5648 | 0.700281618 |
| SL1344_RS21135_ | 0.75750558 | 0.778462714 | 0.740551837 | -0.035435912 | 0.140099726 | -0.235778718 | 0.390000004 | 0.519487915 | 0.661781934 | 0.6292 | 0.737691267 |
| SL1344_RS21140_ | 1.075466253 | 1.135953753 | 1.002392376 | -0.136260539 | 0.139216624 | -0.892488201 | 0.156000024 | 0.235333192 | 0.399616538 | 0.3214 | 0.475341409 |
| SL1344_RS21145_ | -0.901898132 | -0.870261331 | -0.946440432 | -0.059721009 | 0.317648562 | -0.183438297 | 0.423051302 | 0.617623432 | 0.746004279 | 0.723 | 0.815174947 |
| SL1344_RS21150_ | 0.190014509 | 0.087031544 | 0.466892157 | 0.390135441 | 0.327152868 | 1.121694633 | 0.122000033 | 0.176347972 | 0.356000705 | 0.2628 | 0.433303269 |
| SL1344_RS21155_ | 3.471450549 | 3.151263135 | 3.710986163 | 0.543278352 | 0.091499401 | 5.839756768 | 4.68E-05 | 0.004457547 | 0.042533453 | 0.1 | 0.243021319 |
| SL1344_RS21160_ | 4.010415735 | 3.940817521 | 4.058421993 | 0.126410968 | 0.086141466 | 1.488623107 | 0.03530992 | 0.087592781 | 0.232293226 | 0.1387 | 0.293205819 |
| SL1344_RS21165_ | 0.560219443 | 0.607271214 | 0.512771446 | -0.096604012 | 0.145722166 | -0.619092904 | 0.246502344 | 0.368251219 | 0.5224107 | 0.4574 | 0.592988557 |
| SL1344_RS21170_ | -0.555980014 | -0.631562415 | -0.47283765 | 0.214930719 | 0.351380657 | 0.584565973 | 0.254000012 | 0.377269825 | 0.551111741 | 0.5205 | 0.652311503 |
| SL1344_RS21175_ | 0.040370443 | 0.064039866 | -0.013409485 | -0.046126267 | 0.261451346 | -0.165135285 | 0.428666669 | 0.661341518 | 0.785067607 | 0.7716 | 0.857717408 |
| SL1344_RS21180_ | 1.311304076 | 1.349033072 | 1.270028595 | -0.093422967 | 0.141825435 | -0.632898985 | 0.247333346 | 0.364763552 | 0.530467328 | 0.4745 | 0.609260535 |
| SL1344_RS21185_ | 2.96833666 | 2.96331809 | 2.975847189 | 0.006800936 | 0.08222245 | 0.071456311 | 0.472000001 | 0.631432652 | 0.758360089 | 0.7355 | 0.826989209 |
| SL1344_RS21195_ | 2.307270126 | 2.258747362 | 2.425150477 | 0.148473683 | 0.168228463 | 0.900531494 | 0.1798801 | 0.246458181 | 0.433538186 | 0.346 | 0.524761721 |
| SL1344_RS21200_ | 1.138353468 | 1.11044649 | 1.1588701 | 0.04702976 | 0.146372351 | 0.300745958 | 0.370000005 | 0.538468229 | 0.680927416 | 0.6542 | 0.759851064 |
| SL1344_RS21205_ | 2.053208901 | 1.867677113 | 2.27460254 | 0.403573284 | 0.09931068 | 3.999705182 | 4.68E-05 | 0.007940807 | 0.055682569 | 0.1 | 0.243021319 |
| SL1344_RS21210_ | 2.953337657 | 2.820997538 | 3.131646988 | 0.312652077 | 0.090641568 | 3.356788271 | 4.68E-05 | 0.011512055 | 0.07060571 | 0.1 | 0.243021319 |
| SL1344_RS21215_ | 2.9589164 | 2.743589143 | 3.179345553 | 0.422495915 | 0.119868551 | 3.558723662 | 4.68E-05 | 0.007829943 | 0.058829776 | 0.1 | 0.243021319 |
| SL1344_RS21220_ | -2.723307452 | -3.146306113 | -2.425915274 | 0.891708607 | 0.972284601 | 0.930638689 | 0.149900092 | 0.220187586 | 0.406454461 | 0.3094 | 0.480693804 |
| SL1344_RS21225_ | 1.493426674 | 1.368104095 | 1.616841561 | 0.215088546 | 0.211765034 | 0.968390459 | 0.141333361 | 0.203849178 | 0.388081299 | 0.2748 | 0.452806846 |
| SL1344_RS21230_ | -1.251423437 | -1.215739045 | -1.338672431 | -0.121825155 | 0.419845954 | -0.264951625 | 0.390666671 | 0.607629055 | 0.744350482 | 0.7258 | 0.823327306 |
| SL1344_RS21235_ | -1.038522129 | -0.945239728 | -1.253362199 | -0.242387671 | 0.56520064 | -0.262557051 | 0.381745508 | 0.853198681 | 0.918154514 | 0.7846 | 0.877561015 |
| SL1344_RS21245_ | 1.550582882 | 1.503107499 | 1.610344719 | 0.108035751 | 0.116795151 | 0.863692398 | 0.178547655 | 0.263277866 | 0.428096927 | 0.3593 | 0.511179667 |
| SL1344_RS21250_ | 1.958509476 | 1.865975666 | 2.036869403 | 0.135207101 | 0.172334522 | 0.798998568 | 0.189873436 | 0.284516104 | 0.471694096 | 0.3662 | 0.538939736 |
| SL1344_RS21255_ | 2.403607452 | 2.112697347 | 2.651440299 | 0.557551096 | 0.126216416 | 4.401756763 | 4.68E-05 | 0.004947405 | 0.045460111 | 0.1 | 0.243021319 |
| SL1344_RS21260_ | -7.247378396 | -6.845805899 | -7.62336977 | -0.63896278 | 3.040301251 | -0.172994479 | 0.395069957 | 0.574727472 | 0.715429652 | 0.6793 | 0.783220157 |
| SL1344_RS21265_ | 3.876584836 | 3.685682185 | 3.962698962 | 0.256363368 | 0.205908674 | 1.19959614 | 0.114590308 | 0.141225931 | 0.320872512 | 0.2157 | 0.398554003 |
| SL1344_RS21270_ | 4.794416144 | 4.627787373 | 4.922539989 | 0.225400026 | 0.183633832 | 1.269519665 | 0.115333368 | 0.126621404 | 0.30114318 | 0.1987 | 0.381521387 |

| | | | | | | | | | | | |
|---|---|---|---|---|---|---|---|---|---|---|---|
| SL1344_RS21275_ | 6.136072666 | 6.018165179 | 6.254057749 | 0.233277317 | 0.167808746 | 1.261133449 | 0.092000045 | 0.137396901 | 0.318102058 | 0.1869 | 0.365044499 |
| SL1344_RS21280_ | 5.248330941 | 5.192748473 | 5.297543422 | 0.132031516 | 0.15079051 | 0.840181878 | 0.18000002 | 0.23781895 | 0.439667642 | 0.3374 | 0.52261785 |
| SL1344_RS21285_ | 8.111587308 | 8.141774164 | 8.016487294 | -0.125841202 | 0.106486424 | -0.747792858 | 0.320666674 | 0.376426344 | 0.579248221 | 0.6652 | 0.802783387 |
| SL1344_RS21290_ | 4.922514654 | 4.923544502 | 4.921006306 | -0.003480391 | 0.101344717 | -0.038038403 | 0.475333334 | 0.814726336 | 0.893965308 | 0.8434 | 0.90429868 |
| SL1344_RS21300_ | 6.74473007 | 6.63918934 | 6.906133896 | 0.263987217 | 0.068719905 | 3.690243579 | 4.68E-05 | 0.018858643 | 0.098700963 | 0.1 | 0.243021319 |
| SL1344_RS21305_ | 7.90068659 | 7.804800514 | 8.05224098 | 0.247966486 | 0.064770402 | 3.936953028 | 4.68E-05 | 0.022634436 | 0.110168 | 0.1 | 0.243021319 |
| SL1344_RS21310_ | 8.452152736 | 8.299936931 | 8.526360123 | 0.22269934 | 0.104081778 | 2.368861976 | 4.68E-05 | 0.014929433 | 0.087913677 | 0.1 | 0.243021319 |
| SL1344_RS21315_ | 9.618822693 | 9.518000056 | 9.688004727 | 0.166546542 | 0.08555565 | 1.803311765 | 0.00067292 | 0.049309118 | 0.172486555 | 0.1007 | 0.244015794 |
| SL1344_RS21320_ | 7.504977913 | 7.498102992 | 7.594067283 | 0.098493518 | 0.127298095 | 0.855124095 | 0.217854779 | 0.263252274 | 0.469679557 | 0.3866 | 0.583880053 |
| SL1344_RS21325_ | 7.740938163 | 7.711029297 | 7.800174661 | 0.086898756 | 0.127250006 | 0.57589348 | 0.325782819 | 0.39945102 | 0.599562277 | 0.6822 | 0.815216081 |
| SL1344_RS21330_ | -0.063160453 | -0.148498266 | 0.009336987 | 0.15714786 | 0.257735674 | 0.581542048 | 0.263824128 | 0.417414631 | 0.581425514 | 0.5406 | 0.672445166 |
| SL1344_RS21335_ | -2.073970266 | -2.453920929 | -1.747946682 | 0.818969555 | 0.614298421 | 1.291421944 | 0.055962769 | 0.115690865 | 0.271624575 | 0.1699 | 0.327519352 |
| SL1344_RS21340_ | 0.576593489 | 0.494483233 | 0.773974029 | 0.256640756 | 0.197920149 | 1.162344783 | 0.135909422 | 0.186270843 | 0.356689796 | 0.2899 | 0.455487195 |
| SL1344_RS21345_ | -1.32962655 | -1.577720251 | -1.114100188 | 0.472614728 | 0.319036415 | 1.383951962 | 0.047333426 | 0.108021993 | 0.248583862 | 0.1707 | 0.323546262 |
| SL1344_RS21350_ | 0.178652778 | 0.082296017 | 0.251770481 | 0.203885162 | 0.281233809 | 0.699214809 | 0.173884098 | 0.285109392 | 0.476151321 | 0.3578 | 0.509202321 |
| SL1344_RS21355_ | -1.31245419 | -1.495207636 | -0.799219134 | 0.637774878 | 0.430041543 | 1.320195299 | 0.114666702 | 0.148467333 | 0.313547792 | 0.2476 | 0.412783618 |
| SL1344_RS21360_ | -2.925636335 | -3.258993251 | -2.679938647 | 0.622998749 | 0.714044784 | 0.819326352 | 0.157333357 | 0.252129404 | 0.434384629 | 0.319 | 0.471701877 |
| SL1344_RS21365_ | -1.11463927 | -1.281429544 | -1.01861144 | 0.280323907 | 0.367322659 | 0.692376752 | 0.208666683 | 0.307823968 | 0.485367786 | 0.4172 | 0.563783391 |
| SL1344_RS21370_ | -1.78260194 | -1.867983194 | -1.535942619 | 0.293266996 | 0.489371182 | 0.512851838 | 0.310459702 | 0.549871082 | 0.695092751 | 0.636 | 0.76226622 |
| SL1344_RS21375_ | 0.015460303 | -0.059882684 | 0.080818173 | 0.128149361 | 0.317477375 | 0.389731153 | 0.319333341 | 0.54233481 | 0.695247503 | 0.6631 | 0.775853985 |
| SL1344_RS21380_ | -0.876066293 | -0.683103982 | -1.038490299 | -0.356113647 | 0.26269142 | -1.27847451 | 0.084610309 | 0.140762722 | 0.28720313 | 0.2168 | 0.371905834 |
| SL1344_RS21390_ | 2.503756226 | 2.345913135 | 2.572242204 | 0.21871047 | 0.113208792 | 1.775592612 | 0.025982851 | 0.080554273 | 0.219277595 | 0.1263 | 0.27701879 |
| SL1344_RS21395_ | 1.827974627 | 1.768873019 | 1.924922461 | 0.15421715 | 0.109415223 | 1.318690966 | 0.073950757 | 0.1343489 | 0.284500834 | 0.2072 | 0.362760025 |
| SL1344_RS21400_ | 2.2987671 | 2.239536818 | 2.355059097 | 0.124512063 | 0.112607361 | 1.055491958 | 0.122666699 | 0.185224576 | 0.353512071 | 0.2696 | 0.43077975 |
| SL1344_RS21420_ | 0.983640156 | 0.862138247 | 1.049216618 | 0.181060105 | 0.170775265 | 0.985747285 | 0.151232537 | 0.228107122 | 0.399816803 | 0.3264 | 0.488290589 |
| SL1344_RS21425_ | 1.339488378 | 1.30023133 | 1.382228935 | 0.089793566 | 0.159390139 | 0.526945438 | 0.269333344 | 0.42517696 | 0.58865941 | 0.5564 | 0.682486061 |
| SL1344_RS21430_ | 1.225519631 | 1.207078092 | 1.246592908 | 0.045817734 | 0.175494156 | 0.257634015 | 0.386000004 | 0.592131464 | 0.729011567 | 0.7182 | 0.81084581 |
| SL1344_RS21435_ | 1.301746045 | 1.289499366 | 1.315048424 | 0.022879086 | 0.117088175 | 0.202902533 | 0.408000004 | 0.515308261 | 0.658569957 | 0.6275 | 0.73577061 |
| SL1344_RS21440_ | -1.554082658 | -1.614975337 | -1.427707483 | 0.225207822 | 0.595171813 | 0.364145043 | 0.340666673 | 0.497556776 | 0.66771842 | 0.6775 | 0.785053122 |
| SL1344_RS21445_ | -3.932982747 | -4.095543204 | -3.790392599 | 0.345214312 | 0.859179591 | 0.371450545 | 0.332666674 | 0.490563566 | 0.644123968 | 0.6147 | 0.727472315 |
| SL1344_RS21450_ | 0.412062169 | 0.249113721 | 0.551049175 | 0.314360657 | 0.478222318 | 0.608477194 | 0.263333345 | 0.365008007 | 0.560643967 | 0.5044 | 0.666839552 |
| SL1344_RS21460_ | -1.87381624 | -2.164182383 | -1.617645699 | 0.533727983 | 0.469186255 | 1.065597638 | 0.114000035 | 0.180597731 | 0.346675868 | 0.2609 | 0.424227272 |
| SL1344_RS21465_ | 0.831289586 | 0.867971757 | 0.790599654 | -0.10110762 | 0.326627522 | -0.310740615 | 0.346000006 | 0.542185297 | 0.704564145 | 0.6998 | 0.8047255 |
| SL1344_RS21475_ | -1.221896341 | -1.15549541 | -1.292531738 | -0.135463449 | 0.26144109 | -0.49234524 | 0.299333342 | 0.421761005 | 0.573339982 | 0.5282 | 0.655448148 |
| SL1344_RS21480_ | -0.383134459 | -0.421440186 | -0.335355737 | 0.100405852 | 0.254945165 | 0.361842997 | 0.341105936 | 0.505884584 | 0.655631261 | 0.6194 | 0.733173535 |
| SL1344_RS21485_ | -0.27333403 | -0.19772221 | -0.333966457 | -0.14494153 | 0.262746739 | -0.535906544 | 0.268000011 | 0.419839575 | 0.581442624 | 0.5307 | 0.659019023 |
| SL1344_RS21490_ | 0.979045936 | 0.949704441 | 1.037803764 | 0.097042227 | 0.17332475 | 0.506145227 | 0.290666676 | 0.46012771 | 0.622001473 | 0.5833 | 0.708363963 |
| SL1344_RS21495_ | 1.949389277 | 1.941368951 | 1.955652638 | 0.021577367 | 0.156272122 | 0.125259254 | 0.444000002 | 0.675160129 | 0.794236252 | 0.7799 | 0.857962109 |
| SL1344_RS21505_ | 1.394669846 | 1.334397636 | 1.454109469 | 0.114957594 | 0.143008241 | 0.755951102 | 0.193333352 | 0.305807387 | 0.477709855 | 0.4088 | 0.560953779 |
| SL1344_RS21510_ | 0.036745512 | 0.237006454 | -0.164513326 | -0.403223141 | 0.19843734 | -1.994184205 | 0.014666978 | 0.04948324 | 0.15704616 | 0.1136 | 0.260202577 |
| SL1344_RS21515_ | -3.526153319 | -3.67794908 | -3.396830969 | 0.260367519 | 0.684121363 | 0.364584438 | 0.348000006 | 0.494146159 | 0.644172092 | 0.6069 | 0.723967182 |
| SL1344_RS21520_ | 2.272684439 | 2.204079006 | 2.40385348 | 0.19789813 | 0.129926738 | 1.421095692 | 0.069953426 | 0.106444757 | 0.258000566 | 0.1775 | 0.338902971 |
| SL1344_RS21525_ | 2.630992689 | 2.435225618 | 2.842930681 | 0.396621933 | 0.106652324 | 3.630115338 | 4.68E-05 | 0.008541271 | 0.059850117 | 0.1 | 0.243021319 |
| SL1344_RS21530_ | 0.483756896 | 0.258507966 | 0.6455469 | 0.396978766 | 0.260067273 | 1.462790547 | 0.052000084 | 0.092723844 | 0.238395075 | 0.154 | 0.312363074 |
| SL1344_RS21535_ | -1.08967859 | -1.232852462 | -0.953936327 | 0.288265117 | 0.319724303 | 0.813391163 | 0.176000021 | 0.266000985 | 0.433847426 | 0.3609 | 0.510743414 |
| SL1344_RS21540_ | -3.140019748 | -3.453077316 | -2.661725936 | 0.757066028 | 0.629065601 | 1.12760464 | 0.125916087 | 0.1866523 | 0.346564519 | 0.2716 | 0.43269738 |
| SL1344_RS21545_ | -3.675810873 | -3.810235455 | -3.571332871 | 0.230518263 | 0.686057722 | 0.32867851 | 0.363091278 | 0.509623794 | 0.657481912 | 0.627 | 0.737879934 |
| SL1344_RS21550_ | -1.551498051 | -1.627410161 | -1.447809838 | 0.19499309 | 0.365703667 | 0.497372499 | 0.293333343 | 0.42671495 | 0.580681787 | 0.5363 | 0.659902515 |
| SL1344_RS21555_ | -2.937124315 | -2.982052418 | -2.882655154 | 0.10274911 | 0.544858912 | 0.173689475 | 0.424383747 | 0.570989951 | 0.707078102 | 0.6773 | 0.778369753 |
| SL1344_RS21560_ | -2.053998233 | -2.077858862 | -2.006681624 | 0.134016124 | 0.519186079 | 0.216540225 | 0.40373085 | 0.58975609 | 0.728540841 | 0.7249 | 0.816140063 |
| SL1344_RS21565_ | -4.020503832 | -4.438334917 | -3.602896318 | 0.825788155 | 0.765235365 | 1.041201364 | 0.130666697 | 0.204554562 | 0.361948093 | 0.2878 | 0.44257092 |
| SL1344_RS21570_ | -2.743062516 | -2.915436191 | -2.643557416 | 0.291094954 | 0.700353073 | 0.371526872 | 0.332445044 | 0.495592352 | 0.655799348 | 0.6422 | 0.755040413 |
| SL1344_RS21575_ | -4.301072322 | -4.28394032 | -4.311889911 | 0.073424794 | 1.205952311 | 0.060470767 | 0.473333334 | 0.641841634 | 0.770702922 | 0.7632 | 0.846625309 |

| | | | | | | | | | | | |
|---|---|---|---|---|---|---|---|---|---|---|---|
| SL1344_RS21580_ | 1.389064616 | 1.186941713 | 1.628731312 | 0.437500254 | 0.153143906 | 2.786284925 | 0.00067292 | 0.018821379 | 0.092619177 | 0.1004 | 0.2435842 |
| SL1344_RS21585_ | 0.002739747 | -0.306019124 | 0.32721585 | 0.632049122 | 0.213350617 | 2.907571529 | 0.001335878 | 0.018295942 | 0.089058 | 0.1006 | 0.243869875 |
| SL1344_RS21590_ | -0.80187073 | -1.317146085 | -0.602856611 | 0.686494523 | 0.369773833 | 1.697354787 | 0.03200014 | 0.087101672 | 0.230341672 | 0.1407 | 0.294061447 |
| SL1344_RS21595_ | -1.539892554 | -1.595314539 | -1.514259627 | 0.078508804 | 0.501921947 | 0.143014509 | 0.451333335 | 0.703587857 | 0.817171907 | 0.8108 | 0.885494722 |
| SL1344_RS21600_ | -2.367591981 | -2.461948981 | -2.243119451 | 0.207863648 | 0.536334507 | 0.355233674 | 0.349766828 | 0.552704491 | 0.69497392 | 0.6494 | 0.758151341 |
| SL1344_RS21605_ | -4.214618182 | -3.684120648 | -5.051403413 | -1.37325237 | 0.895444301 | -1.443070255 | 0.039307241 | 0.102293139 | 0.239441424 | 0.1533 | 0.305677796 |
| SL1344_RS21610_ | -2.676722627 | -2.542430602 | -2.841395632 | -0.292542591 | 0.475496176 | -0.582475529 | 0.24266668 | 0.374392617 | 0.533365905 | 0.485 | 0.615786935 |
| SL1344_RS21615_ | -1.633410324 | -1.653408285 | -1.618063827 | 0.011373508 | 0.479328839 | 0.028459118 | 0.490339774 | 0.679394914 | 0.796707854 | 0.7828 | 0.860023926 |
| SL1344_RS21620_ | -2.360630777 | -2.239501504 | -2.469945258 | -0.234212754 | 0.4623733 | -0.46546348 | 0.300666675 | 0.440542315 | 0.594944912 | 0.54 | 0.666900278 |
| SL1344_RS21625_ | -2.702778594 | -2.561238741 | -2.828346188 | -0.269062365 | 0.470556788 | -0.569619759 | 0.268666678 | 0.394386497 | 0.549459067 | 0.4994 | 0.629091876 |
| SL1344_RS21630_ | -1.418694168 | -1.521076231 | -1.372501967 | 0.110176863 | 0.466504788 | 0.223349264 | 0.413057965 | 0.66728652 | 0.789503175 | 0.7796 | 0.862149427 |
| SL1344_RS21635_ | -5.301070268 | -5.66424772 | -4.945647784 | 0.654861115 | 1.270692244 | 0.478637684 | 0.280666677 | 0.427807487 | 0.584077788 | 0.5373 | 0.664173358 |
| SL1344_RS21640_ | -5.578708548 | -5.627958695 | -5.500166135 | 0.110601354 | 1.27196474 | 0.078617744 | 0.468354432 | 0.536450748 | 0.675093185 | 0.6504 | 0.75337516 |
| SL1344_RS21645_ | -3.938668977 | -3.850835187 | -4.193816314 | -0.334652847 | 1.145573143 | -0.245205285 | 0.40373085 | 0.626611703 | 0.761985172 | 0.7659 | 0.857032533 |
| SL1344_RS21650_ | -3.342288009 | -3.28103987 | -3.415656201 | -0.107011135 | 0.594012164 | -0.167464664 | 0.41666667 | 0.527741673 | 0.672088319 | 0.634 | 0.745264328 |
| SL1344_RS21655_ | -3.224817957 | -3.019367009 | -3.403397091 | -0.37309936 | 0.759825634 | -0.46003581 | 0.294470362 | 0.472719259 | 0.627727153 | 0.5671 | 0.694076958 |
| SL1344_RS21660_ | -0.739584094 | -0.80152319 | -0.599389239 | 0.150719661 | 0.41614308 | 0.291062093 | 0.390666671 | 0.762679064 | 0.85822544 | 0.7778 | 0.87031745 |
| SL1344_RS21665_ | 1.384657713 | 1.308636282 | 1.491381191 | 0.183508539 | 0.115313785 | 1.538170501 | 0.050666753 | 0.097306551 | 0.229892068 | 0.1642 | 0.31802001 |
| SL1344_RS21670_ | 4.255997906 | 4.072175454 | 4.489502734 | 0.415598752 | 0.10549531 | 3.973231834 | 4.68E-05 | 0.015963018 | 0.088837501 | 0.1 | 0.243021319 |
| SL1344_RS21675_ | -0.572130556 | -0.567005028 | -0.576922644 | -0.069983045 | 0.462659204 | -0.152667842 | 0.435043307 | 0.715825849 | 0.826318065 | 0.8508 | 0.907315224 |
| SL1344_RS21680_ | -2.517057037 | -2.424339815 | -2.597860852 | -0.177067564 | 0.444186992 | -0.372893065 | 0.333111266 | 0.474518406 | 0.62424055 | 0.582 | 0.702023926 |
| SL1344_RS21685_ | -3.190927181 | -3.456447367 | -3.050134804 | 0.367989575 | 0.837879608 | 0.422416969 | 0.317121927 | 0.493900278 | 0.655884342 | 0.6311 | 0.749607556 |
| SL1344_RS21690_ | 0.452269725 | 0.308340837 | 0.58942154 | 0.284548501 | 0.16114418 | 1.711694178 | 0.03200014 | 0.070408369 | 0.188788293 | 0.1395 | 0.29025667 |
| SL1344_RS21695_ | -0.980484085 | -0.987003995 | -0.973216618 | 0.012265271 | 0.271686202 | 0.045233707 | 0.476000001 | 0.559972126 | 0.69759294 | 0.6712 | 0.773389909 |
| SL1344_RS21710_ | -1.695893586 | -1.802059249 | -1.611239415 | 0.17479819 | 0.409664132 | 0.416151309 | 0.326000007 | 0.501955853 | 0.653391193 | 0.6174 | 0.734331656 |
| SL1344_RS21715_ | -2.921217835 | -3.093709722 | -2.775839964 | 0.317359636 | 0.528120756 | 0.561319206 | 0.262000011 | 0.397007025 | 0.556453491 | 0.5011 | 0.633559936 |
| SL1344_RS21725_ | -1.420731211 | -1.442381537 | -1.386897208 | 0.070753946 | 0.354916221 | 0.196255684 | 0.418000003 | 0.58075453 | 0.717270315 | 0.6885 | 0.788049638 |
| SL1344_RS21730_ | 1.224208956 | 1.026980359 | 1.531628829 | 0.445630614 | 0.29199682 | 1.562309768 | 0.050000088 | 0.072826926 | 0.210379452 | 0.1454 | 0.305377234 |
| SL1344_RS21735_ | 1.157983305 | 0.975222739 | 1.316964426 | 0.313715411 | 0.185487412 | 1.655919793 | 0.03533346 | 0.071607674 | 0.202363539 | 0.1399 | 0.296125443 |
| SL1344_RS21740_ | 4.869675911 | 4.761314971 | 5.013536147 | 0.254730387 | 0.06863677 | 3.641335678 | 4.68E-05 | 0.018085678 | 0.093794639 | 0.1 | 0.243021319 |
| SL1344_RS21745_ | 0.958665604 | 0.752010991 | 1.247583453 | 0.502089057 | 0.209352 | 2.340931425 | 0.002000966 | 0.030725136 | 0.123236702 | 0.1019 | 0.24561994 |
| SL1344_RS21755_ | 2.018576865 | 1.929026792 | 2.142907684 | 0.166294166 | 0.19426483 | 0.834605774 | 0.176666687 | 0.266680399 | 0.457941478 | 0.345 | 0.521223261 |
| SL1344_RS21760_ | -0.121132132 | -0.284485332 | -0.0074765 | 0.272743842 | 0.248389621 | 1.045871176 | 0.129333364 | 0.197006699 | 0.365109706 | 0.2806 | 0.439222764 |
| SL1344_RS21765_ | 2.545630455 | 2.382041464 | 2.707907903 | 0.321609298 | 0.123015442 | 2.579761482 | 0.00067292 | 0.022809685 | 0.104275072 | 0.1018 | 0.245541841 |
| SL1344_RS21770_ | 1.468880997 | 1.401804209 | 1.550281225 | 0.149489014 | 0.119984321 | 1.182957826 | 0.091938753 | 0.162874339 | 0.314324895 | 0.2374 | 0.391820503 |
| SL1344_RS21775_ | 0.043087132 | 0.201702398 | -0.142974687 | -0.354272368 | 0.197395582 | -1.691451628 | 0.027315288 | 0.074810577 | 0.198244129 | 0.1327 | 0.283031096 |
| SL1344_RS21780_ | 3.468934571 | 3.459516164 | 3.480627949 | 0.003303697 | 0.085820156 | 0.038493218 | 0.485009994 | 0.703265935 | 0.815216891 | 0.8186 | 0.884613583 |
| SL1344_RS21785_ | -0.494079951 | -0.720456909 | -0.232589005 | 0.489772502 | 0.283903516 | 1.644520384 | 0.03397748 | 0.074110724 | 0.201787788 | 0.1354 | 0.287823627 |
| SL1344_RS21790_ | 0.414702688 | 0.444464345 | 0.391046375 | -0.059673297 | 0.185074659 | -0.287921351 | 0.375333338 | 0.543062081 | 0.683336015 | 0.6594 | 0.762006428 |
| SL1344_RS21795_ | 3.187456127 | 3.099795271 | 3.305978392 | 0.204799657 | 0.098667166 | 1.975730793 | 0.011333737 | 0.05008588 | 0.16465635 | 0.1127 | 0.26024066 |
| SL1344_RS21800_ | 1.979257681 | 1.953084316 | 2.056023389 | 0.107661697 | 0.188732048 | 0.533196046 | 0.294666676 | 0.425650598 | 0.608676479 | 0.6023 | 0.729810065 |
| SL1344_RS21805_ | 1.347512261 | 1.159748132 | 1.621163951 | 0.452273095 | 0.132459459 | 3.343650814 | 4.68E-05 | 0.014930528 | 0.079018348 | 0.1 | 0.243021319 |
| SL1344_RS21810_ | 1.852841805 | 1.919016125 | 1.783721561 | -0.126922035 | 0.12513754 | -0.977939637 | 0.139333361 | 0.214501369 | 0.382967778 | 0.2993 | 0.460725734 |
| SL1344_RS21815_ | 1.555289984 | 1.421280579 | 1.697600183 | 0.301227003 | 0.171890468 | 1.77858324 | 0.004667653 | 0.071333973 | 0.205170643 | 0.1074 | 0.252864937 |
| SL1344_RS21820_ | 0.14849832 | -0.039780982 | 0.220969264 | 0.255755434 | 0.25711164 | 0.877024146 | 0.17800002 | 0.260723327 | 0.441991069 | 0.356 | 0.515865357 |
| SL1344_RS21825_ | -3.353917446 | -3.752899784 | -2.953089899 | 0.780060526 | 0.731499364 | 1.013027976 | 0.144570313 | 0.213012434 | 0.380217302 | 0.294 | 0.456184756 |
| SL1344_RS21830_ | -3.467330206 | -3.442976284 | -3.510476273 | -0.136641657 | 0.77732713 | -0.14877802 | 0.42466667 | 0.579089791 | 0.718951997 | 0.7036 | 0.798674796 |
| SL1344_RS21835_ | 3.457176098 | 3.508837859 | 3.40437824 | -0.098125091 | 0.072252402 | -1.283043516 | 0.084610309 | 0.133355336 | 0.288519828 | 0.2025 | 0.365697715 |
| SL1344_RS21840_ | -3.385783767 | -3.235430815 | -3.598494389 | -0.362013494 | 0.647376373 | -0.527610031 | 0.281145913 | 0.414862008 | 0.571542391 | 0.5216 | 0.649708616 |
| SL1344_RS21845_ | 4.118711792 | 4.101152639 | 4.146897818 | 0.050058369 | 0.071849259 | 0.673009086 | 0.23866668 | 0.360821531 | 0.540489041 | 0.483 | 0.630306103 |
| SL1344_RS21880_ | 1.391047185 | 1.426101809 | 1.371141104 | -0.059668365 | 0.146798855 | -0.378110387 | 0.348434383 | 0.532756734 | 0.679038209 | 0.651 | 0.760566761 |
| SL1344_RS21885_ | 0.227151223 | 0.073003858 | 0.365044552 | 0.356612659 | 0.303518157 | 1.144191282 | 0.061333404 | 0.141080817 | 0.309598359 | 0.1884 | 0.346405538 |
| SL1344_RS21890_ | -0.36948865 | -0.075489633 | -0.5887981 | -0.456686362 | 0.404456821 | -1.050429167 | 0.134666696 | 0.188088501 | 0.368049654 | 0.2664 | 0.441580942 |

| | | | | | | | | | | | |
|---|---|---|---|---|---|---|---|---|---|---|---|
| SL1344_RS21895_ | -0.218172849 | -0.14415801 | -0.289630641 | -0.140882992 | 0.209197361 | -0.626783029 | 0.244666679 | 0.370556784 | 0.528201684 | 0.4728 | 0.608203694 |
| SL1344_RS21900_ | -1.064344697 | -1.103966269 | -0.982696522 | 0.119903785 | 0.403496107 | 0.277006272 | 0.395333337 | 0.620666478 | 0.752836843 | 0.7411 | 0.832344153 |
| SL1344_RS21905_ | 2.766682578 | 2.721504577 | 2.830087179 | 0.112806821 | 0.094714561 | 1.128344569 | 0.093271197 | 0.164493969 | 0.33203391 | 0.2201 | 0.379028804 |
| SL1344_RS21910_ | 1.217035671 | 1.307480511 | 1.136361037 | -0.173139772 | 0.16879989 | -0.974389284 | 0.142666694 | 0.209335514 | 0.38388534 | 0.2951 | 0.461962163 |
| SL1344_RS21915_ | 1.625626191 | 1.575894261 | 1.685580783 | 0.118742472 | 0.155378619 | 0.708256967 | 0.202531663 | 0.30919173 | 0.487216733 | 0.4134 | 0.568678056 |
| SL1344_RS21920_ | -1.013080395 | -1.082703648 | -0.943655396 | 0.135547009 | 0.25436794 | 0.530169695 | 0.284477025 | 0.415632666 | 0.567989859 | 0.5205 | 0.64572502 |
| SL1344_RS21925_ | -0.554653579 | -0.707305081 | -0.327798733 | 0.380773632 | 0.262056832 | 1.372089192 | 0.066622316 | 0.113910298 | 0.260309265 | 0.1752 | 0.331220722 |
| SL1344_RS21930_ | -1.195432587 | -1.166769428 | -1.224220888 | -0.060082257 | 0.261644223 | -0.202980137 | 0.409726852 | 0.498497436 | 0.644112679 | 0.6092 | 0.722972314 |
| SL1344_RS21935_ | -3.91354976 | -4.001323746 | -3.802498046 | 0.169794201 | 1.208767587 | 0.122250588 | 0.431712194 | 0.695341174 | 0.809582934 | 0.7975 | 0.871563745 |
| SL1344_RS21940_ | -1.176592813 | -0.810061909 | -1.86606011 | -1.037747655 | 0.417411406 | -2.335470088 | 0.002666616 | 0.039799412 | 0.142522095 | 0.1025 | 0.246537108 |
| SL1344_RS21945_ | -1.484728616 | -1.280534005 | -1.715589672 | -0.473646335 | 0.53625548 | -0.856388412 | 0.140000028 | 0.238991913 | 0.426691508 | 0.3007 | 0.465718692 |
| SL1344_RS21950_ | -1.418443245 | -1.580842118 | -1.333384036 | 0.192129099 | 0.535409902 | 0.329686564 | 0.367333339 | 0.63275912 | 0.766115616 | 0.7102 | 0.817481338 |
| SL1344_RS21965_ | -1.643559867 | -1.286571698 | -2.043342809 | -0.759900071 | 0.328140001 | -2.262870414 | 0.005334196 | 0.034826737 | 0.12658152 | 0.107 | 0.251921011 |
| SL1344_RS21970_ | -1.322508312 | -1.204394477 | -1.460083536 | -0.252180142 | 0.312518345 | -0.770416788 | 0.199333351 | 0.293852594 | 0.454805326 | 0.3839 | 0.530044598 |
| SL1344_RS21980_ | 3.853098157 | 3.820732679 | 3.877548777 | 0.06118629 | 0.080667098 | 0.728823708 | 0.200666684 | 0.304866064 | 0.486631783 | 0.4056 | 0.564809489 |
| SL1344_RS21985_ | 0.733936926 | 0.071026303 | 1.302722343 | 1.227867633 | 0.16945707 | 7.180261678 | 4.68E-05 | 0.001865788 | 0.026149853 | 0.1 | 0.243021319 |
| SL1344_RS21990_ | 0.89675679 | 0.976660318 | 0.80261341 | -0.210597035 | 0.242695195 | -0.852453011 | 0.148000026 | 0.234542611 | 0.421314359 | 0.3059 | 0.471513445 |
| SL1344_RS21995_ | -2.872766467 | -2.863873766 | -2.881264194 | -0.046800635 | 0.668065808 | -0.064622005 | 0.468666668 | 0.640949221 | 0.767674725 | 0.753 | 0.840319092 |
| SL1344_RS22000_ | 1.643989353 | 1.629099232 | 1.663170927 | 0.032503707 | 0.150711812 | 0.200070207 | 0.416389077 | 0.645062186 | 0.770221565 | 0.7667 | 0.846684136 |
| SL1344_RS22005_ | -1.594982678 | -1.43212411 | -1.795416417 | -0.355023086 | 0.33769763 | -0.991077585 | 0.127333364 | 0.210426393 | 0.367694443 | 0.3032 | 0.456515462 |
| SL1344_RS22010_ | -0.988487315 | -1.130005333 | -0.820491071 | 0.309139926 | 0.258580686 | 1.155364031 | 0.110592975 | 0.174146838 | 0.326917446 | 0.2536 | 0.409955744 |
| SL1344_RS22015_ | 3.10138692 | 3.021500172 | 3.203975856 | 0.179401207 | 0.087535093 | 2.017007435 | 0.012000381 | 0.045804872 | 0.153244946 | 0.1131 | 0.260574514 |
| SL1344_RS22020_ | 2.557079484 | 2.678733324 | 2.396577829 | -0.302409853 | 0.145675881 | -2.103266499 | 0.00067292 | 0.059962081 | 0.187230573 | 0.1003 | 0.243443111 |
| SL1344_RS22030_ | 2.41344513 | 1.931293346 | 2.79394731 | 0.859239652 | 0.096473908 | 8.79710818 | 4.68E-05 | 0.001353021 | 0.022343724 | 0.1 | 0.243021319 |
| SL1344_RS22035_ | 1.96413537 | 1.488718404 | 2.453716926 | 0.968303786 | 0.137614949 | 7.02250167 | 4.68E-05 | 0.001892091 | 0.027746868 | 0.1 | 0.243021319 |
| SL1344_RS22040_ | 3.174909785 | 2.449211881 | 3.722117569 | 1.267158258 | 0.119887168 | 10.43306078 | 4.68E-05 | 0.000898576 | 0.019521133 | 0.1 | 0.243021319 |
| SL1344_RS22045_ | -0.389802652 | -0.531431369 | -0.277184439 | 0.262952739 | 0.246076152 | 1.034118326 | 0.122666699 | 0.201387821 | 0.364108263 | 0.2789 | 0.434302566 |
| SL1344_RS22050_ | -0.208950213 | -0.215155214 | -0.198094644 | -0.043010276 | 0.329322405 | -0.118440392 | 0.459027317 | 0.674525808 | 0.794994443 | 0.8212 | 0.885258336 |
| SL1344_RS22055_ | -0.864425968 | -0.839383745 | -0.897668525 | -0.055966291 | 0.468958382 | -0.109571332 | 0.452000002 | 0.718244599 | 0.828233368 | 0.843 | 0.903049925 |
| SL1344_RS22060_ | -1.197220656 | -1.275792196 | -1.082199502 | 0.215159857 | 0.395844396 | 0.509115844 | 0.303131254 | 0.459629655 | 0.621416925 | 0.5852 | 0.709703453 |
| SL1344_RS22065_ | -1.747984826 | -1.6675299 | -1.893582074 | -0.262254585 | 0.472403408 | -0.504187905 | 0.287141915 | 0.421464157 | 0.590041003 | 0.5591 | 0.68222082 |
| SL1344_RS22070_ | -2.239530425 | -2.141899573 | -2.328360404 | -0.266194708 | 0.726783016 | -0.371349453 | 0.352431718 | 0.536276641 | 0.690467516 | 0.6675 | 0.77656347 |
| SL1344_RS22075_ | -1.761039907 | -1.80618554 | -1.715823645 | 0.096194959 | 0.394513953 | 0.21782626 | 0.394000004 | 0.538479086 | 0.682429735 | 0.6454 | 0.756251949 |
| SL1344_RS22080_ | 0.504275459 | 0.501200869 | 0.509762279 | 0.007569143 | 0.185353037 | 0.04628028 | 0.484666667 | 0.617510813 | 0.747632633 | 0.7227 | 0.8144813 |
| SL1344_RS22085_ | -0.975241361 | -0.876209611 | -1.08176408 | -0.20612651 | 0.293659376 | -0.651144594 | 0.217854779 | 0.351290259 | 0.511093245 | 0.4538 | 0.588508201 |
| SL1344_RS22090_ | -2.904959873 | -2.543240867 | -3.379981708 | -0.82666199 | 0.500564387 | -1.577489415 | 0.034666795 | 0.087203816 | 0.213170898 | 0.1476 | 0.298717496 |
| SL1344_RS22095_ | 0.781155588 | 0.713495051 | 0.847492503 | 0.131723277 | 0.131379215 | 0.948471955 | 0.146000026 | 0.235557404 | 0.388675069 | 0.3168 | 0.466696596 |
| SL1344_RS22100_ | 1.263504711 | 1.193650225 | 1.331136366 | 0.138296149 | 0.134144013 | 0.989807931 | 0.133333363 | 0.209179692 | 0.370664944 | 0.3012 | 0.455953825 |
| SL1344_RS22105_ | -1.593162419 | -1.552631365 | -1.66229424 | -0.159154745 | 0.452443593 | -0.321257568 | 0.356000006 | 0.512220041 | 0.670265939 | 0.6701 | 0.771327702 |
| SL1344_RS22110_ | -2.91702304 | -3.135848315 | -2.777570391 | 0.402277938 | 0.654940675 | 0.556819065 | 0.23666668 | 0.374491112 | 0.545658195 | 0.4843 | 0.616653545 |
| SL1344_RS22115_ | -2.699099004 | -2.746452334 | -2.6573814 | 0.080288071 | 0.463312844 | 0.151260342 | 0.430379749 | 0.512347222 | 0.655610073 | 0.6172 | 0.730233023 |
| SL1344_RS22120_ | -1.387172029 | -1.250791745 | -1.590658703 | -0.331490717 | 0.332294262 | -0.928448219 | 0.156000024 | 0.237651714 | 0.399712177 | 0.3321 | 0.486167828 |
| SL1344_RS22125_ | -2.765428035 | -2.951642415 | -2.543270234 | 0.505646539 | 0.734887898 | 0.649925462 | 0.232666681 | 0.351126575 | 0.528046466 | 0.4885 | 0.628891458 |
| SL1344_RS22130_ | -0.946403135 | -0.383069946 | -1.766559383 | -1.39261851 | 0.363703724 | -3.824819079 | 4.68E-05 | 0.01072919 | 0.065608028 | 0.1 | 0.243021319 |
| SL1344_RS22155_ | -6.529243021 | -6.297678953 | -6.777124848 | -0.480951665 | 1.824220324 | -0.239054559 | 0.388000004 | 0.527919307 | 0.671009762 | 0.6286 | 0.740711927 |
| SL1344_RS22160_ | 0.13264218 | 0.190728649 | 0.06007189 | -0.123083399 | 0.225194084 | -0.505283287 | 0.299333342 | 0.467234216 | 0.623128416 | 0.5716 | 0.697917275 |
| SL1344_RS22165_ | 0.873284628 | 0.906464475 | 0.84119732 | -0.073431356 | 0.168572538 | -0.421454564 | 0.325333341 | 0.483408193 | 0.636895669 | 0.6129 | 0.725617908 |
| SL1344_RS22170_ | 1.791083695 | 1.66741342 | 1.886494734 | 0.191881485 | 0.154191239 | 1.19056775 | 0.10926053 | 0.152912686 | 0.320061583 | 0.228 | 0.397878832 |
| SL1344_RS22175_ | -2.562017456 | -2.626028847 | -2.472745389 | 0.156651036 | 0.498279986 | 0.277417233 | 0.375749505 | 0.53293749 | 0.675424462 | 0.6401 | 0.74873678 |
| SL1344_RS22180_ | -0.148874842 | -0.184296474 | -0.105044814 | 0.046638182 | 0.417898043 | 0.113166142 | 0.440000002 | 0.773066773 | 0.865004049 | 0.7969 | 0.874558404 |
| SL1344_RS22185_ | -0.288558728 | -0.464112987 | -0.138049379 | 0.345327947 | 0.247336847 | 1.315401695 | 0.050666753 | 0.117226956 | 0.264579724 | 0.1682 | 0.322914126 |
| SL1344_RS22190_ | 0.622686657 | 0.638421457 | 0.599794042 | -0.035474954 | 0.223578129 | -0.154040023 | 0.434000002 | 0.654652676 | 0.779459788 | 0.7668 | 0.850106511 |
| SL1344_RS22195_ | 0.890178926 | 0.885578452 | 0.896986608 | 0.014256091 | 0.140231138 | 0.0973949 | 0.457028649 | 0.547707812 | 0.691170983 | 0.6661 | 0.771268022 |

| | | | | | | | | | | | |
|---|---|---|---|---|---|---|---|---|---|---|---|
| SL1344_RS22200_ | 0.085508069 | 0.021596237 | 0.136834603 | 0.172048629 | 0.296811871 | 0.556361511 | 0.28000001 | 0.432276783 | 0.604851019 | 0.5916 | 0.715053301 |
| SL1344_RS22205_ | 0.733183526 | 0.6253663 | 0.789519734 | 0.153823811 | 0.253603237 | 0.600595743 | 0.275333344 | 0.433780856 | 0.607270589 | 0.5383 | 0.684294324 |
| SL1344_RS22210_ | 3.5281423 | 3.783765543 | 3.234985926 | -0.547343137 | 0.05817148 | -9.401020371 | 4.68E-05 | 0.000835322 | 0.017914684 | 0.1 | 0.243021319 |
| SL1344_RS22215_ | 0.57679089 | 0.729197388 | 0.39767495 | -0.335513611 | 0.180205237 | -1.818025211 | 0.021985549 | 0.065940348 | 0.187159209 | 0.124 | 0.272644171 |
| SL1344_RS22220_ | 2.834133964 | 3.165921777 | 2.789312668 | -0.372864974 | 0.187476872 | -1.245376062 | 0.108666704 | 0.153068655 | 0.336904426 | 0.2397 | 0.41280426 |
| SL1344_RS22225_ | 0.389476236 | 0.172562343 | 0.568082135 | 0.403753793 | 0.208337183 | 1.928148053 | 0.006667356 | 0.057790993 | 0.175722574 | 0.1078 | 0.253391714 |
| SL1344_RS22235_ | 7.971384734 | 8.017046659 | 7.96537596 | -0.048437073 | 0.033177065 | -0.626024801 | 0.332445044 | 0.521326455 | 0.695525192 | 0.6955 | 0.826126019 |
| SL1344_RS22240_ | 3.346241523 | 3.378883296 | 3.319560081 | -0.055350184 | 0.066466445 | -0.77167875 | 0.193333352 | 0.297519233 | 0.46429068 | 0.3985 | 0.547826707 |
| SL1344_RS22245_ | -0.127181602 | -0.332603825 | 0.056466004 | 0.394156706 | 0.23596054 | 1.624100557 | 0.025982851 | 0.079429172 | 0.210854604 | 0.1328 | 0.283842544 |
| SL1344_RS22250_ | 2.002812542 | 1.735427689 | 2.231864802 | 0.496540861 | 0.096261824 | 5.08741 | 4.68E-05 | 0.004523349 | 0.04102144 | 0.1 | 0.243021319 |
| SL1344_RS22255_ | 4.566717359 | 4.580376471 | 4.530680036 | -0.045303427 | 0.096234908 | -0.423981853 | 0.338000007 | 0.646556723 | 0.78132664 | 0.6992 | 0.817095492 |
| SL1344_RS22260_ | -0.208398328 | -0.387007551 | 0.057102572 | 0.436258012 | 0.208516648 | 1.99076154 | 0.01866691 | 0.055436746 | 0.166284228 | 0.1254 | 0.274763214 |
| SL1344_RS22265_ | 0.993516052 | 0.959662263 | 1.013167273 | 0.048242302 | 0.14884138 | 0.304194535 | 0.376666672 | 0.552686931 | 0.691394784 | 0.6474 | 0.758759326 |
| SL1344_RS22275_ | 0.380863832 | 0.201335355 | 0.514465973 | 0.302304712 | 0.2028794 | 1.372480205 | 0.067288538 | 0.113062283 | 0.25976532 | 0.1849 | 0.345230044 |
| SL1344_RS22280_ | -2.507665829 | -2.598870748 | -2.318281789 | 0.260393331 | 0.602442999 | 0.390095981 | 0.337774823 | 0.555276333 | 0.696541962 | 0.6581 | 0.767714911 |
| SL1344_RS22285_ | -0.649469616 | -0.575615089 | -0.738707524 | -0.166577718 | 0.225466927 | -0.661178046 | 0.22918056 | 0.336085474 | 0.493305662 | 0.435 | 0.573714847 |
| SL1344_RS22290_ | -1.75517358 | -1.600343099 | -1.950791872 | -0.339499882 | 0.545705491 | -0.593991842 | 0.248000013 | 0.400264676 | 0.573806245 | 0.5155 | 0.657718193 |
| SL1344_RS22295_ | -0.232185601 | -0.117703889 | -0.360087051 | -0.242040616 | 0.18904109 | -1.236063903 | 0.086608976 | 0.143494846 | 0.284157483 | 0.2249 | 0.37728925 |
| SL1344_RS22300_ | 1.079459566 | 1.127352574 | 1.016492581 | -0.114678648 | 0.133220589 | -0.822381108 | 0.17800002 | 0.281781605 | 0.443561449 | 0.3795 | 0.521704478 |
| SL1344_RS22305_ | 2.650857117 | 2.569525118 | 2.835688418 | 0.268082491 | 0.146447371 | 1.726990463 | 0.012658588 | 0.089381478 | 0.239446454 | 0.1135 | 0.260862565 |
| SL1344_RS22310_ | -0.391408244 | -0.871725051 | -0.027045118 | 0.844142614 | 0.251744136 | 3.383276763 | 4.68E-05 | 0.018924821 | 0.089670918 | 0.1 | 0.243021319 |
| SL1344_RS22315_ | 0.23769241 | 0.121814237 | 0.367307682 | 0.263412685 | 0.222721242 | 1.129769456 | 0.093271197 | 0.163701896 | 0.329191983 | 0.2229 | 0.382623922 |
| SL1344_RS22320_ | 2.948584106 | 2.800062978 | 3.123123959 | 0.321110765 | 0.09781152 | 3.187046074 | 4.68E-05 | 0.013977275 | 0.078378344 | 0.1 | 0.243021319 |
| SL1344_RS22325_ | 1.70542129 | 1.546205726 | 1.851779436 | 0.306980089 | 0.113941435 | 2.672814783 | 0.001335878 | 0.025333394 | 0.106868844 | 0.1009 | 0.244291382 |
| SL1344_RS22330_ | 2.374182472 | 2.176188209 | 2.430625072 | 0.233603072 | 0.149618516 | 1.279814867 | 0.104000039 | 0.14285641 | 0.315691595 | 0.2346 | 0.40478561 |
| SL1344_RS22335_ | 1.54790282 | 1.49789031 | 1.594486558 | 0.015378991 | 0.303886426 | 0.051441113 | 0.475016656 | 0.827291426 | 0.901725158 | 0.9178 | 0.952553913 |
| SL1344_RS22340_ | 0.229194535 | 0.291295576 | 0.155335048 | -0.155175412 | 0.251143166 | -0.553143199 | 0.263333345 | 0.390798943 | 0.564091402 | 0.5191 | 0.65549514 |
| SL1344_RS22345_ | 5.500806815 | 5.207341202 | 5.767598022 | 0.501261485 | 0.187716491 | 2.639041428 | 4.68E-05 | 0.012079577 | 0.078396398 | 0.1 | 0.243021319 |
| SL1344_RS22350_ | 4.331203434 | 4.20112264 | 4.405502547 | 0.217957623 | 0.115885112 | 2.025124229 | 4.68E-05 | 0.040901689 | 0.15128796 | 0.1002 | 0.243306537 |
| SL1344_RS22355_ | 1.479545972 | 1.445224213 | 1.576779613 | 0.121990115 | 0.167001103 | 0.612096315 | 0.275816133 | 0.436221261 | 0.60699582 | 0.5765 | 0.711268357 |
| SL1344_RS22360_ | 1.10444126 | 1.055655425 | 1.16320411 | 0.108938412 | 0.131282474 | 0.778119744 | 0.211858777 | 0.300050704 | 0.465169697 | 0.4029 | 0.550224983 |
| SL1344_RS22365_ | 0.924376601 | 0.948578817 | 0.903645446 | -0.045770761 | 0.131535064 | -0.29945691 | 0.365333339 | 0.488547762 | 0.634153398 | 0.5914 | 0.70818398 |
| SL1344_RS22370_ | -0.298650382 | -0.287659346 | -0.319458401 | -0.034960326 | 0.230641517 | -0.138078315 | 0.446369089 | 0.598728014 | 0.733041141 | 0.7105 | 0.807606015 |
| SL1344_RS22375_ | 2.172230229 | 2.111983537 | 2.254382082 | 0.156196608 | 0.116401087 | 1.27845922 | 0.061958764 | 0.136802236 | 0.296343891 | 0.1802 | 0.332406419 |
| SL1344_RS22380_ | 2.718953369 | 2.634517536 | 2.819187157 | 0.18608031 | 0.07419669 | 2.427336477 | 0.002668394 | 0.029993486 | 0.116910354 | 0.1036 | 0.247694025 |
| SL1344_RS22385_ | 4.335465559 | 4.249005323 | 4.380172271 | 0.100648618 | 0.103488702 | 0.959722141 | 0.161225873 | 0.219382847 | 0.410984155 | 0.2945 | 0.47844872 |
| SL1344_RS22395_ | 4.534638424 | 4.526753945 | 4.551639207 | 0.024610957 | 0.068161976 | 0.30885396 | 0.380413063 | 0.625509567 | 0.75847512 | 0.7404 | 0.833159167 |
| SL1344_RS22400_ | 3.979487683 | 3.939547148 | 4.041125917 | 0.090336758 | 0.106896742 | 0.801646825 | 0.203197885 | 0.286569696 | 0.480495051 | 0.3814 | 0.554559783 |
| SL1344_RS22405_ | 4.831271438 | 4.781596769 | 4.892712631 | 0.114000451 | 0.050641907 | 2.249722547 | 0.004664544 | 0.038412564 | 0.139668845 | 0.1027 | 0.246743379 |
| SL1344_RS22410_ | 0.032169094 | -0.170099652 | 0.224344105 | 0.39314987 | 0.185205598 | 2.036301594 | 0.011992386 | 0.041841236 | 0.140674981 | 0.1103 | 0.256436545 |
| SL1344_RS22415_ | 6.208512297 | 6.180698863 | 6.25503729 | 0.078318021 | 0.084501808 | 0.932428597 | 0.14533336 | 0.205311694 | 0.400711271 | 0.2656 | 0.451240223 |
| SL1344_RS22420_ | 1.633589479 | 1.542082683 | 1.703115948 | 0.161858126 | 0.15209059 | 1.060710599 | 0.136575645 | 0.195090619 | 0.36743842 | 0.2803 | 0.446768454 |
| SL1344_RS22425_ | 4.020921512 | 3.98729255 | 4.067852405 | 0.089363901 | 0.085966386 | 1.046093022 | 0.115922753 | 0.171369173 | 0.346754071 | 0.2526 | 0.4205764 |
| SL1344_RS22430_ | 3.367124281 | 3.242932112 | 3.466425478 | 0.231873945 | 0.086100054 | 2.732086633 | 4.68E-05 | 0.031449698 | 0.125154868 | 0.1001 | 0.243162854 |
| SL1344_RS22435_ | -4.437141963 | -4.508045272 | -4.366202089 | 0.167626023 | 0.881604043 | 0.166698009 | 0.423333336 | 0.531420993 | 0.674552433 | 0.6335 | 0.744368778 |
| SL1344_RS22440_ | -2.280194689 | -2.438375809 | -2.120391727 | 0.334811525 | 0.39361762 | 0.751318743 | 0.195333351 | 0.294436954 | 0.452441011 | 0.3889 | 0.533797193 |
| SL1344_RS22445_ | -1.982692189 | -1.998733876 | -1.968354109 | 0.073378796 | 0.533279172 | 0.131959901 | 0.442371754 | 0.647644918 | 0.77364952 | 0.7544 | 0.841457861 |
| SL1344_RS22450_ | -3.302652578 | -3.551972564 | -3.007570782 | 0.551276437 | 0.736774318 | 0.712439542 | 0.207861442 | 0.319118514 | 0.489786528 | 0.4236 | 0.56975716 |
| SL1344_RS22455_ | -3.80196189 | -3.899535393 | -3.732443843 | 0.170928608 | 0.814411912 | 0.209049214 | 0.40466667 | 0.554978756 | 0.69763494 | 0.6766 | 0.777886917 |
| SL1344_RS22460_ | -2.612524463 | -2.50408799 | -2.692987867 | -0.216478041 | 0.50805114 | -0.396407425 | 0.343770826 | 0.495116421 | 0.645588932 | 0.6128 | 0.728069689 |
| SL1344_RS22465_ | -0.789168416 | -0.851075696 | -0.727918261 | 0.128332398 | 0.250700527 | 0.474687576 | 0.29913392 | 0.433806916 | 0.587182878 | 0.5498 | 0.67252035 |
| SL1344_RS22470_ | -0.993058319 | -1.09555759 | -0.87643091 | 0.22525414 | 0.297652512 | 0.683402872 | 0.219853446 | 0.334674441 | 0.499454901 | 0.4444 | 0.58428151 |
| SL1344_RS22475_ | -1.115921142 | -1.113333466 | -1.117598379 | 0.009622606 | 0.481933165 | 0.015534869 | 0.490339774 | 0.736575351 | 0.839345464 | 0.8362 | 0.8984854 |

| | | | | | | | | | | | |
|---|---|---|---|---|---|---|---|---|---|---|---|
| SL1344_RS22480_ | -0.592938638 | -0.900510123 | -0.471190129 | 0.40643341 | 0.39210938 | 0.986720155 | 0.165223207 | 0.214951062 | 0.398764282 | 0.3183 | 0.49282906 |
| SL1344_RS22485_ | -0.079920661 | 0.069202688 | -0.237816054 | -0.298913167 | 0.181442751 | -1.587015871 | 0.041972124 | 0.08446141 | 0.210467912 | 0.1565 | 0.309363897 |
| SL1344_RS22490_ | 1.131995756 | 0.995288629 | 1.257490025 | 0.159766902 | 0.273971826 | 0.625222139 | 0.238507675 | 0.39730545 | 0.580550652 | 0.484 | 0.63694192 |
| SL1344_RS22495_ | -0.967087561 | -1.061692109 | -0.850150989 | 0.162013969 | 0.617840513 | 0.267733452 | 0.375083283 | 0.680836454 | 0.802912327 | 0.764 | 0.85292299 |
| SL1344_RS22500_ | -2.914249754 | -2.758806661 | -3.08197239 | -0.323025232 | 0.514947495 | -0.591435055 | 0.249167234 | 0.379170619 | 0.536315217 | 0.4773 | 0.613741014 |
| SL1344_RS22505_ | -4.06530623 | -3.892268092 | -4.304837653 | -0.560082188 | 1.280990525 | -0.365681026 | 0.314457037 | 0.459850187 | 0.632778391 | 0.6029 | 0.722410104 |
| SL1344_RS22510_ | -3.436289489 | -3.131077502 | -3.848089503 | -0.719925301 | 0.580055121 | -1.152158676 | 0.08933338 | 0.157724934 | 0.306859278 | 0.2276 | 0.382188414 |
| SL1344_RS22520_ | -1.9441693 | -1.700906288 | -2.295030924 | -0.615977355 | 0.430417343 | -1.355099747 | 0.045303228 | 0.127638451 | 0.279105587 | 0.1672 | 0.319239458 |
| SL1344_RS22525_ | -1.595746702 | -1.476395328 | -1.766696911 | -0.311091112 | 0.411409647 | -0.736458497 | 0.186666686 | 0.296471687 | 0.471559873 | 0.395 | 0.540026162 |
| SL1344_RS22530_ | -1.227607557 | -1.273628007 | -1.178643714 | 0.103304066 | 0.32041931 | 0.278154378 | 0.373750838 | 0.525091301 | 0.667765973 | 0.6342 | 0.743611414 |
| SL1344_RS22535_ | 4.419847032 | 4.298799072 | 4.710763817 | 0.292878312 | 0.210453123 | 1.31324509 | 0.115256531 | 0.136874991 | 0.316173367 | 0.2017 | 0.38487149 |
| SL1344_RS22540_ | 6.327890518 | 6.221500901 | 6.548103914 | 0.330221324 | 0.092484429 | 3.396640869 | 4.68E-05 | 0.021551589 | 0.10678589 | 0.1 | 0.243021319 |
| SL1344_RS22545_ | 6.711757854 | 6.53153745 | 6.778408048 | 0.212959711 | 0.113468977 | 1.69611234 | 0.015989625 | 0.066495299 | 0.205064651 | 0.1148 | 0.263770137 |
| SL1344_RS22550_ | 5.808060463 | 5.722483993 | 5.942128166 | 0.211726093 | 0.069721805 | 2.844610437 | 4.68E-05 | 0.02680679 | 0.11852673 | 0.1 | 0.243021319 |
| SL1344_RS22555_ | 7.277233071 | 7.116350633 | 7.424017715 | 0.323253013 | 0.050363792 | 6.553586726 | 4.68E-05 | 0.000722384 | 0.019422986 | 0.1 | 0.243021319 |
| SL1344_RS22560_ | -0.732685229 | -1.075792842 | -0.406622554 | 0.663836105 | 0.231192735 | 2.830469726 | 0.001336769 | 0.018065091 | 0.086655796 | 0.1004 | 0.243581003 |
| SL1344_RS22565_ | 1.14664994 | 1.049786749 | 1.248762241 | 0.200881131 | 0.117211358 | 1.629771563 | 0.036000124 | 0.080950456 | 0.204077747 | 0.1546 | 0.30544255 |
| SL1344_RS22570_ | 2.270625591 | 2.297552723 | 1.799637138 | -0.500494199 | 0.668654458 | -0.722128921 | 0.33333334 | 0.386336339 | 0.587898608 | 0.7 | 0.829598552 |
| SL1344_RS22575_ | 2.71035416 | 2.603112602 | 2.914817252 | 0.317367803 | 0.146009693 | 2.158331424 | 0.00067292 | 0.0623794 | 0.193539852 | 0.1006 | 0.243859906 |
| SL1344_RS22580_ | -1.325361814 | -1.334653719 | -1.320592043 | -0.013940476 | 0.434030787 | -0.030374763 | 0.487333334 | 0.696642401 | 0.811044349 | 0.7943 | 0.872967905 |
| SL1344_RS22585_ | 0.061752004 | 0.328419373 | -0.098123319 | -0.412780915 | 0.312921049 | -1.255422585 | 0.085942754 | 0.127951662 | 0.290927713 | 0.2048 | 0.374319328 |
| SL1344_RS22590_ | -2.515523393 | -2.431672235 | -2.576554693 | -0.075338021 | 0.710684567 | -0.102911774 | 0.461333335 | 0.712139546 | 0.82191857 | 0.819 | 0.886529971 |
| SL1344_RS22595_ | -0.903498556 | -0.830883633 | -0.952833135 | -0.116044568 | 0.279812694 | -0.385960609 | 0.336442379 | 0.509443695 | 0.656147687 | 0.6134 | 0.728608848 |
| SL1344_RS22600_ | 0.453708921 | 0.473919831 | 0.42584325 | -0.050674638 | 0.172974506 | -0.279366986 | 0.381333338 | 0.526617775 | 0.670552847 | 0.6502 | 0.754609725 |
| SL1344_RS22610_ | -0.437668227 | -0.520478354 | -0.374520281 | 0.1479498 | 0.238936763 | 0.55180721 | 0.251333346 | 0.382261189 | 0.542936669 | 0.4881 | 0.619657601 |
| SL1344_RS22615_ | 3.614644548 | 4.037499057 | 3.298031212 | -0.696506911 | 0.227967349 | -3.216264791 | 4.68E-05 | 0.006888349 | 0.057836347 | 0.1 | 0.243021319 |
| SL1344_RS22620_ | 3.270205483 | 3.189351315 | 3.348457457 | 0.217630863 | 0.141265216 | 1.551045413 | 0.007329074 | 0.06948817 | 0.204852011 | 0.1086 | 0.254458333 |
| SL1344_RS22625_ | 0.854072924 | 1.182513931 | 0.681077625 | -0.561315465 | 0.368428575 | -1.504068356 | 0.03000015 | 0.078389573 | 0.222202689 | 0.1302 | 0.283710663 |
| SL1344_RS22630_ | 2.205138782 | 2.014500679 | 2.563036524 | 0.519493907 | 0.171611826 | 2.730573234 | 4.68E-05 | 0.027511794 | 0.118830723 | 0.1 | 0.243021319 |
| SL1344_RS22635_ | 3.98117943 | 3.80169647 | 4.175944392 | 0.375492528 | 0.052936304 | 7.068221416 | 4.68E-05 | 0.001635629 | 0.025141789 | 0.1 | 0.243021319 |
| SL1344_RS22640_ | 4.139361726 | 4.172484514 | 3.990997947 | -0.150785251 | 0.125685806 | -0.912335457 | 0.229846783 | 0.270261228 | 0.470449984 | 0.4407 | 0.610016644 |
| SL1344_RS22645_ | -1.629019813 | -1.590209678 | -1.664595101 | -0.063702141 | 0.35850925 | -0.156945478 | 0.433333336 | 0.585420702 | 0.721208546 | 0.687 | 0.788258763 |
| SL1344_RS22650_ | -2.251436365 | -1.602325923 | -2.78395185 | -1.219976293 | 0.645880343 | -1.860890881 | 0.015989625 | 0.055052465 | 0.173164365 | 0.1128 | 0.260488637 |
| SL1344_RS22655_ | 5.087847511 | 5.032365611 | 5.174633761 | 0.150602182 | 0.082636737 | 1.78254425 | 0.008000574 | 0.063810441 | 0.197138442 | 0.1057 | 0.250811766 |
| SL1344_RS22660_ | 2.393690031 | 2.423673573 | 2.346271415 | -0.084883523 | 0.099750689 | -0.797574388 | 0.170000022 | 0.280915721 | 0.454805259 | 0.3759 | 0.520137879 |
| SL1344_RS22665_ | 3.082506722 | 3.046635731 | 3.214621522 | 0.148908587 | 0.135355199 | 0.880221865 | 0.214000016 | 0.281108855 | 0.476641514 | 0.4172 | 0.585848129 |
| SL1344_RS22670_ | -3.064047394 | -2.837903987 | -3.254239367 | -0.412558308 | 0.658196216 | -0.576110976 | 0.265333344 | 0.414299101 | 0.575167446 | 0.5181 | 0.654847661 |
| SL1344_RS22675_ | 2.422257689 | 2.293304587 | 2.499481816 | 0.207335673 | 0.099107252 | 1.950787291 | 0.017333596 | 0.0577581 | 0.175409794 | 0.1168 | 0.264777475 |
| SL1344_RS22680_ | -0.564434733 | -0.604009982 | -0.531275172 | 0.075385683 | 0.225514704 | 0.305376112 | 0.358427721 | 0.501558102 | 0.648907503 | 0.6125 | 0.727160097 |
| SL1344_RS22685_ | -5.185593345 | -5.982842056 | -4.813873754 | 0.905958372 | 2.60919806 | 0.328625304 | 0.37974684 | 0.631594096 | 0.770600431 | 0.7582 | 0.854437111 |
| SL1344_RS22690_ | -0.455464878 | -0.355596844 | -0.622818431 | -0.269462247 | 0.261627432 | -0.932146711 | 0.148666692 | 0.23494125 | 0.404947202 | 0.333 | 0.488425031 |
| SL1344_RS22695_ | -0.788568087 | -0.695542181 | -0.835189672 | -0.186297394 | 0.485001423 | -0.36162035 | 0.36642239 | 0.59633038 | 0.739896261 | 0.7342 | 0.829831357 |
| SL1344_RS22700_ | -1.569574997 | -1.604668583 | -1.527277904 | 0.087887475 | 0.372431278 | 0.218030013 | 0.40373085 | 0.567497552 | 0.70529834 | 0.6803 | 0.779727263 |
| SL1344_RS22705_ | -3.811850767 | -3.632668192 | -3.942115072 | -0.326379938 | 0.992200835 | -0.309925208 | 0.369753503 | 0.592038677 | 0.7272865 | 0.7088 | 0.803710342 |
| SL1344_RS22710_ | -4.334463468 | -4.71105684 | -3.970406718 | 0.750989987 | 0.868579726 | 0.805720475 | 0.181878767 | 0.275872371 | 0.436813995 | 0.3637 | 0.513531401 |
| SL1344_RS22715_ | -1.494483947 | -1.494539982 | -1.494429798 | -0.012602942 | 0.316140141 | -0.033899972 | 0.488666667 | 0.553042227 | 0.692260117 | 0.6583 | 0.764232103 |
| SL1344_RS22720_ | -0.882574175 | -0.93403652 | -0.84797953 | 0.071789261 | 0.337513416 | 0.200219594 | 0.409726852 | 0.626372579 | 0.756170501 | 0.7375 | 0.829358508 |
| SL1344_RS22725_ | -0.806408585 | -0.77429562 | -0.855726577 | -0.112648757 | 0.351896864 | -0.27611364 | 0.36908728 | 0.522081961 | 0.678519736 | 0.6702 | 0.770685888 |
| SL1344_RS22730_ | -2.069193398 | -1.964392121 | -2.14577097 | -0.243767549 | 0.538031214 | -0.418080974 | 0.322000008 | 0.48574925 | 0.644006423 | 0.6233 | 0.7372447 |
| SL1344_RS22735_ | -0.969420362 | -0.857080909 | -1.164904482 | -0.292646471 | 0.393093126 | -0.700263726 | 0.250666679 | 0.364452058 | 0.542232652 | 0.4834 | 0.630022111 |
| SL1344_RS22740_ | -1.907864862 | -1.733518985 | -2.153160212 | -0.437273388 | 0.416264696 | -0.998880056 | 0.125333365 | 0.212653857 | 0.379454091 | 0.289 | 0.441341722 |
| SL1344_RS22745_ | -1.4479294 | -1.640834591 | -1.014686297 | 0.647225141 | 0.459127646 | 1.311275434 | 0.086000049 | 0.13937189 | 0.303438695 | 0.202 | 0.362432321 |
| SL1344_RS22750_ | -0.929168126 | -0.759683226 | -1.113787726 | -0.350142419 | 0.242212989 | -1.408311626 | 0.066000065 | 0.115865779 | 0.253584628 | 0.1851 | 0.33958147 |

| | | | | | | | | | | | |
|---|---|---|---|---|---|---|---|---|---|---|---|
| SL1344_RS22755_ | -3.957780692 | -3.352598126 | -4.229248274 | -0.750604787 | 1.345692277 | -0.501561927 | 0.28600001 | 0.528303014 | 0.681107676 | 0.5894 | 0.728096544 |
| SL1344_RS22760_ | -2.469136003 | -2.2426602 | -2.607133584 | -0.341794914 | 0.667049805 | -0.468503635 | 0.309793479 | 0.51112286 | 0.66519933 | 0.6291 | 0.74680562 |
| SL1344_RS22765_ | 2.048734316 | 1.880994364 | 2.172891634 | 0.294900025 | 0.126628035 | 2.248496204 | 0.005330642 | 0.044339673 | 0.151993886 | 0.1048 | 0.249507554 |
| SL1344_RS22770_ | 2.712395629 | 2.67696518 | 2.72579032 | 0.046034909 | 0.118472772 | 0.343090583 | 0.381333338 | 0.651196689 | 0.778906064 | 0.7571 | 0.850121412 |
| SL1344_RS22775_ | 3.681474133 | 3.765951933 | 3.610949847 | -0.164650445 | 0.100176869 | -1.702296896 | 0.012658588 | 0.071771347 | 0.206356198 | 0.1118 | 0.258902676 |
| SL1344_RS22780_ | -4.94412919 | -4.785394332 | -5.112835829 | -0.308731302 | 1.194053462 | -0.233051109 | 0.389740178 | 0.560181396 | 0.699156573 | 0.671 | 0.773571744 |
| SL1344_RS22785_ | -3.757390304 | -3.673524701 | -3.860106962 | -0.276577919 | 1.014333005 | -0.255220682 | 0.375333338 | 0.559612173 | 0.70619514 | 0.7106 | 0.803955196 |
| SL1344_RS22790_ | -4.727233628 | -4.592136235 | -4.921333118 | -0.317315691 | 1.099711038 | -0.260006709 | 0.374000005 | 0.534304306 | 0.679555072 | 0.6529 | 0.759390579 |
| SL1344_RS22795_ | -3.333130592 | -3.279487756 | -3.373620486 | -0.084250523 | 0.575621565 | -0.135614487 | 0.434377084 | 0.509814419 | 0.657541021 | 0.6291 | 0.739588642 |
| SL1344_RS22800_ | -3.436689592 | -3.203323071 | -3.675511251 | -0.472827947 | 1.071703505 | -0.400482454 | 0.292000009 | 0.479885193 | 0.646025985 | 0.5831 | 0.714229519 |
| SL1344_RS22805_ | -2.286763894 | -2.268243346 | -2.311349378 | -0.082293229 | 0.509709616 | -0.140229296 | 0.441039309 | 0.608083925 | 0.741490953 | 0.7256 | 0.817177982 |
| SL1344_RS22810_ | -2.623806211 | -2.787195514 | -2.488076892 | 0.278451656 | 0.528755757 | 0.470438153 | 0.295136585 | 0.454439876 | 0.608129196 | 0.5647 | 0.689166259 |
| SL1344_RS22815_ | -2.470195761 | -2.344903386 | -2.628707414 | -0.227696116 | 0.635645909 | -0.337271807 | 0.349766828 | 0.571392349 | 0.714777995 | 0.68 | 0.787606113 |
| SL1344_RS22820_ | 0.634531663 | 0.600342495 | 0.681279142 | 0.080827442 | 0.149997535 | 0.504221 | 0.280666677 | 0.414487453 | 0.568330145 | 0.5246 | 0.651713956 |
| SL1344_RS22825_ | 0.323809281 | 0.313965225 | 0.334563111 | 0.033622668 | 0.216916868 | 0.143752594 | 0.439706864 | 0.622759019 | 0.751757048 | 0.7498 | 0.833142146 |
| SL1344_RS22830_ | 0.131319222 | 0.174127966 | 0.107512364 | -0.041441374 | 0.248408154 | -0.154214948 | 0.427714859 | 0.657106268 | 0.779627666 | 0.7507 | 0.840585932 |
| SL1344_RS22835_ | -2.751169026 | -2.913756997 | -2.70414818 | 0.17888818 | 0.895394332 | 0.167638939 | 0.440373087 | 0.76989024 | 0.864236323 | 0.8292 | 0.901543654 |
| SL1344_RS22840_ | -0.560819743 | -0.794748479 | -0.384863389 | 0.353048653 | 0.44011092 | 0.807723078 | 0.200666684 | 0.292682364 | 0.481132285 | 0.3752 | 0.548256509 |
| SL1344_RS22845_ | -6.190598277 | -6.314147847 | -6.078117898 | 0.378391951 | 2.482830001 | 0.118602098 | 0.415722855 | 0.578284462 | 0.723357915 | 0.7198 | 0.81156391 |
| SL1344_RS22850_ | -1.634595232 | -1.747314568 | -1.498460462 | 0.238679863 | 0.412451302 | 0.539377424 | 0.285143248 | 0.450559393 | 0.605902954 | 0.5565 | 0.682637572 |
| SL1344_RS22855_ | -1.128661323 | -1.253730373 | -0.974968422 | 0.279220175 | 0.377219566 | 0.709930988 | 0.207861442 | 0.325938884 | 0.498910037 | 0.422 | 0.574648505 |
| SL1344_RS22860_ | -0.362616661 | -0.443643462 | -0.275880773 | 0.16589535 | 0.223275096 | 0.715043726 | 0.224000015 | 0.324915623 | 0.490285299 | 0.4299 | 0.574181421 |
| SL1344_RS22865_ | 1.472582932 | 1.306027218 | 1.780873497 | 0.463084461 | 0.180013196 | 2.417720555 | 0.003332494 | 0.043536637 | 0.153796697 | 0.1021 | 0.245981102 |
| SL1344_RS22870_ | 1.413137234 | 1.422383001 | 1.39843839 | -0.023131186 | 0.205023827 | -0.091540962 | 0.461692207 | 0.742776505 | 0.845339364 | 0.8475 | 0.907391272 |
| SL1344_RS22875_ | 4.750437981 | 4.688050408 | 4.780176859 | 0.077627919 | 0.068251405 | 1.056548524 | 0.127333364 | 0.18896506 | 0.368946826 | 0.2728 | 0.44709708 |
| SL1344_RS22880_ | -0.962686981 | -0.867015195 | -1.043050717 | -0.168791262 | 0.262688454 | -0.626466326 | 0.24266668 | 0.373599652 | 0.530007189 | 0.4823 | 0.61567928 |
| SL1344_RS22885_ | -0.087933804 | -0.06782977 | -0.106636017 | -0.079443486 | 0.31381019 | -0.233598284 | 0.402398405 | 0.623187463 | 0.755024172 | 0.7501 | 0.835505281 |
| SL1344_RS22890_ | -2.519566571 | -2.351792769 | -2.704770505 | -0.335134712 | 0.553777735 | -0.582607373 | 0.258000012 | 0.396165007 | 0.560081214 | 0.5054 | 0.64322656 |
| SL1344_RS22895_ | -3.436381833 | -3.494583785 | -3.413906747 | 0.058774633 | 0.969996082 | 0.052047945 | 0.477333334 | 0.709319696 | 0.820185416 | 0.8189 | 0.887984373 |
| SL1344_RS22900_ | -0.995850681 | -1.196806715 | -0.763765227 | 0.458504991 | 0.333218772 | 1.339520515 | 0.066622316 | 0.120377063 | 0.27064615 | 0.1842 | 0.342972416 |
| SL1344_RS22905_ | -3.16889087 | -3.433031528 | -2.932657658 | 0.172182627 | 1.445846554 | 0.112916968 | 0.452365092 | 0.791286462 | 0.878908584 | 0.913 | 0.949130264 |
| SL1344_RS22910_ | -4.066622754 | -4.202460921 | -3.936822497 | 0.371692246 | 1.019021261 | 0.334068834 | 0.356000006 | 0.543276922 | 0.688941801 | 0.6793 | 0.780590502 |
| SL1344_RS22915_ | -3.102109447 | -3.124185115 | -3.079094655 | 0.197492756 | 1.089489998 | 0.170864386 | 0.431333336 | 0.681966696 | 0.8021338 | 0.8379 | 0.898579237 |
| SL1344_RS22920_ | -0.75867164 | -0.724573481 | -0.774822029 | -0.053847381 | 0.307378779 | -0.157177937 | 0.433044639 | 0.623071715 | 0.752738836 | 0.7372 | 0.828058972 |
| SL1344_RS22925_ | -0.886899101 | -0.749907244 | -1.001101106 | -0.302151546 | 0.460790155 | -0.636283885 | 0.23451034 | 0.349319568 | 0.537120513 | 0.4722 | 0.622451123 |
| SL1344_RS22930_ | 1.906012621 | 1.949180936 | 1.84348171 | -0.113989294 | 0.130642207 | -0.791024416 | 0.187874769 | 0.283457328 | 0.461266445 | 0.3849 | 0.536366136 |
| SL1344_RS22935_ | 1.457232079 | 1.3016256 | 1.569073019 | 0.265156375 | 0.120053736 | 2.149286988 | 0.010659989 | 0.042086485 | 0.140932162 | 0.1104 | 0.256337112 |
| SL1344_RS22940_ | -0.628152418 | -0.903695513 | -0.41126652 | 0.48576305 | 0.266323243 | 1.766018775 | 0.028000161 | 0.065960999 | 0.186969403 | 0.1291 | 0.280242724 |
| SL1344_RS22945_ | 0.879819717 | 0.688218077 | 1.114712158 | 0.435446265 | 0.192796189 | 2.21839279 | 0.003332494 | 0.037517808 | 0.137121752 | 0.1039 | 0.248417514 |
| SL1344_RS22950_ | -0.35419118 | -0.431748942 | -0.284965025 | 0.138752887 | 0.355420366 | 0.364049061 | 0.344000006 | 0.562773142 | 0.711369629 | 0.6827 | 0.78941902 |
| SL1344_RS22955_ | 5.467944342 | 5.369213666 | 5.587179747 | 0.209700059 | 0.051493759 | 3.969703534 | 4.68E-05 | 0.007023376 | 0.055187913 | 0.1 | 0.243021319 |
| SL1344_RS22960_ | 2.580765681 | 2.568882915 | 2.617019626 | 0.048137916 | 0.153544098 | 0.286494883 | 0.40266667 | 0.691890026 | 0.810250467 | 0.7938 | 0.878047123 |
| SL1344_RS22965_ | 4.014182379 | 4.044815134 | 3.905751632 | -0.124585029 | 0.096945352 | -1.088543688 | 0.165223207 | 0.194553585 | 0.381579545 | 0.3088 | 0.486774777 |
| SL1344_RS22970_ | 2.471912138 | 2.313831303 | 2.533832217 | 0.210278351 | 0.113073519 | 1.688722711 | 0.043333435 | 0.088498836 | 0.230114565 | 0.1491 | 0.303684617 |
| SL1344_RS22975_ | 2.729826089 | 2.715022482 | 2.781125613 | 0.065339185 | 0.164635366 | 0.304444653 | 0.384000005 | 0.796277605 | 0.882413655 | 0.7819 | 0.876103295 |
| SL1344_RS22980_ | 2.49355917 | 2.479496387 | 2.538922289 | 0.061056623 | 0.215174551 | 0.244815495 | 0.393333337 | 0.811953192 | 0.89217324 | 0.7936 | 0.882340966 |
| SL1344_RS22985_ | -2.302166367 | -2.115642646 | -2.499261798 | -0.390148545 | 0.486115272 | -0.762382322 | 0.201199218 | 0.30304789 | 0.470556027 | 0.4032 | 0.550423199 |
| SL1344_RS22995_ | -3.067080156 | -3.261379101 | -2.828654644 | 0.418415645 | 0.632798845 | 0.623602087 | 0.251333346 | 0.403994791 | 0.562907884 | 0.5013 | 0.637652345 |
| SL1344_RS23000_ | -4.096082084 | -4.410778756 | -3.809354477 | 0.83225038 | 1.22894612 | 0.621407334 | 0.244666679 | 0.349083472 | 0.52688617 | 0.4767 | 0.620338177 |
| SL1344_RS23010_ | -0.486344882 | -0.308171755 | -0.671680513 | -0.357034868 | 0.222384265 | -1.575039637 | 0.046000096 | 0.089948 | 0.217701743 | 0.1576 | 0.310453811 |
| SL1344_RS23015_ | 1.605343665 | 1.231855892 | 1.752627943 | 0.536786772 | 0.302194971 | 1.724102159 | 0.005334196 | 0.083280002 | 0.231876357 | 0.1059 | 0.251020334 |
| SL1344_RS23020_ | -2.502465725 | -3.033738497 | -2.057927542 | 0.976342362 | 0.434489098 | 2.185155817 | 0.006662914 | 0.03838861 | 0.132982743 | 0.1091 | 0.254531451 |
| SL1344_RS23025_ | 1.616989885 | 1.679070769 | 1.525311117 | -0.139734365 | 0.208826167 | -0.663560547 | 0.228666681 | 0.370527911 | 0.554111002 | 0.4818 | 0.64105397 |

| | | | | | | | | | | | |
|---|---|---|---|---|---|---|---|---|---|---|---|
| SL1344_RS23030_ | 1.302363236 | 1.540247048 | 1.017370088 | -0.525132535 | 0.127526944 | -4.040595447 | 4.68E-05 | 0.008428637 | 0.057084308 | 0.1 | 0.243021319 |
| SL1344_RS23035_ | 3.074673561 | 3.027298024 | 3.127081236 | 0.093434295 | 0.095751441 | 0.979511836 | 0.146666693 | 0.22142727 | 0.397013449 | 0.3045 | 0.471619237 |
| SL1344_RS23040_ | 3.559047163 | 3.495993296 | 3.613307721 | 0.117410676 | 0.088090754 | 1.296017928 | 0.074666724 | 0.118303939 | 0.275377648 | 0.1863 | 0.350581071 |
| SL1344_RS23045_ | 2.202447929 | 2.203165486 | 2.202037382 | -0.000743986 | 0.130331194 | -0.005586861 | 0.499333333 | 0.682904484 | 0.799331568 | 0.7771 | 0.857831412 |
| SL1344_RS23050_ | 2.772931174 | 2.715754328 | 2.814532104 | 0.096634236 | 0.096439108 | 0.969993773 | 0.145902758 | 0.227476642 | 0.402555103 | 0.3138 | 0.474763451 |
| SL1344_RS23055_ | 3.307800273 | 3.248343685 | 3.363009465 | 0.115157773 | 0.06849792 | 1.634227067 | 0.026000173 | 0.071432065 | 0.195804445 | 0.1335 | 0.283663214 |
| SL1344_RS23060_ | 4.037362893 | 3.904843246 | 4.162045496 | 0.260346338 | 0.075854425 | 3.442778801 | 4.68E-05 | 0.01092663 | 0.069476555 | 0.1001 | 0.243158931 |
| SL1344_RS23065_ | 1.791254194 | 1.61843375 | 1.846898347 | 0.187593184 | 0.2319123 | 0.67561727 | 0.285143248 | 0.416780653 | 0.604483895 | 0.582 | 0.729088539 |
| SL1344_RS23070_ | 1.726416462 | 1.563390475 | 1.848126927 | 0.246516357 | 0.204723421 | 1.156584956 | 0.101333374 | 0.156561904 | 0.331387568 | 0.2223 | 0.398313242 |
| SL1344_RS23075_ | -3.565206637 | -3.82526412 | -3.371478969 | 0.507360613 | 0.807002752 | 0.596684507 | 0.244000013 | 0.367925113 | 0.539212944 | 0.4779 | 0.614625321 |
| SL1344_RS23080_ | 2.396165399 | 2.425192723 | 2.380548206 | -0.038793911 | 0.101445619 | -0.343328616 | 0.36109261 | 0.575226381 | 0.712754575 | 0.6704 | 0.778756203 |
| SL1344_RS23085_ | -0.162876305 | -0.09777832 | -0.213970388 | -0.120836058 | 0.214569625 | -0.570707147 | 0.265822796 | 0.398615897 | 0.559479876 | 0.5119 | 0.643357439 |
| SL1344_RS23090_ | -3.026998138 | -3.188563165 | -2.858091356 | 0.422662715 | 0.788941747 | 0.49918744 | 0.292000009 | 0.434648938 | 0.601724497 | 0.5699 | 0.695453836 |
| SL1344_RS23095_ | -3.504643279 | -3.367308219 | -3.959063438 | -0.526351379 | 0.908870682 | -0.48661011 | 0.310000008 | 0.469233853 | 0.636521651 | 0.6214 | 0.748964128 |
| SL1344_RS23100_ | 0.712371144 | 0.633565309 | 0.750407394 | 0.106486046 | 0.201513097 | 0.492210826 | 0.298666676 | 0.499771964 | 0.653268046 | 0.6164 | 0.736752473 |
| SL1344_RS23105_ | -2.439855386 | -2.380495289 | -2.485806025 | -0.117249174 | 0.472560053 | -0.230158149 | 0.405729517 | 0.559399173 | 0.695818898 | 0.6582 | 0.76391729 |
| SL1344_RS23110_ | -1.909377448 | -2.034494636 | -1.833163936 | 0.221762408 | 0.440631882 | 0.467151889 | 0.312000008 | 0.46407405 | 0.61835681 | 0.585 | 0.704777059 |
| SL1344_RS23115_ | -0.335806901 | -0.381002449 | -0.289803437 | 0.093364779 | 0.192075263 | 0.452622225 | 0.313790814 | 0.425642803 | 0.577153938 | 0.5268 | 0.652213027 |
| SL1344_RS23120_ | 2.231281876 | 2.224791963 | 2.23925316 | 0.017388792 | 0.240617342 | 0.075531537 | 0.463690874 | 0.823303762 | 0.899071289 | 0.8992 | 0.939737987 |
| SL1344_RS23125_ | -1.822898033 | -1.858276394 | -1.796786048 | 0.083524609 | 0.448031 | 0.170818043 | 0.431045972 | 0.604618016 | 0.738408307 | 0.7291 | 0.818772236 |
| SL1344_RS23130_ | -4.667385678 | -4.679480545 | -4.663721559 | 0.026172594 | 1.005645465 | 0.027622812 | 0.489333334 | 0.584208734 | 0.720743912 | 0.7053 | 0.801275646 |
| SL1344_RS23140_ | -1.498807149 | -1.393468882 | -1.723539139 | -0.351830054 | 0.486229235 | -0.687961773 | 0.223850781 | 0.330676119 | 0.517881975 | 0.4551 | 0.601206979 |
| SL1344_RS23145_ | 0.229547543 | 0.294129206 | 0.157763974 | -0.128772018 | 0.20023211 | -0.583585195 | 0.259826794 | 0.409639472 | 0.572509984 | 0.5209 | 0.655318641 |
| SL1344_RS23150_ | 1.136610413 | 1.086225771 | 1.229679219 | 0.146035664 | 0.159382425 | 0.860849133 | 0.175333354 | 0.262334252 | 0.436253437 | 0.3565 | 0.509559748 |
| SL1344_RS23155_ | -2.89978931 | -2.920105207 | -2.871502575 | 0.021734542 | 0.79993056 | 0.030122076 | 0.485009994 | 0.712751751 | 0.822380208 | 0.8088 | 0.880116082 |
| SL1344_RS23160_ | -4.157085227 | -4.3795882 | -3.949102613 | 0.451146737 | 0.840440002 | 0.495671199 | 0.28600001 | 0.412026404 | 0.569161811 | 0.5149 | 0.645530077 |
| SL1344_RS23165_ | -5.066039977 | -4.480635605 | -5.66487086 | -1.262874582 | 1.292265189 | -0.912332877 | 0.149333359 | 0.23456081 | 0.398597305 | 0.3268 | 0.478507545 |
| SL1344_RS23170_ | -0.882601352 | -0.796224553 | -0.991397997 | -0.262024013 | 0.519477354 | -0.503292806 | 0.291805472 | 0.425029433 | 0.604773041 | 0.5618 | 0.699576792 |
| SL1344_RS23175_ | -0.405786811 | -0.692468102 | -0.146399928 | 0.544357532 | 0.211929954 | 2.497439044 | 0.0020023 | 0.026463479 | 0.107035427 | 0.1031 | 0.247135699 |
| SL1344_RS23180_ | -0.089575625 | -0.087393104 | -0.094301406 | 0.003217847 | 0.256757085 | 0.013018139 | 0.495669554 | 0.667815502 | 0.78911132 | 0.7737 | 0.855690473 |
| SL1344_RS23185_ | -2.015046346 | -1.864781335 | -2.139060795 | -0.327552388 | 0.629389718 | -0.511609567 | 0.298666676 | 0.458774746 | 0.625714475 | 0.5869 | 0.715955452 |
| SL1344_RS23190_ | -0.821074938 | -0.812982484 | -0.831793552 | -0.005913983 | 0.284046281 | -0.020336624 | 0.493333334 | 0.609218705 | 0.739783195 | 0.7109 | 0.807854802 |
| SL1344_RS23195_ | -5.121633803 | -4.790840688 | -5.566389094 | -0.753994544 | 1.153361928 | -0.608546486 | 0.238000013 | 0.351143484 | 0.514782898 | 0.4549 | 0.592336347 |
| SL1344_RS23200_ | -4.820120924 | -4.744333861 | -4.901008656 | -0.082216264 | 1.123221149 | -0.061137413 | 0.472000001 | 0.601395084 | 0.734055009 | 0.7181 | 0.809412404 |
| SL1344_RS23205_ | -0.813959403 | -0.615906787 | -1.036732086 | -0.437589005 | 0.352017113 | -1.239317156 | 0.085942754 | 0.14696385 | 0.308866474 | 0.2128 | 0.374497292 |
| SL1344_RS23210_ | -3.122614996 | -2.886878119 | -3.351386012 | -0.473047682 | 0.591894738 | -0.741120682 | 0.198666684 | 0.303727062 | 0.467803472 | 0.3994 | 0.545336178 |
| SL1344_RS23215_ | 1.665740292 | 1.60510086 | 1.721488821 | 0.116911898 | 0.147703979 | 0.752150927 | 0.206528997 | 0.309107974 | 0.485716339 | 0.4187 | 0.571579458 |
| SL1344_RS23220_ | 2.034152537 | 2.005697457 | 2.082335303 | 0.076702384 | 0.132956673 | 0.504549422 | 0.289806805 | 0.467181054 | 0.62769456 | 0.5801 | 0.710329661 |
| SL1344_RS23225_ | 2.0079967 | 1.970888159 | 2.032222621 | 0.063868379 | 0.190010634 | 0.322167917 | 0.348434383 | 0.621581116 | 0.757798321 | 0.7002 | 0.807323208 |
| SL1344_RS23230_ | 0.321958114 | 0.436924961 | 0.168585339 | -0.292731358 | 0.355885797 | -0.81841817 | 0.152564982 | 0.254933802 | 0.451167846 | 0.3115 | 0.480022257 |
| SL1344_RS23235_ | -0.331426264 | -0.18002083 | -0.418650984 | -0.248781158 | 0.289151458 | -0.814142198 | 0.182666686 | 0.274519548 | 0.454752696 | 0.3707 | 0.524289957 |
| SL1344_RS23240_ | -0.121336837 | 0.009246973 | -0.219167738 | -0.237340945 | 0.250834884 | -0.896674496 | 0.170552986 | 0.249171288 | 0.422236993 | 0.3333 | 0.492674355 |
| SL1344_RS23245_ | 1.70547564 | 1.66512971 | 1.921158611 | 0.229932374 | 0.191672858 | 0.904076521 | 0.229333348 | 0.285749302 | 0.479627484 | 0.4614 | 0.626952253 |
| SL1344_RS23250_ | 0.013721485 | -0.071206437 | 0.106642897 | 0.181751396 | 0.178820263 | 0.942776497 | 0.148567647 | 0.22413581 | 0.380850469 | 0.3147 | 0.466657914 |
| SL1344_RS23265_ | 1.338709519 | 1.86143451 | 0.778542431 | -1.089521975 | 0.150337644 | -7.179003978 | 4.68E-05 | 0.001571314 | 0.025271759 | 0.1 | 0.243021319 |
| SL1344_RS23270_ | -3.859867351 | -3.624962669 | -4.392247126 | -0.749907109 | 1.025233899 | -0.640662906 | 0.247168567 | 0.356729412 | 0.534740262 | 0.4902 | 0.632199901 |
| SL1344_RS23275_ | -5.144179768 | -5.117640798 | -5.174652429 | -0.066527157 | 1.441872679 | -0.044361008 | 0.479680214 | 0.62634836 | 0.757237953 | 0.7464 | 0.833686797 |
| SL1344_RS23280_ | -4.335866843 | -4.208504353 | -4.440206231 | -0.286291019 | 1.043782435 | -0.265938098 | 0.384000005 | 0.564656348 | 0.704911715 | 0.6762 | 0.779669021 |
| SL1344_RS23285_ | -3.604902664 | -3.55123656 | -3.660023246 | -0.109524315 | 0.656551836 | -0.164117675 | 0.430666669 | 0.547272414 | 0.68597391 | 0.6585 | 0.76115266 |
| SL1344_RS23290_ | -3.470782442 | -3.167508033 | -4.043811061 | -0.91044493 | 0.957943233 | -0.91831402 | 0.1206667 | 0.225069324 | 0.408929303 | 0.2885 | 0.448666942 |
| SL1344_RS23295_ | -0.675366029 | -0.549409043 | -0.776069738 | -0.219580578 | 0.351458125 | -0.580149173 | 0.274000011 | 0.421964865 | 0.589236577 | 0.5123 | 0.656408711 |
| SL1344_RS23300_ | 4.09841036 | 3.881183423 | 4.319984693 | 0.476784249 | 0.100400121 | 4.915941103 | 4.68E-05 | 0.002411113 | 0.03303838 | 0.1 | 0.243021319 |
| SL1344_RS23305_ | 0.931420163 | 0.759455026 | 1.129653776 | 0.372555009 | 0.149172188 | 2.485612845 | 0.003334715 | 0.027523692 | 0.111026731 | 0.1013 | 0.244776273 |

| | | | | | | | | | | | |
|---|---|---|---|---|---|---|---|---|---|---|---|
| SL1344_RS23310_ | 2.015454081 | 1.909006763 | 2.148570069 | 0.240633659 | 0.104578855 | 2.213996221 | 0.005996769 | 0.039159087 | 0.137259919 | 0.1072 | 0.25224083 |
| SL1344_RS23315_ | 1.285664678 | 1.218553712 | 1.339855661 | 0.124860154 | 0.148973342 | 0.842397174 | 0.176000021 | 0.277182504 | 0.44821878 | 0.3768 | 0.526977212 |
| SL1344_RS23320_ | -1.54738719 | -1.636076765 | -1.464794475 | 0.146464601 | 0.383977719 | 0.381393668 | 0.338000007 | 0.53137539 | 0.676795889 | 0.6414 | 0.751794992 |
| SL1344_RS23325_ | -1.159253507 | -1.155824726 | -1.162935526 | 2.10E-05 | 0.291447491 | 0.000257057 | 0.5 | 0.574333713 | 0.711825549 | 0.6916 | 0.790273619 |
| SL1344_RS23330_ | -5.491572109 | -5.056833615 | -5.814390455 | -0.749094912 | 1.489113311 | -0.440453024 | 0.309333342 | 0.487921899 | 0.635804833 | 0.5843 | 0.704386681 |
| SL1344_RS23335_ | -4.903874208 | -4.750178339 | -5.08561561 | -0.339987653 | 1.154455577 | -0.26990369 | 0.37974684 | 0.531931886 | 0.677338554 | 0.6484 | 0.756616231 |
| SL1344_RS23340_ | 1.812103657 | 1.61994268 | 1.989832386 | 0.394517291 | 0.162173802 | 2.429177235 | 0.00067292 | 0.025518318 | 0.111124647 | 0.1004 | 0.243572544 |
| SL1344_RS23345_ | 2.975120266 | 2.81958724 | 3.326926078 | 0.494867376 | 0.112141146 | 3.839239947 | 4.68E-05 | 0.020466312 | 0.101160913 | 0.1 | 0.243021319 |
| SL1344_RS23350_ | 1.112161203 | 0.934523375 | 1.35275571 | 0.417529944 | 0.149619184 | 2.733575865 | 0.002000966 | 0.023520626 | 0.102763779 | 0.1012 | 0.244626195 |
| SL1344_RS23355_ | -1.840283572 | -2.439827077 | -1.479716731 | 0.92325128 | 1.172604341 | 0.743314675 | 0.200666684 | 0.296987808 | 0.497074399 | 0.3903 | 0.569197017 |
| SL1344_RS23360_ | 5.020696928 | 4.973853718 | 5.089711681 | 0.116608218 | 0.108746529 | 1.020834271 | 0.098600974 | 0.264647872 | 0.47027014 | 0.2026 | 0.378205321 |
| SL1344_RS23365_ | 6.398995227 | 6.353541378 | 6.411968911 | 0.059113973 | 0.073342754 | 0.770944687 | 0.230000014 | 0.321440323 | 0.522299248 | 0.4198 | 0.609081593 |
| SL1344_RS23370_ | 4.985714997 | 4.931466574 | 5.02540871 | 0.096325147 | 0.046853842 | 1.950966088 | 0.015323415 | 0.05179688 | 0.166109923 | 0.1117 | 0.258572288 |
| SL1344_RS23375_ | 2.692481995 | 2.418981982 | 2.96453673 | 0.549220616 | 0.080873432 | 6.673091847 | 4.68E-05 | 0.001891369 | 0.026620408 | 0.1 | 0.243021319 |
| SL1344_RS23380_ | 2.113031154 | 2.019534382 | 2.203887643 | 0.185562598 | 0.171495942 | 1.057613496 | 0.10926053 | 0.173957917 | 0.3532825 | 0.2251 | 0.395633487 |
| SL1344_RS23385_ | 1.382426717 | 1.356058286 | 1.413323983 | 0.069900758 | 0.181893782 | 0.381613826 | 0.346666673 | 0.527777717 | 0.680603665 | 0.66 | 0.769715275 |
| SL1344_RS23390_ | 2.226285228 | 2.103994164 | 2.327737275 | 0.227604369 | 0.101509479 | 2.184012603 | 0.005334196 | 0.042744838 | 0.146326151 | 0.1042 | 0.248639987 |
| SL1344_RS23395_ | 4.210913297 | 4.119374108 | 4.319246037 | 0.207038335 | 0.086541312 | 2.373909722 | 0.000673369 | 0.039690824 | 0.147021733 | 0.1002 | 0.243303969 |
| SL1344_RS23400_ | 0.760776527 | 0.957325824 | 0.574968977 | -0.386407312 | 0.166372715 | -2.267473262 | 0.003334715 | 0.035947606 | 0.130996716 | 0.1059 | 0.250827568 |
| SL1344_RS23410_ | 0.678355511 | 0.736228772 | 0.615985241 | -0.118286165 | 0.136090043 | -0.833017786 | 0.169886764 | 0.274455473 | 0.427082136 | 0.3642 | 0.507746206 |
| SL1344_RS23415_ | 0.214672158 | 0.16260923 | 0.306137405 | 0.131507941 | 0.247825162 | 0.49428096 | 0.294470362 | 0.490462974 | 0.644726141 | 0.5998 | 0.723192726 |
| SL1344_RS23420_ | -2.929223217 | -2.962959626 | -2.891196163 | 0.059278705 | 0.566065693 | 0.089645774 | 0.457028649 | 0.586541403 | 0.721332568 | 0.6856 | 0.787817837 |
| SL1344_RS23425_ | -2.488732046 | -2.356672613 | -2.597082696 | -0.231587916 | 0.487931174 | -0.442977858 | 0.313333341 | 0.474329438 | 0.620498291 | 0.5709 | 0.691554218 |
| SL1344_RS23430_ | 0.877167278 | 0.702995183 | 1.106426704 | 0.390744847 | 0.162213808 | 2.346257972 | 0.005996769 | 0.033681108 | 0.126734798 | 0.1061 | 0.251290982 |
| SL1344_RS23435_ | 1.026495044 | 0.864799111 | 1.126238311 | 0.262062446 | 0.159602199 | 1.573526364 | 0.032666804 | 0.088377109 | 0.222662943 | 0.1484 | 0.301171233 |
| SL1344_RS23440_ | 3.520593917 | 3.597977066 | 3.329765018 | -0.244314205 | 0.185959295 | -1.243788793 | 0.114590308 | 0.136029428 | 0.312067127 | 0.2111 | 0.390713641 |
| SL1344_RS23445_ | 3.872483137 | 3.927146657 | 3.736449279 | -0.193448986 | 0.127268074 | -1.480357008 | 0.004667653 | 0.157673516 | 0.343651533 | 0.1045 | 0.249172229 |
| SL1344_RS23450_ | -0.480322207 | -0.71011305 | -0.373070012 | 0.304134654 | 0.320904965 | 0.865431058 | 0.18200002 | 0.262285258 | 0.444768372 | 0.3647 | 0.530575011 |
| SL1344_RS23455_ | -0.122562057 | -0.302138072 | 0.029619767 | 0.330329116 | 0.208348879 | 1.531170471 | 0.043304566 | 0.092147074 | 0.226935526 | 0.1519 | 0.305512431 |
| SL1344_RS23460_ | -3.199964495 | -3.650810004 | -2.87595009 | 0.779480544 | 0.588148856 | 1.258445651 | 0.090606308 | 0.148175222 | 0.300515419 | 0.2175 | 0.37333442 |
| SL1344_RS23465_ | -2.0016319 | -2.140951382 | -1.876020739 | 0.28343345 | 0.461408431 | 0.570280972 | 0.257333345 | 0.390020673 | 0.553717791 | 0.5046 | 0.637736378 |
| SL1344_RS23470_ | -2.216339049 | -2.471980808 | -1.974588413 | 0.500222293 | 0.435092604 | 1.071382794 | 0.114590308 | 0.183841838 | 0.33967655 | 0.2662 | 0.42225513 |
| SL1344_RS23480_ | 0.071437155 | -0.018090217 | 0.16131574 | 0.179078414 | 0.196632733 | 0.850126506 | 0.167221874 | 0.265861002 | 0.428825799 | 0.3594 | 0.508155959 |
| SL1344_RS23485_ | 2.152322983 | 2.067087243 | 2.280054027 | 0.219312009 | 0.165266164 | 1.311330481 | 0.083333384 | 0.124038701 | 0.285040921 | 0.1931 | 0.359863666 |
| SL1344_RS23495_ | 2.732204314 | 2.701253454 | 2.767339663 | 0.06025634 | 0.108353697 | 0.5523221 | 0.274666677 | 0.429961647 | 0.59882893 | 0.5415 | 0.678310886 |
| SL1344_RS23500_ | 5.377308425 | 5.361738318 | 5.394932692 | 0.024581941 | 0.153877818 | 0.145246019 | 0.444370422 | 0.845514012 | 0.914277668 | 0.9679 | 0.981257478 |
| SL1344_RS23505_ | 1.505499026 | 1.432140968 | 1.556109525 | 0.122896837 | 0.117594604 | 0.991295922 | 0.144666693 | 0.2220079 | 0.384389244 | 0.316 | 0.470488536 |
| SL1344_RS23510_ | 0.82403233 | 0.734552691 | 1.037653821 | 0.287871899 | 0.204249576 | 1.276065749 | 0.097934752 | 0.147478031 | 0.315030477 | 0.2242 | 0.389643228 |
| SL1344_RS23515_ | 0.434997915 | 0.346472745 | 0.50599679 | 0.120484812 | 0.236702123 | 0.492426242 | 0.305129922 | 0.488577028 | 0.648163429 | 0.5943 | 0.721220915 |
| SL1344_RS23520_ | 2.078868232 | 2.067054017 | 2.094717412 | 0.045657512 | 0.146054695 | 0.28900753 | 0.372000005 | 0.606581473 | 0.743074646 | 0.7451 | 0.832219761 |
| SL1344_RS23525_ | 3.459973978 | 3.392554565 | 3.586410438 | 0.185864086 | 0.080290693 | 2.181899608 | 0.012667027 | 0.042525661 | 0.147703469 | 0.1099 | 0.25625109 |
| SL1344_RS23530_ | 1.472042831 | 1.604742268 | 1.390667184 | -0.208704462 | 0.124348758 | -1.571186269 | 0.054630326 | 0.095092201 | 0.232755534 | 0.1647 | 0.320676095 |
| SL1344_RS23535_ | 2.021279872 | 2.061276523 | 1.954671064 | -0.119307079 | 0.171345351 | -0.708890695 | 0.214000016 | 0.313603553 | 0.501078875 | 0.4187 | 0.583132022 |
| SL1344_RS23540_ | 1.614189666 | 1.617689924 | 1.608782358 | -0.00925131 | 0.107349788 | -0.080238842 | 0.458361094 | 0.533501529 | 0.673857342 | 0.6452 | 0.750934361 |
| SL1344_RS23545_ | -0.425106994 | -0.52329337 | -0.344381785 | 0.17512374 | 0.22785683 | 0.725173787 | 0.213191222 | 0.319361599 | 0.480677842 | 0.4131 | 0.556718025 |
| SL1344_RS23550_ | -1.739624222 | -1.863206099 | -1.584755227 | 0.275859083 | 0.363467582 | 0.723220362 | 0.220519669 | 0.323800294 | 0.482972996 | 0.4271 | 0.567023521 |
| SL1344_RS23555_ | -3.958252418 | -4.089810236 | -3.806615809 | 0.25278265 | 1.068681596 | 0.214775451 | 0.402000004 | 0.636576888 | 0.762534029 | 0.7378 | 0.827019166 |
| SL1344_RS23560_ | -1.322786215 | -1.459635174 | -1.13813015 | 0.346715099 | 0.420846412 | 0.794908623 | 0.189333352 | 0.28136484 | 0.460738063 | 0.3867 | 0.541155967 |
| SL1344_RS23565_ | -2.998696422 | -3.481486059 | -2.657029133 | 0.83479501 | 0.601849544 | 1.310361338 | 0.057295212 | 0.12778244 | 0.27864882 | 0.1707 | 0.323179124 |
| SL1344_RS23570_ | -1.893658825 | -2.090959586 | -1.598196621 | 0.477020993 | 0.427673347 | 1.031586401 | 0.139906757 | 0.205387774 | 0.370329186 | 0.2922 | 0.452238111 |
| SL1344_RS23575_ | 0.659478538 | 0.571629199 | 0.755395835 | 0.178959975 | 0.168992872 | 0.998485443 | 0.12658231 | 0.20584688 | 0.367095145 | 0.2891 | 0.447740263 |
| SL1344_RS23580_ | 0.197981898 | -0.148752005 | 0.540768518 | 0.723789795 | 0.266542865 | 2.836430434 | 4.68E-05 | 0.027484944 | 0.117299772 | 0.1 | 0.243021319 |
| SL1344_RS23590_ | -5.553064095 | -5.936182434 | -5.142542369 | 0.745464595 | 1.571024934 | 0.427071556 | 0.292000009 | 0.474531253 | 0.62927394 | 0.5694 | 0.695474418 |

| | | | | | | | | | | | |
|---|---|---|---|---|---|---|---|---|---|---|---|
| SL1344_RS23595_ | -0.525236618 | -0.586827308 | -0.414311897 | 0.132781388 | 0.309607389 | 0.384673739 | 0.34533334 | 0.589449948 | 0.724660562 | 0.6755 | 0.786569481 |
| SL1344_RS23640_ | 0.482762104 | 0.602736084 | 0.358277023 | -0.271354823 | 0.236963359 | -1.119076059 | 0.084666716 | 0.170058452 | 0.342834699 | 0.2208 | 0.382454383 |
| SL1344_RS23645_ | -0.650308512 | -0.694125775 | -0.619023433 | 0.060912305 | 0.361548695 | 0.16058811 | 0.431333336 | 0.664641675 | 0.786886133 | 0.7682 | 0.851862972 |
| SL1344_RS23650_ | 1.48405825 | 1.549523381 | 1.404170343 | -0.154087333 | 0.144961802 | -1.021506835 | 0.115256531 | 0.189980391 | 0.358968569 | 0.2528 | 0.409730939 |
| SL1344_RS23655_ | 0.634401548 | 0.636672573 | 0.63148764 | -0.006346199 | 0.175352012 | -0.037971278 | 0.484666667 | 0.620554677 | 0.74932184 | 0.7365 | 0.82514806 |
| SL1344_RS23660_ | 1.539987045 | 1.556297334 | 1.518446398 | -0.037166071 | 0.111737243 | -0.307255326 | 0.366000005 | 0.520930034 | 0.663087892 | 0.6237 | 0.736353418 |
| SL1344_RS23665_ | -1.080460985 | -0.758541932 | -1.396491298 | -0.65576749 | 0.320813599 | -1.944885044 | 0.015333631 | 0.049721743 | 0.158924599 | 0.1139 | 0.260702225 |
| SL1344_RS23670_ | 3.467748264 | 3.543403639 | 3.36684196 | -0.187788026 | 0.121782718 | -1.534677197 | 0.013990999 | 0.122272253 | 0.292424232 | 0.1112 | 0.258204602 |
| SL1344_RS23675_ | 1.584226634 | 1.633029281 | 1.549797341 | -0.074384477 | 0.155386039 | -0.457641091 | 0.313333341 | 0.495203318 | 0.652293176 | 0.6075 | 0.729037287 |
| SL1344_RS23680_ | 1.3993916 | 1.367363324 | 1.414562409 | 0.047761044 | 0.1415223 | 0.305857381 | 0.372666672 | 0.567113574 | 0.707384234 | 0.6776 | 0.782358218 |
| SL1344_RS23685_ | -1.721027329 | -1.804515586 | -1.629979617 | 0.155861874 | 0.386669743 | 0.384532683 | 0.331112599 | 0.506650942 | 0.653488221 | 0.6179 | 0.731609503 |
| SL1344_RS23690_ | -2.586551823 | -2.80559556 | -2.450219976 | 0.320206454 | 0.5688413 | 0.528898399 | 0.282666677 | 0.43865161 | 0.601583153 | 0.5595 | 0.690623525 |
| SL1344_RS23695_ | -3.32371395 | -3.686618388 | -3.027670052 | 0.655634393 | 0.816662817 | 0.753256841 | 0.183333353 | 0.294160016 | 0.469967629 | 0.3943 | 0.547723049 |
| SL1344_RS23700_ | -2.133035424 | -1.996606345 | -2.279839462 | -0.286097871 | 0.431724216 | -0.623760721 | 0.233177895 | 0.356038011 | 0.519073318 | 0.4675 | 0.602791611 |
| SL1344_RS23705_ | -3.593381872 | -3.401630324 | -3.752565108 | -0.371990444 | 0.653301131 | -0.51041823 | 0.280666677 | 0.430257906 | 0.580787007 | 0.5323 | 0.657136232 |
| SL1344_RS23710_ | -1.025084527 | -1.036131152 | -1.007674541 | 0.021003077 | 0.36039988 | 0.055730192 | 0.476000001 | 0.653304995 | 0.775810578 | 0.763 | 0.844680584 |
| SL1344_RS23715_ | -2.493571634 | -2.359867838 | -2.675608998 | -0.348447858 | 0.66306773 | -0.499505217 | 0.27514991 | 0.42019056 | 0.591633876 | 0.5445 | 0.675482174 |
| SL1344_RS23720_ | -2.321432372 | -2.187342973 | -2.462233572 | -0.274170128 | 0.400921685 | -0.626747865 | 0.232511673 | 0.358908356 | 0.514246603 | 0.4605 | 0.593994427 |
| SL1344_RS23725_ | -3.096645378 | -2.890705526 | -3.332098907 | -0.452820176 | 0.601915829 | -0.705203871 | 0.20933335 | 0.325869927 | 0.490497029 | 0.4216 | 0.565905055 |
| SL1344_RS23730_ | -2.639374064 | -2.840459769 | -2.462484203 | 0.386210446 | 0.521086165 | 0.689219106 | 0.217188556 | 0.330046698 | 0.493413754 | 0.4327 | 0.574597233 |
| SL1344_RS23735_ | -3.090019006 | -3.081198119 | -3.101737489 | 0.008168411 | 0.628674653 | 0.009300198 | 0.495003331 | 0.59041457 | 0.726811176 | 0.7043 | 0.801763691 |
| SL1344_RS23740_ | -3.253069082 | -3.323098832 | -3.166114956 | 0.154859641 | 0.62421629 | 0.224174244 | 0.399733515 | 0.538758863 | 0.680508052 | 0.644 | 0.754195643 |
| SL1344_RS23745_ | -0.835880605 | -0.806819422 | -0.862842919 | -0.060646048 | 0.293512392 | -0.191745595 | 0.415722855 | 0.597098688 | 0.729538553 | 0.7099 | 0.803507485 |
| SL1344_RS23750_ | -4.462983914 | -4.285701891 | -4.588711211 | -0.277652338 | 1.101348974 | -0.219266282 | 0.40066667 | 0.615820089 | 0.744604357 | 0.7221 | 0.814800894 |
| SL1344_RS23755_ | -3.972909867 | -3.976258605 | -3.968936529 | 0.040705989 | 0.828286852 | 0.050697425 | 0.479680214 | 0.592251354 | 0.725211991 | 0.7013 | 0.795350053 |
| SL1344_RS23760_ | -4.891284063 | -4.883413784 | -4.906178688 | -0.043045256 | 1.059068319 | -0.039050902 | 0.483677549 | 0.564101984 | 0.70203212 | 0.6802 | 0.780202365 |
| SL1344_RS23765_ | -2.46063808 | -2.266346504 | -2.771883827 | -0.469243589 | 0.477937841 | -0.935583955 | 0.14656898 | 0.238013614 | 0.401108349 | 0.3229 | 0.4753522 |
| SL1344_RS23770_ | -3.475668924 | -3.313722065 | -3.654084157 | -0.341623211 | 0.643239822 | -0.497918366 | 0.281812135 | 0.422541883 | 0.575753885 | 0.5289 | 0.653979076 |
| SL1344_RS23775_ | -2.663230691 | -2.326267756 | -2.94896999 | -0.592911388 | 0.612002608 | -0.925415565 | 0.16322454 | 0.249779722 | 0.420146155 | 0.3294 | 0.491789447 |
| SL1344_RS23780_ | -2.950089728 | -2.528646539 | -3.257244977 | -0.719718224 | 0.662694548 | -1.02931947 | 0.13200003 | 0.197052865 | 0.36655824 | 0.2787 | 0.441883558 |
| SL1344_RS23785_ | -2.652169651 | -2.714842322 | -2.596041251 | 0.097042488 | 0.561745402 | 0.168189996 | 0.425716192 | 0.589645904 | 0.725545353 | 0.7029 | 0.800532847 |
| SL1344_RS23790_ | -4.374096613 | -4.323545258 | -4.4311651 | -0.118321824 | 0.954729256 | -0.109841941 | 0.449033979 | 0.582878892 | 0.719164687 | 0.6906 | 0.789433456 |
| SL1344_RS23795_ | -3.587278805 | -3.461873233 | -3.766452957 | -0.352552193 | 0.796721996 | -0.419830122 | 0.305796144 | 0.455309643 | 0.614243631 | 0.5815 | 0.698213841 |
| SL1344_RS23800_ | -3.735583472 | -3.780783811 | -3.684353147 | 0.096744032 | 0.770896559 | 0.126085031 | 0.440666669 | 0.576408086 | 0.713670072 | 0.6884 | 0.789306328 |
| SL1344_RS23805_ | -3.501321669 | -3.31094362 | -3.688767664 | -0.373294723 | 0.602047041 | -0.549563248 | 0.262666678 | 0.390478312 | 0.542137465 | 0.4872 | 0.617808287 |
| SL1344_RS23810_ | -1.003921664 | -0.701455489 | -1.213661459 | -0.500174267 | 0.305644058 | -1.546481444 | 0.048666757 | 0.088772586 | 0.222889515 | 0.1544 | 0.309626821 |
| SL1344_RS23815_ | -1.820214109 | -1.905017677 | -1.776279584 | 0.125722027 | 0.445974381 | 0.255788018 | 0.401333337 | 0.595126781 | 0.731016586 | 0.71 | 0.807748349 |
| SL1344_RS23820_ | -3.581498848 | -3.601195199 | -3.542624396 | 0.038393854 | 0.728454922 | 0.050273037 | 0.482345104 | 0.588578591 | 0.724264779 | 0.6993 | 0.797489782 |
| SL1344_RS23825_ | -2.631611169 | -2.740746431 | -2.496368753 | 0.135878294 | 0.718374222 | 0.184877011 | 0.415333337 | 0.68568897 | 0.801649494 | 0.7757 | 0.858341789 |
| SL1344_RS23830_ | -2.669004388 | -2.400915437 | -3.267507401 | -0.842893386 | 0.698737289 | -1.110851651 | 0.11792142 | 0.176774236 | 0.351662883 | 0.2525 | 0.416987159 |
| SL1344_RS23840_ | -3.11305567 | -3.083281113 | -3.141313004 | -0.06406585 | 0.739634861 | -0.080263387 | 0.465023319 | 0.654360225 | 0.777500155 | 0.7506 | 0.836642645 |
| SL1344_RS23845_ | -5.031625186 | -4.261816264 | -5.700667536 | -1.455994582 | 1.533266026 | -0.864072466 | 0.150666692 | 0.242507274 | 0.418560967 | 0.3065 | 0.471820696 |
| SL1344_RS23850_ | -4.650290241 | -4.534719251 | -4.739454053 | -0.221837762 | 1.10569093 | -0.17496053 | 0.416000003 | 0.587422772 | 0.721650153 | 0.7019 | 0.79716786 |
| SL1344_RS23855_ | -4.77418008 | -4.789872548 | -4.73840586 | 0.021477222 | 0.981228944 | 0.025587589 | 0.490666667 | 0.563114371 | 0.701756399 | 0.6789 | 0.780431309 |
| SL1344_RS23860_ | -0.75041728 | -0.646921742 | -0.83965652 | -0.220042109 | 0.315217578 | -0.647872044 | 0.223184558 | 0.335827133 | 0.50540902 | 0.4577 | 0.59641624 |
| SL1344_RS23865_ | -3.393915675 | -3.076805608 | -3.735502464 | -0.652678078 | 0.566338881 | -1.114123078 | 0.106666705 | 0.180142221 | 0.327211123 | 0.2555 | 0.408150719 |
| SL1344_RS23870_ | -4.026623434 | -3.693252271 | -4.533860528 | -0.855897059 | 0.962255487 | -0.822957468 | 0.147901425 | 0.239337691 | 0.414103313 | 0.3116 | 0.464403446 |
| SL1344_RS23875_ | -3.061387489 | -2.74164487 | -3.417245477 | -0.676577495 | 0.660506847 | -0.968495067 | 0.143904091 | 0.209807504 | 0.379634104 | 0.2925 | 0.453988546 |
| SL1344_RS23880_ | -6.986792685 | -9.514292579 | -5.536403021 | 3.940752969 | 2.792143748 | 1.36632254 | 0.03597614 | 0.107804942 | 0.250419767 | 0.1418 | 0.293353305 |
| SL1344_RS23885_ | -6.548896486 | -6.128864678 | -7.226536547 | -1.139104416 | 2.115435015 | -0.441358594 | 0.27800001 | 0.39004541 | 0.563694694 | 0.498 | 0.633735994 |
| SL1344_RS23890_ | -6.416758335 | -6.6010449 | -6.224186206 | 0.394945625 | 1.739820709 | 0.209966778 | 0.39573618 | 0.494333197 | 0.644423205 | 0.6184 | 0.730336697 |
| SL1344_RS23895_ | -5.225346219 | -5.500542693 | -5.001923265 | 0.427337437 | 1.618455997 | 0.24833857 | 0.386666671 | 0.585850375 | 0.72517306 | 0.705 | 0.801777172 |
| SL1344_RS23900_ | -5.400572386 | -5.249856596 | -5.53904481 | -0.292118944 | 1.280325635 | -0.209435129 | 0.394666671 | 0.531518664 | 0.674843819 | 0.6444 | 0.750967788 |

| | | | | | | | | | | | |
|---|---|---|---|---|---|---|---|---|---|---|---|
| SL1344_RS23905_ | -7.856696665 | -7.333091779 | -8.423668503 | -0.937557681 | 3.58235598 | -0.214725862 | 0.37441706 | 0.530231605 | 0.67744167 | 0.6446 | 0.754526329 |
| SL1344_RS23910_ | -0.529953387 | -0.470450114 | -0.580114645 | -0.110030136 | 0.23627643 | -0.440172221 | 0.315789482 | 0.468757777 | 0.619690013 | 0.5778 | 0.698899522 |
| SL1344_RS23915_ | 0.437055465 | 0.525096415 | 0.337578852 | -0.189663724 | 0.166164413 | -1.089768716 | 0.118000034 | 0.188479837 | 0.342583897 | 0.2642 | 0.418023156 |
| SL1344_RS23920_ | 3.348363797 | 3.335516719 | 3.36869036 | 0.026642515 | 0.076650055 | 0.324918299 | 0.356000006 | 0.582036497 | 0.717710355 | 0.6804 | 0.785906718 |
| SL1344_RS23925_ | 2.722933993 | 2.672813998 | 2.785973673 | 0.11547073 | 0.109034091 | 1.030443707 | 0.120000033 | 0.194179685 | 0.370481491 | 0.2579 | 0.423465334 |
| SL1344_RS23930_ | 3.581406899 | 3.62027934 | 3.534640502 | -0.118334174 | 0.150836694 | -0.790136247 | 0.170000022 | 0.259987873 | 0.460880628 | 0.3188 | 0.493490878 |
| SL1344_RS23935_ | 1.314561156 | 1.215013447 | 1.446617128 | 0.22721034 | 0.131429809 | 1.639152317 | 0.039307241 | 0.081580495 | 0.210844272 | 0.1495 | 0.302937231 |
| SL1344_RS23940_ | 0.542380862 | 0.61801869 | 0.467867191 | -0.157682856 | 0.17427451 | -0.85309213 | 0.175216543 | 0.260510176 | 0.42831538 | 0.3501 | 0.503035581 |
| SL1344_RS23945_ | 0.677700525 | 0.764845377 | 0.55687154 | -0.216102258 | 0.170788237 | -1.217402312 | 0.077281867 | 0.155277557 | 0.313931361 | 0.2123 | 0.3663464 |
| SL1344_RS23950_ | 2.43360289 | 2.623840203 | 2.382461774 | -0.219443078 | 0.14368533 | -1.271229377 | 0.129913421 | 0.156098876 | 0.331720675 | 0.2602 | 0.431828447 |
| SL1344_RS23955_ | 0.498694414 | 0.739920337 | 0.394890341 | -0.323054484 | 0.24473949 | -1.173089744 | 0.117255197 | 0.158624412 | 0.330141792 | 0.2353 | 0.403934799 |
| SL1344_RS23960_ | 2.897114255 | 2.903119107 | 2.890686212 | -0.024555854 | 0.12884923 | -0.190774551 | 0.410393075 | 0.657359671 | 0.783723695 | 0.7853 | 0.862924187 |
| SL1344_RS23965_ | 2.21317925 | 2.274892432 | 2.112183545 | -0.125395561 | 0.17373958 | -0.685783659 | 0.249167234 | 0.377743362 | 0.56381788 | 0.4896 | 0.645565037 |
| SL1344_RS23970_ | 2.737797336 | 2.776098865 | 2.716481229 | -0.0553178 | 0.105431414 | -0.495968241 | 0.300466365 | 0.492243815 | 0.649347557 | 0.5933 | 0.723212458 |
| SL1344_RS23975_ | 1.449000818 | 1.427625219 | 1.476139481 | 0.035240548 | 0.201071081 | 0.175115013 | 0.419333336 | 0.702921176 | 0.814721395 | 0.7603 | 0.84904765 |
| SL1344_RS23980_ | -1.665277379 | -1.993690797 | -1.407224074 | 0.599969614 | 0.392548991 | 1.46370083 | 0.043304566 | 0.101677786 | 0.242171124 | 0.1591 | 0.311685935 |
| SL1344_RS23985_ | -3.973287212 | -4.307062593 | -3.774584332 | 0.458168076 | 1.007067069 | 0.405388306 | 0.325333341 | 0.50870621 | 0.66274445 | 0.6306 | 0.747602278 |
| SL1344_RS23990_ | -5.278056672 | -5.46761778 | -5.058949617 | 0.443509708 | 1.351737319 | 0.303118967 | 0.361333339 | 0.535965941 | 0.680733879 | 0.6576 | 0.761919481 |
| SL1344_RS23995_ | -3.618642118 | -3.554721537 | -3.677515563 | -0.133295477 | 0.772110419 | -0.159786675 | 0.429713527 | 0.598011508 | 0.730013474 | 0.7013 | 0.798114604 |
| SL1344_RS24000_ | -1.734118324 | -1.55038218 | -1.871998226 | -0.325439016 | 0.374744082 | -0.81410269 | 0.183211212 | 0.285997821 | 0.448623431 | 0.3867 | 0.532768467 |
| SL1344_RS24005_ | -6.113542584 | -5.890381369 | -6.411377887 | -0.402564321 | 1.953470901 | -0.171754611 | 0.41866667 | 0.578296716 | 0.718113537 | 0.7039 | 0.800789615 |
| SL1344_RS24010_ | -4.850342428 | -4.800726822 | -4.888912481 | 0.024338543 | 1.309257177 | 0.013370401 | 0.492 | 0.624892897 | 0.756066877 | 0.7432 | 0.83059669 |
| SL1344_RS24015_ | -0.846436651 | -0.911995936 | -0.771573664 | 0.142189335 | 0.250389934 | 0.517030112 | 0.28600001 | 0.417286915 | 0.569698899 | 0.5209 | 0.646787579 |
| SL1344_RS24020_ | 4.031757159 | 4.012924084 | 4.143697581 | 0.115854887 | 0.112465854 | 0.720546217 | 0.289806805 | 0.413102676 | 0.603319945 | 0.5947 | 0.74677611 |
| SL1344_RS24025_ | -5.225526511 | -5.069095556 | -5.407762737 | -0.384959624 | 1.549790185 | -0.235203711 | 0.397333337 | 0.580863897 | 0.7211772 | 0.7011 | 0.800361169 |
| SL1344_RS24030_ | -3.691998814 | -3.722005613 | -3.649867889 | 0.06459676 | 0.619645329 | 0.106411545 | 0.449700202 | 0.515720653 | 0.661783218 | 0.6272 | 0.740363187 |
| SL1344_RS24035_ | -1.949357344 | -1.78799639 | -2.175273962 | -0.382566883 | 0.450820482 | -0.80947175 | 0.183877435 | 0.290199387 | 0.460203611 | 0.3875 | 0.539180539 |
| SL1344_RS24040_ | -0.901238388 | -0.936309985 | -0.871746522 | 0.033963653 | 0.330557454 | 0.097855913 | 0.456362427 | 0.63922744 | 0.765609671 | 0.7518 | 0.838225384 |
| SL1344_RS24045_ | 0.708389997 | 0.563637425 | 0.848421157 | 0.238980454 | 0.263882053 | 0.894705255 | 0.163333356 | 0.249638995 | 0.434144464 | 0.3247 | 0.498058771 |
| SL1344_RS24050_ | 1.524977279 | 1.462219584 | 1.590190593 | 0.140335147 | 0.161126045 | 0.836517827 | 0.188666685 | 0.266477625 | 0.444429643 | 0.3583 | 0.517366575 |
| SL1344_RS24055_ | 1.441260778 | 1.466994805 | 1.411381764 | -0.055448362 | 0.109108769 | -0.465134457 | 0.28600001 | 0.414331118 | 0.565401813 | 0.5219 | 0.646377201 |
| SL1344_RS24060_ | 1.310597039 | 1.27518347 | 1.342900186 | 0.075787287 | 0.146315385 | 0.501669526 | 0.290000009 | 0.450669631 | 0.61010098 | 0.5758 | 0.69870521 |
| SL1344_RS24065_ | 1.418728897 | 1.33721655 | 1.487335989 | 0.153308317 | 0.134026239 | 1.110445299 | 0.098000042 | 0.16937014 | 0.330298472 | 0.2349 | 0.392201632 |
| SL1344_RS24070_ | 1.239631597 | 1.151277534 | 1.273658088 | 0.102089404 | 0.167682797 | 0.535046473 | 0.300000009 | 0.485520324 | 0.644823279 | 0.5981 | 0.730704717 |
| SL1344_RS24075_ | 0.83455748 | 0.910839064 | 0.723715512 | -0.206001108 | 0.208317938 | -0.944801815 | 0.1206667 | 0.219449368 | 0.401442047 | 0.2817 | 0.439755629 |
| SL1344_RS24080_ | -1.888339068 | -2.110484119 | -1.711086952 | 0.509473303 | 0.646149048 | 0.743603011 | 0.206000017 | 0.301316133 | 0.486785144 | 0.4257 | 0.582571126 |
| SL1344_RS24085_ | 4.250382726 | 4.194558679 | 4.295746474 | 0.103984731 | 0.063665058 | 1.590034687 | 0.038000117 | 0.079508889 | 0.215185456 | 0.14 | 0.293720201 |
| SL1344_RS24090_ | -2.235954149 | -2.208257383 | -2.260955157 | -0.03663091 | 0.443342139 | -0.076161405 | 0.464666668 | 0.581860445 | 0.718860862 | 0.6976 | 0.796616121 |
| SL1344_RS24095_ | -1.588552594 | -1.733261743 | -1.393227592 | 0.336872985 | 0.341962344 | 0.934806954 | 0.148666692 | 0.242422572 | 0.403393212 | 0.33 | 0.480010742 |
| SL1344_RS24100_ | -3.561298098 | -3.842240891 | -3.375965336 | 0.490656664 | 0.795378879 | 0.541606659 | 0.23984012 | 0.366084608 | 0.538904667 | 0.4801 | 0.616221955 |
| SL1344_RS24105_ | 0.421317499 | 0.242389442 | 0.556816179 | 0.305124695 | 0.190484285 | 1.522262651 | 0.052000084 | 0.095829512 | 0.233408716 | 0.1623 | 0.319666207 |
| SL1344_RS24110_ | -1.812057627 | -1.811368278 | -1.813954359 | 0.014088595 | 0.408895973 | 0.026194408 | 0.489007329 | 0.618803844 | 0.748629678 | 0.733 | 0.822635838 |
| SL1344_RS24115_ | -2.584998682 | -2.590112216 | -2.578425978 | 0.004598929 | 0.504268964 | 0.008293237 | 0.498 | 0.593879046 | 0.72464693 | 0.7043 | 0.798770992 |
| SL1344_RS24120_ | -1.813382039 | -1.854434125 | -1.751170247 | 0.110099034 | 0.432191242 | 0.249134846 | 0.39307129 | 0.567613841 | 0.70490319 | 0.6856 | 0.7842892 |
| SL1344_RS24125_ | 3.628187014 | 3.518875502 | 3.903542879 | 0.384333362 | 0.156861401 | 2.318895636 | 4.68E-05 | 0.062235175 | 0.196209177 | 0.1001 | 0.243162813 |
| SL1344_RS24130_ | -0.331801149 | -0.571382819 | -0.135968947 | 0.426106221 | 0.257476868 | 1.612360446 | 0.03397748 | 0.082234977 | 0.215315264 | 0.1448 | 0.298723201 |
| SL1344_RS24135_ | 0.10857652 | -0.19956763 | 0.346703364 | 0.555290735 | 0.204197721 | 2.697240816 | 0.00067292 | 0.028413559 | 0.113676771 | 0.1004 | 0.243578174 |
| SL1344_RS24140_ | -1.201648615 | -1.505447851 | -1.041267402 | 0.430624217 | 0.404158235 | 0.966143568 | 0.135909422 | 0.212572284 | 0.387485801 | 0.2984 | 0.463537467 |
| SL1344_RS24145_ | -1.520948762 | -1.803020004 | -1.275992722 | 0.530758032 | 0.351018698 | 1.463083566 | 0.043333435 | 0.094940126 | 0.229225657 | 0.1587 | 0.311747572 |
| SL1344_RS24150_ | -0.518233711 | -0.523279824 | -0.508607738 | 0.032990783 | 0.354327886 | 0.089219577 | 0.468354432 | 0.729011255 | 0.8337126 | 0.8317 | 0.896175405 |
| SL1344_RS24155_ | -0.037085833 | -0.02386349 | -0.048862397 | -0.021723635 | 0.17210728 | -0.118895227 | 0.449033979 | 0.515904389 | 0.656887286 | 0.6196 | 0.730352949 |
| SL1344_RS24160_ | -5.778963923 | -5.332272194 | -6.088624111 | -0.456407609 | 2.553282777 | -0.120870363 | 0.407728185 | 0.672174798 | 0.794883317 | 0.7672 | 0.854083925 |
| SL1344_RS24165_ | 2.635779685 | 2.693509962 | 2.616195519 | -0.064156351 | 0.119037531 | -0.422272575 | 0.34533334 | 0.637236402 | 0.768407469 | 0.6982 | 0.817155449 |

| | | | | | | | | | | | |
|---|---|---|---|---|---|---|---|---|---|---|---|
| SL1344_RS24170_ | 2.282185477 | 2.325723917 | 2.233339115 | -0.10111696 | 0.144640376 | -0.679892018 | 0.227333348 | 0.345398803 | 0.530531073 | 0.4658 | 0.618598075 |
| SL1344_RS24175_ | 4.317371724 | 4.292601629 | 4.347324568 | 0.048161261 | 0.186183569 | 0.25458611 | 0.385333338 | 0.670065235 | 0.802687342 | 0.8305 | 0.896170158 |
| SL1344_RS24180_ | 3.057175296 | 2.977462828 | 3.247127358 | 0.266630588 | 0.121461249 | 2.038835313 | 0.003998486 | 0.069826847 | 0.207000174 | 0.1027 | 0.246684984 |
| SL1344_RS24185_ | -0.388371246 | -0.378402394 | -0.398168452 | -0.028275563 | 0.2851295 | -0.086228952 | 0.464000001 | 0.651994749 | 0.775739862 | 0.7654 | 0.847081443 |
| SL1344_RS24190_ | 0.478492282 | 0.258139253 | 0.6646886 | 0.399309379 | 0.187669306 | 2.038731678 | 0.013990999 | 0.046555197 | 0.153330876 | 0.1142 | 0.261584187 |
| SL1344_RS24195_ | -0.378569747 | -0.348272859 | -0.403343624 | -0.095296564 | 0.346530055 | -0.249985989 | 0.37974684 | 0.59993969 | 0.7395996 | 0.73 | 0.822251887 |
| SL1344_RS24200_ | -5.299764371 | -4.972077698 | -5.917652644 | -0.925053992 | 1.579179935 | -0.519971951 | 0.266489018 | 0.397320236 | 0.569917001 | 0.523 | 0.66011159 |
| SL1344_RS24205_ | 3.348097559 | 3.258461515 | 3.424232823 | 0.215051025 | 0.134001326 | 1.638646524 | 0.007329074 | 0.062304889 | 0.19132744 | 0.1073 | 0.252715109 |
| SL1344_RS24210_ | 0.118664894 | 0.104843638 | 0.146975422 | 0.027942764 | 0.230168339 | 0.097333359 | 0.451698869 | 0.661146954 | 0.782837088 | 0.7546 | 0.843771419 |
| SL1344_RS24215_ | 0.330640472 | 0.421069741 | 0.232806501 | -0.202465161 | 0.21763211 | -0.88699541 | 0.146000026 | 0.233038329 | 0.404672929 | 0.3264 | 0.483148523 |
| SL1344_RS24220_ | 1.900353977 | 1.933676127 | 1.866696107 | -0.066648235 | 0.094228566 | -0.655501321 | 0.240000013 | 0.354831527 | 0.509236586 | 0.4549 | 0.589327201 |
| SL1344_RS24225_ | 0.422871505 | 0.403690807 | 0.438110583 | -0.012339374 | 0.27181547 | -0.056380343 | 0.483011326 | 0.715592841 | 0.824533246 | 0.8283 | 0.892938536 |
| SL1344_RS24230_ | -0.854594581 | -0.921838206 | -0.784717665 | 0.153759421 | 0.29114914 | 0.498603966 | 0.302000009 | 0.444510369 | 0.60235869 | 0.5716 | 0.690663088 |
| SL1344_RS24235_ | -0.01386327 | -0.060723527 | 0.037951009 | 0.103629736 | 0.236051689 | 0.415822849 | 0.327781486 | 0.501049631 | 0.651318522 | 0.6111 | 0.728883332 |
| SL1344_RS24240_ | 0.558603313 | 0.488049223 | 0.64747509 | 0.154057073 | 0.150590513 | 0.958245678 | 0.139240534 | 0.228611624 | 0.387366876 | 0.3147 | 0.467969599 |
| SL1344_RS24245_ | 0.813113522 | 0.869880226 | 0.728258834 | -0.142271943 | 0.173911588 | -0.763065662 | 0.203864107 | 0.30229588 | 0.473457374 | 0.404 | 0.551433937 |
| SL1344_RS24250_ | 3.067654136 | 3.253777557 | 2.84878016 | -0.404944576 | 0.129908153 | -3.088027491 | 4.68E-05 | 0.012138735 | 0.075155856 | 0.1 | 0.243021319 |
| SL1344_RS24255_ | -0.030584473 | -0.006558185 | -0.051699771 | -0.037530242 | 0.233489184 | -0.150491335 | 0.437041974 | 0.624830034 | 0.753703998 | 0.7432 | 0.830506207 |
| SL1344_RS24260_ | -0.156669131 | -0.16690604 | -0.144976424 | 0.015127587 | 0.211306872 | 0.070825355 | 0.472000001 | 0.575485209 | 0.712706498 | 0.6862 | 0.785830185 |
| SL1344_RS24265_ | 0.200763903 | 0.14705884 | 0.264985562 | 0.118556631 | 0.283384706 | 0.387881502 | 0.325782819 | 0.526301221 | 0.679253179 | 0.6147 | 0.739061289 |
| SL1344_RS24270_ | -1.012596849 | -1.130321922 | -0.898459252 | 0.229107652 | 0.246455561 | 0.868569466 | 0.16066669 | 0.255041657 | 0.407282923 | 0.3448 | 0.492212538 |
| SL1344_RS24275_ | -1.648509753 | -1.65624927 | -1.627919401 | 0.020121524 | 0.417909714 | 0.047006276 | 0.482345104 | 0.651096734 | 0.774276998 | 0.7576 | 0.843081178 |
| SL1344_RS24280_ | -1.031547057 | -1.068992524 | -1.001861328 | 0.088434506 | 0.346411968 | 0.231178382 | 0.398000004 | 0.57642147 | 0.715660969 | 0.6938 | 0.790616948 |
| SL1344_RS24285_ | 2.012142061 | 2.037085155 | 1.99549862 | -0.047224629 | 0.13133265 | -0.355296677 | 0.355762831 | 0.550001785 | 0.695274367 | 0.6742 | 0.778046549 |
| SL1344_RS24290_ | 1.94850299 | 2.083728235 | 1.8170089 | -0.228404971 | 0.170468008 | -1.313275626 | 0.088666714 | 0.121423303 | 0.281697643 | 0.1888 | 0.359272131 |
| SL1344_RS24295_ | 2.49763064 | 2.529489987 | 2.472890408 | -0.058008311 | 0.079695673 | -0.699517125 | 0.224000015 | 0.346446702 | 0.50410141 | 0.4497 | 0.588571247 |
| SL1344_RS24300_ | 1.798748104 | 1.824764722 | 1.753482371 | -0.072769681 | 0.128340411 | -0.514820605 | 0.282478358 | 0.442553147 | 0.603150361 | 0.5611 | 0.688657652 |
| SL1344_RS24305_ | -0.396221809 | -0.26747897 | -0.578511305 | -0.307153773 | 0.25750937 | -1.152236875 | 0.10200004 | 0.172496555 | 0.333688521 | 0.2524 | 0.412434816 |
| SL1344_RS24310_ | -1.869284011 | -1.625554745 | -2.182950348 | -0.55872722 | 0.371763117 | -1.4041838 | 0.066622316 | 0.107599059 | 0.248302446 | 0.1716 | 0.326911039 |
| SL1344_RS24315_ | -4.989837368 | -4.90866028 | -5.056591178 | -0.129027695 | 1.102801179 | -0.107422447 | 0.445036644 | 0.543139471 | 0.686508419 | 0.6528 | 0.761059462 |
| SL1344_RS24320_ | -6.985148389 | -8.43062635 | -6.055063649 | 2.328462691 | 2.68719197 | 0.798176225 | 0.14333336 | 0.24826991 | 0.421008394 | 0.3095 | 0.465808907 |
| SL1344_RS24325_ | -4.555753429 | -4.17662283 | -5.12866401 | -0.976562506 | 0.978870319 | -0.899390101 | 0.151898759 | 0.237559977 | 0.401641632 | 0.3176 | 0.471454521 |
| SL1344_RS24330_ | -3.338487447 | -2.988153044 | -3.746015482 | -0.803062743 | 0.974773817 | -0.778454479 | 0.161892095 | 0.254385778 | 0.439654318 | 0.328 | 0.48987071 |
| SL1344_RS24335_ | -4.815667407 | -4.410545078 | -5.253900879 | -0.890775201 | 1.122493883 | -0.733296463 | 0.181878767 | 0.28917701 | 0.458739697 | 0.3841 | 0.53103276 |
| SL1344_RS24340_ | -4.17195147 | -3.851277909 | -4.685665618 | -0.804134974 | 1.1402714 | -0.684537911 | 0.226666681 | 0.357066999 | 0.532628922 | 0.4707 | 0.621251249 |
| SL1344_RS24345_ | -4.032073005 | -3.845093939 | -4.416949946 | -0.581622703 | 0.944466074 | -0.534214979 | 0.264000011 | 0.38919426 | 0.559143543 | 0.5074 | 0.643240676 |
| SL1344_RS24350_ | -4.922631702 | -4.498986684 | -5.383736781 | -0.89980998 | 0.995716114 | -0.831594607 | 0.171219209 | 0.266935652 | 0.424808227 | 0.3512 | 0.497392064 |
| SL1344_RS24355_ | -5.718748492 | -5.340297578 | -6.437575624 | -1.038981052 | 1.550678654 | -0.584550446 | 0.243837455 | 0.352000991 | 0.525865231 | 0.4674 | 0.60931357 |
| SL1344_RS24360_ | -4.750798559 | -4.439147294 | -4.968072133 | -0.569894102 | 1.130483271 | -0.485511728 | 0.296666676 | 0.449597454 | 0.604615241 | 0.5713 | 0.691061961 |
| SL1344_RS24365_ | -5.161350985 | -5.093360989 | -5.302516485 | -0.249870528 | 1.843518971 | -0.113109573 | 0.441705532 | 0.666326811 | 0.78926562 | 0.8005 | 0.872876489 |
| SL1344_RS24370_ | -1.746041406 | -1.748015023 | -1.74486069 | -0.007110293 | 0.387576431 | -0.016473512 | 0.490666667 | 0.603079374 | 0.734893933 | 0.7124 | 0.807025699 |
| SL1344_RS24380_ | -1.901631877 | -1.904474278 | -1.899104189 | 0.006383637 | 0.372571593 | 0.014555166 | 0.495333334 | 0.58225574 | 0.715331555 | 0.6923 | 0.792013608 |
| SL1344_RS24385_ | -1.68527325 | -1.593406795 | -1.74963502 | -0.158265099 | 0.429968116 | -0.343018795 | 0.348666673 | 0.549037314 | 0.691222227 | 0.6642 | 0.769851263 |
| SL1344_RS24390_ | -5.826514 | -5.994948956 | -5.560302619 | 0.435212304 | 1.754089085 | 0.202305276 | 0.40639574 | 0.607817098 | 0.740314285 | 0.7152 | 0.810213617 |
| SL1344_RS24395_ | -2.698256742 | -2.585858368 | -2.784207213 | -0.221270214 | 0.498456411 | -0.403871539 | 0.328666674 | 0.481952416 | 0.629480362 | 0.5901 | 0.705342401 |
| SL1344_RS24400_ | -4.640567182 | -4.553163809 | -4.714846925 | -0.166363688 | 1.379923071 | -0.107259666 | 0.453333335 | 0.660704894 | 0.783239612 | 0.7742 | 0.854793492 |
| SL1344_RS24405_ | -4.427446769 | -4.214010263 | -4.683459872 | -0.514163209 | 0.859257491 | -0.543637043 | 0.262666678 | 0.378781897 | 0.539013552 | 0.4825 | 0.61453637 |
| SL1344_RS24410_ | 0.559503372 | 0.603899027 | 0.492633648 | -0.128121776 | 0.210485898 | -0.550951077 | 0.265156573 | 0.399122102 | 0.567876738 | 0.5245 | 0.656048403 |
| SL1344_RS24415_ | -3.55404523 | -3.540597949 | -3.574162611 | -0.019736331 | 0.842114384 | -0.022705017 | 0.490339774 | 0.656434288 | 0.778364477 | 0.7702 | 0.853762755 |
| SL1344_RS24420_ | -0.782979305 | -0.79928449 | -0.764614635 | 0.03318556 | 0.255180738 | 0.113380608 | 0.450000002 | 0.574170344 | 0.711242198 | 0.6806 | 0.782401654 |
| SL1344_RS24425_ | -4.038902567 | -3.654741733 | -4.292093202 | -0.635095288 | 0.918103666 | -0.651599708 | 0.231333347 | 0.363085091 | 0.526693037 | 0.4673 | 0.608709043 |
| SL1344_RS24430_ | 2.49564397 | 2.664259419 | 2.322969718 | -0.335632963 | 0.13794939 | -2.423075489 | 0.001336769 | 0.030041735 | 0.123003902 | 0.1011 | 0.244577541 |
| SL1344_RS24435_ | 0.516402967 | 0.617613143 | 0.405504935 | -0.208639796 | 0.156424643 | -1.265596215 | 0.073284535 | 0.131927053 | 0.2751435 | 0.2036 | 0.358102816 |

| | | | | | | | | | | | |
|---|---|---|---|---|---|---|---|---|---|---|---|
| SL1344_RS24440_ | 2.856297736 | 2.863637656 | 2.849727388 | -0.012057853 | 0.112159485 | -0.096028951 | 0.457028649 | 0.705067088 | 0.817092492 | 0.8153 | 0.883063605 |
| SL1344_RS24445_ | 0.402805553 | 0.364282825 | 0.49076805 | 0.130296886 | 0.273269183 | 0.449125301 | 0.313333341 | 0.469875422 | 0.638962809 | 0.6294 | 0.746123974 |
| SL1344_RS24450_ | -1.486570049 | -1.596982136 | -1.376702896 | 0.220871973 | 0.299800352 | 0.683146776 | 0.219333349 | 0.322381045 | 0.480637926 | 0.4258 | 0.56466875 |
| SL1344_RS24455_ | -2.404224772 | -2.420013797 | -2.39265753 | 0.032705536 | 0.454925747 | 0.069667659 | 0.467333335 | 0.582205534 | 0.717187635 | 0.6991 | 0.795554146 |
| SL1344_RS24460_ | 7.966752331 | 7.971820663 | 7.961432859 | -0.008720246 | 0.082409082 | -0.092603934 | 0.468354432 | 0.910417026 | 0.952107372 | 0.9925 | 0.995741565 |
| SL1344_RS24465_ | 0.020804501 | -0.019237295 | 0.07247368 | 0.084851052 | 0.316814188 | 0.259210271 | 0.386000004 | 0.622437664 | 0.755069038 | 0.7378 | 0.829223998 |
| SL1344_RS24470_ | 6.138479307 | 6.129167987 | 6.146917655 | 0.018277542 | 0.174673207 | 0.093956549 | 0.462000001 | 0.874774041 | 0.932296202 | 0.9988 | 0.999320172 |
| SL1344_RS24475_ | 1.224295824 | 1.344531904 | 1.131724202 | -0.167450707 | 0.216451958 | -0.751958845 | 0.20733335 | 0.323616538 | 0.507898083 | 0.411 | 0.576284423 |
| SL1344_RS24480_ | 3.20061092 | 3.22449619 | 3.112362757 | -0.104812844 | 0.097311788 | -0.880490292 | 0.210666683 | 0.294837582 | 0.482491556 | 0.4474 | 0.605612253 |
| SL1344_RS24485_ | -0.3062648 | -0.345574143 | -0.260236611 | 0.031602592 | 0.324850121 | 0.101150868 | 0.456000002 | 0.703809094 | 0.816136071 | 0.8243 | 0.888687144 |
| SL1344_RS24490_ | -2.315307267 | -2.206948219 | -2.503464037 | -0.27834695 | 0.715477757 | -0.379164179 | 0.353097941 | 0.584079433 | 0.730783306 | 0.7088 | 0.813223442 |
| SL1344_RS24495_ | -2.15231256 | -2.228011463 | -2.103292969 | 0.124088387 | 0.443845852 | 0.260446033 | 0.389073955 | 0.573116893 | 0.711586517 | 0.6841 | 0.78768053 |
| SL1344_RS24500_ | -0.09800595 | -0.172957123 | -0.010680966 | 0.158106332 | 0.20184409 | 0.750772576 | 0.204666683 | 0.314790187 | 0.477310564 | 0.4098 | 0.555504251 |
| SL1344_RS24505_ | -3.181188916 | -3.100889682 | -3.323885131 | -0.24217093 | 0.702807906 | -0.315206273 | 0.355333339 | 0.49910581 | 0.653241128 | 0.6339 | 0.742949935 |
| SL1344_RS24510_ | -6.521652158 | -6.331574124 | -6.912432865 | -0.598161667 | 2.61307353 | -0.180128879 | 0.400000004 | 0.551774625 | 0.704256099 | 0.7165 | 0.813692443 |
| SL1344_RS24515_ | -3.946347304 | -3.833172667 | -4.033237756 | -0.223833747 | 0.761760547 | -0.272488022 | 0.384410398 | 0.53513181 | 0.677065278 | 0.6398 | 0.749703402 |
| SL1344_RS24520_ | -3.303212639 | -3.369033474 | -3.229873098 | 0.171620134 | 0.612963164 | 0.255700781 | 0.386000004 | 0.527281732 | 0.670313436 | 0.6305 | 0.742894662 |
| SL1344_RS24525_ | -1.703455249 | -1.82806313 | -1.589904574 | 0.249313784 | 0.343937813 | 0.677906319 | 0.214000016 | 0.336024661 | 0.495047469 | 0.4348 | 0.574787414 |
| SL1344_RS24530_ | -2.444024807 | -2.48698555 | -2.387666255 | 0.092871539 | 0.45201508 | 0.194206876 | 0.419053967 | 0.550015685 | 0.689765204 | 0.6604 | 0.764705688 |
| SL1344_RS24535_ | -2.822628736 | -2.512543222 | -3.24282749 | -0.757925303 | 0.61573827 | -1.18006356 | 0.079333387 | 0.156757195 | 0.318452105 | 0.2141 | 0.371327494 |
| SL1344_RS24540_ | -2.892405329 | -2.887309102 | -2.895835339 | 0.014002447 | 0.604028683 | 0.022615991 | 0.488666667 | 0.602276941 | 0.733966294 | 0.7124 | 0.806995445 |
| SL1344_RS24545_ | -4.121966401 | -4.001943015 | -4.25333532 | -0.298007318 | 0.89811949 | -0.30493 | 0.360666672 | 0.537209298 | 0.678663956 | 0.6584 | 0.761693905 |
| SL1344_RS24550_ | -0.993063292 | -1.08802502 | -0.89068688 | 0.196960087 | 0.280451021 | 0.666124462 | 0.24250501 | 0.36197858 | 0.521864376 | 0.4604 | 0.598835619 |
| SL1344_RS24555_ | -2.500194275 | -2.382891207 | -2.629223028 | -0.243922131 | 0.436469568 | -0.522051904 | 0.277333344 | 0.407861874 | 0.561869141 | 0.5141 | 0.642253773 |
| SL1344_RS24560_ | -2.038307405 | -2.022774653 | -2.052317048 | -0.069341286 | 0.502558334 | -0.123540661 | 0.442666669 | 0.621523255 | 0.75190054 | 0.7537 | 0.83628656 |
| SL1344_RS24565_ | -2.141687115 | -2.251825014 | -1.95170272 | 0.323856124 | 0.545217355 | 0.540483903 | 0.2778148 | 0.431316693 | 0.59693199 | 0.5547 | 0.683610688 |
| SL1344_RS24570_ | -2.877504309 | -3.107323308 | -2.627095934 | 0.479211558 | 0.500116429 | 0.93074553 | 0.150666692 | 0.245412308 | 0.400356857 | 0.3335 | 0.482624362 |
| SL1344_RS24575_ | -2.067171679 | -1.69753346 | -2.718007151 | -1.026829281 | 0.472768459 | -2.111903793 | 0.004664544 | 0.052529658 | 0.163656543 | 0.109 | 0.254141491 |
| SL1344_RS24580_ | 1.524403395 | 2.12818127 | 0.69579905 | -1.428800318 | 0.137905266 | -10.281803 | 4.68E-05 | 0.00133198 | 0.021893522 | 0.1 | 0.243021319 |
| SL1344_RS24585_ | -0.097714481 | 0.38388859 | -0.649586245 | -1.021139453 | 0.234176143 | -4.339639938 | 4.68E-05 | 0.006780228 | 0.051006241 | 0.1 | 0.243021319 |
| SL1344_RS24590_ | 0.558279779 | 0.489552053 | 0.647468187 | 0.167589671 | 0.187491093 | 0.834034194 | 0.171885431 | 0.262850738 | 0.433464972 | 0.3501 | 0.503499035 |
| SL1344_RS24595_ | -5.803802214 | -6.526526559 | -5.05632123 | 1.483180117 | 1.539142101 | 0.883737059 | 0.142000027 | 0.239333485 | 0.40446178 | 0.3242 | 0.47679604 |
| SL1344_RS24600_ | -2.844696995 | -2.716583171 | -3.068098733 | -0.387169891 | 0.631406756 | -0.549059729 | 0.257333345 | 0.396438122 | 0.566880896 | 0.5138 | 0.647464272 |
| SL1344_RS24605_ | -2.853631178 | -2.856018519 | -2.850967753 | -0.052864823 | 0.731845589 | -0.064247756 | 0.470353099 | 0.681899558 | 0.799675747 | 0.7996 | 0.872646396 |
| SL1344_RS24610_ | -1.053349187 | -1.33340699 | -0.852849155 | 0.479771923 | 0.299397924 | 1.532261433 | 0.041972124 | 0.09365538 | 0.229212986 | 0.1549 | 0.306899249 |
| SL1344_RS24615_ | -2.709001766 | -2.559601429 | -2.881506484 | -0.310749158 | 0.529800082 | -0.533061348 | 0.270666677 | 0.404444854 | 0.562961209 | 0.5109 | 0.643578936 |
| SL1344_RS24620_ | -3.030549373 | -3.083409814 | -2.947729403 | 0.167075217 | 0.587193087 | 0.252575045 | 0.38774151 | 0.555341228 | 0.694914272 | 0.6738 | 0.77469638 |
| SL1344_RS24625_ | -5.432683917 | -5.591051471 | -5.280323085 | 0.315385151 | 1.155525957 | 0.240844207 | 0.385742843 | 0.501496519 | 0.648082751 | 0.6002 | 0.71609726 |
| SL1344_RS24630_ | -1.365610236 | -1.28707565 | -1.473394662 | -0.167876679 | 0.426797703 | -0.352465028 | 0.340439713 | 0.537635774 | 0.6887628 | 0.6659 | 0.776869621 |
| SL1344_RS24635_ | -1.995360146 | -1.924936085 | -2.033750629 | -0.069055558 | 0.695500658 | -0.089171024 | 0.462000001 | 0.756958696 | 0.855012251 | 0.8402 | 0.903286115 |
| SL1344_RS24640_ | -5.595652001 | -5.914252317 | -5.022281001 | 0.817577007 | 1.940010797 | 0.367104814 | 0.316666675 | 0.576103032 | 0.713964357 | 0.6533 | 0.767693013 |
| SL1344_RS24645_ | -4.150704751 | -4.347176652 | -3.941486138 | 0.415804197 | 0.894715043 | 0.424323249 | 0.308666675 | 0.460536858 | 0.615077347 | 0.5741 | 0.695047999 |
| SL1344_RS24650_ | -3.157542599 | -3.340816392 | -3.009581813 | 0.321226175 | 0.609246818 | 0.499596565 | 0.290473027 | 0.434104481 | 0.590744997 | 0.5446 | 0.67050723 |
| SL1344_RS24655_ | -2.558520232 | -2.807245051 | -2.27383123 | 0.471470109 | 0.614624432 | 0.752294167 | 0.214000016 | 0.333572013 | 0.504265793 | 0.4236 | 0.576528947 |
| SL1344_RS24660_ | 0.741133682 | 0.754080754 | 0.726100395 | -0.028425633 | 0.156584357 | -0.186430298 | 0.421718857 | 0.575126837 | 0.712664483 | 0.6959 | 0.793287162 |
| SL1344_RS24665_ | 0.236179106 | 0.22888092 | 0.244308174 | 0.017918439 | 0.390646125 | 0.049591698 | 0.475682879 | 0.793059175 | 0.878373842 | 0.8506 | 0.908639482 |
| SL1344_RS24670_ | -2.406168563 | -2.481259357 | -2.337322237 | 0.135686636 | 0.407723761 | 0.326167898 | 0.363091278 | 0.482044225 | 0.626847069 | 0.5908 | 0.704405524 |
| SL1344_RS24675_ | -1.411220628 | -1.259039555 | -1.484075042 | -0.206957485 | 0.376090289 | -0.5183979 | 0.29380414 | 0.476434108 | 0.631760187 | 0.5851 | 0.711683501 |
| SL1344_RS24690_ | 6.448518123 | 6.599700956 | 6.221557731 | -0.380217808 | 0.132954185 | -3.131457447 | 4.68E-05 | 0.008808906 | 0.066087853 | 0.1 | 0.243021319 |
| SL1344_RS24695_ | 3.154313982 | 3.718378658 | 2.572006456 | -1.14507897 | 0.127504459 | -8.963596752 | 4.68E-05 | 0.000486603 | 0.015782944 | 0.1 | 0.243021319 |
| SL1344_RS24700_ | 3.391146397 | 3.970362108 | 2.848051222 | -1.121484362 | 0.105939593 | -10.62738386 | 4.68E-05 | 0.00062042 | 0.016859288 | 0.1 | 0.243021319 |
| SL1344_RS24705_ | 3.208606094 | 3.623647838 | 2.64622589 | -0.975342453 | 0.136354435 | -7.147782329 | 4.68E-05 | 0.005203584 | 0.048581417 | 0.1 | 0.243021319 |
| SL1344_RS24710_ | 2.83155763 | 3.206583496 | 2.280062724 | -0.903153305 | 0.143558002 | -6.121462097 | 4.68E-05 | 0.004738098 | 0.044763485 | 0.1 | 0.243021319 |

| | | | | | | | | | | | |
|---|---|---|---|---|---|---|---|---|---|---|---|
| SL1344_RS24730_ | 5.666910508 | 5.924730986 | 5.517336015 | -0.412195757 | 0.103813583 | -3.946286356 | 4.68E-05 | 0.011120552 | 0.074007918 | 0.1 | 0.243021319 |
| SL1344_RS24735_ | 6.359245903 | 6.55473375 | 6.156566625 | -0.42109238 | 0.103837757 | -3.895012441 | 4.68E-05 | 0.003373549 | 0.040833005 | 0.1 | 0.243021319 |
| SL1344_RS24740_ | -3.456754057 | -3.24521444 | -3.762602154 | -0.516606089 | 0.739306603 | -0.614440674 | 0.245333346 | 0.375017735 | 0.541384121 | 0.4903 | 0.629791125 |
| SL1344_RS24745_ | -6.969349626 | -6.902866942 | -7.072667068 | -0.251363532 | 2.564135315 | -0.08014552 | 0.456666668 | 0.577582589 | 0.720201695 | 0.724 | 0.815246069 |
| SL1344_RS24750_ | 0.137322233 | -0.029957111 | 0.263260575 | 0.332756605 | 0.401717345 | 0.843011012 | 0.167333355 | 0.251702731 | 0.444824589 | 0.3454 | 0.518806352 |
| SL1344_RS24755_ | 1.082961567 | 0.993422311 | 1.170948952 | 0.177889076 | 0.12270716 | 1.400855697 | 0.064666733 | 0.116050518 | 0.249148945 | 0.1871 | 0.339526717 |
| SL1344_RS24760_ | 0.079201695 | -0.218757095 | 0.340521801 | 0.567527013 | 0.23085595 | 2.414796244 | 0.0020023 | 0.026953206 | 0.112136588 | 0.1018 | 0.245527461 |
| SL1344_RS24780_ | -2.26941468 | -2.168215052 | -2.582850203 | -0.36503425 | 0.670373573 | -0.488160879 | 0.318666674 | 0.485511416 | 0.652702851 | 0.6375 | 0.761936591 |
| SL1344_RS24785_ | 6.248441633 | 6.124437747 | 6.423362337 | 0.31505004 | 0.113399793 | 2.682603243 | 4.68E-05 | 0.031077913 | 0.131539445 | 0.1 | 0.243021319 |
| SL1344_RS24790_ | 6.01941948 | 5.917134523 | 6.050274128 | 0.133381263 | 0.124568025 | 1.124010915 | 0.142666694 | 0.141166447 | 0.321494782 | 0.2476 | 0.432804464 |
| SL1344_RS24795_ | 0.146635559 | 0.024941305 | 0.257046116 | 0.244771329 | 0.206447592 | 1.130251454 | 0.098600974 | 0.166333562 | 0.325944857 | 0.2419 | 0.397557786 |
| SL1344_RS24800_ | -3.729886916 | -3.744580842 | -3.710726185 | 0.088006647 | 1.218819368 | 0.069238184 | 0.464666668 | 0.701735572 | 0.815314859 | 0.7959 | 0.87157304 |
| SL1344_RS24805_ | 0.451523077 | 0.222552861 | 0.602596718 | 0.366646065 | 0.241562469 | 1.503832945 | 0.046635671 | 0.086395955 | 0.224411515 | 0.1497 | 0.305073039 |
| SL1344_RS24810_ | 1.322681199 | 0.958545597 | 1.525339749 | 0.54980433 | 0.197209409 | 2.579463063 | 0.000673369 | 0.034087187 | 0.132782354 | 0.1008 | 0.244144027 |
| SL1344_RS24820_ | 3.510508891 | 4.160676767 | 2.909135295 | -1.20724756 | 0.174522804 | -6.817833521 | 4.68E-05 | 0.001728603 | 0.028618604 | 0.1 | 0.243021319 |
| SL1344_RS24825_ | 5.451665485 | 5.473464531 | 5.407691593 | -0.06007213 | 0.194764522 | -0.372534808 | 0.23666668 | 0.572410366 | 0.735535768 | 0.4447 | 0.631924346 |
| SL1344_RS24830_ | 1.484826278 | 1.60362009 | 1.344865671 | -0.271225776 | 0.156142241 | -1.698770848 | 0.011992386 | 0.072166847 | 0.201189429 | 0.1197 | 0.267376923 |
| SL1344_RS24835_ | -1.365342969 | -1.291280934 | -1.437394597 | -0.145556954 | 0.406735749 | -0.346911414 | 0.351099273 | 0.548711865 | 0.69242862 | 0.6538 | 0.762573884 |
| SL1344_RS24840_ | -3.129813291 | -3.139528581 | -3.123607985 | 0.05709318 | 0.666268948 | 0.080777502 | 0.464666668 | 0.611360009 | 0.744753334 | 0.7204 | 0.814424562 |
| SL1344_RS24845_ | -3.910480248 | -4.313375334 | -3.772221895 | 0.539963137 | 1.447315071 | 0.340857476 | 0.359760166 | 0.507808897 | 0.676420327 | 0.7104 | 0.81488178 |
| SL1344_RS24850_ | -5.055042126 | -5.123802715 | -4.973334084 | 0.160255386 | 1.175738228 | 0.125989335 | 0.440000002 | 0.563457315 | 0.701342965 | 0.6756 | 0.777280406 |
| SL1344_RS24860_ | 2.727430673 | 2.310152642 | 2.953282427 | 0.628601982 | 0.129783258 | 4.524838564 | 4.68E-05 | 0.010445362 | 0.068474927 | 0.1 | 0.243021319 |
| SL1344_RS24865_ | -6.013357527 | -6.185018226 | -5.883483211 | 0.34907964 | 1.661590168 | 0.182703332 | 0.417333336 | 0.554668993 | 0.695887577 | 0.6811 | 0.78060824 |
| SL1344_RS24875_ | -5.121000713 | -5.270083359 | -4.928276327 | 0.362235726 | 1.46773062 | 0.215482253 | 0.40466667 | 0.609576214 | 0.740679358 | 0.7127 | 0.809321432 |
| SL1344_RS24885_ | 0.885882828 | 0.464537833 | 1.149490844 | 0.684990636 | 0.210243061 | 3.162143128 | 4.68E-05 | 0.017238296 | 0.087758556 | 0.1 | 0.243021319 |
| SLnc0002_ | -3.321865955 | -3.35691675 | -3.293395728 | 0.080607814 | 0.785755673 | 0.093425467 | 0.457028649 | 0.647655239 | 0.772423754 | 0.7688 | 0.849024453 |
| SLnc0010_ | -2.943970915 | -2.911514878 | -3.006210302 | -0.061089823 | 0.619189102 | -0.090734281 | 0.457333335 | 0.629478074 | 0.759724984 | 0.7392 | 0.831362605 |
| SLnc0010a_ | -0.909011532 | -0.990621078 | -0.789482954 | 0.255259467 | 0.447053729 | 0.564089649 | 0.275333344 | 0.42961982 | 0.60293112 | 0.5607 | 0.69662057 |
| SLnc0014_ | 0.789820856 | 1.30706173 | 0.084169824 | -1.220894307 | 0.194569553 | -6.259811614 | 4.68E-05 | 0.005634704 | 0.046883662 | 0.1 | 0.243021319 |
| SLnc0022_ | -0.626341403 | -0.876013268 | -0.444789149 | 0.439974741 | 0.271498109 | 1.550836502 | 0.029333486 | 0.091021186 | 0.229170945 | 0.1423 | 0.293984671 |
| SLnc0030_ | 3.287872053 | 3.068575527 | 3.527080738 | 0.468667022 | 0.139226808 | 3.380837149 | 4.68E-05 | 0.013882473 | 0.081668979 | 0.1 | 0.243021319 |
| SLnc0030a_ | -1.124045934 | -1.161282577 | -1.098408705 | 0.061775184 | 0.302126212 | 0.188747288 | 0.416389077 | 0.579716622 | 0.716393502 | 0.6903 | 0.791195787 |
| SLnc0031_ | -3.402548674 | -3.590545539 | -3.203272862 | 0.431276825 | 0.750629768 | 0.567460263 | 0.274483688 | 0.415033032 | 0.575758683 | 0.5301 | 0.660242431 |
| SLnc0035_ | 3.877096045 | 3.769908728 | 3.972139982 | 0.182909247 | 0.094663504 | 1.941126925 | 0.015323415 | 0.04420907 | 0.155054237 | 0.1146 | 0.263540742 |
| SLnc0039_ | -0.108499992 | -0.325766181 | 0.273505003 | 0.626406599 | 0.375486782 | 1.623849991 | 0.018654474 | 0.093736078 | 0.245072798 | 0.1201 | 0.269450602 |
| SLnc0048_ | -1.867487072 | -1.907763448 | -1.842322149 | 0.107994525 | 0.594255396 | 0.178351033 | 0.426666669 | 0.654680718 | 0.781244139 | 0.7876 | 0.86398346 |
| SLnc1001_ | -8.99487492 | -9.540118103 | -8.461232281 | 0.947534678 | 4.13969924 | 0.189795564 | 0.392405067 | 0.506305612 | 0.652794238 | 0.6159 | 0.727545137 |
| SLnc1002_ | -1.19186119 | -1.211863566 | -1.169966059 | 0.034925181 | 0.291148531 | 0.113616148 | 0.449700202 | 0.574057963 | 0.708260038 | 0.6748 | 0.776569921 |
| SLnc1006_ | 1.030160093 | 1.057298597 | 0.993646063 | -0.083898055 | 0.220064128 | -0.360121598 | 0.322000008 | 0.474698549 | 0.6435744 | 0.6344 | 0.745300666 |
| SLnc1008_ | -5.242306965 | -5.753127695 | -4.600148524 | 1.10206739 | 1.338165392 | 0.758095152 | 0.185333353 | 0.315088523 | 0.47949325 | 0.402 | 0.55385092 |
| SLnc1013_ | -1.95686196 | -1.988687932 | -1.916378651 | 0.035464847 | 0.483454307 | 0.064232741 | 0.470353099 | 0.660282369 | 0.782303171 | 0.7715 | 0.851686718 |
| SLnc1014_ | -3.253958839 | -3.273114916 | -3.24513798 | 0.024064019 | 0.854150146 | 0.022316261 | 0.49 | 0.688683406 | 0.804715483 | 0.7921 | 0.871913851 |
| SLnc1015_ | -2.824403958 | -2.989998218 | -2.49234146 | 0.438092749 | 0.599835331 | 0.629287674 | 0.259160571 | 0.421693925 | 0.581229093 | 0.5199 | 0.660143362 |
| SLnc1016_ | -2.5073985 | -2.673303665 | -2.368518335 | 0.311852916 | 0.487065781 | 0.612545 | 0.256000012 | 0.37305388 | 0.532774604 | 0.4755 | 0.612378587 |
| SLnc1017_ | 0.165556032 | -0.162699175 | 0.539776102 | 0.710509879 | 0.261327956 | 2.688016681 | 0.001335878 | 0.021614531 | 0.101852945 | 0.1004 | 0.243585946 |
| SLnc1018_ | -3.381932889 | -3.593325799 | -3.253941567 | 0.367443161 | 0.770291717 | 0.462007696 | 0.304666675 | 0.455412766 | 0.617162437 | 0.5751 | 0.69496039 |
| SLnc1019_ | -2.541414172 | -2.556225715 | -2.52469681 | 0.036408689 | 0.497239046 | 0.068964809 | 0.467333335 | 0.590300046 | 0.725665718 | 0.6952 | 0.794890657 |
| SLnc1021_ | -1.456666927 | -1.552393087 | -1.286824212 | 0.222557363 | 0.421603456 | 0.463207772 | 0.316000008 | 0.528126257 | 0.676156381 | 0.6252 | 0.746441086 |
| SLnc1022_ | -4.281109891 | -4.636925941 | -3.95252037 | 0.691136415 | 0.849108651 | 0.744423717 | 0.196000018 | 0.300930662 | 0.460965022 | 0.3922 | 0.535674953 |
| SLnc1024_ | -1.99321967 | -2.384202144 | -1.478981041 | 0.879994988 | 0.402104353 | 2.110701984 | 0.012000381 | 0.042058575 | 0.142320148 | 0.1128 | 0.259485767 |
| SLnc1025_ | -2.221538424 | -2.602041055 | -1.768298197 | 0.834214175 | 0.502307792 | 1.615364272 | 0.03730858 | 0.080283444 | 0.212961719 | 0.1386 | 0.291691959 |
| SLnc1026_ | -3.093208869 | -3.00377915 | -3.213334845 | -0.238538127 | 0.689226011 | -0.320163693 | 0.355096608 | 0.535009399 | 0.6810717 | 0.6517 | 0.759377892 |
| SLnc1027_ | -2.595475678 | -2.771569462 | -2.375241653 | 0.398848407 | 0.51663903 | 0.719934317 | 0.211333349 | 0.326073523 | 0.490266141 | 0.4356 | 0.578962798 |

| | | | | | | | | | | | |
|---|---|---|---|---|---|---|---|---|---|---|---|
| SLnc1028_ | -2.060805845 | -2.331575219 | -1.912397156 | 0.404292875 | 0.464735483 | 0.821948955 | 0.181212545 | 0.279353821 | 0.450904518 | 0.3737 | 0.525318835 |
| SLnc1029_ | -4.292818607 | -4.702725951 | -3.872253114 | 0.815981349 | 0.956205327 | 0.784037613 | 0.188540992 | 0.289743062 | 0.458317506 | 0.3789 | 0.534029785 |
| sRNA1_ | 3.590201364 | 3.577120153 | 3.609938146 | 0.050045947 | 0.096631876 | 0.500472078 | 0.29113925 | 0.415367704 | 0.594910082 | 0.5869 | 0.708449496 |
| sRNA10_ | 1.329180459 | 1.298557492 | 1.426946515 | 0.107984664 | 0.160134477 | 0.572775234 | 0.295136585 | 0.470097682 | 0.632973226 | 0.5784 | 0.716365445 |
| sRNA11_ | -2.521588959 | -2.784839025 | -2.294045555 | 0.512047469 | 0.907764284 | 0.526963616 | 0.259333345 | 0.402853002 | 0.585466453 | 0.5172 | 0.664570157 |
| sRNA12_ | -6.855856439 | -6.876404036 | -6.797870272 | 0.098217342 | 2.74899972 | 0.035418695 | 0.485333334 | 0.631273776 | 0.763063103 | 0.7719 | 0.854501722 |
| sRNA15_ | -7.557664286 | -7.304836421 | -7.959438487 | -0.374586267 | 3.951246727 | -0.083695004 | 0.453333335 | 0.627677473 | 0.760790006 | 0.7802 | 0.859908535 |
| sRNA16_ | -4.043837536 | -4.143220697 | -3.98547167 | 0.106874414 | 0.970557195 | 0.093019413 | 0.462358429 | 0.630588368 | 0.760936721 | 0.7535 | 0.841586104 |
| sRNA18_ | -5.424368263 | -6.296877977 | -4.793200657 | 1.50646614 | 1.214839827 | 1.166683055 | 0.088666714 | 0.156791194 | 0.304810116 | 0.2266 | 0.380490871 |
| sRNA19_ | -8.934687403 | -8.424218659 | -9.396865172 | -0.803931874 | 4.065508544 | -0.170907868 | 0.40106596 | 0.525001245 | 0.66915664 | 0.6403 | 0.746689828 |
| sRNA2_ | 2.850161829 | 2.664251411 | 3.16265937 | 0.50156421 | 0.193359301 | 2.596037342 | 4.68E-05 | 0.043812777 | 0.159131037 | 0.1 | 0.243021319 |
| sRNA21_ | -1.516526544 | -1.219966593 | -1.791344895 | -0.568494406 | 0.355609851 | -1.553166823 | 0.039333446 | 0.084561178 | 0.215096854 | 0.1476 | 0.300040764 |
| sRNA22_ | -2.06719778 | -1.978473756 | -2.17069013 | -0.278669383 | 0.572357462 | -0.453347629 | 0.311125924 | 0.455437186 | 0.621942384 | 0.6131 | 0.72648377 |
| sRNA23_ | -3.545182872 | -3.564327853 | -3.531899147 | 0.093334332 | 1.1848602 | 0.071049143 | 0.458000002 | 0.708617211 | 0.820104549 | 0.8189 | 0.886747534 |
| sRNA24_ | -4.808280038 | -4.489128561 | -5.089465476 | -0.607884298 | 1.095212595 | -0.532675289 | 0.272485021 | 0.40424441 | 0.563502885 | 0.5189 | 0.647292684 |
| sRNA25_ | -6.091604365 | -5.727065255 | -6.252193279 | -0.654129229 | 3.593491096 | -0.15298357 | 0.425716192 | 0.708412444 | 0.819170968 | 0.8096 | 0.883458105 |
| sRNA26_ | -5.488314843 | -5.573087424 | -5.404696663 | 0.254996376 | 1.597266083 | 0.132094079 | 0.434377084 | 0.593745042 | 0.730735127 | 0.7216 | 0.81351803 |
| sRNA28_ | -5.136453797 | -4.488576071 | -5.902660828 | -1.421593771 | 1.272439473 | -1.044061955 | 0.109926752 | 0.183671042 | 0.342423985 | 0.2623 | 0.420220987 |
| sRNA29_ | -5.353183677 | -5.705242303 | -5.156451102 | 0.55570372 | 1.49123177 | 0.319682428 | 0.346000006 | 0.487267379 | 0.644889946 | 0.6187 | 0.733956531 |
| sRNA31_ | -4.86569491 | -4.90390655 | -4.802564398 | 0.265974183 | 1.516534981 | 0.163578777 | 0.437041974 | 0.639201494 | 0.766622333 | 0.7596 | 0.845399607 |
| sRNA32_ | -4.405811299 | -4.984981999 | -3.757187298 | 1.227163346 | 0.968050064 | 1.202718863 | 0.100000041 | 0.157981577 | 0.313974313 | 0.2315 | 0.390870284 |
| sRNA33_ | -2.651810811 | -2.741041074 | -2.557046302 | 0.18788325 | 0.585166259 | 0.286794919 | 0.372418393 | 0.570915156 | 0.709865447 | 0.6763 | 0.780837565 |
| sRNA5_ | -2.452503907 | -2.532919342 | -2.35140932 | 0.079077107 | 0.713287966 | 0.101810602 | 0.459027317 | 0.724651168 | 0.831145584 | 0.8285 | 0.89439716 |
| sRNA7_ | -0.533033206 | -0.701783508 | -0.384355494 | 0.320341035 | 0.203026611 | 1.523817329 | 0.047333426 | 0.098662707 | 0.227257893 | 0.1706 | 0.322228296 |
| STnc10_ | -5.915628254 | -6.187582503 | -5.639769514 | 0.462114208 | 1.83459023 | 0.229606471 | 0.38774151 | 0.571512762 | 0.711178736 | 0.6802 | 0.782526689 |
| STnc1010_ | -3.045289353 | -3.134759561 | -2.944834467 | 0.154544222 | 0.801090103 | 0.175037182 | 0.41866667 | 0.653087684 | 0.7782061 | 0.7655 | 0.849524372 |
| STnc1030_ | -2.424944089 | -2.331847554 | -2.560108657 | -0.254699114 | 0.710102972 | -0.348188494 | 0.339773491 | 0.562786204 | 0.71027963 | 0.6798 | 0.788683908 |
| STnc1040_ | -4.242229563 | -4.399912032 | -4.114571776 | 0.266437855 | 0.872921322 | 0.287591594 | 0.366000005 | 0.504331845 | 0.651973232 | 0.6169 | 0.730263987 |
| STnc1065_ | 3.646791026 | 3.533192502 | 3.752945529 | 0.233026456 | 0.103844583 | 2.101373208 | 4.68E-05 | 0.093976536 | 0.250805184 | 0.1 | 0.243021319 |
| STnc1070_ | -1.448991636 | -1.564875772 | -1.402060732 | 0.145691217 | 0.464627537 | 0.282525558 | 0.389073955 | 0.617541462 | 0.753572938 | 0.747 | 0.839588541 |
| STnc1080_ | -2.983188558 | -2.928802625 | -3.032442212 | -0.130917821 | 0.644400453 | -0.184468411 | 0.423051302 | 0.590511238 | 0.72804788 | 0.7098 | 0.805381791 |
| STnc1090_ | -0.919225775 | -0.869328658 | -0.968838396 | -0.086345104 | 0.248045439 | -0.333863694 | 0.345103271 | 0.486242693 | 0.631981252 | 0.5848 | 0.702980685 |
| STnc1100_ | -3.011755629 | -3.286495601 | -2.787410336 | 0.498308739 | 0.581449555 | 0.800805707 | 0.190000019 | 0.290733251 | 0.452795576 | 0.3845 | 0.530873361 |
| STnc1130_ | -0.484395067 | -0.177964701 | -0.914013222 | -0.736682691 | 0.242744409 | -2.939434008 | 0.000673369 | 0.022027438 | 0.09728465 | 0.1007 | 0.24393981 |
| STnc1150_ | 0.653342282 | 0.604004836 | 0.709102237 | 0.109972564 | 0.174462686 | 0.58771766 | 0.249833457 | 0.399141842 | 0.557897982 | 0.5098 | 0.640442252 |
| STnc1160_ | 1.916752143 | 1.766415433 | 2.138505826 | 0.344223536 | 0.153260629 | 2.167339775 | 0.008661422 | 0.037392207 | 0.137934752 | 0.1093 | 0.255639011 |
| STnc1170_ | -4.224110857 | -4.312580201 | -4.100300271 | 0.3428727 | 1.055460038 | 0.298587007 | 0.386666671 | 0.555128175 | 0.698794641 | 0.6851 | 0.78620096 |
| STnc1180_ | -2.673505762 | -2.542904313 | -2.841093034 | -0.286967017 | 0.57201474 | -0.46880486 | 0.305333342 | 0.464505447 | 0.620402945 | 0.5738 | 0.700012024 |
| STnc1190_ | 1.177529001 | 1.109500335 | 1.217936813 | 0.102601869 | 0.137585895 | 0.666551208 | 0.235842785 | 0.368665486 | 0.531822921 | 0.4809 | 0.621804273 |
| STnc1200_ | -1.940295456 | -2.016900856 | -1.875956476 | 0.132844644 | 0.383420408 | 0.325380785 | 0.356429053 | 0.505883165 | 0.648919997 | 0.61 | 0.721390158 |
| STnc1210_ | -4.789433068 | -4.909930056 | -4.701813773 | 0.238952646 | 1.228311375 | 0.175429907 | 0.413724187 | 0.558514189 | 0.703089157 | 0.686 | 0.785917865 |
| STnc1220_ | -8.516268561 | -8.57904206 | -8.455622479 | 0.017657278 | 3.914192656 | 0.004946864 | 0.497333333 | 0.569033632 | 0.711122839 | 0.6889 | 0.788946465 |
| STnc1275_ | -0.798137589 | -0.646424714 | -0.970963718 | -0.362088744 | 0.359907364 | -0.971516332 | 0.115256531 | 0.19279342 | 0.365560432 | 0.2753 | 0.433079785 |
| STnc1280_ | -4.529522161 | -4.326445137 | -4.821413101 | -0.498160044 | 0.902199819 | -0.503100862 | 0.273817466 | 0.403809024 | 0.563277486 | 0.5121 | 0.643186616 |
| STnc1285_ | 2.531325255 | 3.610690248 | 1.414031333 | -2.186225905 | 0.2884833 | -7.563495233 | 4.68E-05 | 0.004525527 | 0.046021047 | 0.1 | 0.243021319 |
| STnc1290_ | 1.524754902 | 1.257519965 | 1.863901328 | 0.602564556 | 0.140966658 | 4.279255389 | 4.68E-05 | 0.006998175 | 0.053826477 | 0.1 | 0.243021319 |
| STnc130_ | -4.328108892 | -4.351403713 | -4.302651458 | 0.074579343 | 1.001568955 | 0.066877572 | 0.469020654 | 0.621769952 | 0.749824391 | 0.7255 | 0.816716598 |
| STnc1320_ | -3.728513572 | -4.027509651 | -3.520314321 | 0.528694135 | 0.759650236 | 0.639126361 | 0.239173897 | 0.348954257 | 0.51575682 | 0.4551 | 0.595443931 |
| STnc1340_ | -0.720927443 | -1.116646477 | -0.387109097 | 0.734745891 | 0.262424082 | 2.780257698 | 0.000673369 | 0.022768762 | 0.099425806 | 0.1006 | 0.24386408 |
| STnc1380_ | -3.020460913 | -3.179880965 | -2.890378374 | 0.28904669 | 0.539170481 | 0.505798724 | 0.28200001 | 0.406476031 | 0.55969343 | 0.5122 | 0.640768445 |
| STnc1385_ | 3.075561577 | 2.908780466 | 3.478754662 | 0.576413701 | 0.241800754 | 2.398783157 | 4.68E-05 | 0.025701882 | 0.117060258 | 0.1 | 0.243021319 |
| STnc1390_ | -2.926424374 | -3.162831654 | -2.482732072 | 0.592738998 | 0.768763351 | 0.708375447 | 0.221185891 | 0.349434951 | 0.526590728 | 0.4343 | 0.598316191 |
| STnc140_ | -6.284645747 | -6.507809104 | -6.089895407 | 0.305230208 | 2.073868076 | 0.135195226 | 0.439333336 | 0.596502406 | 0.733394516 | 0.7253 | 0.817892784 |

| | | | | | | | | | | | |
|---|---|---|---|---|---|---|---|---|---|---|---|
| STnc1400_ | 0.741635182 | 0.577217943 | 0.867583879 | 0.284541899 | 0.176666449 | 1.563154581 | 0.045969449 | 0.085626789 | 0.219196173 | 0.1513 | 0.306704493 |
| STnc1405_ | 0.117124101 | 1.05995444 | -0.918413438 | -2.017530243 | 0.264068823 | -7.752706736 | 4.68E-05 | 0.001072094 | 0.021622798 | 0.1 | 0.243021319 |
| STnc1410_ | -2.360907715 | -2.480634587 | -2.157822869 | 0.303569737 | 0.475309128 | 0.561938207 | 0.269820131 | 0.438734873 | 0.591752672 | 0.5365 | 0.665953451 |
| STnc1460_ | 6.618741315 | 6.378658671 | 6.868497976 | 0.505650595 | 0.08034122 | 6.488918286 | 4.68E-05 | 0.011390491 | 0.075940174 | 0.1 | 0.243021319 |
| STnc1490_ | 0.298135259 | 0.004916726 | 0.574246907 | 0.57146298 | 0.297547783 | 1.866304127 | 0.019333568 | 0.050964253 | 0.168353444 | 0.1143 | 0.262676067 |
| STnc150_ | -0.60214298 | -0.836929869 | -0.274108691 | 0.6009638 | 0.400734204 | 1.471997694 | 0.033333467 | 0.083284712 | 0.224542181 | 0.135 | 0.288004832 |
| STnc1500_ | -2.993644318 | -3.032598909 | -2.953628402 | 0.078811562 | 0.562853927 | 0.125888338 | 0.441333336 | 0.562511751 | 0.701236096 | 0.6778 | 0.778567245 |
| STnc1510_ | -0.93421316 | -0.969066523 | -0.893030593 | 0.074222689 | 0.263813224 | 0.272153435 | 0.380000005 | 0.531535847 | 0.673014973 | 0.6378 | 0.74657992 |
| STnc1530_ | -5.530075747 | -6.097322527 | -5.059909801 | 1.041142433 | 1.22342757 | 0.803287368 | 0.169333355 | 0.281938882 | 0.43949789 | 0.368 | 0.51311968 |
| STnc1540_ | -0.617881387 | -0.934349965 | -0.176227528 | 0.746407649 | 0.295170034 | 2.491365484 | 0.002668394 | 0.028710796 | 0.116964895 | 0.102 | 0.24583726 |
| STnc1545_ | 1.28772533 | 1.042126468 | 1.413194165 | 0.355661195 | 0.156422056 | 2.216265547 | 0.006000767 | 0.047196168 | 0.156536427 | 0.1067 | 0.251695647 |
| STnc1550_ | -0.477289162 | -0.262004722 | -0.773054378 | -0.474805567 | 0.31716648 | -1.413002223 | 0.066666731 | 0.107536431 | 0.257477678 | 0.1736 | 0.336232914 |
| STnc1560_ | 4.537161214 | 4.404461535 | 4.647984058 | 0.273769341 | 0.105501375 | 2.470068542 | 4.68E-05 | 0.157115003 | 0.345206953 | 0.1 | 0.243021319 |
| STnc1565_ | -0.60370353 | -0.680665073 | -0.525246666 | 0.245557617 | 0.412907026 | 0.551241213 | 0.258000012 | 0.372035811 | 0.55510581 | 0.5424 | 0.667369922 |
| STnc1580_ | -0.738762282 | -1.063514544 | -0.593889388 | 0.447186633 | 0.333115519 | 1.228540441 | 0.112666703 | 0.167446749 | 0.337149146 | 0.2591 | 0.423678328 |
| STnc1590_ | 3.006106467 | 2.627346359 | 3.271219988 | 0.670455575 | 0.242886068 | 2.995960669 | 4.68E-05 | 0.016765049 | 0.092300971 | 0.1 | 0.243021319 |
| STnc1600_ | -2.62840302 | -2.774239711 | -2.482472988 | 0.06879518 | 0.833641365 | 0.091230673 | 0.462000001 | 0.723126256 | 0.830450911 | 0.8522 | 0.907950415 |
| STnc1640_ | -2.757673062 | -2.988226572 | -2.506454084 | 0.47605098 | 0.485164579 | 0.915378113 | 0.140572979 | 0.225668089 | 0.385759074 | 0.3167 | 0.468590442 |
| STnc1650_ | -4.294822952 | -4.798438533 | -3.935534817 | 0.900291295 | 1.019844722 | 0.804631133 | 0.161333357 | 0.259531382 | 0.432358984 | 0.3437 | 0.496982583 |
| STnc1660_ | -0.18577386 | -0.282383586 | -0.126132167 | 0.116790538 | 0.333169999 | 0.342946748 | 0.363091278 | 0.618561124 | 0.753287647 | 0.7093 | 0.81378609 |
| STnc1670_ | -3.884260421 | -4.397448706 | -3.447885184 | 1.07375884 | 1.458838117 | 0.704695126 | 0.130579644 | 0.275934549 | 0.47229548 | 0.2902 | 0.455149136 |
| STnc1675_ | 7.532527581 | 7.459710238 | 7.595731233 | 0.156650126 | 0.073029338 | 1.97690623 | 4.68E-05 | 0.091996176 | 0.249689399 | 0.1 | 0.243021319 |
| STnc1680_ | 1.426542147 | 1.211603141 | 1.77150765 | 0.566363782 | 0.214019528 | 2.584867715 | 4.68E-05 | 0.038547975 | 0.14530591 | 0.1004 | 0.243517657 |
| STnc1685_ | 5.650500655 | 3.423080944 | 5.699334199 | 2.264971172 | 0.243935617 | 0.84175511 | 0.33333334 | 0.277289563 | 0.484850822 | 0.7 | 0.829598552 |
| STnc1690_ | -2.886126405 | -2.422514256 | -3.514047838 | -1.095424579 | 0.511483689 | -2.067595152 | 0.007995244 | 0.046439325 | 0.147250528 | 0.1112 | 0.257168095 |
| STnc170_ | -3.63130865 | -3.010575455 | -4.760521456 | -1.789206042 | 1.266637673 | -1.473977228 | 0.008000574 | 0.135176798 | 0.302456567 | 0.1079 | 0.253055196 |
| STnc1700_ | -3.037386259 | -3.11081288 | -2.952318598 | 0.152964846 | 0.656409758 | 0.215358176 | 0.409333337 | 0.581565536 | 0.718045011 | 0.6932 | 0.79229438 |
| STnc1710_ | 0.758943151 | 0.647182821 | 0.991390923 | 0.332197597 | 0.210998058 | 1.447245516 | 0.066000065 | 0.116989036 | 0.273586146 | 0.184 | 0.343802438 |
| STnc1760_ | -2.901706251 | -2.96790822 | -2.804817051 | 0.118707602 | 0.63898084 | 0.166576274 | 0.42466667 | 0.619975952 | 0.751629326 | 0.7211 | 0.816748988 |
| STnc1820_ | -0.350507224 | -0.672271154 | -0.025768787 | 0.650316783 | 0.249166183 | 2.570359809 | 0.004667653 | 0.025062428 | 0.106486725 | 0.1025 | 0.246407884 |
| STnc1830_ | -2.662370678 | -2.427832335 | -3.140998787 | -0.683252245 | 0.629678739 | -1.054277439 | 0.138000028 | 0.20830383 | 0.38173071 | 0.3031 | 0.465153357 |
| STnc1870_ | -5.380712108 | -5.282434359 | -5.471202625 | -0.172416941 | 1.575237373 | -0.100563365 | 0.450666669 | 0.618409204 | 0.748886108 | 0.7369 | 0.824857851 |
| STnc1880_ | -1.710769253 | -1.762538299 | -1.635559669 | 0.182661386 | 0.458756064 | 0.365587102 | 0.345103271 | 0.514351752 | 0.668130409 | 0.6649 | 0.768889902 |
| STnc1890_ | -5.82943921 | -5.908277209 | -5.748486229 | 0.098684639 | 3.795474585 | 0.019941173 | 0.484343771 | 0.687011031 | 0.807345867 | 0.7999 | 0.874434174 |
| STnc190_ | -2.040139836 | -1.502682907 | -2.693797632 | -1.198353592 | 0.393612832 | -3.036330799 | 0.000673369 | 0.017859837 | 0.085633488 | 0.1004 | 0.243589849 |
| STnc1920_ | -0.729106199 | -1.043549681 | -0.456762038 | 0.591133162 | 0.246575639 | 2.324287599 | 0.002666616 | 0.031672101 | 0.118903048 | 0.1026 | 0.246599942 |
| STnc1990_ | -0.703106361 | -0.977888345 | -0.470328836 | 0.507346 | 0.231627118 | 2.14560241 | 0.008667196 | 0.039099526 | 0.132336535 | 0.1099 | 0.255572491 |
| STnc20_ | -2.558274603 | -2.515707114 | -2.588150803 | -0.12941765 | 0.639866757 | -0.183800638 | 0.418387745 | 0.626826047 | 0.756540172 | 0.7382 | 0.827833427 |
| STnc200_ | -0.435684672 | -0.169604385 | -0.710252781 | -0.567118939 | 0.319685057 | -1.790164133 | 0.015333631 | 0.066016579 | 0.193006634 | 0.114 | 0.261195165 |
| STnc2000_ | 1.218339794 | 1.291720609 | 1.137122414 | -0.192047891 | 0.210159516 | -0.904801266 | 0.152666692 | 0.220628389 | 0.40283521 | 0.337 | 0.499235172 |
| STnc2010_ | -0.976707512 | -1.082724221 | -0.886194869 | 0.204336828 | 0.285189237 | 0.671259066 | 0.221852113 | 0.335341396 | 0.498711492 | 0.4359 | 0.576416274 |
| STnc2030_ | -4.435098179 | -4.510027036 | -4.372241733 | 0.153607864 | 0.970206647 | 0.154203562 | 0.431045972 | 0.562364333 | 0.70339001 | 0.6736 | 0.77679696 |
| STnc2040_ | -6.176463551 | -4.722984266 | -8.46006181 | -3.733438148 | 2.478285939 | -1.431834259 | 0.025316634 | 0.105254842 | 0.250231616 | 0.1314 | 0.281956894 |
| STnc205_ | 4.775777624 | 5.957002726 | 3.535626054 | -2.420647036 | 0.061165884 | -39.59002533 | 4.68E-05 | 1.51E-05 | 0.003438448 | 0.1 | 0.243021319 |
| STnc2070_ | -0.025473918 | -0.21327739 | 0.144679255 | 0.377737992 | 0.251966901 | 1.460235495 | 0.041972124 | 0.099024231 | 0.243778729 | 0.1533 | 0.306900487 |
| STnc2080_ | 0.334682913 | -0.015187789 | 0.62294365 | 0.632428857 | 0.180342677 | 3.499583617 | 4.68E-05 | 0.013761385 | 0.075204856 | 0.1 | 0.243021319 |
| STnc2090_ | -0.272944209 | -0.011972749 | -0.598048728 | -0.624097239 | 0.460615675 | -1.336341985 | 0.065289873 | 0.105101627 | 0.262026971 | 0.1653 | 0.329134929 |
| STnc210_ | -6.130887003 | -5.957466065 | -6.305328372 | -0.231687673 | 3.626655824 | -0.053881885 | 0.460666668 | 0.669809032 | 0.794103253 | 0.8134 | 0.881602199 |
| STnc2100_ | -1.151853984 | -1.195966427 | -1.120606193 | 0.057542273 | 0.322882892 | 0.165873783 | 0.418387745 | 0.588951289 | 0.724164025 | 0.6987 | 0.797875114 |
| STnc2110_ | -1.229540921 | -0.978861561 | -1.543231634 | -0.626850243 | 0.525765453 | -1.17707602 | 0.082000051 | 0.145149438 | 0.314953749 | 0.2002 | 0.363699025 |
| STnc2130_ | -4.601913445 | -4.506821945 | -4.712493936 | -0.210621072 | 0.84559718 | -0.242211377 | 0.402398405 | 0.509820232 | 0.653356435 | 0.6104 | 0.724049668 |
| STnc2160_ | -5.273927484 | -5.306130976 | -5.247315562 | -0.490878084 | 2.956698057 | -0.163533748 | 0.439706864 | 0.685617918 | 0.802939438 | 0.854 | 0.907292728 |
| STnc2170_ | -1.985338439 | -2.166032684 | -1.838967668 | 0.319854083 | 0.430017539 | 0.720837534 | 0.210666683 | 0.321909133 | 0.487537165 | 0.4179 | 0.564654399 |

| | | | | | | | | | | | |
|---|---|---|---|---|---|---|---|---|---|---|---|
| STnc2175_ | -2.640509644 | -2.790224576 | -2.530633464 | 0.339796888 | 1.417145508 | 0.268987061 | 0.353097941 | 0.685579948 | 0.809256804 | 0.7189 | 0.820650127 |
| STnc2180_ | 3.855318096 | 4.119034789 | 3.443646231 | -0.650473305 | 0.14382596 | -4.492494119 | 4.68E-05 | 0.00381841 | 0.041609022 | 0.1 | 0.243021319 |
| STnc220_ | -2.958311403 | -3.676645065 | -2.400834673 | 1.194749629 | 0.773102654 | 1.47933037 | 0.046000096 | 0.094416246 | 0.240086032 | 0.1521 | 0.311211296 |
| STnc230_ | -7.819187095 | -8.633492261 | -7.1742063 | 1.305072305 | 3.614544462 | 0.300466866 | 0.333777489 | 0.490528241 | 0.644937235 | 0.602 | 0.720345091 |
| STnc240_ | 0.2489468 | -0.385403505 | 0.696520616 | 1.010450855 | 0.321407513 | 2.994344103 | 4.68E-05 | 0.017410102 | 0.090377995 | 0.1001 | 0.243161833 |
| STnc260_ | -6.274915822 | -6.074086455 | -6.575069029 | -0.488959719 | 1.753952978 | -0.240009479 | 0.386000004 | 0.507625074 | 0.656472187 | 0.6258 | 0.738522529 |
| STnc280_ | -5.440015525 | -5.741976711 | -5.252356414 | 0.553139303 | 1.352312289 | 0.34722388 | 0.333777489 | 0.466706835 | 0.624937737 | 0.5833 | 0.700619165 |
| STnc290_ | 0.417965658 | 0.449697281 | 0.383162591 | -0.071956326 | 0.183072286 | -0.377783624 | 0.343104603 | 0.500950497 | 0.647425705 | 0.6192 | 0.729710974 |
| STnc30_ | 1.092593543 | 0.878132736 | 1.25403121 | 0.451434289 | 0.300574915 | 1.527303837 | 0.016655836 | 0.077020685 | 0.21823416 | 0.1161 | 0.264941628 |
| STnc300_ | -6.883301577 | -7.019907805 | -6.799229819 | 0.150197563 | 3.178289852 | 0.039483543 | 0.469333334 | 0.619358689 | 0.754038823 | 0.7472 | 0.833699841 |
| STnc3000_ | -5.646157951 | -5.588659208 | -5.687266745 | -0.04575919 | 1.599622956 | -0.024922084 | 0.487674884 | 0.604334793 | 0.738009713 | 0.7226 | 0.815311458 |
| STnc3020_ | -4.304202712 | -4.951726068 | -3.69227545 | 1.306333575 | 1.07508988 | 1.136438835 | 0.093271197 | 0.152609584 | 0.312412675 | 0.2148 | 0.373689495 |
| STnc3030_ | -4.010478446 | -4.068738722 | -3.926883654 | 0.198577025 | 0.914048077 | 0.208163377 | 0.407728185 | 0.586473162 | 0.721625881 | 0.7 | 0.798215659 |
| STnc3040_ | -4.885966843 | -5.128134268 | -4.679127815 | 0.45552655 | 1.204730728 | 0.344429619 | 0.348666673 | 0.514322765 | 0.662546175 | 0.6323 | 0.744286908 |
| STnc3060_ | -2.957378534 | -3.179136084 | -2.758980331 | 0.43038791 | 0.526325003 | 0.780983638 | 0.189873436 | 0.287602767 | 0.449375155 | 0.3794 | 0.528188837 |
| STnc3070_ | -2.276747202 | -2.230425284 | -2.326718382 | -0.102484726 | 0.488708713 | -0.193111006 | 0.418387745 | 0.586915376 | 0.721033604 | 0.7081 | 0.802112213 |
| STnc3080_ | -3.089755967 | -3.68133972 | -2.937348293 | 0.656050136 | 0.901900997 | 0.631813194 | 0.29646903 | 0.446148734 | 0.619487033 | 0.6104 | 0.745084527 |
| STnc3090_ | -4.963232363 | -5.150153651 | -4.882290504 | 0.197894968 | 1.460998173 | 0.118891012 | 0.447333335 | 0.634868633 | 0.766310467 | 0.7652 | 0.851889755 |
| STnc310_ | -4.235139668 | -4.373226699 | -4.090697428 | 0.292996788 | 0.806381552 | 0.344655941 | 0.34733334 | 0.479938779 | 0.62535595 | 0.585 | 0.700931368 |
| STnc3100_ | 0.573853776 | 0.416215068 | 0.726547375 | 0.304574056 | 0.173222731 | 1.683850061 | 0.038000117 | 0.069608475 | 0.191348755 | 0.1373 | 0.288945474 |
| STnc3110_ | -0.587482225 | -0.554615742 | -0.61766004 | -0.067724859 | 0.272041569 | -0.238544847 | 0.396402402 | 0.580143812 | 0.718260415 | 0.6918 | 0.793492896 |
| STnc3120_ | -0.974111736 | -1.135947322 | -0.853405943 | 0.289944426 | 0.342264808 | 0.797049879 | 0.180666687 | 0.276766584 | 0.451958744 | 0.3781 | 0.532648967 |
| STnc3150_ | -0.023278986 | -0.07887838 | 0.045629328 | 0.007008258 | 0.379585419 | 0.023535712 | 0.490666667 | 0.767353548 | 0.860899744 | 0.895 | 0.935276882 |
| STnc3160_ | -3.328726943 | -3.289200516 | -3.3801074 | -0.073700287 | 0.859903363 | -0.0813459 | 0.471333334 | 0.680648378 | 0.798641201 | 0.7948 | 0.869680437 |
| STnc3170_ | -3.144748228 | -3.005574987 | -3.321779829 | -0.312423147 | 0.665116812 | -0.443499127 | 0.318454371 | 0.487254849 | 0.637975347 | 0.6045 | 0.721348139 |
| STnc3180_ | -4.301570883 | -4.442020481 | -4.135300328 | 0.330787131 | 0.812418945 | 0.384455064 | 0.333777489 | 0.468746162 | 0.615090948 | 0.5736 | 0.692144233 |
| STnc320_ | -2.38284303 | -2.411653307 | -2.356183732 | 0.06555557 | 0.442442558 | 0.14162017 | 0.441039309 | 0.558847707 | 0.697328087 | 0.6675 | 0.772232955 |
| STnc3210_ | -5.1610399 | -4.732166106 | -5.53954797 | -0.800017108 | 1.186530927 | -0.635731525 | 0.233333347 | 0.366761707 | 0.52230031 | 0.4603 | 0.596741331 |
| STnc3220_ | -0.365435008 | -0.39428742 | -0.32763944 | 0.072437618 | 0.339795483 | 0.200220075 | 0.422000003 | 0.677237543 | 0.795737738 | 0.7868 | 0.86407029 |
| STnc3230_ | -3.943569671 | -4.243227022 | -3.633102702 | 0.582288193 | 0.744279886 | 0.734937258 | 0.20333335 | 0.311796832 | 0.470810503 | 0.4051 | 0.548317932 |
| STnc3270_ | -3.143538436 | -3.194941101 | -3.076294947 | 0.111911924 | 0.649101853 | 0.154902221 | 0.427333336 | 0.5958279 | 0.729001294 | 0.7041 | 0.80130457 |
| STnc3280_ | -3.466775204 | -3.8074831 | -3.134429932 | 0.694798437 | 0.678174541 | 0.966500463 | 0.133244533 | 0.216768748 | 0.381278224 | 0.3027 | 0.45858761 |
| STnc3290_ | -2.094840549 | -2.573234341 | -1.716662862 | 0.85486992 | 0.388334243 | 2.130879403 | 0.008000574 | 0.044382518 | 0.145344264 | 0.1081 | 0.253438576 |
| STnc3300_ | -4.019165721 | -4.576695756 | -3.548087727 | 1.0300941 | 0.821257882 | 1.205340668 | 0.080612977 | 0.149933998 | 0.303362871 | 0.2151 | 0.370460816 |
| STnc3310_ | 3.095915768 | 3.099947101 | 3.091149761 | -0.000896044 | 0.088973525 | -0.010674856 | 0.493333334 | 0.662569378 | 0.784034751 | 0.7748 | 0.854268044 |
| STnc3330_ | -6.575899522 | -6.89664257 | -6.441004003 | 0.408096498 | 2.717479107 | 0.123493992 | 0.434666669 | 0.602506241 | 0.742531709 | 0.7588 | 0.846228035 |
| STnc3340_ | -2.712211068 | -2.546009682 | -2.936588253 | -0.413803869 | 0.540767282 | -0.71221578 | 0.206666683 | 0.329755984 | 0.494099147 | 0.4366 | 0.576378426 |
| STnc3350_ | -2.201287312 | -2.170289306 | -2.22063572 | -0.021411323 | 0.475102687 | -0.045116314 | 0.481333334 | 0.629895088 | 0.75690419 | 0.7285 | 0.821523614 |
| STnc340_ | -1.470444083 | -1.347054408 | -1.635889843 | -0.275820375 | 0.388823414 | -0.652648598 | 0.227181893 | 0.343720951 | 0.513758126 | 0.4485 | 0.593280738 |
| STnc3400_ | -4.341723258 | -4.443325745 | -4.239947577 | 0.1466725 | 1.108394355 | 0.121635719 | 0.435043307 | 0.637436395 | 0.763967251 | 0.7429 | 0.831187483 |
| STnc3410_ | 0.230650029 | 0.148399018 | 0.355863796 | 0.188147984 | 0.291449244 | 0.61442619 | 0.247168567 | 0.388113719 | 0.566571024 | 0.4899 | 0.643569454 |
| STnc3420_ | -2.985630568 | -3.175794786 | -2.765353486 | 0.460929073 | 0.631444414 | 0.664611409 | 0.236509007 | 0.343098366 | 0.50921287 | 0.4517 | 0.591502879 |
| STnc3430_ | -1.254960091 | -1.553154357 | -0.871839925 | 0.671658363 | 0.31774039 | 2.004723876 | 0.013333676 | 0.047314159 | 0.152545687 | 0.1187 | 0.266322348 |
| STnc3440_ | -2.567073553 | -2.692361802 | -2.375558821 | 0.302426474 | 0.65848761 | 0.435410318 | 0.324450374 | 0.535203062 | 0.683844238 | 0.6467 | 0.760405368 |
| STnc3460_ | 1.992567631 | 1.773544493 | 2.142432438 | 0.372017932 | 0.104705994 | 3.507753365 | 4.68E-05 | 0.014322045 | 0.076307973 | 0.1 | 0.243021319 |
| STnc3480_ | -1.248781577 | -0.480545065 | -2.198794665 | -1.748002585 | 0.385357013 | -4.480641052 | 4.68E-05 | 0.004826726 | 0.043859897 | 0.1 | 0.243021319 |
| STnc3490_ | -3.617822202 | -3.730578192 | -3.506435074 | 0.050013711 | 1.090083507 | 0.043732697 | 0.479333334 | 0.700896693 | 0.81535295 | 0.8341 | 0.895042699 |
| STnc350_ | -2.592795597 | -3.541683398 | -2.150236328 | 1.31369291 | 0.88237013 | 1.381545012 | 0.062624986 | 0.116296287 | 0.277611921 | 0.1643 | 0.326501304 |
| STnc3500_ | -4.224067224 | -4.224598052 | -4.222348467 | -0.024486441 | 0.844232863 | -0.026328448 | 0.491672219 | 0.567200079 | 0.706152406 | 0.6812 | 0.781579111 |
| STnc3510_ | -0.806897483 | -0.866714073 | -0.754178162 | 0.113034155 | 0.277954449 | 0.374825569 | 0.339107268 | 0.492996179 | 0.64227322 | 0.6014 | 0.717897521 |
| STnc3540_ | 2.420380867 | 2.717716305 | 2.025748295 | -0.689247081 | 0.127305618 | -5.39536266 | 4.68E-05 | 0.004688963 | 0.043397168 | 0.1 | 0.243021319 |
| STnc3550_ | -3.38562006 | -3.97790836 | -2.85635992 | 1.123046968 | 0.853599455 | 1.258626965 | 0.081279199 | 0.126065718 | 0.281608896 | 0.1977 | 0.361024445 |
| STnc3580_ | -4.675337885 | -4.504160112 | -4.797652033 | -0.24225744 | 1.248892137 | -0.179833696 | 0.419333336 | 0.627610834 | 0.755617814 | 0.7256 | 0.819312523 |

| | | | | | | | | | | | |
|---|---|---|---|---|---|---|---|---|---|---|---|
| STnc360_ | -5.444082474 | -5.740036447 | -5.06136321 | 0.694462873 | 1.281525196 | 0.512351518 | 0.284477025 | 0.439953437 | 0.591569639 | 0.5386 | 0.665355445 |
| STnc3600_ | -1.202719442 | -1.16089153 | -1.249386807 | -0.123720276 | 0.435062573 | -0.266402181 | 0.395069957 | 0.625622578 | 0.758220934 | 0.7483 | 0.834107779 |
| STnc3610_ | -0.110646328 | 0.079742702 | -0.28608581 | -0.413447073 | 0.290437425 | -1.418558873 | 0.036666788 | 0.105407709 | 0.256096278 | 0.146 | 0.298021002 |
| STnc3620_ | -3.787835509 | -3.901913829 | -3.697199329 | 0.250083012 | 0.839120432 | 0.267899059 | 0.370666672 | 0.544064588 | 0.688462625 | 0.661 | 0.765865473 |
| STnc3630_ | -3.422349295 | -3.364442703 | -3.494713877 | -0.15151868 | 0.632198131 | -0.218306084 | 0.40373085 | 0.528569307 | 0.672917826 | 0.6427 | 0.750712215 |
| STnc3640_ | -2.317616766 | -2.241370449 | -2.437624481 | -0.231270106 | 0.591501275 | -0.355923471 | 0.34133334 | 0.52026153 | 0.671877402 | 0.659 | 0.765097117 |
| STnc3680_ | 1.99222491 | 1.703027307 | 2.245591374 | 0.547762192 | 0.145320251 | 3.776534175 | 4.68E-05 | 0.007933082 | 0.058305847 | 0.1 | 0.243021319 |
| STnc370_ | -4.9699178 | -5.378909731 | -4.658626454 | 0.739652465 | 1.135856804 | 0.603616171 | 0.240000013 | 0.358317119 | 0.522923594 | 0.4648 | 0.601371014 |
| STnc3700_ | -2.743137504 | -2.910684938 | -2.622452539 | 0.274844952 | 0.555642339 | 0.451448496 | 0.304463699 | 0.468714189 | 0.621679838 | 0.5845 | 0.705492447 |
| STnc3720_ | -4.183026862 | -4.426697179 | -3.44383411 | 0.879325402 | 1.068665708 | 0.680944654 | 0.263824128 | 0.438224264 | 0.600379666 | 0.5338 | 0.678628712 |
| STnc3730_ | -1.04604504 | -1.020556853 | -1.071701756 | -0.051667311 | 0.290433121 | -0.176216313 | 0.434000002 | 0.590231581 | 0.723379036 | 0.6933 | 0.792036244 |
| STnc3750_ | -3.641778249 | -3.986261211 | -3.283643632 | 0.676319825 | 0.868010044 | 0.723595977 | 0.205862775 | 0.317058725 | 0.490931978 | 0.4164 | 0.569699045 |
| STnc3770_ | -2.939446414 | -3.126472775 | -2.803375153 | 0.357156387 | 0.55786053 | 0.605462214 | 0.244503677 | 0.374752568 | 0.536354531 | 0.4793 | 0.612448599 |
| STnc3790_ | -4.68821571 | -4.717253371 | -4.668856816 | -0.139862896 | 1.540130075 | -0.094462086 | 0.466355764 | 0.681322476 | 0.798659548 | 0.7794 | 0.860092029 |
| STnc380_ | -3.45880714 | -3.298418053 | -3.64026686 | -0.334984024 | 0.687806042 | -0.440412774 | 0.311125924 | 0.460873289 | 0.613968535 | 0.5681 | 0.691841467 |
| STnc3800_ | -0.695439821 | -1.027793154 | -0.372958843 | 0.666637019 | 0.281384867 | 2.359920919 | 0.003332494 | 0.03226118 | 0.123071778 | 0.1033 | 0.247273869 |
| STnc3830_ | -3.399157096 | -3.516318007 | -3.316768865 | 0.121199892 | 0.827709601 | 0.137119923 | 0.438000002 | 0.644271696 | 0.77075401 | 0.766 | 0.848144712 |
| STnc3850_ | -3.524617972 | -3.804211596 | -3.33620518 | 0.49058348 | 0.832929278 | 0.535653946 | 0.255333345 | 0.395037399 | 0.567490995 | 0.5177 | 0.65012599 |
| STnc3860_ | -3.5347729 | -3.589674062 | -3.483744724 | 0.054794309 | 1.042313503 | 0.046421076 | 0.476666668 | 0.699237886 | 0.812913735 | 0.8164 | 0.883706021 |
| STnc3870_ | -2.233741958 | -2.279831348 | -2.185584567 | 0.081407923 | 0.442786312 | 0.164601225 | 0.426382415 | 0.571002067 | 0.70707995 | 0.6815 | 0.782036646 |
| STnc3880_ | -5.810413957 | -5.829522499 | -5.77436421 | 0.065798416 | 1.610647449 | 0.041104398 | 0.483333334 | 0.603801439 | 0.736276081 | 0.7131 | 0.807514216 |
| STnc3890_ | -3.57805138 | -3.663195532 | -3.510974845 | 0.182680611 | 0.761780641 | 0.221865082 | 0.392405067 | 0.556411513 | 0.698610972 | 0.6735 | 0.774922106 |
| STnc390_ | 1.459669298 | 1.394078625 | 1.53975197 | 0.178442595 | 0.240553246 | 0.790088413 | 0.152000025 | 0.264322108 | 0.459694117 | 0.3378 | 0.498483445 |
| STnc3920_ | -5.955120399 | -6.059544325 | -5.769766775 | 0.116075738 | 2.139596243 | 0.036683213 | 0.473684212 | 0.630556086 | 0.763354658 | 0.757 | 0.845321864 |
| STnc3950_ | -1.422516892 | -1.303358351 | -1.634673398 | -0.326346932 | 0.44136334 | -0.706791212 | 0.221852113 | 0.347912104 | 0.526678239 | 0.46 | 0.612447435 |
| STnc3960_ | -0.521092498 | -0.496571947 | -0.552771791 | -0.063232673 | 0.242281675 | -0.23861149 | 0.395333337 | 0.54141831 | 0.682409382 | 0.653 | 0.760035008 |
| STnc3970_ | -4.421006344 | -4.869166499 | -4.037783885 | 0.8269524 | 0.968672765 | 0.809057868 | 0.178547655 | 0.28339626 | 0.451421937 | 0.3715 | 0.524483631 |
| STnc3990_ | -0.757572776 | -0.692638388 | -0.795281342 | -0.025440892 | 0.436549621 | -0.062778928 | 0.477333334 | 0.742200971 | 0.84526536 | 0.8781 | 0.926398372 |
| STnc40_ | 1.018100262 | 0.933504957 | 1.121844963 | 0.204820867 | 0.161309183 | 1.18230023 | 0.092666712 | 0.149035214 | 0.308720302 | 0.2182 | 0.376896156 |
| STnc4000_ | -2.631609321 | -2.812202036 | -2.464233422 | 0.357750707 | 0.475587735 | 0.723014897 | 0.208000016 | 0.311135199 | 0.473307786 | 0.4165 | 0.557021881 |
| STnc4010_ | -6.268839977 | -5.788325078 | -6.847006188 | -0.673296935 | 3.750331111 | -0.168629876 | 0.40106596 | 0.652026592 | 0.780308414 | 0.782 | 0.861433124 |
| STnc4020_ | -4.941330693 | -4.929716073 | -4.95732147 | 0.143435439 | 2.359352961 | 0.056095464 | 0.470353099 | 0.719599849 | 0.828938238 | 0.8302 | 0.893902214 |
| STnc4040_ | -1.2883862 | -1.409280659 | -1.175763259 | 0.238206122 | 0.296263935 | 0.744868304 | 0.209860109 | 0.311920156 | 0.473992173 | 0.4086 | 0.551895003 |
| STnc4070_ | -3.330536931 | -3.408500505 | -3.184211534 | 0.228824688 | 0.730205922 | 0.285695265 | 0.378000005 | 0.583410615 | 0.717307411 | 0.6904 | 0.791249695 |
| STnc4080_ | -2.501214617 | -2.645294616 | -2.199735435 | 0.368092928 | 0.621708241 | 0.531145795 | 0.302465032 | 0.499192757 | 0.654340161 | 0.5965 | 0.727280252 |
| STnc410_ | -4.515596647 | -4.644517472 | -4.366661147 | 0.27355761 | 0.917932805 | 0.267249606 | 0.386409065 | 0.536040146 | 0.676281457 | 0.6268 | 0.739181613 |
| STnc4100_ | -3.110116994 | -2.512572914 | -3.761301554 | -1.383944307 | 0.81408638 | -1.756105116 | 0.013333676 | 0.066656706 | 0.194528996 | 0.1148 | 0.262368378 |
| STnc4110_ | -3.846800971 | -3.922096672 | -3.762381086 | 0.154654561 | 0.867273599 | 0.161198829 | 0.425716192 | 0.592798535 | 0.727343506 | 0.6933 | 0.792472797 |
| STnc4120_ | -1.804553687 | -1.956758095 | -1.67072621 | 0.284686354 | 0.361258601 | 0.748561549 | 0.206666683 | 0.307360279 | 0.469872915 | 0.4063 | 0.550483127 |
| STnc4130_ | -2.872414955 | -2.816439514 | -2.976092693 | -0.169514172 | 0.65085704 | -0.239630453 | 0.389073955 | 0.579723777 | 0.721778224 | 0.7043 | 0.804614975 |
| STnc4140_ | -1.636678031 | -1.737426256 | -1.543209788 | 0.220864889 | 0.439550405 | 0.472476474 | 0.298467697 | 0.452456154 | 0.612719908 | 0.588 | 0.706517934 |
| STnc4150_ | -7.03177535 | -6.855677296 | -7.258306548 | -0.44492786 | 2.44037713 | -0.143446702 | 0.423333336 | 0.545593136 | 0.693374905 | 0.6776 | 0.77947499 |
| STnc4160_ | -2.68750794 | -2.608374369 | -2.788408174 | -0.175662006 | 0.508604444 | -0.32626392 | 0.348666673 | 0.502455077 | 0.648782533 | 0.5974 | 0.714946838 |
| STnc4170_ | -3.498809207 | -3.652826485 | -3.304861879 | 0.33570677 | 0.75249583 | 0.407773716 | 0.327781486 | 0.506649089 | 0.652245384 | 0.6098 | 0.726383397 |
| STnc4190_ | -4.747972708 | -4.858804835 | -4.593578858 | 0.254058135 | 1.003077001 | 0.244812525 | 0.398666671 | 0.540403991 | 0.680912372 | 0.6402 | 0.750783453 |
| STnc420_ | -2.004871439 | -1.978750716 | -2.02816774 | -0.039560801 | 0.384189237 | -0.105098668 | 0.457333335 | 0.562086819 | 0.701982809 | 0.6737 | 0.776672089 |
| STnc4200_ | -2.196539995 | -2.342108961 | -2.045585616 | 0.282336845 | 0.406291692 | 0.661095441 | 0.23466668 | 0.355051278 | 0.511397657 | 0.4547 | 0.590904713 |
| STnc4210_ | -3.133413628 | -2.303895309 | -4.589143705 | -2.292145158 | 0.803842451 | -2.821461158 | 4.68E-05 | 0.030999026 | 0.120057813 | 0.1 | 0.243021319 |
| STnc4220_ | 1.129289349 | 0.738849739 | 1.309661112 | 0.551666957 | 0.198768027 | 2.490434586 | 0.000673369 | 0.039239273 | 0.144131011 | 0.1012 | 0.244582173 |
| STnc4230_ | -0.553230883 | -0.677446923 | -0.487892231 | 0.125225345 | 0.431299935 | 0.264471235 | 0.399733515 | 0.723074275 | 0.831110366 | 0.7914 | 0.875403757 |
| STnc4240_ | 1.553284807 | 1.327953944 | 1.821613601 | 0.485153291 | 0.13052943 | 3.658404585 | 4.68E-05 | 0.010518106 | 0.066274005 | 0.1 | 0.243021319 |
| STnc4250_ | 4.451412394 | 4.295083537 | 4.628741107 | 0.293776768 | 0.11142205 | 2.72960351 | 4.68E-05 | 0.014286528 | 0.083372476 | 0.1001 | 0.243157978 |
| STnc4260_ | -4.258051926 | -4.065044728 | -4.516738641 | -0.4525726 | 1.261081836 | -0.329529802 | 0.342438381 | 0.532976678 | 0.687431059 | 0.664 | 0.773905026 |

| | | | | | | | | | | | |
|---|---|---|---|---|---|---|---|---|---|---|---|
| STnc430_ | -4.795059366 | -4.715144399 | -4.889930088 | -0.211169876 | 0.917536262 | -0.201558047 | 0.416389077 | 0.500146477 | 0.645049731 | 0.6185 | 0.727855976 |
| STnc440_ | -2.767816286 | -2.742568109 | -2.792668262 | -0.051741131 | 0.580707896 | -0.085221451 | 0.466000001 | 0.611237105 | 0.742940793 | 0.7201 | 0.81358258 |
| STnc450_ | 0.013353744 | -0.054416007 | 0.092022677 | 0.156238838 | 0.246331561 | 0.61515924 | 0.257333345 | 0.407108098 | 0.57494864 | 0.5281 | 0.661969451 |
| STnc460_ | -6.716677199 | -6.634753684 | -6.855181327 | -0.246405668 | 2.79164991 | -0.071707736 | 0.462358429 | 0.615338237 | 0.752132068 | 0.7724 | 0.853530893 |
| STnc470_ | -2.428285086 | -2.400578494 | -2.456638446 | -0.056525831 | 0.391680157 | -0.137172359 | 0.430666669 | 0.519321593 | 0.66366294 | 0.6281 | 0.739692949 |
| STnc475_ | 5.005459118 | 5.019363238 | 4.989478091 | -0.043248408 | 0.081863767 | -0.481908302 | 0.248666679 | 0.374760244 | 0.575687852 | 0.5138 | 0.656530006 |
| STnc480_ | -3.63513735 | -3.60143347 | -3.656111208 | -0.057908105 | 0.677735383 | -0.087972859 | 0.463024652 | 0.561024917 | 0.699348918 | 0.6763 | 0.776538625 |
| STnc50_ | -4.786743799 | -5.100789639 | -4.519105467 | 0.503642035 | 1.511730099 | 0.29027734 | 0.358000006 | 0.561031629 | 0.706404796 | 0.6719 | 0.778647606 |
| STnc500_ | -4.163146518 | -4.2834694 | -4.064976913 | 0.190363387 | 0.983008885 | 0.174593493 | 0.415333337 | 0.598691249 | 0.735210074 | 0.7197 | 0.81236518 |
| STnc510_ | -2.381518518 | -2.385983411 | -2.374603042 | -0.045487684 | 0.597083276 | -0.070643341 | 0.474666668 | 0.663971414 | 0.783924005 | 0.7767 | 0.854476653 |
| STnc520_ | -1.579644539 | -1.74018378 | -1.41884943 | 0.319914071 | 0.330222446 | 0.905619551 | 0.155333358 | 0.24623546 | 0.406328455 | 0.3428 | 0.493170525 |
| STnc530_ | -5.144401406 | -5.104250591 | -5.203963215 | -0.0676282 | 1.301357998 | -0.047644538 | 0.480346436 | 0.602887926 | 0.7382033 | 0.7307 | 0.822356904 |
| STnc540_ | 4.012526599 | 4.028319977 | 3.856584134 | -0.163766045 | 0.084340875 | -0.982542096 | 0.223184558 | 0.241422532 | 0.439427415 | 0.4725 | 0.630781231 |
| STnc590_ | -1.57501062 | -1.298457937 | -2.236728255 | -0.918125592 | 0.503921846 | -1.661053566 | 0.027315288 | 0.087164944 | 0.229964128 | 0.1288 | 0.280151608 |
| STnc60_ | -7.223823873 | -6.830159932 | -7.541030675 | -0.546786281 | 3.081137312 | -0.13414458 | 0.41866667 | 0.594905321 | 0.728509075 | 0.6854 | 0.786113019 |
| STnc600_ | 0.995777 | 1.651475766 | -0.151716553 | -1.816017867 | 0.296942055 | -5.934040542 | 4.68E-05 | 0.004611735 | 0.043598535 | 0.1 | 0.243021319 |
| STnc610_ | 3.356275574 | 3.349864954 | 3.361910332 | 0.014790438 | 0.227301357 | 0.070123338 | 0.444370422 | 0.845465442 | 0.913397371 | 0.7819 | 0.864505874 |
| STnc620_ | 2.25574128 | 2.279566144 | 2.220704477 | -0.014973623 | 0.204141958 | -0.061574686 | 0.472666668 | 0.770299132 | 0.864676325 | 0.8971 | 0.94010022 |
| STnc640_ | -2.162923031 | -2.084583705 | -2.339376746 | -0.231431955 | 0.555860081 | -0.355056027 | 0.360426388 | 0.57074094 | 0.715806063 | 0.6935 | 0.800102413 |
| STnc650_ | 4.859098661 | 4.788109316 | 5.016539122 | 0.198878714 | 0.303025572 | 0.69237091 | 0.223850781 | 0.368410135 | 0.571276032 | 0.4024 | 0.600327269 |
| STnc660_ | 1.736261723 | 1.835541733 | 1.617977834 | -0.200889141 | 0.148208843 | -1.294652143 | 0.081333385 | 0.123669592 | 0.279302357 | 0.1967 | 0.361318436 |
| STnc670_ | 0.361916086 | 0.858549207 | -0.269924852 | -1.13563339 | 0.263961738 | -4.340655035 | 4.68E-05 | 0.015686101 | 0.084700795 | 0.1 | 0.243021319 |
| STnc680_ | 2.194332259 | 2.134789032 | 2.298906542 | 0.161356922 | 0.176735377 | 0.846257475 | 0.17721521 | 0.258006949 | 0.45061127 | 0.3497 | 0.521557482 |
| STnc690_ | 1.22170049 | 0.641535309 | 1.573905791 | 0.917875133 | 0.176539068 | 4.965760743 | 4.68E-05 | 0.008036467 | 0.057771844 | 0.1 | 0.243021319 |
| STnc695_ | -2.017552408 | -2.275667354 | -1.808526834 | 0.480957599 | 0.418083707 | 1.086253144 | 0.109926752 | 0.181965648 | 0.342492946 | 0.2579 | 0.415587189 |
| STnc70_ | 2.576675076 | 2.409302699 | 2.782492458 | 0.311982735 | 0.173633587 | 1.850831157 | 0.026649069 | 0.049769401 | 0.167539809 | 0.1243 | 0.276937819 |
| STnc720_ | -2.527044061 | -2.710119108 | -2.385314147 | 0.34103185 | 0.472254805 | 0.659354283 | 0.23184545 | 0.350895574 | 0.512189142 | 0.4488 | 0.588481292 |
| STnc730_ | 2.042578225 | 2.02275992 | 2.075293735 | 0.046106873 | 0.351576497 | 0.124010784 | 0.448367757 | 0.826037917 | 0.902068549 | 0.9244 | 0.957043985 |
| STnc740_ | -6.461426048 | -6.124916109 | -6.766187668 | -0.599021925 | 1.943597574 | -0.274969181 | 0.38241173 | 0.512611552 | 0.659845373 | 0.6189 | 0.733540073 |
| STnc750_ | 0.589938925 | 0.50920884 | 0.880119693 | 0.358810835 | 0.337570737 | 0.930110776 | 0.187874769 | 0.249283218 | 0.444945861 | 0.3643 | 0.53765875 |
| STnc760_ | -1.286045715 | -1.400132388 | -1.246257646 | 0.135423609 | 0.401093725 | 0.299205284 | 0.373084615 | 0.587478573 | 0.727592225 | 0.7015 | 0.805626218 |
| STnc770_ | 2.272450139 | 2.303338817 | 2.239911327 | -0.065957785 | 0.28441032 | -0.210025991 | 0.410000003 | 0.629879607 | 0.773650705 | 0.8657 | 0.918594335 |
| STnc780_ | 1.417658133 | 1.421103403 | 1.411922336 | 0.009509323 | 0.170156643 | 0.05246444 | 0.485009994 | 0.683137798 | 0.801231149 | 0.7903 | 0.86713124 |
| STnc790_ | -1.05825617 | -1.063715702 | -1.04443014 | -0.010562564 | 0.387223748 | -0.028517883 | 0.488341106 | 0.674534911 | 0.794305285 | 0.7756 | 0.859169909 |
| STnc800_ | 3.716194447 | 3.447635761 | 4.007250295 | 0.572611839 | 0.103487485 | 5.613057622 | 4.68E-05 | 0.003459814 | 0.038583905 | 0.1 | 0.243021319 |
| STnc810_ | -4.650272546 | -4.935614647 | -4.477464921 | 0.401919219 | 1.222812925 | 0.322149595 | 0.364000005 | 0.535932395 | 0.685118024 | 0.662 | 0.770121556 |
| STnc835_ | 3.24298529 | 3.273658102 | 3.205986263 | -0.023974362 | 0.17648406 | -0.129346102 | 0.441333336 | 0.837570367 | 0.90811373 | 0.8845 | 0.933838199 |
| STnc840_ | 1.959239357 | 2.558733575 | 1.272219215 | -1.277608432 | 0.169340343 | -7.611889864 | 4.68E-05 | 0.000873577 | 0.019827995 | 0.1 | 0.243021319 |
| STnc850_ | -3.296187617 | -3.306012476 | -3.280067287 | 0.003929304 | 0.697028622 | 0.004442087 | 0.498666667 | 0.626004731 | 0.754925022 | 0.7385 | 0.827690892 |
| STnc870_ | -0.821921242 | -0.684667669 | -0.921828547 | -0.2663424 | 0.34531431 | -0.746273608 | 0.19920055 | 0.308845724 | 0.485257824 | 0.4101 | 0.557676405 |
| STnc880_ | -1.206041258 | -1.344872693 | -0.97993259 | 0.352532815 | 0.38083393 | 0.875648086 | 0.177333354 | 0.267424365 | 0.443334895 | 0.3572 | 0.520259217 |
| STnc890_ | -0.14826211 | -0.186479157 | -0.05466532 | 0.131003939 | 0.316589388 | 0.366920797 | 0.363091278 | 0.601476395 | 0.740358906 | 0.7196 | 0.820141204 |
| STnc920_ | 1.080411773 | 0.879438123 | 1.279400318 | 0.401978934 | 0.158749273 | 2.493441996 | 0.003332494 | 0.028039565 | 0.114471061 | 0.1029 | 0.246870827 |
| STnc925_ | -0.362473976 | -0.278033017 | -0.425615781 | -0.128969643 | 0.2346251 | -0.500727413 | 0.28580947 | 0.453138153 | 0.608376163 | 0.5549 | 0.683001597 |
| STnc930_ | -5.235114723 | -4.944324562 | -5.809898063 | -0.942145909 | 1.909188384 | -0.414546485 | 0.239333347 | 0.379337244 | 0.566365134 | 0.4939 | 0.629154043 |
| STnc940_ | -4.194772529 | -3.965928908 | -4.466728513 | -0.444583576 | 1.11671108 | -0.374487558 | 0.338441046 | 0.528222185 | 0.678000449 | 0.6377 | 0.752250043 |
| STnc950_ | -4.141397198 | -3.851684635 | -4.364030767 | -0.466002693 | 0.898034383 | -0.486667239 | 0.290473027 | 0.466784947 | 0.615450182 | 0.5655 | 0.688013176 |
| STnc970_ | 0.456234322 | 0.591733404 | 0.033538246 | -0.544087629 | 0.303261578 | -1.557993814 | 0.034666795 | 0.109168127 | 0.26901545 | 0.1379 | 0.291484801 |
| StyR.103a_ | -8.937118587 | -8.420785839 | -9.42601503 | -0.885326306 | 3.973763447 | -0.193218493 | 0.394666671 | 0.507618099 | 0.655308807 | 0.6199 | 0.731477791 |
| StyR.103b_ | -8.928472666 | -9.569356085 | -8.102485612 | 1.465280547 | 4.139600999 | 0.309367604 | 0.348000006 | 0.505183003 | 0.651055552 | 0.6178 | 0.729432563 |
| StyR.103c_ | -7.381441897 | -7.119435899 | -7.997062996 | -0.718577822 | 3.235907609 | -0.19004352 | 0.41666667 | 0.536875276 | 0.690313607 | 0.6974 | 0.802031229 |
| StyR.103e_ | -8.931174027 | -9.523627098 | -8.116689678 | 1.458158821 | 4.211170888 | 0.287549147 | 0.349333339 | 0.50133197 | 0.648693213 | 0.612 | 0.724286283 |
| StyR.103g_ | -8.486859301 | -8.552332262 | -8.415299855 | 0.129629772 | 3.858294906 | 0.032101939 | 0.481333334 | 0.556980152 | 0.698546778 | 0.6858 | 0.784013079 |

| | | | | | | | | | | | |
|---|---|---|---|---|---|---|---|---|---|---|---|
| StyR.122_ | -4.772899912 | -5.044654758 | -4.188057709 | 0.750542915 | 1.268251577 | 0.51429413 | 0.294000009 | 0.499556113 | 0.647961971 | 0.5825 | 0.71364999 |
| StyR.126_ | -1.372886051 | -1.281913067 | -1.443176554 | -0.220457245 | 0.824547519 | -0.260745153 | 0.358000006 | 0.600584703 | 0.748464996 | 0.7111 | 0.815441529 |
| StyR.128_ | -4.267249106 | -4.039648517 | -4.552250205 | -0.551457958 | 0.967512196 | -0.524752429 | 0.27600001 | 0.41455199 | 0.575676642 | 0.5285 | 0.656686447 |
| StyR.134_ | 0.551308446 | 0.530024535 | 0.603008142 | 0.081703434 | 0.233647209 | 0.332054247 | 0.375333338 | 0.593173069 | 0.730536505 | 0.7144 | 0.810729326 |
| StyR.150_ | -2.273253082 | -2.238151467 | -2.348820308 | -0.062840621 | 0.510361589 | -0.113522591 | 0.455696204 | 0.655346705 | 0.777821122 | 0.7551 | 0.844287731 |
| StyR.169_ | -0.623563489 | -0.944338959 | -0.29843477 | 0.634954124 | 0.256814156 | 2.421063796 | 0.005996769 | 0.02958766 | 0.116420057 | 0.1051 | 0.249665506 |
| StyR.172_ | -8.925472176 | -8.406499839 | -9.445777122 | -1.04448274 | 4.006124075 | -0.22666274 | 0.381745508 | 0.502332793 | 0.651089922 | 0.6102 | 0.723099308 |
| StyR.178_ | -8.486504484 | -8.581600744 | -8.386093039 | 0.06058065 | 3.917704918 | 0.012283909 | 0.490339774 | 0.565451458 | 0.707806546 | 0.6966 | 0.793934462 |
| StyR.195_ | -3.157819573 | -3.050386715 | -3.371462783 | -0.255358184 | 0.773674333 | -0.307822268 | 0.384000005 | 0.614394092 | 0.749475921 | 0.7261 | 0.82646741 |
| StyR.196_ | -6.578824828 | -6.623992066 | -6.538562868 | 0.086846732 | 2.061794579 | 0.039771706 | 0.480666667 | 0.573613321 | 0.712432992 | 0.6919 | 0.789916329 |
| StyR.199_ | -3.214219621 | -3.344823576 | -3.037893085 | 0.316279774 | 0.615747499 | 0.48206182 | 0.295136585 | 0.431312223 | 0.588085907 | 0.5542 | 0.677028861 |
| StyR.215a_ | 4.601342254 | 4.415316462 | 4.719488067 | 0.491103235 | 0.29732678 | 1.790806468 | 4.68E-05 | 0.042188263 | 0.157088415 | 0.1 | 0.243021319 |
| StyR.215b_ | 4.115958561 | 3.769861062 | 4.343650052 | 0.610815957 | 0.207098153 | 3.120763106 | 4.68E-05 | 0.013587662 | 0.082804764 | 0.1 | 0.243021319 |
| StyR.215c_ | 1.400790099 | 1.122140646 | 1.647243565 | 0.564124559 | 0.251559599 | 2.273478628 | 0.001336769 | 0.028107746 | 0.12045546 | 0.1007 | 0.243991182 |
| StyR.215d_ | 2.116992846 | 1.967580273 | 2.210325425 | 0.329517018 | 0.282458028 | 1.274888383 | 0.030646382 | 0.113903353 | 0.279652222 | 0.1346 | 0.28827187 |
| StyR.215e_ | 1.076357912 | 0.953233133 | 1.331119376 | 0.649441946 | 0.569710157 | 1.169485807 | 0.061292542 | 0.128244998 | 0.302530312 | 0.1595 | 0.319804487 |
| StyR.215f_ | 3.074026629 | 2.767027282 | 3.347950654 | 0.544186454 | 0.123570595 | 4.413068949 | 4.68E-05 | 0.004110538 | 0.042555106 | 0.1 | 0.243021319 |
| StyR.215g_ | 1.061196224 | 0.578335634 | 1.21509497 | 0.591825099 | 0.279890052 | 1.636247096 | 0.030666813 | 0.091096569 | 0.243388071 | 0.1328 | 0.28654792 |
| StyR.219_ | -3.976563367 | -3.994669367 | -3.951466714 | 0.003751806 | 0.809863488 | 0.006132508 | 0.496335776 | 0.595494153 | 0.731510131 | 0.7101 | 0.806733664 |
| StyR.222_ | -6.937782532 | -6.803988136 | -7.10651508 | -0.372190065 | 2.477365496 | -0.111310907 | 0.434666669 | 0.550869765 | 0.697859492 | 0.6802 | 0.780417791 |
| StyR.234_ | -3.97546796 | -3.88060452 | -4.037726175 | -0.103088604 | 0.922561792 | -0.110063681 | 0.449700202 | 0.648413358 | 0.771303178 | 0.7393 | 0.830349301 |
| StyR.243_ | -0.300450854 | -0.036479618 | -0.661941846 | -0.621288817 | 0.210329811 | -2.899031232 | 0.001335878 | 0.021401116 | 0.094641455 | 0.1007 | 0.244002225 |
| StyR.248_ | -3.079915274 | -3.185883827 | -2.998195204 | 0.173119759 | 0.593076637 | 0.26956039 | 0.384666671 | 0.532579558 | 0.677829056 | 0.6435 | 0.755024786 |
| StyR.24a_ | -4.947988796 | -4.338963561 | -5.673903591 | -1.32816692 | 1.009421259 | -1.233673682 | 0.079280533 | 0.141328715 | 0.28481472 | 0.2133 | 0.367794944 |
| StyR.24b_ | -2.729831258 | -1.388052186 | -5.685459188 | -4.30429902 | 1.13898653 | -3.906147904 | 4.68E-05 | 0.026562243 | 0.109462139 | 0.1 | 0.243021319 |
| StyR.24c_ | -3.377242674 | -3.391545904 | -3.365908183 | -0.005868978 | 0.744661749 | -0.009513207 | 0.497001999 | 0.626808699 | 0.756108613 | 0.7442 | 0.833155229 |
| StyR.24d_ | -4.561132139 | -5.488152024 | -3.965636953 | 1.533631624 | 1.019851632 | 1.445225977 | 0.043970787 | 0.105683697 | 0.244378807 | 0.155 | 0.305794001 |
| StyR.24e_ | -5.472872883 | -4.674560516 | -6.484489222 | -1.829662505 | 1.357480416 | -1.30551731 | 0.066000065 | 0.131979664 | 0.276919188 | 0.1901 | 0.34345789 |
| StyR.24f_ | -2.585619686 | -2.281473981 | -3.105126495 | -0.790856728 | 0.791948656 | -0.973723288 | 0.151333358 | 0.216024834 | 0.399977919 | 0.3005 | 0.475322591 |
| StyR.24g_ | -3.656032556 | -2.557662933 | -4.974307795 | -2.39862752 | 0.730863298 | -3.195082275 | 0.000673369 | 0.015094138 | 0.07762492 | 0.1003 | 0.243370522 |
| StyR.250_ | -3.734924334 | -3.976847115 | -3.38512089 | 0.554556819 | 0.719551918 | 0.720547776 | 0.227181893 | 0.347547076 | 0.507491255 | 0.4393 | 0.584497777 |
| StyR.252_ | -1.19165534 | -0.982644329 | -1.355672304 | -0.354561548 | 0.369122979 | -0.916367993 | 0.161333357 | 0.247824884 | 0.420802397 | 0.3375 | 0.499469692 |
| StyR.254_ | -6.42834008 | -5.763806517 | -7.213651178 | -1.417688042 | 2.474689358 | -0.49100086 | 0.244503677 | 0.395437733 | 0.565030523 | 0.4841 | 0.625976357 |
| StyR.262_ | -1.199785992 | -1.232177647 | -1.156980227 | 0.087356445 | 0.318100359 | 0.252601753 | 0.399333337 | 0.570449809 | 0.707566148 | 0.6841 | 0.78307176 |
| StyR.264_ | -1.493935202 | -1.784182951 | -1.070240941 | 0.743644967 | 0.473486522 | 1.499943393 | 0.050000088 | 0.094958423 | 0.240528328 | 0.1524 | 0.309275692 |
| StyR.27_ | -4.23505015 | -4.206476318 | -4.270204843 | -0.094482584 | 0.975780385 | -0.080959195 | 0.461333335 | 0.592999947 | 0.728206498 | 0.7088 | 0.805442935 |
| StyR.277_ | -0.122166408 | -0.191086898 | -0.054311132 | 0.190106028 | 0.352069687 | 0.551588434 | 0.273817466 | 0.407287482 | 0.585403024 | 0.5636 | 0.693291242 |
| StyR.281_ | -4.821811174 | -4.538356361 | -5.297844745 | -0.828407134 | 1.424690838 | -0.523879469 | 0.257828126 | 0.375569653 | 0.552676066 | 0.4966 | 0.631731042 |
| StyR.285_ | -3.845175516 | -4.28737546 | -3.46550561 | 0.814920519 | 0.743014031 | 1.028204753 | 0.119333367 | 0.195106344 | 0.35095743 | 0.2716 | 0.425791571 |
| StyR.287_ | -1.696703939 | -1.616635453 | -1.783097048 | -0.166458242 | 0.395110803 | -0.389911205 | 0.324450374 | 0.49024793 | 0.641852023 | 0.6062 | 0.721641717 |
| StyR.29_ | 5.670434233 | 5.659330294 | 5.938808347 | 0.276807065 | 0.123785994 | 0.88572592 | 0.247168567 | 0.257337982 | 0.463111437 | 0.4696 | 0.652022427 |
| StyR.291_ | -1.878177528 | -1.63117535 | -2.46942628 | -0.827335155 | 0.530440033 | -1.406443096 | 0.045333431 | 0.120639914 | 0.279308953 | 0.1611 | 0.315862375 |
| StyR.292_ | -5.485900262 | -5.783510831 | -5.263706349 | 0.529785449 | 1.224426065 | 0.392079674 | 0.315333341 | 0.440440118 | 0.592646378 | 0.5507 | 0.672405921 |
| StyR.293_ | -3.802382672 | -3.803353777 | -3.802001933 | -0.008694731 | 0.665528953 | -0.011164777 | 0.494666667 | 0.537666967 | 0.678878523 | 0.6441 | 0.752008812 |
| StyR.295_ | -3.789217981 | -4.009381336 | -3.574070289 | 0.434174705 | 0.674014114 | 0.617607201 | 0.239173897 | 0.368167457 | 0.522770911 | 0.4622 | 0.597437924 |
| StyR.296_ | 3.753545826 | 3.705651621 | 3.796506672 | 0.091139391 | 0.476805208 | 0.20391902 | 0.451333335 | 0.759922232 | 0.864602662 | 0.9943 | 0.996777895 |
| StyR.3_ | -1.092434404 | -0.861288071 | -1.25840621 | -0.386063318 | 0.316083194 | -1.154112397 | 0.108000038 | 0.159317166 | 0.316125353 | 0.2382 | 0.398986283 |
| StyR.302_ | -7.064849501 | -7.098110425 | -7.010260803 | 0.081937482 | 3.236260289 | 0.01629843 | 0.486000001 | 0.616010502 | 0.752188871 | 0.7617 | 0.845953077 |
| StyR.305_ | -1.037292468 | -0.292261718 | -1.887203065 | -1.594256964 | 0.284190631 | -5.578246205 | 4.68E-05 | 0.004018854 | 0.037661378 | 0.1 | 0.243021319 |
| StyR.306_ | 0.898320845 | 0.775018503 | 1.10788178 | 0.305028993 | 0.377754113 | 0.684104781 | 0.245333346 | 0.351411923 | 0.547911965 | 0.4677 | 0.629285437 |
| StyR.315_ | -7.216722008 | -6.835536217 | -7.620802242 | -0.725031569 | 2.980699101 | -0.198882433 | 0.384666671 | 0.571383009 | 0.712493795 | 0.6759 | 0.780227501 |
| StyR.322_ | -3.909504112 | -4.398210123 | -3.473552449 | 0.909933247 | 0.719172967 | 1.204528322 | 0.088666714 | 0.14393836 | 0.289651691 | 0.2188 | 0.373105186 |
| StyR.327_ | -4.548044797 | -4.802124983 | -4.284187331 | 0.424254134 | 1.218263239 | 0.329973629 | 0.357761498 | 0.57015742 | 0.711199036 | 0.6722 | 0.778462743 |

| | | | | | | | | | | | |
|---|---|---|---|---|---|---|---|---|---|---|---|
| StyR.328_ | -6.610074805 | -7.122222897 | -6.16124207 | 0.618961345 | 3.40805118 | 0.158601294 | 0.401732182 | 0.61287764 | 0.748592593 | 0.7222 | 0.817379965 |
| StyR.329a_ | -0.446526965 | -0.597175183 | -0.275502399 | 0.323209713 | 0.21983089 | 1.417178805 | 0.048666757 | 0.108292898 | 0.24708404 | 0.1726 | 0.32641146 |
| StyR.329b_ | -4.010358657 | -4.044675067 | -3.967495847 | -0.008591234 | 1.2201494 | -0.00616575 | 0.496335776 | 0.695835792 | 0.811648513 | 0.827 | 0.891822522 |
| StyR.336_ | -3.864444395 | -3.686847294 | -4.265640438 | -0.52768327 | 0.933866966 | -0.507809747 | 0.305333342 | 0.45981597 | 0.62191885 | 0.5844 | 0.713002081 |
| StyR.341_ | -4.390565502 | -4.18199077 | -4.62180155 | -0.572583735 | 1.252124556 | -0.411903898 | 0.304666675 | 0.438548109 | 0.608035594 | 0.5866 | 0.706166 |
| StyR.354_ | -1.750607192 | -2.428431364 | -1.18224839 | 1.496839356 | 0.775446939 | 1.986138811 | 0.001335878 | 0.040710997 | 0.148302475 | 0.1006 | 0.243865404 |
| StyR.358_ | -3.742021584 | -3.934507066 | -3.545479766 | 0.246800638 | 1.097604454 | 0.210771571 | 0.410393075 | 0.641386729 | 0.769491966 | 0.7583 | 0.842820799 |
| StyR.362_ | -4.055866519 | -3.922971633 | -4.244080574 | -0.325034534 | 0.836506386 | -0.381264039 | 0.331112599 | 0.478848988 | 0.629097071 | 0.5798 | 0.696565748 |
| StyR.369_ | -5.008900036 | -4.883833766 | -5.194697674 | -0.333740141 | 1.845535285 | -0.171108642 | 0.407333337 | 0.62484691 | 0.759978193 | 0.7843 | 0.860248863 |
| StyR.377_ | 1.687346949 | 1.731258834 | 1.526426903 | -0.223494874 | 0.362366866 | -0.592098172 | 0.274483688 | 0.409592883 | 0.603995699 | 0.5434 | 0.706540511 |
| StyR.378_ | -7.099525007 | -6.321492024 | -8.127498587 | -2.993565668 | 4.429600568 | -0.613151332 | 0.243837455 | 0.333613079 | 0.51766991 | 0.4831 | 0.630364138 |
| StyR.44a_ | 0.43729314 | 0.163010686 | 0.706235599 | 0.597067518 | 0.326255319 | 1.895084847 | 0.002000966 | 0.075342504 | 0.217049298 | 0.1012 | 0.244721245 |
| StyR.44b_ | 2.140655406 | 1.614359552 | 2.37067341 | 0.738203857 | 0.194919251 | 3.254762577 | 4.68E-05 | 0.02298478 | 0.108243768 | 0.1 | 0.243021319 |
| StyR.44c_ | -0.422673063 | -0.943018921 | 0.09250368 | 1.011140809 | 0.303326634 | 3.282658392 | 4.68E-05 | 0.012503134 | 0.073522922 | 0.1 | 0.243021319 |
| StyR.44d_ | -0.646624293 | -1.077270463 | -0.328762133 | 0.762923484 | 0.345526219 | 2.181309053 | 0.002668394 | 0.04959437 | 0.162670797 | 0.1025 | 0.246382352 |
| StyR.44e_ | -2.1983128 | -2.416551116 | -2.012568378 | 0.441887587 | 0.588980943 | 0.71411516 | 0.213191222 | 0.332007065 | 0.510500596 | 0.4409 | 0.594118758 |
| StyR.44f_ | -3.114841725 | -3.487005807 | -2.632567666 | 0.797598861 | 0.711173247 | 1.078986822 | 0.120000033 | 0.186438118 | 0.353561804 | 0.2626 | 0.427864204 |
| StyR.44g_ | -2.930854566 | -3.156123771 | -2.749800643 | 0.402869445 | 0.555510082 | 0.677138135 | 0.220000015 | 0.334708343 | 0.499911083 | 0.4291 | 0.573081851 |
| StyR.51_ | 3.117861763 | 3.00709668 | 3.284451846 | 0.284786122 | 0.094811395 | 2.98348079 | 0.00067292 | 0.026188916 | 0.113207504 | 0.1 | 0.243021319 |
| StyR.55_ | 5.832309738 | 5.652694069 | 5.97896383 | 0.30876508 | 0.106465814 | 2.903857241 | 4.68E-05 | 0.009647455 | 0.06904727 | 0.1 | 0.243021319 |
| StyR.8_ | 5.022912539 | 4.882716916 | 5.096510969 | 0.209739291 | 0.101108817 | 2.20265942 | 4.68E-05 | 0.023833377 | 0.11108024 | 0.1001 | 0.243158537 |
| StyR.88_ | 1.574061477 | 1.503290715 | 1.62438306 | 0.123709451 | 0.192388709 | 0.626143874 | 0.250666679 | 0.377614088 | 0.559566167 | 0.4919 | 0.645380587 |
| StyR.90_ | -3.161072371 | -3.140136646 | -3.186986497 | -0.052779735 | 0.534209076 | -0.091830701 | 0.456000002 | 0.53218034 | 0.674239734 | 0.6427 | 0.749545505 |
| StyR.99_ | -0.692440413 | -0.760909049 | -0.502002482 | 0.218983436 | 0.359642867 | 0.546415014 | 0.300000009 | 0.512609325 | 0.668594812 | 0.607 | 0.741491295 |
| tp2_ | -5.291902331 | -5.860997268 | -4.71902514 | 1.250783075 | 1.538065766 | 0.755413374 | 0.181878767 | 0.272833386 | 0.448584349 | 0.3633 | 0.515717532 |
| tp28_ | 1.026656364 | 1.068592575 | 0.980590329 | -0.104689466 | 0.242002258 | -0.402480065 | 0.319786816 | 0.485518972 | 0.652549436 | 0.6018 | 0.729752865 |
| tpk3_ | 2.043718791 | 1.490363059 | 2.278971006 | 0.779992175 | 0.203099836 | 3.354247357 | 4.68E-05 | 0.032685694 | 0.133911874 | 0.1 | 0.243021319 |
| tpke11_ | 4.468192515 | 4.538086911 | 4.391341407 | -0.019882379 | 0.367786508 | -0.057891847 | 0.480000001 | 0.916288515 | 0.955646756 | 1 | 1 |
| tpke70_ | -2.10768045 | -2.187472413 | -2.07294305 | 0.084426194 | 0.637851283 | 0.115877714 | 0.451698869 | 0.722311974 | 0.830629873 | 0.8314 | 0.898735678 |
| zipA.leader_ | 2.110590729 | 1.939510624 | 2.365024691 | 0.417373733 | 0.222273197 | 1.829349201 | 0.011992386 | 0.068129374 | 0.204428529 | 0.1116 | 0.259190917 |

# ME

| | rab.all | rab.win.tnpA | rab.win.WT | diff.btw | diff.win | effect | overlap | we.ep | we.eBH | wi.ep | wi.eBH |
|---|---|---|---|---|---|---|---|---|---|---|---|
| aceF_aceF | 5.419365635 | 4.969867185 | 5.47514905 | 0.504427072 | 0.440887393 | 1.003080487 | 0.205862775 | 0.184115567 | 0.483619311 | 0.378 | 0.657896149 |
| SL1344_RS11990_ackA | 3.093396384 | 2.922198798 | 3.530643206 | 0.612183035 | 0.171274521 | 2.815617189 | 4.68E-05 | 0.044952758 | 0.251425797 | 0.1 | 0.330257709 |
| SL1344_RS08180_acpD | 0.538335685 | 0.784858053 | 0.242884774 | -0.533588582 | 0.265494908 | -1.962486859 | 0.005996769 | 0.048680509 | 0.249489716 | 0.1067 | 0.339919425 |
| acpS_acpS | 0.719357251 | 0.662287698 | 0.792233357 | 0.163079471 | 0.227222059 | 0.702603832 | 0.236000014 | 0.332523239 | 0.601732455 | 0.4637 | 0.677887949 |
| adrA_adrA | -0.504505977 | -0.518853412 | -0.487540519 | 0.057784484 | 0.264832521 | 0.206553107 | 0.410000003 | 0.609743079 | 0.797905654 | 0.7377 | 0.857425218 |
| SL1344_RS16630_aer | 3.732620386 | 4.203982923 | 3.297698221 | -1.140143446 | 0.317961 | -3.6495279 | 4.68E-05 | 0.003047353 | 0.07468403 | 0.1 | 0.330257709 |
| SL1344_RS13055_alpha.beta.hydrolase | -2.111425961 | -2.134926402 | -2.075102229 | 0.049396557 | 0.457218273 | 0.1099409 | 0.459333335 | 0.664232785 | 0.833677296 | 0.7667 | 0.881341731 |
| AmgR_amgR | -4.74733566 | -5.072060314 | -4.468172815 | 0.608880958 | 0.798137363 | 0.696711847 | 0.201199218 | 0.3149222 | 0.548267211 | 0.4132 | 0.618872671 |
| amiC_amiC | 1.852195575 | 1.934491116 | 1.720409501 | -0.212696576 | 0.103254219 | -2.040415649 | 0.012667027 | 0.056310052 | 0.254196677 | 0.1127 | 0.346200012 |
| SL1344_RS04710_amino.acid.permease | 0.221635565 | -0.115154764 | 0.525302726 | 0.674558734 | 0.255283464 | 2.778817514 | 4.68E-05 | 0.027417558 | 0.19166357 | 0.1 | 0.330257709 |
| SL1344_RS01920_ampH | 0.04068258 | -0.680032968 | 0.399531657 | 1.074498123 | 0.330912041 | 3.024833339 | 4.68E-05 | 0.028028657 | 0.194754196 | 0.1 | 0.330257709 |
| SL1344_RS23585_arcA | 4.531393585 | 4.430517357 | 4.543246224 | 0.106108935 | 0.065802602 | 0.974822505 | 0.216666682 | 0.24236818 | 0.537901924 | 0.4456 | 0.676648432 |
| ArcZ_arcZ | 1.914219391 | 1.748290489 | 2.16049271 | 0.47299007 | 0.298989897 | 1.486533704 | 0.00067292 | 0.167491172 | 0.461989536 | 0.1003 | 0.33069165 |
| STnc490a_art200 | -6.843268673 | -7.225268377 | -6.635727664 | 0.619432246 | 2.509809343 | 0.195100131 | 0.391072622 | 0.538234085 | 0.75772098 | 0.6881 | 0.83151223 |
| STnc490b_art200 | -6.981902256 | -9.91687325 | -5.54534575 | 4.363257024 | 2.662672819 | 1.565714716 | 0.018000252 | 0.089666931 | 0.310987848 | 0.1178 | 0.352438475 |
| STnc490c_art200 | -7.789243435 | -9.949571991 | -6.381984653 | 3.4353727 | 2.955920673 | 1.091985335 | 0.078614311 | 0.160802123 | 0.409362845 | 0.2088 | 0.449425424 |
| STnc490d_art200 | -9.36069959 | -9.954994014 | -8.747640657 | 1.038087746 | 3.99192634 | 0.20983806 | 0.375333338 | 0.497018432 | 0.708702361 | 0.6063 | 0.767443862 |
| STnc490e_art200 | -9.359174119 | -9.898934964 | -8.823450938 | 1.081524643 | 4.179914169 | 0.21495206 | 0.38241173 | 0.4876135 | 0.705153371 | 0.6026 | 0.765979937 |
| STnc490f_art200 | -7.902926957 | -9.883230502 | -6.62663971 | 3.157387754 | 2.866146584 | 0.998101918 | 0.093271197 | 0.187431416 | 0.432257793 | 0.2353 | 0.46837408 |
| STnc490g_art200 | -9.309075445 | -9.986352524 | -8.314919797 | 1.732628625 | 4.318156619 | 0.357605111 | 0.327115264 | 0.491621149 | 0.709309427 | 0.5993 | 0.767264305 |
| artM_artM | 1.935608026 | 1.814589163 | 2.022423623 | 0.202213719 | 0.122012593 | 1.574800906 | 0.03597614 | 0.078914649 | 0.303323232 | 0.1436 | 0.386608461 |
| artP_artP | 2.679064248 | 3.162090931 | 2.55032625 | -0.434273115 | 0.47023853 | -0.467625081 | 0.33333334 | 0.817037069 | 0.929455518 | 0.7 | 0.872542119 |
| asarfA_asarfA | 0.952947801 | 0.356764929 | 1.016051669 | 0.637336289 | 0.512842615 | 0.823342839 | 0.255333345 | 0.325825149 | 0.62352196 | 0.4843 | 0.734369315 |
| asnB_asnB | 2.223800503 | 2.184051983 | 2.248853084 | 0.105143652 | 0.186877099 | 0.53727505 | 0.303131254 | 0.429449707 | 0.692956108 | 0.6017 | 0.789537827 |
| SL1344_RS13060_asrA | -3.610163042 | -3.698860254 | -3.51963163 | 0.179879334 | 0.540433012 | 0.303549176 | 0.360426388 | 0.478982583 | 0.695790612 | 0.5836 | 0.753747584 |
| SL1344_RS13065_asrB | -2.395116618 | -2.464290992 | -2.280861501 | 0.116281165 | 0.54961136 | 0.17561456 | 0.41866667 | 0.693068601 | 0.85264218 | 0.7694 | 0.887243339 |
| SL1344_RS13070_asrC | -1.639731669 | -1.540489532 | -1.746023939 | 0.003346629 | 0.655500608 | 0.006842449 | 0.498 | 0.787999944 | 0.908565197 | 0.9255 | 0.966410008 |
| asyhbQ_asyhbQ | 0.579389282 | 0.472166109 | 0.686207479 | 0.169689288 | 0.603389603 | 0.253248031 | 0.431712194 | 0.808783077 | 0.923109859 | 0.9556 | 0.981084128 |
| atpC_atpC | 6.096507635 | 5.742005958 | 6.292370322 | 0.494701085 | 0.105713828 | 3.842234761 | 4.68E-05 | 0.019233693 | 0.171331741 | 0.1 | 0.330257709 |
| SL1344_RS15305_barA | 1.765788837 | 1.709082029 | 1.813355299 | 0.113614469 | 0.108684045 | 1.031509888 | 0.125916087 | 0.198342017 | 0.455346708 | 0.2798 | 0.514030579 |
| bcsA_bcsA | 1.589051983 | 1.175274777 | 2.00260045 | 0.873319503 | 0.193324545 | 4.590252358 | 4.68E-05 | 0.004360468 | 0.082476779 | 0.1 | 0.330257709 |
| SL1344_RS17950_bigA | 0.407625167 | 0.515702584 | 0.277943443 | -0.245843186 | 0.162656443 | -1.399623094 | 0.054630326 | 0.107320662 | 0.338321613 | 0.169 | 0.40865231 |
| bssR_bssR | -1.314009589 | -1.491552169 | -1.132438338 | 0.362558989 | 0.283421743 | 1.232866981 | 0.08933338 | 0.143708801 | 0.385080177 | 0.2154 | 0.455727951 |
| bssS_bssS | 1.763800967 | 1.609811233 | 2.066214807 | 0.547082309 | 0.307961119 | 1.885018865 | 0.00067292 | 0.041572491 | 0.23936076 | 0.1004 | 0.330863898 |
| cadB_cadB | 1.606309367 | 1.35300948 | 2.582674402 | 1.001087684 | 0.660038003 | 1.403337521 | 0.107261863 | 0.108713256 | 0.378315242 | 0.1938 | 0.466266212 |
| cbiC_cbiC | -1.259359155 | -1.458449731 | -1.125640687 | 0.39269523 | 0.461775451 | 0.840300231 | 0.129247199 | 0.240327134 | 0.528317107 | 0.2898 | 0.526390547 |
| cbiD_cbiD | -0.348135429 | -0.498846648 | -0.107806486 | 0.416175417 | 0.526793591 | 0.83946061 | 0.145236535 | 0.250753344 | 0.549591617 | 0.2738 | 0.540204362 |
| SL1344_RS11580_ccmA | -2.574737583 | -1.88824629 | -2.91595956 | -0.767060853 | 1.281361156 | -0.429815502 | 0.346000006 | 0.739695822 | 0.886242574 | 0.7369 | 0.886116511 |
| SL1344_RS11575_ccmB | -4.024239199 | -4.047203431 | -4.003530542 | 0.054835172 | 0.712160035 | 0.078303833 | 0.462666668 | 0.571474796 | 0.769492372 | 0.6821 | 0.823326671 |
| cdsA_cdsA | 2.593433662 | 2.581971836 | 2.601599631 | 0.01237782 | 0.21053125 | 0.056472053 | 0.482666667 | 0.84696866 | 0.939312717 | 0.9589 | 0.982387332 |
| SL1344_RS02340_cdsH | -0.42101963 | -0.411634075 | -0.426515278 | -0.054076156 | 0.270525975 | -0.170164706 | 0.41972019 | 0.625721495 | 0.810533691 | 0.7547 | 0.871681562 |
| chaB_chaB | -0.901637153 | -0.566981324 | -1.232855359 | -0.673477137 | 0.335328561 | -1.983295623 | 0.011992386 | 0.048505479 | 0.24041782 | 0.1102 | 0.344061177 |
| SL1344_RS09610_cheA | 6.314550582 | 7.354236963 | 5.352782889 | -1.999182848 | 0.183718037 | -10.97667437 | 4.68E-05 | 0.00133413 | 0.052586083 | 0.1 | 0.330257709 |
| SL1344_RS09600_cheM | 5.313096221 | 6.327482473 | 3.945069949 | -2.293831679 | 0.282546149 | -8.058211331 | 4.68E-05 | 0.000849805 | 0.044105446 | 0.1 | 0.330257709 |
| SL1344_RS09595_chemotaxis.protein.methyltransferase | 4.276339117 | 5.24220378 | 3.223302266 | -1.911027479 | 0.216183871 | -8.968186724 | 4.68E-05 | 0.000103417 | 0.021658227 | 0.1 | 0.330257709 |
| SL1344_RS09585_chemotaxis.regulator | 3.775865853 | 4.703330273 | 3.019944603 | -1.692366119 | 0.271404975 | -6.284120138 | 4.68E-05 | 0.003846505 | 0.082530603 | 0.1 | 0.330257709 |
| SL1344_RS09590_chemotaxis.response.regulator.protein.glutamate.methylesterase | 3.81034308 | 4.757886144 | 2.943074892 | -1.817268639 | 0.135259953 | -13.47054511 | 4.68E-05 | 0.001212805 | 0.04948947 | 0.1 | 0.330257709 |
| SL1344_RS11875_cheV | 4.742831439 | 5.329635715 | 4.140488393 | -1.206759925 | 0.314637779 | -3.430115426 | 4.68E-05 | 0.00600962 | 0.10171457 | 0.1 | 0.330257709 |
| SL1344_RS09605_CheW | 6.259060363 | 7.397403443 | 5.362645786 | -2.029843943 | 0.136003083 | -14.49714706 | 4.68E-05 | 0.000836607 | 0.043913517 | 0.1 | 0.330257709 |
| SL1344_RS09580_cheZ | 5.267001926 | 6.214357267 | 4.312117338 | -1.915775088 | 0.089085338 | -21.54586406 | 4.68E-05 | 3.95E-05 | 0.016101994 | 0.1 | 0.330257709 |
| chiP_chiP | 2.748201971 | 2.202116983 | 2.897570408 | 0.696794413 | 0.463054124 | 1.241697594 | 0.114000035 | 0.146562556 | 0.435019651 | 0.208 | 0.484474315 |
| ChiX_chiX | 1.667894079 | 2.14794784 | 1.173187515 | -0.566696861 | 1.448536713 | -0.389470243 | 0.444666669 | 0.664120002 | 0.854645044 | 0.9976 | 0.998968513 |

| | | | | | | | | | | | |
|---|---|---|---|---|---|---|---|---|---|---|---|
| SL1344_RS02290_clpP | 3.140442359 | 3.161544239 | 3.129637996 | -0.028219735 | 0.138439284 | -0.152051317 | 0.428381082 | 0.830986222 | 0.931528611 | 0.8593 | 0.940129301 |
| SL1344_RS02295_clpX | 5.354333684 | 5.35256607 | 5.35598015 | 0.041318265 | 0.128812223 | 0.319018512 | 0.378666671 | 0.661606836 | 0.849225245 | 0.8344 | 0.928697485 |
| SL1344_RS09530_cmoA | 2.107624824 | 2.521645695 | 1.664717269 | -0.871117121 | 0.136442111 | -6.276624248 | 4.68E-05 | 0.001817327 | 0.055069971 | 0.1 | 0.330257709 |
| cobS_cobS | -0.26272592 | -0.287962658 | -0.240653146 | 0.061151994 | 0.230404077 | 0.253579003 | 0.398000004 | 0.579843368 | 0.774897011 | 0.6957 | 0.830316719 |
| copA_copA | 2.976905327 | 2.920243091 | 3.3552097 | 0.416595271 | 0.497172272 | 0.795024631 | 0.232000014 | 0.332489442 | 0.629930005 | 0.4039 | 0.682816585 |
| SL1344_RS20320_corA | 1.898480986 | 2.044074484 | 1.862494733 | -0.203452943 | 0.34263145 | -0.652854987 | 0.218666682 | 0.334248055 | 0.625149229 | 0.4076 | 0.682167769 |
| SL1344_RS20860_cpxA | 3.233490437 | 3.131303808 | 3.353164071 | 0.202357475 | 0.150189249 | 1.315250777 | 0.088666714 | 0.107457879 | 0.368620319 | 0.1796 | 0.443097021 |
| cpxP_cpxP | 1.192055382 | 0.968589474 | 1.7674024 | 0.773826204 | 0.357886269 | 2.237338445 | 0.00067292 | 0.027767169 | 0.19857211 | 0.1001 | 0.330406553 |
| SL1344_RS20865_cpxR | 1.934826343 | 1.755557958 | 2.020895932 | 0.263615191 | 0.175409797 | 1.388071676 | 0.046666761 | 0.124728559 | 0.394097175 | 0.1502 | 0.396990742 |
| SL1344_RS17890_crp | 5.26454896 | 5.224621296 | 5.394965127 | 0.195864939 | 0.130156305 | 1.609358797 | 0.005330642 | 0.0566863 | 0.278016869 | 0.1042 | 0.335881827 |
| CrpT_crpT | -0.143596383 | -0.273446988 | -0.106695471 | 0.16471266 | 0.563086734 | 0.284807727 | 0.376415728 | 0.624286976 | 0.826921381 | 0.781 | 0.906571233 |
| csgB_csgB | -5.693409428 | -5.730932975 | -5.640451514 | 0.094024011 | 1.345288896 | 0.071297666 | 0.468354432 | 0.580062002 | 0.776063872 | 0.7035 | 0.838725863 |
| SL1344_RS05640_csgC | 1.038960674 | 1.443819517 | 0.330740808 | -1.278959894 | 0.652866296 | -2.2336046 | 4.68E-05 | 0.025754767 | 0.195438687 | 0.1 | 0.330257709 |
| SL1344_RS18810_cspA | 5.071192534 | 4.447459897 | 5.846390665 | 1.19300838 | 0.371577796 | 3.716275878 | 4.68E-05 | 0.002925771 | 0.073840171 | 0.1 | 0.330257709 |
| SL1344_RS09980_cspB | -0.554510842 | -1.03619055 | -0.122802823 | 0.929934479 | 0.238730201 | 3.921476674 | 4.68E-05 | 0.00907512 | 0.108267322 | 0.1 | 0.330257709 |
| SL1344_RS06145_cspH | -2.248013259 | -2.263064582 | -2.241347978 | 0.165122623 | 0.71178824 | 0.21254786 | 0.405063295 | 0.647829613 | 0.830716063 | 0.8234 | 0.912520515 |
| SL1344_RS14635_csrA | 4.435482441 | 4.396479217 | 4.468338487 | 0.035405478 | 0.180564282 | 0.220953119 | 0.381333338 | 0.778098252 | 0.907814563 | 0.8047 | 0.910379155 |
| CsrB_csrB | 9.214744723 | 9.294538207 | 9.057056999 | -0.236800494 | 0.122440064 | -2.048069645 | 4.68E-05 | 0.040819567 | 0.242074566 | 0.1 | 0.330257709 |
| SL1344_RS15345_csrB | 8.873690578 | 9.109255972 | 8.798378332 | -0.206129371 | 0.308906995 | -0.728150507 | 0.223184558 | 0.381797947 | 0.671412941 | 0.4012 | 0.681130087 |
| CsrC_csrC | 7.608217439 | 7.848052746 | 7.157306915 | -0.691767519 | 0.192017941 | -3.654211466 | 4.68E-05 | 0.005544118 | 0.098108906 | 0.1 | 0.330257709 |
| SL1344_RS17435_csrD | 2.762178959 | 3.117396029 | 2.489578218 | -0.62481026 | 0.207758252 | -3.100655416 | 4.68E-05 | 0.00808571 | 0.112480245 | 0.1 | 0.330257709 |
| CyaR_cyaR | 4.459787362 | 4.383747642 | 4.523146186 | 0.126991664 | 0.268826999 | 0.444570366 | 0.325782819 | 0.505496953 | 0.758276817 | 0.6799 | 0.859188795 |
| SL1344_RS12535_cysA | 1.003220983 | 1.138762474 | 0.952120559 | -0.146543482 | 0.256048483 | -0.398551033 | 0.350666673 | 0.782147037 | 0.907169458 | 0.7411 | 0.88893756 |
| SL1344_RS08565_cysB | 1.780383274 | 1.932235248 | 1.54545822 | -0.375278772 | 0.186331362 | -1.903059292 | 0.009327606 | 0.062153676 | 0.282551833 | 0.107 | 0.340387642 |
| SL1344_RS15175_cysC | 0.526279854 | 0.683894832 | 0.476311485 | -0.210307255 | 0.3746102 | -0.601213535 | 0.289140582 | 0.410280774 | 0.68469035 | 0.5923 | 0.789375186 |
| SL1344_RS15185_cysD | 1.396299897 | 1.294110095 | 1.497198506 | 0.234098921 | 0.875005631 | 0.240722587 | 0.398000004 | 0.599004433 | 0.817603689 | 0.9034 | 0.959138626 |
| SL1344_RS19060_cysE | 3.298100645 | 3.276180735 | 3.313408843 | 0.026551863 | 0.135743428 | 0.182385765 | 0.420000003 | 0.725247974 | 0.877044712 | 0.8644 | 0.935482549 |
| SL1344_RS17945_cysG | 2.230776235 | 2.091753452 | 2.277373679 | 0.174352126 | 0.150217889 | 1.063785929 | 0.148567647 | 0.184517383 | 0.464199826 | 0.2834 | 0.546225731 |
| SL1344_RS15240_cysH | 0.084042403 | 0.376873877 | -0.011944742 | -0.398063873 | 0.294082198 | -1.169929274 | 0.100666708 | 0.171346099 | 0.454418335 | 0.2261 | 0.474487919 |
| SL1344_RS15245_cysI | 2.311917279 | 2.586764382 | 2.281848514 | -0.286535445 | 0.17718124 | -0.433550885 | 0.33333334 | 0.824163577 | 0.930887699 | 0.7 | 0.872542119 |
| SL1344_RS15250_cysJ | 1.868725922 | 1.658042121 | 2.081233504 | 0.101021545 | 0.649110279 | 0.184037918 | 0.394403735 | 0.931451294 | 0.975289295 | 0.8371 | 0.930847414 |
| SL1344_RS12475_cysK | 3.948853948 | 3.712694704 | 3.96800327 | 0.243675763 | 0.25596017 | 0.736327306 | 0.232511673 | 0.352853842 | 0.646603539 | 0.4318 | 0.700717331 |
| SL1344_RS12530_cysM | 1.446950274 | 1.191015573 | 1.580323979 | 0.350117778 | 0.199528103 | 1.593480745 | 0.048666757 | 0.08796126 | 0.331487438 | 0.1457 | 0.394830016 |
| SL1344_RS15180_cysN | 1.479908253 | 1.263400317 | 1.657370534 | 0.096021029 | 0.548027835 | 0.234972571 | 0.404000004 | 0.901953872 | 0.963451599 | 0.8491 | 0.935792149 |
| SL1344_RS12550_cysP | 1.820366696 | 1.761940619 | 1.876223484 | 0.321655752 | 0.766412815 | 0.38669318 | 0.320000008 | 0.392684369 | 0.679380464 | 0.6502 | 0.837885787 |
| SL1344_RS22605_cysQ | 1.710974071 | 1.708532781 | 1.71335325 | 0.010796887 | 0.307653783 | 0.026469736 | 0.491005996 | 0.86350357 | 0.946739134 | 0.9649 | 0.984913491 |
| SL1344_RS02725_cysS | 3.163314724 | 3.074496136 | 3.328864488 | 0.242018427 | 0.099833947 | 2.228311196 | 0.005996769 | 0.047655004 | 0.247538784 | 0.1057 | 0.33861236 |
| SL1344_RS09720_cysT | 3.976917242 | 5.133084708 | 3.696135243 | -1.344479095 | 0.398730579 | -1.531676806 | 4.68E-05 | 0.092253497 | 0.351865609 | 0.1 | 0.330257709 |
| SL1344_RS12545_cysU | 0.053394439 | -0.17315743 | 0.223769579 | 0.207894186 | 0.585368444 | 0.364556859 | 0.414000003 | 0.702544534 | 0.869241549 | 0.8789 | 0.947766302 |
| SL1344_RS12540_cysW | 0.680384413 | 0.515698114 | 0.848154711 | 0.113462798 | 0.599927689 | 0.236768811 | 0.426000003 | 0.795760679 | 0.915885856 | 0.8839 | 0.949643196 |
| SL1344_RS12470_cysZ | 0.727167574 | 0.421737803 | 0.931086949 | 0.508515782 | 0.223729204 | 2.228126137 | 0.000673369 | 0.030633668 | 0.200274553 | 0.1015 | 0.332435973 |
| SL1344_RS16735_cyuA | 0.282879997 | 0.10879337 | 0.407455998 | 0.298821215 | 0.157437304 | 1.802091454 | 0.028000161 | 0.06223358 | 0.259803957 | 0.126 | 0.362297936 |
| SL1344_RS16740_cyuP | -1.100293566 | -1.167351597 | -0.884710809 | 0.255503354 | 0.319080967 | 0.673752986 | 0.274000011 | 0.428878532 | 0.673568819 | 0.5553 | 0.754358277 |
| SL1344_RS10610_dacD | -1.866533358 | -1.94121132 | -1.803301725 | 0.146846704 | 0.315950366 | 0.415514785 | 0.315333341 | 0.471986447 | 0.68882021 | 0.5802 | 0.747493771 |
| SL1344_RS17980_dam | 2.655142624 | 2.501870491 | 2.734179638 | 0.233245425 | 0.1399131 | 1.616201092 | 0.011326186 | 0.101940244 | 0.360287616 | 0.1125 | 0.347640737 |
| DapZ_dapZ | 0.986278558 | 1.070248351 | 0.934719033 | -0.146367632 | 0.311471765 | -0.428794112 | 0.339773491 | 0.515495417 | 0.755519322 | 0.6823 | 0.8421689 |
| ddl_ddl | 2.857642708 | 2.866958224 | 2.837055852 | -0.022091182 | 0.08315567 | -0.236691878 | 0.40466667 | 0.660919428 | 0.832920233 | 0.7484 | 0.876879323 |
| SL1344_RS09505_dihydroneopterin.triphosphate.diphosphatase | 2.596885743 | 3.006686913 | 2.140347022 | -0.861307901 | 0.098552379 | -8.856241718 | 4.68E-05 | 0.00097846 | 0.040044957 | 0.1 | 0.330257709 |
| dinG_dinG | 2.124603088 | 2.13903916 | 2.090287399 | -0.051258288 | 0.135084716 | -0.353603237 | 0.359333339 | 0.555890963 | 0.772102441 | 0.7157 | 0.853993263 |
| SL1344_RS05725_dinI | 2.074533539 | 2.258266301 | 1.907021375 | -0.199117982 | 0.293948289 | -0.922235927 | 0.154000025 | 0.261476411 | 0.55865881 | 0.2689 | 0.542168607 |
| SL1344_RS04685_dmsA | -1.121188962 | -1.106649571 | -1.135999236 | -0.025230024 | 0.236059463 | -0.111405853 | 0.451698869 | 0.532594961 | 0.735820168 | 0.6448 | 0.79532183 |
| SL1344_RS04690_dmsB | -4.484726439 | -4.968955832 | -4.101375804 | 0.902396144 | 0.984793638 | 0.877041557 | 0.138000028 | 0.234019423 | 0.496853782 | 0.3001 | 0.530448124 |
| SL1344_RS04695_dmsC | -2.548835028 | -2.444015801 | -2.671082966 | -0.069571792 | 0.749928206 | -0.087355666 | 0.463024652 | 0.741000348 | 0.883000334 | 0.851 | 0.92899906 |

| | | | | | | | | | | | |
|---|---|---|---|---|---|---|---|---|---|---|---|
| SL1344_RS06610_DNA.RNA.non.specific.endonuclease | -0.3243512 | -0.193406106 | -0.370035154 | -0.169851095 | 0.327336348 | -0.442694096 | 0.33733334 | 0.559746563 | 0.773411795 | 0.6951 | 0.846133832 |
| dnaA_dnaA | 3.20253934 | 3.209769333 | 3.199510311 | -0.010392453 | 0.1451457 | -0.053119317 | 0.473684212 | 0.789770023 | 0.911008509 | 0.8861 | 0.95126231 |
| SL1344_RS00060_dnaK | 6.995456399 | 6.963758784 | 7.024466777 | -0.176378058 | 0.409778594 | -0.430627943 | 0.439706864 | 0.528374781 | 0.774446156 | 1 | 1 |
| dpiA_dpiA | -1.400151122 | -1.401073343 | -1.398845585 | 0.03543278 | 0.353947476 | 0.092425604 | 0.458666668 | 0.62605666 | 0.80881066 | 0.7541 | 0.869284178 |
| dpiA_dpiA.1 | -1.139859653 | -1.054046487 | -1.256314748 | -0.137633846 | 0.409278109 | -0.324332765 | 0.368666672 | 0.613614243 | 0.804735609 | 0.7339 | 0.861586728 |
| dpiB_dpiB | -0.882363956 | -0.747465823 | -1.081757418 | -0.331643651 | 0.332180371 | -0.975990699 | 0.139240534 | 0.210898746 | 0.48312283 | 0.2973 | 0.547197212 |
| DsrA_dsrA | -4.042365729 | -4.887412114 | -2.799468057 | 2.006205805 | 1.058468269 | 1.836782178 | 0.01798826 | 0.059918287 | 0.269337822 | 0.1229 | 0.360774915 |
| SL1344_RS09330_DUF1441.domain.containing.protein | -4.141580187 | -3.289215262 | -5.073130714 | -1.764617648 | 0.759615195 | -2.272873924 | 0.003998486 | 0.032597513 | 0.193673557 | 0.105 | 0.337115999 |
| SL1344_RS06460_DUF1496.domain.containing.protein | 2.618099075 | 3.210950271 | 1.910308729 | -1.324919479 | 0.336667701 | -3.706358759 | 4.68E-05 | 0.004486482 | 0.08862967 | 0.1 | 0.330257709 |
| SL1344_RS16295_DUF2623.domain.containing.protein | 1.201114498 | 2.194144178 | 0.600189779 | -1.538691916 | 0.299107897 | -4.679464217 | 4.68E-05 | 0.010179712 | 0.125216639 | 0.1 | 0.330257709 |
| emrB_emrB | 1.424580561 | 1.345309691 | 1.533447034 | 0.187688165 | 0.144774998 | 1.263876695 | 0.074616979 | 0.133098108 | 0.384497399 | 0.2025 | 0.453112309 |
| emrD_emrD | 1.692475222 | 1.70590005 | 1.678972006 | -0.060936088 | 0.288018751 | -0.224024807 | 0.409333337 | 0.588001925 | 0.80390853 | 0.8472 | 0.927493687 |
| emtA_emtA | 0.787563296 | 0.942282905 | 0.67591599 | -0.250850107 | 0.213722403 | -1.135381622 | 0.10933337 | 0.157637381 | 0.427583528 | 0.229 | 0.487735322 |
| SL1344_RS03015_entB | -0.487859599 | -0.209440904 | -0.729798695 | -0.499696493 | 0.319929843 | -1.498410774 | 0.045333431 | 0.085031563 | 0.319190035 | 0.1484 | 0.395361849 |
| entE_entE | -0.023400624 | 0.155895888 | -0.277560902 | -0.384280693 | 0.338684303 | -1.141582502 | 0.115922753 | 0.156542565 | 0.43048138 | 0.2303 | 0.495910145 |
| entF_entF | 0.784738273 | 0.90435794 | 0.39538929 | -0.506843307 | 0.373201244 | -1.419128658 | 0.052000084 | 0.081603997 | 0.324339119 | 0.1533 | 0.400118499 |
| SL1344_RS18065_envZ | 2.702488635 | 2.558138366 | 2.730545299 | 0.177192251 | 0.249002524 | 0.639657865 | 0.287141915 | 0.471731048 | 0.733101106 | 0.5719 | 0.789928451 |
| eutA_eutA | -2.124900315 | -2.403964454 | -1.940175449 | 0.50360573 | 0.48762263 | 0.988489096 | 0.107261863 | 0.199490268 | 0.473083498 | 0.2417 | 0.479292259 |
| SL1344_RS12680_eutS | -5.687572684 | -5.915200525 | -5.480487684 | 0.35906336 | 1.526481399 | 0.207313666 | 0.386000004 | 0.579635054 | 0.777483143 | 0.6982 | 0.836034848 |
| fabG_fabG | 3.99072291 | 3.990753954 | 3.990685982 | 0.001791052 | 0.093903532 | 0.013551534 | 0.491005996 | 0.810329341 | 0.920565824 | 0.868 | 0.943509276 |
| fadA_fadA | 2.15378806 | 2.689081889 | 2.073488864 | -0.544273044 | 0.263792623 | -0.974117608 | 0.23451034 | 0.234852295 | 0.537928779 | 0.4534 | 0.693262348 |
| SL1344_RS09080_fadD | 2.097491122 | 2.44557631 | 1.417435747 | -1.032997988 | 0.318013379 | -3.245387587 | 4.68E-05 | 0.011742024 | 0.133766983 | 0.1 | 0.330257709 |
| fadE_fadE | 2.276485837 | 2.300027551 | 2.261069838 | -0.055025975 | 0.429571921 | -0.129544316 | 0.434000002 | 0.889256374 | 0.958321054 | 0.8887 | 0.952656807 |
| fadI_fadI | 0.047400945 | -0.411585281 | 0.154692668 | 0.535269629 | 0.430589366 | 1.195754287 | 0.151333358 | 0.156639138 | 0.438988171 | 0.275 | 0.544747206 |
| SL1344_RS13945_Fels.2.prophage.protein | -3.212799732 | -3.441749565 | -2.925518967 | 0.457487602 | 1.195048381 | 0.388396887 | 0.310000008 | 0.575389053 | 0.785328008 | 0.6433 | 0.816622446 |
| feoA_feoA | 1.263557882 | 1.122426411 | 1.318184566 | 0.196190427 | 0.323669684 | 0.567404535 | 0.273333344 | 0.382692121 | 0.666621354 | 0.562 | 0.76842154 |
| ffs_ffs | 3.307673281 | 2.926593816 | 3.667852666 | -0.143199697 | 1.53518194 | -0.096504498 | 0.449033979 | 0.967892082 | 0.988805573 | 1 | 1 |
| fimA_fimA | -2.016317309 | -1.986880657 | -2.059248196 | -0.102491435 | 0.463365619 | -0.205097569 | 0.40066667 | 0.579544591 | 0.781116724 | 0.7188 | 0.846150701 |
| fimF_fimF | -2.049404922 | -2.064074643 | -2.032345017 | 0.0419585 | 0.37525369 | 0.095485316 | 0.460666668 | 0.592665696 | 0.784390277 | 0.7054 | 0.840347927 |
| SL1344_RS02785_fimZ | -3.85663592 | -3.765527699 | -4.011680292 | -0.228995435 | 1.865875459 | -0.12050004 | 0.454666668 | 0.819572223 | 0.926343889 | 0.8851 | 0.951130181 |
| fis_fis | 2.053468055 | 1.875108317 | 2.140115757 | 0.256302525 | 0.110634489 | 2.167818292 | 0.009327606 | 0.052551252 | 0.253600687 | 0.109 | 0.342403184 |
| fixB_fixB | -1.125163101 | -1.139935717 | -1.087599377 | 0.04153969 | 0.414356732 | 0.095301281 | 0.466000001 | 0.752118322 | 0.889324769 | 0.8278 | 0.922809245 |
| flgD_flgD | 4.029160576 | 4.541567768 | 3.543110345 | -1.069428666 | 0.155134549 | -6.937712476 | 4.68E-05 | 0.00041615 | 0.033432083 | 0.1 | 0.330257709 |
| flgF_flgF | 4.636435777 | 5.073516494 | 4.173450323 | -0.919164945 | 0.071991538 | -12.8114279 | 4.68E-05 | 0.000150863 | 0.02203621 | 0.1 | 0.330257709 |
| flgI_flgI | 3.201675703 | 3.730628479 | 2.765864997 | -0.947825536 | 0.100401138 | -9.39617646 | 4.68E-05 | 0.001119277 | 0.044279365 | 0.1 | 0.330257709 |
| SL1344_RS05830_flgK | 5.499988746 | 6.130492834 | 4.748293183 | -1.381092731 | 0.266639678 | -5.3729771 | 4.68E-05 | 0.000995286 | 0.047211767 | 0.1 | 0.330257709 |
| flgL_flgL | 5.305995048 | 5.930387622 | 4.738390185 | -1.150473643 | 0.241562085 | -4.816603858 | 4.68E-05 | 0.002492823 | 0.068600187 | 0.1 | 0.330257709 |
| SL1344_RS05775_flgM | 4.207852145 | 5.107370046 | 3.749458613 | -1.356801485 | 0.333545963 | -3.537100254 | 4.68E-05 | 0.029692571 | 0.210148746 | 0.1 | 0.330257709 |
| SL1344_RS05770_flgN | 4.785938376 | 5.343270624 | 4.021441564 | -1.364324443 | 0.310106731 | -4.510186073 | 4.68E-05 | 0.001350948 | 0.052992589 | 0.1 | 0.330257709 |
| SL1344_RS09570_flhA | 3.736032414 | 4.679714605 | 2.865566404 | -1.813448134 | 0.172153993 | -10.54566505 | 4.68E-05 | 0.001721401 | 0.05746736 | 0.1 | 0.330257709 |
| SL1344_RS09575_flhB | 3.40364242 | 4.335952996 | 2.400015285 | -2.002389768 | 0.322813581 | -6.548941197 | 4.68E-05 | 0.000828281 | 0.043415737 | 0.1 | 0.330257709 |
| SL1344_RS09625_flhC | 4.085721547 | 4.735497711 | 3.383129796 | -1.358873422 | 0.084598008 | -16.54925921 | 4.68E-05 | 0.001319364 | 0.05009393 | 0.1 | 0.330257709 |
| SL1344_RS09630_flhD | 2.642637942 | 2.963263132 | 2.385260238 | -0.635717832 | 0.230681707 | -2.713686827 | 4.68E-05 | 0.014487033 | 0.14839584 | 0.1 | 0.330257709 |
| SL1344_RS09565_flhE | 1.717458529 | 2.48486614 | 0.804378682 | -1.713564252 | 0.260207017 | -6.698965099 | 4.68E-05 | 0.003120995 | 0.072869847 | 0.1 | 0.330257709 |
| SL1344_RS09785_fliA | 5.375667337 | 6.38016751 | 4.465469404 | -1.970516075 | 0.239510393 | -8.227138263 | 4.68E-05 | 0.000572314 | 0.037943293 | 0.1 | 0.330257709 |
| SL1344_RS09800_fliC | 7.479315537 | 9.915510019 | 6.311191609 | -3.599010627 | 0.141933698 | -23.006054 | 4.68E-05 | 0.018712925 | 0.170334379 | 0.1 | 0.330257709 |
| SL1344_RS09845_fliE | 2.05624557 | 2.404796638 | 1.646892631 | -0.82468613 | 0.234622554 | -3.626154178 | 4.68E-05 | 0.008858384 | 0.117251724 | 0.1 | 0.330257709 |
| fliI_fliI | 3.507151243 | 3.909650299 | 3.22139424 | -0.688824269 | 0.158481208 | -4.568465616 | 4.68E-05 | 0.01547054 | 0.153565019 | 0.1 | 0.330257709 |
| fliJ_fliJ | 2.418608741 | 2.628753667 | 1.935380175 | -0.694316694 | 0.183700889 | -3.514652886 | 4.68E-05 | 0.028834146 | 0.202699153 | 0.1 | 0.330257709 |
| fliL_fliL | 3.222550044 | 3.569507745 | 2.890181821 | -0.730759644 | 0.139927898 | -5.311380486 | 4.68E-05 | 0.001565212 | 0.05376875 | 0.1 | 0.330257709 |
| fliQ_fliQ | 1.487871224 | 1.808438868 | 1.065028101 | -0.742827892 | 0.099591885 | -7.36352074 | 4.68E-05 | 0.00233478 | 0.055276079 | 0.1 | 0.330257709 |
| SL1344_RS09815_fliT | 4.913441487 | 5.393659712 | 4.684423543 | -0.69139139 | 0.234742903 | -2.491086216 | 4.68E-05 | 0.030076217 | 0.210688796 | 0.1 | 0.330257709 |
| SL1344_RS09775_fliY | 3.653078021 | 4.628630272 | 2.838198685 | -1.790975689 | 0.236257865 | -7.650325424 | 4.68E-05 | 0.00453284 | 0.089035058 | 0.1 | 0.330257709 |

| | | | | | | | | | | | |
|---|---|---|---|---|---|---|---|---|---|---|---|
| SL1344_RS09780_fliZ | 4.662276312 | 5.54323743 | 3.665553667 | -1.953904365 | 0.210500175 | -9.405472143 | 4.68E-05 | 8.34E-05 | 0.020573912 | 0.1 | 0.330257709 |
| SL1344_RS14340_fljA | 3.069929903 | -1.00703497 | 3.135786205 | 4.141518822 | 0.57393665 | 1.026423247 | 0.188540992 | 0.186768624 | 0.486920881 | 0.3417 | 0.620133264 |
| flk_flk | 0.336116621 | 0.085403186 | 0.571767641 | 0.533030463 | 0.317304384 | 1.631846124 | 0.020653117 | 0.065368788 | 0.289130592 | 0.12 | 0.358479809 |
| SL1344_RS08285_fnr | 3.821299754 | 4.189695159 | 3.607790233 | -0.504971473 | 0.227709169 | -2.07109641 | 4.68E-05 | 0.043153268 | 0.246772332 | 0.1006 | 0.331134655 |
| FnrS_fnrS | 0.453852002 | 0.327821433 | 1.030979818 | 0.610668047 | 0.472079506 | 0.693332266 | 0.323117929 | 0.448711819 | 0.717539568 | 0.675 | 0.856401551 |
| SL1344_RS04735_focA | 2.342938033 | 2.1855398 | 2.489935623 | 0.146012419 | 0.510199986 | 0.261596009 | 0.447035312 | 0.75749165 | 0.90056469 | 0.9982 | 0.999233431 |
| folA_folA | 1.101028511 | 0.875599003 | 1.151468112 | 0.246039428 | 0.220272603 | 0.854755137 | 0.244000013 | 0.304528121 | 0.593359846 | 0.4803 | 0.710470582 |
| frsA_frsA | 1.97473313 | 1.917983555 | 2.026751905 | 0.122839058 | 0.129783163 | 0.944976715 | 0.110592975 | 0.210726961 | 0.490010052 | 0.2594 | 0.499228401 |
| SL1344_RS09665_ftnB | -0.230819674 | 0.327840644 | -0.385385063 | -0.691688701 | 0.416726985 | -1.379879267 | 0.079280533 | 0.117324061 | 0.383643354 | 0.1843 | 0.437097661 |
| SL1344_RS06895_fumD | 0.147447429 | 0.050527716 | 0.267455023 | 0.239721265 | 0.276553068 | 0.865829961 | 0.141905424 | 0.227675199 | 0.50836697 | 0.3055 | 0.552250553 |
| SL1344_RS03500_fur | 4.17512914 | 4.193071447 | 4.152115051 | -0.11294032 | 0.132218162 | -0.828685215 | 0.198666684 | 0.236943133 | 0.532474125 | 0.3989 | 0.626951685 |
| gapA_gapA | 2.765772535 | 2.637085667 | 2.795891775 | 0.153152269 | 0.136654113 | 0.962474702 | 0.183877435 | 0.238908304 | 0.531918966 | 0.3685 | 0.628273064 |
| GcvB_gcvB | 7.091755256 | 7.041498266 | 7.150557445 | 0.018959187 | 0.64376384 | 0.039071561 | 0.457333335 | 0.963612234 | 0.987338446 | 1 | 1 |
| gcvT_gcvT | 4.817280041 | 4.859609079 | 4.724584653 | -0.136403758 | 0.064476293 | -2.037318911 | 0.0020023 | 0.070292223 | 0.30166552 | 0.1017 | 0.332466365 |
| gidB_gidB | 0.722925202 | 0.574641201 | 0.868542009 | 0.355471748 | 0.233927904 | 1.516717375 | 0.015989625 | 0.078433902 | 0.310075184 | 0.1199 | 0.356047373 |
| gldA_gldA | -0.159107471 | -0.15415476 | -0.160396148 | 0.012882692 | 0.265300032 | 0.042458802 | 0.480666667 | 0.711122594 | 0.863577074 | 0.8133 | 0.909438126 |
| gldA_gldA.1 | 1.976442744 | 1.909968418 | 2.096046661 | 0.181139974 | 0.141009231 | 1.291953736 | 0.082611643 | 0.125161853 | 0.381791356 | 0.2001 | 0.454195922 |
| GlmY_glmY | 3.22472446 | 3.211392319 | 3.237792207 | 0.029998726 | 0.877840692 | 0.035177203 | 0.458361094 | 0.952427835 | 0.983220855 | 0.9586 | 0.982315826 |
| GlmZ_glmZ | 5.197551135 | 5.13948033 | 5.245680715 | -0.188244648 | 0.439880743 | -0.416895097 | 0.439706864 | 0.663422091 | 0.854353305 | 1 | 1 |
| glnQ_glnQ | 2.897964501 | 2.884219542 | 2.909221206 | 0.024411816 | 0.306391213 | 0.068554752 | 0.459333335 | 0.616270558 | 0.826442881 | 0.9343 | 0.971380906 |
| SL1344_RS13690_gluconolactonase | -1.414579684 | -1.642379872 | -1.155222945 | 0.564020906 | 0.565025652 | 1.0503337 | 0.08933338 | 0.189391036 | 0.475759989 | 0.2241 | 0.475389522 |
| SL1344_RS09560_glycoside.hydrolase.family.105.protein | 1.050008137 | 1.645332862 | 0.239598326 | -1.427258739 | 0.189861285 | -7.644935239 | 4.68E-05 | 0.000965693 | 0.041549685 | 0.1 | 0.330257709 |
| gntK_gntK | -1.86592154 | -2.057786887 | -1.676767011 | 0.384565386 | 0.390421271 | 0.898570661 | 0.152000025 | 0.247715986 | 0.507742213 | 0.3391 | 0.574059594 |
| SL1344_RS18265_gntR | 1.433827282 | 1.448577617 | 1.424705763 | -0.048735858 | 0.182122058 | -0.275329583 | 0.388000004 | 0.602646113 | 0.800758198 | 0.7687 | 0.881080994 |
| gpmA_gpmA | 5.625476874 | 5.605200125 | 5.646227683 | 0.027246252 | 0.232184574 | 0.104529076 | 0.452666668 | 0.951549604 | 0.982807952 | 1 | 1 |
| GraL_graL | 1.75189443 | 1.692297966 | 1.805252926 | 0.036355222 | 0.55675162 | 0.067873061 | 0.466000001 | 0.885623667 | 0.956676248 | 0.9763 | 0.98944591 |
| groES_groES | 5.915809826 | 6.035848061 | 5.82842844 | -0.202963547 | 0.234374171 | -0.627282347 | 0.264666678 | 0.372141284 | 0.663636397 | 0.5472 | 0.757374174 |
| SL1344_RS14595_gshA | 3.098547871 | 2.961783692 | 3.155640112 | 0.196101981 | 0.118602096 | 1.628347892 | 0.03597614 | 0.082868825 | 0.323636505 | 0.136 | 0.378040431 |
| SL1344_RS16010_gshB | 2.504778715 | 2.424787986 | 2.599106918 | 0.183900397 | 0.102348574 | 1.768299899 | 0.013333676 | 0.067314565 | 0.284210416 | 0.1146 | 0.349870252 |
| SL1344_RS05170_gtgD | -3.693606683 | -3.707475254 | -3.683075251 | 0.010964515 | 0.595706949 | 0.019761496 | 0.490666667 | 0.57679391 | 0.770674993 | 0.6854 | 0.824454312 |
| SL1344_RS05180_gtgE | 0.550337511 | 0.67401961 | 0.424209086 | -0.220496559 | 0.233772052 | -0.922946174 | 0.159227205 | 0.231151159 | 0.50894116 | 0.3091 | 0.563599937 |
| SL1344_RS05185_gtgF | -4.059219578 | -4.138994289 | -3.992036457 | 0.177867099 | 0.719216275 | 0.227886305 | 0.392666671 | 0.546567456 | 0.747309601 | 0.6546 | 0.802141416 |
| SL1344_RS11665_gyrA | 5.199442465 | 5.059635074 | 5.357495687 | 0.305526832 | 0.064302282 | 4.792491126 | 4.68E-05 | 0.002879114 | 0.069201901 | 0.1 | 0.330257709 |
| SL1344_RS13995_head.completion.stabilization.protein | -4.978910404 | -4.764533188 | -5.07530198 | -0.151423068 | 1.667574512 | -0.063160481 | 0.466000001 | 0.696865762 | 0.860473472 | 0.8098 | 0.913737221 |
| helD_helD | 1.948244255 | 2.015992872 | 1.899380056 | -0.102508937 | 0.231519899 | -0.413166526 | 0.358427721 | 0.666134529 | 0.846186683 | 0.7792 | 0.899089795 |
| hemG_hemG | 0.833510395 | 0.79399541 | 0.850612732 | 0.034876561 | 0.184230581 | 0.15872867 | 0.434377084 | 0.721203889 | 0.871349789 | 0.8 | 0.908094712 |
| hflB_hflB | 6.112110008 | 6.058661557 | 6.192203643 | 0.136981738 | 0.050331893 | 2.787970726 | 4.68E-05 | 0.019573712 | 0.166425674 | 0.1 | 0.330257709 |
| SL1344_RS22390_hfq | 4.850162917 | 4.777576076 | 4.879929142 | 0.105663033 | 0.108319913 | 1.01707069 | 0.137241867 | 0.169296171 | 0.457371416 | 0.2608 | 0.532069508 |
| SL1344_RS02410_hha | 1.993394305 | 1.683967917 | 2.452026036 | 0.769158797 | 0.232815172 | 3.431270067 | 4.68E-05 | 0.006534694 | 0.100646237 | 0.1 | 0.330257709 |
| SL1344_RS02415_Hha.toxicity.attenuator | 1.991254558 | 1.690923027 | 2.403680071 | 0.872065888 | 0.348775832 | 2.565139599 | 4.68E-05 | 0.014660952 | 0.150099186 | 0.1 | 0.330257709 |
| SL1344_RS14890_hilA | 1.262688343 | 2.359693773 | 0.193604829 | -2.108941992 | 0.237274655 | -8.911363504 | 4.68E-05 | 0.000770375 | 0.03864478 | 0.1 | 0.330257709 |
| SL1344_RS14845_hilC | 2.965353185 | 3.479836123 | 2.487359494 | -0.973526892 | 0.162348361 | -5.925494815 | 4.68E-05 | 0.001987027 | 0.059569728 | 0.1 | 0.330257709 |
| SL1344_RS14885_hilD | 2.615705401 | 3.279793231 | 2.068069816 | -1.183421267 | 0.261305544 | -4.399957422 | 4.68E-05 | 0.00590635 | 0.098847727 | 0.1 | 0.330257709 |
| SL1344_RS23135_hilE | -1.385828664 | -1.348706878 | -1.402721196 | -0.067660001 | 0.429796326 | -0.139822448 | 0.442666669 | 0.705521278 | 0.862075814 | 0.8119 | 0.911075803 |
| SL1344_RS21865_HlyD.family.type.I.secretion.periplasmic.adaptor.subunit | -1.179081849 | -0.503832008 | -2.282951933 | -1.781404087 | 0.388413706 | -4.519230738 | 4.68E-05 | 0.008903831 | 0.106220281 | 0.1 | 0.330257709 |
| SL1344_RS08745_hns | 6.939148629 | 6.965082976 | 6.84022222 | -0.114140132 | 0.179758835 | -0.727275498 | 0.219853446 | 0.332001107 | 0.630152089 | 0.4 | 0.680350104 |
| hscB_hscB | 1.787534337 | 1.788775576 | 1.786460927 | -0.007398134 | 0.295197912 | -0.022310422 | 0.486342439 | 0.848445293 | 0.939457311 | 0.9121 | 0.960548721 |
| SL1344_RS08210_hslJ | -0.492216775 | -0.115201946 | -0.634969442 | -0.401829453 | 0.456829321 | -0.816863593 | 0.230666681 | 0.330217154 | 0.6144939 | 0.4502 | 0.689016221 |
| SL1344_RS21405_hupA | 7.063823903 | 7.156197261 | 7.028585023 | -0.094880546 | 0.231587551 | -0.315162587 | 0.33333334 | 0.926933614 | 0.974134405 | 0.7 | 0.872542119 |
| SL1344_RS21410_hupB | 4.3883231 | 4.391819209 | 4.385040224 | 0.01366983 | 0.241193889 | 0.072912226 | 0.432000003 | 0.885476157 | 0.956362303 | 0.8875 | 0.947996753 |
| hycD_hycD | -2.729863865 | -2.784203556 | -2.668210825 | 0.057551721 | 0.638806945 | 0.09175608 | 0.460666668 | 0.69303209 | 0.853799959 | 0.7899 | 0.896225898 |
| hycE_hycE | -1.507774691 | -1.604979536 | -1.431647987 | 0.159072517 | 0.326388012 | 0.43943284 | 0.315789482 | 0.470717485 | 0.695295243 | 0.579 | 0.755946729 |
| SL1344_RS06065_hypothetical.protein | 0.332317142 | 1.160606621 | -0.609470806 | -1.780848326 | 0.171435463 | -10.46275179 | 4.68E-05 | 0.000539052 | 0.031454413 | 0.1 | 0.330257709 |

| | | | | | | | | | | |
|---|---|---|---|---|---|---|---|---|---|---|
| SL1344_RS07940_hypothetical.protein | 3.225944222 | 4.108031717 | 2.474355735 | -1.565700497 | 0.206909881 | -7.389563634 | 4.68E-05 | 0.000960841 | 0.044176783 | 0.1 | 0.330257709 |
| SL1344_RS09195_hypothetical.protein | -0.253718447 | 0.392319664 | -0.593034431 | -1.029746843 | 0.517831205 | -1.892666814 | 0.003332494 | 0.047918391 | 0.255169781 | 0.1028 | 0.334371665 |
| SL1344_RS09675_hypothetical.protein | 1.711741766 | 2.906395905 | 0.397263311 | -2.50262221 | 0.13858671 | -18.03988309 | 4.68E-05 | 0.000254817 | 0.022712892 | 0.1 | 0.330257709 |
| SL1344_RS09745_hypothetical.protein | 0.440918982 | 0.834802958 | 0.272918701 | -0.655303992 | 0.407041351 | -1.600323285 | 0.022651765 | 0.074449655 | 0.311938841 | 0.1198 | 0.356618629 |
| SL1344_RS22135_hypothetical.protein | -1.57333194 | -0.602178725 | -2.307578911 | -1.646892317 | 0.403641021 | -3.950951621 | 4.68E-05 | 0.007644313 | 0.103398201 | 0.1 | 0.330257709 |
| iacP_iacP | 0.067269163 | 0.400670646 | -0.274163734 | -0.969032867 | 0.569762362 | -1.777141104 | 4.68E-05 | 0.053791982 | 0.268282528 | 0.1 | 0.330257709 |
| SL1344_RS14895_iagB | 0.963783237 | 1.926038301 | -0.129471078 | -2.005583187 | 0.264220148 | -7.554433999 | 4.68E-05 | 0.000692979 | 0.037514827 | 0.1 | 0.330257709 |
| lamB_iamB | 4.12261369 | 3.797204968 | 4.634304793 | 1.036724453 | 0.465447633 | 2.594685234 | 4.68E-05 | 0.016280952 | 0.159082074 | 0.1 | 0.330257709 |
| idnK_idnK | -1.619671055 | -1.739357049 | -1.17519936 | 0.486051172 | 0.512621221 | 0.798348318 | 0.238507675 | 0.340561323 | 0.615510359 | 0.4824 | 0.709327189 |
| SL1344_RS22990_idnT | -0.318290045 | -0.50378047 | -0.147320042 | 0.402829868 | 0.289248909 | 1.334894772 | 0.036000124 | 0.111952103 | 0.363932099 | 0.146 | 0.385035172 |
| SL1344_RS06655_ihfA | 4.53670781 | 4.542365473 | 4.531617537 | 0.024155637 | 0.218886865 | 0.115815752 | 0.431333336 | 0.816885682 | 0.927501795 | 0.9244 | 0.967829901 |
| SL1344_RS04775_ihfB | 2.960973929 | 2.986542437 | 2.937465514 | -0.049936911 | 0.199463742 | -0.219948226 | 0.423333336 | 0.710764035 | 0.87239476 | 0.9095 | 0.959477001 |
| lldD_ildD | 3.926485494 | 4.456687945 | 3.823314303 | -0.468569848 | 0.319688707 | -1.021431353 | 0.207861442 | 0.223505884 | 0.52716289 | 0.3641 | 0.640725012 |
| ilvM_ilvM | 0.306623229 | 0.071482011 | 0.408553123 | 0.309772873 | 0.32745063 | 0.856297393 | 0.180666687 | 0.258067037 | 0.550195325 | 0.3487 | 0.604533654 |
| SL1344_RS14990_invA | 1.202701029 | 2.347211476 | -0.230979315 | -2.601637527 | 0.160703753 | -16.14451736 | 4.68E-05 | 0.000145356 | 0.020379878 | 0.1 | 0.330257709 |
| SL1344_RS14980_invC | 0.169792706 | 1.188272753 | -0.955937495 | -2.155684713 | 0.209955296 | -10.22857523 | 4.68E-05 | 0.000456591 | 0.030325846 | 0.1 | 0.330257709 |
| SL1344_RS14995_invE | 0.985765192 | 2.067016902 | -0.528145204 | -2.597086472 | 0.182858837 | -14.04864359 | 4.68E-05 | 0.001169652 | 0.042809879 | 0.1 | 0.330257709 |
| SL1344_RS15005_invF | 1.552501907 | 2.683617203 | 0.111887234 | -2.572675138 | 0.153054626 | -16.62891534 | 4.68E-05 | 0.000709059 | 0.035732696 | 0.1 | 0.330257709 |
| SL1344_RS15000_invG | 1.282412599 | 2.498852504 | -0.128884332 | -2.627483174 | 0.216574572 | -12.12511595 | 4.68E-05 | 0.000219397 | 0.024683533 | 0.1 | 0.330257709 |
| InvR_invR | 3.230376951 | 3.82055817 | 2.390460812 | -1.457744573 | 0.32847607 | -4.242615381 | 4.68E-05 | 0.014477453 | 0.150331262 | 0.1 | 0.330257709 |
| SL1344_RS14355_iroB | -1.419867489 | -0.166491533 | -2.139498024 | -1.820453133 | 0.651798787 | -2.7294638 | 4.68E-05 | 0.019804207 | 0.166835762 | 0.1 | 0.330257709 |
| SL1344_RS24725_IS91.family.transposase | 0.695120662 | 1.890037449 | -0.419411026 | -2.324562764 | 0.437625959 | -5.283311875 | 4.68E-05 | 0.004008147 | 0.082055852 | 0.1 | 0.330257709 |
| IsrA_isrA | -6.504826857 | -6.665527118 | -6.365427506 | 0.302299059 | 1.613553417 | 0.167596911 | 0.420386412 | 0.535510324 | 0.739310251 | 0.6331 | 0.790161113 |
| IsrB_isrB | -2.705367775 | -2.837228387 | -2.471781108 | 0.334363222 | 0.834431895 | 0.346043374 | 0.361333339 | 0.651590418 | 0.829599105 | 0.7642 | 0.88166535 |
| IsrC_isrC | -2.427886211 | -2.478720653 | -2.367167551 | -0.011487769 | 0.746234358 | -0.01170717 | 0.492666667 | 0.753691599 | 0.89099714 | 0.8788 | 0.943405231 |
| IsrD_isrD | -9.258072135 | -9.874257822 | -8.459746143 | 1.382394808 | 4.295302446 | 0.275781698 | 0.354000006 | 0.513752778 | 0.723888498 | 0.6278 | 0.783412438 |
| IsrE_isrE | 0.081126683 | 0.405589059 | -0.202874209 | -0.069480439 | 0.92196279 | -0.096960994 | 0.454000002 | 0.74291849 | 0.891481006 | 0.9568 | 0.98021626 |
| IsrF_isrF | -0.263951271 | 0.492125395 | -0.98788143 | -1.441854932 | 0.2895914 | -4.893791396 | 4.68E-05 | 0.003301595 | 0.070814271 | 0.1 | 0.330257709 |
| IsrG_isrG | -2.606343963 | -2.102268605 | -2.780682742 | -0.633042498 | 0.751302716 | -0.731921061 | 0.248000013 | 0.375735971 | 0.644822896 | 0.5089 | 0.726490761 |
| IsrH.1_isrH | -3.602585656 | -3.419906501 | -3.789228203 | -0.36188101 | 0.647009215 | -0.537689088 | 0.271333344 | 0.408508567 | 0.643794341 | 0.5201 | 0.709064934 |
| IsrH.2_isrH2 | -3.662569577 | -3.419282022 | -3.931798541 | -0.5333829 | 0.736474108 | -0.668297741 | 0.233844118 | 0.354860001 | 0.605541005 | 0.4664 | 0.673665024 |
| IsrI_isrI | -2.877758634 | -2.901425637 | -2.853749492 | 0.056047519 | 0.403125458 | 0.1310827 | 0.441705532 | 0.507645438 | 0.719002274 | 0.6149 | 0.776603891 |
| IsrJ_isrJ | -1.904805062 | -1.964989995 | -1.881247197 | 0.043143386 | 0.472730893 | 0.084089641 | 0.467021987 | 0.702189834 | 0.85989007 | 0.8053 | 0.907379138 |
| IsrK_isrK | 1.014770199 | 1.184289838 | 0.853711074 | -0.145851183 | 0.81481711 | -0.197140466 | 0.432000003 | 0.839968279 | 0.937048935 | 0.9493 | 0.978031536 |
| IsrL_isrL | -2.013780265 | -1.929818533 | -2.096832378 | -0.089095624 | 0.499875569 | -0.168472884 | 0.424000003 | 0.673230102 | 0.840401437 | 0.7754 | 0.886416333 |
| IsrM_isrM | -4.706836345 | -4.917601977 | -4.402750409 | 0.442324056 | 1.084859361 | 0.389470934 | 0.325333341 | 0.533270537 | 0.74248833 | 0.6398 | 0.80020483 |
| IsrN_isrN | -7.877080763 | -8.435681976 | -7.312379643 | 1.136746895 | 3.772156289 | 0.255281998 | 0.352666673 | 0.543140063 | 0.754362124 | 0.6677 | 0.817942355 |
| IsrO_isrO | -0.54910068 | -0.753920663 | -0.314963958 | 0.320092219 | 0.60080114 | 0.531777167 | 0.299333342 | 0.46979153 | 0.725650373 | 0.6079 | 0.797283648 |
| IsrP_isrP | -1.781114434 | -1.749191033 | -1.796938093 | -0.009778101 | 0.460389289 | -0.014917931 | 0.492 | 0.682076938 | 0.848347537 | 0.7956 | 0.901967476 |
| IsrQ_isrQ | -6.491332068 | -6.175524875 | -6.839274981 | -0.680040285 | 1.630922722 | -0.374208233 | 0.341105936 | 0.479009957 | 0.694905004 | 0.5867 | 0.753432497 |
| IstR.1_istR1 | -2.984914012 | -3.156323115 | -2.664007427 | 0.479266506 | 0.700836904 | 0.655310829 | 0.257161904 | 0.405976061 | 0.656237239 | 0.5261 | 0.725347444 |
| IstR.2_istR2 | -2.6682617 | -2.802486132 | -2.48117318 | 0.279923822 | 0.767849113 | 0.342062682 | 0.351765496 | 0.598695126 | 0.796363367 | 0.7178 | 0.853943357 |
| ksgA_ksgA | 2.827677197 | 2.604620784 | 2.897404874 | 0.290385707 | 0.160602583 | 1.625136652 | 0.018654474 | 0.099595256 | 0.358010098 | 0.1232 | 0.361056054 |
| SL1344_RS08205_ldhA | 1.470520268 | 1.768724215 | 1.224793164 | -0.528709396 | 0.248321802 | -2.055200201 | 0.005334196 | 0.04151822 | 0.236767588 | 0.1037 | 0.335691736 |
| SL1344_RS00585_leuO | -4.117387949 | -4.152664746 | -4.050592342 | 0.07177985 | 1.117831948 | 0.057413342 | 0.478000001 | 0.723795027 | 0.872621076 | 0.8191 | 0.916404277 |
| SL1344_RS21750_lexA | 3.978740154 | 4.123108386 | 3.809064362 | -0.240940131 | 0.202038636 | -1.332799494 | 0.116666701 | 0.110398624 | 0.378948446 | 0.1975 | 0.472766282 |
| ligA_ligA | 3.165271264 | 2.86707111 | 3.351220314 | 0.499251835 | 0.145591645 | 3.471255082 | 4.68E-05 | 0.020077767 | 0.169847828 | 0.1 | 0.330257709 |
| livF_livF | -1.412425464 | -1.318607226 | -1.467001091 | -0.15559046 | 0.379045774 | -0.387075551 | 0.353333339 | 0.539094421 | 0.755007605 | 0.6619 | 0.816653929 |
| lolA_lolA | 2.354283046 | 2.147974707 | 2.422816055 | 0.269563196 | 0.259074895 | 0.885913368 | 0.199333351 | 0.266745695 | 0.567337931 | 0.3628 | 0.62479776 |
| lolB_lolB | 0.455158768 | 0.524161072 | 0.41836949 | -0.125289083 | 0.26115805 | -0.429847405 | 0.338000007 | 0.523273744 | 0.750867579 | 0.6757 | 0.827284758 |
| lplA_lplA | 1.33873071 | 1.339360518 | 1.3381654 | -0.042318533 | 0.190659732 | -0.228365391 | 0.419053967 | 0.665824283 | 0.837958488 | 0.8237 | 0.910468631 |
| SL1344_RS06600_lpxR | -0.077360988 | 0.977654749 | -0.93855733 | -1.782080668 | 0.442853076 | -3.956001143 | 4.68E-05 | 0.006302271 | 0.098439933 | 0.1 | 0.330257709 |
| SL1344_RS11955_lrhA | 1.266557014 | 0.896065473 | 1.559627175 | 0.662816721 | 0.162766564 | 4.04172197 | 4.68E-05 | 0.007023707 | 0.099002291 | 0.1 | 0.330257709 |

| | | | | | | | | | | |
|---|---|---|---|---|---|---|---|---|---|---|
| SL1344_RS04660_lrp | 4.500501031 | 4.38783431 | 4.51428148 | 0.128576889 | 0.207696111 | 0.605411942 | 0.33333334 | 0.489938599 | 0.747452402 | 0.7003 | 0.872601817 |
| SL1344_RS09120_L.serine.ammonia.lyase | 4.444281957 | 4.767240668 | 4.076691588 | -0.696703271 | 0.07430394 | -9.449236437 | 4.68E-05 | 0.001851576 | 0.055997795 | 0.1 | 0.330257709 |
| SL1344_RS20930_lsrR | 1.093024778 | 1.060201191 | 1.210047863 | 0.126066024 | 0.193677168 | 0.486175757 | 0.339773491 | 0.621178357 | 0.813813647 | 0.7024 | 0.862737087 |
| SL1344_RS14590_luxS | 3.26949231 | 3.201881746 | 3.338834128 | 0.185650479 | 0.129948378 | 1.442449979 | 0.023333527 | 0.083604526 | 0.325022439 | 0.1201 | 0.357806787 |
| SL1344_RS09795_lysine.N.methylase | 2.885362867 | 3.366982967 | 2.338474768 | -1.070255565 | 0.176355514 | -6.251847084 | 4.68E-05 | 0.00155704 | 0.0532943 | 0.1 | 0.330257709 |
| malF_malF | 2.667047386 | 2.436804922 | 2.897525994 | 0.488868818 | 0.161412249 | 3.093944969 | 4.68E-05 | 0.01709537 | 0.155324054 | 0.1 | 0.330257709 |
| malG_malG | 1.633632882 | 1.176623966 | 1.996685556 | 0.848543904 | 0.173917183 | 4.944933686 | 4.68E-05 | 0.003170575 | 0.07120531 | 0.1 | 0.330257709 |
| SL1344_RS18135_malT | 4.123293891 | 3.924622545 | 4.259745343 | 0.324789175 | 0.181370629 | 1.675340358 | 0.021333546 | 0.063443197 | 0.293646402 | 0.1176 | 0.355925099 |
| SL1344_RS07280_malY. | -1.226783122 | -1.196204837 | -1.261602955 | -0.063821943 | 0.277723726 | -0.201689606 | 0.40373085 | 0.559347396 | 0.76117872 | 0.6685 | 0.81707772 |
| mdoD_mdoD | 2.949313461 | 3.070173715 | 2.821852765 | -0.245887821 | 0.070967175 | -3.398321811 | 4.68E-05 | 0.012274401 | 0.12449342 | 0.1 | 0.330257709 |
| mdoG_mdoG | 3.887895841 | 3.860562647 | 3.914347735 | 0.111976154 | 0.144340362 | 0.767621991 | 0.152666692 | 0.26873133 | 0.569927903 | 0.322 | 0.558423043 |
| SL1344_RS09375_membrane.protein | 2.724861149 | 3.112157031 | 2.208119129 | -0.891786577 | 0.125531025 | -6.941889321 | 4.68E-05 | 0.002186475 | 0.057811878 | 0.1 | 0.330257709 |
| SL1344_RS21870_membrane.protein | 3.295893311 | 3.49154478 | 3.104777591 | -0.43358976 | 0.153719677 | -2.995795065 | 4.68E-05 | 0.013597345 | 0.141734657 | 0.1 | 0.330257709 |
| SL1344_RS11835_menC | 1.37580591 | 1.199356039 | 1.628288269 | 0.391486043 | 0.209398525 | 1.900157731 | 0.014657206 | 0.046119614 | 0.244551652 | 0.1141 | 0.350533008 |
| SL1344_RS21470_metA | -0.662304319 | -0.598188293 | -0.728109205 | -0.124151138 | 0.248592958 | -0.494335261 | 0.293333343 | 0.449594404 | 0.677034012 | 0.5487 | 0.732258152 |
| SL1344_RS21065_metB | 0.965154698 | 0.985551317 | 0.933543251 | -0.056817918 | 0.152846872 | -0.344062494 | 0.358000006 | 0.549813183 | 0.758260126 | 0.6676 | 0.818436995 |
| SL1344_RS16345_metC | 0.575008122 | 0.721053619 | 0.478664166 | -0.204958695 | 0.206908151 | -0.904582613 | 0.185876102 | 0.258645914 | 0.530799364 | 0.3541 | 0.598721455 |
| SL1344_RS20385_metE | 0.740964416 | 0.880473536 | 0.579148468 | -0.302174177 | 0.154387653 | -1.923567645 | 0.01666694 | 0.053861642 | 0.245953018 | 0.1169 | 0.352042299 |
| SL1344_RS21095_metF | -0.465180864 | -0.219968725 | -0.747947134 | -0.569765803 | 0.292607574 | -1.941800901 | 0.005996769 | 0.048047005 | 0.240067527 | 0.1069 | 0.339718331 |
| metG_metG | 3.206282271 | 3.006793412 | 3.341315078 | 0.356989881 | 0.120453949 | 3.067926047 | 4.68E-05 | 0.027317499 | 0.194520508 | 0.1 | 0.330257709 |
| SL1344_RS21500_metH | 2.682929278 | 2.710230072 | 2.65544145 | -0.027775497 | 0.251433335 | -0.101735544 | 0.454000002 | 0.852514683 | 0.941348714 | 0.9379 | 0.973017945 |
| SL1344_RS18435_methyl.accepting.chemotaxis.citrate.transducer | 4.127337122 | 5.179616985 | 3.42246553 | -1.760193306 | 0.29832397 | -4.999280425 | 4.68E-05 | 0.010955235 | 0.133510326 | 0.1 | 0.330257709 |
| SL1344_RS08105_methyl.accepting.chemotaxis.protein | 4.619985128 | 5.438066304 | 3.763205473 | -1.679924737 | 0.132878134 | -12.71230352 | 4.68E-05 | 0.002692904 | 0.070619908 | 0.1 | 0.330257709 |
| SL1344_RS16230_methyl.accepting.chemotaxis.protein | 3.200388158 | 3.461647324 | 3.114581395 | -0.337336656 | 0.187708524 | -1.345767527 | 0.065333399 | 0.129876215 | 0.408615801 | 0.1649 | 0.421383838 |
| SL1344_RS16300_methyl.accepting.chemotaxis.protein | 3.866711648 | 4.699414803 | 3.159928376 | -1.388319244 | 0.309365566 | -4.905674559 | 4.68E-05 | 0.001093369 | 0.048569076 | 0.1 | 0.330257709 |
| SL1344_RS16625_methyl.accepting.chemotaxis.protein | 5.546734684 | 6.114300215 | 4.741203605 | -1.425790743 | 0.326122421 | -4.678756646 | 4.68E-05 | 0.000967909 | 0.04676688 | 0.1 | 0.330257709 |
| SL1344_RS21060_metJ | 2.664062294 | 2.753877367 | 2.455163022 | -0.305875557 | 0.159466976 | -1.96413786 | 0.004001152 | 0.039006357 | 0.229108152 | 0.1037 | 0.335291034 |
| SL1344_RS21070_metL | 2.555369259 | 2.397368521 | 2.621609064 | 0.165330644 | 0.172832105 | 0.926864009 | 0.189873436 | 0.269501184 | 0.562575934 | 0.3628 | 0.618232575 |
| metQ_metQ | 3.019219742 | 3.031231281 | 3.013031155 | -0.004287896 | 0.109546809 | -0.024377801 | 0.480346436 | 0.602902276 | 0.813287243 | 0.8245 | 0.918394618 |
| SL1344_RS20380_metR | -1.334300408 | -1.133375597 | -1.471554571 | -0.360193961 | 0.344042168 | -0.980203529 | 0.144570313 | 0.220251657 | 0.487800355 | 0.3131 | 0.551181731 |
| MgrR_mgrR | 1.75338813 | 1.839949904 | 1.665807321 | -0.059035674 | 0.789337536 | -0.073100594 | 0.465689542 | 0.932383089 | 0.975726239 | 0.9964 | 0.998457809 |
| MicA_micA | -2.057116462 | -2.001157141 | -2.105516475 | -0.104176876 | 0.782441683 | -0.138060976 | 0.441039309 | 0.761102821 | 0.895117259 | 0.8435 | 0.925564194 |
| MicC_micC | -2.657479607 | -2.598409798 | -2.725536787 | -0.121295473 | 0.439737269 | -0.254480215 | 0.378666671 | 0.513348265 | 0.723892339 | 0.6296 | 0.78512684 |
| MicF_micF | 0.607737408 | -0.147760684 | 1.518261979 | 1.58547502 | 0.404427957 | 3.869654027 | 4.68E-05 | 0.008785199 | 0.116839248 | 0.1 | 0.330257709 |
| SL1344_RS07395_mlc | 1.34735704 | 1.349267857 | 1.344892791 | 0.013437601 | 0.215206164 | 0.061562553 | 0.476000001 | 0.776030472 | 0.902419619 | 0.8745 | 0.941733006 |
| moaA_moaA | 1.010322836 | 1.034148138 | 0.992024264 | -0.045476142 | 0.159798134 | -0.278854014 | 0.376415728 | 0.586729145 | 0.783761787 | 0.6908 | 0.833424117 |
| moaD_moaD | -0.514685507 | -0.568564506 | -0.44593204 | 0.154770844 | 0.337541531 | 0.432544584 | 0.315123259 | 0.5200142 | 0.743141751 | 0.6562 | 0.813410677 |
| moaE_moaE | 0.249283441 | 0.191200269 | 0.308201196 | 0.143231993 | 0.245043965 | 0.568453867 | 0.271333344 | 0.411730169 | 0.664506781 | 0.5408 | 0.735225967 |
| mobA_mobA | 0.501540898 | 0.493513227 | 0.513213308 | 0.041812392 | 0.201427861 | 0.207114816 | 0.413724187 | 0.60517126 | 0.796779936 | 0.7368 | 0.860532145 |
| SL1344_RS24715_mobA | 4.149796836 | 5.138473469 | 2.807088799 | -2.282819124 | 0.279875515 | -8.184517919 | 4.68E-05 | 0.000621197 | 0.038751025 | 0.1 | 0.330257709 |
| SL1344_RS24720_mobC | 1.898931803 | 3.215964494 | 0.844156757 | -2.373247759 | 0.221265872 | -9.972189194 | 4.68E-05 | 0.004132229 | 0.084646154 | 0.1 | 0.330257709 |
| SL1344_RS03955_modA | 3.510761186 | 4.254995185 | 2.610076516 | -1.424749092 | 0.320405632 | -4.533667447 | 4.68E-05 | 0.001922778 | 0.060446146 | 0.1 | 0.330257709 |
| modB_modB | 3.371650101 | 4.024204007 | 2.560389385 | -1.401707644 | 0.231003877 | -5.584431998 | 4.68E-05 | 0.004258333 | 0.084713631 | 0.1 | 0.330257709 |
| SL1344_RS03965_modC | 3.523633315 | 4.022957428 | 3.051362679 | -1.122358442 | 0.350158066 | -3.176718914 | 4.68E-05 | 0.006300887 | 0.103454608 | 0.1 | 0.330257709 |
| SL1344_RS09620_motA | 5.067574502 | 6.148330881 | 3.610429536 | -2.529978836 | 0.353937917 | -7.19106547 | 4.68E-05 | 0.000389934 | 0.033232465 | 0.1 | 0.330257709 |
| motB_motB | 5.175884004 | 6.225983125 | 4.008267648 | -2.113708605 | 0.234824888 | -9.234705873 | 4.68E-05 | 7.41E-05 | 0.020176911 | 0.1 | 0.330257709 |
| SL1344_RS09450_msbB | 2.109463589 | 2.461189354 | 1.69135083 | -0.770267821 | 0.080424678 | -9.48914035 | 4.68E-05 | 0.000843545 | 0.036631429 | 0.1 | 0.330257709 |
| Msr_Msr | -0.592195887 | -0.63142567 | -0.549098725 | 0.096363065 | 1.049850811 | 0.105016349 | 0.453697537 | 0.862702959 | 0.946368119 | 0.9394 | 0.973095248 |
| mtlR_mtlR | 1.470214451 | 1.482678422 | 1.457769657 | -0.02252121 | 0.099135219 | -0.215834359 | 0.393737512 | 0.534994799 | 0.738021398 | 0.6476 | 0.798286018 |
| mukB_mukB | 5.174848805 | 5.383637571 | 5.079325671 | -0.303118941 | 0.067751718 | -3.942375341 | 4.68E-05 | 0.024735317 | 0.190300257 | 0.1 | 0.330257709 |
| murD_murD | 3.691656938 | 3.674786702 | 3.771854852 | 0.095897399 | 0.099088801 | 0.808156407 | 0.249333346 | 0.323158858 | 0.611095566 | 0.4991 | 0.724798163 |
| SL1344_RS00625_murE | 3.049421243 | 2.948103693 | 3.134504092 | 0.097070808 | 0.222730363 | 0.493544816 | 0.320000008 | 0.496465447 | 0.747355483 | 0.6599 | 0.822877655 |
| SL1344_RS09280_N.acetyltransferase | 0.405399871 | 0.677628748 | -0.100671035 | -0.828214585 | 0.524482287 | -1.76378264 | 0.0020023 | 0.056236057 | 0.276443715 | 0.1016 | 0.332502975 |

| | | | | | | | | | | | |
|---|---|---|---|---|---|---|---|---|---|---|---|
| napB_napB | -0.776167214 | -0.850945451 | -0.715256157 | 0.139064377 | 0.249887765 | 0.507330352 | 0.280666677 | 0.42409336 | 0.655279861 | 0.5275 | 0.714669709 |
| SL1344_RS08805_narG | 0.581428033 | 0.696166798 | 0.136666766 | -0.572839761 | 0.415660571 | -1.255957694 | 0.08794142 | 0.159655725 | 0.447696359 | 0.2016 | 0.459897543 |
| SL1344_RS08800_narH | -1.3741624 | -1.028789285 | -1.647632769 | -0.611913458 | 0.311264622 | -1.882135238 | 0.019986902 | 0.055308368 | 0.252258665 | 0.1184 | 0.354814657 |
| SL1344_RS08790_narI | -2.143442225 | -1.812727043 | -2.388283323 | -0.576298695 | 0.391118529 | -1.412269165 | 0.051965441 | 0.108117814 | 0.339030647 | 0.1776 | 0.41808351 |
| SL1344_RS08795_narJ | -2.730464792 | -2.333551584 | -3.241576229 | -0.923365784 | 0.604393649 | -1.483108028 | 0.049300555 | 0.094528473 | 0.328677145 | 0.1579 | 0.403825791 |
| SL1344_RS08810_narK | -0.356573525 | -0.223591657 | -0.503589721 | -0.348991688 | 0.377295083 | -0.919983145 | 0.10326453 | 0.21526999 | 0.504160957 | 0.2455 | 0.490212138 |
| SL1344_RS08820_narL | -0.052018179 | 0.27758044 | -0.31938077 | -0.58546831 | 0.261601107 | -2.225680477 | 0.004664544 | 0.035224951 | 0.210002772 | 0.1037 | 0.335689388 |
| nepI_nepI | -1.448728316 | -1.392619914 | -1.578909143 | -0.159747931 | 0.443131063 | -0.294344752 | 0.389073955 | 0.665220321 | 0.838905144 | 0.7653 | 0.889565854 |
| nhaA_nhaA | 1.737609502 | 1.748728559 | 1.727554967 | -0.012722045 | 0.124755107 | -0.098918011 | 0.457028649 | 0.655283384 | 0.826608057 | 0.7547 | 0.870811153 |
| nirD_nirD | -3.215090211 | -3.447872835 | -2.988276769 | 0.45587012 | 0.529814828 | 0.816982406 | 0.177333354 | 0.27292932 | 0.517670438 | 0.3622 | 0.583893821 |
| nlpD_nlpD | 6.03760131 | 6.072375306 | 5.994757282 | -0.057786808 | 0.189261034 | -0.290736325 | 0.411333337 | 0.740653426 | 0.893298337 | 0.9067 | 0.959921965 |
| nrdI_nrdI | -1.816969993 | -1.762041547 | -1.872425013 | -0.156398829 | 0.413902047 | -0.356419006 | 0.353333339 | 0.53492044 | 0.74556969 | 0.6722 | 0.816143324 |
| SL1344_RS21955_nrfB | -0.046488915 | 0.851345065 | -1.073007671 | -1.915449109 | 0.236353621 | -8.066405143 | 4.68E-05 | 0.000944971 | 0.040769193 | 0.1 | 0.330257709 |
| SL1344_RS21960_nrfC | 0.171625414 | 0.979878783 | -0.963846945 | -1.9394245 | 0.233957127 | -8.172947355 | 4.68E-05 | 0.003133545 | 0.065951372 | 0.1 | 0.330257709 |
| SL1344_RS21975_nrfG | -2.535217876 | -2.279352059 | -2.699943241 | -0.383949029 | 0.451374868 | -0.773449232 | 0.211192554 | 0.329563952 | 0.579615206 | 0.4295 | 0.648282178 |
| nudC_nudC | 0.572778655 | 0.476408558 | 0.785470874 | 0.283081119 | 0.175272575 | 1.467735002 | 0.060000072 | 0.109788311 | 0.353196862 | 0.186 | 0.431502846 |
| nusA_nusA | 4.456206641 | 4.355565772 | 4.53986543 | 0.179907142 | 0.108501688 | 1.582322658 | 0.029980163 | 0.064314643 | 0.291921522 | 0.1273 | 0.369969957 |
| nusB_nusB | 3.803443902 | 3.681842587 | 3.988600462 | 0.310757748 | 0.325181454 | 0.864637505 | 0.169220541 | 0.2912541 | 0.59365553 | 0.3101 | 0.581904945 |
| nusG_nusG | 5.001036282 | 5.014283661 | 4.984562935 | 0.019997259 | 0.1359371 | 0.158788453 | 0.454363759 | 0.758564887 | 0.89833515 | 0.9679 | 0.986185971 |
| obgE_obgE | 1.750064297 | 1.79345524 | 1.725101085 | -0.064930064 | 0.104677661 | -0.550890382 | 0.278481023 | 0.451582992 | 0.68023156 | 0.5567 | 0.741454229 |
| ompL_ompL | -4.396235278 | -4.053202058 | -4.59239716 | -0.535781046 | 1.411913749 | -0.321954762 | 0.355333339 | 0.646956213 | 0.824443177 | 0.7238 | 0.861803301 |
| SL1344_RS18070_ompR | 2.345064691 | 2.277404233 | 2.420336189 | 0.133227356 | 0.260168809 | 0.490923471 | 0.307128589 | 0.488213308 | 0.741820813 | 0.6367 | 0.822201855 |
| ompX_ompX | 5.753049985 | 5.502869376 | 6.05107434 | 0.414382282 | 0.273711478 | 1.80083128 | 0.003998486 | 0.043472014 | 0.249106353 | 0.1038 | 0.335813107 |
| OmrA_omrA | -1.668454754 | -1.956718925 | -0.944667792 | 0.816045244 | 0.6877193 | 1.137866308 | 0.126666698 | 0.172079275 | 0.453033549 | 0.2691 | 0.535357058 |
| OmrB_omrB | -2.51935842 | -2.720824 | -2.307064001 | 0.410311456 | 0.471215916 | 0.824305134 | 0.164556985 | 0.262032614 | 0.519772477 | 0.3563 | 0.584411249 |
| oppB_oppB | 1.226175718 | 1.286782574 | 1.198821514 | -0.074199679 | 0.240458266 | -0.155529646 | 0.400399737 | 0.733355224 | 0.886278763 | 0.8194 | 0.923129868 |
| SL1344_RS14860_orgA | 2.082579366 | 3.096607515 | 0.979788748 | -2.122266421 | 0.172422874 | -12.21744399 | 4.68E-05 | 0.000163606 | 0.022920521 | 0.1 | 0.330257709 |
| OxyS_oxyS | -1.217053084 | -1.270992582 | -1.164146693 | -0.04466753 | 0.547893917 | -0.085164782 | 0.467688209 | 0.7334424 | 0.879283806 | 0.9 | 0.952491608 |
| pabB_pabB | 1.944978287 | 2.287733277 | 1.574183892 | -0.716580137 | 0.093686012 | -7.68528507 | 4.68E-05 | 0.002054321 | 0.053014873 | 0.1 | 0.330257709 |
| pagP_pagP | 1.772076277 | 1.563694834 | 1.859419785 | 0.299632939 | 0.30136705 | 0.950062504 | 0.153333358 | 0.221370837 | 0.520902348 | 0.2701 | 0.54329636 |
| SL1344_RS10525_pduA | -4.940597301 | -4.922112587 | -4.947366187 | -0.067355073 | 1.102216078 | -0.050598155 | 0.474666668 | 0.617482352 | 0.80351781 | 0.7394 | 0.864188578 |
| pduC_pduC | -4.045031275 | -4.013722987 | -4.057039457 | 0.00984574 | 1.033868334 | 0.010175964 | 0.498 | 0.701286129 | 0.859938769 | 0.8187 | 0.914947047 |
| pduD_pduD | -5.181180225 | -5.072936741 | -5.289227254 | -0.430680379 | 1.662776873 | -0.238369659 | 0.383077953 | 0.566260696 | 0.776644657 | 0.7313 | 0.853142342 |
| pduE_pduE | -5.212970998 | -5.094480249 | -5.314791535 | -0.180304938 | 1.514064199 | -0.11708668 | 0.448000002 | 0.647253921 | 0.824006827 | 0.7453 | 0.867586066 |
| pdxK_pdxK | -0.512534881 | -0.535881758 | -0.491759206 | 0.045771913 | 0.295695655 | 0.148280249 | 0.425333336 | 0.662707006 | 0.834934554 | 0.7493 | 0.873340758 |
| SL1344_RS06590_pfkB | 2.463359231 | 2.164454824 | 2.768484453 | 0.561709454 | 0.156954189 | 3.639859291 | 4.68E-05 | 0.006430603 | 0.099125251 | 0.1 | 0.330257709 |
| SL1344_RS13960_phage.virion.morphogenesis.protein | -4.642842868 | -5.153141804 | -4.275791346 | 0.884786371 | 0.895728732 | 0.919618062 | 0.125333365 | 0.22303628 | 0.479213661 | 0.2955 | 0.523761516 |
| SL1344_RS02035_phoB | 2.62794293 | 2.472182742 | 2.829194451 | 0.353316724 | 0.109738536 | 3.168702024 | 4.68E-05 | 0.013780558 | 0.13709956 | 0.1 | 0.330257709 |
| SL1344_RS11645_phoE | 7.410947504 | 7.09072434 | 7.756519249 | 0.706886308 | 0.111611473 | 6.398686072 | 4.68E-05 | 0.012279711 | 0.140543031 | 0.1 | 0.330257709 |
| SL1344_RS05535_phoH | 1.531189811 | 1.865442429 | 1.405389938 | -0.46731662 | 0.314819297 | -1.573977147 | 0.023984199 | 0.065030509 | 0.293513906 | 0.1246 | 0.364546732 |
| SL1344_RS06080_phoP | 4.054031296 | 4.030849946 | 4.076634579 | 0.017251191 | 0.139869692 | 0.122538806 | 0.455696204 | 0.81685949 | 0.924614421 | 0.9307 | 0.968854825 |
| SL1344_RS06075_phoQ | 3.824199318 | 3.794802369 | 3.837471974 | 0.059668314 | 0.142167893 | 0.446435124 | 0.304000009 | 0.431380198 | 0.702584042 | 0.5974 | 0.793218027 |
| phoR_phoR | 2.181760511 | 1.857439073 | 2.548241793 | 0.699069018 | 0.133147332 | 5.277338808 | 4.68E-05 | 0.003393162 | 0.071462311 | 0.1 | 0.330257709 |
| SL1344_RS13700_phospho.2.dehydro.3.deoxyheptonate.aldolase.Tyr.sensitive | 0.536683242 | 0.262250045 | 0.80163166 | 0.535860774 | 0.150805724 | 3.473082409 | 4.68E-05 | 0.011265359 | 0.119154813 | 0.1001 | 0.330398689 |
| SL1344_RS10625_phsA | 1.262969459 | 0.996897097 | 1.532233056 | 0.549494155 | 0.18585608 | 3.063728466 | 4.68E-05 | 0.019887659 | 0.162296259 | 0.1 | 0.330257709 |
| SL1344_RS10620_phsB | -0.754595132 | -0.904421607 | -0.456055401 | 0.419887405 | 0.328044385 | 1.220673123 | 0.101333374 | 0.143263866 | 0.402526778 | 0.2242 | 0.479315151 |
| SL1344_RS10615_phsC | 0.40396974 | 0.216089714 | 0.692654033 | 0.499315185 | 0.35592762 | 1.435811483 | 0.038000117 | 0.090400974 | 0.339296619 | 0.1324 | 0.375506753 |
| potG_potG | -0.794782605 | -0.986683138 | -0.649865143 | 0.360617139 | 0.298174663 | 1.156332879 | 0.071952091 | 0.162832847 | 0.431709398 | 0.1927 | 0.435237857 |
| SL1344_RS09215_prc | 5.236206925 | 5.627394593 | 4.822923525 | -0.722324737 | 0.135535178 | -5.556688441 | 4.68E-05 | 0.000678258 | 0.040486777 | 0.1 | 0.330257709 |
| SL1344_RS14880_prgH | 1.702464212 | 2.942434555 | 0.255852773 | -2.759712699 | 0.302378095 | -9.215940202 | 4.68E-05 | 0.000216798 | 0.02566909 | 0.1 | 0.330257709 |
| SL1344_RS14875_prgI | 0.385362699 | 1.876523752 | -1.067738114 | -2.912282041 | 0.392086361 | -7.378251797 | 4.68E-05 | 0.000703168 | 0.037708502 | 0.1 | 0.330257709 |
| SL1344_RS14870_prgJ | 1.310280897 | 2.675676501 | -0.113750943 | -2.770031767 | 0.249815646 | -11.16039759 | 4.68E-05 | 0.000317308 | 0.026857306 | 0.1 | 0.330257709 |
| SL1344_RS14865_prgK | 2.23497102 | 3.394485835 | 1.076568662 | -2.297131968 | 0.187406972 | -12.23103803 | 4.68E-05 | 0.000245624 | 0.025807149 | 0.1 | 0.330257709 |

| | | | | | | | | | | | |
|---|---|---|---|---|---|---|---|---|---|---|---|
| SL1344_RS09220_proQ | 4.717507528 | 5.018024209 | 4.331777298 | -0.666436458 | 0.123546793 | -5.232478887 | 4.68E-05 | 0.003601025 | 0.077845787 | 0.1 | 0.330257709 |
| SL1344_RS09255_prpA.pphA | -1.900920719 | -1.47328141 | -2.586060043 | -1.120273455 | 0.3750262 | -2.966954613 | 4.68E-05 | 0.025862329 | 0.172240314 | 0.1003 | 0.330703955 |
| psiB_psiB | -2.440980146 | -2.430762487 | -2.448164218 | -0.03582979 | 0.482139175 | -0.067171603 | 0.470000001 | 0.641922311 | 0.817161923 | 0.7471 | 0.867381661 |
| psiB_psiB.1 | -2.924009633 | -3.131728942 | -2.732759197 | 0.406817694 | 0.433964543 | 0.903658188 | 0.169333355 | 0.262496733 | 0.50176808 | 0.3507 | 0.568713837 |
| pssA_pssA | 3.523919075 | 3.479452398 | 3.54290511 | 0.079757464 | 0.151079815 | 0.551358838 | 0.273817466 | 0.393122736 | 0.673580022 | 0.5324 | 0.755738908 |
| SL1344_RS11995_pta | 4.933347466 | 4.188379102 | 5.000388865 | 0.712065967 | 0.514283751 | 1.057838612 | 0.222518336 | 0.241534922 | 0.54658941 | 0.4 | 0.680350104 |
| SL1344_RS15255_ptpS | 0.024842651 | -0.640666137 | 0.499410638 | 1.137951937 | 0.400994707 | 2.752709087 | 4.68E-05 | 0.024626154 | 0.182882873 | 0.1 | 0.330257709 |
| SL1344_RS02910_PTS.mannose.fructose.sorbose.transporter.subunit.IIB | -5.643683944 | -5.456529795 | -5.821462315 | -0.369900742 | 1.440719562 | -0.253534987 | 0.379080618 | 0.55650634 | 0.762346467 | 0.677 | 0.821917315 |
| SL1344_RS07105_purR | 1.613622867 | 1.466769907 | 1.664407425 | 0.187293752 | 0.153007594 | 1.085027007 | 0.141905424 | 0.200050601 | 0.481761173 | 0.287 | 0.53884835 |
| putA_putA | 6.140512977 | 6.175297676 | 6.101231572 | 0.069282952 | 0.18054618 | 0.381113071 | 0.440373087 | 0.760946092 | 0.901937125 | 1 | 1 |
| SL1344_RS23405_putative.inner.membrane.protein | 2.017814235 | 2.182409315 | 1.980959731 | -0.217656022 | 0.270087477 | -0.774194956 | 0.228000014 | 0.318091303 | 0.611313496 | 0.4333 | 0.695716635 |
| SL1344_RS09445_pyruvate.kinase.II | 4.05708307 | 4.475212542 | 3.752673434 | -0.683830647 | 0.162003554 | -4.34129839 | 4.68E-05 | 0.002749233 | 0.069844404 | 0.1 | 0.330257709 |
| SL1344_RS16430_qseB | -1.524228738 | -1.475061583 | -1.565534286 | -0.125689156 | 0.472808246 | -0.259840396 | 0.378000005 | 0.601789074 | 0.796923415 | 0.7088 | 0.846524225 |
| SL1344_RS16435_qseC | -0.551352906 | -0.681774739 | -0.440232884 | 0.241219217 | 0.208334933 | 1.0941756 | 0.122666699 | 0.193720064 | 0.441153254 | 0.2678 | 0.506316272 |
| SL1344_RS23490_radA | 2.117505216 | 1.869502031 | 2.280059803 | 0.413531797 | 0.127861559 | 3.089983148 | 4.68E-05 | 0.027536041 | 0.189856197 | 0.1 | 0.330257709 |
| RaoN_raoN | -2.82735553 | -2.981667882 | -2.657146319 | 0.325153123 | 0.497260937 | 0.628636408 | 0.245169899 | 0.363474698 | 0.605612782 | 0.4802 | 0.680285939 |
| SL1344_RS12895_ratA | 1.245906993 | 1.228528458 | 1.26823086 | -0.021800706 | 0.245546005 | -0.077558554 | 0.466355764 | 0.747848558 | 0.887283396 | 0.8748 | 0.941146774 |
| rcsA_rcsA | -1.635945062 | -1.595979254 | -1.664325068 | -0.073856835 | 0.354153539 | -0.200259932 | 0.41172552 | 0.595179153 | 0.784346157 | 0.7059 | 0.841514641 |
| SL1344_RS11655_rcsB | 3.243738232 | 3.070654001 | 3.427503721 | 0.383245413 | 0.118811709 | 3.257705017 | 4.68E-05 | 0.013790828 | 0.141379118 | 0.1 | 0.330257709 |
| SL1344_RS11660_rcsC | 2.194539042 | 1.983899489 | 2.37651726 | 0.423566419 | 0.158430527 | 2.692134243 | 4.68E-05 | 0.023633016 | 0.179009053 | 0.1 | 0.330257709 |
| SL1344_RS11650_rcsD | 3.071640965 | 2.831851813 | 3.290276235 | 0.467445622 | 0.095204407 | 4.904075421 | 4.68E-05 | 0.031633928 | 0.212782455 | 0.1 | 0.330257709 |
| rcsF_rcsF | 1.585211068 | 1.422056973 | 1.799144161 | 0.363566073 | 0.253606292 | 1.454533753 | 0.048634334 | 0.096923376 | 0.350897476 | 0.144 | 0.393110252 |
| RdlD_rdlD | -1.059299822 | -1.083654121 | -1.039892674 | 0.061276822 | 1.122800949 | 0.055863911 | 0.478000001 | 0.894895273 | 0.960494122 | 0.9325 | 0.971304185 |
| RdlE_rdlE | -2.14543687 | -2.024461161 | -2.435581775 | -0.550795142 | 1.384426278 | -0.426862562 | 0.268487686 | 0.416859555 | 0.693780369 | 0.5099 | 0.744049673 |
| SL1344_RS14650_recA | 4.185360015 | 4.298524776 | 4.04333711 | -0.252753242 | 0.229327921 | -1.032388077 | 0.126000031 | 0.190388654 | 0.48896723 | 0.2216 | 0.498626916 |
| SL1344_RS13835_recN | 2.330248025 | 2.427691362 | 2.109850767 | -0.304779157 | 0.132298869 | -2.123225464 | 0.005330642 | 0.049470123 | 0.250898651 | 0.1069 | 0.339578295 |
| SL1344_RS06620_rfc | 1.096211775 | 0.960548222 | 1.415344768 | 0.457062126 | 0.235482771 | 1.763952599 | 0.017322048 | 0.082812656 | 0.325452967 | 0.1143 | 0.349895777 |
| SL1344_RS22270_rflM | 2.150272729 | 2.624204015 | 1.407148995 | -1.25487404 | 0.284159287 | -4.485639881 | 4.68E-05 | 0.002545278 | 0.067069862 | 0.1 | 0.330257709 |
| rhtB_rhtB | -1.785170935 | -1.897369776 | -1.659047589 | 0.144897416 | 0.525019279 | 0.26561117 | 0.388000004 | 0.655533911 | 0.830932191 | 0.7563 | 0.875939909 |
| rimM_rimM | 5.126697983 | 5.099191608 | 5.1406518 | 0.150147995 | 0.240980633 | 0.642002888 | 0.263333345 | 0.317213519 | 0.616741274 | 0.5129 | 0.749932283 |
| rimO_rimO | 1.588941181 | 1.570171079 | 1.600452607 | 0.016968659 | 0.21374523 | 0.072807166 | 0.477015324 | 0.798535368 | 0.914440438 | 0.8693 | 0.941053695 |
| SL1344_RS03220_rlpA | 1.608377575 | 1.570775514 | 1.623988697 | 0.048943413 | 0.18513956 | 0.235781353 | 0.416389077 | 0.707767602 | 0.866392458 | 0.8177 | 0.916976371 |
| SL1344_RS05840_rne | 5.333982126 | 5.219923114 | 5.3894353 | 0.192095889 | 0.108254734 | 1.870650937 | 4.68E-05 | 0.047047411 | 0.254925141 | 0.1 | 0.330257709 |
| rnpB_rnpB | 10.91564679 | 10.92007255 | 10.91111782 | 4.57E-05 | 0.122648685 | 0.000641278 | 0.499666889 | 0.868845896 | 0.951169828 | 0.9856 | 0.993824482 |
| RnpB_RnpB | 10.91581901 | 10.92021647 | 10.91125184 | 0.0004215 | 0.122887331 | 0.003928775 | 0.496666667 | 0.869102227 | 0.951228964 | 0.9859 | 0.993956866 |
| SL1344_RS15130_rpoS | 5.556741395 | 5.628822197 | 5.478459459 | -0.156037008 | 0.240934605 | -0.68442886 | 0.269820131 | 0.320133592 | 0.620078105 | 0.5093 | 0.746334304 |
| rprA_rprA | -1.945698623 | -2.040715774 | -1.830504629 | -0.014944144 | 0.805555767 | -0.016510078 | 0.492666667 | 0.79750878 | 0.914506408 | 0.9328 | 0.969110162 |
| rpsN_rpsN | 5.511277897 | 5.350067166 | 5.597859017 | 0.24447438 | 0.228010365 | 0.640472668 | 0.33333334 | 0.469897286 | 0.735373363 | 0.7 | 0.872542119 |
| rpsV_rpsV | 2.769760082 | 3.019098169 | 2.460476789 | -0.549865979 | 0.159996564 | -3.460060173 | 4.68E-05 | 0.007682979 | 0.108554192 | 0.1 | 0.330257709 |
| RseX_rseX | -5.594039484 | -5.315409632 | -5.947190185 | -0.616730667 | 1.224519229 | -0.468500948 | 0.294000009 | 0.439329562 | 0.668195384 | 0.5467 | 0.730203418 |
| rsmD_rsmD | 0.409760061 | 0.324314555 | 0.608529232 | 0.269279762 | 0.2545093 | 1.030475714 | 0.138000028 | 0.196167721 | 0.472214156 | 0.2906 | 0.551407711 |
| SL1344_RS09240_rsmF.yebU | 1.409490094 | 1.820800427 | 0.969099043 | -0.866863277 | 0.135778869 | -6.385109426 | 4.68E-05 | 0.002569668 | 0.061907258 | 0.1 | 0.330257709 |
| SL1344_RS22150_rtsA | 1.692100934 | 2.596461084 | 1.03813402 | -1.529766779 | 0.27295862 | -5.661124278 | 4.68E-05 | 0.001414132 | 0.051659757 | 0.1 | 0.330257709 |
| SL1344_RS22145_rtsB | 0.160663031 | 0.832831283 | -0.681875157 | -1.504558917 | 0.2385591 | -6.290887347 | 4.68E-05 | 0.002131263 | 0.055877592 | 0.1 | 0.330257709 |
| rumA_rumA | 2.961248863 | 2.858054777 | 3.002354213 | 0.141357919 | 0.124147206 | 1.167843382 | 0.117255197 | 0.152089462 | 0.426179848 | 0.2484 | 0.512871146 |
| SL1344_RS09480_ruvA | 1.516051729 | 1.954842632 | 1.040172887 | -0.935268798 | 0.17006093 | -5.701708144 | 4.68E-05 | 0.008421137 | 0.108889391 | 0.1 | 0.330257709 |
| SL1344_RS09475_ruvB | 2.351928013 | 2.781461259 | 1.855953253 | -0.920136908 | 0.179461958 | -5.126499389 | 4.68E-05 | 0.002023645 | 0.059599869 | 0.1 | 0.330257709 |
| RybA_rybA | 3.174509496 | 3.10722481 | 3.262944614 | -0.0081848 | 0.267335215 | -0.042509011 | 0.486342439 | 0.839734638 | 0.936010984 | 0.9868 | 0.994259875 |
| RybB_rybB | -0.778014788 | -0.944471901 | -0.614758298 | 0.294708115 | 0.813247449 | 0.333394089 | 0.381079285 | 0.599767895 | 0.811985214 | 0.8214 | 0.91663491 |
| RybD_rybD | 3.461811401 | 3.791545632 | 3.090422858 | -0.704566645 | 0.155173409 | -4.632389621 | 4.68E-05 | 0.00329643 | 0.074723072 | 0.1 | 0.330257709 |
| RydB_rydB | -3.165112467 | -3.349420968 | -3.061450551 | 0.145627423 | 0.881989742 | 0.146407639 | 0.442000002 | 0.724345441 | 0.874495173 | 0.8299 | 0.921982042 |
| RydC_rydC | 0.134638197 | 0.366264409 | -0.538952481 | -0.825585075 | 0.607888012 | -1.377584058 | 0.079946755 | 0.099034921 | 0.358874178 | 0.1794 | 0.441793373 |
| RyeF_ryeF | -0.063672368 | 0.443572432 | -0.560324307 | -1.004443245 | 0.223924245 | -4.441145611 | 4.68E-05 | 0.007355069 | 0.097032393 | 0.1 | 0.330257709 |

| | | | | | | | | | | | |
|---|---|---|---|---|---|---|---|---|---|---|---|
| RyfA_ryfA | 2.261210244 | 2.321722651 | 2.157186907 | -0.254639327 | 0.345464873 | -0.773913381 | 0.210000016 | 0.262254853 | 0.563958801 | 0.3799 | 0.641235361 |
| RyfD_ryfD | 3.131311227 | 3.00833696 | 3.506985132 | 0.453399237 | 0.494883205 | 0.90569696 | 0.213857444 | 0.260295126 | 0.564200922 | 0.3957 | 0.675847938 |
| RyhB_ryhB | -0.723297123 | -0.724715752 | -0.722059359 | 0.104933977 | 0.775646442 | 0.134730172 | 0.438374419 | 0.807941552 | 0.920657905 | 0.8869 | 0.950799725 |
| RyjB_ryjB | -1.911436761 | -1.682109817 | -2.80623733 | -1.078168852 | 0.577441639 | -1.517700906 | 0.062624986 | 0.120475846 | 0.385531607 | 0.1711 | 0.416081485 |
| Sar_sar | -0.018498513 | 0.162771504 | -0.242597869 | 0.1901746 | 1.132500545 | 0.172419263 | 0.461025984 | 0.86796428 | 0.94939699 | 0.9997 | 0.999869064 |
| SL1344_RS10635_sbcB | 1.602879192 | 1.3844287 | 1.847983455 | 0.417366528 | 0.184976861 | 2.234571307 | 0.006662914 | 0.030882757 | 0.201605162 | 0.1068 | 0.340206623 |
| SL1344_RS01925_sbmA | -0.266030206 | -0.284057908 | -0.25711538 | 0.006553468 | 0.222541718 | 0.028510919 | 0.486000001 | 0.652182955 | 0.826579237 | 0.7531 | 0.874841192 |
| SL1344_RS10605_sbmC | 3.248474226 | 3.276332378 | 3.198583854 | -0.205652618 | 0.253899283 | -0.813132686 | 0.212524999 | 0.241460935 | 0.541692105 | 0.4045 | 0.655471712 |
| SL1344_RS20885_sbp | -0.625854045 | -0.672496952 | -0.602278845 | 0.063013991 | 0.55118462 | 0.093544569 | 0.462666668 | 0.719524447 | 0.876351434 | 0.9158 | 0.962552929 |
| sdhC_sdhC | 3.95386893 | 3.844753259 | 4.187311953 | 0.216523829 | 0.373871466 | 0.770095656 | 0.205862775 | 0.289232178 | 0.591720414 | 0.379 | 0.658931761 |
| SL1344_RS09755_sdiA | 3.420859021 | 4.787490666 | 2.522366389 | -2.232997147 | 0.296635149 | -6.887854523 | 4.68E-05 | 0.004288536 | 0.086786951 | 0.1 | 0.330257709 |
| SdsR_sdsR | -1.06236242 | -1.106934746 | -1.017227475 | 0.05743907 | 0.784075035 | 0.067783555 | 0.472000001 | 0.821733361 | 0.927082814 | 0.9385 | 0.973047127 |
| sgbH_sgbH | -4.668438329 | -4.38647795 | -4.919919527 | -0.501201386 | 1.198344384 | -0.38569152 | 0.311125924 | 0.514511035 | 0.731963722 | 0.6035 | 0.775516627 |
| SgrS_sgrS | 3.395474911 | 3.317706057 | 3.415531248 | 0.104329064 | 0.301323248 | 0.374047815 | 0.341105936 | 0.619618657 | 0.82811429 | 0.7258 | 0.883552242 |
| SibA_sibA | 1.511087239 | 1.552217038 | 1.475037121 | -0.086208296 | 1.200606262 | -0.064471924 | 0.464000001 | 0.947075 | 0.981234707 | 1 | 1 |
| SibC_sibC | 3.108197073 | 3.499432568 | 2.823209016 | -0.482517363 | 0.846586861 | -0.658816844 | 0.226666681 | 0.343310676 | 0.640221673 | 0.4002 | 0.68033961 |
| SibD_sibD | 3.067984942 | 3.444211331 | 2.668360046 | -0.571899195 | 1.163107248 | -0.426155076 | 0.326449041 | 0.534018721 | 0.778037104 | 0.7 | 0.872542119 |
| SL1344_RS14940_sicA | 0.318794582 | 1.928106023 | -1.544216347 | -3.521955923 | 0.330409238 | -10.72445589 | 4.68E-05 | 0.000344851 | 0.028904491 | 0.1 | 0.330257709 |
| sifA_sifA | -0.508688238 | -1.343960517 | -0.386525795 | 0.92885231 | 0.392908325 | 1.338008872 | 0.070666727 | 0.140278863 | 0.420915871 | 0.183 | 0.427687164 |
| sifB_sifB | -3.812388178 | -3.46083044 | -4.363730842 | -0.896816925 | 0.692518972 | -1.193929113 | 0.076615645 | 0.148579886 | 0.396225989 | 0.2073 | 0.447348326 |
| SL1344_RS05355_sigE | 1.030820806 | 2.848520627 | -0.810625713 | -3.505004703 | 0.434269269 | -8.188222177 | 4.68E-05 | 0.000281616 | 0.028026183 | 0.1 | 0.330257709 |
| SL1344_RS21850_siiA | -2.203841027 | -1.730273239 | -3.171559176 | -1.490553816 | 0.646743119 | -2.256312121 | 0.002668394 | 0.038374716 | 0.219253968 | 0.1025 | 0.33365065 |
| SL1344_RS21855_siiB | -1.328315758 | -0.834153021 | -2.071847504 | -1.264993153 | 0.441859311 | -2.803333862 | 4.68E-05 | 0.027868582 | 0.186593883 | 0.1 | 0.330257709 |
| SL1344_RS21875_siiF | 1.871229417 | 1.842101647 | 1.934775023 | 0.082944501 | 0.233795153 | 0.342713433 | 0.353333339 | 0.719665737 | 0.874128006 | 0.7385 | 0.877672061 |
| SL1344_RS01560_sinR | 0.748479227 | 0.874750398 | 0.604569024 | -0.275646323 | 0.140910295 | -1.878806283 | 0.010667096 | 0.050914227 | 0.239204652 | 0.1121 | 0.346462376 |
| SL1344_RS14920_sipA | 1.18534644 | 2.334105594 | -0.157104952 | -2.543344006 | 0.28757154 | -8.961169503 | 4.68E-05 | 0.000569638 | 0.034807389 | 0.1 | 0.330257709 |
| SL1344_RS14935_sipB | 1.925299485 | 3.471767331 | 0.230556095 | -3.266159246 | 0.221811896 | -14.84577921 | 4.68E-05 | 0.000245589 | 0.025874124 | 0.1 | 0.330257709 |
| SL1344_RS14930_sipC | 2.528367853 | 4.363357329 | 0.584523648 | -3.751749189 | 0.187015584 | -19.903222 | 4.68E-05 | 3.69E-05 | 0.015517636 | 0.1 | 0.330257709 |
| SL1344_RS14925_sipD | -0.060121071 | 1.251552029 | -1.753138958 | -3.093917003 | 0.400110577 | -7.942234095 | 4.68E-05 | 0.000677345 | 0.037385896 | 0.1 | 0.330257709 |
| SL1344_RS09740_sirA | 3.490204479 | 3.686204951 | 3.282446089 | -0.423601703 | 0.10201918 | -4.217584592 | 4.68E-05 | 0.006150217 | 0.096343294 | 0.1 | 0.330257709 |
| SL1344_RS10425_sirohydrochlorin.cobaltochelatase | -0.90437327 | -0.929871736 | -0.875553803 | 0.141324323 | 1.025321924 | 0.140282576 | 0.381079285 | 0.67254797 | 0.853260098 | 0.7651 | 0.884882873 |
| SL1344_RS09840_SL1896 | -1.335638882 | -1.167804787 | -1.558017092 | -0.390433104 | 0.356733156 | -1.065263314 | 0.124666699 | 0.186360855 | 0.449156951 | 0.269 | 0.513317757 |
| SL1344_RS07175_slyA | 3.238072617 | 3.15372492 | 3.260337577 | 0.105440181 | 0.105319927 | 0.938309817 | 0.175216543 | 0.249747434 | 0.541643346 | 0.3585 | 0.595980835 |
| SL1344_RS07110_sodB | 3.904004579 | 3.927821775 | 3.87973315 | -0.007507827 | 0.380026804 | -0.023363464 | 0.480000001 | 0.864641904 | 0.948646213 | 0.9742 | 0.988976353 |
| SL1344_RS10630_sopA | -0.935780218 | 0.053497257 | -2.11292451 | -2.199911481 | 0.307200939 | -7.098623599 | 4.68E-05 | 0.001311986 | 0.046630313 | 0.1 | 0.330257709 |
| sopB_sopB | 1.136603764 | 2.883886376 | -0.76841136 | -3.656526249 | 0.313439593 | -11.66799894 | 4.68E-05 | 0.000241362 | 0.025019864 | 0.1 | 0.330257709 |
| SL1344_RS15235_sopD | -0.895912627 | -0.292433632 | -1.525051039 | -1.206081014 | 0.286302429 | -4.164738386 | 4.68E-05 | 0.00635518 | 0.093487879 | 0.1 | 0.330257709 |
| SL1344_RS13925_sopE | 0.925993217 | 2.770729463 | -1.2051904 | -3.976163808 | 0.417099785 | -9.393196846 | 4.68E-05 | 0.000412503 | 0.032021139 | 0.1 | 0.330257709 |
| SL1344_RS09265_sopE2 | -0.415249403 | 1.487664965 | -3.192625846 | -4.73435865 | 0.486756094 | -9.893082718 | 4.68E-05 | 0.003476235 | 0.069731716 | 0.1 | 0.330257709 |
| spaK_spaK | -1.04209386 | 0.127436083 | -2.512991131 | -2.648766267 | 0.396743003 | -6.665884428 | 4.68E-05 | 0.004367552 | 0.076261503 | 0.1 | 0.330257709 |
| spaM_spaM | -0.597855633 | 0.739884071 | -2.364291728 | -3.013190629 | 0.412308182 | -7.270615877 | 4.68E-05 | 0.00174245 | 0.049991225 | 0.1 | 0.330257709 |
| spaN_spaN | 0.114803244 | 1.169074271 | -1.363757898 | -2.550457082 | 0.255255827 | -9.909438287 | 4.68E-05 | 0.00093969 | 0.038263414 | 0.1 | 0.330257709 |
| SL1344_RS14965_spaO | -0.21400462 | 1.019100782 | -1.640519892 | -2.623494503 | 0.300219854 | -8.715630905 | 4.68E-05 | 0.001077235 | 0.0405136 | 0.1 | 0.330257709 |
| SL1344_RS14960_spaP | -0.383908412 | 0.628480142 | -1.774743193 | -2.425609301 | 0.364503329 | -6.701358035 | 4.68E-05 | 0.008169897 | 0.107225985 | 0.1 | 0.330257709 |
| SL1344_RS14955_spaQ | -0.64574206 | 0.22772356 | -2.029898136 | -2.313336623 | 0.384879881 | -6.050540187 | 4.68E-05 | 0.002248678 | 0.057654803 | 0.1 | 0.330257709 |
| SL1344_RS14950_spaR | -0.897505667 | 0.001271349 | -2.311831161 | -2.323069128 | 0.593901432 | -4.016794148 | 4.68E-05 | 0.025725042 | 0.189023532 | 0.1 | 0.330257709 |
| SL1344_RS14945_spaS | -0.956871728 | -0.120465566 | -2.042608853 | -1.926782718 | 0.318508151 | -5.9830772 | 4.68E-05 | 0.004496073 | 0.074212707 | 0.1 | 0.330257709 |
| Spf_spf | 4.630223338 | 4.597206637 | 5.222157334 | 0.605262506 | 0.499912779 | 0.603118304 | 0.329113931 | 0.415870596 | 0.697027894 | 0.6862 | 0.863682089 |
| SL1344_RS14840_spi4.transcriptional.regulator | 1.080364611 | 1.607890812 | 0.502771993 | -1.193881175 | 0.289586521 | -4.21793726 | 4.68E-05 | 0.003481929 | 0.07602728 | 0.1 | 0.330257709 |
| spiC_spiC | -6.622113817 | -8.451896249 | -5.686004454 | 2.698727036 | 2.756341407 | 0.941482579 | 0.103930752 | 0.198276238 | 0.47448541 | 0.2409 | 0.48498465 |
| spr_spr | 5.760757586 | 5.733222095 | 5.780516546 | 0.100984558 | 0.1991752 | 0.611316672 | 0.282666677 | 0.351795327 | 0.646295892 | 0.5468 | 0.773738643 |
| SraA_sraA | -5.282919001 | -4.994610163 | -6.274891531 | -1.130143205 | 1.804314194 | -0.520706364 | 0.314666675 | 0.435041081 | 0.691271941 | 0.6239 | 0.802663086 |
| SraB_sraB | 0.255354315 | 0.201439992 | 0.286873602 | 0.093084232 | 0.361244424 | 0.227364872 | 0.416000003 | 0.775632854 | 0.903671246 | 0.8545 | 0.935987192 |

| | | | | | | | | | | | |
|---|---|---|---|---|---|---|---|---|---|---|---|
| SraC_sraC | 3.592620052 | 4.19081467 | 3.140345875 | -1.093626073 | 0.275663175 | -4.184611218 | 4.68E-05 | 0.007730812 | 0.112990586 | 0.1 | 0.330257709 |
| SraF_sraF | 1.354357237 | 1.311070334 | 1.440093741 | 0.426292273 | 0.668998099 | 0.622729229 | 0.271818798 | 0.352422083 | 0.64485308 | 0.5329 | 0.761691459 |
| SraG_sraG | -3.123268088 | -3.217447483 | -2.983525234 | 0.232536111 | 0.54331529 | 0.388546467 | 0.341772158 | 0.496140362 | 0.708527261 | 0.5969 | 0.764515427 |
| SraL_sraL | -1.227115994 | -1.203369808 | -1.240543473 | -0.014095437 | 0.387826685 | -0.029775022 | 0.486666667 | 0.710812214 | 0.864110389 | 0.8146 | 0.909322066 |
| SL1344_RS07945_srfB | 3.65595235 | 4.391199128 | 2.893827174 | -1.469222317 | 0.193212322 | -7.481062172 | 4.68E-05 | 0.000523504 | 0.036645274 | 0.1 | 0.330257709 |
| SL1344_RS07950_srfC | 3.256693227 | 4.022870507 | 2.565054643 | -1.427674662 | 0.12106283 | -11.72660473 | 4.68E-05 | 0.000463613 | 0.031488517 | 0.1 | 0.330257709 |
| srlA_srlA | -0.768951191 | -0.846147723 | -0.683761389 | 0.154034058 | 0.276309472 | 0.528977465 | 0.299333342 | 0.469429729 | 0.695854312 | 0.5706 | 0.752627054 |
| srlE_srlE | -0.540040451 | -0.516604441 | -0.565538563 | -0.018652118 | 0.276592601 | -0.065924707 | 0.477333334 | 0.661329388 | 0.831708664 | 0.7737 | 0.884585807 |
| srlR_srlR | 1.537132285 | 1.3941855 | 1.667163531 | 0.281790371 | 0.118434368 | 2.36515135 | 0.003332494 | 0.037286934 | 0.20918236 | 0.1039 | 0.33552801 |
| SroA_sroA | -1.865995313 | -1.981663174 | -1.686178683 | 0.347767177 | 0.534984186 | 0.615241491 | 0.249167234 | 0.374676281 | 0.635360505 | 0.4985 | 0.707147672 |
| SroC_sroC | 7.223040364 | 7.230164183 | 7.14689021 | -0.234687379 | 0.271041035 | -0.844582533 | 0.214523667 | 0.230425184 | 0.535015032 | 0.3882 | 0.668166388 |
| SroD_sroD | -0.462823647 | 0.115344014 | -1.104018869 | -1.20657633 | 0.467946238 | -2.514372391 | 4.68E-05 | 0.022708025 | 0.179531193 | 0.1001 | 0.330407312 |
| SroG_sroG | 4.137398196 | 4.141939876 | 4.132246345 | -0.335383063 | 0.48956068 | -0.549840632 | 0.360426388 | 0.341974577 | 0.638932987 | 0.787 | 0.902419142 |
| SL1344_RS06985_ssaG | -5.713119707 | -6.347411138 | -5.0321773 | 1.300776752 | 2.302801465 | 0.501901143 | 0.221185891 | 0.372799482 | 0.64150619 | 0.4412 | 0.664955867 |
| SL1344_RS07050_ssaR | -2.556017167 | -2.920018936 | -2.202216537 | 0.722966999 | 0.376968782 | 1.798552205 | 0.022651765 | 0.058010211 | 0.243391601 | 0.126 | 0.362033916 |
| SL1344_RS05160_sseI | -1.872147956 | -2.016193528 | -1.685426545 | 0.305976248 | 0.37372519 | 0.777712598 | 0.196000018 | 0.297359146 | 0.553258419 | 0.3962 | 0.619207132 |
| sspA_sspA | 3.753399523 | 3.706370798 | 3.844495538 | 0.136668957 | 0.081689931 | 1.618717758 | 0.033333467 | 0.100398199 | 0.351765198 | 0.136 | 0.378334336 |
| SL1344_RS06915_ssrA | -0.90783757 | -0.919958255 | -0.895580232 | 0.045115693 | 0.413773296 | 0.116864091 | 0.445333335 | 0.707927375 | 0.863006058 | 0.7796 | 0.890560353 |
| SsrA_ssrA | 10.68481446 | 10.67117208 | 10.7673768 | 0.095026747 | 0.094184208 | 0.957792248 | 0.20333335 | 0.196901384 | 0.498195946 | 0.3756 | 0.655515709 |
| SL1344_RS06910_ssrB | -1.94314649 | -2.185325372 | -1.776602774 | 0.419187168 | 0.461318847 | 0.851120767 | 0.159227205 | 0.250579846 | 0.520643286 | 0.331 | 0.565714481 |
| ssrS_ssrS | 8.718666254 | 8.655043487 | 9.093896517 | 0.440321385 | 0.515142503 | 0.627259305 | 0.33333334 | 0.516251669 | 0.766736804 | 0.7 | 0.872542119 |
| SsrS_ssrS | 8.718680751 | 8.654540207 | 9.093780691 | 0.438479794 | 0.514802764 | 0.621738867 | 0.33333334 | 0.516328144 | 0.766836109 | 0.7 | 0.872542119 |
| SL1344_RS16670_sstT | 2.910934905 | 2.903431241 | 2.915278902 | 0.022471402 | 0.493934689 | 0.06130255 | 0.423051302 | 0.910676138 | 0.967048534 | 0.8248 | 0.916098119 |
| SL1344_RS01735_stbB | -5.45519886 | -5.185348766 | -5.886198225 | -0.67505918 | 1.299417952 | -0.481865651 | 0.299800142 | 0.448766433 | 0.681667993 | 0.5723 | 0.749253993 |
| SL1344_RS05145_stfT | -1.814647962 | -1.547663215 | -2.143102702 | -0.594094865 | 0.332945558 | -1.692833664 | 0.027981506 | 0.07139406 | 0.277857147 | 0.1343 | 0.371666144 |
| STnc3140_STnc3140..sopD. | -2.55178837 | -2.53805287 | -2.556149867 | -0.061457896 | 0.729545782 | -0.079038495 | 0.473684212 | 0.73181526 | 0.87834213 | 0.8468 | 0.929314375 |
| STnc3200_STnc3200 | -5.77595873 | -5.435174154 | -6.008847932 | -0.679752565 | 1.50714292 | -0.42876927 | 0.320453039 | 0.479206345 | 0.70015618 | 0.6018 | 0.76606449 |
| SL1344_RS15350_syd | 1.140516452 | 1.237790027 | 1.068052006 | -0.156983035 | 0.33347035 | -0.429374251 | 0.352431718 | 0.528506236 | 0.76650814 | 0.7605 | 0.882165491 |
| SymR_symR | -2.726938837 | -2.853355188 | -2.603920747 | 0.261711824 | 0.571705105 | 0.42950577 | 0.309333342 | 0.494870486 | 0.717126285 | 0.5968 | 0.770626845 |
| SL1344_RS13990_tail.protein | -8.037573213 | -7.986178821 | -8.13745206 | 0.283733728 | 4.161210977 | 0.064232258 | 0.474666668 | 0.62185867 | 0.811270222 | 0.7855 | 0.891801713 |
| tatE_tatE | 0.732849573 | 0.751218389 | 0.713216389 | 0.02023734 | 0.26611115 | 0.068286961 | 0.473017989 | 0.75488288 | 0.891315316 | 0.8838 | 0.946873629 |
| tcuB_tcuB | -1.966255344 | -1.614622448 | -2.120856085 | -0.476286614 | 0.435692067 | -1.004323646 | 0.160000023 | 0.228038406 | 0.492923083 | 0.3294 | 0.568853558 |
| SL1344_RS16770_tdcA | 0.556328581 | 0.221433402 | 0.935634324 | 0.645567344 | 0.520374875 | 1.261974295 | 0.090666713 | 0.113037677 | 0.382047939 | 0.1891 | 0.459493196 |
| SL1344_RS16765_tdcB | -0.095019319 | 0.03622989 | -0.343138493 | -0.404613477 | 0.3446835 | -1.13126543 | 0.106000038 | 0.152471141 | 0.424500668 | 0.23 | 0.488607658 |
| SL1344_RS16760_tdcC | 0.784871087 | 1.360692012 | 0.547120768 | -0.783341876 | 0.339812304 | -2.243642435 | 0.001335878 | 0.035463337 | 0.220810341 | 0.1008 | 0.331425856 |
| SL1344_RS05150_tfaT | -1.718990236 | -1.383556439 | -2.011698151 | -0.60992486 | 0.497434115 | -1.174512791 | 0.106666705 | 0.144105497 | 0.41115874 | 0.2111 | 0.468781715 |
| Tff_tff | 4.278107097 | 4.296701534 | 4.259733979 | -0.031459193 | 0.447001246 | -0.06786537 | 0.471333334 | 0.92906484 | 0.974352408 | 1 | 1 |
| thiP_thiP | 0.901038144 | 0.784045662 | 1.004552884 | 0.174417886 | 0.203911012 | 0.83930685 | 0.177881433 | 0.264190686 | 0.54224283 | 0.3457 | 0.596773786 |
| thiQ_thiQ | -0.301805383 | -0.356954094 | -0.238566164 | 0.143974489 | 0.274817197 | 0.506728262 | 0.302000009 | 0.478316786 | 0.710296777 | 0.5937 | 0.770841519 |
| SL1344_RS06625_thrS | 5.128987027 | 4.921020405 | 5.330360065 | 0.413224909 | 0.040019175 | 10.47252898 | 4.68E-05 | 0.001764309 | 0.052466782 | 0.1 | 0.330257709 |
| SL1344_RS04595_tnpA1 | -3.073291671 | -1.507140318 | -5.523564204 | -3.951890053 | 0.840371438 | -4.705052842 | 4.68E-05 | 0.008955879 | 0.099158738 | 0.1 | 0.330257709 |
| SL1344_RS06060_tnpA2 | -2.887989581 | -2.553727697 | -3.573564368 | -1.042305774 | 0.686083779 | -1.390867144 | 0.048634334 | 0.120496009 | 0.373966578 | 0.1553 | 0.397728104 |
| SL1344_RS09075_tnpA3 | -3.905792744 | -2.897600447 | -5.450047924 | -2.563456194 | 0.774663214 | -3.271041868 | 4.68E-05 | 0.017705572 | 0.13989362 | 0.1001 | 0.330407235 |
| SL1344_RS09790_tnpA4 | -5.713203429 | -4.668374219 | -6.901464516 | -2.206830242 | 1.58524482 | -1.301282015 | 0.049333422 | 0.111295361 | 0.350429894 | 0.166 | 0.410702178 |
| SL1344_RS12685_tnpA5 | -4.3250275 | -4.03724232 | -4.712237517 | -0.677333292 | 0.84281731 | -0.774093282 | 0.191205881 | 0.28233398 | 0.535284668 | 0.3861 | 0.601687625 |
| SL1344_RS17020_tnpA6 | -3.015463358 | -2.643185945 | -3.757574452 | -1.035631719 | 0.912270854 | -1.068964806 | 0.125916087 | 0.182141976 | 0.465483223 | 0.2462 | 0.506725181 |
| SL1344_RS17955_tnpA7 | -3.451603986 | -3.373451762 | -3.501678148 | -0.316281286 | 1.01987878 | -0.28163533 | 0.388407733 | 0.626922152 | 0.814580181 | 0.7888 | 0.892949621 |
| tolA_tolA | 2.049589987 | 1.90088246 | 2.098345139 | 0.20381683 | 0.207756231 | 0.997382473 | 0.095936086 | 0.232031766 | 0.531171594 | 0.2357 | 0.482393336 |
| SL1344_RS16475_tolC | 3.815678038 | 3.710560389 | 4.019966245 | 0.342951051 | 0.170856221 | 2.164027696 | 4.68E-05 | 0.025325077 | 0.191422448 | 0.1 | 0.330257709 |
| SL1344_RS21860_TolC.family.protein | -1.514430172 | -0.808076152 | -2.094843787 | -1.250112829 | 0.38195162 | -3.189274765 | 0.00067292 | 0.014273108 | 0.135796607 | 0.1001 | 0.33040569 |
| tqsA_tqsA | -1.127646236 | -1.182144327 | -1.077237751 | 0.112914711 | 0.261686808 | 0.4046396 | 0.319120594 | 0.469937308 | 0.692259324 | 0.586 | 0.755638402 |
| SL1344_RS14145_transcriptional.regulator | -5.719107855 | -6.253545063 | -5.058757797 | 1.093905766 | 1.820417988 | 0.55217298 | 0.249333346 | 0.419420883 | 0.657494687 | 0.489 | 0.69806739 |
| trkD_trkD | 1.887809579 | 1.779312518 | 2.032619258 | 0.235044166 | 0.130723177 | 1.689093239 | 0.042000105 | 0.070562152 | 0.291447118 | 0.1358 | 0.38041342 |

| | | | | | | | | | | | |
|---|---|---|---|---|---|---|---|---|---|---|---|
| SL1344_RS06175_tRNA.arg | -3.914277184 | -3.031025008 | -5.172013037 | -2.160541823 | 0.822211415 | -2.595204708 | 4.68E-05 | 0.027425647 | 0.176806485 | 0.1003 | 0.330691582 |
| SL1344_RS13600_tRNA.rRNA.methyltransferase | 1.727155397 | 1.407433772 | 2.086310886 | 0.662686621 | 0.12542492 | 5.192010481 | 4.68E-05 | 0.004109208 | 0.076686717 | 0.1 | 0.330257709 |
| truA_truA | 0.797261444 | 0.530475865 | 0.996660436 | 0.471808195 | 0.176781893 | 2.613946524 | 0.001335878 | 0.021464498 | 0.166805404 | 0.101 | 0.331713861 |
| SL1344_RS23260_tsr | 5.931160909 | 6.643374538 | 5.299703161 | -1.21351183 | 0.223794866 | -5.391245521 | 4.68E-05 | 0.001178105 | 0.050116244 | 0.1 | 0.330257709 |
| SL1344_RS06870_ttrA | -1.713331319 | -1.792129306 | -1.627584034 | 0.166684386 | 0.337858722 | 0.468037153 | 0.314457037 | 0.460372996 | 0.685074739 | 0.5634 | 0.74255405 |
| SL1344_RS06880_ttrB | -5.330993521 | -5.429166206 | -5.241784544 | 0.06339326 | 1.41501309 | 0.035452588 | 0.483677549 | 0.639146846 | 0.818131031 | 0.754 | 0.870492803 |
| SL1344_RS06875_ttrC | -4.685869773 | -4.91997752 | -4.367747527 | 0.530234788 | 1.079230663 | 0.439991271 | 0.296000009 | 0.480349233 | 0.704490879 | 0.5767 | 0.755811079 |
| SL1344_RS06890_ttrR | 0.686426279 | 0.606528393 | 0.739869127 | 0.203250724 | 0.280138713 | 0.724227018 | 0.214523667 | 0.303057421 | 0.589103513 | 0.4389 | 0.661295492 |
| SL1344_RS06885_ttrS | 2.530516001 | 2.304884485 | 2.554430438 | 0.236779381 | 0.139192829 | 0.891988023 | 0.265156573 | 0.287582854 | 0.583728589 | 0.5372 | 0.759520033 |
| tyrA_tyrA | 1.134643954 | 0.909788324 | 1.357912812 | 0.449236799 | 0.117468906 | 3.788304486 | 4.68E-05 | 0.008937951 | 0.105612993 | 0.1 | 0.330257709 |
| ubiB_ubiB | 3.508298661 | 3.450968653 | 3.548404795 | 0.140892349 | 0.148842894 | 0.981711396 | 0.105929419 | 0.193477453 | 0.487302587 | 0.226 | 0.481641847 |
| ubiF_ubiF | 1.304372185 | 1.318084437 | 1.297799924 | -0.025253005 | 0.166498527 | -0.136950946 | 0.452000002 | 0.696975274 | 0.856444622 | 0.8051 | 0.906048067 |
| SL1344_RS21190_udhA | 3.848949029 | 3.695506172 | 3.890971708 | 0.193244216 | 0.194365054 | 0.808394236 | 0.267333344 | 0.326508047 | 0.62359229 | 0.5347 | 0.762478579 |
| SL1344_RS03635_UDP.galactopyranose.mutase | -4.058511922 | -3.993996203 | -4.128119794 | -0.145307633 | 0.727033484 | -0.187993745 | 0.424383747 | 0.558203949 | 0.758702703 | 0.6666 | 0.816190501 |
| ugpC_ugpC | -0.312461304 | -0.451279483 | -0.202516208 | 0.27855938 | 0.276849105 | 0.996139353 | 0.10200004 | 0.20273116 | 0.47779009 | 0.2457 | 0.481699688 |
| ulaD_ulaD | -2.497654232 | -2.276679824 | -2.702087501 | -0.426964364 | 0.431885179 | -0.941509736 | 0.148567647 | 0.22685646 | 0.47732786 | 0.3133 | 0.542757499 |
| ushA_ushA | 3.349226386 | 3.278479969 | 3.461047498 | 0.208430601 | 0.163568429 | 1.370108698 | 0.026000173 | 0.114660627 | 0.383403668 | 0.1229 | 0.362666857 |
| SL1344_RS09735_uvrC | 3.240891218 | 3.569222265 | 3.095777563 | -0.47546707 | 0.21448573 | -2.400182267 | 4.68E-05 | 0.064838456 | 0.298016799 | 0.1 | 0.330257709 |
| SL1344_RS11530_virulence.protein.MsgA | -6.326941445 | -6.35178865 | -6.282167483 | 0.089209694 | 2.261867492 | 0.030865739 | 0.481333334 | 0.666938703 | 0.838118831 | 0.7852 | 0.892494998 |
| wecC_wecC | 2.629606484 | 2.510646724 | 2.922596632 | 0.398253571 | 0.119712007 | 2.797515044 | 0.00067292 | 0.035432852 | 0.217982516 | 0.1 | 0.330257709 |
| SL1344_RS10695_wzzB | 2.505508582 | 1.814397904 | 3.276316189 | 1.452513268 | 0.132800096 | 10.88080659 | 4.68E-05 | 0.000562989 | 0.033861131 | 0.1 | 0.330257709 |
| SL1344_RS02345_ybaO | 0.868957395 | 0.868298166 | 0.869335368 | 0.053603252 | 0.344095634 | 0.158858667 | 0.430000003 | 0.776381366 | 0.903641379 | 0.8802 | 0.945706207 |
| SL1344_RS03185_ybeM | -1.616147321 | -1.286438247 | -1.974662956 | -0.797082159 | 0.490787071 | -1.618936806 | 0.012667027 | 0.069220338 | 0.292133249 | 0.1143 | 0.348948228 |
| SL1344_RS04285_ybiK | 2.258727776 | 2.229560678 | 2.29143658 | 0.102420172 | 0.154425557 | 0.634391607 | 0.235176563 | 0.322120345 | 0.604558649 | 0.4718 | 0.683367985 |
| SL1344_RS08990_ycgO | 3.405480739 | 4.01897233 | 2.515181484 | -1.505485775 | 0.241810728 | -6.168496481 | 4.68E-05 | 0.001879654 | 0.059064794 | 0.1 | 0.330257709 |
| SL1344_RS08975_ycgR | 2.67757148 | 3.345178453 | 2.091519381 | -1.155171306 | 0.225749143 | -5.30189738 | 4.68E-05 | 0.001110945 | 0.047839266 | 0.1 | 0.330257709 |
| SL1344_RS08585_yciK | 2.083894221 | 2.12676848 | 2.016390722 | -0.220433877 | 0.250727446 | -0.871500816 | 0.16266669 | 0.218357521 | 0.512193995 | 0.3387 | 0.587662185 |
| SL1344_RS08495_yciW | 0.092825485 | 0.592183324 | 0.037514457 | -0.547691198 | 0.345390059 | -0.869916958 | 0.280666677 | 0.308397833 | 0.603004629 | 0.5657 | 0.776998168 |
| SL1344_RS07545_ydeD | -0.905273112 | -1.105035132 | -0.729164651 | 0.374569718 | 0.260962842 | 1.394621531 | 0.061958764 | 0.109304932 | 0.343394765 | 0.18 | 0.423915437 |
| SL1344_RS06680_ydiV | -0.820584218 | -1.247271259 | -0.404182842 | 0.844895193 | 0.215395339 | 3.903630876 | 4.68E-05 | 0.007968179 | 0.0996177 | 0.1 | 0.330257709 |
| SL1344_RS06595_ydiY | 1.690339552 | 1.616490635 | 2.031102035 | 0.411120333 | 0.378609395 | 1.179468847 | 0.104596974 | 0.116406242 | 0.386260818 | 0.221 | 0.483812049 |
| SL1344_RS06585_ydiZ | -2.091964948 | -2.232579277 | -1.928319845 | 0.296605621 | 0.378198752 | 0.727346453 | 0.214666682 | 0.322210165 | 0.570727689 | 0.4248 | 0.638782909 |
| SL1344_RS06560_ydjN | 2.185361538 | 2.130390448 | 2.211658757 | 0.058818419 | 0.155366075 | 0.16864948 | 0.380413063 | 0.734301984 | 0.887424231 | 0.7831 | 0.907807811 |
| SL1344_RS09415_yebG | -0.29935637 | 0.202614444 | -1.027644774 | -1.281240536 | 0.360953816 | -3.702698093 | 4.68E-05 | 0.007548989 | 0.105832794 | 0.1 | 0.330257709 |
| SL1344_RS09225_yebR | 3.574959191 | 3.972296778 | 3.161261294 | -0.763115374 | 0.151226431 | -5.092635773 | 4.68E-05 | 0.001717766 | 0.056884003 | 0.1 | 0.330257709 |
| SL1344_RS09230_yebS | 1.248995121 | 1.579299967 | 0.892373644 | -0.685661976 | 0.138356837 | -4.972454244 | 4.68E-05 | 0.004980226 | 0.081616512 | 0.1 | 0.330257709 |
| SL1344_RS09760_yecC | 1.188718045 | 1.806004693 | 0.53425333 | -1.288178453 | 0.149530375 | -8.599505296 | 4.68E-05 | 0.000839927 | 0.038884828 | 0.1 | 0.330257709 |
| SL1344_RS09750_yecF | 2.16900388 | 2.419756756 | 1.944021237 | -0.505832 | 0.215166687 | -2.372602643 | 4.68E-05 | 0.024426179 | 0.186610798 | 0.1002 | 0.330552327 |
| SL1344_RS09765_yecS | 0.093730262 | 0.883109439 | -0.571443502 | -1.37932395 | 0.368268441 | -3.77174052 | 4.68E-05 | 0.005323469 | 0.090769515 | 0.1 | 0.330257709 |
| SL1344_RS09825_yedD | 1.36126031 | 1.167576049 | 1.426855525 | 0.206603349 | 0.220181128 | 0.861506388 | 0.210666683 | 0.294476298 | 0.580198534 | 0.4153 | 0.661374969 |
| SL1344_RS09830_yedE | -0.706567507 | -1.072657577 | -0.165073171 | 0.904380396 | 0.2867674 | 3.031840935 | 4.68E-05 | 0.01939908 | 0.159565853 | 0.1 | 0.330257709 |
| SL1344_RS09835_yedF | -1.201071792 | -1.624035315 | -0.868579888 | 0.756338211 | 0.250586692 | 3.036262539 | 0.00067292 | 0.019147819 | 0.147273803 | 0.1004 | 0.330855001 |
| SL1344_RS09770_yedO | 1.257479393 | 1.961696881 | 0.726932597 | -1.199224616 | 0.21285504 | -5.348959545 | 4.68E-05 | 0.006537691 | 0.100007512 | 0.1 | 0.330257709 |
| SL1344_RS10925_yegD | -0.943401469 | -1.241529572 | -0.668350276 | 0.647946337 | 0.366282621 | 1.770313877 | 0.010659989 | 0.057034777 | 0.263547619 | 0.1109 | 0.344518751 |
| SL1344_RS13585_yfiK | -2.260714841 | -2.399387727 | -2.170380575 | 0.219444469 | 0.40218285 | 0.502395561 | 0.275333344 | 0.425413401 | 0.65945962 | 0.5427 | 0.725344649 |
| SL1344_RS17960_yhfL | 1.23462322 | 1.665552388 | 0.832775202 | -0.806189593 | 0.247946387 | -3.24358872 | 4.68E-05 | 0.009463277 | 0.118630119 | 0.1 | 0.330257709 |
| SL1344_RS18605_yhjH | 4.137861738 | 4.864887935 | 3.004500092 | -1.850002685 | 0.263915041 | -6.304789815 | 4.68E-05 | 0.00941251 | 0.124353491 | 0.1 | 0.330257709 |
| SL1344_RS18650_yhjQ | 1.065441556 | 0.362983306 | 1.541201423 | 1.147005123 | 0.270914116 | 4.157608202 | 4.68E-05 | 0.005777209 | 0.092937728 | 0.1 | 0.330257709 |
| SL1344_RS23255_yjiY | 5.617994277 | 5.183856062 | 5.921221734 | 0.755436217 | 0.368333345 | 1.917909979 | 4.68E-05 | 0.094702017 | 0.356214585 | 0.1 | 0.330257709 |
| SL1344_RS05645_ymdA | 0.892069952 | 1.565762107 | 0.24714363 | -1.292494092 | 0.222345118 | -5.776581771 | 4.68E-05 | 0.001830146 | 0.055485426 | 0.1 | 0.330257709 |
| SL1344_RS05655_ymdC | 2.065955743 | 2.413553255 | 1.623064049 | -0.797793756 | 0.141958572 | -5.543740033 | 4.68E-05 | 0.00414078 | 0.076639796 | 0.1 | 0.330257709 |
| yopJ_yopJ | 0.75596586 | 1.333118932 | 0.510238089 | -0.806611224 | 0.263064305 | -2.635002067 | 0.00067292 | 0.043667752 | 0.24457503 | 0.1 | 0.330257709 |
| SL1344_RS15335_yqcB | 0.921111706 | 0.914299774 | 0.929755965 | 0.009799856 | 0.144367632 | 0.062978594 | 0.469020654 | 0.623671865 | 0.805520721 | 0.7273 | 0.856066077 |

| | | | | | | | | | | | |
|---|---|---|---|---|---|---|---|---|---|---|---|
| SL1344_RS15340_yqcC..next.to.csrB. | -0.167660651 | -0.10568615 | -0.24653352 | -0.137797103 | 0.199975261 | -0.631581227 | 0.24266668 | 0.367999902 | 0.609028986 | 0.4649 | 0.670436862 |
| YrlA_yrlA | -4.29928224 | -4.021167798 | -4.601739264 | -0.566227977 | 0.841798325 | -0.643044087 | 0.241172565 | 0.365973206 | 0.607960833 | 0.4658 | 0.668076101 |
| YrlB_yrlB | -3.790696329 | -3.533672748 | -3.91246967 | -0.269602835 | 0.939461489 | -0.217494698 | 0.399333337 | 0.723752111 | 0.871659216 | 0.7717 | 0.89381808 |
| zntA_zntA | 2.321898522 | 2.513056327 | 2.15670761 | -0.281874745 | 0.263491851 | -1.120780014 | 0.118587642 | 0.151339386 | 0.434931595 | 0.2093 | 0.484222681 |
| zntB_zntB | 0.662652243 | 0.87230586 | 0.493033269 | -0.38340012 | 0.165131859 | -2.279289042 | 0.004664544 | 0.036235342 | 0.208545221 | 0.1036 | 0.335096144 |
| zraP_zraP | -2.63431828 | -2.63393449 | -2.63517622 | -0.003886265 | 0.490217382 | -0.01009933 | 0.497333333 | 0.617774973 | 0.802601812 | 0.7367 | 0.861931675 |
| crRNA1_ | 0.057555834 | -0.048288492 | 0.111876388 | 0.161494615 | 0.198431882 | 0.737448175 | 0.220666682 | 0.334558352 | 0.588052208 | 0.4459 | 0.657569684 |
| crRNA2_ | 1.09950291 | 0.924241512 | 1.167111689 | 0.236786058 | 0.173779775 | 1.222220604 | 0.094000044 | 0.156908095 | 0.426224175 | 0.2323 | 0.478224004 |
| ECS021_ | -3.702431582 | -3.652482262 | -3.740425967 | -0.049904582 | 1.028255893 | -0.03931821 | 0.478000001 | 0.713661584 | 0.864663356 | 0.8019 | 0.902273233 |
| ECS173_ | -2.078878835 | -2.000195549 | -2.145811695 | -0.141230611 | 0.366242431 | -0.364278001 | 0.355333339 | 0.508176341 | 0.720633969 | 0.6157 | 0.779483153 |
| Ffs_ | 3.296256011 | 2.929728165 | 3.667197819 | -0.142913303 | 1.523874438 | -0.103562067 | 0.443037977 | 0.965166014 | 0.98789864 | 1 | 1 |
| fimA.trailer_ | -4.389588667 | -4.595279791 | -4.207659295 | 0.440227755 | 0.957723067 | 0.41024114 | 0.308666675 | 0.45548266 | 0.689871497 | 0.5826 | 0.75473425 |
| GO.10702_ | -0.936047495 | -0.799835963 | -1.009269557 | -0.156129336 | 0.362694171 | -0.373311033 | 0.355333339 | 0.634685414 | 0.817940401 | 0.7127 | 0.856393285 |
| GO.10704_ | 2.404930791 | 2.475373633 | 2.338134534 | -0.070588013 | 0.460563402 | -0.1387368 | 0.448000002 | 0.928922372 | 0.97416665 | 0.9997 | 0.999870836 |
| Ig1085_ | -2.080189368 | -2.258852726 | -1.829949044 | 0.442119407 | 0.472022112 | 0.901731411 | 0.173217876 | 0.258048657 | 0.522660911 | 0.3538 | 0.585606229 |
| Ig1140_ | -2.139060596 | -1.698910773 | -2.481501858 | -0.756549567 | 0.386810893 | -1.88311126 | 0.023984199 | 0.05738112 | 0.251756612 | 0.1249 | 0.361774025 |
| Ig2335_ | 0.549436024 | 0.346034328 | 0.829251601 | 0.473747282 | 0.156791111 | 2.92116668 | 0.000673369 | 0.019648846 | 0.153147266 | 0.1012 | 0.331916603 |
| Ig959_ | -9.298020086 | -8.88461939 | -9.765183019 | -0.851286371 | 3.950190686 | -0.180604356 | 0.405063295 | 0.518733971 | 0.730708019 | 0.6413 | 0.792377638 |
| IS042_ | -4.410307719 | -4.252065721 | -4.580662237 | -0.623575774 | 1.284314755 | -0.447676475 | 0.310459702 | 0.456435016 | 0.69964311 | 0.6413 | 0.795631561 |
| lysC.leader_ | -1.086266301 | -1.157870472 | -0.956819005 | 0.230666612 | 0.610544092 | 0.35098126 | 0.347768161 | 0.542463711 | 0.771182674 | 0.716 | 0.859818511 |
| mreB.leader_ | 1.170129456 | 1.311168442 | 0.956780631 | -0.377052102 | 0.19684874 | -1.855483949 | 0.006662914 | 0.070556633 | 0.29658931 | 0.1063 | 0.339033255 |
| pRNA_ | -4.50648736 | -4.656077834 | -4.413441846 | 0.225376425 | 0.904247494 | 0.239912097 | 0.391072622 | 0.549934935 | 0.754673768 | 0.6666 | 0.816716183 |
| RprA_ | -1.946987099 | -2.048695431 | -1.826582517 | 0.002747301 | 0.824463218 | 0.001793173 | 0.498334444 | 0.808631726 | 0.919937101 | 0.9408 | 0.973786898 |
| rpsL.leader_ | 2.968151571 | 2.992921037 | 2.944704597 | -0.04560703 | 0.4544295 | -0.088917274 | 0.456362427 | 0.773992652 | 0.90943938 | 0.9985 | 0.999357826 |
| rpsP.leader_ | 2.933315406 | 2.872158346 | 2.987684011 | 0.10124431 | 0.45539849 | 0.217742403 | 0.42066667 | 0.734561047 | 0.889693124 | 0.9391 | 0.973325846 |
| rrf_ | -6.682959001 | -6.9106196 | -6.429658436 | 0.481534489 | 1.842961104 | 0.231962691 | 0.385742843 | 0.541630351 | 0.746320979 | 0.6555 | 0.804969153 |
| rrf_.1 | -5.738523789 | -6.140467333 | -4.574850273 | 1.247528906 | 2.509554155 | 0.305525086 | 0.327115264 | 0.712186713 | 0.862509338 | 0.6858 | 0.854415558 |
| rrf_.2 | -4.980106618 | -5.829182024 | -4.188837612 | 1.613835818 | 1.098481982 | 1.404638997 | 0.055296548 | 0.106132427 | 0.335662538 | 0.1672 | 0.409894123 |
| rrf_.3 | -6.652796921 | -6.632051416 | -6.669840024 | 0.261318664 | 2.956539663 | 0.066661842 | 0.465023319 | 0.644961914 | 0.829950661 | 0.8134 | 0.908184124 |
| rrf_.4 | -2.644918284 | -3.821113683 | -2.415084665 | 1.354451513 | 1.211327524 | 1.12337567 | 0.16266669 | 0.166905455 | 0.452731383 | 0.2981 | 0.570554805 |
| rrf_.5 | -6.176966865 | -6.827855853 | -5.418779424 | 1.257035137 | 2.217549566 | 0.523864748 | 0.271333344 | 0.458091404 | 0.691245447 | 0.5383 | 0.736355161 |
| rrf_.6 | -7.901753416 | -9.826984803 | -6.832752532 | 2.890857942 | 2.829988432 | 0.943385739 | 0.107333371 | 0.199455104 | 0.450929969 | 0.2553 | 0.486074633 |
| rrf_.7 | -7.276149777 | -7.355806025 | -7.149882457 | 0.549658766 | 3.624214603 | 0.120632608 | 0.432666669 | 0.647038437 | 0.826164204 | 0.7996 | 0.898960401 |
| RUF_107c.1_ | 0.778581278 | 0.663914114 | 0.81312435 | 0.1189119 | 0.219733321 | 0.388207929 | 0.358666672 | 0.724053363 | 0.874783715 | 0.7499 | 0.891576518 |
| RUF_107c.10_ | 0.799737203 | 0.84732894 | 0.75131556 | -0.129429342 | 0.243918051 | -0.534337168 | 0.23866668 | 0.355461243 | 0.637985308 | 0.4749 | 0.68070185 |
| RUF_107c.102_ | -2.781886788 | -2.827138658 | -2.743760379 | 0.100568046 | 0.462263677 | 0.209177112 | 0.409726852 | 0.546140091 | 0.749207508 | 0.6565 | 0.805510833 |
| RUF_107c.105_ | 0.108811424 | 0.089352154 | 0.13040649 | -0.008314671 | 0.249777273 | -0.036575928 | 0.485333334 | 0.715765259 | 0.867085164 | 0.8405 | 0.921850715 |
| RUF_107c.107_ | -1.443327612 | -1.486626007 | -1.416918051 | 0.041251749 | 0.422616863 | 0.105162362 | 0.457694872 | 0.716972352 | 0.86885036 | 0.8202 | 0.913805684 |
| RUF_107c.108_ | 1.712698522 | 1.008494807 | 1.81973301 | 0.785247447 | 0.602899403 | 0.654634025 | 0.33333334 | 0.487407916 | 0.747225116 | 0.7 | 0.872542119 |
| RUF_107c.111_ | 1.843257329 | 1.664029018 | 1.948277479 | 0.204595035 | 0.257239975 | 0.862652883 | 0.163890762 | 0.25579651 | 0.55016017 | 0.3133 | 0.584348652 |
| RUF_107c.112_ | 1.136970623 | 0.904255549 | 1.215161592 | 0.35188411 | 0.364969423 | 0.972149939 | 0.14923387 | 0.197791383 | 0.492304908 | 0.2998 | 0.572060595 |
| RUF_107c.18_ | -2.638532861 | -2.357320062 | -2.838479209 | -0.563259597 | 0.722060076 | -0.731661631 | 0.221333348 | 0.316890249 | 0.591373957 | 0.4405 | 0.664707708 |
| RUF_107c.19_ | 0.038039721 | -0.083716117 | 0.147631825 | 0.223156823 | 0.191314629 | 1.073560029 | 0.122000033 | 0.184935315 | 0.438992214 | 0.2658 | 0.509310309 |
| RUF_107c.2_ | -2.234743932 | -2.253026527 | -2.220923393 | 0.049146485 | 0.467218757 | 0.099805521 | 0.459333335 | 0.647871511 | 0.822613791 | 0.754 | 0.872312853 |
| RUF_107c.23_ | 0.657105852 | 0.76480995 | 0.579393034 | -0.161960982 | 0.359254222 | -0.367895326 | 0.378000005 | 0.64729744 | 0.83557534 | 0.817 | 0.916566706 |
| RUF_107c.24_ | 0.862004996 | 0.731939985 | 0.962518291 | 0.263754009 | 0.204580813 | 1.211861825 | 0.07533339 | 0.129232888 | 0.387977485 | 0.1888 | 0.435972302 |
| RUF_107c.26_ | -4.571037694 | -4.462746613 | -4.671940768 | -0.29807986 | 1.052717139 | -0.263504441 | 0.38241173 | 0.582905183 | 0.780190578 | 0.7153 | 0.8454677 |
| RUF_107c.30_ | -5.736467684 | -5.537707476 | -5.909916891 | -0.401914609 | 1.875494185 | -0.183515894 | 0.403333337 | 0.627517499 | 0.81431894 | 0.76 | 0.876482542 |
| RUF_107c.38_ | -0.957142908 | -1.125568306 | -0.89503228 | 0.18577955 | 0.41988395 | 0.314937341 | 0.380000005 | 0.788458833 | 0.909924751 | 0.7687 | 0.900547289 |
| RUF_107c.47_ | 0.873812771 | 0.44371192 | 1.074055882 | 0.587951075 | 0.564318032 | 0.874059441 | 0.185209879 | 0.276334533 | 0.578172301 | 0.3345 | 0.605811498 |
| RUF_107c.50_ | -0.738468277 | -0.809316416 | -0.711148867 | 0.054489583 | 0.35377719 | 0.120375665 | 0.451698869 | 0.744211798 | 0.884919736 | 0.808 | 0.913450839 |
| RUF_107c.51_ | -0.446723341 | -0.486808808 | -0.395812715 | 0.085297117 | 0.217709145 | 0.358371464 | 0.34733334 | 0.527140598 | 0.731283006 | 0.6315 | 0.78771077 |
| RUF_107c.53_ | 0.854671766 | 0.980248689 | 0.705719541 | -0.321135853 | 0.277889729 | -1.195219896 | 0.042000105 | 0.193130053 | 0.489162444 | 0.1386 | 0.381537193 |

| | | | | | | | | | | | |
|---|---|---|---|---|---|---|---|---|---|---|---|
| RUF_107c.56_ | 0.583038037 | 0.172875537 | 0.710983291 | 0.508909959 | 0.326709859 | 1.428042983 | 0.077281867 | 0.113242856 | 0.376521634 | 0.188 | 0.44492984 |
| RUF_107c.57_ | -1.413721045 | -1.145382652 | -1.514020555 | -0.343921119 | 0.345117297 | -0.882457393 | 0.190666685 | 0.280242929 | 0.543129448 | 0.3942 | 0.624399873 |
| RUF_107c.58_ | 1.582069537 | 1.604473504 | 1.553885754 | -0.09829538 | 0.919496798 | -0.140112033 | 0.366666672 | 0.807092137 | 0.924195439 | 0.7522 | 0.881907061 |
| RUF_107c.59_ | 0.304522927 | 0.051235034 | 0.484049345 | 0.229245822 | 0.717685113 | 0.256993192 | 0.376000005 | 0.920462807 | 0.970805167 | 0.8011 | 0.915504503 |
| RUF_107c.63_ | -0.874611402 | -1.161961151 | -0.760639173 | 0.321715639 | 1.131802719 | 0.246174264 | 0.358666672 | 0.907262762 | 0.965566943 | 0.7477 | 0.892685382 |
| RUF_107c.69_ | -0.114337893 | -0.067308066 | -0.188872295 | -0.095053684 | 0.313704786 | -0.279012547 | 0.380000005 | 0.645595453 | 0.828572742 | 0.7628 | 0.88238503 |
| RUF_107c.70_ | -1.521165256 | -1.794777616 | -1.327126614 | 0.507215481 | 0.423243539 | 1.138853081 | 0.100000041 | 0.156627502 | 0.424290229 | 0.2233 | 0.475080576 |
| RUF_107c.72_ | -1.200475342 | -1.354081595 | -1.118864075 | 0.164396421 | 0.685679634 | 0.221900439 | 0.401333337 | 0.815246881 | 0.923698536 | 0.8083 | 0.916394931 |
| RUF_107c.77_ | -0.621302312 | -0.646667215 | -0.583789154 | 0.110911504 | 0.63333962 | 0.174535107 | 0.392405067 | 0.72627675 | 0.877828866 | 0.7826 | 0.893905257 |
| RUF_107c.8_ | -2.689956696 | -3.112963395 | -2.091159483 | 0.993573406 | 0.53277222 | 1.808718687 | 0.027315288 | 0.062818932 | 0.268714894 | 0.1316 | 0.370672959 |
| RUF_107c.81_ | -3.754762175 | -3.911467087 | -3.65150593 | 0.191955955 | 0.781892356 | 0.227001162 | 0.405729517 | 0.614996884 | 0.799867929 | 0.731 | 0.85543703 |
| RUF_107c.82_ | 2.024197494 | 1.718987796 | 2.104589276 | 0.323245966 | 0.427713229 | 0.624440297 | 0.295333342 | 0.472494414 | 0.735384283 | 0.6097 | 0.814621501 |
| RUF_107c.83_ | -4.243161857 | -4.318066917 | -4.199318786 | 0.007838919 | 1.026807728 | 0.008387577 | 0.496 | 0.688582059 | 0.850217046 | 0.7891 | 0.896221203 |
| RUF_107c.86_ | -0.258135407 | -0.296310521 | -0.195186335 | -0.092938772 | 0.547321197 | -0.179525751 | 0.441705532 | 0.756756599 | 0.893582806 | 0.9218 | 0.96367873 |
| RUF_107c.89_ | 0.861179639 | 0.927015693 | 0.789764238 | -0.1984085 | 0.65943622 | -0.28136133 | 0.344000006 | 0.478762997 | 0.740212122 | 0.7195 | 0.871862482 |
| RUF_107c.90_ | -3.071072975 | -3.452419457 | -2.748043831 | 0.703613564 | 0.495195463 | 1.37066659 | 0.055962769 | 0.117025937 | 0.347917043 | 0.1878 | 0.429210633 |
| RUF_107c.92_ | 1.608971007 | 1.710033665 | 1.580380804 | -0.132484212 | 0.383868756 | -0.33407755 | 0.36109261 | 0.639928116 | 0.836579843 | 0.7465 | 0.89202373 |
| RUF_107c.94_ | -3.716774903 | -3.819496326 | -3.610298894 | 0.183232683 | 0.793434917 | 0.216698952 | 0.40266667 | 0.598407797 | 0.789279854 | 0.701 | 0.83881157 |
| RUF_107c.95_ | 2.198924611 | 2.145123315 | 2.280368461 | 0.160639874 | 0.172526239 | 0.894590967 | 0.159893428 | 0.224240083 | 0.512127874 | 0.3327 | 0.59014527 |
| RUF_107c.96_ | -1.025925521 | -0.93471374 | -1.201493012 | -0.262519315 | 0.465114201 | -0.556187978 | 0.27514991 | 0.442683908 | 0.694791997 | 0.5442 | 0.754640211 |
| RUF_111_ | 5.349948634 | 5.244276431 | 5.45705783 | 0.215241916 | 0.280691811 | 0.628093144 | 0.297135252 | 0.393435025 | 0.679797958 | 0.5864 | 0.798314372 |
| RUF_121_ | -1.691980206 | -1.328580467 | -2.223327293 | -0.991240367 | 0.605198419 | -1.686034619 | 0.013333676 | 0.063012803 | 0.283216667 | 0.1122 | 0.34709584 |
| RUF_125_ | 2.388038256 | 3.339584482 | 1.352205132 | -1.973399189 | 0.198449943 | -9.905683186 | 4.68E-05 | 0.00027133 | 0.026904216 | 0.1 | 0.330257709 |
| RUF_167_ | -1.276343483 | -1.462538747 | -1.173218611 | 0.236931423 | 0.417457134 | 0.538257569 | 0.283333343 | 0.452952583 | 0.695458846 | 0.5798 | 0.768231388 |
| RUF_178_ | 1.193684602 | 0.807943977 | 1.691263082 | 0.912711447 | 0.676775816 | 1.476543436 | 0.013333676 | 0.085336309 | 0.337748394 | 0.1098 | 0.344571009 |
| RUF_219c_ | 0.840537601 | 1.392233954 | 0.416933486 | -0.93822684 | 0.213436958 | -4.247589069 | 4.68E-05 | 0.008319398 | 0.109680392 | 0.1 | 0.330257709 |
| RUF_220c_ | -0.876888624 | -0.150298933 | -1.830335997 | -1.683976774 | 0.299982253 | -5.519969368 | 4.68E-05 | 0.003855561 | 0.07055079 | 0.1 | 0.330257709 |
| RUF_226_ | -2.395269529 | -2.403628268 | -2.390052041 | -0.019783101 | 0.507644616 | -0.035938191 | 0.486000001 | 0.667051351 | 0.834486352 | 0.7674 | 0.88321542 |
| RUF_247c_ | 0.991248011 | 0.868951118 | 1.134171533 | 0.186894528 | 0.325888673 | 0.587177165 | 0.248501012 | 0.406226119 | 0.677817207 | 0.4745 | 0.71009249 |
| RUF_257c_ | 0.829088666 | 0.658608277 | 0.902272947 | 0.205636724 | 0.221403935 | 0.874499638 | 0.197868106 | 0.270362605 | 0.551784423 | 0.3759 | 0.624372202 |
| RUF_279_ | 2.233437202 | 1.83427971 | 2.304451282 | 0.398879561 | 0.275061709 | 0.877054912 | 0.257828126 | 0.297873769 | 0.598666086 | 0.5026 | 0.743960028 |
| RUF_290_ | 2.171554098 | 2.172364026 | 2.171027523 | -0.004766428 | 0.194991265 | -0.017966765 | 0.491672219 | 0.765481041 | 0.899589975 | 0.8968 | 0.955308703 |
| RUF_321c_ | -8.01254293 | -8.058145823 | -7.958932034 | 0.08045369 | 3.012871818 | 0.018584386 | 0.486666667 | 0.568912327 | 0.770932143 | 0.6942 | 0.831820216 |
| RUF_328c_ | -5.917503038 | -5.931830274 | -5.901479558 | 0.140742704 | 1.593787106 | 0.069072946 | 0.462666668 | 0.588217452 | 0.783449379 | 0.6935 | 0.8324784 |
| RUF_341c.10_ | -1.286052986 | -1.316712486 | -1.266241435 | 0.056257331 | 0.303599445 | 0.170875146 | 0.419053967 | 0.585806925 | 0.780474281 | 0.7025 | 0.836515164 |
| RUF_341c.11_ | 0.749543202 | 0.600495196 | 0.896780501 | 0.325596118 | 0.191407468 | 1.698353906 | 0.011992386 | 0.068436806 | 0.288491035 | 0.1156 | 0.350676767 |
| RUF_341c.2_ | -2.80303681 | -2.775543554 | -2.838770496 | 0.079855126 | 0.71107952 | 0.108904938 | 0.459333335 | 0.681755792 | 0.848662881 | 0.8398 | 0.920753576 |
| SL1344_RS00005_ | 4.004341447 | 4.059722143 | 3.946479741 | 0.012907303 | 0.253663023 | 0.063985441 | 0.477681546 | 0.89887872 | 0.962262225 | 0.9943 | 0.997500765 |
| SL1344_RS00010_ | 1.839091649 | 1.841664393 | 1.834023585 | 0.0170111 | 0.264928324 | 0.056666482 | 0.476666668 | 0.845780127 | 0.938048144 | 0.9433 | 0.974677934 |
| SL1344_RS00015_ | 1.547528156 | 1.439948308 | 1.580646754 | 0.120589178 | 0.175726248 | 0.594958797 | 0.301798809 | 0.489852443 | 0.728493538 | 0.5959 | 0.793811816 |
| SL1344_RS00020_ | 2.38616919 | 2.332694425 | 2.471996498 | 0.069429989 | 0.187119683 | 0.367049615 | 0.365089945 | 0.646450047 | 0.83343009 | 0.7495 | 0.879812672 |
| SL1344_RS00025_ | 1.452757169 | 1.456563313 | 1.446458016 | -0.008782496 | 0.107606819 | -0.074695187 | 0.468000001 | 0.576805515 | 0.771728739 | 0.687 | 0.828602352 |
| SL1344_RS00030_ | 1.05328156 | 1.021409688 | 1.135604393 | 0.110592393 | 0.1872541 | 0.51066075 | 0.309333342 | 0.50684274 | 0.734512359 | 0.6331 | 0.806053317 |
| SL1344_RS00035_ | 6.182413525 | 6.109888269 | 6.272770944 | 0.126859871 | 0.095251576 | 1.556476365 | 0.074666724 | 0.073558828 | 0.313607695 | 0.1687 | 0.430666531 |
| SL1344_RS00040_ | 1.895372972 | 1.77419668 | 1.930147447 | 0.156131486 | 0.207183908 | 0.748323976 | 0.220519669 | 0.320562416 | 0.603730266 | 0.4121 | 0.673285984 |
| SL1344_RS00045_ | 0.437550816 | 0.106133298 | 0.613823321 | 0.495810476 | 0.1837629 | 2.546023494 | 0.002000966 | 0.032949747 | 0.199953706 | 0.1023 | 0.333374415 |
| SL1344_RS00050_ | -3.634445275 | -4.014279288 | -3.252250549 | 0.811804637 | 0.716615228 | 1.097689904 | 0.105333372 | 0.173341717 | 0.431638878 | 0.2557 | 0.495049197 |
| SL1344_RS00055_ | -3.308787822 | -3.445915958 | -3.197908289 | 0.216323222 | 0.542882596 | 0.383265422 | 0.331112599 | 0.48867942 | 0.702456171 | 0.5893 | 0.757639005 |
| SL1344_RS00065_ | 3.699676276 | 3.740836565 | 3.659987822 | -0.105111744 | 0.385669371 | -0.32048025 | 0.323333341 | 0.596611529 | 0.815608451 | 0.6724 | 0.852554448 |
| SL1344_RS00070_ | -1.293365194 | -1.394112321 | -1.197240235 | 0.196508583 | 0.322299597 | 0.575373693 | 0.260000012 | 0.401492296 | 0.644140316 | 0.5101 | 0.705561347 |
| SL1344_RS00075_ | -6.273224125 | -5.990202991 | -7.083044137 | -0.780425997 | 3.167144702 | -0.190224838 | 0.41972019 | 0.604510154 | 0.807272532 | 0.8121 | 0.913193134 |
| SL1344_RS00080_ | -3.931068114 | -3.8010802 | -4.247897925 | -0.403533679 | 0.819394848 | -0.427186659 | 0.314457037 | 0.480385469 | 0.711257226 | 0.6069 | 0.782520828 |
| SL1344_RS00085_ | -5.521543306 | -5.499286972 | -5.551497669 | -0.110323449 | 1.230626512 | -0.080472052 | 0.466355764 | 0.563133841 | 0.762058157 | 0.6791 | 0.820923774 |

| | | | | | | | | | | | |
|---|---|---|---|---|---|---|---|---|---|---|---|
| SL1344_RS00090_ | -0.3655661 | -0.301421866 | -0.444536295 | -0.107073993 | 0.281625272 | -0.376468917 | 0.346435716 | 0.578531914 | 0.779960037 | 0.6907 | 0.835856288 |
| SL1344_RS00095_ | -0.88472772 | -0.744211256 | -1.08793274 | -0.365753857 | 0.358626177 | -0.990133809 | 0.127333364 | 0.195209045 | 0.469258438 | 0.279 | 0.528214354 |
| SL1344_RS00100_ | -2.905495602 | -2.838503025 | -2.976015842 | -0.20818148 | 0.650878673 | -0.29566496 | 0.375333338 | 0.556988723 | 0.762809411 | 0.6998 | 0.833680106 |
| SL1344_RS00105_ | -2.988054593 | -3.167277019 | -2.785371605 | 0.396437175 | 0.450981749 | 0.821854656 | 0.184000019 | 0.288257034 | 0.529141684 | 0.3747 | 0.590276397 |
| SL1344_RS00110_ | -3.808093618 | -3.715337912 | -3.924923769 | -0.201036362 | 0.690387652 | -0.256436571 | 0.38241173 | 0.547658797 | 0.751738001 | 0.6689 | 0.814764292 |
| SL1344_RS00115_ | -1.796458083 | -1.714492202 | -1.833872607 | -0.090556558 | 0.405953961 | -0.20078481 | 0.41972019 | 0.651296635 | 0.82593937 | 0.749 | 0.873533539 |
| SL1344_RS00120_ | -2.596182899 | -2.571282235 | -2.625077917 | -0.048862763 | 0.409672016 | -0.115379683 | 0.448367757 | 0.542988219 | 0.747081466 | 0.6605 | 0.807112498 |
| SL1344_RS00125_ | -3.658570954 | -3.738427944 | -3.630667428 | -0.002994203 | 0.842793587 | -0.003392449 | 0.498334444 | 0.682901416 | 0.846242855 | 0.7754 | 0.890762553 |
| SL1344_RS00130_ | -3.621308031 | -3.77920129 | -3.513861513 | 0.258823333 | 0.636826133 | 0.377621564 | 0.331112599 | 0.487694305 | 0.702972739 | 0.6004 | 0.76523918 |
| SL1344_RS00135_ | -2.997240509 | -3.125625048 | -2.892585416 | 0.238919848 | 0.496425358 | 0.448550153 | 0.291805472 | 0.439597479 | 0.666906419 | 0.5449 | 0.723368115 |
| SL1344_RS00140_ | -1.651905222 | -1.596918143 | -1.703545323 | -0.106556416 | 0.298776028 | -0.320914352 | 0.358427721 | 0.485379366 | 0.700282036 | 0.5932 | 0.759979157 |
| SL1344_RS00145_ | -1.887348995 | -1.811963771 | -1.932163719 | -0.078893991 | 0.413138738 | -0.180798451 | 0.42066667 | 0.641198085 | 0.817814286 | 0.7369 | 0.863953253 |
| SL1344_RS00150_ | -5.970328105 | -6.43903408 | -5.488435488 | 0.962500103 | 1.487818032 | 0.57653027 | 0.233333347 | 0.366583823 | 0.612127483 | 0.4635 | 0.664940448 |
| SL1344_RS00155_ | -3.221683585 | -3.13842862 | -3.284445842 | -0.1556917 | 0.600190188 | -0.226643841 | 0.398666671 | 0.588771225 | 0.781551599 | 0.7 | 0.83673049 |
| SL1344_RS00160_ | -3.814973369 | -3.885899928 | -3.75229522 | 0.149859742 | 0.657836672 | 0.212694315 | 0.407333337 | 0.534091549 | 0.739985951 | 0.6501 | 0.800853015 |
| SL1344_RS00165_ | -1.716936742 | -1.815757932 | -1.626407922 | 0.116242928 | 0.48967979 | 0.238354954 | 0.399733515 | 0.664554495 | 0.836072901 | 0.776 | 0.885227153 |
| SL1344_RS00170_ | -0.695028965 | -0.763871192 | -0.614833809 | 0.149265738 | 0.218713994 | 0.620194428 | 0.234000014 | 0.36044787 | 0.595999424 | 0.46 | 0.660814952 |
| SL1344_RS00175_ | -5.522873122 | -5.530612395 | -5.50024485 | 0.140976449 | 1.616383943 | 0.079682476 | 0.465689542 | 0.661911993 | 0.833065811 | 0.7848 | 0.893079834 |
| SL1344_RS00180_ | 0.003384183 | 0.038686058 | -0.022443221 | -0.046494373 | 0.225332733 | -0.185602031 | 0.418387745 | 0.650754544 | 0.82330192 | 0.7436 | 0.867100396 |
| SL1344_RS00185_ | -0.965672189 | -1.027835098 | -0.922165291 | 0.101546879 | 0.309923406 | 0.325615178 | 0.368000005 | 0.551223674 | 0.758062578 | 0.6609 | 0.812945368 |
| SL1344_RS00190_ | -2.00904542 | -2.048444676 | -1.9707942 | 0.072939428 | 0.373684786 | 0.181637483 | 0.41866667 | 0.578393408 | 0.772358169 | 0.69 | 0.828414645 |
| SL1344_RS00195_ | 0.208195304 | 0.092669049 | 0.275926887 | 0.119926436 | 0.294344095 | 0.383697396 | 0.354000006 | 0.608922427 | 0.806325825 | 0.7172 | 0.859343164 |
| SL1344_RS00205_ | 1.573170362 | 1.672317469 | 1.45844969 | -0.219121907 | 0.12472572 | -1.739237325 | 0.025333511 | 0.065346574 | 0.273678463 | 0.1281 | 0.366412069 |
| SL1344_RS00210_ | -0.535052456 | -0.583469429 | -0.483751368 | 0.114751642 | 0.277841148 | 0.397020421 | 0.338441046 | 0.517049183 | 0.735707272 | 0.6341 | 0.795840891 |
| SL1344_RS00215_ | -3.611374935 | -3.968431998 | -3.291740087 | 0.680409449 | 0.587482453 | 1.102208635 | 0.101265863 | 0.174437219 | 0.416269568 | 0.2492 | 0.484385276 |
| SL1344_RS00220_ | 6.729040708 | 6.711259408 | 6.84989149 | 0.140745276 | 0.179122952 | 0.822908766 | 0.195333351 | 0.193991493 | 0.494247206 | 0.3639 | 0.643472947 |
| SL1344_RS00225_ | -1.193167276 | -0.979783305 | -1.424707313 | -0.437870324 | 0.4022118 | -1.054985908 | 0.120586309 | 0.185604954 | 0.458918149 | 0.2549 | 0.508414833 |
| SL1344_RS00230_ | 2.650831626 | 2.635632058 | 2.661569936 | -0.017206043 | 0.14973997 | -0.115900046 | 0.460000001 | 0.754775531 | 0.890243717 | 0.9045 | 0.954979009 |
| SL1344_RS00235_ | 4.727709304 | 4.701622913 | 4.771786165 | 0.095352432 | 0.21266143 | 0.49856522 | 0.251832124 | 0.438949644 | 0.711334755 | 0.4819 | 0.732357203 |
| SL1344_RS00240_ | 2.80549475 | 2.605635353 | 2.861352738 | 0.198120288 | 0.249858311 | 0.652713429 | 0.309333342 | 0.472343923 | 0.733420182 | 0.6333 | 0.829334321 |
| SL1344_RS00245_ | 1.891780633 | 1.833987051 | 1.928884302 | 0.089073833 | 0.115922193 | 0.706674372 | 0.225333348 | 0.35431562 | 0.60670262 | 0.4562 | 0.670884375 |
| SL1344_RS00250_ | 2.863765114 | 2.799693987 | 2.928650207 | 0.124711012 | 0.085297123 | 1.426838552 | 0.055962769 | 0.1048383 | 0.346181695 | 0.1705 | 0.417650366 |
| SL1344_RS00255_ | 1.657155356 | 1.716355802 | 1.531074341 | -0.180352663 | 0.221815649 | -0.802482132 | 0.185876102 | 0.2655849 | 0.555320124 | 0.3521 | 0.616942493 |
| SL1344_RS00260_ | 1.837175311 | 2.233313001 | 1.653826146 | -0.556624041 | 0.156717421 | -3.12851414 | 4.68E-05 | 0.023073311 | 0.177882261 | 0.1 | 0.330257709 |
| SL1344_RS00270_ | -0.585676009 | -0.789579475 | -0.477951523 | 0.292474744 | 0.247985524 | 1.118333379 | 0.116588975 | 0.179909207 | 0.439075459 | 0.2604 | 0.504196795 |
| SL1344_RS00275_ | -4.072099069 | -4.234660446 | -3.927381715 | 0.356178222 | 0.788393048 | 0.420159591 | 0.317333341 | 0.469230738 | 0.69633303 | 0.5888 | 0.758047228 |
| SL1344_RS00280_ | -4.804391634 | -4.875810341 | -4.712047495 | 0.163426177 | 0.964625775 | 0.157114408 | 0.431333336 | 0.586592026 | 0.778328267 | 0.6926 | 0.831395718 |
| SL1344_RS00285_ | -6.468132315 | -6.515150247 | -6.423815356 | 0.106426617 | 1.795898361 | 0.045706529 | 0.475333334 | 0.58870633 | 0.780416792 | 0.71 | 0.840645471 |
| SL1344_RS00290_ | -2.418237054 | -2.337568265 | -2.501544521 | -0.162365263 | 0.428451026 | -0.339905548 | 0.356666672 | 0.50260074 | 0.714545801 | 0.6019 | 0.767492747 |
| SL1344_RS00295_ | -3.12767534 | -3.387818332 | -2.897266185 | 0.483268177 | 0.454813845 | 1.046425359 | 0.122666699 | 0.201848601 | 0.438849754 | 0.2856 | 0.51173994 |
| SL1344_RS00300_ | -5.238772916 | -5.125943829 | -5.344609813 | -0.335264597 | 1.393199293 | -0.241306765 | 0.39573618 | 0.602768434 | 0.794876015 | 0.7206 | 0.850290037 |
| SL1344_RS00305_ | -3.112539627 | -3.08666348 | -3.132833057 | 0.055855717 | 0.800051556 | 0.065596594 | 0.465023319 | 0.695358796 | 0.855947885 | 0.8254 | 0.913246103 |
| SL1344_RS00310_ | -2.251627127 | -2.347280739 | -2.18350002 | 0.140904513 | 0.502455239 | 0.264131489 | 0.38774151 | 0.621605058 | 0.809153332 | 0.7251 | 0.858786082 |
| SL1344_RS00315_ | -2.734207855 | -2.494354301 | -3.065984836 | -0.560314234 | 0.432558865 | -1.234325047 | 0.093937419 | 0.157476886 | 0.39756323 | 0.2344 | 0.471577597 |
| SL1344_RS00320_ | -1.817287036 | -1.766624333 | -1.851018411 | -0.123996771 | 0.453220754 | -0.266161073 | 0.388407733 | 0.596383064 | 0.791465065 | 0.7283 | 0.855642752 |
| SL1344_RS00325_ | 2.258718413 | 2.231283313 | 2.269832716 | 0.047786017 | 0.155068225 | 0.257602947 | 0.370666672 | 0.508384979 | 0.749916904 | 0.7182 | 0.853373004 |
| SL1344_RS00330_ | -3.169758015 | -3.578236369 | -2.822114992 | 0.772300914 | 0.571599288 | 1.331079157 | 0.064000067 | 0.127828608 | 0.368952418 | 0.1897 | 0.433420257 |
| SL1344_RS00335_ | -0.070198608 | -0.305972453 | 0.258619137 | 0.560493321 | 0.210887579 | 2.574383846 | 0.002668394 | 0.027511656 | 0.183516516 | 0.1017 | 0.332643865 |
| SL1344_RS00340_ | 1.871498875 | 1.548831275 | 2.123697611 | 0.56972166 | 0.166052773 | 3.409075102 | 4.68E-05 | 0.009837736 | 0.119578512 | 0.1 | 0.330257709 |
| SL1344_RS00345_ | -0.087361953 | -0.272320854 | 0.056109366 | 0.323878688 | 0.203137874 | 1.544544686 | 0.049333422 | 0.094017959 | 0.319594098 | 0.1603 | 0.403176433 |
| SL1344_RS00350_ | -2.718692284 | -2.537701196 | -2.851178991 | -0.259222939 | 0.698321902 | -0.34434766 | 0.349333339 | 0.612858466 | 0.802182261 | 0.6917 | 0.840775981 |
| SL1344_RS00355_ | -2.100517492 | -2.031756389 | -2.184719354 | -0.125693311 | 0.38307537 | -0.306396488 | 0.367088613 | 0.533944795 | 0.739735594 | 0.6407 | 0.796253994 |
| SL1344_RS00360_ | -1.700837979 | -1.459266477 | -1.952453194 | -0.485494956 | 0.387489039 | -1.217815794 | 0.077948089 | 0.138420045 | 0.388259617 | 0.1963 | 0.442411026 |

| | | | | | | | | | | | |
|---|---|---|---|---|---|---|---|---|---|---|---|
| SL1344_RS00365_ | -1.386556323 | -1.06774452 | -1.579060486 | -0.418865978 | 0.468798487 | -0.875460095 | 0.180666687 | 0.265091147 | 0.54386686 | 0.352 | 0.601121623 |
| SL1344_RS00370_ | -0.566301437 | -0.351132818 | -0.683785436 | -0.325656336 | 0.329769998 | -0.943890008 | 0.16055965 | 0.224283572 | 0.499261022 | 0.3196 | 0.57404124 |
| SL1344_RS00375_ | -3.888627681 | -4.044530726 | -3.740611692 | 0.324901844 | 0.654691891 | 0.449504981 | 0.306000008 | 0.439680393 | 0.665689554 | 0.5514 | 0.727794245 |
| SL1344_RS00380_ | -6.4553382 | -6.49230621 | -6.40593032 | -0.099248692 | 2.506098881 | -0.030506975 | 0.479333334 | 0.628221886 | 0.817598818 | 0.771 | 0.881547123 |
| SL1344_RS00385_ | -2.573086402 | -2.831590997 | -2.243727026 | 0.402774037 | 0.806949037 | 0.47623839 | 0.305796144 | 0.533128553 | 0.756880949 | 0.6311 | 0.804588413 |
| SL1344_RS00390_ | -3.183937885 | -3.006451907 | -3.421758366 | -0.454848322 | 0.61064948 | -0.66986939 | 0.216666682 | 0.331232675 | 0.589208993 | 0.4302 | 0.641268159 |
| SL1344_RS00395_ | -2.969548371 | -3.013892049 | -2.909914264 | 0.106181951 | 0.451380279 | 0.219368091 | 0.397734847 | 0.520722312 | 0.72862913 | 0.6258 | 0.784187846 |
| SL1344_RS00400_ | -3.198245569 | -3.46630898 | -3.004986799 | 0.470148469 | 0.555466823 | 0.843509477 | 0.16066669 | 0.265855567 | 0.514517829 | 0.3478 | 0.566862117 |
| SL1344_RS00405_ | -0.81148708 | -0.848820841 | -0.7767656 | 0.095761432 | 0.348007698 | 0.266637519 | 0.376415728 | 0.59232666 | 0.790615856 | 0.7193 | 0.850411315 |
| SL1344_RS00410_ | -2.157404858 | -2.134620236 | -2.179520221 | 0.055370509 | 0.596689609 | 0.095073016 | 0.462000001 | 0.684731748 | 0.849691386 | 0.8311 | 0.915962984 |
| SL1344_RS00415_ | -0.286491029 | -0.464348679 | -0.229167404 | 0.209239206 | 0.319637007 | 0.574001425 | 0.305796144 | 0.487790349 | 0.727484137 | 0.6138 | 0.802393093 |
| SL1344_RS00420_ | -1.672282589 | -1.916667968 | -1.394826254 | 0.501355733 | 0.392878686 | 1.230812433 | 0.08794142 | 0.134306078 | 0.386237793 | 0.2076 | 0.456638453 |
| SL1344_RS00425_ | -5.826101968 | -5.832383308 | -5.816333891 | 0.137572898 | 1.645206344 | 0.074392723 | 0.462358429 | 0.608023058 | 0.796783136 | 0.7322 | 0.854299141 |
| SL1344_RS00430_ | -0.63911001 | -0.777534699 | -0.530447165 | 0.240439365 | 0.262412601 | 0.878273406 | 0.164556985 | 0.250714443 | 0.512854977 | 0.3402 | 0.57741947 |
| SL1344_RS00435_ | -2.451698906 | -2.507918005 | -2.403083512 | 0.118479503 | 0.372155033 | 0.29820415 | 0.371333338 | 0.491035957 | 0.704958187 | 0.6014 | 0.76513892 |
| SL1344_RS00440_ | -0.143243656 | -0.337433111 | -0.08614977 | 0.218061795 | 0.334899142 | 0.540916163 | 0.324666674 | 0.512985719 | 0.74862374 | 0.6549 | 0.831830306 |
| SL1344_RS00450_ | 2.294860938 | 2.220906668 | 2.324813502 | 0.088524642 | 0.131757953 | 0.634768984 | 0.27514991 | 0.456840499 | 0.705199237 | 0.551 | 0.762052629 |
| SL1344_RS00455_ | 1.908765775 | 1.805431011 | 2.105825802 | 0.268888657 | 0.143732503 | 1.730838224 | 0.038641021 | 0.071929254 | 0.296354828 | 0.1375 | 0.381748652 |
| SL1344_RS00465_ | 2.968217878 | 2.885898072 | 3.067962995 | 0.233335034 | 0.141446743 | 1.699884296 | 0.004664544 | 0.06019948 | 0.279375232 | 0.1043 | 0.336256539 |
| SL1344_RS00470_ | 3.893424846 | 3.731124803 | 4.127149759 | 0.359691593 | 0.118861516 | 3.054261179 | 4.68E-05 | 0.010322274 | 0.124398001 | 0.1 | 0.330257709 |
| SL1344_RS00475_ | 3.843577874 | 3.806435804 | 3.930237235 | 0.105357302 | 0.153659973 | 0.734378899 | 0.207861442 | 0.322975038 | 0.614914076 | 0.4028 | 0.670518623 |
| SL1344_RS00480_ | 1.108680024 | 0.924318652 | 1.269604835 | 0.372843649 | 0.173360869 | 2.14784915 | 0.0020023 | 0.050041676 | 0.249451007 | 0.101 | 0.331627179 |
| SL1344_RS00485_ | 0.644121002 | 0.401707634 | 0.721108228 | 0.253608527 | 0.360994839 | 0.553632623 | 0.337774823 | 0.572443869 | 0.793477774 | 0.6994 | 0.866685398 |
| SL1344_RS00490_ | 1.847971445 | 1.756894442 | 1.879271748 | 0.123667658 | 0.23842534 | 0.445764479 | 0.358000006 | 0.571919488 | 0.794410792 | 0.7415 | 0.885731442 |
| SL1344_RS00495_ | 0.80408245 | 0.80110085 | 0.807100669 | 0.00239143 | 0.152357916 | 0.014081824 | 0.493670886 | 0.619349037 | 0.801088079 | 0.7295 | 0.854130605 |
| SL1344_RS00500_ | -2.095402081 | -2.108382997 | -2.073964447 | -0.025331808 | 0.532511184 | -0.047140225 | 0.481333334 | 0.680378168 | 0.846754133 | 0.7989 | 0.900711769 |
| SL1344_RS00505_ | -3.806436494 | -4.364959161 | -3.564599167 | 0.775630599 | 0.782388863 | 0.885672816 | 0.164556985 | 0.249092525 | 0.519843619 | 0.3535 | 0.587675557 |
| SL1344_RS00510_ | -2.611465592 | -2.474298499 | -2.736601041 | -0.239587145 | 0.447229267 | -0.509913711 | 0.289140582 | 0.435285826 | 0.663222829 | 0.5399 | 0.726433177 |
| SL1344_RS00515_ | -2.055585583 | -2.064810221 | -2.050014315 | 0.019204276 | 0.511063746 | 0.030940859 | 0.485009994 | 0.70033483 | 0.857844624 | 0.8064 | 0.90522264 |
| SL1344_RS00520_ | -2.839289616 | -2.708079567 | -2.985278238 | -0.367937368 | 0.712635224 | -0.483939761 | 0.295136585 | 0.460044466 | 0.701579893 | 0.6051 | 0.780304761 |
| SL1344_RS00525_ | 1.259514941 | 1.260898745 | 1.258628926 | 0.015177557 | 0.299960269 | 0.05066351 | 0.482000001 | 0.810702186 | 0.921742628 | 0.9496 | 0.977793941 |
| SL1344_RS00530_ | 0.938574958 | 0.8928271 | 0.968743545 | 0.044882398 | 0.35734219 | 0.080342855 | 0.444000002 | 0.723689228 | 0.882062031 | 0.8797 | 0.948402573 |
| SL1344_RS00545_ | 0.134873673 | 0.095494433 | 0.210260409 | 0.071471629 | 0.273775055 | 0.226082059 | 0.413057965 | 0.743094655 | 0.883338301 | 0.7852 | 0.902509509 |
| SL1344_RS00550_ | -0.252308382 | -0.223222869 | -0.305876679 | -0.065271803 | 0.238916073 | -0.249994467 | 0.397068625 | 0.619279341 | 0.804959607 | 0.7292 | 0.860726158 |
| SL1344_RS00555_ | 3.288927366 | 3.255005673 | 3.313768737 | 0.093061804 | 0.219740372 | 0.458893412 | 0.290666676 | 0.41293795 | 0.690666636 | 0.6024 | 0.801140151 |
| SL1344_RS00560_ | -0.887091731 | -1.126146843 | -0.640984959 | 0.50492345 | 0.269148343 | 1.816715785 | 0.017322048 | 0.057601169 | 0.257289339 | 0.1174 | 0.352426683 |
| SL1344_RS00565_ | -0.375837447 | -0.453488587 | -0.324205882 | 0.159174685 | 0.297383738 | 0.546474805 | 0.249833457 | 0.375665395 | 0.643368492 | 0.5066 | 0.697681776 |
| SL1344_RS00570_ | -0.280815091 | -0.400220813 | -0.190873574 | 0.21772628 | 0.208778977 | 0.969533435 | 0.134576978 | 0.215483328 | 0.475635564 | 0.2884 | 0.525479678 |
| SL1344_RS00575_ | -0.741735297 | -1.021235695 | -0.660041639 | 0.344618601 | 0.436786665 | 0.705359982 | 0.249167234 | 0.352919371 | 0.632236214 | 0.4974 | 0.726036162 |
| SL1344_RS00580_ | 0.222604961 | 0.483209918 | -0.090761571 | -0.411238192 | 0.703429249 | -0.522497868 | 0.308666675 | 0.456632093 | 0.722564408 | 0.6451 | 0.817539814 |
| SL1344_RS00590_ | 0.175970281 | 0.270805467 | 0.037446073 | -0.245387151 | 0.2131108 | -1.108244016 | 0.099267197 | 0.169876265 | 0.432451983 | 0.2378 | 0.48176204 |
| SL1344_RS00595_ | 0.443711381 | 0.316470299 | 0.497997925 | 0.162516296 | 0.365190189 | 0.4679327 | 0.264666678 | 0.502764899 | 0.743141975 | 0.5265 | 0.754025812 |
| SL1344_RS00600_ | 3.638921723 | 3.352319539 | 3.712244799 | 0.322684467 | 0.139365895 | 1.3860986 | 0.060000072 | 0.118297246 | 0.391224684 | 0.162 | 0.417505625 |
| SL1344_RS00605_ | 4.099508315 | 3.972288475 | 4.118964925 | 0.147357798 | 0.122002461 | 1.067844213 | 0.186666686 | 0.199802394 | 0.494770296 | 0.344 | 0.616667517 |
| SL1344_RS00610_ | 4.148242625 | 4.16404242 | 4.136410373 | -0.012476243 | 0.085086867 | -0.081851843 | 0.447701534 | 0.673464675 | 0.854110192 | 0.8604 | 0.938772272 |
| SL1344_RS00615_ | 3.389583344 | 3.321282783 | 3.415582525 | 0.095853562 | 0.261517711 | 0.373316979 | 0.354000006 | 0.604547026 | 0.8189405 | 0.7495 | 0.893376618 |
| SL1344_RS00620_ | 5.701120481 | 5.752729985 | 5.658921234 | -0.084207375 | 0.289685512 | -0.220254278 | 0.447035312 | 0.849982921 | 0.943644149 | 1 | 1 |
| SL1344_RS00630_ | 3.431825443 | 3.314953256 | 3.535291787 | 0.2196956 | 0.114322207 | 1.924788926 | 0.003998486 | 0.063802059 | 0.288713054 | 0.1035 | 0.335351084 |
| SL1344_RS00635_ | 3.272125657 | 3.151178193 | 3.331158367 | 0.172720122 | 0.154699984 | 1.021045775 | 0.141333361 | 0.198177023 | 0.49213145 | 0.2596 | 0.531280741 |
| SL1344_RS00645_ | 3.370911947 | 3.331036157 | 3.558113839 | 0.190893881 | 0.227146617 | 0.765136342 | 0.257828126 | 0.359782366 | 0.6495842 | 0.4863 | 0.735526167 |
| SL1344_RS00650_ | 3.050070343 | 3.001974641 | 3.096105692 | -0.015810768 | 0.211915846 | -0.089551801 | 0.477681546 | 0.848302043 | 0.939600465 | 0.9853 | 0.993592598 |
| SL1344_RS00655_ | 3.378849227 | 3.396038513 | 3.35940807 | -0.033020514 | 0.168760974 | -0.224694993 | 0.360666672 | 0.723755294 | 0.877153359 | 0.7436 | 0.874697211 |
| SL1344_RS00665_ | 2.500503737 | 2.558496081 | 2.304274112 | -0.206402982 | 0.179977558 | -1.025493783 | 0.170552986 | 0.220504198 | 0.514302207 | 0.3285 | 0.593861512 |

| | | | | | | | | | | | |
|---|---|---|---|---|---|---|---|---|---|---|---|
| SL1344_RS00670_ | 3.640630606 | 3.644526883 | 3.636136299 | 0.006578563 | 0.077127339 | 0.080132148 | 0.470000001 | 0.718745091 | 0.870324251 | 0.8458 | 0.925174512 |
| SL1344_RS00675_ | 4.104893785 | 4.092415408 | 4.141737145 | 0.039986304 | 0.067828671 | 0.481457606 | 0.324450374 | 0.603436042 | 0.800018847 | 0.6855 | 0.85006247 |
| SL1344_RS00680_ | 5.646806248 | 5.699380333 | 5.578364196 | -0.120987447 | 0.155939537 | -0.741921537 | 0.147333359 | 0.270607849 | 0.574156532 | 0.2772 | 0.551909868 |
| SL1344_RS00685_ | 2.556579009 | 2.554893118 | 2.557710581 | 0.051363571 | 0.172662625 | 0.306297993 | 0.399333337 | 0.634679099 | 0.825070155 | 0.8171 | 0.911539216 |
| SL1344_RS00690_ | 4.739896618 | 4.477385709 | 4.885308276 | 0.39868416 | 0.064823717 | 5.408298567 | 4.68E-05 | 0.010310153 | 0.124322561 | 0.1 | 0.330257709 |
| SL1344_RS00695_ | -0.943510671 | -0.988911367 | -0.903485757 | 0.042690634 | 0.566086438 | 0.072214675 | 0.469020654 | 0.813896778 | 0.923044516 | 0.8968 | 0.953947808 |
| SL1344_RS00700_ | 1.094272794 | 1.006142864 | 1.171273864 | 0.178527418 | 0.17206641 | 0.963560197 | 0.132666696 | 0.200067923 | 0.470512775 | 0.2866 | 0.532856251 |
| SL1344_RS00705_ | 1.159824214 | 1.005527419 | 1.219964455 | 0.200044054 | 0.189714662 | 1.01581866 | 0.156562316 | 0.199708451 | 0.477223534 | 0.3036 | 0.562476523 |
| SL1344_RS00710_ | 1.452862902 | 1.396209254 | 1.48528359 | 0.086708381 | 0.132412876 | 0.584363129 | 0.263333345 | 0.408498701 | 0.653564456 | 0.5276 | 0.721117752 |
| SL1344_RS00715_ | 1.493404261 | 1.575687307 | 1.460864373 | -0.110891542 | 0.190904658 | -0.503949664 | 0.306666675 | 0.494110269 | 0.734413911 | 0.6248 | 0.809035906 |
| SL1344_RS00720_ | 4.262754563 | 4.020180841 | 4.462168607 | 0.436065124 | 0.131867046 | 3.006786156 | 4.68E-05 | 0.067364845 | 0.304045867 | 0.1 | 0.330257709 |
| SL1344_RS00725_ | -3.441345372 | -3.270658886 | -3.603118598 | -0.320245661 | 0.55753251 | -0.554164705 | 0.265822796 | 0.409953143 | 0.637737896 | 0.5006 | 0.692516043 |
| SL1344_RS00730_ | -4.126057751 | -4.297057121 | -3.926340237 | 0.351722298 | 0.745421285 | 0.443528431 | 0.317333341 | 0.480858891 | 0.696917727 | 0.5869 | 0.756239965 |
| SL1344_RS00735_ | -6.009064299 | -6.282788258 | -5.750202381 | 0.537473092 | 1.572512222 | 0.305109989 | 0.351333339 | 0.509923466 | 0.727054764 | 0.6217 | 0.782304893 |
| SL1344_RS00740_ | 1.035268357 | 0.899951757 | 1.218541563 | 0.290397994 | 0.235652623 | 1.189933346 | 0.111925419 | 0.151595449 | 0.425604183 | 0.2243 | 0.489309885 |
| SL1344_RS00745_ | 0.620535783 | 0.557803196 | 0.713019906 | 0.100133008 | 0.240009668 | 0.377988015 | 0.362425055 | 0.638475683 | 0.821354494 | 0.7408 | 0.870527934 |
| SL1344_RS00750_ | 1.441775091 | 1.468096797 | 1.429584751 | -0.052955499 | 0.208821653 | -0.217808355 | 0.376666672 | 0.493026744 | 0.742543467 | 0.7161 | 0.848277882 |
| SL1344_RS00755_ | -0.744004203 | -0.84986224 | -0.669404729 | 0.171440702 | 0.265164767 | 0.598862279 | 0.251165902 | 0.411412682 | 0.646655495 | 0.5118 | 0.707689048 |
| SL1344_RS00760_ | -2.303574274 | -2.243889031 | -2.343765886 | -0.087406856 | 0.423871735 | -0.180395986 | 0.425716192 | 0.584556935 | 0.778141528 | 0.6846 | 0.829341578 |
| SL1344_RS00765_ | 0.843844519 | 0.81760436 | 0.863287901 | 0.030387251 | 0.254848597 | 0.112028438 | 0.447701534 | 0.765246052 | 0.896914583 | 0.7851 | 0.897314094 |
| SL1344_RS00770_ | 2.457950083 | 2.446231388 | 2.463829968 | 0.002754316 | 0.139270709 | 0.01129423 | 0.489333334 | 0.613991493 | 0.818849286 | 0.811 | 0.911545271 |
| SL1344_RS00775_ | 5.022197595 | 4.819003469 | 5.133278338 | 0.436448021 | 0.279297531 | 1.564210317 | 4.68E-05 | 0.062584289 | 0.294145823 | 0.1 | 0.330257709 |
| SL1344_RS00785_ | 6.366020622 | 6.340525123 | 6.374361436 | 0.041507419 | 0.148654723 | 0.35675508 | 0.240000013 | 0.438365642 | 0.712883148 | 0.4534 | 0.713183479 |
| SL1344_RS00790_ | -2.625460427 | -2.377948077 | -2.899896975 | -0.548890003 | 0.498180268 | -1.078717194 | 0.10926053 | 0.181441195 | 0.437525327 | 0.2589 | 0.497282018 |
| SL1344_RS00795_ | -3.120503256 | -3.148699157 | -3.09532814 | 0.051547289 | 0.438631527 | 0.104243839 | 0.453031314 | 0.49911399 | 0.709058126 | 0.6026 | 0.766744789 |
| SL1344_RS00800_ | -1.418460763 | -1.499798453 | -1.346639916 | 0.101517722 | 0.380508216 | 0.241496977 | 0.391333338 | 0.620189568 | 0.807735069 | 0.7246 | 0.85707195 |
| SL1344_RS00805_ | 5.1258256 | 4.956976848 | 5.457232975 | 0.494486453 | 0.360285628 | 1.417817718 | 0.074666724 | 0.081472794 | 0.332214096 | 0.1688 | 0.430838614 |
| SL1344_RS00810_ | 1.917027143 | 1.72619828 | 2.056776247 | 0.349940959 | 0.270058864 | 1.328730882 | 0.074666724 | 0.104832228 | 0.366551166 | 0.1695 | 0.428588764 |
| SL1344_RS00815_ | 1.99093991 | 1.966275737 | 2.02855002 | 0.083541514 | 0.184972897 | 0.435787935 | 0.299333342 | 0.475846327 | 0.722147454 | 0.583 | 0.775772026 |
| SL1344_RS00820_ | 3.118052347 | 3.005170419 | 3.15258781 | 0.12041855 | 0.153610088 | 0.18369899 | 0.337774823 | 0.675155313 | 0.860401164 | 0.7102 | 0.876872426 |
| SL1344_RS00825_ | 3.589352104 | 3.523918855 | 3.632537044 | 0.095562259 | 0.204724221 | 0.231372711 | 0.382666671 | 0.901126676 | 0.963337261 | 0.829 | 0.927411419 |
| SL1344_RS00830_ | 3.352241281 | 3.122803368 | 3.373135418 | 0.262037803 | 0.463906252 | 0.578444415 | 0.327115264 | 0.38444416 | 0.673131217 | 0.6787 | 0.858878968 |
| SL1344_RS00835_ | 3.793801811 | 3.580683377 | 3.813818804 | 0.221225317 | 0.096720866 | 1.004436174 | 0.213333349 | 0.215714455 | 0.515137251 | 0.42 | 0.667934867 |
| SL1344_RS00840_ | 1.87934872 | 1.861687776 | 1.899515012 | 0.032604456 | 0.286514252 | 0.11658189 | 0.431333336 | 0.817477532 | 0.924339959 | 0.8536 | 0.931003752 |
| SL1344_RS00845_ | 1.272422921 | 1.267468746 | 1.275589767 | -0.028917204 | 0.270416147 | -0.101682845 | 0.458361094 | 0.777420517 | 0.904137694 | 0.8937 | 0.952031219 |
| SL1344_RS00850_ | 0.304303892 | 0.387083426 | 0.248832262 | -0.086792755 | 0.240729932 | -0.35030013 | 0.356000006 | 0.590065955 | 0.787828274 | 0.6876 | 0.8340251 |
| SL1344_RS00855_ | 1.854059214 | 1.812020889 | 2.211569966 | 0.366566297 | 0.510559071 | 0.777233738 | 0.219187224 | 0.284899881 | 0.585141444 | 0.3997 | 0.679695977 |
| SL1344_RS00860_ | 0.508477064 | 0.548317254 | 0.444932822 | -0.093635145 | 0.16737424 | -0.52224868 | 0.285143248 | 0.441485057 | 0.673987352 | 0.5539 | 0.738241066 |
| SL1344_RS00865_ | 2.276515211 | 2.257426576 | 2.304801046 | 0.048626059 | 0.116263769 | 0.397482505 | 0.333777489 | 0.547028911 | 0.75848392 | 0.6464 | 0.809148213 |
| SL1344_RS00870_ | 4.477729386 | 4.471085782 | 4.487242601 | 0.02685357 | 0.126603944 | 0.23362306 | 0.368666672 | 0.709455107 | 0.870823302 | 0.7568 | 0.882892113 |
| SL1344_RS00875_ | 0.053840436 | 0.008202359 | 0.09325472 | 0.087443986 | 0.213483545 | 0.399152988 | 0.332666674 | 0.510935074 | 0.729688849 | 0.6305 | 0.791901275 |
| SL1344_RS00880_ | 0.702104383 | 0.580750729 | 0.797962251 | 0.193365259 | 0.502010551 | 0.317770515 | 0.413333337 | 0.686285634 | 0.862070591 | 0.9013 | 0.9578931 |
| SL1344_RS00885_ | -1.651231231 | -1.606384397 | -1.698634551 | -0.146873419 | 0.427763841 | -0.35368911 | 0.363091278 | 0.544904926 | 0.758032124 | 0.705 | 0.838346187 |
| SL1344_RS00890_ | -2.079215947 | -2.002359166 | -2.138721721 | -0.117225156 | 0.570451031 | -0.203567699 | 0.402000004 | 0.663765133 | 0.836760215 | 0.7557 | 0.877028949 |
| SL1344_RS00895_ | -5.064723185 | -5.37373965 | -4.812863854 | 0.626168385 | 1.190675123 | 0.455459857 | 0.269820131 | 0.410395551 | 0.656856989 | 0.536 | 0.719384491 |
| SL1344_RS00900_ | -2.97002102 | -3.151205272 | -2.818835057 | 0.352886742 | 0.535672991 | 0.61664319 | 0.239333347 | 0.375742136 | 0.620543143 | 0.4831 | 0.679700213 |
| SL1344_RS00905_ | -2.851749321 | -3.008290495 | -2.740243522 | 0.201682352 | 0.672920717 | 0.298295184 | 0.383077953 | 0.622222995 | 0.808969356 | 0.7267 | 0.859225938 |
| SL1344_RS00910_ | 0.542651561 | 0.540710985 | 0.543638993 | -0.006605155 | 0.177929475 | -0.038931412 | 0.485009994 | 0.642481437 | 0.819526281 | 0.7528 | 0.87198924 |
| SL1344_RS00915_ | 2.418277304 | 2.361157488 | 2.595897463 | 0.206394533 | 0.164967357 | 1.068519777 | 0.160000023 | 0.211277584 | 0.50333016 | 0.314 | 0.575952031 |
| SL1344_RS00920_ | 1.202348903 | 1.073931962 | 1.301515891 | 0.24755855 | 0.17642653 | 1.346718772 | 0.050632998 | 0.110018625 | 0.362526945 | 0.1507 | 0.394142341 |
| SL1344_RS00925_ | 0.639144239 | 0.43736612 | 0.753846049 | 0.254445529 | 0.344857373 | 0.74650048 | 0.202666684 | 0.296926444 | 0.585339904 | 0.3697 | 0.631677625 |
| SL1344_RS00930_ | 1.588772199 | 1.534764249 | 1.631730623 | 0.069249845 | 0.322097729 | 0.191972235 | 0.424000003 | 0.808934759 | 0.921324308 | 0.8929 | 0.952807009 |
| SL1344_RS00935_ | 3.769896636 | 3.69114267 | 3.804108287 | 0.113558384 | 0.175786294 | 0.589793053 | 0.262666678 | 0.398488205 | 0.679760813 | 0.4928 | 0.737003664 |

| | | | | | | | | | | | |
|---|---|---|---|---|---|---|---|---|---|---|---|
| SL1344_RS00940_ | 1.543868848 | 1.532157436 | 1.551251141 | 0.025518605 | 0.205924013 | 0.123529655 | 0.458000002 | 0.723582629 | 0.87550142 | 0.8691 | 0.94023253 |
| SL1344_RS00945_ | 6.160444504 | 6.265744464 | 6.065929269 | -0.144312321 | 0.15987699 | -1.005727982 | 0.111925419 | 0.211865119 | 0.513852259 | 0.2024 | 0.478995064 |
| SL1344_RS00950_ | 0.9446183 | 0.911239753 | 1.100327976 | 0.166916058 | 0.181932375 | 0.692467107 | 0.283333343 | 0.44244032 | 0.69173796 | 0.5793 | 0.781757568 |
| SL1344_RS00955_ | 0.710479589 | 0.731103747 | 0.69593474 | -0.065759295 | 0.302364056 | -0.197789284 | 0.418000003 | 0.713440321 | 0.868952861 | 0.8561 | 0.932572739 |
| SL1344_RS00960_ | 2.319552679 | 2.357062275 | 2.295324636 | -0.076338944 | 0.130989479 | -0.567512521 | 0.267333344 | 0.391616391 | 0.653184161 | 0.5373 | 0.732800538 |
| SL1344_RS00965_ | 3.60554813 | 3.58348064 | 3.631043068 | 0.061895948 | 0.082731032 | 0.721293545 | 0.210000016 | 0.295380059 | 0.570856984 | 0.4293 | 0.650662892 |
| SL1344_RS00970_ | 3.564141644 | 4.029987146 | 3.235119299 | -0.77152341 | 0.329724173 | -2.32149641 | 4.68E-05 | 0.029043277 | 0.206989125 | 0.1 | 0.330257709 |
| SL1344_RS00975_ | 0.960591481 | 1.319208908 | 0.625088254 | -0.707987234 | 0.206073306 | -3.427189672 | 4.68E-05 | 0.009876444 | 0.118252612 | 0.1 | 0.330257709 |
| SL1344_RS00980_ | 1.205076509 | 1.409165222 | 0.945292025 | -0.599049113 | 0.320994621 | -1.963337871 | 4.68E-05 | 0.03520972 | 0.219933722 | 0.1 | 0.330257709 |
| SL1344_RS00985_ | 2.005119844 | 2.226206508 | 1.883492716 | -0.360863915 | 0.373019769 | -0.996673297 | 0.135909422 | 0.178410949 | 0.472300977 | 0.241 | 0.516778943 |
| SL1344_RS00990_ | -3.367711622 | -3.849091016 | -2.884349328 | 0.897423531 | 0.84565905 | 1.010212619 | 0.134666696 | 0.196567349 | 0.467361644 | 0.28 | 0.532038014 |
| SL1344_RS00995_ | -1.87062583 | -1.967309044 | -1.756214274 | 0.255201402 | 0.482921409 | 0.51145383 | 0.280666677 | 0.441798709 | 0.682305404 | 0.5854 | 0.760352287 |
| SL1344_RS01000_ | -3.674520121 | -3.695695798 | -3.62519639 | -0.022940949 | 1.009962004 | -0.019394306 | 0.491672219 | 0.675298898 | 0.847474002 | 0.8073 | 0.907204595 |
| SL1344_RS01005_ | -3.664140567 | -3.597945528 | -3.734084366 | -0.168230173 | 0.715585964 | -0.229657422 | 0.400399737 | 0.575652466 | 0.773213027 | 0.7015 | 0.835264508 |
| SL1344_RS01010_ | -2.820283904 | -2.575952572 | -3.204640457 | -0.599341234 | 0.473739449 | -1.21890797 | 0.092000045 | 0.152876865 | 0.396517241 | 0.2304 | 0.470482315 |
| SL1344_RS01015_ | -3.353101682 | -3.012439836 | -3.817162302 | -0.808513917 | 0.559780739 | -1.388583882 | 0.054630326 | 0.111751244 | 0.34376589 | 0.1741 | 0.414905997 |
| SL1344_RS01020_ | 0.470360785 | 0.358846523 | 0.518450634 | 0.115819427 | 0.315056579 | 0.214189751 | 0.390000004 | 0.835356057 | 0.933662573 | 0.8218 | 0.924269505 |
| SL1344_RS01025_ | 2.614130908 | 2.588999732 | 2.644655219 | 0.077450967 | 0.118764763 | 0.6236103 | 0.243333346 | 0.338131345 | 0.612545003 | 0.4889 | 0.694533553 |
| SL1344_RS01030_ | 0.80536094 | 0.822738889 | 0.787514434 | -0.037982171 | 0.144169001 | -0.246815585 | 0.39573618 | 0.576317659 | 0.771562146 | 0.6804 | 0.82230228 |
| SL1344_RS01035_ | 3.025399816 | 3.220250831 | 2.882984589 | -0.279835305 | 0.253093609 | -1.120544879 | 0.120000033 | 0.146417074 | 0.431216811 | 0.2054 | 0.481273066 |
| SL1344_RS01040_ | 0.759200619 | 0.792632201 | 0.736025706 | -0.002681998 | 0.274380574 | -0.009540329 | 0.494 | 0.676349484 | 0.852018624 | 0.8707 | 0.94118125 |
| SL1344_RS01045_ | 0.967176448 | 1.024070713 | 0.910593108 | -0.13327328 | 0.175372486 | -0.731616273 | 0.206528997 | 0.298218081 | 0.569122111 | 0.416 | 0.635562909 |
| SL1344_RS01050_ | 2.232618991 | 2.304220482 | 2.12540682 | -0.186093401 | 0.125969991 | -1.446417048 | 0.03000015 | 0.113600691 | 0.368565391 | 0.1342 | 0.373757541 |
| SL1344_RS01055_ | 0.435502909 | 0.299945096 | 0.526534693 | 0.244031385 | 0.257162138 | 0.920411312 | 0.150666692 | 0.215031306 | 0.497105044 | 0.3017 | 0.55112921 |
| SL1344_RS01060_ | 2.993393388 | 2.723479106 | 3.295440878 | 0.62962014 | 0.198118335 | 3.314948431 | 4.68E-05 | 0.012304073 | 0.136042309 | 0.1 | 0.330257709 |
| SL1344_RS01065_ | 1.292651063 | 1.345957987 | 1.205709841 | -0.126502893 | 0.317720044 | -0.422815755 | 0.278666677 | 0.529816955 | 0.763933807 | 0.5558 | 0.771425227 |
| SL1344_RS01070_ | 4.822364329 | 4.821539202 | 4.82346651 | -0.085072156 | 0.216531521 | -0.176709348 | 0.39840107 | 0.439025169 | 0.713756701 | 0.8417 | 0.929046501 |
| SL1344_RS01075_ | 0.994558948 | 1.151280807 | 0.926204118 | -0.22868983 | 0.187659371 | -1.117494609 | 0.112591641 | 0.183822982 | 0.459378727 | 0.2532 | 0.499251138 |
| SL1344_RS01080_ | 4.298284997 | 4.148390929 | 4.460409041 | 0.193787322 | 0.178644932 | 1.544956042 | 0.107261863 | 0.086968175 | 0.338688722 | 0.1916 | 0.464239272 |
| SL1344_RS01085_ | 1.617597333 | 1.535439075 | 1.679388784 | 0.15726412 | 0.143002357 | 1.044661066 | 0.114666702 | 0.177388912 | 0.446725792 | 0.2506 | 0.499754308 |
| SL1344_RS01090_ | 3.109925263 | 3.089678645 | 3.122451022 | 0.07042983 | 0.165884265 | 0.414668703 | 0.318454371 | 0.414968086 | 0.688767202 | 0.6499 | 0.815527515 |
| SL1344_RS01095_ | 6.812695755 | 6.86446523 | 6.765122639 | -0.03765203 | 0.131356927 | -0.319930151 | 0.42466667 | 0.719118879 | 0.882116738 | 0.9457 | 0.976916847 |
| SL1344_RS01100_ | 7.370600839 | 7.161199373 | 7.441834744 | 0.259132135 | 0.184499307 | 0.65090659 | 0.33333334 | 0.48874294 | 0.748542779 | 0.7 | 0.872542119 |
| SL1344_RS01105_ | 3.000218244 | 2.953889631 | 3.100917126 | 0.139808625 | 0.122832126 | 1.119018659 | 0.129247199 | 0.162140246 | 0.437667824 | 0.2503 | 0.514132562 |
| SL1344_RS01110_ | 5.355745567 | 5.209325836 | 5.466034239 | 0.213154381 | 0.119764161 | 1.931139028 | 0.005334196 | 0.039217667 | 0.235310305 | 0.106 | 0.338990083 |
| SL1344_RS01115_ | 2.399426417 | 2.364394923 | 2.435571114 | 0.075721161 | 0.10400154 | 0.699093033 | 0.217188556 | 0.329989275 | 0.590510427 | 0.4496 | 0.664034191 |
| SL1344_RS01120_ | 2.415721454 | 2.497367049 | 2.226899322 | -0.219327181 | 0.227480324 | -0.829953068 | 0.206528997 | 0.296775517 | 0.592561579 | 0.4052 | 0.654879545 |
| SL1344_RS01130_ | 4.639207198 | 4.582977043 | 4.657680458 | 0.073074239 | 0.163775415 | 0.437591867 | 0.348000006 | 0.583129773 | 0.805546288 | 0.733 | 0.88631733 |
| SL1344_RS01135_ | 5.562802129 | 5.512811843 | 5.609745647 | 0.131622372 | 0.083985792 | 1.645181288 | 0.00067292 | 0.098735023 | 0.359691293 | 0.1 | 0.330257709 |
| SL1344_RS01140_ | 5.914875542 | 5.893476189 | 5.930840134 | 0.067546048 | 0.185102158 | 0.423237952 | 0.274000011 | 0.482596119 | 0.743171801 | 0.5425 | 0.770256675 |
| SL1344_RS01145_ | 5.376602584 | 5.370363863 | 5.381939243 | 0.00903293 | 0.152927561 | 0.061569856 | 0.472000001 | 0.896674343 | 0.961363223 | 0.9523 | 0.979586173 |
| SL1344_RS01150_ | 2.761356218 | 2.780718396 | 2.733043951 | -0.043452073 | 0.092446716 | -0.4281013 | 0.323784151 | 0.514974147 | 0.735150379 | 0.6172 | 0.789355773 |
| SL1344_RS01155_ | 4.041278127 | 4.083255368 | 3.993392084 | -0.086182064 | 0.123758721 | -0.681390084 | 0.230000014 | 0.331640668 | 0.621326545 | 0.4427 | 0.687822069 |
| SL1344_RS01160_ | 4.089710674 | 4.11141694 | 4.072840087 | -0.034508237 | 0.144391672 | -0.088151472 | 0.434377084 | 0.490117536 | 0.748737373 | 0.9245 | 0.965443434 |
| SL1344_RS01165_ | 2.383118872 | 2.411556237 | 2.358858328 | -0.066689186 | 0.165970744 | -0.376195935 | 0.348000006 | 0.571909348 | 0.787288793 | 0.7151 | 0.857951896 |
| SL1344_RS01170_ | 3.836313468 | 3.866791789 | 3.815810499 | -0.079767205 | 0.103898357 | -0.760156132 | 0.195333351 | 0.266512532 | 0.554877866 | 0.4037 | 0.636221216 |
| SL1344_RS01175_ | 3.37735049 | 3.335215591 | 3.421399968 | 0.122140864 | 0.127259838 | 1.006598344 | 0.104666706 | 0.177788671 | 0.462837276 | 0.2329 | 0.483051564 |
| SL1344_RS01180_ | 0.720371434 | 0.728791637 | 0.710793176 | -0.018533271 | 0.121863643 | -0.144647608 | 0.440666669 | 0.519204321 | 0.726812631 | 0.6231 | 0.782175676 |
| SL1344_RS01185_ | 1.711113177 | 1.587094814 | 1.798577363 | 0.227129441 | 0.149355444 | 1.552517111 | 0.014666978 | 0.120951231 | 0.38662495 | 0.1128 | 0.346785935 |
| SL1344_RS01190_ | 0.240550939 | 0.300384354 | 0.196953571 | -0.108075207 | 0.176877049 | -0.577627552 | 0.257828126 | 0.402114719 | 0.639774024 | 0.5178 | 0.708354201 |
| SL1344_RS01195_ | 1.511808501 | 1.37932456 | 1.561084404 | 0.186167868 | 0.225560632 | 0.849228768 | 0.182666686 | 0.251225612 | 0.539143864 | 0.3505 | 0.619215315 |
| SL1344_RS01200_ | 2.541631274 | 2.595102883 | 2.435256326 | -0.172543552 | 0.603217766 | -0.361906013 | 0.23866668 | 0.534156998 | 0.777210495 | 0.4414 | 0.706329004 |
| SL1344_RS01205_ | 2.159487398 | 2.163965075 | 2.157072632 | -0.019366283 | 0.144607946 | -0.114934231 | 0.442000002 | 0.585068355 | 0.796032369 | 0.754 | 0.876502957 |

| | | | | | | | | | | | |
|---|---|---|---|---|---|---|---|---|---|---|---|
| SL1344_RS01210_ | 0.080063426 | -0.037883816 | 0.177988445 | 0.2170457 | 0.158479294 | 1.301236817 | 0.068666729 | 0.128198299 | 0.354907332 | 0.198 | 0.43419957 |
| SL1344_RS01215_ | 0.271063207 | 0.094901978 | 0.323127059 | 0.197373628 | 0.233679914 | 0.701546967 | 0.266666678 | 0.394312355 | 0.653795681 | 0.5389 | 0.744601742 |
| SL1344_RS01220_ | 1.347904206 | 1.184128137 | 1.404871441 | 0.208598144 | 0.139763419 | 1.29624394 | 0.101932086 | 0.150691289 | 0.411116238 | 0.2458 | 0.489727119 |
| SL1344_RS01225_ | 4.179945558 | 4.098214504 | 4.241756103 | 0.150844773 | 0.069840698 | 2.155885967 | 0.001336769 | 0.051945463 | 0.257577724 | 0.1009 | 0.331551222 |
| SL1344_RS01230_ | 2.121528693 | 2.013303396 | 2.233311297 | 0.240060327 | 0.128677851 | 1.826276684 | 0.006662914 | 0.066774107 | 0.287147855 | 0.1062 | 0.338935377 |
| SL1344_RS01245_ | 0.602703147 | 0.690958929 | 0.557434922 | -0.108462629 | 0.226566031 | -0.448963623 | 0.318000008 | 0.546441186 | 0.760517165 | 0.6423 | 0.811818373 |
| SL1344_RS01250_ | 0.782510515 | 0.778167696 | 0.78689541 | -0.004393891 | 0.302973214 | -0.017556638 | 0.492 | 0.790190357 | 0.910632001 | 0.7954 | 0.902872847 |
| SL1344_RS01255_ | 0.079432145 | -0.124150076 | 0.148605078 | 0.224114491 | 0.319832996 | 0.622955788 | 0.296666676 | 0.464125013 | 0.714590633 | 0.6077 | 0.799164901 |
| SL1344_RS01260_ | 0.090450452 | -0.363910582 | 0.246288658 | 0.728793594 | 0.997756422 | 0.832256716 | 0.153897426 | 0.232738375 | 0.535819103 | 0.2803 | 0.556291504 |
| SL1344_RS01265_ | 3.873738496 | 3.849635334 | 3.912474066 | 0.072364565 | 0.141687486 | 0.522823954 | 0.279333343 | 0.3836245 | 0.666148095 | 0.5434 | 0.757204478 |
| SL1344_RS01270_ | 3.444916977 | 3.449543729 | 3.440444923 | 0.151720896 | 0.270261181 | 0.484739036 | 0.382666671 | 0.38175195 | 0.668007962 | 0.8451 | 0.921686746 |
| SL1344_RS01275_ | 1.17071884 | 1.007655693 | 1.21616668 | 0.244328817 | 0.680094061 | 0.440521216 | 0.233177895 | 0.445998918 | 0.7167903 | 0.4216 | 0.693553471 |
| SL1344_RS01285_ | -0.840060559 | -0.680328382 | -1.017745404 | -0.324078482 | 0.904695027 | -0.30228499 | 0.404397072 | 0.673928004 | 0.854245725 | 0.8876 | 0.950749031 |
| SL1344_RS01290_ | 0.559177524 | 0.636967303 | 0.502235055 | -0.136280636 | 0.221024667 | -0.603442069 | 0.236509007 | 0.396213966 | 0.650712799 | 0.4821 | 0.70035097 |
| SL1344_RS01295_ | -0.712798718 | -0.671627239 | -0.744575966 | -0.122694876 | 0.381868896 | -0.313646136 | 0.359093943 | 0.561271064 | 0.7746776 | 0.7054 | 0.844544475 |
| SL1344_RS01300_ | -1.444636023 | -1.393780579 | -1.495446094 | -0.10764685 | 0.244547966 | -0.411523446 | 0.320666674 | 0.452579095 | 0.667818633 | 0.5524 | 0.726480737 |
| SL1344_RS01305_ | 1.523949842 | 1.423694857 | 1.621718532 | 0.21343022 | 0.14560046 | 1.426905346 | 0.046635671 | 0.095520468 | 0.334363049 | 0.1475 | 0.390017974 |
| SL1344_RS01310_ | 0.033502752 | 0.024022967 | 0.043098059 | 0.03594805 | 0.276443641 | 0.119736668 | 0.441705532 | 0.693438045 | 0.855468958 | 0.8019 | 0.902784409 |
| SL1344_RS01315_ | 4.00711695 | 4.241758882 | 3.816419801 | -0.418529717 | 0.21828895 | -1.781233334 | 0.00067292 | 0.072425502 | 0.313553222 | 0.1008 | 0.331418906 |
| SL1344_RS01320_ | 0.153140107 | 0.06443651 | 0.199985486 | 0.119512697 | 0.272925802 | 0.428010189 | 0.326666674 | 0.523710161 | 0.74973274 | 0.6683 | 0.824612664 |
| SL1344_RS01325_ | -0.429080719 | -0.388877152 | -0.491077644 | -0.045145066 | 0.382517815 | -0.111156723 | 0.458000002 | 0.776315387 | 0.902082504 | 0.8856 | 0.948702696 |
| SL1344_RS01330_ | 0.676344797 | 0.623578609 | 0.739609565 | 0.121626401 | 0.205924944 | 0.574703418 | 0.266666678 | 0.403074668 | 0.657468389 | 0.5128 | 0.719672318 |
| SL1344_RS01335_ | 1.615960825 | 1.563432946 | 1.732164824 | 0.166644814 | 0.147795909 | 1.017199099 | 0.148666692 | 0.207110907 | 0.479576158 | 0.3056 | 0.556868135 |
| SL1344_RS01340_ | 1.03639324 | 1.19665826 | 0.875367961 | -0.322597276 | 0.631700487 | -0.435988644 | 0.352431718 | 0.546803413 | 0.783705651 | 0.7549 | 0.88684477 |
| SL1344_RS01345_ | -2.083413017 | -2.0757333 | -2.088767644 | 0.037726029 | 0.458762949 | 0.069600946 | 0.466000001 | 0.634134969 | 0.817894623 | 0.7521 | 0.872116027 |
| SL1344_RS01350_ | -2.998547431 | -2.903737663 | -3.134462036 | -0.249666374 | 0.561047171 | -0.400783951 | 0.32933334 | 0.501467083 | 0.717733675 | 0.6145 | 0.777259258 |
| SL1344_RS01355_ | -1.740914044 | -1.926997164 | -1.606086751 | 0.308821386 | 0.309348341 | 0.949786831 | 0.157333357 | 0.242445116 | 0.492409554 | 0.3385 | 0.566512323 |
| SL1344_RS01360_ | -3.491302785 | -3.489917972 | -3.493199588 | -0.012284072 | 0.602671339 | -0.017971048 | 0.490666667 | 0.595715772 | 0.786239233 | 0.6924 | 0.835186088 |
| SL1344_RS01365_ | -2.137999709 | -2.16665023 | -2.112638902 | 0.057169329 | 0.368007252 | 0.14801459 | 0.439040642 | 0.572775065 | 0.76697586 | 0.6843 | 0.822397684 |
| SL1344_RS01370_ | -3.18398496 | -3.337311851 | -3.016197255 | 0.419195598 | 0.714435309 | 0.557203865 | 0.268487686 | 0.400803978 | 0.650381751 | 0.5481 | 0.731868624 |
| SL1344_RS01375_ | -8.222257992 | -8.536258584 | -8.033491162 | 0.441429164 | 3.695071489 | 0.098505741 | 0.442666669 | 0.555589501 | 0.765823341 | 0.6955 | 0.835958668 |
| SL1344_RS01380_ | -0.359025685 | -0.437632921 | -0.292244565 | 0.012725318 | 0.507346369 | 0.021624643 | 0.492666667 | 0.81461121 | 0.92356796 | 0.9235 | 0.965880193 |
| SL1344_RS01385_ | -2.512803821 | -2.633644214 | -2.425721707 | 0.210108746 | 0.442710017 | 0.438616504 | 0.306000008 | 0.440332999 | 0.671702568 | 0.5478 | 0.728190136 |
| SL1344_RS01390_ | -1.651895157 | -1.62314058 | -1.699749652 | -0.074642304 | 0.310174993 | -0.2337685 | 0.398666671 | 0.532701847 | 0.741556464 | 0.6453 | 0.800717076 |
| SL1344_RS01395_ | -2.792538553 | -2.843562538 | -2.740477081 | 0.102325872 | 0.425566482 | 0.21818244 | 0.399067292 | 0.519420318 | 0.728428834 | 0.6323 | 0.786973692 |
| SL1344_RS01400_ | -3.80754067 | -3.841980237 | -3.756566335 | 0.076534331 | 0.728718466 | 0.094816846 | 0.460000001 | 0.631314759 | 0.811506446 | 0.7441 | 0.867259588 |
| SL1344_RS01405_ | -3.813280708 | -3.571785751 | -4.114790117 | -0.542171814 | 0.635934061 | -0.78786355 | 0.187333352 | 0.280320616 | 0.527249939 | 0.3706 | 0.589057445 |
| SL1344_RS01410_ | -4.891164191 | -5.323586559 | -4.685535452 | 0.686813592 | 1.364495803 | 0.453989268 | 0.309333342 | 0.452941513 | 0.699102894 | 0.6073 | 0.781913854 |
| SL1344_RS01415_ | -3.818378276 | -3.937014964 | -3.641809587 | 0.342633033 | 0.906939279 | 0.335816663 | 0.359333339 | 0.571089577 | 0.772671648 | 0.6775 | 0.824729806 |
| SL1344_RS01420_ | -4.192006067 | -4.000153548 | -4.30744662 | -0.270437266 | 0.77230099 | -0.322873385 | 0.361758833 | 0.539967011 | 0.74478441 | 0.6472 | 0.802538091 |
| SL1344_RS01425_ | -2.482632629 | -2.554570057 | -2.429239403 | 0.171016087 | 0.502775965 | 0.299830829 | 0.363333339 | 0.541863049 | 0.750083508 | 0.6667 | 0.813065331 |
| SL1344_RS01430_ | -2.029227486 | -1.997252996 | -2.058954871 | -0.074439799 | 0.400998345 | -0.169599895 | 0.432378417 | 0.60935301 | 0.794876917 | 0.7213 | 0.849254391 |
| SL1344_RS01435_ | -3.638170396 | -3.76116048 | -3.427823981 | 0.261792233 | 0.727471217 | 0.332143533 | 0.346000006 | 0.572760756 | 0.769608905 | 0.6685 | 0.820895033 |
| SL1344_RS01440_ | -5.036838173 | -5.062267159 | -5.012348966 | 0.022973811 | 1.1473958 | 0.02104722 | 0.492338441 | 0.629365067 | 0.810412158 | 0.7489 | 0.865382426 |
| SL1344_RS01445_ | -0.353740336 | -0.427201108 | -0.311390622 | 0.104617231 | 0.2944845 | 0.329944729 | 0.359333339 | 0.568013943 | 0.775190069 | 0.6969 | 0.839653125 |
| SL1344_RS01450_ | -2.965330676 | -2.998070508 | -2.923698006 | 0.073596676 | 0.552817752 | 0.1329728 | 0.446369089 | 0.61212717 | 0.794617454 | 0.7166 | 0.845726768 |
| SL1344_RS01455_ | -3.891517725 | -3.711526405 | -3.986388421 | -0.203542499 | 0.820257943 | -0.207685284 | 0.40466667 | 0.666589733 | 0.835356379 | 0.7534 | 0.879914865 |
| SL1344_RS01460_ | -2.398035461 | -2.28162543 | -2.533489255 | -0.249570509 | 0.357461284 | -0.647975209 | 0.233844118 | 0.355025765 | 0.588496484 | 0.4538 | 0.655012213 |
| SL1344_RS01465_ | -1.344633991 | -1.3111817 | -1.439295621 | -0.118913879 | 0.44512868 | -0.222726746 | 0.413333337 | 0.692492874 | 0.855571206 | 0.7988 | 0.907155908 |
| SL1344_RS01470_ | -4.660243929 | -4.790157602 | -4.539923253 | 0.257170409 | 0.906857904 | 0.272911192 | 0.381745508 | 0.554700799 | 0.752847688 | 0.6688 | 0.811803422 |
| SL1344_RS01475_ | 0.578229744 | 0.535115652 | 0.634422893 | 0.111374287 | 0.321880104 | 0.361215223 | 0.324000007 | 0.567750807 | 0.782959095 | 0.6541 | 0.821306331 |
| SL1344_RS01480_ | 0.826038353 | 0.695723599 | 0.918196249 | 0.269286185 | 0.272776396 | 1.032882438 | 0.099267197 | 0.19033812 | 0.478405256 | 0.2178 | 0.470376571 |
| SL1344_RS01485_ | 0.268408271 | 0.329706201 | 0.155418088 | -0.175398806 | 0.204057626 | -0.816383457 | 0.182666686 | 0.279589358 | 0.547768858 | 0.3875 | 0.611600503 |

| | | | | | | | | | | | |
|---|---|---|---|---|---|---|---|---|---|---|---|
| SL1344_RS01490_ | -0.654039263 | -0.373092732 | -0.765609818 | -0.373378037 | 0.334652282 | -1.006567377 | 0.170000022 | 0.230420507 | 0.507832305 | 0.3333 | 0.578414361 |
| SL1344_RS01495_ | -2.180699821 | -2.439065487 | -2.012359822 | 0.433832545 | 0.543839028 | 0.802140231 | 0.205862775 | 0.301551738 | 0.569033855 | 0.3986 | 0.635717124 |
| SL1344_RS01500_ | -6.238922975 | -5.946785719 | -6.446377965 | -0.185902972 | 2.729176031 | -0.050003796 | 0.462666668 | 0.681847369 | 0.85007787 | 0.8113 | 0.909981897 |
| SL1344_RS01505_ | -2.950108391 | -2.951355749 | -2.946313019 | 0.015426635 | 0.408565693 | 0.035690805 | 0.484000001 | 0.521050336 | 0.729669301 | 0.637 | 0.791677116 |
| SL1344_RS01515_ | -3.141914084 | -3.034314928 | -3.203804884 | -0.111404243 | 0.730178582 | -0.148898525 | 0.436666669 | 0.690902377 | 0.850816258 | 0.7881 | 0.895364839 |
| SL1344_RS01520_ | -5.823611839 | -6.027973003 | -5.701514604 | 0.235897793 | 1.885407544 | 0.116304409 | 0.450366424 | 0.63928153 | 0.822044971 | 0.777 | 0.88732455 |
| SL1344_RS01525_ | -1.365434361 | -1.113460775 | -1.555921566 | -0.486619082 | 0.439523085 | -1.123297832 | 0.070666727 | 0.17902198 | 0.460396543 | 0.2007 | 0.442510088 |
| SL1344_RS01530_ | -2.64089098 | -2.764231934 | -2.551383286 | 0.137055143 | 0.587699379 | 0.215015992 | 0.409726852 | 0.630474872 | 0.813411172 | 0.7429 | 0.867645257 |
| SL1344_RS01535_ | -0.07743303 | -0.117588816 | -0.059901764 | 0.041950217 | 0.222459432 | 0.169618167 | 0.434000002 | 0.667563744 | 0.837873949 | 0.7642 | 0.88659097 |
| SL1344_RS01540_ | -2.014876794 | -1.980857204 | -2.050264743 | -0.061092746 | 0.333742115 | -0.184988399 | 0.421718857 | 0.528703479 | 0.735764727 | 0.6371 | 0.793529407 |
| SL1344_RS01545_ | -2.298471086 | -2.375301167 | -2.230151802 | 0.117424264 | 0.456199955 | 0.240153778 | 0.385742843 | 0.573947968 | 0.772554908 | 0.676 | 0.823040632 |
| SL1344_RS01550_ | -3.12420491 | -3.534504867 | -2.853329085 | 0.725414354 | 0.646791582 | 1.051647576 | 0.086608976 | 0.173665844 | 0.435527367 | 0.2154 | 0.455258506 |
| SL1344_RS01565_ | -2.400027483 | -2.277631002 | -2.53122748 | -0.249214506 | 0.415129132 | -0.556064728 | 0.261825461 | 0.419782756 | 0.648845843 | 0.5251 | 0.711815269 |
| SL1344_RS01570_ | 0.069262139 | 0.022079101 | 0.102292386 | 0.106571014 | 0.230691762 | 0.406091535 | 0.302465032 | 0.431453913 | 0.682137488 | 0.57 | 0.74314606 |
| SL1344_RS01575_ | -0.676111202 | -0.951448261 | -0.501329723 | 0.526686974 | 0.49573014 | 1.061288546 | 0.071333393 | 0.199363005 | 0.494348392 | 0.1752 | 0.423332657 |
| SL1344_RS01580_ | 0.19431199 | 0.164158533 | 0.225562497 | -0.035747238 | 0.312826048 | -0.109248953 | 0.461692207 | 0.723653875 | 0.873106696 | 0.9062 | 0.954869345 |
| SL1344_RS01590_ | 2.374142696 | 2.151268287 | 2.494593529 | 0.320536283 | 0.17256193 | 1.823796535 | 0.021319333 | 0.058779245 | 0.276738409 | 0.1189 | 0.357478853 |
| SL1344_RS01595_ | 1.800392409 | 1.766190096 | 1.827160801 | 0.061903 | 0.208514679 | 0.305136135 | 0.37441706 | 0.668516477 | 0.844045296 | 0.7688 | 0.888400291 |
| SL1344_RS01600_ | 2.424844338 | 2.343646305 | 2.470729279 | 0.17644132 | 0.183781502 | 0.921903924 | 0.158560983 | 0.213636686 | 0.505391157 | 0.3144 | 0.574053756 |
| SL1344_RS01605_ | 1.264184022 | 1.303501072 | 1.156028289 | -0.15453426 | 0.222700147 | -0.616854755 | 0.253830791 | 0.356301713 | 0.636626973 | 0.4981 | 0.718231654 |
| SL1344_RS01610_ | -1.063535736 | -1.029423698 | -1.117312107 | -0.097588035 | 0.27686085 | -0.321528993 | 0.365333339 | 0.530232092 | 0.738622207 | 0.6523 | 0.801941658 |
| SL1344_RS01615_ | -0.786327586 | -0.704931398 | -0.838453784 | -0.157942274 | 0.266677443 | -0.544367445 | 0.269820131 | 0.406389861 | 0.649663855 | 0.5253 | 0.711601468 |
| SL1344_RS01620_ | 4.082998332 | 3.885515631 | 4.148912265 | 0.238051999 | 0.236709534 | 1.068094057 | 0.136575645 | 0.150903805 | 0.438672688 | 0.2377 | 0.514128272 |
| SL1344_RS01625_ | 2.745068799 | 2.411474451 | 2.787334994 | 0.363438787 | 0.152342402 | 1.239836004 | 0.11133337 | 0.148253695 | 0.43424827 | 0.2274 | 0.48597383 |
| SL1344_RS01635_ | 2.652997195 | 2.684263056 | 2.496183253 | -0.186189909 | 0.199078763 | -0.753040516 | 0.287141915 | 0.376907743 | 0.660946413 | 0.5625 | 0.782289862 |
| SL1344_RS01640_ | -1.393079187 | -1.488821859 | -1.296681042 | 0.201403922 | 0.246920583 | 0.730458484 | 0.209193887 | 0.313739483 | 0.550368616 | 0.4048 | 0.613822645 |
| SL1344_RS01645_ | 2.161056009 | 2.024670826 | 2.219589341 | 0.205674881 | 0.178331297 | 1.141575198 | 0.104596974 | 0.149354143 | 0.425425887 | 0.2321 | 0.49482574 |
| SL1344_RS01650_ | 3.66897001 | 3.527613935 | 3.776425699 | 0.239185111 | 0.242247534 | 0.882328424 | 0.139906757 | 0.263377878 | 0.566327846 | 0.2441 | 0.518787651 |
| SL1344_RS01655_ | -0.621511263 | -0.336841013 | -0.810091395 | -0.313125076 | 0.5826471 | -0.463232424 | 0.359093943 | 0.650572515 | 0.837375928 | 0.7637 | 0.892886399 |
| SL1344_RS01660_ | -4.398804239 | -4.245255028 | -4.663222857 | -0.176129466 | 1.694368253 | -0.096079769 | 0.463024652 | 0.733857472 | 0.880473417 | 0.8776 | 0.942945087 |
| SL1344_RS01665_ | -5.190701248 | -4.960147872 | -5.424547153 | -0.500491051 | 1.050967777 | -0.447899716 | 0.301798809 | 0.438835042 | 0.668444633 | 0.5603 | 0.736031537 |
| SL1344_RS01670_ | -3.945441528 | -3.867133159 | -4.036367545 | -0.200125914 | 0.812115869 | -0.228621417 | 0.400399737 | 0.556285101 | 0.758883826 | 0.6701 | 0.816129268 |
| SL1344_RS01675_ | 3.881289542 | 3.705111367 | 3.949826597 | 0.236745981 | 0.161683189 | 1.243760473 | 0.103333373 | 0.152902089 | 0.440152162 | 0.201 | 0.470168875 |
| SL1344_RS01680_ | -2.367428368 | -2.394057603 | -2.337786586 | 0.008790488 | 0.555025133 | 0.013408245 | 0.494666667 | 0.683625128 | 0.846822754 | 0.7973 | 0.898776761 |
| SL1344_RS01685_ | -2.684447812 | -3.024256125 | -2.418444459 | 0.606645949 | 0.402779097 | 1.450212144 | 0.047968113 | 0.102294616 | 0.321988835 | 0.1693 | 0.408464192 |
| SL1344_RS01690_ | -3.869702785 | -3.968583769 | -3.67760523 | 0.236235447 | 0.851555834 | 0.238186462 | 0.398666671 | 0.658594842 | 0.829073387 | 0.7397 | 0.868398954 |
| SL1344_RS01695_ | -1.975050069 | -1.812994953 | -2.280088704 | -0.458427965 | 0.432863938 | -0.995434701 | 0.140000028 | 0.210939344 | 0.478415895 | 0.2986 | 0.543258498 |
| SL1344_RS01700_ | -0.724699949 | -0.767475023 | -0.69952961 | 0.048204269 | 0.290174273 | 0.141387353 | 0.432666669 | 0.655026655 | 0.827788955 | 0.7678 | 0.881207817 |
| SL1344_RS01705_ | -0.024966627 | -0.101528252 | 0.105238935 | 0.170697455 | 0.240265742 | 0.668931305 | 0.244503677 | 0.399869606 | 0.648659564 | 0.4993 | 0.707432781 |
| SL1344_RS01710_ | -2.642250166 | -2.892112676 | -2.510009084 | 0.380654801 | 0.531006146 | 0.68392352 | 0.223184558 | 0.341529257 | 0.600647786 | 0.4538 | 0.661633879 |
| SL1344_RS01715_ | -3.587540748 | -3.336717531 | -3.772234839 | -0.401664022 | 0.656225127 | -0.57943809 | 0.266489018 | 0.419000141 | 0.650959295 | 0.515 | 0.708209755 |
| SL1344_RS01720_ | -1.356457097 | -1.401443293 | -1.290432232 | 0.09938162 | 0.308066671 | 0.296535479 | 0.379080618 | 0.547088934 | 0.752238592 | 0.653 | 0.807696141 |
| SL1344_RS01725_ | -2.048418008 | -2.382450157 | -1.879781108 | 0.47380099 | 0.439165081 | 0.993647399 | 0.147235203 | 0.219806435 | 0.488174343 | 0.3159 | 0.557129386 |
| SL1344_RS01730_ | -1.52943903 | -1.476420111 | -1.590071031 | -0.11510772 | 0.321340492 | -0.337108056 | 0.349766828 | 0.523487433 | 0.73182793 | 0.6303 | 0.787677937 |
| SL1344_RS01740_ | -6.172461471 | -6.651630252 | -5.783794924 | 0.861395931 | 1.529808981 | 0.512159322 | 0.266489018 | 0.408831597 | 0.64341629 | 0.5076 | 0.700550976 |
| SL1344_RS01745_ | -0.531224143 | -0.510644802 | -0.571439045 | -0.036832864 | 0.249102836 | -0.141488891 | 0.447035312 | 0.641230717 | 0.819758021 | 0.748 | 0.873093852 |
| SL1344_RS01750_ | -1.677793499 | -1.587908707 | -1.769632587 | -0.188738024 | 0.283810605 | -0.630657504 | 0.240506342 | 0.37040195 | 0.604784742 | 0.4767 | 0.670196942 |
| SL1344_RS01755_ | -0.854255907 | -0.974638184 | -0.792241531 | 0.189415993 | 0.276606749 | 0.620071216 | 0.253333345 | 0.381893924 | 0.632878154 | 0.4994 | 0.697951206 |
| SL1344_RS01760_ | 0.46477508 | 0.518033735 | 0.436922436 | -0.055115535 | 0.206819398 | -0.225885974 | 0.408000004 | 0.680321921 | 0.845499084 | 0.7448 | 0.87828906 |
| SL1344_RS01765_ | -0.381217119 | -0.39154061 | -0.367711672 | 0.031033052 | 0.24584118 | 0.121226789 | 0.450366424 | 0.64780176 | 0.821740891 | 0.7452 | 0.867876457 |
| SL1344_RS01770_ | -4.56603499 | -4.701519124 | -4.499390164 | 0.175014421 | 0.949386828 | 0.158743123 | 0.436375752 | 0.612231344 | 0.800663986 | 0.7219 | 0.854516028 |
| SL1344_RS01775_ | -4.317713174 | -4.948569985 | -3.907117771 | 0.970192079 | 1.308275291 | 0.677594896 | 0.226000015 | 0.339312161 | 0.613899994 | 0.4454 | 0.674366898 |
| SL1344_RS01780_ | -4.083860992 | -3.936801218 | -4.267585234 | -0.33612312 | 0.762744846 | -0.40408934 | 0.317788149 | 0.475840254 | 0.695657563 | 0.5792 | 0.752342939 |

| | | | | | | | | | | | |
|---|---|---|---|---|---|---|---|---|---|---|---|
| SL1344_RS01785_ | -1.680606681 | -1.854912065 | -1.514108563 | 0.343090604 | 0.277454363 | 1.164414647 | 0.103333373 | 0.164197142 | 0.398469632 | 0.241 | 0.474699026 |
| SL1344_RS01790_ | -2.777224827 | -2.832047343 | -2.727758628 | 0.100975908 | 0.49855104 | 0.196023658 | 0.41266667 | 0.588036271 | 0.780832987 | 0.7004 | 0.836702013 |
| SL1344_RS01795_ | -1.92486746 | -1.851338287 | -1.970263778 | -0.066211171 | 0.461255213 | -0.116591573 | 0.444666669 | 0.703124055 | 0.860974167 | 0.7964 | 0.902781426 |
| SL1344_RS01800_ | -4.464837539 | -4.635808026 | -4.336346154 | 0.278294317 | 0.928940821 | 0.283028783 | 0.390666671 | 0.564135104 | 0.765580316 | 0.6866 | 0.83039027 |
| SL1344_RS01805_ | -1.436366518 | -1.465084509 | -1.413629526 | 0.042806217 | 0.300546133 | 0.130382684 | 0.442000002 | 0.59357502 | 0.786173477 | 0.7129 | 0.844326302 |
| SL1344_RS01810_ | -2.393640973 | -2.240286983 | -2.534865136 | -0.290179737 | 0.348087717 | -0.805744475 | 0.187333352 | 0.288530187 | 0.534566714 | 0.3797 | 0.596388498 |
| SL1344_RS01815_ | -1.168516311 | -1.319653576 | -1.012153358 | 0.304463815 | 0.252041075 | 1.162153992 | 0.101333374 | 0.169201853 | 0.408396353 | 0.2489 | 0.483071133 |
| SL1344_RS01820_ | -2.97972482 | -3.321437136 | -2.822317837 | 0.526523061 | 0.767745784 | 0.608572565 | 0.211858777 | 0.326084857 | 0.605013508 | 0.4374 | 0.654563363 |
| SL1344_RS01825_ | 1.169745233 | 0.725904272 | 1.231583272 | 0.513285996 | 0.349775814 | 1.148284465 | 0.152000025 | 0.192568864 | 0.489136199 | 0.2962 | 0.558379675 |
| SL1344_RS01830_ | 1.68960133 | 1.726286073 | 1.670153289 | -0.039035021 | 0.150216981 | -0.230264269 | 0.41466667 | 0.727397442 | 0.874276255 | 0.7961 | 0.906572694 |
| SL1344_RS01835_ | 2.696629515 | 2.625823866 | 2.778660034 | 0.160934204 | 0.090794066 | 1.735310625 | 0.016655836 | 0.064069289 | 0.272797748 | 0.1161 | 0.35165093 |
| SL1344_RS01840_ | -3.184747556 | -3.541890541 | -2.940502542 | 0.612814654 | 0.527790145 | 1.103088395 | 0.101265863 | 0.174837734 | 0.421846447 | 0.2451 | 0.479463724 |
| SL1344_RS01845_ | -1.734665839 | -1.992493772 | -1.513658509 | 0.469110252 | 0.322452204 | 1.39074953 | 0.058000075 | 0.114862339 | 0.343155442 | 0.1821 | 0.420868438 |
| SL1344_RS01850_ | 0.104185366 | 0.154492064 | 0.083388951 | -0.077896243 | 0.405875972 | -0.170000951 | 0.425716192 | 0.649088858 | 0.836797716 | 0.8204 | 0.918087252 |
| SL1344_RS01855_ | -2.383962367 | -2.42077638 | -2.295482421 | 0.112736317 | 0.50337443 | 0.218967352 | 0.421718857 | 0.633450186 | 0.81322007 | 0.733 | 0.862318435 |
| SL1344_RS01860_ | 1.374136741 | 1.506329904 | 1.172794533 | -0.280019004 | 0.209832774 | -1.283778829 | 0.091333379 | 0.124885992 | 0.388884037 | 0.1877 | 0.44888167 |
| SL1344_RS01865_ | -0.274146219 | -0.30442935 | -0.237834989 | 0.06778482 | 0.199891363 | 0.323414717 | 0.357333339 | 0.518810984 | 0.727717483 | 0.6424 | 0.792879728 |
| SL1344_RS01870_ | 1.191124582 | 1.1004489 | 1.401411349 | 0.271052275 | 0.23187805 | 1.09301903 | 0.127333364 | 0.175133838 | 0.45748811 | 0.2599 | 0.526362937 |
| SL1344_RS01875_ | 1.792745448 | 1.747251722 | 1.892613135 | 0.16419306 | 0.27710529 | 0.646505371 | 0.20733335 | 0.325679792 | 0.619885946 | 0.3974 | 0.659208422 |
| SL1344_RS01880_ | 3.093497068 | 2.790415076 | 3.187564284 | 0.381634692 | 0.453305093 | 0.518488608 | 0.33333334 | 0.666383089 | 0.855763928 | 0.7 | 0.872542119 |
| SL1344_RS01885_ | -2.87741235 | -2.561400309 | -3.243104125 | -0.721282575 | 0.569832404 | -1.18688283 | 0.089940086 | 0.150270337 | 0.405755645 | 0.2233 | 0.469527366 |
| SL1344_RS01890_ | -2.556022543 | -2.413776183 | -2.658807785 | -0.274241992 | 0.531926454 | -0.480191156 | 0.281145913 | 0.447094118 | 0.679790185 | 0.5683 | 0.744661776 |
| SL1344_RS01895_ | -1.825740802 | -1.900457314 | -1.651601376 | 0.207138439 | 0.411325964 | 0.441947551 | 0.332445044 | 0.560555161 | 0.76493914 | 0.6603 | 0.820455756 |
| SL1344_RS01900_ | -1.220111779 | -1.20488739 | -1.244996186 | -0.033511241 | 0.348740054 | -0.09128295 | 0.459027317 | 0.674562419 | 0.840681766 | 0.7803 | 0.888546513 |
| SL1344_RS01905_ | 3.117583969 | 3.068868304 | 3.315175525 | 0.242550325 | 0.132239835 | 1.488736401 | 0.055333412 | 0.110904123 | 0.373731424 | 0.1667 | 0.41413739 |
| SL1344_RS01910_ | 1.244083473 | 1.327438985 | 1.182711887 | -0.142001805 | 0.171905599 | -0.782940417 | 0.191872104 | 0.28469041 | 0.558932501 | 0.3919 | 0.624932041 |
| SL1344_RS01915_ | -1.99766162 | -2.157904643 | -1.876241069 | 0.275930577 | 0.378824501 | 0.665512872 | 0.213191222 | 0.340109336 | 0.588991324 | 0.4415 | 0.650213349 |
| SL1344_RS01930_ | -0.252358144 | -0.351023773 | -0.16490513 | 0.180785357 | 0.198633995 | 0.859941443 | 0.165223207 | 0.266370172 | 0.511432952 | 0.3536 | 0.575926487 |
| SL1344_RS01935_ | -2.66815909 | -2.862213677 | -2.435887262 | 0.450402235 | 0.473478377 | 0.918701877 | 0.153231204 | 0.240905797 | 0.497199111 | 0.3345 | 0.560420633 |
| SL1344_RS01940_ | 2.025945739 | 1.86959916 | 2.293057503 | 0.407347014 | 0.3240962 | 1.138707212 | 0.11792142 | 0.183044704 | 0.478039424 | 0.2155 | 0.489840043 |
| SL1344_RS01945_ | 2.594729704 | 2.559119475 | 2.680422555 | 0.107002782 | 0.138889481 | 0.697112494 | 0.243837455 | 0.357710473 | 0.632952799 | 0.4674 | 0.704047921 |
| SL1344_RS01950_ | -2.154401254 | -2.224060156 | -2.127834939 | 0.106709371 | 0.605644967 | 0.15060386 | 0.433044639 | 0.689004558 | 0.854335281 | 0.8166 | 0.914700138 |
| SL1344_RS01955_ | -1.916348248 | -1.877026472 | -1.969830334 | -0.098078337 | 0.306122465 | -0.298073498 | 0.364423723 | 0.491036361 | 0.708832345 | 0.6025 | 0.768688282 |
| SL1344_RS01960_ | 3.910436867 | 3.869634167 | 4.038502196 | 0.183634458 | 0.360272515 | 0.567230186 | 0.221852113 | 0.354810068 | 0.648005632 | 0.4015 | 0.681309749 |
| SL1344_RS01965_ | -0.40560752 | -0.662684473 | -0.181240582 | 0.478941119 | 0.181981331 | 2.531398792 | 0.002666616 | 0.027001396 | 0.174721349 | 0.1028 | 0.334287041 |
| SL1344_RS01975_ | 2.652226338 | 2.623451503 | 2.672057788 | 0.062207796 | 0.118831392 | 0.521181772 | 0.261333345 | 0.386309185 | 0.654985474 | 0.5144 | 0.706991996 |
| SL1344_RS01980_ | -0.039633584 | -0.02508585 | -0.06258121 | -0.021261738 | 0.195899885 | -0.102491363 | 0.454666668 | 0.612978515 | 0.801952495 | 0.7234 | 0.857573547 |
| SL1344_RS01985_ | 1.093402115 | 1.011128883 | 1.175154236 | 0.162034113 | 0.109311988 | 1.427625879 | 0.058000075 | 0.110661412 | 0.332871031 | 0.1798 | 0.418537886 |
| SL1344_RS01990_ | 1.197956289 | 0.992810859 | 1.345115634 | 0.364430873 | 0.140178492 | 2.592902999 | 0.001335878 | 0.034836531 | 0.204462014 | 0.1012 | 0.331915916 |
| SL1344_RS01995_ | 1.10554177 | 1.015035182 | 1.230569775 | 0.242847397 | 0.186104589 | 1.284544806 | 0.067333397 | 0.120906801 | 0.374978201 | 0.1842 | 0.431058568 |
| SL1344_RS02000_ | 2.427781948 | 2.410177813 | 2.441688566 | 0.025634806 | 0.269483006 | 0.082944192 | 0.470353099 | 0.839576268 | 0.93571216 | 0.9838 | 0.99312032 |
| SL1344_RS02005_ | 2.031263509 | 1.96542315 | 2.219810266 | 0.242617777 | 0.177566535 | 1.371408139 | 0.077948089 | 0.105848288 | 0.360545308 | 0.1946 | 0.44927691 |
| SL1344_RS02010_ | 0.284053773 | 0.483803576 | 0.224140319 | -0.223058005 | 0.254050573 | -0.760223139 | 0.23717523 | 0.349630538 | 0.621219673 | 0.4816 | 0.705702013 |
| SL1344_RS02015_ | -0.085316802 | -0.025597526 | -0.141535018 | -0.11604888 | 0.16687797 | -0.6576062 | 0.230666681 | 0.353306897 | 0.587514928 | 0.4462 | 0.647682106 |
| SL1344_RS02020_ | 1.823061631 | 1.856020134 | 1.777659354 | -0.08213719 | 0.186373919 | -0.420377232 | 0.310000008 | 0.519272412 | 0.749199231 | 0.6201 | 0.79829323 |
| SL1344_RS02025_ | 0.393739532 | 0.467863826 | 0.334202683 | -0.136153038 | 0.190987034 | -0.694504735 | 0.221852113 | 0.336334824 | 0.587624624 | 0.4377 | 0.650149956 |
| SL1344_RS02030_ | -5.711144741 | -5.859718084 | -5.47747786 | 0.133638865 | 2.328687374 | 0.039493152 | 0.470353099 | 0.660655336 | 0.840489821 | 0.8143 | 0.912734669 |
| SL1344_RS02045_ | 2.967699189 | 3.232311877 | 2.901764064 | -0.343159362 | 0.244521849 | -1.452847911 | 0.048634334 | 0.087329511 | 0.339018221 | 0.1459 | 0.39244447 |
| SL1344_RS02050_ | 2.159807853 | 2.063374775 | 2.269866695 | 0.217476638 | 0.106161638 | 2.033787896 | 0.0020023 | 0.054715755 | 0.258102666 | 0.1056 | 0.337938208 |
| SL1344_RS02055_ | -0.529230867 | -0.574126688 | -0.446192745 | 0.109662854 | 0.273269142 | 0.355847 | 0.357095276 | 0.60364211 | 0.794437388 | 0.6979 | 0.8441883 |
| SL1344_RS02060_ | 4.501156089 | 4.494694982 | 4.508186223 | 0.018402082 | 0.159313231 | 0.10814675 | 0.436666669 | 0.857095296 | 0.944163593 | 0.9175 | 0.964810469 |
| SL1344_RS02065_ | 2.176562482 | 2.212546468 | 2.13998821 | 0.062901696 | 0.268652911 | 0.237371108 | 0.446000002 | 0.748485415 | 0.890944369 | 0.9718 | 0.987413908 |
| SL1344_RS02070_ | 1.572602431 | 1.583384172 | 1.559977087 | -0.010554207 | 0.142444605 | -0.075041143 | 0.468354432 | 0.702814017 | 0.858908441 | 0.7808 | 0.892253494 |

| | | | | | | | | | | | |
|---|---|---|---|---|---|---|---|---|---|---|---|
| SL1344_RS02075_ | 2.860505249 | 2.691818401 | 2.982331211 | 0.248800011 | 0.142235915 | 1.725135911 | 0.038641021 | 0.062684718 | 0.283974242 | 0.1325 | 0.377288267 |
| SL1344_RS02080_ | 3.296596074 | 3.2903232 | 3.301796009 | 0.037548689 | 0.134943385 | 0.261647414 | 0.394000004 | 0.551215239 | 0.780684048 | 0.8178 | 0.91140571 |
| SL1344_RS02085_ | 4.531679851 | 4.427664902 | 4.635676317 | 0.203136084 | 0.068735135 | 2.93024626 | 4.68E-05 | 0.01590858 | 0.149039414 | 0.1 | 0.330257709 |
| SL1344_RS02090_ | 3.228137967 | 3.271389695 | 3.17709574 | 0.004736956 | 0.235283746 | 0.022022378 | 0.493670886 | 0.876520579 | 0.952515369 | 0.9931 | 0.997010206 |
| SL1344_RS02095_ | -1.018316784 | -1.064273922 | -0.990143225 | 0.064350522 | 0.262833931 | 0.209512379 | 0.411333337 | 0.579353999 | 0.775654346 | 0.6793 | 0.827364359 |
| SL1344_RS02100_ | -0.683105001 | -0.625587941 | -0.740625776 | -0.199663011 | 0.361583612 | -0.550975586 | 0.281145913 | 0.400032103 | 0.657741693 | 0.5773 | 0.74922931 |
| SL1344_RS02105_ | 1.417831878 | 1.442694874 | 1.40039102 | 0.014166584 | 0.303563997 | 0.049139272 | 0.479680214 | 0.812818462 | 0.922390002 | 0.9469 | 0.975464234 |
| SL1344_RS02110_ | 2.407407773 | 2.821995063 | 2.01922745 | -0.879486378 | 0.228701926 | -3.839363134 | 4.68E-05 | 0.003911953 | 0.081973035 | 0.1 | 0.330257709 |
| SL1344_RS02115_ | 4.368165995 | 4.780258371 | 4.350172163 | -0.435942935 | 0.471478118 | -1.019136396 | 0.222000015 | 0.183156074 | 0.482130099 | 0.4 | 0.680350104 |
| SL1344_RS02120_ | -2.01913059 | -1.971485487 | -2.06135137 | -0.157285539 | 0.572489429 | -0.281681707 | 0.390000004 | 0.599931737 | 0.796224472 | 0.737 | 0.860761028 |
| SL1344_RS02125_ | 1.354338862 | 1.249349326 | 1.520585504 | 0.255065561 | 0.392362141 | 0.6515509 | 0.23717523 | 0.365482691 | 0.651596803 | 0.4621 | 0.712926179 |
| SL1344_RS02130_ | 1.855728844 | 1.835283729 | 1.873161574 | 0.05452282 | 0.158847797 | 0.336260392 | 0.353764163 | 0.570310784 | 0.779808557 | 0.6959 | 0.840953316 |
| SL1344_RS02135_ | 1.883671065 | 1.818463711 | 1.929884667 | 0.122758897 | 0.457369388 | 0.268069917 | 0.295333342 | 0.521945053 | 0.767874751 | 0.5947 | 0.793072626 |
| SL1344_RS02145_ | 0.495250609 | 0.400225605 | 0.587601971 | 0.18602072 | 0.144810186 | 1.233720752 | 0.074000058 | 0.146059872 | 0.382848474 | 0.2168 | 0.455617005 |
| SL1344_RS02150_ | 0.283031692 | 0.139389959 | 0.515562013 | 0.381819427 | 0.258096701 | 1.40062141 | 0.046000096 | 0.092062585 | 0.331506183 | 0.1546 | 0.399806731 |
| SL1344_RS02155_ | 0.885771193 | 0.83633424 | 0.961884416 | 0.129059774 | 0.174842437 | 0.683934122 | 0.222518336 | 0.350590234 | 0.610288985 | 0.4604 | 0.677666933 |
| SL1344_RS02160_ | 2.855274869 | 2.824034334 | 2.883711896 | 0.075858019 | 0.114073538 | 0.652107405 | 0.244000013 | 0.367121297 | 0.634235831 | 0.5072 | 0.716962287 |
| SL1344_RS02165_ | 2.556091149 | 2.503988596 | 2.599314071 | 0.096740355 | 0.087232596 | 1.056618275 | 0.108000038 | 0.178120574 | 0.435322516 | 0.2577 | 0.49945065 |
| SL1344_RS02170_ | 1.364558828 | 1.320594088 | 1.44430115 | 0.137197348 | 0.192315822 | 0.710448632 | 0.204666683 | 0.322544353 | 0.599771417 | 0.4211 | 0.663210104 |
| SL1344_RS02175_ | 1.880576549 | 1.826784532 | 2.017388548 | 0.162479163 | 0.15222544 | 0.992430687 | 0.168000022 | 0.237141461 | 0.52004846 | 0.34 | 0.595835079 |
| SL1344_RS02180_ | -1.616804939 | -1.589262539 | -1.637243052 | -0.02474044 | 0.411419846 | -0.057158159 | 0.477015324 | 0.661181077 | 0.832306511 | 0.7734 | 0.884947588 |
| SL1344_RS02185_ | -2.606692896 | -2.520838914 | -2.693426494 | -0.112306893 | 0.610083992 | -0.177827905 | 0.427333336 | 0.669320441 | 0.836876633 | 0.7742 | 0.884088951 |
| SL1344_RS02190_ | -1.705117423 | -1.540232189 | -1.82651467 | -0.270586806 | 0.391660398 | -0.620723796 | 0.256666678 | 0.403898566 | 0.649444098 | 0.5036 | 0.71059357 |
| SL1344_RS02195_ | -1.680831118 | -1.808373512 | -1.527274091 | 0.27848537 | 0.34805069 | 0.760568377 | 0.19920055 | 0.315519465 | 0.563954126 | 0.4176 | 0.633265557 |
| SL1344_RS02200_ | -0.029482982 | -0.068815208 | 0.012441657 | 0.075096066 | 0.200364814 | 0.353898723 | 0.350000006 | 0.52724782 | 0.737100846 | 0.6364 | 0.793355086 |
| SL1344_RS02205_ | -2.539869711 | -2.537016665 | -2.542629762 | 0.012318771 | 0.568704826 | 0.022708391 | 0.489007329 | 0.680841926 | 0.844699867 | 0.7924 | 0.896279547 |
| SL1344_RS02210_ | -2.967851809 | -3.023617575 | -2.90156156 | 0.11623277 | 0.47361973 | 0.225557834 | 0.394403735 | 0.528319464 | 0.735225065 | 0.6415 | 0.795256334 |
| SL1344_RS02215_ | 1.212830707 | 1.078952588 | 1.25530498 | 0.157331351 | 0.244533956 | 0.39821118 | 0.351765496 | 0.798270726 | 0.916003811 | 0.7444 | 0.891332156 |
| SL1344_RS02220_ | 0.288853852 | 0.191545362 | 0.323723606 | 0.11144783 | 0.225491294 | 0.441171017 | 0.338000007 | 0.582446868 | 0.786212406 | 0.6903 | 0.844663038 |
| SL1344_RS02225_ | 2.931518958 | 2.947225552 | 2.909512219 | -0.04220987 | 0.098970702 | -0.396897304 | 0.332666674 | 0.541339647 | 0.757719677 | 0.6766 | 0.826337579 |
| SL1344_RS02230_ | 1.173936196 | 1.194761697 | 1.159166353 | -0.030927724 | 0.121897653 | -0.225849806 | 0.396000004 | 0.561423458 | 0.760754517 | 0.6761 | 0.818206321 |
| SL1344_RS02235_ | -0.94561681 | -1.031663021 | -0.833201545 | 0.193788372 | 0.241494513 | 0.78003774 | 0.194000018 | 0.303462364 | 0.546746731 | 0.4011 | 0.611130916 |
| SL1344_RS02240_ | 0.413632912 | 0.206453896 | 0.55436253 | 0.3451961 | 0.157970504 | 2.102758557 | 0.011326186 | 0.044965944 | 0.226258534 | 0.1121 | 0.345862951 |
| SL1344_RS02245_ | 6.009246976 | 5.966354452 | 6.068727645 | 0.184690736 | 0.137661018 | 1.443320606 | 4.68E-05 | 0.108301751 | 0.377525121 | 0.1001 | 0.330401797 |
| SL1344_RS02250_ | 3.829627297 | 3.673263622 | 3.883694909 | 0.208449543 | 0.100417025 | 1.81486437 | 0.010000458 | 0.083289691 | 0.328786494 | 0.107 | 0.340125848 |
| SL1344_RS02255_ | 4.452964483 | 4.40767857 | 4.588006484 | 0.187897955 | 0.139327566 | 1.51841463 | 0.014000326 | 0.070115813 | 0.304997648 | 0.1141 | 0.34949376 |
| SL1344_RS02260_ | 6.296299898 | 6.267405617 | 6.42820859 | 0.164609568 | 0.109680042 | 1.451038786 | 0.006662914 | 0.10920476 | 0.378926099 | 0.1066 | 0.339245709 |
| SL1344_RS02265_ | 6.478452558 | 6.383968462 | 6.58287853 | 0.264376473 | 0.118361014 | 2.279061632 | 4.68E-05 | 0.034011353 | 0.221896437 | 0.1 | 0.330257709 |
| SL1344_RS02270_ | 2.422215008 | 2.350194686 | 2.519349132 | 0.166447946 | 0.085049418 | 1.875154943 | 0.019986902 | 0.058935275 | 0.255856568 | 0.1284 | 0.366130528 |
| SL1344_RS02275_ | 2.024244506 | 1.982557694 | 2.05358128 | 0.181376801 | 0.296360927 | 0.610641284 | 0.260000012 | 0.314728639 | 0.610906638 | 0.514 | 0.72633745 |
| SL1344_RS02280_ | 2.966877959 | 2.870965871 | 3.003294535 | 0.128607159 | 0.094786805 | 1.201588254 | 0.102666707 | 0.171300346 | 0.446555843 | 0.2479 | 0.498525595 |
| SL1344_RS02285_ | 6.441621874 | 6.306443075 | 6.500061045 | 0.192744559 | 0.087212363 | 2.372447099 | 4.68E-05 | 0.018860304 | 0.16779418 | 0.1 | 0.330257709 |
| SL1344_RS02300_ | 5.376532996 | 5.481399812 | 5.254182152 | -0.234276449 | 0.211350795 | -1.298377453 | 0.072666725 | 0.108644505 | 0.378213214 | 0.1626 | 0.422001336 |
| SL1344_RS02305_ | 7.472643072 | 7.49727568 | 7.289000468 | -0.204713753 | 0.244841303 | -0.877551761 | 0.223184558 | 0.206444216 | 0.508922057 | 0.4 | 0.680350104 |
| SL1344_RS02310_ | 3.544127948 | 3.329394234 | 3.565158358 | 0.225346042 | 0.103899897 | 1.058262233 | 0.176666687 | 0.198369675 | 0.494230415 | 0.374 | 0.61739473 |
| SL1344_RS02315_ | -2.901480105 | -2.672774095 | -3.177455598 | -0.312554086 | 0.837099519 | -0.365600846 | 0.355096608 | 0.592032527 | 0.792931322 | 0.707 | 0.844611146 |
| SL1344_RS02320_ | -0.998678285 | -1.020683946 | -0.97789254 | 0.0499839 | 0.275505526 | 0.163590313 | 0.428666669 | 0.600643597 | 0.786798402 | 0.7094 | 0.839987608 |
| SL1344_RS02325_ | 1.451763169 | 1.517511629 | 1.420233236 | -0.107040979 | 0.360213848 | -0.279518949 | 0.360426388 | 0.537923045 | 0.775728413 | 0.7379 | 0.877607748 |
| SL1344_RS02330_ | 1.248786253 | 1.034337941 | 1.314245722 | 0.219178905 | 0.287155779 | 0.366893395 | 0.338441046 | 0.906078029 | 0.965208399 | 0.7126 | 0.877892365 |
| SL1344_RS02335_ | -0.70488905 | -0.730252009 | -0.669475292 | 0.032581508 | 0.285639497 | 0.110174083 | 0.447035312 | 0.668324531 | 0.837092 | 0.7798 | 0.888781871 |
| SL1344_RS02350_ | 1.393074156 | 1.290844271 | 1.428034312 | 0.12184543 | 0.171050868 | 0.632700926 | 0.272000011 | 0.42410296 | 0.677249096 | 0.5566 | 0.758597819 |
| SL1344_RS02355_ | 1.217328855 | 1.222452141 | 1.213071868 | -0.016339823 | 0.127152188 | -0.116681739 | 0.443704199 | 0.604768533 | 0.791102595 | 0.7253 | 0.848751545 |
| SL1344_RS02360_ | -4.313231142 | -4.391590009 | -4.251376832 | 0.150146782 | 0.772616167 | 0.178658949 | 0.42238508 | 0.549915487 | 0.753219377 | 0.6718 | 0.815659013 |

| | | | | | | | | | | | |
|---|---|---|---|---|---|---|---|---|---|---|---|
| SL1344_RS02365_ | -0.888550637 | -0.765237671 | -0.95890156 | -0.212640384 | 0.3153646 | -0.633839283 | 0.246666679 | 0.371938266 | 0.62980849 | 0.5008 | 0.698862527 |
| SL1344_RS02370_ | 1.087808272 | 0.966036417 | 1.141344894 | 0.150254781 | 0.171388621 | 0.784432447 | 0.217333349 | 0.322754629 | 0.593166735 | 0.4395 | 0.668284423 |
| SL1344_RS02375_ | 1.102702982 | 1.262561114 | 1.051216041 | -0.180254881 | 0.263207135 | -0.46838774 | 0.344437048 | 0.706864519 | 0.868078112 | 0.7307 | 0.883861212 |
| SL1344_RS02380_ | -0.153535151 | -0.165022243 | -0.141691125 | 0.029590929 | 0.175985597 | 0.152225508 | 0.432666669 | 0.546416754 | 0.745927087 | 0.652 | 0.801443786 |
| SL1344_RS02385_ | 1.148262606 | 1.115844674 | 1.177676534 | 0.072196921 | 0.175308739 | 0.403595177 | 0.322666674 | 0.503915535 | 0.732496495 | 0.6356 | 0.797546605 |
| SL1344_RS02390_ | -1.989590145 | -1.877567182 | -2.069682989 | -0.127353482 | 0.470686912 | -0.240467773 | 0.404397072 | 0.657934237 | 0.830444191 | 0.7544 | 0.876450814 |
| SL1344_RS02395_ | -0.339952772 | -0.328641014 | -0.354515554 | -0.032894175 | 0.206568741 | -0.153969031 | 0.435333336 | 0.579238271 | 0.773814304 | 0.6959 | 0.831714363 |
| SL1344_RS02400_ | 3.322716781 | 3.415920724 | 3.047891787 | -0.365676184 | 0.226786285 | -1.689928562 | 0.001336769 | 0.113855353 | 0.385914812 | 0.101 | 0.331720728 |
| SL1344_RS02405_ | 2.267640444 | 1.941157738 | 2.56234972 | 0.627270643 | 0.112955948 | 5.47526141 | 4.68E-05 | 0.002727234 | 0.063988668 | 0.1 | 0.330257709 |
| SL1344_RS02420_ | 4.530056906 | 4.369194464 | 4.696035295 | 0.328071817 | 0.049389019 | 6.652997147 | 4.68E-05 | 0.002370159 | 0.059900482 | 0.1 | 0.330257709 |
| SL1344_RS02425_ | 3.295907184 | 3.023981466 | 3.427612149 | 0.401054981 | 0.153883278 | 2.550704989 | 4.68E-05 | 0.028123488 | 0.198677573 | 0.1 | 0.330257709 |
| SL1344_RS02430_ | 1.593509888 | 1.546189317 | 1.635823161 | 0.039039628 | 0.181421656 | 0.230536229 | 0.41666667 | 0.686261863 | 0.851488309 | 0.811 | 0.905581752 |
| SL1344_RS02435_ | 2.830879029 | 2.799995363 | 2.862614282 | 0.0978591 | 0.136972196 | 0.719500829 | 0.222666682 | 0.29716265 | 0.579781356 | 0.4551 | 0.6753777 |
| SL1344_RS02440_ | 1.441566278 | 1.478400337 | 1.404494168 | -0.077920149 | 0.132503264 | -0.564592201 | 0.266000011 | 0.40652845 | 0.647677778 | 0.5251 | 0.715677809 |
| SL1344_RS02445_ | -0.327711189 | -0.251470576 | -0.42202558 | -0.193613093 | 0.270402182 | -0.673676253 | 0.198666684 | 0.313305323 | 0.584200534 | 0.4029 | 0.625300119 |
| SL1344_RS02450_ | 0.400061551 | 0.423405629 | 0.368803808 | -0.050117394 | 0.171136577 | -0.279077998 | 0.384000005 | 0.548989221 | 0.75464904 | 0.6674 | 0.815248141 |
| SL1344_RS02455_ | 0.392295253 | 0.554253317 | 0.339509979 | -0.203564881 | 0.218275207 | -0.828952965 | 0.220666682 | 0.322995171 | 0.595083447 | 0.449 | 0.673746519 |
| SL1344_RS02460_ | 2.153081159 | 2.15226478 | 2.153570097 | 0.01429125 | 0.113798889 | 0.099427276 | 0.453333335 | 0.653891012 | 0.828366625 | 0.7607 | 0.877443943 |
| SL1344_RS02465_ | 3.392843199 | 3.415158341 | 3.375685207 | -0.035407865 | 0.151050859 | -0.201470436 | 0.436375752 | 0.762405099 | 0.898035135 | 0.9101 | 0.960450425 |
| SL1344_RS02470_ | 3.346902877 | 3.155863477 | 3.490378188 | 0.310795548 | 0.166944335 | 1.934449798 | 0.005996769 | 0.042308542 | 0.24099336 | 0.1062 | 0.339325813 |
| SL1344_RS02475_ | 3.841260077 | 3.808610412 | 3.992310385 | 0.151217329 | 0.247075914 | 0.681838862 | 0.223850781 | 0.36477743 | 0.653484197 | 0.4131 | 0.688610202 |
| SL1344_RS02480_ | 5.950217676 | 6.267927742 | 5.9057484 | -0.303327853 | 0.341412783 | -0.712059231 | 0.23184545 | 0.37477454 | 0.665761702 | 0.4258 | 0.696830241 |
| SL1344_RS02485_ | 4.212954425 | 4.077283896 | 4.335077287 | 0.285447524 | 0.100222324 | 2.820610418 | 4.68E-05 | 0.038570467 | 0.232283776 | 0.1 | 0.330257709 |
| SL1344_RS02490_ | 1.064344939 | 1.144270934 | 1.027859402 | -0.094585919 | 0.21613229 | -0.373995895 | 0.359760166 | 0.665408941 | 0.839553402 | 0.7329 | 0.874496265 |
| SL1344_RS02495_ | -1.057593816 | -0.986017388 | -1.131856132 | -0.135568864 | 0.387363775 | -0.337409042 | 0.360666672 | 0.567415213 | 0.777128478 | 0.6706 | 0.827310085 |
| SL1344_RS02500_ | 0.82430003 | 0.797806473 | 0.847330827 | 0.058502175 | 0.167782035 | 0.308251377 | 0.357761498 | 0.545998709 | 0.752726797 | 0.657 | 0.807699249 |
| SL1344_RS02505_ | 2.380190332 | 2.584521822 | 2.177796489 | -0.406448155 | 0.099691081 | -4.01862757 | 4.68E-05 | 0.008600699 | 0.106314591 | 0.1 | 0.330257709 |
| SL1344_RS02510_ | 1.021366298 | 0.926924156 | 1.261185932 | 0.316108823 | 0.169798255 | 1.671246957 | 0.046000096 | 0.086482674 | 0.320140628 | 0.1529 | 0.396301346 |
| SL1344_RS02520_ | 0.87673684 | 0.813810733 | 0.950117076 | 0.133963467 | 0.118390332 | 1.065008971 | 0.121918754 | 0.194448374 | 0.431404126 | 0.2804 | 0.512185338 |
| SL1344_RS02525_ | 0.549385897 | 0.655449534 | 0.513159657 | -0.128769191 | 0.179139296 | -0.616598768 | 0.262491684 | 0.421328982 | 0.668197233 | 0.5498 | 0.745249709 |
| SL1344_RS02530_ | -0.974985091 | -1.228699205 | -0.646340847 | 0.627757402 | 0.347375368 | 1.793149754 | 0.009327606 | 0.073815779 | 0.300365989 | 0.1088 | 0.34183408 |
| SL1344_RS02540_ | 0.245202238 | 0.121382352 | 0.306022959 | 0.150470419 | 0.247116394 | 0.564719458 | 0.294666676 | 0.452712749 | 0.694049272 | 0.573 | 0.76184391 |
| SL1344_RS02545_ | 0.59398501 | 0.829029668 | 0.247331829 | -0.5869202 | 0.194141881 | -2.926990686 | 0.001335878 | 0.019262874 | 0.156099238 | 0.1005 | 0.330889407 |
| SL1344_RS02550_ | 0.30455826 | 0.612387866 | 0.110014412 | -0.509102507 | 0.219076923 | -2.314591177 | 0.003332494 | 0.046157054 | 0.241638385 | 0.1026 | 0.333622807 |
| SL1344_RS02555_ | -0.569097247 | -0.632079326 | -0.495356706 | 0.143398395 | 0.381122062 | 0.372021975 | 0.334666674 | 0.552941918 | 0.7682251 | 0.6668 | 0.823700435 |
| SL1344_RS02560_ | -0.174247591 | -0.377000941 | -0.03563298 | 0.303997383 | 0.276671834 | 1.067017312 | 0.125249865 | 0.184937585 | 0.454944475 | 0.2675 | 0.523657358 |
| SL1344_RS02565_ | 3.37844222 | 3.191430734 | 3.456943367 | 0.221337464 | 0.222569649 | 0.513101273 | 0.33333334 | 0.685822523 | 0.864831734 | 0.703 | 0.873797538 |
| SL1344_RS02570_ | 0.958705105 | 0.885213423 | 1.002797528 | 0.104737063 | 0.180796748 | 0.532202491 | 0.287333343 | 0.455061929 | 0.696769901 | 0.5693 | 0.76213067 |
| SL1344_RS02575_ | 0.421488747 | 0.419187711 | 0.42360223 | 0.055780025 | 0.251775498 | 0.208908498 | 0.415056632 | 0.629395149 | 0.816998948 | 0.8155 | 0.906007062 |
| SL1344_RS02580_ | -0.713835251 | -0.693335873 | -0.721862982 | -0.013298988 | 0.355317032 | -0.032787871 | 0.492338441 | 0.740070184 | 0.882382094 | 0.8437 | 0.928338703 |
| SL1344_RS02585_ | 1.204737488 | 1.198850675 | 1.208146233 | 0.005627132 | 0.132416759 | 0.034862169 | 0.483677549 | 0.632234742 | 0.813261978 | 0.7448 | 0.867493342 |
| SL1344_RS02590_ | 0.100663212 | 0.142764583 | 0.066662542 | -0.079524174 | 0.207362596 | -0.344604027 | 0.350666673 | 0.55365782 | 0.759190359 | 0.6646 | 0.814935284 |
| SL1344_RS02595_ | -0.497402911 | -0.493761963 | -0.500291882 | -0.062812641 | 0.377926779 | -0.132603029 | 0.427333336 | 0.552742273 | 0.777915668 | 0.7789 | 0.884611195 |
| SL1344_RS02600_ | -0.587244778 | -0.416988776 | -0.746061145 | -0.326529635 | 0.199871443 | -1.569639665 | 0.042000105 | 0.085749973 | 0.299124185 | 0.1527 | 0.392073364 |
| SL1344_RS02605_ | -0.14026623 | -0.155635197 | -0.1247569 | 0.008127045 | 0.343879661 | 0.018792513 | 0.487333334 | 0.771201302 | 0.899207808 | 0.8334 | 0.921783999 |
| SL1344_RS02610_ | 1.684609703 | 1.658782551 | 1.701710553 | 0.06575272 | 0.144094958 | 0.416444667 | 0.317333341 | 0.462513015 | 0.703734716 | 0.6283 | 0.786541885 |
| SL1344_RS02615_ | -1.896333201 | -1.832743703 | -1.932494167 | -0.095506648 | 0.39997766 | -0.224847647 | 0.398666671 | 0.587706715 | 0.780766791 | 0.6944 | 0.833498462 |
| SL1344_RS02620_ | -1.054034856 | -1.081130619 | -1.021582413 | 0.056257774 | 0.269905225 | 0.189558385 | 0.409333337 | 0.560774872 | 0.758702439 | 0.6656 | 0.812252946 |
| SL1344_RS02625_ | 1.986611529 | 2.057222539 | 1.887299519 | -0.147644807 | 0.287053447 | -0.535286385 | 0.263824128 | 0.424933735 | 0.696462219 | 0.5018 | 0.737530857 |
| SL1344_RS02630_ | -2.599035369 | -2.729850436 | -2.458769356 | 0.302449294 | 0.510755019 | 0.570239715 | 0.271333344 | 0.414127865 | 0.649474893 | 0.5251 | 0.71143572 |
| SL1344_RS02635_ | -3.219247712 | -3.326423214 | -3.124513594 | 0.186479225 | 0.481186509 | 0.359807151 | 0.349766828 | 0.468647844 | 0.68784571 | 0.5828 | 0.750971217 |
| SL1344_RS02640_ | -2.800828561 | -2.801924668 | -2.800136617 | 0.017015559 | 0.409143916 | 0.042859475 | 0.484343771 | 0.541849342 | 0.746164074 | 0.6556 | 0.805051813 |
| SL1344_RS02645_ | -2.715991712 | -2.848750769 | -2.580394955 | 0.265028831 | 0.415091701 | 0.610296939 | 0.251333346 | 0.374907273 | 0.610986026 | 0.4769 | 0.674040358 |

| | | | | | | | | | | | |
|---|---|---|---|---|---|---|---|---|---|---|---|
| SL1344_RS02650_ | -3.520726195 | -4.302135072 | -2.988912081 | 1.35482599 | 0.821381778 | 1.59440379 | 0.017333596 | 0.095751996 | 0.334528866 | 0.1155 | 0.350979981 |
| SL1344_RS02655_ | -1.870780928 | -1.8702911 | -1.871039513 | -0.104734386 | 0.53603882 | -0.189707121 | 0.41266667 | 0.649088952 | 0.826596098 | 0.7991 | 0.896560962 |
| SL1344_RS02660_ | -3.51231562 | -3.36174055 | -3.674815341 | -0.314073541 | 0.491215593 | -0.581490244 | 0.252666679 | 0.361054702 | 0.592915973 | 0.4586 | 0.656228598 |
| SL1344_RS02665_ | -2.558557521 | -2.69768321 | -2.308008578 | 0.35831505 | 0.524436551 | 0.648386474 | 0.256000012 | 0.409965176 | 0.65136675 | 0.5152 | 0.71745122 |
| SL1344_RS02670_ | 0.110409289 | -0.046790672 | 0.247901038 | 0.304461646 | 0.228396537 | 1.304185548 | 0.076666722 | 0.127038989 | 0.378692253 | 0.196 | 0.445873046 |
| SL1344_RS02675_ | -2.514123358 | -2.633983197 | -2.359535306 | 0.26875719 | 0.432847536 | 0.561236002 | 0.267155241 | 0.416041444 | 0.650804363 | 0.5178 | 0.713142767 |
| SL1344_RS02680_ | -2.717001913 | -2.670801135 | -2.769037681 | -0.102989739 | 0.444575886 | -0.207276849 | 0.40266667 | 0.546060872 | 0.745726621 | 0.6522 | 0.801605718 |
| SL1344_RS02685_ | -2.388541463 | -2.394136689 | -2.385710636 | 0.000514459 | 0.472599048 | 0.001467612 | 0.499666889 | 0.637989494 | 0.815113786 | 0.7449 | 0.867001243 |
| SL1344_RS02690_ | -3.232201354 | -3.301341947 | -3.130562638 | 0.239789216 | 0.667284647 | 0.344981485 | 0.363091278 | 0.531265731 | 0.743396329 | 0.6627 | 0.806333096 |
| SL1344_RS02695_ | -3.646865976 | -3.713971308 | -3.567312225 | 0.161361644 | 0.708084293 | 0.234141852 | 0.413057965 | 0.583750909 | 0.776906399 | 0.6971 | 0.833873111 |
| SL1344_RS02700_ | -1.794046494 | -1.645817753 | -1.992106844 | -0.374609112 | 0.403454381 | -0.851639383 | 0.154563649 | 0.244355291 | 0.508708501 | 0.3324 | 0.563825947 |
| SL1344_RS02705_ | 1.069639634 | 1.044650453 | 1.090240666 | 0.011061038 | 0.182766161 | 0.05451209 | 0.473333334 | 0.71751173 | 0.869266805 | 0.8332 | 0.919480699 |
| SL1344_RS02710_ | -0.07431008 | -0.030824135 | -0.098142845 | -0.094470905 | 0.307328642 | -0.304930319 | 0.372418393 | 0.573433371 | 0.783251394 | 0.7335 | 0.860397569 |
| SL1344_RS02715_ | 2.765482989 | 2.623217682 | 2.852042472 | 0.131589142 | 0.295544288 | 0.566144176 | 0.238000013 | 0.42773448 | 0.700961559 | 0.4603 | 0.71414189 |
| SL1344_RS02720_ | 4.650214557 | 4.577734103 | 4.759644003 | 0.320841109 | 0.226329445 | 1.593674706 | 4.68E-05 | 0.071756181 | 0.311811742 | 0.1001 | 0.330406921 |
| SL1344_RS02730_ | -0.918238474 | -0.757143723 | -0.988040213 | -0.190238134 | 0.33687839 | -0.489675533 | 0.310459702 | 0.53954355 | 0.754366843 | 0.636 | 0.80970845 |
| SL1344_RS02735_ | -1.716095152 | -1.668806871 | -1.77408266 | -0.108174132 | 0.295527453 | -0.337043038 | 0.354430386 | 0.497781835 | 0.710950373 | 0.6025 | 0.768936438 |
| SL1344_RS02740_ | -2.233484506 | -2.24171779 | -2.223920342 | 0.015124556 | 0.37341928 | 0.036373908 | 0.483677549 | 0.584751129 | 0.777300199 | 0.6928 | 0.830632826 |
| SL1344_RS02745_ | 1.893383235 | 1.99922393 | 1.754541936 | -0.238155505 | 0.111149183 | -2.053350384 | 0.012667027 | 0.043983552 | 0.229263149 | 0.1119 | 0.346820369 |
| SL1344_RS02750_ | 2.208905277 | 2.185879348 | 2.241658947 | 0.025568017 | 0.150215294 | 0.166262475 | 0.431045972 | 0.742767768 | 0.884270261 | 0.8396 | 0.924723913 |
| SL1344_RS02760_ | -4.111151975 | -4.029525886 | -4.215445532 | -0.178863259 | 0.946663219 | -0.181148939 | 0.415722855 | 0.623205636 | 0.807274397 | 0.7359 | 0.861749001 |
| SL1344_RS02765_ | -4.224477445 | -4.243842714 | -4.214388826 | -0.018215529 | 0.845642337 | -0.020880355 | 0.490339774 | 0.620632008 | 0.804693777 | 0.7374 | 0.86284295 |
| SL1344_RS02770_ | -1.972946589 | -1.888740481 | -2.023527092 | -0.080899118 | 0.483160034 | -0.150482144 | 0.438000002 | 0.691801759 | 0.852578315 | 0.7672 | 0.885466816 |
| SL1344_RS02775_ | -2.215568064 | -2.167028029 | -2.262798885 | -0.089178925 | 0.353468327 | -0.231203419 | 0.404000004 | 0.528596953 | 0.730497123 | 0.6341 | 0.787442483 |
| SL1344_RS02790_ | -4.904573445 | -5.210205582 | -4.227362475 | 1.124901103 | 1.535465645 | 0.730562267 | 0.217188556 | 0.331230233 | 0.606444647 | 0.4432 | 0.658341335 |
| SL1344_RS02795_ | -6.309897488 | -6.890558118 | -5.803641909 | 1.042776693 | 1.554361263 | 0.596862918 | 0.232666681 | 0.362969224 | 0.599039687 | 0.4571 | 0.658964743 |
| SL1344_RS02800_ | -2.630209396 | -2.682339638 | -2.525445165 | 0.110052768 | 0.680395914 | 0.136914889 | 0.438000002 | 0.716044147 | 0.866942641 | 0.8036 | 0.906220902 |
| SL1344_RS02805_ | -0.939557816 | -0.647563353 | -1.296434854 | -0.706844595 | 0.921913374 | -0.732330685 | 0.241333346 | 0.304074231 | 0.601752367 | 0.4405 | 0.689566223 |
| SL1344_RS02815_ | -0.958572929 | -0.876707763 | -1.053638296 | -0.321310294 | 0.739839874 | -0.458260957 | 0.320666674 | 0.44064832 | 0.706353993 | 0.649 | 0.820948521 |
| SL1344_RS02820_ | -1.365347539 | -0.824317728 | -1.665551375 | -0.697550318 | 0.610340796 | -1.128974642 | 0.1352432 | 0.16615065 | 0.446391819 | 0.2431 | 0.51148861 |
| SL1344_RS02825_ | -2.856337672 | -2.578047298 | -3.174256056 | -0.889702979 | 0.986278735 | -0.916854737 | 0.14533336 | 0.208092903 | 0.491858638 | 0.3041 | 0.544858828 |
| SL1344_RS02830_ | -4.330188808 | -4.705142129 | -4.128089417 | 0.603298673 | 1.044919904 | 0.537506301 | 0.248666679 | 0.377615363 | 0.639573001 | 0.5046 | 0.698484915 |
| SL1344_RS02835_ | -5.459080724 | -5.289889632 | -5.593658537 | -0.108838598 | 1.838603424 | -0.042889997 | 0.471019322 | 0.690475949 | 0.85249395 | 0.8035 | 0.902991179 |
| SL1344_RS02840_ | -0.691960898 | -0.770522008 | -0.598690528 | 0.139642339 | 0.441773251 | 0.287506223 | 0.381079285 | 0.664101216 | 0.838755746 | 0.7954 | 0.899660891 |
| SL1344_RS02845_ | -1.397103808 | -1.337218637 | -1.478536726 | -0.118574305 | 0.356859043 | -0.314609327 | 0.361333339 | 0.580305343 | 0.778539895 | 0.6953 | 0.83654624 |
| SL1344_RS02850_ | -1.634251234 | -1.585105248 | -1.673630838 | -0.100098939 | 0.350433845 | -0.265022817 | 0.381333338 | 0.566679121 | 0.764912858 | 0.6738 | 0.820258472 |
| SL1344_RS02855_ | -5.793186169 | -5.759622231 | -5.842337872 | -0.172767049 | 1.607569923 | -0.094215723 | 0.452365092 | 0.611966617 | 0.800434463 | 0.7225 | 0.851854601 |
| SL1344_RS02860_ | -3.627535987 | -3.505089537 | -3.741649939 | -0.22262003 | 0.556060935 | -0.383692239 | 0.321333341 | 0.457024115 | 0.675158964 | 0.5547 | 0.730200608 |
| SL1344_RS02865_ | 2.343923781 | 2.704375412 | 1.820042042 | -0.994583203 | 0.403391548 | -2.432895623 | 4.68E-05 | 0.016423927 | 0.158873132 | 0.1 | 0.330257709 |
| SL1344_RS02870_ | 1.356078715 | 1.431735092 | 1.186094499 | -0.233570881 | 0.151499948 | -1.40960434 | 0.069287204 | 0.115754527 | 0.368009757 | 0.1802 | 0.427877852 |
| SL1344_RS02875_ | 0.710988375 | 0.49181459 | 1.111272887 | 0.567626225 | 0.255142221 | 2.110936935 | 0.009327606 | 0.041129528 | 0.231754984 | 0.1065 | 0.33978351 |
| SL1344_RS02880_ | -0.108623089 | -0.059197984 | -0.159727542 | -0.031951478 | 0.376020399 | -0.090839188 | 0.463333335 | 0.774897462 | 0.90226899 | 0.8718 | 0.939969381 |
| SL1344_RS02885_ | 1.498872573 | 1.426637758 | 1.562820229 | 0.049847672 | 0.316634791 | 0.160031625 | 0.442000002 | 0.852485065 | 0.941140671 | 0.8977 | 0.956416706 |
| SL1344_RS02890_ | -2.97314566 | -3.00962371 | -2.924154089 | 0.088283339 | 0.433565812 | 0.182784271 | 0.415333337 | 0.51531982 | 0.718447653 | 0.6212 | 0.777316249 |
| SL1344_RS02895_ | -3.556265045 | -3.590937841 | -3.524712524 | -0.069166519 | 0.846282192 | -0.076052489 | 0.466355764 | 0.682311587 | 0.846071412 | 0.8043 | 0.900873876 |
| SL1344_RS02900_ | -4.083587124 | -3.844672886 | -4.265232236 | -0.214061837 | 1.284902674 | -0.149988235 | 0.433044639 | 0.729263232 | 0.87626334 | 0.8179 | 0.913891274 |
| SL1344_RS02905_ | -4.52312059 | -4.272776268 | -4.775499492 | -0.507034192 | 0.899588957 | -0.510938116 | 0.276482355 | 0.415111997 | 0.65244453 | 0.5283 | 0.715933745 |
| SL1344_RS02915_ | -5.210924026 | -4.999232988 | -5.615861949 | -0.747316684 | 1.902091463 | -0.338476406 | 0.284477025 | 0.398054218 | 0.671792382 | 0.5299 | 0.712416426 |
| SL1344_RS02920_ | 2.25888651 | 2.186292107 | 2.334757293 | 0.15519695 | 0.154369877 | 0.983829845 | 0.1352432 | 0.197976881 | 0.482879835 | 0.26 | 0.520266631 |
| SL1344_RS02925_ | -0.550598037 | -0.531664833 | -0.563351671 | -0.043428275 | 0.220702244 | -0.177006436 | 0.421333336 | 0.571683763 | 0.769547905 | 0.6917 | 0.831446002 |
| SL1344_RS02930_ | 0.606136488 | 0.561265344 | 0.703083089 | 0.101301988 | 0.233173625 | 0.363172834 | 0.362425055 | 0.68473636 | 0.851171611 | 0.7436 | 0.882206854 |
| SL1344_RS02935_ | -1.43497592 | -1.381548703 | -1.495366457 | -0.114418552 | 0.268803708 | -0.385412789 | 0.33533334 | 0.472279218 | 0.691618105 | 0.5772 | 0.750559294 |
| SL1344_RS02940_ | -1.239839101 | -1.358390707 | -1.135617564 | 0.203664656 | 0.337751399 | 0.569262684 | 0.264000011 | 0.43000347 | 0.669324408 | 0.5472 | 0.735943485 |

| | | | | | | | | | | | |
|---|---|---|---|---|---|---|---|---|---|---|---|
| SL1344_RS02945_ | -1.865360104 | -1.835055812 | -1.911508132 | -0.060833083 | 0.364441317 | -0.152062323 | 0.42504997 | 0.60651928 | 0.794638778 | 0.7209 | 0.852440004 |
| SL1344_RS02950_ | -3.485201665 | -3.286850292 | -3.641358445 | -0.412690615 | 0.665558658 | -0.585492147 | 0.266666678 | 0.401535241 | 0.642725766 | 0.5138 | 0.703701463 |
| SL1344_RS02955_ | 0.248198044 | 0.53519292 | 0.080603279 | -0.388299764 | 0.356131618 | -1.051246166 | 0.143904091 | 0.202867951 | 0.490975944 | 0.2751 | 0.541556721 |
| SL1344_RS02960_ | 0.176384977 | 0.354441979 | -0.236811878 | -0.746631654 | 0.515908834 | -1.46953051 | 0.020653117 | 0.082482297 | 0.327658686 | 0.1236 | 0.362094352 |
| SL1344_RS02965_ | -3.540165941 | -2.90215416 | -4.072498123 | -1.109033023 | 0.936223309 | -1.146866442 | 0.108594308 | 0.161878785 | 0.430449099 | 0.2369 | 0.492733921 |
| SL1344_RS02975_ | 1.604789477 | 1.410234691 | 1.811188713 | 0.415985547 | 0.137760482 | 3.063820204 | 4.68E-05 | 0.021323667 | 0.16703347 | 0.1 | 0.330257709 |
| SL1344_RS02980_ | -0.742617162 | -0.235672329 | -1.1167009 | -0.909983863 | 0.338313769 | -2.645785283 | 0.00067292 | 0.022197333 | 0.171359949 | 0.1002 | 0.330551659 |
| SL1344_RS02985_ | 0.525537247 | 0.947305417 | -0.066027534 | -0.986200706 | 0.437297772 | -2.172458088 | 4.68E-05 | 0.059303518 | 0.282907802 | 0.1002 | 0.330557578 |
| SL1344_RS02990_ | 0.216979476 | 0.601829964 | -0.273389534 | -0.93951418 | 0.276958971 | -3.469559962 | 4.68E-05 | 0.008695909 | 0.110862453 | 0.1 | 0.330257709 |
| SL1344_RS02995_ | 0.360699355 | 0.757635505 | -0.237660076 | -0.989397463 | 0.189889556 | -5.130889464 | 4.68E-05 | 0.007037052 | 0.094719051 | 0.1 | 0.330257709 |
| SL1344_RS03000_ | -0.27964309 | -0.113418085 | -0.415496601 | -0.345138635 | 0.355246665 | -0.957222259 | 0.134000029 | 0.207123735 | 0.491309774 | 0.28 | 0.535880505 |
| SL1344_RS03005_ | -0.095573279 | 0.317146623 | -0.342404474 | -0.625378637 | 0.264205566 | -2.211284512 | 0.003332494 | 0.035177235 | 0.211030075 | 0.1033 | 0.334972203 |
| SL1344_RS03020_ | -0.71667788 | -0.475857292 | -1.051987338 | -0.596465364 | 0.298978412 | -1.94600143 | 0.008661422 | 0.059892824 | 0.266911367 | 0.108 | 0.340999935 |
| SL1344_RS03025_ | -1.576389425 | -1.402585042 | -1.776414044 | -0.371655348 | 0.336892839 | -1.023506648 | 0.139906757 | 0.213054787 | 0.466888952 | 0.3017 | 0.537477878 |
| SL1344_RS03030_ | 4.026832894 | 4.285762759 | 3.891136518 | -0.17069322 | 0.342254089 | -0.722740173 | 0.311792147 | 0.437732225 | 0.711856022 | 0.6455 | 0.836517721 |
| SL1344_RS03035_ | 0.728971792 | 0.824393819 | 0.67350518 | -0.111052105 | 0.320710734 | -0.290081713 | 0.39307129 | 0.765856165 | 0.89815619 | 0.8339 | 0.926953816 |
| SL1344_RS03040_ | -0.840402007 | -0.690525362 | -0.92775741 | -0.271493257 | 0.350132583 | -0.731382566 | 0.228514338 | 0.339766911 | 0.605603076 | 0.4607 | 0.678731729 |
| SL1344_RS03045_ | -0.303854019 | -0.18497319 | -0.359551822 | -0.170932961 | 0.295136631 | -0.535319024 | 0.285333343 | 0.461357084 | 0.701919514 | 0.5716 | 0.763384964 |
| SL1344_RS03050_ | -1.024714944 | -1.10234328 | -0.890823122 | 0.212459929 | 0.297589701 | 0.668752165 | 0.244000013 | 0.37529173 | 0.624121262 | 0.493 | 0.694747719 |
| SL1344_RS03055_ | -0.727541499 | -1.035945099 | -0.553169727 | 0.472036556 | 0.301563126 | 1.483898648 | 0.058627656 | 0.102096634 | 0.343619435 | 0.1648 | 0.41081231 |
| SL1344_RS03060_ | -0.16851888 | -0.356352903 | -0.06503947 | 0.322332526 | 0.361549039 | 0.841912363 | 0.167888097 | 0.245605087 | 0.537532251 | 0.3289 | 0.583057447 |
| SL1344_RS03065_ | 1.471512213 | 1.398117923 | 1.5551797 | 0.162605702 | 0.114044774 | 1.361150998 | 0.070619647 | 0.117691515 | 0.354718193 | 0.1831 | 0.424659754 |
| SL1344_RS03070_ | 4.621551501 | 4.583076726 | 4.661695396 | 0.068962269 | 0.189621546 | 0.276333424 | 0.445036644 | 0.723928368 | 0.883568996 | 0.9886 | 0.995064279 |
| SL1344_RS03075_ | 2.378980429 | 2.416249062 | 2.346903065 | -0.08276541 | 0.142483335 | -0.589104176 | 0.269333344 | 0.408625175 | 0.670502941 | 0.5349 | 0.738514438 |
| SL1344_RS03080_ | -5.074900973 | -5.214913653 | -4.932779048 | 0.585082222 | 1.612651785 | 0.355179532 | 0.359093943 | 0.550774584 | 0.762216735 | 0.7245 | 0.848439989 |
| SL1344_RS03085_ | -1.176900939 | -1.324351027 | -1.000274345 | 0.319988536 | 0.255574317 | 1.165496254 | 0.098000042 | 0.156875122 | 0.398249501 | 0.2341 | 0.472890918 |
| SL1344_RS03090_ | -2.604010956 | -2.484142996 | -2.702890989 | -0.282227668 | 0.662868086 | -0.413350354 | 0.33933334 | 0.534250015 | 0.753341663 | 0.6718 | 0.822829891 |
| SL1344_RS03095_ | 1.04296137 | 1.116497214 | 0.982854109 | -0.125966033 | 0.136461664 | -0.883193666 | 0.169886764 | 0.257263848 | 0.514283606 | 0.3532 | 0.582468448 |
| SL1344_RS03100_ | 4.552457922 | 4.578182984 | 4.522805547 | -0.059503863 | 0.183956371 | -0.29296541 | 0.389333338 | 0.678320141 | 0.859905863 | 0.85 | 0.935532027 |
| SL1344_RS03105_ | -1.76639867 | -1.889306916 | -1.671974757 | 0.217870231 | 0.323568365 | 0.618791401 | 0.244000013 | 0.369606212 | 0.607698853 | 0.4698 | 0.6690061 |
| SL1344_RS03110_ | 0.975238629 | 1.186557647 | 0.913708728 | -0.272236959 | 0.224107667 | -1.03833142 | 0.16055965 | 0.211151507 | 0.498239597 | 0.3159 | 0.571099159 |
| SL1344_RS03115_ | 2.683423609 | 2.688152461 | 2.680261682 | -0.001185601 | 0.307938526 | -0.004992384 | 0.498334444 | 0.838360539 | 0.935969048 | 0.9682 | 0.986342485 |
| SL1344_RS03120_ | 0.31361976 | 0.632246866 | 0.123569951 | -0.490272873 | 0.210531052 | -2.187755819 | 0.005330642 | 0.038650926 | 0.218088147 | 0.1068 | 0.339602129 |
| SL1344_RS03125_ | -5.12249139 | -5.090450495 | -5.149529722 | -0.002481583 | 1.301823321 | -0.001672922 | 0.498666667 | 0.636044639 | 0.816701463 | 0.7467 | 0.86837917 |
| SL1344_RS03130_ | -4.815421202 | -4.249400553 | -5.258217367 | -0.919346891 | 1.061934358 | -0.819177453 | 0.170552986 | 0.287825436 | 0.539390958 | 0.3731 | 0.603116288 |
| SL1344_RS03135_ | -3.799430127 | -4.1100912 | -3.512598492 | 0.645666175 | 0.797925743 | 0.737015619 | 0.200000017 | 0.300270507 | 0.56119021 | 0.4003 | 0.623956593 |
| SL1344_RS03140_ | -4.508310779 | -5.097439291 | -4.015419662 | 0.990576863 | 1.124960779 | 0.828858524 | 0.161333357 | 0.260089383 | 0.530057861 | 0.336 | 0.576090427 |
| SL1344_RS03145_ | -6.446253873 | -6.124976258 | -7.237927723 | -1.102307 | 2.233651962 | -0.420706678 | 0.290473027 | 0.406522182 | 0.664866768 | 0.5494 | 0.736964198 |
| SL1344_RS03150_ | -4.638480387 | -4.63509274 | -4.643631052 | 0.168791263 | 1.378824266 | 0.118553363 | 0.460000001 | 0.696219631 | 0.856360198 | 0.8201 | 0.911244607 |
| SL1344_RS03165_ | -1.739947065 | -1.816376459 | -1.672318973 | 0.146778646 | 0.278317847 | 0.492042764 | 0.286666676 | 0.404663729 | 0.627498844 | 0.5019 | 0.688891852 |
| SL1344_RS03175_ | 7.533819551 | 7.480320537 | 7.702759269 | 0.222220997 | 0.166670718 | 1.338085661 | 0.006000767 | 0.12835435 | 0.409850462 | 0.1059 | 0.338902259 |
| SL1344_RS03180_ | -0.914216307 | -0.842443809 | -0.999804142 | -0.149020417 | 0.245944653 | -0.574598463 | 0.262000011 | 0.406040137 | 0.63687182 | 0.5133 | 0.703841113 |
| SL1344_RS03195_ | 3.335207934 | 3.204563958 | 3.464521936 | 0.264530928 | 0.070869556 | 3.795646953 | 4.68E-05 | 0.010491953 | 0.116986091 | 0.1 | 0.330257709 |
| SL1344_RS03200_ | -0.470624179 | -0.914112225 | -0.03928594 | 0.878441432 | 0.22310254 | 3.913349553 | 4.68E-05 | 0.007777444 | 0.101370727 | 0.1 | 0.330257709 |
| SL1344_RS03205_ | 1.397250953 | 1.580543799 | 1.339675713 | -0.179114192 | 0.24118229 | -0.622848671 | 0.29113925 | 0.470313566 | 0.723219593 | 0.6004 | 0.802295003 |
| SL1344_RS03210_ | 3.52043342 | 3.538686406 | 3.500823125 | -0.032867426 | 0.435168825 | -0.070548165 | 0.468000001 | 0.885909645 | 0.957672782 | 0.9955 | 0.998094132 |
| SL1344_RS03215_ | 3.203008805 | 3.029981673 | 3.227905184 | 0.183274772 | 0.126355617 | 1.016631346 | 0.224000015 | 0.248124342 | 0.545312814 | 0.4258 | 0.68105266 |
| SL1344_RS03225_ | 1.157980553 | 1.144183417 | 1.185042397 | 0.04490661 | 0.150903655 | 0.280288099 | 0.381079285 | 0.591914605 | 0.786710451 | 0.7126 | 0.848553674 |
| SL1344_RS03230_ | 2.275426578 | 2.22088968 | 2.30979468 | 0.094848034 | 0.096303927 | 0.921061712 | 0.149333359 | 0.243696417 | 0.511446953 | 0.3267 | 0.559220669 |
| SL1344_RS03235_ | 2.145454695 | 2.097520803 | 2.260938353 | 0.147679605 | 0.194188135 | 0.64658782 | 0.28847436 | 0.414791775 | 0.684589149 | 0.5851 | 0.781865702 |
| SL1344_RS03240_ | 1.729352102 | 1.708711945 | 1.753310995 | 0.102895819 | 0.287621197 | 0.351367096 | 0.363333339 | 0.588313763 | 0.802014998 | 0.7569 | 0.886654237 |
| SL1344_RS03245_ | 0.073779797 | -0.032730406 | 0.208200749 | 0.18457947 | 0.329214455 | 0.540931057 | 0.281145913 | 0.473618405 | 0.718237008 | 0.5785 | 0.771008236 |
| SL1344_RS03250_ | -0.272741498 | -0.302406901 | -0.244221073 | 0.062683839 | 0.187169432 | 0.315433057 | 0.358427721 | 0.498200071 | 0.710471461 | 0.608 | 0.771146733 |

| | | | | | | | | | | |
|---|---|---|---|---|---|---|---|---|---|---|
| SL1344_RS03255_ | 0.240937524 | 0.254060952 | 0.233827294 | -0.049524949 | 0.232290582 | -0.179779482 | 0.415333337 | 0.612016182 | 0.803597658 | 0.7398 | 0.865202086 |
| SL1344_RS03260_ | 0.648504049 | 0.623665265 | 0.681573784 | 0.070376726 | 0.167042535 | 0.399959828 | 0.327333341 | 0.488729612 | 0.713192531 | 0.6237 | 0.781529507 |
| SL1344_RS03265_ | 2.466456086 | 2.428013575 | 2.506597459 | 0.060199825 | 0.162344618 | 0.375435354 | 0.341105936 | 0.569782488 | 0.784111536 | 0.6938 | 0.845596869 |
| SL1344_RS03270_ | 2.801402989 | 2.709755151 | 2.868938346 | 0.147679285 | 0.118650956 | 1.211273456 | 0.09726853 | 0.133902979 | 0.397194267 | 0.2034 | 0.461575293 |
| SL1344_RS03275_ | 4.493184937 | 4.349567849 | 4.527051669 | 0.168349907 | 0.091498525 | 1.561975271 | 0.045303228 | 0.090752672 | 0.342610014 | 0.1493 | 0.394331702 |
| SL1344_RS03280_ | -5.017919986 | -5.23601136 | -4.786853612 | 0.511959101 | 1.294921913 | 0.367938266 | 0.34133334 | 0.515651086 | 0.732604693 | 0.6285 | 0.789977228 |
| SL1344_RS03285_ | -2.301448774 | -2.476530015 | -2.212383372 | 0.231455245 | 0.512020989 | 0.410403065 | 0.33933334 | 0.551324983 | 0.762610337 | 0.6635 | 0.824265256 |
| SL1344_RS03290_ | -1.358782669 | -1.250422689 | -1.527375814 | -0.28193205 | 0.368850075 | -0.709515673 | 0.213333349 | 0.318751338 | 0.583294647 | 0.4255 | 0.649102612 |
| SL1344_RS03295_ | 1.10205722 | 1.147556485 | 1.045608996 | -0.102510061 | 0.110937254 | -0.887846492 | 0.171333355 | 0.252688038 | 0.498318067 | 0.342 | 0.565760156 |
| SL1344_RS03300_ | 0.95915965 | 0.766858241 | 1.179291802 | 0.281352685 | 0.338490191 | 0.909902922 | 0.148666692 | 0.22882249 | 0.521954477 | 0.2678 | 0.538944378 |
| SL1344_RS03305_ | -2.028390053 | -2.028462532 | -2.027767755 | -0.009366878 | 0.357999924 | -0.023996503 | 0.491005996 | 0.580403301 | 0.773073745 | 0.6915 | 0.829084656 |
| SL1344_RS03310_ | -3.784772189 | -3.869339613 | -3.687817644 | 0.163169286 | 0.659629332 | 0.222624701 | 0.396000004 | 0.558579052 | 0.75563492 | 0.6702 | 0.814425844 |
| SL1344_RS03315_ | -2.235132248 | -2.354351163 | -2.147856775 | 0.191317504 | 0.468424421 | 0.378254371 | 0.335776156 | 0.518541851 | 0.735431319 | 0.6329 | 0.795875185 |
| SL1344_RS03320_ | -3.294806417 | -3.314912665 | -3.275818909 | 0.024848128 | 0.50832381 | 0.042141107 | 0.482666667 | 0.552550617 | 0.75128703 | 0.6542 | 0.801560684 |
| SL1344_RS03325_ | -1.147878123 | -1.198196471 | -1.121718575 | 0.067151887 | 0.348285353 | 0.173331847 | 0.426382415 | 0.652284664 | 0.82634643 | 0.7619 | 0.879379864 |
| SL1344_RS03330_ | 0.228537078 | 0.04025399 | 0.287459681 | 0.252719249 | 0.382010136 | 0.616228653 | 0.271818798 | 0.41754386 | 0.686729227 | 0.5447 | 0.76385453 |
| SL1344_RS03335_ | 2.003185957 | 2.024130052 | 1.984370553 | -0.033427384 | 0.144788074 | -0.223178651 | 0.404000004 | 0.664481661 | 0.83720527 | 0.7779 | 0.88646843 |
| SL1344_RS03340_ | 1.544062853 | 1.561026512 | 1.521916263 | -0.041432058 | 0.116045748 | -0.325202409 | 0.350666673 | 0.521362072 | 0.733160981 | 0.6319 | 0.789735245 |
| SL1344_RS03350_ | 1.987058162 | 2.121497549 | 1.940087896 | -0.182436576 | 0.555800903 | -0.263643639 | 0.351099273 | 0.930483921 | 0.97486374 | 0.7402 | 0.889608345 |
| SL1344_RS03355_ | 2.351824755 | 2.384848957 | 2.32128123 | -0.061797193 | 0.610937566 | -0.089866404 | 0.460359762 | 0.910649669 | 0.967422885 | 0.9997 | 0.999883855 |
| SL1344_RS03360_ | 7.093560186 | 7.095921953 | 7.090943452 | -0.171051174 | 0.329219884 | -0.336043071 | 0.377748173 | 0.415155068 | 0.696603092 | 0.7954 | 0.912267383 |
| SL1344_RS03365_ | 3.131583629 | 3.098566147 | 3.160229364 | 0.075288529 | 0.10796368 | 0.675598979 | 0.219333349 | 0.324729492 | 0.605642939 | 0.4324 | 0.660903348 |
| SL1344_RS03370_ | 3.289094198 | 3.278429735 | 3.300310692 | 0.028795712 | 0.218291613 | 0.129582057 | 0.444000002 | 0.793147509 | 0.914699467 | 0.9193 | 0.963709891 |
| SL1344_RS03375_ | 2.312213417 | 2.276631105 | 2.345060625 | 0.054713523 | 0.205188955 | 0.261242935 | 0.409726852 | 0.719029041 | 0.874700719 | 0.8699 | 0.940129048 |
| SL1344_RS03380_ | 2.835211305 | 2.813284242 | 2.858400717 | 0.054563089 | 0.274361652 | 0.181619098 | 0.438374419 | 0.70101187 | 0.870678558 | 0.9469 | 0.976690429 |
| SL1344_RS03385_ | 3.291485268 | 3.187638065 | 3.376945446 | 0.197754532 | 0.199022069 | 1.008976537 | 0.111925419 | 0.192887873 | 0.490225937 | 0.212 | 0.482050431 |
| SL1344_RS03395_ | -1.638205929 | -1.595747396 | -1.659162277 | -0.013836926 | 0.475323243 | -0.023313918 | 0.489673551 | 0.702423124 | 0.861715161 | 0.8134 | 0.912013953 |
| SL1344_RS03400_ | 4.269806589 | 4.277478255 | 4.261077488 | -0.020945982 | 0.764021611 | -0.027154947 | 0.473017989 | 0.865997822 | 0.950613536 | 1 | 1 |
| SL1344_RS03405_ | 3.563134434 | 3.776935067 | 3.543982965 | -0.23304798 | 0.216146041 | -0.435386413 | 0.33333334 | 0.783733933 | 0.913714543 | 0.7 | 0.872542119 |
| SL1344_RS03410_ | 3.666049802 | 3.653847432 | 3.678258565 | -0.011855802 | 0.28304116 | -0.052103975 | 0.471685544 | 0.858532309 | 0.945242864 | 0.9577 | 0.982005004 |
| SL1344_RS03415_ | 4.729419377 | 4.721924199 | 4.734675763 | 0.009015033 | 0.578652432 | 0.014654874 | 0.478347769 | 0.792084192 | 0.91867836 | 0.982 | 0.992314621 |
| SL1344_RS03420_ | 5.543017221 | 5.621734638 | 5.471496759 | -0.146575682 | 0.317794833 | -0.324319483 | 0.391738845 | 0.773266956 | 0.909917241 | 0.8443 | 0.933785138 |
| SL1344_RS03425_ | 5.556589872 | 5.628914753 | 5.495283083 | -0.044687632 | 0.161059126 | -0.242529026 | 0.387333338 | 0.816306053 | 0.929069747 | 0.8302 | 0.927866444 |
| SL1344_RS03430_ | 4.890231223 | 4.989266305 | 4.879578627 | -0.10968492 | 0.326364879 | -0.274608998 | 0.33333334 | 0.951450758 | 0.982893427 | 0.7 | 0.872542119 |
| SL1344_RS03440_ | 2.223881667 | 2.229280689 | 2.219363523 | 0.008688065 | 0.341023785 | 0.031053358 | 0.485333334 | 0.879559622 | 0.953554054 | 0.919 | 0.962977347 |
| SL1344_RS03445_ | 2.010388905 | 2.008827646 | 2.011251638 | -0.010740162 | 0.170443479 | -0.061671219 | 0.478347769 | 0.769418227 | 0.899022102 | 0.8802 | 0.946583022 |
| SL1344_RS03450_ | 2.514186515 | 2.57487418 | 2.462501145 | -0.104596498 | 0.269993814 | -0.28304424 | 0.39840107 | 0.832306734 | 0.932872031 | 0.8824 | 0.949904123 |
| SL1344_RS03455_ | 1.597308341 | 1.462296833 | 1.643613439 | 0.165637519 | 0.255762672 | 0.374888002 | 0.351333339 | 0.84372194 | 0.938064372 | 0.7498 | 0.893699965 |
| SL1344_RS03460_ | 4.312783645 | 4.237361747 | 4.378133122 | 0.136708217 | 0.190811437 | 0.585984612 | 0.242000013 | 0.359183906 | 0.651503986 | 0.4741 | 0.705002955 |
| SL1344_RS03465_ | 3.751425353 | 3.782520604 | 3.737488406 | -0.051050286 | 0.080136383 | -0.609916208 | 0.260000012 | 0.378190195 | 0.651576679 | 0.5336 | 0.729967572 |
| SL1344_RS03475_ | 0.059277555 | -0.252970824 | 0.584352486 | 0.827019949 | 0.253886462 | 3.097636979 | 4.68E-05 | 0.0202419 | 0.165142595 | 0.1 | 0.330257709 |
| SL1344_RS03480_ | -0.809343327 | -0.810292269 | -0.808741119 | 0.00033861 | 0.217337639 | 0.001277609 | 0.498666667 | 0.558096595 | 0.756986213 | 0.6737 | 0.815692981 |
| SL1344_RS03490_ | -1.23597035 | -1.262437047 | -1.221749801 | 0.007250323 | 0.328967322 | 0.019675353 | 0.491672219 | 0.659694332 | 0.831388672 | 0.7586 | 0.879417936 |
| SL1344_RS03495_ | 0.512907003 | 0.475632425 | 0.568084405 | 0.094789075 | 0.176541631 | 0.525578509 | 0.297333342 | 0.453765529 | 0.684467971 | 0.5704 | 0.751573853 |
| SL1344_RS03505_ | 3.647053177 | 3.623921519 | 3.780226904 | 0.148337839 | 0.234965636 | 0.695616135 | 0.220000015 | 0.309573877 | 0.60668395 | 0.3987 | 0.678936796 |
| SL1344_RS03510_ | 0.642672787 | 0.819246958 | 0.471781679 | 0.023274847 | 0.669995452 | 0.034319495 | 0.484666667 | 0.863447303 | 0.946887239 | 0.9856 | 0.993555559 |
| SL1344_RS03515_ | 1.460516314 | 1.372494826 | 1.631401922 | 0.251816426 | 0.130228591 | 1.78913146 | 0.0346437 | 0.069644147 | 0.284543004 | 0.133 | 0.372385397 |
| SL1344_RS03520_ | 2.532261517 | 2.452595473 | 2.611916751 | 0.192687555 | 0.147488337 | 1.309537581 | 0.031978821 | 0.123534614 | 0.390478431 | 0.139 | 0.380465527 |
| SL1344_RS03525_ | 4.750246437 | 4.608908751 | 4.82401098 | 0.208276346 | 0.064378578 | 2.944770803 | 4.68E-05 | 0.024565969 | 0.182756713 | 0.1 | 0.330257709 |
| SL1344_RS03530_ | -1.079745846 | -1.582248481 | -0.594606368 | 1.014653575 | 0.307872609 | 3.302697637 | 4.68E-05 | 0.011971611 | 0.125358482 | 0.1 | 0.330257709 |
| SL1344_RS03535_ | -0.304977369 | -1.327448174 | 0.361756518 | 1.675377535 | 0.308030384 | 5.247457011 | 4.68E-05 | 0.009206195 | 0.112651715 | 0.1 | 0.330257709 |
| SL1344_RS03540_ | 0.577001654 | -0.276323217 | 1.386343822 | 1.655232862 | 0.171122298 | 9.701576847 | 4.68E-05 | 0.000715138 | 0.035170415 | 0.1 | 0.330257709 |
| SL1344_RS03545_ | -5.958907396 | -5.835953919 | -6.23141465 | -0.51872295 | 2.13470043 | -0.170348392 | 0.380666671 | 0.521717046 | 0.753003142 | 0.7074 | 0.8433138 |

| | | | | | | | | | | | |
|---|---|---|---|---|---|---|---|---|---|---|---|
| SL1344_RS03550_ | 0.28342614 | 0.319946287 | 0.174157126 | -0.152352793 | 0.297179399 | -0.477164888 | 0.313124592 | 0.426974122 | 0.694622328 | 0.6246 | 0.798909029 |
| SL1344_RS03555_ | 0.904242847 | 0.975113683 | 0.828762868 | -0.132681707 | 0.168166553 | -0.72908811 | 0.214523667 | 0.329207611 | 0.591798732 | 0.4274 | 0.654278472 |
| SL1344_RS03560_ | -1.825301792 | -1.677145773 | -1.957511722 | -0.289907315 | 0.36009238 | -0.762181595 | 0.191205881 | 0.302468595 | 0.550902768 | 0.4017 | 0.617050497 |
| SL1344_RS03565_ | -2.498747176 | -2.324487881 | -2.640983322 | -0.313898743 | 0.382814817 | -0.744426765 | 0.208666683 | 0.316514482 | 0.561565675 | 0.4141 | 0.627145358 |
| SL1344_RS03570_ | -3.648152408 | -3.482747194 | -3.776884756 | -0.263612321 | 0.630519462 | -0.393952404 | 0.330000007 | 0.492906114 | 0.708137258 | 0.6016 | 0.769379407 |
| SL1344_RS03575_ | -7.844058325 | -8.476867698 | -7.361074966 | 0.977455396 | 3.775756894 | 0.223679725 | 0.37441706 | 0.558537242 | 0.766546693 | 0.6829 | 0.826222777 |
| SL1344_RS03580_ | 0.440651384 | 0.50840694 | 0.338487127 | -0.172439286 | 0.431044394 | -0.386202472 | 0.33533334 | 0.589349941 | 0.803613331 | 0.6839 | 0.846002883 |
| SL1344_RS03585_ | -0.3411196 | -0.577242913 | -0.163393692 | 0.417138406 | 0.206438422 | 2.00906607 | 0.012658588 | 0.056045563 | 0.249719748 | 0.1158 | 0.349990027 |
| SL1344_RS03590_ | -0.876540245 | -0.794086366 | -0.979030965 | -0.174547347 | 0.249310861 | -0.644323835 | 0.232511673 | 0.349463449 | 0.595687803 | 0.4596 | 0.663670073 |
| SL1344_RS03595_ | 1.313898791 | 1.213035537 | 1.410041764 | 0.096816915 | 0.270057233 | 0.385055629 | 0.354000006 | 0.586653928 | 0.797144136 | 0.7229 | 0.859645075 |
| SL1344_RS03600_ | 1.008778725 | 0.916783389 | 1.039209383 | 0.115078223 | 0.198419899 | 0.517550103 | 0.318454371 | 0.492520119 | 0.729218725 | 0.6471 | 0.816670328 |
| SL1344_RS03605_ | 1.393609777 | 1.259617133 | 1.427794079 | 0.165136495 | 0.25017611 | 0.586235734 | 0.314000008 | 0.440826542 | 0.705932241 | 0.645 | 0.825083803 |
| SL1344_RS03610_ | 1.500113041 | 1.503514314 | 1.49810463 | 0.013884622 | 0.161394183 | 0.078699003 | 0.463333335 | 0.654684683 | 0.834145548 | 0.7881 | 0.895511838 |
| SL1344_RS03615_ | -4.625579495 | -5.034391751 | -4.326707849 | 0.723941314 | 0.928113882 | 0.708194026 | 0.192666685 | 0.31152764 | 0.5642256 | 0.4106 | 0.624290359 |
| SL1344_RS03620_ | -0.47659621 | -0.67777321 | -0.327492973 | 0.350460814 | 0.201914211 | 1.65559257 | 0.027333498 | 0.080944424 | 0.293647153 | 0.1394 | 0.377444056 |
| SL1344_RS03625_ | -4.905327875 | -5.304761786 | -4.54264352 | 0.768309562 | 0.88098968 | 0.831215804 | 0.166666689 | 0.269492062 | 0.506274162 | 0.3602 | 0.575943433 |
| SL1344_RS03630_ | -3.052717709 | -2.856390992 | -3.22096061 | -0.386424298 | 0.474750594 | -0.754379981 | 0.199866773 | 0.314820865 | 0.555621667 | 0.4074 | 0.616887709 |
| SL1344_RS03640_ | -4.275685151 | -4.225638482 | -4.322692035 | -0.081026099 | 0.686861895 | -0.1075855 | 0.450000002 | 0.525486058 | 0.732301369 | 0.6431 | 0.795183043 |
| SL1344_RS03645_ | -3.791991535 | -3.692832112 | -3.886967211 | -0.199623096 | 0.690153372 | -0.270215214 | 0.376000005 | 0.541691778 | 0.747161365 | 0.6521 | 0.803152235 |
| SL1344_RS03650_ | -4.272254922 | -4.263207039 | -4.276420233 | 0.01991438 | 0.848216406 | 0.025467147 | 0.485676216 | 0.603491297 | 0.793522762 | 0.7199 | 0.850650142 |
| SL1344_RS03655_ | -4.403942047 | -4.472113002 | -4.270171963 | 0.212585177 | 0.983075339 | 0.18346329 | 0.41439041 | 0.635415506 | 0.814307017 | 0.7376 | 0.8634488 |
| SL1344_RS03660_ | -2.882902836 | -2.925928672 | -2.853228863 | 0.053668735 | 0.58184069 | 0.082107916 | 0.463690874 | 0.636609889 | 0.818498888 | 0.76 | 0.8777192 |
| SL1344_RS03665_ | -4.454568922 | -4.195445669 | -4.65026531 | -0.469462671 | 0.971431816 | -0.449304293 | 0.310666675 | 0.489282226 | 0.7088711 | 0.6013 | 0.768464867 |
| SL1344_RS03670_ | -2.757032334 | -2.57641887 | -2.935591068 | -0.347958589 | 0.430711189 | -0.78216467 | 0.189873436 | 0.284827839 | 0.52726237 | 0.3783 | 0.595743835 |
| SL1344_RS03675_ | -1.844230677 | -2.175276383 | -1.592174785 | 0.316193678 | 0.771287927 | 0.417730684 | 0.342000007 | 0.548653054 | 0.772553891 | 0.6806 | 0.833486126 |
| SL1344_RS03680_ | 0.984908986 | 0.992205057 | 0.973549118 | -0.016832234 | 0.137552459 | -0.107046761 | 0.453031314 | 0.607249863 | 0.793511064 | 0.7064 | 0.844132701 |
| SL1344_RS03685_ | 0.873981464 | 0.874317556 | 0.873686626 | -0.016952849 | 0.259350642 | -0.06033235 | 0.474666668 | 0.672502664 | 0.848302704 | 0.8408 | 0.925415953 |
| SL1344_RS03690_ | 6.285955232 | 6.297811103 | 6.199248466 | -0.104844362 | 0.165887167 | -0.677030004 | 0.220519669 | 0.284862385 | 0.587331972 | 0.4 | 0.680350104 |
| SL1344_RS03695_ | -3.461555439 | -3.41202362 | -3.503599644 | -0.033561694 | 1.411138503 | -0.026933478 | 0.485676216 | 0.773554934 | 0.901560056 | 0.8096 | 0.909594469 |
| SL1344_RS03705_ | 3.042650242 | 2.993064328 | 3.090388217 | 0.092587687 | 0.086960981 | 0.989222556 | 0.133244533 | 0.199864652 | 0.469002478 | 0.2846 | 0.535806012 |
| SL1344_RS03710_ | 4.90986462 | 5.156208248 | 4.831961581 | -0.222320144 | 0.421225334 | -0.464214776 | 0.33333334 | 0.656919923 | 0.850888126 | 0.7 | 0.872542119 |
| SL1344_RS03715_ | 5.635099551 | 5.594331024 | 5.679995461 | 0.080969655 | 0.452591478 | 0.105456919 | 0.446000002 | 0.753669401 | 0.900829701 | 1 | 1 |
| SL1344_RS03720_ | 6.718549754 | 6.785899616 | 6.617500271 | -0.179569628 | 0.138903037 | -1.448183287 | 0.001336769 | 0.112572677 | 0.385248703 | 0.1008 | 0.331419535 |
| SL1344_RS03725_ | 6.537450856 | 6.687522134 | 6.408504961 | -0.301594759 | 0.143185253 | -2.221469877 | 4.68E-05 | 0.022888044 | 0.185728083 | 0.1 | 0.330257709 |
| SL1344_RS03730_ | 5.873281175 | 6.111835788 | 5.658099427 | -0.444609227 | 0.211128588 | -1.930146432 | 4.68E-05 | 0.041160053 | 0.242870548 | 0.1 | 0.330257709 |
| SL1344_RS03735_ | 5.751913614 | 6.371266028 | 5.579855552 | -0.761888196 | 0.306469074 | -1.988819976 | 4.68E-05 | 0.04960913 | 0.264685347 | 0.1 | 0.330257709 |
| SL1344_RS03740_ | 2.696660629 | 3.036765014 | 2.276826135 | -0.679923666 | 0.242713562 | -2.93184016 | 4.68E-05 | 0.008815005 | 0.118304031 | 0.1 | 0.330257709 |
| SL1344_RS03745_ | 4.443627714 | 4.431605857 | 4.45077199 | 0.023328533 | 0.204343568 | 0.097592345 | 0.440000002 | 0.919499166 | 0.970694618 | 0.8893 | 0.952912917 |
| SL1344_RS03750_ | 3.156110795 | 3.125591697 | 3.186565986 | -0.085146457 | 0.376404293 | -0.221542491 | 0.431333336 | 0.792163855 | 0.916584015 | 0.9697 | 0.987112035 |
| SL1344_RS03755_ | 2.511693489 | 2.517567038 | 2.506253702 | 0.138852554 | 0.312545396 | 0.343586778 | 0.391738845 | 0.437811214 | 0.710776157 | 0.8434 | 0.925156573 |
| SL1344_RS03760_ | 2.965479151 | 2.928578065 | 3.105989192 | 0.144936242 | 0.173068604 | 0.615614133 | 0.320666674 | 0.528541225 | 0.767868071 | 0.6617 | 0.847388116 |
| SL1344_RS03765_ | 2.795171126 | 3.100626704 | 2.640544545 | -0.35918605 | 0.278620927 | -1.269982683 | 0.117255197 | 0.140572455 | 0.423579356 | 0.205 | 0.480885395 |
| SL1344_RS03770_ | 1.602493378 | 1.567634661 | 1.699241885 | 0.145810121 | 0.219950932 | 0.609270136 | 0.273333344 | 0.396455586 | 0.668553855 | 0.5306 | 0.750087104 |
| SL1344_RS03775_ | 2.635560797 | 2.52011535 | 2.814515832 | 0.301143857 | 0.154539945 | 1.987627768 | 0.004664544 | 0.040778812 | 0.233001082 | 0.1031 | 0.334757442 |
| SL1344_RS03780_ | 1.989937522 | 1.852155658 | 2.077609476 | 0.23810946 | 0.179962443 | 1.334480923 | 0.057961434 | 0.104470165 | 0.358315491 | 0.1615 | 0.411667783 |
| SL1344_RS03790_ | 4.721634814 | 4.705309784 | 4.766650919 | 0.137226585 | 0.221708544 | 0.622361563 | 0.243837455 | 0.302217404 | 0.60295899 | 0.4314 | 0.696340623 |
| SL1344_RS03795_ | 5.958549803 | 5.942307662 | 5.974112965 | 0.048904559 | 0.143134896 | 0.428208961 | 0.329113931 | 0.480930542 | 0.741139561 | 0.7072 | 0.873165755 |
| SL1344_RS03800_ | 5.215321265 | 5.216245195 | 5.214358902 | -0.028092437 | 0.181474049 | -0.187720373 | 0.409333337 | 0.650720906 | 0.845660968 | 0.8776 | 0.947835307 |
| SL1344_RS03805_ | 3.631148255 | 3.912445629 | 3.376846578 | -0.229192832 | 0.577204796 | -0.433609536 | 0.371333338 | 0.521673605 | 0.769436653 | 0.8089 | 0.917295553 |
| SL1344_RS03810_ | 4.129923052 | 4.274313071 | 3.981210812 | -0.142861305 | 0.380094042 | -0.314676989 | 0.430379749 | 0.943041956 | 0.979780552 | 0.9787 | 0.990914572 |
| SL1344_RS03815_ | 3.209828303 | 3.352500025 | 3.087644708 | -0.106932679 | 0.431280838 | -0.366601999 | 0.355333339 | 0.776743825 | 0.909864364 | 0.7606 | 0.897956127 |
| SL1344_RS03820_ | 3.264628443 | 3.369150793 | 3.153137008 | -0.065263556 | 0.451765598 | -0.180489752 | 0.399067292 | 0.734128557 | 0.889355014 | 0.8545 | 0.936483064 |
| SL1344_RS03825_ | 2.489016911 | 2.531090802 | 2.443538114 | -0.082265852 | 0.572952205 | -0.140176314 | 0.456666668 | 0.817480132 | 0.928709418 | 0.9949 | 0.997843339 |

| | | | | | | | | | | | |
|---|---|---|---|---|---|---|---|---|---|---|---|
| SL1344_RS03830_ | -1.693940112 | -1.77152617 | -1.622344869 | 0.123891105 | 0.381921861 | 0.30914794 | 0.37175217 | 0.575684072 | 0.774476311 | 0.6871 | 0.829180479 |
| SL1344_RS03835_ | -0.671798228 | -0.680776136 | -0.655364052 | 0.01286506 | 0.270503922 | 0.042684464 | 0.484666667 | 0.662264785 | 0.83103302 | 0.7713 | 0.883461963 |
| SL1344_RS03840_ | -0.452149402 | -0.527911207 | -0.365730465 | 0.168915192 | 0.241551088 | 0.647419726 | 0.228000014 | 0.358399299 | 0.610673439 | 0.4708 | 0.677770935 |
| SL1344_RS03845_ | -2.030966985 | -2.196320839 | -1.817830312 | 0.372364875 | 0.349863921 | 0.984215637 | 0.136666695 | 0.223111813 | 0.465433977 | 0.3075 | 0.539812612 |
| SL1344_RS03850_ | 3.327217918 | 3.038755383 | 3.499919741 | 0.375702809 | 0.253894406 | 1.648013455 | 0.038000117 | 0.060344739 | 0.286397196 | 0.1394 | 0.387889433 |
| SL1344_RS03855_ | -1.016090385 | -1.136764842 | -0.917083164 | 0.218000636 | 0.254456705 | 0.829248129 | 0.174550321 | 0.28187861 | 0.533058087 | 0.3766 | 0.596225492 |
| SL1344_RS03860_ | -1.181250887 | -1.217573586 | -1.141655271 | 0.135470365 | 0.378205572 | 0.333491053 | 0.36908728 | 0.574441744 | 0.779922926 | 0.722 | 0.850982718 |
| SL1344_RS03865_ | 0.367411969 | 0.316131522 | 0.401963709 | 0.100672759 | 0.190161613 | 0.505532906 | 0.28047969 | 0.41897429 | 0.663155329 | 0.5465 | 0.724644164 |
| SL1344_RS03870_ | -2.385615767 | -2.544180133 | -2.235212171 | 0.319988253 | 0.374760379 | 0.797149751 | 0.180666687 | 0.282913814 | 0.528634138 | 0.3822 | 0.599331937 |
| SL1344_RS03875_ | -4.007002132 | -3.945343651 | -4.14886295 | -0.247479476 | 0.987200765 | -0.210832993 | 0.40106596 | 0.606209986 | 0.802037415 | 0.746 | 0.8718141 |
| SL1344_RS03880_ | -6.111189424 | -5.826852883 | -6.389779683 | -0.543294638 | 1.392283969 | -0.34182967 | 0.345769493 | 0.484071748 | 0.698764484 | 0.5819 | 0.750871222 |
| SL1344_RS03885_ | -2.679897319 | -2.865446865 | -2.519425272 | 0.355107285 | 0.415343188 | 0.827531961 | 0.174550321 | 0.26940593 | 0.515304708 | 0.3677 | 0.583248627 |
| SL1344_RS03890_ | -2.585028985 | -2.85592225 | -2.371881747 | 0.481847001 | 0.443757865 | 1.026803727 | 0.125333365 | 0.196650171 | 0.448799681 | 0.2791 | 0.517704472 |
| SL1344_RS03895_ | -0.78187003 | -0.786513261 | -0.77666901 | 0.012210972 | 0.336246479 | 0.030732891 | 0.484000001 | 0.698283752 | 0.856739403 | 0.8107 | 0.90663722 |
| SL1344_RS03900_ | -2.067666517 | -2.115559002 | -1.998681593 | 0.117190466 | 0.45271643 | 0.245603042 | 0.399733515 | 0.608045959 | 0.797019454 | 0.7269 | 0.856327526 |
| SL1344_RS03905_ | -2.625461577 | -2.672045136 | -2.56035073 | 0.099304354 | 0.498081611 | 0.181594159 | 0.420386412 | 0.613231351 | 0.799308074 | 0.7188 | 0.849454926 |
| SL1344_RS03915_ | 3.249775239 | 3.53750366 | 3.224359275 | -0.321011789 | 0.24690374 | -0.845044055 | 0.268487686 | 0.318026777 | 0.617137806 | 0.5118 | 0.751798969 |
| SL1344_RS03920_ | 2.933075119 | 3.0251535 | 2.769012366 | -0.274091153 | 0.135838227 | -2.087563767 | 0.000673369 | 0.034056624 | 0.213454207 | 0.1017 | 0.332581102 |
| SL1344_RS03925_ | 2.500054359 | 2.632582407 | 2.283166478 | -0.378000461 | 0.167205653 | -2.404022532 | 4.68E-05 | 0.026214944 | 0.190511752 | 0.1 | 0.330257709 |
| SL1344_RS03930_ | 2.252753026 | 2.4997593 | 2.186432472 | -0.301475323 | 0.510767008 | -0.577905346 | 0.334443711 | 0.498535497 | 0.753518303 | 0.6997 | 0.872160378 |
| SL1344_RS03935_ | -2.590255536 | -2.495587837 | -2.684339121 | -0.16650275 | 0.482590336 | -0.33344337 | 0.359093943 | 0.545006033 | 0.749448639 | 0.6581 | 0.808780054 |
| SL1344_RS03940_ | 1.960634782 | 1.99927916 | 1.930883143 | 0.012844183 | 0.224453079 | 0.064334107 | 0.480346436 | 0.799678001 | 0.915378927 | 0.9506 | 0.977469936 |
| SL1344_RS03945_ | 1.027566757 | 0.991706269 | 1.058191727 | 0.085187278 | 0.179497064 | 0.438551916 | 0.304666675 | 0.455507683 | 0.695405802 | 0.6117 | 0.776344611 |
| SL1344_RS03950_ | 4.26375608 | 4.416358061 | 4.112597167 | -0.132900799 | 0.50646942 | -0.267200987 | 0.359093943 | 0.655002437 | 0.849832467 | 0.7723 | 0.903090094 |
| SL1344_RS03970_ | 0.841987921 | 0.682147363 | 0.978454903 | 0.297098138 | 0.154338868 | 1.863257043 | 0.021985549 | 0.057646935 | 0.25678113 | 0.1237 | 0.361009243 |
| SL1344_RS03975_ | 1.452182335 | 1.466025592 | 1.44727297 | -0.022308857 | 0.168259085 | -0.102358534 | 0.460666668 | 0.732790954 | 0.878296311 | 0.8282 | 0.921375111 |
| SL1344_RS03980_ | 3.622876812 | 3.402481925 | 3.858529651 | 0.391558148 | 0.140323553 | 2.945731056 | 4.68E-05 | 0.00953229 | 0.121110053 | 0.1 | 0.330257709 |
| SL1344_RS03985_ | 1.711490527 | 1.460701176 | 1.755898919 | 0.281146428 | 0.258506509 | 0.789019214 | 0.281145913 | 0.354118262 | 0.643157062 | 0.5453 | 0.770832822 |
| SL1344_RS03990_ | 1.251778391 | 1.219648487 | 1.267805384 | 0.057906028 | 0.471287704 | 0.111482519 | 0.435709529 | 0.907860761 | 0.965712466 | 0.8872 | 0.951903078 |
| SL1344_RS03995_ | 0.831260816 | 0.735527797 | 0.859765165 | 0.12569863 | 0.481195544 | 0.255835192 | 0.379080618 | 0.71472627 | 0.876891669 | 0.7855 | 0.908745515 |
| SL1344_RS04000_ | 1.644643215 | 1.629071048 | 1.653207522 | 0.043112334 | 0.469936979 | 0.092927541 | 0.446369089 | 0.66216528 | 0.851125069 | 0.8902 | 0.951016621 |
| SL1344_RS04005_ | 1.872776514 | 1.908679175 | 1.844206791 | -0.060056738 | 0.437756679 | -0.100270475 | 0.449033979 | 0.816247787 | 0.927540073 | 0.9685 | 0.986710594 |
| SL1344_RS04010_ | 0.672070013 | 0.491843865 | 0.725291979 | 0.200630413 | 0.233570307 | 0.71987463 | 0.247333346 | 0.364886641 | 0.636424944 | 0.5009 | 0.725658048 |
| SL1344_RS04015_ | -2.156774606 | -2.127078426 | -2.193945505 | -0.046391044 | 0.36916796 | -0.116733872 | 0.447333335 | 0.551632709 | 0.754568862 | 0.6677 | 0.814400376 |
| SL1344_RS04020_ | -2.371891924 | -2.202897958 | -2.458867685 | -0.198078238 | 0.499646031 | -0.356926942 | 0.348666673 | 0.582830576 | 0.78097387 | 0.6644 | 0.824180164 |
| SL1344_RS04025_ | -2.527327035 | -2.706013725 | -2.375834223 | 0.296195712 | 0.48917003 | 0.558478471 | 0.266666678 | 0.414969943 | 0.653891239 | 0.5215 | 0.717233999 |
| SL1344_RS04030_ | -2.689554084 | -2.621438392 | -2.768671968 | -0.297037083 | 1.010134571 | -0.313767756 | 0.355096608 | 0.499885266 | 0.742216076 | 0.6804 | 0.83124592 |
| SL1344_RS04035_ | -3.40565418 | -3.435906388 | -3.39145529 | 0.087404076 | 0.613517047 | 0.136938649 | 0.436375752 | 0.585987507 | 0.780665157 | 0.7038 | 0.838050731 |
| SL1344_RS04040_ | 2.424088561 | 2.642692343 | 2.333720067 | -0.295664329 | 0.15473036 | -1.823718755 | 0.013324793 | 0.049515543 | 0.254262833 | 0.1136 | 0.349010237 |
| SL1344_RS04045_ | -3.201138376 | -2.925422861 | -3.775579212 | -0.852983354 | 0.691121673 | -1.132359514 | 0.099933419 | 0.170626367 | 0.434372539 | 0.2322 | 0.472894701 |
| SL1344_RS04050_ | 1.451035756 | 1.45687982 | 1.442898099 | 0.030210937 | 0.190742392 | 0.16441077 | 0.431333336 | 0.683540748 | 0.847889608 | 0.8288 | 0.914078274 |
| SL1344_RS04055_ | 0.801923461 | 0.620116987 | 0.917592717 | 0.287849112 | 0.180125666 | 1.557125986 | 0.046000096 | 0.089955594 | 0.323566272 | 0.1555 | 0.399802613 |
| SL1344_RS04065_ | 0.457034975 | 0.429836649 | 0.509885515 | 0.071272261 | 0.196837288 | 0.30252641 | 0.372666672 | 0.603050658 | 0.794358148 | 0.7058 | 0.845348049 |
| SL1344_RS04070_ | 0.173853053 | 0.166032572 | 0.183323297 | 0.016564014 | 0.18276947 | 0.076579795 | 0.464000001 | 0.611985062 | 0.797694762 | 0.7153 | 0.848214118 |
| SL1344_RS04085_ | 3.303111052 | 3.092318732 | 3.330026976 | 0.227084709 | 0.1535818 | 0.625941063 | 0.33333334 | 0.531836633 | 0.774439072 | 0.7 | 0.872542119 |
| SL1344_RS04090_ | -4.62776182 | -4.736456702 | -4.525835413 | 0.201857172 | 0.788458763 | 0.221980856 | 0.401333337 | 0.51146183 | 0.721474229 | 0.6166 | 0.777075228 |
| SL1344_RS04095_ | -4.005652946 | -3.747167561 | -4.222846444 | -0.449160817 | 0.754338841 | -0.535201233 | 0.267821463 | 0.432282998 | 0.660360479 | 0.5239 | 0.715742467 |
| SL1344_RS04100_ | -3.878192341 | -3.72737804 | -4.05509961 | -0.349052861 | 0.609961097 | -0.510300308 | 0.273151243 | 0.397836966 | 0.63062395 | 0.5051 | 0.692065048 |
| SL1344_RS04105_ | -1.21707825 | -1.259572098 | -1.200325655 | 0.045863243 | 0.496040741 | 0.080865633 | 0.471333334 | 0.781627178 | 0.9055064 | 0.8556 | 0.937370222 |
| SL1344_RS04110_ | -2.200872564 | -2.788399339 | -2.026962928 | 0.713949381 | 0.544717483 | 1.112701676 | 0.124000032 | 0.180178187 | 0.456241362 | 0.2782 | 0.524511473 |
| SL1344_RS04115_ | -2.964301475 | -2.834786664 | -3.189503285 | -0.271003561 | 0.721442864 | -0.354824948 | 0.358000006 | 0.584572422 | 0.78481612 | 0.7035 | 0.844048009 |
| SL1344_RS04120_ | 3.13546119 | 3.095231903 | 3.340619902 | 0.243949617 | 0.216899105 | 0.935666072 | 0.220519669 | 0.257328618 | 0.55887415 | 0.4043 | 0.670934042 |
| SL1344_RS04125_ | 0.144749669 | 0.059726144 | 0.273112812 | 0.213151964 | 0.195131212 | 1.021472705 | 0.144570313 | 0.220453269 | 0.481062493 | 0.3063 | 0.542051404 |

| | | | | | | | | | | |
|---|---|---|---|---|---|---|---|---|---|---|
| SL1344_RS04130_ | 0.391058351 | 0.274310583 | 0.451085221 | 0.192745629 | 0.237744742 | 0.765657372 | 0.194000018 | 0.286424672 | 0.567884597 | 0.3846 | 0.612143818 |
| SL1344_RS04135_ | 1.022617508 | 0.914838334 | 1.080237054 | 0.168111217 | 0.340636092 | 0.418949303 | 0.358427721 | 0.574618293 | 0.796305573 | 0.7618 | 0.891885642 |
| SL1344_RS04140_ | 0.163117807 | 0.152148726 | 0.16991868 | -0.015390185 | 0.393717188 | -0.0372559 | 0.480000001 | 0.809370995 | 0.920689413 | 0.9206 | 0.964633374 |
| SL1344_RS04145_ | 0.021458768 | -0.096815395 | 0.135060942 | 0.243916038 | 0.232920536 | 0.987980234 | 0.124000032 | 0.196042515 | 0.462704993 | 0.2686 | 0.515963204 |
| SL1344_RS04150_ | -0.142873102 | -0.137499799 | -0.145593911 | 0.010995556 | 0.315332694 | 0.032603045 | 0.489007329 | 0.715390384 | 0.869863505 | 0.8291 | 0.922325428 |
| SL1344_RS04160_ | 1.029664035 | 0.968353622 | 1.103885623 | 0.138397724 | 0.149245461 | 0.894848497 | 0.164000023 | 0.253385263 | 0.514692796 | 0.3351 | 0.572530654 |
| SL1344_RS04165_ | -1.86086098 | -1.885956483 | -1.841928209 | 0.053577002 | 0.665085006 | 0.080297669 | 0.459693539 | 0.731978006 | 0.877505268 | 0.8171 | 0.911017143 |
| SL1344_RS04170_ | -3.781140898 | -3.637585307 | -3.937530574 | -0.310868433 | 0.567246204 | -0.514952155 | 0.283810803 | 0.407965487 | 0.632613783 | 0.5064 | 0.691190782 |
| SL1344_RS04175_ | -1.528509306 | -1.528865843 | -1.528204608 | -0.006055833 | 0.341435576 | -0.021903673 | 0.488 | 0.626607735 | 0.809803893 | 0.7332 | 0.860879104 |
| SL1344_RS04180_ | -0.062474866 | -0.119829643 | -0.015435372 | 0.101484804 | 0.274324835 | 0.361817548 | 0.339107268 | 0.567397844 | 0.772924643 | 0.6764 | 0.827706526 |
| SL1344_RS04185_ | -0.375037113 | -0.365834173 | -0.388578277 | -0.021037906 | 0.198684842 | -0.089814526 | 0.453333335 | 0.550223457 | 0.750324966 | 0.6609 | 0.807312097 |
| SL1344_RS04195_ | 1.814289676 | 1.736938431 | 1.911086874 | -0.00181726 | 0.399942821 | -0.00391012 | 0.498 | 0.882003232 | 0.954834944 | 0.9712 | 0.987612423 |
| SL1344_RS04200_ | 4.440297594 | 4.025878438 | 4.526519069 | 0.506762711 | 0.346691732 | 1.599168161 | 4.68E-05 | 0.113716626 | 0.387298811 | 0.1 | 0.330257709 |
| SL1344_RS04205_ | 2.6215046 | 2.588413907 | 2.655409432 | 0.086195584 | 0.262883565 | 0.290623896 | 0.350433051 | 0.462538783 | 0.728620343 | 0.7266 | 0.869307232 |
| SL1344_RS04210_ | 0.07585145 | -0.021432381 | 0.182465202 | 0.204089728 | 0.160633105 | 1.220190977 | 0.088607642 | 0.14772314 | 0.381139271 | 0.2228 | 0.460285985 |
| SL1344_RS04220_ | 0.884910963 | 0.573015748 | 1.031904788 | 0.437778469 | 0.189793945 | 2.117302515 | 0.007995244 | 0.048571579 | 0.245285402 | 0.1058 | 0.338498288 |
| SL1344_RS04225_ | -0.077281253 | 0.000429706 | -0.143980353 | -0.140686081 | 0.199305708 | -0.653052715 | 0.239333347 | 0.361926387 | 0.60914979 | 0.4724 | 0.678981728 |
| SL1344_RS04230_ | 0.578955026 | 0.710471523 | 0.432320314 | -0.298399735 | 0.192790663 | -1.536777804 | 0.025316634 | 0.08620133 | 0.318235506 | 0.1344 | 0.373876933 |
| SL1344_RS04235_ | 2.080447192 | 2.009162068 | 2.135851069 | 0.109904051 | 0.19476384 | 0.542967007 | 0.272485021 | 0.416929937 | 0.68585971 | 0.5595 | 0.760593167 |
| SL1344_RS04240_ | 2.137745033 | 2.181384191 | 2.055615694 | -0.121301143 | 0.150834201 | -0.79251051 | 0.202666684 | 0.288719683 | 0.569471391 | 0.3814 | 0.63288796 |
| SL1344_RS04245_ | -2.850085582 | -2.638684805 | -3.099701215 | -0.457675362 | 0.458449772 | -0.957321384 | 0.140666694 | 0.226770828 | 0.474047006 | 0.3107 | 0.539850842 |
| SL1344_RS04250_ | -5.577324966 | -5.833721489 | -5.329381167 | 0.503394224 | 1.103123723 | 0.411702405 | 0.319333341 | 0.430650335 | 0.654005368 | 0.527 | 0.710946935 |
| SL1344_RS04255_ | -1.341883153 | -1.326814696 | -1.359368548 | -0.03026474 | 0.283094273 | -0.090041776 | 0.453031314 | 0.574913753 | 0.769187738 | 0.6856 | 0.825143402 |
| SL1344_RS04260_ | 1.585587538 | 1.51132788 | 1.764095712 | 0.217411902 | 0.167165659 | 1.206143571 | 0.125333365 | 0.167364654 | 0.439812023 | 0.2542 | 0.512697701 |
| SL1344_RS04265_ | -2.484177725 | -2.564471854 | -2.293094394 | 0.235286675 | 0.509809051 | 0.388614147 | 0.33133334 | 0.54783278 | 0.754627462 | 0.6494 | 0.812037343 |
| SL1344_RS04270_ | -3.77371799 | -3.90733937 | -3.677446812 | 0.220519262 | 0.71454472 | 0.291543549 | 0.376415728 | 0.547546009 | 0.752677608 | 0.6546 | 0.808199353 |
| SL1344_RS04275_ | 0.959114989 | 0.91780598 | 1.041381297 | 0.118649167 | 0.132934712 | 0.811375599 | 0.20333335 | 0.328286869 | 0.575992013 | 0.4281 | 0.646916058 |
| SL1344_RS04280_ | 0.914077891 | 0.868921384 | 0.937504232 | 0.05439489 | 0.171291589 | 0.287349013 | 0.387333338 | 0.61352652 | 0.803508219 | 0.7279 | 0.859092066 |
| SL1344_RS04290_ | 2.671954236 | 2.54842384 | 2.702912177 | 0.148560654 | 0.221375458 | 0.606376665 | 0.292000009 | 0.387641826 | 0.670619969 | 0.607 | 0.803177381 |
| SL1344_RS04295_ | 1.761257919 | 1.430452274 | 1.820717417 | 0.356965756 | 0.205183149 | 1.220439651 | 0.137333362 | 0.164202899 | 0.450586962 | 0.2781 | 0.541381382 |
| SL1344_RS04300_ | 1.553302352 | 1.488165195 | 1.619897051 | 0.171769826 | 0.193657232 | 0.882825148 | 0.145236535 | 0.217468645 | 0.503486468 | 0.3088 | 0.556428988 |
| SL1344_RS04305_ | 1.095656259 | 1.020185091 | 1.162028633 | 0.131160901 | 0.188259545 | 0.672722201 | 0.228000014 | 0.357398633 | 0.619196786 | 0.458 | 0.678642878 |
| SL1344_RS04320_ | -3.732494537 | -3.648541936 | -3.813966189 | -0.176827476 | 0.790086026 | -0.208892659 | 0.399333337 | 0.608010887 | 0.794324921 | 0.7102 | 0.84353391 |
| SL1344_RS04325_ | -3.095948356 | -3.324689486 | -2.756000872 | 0.487670168 | 0.696218382 | 0.659700923 | 0.230513005 | 0.37486932 | 0.626413393 | 0.4687 | 0.685151089 |
| SL1344_RS04330_ | -2.709678079 | -2.833146371 | -2.51063367 | 0.343941792 | 0.49680377 | 0.634691786 | 0.24266668 | 0.376214421 | 0.619033385 | 0.4926 | 0.682221912 |
| SL1344_RS04335_ | -4.595859743 | -4.79710271 | -4.477219797 | 0.448200034 | 1.231934741 | 0.299209091 | 0.342000007 | 0.500166069 | 0.731657736 | 0.652 | 0.801178011 |
| SL1344_RS04340_ | -1.601992457 | -1.618642364 | -1.586813117 | 0.032892586 | 0.309920209 | 0.099279766 | 0.457694872 | 0.573286974 | 0.772141698 | 0.6874 | 0.827911351 |
| SL1344_RS04345_ | -1.983280102 | -2.054992143 | -1.918327678 | 0.189690677 | 0.493736889 | 0.375832487 | 0.354666673 | 0.544936932 | 0.758884081 | 0.6801 | 0.828689762 |
| SL1344_RS04350_ | -1.074624472 | -1.221914636 | -0.942846318 | 0.335531065 | 0.398962524 | 0.816814234 | 0.175216543 | 0.260014381 | 0.540468799 | 0.3614 | 0.598640551 |
| SL1344_RS04355_ | -0.397758473 | -0.494379953 | -0.293045617 | 0.21175284 | 0.364736534 | 0.578040013 | 0.270486353 | 0.412773043 | 0.672799684 | 0.5281 | 0.737534845 |
| SL1344_RS04360_ | -1.607593386 | -1.8312912 | -1.429770421 | 0.394447277 | 0.295577002 | 1.259868583 | 0.079946755 | 0.141324346 | 0.378828888 | 0.2166 | 0.454556762 |
| SL1344_RS04365_ | 1.345005124 | 1.433917972 | 1.293525471 | -0.130603177 | 0.143792837 | -0.858316425 | 0.187208547 | 0.28778516 | 0.552920148 | 0.3832 | 0.618647007 |
| SL1344_RS04370_ | 1.71169995 | 1.656089065 | 1.795788525 | 0.174523022 | 0.220156519 | 0.834534184 | 0.129333364 | 0.238930936 | 0.532857096 | 0.2865 | 0.540695885 |
| SL1344_RS04375_ | 2.418842215 | 2.381921802 | 2.465107331 | 0.103978761 | 0.142353734 | 0.730988529 | 0.183211212 | 0.278603129 | 0.565452005 | 0.3712 | 0.609360646 |
| SL1344_RS04380_ | 1.016718219 | 0.658612708 | 1.350686034 | 0.692908873 | 0.121159773 | 5.703656482 | 4.68E-05 | 0.002878083 | 0.062955359 | 0.1 | 0.330257709 |
| SL1344_RS04385_ | 0.297898627 | 0.338049318 | 0.278016161 | -0.041465499 | 0.29529622 | -0.1260502 | 0.450366424 | 0.780990729 | 0.9052863 | 0.8583 | 0.936863154 |
| SL1344_RS04390_ | 0.601935219 | 0.507580792 | 0.742336531 | 0.249549147 | 0.202817861 | 1.146029386 | 0.090000046 | 0.157693116 | 0.424421802 | 0.223 | 0.471461204 |
| SL1344_RS04395_ | 0.868346483 | 0.804219223 | 0.960463095 | 0.151273636 | 0.154972719 | 0.929411404 | 0.167221874 | 0.250027164 | 0.514126823 | 0.3428 | 0.580241414 |
| SL1344_RS04400_ | -1.846770316 | -1.807684101 | -1.92123294 | -0.081355136 | 0.493229893 | -0.137590593 | 0.451333335 | 0.72301387 | 0.872637288 | 0.8073 | 0.910888301 |
| SL1344_RS04405_ | 1.622743972 | 1.248786397 | 2.024774459 | 0.694284646 | 0.285221362 | 2.463226321 | 4.68E-05 | 0.019219083 | 0.167695688 | 0.1001 | 0.330407687 |
| SL1344_RS04410_ | -2.862515663 | -2.964165673 | -2.74389237 | 0.287577853 | 0.649574307 | 0.409209977 | 0.324666674 | 0.504458577 | 0.728546498 | 0.6414 | 0.797998616 |
| SL1344_RS04415_ | 3.618855639 | 3.690579784 | 3.537536222 | -0.149579143 | 0.194090745 | -0.664616566 | 0.186542324 | 0.305930595 | 0.605318413 | 0.3453 | 0.608393898 |
| SL1344_RS04420_ | -1.232003669 | -1.198245385 | -1.277547364 | -0.085891333 | 0.3087397 | -0.271417989 | 0.388666671 | 0.553024694 | 0.757018564 | 0.6695 | 0.816099377 |

| | | | | | | | | | | | |
|---|---|---|---|---|---|---|---|---|---|---|---|
| SL1344_RS04425_ | -0.279996777 | -0.372843493 | -0.198874045 | 0.179145222 | 0.238220066 | 0.692062194 | 0.221185891 | 0.327841481 | 0.586849681 | 0.4352 | 0.651395342 |
| SL1344_RS04430_ | 0.615274831 | 0.626875433 | 0.585467937 | -0.04229055 | 0.252273237 | -0.15295282 | 0.441333336 | 0.725033055 | 0.875314999 | 0.8429 | 0.929110626 |
| SL1344_RS04435_ | 1.337321101 | 1.261566378 | 1.393975066 | 0.128610977 | 0.263880314 | 0.48967709 | 0.281812135 | 0.474479038 | 0.725237768 | 0.5666 | 0.771447353 |
| SL1344_RS04440_ | -0.423389204 | -0.599862253 | -0.372397527 | 0.19388513 | 0.318250333 | 0.512884585 | 0.326449041 | 0.54428469 | 0.764557949 | 0.6702 | 0.837743838 |
| SL1344_RS04450_ | -1.564794074 | -1.612159499 | -1.528062572 | 0.113831896 | 0.39803698 | 0.264952572 | 0.389333338 | 0.584764719 | 0.78121214 | 0.7219 | 0.848712203 |
| SL1344_RS04455_ | -0.821870186 | -0.709508422 | -0.891055392 | -0.172708738 | 0.425674937 | -0.38877515 | 0.316000008 | 0.546820863 | 0.764170302 | 0.6168 | 0.797872994 |
| SL1344_RS04460_ | -0.266177685 | -0.111796512 | -0.450229019 | -0.345829682 | 0.195705881 | -1.65757417 | 0.028000161 | 0.072827392 | 0.276798582 | 0.1319 | 0.369570103 |
| SL1344_RS04465_ | 0.841979558 | 0.789522782 | 0.985028615 | 0.166943094 | 0.217405462 | 0.665603404 | 0.275333344 | 0.428735692 | 0.683495768 | 0.5601 | 0.76297608 |
| SL1344_RS04470_ | -3.434693104 | -3.736667604 | -3.260924289 | 0.446696568 | 0.642468653 | 0.656426099 | 0.245836122 | 0.356908048 | 0.609253297 | 0.4713 | 0.678698734 |
| SL1344_RS04475_ | -4.376559027 | -4.812878522 | -4.092553314 | 0.802018784 | 1.113729538 | 0.684748511 | 0.174000021 | 0.294930646 | 0.572796279 | 0.3685 | 0.587765302 |
| SL1344_RS04480_ | -1.039028652 | -1.045183011 | -1.032695524 | 0.017023483 | 0.22396457 | 0.065690766 | 0.472000001 | 0.529804988 | 0.734822902 | 0.6463 | 0.796411547 |
| SL1344_RS04485_ | -0.971572792 | -1.2047275 | -0.707122426 | 0.504852122 | 0.245987511 | 1.987029911 | 0.013324793 | 0.048368084 | 0.230247481 | 0.1168 | 0.351132863 |
| SL1344_RS04495_ | 1.768715822 | 1.71607856 | 1.855802794 | 0.171967371 | 0.19120941 | 0.897154039 | 0.154000025 | 0.217843677 | 0.503365658 | 0.316 | 0.57228335 |
| SL1344_RS04500_ | 2.387959455 | 2.337517004 | 2.532099296 | 0.20376446 | 0.16593876 | 1.189018819 | 0.092604975 | 0.172356293 | 0.459864691 | 0.2169 | 0.469621516 |
| SL1344_RS04505_ | 2.097331993 | 2.016173874 | 2.140072939 | 0.131502294 | 0.119377723 | 1.069914932 | 0.105263197 | 0.193501988 | 0.472934791 | 0.2353 | 0.479100676 |
| SL1344_RS04510_ | 1.642563433 | 1.599221169 | 1.758982041 | 0.201245992 | 0.369658841 | 0.600496167 | 0.230000014 | 0.339865716 | 0.631768784 | 0.4276 | 0.690561844 |
| SL1344_RS04515_ | 0.017007383 | 0.027973234 | 0.010586884 | -0.05791157 | 0.265798169 | -0.203912278 | 0.42504997 | 0.660159651 | 0.833893849 | 0.7955 | 0.897850292 |
| SL1344_RS04520_ | 1.078561645 | 1.097171547 | 1.056981316 | -0.042442502 | 0.132526447 | -0.296888992 | 0.370000005 | 0.542316897 | 0.747936169 | 0.6579 | 0.808036956 |
| SL1344_RS04525_ | 1.377801003 | 1.431233522 | 1.336016531 | -0.09751009 | 0.194252666 | -0.475183688 | 0.300466365 | 0.480449555 | 0.720771568 | 0.5937 | 0.781693279 |
| SL1344_RS04530_ | 1.16202017 | 1.190649281 | 1.117062564 | -0.084264073 | 0.292531872 | -0.298873475 | 0.34333334 | 0.634654013 | 0.825931319 | 0.6949 | 0.847978013 |
| SL1344_RS04535_ | 1.18305481 | 1.162082607 | 1.201248662 | 0.031484968 | 0.189391998 | 0.170720648 | 0.423333336 | 0.680195188 | 0.845129515 | 0.7899 | 0.893738268 |
| SL1344_RS04540_ | -0.562845204 | -0.629271492 | -0.510931867 | 0.097454168 | 0.381821161 | 0.25015139 | 0.386000004 | 0.65257291 | 0.831280658 | 0.7687 | 0.883867169 |
| SL1344_RS04545_ | -0.350584548 | -0.310313198 | -0.382416151 | -0.068319864 | 0.198308919 | -0.313380372 | 0.357095276 | 0.505451946 | 0.716890202 | 0.6189 | 0.77717262 |
| SL1344_RS04550_ | -0.314057761 | -0.303741742 | -0.331319679 | -0.059519283 | 0.254528651 | -0.210343005 | 0.410393075 | 0.597960033 | 0.790230494 | 0.7356 | 0.858413714 |
| SL1344_RS04555_ | 1.434953031 | 1.462855796 | 1.403575281 | -0.025600467 | 0.215308473 | -0.107564284 | 0.455696204 | 0.791240836 | 0.910600007 | 0.8583 | 0.936746181 |
| SL1344_RS04560_ | 1.188217538 | 1.287641669 | 1.118713952 | -0.182475534 | 0.182653059 | -0.968620934 | 0.136000029 | 0.19825075 | 0.475638828 | 0.2834 | 0.537443892 |
| SL1344_RS04565_ | 3.521315365 | 3.519978761 | 3.522671138 | 0.066952665 | 0.186283178 | 0.293250884 | 0.364000005 | 0.443592168 | 0.714213965 | 0.7949 | 0.898350142 |
| SL1344_RS04570_ | 1.429691204 | 1.416428863 | 1.454008633 | 0.06535967 | 0.279685013 | 0.248792455 | 0.361758833 | 0.701324273 | 0.863881592 | 0.7473 | 0.875427365 |
| SL1344_RS04575_ | 1.673368896 | 1.64738276 | 1.699981293 | 0.096307446 | 0.200897029 | 0.45642488 | 0.317121927 | 0.459086442 | 0.711296941 | 0.649 | 0.812999383 |
| SL1344_RS04580_ | 3.98469424 | 4.340509189 | 3.661560133 | -0.687161245 | 0.173028883 | -4.090757001 | 4.68E-05 | 0.009553195 | 0.123332443 | 0.1 | 0.330257709 |
| SL1344_RS04585_ | 1.831025593 | 1.888583768 | 1.794131716 | -0.048300632 | 0.210244466 | -0.200210449 | 0.417333336 | 0.812987054 | 0.922355659 | 0.8371 | 0.92886777 |
| SL1344_RS04590_ | 4.066928226 | 4.251870283 | 4.010441547 | -0.15587879 | 0.163559234 | -0.4149476 | 0.33333334 | 0.923017392 | 0.972122711 | 0.7 | 0.872542119 |
| SL1344_RS04600_ | -2.372982319 | -2.474143981 | -2.258254137 | 0.233962691 | 0.451825708 | 0.483549814 | 0.313790814 | 0.466421235 | 0.692970383 | 0.5745 | 0.752356174 |
| SL1344_RS04605_ | -2.088904835 | -2.021020359 | -2.148821085 | -0.138032197 | 0.500144306 | -0.248346312 | 0.389333338 | 0.601754016 | 0.796495715 | 0.709 | 0.846011847 |
| SL1344_RS04610_ | -0.261154684 | 0.270031007 | -0.342552563 | -0.637236915 | 0.896032725 | -0.678249807 | 0.318666674 | 0.43883529 | 0.711066836 | 0.6655 | 0.850304419 |
| SL1344_RS04615_ | -0.888849819 | -0.849331344 | -0.911745526 | -0.076796089 | 0.355727991 | -0.195607222 | 0.417333336 | 0.66983602 | 0.839227734 | 0.7909 | 0.895627018 |
| SL1344_RS04620_ | -2.677243923 | -2.45675726 | -2.840717213 | -0.366158369 | 0.455133557 | -0.742280117 | 0.206528997 | 0.325903723 | 0.572317555 | 0.4246 | 0.640139917 |
| SL1344_RS04625_ | -1.895774486 | -1.747119039 | -2.102743981 | -0.428194603 | 0.473417898 | -0.849159018 | 0.150566314 | 0.247382175 | 0.528689154 | 0.3239 | 0.554979027 |
| SL1344_RS04630_ | 4.734115251 | 4.704150093 | 4.758882669 | -0.04311484 | 0.180920467 | -0.20920753 | 0.448367757 | 0.655559494 | 0.847228759 | 0.9922 | 0.99670693 |
| SL1344_RS04635_ | 4.19885543 | 4.19536807 | 4.202660702 | -0.032760394 | 0.512858809 | -0.084585178 | 0.427333336 | 0.840687287 | 0.939482595 | 0.9181 | 0.964649126 |
| SL1344_RS04640_ | 1.211062185 | 1.084853996 | 1.285787873 | 0.15052009 | 0.215365139 | 0.687957959 | 0.244666679 | 0.38184769 | 0.652219245 | 0.4834 | 0.70845036 |
| SL1344_RS04645_ | 1.981221094 | 1.848348497 | 2.022157272 | 0.13800445 | 0.242348514 | 0.470620754 | 0.32933334 | 0.649785571 | 0.839594973 | 0.6895 | 0.8612012 |
| SL1344_RS04650_ | 2.348404078 | 2.29430508 | 2.392316885 | 0.043880981 | 0.212690594 | 0.216417088 | 0.416000003 | 0.720161456 | 0.874502027 | 0.8607 | 0.933147214 |
| SL1344_RS04655_ | 4.038718949 | 4.034294872 | 4.043444187 | 0.011969453 | 0.149228585 | 0.081538094 | 0.474350434 | 0.710782108 | 0.874437223 | 0.9544 | 0.980252241 |
| SL1344_RS04665_ | 4.513188409 | 4.498890172 | 4.627156539 | 0.120759846 | 0.155848471 | 0.756492041 | 0.235176563 | 0.332680989 | 0.626141898 | 0.4204 | 0.693180127 |
| SL1344_RS04675_ | 2.82828664 | 2.737436108 | 2.88225103 | 0.143203552 | 0.090575465 | 1.528255267 | 0.057295212 | 0.091939236 | 0.329587365 | 0.1602 | 0.407267119 |
| SL1344_RS04680_ | 4.88207326 | 4.798511862 | 4.977424236 | 0.227875101 | 0.115007101 | 2.087370856 | 4.68E-05 | 0.047312672 | 0.255914144 | 0.1 | 0.330257709 |
| SL1344_RS04700_ | -4.275425545 | -4.153008757 | -4.348767546 | -0.180809886 | 0.886775682 | -0.178875367 | 0.415333337 | 0.609979318 | 0.797746025 | 0.715 | 0.848168277 |
| SL1344_RS04705_ | -0.269618199 | -0.343409833 | -0.117177387 | 0.184730034 | 0.387458705 | 0.377683286 | 0.372666672 | 0.679015669 | 0.849742765 | 0.7772 | 0.898112382 |
| SL1344_RS04715_ | 3.122257165 | 3.033618057 | 3.193527763 | 0.167533437 | 0.170785444 | 0.975846795 | 0.134666696 | 0.198781011 | 0.492743948 | 0.2551 | 0.52538873 |
| SL1344_RS04720_ | -1.651707268 | -1.489644638 | -1.868694455 | -0.193722845 | 0.68961303 | -0.265766841 | 0.415056632 | 0.74973573 | 0.889617053 | 0.8619 | 0.938070381 |
| SL1344_RS04725_ | 0.24752418 | 0.1084892 | 0.414489966 | 0.311200849 | 0.162696648 | 1.86114024 | 0.024000188 | 0.063015366 | 0.263785062 | 0.1245 | 0.361439732 |
| SL1344_RS04730_ | 3.809341753 | 3.532016766 | 4.035862676 | 0.502433852 | 0.302172518 | 1.507757143 | 0.062624986 | 0.09154991 | 0.34966545 | 0.1531 | 0.407738601 |

| | | | | | | | | | | | |
|---|---|---|---|---|---|---|---|---|---|---|---|
| SL1344_RS04740_ | 1.413285489 | 1.157972294 | 1.467174056 | 0.279708058 | 0.32691615 | 0.908586159 | 0.205196552 | 0.24105475 | 0.537866795 | 0.3662 | 0.64364654 |
| SL1344_RS04745_ | -1.762778268 | -1.966103602 | -1.617084224 | 0.304069498 | 0.427980572 | 0.676294059 | 0.252498347 | 0.379935164 | 0.634622718 | 0.4903 | 0.699909132 |
| SL1344_RS04750_ | 2.938362433 | 2.787010926 | 3.09118104 | 0.210364972 | 0.257075244 | 0.947983486 | 0.134666696 | 0.217667458 | 0.515828425 | 0.2363 | 0.512542131 |
| SL1344_RS04755_ | 2.380416834 | 2.292577681 | 2.433946531 | 0.171811775 | 0.224526701 | 0.753144744 | 0.224000015 | 0.288761291 | 0.582729068 | 0.4174 | 0.672524227 |
| SL1344_RS04760_ | 0.158058064 | 0.092319751 | 0.255632408 | 0.154697854 | 0.243038853 | 0.618359991 | 0.252498347 | 0.409681897 | 0.658301995 | 0.5077 | 0.717077379 |
| SL1344_RS04765_ | 2.922132711 | 2.864209621 | 2.97480196 | 0.109412106 | 0.270255175 | 0.327741956 | 0.400399737 | 0.760093368 | 0.900563781 | 0.8992 | 0.957087203 |
| SL1344_RS04770_ | 7.313473785 | 7.286997273 | 7.381224229 | 0.091985493 | 0.094036287 | 0.938283202 | 0.133333363 | 0.244696156 | 0.548532501 | 0.2492 | 0.521927655 |
| SL1344_RS04780_ | -0.866391318 | -0.715871595 | -1.012028087 | -0.319139526 | 0.284801633 | -1.084833578 | 0.115256531 | 0.178028858 | 0.436376332 | 0.2576 | 0.492769089 |
| SL1344_RS04785_ | 2.781330264 | 2.793346542 | 2.767820544 | 0.001738541 | 0.173828917 | 0.008551996 | 0.496 | 0.803695771 | 0.917831686 | 0.9031 | 0.95679772 |
| SL1344_RS04790_ | 2.10322316 | 2.179729129 | 2.082241326 | -0.083886614 | 0.138978693 | -0.459846064 | 0.33733334 | 0.609766332 | 0.808227792 | 0.7003 | 0.86143073 |
| SL1344_RS04795_ | -0.343254536 | -0.266230292 | -0.395460961 | -0.138992688 | 0.295728674 | -0.45484913 | 0.311125924 | 0.503975536 | 0.72951184 | 0.6355 | 0.798444126 |
| SL1344_RS04800_ | -1.700259417 | -1.66566367 | -1.735270647 | 0.011089909 | 0.629432458 | 0.016914873 | 0.493004664 | 0.771253687 | 0.900013575 | 0.8893 | 0.948743084 |
| SL1344_RS04805_ | 1.308984502 | 1.275916359 | 1.337485121 | 0.053855511 | 0.301215236 | 0.168944165 | 0.426666669 | 0.768757173 | 0.900344187 | 0.8699 | 0.94088237 |
| SL1344_RS04810_ | -0.423546717 | -0.413903904 | -0.434821007 | 0.031937622 | 0.409831191 | 0.082022467 | 0.468354432 | 0.756133739 | 0.890580537 | 0.8648 | 0.935788744 |
| SL1344_RS04815_ | -0.629766991 | -0.768968976 | -0.565753927 | 0.165125639 | 0.301673469 | 0.506732786 | 0.309127257 | 0.510710333 | 0.733814202 | 0.6223 | 0.797473708 |
| SL1344_RS04820_ | 1.458362919 | 1.827012218 | 1.350773718 | -0.46313996 | 0.35468624 | -1.404367386 | 0.056666743 | 0.082670776 | 0.328125761 | 0.1584 | 0.40789871 |
| SL1344_RS04825_ | 2.428020858 | 2.751523189 | 2.211409896 | -0.481179676 | 0.232603508 | -2.151966994 | 0.00067292 | 0.027476685 | 0.197708055 | 0.101 | 0.33171998 |
| SL1344_RS04830_ | 2.618696325 | 2.97864936 | 2.571464506 | -0.389908722 | 0.186552674 | -1.261176194 | 0.10926053 | 0.135055035 | 0.415503968 | 0.2244 | 0.486192035 |
| SL1344_RS04840_ | 3.225966487 | 3.356830685 | 3.081120172 | -0.237309609 | 0.149101296 | -1.578639415 | 0.050000088 | 0.073872019 | 0.310063535 | 0.146 | 0.397326969 |
| SL1344_RS04845_ | 2.395465596 | 2.414326239 | 2.371886324 | 0.052235759 | 0.208593821 | 0.259521328 | 0.433710862 | 0.687348553 | 0.856091432 | 0.9393 | 0.971461142 |
| SL1344_RS04850_ | 3.001747427 | 2.912485058 | 3.089916641 | 0.211604438 | 0.141630398 | 1.604860929 | 0.004664544 | 0.095756919 | 0.350679197 | 0.1043 | 0.336265209 |
| SL1344_RS04855_ | 4.36480063 | 4.156814197 | 4.601365652 | 0.405454038 | 0.122737272 | 3.315837318 | 4.68E-05 | 0.008409821 | 0.115304698 | 0.1 | 0.330257709 |
| SL1344_RS04860_ | 5.113717848 | 5.614019279 | 4.602388812 | -0.755099857 | 0.382241775 | -2.457833567 | 4.68E-05 | 0.01565656 | 0.15670695 | 0.1 | 0.330257709 |
| SL1344_RS04865_ | 4.896351731 | 4.772382435 | 5.034019698 | 0.251037389 | 0.0736035 | 3.376739361 | 4.68E-05 | 0.008765464 | 0.114668766 | 0.1 | 0.330257709 |
| SL1344_RS04870_ | -0.326980765 | -0.375972567 | -0.288274889 | 0.117516266 | 0.301985386 | 0.368083139 | 0.32933334 | 0.513936464 | 0.740181361 | 0.6572 | 0.811413469 |
| SL1344_RS04875_ | -2.842263784 | -2.997822779 | -2.674428251 | 0.31088682 | 0.553883412 | 0.51862642 | 0.269820131 | 0.413039095 | 0.652507284 | 0.5355 | 0.72054443 |
| SL1344_RS04880_ | -2.031052403 | -1.908620018 | -2.117135555 | -0.188645167 | 0.470243313 | -0.364437731 | 0.342000007 | 0.543387652 | 0.75346267 | 0.6622 | 0.815916503 |
| SL1344_RS04885_ | -3.396755204 | -3.293808972 | -3.520775437 | -0.244981155 | 0.742413826 | -0.316274998 | 0.362000006 | 0.554832417 | 0.762200385 | 0.6727 | 0.820696234 |
| SL1344_RS04890_ | 1.747104774 | 1.751117408 | 1.737722502 | -0.012423889 | 0.146988801 | -0.072468373 | 0.473017989 | 0.722598564 | 0.871831086 | 0.8124 | 0.913992415 |
| SL1344_RS04895_ | 2.24071847 | 2.240019244 | 2.240995071 | -0.001760678 | 0.106689201 | -0.011556672 | 0.494666667 | 0.670433786 | 0.83849729 | 0.7651 | 0.880705469 |
| SL1344_RS04900_ | -3.974363932 | -4.127264475 | -3.830536107 | 0.305047749 | 0.712439823 | 0.40447431 | 0.329113931 | 0.470086483 | 0.690899537 | 0.5811 | 0.752502616 |
| SL1344_RS04905_ | -5.813794873 | -5.892191003 | -5.703241336 | 0.203924715 | 1.557692759 | 0.124945202 | 0.443037977 | 0.60731343 | 0.797185052 | 0.7287 | 0.855790104 |
| SL1344_RS04910_ | -4.020732643 | -3.664180383 | -4.536492752 | -0.689723633 | 1.118985442 | -0.595445219 | 0.264000011 | 0.41222553 | 0.665226739 | 0.5241 | 0.726848502 |
| SL1344_RS04915_ | -2.843642329 | -2.974976549 | -2.654819223 | 0.371886524 | 0.52066274 | 0.70045574 | 0.222666682 | 0.347137018 | 0.597717793 | 0.4536 | 0.655023248 |
| SL1344_RS04920_ | -4.968360578 | -5.839370311 | -4.123794128 | 1.677546957 | 1.100826249 | 1.45727769 | 0.045303228 | 0.101487035 | 0.333035732 | 0.1516 | 0.393191054 |
| SL1344_RS04925_ | -5.750117754 | -6.088240236 | -5.44840116 | 0.652046139 | 1.243719788 | 0.497350521 | 0.280666677 | 0.406994726 | 0.638087137 | 0.5197 | 0.704374456 |
| SL1344_RS04930_ | 2.583691314 | 2.611354792 | 2.50184942 | -0.105561038 | 0.176585357 | -0.621635142 | 0.246502344 | 0.421259062 | 0.688260858 | 0.4721 | 0.721692547 |
| SL1344_RS04935_ | -3.832920318 | -3.86576753 | -3.815010956 | 0.133253448 | 0.878730511 | 0.12778079 | 0.442371754 | 0.616972893 | 0.808273466 | 0.7567 | 0.873110189 |
| SL1344_RS04940_ | -3.613676629 | -3.473081641 | -3.789138261 | -0.292589671 | 0.718713853 | -0.393521324 | 0.328666674 | 0.510433468 | 0.72428642 | 0.6194 | 0.783144594 |
| SL1344_RS04945_ | -2.180858033 | -2.166610429 | -2.196763535 | -0.025757542 | 0.351072996 | -0.078661207 | 0.469333334 | 0.561189662 | 0.760298445 | 0.6662 | 0.81505321 |
| SL1344_RS04950_ | -4.345656578 | -4.459915376 | -4.187796575 | 0.239675805 | 1.072923715 | 0.212992074 | 0.392666671 | 0.631471787 | 0.81310657 | 0.7403 | 0.864848042 |
| SL1344_RS04955_ | -4.998422518 | -5.004948568 | -4.991512287 | 0.037371241 | 0.957193197 | 0.038798385 | 0.481678881 | 0.572672774 | 0.765569845 | 0.6806 | 0.821124513 |
| SL1344_RS04960_ | -1.589888315 | -1.899034373 | -1.365854608 | 0.545293443 | 0.336977159 | 1.563097765 | 0.029980163 | 0.089829613 | 0.316351357 | 0.1365 | 0.375054577 |
| SL1344_RS04965_ | 2.320203396 | 2.149990678 | 2.428802211 | 0.263999367 | 0.133852374 | 1.964557061 | 0.017322048 | 0.048115714 | 0.246541328 | 0.1148 | 0.35098589 |
| SL1344_RS04970_ | -0.576582708 | -0.705769651 | -0.501913378 | 0.18501354 | 0.268280054 | 0.670018653 | 0.252498347 | 0.376806506 | 0.628892231 | 0.4858 | 0.696194951 |
| SL1344_RS04975_ | 0.027811908 | 0.030517906 | 0.026233328 | 0.005337326 | 0.206394975 | 0.022384498 | 0.488666667 | 0.639063018 | 0.818457208 | 0.7492 | 0.869081936 |
| SL1344_RS04980_ | -1.070419671 | -1.181977039 | -0.995904294 | 0.255060247 | 0.517073521 | 0.440068229 | 0.296666676 | 0.422737333 | 0.689873681 | 0.5959 | 0.778197008 |
| SL1344_RS04985_ | -6.11116828 | -5.834126231 | -7.071493054 | -1.179756702 | 2.506548009 | -0.406703373 | 0.311333341 | 0.434821611 | 0.691042976 | 0.6242 | 0.79599948 |
| SL1344_RS04990_ | -4.889886004 | -4.946123997 | -4.853013608 | 0.014167865 | 1.108332122 | 0.01182942 | 0.494 | 0.628936839 | 0.8114456 | 0.7487 | 0.869148478 |
| SL1344_RS04995_ | -6.524373144 | -6.880513231 | -6.05030927 | 0.749377534 | 1.76479846 | 0.371302191 | 0.324450374 | 0.514055418 | 0.720850876 | 0.6004 | 0.767838655 |
| SL1344_RS05000_ | -3.304872529 | -3.155075904 | -3.568894178 | -0.401247809 | 0.892104428 | -0.395311825 | 0.328666674 | 0.502245936 | 0.734557897 | 0.644 | 0.804375952 |
| SL1344_RS05005_ | 0.313176484 | 0.314668204 | 0.311588945 | 0.002648517 | 0.175080164 | 0.009964657 | 0.495003331 | 0.635329925 | 0.814910291 | 0.7445 | 0.867284507 |
| SL1344_RS05010_ | -6.691139694 | -6.522225938 | -6.84559987 | -0.279404475 | 1.816065186 | -0.145350123 | 0.429713527 | 0.539104034 | 0.741603608 | 0.6352 | 0.791190547 |

| | | | | | | | | | | | |
|---|---|---|---|---|---|---|---|---|---|---|---|
| SL1344_RS05015_ | -1.308405584 | -1.458425163 | -1.150772609 | 0.293433012 | 0.327688448 | 0.845213815 | 0.203864107 | 0.285333271 | 0.547370019 | 0.3823 | 0.612778645 |
| SL1344_RS05020_ | -1.212099368 | -1.326302002 | -0.971361925 | 0.353591718 | 0.371349505 | 0.892497 | 0.184666686 | 0.265461901 | 0.539041647 | 0.3793 | 0.61517506 |
| SL1344_RS05025_ | -7.708971039 | -8.371967579 | -7.304465515 | 0.504760701 | 4.297789312 | 0.094015521 | 0.455696204 | 0.657257012 | 0.836180123 | 0.8298 | 0.919804601 |
| SL1344_RS05030_ | -3.040889996 | -3.322251462 | -2.735459103 | 0.580910309 | 0.465160999 | 1.178418958 | 0.099933419 | 0.160830956 | 0.396636338 | 0.2435 | 0.480198558 |
| SL1344_RS05035_ | -4.418969325 | -4.295159733 | -4.53263615 | -0.195385731 | 0.906259827 | -0.190744418 | 0.41172552 | 0.579702066 | 0.777843016 | 0.6959 | 0.8364586 |
| SL1344_RS05040_ | -2.356904761 | -2.536259936 | -2.178959116 | 0.333267385 | 0.461365628 | 0.670239221 | 0.232511673 | 0.374665785 | 0.621241868 | 0.4759 | 0.681812948 |
| SL1344_RS05045_ | -1.472667548 | -1.304601037 | -1.642689726 | -0.337306773 | 0.280862501 | -1.156118764 | 0.101932086 | 0.171713219 | 0.414371457 | 0.2494 | 0.484823899 |
| SL1344_RS05050_ | -5.387355529 | -5.214487534 | -5.600869486 | -0.376846555 | 1.179902997 | -0.29456002 | 0.370000005 | 0.5009802 | 0.722114742 | 0.6281 | 0.788033013 |
| SL1344_RS05055_ | -1.646140911 | -1.6077783 | -1.673554811 | -0.045833783 | 0.421223074 | -0.099075731 | 0.454666668 | 0.680075562 | 0.844506322 | 0.7772 | 0.888901496 |
| SL1344_RS05060_ | -1.110075878 | -1.037618174 | -1.289658566 | -0.228382051 | 0.319041647 | -0.644390165 | 0.250666679 | 0.380257817 | 0.636692379 | 0.5092 | 0.713279358 |
| SL1344_RS05065_ | -4.658331918 | -4.458456807 | -4.85044544 | -0.367546966 | 0.952668327 | -0.348621544 | 0.349766828 | 0.525088539 | 0.732773623 | 0.6201 | 0.782470136 |
| SL1344_RS05070_ | -5.262572897 | -4.976136765 | -5.660008614 | -0.681216792 | 1.052544186 | -0.608070214 | 0.247333346 | 0.366714508 | 0.606766983 | 0.4647 | 0.664649339 |
| SL1344_RS05075_ | -3.868874638 | -3.672964746 | -4.010924079 | -0.328549272 | 0.670663028 | -0.460486183 | 0.306462367 | 0.463492764 | 0.684829204 | 0.5643 | 0.744862839 |
| SL1344_RS05080_ | -3.955764143 | -3.541385 | -4.713780887 | -1.175615146 | 0.768554766 | -1.386506434 | 0.059333406 | 0.113627008 | 0.351941764 | 0.1761 | 0.41598767 |
| SL1344_RS05085_ | -2.263391777 | -2.246857438 | -2.284764212 | -0.039824736 | 0.408784212 | -0.09569938 | 0.458361094 | 0.604065453 | 0.791715007 | 0.7114 | 0.845296185 |
| SL1344_RS05090_ | -2.632371091 | -2.768230156 | -2.533552864 | 0.207255804 | 0.476210436 | 0.385682893 | 0.348666673 | 0.529084029 | 0.738163258 | 0.645 | 0.800425314 |
| SL1344_RS05095_ | -3.305089986 | -3.588973267 | -3.138185942 | 0.424485979 | 0.738448003 | 0.556601564 | 0.279147245 | 0.444380252 | 0.683899059 | 0.555 | 0.747595401 |
| SL1344_RS05100_ | -0.421307323 | -0.316799714 | -0.512297098 | -0.202122428 | 0.195298009 | -0.981451212 | 0.137333362 | 0.218423595 | 0.46427812 | 0.3005 | 0.527310991 |
| SL1344_RS05105_ | -3.766191526 | -3.853894202 | -3.689092609 | 0.152866395 | 0.691679709 | 0.209537916 | 0.404397072 | 0.560437796 | 0.761235539 | 0.6741 | 0.820182402 |
| SL1344_RS05110_ | 0.550048579 | 0.663886607 | 0.446820696 | -0.177693628 | 0.318891034 | -0.527670838 | 0.271333344 | 0.445454681 | 0.70234813 | 0.5552 | 0.76122347 |
| SL1344_RS05115_ | 1.915906774 | 2.159427569 | 1.836230296 | -0.238123411 | 0.305537997 | -0.429149153 | 0.333777489 | 0.862188169 | 0.94682371 | 0.7027 | 0.87368088 |
| SL1344_RS05120_ | -1.30635415 | -1.401024359 | -1.254216653 | 0.144855975 | 0.398599878 | 0.329022682 | 0.36109261 | 0.564417326 | 0.774136365 | 0.6971 | 0.840578374 |
| SL1344_RS05125_ | -2.043101874 | -2.185915899 | -1.908626477 | 0.279135366 | 0.321997947 | 0.795903462 | 0.185209879 | 0.277671388 | 0.51825448 | 0.3751 | 0.591467406 |
| SL1344_RS05130_ | -1.717568986 | -1.815468397 | -1.57309201 | 0.250892553 | 0.354449603 | 0.651716189 | 0.248666679 | 0.383394321 | 0.62666184 | 0.4906 | 0.692801398 |
| SL1344_RS05135_ | -0.269046235 | -0.199964594 | -0.346968405 | -0.147828915 | 0.175111197 | -0.802450966 | 0.186666686 | 0.290570611 | 0.529710938 | 0.3886 | 0.602180618 |
| SL1344_RS05140_ | -3.140493593 | -2.803899628 | -3.371409031 | -0.578757612 | 0.562318601 | -0.951321674 | 0.150666692 | 0.234280392 | 0.488423872 | 0.3208 | 0.55314663 |
| SL1344_RS05165_ | -4.508370563 | -4.638418306 | -4.293370347 | 0.309005624 | 0.934876714 | 0.299413592 | 0.36642239 | 0.560789217 | 0.756901741 | 0.6522 | 0.806185955 |
| SL1344_RS05175_ | -0.997879298 | -0.784138365 | -1.221476678 | -0.490800438 | 0.349049312 | -1.394063966 | 0.049966777 | 0.107654903 | 0.353559258 | 0.1585 | 0.398095678 |
| SL1344_RS05190_ | 3.601350771 | 3.579884356 | 3.622909756 | -0.010605368 | 0.158508054 | -0.053860405 | 0.475682879 | 0.652571134 | 0.843647579 | 0.9568 | 0.980726941 |
| SL1344_RS05195_ | 0.356673755 | 0.263054843 | 0.405674496 | 0.182625071 | 0.297802616 | 0.570885349 | 0.272485021 | 0.390106086 | 0.659136761 | 0.5367 | 0.745509106 |
| SL1344_RS05200_ | 1.856911223 | 1.659793547 | 2.082273127 | 0.409309443 | 0.196394882 | 2.045449754 | 0.005996769 | 0.033988423 | 0.214102024 | 0.1041 | 0.336231552 |
| SL1344_RS05205_ | 0.014996005 | 0.074525495 | -0.023334472 | -0.079865092 | 0.244173584 | -0.312641365 | 0.378666671 | 0.623550669 | 0.809797479 | 0.7154 | 0.8518657 |
| SL1344_RS05210_ | 2.265221222 | 2.291356067 | 2.222587034 | -0.066658351 | 0.145299102 | -0.46608353 | 0.298000009 | 0.510120987 | 0.740036344 | 0.5837 | 0.778887037 |
| SL1344_RS05215_ | 2.675956976 | 2.631128429 | 2.726118293 | 0.036851033 | 0.220543686 | 0.192609807 | 0.412391742 | 0.777362686 | 0.905841908 | 0.8554 | 0.934497535 |
| SL1344_RS05220_ | 1.224790394 | 1.272084512 | 1.191839517 | -0.080166645 | 0.136057854 | -0.536612388 | 0.272666677 | 0.436059766 | 0.672014705 | 0.5379 | 0.730696899 |
| SL1344_RS05225_ | 1.590850493 | 1.566868915 | 1.617316434 | 0.051011756 | 0.09259309 | 0.526460809 | 0.28847436 | 0.406127849 | 0.63485693 | 0.5138 | 0.701572995 |
| SL1344_RS05230_ | 1.458900245 | 1.33754531 | 1.529102897 | 0.184297264 | 0.128630668 | 1.394942158 | 0.074616979 | 0.122702843 | 0.368903973 | 0.2015 | 0.447560106 |
| SL1344_RS05235_ | 2.33352118 | 2.435189769 | 2.21830737 | -0.270703337 | 0.188461498 | -1.59343008 | 0.003332494 | 0.094509094 | 0.349478671 | 0.1035 | 0.335307526 |
| SL1344_RS05240_ | 3.863755498 | 4.169563655 | 3.672321397 | -0.359363988 | 0.271018355 | -1.463640027 | 0.107928086 | 0.09616845 | 0.356662962 | 0.1934 | 0.466886128 |
| SL1344_RS05245_ | 2.632235684 | 2.633490901 | 2.630980905 | 0.031311202 | 0.125263113 | 0.237875721 | 0.410000003 | 0.639705875 | 0.824421635 | 0.8315 | 0.914978965 |
| SL1344_RS05250_ | 0.609389252 | 0.695735487 | 0.575977246 | -0.094144805 | 0.214315364 | -0.36167231 | 0.362666672 | 0.683536471 | 0.84911092 | 0.7365 | 0.877536105 |
| SL1344_RS05255_ | 8.013762348 | 7.725733916 | 8.251396665 | 0.451575813 | 0.170743643 | 2.882421526 | 4.68E-05 | 0.012365956 | 0.140915136 | 0.1 | 0.330257709 |
| SL1344_RS05260_ | 1.186824969 | 1.505390729 | 1.101244687 | -0.367189706 | 0.221398814 | -1.288976677 | 0.109926752 | 0.144543898 | 0.419420975 | 0.2368 | 0.493387031 |
| SL1344_RS05265_ | -0.316116632 | -0.244880413 | -0.406857592 | -0.173012099 | 0.232046759 | -0.687255412 | 0.210526332 | 0.31159246 | 0.572098377 | 0.4186 | 0.632786127 |
| SL1344_RS05270_ | 1.008198745 | 0.988429456 | 1.031456004 | 0.043164926 | 0.113221774 | 0.350284136 | 0.348434383 | 0.476135171 | 0.695325288 | 0.5886 | 0.759095937 |
| SL1344_RS05275_ | -0.898750567 | -0.889831075 | -0.90446883 | -0.025912715 | 0.269491895 | -0.094118275 | 0.463333335 | 0.622320013 | 0.805385108 | 0.7379 | 0.86221233 |
| SL1344_RS05285_ | 2.209301104 | 2.137073478 | 2.319899749 | 0.212265166 | 0.169351239 | 1.26651925 | 0.060626321 | 0.115659463 | 0.375697723 | 0.1697 | 0.417751243 |
| SL1344_RS05290_ | -2.51794038 | -2.57930176 | -2.471293341 | 0.139709321 | 0.485345048 | 0.272497151 | 0.385742843 | 0.553594109 | 0.759042189 | 0.6681 | 0.816160993 |
| SL1344_RS05295_ | 1.780073825 | 1.787751218 | 1.774578627 | -0.037821028 | 0.521928429 | -0.072347581 | 0.453697537 | 0.620788405 | 0.828961124 | 0.9151 | 0.961713978 |
| SL1344_RS05300_ | 2.350888471 | 2.312504935 | 2.388429518 | 0.021438435 | 0.268195204 | 0.050528971 | 0.469686877 | 0.700245493 | 0.870593134 | 0.9322 | 0.971020059 |
| SL1344_RS05305_ | 0.81137981 | 0.789334782 | 0.830220592 | 0.05919723 | 0.225315489 | 0.258065945 | 0.375333338 | 0.637297391 | 0.821064342 | 0.7348 | 0.864727847 |
| SL1344_RS05310_ | -1.367862033 | -1.738418135 | -1.159435083 | 0.546612381 | 0.34847411 | 1.470661091 | 0.056628991 | 0.10479453 | 0.342917765 | 0.1705 | 0.415259568 |
| SL1344_RS05315_ | -0.388584701 | -0.680859523 | -0.078063785 | 0.634368694 | 0.292873486 | 2.175170733 | 0.002666616 | 0.044886698 | 0.235652034 | 0.1022 | 0.333303319 |

| | | | | | | | | | | |
|---|---|---|---|---|---|---|---|---|---|---|
| SL1344_RS05320_ | -2.519789734 | -2.618468181 | -2.337107253 | 0.241177104 | 0.472721327 | 0.446569456 | 0.324450374 | 0.526400686 | 0.733744218 | 0.6182 | 0.787810292 |
| SL1344_RS05325_ | 2.331451714 | 2.338125356 | 2.326362146 | -0.003278981 | 0.134892285 | -0.021860234 | 0.49 | 0.756484023 | 0.891831909 | 0.8436 | 0.928165761 |
| SL1344_RS05330_ | 2.597644004 | 2.652173973 | 2.527631466 | -0.016381969 | 0.348546404 | -0.038337938 | 0.477015324 | 0.71335325 | 0.879174805 | 0.9484 | 0.978060588 |
| SL1344_RS05335_ | 2.919322123 | 3.116529862 | 2.74899767 | -0.375183432 | 0.872540684 | -0.472913446 | 0.264490351 | 0.484641013 | 0.74541434 | 0.5185 | 0.754823415 |
| SL1344_RS05340_ | -1.679776269 | -1.688769503 | -1.67056526 | 0.082682114 | 0.447639888 | 0.176349488 | 0.432378417 | 0.63866959 | 0.820198922 | 0.7831 | 0.887821833 |
| SL1344_RS05345_ | -2.549415561 | -2.130801118 | -3.414408483 | -1.264792691 | 0.633585293 | -1.886397251 | 0.013333676 | 0.059820293 | 0.267533105 | 0.1155 | 0.350982382 |
| SL1344_RS05350_ | -6.473162477 | -5.815953122 | -7.395745655 | -1.679161245 | 2.516059587 | -0.63045608 | 0.217854779 | 0.334344796 | 0.604412455 | 0.4385 | 0.656971469 |
| SL1344_RS05365_ | 1.0294569 | 0.715467563 | 1.138583037 | 0.348154522 | 0.361013552 | 0.857903381 | 0.200666684 | 0.285257291 | 0.582009228 | 0.3967 | 0.646744727 |
| SL1344_RS05370_ | -4.165918595 | -4.353230319 | -3.936109729 | 0.411280964 | 1.054521518 | 0.373013288 | 0.323784151 | 0.528084115 | 0.746447815 | 0.6372 | 0.801912044 |
| SL1344_RS05375_ | 0.575725152 | 0.348200133 | 0.772979574 | 0.406476527 | 0.201667649 | 1.974940193 | 0.013324793 | 0.046311724 | 0.237907107 | 0.1144 | 0.350686698 |
| SL1344_RS05380_ | -0.112408612 | -0.176533909 | -0.055961339 | 0.120263366 | 0.198425342 | 0.561036359 | 0.252000012 | 0.415133632 | 0.651359025 | 0.5241 | 0.71526161 |
| SL1344_RS05385_ | -0.111476533 | -0.1305647 | -0.093003642 | 0.041690507 | 0.191763706 | 0.196084257 | 0.41266667 | 0.561073598 | 0.762016592 | 0.677 | 0.820306209 |
| SL1344_RS05390_ | -0.677704696 | -0.805965595 | -0.573462034 | 0.236425793 | 0.216972437 | 1.033559997 | 0.124583643 | 0.202508835 | 0.445714239 | 0.2887 | 0.518837411 |
| SL1344_RS05395_ | -1.216448045 | -1.161693629 | -1.292099714 | -0.189400751 | 0.455417215 | -0.383351745 | 0.297801475 | 0.412832115 | 0.678196645 | 0.5954 | 0.763538084 |
| SL1344_RS05400_ | -2.407880966 | -2.33696166 | -2.463794867 | -0.068203648 | 0.549201982 | -0.108405707 | 0.450000002 | 0.676853398 | 0.841931751 | 0.7743 | 0.886777452 |
| SL1344_RS05405_ | -1.325499199 | -1.015910762 | -1.468600689 | -0.414254987 | 0.347324334 | -1.124303615 | 0.127248532 | 0.192910305 | 0.454286627 | 0.2895 | 0.533028447 |
| SL1344_RS05410_ | -0.307037018 | -0.467939101 | -0.247630352 | 0.184463449 | 0.385908368 | 0.37356068 | 0.366000005 | 0.730262854 | 0.87959534 | 0.7637 | 0.896897114 |
| SL1344_RS05415_ | -0.420887896 | -0.375352355 | -0.452710014 | -0.028660767 | 0.402709755 | -0.064995378 | 0.461025984 | 0.778827849 | 0.903610119 | 0.8271 | 0.920924566 |
| SL1344_RS05420_ | -1.302159144 | -1.165781737 | -1.438969562 | -0.3222877 | 0.379319828 | -0.870061958 | 0.170000022 | 0.250664575 | 0.522746963 | 0.3536 | 0.580915341 |
| SL1344_RS05425_ | -1.556332966 | -1.377052061 | -1.674046472 | -0.345723844 | 0.402225734 | -0.856933285 | 0.158000024 | 0.263370035 | 0.536729528 | 0.3493 | 0.575355778 |
| SL1344_RS05430_ | -1.620769399 | -1.645403584 | -1.587310655 | 0.01072121 | 0.429156673 | 0.024799132 | 0.491005996 | 0.685693004 | 0.84843484 | 0.7987 | 0.899378398 |
| SL1344_RS05435_ | -1.74541111 | -1.663270199 | -1.820666682 | -0.221012173 | 0.533858029 | -0.390222582 | 0.304666675 | 0.459952714 | 0.708533478 | 0.6066 | 0.778394581 |
| SL1344_RS05440_ | -1.582018123 | -1.360373803 | -1.728680296 | -0.347243257 | 0.381255304 | -0.85350091 | 0.175333354 | 0.264572641 | 0.527138644 | 0.354 | 0.592881237 |
| SL1344_RS05445_ | -2.078521571 | -2.022676747 | -2.113704709 | -0.070544125 | 0.401424245 | -0.171238009 | 0.427048637 | 0.605768945 | 0.795446057 | 0.7182 | 0.850289724 |
| SL1344_RS05450_ | -0.575489271 | -0.512517362 | -0.671731931 | -0.172302488 | 0.25182846 | -0.610744859 | 0.226515671 | 0.348604796 | 0.602357969 | 0.4686 | 0.669250158 |
| SL1344_RS05455_ | -3.608718942 | -4.404820786 | -3.051278588 | 1.349546539 | 0.610374201 | 2.193357997 | 0.008000574 | 0.041378113 | 0.211443947 | 0.1096 | 0.342235493 |
| SL1344_RS05460_ | 0.930525357 | 1.085735194 | 0.885536141 | -0.201360856 | 0.30798639 | -0.689488128 | 0.227848116 | 0.356706484 | 0.636872156 | 0.4315 | 0.692325265 |
| SL1344_RS05465_ | 1.467691916 | 1.309699433 | 1.506438395 | 0.186987934 | 0.142227307 | 1.016398205 | 0.191333352 | 0.248019449 | 0.527953279 | 0.3809 | 0.619098693 |
| SL1344_RS05470_ | 0.660906381 | 0.723583194 | 0.633013557 | -0.062359112 | 0.229510348 | -0.224972701 | 0.407728185 | 0.768785277 | 0.898339868 | 0.8113 | 0.917150938 |
| SL1344_RS05475_ | -0.93991856 | -0.893730397 | -0.981726237 | -0.04932113 | 0.362626909 | -0.125316587 | 0.445333335 | 0.707784125 | 0.86283922 | 0.8074 | 0.905650214 |
| SL1344_RS05480_ | -3.06845406 | -3.182681582 | -3.007962611 | 0.172794702 | 0.559181551 | 0.281039498 | 0.375083283 | 0.550575376 | 0.756178407 | 0.6678 | 0.81643522 |
| SL1344_RS05485_ | -1.737168947 | -1.657683473 | -1.8260654 | -0.262337109 | 0.506924927 | -0.473104714 | 0.298000009 | 0.442728618 | 0.690670926 | 0.6096 | 0.780305971 |
| SL1344_RS05490_ | 2.497905304 | 2.772583749 | 2.467282776 | -0.29384081 | 0.164284514 | -0.958646437 | 0.230666681 | 0.242804213 | 0.542489774 | 0.4614 | 0.697777544 |
| SL1344_RS05495_ | 1.594215856 | 1.522547005 | 1.673475778 | 0.153410768 | 0.434531701 | 0.337932825 | 0.364666672 | 0.541131457 | 0.779363505 | 0.7825 | 0.902922647 |
| SL1344_RS05500_ | 0.28743275 | 0.224344073 | 0.356158172 | 0.165015813 | 0.292171217 | 0.522405099 | 0.245169899 | 0.363064549 | 0.642436004 | 0.4803 | 0.693371738 |
| SL1344_RS05505_ | -0.92286909 | -0.544678066 | -1.040942 | -0.452609391 | 0.327912686 | -1.202175657 | 0.131333363 | 0.175790884 | 0.443834177 | 0.2814 | 0.528434641 |
| SL1344_RS05510_ | -0.523593425 | -0.416183268 | -0.643178887 | -0.226338115 | 0.200361637 | -1.092708481 | 0.112591641 | 0.191111274 | 0.434206276 | 0.2678 | 0.500710944 |
| SL1344_RS05515_ | 3.200296692 | 3.232279 | 3.18452391 | -0.047107787 | 0.136175182 | -0.302842063 | 0.396000004 | 0.679785223 | 0.85349418 | 0.8199 | 0.91998703 |
| SL1344_RS05525_ | 3.846652373 | 3.796276121 | 3.89348001 | 0.16743914 | 0.343722505 | 0.565601202 | 0.291333343 | 0.353431128 | 0.64760188 | 0.6137 | 0.815277246 |
| SL1344_RS05530_ | 1.245918226 | 1.554605133 | 1.202070472 | -0.334258817 | 0.207673415 | -1.061372398 | 0.17721521 | 0.208713904 | 0.500810339 | 0.3587 | 0.607181973 |
| SL1344_RS05540_ | 2.306863598 | 2.273375923 | 2.435341293 | 0.166708549 | 0.209369232 | 0.834594883 | 0.184543657 | 0.232233842 | 0.523033391 | 0.3425 | 0.615479332 |
| SL1344_RS05545_ | 0.404245106 | 0.459908091 | 0.350440734 | -0.126548747 | 0.892912166 | -0.127362875 | 0.446666669 | 0.863913197 | 0.947724483 | 0.9715 | 0.987827573 |
| SL1344_RS05550_ | 1.214420485 | 1.082276926 | 1.330405136 | 0.192609532 | 1.435661762 | 0.119120173 | 0.447035312 | 0.93708114 | 0.977625774 | 1 | 1 |
| SL1344_RS05555_ | -0.250557145 | -0.296773339 | -0.213040188 | -0.230396404 | 1.122249719 | -0.207989145 | 0.415722855 | 0.740368891 | 0.891586472 | 0.9133 | 0.961844942 |
| SL1344_RS05560_ | -1.070044385 | -0.890462648 | -1.27206555 | 0.066856176 | 1.168701816 | 0.065179766 | 0.476000001 | 0.890084329 | 0.958354775 | 0.9856 | 0.993867346 |
| SL1344_RS05565_ | -2.058279644 | -1.998961875 | -2.097901137 | -0.197450782 | 0.726465904 | -0.260258654 | 0.392666671 | 0.659413738 | 0.836650821 | 0.7936 | 0.898162623 |
| SL1344_RS05570_ | -0.077387341 | -0.034139369 | -0.110700016 | -0.081746301 | 0.221752218 | -0.338343091 | 0.355333339 | 0.54872086 | 0.755433734 | 0.6715 | 0.817546908 |
| SL1344_RS05580_ | 1.699618386 | 2.402468609 | 1.659478433 | -0.742057845 | 0.290538396 | -0.644942808 | 0.33333334 | 0.475277896 | 0.738381375 | 0.7 | 0.872542119 |
| SL1344_RS05585_ | -5.494847944 | -5.425729533 | -5.624401617 | -0.204959247 | 1.527119259 | -0.107216517 | 0.444000002 | 0.616644721 | 0.809008801 | 0.7625 | 0.87848967 |
| SL1344_RS05590_ | 1.752655467 | 1.787945301 | 1.682512934 | -0.094796645 | 0.125056513 | -0.723999408 | 0.233844118 | 0.360287999 | 0.613662412 | 0.4681 | 0.684819226 |
| SL1344_RS05595_ | 1.522825469 | 1.532211645 | 1.515372868 | 0.00789974 | 0.164833395 | 0.033402616 | 0.485333334 | 0.665098615 | 0.841525957 | 0.8219 | 0.913170407 |
| SL1344_RS05600_ | 0.745905959 | 0.682800476 | 0.786255775 | 0.064268521 | 0.205348824 | 0.276096517 | 0.388666671 | 0.627286211 | 0.814667988 | 0.7355 | 0.864802108 |
| SL1344_RS05605_ | 1.283803191 | 1.358501456 | 1.158482124 | -0.192686206 | 0.189349781 | -0.974198613 | 0.16066669 | 0.224377514 | 0.50557591 | 0.3134 | 0.568112127 |

| | | | | | | | | | | | |
|---|---|---|---|---|---|---|---|---|---|---|---|
| SL1344_RS05610_ | -1.88994354 | -1.858014134 | -1.924688886 | -0.068100464 | 0.287824202 | -0.214428206 | 0.402000004 | 0.497621824 | 0.708509313 | 0.6088 | 0.770205814 |
| SL1344_RS05615_ | -4.288119486 | -4.494890549 | -4.110433241 | 0.387194769 | 0.766541672 | 0.496159944 | 0.301132587 | 0.433423886 | 0.660196698 | 0.5419 | 0.722455907 |
| SL1344_RS05620_ | -4.09702652 | -4.268546238 | -3.919414615 | 0.334085922 | 0.750253574 | 0.411096572 | 0.322000008 | 0.502790228 | 0.714875423 | 0.6067 | 0.771862644 |
| SL1344_RS05625_ | -2.97228703 | -3.149787064 | -2.745061586 | 0.37571501 | 0.529866276 | 0.655044696 | 0.229333348 | 0.371147139 | 0.607455721 | 0.4743 | 0.671123993 |
| SL1344_RS05635_ | -3.542547376 | -3.350853486 | -3.801949298 | -0.461298334 | 0.642311678 | -0.661151989 | 0.212000016 | 0.326561498 | 0.577911941 | 0.4248 | 0.632234661 |
| SL1344_RS05650_ | 0.993414749 | 1.502857647 | 0.447981655 | -1.058859616 | 0.138371296 | -7.665170385 | 4.68E-05 | 0.002263604 | 0.054736877 | 0.1 | 0.330257709 |
| SL1344_RS05660_ | 0.160172633 | 3.40E-05 | 0.405234486 | 0.383572332 | 0.224085122 | 1.6136506 | 0.043304566 | 0.077061588 | 0.299370964 | 0.1481 | 0.392854257 |
| SL1344_RS05670_ | 4.181484192 | 4.173516211 | 4.187445776 | 0.011398854 | 0.079510513 | 0.136611817 | 0.450666669 | 0.741902975 | 0.883436447 | 0.8612 | 0.934444994 |
| SL1344_RS05675_ | -1.53779331 | -1.521852162 | -1.553403691 | -0.074862123 | 0.398579711 | -0.173778474 | 0.42066667 | 0.642571425 | 0.819908102 | 0.7758 | 0.883835031 |
| SL1344_RS05680_ | 0.350596062 | 0.327083351 | 0.374834004 | 0.136620859 | 0.293978835 | 0.449472091 | 0.332666674 | 0.478228605 | 0.722958052 | 0.6895 | 0.83329132 |
| SL1344_RS05685_ | -0.800747509 | -0.896863277 | -0.677314018 | 0.216194911 | 0.228034205 | 0.917688244 | 0.16322454 | 0.256446859 | 0.502811801 | 0.3466 | 0.56980938 |
| SL1344_RS05690_ | 1.991236864 | 2.007114068 | 1.955961312 | -0.032787295 | 0.127071663 | -0.227606659 | 0.418387745 | 0.732899432 | 0.877947575 | 0.7841 | 0.90325516 |
| SL1344_RS05695_ | 1.664801536 | 1.699924708 | 1.638091424 | -0.056716864 | 0.220898061 | -0.243452988 | 0.394666671 | 0.637213012 | 0.827392894 | 0.7968 | 0.900316319 |
| SL1344_RS05700_ | -0.798519486 | -0.728314045 | -0.855352618 | -0.123055736 | 0.215511758 | -0.546000279 | 0.266000011 | 0.400983165 | 0.634121392 | 0.502 | 0.696110624 |
| SL1344_RS05705_ | -2.243883674 | -2.308213742 | -2.207010596 | 0.156196889 | 0.513303567 | 0.274383061 | 0.369333339 | 0.529175485 | 0.750513783 | 0.6727 | 0.818315787 |
| SL1344_RS05710_ | -3.364605582 | -3.80607195 | -3.029024402 | 0.77018094 | 0.523334356 | 1.413870945 | 0.055962769 | 0.112743421 | 0.338808211 | 0.1779 | 0.417376468 |
| SL1344_RS05715_ | 1.963055621 | 1.807253174 | 2.279527849 | 0.442824006 | 0.156707827 | 2.562222417 | 0.00067292 | 0.031237558 | 0.204611076 | 0.1007 | 0.3312952 |
| SL1344_RS05730_ | 2.900750442 | 2.901328345 | 2.900130521 | 0.03444074 | 0.151044058 | 0.205275865 | 0.40466667 | 0.531758145 | 0.766663779 | 0.8242 | 0.911319 |
| SL1344_RS05735_ | 2.361226579 | 2.288531198 | 2.533244406 | 0.17746968 | 0.185984465 | 0.957407769 | 0.186666686 | 0.253271985 | 0.544430632 | 0.3386 | 0.600502476 |
| SL1344_RS05740_ | 1.666704232 | 1.857247507 | 1.56379399 | -0.19996617 | 0.24215291 | -0.897578302 | 0.178547655 | 0.276058702 | 0.565212258 | 0.3467 | 0.60273573 |
| SL1344_RS05745_ | 0.156553131 | 0.29298425 | 0.042696397 | -0.262761667 | 0.185751472 | -1.383228279 | 0.055296548 | 0.116432719 | 0.35351638 | 0.1728 | 0.412158319 |
| SL1344_RS05750_ | 1.757053958 | 1.690614509 | 1.795834243 | 0.124038982 | 0.167156696 | 0.705092267 | 0.222518336 | 0.319744462 | 0.597987044 | 0.4521 | 0.67464464 |
| SL1344_RS05755_ | 2.230068445 | 2.125879502 | 2.251978937 | 0.124170911 | 0.252328675 | 0.477864186 | 0.307794812 | 0.412365161 | 0.690830492 | 0.6497 | 0.831121689 |
| SL1344_RS05760_ | 2.536030416 | 2.57084382 | 2.522997071 | -0.039652278 | 0.12368606 | -0.122723753 | 0.397068625 | 0.649678709 | 0.842958423 | 0.7981 | 0.913698667 |
| SL1344_RS05765_ | 1.209876413 | 1.239535773 | 1.193868997 | -0.032520816 | 0.203651674 | -0.126026981 | 0.445702867 | 0.780638139 | 0.905120347 | 0.855 | 0.935509574 |
| SL1344_RS05780_ | 3.952081188 | 4.408167008 | 3.481668525 | -0.937998174 | 0.091451555 | -10.28475682 | 4.68E-05 | 0.000357337 | 0.029151511 | 0.1 | 0.330257709 |
| SL1344_RS05785_ | 3.870752099 | 4.376561284 | 3.316239735 | -1.072605181 | 0.085218576 | -12.66204721 | 4.68E-05 | 0.00030718 | 0.027262654 | 0.1 | 0.330257709 |
| SL1344_RS05790_ | 3.390480239 | 3.886099566 | 2.936158118 | -0.955782382 | 0.118645492 | -8.110650457 | 4.68E-05 | 0.003452979 | 0.075780581 | 0.1 | 0.330257709 |
| SL1344_RS05800_ | 4.91579576 | 5.420267986 | 4.483788545 | -0.935100577 | 0.083520027 | -10.89592608 | 4.68E-05 | 0.001335904 | 0.050492464 | 0.1 | 0.330257709 |
| SL1344_RS05810_ | 5.38302617 | 5.848500219 | 4.907530992 | -0.873303084 | 0.113048748 | -7.942332056 | 4.68E-05 | 0.00020558 | 0.026011235 | 0.1 | 0.330257709 |
| SL1344_RS05815_ | 3.326181747 | 3.867106028 | 2.858554485 | -0.997718625 | 0.11426721 | -8.501117418 | 4.68E-05 | 0.001117785 | 0.044886287 | 0.1 | 0.330257709 |
| SL1344_RS05825_ | 3.081914289 | 3.513187793 | 2.541073595 | -0.984859828 | 0.14104475 | -7.11590554 | 4.68E-05 | 0.000915042 | 0.042428778 | 0.1 | 0.330257709 |
| SL1344_RS05845_ | -5.854756015 | -6.438195465 | -5.286889291 | 1.153279101 | 1.261862582 | 0.83824588 | 0.169220541 | 0.26760034 | 0.508936532 | 0.3507 | 0.570651587 |
| SL1344_RS05850_ | 2.129274777 | 2.109719118 | 2.170153958 | 0.056491625 | 0.117632324 | 0.443699301 | 0.326000007 | 0.53384679 | 0.753401369 | 0.6575 | 0.819648469 |
| SL1344_RS05855_ | 0.704821505 | 0.676614887 | 0.724701049 | 0.091074699 | 0.258962408 | 0.351838444 | 0.351765496 | 0.589526444 | 0.794495416 | 0.7198 | 0.85576523 |
| SL1344_RS05860_ | -1.634483565 | -1.712501075 | -1.55844991 | 0.207857646 | 0.415481915 | 0.474518521 | 0.294000009 | 0.446359004 | 0.684829106 | 0.5982 | 0.763837179 |
| SL1344_RS05865_ | 0.73718483 | 0.94484003 | 0.651880335 | -0.282853752 | 0.21791024 | -1.16906213 | 0.127333364 | 0.178453456 | 0.455782058 | 0.2616 | 0.51706037 |
| SL1344_RS05870_ | 5.29781845 | 5.232609891 | 5.364640446 | 0.071262122 | 0.172115985 | 0.41610878 | 0.386666671 | 0.571961835 | 0.799736931 | 0.8426 | 0.929291202 |
| SL1344_RS05875_ | 5.595183319 | 5.496394579 | 5.745926498 | 0.294665957 | 0.163195064 | 2.029855607 | 4.68E-05 | 0.045536347 | 0.253526153 | 0.1 | 0.330257709 |
| SL1344_RS05880_ | 4.434709689 | 4.265332355 | 4.555020868 | 0.133614669 | 0.304681305 | 0.367801609 | 0.339107268 | 0.964280016 | 0.987518237 | 0.7153 | 0.878971698 |
| SL1344_RS05885_ | 3.073420066 | 3.029621737 | 3.085292093 | 0.055134155 | 0.111314171 | 0.430034187 | 0.340439713 | 0.543555133 | 0.76958264 | 0.7128 | 0.862756181 |
| SL1344_RS05890_ | 3.301774619 | 3.338891185 | 3.279328244 | -0.058058741 | 0.077705995 | -0.708703936 | 0.229333348 | 0.350401726 | 0.61377889 | 0.4604 | 0.679653962 |
| SL1344_RS05900_ | 6.987871595 | 7.052729605 | 6.914869112 | 0.003368709 | 0.211578769 | 0.022837618 | 0.488341106 | 0.81091902 | 0.926614268 | 1 | 1 |
| SL1344_RS05905_ | 6.180984355 | 6.124392381 | 6.20788957 | 0.084494082 | 0.115879351 | 0.556084671 | 0.337774823 | 0.529849313 | 0.77386141 | 0.7087 | 0.874461898 |
| SL1344_RS05910_ | 0.699337341 | 0.749097065 | 0.661455726 | -0.079998564 | 0.204229477 | -0.368639498 | 0.348666673 | 0.585914806 | 0.785772731 | 0.7102 | 0.846534443 |
| SL1344_RS05915_ | 1.208817856 | 1.345155335 | 1.172277501 | -0.160213686 | 0.161839846 | -0.837685088 | 0.211192554 | 0.307129027 | 0.580660106 | 0.4525 | 0.679520305 |
| SL1344_RS05920_ | 1.311548656 | 1.391494763 | 1.264682216 | -0.121997136 | 0.131155496 | -0.850716379 | 0.187874769 | 0.277602973 | 0.539254434 | 0.3801 | 0.612801701 |
| SL1344_RS05925_ | 1.529428853 | 1.571165705 | 1.482373136 | -0.094164415 | 0.125503898 | -0.727359178 | 0.209860109 | 0.315473791 | 0.574929202 | 0.4149 | 0.633440594 |
| SL1344_RS05930_ | 1.388410915 | 1.393884717 | 1.384370739 | 0.053493236 | 0.237840524 | 0.223462109 | 0.42066667 | 0.587791319 | 0.7965628 | 0.8399 | 0.917271411 |
| SL1344_RS05935_ | 4.319963959 | 4.441041696 | 4.250680701 | -0.192181428 | 0.098802516 | -1.937549074 | 0.000673369 | 0.07592633 | 0.317105466 | 0.1007 | 0.331274541 |
| SL1344_RS05940_ | -1.175423997 | -0.833825713 | -1.512110105 | -0.704753446 | 0.354248282 | -1.979527536 | 0.009333825 | 0.049089841 | 0.244100921 | 0.1087 | 0.342034749 |
| SL1344_RS05945_ | 1.882791886 | 1.777944904 | 1.908572269 | 0.115035675 | 0.15471302 | 0.465232 | 0.344666673 | 0.703566299 | 0.866580901 | 0.7189 | 0.878295234 |
| SL1344_RS05950_ | 2.01241072 | 1.878766544 | 2.067195991 | 0.146901141 | 0.16637501 | 0.803070588 | 0.220000015 | 0.319618644 | 0.603496246 | 0.4267 | 0.667726384 |

| | | | | | | | | | | | |
|---|---|---|---|---|---|---|---|---|---|---|---|
| SL1344_RS05955_ | 1.522102825 | 1.434573896 | 1.594026381 | 0.044871164 | 0.266897505 | 0.195229222 | 0.437041974 | 0.751572482 | 0.890556997 | 0.8687 | 0.938739694 |
| SL1344_RS05960_ | 1.178758589 | 1.28513902 | 1.095474182 | -0.175738955 | 0.17618209 | -0.965409624 | 0.151333358 | 0.21965834 | 0.497562578 | 0.3043 | 0.560006792 |
| SL1344_RS05965_ | 1.293659215 | 1.304289271 | 1.284602393 | -0.062509754 | 0.233053421 | -0.265923933 | 0.390666671 | 0.618337396 | 0.81443388 | 0.7944 | 0.898069287 |
| SL1344_RS05970_ | 2.538207307 | 2.585368866 | 2.520310541 | -0.063262215 | 0.166378464 | -0.32696602 | 0.383077953 | 0.713820838 | 0.871511216 | 0.7915 | 0.909092695 |
| SL1344_RS05975_ | 1.052680413 | 1.499835876 | 0.823704195 | -0.677101473 | 0.426232977 | -1.630477125 | 0.023984199 | 0.060867089 | 0.286207784 | 0.1208 | 0.360514043 |
| SL1344_RS05980_ | -1.830632241 | -1.955697843 | -1.64177493 | 0.208637871 | 0.522042625 | 0.370883768 | 0.350666673 | 0.612598951 | 0.803944105 | 0.7133 | 0.853309613 |
| SL1344_RS05985_ | 2.237060869 | 2.323435882 | 2.192359358 | -0.160382214 | 0.38476186 | -0.464613464 | 0.254000012 | 0.468972145 | 0.730520927 | 0.4891 | 0.735632984 |
| SL1344_RS05990_ | 0.161180299 | 0.535703943 | -0.15392552 | -0.682167413 | 0.180971552 | -3.710296732 | 0.00067292 | 0.009910896 | 0.112431724 | 0.1 | 0.330257709 |
| SL1344_RS05995_ | -0.230165008 | -0.307902081 | -0.195863104 | 0.103987347 | 0.294960405 | 0.329611176 | 0.372666672 | 0.595510213 | 0.796250121 | 0.7367 | 0.867140961 |
| SL1344_RS06000_ | 2.931303472 | 2.906212527 | 2.962927299 | 0.053749747 | 0.087632001 | 0.56945721 | 0.263157906 | 0.420263842 | 0.663582511 | 0.527 | 0.724842522 |
| SL1344_RS06005_ | 1.997250785 | 2.022471022 | 1.975237044 | -0.047437845 | 0.086097954 | -0.532425857 | 0.28047969 | 0.413561426 | 0.644499462 | 0.5202 | 0.706989444 |
| SL1344_RS06010_ | 1.243305406 | 1.263030699 | 1.207726354 | -0.042403231 | 0.239318931 | -0.158554451 | 0.439040642 | 0.72335844 | 0.87539749 | 0.8655 | 0.938402389 |
| SL1344_RS06015_ | 1.653939786 | 1.703444842 | 1.617324799 | -0.083577209 | 0.110617658 | -0.692932187 | 0.225183226 | 0.34262729 | 0.590666271 | 0.4453 | 0.655941225 |
| SL1344_RS06020_ | 2.090758418 | 2.092167324 | 2.089238571 | -0.05213463 | 0.237082048 | -0.185748054 | 0.405729517 | 0.499809801 | 0.752288165 | 0.8443 | 0.923735589 |
| SL1344_RS06025_ | 2.141788059 | 2.189681658 | 2.108475982 | -0.094167536 | 0.123180127 | -0.747948621 | 0.158000024 | 0.277591703 | 0.560089908 | 0.3524 | 0.576572056 |
| SL1344_RS06030_ | 2.091462941 | 1.964580363 | 2.225124675 | 0.289151695 | 0.162231437 | 1.780569579 | 0.009327606 | 0.058009647 | 0.272835698 | 0.1061 | 0.338900091 |
| SL1344_RS06035_ | 0.908547482 | 0.6497912 | 1.025460294 | 0.312810802 | 0.234203646 | 1.245849872 | 0.105929419 | 0.145949189 | 0.418724116 | 0.2142 | 0.477518036 |
| SL1344_RS06045_ | 0.851179307 | 0.970145315 | 0.800859228 | -0.130331761 | 0.34901017 | -0.295484618 | 0.355096608 | 0.834179609 | 0.932561948 | 0.739 | 0.88708291 |
| SL1344_RS06050_ | 1.420911878 | 1.49939102 | 1.360925884 | -0.124374207 | 0.142288092 | -0.808105553 | 0.205862775 | 0.313433437 | 0.577339344 | 0.407 | 0.639352953 |
| SL1344_RS06055_ | 0.424882508 | 0.416487058 | 0.427402989 | 0.008113183 | 0.275377659 | 0.030376448 | 0.488666667 | 0.778176783 | 0.904334792 | 0.8418 | 0.931867379 |
| SL1344_RS06070_ | 2.540430096 | 2.25967344 | 2.717524034 | 0.433904113 | 0.162911655 | 2.470035911 | 0.00067292 | 0.037166251 | 0.225541716 | 0.1002 | 0.330551488 |
| SL1344_RS06085_ | 2.18680803 | 1.736964163 | 2.349589442 | 0.612566562 | 0.227412768 | 2.504259541 | 4.68E-05 | 0.053258982 | 0.27024818 | 0.1 | 0.330257709 |
| SL1344_RS06090_ | 1.60332665 | 1.308590828 | 1.707628396 | 0.400832009 | 0.209509303 | 1.695433591 | 0.019333568 | 0.083695457 | 0.327855772 | 0.1213 | 0.358696972 |
| SL1344_RS06095_ | 3.171419544 | 3.039324894 | 3.313232867 | 0.290765354 | 0.094932458 | 3.093413284 | 4.68E-05 | 0.014450111 | 0.141453009 | 0.1 | 0.330257709 |
| SL1344_RS06100_ | -0.468294248 | -0.688008222 | -0.265935068 | 0.417518592 | 0.179980815 | 2.261394323 | 0.005330642 | 0.036885656 | 0.19903326 | 0.1099 | 0.34261797 |
| SL1344_RS06105_ | -0.599285015 | -0.858243959 | -0.347243418 | 0.511666213 | 0.210251268 | 2.376110928 | 0.005334196 | 0.033696355 | 0.194178156 | 0.1047 | 0.336638823 |
| SL1344_RS06110_ | -0.065846278 | -0.194542647 | 0.026841805 | 0.243418281 | 0.235868427 | 1.008937624 | 0.092000045 | 0.193259392 | 0.466313239 | 0.2401 | 0.475032029 |
| SL1344_RS06115_ | 5.954026401 | 5.99138887 | 5.918689909 | 0.03449407 | 0.170423804 | 0.218166979 | 0.445333335 | 0.905917994 | 0.964978198 | 1 | 1 |
| SL1344_RS06120_ | 0.382328617 | 0.498703318 | 0.288962861 | -0.124642486 | 0.438130926 | -0.299739907 | 0.377333338 | 0.757424129 | 0.894477909 | 0.793 | 0.905641321 |
| SL1344_RS06125_ | -4.943010253 | -2.799320249 | -5.954554307 | -2.569322003 | 2.074041474 | -1.255399581 | 0.10933337 | 0.136841235 | 0.402619072 | 0.2309 | 0.492942261 |
| SL1344_RS06130_ | -3.106843321 | -3.148848106 | -3.06596342 | 0.135573574 | 0.664804239 | 0.197361007 | 0.41439041 | 0.604282032 | 0.797789627 | 0.7283 | 0.854795662 |
| SL1344_RS06135_ | 2.070548393 | 2.114426721 | 2.006811164 | -0.140240846 | 0.221806854 | -0.614721787 | 0.23717523 | 0.339947907 | 0.629907439 | 0.4645 | 0.698797945 |
| SL1344_RS06140_ | -0.821061284 | -1.087594984 | -0.519939103 | 0.572791948 | 0.228654661 | 2.390506154 | 0.003334715 | 0.030276123 | 0.187444689 | 0.1037 | 0.335230633 |
| SL1344_RS06150_ | -4.670885569 | -4.600070445 | -4.791501258 | -0.286054513 | 1.226604389 | -0.207843378 | 0.393333337 | 0.54744541 | 0.765095476 | 0.6999 | 0.8349536 |
| SL1344_RS06155_ | -9.38020152 | -8.892850922 | -9.857135242 | -0.814377552 | 4.021659128 | -0.16691101 | 0.411059297 | 0.50357286 | 0.720611764 | 0.6201 | 0.779460375 |
| SL1344_RS06160_ | -3.340192573 | -3.431807772 | -3.249447554 | 0.128184057 | 0.728825833 | 0.160799109 | 0.430000003 | 0.633581317 | 0.814730862 | 0.7418 | 0.866027687 |
| SL1344_RS06165_ | 0.22405274 | 0.178624041 | 0.270112001 | 0.005307273 | 0.329235525 | 0.013866683 | 0.495003331 | 0.769283924 | 0.899095561 | 0.9044 | 0.955687006 |
| SL1344_RS06170_ | -2.120181412 | -2.238992441 | -2.039543776 | 0.155953508 | 0.429563092 | 0.336696605 | 0.355762831 | 0.556943653 | 0.760860613 | 0.6665 | 0.816895605 |
| SL1344_RS06180_ | 0.848649539 | 0.627764157 | 1.04161868 | 0.431942931 | 0.177886157 | 2.418942434 | 0.002000966 | 0.030179336 | 0.194070659 | 0.1007 | 0.331272355 |
| SL1344_RS06185_ | -4.312989017 | -4.308174322 | -4.317174593 | 0.008339245 | 0.861396335 | 0.00776946 | 0.497001999 | 0.619868393 | 0.801216401 | 0.7228 | 0.851572969 |
| SL1344_RS06190_ | -3.451570899 | -3.250214938 | -3.58927465 | -0.311222269 | 0.761859141 | -0.395745939 | 0.328666674 | 0.549154133 | 0.753316215 | 0.6539 | 0.807051874 |
| SL1344_RS06195_ | 0.144142759 | 0.136799669 | 0.15563046 | 0.087289748 | 0.436642126 | 0.201541773 | 0.41972019 | 0.736924691 | 0.883173727 | 0.8594 | 0.934603836 |
| SL1344_RS06200_ | -2.03754423 | -2.532719339 | -1.844315989 | 0.625066993 | 0.593134474 | 0.945493377 | 0.166000022 | 0.232287225 | 0.516378653 | 0.3368 | 0.58828773 |
| SL1344_RS06205_ | 2.399063566 | 2.631195503 | 2.264222713 | -0.396560272 | 0.594553006 | -0.774927123 | 0.174000021 | 0.250108219 | 0.554223058 | 0.3155 | 0.59189453 |
| SL1344_RS06210_ | -3.572688728 | -3.355304322 | -3.801194298 | -0.46567999 | 0.619209682 | -0.72247332 | 0.202531663 | 0.325796897 | 0.572174876 | 0.431 | 0.63889137 |
| SL1344_RS06215_ | -0.598481613 | -0.690817963 | -0.539187962 | 0.143238274 | 0.248917281 | 0.538712223 | 0.289806805 | 0.449542997 | 0.681700883 | 0.5533 | 0.74033192 |
| SL1344_RS06220_ | 0.774707239 | 0.244575904 | 1.032769474 | 0.786693355 | 0.300362964 | 2.485394982 | 4.68E-05 | 0.045590922 | 0.247917044 | 0.1 | 0.330257709 |
| SL1344_RS06225_ | -0.807742876 | -1.164841923 | -0.394526853 | 0.742583301 | 0.288538742 | 2.520566527 | 0.003334715 | 0.025687971 | 0.177659007 | 0.1036 | 0.335235379 |
| SL1344_RS06230_ | 2.083033467 | 1.956802759 | 2.198624273 | 0.244681657 | 0.092527429 | 2.573078841 | 0.0020023 | 0.025749501 | 0.175469362 | 0.1024 | 0.333751298 |
| SL1344_RS06235_ | 0.225657125 | 0.113901898 | 0.289500165 | 0.164072406 | 0.273500943 | 0.535000824 | 0.290666676 | 0.442078211 | 0.693635786 | 0.5817 | 0.769688708 |
| SL1344_RS06240_ | -1.507218182 | -1.634399721 | -1.44619962 | 0.170488838 | 0.369648162 | 0.416840278 | 0.337774823 | 0.538514134 | 0.751272377 | 0.6607 | 0.818534125 |
| SL1344_RS06245_ | -1.112000537 | -1.151199526 | -1.070798753 | 0.079046032 | 0.308530738 | 0.263682729 | 0.384666671 | 0.577978177 | 0.775459292 | 0.6872 | 0.829908213 |
| SL1344_RS06250_ | -1.171249642 | -1.142368306 | -1.191364813 | -0.031793211 | 0.385350538 | -0.077040267 | 0.465333335 | 0.709694823 | 0.863427006 | 0.796 | 0.899698668 |

| | | | | | | | | | | | |
|---|---|---|---|---|---|---|---|---|---|---|---|
| SL1344_RS06255_ | -0.924478072 | -1.228870643 | -0.687794806 | 0.537589808 | 0.241832835 | 2.188688711 | 0.007995244 | 0.041002641 | 0.213805561 | 0.1096 | 0.342689597 |
| SL1344_RS06260_ | -1.376586723 | -1.855112152 | -0.931318656 | 0.962717679 | 0.450154779 | 2.175372031 | 4.68E-05 | 0.062964754 | 0.281566852 | 0.1002 | 0.330539174 |
| SL1344_RS06265_ | -2.648952227 | -3.25425489 | -2.183732826 | 1.061248031 | 0.596523539 | 1.744979897 | 0.014000326 | 0.072123693 | 0.290754094 | 0.1152 | 0.350942553 |
| SL1344_RS06270_ | -0.73772405 | -1.000725945 | -0.558448111 | 0.438270989 | 0.251654381 | 1.669201477 | 0.029980163 | 0.076239661 | 0.295449088 | 0.1351 | 0.375615204 |
| SL1344_RS06275_ | -0.017211894 | -0.143643487 | 0.075574937 | 0.220147217 | 0.177546367 | 1.135028806 | 0.085333383 | 0.160105433 | 0.402203874 | 0.2321 | 0.467246483 |
| SL1344_RS06280_ | -3.10342659 | -3.363029208 | -2.708934522 | 0.634198424 | 0.515859981 | 1.150373483 | 0.118587642 | 0.180436507 | 0.423845029 | 0.2641 | 0.505446254 |
| SL1344_RS06285_ | 0.705059842 | 0.451504116 | 0.89319289 | 0.46311042 | 0.192646863 | 2.407306386 | 0.001335878 | 0.046244863 | 0.240096842 | 0.1001 | 0.330408565 |
| SL1344_RS06290_ | 0.449645605 | 0.010960444 | 0.867889767 | 0.900381244 | 0.539766384 | 1.698997466 | 0.00067292 | 0.084361796 | 0.334191393 | 0.1004 | 0.330825568 |
| SL1344_RS06295_ | -5.653855165 | -6.082733982 | -5.055642413 | 1.063020111 | 1.379399113 | 0.701457291 | 0.212000016 | 0.335964456 | 0.580245612 | 0.4337 | 0.642452659 |
| SL1344_RS06300_ | 0.603177182 | 0.329500264 | 0.825466043 | 0.571625663 | 0.366953242 | 1.573246874 | 0.003334715 | 0.09624013 | 0.352566541 | 0.1043 | 0.33644027 |
| SL1344_RS06305_ | 1.1822478 | 0.975369479 | 1.380711101 | 0.402605532 | 0.133953398 | 2.957073745 | 4.68E-05 | 0.017284582 | 0.146449697 | 0.1003 | 0.330693984 |
| SL1344_RS06310_ | -1.574823966 | -1.540061421 | -1.607456614 | -0.079317568 | 0.336520399 | -0.223476155 | 0.399733515 | 0.581715935 | 0.77619785 | 0.695 | 0.832751822 |
| SL1344_RS06315_ | 0.25244934 | 0.09749616 | 0.406326124 | 0.306375175 | 0.249206637 | 1.20999154 | 0.097934752 | 0.14159057 | 0.404159159 | 0.2096 | 0.46422763 |
| SL1344_RS06320_ | -2.093774729 | -2.18302153 | -1.986809511 | 0.201581759 | 0.345876344 | 0.530947964 | 0.270486353 | 0.408506644 | 0.635923508 | 0.5192 | 0.703306265 |
| SL1344_RS06325_ | -2.460948812 | -2.787158528 | -2.32551395 | 0.442656182 | 0.562001485 | 0.723622341 | 0.220666682 | 0.317019984 | 0.587894393 | 0.4456 | 0.665307611 |
| SL1344_RS06330_ | 0.632442068 | 0.420627018 | 0.781293988 | 0.332195248 | 0.34169437 | 0.900879488 | 0.159227205 | 0.237033911 | 0.532136582 | 0.3022 | 0.566680503 |
| SL1344_RS06335_ | -0.031999765 | -0.527880267 | 0.115618502 | 0.624047708 | 0.279632684 | 1.832771183 | 0.016655836 | 0.083930914 | 0.323335769 | 0.1205 | 0.357309741 |
| SL1344_RS06340_ | 1.306164147 | 0.97246077 | 1.609241774 | 0.580480686 | 0.216462329 | 2.678785758 | 4.68E-05 | 0.018404462 | 0.160188154 | 0.1002 | 0.3305455 |
| SL1344_RS06345_ | 0.275411388 | 0.116835733 | 0.571596844 | 0.455247041 | 0.229865935 | 1.814886021 | 0.013324793 | 0.065096535 | 0.282936582 | 0.1144 | 0.349584246 |
| SL1344_RS06350_ | -1.396215831 | -1.464340068 | -1.271928985 | 0.170033903 | 0.332245648 | 0.45096085 | 0.318000008 | 0.524492882 | 0.736168645 | 0.6151 | 0.789338899 |
| SL1344_RS06355_ | -0.560171415 | -0.563361545 | -0.558547622 | 0.002215666 | 0.270136118 | 0.006720904 | 0.498 | 0.659006792 | 0.828713992 | 0.7649 | 0.877629349 |
| SL1344_RS06360_ | -1.238031643 | -1.268513531 | -1.218216888 | 0.090808253 | 0.372873228 | 0.221693808 | 0.383077953 | 0.549840789 | 0.766057563 | 0.6968 | 0.833868842 |
| SL1344_RS06365_ | -4.884340446 | -5.358416655 | -4.562773561 | 0.779873954 | 1.091682024 | 0.643439374 | 0.222666682 | 0.341134989 | 0.594285522 | 0.4507 | 0.65845646 |
| SL1344_RS06370_ | -0.96234496 | -1.334220769 | -0.57621555 | 0.755369404 | 0.250961262 | 2.946128543 | 0.001335878 | 0.017419022 | 0.144482107 | 0.1007 | 0.331258149 |
| SL1344_RS06375_ | -0.879995117 | -1.254592382 | -0.612881476 | 0.627181035 | 0.325696377 | 1.886432154 | 0.014000326 | 0.062298165 | 0.271675612 | 0.1154 | 0.351388626 |
| SL1344_RS06380_ | 0.8729119 | 0.822831943 | 0.996002245 | 0.161044112 | 0.14927714 | 0.995962067 | 0.164666689 | 0.247141672 | 0.512023543 | 0.3535 | 0.588120259 |
| SL1344_RS06385_ | 1.057542784 | 0.775536317 | 1.293447357 | 0.515468146 | 0.11405171 | 4.46102904 | 4.68E-05 | 0.006346481 | 0.089850778 | 0.1 | 0.330257709 |
| SL1344_RS06390_ | 3.979818099 | 3.977703226 | 3.983204096 | 0.021581356 | 0.092839707 | 0.235880926 | 0.391333338 | 0.552378587 | 0.775880461 | 0.7756 | 0.883344001 |
| SL1344_RS06395_ | -1.907020512 | -1.956717352 | -1.829275677 | 0.170880252 | 0.43577424 | 0.358273404 | 0.331778821 | 0.503508329 | 0.727418752 | 0.6336 | 0.791443542 |
| SL1344_RS06400_ | 0.511822087 | 0.402352331 | 0.54835497 | 0.13918246 | 0.205703904 | 0.597392465 | 0.273333344 | 0.416284664 | 0.669573247 | 0.5562 | 0.74660827 |
| SL1344_RS06405_ | 1.111615064 | 1.153504719 | 1.093158007 | -0.052131765 | 0.195007868 | -0.221566545 | 0.410393075 | 0.712972589 | 0.866181805 | 0.809 | 0.910455538 |
| SL1344_RS06410_ | 6.62969594 | 6.587499063 | 6.707178766 | 0.121673362 | 0.120450435 | 1.039815966 | 0.106595641 | 0.21678354 | 0.520037914 | 0.1943 | 0.467184174 |
| SL1344_RS06415_ | 2.656317303 | 2.544921539 | 2.97733494 | 0.417156791 | 0.276450985 | 1.657491742 | 0.010000458 | 0.052813958 | 0.268552015 | 0.1109 | 0.345984835 |
| SL1344_RS06420_ | 2.987918723 | 2.880223138 | 3.088888886 | 0.142546431 | 0.15496768 | 1.039140575 | 0.127333364 | 0.191883786 | 0.47970303 | 0.2391 | 0.51157103 |
| SL1344_RS06425_ | 0.426254593 | 0.366871477 | 0.50164877 | 0.156819163 | 0.196894362 | 0.773544198 | 0.18254499 | 0.273039977 | 0.545854164 | 0.372 | 0.594388971 |
| SL1344_RS06430_ | 0.479922977 | 0.238125685 | 0.638256453 | 0.398532795 | 0.166945124 | 2.26804598 | 0.004001152 | 0.041627501 | 0.21972351 | 0.1048 | 0.336977582 |
| SL1344_RS06435_ | 2.026668684 | 1.835866418 | 2.218508727 | 0.394455648 | 0.131324896 | 3.005706547 | 4.68E-05 | 0.016785014 | 0.149665753 | 0.1001 | 0.330399519 |
| SL1344_RS06440_ | 1.156193887 | 0.817542295 | 1.202573969 | 0.368202206 | 0.195723949 | 1.099185809 | 0.165223207 | 0.198598416 | 0.490132908 | 0.3648 | 0.604181959 |
| SL1344_RS06445_ | 2.021980864 | 1.88791725 | 2.119288504 | 0.237694876 | 0.141030891 | 1.631473973 | 0.029333486 | 0.075399131 | 0.303562981 | 0.1272 | 0.365814963 |
| SL1344_RS06450_ | 2.957490453 | 2.81868199 | 3.078075206 | 0.237079459 | 0.138448991 | 1.734971877 | 0.028647725 | 0.056834393 | 0.272028026 | 0.1241 | 0.365234428 |
| SL1344_RS06455_ | 1.434372813 | 1.376969546 | 1.493891691 | 0.145319452 | 0.173220632 | 0.82061659 | 0.155229871 | 0.245988428 | 0.52766486 | 0.3369 | 0.574150023 |
| SL1344_RS06465_ | -1.560035521 | -2.133063214 | -1.05764152 | 1.064062945 | 0.342848087 | 3.043378263 | 4.68E-05 | 0.016453513 | 0.142208611 | 0.1005 | 0.330991751 |
| SL1344_RS06470_ | 1.62356954 | 1.385738228 | 1.773537943 | 0.503646691 | 0.311289046 | 1.660170981 | 0.005996769 | 0.057197174 | 0.276036143 | 0.1056 | 0.338231541 |
| SL1344_RS06475_ | -0.52403036 | -0.81619574 | -0.243153771 | 0.699007457 | 0.405934337 | 1.709658637 | 0.003332494 | 0.054611111 | 0.263327551 | 0.1044 | 0.336076498 |
| SL1344_RS06480_ | -0.9495209 | -1.200680647 | -0.674528592 | 0.698234193 | 0.562324227 | 1.208686789 | 0.060626321 | 0.1202414 | 0.38526143 | 0.1734 | 0.422221903 |
| SL1344_RS06485_ | -0.184837035 | -0.237058272 | -0.121037501 | 0.19588477 | 0.37259155 | 0.517844454 | 0.278481023 | 0.398873651 | 0.666228115 | 0.5658 | 0.754515597 |
| SL1344_RS06490_ | -0.648890981 | -0.767494414 | -0.46340226 | 0.332558099 | 0.430205904 | 0.780409422 | 0.182666686 | 0.256419863 | 0.542483629 | 0.338 | 0.593272363 |
| SL1344_RS06495_ | -0.912533899 | -1.020403059 | -0.874022131 | 0.151571657 | 0.38180327 | 0.332300586 | 0.373750838 | 0.601951663 | 0.800983745 | 0.7389 | 0.872247704 |
| SL1344_RS06500_ | -0.228693178 | -0.355405593 | -0.060041211 | 0.3031004 | 0.203466521 | 1.472406719 | 0.050000088 | 0.110963492 | 0.34547056 | 0.1633 | 0.4022667 |
| SL1344_RS06505_ | 0.261518063 | 0.314326905 | 0.22169001 | -0.092764594 | 0.17500951 | -0.482349016 | 0.301132587 | 0.458558174 | 0.687030852 | 0.5675 | 0.745904903 |
| SL1344_RS06510_ | 2.309181021 | 2.066644568 | 2.663149662 | 0.562121483 | 0.257932805 | 2.155002237 | 4.68E-05 | 0.027969449 | 0.200038341 | 0.1002 | 0.330549053 |
| SL1344_RS06515_ | 2.419650864 | 2.285857966 | 2.718039575 | 0.430685527 | 0.332852387 | 1.432310865 | 0.014657206 | 0.107893155 | 0.375768581 | 0.1127 | 0.348723595 |
| SL1344_RS06520_ | 0.789571548 | 0.588623221 | 1.164589938 | 0.576456394 | 0.430972332 | 1.384146381 | 0.063291207 | 0.092124629 | 0.345625196 | 0.1537 | 0.405931717 |

| | | | | | | | | | | | |
|---|---|---|---|---|---|---|---|---|---|---|---|
| SL1344_RS06525_ | 0.455822828 | -0.379392416 | 0.604234356 | 0.960707632 | 0.307288996 | 1.69427973 | 0.006000767 | 0.100350424 | 0.362069774 | 0.1062 | 0.338891472 |
| SL1344_RS06530_ | -1.127015399 | -1.378671522 | -1.051733952 | 0.284773545 | 0.424910288 | 0.589828848 | 0.301333342 | 0.475888121 | 0.719017378 | 0.6225 | 0.802877079 |
| SL1344_RS06535_ | -0.309342798 | -0.586349795 | -0.121448073 | 0.467003898 | 0.224511239 | 2.072422418 | 0.011992386 | 0.051391377 | 0.244153423 | 0.112 | 0.346414972 |
| SL1344_RS06540_ | -1.666226853 | -1.872148795 | -1.466357644 | 0.39950496 | 0.325142729 | 1.170161863 | 0.106000038 | 0.161060116 | 0.408381121 | 0.2375 | 0.481723879 |
| SL1344_RS06545_ | -0.054162022 | -0.166480627 | 0.149123383 | 0.310051558 | 0.275038405 | 1.074105652 | 0.131245866 | 0.1806825 | 0.453086058 | 0.2678 | 0.523431111 |
| SL1344_RS06550_ | 0.693688683 | 0.519919472 | 0.869741328 | 0.32377551 | 0.558997896 | 0.579525103 | 0.261825461 | 0.388062505 | 0.671649682 | 0.5061 | 0.741769257 |
| SL1344_RS06555_ | -0.651326536 | -1.055125735 | -0.368412504 | 0.680900551 | 0.210327539 | 3.227710148 | 0.00067292 | 0.015286058 | 0.131451456 | 0.1004 | 0.330754667 |
| SL1344_RS06565_ | -0.945772178 | -1.034948625 | -0.884056618 | 0.087153882 | 0.739113077 | 0.117436849 | 0.437333336 | 0.846941774 | 0.938845602 | 0.8221 | 0.921811132 |
| SL1344_RS06570_ | 1.797955419 | 1.638696336 | 1.947866517 | 0.360839115 | 0.177056846 | 2.041237338 | 4.68E-05 | 0.043156994 | 0.237308594 | 0.1001 | 0.33040342 |
| SL1344_RS06575_ | 0.097085343 | -0.178853809 | 0.348254321 | 0.515422993 | 0.203485116 | 2.475276747 | 0.003334715 | 0.026596259 | 0.180376654 | 0.1024 | 0.33362003 |
| SL1344_RS06580_ | 1.206411981 | 1.085936287 | 1.35453642 | 0.281167704 | 0.148891658 | 1.8030536 | 0.019986902 | 0.05995681 | 0.266736903 | 0.1186 | 0.355207962 |
| SL1344_RS06605_ | -3.099217329 | -2.916144678 | -3.211867707 | -0.28908582 | 0.809825928 | -0.322433456 | 0.366000005 | 0.61679188 | 0.807074091 | 0.7362 | 0.864998288 |
| SL1344_RS06615_ | -4.336615928 | -4.54329026 | -4.147155781 | 0.397619641 | 0.836703933 | 0.444459057 | 0.307794812 | 0.454419719 | 0.68287938 | 0.5642 | 0.74156404 |
| SL1344_RS06630_ | 7.565960073 | 7.526781998 | 7.620048092 | 0.137481213 | 0.094351609 | 1.574827314 | 4.68E-05 | 0.063996734 | 0.296187624 | 0.1 | 0.330257709 |
| SL1344_RS06635_ | 7.846952823 | 7.571271446 | 7.856099359 | 0.285050341 | 0.097834725 | 0.623621792 | 0.33333334 | 0.498844158 | 0.755273347 | 0.7 | 0.872542119 |
| SL1344_RS06640_ | 7.821995601 | 7.822626659 | 7.82147927 | -0.01365218 | 0.204123285 | -0.075484296 | 0.409726852 | 0.90644121 | 0.966445495 | 0.8707 | 0.945036144 |
| SL1344_RS06645_ | 2.377367922 | 2.237013046 | 2.591819814 | 0.358970797 | 0.183636889 | 2.035834968 | 0.001335878 | 0.042784478 | 0.239069097 | 0.1027 | 0.334215784 |
| SL1344_RS06650_ | 3.779526925 | 3.596771676 | 3.850974392 | 0.194186522 | 0.212127149 | 1.095097448 | 0.133333363 | 0.159996417 | 0.449168963 | 0.2476 | 0.524853781 |
| SL1344_RS06660_ | 0.891206905 | 0.722474216 | 1.056205651 | 0.343213242 | 0.208835956 | 1.587134738 | 0.026666836 | 0.07214419 | 0.296423008 | 0.1319 | 0.373587961 |
| SL1344_RS06665_ | -0.39854742 | -0.431970896 | -0.365162758 | 0.076677489 | 0.284965819 | 0.259977524 | 0.388000004 | 0.634671627 | 0.817146175 | 0.7237 | 0.857637007 |
| SL1344_RS06670_ | -0.378456934 | -0.577501406 | -0.175410445 | 0.303304587 | 0.410224629 | 0.716821953 | 0.200000017 | 0.314415391 | 0.59751036 | 0.4005 | 0.647819331 |
| SL1344_RS06675_ | 0.163495989 | -0.309713805 | 0.543307181 | 0.867699512 | 0.224388665 | 3.955625752 | 4.68E-05 | 0.016438483 | 0.144772689 | 0.1 | 0.330257709 |
| SL1344_RS06685_ | 1.078817101 | 1.026430536 | 1.109468711 | 0.101871079 | 0.201527379 | 0.468252919 | 0.269820131 | 0.381525104 | 0.656475306 | 0.5175 | 0.708226203 |
| SL1344_RS06690_ | 0.768126215 | 0.892054806 | 0.678181419 | -0.133216177 | 0.433761155 | -0.263405038 | 0.381079285 | 0.86265112 | 0.945935482 | 0.8023 | 0.915625148 |
| SL1344_RS06695_ | 1.405942439 | 1.335683487 | 1.614579895 | 0.277357826 | 0.179477906 | 1.394142666 | 0.058627656 | 0.135812891 | 0.407284438 | 0.1754 | 0.418658382 |
| SL1344_RS06700_ | 0.221432851 | 0.126050692 | 0.374554953 | 0.261784259 | 0.204138785 | 1.192382506 | 0.092604975 | 0.158515757 | 0.421572198 | 0.2245 | 0.466773709 |
| SL1344_RS06705_ | 5.174493712 | 5.063586784 | 5.329348371 | 0.237884359 | 0.110110912 | 2.096460247 | 0.001335878 | 0.036312438 | 0.226535155 | 0.1006 | 0.331122153 |
| SL1344_RS06710_ | -1.614577129 | -1.650415641 | -1.583423306 | 0.067611314 | 0.359254466 | 0.184145147 | 0.42466667 | 0.619948258 | 0.805355716 | 0.724 | 0.854295543 |
| SL1344_RS06715_ | -4.563045194 | -4.592046606 | -4.517335752 | 0.098271605 | 0.900444442 | 0.098842739 | 0.457333335 | 0.586500626 | 0.780472911 | 0.7036 | 0.840564185 |
| SL1344_RS06720_ | -2.755221205 | -2.912928402 | -2.65189554 | 0.256830774 | 0.471857757 | 0.509487664 | 0.280666677 | 0.422687351 | 0.652269809 | 0.5355 | 0.718903727 |
| SL1344_RS06730_ | -4.453240406 | -4.802706358 | -3.943749166 | 0.815335328 | 0.910816762 | 0.813423258 | 0.195203216 | 0.313578735 | 0.557374496 | 0.4021 | 0.618498266 |
| SL1344_RS06735_ | -3.17585872 | -3.600795711 | -2.739166473 | 0.855536161 | 0.567964304 | 1.44544607 | 0.058000075 | 0.106381306 | 0.33370804 | 0.1766 | 0.416598266 |
| SL1344_RS06740_ | -0.80546488 | -0.949957138 | -0.674110312 | 0.296338642 | 0.262065354 | 1.088383471 | 0.105333372 | 0.18006584 | 0.435358759 | 0.2604 | 0.49733956 |
| SL1344_RS06745_ | -2.605120375 | -2.795220478 | -2.485773917 | 0.333275478 | 0.64877144 | 0.488345135 | 0.284666676 | 0.410114512 | 0.669793694 | 0.5525 | 0.739122693 |
| SL1344_RS06750_ | 0.782461314 | 0.521697823 | 1.014592362 | 0.59569275 | 0.306337705 | 1.962200369 | 0.000673369 | 0.042583818 | 0.237187951 | 0.1007 | 0.331264014 |
| SL1344_RS06755_ | -0.575252929 | -0.945528048 | -0.438535562 | 0.48383227 | 0.460884563 | 0.913971436 | 0.157894761 | 0.226938031 | 0.520728948 | 0.2965 | 0.555489837 |
| SL1344_RS06760_ | -2.085500283 | -2.224852253 | -1.98936286 | 0.222033021 | 0.770418563 | 0.260740619 | 0.390000004 | 0.603261007 | 0.806163655 | 0.7837 | 0.891316198 |
| SL1344_RS06765_ | -3.753807839 | -3.689135866 | -3.840437972 | -0.160578416 | 0.599595236 | -0.257942797 | 0.392405067 | 0.506524993 | 0.71853675 | 0.6169 | 0.777524864 |
| SL1344_RS06770_ | -4.270165974 | -4.692127502 | -3.695138919 | 0.982743883 | 0.807499151 | 1.177550674 | 0.101333374 | 0.166628794 | 0.412925505 | 0.2462 | 0.488541648 |
| SL1344_RS06775_ | 1.103503478 | 0.92287298 | 1.259570477 | 0.346978093 | 0.142687691 | 2.379957129 | 0.002668394 | 0.035275685 | 0.206302484 | 0.1013 | 0.332106993 |
| SL1344_RS06780_ | 2.222743673 | 2.051736087 | 2.290463974 | 0.230192864 | 0.114357634 | 1.769892097 | 0.027333498 | 0.080311213 | 0.311502836 | 0.1342 | 0.373843283 |
| SL1344_RS06785_ | -0.297961036 | -0.419286552 | -0.237912189 | 0.178648554 | 0.264752445 | 0.627530554 | 0.255829459 | 0.395602921 | 0.650116355 | 0.5252 | 0.726394251 |
| SL1344_RS06790_ | 1.209737969 | 1.187613889 | 1.267890464 | 0.062213678 | 0.178456722 | 0.272903543 | 0.389740178 | 0.763587387 | 0.895286602 | 0.7901 | 0.907345255 |
| SL1344_RS06795_ | -0.368866081 | -0.735783957 | 0.206247296 | 0.896033258 | 0.327105238 | 2.698916238 | 4.68E-05 | 0.019983299 | 0.162734245 | 0.1002 | 0.330545993 |
| SL1344_RS06800_ | -2.143969477 | -1.749253042 | -2.649151354 | -0.897151853 | 0.381670637 | -2.308503066 | 0.006000767 | 0.032962803 | 0.194108767 | 0.1042 | 0.336169276 |
| SL1344_RS06805_ | 0.025464336 | 0.084523701 | -0.022992642 | -0.113561883 | 0.26118708 | -0.404981868 | 0.336442379 | 0.542694389 | 0.756843092 | 0.6654 | 0.818943781 |
| SL1344_RS06810_ | 0.352801868 | 0.403676744 | 0.270951442 | -0.149055856 | 0.201555923 | -0.679347221 | 0.206528997 | 0.317357009 | 0.584222212 | 0.4154 | 0.627150479 |
| SL1344_RS06815_ | 0.442928764 | 0.608372762 | 0.359143995 | -0.220069234 | 0.206640025 | -1.010590186 | 0.145902758 | 0.222627185 | 0.492512423 | 0.318 | 0.565920512 |
| SL1344_RS06820_ | 0.184290829 | 0.293124987 | 0.031605922 | -0.214432282 | 0.283780903 | -0.706590496 | 0.217854779 | 0.333841296 | 0.608659816 | 0.4433 | 0.675487329 |
| SL1344_RS06825_ | -1.172265758 | -1.092476902 | -1.245462583 | -0.153139734 | 0.237537477 | -0.612579557 | 0.247834789 | 0.372631456 | 0.603817855 | 0.4706 | 0.663933654 |
| SL1344_RS06830_ | 0.750626144 | 0.62852765 | 0.862379901 | 0.257695906 | 0.208263416 | 1.231122339 | 0.061958764 | 0.134866644 | 0.398370753 | 0.1712 | 0.416942781 |
| SL1344_RS06835_ | 0.84359621 | 0.816009192 | 0.869700024 | 0.120631658 | 0.296997914 | 0.339566913 | 0.317788149 | 0.412331349 | 0.687484492 | 0.6461 | 0.802145264 |
| SL1344_RS06840_ | 10.38917371 | 10.37262468 | 10.49710605 | 0.123752692 | 0.067509064 | 0.717998207 | 0.33333334 | 0.431541718 | 0.708280041 | 0.7 | 0.872542119 |

| | | | | | | | | | | | |
|---|---|---|---|---|---|---|---|---|---|---|---|
| SL1344_RS06845_ | 3.112110424 | 2.857717523 | 3.426055621 | 0.546825432 | 0.107172092 | 5.107162071 | 4.68E-05 | 0.002811559 | 0.067279546 | 0.1 | 0.330257709 |
| SL1344_RS06850_ | -1.299579514 | -1.373163629 | -1.23148122 | 0.15331073 | 0.318565981 | 0.454766732 | 0.309793479 | 0.463265194 | 0.692029976 | 0.5859 | 0.755640322 |
| SL1344_RS06855_ | -3.155580517 | -3.481186309 | -2.754495569 | 0.617519818 | 0.777521067 | 0.735470882 | 0.215856111 | 0.331735867 | 0.597669738 | 0.4254 | 0.656585542 |
| SL1344_RS06860_ | -1.877845008 | -2.051854846 | -1.498597976 | 0.490010462 | 0.499502628 | 0.909318817 | 0.186542324 | 0.265187012 | 0.540921585 | 0.3649 | 0.614865959 |
| SL1344_RS06865_ | -1.798422894 | -2.329267866 | -1.438314309 | 0.777605307 | 0.556448678 | 1.35463435 | 0.064623651 | 0.111563335 | 0.366626743 | 0.1771 | 0.433417955 |
| SL1344_RS06900_ | 2.006766634 | 1.749309196 | 2.225618499 | 0.435517976 | 0.175234382 | 2.47782361 | 0.001335878 | 0.020961541 | 0.170680021 | 0.101 | 0.331720975 |
| SL1344_RS06905_ | -0.433409851 | -0.507844742 | -0.360124811 | 0.148391474 | 0.203133562 | 0.680162711 | 0.231179228 | 0.343346624 | 0.58703275 | 0.4472 | 0.652638209 |
| SL1344_RS06925_ | -3.459895474 | -3.994985993 | -3.141620503 | 1.004438824 | 1.057408289 | 0.918367255 | 0.14923387 | 0.211834175 | 0.492279313 | 0.3127 | 0.564312321 |
| SL1344_RS06930_ | -3.318880672 | -3.454574602 | -3.187981538 | 0.434376767 | 0.919931249 | 0.459909522 | 0.313124592 | 0.476276939 | 0.717750222 | 0.6382 | 0.802009019 |
| SL1344_RS06935_ | -3.849697396 | -4.493798672 | -3.307070346 | 1.213902777 | 0.766070106 | 1.514661271 | 0.0440001 | 0.093230787 | 0.319189624 | 0.1478 | 0.387002302 |
| SL1344_RS06940_ | -3.623759782 | -3.682301664 | -3.546990461 | 0.144773038 | 0.584696575 | 0.21961142 | 0.39840107 | 0.533429761 | 0.738667185 | 0.6468 | 0.798661013 |
| SL1344_RS06945_ | -2.779139637 | -3.182829095 | -2.472499895 | 0.70829877 | 0.43200493 | 1.571071739 | 0.034000131 | 0.087025148 | 0.300098861 | 0.147 | 0.384856752 |
| SL1344_RS06950_ | -3.000632315 | -3.091907016 | -2.879487241 | 0.153036995 | 1.172790278 | 0.132905527 | 0.41466667 | 0.756034193 | 0.890846088 | 0.7805 | 0.893214968 |
| SL1344_RS06955_ | -1.835299201 | -2.050348598 | -1.723879735 | 0.317430856 | 0.473876276 | 0.61379173 | 0.253830791 | 0.388785979 | 0.648284901 | 0.5151 | 0.720655549 |
| SL1344_RS06960_ | -3.616076076 | -3.5530481 | -3.694057484 | -0.129828653 | 0.699167403 | -0.169865124 | 0.420386412 | 0.591115107 | 0.780230101 | 0.6935 | 0.83007462 |
| SL1344_RS06965_ | -3.357619462 | -3.768252969 | -3.062469217 | 0.701671445 | 0.538511982 | 1.24735561 | 0.077281867 | 0.1416514 | 0.376721771 | 0.2148 | 0.451295346 |
| SL1344_RS06970_ | -2.899834471 | -3.135199572 | -2.675835208 | 0.475055022 | 0.471172934 | 0.983919129 | 0.14333336 | 0.223093415 | 0.469752708 | 0.3101 | 0.539593684 |
| SL1344_RS06975_ | -2.952790101 | -3.540481422 | -2.52416035 | 1.023027486 | 0.686208321 | 1.417430309 | 0.0440001 | 0.106414299 | 0.352817999 | 0.1386 | 0.380312581 |
| SL1344_RS06980_ | -1.898079652 | -1.90380408 | -1.88977307 | 0.03715384 | 0.377126359 | 0.094933383 | 0.460000001 | 0.605374705 | 0.793680055 | 0.7091 | 0.843097187 |
| SL1344_RS06990_ | -5.544963194 | -5.830105165 | -5.337298337 | 0.473894366 | 1.33855531 | 0.315124762 | 0.349766828 | 0.499744202 | 0.721656029 | 0.6208 | 0.784225346 |
| SL1344_RS06995_ | -5.755523403 | -5.757239137 | -5.753514789 | -0.236058996 | 2.772178481 | -0.064655751 | 0.454000002 | 0.656102916 | 0.838517045 | 0.8384 | 0.921555701 |
| SL1344_RS07000_ | -3.647268207 | -3.977636186 | -3.341580338 | 0.629208268 | 0.608004188 | 0.998085155 | 0.131245866 | 0.206349384 | 0.45295074 | 0.2868 | 0.518410991 |
| SL1344_RS07005_ | -3.199664442 | -3.503499483 | -3.059294559 | 0.378068147 | 0.807300678 | 0.42584539 | 0.334443711 | 0.535760918 | 0.753988443 | 0.6551 | 0.815755818 |
| SL1344_RS07010_ | -4.312269456 | -4.426259808 | -4.153987731 | 0.282331972 | 0.777908591 | 0.333657991 | 0.356000006 | 0.503154938 | 0.714203154 | 0.6175 | 0.775927756 |
| SL1344_RS07015_ | -2.598087186 | -2.701472275 | -2.50211135 | 0.31499029 | 0.716158859 | 0.416261198 | 0.324666674 | 0.491647514 | 0.726088391 | 0.6358 | 0.800677209 |
| SL1344_RS07020_ | -5.07974304 | -5.051391581 | -5.101855812 | -0.009373587 | 1.234921145 | -0.005919407 | 0.497001999 | 0.623454857 | 0.806523566 | 0.7261 | 0.853477119 |
| SL1344_RS07025_ | -1.968980894 | -2.210700616 | -1.856266644 | 0.35604208 | 0.476198452 | 0.703287375 | 0.197333351 | 0.303640438 | 0.579086383 | 0.414 | 0.634136934 |
| SL1344_RS07030_ | -1.27385633 | -1.517631179 | -1.101149227 | 0.426753838 | 0.294034203 | 1.40415998 | 0.058000075 | 0.121039138 | 0.360754376 | 0.1745 | 0.41231045 |
| SL1344_RS07035_ | -2.843418635 | -3.246379937 | -2.397598065 | 0.847080904 | 0.552636519 | 1.475288811 | 0.046666761 | 0.096643941 | 0.327752366 | 0.1654 | 0.409146899 |
| SL1344_RS07040_ | -2.886889769 | -3.202693755 | -2.596378719 | 0.589345664 | 0.585814807 | 0.999790613 | 0.147901425 | 0.212068557 | 0.474395369 | 0.2996 | 0.543039878 |
| SL1344_RS07045_ | -1.401394246 | -1.626729366 | -1.129640371 | 0.519511037 | 0.322131604 | 1.562464075 | 0.034666795 | 0.086682848 | 0.309622595 | 0.145 | 0.384560405 |
| SL1344_RS07055_ | -4.578807268 | -4.561693461 | -4.599760067 | -0.054639047 | 1.044035534 | -0.048843738 | 0.476000001 | 0.627732054 | 0.810332344 | 0.7524 | 0.87210348 |
| SL1344_RS07060_ | -3.450628229 | -3.888876723 | -2.971858306 | 0.905924787 | 0.617495849 | 1.409475046 | 0.063957429 | 0.109290982 | 0.34021283 | 0.1805 | 0.423612099 |
| SL1344_RS07065_ | -2.950145337 | -3.022284923 | -2.867233564 | 0.148277019 | 0.445985252 | 0.330224597 | 0.356429053 | 0.467619533 | 0.683727142 | 0.5741 | 0.745161339 |
| SL1344_RS07070_ | 0.312623087 | 0.613614628 | 0.247255038 | -0.368182194 | 0.578211874 | -0.405272099 | 0.335109934 | 0.857113434 | 0.944458443 | 0.7039 | 0.874164187 |
| SL1344_RS07075_ | 0.635360861 | 1.322000664 | 0.550572467 | -0.686928212 | 0.358101839 | -0.634694995 | 0.33333334 | 0.498312619 | 0.752413511 | 0.7 | 0.872542119 |
| SL1344_RS07080_ | 1.456714277 | 1.353572425 | 1.557905092 | 0.287338724 | 0.227969954 | 1.272200153 | 0.031312601 | 0.109316963 | 0.367568301 | 0.139 | 0.37701468 |
| SL1344_RS07085_ | 0.944854613 | 0.728744425 | 1.008733403 | 0.27009489 | 0.210603693 | 1.063066104 | 0.140666694 | 0.195366815 | 0.47823478 | 0.3001 | 0.55321401 |
| SL1344_RS07090_ | 2.369780349 | 2.408296318 | 2.347522206 | -0.057276296 | 0.119907658 | -0.437171133 | 0.320000008 | 0.53162931 | 0.752361534 | 0.6438 | 0.812546391 |
| SL1344_RS07095_ | -0.821460304 | -0.93196473 | -0.734689128 | 0.192288129 | 0.3556881 | 0.534859823 | 0.28580947 | 0.453837437 | 0.694355729 | 0.5634 | 0.751769963 |
| SL1344_RS07100_ | -1.37981344 | -1.61697812 | -1.31836397 | 0.277530989 | 0.482789175 | 0.451579152 | 0.360666672 | 0.626752001 | 0.820641764 | 0.7395 | 0.884457021 |
| SL1344_RS07115_ | 1.985415202 | 1.69679683 | 2.307803813 | 0.619944506 | 0.137181964 | 4.539467886 | 4.68E-05 | 0.005866237 | 0.092009734 | 0.1 | 0.330257709 |
| SL1344_RS07120_ | 4.397589905 | 4.298484459 | 4.509355422 | 0.216505547 | 0.061626248 | 3.506565042 | 4.68E-05 | 0.013601704 | 0.136476281 | 0.1 | 0.330257709 |
| SL1344_RS07125_ | 1.146954138 | 0.93048589 | 1.397902938 | 0.473118551 | 0.160831296 | 2.970327529 | 4.68E-05 | 0.017055628 | 0.148721428 | 0.1001 | 0.330405227 |
| SL1344_RS07130_ | 1.785388546 | 1.651720147 | 1.909047395 | 0.261819301 | 0.108008031 | 2.399645798 | 0.003334715 | 0.028747582 | 0.186107945 | 0.1033 | 0.33486133 |
| SL1344_RS07135_ | 1.004192807 | 0.851274074 | 1.077585762 | 0.225958618 | 0.195168125 | 1.08235517 | 0.121333366 | 0.182745228 | 0.460882946 | 0.2585 | 0.508564916 |
| SL1344_RS07140_ | -0.32255295 | -0.375514228 | -0.265466257 | 0.111180534 | 0.177095443 | 0.575420259 | 0.260000012 | 0.38511821 | 0.615005056 | 0.4925 | 0.682830914 |
| SL1344_RS07145_ | -2.39927589 | -2.463666045 | -2.325026932 | 0.144423033 | 0.375321844 | 0.354405864 | 0.346666673 | 0.478168377 | 0.694737448 | 0.596 | 0.758508311 |
| SL1344_RS07150_ | 1.28358776 | 0.979457382 | 1.514027467 | 0.548821651 | 0.163161266 | 3.285999776 | 4.68E-05 | 0.011919179 | 0.126961839 | 0.1 | 0.330257709 |
| SL1344_RS07155_ | 0.541321998 | 0.085072037 | 0.615555406 | 0.492197591 | 0.277730783 | 1.139366005 | 0.156000024 | 0.189646752 | 0.479849991 | 0.3262 | 0.577162604 |
| SL1344_RS07160_ | 0.091554625 | -0.25930307 | 0.378245516 | 0.639219409 | 0.242451756 | 2.544714431 | 0.000673369 | 0.026859024 | 0.184761153 | 0.1007 | 0.331281063 |
| SL1344_RS07165_ | -0.431821885 | -0.582809859 | -0.274300292 | 0.304231063 | 0.235464217 | 1.248923049 | 0.076615645 | 0.135628139 | 0.380100981 | 0.2044 | 0.448676845 |
| SL1344_RS07170_ | -1.961065404 | -2.20770584 | -1.703394038 | 0.510534828 | 0.329976088 | 1.480955192 | 0.053297883 | 0.097533228 | 0.316096353 | 0.1678 | 0.40603125 |

| | | | | | | | | | | | |
|---|---|---|---|---|---|---|---|---|---|---|---|
| SL1344_RS07180_ | 3.065703522 | 2.700064839 | 3.431138634 | 0.765537907 | 0.23500631 | 3.309454332 | 4.68E-05 | 0.00858193 | 0.117309139 | 0.1 | 0.330257709 |
| SL1344_RS07185_ | 0.272263267 | 0.291484218 | 0.25824586 | -0.027784688 | 0.208799722 | -0.12659606 | 0.444370422 | 0.664739453 | 0.834578846 | 0.7772 | 0.885928027 |
| SL1344_RS07190_ | 0.30304745 | 0.286479121 | 0.319275307 | 0.031554094 | 0.192735834 | 0.155303244 | 0.432666669 | 0.631084664 | 0.809607002 | 0.7422 | 0.862845851 |
| SL1344_RS07195_ | 1.414128583 | 1.134953442 | 1.567993645 | 0.404485029 | 0.201629813 | 1.895544259 | 0.01866691 | 0.061259359 | 0.279793139 | 0.1129 | 0.348989867 |
| SL1344_RS07200_ | 4.06266504 | 3.795914669 | 4.235638916 | 0.438238082 | 0.126095387 | 3.62302499 | 4.68E-05 | 0.004629664 | 0.087703439 | 0.1 | 0.330257709 |
| SL1344_RS07205_ | 1.976235525 | 1.81973332 | 2.127935687 | 0.355319365 | 0.174774778 | 2.1018216 | 0.00067292 | 0.044949291 | 0.242545745 | 0.1003 | 0.33069854 |
| SL1344_RS07210_ | 1.583535165 | 1.548965632 | 1.603885443 | 0.058322032 | 0.172627302 | 0.317740216 | 0.361758833 | 0.545141758 | 0.767795194 | 0.7194 | 0.852478402 |
| SL1344_RS07215_ | 3.251553846 | 2.977771872 | 3.4848588 | 0.505881534 | 0.071781039 | 7.030824642 | 4.68E-05 | 0.002129192 | 0.055840303 | 0.1 | 0.330257709 |
| SL1344_RS07220_ | 1.001177527 | 0.765443728 | 1.163119073 | 0.241821477 | 0.428621279 | 0.654272874 | 0.228514338 | 0.359630584 | 0.646115279 | 0.4147 | 0.667367283 |
| SL1344_RS07225_ | 0.805034779 | 0.703172907 | 1.003044926 | 0.19819895 | 0.394898985 | 0.524433043 | 0.294000009 | 0.537449039 | 0.770984764 | 0.5908 | 0.796331302 |
| SL1344_RS07230_ | 0.398173134 | 0.285177936 | 0.705713771 | 0.369786934 | 0.562682403 | 0.716621466 | 0.219333349 | 0.332752863 | 0.6239981 | 0.4001 | 0.673728996 |
| SL1344_RS07235_ | 0.671233694 | 0.50986112 | 0.738272575 | 0.325522611 | 0.383426677 | 0.851403304 | 0.195203216 | 0.238806383 | 0.53354165 | 0.3712 | 0.629446048 |
| SL1344_RS07240_ | 2.278390626 | 2.000037943 | 2.337867292 | 0.33561271 | 0.188413611 | 1.387497383 | 0.058000075 | 0.132334675 | 0.40964888 | 0.1652 | 0.411963452 |
| SL1344_RS07245_ | 0.936142921 | 0.881497528 | 0.980111979 | 0.095307538 | 0.129889219 | 0.684701564 | 0.222000015 | 0.346285651 | 0.590362249 | 0.448 | 0.656080441 |
| SL1344_RS07250_ | 2.021988666 | 2.010523762 | 2.035434736 | 0.029770194 | 0.170916899 | 0.177896724 | 0.397068625 | 0.698983545 | 0.858086819 | 0.7394 | 0.869280552 |
| SL1344_RS07255_ | 2.107206493 | 1.949153205 | 2.23942379 | 0.236986728 | 0.262701776 | 0.913848269 | 0.150566314 | 0.231182516 | 0.528874283 | 0.2893 | 0.562698385 |
| SL1344_RS07260_ | 1.020421394 | 0.902937507 | 1.251090357 | 0.344490496 | 0.524261775 | 0.70521995 | 0.188666685 | 0.279798553 | 0.581016289 | 0.3313 | 0.60687072 |
| SL1344_RS07265_ | 0.692892452 | 0.440055055 | 0.807157108 | 0.347134895 | 0.229356097 | 1.380014517 | 0.061333404 | 0.122074911 | 0.386284625 | 0.1689 | 0.419649986 |
| SL1344_RS07270_ | 1.795643023 | 1.507314275 | 2.050354471 | 0.567752491 | 0.18262801 | 3.219684686 | 4.68E-05 | 0.021418699 | 0.172003721 | 0.1 | 0.330257709 |
| SL1344_RS07275_ | 1.485083551 | 1.409729479 | 1.604240326 | 0.189602653 | 0.140827019 | 1.25441564 | 0.08933338 | 0.141699356 | 0.395584054 | 0.2171 | 0.465781117 |
| SL1344_RS07285_ | 2.528163085 | 2.392080642 | 2.684040909 | 0.295243857 | 0.171052709 | 1.71415383 | 0.015333631 | 0.067352596 | 0.295497379 | 0.1151 | 0.351987565 |
| SL1344_RS07290_ | 1.878780056 | 1.712028707 | 2.09637462 | 0.420629238 | 0.187229894 | 2.255255494 | 0.00067292 | 0.029704636 | 0.199767839 | 0.1004 | 0.330841338 |
| SL1344_RS07295_ | 4.7649893 | 4.640633465 | 4.883926152 | 0.26738522 | 0.262169764 | 1.170356406 | 0.116588975 | 0.145682003 | 0.433457843 | 0.1988 | 0.474711584 |
| SL1344_RS07300_ | 3.017176608 | 2.977678225 | 3.054889102 | 0.151393315 | 0.183299138 | 0.855065633 | 0.116666701 | 0.279054756 | 0.58104755 | 0.2755 | 0.513340981 |
| SL1344_RS07305_ | -1.14739438 | -1.269969775 | -0.92095114 | 0.333648813 | 0.283188612 | 1.058133456 | 0.144000027 | 0.207804562 | 0.466886382 | 0.2986 | 0.537647059 |
| SL1344_RS07310_ | -0.570332297 | -0.695996234 | -0.351575721 | 0.338955507 | 0.329535413 | 1.010475497 | 0.149333359 | 0.217332964 | 0.492407421 | 0.3109 | 0.562306325 |
| SL1344_RS07315_ | -3.762288168 | -3.684649608 | -3.826396591 | -0.148337917 | 0.652293927 | -0.2090405 | 0.401333337 | 0.548075089 | 0.75130446 | 0.6657 | 0.81344485 |
| SL1344_RS07320_ | 0.018293306 | -0.081381367 | 0.082846952 | 0.144766215 | 0.215518343 | 0.641300824 | 0.242000013 | 0.391805881 | 0.63939737 | 0.499 | 0.704035863 |
| SL1344_RS07325_ | -4.705522334 | -4.767127554 | -4.66800408 | 0.11148234 | 1.068548072 | 0.098544019 | 0.456000002 | 0.614657674 | 0.799955524 | 0.7189 | 0.848985705 |
| SL1344_RS07330_ | 3.145863978 | 3.040963899 | 3.233077809 | 0.204514338 | 0.130528289 | 1.563789675 | 0.027981506 | 0.086975711 | 0.332744363 | 0.1246 | 0.365140874 |
| SL1344_RS07335_ | -0.841146673 | -0.982429679 | -0.779218399 | 0.18264342 | 0.317089512 | 0.510273774 | 0.29646903 | 0.468469711 | 0.706639041 | 0.584 | 0.772070628 |
| SL1344_RS07340_ | -0.445614315 | -0.703722994 | -0.277822166 | 0.512257901 | 0.388222158 | 1.298470741 | 0.049333422 | 0.106068813 | 0.359730586 | 0.1612 | 0.405815797 |
| SL1344_RS07345_ | 3.292936338 | 3.073543948 | 3.48708097 | 0.27605075 | 0.256598822 | 1.395770336 | 0.098000042 | 0.099825393 | 0.36086473 | 0.1943 | 0.46819447 |
| SL1344_RS07350_ | 2.825711716 | 2.490897101 | 3.106746066 | 0.58352708 | 0.111898193 | 5.119573734 | 4.68E-05 | 0.003340775 | 0.070757842 | 0.1 | 0.330257709 |
| SL1344_RS07355_ | 2.986079043 | 2.735729118 | 3.221992398 | 0.491190128 | 0.128194411 | 3.838513741 | 4.68E-05 | 0.006713919 | 0.100177624 | 0.1 | 0.330257709 |
| SL1344_RS07365_ | -0.770229693 | -1.763793111 | -0.188550058 | 1.559145587 | 0.309821383 | 4.877041138 | 4.68E-05 | 0.009837278 | 0.112242925 | 0.1 | 0.330257709 |
| SL1344_RS07370_ | -0.462212234 | -1.434897171 | 0.154663885 | 1.602502102 | 0.357184598 | 4.498303238 | 4.68E-05 | 0.011223148 | 0.121778566 | 0.1 | 0.330257709 |
| SL1344_RS07375_ | -0.131840786 | -0.211529368 | -0.065194028 | 0.15526635 | 0.211783335 | 0.691964745 | 0.227333348 | 0.347662696 | 0.596858593 | 0.4611 | 0.66618707 |
| SL1344_RS07380_ | -0.645725369 | -1.370411672 | -0.396248407 | 0.922856072 | 0.436223982 | 1.802460899 | 0.01666694 | 0.071941856 | 0.304287391 | 0.1176 | 0.354494024 |
| SL1344_RS07385_ | -0.077222209 | -0.304154541 | 0.24542045 | 0.538954198 | 0.18116953 | 2.869127125 | 0.001336769 | 0.02164398 | 0.160534681 | 0.1017 | 0.332671113 |
| SL1344_RS07390_ | -1.514177193 | -1.580286216 | -1.422726992 | 0.127610175 | 0.339028771 | 0.350840268 | 0.352431718 | 0.540890791 | 0.748306742 | 0.6475 | 0.804201418 |
| SL1344_RS07400_ | -0.11683317 | -0.57579266 | -0.035307731 | 0.52479149 | 0.484644364 | 0.876292616 | 0.239173897 | 0.282828775 | 0.580277212 | 0.4524 | 0.700476634 |
| SL1344_RS07405_ | -0.692306571 | -0.712301352 | -0.675649696 | 0.108382705 | 0.427268057 | 0.246544792 | 0.405333337 | 0.637558939 | 0.825025468 | 0.8112 | 0.90649505 |
| SL1344_RS07410_ | -0.935374035 | -0.988900488 | -0.886287678 | 0.110453777 | 0.253618894 | 0.39328395 | 0.317788149 | 0.469156542 | 0.69168583 | 0.5841 | 0.75348853 |
| SL1344_RS07415_ | -2.836941509 | -2.879382381 | -2.782910294 | 0.121968072 | 0.540631246 | 0.216791788 | 0.407061962 | 0.578083129 | 0.775064998 | 0.7044 | 0.838437838 |
| SL1344_RS07420_ | -3.528565556 | -3.442462725 | -3.640504853 | -0.21430885 | 0.626061832 | -0.311095339 | 0.363333339 | 0.499768674 | 0.714481886 | 0.6093 | 0.77227057 |
| SL1344_RS07425_ | -3.753599814 | -4.009702965 | -3.559588346 | 0.542139553 | 0.990771525 | 0.497337918 | 0.260666678 | 0.392847608 | 0.653960518 | 0.5146 | 0.710980961 |
| SL1344_RS07430_ | 0.888232558 | 0.83404984 | 1.058576113 | 0.202460477 | 0.248483227 | 0.769124832 | 0.225333348 | 0.33458759 | 0.612813278 | 0.4383 | 0.685992776 |
| SL1344_RS07435_ | 0.193759531 | 0.213377527 | 0.177713747 | 0.008496774 | 0.254237539 | 0.032941302 | 0.489333334 | 0.722525518 | 0.870954798 | 0.8324 | 0.918275705 |
| SL1344_RS07440_ | -4.591467438 | -4.469473482 | -4.684591692 | -0.148997511 | 1.119064014 | -0.126492058 | 0.437333336 | 0.652161543 | 0.826910513 | 0.758 | 0.87592839 |
| SL1344_RS07445_ | -2.087764188 | -2.442560379 | -1.70863868 | 0.742771211 | 0.521274438 | 1.368130981 | 0.064000067 | 0.111400494 | 0.35740012 | 0.1778 | 0.425966169 |
| SL1344_RS07450_ | -1.336475575 | -1.424099807 | -1.268382494 | 0.143073799 | 0.385518386 | 0.356126778 | 0.357095276 | 0.58389672 | 0.782761252 | 0.6916 | 0.83639429 |
| SL1344_RS07455_ | 0.708980841 | 0.632197114 | 0.785049881 | 0.173627564 | 0.260872889 | 0.677021686 | 0.202531663 | 0.318756628 | 0.603621385 | 0.417 | 0.655086513 |

| | | | | | | | | | | | |
|---|---|---|---|---|---|---|---|---|---|---|---|
| SL1344_RS07460_ | -0.609347434 | -0.487825264 | -0.707639123 | -0.217383545 | 0.248589703 | -0.839225815 | 0.180546322 | 0.273635996 | 0.534666908 | 0.3822 | 0.607419067 |
| SL1344_RS07465_ | 2.460887968 | 2.720787232 | 2.434426883 | -0.266929422 | 0.145947166 | -0.780103081 | 0.309127257 | 0.362347386 | 0.650128905 | 0.635 | 0.82976267 |
| SL1344_RS07470_ | 2.312793902 | 2.258215448 | 2.346401397 | 0.102125618 | 0.267338914 | 0.377790984 | 0.347101938 | 0.515732827 | 0.761279002 | 0.7189 | 0.864413064 |
| SL1344_RS07475_ | -1.089674336 | -0.980820633 | -1.213382954 | -0.228805273 | 0.275709948 | -0.794683077 | 0.187874769 | 0.300256 | 0.552437172 | 0.3962 | 0.617227914 |
| SL1344_RS07480_ | 0.230995678 | 0.371897581 | -0.107722449 | -0.466013623 | 0.225210943 | -1.857902986 | 0.02466685 | 0.066351924 | 0.28189956 | 0.1259 | 0.36352911 |
| SL1344_RS07485_ | -0.41817604 | -0.213302493 | -0.811970191 | -0.585982511 | 0.251965335 | -2.219306215 | 0.004664544 | 0.04831579 | 0.237996476 | 0.1058 | 0.33814345 |
| SL1344_RS07490_ | -0.086695699 | 0.206710676 | -0.316467949 | -0.446909517 | 0.325300123 | -1.312835051 | 0.074666724 | 0.113736438 | 0.369789408 | 0.1815 | 0.440218885 |
| SL1344_RS07495_ | 0.363403078 | 0.413418486 | 0.338760656 | -0.049421441 | 0.239451374 | -0.171224465 | 0.427333336 | 0.752457998 | 0.889853691 | 0.7991 | 0.910329634 |
| SL1344_RS07500_ | -0.607113994 | -0.779935854 | -0.13868169 | 0.527865932 | 0.440628347 | 1.095884561 | 0.156000024 | 0.194081556 | 0.47640097 | 0.3017 | 0.5598409 |
| SL1344_RS07505_ | 1.33244368 | 1.211334379 | 1.500640582 | 0.296147235 | 0.136833977 | 2.082932361 | 0.004664544 | 0.05010965 | 0.243912237 | 0.1065 | 0.339025404 |
| SL1344_RS07510_ | 2.553954151 | 2.47004575 | 2.706795435 | 0.242968984 | 0.128718815 | 1.820678269 | 0.012658588 | 0.05448744 | 0.26251815 | 0.1154 | 0.351303532 |
| SL1344_RS07515_ | 1.221352166 | 1.13198293 | 1.26416919 | 0.203702254 | 0.307820939 | 0.660687079 | 0.251333346 | 0.33871594 | 0.625249082 | 0.4906 | 0.724402448 |
| SL1344_RS07520_ | -3.468997933 | -3.262743326 | -3.590646785 | -0.262389768 | 0.861228153 | -0.270990672 | 0.385333338 | 0.65176201 | 0.825487781 | 0.7368 | 0.866091319 |
| SL1344_RS07525_ | -3.023542046 | -3.020430115 | -3.025211963 | 0.017147553 | 0.527360127 | 0.03157248 | 0.482000001 | 0.605974161 | 0.794058093 | 0.7196 | 0.849028611 |
| SL1344_RS07530_ | -2.179531422 | -2.313825382 | -2.042583228 | 0.285640555 | 0.410255145 | 0.658921504 | 0.225183226 | 0.342698078 | 0.589833514 | 0.4521 | 0.653954727 |
| SL1344_RS07535_ | 1.411889207 | 1.302163248 | 1.523990765 | 0.141862776 | 0.344028802 | 0.416433079 | 0.345103271 | 0.555830899 | 0.784465219 | 0.6965 | 0.854343966 |
| SL1344_RS07540_ | -0.939651949 | -0.863959827 | -1.044357799 | -0.18778927 | 0.248528848 | -0.698734673 | 0.219333349 | 0.335718145 | 0.583976741 | 0.4402 | 0.647888937 |
| SL1344_RS07550_ | -1.14218786 | -1.301611162 | -0.944153087 | 0.351922253 | 0.255421306 | 1.305190494 | 0.085942754 | 0.139599666 | 0.374720463 | 0.2197 | 0.46077525 |
| SL1344_RS07555_ | -0.80175191 | -0.980614 | -0.650136353 | 0.332165874 | 0.203959445 | 1.56892597 | 0.047968113 | 0.094031125 | 0.311914212 | 0.1552 | 0.39420307 |
| SL1344_RS07560_ | -0.623430121 | -0.673342938 | -0.566846649 | 0.106417699 | 0.22384512 | 0.444695536 | 0.312458369 | 0.466533337 | 0.688405704 | 0.5635 | 0.742815352 |
| SL1344_RS07565_ | 0.227702675 | 0.191270529 | 0.260557615 | 0.079984941 | 0.239299012 | 0.328527642 | 0.358427721 | 0.582730244 | 0.783572804 | 0.6978 | 0.83877288 |
| SL1344_RS07570_ | -0.58566599 | -0.461339021 | -0.7183935 | -0.259584708 | 0.22188277 | -1.127187439 | 0.099267197 | 0.169551486 | 0.414256959 | 0.2422 | 0.480165082 |
| SL1344_RS07575_ | -1.094522843 | -1.158773134 | -1.022156422 | 0.137537187 | 0.275102552 | 0.455898036 | 0.301333342 | 0.467165077 | 0.6936222 | 0.5648 | 0.747328893 |
| SL1344_RS07580_ | -0.767615632 | -0.694027079 | -0.814391024 | -0.136033945 | 0.271007702 | -0.474841073 | 0.302000009 | 0.451235313 | 0.683227654 | 0.5721 | 0.74734501 |
| SL1344_RS07585_ | -0.296327859 | -0.259338724 | -0.378314138 | -0.096126609 | 0.261929142 | -0.326884537 | 0.367088613 | 0.618438884 | 0.806345483 | 0.7125 | 0.854779747 |
| SL1344_RS07590_ | -0.902254053 | -1.029877354 | -0.824385277 | 0.114383506 | 0.388273707 | 0.286537169 | 0.390000004 | 0.646329631 | 0.826523242 | 0.7536 | 0.876832053 |
| SL1344_RS07595_ | 0.066679132 | -0.208783976 | 0.54013484 | 0.546425247 | 0.545215944 | 1.080614665 | 0.133333363 | 0.191962304 | 0.484639995 | 0.25 | 0.52514724 |
| SL1344_RS07600_ | -1.894691603 | -1.332158741 | -2.775576202 | -1.490971708 | 0.514138595 | -2.80593087 | 0.00067292 | 0.018504794 | 0.153947084 | 0.1001 | 0.330405725 |
| SL1344_RS07605_ | 0.165997197 | 0.380480308 | 0.122567487 | -0.259921501 | 0.387163058 | -0.606850765 | 0.323333341 | 0.481908959 | 0.73383787 | 0.6679 | 0.844720552 |
| SL1344_RS07610_ | -2.258054744 | -2.201351456 | -2.346299548 | -0.12389056 | 0.469231662 | -0.247394819 | 0.398666671 | 0.599986094 | 0.793264001 | 0.7172 | 0.851350396 |
| SL1344_RS07615_ | -0.717553187 | -0.686929812 | -0.745544204 | -0.056235395 | 0.200549429 | -0.285144415 | 0.38774151 | 0.499060038 | 0.711752848 | 0.6127 | 0.773866773 |
| SL1344_RS07620_ | -0.178018886 | -0.20368267 | -0.133717293 | 0.065975858 | 0.197805265 | 0.300687668 | 0.370419725 | 0.546662208 | 0.749295179 | 0.6546 | 0.806129463 |
| SL1344_RS07625_ | -1.593185097 | -1.966253431 | -1.437957381 | 0.443250067 | 0.549099969 | 0.736604217 | 0.232000014 | 0.335356768 | 0.614186823 | 0.46 | 0.689389614 |
| SL1344_RS07630_ | -1.814380497 | -1.734152587 | -1.853961452 | -0.143695804 | 0.452010344 | -0.291503946 | 0.386000004 | 0.619473834 | 0.804410399 | 0.7392 | 0.862616767 |
| SL1344_RS07635_ | -0.29383799 | -0.304138142 | -0.279282456 | 0.013773905 | 0.233585889 | 0.052617723 | 0.477015324 | 0.628628362 | 0.811679075 | 0.7391 | 0.863264987 |
| SL1344_RS07640_ | -0.191198052 | -0.310248017 | -0.138742362 | 0.153261076 | 0.216580729 | 0.61771837 | 0.267821463 | 0.425456588 | 0.663437825 | 0.5393 | 0.734400553 |
| SL1344_RS07645_ | 0.20610169 | -0.047806039 | 0.444795008 | 0.495367953 | 0.185469205 | 2.658621424 | 0.001335878 | 0.023676908 | 0.168745961 | 0.1006 | 0.331119465 |
| SL1344_RS07650_ | -0.844574159 | -0.959471119 | -0.720816051 | 0.254273494 | 0.323925332 | 0.739481075 | 0.200666684 | 0.303744519 | 0.569064788 | 0.4149 | 0.638769731 |
| SL1344_RS07655_ | -0.470655884 | -0.923122955 | -0.176153359 | 0.761275348 | 0.270299286 | 2.7629137 | 4.68E-05 | 0.0315491 | 0.195530915 | 0.1002 | 0.330547938 |
| SL1344_RS07660_ | 1.121251495 | 1.167905252 | 1.077032237 | -0.120847981 | 0.190480581 | -0.617803847 | 0.225333348 | 0.333871869 | 0.608360853 | 0.4616 | 0.672804358 |
| SL1344_RS07665_ | 0.309999048 | -0.134588181 | 0.462263257 | 0.577194286 | 0.291528321 | 1.619338092 | 0.042000105 | 0.102182513 | 0.358680667 | 0.139 | 0.38364111 |
| SL1344_RS07670_ | -0.702562969 | -1.09417088 | -0.388526782 | 0.742663561 | 0.283907601 | 2.515816684 | 0.001336769 | 0.026596365 | 0.181787361 | 0.1006 | 0.331141823 |
| SL1344_RS07675_ | -2.5071693 | -2.29614963 | -2.596868985 | -0.258287684 | 0.503476847 | -0.439082604 | 0.325782819 | 0.562165042 | 0.764291602 | 0.6523 | 0.816538917 |
| SL1344_RS07680_ | -1.933449571 | -1.845984977 | -1.981262619 | -0.100828287 | 0.441972915 | -0.216258405 | 0.41466667 | 0.658304719 | 0.829735001 | 0.7458 | 0.871638735 |
| SL1344_RS07685_ | -2.136430369 | -2.073574081 | -2.295000021 | -0.184302131 | 0.485868699 | -0.337246284 | 0.371333338 | 0.610048862 | 0.801903654 | 0.7108 | 0.856045789 |
| SL1344_RS07690_ | 0.377071512 | 0.469964952 | 0.299039807 | -0.191954008 | 0.182210519 | -1.035555303 | 0.120586309 | 0.200594125 | 0.463520375 | 0.27 | 0.506307381 |
| SL1344_RS07695_ | -0.76249657 | -0.466207141 | -1.015488859 | -0.539061795 | 0.261188763 | -1.974426134 | 0.014000326 | 0.04771909 | 0.234920504 | 0.1172 | 0.353099707 |
| SL1344_RS07700_ | -0.137287855 | 0.037933165 | -0.307479765 | -0.419724161 | 0.312145049 | -1.35572252 | 0.043970787 | 0.115405554 | 0.372866117 | 0.1541 | 0.395426744 |
| SL1344_RS07705_ | 0.583490788 | 0.552342028 | 0.610277445 | 0.105568901 | 0.249084382 | 0.412850642 | 0.321333341 | 0.452972038 | 0.705625903 | 0.6497 | 0.803669181 |
| SL1344_RS07710_ | -0.017516436 | 0.00151697 | -0.032009069 | 0.014700539 | 0.368742342 | 0.036122501 | 0.487008661 | 0.724317463 | 0.875679138 | 0.873 | 0.94147044 |
| SL1344_RS07715_ | -1.980463743 | -1.982633157 | -1.97835443 | 0.071275788 | 0.448658112 | 0.144391652 | 0.450000002 | 0.65551248 | 0.828288461 | 0.7837 | 0.887411493 |
| SL1344_RS07720_ | -2.065682124 | -2.057260051 | -2.075456978 | -0.022956936 | 0.37825392 | -0.059942956 | 0.470353099 | 0.591365495 | 0.783274173 | 0.7007 | 0.835807201 |
| SL1344_RS07725_ | -1.004954971 | -0.89270687 | -1.286375407 | -0.367228979 | 0.323832446 | -1.062207391 | 0.140666694 | 0.214667542 | 0.484162368 | 0.3148 | 0.556219121 |

| | | | | | | | | | | | |
|---|---|---|---|---|---|---|---|---|---|---|---|
| SL1344_RS07735_ | -0.328909324 | 0.403810786 | -1.066428147 | -1.43056219 | 0.287519171 | -4.992321621 | 4.68E-05 | 0.00340434 | 0.071000764 | 0.1 | 0.330257709 |
| SL1344_RS07740_ | 1.956174076 | 2.930766438 | 1.288501236 | -1.620717211 | 0.239616839 | -6.052155087 | 4.68E-05 | 0.008312456 | 0.115446527 | 0.1 | 0.330257709 |
| SL1344_RS07745_ | -0.3677649 | -0.228162371 | -0.536912036 | -0.315605081 | 0.193400134 | -1.574121153 | 0.039307241 | 0.09070979 | 0.306482519 | 0.1511 | 0.388522841 |
| SL1344_RS07750_ | -1.050679561 | -0.593433773 | -1.210654216 | -0.590234747 | 0.409883784 | -1.330055165 | 0.105929419 | 0.138839496 | 0.403878912 | 0.2283 | 0.48338484 |
| SL1344_RS07755_ | -0.040552282 | 0.207302802 | -0.417119793 | -0.620999663 | 0.197844318 | -3.079102119 | 0.00067292 | 0.020472784 | 0.155970008 | 0.1004 | 0.330853289 |
| SL1344_RS07760_ | 0.161690569 | 0.272505967 | -0.138698485 | -0.396031015 | 0.240585474 | -1.476909329 | 0.061958764 | 0.116042518 | 0.369577604 | 0.1726 | 0.416729378 |
| SL1344_RS07765_ | -0.629473875 | -0.556884934 | -0.694749495 | -0.135400699 | 0.201822965 | -0.62631428 | 0.235176563 | 0.365960197 | 0.595641741 | 0.4684 | 0.663464666 |
| SL1344_RS07770_ | -1.04855096 | -1.070885558 | -1.016835144 | 0.066168191 | 0.252717276 | 0.244390912 | 0.388000004 | 0.553548281 | 0.75446842 | 0.661 | 0.808389361 |
| SL1344_RS07775_ | -2.66345662 | -2.735788422 | -2.547110677 | 0.197529392 | 0.475145353 | 0.369708998 | 0.343104603 | 0.52775489 | 0.734922896 | 0.6335 | 0.791883145 |
| SL1344_RS07780_ | 0.624438705 | 0.847882922 | 0.368536262 | -0.503478282 | 0.257663766 | -1.96683301 | 0.006667356 | 0.045420916 | 0.241718951 | 0.1053 | 0.337875647 |
| SL1344_RS07785_ | -1.109718649 | -0.891566874 | -1.263752701 | -0.34320725 | 0.310231076 | -1.057238461 | 0.137241867 | 0.202338676 | 0.462689117 | 0.2876 | 0.533799378 |
| SL1344_RS07790_ | -2.397926637 | -2.36399991 | -2.43476188 | -0.06881666 | 0.406175784 | -0.164945891 | 0.433044639 | 0.553286385 | 0.755839036 | 0.6575 | 0.809249289 |
| SL1344_RS07800_ | 3.794326766 | 3.915334434 | 3.72927632 | -0.189626821 | 0.109325779 | -1.75486624 | 0.009333825 | 0.051056131 | 0.258911782 | 0.1121 | 0.347200459 |
| SL1344_RS07805_ | 1.803171876 | 2.13744365 | 1.430425007 | -0.710558884 | 0.117134512 | -6.007848667 | 4.68E-05 | 0.002431944 | 0.059510739 | 0.1 | 0.330257709 |
| SL1344_RS07810_ | 2.762406184 | 2.977003087 | 2.350882791 | -0.593172029 | 0.168914522 | -3.129946839 | 4.68E-05 | 0.019542419 | 0.167281519 | 0.1 | 0.330257709 |
| SL1344_RS07815_ | -1.47255295 | -1.380544683 | -1.596120507 | -0.207275844 | 0.291267421 | -0.687059349 | 0.23466668 | 0.34321235 | 0.582009157 | 0.435 | 0.643264234 |
| SL1344_RS07820_ | 0.540763561 | 0.589540332 | 0.480031278 | -0.131429789 | 0.232705861 | -0.551660562 | 0.264000011 | 0.395713292 | 0.657756203 | 0.5309 | 0.728396207 |
| SL1344_RS07825_ | 0.266957165 | 0.361782865 | 0.192081011 | -0.17520215 | 0.171872313 | -0.952424794 | 0.143904091 | 0.223227216 | 0.47874701 | 0.31 | 0.539461518 |
| SL1344_RS07830_ | -1.633712464 | -1.380967667 | -1.837603501 | -0.461443648 | 0.297501495 | -1.432554651 | 0.060666738 | 0.101625469 | 0.324220808 | 0.1725 | 0.414790378 |
| SL1344_RS07835_ | 8.429448397 | 8.663306526 | 8.419731857 | -0.238334393 | 0.039689624 | -0.910283567 | 0.327333341 | 0.23384435 | 0.538502964 | 0.6815 | 0.860416594 |
| SL1344_RS07840_ | 0.997295255 | 1.201077163 | 0.723929765 | -0.513948397 | 0.202379716 | -2.538392514 | 0.00067292 | 0.021844726 | 0.171061772 | 0.1001 | 0.330400895 |
| SL1344_RS07845_ | 1.672836467 | 2.151628741 | 1.24364662 | -0.898165888 | 1.457936943 | -0.631309188 | 0.228514338 | 0.384887581 | 0.673462766 | 0.4237 | 0.695548594 |
| SL1344_RS07850_ | 0.918201537 | 1.220570054 | 0.692293019 | -0.708680419 | 0.649268594 | -1.031419398 | 0.110000037 | 0.158395669 | 0.448621608 | 0.2167 | 0.491406186 |
| SL1344_RS07855_ | -2.106141315 | -2.025042305 | -2.151588907 | -0.111148189 | 0.427536775 | -0.246172836 | 0.396402402 | 0.606450363 | 0.79442557 | 0.7163 | 0.848414525 |
| SL1344_RS07860_ | -0.527432852 | -0.443020832 | -0.613868422 | -0.201544956 | 0.258727411 | -0.735152297 | 0.20186544 | 0.306636796 | 0.568479846 | 0.4213 | 0.634611567 |
| SL1344_RS07865_ | -1.756350343 | -1.642581071 | -1.889237831 | -0.283946231 | 0.429394543 | -0.631993602 | 0.23666668 | 0.356086109 | 0.614860139 | 0.4873 | 0.687330462 |
| SL1344_RS07870_ | -3.385547482 | -2.960401685 | -3.627329916 | -0.590295084 | 0.702191588 | -0.787938884 | 0.209860109 | 0.316021524 | 0.576154688 | 0.413 | 0.640673022 |
| SL1344_RS07875_ | -2.270526428 | -1.984450161 | -2.464837666 | -0.444075537 | 0.587380902 | -0.751196117 | 0.20453033 | 0.31690747 | 0.58549364 | 0.407 | 0.644373443 |
| SL1344_RS07880_ | 0.065129778 | 0.359001544 | -0.131745846 | -0.491199645 | 0.220651192 | -2.141797454 | 0.007329074 | 0.043662786 | 0.231075084 | 0.1066 | 0.338975925 |
| SL1344_RS07885_ | 0.157548988 | 0.312270514 | 0.000421879 | -0.309332603 | 0.18462296 | -1.626625946 | 0.028666823 | 0.072906227 | 0.284769056 | 0.1324 | 0.371340999 |
| SL1344_RS07890_ | 2.018356824 | 2.333159564 | 1.877381184 | -0.42027278 | 0.183608574 | -2.048593486 | 0.006000767 | 0.048043822 | 0.252041856 | 0.1046 | 0.336962384 |
| SL1344_RS07895_ | 1.036041919 | 1.135953763 | 0.893839407 | -0.271776768 | 0.217939079 | -1.205223703 | 0.091333379 | 0.130871797 | 0.393713703 | 0.2006 | 0.454684194 |
| SL1344_RS07900_ | -2.927205519 | -2.711944846 | -3.201875876 | -0.494852079 | 0.436194488 | -1.055937444 | 0.119920087 | 0.200815302 | 0.440303476 | 0.2827 | 0.511538222 |
| SL1344_RS07905_ | 1.562747838 | 1.855778426 | 1.326597829 | -0.643925617 | 0.286054212 | -2.363751531 | 4.68E-05 | 0.022954502 | 0.180823997 | 0.1 | 0.330257709 |
| SL1344_RS07910_ | -0.020143925 | 0.070615477 | -0.147325947 | -0.217911821 | 0.184167305 | -1.095976527 | 0.118000034 | 0.185580912 | 0.434444871 | 0.2687 | 0.50718591 |
| SL1344_RS07915_ | -1.458937697 | -1.835696515 | -1.132135466 | 0.710381851 | 0.281886343 | 2.446611587 | 0.002000966 | 0.026604456 | 0.171824058 | 0.1032 | 0.334490703 |
| SL1344_RS07920_ | 0.938367833 | 1.083484768 | 0.695871688 | -0.451255844 | 0.287989554 | -1.616890942 | 0.017322048 | 0.06492466 | 0.289368984 | 0.1139 | 0.349347661 |
| SL1344_RS07925_ | 0.422745655 | 0.71640779 | 0.284480142 | -0.467463348 | 0.319614531 | -1.44174656 | 0.040000111 | 0.085516462 | 0.328419021 | 0.1455 | 0.390304309 |
| SL1344_RS07930_ | 0.489848327 | 0.646841291 | 0.205084132 | -0.506144564 | 0.377026057 | -1.54947034 | 0.010667096 | 0.096621264 | 0.352646728 | 0.1108 | 0.344831329 |
| SL1344_RS07935_ | -1.368306813 | -1.137845323 | -1.457491847 | -0.303829359 | 0.426186207 | -0.609022072 | 0.277148578 | 0.442138245 | 0.689698916 | 0.5664 | 0.760272173 |
| SL1344_RS07955_ | -1.300081506 | -1.424548506 | -1.168130866 | 0.242639244 | 0.32277625 | 0.719453041 | 0.218666682 | 0.331556848 | 0.586667235 | 0.4371 | 0.654690122 |
| SL1344_RS07960_ | 1.083690474 | 1.162370662 | 0.971431283 | -0.197746482 | 0.175684875 | -1.123606064 | 0.116000035 | 0.171577342 | 0.441624851 | 0.2455 | 0.499650205 |
| SL1344_RS07965_ | 0.874762721 | 1.077825525 | 0.757517965 | -0.321815566 | 0.213234428 | -1.39547965 | 0.054630326 | 0.108514668 | 0.36092085 | 0.1613 | 0.409417739 |
| SL1344_RS07970_ | 0.145700931 | 0.577976256 | 0.040848198 | -0.480198886 | 0.337916262 | -1.09299083 | 0.156666691 | 0.19826823 | 0.488226585 | 0.3198 | 0.57781514 |
| SL1344_RS07975_ | -3.270079419 | -2.905012757 | -4.033886743 | -1.074516741 | 0.964688792 | -1.00678355 | 0.121252531 | 0.195380623 | 0.479276626 | 0.2552 | 0.510536702 |
| SL1344_RS07980_ | -2.476831788 | -2.279032416 | -2.737085515 | -0.455618855 | 0.421910535 | -1.016736701 | 0.122584976 | 0.198834316 | 0.447711397 | 0.2795 | 0.512038068 |
| SL1344_RS07990_ | -2.273137977 | -1.607274109 | -2.860290986 | -1.224476206 | 0.438041678 | -2.721851319 | 0.002000966 | 0.020830343 | 0.160340725 | 0.1011 | 0.331844168 |
| SL1344_RS07995_ | 1.890342531 | 1.997174378 | 1.74807309 | -0.283211639 | 0.181805759 | -1.532808668 | 0.029980163 | 0.075016018 | 0.306849502 | 0.1304 | 0.370800617 |
| SL1344_RS08000_ | 1.042649392 | 1.106489645 | 1.015679526 | -0.078792141 | 0.174599408 | -0.392000816 | 0.349333339 | 0.588556701 | 0.788544811 | 0.6976 | 0.845459357 |
| SL1344_RS08005_ | 0.781655449 | 0.878135818 | 0.657234655 | -0.111519722 | 0.324043456 | -0.358787917 | 0.369333339 | 0.62510671 | 0.821126088 | 0.7744 | 0.889484361 |
| SL1344_RS08010_ | 2.987048804 | 3.216495162 | 2.94443765 | -0.258588026 | 0.126951588 | -1.283209542 | 0.100000041 | 0.141893714 | 0.421138962 | 0.2217 | 0.477091619 |
| SL1344_RS08015_ | 1.337830626 | 1.510449309 | 1.267328008 | -0.231752198 | 0.135085139 | -1.528355765 | 0.062666736 | 0.107552324 | 0.351349612 | 0.1788 | 0.422967325 |
| SL1344_RS08020_ | 0.893293478 | 1.010266117 | 0.6870544 | -0.328820218 | 0.173152777 | -1.799431756 | 0.013333676 | 0.074229171 | 0.296658244 | 0.1145 | 0.347859697 |

| | | | | | | | | | | | |
|---|---|---|---|---|---|---|---|---|---|---|---|
| SL1344_RS08025_ | -1.953881069 | -1.705723647 | -2.135310317 | -0.459651823 | 0.459029165 | -0.966369971 | 0.148000026 | 0.225813486 | 0.49568918 | 0.3189 | 0.556357205 |
| SL1344_RS08030_ | -1.178274485 | -0.986273098 | -1.37193453 | -0.44160471 | 0.406958856 | -1.110245242 | 0.089273864 | 0.165070463 | 0.436830353 | 0.2132 | 0.459646471 |
| SL1344_RS08035_ | -2.246527269 | -1.993832258 | -2.59121373 | -0.589121166 | 0.397650756 | -1.402305737 | 0.053964105 | 0.108465451 | 0.337389865 | 0.1779 | 0.421821637 |
| SL1344_RS08040_ | -4.257612818 | -3.814104259 | -4.637438141 | -0.806159657 | 0.812799722 | -0.920873993 | 0.16266669 | 0.240909319 | 0.492126339 | 0.3194 | 0.5542143 |
| SL1344_RS08045_ | -1.83841975 | -1.500400426 | -2.222186903 | -0.789458947 | 0.499771146 | -1.628974772 | 0.013990999 | 0.077991525 | 0.307309997 | 0.1167 | 0.351824533 |
| SL1344_RS08050_ | -2.130843266 | -1.633760462 | -2.713093413 | -1.048850185 | 0.468938313 | -2.208349145 | 0.006667356 | 0.037657959 | 0.214596712 | 0.1068 | 0.339888474 |
| SL1344_RS08055_ | -2.906807742 | -2.66495048 | -3.07863413 | -0.303478615 | 0.702698865 | -0.399988241 | 0.327781486 | 0.57201613 | 0.774463358 | 0.6507 | 0.813706716 |
| SL1344_RS08060_ | -2.471424776 | -2.308666937 | -2.64624801 | -0.353686766 | 0.426354063 | -0.791391383 | 0.189873436 | 0.298554104 | 0.547279954 | 0.4018 | 0.615137697 |
| SL1344_RS08065_ | -1.497930969 | -1.189383797 | -1.746139156 | -0.566699834 | 0.349206013 | -1.58948391 | 0.046666761 | 0.090560354 | 0.317271521 | 0.1534 | 0.396072278 |
| SL1344_RS08070_ | -1.560005245 | -1.280257934 | -1.9230555 | -0.642099743 | 0.303630029 | -1.983223307 | 0.016000285 | 0.050081218 | 0.233961514 | 0.1169 | 0.351960538 |
| SL1344_RS08075_ | -1.066458338 | -1.05028564 | -1.075717387 | 0.017637085 | 0.373357324 | 0.042211057 | 0.477015324 | 0.666639278 | 0.840117783 | 0.795 | 0.900132457 |
| SL1344_RS08080_ | -3.227229899 | -3.177703695 | -3.280386421 | -0.090578168 | 0.558150526 | -0.157203369 | 0.436000002 | 0.570387266 | 0.766579848 | 0.6798 | 0.821898681 |
| SL1344_RS08090_ | 0.925583803 | 0.977735713 | 0.892458176 | -0.158812462 | 0.269259237 | -0.576552591 | 0.27800001 | 0.399932839 | 0.668392365 | 0.5825 | 0.772112352 |
| SL1344_RS08095_ | 1.377270422 | 1.353938227 | 1.412141735 | 0.036874861 | 0.208442299 | 0.158752162 | 0.440000002 | 0.745388046 | 0.88615091 | 0.8684 | 0.93710165 |
| SL1344_RS08100_ | 2.857578674 | 2.666235234 | 3.03623504 | -0.141337763 | 0.839143836 | -0.180491109 | 0.448367757 | 0.957512778 | 0.985028961 | 1 | 1 |
| SL1344_RS08110_ | -0.049165822 | 0.051659468 | -0.147205131 | -0.221843506 | 0.272858603 | -0.804638534 | 0.172666688 | 0.269903861 | 0.547572171 | 0.358 | 0.595335629 |
| SL1344_RS08115_ | -1.805566465 | -1.913549752 | -1.714207449 | 0.194775708 | 0.299134153 | 0.612629166 | 0.260666678 | 0.377580365 | 0.609462362 | 0.4848 | 0.677180759 |
| SL1344_RS08120_ | -4.867022579 | -4.471702361 | -5.564060443 | -1.123929182 | 1.124473413 | -0.915983166 | 0.130579644 | 0.217061491 | 0.479966126 | 0.2848 | 0.51853437 |
| SL1344_RS08125_ | -3.926880269 | -3.562724976 | -4.203542755 | -0.721952433 | 1.081776736 | -0.633105889 | 0.230000014 | 0.362732404 | 0.625103165 | 0.4755 | 0.690836553 |
| SL1344_RS08130_ | -4.023373713 | -4.024388608 | -4.021878926 | 0.000761287 | 0.732955497 | 0.001450804 | 0.499000666 | 0.586803824 | 0.779178335 | 0.698 | 0.834522786 |
| SL1344_RS08135_ | -1.697342568 | -1.439344082 | -1.964560927 | -0.521667268 | 0.496419791 | -0.985808872 | 0.139240534 | 0.196673307 | 0.470325842 | 0.2816 | 0.536709016 |
| SL1344_RS08140_ | -2.211849874 | -1.777287 | -2.581628872 | -0.806477864 | 0.412958315 | -1.836825397 | 0.019333568 | 0.056553379 | 0.254112271 | 0.1208 | 0.357542391 |
| SL1344_RS08145_ | -2.955071399 | -2.715536434 | -3.876261122 | -1.147512424 | 0.968524416 | -1.088177244 | 0.154666691 | 0.182401537 | 0.467469739 | 0.292 | 0.559851417 |
| SL1344_RS08150_ | -4.328910423 | -3.862152126 | -4.75190373 | -0.867980848 | 0.921363125 | -0.873084383 | 0.159893428 | 0.254725231 | 0.513246534 | 0.3501 | 0.582715231 |
| SL1344_RS08155_ | -1.846671634 | -1.556666893 | -2.110244053 | -0.620951179 | 0.473512817 | -1.338988061 | 0.055333412 | 0.123265069 | 0.377149988 | 0.1696 | 0.413827364 |
| SL1344_RS08160_ | 1.814776415 | 2.502197066 | 1.603988694 | -0.823123549 | 0.248359608 | -1.690851428 | 0.009327606 | 0.084977502 | 0.336790222 | 0.1081 | 0.342090374 |
| SL1344_RS08165_ | 0.888349544 | 1.060613144 | 0.840558437 | -0.199846859 | 0.186675702 | -0.92769837 | 0.197868106 | 0.278261228 | 0.558452639 | 0.4098 | 0.645138688 |
| SL1344_RS08170_ | -0.172149146 | 0.199024982 | -0.456884226 | -0.637534135 | 0.193317941 | -3.166789258 | 4.68E-05 | 0.015398253 | 0.137377629 | 0.1002 | 0.330559573 |
| SL1344_RS08175_ | 2.379688705 | 2.639998834 | 2.261173366 | -0.376863279 | 0.141174068 | -2.441612024 | 0.000673369 | 0.048482846 | 0.254306824 | 0.1002 | 0.330542564 |
| SL1344_RS08185_ | -0.953839777 | -0.780325686 | -1.088049885 | -0.340785745 | 0.318863178 | -1.082299315 | 0.092666712 | 0.182441355 | 0.449819105 | 0.2379 | 0.477464037 |
| SL1344_RS08190_ | -1.576410679 | -1.337896622 | -1.683189608 | -0.323082853 | 0.358167185 | -0.841146695 | 0.208000016 | 0.311277466 | 0.568950149 | 0.4163 | 0.642209354 |
| SL1344_RS08195_ | -1.323977123 | -1.148526191 | -1.601837629 | -0.498023894 | 0.411742484 | -1.135572764 | 0.070619647 | 0.166514974 | 0.440319272 | 0.1827 | 0.424199603 |
| SL1344_RS08200_ | 1.471348779 | 1.666341207 | 1.269256885 | -0.436660449 | 0.221558562 | -2.082116222 | 4.68E-05 | 0.057630409 | 0.274049164 | 0.1004 | 0.330832827 |
| SL1344_RS08215_ | 1.647575993 | 1.777126624 | 1.571326224 | -0.246575658 | 0.30437675 | -0.879125795 | 0.140000028 | 0.218328986 | 0.514979766 | 0.2695 | 0.53784214 |
| SL1344_RS08220_ | -3.752217047 | -3.733797396 | -3.772941121 | -0.006744735 | 0.711190655 | -0.007182787 | 0.497001999 | 0.626444989 | 0.807237835 | 0.7385 | 0.862070596 |
| SL1344_RS08225_ | 0.879530897 | 0.956939269 | 0.769521947 | -0.19607871 | 0.153233427 | -1.221482685 | 0.066000065 | 0.146535327 | 0.401531193 | 0.1908 | 0.43192872 |
| SL1344_RS08230_ | -2.562989594 | -2.522457189 | -2.60421045 | -0.09060245 | 0.427574358 | -0.185545672 | 0.419333336 | 0.542544125 | 0.746720733 | 0.6581 | 0.805321788 |
| SL1344_RS08235_ | 3.655901903 | 4.244817849 | 3.247467238 | -0.897100151 | 0.198524448 | -4.052243365 | 4.68E-05 | 0.007385811 | 0.109387971 | 0.1 | 0.330257709 |
| SL1344_RS08240_ | 0.519307741 | 0.440427137 | 0.872802247 | 0.363317483 | 0.317969723 | 0.800810414 | 0.260000012 | 0.343140311 | 0.627966509 | 0.5303 | 0.752098161 |
| SL1344_RS08245_ | -1.173077903 | -1.266868239 | -0.981518461 | 0.25635892 | 0.340060222 | 0.703178522 | 0.242000013 | 0.394972708 | 0.644028999 | 0.4991 | 0.711203185 |
| SL1344_RS08250_ | -0.997545915 | -0.683721067 | -1.132606472 | -0.446840276 | 0.402984004 | -1.008052266 | 0.155896093 | 0.212960742 | 0.494712084 | 0.3061 | 0.559132446 |
| SL1344_RS08255_ | 2.023821215 | 2.086307832 | 1.941665093 | -0.152330909 | 0.151135421 | -0.951113196 | 0.129247199 | 0.208064949 | 0.491631239 | 0.2782 | 0.533968352 |
| SL1344_RS08260_ | 0.443153184 | 0.609870243 | 0.248714504 | -0.352311923 | 0.166359126 | -2.049437408 | 0.011326186 | 0.045258347 | 0.228876062 | 0.1113 | 0.345851275 |
| SL1344_RS08270_ | 0.679469963 | 0.910248405 | 0.399194812 | -0.511063989 | 0.125978449 | -3.9943629 | 4.68E-05 | 0.008366916 | 0.10083974 | 0.1 | 0.330257709 |
| SL1344_RS08275_ | 1.618275263 | 1.889068521 | 1.424439207 | -0.464387436 | 0.133060209 | -3.406017105 | 4.68E-05 | 0.016556947 | 0.148573828 | 0.1 | 0.330257709 |
| SL1344_RS08280_ | 0.626012989 | 0.768272448 | 0.500747225 | -0.294714121 | 0.208665585 | -1.432336965 | 0.0379748 | 0.112337277 | 0.363415513 | 0.1413 | 0.380380477 |
| SL1344_RS08290_ | 2.951563198 | 3.12720919 | 2.772943985 | -0.360478481 | 0.085369671 | -4.210285466 | 4.68E-05 | 0.011206496 | 0.120659747 | 0.1 | 0.330257709 |
| SL1344_RS08295_ | 2.517698506 | 2.866603125 | 2.247165539 | -0.611747507 | 0.291824074 | -1.931969725 | 4.68E-05 | 0.074133816 | 0.317020734 | 0.1 | 0.330257709 |
| SL1344_RS08300_ | -0.851772178 | -0.821855162 | -0.900427963 | -0.056839008 | 0.321587347 | -0.167379886 | 0.423333336 | 0.652672573 | 0.828652389 | 0.7587 | 0.877513406 |
| SL1344_RS08305_ | 1.88E-05 | 0.077520039 | -0.079621009 | -0.149408083 | 0.203796142 | -0.694755917 | 0.217188556 | 0.335847193 | 0.5869184 | 0.4302 | 0.645134786 |
| SL1344_RS08310_ | -0.724499744 | -0.789363127 | -0.64252792 | 0.197310557 | 0.405294604 | 0.461847527 | 0.300000009 | 0.477050751 | 0.718386451 | 0.6104 | 0.788352205 |
| SL1344_RS08315_ | -1.541530493 | -1.774490257 | -1.357734568 | 0.436111709 | 0.379943937 | 1.083193714 | 0.10326453 | 0.169822225 | 0.429950876 | 0.2365 | 0.482095851 |
| SL1344_RS08320_ | -3.954093966 | -3.802732015 | -4.134563003 | -0.334817121 | 0.683592939 | -0.443382807 | 0.299800142 | 0.446973545 | 0.672558297 | 0.56 | 0.737516805 |

| | | | | | | | | | | | |
|---|---|---|---|---|---|---|---|---|---|---|---|
| SL1344_RS08325_ | -4.59778257 | -4.304648526 | -4.888202166 | -0.295959986 | 1.857294146 | -0.146665817 | 0.424000003 | 0.733811822 | 0.878450317 | 0.8202 | 0.912948837 |
| SL1344_RS08330_ | -2.991481109 | -2.743992744 | -3.37313271 | -0.648516702 | 0.558519142 | -1.072163963 | 0.100666708 | 0.181626125 | 0.441588091 | 0.246 | 0.484184412 |
| SL1344_RS08335_ | -4.237851548 | -3.791734856 | -4.798667897 | -1.000382431 | 0.713804268 | -1.340442309 | 0.066666731 | 0.12180517 | 0.347915073 | 0.1866 | 0.424413777 |
| SL1344_RS08340_ | -4.621125093 | -4.408901463 | -4.787101695 | -0.35231604 | 1.082594968 | -0.282575261 | 0.372666672 | 0.590716036 | 0.783427996 | 0.6868 | 0.830166613 |
| SL1344_RS08345_ | 2.15561443 | 2.559100276 | 2.046252325 | -0.419467031 | 0.348374727 | -0.753531384 | 0.319786816 | 0.395966727 | 0.679824756 | 0.6702 | 0.852831207 |
| SL1344_RS08350_ | -1.055686115 | -0.869780921 | -1.204546058 | -0.360609877 | 0.312490783 | -1.094847741 | 0.106595641 | 0.179250408 | 0.439014205 | 0.2541 | 0.495304585 |
| SL1344_RS08355_ | -1.859287867 | -1.498576386 | -2.263793602 | -0.779767505 | 0.360739562 | -2.093044791 | 0.008000574 | 0.041774087 | 0.217291476 | 0.1077 | 0.340554299 |
| SL1344_RS08360_ | -0.154956388 | -0.363400787 | -0.06968341 | 0.237144591 | 0.616869801 | 0.302782801 | 0.358666672 | 0.855879137 | 0.943419999 | 0.7426 | 0.890661157 |
| SL1344_RS08365_ | -0.054136506 | -0.126335104 | -0.008595676 | 0.110424248 | 0.186026718 | 0.558534242 | 0.271818798 | 0.427125628 | 0.656029805 | 0.5322 | 0.720043653 |
| SL1344_RS08370_ | -2.013436002 | -1.858519713 | -2.27833185 | -0.410800544 | 0.496846901 | -0.773093513 | 0.206666683 | 0.29953645 | 0.569205559 | 0.4004 | 0.636281271 |
| SL1344_RS08375_ | -3.187998168 | -3.125213267 | -3.274379019 | -0.067583031 | 0.738809048 | -0.085003169 | 0.464000001 | 0.685306365 | 0.849561043 | 0.802 | 0.903966136 |
| SL1344_RS08380_ | 2.377819895 | 2.557790915 | 2.282240394 | -0.272134812 | 0.133086592 | -1.95303407 | 0.011992386 | 0.047339369 | 0.245595914 | 0.1107 | 0.344611527 |
| SL1344_RS08385_ | -0.075939024 | 0.187034146 | -0.371588579 | -0.558527308 | 0.244935269 | -2.258487575 | 0.003998486 | 0.034823597 | 0.20768002 | 0.1024 | 0.333659757 |
| SL1344_RS08390_ | 0.087856262 | 0.401207797 | -0.16778168 | -0.562644185 | 0.230858092 | -2.382541488 | 0.002000966 | 0.03104598 | 0.197349293 | 0.1033 | 0.3350409 |
| SL1344_RS08395_ | 4.957048408 | 4.970979422 | 4.76081724 | -0.432336582 | 0.462893315 | -0.948437119 | 0.166666689 | 0.157352967 | 0.449975673 | 0.3154 | 0.589586712 |
| SL1344_RS08400_ | 1.631038763 | 1.734930795 | 1.510726716 | -0.232739705 | 0.127692772 | -1.777705555 | 0.015323415 | 0.069654716 | 0.286489309 | 0.117 | 0.352664677 |
| SL1344_RS08405_ | 1.361715603 | 1.598170844 | 1.109199324 | -0.458520728 | 0.173254338 | -2.633183017 | 0.000673369 | 0.020465911 | 0.163874142 | 0.1011 | 0.331848777 |
| SL1344_RS08410_ | 1.166344844 | 1.320790815 | 1.020312374 | -0.324593648 | 0.244768151 | -1.323176606 | 0.062666736 | 0.121761898 | 0.388014753 | 0.1621 | 0.413103385 |
| SL1344_RS08415_ | 2.2129128 | 2.504461033 | 2.103106517 | -0.418564701 | 0.295845309 | -1.494765397 | 0.032666804 | 0.071546604 | 0.307914082 | 0.1272 | 0.368466093 |
| SL1344_RS08420_ | -1.141354415 | -0.886756013 | -1.335481684 | -0.451504238 | 0.300777557 | -1.457462718 | 0.054000081 | 0.103850291 | 0.336931407 | 0.167 | 0.409292098 |
| SL1344_RS08425_ | -0.740511209 | -0.523202621 | -1.081220442 | -0.557528316 | 0.309973769 | -1.729586033 | 0.025316634 | 0.068386418 | 0.283502775 | 0.1296 | 0.369113354 |
| SL1344_RS08430_ | -1.650165875 | -1.455554846 | -1.776729391 | -0.282670579 | 0.409544188 | -0.664648539 | 0.23184545 | 0.37510272 | 0.628102802 | 0.4789 | 0.692292439 |
| SL1344_RS08435_ | 1.999471456 | 2.076386748 | 1.914877031 | -0.161513598 | 0.095859752 | -1.651247023 | 0.030646382 | 0.075018253 | 0.287438188 | 0.137 | 0.375098403 |
| SL1344_RS08440_ | -0.336330173 | -0.201802307 | -0.438902611 | -0.251583587 | 0.330297348 | -0.717411637 | 0.212000016 | 0.316421525 | 0.591279143 | 0.432 | 0.664646581 |
| SL1344_RS08445_ | 1.593227923 | 1.686257053 | 1.441631939 | -0.240988225 | 0.104632906 | -2.220500232 | 0.009993796 | 0.045300771 | 0.22584934 | 0.1101 | 0.34354805 |
| SL1344_RS08450_ | 0.556008083 | 0.56271587 | 0.548732442 | -0.034896314 | 0.193764191 | -0.175609756 | 0.427333336 | 0.631524599 | 0.81271331 | 0.7612 | 0.873576787 |
| SL1344_RS08455_ | 0.512865245 | 0.62290127 | 0.38699921 | -0.23866561 | 0.142605919 | -1.610418221 | 0.0379748 | 0.082501891 | 0.297440451 | 0.1457 | 0.385619619 |
| SL1344_RS08460_ | 0.704049613 | 0.744393848 | 0.654729908 | -0.089628212 | 0.1313436 | -0.639145995 | 0.23984012 | 0.346227625 | 0.587636981 | 0.4588 | 0.659418011 |
| SL1344_RS08465_ | 0.177654066 | 0.328528922 | 0.048688905 | -0.294528482 | 0.190047742 | -1.522879641 | 0.040000111 | 0.095249106 | 0.324994997 | 0.1411 | 0.378647811 |
| SL1344_RS08470_ | -0.788098436 | -0.763069199 | -0.823254948 | -0.046560839 | 0.263903465 | -0.174153516 | 0.418000003 | 0.592411691 | 0.784900608 | 0.698 | 0.837236069 |
| SL1344_RS08475_ | -2.187915065 | -2.01021301 | -2.611494127 | -0.554645379 | 0.47842255 | -1.075693832 | 0.140572979 | 0.203341893 | 0.475323851 | 0.3011 | 0.545411596 |
| SL1344_RS08480_ | -6.373814631 | -6.232821389 | -6.578275677 | -0.445393714 | 2.199563338 | -0.151633424 | 0.398000004 | 0.525246247 | 0.75170472 | 0.682 | 0.825799789 |
| SL1344_RS08485_ | -1.260756033 | -1.007990438 | -1.512061037 | -0.543492567 | 0.603363228 | -0.883763848 | 0.161892095 | 0.224374242 | 0.51524245 | 0.2995 | 0.562406859 |
| SL1344_RS08490_ | 5.074389358 | 5.184354593 | 4.830818933 | -0.351909898 | 0.087739762 | -3.27747344 | 4.68E-05 | 0.036744747 | 0.229162199 | 0.1 | 0.330257709 |
| SL1344_RS08500_ | 3.734399158 | 3.846601361 | 3.711772498 | -0.139016488 | 0.136355024 | -0.950104537 | 0.196666684 | 0.236168866 | 0.533887569 | 0.3627 | 0.634998756 |
| SL1344_RS08505_ | 0.777412063 | 0.956370789 | 0.666808674 | -0.280620858 | 0.216125542 | -1.240161091 | 0.093271197 | 0.129702641 | 0.387990062 | 0.206 | 0.462225193 |
| SL1344_RS08510_ | 1.318309552 | 1.433635846 | 1.226692961 | -0.172439741 | 0.269778081 | -0.623786733 | 0.233177895 | 0.360780724 | 0.6411229 | 0.4481 | 0.688127448 |
| SL1344_RS08515_ | 1.598021536 | 1.682798847 | 1.533322058 | -0.296026508 | 0.326803689 | -0.963993079 | 0.105263197 | 0.185194582 | 0.47733383 | 0.2404 | 0.490879575 |
| SL1344_RS08520_ | 1.032430451 | 1.296679173 | 0.934388718 | -0.363123589 | 0.282139194 | -1.357854697 | 0.064000067 | 0.099829637 | 0.352187766 | 0.1732 | 0.423923531 |
| SL1344_RS08525_ | -0.121229057 | -0.010613556 | -0.278598249 | -0.246840491 | 0.306950311 | -0.778145056 | 0.19920055 | 0.3063675 | 0.583606238 | 0.3982 | 0.643303211 |
| SL1344_RS08530_ | 0.20465696 | 0.315404749 | 0.01648021 | -0.319046351 | 0.26703756 | -1.125101811 | 0.104000039 | 0.160292412 | 0.434347346 | 0.2243 | 0.479492556 |
| SL1344_RS08535_ | 2.30200401 | 2.553740933 | 1.90053653 | -0.632646906 | 0.124246888 | -4.864010014 | 4.68E-05 | 0.006251662 | 0.092953269 | 0.1 | 0.330257709 |
| SL1344_RS08540_ | 2.233723082 | 2.494080158 | 1.941908824 | -0.556886363 | 0.109103407 | -5.108200848 | 4.68E-05 | 0.004011215 | 0.075080449 | 0.1 | 0.330257709 |
| SL1344_RS08545_ | -0.690270872 | -0.557061991 | -0.781958796 | -0.261246445 | 0.29704811 | -0.87888669 | 0.154563649 | 0.252927863 | 0.526218066 | 0.3243 | 0.553007461 |
| SL1344_RS08550_ | 2.026466835 | 2.209612643 | 1.872028753 | -0.212737586 | 0.272338814 | -0.937162735 | 0.143237868 | 0.230587212 | 0.526626807 | 0.2561 | 0.530669272 |
| SL1344_RS08555_ | 1.970493367 | 2.395121041 | 1.621518143 | -0.763080396 | 0.16462311 | -4.707679861 | 4.68E-05 | 0.003260587 | 0.072172039 | 0.1 | 0.330257709 |
| SL1344_RS08560_ | 1.437338459 | 1.399219066 | 1.55370562 | 0.144598619 | 0.20746714 | 0.595576958 | 0.300666675 | 0.451250542 | 0.709357949 | 0.6254 | 0.810288304 |
| SL1344_RS08570_ | 4.076486324 | 4.210773761 | 3.961522903 | -0.246232856 | 0.052254077 | -4.618126472 | 4.68E-05 | 0.005465811 | 0.086617419 | 0.1 | 0.330257709 |
| SL1344_RS08575_ | 3.092754734 | 3.162389952 | 2.930330758 | -0.236223625 | 0.116032253 | -1.931428623 | 0.006000767 | 0.076976385 | 0.314607902 | 0.1053 | 0.337971452 |
| SL1344_RS08580_ | 1.050674362 | 1.385169034 | 0.962604325 | -0.397429321 | 0.249314957 | -1.381605484 | 0.076666722 | 0.125573058 | 0.394813692 | 0.197 | 0.452656272 |
| SL1344_RS08590_ | 2.035095115 | 2.23616556 | 1.920109283 | -0.332254701 | 0.321956169 | -1.053022854 | 0.115922753 | 0.169357911 | 0.461088162 | 0.2259 | 0.499134481 |
| SL1344_RS08595_ | 1.887929747 | 2.053481146 | 1.669445159 | -0.409317432 | 0.17480581 | -2.349547386 | 4.68E-05 | 0.02619804 | 0.187701073 | 0.1007 | 0.331268856 |
| SL1344_RS08600_ | 1.579678909 | 1.776557772 | 1.388475992 | -0.391089013 | 0.184068171 | -2.113215126 | 0.003334715 | 0.042809439 | 0.234917003 | 0.1039 | 0.335991557 |

| | | | | | | | | | | | |
|---|---|---|---|---|---|---|---|---|---|---|---|
| SL1344_RS08605_ | 1.425208209 | 1.607808011 | 1.274886498 | -0.380154417 | 0.213897617 | -1.775858602 | 0.005996769 | 0.053589746 | 0.262602941 | 0.1058 | 0.338373905 |
| SL1344_RS08610_ | -2.053182958 | -1.955814496 | -2.208340715 | -0.257461083 | 0.399782196 | -0.577890414 | 0.243333346 | 0.365865488 | 0.613545185 | 0.4845 | 0.678668646 |
| SL1344_RS08615_ | -1.068331351 | -0.898438126 | -1.313733681 | -0.420264067 | 0.257556299 | -1.560586931 | 0.04266677 | 0.092470543 | 0.31597455 | 0.147 | 0.385641596 |
| SL1344_RS08620_ | -0.142915756 | -0.001558401 | -0.255758911 | -0.286117046 | 0.32493324 | -0.892821874 | 0.137908089 | 0.23396521 | 0.521974908 | 0.2898 | 0.535948752 |
| SL1344_RS08625_ | 0.145639276 | 0.354012424 | -0.300558976 | -0.596433717 | 0.289367695 | -1.866718768 | 0.025316634 | 0.067917356 | 0.294019683 | 0.1226 | 0.362756408 |
| SL1344_RS08630_ | 0.184672749 | 0.470832995 | -0.000232908 | -0.429918643 | 0.326489443 | -1.281559811 | 0.091272531 | 0.127770308 | 0.393074728 | 0.1965 | 0.456521124 |
| SL1344_RS08635_ | -3.903751783 | -3.980787231 | -3.827101843 | 0.157900764 | 0.607670425 | 0.254853363 | 0.381333338 | 0.493266725 | 0.708873736 | 0.605 | 0.76875911 |
| SL1344_RS08640_ | -4.737133086 | -4.786820777 | -4.681038012 | 0.11841481 | 0.83381218 | 0.131967104 | 0.437333336 | 0.549523172 | 0.749283591 | 0.6477 | 0.798327815 |
| SL1344_RS08645_ | -5.294858913 | -4.890059867 | -5.668107993 | -0.935818328 | 1.372799016 | -0.63778472 | 0.245333346 | 0.361716064 | 0.613936162 | 0.4919 | 0.69083534 |
| SL1344_RS08650_ | -2.264241775 | -2.055352824 | -2.419088565 | -0.355592576 | 0.414529163 | -0.791034592 | 0.180666687 | 0.289605805 | 0.542617088 | 0.3958 | 0.614755396 |
| SL1344_RS08655_ | 1.344627261 | 1.397472106 | 1.237524016 | -0.177609296 | 0.198165717 | -0.895675058 | 0.171333355 | 0.240309319 | 0.524381285 | 0.3429 | 0.598041911 |
| SL1344_RS08660_ | 0.19268864 | 0.157504925 | 0.242818487 | 0.086100159 | 0.183472422 | 0.440685711 | 0.322666674 | 0.501102091 | 0.717223629 | 0.6112 | 0.776723988 |
| SL1344_RS08665_ | 2.562031692 | 2.621651485 | 2.500348502 | -0.139622093 | 0.145721519 | -0.914963707 | 0.112666703 | 0.212910629 | 0.505138986 | 0.2377 | 0.491637732 |
| SL1344_RS08670_ | 2.637305351 | 2.716828631 | 2.553764864 | -0.179369527 | 0.108332982 | -1.611175056 | 0.018000252 | 0.073197094 | 0.297712501 | 0.1203 | 0.357274187 |
| SL1344_RS08675_ | 1.523634225 | 1.587643757 | 1.454285753 | -0.173161619 | 0.204286002 | -0.842830997 | 0.176548988 | 0.255135657 | 0.544619785 | 0.3541 | 0.598565058 |
| SL1344_RS08680_ | 2.340788426 | 2.809404827 | 2.006740938 | -0.756857781 | 0.179109679 | -4.078141174 | 4.68E-05 | 0.006542906 | 0.099597306 | 0.1 | 0.330257709 |
| SL1344_RS08685_ | 1.025202046 | 1.317063125 | 0.858433809 | -0.508123349 | 0.385058386 | -1.386755734 | 0.019333568 | 0.120888134 | 0.393327078 | 0.1201 | 0.358312154 |
| SL1344_RS08690_ | 0.783134078 | 1.110808081 | 0.393549968 | -0.717275017 | 0.128382557 | -5.576133815 | 4.68E-05 | 0.003938512 | 0.070311255 | 0.1 | 0.330257709 |
| SL1344_RS08695_ | 1.463696573 | 1.839107471 | 1.100307922 | -0.715746178 | 0.217148846 | -3.389337871 | 4.68E-05 | 0.00939253 | 0.115910588 | 0.1 | 0.330257709 |
| SL1344_RS08700_ | -0.076887853 | 0.119652079 | -0.176327565 | -0.27413673 | 0.239380431 | -1.095810932 | 0.12391742 | 0.186200078 | 0.452145213 | 0.28 | 0.529173113 |
| SL1344_RS08705_ | 1.897398508 | 2.211185347 | 1.773557643 | -0.420131442 | 0.352484857 | -1.23982915 | 0.086666715 | 0.108732495 | 0.374602104 | 0.1842 | 0.447142647 |
| SL1344_RS08710_ | 1.525482261 | 1.650645826 | 1.442571015 | -0.120517281 | 0.30480697 | -0.32491033 | 0.384410398 | 0.817416211 | 0.92494158 | 0.8237 | 0.924480117 |
| SL1344_RS08715_ | 0.991060379 | 0.945487602 | 1.023002011 | 0.054238785 | 0.474327851 | 0.103040601 | 0.461333335 | 0.804822795 | 0.920971301 | 0.9703 | 0.987334576 |
| SL1344_RS08725_ | 3.158678208 | 3.263861348 | 3.058480871 | -0.043577813 | 0.294566076 | -0.199853212 | 0.428381082 | 0.718701892 | 0.87897486 | 0.8932 | 0.952490052 |
| SL1344_RS08730_ | -0.024591937 | 0.036518847 | -0.097859545 | -0.140699197 | 0.234458129 | -0.551506384 | 0.262666678 | 0.41129423 | 0.658968366 | 0.5211 | 0.720002401 |
| SL1344_RS08735_ | 4.006204382 | 4.099763491 | 3.950019374 | -0.299719934 | 0.255427023 | -1.259066499 | 0.009333825 | 0.116049507 | 0.388672261 | 0.1071 | 0.340283637 |
| SL1344_RS08740_ | 0.741359145 | 0.902859595 | 0.556952253 | -0.205616632 | 0.326103673 | -0.730205193 | 0.209860109 | 0.342593185 | 0.624744393 | 0.4091 | 0.65199266 |
| SL1344_RS08750_ | 3.935607658 | 4.100961001 | 3.812266466 | -0.318959938 | 0.142674964 | -2.209246182 | 4.68E-05 | 0.026747539 | 0.196276889 | 0.1 | 0.330257709 |
| SL1344_RS08755_ | 1.634373895 | 1.955713751 | 1.480116484 | -0.453355337 | 0.149334889 | -2.764621957 | 4.68E-05 | 0.030900525 | 0.202332154 | 0.1002 | 0.330548947 |
| SL1344_RS08760_ | 1.835012435 | 2.073447513 | 1.590750151 | -0.476060113 | 0.139456844 | -3.42404005 | 4.68E-05 | 0.010909745 | 0.122058985 | 0.1 | 0.330257709 |
| SL1344_RS08765_ | 0.703379908 | 1.128235857 | 0.497009513 | -0.604285782 | 0.215326546 | -2.555061191 | 0.001335878 | 0.033508875 | 0.208613206 | 0.101 | 0.331710938 |
| SL1344_RS08770_ | 2.251778178 | 2.631010817 | 1.859643933 | -0.81237922 | 0.144545643 | -5.550466047 | 4.68E-05 | 0.002127282 | 0.059270619 | 0.1 | 0.330257709 |
| SL1344_RS08775_ | 2.592824643 | 3.125197075 | 2.392791769 | -0.71458435 | 0.57784525 | -1.275454284 | 0.11133337 | 0.121831544 | 0.398972216 | 0.2039 | 0.480310493 |
| SL1344_RS08780_ | 1.267649691 | 1.869476569 | 0.905451135 | -0.632793534 | 0.75948046 | -1.092898043 | 0.119253864 | 0.190912147 | 0.489283296 | 0.2114 | 0.48819958 |
| SL1344_RS08785_ | -0.437427428 | -0.308589884 | -0.556157841 | -0.289625901 | 0.271550591 | -1.022121265 | 0.115333368 | 0.1930589 | 0.464170172 | 0.2622 | 0.500958104 |
| SL1344_RS08815_ | 0.960878354 | 1.219766004 | 0.696883659 | -0.518020635 | 0.153751805 | -3.271609891 | 4.68E-05 | 0.01234655 | 0.126695873 | 0.1 | 0.330257709 |
| SL1344_RS08825_ | 0.440762438 | 0.714072741 | 0.299497592 | -0.460185296 | 0.323596472 | -1.43507493 | 0.039333446 | 0.088599512 | 0.333364002 | 0.1477 | 0.391967534 |
| SL1344_RS08830_ | 2.157978793 | 2.366183466 | 1.987407765 | -0.364922225 | 0.122338573 | -2.912456121 | 0.00067292 | 0.015299855 | 0.143486283 | 0.1002 | 0.33054635 |
| SL1344_RS08840_ | -0.191514597 | -0.209182453 | -0.184701343 | 0.051962198 | 0.302367595 | 0.141591463 | 0.421718857 | 0.57111998 | 0.787244531 | 0.7542 | 0.874261934 |
| SL1344_RS08845_ | 3.658524442 | 3.809061816 | 3.460170486 | -0.341298223 | 0.06616138 | -5.106585672 | 4.68E-05 | 0.00436717 | 0.077845175 | 0.1 | 0.330257709 |
| SL1344_RS08850_ | 1.754233298 | 1.960357961 | 1.641165888 | -0.301151996 | 0.128255884 | -2.20574797 | 0.008667196 | 0.041741966 | 0.226338301 | 0.1081 | 0.341140666 |
| SL1344_RS08855_ | 0.272384157 | 0.396759865 | 0.171507375 | -0.250696824 | 0.215865332 | -1.11040865 | 0.083944087 | 0.162176653 | 0.429842742 | 0.2063 | 0.448603999 |
| SL1344_RS08860_ | 1.513415795 | 1.562271989 | 1.444237963 | -0.099326193 | 0.197521347 | -0.483911436 | 0.305129922 | 0.47666216 | 0.720809785 | 0.6131 | 0.792505671 |
| SL1344_RS08865_ | 2.299833467 | 2.360916219 | 2.197337386 | -0.157423477 | 0.171409563 | -0.907551861 | 0.178547655 | 0.239606225 | 0.530350115 | 0.3281 | 0.593837948 |
| SL1344_RS08870_ | 3.346523835 | 3.487235509 | 3.178677194 | -0.31582678 | 0.079592691 | -4.061248969 | 4.68E-05 | 0.021684508 | 0.172573863 | 0.1 | 0.330257709 |
| SL1344_RS08880_ | 4.306302032 | 4.583686654 | 4.098868819 | -0.473812154 | 0.253448131 | -1.928689005 | 4.68E-05 | 0.037307913 | 0.231424623 | 0.1 | 0.330257709 |
| SL1344_RS08885_ | 5.466232434 | 5.642080816 | 5.375672069 | -0.2696485 | 0.136531905 | -1.747549763 | 4.68E-05 | 0.13252321 | 0.415769919 | 0.1 | 0.330257709 |
| SL1344_RS08890_ | 1.473775123 | 1.667818063 | 1.263235675 | -0.432293099 | 0.173133896 | -2.511972199 | 4.68E-05 | 0.031275981 | 0.203933266 | 0.1001 | 0.330404651 |
| SL1344_RS08895_ | 0.030820928 | 0.2288078 | -0.356205866 | -0.404846397 | 0.775062981 | -0.566749748 | 0.259333345 | 0.461564763 | 0.72579701 | 0.4979 | 0.741476501 |
| SL1344_RS08900_ | 0.591366606 | 0.77444213 | 0.455801277 | -0.366906543 | 0.280861403 | -1.313703534 | 0.029333486 | 0.12066288 | 0.386218136 | 0.1373 | 0.378059575 |
| SL1344_RS08905_ | 3.63831336 | 3.763797193 | 3.611593012 | -0.132202499 | 0.100442425 | -1.023644136 | 0.189207214 | 0.22364975 | 0.516603336 | 0.3658 | 0.618444537 |
| SL1344_RS08910_ | 0.752632129 | 0.919646555 | 0.511513314 | -0.402436328 | 0.13683896 | -2.873526257 | 0.00067292 | 0.022228181 | 0.162491902 | 0.1002 | 0.330556482 |
| SL1344_RS08915_ | -5.005334858 | -4.869491931 | -5.291780894 | -0.493769705 | 1.437751108 | -0.297501085 | 0.33733334 | 0.489075963 | 0.725189469 | 0.6413 | 0.79384115 |

| | | | | | | | | | | | |
|---|---|---|---|---|---|---|---|---|---|---|---|
| SL1344_RS08920_ | -1.858811997 | -1.634507459 | -2.187981822 | -0.554147051 | 0.331053461 | -1.599096019 | 0.04266677 | 0.088154703 | 0.308266936 | 0.1487 | 0.386666151 |
| SL1344_RS08925_ | -4.950298848 | -4.180810669 | -5.579187914 | -1.337855919 | 1.125124068 | -1.121259538 | 0.10933337 | 0.174960733 | 0.428322508 | 0.2544 | 0.49877693 |
| SL1344_RS08930_ | -4.72006749 | -3.800538354 | -5.451235518 | -1.561787078 | 0.937987286 | -1.580958859 | 0.054630326 | 0.092577791 | 0.311684463 | 0.1629 | 0.404546777 |
| SL1344_RS08935_ | -5.179734648 | -4.529893651 | -5.855499887 | -1.211729253 | 1.419959568 | -0.806909136 | 0.175333354 | 0.271348758 | 0.539687516 | 0.3517 | 0.593104989 |
| SL1344_RS08940_ | -3.744823424 | -3.385445426 | -4.120324374 | -0.738044549 | 0.599929336 | -1.175648703 | 0.089940086 | 0.154534479 | 0.39040492 | 0.228 | 0.463227625 |
| SL1344_RS08945_ | -2.556780901 | -2.251571175 | -2.850534661 | -0.610441999 | 0.433888873 | -1.336163094 | 0.070666727 | 0.120047728 | 0.353806481 | 0.1887 | 0.430979381 |
| SL1344_RS08950_ | -2.543159126 | -2.489252492 | -2.609398172 | -0.149123567 | 0.4334298 | -0.306697611 | 0.362666672 | 0.513873344 | 0.724405622 | 0.6249 | 0.781220725 |
| SL1344_RS08955_ | -2.610424398 | -2.34523895 | -2.91477712 | -0.581327811 | 0.475810609 | -1.157064865 | 0.091333379 | 0.161232272 | 0.411978467 | 0.2242 | 0.466629789 |
| SL1344_RS08960_ | 5.875783392 | 6.132781091 | 5.663131289 | -0.501364002 | 0.131950636 | -3.725544004 | 4.68E-05 | 0.018273651 | 0.16741119 | 0.1 | 0.330257709 |
| SL1344_RS08965_ | 1.877436024 | 2.053987299 | 1.568918096 | -0.511603534 | 0.221578854 | -2.316071637 | 4.68E-05 | 0.043045607 | 0.241190586 | 0.1 | 0.330257709 |
| SL1344_RS08970_ | -1.603912204 | -1.379955476 | -1.848826893 | -0.46758731 | 0.296089681 | -1.51139365 | 0.045969449 | 0.094324936 | 0.309930614 | 0.1613 | 0.399973047 |
| SL1344_RS08985_ | 0.129202546 | 0.297771719 | -0.122493541 | -0.426994395 | 0.188809471 | -2.214490294 | 0.003998486 | 0.04523149 | 0.226618551 | 0.1052 | 0.337233047 |
| SL1344_RS08995_ | 3.185517162 | 4.011851473 | 2.997124231 | -1.013809018 | 0.591311492 | -0.956748614 | 0.28847436 | 0.268470588 | 0.573380706 | 0.5781 | 0.791662139 |
| SL1344_RS09000_ | 3.211203369 | 3.824116212 | 2.85193107 | -0.792058348 | 0.419959998 | -1.645383636 | 0.040000111 | 0.071602605 | 0.312647585 | 0.1345 | 0.380748764 |
| SL1344_RS09005_ | 0.204554239 | 0.332976578 | 0.129755622 | -0.196394402 | 0.204846009 | -0.902350211 | 0.166555652 | 0.253175778 | 0.521043495 | 0.3548 | 0.59109096 |
| SL1344_RS09010_ | 2.350661689 | 2.565492846 | 2.277005148 | -0.271443583 | 0.144531593 | -1.639814801 | 0.03331126 | 0.081833455 | 0.320388878 | 0.1416 | 0.384484207 |
| SL1344_RS09015_ | 1.719550984 | 1.774741977 | 1.69312768 | -0.063204758 | 0.148076084 | -0.395758944 | 0.337774823 | 0.596559057 | 0.79343908 | 0.6771 | 0.8362633 |
| SL1344_RS09020_ | 2.858874174 | 3.166725704 | 2.538172698 | -0.597681219 | 0.110433534 | -5.384071553 | 4.68E-05 | 0.002179109 | 0.05964986 | 0.1 | 0.330257709 |
| SL1344_RS09025_ | -2.117155616 | -1.854750797 | -2.559592091 | -0.695618287 | 0.381585304 | -1.729394564 | 0.027333498 | 0.075971068 | 0.287493539 | 0.1351 | 0.371715321 |
| SL1344_RS09030_ | 2.007863385 | 2.048100038 | 1.991747726 | -0.06349869 | 0.172081391 | -0.311340292 | 0.318666674 | 0.439265549 | 0.707558848 | 0.6572 | 0.816930674 |
| SL1344_RS09035_ | -1.313037463 | -1.159557995 | -1.593013603 | -0.433093047 | 0.424463606 | -0.98468506 | 0.139240534 | 0.210382068 | 0.48408586 | 0.3044 | 0.557248443 |
| SL1344_RS09040_ | 1.351390318 | 1.483351791 | 1.197244358 | -0.303474644 | 0.177434814 | -1.723068388 | 0.018000252 | 0.065743922 | 0.285033827 | 0.1189 | 0.355904916 |
| SL1344_RS09045_ | 2.147622484 | 2.034140796 | 2.233352503 | 0.139681857 | 0.253375585 | 0.516577538 | 0.297801475 | 0.470990849 | 0.728252325 | 0.6258 | 0.805427927 |
| SL1344_RS09050_ | 0.950596523 | 0.939841009 | 0.963503562 | 0.020505795 | 0.256981606 | 0.073081372 | 0.470000001 | 0.77663973 | 0.903181997 | 0.9 | 0.9548384 |
| SL1344_RS09055_ | 3.620467554 | 3.731506158 | 3.488548104 | -0.253958397 | 0.088143397 | -3.025572441 | 4.68E-05 | 0.030858202 | 0.202898605 | 0.1 | 0.330257709 |
| SL1344_RS09060_ | 4.398209057 | 4.504991832 | 4.277935649 | -0.251269825 | 0.088966417 | -2.869117645 | 4.68E-05 | 0.013414821 | 0.140353954 | 0.1 | 0.330257709 |
| SL1344_RS09065_ | 4.257030038 | 4.393101402 | 4.01124518 | -0.374630021 | 0.070513493 | -4.944619086 | 4.68E-05 | 0.010467624 | 0.123495706 | 0.1 | 0.330257709 |
| SL1344_RS09070_ | -0.435285047 | 0.177934435 | -1.009015544 | -1.153343253 | 0.376353447 | -3.090716518 | 4.68E-05 | 0.01426231 | 0.138769756 | 0.1 | 0.330257709 |
| SL1344_RS09085_ | 1.551432232 | 2.119096564 | 1.007897498 | -1.098668021 | 0.122796331 | -8.907509162 | 4.68E-05 | 0.000942632 | 0.040189327 | 0.1 | 0.330257709 |
| SL1344_RS09090_ | 2.034847314 | 2.518276502 | 1.439331725 | -1.080844425 | 0.094689274 | -11.27655577 | 4.68E-05 | 0.000557841 | 0.031072277 | 0.1 | 0.330257709 |
| SL1344_RS09095_ | 2.006444168 | 2.49769367 | 1.477093564 | -1.031286926 | 0.104453763 | -9.782567894 | 4.68E-05 | 0.000819439 | 0.03769026 | 0.1 | 0.330257709 |
| SL1344_RS09100_ | 3.353336304 | 3.922846788 | 2.887743084 | -1.029059057 | 0.208427832 | -5.091116143 | 4.68E-05 | 0.001199913 | 0.049255693 | 0.1 | 0.330257709 |
| SL1344_RS09105_ | 0.899394919 | 1.180282421 | 0.57466885 | -0.801318066 | 0.37866021 | -2.262849225 | 4.68E-05 | 0.025231781 | 0.190012142 | 0.1 | 0.330257709 |
| SL1344_RS09115_ | 1.02683614 | 1.481140906 | 0.637816339 | -0.841710208 | 0.203870761 | -4.16256105 | 4.68E-05 | 0.005576752 | 0.091060164 | 0.1 | 0.330257709 |
| SL1344_RS09125_ | -1.758941021 | -1.227061923 | -2.120606452 | -0.91135711 | 0.581899133 | -1.550634394 | 0.043304566 | 0.07804314 | 0.3103601 | 0.1387 | 0.382376585 |
| SL1344_RS09130_ | 0.734301212 | 1.272598274 | 0.375257948 | -0.831979997 | 0.27223977 | -2.876859148 | 4.68E-05 | 0.020214033 | 0.168709488 | 0.1 | 0.330257709 |
| SL1344_RS09135_ | -0.500227347 | -0.134258021 | -0.633594045 | -0.504297036 | 0.329246651 | -1.367725959 | 0.062624986 | 0.1359014 | 0.40470434 | 0.1841 | 0.428710949 |
| SL1344_RS09140_ | 3.796906423 | 4.345512283 | 3.370511617 | -0.953401014 | 0.115156963 | -7.987589355 | 4.68E-05 | 0.001592758 | 0.053270424 | 0.1 | 0.330257709 |
| SL1344_RS09145_ | 3.197004119 | 3.604555697 | 2.622981303 | -1.000999235 | 0.219631153 | -4.662344399 | 4.68E-05 | 0.002921697 | 0.0716688 | 0.1 | 0.330257709 |
| SL1344_RS09150_ | 4.489119376 | 5.024502959 | 3.910743997 | -1.189723074 | 0.192923332 | -6.202490913 | 4.68E-05 | 0.001822454 | 0.058935214 | 0.1 | 0.330257709 |
| SL1344_RS09155_ | -0.090991487 | 0.367404166 | -0.549064473 | -0.809481916 | 0.388528961 | -2.054875441 | 0.008000574 | 0.034460267 | 0.216287314 | 0.1071 | 0.340625958 |
| SL1344_RS09160_ | 0.832668492 | 0.826830872 | 0.835547389 | 0.008212939 | 0.212137561 | 0.033660781 | 0.487008661 | 0.743903568 | 0.884019917 | 0.8382 | 0.925395454 |
| SL1344_RS09165_ | 0.675828919 | 1.069719191 | 0.032098229 | -1.008783888 | 0.248814923 | -3.996028603 | 4.68E-05 | 0.008310146 | 0.107340219 | 0.1 | 0.330257709 |
| SL1344_RS09170_ | 0.75666024 | 1.160400995 | 0.26381867 | -0.90555307 | 0.165296234 | -5.544704608 | 4.68E-05 | 0.004484622 | 0.07655703 | 0.1 | 0.330257709 |
| SL1344_RS09175_ | 7.190891479 | 7.380478077 | 6.970432523 | -0.418382452 | 0.070591682 | -5.823646519 | 4.68E-05 | 0.008513706 | 0.117980491 | 0.1 | 0.330257709 |
| SL1344_RS09180_ | 6.14127983 | 6.657838297 | 6.058733704 | -0.598468769 | 0.498757561 | -1.342601097 | 0.021319333 | 0.08227453 | 0.333955854 | 0.1205 | 0.36021244 |
| SL1344_RS09185_ | 1.785912801 | 1.987076865 | 1.4061917 | -0.637896425 | 0.270754313 | -2.5448903 | 4.68E-05 | 0.018156778 | 0.163297065 | 0.1 | 0.330257709 |
| SL1344_RS09190_ | 0.363931924 | 0.480093562 | 0.283018857 | -0.223127324 | 0.255295496 | -0.921249424 | 0.110592975 | 0.294322395 | 0.592620597 | 0.2691 | 0.507969636 |
| SL1344_RS09200_ | 3.185419629 | 3.344709072 | 3.017946808 | -0.301703348 | 0.095604333 | -3.117707941 | 4.68E-05 | 0.011587489 | 0.127021358 | 0.1 | 0.330257709 |
| SL1344_RS09205_ | -0.694494558 | -0.283437709 | -1.106670246 | -0.823462196 | 0.283275847 | -2.934307956 | 4.68E-05 | 0.020885168 | 0.161615414 | 0.1 | 0.330257709 |
| SL1344_RS09210_ | 3.014097699 | 3.381923936 | 2.854229749 | -0.487092086 | 0.174374795 | -2.137415321 | 0.002668394 | 0.053982458 | 0.271713119 | 0.1014 | 0.332307412 |
| SL1344_RS09235_ | 2.591013703 | 2.960188041 | 2.218029631 | -0.738541672 | 0.083911334 | -8.757967088 | 4.68E-05 | 0.000851708 | 0.038162555 | 0.1 | 0.330257709 |
| SL1344_RS09245_ | 0.399851891 | 0.959019304 | -0.118876633 | -0.969096644 | 0.381696388 | -2.542322418 | 0.00067292 | 0.018766902 | 0.164627408 | 0.1003 | 0.330689776 |

| | | | | | | | | | | | |
|---|---|---|---|---|---|---|---|---|---|---|---|
| SL1344_RS09250_ | -1.619075141 | -1.330423169 | -2.013024074 | -0.749758872 | 0.448232787 | -1.644811989 | 0.020666886 | 0.069471653 | 0.288008081 | 0.1265 | 0.36429335 |
| SL1344_RS09260_ | -0.107757424 | 0.33470216 | -1.069379963 | -1.373438454 | 0.346316795 | -3.530626388 | 4.68E-05 | 0.024635715 | 0.184642193 | 0.1 | 0.330257709 |
| SL1344_RS09270_ | -3.586888245 | -3.540160771 | -3.633251044 | -0.111691343 | 0.669972152 | -0.156966829 | 0.431333336 | 0.590809098 | 0.78281729 | 0.7068 | 0.839438909 |
| SL1344_RS09275_ | 0.589735998 | 1.147616856 | 0.288873406 | -0.741874993 | 0.503761783 | -1.58806349 | 0.048000092 | 0.06326143 | 0.29071744 | 0.1396 | 0.388013713 |
| SL1344_RS09285_ | -1.642113411 | -1.335716454 | -1.940279046 | -0.626920651 | 0.399488176 | -1.510978765 | 0.040000111 | 0.091647196 | 0.323339029 | 0.1416 | 0.383191236 |
| SL1344_RS09290_ | -1.058427614 | -0.302578506 | -1.206769996 | -0.844054313 | 0.45669838 | -1.164496529 | 0.157333357 | 0.181330432 | 0.469944606 | 0.3054 | 0.558594563 |
| SL1344_RS09295_ | -2.71288208 | -2.182842773 | -3.135698878 | -0.918841101 | 0.58407005 | -1.49066879 | 0.059333406 | 0.09225966 | 0.325553013 | 0.1568 | 0.403767967 |
| SL1344_RS09300_ | -3.763737393 | -3.341054348 | -4.668909537 | -1.317954384 | 0.985947725 | -1.243456644 | 0.081333385 | 0.12972513 | 0.389026955 | 0.2016 | 0.453796861 |
| SL1344_RS09305_ | -1.171916595 | -0.801426292 | -1.578659104 | -0.775559073 | 0.254255138 | -3.034953806 | 0.000673369 | 0.017020548 | 0.141650331 | 0.1007 | 0.331270188 |
| SL1344_RS09310_ | -4.679129478 | -4.420529544 | -5.048892125 | -0.697299729 | 1.041688892 | -0.592461919 | 0.247168567 | 0.368818481 | 0.622524497 | 0.4871 | 0.684163173 |
| SL1344_RS09315_ | -2.624877708 | -2.429842656 | -2.856805808 | -0.430304491 | 0.422044338 | -0.996325544 | 0.133910755 | 0.219405001 | 0.463617102 | 0.3024 | 0.530200133 |
| SL1344_RS09320_ | -0.22873314 | -0.10009317 | -0.300968949 | -0.177980556 | 0.640670668 | -0.304673744 | 0.318454371 | 0.676439139 | 0.850911903 | 0.6465 | 0.824190313 |
| SL1344_RS09325_ | -3.318426431 | -2.814351664 | -4.121569769 | -1.300503827 | 0.600082971 | -2.117741962 | 0.00733396 | 0.046154572 | 0.228632511 | 0.1076 | 0.340378411 |
| SL1344_RS09335_ | -3.986326891 | -4.103468898 | -3.853181521 | 0.305733878 | 1.026650133 | 0.284643539 | 0.354666673 | 0.546261508 | 0.762036763 | 0.6722 | 0.819972574 |
| SL1344_RS09340_ | 0.590527373 | 1.712481491 | -0.196000424 | -1.591730584 | 0.602900215 | -2.591880396 | 4.68E-05 | 0.018986719 | 0.168910953 | 0.1 | 0.330257709 |
| SL1344_RS09345_ | -4.396455591 | -3.817613005 | -5.818408683 | -1.97654067 | 1.163745401 | -1.585442237 | 0.026649069 | 0.095813396 | 0.332048079 | 0.1336 | 0.371646362 |
| SL1344_RS09350_ | -4.687386066 | -4.000972582 | -5.768615975 | -1.755788803 | 0.878699097 | -1.962058248 | 0.012658588 | 0.056905523 | 0.246842851 | 0.1146 | 0.34843614 |
| SL1344_RS09355_ | -3.662977987 | -3.159622073 | -4.008544017 | -0.803559215 | 0.725933733 | -1.03913234 | 0.136575645 | 0.202276333 | 0.462931097 | 0.2903 | 0.53462399 |
| SL1344_RS09360_ | -2.593728452 | -2.16822073 | -3.063203439 | -0.913060025 | 0.480724686 | -1.804208211 | 0.021319333 | 0.058299851 | 0.255690213 | 0.1244 | 0.361159865 |
| SL1344_RS09365_ | -0.578813769 | -0.678873866 | -0.48532785 | 0.128508149 | 0.772142981 | 0.161711807 | 0.450000002 | 0.867752292 | 0.948466353 | 0.9688 | 0.986519726 |
| SL1344_RS09370_ | 2.725107918 | 3.184545544 | 2.323026744 | -0.755209664 | 0.217513552 | -3.565639705 | 4.68E-05 | 0.00610817 | 0.099419289 | 0.1 | 0.330257709 |
| SL1344_RS09380_ | 2.372363317 | 2.778919553 | 2.197118547 | -0.589420281 | 0.270820036 | -2.208464049 | 4.68E-05 | 0.04865863 | 0.259809136 | 0.1 | 0.330257709 |
| SL1344_RS09385_ | 0.586107396 | 1.126923739 | 0.188962073 | -0.921854945 | 0.424186821 | -2.08875199 | 0.001335878 | 0.030838239 | 0.208848413 | 0.1007 | 0.331258917 |
| SL1344_RS09390_ | 0.65922122 | 0.861750948 | 0.327298719 | -0.528569428 | 0.181182808 | -2.747919429 | 4.68E-05 | 0.025217043 | 0.177593984 | 0.1007 | 0.331310227 |
| SL1344_RS09395_ | 1.373694526 | 1.736743409 | 0.943940601 | -0.794596283 | 0.121847147 | -6.499944723 | 4.68E-05 | 0.002588681 | 0.05900801 | 0.1 | 0.330257709 |
| SL1344_RS09400_ | 1.606474981 | 2.031992329 | 1.186798617 | -0.867556124 | 0.154641477 | -5.567035997 | 4.68E-05 | 0.002160951 | 0.059418407 | 0.1 | 0.330257709 |
| SL1344_RS09405_ | 0.519241273 | 1.000793647 | -0.159971178 | -1.176643112 | 0.222939719 | -5.371764534 | 4.68E-05 | 0.008170054 | 0.10458182 | 0.1 | 0.330257709 |
| SL1344_RS09410_ | 1.289524313 | 1.872132599 | 0.682646126 | -1.188610595 | 0.118382805 | -10.07590544 | 4.68E-05 | 0.00066338 | 0.03386129 | 0.1 | 0.330257709 |
| SL1344_RS09420_ | 0.181007021 | 0.622855703 | -0.360803829 | -1.025953191 | 0.261024096 | -3.953421999 | 4.68E-05 | 0.007560332 | 0.102448697 | 0.1 | 0.330257709 |
| SL1344_RS09425_ | 2.351918705 | 2.779061701 | 2.040966854 | -0.727336081 | 0.152368505 | -4.576605295 | 4.68E-05 | 0.011245769 | 0.129129854 | 0.1 | 0.330257709 |
| SL1344_RS09430_ | 1.398185267 | 1.677064806 | 1.10134642 | -0.57771623 | 0.128971481 | -4.449472624 | 4.68E-05 | 0.008534289 | 0.104456041 | 0.1 | 0.330257709 |
| SL1344_RS09435_ | 3.948418767 | 4.32322626 | 3.459321459 | -0.906820054 | 0.233882725 | -3.803943276 | 4.68E-05 | 0.01460011 | 0.151345765 | 0.1 | 0.330257709 |
| SL1344_RS09440_ | 2.055906365 | 2.283035568 | 1.88121797 | -0.393875301 | 0.141744758 | -2.795642886 | 4.68E-05 | 0.016799346 | 0.150862245 | 0.1001 | 0.330408442 |
| SL1344_RS09455_ | 2.755101115 | 3.070805109 | 2.291002196 | -0.768602002 | 0.124338113 | -5.989318488 | 4.68E-05 | 0.002606737 | 0.063541165 | 0.1 | 0.330257709 |
| SL1344_RS09460_ | 1.784470763 | 2.160528779 | 1.307525432 | -0.817516169 | 0.164481962 | -4.849145143 | 4.68E-05 | 0.004166588 | 0.077487261 | 0.1 | 0.330257709 |
| SL1344_RS09465_ | 0.72758995 | 0.977474192 | 0.313627998 | -0.672767843 | 0.204757794 | -3.30862214 | 4.68E-05 | 0.025549242 | 0.181878447 | 0.1 | 0.330257709 |
| SL1344_RS09470_ | 0.956972374 | 1.361045287 | 0.529640013 | -0.856308819 | 0.184375666 | -4.642290268 | 4.68E-05 | 0.004579122 | 0.082582511 | 0.1 | 0.330257709 |
| SL1344_RS09485_ | -2.660226934 | -2.309634757 | -3.171460049 | -0.862553295 | 0.423300916 | -1.958935924 | 0.016000285 | 0.055796828 | 0.241439181 | 0.1183 | 0.352603021 |
| SL1344_RS09490_ | -2.152015536 | -1.551050604 | -2.690401573 | -1.203682661 | 0.473631951 | -2.495720835 | 4.68E-05 | 0.031961865 | 0.20131687 | 0.1009 | 0.331480731 |
| SL1344_RS09495_ | 2.866299225 | 3.354367328 | 2.42856487 | -0.902560154 | 0.159799556 | -5.612252951 | 4.68E-05 | 0.001636221 | 0.054381731 | 0.1 | 0.330257709 |
| SL1344_RS09500_ | 2.630015961 | 3.065995694 | 2.063082892 | -0.954098433 | 0.198623877 | -4.811011224 | 4.68E-05 | 0.002672331 | 0.067447795 | 0.1 | 0.330257709 |
| SL1344_RS09510_ | 4.127384055 | 4.50065619 | 3.702641487 | -0.796196008 | 0.057562514 | -13.82121388 | 4.68E-05 | 0.000338857 | 0.026288216 | 0.1 | 0.330257709 |
| SL1344_RS09515_ | 0.708106909 | 1.11605019 | 0.47296604 | -0.634897806 | 0.187290256 | -3.20251238 | 4.68E-05 | 0.020585761 | 0.165126171 | 0.1 | 0.330257709 |
| SL1344_RS09520_ | 0.621858909 | 0.95042595 | 0.293939399 | -0.676337432 | 0.190897066 | -3.547333018 | 4.68E-05 | 0.008975333 | 0.111204894 | 0.1 | 0.330257709 |
| SL1344_RS09525_ | 2.053138762 | 2.517165931 | 1.594606913 | -0.846303774 | 0.234474638 | -3.64914229 | 4.68E-05 | 0.004824682 | 0.088457692 | 0.1 | 0.330257709 |
| SL1344_RS09535_ | 2.258045674 | 2.587714008 | 1.919951956 | -0.685856933 | 0.133558092 | -5.10385488 | 4.68E-05 | 0.004862328 | 0.082936633 | 0.1 | 0.330257709 |
| SL1344_RS09540_ | 0.728549779 | 1.208715338 | 0.194479208 | -1.022411785 | 0.169028065 | -6.062111203 | 4.68E-05 | 0.003999249 | 0.072300266 | 0.1 | 0.330257709 |
| SL1344_RS09545_ | 0.069818893 | 0.594657965 | -0.433684166 | -1.028787738 | 0.199654426 | -5.109828741 | 4.68E-05 | 0.003986234 | 0.07576173 | 0.1 | 0.330257709 |
| SL1344_RS09550_ | 3.257556063 | 3.694634807 | 2.796115926 | -0.8857747 | 0.204787389 | -4.293232695 | 4.68E-05 | 0.003879164 | 0.081776619 | 0.1 | 0.330257709 |
| SL1344_RS09555_ | -1.116661856 | -0.573116385 | -1.565301493 | -0.979240406 | 0.350723991 | -2.692877339 | 0.00067292 | 0.020617614 | 0.164673928 | 0.1002 | 0.330548971 |
| SL1344_RS09635_ | -1.064519687 | -0.589486533 | -1.38222814 | -0.760361795 | 0.318947741 | -2.266153136 | 0.008000574 | 0.035794208 | 0.209514354 | 0.1076 | 0.340954865 |
| SL1344_RS09640_ | 0.848548407 | 1.055429556 | 0.505593836 | -0.649016547 | 0.335048598 | -2.006412409 | 0.000673369 | 0.038518013 | 0.227522348 | 0.1002 | 0.330555887 |
| SL1344_RS09645_ | 0.015221612 | 0.551645771 | -0.261652672 | -0.783350934 | 0.235586789 | -3.088228572 | 0.00067292 | 0.023667945 | 0.174415252 | 0.1 | 0.330257709 |

| | | | | | | | | | | | |
|---|---|---|---|---|---|---|---|---|---|---|---|
| SL1344_RS09650_ | -1.332715785 | -0.899067991 | -1.741079011 | -0.838407269 | 0.25233109 | -3.269962075 | 0.00067292 | 0.013632097 | 0.126569674 | 0.1001 | 0.33040795 |
| SL1344_RS09655_ | -7.117178866 | -7.480833543 | -6.549636411 | 0.731866515 | 2.512755192 | 0.232181962 | 0.369333339 | 0.604375068 | 0.788508167 | 0.6829 | 0.827916785 |
| SL1344_RS09660_ | -1.88281195 | -1.361642957 | -2.405783772 | -0.956739619 | 0.603895566 | -1.572389496 | 0.048634334 | 0.075358397 | 0.305431646 | 0.1451 | 0.393329028 |
| SL1344_RS09670_ | -1.21397728 | -1.12989104 | -1.271036451 | -0.182220276 | 0.560165171 | -0.322461906 | 0.375083283 | 0.633309561 | 0.821148694 | 0.7577 | 0.879530176 |
| SL1344_RS09680_ | 2.595976529 | 2.712207451 | 2.498019447 | -0.198507562 | 0.098908019 | -1.925701323 | 0.018000252 | 0.050433455 | 0.248484871 | 0.1224 | 0.362143282 |
| SL1344_RS09685_ | -3.195341612 | -2.825719763 | -3.635204617 | -0.804415495 | 0.558789202 | -1.396837827 | 0.059960099 | 0.107862661 | 0.337401727 | 0.1736 | 0.414909288 |
| SL1344_RS09690_ | 0.589843881 | 0.86512664 | 0.410548584 | -0.367489832 | 0.357838174 | -1.087793367 | 0.125333365 | 0.165610491 | 0.448746099 | 0.2356 | 0.508758158 |
| SL1344_RS09695_ | 2.395548584 | 2.862613567 | 2.024983352 | -0.80165125 | 0.206485302 | -3.68895774 | 4.68E-05 | 0.011082532 | 0.12953939 | 0.1 | 0.330257709 |
| SL1344_RS09700_ | 1.988483484 | 2.131535052 | 1.523983249 | -0.596818024 | 0.159169663 | -2.5623751 | 4.68E-05 | 0.051229393 | 0.263124786 | 0.1 | 0.330257709 |
| SL1344_RS09705_ | 1.277687747 | 1.413614716 | 1.154541818 | -0.237507341 | 0.16450548 | -1.371525002 | 0.071285869 | 0.109991575 | 0.357198298 | 0.1766 | 0.43038812 |
| SL1344_RS09710_ | 0.162944828 | 0.326811759 | 0.001538992 | -0.329999549 | 0.211038926 | -1.537815532 | 0.036666788 | 0.086834209 | 0.313502806 | 0.1477 | 0.389707223 |
| SL1344_RS09715_ | 3.850021003 | 4.976067222 | 3.486043763 | -1.142798014 | 0.930440097 | -1.562324429 | 0.088607642 | 0.076012069 | 0.322132449 | 0.1837 | 0.452589765 |
| SL1344_RS09725_ | 3.70935733 | 4.272989852 | 3.159583692 | -1.057441358 | 0.28137732 | -3.714639191 | 4.68E-05 | 0.00492821 | 0.092660457 | 0.1 | 0.330257709 |
| SL1344_RS09730_ | 2.578576006 | 3.014745656 | 2.226476606 | -0.773950817 | 0.123773522 | -6.153539522 | 4.68E-05 | 0.002256644 | 0.060040998 | 0.1 | 0.330257709 |
| SL1344_RS09805_ | 5.954490022 | 6.539914174 | 5.747000703 | -0.793792481 | 0.123233438 | -4.158308347 | 4.68E-05 | 0.044929329 | 0.253121787 | 0.1 | 0.330257709 |
| SL1344_RS09810_ | 4.693002112 | 5.124096266 | 4.389171823 | -0.728308954 | 0.124598243 | -5.197367 | 4.68E-05 | 0.01125061 | 0.133585308 | 0.1 | 0.330257709 |
| SL1344_RS09820_ | 1.516605335 | 1.850046425 | 0.975756524 | -0.844008416 | 0.457873194 | -1.969150238 | 0.001335878 | 0.035791376 | 0.226190718 | 0.1008 | 0.331437147 |
| SL1344_RS09850_ | 5.088446793 | 5.355561592 | 4.800739104 | -0.568747943 | 0.091024095 | -6.292049579 | 4.68E-05 | 0.000728177 | 0.040871463 | 0.1 | 0.330257709 |
| SL1344_RS09855_ | 4.066475627 | 4.328918082 | 3.775821756 | -0.598844485 | 0.147091119 | -4.216255459 | 4.68E-05 | 0.003239981 | 0.075359415 | 0.1 | 0.330257709 |
| SL1344_RS09860_ | 3.764187556 | 4.04578021 | 3.403447969 | -0.631268763 | 0.115803947 | -5.360643355 | 4.68E-05 | 0.001438916 | 0.052386495 | 0.1 | 0.330257709 |
| SL1344_RS09875_ | 3.566213591 | 3.827458217 | 3.256594291 | -0.582103261 | 0.104181384 | -5.696363335 | 4.68E-05 | 0.001899249 | 0.057094137 | 0.1 | 0.330257709 |
| SL1344_RS09885_ | 4.140669224 | 4.459880008 | 3.805510412 | -0.655038476 | 0.04707893 | -13.90129551 | 4.68E-05 | 0.000645056 | 0.033153262 | 0.1 | 0.330257709 |
| SL1344_RS09890_ | 3.144108654 | 3.485755003 | 2.849284837 | -0.644791855 | 0.132967769 | -4.970184937 | 4.68E-05 | 0.008574395 | 0.114758375 | 0.1 | 0.330257709 |
| SL1344_RS09895_ | 2.812805817 | 2.989649786 | 2.55608179 | -0.554068084 | 0.30918723 | -1.914558224 | 4.68E-05 | 0.049485568 | 0.262615018 | 0.1 | 0.330257709 |
| SL1344_RS09900_ | 3.182105078 | 3.479964138 | 2.880479031 | -0.61424717 | 0.081396712 | -7.522337156 | 4.68E-05 | 0.001339903 | 0.046984881 | 0.1 | 0.330257709 |
| SL1344_RS09910_ | 1.341628773 | 1.743371975 | 1.073867264 | -0.64048153 | 0.20720235 | -3.021289654 | 4.68E-05 | 0.017128391 | 0.154607348 | 0.1001 | 0.330402907 |
| SL1344_RS09920_ | 1.477948806 | 1.207392171 | 1.680540909 | 0.446188489 | 0.338131507 | 1.225128711 | 0.106000038 | 0.152537244 | 0.438469865 | 0.1953 | 0.466567137 |
| SL1344_RS09925_ | -2.320517657 | -2.473923879 | -2.019009185 | 0.411206484 | 0.510053595 | 0.738095431 | 0.217854779 | 0.333011017 | 0.593780351 | 0.432 | 0.661926313 |
| SL1344_RS09930_ | -2.170632854 | -2.33235948 | -1.971142939 | 0.329806167 | 0.422418196 | 0.730218076 | 0.206528997 | 0.332793041 | 0.582240597 | 0.4282 | 0.644135157 |
| SL1344_RS09935_ | 0.127901733 | -0.119608408 | 0.298907377 | 0.503403906 | 0.343308943 | 1.471252655 | 0.020666886 | 0.083773104 | 0.323677458 | 0.1248 | 0.363741406 |
| SL1344_RS09940_ | 2.198271022 | 2.038917888 | 2.454462582 | 0.415077627 | 0.199867782 | 2.049552551 | 0.00067292 | 0.063046539 | 0.290319774 | 0.1003 | 0.330699013 |
| SL1344_RS09945_ | 0.186613808 | 0.031276508 | 0.302519704 | 0.266797818 | 0.179733894 | 1.397121113 | 0.057333409 | 0.112146699 | 0.344606888 | 0.1804 | 0.421790846 |
| SL1344_RS09950_ | -1.211348498 | -1.450710807 | -1.027310823 | 0.439654539 | 0.285241148 | 1.471382563 | 0.041972124 | 0.109900637 | 0.343546463 | 0.1616 | 0.399385288 |
| SL1344_RS09955_ | -0.174978147 | -0.316620437 | -0.065275987 | 0.267755313 | 0.280475957 | 0.917083294 | 0.137908089 | 0.218078211 | 0.494814035 | 0.3037 | 0.548777211 |
| SL1344_RS09960_ | 1.073416741 | 0.895990198 | 1.174350482 | 0.24264645 | 0.207594272 | 1.128863615 | 0.12391742 | 0.167552236 | 0.443167866 | 0.2514 | 0.51578453 |
| SL1344_RS09965_ | -1.160466037 | -1.270642888 | -1.022290749 | 0.255565998 | 0.27912531 | 0.859572589 | 0.169333355 | 0.271478265 | 0.522268015 | 0.3678 | 0.589057213 |
| SL1344_RS09970_ | -3.974862411 | -3.842624238 | -4.065430907 | -0.229763045 | 0.907546346 | -0.227031722 | 0.397333337 | 0.601576367 | 0.792353324 | 0.7073 | 0.843850844 |
| SL1344_RS09975_ | 0.036208937 | 0.086230903 | 0.002236756 | -0.053819175 | 0.239945029 | -0.212387864 | 0.41439041 | 0.67562631 | 0.841322242 | 0.7816 | 0.891703278 |
| SL1344_RS09985_ | -0.345111894 | -0.473551347 | -0.21457887 | 0.263485289 | 0.22699187 | 1.12762811 | 0.105263197 | 0.166409332 | 0.419684377 | 0.2369 | 0.481086149 |
| SL1344_RS09990_ | -2.014095632 | -2.270320221 | -1.804910794 | 0.475389071 | 0.353528213 | 1.312924072 | 0.077333388 | 0.133661642 | 0.369738869 | 0.2041 | 0.442411126 |
| SL1344_RS09995_ | -4.611647028 | -4.72500413 | -4.488909902 | 0.226117174 | 0.737951446 | 0.268845228 | 0.377333338 | 0.487509674 | 0.698995984 | 0.5937 | 0.758450346 |
| SL1344_RS10000_ | -2.794575235 | -2.855721127 | -2.747671625 | 0.122328981 | 0.610241885 | 0.186735991 | 0.417333336 | 0.6212141 | 0.806555789 | 0.7352 | 0.85991092 |
| SL1344_RS10005_ | -2.07555387 | -2.043616658 | -2.095630492 | -0.068181137 | 0.481377154 | -0.140884606 | 0.441705532 | 0.663435712 | 0.833026073 | 0.7777 | 0.886739385 |
| SL1344_RS10010_ | -2.187559109 | -1.984934761 | -2.562390816 | -0.531833389 | 0.488480033 | -1.02694436 | 0.134666696 | 0.201878954 | 0.472429843 | 0.2894 | 0.537748783 |
| SL1344_RS10015_ | -4.176410105 | -4.069891668 | -4.31859584 | -0.318210392 | 1.120305639 | -0.267277779 | 0.379333338 | 0.595640181 | 0.793406746 | 0.7176 | 0.852134918 |
| SL1344_RS10020_ | -3.493398404 | -3.326376029 | -3.710161683 | -0.469233831 | 0.791900711 | -0.560739001 | 0.248000013 | 0.388219782 | 0.642012931 | 0.5167 | 0.710667094 |
| SL1344_RS10025_ | -3.724294673 | -3.585243173 | -3.950830345 | -0.347427333 | 0.676778721 | -0.474742375 | 0.29646903 | 0.45685945 | 0.682258493 | 0.5704 | 0.748693349 |
| SL1344_RS10030_ | -4.413879826 | -3.851302886 | -5.090073825 | -1.258155913 | 0.851279964 | -1.427499327 | 0.056000078 | 0.102347763 | 0.328869855 | 0.1646 | 0.402485587 |
| SL1344_RS10035_ | -2.563460981 | -2.587602771 | -2.542822044 | 0.036449078 | 0.414553013 | 0.081379219 | 0.463024652 | 0.560717625 | 0.761553232 | 0.6803 | 0.82257101 |
| SL1344_RS10040_ | -1.794596409 | -1.808425074 | -1.775921727 | 0.031984432 | 0.45921426 | 0.064881288 | 0.471685544 | 0.69053674 | 0.849736374 | 0.793 | 0.895712078 |
| SL1344_RS10045_ | -2.044335081 | -2.01170604 | -2.094589199 | -0.046752021 | 0.436063778 | -0.10038346 | 0.456666668 | 0.642504445 | 0.818246261 | 0.7533 | 0.869948409 |
| SL1344_RS10050_ | -0.430073889 | -0.390239127 | -0.457457828 | -0.063095576 | 0.221756177 | -0.255918958 | 0.395069957 | 0.56255141 | 0.762045391 | 0.6664 | 0.815215043 |
| SL1344_RS10055_ | -2.808758771 | -2.818546353 | -2.800179252 | -0.004088053 | 0.474494773 | -0.00814865 | 0.493004664 | 0.597128496 | 0.78271452 | 0.7082 | 0.840492283 |

| | | | | | | | | | | | |
|---|---|---|---|---|---|---|---|---|---|---|---|
| SL1344_RS10060_ | -2.238100136 | -2.368266451 | -2.108146335 | 0.257772745 | 0.34320965 | 0.718168633 | 0.202666684 | 0.314812228 | 0.556802494 | 0.4167 | 0.626641024 |
| SL1344_RS10065_ | -1.834637293 | -1.562999468 | -1.969054305 | -0.393015902 | 0.383533644 | -0.951876237 | 0.163333356 | 0.241416906 | 0.507187786 | 0.3452 | 0.580860779 |
| SL1344_RS10070_ | -3.420622551 | -3.239486008 | -3.620278473 | -0.372420075 | 0.49548589 | -0.700019969 | 0.219333349 | 0.329463196 | 0.560838053 | 0.422 | 0.623793384 |
| SL1344_RS10075_ | -3.339665801 | -3.056183115 | -3.844688605 | -0.781974435 | 0.661455354 | -1.086891014 | 0.104666706 | 0.179006725 | 0.439118771 | 0.2555 | 0.496688648 |
| SL1344_RS10080_ | -3.874140825 | -3.748585142 | -3.944333623 | -0.156270785 | 0.714164916 | -0.194044416 | 0.417333336 | 0.597279107 | 0.782433951 | 0.6827 | 0.825727769 |
| SL1344_RS10085_ | -3.656837659 | -3.619831903 | -3.685788226 | -0.104540187 | 0.88218061 | -0.105966447 | 0.446000002 | 0.676171619 | 0.843100034 | 0.79 | 0.894500474 |
| SL1344_RS10090_ | -3.881786831 | -3.936958368 | -3.823201938 | 0.129294996 | 0.86629812 | 0.138438061 | 0.440000002 | 0.633176429 | 0.81572011 | 0.7403 | 0.866006668 |
| SL1344_RS10095_ | -1.932640676 | -1.625799913 | -2.205602866 | -0.573042774 | 0.368767411 | -1.536897027 | 0.04266677 | 0.089097522 | 0.309219379 | 0.154 | 0.394230639 |
| SL1344_RS10100_ | -3.58354472 | -3.114060342 | -3.836389168 | -0.673959229 | 0.695217979 | -0.892705172 | 0.175882766 | 0.26156195 | 0.523476665 | 0.3617 | 0.594481525 |
| SL1344_RS10105_ | -2.450009236 | -2.178833771 | -2.81272331 | -0.641588015 | 0.494919142 | -1.271349415 | 0.076615645 | 0.142685846 | 0.394993112 | 0.1964 | 0.439698358 |
| SL1344_RS10110_ | -5.255744737 | -5.39467326 | -5.119664493 | 0.295123741 | 1.15975321 | 0.226906845 | 0.395333337 | 0.524938874 | 0.735873899 | 0.6497 | 0.799858091 |
| SL1344_RS10115_ | -1.524339676 | -1.459024886 | -1.559322317 | -0.075068649 | 0.366739075 | -0.192695558 | 0.424383747 | 0.651493084 | 0.827028477 | 0.7551 | 0.877946899 |
| SL1344_RS10120_ | -2.979562458 | -2.928869687 | -3.062390386 | -0.132299941 | 0.530660023 | -0.224038154 | 0.40906063 | 0.577306232 | 0.774966347 | 0.693 | 0.83361648 |
| SL1344_RS10130_ | -1.722542369 | -1.610855255 | -1.827286024 | -0.20919659 | 0.402530142 | -0.479015003 | 0.306666675 | 0.489480036 | 0.714720491 | 0.6029 | 0.774209013 |
| SL1344_RS10135_ | -3.414633446 | -3.221303057 | -3.676060129 | -0.451930482 | 0.565689685 | -0.752702929 | 0.188000019 | 0.290216227 | 0.538330011 | 0.3941 | 0.607616862 |
| SL1344_RS10140_ | -2.64799824 | -2.467985116 | -2.864462724 | -0.408035404 | 0.423818704 | -0.96798437 | 0.153231204 | 0.238852279 | 0.486367128 | 0.3201 | 0.54839823 |
| SL1344_RS10145_ | -2.941779415 | -2.884523603 | -3.003504957 | -0.127542642 | 0.482105905 | -0.23137366 | 0.403333337 | 0.540305056 | 0.74373792 | 0.6512 | 0.800896751 |
| SL1344_RS10150_ | -3.871628495 | -3.473292422 | -4.192414692 | -0.671924271 | 0.781019539 | -0.803432723 | 0.181333353 | 0.282000821 | 0.53807192 | 0.3721 | 0.603040387 |
| SL1344_RS10155_ | -1.533861155 | -1.528738587 | -1.53923925 | 0.114927101 | 0.485716863 | 0.222130078 | 0.418387745 | 0.632435916 | 0.817057109 | 0.8119 | 0.901758525 |
| SL1344_RS10160_ | -0.941737216 | -0.832481304 | -1.098129737 | -0.288326424 | 0.3528029 | -0.772819012 | 0.167333355 | 0.270994795 | 0.549911802 | 0.3662 | 0.593078224 |
| SL1344_RS10165_ | -1.923404946 | -1.712409772 | -2.031755456 | -0.293394854 | 0.383997052 | -0.67080755 | 0.246666679 | 0.381553685 | 0.624643058 | 0.4869 | 0.693209597 |
| SL1344_RS10170_ | -2.160490016 | -2.025498908 | -2.404876195 | -0.334241616 | 0.4891592 | -0.622162534 | 0.255163236 | 0.396816321 | 0.647740292 | 0.5131 | 0.71632257 |
| SL1344_RS10175_ | -3.303310282 | -3.307846304 | -3.298877608 | 0.068618038 | 0.704568239 | 0.087835601 | 0.464000001 | 0.65106553 | 0.825758753 | 0.7768 | 0.884498915 |
| SL1344_RS10180_ | -2.535036936 | -2.532015033 | -2.537176013 | -0.005732026 | 0.391442251 | -0.016586964 | 0.493004664 | 0.551690271 | 0.752409515 | 0.6558 | 0.806057499 |
| SL1344_RS10185_ | -2.743009192 | -2.820117994 | -2.67017589 | 0.145120873 | 0.423626255 | 0.333656002 | 0.364000005 | 0.487874514 | 0.703535881 | 0.5953 | 0.761524225 |
| SL1344_RS10190_ | -1.900407511 | -1.876670954 | -1.930025776 | -0.057053367 | 0.336475834 | -0.144691933 | 0.433333336 | 0.561102855 | 0.759735127 | 0.6625 | 0.811336012 |
| SL1344_RS10195_ | -3.648887931 | -3.559411718 | -3.743153035 | -0.181744094 | 0.576398017 | -0.297107369 | 0.368666672 | 0.502727664 | 0.716737286 | 0.609 | 0.773317426 |
| SL1344_RS10200_ | 1.978055246 | 2.070748423 | 1.862160488 | -0.066198918 | 0.446569852 | -0.131207309 | 0.454000002 | 0.893657295 | 0.959946767 | 0.9874 | 0.994639215 |
| SL1344_RS10205_ | -1.7950565 | -1.711552361 | -1.900654232 | -0.173740137 | 0.347657976 | -0.445706398 | 0.300666675 | 0.471950628 | 0.694339545 | 0.5753 | 0.751688236 |
| SL1344_RS10210_ | -2.659069942 | -2.378955404 | -3.018047192 | -0.632865252 | 0.46565347 | -1.28740571 | 0.078614311 | 0.138309682 | 0.376276098 | 0.2105 | 0.451198424 |
| SL1344_RS10215_ | 3.2991669 | 3.276620156 | 3.310300529 | 0.051617727 | 0.193289548 | 0.274083074 | 0.359333339 | 0.636566354 | 0.833327192 | 0.7381 | 0.872705587 |
| SL1344_RS10225_ | -1.074385762 | -1.183033868 | -0.946168842 | 0.230914968 | 0.262670934 | 0.842946771 | 0.175333354 | 0.26692116 | 0.517405397 | 0.3534 | 0.583012247 |
| SL1344_RS10230_ | 0.643014885 | 0.643918178 | 0.642745207 | -0.064876754 | 0.284435107 | -0.218982805 | 0.423051302 | 0.693055215 | 0.856104298 | 0.8318 | 0.917734435 |
| SL1344_RS10235_ | -3.399554114 | -3.221465129 | -3.776048689 | -0.535375795 | 0.887074057 | -0.560984533 | 0.2778148 | 0.410946768 | 0.669050704 | 0.5513 | 0.747014694 |
| SL1344_RS10240_ | -4.194882989 | -4.360220725 | -4.138916458 | 0.191327786 | 1.366355638 | 0.122068598 | 0.445702867 | 0.715825341 | 0.871237288 | 0.8407 | 0.92918937 |
| SL1344_RS10245_ | -3.678926202 | -3.881698043 | -3.497151531 | 0.41974359 | 0.812754804 | 0.509929096 | 0.302666675 | 0.465416116 | 0.695313695 | 0.5777 | 0.758575674 |
| SL1344_RS10250_ | -3.287595309 | -3.205415399 | -3.332667942 | -0.196807755 | 0.806634539 | -0.224167378 | 0.405333337 | 0.597153553 | 0.793752772 | 0.7263 | 0.856595053 |
| SL1344_RS10255_ | -1.675490482 | -1.64439593 | -1.707444421 | -0.061573124 | 0.282251743 | -0.192046612 | 0.42238508 | 0.516143053 | 0.725569962 | 0.6274 | 0.783118068 |
| SL1344_RS10260_ | -1.751902387 | -1.881047208 | -1.625845959 | 0.254809381 | 0.346286439 | 0.697704457 | 0.213857444 | 0.348132872 | 0.594753337 | 0.4522 | 0.659398692 |
| SL1344_RS10265_ | -2.498171511 | -2.424056719 | -2.541727704 | -0.131961041 | 0.464558989 | -0.264805991 | 0.390666671 | 0.565180706 | 0.761785296 | 0.6725 | 0.816695364 |
| SL1344_RS10270_ | -1.329023482 | -1.476674879 | -0.998156678 | 0.440556455 | 0.373024801 | 1.072328944 | 0.140666694 | 0.203008132 | 0.474426698 | 0.2958 | 0.546015313 |
| SL1344_RS10275_ | 1.469175955 | 1.36573778 | 2.111410857 | 0.720196832 | 0.176584126 | 1.513051789 | 0.020653117 | 0.109846229 | 0.377976218 | 0.1219 | 0.359058638 |
| SL1344_RS10280_ | 2.073181241 | 1.78640747 | 2.216595087 | 0.361994693 | 0.294808846 | 1.152790485 | 0.118000034 | 0.16164801 | 0.451052448 | 0.2184 | 0.493960866 |
| SL1344_RS10285_ | -0.026603627 | 0.157839054 | -0.166288989 | -0.305522405 | 0.678960824 | -0.343836686 | 0.396666671 | 0.685164201 | 0.8614106 | 0.868 | 0.943696378 |
| SL1344_RS10290_ | -4.382889986 | -4.636586902 | -4.146738829 | 0.506016495 | 0.804059303 | 0.591784216 | 0.250000012 | 0.381486261 | 0.61941704 | 0.488 | 0.683381508 |
| SL1344_RS10295_ | -6.111704596 | -5.491647623 | -6.911653634 | -1.406189979 | 1.583526232 | -0.841492539 | 0.16066669 | 0.257403586 | 0.516973579 | 0.3386 | 0.568074923 |
| SL1344_RS10300_ | -0.57950983 | -0.876885787 | -0.49652747 | 0.423814435 | 0.684433219 | 0.587308251 | 0.287141915 | 0.431095593 | 0.703215957 | 0.5891 | 0.797514597 |
| SL1344_RS10305_ | -1.316131239 | -1.420901955 | -1.115807793 | 0.340243898 | 0.413436433 | 0.732509072 | 0.216000016 | 0.321578492 | 0.596381854 | 0.443 | 0.667603301 |
| SL1344_RS10310_ | -4.645678719 | -4.474287294 | -4.902532227 | -0.437386287 | 1.016084809 | -0.381040795 | 0.330666674 | 0.4987826 | 0.718952988 | 0.613 | 0.776299369 |
| SL1344_RS10315_ | -1.965706554 | -2.280541763 | -1.733584604 | 0.528455501 | 0.412877053 | 1.223069296 | 0.088666714 | 0.145226718 | 0.393549031 | 0.2159 | 0.463246768 |
| SL1344_RS10320_ | 2.074967606 | 1.78299677 | 2.186497421 | 0.302975164 | 0.376940464 | 0.941594728 | 0.184000019 | 0.226630954 | 0.526375236 | 0.3128 | 0.591264829 |
| SL1344_RS10325_ | -3.051224011 | -3.143603155 | -2.929089253 | 0.226218132 | 0.501063565 | 0.433161848 | 0.324666674 | 0.488895935 | 0.699815448 | 0.5918 | 0.757109345 |
| SL1344_RS10330_ | 2.518024424 | 2.369812003 | 2.549807841 | 0.197709965 | 0.215046461 | 0.905184595 | 0.197201883 | 0.235434298 | 0.533090562 | 0.3631 | 0.637141679 |

| | | | | | | | | | | | |
|---|---|---|---|---|---|---|---|---|---|---|---|
| SL1344_RS10335_ | -2.367236005 | -2.680690728 | -2.219937495 | 0.434228102 | 0.488983569 | 0.810650315 | 0.20333335 | 0.308181297 | 0.572546303 | 0.4143 | 0.640690921 |
| SL1344_RS10340_ | -0.604682912 | -0.781377053 | -0.50502722 | 0.290832999 | 0.281027009 | 0.947766226 | 0.125249865 | 0.212951463 | 0.487102693 | 0.2848 | 0.521712743 |
| SL1344_RS10345_ | -0.331827019 | -0.550423157 | -0.241426896 | 0.300299522 | 0.281316315 | 0.956258141 | 0.155229871 | 0.226114636 | 0.501163001 | 0.3225 | 0.565802098 |
| SL1344_RS10350_ | -0.435108052 | -0.620390106 | -0.238945726 | 0.426085106 | 0.315688268 | 1.365962853 | 0.045303228 | 0.121108737 | 0.379428523 | 0.1551 | 0.398177369 |
| SL1344_RS10355_ | 1.997111566 | 1.825416767 | 2.176573825 | 0.336839085 | 0.817610778 | 0.316721918 | 0.440000002 | 0.741637752 | 0.894418497 | 0.9949 | 0.997796632 |
| SL1344_RS10360_ | -0.933123368 | -1.316625428 | -0.620062814 | 0.63826059 | 0.40078732 | 1.553484036 | 0.048634334 | 0.086427967 | 0.32306613 | 0.1491 | 0.39748852 |
| SL1344_RS10365_ | -0.053508987 | 0.104768338 | -0.094786294 | -0.178810153 | 0.280882706 | -0.28362816 | 0.354430386 | 0.839064515 | 0.934984863 | 0.7402 | 0.889483458 |
| SL1344_RS10370_ | -4.47623345 | -4.50457728 | -4.451371171 | 0.077192218 | 0.884293029 | 0.082342529 | 0.465689542 | 0.595654006 | 0.784368993 | 0.6989 | 0.833894177 |
| SL1344_RS10375_ | -0.232080871 | -0.252360435 | -0.217716362 | 0.011361006 | 0.249468867 | 0.048549466 | 0.482000001 | 0.676395223 | 0.843525993 | 0.778 | 0.889836579 |
| SL1344_RS10380_ | 0.127551115 | 0.169834491 | 0.104367113 | -0.046128919 | 0.22199023 | -0.182043048 | 0.4170553 | 0.667049398 | 0.83672631 | 0.7557 | 0.878556074 |
| SL1344_RS10390_ | -1.440771934 | -1.357916177 | -1.521401625 | -0.129152913 | 0.461216088 | -0.271074275 | 0.374666672 | 0.620139299 | 0.806703522 | 0.7196 | 0.85228131 |
| SL1344_RS10395_ | -0.03440796 | 0.038544956 | -0.113145943 | -0.16460928 | 0.220278495 | -0.724392479 | 0.197868106 | 0.317600741 | 0.576372552 | 0.431 | 0.640923854 |
| SL1344_RS10400_ | -0.719806755 | -0.738495478 | -0.705538214 | 0.138181494 | 0.528981922 | 0.27667157 | 0.388000004 | 0.654064387 | 0.836877737 | 0.8079 | 0.906802339 |
| SL1344_RS10405_ | -1.203994499 | -1.294432388 | -1.115308919 | 0.163325135 | 0.766236249 | 0.209553856 | 0.402000004 | 0.71567336 | 0.872946389 | 0.8486 | 0.931186434 |
| SL1344_RS10410_ | -2.91480357 | -2.819829974 | -2.989683727 | -0.131502755 | 0.817824242 | -0.154102023 | 0.435333336 | 0.700709095 | 0.858138379 | 0.8185 | 0.909354402 |
| SL1344_RS10415_ | -1.183715305 | -1.101429155 | -1.281808892 | -0.083812605 | 0.850083666 | -0.106698536 | 0.436000002 | 0.811893613 | 0.922075319 | 0.82 | 0.914487073 |
| SL1344_RS10420_ | -1.056577568 | -1.070491592 | -1.039749395 | 0.324243357 | 0.806293254 | 0.404339048 | 0.361758833 | 0.505593057 | 0.752258219 | 0.7728 | 0.889294864 |
| SL1344_RS10430_ | -0.434076795 | -0.558205887 | -0.186330504 | 0.321075348 | 0.612866268 | 0.468748078 | 0.327781486 | 0.611036927 | 0.816487663 | 0.6765 | 0.8486868 |
| SL1344_RS10435_ | -0.269751887 | -0.347738544 | -0.070506614 | 0.302521576 | 0.78441845 | 0.443582098 | 0.237333347 | 0.490231927 | 0.743431357 | 0.4571 | 0.714603771 |
| SL1344_RS10440_ | -0.684149871 | -1.044258091 | -0.393998967 | 0.537657125 | 0.652610088 | 0.793350905 | 0.18254499 | 0.276121277 | 0.572563795 | 0.3351 | 0.598994236 |
| SL1344_RS10445_ | -1.503096291 | -1.632589581 | -1.403094077 | 0.288624058 | 0.548417173 | 0.50265545 | 0.290000009 | 0.442600221 | 0.697748484 | 0.5994 | 0.782478231 |
| SL1344_RS10450_ | -1.322004967 | -1.64908626 | -1.126617063 | 0.461704521 | 0.471590454 | 0.96861104 | 0.159333357 | 0.231659576 | 0.5126939 | 0.3132 | 0.57093745 |
| SL1344_RS10455_ | -1.154387147 | -1.319127065 | -1.021209164 | 0.345410704 | 0.457285764 | 0.772233154 | 0.15866669 | 0.280256818 | 0.568791131 | 0.3342 | 0.568316549 |
| SL1344_RS10470_ | -0.272636908 | -0.447680797 | -0.20478071 | 0.250941701 | 0.370941723 | 0.706310654 | 0.187208547 | 0.31371667 | 0.605682725 | 0.4037 | 0.630186651 |
| SL1344_RS10475_ | -0.193661488 | -0.31167289 | -0.057303453 | 0.25354274 | 0.382947678 | 0.623610355 | 0.248666679 | 0.361173284 | 0.638029141 | 0.4858 | 0.707098199 |
| SL1344_RS10480_ | 0.047708869 | -0.05949791 | 0.159626574 | 0.290382451 | 0.32593285 | 0.878622609 | 0.133333363 | 0.224272323 | 0.514424588 | 0.2821 | 0.527671771 |
| SL1344_RS10485_ | -1.705684039 | -1.769930134 | -1.629911671 | 0.142863082 | 0.292781716 | 0.453841736 | 0.303333342 | 0.441481771 | 0.665416452 | 0.554 | 0.730385145 |
| SL1344_RS10490_ | -6.736516656 | -6.961608164 | -6.596454787 | 0.366726598 | 1.722284041 | 0.195584083 | 0.40866667 | 0.526795126 | 0.736277516 | 0.6456 | 0.79814339 |
| SL1344_RS10495_ | -4.542691809 | -4.353189292 | -4.689026759 | -0.30613229 | 0.995596021 | -0.299568271 | 0.373750838 | 0.568794697 | 0.766473521 | 0.6751 | 0.820619106 |
| SL1344_RS10515_ | -3.108343118 | -3.006745298 | -3.192062347 | -0.167968766 | 0.531398525 | -0.301149923 | 0.370666672 | 0.561883869 | 0.757517337 | 0.671 | 0.815333192 |
| SL1344_RS10520_ | -5.726703602 | -5.710931617 | -5.74400277 | -0.076578869 | 1.569072529 | -0.045636362 | 0.476000001 | 0.618127843 | 0.805919342 | 0.7268 | 0.854422313 |
| SL1344_RS10530_ | -4.345348598 | -4.356362755 | -4.330527111 | 0.049915932 | 0.84154746 | 0.055660108 | 0.473333334 | 0.588184761 | 0.780978432 | 0.7009 | 0.836471255 |
| SL1344_RS10535_ | -4.133881161 | -4.003922802 | -4.274061686 | -0.268745122 | 0.664369277 | -0.362590079 | 0.33933334 | 0.464009684 | 0.682046549 | 0.5669 | 0.741051868 |
| SL1344_RS10540_ | -3.89447534 | -3.717488809 | -4.071632189 | -0.426101206 | 0.877284876 | -0.488011018 | 0.292666676 | 0.447537261 | 0.679638087 | 0.5779 | 0.748532183 |
| SL1344_RS10545_ | -4.634319326 | -4.337564567 | -5.065526271 | -0.734872799 | 0.860353974 | -0.787727915 | 0.166666689 | 0.266505714 | 0.518854321 | 0.3553 | 0.577535327 |
| SL1344_RS10550_ | -4.053984318 | -3.610082283 | -4.241949731 | -0.565437585 | 0.912191824 | -0.563753406 | 0.287141915 | 0.484215413 | 0.707354995 | 0.578 | 0.763764155 |
| SL1344_RS10555_ | -3.420567875 | -3.2677498 | -3.531250688 | -0.177159328 | 0.835981713 | -0.182049098 | 0.415722855 | 0.688356077 | 0.849635453 | 0.7583 | 0.88125485 |
| SL1344_RS10560_ | -3.291575494 | -3.154334984 | -3.602020949 | -0.39839011 | 0.802217183 | -0.449859213 | 0.350433051 | 0.538725567 | 0.757700507 | 0.6807 | 0.832452085 |
| SL1344_RS10565_ | -2.878617208 | -2.478911192 | -3.522748439 | -1.033866 | 0.492555785 | -2.012336499 | 0.010667096 | 0.054204443 | 0.244630745 | 0.1158 | 0.350286913 |
| SL1344_RS10570_ | -4.113709439 | -3.891966368 | -4.373661785 | -0.467843898 | 0.743335586 | -0.596889788 | 0.266666678 | 0.398842501 | 0.630768428 | 0.5019 | 0.694339245 |
| SL1344_RS10575_ | -5.015482249 | -4.853656858 | -5.092172924 | -0.204629159 | 1.23982339 | -0.143360707 | 0.427333336 | 0.650166401 | 0.824462828 | 0.7597 | 0.877042245 |
| SL1344_RS10580_ | -4.8994275 | -5.000773124 | -4.793652146 | 0.1886378 | 0.877229754 | 0.199807833 | 0.418387745 | 0.523653878 | 0.729771866 | 0.6265 | 0.782869597 |
| SL1344_RS10585_ | -2.115511996 | -2.009791534 | -2.258184464 | -0.244964778 | 0.360914286 | -0.616429308 | 0.239333347 | 0.35955911 | 0.601231063 | 0.4677 | 0.663714893 |
| SL1344_RS10590_ | -1.172152181 | -1.316825238 | -1.039807741 | 0.30983183 | 0.341618888 | 0.855111549 | 0.154666691 | 0.245624659 | 0.514667477 | 0.3417 | 0.569273767 |
| SL1344_RS10595_ | 3.645828673 | 3.466552921 | 4.032688525 | 0.511023824 | 0.297493627 | 1.969619321 | 4.68E-05 | 0.030471577 | 0.210688551 | 0.1 | 0.330257709 |
| SL1344_RS10600_ | 1.392290161 | 1.45286999 | 1.255857979 | -0.188799261 | 0.135201414 | -1.247705108 | 0.105263197 | 0.155262096 | 0.415895944 | 0.2369 | 0.485639294 |
| SL1344_RS10640_ | 3.278040534 | 2.740969599 | 3.412640719 | 0.682460648 | 0.321618446 | 1.994697937 | 4.68E-05 | 0.076497573 | 0.322274192 | 0.1 | 0.330257709 |
| SL1344_RS10645_ | 0.319457775 | 0.118517698 | 0.396143129 | 0.254833823 | 0.226353612 | 1.005534805 | 0.148000026 | 0.218359737 | 0.496934495 | 0.3167 | 0.563676735 |
| SL1344_RS10650_ | 1.744823526 | 1.616161082 | 1.890910002 | 0.310485872 | 0.194964942 | 1.592908147 | 0.017322048 | 0.079809686 | 0.317484701 | 0.1198 | 0.357206838 |
| SL1344_RS10655_ | -0.033121566 | -0.529920739 | 0.476423832 | 0.97749082 | 0.301051094 | 3.251397219 | 4.68E-05 | 0.012359584 | 0.130112368 | 0.1 | 0.330257709 |
| SL1344_RS10660_ | -0.005924386 | -0.381641448 | 0.408492748 | 0.807499453 | 0.237801313 | 3.373660324 | 4.68E-05 | 0.011507438 | 0.12428203 | 0.1 | 0.330257709 |
| SL1344_RS10665_ | 0.095707065 | -0.335420667 | 0.49026729 | 0.826533982 | 0.215014283 | 3.946705299 | 4.68E-05 | 0.010282445 | 0.11471223 | 0.1 | 0.330257709 |
| SL1344_RS10670_ | 1.061141036 | 0.635765847 | 1.411186706 | 0.775840528 | 0.11180897 | 6.829084373 | 4.68E-05 | 0.002098674 | 0.053488624 | 0.1 | 0.330257709 |

| | | | | | | | | | | | |
|---|---|---|---|---|---|---|---|---|---|---|---|
| SL1344_RS10675_ | -0.073418152 | -0.5355448 | 0.283595925 | 0.82581965 | 0.272715873 | 2.94517745 | 4.68E-05 | 0.017808828 | 0.152441769 | 0.1 | 0.330257709 |
| SL1344_RS10680_ | 0.495219841 | 0.166286892 | 0.860615345 | 0.693219105 | 0.1511467 | 4.5649736 | 4.68E-05 | 0.00543154 | 0.08360023 | 0.1 | 0.330257709 |
| SL1344_RS10685_ | 0.968698708 | 0.664439944 | 1.356539465 | 0.691767293 | 0.186711297 | 3.734052364 | 4.68E-05 | 0.008644557 | 0.108993371 | 0.1 | 0.330257709 |
| SL1344_RS10690_ | 0.560535271 | 0.166624914 | 1.143233939 | 0.936094386 | 0.277064653 | 3.294626949 | 4.68E-05 | 0.011502275 | 0.128588214 | 0.1 | 0.330257709 |
| SL1344_RS10700_ | 1.671387868 | 1.139086304 | 2.149043807 | 1.001850973 | 0.189596832 | 5.182941724 | 4.68E-05 | 0.00235796 | 0.062280453 | 0.1 | 0.330257709 |
| SL1344_RS10705_ | 2.963320177 | 2.454957664 | 3.236351594 | 0.769917844 | 0.167460113 | 4.068755045 | 4.68E-05 | 0.010952458 | 0.128725575 | 0.1 | 0.330257709 |
| SL1344_RS10710_ | 2.194235097 | 1.870843918 | 2.499591099 | 0.492910089 | 0.30703275 | 1.819177937 | 0.022000206 | 0.044805106 | 0.248737127 | 0.1202 | 0.359772766 |
| SL1344_RS10715_ | 2.202126081 | 1.760053454 | 2.380304933 | 0.591648487 | 0.187180026 | 2.533511691 | 4.68E-05 | 0.03819497 | 0.229289437 | 0.1 | 0.330257709 |
| SL1344_RS10720_ | 2.874637427 | 2.507682762 | 3.070890901 | 0.531930443 | 0.166307468 | 3.007944551 | 4.68E-05 | 0.015931778 | 0.151877087 | 0.1 | 0.330257709 |
| SL1344_RS10725_ | 2.247121121 | 1.819126381 | 2.452727004 | 0.625707061 | 0.220962991 | 2.729190802 | 4.68E-05 | 0.020342798 | 0.170618851 | 0.1 | 0.330257709 |
| SL1344_RS10730_ | 2.153497236 | 1.901157279 | 2.382072757 | 0.593236015 | 0.242761132 | 2.535721348 | 4.68E-05 | 0.01658422 | 0.15615669 | 0.1 | 0.330257709 |
| SL1344_RS10735_ | 2.554620956 | 2.334075124 | 2.785198761 | 0.50877573 | 0.16343985 | 3.17256006 | 4.68E-05 | 0.010990976 | 0.126810567 | 0.1 | 0.330257709 |
| SL1344_RS10740_ | 2.381590601 | 2.091433817 | 2.690326847 | 0.613188407 | 0.120198855 | 5.177061201 | 4.68E-05 | 0.006465861 | 0.09703552 | 0.1 | 0.330257709 |
| SL1344_RS10745_ | 3.39574856 | 3.107384415 | 3.789741997 | 0.654929762 | 0.163275148 | 4.155180802 | 4.68E-05 | 0.002969552 | 0.072144263 | 0.1 | 0.330257709 |
| SL1344_RS10750_ | 4.621607309 | 4.133222318 | 4.851278317 | 0.711687103 | 0.171779669 | 3.434083369 | 4.68E-05 | 0.016687947 | 0.160239153 | 0.1 | 0.330257709 |
| SL1344_RS10755_ | 2.839685663 | 2.368075632 | 3.163672411 | 0.593017368 | 0.333906039 | 1.917646772 | 0.007995244 | 0.04276281 | 0.245420524 | 0.1061 | 0.339252089 |
| SL1344_RS10760_ | 2.102876773 | 1.684889561 | 2.32760345 | 0.593635481 | 0.188898329 | 2.68510493 | 4.68E-05 | 0.029811314 | 0.203444242 | 0.1 | 0.330257709 |
| SL1344_RS10765_ | 2.368899286 | 2.059575619 | 2.697983694 | 0.62538491 | 0.23170455 | 2.655706627 | 4.68E-05 | 0.016561539 | 0.156147553 | 0.1 | 0.330257709 |
| SL1344_RS10770_ | 1.916431321 | 1.558035297 | 2.274202692 | 0.670767456 | 0.195965016 | 3.40946098 | 4.68E-05 | 0.009761049 | 0.119543186 | 0.1 | 0.330257709 |
| SL1344_RS10775_ | 1.725001202 | 1.495857581 | 1.95353081 | 0.515696493 | 0.305399631 | 1.627972843 | 0.010000458 | 0.05556871 | 0.273764342 | 0.106 | 0.339055375 |
| SL1344_RS10780_ | 2.280794095 | 2.001370733 | 2.61973546 | 0.62312735 | 0.13472714 | 4.582377812 | 4.68E-05 | 0.003671085 | 0.075762039 | 0.1 | 0.330257709 |
| SL1344_RS10785_ | 2.614921932 | 2.325527919 | 2.923964946 | 0.606261819 | 0.198589791 | 3.151307321 | 4.68E-05 | 0.013241405 | 0.1395405 | 0.1 | 0.330257709 |
| SL1344_RS10790_ | 3.591225951 | 3.474533972 | 3.765530107 | 0.296757064 | 0.077000947 | 3.851857207 | 4.68E-05 | 0.007008748 | 0.09870057 | 0.1 | 0.330257709 |
| SL1344_RS10795_ | -2.530196199 | -2.455651293 | -2.633154838 | -0.20047331 | 0.477155287 | -0.372891192 | 0.325782819 | 0.488881198 | 0.713248302 | 0.6129 | 0.774031579 |
| SL1344_RS10800_ | -2.850781148 | -3.056747468 | -2.66343341 | 0.453532578 | 0.677295086 | 0.629225473 | 0.223184558 | 0.34213011 | 0.612028722 | 0.4433 | 0.664536234 |
| SL1344_RS10805_ | -2.988930038 | -3.096857329 | -2.896884631 | 0.181608593 | 0.527283599 | 0.324557333 | 0.354666673 | 0.509357695 | 0.725777363 | 0.6173 | 0.781342378 |
| SL1344_RS10810_ | -2.735692566 | -2.845457868 | -2.649778495 | 0.19511907 | 0.449715517 | 0.410346075 | 0.322666674 | 0.461220128 | 0.683203188 | 0.574 | 0.744827182 |
| SL1344_RS10815_ | -3.087761775 | -2.955370214 | -3.198808391 | -0.227824629 | 0.623189328 | -0.34487917 | 0.353097941 | 0.545010322 | 0.752362694 | 0.6554 | 0.809800951 |
| SL1344_RS10820_ | -3.64765674 | -3.296503953 | -3.815406475 | -0.466334567 | 0.724789145 | -0.56721238 | 0.286475693 | 0.471595884 | 0.694972664 | 0.5569 | 0.748178118 |
| SL1344_RS10825_ | -2.722746675 | -2.756129195 | -2.691348185 | 0.062879118 | 0.376406428 | 0.158975284 | 0.42504997 | 0.48812416 | 0.703383635 | 0.5975 | 0.762732223 |
| SL1344_RS10830_ | -2.375642415 | -2.227858398 | -2.580476106 | -0.356742775 | 0.363956874 | -0.918908744 | 0.164666689 | 0.247862253 | 0.48801048 | 0.3385 | 0.558469669 |
| SL1344_RS10835_ | -3.860691018 | -3.907051296 | -3.820214181 | -0.149007189 | 1.155054313 | -0.126087289 | 0.451698869 | 0.671828088 | 0.841608533 | 0.8055 | 0.904048093 |
| SL1344_RS10840_ | -2.867202964 | -3.037660526 | -2.785001104 | 0.207961931 | 0.597747807 | 0.311965891 | 0.364000005 | 0.572270534 | 0.774314494 | 0.6918 | 0.83488804 |
| SL1344_RS10845_ | -3.217231196 | -3.413410912 | -3.055715341 | 0.368714702 | 0.527387099 | 0.639877895 | 0.23466668 | 0.344147012 | 0.586861594 | 0.4417 | 0.648976502 |
| SL1344_RS10850_ | -5.162647228 | -5.27160237 | -5.069902326 | 0.245860294 | 1.077900016 | 0.203567845 | 0.406000004 | 0.547018521 | 0.752924936 | 0.6691 | 0.814009888 |
| SL1344_RS10855_ | -3.90878195 | -4.207198712 | -3.628846455 | 0.485276925 | 0.974937473 | 0.472628993 | 0.298666676 | 0.480390807 | 0.713394764 | 0.584 | 0.767340067 |
| SL1344_RS10860_ | -2.711756093 | -2.864033347 | -2.624060768 | 0.183492235 | 0.623607688 | 0.28822515 | 0.384000005 | 0.635150661 | 0.819283332 | 0.7278 | 0.865490943 |
| SL1344_RS10865_ | -2.828025096 | -2.931916914 | -2.654501807 | 0.331039535 | 0.705356392 | 0.427625133 | 0.328666674 | 0.505768152 | 0.731277363 | 0.6477 | 0.806225731 |
| SL1344_RS10870_ | -4.068501108 | -3.909554646 | -4.239707135 | -0.338406449 | 0.647115751 | -0.491365081 | 0.292666676 | 0.43074312 | 0.651425627 | 0.5314 | 0.71008112 |
| SL1344_RS10875_ | -3.518178222 | -3.327621208 | -3.744585708 | -0.42497506 | 0.562260631 | -0.699183382 | 0.220519669 | 0.331095925 | 0.57380263 | 0.4323 | 0.638264268 |
| SL1344_RS10880_ | -2.289576491 | -2.279767451 | -2.305395545 | -0.011862336 | 0.363632586 | -0.034721732 | 0.486000001 | 0.558912964 | 0.758233009 | 0.669 | 0.813911787 |
| SL1344_RS10885_ | -5.04837029 | -5.304084617 | -4.81606868 | 0.480787104 | 0.938978615 | 0.472620378 | 0.288666676 | 0.416192054 | 0.644653238 | 0.5225 | 0.708259801 |
| SL1344_RS10890_ | -4.293851629 | -4.351820681 | -4.235784156 | 0.141923017 | 0.832937442 | 0.164717204 | 0.429333336 | 0.58213964 | 0.776709377 | 0.6996 | 0.835940328 |
| SL1344_RS10895_ | 0.682553123 | 0.54377485 | 0.819466749 | 0.345636014 | 0.252600921 | 1.359362221 | 0.03331126 | 0.099311697 | 0.346919998 | 0.1383 | 0.378293554 |
| SL1344_RS10900_ | 3.44287302 | 3.070336925 | 3.588634759 | 0.511503069 | 0.107847957 | 4.22542798 | 4.68E-05 | 0.019744719 | 0.169320912 | 0.1 | 0.330257709 |
| SL1344_RS10905_ | 1.85835686 | 1.610178473 | 2.104673615 | 0.410125328 | 0.214275301 | 1.982579555 | 0.020653117 | 0.041433675 | 0.236204874 | 0.1161 | 0.35385548 |
| SL1344_RS10910_ | 1.788285188 | 1.481697595 | 1.924036831 | 0.466948568 | 0.248814886 | 1.846385298 | 0.002666616 | 0.064704017 | 0.292771418 | 0.1013 | 0.33206334 |
| SL1344_RS10915_ | 1.349156773 | 1.120528104 | 1.569267067 | 0.475199356 | 0.181408981 | 2.583744998 | 0.00067292 | 0.025096306 | 0.183502275 | 0.1001 | 0.330408442 |
| SL1344_RS10920_ | -2.871521247 | -2.974904491 | -2.749913944 | 0.260521098 | 0.592502631 | 0.451963586 | 0.323784151 | 0.487939993 | 0.710760884 | 0.6097 | 0.77414399 |
| SL1344_RS10930_ | -1.506683628 | -1.610011011 | -1.415766931 | 0.206316634 | 0.337071305 | 0.565014793 | 0.256666678 | 0.392686851 | 0.638390188 | 0.5134 | 0.702532067 |
| SL1344_RS10935_ | -0.852436908 | -1.050383713 | -0.756825832 | 0.277553009 | 0.293631275 | 0.88387998 | 0.164556985 | 0.246328874 | 0.516316681 | 0.3433 | 0.580629374 |
| SL1344_RS10940_ | -0.782730237 | -0.975612136 | -0.667939462 | 0.29933695 | 0.269078305 | 1.041278901 | 0.136000029 | 0.201818523 | 0.46611494 | 0.2865 | 0.530782544 |
| SL1344_RS10945_ | -0.7189607 | -0.901091449 | -0.607960054 | 0.295210495 | 0.268183656 | 1.04101112 | 0.123333366 | 0.192877608 | 0.459690987 | 0.2571 | 0.50046007 |

| | | | | | | | | | | | |
|---|---|---|---|---|---|---|---|---|---|---|---|
| SL1344_RS10950_ | -0.15109388 | -0.227273122 | -0.014043234 | 0.146265575 | 0.322752907 | 0.415465102 | 0.357333339 | 0.631100767 | 0.817471308 | 0.7172 | 0.861611485 |
| SL1344_RS10955_ | 0.630580312 | 0.602213881 | 0.701289797 | 0.085807916 | 0.236447093 | 0.310823947 | 0.384666671 | 0.687890399 | 0.851551987 | 0.7691 | 0.891136859 |
| SL1344_RS10960_ | -3.467505902 | -4.298031727 | -3.082958041 | 1.204747435 | 0.75008449 | 1.492053561 | 0.03664236 | 0.106336165 | 0.345812765 | 0.1487 | 0.387194211 |
| SL1344_RS10965_ | -3.656339707 | -3.753178932 | -3.571715109 | 0.205953954 | 0.605433096 | 0.304030987 | 0.357333339 | 0.496990931 | 0.712263321 | 0.6048 | 0.767040313 |
| SL1344_RS10970_ | -1.264657707 | -1.591832317 | -0.972034123 | 0.608605526 | 0.243003331 | 2.43644873 | 0.003334715 | 0.026584277 | 0.173205314 | 0.1027 | 0.333946651 |
| SL1344_RS10975_ | -0.348411439 | -0.421306108 | -0.243507642 | 0.203084002 | 0.325669924 | 0.603956922 | 0.230666681 | 0.36788926 | 0.634675658 | 0.4691 | 0.692527494 |
| SL1344_RS10980_ | -3.947916685 | -3.869921647 | -3.991357234 | -0.007605836 | 1.230083739 | -0.004025589 | 0.495333334 | 0.706809718 | 0.864202873 | 0.8164 | 0.912601539 |
| SL1344_RS10985_ | -2.898878992 | -2.944275502 | -2.829610526 | 0.040934932 | 0.671110036 | 0.052760883 | 0.475682879 | 0.694244221 | 0.854747298 | 0.7941 | 0.900424815 |
| SL1344_RS10990_ | -2.332575328 | -2.276211148 | -2.378870038 | -0.087824076 | 0.393524077 | -0.216078954 | 0.408394407 | 0.540405832 | 0.744065918 | 0.6543 | 0.803586793 |
| SL1344_RS10995_ | -4.930629561 | -4.758196302 | -5.090562017 | -0.305347345 | 1.125149767 | -0.256241858 | 0.386666671 | 0.582231371 | 0.777698283 | 0.686 | 0.827338889 |
| SL1344_RS11000_ | -0.447990115 | -0.786171065 | -0.165970087 | 0.625518276 | 0.201314697 | 3.064756586 | 4.68E-05 | 0.017332274 | 0.141366665 | 0.1001 | 0.330403419 |
| SL1344_RS11005_ | 0.193491732 | 0.149525661 | 0.234797461 | 0.080178154 | 0.205333016 | 0.366476139 | 0.345769493 | 0.532256648 | 0.743940213 | 0.6394 | 0.799768526 |
| SL1344_RS11010_ | -0.118845612 | -0.040799871 | -0.170794541 | -0.100760697 | 0.276343441 | -0.360944637 | 0.359333339 | 0.619598451 | 0.804910872 | 0.7139 | 0.852357761 |
| SL1344_RS11015_ | 0.029685926 | 0.035489884 | 0.025427419 | -0.00926169 | 0.172901547 | -0.051638642 | 0.478000001 | 0.581659334 | 0.773996736 | 0.6946 | 0.830003295 |
| SL1344_RS11020_ | -0.150144415 | -0.10490624 | -0.218716585 | -0.10155784 | 0.194241166 | -0.482783788 | 0.28400001 | 0.452252862 | 0.677509768 | 0.5654 | 0.741543229 |
| SL1344_RS11025_ | 0.639404554 | 0.52078785 | 0.812940244 | 0.284320227 | 0.155182185 | 1.745467477 | 0.037333453 | 0.071437069 | 0.281790656 | 0.1387 | 0.379321739 |
| SL1344_RS11030_ | -1.121636023 | -1.34353934 | -0.887758945 | 0.538863063 | 0.375362657 | 1.396928449 | 0.041333441 | 0.101181249 | 0.343798746 | 0.1499 | 0.391509119 |
| SL1344_RS11035_ | -1.044398719 | -1.109323321 | -0.957726435 | 0.258346702 | 0.453037714 | 0.556092817 | 0.275333344 | 0.390720749 | 0.656788622 | 0.5581 | 0.745771991 |
| SL1344_RS11040_ | 1.074425972 | 0.715710774 | 1.392427569 | 0.648310369 | 0.264421011 | 2.408640844 | 0.000673369 | 0.031737511 | 0.207124434 | 0.1005 | 0.330989127 |
| SL1344_RS11045_ | -3.243633886 | -3.519665735 | -3.096771204 | 0.340714882 | 0.783089141 | 0.416825005 | 0.346000006 | 0.542162485 | 0.760775675 | 0.6698 | 0.827096574 |
| SL1344_RS11050_ | -2.672746416 | -2.67479455 | -2.670853536 | 0.050859716 | 0.539942392 | 0.088923615 | 0.463690874 | 0.643557677 | 0.821208336 | 0.7591 | 0.875192862 |
| SL1344_RS11055_ | -5.161676041 | -5.439895531 | -4.915656386 | 0.57417093 | 1.636188081 | 0.313852895 | 0.351765496 | 0.553650326 | 0.766127128 | 0.6839 | 0.83229637 |
| SL1344_RS11060_ | -4.523350505 | -4.675104849 | -4.281377761 | 0.439816537 | 1.02022059 | 0.412472343 | 0.33933334 | 0.526215453 | 0.737506933 | 0.6472 | 0.801910557 |
| SL1344_RS11065_ | -0.597278769 | -0.73576966 | -0.511200505 | 0.267600911 | 0.358124677 | 0.709384573 | 0.202666684 | 0.300997214 | 0.586796892 | 0.4091 | 0.637366079 |
| SL1344_RS11070_ | 3.017347568 | 2.837466289 | 3.245618443 | 0.378383657 | 0.112740981 | 3.353570105 | 4.68E-05 | 0.010020984 | 0.12027436 | 0.1 | 0.330257709 |
| SL1344_RS11080_ | -1.504878977 | -1.594844791 | -1.421204819 | 0.238466185 | 0.515952081 | 0.426624238 | 0.318666674 | 0.484776908 | 0.722181452 | 0.631 | 0.799625272 |
| SL1344_RS11085_ | -2.645336335 | -3.316135352 | -2.476030053 | 0.805822588 | 0.721841008 | 0.949579321 | 0.157228538 | 0.229016333 | 0.517954137 | 0.3337 | 0.575720697 |
| SL1344_RS11090_ | 1.450693632 | 1.213206832 | 1.594631627 | 0.372069259 | 0.132661088 | 2.64406445 | 0.001335878 | 0.029499509 | 0.191342978 | 0.1008 | 0.331434341 |
| SL1344_RS11095_ | 1.255289355 | 1.0168764 | 1.612686661 | 0.576167558 | 0.14805559 | 3.820323159 | 4.68E-05 | 0.010747967 | 0.120445769 | 0.1 | 0.330257709 |
| SL1344_RS11100_ | 2.298676974 | 2.12074203 | 2.616485259 | 0.401604011 | 0.293307874 | 1.479086342 | 0.085333383 | 0.093515619 | 0.349521751 | 0.1794 | 0.44577892 |
| SL1344_RS11105_ | -1.546142816 | -2.030129428 | -1.059069928 | 0.971531186 | 0.298954946 | 3.1420211 | 0.00067292 | 0.013491676 | 0.127968267 | 0.1002 | 0.330562441 |
| SL1344_RS11110_ | 0.420242712 | 0.242102586 | 0.606556993 | 0.411871259 | 0.301736317 | 1.363944995 | 0.022651765 | 0.12877065 | 0.40153474 | 0.1188 | 0.355880668 |
| SL1344_RS11115_ | -1.274274653 | -1.662325672 | -0.771005287 | 0.903665968 | 0.326965611 | 2.680488884 | 0.00067292 | 0.025601107 | 0.177646945 | 0.1004 | 0.330855409 |
| SL1344_RS11120_ | -0.509314904 | -0.784149063 | -0.26777002 | 0.530541444 | 0.250068187 | 2.129008828 | 0.003334715 | 0.043932398 | 0.227132516 | 0.1041 | 0.335977921 |
| SL1344_RS11125_ | -0.058870166 | -0.361765384 | 0.198525018 | 0.593100144 | 0.255986225 | 2.334438667 | 0.001335878 | 0.030804489 | 0.196606328 | 0.1014 | 0.332317761 |
| SL1344_RS11130_ | -0.139063092 | -0.458396184 | 0.135064757 | 0.58760514 | 0.171022385 | 3.416780839 | 0.000673369 | 0.012447849 | 0.121444585 | 0.1001 | 0.330406963 |
| SL1344_RS11135_ | 1.733111905 | 1.418413374 | 2.031541368 | 0.610213511 | 0.113947025 | 5.227353168 | 4.68E-05 | 0.004010895 | 0.074240166 | 0.1 | 0.330257709 |
| SL1344_RS11140_ | 1.691276136 | 1.456967625 | 1.960236919 | 0.51392811 | 0.172860915 | 2.912761338 | 4.68E-05 | 0.013801523 | 0.138926301 | 0.1 | 0.330257709 |
| SL1344_RS11145_ | 1.96536837 | 1.606594694 | 2.43706107 | 0.81529319 | 0.178203448 | 4.504741217 | 4.68E-05 | 0.006178766 | 0.09679561 | 0.1 | 0.330257709 |
| SL1344_RS11150_ | -3.108223883 | -3.154307905 | -3.064125239 | 0.099330897 | 0.504889635 | 0.183871689 | 0.419053967 | 0.541455066 | 0.745266376 | 0.6641 | 0.807596874 |
| SL1344_RS11155_ | -2.433636084 | -2.692544762 | -2.201258761 | 0.510691986 | 0.425301879 | 1.156798564 | 0.088000048 | 0.154599764 | 0.40078028 | 0.2221 | 0.461997735 |
| SL1344_RS11160_ | -1.852144976 | -2.149962111 | -1.616297502 | 0.528651793 | 0.30043997 | 1.663703468 | 0.03664236 | 0.074134376 | 0.280277489 | 0.1406 | 0.37942284 |
| SL1344_RS11165_ | -2.172663083 | -2.385705605 | -1.982342476 | 0.393385267 | 0.408164855 | 0.91605114 | 0.164000023 | 0.243465031 | 0.503586706 | 0.3344 | 0.568966807 |
| SL1344_RS11170_ | 0.094353838 | -0.126240799 | 0.179823122 | 0.285519651 | 0.238970093 | 1.080215707 | 0.143237868 | 0.196001469 | 0.468185274 | 0.2919 | 0.542054951 |
| SL1344_RS11175_ | -0.435350563 | -0.544840745 | -0.351138938 | 0.189541683 | 0.202350376 | 0.884363225 | 0.172551653 | 0.259579995 | 0.507227801 | 0.3541 | 0.576216716 |
| SL1344_RS11180_ | -2.117423284 | -2.31516926 | -2.015745828 | 0.32519605 | 0.472873009 | 0.611701091 | 0.268000011 | 0.395804056 | 0.646913888 | 0.517 | 0.71290084 |
| SL1344_RS11185_ | -2.560764774 | -2.549930042 | -2.579730315 | -0.012601736 | 0.56589291 | -0.023149034 | 0.490666667 | 0.675632578 | 0.841788875 | 0.7783 | 0.888428473 |
| SL1344_RS11190_ | -2.797577842 | -2.855529024 | -2.746801144 | 0.092251551 | 0.599734428 | 0.148612829 | 0.435043307 | 0.641280398 | 0.820160876 | 0.7492 | 0.870565755 |
| SL1344_RS11195_ | -2.44872106 | -2.482088689 | -2.432982801 | 0.01473068 | 0.732997106 | 0.018659702 | 0.494666667 | 0.773002536 | 0.900318557 | 0.8633 | 0.937811099 |
| SL1344_RS11200_ | 0.047391777 | -0.122955827 | 0.133798682 | 0.238352126 | 0.20935972 | 1.089873747 | 0.139240534 | 0.207192608 | 0.472122399 | 0.3039 | 0.54545385 |
| SL1344_RS11205_ | -1.623982515 | -1.733873369 | -1.487096354 | 0.243600431 | 0.309885309 | 0.744955657 | 0.199333351 | 0.325754856 | 0.565903987 | 0.4267 | 0.631328049 |
| SL1344_RS11210_ | -1.006757771 | -1.123648213 | -0.863551014 | 0.255152651 | 0.251686934 | 0.988064458 | 0.143237868 | 0.2219761 | 0.468632963 | 0.3029 | 0.536458182 |
| SL1344_RS11215_ | 1.226571229 | 1.262781035 | 1.124357324 | -0.139095928 | 0.158199766 | -0.789056806 | 0.212666683 | 0.304742973 | 0.57907919 | 0.4348 | 0.654057015 |

| | | | | | | | | | | | |
|---|---|---|---|---|---|---|---|---|---|---|---|
| SL1344_RS11220_ | 1.572940783 | 1.281498944 | 1.745646266 | 0.451049284 | 0.150976234 | 2.837503275 | 4.68E-05 | 0.024108554 | 0.177524893 | 0.1002 | 0.330540769 |
| SL1344_RS11225_ | 0.123391354 | -0.236252766 | 0.373872189 | 0.598779775 | 0.256403286 | 2.2752714 | 0.003332494 | 0.040052236 | 0.22518042 | 0.1028 | 0.334353916 |
| SL1344_RS11230_ | -0.860335753 | -1.359853681 | -0.43130074 | 0.925507895 | 0.249202997 | 3.733092377 | 4.68E-05 | 0.009645007 | 0.109702563 | 0.1 | 0.330257709 |
| SL1344_RS11235_ | 0.224425474 | -0.265478659 | 0.547399224 | 0.819643985 | 0.209413074 | 3.852683609 | 4.68E-05 | 0.010185288 | 0.115123103 | 0.1 | 0.330257709 |
| SL1344_RS11240_ | 3.271020781 | 3.264615577 | 3.276622327 | -0.057228122 | 0.288623842 | -0.218364111 | 0.410000003 | 0.731239793 | 0.885682538 | 0.8854 | 0.949915586 |
| SL1344_RS11245_ | 5.018241993 | 5.07684885 | 4.947585582 | -0.037782187 | 0.308730159 | -0.122404895 | 0.457028649 | 0.934692575 | 0.976628764 | 1 | 1 |
| SL1344_RS11250_ | 6.713623496 | 6.731222384 | 6.695154592 | -0.037344444 | 0.276355134 | -0.128659272 | 0.444370422 | 0.898608883 | 0.963516045 | 1 | 1 |
| SL1344_RS11255_ | 2.61836582 | 2.54223609 | 2.697745925 | 0.153595516 | 0.424141979 | 0.305066984 | 0.440000002 | 0.74843852 | 0.896408747 | 0.9895 | 0.995557375 |
| SL1344_RS11260_ | 1.194411576 | 1.049420388 | 1.377618126 | 0.375755283 | 0.211793756 | 1.797890274 | 0.009333825 | 0.053560445 | 0.260030386 | 0.1099 | 0.343424792 |
| SL1344_RS11265_ | 1.20521431 | 0.970495435 | 1.480519758 | 0.452168467 | 0.230709257 | 1.945505937 | 0.01798826 | 0.040007418 | 0.228252321 | 0.1133 | 0.349451645 |
| SL1344_RS11270_ | 3.083784483 | 2.801397451 | 3.218181501 | 0.40967037 | 0.111803972 | 3.354204556 | 4.68E-05 | 0.02361265 | 0.181016677 | 0.1 | 0.330257709 |
| SL1344_RS11275_ | -1.852013011 | -1.935504101 | -1.747224485 | 0.186519957 | 0.470417712 | 0.386345276 | 0.340666673 | 0.550270932 | 0.758527667 | 0.6647 | 0.816611565 |
| SL1344_RS11280_ | -0.805626712 | -0.87836351 | -0.709359672 | 0.147137576 | 0.26006615 | 0.526844322 | 0.280666677 | 0.462537061 | 0.688535792 | 0.5692 | 0.751284813 |
| SL1344_RS11285_ | -4.891264688 | -5.365803561 | -4.652897324 | 0.580776996 | 1.360093986 | 0.386451724 | 0.343770826 | 0.52651456 | 0.74794584 | 0.6558 | 0.815778952 |
| SL1344_RS11290_ | -2.639411542 | -2.506038187 | -2.726744792 | -0.158594118 | 0.608108651 | -0.230866971 | 0.394000004 | 0.65020794 | 0.824978535 | 0.733 | 0.863374545 |
| SL1344_RS11295_ | -0.818065707 | -0.53556466 | -1.059735274 | -0.571470812 | 0.307043948 | -1.829384238 | 0.011326186 | 0.066031778 | 0.28142838 | 0.109 | 0.342128689 |
| SL1344_RS11300_ | 2.156396927 | 2.165231476 | 2.145600564 | -0.01533271 | 0.185560865 | -0.081379409 | 0.463690874 | 0.78889931 | 0.910026137 | 0.8984 | 0.953716438 |
| SL1344_RS11305_ | 0.863534491 | 1.060816826 | 0.792135082 | -0.292149755 | 0.285487002 | -0.985866273 | 0.166000022 | 0.201544426 | 0.488941485 | 0.3012 | 0.569323982 |
| SL1344_RS11310_ | 0.050617425 | -0.064461324 | 0.275937918 | 0.339840647 | 0.356583331 | 0.993894412 | 0.139906757 | 0.194094447 | 0.477534656 | 0.2745 | 0.540661965 |
| SL1344_RS11315_ | 1.452526468 | 1.129214298 | 1.665708819 | 0.532075168 | 0.146382775 | 3.503816699 | 4.68E-05 | 0.015250663 | 0.140220277 | 0.1001 | 0.330412106 |
| SL1344_RS11320_ | -0.68495391 | -0.746138076 | -0.608278211 | 0.170094295 | 0.303007635 | 0.539518792 | 0.260666678 | 0.404711624 | 0.654522767 | 0.5441 | 0.725736758 |
| SL1344_RS11325_ | -1.984329295 | -2.047876852 | -1.901764506 | 0.168152768 | 0.405707468 | 0.385948877 | 0.339107268 | 0.511421159 | 0.726527134 | 0.6366 | 0.792888679 |
| SL1344_RS11330_ | -2.037733636 | -2.021624424 | -2.055169127 | -0.06314246 | 0.440006375 | -0.1342939 | 0.446666669 | 0.619903575 | 0.803144786 | 0.7391 | 0.860476269 |
| SL1344_RS11335_ | -2.029154925 | -2.079039513 | -1.974530046 | 0.098757357 | 0.34908027 | 0.272513743 | 0.375749505 | 0.529899335 | 0.732723808 | 0.6343 | 0.788328904 |
| SL1344_RS11340_ | -2.432973873 | -2.510582488 | -2.344440362 | 0.168630597 | 0.42795108 | 0.380074445 | 0.338000007 | 0.503192628 | 0.717956225 | 0.6171 | 0.777333593 |
| SL1344_RS11345_ | -4.628437876 | -4.591537441 | -4.667273658 | -0.047405833 | 1.127893804 | -0.032318474 | 0.486342439 | 0.650549253 | 0.828809407 | 0.7637 | 0.881453603 |
| SL1344_RS11350_ | -2.16952062 | -2.285612623 | -2.019796006 | 0.294210372 | 0.421509072 | 0.673845635 | 0.228666681 | 0.344458647 | 0.592975194 | 0.4592 | 0.662498507 |
| SL1344_RS11355_ | 3.01536124 | 2.498757708 | 3.277783523 | 0.770010048 | 0.121585897 | 5.295719938 | 4.68E-05 | 0.015028832 | 0.149990451 | 0.1 | 0.330257709 |
| SL1344_RS11360_ | 0.043610571 | -0.281629622 | 0.361788583 | 0.636468286 | 0.214417662 | 2.890256765 | 0.000673369 | 0.017636984 | 0.149850739 | 0.1004 | 0.330833529 |
| SL1344_RS11365_ | 1.879786457 | 1.490304868 | 2.051914147 | 0.543752482 | 0.222976666 | 2.11602981 | 0.001335878 | 0.052717007 | 0.266072698 | 0.1018 | 0.33285242 |
| SL1344_RS11375_ | 1.253402131 | 1.160494843 | 1.498615949 | 0.319478748 | 0.153184949 | 1.874496877 | 0.030666813 | 0.07312936 | 0.297839721 | 0.13 | 0.368013264 |
| SL1344_RS11380_ | 1.078002265 | 0.956935751 | 1.249361283 | 0.301198282 | 0.177236159 | 1.661889783 | 0.022000206 | 0.069786566 | 0.288754098 | 0.1289 | 0.367895819 |
| SL1344_RS11385_ | 0.778893553 | 0.52558016 | 0.939770023 | 0.420048388 | 0.245480727 | 1.710577422 | 0.022651765 | 0.056532187 | 0.268258593 | 0.1169 | 0.353825503 |
| SL1344_RS11390_ | 0.369329996 | 0.183678647 | 0.560010161 | 0.372320975 | 0.167337066 | 2.155227716 | 0.009993796 | 0.040766977 | 0.214986358 | 0.1113 | 0.345351141 |
| SL1344_RS11395_ | 1.26020767 | 1.048738177 | 1.648641992 | 0.589985585 | 0.147389825 | 3.848558525 | 4.68E-05 | 0.01477233 | 0.138733444 | 0.1 | 0.330257709 |
| SL1344_RS11400_ | 1.465742135 | 1.307669913 | 1.806924978 | 0.473865255 | 0.165101416 | 2.490939338 | 0.000673369 | 0.040823804 | 0.232022974 | 0.1004 | 0.330829651 |
| SL1344_RS11405_ | -2.802932565 | -2.918310835 | -2.672843581 | 0.23975463 | 0.438668668 | 0.506165108 | 0.294666676 | 0.421484011 | 0.645085803 | 0.5161 | 0.70489653 |
| SL1344_RS11410_ | 1.050706181 | 0.840185844 | 1.348259277 | 0.514396091 | 0.195234446 | 2.6125435 | 0.000673369 | 0.025467544 | 0.18345383 | 0.1005 | 0.330976427 |
| SL1344_RS11415_ | 1.017069298 | 0.788714796 | 1.295332917 | 0.508293983 | 0.188175941 | 2.703594063 | 0.00067292 | 0.02142463 | 0.167341909 | 0.1008 | 0.331342836 |
| SL1344_RS11420_ | 0.886240938 | 0.671034887 | 1.170571913 | 0.483582749 | 0.161596384 | 2.891404847 | 0.00067292 | 0.019309324 | 0.157130523 | 0.1008 | 0.331436618 |
| SL1344_RS11425_ | 4.90430317 | 4.688621832 | 5.147466068 | 0.43921599 | 0.131458479 | 3.247670735 | 4.68E-05 | 0.007745058 | 0.112907694 | 0.1 | 0.330257709 |
| SL1344_RS11430_ | 0.68757147 | 0.473885884 | 0.785509053 | 0.298120621 | 0.221018552 | 1.274551397 | 0.08794142 | 0.13697124 | 0.404263717 | 0.204 | 0.459158332 |
| SL1344_RS11435_ | 1.268472101 | 1.06550985 | 1.504438889 | 0.414307052 | 0.167677263 | 2.417912089 | 0.003334715 | 0.027220819 | 0.187459365 | 0.103 | 0.334630136 |
| SL1344_RS11440_ | 1.437130581 | 1.162714886 | 1.546760419 | 0.380077249 | 0.260200388 | 1.382520965 | 0.027981506 | 0.128633442 | 0.404004062 | 0.1292 | 0.36990075 |
| SL1344_RS11445_ | 2.90284929 | 2.789939206 | 3.09951224 | 0.348787119 | 0.201704007 | 1.886162692 | 0.00067292 | 0.043988764 | 0.245816413 | 0.1004 | 0.330743091 |
| SL1344_RS11450_ | 0.615626587 | 1.253136841 | 0.188046054 | -0.813312608 | 0.899774365 | -0.564519209 | 0.33333334 | 0.538986082 | 0.779902402 | 0.7 | 0.872542119 |
| SL1344_RS11455_ | -3.772442583 | -4.007934926 | -3.550368508 | 0.415417911 | 0.708369485 | 0.545159394 | 0.267333344 | 0.417955393 | 0.653960289 | 0.5209 | 0.713275046 |
| SL1344_RS11460_ | -5.946783188 | -5.942566638 | -5.949714307 | 0.055490661 | 1.665915352 | 0.029338127 | 0.485333334 | 0.608909086 | 0.798889078 | 0.732 | 0.856968717 |
| SL1344_RS11465_ | 2.178458283 | 1.891427221 | 2.483647216 | 0.590698818 | 0.166214503 | 3.52850206 | 4.68E-05 | 0.007693436 | 0.107047531 | 0.1 | 0.330257709 |
| SL1344_RS11470_ | -3.322449691 | -3.666278276 | -3.001982871 | 0.655911987 | 0.499780352 | 1.241376799 | 0.077948089 | 0.139446597 | 0.368578742 | 0.2089 | 0.444832103 |
| SL1344_RS11475_ | -3.094743233 | -3.140383075 | -3.065220113 | 0.079043943 | 0.601664728 | 0.128787593 | 0.444666669 | 0.615264787 | 0.801674767 | 0.7282 | 0.85640056 |
| SL1344_RS11480_ | -2.738113252 | -3.02454454 | -2.534768716 | 0.504646085 | 0.552646649 | 0.867444844 | 0.145902758 | 0.239686799 | 0.508298907 | 0.3155 | 0.54929517 |
| SL1344_RS11485_ | -4.947365115 | -5.213731775 | -4.206702824 | 0.875140935 | 1.208793846 | 0.591836476 | 0.278481023 | 0.494831955 | 0.711361756 | 0.5607 | 0.759094483 |

| | | | | | | | | | | | |
|---|---|---|---|---|---|---|---|---|---|---|---|
| SL1344_RS11490_ | -5.234353876 | -5.580675099 | -5.075447596 | 0.567434584 | 1.553571085 | 0.295143168 | 0.359333339 | 0.517421245 | 0.747893214 | 0.6944 | 0.838178233 |
| SL1344_RS11495_ | 2.273520654 | 1.942254775 | 2.669095356 | 0.774703666 | 0.230068595 | 3.361966864 | 4.68E-05 | 0.033407354 | 0.218295792 | 0.1 | 0.330257709 |
| SL1344_RS11500_ | -2.821784615 | -2.803655305 | -2.831284717 | -0.037102008 | 0.632234083 | -0.060897871 | 0.477015324 | 0.681534383 | 0.84522467 | 0.7932 | 0.896301891 |
| SL1344_RS11505_ | -2.613128104 | -2.853802202 | -2.371345415 | 0.468389262 | 0.408958436 | 1.136255329 | 0.119333367 | 0.188156727 | 0.428336017 | 0.2659 | 0.500500223 |
| SL1344_RS11510_ | -3.469568373 | -3.441392023 | -3.517642428 | -0.083454703 | 0.667386767 | -0.122602183 | 0.454363759 | 0.616493339 | 0.803319025 | 0.7392 | 0.864502711 |
| SL1344_RS11515_ | 0.593073813 | 0.515262975 | 0.64779194 | 0.148797499 | 0.191550895 | 0.764753212 | 0.172666688 | 0.285701142 | 0.558046458 | 0.3816 | 0.600700557 |
| SL1344_RS11520_ | -6.043155246 | -6.506188975 | -5.765087376 | 0.779873662 | 1.550525403 | 0.432292389 | 0.290666676 | 0.425294247 | 0.663789839 | 0.549 | 0.73225598 |
| SL1344_RS11525_ | -3.049096568 | -3.528302582 | -2.65924298 | 0.86751073 | 0.437873093 | 1.927269664 | 0.020000227 | 0.052516438 | 0.236978807 | 0.1201 | 0.354736864 |
| SL1344_RS11535_ | -4.649963993 | -5.003560383 | -4.347633754 | 0.662990992 | 0.811300146 | 0.778710297 | 0.188540992 | 0.283007799 | 0.527416892 | 0.3786 | 0.592046291 |
| SL1344_RS11540_ | 0.322977932 | 0.083404642 | 0.491462178 | 0.402985968 | 0.162641155 | 2.38516915 | 0.005334196 | 0.034786005 | 0.200313898 | 0.1046 | 0.336626796 |
| SL1344_RS11545_ | -0.421126777 | -0.685048815 | -0.182534387 | 0.515353661 | 0.20914757 | 2.398966746 | 0.002668394 | 0.029262948 | 0.18526952 | 0.103 | 0.334461285 |
| SL1344_RS11550_ | -3.911308483 | -4.364376444 | -3.755253581 | 0.559033371 | 1.191982284 | 0.409246514 | 0.360000006 | 0.565877979 | 0.781940975 | 0.7278 | 0.86956273 |
| SL1344_RS11555_ | -2.134528027 | -2.327340637 | -1.893548343 | 0.448634856 | 0.379231245 | 1.103974707 | 0.112666703 | 0.180980002 | 0.432011291 | 0.2547 | 0.49225592 |
| SL1344_RS11560_ | -2.413439346 | -2.917788177 | -2.049745436 | 0.878033243 | 0.39638725 | 2.090451921 | 0.009333825 | 0.046399163 | 0.22329216 | 0.1113 | 0.344847033 |
| SL1344_RS11565_ | -5.472924435 | -5.926376063 | -5.052216195 | 0.895786741 | 1.462446258 | 0.589610023 | 0.23984012 | 0.386182364 | 0.630906221 | 0.4912 | 0.693997081 |
| SL1344_RS11570_ | -3.366110818 | -3.583456067 | -3.164789454 | 0.422511989 | 0.65968383 | 0.623336996 | 0.241838787 | 0.374056878 | 0.62175551 | 0.4863 | 0.689653066 |
| SL1344_RS11585_ | -0.324190641 | -0.59097967 | -0.176568964 | 0.40523369 | 0.233799086 | 1.606102613 | 0.050000088 | 0.090118883 | 0.317453552 | 0.1547 | 0.396490338 |
| SL1344_RS11595_ | 1.113312069 | 0.923893384 | 1.29543268 | 0.388513549 | 0.182906358 | 2.17754133 | 0.002000966 | 0.054079551 | 0.261343593 | 0.1015 | 0.332366714 |
| SL1344_RS11600_ | 0.256310753 | -0.019573626 | 0.320483065 | 0.333981291 | 0.265365608 | 1.068827625 | 0.150666692 | 0.210440498 | 0.497081911 | 0.3161 | 0.555912562 |
| SL1344_RS11605_ | 2.445971926 | 2.174514764 | 2.639341064 | 0.468337006 | 0.137987578 | 3.448863935 | 4.68E-05 | 0.007920956 | 0.10829578 | 0.1 | 0.330257709 |
| SL1344_RS11610_ | -0.615831416 | -0.86439005 | -0.104633148 | 0.717825269 | 0.275626754 | 2.412477587 | 0.006662914 | 0.03892169 | 0.217723378 | 0.1042 | 0.336276978 |
| SL1344_RS11615_ | 1.069918402 | 0.646085785 | 1.407019047 | 0.789001256 | 0.212157232 | 3.761414755 | 4.68E-05 | 0.010431062 | 0.122376654 | 0.1 | 0.330257709 |
| SL1344_RS11620_ | 0.658182927 | -0.146711531 | 1.423376305 | 1.59577532 | 0.224861097 | 7.159589847 | 4.68E-05 | 0.001125599 | 0.044983876 | 0.1 | 0.330257709 |
| SL1344_RS11625_ | 1.181038943 | 1.228026862 | 1.116052028 | -0.106950699 | 0.123359899 | -0.802690199 | 0.195203216 | 0.294714813 | 0.546263612 | 0.3939 | 0.615256253 |
| SL1344_RS11630_ | -0.54672465 | -0.492716852 | -0.602443585 | -0.161864754 | 0.31568694 | -0.489998207 | 0.292000009 | 0.436039259 | 0.680652084 | 0.5941 | 0.765049038 |
| SL1344_RS11635_ | 0.521467458 | 0.543506907 | 0.506752771 | -0.076816826 | 0.265789963 | -0.269253119 | 0.368000005 | 0.53021973 | 0.758433115 | 0.7303 | 0.855014127 |
| SL1344_RS11640_ | -0.07954779 | 0.007739273 | -0.185058912 | -0.190044995 | 0.178346037 | -1.041571374 | 0.129913421 | 0.203350425 | 0.45165208 | 0.2895 | 0.523394496 |
| SL1344_RS11670_ | -1.351950243 | -1.422931788 | -1.253657614 | 0.222966046 | 0.429975379 | 0.521367771 | 0.290000009 | 0.439611906 | 0.688029962 | 0.5912 | 0.765452263 |
| SL1344_RS11675_ | -3.326516749 | -3.055140728 | -3.499822597 | -0.49402193 | 1.403117932 | -0.318248977 | 0.37708195 | 0.655676819 | 0.837234788 | 0.7817 | 0.895655916 |
| SL1344_RS11680_ | -2.007214308 | -1.656273862 | -2.420528249 | -0.756796237 | 0.377254361 | -1.950136568 | 0.019986902 | 0.053357855 | 0.242663449 | 0.1196 | 0.355107136 |
| SL1344_RS11685_ | 2.0599693 | 2.013259873 | 2.099873361 | 0.138120196 | 0.194349851 | 0.724777467 | 0.195333351 | 0.279206486 | 0.571792599 | 0.4015 | 0.629859938 |
| SL1344_RS11690_ | 4.304318669 | 4.214846549 | 4.466975589 | 0.26912465 | 0.128696605 | 2.303809577 | 4.68E-05 | 0.027942125 | 0.199457268 | 0.1 | 0.330257709 |
| SL1344_RS11695_ | 3.036955725 | 2.752474128 | 3.227502828 | 0.476487348 | 0.168886305 | 2.83041329 | 4.68E-05 | 0.054076097 | 0.27341453 | 0.1 | 0.330257709 |
| SL1344_RS11700_ | 2.780768197 | 2.544527855 | 2.971028244 | 0.419588438 | 0.085316814 | 4.817315493 | 4.68E-05 | 0.006135326 | 0.090907319 | 0.1 | 0.330257709 |
| SL1344_RS11705_ | -0.28489566 | -0.351234744 | -0.015449549 | 0.280684192 | 0.333660355 | 0.553587864 | 0.323333341 | 0.567417449 | 0.783963682 | 0.6688 | 0.846224658 |
| SL1344_RS11710_ | 0.735377242 | 0.704403303 | 0.750410953 | 0.014682239 | 0.251156351 | 0.054187292 | 0.477681546 | 0.759892634 | 0.893804448 | 0.8529 | 0.934733702 |
| SL1344_RS11715_ | 7.246094288 | 7.280405296 | 7.207134806 | 0.038051333 | 0.236592462 | 0.170496112 | 0.451032647 | 0.781533724 | 0.913438048 | 1 | 1 |
| SL1344_RS11720_ | 6.743431523 | 6.54190845 | 6.838610668 | 0.232802456 | 0.204507585 | 1.492751094 | 0.091272531 | 0.07493711 | 0.319492496 | 0.1866 | 0.456941418 |
| SL1344_RS11725_ | 3.642496022 | 3.410745927 | 3.949077254 | 0.534310903 | 0.150280331 | 3.651606368 | 4.68E-05 | 0.020895702 | 0.176262064 | 0.1 | 0.330257709 |
| SL1344_RS11730_ | 3.299479161 | 3.015217154 | 3.492076586 | 0.488237643 | 0.148224081 | 3.280872513 | 4.68E-05 | 0.036247493 | 0.226966684 | 0.1 | 0.330257709 |
| SL1344_RS11735_ | 3.485915654 | 3.353490301 | 3.775406653 | 0.43470368 | 0.288934665 | 1.719438085 | 4.68E-05 | 0.062362088 | 0.291971449 | 0.1002 | 0.330547041 |
| SL1344_RS11740_ | -5.683825638 | -6.210269481 | -5.338755614 | 0.88008476 | 2.699527853 | 0.263249169 | 0.351333339 | 0.517249534 | 0.752537214 | 0.7013 | 0.844363206 |
| SL1344_RS11745_ | -2.705888943 | -2.911283159 | -2.214638547 | 0.693337649 | 0.568656253 | 1.106301263 | 0.1352432 | 0.198648586 | 0.468384176 | 0.2963 | 0.538741863 |
| SL1344_RS11750_ | -1.076706753 | -1.224311086 | -0.829519883 | 0.385536398 | 0.289458027 | 1.245781401 | 0.091272531 | 0.150043099 | 0.400028543 | 0.2338 | 0.476188389 |
| SL1344_RS11755_ | -1.980561941 | -2.225008902 | -1.70659909 | 0.5112717 | 0.426604959 | 1.149333747 | 0.106666705 | 0.1634147 | 0.42489508 | 0.2384 | 0.488366678 |
| SL1344_RS11760_ | -1.161181064 | -1.496408172 | -0.877738169 | 0.61918603 | 0.238120208 | 2.5397403 | 0.003332494 | 0.027199497 | 0.175873401 | 0.1026 | 0.333907304 |
| SL1344_RS11765_ | -1.956035216 | -2.093079153 | -1.784297571 | 0.383771498 | 0.681680061 | 0.518409233 | 0.287808137 | 0.404945844 | 0.672055074 | 0.5673 | 0.757506447 |
| SL1344_RS11770_ | 1.433973842 | 1.226402499 | 1.590301844 | 0.409380992 | 0.211715413 | 1.92365817 | 0.003998486 | 0.046333334 | 0.246007776 | 0.1034 | 0.335047239 |
| SL1344_RS11775_ | 1.113591312 | 0.943480545 | 1.268125871 | 0.329301686 | 0.141369099 | 2.305413326 | 0.004001152 | 0.032856841 | 0.19908498 | 0.1042 | 0.336165558 |
| SL1344_RS11780_ | 0.705444634 | 0.477049675 | 0.945760609 | 0.468922768 | 0.125703304 | 3.699322358 | 4.68E-05 | 0.009632657 | 0.109109482 | 0.1 | 0.330257709 |
| SL1344_RS11785_ | 1.448219914 | 1.120061827 | 1.807755333 | 0.674144454 | 0.130371453 | 5.118184109 | 4.68E-05 | 0.003523197 | 0.07161592 | 0.1 | 0.330257709 |
| SL1344_RS11790_ | 2.006426078 | 1.781716464 | 2.311892899 | 0.562045456 | 0.158439647 | 3.568461393 | 4.68E-05 | 0.007645182 | 0.106192975 | 0.1 | 0.330257709 |
| SL1344_RS11795_ | 2.135626027 | 1.811708678 | 2.366803001 | 0.572025965 | 0.143404893 | 4.087092632 | 4.68E-05 | 0.021234606 | 0.171999983 | 0.1 | 0.330257709 |

| | | | | | | | | | | | |
|---|---|---|---|---|---|---|---|---|---|---|---|
| SL1344_RS11800_ | 3.23735055 | 2.819952894 | 3.577826888 | 0.756170069 | 0.154496116 | 4.99916556 | 4.68E-05 | 0.021244878 | 0.178827479 | 0.1 | 0.330257709 |
| SL1344_RS11805_ | 2.00250688 | 1.463630711 | 2.367850922 | 0.90331779 | 0.243927775 | 3.73280928 | 4.68E-05 | 0.021357932 | 0.17692351 | 0.1 | 0.330257709 |
| SL1344_RS11810_ | 1.934991706 | 1.612780127 | 2.172927018 | 0.564045234 | 0.139804079 | 4.02677292 | 4.68E-05 | 0.017117142 | 0.153055054 | 0.1 | 0.330257709 |
| SL1344_RS11815_ | 0.429002078 | 0.228490913 | 0.622913137 | 0.476715864 | 0.298088465 | 1.640439616 | 0.006667356 | 0.070057054 | 0.298178811 | 0.1054 | 0.337739057 |
| SL1344_RS11820_ | -0.326620353 | -0.510297511 | -0.175669557 | 0.345314974 | 0.273852141 | 1.219004175 | 0.073284535 | 0.140410867 | 0.400425459 | 0.1923 | 0.438255592 |
| SL1344_RS11825_ | 1.578502322 | 1.608787862 | 1.56503711 | -0.022582768 | 0.15733539 | -0.139658225 | 0.448000002 | 0.770671889 | 0.899097448 | 0.8388 | 0.929502679 |
| SL1344_RS11830_ | 1.765724542 | 1.525144946 | 1.96810492 | 0.445117039 | 0.10844804 | 4.054796583 | 4.68E-05 | 0.00776425 | 0.100737556 | 0.1 | 0.330257709 |
| SL1344_RS11840_ | 1.070012648 | 0.856729418 | 1.300347519 | 0.436075504 | 0.140012224 | 3.032696176 | 0.00067292 | 0.01604993 | 0.142613273 | 0.1004 | 0.330824671 |
| SL1344_RS11845_ | 0.057805754 | -0.151245958 | 0.192669844 | 0.350606325 | 0.230548185 | 1.452303564 | 0.050632998 | 0.104494327 | 0.348262989 | 0.1566 | 0.401654541 |
| SL1344_RS11850_ | 1.381074118 | 1.129929277 | 1.589508607 | 0.452948195 | 0.125234253 | 3.48322162 | 4.68E-05 | 0.011518516 | 0.122087234 | 0.1 | 0.330257709 |
| SL1344_RS11855_ | 1.250275932 | 1.102997383 | 1.299706006 | 0.180494276 | 0.148780771 | 1.04338915 | 0.146000026 | 0.210561343 | 0.485907936 | 0.3167 | 0.558651261 |
| SL1344_RS11860_ | 2.362126674 | 2.129778306 | 2.714035393 | 0.581097444 | 0.162049466 | 3.438647447 | 4.68E-05 | 0.015400936 | 0.1488037 | 0.1 | 0.330257709 |
| SL1344_RS11865_ | -1.184052013 | -1.395795105 | -0.958590005 | 0.257948265 | 0.523078498 | 0.531445761 | 0.29913392 | 0.498376957 | 0.735363492 | 0.6011 | 0.782931012 |
| SL1344_RS11870_ | 0.434927808 | 0.380115314 | 0.487981325 | 0.107156801 | 0.162960033 | 0.634433242 | 0.23666668 | 0.371112155 | 0.614892893 | 0.4767 | 0.68002549 |
| SL1344_RS11880_ | -0.731709052 | -0.791579663 | -0.660031092 | 0.154387149 | 0.266046036 | 0.548840705 | 0.262000011 | 0.403262424 | 0.645154213 | 0.5305 | 0.714472932 |
| SL1344_RS11885_ | 3.822645065 | 3.623489336 | 3.90572716 | 0.254514692 | 0.24986456 | 1.040536589 | 0.137908089 | 0.164625455 | 0.456512227 | 0.2449 | 0.522014872 |
| SL1344_RS11890_ | 3.089081777 | 3.059263882 | 3.264745365 | 0.259388853 | 0.280166199 | 0.981523709 | 0.193333352 | 0.189947628 | 0.48672815 | 0.3521 | 0.630249551 |
| SL1344_RS11895_ | 4.559080933 | 4.335704984 | 4.764709524 | 0.428540713 | 0.040177034 | 10.52887269 | 4.68E-05 | 0.000885649 | 0.037464417 | 0.1 | 0.330257709 |
| SL1344_RS11900_ | 2.162668189 | 1.983146728 | 2.395277142 | 0.412814075 | 0.150167957 | 2.784574844 | 4.68E-05 | 0.017298654 | 0.154432076 | 0.1 | 0.330257709 |
| SL1344_RS11905_ | 3.708240965 | 3.278121166 | 3.807720165 | 0.532659686 | 0.18774006 | 1.789154204 | 0.001335878 | 0.084953248 | 0.337595701 | 0.1004 | 0.33084707 |
| SL1344_RS11910_ | 3.172347342 | 2.930440065 | 3.44190203 | 0.500917079 | 0.098908718 | 4.978787972 | 4.68E-05 | 0.003880693 | 0.076938051 | 0.1 | 0.330257709 |
| SL1344_RS11915_ | 3.933320828 | 3.736981815 | 4.116648829 | 0.385912328 | 0.059268277 | 6.526828222 | 4.68E-05 | 0.001879267 | 0.053900308 | 0.1 | 0.330257709 |
| SL1344_RS11920_ | 5.677929276 | 5.379825115 | 5.896899024 | 0.506497953 | 0.063574601 | 7.534587416 | 4.68E-05 | 0.002956168 | 0.070417956 | 0.1 | 0.330257709 |
| SL1344_RS11925_ | 4.751922311 | 4.410004683 | 4.989907095 | 0.575220848 | 0.069677029 | 7.877899667 | 4.68E-05 | 0.004096419 | 0.081061689 | 0.1 | 0.330257709 |
| SL1344_RS11930_ | 3.523943407 | 3.204608414 | 3.78637113 | 0.580476551 | 0.058021314 | 9.966371698 | 4.68E-05 | 0.000826479 | 0.036842375 | 0.1 | 0.330257709 |
| SL1344_RS11935_ | 5.208678558 | 4.909835634 | 5.474871294 | 0.555292421 | 0.072908209 | 7.627843437 | 4.68E-05 | 0.000443797 | 0.033038722 | 0.1 | 0.330257709 |
| SL1344_RS11940_ | 4.146877162 | 3.933770262 | 4.458542952 | 0.555640553 | 0.168355664 | 3.601200908 | 4.68E-05 | 0.004865798 | 0.090419194 | 0.1 | 0.330257709 |
| SL1344_RS11945_ | 4.078123435 | 3.83856708 | 4.351018578 | 0.53177789 | 0.089421417 | 5.909187098 | 4.68E-05 | 0.001654421 | 0.054002763 | 0.1 | 0.330257709 |
| SL1344_RS11950_ | -2.902982109 | -3.325813662 | -2.737382108 | 0.581052915 | 0.714897899 | 0.711334649 | 0.214666682 | 0.322134377 | 0.592225602 | 0.4439 | 0.663780907 |
| SL1344_RS11960_ | 3.238861671 | 2.859201171 | 3.556983099 | 0.700147725 | 0.105162419 | 6.663354237 | 4.68E-05 | 0.005482444 | 0.091339099 | 0.1 | 0.330257709 |
| SL1344_RS11965_ | -0.276256783 | -0.596526307 | 0.061272841 | 0.617068822 | 0.304816423 | 1.972119899 | 0.017322048 | 0.044427839 | 0.23554844 | 0.1138 | 0.350149421 |
| SL1344_RS11970_ | 1.410396525 | 1.349117822 | 1.440435155 | 0.13614134 | 0.324812637 | 0.467274631 | 0.311125924 | 0.459936347 | 0.722766803 | 0.636 | 0.816441049 |
| SL1344_RS11975_ | 2.737224134 | 2.563114932 | 2.898560693 | 0.351207138 | 0.123801643 | 2.949596428 | 4.68E-05 | 0.028393885 | 0.196088879 | 0.1 | 0.330257709 |
| SL1344_RS11980_ | 1.454957865 | 1.203701525 | 1.6244513 | 0.428204734 | 0.201141544 | 2.064737183 | 0.003334715 | 0.04587732 | 0.245225906 | 0.1023 | 0.333526718 |
| SL1344_RS11985_ | 0.994835319 | 0.851445922 | 1.153787374 | 0.301730237 | 0.130840779 | 2.225377287 | 0.009327606 | 0.03643372 | 0.202858592 | 0.1071 | 0.339792592 |
| SL1344_RS12000_ | -1.598181784 | -1.71932874 | -1.532680272 | 0.192565116 | 0.351317079 | 0.499845376 | 0.288666676 | 0.438289236 | 0.674421529 | 0.5572 | 0.739704096 |
| SL1344_RS12005_ | 5.30309026 | 5.257819156 | 5.359394861 | 0.302986148 | 0.515273221 | 0.642448049 | 0.328666674 | 0.32896369 | 0.628382482 | 0.6928 | 0.867897588 |
| SL1344_RS12010_ | 4.054728251 | 3.942769174 | 4.133938169 | 0.426624248 | 0.375075174 | 1.21093141 | 0.017333596 | 0.125511096 | 0.404645285 | 0.1169 | 0.354737856 |
| SL1344_RS12015_ | 4.9304232 | 4.745826117 | 5.078734799 | 0.440994853 | 0.249812385 | 2.033423663 | 4.68E-05 | 0.037040922 | 0.231222727 | 0.1 | 0.330257709 |
| SL1344_RS12020_ | 2.80436604 | 2.659701438 | 2.931030972 | 0.502521469 | 0.360658267 | 1.481831701 | 0.00067292 | 0.095451366 | 0.355582265 | 0.1 | 0.330257709 |
| SL1344_RS12025_ | 3.02240696 | 2.791816168 | 3.086213413 | 0.356597602 | 0.412900637 | 0.927912261 | 0.184666686 | 0.200372024 | 0.501176186 | 0.3265 | 0.605457018 |
| SL1344_RS12030_ | -5.91552983 | -6.594008124 | -5.654290909 | 0.916217927 | 1.837114341 | 0.438839505 | 0.306666675 | 0.461451199 | 0.702228435 | 0.6056 | 0.782073766 |
| SL1344_RS12035_ | 1.778492407 | 1.502567708 | 1.967128928 | 0.413010615 | 0.298238127 | 1.375751494 | 0.081333385 | 0.093696492 | 0.348825491 | 0.1737 | 0.437582838 |
| SL1344_RS12040_ | 2.781300104 | 2.697613932 | 2.887944563 | 0.199254932 | 0.10474902 | 1.925026425 | 0.001336769 | 0.078630608 | 0.31705598 | 0.1014 | 0.332308005 |
| SL1344_RS12045_ | 1.321203414 | 1.283449989 | 1.523852767 | 0.222873358 | 0.166385319 | 1.014576724 | 0.202666684 | 0.241161739 | 0.526321247 | 0.4004 | 0.635065899 |
| SL1344_RS12050_ | 0.075712295 | 0.002973193 | 0.133098433 | 0.132307127 | 0.250332096 | 0.503811377 | 0.298666676 | 0.466870908 | 0.702882369 | 0.5857 | 0.766351349 |
| SL1344_RS12055_ | -3.194728599 | -3.563690873 | -2.86578275 | 0.703499827 | 0.505488329 | 1.306772415 | 0.067288538 | 0.125131189 | 0.355010701 | 0.1913 | 0.428934817 |
| SL1344_RS12060_ | 0.34675915 | 0.112707969 | 0.475693521 | 0.357492466 | 0.370372546 | 0.893674623 | 0.164666689 | 0.236847166 | 0.530231737 | 0.3127 | 0.576179224 |
| SL1344_RS12065_ | -0.523316246 | -0.639946442 | -0.384294615 | 0.297387358 | 0.408718214 | 0.740814015 | 0.179333354 | 0.280980067 | 0.566957541 | 0.36 | 0.603762697 |
| SL1344_RS12070_ | -0.648471405 | -0.730097428 | -0.589770512 | 0.159265769 | 0.26439399 | 0.579301076 | 0.256495681 | 0.386485447 | 0.638467536 | 0.5059 | 0.69463289 |
| SL1344_RS12075_ | -0.343212001 | -0.452385394 | -0.289513477 | 0.169695938 | 0.530146002 | 0.296317946 | 0.396000004 | 0.679128397 | 0.85526511 | 0.8362 | 0.928370784 |
| SL1344_RS12080_ | 2.785648673 | 2.485950377 | 2.955257396 | 0.470800277 | 0.192532736 | 2.366355999 | 4.68E-05 | 0.087410617 | 0.342062632 | 0.1 | 0.330257709 |
| SL1344_RS12085_ | -0.058469453 | -0.33964414 | 0.167486411 | 0.555963789 | 0.458942143 | 1.201966796 | 0.088000048 | 0.13039903 | 0.403219446 | 0.1921 | 0.453891336 |

| | | | | | | | | | | |
|---|---|---|---|---|---|---|---|---|---|---|
| SL1344_RS12090_ | 1.55272226 | 1.523948453 | 1.607020259 | 0.088277106 | 0.244833502 | 0.380334323 | 0.295136585 | 0.559464055 | 0.778794355 | 0.5862 | 0.785067534 |
| SL1344_RS12095_ | 1.321895136 | 1.131483597 | 1.422620814 | 0.289896927 | 0.171970488 | 1.563695782 | 0.029333486 | 0.099150385 | 0.348616844 | 0.1304 | 0.369386912 |
| SL1344_RS12100_ | -0.516163235 | -0.750372555 | -0.286599916 | 0.475219472 | 0.198145892 | 2.267542152 | 0.005996769 | 0.035094382 | 0.196648595 | 0.106 | 0.338446112 |
| SL1344_RS12105_ | -1.30598971 | -1.516011038 | -1.128438026 | 0.407651219 | 0.357171484 | 1.083904752 | 0.116000035 | 0.175739465 | 0.437390552 | 0.2522 | 0.498598545 |
| SL1344_RS12110_ | -1.707299634 | -1.780580265 | -1.658081858 | 0.128040303 | 0.470386419 | 0.263976471 | 0.389073955 | 0.622338049 | 0.812173489 | 0.7418 | 0.870084568 |
| SL1344_RS12115_ | 2.088732842 | 1.972272516 | 2.283514603 | 0.305477297 | 0.1275561 | 2.308392565 | 0.002666616 | 0.031703123 | 0.202797645 | 0.1019 | 0.332982834 |
| SL1344_RS12120_ | 2.437687446 | 2.114640963 | 2.471543586 | 0.361375219 | 0.227108959 | 1.071675086 | 0.202666684 | 0.205821064 | 0.505226105 | 0.358 | 0.626078777 |
| SL1344_RS12125_ | 1.350148297 | 1.191735283 | 1.406469539 | 0.205922145 | 0.254125296 | 0.812634003 | 0.200532995 | 0.265982503 | 0.555396097 | 0.3664 | 0.633821685 |
| SL1344_RS12130_ | 2.411124155 | 2.299425147 | 2.503790953 | 0.208935873 | 0.110768866 | 1.866585564 | 0.015323415 | 0.053293114 | 0.254873494 | 0.1162 | 0.352501971 |
| SL1344_RS12135_ | 2.553012532 | 2.409972401 | 2.719904126 | 0.30820158 | 0.087487553 | 3.494856073 | 4.68E-05 | 0.012177957 | 0.125993964 | 0.1 | 0.330257709 |
| SL1344_RS12140_ | 4.467781575 | 4.301194114 | 4.718512029 | 0.421587402 | 0.161154186 | 2.819484178 | 4.68E-05 | 0.010894112 | 0.130536913 | 0.1 | 0.330257709 |
| SL1344_RS12145_ | 0.527362697 | 0.290671253 | 0.756632506 | 0.560951105 | 0.282478973 | 2.03243418 | 0.00067292 | 0.037702548 | 0.222374485 | 0.1012 | 0.331831609 |
| SL1344_RS12155_ | 1.809283487 | 1.681728443 | 1.989136088 | 0.29855193 | 0.107426906 | 2.658898521 | 0.002666616 | 0.024139554 | 0.171501236 | 0.1017 | 0.332566573 |
| SL1344_RS12160_ | 2.952216113 | 2.779317645 | 3.11659169 | 0.381265827 | 0.12407561 | 3.12565837 | 4.68E-05 | 0.012675369 | 0.133983726 | 0.1 | 0.330257709 |
| SL1344_RS12170_ | 1.372369546 | 1.276224468 | 1.408795962 | 0.129597495 | 0.314328376 | 0.439322985 | 0.271333344 | 0.543004552 | 0.773977832 | 0.541 | 0.764425474 |
| SL1344_RS12175_ | -4.482968233 | -4.694288191 | -4.211182033 | 0.427373279 | 1.036574933 | 0.379924448 | 0.323117929 | 0.523504357 | 0.735335853 | 0.6219 | 0.787551202 |
| SL1344_RS12180_ | -5.486078731 | -5.611270801 | -5.368671643 | 0.237094112 | 1.155265488 | 0.194855182 | 0.420000003 | 0.537081391 | 0.743568477 | 0.6519 | 0.803343186 |
| SL1344_RS12185_ | -5.186887502 | -5.679743096 | -4.849937958 | 0.852280108 | 1.349723236 | 0.575659988 | 0.212000016 | 0.339736028 | 0.604769009 | 0.4314 | 0.639649296 |
| SL1344_RS12190_ | -4.389936414 | -4.641819733 | -4.202931641 | 0.346285362 | 1.055965332 | 0.299142291 | 0.370419725 | 0.578537081 | 0.774865002 | 0.6819 | 0.826953148 |
| SL1344_RS12195_ | -0.94213643 | -0.822549684 | -1.066543624 | -0.247816065 | 0.208878904 | -1.096091373 | 0.119333367 | 0.189503776 | 0.422528857 | 0.2766 | 0.506015664 |
| SL1344_RS12200_ | 5.14122516 | 4.991994764 | 5.149300899 | 0.157219163 | 0.169114875 | 0.587981319 | 0.33333334 | 0.637282598 | 0.839302926 | 0.7 | 0.872542119 |
| SL1344_RS12205_ | 1.30342459 | 1.254768114 | 1.346864314 | 0.087594435 | 0.148294863 | 0.580220393 | 0.279147245 | 0.429326278 | 0.674938497 | 0.5399 | 0.735982536 |
| SL1344_RS12210_ | 1.874750726 | 1.499629049 | 2.189588281 | 0.696788295 | 0.134724781 | 5.14580319 | 4.68E-05 | 0.004015013 | 0.075779912 | 0.1 | 0.330257709 |
| SL1344_RS12215_ | 0.482989387 | 0.239604438 | 0.740396069 | 0.58297278 | 0.289929195 | 2.052348115 | 0.000673369 | 0.035937928 | 0.218385763 | 0.1009 | 0.331496676 |
| SL1344_RS12220_ | 0.673261304 | 0.499837075 | 0.882152812 | 0.406932538 | 0.193599906 | 2.08617028 | 0.003998486 | 0.046565313 | 0.241410979 | 0.1032 | 0.334762261 |
| SL1344_RS12225_ | 1.050777247 | 0.862782038 | 1.227197813 | 0.402307837 | 0.196192481 | 2.140827419 | 0.001336769 | 0.046303532 | 0.242798702 | 0.1016 | 0.332530356 |
| SL1344_RS12230_ | 1.728539024 | 1.566169442 | 1.920865193 | 0.331044015 | 0.168489583 | 1.959253377 | 0.017322048 | 0.046477239 | 0.245134354 | 0.1156 | 0.352783869 |
| SL1344_RS12235_ | 1.784019587 | 1.54785142 | 1.948701543 | 0.410954715 | 0.137305702 | 2.949528376 | 4.68E-05 | 0.029218359 | 0.194630564 | 0.1 | 0.330257709 |
| SL1344_RS12240_ | -0.424286262 | -0.587152181 | -0.261368031 | 0.32732135 | 0.223500301 | 1.391794337 | 0.059293877 | 0.107662274 | 0.341600119 | 0.1726 | 0.416240616 |
| SL1344_RS12245_ | 3.193645246 | 2.941815449 | 3.245714516 | 0.263883188 | 0.216982954 | 0.612584544 | 0.33333334 | 0.555778391 | 0.790343628 | 0.7 | 0.872542119 |
| SL1344_RS12250_ | 1.65130074 | 1.343180634 | 1.723000545 | 0.389109541 | 0.317097629 | 1.250628159 | 0.110592975 | 0.122024412 | 0.393817366 | 0.2246 | 0.488200066 |
| SL1344_RS12260_ | 1.576341897 | 1.49754862 | 1.674004203 | 0.305807675 | 0.34074353 | 0.929770537 | 0.096602308 | 0.1987544 | 0.494491013 | 0.218 | 0.468602279 |
| SL1344_RS12265_ | 3.44580158 | 3.055194047 | 3.582587981 | 0.512327876 | 0.200368823 | 2.313391458 | 4.68E-05 | 0.032689576 | 0.215849421 | 0.1 | 0.330257709 |
| SL1344_RS12270_ | 3.7579563 | 3.551689896 | 3.905465426 | 0.261936003 | 0.175548674 | 1.697619723 | 0.059333406 | 0.064809743 | 0.294840339 | 0.1528 | 0.407505947 |
| SL1344_RS12275_ | -1.565323845 | -2.044136133 | -1.250857029 | 0.653157034 | 0.52334846 | 1.226477292 | 0.101932086 | 0.152542839 | 0.424809335 | 0.217 | 0.479698513 |
| SL1344_RS12280_ | -1.602629475 | -1.641103965 | -1.563718933 | 0.003171284 | 0.480825428 | 0.005755682 | 0.497668221 | 0.715399527 | 0.867479187 | 0.8384 | 0.920544322 |
| SL1344_RS12285_ | -1.289208627 | -1.723035159 | -1.117610798 | 0.579464922 | 0.367603953 | 1.416570512 | 0.069333395 | 0.116878972 | 0.368931738 | 0.1826 | 0.428702643 |
| SL1344_RS12290_ | 0.81320991 | 0.584084169 | 1.025957069 | 0.449174178 | 0.175606323 | 2.567513498 | 0.001336769 | 0.027591324 | 0.185899201 | 0.1006 | 0.331116898 |
| SL1344_RS12295_ | 0.71242324 | 0.447210916 | 0.814298088 | 0.353270083 | 0.168784099 | 1.88906938 | 0.026000173 | 0.072861895 | 0.292690045 | 0.1287 | 0.366646501 |
| SL1344_RS12300_ | -0.672770573 | -1.104974934 | -0.416500087 | 0.73064183 | 0.377428563 | 1.877121629 | 0.00733396 | 0.058044832 | 0.269399733 | 0.1078 | 0.340870142 |
| SL1344_RS12305_ | -2.809528385 | -3.258010752 | -2.436473647 | 0.861882379 | 0.598973267 | 1.39014443 | 0.057333409 | 0.103752965 | 0.341247412 | 0.1661 | 0.410919894 |
| SL1344_RS12310_ | -2.849941071 | -2.949145454 | -2.752551019 | 0.189529084 | 0.419293958 | 0.425943196 | 0.316455704 | 0.452879833 | 0.671442302 | 0.5618 | 0.733701988 |
| SL1344_RS12315_ | -0.610182321 | -0.895549114 | -0.162968767 | 0.56103665 | 0.654086833 | 0.917344954 | 0.157894761 | 0.23046764 | 0.526202509 | 0.2788 | 0.553888913 |
| SL1344_RS12320_ | -6.491451714 | -6.257066029 | -6.871342672 | -0.699959795 | 2.17349574 | -0.243074008 | 0.342666673 | 0.47888078 | 0.716851976 | 0.6183 | 0.781405949 |
| SL1344_RS12325_ | -1.059444278 | -1.066079108 | -1.047705602 | 0.008973624 | 0.25237255 | 0.030659358 | 0.483011326 | 0.585069418 | 0.777001229 | 0.685 | 0.826185097 |
| SL1344_RS12330_ | 2.578422555 | 2.565314718 | 2.598449743 | 0.067852237 | 0.16579394 | 0.387017516 | 0.343104603 | 0.555343855 | 0.776114397 | 0.7061 | 0.851417473 |
| SL1344_RS12335_ | 2.472089287 | 2.147018811 | 2.702766945 | 0.553988754 | 0.142200565 | 3.846579752 | 4.68E-05 | 0.015104247 | 0.145642467 | 0.1 | 0.330257709 |
| SL1344_RS12340_ | -0.166792306 | -0.33975647 | -0.003768803 | 0.353698184 | 0.243860874 | 1.442801792 | 0.046000096 | 0.103305591 | 0.342614873 | 0.1581 | 0.400679792 |
| SL1344_RS12345_ | 0.001378976 | -0.125553361 | 0.084896502 | 0.283888389 | 0.359336183 | 0.782831315 | 0.210526332 | 0.275949858 | 0.562573946 | 0.3981 | 0.641453133 |
| SL1344_RS12350_ | 0.787502867 | 0.687422345 | 0.884452797 | 0.200181166 | 0.122965306 | 1.574039377 | 0.046635671 | 0.085517991 | 0.295737455 | 0.1511 | 0.389301635 |
| SL1344_RS12355_ | 0.251099212 | 0.172380021 | 0.466459413 | 0.27440663 | 0.195055682 | 1.262243365 | 0.11133337 | 0.155565969 | 0.413948007 | 0.2499 | 0.493936151 |
| SL1344_RS12360_ | 0.080372073 | -0.161920989 | 0.291390186 | 0.453681351 | 0.161882879 | 2.754927873 | 0.0020023 | 0.021912174 | 0.158039518 | 0.1022 | 0.333047763 |
| SL1344_RS12365_ | 2.559827788 | 2.417720909 | 2.665413507 | 0.227231969 | 0.143523527 | 1.545139615 | 0.050000088 | 0.079826656 | 0.315805499 | 0.1476 | 0.397191436 |

| | | | | | | | | | | | |
|---|---|---|---|---|---|---|---|---|---|---|---|
| SL1344_RS12370_ | 1.589646873 | 1.459146942 | 1.68316604 | 0.222614986 | 0.111595422 | 1.92696691 | 0.020000227 | 0.053826123 | 0.246944131 | 0.1191 | 0.354738475 |
| SL1344_RS12375_ | 3.184959361 | 3.817895144 | 2.965448696 | -0.856963768 | 0.964739054 | -0.56704631 | 0.33333334 | 0.570338531 | 0.800829526 | 0.7 | 0.872542119 |
| SL1344_RS12380_ | 5.162980389 | 5.594219452 | 4.71481223 | -0.295020715 | 1.039670223 | -0.37599648 | 0.319120594 | 0.484657522 | 0.745876849 | 0.655 | 0.843686468 |
| SL1344_RS12385_ | 1.714158657 | 1.365631754 | 1.902125865 | 0.531161113 | 0.433215954 | 1.2481327 | 0.090606308 | 0.138007041 | 0.421389265 | 0.1834 | 0.450681481 |
| SL1344_RS12390_ | 2.378199738 | 2.320030785 | 2.530343335 | 0.304711284 | 0.273168454 | 1.164723376 | 0.078614311 | 0.125361843 | 0.399537068 | 0.1795 | 0.431272804 |
| SL1344_RS12395_ | 3.747478371 | 3.520696072 | 3.960821283 | 0.439172121 | 0.100422173 | 4.429252262 | 4.68E-05 | 0.009822688 | 0.120197029 | 0.1 | 0.330257709 |
| SL1344_RS12400_ | 4.422865361 | 4.554323274 | 4.354305005 | -0.196781419 | 0.295669111 | -0.270877659 | 0.336442379 | 0.905936743 | 0.966167237 | 0.7042 | 0.874297381 |
| SL1344_RS12405_ | 4.185222806 | 4.134886448 | 4.231377527 | 0.088891223 | 0.828268304 | 0.076935444 | 0.447701534 | 0.877801678 | 0.955405972 | 1 | 1 |
| SL1344_RS12410_ | 4.719701612 | 4.843016042 | 4.605224653 | -0.228325973 | 0.567206838 | -0.20297759 | 0.441333336 | 0.946629875 | 0.981346983 | 1 | 1 |
| SL1344_RS12415_ | 3.617511434 | 3.725867034 | 3.503423302 | -0.038198726 | 0.649543602 | -0.06051281 | 0.462666668 | 0.890526708 | 0.959999732 | 1 | 1 |
| SL1344_RS12420_ | -2.317956741 | -2.374330078 | -2.223208092 | 0.122117419 | 0.458567765 | 0.249191506 | 0.39573618 | 0.626223733 | 0.807582209 | 0.7127 | 0.849795744 |
| SL1344_RS12425_ | -5.938099525 | -5.769619297 | -6.149291972 | -0.34891492 | 1.579468973 | -0.20093035 | 0.407333337 | 0.582738663 | 0.778309685 | 0.703 | 0.839819485 |
| SL1344_RS12430_ | -2.023523105 | -1.94635532 | -2.067346341 | -0.12206478 | 0.439500563 | -0.238801254 | 0.397734847 | 0.617137719 | 0.803805165 | 0.7176 | 0.85427617 |
| SL1344_RS12435_ | -5.230086473 | -5.470968111 | -5.075349065 | 0.439269014 | 1.206848669 | 0.339575873 | 0.354666673 | 0.494198628 | 0.717560471 | 0.6143 | 0.775964502 |
| SL1344_RS12440_ | -2.843670086 | -3.048570108 | -2.586534162 | 0.467244117 | 0.565938041 | 0.778655629 | 0.196000018 | 0.305598885 | 0.562425279 | 0.4053 | 0.627842535 |
| SL1344_RS12445_ | -0.993179356 | -1.082844265 | -0.896052626 | 0.190906453 | 0.323320929 | 0.55367903 | 0.253830791 | 0.413746281 | 0.659103914 | 0.5173 | 0.718282473 |
| SL1344_RS12450_ | 0.662746189 | 0.496062755 | 0.787538695 | 0.292776966 | 0.162501779 | 1.723898772 | 0.018000252 | 0.070748462 | 0.286145946 | 0.1225 | 0.358779607 |
| SL1344_RS12455_ | 0.697237325 | 0.389124065 | 0.930252687 | 0.535764781 | 0.158943 | 3.31651153 | 4.68E-05 | 0.01503928 | 0.136637998 | 0.1002 | 0.330548519 |
| SL1344_RS12465_ | 3.981294778 | 3.69564067 | 4.2374097 | 0.497480508 | 0.109288947 | 4.648451923 | 4.68E-05 | 0.002603583 | 0.066753231 | 0.1 | 0.330257709 |
| SL1344_RS12480_ | -5.553661892 | -6.037260491 | -5.343530614 | 0.518555208 | 1.85004074 | 0.254361932 | 0.392666671 | 0.603788754 | 0.802369326 | 0.7605 | 0.881174039 |
| SL1344_RS12485_ | 3.566600256 | 3.470445702 | 3.682477514 | 0.301786044 | 0.223921286 | 1.399448638 | 0.003332494 | 0.125668562 | 0.404032521 | 0.1013 | 0.332151868 |
| SL1344_RS12490_ | 6.450844093 | 6.357451543 | 6.564854657 | 0.294178516 | 0.128516575 | 2.39989011 | 4.68E-05 | 0.015980022 | 0.157328089 | 0.1 | 0.330257709 |
| SL1344_RS12495_ | 4.890044796 | 4.758947943 | 4.9568546 | 0.130132356 | 0.161644087 | 0.819500709 | 0.201333351 | 0.305843371 | 0.605034525 | 0.3823 | 0.635956032 |
| SL1344_RS12500_ | -2.040202609 | -1.911212879 | -2.197668609 | -0.248468371 | 0.456306821 | -0.490086062 | 0.29380414 | 0.4650828 | 0.696956545 | 0.579 | 0.759316974 |
| SL1344_RS12510_ | -0.636320268 | -0.622441044 | -0.666049606 | -0.022761848 | 0.319754342 | -0.066686063 | 0.475682879 | 0.714805698 | 0.866579599 | 0.8188 | 0.913371476 |
| SL1344_RS12515_ | -1.209136336 | -1.314031165 | -1.092334247 | 0.217615314 | 0.259764801 | 0.809095708 | 0.184543657 | 0.290108094 | 0.534466913 | 0.3816 | 0.598601765 |
| SL1344_RS12520_ | -2.150353415 | -2.227352395 | -2.094492176 | 0.114663828 | 0.461216285 | 0.245112867 | 0.393333337 | 0.619874141 | 0.803066981 | 0.717 | 0.850121572 |
| SL1344_RS12525_ | -1.441117687 | -1.496582966 | -1.40622266 | 0.064852976 | 0.314705002 | 0.197134145 | 0.42066667 | 0.620179291 | 0.804536778 | 0.7221 | 0.856548013 |
| SL1344_RS12555_ | 3.55309543 | 3.288766594 | 3.633763379 | 0.356822628 | 0.192831804 | 1.852794007 | 0.00067292 | 0.051254735 | 0.265513096 | 0.1003 | 0.330700982 |
| SL1344_RS12560_ | 1.778163613 | 1.557557628 | 2.070932234 | 0.498096908 | 0.17949919 | 2.661271843 | 0.00067292 | 0.025855248 | 0.187332258 | 0.1003 | 0.330687574 |
| SL1344_RS12565_ | 0.780270479 | 0.62694868 | 0.93550474 | 0.308415838 | 0.126632394 | 2.350844967 | 0.006667356 | 0.032676067 | 0.192888948 | 0.1057 | 0.337837176 |
| SL1344_RS12570_ | 0.80893961 | 0.586378085 | 0.947005855 | 0.367928698 | 0.254631775 | 1.43453954 | 0.047301892 | 0.086349967 | 0.326155304 | 0.1447 | 0.390370411 |
| SL1344_RS12575_ | 1.441282344 | 1.234187958 | 1.505548843 | 0.28028403 | 0.288232362 | 1.035860826 | 0.136575645 | 0.166387538 | 0.449632671 | 0.2685 | 0.53697566 |
| SL1344_RS12580_ | 0.583773222 | 0.27788335 | 0.894032561 | 0.614126508 | 0.138457861 | 4.437581731 | 4.68E-05 | 0.006056931 | 0.087303476 | 0.1 | 0.330257709 |
| SL1344_RS12585_ | 0.057925769 | -0.030844812 | 0.188572841 | 0.1770744 | 0.27369231 | 0.618705272 | 0.253830791 | 0.406134213 | 0.6598099 | 0.4988 | 0.711900239 |
| SL1344_RS12590_ | -1.626786353 | -1.874981057 | -1.350570358 | 0.523860263 | 0.304958758 | 1.624167865 | 0.03200014 | 0.079084142 | 0.288710622 | 0.1468 | 0.385461282 |
| SL1344_RS12595_ | -0.133318308 | -0.473588645 | 0.169174245 | 0.621140993 | 0.257469963 | 2.37415237 | 0.004001152 | 0.03309078 | 0.203297714 | 0.103 | 0.334540668 |
| SL1344_RS12600_ | -1.974802147 | -2.09052596 | -1.927033163 | 0.115897542 | 0.466566127 | 0.202896139 | 0.422000003 | 0.709307968 | 0.863736205 | 0.7782 | 0.897655863 |
| SL1344_RS12605_ | -1.620213597 | -1.67015292 | -1.563825926 | 0.107767606 | 0.310150519 | 0.315079463 | 0.356000006 | 0.507594601 | 0.721512654 | 0.6186 | 0.778730117 |
| SL1344_RS12610_ | -1.575927907 | -1.519882097 | -1.635595413 | 0.065602681 | 0.585093891 | 0.106031094 | 0.455029982 | 0.730283377 | 0.876570981 | 0.8881 | 0.945350667 |
| SL1344_RS12615_ | -3.433247068 | -3.646731101 | -3.255997038 | 0.436370792 | 0.698235139 | 0.58275783 | 0.235176563 | 0.371520807 | 0.624523074 | 0.4846 | 0.683454451 |
| SL1344_RS12620_ | -2.757594594 | -2.962539762 | -2.486531325 | 0.450881084 | 0.491320667 | 0.857408005 | 0.177333354 | 0.275369775 | 0.533734592 | 0.3669 | 0.595777966 |
| SL1344_RS12630_ | -2.49163276 | -2.654734027 | -2.276202763 | 0.412096465 | 0.625853008 | 0.615246626 | 0.250499679 | 0.383747448 | 0.64062145 | 0.4986 | 0.705231708 |
| SL1344_RS12635_ | -3.763698063 | -3.918112034 | -3.468675279 | 0.39543496 | 0.78716313 | 0.422645597 | 0.325782819 | 0.561018781 | 0.75864458 | 0.6543 | 0.813547475 |
| SL1344_RS12640_ | -2.905910566 | -2.99314258 | -2.841928862 | 0.130266958 | 0.480796282 | 0.251540157 | 0.388000004 | 0.544514299 | 0.748311678 | 0.6531 | 0.804336818 |
| SL1344_RS12645_ | -2.473689012 | -2.49560489 | -2.444941072 | 0.13919261 | 0.59318283 | 0.229922901 | 0.40906063 | 0.624695286 | 0.811356667 | 0.7532 | 0.872771314 |
| SL1344_RS12650_ | -4.693461051 | -4.539139971 | -4.912042416 | -0.370484184 | 1.069904945 | -0.30550015 | 0.364666672 | 0.553450071 | 0.761616887 | 0.6725 | 0.823250019 |
| SL1344_RS12655_ | -3.589155747 | -3.583443264 | -3.594279095 | -0.006639043 | 0.538501095 | -0.011324695 | 0.495333334 | 0.527121482 | 0.731331656 | 0.648 | 0.796678211 |
| SL1344_RS12660_ | -3.940814223 | -4.147728241 | -3.714476844 | 0.433382461 | 0.713227883 | 0.580134796 | 0.253830791 | 0.386404137 | 0.623839916 | 0.4842 | 0.68257087 |
| SL1344_RS12665_ | -5.279516654 | -6.16075955 | -4.845277149 | 1.289812602 | 1.285707452 | 0.894010482 | 0.16066669 | 0.248086882 | 0.513951342 | 0.3375 | 0.570703177 |
| SL1344_RS12670_ | -4.114232509 | -4.332980956 | -3.933145039 | 0.375104832 | 0.849021624 | 0.418495924 | 0.323117929 | 0.498720575 | 0.718102475 | 0.6032 | 0.775117772 |
| SL1344_RS12675_ | -4.880865489 | -4.803753467 | -4.98893896 | -0.176816605 | 0.959097422 | -0.168865822 | 0.421052635 | 0.546594134 | 0.751445969 | 0.6615 | 0.81206694 |
| SL1344_RS12690_ | 4.850179795 | 4.853608592 | 4.848214593 | 0.061886423 | 0.130755708 | 0.475510071 | 0.390666671 | 0.49855452 | 0.749275156 | 0.8562 | 0.931991476 |

| | | | | | | | | | | | |
|---|---|---|---|---|---|---|---|---|---|---|---|
| SL1344_RS12695_ | -0.218011299 | -0.44559505 | -0.068389933 | 0.392576358 | 0.260193516 | 1.428419657 | 0.037333453 | 0.110998628 | 0.36157761 | 0.1466 | 0.387213409 |
| SL1344_RS12700_ | 0.658460841 | 0.553762346 | 0.769589695 | 0.223096385 | 0.171968226 | 1.267308689 | 0.079280533 | 0.136536714 | 0.389537991 | 0.1988 | 0.445315274 |
| SL1344_RS12705_ | 0.500664902 | 0.222781007 | 0.564608448 | 0.31130507 | 0.332368022 | 0.898939313 | 0.206528997 | 0.262396668 | 0.553596603 | 0.3821 | 0.649588475 |
| SL1344_RS12710_ | -0.754785981 | -0.897603309 | -0.66447957 | 0.236504286 | 0.36090566 | 0.656248976 | 0.236509007 | 0.365115604 | 0.62757364 | 0.4619 | 0.688449423 |
| SL1344_RS12715_ | 0.556646411 | 0.471033508 | 0.660557795 | 0.163506143 | 0.233649089 | 0.673323153 | 0.23984012 | 0.36709668 | 0.63238517 | 0.4653 | 0.68777381 |
| SL1344_RS12720_ | -1.411501495 | -1.624879367 | -1.239296438 | 0.386914443 | 0.291738971 | 1.254984984 | 0.080612977 | 0.139794426 | 0.380872577 | 0.208 | 0.447652085 |
| SL1344_RS12725_ | -1.806311774 | -1.74598934 | -1.852042923 | -0.083199334 | 0.405419433 | -0.189165192 | 0.419333336 | 0.639152556 | 0.817834729 | 0.7504 | 0.870949679 |
| SL1344_RS12730_ | -1.084517664 | -1.263275071 | -0.791641595 | 0.467252812 | 0.338015025 | 1.335571823 | 0.067288538 | 0.115597191 | 0.362871896 | 0.19 | 0.439067716 |
| SL1344_RS12735_ | 0.514999544 | 0.491247309 | 0.545368068 | 0.056936751 | 0.132238154 | 0.417168272 | 0.323784151 | 0.456141258 | 0.676417276 | 0.5702 | 0.741354826 |
| SL1344_RS12740_ | 0.40959095 | 0.158718774 | 0.721936814 | 0.64840394 | 0.327769966 | 2.021953323 | 0.001336769 | 0.041011689 | 0.232653123 | 0.1004 | 0.330854151 |
| SL1344_RS12745_ | 2.321046795 | 2.016622031 | 2.591956159 | 0.57266522 | 0.101519343 | 5.655432466 | 4.68E-05 | 0.004054675 | 0.075030014 | 0.1 | 0.330257709 |
| SL1344_RS12750_ | 1.511819217 | 1.233231483 | 1.799297782 | 0.56716829 | 0.128925727 | 4.355324151 | 4.68E-05 | 0.006301869 | 0.092629773 | 0.1 | 0.330257709 |
| SL1344_RS12755_ | 1.442448245 | 1.409291514 | 1.456965494 | 0.033274472 | 0.166178192 | 0.151913501 | 0.442000002 | 0.751135227 | 0.889061699 | 0.8107 | 0.916243598 |
| SL1344_RS12760_ | 0.859797321 | 0.847380527 | 0.868521319 | 0.024015905 | 0.138940033 | 0.156006561 | 0.427048637 | 0.589519634 | 0.783066152 | 0.7023 | 0.838771389 |
| SL1344_RS12765_ | 1.239887936 | 1.216306343 | 1.272564466 | 0.036886192 | 0.165723703 | 0.210813625 | 0.416000003 | 0.695534848 | 0.855328409 | 0.7712 | 0.889247443 |
| SL1344_RS12770_ | 5.037435195 | 4.883477338 | 5.155274595 | 0.268307652 | 0.067484888 | 3.903155069 | 4.68E-05 | 0.004980476 | 0.088553533 | 0.1 | 0.330257709 |
| SL1344_RS12775_ | 3.96594389 | 3.92209996 | 4.13309546 | 0.213321623 | 0.098276653 | 1.739576387 | 0.017333596 | 0.101727797 | 0.36288932 | 0.1145 | 0.350043926 |
| SL1344_RS12780_ | 2.331834533 | 2.274381585 | 2.517040991 | 0.219930429 | 0.139604277 | 1.292888078 | 0.111259197 | 0.143393701 | 0.415324514 | 0.2277 | 0.48374489 |
| SL1344_RS12785_ | 3.961813836 | 3.817057607 | 4.181351045 | 0.333521241 | 0.210160341 | 1.702338354 | 0.020653117 | 0.053204819 | 0.270776369 | 0.1175 | 0.355801018 |
| SL1344_RS12790_ | -2.483260997 | -2.443890221 | -2.525713165 | -0.076041452 | 0.391722815 | -0.18380798 | 0.408000004 | 0.523891362 | 0.73291323 | 0.633 | 0.789659288 |
| SL1344_RS12795_ | 0.792540769 | 0.931705162 | 0.678815922 | -0.279915852 | 0.211117356 | -1.263314565 | 0.074616979 | 0.124603391 | 0.382430659 | 0.1859 | 0.435646489 |
| SL1344_RS12800_ | 1.768042043 | 1.694367664 | 1.791360853 | 0.090904105 | 0.146673893 | 0.530608897 | 0.314457037 | 0.480143218 | 0.722114307 | 0.6442 | 0.815658888 |
| SL1344_RS12805_ | 1.650579019 | 1.524188941 | 1.743526563 | 0.190340308 | 0.179502043 | 1.020221951 | 0.140666694 | 0.201778472 | 0.479345854 | 0.2775 | 0.53932345 |
| SL1344_RS12810_ | 2.25486718 | 2.354628915 | 2.227698088 | -0.114657591 | 0.176090132 | -0.499042342 | 0.334000007 | 0.602784101 | 0.808930568 | 0.7055 | 0.869179249 |
| SL1344_RS12815_ | 1.187772738 | 0.886469839 | 1.406397324 | 0.538134569 | 0.196754275 | 2.718136805 | 4.68E-05 | 0.04616064 | 0.246941499 | 0.1 | 0.330257709 |
| SL1344_RS12820_ | 2.776785768 | 2.654413328 | 2.808702195 | 0.150543605 | 0.189417014 | 0.652584161 | 0.316000008 | 0.439099897 | 0.707985064 | 0.647 | 0.834745712 |
| SL1344_RS12825_ | 1.377578693 | 1.447004688 | 1.297785354 | -0.1611588 | 0.178202053 | -0.910401672 | 0.165223207 | 0.243652782 | 0.522958214 | 0.3356 | 0.582444635 |
| SL1344_RS12830_ | 1.294127791 | 1.343793692 | 1.249654224 | -0.091180143 | 0.14021822 | -0.63671186 | 0.23717523 | 0.363869675 | 0.616859733 | 0.4746 | 0.682547306 |
| SL1344_RS12835_ | 2.748288693 | 2.371577489 | 2.97122723 | 0.578154233 | 0.142352213 | 3.600339444 | 4.68E-05 | 0.017757701 | 0.159196768 | 0.1 | 0.330257709 |
| SL1344_RS12840_ | 2.658828833 | 2.354899831 | 3.005450372 | 0.639919123 | 0.195653038 | 3.299727492 | 4.68E-05 | 0.009385493 | 0.119600754 | 0.1 | 0.330257709 |
| SL1344_RS12845_ | -2.090816907 | -2.086482315 | -2.093332739 | -0.042386534 | 0.434640835 | -0.092739224 | 0.457694872 | 0.637784893 | 0.816213961 | 0.7452 | 0.867117352 |
| SL1344_RS12850_ | -3.024597553 | -3.114006259 | -2.960893513 | 0.188089231 | 0.535735153 | 0.296780835 | 0.356000006 | 0.490649095 | 0.711967214 | 0.607 | 0.769888268 |
| SL1344_RS12855_ | 1.852667517 | 1.940655794 | 1.809000006 | -0.127788005 | 0.29741403 | -0.323745483 | 0.36642239 | 0.860076569 | 0.945267203 | 0.7822 | 0.907384099 |
| SL1344_RS12860_ | -2.726969517 | -2.783083503 | -2.678951088 | 0.142669361 | 0.56399521 | 0.257976148 | 0.390666671 | 0.572443065 | 0.774094979 | 0.69 | 0.830183079 |
| SL1344_RS12865_ | -3.598249664 | -3.821532206 | -3.384704124 | 0.451273022 | 0.573243066 | 0.750740249 | 0.205196552 | 0.317447633 | 0.550931346 | 0.4116 | 0.61781528 |
| SL1344_RS12870_ | 3.212383729 | 2.853837429 | 3.289441838 | 0.379997593 | 0.356265581 | 1.133905465 | 0.157894761 | 0.162557903 | 0.454674373 | 0.2946 | 0.572942027 |
| SL1344_RS12875_ | 1.348303265 | 1.59963745 | 1.220926869 | -0.327753382 | 0.210388009 | -1.483178374 | 0.072618313 | 0.093020425 | 0.338511081 | 0.1687 | 0.423392262 |
| SL1344_RS12880_ | 1.514908296 | 1.490246713 | 1.533421337 | 0.029459483 | 0.288123548 | 0.106911688 | 0.446666669 | 0.789241197 | 0.910067915 | 0.853 | 0.931353553 |
| SL1344_RS12885_ | 0.586028358 | 0.578894849 | 0.59245777 | 0.016117156 | 0.137494295 | 0.104156689 | 0.452000002 | 0.568509321 | 0.764118556 | 0.6705 | 0.816710306 |
| SL1344_RS12890_ | 2.345608489 | 2.3118905 | 2.368023989 | 0.054915601 | 0.117089815 | 0.436133027 | 0.330000007 | 0.514617175 | 0.738509002 | 0.639 | 0.802053396 |
| SL1344_RS12900_ | -1.82221093 | -1.732569412 | -1.911884005 | -0.169673422 | 0.385033234 | -0.417830264 | 0.319786816 | 0.490146579 | 0.711926309 | 0.595 | 0.767109927 |
| SL1344_RS12905_ | -2.563973481 | -2.402005871 | -2.692236881 | -0.317878858 | 0.515753316 | -0.588001959 | 0.257828126 | 0.403813703 | 0.645151712 | 0.5269 | 0.713680466 |
| SL1344_RS12910_ | -0.247080126 | -0.526585794 | -0.169748006 | 0.352907985 | 0.315448477 | 0.960845218 | 0.165223207 | 0.230209168 | 0.519014109 | 0.341 | 0.583827208 |
| SL1344_RS12915_ | 2.229017906 | 2.125053927 | 2.35561389 | 0.243729389 | 0.147232588 | 1.679210972 | 0.010000458 | 0.072967177 | 0.301657408 | 0.1106 | 0.344702064 |
| SL1344_RS12920_ | 3.048064491 | 3.004608087 | 3.0671132 | 0.046845756 | 0.116001529 | 0.323155035 | 0.369333339 | 0.766674584 | 0.898006186 | 0.7696 | 0.899809923 |
| SL1344_RS12925_ | 2.435095609 | 2.371070606 | 2.474977566 | 0.149848819 | 0.166688109 | 0.900912682 | 0.165223207 | 0.213048904 | 0.501666516 | 0.3178 | 0.569128817 |
| SL1344_RS12930_ | 3.123998875 | 3.09587511 | 3.240205255 | 0.126078648 | 0.106411228 | 1.000523684 | 0.189207214 | 0.242652042 | 0.529907326 | 0.3819 | 0.631288054 |
| SL1344_RS12935_ | 3.303006275 | 3.197381705 | 3.322917869 | 0.128613169 | 0.240377868 | 0.516337354 | 0.335776156 | 0.540382762 | 0.779409108 | 0.7024 | 0.873084472 |
| SL1344_RS12940_ | 2.576729173 | 2.586395442 | 2.565462639 | -0.029553037 | 0.091363419 | -0.304467484 | 0.367754835 | 0.53886288 | 0.75029157 | 0.6845 | 0.823275561 |
| SL1344_RS12945_ | 2.530616157 | 2.470677301 | 2.594707643 | 0.093937225 | 0.254688117 | 0.356793809 | 0.338441046 | 0.61141991 | 0.818951815 | 0.7421 | 0.882593634 |
| SL1344_RS12950_ | 4.58169153 | 4.5824126 | 4.580821587 | -0.172186848 | 0.372669391 | -0.459220266 | 0.362425055 | 0.53363064 | 0.776723384 | 0.7927 | 0.899885509 |
| SL1344_RS12955_ | -2.689889944 | -2.823197914 | -2.576462814 | 0.256496069 | 0.471201556 | 0.504731636 | 0.274483688 | 0.425971151 | 0.657756938 | 0.5377 | 0.719803095 |
| SL1344_RS12960_ | -2.158899093 | -2.296432806 | -1.97279812 | 0.326996989 | 0.395418548 | 0.756101667 | 0.200532995 | 0.319521936 | 0.571524602 | 0.4193 | 0.637473806 |

| | | | | | | | | | | | |
|---|---|---|---|---|---|---|---|---|---|---|---|
| SL1344_RS12965_ | -2.980453494 | -2.873033557 | -3.254638873 | -0.401869734 | 0.79564093 | -0.469323883 | 0.296666676 | 0.442689196 | 0.697179243 | 0.6042 | 0.781053317 |
| SL1344_RS12970_ | -1.053125155 | -1.030944571 | -1.079361062 | -0.015471509 | 0.377552763 | -0.039648332 | 0.482345104 | 0.717162258 | 0.868749868 | 0.804 | 0.904642291 |
| SL1344_RS12975_ | 1.287290424 | 1.236466541 | 1.317289281 | 0.052255525 | 0.246612185 | 0.170410021 | 0.4170553 | 0.816488866 | 0.923973003 | 0.8071 | 0.916710774 |
| SL1344_RS12980_ | 3.50505973 | 3.326447879 | 3.548342567 | 0.176904652 | 0.14990453 | 0.916187377 | 0.224517003 | 0.271425664 | 0.569877547 | 0.4223 | 0.677042882 |
| SL1344_RS12985_ | 2.232626877 | 2.196558525 | 2.254111729 | 0.058079873 | 0.093663958 | 0.599381648 | 0.260666678 | 0.387616672 | 0.637000946 | 0.5057 | 0.703403072 |
| SL1344_RS12990_ | -1.269814247 | -1.496688773 | -1.188107673 | 0.250969527 | 0.511394098 | 0.28966204 | 0.365333339 | 0.853640702 | 0.941778265 | 0.7582 | 0.8970765 |
| SL1344_RS12995_ | 1.634009338 | 1.75312972 | 1.5911692 | -0.187778716 | 0.298878983 | -0.652487028 | 0.217333349 | 0.310768986 | 0.603959855 | 0.4032 | 0.667832391 |
| SL1344_RS13000_ | 2.377275856 | 2.503585639 | 2.242874655 | -0.248778781 | 0.102503584 | -2.408685669 | 0.004664544 | 0.030580854 | 0.19636201 | 0.1045 | 0.336728781 |
| SL1344_RS13005_ | 2.247297783 | 2.112223006 | 2.425688601 | 0.293449967 | 0.236134428 | 1.161185901 | 0.114590308 | 0.170228544 | 0.460429936 | 0.2056 | 0.478365423 |
| SL1344_RS13010_ | 2.563381106 | 2.466503932 | 2.617370685 | 0.169155995 | 0.137250598 | 1.234354016 | 0.077948089 | 0.13545301 | 0.404760342 | 0.19 | 0.442589462 |
| SL1344_RS13015_ | 3.344909454 | 3.307473566 | 3.372466157 | 0.068557763 | 0.070263151 | 0.941106472 | 0.148000026 | 0.222746389 | 0.489879303 | 0.3091 | 0.546524543 |
| SL1344_RS13025_ | 3.578938286 | 3.609246851 | 3.546990918 | -0.235468663 | 0.24631342 | -0.955475384 | 0.129913421 | 0.197714963 | 0.494554581 | 0.2866 | 0.526208143 |
| SL1344_RS13030_ | 2.84044303 | 2.828156795 | 2.848679587 | 0.001841326 | 0.273975035 | 0.00649074 | 0.496666667 | 0.85452221 | 0.943162146 | 0.979 | 0.990998645 |
| SL1344_RS13035_ | 4.163167702 | 4.333390801 | 4.110882582 | -0.20786433 | 0.079951424 | -1.788555587 | 0.019333568 | 0.076095838 | 0.315477524 | 0.1172 | 0.35481573 |
| SL1344_RS13040_ | 2.711538013 | 2.854908216 | 2.592072093 | -0.24999009 | 0.11249064 | -2.120894335 | 0.008000574 | 0.034933452 | 0.212261532 | 0.1051 | 0.337646766 |
| SL1344_RS13045_ | 2.055761235 | 1.953340576 | 2.262935027 | 0.291705228 | 0.116648285 | 2.32157107 | 0.004667653 | 0.044374801 | 0.235995603 | 0.106 | 0.338889716 |
| SL1344_RS13050_ | 2.755435375 | 2.775342116 | 2.734388604 | -0.044865995 | 0.527505976 | -0.08548906 | 0.466000001 | 0.942797052 | 0.979553955 | 1 | 1 |
| SL1344_RS13075_ | -0.183143348 | -0.13854984 | -0.214291013 | -0.069994821 | 0.35216605 | -0.188588581 | 0.402398405 | 0.707371608 | 0.863915425 | 0.7562 | 0.881021198 |
| SL1344_RS13080_ | -1.01205434 | -0.984634931 | -1.045268038 | -0.078002628 | 0.289103137 | -0.246897133 | 0.396666671 | 0.592401371 | 0.786814659 | 0.7106 | 0.843539988 |
| SL1344_RS13085_ | 0.027043031 | -0.677773236 | 0.65075374 | 1.237902794 | 0.374759962 | 3.313453025 | 4.68E-05 | 0.01004796 | 0.120203143 | 0.1 | 0.330257709 |
| SL1344_RS13090_ | 1.357301533 | 1.411063128 | 1.30088687 | -0.050761473 | 0.19880684 | -0.258184063 | 0.391072622 | 0.662129724 | 0.835431301 | 0.7752 | 0.883564234 |
| SL1344_RS13095_ | 5.227300906 | 5.142405274 | 5.360720576 | 0.221068959 | 0.059406179 | 3.656225147 | 4.68E-05 | 0.014424185 | 0.143823908 | 0.1 | 0.330257709 |
| SL1344_RS13100_ | 0.844408204 | 0.885378689 | 0.821220208 | -0.060010878 | 0.178503724 | -0.315623887 | 0.372000005 | 0.598248528 | 0.791244046 | 0.7026 | 0.84418662 |
| SL1344_RS13105_ | 1.467793958 | 1.355949414 | 1.56574424 | 0.212693672 | 0.108264303 | 1.918815226 | 0.011992386 | 0.055457905 | 0.246028616 | 0.1164 | 0.3502422 |
| SL1344_RS13115_ | 2.243641139 | 2.23469027 | 2.257878667 | 0.029185049 | 0.280324377 | 0.118594091 | 0.423717525 | 0.813281967 | 0.922667713 | 0.8212 | 0.915109214 |
| SL1344_RS13120_ | -0.404027925 | -0.546016631 | -0.240658475 | 0.346219743 | 0.296181983 | 1.153073075 | 0.092666712 | 0.151172183 | 0.414049465 | 0.2293 | 0.475618379 |
| SL1344_RS13125_ | 2.574787963 | 2.376444312 | 2.601340989 | 0.224243701 | 0.188453515 | 1.006501066 | 0.198666684 | 0.23539867 | 0.533390673 | 0.398 | 0.634169216 |
| SL1344_RS13130_ | 2.282612835 | 2.179658796 | 2.397523709 | 0.202208378 | 0.136767667 | 1.43420949 | 0.062666736 | 0.099966862 | 0.347314195 | 0.1674 | 0.420381512 |
| SL1344_RS13135_ | 1.033428387 | 1.029135946 | 1.039603564 | 0.014062092 | 0.131171607 | 0.101466772 | 0.463333335 | 0.615074516 | 0.798097525 | 0.719 | 0.849274835 |
| SL1344_RS13140_ | 1.351092635 | 1.461479982 | 1.296015447 | -0.149445764 | 0.168652791 | -0.830223871 | 0.196000018 | 0.290434603 | 0.56543064 | 0.3937 | 0.638459066 |
| SL1344_RS13145_ | 1.895103849 | 1.968985943 | 1.800708461 | -0.15032714 | 0.1585598 | -0.935221502 | 0.156000024 | 0.230674539 | 0.514085732 | 0.3157 | 0.575845027 |
| SL1344_RS13150_ | -1.58806825 | -1.634833939 | -1.548734042 | -0.121959287 | 0.688576211 | -0.167243082 | 0.430379749 | 0.712995532 | 0.869106332 | 0.8882 | 0.947257382 |
| SL1344_RS13155_ | 1.130587697 | 1.110743713 | 1.147091204 | 0.026025112 | 0.174052638 | 0.134225069 | 0.436375752 | 0.690193004 | 0.850033423 | 0.7974 | 0.898429628 |
| SL1344_RS13160_ | 0.938767753 | 0.687274154 | 0.993231345 | 0.302353203 | 0.301702177 | 0.847558084 | 0.221333348 | 0.290274917 | 0.585594863 | 0.442 | 0.689272965 |
| SL1344_RS13165_ | 1.361545578 | 1.218318047 | 1.459420105 | 0.286540873 | 0.232178753 | 1.241681411 | 0.034666795 | 0.159003719 | 0.444112619 | 0.1396 | 0.380518481 |
| SL1344_RS13170_ | 0.998918072 | 0.83688053 | 1.124503885 | 0.486974972 | 0.449017894 | 1.166433529 | 0.034000131 | 0.13492942 | 0.413904379 | 0.1271 | 0.366722101 |
| SL1344_RS13175_ | 0.743075141 | 0.270402705 | 0.831537508 | 0.581233406 | 0.439177948 | 1.192479498 | 0.085942754 | 0.169039446 | 0.460939531 | 0.2048 | 0.460585715 |
| SL1344_RS13180_ | -0.474887978 | -0.514646294 | -0.404922257 | 0.154875022 | 0.384323585 | 0.38067387 | 0.321119261 | 0.448739978 | 0.705997075 | 0.6406 | 0.7986262 |
| SL1344_RS13185_ | -0.381788321 | -0.371644518 | -0.39416507 | -0.026404173 | 0.201886596 | -0.113101259 | 0.447035312 | 0.587319687 | 0.781397112 | 0.6958 | 0.834774288 |
| SL1344_RS13190_ | -4.121698779 | -4.371181672 | -3.845193332 | 0.434281694 | 1.367929991 | 0.30104782 | 0.365089945 | 0.639227256 | 0.820491936 | 0.7401 | 0.866745476 |
| SL1344_RS13195_ | -1.618744457 | -1.897899012 | -1.347080333 | 0.575116222 | 0.403652496 | 1.370853858 | 0.053333415 | 0.109460171 | 0.352549564 | 0.1757 | 0.421562522 |
| SL1344_RS13200_ | -1.222134568 | -1.178099688 | -1.28080609 | -0.089177899 | 0.297939433 | -0.284088214 | 0.378666671 | 0.562911657 | 0.765001802 | 0.6794 | 0.824288795 |
| SL1344_RS13210_ | 1.01309699 | 0.917742549 | 1.132194216 | 0.205227638 | 0.193618234 | 1.0372528 | 0.133910755 | 0.188778529 | 0.462185112 | 0.267 | 0.523741848 |
| SL1344_RS13215_ | 2.002882458 | 1.697544704 | 2.058409517 | 0.363447184 | 0.223201788 | 1.21134513 | 0.125249865 | 0.174041661 | 0.465618237 | 0.2588 | 0.517219309 |
| SL1344_RS13220_ | 2.801521251 | 2.60512228 | 2.919853723 | 0.341719642 | 0.151673524 | 2.283394856 | 0.00067292 | 0.030269427 | 0.203301707 | 0.1005 | 0.330901515 |
| SL1344_RS13225_ | 2.542991231 | 2.368001905 | 2.706330599 | 0.344513344 | 0.096960269 | 3.560178074 | 4.68E-05 | 0.010769757 | 0.120004776 | 0.1 | 0.330257709 |
| SL1344_RS13230_ | 3.817964823 | 3.617636955 | 3.835395993 | 0.213240936 | 0.104514476 | 1.037818901 | 0.20186544 | 0.213420668 | 0.512629621 | 0.4018 | 0.640576571 |
| SL1344_RS13235_ | 3.732883384 | 3.598768917 | 3.893418554 | 0.284235315 | 0.081436964 | 3.449701585 | 4.68E-05 | 0.008632523 | 0.111510449 | 0.1 | 0.330257709 |
| SL1344_RS13240_ | 1.09554727 | 0.997510245 | 1.162717078 | 0.162705748 | 0.121421327 | 1.215744223 | 0.100000041 | 0.152791239 | 0.39729054 | 0.2328 | 0.474356488 |
| SL1344_RS13245_ | -0.537069125 | -0.660530231 | -0.386108993 | 0.238596797 | 0.301435479 | 0.730519902 | 0.218521001 | 0.322867926 | 0.58898171 | 0.4233 | 0.653934603 |
| SL1344_RS13250_ | -5.427472669 | -4.990624694 | -5.887060904 | -0.923595992 | 1.250632525 | -0.699676796 | 0.20186544 | 0.313566174 | 0.565042076 | 0.4224 | 0.632062705 |
| SL1344_RS13255_ | -4.533532601 | -4.280979425 | -5.296650515 | -0.910687835 | 1.356668594 | -0.571292923 | 0.257333345 | 0.374624518 | 0.644346418 | 0.5289 | 0.732545046 |
| SL1344_RS13260_ | -4.045030898 | -3.791402706 | -4.378921573 | -0.59860019 | 0.691105598 | -0.807843082 | 0.190666685 | 0.289276849 | 0.534094833 | 0.3825 | 0.601084962 |

| | | | | | | | | | | | |
|---|---|---|---|---|---|---|---|---|---|---|---|
| SL1344_RS13265_ | -2.043316915 | -1.998824204 | -2.088365455 | -0.112500764 | 0.424157346 | -0.246727819 | 0.396666671 | 0.585102101 | 0.779597649 | 0.6953 | 0.835316012 |
| SL1344_RS13270_ | -4.03923588 | -3.660991305 | -4.50089457 | -0.832055088 | 0.702598869 | -1.143077437 | 0.096666709 | 0.172986375 | 0.415577085 | 0.2469 | 0.481615503 |
| SL1344_RS13275_ | -1.748584318 | -1.905285183 | -1.630283564 | 0.251371508 | 0.317462511 | 0.76280051 | 0.200000017 | 0.310939312 | 0.555207364 | 0.4114 | 0.62327398 |
| SL1344_RS13280_ | -2.1756825 | -2.14078759 | -2.224684435 | -0.049410233 | 0.480511447 | -0.092698548 | 0.460000001 | 0.663385704 | 0.833104226 | 0.7656 | 0.880605348 |
| SL1344_RS13285_ | -2.776225671 | -2.740204866 | -2.819402579 | -0.085816862 | 0.443371638 | -0.17342296 | 0.42266667 | 0.544878208 | 0.74746088 | 0.6535 | 0.80411832 |
| SL1344_RS13290_ | -2.177282078 | -2.354633792 | -2.009307375 | 0.349030966 | 0.346474733 | 0.979323282 | 0.143904091 | 0.222901946 | 0.468722788 | 0.3045 | 0.533889122 |
| SL1344_RS13295_ | -4.158114349 | -4.039547709 | -4.267822144 | -0.230909975 | 0.794713934 | -0.271085169 | 0.369333339 | 0.552158547 | 0.750072875 | 0.6609 | 0.807697189 |
| SL1344_RS13300_ | -0.351395191 | -0.389469107 | -0.31110757 | 0.080361811 | 0.199712719 | 0.371390976 | 0.329113931 | 0.492294684 | 0.709349494 | 0.6142 | 0.776455395 |
| SL1344_RS13305_ | -4.316533562 | -4.45836524 | -4.132548637 | 0.312197686 | 0.755939681 | 0.373668734 | 0.328666674 | 0.480051861 | 0.694633077 | 0.5744 | 0.74764595 |
| SL1344_RS13310_ | -4.891805338 | -4.789898538 | -4.97654885 | -0.186608906 | 1.102131299 | -0.143969175 | 0.434377084 | 0.623710413 | 0.806641646 | 0.7265 | 0.857064533 |
| SL1344_RS13315_ | -0.852945081 | -0.883265816 | -0.810978983 | 0.068765607 | 0.280463581 | 0.218672698 | 0.40466667 | 0.606294769 | 0.794015985 | 0.7151 | 0.847923179 |
| SL1344_RS13320_ | -2.89280345 | -2.926583762 | -2.85797752 | 0.082130367 | 0.46606263 | 0.174258722 | 0.428000003 | 0.554472509 | 0.754257627 | 0.6674 | 0.811675337 |
| SL1344_RS13325_ | 4.489652354 | 4.453728024 | 4.521643667 | 0.060385942 | 0.305688782 | 0.171859331 | 0.443037977 | 0.845900104 | 0.941315901 | 0.9949 | 0.997826279 |
| SL1344_RS13330_ | -2.768087473 | -2.576155332 | -2.982988412 | -0.365991125 | 0.582296704 | -0.613754314 | 0.251832124 | 0.398615874 | 0.64610907 | 0.4982 | 0.702721283 |
| SL1344_RS13335_ | -3.368829876 | -3.35678389 | -3.386463768 | -0.035039423 | 0.559888249 | -0.058749267 | 0.470000001 | 0.568708744 | 0.762880727 | 0.6816 | 0.825087977 |
| SL1344_RS13340_ | -3.726134505 | -3.437364105 | -3.884631514 | -0.366733902 | 0.868462344 | -0.382358326 | 0.339773491 | 0.578741142 | 0.775709526 | 0.6598 | 0.81952229 |
| SL1344_RS13345_ | -2.188623194 | -2.107903095 | -2.310035284 | -0.210850145 | 0.50821925 | -0.415730017 | 0.33533334 | 0.521031635 | 0.738146629 | 0.6435 | 0.798637419 |
| SL1344_RS13350_ | -3.00990498 | -3.024823367 | -2.991010717 | 0.004933482 | 0.614005758 | 0.007949451 | 0.495333334 | 0.658250895 | 0.830418982 | 0.7643 | 0.879416337 |
| SL1344_RS13355_ | -1.951111912 | -2.047783206 | -1.844926877 | 0.189641967 | 0.425641974 | 0.428672707 | 0.319333341 | 0.499271155 | 0.72126966 | 0.6133 | 0.782168176 |
| SL1344_RS13360_ | -1.448725268 | -1.509490888 | -1.390220576 | 0.108396284 | 0.307091406 | 0.325005495 | 0.37175217 | 0.542401002 | 0.7472564 | 0.6489 | 0.801383644 |
| SL1344_RS13365_ | -3.857885885 | -3.93309715 | -3.741799578 | 0.111487922 | 0.901554906 | 0.11397566 | 0.442371754 | 0.664950582 | 0.834987618 | 0.7695 | 0.883152971 |
| SL1344_RS13370_ | -1.902446465 | -1.977791944 | -1.84593514 | 0.135277809 | 0.3632538 | 0.341857594 | 0.346435716 | 0.501914669 | 0.716886524 | 0.6154 | 0.77559516 |
| SL1344_RS13375_ | -4.790894137 | -4.845559082 | -4.756903483 | 0.406211588 | 1.724650142 | 0.206443088 | 0.395333337 | 0.584563422 | 0.793546016 | 0.7933 | 0.893306319 |
| SL1344_RS13380_ | -1.708812921 | -1.983453042 | -1.438506613 | 0.558125241 | 0.365946148 | 1.415027358 | 0.056628991 | 0.104374359 | 0.341301702 | 0.1656 | 0.409789036 |
| SL1344_RS13385_ | -3.381364632 | -3.482342749 | -3.303830391 | 0.175238824 | 0.600900457 | 0.26831276 | 0.383333338 | 0.540820881 | 0.746255423 | 0.6518 | 0.806081563 |
| SL1344_RS13390_ | -3.312069595 | -3.185062831 | -3.472803691 | -0.122088522 | 1.020887139 | -0.116487441 | 0.458361094 | 0.750042968 | 0.889045694 | 0.8363 | 0.923553979 |
| SL1344_RS13395_ | -6.643709221 | -7.634650243 | -5.914794819 | 1.719785873 | 2.115543089 | 0.738495951 | 0.153897426 | 0.265776458 | 0.532756955 | 0.331 | 0.561479065 |
| SL1344_RS13400_ | 0.199144309 | 0.198053708 | 0.199787058 | 0.014090947 | 0.26012567 | 0.041055161 | 0.479680214 | 0.704622858 | 0.862151951 | 0.8205 | 0.91558911 |
| SL1344_RS13405_ | -2.997226152 | -3.02152148 | -2.966018749 | 0.051338278 | 0.440330801 | 0.108957244 | 0.458361094 | 0.54864029 | 0.74793634 | 0.6573 | 0.803793325 |
| SL1344_RS13410_ | -2.514512497 | -2.718365713 | -2.294461606 | 0.394298551 | 0.530006555 | 0.727684784 | 0.209193887 | 0.329409077 | 0.591157802 | 0.4301 | 0.654063344 |
| SL1344_RS13415_ | -0.519877099 | -0.461995225 | -0.592473235 | -0.146188003 | 0.281901496 | -0.480356894 | 0.290000009 | 0.452073103 | 0.691703979 | 0.5854 | 0.762483264 |
| SL1344_RS13420_ | -3.537505662 | -3.425081457 | -3.633886467 | -0.198291292 | 0.674322707 | -0.277483677 | 0.373084615 | 0.555260902 | 0.754095789 | 0.6548 | 0.803792263 |
| SL1344_RS13425_ | -7.080587263 | -7.493384559 | -6.79442539 | 0.835457809 | 2.690090374 | 0.246348624 | 0.377748173 | 0.531239921 | 0.749506669 | 0.6657 | 0.815842012 |
| SL1344_RS13430_ | -4.748795582 | -5.054550336 | -4.064099516 | 0.910279054 | 1.140912893 | 0.698700811 | 0.241172565 | 0.382874178 | 0.631219659 | 0.4866 | 0.699779066 |
| SL1344_RS13435_ | -6.549930359 | -6.352099136 | -6.735860026 | -0.313126818 | 2.130500779 | -0.134800203 | 0.431712194 | 0.596930384 | 0.792290616 | 0.7234 | 0.854432535 |
| SL1344_RS13440_ | 0.597186758 | 0.651092128 | 0.532473588 | 0.015717607 | 0.408372745 | 0.036057103 | 0.487674884 | 0.829666522 | 0.929912715 | 0.9547 | 0.98025546 |
| SL1344_RS13450_ | -0.522334213 | -0.466873282 | -0.554101399 | -0.048615976 | 0.298236988 | -0.137545653 | 0.432000003 | 0.71342785 | 0.865641804 | 0.7824 | 0.897068782 |
| SL1344_RS13455_ | 0.693083549 | 0.264840406 | 0.997257014 | 0.720000102 | 0.285132533 | 2.445501855 | 4.68E-05 | 0.038116214 | 0.224939835 | 0.1004 | 0.330833785 |
| SL1344_RS13460_ | -2.916038568 | -2.879208956 | -2.949420955 | -0.067572149 | 0.453003937 | -0.131160567 | 0.436666669 | 0.519061605 | 0.728817235 | 0.6267 | 0.786513762 |
| SL1344_RS13465_ | -1.882185755 | -1.842097782 | -1.943206314 | -0.109711571 | 0.386562504 | -0.259429869 | 0.38774151 | 0.551189976 | 0.757048467 | 0.6589 | 0.809632798 |
| SL1344_RS13470_ | -3.389517229 | -3.411816506 | -3.363810631 | 0.03733437 | 0.578481941 | 0.05989529 | 0.474000001 | 0.578034323 | 0.77412102 | 0.6866 | 0.828648894 |
| SL1344_RS13475_ | -2.839967223 | -2.943072602 | -2.719767602 | 0.224622669 | 0.453937665 | 0.484236662 | 0.307333342 | 0.440572695 | 0.667034468 | 0.5609 | 0.736256793 |
| SL1344_RS13480_ | -3.601331605 | -3.679960804 | -3.515776629 | 0.180583923 | 0.631592198 | 0.270666109 | 0.384410398 | 0.537645589 | 0.743370046 | 0.6488 | 0.799735678 |
| SL1344_RS13485_ | -2.475817855 | -2.441892963 | -2.532698159 | -0.159191891 | 0.566671169 | -0.238802023 | 0.391333338 | 0.561714046 | 0.77252776 | 0.7196 | 0.849026395 |
| SL1344_RS13490_ | -2.994375908 | -2.941400431 | -3.030239732 | -0.096739634 | 0.536701514 | -0.177129292 | 0.424000003 | 0.592341527 | 0.781837754 | 0.6974 | 0.835253662 |
| SL1344_RS13495_ | -3.97352989 | -3.79876093 | -4.162842993 | -0.357742256 | 0.627404361 | -0.543699798 | 0.266489018 | 0.390547119 | 0.621099107 | 0.4914 | 0.683661927 |
| SL1344_RS13500_ | 2.727304697 | 2.653997892 | 2.779307147 | 0.158699059 | 0.193215387 | 0.795866531 | 0.20733335 | 0.269114128 | 0.564605046 | 0.3791 | 0.637790855 |
| SL1344_RS13505_ | 4.870346419 | 4.784937184 | 5.022512768 | 0.282051289 | 0.265306612 | 1.250123803 | 0.01866691 | 0.159341025 | 0.452406393 | 0.121 | 0.36102638 |
| SL1344_RS13510_ | -4.456179656 | -3.935778595 | -4.698107989 | -0.538385285 | 1.269270233 | -0.366428805 | 0.34533334 | 0.654280403 | 0.827094773 | 0.7047 | 0.853559035 |
| SL1344_RS13515_ | -3.536330261 | -3.344337392 | -3.750663008 | -0.42411361 | 0.526485556 | -0.749584567 | 0.204000017 | 0.30767914 | 0.550349731 | 0.4071 | 0.618596468 |
| SL1344_RS13520_ | -1.455687395 | -1.568444349 | -1.380657378 | 0.185110023 | 0.301008114 | 0.563793742 | 0.257333345 | 0.398060154 | 0.638094445 | 0.5133 | 0.705488223 |
| SL1344_RS13525_ | -3.379666481 | -3.295560194 | -3.551007091 | -0.219153963 | 0.660943613 | -0.309650838 | 0.381333338 | 0.570391926 | 0.770172848 | 0.6754 | 0.824235827 |
| SL1344_RS13530_ | -5.558036719 | -5.236264106 | -5.782952054 | -0.389528039 | 1.776369206 | -0.181976663 | 0.401732182 | 0.663309607 | 0.83212742 | 0.7445 | 0.869685091 |

| | | | | | | | | | | | |
|---|---|---|---|---|---|---|---|---|---|---|---|
| SL1344_RS13535_ | -2.878335228 | -2.812987855 | -2.942033181 | -0.117452256 | 0.477522392 | -0.233523238 | 0.386409065 | 0.532387254 | 0.733894422 | 0.6384 | 0.790177987 |
| SL1344_RS13540_ | -0.050713907 | -0.199291066 | 0.065207443 | 0.266576935 | 0.179365036 | 1.41167363 | 0.068000063 | 0.114094944 | 0.345244117 | 0.1816 | 0.422951919 |
| SL1344_RS13545_ | 1.249189295 | 1.023718409 | 1.291970101 | 0.251986015 | 0.266384118 | 0.57290469 | 0.332666674 | 0.599333371 | 0.812161904 | 0.7012 | 0.872092875 |
| SL1344_RS13550_ | 2.245467003 | 2.247169537 | 2.243076599 | 0.047081489 | 0.216929026 | 0.221163713 | 0.409333337 | 0.738481653 | 0.883941396 | 0.8665 | 0.938314319 |
| SL1344_RS13555_ | 2.326110763 | 2.365775117 | 2.281684857 | -0.077115506 | 0.246800716 | -0.312809995 | 0.346666673 | 0.610401006 | 0.81648233 | 0.7218 | 0.86189229 |
| SL1344_RS13560_ | 2.501627801 | 2.505117483 | 2.497116057 | 0.035577647 | 0.230266722 | 0.163011984 | 0.420386412 | 0.728092544 | 0.879434219 | 0.8593 | 0.932498712 |
| SL1344_RS13565_ | -1.17365671 | -1.047377736 | -1.325475465 | -0.27671062 | 0.305099007 | -0.878901722 | 0.170552986 | 0.251156621 | 0.510558699 | 0.3374 | 0.571103895 |
| SL1344_RS13570_ | 1.475495939 | 1.384349665 | 1.523794224 | 0.138905302 | 0.128302012 | 1.006423384 | 0.134666696 | 0.212591816 | 0.48068173 | 0.3001 | 0.540886281 |
| SL1344_RS13575_ | 3.200074951 | 3.003062092 | 3.367128281 | 0.361146022 | 0.191015329 | 1.836993392 | 0.004664544 | 0.06307769 | 0.291295696 | 0.1056 | 0.338428014 |
| SL1344_RS13580_ | -0.539107816 | -0.458002144 | -0.701490827 | -0.231658561 | 0.298960391 | -0.743522472 | 0.211858777 | 0.319842115 | 0.592648633 | 0.4305 | 0.654838215 |
| SL1344_RS13590_ | 2.049746251 | 2.098225284 | 1.707084554 | -0.361182432 | 0.363596901 | -0.544327606 | 0.33333334 | 0.572159636 | 0.800217603 | 0.6991 | 0.871966946 |
| SL1344_RS13595_ | 1.824058984 | 1.865942514 | 1.782231979 | 0.007325391 | 0.268265046 | 0.021817127 | 0.493333334 | 0.761388222 | 0.896229204 | 0.9526 | 0.978858411 |
| SL1344_RS13605_ | 2.177681513 | 2.027898041 | 2.204117844 | 0.158956004 | 0.149453974 | 0.764462237 | 0.28600001 | 0.384306752 | 0.660649355 | 0.5857 | 0.791156319 |
| SL1344_RS13610_ | 0.724615231 | 0.736893034 | 0.716764084 | 0.004599194 | 0.200229732 | 0.019753074 | 0.489333334 | 0.691113986 | 0.852758347 | 0.7946 | 0.900380159 |
| SL1344_RS13615_ | 2.954585005 | 2.885002238 | 2.983129958 | 0.089248556 | 0.140032367 | 0.678866618 | 0.233177895 | 0.375001729 | 0.651332219 | 0.4491 | 0.701236761 |
| SL1344_RS13625_ | 0.149956076 | 0.124424776 | 0.160015202 | 0.028196571 | 0.239194378 | 0.103666724 | 0.460000001 | 0.71586991 | 0.867667187 | 0.8078 | 0.909951158 |
| SL1344_RS13630_ | 0.318926671 | 0.293586606 | 0.341359934 | 0.058032611 | 0.264077586 | 0.236976707 | 0.40466667 | 0.632367398 | 0.817768892 | 0.745 | 0.868046159 |
| SL1344_RS13635_ | -3.717495946 | -3.929950746 | -3.501412699 | 0.46307274 | 0.70839239 | 0.634045103 | 0.250499679 | 0.373507451 | 0.614962463 | 0.4859 | 0.680650543 |
| SL1344_RS13645_ | 3.733604332 | 3.67498018 | 3.875339189 | 0.202417091 | 0.648079772 | 0.380604352 | 0.333777489 | 0.543289509 | 0.78369086 | 0.7015 | 0.873065728 |
| SL1344_RS13650_ | -0.015262737 | -0.235842019 | 0.322609365 | 0.559380939 | 0.463718266 | 1.186902532 | 0.070666727 | 0.153268679 | 0.436878015 | 0.1759 | 0.432749165 |
| SL1344_RS13655_ | 0.667627214 | -0.09868087 | 0.759690027 | 0.957564017 | 1.295320955 | 0.808236982 | 0.220666682 | 0.201954385 | 0.503003531 | 0.3907 | 0.670822405 |
| SL1344_RS13660_ | 4.605342132 | 4.688650833 | 4.514228947 | -0.189668865 | 0.420044108 | -0.626117285 | 0.304463699 | 0.345155829 | 0.641458621 | 0.632 | 0.827698558 |
| SL1344_RS13665_ | 2.384339873 | 2.346004334 | 2.503864502 | 0.17336681 | 0.199190954 | 0.890791225 | 0.173217876 | 0.252458864 | 0.551017472 | 0.3331 | 0.594536776 |
| SL1344_RS13670_ | 2.805011032 | 2.722223804 | 2.891408362 | 0.179838242 | 0.093921182 | 1.931916252 | 0.004667653 | 0.056046578 | 0.261985513 | 0.1049 | 0.337195464 |
| SL1344_RS13675_ | 3.07953401 | 2.955262254 | 3.234058713 | 0.300576587 | 0.138538673 | 2.1979693 | 4.68E-05 | 0.05128348 | 0.262248367 | 0.1 | 0.330257709 |
| SL1344_RS13680_ | 4.986627428 | 4.743305703 | 5.170163107 | 0.227683561 | 0.419664973 | 0.756535689 | 0.166666689 | 0.333219993 | 0.631675003 | 0.2996 | 0.576359256 |
| SL1344_RS13685_ | 2.344970365 | 2.290597688 | 2.467733027 | 0.191459123 | 0.173649161 | 1.10889275 | 0.122000033 | 0.153918145 | 0.431507319 | 0.2416 | 0.504725383 |
| SL1344_RS13705_ | -1.557856191 | -1.865653603 | -1.162672416 | 0.718038737 | 0.38219516 | 1.795106042 | 0.017333596 | 0.062855775 | 0.272261147 | 0.1181 | 0.354095743 |
| SL1344_RS13710_ | -0.331072787 | -0.599488539 | -0.001428909 | 0.595285904 | 0.222748874 | 2.653982472 | 0.000673369 | 0.023526312 | 0.16992838 | 0.1007 | 0.33129397 |
| SL1344_RS13715_ | 2.752626728 | 2.624610819 | 2.798705483 | 0.147681296 | 0.132383237 | 1.010716317 | 0.154563649 | 0.209838448 | 0.492638708 | 0.3025 | 0.563411385 |
| SL1344_RS13720_ | 2.765127491 | 2.789173215 | 2.745206113 | 0.022083294 | 0.23542667 | 0.08448681 | 0.465333335 | 0.647693063 | 0.842238183 | 0.9658 | 0.984553618 |
| SL1344_RS13725_ | 2.06710151 | 2.081102677 | 2.056491999 | -0.00700266 | 0.194926138 | -0.036572067 | 0.488 | 0.809070075 | 0.920280217 | 0.9236 | 0.96573903 |
| SL1344_RS13730_ | -6.240974306 | -6.0801564 | -6.414194835 | -0.294106783 | 1.382669361 | -0.193908807 | 0.411059297 | 0.521283254 | 0.728990178 | 0.6274 | 0.786058966 |
| SL1344_RS13735_ | -5.032908574 | -4.676317813 | -5.523491873 | -0.777142093 | 1.347898255 | -0.545826692 | 0.262491684 | 0.412712013 | 0.661298103 | 0.5161 | 0.715872744 |
| SL1344_RS13745_ | -5.36287474 | -5.440028612 | -5.295081212 | 0.158957567 | 1.210371641 | 0.129664799 | 0.443704199 | 0.580676686 | 0.776124518 | 0.7014 | 0.834674239 |
| SL1344_RS13750_ | -6.101824741 | -5.850654058 | -6.443094679 | -0.593431578 | 1.552099954 | -0.348222637 | 0.342666673 | 0.471149723 | 0.699229483 | 0.6023 | 0.768237839 |
| SL1344_RS13755_ | -6.520111888 | -6.705270122 | -6.275478794 | 0.457528215 | 1.851165543 | 0.213943495 | 0.405063295 | 0.559748178 | 0.762081725 | 0.676 | 0.820982129 |
| SL1344_RS13760_ | -5.784050384 | -5.863785899 | -5.653945929 | 0.098247844 | 1.620166458 | 0.05706445 | 0.471333334 | 0.646749151 | 0.823739631 | 0.7537 | 0.87397113 |
| SL1344_RS13765_ | -3.527538706 | -3.591028449 | -3.448667369 | 0.182309846 | 0.957251604 | 0.18044217 | 0.42238508 | 0.685590458 | 0.849762918 | 0.7857 | 0.893254766 |
| SL1344_RS13770_ | -3.276235555 | -3.12203778 | -3.479054816 | -0.381731281 | 0.607318026 | -0.574120552 | 0.264666678 | 0.402257143 | 0.645116602 | 0.5197 | 0.710928886 |
| SL1344_RS13775_ | -1.4007363 | -1.455192841 | -1.309070334 | 0.191455163 | 0.382082695 | 0.487487479 | 0.301798809 | 0.432164394 | 0.679272535 | 0.5704 | 0.744634017 |
| SL1344_RS13780_ | 3.053997828 | 2.893343457 | 3.280046522 | 0.328813836 | 0.213991479 | 1.608060741 | 0.050666753 | 0.064619977 | 0.294984116 | 0.1451 | 0.396230726 |
| SL1344_RS13785_ | 5.65158175 | 5.719258833 | 5.62403761 | -0.060060616 | 0.096172093 | -0.448470657 | 0.339773491 | 0.761370656 | 0.901234118 | 0.7159 | 0.879350419 |
| SL1344_RS13790_ | 5.698058085 | 5.636460692 | 5.753466782 | 0.07974409 | 0.085727626 | 1.124210911 | 0.1206667 | 0.171825829 | 0.461366856 | 0.2119 | 0.487786265 |
| SL1344_RS13800_ | 4.498590584 | 4.342965311 | 4.616586749 | 0.180977565 | 0.255080445 | 0.797456643 | 0.147333359 | 0.292767398 | 0.594660824 | 0.2747 | 0.549902994 |
| SL1344_RS13805_ | 2.816226228 | 2.650019301 | 2.834919696 | 0.176476643 | 0.134616096 | 0.597408557 | 0.333111266 | 0.568621522 | 0.793837738 | 0.7012 | 0.873036452 |
| SL1344_RS13810_ | 1.812436062 | 1.784253284 | 1.839951074 | 0.071018867 | 0.121905309 | 0.534277414 | 0.281333343 | 0.41742318 | 0.661713716 | 0.5511 | 0.731974412 |
| SL1344_RS13815_ | 1.781708192 | 1.68641033 | 1.828113093 | 0.116550554 | 0.176433469 | 0.619993682 | 0.268666678 | 0.412877083 | 0.67456746 | 0.5359 | 0.745924434 |
| SL1344_RS13820_ | -0.912518508 | -0.891950682 | -0.940882572 | -0.061371008 | 0.259280923 | -0.204378053 | 0.407061962 | 0.566108178 | 0.765988234 | 0.684 | 0.82501437 |
| SL1344_RS13825_ | 5.218631792 | 5.330917926 | 5.129762629 | -0.178458191 | 0.18216744 | -0.927173324 | 0.129247199 | 0.281269103 | 0.583878035 | 0.2415 | 0.51783725 |
| SL1344_RS13830_ | 1.048968764 | 1.053831341 | 1.044512924 | 0.001590173 | 0.198547235 | 0.009134127 | 0.494666667 | 0.726041464 | 0.875201403 | 0.8494 | 0.928345714 |
| SL1344_RS13840_ | 2.836269796 | 2.840269587 | 2.83320237 | -0.010483186 | 0.210065763 | -0.043993704 | 0.482345104 | 0.856316609 | 0.943258673 | 0.9349 | 0.971863663 |
| SL1344_RS13845_ | 1.391538633 | 1.464655887 | 1.357508418 | -0.135048096 | 0.216940546 | -0.605123323 | 0.270666677 | 0.372193968 | 0.649387028 | 0.5367 | 0.740626263 |

| | | | | | | | | | | | |
|---|---|---|---|---|---|---|---|---|---|---|---|
| SL1344_RS13850_ | 2.306867898 | 2.262173356 | 2.36355158 | 0.009366316 | 0.278622591 | 0.031668316 | 0.488341106 | 0.851276988 | 0.940914438 | 0.9508 | 0.978119573 |
| SL1344_RS13855_ | 3.124005369 | 3.013658502 | 3.182716352 | 0.227308779 | 0.214236579 | 1.129695003 | 0.064623651 | 0.142320695 | 0.424842529 | 0.1658 | 0.417127513 |
| SL1344_RS13860_ | 0.083059481 | 0.090290037 | 0.075915023 | -0.014263029 | 0.178228389 | -0.075675753 | 0.468354432 | 0.596763598 | 0.789967921 | 0.7074 | 0.844683872 |
| SL1344_RS13865_ | -2.945901449 | -3.197447693 | -2.795047532 | 0.402769386 | 0.542298671 | 0.674917083 | 0.213857444 | 0.33807564 | 0.590169486 | 0.444 | 0.651178362 |
| SL1344_RS13870_ | -2.353361681 | -2.519449699 | -2.197920317 | 0.317575238 | 0.50909353 | 0.572681963 | 0.251165902 | 0.409816218 | 0.655692632 | 0.5233 | 0.721542938 |
| SL1344_RS13875_ | -2.694703807 | -2.637137921 | -2.812150339 | -0.077335553 | 0.659360845 | -0.116154651 | 0.456362427 | 0.716170073 | 0.86694767 | 0.8117 | 0.911571143 |
| SL1344_RS13880_ | -2.762241221 | -2.753616582 | -2.767069411 | -0.032910619 | 0.477898083 | -0.063287769 | 0.477333334 | 0.591822375 | 0.785326499 | 0.7086 | 0.842920477 |
| SL1344_RS13885_ | -2.427469829 | -2.235022748 | -2.728592915 | -0.498771889 | 0.602683546 | -0.807729697 | 0.186666686 | 0.281894373 | 0.553607897 | 0.3799 | 0.622391214 |
| SL1344_RS13890_ | -3.899214008 | -3.874507369 | -3.921388771 | -0.051698717 | 0.602245557 | -0.082544132 | 0.466000001 | 0.527292215 | 0.733728161 | 0.6374 | 0.793375131 |
| SL1344_RS13895_ | -1.19885293 | -1.245526993 | -1.149265286 | 0.102094923 | 0.258715486 | 0.349722463 | 0.343104603 | 0.497563414 | 0.710454977 | 0.6085 | 0.769765337 |
| SL1344_RS13900_ | -4.286316849 | -4.383840102 | -4.156478113 | 0.210924038 | 0.808102388 | 0.229516809 | 0.396402402 | 0.553470578 | 0.756132867 | 0.6568 | 0.808881592 |
| SL1344_RS13905_ | -4.43522037 | -4.329827048 | -4.508016772 | -0.38728806 | 1.20197346 | -0.307284425 | 0.374000005 | 0.572923141 | 0.775814463 | 0.722 | 0.850914584 |
| SL1344_RS13910_ | -4.332800668 | -4.44576543 | -4.165350717 | 0.232184828 | 0.917278916 | 0.238153649 | 0.394000004 | 0.587507952 | 0.780098868 | 0.6862 | 0.828582815 |
| SL1344_RS13915_ | -2.798742828 | -2.557148836 | -3.033710569 | -0.477488903 | 0.452005515 | -0.968745365 | 0.1352432 | 0.208042694 | 0.457008873 | 0.2893 | 0.526240952 |
| SL1344_RS13920_ | -5.149662056 | -5.013332517 | -5.325656929 | -0.356961796 | 0.979808631 | -0.33432673 | 0.351333339 | 0.48569066 | 0.69814093 | 0.6062 | 0.76420035 |
| SL1344_RS13930_ | -1.648450412 | -1.768610755 | -1.544368252 | 0.25696494 | 0.418044706 | 0.560716837 | 0.249333346 | 0.385199524 | 0.642409664 | 0.5092 | 0.700949858 |
| SL1344_RS13935_ | -2.057755804 | -2.149415451 | -1.976714472 | 0.169620219 | 0.330201181 | 0.484190532 | 0.28800001 | 0.421515366 | 0.651467622 | 0.5313 | 0.714192793 |
| SL1344_RS13940_ | -4.522875878 | -4.62118252 | -4.38745242 | 0.301905089 | 1.06641461 | 0.271472458 | 0.371333338 | 0.582282415 | 0.778186023 | 0.6983 | 0.836650871 |
| SL1344_RS13950_ | -6.277374523 | -6.434698056 | -5.969102171 | 0.52690587 | 1.783478459 | 0.25333993 | 0.389073955 | 0.605404903 | 0.790692202 | 0.7153 | 0.846943709 |
| SL1344_RS13955_ | -5.197244656 | -5.580209444 | -4.432738043 | 1.013319482 | 1.315541021 | 0.670634529 | 0.249833457 | 0.423104885 | 0.65430523 | 0.5078 | 0.713475094 |
| SL1344_RS13965_ | -4.366012272 | -4.4589383 | -4.247708097 | 0.279321899 | 1.052534963 | 0.241148738 | 0.384410398 | 0.578084878 | 0.77962021 | 0.7155 | 0.847075359 |
| SL1344_RS13970_ | -4.505851478 | -4.42583175 | -4.611756834 | -0.183486077 | 0.8956637 | -0.191983728 | 0.418000003 | 0.574607303 | 0.773154846 | 0.6964 | 0.832925371 |
| SL1344_RS13975_ | -5.401085154 | -5.695929993 | -4.79563241 | 0.74758646 | 1.408942634 | 0.463935374 | 0.305129922 | 0.542615432 | 0.747010362 | 0.6127 | 0.788339517 |
| SL1344_RS13980_ | -5.034878671 | -5.597223609 | -4.44512312 | 1.116093881 | 1.082957185 | 0.965121676 | 0.135909422 | 0.215852835 | 0.470941089 | 0.2919 | 0.532704871 |
| SL1344_RS13985_ | -7.012657637 | -8.409352576 | -5.743959602 | 1.928126722 | 4.27484937 | 0.417925459 | 0.274483688 | 0.493789988 | 0.719201921 | 0.5362 | 0.736882139 |
| SL1344_RS14000_ | -4.446534715 | -4.153869103 | -4.770305773 | -0.95263058 | 1.520466549 | -0.628255326 | 0.253830791 | 0.355340564 | 0.628296889 | 0.5015 | 0.70888497 |
| SL1344_RS14005_ | -3.484657981 | -3.670638568 | -3.232226645 | 0.39274461 | 0.779221281 | 0.471881991 | 0.307333342 | 0.497905679 | 0.721081721 | 0.6007 | 0.776376746 |
| SL1344_RS14010_ | -4.989926497 | -4.837382958 | -5.07070054 | -0.057444445 | 1.663309596 | -0.025579719 | 0.486000001 | 0.676702026 | 0.848122272 | 0.7994 | 0.903566111 |
| SL1344_RS14015_ | -3.555359265 | -3.658033695 | -3.4875839 | 0.17704925 | 0.662541584 | 0.245785205 | 0.393333337 | 0.564867244 | 0.76415025 | 0.6696 | 0.818324685 |
| SL1344_RS14020_ | -4.103578899 | -4.185713304 | -4.001110098 | 0.126302529 | 0.927088984 | 0.117207149 | 0.442666669 | 0.636273788 | 0.81691171 | 0.7441 | 0.868041872 |
| SL1344_RS14025_ | 2.576972917 | 2.326641252 | 2.798506602 | 0.356471428 | 0.329683425 | 1.204477206 | 0.118000034 | 0.130457356 | 0.408796967 | 0.1995 | 0.475582485 |
| SL1344_RS14030_ | 2.295730272 | 2.149325965 | 2.65842048 | 0.496594775 | 0.219416667 | 1.946203579 | 0.003332494 | 0.057997324 | 0.279787818 | 0.1032 | 0.33494393 |
| SL1344_RS14035_ | 0.662218545 | 0.667100381 | 0.656531012 | -0.044613705 | 0.493332295 | -0.093967234 | 0.443037977 | 0.750630702 | 0.893427569 | 0.8416 | 0.925528499 |
| SL1344_RS14040_ | 2.896126078 | 2.822862117 | 2.941633683 | 0.088145619 | 0.170404486 | 0.554583479 | 0.264490351 | 0.472881573 | 0.726047453 | 0.539 | 0.759063731 |
| SL1344_RS14045_ | 2.184762195 | 2.15237087 | 2.231824548 | 0.089017422 | 0.180440131 | 0.485246991 | 0.275816133 | 0.45019774 | 0.705564928 | 0.5483 | 0.755876781 |
| SL1344_RS14050_ | -0.563255068 | -0.435463611 | -0.632662239 | -0.150281778 | 0.506462922 | -0.296007129 | 0.357095276 | 0.745583073 | 0.886469937 | 0.738 | 0.879473611 |
| SL1344_RS14055_ | -1.12996873 | -0.98032345 | -1.281238124 | -0.313513598 | 0.282378164 | -1.033059675 | 0.113333369 | 0.192349577 | 0.447337151 | 0.274 | 0.510116413 |
| SL1344_RS14060_ | -1.577304172 | -1.47114429 | -1.686543813 | -0.243737046 | 0.410051704 | -0.566607686 | 0.266000011 | 0.405746937 | 0.653938313 | 0.5182 | 0.71554225 |
| SL1344_RS14065_ | -2.976252698 | -2.977218565 | -2.97452305 | -0.069293071 | 0.848261991 | -0.080884629 | 0.464357097 | 0.67108368 | 0.846030775 | 0.8272 | 0.918610867 |
| SL1344_RS14070_ | -5.773494189 | -5.495688655 | -6.06075338 | -0.513250679 | 1.320067801 | -0.380174817 | 0.321119261 | 0.484208611 | 0.703506019 | 0.5945 | 0.762693626 |
| SL1344_RS14075_ | -5.775250454 | -5.547064979 | -6.035973779 | -0.437269474 | 1.577501324 | -0.236838019 | 0.391072622 | 0.574266246 | 0.773035199 | 0.6825 | 0.824747944 |
| SL1344_RS14080_ | -5.194023585 | -5.504959317 | -4.880116462 | 0.624550177 | 1.115611987 | 0.495357321 | 0.278481023 | 0.446300644 | 0.670523095 | 0.5564 | 0.733060804 |
| SL1344_RS14085_ | 0.079236192 | 0.332401564 | -0.119282694 | -0.188303582 | 0.591455284 | -0.373323958 | 0.372418393 | 0.709360056 | 0.871019395 | 0.7882 | 0.906194813 |
| SL1344_RS14090_ | -4.471284059 | -4.459928057 | -4.48784386 | 0.008998253 | 0.824504803 | 0.011093592 | 0.494337109 | 0.578156563 | 0.774648102 | 0.6861 | 0.829811533 |
| SL1344_RS14095_ | -4.563584597 | -4.122246305 | -6.041369453 | -2.005968094 | 1.732137277 | -1.089376746 | 0.133244533 | 0.169619505 | 0.451644147 | 0.2756 | 0.540067348 |
| SL1344_RS14100_ | 0.327663262 | 0.216735532 | 0.361367985 | 0.144292119 | 0.291144612 | 0.402995468 | 0.366666672 | 0.663474922 | 0.841386446 | 0.767 | 0.896473574 |
| SL1344_RS14105_ | 1.177587579 | 0.990725648 | 1.210092313 | 0.202381642 | 0.19622488 | 0.746666957 | 0.285333343 | 0.391597345 | 0.663418187 | 0.5908 | 0.793249714 |
| SL1344_RS14110_ | 1.0919613 | 0.937173342 | 1.32874867 | 0.324035056 | 0.232060368 | 1.406018795 | 0.073950757 | 0.102686626 | 0.354409685 | 0.1743 | 0.433609511 |
| SL1344_RS14115_ | -0.803572612 | -0.793991998 | -0.813810468 | -0.070375016 | 0.347924104 | -0.197561105 | 0.426000003 | 0.609835543 | 0.804327983 | 0.7667 | 0.877140232 |
| SL1344_RS14120_ | 2.854726057 | 2.79978285 | 2.91476574 | 0.069139761 | 0.259805987 | 0.2793595 | 0.375333338 | 0.70111875 | 0.869225462 | 0.8174 | 0.918534884 |
| SL1344_RS14125_ | -0.439303304 | -0.414229926 | -0.453583109 | -0.004879298 | 0.283942773 | -0.014982637 | 0.493670886 | 0.698460937 | 0.857214623 | 0.7969 | 0.902184533 |
| SL1344_RS14130_ | 0.514474724 | 0.462383377 | 0.612291642 | 0.166527681 | 0.24830218 | 0.628248179 | 0.247333346 | 0.383756924 | 0.64779711 | 0.5122 | 0.724203112 |
| SL1344_RS14135_ | 1.438981537 | 1.393684626 | 1.536410659 | 0.242887112 | 0.350641594 | 0.723889984 | 0.23717523 | 0.306738878 | 0.601317091 | 0.4645 | 0.709744891 |

| | | | | | | | | | | | |
|---|---|---|---|---|---|---|---|---|---|---|---|
| SL1344_RS14140_ | -4.433278902 | -5.069134483 | -3.928180397 | 1.127206701 | 0.781185002 | 1.388226149 | 0.059333406 | 0.116245555 | 0.343418187 | 0.1783 | 0.417065194 |
| SL1344_RS14150_ | -3.399184136 | -3.334949799 | -3.489211014 | -0.135006356 | 0.606470616 | -0.19879273 | 0.41066667 | 0.569973959 | 0.767799422 | 0.6777 | 0.823239059 |
| SL1344_RS14155_ | -3.744247699 | -3.630796413 | -3.902792593 | -0.288489678 | 0.672601551 | -0.369091022 | 0.332000007 | 0.484799135 | 0.70623271 | 0.6078 | 0.770562031 |
| SL1344_RS14165_ | -4.186101885 | -4.942788685 | -3.915064641 | 0.981090487 | 0.951298942 | 0.927351797 | 0.165333356 | 0.243851627 | 0.515688052 | 0.3545 | 0.587997306 |
| SL1344_RS14170_ | -3.358785158 | -3.385936256 | -3.339505313 | 0.074497266 | 0.590113688 | 0.116364134 | 0.443333335 | 0.573443802 | 0.770527647 | 0.6838 | 0.826173428 |
| SL1344_RS14175_ | 0.320609615 | 0.265355441 | 0.402960507 | 0.140214174 | 0.162670037 | 0.823034398 | 0.179333354 | 0.284415153 | 0.535752389 | 0.389 | 0.60352149 |
| SL1344_RS14180_ | -0.826395719 | -0.797886626 | -0.840536367 | -0.054605841 | 0.275268005 | -0.185556726 | 0.431045972 | 0.610110513 | 0.796577338 | 0.7207 | 0.851023368 |
| SL1344_RS14190_ | -2.355761251 | -2.330368822 | -2.412591211 | -0.037773594 | 0.484659503 | -0.064308865 | 0.472351767 | 0.650806742 | 0.826755097 | 0.7585 | 0.879959439 |
| SL1344_RS14195_ | 1.555970953 | 1.65758324 | 1.496038983 | -0.138843635 | 0.141648644 | -0.915827042 | 0.175216543 | 0.255291222 | 0.52367853 | 0.3447 | 0.586623309 |
| SL1344_RS14200_ | 0.900100646 | 0.860760876 | 0.934418639 | 0.110203586 | 0.232274904 | 0.448630799 | 0.291805472 | 0.417943031 | 0.680827433 | 0.5795 | 0.760588463 |
| SL1344_RS14205_ | -5.149555151 | -5.171626264 | -5.12892122 | 0.116222667 | 1.381251143 | 0.079995835 | 0.457028649 | 0.636296882 | 0.818764081 | 0.746 | 0.868575777 |
| SL1344_RS14210_ | -2.607700764 | -2.705937589 | -2.414140167 | 0.355207042 | 0.722686559 | 0.450649356 | 0.322666674 | 0.475345863 | 0.717841283 | 0.6393 | 0.804042026 |
| SL1344_RS14215_ | 2.198810083 | 2.039308048 | 2.357628442 | 0.364335631 | 0.198648328 | 2.0910935 | 4.68E-05 | 0.052064891 | 0.264359273 | 0.1001 | 0.330407704 |
| SL1344_RS14220_ | 1.166571873 | 0.959932014 | 1.42831903 | 0.480388025 | 0.173087252 | 2.802691352 | 4.68E-05 | 0.029582101 | 0.198590821 | 0.1 | 0.330257709 |
| SL1344_RS14225_ | 1.372393477 | 1.342169437 | 1.396627641 | 0.068394162 | 0.271598712 | 0.25155787 | 0.362425055 | 0.68541178 | 0.853952 | 0.7516 | 0.878270485 |
| SL1344_RS14230_ | -4.300243076 | -3.999528744 | -4.708072022 | -0.666313592 | 0.930737351 | -0.704982225 | 0.228000014 | 0.356854478 | 0.608460294 | 0.4578 | 0.671191261 |
| SL1344_RS14235_ | -5.095987266 | -5.401792493 | -4.638089963 | 0.754137756 | 1.113479652 | 0.640730033 | 0.231179228 | 0.367765077 | 0.60904433 | 0.4703 | 0.671062602 |
| SL1344_RS14240_ | -5.338871973 | -5.190075333 | -5.539190525 | -0.285686989 | 1.348366809 | -0.191330044 | 0.41666667 | 0.579631636 | 0.780133011 | 0.7147 | 0.848062752 |
| SL1344_RS14245_ | -4.029290851 | -4.001191901 | -4.063886622 | -0.064434086 | 0.652255419 | -0.100209554 | 0.455696204 | 0.550797789 | 0.750344841 | 0.6661 | 0.809902334 |
| SL1344_RS14250_ | -2.915623649 | -2.868603425 | -3.041125742 | -0.149624172 | 0.855080307 | -0.148371917 | 0.442666669 | 0.737098428 | 0.882299453 | 0.8226 | 0.922004078 |
| SL1344_RS14255_ | 1.436026395 | 1.38583387 | 1.507998893 | 0.140127505 | 0.298423104 | 0.485050853 | 0.267821463 | 0.453348762 | 0.71385518 | 0.5345 | 0.75501503 |
| SL1344_RS14260_ | -2.861049074 | -2.614660501 | -3.155657645 | -0.535847372 | 0.535128073 | -0.946952073 | 0.16066669 | 0.228042948 | 0.487014777 | 0.3152 | 0.551162683 |
| SL1344_RS14265_ | -4.294576801 | -4.312082077 | -4.288585923 | -0.060751681 | 1.10377839 | -0.050443305 | 0.476349101 | 0.693910124 | 0.853460823 | 0.8053 | 0.90453265 |
| SL1344_RS14270_ | -3.385476638 | -3.861286822 | -2.932530989 | 0.722362645 | 1.222675542 | 0.55507345 | 0.271333344 | 0.429299786 | 0.690052475 | 0.5419 | 0.747497531 |
| SL1344_RS14275_ | -3.585543961 | -3.705796409 | -3.499148861 | 0.217624732 | 0.644134509 | 0.314864932 | 0.3637575 | 0.52060142 | 0.733406484 | 0.63 | 0.791557415 |
| SL1344_RS14280_ | -2.504085118 | -2.79156779 | -2.249317629 | 0.542936673 | 0.387517037 | 1.337416784 | 0.066000065 | 0.122367388 | 0.352442384 | 0.1938 | 0.433705003 |
| SL1344_RS14285_ | -5.968510421 | -5.999319199 | -5.955291649 | -0.019787684 | 1.843038036 | -0.008549905 | 0.495333334 | 0.651319266 | 0.828246136 | 0.7776 | 0.889406172 |
| SL1344_RS14290_ | -3.015577593 | -2.822933493 | -3.277988222 | -0.4465366 | 0.49031054 | -0.884959359 | 0.172000021 | 0.267295758 | 0.519454485 | 0.3623 | 0.58569292 |
| SL1344_RS14295_ | -0.930472773 | -1.069437107 | -0.802410528 | 0.283663639 | 0.414951714 | 0.66794223 | 0.212666683 | 0.321307791 | 0.600559481 | 0.4229 | 0.65513603 |
| SL1344_RS14300_ | -0.392360737 | -0.442600975 | -0.334959458 | 0.130598413 | 0.283317254 | 0.452291464 | 0.316455704 | 0.496376106 | 0.725522184 | 0.6301 | 0.796036412 |
| SL1344_RS14305_ | 0.445636535 | 0.211143862 | 0.681489452 | 0.254236231 | 0.55360778 | 0.555833365 | 0.260666678 | 0.439030435 | 0.707204251 | 0.5285 | 0.748186376 |
| SL1344_RS14310_ | -4.192814422 | -3.82677468 | -4.589044613 | -0.621682568 | 1.074482689 | -0.552378955 | 0.270486353 | 0.434759544 | 0.677418707 | 0.5426 | 0.737962145 |
| SL1344_RS14315_ | -3.449726408 | -3.383891849 | -3.502758096 | -0.249910211 | 0.781411403 | -0.282117663 | 0.388000004 | 0.578176138 | 0.778660122 | 0.7298 | 0.851927647 |
| SL1344_RS14320_ | 1.778980163 | 1.779163806 | 1.777869647 | 0.075734108 | 0.216100984 | 0.362804686 | 0.384666671 | 0.569922223 | 0.783764375 | 0.8156 | 0.904492264 |
| SL1344_RS14325_ | 3.492037467 | 3.393365679 | 3.542594982 | 0.109983049 | 0.136577311 | 0.765795763 | 0.225333348 | 0.32775922 | 0.616593319 | 0.436 | 0.676285066 |
| SL1344_RS14330_ | -1.951941875 | -1.984575003 | -1.926023338 | 0.032734924 | 0.438533918 | 0.064329853 | 0.466000001 | 0.665593598 | 0.836728971 | 0.7757 | 0.88764053 |
| SL1344_RS14335_ | -2.446437759 | -2.428708937 | -2.465262502 | -0.035674005 | 0.432144381 | -0.077622173 | 0.465333335 | 0.574107283 | 0.770440686 | 0.687 | 0.828316632 |
| SL1344_RS14345_ | 9.05301794 | 5.408789596 | 9.087829503 | 3.682768099 | 0.32537807 | 0.975142559 | 0.33333334 | 0.19833289 | 0.500202919 | 0.7 | 0.872542119 |
| SL1344_RS14350_ | -2.476983886 | -2.824305591 | -2.014408971 | 0.804220913 | 0.552344197 | 1.40386143 | 0.055333412 | 0.104629132 | 0.34475545 | 0.1707 | 0.418467957 |
| SL1344_RS14360_ | -0.01735939 | 0.636241241 | -0.386001422 | -1.014527872 | 0.305446049 | -3.096174186 | 4.68E-05 | 0.024976258 | 0.185935354 | 0.1 | 0.330257709 |
| SL1344_RS14365_ | -1.842020164 | -1.575174232 | -2.113100619 | -0.480567987 | 0.472350964 | -0.99385371 | 0.147333359 | 0.219844435 | 0.491229582 | 0.2979 | 0.549345071 |
| SL1344_RS14370_ | -1.971079185 | -1.57434578 | -2.0644098 | -0.430449953 | 0.519224862 | -0.621287416 | 0.306462367 | 0.523718112 | 0.748592651 | 0.6358 | 0.820441658 |
| SL1344_RS14375_ | -0.138453919 | 0.033055913 | -0.248776471 | -0.276946444 | 0.19568973 | -1.332971357 | 0.079946755 | 0.126367796 | 0.36761887 | 0.2023 | 0.448756975 |
| SL1344_RS14380_ | -2.376085678 | -2.173019824 | -2.575362779 | -0.408753018 | 0.415397249 | -0.956329372 | 0.14923387 | 0.230255283 | 0.481964167 | 0.3151 | 0.545392689 |
| SL1344_RS14385_ | -1.141347304 | -1.186638179 | -1.067926985 | 0.132815565 | 0.340725472 | 0.340274153 | 0.358427721 | 0.563722035 | 0.76675925 | 0.6776 | 0.824543447 |
| SL1344_RS14390_ | 2.796119538 | 2.759502562 | 2.822543576 | 0.080653867 | 0.239700655 | 0.332218252 | 0.341105936 | 0.469707877 | 0.732728757 | 0.6959 | 0.852344691 |
| SL1344_RS14395_ | 3.251087104 | 3.231838183 | 3.26580226 | 0.141828573 | 0.194314337 | 0.720607408 | 0.273151243 | 0.30798497 | 0.601901595 | 0.5864 | 0.770300454 |
| SL1344_RS14400_ | 0.078362801 | -0.070907923 | 0.136512289 | 0.195722331 | 0.365046645 | 0.524789068 | 0.292000009 | 0.498328013 | 0.740119348 | 0.5652 | 0.777397846 |
| SL1344_RS14405_ | 0.661290229 | 0.70451642 | 0.632143535 | -0.051478974 | 0.313570229 | -0.155304645 | 0.447333335 | 0.790573811 | 0.910834449 | 0.9091 | 0.959673513 |
| SL1344_RS14410_ | 1.814936386 | 1.913995322 | 1.725503915 | -0.168376793 | 0.544227403 | -0.23019476 | 0.452000002 | 0.827389147 | 0.932243074 | 0.9994 | 0.999752194 |
| SL1344_RS14415_ | 1.720002606 | 1.73306857 | 1.706621312 | -0.059568424 | 0.647624692 | -0.108813224 | 0.407333337 | 0.860973731 | 0.946550321 | 0.8542 | 0.93662759 |
| SL1344_RS14420_ | 0.414870868 | 0.491743197 | 0.350798276 | -0.118468458 | 0.6400249 | -0.155544335 | 0.452000002 | 0.799173229 | 0.918155338 | 0.9874 | 0.994566223 |
| SL1344_RS14425_ | 2.634427839 | 2.660704921 | 2.516352521 | -0.397323673 | 0.706945089 | -0.601841254 | 0.234000014 | 0.355209959 | 0.648927847 | 0.4141 | 0.688825163 |

| | | | | | | | | | | | |
|---|---|---|---|---|---|---|---|---|---|---|---|
| SL1344_RS14430_ | 0.888588066 | 0.957247434 | 0.69666388 | -0.343697506 | 0.620514907 | -0.626127065 | 0.23184545 | 0.344790372 | 0.637202815 | 0.4322 | 0.699419933 |
| SL1344_RS14435_ | 2.705440283 | 3.131476367 | 2.50805483 | -0.58663498 | 0.578887324 | -0.877872991 | 0.184000019 | 0.272580536 | 0.576400766 | 0.336 | 0.614473441 |
| SL1344_RS14440_ | -1.557820787 | -1.426530848 | -1.878553275 | -0.364319516 | 0.72013345 | -0.499009095 | 0.304463699 | 0.562149721 | 0.7852076 | 0.6202 | 0.814147507 |
| SL1344_RS14445_ | -1.34479891 | -1.084676833 | -1.478218382 | -0.421786252 | 0.722699462 | -0.579898878 | 0.249167234 | 0.418046694 | 0.687008073 | 0.4946 | 0.733409318 |
| SL1344_RS14450_ | -0.971650251 | -0.39245484 | -1.31692392 | -0.888957259 | 0.414677556 | -2.059492172 | 0.008000574 | 0.037173842 | 0.220250648 | 0.1049 | 0.337121414 |
| SL1344_RS14455_ | -0.302182158 | 0.083503224 | -0.817192498 | -0.90719109 | 0.233974673 | -3.852731511 | 4.68E-05 | 0.011572906 | 0.119936628 | 0.1 | 0.330257709 |
| SL1344_RS14460_ | -0.892951459 | -0.721034419 | -0.994818177 | -0.266471437 | 0.363808791 | -0.705078514 | 0.236509007 | 0.354501794 | 0.621558944 | 0.4776 | 0.693013249 |
| SL1344_RS14465_ | -1.599744138 | -1.425001058 | -1.828106989 | -0.379029416 | 0.399995843 | -0.871434649 | 0.181212545 | 0.260174831 | 0.529298734 | 0.3573 | 0.597205079 |
| SL1344_RS14470_ | -1.595831017 | -1.719271553 | -1.386986963 | 0.295679162 | 0.374642342 | 0.763312636 | 0.210526332 | 0.332877895 | 0.589068469 | 0.4298 | 0.653713468 |
| SL1344_RS14475_ | -2.28164821 | -2.255648346 | -2.303314858 | -0.046070455 | 0.35455899 | -0.113288081 | 0.437708197 | 0.549532079 | 0.747997793 | 0.6628 | 0.807535428 |
| SL1344_RS14480_ | -2.240663759 | -2.337254309 | -2.165782114 | 0.185364851 | 0.3927421 | 0.436621961 | 0.320453039 | 0.466992196 | 0.69091808 | 0.581 | 0.752316208 |
| SL1344_RS14485_ | -0.982344802 | -0.889092845 | -1.126904413 | -0.230386297 | 0.263276263 | -0.796316838 | 0.177881433 | 0.28042065 | 0.533805299 | 0.3762 | 0.59898526 |
| SL1344_RS14490_ | 0.656013463 | 0.795648403 | 0.542747144 | -0.290357393 | 0.228083413 | -1.294339903 | 0.052000084 | 0.129647673 | 0.39267577 | 0.164 | 0.40747419 |
| SL1344_RS14495_ | 3.646948595 | 3.495050234 | 3.743481393 | 0.252285812 | 0.117680635 | 2.233137279 | 4.68E-05 | 0.062544296 | 0.289316155 | 0.1 | 0.330257709 |
| SL1344_RS14500_ | 1.442722586 | 0.722395316 | 2.231461782 | 1.484117725 | 0.512416176 | 2.822266988 | 4.68E-05 | 0.018845407 | 0.16790967 | 0.1 | 0.330257709 |
| SL1344_RS14505_ | -1.887917514 | -1.879433181 | -1.896700716 | -0.013150077 | 0.316154771 | -0.03804821 | 0.480346436 | 0.553556183 | 0.754647978 | 0.667 | 0.815035095 |
| SL1344_RS14510_ | -0.193778484 | -0.151471163 | -0.249641199 | -0.083811215 | 0.361357267 | -0.228712355 | 0.412391742 | 0.702009479 | 0.861131511 | 0.8253 | 0.915532175 |
| SL1344_RS14515_ | 0.346713862 | 0.315385212 | 0.363210285 | 0.037850608 | 0.200422111 | 0.175134437 | 0.425333336 | 0.657646663 | 0.829228432 | 0.7555 | 0.876306825 |
| SL1344_RS14520_ | -2.25607023 | -2.442303842 | -2.155319765 | 0.283626928 | 0.423095088 | 0.610053962 | 0.24250501 | 0.376489772 | 0.623429204 | 0.4885 | 0.686827726 |
| SL1344_RS14525_ | -4.10580281 | -4.433604217 | -3.83202442 | 0.609715228 | 0.793853556 | 0.719104656 | 0.205862775 | 0.32362996 | 0.571814704 | 0.4213 | 0.631939091 |
| SL1344_RS14530_ | -2.111594293 | -2.137334638 | -2.090182497 | 0.131911528 | 0.516184703 | 0.228358439 | 0.40266667 | 0.593194292 | 0.7896564 | 0.7425 | 0.860126025 |
| SL1344_RS14540_ | -0.178625042 | -0.235737739 | -0.123616022 | 0.133798865 | 0.311487844 | 0.423018468 | 0.312000008 | 0.49630959 | 0.729712251 | 0.6107 | 0.787261821 |
| SL1344_RS14545_ | -0.417431863 | -0.385319061 | -0.445633139 | -0.017039384 | 0.303401393 | -0.053333659 | 0.477333334 | 0.703526005 | 0.860239771 | 0.8268 | 0.914290013 |
| SL1344_RS14550_ | 0.10834026 | 0.072419248 | 0.137189326 | 0.136206153 | 0.343641356 | 0.381012012 | 0.333777489 | 0.482275824 | 0.727146835 | 0.6755 | 0.82445016 |
| SL1344_RS14555_ | -0.788299977 | -0.963487735 | -0.601190615 | -0.04531986 | 0.745315378 | -0.068227851 | 0.474350434 | 0.849908245 | 0.940028055 | 0.9791 | 0.990455172 |
| SL1344_RS14560_ | 0.114071941 | 0.090688107 | 0.138537966 | 0.04540282 | 0.165967167 | 0.265507599 | 0.388407733 | 0.530874034 | 0.734306414 | 0.6362 | 0.790538061 |
| SL1344_RS14565_ | -0.694191599 | -0.767710436 | -0.607221365 | 0.215240833 | 0.347996079 | 0.589950122 | 0.252498347 | 0.3716499 | 0.633602944 | 0.5281 | 0.721626085 |
| SL1344_RS14570_ | 2.351831919 | 2.177658037 | 2.526993867 | 0.155970755 | 0.334602513 | 0.722452633 | 0.202666684 | 0.363037212 | 0.650789532 | 0.3743 | 0.626846118 |
| SL1344_RS14575_ | 1.810179758 | 1.70072568 | 1.886810695 | 0.148203236 | 0.16353332 | 0.883634453 | 0.175882766 | 0.259084701 | 0.540458187 | 0.3379 | 0.591214953 |
| SL1344_RS14585_ | -1.711104847 | -1.90253929 | -1.582841593 | 0.324779681 | 0.389396602 | 0.78981768 | 0.183333353 | 0.281338589 | 0.54345458 | 0.3788 | 0.60272104 |
| SL1344_RS14600_ | 1.929715989 | 1.910705148 | 1.952083877 | 0.031045947 | 0.126242381 | 0.232993313 | 0.40666667 | 0.637333972 | 0.81673019 | 0.7438 | 0.867775337 |
| SL1344_RS14605_ | 2.328006368 | 2.10920921 | 2.360271786 | 0.229854227 | 0.170304364 | 0.772889124 | 0.294470362 | 0.365878154 | 0.652545337 | 0.601 | 0.807298877 |
| SL1344_RS14610_ | 1.51296619 | 1.733619086 | 1.297948649 | -0.337959335 | 1.015858139 | -0.295230512 | 0.417333336 | 0.703765182 | 0.874955388 | 0.9205 | 0.966126279 |
| SL1344_RS14615_ | 3.312961372 | 3.567537343 | 3.048990116 | -0.445836954 | 0.834828298 | -0.443668809 | 0.444000002 | 0.648788005 | 0.846356067 | 0.9889 | 0.99524298 |
| SL1344_RS14620_ | 4.342280405 | 4.488521919 | 4.19692867 | -0.295191678 | 0.960592999 | -0.303435675 | 0.440000002 | 0.742786938 | 0.895715151 | 1 | 1 |
| SL1344_RS14625_ | 4.803651505 | 4.961700346 | 4.628401887 | -0.292110797 | 0.497544661 | -0.396868454 | 0.451698869 | 0.624645525 | 0.832876699 | 0.9997 | 0.999868867 |
| SL1344_RS14630_ | 6.425557107 | 6.567828567 | 6.325480382 | -0.212946296 | 0.287002479 | -0.501346612 | 0.333777489 | 0.638366296 | 0.840713216 | 0.7 | 0.872542119 |
| SL1344_RS14640_ | 4.470986909 | 4.401298424 | 4.641810849 | 0.199695274 | 0.22879744 | 0.954651867 | 0.163333356 | 0.197864664 | 0.497190879 | 0.2918 | 0.570053986 |
| SL1344_RS14645_ | -0.786853867 | -0.818882806 | -0.768264572 | 0.039796639 | 0.288169411 | 0.122338891 | 0.443037977 | 0.65016242 | 0.824323719 | 0.7583 | 0.87723886 |
| SL1344_RS14655_ | 1.305066125 | 1.291981481 | 1.316869127 | 0.035737605 | 0.1952558 | 0.164662782 | 0.428000003 | 0.699292652 | 0.860122441 | 0.8286 | 0.917452079 |
| SL1344_RS14660_ | 1.55505232 | 1.424250342 | 1.704443068 | 0.256497388 | 0.148416603 | 1.683397856 | 0.036666788 | 0.069627348 | 0.29037414 | 0.1381 | 0.383738185 |
| SL1344_RS14675_ | -1.634488421 | -1.597527292 | -1.665009507 | -0.073902914 | 0.330838047 | -0.21737881 | 0.409726852 | 0.581197603 | 0.773445851 | 0.6949 | 0.831186372 |
| SL1344_RS14680_ | 0.7232566 | 0.787875716 | 0.645670185 | -0.143477882 | 0.136668007 | -0.979844685 | 0.143237868 | 0.212808516 | 0.464841838 | 0.2917 | 0.523129837 |
| SL1344_RS14685_ | -0.101262647 | -0.171385595 | -0.001053586 | 0.192953447 | 0.258822129 | 0.708756514 | 0.217854779 | 0.320058056 | 0.590682664 | 0.4302 | 0.646884677 |
| SL1344_RS14690_ | -0.25111128 | -0.450043089 | -0.066728503 | 0.337315847 | 0.286225751 | 1.157417597 | 0.109926752 | 0.165614199 | 0.431905093 | 0.2373 | 0.495955795 |
| SL1344_RS14700_ | 1.779689168 | 1.735986333 | 1.798496459 | 0.071040826 | 0.289613599 | 0.228807527 | 0.416389077 | 0.858974225 | 0.944194597 | 0.8629 | 0.941589088 |
| SL1344_RS14705_ | -0.081875076 | -0.039153508 | -0.106075459 | -0.037230224 | 0.258423404 | -0.144250102 | 0.438374419 | 0.701343741 | 0.859597323 | 0.8126 | 0.908822187 |
| SL1344_RS14710_ | -2.36284139 | -2.280164342 | -2.556213349 | -0.290963389 | 0.616070699 | -0.445790712 | 0.313124592 | 0.455885091 | 0.704015002 | 0.6202 | 0.789278976 |
| SL1344_RS14715_ | -2.082930544 | -2.012723989 | -2.154516833 | -0.146425745 | 0.330345114 | -0.411763752 | 0.318666674 | 0.456761851 | 0.679155254 | 0.5672 | 0.740340329 |
| SL1344_RS14720_ | 1.241143084 | 1.190721443 | 1.280121776 | 0.091718676 | 0.167066521 | 0.503514723 | 0.291805472 | 0.453499931 | 0.694947268 | 0.5809 | 0.762565581 |
| SL1344_RS14725_ | -3.343261242 | -3.319260956 | -3.365885341 | -0.040467969 | 0.5380094 | -0.074831691 | 0.467688209 | 0.550330382 | 0.751188756 | 0.6619 | 0.807994758 |
| SL1344_RS14730_ | -1.224691692 | -1.169534217 | -1.278300983 | -0.084096602 | 0.391563608 | -0.208014875 | 0.405063295 | 0.645715019 | 0.823479943 | 0.7502 | 0.871876234 |
| SL1344_RS14735_ | -2.08746723 | -1.930532482 | -2.152572256 | -0.203348058 | 0.477309508 | -0.388081005 | 0.34133334 | 0.57820139 | 0.775902966 | 0.6931 | 0.835842547 |

| | | | | | | | | | | | |
|---|---|---|---|---|---|---|---|---|---|---|---|
| SL1344_RS14740_ | -2.243634959 | -2.080203848 | -2.472231995 | -0.380037295 | 0.447774507 | -0.822332524 | 0.184543657 | 0.285436635 | 0.543936328 | 0.3846 | 0.608301134 |
| SL1344_RS14745_ | -1.934524141 | -1.960781793 | -1.903802282 | 0.049909535 | 0.330924727 | 0.150611678 | 0.432666669 | 0.52414827 | 0.730958417 | 0.6376 | 0.789652224 |
| SL1344_RS14750_ | -2.485415424 | -2.653037071 | -2.329759124 | 0.319111045 | 0.466420769 | 0.647659271 | 0.241838787 | 0.362861476 | 0.608310172 | 0.4671 | 0.673762306 |
| SL1344_RS14765_ | -1.698849736 | -1.714056765 | -1.672162548 | 0.034277126 | 0.412060845 | 0.067750862 | 0.470666668 | 0.68597026 | 0.848152062 | 0.8003 | 0.902010843 |
| SL1344_RS14770_ | -4.238955092 | -4.497648321 | -4.109273207 | 0.356146563 | 1.047515243 | 0.311946423 | 0.366000005 | 0.562258431 | 0.771262712 | 0.7012 | 0.842373986 |
| SL1344_RS14775_ | -4.204436316 | -4.223832786 | -4.184676643 | 0.048848144 | 0.748949367 | 0.064823898 | 0.467688209 | 0.559419589 | 0.760040563 | 0.6767 | 0.819739744 |
| SL1344_RS14780_ | 0.215130513 | -0.043534233 | 0.295779304 | 0.340014479 | 0.415099502 | 0.847381666 | 0.192538326 | 0.240610945 | 0.53320902 | 0.3475 | 0.620036562 |
| SL1344_RS14785_ | -0.069392847 | -0.186629845 | 0.125647462 | 0.30272811 | 0.216910606 | 1.278927131 | 0.084610309 | 0.139114419 | 0.389415089 | 0.2151 | 0.463275881 |
| SL1344_RS14790_ | -1.330381317 | -1.301154755 | -1.357678013 | -0.047296489 | 0.296087826 | -0.149511583 | 0.440666669 | 0.596247401 | 0.785236147 | 0.7017 | 0.837567702 |
| SL1344_RS14795_ | 0.207138503 | 0.173845563 | 0.244106541 | 0.062828057 | 0.185655565 | 0.313498147 | 0.359760166 | 0.5502042 | 0.754298059 | 0.6531 | 0.807325652 |
| SL1344_RS14800_ | 0.479677402 | 0.364132059 | 0.515988109 | 0.130230039 | 0.254188524 | 0.375521669 | 0.367088613 | 0.717960619 | 0.871685612 | 0.7661 | 0.898373965 |
| SL1344_RS14805_ | 0.256538059 | 0.14839275 | 0.369977046 | 0.220264289 | 0.137744966 | 1.51220983 | 0.054666746 | 0.096889207 | 0.308362753 | 0.1657 | 0.402226623 |
| SL1344_RS14810_ | -2.109757765 | -1.913752454 | -2.222241192 | -0.264475707 | 0.513658508 | -0.46328852 | 0.326000007 | 0.539230428 | 0.755151158 | 0.6372 | 0.80596911 |
| SL1344_RS14815_ | -0.262001039 | 0.186798213 | -0.37297044 | -0.499620053 | 0.360557486 | -1.110633355 | 0.155229871 | 0.196633464 | 0.483506871 | 0.3063 | 0.561709184 |
| SL1344_RS14820_ | -0.986747445 | -0.635200881 | -1.292618328 | -0.658763885 | 0.242259781 | -2.638786426 | 0.003332494 | 0.022937869 | 0.164860052 | 0.1018 | 0.332576155 |
| SL1344_RS14825_ | -1.704653571 | -1.385484168 | -2.095559721 | -0.735348859 | 0.369564965 | -1.929380466 | 0.010667096 | 0.048311794 | 0.236780154 | 0.1111 | 0.345645864 |
| SL1344_RS14830_ | -0.194275235 | -0.13520965 | -0.276273202 | -0.172351702 | 0.287040209 | -0.574954142 | 0.230666681 | 0.34898005 | 0.626953959 | 0.4685 | 0.674347632 |
| SL1344_RS14850_ | 2.24021269 | 2.510492492 | 1.977001125 | -0.426955404 | 0.304840082 | -1.532632166 | 0.075283201 | 0.072875663 | 0.311580714 | 0.164 | 0.42387532 |
| SL1344_RS14855_ | 2.179738697 | 2.632924985 | 1.917758073 | -0.678758293 | 0.23563992 | -3.049571791 | 4.68E-05 | 0.009976257 | 0.123626413 | 0.1 | 0.330257709 |
| SL1344_RS14900_ | 2.351615109 | 2.313751828 | 2.366385563 | 0.038210906 | 0.126166661 | 0.19654451 | 0.40266667 | 0.797944982 | 0.914363303 | 0.805 | 0.916487631 |
| SL1344_RS14905_ | 0.724047762 | 0.820963432 | 0.556934334 | -0.239237773 | 0.227482768 | -0.99554858 | 0.153897426 | 0.208950668 | 0.487325809 | 0.2948 | 0.552206928 |
| SL1344_RS14910_ | 0.961277804 | 1.144271618 | 0.889141496 | -0.249452951 | 0.18427801 | -1.214429513 | 0.102666707 | 0.159111565 | 0.431745363 | 0.2278 | 0.476943598 |
| SL1344_RS15010_ | 0.524712108 | 1.111125283 | 0.425329471 | -0.67304442 | 0.598030936 | -1.204529226 | 0.112000036 | 0.112003572 | 0.379390131 | 0.231 | 0.493930943 |
| SL1344_RS15015_ | 1.645801526 | 1.580333265 | 1.690058306 | 0.096750619 | 0.255425385 | 0.329988055 | 0.395333337 | 0.702007825 | 0.866667742 | 0.8431 | 0.931329533 |
| SL1344_RS15020_ | 0.008452066 | 0.044211102 | -0.015179333 | -0.03398767 | 0.333121389 | -0.093128565 | 0.461025984 | 0.776209948 | 0.903224812 | 0.8852 | 0.947837052 |
| SL1344_RS15025_ | -1.61117861 | -1.686007226 | -1.518718024 | 0.241278789 | 0.477310752 | 0.512869959 | 0.301798809 | 0.420597411 | 0.673524536 | 0.5927 | 0.76357103 |
| SL1344_RS15030_ | 0.93276303 | 0.910989405 | 0.964349555 | 0.114518635 | 0.365094015 | 0.308454861 | 0.352431718 | 0.525221046 | 0.765451273 | 0.7373 | 0.870579837 |
| SL1344_RS15035_ | 1.354901534 | 1.336518195 | 1.369038072 | 0.05322391 | 0.186314921 | 0.238487587 | 0.356666672 | 0.471926292 | 0.724473453 | 0.6678 | 0.813093321 |
| SL1344_RS15040_ | -5.079412186 | -5.484538203 | -4.556353113 | 0.904445923 | 0.981374253 | 0.85607263 | 0.176548988 | 0.282012915 | 0.52550361 | 0.3691 | 0.593245695 |
| SL1344_RS15045_ | -1.763938532 | -2.098472724 | -1.443649707 | 0.662911416 | 0.30277117 | 2.125592884 | 0.011333737 | 0.040955014 | 0.212591278 | 0.1104 | 0.343718898 |
| SL1344_RS15050_ | -4.424951892 | -4.692148314 | -4.173843042 | 0.509071288 | 0.772738271 | 0.611220986 | 0.23666668 | 0.372367486 | 0.608669779 | 0.4775 | 0.671208444 |
| SL1344_RS15055_ | 2.889767487 | 2.843773634 | 2.9492976 | 0.105313057 | 0.232798259 | 0.448738777 | 0.297135252 | 0.485496053 | 0.740150344 | 0.6092 | 0.807041466 |
| SL1344_RS15060_ | -2.449597775 | -2.522159496 | -2.362504029 | 0.203227605 | 0.46999059 | 0.394838482 | 0.33733334 | 0.507573513 | 0.72354896 | 0.6335 | 0.788003167 |
| SL1344_RS15065_ | -2.041813059 | -2.04849219 | -2.028548122 | 0.004218083 | 0.366233476 | 0.009030515 | 0.495333334 | 0.617019238 | 0.801226327 | 0.7211 | 0.851888821 |
| SL1344_RS15070_ | -1.428179322 | -1.432152417 | -1.423350207 | 0.016025258 | 0.370858113 | 0.038118808 | 0.486666667 | 0.668801612 | 0.837054762 | 0.7795 | 0.890418591 |
| SL1344_RS15075_ | -0.011863057 | 0.106657189 | -0.129738715 | -0.238062916 | 0.241190411 | -0.950748077 | 0.149333359 | 0.215986934 | 0.484798731 | 0.2972 | 0.543979757 |
| SL1344_RS15080_ | -1.638235331 | -1.526027935 | -1.750241477 | -0.225332914 | 0.293424577 | -0.76162164 | 0.215856111 | 0.328367813 | 0.566703678 | 0.4294 | 0.633801789 |
| SL1344_RS15085_ | -1.386436192 | -1.431953787 | -1.352151663 | 0.037327127 | 0.368889744 | 0.096646928 | 0.460359762 | 0.666998896 | 0.836738519 | 0.7817 | 0.89004396 |
| SL1344_RS15090_ | -1.13417419 | -1.182148003 | -1.108381427 | 0.059286907 | 0.337584177 | 0.156775047 | 0.446369089 | 0.654956636 | 0.828938375 | 0.7639 | 0.881695444 |
| SL1344_RS15095_ | -0.390091416 | -0.294620103 | -0.433929802 | -0.152010365 | 0.309900536 | -0.463757335 | 0.322000008 | 0.489314552 | 0.727980731 | 0.6485 | 0.811015665 |
| SL1344_RS15100_ | -0.493880924 | -0.532597073 | -0.479073365 | 0.065040932 | 0.321977652 | 0.171484131 | 0.436666669 | 0.684164844 | 0.849657075 | 0.7997 | 0.904378523 |
| SL1344_RS15105_ | 0.755947086 | 0.649043977 | 0.878789263 | 0.216357972 | 0.167128047 | 1.230538364 | 0.102666707 | 0.152387962 | 0.410203216 | 0.2251 | 0.478393096 |
| SL1344_RS15110_ | 0.129662489 | 0.058046747 | 0.216381863 | 0.164337558 | 0.210469151 | 0.725265757 | 0.200532995 | 0.315374999 | 0.572500269 | 0.4091 | 0.631501763 |
| SL1344_RS15115_ | -4.33969984 | -3.912589466 | -4.795123227 | -0.874181069 | 0.74023665 | -1.115648102 | 0.10326453 | 0.173047644 | 0.410244205 | 0.2409 | 0.477403659 |
| SL1344_RS15120_ | -1.933223058 | -1.936380893 | -1.925688785 | -0.017537693 | 0.369240047 | -0.039626959 | 0.481012659 | 0.598085151 | 0.788573921 | 0.7101 | 0.842015617 |
| SL1344_RS15125_ | -3.272840781 | -3.272269564 | -3.273253674 | 0.021940983 | 0.588266666 | 0.028976799 | 0.482345104 | 0.621560589 | 0.803203718 | 0.732 | 0.857734331 |
| SL1344_RS15140_ | 2.286099326 | 2.238328547 | 2.319168825 | 0.08155104 | 0.084603039 | 0.92312662 | 0.145902758 | 0.241610821 | 0.496614327 | 0.3322 | 0.556643433 |
| SL1344_RS15145_ | 1.90186754 | 1.791960945 | 1.976140595 | 0.208366935 | 0.155575523 | 1.285072449 | 0.058000075 | 0.119408696 | 0.376327451 | 0.173 | 0.419815628 |
| SL1344_RS15150_ | 2.169268721 | 1.997088688 | 2.246900065 | 0.252870866 | 0.159940125 | 1.533057296 | 0.019986902 | 0.113166498 | 0.377555463 | 0.1176 | 0.354054621 |
| SL1344_RS15155_ | 0.691469007 | 0.653964344 | 0.716012163 | 0.077003804 | 0.178789703 | 0.412765451 | 0.324000007 | 0.475350906 | 0.708862367 | 0.6287 | 0.782053613 |
| SL1344_RS15160_ | 1.287065435 | 1.198316961 | 1.403631489 | 0.200953852 | 0.124581837 | 1.501553012 | 0.050632998 | 0.093000038 | 0.320014926 | 0.1615 | 0.405491389 |
| SL1344_RS15165_ | 1.865651667 | 1.834299374 | 1.918167742 | 0.09110128 | 0.117886457 | 0.712903145 | 0.220000015 | 0.31834997 | 0.579714109 | 0.4292 | 0.644070329 |
| SL1344_RS15170_ | 0.505763581 | 0.502063276 | 0.513469485 | 0.007855414 | 0.173008328 | 0.042295763 | 0.481678881 | 0.644375421 | 0.818577378 | 0.7482 | 0.869166673 |

| | | | | | | | | | | | |
|---|---|---|---|---|---|---|---|---|---|---|---|
| SL1344_RS15190_ | 0.783578828 | 0.602065139 | 1.040527825 | 0.375116799 | 0.267693021 | 1.411663731 | 0.085333383 | 0.10289602 | 0.35680141 | 0.1772 | 0.437706855 |
| SL1344_RS15195_ | -1.937631177 | -1.749890365 | -2.097016268 | -0.352479547 | 0.415101733 | -0.812894048 | 0.187208547 | 0.288919241 | 0.550630831 | 0.3872 | 0.616063428 |
| SL1344_RS15200_ | -0.245972385 | -0.151630395 | -0.315743108 | -0.170652966 | 0.197439035 | -0.812185496 | 0.174000021 | 0.282478272 | 0.531321914 | 0.3737 | 0.58875611 |
| SL1344_RS15205_ | -1.057026766 | -0.97928444 | -1.149826683 | -0.164032721 | 0.288865031 | -0.552809322 | 0.270000011 | 0.427600148 | 0.660439979 | 0.54 | 0.72604957 |
| SL1344_RS15210_ | -1.282832003 | -1.158953463 | -1.335008412 | -0.13362353 | 0.365802678 | -0.275592758 | 0.389333338 | 0.708200665 | 0.862394939 | 0.7559 | 0.886540272 |
| SL1344_RS15215_ | -0.869893581 | -0.834423132 | -0.900681994 | -0.011194799 | 0.385824032 | -0.029460084 | 0.486666667 | 0.734575273 | 0.879409902 | 0.8422 | 0.92524694 |
| SL1344_RS15220_ | -2.445030534 | -2.529348081 | -2.383291546 | 0.110800364 | 0.45158385 | 0.231989144 | 0.405729517 | 0.573685198 | 0.770487342 | 0.6716 | 0.819119985 |
| SL1344_RS15225_ | -1.068193636 | -1.015021627 | -1.119608184 | -0.098687541 | 0.220166915 | -0.423156436 | 0.325333341 | 0.435681126 | 0.657196238 | 0.5342 | 0.714738931 |
| SL1344_RS15230_ | -1.368874571 | -1.308544676 | -1.466686876 | -0.19599881 | 0.415933513 | -0.446392735 | 0.317121927 | 0.498980749 | 0.727087752 | 0.6377 | 0.798219725 |
| SL1344_RS15260_ | -0.327460376 | -0.232626096 | -0.400767745 | -0.194113379 | 0.277141107 | -0.660670919 | 0.227181893 | 0.337265067 | 0.603178816 | 0.4628 | 0.674422683 |
| SL1344_RS15265_ | 2.039651235 | 2.003043609 | 2.126677296 | 0.099655108 | 0.128731039 | 0.69959316 | 0.254000012 | 0.384595853 | 0.645536773 | 0.4976 | 0.714242985 |
| SL1344_RS15270_ | 5.787325077 | 5.599322339 | 5.873928931 | 0.284761473 | 0.121658359 | 2.604856045 | 4.68E-05 | 0.012094173 | 0.137832135 | 0.1 | 0.330257709 |
| SL1344_RS15275_ | 4.15327527 | 4.051411819 | 4.301385661 | 0.257220832 | 0.083699589 | 3.047090401 | 4.68E-05 | 0.01936552 | 0.164240277 | 0.1 | 0.330257709 |
| SL1344_RS15280_ | 1.288761532 | 1.20241913 | 1.367237848 | 0.16592669 | 0.121368729 | 1.303519983 | 0.072666725 | 0.131465912 | 0.368563556 | 0.1989 | 0.440822489 |
| SL1344_RS15285_ | 1.126547565 | 1.213217223 | 1.082680594 | -0.104842957 | 0.167044368 | -0.592163676 | 0.275333344 | 0.45623409 | 0.691971655 | 0.5592 | 0.751964203 |
| SL1344_RS15290_ | 1.856402355 | 1.890255466 | 1.831367251 | -0.035328339 | 0.177733652 | -0.180693691 | 0.426000003 | 0.761900102 | 0.894348859 | 0.8369 | 0.92518953 |
| SL1344_RS15295_ | 3.983839098 | 3.954526778 | 4.049086484 | 0.082180576 | 0.071463985 | 1.065548419 | 0.141905424 | 0.201547861 | 0.47840259 | 0.2897 | 0.544923475 |
| SL1344_RS15310_ | -0.433294789 | -0.520771872 | -0.351111517 | 0.156614329 | 0.331588873 | 0.448840297 | 0.304000009 | 0.494076672 | 0.727184844 | 0.5992 | 0.781374567 |
| SL1344_RS15315_ | 0.232466808 | 0.141060703 | 0.376876966 | 0.256179256 | 0.246130805 | 1.068784378 | 0.087275198 | 0.192281252 | 0.473377121 | 0.2206 | 0.459371655 |
| SL1344_RS15320_ | -0.39140634 | -0.590083275 | -0.171463828 | 0.494348828 | 0.413087382 | 1.277593908 | 0.041333441 | 0.127973178 | 0.397020545 | 0.1446 | 0.387814343 |
| SL1344_RS15325_ | -0.66533164 | -1.05173018 | -0.070711588 | 0.947447785 | 0.690476769 | 1.371538989 | 0.069953426 | 0.094616129 | 0.350010841 | 0.1694 | 0.428828991 |
| SL1344_RS15330_ | 1.886777843 | 1.938432611 | 1.806064458 | -0.148571015 | 0.148737184 | -0.974843897 | 0.131333363 | 0.203191665 | 0.48301462 | 0.2808 | 0.526285669 |
| SL1344_RS15355_ | 1.033771057 | 0.867207276 | 1.082657711 | 0.20573529 | 0.226067156 | 0.788099004 | 0.230000014 | 0.317812591 | 0.60326727 | 0.4637 | 0.693043089 |
| SL1344_RS15360_ | 4.462725848 | 4.261719401 | 4.616088216 | 0.373613988 | 0.146087284 | 2.532055815 | 4.68E-05 | 0.046360543 | 0.255331386 | 0.1 | 0.330257709 |
| SL1344_RS15365_ | 6.489290329 | 6.338149257 | 6.550654919 | 0.110928031 | 0.154784371 | 0.70056338 | 0.33333334 | 0.51055326 | 0.762305665 | 0.7 | 0.872542119 |
| SL1344_RS15370_ | 6.134702069 | 6.004247697 | 6.174846235 | 0.16326737 | 0.126779221 | 0.814759078 | 0.321333341 | 0.358840839 | 0.652128989 | 0.6729 | 0.854992422 |
| SL1344_RS15375_ | 0.917297136 | 0.852614982 | 0.943523857 | 0.067967061 | 0.184293334 | 0.297079845 | 0.385333338 | 0.705850507 | 0.861593007 | 0.7607 | 0.89126584 |
| SL1344_RS15380_ | -1.67619642 | -1.368056242 | -1.857465267 | -0.345427016 | 0.615040547 | -0.537301361 | 0.287141915 | 0.479697459 | 0.725096866 | 0.6016 | 0.786906266 |
| SL1344_RS15385_ | -2.863895342 | -3.166941764 | -2.764880577 | 0.358224035 | 0.701771304 | 0.418730855 | 0.360000006 | 0.591195239 | 0.793866619 | 0.7182 | 0.866170119 |
| SL1344_RS15390_ | -1.527330209 | -2.162376369 | -1.375054022 | 0.75002394 | 0.506885702 | 1.223541802 | 0.10933337 | 0.167711224 | 0.449292518 | 0.2418 | 0.49252925 |
| SL1344_RS15395_ | -1.009222093 | -1.055993989 | -0.940427664 | 0.115468087 | 0.28550488 | 0.365375093 | 0.341105936 | 0.524581971 | 0.73588467 | 0.633 | 0.793800152 |
| SL1344_RS15400_ | -1.545601671 | -1.806514763 | -1.357898505 | 0.4517568 | 0.319162015 | 1.319271569 | 0.070619647 | 0.127329593 | 0.36459507 | 0.192 | 0.432587345 |
| SL1344_RS15405_ | -1.387444896 | -1.30250876 | -1.468707609 | -0.161776154 | 0.309941501 | -0.501796342 | 0.292666676 | 0.452594249 | 0.679536548 | 0.5645 | 0.743526966 |
| SL1344_RS15410_ | 1.879337016 | 1.772577686 | 1.962216075 | 0.21145455 | 0.18444858 | 1.169073591 | 0.033333467 | 0.246695035 | 0.549375903 | 0.1359 | 0.37590174 |
| SL1344_RS15415_ | 2.015807808 | 1.959581521 | 2.072434845 | 0.114418256 | 0.100256058 | 1.102053514 | 0.103333373 | 0.173417176 | 0.427043865 | 0.2511 | 0.491712487 |
| SL1344_RS15420_ | 0.041341924 | 0.017822796 | 0.07446236 | 0.071427408 | 0.235512076 | 0.293015453 | 0.37708195 | 0.569869045 | 0.772559936 | 0.7107 | 0.84263664 |
| SL1344_RS15425_ | 0.854293808 | 0.873390142 | 0.835824853 | -0.079072616 | 0.235369489 | -0.3163399 | 0.367333339 | 0.588428492 | 0.790630827 | 0.7408 | 0.865176897 |
| SL1344_RS15430_ | 1.488119792 | 1.50829876 | 1.465666686 | 0.125164661 | 0.324862046 | 0.381745532 | 0.409726852 | 0.575683175 | 0.794025675 | 0.9113 | 0.957942221 |
| SL1344_RS15435_ | 0.934369978 | 0.837573116 | 1.067987258 | 0.229152988 | 0.133733289 | 1.63017453 | 0.046000096 | 0.090253333 | 0.313108688 | 0.1547 | 0.395398647 |
| SL1344_RS15440_ | 0.246284792 | 0.156213616 | 0.369682224 | 0.236652018 | 0.197312563 | 1.15727388 | 0.081279199 | 0.168421884 | 0.433185749 | 0.2179 | 0.455848891 |
| SL1344_RS15445_ | -1.78640264 | -1.938441146 | -1.355180663 | 0.529981893 | 0.409438265 | 1.166654063 | 0.129333364 | 0.185947217 | 0.450143091 | 0.2937 | 0.536677145 |
| SL1344_RS15450_ | 1.330954757 | 1.375315385 | 1.29699712 | -0.034361925 | 0.271462508 | -0.110675474 | 0.448367757 | 0.807395953 | 0.919483191 | 0.8639 | 0.940508691 |
| SL1344_RS15455_ | 2.563094482 | 2.575849099 | 2.538244182 | -0.050459917 | 0.132512916 | -0.356683435 | 0.351099273 | 0.569073355 | 0.781275797 | 0.7139 | 0.853262969 |
| SL1344_RS15460_ | 2.353962004 | 2.716114845 | 2.32309457 | -0.393953282 | 0.248600762 | -0.513242047 | 0.33333334 | 0.692943766 | 0.868953178 | 0.7 | 0.872542119 |
| SL1344_RS15465_ | 4.133408816 | 4.042691856 | 4.155520382 | 0.117442607 | 0.81374388 | 0.163488932 | 0.335109934 | 0.894602013 | 0.962090103 | 0.7045 | 0.874430761 |
| SL1344_RS15475_ | 0.413550499 | 0.205986368 | 0.477994177 | 0.249492239 | 0.207669387 | 0.967894565 | 0.167221874 | 0.232065278 | 0.508086427 | 0.3546 | 0.59533948 |
| SL1344_RS15480_ | 1.222902546 | 1.115190378 | 1.360532778 | 0.237336423 | 0.153732546 | 1.491855766 | 0.056628991 | 0.093431953 | 0.32872987 | 0.1572 | 0.403604249 |
| SL1344_RS15485_ | 2.480434427 | 2.422996937 | 2.605791172 | 0.175948036 | 0.099249602 | 1.668579659 | 0.031978821 | 0.092170807 | 0.330999005 | 0.1398 | 0.379468664 |
| SL1344_RS15490_ | 2.207199871 | 2.083464949 | 2.325752356 | 0.245419688 | 0.092765838 | 2.639285737 | 0.001335878 | 0.023231969 | 0.167113616 | 0.1006 | 0.331142832 |
| SL1344_RS15495_ | 2.2084435 | 2.127311433 | 2.329588528 | 0.250646127 | 0.173898651 | 1.43509465 | 0.03000015 | 0.087050385 | 0.330103392 | 0.1333 | 0.374451976 |
| SL1344_RS15500_ | -2.192541337 | -2.256976593 | -2.125761573 | 0.134758438 | 0.379237476 | 0.330363041 | 0.350000006 | 0.505811339 | 0.718151626 | 0.6135 | 0.776104756 |
| SL1344_RS15505_ | -2.776644998 | -2.815987663 | -2.714737466 | 0.105297329 | 0.481743903 | 0.193911572 | 0.416000003 | 0.572518199 | 0.767030845 | 0.6765 | 0.819943813 |
| SL1344_RS15510_ | -2.52112711 | -2.558412275 | -2.480062654 | 0.06975051 | 0.407428435 | 0.171220904 | 0.429713527 | 0.543457059 | 0.745713142 | 0.643 | 0.796231224 |

| | | | | | | | | | | | |
|---|---|---|---|---|---|---|---|---|---|---|---|
| SL1344_RS15515_ | -3.971501237 | -3.672453137 | -4.25777885 | -0.602361534 | 0.740208996 | -0.760956422 | 0.195869438 | 0.292617019 | 0.540884913 | 0.394 | 0.607501665 |
| SL1344_RS15520_ | -4.855474584 | -4.76077736 | -4.994399852 | -0.260569796 | 0.994608748 | -0.244566426 | 0.399067292 | 0.538548119 | 0.746112849 | 0.6604 | 0.809630464 |
| SL1344_RS15525_ | 2.263962563 | 2.191969736 | 2.496654534 | 0.322807628 | 0.205372226 | 1.451520106 | 0.039973462 | 0.08580518 | 0.33203341 | 0.1427 | 0.385750404 |
| SL1344_RS15530_ | 1.686085542 | 1.580709239 | 1.715884673 | 0.138727827 | 0.206421516 | 0.68201077 | 0.230513005 | 0.332263593 | 0.622139389 | 0.4591 | 0.677429283 |
| SL1344_RS15535_ | 3.479677519 | 3.583128139 | 3.441070954 | -0.114896196 | 0.151348188 | -0.661855063 | 0.287808137 | 0.432498395 | 0.700835304 | 0.578 | 0.789389117 |
| SL1344_RS15540_ | 2.753639765 | 2.706925093 | 2.775138902 | 0.079510237 | 0.260038042 | 0.310775823 | 0.361333339 | 0.724188224 | 0.88081938 | 0.7483 | 0.884013433 |
| SL1344_RS15545_ | -3.12638939 | -3.392898198 | -2.647810599 | 0.706461553 | 0.599601466 | 1.103564705 | 0.135333362 | 0.200973535 | 0.457518042 | 0.29 | 0.532329344 |
| SL1344_RS15550_ | 2.01023831 | 1.932630631 | 2.031147429 | 0.085736847 | 0.128115961 | 0.512221415 | 0.334443711 | 0.601137254 | 0.8009046 | 0.693 | 0.857210496 |
| SL1344_RS15555_ | 2.136452839 | 2.166823251 | 2.108850399 | -0.07426267 | 0.137899416 | -0.550164807 | 0.253333345 | 0.373406699 | 0.642691754 | 0.4994 | 0.699407989 |
| SL1344_RS15560_ | -0.666514339 | -0.786367828 | -0.49117373 | 0.314839058 | 0.348497077 | 0.898214777 | 0.155896093 | 0.224205295 | 0.503035687 | 0.3138 | 0.56760206 |
| SL1344_RS15565_ | 1.046093109 | 0.946698469 | 1.121782399 | 0.188040854 | 0.184120685 | 0.990883088 | 0.131333363 | 0.192018567 | 0.464182906 | 0.2787 | 0.526936631 |
| SL1344_RS15570_ | 1.360810917 | 1.307932608 | 1.395249406 | 0.047328085 | 0.250997066 | 0.179015743 | 0.434377084 | 0.744149384 | 0.8868668 | 0.8939 | 0.950879376 |
| SL1344_RS15575_ | 2.477125365 | 2.476471702 | 2.477415193 | 0.001517702 | 0.222159182 | 0.007487641 | 0.496666667 | 0.836502434 | 0.933727028 | 0.8983 | 0.953776349 |
| SL1344_RS15580_ | 1.961807437 | 1.816078013 | 2.082244164 | 0.274693291 | 0.120628142 | 2.286314646 | 0.001335878 | 0.044681297 | 0.233989266 | 0.1007 | 0.331287425 |
| SL1344_RS15585_ | 1.687240415 | 1.477590839 | 1.721641463 | 0.245087391 | 0.246742822 | 0.773261875 | 0.289333343 | 0.368129791 | 0.653236091 | 0.5807 | 0.791536045 |
| SL1344_RS15590_ | -1.91898487 | -1.489439777 | -2.458065635 | -1.059252497 | 0.656968047 | -1.649443678 | 0.003332494 | 0.097059555 | 0.347806862 | 0.103 | 0.334389285 |
| SL1344_RS15595_ | -0.673930742 | -0.675676966 | -0.672546608 | -0.026030286 | 0.305820869 | -0.087461893 | 0.462666668 | 0.665098163 | 0.835504879 | 0.7963 | 0.896693116 |
| SL1344_RS15600_ | 0.836155067 | 0.762226824 | 0.873995189 | 0.119432383 | 0.281796054 | 0.421058046 | 0.292666676 | 0.529699231 | 0.758017721 | 0.6018 | 0.793828031 |
| SL1344_RS15605_ | -0.860727728 | -1.069960426 | -0.704100316 | 0.342937329 | 0.282894817 | 1.1415227 | 0.114000035 | 0.173566631 | 0.431729169 | 0.2545 | 0.502082227 |
| SL1344_RS15610_ | -1.356316961 | -1.439850382 | -1.248519539 | 0.194348321 | 0.329872495 | 0.546613425 | 0.28400001 | 0.437549572 | 0.672151586 | 0.5391 | 0.731014054 |
| SL1344_RS15615_ | -2.425065827 | -2.537925854 | -2.258755128 | 0.29872189 | 0.455064188 | 0.641058891 | 0.251832124 | 0.385295962 | 0.625225696 | 0.4992 | 0.691534723 |
| SL1344_RS15620_ | 2.8167734 | 2.835965227 | 2.794065279 | -0.048478195 | 0.532298495 | -0.0795349 | 0.463333335 | 0.866110043 | 0.949828053 | 1 | 1 |
| SL1344_RS15625_ | 0.400739059 | 0.58729431 | 0.288283283 | -0.304233618 | 0.29032551 | -0.984529061 | 0.139333361 | 0.207035453 | 0.494492049 | 0.2822 | 0.542868702 |
| SL1344_RS15630_ | -1.333744558 | -1.423581821 | -1.251545751 | 0.165001142 | 0.382278124 | 0.414293855 | 0.330446376 | 0.523175049 | 0.741425617 | 0.6343 | 0.797836056 |
| SL1344_RS15635_ | 0.473072595 | 0.166532398 | 0.781814647 | 0.644651144 | 0.228445335 | 2.845116747 | 4.68E-05 | 0.021422457 | 0.169363305 | 0.1 | 0.330257709 |
| SL1344_RS15640_ | -0.230425077 | -0.147254579 | -0.302938996 | -0.133888727 | 0.315017356 | -0.419666714 | 0.334000007 | 0.559564568 | 0.771379825 | 0.6727 | 0.827447114 |
| SL1344_RS15645_ | -1.472309689 | -1.565080865 | -1.404548734 | 0.154186787 | 0.395507786 | 0.365864798 | 0.339773491 | 0.528845186 | 0.74688694 | 0.6495 | 0.808227554 |
| SL1344_RS15650_ | -2.004454592 | -1.99419324 | -2.019300603 | -0.016477215 | 0.352531963 | -0.04197583 | 0.482666667 | 0.582398064 | 0.774827793 | 0.6894 | 0.828327344 |
| SL1344_RS15655_ | -4.610821758 | -4.681662533 | -4.518695601 | 0.211556314 | 0.934671805 | 0.211093464 | 0.404000004 | 0.563403269 | 0.76153692 | 0.6792 | 0.821995358 |
| SL1344_RS15660_ | -2.733927465 | -2.6684568 | -2.791415983 | -0.150391189 | 0.794745659 | -0.182374475 | 0.433044639 | 0.715952575 | 0.868135159 | 0.8344 | 0.919597154 |
| SL1344_RS15665_ | -3.40845525 | -3.651658342 | -3.283997332 | 0.39707298 | 0.77836434 | 0.48082081 | 0.298000009 | 0.43690188 | 0.683938064 | 0.5728 | 0.750783969 |
| SL1344_RS15670_ | -2.501696968 | -2.624296394 | -2.38136264 | 0.235522956 | 0.464257919 | 0.476208497 | 0.305796144 | 0.466679627 | 0.692961888 | 0.5802 | 0.754689103 |
| SL1344_RS15675_ | -4.497380731 | -4.386761372 | -4.601733676 | -0.193452233 | 0.827890178 | -0.214030834 | 0.402398405 | 0.55609907 | 0.757111507 | 0.6693 | 0.815377045 |
| SL1344_RS15680_ | -0.051488796 | -0.000786696 | -0.099248504 | -0.060859293 | 0.374877156 | -0.162000714 | 0.433333336 | 0.747257429 | 0.886158064 | 0.855 | 0.930918836 |
| SL1344_RS15685_ | -3.406144911 | -3.363174211 | -3.471505011 | -0.081075091 | 0.731961552 | -0.109834627 | 0.455696204 | 0.661156112 | 0.833293497 | 0.7765 | 0.887306413 |
| SL1344_RS15690_ | 1.850456944 | 1.75665185 | 1.923500217 | 0.086384925 | 0.276968238 | 0.346620456 | 0.351765496 | 0.630523259 | 0.826436891 | 0.7318 | 0.866380621 |
| SL1344_RS15695_ | 1.527103263 | 1.274878545 | 1.690321899 | 0.304282525 | 0.385832249 | 0.842167988 | 0.165333356 | 0.255743193 | 0.55453945 | 0.3015 | 0.575002526 |
| SL1344_RS15700_ | -5.653367509 | -5.423710118 | -5.896409025 | -0.473738891 | 1.224688973 | -0.343432416 | 0.338666673 | 0.506580944 | 0.71567067 | 0.6076 | 0.769521016 |
| SL1344_RS15705_ | -2.550385909 | -2.831412512 | -2.305926101 | 0.532515947 | 0.390313115 | 1.328407839 | 0.070619647 | 0.130596011 | 0.360226562 | 0.2027 | 0.439171893 |
| SL1344_RS15710_ | 1.871461863 | 2.009426809 | 1.644293258 | -0.356000065 | 0.452662107 | -0.621615743 | 0.242000013 | 0.356845979 | 0.649735798 | 0.4929 | 0.716299094 |
| SL1344_RS15715_ | 1.493337507 | 1.536657982 | 1.432599229 | -0.106010812 | 0.129720113 | -0.761201119 | 0.200532995 | 0.29897641 | 0.560728307 | 0.3973 | 0.623123648 |
| SL1344_RS15720_ | 2.004210963 | 1.93766305 | 2.049240126 | 0.111117083 | 0.152646586 | 0.685778205 | 0.222666682 | 0.343131116 | 0.615328575 | 0.4508 | 0.680317633 |
| SL1344_RS15725_ | 4.384496973 | 4.373223888 | 4.394515884 | -0.011607899 | 0.199957916 | -0.054052711 | 0.473017989 | 0.890776574 | 0.958694546 | 0.9538 | 0.980395512 |
| SL1344_RS15730_ | 3.939833628 | 3.886199448 | 3.978822135 | 0.089451871 | 0.167599076 | 0.441599375 | 0.353333339 | 0.46985814 | 0.732100312 | 0.7664 | 0.885934777 |
| SL1344_RS15735_ | 3.205514939 | 3.059253305 | 3.307343088 | 0.248989397 | 0.124890801 | 1.983107699 | 0.00067292 | 0.067029195 | 0.296484142 | 0.1015 | 0.332405523 |
| SL1344_RS15740_ | 2.333021011 | 2.130740494 | 2.359860573 | 0.224282711 | 0.196758872 | 0.77397239 | 0.311333341 | 0.383384553 | 0.666693898 | 0.6461 | 0.836082745 |
| SL1344_RS15745_ | 1.573549757 | 1.456444005 | 1.644541482 | 0.121160416 | 0.214062025 | 0.560647038 | 0.309793479 | 0.484847954 | 0.729456207 | 0.6066 | 0.791554376 |
| SL1344_RS15750_ | 1.438696518 | 1.489048421 | 1.386849961 | -0.051976728 | 0.191057136 | -0.284222136 | 0.393333337 | 0.665212516 | 0.839251984 | 0.7842 | 0.891341703 |
| SL1344_RS15755_ | 1.96411377 | 1.947831504 | 1.97541561 | 0.050834555 | 0.226397889 | 0.226729193 | 0.389740178 | 0.728540235 | 0.878978398 | 0.8062 | 0.90727499 |
| SL1344_RS15760_ | 0.926916511 | 0.922955151 | 0.929356276 | 0.054214471 | 0.27994897 | 0.172437725 | 0.422000003 | 0.617648341 | 0.818140877 | 0.8489 | 0.927332401 |
| SL1344_RS15765_ | -1.374382188 | -1.379868669 | -1.367466705 | -0.02386398 | 0.396648231 | -0.053833315 | 0.474666668 | 0.684401585 | 0.847484728 | 0.8027 | 0.900233509 |
| SL1344_RS15770_ | 2.629823894 | 2.535156505 | 2.761347028 | 0.17121901 | 0.175037235 | 1.009356631 | 0.141333361 | 0.205847019 | 0.495866522 | 0.2625 | 0.532445018 |
| SL1344_RS15775_ | 2.067963441 | 2.044281781 | 2.119294697 | 0.069866384 | 0.335920852 | 0.226960071 | 0.33933334 | 0.759088478 | 0.896650167 | 0.6841 | 0.846116704 |

| | | | | | | | | | | | |
|---|---|---|---|---|---|---|---|---|---|---|---|
| SL1344_RS15780_ | 0.392086444 | 0.277480011 | 0.467471384 | 0.128950819 | 0.338479913 | 0.357231499 | 0.373084615 | 0.662679705 | 0.841135213 | 0.7609 | 0.886274797 |
| SL1344_RS15785_ | 0.777958633 | 0.530337389 | 0.833533623 | 0.271213168 | 0.252718499 | 0.796864179 | 0.254666679 | 0.335055158 | 0.618694113 | 0.514 | 0.736390364 |
| SL1344_RS15790_ | -1.867337477 | -2.03834867 | -1.707068832 | 0.329902876 | 0.346293894 | 0.902993469 | 0.161333357 | 0.256217424 | 0.506276765 | 0.3424 | 0.569379725 |
| SL1344_RS15795_ | 5.946771856 | 5.963137672 | 5.928175609 | 0.009375353 | 0.102014086 | 0.102221731 | 0.469686877 | 0.859343909 | 0.944865251 | 0.9907 | 0.996063451 |
| SL1344_RS15800_ | 4.646632903 | 4.7059875 | 4.61564131 | -0.061009848 | 0.079651855 | -0.78446171 | 0.204666683 | 0.315234243 | 0.599348967 | 0.3936 | 0.64096883 |
| SL1344_RS15810_ | 2.007754836 | 1.971062399 | 2.06845245 | 0.073972883 | 0.139536457 | 0.490104263 | 0.303131254 | 0.503201182 | 0.731899209 | 0.6023 | 0.785302839 |
| SL1344_RS15815_ | 2.188516772 | 2.213152444 | 2.172269835 | -0.038579921 | 0.112828712 | -0.312079044 | 0.366666672 | 0.593313332 | 0.790418969 | 0.7071 | 0.847835411 |
| SL1344_RS15820_ | 2.649401472 | 2.573608636 | 2.67669369 | 0.091749203 | 0.099419419 | 0.81645074 | 0.216000016 | 0.313486644 | 0.586133798 | 0.4389 | 0.669056312 |
| SL1344_RS15825_ | 2.468973603 | 2.310064974 | 2.557670324 | 0.226785087 | 0.172592873 | 1.326769142 | 0.084666716 | 0.108145414 | 0.366393286 | 0.1837 | 0.443903984 |
| SL1344_RS15830_ | 3.550500232 | 3.511150403 | 3.572350509 | 0.069331185 | 0.12084632 | 0.536022948 | 0.227848116 | 0.353880885 | 0.643720759 | 0.4686 | 0.691255114 |
| SL1344_RS15835_ | 1.362079574 | 1.204581983 | 1.451371244 | 0.190328016 | 0.232375918 | 0.845853964 | 0.187333352 | 0.28331751 | 0.570175395 | 0.367 | 0.62333113 |
| SL1344_RS15840_ | 1.815553409 | 1.93337153 | 1.674597359 | 0.050456922 | 0.4927039 | 0.100794888 | 0.461692207 | 0.835970927 | 0.935279707 | 0.9922 | 0.996690608 |
| SL1344_RS15845_ | 3.012659369 | 2.846014622 | 3.205749449 | 0.362709746 | 0.077508827 | 4.600149371 | 4.68E-05 | 0.007555253 | 0.100858494 | 0.1 | 0.330257709 |
| SL1344_RS15850_ | -0.17004141 | -0.190270895 | -0.142246929 | 0.099445993 | 0.299389406 | 0.300364881 | 0.37974684 | 0.551663882 | 0.766366385 | 0.7233 | 0.850050152 |
| SL1344_RS15855_ | -3.144783829 | -3.340275769 | -2.986471998 | 0.122272923 | 1.124991844 | 0.101336538 | 0.452365092 | 0.754847081 | 0.891175713 | 0.8658 | 0.936118351 |
| SL1344_RS15860_ | 1.587076093 | 1.434665445 | 1.734857419 | 0.332426695 | 0.153670924 | 2.133403941 | 0.004001152 | 0.035265881 | 0.210612889 | 0.1032 | 0.334006097 |
| SL1344_RS15865_ | -1.40110088 | -1.556464839 | -1.14644089 | 0.426543095 | 0.393735477 | 1.036651181 | 0.127914754 | 0.204491512 | 0.474439626 | 0.2903 | 0.53334371 |
| SL1344_RS15870_ | 1.520196182 | 1.398055065 | 1.687552012 | 0.293445915 | 0.142416736 | 2.007074662 | 0.011333737 | 0.048027929 | 0.240578171 | 0.1093 | 0.342544082 |
| SL1344_RS15875_ | 5.894896128 | 5.691189494 | 5.981263018 | 0.292670981 | 0.107153862 | 2.84325521 | 4.68E-05 | 0.01245311 | 0.139232134 | 0.1 | 0.330257709 |
| SL1344_RS15880_ | 3.97153798 | 3.947457565 | 4.012963394 | 0.056892241 | 0.203955593 | 0.274273266 | 0.371085948 | 0.817995202 | 0.926948208 | 0.7945 | 0.91115444 |
| SL1344_RS15890_ | -2.676809933 | -2.822345717 | -2.444368074 | 0.39801328 | 0.708249156 | 0.530739817 | 0.273817466 | 0.446119617 | 0.690350868 | 0.5534 | 0.751211391 |
| SL1344_RS15895_ | -2.747949456 | -2.768270517 | -2.735994616 | 0.089674806 | 0.554179595 | 0.146897772 | 0.440000002 | 0.605529222 | 0.795560751 | 0.732 | 0.855761412 |
| SL1344_RS15900_ | -1.921946222 | -2.037552537 | -1.791319916 | 0.251575079 | 0.328537633 | 0.703899734 | 0.207861442 | 0.314940088 | 0.560353039 | 0.4169 | 0.627875957 |
| SL1344_RS15905_ | -1.180953544 | -1.200376737 | -1.160131625 | 0.04828276 | 0.293942564 | 0.150705338 | 0.436666669 | 0.608164138 | 0.796750088 | 0.7198 | 0.850398249 |
| SL1344_RS15910_ | -2.710006453 | -2.812289754 | -2.621723072 | 0.147900411 | 0.574577855 | 0.24806516 | 0.394666671 | 0.600689865 | 0.793232979 | 0.7181 | 0.848539211 |
| SL1344_RS15915_ | 5.77106688 | 5.5490387 | 5.904428211 | 0.344125147 | 0.107560203 | 2.972550096 | 4.68E-05 | 0.019955971 | 0.173104027 | 0.1 | 0.330257709 |
| SL1344_RS15920_ | 1.184273595 | 1.069706701 | 1.286566235 | 0.229220335 | 0.162545465 | 1.345960063 | 0.056628991 | 0.109352583 | 0.352038613 | 0.1712 | 0.415444754 |
| SL1344_RS15925_ | 4.096275439 | 3.824765724 | 4.183329085 | 0.250132855 | 0.268224464 | 0.752991019 | 0.321333341 | 0.399742454 | 0.68416394 | 0.6694 | 0.852917984 |
| SL1344_RS15930_ | -1.886103585 | -1.827739417 | -1.961687938 | -0.143295788 | 0.330611893 | -0.39930294 | 0.319120594 | 0.458173332 | 0.679480139 | 0.5704 | 0.742455543 |
| SL1344_RS15935_ | -2.389249961 | -2.292200785 | -2.508317974 | -0.211972662 | 0.442798751 | -0.457285021 | 0.305129922 | 0.487266717 | 0.706922524 | 0.6033 | 0.771555833 |
| SL1344_RS15940_ | -1.14791299 | -1.109327205 | -1.17289136 | -0.082106199 | 0.31680808 | -0.241197192 | 0.385333338 | 0.572987679 | 0.772070712 | 0.6871 | 0.82872885 |
| SL1344_RS15945_ | -0.850534479 | -0.660789373 | -1.038905464 | -0.382980823 | 0.229243359 | -1.656529554 | 0.034666795 | 0.076703873 | 0.286128271 | 0.1409 | 0.379341406 |
| SL1344_RS15950_ | -2.363307689 | -2.252458916 | -2.437078249 | -0.145208382 | 0.523288906 | -0.25074243 | 0.390666671 | 0.634659945 | 0.81580461 | 0.7416 | 0.867211144 |
| SL1344_RS15955_ | -2.938404643 | -2.740039904 | -3.13649758 | -0.38688173 | 0.443408701 | -0.853855584 | 0.168554319 | 0.263618318 | 0.506581851 | 0.3603 | 0.5788642 |
| SL1344_RS15960_ | 2.010012299 | 1.80818971 | 2.044970214 | 0.23433549 | 0.310213289 | 0.615280688 | 0.332000007 | 0.467061435 | 0.73032027 | 0.6942 | 0.867878817 |
| SL1344_RS15965_ | -2.673537115 | -2.747962121 | -2.558370623 | 0.188269468 | 0.530574215 | 0.312314947 | 0.358000006 | 0.571053619 | 0.768595999 | 0.6715 | 0.82223043 |
| SL1344_RS15970_ | 3.735751737 | 3.568956775 | 3.751162161 | 0.183658369 | 0.17161434 | 0.944985674 | 0.238000013 | 0.258652301 | 0.560923718 | 0.4635 | 0.695199135 |
| SL1344_RS15975_ | 0.89640566 | 0.924686426 | 0.872771054 | 0.052517314 | 0.39350895 | 0.144375061 | 0.443037977 | 0.780857129 | 0.90685162 | 0.925 | 0.966545041 |
| SL1344_RS15985_ | 3.464239812 | 3.44436916 | 3.485373339 | -0.060985524 | 0.212126821 | -0.211616344 | 0.433044639 | 0.505821649 | 0.757342842 | 0.9572 | 0.979610023 |
| SL1344_RS15990_ | 4.373092081 | 4.708227945 | 4.027184883 | -0.688417788 | 0.067076121 | -10.64177672 | 4.68E-05 | 0.023470678 | 0.18857992 | 0.1 | 0.330257709 |
| SL1344_RS15995_ | -0.093212144 | 0.009135564 | -0.256691143 | -0.177052203 | 0.361407729 | -0.488501937 | 0.320666674 | 0.536219044 | 0.763455326 | 0.6642 | 0.827112668 |
| SL1344_RS16000_ | 0.261402329 | -0.079215131 | 0.523478819 | 0.602761315 | 0.17285024 | 3.401242981 | 4.68E-05 | 0.01376253 | 0.129691723 | 0.1 | 0.330257709 |
| SL1344_RS16005_ | 1.633620726 | 1.525650011 | 1.738831616 | 0.228203325 | 0.130197836 | 1.715769979 | 0.021319333 | 0.067058434 | 0.281519716 | 0.1215 | 0.358081863 |
| SL1344_RS16015_ | 1.174863437 | 1.159961813 | 1.19134224 | -0.01486087 | 0.196411377 | -0.078122961 | 0.469686877 | 0.711456308 | 0.865329548 | 0.86 | 0.930861516 |
| SL1344_RS16020_ | 1.071895683 | 1.029854916 | 1.153818569 | 0.115041925 | 0.276473551 | 0.434140668 | 0.289333343 | 0.538932682 | 0.764425872 | 0.5573 | 0.769180671 |
| SL1344_RS16025_ | -0.701005492 | -0.699146487 | -0.70516394 | -0.01434448 | 0.251608997 | -0.055193431 | 0.477333334 | 0.634951082 | 0.814723055 | 0.7401 | 0.864404249 |
| SL1344_RS16030_ | -2.005308154 | -1.664027166 | -2.452766128 | -0.806835097 | 0.393723119 | -1.944425796 | 0.009333825 | 0.059830685 | 0.25980047 | 0.1105 | 0.343167943 |
| SL1344_RS16035_ | 1.203585373 | 1.191212517 | 1.216768053 | 0.076568733 | 0.235426391 | 0.301629752 | 0.366000005 | 0.545905921 | 0.771736777 | 0.7572 | 0.876150346 |
| SL1344_RS16040_ | 1.274971911 | 1.257506039 | 1.280592416 | 0.023672378 | 0.162068152 | 0.137384924 | 0.446369089 | 0.696881834 | 0.856393076 | 0.788 | 0.899691512 |
| SL1344_RS16045_ | 0.797431568 | 0.803175508 | 0.793286602 | 0.003146373 | 0.187427049 | 0.01692143 | 0.493670886 | 0.690144381 | 0.853474654 | 0.8091 | 0.907634369 |
| SL1344_RS16050_ | 1.402919997 | 1.259473854 | 1.507166136 | 0.258235752 | 0.154161925 | 1.72627754 | 0.008661422 | 0.092298418 | 0.334773807 | 0.1084 | 0.341147878 |
| SL1344_RS16055_ | 2.41202995 | 2.339946025 | 2.570780607 | 0.224759373 | 0.115379348 | 1.782981324 | 0.022666866 | 0.073159183 | 0.299865714 | 0.1254 | 0.363347482 |
| SL1344_RS16060_ | 0.307930991 | 0.355322376 | 0.26315922 | -0.003442182 | 0.576119521 | -0.006794806 | 0.494337109 | 0.859579713 | 0.944667235 | 0.904 | 0.956856543 |

| | | | | | | | | | | | |
|---|---|---|---|---|---|---|---|---|---|---|---|
| SL1344_RS16065_ | 5.635915077 | 5.655846463 | 5.540586658 | -0.110011449 | 0.119899101 | -0.439091757 | 0.33333334 | 0.82177343 | 0.931078915 | 0.7003 | 0.872669503 |
| SL1344_RS16070_ | 1.559287837 | 1.56998843 | 1.545178232 | 0.061714637 | 0.218385201 | 0.286686238 | 0.414000003 | 0.629068215 | 0.819701514 | 0.8776 | 0.938743149 |
| SL1344_RS16075_ | 1.965900297 | 1.949718208 | 1.984318037 | 0.091837915 | 0.182112807 | 0.501851255 | 0.325782819 | 0.448385412 | 0.703311378 | 0.6729 | 0.823436879 |
| SL1344_RS16080_ | 1.98048762 | 1.942943541 | 1.996439824 | 0.188866048 | 0.340426976 | 0.579671336 | 0.303333342 | 0.409949549 | 0.686313754 | 0.6278 | 0.811532739 |
| SL1344_RS16085_ | 0.969088315 | 0.857957274 | 1.120947972 | 0.264705699 | 0.134740971 | 1.902945688 | 0.018000252 | 0.056859034 | 0.253740696 | 0.1204 | 0.356876566 |
| SL1344_RS16090_ | 1.279662626 | 1.006746107 | 1.347928829 | 0.312226139 | 0.260891663 | 0.988386416 | 0.194000018 | 0.232269328 | 0.527893611 | 0.3535 | 0.616578586 |
| SL1344_RS16095_ | 1.44481709 | 1.355285215 | 1.530309083 | 0.29614556 | 0.283410571 | 1.083282155 | 0.071952091 | 0.154384737 | 0.43563501 | 0.1851 | 0.430162629 |
| SL1344_RS16100_ | 2.749315575 | 2.77595989 | 2.722118158 | -0.043435555 | 0.155058708 | -0.281745245 | 0.383744175 | 0.674357963 | 0.846676184 | 0.7905 | 0.896938299 |
| SL1344_RS16105_ | 0.816340835 | 0.730480786 | 0.985637493 | 0.234326261 | 0.196536104 | 1.110531523 | 0.1206667 | 0.173055573 | 0.442061141 | 0.2632 | 0.519249066 |
| SL1344_RS16110_ | -4.914313128 | -5.507662248 | -4.636682159 | 0.863495453 | 1.218390069 | 0.631466688 | 0.238000013 | 0.351702714 | 0.614366941 | 0.4705 | 0.682186458 |
| SL1344_RS16115_ | -1.64669197 | -1.83628345 | -1.514338588 | 0.321400208 | 0.296389569 | 1.025672018 | 0.120000033 | 0.20776826 | 0.457859699 | 0.2892 | 0.520285157 |
| SL1344_RS16120_ | -0.128417323 | 0.282770111 | -0.293487573 | -0.571510108 | 0.510575533 | -1.030637705 | 0.13200003 | 0.216114209 | 0.512886121 | 0.2747 | 0.539571125 |
| SL1344_RS16125_ | -1.87930757 | -1.916056589 | -1.82879716 | 0.063858017 | 0.369479775 | 0.167877493 | 0.431045972 | 0.587753633 | 0.780972269 | 0.689 | 0.831691051 |
| SL1344_RS16130_ | -0.236678859 | -0.602950378 | -0.093008763 | 0.481501368 | 0.253561402 | 1.73764066 | 0.03533346 | 0.086423255 | 0.319428687 | 0.1416 | 0.381761541 |
| SL1344_RS16135_ | -1.682599923 | -1.904937914 | -1.299523873 | 0.577940416 | 0.433374628 | 1.259412369 | 0.100599641 | 0.14215024 | 0.398805811 | 0.2184 | 0.469021182 |
| SL1344_RS16140_ | -0.560028636 | -0.667979434 | -0.420028773 | 0.297406936 | 0.343718553 | 0.833974071 | 0.174000021 | 0.256703329 | 0.53660286 | 0.3428 | 0.588403414 |
| SL1344_RS16145_ | 0.481887494 | 0.293747371 | 0.624040951 | 0.33654315 | 0.252183682 | 1.297220099 | 0.081945421 | 0.12170869 | 0.379687492 | 0.1937 | 0.447992237 |
| SL1344_RS16150_ | 0.092502391 | 0.108066702 | 0.08302205 | -0.038513624 | 0.227070224 | -0.155394492 | 0.430379749 | 0.642653557 | 0.821593935 | 0.749 | 0.870349261 |
| SL1344_RS16155_ | -3.155818381 | -3.208243273 | -3.100771744 | 0.116107959 | 0.480719262 | 0.225120216 | 0.405333337 | 0.525833719 | 0.730299379 | 0.6312 | 0.786792773 |
| SL1344_RS16160_ | -0.151288821 | -0.333915795 | 0.311247389 | 0.81498012 | 0.760993992 | 1.035748207 | 0.142666694 | 0.171211846 | 0.463826486 | 0.269 | 0.544328733 |
| SL1344_RS16165_ | -1.201653482 | -1.263521545 | -1.130704475 | 0.137298801 | 0.274850411 | 0.473869487 | 0.310666675 | 0.461327777 | 0.684915182 | 0.5633 | 0.741818169 |
| SL1344_RS16170_ | -0.454298914 | -0.45013817 | -0.459578969 | -0.011176585 | 0.234269899 | -0.042391988 | 0.482345104 | 0.636378046 | 0.815756681 | 0.7482 | 0.869795275 |
| SL1344_RS16175_ | -1.582714114 | -1.707562855 | -1.452649001 | 0.254787071 | 0.297410758 | 0.78118003 | 0.180546322 | 0.290649792 | 0.534135438 | 0.382 | 0.598086535 |
| SL1344_RS16180_ | -0.378573706 | -0.446042551 | -0.308113303 | 0.134557001 | 0.18461351 | 0.692538478 | 0.209193887 | 0.322540598 | 0.562903174 | 0.4174 | 0.628750412 |
| SL1344_RS16185_ | -0.429780643 | -0.407149039 | -0.458593899 | -0.055868833 | 0.387048489 | -0.14823652 | 0.438000002 | 0.697226273 | 0.860197066 | 0.8296 | 0.917900435 |
| SL1344_RS16190_ | -2.131297167 | -2.016570074 | -2.252611591 | -0.240745928 | 0.325157025 | -0.683081226 | 0.230666681 | 0.34780756 | 0.584883226 | 0.4524 | 0.652168819 |
| SL1344_RS16195_ | -1.966053731 | -1.870992634 | -2.065951779 | -0.185078992 | 0.355249178 | -0.501513729 | 0.302666675 | 0.44578848 | 0.671128571 | 0.5636 | 0.737964094 |
| SL1344_RS16200_ | -1.920858402 | -1.914306008 | -1.927321921 | -0.000789087 | 0.358533505 | -0.001614342 | 0.498666667 | 0.602537536 | 0.791643937 | 0.7121 | 0.845944016 |
| SL1344_RS16205_ | -3.101274902 | -3.470237383 | -2.986630172 | 0.448502616 | 0.71263476 | 0.571441435 | 0.29113925 | 0.432293982 | 0.680575123 | 0.5811 | 0.76669359 |
| SL1344_RS16210_ | 2.271994079 | 1.858992925 | 2.346230899 | 0.474911124 | 0.441309141 | 0.773812787 | 0.331112599 | 0.380002908 | 0.66883986 | 0.6964 | 0.870206943 |
| SL1344_RS16215_ | 3.726644923 | 3.346989237 | 3.817001085 | 0.467288884 | 0.37961555 | 1.247679433 | 0.005330642 | 0.169098773 | 0.464858597 | 0.1063 | 0.33942548 |
| SL1344_RS16220_ | 2.723347867 | 2.53404568 | 2.761134008 | 0.239105413 | 0.356070728 | 0.655517623 | 0.230666681 | 0.328928575 | 0.627355863 | 0.4265 | 0.692042719 |
| SL1344_RS16225_ | 2.991939439 | 2.99357247 | 2.989296571 | 0.036329522 | 0.205451967 | 0.08585798 | 0.409333337 | 0.478791219 | 0.740713346 | 0.8276 | 0.916763614 |
| SL1344_RS16235_ | 2.009117114 | 1.884287525 | 2.177271977 | 0.303698085 | 0.161373083 | 1.923304145 | 0.003334715 | 0.056214345 | 0.268256011 | 0.1038 | 0.335530219 |
| SL1344_RS16240_ | 0.555504916 | 0.48649368 | 0.650408236 | 0.119262506 | 0.234156727 | 0.474998331 | 0.304463699 | 0.516709397 | 0.743035334 | 0.6312 | 0.80226517 |
| SL1344_RS16245_ | -1.745033033 | -1.581192351 | -1.896192887 | -0.325394304 | 0.381149838 | -0.826469855 | 0.186000019 | 0.279183313 | 0.538394647 | 0.3804 | 0.607058376 |
| SL1344_RS16250_ | -2.020077456 | -1.686329851 | -2.111853971 | -0.379410707 | 0.516382915 | -0.61907238 | 0.293333343 | 0.486246536 | 0.721482463 | 0.6085 | 0.795730973 |
| SL1344_RS16255_ | 0.714222076 | 0.947707193 | 0.567880515 | -0.384954251 | 0.178278525 | -2.069180242 | 0.007995244 | 0.052878057 | 0.252876042 | 0.1091 | 0.342146183 |
| SL1344_RS16260_ | -0.221385991 | 0.022129901 | -0.369671104 | -0.424721905 | 0.299196036 | -1.34723845 | 0.048634334 | 0.12442645 | 0.387648605 | 0.1506 | 0.392466411 |
| SL1344_RS16265_ | -0.046907715 | 0.252995273 | -0.234077516 | -0.481146339 | 0.249989767 | -1.87264374 | 0.018654474 | 0.053351038 | 0.25315367 | 0.1171 | 0.353177274 |
| SL1344_RS16270_ | 1.005588547 | 1.181685839 | 0.904145594 | -0.309876837 | 0.235351544 | -1.311046791 | 0.054630326 | 0.119435444 | 0.380973759 | 0.169 | 0.415532895 |
| SL1344_RS16275_ | 2.278639031 | 2.511785544 | 2.182365287 | -0.321530604 | 0.139511583 | -2.041134247 | 0.008000574 | 0.064777741 | 0.289347654 | 0.106 | 0.338815303 |
| SL1344_RS16280_ | 0.762439981 | 1.141277611 | 0.658473436 | -0.439600021 | 0.28591377 | -1.252452304 | 0.108594308 | 0.14968813 | 0.430032695 | 0.2331 | 0.494904111 |
| SL1344_RS16285_ | 0.14477744 | 0.761330191 | -0.086930582 | -0.741553671 | 0.437952786 | -1.561552987 | 0.048634334 | 0.080947254 | 0.323747462 | 0.1446 | 0.393292639 |
| SL1344_RS16290_ | 1.308161004 | 1.761709391 | 1.192764093 | -0.486098844 | 0.359481351 | -1.084545724 | 0.156000024 | 0.185162483 | 0.479008956 | 0.2875 | 0.559883415 |
| SL1344_RS16305_ | 2.40999167 | 2.659747431 | 2.226447123 | -0.431268546 | 0.081038911 | -5.248623823 | 4.68E-05 | 0.004439412 | 0.076484745 | 0.1 | 0.330257709 |
| SL1344_RS16310_ | 3.11471606 | 3.649213296 | 2.617607578 | -0.972374851 | 0.305418667 | -3.097076208 | 4.68E-05 | 0.007528863 | 0.111847543 | 0.1 | 0.330257709 |
| SL1344_RS16315_ | 1.532998576 | 2.362586547 | 1.222933303 | -1.115333989 | 0.158935345 | -5.050988434 | 4.68E-05 | 0.03039587 | 0.210217203 | 0.1 | 0.330257709 |
| SL1344_RS16320_ | 1.929582134 | 2.528492368 | 1.485073884 | -1.048964757 | 0.19535304 | -5.344581285 | 4.68E-05 | 0.005205046 | 0.090272388 | 0.1 | 0.330257709 |
| SL1344_RS16325_ | 1.966816531 | 2.332940801 | 1.585212987 | -0.742723119 | 0.102310972 | -7.224384971 | 4.68E-05 | 0.001487864 | 0.047979431 | 0.1 | 0.330257709 |
| SL1344_RS16330_ | 2.295388607 | 2.420317858 | 2.209032698 | -0.24588795 | 0.21316254 | -1.148681921 | 0.104596974 | 0.139703888 | 0.416512652 | 0.2075 | 0.470124234 |
| SL1344_RS16335_ | 2.903225804 | 3.04943328 | 2.778696863 | -0.164111797 | 0.2445799 | -0.892389123 | 0.169220541 | 0.281300806 | 0.578450936 | 0.2963 | 0.567126458 |
| SL1344_RS16340_ | 1.766759345 | 1.784247939 | 1.744541517 | -0.026499067 | 0.431918701 | -0.05538715 | 0.485333334 | 0.901750415 | 0.963149888 | 0.9952 | 0.997929714 |

| | | | | | | | | | | | |
|---|---|---|---|---|---|---|---|---|---|---|---|
| SL1344_RS16350_ | 1.69934216 | 1.544400639 | 1.81541878 | 0.256734143 | 0.319126615 | 0.767052839 | 0.20933335 | 0.296581679 | 0.592892273 | 0.396 | 0.655012472 |
| SL1344_RS16355_ | 0.521426276 | 0.528055076 | 0.513564005 | -0.012221617 | 0.184973855 | -0.063817883 | 0.476000001 | 0.659314937 | 0.830868081 | 0.7667 | 0.880466871 |
| SL1344_RS16360_ | 0.975983773 | 0.896531725 | 1.101945336 | 0.192948318 | 0.160975036 | 1.140132834 | 0.10933337 | 0.164417175 | 0.424877333 | 0.2379 | 0.485901609 |
| SL1344_RS16365_ | 0.765400695 | 0.698136275 | 0.828478789 | 0.085561449 | 0.239406921 | 0.333682339 | 0.350433051 | 0.605796741 | 0.802579051 | 0.7201 | 0.855684977 |
| SL1344_RS16370_ | -1.371199133 | -1.515208443 | -1.239028096 | 0.275569787 | 0.267879575 | 0.978824514 | 0.145236535 | 0.222262746 | 0.463579396 | 0.3128 | 0.53767824 |
| SL1344_RS16375_ | 1.034176431 | 0.813985111 | 1.092341467 | 0.259853642 | 0.225473481 | 0.943536435 | 0.200532995 | 0.255789525 | 0.545491852 | 0.3937 | 0.640648424 |
| SL1344_RS16380_ | -2.037125783 | -1.986951982 | -2.102770088 | -0.257129848 | 0.774156548 | -0.349097976 | 0.360426388 | 0.545447701 | 0.769887414 | 0.7256 | 0.858631116 |
| SL1344_RS16385_ | 2.32350721 | 2.096878328 | 2.403533986 | 0.265350737 | 0.209129355 | 1.270795613 | 0.121333366 | 0.128727204 | 0.401041411 | 0.2191 | 0.486978862 |
| SL1344_RS16390_ | -1.635179338 | -1.488577422 | -1.795025062 | -0.315426256 | 0.35359371 | -0.853231403 | 0.163333356 | 0.257959382 | 0.518716412 | 0.3455 | 0.572647517 |
| SL1344_RS16395_ | -3.875006041 | -3.905574224 | -3.833491562 | 0.083902764 | 0.730407887 | 0.113548907 | 0.452365092 | 0.596885555 | 0.785412917 | 0.7104 | 0.84115543 |
| SL1344_RS16400_ | -1.332153149 | -1.330951411 | -1.333573438 | -0.002216668 | 0.297961898 | -0.009301062 | 0.496335776 | 0.602099821 | 0.790725855 | 0.6999 | 0.838917106 |
| SL1344_RS16405_ | 2.081834904 | 1.92191084 | 2.218318794 | 0.29137395 | 0.235357081 | 1.174067523 | 0.096602308 | 0.143821792 | 0.424318275 | 0.1938 | 0.460218042 |
| SL1344_RS16410_ | 1.034164674 | 0.815364527 | 1.274251928 | 0.457428905 | 0.126280664 | 3.520181394 | 4.68E-05 | 0.010548234 | 0.115391103 | 0.1 | 0.330257709 |
| SL1344_RS16415_ | 3.317071531 | 3.303535402 | 3.326750437 | 0.0086911 | 0.111570699 | 0.068419864 | 0.467688209 | 0.782960901 | 0.906057452 | 0.8522 | 0.93293612 |
| SL1344_RS16420_ | 1.844740622 | 1.915214117 | 1.740369211 | -0.258084761 | 0.357735021 | -0.698264332 | 0.236000014 | 0.291150047 | 0.589229221 | 0.4497 | 0.699512534 |
| SL1344_RS16425_ | 2.402461555 | 2.436496236 | 2.298457466 | -0.187350754 | 0.288165449 | -0.696706743 | 0.228000014 | 0.318915247 | 0.612684626 | 0.4294 | 0.694183414 |
| SL1344_RS16440_ | 0.129432687 | 0.140980313 | 0.109197427 | -0.066319124 | 0.300928619 | -0.212001065 | 0.405729517 | 0.630877167 | 0.820353587 | 0.808 | 0.904224143 |
| SL1344_RS16445_ | 3.164423894 | 3.047087742 | 3.271324484 | 0.18182463 | 0.298310709 | 0.512175487 | 0.349766828 | 0.500934846 | 0.75409125 | 0.7403 | 0.876657537 |
| SL1344_RS16450_ | 4.046840553 | 3.881379325 | 4.172685736 | 0.289439842 | 0.161941947 | 1.763277265 | 0.004664544 | 0.070326341 | 0.30738924 | 0.1039 | 0.33593354 |
| SL1344_RS16455_ | 2.687581368 | 2.570510345 | 2.716709219 | 0.137405011 | 0.171385676 | 0.822467656 | 0.215333349 | 0.292577437 | 0.582759449 | 0.3983 | 0.668546863 |
| SL1344_RS16460_ | 3.453832069 | 3.371538333 | 3.53174442 | 0.170342224 | 0.09575514 | 1.782972623 | 0.005996769 | 0.07385243 | 0.307123989 | 0.1052 | 0.337652215 |
| SL1344_RS16465_ | 1.645326621 | 1.559464816 | 1.748767046 | 0.342049323 | 0.329927747 | 1.063162892 | 0.054666746 | 0.153386445 | 0.437952872 | 0.1663 | 0.410059291 |
| SL1344_RS16470_ | 0.349678533 | 0.346244296 | 0.35789594 | 0.034879973 | 0.24848803 | 0.129544524 | 0.444000002 | 0.701228747 | 0.859881163 | 0.8312 | 0.917707922 |
| SL1344_RS16480_ | 2.406030042 | 2.170449762 | 2.653129111 | 0.316696312 | 0.32044721 | 1.28176135 | 0.105333372 | 0.129831778 | 0.40683137 | 0.1989 | 0.474528592 |
| SL1344_RS16485_ | 3.480813672 | 3.254548339 | 3.596769384 | 0.342431773 | 0.113099419 | 2.943101795 | 4.68E-05 | 0.035888253 | 0.222878641 | 0.1 | 0.330257709 |
| SL1344_RS16490_ | 1.006761532 | 1.15671506 | 0.843618143 | -0.309179505 | 0.128922425 | -2.34851683 | 0.004664544 | 0.032578947 | 0.192615016 | 0.1058 | 0.337856347 |
| SL1344_RS16495_ | 1.190433424 | 1.178720799 | 1.19611775 | 0.075486014 | 0.270300317 | 0.282692108 | 0.382000005 | 0.63705486 | 0.825990782 | 0.7972 | 0.900030124 |
| SL1344_RS16500_ | -2.708281333 | -2.962185135 | -2.159986634 | 0.597663867 | 0.826025778 | 0.671641159 | 0.248501012 | 0.408832097 | 0.666099587 | 0.4969 | 0.718443516 |
| SL1344_RS16505_ | -1.411012465 | -1.469705722 | -1.358853151 | 0.130501557 | 0.313071153 | 0.398189675 | 0.318454371 | 0.482083347 | 0.703938886 | 0.6076 | 0.76952617 |
| SL1344_RS16510_ | -2.751988201 | -2.918076053 | -2.483389795 | 0.4827179 | 0.584208038 | 0.752893555 | 0.203197885 | 0.311065096 | 0.570876066 | 0.4229 | 0.637900627 |
| SL1344_RS16515_ | -0.463526462 | -0.430478 | -0.52585367 | -0.06501165 | 0.378113964 | -0.167514205 | 0.440000002 | 0.743999177 | 0.886439777 | 0.8514 | 0.933414587 |
| SL1344_RS16520_ | 4.71591481 | 4.795458945 | 4.669442205 | -0.344111381 | 0.355635121 | -0.958951444 | 0.150566314 | 0.203378576 | 0.505024729 | 0.2734 | 0.549221878 |
| SL1344_RS16525_ | 2.414516537 | 2.136513858 | 2.54889746 | 0.414125621 | 0.191080852 | 2.185085077 | 4.68E-05 | 0.061548444 | 0.286658792 | 0.1003 | 0.330692807 |
| SL1344_RS16530_ | 0.076590836 | -0.195764044 | 0.310676683 | 0.494677025 | 0.532254565 | 0.861600136 | 0.152564982 | 0.236527122 | 0.534632866 | 0.28 | 0.550119062 |
| SL1344_RS16535_ | -3.026547452 | -2.946828547 | -3.08897488 | -0.176740653 | 0.637206238 | -0.269004992 | 0.38241173 | 0.596990226 | 0.79155846 | 0.7192 | 0.851178065 |
| SL1344_RS16540_ | 0.064073875 | 0.051172409 | 0.082940787 | -0.002152945 | 0.246440921 | -0.008341171 | 0.496 | 0.693275425 | 0.853463073 | 0.8209 | 0.911075179 |
| SL1344_RS16545_ | 2.023514441 | 1.893527133 | 2.149618129 | 0.28154237 | 0.132452921 | 2.119626768 | 0.003332494 | 0.039278763 | 0.223223535 | 0.1011 | 0.331849679 |
| SL1344_RS16550_ | 2.638626373 | 2.562822055 | 2.690426995 | 0.136576541 | 0.124678957 | 1.114278185 | 0.054000081 | 0.198627901 | 0.491518463 | 0.1669 | 0.410177143 |
| SL1344_RS16555_ | 1.681867681 | 1.586192359 | 1.830577916 | 0.235874303 | 0.151058737 | 1.522986869 | 0.045303228 | 0.083948827 | 0.316977211 | 0.152 | 0.398048672 |
| SL1344_RS16560_ | 0.863415028 | 0.81341551 | 0.904650165 | 0.091917621 | 0.171293817 | 0.508997082 | 0.27800001 | 0.443155839 | 0.682100364 | 0.5574 | 0.743524841 |
| SL1344_RS16565_ | 1.948303099 | 1.905757111 | 1.971492034 | 0.072357899 | 0.275643999 | 0.230727865 | 0.41066667 | 0.855141361 | 0.942811828 | 0.8617 | 0.941029569 |
| SL1344_RS16570_ | 1.250660587 | 1.161794692 | 1.364929046 | 0.219562423 | 0.203544448 | 1.032517257 | 0.119253864 | 0.176137822 | 0.45307314 | 0.248 | 0.505547731 |
| SL1344_RS16575_ | -0.621884384 | -0.714970978 | -0.562004885 | 0.156800279 | 0.251183463 | 0.589259992 | 0.265333344 | 0.394921578 | 0.639812772 | 0.5155 | 0.704600434 |
| SL1344_RS16580_ | 1.748564079 | 1.725797711 | 1.77221183 | 0.046458219 | 0.10399971 | 0.407512045 | 0.322000008 | 0.488989363 | 0.708473816 | 0.5955 | 0.765522936 |
| SL1344_RS16585_ | 1.862751255 | 1.691427907 | 2.008382172 | 0.267660129 | 0.164282541 | 1.639007408 | 0.054666746 | 0.076466586 | 0.306465867 | 0.1507 | 0.40228869 |
| SL1344_RS16590_ | 6.723176697 | 6.785426369 | 6.651900941 | -0.131445942 | 0.177564525 | -0.617927977 | 0.316000008 | 0.397142716 | 0.682496103 | 0.6412 | 0.834953661 |
| SL1344_RS16595_ | 3.957656719 | 3.898548966 | 4.013456163 | 0.087683777 | 0.306804193 | 0.22018643 | 0.442000002 | 0.901796014 | 0.963580161 | 1 | 1 |
| SL1344_RS16600_ | 4.398977132 | 4.448860855 | 4.361914285 | -0.082689578 | 0.148295397 | -0.504219796 | 0.279813468 | 0.392043106 | 0.675469492 | 0.5713 | 0.773823076 |
| SL1344_RS16605_ | -1.379130511 | -1.2853206 | -1.431325208 | -0.181661078 | 0.492749137 | -0.324561666 | 0.374000005 | 0.577851014 | 0.787478299 | 0.7382 | 0.866627526 |
| SL1344_RS16610_ | -0.575599839 | -0.015828859 | -0.767182827 | -0.547174686 | 0.629670646 | -0.484874682 | 0.333777489 | 0.76812102 | 0.903028047 | 0.7015 | 0.87317403 |
| SL1344_RS16615_ | 0.543572473 | 0.675613441 | 0.385630592 | -0.291910384 | 0.158225947 | -1.809501744 | 0.016000285 | 0.064045542 | 0.267895235 | 0.1225 | 0.357930512 |
| SL1344_RS16620_ | -0.020886589 | -0.192098297 | 0.199165184 | 0.38840578 | 0.207143041 | 1.797832108 | 0.022651765 | 0.061268301 | 0.262861952 | 0.1268 | 0.364858397 |
| SL1344_RS16635_ | -1.03822682 | -1.006440358 | -1.116277376 | -0.126173741 | 0.321826655 | -0.345638544 | 0.355762831 | 0.544741102 | 0.756943677 | 0.676 | 0.823198846 |

| | | | | | | | | | | | |
|---|---|---|---|---|---|---|---|---|---|---|---|
| SL1344_RS16640_ | 0.80995685 | 0.600833159 | 0.866816976 | 0.228403317 | 0.353478 | 0.420361664 | 0.340439713 | 0.799664009 | 0.917563671 | 0.7168 | 0.879543219 |
| SL1344_RS16645_ | 0.956090589 | 0.865498324 | 1.018104914 | 0.144497412 | 0.167425917 | 0.843016746 | 0.185209879 | 0.275976064 | 0.54380091 | 0.3726 | 0.608873766 |
| SL1344_RS16650_ | 0.302936831 | 0.219892907 | 0.378516138 | 0.179075677 | 0.202453152 | 0.84259256 | 0.162000023 | 0.247035759 | 0.515003947 | 0.3488 | 0.57552866 |
| SL1344_RS16655_ | -1.290364815 | -1.291743037 | -1.28940914 | -0.018526342 | 0.359808496 | -0.048655438 | 0.476349101 | 0.663753427 | 0.834429196 | 0.7659 | 0.879772769 |
| SL1344_RS16660_ | 1.921193914 | 1.809965383 | 2.110875401 | 0.315073548 | 0.154974653 | 2.003310864 | 0.006667356 | 0.044018471 | 0.238212697 | 0.1063 | 0.338886654 |
| SL1344_RS16665_ | 0.233331955 | 0.288040543 | 0.191583261 | -0.149168252 | 0.31331662 | -0.486766279 | 0.290666676 | 0.419200926 | 0.683051026 | 0.6056 | 0.778625499 |
| SL1344_RS16675_ | 2.094409801 | 2.055003703 | 2.254877969 | 0.181729668 | 0.148467102 | 1.002238608 | 0.181878767 | 0.225873348 | 0.511806457 | 0.3607 | 0.611220347 |
| SL1344_RS16680_ | 1.349352358 | 1.398575158 | 1.325665942 | -0.054105979 | 0.221006093 | -0.210314377 | 0.41666667 | 0.811443582 | 0.921123724 | 0.8395 | 0.930499105 |
| SL1344_RS16685_ | 2.812744645 | 2.63218066 | 2.974507781 | 0.337605799 | 0.213227854 | 1.585709766 | 0.009993796 | 0.088523904 | 0.340793375 | 0.1103 | 0.345349988 |
| SL1344_RS16690_ | 2.291851576 | 2.077313008 | 2.361286184 | 0.293014601 | 0.215266424 | 1.291198499 | 0.071952091 | 0.145302843 | 0.428124055 | 0.1754 | 0.428579909 |
| SL1344_RS16695_ | 2.764601614 | 2.570108745 | 2.871783166 | 0.294711527 | 0.083334461 | 3.34161707 | 4.68E-05 | 0.021458086 | 0.166180186 | 0.1 | 0.330257709 |
| SL1344_RS16700_ | 2.895897185 | 2.837092762 | 2.919407841 | 0.154516837 | 0.240473781 | 0.652751222 | 0.249167234 | 0.345957634 | 0.636083846 | 0.4927 | 0.734208918 |
| SL1344_RS16705_ | -3.534410295 | -3.29381432 | -3.761733206 | -0.398365809 | 0.766707232 | -0.474382061 | 0.302666675 | 0.477724747 | 0.705757396 | 0.5757 | 0.759415875 |
| SL1344_RS16710_ | -0.241517404 | -0.31915952 | -0.167846102 | 0.199537083 | 0.271007925 | 0.710393223 | 0.222666682 | 0.321908925 | 0.591092272 | 0.4611 | 0.672016865 |
| SL1344_RS16715_ | 2.010318454 | 1.845710282 | 2.365286841 | 0.497193784 | 0.185664905 | 2.608426326 | 0.00067292 | 0.023185301 | 0.179779262 | 0.1001 | 0.330401606 |
| SL1344_RS16720_ | 0.154855343 | 0.267110982 | 0.076035579 | -0.245998461 | 0.332426165 | -0.703560893 | 0.225849448 | 0.311113473 | 0.59381349 | 0.4457 | 0.680209732 |
| SL1344_RS16725_ | -0.682355169 | -0.763114145 | -0.615871271 | 0.10722655 | 0.393040739 | 0.269209018 | 0.38774151 | 0.676611571 | 0.846974737 | 0.7678 | 0.886199826 |
| SL1344_RS16730_ | 0.885450401 | 0.700848201 | 1.014173692 | 0.321356136 | 0.181743473 | 1.731196708 | 0.025982851 | 0.065372759 | 0.279546261 | 0.13 | 0.36885611 |
| SL1344_RS16745_ | -0.041319598 | 0.19078927 | -0.473294173 | -0.648716668 | 0.235076003 | -2.532108944 | 0.001335878 | 0.035909934 | 0.210088871 | 0.102 | 0.332852999 |
| SL1344_RS16750_ | 0.221923603 | 0.670051352 | 0.01049969 | -0.654591627 | 0.266669526 | -2.255568911 | 0.003334715 | 0.051157299 | 0.257568357 | 0.1033 | 0.334925095 |
| SL1344_RS16755_ | -0.61701239 | -0.01923198 | -0.843840933 | -0.781907814 | 0.299828599 | -2.171959223 | 0.010659989 | 0.056564127 | 0.265386707 | 0.1094 | 0.342751378 |
| SL1344_RS16775_ | -0.549456119 | -0.743147445 | -0.380063347 | 0.406939581 | 0.357861382 | 1.191924786 | 0.082611643 | 0.147631348 | 0.417887759 | 0.1958 | 0.445902366 |
| SL1344_RS16780_ | 0.218759532 | -0.384474897 | 0.504579922 | 0.890743727 | 0.363643592 | 2.491212764 | 4.68E-05 | 0.020314508 | 0.168544804 | 0.1 | 0.330257709 |
| SL1344_RS16785_ | -0.697735668 | -1.68322599 | -0.487677163 | 1.238059855 | 0.795550155 | 1.522460198 | 0.029313944 | 0.087703268 | 0.339551115 | 0.1305 | 0.371157338 |
| SL1344_RS16790_ | -0.523910331 | -0.805450981 | -0.1782722 | 0.63066543 | 0.252923598 | 2.430296189 | 0.002000966 | 0.029852846 | 0.192364828 | 0.1019 | 0.332968358 |
| SL1344_RS16795_ | 1.11663089 | 1.089997583 | 1.17296032 | 0.068919193 | 0.13960643 | 0.468787604 | 0.304000009 | 0.504176525 | 0.723270177 | 0.6157 | 0.785539157 |
| SL1344_RS16800_ | 1.55873271 | 1.404569045 | 1.607719677 | 0.210179182 | 0.232064585 | 0.863478474 | 0.189333352 | 0.26201806 | 0.555274814 | 0.3762 | 0.636908246 |
| SL1344_RS16805_ | -2.617851788 | -2.256873952 | -2.967684836 | -0.743672698 | 0.670905912 | -1.058875363 | 0.111925419 | 0.170174019 | 0.439559092 | 0.241 | 0.493815472 |
| SL1344_RS16810_ | -3.700513767 | -3.79396422 | -3.534505514 | 0.209851671 | 0.781454583 | 0.236905937 | 0.397068625 | 0.640836855 | 0.81571143 | 0.7368 | 0.862035464 |
| SL1344_RS16815_ | -2.962789027 | -3.012423685 | -2.890437949 | 0.120727915 | 0.520048931 | 0.201403459 | 0.417721522 | 0.579686758 | 0.77379718 | 0.6833 | 0.826936507 |
| SL1344_RS16820_ | -3.315879195 | -3.66976468 | -2.993008617 | 0.663227929 | 0.471253041 | 1.347704307 | 0.058000075 | 0.117671269 | 0.335570172 | 0.1876 | 0.423710844 |
| SL1344_RS16825_ | -2.853590735 | -2.681407392 | -2.983890744 | -0.324766445 | 0.669650979 | -0.468889536 | 0.295333342 | 0.48643405 | 0.714225524 | 0.6043 | 0.778227905 |
| SL1344_RS16830_ | -4.40136312 | -4.291932667 | -4.513659386 | -0.227901236 | 0.723462338 | -0.27509774 | 0.376666672 | 0.487980537 | 0.701683528 | 0.5941 | 0.759040234 |
| SL1344_RS16835_ | -4.107832002 | -3.911977854 | -4.352041288 | -0.404964985 | 0.785472687 | -0.488836719 | 0.292666676 | 0.427793945 | 0.661387075 | 0.5318 | 0.719613491 |
| SL1344_RS16840_ | -1.471998037 | -1.289845152 | -1.6097281 | -0.300014947 | 0.325154545 | -0.850272026 | 0.178547655 | 0.283833759 | 0.538878549 | 0.383 | 0.611467092 |
| SL1344_RS16845_ | -3.598296935 | -3.606733593 | -3.587447708 | 0.001527017 | 0.601106705 | 0.001768317 | 0.498334444 | 0.566124549 | 0.762054765 | 0.6817 | 0.820133579 |
| SL1344_RS16850_ | 1.453009665 | 1.400521914 | 1.501254514 | 0.117231155 | 0.158115002 | 0.71320162 | 0.203864107 | 0.308045448 | 0.583297544 | 0.4094 | 0.635171411 |
| SL1344_RS16855_ | 1.041270962 | 0.959933681 | 1.132336203 | 0.152263143 | 0.164010174 | 0.870649374 | 0.177333354 | 0.271942564 | 0.539750161 | 0.3634 | 0.602920633 |
| SL1344_RS16860_ | 3.053103878 | 3.03130326 | 3.180036823 | 0.142114329 | 0.106250249 | 0.997709877 | 0.211192554 | 0.250106072 | 0.540604031 | 0.4255 | 0.657721668 |
| SL1344_RS16865_ | 2.000265877 | 1.918397347 | 2.070791992 | 0.117612236 | 0.181361777 | 0.607382687 | 0.250000012 | 0.393400288 | 0.662334806 | 0.4978 | 0.718244023 |
| SL1344_RS16870_ | 1.493354422 | 1.464391565 | 1.503875521 | 0.037581183 | 0.222169023 | 0.150870474 | 0.444000002 | 0.715602372 | 0.872038732 | 0.8499 | 0.934033083 |
| SL1344_RS16875_ | 2.547827466 | 2.497237147 | 2.65930639 | 0.157137892 | 0.14672863 | 1.024508982 | 0.152000025 | 0.214653782 | 0.504045951 | 0.2901 | 0.546131173 |
| SL1344_RS16880_ | -1.412016744 | -1.607524913 | -1.331418922 | 0.263601169 | 0.429780206 | 0.541603581 | 0.289333343 | 0.443898418 | 0.691230759 | 0.5848 | 0.769073665 |
| SL1344_RS16885_ | -1.593777087 | -1.928755897 | -1.456741568 | 0.457290805 | 0.378158268 | 1.094430576 | 0.13000003 | 0.189269222 | 0.456766407 | 0.2793 | 0.521877991 |
| SL1344_RS16890_ | -1.056638208 | -0.885030901 | -1.238093242 | -0.356084839 | 0.318666821 | -1.070972525 | 0.107928086 | 0.180309773 | 0.444681499 | 0.2497 | 0.494824981 |
| SL1344_RS16895_ | -0.926287986 | -0.941599832 | -0.914752526 | 0.020124977 | 0.278646474 | 0.061649646 | 0.477681546 | 0.650006752 | 0.822011418 | 0.7544 | 0.873175412 |
| SL1344_RS16900_ | 1.878016762 | 1.826965176 | 1.92005014 | 0.073749928 | 0.36863025 | 0.194206771 | 0.42066667 | 0.699428689 | 0.868134364 | 0.8887 | 0.948494648 |
| SL1344_RS16905_ | 1.70884578 | 1.631622305 | 1.774758394 | 0.209886527 | 0.23884153 | 0.880214477 | 0.177333354 | 0.219243515 | 0.508906863 | 0.3407 | 0.59333804 |
| SL1344_RS16910_ | -1.862370747 | -1.996649294 | -1.747742033 | 0.277756118 | 0.473013663 | 0.553162673 | 0.267155241 | 0.408954667 | 0.66327863 | 0.5308 | 0.731841846 |
| SL1344_RS16915_ | -2.899966289 | -3.261764431 | -2.543525496 | 0.709052821 | 0.437375874 | 1.56293017 | 0.03664236 | 0.0850575 | 0.294769642 | 0.1513 | 0.388867586 |
| SL1344_RS16920_ | -0.00384153 | -0.284516288 | 0.203442038 | 0.498456339 | 0.204154338 | 2.381517991 | 0.001335878 | 0.040059164 | 0.216745626 | 0.101 | 0.331628623 |
| SL1344_RS16925_ | -3.350712956 | -3.500594096 | -3.239785078 | 0.278612999 | 0.580093896 | 0.433324898 | 0.318454371 | 0.459121768 | 0.685203424 | 0.564 | 0.741818231 |
| SL1344_RS16930_ | -4.660228605 | -4.640362598 | -4.69563434 | -0.039240235 | 0.824333985 | -0.041854572 | 0.479333334 | 0.557107311 | 0.756914624 | 0.6717 | 0.81547818 |

| | | | | | | | | | | | |
|---|---|---|---|---|---|---|---|---|---|---|---|
| SL1344_RS16935_ | -0.561397696 | -0.664315585 | -0.458692366 | 0.220254233 | 0.323764106 | 0.650380403 | 0.219187224 | 0.322875631 | 0.595779091 | 0.4274 | 0.650299606 |
| SL1344_RS16940_ | 2.165432968 | 2.215947445 | 1.896435004 | -0.320195986 | 0.286786894 | -1.041489778 | 0.108594308 | 0.202921678 | 0.502098577 | 0.2397 | 0.495403263 |
| SL1344_RS16945_ | 5.558779508 | 5.579324591 | 5.514629965 | -0.154525788 | 0.152522419 | -1.033012262 | 0.083277865 | 0.160091331 | 0.450358513 | 0.2003 | 0.456136465 |
| SL1344_RS16950_ | 4.968359845 | 4.936463689 | 5.002300965 | 0.050346033 | 0.068950995 | 0.73681949 | 0.213191222 | 0.324216112 | 0.608007204 | 0.4093 | 0.654530468 |
| SL1344_RS16955_ | 6.327932392 | 6.181635913 | 6.351216796 | 0.169071923 | 0.189662457 | 0.775281152 | 0.256495681 | 0.333868105 | 0.632316468 | 0.4957 | 0.741693486 |
| SL1344_RS16960_ | 2.637271096 | 2.596018119 | 2.724513853 | 0.133273967 | 0.140328942 | 0.899666364 | 0.175333354 | 0.246932822 | 0.538925906 | 0.3287 | 0.580437082 |
| SL1344_RS16965_ | 1.711022614 | 1.646474658 | 1.781022311 | 0.134208578 | 0.099445165 | 1.343767984 | 0.081279199 | 0.138124902 | 0.373571991 | 0.2067 | 0.447292017 |
| SL1344_RS16970_ | 5.317434081 | 5.243724349 | 5.406020857 | 0.119382506 | 0.093121611 | 1.492795089 | 0.089940086 | 0.087390601 | 0.337130404 | 0.1766 | 0.442240209 |
| SL1344_RS16980_ | 2.574487371 | 2.534752174 | 2.651042711 | 0.113830299 | 0.092856056 | 1.14291898 | 0.131245866 | 0.1859099 | 0.455236604 | 0.2746 | 0.523990528 |
| SL1344_RS16985_ | 3.474869849 | 4.096257321 | 3.209932585 | -0.664252646 | 0.65408396 | -0.781160668 | 0.320453039 | 0.363158694 | 0.656513452 | 0.6631 | 0.848890235 |
| SL1344_RS16990_ | 0.222561582 | -0.185852724 | 0.524926972 | 0.699086174 | 0.182233608 | 3.742659899 | 4.68E-05 | 0.013395629 | 0.128751827 | 0.1 | 0.330257709 |
| SL1344_RS16995_ | 0.625756304 | 0.820397316 | 0.292416017 | -0.560811114 | 0.279339095 | -2.096244504 | 0.001336769 | 0.034647638 | 0.215079041 | 0.1016 | 0.3324896 |
| SL1344_RS17000_ | 3.858674415 | 4.225279272 | 3.838932757 | -0.387204989 | 0.632354108 | -0.709729986 | 0.218521001 | 0.281250349 | 0.58546082 | 0.4 | 0.680350104 |
| SL1344_RS17005_ | 5.961988988 | 6.339925758 | 5.447971496 | -0.343176725 | 0.651848305 | -1.174711013 | 0.104596974 | 0.248756202 | 0.553993046 | 0.2 | 0.476573497 |
| SL1344_RS17010_ | 3.164017889 | 3.225327361 | 3.104757557 | -0.064250118 | 0.170228585 | -0.407959814 | 0.334443711 | 0.563936304 | 0.786305616 | 0.6866 | 0.842649568 |
| SL1344_RS17015_ | 1.011791737 | 1.080946949 | 0.76058464 | -0.304562095 | 0.257216139 | -1.115162784 | 0.150000025 | 0.174781 | 0.459507036 | 0.2878 | 0.555087666 |
| SL1344_RS17030_ | 3.393610364 | 3.36996837 | 3.474644212 | 0.118715446 | 0.188499248 | 0.664202547 | 0.235176563 | 0.38010459 | 0.664155835 | 0.4364 | 0.70286104 |
| SL1344_RS17035_ | 3.60647769 | 3.55186207 | 3.741061514 | 0.19182675 | 0.121319858 | 1.686027992 | 0.018654474 | 0.0583267 | 0.275541562 | 0.1192 | 0.355857295 |
| SL1344_RS17040_ | 2.727409068 | 2.744245588 | 2.714820067 | 0.010767998 | 0.171779861 | 0.067928739 | 0.475016656 | 0.766293675 | 0.899006146 | 0.9298 | 0.96891028 |
| SL1344_RS17045_ | 1.896963837 | 1.803119836 | 1.973255786 | 0.12656428 | 0.442943108 | 0.245636753 | 0.444370422 | 0.752662962 | 0.896699115 | 0.9841 | 0.993089557 |
| SL1344_RS17050_ | 2.369258786 | 2.37649144 | 2.356788465 | -0.019817528 | 0.087791996 | -0.194107042 | 0.418000003 | 0.5939521 | 0.78648885 | 0.7069 | 0.844188931 |
| SL1344_RS17060_ | 1.293864746 | 1.239485536 | 1.338328366 | 0.118318663 | 0.151587238 | 0.730578646 | 0.181212545 | 0.285774343 | 0.560535462 | 0.3764 | 0.600284773 |
| SL1344_RS17065_ | 4.730705949 | 4.714841517 | 4.74842419 | 0.048037991 | 0.22284047 | 0.273545434 | 0.349766828 | 0.616758523 | 0.826818995 | 0.7411 | 0.879781467 |
| SL1344_RS17070_ | 5.247333333 | 5.271379535 | 5.059378647 | -0.18581612 | 0.088008335 | -0.645249547 | 0.33333334 | 0.483408145 | 0.743630538 | 0.7 | 0.872542119 |
| SL1344_RS17075_ | 3.800229993 | 3.735491205 | 4.073928217 | 0.31901003 | 0.113856814 | 1.68375286 | 0.012000381 | 0.082396235 | 0.330183768 | 0.1098 | 0.344213456 |
| SL1344_RS17080_ | -1.334993432 | -1.321363801 | -1.362524978 | -0.027906088 | 0.315285537 | -0.088288918 | 0.462000001 | 0.63457569 | 0.81524114 | 0.7365 | 0.865542279 |
| SL1344_RS17085_ | 3.256821439 | 3.181171117 | 3.366633872 | 0.16032391 | 0.12507609 | 1.298228416 | 0.100599641 | 0.123290717 | 0.388848372 | 0.1973 | 0.462528051 |
| SL1344_RS17090_ | 0.296639986 | 0.274501911 | 0.333360187 | 0.04815333 | 0.211376171 | 0.212932902 | 0.413724187 | 0.66424684 | 0.835803855 | 0.7765 | 0.888869153 |
| SL1344_RS17095_ | 2.530473333 | 2.287001539 | 2.742622103 | 0.440373156 | 0.119064545 | 3.641753821 | 4.68E-05 | 0.007980435 | 0.1077249 | 0.1 | 0.330257709 |
| SL1344_RS17100_ | 2.017838397 | 1.888557348 | 2.198344943 | 0.335457615 | 0.197415941 | 1.803492188 | 0.001336769 | 0.06436396 | 0.289459535 | 0.1034 | 0.335240826 |
| SL1344_RS17105_ | 0.73429161 | 0.698400618 | 0.752500236 | 0.055621924 | 0.20218424 | 0.254816496 | 0.391333338 | 0.60138949 | 0.797186511 | 0.7394 | 0.864457683 |
| SL1344_RS17110_ | 1.027136942 | 1.015728859 | 1.034243012 | -0.007938922 | 0.215537352 | -0.031115669 | 0.484343771 | 0.753454445 | 0.8888397 | 0.869 | 0.937261791 |
| SL1344_RS17115_ | 1.662732482 | 1.566699326 | 1.726492474 | 0.101934573 | 0.226885572 | 0.43544139 | 0.339107268 | 0.623412725 | 0.818844384 | 0.707 | 0.859227183 |
| SL1344_RS17120_ | 2.281833959 | 2.314220252 | 2.236193206 | -0.080714765 | 0.090460839 | -0.866399982 | 0.170552986 | 0.266952376 | 0.526655231 | 0.36 | 0.586071312 |
| SL1344_RS17125_ | 2.334972 | 2.282921783 | 2.392683049 | 0.111809456 | 0.089962844 | 1.197135678 | 0.079280533 | 0.148366164 | 0.398544221 | 0.2174 | 0.459395646 |
| SL1344_RS17130_ | 2.83388017 | 2.775572522 | 2.88341594 | 0.098061107 | 0.112527634 | 0.854375669 | 0.166555652 | 0.246996738 | 0.526523401 | 0.3357 | 0.587749888 |
| SL1344_RS17135_ | 2.476724621 | 2.410444219 | 2.543153884 | 0.198462714 | 0.17194745 | 1.211392284 | 0.049300555 | 0.125105587 | 0.39117794 | 0.1657 | 0.407470031 |
| SL1344_RS17140_ | 2.615402629 | 2.446069063 | 2.734474076 | 0.321954366 | 0.244012347 | 1.375793245 | 0.028000161 | 0.106705307 | 0.371907362 | 0.1239 | 0.364923131 |
| SL1344_RS17145_ | 2.638362964 | 2.532440922 | 2.803804641 | 0.281045138 | 0.12997003 | 2.177196384 | 0.000673369 | 0.033066636 | 0.209139156 | 0.1016 | 0.33252665 |
| SL1344_RS17150_ | 3.878990147 | 3.700344915 | 3.932346437 | 0.217037584 | 0.125619361 | 1.589971203 | 0.03533346 | 0.079820545 | 0.32296609 | 0.1353 | 0.379043036 |
| SL1344_RS17155_ | 2.570923027 | 2.634906318 | 2.546649354 | -0.071821654 | 0.146323132 | -0.245571332 | 0.362425055 | 0.829768522 | 0.931563737 | 0.751 | 0.894130783 |
| SL1344_RS17160_ | 2.414561839 | 2.323658039 | 2.49306664 | 0.181097325 | 0.108051008 | 1.633516775 | 0.014000326 | 0.079992376 | 0.309238313 | 0.12 | 0.35607073 |
| SL1344_RS17165_ | 3.273456036 | 3.207842799 | 3.427789539 | 0.238363373 | 0.172485641 | 1.504879502 | 0.022000206 | 0.074092315 | 0.310943232 | 0.1205 | 0.35852689 |
| SL1344_RS17170_ | 1.541553704 | 1.49982797 | 1.65277225 | 0.153940557 | 0.14921334 | 0.945707302 | 0.147333359 | 0.242916721 | 0.522729636 | 0.3398 | 0.575262489 |
| SL1344_RS17175_ | -2.01355224 | -2.136990014 | -1.889688503 | 0.229071837 | 0.348200112 | 0.623681235 | 0.241838787 | 0.380367679 | 0.61603435 | 0.476 | 0.6764267 |
| SL1344_RS17180_ | 2.132951611 | 2.589359488 | 1.803161056 | -0.840616315 | 0.36476798 | -2.47212673 | 4.68E-05 | 0.018405046 | 0.166075064 | 0.1 | 0.330257709 |
| SL1344_RS17185_ | 2.054711306 | 2.299508907 | 1.801214134 | -0.59466858 | 0.269377949 | -2.412260134 | 4.68E-05 | 0.026730764 | 0.195999659 | 0.1 | 0.330257709 |
| SL1344_RS17190_ | 2.803841176 | 3.023845899 | 2.753050633 | -0.253679451 | 0.136336464 | -1.346260365 | 0.092000045 | 0.140322009 | 0.419346554 | 0.2081 | 0.464510483 |
| SL1344_RS17195_ | -1.487185682 | -1.555285652 | -1.42677725 | 0.13532057 | 0.283367912 | 0.433702489 | 0.306666675 | 0.441211986 | 0.663948009 | 0.5559 | 0.730731894 |
| SL1344_RS17200_ | -4.581187486 | -4.625052521 | -4.557307674 | -0.007571689 | 1.000547662 | -0.01061857 | 0.496335776 | 0.641446383 | 0.819100952 | 0.7466 | 0.871251668 |
| SL1344_RS17205_ | 2.318902031 | 2.056357867 | 2.561440984 | 0.60040533 | 0.293725329 | 2.08844174 | 4.68E-05 | 0.029748266 | 0.207234175 | 0.1 | 0.330257709 |
| SL1344_RS17210_ | 1.180705391 | 0.887365275 | 1.416272434 | 0.540927465 | 0.168539386 | 3.210412619 | 4.68E-05 | 0.017921505 | 0.151859331 | 0.1 | 0.330257709 |
| SL1344_RS17215_ | -2.514318238 | -2.380574421 | -2.819201347 | -0.40336808 | 0.723993102 | -0.517656709 | 0.309127257 | 0.463355651 | 0.713223087 | 0.6199 | 0.798334356 |

| | | | | | | | | | | | |
|---|---|---|---|---|---|---|---|---|---|---|---|
| SL1344_RS17220_ | -0.569313763 | -0.624112934 | -0.548319457 | 0.090713171 | 0.380115177 | 0.2293139 | 0.419333336 | 0.616451614 | 0.814258173 | 0.7815 | 0.896412388 |
| SL1344_RS17225_ | 1.285435559 | 1.237274077 | 1.352497106 | 0.127139222 | 0.140253782 | 0.904007918 | 0.144666693 | 0.234458302 | 0.505622696 | 0.3129 | 0.542144351 |
| SL1344_RS17230_ | 2.924706578 | 3.019110083 | 2.827158986 | -0.157356024 | 0.351813433 | -0.428799758 | 0.301333342 | 0.509678791 | 0.759967204 | 0.6086 | 0.81282815 |
| SL1344_RS17235_ | 2.371226294 | 2.55261461 | 2.206449817 | -0.257867389 | 0.387736254 | -0.654904211 | 0.238000013 | 0.358537306 | 0.649417438 | 0.4386 | 0.691829332 |
| SL1344_RS17240_ | 2.392311799 | 2.449443396 | 2.327849828 | -0.145521852 | 0.390580687 | -0.370422832 | 0.341772158 | 0.58100597 | 0.804139283 | 0.7417 | 0.888930109 |
| SL1344_RS17245_ | 3.475190983 | 3.569251676 | 3.174639574 | -0.314661549 | 0.596518886 | -0.610966927 | 0.224000015 | 0.429365956 | 0.706561238 | 0.4 | 0.680350104 |
| SL1344_RS17250_ | 4.511663881 | 4.61798776 | 4.385108274 | -0.192358386 | 0.522618539 | -0.36372368 | 0.311125924 | 0.622164002 | 0.831413772 | 0.6295 | 0.818353119 |
| SL1344_RS17255_ | 2.571353919 | 2.599145691 | 2.525578598 | -0.085660449 | 0.1341754 | -0.616881522 | 0.250666679 | 0.375981147 | 0.645544846 | 0.4949 | 0.71706096 |
| SL1344_RS17260_ | 4.532498594 | 4.347663281 | 4.552442985 | 0.204482447 | 0.120284764 | 0.682066529 | 0.33333334 | 0.45221408 | 0.721864412 | 0.7 | 0.872542119 |
| SL1344_RS17270_ | -0.231937446 | -0.174874107 | -0.253519954 | -0.073484213 | 0.393369386 | -0.188351685 | 0.427333336 | 0.736412366 | 0.881523525 | 0.8317 | 0.925835609 |
| SL1344_RS17275_ | 6.566950182 | 6.552943654 | 6.57758574 | 0.061325642 | 0.115389309 | 0.415382745 | 0.282478358 | 0.363669559 | 0.656517721 | 0.5746 | 0.787803311 |
| SL1344_RS17280_ | 5.395609719 | 5.326052508 | 5.455217355 | 0.125903987 | 0.19402112 | 0.415640157 | 0.37175217 | 0.686260043 | 0.865873283 | 0.7969 | 0.913645741 |
| SL1344_RS17285_ | 1.870716622 | 1.833953299 | 1.92364276 | 0.060329129 | 0.152318667 | 0.398894005 | 0.352000006 | 0.606766367 | 0.801908757 | 0.715 | 0.853627377 |
| SL1344_RS17290_ | 3.590957991 | 3.560841209 | 3.620689249 | 0.06494364 | 0.216005132 | 0.294672126 | 0.39307129 | 0.569871305 | 0.796909593 | 0.8302 | 0.923253595 |
| SL1344_RS17295_ | 2.975545656 | 2.98019054 | 2.96926703 | 0.184627771 | 0.293986356 | 0.60504995 | 0.391072622 | 0.385883993 | 0.669731999 | 0.8549 | 0.927171316 |
| SL1344_RS17300_ | 2.068414604 | 2.024573644 | 2.223743669 | 0.187794978 | 0.122078014 | 1.245773769 | 0.119920087 | 0.16097493 | 0.430427769 | 0.257 | 0.500595132 |
| SL1344_RS17305_ | -3.141992308 | -3.377414548 | -2.963990444 | 0.422139497 | 0.493465651 | 0.79325934 | 0.189333352 | 0.281214279 | 0.525630484 | 0.3869 | 0.599720728 |
| SL1344_RS17310_ | -1.664443912 | -1.922812884 | -1.454069554 | 0.471608583 | 0.290765181 | 1.542446584 | 0.052631662 | 0.090872844 | 0.305999184 | 0.1591 | 0.398099779 |
| SL1344_RS17315_ | -1.010006515 | -1.116142692 | -0.935194848 | 0.180880724 | 0.312260565 | 0.558572598 | 0.270486353 | 0.414220061 | 0.656325857 | 0.5249 | 0.715889651 |
| SL1344_RS17320_ | -5.175001419 | -5.321704221 | -5.091480095 | 0.289186504 | 1.580214512 | 0.159508032 | 0.417333336 | 0.586470989 | 0.790819197 | 0.7466 | 0.866364064 |
| SL1344_RS17325_ | -2.112120171 | -2.544243324 | -1.861332264 | 0.639379244 | 0.467020146 | 1.269621793 | 0.083333384 | 0.13506795 | 0.388377812 | 0.2076 | 0.456370004 |
| SL1344_RS17330_ | -2.153436108 | -2.337141296 | -2.039764811 | 0.307933107 | 0.516102568 | 0.563678941 | 0.261159239 | 0.392432453 | 0.64744809 | 0.5211 | 0.716107656 |
| SL1344_RS17335_ | -2.872622712 | -3.036423909 | -2.762020491 | 0.341998936 | 0.641944117 | 0.475479968 | 0.301333342 | 0.455958144 | 0.697315513 | 0.5915 | 0.764159676 |
| SL1344_RS17340_ | 1.169778047 | 1.126852955 | 1.287659066 | 0.196482099 | 0.333709712 | 0.627042184 | 0.244503677 | 0.366631493 | 0.647943628 | 0.4574 | 0.709402342 |
| SL1344_RS17345_ | 1.45198419 | 1.371167263 | 1.557989886 | 0.211529894 | 0.185199054 | 1.134293959 | 0.079946755 | 0.149480926 | 0.420854793 | 0.1989 | 0.449199956 |
| SL1344_RS17350_ | 5.788556447 | 5.677707239 | 5.802083209 | 0.148691678 | 0.150540325 | 1.067994445 | 0.167221874 | 0.140748047 | 0.425426315 | 0.3034 | 0.577078155 |
| SL1344_RS17355_ | 2.211220702 | 2.101123946 | 2.262006578 | 0.166391439 | 0.135022485 | 1.207391055 | 0.069953426 | 0.169677585 | 0.452182749 | 0.1856 | 0.430081481 |
| SL1344_RS17360_ | -2.335440279 | -2.573469683 | -2.152795194 | 0.425722518 | 0.406413867 | 1.005181223 | 0.133333363 | 0.211194503 | 0.464365369 | 0.2921 | 0.52161527 |
| SL1344_RS17365_ | -0.669856772 | -0.6460418 | -0.68644467 | -0.023173003 | 0.289196073 | -0.08242023 | 0.462000001 | 0.67278558 | 0.841440761 | 0.7871 | 0.892640401 |
| SL1344_RS17370_ | -1.195600122 | -1.417193876 | -0.960055598 | 0.472942135 | 0.305923199 | 1.508739349 | 0.043333435 | 0.091444358 | 0.316998615 | 0.1471 | 0.385808336 |
| SL1344_RS17375_ | -0.347712234 | -0.261382481 | -0.38617037 | -0.095538875 | 0.25572007 | -0.331107281 | 0.376000005 | 0.653685005 | 0.82685072 | 0.7326 | 0.868290333 |
| SL1344_RS17380_ | -1.315519401 | -1.425643819 | -1.235401482 | 0.185522494 | 0.300489468 | 0.581073262 | 0.246000013 | 0.398618158 | 0.635000284 | 0.4978 | 0.696121192 |
| SL1344_RS17385_ | -4.450024295 | -4.418202745 | -4.501321103 | -0.089091087 | 0.843002897 | -0.104118943 | 0.455029982 | 0.582150703 | 0.775523428 | 0.6997 | 0.835993356 |
| SL1344_RS17390_ | 0.236933728 | 0.227032974 | 0.245417788 | 0.061390829 | 0.244366718 | 0.228269347 | 0.404000004 | 0.617658099 | 0.806829821 | 0.7664 | 0.877779561 |
| SL1344_RS17395_ | 2.331577237 | 2.196294386 | 2.396148435 | 0.293239323 | 0.30334056 | 1.080525601 | 0.067333397 | 0.18081428 | 0.475977849 | 0.1762 | 0.426320856 |
| SL1344_RS17400_ | 3.319935869 | 3.23674186 | 3.389325031 | 0.157948375 | 0.113638545 | 1.379338869 | 0.056000078 | 0.1222722 | 0.38895466 | 0.1478 | 0.395012305 |
| SL1344_RS17405_ | 3.199880538 | 3.188271826 | 3.212039351 | 0.063397156 | 0.114423748 | 0.538480112 | 0.292666676 | 0.412641084 | 0.674184226 | 0.6388 | 0.797654201 |
| SL1344_RS17410_ | 2.471833811 | 2.459227533 | 2.497391537 | 0.049402022 | 0.130403866 | 0.372713806 | 0.318000008 | 0.451807094 | 0.707881142 | 0.629 | 0.790453367 |
| SL1344_RS17415_ | 2.030077645 | 2.059498807 | 2.01452184 | -0.043147486 | 0.172800631 | -0.215147434 | 0.41972019 | 0.743896754 | 0.88476911 | 0.8565 | 0.935376271 |
| SL1344_RS17420_ | 2.94056121 | 2.952719727 | 2.93550768 | -0.01498433 | 0.075899192 | -0.164861477 | 0.423333336 | 0.638735943 | 0.816399482 | 0.741 | 0.865591578 |
| SL1344_RS17425_ | 4.493806978 | 4.442381703 | 4.534447714 | -0.021092162 | 0.196110312 | -0.11380092 | 0.460000001 | 0.897511393 | 0.961238525 | 0.9934 | 0.997225398 |
| SL1344_RS17430_ | -5.034627929 | -4.698944729 | -5.516740051 | -0.823384752 | 1.337042408 | -0.564073339 | 0.257161904 | 0.391694609 | 0.641985214 | 0.5083 | 0.706594943 |
| SL1344_RS17440_ | 1.553673817 | 1.555109553 | 1.551844682 | 0.017335417 | 0.143335714 | 0.115234744 | 0.452666668 | 0.663456476 | 0.834069079 | 0.7885 | 0.890370945 |
| SL1344_RS17445_ | 0.757211519 | 0.606493768 | 0.816697714 | 0.249853185 | 0.340506197 | 0.755412187 | 0.219187224 | 0.29538436 | 0.584826821 | 0.4177 | 0.672969168 |
| SL1344_RS17450_ | 0.049504834 | -0.034274575 | 0.115169678 | 0.096966469 | 0.261237205 | 0.340830289 | 0.351333339 | 0.578984756 | 0.780271563 | 0.6991 | 0.837773755 |
| SL1344_RS17455_ | 4.323552144 | 4.138977985 | 4.491414651 | 0.405363414 | 0.120159249 | 3.484847538 | 4.68E-05 | 0.005944746 | 0.097937127 | 0.1 | 0.330257709 |
| SL1344_RS17460_ | 5.20258004 | 5.019285775 | 5.443944947 | 0.424992989 | 0.052405292 | 7.94272686 | 4.68E-05 | 0.003487534 | 0.075398718 | 0.1 | 0.330257709 |
| SL1344_RS17465_ | -2.387373551 | -2.381470365 | -2.393225093 | -0.004847445 | 0.471568346 | -0.009741369 | 0.496 | 0.650951044 | 0.822699859 | 0.7533 | 0.869284527 |
| SL1344_RS17470_ | 0.729773219 | 0.655571054 | 0.773044724 | 0.103051761 | 0.183145049 | 0.544743453 | 0.290666676 | 0.475600418 | 0.709036173 | 0.5853 | 0.770027952 |
| SL1344_RS17475_ | 1.488879077 | 1.290199928 | 1.522118213 | 0.218790451 | 0.186280356 | 0.859866977 | 0.247168567 | 0.29290575 | 0.582275051 | 0.4752 | 0.705527335 |
| SL1344_RS17480_ | 3.315613604 | 3.278850326 | 3.352569932 | 0.007494086 | 0.179427501 | 0.043338737 | 0.485676216 | 0.843313385 | 0.936788791 | 0.9529 | 0.979468007 |
| SL1344_RS17490_ | -0.936166564 | -1.036177431 | -0.875966869 | 0.154980862 | 0.341574585 | 0.451046781 | 0.310000008 | 0.493546224 | 0.724084492 | 0.6271 | 0.796636929 |
| SL1344_RS17495_ | 0.574869031 | 0.545640242 | 0.630189361 | 0.055709414 | 0.240175759 | 0.217865378 | 0.410393075 | 0.726898657 | 0.873812411 | 0.765 | 0.890104026 |

| | | | | | | | | | | | |
|---|---|---|---|---|---|---|---|---|---|---|---|
| SL1344_RS17500_ | 0.466760408 | 0.365267478 | 0.525538592 | 0.163042037 | 0.172103427 | 0.893488664 | 0.160000023 | 0.245511442 | 0.507940851 | 0.3318 | 0.562401918 |
| SL1344_RS17505_ | -3.528061616 | -3.94967537 | -3.086992933 | 0.856275723 | 0.56380739 | 1.456643486 | 0.058666741 | 0.106664885 | 0.329550785 | 0.1751 | 0.415263853 |
| SL1344_RS17510_ | -3.655060688 | -3.806161755 | -3.502923964 | 0.316166577 | 0.554813185 | 0.536431015 | 0.272485021 | 0.404981473 | 0.634979563 | 0.509 | 0.698245353 |
| SL1344_RS17515_ | -0.734348125 | -0.655373291 | -0.810885896 | -0.165035624 | 0.231292673 | -0.665311972 | 0.233333347 | 0.350969318 | 0.595681102 | 0.4465 | 0.654541062 |
| SL1344_RS17520_ | -0.9624597 | -1.294940922 | -0.770380824 | 0.484195128 | 0.335617037 | 1.371155617 | 0.086000049 | 0.121649992 | 0.373079989 | 0.1934 | 0.44545241 |
| SL1344_RS17530_ | -0.933428932 | -1.080520108 | -0.733050143 | 0.327653989 | 0.749758694 | 0.362971288 | 0.381745508 | 0.731703699 | 0.882122233 | 0.8315 | 0.924636305 |
| SL1344_RS17540_ | 2.06035153 | 1.854227215 | 2.230145966 | 0.384310585 | 0.849535847 | 0.460245464 | 0.273151243 | 0.421139159 | 0.700486015 | 0.5347 | 0.766149512 |
| SL1344_RS17545_ | -1.17751339 | -1.254792553 | -1.090722621 | 0.234537608 | 0.546126658 | 0.378755107 | 0.305796144 | 0.428446695 | 0.697578713 | 0.6288 | 0.792938772 |
| SL1344_RS17550_ | -0.070139506 | -0.404095523 | 0.502488998 | 0.927652543 | 0.797417926 | 1.119824638 | 0.084666716 | 0.187542799 | 0.485900582 | 0.1782 | 0.441282043 |
| SL1344_RS17555_ | 1.230150799 | 1.193338254 | 1.264088665 | 0.056075288 | 0.158601556 | 0.326169548 | 0.358666672 | 0.603047046 | 0.795729586 | 0.7142 | 0.850401728 |
| SL1344_RS17560_ | 1.128854288 | 0.802121025 | 1.18732994 | 0.379641182 | 0.226608166 | 1.261134635 | 0.114666702 | 0.170906436 | 0.457983191 | 0.2528 | 0.496162947 |
| SL1344_RS17565_ | 2.062117369 | 1.727828905 | 2.133939908 | 0.399696511 | 0.239832966 | 1.266526945 | 0.085276532 | 0.15374179 | 0.440386586 | 0.2014 | 0.458887614 |
| SL1344_RS17570_ | 1.505237556 | 1.466368982 | 1.552494531 | 0.089913632 | 0.211076134 | 0.436682016 | 0.321333341 | 0.509064722 | 0.745147088 | 0.6421 | 0.811547093 |
| SL1344_RS17575_ | 3.498046435 | 3.447725313 | 3.525951045 | 0.087889996 | 0.311032769 | 0.280776327 | 0.36642239 | 0.675630422 | 0.85925028 | 0.7906 | 0.910961204 |
| SL1344_RS17580_ | 3.806352658 | 3.785276085 | 3.817297159 | 0.03971243 | 0.123227629 | 0.28345492 | 0.363091278 | 0.481651874 | 0.738751692 | 0.7347 | 0.868162128 |
| SL1344_RS17585_ | 1.616260675 | 1.443639061 | 1.705323622 | 0.184129765 | 0.261746613 | 0.724251994 | 0.224666681 | 0.336843288 | 0.623433551 | 0.4228 | 0.67398807 |
| SL1344_RS17590_ | 2.864901683 | 2.771021574 | 3.00883453 | 0.174517228 | 0.223809878 | 0.8227217 | 0.160000023 | 0.266270129 | 0.564055364 | 0.3066 | 0.579419365 |
| SL1344_RS17595_ | 3.353773414 | 3.324389912 | 3.388793925 | 0.087744729 | 0.114083038 | 0.759324074 | 0.211333349 | 0.292246694 | 0.577414223 | 0.4227 | 0.659984319 |
| SL1344_RS17600_ | 0.325596845 | 0.302930975 | 0.349241656 | 0.151568645 | 0.31395817 | 0.461970283 | 0.325782819 | 0.439447486 | 0.695926051 | 0.695 | 0.831264886 |
| SL1344_RS17605_ | 0.860390588 | 0.456741552 | 0.919449797 | 0.460961949 | 0.297053471 | 1.09972333 | 0.190666685 | 0.199197833 | 0.49372664 | 0.3377 | 0.597888574 |
| SL1344_RS17610_ | 2.880081026 | 2.645384762 | 3.040844018 | 0.383201135 | 0.132055009 | 2.868221799 | 4.68E-05 | 0.029221137 | 0.199688935 | 0.1001 | 0.330403578 |
| SL1344_RS17615_ | 0.919987868 | 1.024158677 | 0.779215757 | -0.230757799 | 0.382889259 | -0.553582388 | 0.29646903 | 0.447130117 | 0.713003851 | 0.6026 | 0.797421081 |
| SL1344_RS17620_ | 0.915385841 | 0.962417575 | 0.825717701 | -0.131482087 | 0.191651317 | -0.631473067 | 0.248666679 | 0.385485933 | 0.645000563 | 0.5077 | 0.715400182 |
| SL1344_RS17625_ | -0.123155104 | -0.100392689 | -0.133107904 | -0.012744525 | 0.278540958 | -0.040648345 | 0.483333334 | 0.718849784 | 0.870450148 | 0.8166 | 0.913995628 |
| SL1344_RS17630_ | 6.196373444 | 5.935594852 | 6.348373464 | 0.304568696 | 0.240416097 | 1.731326012 | 0.013333676 | 0.047944928 | 0.260337971 | 0.1116 | 0.34727999 |
| SL1344_RS17635_ | 7.249634592 | 7.106608211 | 7.295426047 | 0.161978147 | 0.151486098 | 0.983159247 | 0.154666691 | 0.218467131 | 0.52165289 | 0.2832 | 0.561353476 |
| SL1344_RS17640_ | 6.746031446 | 6.664748181 | 6.828855782 | 0.159006647 | 0.372981131 | 0.342554352 | 0.441705532 | 0.808495998 | 0.926146229 | 1 | 1 |
| SL1344_RS17645_ | 4.397317305 | 4.37782378 | 4.41631695 | -0.137000735 | 0.401455125 | -0.321781081 | 0.429047304 | 0.53503582 | 0.778144968 | 0.9712 | 0.987756119 |
| SL1344_RS17650_ | 5.438905575 | 5.3485239 | 5.530615095 | 0.014427402 | 0.339524901 | 0.044456316 | 0.478666667 | 0.961070563 | 0.986285052 | 1 | 1 |
| SL1344_RS17655_ | 5.567924571 | 5.430452881 | 5.646828364 | 0.214026004 | 0.254956065 | 0.674563022 | 0.238507675 | 0.364059486 | 0.656905623 | 0.4487 | 0.711411429 |
| SL1344_RS17660_ | 6.9894641 | 6.906985473 | 7.025418269 | 0.06223127 | 0.162002975 | 0.314959035 | 0.33333334 | 0.962304807 | 0.986792358 | 0.7 | 0.872542119 |
| SL1344_RS17665_ | 5.028387228 | 5.012172087 | 5.045689089 | 0.024474772 | 0.13724006 | 0.159398175 | 0.451698869 | 0.905368164 | 0.964634017 | 0.997 | 0.998705552 |
| SL1344_RS17670_ | 3.88111224 | 3.850247353 | 3.907278459 | 0.017759171 | 0.102725351 | 0.195384503 | 0.438374419 | 0.736416921 | 0.881621438 | 0.8644 | 0.934756399 |
| SL1344_RS17675_ | 5.065933004 | 5.024367608 | 5.104817603 | 0.065271103 | 0.13232746 | 0.431894616 | 0.364666672 | 0.555001443 | 0.78763096 | 0.7617 | 0.889380802 |
| SL1344_RS17680_ | 4.982312577 | 4.799695381 | 5.087562267 | 0.214633732 | 0.263003558 | 0.65910141 | 0.332666674 | 0.454177535 | 0.724217968 | 0.6982 | 0.870853311 |
| SL1344_RS17685_ | 5.65990698 | 5.493913627 | 5.803594326 | 0.157016238 | 0.336594481 | 0.525993046 | 0.313124592 | 0.451822394 | 0.722700362 | 0.6474 | 0.827752363 |
| SL1344_RS17690_ | 5.614367713 | 5.381782384 | 5.749421372 | 0.101415577 | 0.320546054 | 0.926450887 | 0.210000016 | 0.390857325 | 0.677934787 | 0.4121 | 0.660416763 |
| SL1344_RS17700_ | 6.340035925 | 6.174692746 | 6.386638408 | 0.166272546 | 0.118812434 | 0.64779819 | 0.33333334 | 0.523170124 | 0.770598734 | 0.7 | 0.872542119 |
| SL1344_RS17705_ | 5.947486856 | 5.880728053 | 6.001061528 | 0.040305988 | 0.164508912 | 0.235185319 | 0.432378417 | 0.791569873 | 0.916851555 | 0.9589 | 0.982528633 |
| SL1344_RS17710_ | 6.19307101 | 6.156539854 | 6.228342932 | 0.071326558 | 0.158786871 | 0.381414533 | 0.377748173 | 0.586586142 | 0.80941831 | 0.8116 | 0.919781801 |
| SL1344_RS17715_ | 6.699467174 | 6.575387634 | 6.824690286 | 0.181582798 | 0.468743677 | 0.247783313 | 0.438374419 | 0.942742048 | 0.979974592 | 1 | 1 |
| SL1344_RS17720_ | 5.676678929 | 5.608483698 | 5.747207501 | 0.139609496 | 0.25105507 | 0.259868269 | 0.392000004 | 0.972994221 | 0.990532362 | 0.8647 | 0.94227761 |
| SL1344_RS17725_ | 6.847073533 | 6.703343163 | 6.905659296 | 0.202353905 | 0.287364251 | 0.414018381 | 0.33333334 | 0.806017077 | 0.925080127 | 0.7 | 0.872542119 |
| SL1344_RS17730_ | 6.397446219 | 6.293713288 | 6.424295869 | 0.113386077 | 0.152415532 | 0.557321501 | 0.332445044 | 0.544782397 | 0.783975884 | 0.6967 | 0.870421991 |
| SL1344_RS17735_ | 5.487896701 | 5.490982864 | 5.484431788 | 0.08295138 | 0.127429536 | 0.618023241 | 0.387333338 | 0.405240018 | 0.684236131 | 0.8612 | 0.930758116 |
| SL1344_RS17740_ | 5.330897557 | 5.305043742 | 5.345826207 | 0.040413818 | 0.193491842 | 0.201854577 | 0.38241173 | 0.917733367 | 0.970070189 | 0.8149 | 0.921301466 |
| SL1344_RS17745_ | 5.786773799 | 5.773292805 | 5.797848964 | 0.007082912 | 0.078936425 | 0.045070162 | 0.467021987 | 0.629356625 | 0.83375245 | 0.9628 | 0.983945312 |
| SL1344_RS17750_ | 6.082986643 | 6.004676551 | 6.10338992 | 0.102381148 | 0.269481912 | 0.346659715 | 0.33333334 | 0.958075373 | 0.985349685 | 0.7006 | 0.872793903 |
| SL1344_RS17755_ | 5.872264084 | 5.829233703 | 5.914570552 | 0.087095962 | 0.195463517 | 0.299803221 | 0.432378417 | 0.829152736 | 0.934454986 | 0.9718 | 0.987996596 |
| SL1344_RS17760_ | 6.11120995 | 5.928592501 | 6.243270636 | 0.123557763 | 0.297545357 | 0.50954021 | 0.336666673 | 0.720532166 | 0.884398442 | 0.7069 | 0.875470085 |
| SL1344_RS17765_ | 5.035282108 | 4.824114221 | 5.086242805 | 0.260901567 | 0.295162838 | 0.487635897 | 0.33333334 | 0.726179186 | 0.88722027 | 0.7 | 0.872542119 |
| SL1344_RS17770_ | -3.865140066 | -3.828038672 | -3.90437007 | -0.123962883 | 1.230069984 | -0.088817909 | 0.449333335 | 0.712152872 | 0.866060089 | 0.8035 | 0.903367709 |
| SL1344_RS17775_ | 3.04380852 | 3.154863493 | 2.87505215 | -0.262210942 | 0.208436311 | -1.178592991 | 0.110592975 | 0.141510117 | 0.422810698 | 0.2004 | 0.472763012 |

| | | | | | | | | | | | |
|---|---|---|---|---|---|---|---|---|---|---|---|
| SL1344_RS17780_ | 2.858862218 | 3.347289448 | 2.607964564 | -0.469777894 | 0.709994836 | -0.925127007 | 0.153333358 | 0.247538495 | 0.551964868 | 0.2904 | 0.56860753 |
| SL1344_RS17785_ | 7.495367622 | 7.263552593 | 7.516373844 | 0.250271699 | 0.352628186 | 0.37811646 | 0.33333334 | 0.740525964 | 0.894528333 | 0.7 | 0.872542119 |
| SL1344_RS17790_ | 7.764002225 | 7.752610732 | 7.818991391 | 0.10384502 | 0.107156168 | 1.021152255 | 0.080666719 | 0.186453932 | 0.485571457 | 0.1945 | 0.45182605 |
| SL1344_RS17795_ | 5.483840224 | 5.337693284 | 5.521425024 | 0.171764947 | 0.217213375 | 0.504092884 | 0.33333334 | 0.660765469 | 0.852705819 | 0.7 | 0.872542119 |
| SL1344_RS17800_ | 4.982534561 | 4.884007703 | 4.999767664 | 0.11432018 | 0.223733175 | 0.35580018 | 0.33333334 | 0.868378656 | 0.950849605 | 0.7 | 0.872542119 |
| SL1344_RS17805_ | 1.790166644 | 1.833680936 | 1.751336277 | -0.097788712 | 0.201653668 | -0.439456376 | 0.252666679 | 0.385900214 | 0.670151262 | 0.5237 | 0.722579752 |
| SL1344_RS17810_ | 1.542962621 | 1.389787225 | 1.626495556 | 0.246618953 | 0.215364694 | 1.160026193 | 0.106666705 | 0.147899172 | 0.422377659 | 0.2153 | 0.477316233 |
| SL1344_RS17815_ | 1.581504515 | 1.47311733 | 1.752627228 | 0.27399463 | 0.104899668 | 2.488269219 | 0.004667653 | 0.0330973 | 0.195461751 | 0.1053 | 0.337455028 |
| SL1344_RS17820_ | 2.533382045 | 2.398517557 | 2.631914118 | 0.23529121 | 0.150737378 | 1.49213243 | 0.027333498 | 0.102783481 | 0.360590057 | 0.1271 | 0.368951978 |
| SL1344_RS17825_ | 5.117344572 | 5.002281637 | 5.190322702 | 0.178094858 | 0.087056679 | 1.91086398 | 0.007995244 | 0.063809963 | 0.291921226 | 0.1075 | 0.341203221 |
| SL1344_RS17830_ | 0.903071645 | 1.104991045 | 0.813487451 | -0.226943545 | 0.262301298 | -0.878052465 | 0.194000018 | 0.270291778 | 0.556247173 | 0.3582 | 0.619429984 |
| SL1344_RS17835_ | 4.504084915 | 4.287280587 | 4.569394752 | 0.274527889 | 0.119466802 | 1.942122328 | 0.002668394 | 0.056868549 | 0.277777105 | 0.1016 | 0.332612448 |
| SL1344_RS17840_ | 0.460080504 | 0.428974188 | 0.481454908 | 0.077956101 | 0.269277626 | 0.234588714 | 0.350000006 | 0.461225574 | 0.721164547 | 0.6637 | 0.812169009 |
| SL1344_RS17845_ | 2.099084432 | 2.024122507 | 2.152586826 | 0.116012334 | 0.392070007 | 0.235321949 | 0.40373085 | 0.580388074 | 0.804711623 | 0.8785 | 0.947980052 |
| SL1344_RS17850_ | 0.196284926 | -0.130772681 | 0.295595155 | 0.36720888 | 0.461288152 | 0.642038149 | 0.298666676 | 0.444579066 | 0.711607118 | 0.6123 | 0.812813341 |
| SL1344_RS17855_ | 2.628621347 | 2.509753664 | 2.752483539 | 0.252414172 | 0.090795402 | 2.807867203 | 4.68E-05 | 0.022912803 | 0.171090124 | 0.1002 | 0.330546916 |
| SL1344_RS17860_ | -2.097212224 | -2.134500637 | -2.053974485 | 0.076846229 | 0.44680477 | 0.169186731 | 0.425333336 | 0.618861178 | 0.805197422 | 0.7261 | 0.857491651 |
| SL1344_RS17865_ | 0.666095726 | 0.509250389 | 0.741845281 | 0.181061111 | 0.34738618 | 0.482654256 | 0.332666674 | 0.60298927 | 0.810356842 | 0.6985 | 0.858517252 |
| SL1344_RS17870_ | 1.561653417 | 1.564663639 | 1.558830752 | -0.000863762 | 0.115334788 | -0.008317818 | 0.497668221 | 0.62343067 | 0.805664884 | 0.7401 | 0.861783017 |
| SL1344_RS17875_ | -0.354321146 | -0.308505468 | -0.378765001 | -0.029640357 | 0.326115188 | -0.07286377 | 0.466666668 | 0.738596082 | 0.882031339 | 0.8276 | 0.920733286 |
| SL1344_RS17880_ | 1.480538873 | 1.44176283 | 1.519627607 | 0.106039279 | 0.236267295 | 0.445730466 | 0.312000008 | 0.495718647 | 0.737046883 | 0.651 | 0.819042759 |
| SL1344_RS17885_ | 2.452055204 | 2.376195912 | 2.52347349 | 0.1241925 | 0.280377789 | 0.397530079 | 0.367088613 | 0.57679101 | 0.799751114 | 0.7878 | 0.902932366 |
| SL1344_RS17895_ | 3.465923107 | 3.423485231 | 3.538378042 | 0.114073386 | 0.119986 | 0.888788932 | 0.192000018 | 0.250748824 | 0.543426258 | 0.3434 | 0.602304785 |
| SL1344_RS17900_ | 0.821236489 | 0.776840075 | 0.856920488 | 0.072755943 | 0.151100694 | 0.445863306 | 0.315333341 | 0.487196228 | 0.706861433 | 0.595 | 0.766438614 |
| SL1344_RS17905_ | 0.323564245 | 0.285976097 | 0.345476218 | 0.055848833 | 0.180388878 | 0.287518095 | 0.384410398 | 0.560243966 | 0.764431759 | 0.6813 | 0.826020193 |
| SL1344_RS17910_ | -1.265119981 | -1.370983469 | -1.154010915 | 0.238143034 | 0.336860475 | 0.703657079 | 0.226666681 | 0.340387626 | 0.599365113 | 0.4548 | 0.666888195 |
| SL1344_RS17915_ | -1.820563038 | -1.950061009 | -1.654622498 | 0.296721869 | 0.319541447 | 0.856366257 | 0.181212545 | 0.279749498 | 0.524908096 | 0.3773 | 0.595200346 |
| SL1344_RS17920_ | 2.205234292 | 1.856452227 | 2.522458879 | 0.68257731 | 0.167014329 | 4.119471796 | 4.68E-05 | 0.004575011 | 0.085330518 | 0.1 | 0.330257709 |
| SL1344_RS17925_ | -0.828981504 | -0.858364765 | -0.791763882 | 0.057955647 | 0.261524878 | 0.197801473 | 0.410000003 | 0.599590267 | 0.78977207 | 0.7049 | 0.840670803 |
| SL1344_RS17930_ | -0.255423875 | -0.435043662 | 0.361000087 | 0.656059156 | 0.665241816 | 0.719903335 | 0.298000009 | 0.416290348 | 0.692513116 | 0.606 | 0.811525349 |
| SL1344_RS17940_ | -0.335454923 | -0.523418111 | -0.138791899 | 0.430350242 | 0.306949385 | 1.368976321 | 0.053297883 | 0.104799716 | 0.352150974 | 0.1592 | 0.404320293 |
| SL1344_RS17965_ | 3.665496278 | 3.578674087 | 3.722749132 | 0.17489266 | 0.111822465 | 1.578342036 | 0.019986902 | 0.074602446 | 0.309394331 | 0.1195 | 0.357223582 |
| SL1344_RS17970_ | 2.439824405 | 2.274987513 | 2.563006104 | 0.274713442 | 0.103142812 | 2.63850104 | 0.002000966 | 0.027821301 | 0.187426698 | 0.1013 | 0.332156215 |
| SL1344_RS17975_ | 1.916846229 | 1.76392319 | 2.088945484 | 0.355034358 | 0.157611501 | 2.33374052 | 4.68E-05 | 0.03502329 | 0.213862469 | 0.1006 | 0.331121178 |
| SL1344_RS17985_ | 3.82733868 | 3.770164049 | 4.04561128 | 0.266885102 | 0.074082703 | 2.217920245 | 0.001336769 | 0.067758378 | 0.299451049 | 0.1011 | 0.331874815 |
| SL1344_RS17990_ | 3.576704011 | 3.516243244 | 3.600667807 | 0.064853045 | 0.145452796 | 0.346902126 | 0.348434383 | 0.82431006 | 0.928530423 | 0.7333 | 0.886180221 |
| SL1344_RS17995_ | 3.315368312 | 3.332723506 | 3.300620538 | -0.048028006 | 0.125459344 | -0.374148767 | 0.334666674 | 0.52013492 | 0.754754246 | 0.7114 | 0.851514658 |
| SL1344_RS18000_ | -0.979586264 | -0.913158307 | -1.053261648 | -0.101934303 | 0.31708303 | -0.305801914 | 0.367333339 | 0.602901994 | 0.794517147 | 0.7156 | 0.849994752 |
| SL1344_RS18005_ | -2.958688815 | -2.755078892 | -3.105909135 | -0.308777303 | 0.73421569 | -0.403658618 | 0.319333341 | 0.542470909 | 0.754685719 | 0.6234 | 0.797135057 |
| SL1344_RS18010_ | -3.085567885 | -3.008184494 | -3.138614226 | -0.16202286 | 0.545889944 | -0.274055182 | 0.380000005 | 0.559052516 | 0.756291009 | 0.6646 | 0.809663609 |
| SL1344_RS18015_ | -3.154984665 | -2.984172973 | -3.318471311 | -0.348341588 | 0.633243964 | -0.528125908 | 0.290666676 | 0.455919366 | 0.687396204 | 0.5664 | 0.748128177 |
| SL1344_RS18020_ | -2.57414053 | -2.310240761 | -2.825629853 | -0.521966503 | 0.421659403 | -1.176254723 | 0.098666708 | 0.162412933 | 0.408601365 | 0.2367 | 0.474736302 |
| SL1344_RS18025_ | 2.362513428 | 2.371908505 | 2.348916118 | -0.001455305 | 0.195393121 | -0.007643464 | 0.495333334 | 0.816120411 | 0.924035322 | 0.8911 | 0.950210255 |
| SL1344_RS18030_ | 1.316310963 | 1.102589262 | 1.448150619 | 0.351960496 | 0.251741083 | 1.406459371 | 0.021333546 | 0.152582293 | 0.439377385 | 0.1181 | 0.355916342 |
| SL1344_RS18035_ | 3.558323288 | 3.594719486 | 3.52069398 | -0.087815065 | 0.077865736 | -1.126210515 | 0.085942754 | 0.159947669 | 0.428044611 | 0.2165 | 0.459647552 |
| SL1344_RS18040_ | 1.085095739 | 0.97685614 | 1.117753305 | 0.118718053 | 0.230291211 | 0.394050985 | 0.366000005 | 0.748014557 | 0.889481676 | 0.7631 | 0.898261344 |
| SL1344_RS18045_ | 0.621757476 | 0.544278321 | 0.67852251 | 0.064759799 | 0.346434141 | 0.157602789 | 0.439706864 | 0.833036998 | 0.931930485 | 0.9055 | 0.959557007 |
| SL1344_RS18050_ | 1.628098099 | 1.47513391 | 1.759148161 | 0.268464136 | 0.560700847 | 0.381972483 | 0.400399737 | 0.651050049 | 0.84593747 | 0.8812 | 0.949488617 |
| SL1344_RS18055_ | 0.526134099 | 0.516509236 | 0.537717174 | -0.199746455 | 0.433513213 | -0.468257256 | 0.395333337 | 0.501230051 | 0.744755023 | 0.8466 | 0.921575252 |
| SL1344_RS18060_ | 5.402427179 | 5.349703418 | 5.432974825 | 0.081875052 | 0.054791197 | 1.436215976 | 0.061292542 | 0.110372178 | 0.368155901 | 0.1611 | 0.41187776 |
| SL1344_RS18075_ | 0.676415373 | 0.599653802 | 0.786722011 | 0.178725356 | 0.181998809 | 0.926267418 | 0.162000023 | 0.23922718 | 0.504433568 | 0.3351 | 0.576512259 |
| SL1344_RS18080_ | 3.428531746 | 3.239626136 | 3.464603862 | 0.206407655 | 0.215317811 | 0.916632168 | 0.227181893 | 0.279299583 | 0.579420406 | 0.4007 | 0.677282774 |
| SL1344_RS18090_ | 3.391490171 | 3.187424192 | 3.457237629 | 0.30243766 | 0.248690251 | 1.292524234 | 0.073284535 | 0.098329052 | 0.35909139 | 0.1658 | 0.421347008 |

| | | | | | | | | | | | |
|---|---|---|---|---|---|---|---|---|---|---|---|
| SL1344_RS18095_ | 2.303379257 | 2.02862344 | 2.379398099 | 0.301103914 | 0.288400204 | 1.100272368 | 0.151232537 | 0.159835648 | 0.44650056 | 0.2677 | 0.543518737 |
| SL1344_RS18100_ | -0.27295943 | -0.207344936 | -0.338962418 | -0.148323486 | 0.269589661 | -0.538686314 | 0.261825461 | 0.401712072 | 0.649282926 | 0.5434 | 0.730547924 |
| SL1344_RS18105_ | 0.13358071 | 0.173738939 | 0.104233172 | -0.063217587 | 0.46248864 | -0.129398364 | 0.444370422 | 0.822356669 | 0.926737006 | 0.889 | 0.94979798 |
| SL1344_RS18110_ | -0.078442095 | -0.209021538 | 0.032526084 | 0.100371039 | 0.589477569 | 0.16873018 | 0.445036644 | 0.784544518 | 0.909371242 | 0.9382 | 0.972242304 |
| SL1344_RS18115_ | 3.504307695 | 3.507616613 | 3.497983996 | -0.043196435 | 0.17626539 | -0.271192987 | 0.386000004 | 0.606213973 | 0.815462701 | 0.8036 | 0.904990562 |
| SL1344_RS18120_ | 0.796796298 | 0.568484319 | 0.838531557 | 0.245692565 | 0.232478323 | 0.730778631 | 0.294000009 | 0.420759347 | 0.686862347 | 0.6114 | 0.808342337 |
| SL1344_RS18125_ | 1.936425742 | 1.804735197 | 2.301410826 | 0.584566969 | 0.401267452 | 1.490404452 | 0.01666694 | 0.07415065 | 0.315742104 | 0.112 | 0.347546812 |
| SL1344_RS18130_ | 2.658323054 | 2.28469642 | 2.966485919 | 0.689086333 | 0.120145182 | 5.707834196 | 4.68E-05 | 0.003183667 | 0.069318732 | 0.1 | 0.330257709 |
| SL1344_RS18140_ | 0.894060355 | 0.854714789 | 0.928900302 | 0.14257769 | 0.30872068 | 0.448302013 | 0.292666676 | 0.418148965 | 0.689537279 | 0.6376 | 0.804645865 |
| SL1344_RS18145_ | 1.673325001 | 1.631563001 | 1.707157586 | 0.133529943 | 0.344627249 | 0.42899256 | 0.336000007 | 0.500365074 | 0.751104705 | 0.7051 | 0.858509169 |
| SL1344_RS18150_ | 0.166574396 | 0.495677294 | -0.03539608 | -0.525341353 | 0.212382098 | -2.421970754 | 0.002666616 | 0.038792153 | 0.219570247 | 0.1028 | 0.334159277 |
| SL1344_RS18155_ | 0.394115637 | 0.506551484 | 0.149062962 | -0.351149332 | 0.204270395 | -1.597523159 | 0.033333467 | 0.093408751 | 0.332428418 | 0.1344 | 0.373220968 |
| SL1344_RS18160_ | -0.79253982 | -0.671833257 | -0.93149336 | -0.257534856 | 0.220099152 | -1.116575352 | 0.10326453 | 0.175655065 | 0.414536694 | 0.2518 | 0.484663429 |
| SL1344_RS18165_ | 0.690415834 | 0.643510266 | 0.704838013 | 0.049655274 | 0.237511536 | 0.18441928 | 0.432666669 | 0.736115785 | 0.880482063 | 0.8272 | 0.923163731 |
| SL1344_RS18170_ | 1.986505306 | 1.974932049 | 1.995091274 | 0.015532741 | 0.311724183 | 0.052468252 | 0.462666668 | 0.83011725 | 0.930962083 | 0.8845 | 0.946769603 |
| SL1344_RS18175_ | 2.751001567 | 2.73647392 | 2.764220914 | 0.094368387 | 0.175272851 | 0.527075326 | 0.29913392 | 0.387289594 | 0.664497355 | 0.6491 | 0.805588351 |
| SL1344_RS18180_ | 1.286414034 | 1.227273027 | 1.363084769 | 0.149008991 | 0.142488201 | 1.043244806 | 0.108666704 | 0.194505196 | 0.463289575 | 0.2473 | 0.48630142 |
| SL1344_RS18185_ | 7.09461486 | 7.089606375 | 7.097840632 | -0.037443453 | 0.116861245 | -0.354169856 | 0.40866667 | 0.619239694 | 0.828059943 | 0.9103 | 0.958153785 |
| SL1344_RS18190_ | -1.40297536 | -1.357572528 | -1.4725351 | -0.112863944 | 0.329993841 | -0.319595584 | 0.364423723 | 0.539724457 | 0.746683772 | 0.6578 | 0.808023366 |
| SL1344_RS18195_ | 0.595754095 | 0.634526317 | 0.526919521 | -0.080026257 | 0.279212769 | -0.272381042 | 0.403333337 | 0.73938404 | 0.883008044 | 0.8122 | 0.913761057 |
| SL1344_RS18205_ | -2.974362093 | -2.868189419 | -3.057334951 | -0.153305991 | 0.60710124 | -0.241602169 | 0.40066667 | 0.614189973 | 0.80006756 | 0.719 | 0.85101234 |
| SL1344_RS18210_ | -2.195629626 | -2.231896086 | -2.155141638 | 0.093227197 | 0.428031402 | 0.212306077 | 0.400399737 | 0.56392076 | 0.764719675 | 0.6851 | 0.825852997 |
| SL1344_RS18215_ | -1.58962762 | -1.635014409 | -1.529588871 | 0.099397745 | 0.296233736 | 0.311058035 | 0.355333339 | 0.51428428 | 0.724773721 | 0.6306 | 0.788695487 |
| SL1344_RS18220_ | 2.087885283 | 2.087733671 | 2.088102561 | -0.002569825 | 0.094322863 | -0.023793252 | 0.491672219 | 0.629566475 | 0.80884418 | 0.7327 | 0.858685873 |
| SL1344_RS18225_ | 3.288270734 | 3.293213972 | 3.284934779 | 0.005919293 | 0.15912329 | 0.031478338 | 0.484000001 | 0.65879712 | 0.847235116 | 0.9295 | 0.968978934 |
| SL1344_RS18230_ | 1.680675957 | 1.727085374 | 1.655823595 | -0.057532001 | 0.200993497 | -0.277041891 | 0.390666671 | 0.743427013 | 0.886306936 | 0.8007 | 0.909201566 |
| SL1344_RS18235_ | 1.699142244 | 1.716470809 | 1.686194723 | -0.024242374 | 0.201096132 | -0.114494065 | 0.446000002 | 0.764829929 | 0.895977403 | 0.8713 | 0.939765755 |
| SL1344_RS18240_ | 2.22066994 | 2.222775676 | 2.219084351 | 0.093463011 | 0.21374297 | 0.427418186 | 0.373750838 | 0.49578174 | 0.740088823 | 0.8114 | 0.902079115 |
| SL1344_RS18245_ | 2.400150407 | 2.411857316 | 2.393195136 | -0.004918889 | 0.115980948 | -0.032766775 | 0.480666667 | 0.641642055 | 0.831297982 | 0.7935 | 0.903040175 |
| SL1344_RS18250_ | 2.838680061 | 2.872260109 | 2.805318526 | -0.115124908 | 0.14981127 | -0.728823487 | 0.204000017 | 0.275849205 | 0.567237924 | 0.4142 | 0.643733939 |
| SL1344_RS18255_ | -0.077178591 | -0.096080489 | -0.054724004 | 0.005007498 | 0.245910797 | 0.020044181 | 0.492666667 | 0.681616283 | 0.846621927 | 0.8108 | 0.904489649 |
| SL1344_RS18270_ | -0.441286633 | -0.327113567 | -0.549911295 | -0.125872902 | 0.399146546 | -0.314268567 | 0.387333338 | 0.657424065 | 0.834911591 | 0.7667 | 0.882491232 |
| SL1344_RS18275_ | 0.539259201 | 0.526516836 | 0.549298519 | 0.010987264 | 0.27606961 | 0.034710488 | 0.487674884 | 0.764192812 | 0.896381577 | 0.8766 | 0.942798038 |
| SL1344_RS18280_ | 0.139246001 | 0.115972709 | 0.170620737 | 0.065275993 | 0.38821901 | 0.144989382 | 0.440666669 | 0.765620461 | 0.897664249 | 0.884 | 0.946414323 |
| SL1344_RS18285_ | -3.258689613 | -3.454067751 | -3.133804068 | 0.309107686 | 0.712998054 | 0.41321871 | 0.326000007 | 0.512904533 | 0.734067981 | 0.6259 | 0.792544073 |
| SL1344_RS18290_ | -4.243557668 | -4.228912477 | -4.251506103 | 0.074840677 | 1.210796238 | 0.055191483 | 0.469686877 | 0.643459284 | 0.826143183 | 0.789 | 0.892494133 |
| SL1344_RS18295_ | -4.382156034 | -4.868117642 | -3.985949768 | 0.868736306 | 0.747397769 | 1.088616772 | 0.117333368 | 0.182153786 | 0.42406348 | 0.2528 | 0.487357636 |
| SL1344_RS18300_ | -1.841737973 | -1.743649434 | -1.949776339 | -0.205892809 | 0.308435573 | -0.615352064 | 0.249167234 | 0.364579306 | 0.597781398 | 0.4669 | 0.662065402 |
| SL1344_RS18305_ | -0.291738375 | -0.222226426 | -0.378013717 | -0.156397424 | 0.223930425 | -0.654051825 | 0.235842785 | 0.36159623 | 0.609321802 | 0.4758 | 0.678620733 |
| SL1344_RS18310_ | -0.248676558 | -0.103919062 | -0.381037858 | -0.174331222 | 0.459325805 | -0.374835661 | 0.362000006 | 0.633838109 | 0.824703121 | 0.7457 | 0.874626994 |
| SL1344_RS18315_ | 0.239542074 | 0.250272818 | 0.229358201 | -0.044430488 | 0.362066828 | -0.118431192 | 0.437041974 | 0.726400472 | 0.877527859 | 0.8198 | 0.913220591 |
| SL1344_RS18325_ | -0.372965372 | -0.82774677 | -0.116496945 | 0.699079572 | 0.337746322 | 1.944399065 | 0.008661422 | 0.057242977 | 0.268712386 | 0.1095 | 0.343602008 |
| SL1344_RS18330_ | -0.132436576 | -0.426288088 | -0.024494478 | 0.383053523 | 0.235543028 | 1.460968897 | 0.077281867 | 0.12140984 | 0.373125176 | 0.1975 | 0.442184351 |
| SL1344_RS18335_ | 2.066286736 | 1.944346476 | 2.19918783 | 0.279501263 | 0.163691625 | 1.695092865 | 0.008661422 | 0.090046236 | 0.33898216 | 0.108 | 0.341134867 |
| SL1344_RS18340_ | -1.758635746 | -1.848827071 | -1.621170093 | 0.236169243 | 0.43076409 | 0.517626615 | 0.290473027 | 0.48042945 | 0.70959174 | 0.5917 | 0.76992841 |
| SL1344_RS18345_ | -1.902449989 | -1.829107737 | -1.936826323 | -0.082801587 | 0.554165204 | -0.130598166 | 0.448000002 | 0.757806029 | 0.893238189 | 0.8356 | 0.92759051 |
| SL1344_RS18355_ | -1.380634287 | -1.43449462 | -1.336199195 | 0.071910352 | 0.366498117 | 0.199454152 | 0.42238508 | 0.629258175 | 0.81114287 | 0.7331 | 0.860243755 |
| SL1344_RS18360_ | -1.037507746 | -1.086200817 | -1.020817499 | 0.076150122 | 0.440836183 | 0.148357997 | 0.436666669 | 0.660271693 | 0.839164845 | 0.8031 | 0.90710824 |
| SL1344_RS18365_ | -1.748436897 | -1.584670294 | -1.962068507 | -0.345348404 | 0.380292565 | -0.874829899 | 0.175333354 | 0.261801086 | 0.523965864 | 0.3535 | 0.589184702 |
| SL1344_RS18370_ | -1.053487878 | -0.970226194 | -1.152131029 | -0.179149531 | 0.223065384 | -0.736782024 | 0.206528997 | 0.308280505 | 0.546068436 | 0.4001 | 0.611439162 |
| SL1344_RS18375_ | 0.685419297 | 0.598984886 | 0.748989359 | 0.181156928 | 0.25253746 | 0.691726419 | 0.214666682 | 0.310660344 | 0.593347604 | 0.423 | 0.651576892 |
| SL1344_RS18380_ | 1.246331238 | 0.972477024 | 1.321009361 | 0.292288264 | 0.432316645 | 0.413300987 | 0.335776156 | 0.824288107 | 0.930871037 | 0.7069 | 0.875453212 |
| SL1344_RS18385_ | 0.333498496 | 0.775798055 | 0.236868779 | -0.528217444 | 0.40473096 | -1.089814055 | 0.150000025 | 0.193837243 | 0.488304878 | 0.2881 | 0.553405064 |

| | | | | | | | | | | | |
|---|---|---|---|---|---|---|---|---|---|---|---|
| SL1344_RS18390_ | 5.001597548 | 5.069733165 | 4.968944465 | -0.183775437 | 0.239659783 | -0.843065394 | 0.181333353 | 0.232106632 | 0.535722424 | 0.3446 | 0.618895612 |
| SL1344_RS18395_ | 2.958437071 | 2.784172818 | 3.143498951 | 0.220622858 | 0.224661898 | 1.353382576 | 0.106666705 | 0.120006677 | 0.390317808 | 0.1974 | 0.47230236 |
| SL1344_RS18400_ | 1.953520217 | 1.890075596 | 2.002515527 | 0.113210276 | 0.084929963 | 1.255384005 | 0.094000044 | 0.15144264 | 0.384853949 | 0.2319 | 0.467167757 |
| SL1344_RS18405_ | 2.627592471 | 2.58826073 | 2.760340624 | 0.164226629 | 0.220588486 | 0.804984058 | 0.208527665 | 0.289035401 | 0.583796886 | 0.3868 | 0.660735936 |
| SL1344_RS18415_ | 0.6394368 | 0.518951022 | 0.981035571 | 0.441658942 | 0.196723604 | 1.902591672 | 0.018654474 | 0.072989473 | 0.298695889 | 0.1235 | 0.360135522 |
| SL1344_RS18420_ | 0.018282924 | 0.027153776 | 0.012706999 | -0.008542858 | 0.208261736 | -0.035970915 | 0.484000001 | 0.650429582 | 0.823504753 | 0.7579 | 0.873461594 |
| SL1344_RS18425_ | 1.155638982 | 1.246325948 | 1.069360524 | -0.102990975 | 0.346798336 | -0.27891418 | 0.387333338 | 0.682305508 | 0.855512086 | 0.8268 | 0.916624814 |
| SL1344_RS18440_ | 0.511477248 | 0.523725188 | 0.502862706 | 0.072152962 | 0.309702488 | 0.225147383 | 0.418000003 | 0.670813785 | 0.844467719 | 0.8515 | 0.927195253 |
| SL1344_RS18445_ | -0.23187325 | 0.031505679 | -0.371924969 | -0.361685789 | 0.428697755 | -0.814697565 | 0.197868106 | 0.292641316 | 0.580730694 | 0.3837 | 0.64192742 |
| SL1344_RS18450_ | 3.208034089 | 3.206687917 | 3.209577196 | 0.041155528 | 0.19269228 | 0.19455614 | 0.380666671 | 0.540791772 | 0.778415066 | 0.8006 | 0.902638029 |
| SL1344_RS18455_ | 0.950106965 | 0.963003439 | 0.93690934 | 0.049572419 | 0.295993064 | 0.162744138 | 0.437333336 | 0.653924245 | 0.837647222 | 0.8807 | 0.943007215 |
| SL1344_RS18460_ | -0.863742884 | -0.885862322 | -0.847042873 | 0.045082637 | 0.258009691 | 0.168568025 | 0.424000003 | 0.572957122 | 0.771905175 | 0.6841 | 0.82703818 |
| SL1344_RS18465_ | 0.967162244 | 0.983460523 | 0.947707552 | -0.008318258 | 0.221562105 | -0.035325103 | 0.482000001 | 0.745208756 | 0.884974708 | 0.8298 | 0.919154935 |
| SL1344_RS18470_ | 0.795948642 | 0.730969731 | 0.848932386 | 0.060387781 | 0.307326623 | 0.195653634 | 0.40866667 | 0.731230449 | 0.878177614 | 0.8465 | 0.926080022 |
| SL1344_RS18475_ | -0.087931523 | -0.082133452 | -0.090140401 | 0.008171654 | 0.278901123 | 0.028424499 | 0.489673551 | 0.721098679 | 0.870761507 | 0.8417 | 0.924763608 |
| SL1344_RS18480_ | 1.686391944 | 1.821099702 | 1.508692628 | -0.241291869 | 0.230385034 | -1.082621836 | 0.123333366 | 0.172641055 | 0.4562629 | 0.2307 | 0.503131987 |
| SL1344_RS18485_ | 0.186921243 | 0.361743796 | -0.055203326 | -0.435106371 | 0.344039619 | -1.250351285 | 0.084666716 | 0.123133639 | 0.38815267 | 0.1936 | 0.45292917 |
| SL1344_RS18490_ | 1.250083252 | 1.15692795 | 1.34496358 | 0.191998404 | 0.126014709 | 1.456532673 | 0.050000088 | 0.09884242 | 0.32962163 | 0.1625 | 0.405030168 |
| SL1344_RS18495_ | 3.735861979 | 3.580226759 | 3.792571003 | 0.144449895 | 0.216365158 | 0.8445916 | 0.204000017 | 0.280122554 | 0.57982412 | 0.3672 | 0.645445533 |
| SL1344_RS18500_ | 0.700042935 | 0.719294616 | 0.669007795 | -0.049058319 | 0.153448182 | -0.306791296 | 0.377333338 | 0.553616043 | 0.754421029 | 0.6686 | 0.813758577 |
| SL1344_RS18505_ | 5.308352258 | 5.329416537 | 5.29620455 | -0.035289311 | 0.208095682 | -0.198090409 | 0.386666671 | 0.829691632 | 0.934136753 | 0.8224 | 0.924613066 |
| SL1344_RS18510_ | 2.92017768 | 2.62797244 | 3.134143676 | 0.541941455 | 0.132315418 | 4.166321682 | 4.68E-05 | 0.00398432 | 0.079372164 | 0.1 | 0.330257709 |
| SL1344_RS18515_ | 0.995468955 | 1.065922151 | 0.930311235 | -0.123884534 | 0.182625332 | -0.653173229 | 0.245169899 | 0.384225431 | 0.641249686 | 0.4805 | 0.697432283 |
| SL1344_RS18520_ | 2.421190825 | 2.435232779 | 2.407657386 | -0.009932327 | 0.248048774 | -0.040120099 | 0.475682879 | 0.843928956 | 0.937229815 | 0.8515 | 0.930482173 |
| SL1344_RS18525_ | 1.233804184 | 1.42779651 | 1.142325852 | -0.24941086 | 0.255121374 | -0.978630951 | 0.14533336 | 0.202339102 | 0.48969527 | 0.2919 | 0.562824992 |
| SL1344_RS18530_ | 1.217108569 | 1.181269509 | 1.246146386 | 0.079435399 | 0.155735309 | 0.477581019 | 0.300666675 | 0.435193664 | 0.679217458 | 0.5817 | 0.755326652 |
| SL1344_RS18535_ | 2.861463709 | 2.849131692 | 2.871595024 | 0.023010199 | 0.158104831 | 0.156161576 | 0.425716192 | 0.733169465 | 0.880576606 | 0.8035 | 0.905134098 |
| SL1344_RS18540_ | -0.658638139 | -0.629943178 | -0.700786282 | -0.079353635 | 0.254275126 | -0.287883316 | 0.370419725 | 0.541332463 | 0.749414541 | 0.6764 | 0.815958131 |
| SL1344_RS18545_ | -0.630139672 | -0.599783231 | -0.678881051 | -0.056250073 | 0.314108053 | -0.172373948 | 0.428000003 | 0.670753047 | 0.840729453 | 0.7869 | 0.893387489 |
| SL1344_RS18550_ | -1.546034814 | -1.576886342 | -1.509897265 | 0.068730839 | 0.456886982 | 0.143970727 | 0.438374419 | 0.683372142 | 0.846594846 | 0.7995 | 0.899939926 |
| SL1344_RS18555_ | 0.027029929 | 0.058718431 | -0.001876426 | -0.004789072 | 0.285727992 | -0.01482756 | 0.494666667 | 0.739508109 | 0.882097618 | 0.8684 | 0.93821415 |
| SL1344_RS18560_ | 0.456406284 | 0.375489605 | 0.595801525 | 0.163809638 | 0.290150063 | 0.572096642 | 0.293137917 | 0.504563747 | 0.738800427 | 0.5946 | 0.786962986 |
| SL1344_RS18565_ | 0.602783391 | 0.610948967 | 0.582809386 | -0.009715011 | 0.203640077 | -0.045175035 | 0.483677549 | 0.725558531 | 0.873413562 | 0.8117 | 0.913440875 |
| SL1344_RS18570_ | 4.186949733 | 4.399734771 | 3.947735532 | -0.46466143 | 0.08851439 | -5.409803543 | 4.68E-05 | 0.027407859 | 0.201069875 | 0.1 | 0.330257709 |
| SL1344_RS18575_ | -3.513233768 | -3.258702659 | -3.653288089 | -0.326942141 | 0.753523108 | -0.385887405 | 0.339107268 | 0.583519105 | 0.779096372 | 0.6758 | 0.82872165 |
| SL1344_RS18580_ | -2.452729404 | -2.612895006 | -2.332959887 | 0.278650843 | 0.429954087 | 0.614872362 | 0.240000013 | 0.366027767 | 0.609590338 | 0.471 | 0.671943191 |
| SL1344_RS18585_ | -0.278393579 | -0.313803974 | -0.241395531 | 0.15118197 | 0.318113658 | 0.445030881 | 0.314000008 | 0.467545041 | 0.708043258 | 0.6718 | 0.815660528 |
| SL1344_RS18590_ | 0.681606201 | 0.603031765 | 0.744691753 | 0.149476078 | 0.217881605 | 0.65324262 | 0.243333346 | 0.361771752 | 0.62713981 | 0.4698 | 0.684732791 |
| SL1344_RS18595_ | -0.263230369 | -0.202493351 | -0.33896679 | -0.165376513 | 0.279277144 | -0.534268601 | 0.267155241 | 0.402159877 | 0.659136326 | 0.5342 | 0.728680484 |
| SL1344_RS18600_ | 2.29429341 | 2.788267498 | 1.641232274 | -1.151121056 | 0.2001965 | -5.901828969 | 4.68E-05 | 0.013255836 | 0.143127972 | 0.1 | 0.330257709 |
| SL1344_RS18610_ | 0.587645202 | 0.519526431 | 0.934025681 | 0.391381135 | 0.204226084 | 1.31578292 | 0.111259197 | 0.150791714 | 0.424358099 | 0.2527 | 0.496761803 |
| SL1344_RS18615_ | 2.259404732 | 2.22878505 | 2.276278003 | 0.050185392 | 0.095011204 | 0.470501489 | 0.303333342 | 0.445417654 | 0.681832533 | 0.5689 | 0.744503657 |
| SL1344_RS18620_ | 4.752195832 | 4.70346651 | 4.796144003 | 0.05226747 | 0.316038723 | 0.146464568 | 0.390000004 | 0.821702203 | 0.931100271 | 0.847 | 0.93495304 |
| SL1344_RS18625_ | 1.906549511 | 1.951933164 | 1.883563341 | -0.046786065 | 0.131785132 | -0.325823508 | 0.376415728 | 0.666884104 | 0.838089553 | 0.7428 | 0.876508552 |
| SL1344_RS18630_ | 2.552267376 | 2.46698863 | 2.811418547 | 0.326198965 | 0.157560711 | 1.825941599 | 0.01798826 | 0.064641977 | 0.290267194 | 0.1133 | 0.348788076 |
| SL1344_RS18635_ | 1.102435863 | 0.859375654 | 1.463662387 | 0.635656987 | 0.250080984 | 2.56291278 | 4.68E-05 | 0.018468373 | 0.161568645 | 0.1 | 0.330257709 |
| SL1344_RS18640_ | 1.573748023 | 1.189012483 | 1.857684996 | 0.67982445 | 0.167234612 | 4.090997971 | 4.68E-05 | 0.017259396 | 0.154025788 | 0.1 | 0.330257709 |
| SL1344_RS18655_ | -0.622137438 | -1.229844031 | -0.148132861 | 1.080018309 | 0.207047341 | 5.175580821 | 4.68E-05 | 0.00460413 | 0.075422512 | 0.1 | 0.330257709 |
| SL1344_RS18660_ | 1.203865054 | 0.706747591 | 1.590448724 | 0.873803067 | 0.142418401 | 6.044009985 | 4.68E-05 | 0.005400132 | 0.085158678 | 0.1 | 0.330257709 |
| SL1344_RS18665_ | -0.88995996 | -1.285118612 | -0.462878205 | 0.81577769 | 0.284111645 | 2.784651652 | 0.001336769 | 0.019508682 | 0.15519069 | 0.1007 | 0.331285063 |
| SL1344_RS18670_ | 1.462291404 | 1.125213869 | 1.765675951 | 0.635793868 | 0.156783517 | 3.967795678 | 4.68E-05 | 0.006824047 | 0.098693452 | 0.1 | 0.330257709 |
| SL1344_RS18675_ | 2.984959952 | 2.93446401 | 3.037127695 | 0.074186892 | 0.435989139 | 0.153354901 | 0.447333335 | 0.933531486 | 0.976041828 | 0.9997 | 0.999871231 |
| SL1344_RS18680_ | -2.369630756 | -2.068990153 | -2.545814004 | -0.422055555 | 0.608877543 | -0.661013042 | 0.251333346 | 0.384696957 | 0.644633923 | 0.5006 | 0.713360004 |

| | | | | | | | | | | | |
|---|---|---|---|---|---|---|---|---|---|---|---|
| SL1344_RS18685_ | -0.240648338 | -0.498017931 | -0.080102778 | 0.424043409 | 0.337937573 | 1.251078068 | 0.091938753 | 0.129729944 | 0.392594355 | 0.2019 | 0.459373089 |
| SL1344_RS18690_ | 0.973540036 | 1.092034953 | 0.868875414 | -0.180308351 | 0.18826715 | -0.918248626 | 0.153231204 | 0.228636117 | 0.50683958 | 0.3061 | 0.562807178 |
| SL1344_RS18695_ | 0.886203034 | 0.908218623 | 0.875081176 | -0.034225602 | 0.178076684 | -0.18150278 | 0.424383747 | 0.671797437 | 0.838619776 | 0.7822 | 0.891949848 |
| SL1344_RS18700_ | 1.069723436 | 1.058408985 | 1.075757615 | 0.015328035 | 0.138272021 | 0.109492005 | 0.452365092 | 0.611353554 | 0.798038555 | 0.7179 | 0.850246269 |
| SL1344_RS18705_ | 0.420853696 | 0.505409021 | 0.383617277 | -0.101011378 | 0.195654817 | -0.458510704 | 0.329780154 | 0.555343673 | 0.76429697 | 0.6595 | 0.82157735 |
| SL1344_RS18710_ | 2.763692184 | 2.756612262 | 2.772430048 | 0.081276158 | 0.195015741 | 0.423098944 | 0.333111266 | 0.421215539 | 0.691849976 | 0.7259 | 0.850716326 |
| SL1344_RS18715_ | -2.083404149 | -2.202157952 | -1.982956282 | 0.21979341 | 0.38209825 | 0.543051953 | 0.278481023 | 0.411279589 | 0.649311048 | 0.5253 | 0.714440554 |
| SL1344_RS18720_ | -1.34751717 | -1.427893446 | -1.185702109 | 0.222408946 | 0.401538294 | 0.518002104 | 0.298000009 | 0.505373606 | 0.729360195 | 0.5962 | 0.781759422 |
| SL1344_RS18725_ | -1.002134531 | -1.04203181 | -0.977843449 | 0.074385317 | 0.276210915 | 0.237964598 | 0.386666671 | 0.563636816 | 0.767002304 | 0.6841 | 0.826138397 |
| SL1344_RS18735_ | 1.162772083 | 1.65662138 | 1.071799993 | -0.576601559 | 0.766848364 | -0.607614231 | 0.33333334 | 0.525557363 | 0.771569474 | 0.7 | 0.872542119 |
| SL1344_RS18740_ | 1.452568728 | 1.369553712 | 1.546001804 | 0.190142009 | 0.137407387 | 1.387993673 | 0.044666765 | 0.115222456 | 0.364248695 | 0.1539 | 0.394766873 |
| SL1344_RS18745_ | -1.829861016 | -1.980622416 | -1.709486038 | 0.265147969 | 0.334562547 | 0.74316817 | 0.20733335 | 0.30723049 | 0.557183257 | 0.4028 | 0.617266999 |
| SL1344_RS18750_ | -1.498272559 | -1.47393189 | -1.513605838 | -0.004875931 | 0.357821772 | -0.014289497 | 0.495003331 | 0.649772841 | 0.824915068 | 0.7567 | 0.874550937 |
| SL1344_RS18755_ | -1.618870134 | -1.568766246 | -1.661464401 | -0.083357659 | 0.329408943 | -0.23706104 | 0.39840107 | 0.567824826 | 0.765191858 | 0.6661 | 0.815286997 |
| SL1344_RS18760_ | -4.663182797 | -4.352078376 | -4.912299191 | -0.413531054 | 1.15333644 | -0.335624691 | 0.349333339 | 0.58494761 | 0.77849512 | 0.659 | 0.817659774 |
| SL1344_RS18765_ | -3.682500484 | -3.511738514 | -3.882797302 | -0.200621662 | 1.091116149 | -0.174328653 | 0.41666667 | 0.706992485 | 0.862555943 | 0.8005 | 0.900836823 |
| SL1344_RS18770_ | -1.384776916 | -1.582634815 | -1.077393199 | 0.515446622 | 0.365642933 | 1.340511919 | 0.063333401 | 0.120019093 | 0.367349056 | 0.1774 | 0.422434295 |
| SL1344_RS18775_ | 1.656688837 | 1.628981845 | 1.720653953 | 0.070826507 | 0.176542993 | 0.342189459 | 0.362000006 | 0.683434306 | 0.850190097 | 0.7266 | 0.871276285 |
| SL1344_RS18780_ | 1.089738853 | 1.081317499 | 1.097796902 | 0.05133745 | 0.257911739 | 0.179036993 | 0.40666667 | 0.625164341 | 0.819453433 | 0.7715 | 0.886819153 |
| SL1344_RS18785_ | 2.345997169 | 2.26986515 | 2.384616702 | 0.108457612 | 0.093273874 | 1.092663904 | 0.12658231 | 0.189196662 | 0.449661662 | 0.2827 | 0.525559247 |
| SL1344_RS18790_ | 0.928879212 | 0.803580137 | 1.136384325 | 0.334526849 | 0.157118201 | 2.031158955 | 0.013324793 | 0.054819562 | 0.255601697 | 0.1119 | 0.34607545 |
| SL1344_RS18795_ | 1.762081305 | 1.743286074 | 1.775826414 | 0.127026221 | 0.270564125 | 0.47171137 | 0.340666673 | 0.450765082 | 0.71273925 | 0.7161 | 0.856281427 |
| SL1344_RS18800_ | 3.543352419 | 3.457502993 | 3.698044927 | 0.243850391 | 0.106162277 | 2.152360038 | 0.001335878 | 0.050360298 | 0.258238489 | 0.1011 | 0.331867025 |
| SL1344_RS18805_ | 0.497975114 | 0.245063324 | 0.727321568 | 0.497524875 | 0.359011267 | 1.338918073 | 0.019986902 | 0.148487196 | 0.433151287 | 0.1219 | 0.361235598 |
| SL1344_RS18815_ | -0.206630138 | -0.544190709 | 0.352518018 | 0.787356732 | 0.570657732 | 1.253468105 | 0.105929419 | 0.134393455 | 0.411404119 | 0.2015 | 0.471929789 |
| SL1344_RS18820_ | 0.709297303 | 0.703151375 | 0.722492563 | 0.021651637 | 0.218079473 | 0.094711338 | 0.459027317 | 0.733770582 | 0.877872796 | 0.8428 | 0.925084299 |
| SL1344_RS18825_ | 0.627981199 | 0.662031159 | 0.582295349 | -0.087198724 | 0.252471835 | -0.3483099 | 0.345769493 | 0.588814752 | 0.793425276 | 0.6651 | 0.828789416 |
| SL1344_RS18830_ | -1.953623804 | -2.22619724 | -1.544751473 | 0.616096325 | 0.474321282 | 1.209357837 | 0.100599641 | 0.141146692 | 0.400513284 | 0.2192 | 0.474546035 |
| SL1344_RS18835_ | 3.987353522 | 3.872443189 | 4.150987325 | 0.269425228 | 0.066086239 | 3.914955068 | 4.68E-05 | 0.010848248 | 0.122114875 | 0.1 | 0.330257709 |
| SL1344_RS18840_ | 2.419272179 | 2.383967904 | 2.49529909 | 0.109922185 | 0.115255997 | 0.880789842 | 0.171219209 | 0.259322418 | 0.540216254 | 0.3511 | 0.590949589 |
| SL1344_RS18845_ | -0.893441295 | -1.035445441 | -0.742495202 | 0.339143928 | 0.325210462 | 1.000458157 | 0.119253864 | 0.195827038 | 0.463046631 | 0.2825 | 0.522076753 |
| SL1344_RS18850_ | -0.830745145 | -0.718726058 | -0.950261832 | -0.228282589 | 0.274374448 | -0.801609041 | 0.183211212 | 0.278689861 | 0.53847266 | 0.3692 | 0.597686885 |
| SL1344_RS18855_ | -2.899924733 | -2.899924733 | -2.899648245 | 0.03028001 | 0.583935643 | 0.047074571 | 0.476666668 | 0.632166808 | 0.812468321 | 0.7528 | 0.871129238 |
| SL1344_RS18860_ | 0.277915817 | 0.267541316 | 0.288628095 | 0.020185281 | 0.16090492 | 0.121418654 | 0.444370422 | 0.57018712 | 0.766686567 | 0.6875 | 0.828544921 |
| SL1344_RS18865_ | 0.648349521 | 0.713222671 | 0.516052236 | -0.183655361 | 0.172593153 | -0.993671617 | 0.162000023 | 0.227053722 | 0.494027456 | 0.3224 | 0.567227495 |
| SL1344_RS18870_ | 0.152188265 | -0.47071665 | 0.449034922 | 0.905567278 | 0.325819793 | 2.664584175 | 0.00067292 | 0.024879183 | 0.182884279 | 0.1001 | 0.330408033 |
| SL1344_RS18875_ | 3.043646574 | 3.022854329 | 3.05585628 | 0.020518015 | 0.120961455 | 0.126973423 | 0.435333336 | 0.756946232 | 0.894799138 | 0.834 | 0.928170682 |
| SL1344_RS18880_ | -2.426413992 | -2.680571026 | -2.255090239 | 0.369037358 | 0.562777371 | 0.633268314 | 0.256666678 | 0.401017136 | 0.653019511 | 0.5121 | 0.715631947 |
| SL1344_RS18885_ | -0.676458843 | -0.964416229 | -0.556899474 | 0.388551107 | 0.299836822 | 1.171510209 | 0.114666702 | 0.165002475 | 0.430825967 | 0.2498 | 0.500525064 |
| SL1344_RS18890_ | 1.528860049 | 1.639844368 | 1.267784361 | -0.374017402 | 0.232354037 | -1.671871585 | 0.026000173 | 0.058987937 | 0.277460382 | 0.1206 | 0.359017238 |
| SL1344_RS18895_ | 0.475731108 | 0.483081682 | 0.463341637 | -0.025336074 | 0.160419854 | -0.1374825 | 0.440000002 | 0.602269716 | 0.790396035 | 0.7187 | 0.846885012 |
| SL1344_RS18900_ | 0.727280577 | 0.741708681 | 0.709582206 | -0.042069033 | 0.23173666 | -0.170510951 | 0.42238508 | 0.69322047 | 0.854404134 | 0.8112 | 0.906618185 |
| SL1344_RS18905_ | -1.953948146 | -1.903946605 | -2.006678185 | -0.10574708 | 0.364993256 | -0.282730947 | 0.377333338 | 0.516420918 | 0.727145911 | 0.6323 | 0.788552672 |
| SL1344_RS18910_ | -3.106404444 | -2.992355457 | -3.216230998 | -0.054458207 | 0.801980421 | -0.074502269 | 0.472666668 | 0.730656004 | 0.876398103 | 0.851 | 0.927727706 |
| SL1344_RS18915_ | -2.780610419 | -3.098024102 | -2.484204388 | 0.642537266 | 0.537870786 | 1.151745786 | 0.087275198 | 0.154946189 | 0.406293599 | 0.2276 | 0.471601273 |
| SL1344_RS18920_ | -6.089911884 | -6.308803811 | -5.842000037 | 0.402239184 | 1.80397216 | 0.198990511 | 0.399733515 | 0.601291613 | 0.790345978 | 0.6958 | 0.836784494 |
| SL1344_RS18925_ | -4.178750624 | -3.920140173 | -4.37739905 | -0.430472921 | 0.935304497 | -0.424277707 | 0.323117929 | 0.515463027 | 0.730820083 | 0.6134 | 0.783620542 |
| SL1344_RS18930_ | -3.948978963 | -3.823052615 | -4.064610488 | -0.19256733 | 0.805085923 | -0.219255421 | 0.403064627 | 0.590015225 | 0.780873499 | 0.6975 | 0.835160476 |
| SL1344_RS18935_ | -2.296067631 | -2.368411601 | -2.122355941 | 0.174577261 | 0.569808862 | 0.214845068 | 0.399733515 | 0.760433451 | 0.891985675 | 0.7745 | 0.899366372 |
| SL1344_RS18945_ | -3.794750416 | -3.25153974 | -4.145077991 | -0.907372327 | 1.011512918 | -0.858405614 | 0.161225873 | 0.242472494 | 0.51920136 | 0.3266 | 0.576204836 |
| SL1344_RS18950_ | -3.161371815 | -3.108340243 | -3.293921304 | -0.175043832 | 0.690907148 | -0.225119551 | 0.410000003 | 0.596111226 | 0.795246056 | 0.7286 | 0.861309255 |
| SL1344_RS18955_ | 0.008229121 | -0.300714424 | 0.309099389 | 0.607010196 | 0.190733784 | 3.191346525 | 4.68E-05 | 0.013788523 | 0.131335067 | 0.1002 | 0.330543941 |
| SL1344_RS18960_ | -0.421402664 | -0.306068351 | -0.511109663 | -0.209361105 | 0.33163806 | -0.603773469 | 0.254000012 | 0.38625489 | 0.649884418 | 0.5063 | 0.720235189 |

| | | | | | | | | | | | |
|---|---|---|---|---|---|---|---|---|---|---|---|
| SL1344_RS18965_ | -0.102816396 | -0.030541266 | -0.158940604 | -0.093099991 | 0.395479464 | -0.235893071 | 0.38507662 | 0.715988789 | 0.869349367 | 0.7756 | 0.891347712 |
| SL1344_RS18970_ | -0.704419803 | -0.775582452 | -0.613911892 | 0.172817811 | 0.243939935 | 0.675715813 | 0.230666681 | 0.35716362 | 0.604127207 | 0.4615 | 0.66049045 |
| SL1344_RS18975_ | 2.146098709 | 2.101173713 | 2.191213549 | 0.097676767 | 0.108678446 | 0.865099937 | 0.165889429 | 0.259198565 | 0.524278589 | 0.3462 | 0.579496317 |
| SL1344_RS18980_ | 2.375853943 | 2.293209322 | 2.502896442 | 0.173876702 | 0.219007404 | 0.71110137 | 0.23984012 | 0.354596539 | 0.640631922 | 0.4635 | 0.699008649 |
| SL1344_RS18985_ | 1.437201341 | 1.6164247 | 1.403251451 | -0.179508401 | 0.193579727 | -0.51210276 | 0.336442379 | 0.677119574 | 0.85277239 | 0.7078 | 0.875329091 |
| SL1344_RS18990_ | 3.024912165 | 2.901272215 | 3.051644431 | 0.157208501 | 0.171205091 | 0.904086993 | 0.213857444 | 0.262789946 | 0.558676433 | 0.3837 | 0.656233014 |
| SL1344_RS18995_ | 2.069631862 | 2.088740465 | 2.052472667 | -0.049486608 | 0.1264428 | -0.362742039 | 0.33333334 | 0.540582133 | 0.753898877 | 0.6798 | 0.824647061 |
| SL1344_RS19005_ | -3.181952581 | -3.595913883 | -2.806853304 | 0.764190964 | 0.631117612 | 1.160244209 | 0.104596974 | 0.163611008 | 0.416963539 | 0.2332 | 0.481003185 |
| SL1344_RS19010_ | 2.488989875 | 2.144052307 | 2.517890903 | 0.379037924 | 0.392070488 | 0.779001353 | 0.326000007 | 0.351134545 | 0.645459882 | 0.6782 | 0.858458616 |
| SL1344_RS19015_ | -5.440445563 | -5.564240965 | -5.288266931 | 0.312094182 | 1.211660333 | 0.236063045 | 0.393737512 | 0.545159414 | 0.74997108 | 0.6636 | 0.811083203 |
| SL1344_RS19020_ | -0.368222131 | -0.433387413 | -0.300967711 | 0.134815036 | 0.239205234 | 0.519760676 | 0.283333343 | 0.455762208 | 0.687304435 | 0.5528 | 0.740241205 |
| SL1344_RS19025_ | 4.070026464 | 4.004732425 | 4.146146493 | 0.171921099 | 0.694580139 | 0.246353402 | 0.33133334 | 0.693822935 | 0.870681948 | 0.7 | 0.872542119 |
| SL1344_RS19030_ | 3.04626631 | 3.226757839 | 2.844894467 | 0.015291776 | 0.791624097 | 0.020293238 | 0.483011326 | 0.891908367 | 0.960445652 | 1 | 1 |
| SL1344_RS19040_ | 1.247013078 | 1.569413994 | 1.144023118 | -0.295340011 | 0.33770829 | -0.751401634 | 0.287333343 | 0.390388323 | 0.671248451 | 0.5814 | 0.788699997 |
| SL1344_RS19045_ | 0.627844473 | 0.516351642 | 0.664656029 | 0.130645964 | 0.198580898 | 0.564009304 | 0.303131254 | 0.459874827 | 0.704290902 | 0.5938 | 0.782872795 |
| SL1344_RS19050_ | 0.987747868 | 0.681880721 | 1.12556432 | 0.427193941 | 0.172075043 | 2.195709839 | 0.005330642 | 0.049471759 | 0.2476793 | 0.1067 | 0.339256372 |
| SL1344_RS19055_ | -0.262424771 | -0.447862033 | -0.156771163 | 0.310035075 | 0.279400553 | 1.091347143 | 0.078000054 | 0.179190297 | 0.457183369 | 0.2051 | 0.448932325 |
| SL1344_RS19065_ | 4.61067147 | 4.577875497 | 4.644284665 | -0.075184721 | 0.260577963 | -0.285761314 | 0.446369089 | 0.772504964 | 0.907631299 | 0.997 | 0.998721826 |
| SL1344_RS19070_ | 4.735709507 | 4.733313055 | 4.738510329 | 0.015082385 | 0.132519371 | 0.116587775 | 0.397068625 | 0.628238509 | 0.830682983 | 0.7522 | 0.878387073 |
| SL1344_RS19075_ | 3.078471605 | 3.037181294 | 3.23547947 | 0.21232301 | 0.210199503 | 1.120430104 | 0.094603641 | 0.142964658 | 0.424439723 | 0.215 | 0.474136134 |
| SL1344_RS19080_ | 3.254637418 | 3.215968839 | 3.363944791 | 0.188474806 | 0.28552723 | 0.692962126 | 0.216522334 | 0.291563162 | 0.591225847 | 0.4034 | 0.678182927 |
| SL1344_RS19085_ | 1.616431911 | 1.55281718 | 1.663082041 | 0.133755371 | 0.238291981 | 0.555555747 | 0.28314458 | 0.439131365 | 0.700606743 | 0.5809 | 0.780426734 |
| SL1344_RS19090_ | 3.04456661 | 3.022177584 | 3.131218922 | 0.097454377 | 0.092507139 | 0.912786831 | 0.216666682 | 0.288883181 | 0.569002809 | 0.4307 | 0.667525723 |
| SL1344_RS19095_ | 1.855040853 | 1.728045702 | 1.954179875 | 0.22656272 | 0.098787311 | 2.22597789 | 0.007329074 | 0.038941551 | 0.211950835 | 0.1091 | 0.342400665 |
| SL1344_RS19100_ | -0.627354645 | -0.955875679 | -0.233755599 | 0.712607719 | 0.290676036 | 2.396011821 | 0.001335878 | 0.030352256 | 0.194357718 | 0.1018 | 0.33289735 |
| SL1344_RS19105_ | 5.036427724 | 4.877740628 | 5.255428093 | 0.328829291 | 0.187442749 | 1.83351167 | 0.004001152 | 0.041676325 | 0.24349678 | 0.1023 | 0.333596928 |
| SL1344_RS19110_ | 4.396482387 | 4.081699489 | 4.470853269 | 0.376393555 | 0.171809026 | 1.383697271 | 0.059293877 | 0.114576494 | 0.387444331 | 0.1555 | 0.410012566 |
| SL1344_RS19115_ | 2.94590275 | 2.870908199 | 2.982092529 | 0.10885379 | 0.174998341 | 0.56339165 | 0.250499679 | 0.356660532 | 0.647280576 | 0.4891 | 0.709616835 |
| SL1344_RS19120_ | 2.906930736 | 2.88160781 | 2.953519721 | 0.067179631 | 0.072521606 | 0.834813104 | 0.187208547 | 0.289137862 | 0.546321994 | 0.399 | 0.6253449 |
| SL1344_RS19125_ | 2.474819073 | 2.413505339 | 2.529603798 | 0.129473184 | 0.104961147 | 1.206193145 | 0.06795476 | 0.151405075 | 0.418784638 | 0.1802 | 0.423573932 |
| SL1344_RS19130_ | 2.830714439 | 2.613871402 | 3.007685232 | 0.387562736 | 0.212119027 | 1.808031415 | 0.008000574 | 0.051147404 | 0.263448136 | 0.1074 | 0.341134642 |
| SL1344_RS19135_ | 1.76574667 | 1.74912987 | 1.77670107 | 0.008455597 | 0.178454291 | 0.02661266 | 0.480346436 | 0.603168954 | 0.814954615 | 0.83 | 0.921991537 |
| SL1344_RS19140_ | 1.332826109 | 1.240551873 | 1.60242163 | 0.339016798 | 0.230915526 | 1.40564547 | 0.069333395 | 0.108352592 | 0.365513824 | 0.1884 | 0.44288277 |
| SL1344_RS19145_ | 0.813834388 | 0.860468755 | 0.682081202 | -0.210814873 | 0.322102637 | -0.663229348 | 0.23984012 | 0.362811054 | 0.643625138 | 0.4733 | 0.716709956 |
| SL1344_RS19150_ | 1.545143537 | 1.570304029 | 1.511293779 | -0.01974648 | 0.212706829 | -0.095307553 | 0.459333335 | 0.773210927 | 0.901200001 | 0.8907 | 0.949475461 |
| SL1344_RS19155_ | 1.907694206 | 1.974575608 | 1.830977595 | -0.04708709 | 0.306472778 | -0.165113898 | 0.430379749 | 0.824364224 | 0.928059983 | 0.9085 | 0.959357522 |
| SL1344_RS19160_ | 1.977938751 | 1.964764149 | 1.991031999 | 0.054345255 | 0.28068465 | 0.193145284 | 0.387333338 | 0.73836688 | 0.885188238 | 0.7999 | 0.905169817 |
| SL1344_RS19165_ | 0.00719852 | -0.089844142 | 0.080099483 | 0.172976063 | 0.172828883 | 0.921037541 | 0.152000025 | 0.249982491 | 0.501555174 | 0.3405 | 0.566995187 |
| SL1344_RS19170_ | 1.916893178 | 1.900671245 | 1.940267278 | 0.079485409 | 0.405622691 | 0.211030617 | 0.351765496 | 0.624688965 | 0.828628946 | 0.7507 | 0.879011728 |
| SL1344_RS19175_ | 2.345171391 | 2.325679659 | 2.366511543 | 0.062876521 | 0.295433159 | 0.211626078 | 0.356000006 | 0.556841291 | 0.787743781 | 0.7238 | 0.863253263 |
| SL1344_RS19180_ | 2.26616956 | 2.338690593 | 2.236565941 | -0.088783737 | 0.21700798 | -0.31585727 | 0.3637575 | 0.82811567 | 0.929997201 | 0.7645 | 0.898681102 |
| SL1344_RS19185_ | 3.268607071 | 3.044978923 | 3.34886267 | 0.304008059 | 0.14517116 | 1.928306295 | 0.002668394 | 0.066899726 | 0.298602707 | 0.1025 | 0.333776915 |
| SL1344_RS19190_ | 2.166974604 | 2.0601895 | 2.286241565 | 0.240869149 | 0.141154292 | 1.676195675 | 0.022651765 | 0.065751051 | 0.2857038 | 0.122 | 0.360520104 |
| SL1344_RS19195_ | 0.232972051 | 0.314943474 | 0.164126661 | -0.098743456 | 0.584463282 | -0.154743799 | 0.440666669 | 0.819507009 | 0.926068744 | 0.9406 | 0.973399028 |
| SL1344_RS19200_ | 6.537425626 | 6.545734472 | 6.512436257 | -0.035290896 | 0.101043965 | -0.269347366 | 0.351333339 | 0.924738027 | 0.972836771 | 0.739 | 0.889068922 |
| SL1344_RS19205_ | 6.115201937 | 6.081814571 | 6.158388934 | 0.082704344 | 0.41899758 | 0.211116795 | 0.28580947 | 0.725359272 | 0.886779693 | 0.5506 | 0.776702895 |
| SL1344_RS19210_ | 1.123375332 | 1.082250559 | 1.22706252 | 0.155928118 | 0.268044658 | 0.598293445 | 0.254000012 | 0.432880936 | 0.696029253 | 0.5214 | 0.748703178 |
| SL1344_RS19215_ | 2.525251086 | 2.525429567 | 2.524918283 | -0.000197008 | 0.082908772 | -0.001759854 | 0.498334444 | 0.628388213 | 0.808376791 | 0.7409 | 0.862945301 |
| SL1344_RS19220_ | 2.133605212 | 2.110665591 | 2.154902629 | 0.029802887 | 0.205704268 | 0.140724073 | 0.445333335 | 0.798948912 | 0.915358605 | 0.9186 | 0.963219905 |
| SL1344_RS19225_ | 2.236745361 | 2.223824636 | 2.24714285 | 0.035709116 | 0.137148527 | 0.238728193 | 0.3904064 | 0.639376613 | 0.822878327 | 0.763 | 0.879626885 |
| SL1344_RS19230_ | 0.628033311 | 0.427206134 | 0.895417761 | 0.381217484 | 0.441560969 | 0.869729085 | 0.15866669 | 0.241739117 | 0.539251322 | 0.2813 | 0.554809738 |
| SL1344_RS19235_ | 0.413950095 | 0.436941358 | 0.377926759 | -0.048431864 | 0.335167178 | -0.121610603 | 0.458000002 | 0.796405808 | 0.913873821 | 0.9016 | 0.957365568 |
| SL1344_RS19240_ | 2.658269914 | 2.616673423 | 2.685351995 | 0.071187402 | 0.11641928 | 0.548349238 | 0.255163236 | 0.389951456 | 0.654001951 | 0.5168 | 0.72034086 |

| | | | | | | | | | | | |
|---|---|---|---|---|---|---|---|---|---|---|---|
| SL1344_RS19245_ | -0.23566308 | -0.272701919 | -0.201825761 | 0.07386085 | 0.193693945 | 0.375277275 | 0.334666674 | 0.488352932 | 0.7050147 | 0.6062 | 0.769052962 |
| SL1344_RS19250_ | -1.977382753 | -1.973780639 | -1.981625412 | 0.006829109 | 0.373241191 | 0.021252508 | 0.492338441 | 0.606275245 | 0.793136808 | 0.7266 | 0.851772559 |
| SL1344_RS19255_ | 1.269798152 | 1.227224268 | 1.343407053 | 0.155579371 | 0.483816235 | 0.368305698 | 0.282478358 | 0.494456994 | 0.747719171 | 0.5756 | 0.784014236 |
| SL1344_RS19260_ | -0.537347268 | -0.583753674 | -0.465307326 | 0.113259575 | 0.211409018 | 0.495347242 | 0.285333343 | 0.449749201 | 0.676454661 | 0.5566 | 0.738753208 |
| SL1344_RS19265_ | 2.238909392 | 2.208226164 | 2.256070177 | 0.042737099 | 0.211785399 | 0.179660157 | 0.428666669 | 0.622712348 | 0.824621477 | 0.8795 | 0.944657909 |
| SL1344_RS19270_ | 2.864547556 | 2.820739738 | 2.885347583 | 0.05811243 | 0.150962177 | 0.356909864 | 0.375333338 | 0.621879188 | 0.820418615 | 0.7787 | 0.898196595 |
| SL1344_RS19275_ | 4.39410589 | 4.342677248 | 4.460131798 | 0.129669295 | 0.080204225 | 1.616759009 | 0.015323415 | 0.067960613 | 0.295035272 | 0.1165 | 0.353448132 |
| SL1344_RS19280_ | 2.391661 | 2.144387013 | 2.424946058 | 0.267343154 | 0.137561508 | 1.150578631 | 0.153333358 | 0.183811171 | 0.474962349 | 0.3197 | 0.567241593 |
| SL1344_RS19285_ | 2.636982801 | 2.434796478 | 2.669002454 | 0.215623902 | 0.135683596 | 1.06623823 | 0.191205881 | 0.206955591 | 0.500580182 | 0.3678 | 0.618049944 |
| SL1344_RS19290_ | 1.675594306 | 1.524812653 | 1.713277243 | 0.180543569 | 0.160556956 | 0.899507011 | 0.202000017 | 0.275584898 | 0.560789084 | 0.4242 | 0.656641818 |
| SL1344_RS19295_ | 1.69058058 | 1.591175146 | 1.754914738 | 0.15287667 | 0.123831377 | 1.118518558 | 0.118587642 | 0.173425221 | 0.435287868 | 0.2516 | 0.501998477 |
| SL1344_RS19300_ | -0.287589366 | -0.444814373 | -0.22028273 | 0.229272781 | 0.388774993 | 0.600015142 | 0.263824128 | 0.416130581 | 0.678189353 | 0.5029 | 0.731011956 |
| SL1344_RS19305_ | 2.604870885 | 2.566151842 | 2.627987884 | 0.059213122 | 0.273205616 | 0.254650026 | 0.325333341 | 0.711558396 | 0.872870285 | 0.6229 | 0.808865365 |
| SL1344_RS19310_ | -0.555299902 | -0.626554006 | -0.475109231 | 0.153145126 | 0.206362154 | 0.717674848 | 0.214523667 | 0.333323056 | 0.570384447 | 0.4377 | 0.638565898 |
| SL1344_RS19315_ | -1.708126573 | -1.879460956 | -1.5535995 | 0.319641215 | 0.527882171 | 0.590560189 | 0.269333344 | 0.41307437 | 0.671467499 | 0.5397 | 0.742200849 |
| SL1344_RS19320_ | -0.09068396 | 0.058222391 | -0.264790496 | -0.308711813 | 0.763613872 | -0.379802144 | 0.361758833 | 0.608700987 | 0.819418655 | 0.7772 | 0.902269628 |
| SL1344_RS19325_ | -3.150166963 | -3.177307016 | -3.117535865 | 0.080418509 | 0.523054016 | 0.138021709 | 0.442371754 | 0.577021294 | 0.769485367 | 0.687 | 0.827894498 |
| SL1344_RS19330_ | 1.872104889 | 1.848244878 | 1.902565272 | 0.076182005 | 0.2318888 | 0.324207044 | 0.356000006 | 0.60879093 | 0.811225024 | 0.7492 | 0.879530839 |
| SL1344_RS19335_ | 0.105917424 | 0.192929385 | -0.047463315 | -0.238653712 | 0.217284952 | -1.016403702 | 0.129333364 | 0.202491595 | 0.469434115 | 0.2845 | 0.525730467 |
| SL1344_RS19340_ | 1.596707786 | 1.575788706 | 1.621371244 | 0.041228603 | 0.195468116 | 0.203318885 | 0.40373085 | 0.675620331 | 0.846677583 | 0.772 | 0.886558696 |
| SL1344_RS19345_ | -5.030772495 | -5.270445279 | -4.755773377 | 0.398723709 | 1.252647401 | 0.281963319 | 0.363333339 | 0.567418003 | 0.766865643 | 0.6664 | 0.817713354 |
| SL1344_RS19350_ | -0.526247305 | -0.41333975 | -0.59801759 | -0.198275071 | 0.259255655 | -0.730434039 | 0.20453033 | 0.314481791 | 0.576196065 | 0.4213 | 0.634716877 |
| SL1344_RS19355_ | -1.793280121 | -2.141226592 | -1.480498368 | 0.65890065 | 0.307525299 | 2.090481696 | 0.010000458 | 0.042858214 | 0.219318723 | 0.1099 | 0.343539451 |
| SL1344_RS19360_ | -1.602749071 | -1.953521507 | -1.306310331 | 0.653734525 | 0.331909079 | 1.943046001 | 0.014666978 | 0.052770393 | 0.243058625 | 0.1152 | 0.349591073 |
| SL1344_RS19365_ | -0.888752167 | -1.092746418 | -0.606956299 | 0.484999571 | 0.25341643 | 1.809152434 | 0.020666886 | 0.061663522 | 0.259315257 | 0.1286 | 0.366024139 |
| SL1344_RS19370_ | -0.121877799 | -0.197363719 | -0.033514419 | 0.169834585 | 0.203206662 | 0.771529622 | 0.189207214 | 0.282337453 | 0.536797285 | 0.378 | 0.600563581 |
| SL1344_RS19375_ | 0.462788079 | 0.618607077 | 0.272930431 | -0.350434746 | 0.151842335 | -2.273265365 | 0.004664544 | 0.034004307 | 0.196617648 | 0.1031 | 0.334538011 |
| SL1344_RS19380_ | -1.774678686 | -1.449884958 | -1.932146415 | -0.458862687 | 0.493399319 | -0.882297516 | 0.183211212 | 0.267572794 | 0.540760171 | 0.3658 | 0.611110973 |
| SL1344_RS19385_ | 0.613350836 | 0.563439544 | 0.70504817 | 0.151657203 | 0.25120232 | 0.562363744 | 0.248501012 | 0.40258613 | 0.662336497 | 0.479 | 0.708901232 |
| SL1344_RS19390_ | -0.977057785 | -0.965052426 | -0.991510158 | 0.014434795 | 0.301924954 | 0.037432906 | 0.484000001 | 0.67029088 | 0.838121106 | 0.7884 | 0.892070251 |
| SL1344_RS19395_ | -0.625146012 | -0.738933701 | -0.577692196 | 0.136688948 | 0.502922909 | 0.232548948 | 0.40106596 | 0.843390228 | 0.93725711 | 0.8272 | 0.926527976 |
| SL1344_RS19400_ | -1.190197721 | -1.195700844 | -1.181451101 | -0.013721162 | 0.345136186 | -0.041086316 | 0.482000001 | 0.672151193 | 0.839509603 | 0.7721 | 0.884629303 |
| SL1344_RS19405_ | -2.268451389 | -2.266919412 | -2.269458677 | -0.031470602 | 0.430071967 | -0.064255933 | 0.469686877 | 0.608391851 | 0.796461679 | 0.7303 | 0.856161904 |
| SL1344_RS19410_ | -2.217006996 | -2.182857268 | -2.23915569 | -0.035986233 | 0.417486271 | -0.071559888 | 0.470353099 | 0.63000158 | 0.811391541 | 0.7385 | 0.863254023 |
| SL1344_RS19415_ | -2.915364399 | -3.391321214 | -2.608216021 | 0.775717186 | 0.637544287 | 1.138559493 | 0.086666715 | 0.160496683 | 0.424010243 | 0.2162 | 0.463047864 |
| SL1344_RS19420_ | -3.536582354 | -3.783760525 | -3.309979753 | 0.358617806 | 0.92890132 | 0.373927907 | 0.34333334 | 0.552881742 | 0.765645589 | 0.667 | 0.822027028 |
| SL1344_RS19425_ | 2.444860274 | 2.392568623 | 2.470895665 | 0.058237837 | 0.148316522 | 0.319945977 | 0.381745508 | 0.771584544 | 0.900813879 | 0.789 | 0.907739003 |
| SL1344_RS19430_ | -3.024668637 | -3.33558822 | -2.829282501 | 0.538146635 | 0.791309715 | 0.648793413 | 0.236509007 | 0.363837216 | 0.630618132 | 0.4809 | 0.701267757 |
| SL1344_RS19435_ | 1.388374843 | 1.646420335 | 1.199860361 | -0.360724566 | 0.254412007 | -1.497295077 | 0.065333399 | 0.084083958 | 0.326322618 | 0.1619 | 0.419573355 |
| SL1344_RS19445_ | -0.685741504 | -0.78165854 | -0.59807227 | 0.19171517 | 0.1930701 | 0.909033474 | 0.156562316 | 0.238320573 | 0.475344338 | 0.3289 | 0.550894304 |
| SL1344_RS19450_ | -0.389217091 | -0.449138361 | -0.332305677 | 0.120336193 | 0.192875962 | 0.58528167 | 0.258494349 | 0.391087985 | 0.625526132 | 0.4959 | 0.690021345 |
| SL1344_RS19455_ | -2.201203216 | -2.377586107 | -2.011143102 | 0.366614454 | 0.402444852 | 0.870687357 | 0.171333355 | 0.259003655 | 0.512945743 | 0.3448 | 0.573926901 |
| SL1344_RS19460_ | -2.914218764 | -3.168158213 | -2.733984694 | 0.338123512 | 0.781153043 | 0.411948222 | 0.327333341 | 0.529376851 | 0.749405075 | 0.6489 | 0.808214695 |
| SL1344_RS19465_ | 0.456410467 | 0.271333429 | 0.620706358 | 0.40215194 | 0.250453503 | 1.569077424 | 0.020653117 | 0.075498439 | 0.30400555 | 0.1202 | 0.356121121 |
| SL1344_RS19470_ | 0.042364108 | -0.471617513 | 0.335770956 | 0.782551423 | 0.393717834 | 1.819391009 | 0.008000574 | 0.072387743 | 0.30727829 | 0.1077 | 0.341415221 |
| SL1344_RS19475_ | 0.525419651 | 0.449473389 | 0.583455917 | 0.070882676 | 0.320438002 | 0.226092822 | 0.413724187 | 0.70870183 | 0.866433929 | 0.829 | 0.916516603 |
| SL1344_RS19480_ | 0.97010986 | 0.799598556 | 1.172342656 | 0.385189884 | 0.243708804 | 1.493576549 | 0.036666788 | 0.097260591 | 0.348551521 | 0.1364 | 0.380795629 |
| SL1344_RS19485_ | 2.097292206 | 1.856346315 | 2.561196843 | 0.709962242 | 0.277632822 | 2.571215859 | 4.68E-05 | 0.039130372 | 0.235061974 | 0.1 | 0.330257709 |
| SL1344_RS19490_ | 0.632135761 | 0.590071318 | 0.661492185 | 0.089956183 | 0.195895192 | 0.478632193 | 0.29913392 | 0.44858818 | 0.691355578 | 0.5812 | 0.756174864 |
| SL1344_RS19495_ | -1.086310669 | -1.182522203 | -1.015387476 | 0.163412448 | 0.277327587 | 0.573161612 | 0.265822796 | 0.402572834 | 0.638960716 | 0.5077 | 0.699091571 |
| SL1344_RS19500_ | -0.094843619 | -0.186490004 | -0.030653821 | 0.154015788 | 0.308880621 | 0.476262188 | 0.314000008 | 0.480865465 | 0.723545232 | 0.631 | 0.801383151 |
| SL1344_RS19505_ | 0.532663694 | 0.474351685 | 0.605826886 | 0.135417891 | 0.217819938 | 0.602572001 | 0.24250501 | 0.401445386 | 0.654252161 | 0.4975 | 0.708718586 |
| SL1344_RS19510_ | 0.030757452 | 0.112906172 | -0.021977704 | -0.088717709 | 0.303276688 | -0.284863116 | 0.376666672 | 0.679146505 | 0.846889274 | 0.7487 | 0.877821555 |

| | | | | | | | | | | |
|---|---|---|---|---|---|---|---|---|---|---|
| SL1344_RS19515_ | 0.353564152 | 0.349269187 | 0.357659385 | 0.002633437 | 0.196198125 | 0.011494912 | 0.495333334 | 0.663096385 | 0.833744839 | 0.7703 | 0.882524169 |
| SL1344_RS19520_ | -0.398001886 | -0.472567471 | -0.274857135 | 0.166621404 | 0.344079085 | 0.476829631 | 0.287141915 | 0.519024962 | 0.741777903 | 0.5864 | 0.777179556 |
| SL1344_RS19525_ | -0.347508019 | -0.370542172 | -0.326088442 | 0.050244738 | 0.185478172 | 0.240832716 | 0.392405067 | 0.514005337 | 0.725060372 | 0.6309 | 0.786431626 |
| SL1344_RS19530_ | -0.481222872 | -0.559514036 | -0.389706944 | 0.165465323 | 0.20627916 | 0.754073639 | 0.205862775 | 0.323550815 | 0.56371495 | 0.4183 | 0.630710162 |
| SL1344_RS19535_ | -1.153504757 | -0.968745988 | -1.281982422 | -0.345394561 | 0.340021782 | -0.9948946 | 0.134666696 | 0.211099478 | 0.483624903 | 0.2821 | 0.52068194 |
| SL1344_RS19540_ | 0.988111165 | 1.210867748 | 0.870856015 | -0.315285878 | 0.191459401 | -1.55948486 | 0.055333412 | 0.085876083 | 0.320399287 | 0.1548 | 0.402311818 |
| SL1344_RS19545_ | 3.587544247 | 4.143291301 | 3.455055938 | -0.562904632 | 0.31504965 | -0.859920137 | 0.324666674 | 0.303010991 | 0.605249725 | 0.6831 | 0.86156801 |
| SL1344_RS19550_ | 0.901747379 | 0.865522602 | 0.924348076 | 0.059583318 | 0.146222626 | 0.383429597 | 0.333111266 | 0.501771889 | 0.721417904 | 0.6197 | 0.783888884 |
| SL1344_RS19555_ | -0.947659747 | -1.014849334 | -0.884299748 | 0.06567575 | 0.362454203 | 0.176617828 | 0.433710862 | 0.685560754 | 0.850191307 | 0.7941 | 0.898969521 |
| SL1344_RS19565_ | 1.591900462 | 1.622322263 | 1.573613271 | -0.055426521 | 0.300448779 | -0.140922334 | 0.427048637 | 0.560161649 | 0.791060765 | 0.8758 | 0.943811241 |
| SL1344_RS19570_ | 1.081863999 | 1.198862824 | 0.92136094 | -0.198561767 | 0.26071948 | -0.767957066 | 0.20186544 | 0.309600533 | 0.593933867 | 0.3888 | 0.636803914 |
| SL1344_RS19575_ | 3.41070082 | 3.636970435 | 2.681814442 | -0.927415443 | 0.207216028 | -2.937158102 | 4.68E-05 | 0.038053717 | 0.233720526 | 0.1 | 0.330257709 |
| SL1344_RS19580_ | 4.355596904 | 4.928237936 | 3.765182324 | -0.981063162 | 0.410874745 | -2.442761885 | 4.68E-05 | 0.014719026 | 0.152478016 | 0.1 | 0.330257709 |
| SL1344_RS19585_ | 1.450935425 | 1.366753524 | 1.486812578 | 0.124372393 | 0.183757032 | 0.654585988 | 0.227848116 | 0.341289757 | 0.625463544 | 0.4587 | 0.674615541 |
| SL1344_RS19590_ | 0.582612924 | 0.575737201 | 0.589941633 | 0.016516315 | 0.190508556 | 0.079357297 | 0.468354432 | 0.692092068 | 0.851118048 | 0.7918 | 0.894912562 |
| SL1344_RS19595_ | 0.536126532 | 0.568866036 | 0.502135622 | 0.092762588 | 0.406542785 | 0.239660181 | 0.424000003 | 0.622489542 | 0.820723065 | 0.9239 | 0.962705779 |
| SL1344_RS19600_ | 0.876657219 | 1.098856098 | 0.840256912 | -0.235177433 | 0.220772571 | -0.725584534 | 0.29646903 | 0.416109827 | 0.682814407 | 0.6024 | 0.803355618 |
| SL1344_RS19605_ | 1.961762287 | 2.047518642 | 1.701241161 | -0.45587262 | 0.832745762 | -0.627899926 | 0.225333348 | 0.311392949 | 0.612312746 | 0.4057 | 0.683807279 |
| SL1344_RS19610_ | 1.801636289 | 1.862948482 | 1.734581079 | -0.133621519 | 0.639153395 | -0.15948364 | 0.445333335 | 0.669266223 | 0.856851311 | 0.9865 | 0.994234504 |
| SL1344_RS19615_ | 1.727337072 | 1.628316823 | 1.793989546 | 0.201228562 | 0.207134111 | 1.031695313 | 0.103333373 | 0.194947598 | 0.486598366 | 0.221 | 0.475353542 |
| SL1344_RS19620_ | 0.603879929 | 0.641778465 | 0.516981647 | -0.115581215 | 0.208046977 | -0.466217084 | 0.325782819 | 0.550623318 | 0.764898713 | 0.6637 | 0.826855218 |
| SL1344_RS19625_ | -1.596503133 | -1.62624231 | -1.577857368 | -0.054314625 | 0.519394682 | -0.106781925 | 0.464000001 | 0.713788359 | 0.865108492 | 0.852 | 0.926908642 |
| SL1344_RS19630_ | -4.289164873 | -4.27506156 | -4.313102625 | -0.024810472 | 0.778123601 | -0.025552357 | 0.487333334 | 0.590316685 | 0.781432269 | 0.6968 | 0.835125349 |
| SL1344_RS19635_ | -2.456679603 | -2.789781581 | -2.153539682 | 0.544705383 | 0.657554429 | 0.79912722 | 0.194536993 | 0.287178833 | 0.564221275 | 0.3739 | 0.618585814 |
| SL1344_RS19640_ | -3.83348025 | -3.868229088 | -3.790433393 | 0.124624786 | 0.748415736 | 0.150529684 | 0.434377084 | 0.600077854 | 0.79147192 | 0.7166 | 0.84987759 |
| SL1344_RS19645_ | -5.892169198 | -6.731576987 | -4.597938551 | 1.959397581 | 2.570766868 | 0.742199605 | 0.155229871 | 0.269607823 | 0.551118695 | 0.305 | 0.565869039 |
| SL1344_RS19650_ | -3.409259691 | -3.998385349 | -3.030616468 | 0.971631577 | 0.773814743 | 1.22047924 | 0.078614311 | 0.137121529 | 0.392612213 | 0.2062 | 0.456175129 |
| SL1344_RS19655_ | -3.789054309 | -3.913960128 | -3.724509688 | 0.176753415 | 0.80035633 | 0.205982276 | 0.40373085 | 0.586711044 | 0.784325634 | 0.7105 | 0.845518611 |
| SL1344_RS19660_ | -4.370647129 | -4.241509903 | -4.495143905 | -0.029907185 | 1.171911029 | -0.024808808 | 0.488341106 | 0.699194886 | 0.856594602 | 0.8237 | 0.912216798 |
| SL1344_RS19665_ | -1.938063352 | -1.891990902 | -2.000788428 | -0.097618157 | 0.483662943 | -0.189596603 | 0.421718857 | 0.65199475 | 0.829073993 | 0.7696 | 0.883124483 |
| SL1344_RS19670_ | -2.956540426 | -2.793795048 | -3.169406727 | -0.386503842 | 0.571228559 | -0.635998133 | 0.244503677 | 0.371488368 | 0.620057122 | 0.4817 | 0.680595184 |
| SL1344_RS19675_ | -0.884888056 | -0.873682012 | -0.900094698 | -0.032795591 | 0.238693348 | -0.129837631 | 0.444000002 | 0.554776023 | 0.754836858 | 0.6675 | 0.814813934 |
| SL1344_RS19680_ | -2.721561959 | -2.767030182 | -2.660093998 | 0.095186311 | 0.55529357 | 0.155932915 | 0.431045972 | 0.633213142 | 0.81458405 | 0.7413 | 0.865928485 |
| SL1344_RS19685_ | 1.969577003 | 1.910501584 | 2.015419398 | 0.087048039 | 0.187291084 | 0.436070349 | 0.310459702 | 0.530142983 | 0.757205707 | 0.6079 | 0.794015905 |
| SL1344_RS19690_ | -0.861990563 | -0.959645899 | -0.787878449 | 0.17330593 | 0.279455633 | 0.588962647 | 0.260493016 | 0.392140377 | 0.637283798 | 0.5061 | 0.702802281 |
| SL1344_RS19695_ | 0.409974643 | 0.333838569 | 0.547659783 | 0.20640716 | 0.20647234 | 0.935605713 | 0.169333355 | 0.246866737 | 0.516665142 | 0.3455 | 0.589163302 |
| SL1344_RS19700_ | -1.386626654 | -1.348091954 | -1.460006551 | -0.123657328 | 0.434384653 | -0.275274709 | 0.386409065 | 0.61936979 | 0.809198058 | 0.7547 | 0.876210247 |
| SL1344_RS19705_ | -1.113064425 | -1.108184794 | -1.116685174 | -0.060828076 | 0.395323063 | -0.152079095 | 0.442000002 | 0.672422991 | 0.841488464 | 0.7829 | 0.890882986 |
| SL1344_RS19710_ | -2.243175578 | -2.551596868 | -2.052576513 | 0.393818508 | 0.676155331 | 0.553045604 | 0.273817466 | 0.44822381 | 0.697859044 | 0.5545 | 0.75722949 |
| SL1344_RS19715_ | -0.012125817 | 0.03878137 | -0.042814121 | -0.062082263 | 0.295185656 | -0.201971394 | 0.420000003 | 0.720970265 | 0.870704655 | 0.8032 | 0.904389544 |
| SL1344_RS19720_ | -0.591223336 | -0.49391124 | -0.67703443 | -0.129648467 | 0.301390743 | -0.415500836 | 0.325333341 | 0.532746011 | 0.747096079 | 0.6384 | 0.802750577 |
| SL1344_RS19725_ | 1.060219075 | 0.709964503 | 1.223872989 | 0.383633005 | 0.381284942 | 1.122140047 | 0.133910755 | 0.165139243 | 0.45164244 | 0.238 | 0.513468919 |
| SL1344_RS19730_ | -3.020004642 | -3.43566984 | -2.676476082 | 0.746477397 | 0.452233821 | 1.610586198 | 0.0379748 | 0.082523304 | 0.291680463 | 0.1497 | 0.386376561 |
| SL1344_RS19735_ | -1.591337527 | -1.591611162 | -1.590963042 | -0.010580641 | 0.306218906 | -0.027613546 | 0.487333334 | 0.581495383 | 0.775305076 | 0.6989 | 0.833837865 |
| SL1344_RS19740_ | 1.584220546 | 1.131067459 | 1.75093184 | 0.493758642 | 0.356613157 | 1.318093734 | 0.1186667 | 0.129460513 | 0.407330382 | 0.2093 | 0.483073185 |
| SL1344_RS19745_ | 4.517586078 | 4.464783098 | 4.555213588 | 0.095067591 | 0.130269094 | 0.723874305 | 0.225849448 | 0.301608156 | 0.597836033 | 0.4173 | 0.673937238 |
| SL1344_RS19750_ | 2.545706929 | 2.401396348 | 2.581516492 | 0.165103885 | 0.222120924 | 0.78082565 | 0.220519669 | 0.296377794 | 0.5905682 | 0.3895 | 0.666568935 |
| SL1344_RS19755_ | 3.486785134 | 3.446265836 | 3.53785903 | 0.110036473 | 0.093602952 | 1.15621458 | 0.074616979 | 0.150680148 | 0.423048671 | 0.1923 | 0.439531983 |
| SL1344_RS19765_ | 5.494381985 | 5.284543016 | 5.505938883 | 0.217739565 | 0.229143484 | 0.690544819 | 0.235176563 | 0.384448011 | 0.672832731 | 0.4402 | 0.706083349 |
| SL1344_RS19770_ | 3.819663288 | 3.888418817 | 3.773766207 | -0.09843079 | 0.169184342 | -0.333815731 | 0.368000005 | 0.883648501 | 0.955749798 | 0.8065 | 0.91772923 |
| SL1344_RS19775_ | 2.436835349 | 2.394711497 | 2.478619402 | 0.079976306 | 0.26518715 | 0.284834762 | 0.364423723 | 0.666535935 | 0.850201913 | 0.7708 | 0.898962279 |
| SL1344_RS19780_ | 5.110636901 | 5.005511699 | 5.126212431 | 0.152865518 | 0.181680609 | 0.935191932 | 0.213333349 | 0.207996639 | 0.50863121 | 0.3892 | 0.669263222 |
| SL1344_RS19785_ | 1.034889798 | 0.875038345 | 1.222544777 | 0.333220995 | 0.159416577 | 2.022426732 | 0.01666694 | 0.045166936 | 0.233557893 | 0.1147 | 0.350704987 |

| | | | | | | | | | | | |
|---|---|---|---|---|---|---|---|---|---|---|---|
| SL1344_RS19790_ | -2.926762169 | -2.697941511 | -3.164006917 | -0.44611633 | 0.479223974 | -0.894685628 | 0.143237868 | 0.243741199 | 0.487657324 | 0.3435 | 0.565117578 |
| SL1344_RS19795_ | -2.805946143 | -2.491424821 | -3.136576875 | -0.645658251 | 0.551657795 | -1.104201655 | 0.105333372 | 0.172869981 | 0.430085255 | 0.241 | 0.488586686 |
| SL1344_RS19800_ | 2.18775836 | 2.104133772 | 2.263772935 | 0.14372569 | 0.265964744 | 0.508347493 | 0.300000009 | 0.43623511 | 0.705305206 | 0.6072 | 0.798642157 |
| SL1344_RS19805_ | 1.719862048 | 1.599176505 | 1.815584999 | 0.269146855 | 0.219172794 | 1.234676586 | 0.063957429 | 0.123151502 | 0.390089922 | 0.1762 | 0.428680728 |
| SL1344_RS19810_ | -2.043002117 | -2.021565272 | -2.067294142 | -0.049140741 | 0.314050492 | -0.144419746 | 0.437708197 | 0.529674554 | 0.732903079 | 0.6349 | 0.788697745 |
| SL1344_RS19815_ | -0.470914338 | -0.494723542 | -0.440961331 | 0.044486504 | 0.22671114 | 0.176807141 | 0.420000003 | 0.608822574 | 0.794734643 | 0.7074 | 0.844119147 |
| SL1344_RS19820_ | -0.64693918 | -0.903269259 | -0.403593273 | 0.510325477 | 0.254052137 | 1.910296468 | 0.014657206 | 0.05210588 | 0.246264783 | 0.1154 | 0.351007348 |
| SL1344_RS19825_ | 2.021178114 | 1.817133503 | 2.238344114 | 0.462588044 | 0.187233872 | 2.51733312 | 4.68E-05 | 0.042031427 | 0.238342208 | 0.1 | 0.330257709 |
| SL1344_RS19830_ | 1.494365117 | 1.491302511 | 1.497827354 | 0.018124665 | 0.220091895 | 0.078975444 | 0.469333334 | 0.789104259 | 0.909847886 | 0.913 | 0.961396059 |
| SL1344_RS19835_ | 1.418298897 | 1.481637966 | 1.258018419 | -0.254160058 | 0.339900988 | -0.894584603 | 0.157333357 | 0.261470343 | 0.562658807 | 0.2992 | 0.567982813 |
| SL1344_RS19840_ | 3.319080811 | 2.506755812 | 4.142899516 | 1.179927503 | 0.596512553 | 2.614655154 | 4.68E-05 | 0.012441764 | 0.141316256 | 0.1 | 0.330257709 |
| SL1344_RS19845_ | 2.317507894 | 1.756224065 | 3.022754617 | 1.265063914 | 0.629284324 | 2.13733452 | 4.68E-05 | 0.037556353 | 0.232165354 | 0.1 | 0.330257709 |
| SL1344_RS19850_ | 2.02991098 | 1.37406749 | 2.512584667 | 1.2088028 | 0.304712422 | 4.006203148 | 4.68E-05 | 0.013722481 | 0.143962222 | 0.1 | 0.330257709 |
| SL1344_RS19855_ | 2.332129405 | 1.744440369 | 2.864642443 | 1.100581797 | 0.267780293 | 3.831450768 | 4.68E-05 | 0.032434485 | 0.216957258 | 0.1 | 0.330257709 |
| SL1344_RS19860_ | 3.446890316 | 2.74094649 | 3.973688431 | 1.233388181 | 0.298470861 | 4.00157031 | 4.68E-05 | 0.018908841 | 0.170093609 | 0.1 | 0.330257709 |
| SL1344_RS19865_ | -1.128951952 | -1.141733718 | -1.108721101 | 0.061094457 | 0.401338433 | 0.14071695 | 0.438374419 | 0.667192253 | 0.837419142 | 0.7846 | 0.891429782 |
| SL1344_RS19870_ | -3.2382924 | -3.435323656 | -3.065575436 | 0.528548416 | 1.060331878 | 0.460866989 | 0.272666677 | 0.384098696 | 0.6612724 | 0.5366 | 0.723514744 |
| SL1344_RS19875_ | 0.446432977 | 0.329249766 | 0.610077525 | 0.299513109 | 0.284894312 | 1.029592841 | 0.130579644 | 0.197196153 | 0.48050687 | 0.267 | 0.522015102 |
| SL1344_RS19880_ | 4.948237806 | 4.926126697 | 5.057945763 | 0.137915221 | 0.102777162 | 1.279638811 | 0.080666719 | 0.133295269 | 0.411975502 | 0.1999 | 0.456315284 |
| SL1344_RS19885_ | 4.781043015 | 4.687355405 | 4.914807144 | 0.181798716 | 0.117859314 | 1.692665316 | 0.041333441 | 0.056812611 | 0.276409334 | 0.1373 | 0.384780659 |
| SL1344_RS19890_ | -0.462618425 | -0.429129417 | -0.504867059 | -0.073305497 | 0.216503704 | -0.308378585 | 0.347768161 | 0.528500004 | 0.735627669 | 0.6359 | 0.793188465 |
| SL1344_RS19900_ | 6.346790148 | 6.129405602 | 6.563607502 | 0.431564944 | 0.055003588 | 7.857561746 | 4.68E-05 | 0.000904507 | 0.042365104 | 0.1 | 0.330257709 |
| SL1344_RS19905_ | 5.973548041 | 5.67628363 | 6.137443921 | 0.454404247 | 0.085266287 | 4.212877421 | 4.68E-05 | 0.010962793 | 0.131547281 | 0.1 | 0.330257709 |
| SL1344_RS19910_ | 7.312741045 | 7.077568881 | 7.526464957 | 0.462416094 | 0.069813891 | 6.824082165 | 4.68E-05 | 0.00118754 | 0.048754037 | 0.1 | 0.330257709 |
| SL1344_RS19915_ | 5.836150077 | 5.642475082 | 6.121687911 | 0.478223219 | 0.05597911 | 7.96171754 | 4.68E-05 | 0.007423492 | 0.110197855 | 0.1 | 0.330257709 |
| SL1344_RS19920_ | 6.382093 | 6.019847388 | 6.477426349 | 0.452251925 | 0.097185069 | 2.564611249 | 4.68E-05 | 0.062416944 | 0.294026578 | 0.1 | 0.330257709 |
| SL1344_RS19925_ | 5.187670754 | 5.003209345 | 5.365548397 | 0.371924624 | 0.074931301 | 5.008176797 | 4.68E-05 | 0.008796061 | 0.116738771 | 0.1 | 0.330257709 |
| SL1344_RS19930_ | 5.226000582 | 5.016137344 | 5.383703097 | 0.363402316 | 0.127308065 | 2.758060842 | 4.68E-05 | 0.027446152 | 0.2003902 | 0.1 | 0.330257709 |
| SL1344_RS19935_ | 3.957446507 | 3.909097256 | 4.15264516 | 0.238188272 | 0.232654447 | 1.028608953 | 0.178547655 | 0.189493359 | 0.487265917 | 0.3299 | 0.608799071 |
| SL1344_RS19945_ | 2.384532486 | 2.241977147 | 2.446215794 | 0.205644718 | 0.126276825 | 1.541369912 | 0.025982851 | 0.110045892 | 0.367736914 | 0.1334 | 0.372362081 |
| SL1344_RS19950_ | 3.702425271 | 3.779590931 | 3.665375053 | -0.07966218 | 0.244709463 | -0.37529237 | 0.284477025 | 0.63826135 | 0.836231674 | 0.5674 | 0.783473066 |
| SL1344_RS19955_ | -0.099578344 | -0.11540304 | -0.092851931 | 0.027556523 | 0.362506601 | 0.05781192 | 0.469333334 | 0.642306471 | 0.833226588 | 0.8265 | 0.920671292 |
| SL1344_RS19960_ | -0.433384453 | -0.492807449 | -0.371066121 | 0.148397975 | 0.378818761 | 0.367997617 | 0.323784151 | 0.519420433 | 0.749758885 | 0.642 | 0.808394981 |
| SL1344_RS19965_ | 1.243423305 | 1.172072659 | 1.329386306 | 0.181372866 | 0.168990745 | 1.058082014 | 0.08794142 | 0.187000364 | 0.46442384 | 0.2223 | 0.460756271 |
| SL1344_RS19970_ | 1.988243658 | 1.929292523 | 2.059591184 | 0.136965671 | 0.154073735 | 0.86703164 | 0.176666687 | 0.250837289 | 0.530868185 | 0.3411 | 0.592069632 |
| SL1344_RS19980_ | 1.498928571 | 1.742246879 | 1.401473494 | -0.326005536 | 0.457104414 | -0.660266142 | 0.248000013 | 0.341332463 | 0.636128371 | 0.4853 | 0.717515242 |
| SL1344_RS19985_ | -0.297452039 | -0.162742259 | -0.397980698 | -0.276799549 | 0.316582348 | -0.855750439 | 0.179213878 | 0.248235982 | 0.527067393 | 0.3538 | 0.593625177 |
| SL1344_RS19990_ | 0.972516065 | 1.244743425 | 0.744284816 | -0.494550584 | 0.184915634 | -2.599508437 | 4.68E-05 | 0.021368506 | 0.16736052 | 0.1007 | 0.331275265 |
| SL1344_RS19995_ | 4.529465066 | 4.750038676 | 4.428998569 | -0.350112484 | 0.179075464 | -2.042106673 | 4.68E-05 | 0.034040523 | 0.221599309 | 0.1 | 0.330257709 |
| SL1344_RS20000_ | 2.32967445 | 2.142372149 | 2.371260707 | 0.214747841 | 0.226443575 | 0.805644178 | 0.229333348 | 0.306613769 | 0.60165533 | 0.4368 | 0.696840468 |
| SL1344_RS20005_ | 2.092752257 | 1.86531862 | 2.129909872 | 0.251878248 | 0.191518246 | 0.89247639 | 0.245333346 | 0.278145724 | 0.573863619 | 0.4699 | 0.707477412 |
| SL1344_RS20020_ | 1.07070179 | 0.883154332 | 1.242466812 | 0.357222093 | 0.129329197 | 2.721105806 | 0.001335878 | 0.022235927 | 0.163707762 | 0.1013 | 0.332051314 |
| SL1344_RS20025_ | -1.55133336 | -2.198269468 | -1.327527654 | 1.192264671 | 1.1857399 | 1.075426207 | 0.113333369 | 0.156183231 | 0.442740331 | 0.2285 | 0.494207243 |
| SL1344_RS20030_ | -0.999055676 | -1.130102322 | -0.806959484 | 0.448749976 | 0.607882551 | 0.724193033 | 0.236000014 | 0.310229257 | 0.599160375 | 0.4489 | 0.684622397 |
| SL1344_RS20035_ | 1.653958504 | 1.241043237 | 1.749418543 | 0.561476511 | 0.931719313 | 0.687393734 | 0.224517003 | 0.272175027 | 0.575864767 | 0.3994 | 0.679730303 |
| SL1344_RS20045_ | 3.067820815 | 3.46621771 | 3.04595107 | -0.422716578 | 0.308724336 | -0.56285294 | 0.33333334 | 0.633138321 | 0.837400016 | 0.7 | 0.872542119 |
| SL1344_RS20050_ | 4.150022218 | 4.481229546 | 3.951163203 | -0.522723326 | 0.47263678 | -0.428034585 | 0.33333334 | 0.78421467 | 0.915118831 | 0.7 | 0.872542119 |
| SL1344_RS20055_ | 1.446051397 | 1.581505382 | 1.290091526 | -0.035578333 | 0.564080784 | -0.079406105 | 0.467333335 | 0.906097792 | 0.964966205 | 0.9355 | 0.972503469 |
| SL1344_RS20060_ | 3.894874712 | 3.938505129 | 3.852447053 | 0.051434906 | 0.29682243 | 0.195584999 | 0.456000002 | 0.834071815 | 0.93490159 | 0.9973 | 0.998863976 |
| SL1344_RS20065_ | -1.086587367 | -0.922377124 | -1.285632515 | -0.354368391 | 0.30379444 | -1.138025539 | 0.104000039 | 0.163789372 | 0.422608802 | 0.2378 | 0.484660124 |
| SL1344_RS20070_ | 2.872998567 | 2.937501193 | 2.800715146 | -0.219166611 | 1.224011792 | -0.281966488 | 0.324000007 | 0.676382787 | 0.861539349 | 0.6985 | 0.871114073 |
| SL1344_RS20075_ | 0.494426714 | 0.819938946 | 0.309912603 | -0.268252086 | 0.486759037 | -0.746296346 | 0.240506342 | 0.39261651 | 0.671198843 | 0.4758 | 0.709404413 |
| SL1344_RS20085_ | 1.511401916 | 1.442499789 | 1.652397903 | 0.216414971 | 0.181515988 | 1.180708421 | 0.073333391 | 0.171235729 | 0.454640019 | 0.1892 | 0.436290608 |

| | | | | | | | | | | | |
|---|---|---|---|---|---|---|---|---|---|---|---|
| SL1344_RS20090_ | 2.354933171 | 2.316714772 | 2.382263353 | 0.216540187 | 0.339257123 | 0.654253462 | 0.280666677 | 0.325259322 | 0.620267017 | 0.5706 | 0.7735719 |
| SL1344_RS20095_ | 2.881571003 | 2.86088606 | 2.900936989 | 0.042270332 | 0.233238825 | 0.169988725 | 0.446369089 | 0.75963204 | 0.898841826 | 0.9439 | 0.97580244 |
| SL1344_RS20100_ | -0.173218174 | -0.223653784 | -0.13867239 | 0.072124883 | 0.208263569 | 0.32692722 | 0.362425055 | 0.54358351 | 0.75054164 | 0.6501 | 0.806368446 |
| SL1344_RS20105_ | -0.063144835 | -0.121836444 | 0.028180005 | 0.146089801 | 0.242590866 | 0.553703906 | 0.273817466 | 0.441820911 | 0.68067357 | 0.5626 | 0.7482069 |
| SL1344_RS20110_ | 0.152379038 | 0.147813091 | 0.156756203 | 0.007326817 | 0.251545043 | 0.025801844 | 0.486000001 | 0.714015977 | 0.867072248 | 0.7974 | 0.900741217 |
| SL1344_RS20115_ | 2.789881952 | 3.051825131 | 2.644182211 | -0.435924822 | 0.346662162 | -1.204601643 | 0.104596974 | 0.122179688 | 0.398616551 | 0.1987 | 0.472400453 |
| SL1344_RS20120_ | 2.243035917 | 2.277977535 | 2.170587337 | -0.128295478 | 0.416082233 | -0.365001969 | 0.266000011 | 0.493824757 | 0.749025883 | 0.484 | 0.731084183 |
| SL1344_RS20125_ | -0.722469203 | -0.882656716 | -0.597359167 | 0.295883667 | 0.314831851 | 0.93021053 | 0.151232537 | 0.232336058 | 0.506369174 | 0.3207 | 0.562008912 |
| SL1344_RS20130_ | 1.882723106 | 1.869004991 | 1.894715538 | 0.052931455 | 0.180331017 | 0.285951965 | 0.3904064 | 0.641477082 | 0.826221835 | 0.7744 | 0.887795959 |
| SL1344_RS20135_ | 3.854815098 | 3.696535699 | 3.974537383 | 0.244518023 | 0.199824685 | 1.255028533 | 0.110000037 | 0.123135986 | 0.398368109 | 0.1986 | 0.473353634 |
| SL1344_RS20140_ | 3.739957281 | 3.686755293 | 3.939111796 | 0.224463599 | 0.147591765 | 1.213251174 | 0.124583643 | 0.153206902 | 0.438600171 | 0.2388 | 0.509671778 |
| SL1344_RS20145_ | 5.620405498 | 5.514746175 | 5.719757419 | 0.192270879 | 0.089016202 | 2.112234256 | 0.000673369 | 0.031401285 | 0.211783723 | 0.1009 | 0.331545218 |
| SL1344_RS20150_ | 5.742091603 | 5.620859003 | 5.95120078 | 0.280655541 | 0.159540848 | 1.566664906 | 0.072618313 | 0.081256522 | 0.330880414 | 0.1644 | 0.424400458 |
| SL1344_RS20155_ | 2.82200478 | 2.739818976 | 2.859357882 | 0.147323435 | 0.147505729 | 0.961796978 | 0.148567647 | 0.196790572 | 0.484940213 | 0.2925 | 0.553461868 |
| SL1344_RS20160_ | 3.367911426 | 3.282823651 | 3.44077386 | 0.160742782 | 0.058212936 | 2.709528979 | 0.0020023 | 0.025617609 | 0.174315519 | 0.1005 | 0.330989593 |
| SL1344_RS20165_ | 2.841854659 | 2.710388653 | 2.987148943 | 0.277236406 | 0.06687172 | 4.113660416 | 4.68E-05 | 0.007707446 | 0.099318804 | 0.1 | 0.330257709 |
| SL1344_RS20175_ | 2.236689662 | 2.065649384 | 2.391153367 | 0.342049698 | 0.145194089 | 2.253564247 | 4.68E-05 | 0.07826628 | 0.321262783 | 0.1 | 0.330257709 |
| SL1344_RS20180_ | 0.76211231 | 0.545447905 | 0.951795516 | 0.404740659 | 0.150105915 | 2.666287638 | 0.002666616 | 0.025966474 | 0.174657248 | 0.1019 | 0.333006149 |
| SL1344_RS20185_ | 1.331185878 | 1.20797717 | 1.463932852 | 0.331383913 | 0.222545858 | 1.50558022 | 0.016655836 | 0.075683131 | 0.308805152 | 0.1146 | 0.349958203 |
| SL1344_RS20190_ | 1.907505561 | 1.672382162 | 2.045479369 | 0.401238373 | 0.21655185 | 1.882970274 | 0.001335878 | 0.06421065 | 0.291394167 | 0.1019 | 0.332863757 |
| SL1344_RS20195_ | 1.793558706 | 1.595216872 | 1.977261411 | 0.381734036 | 0.100389887 | 3.83414646 | 4.68E-05 | 0.009119386 | 0.10728674 | 0.1 | 0.330257709 |
| SL1344_RS20200_ | 1.246540908 | 1.034133606 | 1.383155123 | 0.302704017 | 0.241641194 | 1.194485531 | 0.101932086 | 0.139818401 | 0.413175332 | 0.2047 | 0.470853756 |
| SL1344_RS20205_ | 1.839773493 | 1.717911533 | 1.991935035 | 0.27417608 | 0.168499864 | 1.571157516 | 0.038000117 | 0.077517223 | 0.31069595 | 0.135 | 0.378710409 |
| SL1344_RS20210_ | 1.180023794 | 1.106575643 | 1.293772789 | 0.186146673 | 0.155718687 | 1.132118961 | 0.103333373 | 0.163613969 | 0.431252854 | 0.2306 | 0.480603656 |
| SL1344_RS20215_ | 2.2810259 | 2.008855165 | 2.483169935 | 0.473883093 | 0.091267031 | 5.150832372 | 4.68E-05 | 0.005321971 | 0.082855215 | 0.1 | 0.330257709 |
| SL1344_RS20220_ | 3.396597922 | 3.569622186 | 3.236475496 | -0.335412681 | 0.612249329 | -0.407336081 | 0.388666671 | 0.551266697 | 0.788605394 | 0.8389 | 0.930726474 |
| SL1344_RS20225_ | 3.86503955 | 4.254452696 | 3.561294635 | -0.163642195 | 0.787556202 | -0.504950272 | 0.33333334 | 0.635813899 | 0.839241145 | 0.7006 | 0.872781673 |
| SL1344_RS20230_ | 4.842130492 | 5.153372748 | 4.537075906 | -0.072374883 | 0.716575032 | -0.20087515 | 0.357333339 | 0.611034371 | 0.825138847 | 0.7654 | 0.900265843 |
| SL1344_RS20235_ | 4.494536729 | 4.642120688 | 4.30449886 | -0.339145318 | 0.907018982 | -0.30704104 | 0.446369089 | 0.814695192 | 0.929052099 | 1 | 1 |
| SL1344_RS20240_ | 3.037620804 | 2.854410215 | 3.128578864 | 0.257433509 | 0.144692263 | 1.757092983 | 0.017333596 | 0.051983235 | 0.261528398 | 0.1174 | 0.354968101 |
| SL1344_RS20245_ | 2.686776743 | 2.605566554 | 2.845650359 | 0.216798009 | 0.166465086 | 1.328816966 | 0.086666715 | 0.109570941 | 0.369163411 | 0.1905 | 0.452311542 |
| SL1344_RS20250_ | 2.681323105 | 2.500728767 | 2.709185325 | 0.207167496 | 0.185427478 | 0.75369443 | 0.296666676 | 0.388173006 | 0.671125281 | 0.5914 | 0.802556211 |
| SL1344_RS20255_ | 2.817678789 | 2.769235914 | 2.970314469 | 0.205884691 | 0.163572087 | 1.277557816 | 0.11133337 | 0.129217242 | 0.400845383 | 0.2216 | 0.483959117 |
| SL1344_RS20260_ | 3.901048039 | 3.876569017 | 3.933167605 | 0.067740058 | 0.246535972 | 0.323828958 | 0.350433051 | 0.620295996 | 0.827830746 | 0.7315 | 0.871023686 |
| SL1344_RS20265_ | -2.895944567 | -2.891706339 | -2.897813176 | -0.046704538 | 0.51495596 | -0.081848934 | 0.465689542 | 0.600315698 | 0.788413565 | 0.7042 | 0.838963219 |
| SL1344_RS20270_ | -3.163997544 | -3.343581009 | -2.978224283 | 0.365839943 | 0.573075867 | 0.61285816 | 0.249333346 | 0.388045023 | 0.629836266 | 0.4942 | 0.692616236 |
| SL1344_RS20275_ | -3.065963226 | -3.480381293 | -2.84230073 | 0.551658095 | 0.761863191 | 0.676132867 | 0.232666681 | 0.347086066 | 0.615195127 | 0.4615 | 0.682735608 |
| SL1344_RS20280_ | 1.547532997 | 1.603090443 | 1.514934245 | -0.08778935 | 0.219668565 | -0.353526362 | 0.367333339 | 0.568860215 | 0.788040055 | 0.7648 | 0.886783428 |
| SL1344_RS20285_ | -3.81287706 | -4.418002483 | -3.483798406 | 0.944410029 | 0.843736673 | 1.052708892 | 0.11792142 | 0.188519749 | 0.457205143 | 0.2587 | 0.501072788 |
| SL1344_RS20290_ | 1.409379887 | 0.990791392 | 1.527622499 | 0.420517823 | 0.466051569 | 1.014559542 | 0.16322454 | 0.193771155 | 0.490131895 | 0.2966 | 0.574729211 |
| SL1344_RS20295_ | 2.70141962 | 2.640354066 | 2.80086346 | 0.167646198 | 0.123598542 | 1.390987765 | 0.057961434 | 0.102124828 | 0.35078462 | 0.1556 | 0.402351781 |
| SL1344_RS20300_ | 2.388606419 | 2.353787383 | 2.42910436 | 0.200787503 | 0.335184627 | 0.612414683 | 0.270666677 | 0.329567034 | 0.624014793 | 0.5539 | 0.76423246 |
| SL1344_RS20305_ | 2.391031404 | 2.346235939 | 2.514802464 | 0.200584624 | 0.207121029 | 1.021816874 | 0.136666695 | 0.170572066 | 0.456067889 | 0.2646 | 0.530269204 |
| SL1344_RS20310_ | 1.366966924 | 1.298059267 | 1.514679153 | 0.214523864 | 0.157846878 | 1.172586204 | 0.118587642 | 0.16457943 | 0.433762155 | 0.2413 | 0.491926319 |
| SL1344_RS20315_ | 3.204222949 | 3.2345136 | 3.175738701 | -0.078326455 | 0.110293532 | -0.693159768 | 0.237333347 | 0.321435458 | 0.600452134 | 0.4609 | 0.686915769 |
| SL1344_RS20325_ | -2.96169466 | -3.380738263 | -2.507464008 | 0.821629168 | 0.621164048 | 1.254828582 | 0.088000048 | 0.138552922 | 0.390538947 | 0.2142 | 0.464374713 |
| SL1344_RS20330_ | -4.319777366 | -4.525601753 | -4.141639519 | 0.339967225 | 1.045069145 | 0.302706325 | 0.362000006 | 0.565107111 | 0.767546101 | 0.6769 | 0.822329144 |
| SL1344_RS20335_ | 0.432082595 | 0.691364664 | 0.375683057 | -0.291065544 | 0.335726093 | -0.564262411 | 0.325782819 | 0.58477432 | 0.800663884 | 0.6881 | 0.862629409 |
| SL1344_RS20340_ | -1.241141011 | -1.107681979 | -1.347638597 | -0.167693348 | 0.784429253 | -0.166992571 | 0.445333335 | 0.863647026 | 0.946591792 | 0.9592 | 0.98268624 |
| SL1344_RS20345_ | 2.931094771 | 2.90267043 | 2.955278734 | 0.078317935 | 0.11356798 | 0.689289169 | 0.203197885 | 0.305480793 | 0.588884288 | 0.4345 | 0.648219854 |
| SL1344_RS20350_ | 2.250418516 | 2.221559701 | 2.281654729 | 0.061805234 | 0.0828466 | 0.695057701 | 0.221333348 | 0.337625592 | 0.581465723 | 0.4373 | 0.645196192 |
| SL1344_RS20355_ | 0.375563563 | 0.297977478 | 0.441800351 | 0.142472566 | 0.150532219 | 0.892394679 | 0.159893428 | 0.253068576 | 0.499947382 | 0.3494 | 0.575083604 |
| SL1344_RS20365_ | 1.895641539 | 1.902575672 | 1.889046888 | 0.018350478 | 0.150979092 | 0.122182774 | 0.450000002 | 0.698707235 | 0.856904849 | 0.8435 | 0.922488821 |

| | | | | | | | | | | | |
|---|---|---|---|---|---|---|---|---|---|---|---|
| SL1344_RS20370_ | 1.341288746 | 1.299296513 | 1.372805592 | 0.081272029 | 0.133638162 | 0.579265635 | 0.253164569 | 0.385145891 | 0.634343245 | 0.4986 | 0.694934357 |
| SL1344_RS20375_ | -0.534824921 | -0.513818355 | -0.552925745 | -0.049410479 | 0.239102203 | -0.199842658 | 0.406000004 | 0.588608693 | 0.783950422 | 0.7022 | 0.838392539 |
| SL1344_RS20390_ | -0.852105097 | -0.923045237 | -0.770546617 | 0.153588791 | 0.231234061 | 0.600906414 | 0.250000012 | 0.382558409 | 0.615782088 | 0.4819 | 0.678021716 |
| SL1344_RS20395_ | 0.883913237 | 0.912107346 | 0.849601971 | -0.070358157 | 0.178485249 | -0.37658069 | 0.33133334 | 0.527154493 | 0.744802482 | 0.6536 | 0.807421119 |
| SL1344_RS20400_ | 3.452171956 | 3.677279669 | 3.241495833 | -0.429541853 | 0.277395125 | -1.351105194 | 0.053297883 | 0.126725453 | 0.406233712 | 0.1483 | 0.401010453 |
| SL1344_RS20405_ | 1.545436485 | 1.503061859 | 1.630496228 | 0.113919055 | 0.238831453 | 0.428120608 | 0.346000006 | 0.62174635 | 0.819519614 | 0.7364 | 0.877518541 |
| SL1344_RS20410_ | 2.533295933 | 2.512842033 | 2.551256527 | 0.051740107 | 0.130372131 | 0.393281203 | 0.324000007 | 0.515781752 | 0.744553748 | 0.6583 | 0.818925664 |
| SL1344_RS20415_ | 2.114259797 | 2.092579221 | 2.125258557 | 0.042060243 | 0.12433873 | 0.310269262 | 0.372000005 | 0.541445042 | 0.759600949 | 0.69 | 0.829742446 |
| SL1344_RS20425_ | 3.564400074 | 3.56006477 | 3.568405931 | 0.021201901 | 0.121361104 | 0.141597025 | 0.408000004 | 0.572514511 | 0.793915724 | 0.7738 | 0.887316985 |
| SL1344_RS20430_ | 3.650051657 | 3.514249886 | 3.686967326 | 0.16497557 | 0.13744519 | 1.229045981 | 0.105263197 | 0.118227698 | 0.385499563 | 0.2241 | 0.487529717 |
| SL1344_RS20435_ | 3.078017861 | 3.01666983 | 3.115071585 | 0.094669635 | 0.077833037 | 1.135424812 | 0.1206667 | 0.179491449 | 0.44553032 | 0.259 | 0.508093827 |
| SL1344_RS20440_ | 0.088102198 | 0.070980113 | 0.109189388 | 0.045879191 | 0.223368975 | 0.191434086 | 0.42238508 | 0.641216531 | 0.820549572 | 0.7522 | 0.871394937 |
| SL1344_RS20445_ | 1.606021194 | 1.476695259 | 1.739103075 | 0.263508636 | 0.160414869 | 1.616504095 | 0.028666823 | 0.069416769 | 0.292317486 | 0.1295 | 0.370572153 |
| SL1344_RS20450_ | 4.115771234 | 4.047739251 | 4.178616274 | 0.11889279 | 0.076880976 | 1.479544537 | 0.056666743 | 0.092451284 | 0.336732091 | 0.1538 | 0.405949118 |
| SL1344_RS20455_ | 3.22517164 | 3.098115717 | 3.259944086 | 0.139430312 | 0.127022617 | 0.916762493 | 0.214000016 | 0.268035093 | 0.560753631 | 0.4034 | 0.654067404 |
| SL1344_RS20460_ | 0.356550563 | 0.298423993 | 0.445025779 | 0.137501883 | 0.173771553 | 0.73475149 | 0.217188556 | 0.340878049 | 0.591760263 | 0.441 | 0.657455663 |
| SL1344_RS20470_ | 2.97141094 | 3.291721292 | 2.77969683 | -0.351112519 | 0.494224583 | -0.691598314 | 0.295802807 | 0.424858713 | 0.702701536 | 0.6024 | 0.809641728 |
| SL1344_RS20475_ | 3.037870552 | 2.920093636 | 3.133012702 | 0.174526797 | 0.189444615 | 0.890240424 | 0.153333358 | 0.244005402 | 0.541531551 | 0.2876 | 0.558819659 |
| SL1344_RS20480_ | 2.48960232 | 2.409898949 | 2.610183162 | 0.141563143 | 0.184168096 | 0.834357216 | 0.182666686 | 0.287007714 | 0.577223242 | 0.3524 | 0.614676991 |
| SL1344_RS20485_ | 1.9053377 | 1.945618761 | 1.883754187 | -0.03463754 | 0.165656623 | -0.175349159 | 0.435333336 | 0.786208067 | 0.908101859 | 0.846 | 0.932658246 |
| SL1344_RS20495_ | -1.610702099 | -1.27589688 | -1.990480635 | -0.070307042 | 1.145644973 | -0.076028694 | 0.472666668 | 0.775147983 | 0.904314662 | 0.9381 | 0.971504984 |
| SL1344_RS20500_ | -0.15836502 | -0.355845059 | 0.127546857 | 0.54610654 | 0.352006315 | 1.56110872 | 0.020653117 | 0.074834916 | 0.305966913 | 0.1212 | 0.358960268 |
| SL1344_RS20505_ | -0.920499232 | -1.167505755 | -0.42825136 | 0.696361735 | 0.392241838 | 1.700926312 | 0.040639683 | 0.068830418 | 0.29046737 | 0.1378 | 0.380934279 |
| SL1344_RS20510_ | -0.67140724 | -1.178857553 | -0.578168778 | 0.65853098 | 1.381465134 | 0.540838449 | 0.225333348 | 0.379625306 | 0.666821269 | 0.4021 | 0.681707061 |
| SL1344_RS20520_ | 0.06683601 | 0.124445646 | -0.013609197 | -0.046212088 | 0.379777697 | -0.117471942 | 0.460666668 | 0.787775322 | 0.909199429 | 0.9035 | 0.955631665 |
| SL1344_RS20530_ | 1.237437775 | 0.993579887 | 1.55340901 | 0.453213672 | 0.272984584 | 1.754194235 | 0.04266677 | 0.058360256 | 0.275285502 | 0.1372 | 0.384726399 |
| SL1344_RS20535_ | 2.762120003 | 2.648616284 | 2.80378644 | 0.147595295 | 0.253956995 | 0.637609239 | 0.244503677 | 0.376381499 | 0.660255542 | 0.431 | 0.698633695 |
| SL1344_RS20540_ | 2.320565486 | 2.181487443 | 2.446968471 | 0.277424559 | 0.14035758 | 2.005792654 | 0.003334715 | 0.063005664 | 0.283986271 | 0.1024 | 0.333676473 |
| SL1344_RS20545_ | 2.27406362 | 2.237141941 | 2.329983668 | 0.069029402 | 0.262949557 | 0.269644867 | 0.344000006 | 0.716541589 | 0.874396074 | 0.7042 | 0.856982072 |
| SL1344_RS20550_ | -2.113215664 | -2.19531765 | -2.020849127 | 0.194370195 | 0.509368538 | 0.349862972 | 0.342000007 | 0.546028486 | 0.758062605 | 0.6608 | 0.814892576 |
| SL1344_RS20555_ | 3.181011873 | 3.115885072 | 3.242694081 | 0.190326552 | 0.180854145 | 1.163876145 | 0.042638345 | 0.157833192 | 0.445135769 | 0.1471 | 0.390651968 |
| SL1344_RS20560_ | 2.315267002 | 2.287753755 | 2.346727565 | 0.066363714 | 0.126558031 | 0.510166216 | 0.294666676 | 0.456018451 | 0.699154571 | 0.59 | 0.771368986 |
| SL1344_RS20565_ | 7.607654509 | 7.84541464 | 7.155513665 | -0.688712138 | 0.193549973 | -3.527364659 | 4.68E-05 | 0.005727741 | 0.099469432 | 0.1 | 0.330257709 |
| SL1344_RS20570_ | 2.542885519 | 2.437676252 | 2.678241154 | 0.249174236 | 0.137377044 | 1.823752405 | 0.015333631 | 0.052000276 | 0.25885422 | 0.1111 | 0.345886457 |
| SL1344_RS20575_ | 2.726288645 | 2.661391058 | 2.766450365 | 0.100066587 | 0.084174596 | 1.128919412 | 0.119920087 | 0.177193124 | 0.436645385 | 0.2602 | 0.506919615 |
| SL1344_RS20580_ | 0.833939498 | 0.856562136 | 0.803524196 | -0.010500898 | 0.251402346 | -0.041961079 | 0.482345104 | 0.765780668 | 0.896546653 | 0.8922 | 0.949957279 |
| SL1344_RS20585_ | 0.118474427 | 0.269535503 | -0.020510023 | -0.310386012 | 0.267239832 | -1.11260062 | 0.123251198 | 0.164245704 | 0.433488923 | 0.2456 | 0.498801211 |
| SL1344_RS20590_ | 3.571773674 | 3.548287976 | 3.593777528 | -0.052327191 | 0.250283214 | -0.206492179 | 0.444666669 | 0.785230988 | 0.911855622 | 0.9607 | 0.983350049 |
| SL1344_RS20595_ | 4.37707466 | 4.33449496 | 4.412454401 | 0.045749055 | 0.224475238 | 0.216053589 | 0.393333337 | 0.88087431 | 0.954832975 | 0.8551 | 0.9382136 |
| SL1344_RS20600_ | -0.969460649 | -0.789485006 | -1.144815934 | -0.351896997 | 0.226900966 | -1.475252026 | 0.052000084 | 0.105378925 | 0.323815278 | 0.1772 | 0.416624183 |
| SL1344_RS20605_ | -2.779422664 | -2.917552046 | -2.59115862 | 0.320686434 | 0.483504022 | 0.598450998 | 0.255163236 | 0.384568464 | 0.620123909 | 0.4862 | 0.682334899 |
| SL1344_RS20610_ | -2.936007253 | -2.91038971 | -2.947700089 | -0.123353344 | 0.716549289 | -0.153238979 | 0.436000002 | 0.641811697 | 0.82169124 | 0.76 | 0.877832826 |
| SL1344_RS20615_ | -2.586013952 | -2.751851665 | -2.36320413 | 0.378578621 | 0.461688307 | 0.758075751 | 0.20533335 | 0.316861184 | 0.57025414 | 0.4118 | 0.631972573 |
| SL1344_RS20620_ | -2.68388374 | -2.558799402 | -2.792597787 | -0.270616304 | 0.54899369 | -0.487419596 | 0.295333342 | 0.449557697 | 0.681692288 | 0.5741 | 0.749368544 |
| SL1344_RS20625_ | -1.982704084 | -1.961491509 | -2.006471339 | -0.053984259 | 0.347658597 | -0.152586803 | 0.430379749 | 0.567322378 | 0.762738139 | 0.6794 | 0.819797969 |
| SL1344_RS20635_ | -1.440174422 | -1.162443115 | -1.57102285 | -0.380522248 | 0.343770319 | -0.993945569 | 0.144570313 | 0.225976029 | 0.490648024 | 0.3337 | 0.568615412 |
| SL1344_RS20640_ | -0.140068227 | -0.196170916 | -0.071575892 | 0.118292084 | 0.193699019 | 0.583241368 | 0.249833457 | 0.38730928 | 0.624547436 | 0.4936 | 0.690473464 |
| SL1344_RS20645_ | -0.577708229 | -0.370991489 | -0.647218698 | -0.237851669 | 0.305121987 | -0.676314815 | 0.270666677 | 0.43196585 | 0.681011349 | 0.5517 | 0.757081122 |
| SL1344_RS20650_ | -1.250106665 | -1.04390727 | -1.51777432 | -0.510429479 | 0.383976182 | -1.328546871 | 0.040666776 | 0.129966978 | 0.38909898 | 0.1505 | 0.390768728 |
| SL1344_RS20655_ | -1.304619243 | -1.191857108 | -1.44209329 | -0.247852782 | 0.257050994 | -0.90873045 | 0.148666692 | 0.245772755 | 0.487806553 | 0.3386 | 0.559936355 |
| SL1344_RS20660_ | -2.331759338 | -2.372945112 | -2.27168111 | 0.044481603 | 0.5134779 | 0.079548499 | 0.459333335 | 0.655030555 | 0.828658167 | 0.7652 | 0.878899044 |
| SL1344_RS20665_ | -3.046871839 | -3.068810904 | -3.035159658 | -0.064162529 | 0.686400759 | -0.085078852 | 0.467333335 | 0.666098512 | 0.836518373 | 0.7788 | 0.889226154 |
| SL1344_RS20670_ | -2.270777006 | -2.094122156 | -2.524327998 | -0.445049979 | 0.439117173 | -0.936084831 | 0.1352432 | 0.217759021 | 0.476273727 | 0.2911 | 0.525212512 |

| | | | | | | | | | | |
|---|---|---|---|---|---|---|---|---|---|---|
| SL1344_RS20675_ | 1.54546832 | 1.470633966 | 1.615724613 | 0.159590434 | 0.145139637 | 1.05774575 | 0.118587642 | 0.177661942 | 0.446066201 | 0.2454 | 0.490669731 |
| SL1344_RS20680_ | 1.922094059 | 1.941821213 | 1.903792961 | -0.059539223 | 0.136888454 | -0.440664474 | 0.327333341 | 0.483278423 | 0.716441302 | 0.6565 | 0.80575514 |
| SL1344_RS20685_ | 1.996681028 | 1.973923359 | 2.011724208 | 0.036333284 | 0.08537615 | 0.390846611 | 0.342000007 | 0.494988952 | 0.709687879 | 0.6022 | 0.767686741 |
| SL1344_RS20690_ | 1.610828644 | 1.540989725 | 1.660206326 | 0.123102781 | 0.114679853 | 0.977485115 | 0.135909422 | 0.206611477 | 0.467698018 | 0.2873 | 0.523353788 |
| SL1344_RS20695_ | 2.593839329 | 2.526334964 | 2.637208314 | 0.120117837 | 0.107205099 | 1.122236217 | 0.064000067 | 0.192459353 | 0.476831179 | 0.1931 | 0.431552872 |
| SL1344_RS20700_ | 0.720403455 | 0.760030092 | 0.695638384 | -0.075974784 | 0.177025535 | -0.400576944 | 0.337108601 | 0.508527379 | 0.72952901 | 0.6373 | 0.793495378 |
| SL1344_RS20705_ | 0.515066421 | 0.617721338 | 0.388931413 | -0.22896686 | 0.140272519 | -1.55934374 | 0.036000124 | 0.086367821 | 0.299768291 | 0.1493 | 0.387852054 |
| SL1344_RS20710_ | 0.178107863 | 0.234751147 | 0.091576138 | -0.167515514 | 0.29406208 | -0.52618465 | 0.286475693 | 0.413948293 | 0.675567174 | 0.5674 | 0.758163119 |
| SL1344_RS20715_ | -0.79816439 | -0.722595083 | -1.043662159 | -0.28912262 | 0.33250737 | -0.766457937 | 0.245333346 | 0.358252329 | 0.623137081 | 0.4936 | 0.71036502 |
| SL1344_RS20720_ | -0.877320726 | -0.853567061 | -0.902035239 | -0.040611777 | 0.250168131 | -0.14186955 | 0.437708197 | 0.59473063 | 0.784589631 | 0.7008 | 0.836115635 |
| SL1344_RS20725_ | 3.387830058 | 3.265167168 | 3.504811105 | 0.229937857 | 0.176766052 | 1.213927031 | 0.084666716 | 0.13326871 | 0.41120905 | 0.1792 | 0.444317292 |
| SL1344_RS20730_ | 3.265610248 | 3.082859988 | 3.416296999 | 0.301390847 | 0.120472129 | 2.520959576 | 0.000673369 | 0.020129149 | 0.167897813 | 0.1005 | 0.330994337 |
| SL1344_RS20735_ | 3.261287588 | 3.062517984 | 3.434917394 | 0.374117328 | 0.101594111 | 3.672144282 | 4.68E-05 | 0.014542127 | 0.142476719 | 0.1 | 0.330257709 |
| SL1344_RS20740_ | 4.651721275 | 4.500035321 | 4.775875852 | 0.284440475 | 0.085891198 | 3.274184247 | 4.68E-05 | 0.011527683 | 0.130483809 | 0.1 | 0.330257709 |
| SL1344_RS20745_ | 4.199776649 | 4.041252695 | 4.269886193 | 0.229690506 | 0.109992865 | 2.075504655 | 0.000673369 | 0.069241883 | 0.304025624 | 0.1002 | 0.33054776 |
| SL1344_RS20750_ | 1.106397111 | 1.140198198 | 1.046644166 | -0.079855596 | 0.196418521 | -0.402654774 | 0.332666674 | 0.58466359 | 0.787141108 | 0.6588 | 0.825403089 |
| SL1344_RS20755_ | -1.980510455 | -2.120168745 | -1.826394891 | 0.299354195 | 0.406693574 | 0.708069108 | 0.215189889 | 0.33950126 | 0.591351691 | 0.4396 | 0.653343915 |
| SL1344_RS20760_ | -2.930076847 | -2.934931885 | -2.925540609 | 0.01890457 | 0.462397542 | 0.034620709 | 0.482345104 | 0.564863979 | 0.759895059 | 0.6753 | 0.817322479 |
| SL1344_RS20765_ | -0.417812801 | -0.363342526 | -0.473386615 | -0.110808098 | 0.196956932 | -0.525819084 | 0.27514991 | 0.411009931 | 0.642264414 | 0.5236 | 0.708354824 |
| SL1344_RS20770_ | 0.254971554 | 0.39191323 | 0.04762593 | -0.308094725 | 0.2673263 | -1.13687339 | 0.122584976 | 0.161850905 | 0.431625076 | 0.2355 | 0.494271029 |
| SL1344_RS20775_ | 1.098159963 | 1.069767807 | 1.129908152 | 0.047130979 | 0.190992708 | 0.230147606 | 0.384410398 | 0.657354512 | 0.832138157 | 0.7504 | 0.873126783 |
| SL1344_RS20780_ | 0.002100897 | 0.177568147 | -0.189644136 | -0.38283536 | 0.209918026 | -1.802156853 | 0.016000285 | 0.066468346 | 0.27725377 | 0.1182 | 0.35379064 |
| SL1344_RS20785_ | -0.103887818 | -0.144379732 | -0.067205826 | 0.108137769 | 0.222743032 | 0.445450714 | 0.316666675 | 0.466010336 | 0.698671598 | 0.6052 | 0.772252556 |
| SL1344_RS20790_ | -1.818084391 | -1.898656267 | -1.720907117 | 0.180778904 | 0.349904655 | 0.478349143 | 0.297333342 | 0.462263787 | 0.686016914 | 0.5677 | 0.746248751 |
| SL1344_RS20795_ | -2.618110405 | -2.68630818 | -2.536881167 | 0.129392161 | 0.600413942 | 0.197000677 | 0.410393075 | 0.650513233 | 0.826058056 | 0.7561 | 0.875223675 |
| SL1344_RS20800_ | -2.965465262 | -2.758322415 | -3.173186302 | -0.472530998 | 0.623574225 | -0.73478925 | 0.197201883 | 0.296510542 | 0.560712993 | 0.4123 | 0.629596896 |
| SL1344_RS20805_ | -0.632792391 | -0.356653639 | -0.84998362 | -0.509387988 | 0.24072429 | -2.034658087 | 0.009993796 | 0.049350233 | 0.240201048 | 0.11 | 0.343246036 |
| SL1344_RS20810_ | -0.75734704 | -0.810585079 | -0.68426884 | 0.092652672 | 0.391646796 | 0.218337755 | 0.40106596 | 0.714631873 | 0.866089417 | 0.7451 | 0.87646652 |
| SL1344_RS20815_ | -1.824566585 | -1.917306601 | -1.763353025 | 0.123709743 | 0.407121942 | 0.285036032 | 0.378000005 | 0.591882563 | 0.786538172 | 0.7015 | 0.841284828 |
| SL1344_RS20820_ | -6.117115913 | -6.254541228 | -5.86277577 | 0.321504257 | 1.769366886 | 0.163598376 | 0.424383747 | 0.611872405 | 0.795722001 | 0.713 | 0.84770141 |
| SL1344_RS20825_ | -3.740486369 | -3.705918459 | -3.776523917 | -0.076467869 | 0.642391003 | -0.11507453 | 0.447333335 | 0.565274678 | 0.763091966 | 0.6777 | 0.820939238 |
| SL1344_RS20830_ | -1.436122036 | -1.524087614 | -1.313036415 | 0.168978334 | 0.372320391 | 0.425235136 | 0.327781486 | 0.537371148 | 0.748124045 | 0.6314 | 0.796987937 |
| SL1344_RS20835_ | -4.662278681 | -5.458735693 | -4.128777747 | 1.346513895 | 1.30074589 | 0.965910312 | 0.125916087 | 0.204258448 | 0.479536753 | 0.2705 | 0.520995973 |
| SL1344_RS20840_ | -3.537814715 | -3.533034056 | -3.546465408 | -0.012887382 | 0.759421636 | -0.013643787 | 0.494337109 | 0.654397249 | 0.827169995 | 0.7646 | 0.878288164 |
| SL1344_RS20845_ | 6.206634822 | 6.176483727 | 6.243024695 | 0.138187283 | 0.141594137 | 1.005549245 | 0.09466671 | 0.165330143 | 0.458264417 | 0.191 | 0.453385338 |
| SL1344_RS20850_ | 0.608993928 | 0.580804613 | 0.623755976 | 0.04318599 | 0.203599627 | 0.199761172 | 0.417721522 | 0.673013072 | 0.841705446 | 0.7773 | 0.890323405 |
| SL1344_RS20855_ | 0.79175514 | 0.871611504 | 0.671085127 | -0.233823999 | 0.335439478 | -0.754514374 | 0.156562316 | 0.276387544 | 0.574876094 | 0.3057 | 0.566211339 |
| SL1344_RS20875_ | 1.246661868 | 1.307824925 | 1.200233202 | -0.127421792 | 0.168954279 | -0.755497436 | 0.189333352 | 0.291532402 | 0.572112113 | 0.3823 | 0.606623479 |
| SL1344_RS20880_ | 1.814901626 | 1.724196369 | 1.91279402 | 0.174142056 | 0.461661385 | 0.297566711 | 0.436666669 | 0.715391429 | 0.878901503 | 0.9577 | 0.981998429 |
| SL1344_RS20890_ | 2.902668019 | 2.849936745 | 2.938211436 | 0.077334459 | 0.165767038 | 0.388876676 | 0.367088613 | 0.556552166 | 0.78300195 | 0.7812 | 0.893658981 |
| SL1344_RS20895_ | 0.181652488 | -0.23765393 | 0.377426171 | 0.592091906 | 0.217348257 | 2.482698459 | 0.002666616 | 0.042272809 | 0.228207214 | 0.102 | 0.333097558 |
| SL1344_RS20900_ | -0.621336861 | -0.780967194 | -0.272260842 | 0.524022865 | 0.388942901 | 1.286814756 | 0.071333393 | 0.135440498 | 0.405520422 | 0.187 | 0.438585123 |
| SL1344_RS20905_ | 1.908844891 | 1.850914955 | 1.968210726 | 0.163330515 | 0.16061048 | 1.008720687 | 0.122000033 | 0.179179298 | 0.455958308 | 0.2675 | 0.509283261 |
| SL1344_RS20910_ | 1.598667517 | 1.584335438 | 1.614093461 | 0.054175968 | 0.147621159 | 0.357545072 | 0.34733334 | 0.550332312 | 0.760715405 | 0.6958 | 0.832632661 |
| SL1344_RS20915_ | -2.992474873 | -2.52296443 | -3.570674312 | -1.067531026 | 0.555244259 | -1.857188376 | 0.013990999 | 0.062284444 | 0.264177232 | 0.1142 | 0.348913368 |
| SL1344_RS20920_ | -0.99545595 | -0.819422506 | -1.217735827 | -0.31181038 | 0.418124782 | -0.726533073 | 0.215189889 | 0.332711718 | 0.604422387 | 0.4312 | 0.665284435 |
| SL1344_RS20925_ | 0.715240388 | 0.723557009 | 0.708993058 | -0.018515807 | 0.369635209 | -0.044309225 | 0.480346436 | 0.676769605 | 0.855841549 | 0.909 | 0.959342587 |
| SL1344_RS20935_ | 0.072258616 | 0.13179723 | -0.074966057 | -0.194756885 | 0.385464665 | -0.499258728 | 0.276482355 | 0.507523536 | 0.746000103 | 0.5527 | 0.769555417 |
| SL1344_RS20940_ | -0.437798123 | -0.355600087 | -0.516570573 | -0.196785571 | 0.484546487 | -0.381497744 | 0.315789482 | 0.441667805 | 0.705163802 | 0.6173 | 0.79208213 |
| SL1344_RS20945_ | -0.804323424 | -0.311490219 | -0.867909213 | -0.515498978 | 0.381711521 | -0.760986707 | 0.300466365 | 0.402058313 | 0.674225632 | 0.6196 | 0.815767313 |
| SL1344_RS20950_ | 0.0994833 | 0.422869773 | -0.064417568 | -0.478484388 | 0.319567941 | -1.403275683 | 0.057333409 | 0.112823883 | 0.372800406 | 0.168 | 0.419968725 |
| SL1344_RS20955_ | 0.318413539 | 0.619471485 | -0.128654159 | -0.81473773 | 0.366224314 | -2.276620049 | 0.00067292 | 0.024615189 | 0.185467748 | 0.1 | 0.330257709 |
| SL1344_RS20960_ | -1.321151049 | -1.160367107 | -1.452446262 | -0.317088605 | 0.341919617 | -0.935647387 | 0.150566314 | 0.232713986 | 0.497302915 | 0.3202 | 0.554576624 |

| | | | | | | | | | | | |
|---|---|---|---|---|---|---|---|---|---|---|---|
| SL1344_RS20965_ | 0.402053752 | 0.584863089 | 0.345496687 | -0.240891902 | 0.277275589 | -0.817791574 | 0.187874769 | 0.284595301 | 0.574687396 | 0.4019 | 0.634762921 |
| SL1344_RS20970_ | 4.234134533 | 4.203651634 | 4.362068873 | 0.153382347 | 0.082662105 | 1.400938047 | 0.068620982 | 0.137028157 | 0.413827413 | 0.1893 | 0.438234207 |
| SL1344_RS20975_ | 0.380746289 | 0.329708794 | 0.437800496 | 0.107797111 | 0.146578633 | 0.698010289 | 0.223850781 | 0.331519513 | 0.569589663 | 0.4346 | 0.636051856 |
| SL1344_RS20980_ | -0.124995282 | -0.181669722 | -0.10081275 | 0.062695492 | 0.249431635 | 0.220116474 | 0.415056632 | 0.675472653 | 0.843688378 | 0.7745 | 0.891249605 |
| SL1344_RS20985_ | 1.94715886 | 1.9825814 | 1.923572446 | -0.036226212 | 0.242406346 | -0.127345293 | 0.41972019 | 0.816416002 | 0.924826278 | 0.817 | 0.920818902 |
| SL1344_RS20990_ | 3.285707078 | 3.508708889 | 3.182619306 | -0.228600033 | 0.27541481 | -0.59216008 | 0.333777489 | 0.561969807 | 0.793800313 | 0.7003 | 0.872604954 |
| SL1344_RS20995_ | 7.920744668 | 7.937609507 | 7.875263258 | -0.053943863 | 0.179205403 | -0.307092772 | 0.329113931 | 0.733795548 | 0.890746741 | 0.6811 | 0.859527151 |
| SL1344_RS21000_ | 6.20400325 | 6.162234692 | 6.244612464 | -0.033652794 | 0.255895149 | -0.151921534 | 0.450666669 | 0.880368484 | 0.955643625 | 1 | 1 |
| SL1344_RS21005_ | 4.043135528 | 3.697760228 | 4.06016652 | 0.35545796 | 0.086130393 | 1.00757127 | 0.198666684 | 0.196363488 | 0.496262934 | 0.4072 | 0.645324727 |
| SL1344_RS21010_ | 3.114797659 | 3.26350964 | 2.948138927 | -0.304285708 | 0.165058248 | -1.802164247 | 0.013324793 | 0.054387404 | 0.270599153 | 0.1094 | 0.344027972 |
| SL1344_RS21015_ | 0.756066402 | 0.687163084 | 0.831599627 | 0.077942973 | 0.291007448 | 0.252223948 | 0.391738845 | 0.686158552 | 0.854038483 | 0.7892 | 0.896735077 |
| SL1344_RS21020_ | 4.323406672 | 4.240066752 | 4.394785261 | 0.14832635 | 0.529241112 | 0.210997933 | 0.446666669 | 0.875801771 | 0.954321706 | 1 | 1 |
| SL1344_RS21025_ | 2.164469232 | 2.168879298 | 2.160187498 | -0.018085874 | 0.525199037 | -0.036498367 | 0.484666667 | 0.854401121 | 0.944187993 | 0.9973 | 0.998847672 |
| SL1344_RS21030_ | 3.152057716 | 3.010453438 | 3.254823507 | 0.257019716 | 0.102771545 | 2.504472537 | 4.68E-05 | 0.039559448 | 0.227763713 | 0.1002 | 0.330558874 |
| SL1344_RS21035_ | 2.944112101 | 3.017502174 | 2.896256413 | -0.075121718 | 0.16236033 | -0.349596521 | 0.378000005 | 0.846172871 | 0.938593555 | 0.7996 | 0.914797939 |
| SL1344_RS21040_ | 1.972438342 | 1.963000116 | 1.9818279 | 0.074158154 | 0.17255645 | 0.42447044 | 0.341772158 | 0.485569741 | 0.726652971 | 0.7331 | 0.853253716 |
| SL1344_RS21045_ | 4.877350768 | 4.685821577 | 4.908000518 | 0.207738238 | 0.44829513 | 0.484064446 | 0.223184558 | 0.521606588 | 0.769705071 | 0.4018 | 0.681507577 |
| SL1344_RS21050_ | -1.055221553 | -1.157434463 | -0.99798025 | 0.122123398 | 0.346551692 | 0.332203429 | 0.375333338 | 0.609918701 | 0.800526516 | 0.714 | 0.85328277 |
| SL1344_RS21055_ | -0.587877417 | -0.740880974 | -0.490491363 | 0.244389497 | 0.235052362 | 0.971956445 | 0.152666692 | 0.22786147 | 0.487277372 | 0.3226 | 0.556697834 |
| SL1344_RS21075_ | -3.871148708 | -4.018367215 | -3.654789242 | 0.331660141 | 0.758235248 | 0.401267677 | 0.325333341 | 0.524814702 | 0.734877033 | 0.6125 | 0.783683674 |
| SL1344_RS21080_ | -3.914685373 | -3.9134449 | -3.915885461 | -0.015219675 | 0.64010689 | -0.018173637 | 0.489007329 | 0.549152871 | 0.754067709 | 0.6547 | 0.80838783 |
| SL1344_RS21085_ | 0.899376352 | 0.880325802 | 0.921121104 | 0.157814036 | 0.334052935 | 0.465450703 | 0.332000007 | 0.480190149 | 0.729683625 | 0.7136 | 0.854961758 |
| SL1344_RS21090_ | -0.059705967 | -0.066404283 | -0.049642772 | 0.020525116 | 0.180731063 | 0.10158077 | 0.449333335 | 0.581535114 | 0.775633842 | 0.6879 | 0.829420751 |
| SL1344_RS21100_ | 2.446275431 | 2.490979884 | 2.361837054 | -0.141996585 | 0.140041758 | -0.961624503 | 0.139906757 | 0.201481642 | 0.487294797 | 0.285 | 0.545617871 |
| SL1344_RS21105_ | -2.049892442 | -2.046579614 | -2.053080463 | -0.008009141 | 0.453189224 | -0.014069992 | 0.492338441 | 0.669295993 | 0.839008015 | 0.7813 | 0.889872478 |
| SL1344_RS21115_ | 0.901217555 | 0.630224474 | 0.959071139 | 0.302735611 | 0.218159086 | 1.059544479 | 0.178666687 | 0.216120388 | 0.504285755 | 0.357 | 0.60458157 |
| SL1344_RS21120_ | 0.867351039 | 0.663230874 | 0.908315155 | 0.225705409 | 0.21812964 | 0.831883401 | 0.252498347 | 0.325949904 | 0.608270253 | 0.5075 | 0.727668059 |
| SL1344_RS21125_ | -1.48363513 | -1.60654603 | -1.367077314 | 0.233263544 | 0.284011194 | 0.770082913 | 0.184000019 | 0.289265828 | 0.533804233 | 0.3866 | 0.601728922 |
| SL1344_RS21130_ | -0.956802452 | -0.779028088 | -1.03281919 | -0.20238029 | 0.422188555 | -0.406341897 | 0.344000006 | 0.664848694 | 0.837209634 | 0.6971 | 0.854927121 |
| SL1344_RS21135_ | 0.911601035 | 0.923594335 | 0.905138757 | -0.010435078 | 0.180008991 | -0.06039027 | 0.476000001 | 0.699909919 | 0.857780512 | 0.8 | 0.902029029 |
| SL1344_RS21140_ | 1.025140171 | 1.051306708 | 0.983914112 | -0.066451494 | 0.269721095 | -0.250966778 | 0.368421058 | 0.665855975 | 0.84141233 | 0.689 | 0.844878483 |
| SL1344_RS21145_ | -0.996228224 | -0.846292738 | -1.067153326 | -0.237687113 | 0.313150366 | -0.702325165 | 0.232666681 | 0.342588378 | 0.605164939 | 0.4606 | 0.668917555 |
| SL1344_RS21150_ | -0.240100637 | -0.257921094 | -0.218127661 | 0.044140074 | 0.182265469 | 0.21903021 | 0.399333337 | 0.544069552 | 0.746349665 | 0.6562 | 0.80386645 |
| SL1344_RS21155_ | 3.072521452 | 2.892474933 | 3.325488892 | 0.429088558 | 0.10409585 | 4.131398513 | 4.68E-05 | 0.005192767 | 0.088010507 | 0.1 | 0.330257709 |
| SL1344_RS21160_ | 3.835588403 | 3.795290507 | 4.034086831 | 0.291217341 | 0.281053956 | 1.119625407 | 0.090666713 | 0.120488595 | 0.395416783 | 0.2098 | 0.470642613 |
| SL1344_RS21165_ | 1.340185846 | 1.16227825 | 1.412824057 | 0.19009374 | 0.233397927 | 0.746290879 | 0.238000013 | 0.355206501 | 0.632944919 | 0.4682 | 0.696276515 |
| SL1344_RS21170_ | -0.552854065 | -0.767712906 | -0.371366761 | 0.542445021 | 0.442892589 | 1.216269882 | 0.057333409 | 0.12321005 | 0.38714692 | 0.1663 | 0.409979598 |
| SL1344_RS21175_ | -0.101009013 | -0.289123037 | 0.185900768 | 0.486770446 | 0.307331082 | 1.538028111 | 0.043304566 | 0.089595376 | 0.330027411 | 0.1435 | 0.387839128 |
| SL1344_RS21180_ | 1.520372676 | 1.242469749 | 1.780617142 | 0.477060949 | 0.218538838 | 2.147254378 | 0.006662914 | 0.031591344 | 0.206310437 | 0.1064 | 0.33960713 |
| SL1344_RS21185_ | 2.843592768 | 2.892835733 | 2.769741175 | -0.063154125 | 0.160558059 | -0.403816507 | 0.347101938 | 0.616039742 | 0.815141979 | 0.746 | 0.877156507 |
| SL1344_RS21195_ | 2.654461924 | 2.66131165 | 2.648621094 | 0.047959917 | 0.230463976 | 0.185853027 | 0.42266667 | 0.558140756 | 0.787546687 | 0.8751 | 0.940470586 |
| SL1344_RS21200_ | 1.981604043 | 1.975007081 | 1.988384223 | 0.022748122 | 0.222201411 | 0.106155189 | 0.431712194 | 0.758007215 | 0.893765984 | 0.7969 | 0.902601533 |
| SL1344_RS21205_ | 0.511657384 | 0.433166186 | 0.621790831 | 0.189664986 | 0.242866709 | 0.734135055 | 0.212524999 | 0.315942388 | 0.591847066 | 0.4141 | 0.653558676 |
| SL1344_RS21210_ | 3.356517975 | 3.220978569 | 3.499412391 | 0.275166556 | 0.117985489 | 2.299211604 | 0.002000966 | 0.027986037 | 0.195647157 | 0.1015 | 0.332467437 |
| SL1344_RS21215_ | 2.930693203 | 2.723706407 | 3.008323699 | 0.187863399 | 0.347879078 | 0.634463385 | 0.245333346 | 0.430560429 | 0.704723469 | 0.4789 | 0.730775447 |
| SL1344_RS21220_ | -2.65022035 | -2.809054262 | -2.43374153 | 0.416582538 | 1.259260095 | 0.342454947 | 0.375749505 | 0.579797702 | 0.795918167 | 0.7773 | 0.893535206 |
| SL1344_RS21225_ | 0.710304501 | 0.506642816 | 0.999070057 | 0.487518119 | 0.325910838 | 1.486258439 | 0.038000117 | 0.107743442 | 0.370305808 | 0.1357 | 0.381908155 |
| SL1344_RS21230_ | -1.550921601 | -1.620484231 | -1.489002977 | 0.099882729 | 0.521388911 | 0.180541332 | 0.414000003 | 0.695044806 | 0.856225366 | 0.8037 | 0.903676541 |
| SL1344_RS21235_ | -1.254188993 | -1.340844109 | -1.178612153 | 0.411321022 | 0.986256212 | 0.420284175 | 0.336442379 | 0.502855461 | 0.751680176 | 0.7037 | 0.858552524 |
| SL1344_RS21245_ | 1.229034876 | 1.142631244 | 1.277857273 | 0.104328524 | 0.196272939 | 0.530078181 | 0.29913392 | 0.497795531 | 0.731859862 | 0.5915 | 0.782422518 |
| SL1344_RS21250_ | 1.624934608 | 1.644799562 | 1.606770775 | -0.063940321 | 0.159887087 | -0.39272345 | 0.334000007 | 0.51190948 | 0.739248507 | 0.6687 | 0.817725018 |
| SL1344_RS21255_ | 2.459478073 | 2.221927011 | 2.714894109 | 0.437443097 | 0.214008781 | 2.065172657 | 0.006000767 | 0.037054774 | 0.226794793 | 0.1054 | 0.338092547 |
| SL1344_RS21260_ | -7.269073615 | -7.049538602 | -7.565890651 | -0.591267437 | 2.465746826 | -0.190060663 | 0.38774151 | 0.533077406 | 0.746943974 | 0.6599 | 0.807895557 |

| | | | | | | | | | | | |
|---|---|---|---|---|---|---|---|---|---|---|---|
| SL1344_RS21265_ | 2.430532636 | 2.95762813 | 2.402057023 | -0.536359901 | 0.17096158 | -0.883096608 | 0.292666676 | 0.258653351 | 0.562124235 | 0.5958 | 0.803685604 |
| SL1344_RS21270_ | 3.493589026 | 3.897376414 | 3.415146702 | -0.368169352 | 0.456138525 | -0.714615979 | 0.244503677 | 0.379801507 | 0.668954526 | 0.4663 | 0.722710848 |
| SL1344_RS21275_ | 5.526769555 | 5.523466611 | 5.529210518 | 0.004907486 | 0.551074567 | 0.006651706 | 0.488 | 0.698093191 | 0.873172925 | 0.985 | 0.993614608 |
| SL1344_RS21280_ | 4.384924847 | 4.621737814 | 4.298910618 | -0.32460435 | 0.452419775 | -0.622652363 | 0.309793479 | 0.366998852 | 0.659616511 | 0.635 | 0.830860621 |
| SL1344_RS21285_ | 7.545508465 | 7.570187355 | 7.520734444 | -0.049477458 | 0.161742595 | -0.298661493 | 0.366000005 | 0.665941445 | 0.855564828 | 0.7696 | 0.902143675 |
| SL1344_RS21290_ | 4.034821618 | 4.056611488 | 4.008780689 | -0.016074752 | 0.157299057 | -0.105035919 | 0.474000001 | 0.871185197 | 0.950222319 | 0.991 | 0.996170883 |
| SL1344_RS21300_ | 5.134785927 | 4.918905584 | 5.246169856 | 0.144089682 | 0.394334064 | 0.582734327 | 0.323117929 | 0.534646661 | 0.778040819 | 0.6688 | 0.852439495 |
| SL1344_RS21305_ | 6.329441548 | 6.328009742 | 6.330983405 | 0.041413903 | 0.223637308 | 0.206288817 | 0.375749505 | 0.674765304 | 0.860233895 | 0.835 | 0.929791535 |
| SL1344_RS21310_ | 7.174320986 | 7.112399227 | 7.351817233 | 0.242590844 | 0.117866251 | 2.126373602 | 4.68E-05 | 0.030400144 | 0.211164911 | 0.1 | 0.330257709 |
| SL1344_RS21315_ | 8.365712374 | 8.284995244 | 8.482958223 | 0.078529265 | 0.302199089 | 0.49628269 | 0.236000014 | 0.547546368 | 0.786500973 | 0.4492 | 0.711794528 |
| SL1344_RS21320_ | 6.393706194 | 6.326580268 | 6.417225721 | 0.066519567 | 0.075043398 | 0.638109663 | 0.330000007 | 0.496211425 | 0.750194147 | 0.6839 | 0.861960554 |
| SL1344_RS21325_ | 6.71518984 | 6.640458164 | 6.722825118 | 0.07984002 | 0.153192844 | 0.392260027 | 0.332000007 | 0.668990637 | 0.856632165 | 0.6952 | 0.869485537 |
| SL1344_RS21330_ | -2.354179715 | -1.948158399 | -2.570464045 | -0.604337451 | 0.461824688 | -1.227225244 | 0.100666708 | 0.161471795 | 0.417588339 | 0.2452 | 0.487104727 |
| SL1344_RS21335_ | -4.469628476 | -4.670050558 | -4.267907262 | 0.426072006 | 0.770032784 | 0.511765614 | 0.280666677 | 0.41475335 | 0.639126425 | 0.5172 | 0.703027585 |
| SL1344_RS21340_ | 0.195241927 | -0.040345354 | 0.324159087 | 0.34699779 | 0.228431962 | 1.449191344 | 0.063333401 | 0.102284908 | 0.344268319 | 0.1719 | 0.420573092 |
| SL1344_RS21345_ | -2.297000324 | -2.619394738 | -2.179485175 | 0.352998363 | 0.616309005 | 0.506637335 | 0.333777489 | 0.556017914 | 0.773147548 | 0.6869 | 0.846571196 |
| SL1344_RS21350_ | -0.198115963 | -0.126897065 | -0.242290841 | -0.065145335 | 0.260523679 | -0.231673379 | 0.407333337 | 0.688646259 | 0.850585657 | 0.7759 | 0.890880349 |
| SL1344_RS21355_ | -1.589215734 | -1.579230519 | -1.598261351 | -0.021346274 | 0.289756582 | -0.069763759 | 0.468354432 | 0.573358397 | 0.771136974 | 0.692 | 0.831699089 |
| SL1344_RS21360_ | -3.135046173 | -2.906744183 | -3.280482736 | -0.360107936 | 0.546582748 | -0.601497319 | 0.247333346 | 0.407121588 | 0.644251077 | 0.5085 | 0.704572669 |
| SL1344_RS21365_ | -1.375939349 | -1.530974605 | -1.311828996 | 0.185468505 | 0.448572944 | 0.378307702 | 0.360000006 | 0.590421757 | 0.793748063 | 0.7185 | 0.859397925 |
| SL1344_RS21370_ | -2.113786054 | -2.129288168 | -2.105143481 | 0.004200542 | 0.4897798 | 0.007452348 | 0.495669554 | 0.677081884 | 0.843343975 | 0.7857 | 0.89244628 |
| SL1344_RS21375_ | -0.502299071 | -0.479769779 | -0.519060454 | -0.0200725 | 0.237314289 | -0.078193513 | 0.465333335 | 0.639440112 | 0.815302394 | 0.7467 | 0.867099463 |
| SL1344_RS21380_ | 0.403057813 | 0.384365831 | 0.422988479 | 0.057177287 | 0.323538125 | 0.173462938 | 0.399333337 | 0.69029927 | 0.85644575 | 0.7558 | 0.880387404 |
| SL1344_RS21390_ | 2.248582472 | 2.187323473 | 2.320668709 | 0.147161721 | 0.117012023 | 1.274781956 | 0.06200007 | 0.145406243 | 0.411589039 | 0.1763 | 0.41837845 |
| SL1344_RS21395_ | 1.689127323 | 1.553593152 | 1.742886034 | 0.186112328 | 0.152350508 | 1.108845049 | 0.130666697 | 0.189672348 | 0.469199227 | 0.2747 | 0.531554452 |
| SL1344_RS21400_ | 1.800005078 | 1.780725706 | 1.823788801 | 0.066594218 | 0.159327469 | 0.398548395 | 0.305129922 | 0.461344062 | 0.704087733 | 0.6404 | 0.795772133 |
| SL1344_RS21420_ | 1.190387385 | 1.178934112 | 1.198766654 | 0.043026971 | 0.174354696 | 0.249969179 | 0.388666671 | 0.592986128 | 0.793693296 | 0.7359 | 0.862156717 |
| SL1344_RS21425_ | 1.44658881 | 1.453441128 | 1.441569927 | -0.011058204 | 0.30515214 | -0.03043986 | 0.488341106 | 0.854127959 | 0.941925439 | 0.9508 | 0.97895009 |
| SL1344_RS21430_ | 0.319011701 | 0.346223258 | 0.304076532 | -0.060305174 | 0.269542236 | -0.195762944 | 0.425716192 | 0.689069661 | 0.85324832 | 0.8131 | 0.911153548 |
| SL1344_RS21435_ | 0.529776512 | 0.565015722 | 0.507059726 | -0.049746369 | 0.203992179 | -0.242893512 | 0.402000004 | 0.636739812 | 0.817289692 | 0.7461 | 0.86833209 |
| SL1344_RS21440_ | -1.951243857 | -2.122891679 | -1.387190726 | 0.757223665 | 0.838036479 | 0.90613307 | 0.161225873 | 0.250025883 | 0.547368811 | 0.3273 | 0.575273586 |
| SL1344_RS21445_ | -4.190992467 | -4.444488714 | -4.025531587 | 0.454517455 | 0.836723043 | 0.501559886 | 0.287808137 | 0.430913913 | 0.666374988 | 0.5537 | 0.731631269 |
| SL1344_RS21450_ | 0.028301733 | -0.025182483 | 0.065110145 | 0.537732253 | 0.909931623 | 0.587919643 | 0.326449041 | 0.386991102 | 0.671569617 | 0.6811 | 0.844860377 |
| SL1344_RS21460_ | -3.722698433 | -3.271615502 | -4.107136115 | -0.819492336 | 0.634145908 | -1.245668376 | 0.087333381 | 0.141845775 | 0.380828611 | 0.2158 | 0.455491539 |
| SL1344_RS21465_ | 0.691948929 | 0.619143391 | 0.740625762 | 0.106084553 | 0.389180009 | 0.295152177 | 0.325333341 | 0.661594029 | 0.842118162 | 0.6357 | 0.815851504 |
| SL1344_RS21475_ | -0.667537455 | -0.85794772 | -0.462641448 | 0.382717286 | 0.26120525 | 1.395455838 | 0.068000063 | 0.112794512 | 0.350716444 | 0.1863 | 0.431902918 |
| SL1344_RS21480_ | 0.050098515 | -0.008527118 | 0.107026261 | 0.116666411 | 0.149567388 | 0.721289164 | 0.209193887 | 0.317014382 | 0.553898931 | 0.4171 | 0.622354189 |
| SL1344_RS21485_ | 0.051796207 | 0.062842132 | 0.03304089 | -0.013461345 | 0.201183934 | -0.060518245 | 0.471019322 | 0.651352028 | 0.82530553 | 0.7553 | 0.876186571 |
| SL1344_RS21490_ | 1.120441883 | 0.977262507 | 1.203643919 | 0.237394663 | 0.165785237 | 1.376428116 | 0.042638345 | 0.126375091 | 0.384293868 | 0.1558 | 0.395468632 |
| SL1344_RS21495_ | 2.050832369 | 1.835140623 | 2.233950272 | 0.393993425 | 0.144563148 | 2.702130444 | 4.68E-05 | 0.018973393 | 0.160965033 | 0.1 | 0.330257709 |
| SL1344_RS21505_ | 1.227585228 | 1.224406343 | 1.229007159 | -0.026424829 | 0.213565651 | -0.105151623 | 0.451333335 | 0.634876102 | 0.824458085 | 0.8133 | 0.908168442 |
| SL1344_RS21510_ | 1.745107317 | 1.786714114 | 1.708962735 | -0.049315913 | 0.42508656 | -0.136057226 | 0.420000003 | 0.872933727 | 0.950609016 | 0.8914 | 0.951755441 |
| SL1344_RS21515_ | -3.41125524 | -3.91038952 | -3.092409904 | 0.902479409 | 1.032603118 | 0.814772248 | 0.184543657 | 0.261872249 | 0.54704954 | 0.3601 | 0.610632251 |
| SL1344_RS21520_ | 1.684333861 | 1.509163698 | 1.864354524 | 0.398839006 | 0.185344772 | 2.321318329 | 4.68E-05 | 0.046096446 | 0.24745961 | 0.1001 | 0.330401723 |
| SL1344_RS21525_ | 2.089095281 | 1.877217662 | 2.265082344 | 0.393372732 | 0.111480267 | 3.486667513 | 4.68E-05 | 0.010162039 | 0.116166818 | 0.1 | 0.330257709 |
| SL1344_RS21530_ | 0.92525874 | 0.783639018 | 1.109644288 | 0.325548638 | 0.137865586 | 2.311702636 | 0.006000767 | 0.036525423 | 0.204887291 | 0.1054 | 0.337864033 |
| SL1344_RS21535_ | -0.689440436 | -0.481752859 | -0.796032917 | -0.252198691 | 0.36585967 | -0.629758043 | 0.260000012 | 0.434494901 | 0.68629586 | 0.5476 | 0.75126523 |
| SL1344_RS21540_ | -3.111096804 | -3.477592974 | -2.832575914 | 0.698399353 | 0.658405457 | 0.965116797 | 0.133333363 | 0.204497683 | 0.475128398 | 0.286 | 0.529134709 |
| SL1344_RS21545_ | -4.092293579 | -4.40272781 | -3.862768561 | 0.569994116 | 0.83349878 | 0.634913317 | 0.222666682 | 0.327663442 | 0.585350365 | 0.427 | 0.633822243 |
| SL1344_RS21550_ | -1.501931756 | -1.646192935 | -1.349011027 | 0.296070985 | 0.293182353 | 0.945586022 | 0.142000027 | 0.226127533 | 0.474413882 | 0.3149 | 0.542887307 |
| SL1344_RS21555_ | -3.27987629 | -3.416042713 | -3.123162464 | 0.390584751 | 0.7382505 | 0.508133088 | 0.286666676 | 0.443643175 | 0.68106533 | 0.5857 | 0.754467355 |
| SL1344_RS21560_ | -2.170941616 | -2.216133878 | -2.126992276 | 0.116710961 | 0.640855635 | 0.175217396 | 0.41466667 | 0.671400848 | 0.841980049 | 0.7544 | 0.876265306 |
| SL1344_RS21565_ | -4.728094006 | -4.809499715 | -4.600983929 | 0.224182154 | 0.953305148 | 0.213363466 | 0.407333337 | 0.56604336 | 0.763530684 | 0.6791 | 0.821777643 |

| | | | | | | | | | | | |
|---|---|---|---|---|---|---|---|---|---|---|---|
| SL1344_RS21570_ | -3.193914874 | -3.442881737 | -3.015081454 | 0.478472779 | 0.665276162 | 0.677622974 | 0.20719522 | 0.320530411 | 0.584679279 | 0.4326 | 0.64001723 |
| SL1344_RS21575_ | -5.126329803 | -5.124499968 | -5.129555037 | 0.023327973 | 1.093557864 | 0.022119457 | 0.489673551 | 0.57276375 | 0.769317193 | 0.6861 | 0.826107645 |
| SL1344_RS21580_ | 0.297171368 | 0.146967987 | 0.419809004 | 0.365057412 | 0.371339655 | 0.986161344 | 0.106666705 | 0.178694442 | 0.466282628 | 0.2242 | 0.47991374 |
| SL1344_RS21585_ | -1.119087853 | -1.421943166 | -0.75734199 | 0.660003121 | 0.290862474 | 2.252753736 | 0.006000767 | 0.035142568 | 0.206200335 | 0.1067 | 0.339617429 |
| SL1344_RS21590_ | -1.584611779 | -1.764966003 | -1.342124378 | 0.399204014 | 0.372669932 | 1.000391674 | 0.152564982 | 0.221336027 | 0.4852745 | 0.3125 | 0.558093777 |
| SL1344_RS21595_ | -1.46562398 | -1.56362393 | -1.351435452 | 0.214041276 | 0.31607039 | 0.633548729 | 0.235176563 | 0.364355845 | 0.608983909 | 0.474 | 0.675207591 |
| SL1344_RS21600_ | -2.224561945 | -2.411913655 | -2.058505252 | 0.35259369 | 0.343393099 | 0.959296396 | 0.146000026 | 0.230465342 | 0.471163552 | 0.3132 | 0.539421345 |
| SL1344_RS21605_ | -3.830619167 | -3.870246446 | -3.766096461 | 0.122890266 | 0.720931074 | 0.165174124 | 0.427048637 | 0.605877309 | 0.793145546 | 0.7155 | 0.848047411 |
| SL1344_RS21610_ | -2.322647412 | -2.378723367 | -2.284471989 | 0.143189754 | 0.49380705 | 0.276062584 | 0.378666671 | 0.563808542 | 0.76735664 | 0.6972 | 0.833661754 |
| SL1344_RS21615_ | -1.81430727 | -1.999636829 | -1.66835696 | 0.338471557 | 0.338973828 | 0.95835169 | 0.132578311 | 0.228396098 | 0.480520689 | 0.3168 | 0.546204224 |
| SL1344_RS21620_ | -1.94601446 | -1.938931852 | -1.955061693 | -4.01E-05 | 0.371285941 | -4.62E-05 | 0.5 | 0.622884085 | 0.807236921 | 0.7288 | 0.858734431 |
| SL1344_RS21625_ | -2.932328172 | -2.831671875 | -3.020412395 | -0.182071835 | 0.44086489 | -0.369606275 | 0.333111266 | 0.489351973 | 0.702038759 | 0.6023 | 0.766628879 |
| SL1344_RS21630_ | -1.844324968 | -1.727884295 | -1.888137962 | -0.101352957 | 0.431766655 | -0.192037536 | 0.413724187 | 0.685416336 | 0.8484215 | 0.7581 | 0.884064701 |
| SL1344_RS21635_ | -5.936026159 | -5.551355024 | -6.244090162 | -0.630557867 | 1.529644466 | -0.384561171 | 0.325116596 | 0.507717569 | 0.721020206 | 0.6117 | 0.776910798 |
| SL1344_RS21640_ | -5.259556641 | -5.104265935 | -5.340951843 | -0.110279352 | 1.719349186 | -0.053287364 | 0.472666668 | 0.683164876 | 0.850184837 | 0.8022 | 0.904855674 |
| SL1344_RS21645_ | -4.29072738 | -4.434596485 | -4.171842575 | 0.238358681 | 0.937597644 | 0.254444634 | 0.386000004 | 0.571105034 | 0.769760941 | 0.6808 | 0.822416725 |
| SL1344_RS21650_ | -3.673428288 | -3.762108587 | -3.582946653 | 0.183679054 | 0.895783287 | 0.177427007 | 0.407728185 | 0.63413566 | 0.817446245 | 0.7356 | 0.862271722 |
| SL1344_RS21655_ | -3.869423712 | -3.893752006 | -3.82424117 | 0.066401697 | 0.786002125 | 0.073669347 | 0.472000001 | 0.632500151 | 0.813074421 | 0.7345 | 0.863480572 |
| SL1344_RS21660_ | -1.2827699 | -1.385231933 | -1.135854628 | 0.25566421 | 0.287913813 | 0.844300508 | 0.176548988 | 0.270323683 | 0.525933201 | 0.3756 | 0.598968302 |
| SL1344_RS21665_ | 1.743762241 | 1.64427395 | 1.81056954 | 0.179262326 | 0.383694749 | 0.484897377 | 0.261159239 | 0.493882567 | 0.747294312 | 0.5352 | 0.764868708 |
| SL1344_RS21670_ | 3.741911235 | 3.609401836 | 3.926561684 | 0.300823589 | 0.082745217 | 3.401902495 | 4.68E-05 | 0.014686963 | 0.143192353 | 0.1 | 0.330257709 |
| SL1344_RS21675_ | -0.497502226 | -0.411193533 | -0.639166379 | -0.221314912 | 0.22496349 | -0.932441542 | 0.152666692 | 0.23410167 | 0.487462802 | 0.3225 | 0.554997923 |
| SL1344_RS21680_ | -3.0519245 | -3.059591457 | -3.040025222 | 0.017185292 | 0.522448818 | 0.029049904 | 0.484666667 | 0.575603467 | 0.770646411 | 0.6918 | 0.829038296 |
| SL1344_RS21685_ | -3.447309273 | -3.19082394 | -3.682136535 | -0.453658397 | 0.813937932 | -0.52629304 | 0.277148578 | 0.458643066 | 0.694254834 | 0.5567 | 0.745967003 |
| SL1344_RS21690_ | 0.474939276 | 0.512459994 | 0.308736871 | -0.177659401 | 0.204373116 | -0.713147646 | 0.264000011 | 0.38681384 | 0.650541757 | 0.5379 | 0.747118397 |
| SL1344_RS21695_ | -1.600658377 | -1.456807287 | -1.77438505 | -0.338896743 | 0.345860887 | -0.935503466 | 0.148567647 | 0.231044438 | 0.489211345 | 0.3069 | 0.540713994 |
| SL1344_RS21710_ | 4.935007906 | 4.369051658 | 5.412539993 | 1.005947189 | 0.142497633 | 7.142363386 | 4.68E-05 | 0.000601759 | 0.037715134 | 0.1 | 0.330257709 |
| SL1344_RS21715_ | 3.567768781 | 3.149871641 | 3.937027714 | 0.778188014 | 0.10496239 | 7.366289947 | 4.68E-05 | 0.002869022 | 0.067173768 | 0.1 | 0.330257709 |
| SL1344_RS21725_ | 2.990249822 | 2.686496321 | 3.372798574 | 0.735266758 | 0.32635233 | 2.550087163 | 4.68E-05 | 0.02600079 | 0.195111879 | 0.1 | 0.330257709 |
| SL1344_RS21730_ | 0.974710726 | 0.640125843 | 1.227673275 | 0.63709279 | 0.234172832 | 2.760831345 | 4.68E-05 | 0.015012729 | 0.14546026 | 0.1 | 0.330257709 |
| SL1344_RS21735_ | 0.87313334 | 0.638504857 | 1.089245392 | 0.471707682 | 0.223047574 | 2.101822911 | 0.003332494 | 0.037163485 | 0.219149316 | 0.1032 | 0.334844095 |
| SL1344_RS21740_ | 4.354978139 | 4.334855011 | 4.387494529 | 0.050517957 | 0.155490079 | 0.350112034 | 0.321333341 | 0.63289552 | 0.831553661 | 0.6424 | 0.823511177 |
| SL1344_RS21745_ | 0.372360648 | 0.227368212 | 0.530195089 | 0.365987583 | 0.268524171 | 1.391466784 | 0.041333441 | 0.105021834 | 0.356478142 | 0.1464 | 0.388205489 |
| SL1344_RS21755_ | 1.123107898 | 1.191979869 | 1.027119744 | -0.166697956 | 0.152262293 | -1.023441928 | 0.111925419 | 0.187067512 | 0.456285916 | 0.2652 | 0.510273293 |
| SL1344_RS21760_ | -0.552848223 | -0.314796199 | -0.61911668 | -0.248297706 | 0.374910074 | -0.456440877 | 0.337108601 | 0.696919424 | 0.858998766 | 0.7038 | 0.868842678 |
| SL1344_RS21765_ | 2.260071521 | 2.351431476 | 2.153260839 | -0.027951609 | 0.319750818 | -0.121584829 | 0.461333335 | 0.850862678 | 0.940866565 | 0.9359 | 0.972510578 |
| SL1344_RS21770_ | 0.928392975 | 0.990775729 | 0.839350086 | -0.166710972 | 0.162515027 | -0.985351445 | 0.112000036 | 0.203706421 | 0.474341991 | 0.2553 | 0.488494004 |
| SL1344_RS21775_ | 0.208701037 | 0.280652177 | 0.155590658 | -0.123101583 | 0.175723757 | -0.688804968 | 0.229333348 | 0.356386926 | 0.598030133 | 0.4609 | 0.664587878 |
| SL1344_RS21780_ | 2.682214314 | 2.807781443 | 2.618429587 | -0.142778326 | 0.1849901 | -0.804703636 | 0.190666685 | 0.281708276 | 0.573775624 | 0.3578 | 0.624508508 |
| SL1344_RS21785_ | -1.685585656 | -1.631715342 | -1.78633017 | -0.126873806 | 0.370658109 | -0.297578986 | 0.378414395 | 0.601751823 | 0.795389942 | 0.71 | 0.851528291 |
| SL1344_RS21790_ | 1.007354125 | 0.967125993 | 1.059074157 | 0.050331393 | 0.234788743 | 0.204253282 | 0.419333336 | 0.711191089 | 0.86712245 | 0.8237 | 0.915800811 |
| SL1344_RS21795_ | 2.66932497 | 2.667326602 | 2.671729795 | 0.002886883 | 0.072215581 | 0.035619502 | 0.482666667 | 0.593340118 | 0.784657474 | 0.7122 | 0.843914817 |
| SL1344_RS21800_ | 1.577668409 | 1.560056133 | 1.596385076 | 0.038701116 | 0.307102345 | 0.132846684 | 0.432666669 | 0.812469089 | 0.922113317 | 0.8884 | 0.949415106 |
| SL1344_RS21805_ | 1.754919601 | 1.566875231 | 1.930241805 | 0.378686548 | 0.178041883 | 2.150244532 | 0.00067292 | 0.052938864 | 0.261549221 | 0.1005 | 0.330998456 |
| SL1344_RS21810_ | 3.445452232 | 3.568678634 | 3.271186024 | -0.326431198 | 0.118608094 | -2.83450859 | 4.68E-05 | 0.014081131 | 0.143101773 | 0.1 | 0.330257709 |
| SL1344_RS21815_ | 1.828544453 | 1.737426559 | 1.904221312 | 0.165726655 | 0.12484677 | 1.261705251 | 0.087333381 | 0.130370029 | 0.382523906 | 0.1985 | 0.449194309 |
| SL1344_RS21820_ | 0.404886278 | 0.33472806 | 0.497811774 | 0.197600533 | 0.298166898 | 0.62239792 | 0.254000012 | 0.362085941 | 0.635162499 | 0.4836 | 0.708331492 |
| SL1344_RS21825_ | -3.792284457 | -3.712646109 | -3.818814004 | -0.150010228 | 1.002383195 | -0.138956367 | 0.446369089 | 0.717923631 | 0.868697344 | 0.8186 | 0.915616721 |
| SL1344_RS21830_ | -5.056954008 | -4.648944032 | -5.350324303 | -0.571221532 | 1.690396278 | -0.309897215 | 0.353764163 | 0.639213751 | 0.816879709 | 0.6859 | 0.837752655 |
| SL1344_RS21835_ | 2.938848052 | 3.044861744 | 2.827834159 | -0.187789298 | 0.156008486 | -1.164168668 | 0.104000039 | 0.143313123 | 0.420787747 | 0.203 | 0.47172959 |
| SL1344_RS21840_ | -3.307616364 | -3.392962411 | -3.27484186 | 0.058877327 | 0.72680527 | 0.076095659 | 0.472000001 | 0.68428985 | 0.849343526 | 0.791 | 0.898880311 |
| SL1344_RS21845_ | 3.435949537 | 3.43957001 | 3.434261816 | 0.000832878 | 0.099625056 | 0.006490313 | 0.496 | 0.686018241 | 0.853060035 | 0.8197 | 0.914651836 |
| SL1344_RS21880_ | -2.231057094 | -2.122549933 | -2.328734943 | -0.22680516 | 0.455073311 | -0.464419186 | 0.297801475 | 0.454897829 | 0.685143342 | 0.5776 | 0.753312271 |

| | | | | | | | | | | | |
|---|---|---|---|---|---|---|---|---|---|---|---|
| SL1344_RS21885_ | -0.229510319 | -0.290969626 | -0.162717752 | 0.12153114 | 0.195400773 | 0.595349894 | 0.269153908 | 0.408552547 | 0.644132546 | 0.5098 | 0.704294937 |
| SL1344_RS21890_ | -1.82044547 | -1.782142387 | -1.849577159 | -0.129172769 | 0.49080785 | -0.265355472 | 0.39307129 | 0.609101042 | 0.800115009 | 0.7155 | 0.849855382 |
| SL1344_RS21895_ | -0.937981314 | -0.848631029 | -1.143759865 | -0.281085719 | 0.289042163 | -0.876736223 | 0.18254499 | 0.269317292 | 0.533824494 | 0.3816 | 0.612772071 |
| SL1344_RS21900_ | -0.661762815 | -0.705025304 | -0.624850161 | 0.049751922 | 0.268174037 | 0.177924143 | 0.4170553 | 0.625724831 | 0.807427551 | 0.7329 | 0.859713041 |
| SL1344_RS21905_ | 1.154792922 | 1.09914837 | 1.183200474 | 0.057899058 | 0.218068714 | 0.199263245 | 0.415333337 | 0.801152745 | 0.916303392 | 0.8095 | 0.917386108 |
| SL1344_RS21910_ | 1.738995051 | 1.834785205 | 1.621007076 | -0.183380719 | 0.252402188 | -0.72054161 | 0.20719522 | 0.306304936 | 0.596884849 | 0.3905 | 0.644792163 |
| SL1344_RS21915_ | 1.231733926 | 1.141351361 | 1.312312372 | 0.171744822 | 0.120660545 | 1.41231432 | 0.053297883 | 0.108329935 | 0.338896856 | 0.1753 | 0.41548876 |
| SL1344_RS21920_ | -1.839950116 | -2.039249132 | -1.716407399 | 0.311609866 | 0.353561805 | 0.821361945 | 0.184000019 | 0.282929474 | 0.535257207 | 0.3839 | 0.605610399 |
| SL1344_RS21925_ | -1.009697094 | -0.981522133 | -1.041430299 | -0.048335431 | 0.285321286 | -0.165809048 | 0.423717525 | 0.604960496 | 0.794631406 | 0.7083 | 0.84452731 |
| SL1344_RS21930_ | 0.450559916 | 0.779894941 | 0.392439054 | -0.336849371 | 0.414724718 | -0.77020591 | 0.247168567 | 0.343974822 | 0.631123804 | 0.4669 | 0.719987888 |
| SL1344_RS21935_ | -1.756604542 | -1.672879418 | -1.791304975 | -0.139307254 | 0.563075585 | -0.227706834 | 0.427048637 | 0.703803845 | 0.861099066 | 0.8107 | 0.91138389 |
| SL1344_RS21940_ | 2.93918884 | 3.083416477 | 2.776228057 | -0.109549204 | 0.570296263 | -0.204302241 | 0.438000002 | 0.849125339 | 0.942618413 | 1 | 1 |
| SL1344_RS21945_ | -0.870675525 | -0.859340432 | -0.889537864 | 0.009903547 | 0.45608761 | 0.022249147 | 0.488666667 | 0.718961747 | 0.871826427 | 0.832 | 0.921811492 |
| SL1344_RS21950_ | -1.100037947 | -1.14735884 | -1.056467954 | 0.016351679 | 0.477118776 | 0.033778335 | 0.484343771 | 0.766634705 | 0.897372977 | 0.8806 | 0.944056055 |
| SL1344_RS21965_ | -1.467899339 | -1.239325788 | -1.673575961 | -0.441117813 | 0.29007693 | -1.460631738 | 0.061333404 | 0.10105425 | 0.328309598 | 0.1635 | 0.405466198 |
| SL1344_RS21970_ | -1.146377398 | -0.968854827 | -1.477621908 | -0.482733678 | 0.308352472 | -1.458231266 | 0.053333415 | 0.106191362 | 0.3438816 | 0.1738 | 0.416895235 |
| SL1344_RS21980_ | 3.423369039 | 3.398784814 | 3.440030411 | 0.016707359 | 0.127929207 | 0.05824284 | 0.459693539 | 0.623355813 | 0.82923313 | 0.9076 | 0.960066183 |
| SL1344_RS21985_ | 1.031728202 | 0.412360698 | 1.446279979 | 1.057186605 | 0.24616617 | 4.242260338 | 4.68E-05 | 0.00732278 | 0.102346205 | 0.1 | 0.330257709 |
| SL1344_RS21990_ | 1.072441045 | 1.140947544 | 0.983946035 | -0.175754649 | 0.163572504 | -1.041230767 | 0.088607642 | 0.189826668 | 0.462381916 | 0.2319 | 0.471719599 |
| SL1344_RS21995_ | -2.611639803 | -2.564632058 | -2.688774533 | -0.16061359 | 0.542735648 | -0.25785725 | 0.374666672 | 0.52521024 | 0.74360287 | 0.6666 | 0.810989572 |
| SL1344_RS22000_ | 1.386054281 | 1.22509147 | 1.58393763 | 0.370246913 | 0.182526599 | 1.94562107 | 0.010000458 | 0.047826439 | 0.246164524 | 0.1095 | 0.343670177 |
| SL1344_RS22005_ | -1.530598322 | -1.543508879 | -1.520157163 | -0.001320472 | 0.657559287 | -0.001300736 | 0.499666889 | 0.787114836 | 0.909646639 | 0.8113 | 0.910306364 |
| SL1344_RS22010_ | -1.530312228 | -1.515237351 | -1.544386906 | -0.025767265 | 0.284306232 | -0.080590039 | 0.466000001 | 0.56619167 | 0.764133213 | 0.6714 | 0.818378473 |
| SL1344_RS22015_ | 1.843356886 | 1.636948166 | 1.936332607 | 0.271596168 | 0.167628134 | 1.495000403 | 0.067288538 | 0.10544893 | 0.360296575 | 0.1655 | 0.416658896 |
| SL1344_RS22020_ | 1.564290677 | 1.373210741 | 1.686482414 | 0.319180551 | 0.18636623 | 1.657234496 | 0.023984199 | 0.077760045 | 0.311175733 | 0.1232 | 0.362680782 |
| SL1344_RS22030_ | 2.122431952 | 1.829565211 | 2.386645118 | 0.586846575 | 0.172180761 | 3.42963206 | 4.68E-05 | 0.02525583 | 0.188326307 | 0.1 | 0.330257709 |
| SL1344_RS22035_ | 1.755884038 | 1.503618579 | 1.992948919 | 0.505303926 | 0.166904991 | 3.014641141 | 4.68E-05 | 0.048657261 | 0.256836275 | 0.1 | 0.330257709 |
| SL1344_RS22040_ | 2.789252135 | 2.43417844 | 3.059189387 | 0.644178065 | 0.180711885 | 3.448671845 | 4.68E-05 | 0.039114316 | 0.235510464 | 0.1 | 0.330257709 |
| SL1344_RS22045_ | -0.738499735 | -0.817360648 | -0.669190627 | 0.142596516 | 0.268047282 | 0.497885819 | 0.294666676 | 0.46996802 | 0.696710696 | 0.5758 | 0.756783764 |
| SL1344_RS22050_ | -0.218496536 | -0.322662359 | -0.153026213 | 0.203561793 | 0.432345263 | 0.455420252 | 0.261825461 | 0.37576758 | 0.656670647 | 0.5226 | 0.726473609 |
| SL1344_RS22055_ | -0.810383213 | -0.900742322 | -0.663647028 | 0.244200369 | 0.346359837 | 0.700707084 | 0.226666681 | 0.344197246 | 0.610139113 | 0.4406 | 0.671975576 |
| SL1344_RS22060_ | -1.179890884 | -1.17879911 | -1.182347697 | -0.022369293 | 0.368870919 | -0.059444651 | 0.480346436 | 0.689333447 | 0.850819774 | 0.8022 | 0.902945372 |
| SL1344_RS22065_ | 1.222083876 | 1.074747577 | 1.327356597 | 0.22077704 | 0.37429391 | 0.53753416 | 0.301798809 | 0.447242561 | 0.714102091 | 0.6294 | 0.813502372 |
| SL1344_RS22070_ | -2.11497965 | -2.290421376 | -1.96265941 | 0.331834846 | 0.394908034 | 0.785009625 | 0.186000019 | 0.2871118 | 0.539234404 | 0.3846 | 0.605411923 |
| SL1344_RS22075_ | -1.944112561 | -1.954898811 | -1.930407885 | 0.02543521 | 0.396681224 | 0.051761565 | 0.475016656 | 0.62988089 | 0.810863197 | 0.7461 | 0.867471334 |
| SL1344_RS22080_ | 1.47115799 | 1.784598178 | 1.41361343 | -0.353304818 | 0.168976397 | -1.334573337 | 0.09726853 | 0.147445446 | 0.425050084 | 0.2355 | 0.481294584 |
| SL1344_RS22085_ | 1.240312887 | 1.352037105 | 1.118955762 | -0.277324719 | 0.209746684 | -1.295737157 | 0.036000124 | 0.110178776 | 0.363711411 | 0.1426 | 0.38370936 |
| SL1344_RS22090_ | 0.564700184 | 0.748895987 | 0.493996573 | -0.237749678 | 0.469303822 | -0.483347285 | 0.284477025 | 0.556829471 | 0.784490379 | 0.5862 | 0.795215192 |
| SL1344_RS22095_ | 1.108258954 | 1.042559867 | 1.138556883 | 0.078003802 | 0.158259286 | 0.463995409 | 0.335776156 | 0.551041565 | 0.761043617 | 0.6629 | 0.82258249 |
| SL1344_RS22100_ | 1.683618205 | 1.559453222 | 1.785599833 | 0.197866188 | 0.150624917 | 1.26194235 | 0.094000044 | 0.126703662 | 0.384888922 | 0.1939 | 0.453075859 |
| SL1344_RS22105_ | -1.183928117 | -1.297081576 | -1.093101054 | 0.24902797 | 0.395068111 | 0.603531329 | 0.207861442 | 0.332763654 | 0.612795457 | 0.4286 | 0.643726813 |
| SL1344_RS22110_ | -2.64036041 | -2.773483195 | -2.504274446 | 0.264613838 | 0.45601986 | 0.53620131 | 0.270486353 | 0.408124539 | 0.643104935 | 0.5081 | 0.698475435 |
| SL1344_RS22115_ | -2.392282226 | -2.524132009 | -2.304790127 | 0.078087645 | 0.684736099 | 0.111085053 | 0.456666668 | 0.731792827 | 0.87831644 | 0.862 | 0.934291909 |
| SL1344_RS22120_ | -0.994617612 | -0.85795504 | -1.116698839 | -0.303870487 | 0.345679376 | -0.847168359 | 0.15866669 | 0.249623154 | 0.518948126 | 0.3555 | 0.583503251 |
| SL1344_RS22125_ | -2.198262604 | -2.263175835 | -2.141710988 | 0.209181807 | 0.636411948 | 0.307982318 | 0.357333339 | 0.534490314 | 0.757674498 | 0.6957 | 0.835703977 |
| SL1344_RS22130_ | -0.937856994 | -0.502302466 | -1.439913984 | -0.92300024 | 0.32137454 | -2.851829612 | 0.00067292 | 0.021256539 | 0.161499901 | 0.1001 | 0.330410035 |
| SL1344_RS22155_ | -6.192867997 | -5.951716319 | -6.563671474 | -0.741476256 | 1.967599749 | -0.305947064 | 0.310666675 | 0.445493152 | 0.695077269 | 0.5833 | 0.75525801 |
| SL1344_RS22160_ | -0.124566549 | -0.267216167 | -0.080760572 | 0.158299223 | 0.271224886 | 0.473327268 | 0.34733334 | 0.619439767 | 0.811946814 | 0.7094 | 0.865156095 |
| SL1344_RS22165_ | 0.690328359 | 0.670219117 | 0.716383577 | 0.07890481 | 0.307032005 | 0.253509069 | 0.372000005 | 0.594658367 | 0.800252094 | 0.7245 | 0.86036601 |
| SL1344_RS22170_ | 1.242204914 | 1.222732226 | 1.258815478 | 0.027514361 | 0.265606855 | 0.101665047 | 0.458361094 | 0.767123183 | 0.898748437 | 0.8892 | 0.947812439 |
| SL1344_RS22175_ | -3.112409546 | -2.990616975 | -3.246451307 | -0.235637095 | 0.582252597 | -0.385576434 | 0.325333341 | 0.493982348 | 0.710968427 | 0.602 | 0.769199675 |
| SL1344_RS22180_ | -0.523100199 | -0.405120395 | -0.643300997 | -0.19777962 | 0.285034545 | -0.652461212 | 0.241838787 | 0.375621301 | 0.627946787 | 0.4749 | 0.687234134 |
| SL1344_RS22185_ | -1.051493371 | -1.134440124 | -1.000995031 | 0.030310439 | 0.550300795 | 0.051670056 | 0.479013991 | 0.765251957 | 0.89718492 | 0.8771 | 0.945609508 |

| | | | | | | | | | | | |
|---|---|---|---|---|---|---|---|---|---|---|---|
| SL1344_RS22190_ | 0.790682966 | 0.696575114 | 0.850435505 | 0.083091864 | 0.257829621 | 0.30523495 | 0.391333338 | 0.700605031 | 0.861286186 | 0.8104 | 0.912470804 |
| SL1344_RS22195_ | 0.900607725 | 0.911181803 | 0.876747158 | -0.014535823 | 0.177729787 | -0.076372702 | 0.471333334 | 0.707227422 | 0.861620622 | 0.7999 | 0.904922341 |
| SL1344_RS22200_ | 0.08405571 | 0.04875392 | 0.127220789 | 0.07526884 | 0.190446833 | 0.362947324 | 0.347768161 | 0.541524369 | 0.748115695 | 0.642 | 0.79962641 |
| SL1344_RS22205_ | 2.182001174 | 2.100574041 | 2.345934279 | 0.239768581 | 0.292118035 | 0.847708633 | 0.159227205 | 0.231814256 | 0.530210938 | 0.3084 | 0.582968799 |
| SL1344_RS22210_ | 7.07160164 | 7.06231008 | 7.161427112 | 0.09673889 | 0.207677743 | 0.496634802 | 0.225333348 | 0.395493456 | 0.681254237 | 0.4 | 0.680350104 |
| SL1344_RS22215_ | 5.331772052 | 5.299419808 | 5.368280518 | 0.205508221 | 0.775622663 | 0.298620101 | 0.259160571 | 0.649894149 | 0.847202812 | 0.5179 | 0.755687537 |
| SL1344_RS22220_ | 2.285970791 | 1.906743451 | 2.415530435 | 0.463614002 | 0.23602169 | 1.500769175 | 0.059960099 | 0.105549548 | 0.370578948 | 0.1542 | 0.40810352 |
| SL1344_RS22225_ | 0.20379737 | 0.30121839 | 0.100613451 | -0.202398894 | 0.147110221 | -1.337455472 | 0.071952091 | 0.127001696 | 0.349914612 | 0.2038 | 0.438151986 |
| SL1344_RS22235_ | 7.886816188 | 8.048100895 | 7.858865233 | -0.18835463 | 0.263806403 | -0.747942457 | 0.216522334 | 0.278528637 | 0.583163408 | 0.4 | 0.680350104 |
| SL1344_RS22240_ | 4.18877046 | 4.242592902 | 4.134392222 | -0.109911853 | 0.288318293 | -0.38615745 | 0.333111266 | 0.549634326 | 0.78692001 | 0.6946 | 0.864960397 |
| SL1344_RS22245_ | -0.709540186 | -0.858418592 | -0.466257261 | 0.368485275 | 0.35941474 | 1.017746268 | 0.128000031 | 0.190572382 | 0.465079938 | 0.2752 | 0.534762357 |
| SL1344_RS22250_ | 1.518469693 | 1.455336803 | 1.620284492 | 0.201063355 | 0.210772425 | 0.920292768 | 0.137908089 | 0.198240548 | 0.483215429 | 0.2686 | 0.524774703 |
| SL1344_RS22255_ | 3.739071533 | 3.71424195 | 3.761034799 | 0.026902981 | 0.093041734 | 0.298706314 | 0.383333338 | 0.663572007 | 0.837833855 | 0.7606 | 0.879580902 |
| SL1344_RS22260_ | -1.215482631 | -1.247755798 | -1.186582052 | 0.012435454 | 0.806574063 | 0.015615296 | 0.485676216 | 0.819051275 | 0.925430602 | 0.8389 | 0.923606604 |
| SL1344_RS22265_ | 2.278876494 | 2.297940713 | 2.262886894 | -0.046718825 | 0.390040928 | -0.129163919 | 0.454666668 | 0.755759309 | 0.898711423 | 0.943 | 0.975550463 |
| SL1344_RS22275_ | -0.250812601 | -0.162785486 | -0.448777883 | -0.303789327 | 0.373018786 | -0.795059445 | 0.207861442 | 0.281643107 | 0.567404808 | 0.3893 | 0.64002217 |
| SL1344_RS22280_ | -0.397684306 | -0.584014565 | -0.307366543 | 0.263963736 | 0.229736385 | 1.052604757 | 0.138666695 | 0.214833883 | 0.477914969 | 0.3078 | 0.548148711 |
| SL1344_RS22285_ | 3.009682231 | 3.076754703 | 2.977497501 | -0.098620694 | 0.172574268 | -0.546881896 | 0.274666677 | 0.457672016 | 0.71886214 | 0.5643 | 0.778926678 |
| SL1344_RS22290_ | 1.898097265 | 1.882351534 | 1.919455787 | -0.013816457 | 0.216023044 | -0.078710362 | 0.468354432 | 0.800538467 | 0.915908751 | 0.9196 | 0.963013894 |
| SL1344_RS22295_ | 3.369722519 | 3.395976486 | 3.350732374 | -0.031270667 | 0.241281279 | -0.147185167 | 0.425716192 | 0.815699859 | 0.925153864 | 0.9094 | 0.958515127 |
| SL1344_RS22300_ | 4.532397174 | 4.533277548 | 4.531508393 | 0.000121542 | 0.178261544 | 0.000311318 | 0.5 | 0.775502532 | 0.907997778 | 0.9121 | 0.960859952 |
| SL1344_RS22305_ | 1.397387514 | 1.304229863 | 1.490271551 | 0.182405146 | 0.119451699 | 1.518802366 | 0.048666757 | 0.095102788 | 0.321238817 | 0.1601 | 0.401700031 |
| SL1344_RS22310_ | -1.281826052 | -1.369406256 | -1.198003395 | 0.165665121 | 0.241870957 | 0.644863973 | 0.226000015 | 0.338472968 | 0.575803658 | 0.4344 | 0.638217902 |
| SL1344_RS22315_ | 0.391546903 | 0.435946166 | 0.356351584 | -0.082737897 | 0.162473212 | -0.473992841 | 0.29913392 | 0.452448682 | 0.676231914 | 0.563 | 0.738618905 |
| SL1344_RS22320_ | 2.545804677 | 2.487064664 | 2.669871978 | 0.176532838 | 0.087110916 | 1.863456842 | 0.022000206 | 0.065303031 | 0.273658579 | 0.1313 | 0.368368554 |
| SL1344_RS22325_ | 1.196497291 | 1.208471304 | 1.186191451 | 0.005199292 | 0.275046864 | 0.015730444 | 0.492 | 0.751191038 | 0.890947168 | 0.829 | 0.921041593 |
| SL1344_RS22330_ | 2.218266443 | 2.184107338 | 2.26697424 | 0.085159716 | 0.110243064 | 0.76563494 | 0.211192554 | 0.320154669 | 0.58380184 | 0.4288 | 0.652341295 |
| SL1344_RS22335_ | 1.277831689 | 1.284185362 | 1.271836783 | -0.083228803 | 0.235268429 | -0.33837607 | 0.375083283 | 0.529249365 | 0.757822866 | 0.7681 | 0.87774548 |
| SL1344_RS22340_ | 0.438761269 | 0.452993538 | 0.425331387 | -0.058345751 | 0.23622262 | -0.240458171 | 0.392666671 | 0.537507436 | 0.761822368 | 0.7117 | 0.84465088 |
| SL1344_RS22345_ | 3.862323759 | 4.059646097 | 3.712420294 | -0.242311673 | 0.46401554 | -0.419099588 | 0.344000006 | 0.713572425 | 0.880532252 | 0.7312 | 0.885838662 |
| SL1344_RS22350_ | 2.925515226 | 3.079770528 | 2.772949281 | -0.131989154 | 0.6177593 | -0.213743541 | 0.437333336 | 0.745623903 | 0.895872172 | 0.9757 | 0.989545632 |
| SL1344_RS22355_ | 0.156113923 | 0.277591896 | 0.031333218 | -0.058811979 | 0.566899701 | -0.118869964 | 0.449700202 | 0.813595057 | 0.922983741 | 0.928 | 0.967515375 |
| SL1344_RS22360_ | 0.216477817 | 0.283195468 | 0.121024049 | -0.198288153 | 0.282456875 | -0.705891441 | 0.213333349 | 0.320282861 | 0.596562933 | 0.4233 | 0.656988955 |
| SL1344_RS22365_ | 1.0422451 | 1.012850586 | 1.067629694 | 0.040552613 | 0.155619351 | 0.254453881 | 0.392666671 | 0.609420557 | 0.798166371 | 0.7236 | 0.85277729 |
| SL1344_RS22370_ | -0.200209512 | -0.411390932 | -0.113941578 | 0.27494782 | 0.341233456 | 0.744658869 | 0.240000013 | 0.338013083 | 0.619049594 | 0.4667 | 0.69668219 |
| SL1344_RS22375_ | 2.190634075 | 2.152468303 | 2.280634156 | 0.152985024 | 0.191176394 | 0.833420122 | 0.183877435 | 0.25005349 | 0.539683495 | 0.3535 | 0.612862926 |
| SL1344_RS22380_ | 2.538603893 | 2.454809439 | 2.587862058 | 0.163754584 | 0.166778917 | 0.96576875 | 0.150666692 | 0.208057945 | 0.497088292 | 0.3152 | 0.577626664 |
| SL1344_RS22385_ | 4.24174519 | 4.132671802 | 4.435506567 | 0.299832399 | 0.180390923 | 1.761433581 | 0.002000966 | 0.054901974 | 0.274664715 | 0.1008 | 0.331451989 |
| SL1344_RS22395_ | 4.54445923 | 4.498025672 | 4.59520667 | 0.115634573 | 0.082566991 | 1.461296285 | 0.005330642 | 0.13468469 | 0.413592332 | 0.1086 | 0.34168643 |
| SL1344_RS22400_ | 3.91651218 | 3.861424043 | 3.97144576 | 0.166700559 | 0.14292054 | 1.284557061 | 0.014000326 | 0.127376708 | 0.402457049 | 0.1134 | 0.348138537 |
| SL1344_RS22405_ | 4.842558596 | 4.663192547 | 4.919166681 | 0.262429163 | 0.146459359 | 1.86050847 | 4.68E-05 | 0.063129843 | 0.293824176 | 0.1 | 0.330257709 |
| SL1344_RS22410_ | -1.066065755 | -1.094834196 | -1.029626274 | 0.022975351 | 0.41064385 | 0.05181676 | 0.480000001 | 0.738742429 | 0.881773659 | 0.8419 | 0.925172742 |
| SL1344_RS22415_ | 5.408319527 | 5.412565167 | 5.404734321 | 0.004052828 | 0.05100668 | 0.078428173 | 0.469333334 | 0.753740585 | 0.890153578 | 0.8786 | 0.942271508 |
| SL1344_RS22420_ | 1.417705877 | 1.34800529 | 1.482220109 | 0.054436214 | 0.265562774 | 0.217387407 | 0.4170553 | 0.725213527 | 0.876055633 | 0.8581 | 0.931876016 |
| SL1344_RS22425_ | 3.997530137 | 4.003602808 | 3.991254811 | 0.037398818 | 0.164019237 | 0.2080776 | 0.411333337 | 0.547435265 | 0.782053455 | 0.8841 | 0.946729868 |
| SL1344_RS22430_ | 3.127425335 | 2.924265509 | 3.220750202 | 0.311668897 | 0.201582569 | 1.74718016 | 0.003332494 | 0.054971364 | 0.272582353 | 0.1018 | 0.332889775 |
| SL1344_RS22435_ | -4.535168119 | -4.473978828 | -4.59780758 | -0.118296769 | 1.072421938 | -0.105416386 | 0.450666669 | 0.642276151 | 0.822798603 | 0.7557 | 0.876227337 |
| SL1344_RS22440_ | -2.415073111 | -2.595643098 | -2.316822012 | 0.227604741 | 0.552272863 | 0.393504117 | 0.351099273 | 0.583646537 | 0.784350849 | 0.6873 | 0.838659802 |
| SL1344_RS22445_ | -2.297427742 | -2.242621432 | -2.35524532 | -0.129066395 | 0.358140796 | -0.343051298 | 0.358666672 | 0.50499985 | 0.715457795 | 0.6167 | 0.774119621 |
| SL1344_RS22450_ | -3.373892875 | -3.401046573 | -3.328554672 | 0.022891161 | 0.599737475 | 0.035366084 | 0.483677549 | 0.596131424 | 0.785275301 | 0.7086 | 0.842612671 |
| SL1344_RS22455_ | -3.877313312 | -3.847670688 | -3.932495493 | -0.074996118 | 0.663899987 | -0.098241151 | 0.460359762 | 0.579311834 | 0.776374298 | 0.6798 | 0.828427788 |
| SL1344_RS22460_ | -2.547313061 | -2.645209626 | -2.46442641 | 0.146069838 | 0.622497838 | 0.216193244 | 0.396000004 | 0.654211297 | 0.830482978 | 0.7613 | 0.879243777 |
| SL1344_RS22465_ | -0.619469809 | -0.699674128 | -0.525286659 | 0.179849873 | 0.2157817 | 0.781174146 | 0.189333352 | 0.296915007 | 0.544196763 | 0.3913 | 0.606945236 |

| | | | | | | | | | | | |
|---|---|---|---|---|---|---|---|---|---|---|---|
| SL1344_RS22470_ | 0.251074074 | 0.007130452 | 0.377501408 | 0.330739004 | 0.311239275 | 0.988761194 | 0.147901425 | 0.216056156 | 0.503304979 | 0.289 | 0.549263984 |
| SL1344_RS22475_ | 1.140975533 | 0.477043542 | 1.926726887 | 1.540329551 | 0.470309015 | 3.106205979 | 4.68E-05 | 0.008985817 | 0.120827263 | 0.1 | 0.330257709 |
| SL1344_RS22480_ | 2.761957325 | 1.371639721 | 3.781948523 | 2.006632485 | 0.687553313 | 3.062784228 | 4.68E-05 | 0.00742559 | 0.111412958 | 0.1 | 0.330257709 |
| SL1344_RS22485_ | 0.096261054 | 0.085414433 | 0.1096817 | 0.074503303 | 0.362074003 | 0.215349661 | 0.418000003 | 0.693021363 | 0.857212244 | 0.832 | 0.91963161 |
| SL1344_RS22490_ | 0.861827189 | 0.819635008 | 0.902858286 | 0.0810607 | 0.203963084 | 0.368723494 | 0.355762831 | 0.580074727 | 0.782496182 | 0.6936 | 0.836862673 |
| SL1344_RS22495_ | -1.158041511 | -0.905103564 | -1.29027649 | -0.355413541 | 0.327421817 | -0.994571289 | 0.135333362 | 0.212605259 | 0.48026739 | 0.3114 | 0.557473071 |
| SL1344_RS22500_ | -2.392117649 | -2.275624188 | -2.561088093 | -0.258183441 | 0.448286815 | -0.53701579 | 0.278666677 | 0.43372654 | 0.66875462 | 0.5463 | 0.734353519 |
| SL1344_RS22505_ | -3.784674219 | -3.987686782 | -3.625992071 | 0.356322712 | 0.719538999 | 0.463995611 | 0.306462367 | 0.465234641 | 0.689402715 | 0.5725 | 0.746240153 |
| SL1344_RS22510_ | -2.919610503 | -2.697545798 | -3.153323874 | -0.458720523 | 0.489461599 | -0.886442592 | 0.155333358 | 0.2494556 | 0.50248562 | 0.3443 | 0.571727478 |
| SL1344_RS22520_ | -1.571564992 | -1.39539687 | -1.951066689 | -0.521553892 | 0.455812788 | -1.053047537 | 0.139333361 | 0.186358562 | 0.455208536 | 0.2742 | 0.527624854 |
| SL1344_RS22525_ | -1.010419318 | -0.834430791 | -1.252973954 | -0.431700717 | 0.414789212 | -1.013043683 | 0.110592975 | 0.193520958 | 0.47489572 | 0.2484 | 0.500742168 |
| SL1344_RS22530_ | -1.663804586 | -1.718270626 | -1.594726488 | 0.132843681 | 0.331689934 | 0.389941487 | 0.336442379 | 0.507355914 | 0.720112042 | 0.6199 | 0.783309547 |
| SL1344_RS22535_ | 2.438513208 | 2.343005934 | 2.507202349 | 0.142899736 | 0.321699447 | 0.199402974 | 0.404397072 | 0.825968112 | 0.932665373 | 0.8947 | 0.955201704 |
| SL1344_RS22540_ | 4.667744593 | 4.637791057 | 4.69533399 | 0.054153607 | 0.245323837 | 0.193078468 | 0.42266667 | 0.908580249 | 0.966214202 | 0.9271 | 0.969020255 |
| SL1344_RS22545_ | 4.838147404 | 4.716165326 | 4.898355694 | 0.090065675 | 0.201007173 | 0.441554071 | 0.340000007 | 0.845005301 | 0.940578035 | 0.7132 | 0.878119686 |
| SL1344_RS22550_ | 4.026605957 | 3.829553694 | 4.092756201 | 0.168904838 | 0.182472218 | 0.598852106 | 0.33333334 | 0.623277225 | 0.830418667 | 0.7 | 0.872542119 |
| SL1344_RS22555_ | 5.266272255 | 5.174928531 | 5.340118733 | 0.157064473 | 0.251443036 | 0.339533327 | 0.357761498 | 0.859306438 | 0.947333473 | 0.7687 | 0.901772127 |
| SL1344_RS22560_ | -1.698229523 | -1.662308204 | -1.726747897 | -0.092970803 | 0.363277341 | -0.224898099 | 0.40866667 | 0.582937949 | 0.779393853 | 0.6948 | 0.834134357 |
| SL1344_RS22565_ | 1.435779851 | 1.33922856 | 1.464739924 | 0.106825671 | 0.167528942 | 0.51187439 | 0.340666673 | 0.606233667 | 0.806286687 | 0.7011 | 0.861842967 |
| SL1344_RS22570_ | 2.108085759 | 2.272078837 | 2.057903593 | -0.445192524 | 1.015283164 | -0.48275988 | 0.225333348 | 0.477382134 | 0.740099076 | 0.4105 | 0.687073899 |
| SL1344_RS22575_ | 2.754907846 | 2.767684964 | 2.743351871 | -0.017440084 | 0.549889612 | -0.028804616 | 0.486666667 | 0.946443418 | 0.980984172 | 0.9958 | 0.998198978 |
| SL1344_RS22580_ | -1.481800594 | -1.281873405 | -1.780880276 | -0.51169158 | 0.3785591 | -1.28595784 | 0.074616979 | 0.122374346 | 0.367751313 | 0.1958 | 0.443850462 |
| SL1344_RS22585_ | -0.221098879 | -0.315538199 | -0.150198325 | 0.080852025 | 0.422281611 | 0.17918663 | 0.433333336 | 0.759522821 | 0.894764115 | 0.8966 | 0.953125611 |
| SL1344_RS22590_ | -2.191831211 | -2.106249215 | -2.267601117 | -0.17259989 | 0.322071946 | -0.499626012 | 0.292666676 | 0.420420638 | 0.647654996 | 0.5297 | 0.713480271 |
| SL1344_RS22595_ | -1.165421162 | -1.109802845 | -1.199532144 | -0.069414967 | 0.360404105 | -0.176434428 | 0.429713527 | 0.697532036 | 0.855291945 | 0.7825 | 0.894464393 |
| SL1344_RS22600_ | 3.11480342 | 3.258189275 | 3.088073276 | -0.161386089 | 0.127454444 | -0.992586337 | 0.213333349 | 0.252426836 | 0.547921594 | 0.4164 | 0.661809326 |
| SL1344_RS22610_ | 0.855335923 | 0.891984255 | 0.837339304 | -0.028379138 | 0.193205284 | -0.142594247 | 0.447035312 | 0.738223329 | 0.88021801 | 0.8022 | 0.91022552 |
| SL1344_RS22615_ | 4.249830034 | 4.452605184 | 3.999719973 | -0.509943113 | 0.242479365 | -2.446799215 | 4.68E-05 | 0.025762571 | 0.195584702 | 0.1 | 0.330257709 |
| SL1344_RS22620_ | 2.43021743 | 2.356863747 | 2.488663263 | 0.130040218 | 0.107061261 | 1.179027864 | 0.103930752 | 0.156762532 | 0.419953136 | 0.2317 | 0.484575739 |
| SL1344_RS22625_ | 1.456291792 | 1.498105446 | 1.421988335 | -0.136628791 | 0.237966598 | -0.576768615 | 0.264000011 | 0.348191025 | 0.631144526 | 0.5352 | 0.731731083 |
| SL1344_RS22630_ | 2.030647133 | 1.744769356 | 2.242570212 | 0.524254561 | 0.18989734 | 2.827963788 | 4.68E-05 | 0.032296594 | 0.210464292 | 0.1 | 0.330257709 |
| SL1344_RS22635_ | 3.758515112 | 3.559631441 | 3.949569063 | 0.412540147 | 0.108093544 | 3.889880408 | 4.68E-05 | 0.013331938 | 0.139159561 | 0.1 | 0.330257709 |
| SL1344_RS22640_ | 3.995303296 | 4.003497109 | 3.985142305 | -0.135695073 | 0.272318137 | -0.250534153 | 0.338666673 | 0.396857574 | 0.682544881 | 0.7204 | 0.867452245 |
| SL1344_RS22645_ | -2.315706286 | -2.31376191 | -2.318197514 | -0.012775091 | 0.407598592 | -0.033178233 | 0.487008661 | 0.60222446 | 0.791018968 | 0.7025 | 0.841071846 |
| SL1344_RS22650_ | -2.423238584 | -1.682054367 | -2.805044727 | -1.128324179 | 0.713176985 | -1.513957179 | 0.047333426 | 0.083957346 | 0.320477161 | 0.1473 | 0.39213743 |
| SL1344_RS22655_ | 4.342058839 | 4.279215663 | 4.409740963 | 0.163874827 | 0.172478587 | 0.955137897 | 0.127914754 | 0.193496353 | 0.490663141 | 0.2486 | 0.522799441 |
| SL1344_RS22660_ | 4.219983021 | 4.091301187 | 4.478596118 | 0.378446847 | 0.084186785 | 3.677838666 | 4.68E-05 | 0.024016987 | 0.185755342 | 0.1 | 0.330257709 |
| SL1344_RS22665_ | 2.766873298 | 2.739621232 | 2.808401492 | 0.098965688 | 0.167447779 | 0.593584099 | 0.230000014 | 0.322835425 | 0.611496897 | 0.4553 | 0.684217843 |
| SL1344_RS22670_ | -2.952096194 | -2.7891681 | -3.094684331 | -0.275009411 | 0.552329864 | -0.444317136 | 0.307333342 | 0.480716405 | 0.702594736 | 0.5911 | 0.762906351 |
| SL1344_RS22675_ | 1.630053804 | 1.588384393 | 1.66475466 | 0.068288271 | 0.18195064 | 0.37250276 | 0.34133334 | 0.583764017 | 0.789590215 | 0.7104 | 0.84914209 |
| SL1344_RS22680_ | -0.967626099 | -0.98744565 | -0.946104853 | 0.041648929 | 0.218870566 | 0.177387926 | 0.422000003 | 0.513466966 | 0.722340432 | 0.6218 | 0.782134795 |
| SL1344_RS22685_ | -6.011772165 | -5.919388926 | -6.054576649 | -0.355305342 | 2.178300447 | -0.14862236 | 0.443037977 | 0.69362981 | 0.853561005 | 0.7995 | 0.902551157 |
| SL1344_RS22690_ | -0.393179487 | -0.331411993 | -0.528128465 | -0.222992432 | 0.365196429 | -0.557117429 | 0.273817466 | 0.385682837 | 0.659017006 | 0.5472 | 0.745051487 |
| SL1344_RS22695_ | 0.079958684 | 0.183584683 | -0.022151928 | -0.125057976 | 0.536144099 | -0.229656605 | 0.400000004 | 0.744638244 | 0.889368245 | 0.8728 | 0.942872336 |
| SL1344_RS22700_ | -0.490850624 | -0.492113834 | -0.487684754 | 0.000791256 | 0.224175294 | 0.004204068 | 0.497668221 | 0.608229388 | 0.793662713 | 0.7126 | 0.844186377 |
| SL1344_RS22705_ | -2.898418363 | -2.960012753 | -2.806906635 | 0.168484566 | 0.915050419 | 0.168385119 | 0.438666669 | 0.745577645 | 0.885185366 | 0.8614 | 0.936056689 |
| SL1344_RS22710_ | -4.387384261 | -4.867340268 | -4.107847268 | 0.608692319 | 1.289680087 | 0.462146307 | 0.301333342 | 0.488377972 | 0.724491266 | 0.615 | 0.789737045 |
| SL1344_RS22715_ | -1.782415757 | -1.839451125 | -1.714692542 | 0.133810956 | 0.356421557 | 0.326356474 | 0.354666673 | 0.511839228 | 0.725130284 | 0.6215 | 0.782334201 |
| SL1344_RS22720_ | -0.948931831 | -0.829494194 | -1.077988773 | -0.248194096 | 0.246209397 | -0.9681204 | 0.142666694 | 0.22091675 | 0.470416736 | 0.3076 | 0.539318114 |
| SL1344_RS22725_ | -0.521064527 | -0.479238076 | -0.561762526 | -0.249343169 | 0.499598008 | -0.501997125 | 0.331778821 | 0.448931731 | 0.707183511 | 0.6951 | 0.839714327 |
| SL1344_RS22730_ | -1.97183125 | -1.897086429 | -2.081990902 | -0.234283151 | 0.47825419 | -0.456488762 | 0.28400001 | 0.423207016 | 0.674692865 | 0.5737 | 0.743841899 |
| SL1344_RS22735_ | -0.467567877 | -0.47133007 | -0.461411854 | 0.013175624 | 0.178506988 | 0.061260499 | 0.474350434 | 0.519360933 | 0.727684955 | 0.6303 | 0.786391649 |
| SL1344_RS22740_ | -1.366093494 | -1.217577866 | -1.706970422 | -0.463956607 | 0.412560239 | -1.06514933 | 0.130579644 | 0.190692386 | 0.463135148 | 0.2856 | 0.539337954 |

| | | | | | | | | | | | |
|---|---|---|---|---|---|---|---|---|---|---|---|
| SL1344_RS22745_ | -2.205684151 | -2.184498449 | -2.225040016 | -0.05184721 | 0.3762738 | -0.12388279 | 0.449333335 | 0.575091017 | 0.768729505 | 0.6858 | 0.824062618 |
| SL1344_RS22750_ | -1.304351866 | -1.118510836 | -1.592800989 | -0.47369434 | 0.284315728 | -1.534662476 | 0.052000084 | 0.100862134 | 0.331553189 | 0.1643 | 0.404417283 |
| SL1344_RS22755_ | -3.325897807 | -3.452878391 | -3.165842703 | 0.280456364 | 0.510572913 | 0.518071355 | 0.270486353 | 0.425294466 | 0.650478888 | 0.5216 | 0.70639399 |
| SL1344_RS22760_ | -2.169593654 | -2.246063197 | -2.089195451 | 0.162421875 | 0.405671497 | 0.387298947 | 0.340439713 | 0.496840334 | 0.712510513 | 0.6088 | 0.771333339 |
| SL1344_RS22765_ | 1.165551529 | 1.127611157 | 1.192306237 | 0.091116458 | 0.190726007 | 0.445610282 | 0.304463699 | 0.426264893 | 0.681126735 | 0.5865 | 0.758515392 |
| SL1344_RS22770_ | 2.729075591 | 2.640866344 | 2.751140873 | 0.106555603 | 0.113043304 | 0.77513333 | 0.249833457 | 0.334201825 | 0.614167658 | 0.495 | 0.714595262 |
| SL1344_RS22775_ | 3.955070552 | 3.979583772 | 3.926964925 | 0.014322163 | 0.161768676 | 0.072846497 | 0.473017989 | 0.660704535 | 0.848903274 | 0.9733 | 0.988655252 |
| SL1344_RS22780_ | -2.457725954 | -2.733651508 | -2.214878049 | 0.522955757 | 0.450320136 | 1.122199391 | 0.101932086 | 0.17739097 | 0.42930403 | 0.2466 | 0.484581329 |
| SL1344_RS22785_ | -1.608460575 | -1.822878695 | -1.194288377 | 0.565966778 | 0.572419748 | 0.945260742 | 0.148567647 | 0.21035185 | 0.491354521 | 0.292 | 0.557586969 |
| SL1344_RS22790_ | -2.182030097 | -2.361838327 | -1.620092639 | 0.646431679 | 0.585252188 | 1.014437873 | 0.174550321 | 0.224135144 | 0.501552307 | 0.3447 | 0.594241147 |
| SL1344_RS22795_ | -1.547340215 | -1.711726312 | -1.347772253 | 0.374344322 | 0.327189759 | 1.10443132 | 0.112666703 | 0.181790766 | 0.431308307 | 0.2599 | 0.497487566 |
| SL1344_RS22800_ | -1.968090695 | -1.967046682 | -1.968541192 | 0.00921745 | 0.466634249 | 0.016475307 | 0.493670886 | 0.681218413 | 0.845577235 | 0.7864 | 0.893888131 |
| SL1344_RS22805_ | -1.310803916 | -1.354201334 | -1.259117093 | 0.215809378 | 0.476217643 | 0.423912378 | 0.331778821 | 0.484838089 | 0.722475115 | 0.6923 | 0.831692102 |
| SL1344_RS22810_ | -1.180223133 | -1.301952198 | -1.043092599 | 0.261624513 | 0.432149517 | 0.59061142 | 0.249833457 | 0.396688154 | 0.657660269 | 0.495 | 0.709071729 |
| SL1344_RS22815_ | -1.375357402 | -1.568653981 | -1.195721579 | 0.377072391 | 0.254132222 | 1.413208723 | 0.05266675 | 0.105704353 | 0.326453586 | 0.1725 | 0.410770519 |
| SL1344_RS22820_ | 1.077905969 | 1.026951662 | 1.135981218 | 0.174489091 | 0.263782729 | 0.658275356 | 0.224517003 | 0.311739448 | 0.599705172 | 0.4669 | 0.681199122 |
| SL1344_RS22825_ | 0.768032764 | 0.565113431 | 0.947302237 | 0.389192136 | 0.179025447 | 2.145270808 | 0.003332494 | 0.04106593 | 0.223323318 | 0.1054 | 0.337532128 |
| SL1344_RS22830_ | 0.575907443 | 0.440796064 | 0.666717423 | 0.229876319 | 0.200088583 | 1.092464792 | 0.111259197 | 0.168129603 | 0.431482306 | 0.2496 | 0.50153362 |
| SL1344_RS22835_ | -3.038184793 | -3.269185191 | -2.504826569 | 0.768603239 | 0.752431866 | 0.950341897 | 0.163890762 | 0.243152625 | 0.519879972 | 0.3415 | 0.585081977 |
| SL1344_RS22840_ | 0.342646943 | -0.080259058 | 0.775630108 | 0.745638174 | 0.329879351 | 2.288673701 | 0.002668394 | 0.026259948 | 0.189015738 | 0.103 | 0.334637639 |
| SL1344_RS22845_ | -5.644893331 | -5.303165679 | -6.068534853 | -0.745846119 | 1.164351672 | -0.593972854 | 0.245333346 | 0.367620996 | 0.601655834 | 0.4658 | 0.663690513 |
| SL1344_RS22850_ | -0.119391763 | -0.324289957 | 0.064587659 | 0.408863294 | 0.22674603 | 1.745761906 | 0.019320688 | 0.065091942 | 0.275714969 | 0.1205 | 0.357223551 |
| SL1344_RS22855_ | 1.463482775 | 1.419713278 | 1.498042166 | 0.191577835 | 0.323474915 | 0.618247588 | 0.284666676 | 0.371890667 | 0.65365871 | 0.5883 | 0.78236599 |
| SL1344_RS22860_ | -0.852168441 | -0.816551472 | -0.916857647 | -0.069675476 | 0.424536258 | -0.154270275 | 0.429333336 | 0.718963313 | 0.87085382 | 0.825 | 0.917197629 |
| SL1344_RS22865_ | 0.089647789 | 0.074783054 | 0.110984721 | 0.015037916 | 0.257005668 | 0.058760478 | 0.475682879 | 0.715285378 | 0.866535936 | 0.7948 | 0.900628695 |
| SL1344_RS22870_ | 1.977058918 | 1.981426369 | 1.972877261 | 0.003192134 | 0.175038146 | 0.015595294 | 0.492666667 | 0.780571697 | 0.904582152 | 0.8417 | 0.924875353 |
| SL1344_RS22875_ | 5.654156344 | 5.57889703 | 5.724934467 | 0.147556631 | 0.030081073 | 4.888338965 | 4.68E-05 | 0.005121743 | 0.083086765 | 0.1 | 0.330257709 |
| SL1344_RS22880_ | -0.86591424 | -0.99045002 | -0.736336554 | 0.248537592 | 0.220713074 | 1.077519561 | 0.113924086 | 0.184252389 | 0.420678122 | 0.2682 | 0.500209949 |
| SL1344_RS22885_ | 0.016328217 | -0.205134568 | 0.073126209 | 0.245276423 | 0.347869616 | 0.551979271 | 0.326666674 | 0.51969007 | 0.755736608 | 0.6748 | 0.847352713 |
| SL1344_RS22890_ | -3.258128586 | -2.836233996 | -3.43974128 | -0.481677144 | 0.822876717 | -0.508426522 | 0.294470362 | 0.534075411 | 0.751712397 | 0.6183 | 0.800217877 |
| SL1344_RS22895_ | -1.788603308 | -1.947399879 | -1.622361732 | 0.299414288 | 0.458259314 | 0.615025558 | 0.242000013 | 0.382827644 | 0.635044514 | 0.485 | 0.695891287 |
| SL1344_RS22900_ | -1.372626792 | -1.274436419 | -1.41474876 | -0.123399681 | 0.392182817 | -0.275469573 | 0.388000004 | 0.664473024 | 0.834274015 | 0.7592 | 0.880646741 |
| SL1344_RS22905_ | -3.719003957 | -3.735127003 | -3.701148106 | 0.355058892 | 1.441792603 | 0.241154201 | 0.407728185 | 0.628190779 | 0.819848763 | 0.8059 | 0.903804219 |
| SL1344_RS22910_ | -4.426153701 | -4.359196471 | -4.453678069 | -0.20647008 | 1.337352726 | -0.130463553 | 0.443333335 | 0.629884667 | 0.818279085 | 0.7589 | 0.876937593 |
| SL1344_RS22915_ | -3.848791093 | -4.004772557 | -3.655174722 | 0.334357909 | 0.804767216 | 0.399777924 | 0.321119261 | 0.510757136 | 0.724562583 | 0.6181 | 0.782367911 |
| SL1344_RS22920_ | -0.199223789 | -0.209236329 | -0.192824789 | 0.041694482 | 0.365508265 | 0.110429599 | 0.451698869 | 0.753702341 | 0.889896974 | 0.8751 | 0.941534496 |
| SL1344_RS22925_ | -0.662677232 | -0.507542197 | -0.806973494 | -0.225120879 | 0.357681473 | -0.62357266 | 0.261333345 | 0.421915408 | 0.67571767 | 0.5254 | 0.731290744 |
| SL1344_RS22930_ | 1.455869843 | 1.539557114 | 1.351284801 | -0.244119362 | 0.266662428 | -1.039091945 | 0.068620982 | 0.190700745 | 0.484503798 | 0.1804 | 0.428555237 |
| SL1344_RS22935_ | 0.7400985 | 0.66294076 | 0.820553014 | 0.162753294 | 0.135994081 | 1.095721929 | 0.105333372 | 0.175065374 | 0.422301716 | 0.2515 | 0.489500998 |
| SL1344_RS22940_ | -0.697426724 | -0.930832766 | -0.498398665 | 0.693076655 | 0.599322479 | 1.219956244 | 0.034666795 | 0.118894803 | 0.38534604 | 0.1438 | 0.385755539 |
| SL1344_RS22945_ | 0.817309029 | 0.584811178 | 0.98969154 | 0.395903529 | 0.196864286 | 1.944536762 | 0.015323415 | 0.04847326 | 0.24402924 | 0.1155 | 0.351858049 |
| SL1344_RS22950_ | -0.925429587 | -0.88755316 | -0.964791254 | -0.0835084 | 0.263380764 | -0.293176869 | 0.376000005 | 0.53571742 | 0.743757865 | 0.6621 | 0.808693331 |
| SL1344_RS22955_ | 4.891641639 | 4.751685805 | 4.980600842 | 0.213310491 | 0.074462417 | 2.644580739 | 4.68E-05 | 0.026919811 | 0.192765668 | 0.1 | 0.330257709 |
| SL1344_RS22960_ | 2.135235857 | 2.092972362 | 2.158597541 | 0.140798854 | 0.232404452 | 0.595702202 | 0.268487686 | 0.353004177 | 0.638436792 | 0.5502 | 0.757630747 |
| SL1344_RS22965_ | 4.205124793 | 4.141950389 | 4.253437969 | 0.048322327 | 0.16771983 | 0.317795973 | 0.41466667 | 0.800906202 | 0.918733991 | 0.8779 | 0.948017202 |
| SL1344_RS22970_ | 2.447688726 | 2.311330771 | 2.477940093 | 0.150933568 | 0.186334377 | 0.297562699 | 0.33533334 | 0.893092925 | 0.959963433 | 0.7087 | 0.876229928 |
| SL1344_RS22975_ | 2.456136561 | 2.481134217 | 2.425914245 | -0.042095287 | 0.197384874 | -0.20171379 | 0.40266667 | 0.713052915 | 0.870954963 | 0.8349 | 0.921446602 |
| SL1344_RS22980_ | 2.084912279 | 2.076724469 | 2.096455731 | -0.006621581 | 0.160599904 | -0.032382346 | 0.487333334 | 0.728034862 | 0.87722693 | 0.8374 | 0.923213705 |
| SL1344_RS22985_ | -1.235479951 | -1.288423708 | -1.158749336 | 0.087438748 | 0.362005493 | 0.221729595 | 0.405333337 | 0.648088021 | 0.824030237 | 0.7482 | 0.87106058 |
| SL1344_RS22995_ | -2.477876762 | -2.585905188 | -2.249920238 | 0.318655527 | 0.488148543 | 0.590474087 | 0.28047969 | 0.427285454 | 0.666424467 | 0.5515 | 0.741332301 |
| SL1344_RS23000_ | -2.752583063 | -2.921339807 | -2.444746452 | 0.53185747 | 0.747490582 | 0.667469493 | 0.247168567 | 0.378395799 | 0.645719147 | 0.5109 | 0.716511053 |
| SL1344_RS23010_ | 0.66439884 | 0.288602257 | 0.745440431 | 0.425119451 | 0.337527562 | 0.949105336 | 0.202666684 | 0.249882456 | 0.547431316 | 0.3914 | 0.645695335 |
| SL1344_RS23015_ | 0.434746977 | 1.730240865 | -0.172057969 | -0.93808535 | 1.702860988 | -0.900076399 | 0.198534328 | 0.299438944 | 0.601286437 | 0.3711 | 0.647798228 |

| | | | | | | | | | | | |
|---|---|---|---|---|---|---|---|---|---|---|---|
| SL1344_RS23020_ | -4.435059653 | -5.026650406 | -4.238039369 | 0.756802235 | 1.072185409 | 0.620893427 | 0.268000011 | 0.391884188 | 0.646617792 | 0.5276 | 0.728799853 |
| SL1344_RS23025_ | 2.154583381 | 2.128764094 | 2.166051423 | 0.044067668 | 0.211158575 | 0.181940165 | 0.435333336 | 0.81796753 | 0.924791033 | 0.8746 | 0.945701165 |
| SL1344_RS23030_ | 2.172681317 | 2.168399766 | 2.174679661 | -0.001637474 | 0.136531491 | -0.011954465 | 0.494 | 0.724824507 | 0.874361977 | 0.8274 | 0.920752533 |
| SL1344_RS23035_ | 3.010233949 | 2.932068055 | 3.07238737 | 0.160025159 | 0.188252698 | 0.837240903 | 0.18000002 | 0.245334396 | 0.543704595 | 0.3363 | 0.602480524 |
| SL1344_RS23040_ | 3.477707553 | 3.420793861 | 3.529510229 | 0.127394797 | 0.102654479 | 1.231449362 | 0.023984199 | 0.175412068 | 0.466436505 | 0.1287 | 0.366447946 |
| SL1344_RS23045_ | 2.139198161 | 2.094815324 | 2.162784469 | 0.084020321 | 0.174824842 | 0.457689147 | 0.316666675 | 0.488510883 | 0.73179084 | 0.603 | 0.793211741 |
| SL1344_RS23050_ | 2.599970701 | 2.60667733 | 2.588926499 | -0.016830826 | 0.097030563 | -0.167742983 | 0.434377084 | 0.662324604 | 0.833813657 | 0.7703 | 0.883766412 |
| SL1344_RS23055_ | 3.123797047 | 2.929348318 | 3.27205049 | 0.286661736 | 0.181038855 | 1.571810648 | 0.063291207 | 0.077498592 | 0.319185494 | 0.1546 | 0.409914252 |
| SL1344_RS23060_ | 3.69349859 | 3.537191596 | 3.821767307 | 0.284120423 | 0.085525367 | 3.335278744 | 4.68E-05 | 0.022014473 | 0.172672662 | 0.1 | 0.330257709 |
| SL1344_RS23065_ | 1.292700533 | 1.242436796 | 1.402088356 | 0.117251439 | 0.223973086 | 0.464436167 | 0.335109934 | 0.61652546 | 0.812346674 | 0.6948 | 0.853567318 |
| SL1344_RS23070_ | 1.405121758 | 1.307419992 | 1.488308447 | 0.19044907 | 0.155810596 | 1.17060396 | 0.105333372 | 0.155263091 | 0.420784659 | 0.2258 | 0.478928965 |
| SL1344_RS23075_ | -5.039564296 | -4.741085463 | -5.34720446 | -0.594954875 | 1.139395769 | -0.487706373 | 0.282478358 | 0.437548886 | 0.667520002 | 0.5408 | 0.724043757 |
| SL1344_RS23080_ | 1.737461993 | 1.83824178 | 1.700870473 | -0.124236619 | 0.180616782 | -0.591328509 | 0.29913392 | 0.500053286 | 0.739303593 | 0.6222 | 0.809609723 |
| SL1344_RS23085_ | 0.610441199 | 0.688960542 | 0.555602059 | -0.13490592 | 0.161702333 | -0.760868216 | 0.213191222 | 0.317046562 | 0.57135506 | 0.4214 | 0.637591203 |
| SL1344_RS23090_ | -0.831486559 | -0.98582427 | -0.737174313 | 0.34156676 | 0.462508745 | 0.738335417 | 0.231333347 | 0.302368905 | 0.58521442 | 0.4506 | 0.677755845 |
| SL1344_RS23095_ | -3.841596231 | -3.924721966 | -3.760654778 | 0.129205207 | 0.788503277 | 0.156200166 | 0.433044639 | 0.616005736 | 0.800864474 | 0.7179 | 0.849308032 |
| SL1344_RS23100_ | 2.42862982 | 2.345109783 | 2.723056502 | 0.368907415 | 0.195448062 | 1.4390623 | 0.068000063 | 0.125622656 | 0.400457809 | 0.1643 | 0.417303479 |
| SL1344_RS23105_ | -2.988856342 | -2.975752985 | -3.000526367 | -0.02523385 | 0.480744784 | -0.053951173 | 0.478666667 | 0.579527313 | 0.773914097 | 0.6929 | 0.830679474 |
| SL1344_RS23110_ | -1.877274743 | -2.004545292 | -1.814335259 | 0.156759628 | 0.418417496 | 0.345045985 | 0.356666672 | 0.574733304 | 0.773186361 | 0.6816 | 0.829571552 |
| SL1344_RS23115_ | -0.580792782 | -0.782392266 | -0.46361733 | 0.279634917 | 0.404937473 | 0.652747706 | 0.244503677 | 0.370295991 | 0.642230787 | 0.4825 | 0.706932406 |
| SL1344_RS23120_ | 2.568054263 | 2.540316708 | 2.62775521 | 0.090523661 | 0.115041971 | 0.747126339 | 0.214523667 | 0.318377281 | 0.595193044 | 0.4427 | 0.664644402 |
| SL1344_RS23125_ | -2.262175429 | -2.270918513 | -2.256592859 | 0.013104672 | 0.377081816 | 0.03677145 | 0.484666667 | 0.57729154 | 0.77283475 | 0.6926 | 0.833501137 |
| SL1344_RS23130_ | -5.78182696 | -5.927680702 | -5.568718999 | 0.397874334 | 1.366657046 | 0.272576079 | 0.376666672 | 0.530774786 | 0.733209318 | 0.6367 | 0.791523429 |
| SL1344_RS23140_ | -1.568378019 | -1.554608441 | -1.578955135 | -0.005169061 | 0.475007489 | -0.012772379 | 0.494666667 | 0.719930249 | 0.870522472 | 0.8388 | 0.92205016 |
| SL1344_RS23145_ | -0.309756566 | -0.174986163 | -0.385220671 | -0.178697633 | 0.290579622 | -0.607645226 | 0.258494349 | 0.433927821 | 0.681494735 | 0.5267 | 0.737408668 |
| SL1344_RS23150_ | 0.767778404 | 0.622252838 | 0.917707168 | 0.287516441 | 0.158555299 | 1.75442559 | 0.030666813 | 0.064260798 | 0.269658583 | 0.1318 | 0.371148281 |
| SL1344_RS23155_ | -3.124438008 | -3.031511264 | -3.198667781 | -0.205782036 | 0.818851989 | -0.218121567 | 0.391738845 | 0.647984513 | 0.827208882 | 0.7439 | 0.868659058 |
| SL1344_RS23160_ | -3.917510327 | -3.894967691 | -3.96688792 | -0.069972065 | 0.875939269 | -0.062895017 | 0.469333334 | 0.654417573 | 0.830931743 | 0.7755 | 0.888766278 |
| SL1344_RS23165_ | -4.506077116 | -4.241973777 | -4.699343792 | -0.577276906 | 0.954695392 | -0.587242466 | 0.257333345 | 0.397873754 | 0.638701628 | 0.5129 | 0.700110961 |
| SL1344_RS23170_ | -0.547025057 | -0.392695702 | -0.621907483 | -0.240819878 | 0.329587653 | -0.688797473 | 0.242000013 | 0.354859231 | 0.625062026 | 0.4853 | 0.700038612 |
| SL1344_RS23175_ | -0.462177631 | -0.616239911 | -0.28008594 | 0.336615604 | 0.210482639 | 1.514249658 | 0.04266677 | 0.092627463 | 0.313347712 | 0.1548 | 0.393503576 |
| SL1344_RS23180_ | -0.214017135 | -0.244843492 | -0.178042528 | 0.020805264 | 0.290072513 | 0.064425656 | 0.473684212 | 0.709232142 | 0.862476173 | 0.8243 | 0.912904717 |
| SL1344_RS23185_ | -1.41110249 | -1.211485508 | -1.566716432 | -0.370883138 | 0.314103578 | -1.107299479 | 0.105929419 | 0.173729255 | 0.428081018 | 0.2502 | 0.489636889 |
| SL1344_RS23190_ | 0.275494086 | -0.10839822 | 0.600656779 | 0.712054242 | 0.141890137 | 4.992481404 | 4.68E-05 | 0.004574349 | 0.076011165 | 0.1 | 0.330257709 |
| SL1344_RS23195_ | -4.899933467 | -4.721011351 | -5.081911405 | -0.387736776 | 1.105647939 | -0.324115416 | 0.365089945 | 0.536495331 | 0.744074187 | 0.6512 | 0.802600015 |
| SL1344_RS23200_ | -5.034671629 | -5.110598233 | -4.940998614 | 0.166767297 | 1.117226443 | 0.13252359 | 0.442666669 | 0.5962533 | 0.787957241 | 0.7035 | 0.840297097 |
| SL1344_RS23205_ | -0.535096548 | -0.143990996 | -0.985882898 | -0.843102758 | 0.25481713 | -3.252583304 | 4.68E-05 | 0.013537528 | 0.129506652 | 0.1 | 0.330257709 |
| SL1344_RS23210_ | -3.248578015 | -3.204241357 | -3.312877332 | -0.119032636 | 0.564913241 | -0.183712203 | 0.420386412 | 0.561234957 | 0.762858256 | 0.6806 | 0.824120187 |
| SL1344_RS23215_ | 0.729142931 | 0.427147166 | 0.799664071 | 0.349695727 | 0.229285769 | 1.213504839 | 0.129333364 | 0.169966246 | 0.451896891 | 0.2733 | 0.526557462 |
| SL1344_RS23220_ | 1.134095485 | 1.095858298 | 1.165961975 | 0.07464565 | 0.144803937 | 0.499936181 | 0.295136585 | 0.456879797 | 0.687671971 | 0.5764 | 0.752544134 |
| SL1344_RS23225_ | 2.105493772 | 2.073603385 | 2.154914583 | 0.090479629 | 0.111034588 | 0.760980332 | 0.193204549 | 0.292009421 | 0.559144669 | 0.3994 | 0.617139671 |
| SL1344_RS23230_ | 0.506454049 | 0.424926605 | 0.581550401 | -0.007069906 | 0.464445125 | -0.012105687 | 0.494666667 | 0.848977936 | 0.939798396 | 0.9655 | 0.985140372 |
| SL1344_RS23235_ | -0.160381997 | -0.225027712 | -0.090230967 | 0.13747473 | 0.233882487 | 0.570777161 | 0.263824128 | 0.408763094 | 0.654219627 | 0.5254 | 0.720818581 |
| SL1344_RS23240_ | -0.257039785 | -0.247630916 | -0.26447876 | -0.020350927 | 0.209694693 | -0.101564894 | 0.460666668 | 0.603502074 | 0.79291032 | 0.7147 | 0.846438146 |
| SL1344_RS23245_ | 3.825834022 | 3.714046941 | 4.206652241 | 0.520515017 | 0.388373907 | 1.471276495 | 0.00067292 | 0.141997537 | 0.429261858 | 0.1 | 0.330257709 |
| SL1344_RS23250_ | 1.416650286 | 0.943084692 | 1.660987798 | 0.696337273 | 0.414534705 | 1.501131871 | 0.049966777 | 0.110417147 | 0.379300723 | 0.1421 | 0.391804599 |
| SL1344_RS23265_ | 1.900318004 | 2.103921618 | 1.675972574 | -0.427248 | 0.132947656 | -3.223045642 | 4.68E-05 | 0.012276614 | 0.129309343 | 0.1 | 0.330257709 |
| SL1344_RS23270_ | -3.131916568 | -3.006541226 | -3.242829612 | -0.461706129 | 0.886211486 | -0.508219953 | 0.292471695 | 0.429833344 | 0.682986256 | 0.6309 | 0.787873269 |
| SL1344_RS23275_ | -4.28436185 | -4.353049674 | -4.236573848 | 0.042600839 | 1.026196965 | 0.040673646 | 0.483677549 | 0.66184497 | 0.833984325 | 0.7654 | 0.882754962 |
| SL1344_RS23280_ | -3.777398238 | -3.718631797 | -3.845665821 | -0.131261172 | 0.656463616 | -0.190177665 | 0.421333336 | 0.560923276 | 0.761202975 | 0.6688 | 0.816305787 |
| SL1344_RS23285_ | -3.500286004 | -3.275366633 | -3.652429155 | -0.367802057 | 0.713347576 | -0.470092604 | 0.312000008 | 0.492907559 | 0.713616811 | 0.5969 | 0.769155148 |
| SL1344_RS23290_ | -3.315546997 | -3.415422179 | -3.212225126 | 0.093484837 | 0.818258106 | 0.111608672 | 0.449333335 | 0.685770759 | 0.850028966 | 0.8021 | 0.901771206 |
| SL1344_RS23295_ | -0.994814383 | -0.957704857 | -1.048632667 | -0.09032975 | 0.259801425 | -0.328125894 | 0.361333339 | 0.526568082 | 0.736614247 | 0.6478 | 0.799181627 |

| | | | | | | | | | | | |
|---|---|---|---|---|---|---|---|---|---|---|---|
| SL1344_RS23300_ | 3.987483323 | 3.875018664 | 4.09395819 | 0.227839335 | 0.070823289 | 3.233711729 | 4.68E-05 | 0.012733196 | 0.132297501 | 0.1 | 0.330257709 |
| SL1344_RS23305_ | 0.445937249 | 0.441283133 | 0.452142181 | 0.023880709 | 0.185273075 | 0.118518652 | 0.444666669 | 0.629594013 | 0.810876751 | 0.748 | 0.866954624 |
| SL1344_RS23310_ | 1.599576688 | 1.541303811 | 1.63951698 | 0.101343147 | 0.153287916 | 0.61008008 | 0.245836122 | 0.378715343 | 0.638243967 | 0.4857 | 0.702083738 |
| SL1344_RS23315_ | 0.944875723 | 0.97296415 | 0.922917464 | -0.038836632 | 0.266909965 | -0.146707447 | 0.435333336 | 0.763627442 | 0.895993638 | 0.837 | 0.924083312 |
| SL1344_RS23320_ | -1.208036138 | -1.243467562 | -1.137539604 | 0.083657725 | 0.315860361 | 0.236900393 | 0.40866667 | 0.630245405 | 0.810669959 | 0.7264 | 0.859784715 |
| SL1344_RS23325_ | -1.138143531 | -1.226702917 | -1.053450486 | 0.167900277 | 0.242202837 | 0.656596461 | 0.227848116 | 0.352002937 | 0.584713571 | 0.4574 | 0.654854035 |
| SL1344_RS23330_ | -6.065516441 | -5.768018858 | -6.404019059 | -0.616576392 | 1.380200792 | -0.421267729 | 0.327115264 | 0.46186073 | 0.677987648 | 0.5594 | 0.732035137 |
| SL1344_RS23335_ | -5.512688598 | -5.520433275 | -5.49563386 | 0.02150285 | 1.33047553 | 0.014137078 | 0.492666667 | 0.621277106 | 0.803590406 | 0.7301 | 0.857771084 |
| SL1344_RS23340_ | 1.767545999 | 1.391668866 | 2.130587864 | 0.746372032 | 0.126684692 | 5.831375645 | 4.68E-05 | 0.003194282 | 0.068816736 | 0.1 | 0.330257709 |
| SL1344_RS23345_ | 2.157325155 | 2.575538873 | 1.862709507 | -0.704737684 | 0.236249686 | -2.998820727 | 4.68E-05 | 0.028719908 | 0.20314612 | 0.1 | 0.330257709 |
| SL1344_RS23350_ | 0.510502334 | 0.310454648 | 0.618704589 | 0.302032294 | 0.172170495 | 1.647201129 | 0.041333441 | 0.084248341 | 0.308059131 | 0.14 | 0.379562456 |
| SL1344_RS23355_ | -2.645389634 | -2.476606517 | -3.394279159 | -0.818478301 | 0.928382294 | -0.538554659 | 0.340666673 | 0.639203189 | 0.833331748 | 0.7174 | 0.879777114 |
| SL1344_RS23360_ | 4.362953473 | 4.368151826 | 4.357135749 | -0.016115963 | 0.575656677 | -0.019625271 | 0.471333334 | 0.585647774 | 0.810158567 | 0.994 | 0.997447282 |
| SL1344_RS23365_ | 5.643571452 | 5.744641349 | 5.345535937 | -0.253803649 | 0.375812105 | -0.963681553 | 0.183333353 | 0.221645561 | 0.525285843 | 0.335 | 0.614091951 |
| SL1344_RS23370_ | 3.80300455 | 4.003942422 | 3.577351099 | -0.093202676 | 0.724565832 | -0.14415999 | 0.447333335 | 0.766236074 | 0.906511268 | 1 | 1 |
| SL1344_RS23375_ | 1.559459256 | 1.400878248 | 1.691859446 | 0.343489216 | 0.215343744 | 1.62713154 | 0.00733396 | 0.081170589 | 0.322347207 | 0.1044 | 0.336248221 |
| SL1344_RS23380_ | 1.290258434 | 1.375323133 | 1.21632248 | -0.130510449 | 0.184821755 | -0.67945384 | 0.221333348 | 0.357377385 | 0.624010568 | 0.4595 | 0.683425873 |
| SL1344_RS23385_ | 0.638654459 | 0.544468918 | 0.759094597 | 0.229520146 | 0.177453651 | 1.227086703 | 0.075949423 | 0.146014561 | 0.403946995 | 0.1985 | 0.440720393 |
| SL1344_RS23390_ | 1.521105944 | 1.477082085 | 1.649181306 | 0.162265833 | 0.162234455 | 0.904319239 | 0.190666685 | 0.263633404 | 0.541934245 | 0.3806 | 0.623151422 |
| SL1344_RS23395_ | 3.099633843 | 2.908909172 | 3.202891792 | 0.292657707 | 0.12958923 | 2.134162201 | 4.68E-05 | 0.054589202 | 0.269323218 | 0.1012 | 0.33204093 |
| SL1344_RS23400_ | 1.566793663 | 1.456130524 | 1.615110783 | 0.164819827 | 0.210513448 | 0.752651753 | 0.194000018 | 0.291072696 | 0.582614544 | 0.3767 | 0.620151721 |
| SL1344_RS23410_ | 0.716097342 | 0.792255665 | 0.664370332 | -0.111935146 | 0.170457601 | -0.650568024 | 0.240000013 | 0.387231557 | 0.634388426 | 0.4891 | 0.697853629 |
| SL1344_RS23415_ | 0.363208402 | 0.353181771 | 0.369540305 | 0.003212961 | 0.280181695 | 0.012833953 | 0.494337109 | 0.76390537 | 0.895653386 | 0.8725 | 0.941473391 |
| SL1344_RS23420_ | -2.572871143 | -2.686052503 | -2.410164576 | 0.256520795 | 0.470253105 | 0.490812064 | 0.293137917 | 0.467077671 | 0.691762268 | 0.5688 | 0.750145064 |
| SL1344_RS23425_ | -1.947526371 | -1.927842463 | -1.967052437 | -0.11096578 | 0.545028726 | -0.19121616 | 0.41266667 | 0.634483402 | 0.819469651 | 0.7911 | 0.892280871 |
| SL1344_RS23430_ | 0.377644897 | 0.425391799 | 0.340166437 | -0.133560452 | 0.258199058 | -0.532261613 | 0.294470362 | 0.41354933 | 0.672859903 | 0.5793 | 0.756701886 |
| SL1344_RS23435_ | 0.642186389 | 0.678687724 | 0.575026718 | -0.091246755 | 0.163480173 | -0.517569455 | 0.292000009 | 0.453417087 | 0.68553626 | 0.5679 | 0.751544649 |
| SL1344_RS23440_ | 2.472772051 | 2.591767981 | 2.288488651 | -0.286320876 | 0.113499895 | -2.406364149 | 0.002000966 | 0.030568575 | 0.19773079 | 0.1017 | 0.332647821 |
| SL1344_RS23445_ | 2.992160905 | 3.131286035 | 2.82989525 | -0.310387051 | 0.144493469 | -2.115176346 | 4.68E-05 | 0.068802864 | 0.302367118 | 0.1001 | 0.330408405 |
| SL1344_RS23450_ | -0.634826477 | -0.704183193 | -0.577911727 | 0.128373518 | 0.209902486 | 0.576339047 | 0.267821463 | 0.414246827 | 0.646112166 | 0.5301 | 0.716552846 |
| SL1344_RS23455_ | -0.694724581 | -0.874561232 | -0.478391505 | 0.405470089 | 0.234024559 | 1.705136999 | 0.029980163 | 0.075901612 | 0.287125069 | 0.1366 | 0.373085015 |
| SL1344_RS23460_ | -4.093744078 | -4.109690382 | -4.078468373 | 0.035455885 | 0.68945905 | 0.048880313 | 0.481333334 | 0.561843821 | 0.762071075 | 0.6753 | 0.822417446 |
| SL1344_RS23465_ | -2.668513476 | -2.600821975 | -2.828013163 | -0.172462904 | 0.588379521 | -0.272614593 | 0.393333337 | 0.629949593 | 0.814810883 | 0.7398 | 0.872022181 |
| SL1344_RS23470_ | -3.230675186 | -3.164353796 | -3.287437185 | -0.122048833 | 0.563806341 | -0.215343904 | 0.401333337 | 0.579841951 | 0.774449906 | 0.6857 | 0.826263453 |
| SL1344_RS23480_ | -0.152832122 | -0.001962091 | -0.274727398 | -0.274352182 | 0.206092325 | -1.282972919 | 0.077333388 | 0.134237411 | 0.376423694 | 0.2014 | 0.442139857 |
| SL1344_RS23485_ | 1.774583103 | 1.694610395 | 1.897833488 | 0.195698559 | 0.113096751 | 1.613044143 | 0.040666776 | 0.076703013 | 0.294542122 | 0.1417 | 0.383599379 |
| SL1344_RS23495_ | 2.355301114 | 2.313825641 | 2.393186202 | 0.086228154 | 0.10224188 | 0.8121677 | 0.159893428 | 0.260893876 | 0.528680477 | 0.3585 | 0.591105603 |
| SL1344_RS23500_ | 5.063700946 | 5.013153032 | 5.090281155 | 0.072865537 | 0.110777987 | 0.532476406 | 0.33733334 | 0.475001001 | 0.734630988 | 0.73 | 0.873273788 |
| SL1344_RS23505_ | 2.232420297 | 1.934464083 | 2.283697029 | 0.326792692 | 0.161543686 | 1.180688665 | 0.128580977 | 0.163914977 | 0.451122824 | 0.2679 | 0.523411916 |
| SL1344_RS23510_ | 0.55636659 | 0.437118856 | 0.616315071 | 0.138462857 | 0.335073063 | 0.338951387 | 0.374000005 | 0.775842395 | 0.904490888 | 0.7621 | 0.89697152 |
| SL1344_RS23515_ | -0.019283312 | -0.004242101 | -0.033771965 | -0.024849806 | 0.176113255 | -0.127509745 | 0.431333336 | 0.568977139 | 0.765669993 | 0.681 | 0.823825475 |
| SL1344_RS23520_ | 2.215172779 | 2.143862491 | 2.352067041 | 0.201749119 | 0.106300547 | 1.750677141 | 0.03664236 | 0.07481943 | 0.298491943 | 0.1369 | 0.377503932 |
| SL1344_RS23525_ | 3.25700474 | 3.22510679 | 3.289143587 | 0.053577122 | 0.159368637 | 0.348938688 | 0.346666673 | 0.590490713 | 0.802296119 | 0.7083 | 0.855410836 |
| SL1344_RS23530_ | 1.528968546 | 1.543206489 | 1.522376172 | -0.008374795 | 0.130307567 | -0.06369354 | 0.478000001 | 0.687998206 | 0.848335995 | 0.7779 | 0.890639653 |
| SL1344_RS23535_ | 2.085565281 | 2.132954655 | 2.024547432 | -0.070138511 | 0.153259736 | -0.442235813 | 0.324000007 | 0.550723066 | 0.7668327 | 0.6682 | 0.821360061 |
| SL1344_RS23540_ | 1.388739537 | 1.424081833 | 1.361767718 | -0.075260731 | 0.2126928 | -0.325400081 | 0.345769493 | 0.515481301 | 0.750056297 | 0.6816 | 0.83230897 |
| SL1344_RS23545_ | -0.520972049 | -0.549651167 | -0.503167559 | 0.027285998 | 0.262185504 | 0.085922051 | 0.463024652 | 0.666668617 | 0.837709937 | 0.7613 | 0.883711475 |
| SL1344_RS23550_ | -2.302019035 | -2.429514286 | -2.133447334 | 0.298365105 | 0.374554266 | 0.728147094 | 0.214000016 | 0.312387657 | 0.550221109 | 0.4034 | 0.613981936 |
| SL1344_RS23555_ | -4.65016376 | -5.090150094 | -4.258883568 | 0.775889971 | 1.153602083 | 0.644473341 | 0.241333346 | 0.361451448 | 0.613380472 | 0.4587 | 0.672100054 |
| SL1344_RS23560_ | -1.847208966 | -2.175028552 | -1.743155212 | 0.386250432 | 0.461628708 | 0.746392921 | 0.236000014 | 0.334876372 | 0.60426392 | 0.4739 | 0.692468813 |
| SL1344_RS23565_ | -3.952444331 | -4.195703687 | -3.594495823 | 0.556074324 | 1.030376154 | 0.497381424 | 0.281333343 | 0.463735616 | 0.698305574 | 0.544 | 0.743526916 |
| SL1344_RS23570_ | -3.299619943 | -3.530819575 | -3.059380934 | 0.473135301 | 0.561481335 | 0.809139357 | 0.185333353 | 0.285563524 | 0.531994839 | 0.3779 | 0.59503957 |
| SL1344_RS23575_ | 0.282176141 | 0.081693431 | 0.409123936 | 0.326780683 | 0.171021543 | 1.807368918 | 0.025333511 | 0.063738131 | 0.266903528 | 0.1229 | 0.35915424 |

| | | | | | | | | | | | |
|---|---|---|---|---|---|---|---|---|---|---|---|
| SL1344_RS23580_ | -0.242418808 | -0.361905943 | -0.126429244 | 0.238948194 | 0.202878335 | 1.083535973 | 0.114590308 | 0.185610136 | 0.43489892 | 0.2644 | 0.502065135 |
| SL1344_RS23590_ | -5.836905421 | -5.866341332 | -5.784451961 | 0.092574608 | 1.328301577 | 0.059570881 | 0.470353099 | 0.567996277 | 0.767760742 | 0.6832 | 0.82581302 |
| SL1344_RS23595_ | -0.511792782 | -0.61713299 | -0.38556795 | 0.247971399 | 0.274136757 | 0.886450739 | 0.161225873 | 0.242466238 | 0.508682806 | 0.341 | 0.572922854 |
| SL1344_RS23640_ | 0.46201205 | 0.574005282 | 0.430446581 | -0.121734425 | 0.200499075 | -0.486687565 | 0.334443711 | 0.59194726 | 0.790789516 | 0.6808 | 0.845147905 |
| SL1344_RS23645_ | -0.509125497 | -0.490147921 | -0.517722393 | -0.014383839 | 0.2878584 | -0.042141083 | 0.484343771 | 0.706892675 | 0.861747145 | 0.7984 | 0.904317585 |
| SL1344_RS23650_ | 1.684615868 | 1.792374044 | 1.64690076 | -0.128121038 | 0.179521876 | -0.670282607 | 0.258666678 | 0.400819788 | 0.66528273 | 0.5076 | 0.735504093 |
| SL1344_RS23655_ | 0.082987043 | 0.116029591 | 0.061009079 | -0.071219512 | 0.388296328 | -0.17499991 | 0.433044639 | 0.780757649 | 0.90529654 | 0.8668 | 0.939169794 |
| SL1344_RS23660_ | 0.972463932 | 1.058916868 | 0.916576316 | -0.129355522 | 0.269892512 | -0.504658765 | 0.27800001 | 0.484469414 | 0.727421796 | 0.5765 | 0.775755347 |
| SL1344_RS23665_ | -0.909619375 | -0.613664468 | -1.262295217 | -0.653683316 | 0.240672086 | -2.705183518 | 0.00067292 | 0.021839726 | 0.158605182 | 0.1016 | 0.332552978 |
| SL1344_RS23670_ | 4.181135473 | 4.127277717 | 4.197057225 | 0.08177961 | 0.245188835 | 0.407479042 | 0.266666678 | 0.590408879 | 0.810604691 | 0.5077 | 0.747294906 |
| SL1344_RS23675_ | 2.367910075 | 2.067091516 | 2.706318599 | 0.41423511 | 0.355147358 | 1.602101135 | 0.082666718 | 0.071320028 | 0.309110298 | 0.1759 | 0.441344487 |
| SL1344_RS23680_ | 1.785063047 | 1.711644891 | 1.878701292 | 0.197407696 | 0.167054192 | 1.157116544 | 0.07533339 | 0.149210099 | 0.421246475 | 0.1953 | 0.441577345 |
| SL1344_RS23685_ | -1.790575091 | -1.943973778 | -1.707231085 | 0.228253959 | 0.438493482 | 0.479320996 | 0.312000008 | 0.474097833 | 0.709319842 | 0.6077 | 0.780942432 |
| SL1344_RS23690_ | -3.180377109 | -3.375417233 | -2.975115742 | 0.407980225 | 0.543078446 | 0.718939559 | 0.204666683 | 0.321001266 | 0.564052879 | 0.4172 | 0.629535929 |
| SL1344_RS23695_ | -3.579082978 | -3.621752918 | -3.518765532 | -0.046111428 | 0.946247738 | -0.042023579 | 0.481012659 | 0.681025531 | 0.847156594 | 0.8219 | 0.91137991 |
| SL1344_RS23700_ | -2.213392183 | -2.114069356 | -2.351354125 | -0.242982487 | 0.387363592 | -0.578838741 | 0.259333345 | 0.385415459 | 0.626069203 | 0.4939 | 0.689368111 |
| SL1344_RS23705_ | -3.533400163 | -3.523467681 | -3.545567459 | -0.027219886 | 0.578205264 | -0.042296425 | 0.481012659 | 0.569496822 | 0.76629329 | 0.6847 | 0.824742451 |
| SL1344_RS23710_ | -0.890289945 | -0.993419596 | -0.811277714 | 0.186095252 | 0.255132702 | 0.658516285 | 0.227181893 | 0.343601674 | 0.587948152 | 0.4476 | 0.650262463 |
| SL1344_RS23715_ | -2.576044948 | -2.5584359 | -2.588546417 | -0.036924202 | 0.458438293 | -0.074146006 | 0.462666668 | 0.59000224 | 0.78214776 | 0.6998 | 0.835321859 |
| SL1344_RS23720_ | -2.401426671 | -2.358557883 | -2.446892778 | -0.073114785 | 0.366937892 | -0.186207226 | 0.41439041 | 0.513730252 | 0.721588485 | 0.6264 | 0.782343992 |
| SL1344_RS23725_ | -3.123013875 | -2.987519568 | -3.295297003 | -0.368121843 | 0.683132402 | -0.502788968 | 0.277333344 | 0.431709622 | 0.675403264 | 0.5487 | 0.735970181 |
| SL1344_RS23730_ | -2.471281481 | -2.665306084 | -2.363070032 | 0.289953618 | 0.469829285 | 0.567319006 | 0.262491684 | 0.425521536 | 0.662400388 | 0.5442 | 0.732406807 |
| SL1344_RS23735_ | -3.237328554 | -3.092420079 | -3.430649226 | -0.35369353 | 0.5176784 | -0.634945353 | 0.24066668 | 0.361721279 | 0.601124805 | 0.4627 | 0.663486079 |
| SL1344_RS23740_ | -3.884314746 | -3.721754427 | -4.142337397 | -0.404424241 | 0.688304483 | -0.54103084 | 0.266000011 | 0.404163179 | 0.642738043 | 0.5065 | 0.701996946 |
| SL1344_RS23745_ | -1.324253859 | -1.351617626 | -1.294948883 | 0.063913539 | 0.306609175 | 0.2058687 | 0.414000003 | 0.584522385 | 0.777328435 | 0.6923 | 0.831113398 |
| SL1344_RS23750_ | -4.849246603 | -4.569601979 | -5.48222335 | -0.756491191 | 1.455376855 | -0.462158892 | 0.307128589 | 0.469142158 | 0.71056996 | 0.6069 | 0.784283794 |
| SL1344_RS23755_ | -4.054084829 | -4.216524732 | -3.919178271 | 0.302516581 | 0.689445211 | 0.419764742 | 0.312000008 | 0.458506377 | 0.681926168 | 0.5792 | 0.748373146 |
| SL1344_RS23760_ | -5.194036379 | -5.189241434 | -5.200828678 | -0.020163758 | 1.272272175 | -0.014511393 | 0.490666667 | 0.636547317 | 0.815953778 | 0.7426 | 0.866479798 |
| SL1344_RS23765_ | -2.34537316 | -2.124093509 | -2.523486019 | -0.392014433 | 0.449080091 | -0.827747544 | 0.181333353 | 0.282533452 | 0.541880037 | 0.3759 | 0.603569617 |
| SL1344_RS23770_ | -3.495719268 | -3.159925494 | -3.782322168 | -0.560638318 | 0.813559944 | -0.653170104 | 0.23866668 | 0.37946056 | 0.631071784 | 0.4639 | 0.681592655 |
| SL1344_RS23775_ | -1.924734007 | -1.477141167 | -2.654820648 | -1.143085149 | 0.541233217 | -2.012224536 | 0.009333825 | 0.048460693 | 0.246151379 | 0.1086 | 0.342584759 |
| SL1344_RS23780_ | -3.147393309 | -3.222226538 | -3.079521518 | 0.160168015 | 0.674290035 | 0.203037419 | 0.404397072 | 0.611473904 | 0.799547849 | 0.7166 | 0.848561092 |
| SL1344_RS23785_ | -2.680902218 | -2.578159593 | -2.876946303 | -0.286196133 | 0.504339292 | -0.514014088 | 0.292666676 | 0.436252149 | 0.670105022 | 0.5479 | 0.735823624 |
| SL1344_RS23790_ | -4.709455559 | -4.670200975 | -4.756017622 | -0.065144417 | 0.836286199 | -0.069575479 | 0.466666668 | 0.55423811 | 0.754584079 | 0.663 | 0.812266608 |
| SL1344_RS23795_ | -4.319484579 | -4.424432403 | -4.189622947 | 0.209583138 | 0.8177832 | 0.236796882 | 0.385333338 | 0.54591479 | 0.751052561 | 0.652 | 0.805124095 |
| SL1344_RS23800_ | -4.481384222 | -4.667946254 | -4.340843878 | 0.254321577 | 1.33526855 | 0.197532997 | 0.405063295 | 0.662097382 | 0.835252302 | 0.7816 | 0.890968867 |
| SL1344_RS23805_ | -3.894133973 | -3.983746479 | -3.809335561 | 0.077049655 | 1.001397921 | 0.080701364 | 0.464666668 | 0.681931243 | 0.846911539 | 0.8011 | 0.89998765 |
| SL1344_RS23810_ | -1.500802841 | -1.086514359 | -1.781878785 | -0.641026178 | 0.595531711 | -1.072608096 | 0.117333368 | 0.169564392 | 0.448600362 | 0.234 | 0.499723067 |
| SL1344_RS23815_ | -1.61986561 | -1.120185965 | -1.795673708 | -0.654140339 | 0.473630803 | -1.213880761 | 0.103333373 | 0.158858871 | 0.431020836 | 0.2401 | 0.491182728 |
| SL1344_RS23820_ | -4.311726397 | -4.185578056 | -4.44516242 | -0.276426181 | 0.657966797 | -0.390057186 | 0.321785484 | 0.443183854 | 0.665135376 | 0.5536 | 0.729413951 |
| SL1344_RS23825_ | -3.271977818 | -3.510658985 | -2.986075388 | 0.507466319 | 0.626529619 | 0.744584062 | 0.206666683 | 0.334445415 | 0.583096031 | 0.428 | 0.644989765 |
| SL1344_RS23830_ | -2.794126776 | -2.692005474 | -2.88543577 | -0.189170139 | 0.44516413 | -0.392120579 | 0.333777489 | 0.479500401 | 0.695315377 | 0.5901 | 0.75819561 |
| SL1344_RS23840_ | -3.032283143 | -2.950261405 | -3.099069395 | -0.114004082 | 0.70369914 | -0.137970242 | 0.425716192 | 0.671714963 | 0.838939191 | 0.7621 | 0.880101886 |
| SL1344_RS23845_ | -4.906117133 | -4.331560803 | -5.665823933 | -1.382047351 | 1.174463334 | -1.120330653 | 0.110592975 | 0.166437804 | 0.422535456 | 0.2404 | 0.484616641 |
| SL1344_RS23850_ | -4.94147413 | -4.672395608 | -5.472257233 | -0.908664581 | 1.939898081 | -0.427176792 | 0.227333348 | 0.362768937 | 0.645414274 | 0.4574 | 0.659878643 |
| SL1344_RS23855_ | -6.024257147 | -6.155401929 | -5.891510187 | 0.230982091 | 1.849865087 | 0.113920575 | 0.449333335 | 0.621946705 | 0.807212375 | 0.7226 | 0.85398359 |
| SL1344_RS23860_ | -1.291068364 | -1.264178175 | -1.325799384 | 0.00847273 | 0.400845515 | 0.022108514 | 0.491333334 | 0.705098719 | 0.862171233 | 0.8315 | 0.917530708 |
| SL1344_RS23865_ | -4.01187743 | -3.950473835 | -4.080172314 | -0.108572398 | 0.815440715 | -0.133311514 | 0.438666669 | 0.612227683 | 0.800641272 | 0.7324 | 0.858826961 |
| SL1344_RS23870_ | -4.168222025 | -3.654730409 | -5.118427898 | -1.507124035 | 1.192511333 | -1.173861277 | 0.065289873 | 0.156117343 | 0.426320358 | 0.1831 | 0.427769764 |
| SL1344_RS23875_ | -2.865645511 | -2.594528309 | -3.236272101 | -0.62817211 | 0.580915821 | -1.003290133 | 0.129247199 | 0.200798145 | 0.463875408 | 0.2825 | 0.522319052 |
| SL1344_RS23880_ | -6.104583028 | -5.422564715 | -6.92968264 | -1.477078731 | 1.684136313 | -0.828210415 | 0.16322454 | 0.260511506 | 0.516573515 | 0.3511 | 0.577970317 |
| SL1344_RS23885_ | -6.202822893 | -6.013837445 | -6.454525192 | -0.389816942 | 1.641474251 | -0.207780498 | 0.395069957 | 0.544984334 | 0.753542625 | 0.6702 | 0.81658453 |
| SL1344_RS23890_ | -7.28906796 | -6.878778674 | -7.62163885 | -0.585331015 | 3.206969069 | -0.151058955 | 0.403064627 | 0.606945898 | 0.797236477 | 0.7147 | 0.847309283 |

| | | | | | | | | | | | |
|---|---|---|---|---|---|---|---|---|---|---|---|
| SL1344_RS23895_ | -5.842049317 | -5.922927798 | -5.732730517 | 0.050023542 | 1.716039906 | 0.027870072 | 0.484343771 | 0.635725306 | 0.817573998 | 0.7572 | 0.873981886 |
| SL1344_RS23900_ | -4.908345776 | -5.083581031 | -4.686117071 | 0.403762756 | 0.987252849 | 0.385771646 | 0.340666673 | 0.487386044 | 0.702658656 | 0.5934 | 0.758938528 |
| SL1344_RS23905_ | -8.224280625 | -8.938534796 | -7.529613238 | 1.235893288 | 3.50582987 | 0.3141432 | 0.321785484 | 0.498416296 | 0.715812888 | 0.6027 | 0.768228233 |
| SL1344_RS23910_ | -0.622777225 | -0.537110042 | -0.7639226 | -0.229400897 | 0.242745685 | -0.889435581 | 0.167221874 | 0.244131514 | 0.507007642 | 0.3364 | 0.566436471 |
| SL1344_RS23915_ | -0.008697441 | -0.002909662 | -0.013248342 | 0.053496109 | 0.339126711 | 0.149263679 | 0.432000003 | 0.69691794 | 0.857300186 | 0.82 | 0.911556228 |
| SL1344_RS23920_ | 4.408069969 | 4.463385915 | 4.383222191 | -0.081524816 | 0.137080481 | -0.34064313 | 0.352431718 | 0.90167808 | 0.963453849 | 0.745 | 0.891700047 |
| SL1344_RS23925_ | 3.686225431 | 3.641168653 | 3.762312473 | 0.118414901 | 0.204699132 | 0.612549221 | 0.227848116 | 0.357610456 | 0.647336273 | 0.4414 | 0.699675595 |
| SL1344_RS23930_ | 3.275511707 | 3.298419291 | 3.178456863 | -0.160833988 | 0.204690907 | -0.82248869 | 0.206000017 | 0.259385723 | 0.558882344 | 0.3888 | 0.663710592 |
| SL1344_RS23935_ | 0.980096257 | 0.982719449 | 0.978028733 | -0.035789244 | 0.23608841 | -0.13827776 | 0.436000002 | 0.723952202 | 0.873961971 | 0.8573 | 0.93186366 |
| SL1344_RS23940_ | 0.227634619 | 0.182214052 | 0.250617764 | 0.066480767 | 0.248966115 | 0.240168669 | 0.402000004 | 0.646386925 | 0.825795518 | 0.7694 | 0.885083321 |
| SL1344_RS23945_ | 0.312975791 | 0.296066422 | 0.321249083 | 0.037094384 | 0.207870124 | 0.163767033 | 0.42238508 | 0.610838172 | 0.801451847 | 0.7278 | 0.858011412 |
| SL1344_RS23950_ | 2.089702247 | 1.99591732 | 2.172233991 | 0.123067254 | 0.274450722 | 0.428742916 | 0.310459702 | 0.51214203 | 0.756262526 | 0.6507 | 0.825420966 |
| SL1344_RS23955_ | -0.033598448 | 0.06400745 | -0.264464489 | -0.295012827 | 0.371498403 | -0.755886576 | 0.222666682 | 0.318727308 | 0.605318913 | 0.446 | 0.68264836 |
| SL1344_RS23960_ | 2.605950526 | 2.640716511 | 2.580336609 | -0.044719203 | 0.227909371 | -0.113123334 | 0.448666669 | 0.74987198 | 0.896189548 | 0.9754 | 0.989496964 |
| SL1344_RS23965_ | 2.11059452 | 2.142876139 | 2.084885946 | -0.099279307 | 0.222016265 | -0.42630472 | 0.315333341 | 0.450809631 | 0.712849875 | 0.6604 | 0.825397699 |
| SL1344_RS23970_ | 2.553291978 | 2.637231197 | 2.497954727 | -0.143804222 | 0.092744598 | -1.51226055 | 0.038666782 | 0.103394181 | 0.346869333 | 0.1434 | 0.383416303 |
| SL1344_RS23975_ | 1.513419379 | 1.592646084 | 1.481904582 | -0.086014119 | 0.202346985 | -0.316751156 | 0.370000005 | 0.812176049 | 0.921537251 | 0.7777 | 0.905001239 |
| SL1344_RS23980_ | -2.128126009 | -2.08215552 | -2.180342037 | -0.103517096 | 0.39149377 | -0.244014943 | 0.38507662 | 0.552291775 | 0.756169229 | 0.6659 | 0.81512643 |
| SL1344_RS23985_ | -4.569577357 | -4.584725276 | -4.55870375 | -0.008435262 | 0.929139171 | -0.007494802 | 0.495333334 | 0.600962526 | 0.791702744 | 0.7237 | 0.852163347 |
| SL1344_RS23990_ | -5.690668439 | -5.738293096 | -5.655330666 | 0.072320338 | 1.255287391 | 0.051569319 | 0.481012659 | 0.57338457 | 0.770978776 | 0.6825 | 0.825454527 |
| SL1344_RS23995_ | -3.999306292 | -4.228745945 | -3.806546859 | 0.443868253 | 0.826936407 | 0.492327174 | 0.266489018 | 0.418854769 | 0.662505151 | 0.5258 | 0.717242779 |
| SL1344_RS24000_ | -1.502386849 | -1.474157043 | -1.540840367 | -0.067249395 | 0.4101535 | -0.154013292 | 0.427714859 | 0.653680609 | 0.828718464 | 0.7696 | 0.883125288 |
| SL1344_RS24005_ | -6.387480088 | -6.112366127 | -6.703761621 | -0.564090292 | 2.717726967 | -0.169495375 | 0.384410398 | 0.569358041 | 0.782210384 | 0.7234 | 0.853242757 |
| SL1344_RS24010_ | -4.68132435 | -4.672719236 | -4.690905841 | -0.045457859 | 0.86395041 | -0.051356135 | 0.478000001 | 0.575280696 | 0.771436053 | 0.6823 | 0.824529265 |
| SL1344_RS24015_ | -1.275872917 | -1.30362857 | -1.261434225 | -0.022667248 | 0.459412544 | -0.053755986 | 0.482666667 | 0.733659697 | 0.878386236 | 0.8457 | 0.927949207 |
| SL1344_RS24020_ | 3.68740548 | 3.672744986 | 3.701749179 | 0.231856288 | 0.471664239 | 0.511917626 | 0.348666673 | 0.452772871 | 0.722605626 | 0.7687 | 0.901419524 |
| SL1344_RS24025_ | -5.828528985 | -6.142623594 | -5.539334653 | 0.624849222 | 1.270126236 | 0.432222786 | 0.319333341 | 0.441054498 | 0.667142388 | 0.5455 | 0.726423382 |
| SL1344_RS24030_ | -4.228502926 | -4.087954149 | -4.533742386 | -0.39485144 | 0.988257034 | -0.39131216 | 0.348434383 | 0.526045459 | 0.746403394 | 0.6564 | 0.815860015 |
| SL1344_RS24035_ | -1.904298735 | -1.778867378 | -2.063447937 | -0.28984171 | 0.32034774 | -0.859210145 | 0.173884098 | 0.267574034 | 0.509974455 | 0.3583 | 0.578354181 |
| SL1344_RS24040_ | -0.92972412 | -0.936664181 | -0.924950553 | 0.006672477 | 0.242499009 | 0.021457332 | 0.490339774 | 0.592851522 | 0.783500483 | 0.7025 | 0.838489265 |
| SL1344_RS24045_ | 0.806870875 | 0.565755141 | 0.839637565 | 0.253932782 | 0.206318476 | 0.370840089 | 0.333777489 | 0.890594449 | 0.958818949 | 0.7018 | 0.873317371 |
| SL1344_RS24050_ | 1.614167236 | 1.561030917 | 1.658544238 | 0.136460298 | 0.200930519 | 0.642240652 | 0.213333349 | 0.306961497 | 0.5948757 | 0.4231 | 0.648499629 |
| SL1344_RS24055_ | 1.456555709 | 1.366606177 | 1.524095722 | 0.155029589 | 0.117784052 | 1.240329278 | 0.089273864 | 0.138983392 | 0.38408121 | 0.2124 | 0.458100036 |
| SL1344_RS24060_ | 1.409035286 | 1.173820427 | 1.440216191 | 0.241820216 | 0.180375738 | 0.815443473 | 0.276482355 | 0.339557252 | 0.624284881 | 0.5721 | 0.779675217 |
| SL1344_RS24065_ | 1.373459347 | 1.283668199 | 1.487727734 | 0.213489174 | 0.139522631 | 1.489043873 | 0.043970787 | 0.094463773 | 0.327852134 | 0.1483 | 0.39011758 |
| SL1344_RS24070_ | 1.299068143 | 1.139890667 | 1.377990407 | 0.241453949 | 0.162110909 | 1.432642875 | 0.054630326 | 0.12112507 | 0.377540989 | 0.1682 | 0.413043574 |
| SL1344_RS24075_ | 0.84284314 | 1.020395973 | 0.788414205 | -0.215102422 | 0.281678469 | -0.708137425 | 0.252666679 | 0.384944905 | 0.657990662 | 0.4956 | 0.730515973 |
| SL1344_RS24080_ | -2.603545724 | -2.817994175 | -2.244521171 | 0.54456958 | 0.513535422 | 1.009520373 | 0.147901425 | 0.219752098 | 0.483324919 | 0.308 | 0.551262898 |
| SL1344_RS24085_ | 3.924144571 | 3.880379009 | 3.956123649 | 0.096247422 | 0.088419084 | 1.041785639 | 0.104596974 | 0.173034601 | 0.455275782 | 0.2348 | 0.488994551 |
| SL1344_RS24090_ | -2.598584361 | -2.430194609 | -2.98925038 | -0.531692625 | 0.529622706 | -0.931442626 | 0.157894761 | 0.24197108 | 0.509865546 | 0.3443 | 0.579895116 |
| SL1344_RS24095_ | -1.930611608 | -2.197988036 | -1.654994506 | 0.569417477 | 0.426513622 | 1.291887745 | 0.067333397 | 0.120361143 | 0.366436685 | 0.1836 | 0.428660022 |
| SL1344_RS24100_ | -5.034414488 | -5.476000923 | -4.744233336 | 0.794606868 | 1.313213953 | 0.541205621 | 0.223850781 | 0.346246249 | 0.613931574 | 0.4477 | 0.651085898 |
| SL1344_RS24105_ | -0.392498929 | -0.629132994 | -0.236045836 | 0.445666513 | 0.376351781 | 1.17826936 | 0.057333409 | 0.161847847 | 0.44322888 | 0.1705 | 0.4172307 |
| SL1344_RS24110_ | -1.989863542 | -2.006218521 | -1.980781071 | 0.057992968 | 0.40115511 | 0.125491474 | 0.452666668 | 0.60560548 | 0.795792037 | 0.7275 | 0.853938954 |
| SL1344_RS24115_ | -2.33588902 | -2.458343236 | -2.225347578 | 0.224780178 | 0.4284531 | 0.483148325 | 0.294470362 | 0.44588384 | 0.674667737 | 0.5498 | 0.734620135 |
| SL1344_RS24120_ | -2.095166292 | -2.098898562 | -2.091131555 | -0.053846024 | 0.548234088 | -0.09174687 | 0.468000001 | 0.695159831 | 0.85509888 | 0.813 | 0.908545007 |
| SL1344_RS24125_ | 2.993359655 | 2.87441448 | 3.252991637 | 0.358793975 | 0.168058146 | 2.157913987 | 0.002668394 | 0.032300093 | 0.212058148 | 0.1008 | 0.331427116 |
| SL1344_RS24130_ | -1.381061765 | -1.468019006 | -1.273754094 | 0.188478082 | 0.253025199 | 0.68764622 | 0.220666682 | 0.32940186 | 0.564660019 | 0.4336 | 0.637808237 |
| SL1344_RS24135_ | -0.773913438 | -0.937504081 | -0.596298411 | 0.356874526 | 0.343323628 | 1.046312208 | 0.095936086 | 0.179023802 | 0.449802941 | 0.2366 | 0.486759614 |
| SL1344_RS24140_ | -2.363468968 | -2.557394092 | -2.000590643 | 0.522810808 | 0.598010702 | 0.855341135 | 0.180666687 | 0.276846791 | 0.549220961 | 0.3646 | 0.614533841 |
| SL1344_RS24145_ | -2.712824573 | -2.907912066 | -2.467912217 | 0.424222318 | 0.465113413 | 0.871381242 | 0.168000022 | 0.264425484 | 0.511599286 | 0.3586 | 0.583545362 |
| SL1344_RS24150_ | -0.964836238 | -1.176361639 | -0.852801756 | 0.313294477 | 0.297472905 | 0.986592122 | 0.153333358 | 0.220360657 | 0.483326157 | 0.3185 | 0.558083876 |
| SL1344_RS24155_ | -0.048346336 | -0.237827621 | 0.093563256 | 0.396021381 | 0.316821155 | 1.216456902 | 0.077333388 | 0.123552035 | 0.384612514 | 0.1852 | 0.433879697 |

| | | | | | | | | | | | |
|---|---|---|---|---|---|---|---|---|---|---|---|
| SL1344_RS24160_ | -5.475634223 | -4.466183011 | -6.069734035 | -1.402684498 | 3.376748596 | -0.408702706 | 0.262491684 | 0.482642495 | 0.725192531 | 0.4883 | 0.723319199 |
| SL1344_RS24165_ | 2.113497473 | 2.348636388 | 1.993700225 | -0.384956714 | 0.422143312 | -0.88599116 | 0.181333353 | 0.235992468 | 0.538357583 | 0.3319 | 0.608779009 |
| SL1344_RS24170_ | 1.924377248 | 1.973609087 | 1.736312809 | -0.276027474 | 0.358604034 | -0.877474076 | 0.18000002 | 0.21557673 | 0.513457153 | 0.322 | 0.596613877 |
| SL1344_RS24175_ | 4.252517029 | 4.357161925 | 4.136828345 | -0.21974906 | 0.365322665 | -0.537935037 | 0.332666674 | 0.450916252 | 0.721816208 | 0.6883 | 0.865095692 |
| SL1344_RS24180_ | 2.88376774 | 2.807839174 | 2.970345728 | 0.139239356 | 0.291790644 | 0.400994508 | 0.396402402 | 0.58544283 | 0.806248873 | 0.8606 | 0.933077503 |
| SL1344_RS24185_ | -0.539345748 | -0.39650437 | -0.658967032 | -0.261960164 | 0.203232031 | -1.231617444 | 0.080666719 | 0.147664856 | 0.38683978 | 0.2176 | 0.456880011 |
| SL1344_RS24190_ | 0.305013678 | -0.155323214 | 0.409030907 | 0.519291432 | 0.299311205 | 1.320208115 | 0.097934752 | 0.143651295 | 0.420534608 | 0.2305 | 0.485782718 |
| SL1344_RS24195_ | -0.466806113 | -0.491253754 | -0.433929221 | 0.01651339 | 0.293218549 | 0.048172042 | 0.483011326 | 0.702232166 | 0.857841364 | 0.8268 | 0.912977241 |
| SL1344_RS24200_ | -6.031060186 | -5.861004295 | -6.239751012 | -0.393869028 | 1.373549565 | -0.260400886 | 0.38774151 | 0.508743254 | 0.721179545 | 0.6205 | 0.780848574 |
| SL1344_RS24205_ | 2.950114356 | 3.216448357 | 2.74095097 | -0.252261853 | 0.757740742 | -0.507956533 | 0.223333348 | 0.485077774 | 0.745263714 | 0.4171 | 0.691261531 |
| SL1344_RS24210_ | 0.01327446 | 0.009291285 | 0.016961641 | 0.006886721 | 0.263574495 | 0.027093659 | 0.49 | 0.708460752 | 0.862353438 | 0.7962 | 0.899044214 |
| SL1344_RS24215_ | 0.098362393 | 0.081512694 | 0.119095061 | 0.04496709 | 0.173643545 | 0.22830891 | 0.397734847 | 0.555018839 | 0.755542335 | 0.6727 | 0.816101013 |
| SL1344_RS24220_ | 1.753957909 | 1.766674171 | 1.737300309 | -0.015020194 | 0.128786978 | -0.109011434 | 0.448000002 | 0.684640865 | 0.848546735 | 0.7868 | 0.894296743 |
| SL1344_RS24225_ | 0.677554775 | 0.715266681 | 0.597615782 | -0.124982688 | 0.194931617 | -0.610960349 | 0.249833457 | 0.37500157 | 0.640748098 | 0.5135 | 0.709264434 |
| SL1344_RS24230_ | -0.640540809 | -0.640698932 | -0.640232219 | 0.04698514 | 0.367660936 | 0.108880124 | 0.450366424 | 0.623534773 | 0.816061985 | 0.8104 | 0.902018711 |
| SL1344_RS24235_ | -0.018666608 | 0.005415718 | -0.045139251 | -0.03658909 | 0.292318247 | -0.12272176 | 0.442000002 | 0.721046452 | 0.871082997 | 0.8173 | 0.910914603 |
| SL1344_RS24240_ | 0.761216683 | 0.784441995 | 0.732387682 | 0.139610229 | 0.426463009 | 0.334411735 | 0.412000003 | 0.619263194 | 0.821635943 | 0.9127 | 0.960405331 |
| SL1344_RS24245_ | 1.178751006 | 1.14841335 | 1.210862988 | 0.082177261 | 0.177197479 | 0.445890339 | 0.313333341 | 0.465854554 | 0.706019353 | 0.6095 | 0.778443331 |
| SL1344_RS24250_ | 3.770008198 | 4.182468247 | 3.552945221 | -0.46937508 | 0.317566268 | -1.74979433 | 0.021319333 | 0.054006912 | 0.273834301 | 0.1197 | 0.359088315 |
| SL1344_RS24255_ | -0.13652432 | 0.072848902 | -0.356673689 | -0.310997628 | 0.404864612 | -0.790051715 | 0.199866773 | 0.306774672 | 0.592082424 | 0.3778 | 0.62996205 |
| SL1344_RS24260_ | -0.640059787 | -0.439746359 | -1.060689694 | -0.608986152 | 0.254805005 | -2.173366483 | 0.007995244 | 0.051912407 | 0.246380762 | 0.1097 | 0.343568436 |
| SL1344_RS24265_ | 0.125027327 | 0.145868006 | 0.097308194 | -0.147321449 | 0.332329435 | -0.430597183 | 0.343104603 | 0.510063458 | 0.745345943 | 0.7229 | 0.855196725 |
| SL1344_RS24270_ | -1.294955234 | -1.319877609 | -1.279708334 | -0.024742379 | 0.464142246 | -0.046824898 | 0.481678881 | 0.723425571 | 0.872720306 | 0.8567 | 0.931846416 |
| SL1344_RS24275_ | -1.370624439 | -1.181260879 | -1.552197391 | -0.38776677 | 0.376841927 | -1.020326852 | 0.137241867 | 0.201145722 | 0.472966054 | 0.2849 | 0.533813134 |
| SL1344_RS24280_ | -0.409318838 | -0.158553771 | -0.645352615 | -0.495458278 | 0.217579226 | -2.215887877 | 0.0020023 | 0.03684486 | 0.20620132 | 0.1049 | 0.3367207 |
| SL1344_RS24285_ | 1.868103809 | 1.8140785 | 1.944731651 | 0.124045898 | 0.107565658 | 1.037608736 | 0.143904091 | 0.200515302 | 0.462242955 | 0.2879 | 0.532248953 |
| SL1344_RS24290_ | 1.784604481 | 1.88297091 | 1.756247252 | -0.126905582 | 0.213598076 | -0.496516301 | 0.343770826 | 0.579034827 | 0.796305515 | 0.7212 | 0.875805056 |
| SL1344_RS24295_ | 2.181128405 | 2.21341941 | 2.150361134 | -0.019533676 | 0.166981659 | -0.118621636 | 0.461333335 | 0.778100918 | 0.903937829 | 0.9016 | 0.955228084 |
| SL1344_RS24300_ | 1.589766679 | 1.608725358 | 1.578914753 | -0.015059411 | 0.126655663 | -0.108631643 | 0.444000002 | 0.663772532 | 0.83373221 | 0.764 | 0.881470985 |
| SL1344_RS24305_ | -0.564984932 | -0.384253301 | -0.632756765 | -0.200754501 | 0.322520947 | -0.583154922 | 0.276666677 | 0.476881177 | 0.71207533 | 0.5711 | 0.770742794 |
| SL1344_RS24310_ | -1.97451661 | -1.809636045 | -2.145847986 | -0.370178824 | 0.413927437 | -0.83480483 | 0.177881433 | 0.272195075 | 0.53405057 | 0.3691 | 0.594813436 |
| SL1344_RS24315_ | -5.593963337 | -5.17440325 | -5.935429346 | -0.578446619 | 1.560817952 | -0.337207521 | 0.344000006 | 0.546803265 | 0.753423957 | 0.6434 | 0.803167984 |
| SL1344_RS24320_ | -6.562635391 | -6.088025516 | -7.109613455 | -0.996089009 | 1.626604825 | -0.55972481 | 0.244000013 | 0.370907661 | 0.609925243 | 0.4748 | 0.670127368 |
| SL1344_RS24325_ | -3.566731953 | -3.427842109 | -3.702091184 | -0.265877755 | 0.561133189 | -0.433182286 | 0.309333342 | 0.45824538 | 0.678380632 | 0.5587 | 0.735700807 |
| SL1344_RS24330_ | -2.853870815 | -2.557470737 | -3.242600455 | -0.690050203 | 0.548965869 | -1.200981513 | 0.097934752 | 0.144934142 | 0.396927122 | 0.2165 | 0.462447741 |
| SL1344_RS24335_ | -4.537054246 | -4.336649054 | -4.81741073 | -0.488561747 | 0.771835244 | -0.580710503 | 0.261333345 | 0.38262587 | 0.618114313 | 0.4876 | 0.680390427 |
| SL1344_RS24340_ | -4.256494369 | -4.333918181 | -4.109152667 | 0.210174954 | 0.894942777 | 0.212757096 | 0.412391742 | 0.614224823 | 0.799225351 | 0.707 | 0.846392433 |
| SL1344_RS24345_ | -3.970220023 | -3.940478538 | -3.998665245 | -0.072909673 | 0.605261023 | -0.109442148 | 0.454000002 | 0.528700781 | 0.733303429 | 0.6375 | 0.790039197 |
| SL1344_RS24350_ | -4.800618426 | -5.33736153 | -4.356859041 | 0.920789018 | 1.206125964 | 0.732081452 | 0.197333351 | 0.317936761 | 0.578880743 | 0.4124 | 0.63876346 |
| SL1344_RS24355_ | -4.808752801 | -4.782435609 | -4.851888943 | 0.024748911 | 1.151376025 | 0.020857952 | 0.491005996 | 0.655907673 | 0.828439095 | 0.7662 | 0.881225702 |
| SL1344_RS24360_ | -5.012790236 | -5.072635302 | -4.920771775 | 0.253381366 | 1.485806386 | 0.154033023 | 0.435333336 | 0.661148151 | 0.832368674 | 0.7776 | 0.885438258 |
| SL1344_RS24365_ | -4.805162677 | -4.67282653 | -4.94764448 | -0.27285898 | 0.872824881 | -0.283963981 | 0.370000005 | 0.510294679 | 0.719031555 | 0.6231 | 0.779640478 |
| SL1344_RS24370_ | -1.648140162 | -1.911266941 | -1.504198349 | 0.391406696 | 0.373630376 | 0.982029854 | 0.14333336 | 0.207868294 | 0.476174392 | 0.2997 | 0.545266159 |
| SL1344_RS24380_ | -1.92768883 | -2.216192581 | -1.667183079 | 0.553024494 | 0.308154323 | 1.706215447 | 0.031312601 | 0.06972411 | 0.266585453 | 0.139 | 0.375546202 |
| SL1344_RS24385_ | -1.33631209 | -1.441366468 | -1.240144094 | 0.191555657 | 0.244343218 | 0.751611692 | 0.200666684 | 0.304498863 | 0.541150849 | 0.3971 | 0.607916351 |
| SL1344_RS24390_ | -5.302328138 | -5.502730979 | -5.080140566 | 0.404462704 | 1.007530135 | 0.35676117 | 0.34133334 | 0.473331731 | 0.692060744 | 0.5861 | 0.756021384 |
| SL1344_RS24395_ | -2.518517605 | -2.464543215 | -2.602643822 | -0.154981015 | 0.471967543 | -0.304546192 | 0.364000005 | 0.547953073 | 0.757026247 | 0.6706 | 0.816398327 |
| SL1344_RS24400_ | -5.152829242 | -4.88261872 | -5.430877876 | -0.519552046 | 0.954539539 | -0.502129113 | 0.28000001 | 0.418890305 | 0.641131292 | 0.5169 | 0.702520046 |
| SL1344_RS24405_ | -4.543425469 | -4.240570147 | -4.951677261 | -0.718159122 | 0.794402226 | -0.841498758 | 0.163890762 | 0.261935008 | 0.507356913 | 0.3495 | 0.571440985 |
| SL1344_RS24410_ | 0.744512039 | 0.728077495 | 0.760703266 | 0.067654605 | 0.220663367 | 0.283732093 | 0.373750838 | 0.565262438 | 0.776627455 | 0.7051 | 0.842484622 |
| SL1344_RS24415_ | -3.285172027 | -3.407423495 | -3.154491068 | 0.322611393 | 0.780140661 | 0.393311025 | 0.328000007 | 0.502655025 | 0.725509882 | 0.6329 | 0.791128943 |
| SL1344_RS24420_ | -0.984925128 | -0.921028229 | -1.064582402 | -0.145475742 | 0.243551344 | -0.556995574 | 0.27600001 | 0.412056481 | 0.641409851 | 0.5186 | 0.703397603 |
| SL1344_RS24425_ | -3.841028718 | -3.647634716 | -4.102712321 | -0.461598356 | 0.623552159 | -0.700225768 | 0.211333349 | 0.320134581 | 0.561257264 | 0.4183 | 0.625199956 |

| | | | | | | | | | | | |
|---|---|---|---|---|---|---|---|---|---|---|---|
| SL1344_RS24430_ | 2.465233472 | 2.588030555 | 2.356630236 | -0.239709693 | 0.325323788 | -0.699554447 | 0.246000013 | 0.337648417 | 0.631822272 | 0.477 | 0.720724233 |
| SL1344_RS24435_ | 0.4577461 | 0.651551013 | 0.334381782 | -0.315720368 | 0.274216459 | -1.133724827 | 0.113333369 | 0.157924257 | 0.43215413 | 0.2387 | 0.499953687 |
| SL1344_RS24440_ | 2.644270278 | 2.830425758 | 2.525410716 | -0.290664949 | 0.10334629 | -2.773107838 | 0.00067292 | 0.021088602 | 0.165673485 | 0.1004 | 0.330848308 |
| SL1344_RS24445_ | 0.289589926 | 0.211968452 | 0.367289868 | 0.214609668 | 0.264453986 | 0.798341014 | 0.165889429 | 0.255520425 | 0.537860221 | 0.3565 | 0.586652588 |
| SL1344_RS24450_ | -1.566787724 | -1.675044715 | -1.44391251 | 0.181759025 | 0.433502153 | 0.386872544 | 0.333111266 | 0.550300734 | 0.7606931 | 0.6485 | 0.808464594 |
| SL1344_RS24455_ | -2.764309377 | -2.901966513 | -2.618478321 | 0.347758151 | 0.578271748 | 0.5642602 | 0.262666678 | 0.392546258 | 0.64036355 | 0.5217 | 0.711695185 |
| SL1344_RS24460_ | 8.159726454 | 8.082665869 | 8.175843363 | 0.094197514 | 0.204519093 | 0.29049934 | 0.33333334 | 0.870481071 | 0.952659043 | 0.7 | 0.872542119 |
| SL1344_RS24465_ | 0.416468896 | 0.404551177 | 0.426580885 | 0.028259377 | 0.526590757 | 0.048432196 | 0.487008661 | 0.879106333 | 0.953030242 | 0.9835 | 0.993025115 |
| SL1344_RS24470_ | 5.99614202 | 6.117784268 | 5.470844066 | -0.64729946 | 0.171445376 | -1.829846708 | 4.68E-05 | 0.074866574 | 0.319954163 | 0.1 | 0.330257709 |
| SL1344_RS24475_ | 1.008992088 | 1.295539441 | 0.972371333 | -0.298960225 | 0.225146234 | -0.583014442 | 0.332000007 | 0.597192047 | 0.809613185 | 0.6961 | 0.869860781 |
| SL1344_RS24480_ | 3.067948013 | 3.0520151 | 3.102524293 | 0.054809223 | 0.127117134 | 0.399137906 | 0.330666674 | 0.500160323 | 0.742083765 | 0.6899 | 0.837802114 |
| SL1344_RS24485_ | -0.550683521 | -0.626471163 | -0.476671826 | 0.142254454 | 0.190042258 | 0.738844617 | 0.220000015 | 0.324267363 | 0.560263791 | 0.429 | 0.63109157 |
| SL1344_RS24490_ | -2.275902612 | -2.313987616 | -2.223111302 | 0.130167972 | 0.484349442 | 0.251609126 | 0.388000004 | 0.604291878 | 0.792034644 | 0.7177 | 0.846031698 |
| SL1344_RS24495_ | -2.173504048 | -2.241777475 | -2.113569106 | 0.106037769 | 0.383707876 | 0.272394584 | 0.380413063 | 0.53849551 | 0.7419531 | 0.6473 | 0.801264913 |
| SL1344_RS24500_ | -0.15836958 | -0.224604579 | -0.047394101 | 0.123148401 | 0.350197492 | 0.31330365 | 0.365756168 | 0.704750815 | 0.862381761 | 0.7271 | 0.871735829 |
| SL1344_RS24505_ | -3.335769463 | -3.613610356 | -3.113269173 | 0.509985405 | 0.501864023 | 0.94952879 | 0.149333359 | 0.234083884 | 0.478020218 | 0.3235 | 0.549136206 |
| SL1344_RS24510_ | -5.513324842 | -5.68892817 | -5.367119216 | 0.354732223 | 1.30500915 | 0.2495019 | 0.3904064 | 0.5596952 | 0.76377077 | 0.6856 | 0.828137826 |
| SL1344_RS24515_ | -4.265394582 | -4.530493236 | -4.03987641 | 0.496182778 | 0.742958189 | 0.646147764 | 0.23666668 | 0.362948791 | 0.597594457 | 0.4571 | 0.656209947 |
| SL1344_RS24520_ | -2.965158852 | -2.87938779 | -3.082388088 | -0.200089865 | 0.486722152 | -0.393858675 | 0.327781486 | 0.484597416 | 0.705267423 | 0.5935 | 0.763016803 |
| SL1344_RS24525_ | -1.538634419 | -1.596162767 | -1.507543649 | 0.038334148 | 0.420217441 | 0.083954072 | 0.472666668 | 0.702969719 | 0.858661473 | 0.8022 | 0.904375083 |
| SL1344_RS24530_ | -2.685698322 | -2.827808542 | -2.506544777 | 0.364214422 | 0.489266925 | 0.694110037 | 0.223333348 | 0.337403956 | 0.587950574 | 0.4514 | 0.6545185 |
| SL1344_RS24535_ | -3.483066896 | -3.127132829 | -3.965070476 | -0.838507576 | 0.680573098 | -1.148270155 | 0.099267197 | 0.158310856 | 0.409327036 | 0.2317 | 0.474728451 |
| SL1344_RS24540_ | -3.308690703 | -3.264397876 | -3.340881071 | -0.062881659 | 0.555481522 | -0.101861019 | 0.447035312 | 0.590145165 | 0.780689871 | 0.6873 | 0.827858043 |
| SL1344_RS24545_ | -3.91975534 | -3.835286462 | -4.013902382 | -0.183118691 | 0.694183146 | -0.238120432 | 0.396666671 | 0.551148896 | 0.755250086 | 0.6649 | 0.812806736 |
| SL1344_RS24550_ | -1.21842855 | -1.193858925 | -1.23986272 | -0.05559821 | 0.390516263 | -0.130904602 | 0.436000002 | 0.671023301 | 0.839129065 | 0.7868 | 0.892058439 |
| SL1344_RS24555_ | -2.578394182 | -2.756596811 | -2.321168625 | 0.50565536 | 0.535471566 | 0.877188828 | 0.153897426 | 0.25008965 | 0.51991205 | 0.3359 | 0.563928508 |
| SL1344_RS24560_ | -2.523852173 | -2.561366162 | -2.472564808 | 0.113662117 | 0.455507402 | 0.225161422 | 0.410393075 | 0.567629393 | 0.767510883 | 0.6899 | 0.829885374 |
| SL1344_RS24565_ | -3.317016326 | -3.018638302 | -3.641133909 | -0.650065889 | 0.634363441 | -0.951262662 | 0.151232537 | 0.230147356 | 0.485544219 | 0.3211 | 0.55628613 |
| SL1344_RS24570_ | -4.080318259 | -4.080318259 | -4.080473052 | -0.065499926 | 0.950149408 | -0.057033277 | 0.471019322 | 0.633210597 | 0.818610506 | 0.7575 | 0.875397435 |
| SL1344_RS24575_ | -2.048200885 | -1.431098683 | -2.642354159 | -1.212462254 | 0.545254903 | -2.230097936 | 0.003334715 | 0.041295221 | 0.22475181 | 0.1027 | 0.334030782 |
| SL1344_RS24580_ | 1.734241024 | 2.62792936 | 1.156637596 | -1.430822693 | 0.203619327 | -6.20172039 | 4.68E-05 | 0.011548039 | 0.134041827 | 0.1 | 0.330257709 |
| SL1344_RS24585_ | -0.157805958 | 0.660767871 | -0.419231502 | -1.035719773 | 0.304682114 | -2.238618381 | 0.001336769 | 0.057734959 | 0.277397776 | 0.1015 | 0.332353254 |
| SL1344_RS24590_ | -0.113076873 | -0.217841582 | -0.024818963 | 0.195053265 | 0.176875077 | 1.044332288 | 0.122666699 | 0.201582693 | 0.451393617 | 0.2847 | 0.51536751 |
| SL1344_RS24595_ | -5.605502855 | -6.020766859 | -5.256304261 | 0.761101041 | 1.270567595 | 0.538424948 | 0.262666678 | 0.387823195 | 0.626670494 | 0.4932 | 0.68860624 |
| SL1344_RS24600_ | -2.679307968 | -2.641139641 | -2.710502287 | -0.082691432 | 0.462629051 | -0.16556787 | 0.434000002 | 0.598383885 | 0.785865323 | 0.7056 | 0.837388412 |
| SL1344_RS24605_ | -3.33319727 | -3.110184617 | -3.454574118 | -0.261232597 | 0.687479684 | -0.343903182 | 0.347768161 | 0.578491476 | 0.773924848 | 0.6668 | 0.822208345 |
| SL1344_RS24610_ | -1.305894887 | -1.384393248 | -1.228343547 | 0.156495507 | 0.587498057 | 0.255382855 | 0.37708195 | 0.665619341 | 0.841677985 | 0.7773 | 0.892445728 |
| SL1344_RS24615_ | -2.564643709 | -2.563272957 | -2.566004016 | 0.010308227 | 0.417693408 | 0.019086104 | 0.488666667 | 0.579485473 | 0.772547208 | 0.6886 | 0.82672092 |
| SL1344_RS24620_ | -2.58396346 | -2.607338381 | -2.555856843 | 0.054837111 | 0.391481007 | 0.123796587 | 0.442000002 | 0.529886168 | 0.734162572 | 0.6368 | 0.789935689 |
| SL1344_RS24625_ | -4.447446699 | -4.485832665 | -4.375649243 | 0.132233109 | 0.962920781 | 0.135046533 | 0.441333336 | 0.609689493 | 0.7983166 | 0.7207 | 0.851006882 |
| SL1344_RS24630_ | -1.527859716 | -1.442379837 | -1.606039096 | -0.177390907 | 0.322869665 | -0.520354656 | 0.279147245 | 0.415532544 | 0.648301087 | 0.5307 | 0.714179254 |
| SL1344_RS24635_ | -2.143533878 | -2.336770328 | -2.04728638 | 0.296287548 | 0.511068353 | 0.519516644 | 0.289140582 | 0.436262576 | 0.683635011 | 0.5675 | 0.751657984 |
| SL1344_RS24640_ | -5.316857991 | -5.318717808 | -5.314717142 | -0.069122773 | 1.258738145 | -0.047924893 | 0.475333334 | 0.604367789 | 0.796245795 | 0.7214 | 0.854788303 |
| SL1344_RS24645_ | -4.275640808 | -4.334305354 | -4.208411325 | 0.080571343 | 0.865263666 | 0.081670349 | 0.459333335 | 0.633955524 | 0.813473214 | 0.7363 | 0.862036255 |
| SL1344_RS24650_ | -3.655978498 | -3.654146527 | -3.657229543 | 0.058992633 | 0.79031877 | 0.069484798 | 0.466666668 | 0.625476674 | 0.809594393 | 0.7482 | 0.866436742 |
| SL1344_RS24655_ | -3.369726373 | -3.446249706 | -3.307588273 | 0.165659976 | 0.625916532 | 0.241317267 | 0.386000004 | 0.546459826 | 0.750868544 | 0.6685 | 0.812470502 |
| SL1344_RS24660_ | 0.579502273 | 0.63848682 | 0.53645696 | -0.036769086 | 0.354372432 | -0.089434341 | 0.457333335 | 0.806801841 | 0.91928182 | 0.9043 | 0.95834508 |
| SL1344_RS24665_ | -0.306119274 | -0.329515454 | -0.28432247 | 0.066123319 | 0.432224096 | 0.148020795 | 0.435709529 | 0.745135357 | 0.88572643 | 0.8435 | 0.926309751 |
| SL1344_RS24670_ | -2.273954717 | -2.192488458 | -2.323312497 | -0.174065313 | 0.472454543 | -0.322445462 | 0.3637575 | 0.541321691 | 0.752012663 | 0.6692 | 0.816947204 |
| SL1344_RS24675_ | -1.693488524 | -1.10446516 | -2.272120478 | -1.13038597 | 0.431028026 | -2.563566409 | 0.003332494 | 0.022293574 | 0.168225928 | 0.1018 | 0.332863826 |
| SL1344_RS24690_ | 6.33748013 | 6.621287725 | 6.038462427 | -0.631278162 | 0.163816393 | -4.227704711 | 4.68E-05 | 0.004175354 | 0.085571097 | 0.1 | 0.330257709 |
| SL1344_RS24695_ | 3.025713482 | 3.567425787 | 2.35205251 | -1.219010362 | 0.123879184 | -9.82383718 | 4.68E-05 | 0.000274979 | 0.027121731 | 0.1 | 0.330257709 |
| SL1344_RS24700_ | 3.301812198 | 3.764186199 | 2.679206789 | -1.084351872 | 0.175946532 | -6.20305346 | 4.68E-05 | 0.008949791 | 0.119490537 | 0.1 | 0.330257709 |

| | | | | | | | | | | | |
|---|---|---|---|---|---|---|---|---|---|---|---|
| SL1344_RS24705_ | 2.888387495 | 3.304812428 | 2.487001599 | -0.848230793 | 0.115518629 | -7.451627064 | 4.68E-05 | 0.000935586 | 0.041937642 | 0.1 | 0.330257709 |
| SL1344_RS24710_ | 2.636862962 | 3.062900426 | 2.214410579 | -0.866523636 | 0.170823883 | -5.103350944 | 4.68E-05 | 0.00276029 | 0.066456155 | 0.1 | 0.330257709 |
| SL1344_RS24730_ | 5.246038119 | 5.531636211 | 4.96847214 | -0.563238798 | 0.057415118 | -9.936420888 | 4.68E-05 | 0.000443483 | 0.031133376 | 0.1 | 0.330257709 |
| SL1344_RS24735_ | 6.129066955 | 6.390545463 | 5.81892834 | -0.561002998 | 0.069193214 | -8.157621556 | 4.68E-05 | 0.000409046 | 0.031866137 | 0.1 | 0.330257709 |
| SL1344_RS24740_ | -3.976974833 | -3.92085336 | -4.033354474 | -0.121850624 | 0.659697352 | -0.172794861 | 0.42504997 | 0.519728189 | 0.727237746 | 0.6319 | 0.788625085 |
| SL1344_RS24745_ | -6.292905872 | -6.709048289 | -5.913561205 | 0.789781212 | 1.542288792 | 0.44857591 | 0.290473027 | 0.441360767 | 0.665603507 | 0.5456 | 0.721747064 |
| SL1344_RS24750_ | -1.104127483 | -1.222819375 | -0.750476347 | 0.441763416 | 0.334511687 | 1.168235668 | 0.137908089 | 0.183756111 | 0.455555787 | 0.2932 | 0.542644015 |
| SL1344_RS24755_ | 0.647299681 | 0.58136979 | 0.696974921 | 0.111825108 | 0.146214843 | 0.738515261 | 0.209193887 | 0.332679889 | 0.57584586 | 0.4329 | 0.640652277 |
| SL1344_RS24760_ | -2.098404 | -2.095929811 | -2.101453293 | 0.086341222 | 0.752434033 | 0.106529216 | 0.441039309 | 0.674533581 | 0.846895075 | 0.7985 | 0.900353799 |
| SL1344_RS24780_ | -1.66012301 | -1.719026843 | -1.594095977 | 0.111408709 | 0.335503677 | 0.317110455 | 0.358000006 | 0.540666028 | 0.745688442 | 0.6505 | 0.8034288 |
| SL1344_RS24785_ | 4.679888663 | 4.828511606 | 4.531128975 | 0.013573586 | 0.560146377 | 0.029081186 | 0.475016656 | 0.865287938 | 0.950152402 | 1 | 1 |
| SL1344_RS24790_ | 4.508182781 | 4.587869259 | 4.418462554 | 0.00480699 | 0.40963242 | 0.011725726 | 0.494 | 0.954946226 | 0.984147054 | 1 | 1 |
| SL1344_RS24795_ | -1.71271877 | -1.782084111 | -1.667639477 | 0.07697186 | 0.538456025 | 0.131896694 | 0.435333336 | 0.676406195 | 0.845089797 | 0.7945 | 0.897474767 |
| SL1344_RS24800_ | -5.789399468 | -6.056874937 | -5.62900137 | 0.147802084 | 1.9569827 | 0.071487721 | 0.470000001 | 0.676627607 | 0.845208459 | 0.8048 | 0.904733759 |
| SL1344_RS24805_ | -2.630822253 | -2.303582594 | -2.849447591 | -0.50103526 | 0.514522184 | -0.911641233 | 0.16322454 | 0.250198584 | 0.510687882 | 0.3448 | 0.582749333 |
| SL1344_RS24810_ | -1.310655259 | -1.068244385 | -1.49585675 | -0.207953663 | 0.733464533 | -0.258869926 | 0.404000004 | 0.80907451 | 0.919283589 | 0.8498 | 0.934318691 |
| SL1344_RS24820_ | 3.41585965 | 3.857721934 | 2.922850061 | -0.953555508 | 0.217124697 | -4.424194981 | 4.68E-05 | 0.029617787 | 0.209489706 | 0.1 | 0.330257709 |
| SL1344_RS24825_ | 3.841361466 | 4.501762768 | 3.766179835 | -0.735664059 | 0.562399754 | -0.879756879 | 0.217188556 | 0.289126569 | 0.592841661 | 0.4012 | 0.681115534 |
| SL1344_RS24830_ | -0.255470218 | -0.095113227 | -0.485994847 | -0.45309817 | 0.50307957 | -0.876475533 | 0.181212545 | 0.225641264 | 0.518744681 | 0.3257 | 0.59118913 |
| SL1344_RS24835_ | -3.605841066 | -3.396314579 | -3.851825695 | -0.456491438 | 0.608329281 | -0.724472503 | 0.212000016 | 0.319938699 | 0.568660341 | 0.4248 | 0.635810334 |
| SL1344_RS24840_ | -5.135046243 | -4.617638154 | -5.896341848 | -1.299578471 | 1.374252045 | -0.847195391 | 0.129333364 | 0.230344407 | 0.501680304 | 0.2851 | 0.520576676 |
| SL1344_RS24845_ | -6.838861161 | -6.803474041 | -6.856218707 | -0.141362484 | 2.058139566 | -0.063375364 | 0.476666668 | 0.606302896 | 0.796421024 | 0.7249 | 0.854677431 |
| SL1344_RS24850_ | -4.365579812 | -4.23076755 | -4.49635495 | -0.261489502 | 0.691001557 | -0.344786552 | 0.352000006 | 0.45632402 | 0.672845578 | 0.5571 | 0.730417043 |
| SL1344_RS24860_ | 2.226312784 | 2.274359119 | 2.17079068 | -0.098372005 | 0.877517644 | -0.117737123 | 0.449033979 | 0.942881391 | 0.979703978 | 1 | 1 |
| SL1344_RS24865_ | -3.382028169 | -3.206612847 | -3.732007966 | -0.500843216 | 0.695298862 | -0.66394925 | 0.222666682 | 0.345223752 | 0.604730547 | 0.4566 | 0.669623636 |
| SL1344_RS24875_ | -4.993159688 | -4.984494254 | -5.003836358 | -0.063588775 | 1.0370194 | -0.06326641 | 0.473017989 | 0.593705403 | 0.78491504 | 0.7029 | 0.840185661 |
| SL1344_RS24885_ | -1.249757905 | -1.157718241 | -1.32913763 | -0.088727659 | 0.702281611 | -0.109538701 | 0.463690874 | 0.833252989 | 0.932006447 | 0.946 | 0.976155221 |
| SLnc0002_ | -3.29386271 | -3.455406959 | -3.016178816 | 0.349755516 | 0.849469724 | 0.367658692 | 0.343104603 | 0.627906252 | 0.811787022 | 0.689 | 0.841988519 |
| SLnc0010_ | -3.15226428 | -3.195150515 | -3.128102602 | 0.07971997 | 0.63342065 | 0.12144358 | 0.450666669 | 0.635156313 | 0.815036886 | 0.7459 | 0.867987853 |
| SLnc0010a_ | -0.434472611 | -0.40364502 | -0.476688242 | 0.025759891 | 0.356008427 | 0.069017726 | 0.474000001 | 0.729551891 | 0.876055921 | 0.8838 | 0.944812946 |
| SLnc0014_ | 1.069689247 | 1.534607061 | 0.940955822 | -0.609971208 | 0.38092521 | -1.493875283 | 0.038000117 | 0.111779293 | 0.380378149 | 0.1434 | 0.387966843 |
| SLnc0022_ | -0.381549829 | -0.310263005 | -0.60228157 | -0.28850292 | 0.307357065 | -0.805736926 | 0.213857444 | 0.300231265 | 0.580885464 | 0.4274 | 0.65660725 |
| SLnc0030_ | 2.605807267 | 2.294271544 | 2.725311709 | 0.245567927 | 0.566344613 | 0.383262696 | 0.334000007 | 0.85988457 | 0.947388286 | 0.7003 | 0.872671677 |
| SLnc0030a_ | -1.073100052 | -1.418044724 | -0.972464976 | 0.390747688 | 0.621724907 | 0.3865975 | 0.346000006 | 0.827981195 | 0.9305238 | 0.7309 | 0.885697127 |
| SLnc0031_ | -3.114305249 | -3.302401499 | -2.975846736 | 0.226258547 | 0.681477932 | 0.325626074 | 0.367088613 | 0.584346652 | 0.784041107 | 0.705 | 0.844074894 |
| SLnc0035_ | 4.686051351 | 4.597978996 | 4.931282368 | 0.37036036 | 0.245591637 | 1.803558248 | 4.68E-05 | 0.046326267 | 0.255589469 | 0.1 | 0.330257709 |
| SLnc0039_ | 3.197840635 | 2.974978609 | 3.299003283 | 0.318545925 | 0.449536296 | 0.636063853 | 0.250000012 | 0.419424258 | 0.699057787 | 0.4738 | 0.727394159 |
| SLnc0048_ | -1.359461159 | -1.475145348 | -1.249624365 | 0.223340797 | 0.263563481 | 0.810708899 | 0.18000002 | 0.290804641 | 0.536240302 | 0.381 | 0.598342755 |
| SLnc1001_ | -5.337317074 | -5.06917117 | -5.737891775 | -0.346435585 | 1.745517288 | -0.178092625 | 0.41972019 | 0.670662184 | 0.839626432 | 0.7943 | 0.89630681 |
| SLnc1002_ | -0.301647486 | -0.211895652 | -0.419349306 | -0.240424405 | 0.366674896 | -0.628538848 | 0.208000016 | 0.316232451 | 0.604157887 | 0.4081 | 0.636461258 |
| SLnc1006_ | -0.073480374 | -0.071736446 | -0.074914382 | -0.044857316 | 0.347153039 | -0.099047656 | 0.449700202 | 0.607658346 | 0.812410187 | 0.7988 | 0.903280535 |
| SLnc1008_ | -3.391296263 | -3.370552026 | -3.443480234 | -0.002294293 | 0.81779033 | -0.002357764 | 0.498666667 | 0.725405311 | 0.872425999 | 0.8151 | 0.913238413 |
| SLnc1013_ | -1.952792001 | -2.35947393 | -1.712707561 | 0.616046802 | 0.504713707 | 1.137665483 | 0.102598308 | 0.164044659 | 0.430082474 | 0.2318 | 0.482784145 |
| SLnc1014_ | -2.172603302 | -2.058465612 | -2.276409925 | -0.150322592 | 0.865548214 | -0.158967676 | 0.432000003 | 0.762792329 | 0.895705495 | 0.8756 | 0.941297605 |
| SLnc1015_ | -2.051387836 | -2.445119886 | -1.934254811 | 0.437421039 | 0.72433555 | 0.38196241 | 0.359760166 | 0.807443157 | 0.920060802 | 0.7513 | 0.894308719 |
| SLnc1016_ | -2.538096257 | -2.742130718 | -2.342701196 | 0.395974263 | 0.381066032 | 0.971136682 | 0.135909422 | 0.221550864 | 0.463040023 | 0.3063 | 0.532111972 |
| SLnc1017_ | -0.673183949 | -0.559279025 | -0.76834036 | -0.197167526 | 1.095750487 | -0.179631031 | 0.4170553 | 0.808546593 | 0.923049643 | 0.9196 | 0.962153394 |
| SLnc1018_ | -3.228644185 | -3.565159112 | -2.839342703 | 0.728729702 | 0.567276876 | 1.242156814 | 0.096666709 | 0.149222221 | 0.393213925 | 0.2244 | 0.463847852 |
| SLnc1019_ | -2.783378407 | -2.727186038 | -2.827876259 | -0.070719527 | 0.597718907 | -0.11136198 | 0.452365092 | 0.658244712 | 0.830033943 | 0.7639 | 0.87895452 |
| SLnc1021_ | -2.992778263 | -2.884616552 | -3.070556575 | -0.192659338 | 0.52932101 | -0.337397554 | 0.351099273 | 0.515613853 | 0.728573585 | 0.6351 | 0.792031586 |
| SLnc1022_ | -2.314141649 | -2.371263588 | -2.226362014 | 0.178390975 | 0.478082845 | 0.357875249 | 0.350433051 | 0.510463464 | 0.729357965 | 0.6374 | 0.793588382 |
| SLnc1024_ | -2.187406004 | -2.788948654 | -2.087861343 | 0.646502215 | 0.621106757 | 0.753741303 | 0.293333343 | 0.385072507 | 0.658315406 | 0.5947 | 0.794588353 |
| SLnc1025_ | -2.102390054 | -2.277569901 | -1.98401758 | 0.259344845 | 0.865176822 | 0.260220249 | 0.383077953 | 0.583953015 | 0.797531059 | 0.8043 | 0.902677677 |

| | | | | | | | | | | | |
|---|---|---|---|---|---|---|---|---|---|---|---|
| SLnc1026_ | -2.912429902 | -2.766714493 | -3.284000531 | -0.437903175 | 0.734172621 | -0.541994809 | 0.299333342 | 0.473072672 | 0.714700627 | 0.5927 | 0.786541412 |
| SLnc1027_ | -1.509526553 | -1.756361584 | -1.4133839 | 0.29527821 | 0.506938127 | 0.495835692 | 0.311125924 | 0.521485547 | 0.749604649 | 0.6471 | 0.820581778 |
| SLnc1028_ | -3.371033522 | -3.030381773 | -3.708760486 | -0.701099208 | 0.672227725 | -0.976001867 | 0.125333365 | 0.207267548 | 0.465385845 | 0.2897 | 0.528632943 |
| SLnc1029_ | -3.56762802 | -3.718648648 | -3.482276968 | 0.227708644 | 0.655883483 | 0.304776743 | 0.365089945 | 0.53117215 | 0.742608948 | 0.6492 | 0.804315359 |
| sRNA1_ | 3.306790106 | 3.009101595 | 3.336854865 | 0.33497752 | 0.288486425 | 0.632408468 | 0.33333334 | 0.51417213 | 0.764582119 | 0.7 | 0.872542119 |
| sRNA10_ | 1.494364898 | 1.513062339 | 1.484582202 | -0.049989327 | 0.181103563 | -0.27190947 | 0.376666672 | 0.51118281 | 0.746419248 | 0.6996 | 0.833787064 |
| sRNA11_ | -2.58036418 | -2.82561438 | -2.36990147 | 0.43657506 | 0.484594067 | 0.855311102 | 0.169220541 | 0.264788373 | 0.523004486 | 0.3617 | 0.592130158 |
| sRNA12_ | -5.388312842 | -5.391555357 | -5.379380676 | -0.012327399 | 1.249045557 | -0.009486536 | 0.496 | 0.599381395 | 0.789177369 | 0.7136 | 0.843874887 |
| sRNA15_ | -6.402921865 | -5.869401173 | -7.115741608 | -1.253929948 | 1.5225692 | -0.777845845 | 0.184543657 | 0.288994377 | 0.535482248 | 0.386 | 0.600214155 |
| sRNA16_ | -2.233470183 | -2.728272292 | -1.929599506 | 0.808464573 | 0.539640137 | 1.41938229 | 0.052631662 | 0.102501832 | 0.347194977 | 0.1587 | 0.404445178 |
| sRNA18_ | -6.354120028 | -6.655021243 | -6.194719483 | 0.392604914 | 1.823191175 | 0.196352341 | 0.410000003 | 0.566807409 | 0.77338143 | 0.7082 | 0.847382378 |
| sRNA19_ | -7.238849398 | -7.189035884 | -7.304344977 | -0.207767046 | 2.607468219 | -0.069821065 | 0.467688209 | 0.583090479 | 0.784167593 | 0.7217 | 0.852688697 |
| sRNA2_ | 3.544477131 | 3.291236156 | 3.581036117 | 0.283306487 | 0.414431413 | 0.711930981 | 0.228000014 | 0.358362008 | 0.651497739 | 0.4009 | 0.680910846 |
| sRNA21_ | -1.52148465 | -1.304271909 | -1.657071023 | -0.341670477 | 0.29754252 | -1.107971354 | 0.134666696 | 0.192915165 | 0.443138325 | 0.2795 | 0.518772771 |
| sRNA22_ | -2.414933707 | -2.302333326 | -2.498940843 | -0.160102677 | 0.526472952 | -0.294968299 | 0.376000005 | 0.607928698 | 0.798586875 | 0.7144 | 0.849079725 |
| sRNA23_ | -4.527965706 | -4.526573685 | -4.529728748 | 0.050664072 | 1.054011711 | 0.040756148 | 0.482000001 | 0.638115482 | 0.817761553 | 0.7687 | 0.879565414 |
| sRNA24_ | -5.409826751 | -5.429486022 | -5.395927618 | 0.025141577 | 1.167751559 | 0.018666381 | 0.492338441 | 0.574249905 | 0.770066375 | 0.6883 | 0.826920148 |
| sRNA25_ | -7.430086429 | -6.614868886 | -8.460387774 | -1.497613824 | 3.932438807 | -0.337392378 | 0.306462367 | 0.521587884 | 0.738261018 | 0.6033 | 0.775761511 |
| sRNA26_ | -5.396404551 | -5.421215812 | -5.358075093 | 0.062531496 | 1.177434182 | 0.048116225 | 0.475682879 | 0.567705709 | 0.766870902 | 0.6911 | 0.828890422 |
| sRNA28_ | -5.151234609 | -4.501283817 | -5.959850715 | -1.413278283 | 1.233520668 | -1.071015374 | 0.107928086 | 0.173996396 | 0.427432544 | 0.248 | 0.489969542 |
| sRNA29_ | -5.421240989 | -5.137167513 | -5.945370665 | -0.813224452 | 1.51412655 | -0.477916711 | 0.275333344 | 0.415213816 | 0.665151416 | 0.5476 | 0.733455983 |
| sRNA31_ | -5.036733404 | -4.950047955 | -5.095026816 | -0.192184864 | 1.135876099 | -0.148291703 | 0.426382415 | 0.610088578 | 0.795734283 | 0.7278 | 0.854909908 |
| sRNA32_ | -4.963945579 | -4.4750015 | -5.254418294 | -0.871608669 | 1.233152707 | -0.630863352 | 0.24266668 | 0.385294117 | 0.631610729 | 0.5019 | 0.69821827 |
| sRNA33_ | -3.091656023 | -2.87748882 | -3.219969318 | -0.310930692 | 0.598635966 | -0.46340608 | 0.310666675 | 0.499245655 | 0.716490047 | 0.5962 | 0.770571792 |
| sRNA5_ | -4.574312792 | -3.875778634 | -5.36984125 | -1.558203696 | 1.097355912 | -1.441816048 | 0.034000131 | 0.090390592 | 0.320464452 | 0.1474 | 0.388307312 |
| sRNA7_ | -1.168266459 | -1.051273088 | -1.275501938 | -0.179077856 | 0.613384825 | -0.276240892 | 0.386666671 | 0.672896821 | 0.846674957 | 0.8094 | 0.908374468 |
| STnc10_ | -5.835611507 | -5.394819407 | -6.300173315 | -0.872435798 | 1.806425021 | -0.433320168 | 0.29113925 | 0.480473853 | 0.709964743 | 0.5879 | 0.766004127 |
| STnc1010_ | -2.476744924 | -2.519474401 | -2.431953257 | 0.097795482 | 0.415179836 | 0.2125236 | 0.40666667 | 0.543275626 | 0.749187916 | 0.6668 | 0.813508505 |
| STnc1030_ | -1.918608592 | -1.993162643 | -1.836443981 | 0.148433316 | 0.334603097 | 0.408151725 | 0.319333341 | 0.482359623 | 0.696781135 | 0.5909 | 0.757186457 |
| STnc1040_ | -5.834141716 | -6.236210565 | -5.437255345 | 0.83813279 | 1.291167815 | 0.586082598 | 0.250666679 | 0.384478249 | 0.617338886 | 0.4812 | 0.677555259 |
| STnc1065_ | 4.475231177 | 4.030775609 | 4.525906385 | 0.437466 | 0.141873017 | 1.016109515 | 0.190666685 | 0.2061586 | 0.508276126 | 0.3618 | 0.619630796 |
| STnc1070_ | -0.80739995 | -0.744474083 | -0.873515708 | -0.1888926 | 0.328796434 | -0.527998314 | 0.281812135 | 0.409481279 | 0.663280942 | 0.5623 | 0.743071686 |
| STnc1080_ | -3.919731975 | -3.778692639 | -4.025621333 | -0.247406286 | 0.676309986 | -0.318663253 | 0.358427721 | 0.525901437 | 0.730952307 | 0.6313 | 0.787599941 |
| STnc1090_ | -1.275492774 | -1.199196782 | -1.391500202 | -0.247741752 | 0.49627721 | -0.459915589 | 0.28400001 | 0.404165095 | 0.670665572 | 0.5635 | 0.75211448 |
| STnc1100_ | -4.282517829 | -3.850267145 | -4.636939064 | -0.824152016 | 0.961296577 | -0.797815477 | 0.183211212 | 0.281456041 | 0.541247834 | 0.3736 | 0.601075305 |
| STnc1130_ | -0.296113693 | 0.071843939 | -0.781047301 | -0.843651599 | 0.246497753 | -3.34039984 | 4.68E-05 | 0.01490797 | 0.137017069 | 0.1002 | 0.330550721 |
| STnc1150_ | 0.524258264 | 0.657760996 | 0.404208054 | -0.0634205 | 0.454388158 | -0.149682373 | 0.445333335 | 0.763769571 | 0.898282902 | 0.9032 | 0.955045017 |
| STnc1160_ | 2.134854209 | 2.001076149 | 2.257471897 | 0.179339628 | 1.238386427 | 0.137866363 | 0.446369089 | 0.965112492 | 0.987864847 | 1 | 1 |
| STnc1170_ | -3.304657777 | -3.615397006 | -2.505457227 | 1.03767328 | 0.69654579 | 1.304755635 | 0.10326453 | 0.141815201 | 0.3981796 | 0.2307 | 0.476603326 |
| STnc1180_ | -2.267847996 | -2.183204863 | -2.444897786 | -0.187158692 | 0.641583714 | -0.247811053 | 0.404397072 | 0.739066795 | 0.882543979 | 0.7904 | 0.907183278 |
| STnc1190_ | 2.375127638 | 2.379704583 | 2.370378805 | -0.020184466 | 0.401030278 | -0.052649481 | 0.480666667 | 0.814634901 | 0.925864781 | 0.9772 | 0.990234673 |
| STnc1200_ | -1.772276148 | -1.828597935 | -1.703239887 | 0.115922705 | 0.364298213 | 0.283598297 | 0.377748173 | 0.574850611 | 0.769814455 | 0.6738 | 0.820201261 |
| STnc1210_ | -4.325369173 | -3.949961523 | -4.518622557 | -0.465398369 | 0.876628169 | -0.484189902 | 0.304666675 | 0.501116872 | 0.713360771 | 0.5838 | 0.763279845 |
| STnc1220_ | -6.856543108 | -6.896159616 | -6.805139945 | 0.073975441 | 1.650535894 | 0.043443447 | 0.480346436 | 0.522432917 | 0.732124885 | 0.6415 | 0.794715599 |
| STnc1275_ | -0.538086941 | -0.292853482 | -0.734197009 | -0.453837726 | 0.306594461 | -1.429963995 | 0.043304566 | 0.098024493 | 0.336638958 | 0.1523 | 0.396272996 |
| STnc1280_ | -1.730186945 | -2.20876948 | -1.374937388 | 0.825730708 | 0.364723227 | 2.220874296 | 0.008667196 | 0.042030627 | 0.220700704 | 0.1089 | 0.342752862 |
| STnc1285_ | 2.48417184 | 3.503262695 | 1.374840762 | -2.132663072 | 0.50385747 | -4.470229326 | 4.68E-05 | 0.003491462 | 0.078376802 | 0.1 | 0.330257709 |
| STnc1290_ | 1.145503205 | 1.185525852 | 1.108281777 | -0.06577797 | 0.614508055 | -0.093747465 | 0.469686877 | 0.881914822 | 0.955133759 | 0.9988 | 0.999491085 |
| STnc130_ | -4.539491815 | -4.779121138 | -4.404496726 | 0.472686359 | 1.198110624 | 0.336656729 | 0.320453039 | 0.451453959 | 0.699097099 | 0.6007 | 0.76749653 |
| STnc1320_ | -4.52034661 | -4.580793203 | -4.451997057 | 0.137071545 | 0.822797558 | 0.153988571 | 0.429333336 | 0.550875754 | 0.753529132 | 0.6605 | 0.80868797 |
| STnc1340_ | -1.260899525 | -1.533954359 | -0.62308594 | 0.939352227 | 0.682639057 | 1.397097879 | 0.040666776 | 0.117789687 | 0.385957472 | 0.1476 | 0.393461898 |
| STnc1380_ | -3.687118892 | -3.519188397 | -3.830971544 | -0.316605444 | 0.599055316 | -0.494777155 | 0.290000009 | 0.425849575 | 0.651205204 | 0.5267 | 0.710902989 |
| STnc1385_ | 3.883197648 | 3.855798719 | 3.910862358 | 0.059916126 | 0.345164808 | 0.144869259 | 0.441333336 | 0.725650753 | 0.886086988 | 0.9967 | 0.9985796 |

| | | | | | | | | | | |
|---|---|---|---|---|---|---|---|---|---|---|
| STnc1390_ | -4.335355577 | -3.966422282 | -4.598734943 | -0.568654158 | 1.221953903 | -0.43128453 | 0.316000008 | 0.537349967 | 0.75129291 | 0.6377 | 0.803447341 |
| STnc140_ | -5.902102483 | -6.503724636 | -5.414596185 | 1.057088078 | 1.39938318 | 0.695965601 | 0.198534328 | 0.305419503 | 0.553472687 | 0.397 | 0.612005794 |
| STnc1400_ | 0.502272692 | 0.497715031 | 0.505057146 | -0.074142946 | 0.538119289 | -0.143627126 | 0.439706864 | 0.730547632 | 0.883311874 | 0.9067 | 0.959460324 |
| STnc1405_ | 0.55831788 | 1.478684853 | -0.602044345 | -2.094301851 | 0.201427769 | -10.41841754 | 4.68E-05 | 0.000746206 | 0.03477911 | 0.1 | 0.330257709 |
| STnc1410_ | -2.697964164 | -2.781022748 | -2.563867633 | 0.203417444 | 0.489367379 | 0.357688077 | 0.34333334 | 0.561848856 | 0.758369887 | 0.6566 | 0.809240278 |
| STnc1460_ | 6.698862551 | 6.392696233 | 7.039903391 | 0.233846522 | 0.779921569 | 0.641615639 | 0.221333348 | 0.446403812 | 0.719288702 | 0.4 | 0.680350104 |
| STnc1490_ | -0.049194072 | -0.153829311 | 0.023600333 | 0.170377606 | 0.201666047 | 0.809331857 | 0.196000018 | 0.295561136 | 0.551707535 | 0.405 | 0.625585292 |
| STnc150_ | -0.075012068 | -0.275599835 | 0.1109245 | 0.439070074 | 0.354759521 | 1.315173089 | 0.030666813 | 0.134615704 | 0.40667041 | 0.1308 | 0.370459884 |
| STnc1500_ | -3.096972889 | -2.737757529 | -3.376186318 | -0.659419831 | 0.550941993 | -1.153289511 | 0.10200004 | 0.169081949 | 0.419146572 | 0.2426 | 0.481434329 |
| STnc1510_ | -0.299387237 | -0.5982186 | -0.138170929 | 0.235337376 | 0.627915461 | 0.276124895 | 0.380413063 | 0.895981933 | 0.960848504 | 0.802 | 0.915788092 |
| STnc1530_ | -5.98188367 | -6.114519973 | -5.84184881 | 0.269113667 | 1.437127694 | 0.178551719 | 0.425716192 | 0.568653779 | 0.762883691 | 0.6665 | 0.81419494 |
| STnc1540_ | -1.384354179 | -1.577777851 | -0.987326695 | 0.562230966 | 0.407113551 | 1.260324431 | 0.096602308 | 0.140468469 | 0.400597385 | 0.223 | 0.477942634 |
| STnc1545_ | 0.730625882 | 0.547155954 | 1.025178349 | 0.469987759 | 0.21076171 | 2.215503105 | 0.004664544 | 0.033168232 | 0.204587158 | 0.1039 | 0.335736598 |
| STnc1550_ | 0.759045368 | 0.685709365 | 0.822948292 | -0.083081142 | 0.494084478 | -0.176582093 | 0.454000002 | 0.81431247 | 0.923177988 | 0.9724 | 0.987838514 |
| STnc1560_ | 3.704312949 | 2.679878786 | 3.767596746 | 1.026842993 | 0.924524714 | 0.455263424 | 0.33333334 | 0.625946797 | 0.833983716 | 0.7 | 0.872542119 |
| STnc1565_ | -0.622522774 | -0.833106971 | -0.522768418 | 0.222411803 | 0.423572718 | 0.472026747 | 0.33733334 | 0.552742798 | 0.775343728 | 0.6977 | 0.850318675 |
| STnc1580_ | -1.190633868 | -1.291592837 | -1.145388309 | 0.111273029 | 0.395635309 | 0.259556755 | 0.40066667 | 0.66670111 | 0.839615415 | 0.7794 | 0.894885064 |
| STnc1590_ | 1.665528462 | 1.475181491 | 1.710869463 | 0.22508915 | 0.467508612 | 0.248917689 | 0.335109934 | 0.90373431 | 0.964572597 | 0.7084 | 0.87615365 |
| STnc1600_ | -1.45998832 | -1.339136089 | -1.629081876 | -0.299530159 | 0.340931247 | -0.828015715 | 0.18000002 | 0.268773013 | 0.530748334 | 0.3639 | 0.592220386 |
| STnc1640_ | -1.540273823 | -1.651026993 | -1.477424588 | 0.141538171 | 0.372560619 | 0.348218674 | 0.362666672 | 0.596959626 | 0.789595688 | 0.6989 | 0.842570153 |
| STnc1650_ | -4.931036708 | -5.072334393 | -4.779537011 | 0.308005234 | 0.923215031 | 0.282424506 | 0.362425055 | 0.497141522 | 0.709985489 | 0.6153 | 0.773180636 |
| STnc1660_ | -0.497248533 | -0.437943301 | -0.543506596 | -0.099090118 | 0.217462818 | -0.434230441 | 0.313124592 | 0.482714894 | 0.699958624 | 0.5752 | 0.751371248 |
| STnc1670_ | -0.54886303 | -0.597377444 | -0.491817042 | 0.106710686 | 1.050745924 | 0.092755346 | 0.442000002 | 0.741194077 | 0.891746279 | 0.8959 | 0.951601281 |
| STnc1675_ | 6.260641609 | 6.127536105 | 6.316722153 | 0.126525062 | 0.161080187 | 0.880308713 | 0.196000018 | 0.285076156 | 0.58776492 | 0.3594 | 0.636556105 |
| STnc1680_ | 1.024931494 | 1.039840489 | 1.009705408 | -0.028493033 | 1.096926962 | -0.025848732 | 0.484000001 | 0.938656115 | 0.978173583 | 0.9958 | 0.998192023 |
| STnc1685_ | 5.987193826 | 3.612146621 | 6.037293627 | 2.421671389 | 0.112095363 | 1.074452251 | 0.0020023 | 0.172776607 | 0.470266864 | 0.1024 | 0.333771085 |
| STnc1690_ | -3.174918602 | -2.735419931 | -3.604432848 | -0.857919172 | 0.555063272 | -1.515662859 | 0.038000117 | 0.08612524 | 0.304745391 | 0.1514 | 0.391348429 |
| STnc170_ | -4.877960329 | -4.127321696 | -5.756274199 | -1.61689054 | 1.101185551 | -1.344677674 | 0.046000096 | 0.109861936 | 0.345440503 | 0.1604 | 0.402835002 |
| STnc1700_ | -3.690053658 | -4.329666212 | -3.305802013 | 0.989108912 | 0.800390106 | 1.129847031 | 0.104000039 | 0.16017429 | 0.417915502 | 0.2345 | 0.481864733 |
| STnc1710_ | 0.257923645 | 0.23646224 | 0.272078895 | 0.047192048 | 0.20033633 | 0.202450133 | 0.405063295 | 0.59316565 | 0.788503651 | 0.7099 | 0.846083054 |
| STnc1760_ | -3.109660947 | -3.490345138 | -2.786100384 | 0.690491853 | 0.48764423 | 1.359094114 | 0.06200007 | 0.118961808 | 0.344145452 | 0.1913 | 0.43147554 |
| STnc1820_ | -1.340887179 | -1.385848983 | -1.27475333 | 0.168367045 | 0.605824854 | 0.27045147 | 0.377333338 | 0.665519784 | 0.840286667 | 0.7632 | 0.883111121 |
| STnc1830_ | -2.577145419 | -2.589225457 | -2.560795805 | 0.040194748 | 0.44029251 | 0.086519168 | 0.464357097 | 0.570993499 | 0.769432677 | 0.6815 | 0.826133587 |
| STnc1870_ | -5.458449306 | -5.480820048 | -5.430373356 | 0.055683821 | 1.006713595 | 0.050192406 | 0.474350434 | 0.52975397 | 0.731174214 | 0.6324 | 0.786042592 |
| STnc1880_ | -2.363308721 | -2.421928731 | -2.25725144 | 0.104416798 | 0.616463071 | 0.146104719 | 0.431045972 | 0.741790955 | 0.882190831 | 0.8067 | 0.909981458 |
| STnc1890_ | -5.838731596 | -6.289357183 | -5.649772784 | 0.608424427 | 1.602619317 | 0.348145143 | 0.356000006 | 0.509024754 | 0.733405357 | 0.6591 | 0.815701695 |
| STnc190_ | -1.499799241 | -1.01841261 | -2.222180294 | -1.215047473 | 0.36272684 | -3.301017237 | 4.68E-05 | 0.015768388 | 0.139919347 | 0.1 | 0.330257709 |
| STnc1920_ | -0.544751691 | -0.64665466 | -0.2813272 | 0.364537297 | 0.304060258 | 1.093159833 | 0.135909422 | 0.19489434 | 0.469874085 | 0.2837 | 0.532728403 |
| STnc1990_ | -1.185653577 | -1.349711065 | -0.942836971 | 0.412061925 | 0.265173185 | 1.456925582 | 0.063333401 | 0.108169336 | 0.337329118 | 0.178 | 0.419485855 |
| STnc20_ | -2.972589507 | -2.9642166 | -2.982768055 | 0.001645613 | 0.584177123 | 0.003295445 | 0.498334444 | 0.645449281 | 0.822138552 | 0.7538 | 0.872806891 |
| STnc200_ | -0.568880236 | -0.247835888 | -1.15157358 | -0.888697789 | 0.47702755 | -1.792345065 | 0.009327606 | 0.080572827 | 0.322091057 | 0.1075 | 0.341070409 |
| STnc2000_ | 0.649765951 | 0.600797449 | 0.705119772 | -0.070888896 | 0.647685331 | -0.128990394 | 0.461333335 | 0.796883327 | 0.916437624 | 0.9601 | 0.982975221 |
| STnc2010_ | -0.927560011 | -1.048082928 | -0.645744432 | 0.36167483 | 0.430261881 | 0.730805128 | 0.241172565 | 0.363301116 | 0.635614789 | 0.4855 | 0.712975055 |
| STnc2030_ | -4.674118302 | -4.473059532 | -4.872292805 | -0.387719682 | 0.813355423 | -0.42093132 | 0.317788149 | 0.442698557 | 0.663024776 | 0.5484 | 0.725722971 |
| STnc2040_ | -6.147223714 | -5.286394308 | -8.494539008 | -3.215075772 | 2.516939305 | -1.210742397 | 0.039973462 | 0.142786814 | 0.406847096 | 0.149 | 0.388277227 |
| STnc205_ | 4.98474965 | 5.999965898 | 3.950127717 | -2.048914648 | 0.117739832 | -17.31534077 | 4.68E-05 | 2.00E-05 | 0.015515064 | 0.1 | 0.330257709 |
| STnc2070_ | 0.363182488 | 0.342636383 | 0.382748794 | -0.02868008 | 1.236448424 | -0.027439584 | 0.481333334 | 0.920899615 | 0.971112727 | 0.9598 | 0.982861185 |
| STnc2080_ | 0.059734531 | 0.013904006 | 0.1323219 | 0.228684087 | 0.526674119 | 0.441175032 | 0.314457037 | 0.453796013 | 0.716731095 | 0.6238 | 0.808217177 |
| STnc2090_ | 4.145180415 | 4.1216717 | 4.175545659 | 0.035334912 | 0.544870063 | 0.075533425 | 0.405729517 | 0.956631961 | 0.984706069 | 0.8749 | 0.946566765 |
| STnc210_ | -6.371376952 | -5.93434569 | -6.899156782 | -1.000793337 | 1.77924501 | -0.506594229 | 0.272485021 | 0.413571862 | 0.648607443 | 0.5252 | 0.710275089 |
| STnc2100_ | -3.078229191 | -3.030150552 | -3.13019801 | -0.026195563 | 0.915515317 | -0.02843472 | 0.486000001 | 0.755841944 | 0.891445518 | 0.8248 | 0.9167328 |
| STnc2110_ | -1.639712714 | -1.105922596 | -1.74181038 | -0.644395881 | 0.736110334 | -0.653984323 | 0.311333341 | 0.455065174 | 0.715731395 | 0.638 | 0.829624679 |
| STnc2130_ | -0.185204882 | -0.38772128 | 0.129242095 | 0.57629624 | 0.351947737 | 1.601542765 | 0.020000227 | 0.073729056 | 0.304097712 | 0.1204 | 0.357488463 |

| | | | | | | | | | | | |
|---|---|---|---|---|---|---|---|---|---|---|---|
| STnc2160_ | -4.571933332 | -4.222581007 | -4.956703175 | -0.726727975 | 0.797329272 | -0.866631764 | 0.171219209 | 0.261857285 | 0.505861553 | 0.3524 | 0.572535161 |
| STnc2170_ | -2.912542191 | -3.183035507 | -2.793421472 | 0.350531645 | 0.69197957 | 0.465431108 | 0.310000008 | 0.500648439 | 0.731004489 | 0.6417 | 0.806692324 |
| STnc2175_ | -2.368547621 | -2.419745317 | -2.31706235 | 0.109386129 | 0.411743066 | 0.25432845 | 0.379080618 | 0.532551749 | 0.739394053 | 0.6459 | 0.798603131 |
| STnc2180_ | 2.638824055 | 3.868460091 | 2.314971383 | -1.218064685 | 1.140781391 | -1.341302089 | 0.104596974 | 0.08838586 | 0.344742013 | 0.1977 | 0.473198594 |
| STnc220_ | -4.517025641 | -4.709380172 | -4.370617213 | 0.355947912 | 0.911508672 | 0.367500814 | 0.339773491 | 0.49357711 | 0.715132115 | 0.6224 | 0.781790356 |
| STnc230_ | -7.755814934 | -8.808380517 | -6.400447755 | 2.117211849 | 3.800468107 | 0.464381185 | 0.248501012 | 0.454827559 | 0.680097824 | 0.4872 | 0.696124731 |
| STnc240_ | -1.679418962 | -2.163872282 | -1.589932432 | 0.574198572 | 0.684308048 | 0.66044737 | 0.308000008 | 0.421748165 | 0.691806818 | 0.6356 | 0.824095057 |
| STnc260_ | -6.50142589 | -7.209947744 | -5.89016648 | 1.31030352 | 1.597867621 | 0.709302225 | 0.194536993 | 0.306269184 | 0.549439005 | 0.3987 | 0.611433655 |
| STnc280_ | -5.062510252 | -5.313142669 | -4.859804353 | 0.462825576 | 0.995688161 | 0.424128781 | 0.314457037 | 0.450956431 | 0.675006586 | 0.5633 | 0.738891807 |
| STnc290_ | 0.675274597 | 0.499232157 | 0.834443128 | 0.029876799 | 0.52639794 | 0.06636894 | 0.479013991 | 0.782485983 | 0.907579699 | 0.9377 | 0.970411287 |
| STnc30_ | 0.502810627 | 0.344823609 | 0.674477835 | 0.170055544 | 0.448403633 | 0.453723845 | 0.316666675 | 0.531886232 | 0.767516902 | 0.6512 | 0.822157597 |
| STnc300_ | -6.404919429 | -6.144063408 | -6.832106701 | -0.752353294 | 2.172581612 | -0.272463166 | 0.333111266 | 0.469404218 | 0.708054063 | 0.6152 | 0.777029717 |
| STnc3000_ | -4.346699458 | -4.197226449 | -4.560644705 | -0.351307337 | 0.81852096 | -0.384812479 | 0.322451706 | 0.477294006 | 0.701431195 | 0.6018 | 0.767399798 |
| STnc3020_ | -5.452534517 | -6.084363814 | -5.072266618 | 1.049399449 | 1.372858739 | 0.674039948 | 0.193870771 | 0.30370065 | 0.568972323 | 0.4031 | 0.619832887 |
| STnc3030_ | -2.452326942 | -3.000692069 | -2.346580867 | 0.599423347 | 0.70730609 | 0.615468815 | 0.327115264 | 0.510281149 | 0.750627958 | 0.6835 | 0.856419897 |
| STnc3040_ | -5.095311119 | -4.804969544 | -5.337779878 | -0.567432152 | 1.289907863 | -0.403893277 | 0.320453039 | 0.50399707 | 0.723750526 | 0.612 | 0.780554171 |
| STnc3060_ | -3.744866728 | -3.542687938 | -3.863459558 | -0.384256961 | 0.742456288 | -0.489169889 | 0.29913392 | 0.470036683 | 0.695101342 | 0.5912 | 0.760430819 |
| STnc3070_ | -1.728495396 | -1.694358984 | -1.776246794 | -0.115076715 | 0.436106714 | -0.250365433 | 0.390666671 | 0.560334738 | 0.769258432 | 0.6878 | 0.832721875 |
| STnc3080_ | -4.471100114 | -4.662164797 | -4.248394174 | 0.393912103 | 0.9970133 | 0.384198062 | 0.328666674 | 0.518459547 | 0.732760352 | 0.6163 | 0.783516722 |
| STnc3090_ | -4.939620297 | -4.65494621 | -5.240036526 | -0.614852448 | 0.883105594 | -0.618499241 | 0.23984012 | 0.364241165 | 0.598815916 | 0.4635 | 0.66227056 |
| STnc310_ | -4.222593585 | -4.096990911 | -4.346224058 | -0.250218573 | 0.758160787 | -0.302171111 | 0.357095276 | 0.504179951 | 0.718439027 | 0.6281 | 0.78416172 |
| STnc3100_ | 0.629030278 | 0.352654849 | 0.99304063 | 0.631836406 | 0.194668942 | 3.232689244 | 0.00067292 | 0.012494855 | 0.128810555 | 0.1 | 0.330257709 |
| STnc3110_ | -0.346196229 | -0.008200304 | -0.430953609 | -0.37403058 | 0.512192621 | -0.687816878 | 0.256495681 | 0.404440697 | 0.676974041 | 0.5009 | 0.741203809 |
| STnc3120_ | -1.160391419 | -1.575035112 | -1.066310281 | 0.486768461 | 0.719625773 | 0.514662862 | 0.341772158 | 0.548190371 | 0.780958491 | 0.7214 | 0.880036313 |
| STnc3150_ | -0.177386313 | 0.259596754 | -0.47602088 | -0.721174235 | 0.262043136 | -2.668467032 | 0.001335878 | 0.024721399 | 0.178125033 | 0.1004 | 0.330847641 |
| STnc3160_ | -2.662920598 | -2.063387066 | -2.800019011 | -0.6745459 | 0.783095091 | -0.563111088 | 0.321119261 | 0.607280983 | 0.808670275 | 0.6789 | 0.854814385 |
| STnc3170_ | -1.728463545 | -1.520613621 | -1.908686747 | -0.398794393 | 0.31761533 | -1.19482646 | 0.088000048 | 0.152408058 | 0.392006891 | 0.2302 | 0.466464653 |
| STnc3180_ | -4.011660184 | -4.319826795 | -3.838022765 | 0.397937321 | 0.983446155 | 0.388011995 | 0.331112599 | 0.544014527 | 0.75545204 | 0.6662 | 0.820412821 |
| STnc320_ | -2.156806089 | -2.410142413 | -1.992309282 | 0.386650197 | 0.604317693 | 0.612659686 | 0.263824128 | 0.404626023 | 0.664602222 | 0.5177 | 0.730320074 |
| STnc3210_ | -5.865119521 | -5.829833766 | -5.924333669 | -0.059358393 | 1.721307142 | -0.029365758 | 0.486666667 | 0.642882577 | 0.822634887 | 0.7806 | 0.889882816 |
| STnc3220_ | -0.558657185 | -0.581648485 | -0.537434272 | 0.140826555 | 0.397279209 | 0.339817282 | 0.363091278 | 0.575617044 | 0.785174849 | 0.7652 | 0.878498694 |
| STnc3230_ | -3.07527808 | -3.453704228 | -2.725253507 | 0.740310257 | 0.500217037 | 1.387327316 | 0.054666746 | 0.106315024 | 0.329193402 | 0.1745 | 0.413010781 |
| STnc3270_ | -2.782555948 | -2.821699609 | -2.698613248 | 0.187193318 | 0.68228641 | 0.26835892 | 0.400000004 | 0.594901074 | 0.79148992 | 0.7346 | 0.857782246 |
| STnc3280_ | -3.371443261 | -3.279689414 | -3.444641496 | -0.162446515 | 0.551131335 | -0.269395245 | 0.380666671 | 0.552956444 | 0.754022405 | 0.6643 | 0.813351554 |
| STnc3290_ | -2.226506508 | -2.306848644 | -2.102667063 | 0.199512923 | 0.61033424 | 0.29465543 | 0.378414395 | 0.651531402 | 0.826960259 | 0.7693 | 0.883796348 |
| STnc3300_ | -2.950950059 | -3.188740556 | -2.726182111 | 0.581270019 | 0.691160143 | 0.825264184 | 0.180546322 | 0.272301729 | 0.541690539 | 0.3991 | 0.619362537 |
| STnc3310_ | 2.646839426 | 2.549317359 | 2.721995795 | 0.151878103 | 0.134425229 | 1.129151775 | 0.111259197 | 0.159203518 | 0.434684757 | 0.2285 | 0.492903608 |
| STnc3330_ | -3.904241748 | -3.935818196 | -3.87574429 | 0.042293501 | 0.743517976 | 0.056969707 | 0.475682879 | 0.605769211 | 0.791991409 | 0.7046 | 0.841624447 |
| STnc3340_ | -3.168350896 | -3.202960553 | -3.128704423 | 0.072869218 | 0.537697638 | 0.127334591 | 0.450000002 | 0.566131226 | 0.762931183 | 0.6777 | 0.818038678 |
| STnc3350_ | -2.857629395 | -2.692646794 | -3.011880944 | -0.301578773 | 0.711269036 | -0.405501686 | 0.331112599 | 0.541114864 | 0.757190789 | 0.6586 | 0.815169255 |
| STnc340_ | -3.287011712 | -2.972320913 | -3.627107675 | -0.650967313 | 0.597251847 | -1.053068954 | 0.109926752 | 0.180782753 | 0.429541001 | 0.2647 | 0.500984953 |
| STnc3400_ | -4.876596635 | -5.764073026 | -4.474700426 | 1.279279102 | 1.155966092 | 1.026429296 | 0.115922753 | 0.194206972 | 0.461443877 | 0.2651 | 0.505471759 |
| STnc3410_ | -0.160165812 | -0.279298856 | -0.065254384 | 0.131006003 | 0.479971117 | 0.289613779 | 0.365089945 | 0.692794606 | 0.860061452 | 0.7466 | 0.877587293 |
| STnc3420_ | -2.165037431 | -2.296662188 | -2.071037654 | 0.207820337 | 0.437047479 | 0.435841233 | 0.325333341 | 0.497782147 | 0.71724553 | 0.6063 | 0.775979803 |
| STnc3430_ | -1.825940586 | -2.271276482 | -1.364275493 | 0.934115259 | 0.397949058 | 2.26955852 | 0.002666616 | 0.032854389 | 0.198654594 | 0.1029 | 0.334419427 |
| STnc3440_ | -2.904126291 | -3.047929068 | -2.780005635 | 0.313095038 | 0.611795559 | 0.482563642 | 0.282478358 | 0.447172705 | 0.681778826 | 0.5664 | 0.744250708 |
| STnc3460_ | 0.762479729 | 0.685188859 | 1.022077696 | 0.354238319 | 0.370592951 | 1.004984283 | 0.130579644 | 0.16620712 | 0.450991351 | 0.2637 | 0.5301199 |
| STnc3480_ | -1.666208223 | -1.050071994 | -2.903186225 | -1.865979943 | 0.461674013 | -4.052846518 | 4.68E-05 | 0.013869777 | 0.130088699 | 0.1 | 0.330257709 |
| STnc3490_ | -4.874131431 | -4.939064151 | -4.80651782 | 0.120588229 | 0.912678553 | 0.112495627 | 0.442666669 | 0.556508497 | 0.75746995 | 0.671 | 0.816173666 |
| STnc350_ | -2.175508984 | -2.41634738 | -1.990053597 | 0.447716619 | 0.481805475 | 0.900777744 | 0.151898759 | 0.231763489 | 0.500383329 | 0.3305 | 0.56714393 |
| STnc3500_ | -4.052163056 | -4.00926174 | -4.090428182 | -0.065865267 | 0.726797067 | -0.090782034 | 0.458666668 | 0.584426811 | 0.776097126 | 0.6794 | 0.823060204 |
| STnc3510_ | -0.508960027 | -0.564933008 | -0.457485775 | 0.131791561 | 0.293790493 | 0.425086563 | 0.324666674 | 0.498321858 | 0.726254264 | 0.6283 | 0.793261559 |
| STnc3540_ | 2.697839807 | 2.920933127 | 2.266839338 | -0.613238022 | 0.198412359 | -2.794736379 | 4.68E-05 | 0.024283507 | 0.185715018 | 0.1 | 0.330257709 |

| | | | | | | | | | | | |
|---|---|---|---|---|---|---|---|---|---|---|---|
| STnc3550_ | -4.0655759 | -4.007209663 | -4.15251905 | -0.228144643 | 1.008330069 | -0.204517277 | 0.396402402 | 0.54940653 | 0.764709043 | 0.6837 | 0.824582264 |
| STnc3580_ | -4.354976005 | -4.638069426 | -3.850233909 | 0.757665886 | 0.931176717 | 0.727462481 | 0.222666682 | 0.351652766 | 0.599587518 | 0.4463 | 0.663698827 |
| STnc360_ | -6.229815647 | -6.717626112 | -5.706310681 | 0.418636413 | 3.140134939 | 0.111938261 | 0.439706864 | 0.707269546 | 0.864228957 | 0.8481 | 0.926358458 |
| STnc3600_ | -1.70078206 | -2.041913404 | -1.358302029 | 0.70670909 | 0.474807454 | 1.455837606 | 0.029313944 | 0.094260984 | 0.333605876 | 0.133 | 0.37354075 |
| STnc3610_ | -0.677627599 | -0.554250763 | -1.231786097 | -0.639940654 | 0.405545326 | -1.261009062 | 0.105929419 | 0.158073518 | 0.439710018 | 0.2465 | 0.499405499 |
| STnc3620_ | -3.663515547 | -3.536795535 | -3.795918511 | -0.266502806 | 0.629021938 | -0.393094604 | 0.330446376 | 0.478723632 | 0.698322892 | 0.5928 | 0.759847299 |
| STnc3630_ | -2.109918018 | -1.497006719 | -2.32030061 | -0.831156795 | 0.763206372 | -0.98640279 | 0.144666693 | 0.214712135 | 0.50700094 | 0.3013 | 0.558250719 |
| STnc3640_ | -2.002219592 | -1.576757532 | -2.548075209 | -0.98933709 | 0.485003207 | -1.973551298 | 0.010659989 | 0.05027041 | 0.245375304 | 0.11 | 0.343984195 |
| STnc3680_ | 1.973808575 | 1.649411966 | 2.100636797 | 0.29709209 | 0.44330698 | 0.862919705 | 0.191333352 | 0.270406344 | 0.57123659 | 0.3536 | 0.632476914 |
| STnc370_ | -6.19039782 | -6.346171469 | -5.958246921 | 0.289660527 | 1.731940174 | 0.14286403 | 0.428666669 | 0.59552229 | 0.785829057 | 0.7074 | 0.842086581 |
| STnc3700_ | -3.639867813 | -3.659261192 | -3.628464095 | 0.046726303 | 0.702239649 | 0.058378314 | 0.471685544 | 0.607879153 | 0.792939499 | 0.7187 | 0.847386438 |
| STnc3720_ | -3.559744633 | -3.908186925 | -3.413367113 | 0.51212177 | 0.818013963 | 0.57694106 | 0.265822796 | 0.386902814 | 0.646467312 | 0.528 | 0.721553622 |
| STnc3730_ | -1.00312075 | -1.128354202 | -0.953816553 | 0.125465659 | 0.424320072 | 0.180035882 | 0.40906063 | 0.785703079 | 0.908361903 | 0.8063 | 0.916060766 |
| STnc3750_ | -3.540426436 | -3.76084812 | -3.363843389 | 0.360621022 | 0.69639979 | 0.482391636 | 0.293137917 | 0.457982572 | 0.68882604 | 0.5634 | 0.74556267 |
| STnc3770_ | -2.636424375 | -2.595590732 | -2.682211861 | -0.146772367 | 0.630659197 | -0.226819414 | 0.40266667 | 0.623799214 | 0.809229208 | 0.7595 | 0.873023658 |
| STnc3790_ | -4.652196207 | -4.432078974 | -4.970334203 | -0.561303744 | 0.963358094 | -0.54924451 | 0.272485021 | 0.402357325 | 0.642364485 | 0.51 | 0.705154034 |
| STnc380_ | -2.931934654 | -3.000230439 | -2.889010384 | 0.17838833 | 0.708154134 | 0.207649648 | 0.397333337 | 0.585288521 | 0.787974711 | 0.7328 | 0.860863513 |
| STnc3800_ | -1.139622496 | -1.45722686 | -0.735538097 | 0.715725908 | 0.300266984 | 2.322098918 | 0.003998486 | 0.03179996 | 0.197938642 | 0.1036 | 0.335071606 |
| STnc3830_ | -2.76756267 | -2.850153289 | -2.686172842 | 0.160432495 | 0.388772254 | 0.386542773 | 0.328447709 | 0.447468633 | 0.664882379 | 0.5372 | 0.717135236 |
| STnc3850_ | -3.160613255 | -3.05041511 | -3.252802791 | -0.203632338 | 0.511852181 | -0.360727371 | 0.328666674 | 0.491458774 | 0.706496093 | 0.5902 | 0.761195983 |
| STnc3860_ | -3.398140578 | -3.428892659 | -3.363406533 | 0.069595449 | 0.5589118 | 0.123026248 | 0.447701534 | 0.553367971 | 0.752414916 | 0.6639 | 0.809380673 |
| STnc3870_ | -1.539078459 | -1.090546746 | -1.703071305 | -0.512139677 | 0.559979347 | -0.886975574 | 0.189333352 | 0.26047717 | 0.545780794 | 0.368 | 0.624869024 |
| STnc3880_ | -2.301449222 | -2.859294426 | -2.105261576 | 0.728487628 | 0.967181217 | 0.758228106 | 0.229846783 | 0.322146963 | 0.611733373 | 0.4244 | 0.683706998 |
| STnc3890_ | -3.433938571 | -3.331280857 | -3.534160283 | -0.176726942 | 0.540919634 | -0.302130496 | 0.354430386 | 0.493105687 | 0.706056567 | 0.6017 | 0.766114233 |
| STnc390_ | 2.818189511 | 2.234347851 | 2.872978464 | 0.642520603 | 0.420664264 | 0.572330685 | 0.33333334 | 0.592154728 | 0.813863396 | 0.7 | 0.872542119 |
| STnc3920_ | -7.235902598 | -6.985289532 | -7.574091101 | -0.667550913 | 2.432100977 | -0.218524095 | 0.384000005 | 0.511193304 | 0.731701031 | 0.6518 | 0.800458169 |
| STnc3950_ | -1.499539171 | -1.337808564 | -1.673895091 | -0.331262963 | 0.310030359 | -1.003635585 | 0.131333363 | 0.211061014 | 0.463363137 | 0.2902 | 0.527891977 |
| STnc3960_ | -0.871456327 | -0.809226398 | -0.923174852 | -0.179970973 | 0.405917317 | -0.421689647 | 0.318000008 | 0.458368489 | 0.703784002 | 0.6303 | 0.792555847 |
| STnc3970_ | -5.324861395 | -5.012763957 | -5.569696782 | -0.389646843 | 1.559382347 | -0.220981433 | 0.396000004 | 0.627033763 | 0.81034899 | 0.7291 | 0.859300646 |
| STnc3990_ | -1.129201413 | -1.307196135 | -1.030947946 | 0.256614525 | 0.341609577 | 0.681520231 | 0.249833457 | 0.37914942 | 0.633601229 | 0.4968 | 0.703375098 |
| STnc40_ | 0.705951938 | 0.596462796 | 0.869297429 | 0.249559014 | 0.204509672 | 1.207700473 | 0.114590308 | 0.157878466 | 0.42077543 | 0.2336 | 0.490638761 |
| STnc4000_ | -3.100500907 | -3.163386885 | -3.061941675 | 0.090932688 | 0.515079321 | 0.164810024 | 0.428000003 | 0.563801027 | 0.761526703 | 0.6756 | 0.819410945 |
| STnc4010_ | -5.378550748 | -5.416538962 | -5.311117165 | -0.048168428 | 1.765323304 | -0.024030698 | 0.487008661 | 0.663301217 | 0.838698294 | 0.7999 | 0.900405607 |
| STnc4020_ | -4.824844404 | -4.72890427 | -4.936610678 | -0.279712356 | 1.113107421 | -0.225801105 | 0.393333337 | 0.568809471 | 0.769145384 | 0.6745 | 0.819534039 |
| STnc4040_ | -0.975222734 | -0.911820812 | -1.068292842 | -0.221971787 | 0.44504878 | -0.487280797 | 0.297333342 | 0.464417177 | 0.71119841 | 0.5981 | 0.782954784 |
| STnc4070_ | -3.442861009 | -3.770268901 | -3.146339735 | 0.621143099 | 0.565863127 | 1.04655254 | 0.1206667 | 0.199218253 | 0.447536025 | 0.2813 | 0.516787685 |
| STnc4080_ | -3.493068224 | -3.428219394 | -3.552765184 | -0.135594389 | 0.708259166 | -0.184869966 | 0.418000003 | 0.610746399 | 0.799227382 | 0.7188 | 0.851215224 |
| STnc410_ | -4.434674238 | -4.345551693 | -4.506012946 | -0.13226012 | 0.77962614 | -0.161667474 | 0.425716192 | 0.559667693 | 0.75627407 | 0.6585 | 0.808213184 |
| STnc4100_ | -3.391815334 | -3.124068944 | -3.884352509 | -0.751691776 | 0.847775733 | -0.810671898 | 0.166555652 | 0.262237668 | 0.539089962 | 0.3516 | 0.588126825 |
| STnc4110_ | -3.60122466 | -3.553478172 | -3.637685225 | -0.07445886 | 0.878135426 | -0.082588068 | 0.459333335 | 0.681733638 | 0.846097372 | 0.7844 | 0.891835826 |
| STnc4120_ | -2.205165495 | -2.308181406 | -2.155113685 | 0.157631736 | 0.524805596 | 0.280660981 | 0.385333338 | 0.577899427 | 0.781167117 | 0.6981 | 0.839862323 |
| STnc4130_ | -2.566783338 | -2.34840213 | -2.707318362 | -0.337890603 | 0.529774314 | -0.579184852 | 0.263157906 | 0.425056718 | 0.665860085 | 0.5342 | 0.72940528 |
| STnc4140_ | -1.695269153 | -1.750794859 | -1.652290594 | 0.046899458 | 0.486053258 | 0.091237895 | 0.457028649 | 0.702917003 | 0.85868742 | 0.8172 | 0.908945586 |
| STnc4150_ | -5.370250746 | -5.184421956 | -5.499021925 | -0.225530388 | 1.35355313 | -0.139982245 | 0.424383747 | 0.637487409 | 0.813623587 | 0.7288 | 0.857354606 |
| STnc4160_ | -2.094671525 | -2.105829962 | -2.087195099 | -0.103873096 | 0.56746627 | -0.173172787 | 0.430000003 | 0.671538291 | 0.841511081 | 0.8193 | 0.90934464 |
| STnc4170_ | -5.308580367 | -4.549075303 | -5.643747376 | -0.862594087 | 2.042631864 | -0.362451785 | 0.320666674 | 0.662032575 | 0.83008844 | 0.6681 | 0.835367069 |
| STnc4190_ | -6.046447246 | -5.960984028 | -6.199534735 | -0.329444357 | 2.293172633 | -0.115931448 | 0.424000003 | 0.574536338 | 0.786297635 | 0.7535 | 0.871672959 |
| STnc420_ | -3.160267802 | -2.965054314 | -3.39060482 | -0.757895037 | 1.209657165 | -0.622543699 | 0.272000011 | 0.362123222 | 0.639391577 | 0.5526 | 0.753908342 |
| STnc4200_ | -2.347756196 | -2.167722254 | -2.589648776 | -0.294218144 | 0.986005732 | -0.285778835 | 0.365756168 | 0.61701075 | 0.81518047 | 0.7451 | 0.869790035 |
| STnc4210_ | -2.58552409 | -1.791329318 | -3.845964622 | -2.058460302 | 0.556399224 | -3.717029196 | 4.68E-05 | 0.01829346 | 0.145425003 | 0.1 | 0.330257709 |
| STnc4220_ | -0.163224737 | -0.248526814 | -0.041436603 | 0.131294528 | 0.369106836 | 0.34095454 | 0.360000006 | 0.637576418 | 0.823414101 | 0.7568 | 0.87775095 |
| STnc4230_ | -1.082356074 | -1.070455459 | -1.08707333 | -0.010041755 | 0.310462611 | -0.029258717 | 0.485676216 | 0.663135367 | 0.833355622 | 0.763 | 0.880556574 |
| STnc4240_ | 0.796837948 | 0.627943964 | 1.041518214 | 0.313474122 | 0.347297308 | 0.918023569 | 0.163890762 | 0.243410776 | 0.536783488 | 0.3004 | 0.568685266 |

| | | | | | | | | | | | |
|---|---|---|---|---|---|---|---|---|---|---|---|
| STnc4250_ | 3.458865271 | 3.370372543 | 3.490824095 | 0.125663436 | 0.270268732 | 0.39495718 | 0.341772158 | 0.820800385 | 0.930178916 | 0.7234 | 0.88251863 |
| STnc4260_ | -4.811376592 | -4.586225777 | -5.017571246 | -0.408500082 | 0.891925981 | -0.4161368 | 0.324000007 | 0.460502006 | 0.676328615 | 0.5645 | 0.737838183 |
| STnc430_ | -5.417664033 | -5.602228122 | -5.19769128 | 0.391024843 | 1.294134847 | 0.277578797 | 0.384410398 | 0.560628363 | 0.761748319 | 0.67 | 0.816309472 |
| STnc440_ | -2.39257233 | -2.22891263 | -2.49623977 | -0.276389254 | 0.813379006 | -0.327441 | 0.352431718 | 0.632228747 | 0.821286961 | 0.701 | 0.849369368 |
| STnc450_ | -0.064602019 | -0.119455833 | -0.030617324 | 0.117754498 | 0.282365157 | 0.387021707 | 0.319120594 | 0.457230068 | 0.706447648 | 0.6262 | 0.787632242 |
| STnc460_ | -5.664695484 | -5.264451838 | -6.181649456 | -0.857345241 | 1.498669026 | -0.524340411 | 0.276482355 | 0.430429409 | 0.664930422 | 0.5447 | 0.730252543 |
| STnc470_ | -2.127191304 | -2.346151493 | -1.977927991 | 0.415281212 | 0.474510909 | 0.827217719 | 0.146666693 | 0.255934298 | 0.530630308 | 0.3294 | 0.558357933 |
| STnc475_ | 5.370630072 | 5.462654787 | 5.270499191 | -0.151461815 | 0.43114724 | -0.291125563 | 0.449333335 | 0.786369716 | 0.916021774 | 1 | 1 |
| STnc480_ | -3.004988156 | -3.425166372 | -2.8738194 | 0.471331391 | 0.865556561 | 0.420248518 | 0.359760166 | 0.674838074 | 0.84905014 | 0.7482 | 0.890800696 |
| STnc50_ | -5.003062947 | -4.910095543 | -5.10214866 | -0.199142279 | 1.077443751 | -0.157358446 | 0.425716192 | 0.571640202 | 0.769583528 | 0.6818 | 0.824853837 |
| STnc500_ | -2.09734745 | -2.038768941 | -2.14666472 | -0.195512616 | 0.797382743 | -0.215408123 | 0.409333337 | 0.649405967 | 0.833160068 | 0.8189 | 0.911586214 |
| STnc510_ | -2.143557204 | -2.23053336 | -2.066018236 | 0.097235491 | 0.556736764 | 0.166195815 | 0.427333336 | 0.687150433 | 0.851749477 | 0.7996 | 0.901261982 |
| STnc520_ | -1.497672147 | -1.566594579 | -1.432279127 | 0.01915818 | 0.736717781 | 0.022855976 | 0.491333334 | 0.823369479 | 0.927210023 | 0.9317 | 0.969749543 |
| STnc530_ | -5.314942264 | -5.010309252 | -5.66028946 | -0.654360624 | 1.264422516 | -0.466467929 | 0.278481023 | 0.433050408 | 0.666891078 | 0.5466 | 0.72909015 |
| STnc540_ | 3.730843742 | 3.694417748 | 3.765882222 | 0.044041333 | 0.216538663 | 0.082152048 | 0.453697537 | 0.644235499 | 0.84341486 | 0.9811 | 0.992008598 |
| STnc590_ | -1.375978465 | -0.937532355 | -2.027273132 | -1.124887381 | 0.440390246 | -2.510509823 | 0.000673369 | 0.021326043 | 0.167565079 | 0.1006 | 0.331138659 |
| STnc60_ | -8.636100643 | -8.905084813 | -8.321540022 | 0.462211649 | 4.076353521 | 0.092743374 | 0.444000002 | 0.564881013 | 0.768948617 | 0.6965 | 0.834428243 |
| STnc600_ | 1.661594428 | 2.49506933 | 0.839949837 | -1.656738803 | 0.18276322 | -9.029311886 | 4.68E-05 | 0.001116126 | 0.04533566 | 0.1 | 0.330257709 |
| STnc610_ | 2.045908614 | 2.755896408 | 1.901434295 | -0.844923219 | 0.658692173 | -0.879791402 | 0.223184558 | 0.271683348 | 0.575706715 | 0.4092 | 0.68582525 |
| STnc620_ | 1.600747933 | 1.572630343 | 1.624404249 | 0.045614162 | 0.371485729 | 0.123532848 | 0.432666669 | 0.840046699 | 0.936280185 | 0.8968 | 0.955669015 |
| STnc640_ | -2.18722368 | -2.053799737 | -2.304959054 | -0.26784949 | 0.405625114 | -0.610565923 | 0.255163236 | 0.382126969 | 0.62109741 | 0.5003 | 0.692918932 |
| STnc650_ | 3.394024217 | 3.321206533 | 3.710818214 | 0.431165834 | 0.527455365 | 0.928827745 | 0.167888097 | 0.176378269 | 0.473332862 | 0.306 | 0.584539331 |
| STnc660_ | 1.670979185 | 1.587214164 | 1.752621683 | 0.006728722 | 0.320058231 | 0.021583497 | 0.491672219 | 0.833210525 | 0.93206702 | 0.9596 | 0.981531463 |
| STnc670_ | 1.653704248 | 1.881586483 | 1.503098034 | -0.446151287 | 0.242287021 | -1.822832766 | 0.001335878 | 0.049524025 | 0.256776937 | 0.102 | 0.33316928 |
| STnc680_ | 1.872541546 | 1.960123063 | 1.826826744 | -0.140802305 | 0.278564747 | -0.507446153 | 0.272000011 | 0.482196708 | 0.735207627 | 0.5451 | 0.768026346 |
| STnc690_ | -0.008209495 | -0.425193307 | 0.066925903 | 0.517064488 | 0.66788674 | 0.635871138 | 0.328447709 | 0.465753305 | 0.729715609 | 0.6844 | 0.861930966 |
| STnc695_ | -1.819905581 | -1.982111346 | -1.66841135 | 0.315436663 | 0.329889172 | 0.908594482 | 0.165223207 | 0.243304989 | 0.495335712 | 0.3336 | 0.563181649 |
| STnc70_ | 2.919324984 | 2.875216531 | 3.142256646 | 0.226362565 | 0.44687855 | 0.395789098 | 0.330446376 | 0.700856579 | 0.872736995 | 0.6934 | 0.868286259 |
| STnc720_ | -2.991420008 | -3.25729024 | -2.78821751 | 0.485843566 | 0.655049987 | 0.694902841 | 0.194666685 | 0.305122138 | 0.571980259 | 0.4025 | 0.623227525 |
| STnc730_ | 1.683174719 | 1.500184313 | 1.838988146 | 0.306538384 | 0.404056953 | 0.595062392 | 0.279813468 | 0.40125189 | 0.682768805 | 0.5843 | 0.78273961 |
| STnc740_ | -5.68525067 | -6.120076819 | -5.325240803 | 0.782655361 | 1.152367234 | 0.642648608 | 0.222666682 | 0.33517094 | 0.577765596 | 0.4397 | 0.64218538 |
| STnc750_ | -0.060322398 | -0.121424795 | 0.149948647 | 0.292115832 | 0.856447954 | 0.390615778 | 0.24250501 | 0.540530235 | 0.776393529 | 0.439 | 0.704219308 |
| STnc760_ | -0.524832134 | -0.690550789 | -0.348826458 | 0.343937509 | 0.227942684 | 1.426342658 | 0.054666746 | 0.099498908 | 0.326957738 | 0.1652 | 0.406805271 |
| STnc770_ | 2.417031911 | 2.459829561 | 2.382801218 | -0.072217729 | 0.237695802 | -0.249043249 | 0.408000004 | 0.807701045 | 0.920364126 | 0.8731 | 0.945072637 |
| STnc780_ | 0.862443347 | 0.804066585 | 0.917791073 | 0.108829674 | 0.155604849 | 0.654885678 | 0.248666679 | 0.366035579 | 0.615367281 | 0.4694 | 0.680486627 |
| STnc790_ | 1.740638156 | 1.905829562 | 1.463880887 | -0.522283013 | 0.268412336 | -2.014894083 | 0.00067292 | 0.033200843 | 0.214836251 | 0.1 | 0.330257709 |
| STnc800_ | 3.441908245 | 3.445356424 | 3.437429203 | -0.090794842 | 0.598077306 | -0.18487049 | 0.393737512 | 0.7876331 | 0.916066553 | 0.8332 | 0.929007247 |
| STnc810_ | 0.014194722 | -0.490941605 | 0.528265052 | 1.158293856 | 0.395916657 | 2.976531628 | 4.68E-05 | 0.010487088 | 0.124771527 | 0.1 | 0.330257709 |
| STnc835_ | 3.676892164 | 3.629909277 | 3.725303324 | 0.083280715 | 0.584192515 | 0.146495175 | 0.441039309 | 0.927392222 | 0.973800407 | 0.9991 | 0.999635048 |
| STnc840_ | 2.421527696 | 2.745795787 | 1.680021638 | -1.072755132 | 0.557813239 | -2.08602664 | 4.68E-05 | 0.02641323 | 0.198148987 | 0.1 | 0.330257709 |
| STnc850_ | -3.754170314 | -3.930106325 | -3.431886193 | 0.460626444 | 0.845833312 | 0.480215098 | 0.310459702 | 0.511333257 | 0.728957257 | 0.6191 | 0.788121157 |
| STnc870_ | -0.912139999 | -1.019049065 | -0.803862711 | 0.149789219 | 0.551327275 | 0.274610008 | 0.370419725 | 0.647856214 | 0.831909429 | 0.7606 | 0.883256805 |
| STnc880_ | 1.222890666 | 1.452788221 | 1.151672673 | -0.296365456 | 0.367820285 | -0.400289014 | 0.338000007 | 0.870795192 | 0.950442528 | 0.7063 | 0.875228672 |
| STnc890_ | -0.510670599 | -0.257917779 | -0.590690192 | -0.307090751 | 0.26775094 | -0.958156593 | 0.183211212 | 0.267911428 | 0.541928398 | 0.3978 | 0.631526243 |
| STnc920_ | 1.035190548 | 0.895087797 | 1.077601898 | 0.176395197 | 0.172336206 | 0.867045938 | 0.180546322 | 0.270533246 | 0.552170209 | 0.3988 | 0.630172726 |
| STnc925_ | -0.773782092 | -0.79536021 | -0.755948053 | 0.019827857 | 0.363109992 | 0.054024277 | 0.472351767 | 0.708906689 | 0.864870534 | 0.8128 | 0.909092071 |
| STnc930_ | -4.177545533 | -3.985421677 | -4.407073025 | -0.418339932 | 0.674685035 | -0.567568947 | 0.256495681 | 0.390521811 | 0.62021423 | 0.4861 | 0.679455173 |
| STnc940_ | -3.519498126 | -3.610815215 | -3.473215409 | 0.155106494 | 0.739782498 | 0.193520672 | 0.420000003 | 0.60507378 | 0.798905658 | 0.738 | 0.864071318 |
| STnc950_ | -3.539230592 | -3.630375695 | -3.464549209 | -0.026988027 | 0.980831933 | -0.025153818 | 0.492 | 0.715091586 | 0.867762719 | 0.8464 | 0.927200382 |
| STnc970_ | -1.205739217 | -1.32942182 | -1.122229623 | 0.218509098 | 0.524434235 | 0.391827837 | 0.318454371 | 0.487642696 | 0.73209445 | 0.6488 | 0.810246578 |
| StyR.103b_ | -8.187868606 | -7.637270681 | -8.816573737 | -1.098239702 | 3.635105084 | -0.260474065 | 0.353097941 | 0.524998038 | 0.735403146 | 0.6183 | 0.780811943 |
| StyR.103c_ | -9.357644036 | -9.828944464 | -8.877968239 | 0.93617716 | 4.105202922 | 0.182570325 | 0.3904064 | 0.505025039 | 0.719278849 | 0.6113 | 0.771176802 |
| StyR.103d_ | -9.331104412 | -8.900092027 | -9.833440703 | -0.775672334 | 4.161497137 | -0.162390476 | 0.40906063 | 0.508841425 | 0.722306652 | 0.6209 | 0.779233493 |

| | | | | | | | | | | | |
|---|---|---|---|---|---|---|---|---|---|---|---|
| StyR.103f_ | -9.27158278 | -8.500613464 | -9.89655063 | -1.608349941 | 4.383983083 | -0.326267786 | 0.349100606 | 0.51166086 | 0.722937686 | 0.6184 | 0.777409881 |
| StyR.103g_ | -8.424789403 | -8.88240071 | -8.027779126 | 0.776543989 | 3.308790634 | 0.18160012 | 0.397734847 | 0.520797032 | 0.734353443 | 0.6373 | 0.790586887 |
| StyR.122_ | -5.040346866 | -4.809967734 | -5.325472277 | -0.074565335 | 1.830997237 | -0.040296792 | 0.482666667 | 0.717688678 | 0.869628099 | 0.8601 | 0.932042736 |
| StyR.126_ | -2.110854949 | -2.145299674 | -2.072481099 | 0.049593288 | 0.488211752 | 0.100683585 | 0.463024652 | 0.676040079 | 0.841998439 | 0.7845 | 0.891187453 |
| StyR.128_ | -5.091157414 | -4.657720721 | -5.394910346 | -0.803726822 | 1.377149375 | -0.520849842 | 0.28800001 | 0.466575475 | 0.698444465 | 0.5839 | 0.763182473 |
| StyR.134_ | 0.232400291 | 0.19868956 | 0.269407368 | 0.079130893 | 0.258525115 | 0.313783834 | 0.365089945 | 0.586249026 | 0.786681287 | 0.7055 | 0.844208967 |
| StyR.137_ | -8.655211837 | -8.356392067 | -8.844119229 | -0.800151057 | 4.42620594 | -0.162671447 | 0.42066667 | 0.578924771 | 0.780049752 | 0.7126 | 0.846944355 |
| StyR.150_ | -3.568600918 | -3.346224161 | -3.777066541 | -0.451226723 | 0.751720791 | -0.566180851 | 0.244503677 | 0.401105375 | 0.644703474 | 0.5022 | 0.700782703 |
| StyR.169_ | -1.301055205 | -1.475401825 | -1.102523914 | 0.398189924 | 0.364430343 | 1.058917558 | 0.1186667 | 0.184460513 | 0.4509388 | 0.2622 | 0.507780157 |
| StyR.172_ | -9.340465126 | -9.879837055 | -8.790299703 | 0.960746205 | 4.039044389 | 0.206947194 | 0.38507662 | 0.50537191 | 0.717185088 | 0.6167 | 0.776678132 |
| StyR.195_ | -3.679554883 | -3.455707153 | -3.930809956 | -0.433786466 | 0.750201728 | -0.548959841 | 0.266000011 | 0.409191825 | 0.651083448 | 0.5117 | 0.709299883 |
| StyR.196_ | -6.653776687 | -6.681595046 | -6.632513046 | 0.149354437 | 2.345630455 | 0.048885939 | 0.463024652 | 0.585575651 | 0.787871902 | 0.7305 | 0.856506736 |
| StyR.199_ | -3.444800513 | -3.426399542 | -3.484646445 | -0.159560825 | 0.773072087 | -0.18675866 | 0.412391742 | 0.617090146 | 0.80721771 | 0.7603 | 0.873987127 |
| StyR.215a_ | 3.153923858 | 3.057306343 | 3.261300611 | 0.238392947 | 0.354932874 | 0.761230431 | 0.171333355 | 0.271503068 | 0.574631574 | 0.3262 | 0.595811854 |
| StyR.215b_ | 2.440647189 | 2.316856393 | 2.517742369 | 0.232804872 | 0.181278832 | 1.222342503 | 0.023317982 | 0.215005128 | 0.516335612 | 0.1246 | 0.362707213 |
| StyR.215c_ | -0.81708319 | -1.208601722 | -0.693379694 | 0.468991573 | 0.39119854 | 1.066881452 | 0.166000022 | 0.200146028 | 0.48406712 | 0.3056 | 0.566218122 |
| StyR.215d_ | 0.573383123 | 0.244110789 | 0.651214731 | 0.33450608 | 0.336187567 | 0.662937454 | 0.324666674 | 0.49458376 | 0.745739062 | 0.6756 | 0.85534909 |
| StyR.215e_ | -0.743015887 | -1.318019022 | -0.541121594 | 0.830612715 | 0.626865446 | 1.30847986 | 0.064666733 | 0.123504795 | 0.3954398 | 0.1691 | 0.421504404 |
| StyR.215f_ | 1.249906783 | 0.929861791 | 1.451949372 | 0.545082492 | 0.29066072 | 1.868263911 | 0.001335878 | 0.074249123 | 0.312787947 | 0.1006 | 0.331115459 |
| StyR.215g_ | -1.648184248 | -1.863570428 | -1.535006445 | 0.279066908 | 0.743610886 | 0.390048237 | 0.301132587 | 0.572364183 | 0.789289456 | 0.6015 | 0.797113938 |
| StyR.219_ | -4.40941791 | -3.957833977 | -5.092615447 | -1.119573514 | 0.962481816 | -1.107314955 | 0.113257864 | 0.178635816 | 0.439302154 | 0.2567 | 0.498914698 |
| StyR.222_ | -6.794081965 | -6.95164505 | -6.578950127 | 0.283500294 | 2.687060902 | 0.077857606 | 0.459333335 | 0.630519827 | 0.815322128 | 0.7729 | 0.883222538 |
| StyR.234_ | -5.269215003 | -5.295743558 | -5.246178531 | 0.204621758 | 1.370281862 | 0.139966598 | 0.441705532 | 0.625249449 | 0.807945319 | 0.7349 | 0.861558021 |
| StyR.243_ | 0.455366244 | 0.102268079 | 0.558340481 | 0.355277092 | 0.375159154 | 0.377763426 | 0.33533334 | 0.918672882 | 0.97018001 | 0.7054 | 0.874873798 |
| StyR.248_ | -1.71050204 | -1.566654584 | -1.827541422 | -0.149999419 | 0.642289433 | -0.23696035 | 0.41439041 | 0.742980364 | 0.884173041 | 0.845 | 0.927055008 |
| StyR.24a_ | -4.199577993 | -4.219481783 | -4.185654505 | 0.01041994 | 0.756787826 | 0.016070991 | 0.495333334 | 0.586864456 | 0.778689561 | 0.7037 | 0.836906967 |
| StyR.24b_ | -2.895914058 | -1.86694976 | -4.464177537 | -2.594924359 | 0.762472542 | -3.464896835 | 4.68E-05 | 0.023790951 | 0.166841043 | 0.1 | 0.330257709 |
| StyR.24c_ | -3.536239108 | -2.618137225 | -4.521479403 | -1.944513848 | 0.709063692 | -2.684013668 | 0.000673369 | 0.020745983 | 0.16025558 | 0.1012 | 0.331915984 |
| StyR.24d_ | -5.416328041 | -6.169311821 | -4.885073417 | 1.311231335 | 1.220157847 | 0.992046443 | 0.119253864 | 0.207498656 | 0.460401493 | 0.2811 | 0.51432389 |
| StyR.24e_ | -5.104283501 | -4.568215297 | -5.868530483 | -1.267347907 | 1.289437619 | -0.9593022 | 0.138574312 | 0.220076311 | 0.485556264 | 0.2974 | 0.539200713 |
| StyR.24f_ | -3.528087302 | -3.09713933 | -4.063424812 | -0.992646473 | 0.751776861 | -1.236838688 | 0.069333395 | 0.131693595 | 0.382710183 | 0.1882 | 0.43226096 |
| StyR.24g_ | -4.000880331 | -2.958420487 | -5.709290766 | -2.894362356 | 1.263551693 | -2.297495759 | 4.68E-05 | 0.057361354 | 0.256209531 | 0.1002 | 0.330546793 |
| StyR.250_ | -4.167153306 | -4.419603531 | -3.967428841 | 0.455837048 | 0.644943137 | 0.654259534 | 0.233844118 | 0.34962432 | 0.586279488 | 0.4501 | 0.650530067 |
| StyR.252_ | -1.689220075 | -1.42613189 | -2.042792055 | -0.663733084 | 0.446034033 | -1.41175477 | 0.044666765 | 0.099792182 | 0.3406798 | 0.1523 | 0.395094879 |
| StyR.254_ | -6.358407473 | -6.101926738 | -6.586394404 | -0.431879267 | 1.62499201 | -0.209161986 | 0.38241173 | 0.561965682 | 0.754945528 | 0.6594 | 0.807477156 |
| StyR.262_ | -1.138917897 | -1.260294784 | -1.027148172 | 0.241933937 | 0.252072906 | 0.893429353 | 0.161225873 | 0.245995629 | 0.495246543 | 0.3347 | 0.558790068 |
| StyR.264_ | -3.008809918 | -3.115967567 | -2.870561332 | 0.214222417 | 0.693411363 | 0.285340754 | 0.369753503 | 0.59937607 | 0.792585969 | 0.6937 | 0.837442904 |
| StyR.27_ | -3.575236381 | -3.396149674 | -3.685810743 | -0.381945429 | 1.191203174 | -0.287060384 | 0.387333338 | 0.644990496 | 0.828155236 | 0.7934 | 0.899107575 |
| StyR.277_ | 0.250215611 | 0.195440652 | 0.302471078 | 0.070694507 | 0.289659915 | 0.231963283 | 0.395069957 | 0.662401869 | 0.836216012 | 0.7701 | 0.883390163 |
| StyR.281_ | -4.735651747 | -4.262537858 | -5.358591692 | -1.12214967 | 1.077488216 | -1.003823485 | 0.127248532 | 0.207417725 | 0.469167055 | 0.2891 | 0.527992734 |
| StyR.285_ | -4.087768756 | -4.716286365 | -3.631531317 | 1.088617479 | 0.822974242 | 1.284637103 | 0.058000075 | 0.128573704 | 0.371931323 | 0.1786 | 0.419063887 |
| StyR.287_ | -1.389239818 | -1.491705252 | -1.254480634 | 0.286759193 | 0.394583931 | 0.688212141 | 0.225333348 | 0.337650551 | 0.60235388 | 0.4591 | 0.670041913 |
| StyR.29_ | 4.048501319 | 3.928296059 | 4.155656249 | 0.220833371 | 0.554884933 | 0.332413327 | 0.445702867 | 0.749727568 | 0.898604551 | 0.9994 | 0.999748423 |
| StyR.291_ | -1.805085409 | -1.30643263 | -1.910967936 | -0.59032929 | 0.590748878 | -0.875704342 | 0.224000015 | 0.296587232 | 0.584910959 | 0.4504 | 0.677824195 |
| StyR.292_ | -8.640500679 | -7.221407477 | -9.862395487 | -2.569770941 | 3.925878574 | -0.572959015 | 0.226515671 | 0.396338051 | 0.627034255 | 0.4767 | 0.674512326 |
| StyR.293_ | -2.982463916 | -2.923680325 | -3.036807464 | -0.344495735 | 0.925676051 | -0.35212709 | 0.368000005 | 0.56875788 | 0.780932642 | 0.7574 | 0.874801201 |
| StyR.295_ | -4.344044964 | -4.39493091 | -4.288498478 | 0.128006414 | 1.050848458 | 0.113683606 | 0.440000002 | 0.630985852 | 0.81292846 | 0.7367 | 0.861210407 |
| StyR.296_ | 3.567092958 | 3.546763678 | 3.617238531 | 0.059997964 | 0.223854922 | 0.316604055 | 0.289140582 | 0.689315757 | 0.862161351 | 0.5752 | 0.78646297 |
| StyR.3_ | -1.726673345 | -1.665101712 | -1.767708248 | -0.141589374 | 0.542026065 | -0.254265796 | 0.40066667 | 0.611125287 | 0.806032363 | 0.7694 | 0.882125847 |
| StyR.302_ | -8.035345188 | -8.635538866 | -7.658935196 | 0.766508272 | 3.751234104 | 0.179271065 | 0.398000004 | 0.569355235 | 0.775007933 | 0.7021 | 0.839826481 |
| StyR.305_ | -1.006414545 | -0.496505078 | -1.789524593 | -1.261332818 | 0.351478974 | -3.470208062 | 4.68E-05 | 0.015407685 | 0.139242468 | 0.1 | 0.330257709 |
| StyR.306_ | 0.778468217 | 0.63347495 | 0.94155024 | 0.482066571 | 0.396279648 | 1.301957617 | 0.021319333 | 0.111515866 | 0.376257512 | 0.121 | 0.358488787 |
| StyR.315_ | -7.422209724 | -8.378043627 | -6.922470618 | 1.071825194 | 4.057488105 | 0.237239246 | 0.370419725 | 0.573448523 | 0.779296659 | 0.7149 | 0.850178947 |

| | | | | | | | | | | | |
|---|---|---|---|---|---|---|---|---|---|---|---|
| StyR.322_ | -4.464962433 | -4.333756499 | -4.692621707 | -0.404435451 | 1.115638162 | -0.327731 | 0.335109934 | 0.491283401 | 0.723867901 | 0.6216 | 0.783956165 |
| StyR.327_ | -5.406428464 | -7.337930259 | -4.582167006 | 2.722644228 | 2.110116081 | 1.204230049 | 0.025333511 | 0.14427335 | 0.410001065 | 0.1341 | 0.374177986 |
| StyR.328_ | -6.452941806 | -6.520280977 | -6.382235041 | 0.135295126 | 1.847080744 | 0.069710259 | 0.465333335 | 0.590065361 | 0.784203577 | 0.7097 | 0.84309228 |
| StyR.329a_ | 0.761502681 | -0.022552135 | 1.639252128 | 1.646608903 | 0.452790736 | 3.538497247 | 4.68E-05 | 0.006005662 | 0.100024583 | 0.1 | 0.330257709 |
| StyR.329b_ | -2.895274068 | -3.280518567 | -2.75709445 | 0.41739878 | 0.794927085 | 0.464595234 | 0.340000007 | 0.582460438 | 0.791211614 | 0.7019 | 0.858271204 |
| StyR.336_ | -3.689085863 | -3.535119509 | -3.895170157 | -0.383304465 | 0.679555896 | -0.5261586 | 0.267821463 | 0.398160204 | 0.641681277 | 0.507 | 0.696298483 |
| StyR.341_ | -5.255875956 | -5.163986788 | -5.354189746 | -0.18677427 | 0.979091619 | -0.171262065 | 0.41466667 | 0.507226574 | 0.718467326 | 0.6146 | 0.775250903 |
| StyR.354_ | 1.135527574 | 0.9273475 | 1.427035444 | 0.681615122 | 0.561515044 | 1.209987832 | 0.007329074 | 0.209461505 | 0.511952846 | 0.1084 | 0.342173913 |
| StyR.358_ | -4.26652584 | -4.234263363 | -4.296526109 | -0.049602555 | 0.854525712 | -0.049920862 | 0.474000001 | 0.622061097 | 0.803789504 | 0.7361 | 0.858124155 |
| StyR.362_ | -5.593801528 | -5.374288427 | -5.879741913 | -0.543113008 | 1.446615238 | -0.333873395 | 0.338000007 | 0.481130176 | 0.710441009 | 0.6095 | 0.772726463 |
| StyR.369_ | -5.611441535 | -5.737308074 | -5.445099057 | 0.156661365 | 1.568684414 | 0.0781541 | 0.457333335 | 0.653928547 | 0.828109635 | 0.7463 | 0.871483072 |
| StyR.377_ | 1.113116134 | 1.508844739 | 0.901174344 | -0.642245133 | 0.39643463 | -1.574272509 | 0.020653117 | 0.083611654 | 0.332478256 | 0.118 | 0.356457939 |
| StyR.378_ | -7.00881533 | -7.205100656 | -6.798822769 | 0.577772791 | 2.642690422 | 0.173256735 | 0.398666671 | 0.552793223 | 0.764123135 | 0.6948 | 0.832725668 |
| StyR.44a_ | -0.39069258 | -0.436175316 | -0.346079306 | 0.032360739 | 1.026831989 | 0.028721118 | 0.481333334 | 0.894608748 | 0.96040762 | 0.9526 | 0.979832162 |
| StyR.44b_ | 0.859670538 | 0.789999844 | 1.300557157 | 0.510335327 | 0.458014952 | 0.953892569 | 0.204000017 | 0.246578318 | 0.547356661 | 0.3897 | 0.654611898 |
| StyR.44c_ | -1.820697214 | -1.975544977 | -1.222967273 | 0.651366042 | 0.778240863 | 0.80388962 | 0.229846783 | 0.32906251 | 0.610132067 | 0.4414 | 0.692602675 |
| StyR.44d_ | -1.391023497 | -1.504844268 | -1.135323906 | 0.402245086 | 0.789817465 | 0.557714307 | 0.240000013 | 0.412466045 | 0.68142117 | 0.4744 | 0.717846714 |
| StyR.44e_ | -3.510125499 | -4.002346469 | -2.980693365 | 1.079411245 | 0.646341992 | 1.601628797 | 0.03397748 | 0.080297722 | 0.297545912 | 0.1388 | 0.377620684 |
| StyR.44f_ | -4.27996876 | -4.501279797 | -3.850680223 | 0.486077144 | 1.038659721 | 0.406778769 | 0.330666674 | 0.581065342 | 0.776475423 | 0.6486 | 0.815214847 |
| StyR.44g_ | -4.030272976 | -4.427964567 | -3.557598926 | 0.856690829 | 0.817915891 | 1.000321178 | 0.134000029 | 0.208795503 | 0.463995398 | 0.295 | 0.532116316 |
| StyR.51_ | 0.415550017 | 0.609665844 | 0.35133321 | -0.266466489 | 0.257260083 | -0.972805569 | 0.148567647 | 0.233221292 | 0.522571944 | 0.3122 | 0.556438054 |
| StyR.55_ | 4.750205077 | 4.701783121 | 4.787543866 | 0.073919817 | 0.289296918 | 0.20216203 | 0.444370422 | 0.900587003 | 0.963526462 | 1 | 1 |
| StyR.8_ | 3.531610356 | 3.450937836 | 3.602168467 | 0.110575151 | 0.26837054 | 0.252576827 | 0.390666671 | 0.929701544 | 0.974620796 | 0.8539 | 0.937733994 |
| StyR.88_ | 0.740500905 | 0.89429379 | 0.666805937 | -0.156308862 | 0.327801834 | -0.384415019 | 0.362666672 | 0.764393147 | 0.897738759 | 0.7601 | 0.896032046 |
| StyR.90_ | -4.672158233 | -4.923171142 | -4.480992342 | 0.417946029 | 1.110672405 | 0.332294426 | 0.338666673 | 0.524895546 | 0.741852935 | 0.6383 | 0.800633159 |
| StyR.99_ | -1.12982003 | -1.36050903 | -1.01506704 | 0.493109547 | 0.58457165 | 0.832502154 | 0.197868106 | 0.251676806 | 0.541695699 | 0.3872 | 0.635233721 |
| tp2_ | -5.767791767 | -5.62918142 | -6.016492171 | -0.368242599 | 1.753002977 | -0.179025811 | 0.415722855 | 0.605924239 | 0.797367596 | 0.7352 | 0.860399805 |
| tp28_ | 0.197887358 | 0.397840613 | 0.134642151 | -0.227590121 | 0.455379054 | -0.489940437 | 0.301132587 | 0.590685874 | 0.802211333 | 0.6111 | 0.810739831 |
| tpk3_ | 0.576869739 | 0.710262332 | 0.391680915 | -0.148668361 | 0.564287051 | -0.262679807 | 0.434377084 | 0.696707128 | 0.866818217 | 0.9234 | 0.964714273 |
| tpke11_ | 4.202037719 | 4.161336167 | 4.234173693 | 0.071168139 | 0.394972111 | 0.081651559 | 0.402000004 | 0.712122769 | 0.880306422 | 0.8746 | 0.946542789 |
| tpke70_ | -1.864112112 | -1.972665279 | -1.75891729 | 0.009940516 | 0.712661804 | 0.014615671 | 0.493670886 | 0.789739111 | 0.909522264 | 0.9068 | 0.958603678 |
| zipA.leader_ | 2.238970434 | 1.95777151 | 2.432910598 | 0.473147554 | 0.140483107 | 3.373158706 | 4.68E-05 | 0.009291327 | 0.1155806 | 0.1 | 0.330257709 |

LE

| | rab.all | rab.win.tnp | rab.win.WT | diff.btw | diff.win | effect | overlap | we.ep | we.eBH | wi.ep | wi.eBH |
|---|---|---|---|---|---|---|---|---|---|---|---|
| aceF_aceF | 5.34687 | 5.395575 | 5.286692 | -0.12409 | 0.069653 | -1.95784 | 4.68E-05 | 0.060274 | 0.186611 | 0.1004 | 0.237123 |
| SL1344_RS11990_ackA | 3.716523 | 3.742125 | 3.664625 | -0.07525 | 0.072983 | -0.94733 | 0.159227 | 0.23056 | 0.400271 | 0.3234 | 0.480262 |
| SL1344_RS08180_acpD | 0.087707 | 0.144498 | 0.028427 | -0.10804 | 0.238083 | -0.43779 | 0.314457 | 0.509758 | 0.65251 | 0.6314 | 0.739698 |
| acpS_acpS | 0.650519 | 0.400776 | 0.907596 | 0.491656 | 0.177351 | 2.670916 | 0.000673 | 0.020281 | 0.101065 | 0.1007 | 0.237534 |
| adrA_adrA | -0.46529 | -0.63724 | -0.26264 | 0.374868 | 0.264711 | 1.381246 | 0.061959 | 0.111648 | 0.253862 | 0.1753 | 0.32605 |
| SL1344_RS16630_aer | 4.124249 | 4.650247 | 3.923611 | -0.71617 | 0.202071 | -2.78246 | 4.68E-05 | 0.021135 | 0.109686 | 0.1 | 0.236592 |
| SL1344_RS13055_alpha.beta.hydrolase | -2.8423 | -2.83287 | -2.87966 | -0.04431 | 0.754507 | -0.05147 | 0.48501 | 0.749914 | 0.843001 | 0.8294 | 0.895852 |
| AmgR_amgR | -5.3602 | -5.3414 | -5.39369 | 0.000479 | 1.618509 | 9.78E-05 | 0.5 | 0.676898 | 0.786715 | 0.777 | 0.852016 |
| amiC_amiC | 1.021203 | 1.311412 | 0.901282 | -0.3876 | 0.204129 | -1.70425 | 0.034644 | 0.074759 | 0.206753 | 0.1368 | 0.282603 |
| SL1344_RS04710_amino.acid.permease | -0.37739 | -0.57493 | -0.20055 | 0.372935 | 0.173722 | 2.118735 | 0.011326 | 0.044055 | 0.144004 | 0.1128 | 0.252344 |
| SL1344_RS01920_ampH | -0.64421 | -1.06436 | -0.26791 | 0.810608 | 0.296223 | 2.712431 | 0.001337 | 0.023345 | 0.107722 | 0.1006 | 0.237348 |
| SL1344_RS23585_arcA | 4.775396 | 4.846645 | 4.750062 | -0.08372 | 0.076903 | -0.96351 | 0.182667 | 0.240254 | 0.419212 | 0.3457 | 0.509449 |
| ArcZ_arcZ | 1.38235 | 1.254461 | 1.489773 | 0.331729 | 0.302464 | 1.179243 | 0.033311 | 0.140156 | 0.304457 | 0.1323 | 0.276114 |
| STnc490a_art200 | -8.711 | -9.87514 | -7.54857 | 2.182346 | 3.725723 | 0.510858 | 0.232667 | 0.387523 | 0.532557 | 0.4631 | 0.590881 |
| STnc490b_art200 | -9.21101 | -9.87736 | -8.43238 | 1.318757 | 4.212685 | 0.277169 | 0.352432 | 0.502638 | 0.638901 | 0.6135 | 0.718945 |
| STnc490c_art200 | -8.67537 | -9.81214 | -7.21926 | 2.470336 | 3.839671 | 0.578275 | 0.227848 | 0.389019 | 0.536779 | 0.47 | 0.601805 |
| STnc490e_art200 | -8.06788 | -8.81602 | -7.2985 | 1.405935 | 3.55456 | 0.342493 | 0.313125 | 0.499614 | 0.637471 | 0.6024 | 0.710439 |
| STnc490f_art200 | -7.72513 | -9.82186 | -6.59871 | 3.220781 | 2.781092 | 1.051079 | 0.069333 | 0.160753 | 0.305936 | 0.2053 | 0.346957 |
| artM_artM | 1.580227 | 1.607233 | 1.526789 | -0.07389 | 0.110854 | -0.60624 | 0.263333 | 0.402901 | 0.553872 | 0.5143 | 0.643591 |
| artP_artP | 0.818775 | 1.015411 | 0.557777 | -0.47329 | 0.180116 | -2.63107 | 0.001336 | 0.029137 | 0.121309 | 0.1004 | 0.237119 |
| asarfA_asarfA | 0.815575 | 0.554569 | 0.980455 | 0.417303 | 0.20293 | 1.995724 | 0.014 | 0.04799 | 0.159864 | 0.1132 | 0.253787 |
| asnB_asnB | 2.879985 | 2.836965 | 3.831769 | 0.993517 | 0.161188 | 1.106262 | 0.090606 | 0.16878 | 0.345052 | 0.2057 | 0.362108 |
| SL1344_RS13060_asrA | -0.41662 | -0.47618 | -0.38829 | 0.070132 | 0.290091 | 0.223074 | 0.418 | 0.669024 | 0.781116 | 0.7714 | 0.851864 |
| SL1344_RS13065_asrB | -0.29284 | -0.17539 | -0.37463 | -0.19886 | 0.207728 | -0.87459 | 0.173218 | 0.257111 | 0.413135 | 0.354 | 0.496949 |
| SL1344_RS13070_asrC | 0.490885 | 0.612721 | 0.393769 | -0.19795 | 0.187733 | -0.96934 | 0.153333 | 0.220134 | 0.382073 | 0.3009 | 0.457105 |
| asyhbQ_asyhbQ | 1.528248 | 1.30686 | 1.854955 | 0.528434 | 0.150373 | 3.420837 | 4.68E-05 | 0.012044 | 0.079579 | 0.1 | 0.236592 |
| atpC_atpC | 5.830014 | 5.597486 | 6.058697 | 0.48378 | 0.098909 | 4.847963 | 4.68E-05 | 0.001854 | 0.040662 | 0.1 | 0.236592 |
| SL1344_RS15305_barA | 2.020502 | 1.935839 | 2.105666 | 0.180209 | 0.123359 | 1.449708 | 0.031979 | 0.100008 | 0.240688 | 0.1418 | 0.28705 |
| bcsA_bcsA | 1.866162 | 1.347579 | 2.22158 | 0.866133 | 0.163862 | 5.206379 | 4.68E-05 | 0.004211 | 0.050763 | 0.1 | 0.236592 |
| SL1344_RS17950_bigA | -0.11938 | -0.08868 | -0.17343 | -0.06417 | 0.241977 | -0.24507 | 0.4004 | 0.633919 | 0.751474 | 0.7327 | 0.820071 |
| bssR_bssR | 1.631216 | 1.395286 | 2.010748 | 0.592689 | 0.157833 | 3.641177 | 4.68E-05 | 0.011124 | 0.07773 | 0.1 | 0.236592 |
| bssS_bssS | 2.338467 | 2.199073 | 2.480649 | 0.277379 | 0.088177 | 3.070137 | 4.68E-05 | 0.014515 | 0.084464 | 0.1006 | 0.237378 |
| cadB_cadB | 1.977807 | 1.651699 | 2.828979 | 1.293409 | 0.689416 | 2.006184 | 4.68E-05 | 0.029252 | 0.129481 | 0.1 | 0.236592 |
| cbiC_cbiC | 0.928616 | 0.402992 | 1.476348 | 1.086375 | 0.141088 | 7.714549 | 4.68E-05 | 0.001414 | 0.034311 | 0.1 | 0.236592 |
| cbiD_cbiD | 1.4825 | 0.924706 | 1.857483 | 0.954323 | 0.241842 | 3.955382 | 4.68E-05 | 0.007384 | 0.065345 | 0.1 | 0.236592 |
| SL1344_RS11580_ccmA | 0.591941 | 0.06791 | 0.812375 | 0.738465 | 0.271022 | 2.492271 | 0.000673 | 0.050469 | 0.168409 | 0.1 | 0.236592 |
| SL1344_RS11575_ccmB | -0.43342 | -0.59945 | -0.19798 | 0.306467 | 0.34684 | 0.887458 | 0.184544 | 0.269816 | 0.441675 | 0.3447 | 0.504235 |
| cdsA_cdsA | 2.402168 | 2.398345 | 2.406293 | -0.04527 | 0.193865 | -0.23668 | 0.402 | 0.665346 | 0.783354 | 0.8462 | 0.900538 |
| SL1344_RS02340_cdsH | -0.75681 | -0.75867 | -0.75584 | -0.00411 | 0.23167 | -0.01405 | 0.493005 | 0.605624 | 0.72693 | 0.7072 | 0.798676 |
| chaB_chaB | 0.224448 | 0.438611 | 0.032024 | -0.38724 | 0.231852 | -1.62961 | 0.049333 | 0.081982 | 0.213815 | 0.1462 | 0.294847 |
| SL1344_RS09610_cheA | 6.520316 | 7.431433 | 5.769992 | -1.66651 | 0.181007 | -9.38992 | 4.68E-05 | 0.0006 | 0.029987 | 0.1 | 0.236592 |
| SL1344_RS09600_cheM | 5.891597 | 6.725627 | 5.17307 | -1.5535 | 0.128287 | -11.8036 | 4.68E-05 | 0.000782 | 0.031673 | 0.1 | 0.236592 |
| SL1344_RS09595_chemotaxis.protein.methyltransferase | 4.343067 | 5.188384 | 3.523798 | -1.66414 | 0.111488 | -14.926 | 4.68E-05 | 8.68E-05 | 0.018204 | 0.1 | 0.236592 |
| SL1344_RS09585_chemotaxis.regulator | 4.351923 | 4.935887 | 3.715895 | -1.29934 | 0.175684 | -7.40268 | 4.68E-05 | 0.003045 | 0.048208 | 0.1 | 0.236592 |

| | | | | | | | | | | |
|---|---|---|---|---|---|---|---|---|---|---|
| SL1344_RS09590_chemotaxis.response.regulator.protein.glutamate.methylesterase | 3.996499 | 4.676904 | 3.251592 | -1.44287 | 0.168937 | -8.65522 | 4.68E-05 | 0.000374 | 0.026482 | 0.1 | 0.236592 |
| SL1344_RS11875_cheV | 5.196218 | 5.863351 | 4.590091 | -1.19826 | 0.139365 | -8.5641 | 4.68E-05 | 0.00045 | 0.027639 | 0.1 | 0.236592 |
| SL1344_RS09605_CheW | 6.623055 | 7.442738 | 5.801331 | -1.71337 | 0.160344 | -10.6825 | 4.68E-05 | 5.54E-05 | 0.01878 | 0.1 | 0.236592 |
| SL1344_RS09580_cheZ | 5.328427 | 6.044429 | 4.650421 | -1.39565 | 0.157961 | -8.67068 | 4.68E-05 | 0.002338 | 0.044167 | 0.1 | 0.236592 |
| chiP_chiP | 2.034927 | 1.660885 | 2.315446 | 0.647111 | 0.143006 | 4.452496 | 4.68E-05 | 0.009487 | 0.071504 | 0.1 | 0.236592 |
| ChiX_chiX | 1.622627 | 1.568144 | 1.69579 | 0.106921 | 1.214168 | 0.088669 | 0.456362 | 0.954145 | 0.974625 | 0.9976 | 0.998577 |
| SL1344_RS02290_clpP | 3.445014 | 3.238813 | 3.532046 | 0.253294 | 0.14615 | 1.669856 | 0.036 | 0.071946 | 0.206718 | 0.1346 | 0.282133 |
| SL1344_RS02295_clpX | 5.011209 | 4.94769 | 5.032442 | 0.078346 | 0.047627 | 1.423847 | 0.078614 | 0.122845 | 0.274254 | 0.2038 | 0.357302 |
| SL1344_RS09530_cmoA | 1.631792 | 1.86464 | 1.302833 | -0.55937 | 0.106914 | -5.1262 | 4.68E-05 | 0.006454 | 0.058164 | 0.1 | 0.236592 |
| cobS_cobS | 0.800156 | 0.398488 | 1.344438 | 0.924031 | 0.190599 | 4.769421 | 4.68E-05 | 0.007949 | 0.066336 | 0.1 | 0.236592 |
| copA_copA | 2.916713 | 2.626689 | 3.200551 | 0.525007 | 0.310899 | 1.60919 | 0.05996 | 0.06665 | 0.20018 | 0.1511 | 0.304727 |
| SL1344_RS20320_corA | 1.623696 | 1.613752 | 1.633528 | 0.059993 | 0.166242 | 0.35431 | 0.367089 | 0.523131 | 0.670421 | 0.7341 | 0.813384 |
| SL1344_RS20860_cpxA | 2.818031 | 2.75749 | 2.868285 | 0.101785 | 0.09032 | 1.075627 | 0.119254 | 0.185357 | 0.344466 | 0.2626 | 0.419769 |
| cpxP_cpxP | 2.985844 | 2.790817 | 3.148898 | 0.14493 | 0.313935 | 0.921412 | 0.140667 | 0.305692 | 0.492743 | 0.258 | 0.428975 |
| SL1344_RS20865_cpxR | 1.874759 | 1.880015 | 1.869845 | -0.01189 | 0.099859 | -0.11327 | 0.454364 | 0.604496 | 0.726467 | 0.7136 | 0.801995 |
| SL1344_RS17890_crp | 5.122984 | 5.108919 | 5.141926 | 0.040202 | 0.050928 | 0.767722 | 0.203864 | 0.287032 | 0.45892 | 0.4143 | 0.56082 |
| CrpT_crpT | -0.95961 | -1.06013 | -0.89835 | 0.082448 | 0.423669 | 0.187434 | 0.428381 | 0.718535 | 0.819057 | 0.8343 | 0.892796 |
| csgB_csgB | -4.96292 | -4.94501 | -4.97241 | -0.11542 | 1.202924 | -0.08145 | 0.46569 | 0.632581 | 0.751359 | 0.7489 | 0.831262 |
| SL1344_RS05640_csgC | 2.140167 | 2.93579 | 1.583748 | -1.35828 | 0.258339 | -5.11144 | 4.68E-05 | 0.005966 | 0.061415 | 0.1 | 0.236592 |
| SL1344_RS18810_cspA | 1.99446 | 1.704228 | 2.026781 | 0.333324 | 0.623786 | 0.631448 | 0.219333 | 0.420537 | 0.60122 | 0.4039 | 0.585459 |
| SL1344_RS09980_cspB | -1.24419 | -1.31772 | -1.05108 | 0.2031 | 0.423267 | 0.388835 | 0.345103 | 0.65731 | 0.772034 | 0.7099 | 0.815895 |
| SL1344_RS06145_cspH | -4.44752 | -4.49413 | -4.38171 | 0.121645 | 0.869228 | 0.13617 | 0.437333 | 0.588818 | 0.715275 | 0.7077 | 0.797006 |
| SL1344_RS14635_csrA | 4.233621 | 4.162343 | 4.466748 | 0.288749 | 0.118481 | 1.703988 | 0.013334 | 0.078619 | 0.218885 | 0.1125 | 0.253222 |
| CsrB_csrB | 9.59326 | 9.498321 | 9.760969 | 0.202459 | 0.182676 | 1.417364 | 0.112 | 0.090414 | 0.238869 | 0.2 | 0.370027 |
| SL1344_RS15345_csrB | 9.392511 | 9.274502 | 9.545862 | 0.221108 | 0.181969 | 1.297159 | 0.115257 | 0.110759 | 0.268692 | 0.2 | 0.370027 |
| CsrC_csrC | 8.448068 | 8.514517 | 8.377275 | 0.013609 | 0.216533 | 0.071517 | 0.453031 | 0.877098 | 0.930488 | 1 | 1 |
| SL1344_RS17435_csrD | 2.572497 | 2.788068 | 2.519627 | -0.27028 | 0.218368 | -1.19247 | 0.128581 | 0.147553 | 0.314585 | 0.2514 | 0.419029 |
| CyaR_cyaR | 5.582104 | 5.344648 | 5.628288 | 0.288287 | 0.258912 | 1.021754 | 0.208 | 0.199842 | 0.38199 | 0.3744 | 0.555124 |
| SL1344_RS12535_cysA | 3.305947 | 4.693057 | 1.58423 | -3.08934 | 0.454829 | -6.58028 | 4.68E-05 | 0.004061 | 0.053781 | 0.1 | 0.236592 |
| SL1344_RS08565_cysB | 2.409409 | 3.455313 | 1.815707 | -1.6369 | 0.28856 | -4.75575 | 4.68E-05 | 0.02122 | 0.110422 | 0.1 | 0.236592 |
| SL1344_RS15175_cysC | 2.649839 | 4.491644 | 0.052628 | -4.33976 | 0.424298 | -9.79408 | 4.68E-05 | 0.003214 | 0.049253 | 0.1 | 0.236592 |
| SL1344_RS15185_cysD | 2.110155 | 4.795192 | -1.05907 | -5.46664 | 0.742856 | -7.77018 | 4.68E-05 | 0.000455 | 0.027523 | 0.1 | 0.236592 |
| SL1344_RS19060_cysE | 2.151589 | 2.184039 | 2.110936 | -0.07583 | 0.087833 | -0.83049 | 0.181879 | 0.279279 | 0.434248 | 0.3783 | 0.516615 |
| SL1344_RS17945_cysG | 2.403386 | 2.27059 | 2.516202 | 0.247544 | 0.087937 | 2.793058 | 4.68E-05 | 0.020379 | 0.10007 | 0.1006 | 0.237396 |
| SL1344_RS15240_cysH | 2.499424 | 4.075874 | -0.337 | -4.43378 | 0.512187 | -7.87148 | 4.68E-05 | 0.012098 | 0.084897 | 0.1 | 0.236592 |
| SL1344_RS15245_cysI | 4.380328 | 6.06424 | 1.989059 | -4.0084 | 0.361921 | -9.00387 | 4.68E-05 | 0.006549 | 0.065154 | 0.1 | 0.236592 |
| SL1344_RS15250_cysJ | 3.234248 | 5.427095 | 0.23531 | -5.12135 | 0.454367 | -10.1552 | 4.68E-05 | 0.00369 | 0.051972 | 0.1 | 0.236592 |
| SL1344_RS12475_cysK | 5.327925 | 7.158138 | 3.226004 | -3.56993 | 0.491526 | -6.95373 | 4.68E-05 | 0.0005 | 0.028858 | 0.1 | 0.236592 |
| SL1344_RS12530_cysM | 1.274891 | 2.171785 | 0.369168 | -1.74795 | 0.215374 | -8.11041 | 4.68E-05 | 0.0006 | 0.028279 | 0.1 | 0.236592 |
| SL1344_RS15180_cysN | 3.439915 | 5.631472 | 0.735726 | -4.69377 | 0.571901 | -7.23945 | 4.68E-05 | 0.001736 | 0.040062 | 0.1 | 0.236592 |
| SL1344_RS12550_cysP | 2.315585 | 5.397266 | -0.93594 | -5.64032 | 0.91948 | -6.70399 | 4.68E-05 | 0.000287 | 0.025723 | 0.1 | 0.236592 |
| SL1344_RS22605_cysQ | 1.938745 | 1.816913 | 2.032689 | 0.216841 | 0.114014 | 1.817833 | 0.019334 | 0.05849 | 0.175083 | 0.124 | 0.266353 |
| SL1344_RS02725_cysS | 2.777493 | 2.756734 | 2.799888 | 0.038115 | 0.074807 | 0.492679 | 0.293138 | 0.455973 | 0.599174 | 0.5612 | 0.678389 |
| SL1344_RS09720_cysT | 2.106792 | 2.199485 | 1.753367 | -0.48239 | 0.332542 | -1.51947 | 0.014657 | 0.084205 | 0.227559 | 0.1162 | 0.257243 |
| SL1344_RS12545_cysU | 1.772297 | 4.010329 | -1.14103 | -4.86458 | 0.746319 | -6.6471 | 4.68E-05 | 0.001409 | 0.036951 | 0.1 | 0.236592 |

| | | | | | | | | | | | |
|---|---|---|---|---|---|---|---|---|---|---|---|
| SL1344_RS12540_cysW | 2.659288 | 4.663417 | 0.177618 | -4.35797 | 0.490092 | -8.24293 | 4.68E-05 | 0.001871 | 0.040642 | 0.1 | 0.236592 |
| SL1344_RS12470_cysZ | 0.475433 | 0.593275 | 0.378481 | -0.21265 | 0.179719 | -1.12147 | 0.109927 | 0.168017 | 0.321018 | 0.2444 | 0.396253 |
| SL1344_RS16735_cyuA | 0.195531 | 0.122369 | 0.322323 | 0.193096 | 0.170783 | 1.073328 | 0.132578 | 0.202339 | 0.355227 | 0.292 | 0.440957 |
| SL1344_RS16740_cyuP | -0.81689 | -0.82143 | -0.81028 | 0.075032 | 0.352907 | 0.175401 | 0.424384 | 0.638348 | 0.758563 | 0.7851 | 0.856311 |
| SL1344_RS10610_dacD | -1.83125 | -2.33347 | -1.4244 | 0.913101 | 0.482049 | 1.82716 | 0.014667 | 0.056311 | 0.173661 | 0.1182 | 0.26 |
| SL1344_RS17980_dam | 1.821405 | 1.724409 | 1.895552 | 0.177007 | 0.124781 | 1.370911 | 0.05463 | 0.107083 | 0.247288 | 0.1717 | 0.321078 |
| DapZ_dapZ | 1.153009 | 1.371274 | 1.034908 | -0.35645 | 0.214129 | -1.59384 | 0.024 | 0.083155 | 0.221386 | 0.1307 | 0.274939 |
| ddl_ddl | 3.041854 | 3.020207 | 3.096168 | 0.061067 | 0.124764 | 0.427262 | 0.338667 | 0.666619 | 0.784093 | 0.7139 | 0.822762 |
| SL1344_RS09505_dihydroneopterin.triphosphate.diphosphatase | 2.525188 | 2.809588 | 2.156385 | -0.65038 | 0.115819 | -5.56593 | 4.68E-05 | 0.007502 | 0.064989 | 0.1 | 0.236592 |
| dinG_dinG | 1.609528 | 1.753182 | 1.481583 | -0.27401 | 0.111935 | -2.38259 | 0.002668 | 0.030988 | 0.123402 | 0.1024 | 0.239699 |
| SL1344_RS05725_dinI | 1.488027 | 1.887749 | 1.183515 | -0.67871 | 0.202928 | -3.32871 | 4.68E-05 | 0.009714 | 0.073907 | 0.1 | 0.236592 |
| SL1344_RS04685_dmsA | 3.647168 | 3.301063 | 3.949756 | 0.692185 | 0.135948 | 5.207872 | 4.68E-05 | 0.003626 | 0.049573 | 0.1 | 0.236592 |
| SL1344_RS04690_dmsB | 0.558936 | 0.186552 | 0.971119 | 0.857887 | 0.35854 | 2.455934 | 4.68E-05 | 0.024103 | 0.113736 | 0.1001 | 0.236725 |
| SL1344_RS04695_dmsC | 1.87732 | 1.39748 | 2.401063 | 0.954327 | 0.273008 | 3.426719 | 4.68E-05 | 0.007656 | 0.06838 | 0.1 | 0.236592 |
| SL1344_RS06610_DNA.RNA.non.specific.endonuclease | 0.886249 | 0.699067 | 0.978921 | 0.273895 | 0.148361 | 1.711187 | 0.039973 | 0.082166 | 0.210377 | 0.1479 | 0.292215 |
| dnaA_dnaA | 2.717386 | 2.78544 | 2.650239 | -0.13543 | 0.069752 | -1.90135 | 0.018654 | 0.053027 | 0.16134 | 0.1201 | 0.260725 |
| SL1344_RS00060_dnaK | 7.003344 | 6.985951 | 7.019811 | 0.021213 | 0.092131 | 0.22386 | 0.417055 | 0.766737 | 0.859713 | 0.9214 | 0.950766 |
| dpiA_dpiA | -1.35077 | -1.37336 | -1.32906 | 0.040563 | 0.384107 | 0.103003 | 0.451699 | 0.687826 | 0.795215 | 0.7841 | 0.857038 |
| dpiA_dpiA.1 | -0.81641 | -0.85468 | -0.78158 | 0.076105 | 0.290043 | 0.245332 | 0.386667 | 0.599883 | 0.723844 | 0.7071 | 0.796776 |
| dpiB_dpiB | -0.74525 | -0.82831 | -0.66631 | 0.15568 | 0.207484 | 0.718687 | 0.203198 | 0.315053 | 0.460658 | 0.405 | 0.540189 |
| DsrA_dsrA | -3.94657 | -4.75818 | -3.69488 | 0.969532 | 1.054285 | 0.806556 | 0.210526 | 0.305769 | 0.479349 | 0.4404 | 0.58896 |
| SL1344_RS09330_DUF1441.domain.containing.protein | -3.62336 | -3.14522 | -4.12085 | -0.97935 | 0.579788 | -1.6155 | 0.037975 | 0.07936 | 0.198585 | 0.1535 | 0.297232 |
| SL1344_RS06460_DUF1496.domain.containing.protein | 3.390193 | 3.841275 | 2.786387 | -1.05192 | 0.185375 | -5.67824 | 4.68E-05 | 0.001926 | 0.040286 | 0.1 | 0.236592 |
| SL1344_RS16295_DUF2623.domain.containing.protein | 0.76472 | 1.71458 | -0.02752 | -1.68376 | 0.269992 | -6.17297 | 4.68E-05 | 0.002337 | 0.04203 | 0.1 | 0.236592 |
| emrB_emrB | 1.179676 | 1.196048 | 1.161731 | -0.03401 | 0.13577 | -0.23533 | 0.401333 | 0.574271 | 0.702933 | 0.6932 | 0.78607 |
| emrD_emrD | 1.224218 | 1.322037 | 1.110948 | -0.21131 | 0.123439 | -1.6719 | 0.028667 | 0.073634 | 0.196419 | 0.1326 | 0.276219 |
| emtA_emtA | 0.121676 | 0.246242 | -0.01447 | -0.2615 | 0.147024 | -1.71139 | 0.034667 | 0.071012 | 0.186867 | 0.1392 | 0.281389 |
| SL1344_RS03015_entB | -2.33819 | -2.43809 | -2.24834 | 0.185562 | 0.373684 | 0.449891 | 0.291805 | 0.426445 | 0.571174 | 0.5475 | 0.662614 |
| entE_entE | -2.25233 | -2.19695 | -2.3057 | -0.18666 | 0.534653 | -0.35572 | 0.357095 | 0.531479 | 0.671628 | 0.7048 | 0.790889 |
| entF_entF | -0.68972 | -0.63502 | -0.75529 | -0.11742 | 0.210671 | -0.52006 | 0.282478 | 0.393678 | 0.538196 | 0.4941 | 0.616221 |
| SL1344_RS18065_envZ | 2.685853 | 2.524971 | 2.868234 | 0.362609 | 0.112442 | 3.332059 | 4.68E-05 | 0.015071 | 0.088008 | 0.1 | 0.236592 |
| eutA_eutA | -1.71886 | -1.84368 | -1.58915 | 0.245339 | 0.333613 | 0.677659 | 0.222667 | 0.335047 | 0.48938 | 0.4408 | 0.575986 |
| SL1344_RS12680_eutS | -4.55669 | -4.41184 | -4.77915 | -0.47032 | 1.141208 | -0.35728 | 0.321119 | 0.450107 | 0.60726 | 0.5867 | 0.697055 |
| fabG_fabG | 3.721861 | 3.680108 | 3.756757 | 0.077655 | 0.060489 | 1.211314 | 0.097269 | 0.152089 | 0.303655 | 0.2238 | 0.376172 |
| fadA_fadA | 0.217448 | 0.539368 | 0.040971 | -0.47552 | 0.225754 | -1.99307 | 0.017322 | 0.048274 | 0.158979 | 0.1165 | 0.257358 |
| SL1344_RS09080_fadD | 1.399017 | 1.929453 | 0.854206 | -1.07622 | 0.141859 | -7.68572 | 4.68E-05 | 0.001033 | 0.031711 | 0.1 | 0.236592 |
| fadE_fadE | 1.323777 | 1.370757 | 1.250202 | -0.12996 | 0.137765 | -0.88854 | 0.156 | 0.244934 | 0.408845 | 0.3332 | 0.477558 |
| fadI_fadI | -1.00438 | -0.82192 | -1.17249 | -0.37093 | 0.283613 | -1.26915 | 0.077948 | 0.137413 | 0.283779 | 0.2079 | 0.356338 |
| SL1344_RS13945_Fels.2.prophage.protein | -3.43542 | -3.41304 | -3.45547 | -0.05328 | 0.531012 | -0.09046 | 0.458361 | 0.546997 | 0.675517 | 0.6546 | 0.751964 |
| feoA_feoA | -1.6018 | -2.18621 | -1.44645 | 0.745357 | 0.73653 | 1.084779 | 0.155896 | 0.17014 | 0.338852 | 0.2913 | 0.461992 |
| ffs_ffs | 3.511637 | 3.526198 | 3.495243 | -0.10012 | 0.189517 | -0.53718 | 0.332667 | 0.412744 | 0.589672 | 0.6783 | 0.788653 |
| fimA_fimA | -1.1503 | -1.53347 | -0.71569 | 0.814407 | 0.259136 | 3.068439 | 4.68E-05 | 0.016936 | 0.090102 | 0.1001 | 0.236725 |
| fimF_fimF | -2.32049 | -2.2219 | -2.38147 | -0.20485 | 0.503765 | -0.3787 | 0.353333 | 0.5341 | 0.669896 | 0.6713 | 0.766175 |
| SL1344_RS02785_fimZ | -3.2014 | -3.27376 | -3.09834 | 0.095193 | 0.766457 | 0.102876 | 0.445037 | 0.694528 | 0.800026 | 0.7846 | 0.860363 |
| fis_fis | 0.951094 | 1.253715 | 0.613809 | -0.75165 | 0.298382 | -2.55033 | 4.68E-05 | 0.025667 | 0.117113 | 0.1 | 0.236592 |

| | | | | | | | | | | | |
|---|---|---|---|---|---|---|---|---|---|---|---|
| fixB_fixB | -1.96702 | -1.90614 | -2.06878 | -0.14758 | 0.412604 | -0.33199 | 0.357095 | 0.551485 | 0.682967 | 0.6643 | 0.763984 |
| flgD_flgD | 4.766925 | 5.077502 | 4.45136 | -0.62995 | 0.102599 | -6.21098 | 4.68E-05 | 0.003866 | 0.051754 | 0.1 | 0.236592 |
| flgF_flgF | 5.257773 | 5.696452 | 4.89938 | -0.77093 | 0.121441 | -6.09905 | 4.68E-05 | 0.002548 | 0.044845 | 0.1 | 0.236592 |
| flgI_flgI | 3.994227 | 4.244327 | 3.626552 | -0.60776 | 0.086078 | -6.7967 | 4.68E-05 | 0.002743 | 0.044211 | 0.1 | 0.236592 |
| SL1344_RS05830_flgK | 5.903171 | 6.509457 | 5.413746 | -1.09548 | 0.156305 | -7.04711 | 4.68E-05 | 0.004581 | 0.056416 | 0.1 | 0.236592 |
| flgL_flgL | 5.807262 | 6.357298 | 5.37926 | -0.98009 | 0.128001 | -7.56601 | 4.68E-05 | 0.003323 | 0.04993 | 0.1 | 0.236592 |
| SL1344_RS05775_flgM | 5.229267 | 5.619885 | 4.729149 | -0.89835 | 0.142276 | -6.26969 | 4.68E-05 | 0.000558 | 0.029319 | 0.1 | 0.236592 |
| SL1344_RS05770_flgN | 5.278533 | 5.744304 | 4.877236 | -0.84764 | 0.169595 | -5.02467 | 4.68E-05 | 0.001405 | 0.037449 | 0.1 | 0.236592 |
| SL1344_RS09570_flhA | 4.388666 | 5.123613 | 3.575599 | -1.5498 | 0.09136 | -16.9336 | 4.68E-05 | 3.91E-05 | 0.015992 | 0.1 | 0.236592 |
| SL1344_RS09575_flhB | 3.752463 | 4.479708 | 2.966941 | -1.51089 | 0.085285 | -17.6954 | 4.68E-05 | 6.57E-05 | 0.01694 | 0.1 | 0.236592 |
| SL1344_RS09625_flhC | 4.54659 | 4.995995 | 4.061095 | -0.95612 | 0.08935 | -10.7192 | 4.68E-05 | 0.003341 | 0.049524 | 0.1 | 0.236592 |
| SL1344_RS09630_flhD | 3.626727 | 4.071918 | 3.099505 | -0.96155 | 0.079832 | -11.9123 | 4.68E-05 | 0.000613 | 0.026541 | 0.1 | 0.236592 |
| SL1344_RS09565_flhE | 2.535433 | 3.211488 | 1.829093 | -1.38104 | 0.110628 | -12.4925 | 4.68E-05 | 0.000267 | 0.022374 | 0.1 | 0.236592 |
| SL1344_RS09785_fliA | 5.69099 | 6.497669 | 5.053939 | -1.4356 | 0.206682 | -6.99115 | 4.68E-05 | 0.000272 | 0.025557 | 0.1 | 0.236592 |
| SL1344_RS09800_fliC | 8.228415 | 10.12967 | 7.088004 | -3.04018 | 0.23412 | -9.82242 | 4.68E-05 | 0.012197 | 0.085449 | 0.1 | 0.236592 |
| SL1344_RS09845_fliE | 2.901454 | 3.245381 | 2.620692 | -0.65122 | 0.204043 | -3.33393 | 4.68E-05 | 0.007848 | 0.068506 | 0.1 | 0.236592 |
| fliI_fliI | 4.037308 | 4.272822 | 3.836237 | -0.44311 | 0.113971 | -4.08624 | 4.68E-05 | 0.006825 | 0.063824 | 0.1 | 0.236592 |
| fliJ_fliJ | 3.022791 | 3.061924 | 2.846779 | -0.20424 | 0.157568 | -1.27709 | 0.124584 | 0.13104 | 0.291245 | 0.2436 | 0.409097 |
| fliL_fliL | 3.847683 | 4.121952 | 3.4518 | -0.66851 | 0.119967 | -5.63368 | 4.68E-05 | 0.001354 | 0.03583 | 0.1 | 0.236592 |
| fliQ_fliQ | 1.674542 | 2.059801 | 1.287556 | -0.78207 | 0.131689 | -6.06842 | 4.68E-05 | 0.006218 | 0.058506 | 0.1 | 0.236592 |
| SL1344_RS09815_fliT | 5.236173 | 5.643594 | 4.756855 | -0.88857 | 0.091728 | -9.80003 | 4.68E-05 | 0.002173 | 0.042258 | 0.1 | 0.236592 |
| SL1344_RS09775_fliY | 4.712991 | 5.37242 | 4.008704 | -1.40944 | 0.158482 | -9.042 | 4.68E-05 | 0.001858 | 0.040708 | 0.1 | 0.236592 |
| SL1344_RS09780_fliZ | 5.40682 | 6.172187 | 4.652751 | -1.58313 | 0.135996 | -11.694 | 4.68E-05 | 6.15E-05 | 0.018876 | 0.1 | 0.236592 |
| SL1344_RS14340_fljA | 2.725421 | -1.27775 | 2.800272 | 4.034293 | 0.371737 | 0.974973 | 0.293138 | 0.197191 | 0.379276 | 0.5927 | 0.735651 |
| flk_flk | -0.21612 | -0.22329 | -0.20317 | 0.039099 | 0.209466 | 0.175147 | 0.431046 | 0.601088 | 0.723446 | 0.7072 | 0.795743 |
| SL1344_RS08285_fnr | 3.365075 | 3.478441 | 3.280583 | -0.16013 | 0.156732 | -1.08447 | 0.126667 | 0.166549 | 0.337351 | 0.228 | 0.398333 |
| FnrS_fnrS | 5.687158 | 5.652912 | 5.721633 | 0.169925 | 0.268787 | 0.600564 | 0.319121 | 0.383127 | 0.568817 | 0.6379 | 0.772409 |
| SL1344_RS04735_focA | 3.468996 | 3.733256 | 3.275987 | -0.44607 | 0.075857 | -5.46824 | 4.68E-05 | 0.007667 | 0.066121 | 0.1 | 0.236592 |
| folA_folA | -0.26775 | -0.0037 | -0.49131 | -0.46889 | 0.227871 | -1.91337 | 0.026 | 0.052474 | 0.163499 | 0.1238 | 0.266947 |
| frsA_frsA | 2.354293 | 2.26837 | 2.405243 | 0.148536 | 0.133004 | 1.091838 | 0.070667 | 0.18798 | 0.360021 | 0.1879 | 0.337026 |
| SL1344_RS09665_ftnB | 1.678046 | 1.772072 | 1.585637 | -0.18821 | 0.086967 | -2.06789 | 0.011326 | 0.040473 | 0.136999 | 0.1119 | 0.251213 |
| SL1344_RS06895_fumD | -0.09779 | -0.40009 | 0.125673 | 0.517306 | 0.214073 | 2.354033 | 0.005331 | 0.033859 | 0.128384 | 0.1043 | 0.242343 |
| SL1344_RS03500_fur | 3.044157 | 3.278341 | 2.889967 | -0.38904 | 0.091878 | -4.03773 | 4.68E-05 | 0.015512 | 0.090973 | 0.1 | 0.236592 |
| gapA_gapA | 2.758305 | 2.614431 | 2.932453 | 0.301856 | 0.113283 | 2.677379 | 0.000673 | 0.018774 | 0.099402 | 0.1004 | 0.237123 |
| GcvB_gcvB | 6.905352 | 6.766956 | 7.036882 | 0.188855 | 0.619622 | 0.33589 | 0.297135 | 0.597502 | 0.744553 | 0.6205 | 0.75852 |
| gcvT_gcvT | 4.932953 | 4.837234 | 4.99022 | 0.146546 | 0.051798 | 2.676846 | 0.001336 | 0.022587 | 0.108372 | 0.1007 | 0.237477 |
| gidB_gidB | 0.370744 | 0.410105 | 0.320492 | -0.08548 | 0.191754 | -0.43402 | 0.315333 | 0.510173 | 0.649735 | 0.6244 | 0.731692 |
| gldA_gldA | -0.56159 | -0.53982 | -0.58688 | -0.05342 | 0.193534 | -0.2474 | 0.390406 | 0.520248 | 0.651753 | 0.627 | 0.729469 |
| gldA_gldA.1 | 3.205746 | 3.089989 | 3.333711 | 0.25545 | 0.118875 | 2.151165 | 0.001336 | 0.039369 | 0.146878 | 0.1013 | 0.238329 |
| GlmY_glmY | 2.996079 | 2.999632 | 2.992118 | 0.091037 | 0.676087 | 0.126747 | 0.399734 | 0.863818 | 0.922462 | 0.8698 | 0.922877 |
| GlmZ_glmZ | 4.815552 | 4.820195 | 4.810974 | -0.11596 | 0.367104 | -0.30036 | 0.365756 | 0.659526 | 0.789035 | 0.8137 | 0.889381 |
| glnQ_glnQ | 1.796152 | 1.99912 | 1.313246 | -0.66823 | 0.176799 | -2.8767 | 4.68E-05 | 0.04059 | 0.150891 | 0.1 | 0.236592 |
| SL1344_RS13690_gluconolactonase | -0.99706 | -1.41869 | -0.6674 | 0.763509 | 0.317576 | 2.395532 | 0.001337 | 0.030076 | 0.121675 | 0.1012 | 0.238203 |
| SL1344_RS09560_glycoside.hydrolase.family.105.protein | 1.130009 | 1.63885 | 0.380711 | -1.25288 | 0.197785 | -6.33205 | 4.68E-05 | 0.005921 | 0.058307 | 0.1 | 0.236592 |
| gntK_gntK | -2.25408 | -2.25289 | -2.2556 | 0.005356 | 0.407942 | 0.012472 | 0.494667 | 0.611966 | 0.73157 | 0.7248 | 0.808198 |

| | | | | | | | | | | | |
|---|---|---|---|---|---|---|---|---|---|---|---|
| SL1344_RS18265_gntR | 1.301884 | 1.348673 | 1.23989 | -0.09992 | 0.146086 | -0.65401 | 0.236 | 0.371353 | 0.529455 | 0.4709 | 0.608139 |
| gpmA_gpmA | 4.750882 | 4.701136 | 4.797198 | -0.00233 | 0.144447 | -0.01783 | 0.492338 | 0.80471 | 0.882708 | 0.9697 | 0.98108 |
| GraL_graL | 0.616526 | 0.950146 | 0.422441 | -0.54792 | 0.280584 | -1.96128 | 0.004001 | 0.04031 | 0.148035 | 0.1037 | 0.241337 |
| groES_groES | 6.15107 | 6.210037 | 6.08328 | -0.12513 | 0.102267 | -1.13496 | 0.112592 | 0.152763 | 0.323531 | 0.2047 | 0.374879 |
| SL1344_RS14595_gshA | 2.747801 | 2.73267 | 2.762291 | 0.055145 | 0.120357 | 0.418246 | 0.324 | 0.458892 | 0.618045 | 0.644 | 0.746823 |
| SL1344_RS16010_gshB | 2.207612 | 2.108815 | 2.340206 | 0.227302 | 0.117214 | 1.904065 | 0.017322 | 0.050305 | 0.164048 | 0.1131 | 0.25337 |
| SL1344_RS05170_gtgD | -4.28008 | -4.0184 | -4.45089 | -0.45392 | 1.017874 | -0.40381 | 0.33511 | 0.547057 | 0.680515 | 0.6545 | 0.758023 |
| SL1344_RS05180_gtgE | 1.515607 | 1.730915 | 1.261936 | -0.44891 | 0.141637 | -3.15139 | 4.68E-05 | 0.013638 | 0.084253 | 0.1001 | 0.236726 |
| SL1344_RS05185_gtgF | -4.49091 | -4.15406 | -4.82499 | -0.68063 | 0.865507 | -0.74951 | 0.198667 | 0.319238 | 0.470294 | 0.4153 | 0.550914 |
| SL1344_RS11665_gyrA | 4.979184 | 4.972155 | 4.984614 | 0.050655 | 0.111949 | 0.461642 | 0.336 | 0.487226 | 0.653013 | 0.7297 | 0.824825 |
| SL1344_RS13995_head.completion.stabilization.protein | -4.97675 | -4.91022 | -5.02651 | -0.13378 | 1.060177 | -0.11968 | 0.446 | 0.587728 | 0.711891 | 0.6975 | 0.788531 |
| helD_helD | 2.175269 | 2.028905 | 2.360528 | 0.312947 | 0.126839 | 2.395618 | 0.002667 | 0.02562 | 0.115229 | 0.102 | 0.239278 |
| hemG_hemG | 0.72495 | 0.716173 | 0.73063 | 0.017993 | 0.11946 | 0.137159 | 0.439333 | 0.521856 | 0.652094 | 0.6371 | 0.735948 |
| hflB_hflB | 5.749705 | 5.692546 | 5.791383 | 0.101218 | 0.034066 | 2.943323 | 4.68E-05 | 0.030335 | 0.125347 | 0.1001 | 0.236725 |
| SL1344_RS22390_hfq | 4.754379 | 4.591611 | 4.966181 | 0.368056 | 0.057785 | 6.236492 | 4.68E-05 | 0.0027 | 0.043383 | 0.1 | 0.236592 |
| SL1344_RS02410_hha | 1.4205 | 1.155662 | 1.698787 | 0.558152 | 0.131213 | 4.276021 | 4.68E-05 | 0.007771 | 0.064394 | 0.1 | 0.236592 |
| SL1344_RS02415_Hha.toxicity.attenuator | 1.668658 | 1.443347 | 2.092272 | 0.628024 | 0.211125 | 2.810645 | 4.68E-05 | 0.021107 | 0.10674 | 0.1 | 0.236592 |
| SL1344_RS14890_hilA | 4.22137 | 4.571456 | 3.971862 | -0.60156 | 0.192558 | -3.33047 | 4.68E-05 | 0.03162 | 0.134595 | 0.1 | 0.236592 |
| SL1344_RS14845_hilC | 4.59042 | 4.782978 | 4.449766 | -0.3836 | 0.233849 | -1.65762 | 0.000673 | 0.055872 | 0.181958 | 0.1004 | 0.237127 |
| SL1344_RS14885_hilD | 5.072855 | 5.34369 | 4.684632 | -0.65591 | 0.108342 | -6.13441 | 4.68E-05 | 0.008821 | 0.073137 | 0.1 | 0.236592 |
| SL1344_RS23135_hilE | -0.25671 | -0.27295 | -0.21313 | 0.035178 | 0.298152 | 0.104193 | 0.463333 | 0.742193 | 0.837451 | 0.8152 | 0.887343 |
| SL1344_RS21865_HlyD.family.type.I.secretion.periplasmic.adaptor.subunit | 1.917744 | 1.335098 | 2.363091 | 1.005485 | 0.221358 | 4.483022 | 4.68E-05 | 0.002738 | 0.045179 | 0.1 | 0.236592 |
| SL1344_RS08745_hns | 6.805141 | 6.934708 | 6.670223 | -0.2879 | 0.055414 | -5.25985 | 4.68E-05 | 0.00409 | 0.05266 | 0.1 | 0.236592 |
| hscB_hscB | 0.335171 | 0.358213 | 0.294482 | -0.05462 | 0.183034 | -0.28121 | 0.381079 | 0.593384 | 0.71857 | 0.6934 | 0.788524 |
| SL1344_RS08210_hslJ | -0.5874 | -0.52295 | -0.66844 | -0.16045 | 0.285569 | -0.55089 | 0.283333 | 0.42654 | 0.580584 | 0.5441 | 0.668502 |
| SL1344_RS21405_hupA | 6.644894 | 6.679363 | 6.584654 | -0.1127 | 0.072223 | -1.62677 | 4.68E-05 | 0.10058 | 0.252899 | 0.1004 | 0.237125 |
| SL1344_RS21410_hupB | 3.676895 | 3.763354 | 3.603944 | -0.18347 | 0.245895 | -0.72053 | 0.246 | 0.315833 | 0.503518 | 0.4772 | 0.638625 |
| hycD_hycD | -2.74393 | -2.7018 | -2.78688 | -0.07489 | 0.427307 | -0.15845 | 0.428667 | 0.535651 | 0.664263 | 0.645 | 0.744341 |
| hycE_hycE | -1.65079 | -1.62552 | -1.67598 | -0.05202 | 0.267705 | -0.19612 | 0.413333 | 0.507771 | 0.640494 | 0.6103 | 0.717905 |
| SL1344_RS06065_hypothetical.protein | 0.64397 | 1.422543 | -0.28 | -1.70532 | 0.131325 | -12.8617 | 4.68E-05 | 0.000552 | 0.024223 | 0.1 | 0.236592 |
| SL1344_RS07940_hypothetical.protein | 3.645149 | 4.243485 | 2.976913 | -1.28172 | 0.102237 | -12.6288 | 4.68E-05 | 0.000368 | 0.024378 | 0.1 | 0.236592 |
| SL1344_RS09195_hypothetical.protein | 1.267522 | 1.521062 | 1.155466 | -0.36121 | 0.1967 | -1.71776 | 0.023318 | 0.074524 | 0.206778 | 0.1256 | 0.268723 |
| SL1344_RS09675_hypothetical.protein | 2.431989 | 3.347787 | 1.465342 | -1.87488 | 0.103025 | -18.1304 | 4.68E-05 | 0.000101 | 0.017415 | 0.1 | 0.236592 |
| SL1344_RS09745_hypothetical.protein | -0.46475 | 0.133484 | -1.2136 | -1.33995 | 0.226194 | -5.86858 | 4.68E-05 | 0.003601 | 0.045897 | 0.1 | 0.236592 |
| SL1344_RS22135_hypothetical.protein | -0.27981 | -0.14403 | -0.38678 | -0.30083 | 0.48273 | -0.60415 | 0.27515 | 0.393436 | 0.56842 | 0.5336 | 0.678462 |
| iacP_iacP | 1.872307 | 2.19364 | 1.593465 | -0.68451 | 0.313637 | -2.34683 | 4.68E-05 | 0.03411 | 0.138822 | 0.1 | 0.236592 |
| SL1344_RS14895_iagB | 3.992779 | 4.349705 | 3.606691 | -0.82486 | 0.175412 | -4.87515 | 4.68E-05 | 0.001873 | 0.040642 | 0.1 | 0.236592 |
| lamB_iamB | 2.992856 | 2.498482 | 3.348536 | 0.846068 | 0.104491 | 8.110626 | 4.68E-05 | 0.002434 | 0.041683 | 0.1 | 0.236592 |
| idnK_idnK | -0.82287 | -1.0452 | -0.67462 | 0.362936 | 0.233163 | 1.495556 | 0.05463 | 0.102506 | 0.23392 | 0.1694 | 0.315558 |
| SL1344_RS22990_idnT | 0.487416 | 0.243226 | 0.693237 | 0.45202 | 0.189728 | 2.388937 | 0.003335 | 0.030902 | 0.123637 | 0.102 | 0.239193 |
| SL1344_RS06655_ihfA | 4.818074 | 4.801671 | 4.957124 | 0.155261 | 0.103356 | 1.148159 | 0.125333 | 0.188362 | 0.36667 | 0.2756 | 0.431082 |
| SL1344_RS04775_ihfB | 3.247551 | 3.192459 | 3.299901 | 0.150511 | 0.140154 | 1.119496 | 0.079333 | 0.15219 | 0.317708 | 0.2047 | 0.358082 |
| lldD_ildD | -0.74939 | -0.56006 | -1.03789 | -0.47242 | 0.238111 | -1.90036 | 0.018667 | 0.060236 | 0.174166 | 0.1231 | 0.264204 |
| ilvM_ilvM | -0.24335 | -0.06623 | -0.30184 | -0.21575 | 0.238601 | -0.77891 | 0.242 | 0.373331 | 0.533963 | 0.4879 | 0.628925 |
| SL1344_RS14990_invA | 3.993422 | 4.50898 | 3.865785 | -0.62344 | 0.100829 | -3.10117 | 4.68E-05 | 0.068664 | 0.204078 | 0.1 | 0.236592 |

| | | | | | | | | | | | |
|---|---|---|---|---|---|---|---|---|---|---|---|
| SL1344_RS14980_invC | 3.337063 | 3.694935 | 3.238562 | -0.45633 | 0.180618 | -2.02394 | 4.68E-05 | 0.086206 | 0.231641 | 0.1003 | 0.236999 |
| SL1344_RS14995_invE | 3.736224 | 4.205242 | 3.542597 | -0.65197 | 0.105991 | -4.44534 | 4.68E-05 | 0.026166 | 0.121882 | 0.1 | 0.236592 |
| SL1344_RS15005_invF | 4.433634 | 4.865307 | 4.192199 | -0.67186 | 0.122657 | -4.6509 | 4.68E-05 | 0.021101 | 0.109884 | 0.1 | 0.236592 |
| SL1344_RS15000_invG | 4.17604 | 4.640814 | 4.07687 | -0.56331 | 0.097233 | -2.24928 | 4.68E-05 | 0.084558 | 0.229673 | 0.1 | 0.236592 |
| InvR_invR | 5.404928 | 5.945875 | 4.805161 | -1.01932 | 0.268534 | -3.85533 | 4.68E-05 | 0.002589 | 0.046066 | 0.1 | 0.236592 |
| SL1344_RS14355_iroB | -2.31465 | -1.59326 | -2.86908 | -1.24941 | 0.634388 | -1.92163 | 0.012667 | 0.048043 | 0.160427 | 0.1124 | 0.25271 |
| SL1344_RS24725_IS91.family.transposase | 0.290979 | 1.14532 | -1.08471 | -2.20694 | 0.374316 | -5.71081 | 4.68E-05 | 0.009522 | 0.073579 | 0.1 | 0.236592 |
| IsrA_isrA | -6.25 | -6.93428 | -5.68561 | 1.376299 | 2.132025 | 0.528547 | 0.219853 | 0.339282 | 0.506557 | 0.4231 | 0.560038 |
| IsrB_isrB | -2.48083 | -2.48689 | -2.47494 | 0.064422 | 0.599179 | 0.116559 | 0.442667 | 0.674649 | 0.784953 | 0.7887 | 0.858543 |
| IsrC_isrC | -2.80138 | -2.54987 | -2.98465 | -0.55994 | 0.787185 | -0.66099 | 0.228514 | 0.344798 | 0.51492 | 0.4805 | 0.622232 |
| IsrD_isrD | -8.42196 | -8.85266 | -8.05106 | 0.846358 | 3.487066 | 0.193303 | 0.390667 | 0.528158 | 0.663811 | 0.6564 | 0.754248 |
| IsrE_isrE | -1.99015 | -1.66909 | -2.16479 | -0.56243 | 0.623317 | -0.89521 | 0.168667 | 0.238318 | 0.41343 | 0.325 | 0.482051 |
| IsrF_isrF | -0.78373 | -0.50664 | -1.19348 | -0.68235 | 0.247766 | -2.67186 | 0.002667 | 0.026357 | 0.111818 | 0.1012 | 0.238189 |
| IsrG_isrG | -3.8868 | -3.44253 | -4.63513 | -1.1868 | 0.780492 | -1.46764 | 0.038667 | 0.105476 | 0.24341 | 0.1425 | 0.285733 |
| IsrH.1_isrH | -4.28389 | -4.20314 | -4.3692 | -0.14928 | 0.701315 | -0.20712 | 0.404 | 0.498455 | 0.632458 | 0.6082 | 0.712968 |
| IsrH.2_isrH2 | -4.73775 | -4.5794 | -4.93082 | -0.35737 | 0.96189 | -0.35023 | 0.353764 | 0.51219 | 0.64806 | 0.6147 | 0.722241 |
| IsrI_isrI | -2.43082 | -2.75743 | -2.11528 | 0.644002 | 0.366203 | 1.708318 | 0.031333 | 0.067328 | 0.181608 | 0.1369 | 0.279262 |
| IsrJ_isrJ | -2.72594 | -2.71597 | -2.74348 | 0.027273 | 0.74412 | 0.031714 | 0.485676 | 0.741931 | 0.835695 | 0.8336 | 0.893854 |
| IsrK_isrK | 0.127152 | 0.164738 | 0.095963 | -0.00271 | 0.361869 | -0.00738 | 0.496336 | 0.780686 | 0.864799 | 0.8795 | 0.924192 |
| IsrL_isrL | -2.13244 | -2.24868 | -2.06142 | 0.160289 | 0.539609 | 0.287282 | 0.381333 | 0.633611 | 0.754489 | 0.7276 | 0.818924 |
| IsrM_isrM | -5.14408 | -5.48785 | -4.73526 | 0.754804 | 1.03334 | 0.704054 | 0.229333 | 0.367325 | 0.513502 | 0.4625 | 0.591012 |
| IsrN_isrN | -7.36248 | -6.88869 | -8.38005 | -1.47991 | 2.90016 | -0.42061 | 0.279147 | 0.408245 | 0.568003 | 0.535 | 0.660159 |
| IsrO_isrO | -0.96442 | -0.87323 | -1.08699 | -0.31495 | 0.420208 | -0.72053 | 0.227182 | 0.309747 | 0.481642 | 0.4544 | 0.593485 |
| IsrP_isrP | -2.3484 | -2.10248 | -2.63458 | -0.53871 | 0.358593 | -1.4654 | 0.044637 | 0.104814 | 0.233663 | 0.1698 | 0.313985 |
| IsrQ_isrQ | -7.4195 | -6.81379 | -8.04636 | -1.31049 | 2.660062 | -0.43273 | 0.292472 | 0.434807 | 0.58369 | 0.545 | 0.661824 |
| IstR.1_istR1 | -4.01944 | -4.04764 | -4.00297 | 0.149447 | 0.980495 | 0.121001 | 0.44437 | 0.611489 | 0.73868 | 0.7673 | 0.843462 |
| IstR.2_istR2 | -3.77449 | -3.74546 | -3.79064 | 0.056306 | 0.947354 | 0.053392 | 0.472667 | 0.649125 | 0.768926 | 0.785 | 0.858925 |
| ksgA_ksgA | 2.169585 | 2.152683 | 2.189809 | 0.038449 | 0.124396 | 0.299802 | 0.357095 | 0.579695 | 0.711109 | 0.6726 | 0.773073 |
| SL1344_RS08205_ldhA | 2.252294 | 2.211045 | 2.295336 | 0.124328 | 0.148677 | 0.833604 | 0.17988 | 0.241219 | 0.415126 | 0.3614 | 0.508654 |
| SL1344_RS00585_leuO | -3.82207 | -3.65149 | -3.99131 | -0.36051 | 0.725442 | -0.4767 | 0.302 | 0.465167 | 0.608619 | 0.5677 | 0.685138 |
| SL1344_RS21750_lexA | 3.86419 | 4.094874 | 3.735303 | -0.30762 | 0.172403 | -1.65274 | 0.043971 | 0.071283 | 0.207006 | 0.14 | 0.289935 |
| ligA_ligA | 2.613945 | 2.514458 | 2.815032 | 0.278679 | 0.145307 | 1.883802 | 0.017334 | 0.052172 | 0.170105 | 0.1184 | 0.260436 |
| livF_livF | -1.95424 | -1.90778 | -1.99514 | -0.08899 | 0.33031 | -0.25407 | 0.38441 | 0.535921 | 0.667595 | 0.6547 | 0.751696 |
| lolA_lolA | 2.470093 | 2.360848 | 2.565896 | 0.153206 | 0.15585 | 1.007947 | 0.140573 | 0.197027 | 0.369228 | 0.2546 | 0.422404 |
| lolB_lolB | 0.385803 | 0.392329 | 0.379582 | -0.01399 | 0.140055 | -0.0996 | 0.456 | 0.536152 | 0.666733 | 0.6396 | 0.741458 |
| lplA_lplA | 1.265917 | 1.241741 | 1.323413 | 0.073701 | 0.140589 | 0.463684 | 0.318667 | 0.516016 | 0.657251 | 0.6254 | 0.740339 |
| SL1344_RS06600_lpxR | 2.920781 | 3.13216 | 2.889335 | -0.22411 | 0.136243 | -1.04916 | 0.189873 | 0.203121 | 0.380364 | 0.3562 | 0.519418 |
| SL1344_RS11955_lrhA | 1.296426 | 1.029094 | 1.620462 | 0.593352 | 0.133019 | 4.402491 | 4.68E-05 | 0.006654 | 0.060524 | 0.1 | 0.236592 |
| SL1344_RS04660_lrp | 4.020425 | 4.000932 | 4.130736 | 0.131204 | 0.144491 | 0.916952 | 0.186 | 0.252451 | 0.437806 | 0.3644 | 0.530904 |
| SL1344_RS09120_L.serine.ammonia.lyase | 4.152881 | 4.315056 | 3.896829 | -0.41322 | 0.071237 | -5.49832 | 4.68E-05 | 0.009843 | 0.074521 | 0.1 | 0.236592 |
| SL1344_RS20930_lsrR | 3.070421 | 3.182326 | 2.990984 | -0.29925 | 0.213871 | -1.44404 | 0.008661 | 0.077184 | 0.215157 | 0.108 | 0.246943 |
| SL1344_RS14590_luxS | 3.169161 | 3.159453 | 3.17445 | 0.0207 | 0.115068 | 0.153522 | 0.445333 | 0.745769 | 0.839422 | 0.8684 | 0.91695 |
| SL1344_RS09795_lysine.N.methylase | 3.81258 | 4.009743 | 3.700502 | -0.32631 | 0.118337 | -2.82473 | 4.68E-05 | 0.012138 | 0.082679 | 0.1 | 0.236592 |
| malF_malF | 0.938605 | 0.570468 | 1.233453 | 0.660704 | 0.122668 | 5.370765 | 4.68E-05 | 0.003968 | 0.047519 | 0.1 | 0.236592 |
| malG_malG | 0.224946 | -0.28616 | 0.638285 | 0.958697 | 0.220758 | 4.406428 | 4.68E-05 | 0.006024 | 0.059093 | 0.1 | 0.236592 |

| | | | | | | | | | | |
|---|---|---|---|---|---|---|---|---|---|---|
| SL1344_RS18135_malT | 4.26231 | 3.935457 | 4.678302 | 0.723515 | 0.083938 | 8.354554 | 4.68E-05 | 0.001889 | 0.039215 | 0.1 | 0.236592 |
| SL1344_RS07280_malY. | -1.49496 | -1.46317 | -1.52195 | -0.10254 | 0.378895 | -0.25425 | 0.397069 | 0.591114 | 0.718194 | 0.7109 | 0.798948 |
| mdoD_mdoD | 2.693377 | 2.67533 | 2.713747 | 0.038046 | 0.065715 | 0.536647 | 0.274667 | 0.40348 | 0.548007 | 0.5048 | 0.628876 |
| mdoG_mdoG | 3.763907 | 3.843467 | 3.730257 | -0.10041 | 0.083931 | -1.10226 | 0.138667 | 0.181104 | 0.347554 | 0.2746 | 0.438064 |
| SL1344_RS09375_membrane.protein | 2.23043 | 2.625555 | 1.845557 | -0.77228 | 0.095695 | -8.07239 | 4.68E-05 | 0.001177 | 0.032466 | 0.1 | 0.236592 |
| SL1344_RS21870_membrane.protein | 4.934048 | 4.413082 | 5.218019 | 0.811184 | 0.216999 | 4.02426 | 4.68E-05 | 0.018392 | 0.102985 | 0.1 | 0.236592 |
| SL1344_RS11835_menC | 2.453996 | 2.190339 | 2.648273 | 0.455731 | 0.142705 | 3.249211 | 4.68E-05 | 0.008973 | 0.071315 | 0.1 | 0.236592 |
| SL1344_RS21470_metA | -0.49685 | -0.65741 | -0.30855 | 0.327664 | 0.279281 | 1.13329 | 0.111925 | 0.166848 | 0.324603 | 0.2423 | 0.400017 |
| SL1344_RS21065_metB | 0.316637 | 0.282338 | 0.389212 | 0.084946 | 0.193927 | 0.383188 | 0.351765 | 0.583614 | 0.712473 | 0.6806 | 0.7854 |
| SL1344_RS16345_metC | 0.527071 | 0.45119 | 0.63076 | 0.20783 | 0.281611 | 0.703665 | 0.218 | 0.309984 | 0.486324 | 0.4305 | 0.584258 |
| SL1344_RS20385_metE | 0.809148 | 0.715612 | 0.846616 | 0.104139 | 0.20291 | 0.456441 | 0.327781 | 0.585922 | 0.717468 | 0.6642 | 0.777553 |
| SL1344_RS21095_metF | -0.94898 | -0.89078 | -0.99823 | -0.10427 | 0.268748 | -0.3897 | 0.336 | 0.511797 | 0.647418 | 0.6268 | 0.730715 |
| metG_metG | 2.803019 | 2.785251 | 2.814347 | 0.034734 | 0.078808 | 0.407842 | 0.313791 | 0.472534 | 0.61656 | 0.5991 | 0.706256 |
| SL1344_RS21500_metH | 2.255838 | 2.19621 | 2.315213 | 0.13915 | 0.120504 | 1.116449 | 0.08994 | 0.15642 | 0.31531 | 0.2153 | 0.366778 |
| SL1344_RS18435_methyl.accepting.chemotaxis.citrate.transducer | 4.93973 | 5.593983 | 4.567686 | -1.02386 | 0.191091 | -4.68093 | 4.68E-05 | 0.006662 | 0.065273 | 0.1 | 0.236592 |
| SL1344_RS08105_methyl.accepting.chemotaxis.protein | 5.156913 | 5.845325 | 4.517491 | -1.35889 | 0.205448 | -6.82203 | 4.68E-05 | 0.000786 | 0.031999 | 0.1 | 0.236592 |
| SL1344_RS16230_methyl.accepting.chemotaxis.protein | 4.985634 | 5.378645 | 4.693923 | -0.65452 | 0.130642 | -4.77973 | 4.68E-05 | 0.004307 | 0.054266 | 0.1 | 0.236592 |
| SL1344_RS16300_methyl.accepting.chemotaxis.protein | 3.726526 | 4.569319 | 3.015596 | -1.55815 | 0.204854 | -7.31505 | 4.68E-05 | 0.00342 | 0.050097 | 0.1 | 0.236592 |
| SL1344_RS16625_methyl.accepting.chemotaxis.protein | 5.617522 | 6.562163 | 4.907299 | -1.6548 | 0.189013 | -7.82379 | 4.68E-05 | 0.005295 | 0.059828 | 0.1 | 0.236592 |
| SL1344_RS21060_metJ | 2.102723 | 2.227061 | 2.018675 | -0.21743 | 0.172866 | -1.25001 | 0.08 | 0.128463 | 0.285788 | 0.1847 | 0.341008 |
| SL1344_RS21070_metL | 2.252257 | 2.211894 | 2.292305 | 0.080612 | 0.084736 | 0.886548 | 0.159893 | 0.247629 | 0.403736 | 0.3362 | 0.482589 |
| metQ_metQ | 3.240077 | 3.233098 | 3.244527 | 0.012101 | 0.058205 | 0.188077 | 0.410393 | 0.562387 | 0.690758 | 0.6752 | 0.768474 |
| SL1344_RS20380_metR | -1.30689 | -1.12973 | -1.50029 | -0.37694 | 0.260294 | -1.38715 | 0.064667 | 0.111061 | 0.242372 | 0.1861 | 0.330593 |
| MgrR_mgrR | 1.743538 | 1.598735 | 1.900937 | 0.283115 | 0.806482 | 0.275299 | 0.444667 | 0.734377 | 0.840171 | 0.9961 | 0.997689 |
| MicA_micA | 0.140418 | -0.42795 | 0.339011 | 0.762849 | 0.483828 | 1.517481 | 0.011992 | 0.120651 | 0.280047 | 0.1095 | 0.249054 |
| MicC_micC | -2.68652 | -2.53703 | -2.81412 | -0.29399 | 0.461188 | -0.58252 | 0.260493 | 0.384854 | 0.532805 | 0.4894 | 0.61455 |
| MicF_micF | 0.825395 | 0.704727 | 1.053175 | 0.32694 | 0.217387 | 1.471144 | 0.065956 | 0.097179 | 0.238296 | 0.1736 | 0.326613 |
| SL1344_RS07395_mlc | 2.203221 | 2.027724 | 2.547323 | 0.50596 | 0.160863 | 2.848353 | 4.68E-05 | 0.030611 | 0.129319 | 0.1 | 0.236592 |
| moaA_moaA | 2.566365 | 2.811854 | 2.354097 | -0.39306 | 0.168169 | -2.45084 | 0.001336 | 0.020932 | 0.106747 | 0.1008 | 0.237654 |
| moaD_moaD | 1.410048 | 1.474051 | 1.372836 | -0.10177 | 0.121585 | -0.78811 | 0.186 | 0.301671 | 0.45817 | 0.3994 | 0.538702 |
| moaE_moaE | 2.013472 | 2.073708 | 1.904005 | -0.15593 | 0.121679 | -1.2069 | 0.118667 | 0.16534 | 0.324212 | 0.2517 | 0.410032 |
| mobA_mobA | 0.38452 | 0.342251 | 0.627787 | 0.280482 | 0.325232 | 0.673885 | 0.314457 | 0.450491 | 0.619454 | 0.6568 | 0.784639 |
| SL1344_RS24715_mobA | 3.69192 | 4.756352 | 2.55973 | -2.23797 | 0.138081 | -16.432 | 4.68E-05 | 0.000262 | 0.02283 | 0.1 | 0.236592 |
| SL1344_RS24720_mobC | 2.242567 | 3.216878 | 0.999972 | -2.19352 | 0.288129 | -7.5449 | 4.68E-05 | 0.003477 | 0.049725 | 0.1 | 0.236592 |
| SL1344_RS03955_modA | 3.682959 | 4.430025 | 3.164602 | -1.25259 | 0.086763 | -13.1026 | 4.68E-05 | 0.0064 | 0.064167 | 0.1 | 0.236592 |
| modB_modB | 3.518414 | 4.203977 | 2.970994 | -1.20937 | 0.13417 | -8.75191 | 4.68E-05 | 0.00108 | 0.033283 | 0.1 | 0.236592 |
| SL1344_RS03965_modC | 3.566851 | 4.16622 | 2.961036 | -1.21568 | 0.127287 | -9.69317 | 4.68E-05 | 0.000993 | 0.032631 | 0.1 | 0.236592 |
| SL1344_RS09620_motA | 5.144831 | 6.158313 | 4.323411 | -1.83975 | 0.234808 | -7.99553 | 4.68E-05 | 0.000245 | 0.025048 | 0.1 | 0.236592 |
| motB_motB | 5.233258 | 6.185771 | 4.350678 | -1.81492 | 0.155293 | -11.8899 | 4.68E-05 | 3.63E-05 | 0.017205 | 0.1 | 0.236592 |
| SL1344_RS09450_msbB | 1.61875 | 1.831985 | 1.360812 | -0.4756 | 0.126951 | -3.77206 | 4.68E-05 | 0.010523 | 0.073356 | 0.1 | 0.236592 |
| Msr_Msr | -1.37486 | -1.53072 | -1.22615 | 0.133208 | 0.720217 | 0.18945 | 0.424667 | 0.741916 | 0.838892 | 0.8732 | 0.920878 |
| mtlR_mtlR | 1.243722 | 1.139298 | 1.350208 | 0.21219 | 0.145833 | 1.453485 | 0.047333 | 0.100426 | 0.237535 | 0.1626 | 0.310827 |
| mukB_mukB | 5.142138 | 5.342938 | 4.876091 | -0.46217 | 0.046429 | -9.58859 | 4.68E-05 | 0.002254 | 0.040532 | 0.1 | 0.236592 |
| murD_murD | 3.337243 | 3.328687 | 3.352959 | 0.013037 | 0.091511 | 0.131967 | 0.441333 | 0.736058 | 0.831586 | 0.8432 | 0.900212 |
| SL1344_RS00625_murE | 2.800915 | 2.701507 | 3.01502 | 0.322804 | 0.153767 | 2.029338 | 0.000673 | 0.051583 | 0.170863 | 0.1015 | 0.23855 |

| | | | | | | | | | | |
|---|---|---|---|---|---|---|---|---|---|---|
| SL1344_RS09280_N.acetyltransferase | -0.58189 | -0.15381 | -1.09917 | -0.94353 | 0.208256 | -4.48508 | 4.68E-05 | 0.007383 | 0.060648 | 0.1 | 0.236592 |
| napB_napB | 3.451856 | 3.131247 | 3.867794 | 0.752186 | 0.174059 | 4.501629 | 4.68E-05 | 0.009151 | 0.07326 | 0.1 | 0.236592 |
| SL1344_RS08805_narG | 4.924709 | 5.720046 | 3.867309 | -1.85657 | 0.361901 | -4.69102 | 4.68E-05 | 0.023196 | 0.115521 | 0.1 | 0.236592 |
| SL1344_RS08800_narH | 4.41476 | 4.927005 | 3.631765 | -1.2986 | 0.308389 | -3.87615 | 4.68E-05 | 0.034828 | 0.14186 | 0.1 | 0.236592 |
| SL1344_RS08790_narI | 3.812495 | 4.04632 | 3.45007 | -0.61316 | 0.203783 | -2.69666 | 4.68E-05 | 0.049186 | 0.169906 | 0.1 | 0.236592 |
| SL1344_RS08795_narJ | 3.121555 | 3.452151 | 2.471037 | -0.98084 | 0.364366 | -2.46062 | 4.68E-05 | 0.072831 | 0.211296 | 0.1 | 0.236592 |
| SL1344_RS08810_narK | 3.958678 | 4.798508 | 2.680012 | -2.11422 | 0.621439 | -3.41845 | 4.68E-05 | 0.032243 | 0.136371 | 0.1 | 0.236592 |
| SL1344_RS08820_narL | 0.891584 | 1.329587 | 0.550016 | -0.76342 | 0.180515 | -4.12003 | 4.68E-05 | 0.008051 | 0.066136 | 0.1 | 0.236592 |
| nepI_nepI | -2.23023 | -2.08088 | -2.38107 | -0.26395 | 0.488915 | -0.51063 | 0.290667 | 0.471466 | 0.61709 | 0.5736 | 0.692746 |
| nhaA_nhaA | 2.112672 | 2.148532 | 2.094036 | -0.04139 | 0.121223 | -0.3109 | 0.38441 | 0.637291 | 0.756328 | 0.7356 | 0.824275 |
| nirD_nirD | 0.474335 | 0.500372 | 0.441955 | -0.06381 | 0.167621 | -0.35358 | 0.33 | 0.503847 | 0.643118 | 0.6211 | 0.7251 |
| nlpD_nlpD | 5.837342 | 5.959142 | 5.648251 | -0.33678 | 0.113488 | -3.19695 | 4.68E-05 | 0.005692 | 0.060944 | 0.1 | 0.236592 |
| nrdI_nrdI | -5.18842 | -5.09902 | -5.24298 | -0.16657 | 1.063831 | -0.15026 | 0.437333 | 0.583262 | 0.707192 | 0.6853 | 0.780692 |
| SL1344_RS21955_nrfB | 1.750272 | 1.662343 | 1.83702 | 0.113404 | 0.243821 | 0.463857 | 0.301799 | 0.487406 | 0.649625 | 0.6201 | 0.741605 |
| SL1344_RS21960_nrfC | 2.012825 | 1.899363 | 2.059952 | 0.167884 | 0.152219 | 1.08148 | 0.103333 | 0.205128 | 0.381065 | 0.2416 | 0.395695 |
| SL1344_RS21975_nrfG | 0.228653 | -0.1961 | 0.36369 | 0.576401 | 0.373579 | 1.470177 | 0.045969 | 0.105528 | 0.256167 | 0.1513 | 0.300027 |
| nudC_nudC | 0.194102 | 0.06439 | 0.282613 | 0.212671 | 0.188186 | 1.093452 | 0.115257 | 0.191609 | 0.34497 | 0.2664 | 0.417753 |
| nusA_nusA | 4.068726 | 4.194501 | 3.950422 | -0.22841 | 0.080988 | -2.81569 | 0.000673 | 0.016165 | 0.093326 | 0.1 | 0.236592 |
| nusB_nusB | 3.763803 | 3.74186 | 3.77393 | 0.021812 | 0.08289 | 0.222477 | 0.406 | 0.779297 | 0.863318 | 0.7949 | 0.875509 |
| nusG_nusG | 4.097131 | 4.39534 | 3.827363 | -0.55359 | 0.110234 | -5.07968 | 4.68E-05 | 0.001672 | 0.038788 | 0.1 | 0.236592 |
| obgE_obgE | 1.687758 | 1.772889 | 1.575945 | -0.18275 | 0.136396 | -1.27636 | 0.075283 | 0.12785 | 0.276606 | 0.1965 | 0.352313 |
| ompL_ompL | -4.56892 | -4.10776 | -4.8444 | -0.75015 | 0.973188 | -0.68413 | 0.226 | 0.339409 | 0.49614 | 0.4506 | 0.585535 |
| SL1344_RS18070_ompR | 2.557313 | 2.344996 | 2.773212 | 0.43643 | 0.097372 | 4.417208 | 4.68E-05 | 0.004951 | 0.05394 | 0.1 | 0.236592 |
| ompX_ompX | 5.099055 | 5.178197 | 5.084911 | -0.08345 | 0.088645 | -0.87642 | 0.226516 | 0.280405 | 0.463506 | 0.4212 | 0.592913 |
| OmrA_omrA | -1.9128 | -1.74247 | -2.06637 | -0.18432 | 0.666464 | -0.29329 | 0.38441 | 0.665455 | 0.781115 | 0.7659 | 0.845547 |
| OmrB_omrB | -3.20822 | -3.13693 | -3.31297 | -0.19507 | 0.607396 | -0.313 | 0.373085 | 0.548668 | 0.681855 | 0.6741 | 0.771286 |
| oppB_oppB | 1.816038 | 1.789788 | 1.838337 | 0.048029 | 0.091291 | 0.490624 | 0.283145 | 0.444247 | 0.586179 | 0.5523 | 0.668392 |
| SL1344_RS14860_orgA | 4.109655 | 4.534921 | 3.807079 | -0.72437 | 0.130012 | -5.37696 | 4.68E-05 | 0.009015 | 0.073596 | 0.1 | 0.236592 |
| OxyS_oxyS | -2.15818 | -1.95971 | -2.34239 | -0.38583 | 0.707051 | -0.51932 | 0.285333 | 0.446548 | 0.606738 | 0.5677 | 0.697184 |
| pabB_pabB | 1.573773 | 1.85706 | 1.24578 | -0.61124 | 0.08702 | -6.94993 | 4.68E-05 | 0.001948 | 0.036597 | 0.1 | 0.236592 |
| pagP_pagP | 1.130174 | 1.115863 | 1.141299 | 0.033669 | 0.14799 | 0.209128 | 0.403731 | 0.582565 | 0.71078 | 0.6926 | 0.7852 |
| SL1344_RS10525_pduA | -4.30529 | -4.82853 | -3.67373 | 1.105654 | 0.908155 | 1.128565 | 0.1 | 0.159812 | 0.309292 | 0.2348 | 0.386991 |
| pduC_pduC | -2.11098 | -3.32392 | -1.28967 | 2.036201 | 0.475646 | 4.189171 | 4.68E-05 | 0.012108 | 0.076583 | 0.1 | 0.236592 |
| pduD_pduD | -3.5738 | -4.63848 | -2.43012 | 2.154789 | 0.864243 | 2.469037 | 0.000673 | 0.028663 | 0.119184 | 0.1004 | 0.237137 |
| pduE_pduE | -3.96463 | -4.47812 | -3.18563 | 1.207946 | 0.829305 | 1.388991 | 0.072 | 0.118465 | 0.260522 | 0.1962 | 0.347993 |
| pdxK_pdxK | -0.89795 | -0.96395 | -0.75829 | 0.160155 | 0.336031 | 0.438286 | 0.33511 | 0.574987 | 0.705596 | 0.6532 | 0.765797 |
| SL1344_RS06590_pfkB | 2.88778 | 2.316838 | 3.481246 | 1.086103 | 0.194319 | 5.649674 | 4.68E-05 | 0.001198 | 0.035089 | 0.1 | 0.236592 |
| SL1344_RS13960_phage.virion.morphogenesis.protein | -4.98034 | -4.46894 | -5.51904 | -1.04874 | 0.995448 | -0.98499 | 0.132 | 0.2115 | 0.362658 | 0.2923 | 0.439506 |
| SL1344_RS02035_phoB | 1.733046 | 1.744788 | 1.714596 | -0.05403 | 0.158436 | -0.29789 | 0.352 | 0.489795 | 0.644575 | 0.6757 | 0.770075 |
| SL1344_RS11645_phoE | 7.723478 | 7.663635 | 7.815773 | 0.246408 | 0.148289 | 1.706998 | 4.68E-05 | 0.067053 | 0.201724 | 0.1 | 0.236592 |
| SL1344_RS05535_phoH | 3.013213 | 3.048561 | 2.904655 | -0.15485 | 0.158754 | -0.98245 | 0.162558 | 0.201722 | 0.377362 | 0.308 | 0.479856 |
| SL1344_RS06080_phoP | 5.200495 | 5.053173 | 5.254159 | 0.200609 | 0.085049 | 2.371699 | 4.68E-05 | 0.066375 | 0.199195 | 0.1 | 0.236592 |
| SL1344_RS06075_phoQ | 4.333462 | 4.190616 | 4.387888 | 0.188177 | 0.085728 | 2.021944 | 0.004665 | 0.04551 | 0.159091 | 0.103 | 0.240571 |
| phoR_phoR | 1.280113 | 1.318077 | 1.223612 | -0.10857 | 0.141667 | -0.74546 | 0.206667 | 0.308245 | 0.472404 | 0.4125 | 0.546092 |
| SL1344_RS13700_phospho.2.dehydro.3.deoxyheptonate.aldolase.Tyr.sensitive | 0.45657 | 0.231642 | 0.676191 | 0.442338 | 0.179897 | 2.384817 | 0.004665 | 0.028783 | 0.119493 | 0.1036 | 0.241321 |

| | | | | | | | | | | | |
|---|---|---|---|---|---|---|---|---|---|---|---|
| SL1344_RS10625_phsA | 1.290928 | -0.79067 | 3.693278 | 4.409865 | 0.322869 | 13.3119 | 4.68E-05 | 0.00143 | 0.03709 | 0.1 | 0.236592 |
| SL1344_RS10620_phsB | -0.43066 | -2.3307 | 1.596571 | 3.794307 | 0.526528 | 7.220972 | 4.68E-05 | 0.00132 | 0.035289 | 0.1 | 0.236592 |
| SL1344_RS10615_phsC | 0.585564 | -1.22025 | 2.538987 | 3.69023 | 0.416199 | 8.957772 | 4.68E-05 | 0.000969 | 0.032475 | 0.1 | 0.236592 |
| potG_potG | -1.46476 | -1.45544 | -1.47094 | -0.06225 | 0.397583 | -0.13823 | 0.449034 | 0.64805 | 0.764542 | 0.7637 | 0.842603 |
| SL1344_RS09215_prc | 4.41582 | 4.812378 | 4.100572 | -0.70281 | 0.065034 | -10.6382 | 4.68E-05 | 0.001413 | 0.035558 | 0.1 | 0.236592 |
| SL1344_RS14880_prgH | 4.611277 | 4.901115 | 4.3717 | -0.52823 | 0.146199 | -3.34395 | 4.68E-05 | 0.031534 | 0.134333 | 0.1 | 0.236592 |
| SL1344_RS14875_prgI | 3.480772 | 3.79443 | 3.262412 | -0.53278 | 0.169015 | -3.37658 | 4.68E-05 | 0.03236 | 0.135607 | 0.1 | 0.236592 |
| SL1344_RS14870_prgJ | 4.20459 | 4.604878 | 3.911441 | -0.6443 | 0.185644 | -3.28386 | 4.68E-05 | 0.012886 | 0.0865 | 0.1 | 0.236592 |
| SL1344_RS14865_prgK | 4.899577 | 5.311694 | 4.67521 | -0.63493 | 0.098007 | -5.27301 | 4.68E-05 | 0.018889 | 0.104098 | 0.1 | 0.236592 |
| SL1344_RS09220_proQ | 3.723874 | 4.16606 | 3.328622 | -0.83496 | 0.088295 | -9.26754 | 4.68E-05 | 0.001771 | 0.038299 | 0.1 | 0.236592 |
| SL1344_RS09255_prpA.pphA | -1.6034 | -1.31892 | -1.79908 | -0.46622 | 0.35251 | -1.21428 | 0.097269 | 0.15189 | 0.298838 | 0.2282 | 0.380107 |
| psiB_psiB | -2.33344 | -2.46255 | -2.20378 | 0.261632 | 0.346969 | 0.714694 | 0.214667 | 0.32466 | 0.472678 | 0.4136 | 0.549515 |
| psiB_psiB.1 | -2.66176 | -2.48995 | -2.86332 | -0.38402 | 0.510022 | -0.72759 | 0.22052 | 0.331903 | 0.489562 | 0.4415 | 0.577444 |
| pssA_pssA | 2.805829 | 2.780059 | 2.83048 | 0.021382 | 0.143289 | 0.152467 | 0.435333 | 0.763326 | 0.852386 | 0.8553 | 0.908557 |
| SL1344_RS11995_pta | 4.881878 | 4.825958 | 4.996608 | 0.180618 | 0.119632 | 1.586618 | 0.014657 | 0.062169 | 0.191057 | 0.1122 | 0.252743 |
| SL1344_RS15255_ptpS | -1.67802 | -1.23517 | -2.27758 | -1.04582 | 0.309616 | -3.34854 | 4.68E-05 | 0.014851 | 0.082283 | 0.1 | 0.236592 |
| SL1344_RS02910_PTS.mannose.fructose.sorbose.transporter.subunit.IIB | -6.04652 | -6.68667 | -5.63685 | 0.891455 | 2.945681 | 0.269137 | 0.337333 | 0.522136 | 0.669057 | 0.6505 | 0.755161 |
| SL1344_RS07105_purR | 1.019255 | 1.096503 | 0.893264 | -0.15906 | 0.209563 | -0.71275 | 0.213857 | 0.336481 | 0.507156 | 0.4317 | 0.58354 |
| putA_putA | 5.851751 | 5.867789 | 5.745029 | -0.11507 | 0.073911 | -0.64379 | 0.333111 | 0.491226 | 0.660139 | 0.7 | 0.822108 |
| SL1344_RS23405_putative.inner.membrane.protein | 3.869321 | 3.706004 | 4.038782 | 0.309978 | 0.108188 | 2.872461 | 4.68E-05 | 0.014493 | 0.089384 | 0.1 | 0.236592 |
| SL1344_RS09445_pyruvate.kinase.II | 4.054011 | 4.313466 | 3.86084 | -0.45474 | 0.090262 | -5.07941 | 4.68E-05 | 0.009743 | 0.074845 | 0.1 | 0.236592 |
| SL1344_RS16430_qseB | -1.84112 | -2.03268 | -1.56198 | 0.466598 | 0.346949 | 1.251528 | 0.086 | 0.143698 | 0.288043 | 0.2132 | 0.360277 |
| SL1344_RS16435_qseC | -0.85147 | -1.01171 | -0.69491 | 0.32027 | 0.218121 | 1.419716 | 0.063333 | 0.117176 | 0.250093 | 0.1871 | 0.333244 |
| SL1344_RS23490_radA | 1.444507 | 1.406693 | 1.474482 | 0.070962 | 0.125251 | 0.537826 | 0.274667 | 0.426303 | 0.575679 | 0.544 | 0.664343 |
| RaoN_raoN | -2.57918 | -2.57137 | -2.59317 | -0.05484 | 0.459134 | -0.09889 | 0.457333 | 0.599835 | 0.723479 | 0.715 | 0.802866 |
| SL1344_RS12895_ratA | 1.212821 | 1.183574 | 1.275322 | 0.078429 | 0.132894 | 0.53825 | 0.286667 | 0.45239 | 0.597814 | 0.5592 | 0.683062 |
| rcsA_rcsA | -1.99509 | -1.72053 | -2.36446 | -0.64881 | 0.375263 | -1.67843 | 0.023318 | 0.080467 | 0.204858 | 0.1297 | 0.271813 |
| SL1344_RS11655_rcsB | 2.742485 | 2.761942 | 2.730634 | -0.03215 | 0.08529 | -0.34264 | 0.361093 | 0.553679 | 0.687746 | 0.666 | 0.767833 |
| SL1344_RS11660_rcsC | 1.729388 | 1.635705 | 1.799423 | 0.165788 | 0.123153 | 1.301124 | 0.081279 | 0.126803 | 0.273435 | 0.1981 | 0.348862 |
| SL1344_RS11650_rcsD | 2.575485 | 2.478979 | 2.660637 | 0.180631 | 0.066701 | 2.647164 | 0.001337 | 0.021845 | 0.100566 | 0.1019 | 0.239071 |
| rcsF_rcsF | 1.204796 | 1.247796 | 1.160396 | -0.09489 | 0.147775 | -0.61964 | 0.251166 | 0.39473 | 0.551746 | 0.5 | 0.632129 |
| RdlD_rdlD | -1.6263 | -1.47951 | -1.72582 | -0.27733 | 0.339952 | -0.76892 | 0.190667 | 0.295057 | 0.456569 | 0.3927 | 0.530019 |
| RdlE_rdlE | -3.10131 | -2.99415 | -3.35277 | -0.22893 | 0.810807 | -0.26742 | 0.403731 | 0.667557 | 0.7823 | 0.7851 | 0.863305 |
| SL1344_RS14650_recA | 4.29801 | 4.216733 | 4.381426 | 0.066552 | 0.168934 | 0.596204 | 0.222667 | 0.406116 | 0.58516 | 0.437 | 0.591829 |
| SL1344_RS13835_recN | 2.2873 | 2.617466 | 1.927892 | -0.69762 | 0.104838 | -6.77617 | 4.68E-05 | 0.00292 | 0.043219 | 0.1 | 0.236592 |
| SL1344_RS06620_rfc | 1.201249 | 1.028439 | 1.398487 | 0.37541 | 0.128963 | 2.879056 | 0.001336 | 0.018577 | 0.095487 | 0.1003 | 0.236997 |
| SL1344_RS22270_rflM | 0.741228 | 1.477708 | 0.006202 | -1.46265 | 0.202271 | -7.24809 | 4.68E-05 | 0.00158 | 0.035343 | 0.1 | 0.236592 |
| rhtB_rhtB | -2.09672 | -2.10277 | -2.0885 | 0.01293 | 0.347018 | 0.037508 | 0.483011 | 0.587204 | 0.710475 | 0.6967 | 0.786922 |
| rimM_rimM | 4.493232 | 4.797497 | 4.201829 | -0.43816 | 0.288357 | -1.83074 | 0.003332 | 0.040324 | 0.152749 | 0.1055 | 0.243936 |
| rimO_rimO | 0.854503 | 0.999938 | 0.747936 | -0.24696 | 0.151172 | -1.53793 | 0.045303 | 0.08821 | 0.218345 | 0.1555 | 0.302241 |
| SL1344_RS03220_rlpA | 1.657735 | 1.612211 | 1.775511 | 0.145152 | 0.197852 | 0.725059 | 0.233333 | 0.348989 | 0.523101 | 0.4373 | 0.599631 |
| SL1344_RS05840_rne | 4.65196 | 4.666748 | 4.633469 | -0.03275 | 0.045843 | -0.6696 | 0.221852 | 0.348865 | 0.508922 | 0.4557 | 0.593402 |
| rnpB_rnpB | 10.77046 | 10.92207 | 10.75316 | -0.1685 | 0.142741 | -1.13668 | 0.05996 | 0.188753 | 0.369354 | 0.1536 | 0.307456 |
| RnpB_RnpB | 10.77046 | 10.92194 | 10.75331 | -0.16868 | 0.142172 | -1.16165 | 0.053298 | 0.188886 | 0.36952 | 0.1523 | 0.305906 |
| SL1344_RS15130_rpoS | 5.570035 | 5.490216 | 5.737316 | 0.156394 | 0.145104 | 1.200663 | 0.122 | 0.163609 | 0.337503 | 0.2053 | 0.375578 |

| | | | | | | | | | | |
|---|---|---|---|---|---|---|---|---|---|---|
| rprA_rprA | -0.37464 | -0.3943 | -0.35402 | 0.156753 | 0.444768 | 0.327469 | 0.357333 | 0.480386 | 0.6428 | 0.7403 | 0.820857 |
| rpsN_rpsN | 5.374153 | 5.485029 | 5.185991 | -0.22652 | 0.191644 | -1.26992 | 0.115257 | 0.149002 | 0.319514 | 0.2006 | 0.370672 |
| rpsV_rpsV | 4.471489 | 4.48747 | 4.41584 | -0.06074 | 0.082144 | -0.59211 | 0.31046 | 0.531879 | 0.685294 | 0.6402 | 0.77027 |
| RseX_rseX | -4.85002 | -4.68982 | -5.09226 | -0.39948 | 0.988801 | -0.36104 | 0.344667 | 0.496222 | 0.63528 | 0.6077 | 0.717111 |
| rsmD_rsmD | 0.072442 | 0.028667 | 0.113297 | 0.134675 | 0.248634 | 0.508804 | 0.286 | 0.440278 | 0.594606 | 0.6088 | 0.71471 |
| SL1344_RS09240_rsmF.yebU | 0.421581 | 0.725731 | -0.07785 | -0.75061 | 0.291958 | -2.41269 | 0.002667 | 0.030713 | 0.12816 | 0.1009 | 0.237783 |
| SL1344_RS22150_rtsA | 4.064717 | 4.325449 | 3.833701 | -0.602 | 0.211485 | -2.9092 | 4.68E-05 | 0.014854 | 0.092576 | 0.1 | 0.236592 |
| SL1344_RS22145_rtsB | 1.982027 | 2.057267 | 1.660662 | -0.43201 | 0.362218 | -1.25852 | 0.08994 | 0.108212 | 0.262846 | 0.1978 | 0.35736 |
| rumA_rumA | 2.976933 | 2.867859 | 3.058544 | 0.200171 | 0.100257 | 1.994077 | 0.006001 | 0.044565 | 0.154464 | 0.1045 | 0.242503 |
| SL1344_RS09480_ruvA | 1.015055 | 1.573814 | 0.38243 | -1.18561 | 0.161992 | -7.17227 | 4.68E-05 | 0.001274 | 0.033542 | 0.1 | 0.236592 |
| SL1344_RS09475_ruvB | 2.003517 | 2.356074 | 1.604922 | -0.75993 | 0.105323 | -7.26703 | 4.68E-05 | 0.001695 | 0.035853 | 0.1 | 0.236592 |
| RybA_rybA | 3.040887 | 3.290532 | 2.747937 | -0.41487 | 0.262043 | -1.76779 | 0.033311 | 0.05471 | 0.178788 | 0.1318 | 0.278981 |
| RybB_rybB | 0.126851 | -0.43783 | 0.218879 | 0.671534 | 0.488243 | 1.175716 | 0.141905 | 0.172928 | 0.347128 | 0.2718 | 0.439541 |
| RybD_rybD | 3.412565 | 3.410618 | 3.415958 | 0.008865 | 0.342263 | 0.023767 | 0.478667 | 0.757963 | 0.855052 | 0.9271 | 0.95526 |
| RydB_rydB | -2.77197 | -2.41082 | -2.98316 | -0.62494 | 0.758345 | -0.78431 | 0.197202 | 0.286347 | 0.456609 | 0.3898 | 0.543269 |
| RydC_rydC | -0.06943 | 0.000499 | -0.36794 | -0.35884 | 0.433221 | -0.74659 | 0.256667 | 0.362933 | 0.542981 | 0.4877 | 0.649087 |
| RyeF_ryeF | 0.367415 | 0.652902 | 0.085124 | -0.60055 | 0.216095 | -2.7738 | 0.000673 | 0.023588 | 0.109855 | 0.1001 | 0.236723 |
| RyfA_ryfA | 3.054799 | 3.124787 | 2.848963 | -0.27048 | 0.104382 | -2.09751 | 0.007329 | 0.063764 | 0.191094 | 0.1061 | 0.244631 |
| RyfD_ryfD | 3.338164 | 3.35957 | 3.315395 | 0.038088 | 0.369313 | 0.114071 | 0.452365 | 0.760572 | 0.856293 | 0.9616 | 0.977154 |
| RyhB_ryhB | -3.12962 | -3.21032 | -3.05295 | 0.17744 | 0.524608 | 0.314124 | 0.368421 | 0.529128 | 0.660844 | 0.6396 | 0.741896 |
| RyjB_ryjB | -2.46219 | -2.3656 | -2.53407 | -0.12764 | 0.528299 | -0.21254 | 0.404667 | 0.652949 | 0.767248 | 0.726 | 0.81742 |
| Sar_sar | -0.44513 | -0.60694 | -0.2997 | 0.319551 | 0.859884 | 0.401593 | 0.306667 | 0.502548 | 0.667398 | 0.6208 | 0.747049 |
| SL1344_RS10635_sbcB | 1.456075 | 1.424374 | 1.533439 | 0.106402 | 0.174004 | 0.551638 | 0.302 | 0.470467 | 0.625288 | 0.596 | 0.721336 |
| SL1344_RS01925_sbmA | -0.25398 | -0.34303 | -0.16666 | 0.170102 | 0.174661 | 0.930815 | 0.140667 | 0.227831 | 0.37303 | 0.3159 | 0.456032 |
| SL1344_RS10605_sbmC | 4.241474 | 4.367377 | 4.209415 | -0.13169 | 0.189729 | -0.58987 | 0.293138 | 0.489858 | 0.658178 | 0.5995 | 0.741492 |
| SL1344_RS20885_sbp | 1.778133 | 5.34087 | -0.60559 | -5.93921 | 0.617006 | -8.89939 | 4.68E-05 | 0.00798 | 0.070575 | 0.1 | 0.236592 |
| sdhC_sdhC | 3.687641 | 3.951685 | 3.299787 | -0.61463 | 0.130967 | -4.64347 | 4.68E-05 | 0.004782 | 0.055504 | 0.1 | 0.236592 |
| SL1344_RS09755_sdiA | 3.076884 | 3.914656 | 2.19999 | -1.7222 | 0.171625 | -10.095 | 4.68E-05 | 0.000783 | 0.030788 | 0.1 | 0.236592 |
| SdsR_sdsR | 0.973629 | 0.69856 | 1.288221 | 0.58227 | 0.477249 | 1.152352 | 0.091273 | 0.165042 | 0.338847 | 0.1817 | 0.3454 |
| sgbH_sgbH | -4.69909 | -4.41302 | -5.01542 | -0.60532 | 0.870112 | -0.61422 | 0.2505 | 0.376284 | 0.526608 | 0.4803 | 0.607979 |
| SgrS_sgrS | 4.452897 | 4.659383 | 4.323846 | -0.37624 | 0.181752 | -2.20625 | 4.68E-05 | 0.023369 | 0.114866 | 0.1 | 0.236592 |
| SibA_sibA | 1.713266 | 1.604778 | 1.820061 | 0.067225 | 1.008329 | 0.069109 | 0.45503 | 0.954784 | 0.97494 | 0.9997 | 0.999824 |
| SibC_sibC | 2.376082 | 2.743541 | 2.059654 | -0.64879 | 0.418368 | -1.37513 | 0.104 | 0.123884 | 0.286206 | 0.198 | 0.367341 |
| SibD_sibD | 3.019179 | 3.115844 | 2.803699 | -0.31489 | 1.088747 | -0.43793 | 0.228 | 0.571915 | 0.725223 | 0.4039 | 0.585454 |
| SL1344_RS14940_sicA | 3.680645 | 4.091554 | 3.252614 | -0.86887 | 0.296334 | -3.29209 | 4.68E-05 | 0.012001 | 0.084141 | 0.1 | 0.236592 |
| sifA_sifA | -0.88512 | -0.84616 | -0.90391 | -0.06206 | 0.388491 | -0.14536 | 0.437708 | 0.711227 | 0.813778 | 0.8042 | 0.87352 |
| sifB_sifB | -4.50409 | -3.8871 | -5.35696 | -1.47137 | 0.934103 | -1.54355 | 0.031313 | 0.092492 | 0.221681 | 0.1441 | 0.287997 |
| SL1344_RS05355_sigE | 4.323093 | 4.76905 | 3.748279 | -1.02319 | 0.222418 | -4.82867 | 4.68E-05 | 0.001921 | 0.041223 | 0.1 | 0.236592 |
| SL1344_RS21850_siiA | 1.028004 | 0.243311 | 1.416856 | 1.164948 | 0.21316 | 5.291348 | 4.68E-05 | 0.010973 | 0.077704 | 0.1 | 0.236592 |
| SL1344_RS21855_siiB | 1.83805 | 1.310739 | 2.296563 | 0.984081 | 0.216673 | 4.552116 | 4.68E-05 | 0.012205 | 0.082414 | 0.1 | 0.236592 |
| SL1344_RS21875_siiF | 1.839943 | 1.888287 | 1.673659 | -0.19505 | 0.161104 | -1.10826 | 0.154667 | 0.200486 | 0.370554 | 0.303 | 0.468923 |
| SL1344_RS01560_sinR | 1.139446 | 1.258808 | 0.830254 | -0.37389 | 0.265599 | -1.20681 | 0.112592 | 0.163198 | 0.33219 | 0.2319 | 0.396646 |
| SL1344_RS14920_sipA | 3.989448 | 4.477475 | 3.734906 | -0.74626 | 0.32095 | -2.45683 | 4.68E-05 | 0.041912 | 0.156074 | 0.1 | 0.236592 |
| SL1344_RS14935_sipB | 4.982012 | 5.492944 | 4.706047 | -0.78606 | 0.317122 | -2.70393 | 4.68E-05 | 0.032184 | 0.136126 | 0.1 | 0.236592 |
| SL1344_RS14930_sipC | 5.56699 | 6.137687 | 5.177914 | -0.96081 | 0.242591 | -3.66156 | 4.68E-05 | 0.02698 | 0.124642 | 0.1 | 0.236592 |

| | | | | | | | | | | |
|---|---|---|---|---|---|---|---|---|---|---|
| SL1344_RS14925_sipD | 2.883503 | 3.431899 | 2.613148 | -0.82147 | 0.311746 | -2.75924 | 4.68E-05 | 0.045214 | 0.162344 | 0.1 | 0.236592 |
| SL1344_RS09740_sirA | 3.194051 | 3.361679 | 2.944466 | -0.41588 | 0.099431 | -4.15218 | 4.68E-05 | 0.005246 | 0.056226 | 0.1 | 0.236592 |
| SL1344_RS10425_sirohydrochlorin.cobaltochelatase | 1.214825 | 0.670379 | 1.726034 | 1.055145 | 0.146313 | 7.226216 | 4.68E-05 | 0.001783 | 0.036441 | 0.1 | 0.236592 |
| SL1344_RS09840_SL1896 | -0.53485 | -0.61354 | -0.49461 | 0.080003 | 0.382054 | 0.178182 | 0.429714 | 0.770423 | 0.857737 | 0.8209 | 0.890844 |
| SL1344_RS07175_slyA | 3.765711 | 3.722647 | 3.827938 | 0.069707 | 0.133018 | 0.535655 | 0.276482 | 0.466395 | 0.631985 | 0.5534 | 0.695397 |
| SL1344_RS07110_sodB | 5.517397 | 5.060082 | 5.879916 | 0.816336 | 0.213065 | 4.01669 | 4.68E-05 | 0.019022 | 0.104715 | 0.1 | 0.236592 |
| SL1344_RS10630_sopA | 1.508671 | 2.252863 | 1.199732 | -1.03025 | 0.170015 | -4.8108 | 4.68E-05 | 0.023977 | 0.115879 | 0.1 | 0.236592 |
| sopB_sopB | 4.293775 | 4.895136 | 3.854953 | -1.03719 | 0.21987 | -4.64807 | 4.68E-05 | 0.01261 | 0.086228 | 0.1 | 0.236592 |
| SL1344_RS15235_sopD | 0.780054 | 1.362463 | 0.317993 | -0.98855 | 0.239357 | -3.97074 | 4.68E-05 | 0.008817 | 0.070248 | 0.1 | 0.236592 |
| SL1344_RS13925_sopE | 4.218407 | 4.920293 | 3.76027 | -1.14795 | 0.310711 | -3.8936 | 4.68E-05 | 0.003737 | 0.05231 | 0.1 | 0.236592 |
| SL1344_RS09265_sopE2 | 2.406072 | 3.363773 | 1.610599 | -1.69648 | 0.211223 | -7.68408 | 4.68E-05 | 0.00223 | 0.042296 | 0.1 | 0.236592 |
| spaK_spaK | 1.981826 | 2.416234 | 1.85638 | -0.51585 | 0.201673 | -1.77072 | 0.012659 | 0.075675 | 0.2132 | 0.1121 | 0.252469 |
| spaM_spaM | 2.403433 | 2.644584 | 2.369628 | -0.28264 | 0.371052 | -0.83189 | 0.215333 | 0.291027 | 0.47836 | 0.4025 | 0.58416 |
| spaN_spaN | 3.212296 | 3.578758 | 3.156789 | -0.42621 | 0.283235 | -1.00719 | 0.213857 | 0.233771 | 0.419617 | 0.3868 | 0.561982 |
| SL1344_RS14965_spaO | 2.91191 | 3.38636 | 2.868693 | -0.52034 | 0.196451 | -1.11264 | 0.164667 | 0.183298 | 0.361929 | 0.3061 | 0.47556 |
| SL1344_RS14960_spaP | 2.332955 | 2.880538 | 2.142769 | -0.71342 | 0.171371 | -2.509 | 4.68E-05 | 0.054334 | 0.178347 | 0.1001 | 0.236725 |
| SL1344_RS14955_spaQ | 2.029127 | 2.450379 | 1.731514 | -0.70771 | 0.202306 | -3.19936 | 4.68E-05 | 0.024395 | 0.116022 | 0.1 | 0.236592 |
| SL1344_RS14950_spaR | 1.537783 | 1.957837 | 1.225438 | -0.71905 | 0.189535 | -3.74754 | 4.68E-05 | 0.007696 | 0.066615 | 0.1 | 0.236592 |
| SL1344_RS14945_spaS | 1.2975 | 1.814188 | 1.161182 | -0.61159 | 0.264314 | -1.75167 | 0.011334 | 0.073951 | 0.209751 | 0.114 | 0.254531 |
| Spf_spf | 3.171525 | 3.649359 | 2.818893 | -0.6195 | 0.33902 | -2.05305 | 0.000673 | 0.033522 | 0.138376 | 0.1003 | 0.23699 |
| SL1344_RS14840_spi4.transcriptional.regulator | 2.879034 | 3.284842 | 2.803997 | -0.46856 | 0.103848 | -1.7566 | 0.006667 | 0.096383 | 0.246375 | 0.1049 | 0.242803 |
| spiC_spiC | -5.30093 | -5.55991 | -5.10384 | 0.435211 | 1.499751 | 0.275092 | 0.382412 | 0.592904 | 0.721134 | 0.703 | 0.796207 |
| spr_spr | 4.871066 | 5.103201 | 4.549721 | -0.54015 | 0.132163 | -4.2293 | 4.68E-05 | 0.002486 | 0.044831 | 0.1 | 0.236592 |
| SraA_sraA | -5.22657 | -4.96565 | -5.35886 | -0.30258 | 1.314598 | -0.1781 | 0.41439 | 0.662999 | 0.773867 | 0.7406 | 0.828797 |
| SraB_sraB | -0.67232 | -0.47645 | -0.88911 | -0.46338 | 0.3455 | -1.38463 | 0.046 | 0.109899 | 0.255405 | 0.1569 | 0.303494 |
| SraC_sraC | 2.330926 | 3.071847 | 1.652613 | -1.4151 | 0.174296 | -7.92879 | 4.68E-05 | 0.001616 | 0.037524 | 0.1 | 0.236592 |
| SraF_sraF | 0.140586 | 0.805856 | -0.26877 | -1.01602 | 0.34224 | -2.89378 | 4.68E-05 | 0.016532 | 0.094687 | 0.1 | 0.236592 |
| SraG_sraG | -3.88518 | -3.56987 | -4.23872 | -0.67169 | 0.581584 | -1.09943 | 0.108594 | 0.177328 | 0.319361 | 0.2531 | 0.396709 |
| SraL_sraL | 0.538718 | 0.121222 | 0.725037 | 0.60693 | 0.264124 | 2.178013 | 0.001336 | 0.06327 | 0.190611 | 0.1016 | 0.23867 |
| SL1344_RS07945_srfB | 4.349648 | 4.7732 | 3.897238 | -0.87877 | 0.045291 | -19.4974 | 4.68E-05 | 0.000293 | 0.021225 | 0.1 | 0.236592 |
| SL1344_RS07950_srfC | 3.881573 | 4.315415 | 3.381772 | -0.93548 | 0.086318 | -10.7352 | 4.68E-05 | 0.000809 | 0.029358 | 0.1 | 0.236592 |
| srlA_srlA | -0.27058 | 0.246544 | -0.33095 | -0.57191 | 0.45405 | -0.98421 | 0.224517 | 0.258274 | 0.442271 | 0.4716 | 0.619397 |
| srlE_srlE | 0.000701 | 0.192549 | -0.13834 | -0.34711 | 0.353228 | -0.93515 | 0.143904 | 0.217643 | 0.39258 | 0.2835 | 0.443715 |
| srlR_srlR | 1.456935 | 1.490113 | 1.430942 | -0.0478 | 0.164125 | -0.2659 | 0.394404 | 0.629978 | 0.750865 | 0.733 | 0.821022 |
| SroA_sroA | -3.011 | -2.92403 | -3.16429 | -0.22587 | 0.768584 | -0.26062 | 0.384667 | 0.60267 | 0.730381 | 0.7384 | 0.82314 |
| SroC_sroC | 7.016526 | 7.230598 | 6.808596 | -0.45237 | 0.08902 | -5.03676 | 4.68E-05 | 0.071393 | 0.20912 | 0.1 | 0.236592 |
| SroD_sroD | -1.14367 | -0.43557 | -1.60758 | -1.13267 | 0.334149 | -3.22185 | 4.68E-05 | 0.01498 | 0.08675 | 0.1 | 0.236592 |
| SroG_sroG | 2.940339 | 2.86755 | 3.119893 | 0.257142 | 0.25774 | 1.125418 | 0.073285 | 0.199896 | 0.381339 | 0.1753 | 0.332461 |
| SL1344_RS06985_ssaG | -4.45174 | -4.1675 | -4.72305 | -0.60608 | 0.906069 | -0.61664 | 0.241839 | 0.370194 | 0.524669 | 0.4747 | 0.602432 |
| SL1344_RS07050_ssaR | -2.80723 | -2.67605 | -2.9197 | -0.25281 | 0.462928 | -0.53177 | 0.285143 | 0.414123 | 0.561617 | 0.5307 | 0.650576 |
| SL1344_RS05160_sseI | -1.85247 | -1.77174 | -1.92658 | -0.13376 | 0.38989 | -0.33173 | 0.367089 | 0.572233 | 0.699875 | 0.6638 | 0.765776 |
| sspA_sspA | 3.213122 | 3.315802 | 3.023683 | -0.28055 | 0.093734 | -2.71386 | 4.68E-05 | 0.028975 | 0.123481 | 0.1002 | 0.236867 |
| SL1344_RS06915_ssrA | 0.25514 | 0.2031 | 0.310832 | 0.10536 | 0.142064 | 0.719214 | 0.202 | 0.319195 | 0.465542 | 0.4153 | 0.545884 |
| SsrA_ssrA | 11.19663 | 11.1597 | 11.23788 | 0.007654 | 0.126101 | 0.072404 | 0.460667 | 0.813271 | 0.891658 | 1 | 1 |
| SL1344_RS06910_ssrB | -0.79499 | -0.87485 | -0.70906 | 0.166844 | 0.207897 | 0.718009 | 0.200667 | 0.311735 | 0.456956 | 0.4052 | 0.536491 |

| | | | | | | | | | | |
|---|---|---|---|---|---|---|---|---|---|---|
| ssrS_ssrS | 9.416566 | 9.386761 | 9.443778 | -0.04014 | 0.274204 | -0.16424 | 0.447702 | 0.894195 | 0.940668 | 1 | 1 |
| SsrS_ssrS | 9.418089 | 9.386681 | 9.443708 | -0.04044 | 0.274691 | -0.17768 | 0.44437 | 0.893338 | 0.94022 | 1 | 1 |
| SL1344_RS16670_sstT | 2.538969 | 2.526367 | 2.564652 | 0.018869 | 0.121945 | 0.131953 | 0.448368 | 0.771689 | 0.85833 | 0.8482 | 0.907497 |
| SL1344_RS01735_stbB | -6.21 | -5.79244 | -6.69488 | -1.24241 | 2.549093 | -0.41668 | 0.286 | 0.41761 | 0.581449 | 0.5682 | 0.687832 |
| SL1344_RS05145_stfT | -1.71614 | -1.57853 | -1.91156 | -0.32981 | 0.397802 | -0.79063 | 0.181879 | 0.278889 | 0.442363 | 0.374 | 0.523441 |
| STnc3140_STnc3140..sopD. | -1.55695 | 0.013151 | -3.55292 | -3.56 | 0.919585 | -4.2078 | 4.68E-05 | 0.0425 | 0.153711 | 0.1 | 0.236592 |
| STnc3200_STnc3200 | -6.08993 | -5.19566 | -8.52115 | -3.26779 | 2.572993 | -1.19938 | 0.034 | 0.138124 | 0.293082 | 0.1389 | 0.281732 |
| SL1344_RS15350_syd | 0.128735 | 0.167794 | 0.074887 | -0.09435 | 0.183572 | -0.46717 | 0.306 | 0.483083 | 0.623255 | 0.5901 | 0.704042 |
| SymR_symR | -2.87317 | -2.41877 | -3.19844 | -0.78994 | 0.5628 | -1.34483 | 0.07062 | 0.123344 | 0.267994 | 0.1925 | 0.342347 |
| SL1344_RS13990_tail.protein | -6.2769 | -6.05035 | -6.64158 | -0.57132 | 2.669749 | -0.17679 | 0.393333 | 0.585575 | 0.721232 | 0.7471 | 0.829561 |
| tatE_tatE | 0.957361 | 1.008176 | 0.821231 | -0.16595 | 0.23048 | -0.71065 | 0.229333 | 0.338432 | 0.510828 | 0.4352 | 0.594542 |
| tcuB_tcuB | -2.34136 | -2.21609 | -2.50302 | -0.30005 | 0.366724 | -0.74153 | 0.209333 | 0.314742 | 0.461441 | 0.4064 | 0.538715 |
| SL1344_RS16770_tdcA | 0.352111 | -0.28758 | 0.90788 | 1.190376 | 0.167298 | 7.1069 | 4.68E-05 | 0.001751 | 0.036067 | 0.1 | 0.236592 |
| SL1344_RS16765_tdcB | 0.458067 | 0.266838 | 0.587211 | 0.329084 | 0.198849 | 1.612004 | 0.033977 | 0.080184 | 0.2102 | 0.136 | 0.279845 |
| SL1344_RS16760_tdcC | 2.032517 | 2.057298 | 2.015895 | -0.02528 | 0.237142 | -0.10252 | 0.442667 | 0.832248 | 0.89968 | 0.847 | 0.905918 |
| SL1344_RS05150_tfaT | -1.69921 | -1.57713 | -1.89193 | -0.33154 | 0.405305 | -0.77945 | 0.193333 | 0.286019 | 0.449762 | 0.3905 | 0.530864 |
| Tff_tff | 3.169035 | 3.398934 | 2.889939 | -0.56817 | 0.192593 | -3.01415 | 4.68E-05 | 0.009123 | 0.073736 | 0.1 | 0.236592 |
| thiP_thiP | 0.407191 | 0.32561 | 0.505619 | 0.174067 | 0.154933 | 1.072621 | 0.124584 | 0.187209 | 0.336804 | 0.2708 | 0.418822 |
| thiQ_thiQ | -1.10335 | -1.13619 | -1.07251 | 0.061509 | 0.312729 | 0.192661 | 0.404667 | 0.605359 | 0.728758 | 0.7197 | 0.806405 |
| SL1344_RS06625_thrS | 5.228745 | 5.023067 | 5.429618 | 0.412778 | 0.0431 | 9.5412 | 4.68E-05 | 0.000515 | 0.0265 | 0.1 | 0.236592 |
| SL1344_RS04595_tnpA1 | -2.41121 | -1.44766 | -4.65407 | -3.17364 | 1.020934 | -2.97236 | 4.68E-05 | 0.037008 | 0.140686 | 0.1 | 0.236592 |
| SL1344_RS06060_tnpA2 | -2.45998 | -1.86737 | -3.20086 | -1.33159 | 0.447001 | -2.92663 | 4.68E-05 | 0.020706 | 0.098712 | 0.1006 | 0.237392 |
| SL1344_RS09075_tnpA3 | -3.72865 | -2.70687 | -4.95543 | -2.21473 | 0.723972 | -3.05084 | 0.001336 | 0.018842 | 0.093881 | 0.1003 | 0.236998 |
| SL1344_RS09790_tnpA4 | -4.85831 | -4.11901 | -5.80556 | -1.69069 | 1.02026 | -1.59978 | 0.038641 | 0.080345 | 0.205094 | 0.1387 | 0.282248 |
| SL1344_RS12685_tnpA5 | -4.37832 | -3.70802 | -4.83406 | -1.13447 | 1.096057 | -0.98162 | 0.135243 | 0.206786 | 0.368099 | 0.2844 | 0.440176 |
| SL1344_RS17020_tnpA6 | -2.85969 | -2.41594 | -3.40661 | -0.99894 | 0.451242 | -2.12554 | 0.01 | 0.041518 | 0.139756 | 0.1084 | 0.247203 |
| SL1344_RS17955_tnpA7 | -3.49046 | -2.88333 | -4.53663 | -1.64324 | 0.641323 | -2.47981 | 0.002001 | 0.038424 | 0.135529 | 0.1014 | 0.23847 |
| tolA_tolA | 1.792843 | 1.617454 | 1.843919 | 0.20304 | 0.18499 | 0.950018 | 0.186 | 0.247166 | 0.426016 | 0.3633 | 0.523283 |
| SL1344_RS16475_tolC | 4.04849 | 3.894868 | 4.310211 | 0.412196 | 0.097038 | 3.969698 | 4.68E-05 | 0.01529 | 0.092147 | 0.1 | 0.236592 |
| SL1344_RS21860_TolC.family.protein | 1.785332 | 1.125651 | 2.16648 | 1.055885 | 0.295048 | 3.729422 | 4.68E-05 | 0.018023 | 0.100119 | 0.1 | 0.236592 |
| tqsA_tqsA | -0.67775 | -0.85767 | -0.55006 | 0.301061 | 0.280709 | 1.02737 | 0.147901 | 0.21213 | 0.371892 | 0.2992 | 0.453222 |
| SL1344_RS14145_transcriptional.regulator | -6.2505 | -5.72129 | -6.9242 | -1.12013 | 1.697127 | -0.62072 | 0.227333 | 0.351862 | 0.505495 | 0.4435 | 0.576576 |
| trkD_trkD | 1.682748 | 1.639222 | 1.752922 | 0.108443 | 0.11167 | 0.890508 | 0.168 | 0.256267 | 0.414439 | 0.3549 | 0.502706 |
| SL1344_RS06175_tRNA.arg | -2.8372 | -2.46359 | -3.23231 | -0.75765 | 0.46002 | -1.60335 | 0.039333 | 0.084629 | 0.206188 | 0.156 | 0.300676 |
| SL1344_RS13600_tRNA.rRNA.methyltransferase | 1.577007 | 1.513647 | 1.690503 | 0.210007 | 0.198911 | 1.058944 | 0.117333 | 0.169215 | 0.335271 | 0.2475 | 0.408917 |
| truA_truA | 0.720869 | 0.546675 | 0.914818 | 0.373063 | 0.137878 | 2.642102 | 0.000673 | 0.024014 | 0.10685 | 0.1011 | 0.238006 |
| SL1344_RS23260_tsr | 5.533486 | 6.412495 | 4.834627 | -1.57573 | 0.15532 | -9.30252 | 4.68E-05 | 0.004002 | 0.053686 | 0.1 | 0.236592 |
| SL1344_RS06870_ttrA | -1.15535 | -1.29745 | -1.04921 | 0.244916 | 0.267592 | 0.882031 | 0.156667 | 0.249457 | 0.402856 | 0.3392 | 0.479937 |
| SL1344_RS06880_ttrB | -4.48518 | -4.56734 | -4.42949 | 0.200803 | 1.036465 | 0.179968 | 0.410393 | 0.592453 | 0.719753 | 0.7158 | 0.80159 |
| SL1344_RS06875_ttrC | -4.13118 | -4.25968 | -3.95874 | 0.329238 | 0.779748 | 0.40754 | 0.338667 | 0.497365 | 0.63165 | 0.6031 | 0.708355 |
| SL1344_RS06890_ttrR | 0.86612 | 0.649782 | 1.390498 | 0.678793 | 0.28041 | 2.04298 | 0.004665 | 0.049401 | 0.166964 | 0.1044 | 0.242455 |
| SL1344_RS06885_ttrS | 2.624492 | 2.317639 | 2.924698 | 0.616944 | 0.090756 | 6.731514 | 4.68E-05 | 0.001687 | 0.036471 | 0.1 | 0.236592 |
| tyrA_tyrA | 0.801928 | 0.636639 | 0.916688 | 0.280803 | 0.171171 | 1.610475 | 0.032 | 0.084165 | 0.215009 | 0.1424 | 0.287834 |
| ubiB_ubiB | 3.109542 | 3.047782 | 3.18236 | 0.135126 | 0.060674 | 2.175361 | 0.007329 | 0.040551 | 0.139062 | 0.1107 | 0.249891 |
| ubiF_ubiF | 1.959287 | 1.731124 | 2.125385 | 0.375231 | 0.168339 | 2.20214 | 0.003998 | 0.031311 | 0.12951 | 0.1035 | 0.241211 |

| | | | | | | | | | | |
|---|---|---|---|---|---|---|---|---|---|---|
| SL1344_RS21190_udhA | 3.392339 | 3.515938 | 3.036945 | -0.47381 | 0.125746 | -2.98605 | 4.68E-05 | 0.036953 | 0.14459 | 0.1 | 0.236592 |
| SL1344_RS03635_UDP.galactopyranose.mutase | -4.2562 | -4.16737 | -4.31224 | -0.15726 | 0.773494 | -0.18855 | 0.411059 | 0.55051 | 0.68158 | 0.6606 | 0.760157 |
| ugpC_ugpC | -0.27842 | -0.55172 | -0.00742 | 0.549995 | 0.205251 | 2.622928 | 0.001336 | 0.022663 | 0.104396 | 0.1018 | 0.238896 |
| ulaD_ulaD | -1.77098 | -1.91705 | -1.58951 | 0.352524 | 0.358497 | 0.940764 | 0.140573 | 0.226416 | 0.381703 | 0.3144 | 0.457362 |
| ushA_ushA | 3.2731 | 3.127348 | 3.442982 | 0.330751 | 0.12453 | 2.684993 | 4.68E-05 | 0.022677 | 0.110814 | 0.1 | 0.236592 |
| SL1344_RS09735_uvrC | 2.93769 | 3.123235 | 2.766029 | -0.36435 | 0.085027 | -4.28737 | 4.68E-05 | 0.005686 | 0.057321 | 0.1 | 0.236592 |
| SL1344_RS11530_virulence.protein.MsgA | -5.50644 | -5.81718 | -5.20827 | 0.604541 | 1.067006 | 0.524733 | 0.272485 | 0.402205 | 0.543703 | 0.4995 | 0.620832 |
| wecC_wecC | 2.369495 | 2.273695 | 2.427581 | 0.160317 | 0.112055 | 1.433115 | 0.036 | 0.118727 | 0.268794 | 0.1446 | 0.289592 |
| SL1344_RS10695_wzzB | 2.46276 | 1.886405 | 2.936531 | 1.050664 | 0.086409 | 12.12891 | 4.68E-05 | 0.000909 | 0.028743 | 0.1 | 0.236592 |
| SL1344_RS02345_ybaO | 1.022651 | 0.985349 | 1.066098 | 0.085249 | 0.122332 | 0.662504 | 0.225183 | 0.341187 | 0.490688 | 0.4521 | 0.579772 |
| SL1344_RS03185_ybeM | -1.13101 | -0.98084 | -1.22353 | -0.24576 | 0.367932 | -0.62786 | 0.251832 | 0.399 | 0.557513 | 0.5081 | 0.641268 |
| SL1344_RS04285_ybiK | 2.461491 | 3.245916 | 1.977779 | -1.26983 | 0.342802 | -3.78476 | 4.68E-05 | 0.006874 | 0.065636 | 0.1 | 0.236592 |
| SL1344_RS08990_ycgO | 3.161766 | 3.726203 | 2.769575 | -0.93016 | 0.124639 | -6.75642 | 4.68E-05 | 0.005806 | 0.060517 | 0.1 | 0.236592 |
| SL1344_RS08975_ycgR | 2.419403 | 3.009826 | 1.818051 | -1.26665 | 0.202475 | -6.44685 | 4.68E-05 | 0.001825 | 0.038754 | 0.1 | 0.236592 |
| SL1344_RS08585_yciK | 1.625703 | 1.541548 | 1.732199 | 0.178574 | 0.130226 | 1.269465 | 0.091333 | 0.133326 | 0.283388 | 0.2106 | 0.36755 |
| SL1344_RS08495_yciW | 0.91287 | 2.672466 | -0.81525 | -3.38611 | 0.467181 | -7.2614 | 4.68E-05 | 0.000631 | 0.029716 | 0.1 | 0.236592 |
| SL1344_RS07545_ydeD | -0.8303 | -1.1841 | -0.55892 | 0.626796 | 0.244365 | 2.534332 | 0.002002 | 0.029829 | 0.119398 | 0.1021 | 0.239125 |
| SL1344_RS06680_ydiV | -1.16696 | -1.43121 | -0.94155 | 0.492384 | 0.254922 | 1.87981 | 0.015334 | 0.055483 | 0.165107 | 0.1187 | 0.259052 |
| SL1344_RS06595_ydiY | 0.292039 | 0.050028 | 0.491405 | 0.443508 | 0.170706 | 2.538285 | 0.001336 | 0.030376 | 0.120804 | 0.1011 | 0.238065 |
| SL1344_RS06585_ydiZ | -1.43567 | -1.94503 | -0.92923 | 1.117307 | 0.47912 | 2.342704 | 0.000673 | 0.026838 | 0.118916 | 0.1004 | 0.237124 |
| SL1344_RS06560_ydjN | 3.367026 | 4.731358 | 1.246728 | -3.44958 | 0.329591 | -8.85201 | 4.68E-05 | 0.007812 | 0.069943 | 0.1 | 0.236592 |
| SL1344_RS09415_yebG | 0.224455 | 0.641193 | -0.16881 | -0.79562 | 0.194974 | -3.94663 | 4.68E-05 | 0.007177 | 0.062234 | 0.1 | 0.236592 |
| SL1344_RS09225_yebR | 2.392314 | 2.76037 | 1.971768 | -0.81147 | 0.12207 | -6.80471 | 4.68E-05 | 0.001925 | 0.038878 | 0.1 | 0.236592 |
| SL1344_RS09230_yebS | 1.389154 | 1.58118 | 1.165771 | -0.42991 | 0.152723 | -2.82312 | 4.68E-05 | 0.027006 | 0.116711 | 0.1 | 0.236592 |
| SL1344_RS09760_yecC | 1.853846 | 2.432362 | 1.164174 | -1.26756 | 0.097997 | -12.9474 | 4.68E-05 | 0.000573 | 0.024991 | 0.1 | 0.236592 |
| SL1344_RS09750_yecF | 1.135943 | 1.839185 | 0.522387 | -1.30814 | 0.17331 | -7.60713 | 4.68E-05 | 0.001789 | 0.037755 | 0.1 | 0.236592 |
| SL1344_RS09765_yecS | 0.873191 | 1.47286 | 0.248745 | -1.23302 | 0.190524 | -6.50803 | 4.68E-05 | 0.002013 | 0.038506 | 0.1 | 0.236592 |
| SL1344_RS09825_yedD | 0.596703 | 0.41657 | 0.649199 | 0.214973 | 0.194845 | 0.925403 | 0.188 | 0.267164 | 0.437027 | 0.4007 | 0.551736 |
| SL1344_RS09830_yedE | -0.89612 | -1.02677 | -0.65344 | 0.387719 | 0.340535 | 1.092094 | 0.103265 | 0.184468 | 0.348351 | 0.246 | 0.395861 |
| SL1344_RS09835_yedF | -1.09687 | -1.32219 | -0.73119 | 0.566099 | 0.291724 | 1.871025 | 0.032645 | 0.06164 | 0.177606 | 0.1301 | 0.272972 |
| SL1344_RS09770_yedO | 1.885323 | 2.459284 | 1.312977 | -1.15898 | 0.126797 | -9.16647 | 4.68E-05 | 0.000976 | 0.031201 | 0.1 | 0.236592 |
| SL1344_RS10925_yegD | -1.54424 | -1.59445 | -1.44013 | 0.156129 | 0.393521 | 0.347571 | 0.359094 | 0.56557 | 0.698407 | 0.6865 | 0.785586 |
| SL1344_RS13585_yfiK | -3.04523 | -3.26424 | -2.96048 | 0.250941 | 0.678114 | 0.343569 | 0.374417 | 0.584594 | 0.715849 | 0.7097 | 0.804829 |
| SL1344_RS17960_yhfL | 0.692181 | 1.302874 | -0.23517 | -1.53246 | 0.233494 | -6.31927 | 4.68E-05 | 0.005838 | 0.058202 | 0.1 | 0.236592 |
| SL1344_RS18605_yhjH | 3.73547 | 4.549282 | 3.274256 | -1.25153 | 0.293119 | -3.94231 | 4.68E-05 | 0.005705 | 0.06125 | 0.1 | 0.236592 |
| SL1344_RS18650_yhjQ | 0.851946 | 0.556661 | 1.144957 | 0.603951 | 0.154119 | 3.851638 | 4.68E-05 | 0.007102 | 0.062138 | 0.1 | 0.236592 |
| SL1344_RS23255_yjiY | -1.02187 | -1.10859 | -0.91284 | 0.20377 | 0.249314 | 0.753398 | 0.199333 | 0.306027 | 0.459255 | 0.4066 | 0.542447 |
| SL1344_RS05645_ymdA | 2.420349 | 2.926079 | 1.825577 | -1.13612 | 0.145468 | -7.93842 | 4.68E-05 | 0.001823 | 0.038884 | 0.1 | 0.236592 |
| SL1344_RS05655_ymdC | 2.963947 | 3.343641 | 2.651454 | -0.68028 | 0.108128 | -6.13974 | 4.68E-05 | 0.003401 | 0.047634 | 0.1 | 0.236592 |
| yopJ_yopJ | 2.725948 | 2.73046 | 2.721338 | -0.00678 | 0.249174 | -0.02496 | 0.491006 | 0.859708 | 0.917645 | 0.9424 | 0.96529 |
| SL1344_RS15335_yqcB | 0.242306 | 0.380265 | 0.125109 | -0.27079 | 0.22408 | -1.16677 | 0.099267 | 0.155916 | 0.31245 | 0.2296 | 0.384137 |
| SL1344_RS15340_yqcC..next.to.csrB. | -0.9888 | -0.87438 | -1.17088 | -0.31405 | 0.288044 | -1.00319 | 0.127915 | 0.205184 | 0.365105 | 0.2875 | 0.435192 |
| YrlA_yrlA | -4.34408 | -4.28125 | -4.41462 | -0.13221 | 0.764791 | -0.15863 | 0.424384 | 0.550289 | 0.679695 | 0.6494 | 0.748971 |
| YrlB_yrlB | -3.80776 | -3.76703 | -3.87396 | -0.08572 | 0.745692 | -0.10417 | 0.452667 | 0.617347 | 0.738638 | 0.7278 | 0.814328 |
| zntA_zntA | 2.139056 | 2.297893 | 1.972152 | -0.31302 | 0.138491 | -2.24353 | 0.003332 | 0.034173 | 0.133853 | 0.1035 | 0.241255 |

| | | | | | | | | | | | |
|---|---|---|---|---|---|---|---|---|---|---|---|
| zntB_zntB | 0.73233 | 0.747503 | 0.715159 | -0.03929 | 0.160889 | -0.22724 | 0.404 | 0.582839 | 0.709137 | 0.7023 | 0.790609 |
| zraP_zraP | -2.42723 | -2.48718 | -2.34025 | 0.142895 | 0.397664 | 0.327657 | 0.36 | 0.503228 | 0.637356 | 0.6086 | 0.716003 |
| crRNA1_ | -0.43273 | -0.3742 | -0.5334 | -0.09803 | 0.3531 | -0.25363 | 0.408394 | 0.720492 | 0.821431 | 0.8328 | 0.894975 |
| crRNA2_ | 0.989005 | 0.803137 | 1.273487 | 0.456639 | 0.145557 | 3.012322 | 4.68E-05 | 0.020566 | 0.101103 | 0.1004 | 0.237131 |
| ECS021_ | -3.44558 | -3.68077 | -2.69946 | 0.871946 | 0.982203 | 0.786117 | 0.21519 | 0.318715 | 0.487906 | 0.423 | 0.58174 |
| ECS173_ | -2.35037 | -2.43344 | -2.28969 | 0.148131 | 0.566983 | 0.241132 | 0.401333 | 0.620694 | 0.743502 | 0.7384 | 0.823088 |
| Ffs_ | 3.51518 | 3.529244 | 3.498316 | -0.10678 | 0.192392 | -0.5592 | 0.317788 | 0.406056 | 0.58435 | 0.6716 | 0.783815 |
| fimA.trailer_ | -2.92155 | -3.41894 | -2.51504 | 0.907939 | 0.444197 | 1.936878 | 0.01599 | 0.050184 | 0.155477 | 0.1141 | 0.253786 |
| GO.10702_ | -1.65436 | -1.59827 | -1.6785 | -0.08005 | 0.426588 | -0.16482 | 0.431046 | 0.665529 | 0.777311 | 0.773 | 0.850104 |
| GO.10704_ | 1.386783 | 1.463099 | 1.345399 | -0.11121 | 0.130231 | -0.76101 | 0.214524 | 0.314721 | 0.475323 | 0.4205 | 0.565326 |
| Ig1085_ | -1.4404 | -1.45233 | -1.40981 | 0.032491 | 0.360958 | 0.081953 | 0.463333 | 0.662912 | 0.775374 | 0.7691 | 0.84759 |
| Ig1140_ | -2.50935 | -2.19541 | -2.9216 | -0.73138 | 0.459943 | -1.57555 | 0.037309 | 0.092648 | 0.222098 | 0.1479 | 0.291208 |
| Ig2335_ | 0.931269 | 0.734305 | 1.06528 | 0.325012 | 0.129027 | 2.438159 | 0.005331 | 0.03231 | 0.123776 | 0.1057 | 0.243669 |
| Ig959_ | -8.88709 | -8.94967 | -8.82839 | 0.165717 | 3.957717 | 0.037285 | 0.48 | 0.568742 | 0.699689 | 0.6879 | 0.782087 |
| IS042_ | -4.28967 | -4.03981 | -4.44217 | -0.47506 | 1.160497 | -0.38362 | 0.348667 | 0.557005 | 0.691114 | 0.6805 | 0.777942 |
| lysC.leader_ | -1.22231 | -1.31437 | -1.12946 | 0.187395 | 0.383518 | 0.44087 | 0.301133 | 0.47866 | 0.627319 | 0.5755 | 0.697774 |
| mreB.leader_ | 0.355636 | 0.559125 | 0.22675 | -0.35382 | 0.217213 | -1.56775 | 0.026667 | 0.098776 | 0.24159 | 0.1274 | 0.269246 |
| pRNA_ | -3.29491 | -2.98275 | -3.67397 | -0.72097 | 0.682859 | -0.9936 | 0.115333 | 0.196094 | 0.354231 | 0.2574 | 0.404578 |
| RprA_ | -0.38367 | -0.40124 | -0.36742 | 0.156367 | 0.449469 | 0.335582 | 0.357761 | 0.485118 | 0.646029 | 0.7427 | 0.820628 |
| rpsL.leader_ | 2.39686 | 2.56906 | 2.022959 | -0.50337 | 0.175648 | -2.18855 | 0.002001 | 0.048485 | 0.166377 | 0.1016 | 0.238712 |
| rpsP.leader_ | 2.154537 | 2.429677 | 1.872429 | -0.50619 | 0.224409 | -2.26238 | 4.68E-05 | 0.026219 | 0.120297 | 0.101 | 0.237924 |
| rrf_ | -7.27117 | -8.93551 | -5.39808 | 3.260965 | 2.944185 | 1.066814 | 0.087941 | 0.17408 | 0.328347 | 0.2307 | 0.384202 |
| rrf_.1 | -5.82603 | -5.983 | -5.62098 | 0.393596 | 3.205159 | 0.094472 | 0.462 | 0.755067 | 0.847477 | 0.8901 | 0.932171 |
| rrf_.2 | -4.85681 | -4.83293 | -4.87486 | 0.320295 | 1.916933 | 0.15417 | 0.424 | 0.704001 | 0.810166 | 0.863 | 0.912454 |
| rrf_.3 | -5.74166 | -5.58062 | -5.96149 | -0.29107 | 1.639948 | -0.17028 | 0.42 | 0.607072 | 0.730927 | 0.7247 | 0.810957 |
| rrf_.4 | -3.24626 | -3.54788 | -2.96301 | 0.629575 | 2.283165 | 0.230685 | 0.410393 | 0.785851 | 0.870119 | 0.9091 | 0.945794 |
| rrf_.5 | -6.41523 | -7.23267 | -5.94711 | 1.428638 | 2.682613 | 0.45737 | 0.278 | 0.411578 | 0.575827 | 0.5377 | 0.668822 |
| rrf_.6 | -7.87412 | -8.57741 | -7.40206 | 0.984408 | 4.013614 | 0.210433 | 0.379747 | 0.576132 | 0.708259 | 0.7013 | 0.793466 |
| rrf_.7 | -8.6618 | -9.92335 | -7.16489 | 2.984118 | 4.219863 | 0.644681 | 0.221333 | 0.370557 | 0.515994 | 0.4566 | 0.585377 |
| RUF_107c.1_ | 0.458573 | 0.282194 | 0.610111 | 0.339771 | 0.22482 | 1.468337 | 0.043333 | 0.090115 | 0.227297 | 0.1483 | 0.296022 |
| RUF_107c.10_ | 0.429977 | 0.745236 | 0.296919 | -0.44753 | 0.276972 | -1.47655 | 0.038667 | 0.106072 | 0.255678 | 0.141 | 0.287254 |
| RUF_107c.102_ | -3.17243 | -3.15176 | -3.18722 | -0.02182 | 0.511185 | -0.03815 | 0.482 | 0.587597 | 0.711515 | 0.6982 | 0.788784 |
| RUF_107c.105_ | -0.35133 | -0.39394 | -0.30469 | 0.081028 | 0.203374 | 0.394358 | 0.331333 | 0.475707 | 0.61431 | 0.5888 | 0.697769 |
| RUF_107c.107_ | -1.30198 | -1.21521 | -1.34418 | -0.06465 | 0.426242 | -0.12821 | 0.440373 | 0.741264 | 0.835639 | 0.8098 | 0.881302 |
| RUF_107c.108_ | 1.313329 | 1.008788 | 1.501759 | 0.485038 | 0.132367 | 3.571136 | 4.68E-05 | 0.014778 | 0.085213 | 0.1 | 0.236592 |
| RUF_107c.111_ | 1.814325 | 1.705616 | 1.890775 | 0.192675 | 0.152024 | 1.235375 | 0.083944 | 0.128882 | 0.282608 | 0.1953 | 0.349959 |
| RUF_107c.112_ | 0.165895 | 0.140168 | 0.234733 | 0.095116 | 0.282812 | 0.311948 | 0.391333 | 0.647143 | 0.767279 | 0.7675 | 0.852946 |
| RUF_107c.18_ | -3.14134 | -2.48698 | -3.475 | -0.7176 | 0.977119 | -0.7189 | 0.224 | 0.349205 | 0.519838 | 0.4414 | 0.593945 |
| RUF_107c.19_ | -0.34665 | -0.34644 | -0.347 | 0.035898 | 0.264255 | 0.130551 | 0.443704 | 0.648646 | 0.763581 | 0.7773 | 0.849206 |
| RUF_107c.2_ | -2.84679 | -2.9686 | -2.65525 | 0.30488 | 0.604743 | 0.48187 | 0.295803 | 0.464956 | 0.61006 | 0.582 | 0.696763 |
| RUF_107c.23_ | -0.03594 | 0.008579 | -0.10928 | -0.11097 | 0.293374 | -0.34 | 0.353333 | 0.610787 | 0.73851 | 0.6965 | 0.797369 |
| RUF_107c.24_ | 0.875438 | 0.688335 | 1.186892 | 0.487122 | 0.162785 | 2.832336 | 0.000673 | 0.023644 | 0.109497 | 0.1005 | 0.237271 |
| RUF_107c.26_ | -4.74619 | -4.55962 | -5.07199 | -0.53868 | 1.030504 | -0.45701 | 0.309127 | 0.449997 | 0.601323 | 0.5747 | 0.69153 |
| RUF_107c.30_ | -6.12719 | -5.601 | -6.84346 | -1.29436 | 2.145661 | -0.48902 | 0.22052 | 0.358082 | 0.52518 | 0.444 | 0.580446 |
| RUF_107c.38_ | -1.1745 | -1.1873 | -1.14721 | 0.05523 | 0.305059 | 0.160713 | 0.422385 | 0.595149 | 0.721169 | 0.7161 | 0.806108 |

| | | | | | | | | | | |
|---|---|---|---|---|---|---|---|---|---|---|
| RUF_107c.47_ | -0.11184 | -0.26641 | -0.04272 | 0.22988 | 0.423141 | 0.550761 | 0.251333 | 0.420997 | 0.589879 | 0.4909 | 0.646474 |
| RUF_107c.50_ | -1.19285 | -1.27427 | -1.06521 | 0.198762 | 0.308652 | 0.623181 | 0.251166 | 0.404753 | 0.558509 | 0.5148 | 0.6442 |
| RUF_107c.51_ | -0.87402 | -1.08559 | -0.69921 | 0.392883 | 0.270725 | 1.405986 | 0.045333 | 0.107613 | 0.24547 | 0.1622 | 0.3073 |
| RUF_107c.53_ | 1.213308 | 1.487196 | 1.099116 | -0.34234 | 0.228603 | -1.31995 | 0.098 | 0.133431 | 0.292456 | 0.1995 | 0.359454 |
| RUF_107c.56_ | 0.257236 | 0.18294 | 0.381521 | 0.211467 | 0.235706 | 0.867799 | 0.166 | 0.260428 | 0.4312 | 0.3537 | 0.500611 |
| RUF_107c.57_ | -1.22063 | -1.44459 | -1.01408 | 0.532783 | 0.451129 | 1.174132 | 0.079333 | 0.145691 | 0.304218 | 0.1991 | 0.348999 |
| RUF_107c.58_ | 1.37747 | 1.255973 | 1.43542 | 0.214207 | 0.345642 | 0.652231 | 0.221333 | 0.318829 | 0.503092 | 0.4239 | 0.590904 |
| RUF_107c.59_ | 0.434614 | 0.31056 | 0.605744 | 0.312167 | 0.233108 | 1.310879 | 0.060667 | 0.13227 | 0.287645 | 0.1625 | 0.310914 |
| RUF_107c.63_ | -0.82419 | -0.89987 | -0.66243 | 0.252609 | 0.394333 | 0.600087 | 0.291139 | 0.419996 | 0.583386 | 0.5618 | 0.688557 |
| RUF_107c.69_ | -0.3127 | -0.39456 | -0.21186 | 0.200499 | 0.236639 | 0.797452 | 0.187333 | 0.287168 | 0.448724 | 0.3884 | 0.528549 |
| RUF_107c.70_ | -2.05654 | -1.96322 | -2.1546 | -0.19212 | 0.352641 | -0.49831 | 0.288 | 0.419448 | 0.565835 | 0.5193 | 0.643026 |
| RUF_107c.72_ | -1.87371 | -1.90169 | -1.81005 | 0.067722 | 0.397544 | 0.156383 | 0.439333 | 0.654856 | 0.767018 | 0.7469 | 0.831208 |
| RUF_107c.77_ | -1.54463 | -1.62248 | -1.41043 | 0.190219 | 0.32838 | 0.527246 | 0.283145 | 0.458723 | 0.601522 | 0.5636 | 0.682501 |
| RUF_107c.8_ | -2.83586 | -3.16547 | -2.69617 | 0.433075 | 0.631784 | 0.640649 | 0.264 | 0.392859 | 0.55413 | 0.5169 | 0.651096 |
| RUF_107c.81_ | -4.45108 | -4.33823 | -4.54998 | -0.20114 | 0.843971 | -0.23037 | 0.407728 | 0.559165 | 0.685734 | 0.6585 | 0.755279 |
| RUF_107c.82_ | 1.745389 | 1.637267 | 1.945549 | 0.300596 | 0.134696 | 2.095224 | 0.009994 | 0.051926 | 0.167038 | 0.109 | 0.247891 |
| RUF_107c.83_ | -4.17439 | -4.12729 | -4.30872 | -0.13914 | 1.044415 | -0.12741 | 0.451699 | 0.671769 | 0.784502 | 0.7822 | 0.860021 |
| RUF_107c.86_ | -0.44376 | -0.54853 | -0.30035 | 0.249722 | 0.235512 | 0.985112 | 0.134 | 0.208915 | 0.366312 | 0.2914 | 0.440772 |
| RUF_107c.89_ | 0.063039 | 0.103388 | 0.042932 | -0.04037 | 0.226117 | -0.14027 | 0.438374 | 0.686365 | 0.794453 | 0.7798 | 0.857517 |
| RUF_107c.90_ | -3.88174 | -3.76921 | -3.99977 | -0.23576 | 0.612343 | -0.35849 | 0.339107 | 0.472069 | 0.610137 | 0.5737 | 0.684435 |
| RUF_107c.92_ | 1.71347 | 1.682437 | 1.830447 | 0.1332 | 0.132694 | 0.819713 | 0.235843 | 0.338629 | 0.506672 | 0.4849 | 0.629721 |
| RUF_107c.94_ | -3.74559 | -3.65302 | -3.81561 | -0.12953 | 0.707211 | -0.16736 | 0.423718 | 0.591091 | 0.714525 | 0.6861 | 0.781661 |
| RUF_107c.95_ | 1.915856 | 1.91894 | 1.914197 | 0.010425 | 0.146977 | 0.06156 | 0.471333 | 0.672981 | 0.786528 | 0.7962 | 0.866763 |
| RUF_107c.96_ | -1.70695 | -2.0352 | -1.38386 | 0.715236 | 0.415886 | 1.680833 | 0.02 | 0.07193 | 0.198332 | 0.1182 | 0.259343 |
| RUF_111_ | 1.175385 | 1.492442 | 1.0604 | -0.42262 | 0.192241 | -1.92117 | 0.013325 | 0.075764 | 0.210376 | 0.1131 | 0.253214 |
| RUF_121_ | -1.81304 | -1.43993 | -2.52701 | -1.09467 | 0.471543 | -2.20543 | 0.002002 | 0.056086 | 0.173068 | 0.1018 | 0.23899 |
| RUF_125_ | 2.953624 | 3.707795 | 2.157361 | -1.58474 | 0.148372 | -10.8599 | 4.68E-05 | 0.000337 | 0.024431 | 0.1 | 0.236592 |
| RUF_167_ | -1.06207 | -1.10171 | -1.03528 | -0.02797 | 0.450494 | -0.05345 | 0.481333 | 0.741042 | 0.836425 | 0.8772 | 0.922824 |
| RUF_178_ | 1.73431 | 1.944035 | 1.66283 | -0.24698 | 0.16398 | -1.31205 | 0.100667 | 0.142386 | 0.302836 | 0.2229 | 0.382688 |
| RUF_219c_ | 2.090933 | 2.41134 | 1.943564 | -0.42817 | 0.237583 | -1.59341 | 0.048667 | 0.089234 | 0.233914 | 0.1475 | 0.299287 |
| RUF_220c_ | 0.64863 | 1.011281 | 0.19922 | -0.82036 | 0.262578 | -3.12668 | 4.68E-05 | 0.011297 | 0.078982 | 0.1 | 0.236592 |
| RUF_226_ | -1.81264 | -1.76504 | -1.86467 | -0.01023 | 0.558602 | -0.01839 | 0.492667 | 0.735047 | 0.831714 | 0.8546 | 0.906794 |
| RUF_247c_ | 0.78837 | 0.760952 | 0.814258 | 0.052022 | 0.145337 | 0.316714 | 0.362 | 0.540195 | 0.672088 | 0.6493 | 0.751281 |
| RUF_257c_ | 0.429017 | 0.285613 | 0.547215 | 0.275664 | 0.187142 | 1.39948 | 0.057333 | 0.102214 | 0.240346 | 0.165 | 0.313111 |
| RUF_279_ | 1.319558 | 1.223568 | 1.404338 | 0.198887 | 0.156497 | 1.242549 | 0.073285 | 0.133261 | 0.28334 | 0.1978 | 0.348181 |
| RUF_290_ | 1.097775 | 1.045324 | 1.172115 | 0.118324 | 0.145923 | 0.736678 | 0.221333 | 0.333328 | 0.49155 | 0.4289 | 0.570787 |
| RUF_321c_ | -7.31993 | -8.04312 | -6.36681 | 1.395251 | 3.303547 | 0.344909 | 0.299333 | 0.537829 | 0.669492 | 0.5946 | 0.712297 |
| RUF_328c_ | -7.85858 | -7.65982 | -8.00819 | -0.31118 | 3.172912 | -0.07789 | 0.451333 | 0.588487 | 0.718075 | 0.7107 | 0.800893 |
| RUF_341c.10_ | 0.147052 | 0.22706 | 0.100267 | -0.16509 | 0.311028 | -0.49965 | 0.292667 | 0.464162 | 0.622237 | 0.6241 | 0.738985 |
| RUF_341c.11_ | 0.567364 | 0.492255 | 0.627896 | 0.14462 | 0.179158 | 0.760445 | 0.194667 | 0.300412 | 0.461902 | 0.3966 | 0.537747 |
| RUF_341c.2_ | -2.77704 | -2.81959 | -2.70173 | 0.14991 | 0.558606 | 0.252424 | 0.395333 | 0.575061 | 0.701634 | 0.692 | 0.782348 |
| SL1344_RS00005_ | 3.759829 | 3.768181 | 3.748414 | -0.01161 | 0.35894 | -0.03168 | 0.48 | 0.936703 | 0.964566 | 0.9826 | 0.989631 |
| SL1344_RS00010_ | 1.39699 | 1.420591 | 1.373172 | -0.05475 | 0.216151 | -0.27967 | 0.374667 | 0.608312 | 0.739815 | 0.7205 | 0.813167 |
| SL1344_RS00015_ | 1.2778 | 1.279072 | 1.27652 | -0.00445 | 0.112525 | -0.0401 | 0.48501 | 0.587941 | 0.712629 | 0.6991 | 0.789692 |
| SL1344_RS00020_ | 2.229483 | 2.169756 | 2.265038 | 0.106804 | 0.146389 | 0.753916 | 0.168554 | 0.279245 | 0.458772 | 0.3401 | 0.487045 |

| | | | | | | | | | | | |
|---|---|---|---|---|---|---|---|---|---|---|---|
| SL1344_RS00025_ | 1.63376 | 1.536738 | 1.741266 | 0.216494 | 0.126921 | 1.699618 | 0.014 | 0.070634 | 0.194765 | 0.119 | 0.259039 |
| SL1344_RS00030_ | 1.190316 | 1.110207 | 1.329292 | 0.210508 | 0.144205 | 1.369472 | 0.078614 | 0.122165 | 0.267117 | 0.1953 | 0.34692 |
| SL1344_RS00035_ | 5.785149 | 5.527273 | 5.982825 | 0.456453 | 0.058977 | 7.627773 | 4.68E-05 | 0.004541 | 0.054925 | 0.1 | 0.236592 |
| SL1344_RS00040_ | 1.88147 | 1.855505 | 1.914293 | 0.075412 | 0.1748 | 0.435714 | 0.304464 | 0.47299 | 0.630906 | 0.5844 | 0.708498 |
| SL1344_RS00045_ | -0.45347 | -0.11438 | -0.93139 | -0.81816 | 0.281268 | -2.80788 | 4.68E-05 | 0.019823 | 0.099661 | 0.1001 | 0.236726 |
| SL1344_RS00050_ | -3.8131 | -3.72817 | -3.92493 | -0.18106 | 0.643128 | -0.26709 | 0.386667 | 0.513635 | 0.64855 | 0.6224 | 0.728388 |
| SL1344_RS00055_ | -3.46852 | -3.46063 | -3.47606 | -0.0197 | 0.660385 | -0.02757 | 0.49 | 0.630144 | 0.748279 | 0.7411 | 0.824056 |
| SL1344_RS00065_ | 3.64827 | 3.628578 | 3.702762 | 0.062858 | 0.157341 | 0.36358 | 0.335776 | 0.716531 | 0.823363 | 0.6865 | 0.804133 |
| SL1344_RS00070_ | -2.18959 | -2.18234 | -2.19802 | 0.000449 | 0.374538 | 0.001037 | 0.499001 | 0.578953 | 0.703363 | 0.6773 | 0.771197 |
| SL1344_RS00075_ | -5.48174 | -5.48984 | -5.47533 | -0.08406 | 1.563849 | -0.04201 | 0.479333 | 0.651272 | 0.769738 | 0.7931 | 0.863748 |
| SL1344_RS00080_ | -4.30447 | -4.05052 | -4.62865 | -0.58378 | 0.838207 | -0.65108 | 0.238 | 0.368624 | 0.522416 | 0.4844 | 0.612788 |
| SL1344_RS00085_ | -5.13224 | -5.08836 | -5.16927 | -0.07141 | 1.006609 | -0.06773 | 0.469333 | 0.554941 | 0.684552 | 0.6692 | 0.766624 |
| SL1344_RS00090_ | -0.48737 | -0.31456 | -0.56733 | -0.24784 | 0.309997 | -0.75959 | 0.215333 | 0.321308 | 0.48801 | 0.4266 | 0.577095 |
| SL1344_RS00095_ | -0.99148 | -0.9278 | -1.04719 | -0.10784 | 0.273534 | -0.37409 | 0.333777 | 0.526918 | 0.6603 | 0.6287 | 0.735951 |
| SL1344_RS00100_ | -1.7539 | -1.47042 | -2.20229 | -0.72743 | 0.332219 | -2.13043 | 0.006663 | 0.045433 | 0.149828 | 0.1067 | 0.244913 |
| SL1344_RS00105_ | -2.0479 | -1.99939 | -2.13321 | -0.11067 | 0.42497 | -0.23915 | 0.407333 | 0.605342 | 0.729576 | 0.7127 | 0.807493 |
| SL1344_RS00110_ | -4.50622 | -4.09538 | -4.80382 | -0.81513 | 1.02197 | -0.78069 | 0.192538 | 0.30367 | 0.463552 | 0.4129 | 0.549326 |
| SL1344_RS00115_ | -1.876 | -1.93603 | -1.79143 | 0.128433 | 0.392714 | 0.302027 | 0.35976 | 0.56246 | 0.691718 | 0.6603 | 0.760828 |
| SL1344_RS00120_ | -2.70116 | -2.5908 | -2.80338 | -0.24454 | 0.605327 | -0.38186 | 0.323784 | 0.509992 | 0.652884 | 0.6254 | 0.734967 |
| SL1344_RS00125_ | -3.8898 | -3.68451 | -4.04848 | -0.35504 | 0.653909 | -0.51168 | 0.287808 | 0.450032 | 0.590058 | 0.558 | 0.673393 |
| SL1344_RS00130_ | -3.45999 | -3.5831 | -3.32943 | 0.263307 | 0.750314 | 0.321779 | 0.347102 | 0.534696 | 0.673852 | 0.6589 | 0.759262 |
| SL1344_RS00135_ | -3.49101 | -3.45563 | -3.51299 | -0.00935 | 0.658121 | -0.01255 | 0.494667 | 0.616688 | 0.738916 | 0.7384 | 0.820967 |
| SL1344_RS00140_ | -2.20572 | -2.25994 | -2.13833 | 0.126796 | 0.348608 | 0.34425 | 0.349101 | 0.491393 | 0.627663 | 0.6002 | 0.706531 |
| SL1344_RS00145_ | -2.40097 | -2.29115 | -2.56913 | -0.27762 | 0.473485 | -0.51725 | 0.271819 | 0.397438 | 0.55223 | 0.5201 | 0.643976 |
| SL1344_RS00150_ | -6.96739 | -7.33506 | -6.80672 | 0.381233 | 2.951057 | 0.11324 | 0.453031 | 0.628232 | 0.753034 | 0.7897 | 0.864145 |
| SL1344_RS00155_ | -3.36003 | -3.28329 | -3.54102 | -0.2774 | 0.717652 | -0.31897 | 0.362425 | 0.530227 | 0.670009 | 0.6672 | 0.768059 |
| SL1344_RS00160_ | -4.4872 | -4.41216 | -4.56487 | -0.16613 | 0.712833 | -0.20872 | 0.410667 | 0.503188 | 0.635715 | 0.6104 | 0.714851 |
| SL1344_RS00165_ | -1.35726 | -1.37207 | -1.34466 | 0.017605 | 0.274923 | 0.054331 | 0.476667 | 0.577891 | 0.703754 | 0.6879 | 0.780825 |
| SL1344_RS00170_ | -0.44267 | -0.58913 | -0.29695 | 0.2957 | 0.177887 | 1.599834 | 0.041306 | 0.085647 | 0.205617 | 0.1525 | 0.295356 |
| SL1344_RS00175_ | -5.70032 | -5.75533 | -5.66209 | 0.062535 | 1.236585 | 0.046723 | 0.477015 | 0.566342 | 0.695658 | 0.6919 | 0.783199 |
| SL1344_RS00180_ | 0.253412 | 0.2644 | 0.242991 | -0.03526 | 0.185633 | -0.1852 | 0.426667 | 0.58795 | 0.712169 | 0.7101 | 0.797237 |
| SL1344_RS00185_ | -1.15447 | -1.1966 | -1.12203 | 0.076194 | 0.269272 | 0.265261 | 0.397735 | 0.554086 | 0.683315 | 0.6636 | 0.761661 |
| SL1344_RS00190_ | -2.37989 | -2.38834 | -2.36999 | 0.010407 | 0.469656 | 0.022559 | 0.490667 | 0.634037 | 0.750614 | 0.7387 | 0.821053 |
| SL1344_RS00195_ | 0.006943 | 0.200585 | -0.05583 | -0.22633 | 0.251763 | -0.78876 | 0.226667 | 0.336159 | 0.504699 | 0.4733 | 0.619946 |
| SL1344_RS00205_ | 2.068139 | 2.149019 | 2.031204 | -0.10816 | 0.098112 | -0.99898 | 0.158561 | 0.229108 | 0.389859 | 0.3305 | 0.481043 |
| SL1344_RS00210_ | -0.50921 | -0.69848 | -0.28972 | 0.408233 | 0.203687 | 1.887429 | 0.015334 | 0.055305 | 0.163049 | 0.122 | 0.261798 |
| SL1344_RS00215_ | -3.0651 | -3.48795 | -2.81672 | 0.633976 | 0.698909 | 0.859287 | 0.168667 | 0.263518 | 0.431353 | 0.3633 | 0.515505 |
| SL1344_RS00220_ | 4.549205 | 5.197778 | 4.092023 | -1.00436 | 0.255725 | -4.0134 | 4.68E-05 | 0.004167 | 0.054318 | 0.1 | 0.236592 |
| SL1344_RS00225_ | -2.31314 | -2.2284 | -2.40057 | -0.17091 | 0.371986 | -0.43912 | 0.312458 | 0.448373 | 0.587084 | 0.5493 | 0.664156 |
| SL1344_RS00230_ | 1.960054 | 2.028503 | 1.933885 | -0.08547 | 0.121976 | -0.61967 | 0.262 | 0.415671 | 0.572616 | 0.5344 | 0.665391 |
| SL1344_RS00235_ | 4.551876 | 4.36985 | 4.666983 | 0.296719 | 0.095591 | 3.315251 | 4.68E-05 | 0.036582 | 0.144137 | 0.1 | 0.236592 |
| SL1344_RS00240_ | 2.475383 | 2.332245 | 2.680652 | 0.34482 | 0.106523 | 3.149866 | 4.68E-05 | 0.019788 | 0.100655 | 0.1001 | 0.236724 |
| SL1344_RS00245_ | 1.458023 | 1.345923 | 1.552663 | 0.197228 | 0.143788 | 1.320923 | 0.080667 | 0.125234 | 0.274167 | 0.1885 | 0.343474 |
| SL1344_RS00250_ | 2.478894 | 2.384708 | 2.545377 | 0.170058 | 0.097673 | 1.732734 | 0.017988 | 0.086173 | 0.220535 | 0.1206 | 0.261395 |
| SL1344_RS00255_ | 1.46323 | 1.601021 | 1.376159 | -0.26354 | 0.237789 | -1.10519 | 0.096 | 0.152333 | 0.316312 | 0.2245 | 0.38509 |

| | | | | | | | | | | | |
|---|---|---|---|---|---|---|---|---|---|---|---|
| SL1344_RS00260_ | 2.226959 | 2.446375 | 2.012253 | -0.43649 | 0.130844 | -3.33332 | 4.68E-05 | 0.012417 | 0.081907 | 0.1 | 0.236592 |
| SL1344_RS00270_ | -0.49293 | -0.62398 | -0.32358 | 0.318903 | 0.241963 | 1.28639 | 0.064 | 0.133779 | 0.281081 | 0.1827 | 0.329398 |
| SL1344_RS00275_ | -4.346 | -4.26922 | -4.41818 | -0.15968 | 0.811839 | -0.1765 | 0.418667 | 0.557868 | 0.68659 | 0.6744 | 0.768386 |
| SL1344_RS00280_ | -4.95489 | -5.11862 | -4.78977 | 0.320512 | 0.860424 | 0.3529 | 0.331779 | 0.443052 | 0.580699 | 0.5521 | 0.662881 |
| SL1344_RS00285_ | -6.91288 | -6.66268 | -7.19096 | -0.5193 | 1.838169 | -0.24903 | 0.381333 | 0.524415 | 0.657478 | 0.6318 | 0.734522 |
| SL1344_RS00290_ | -2.34309 | -2.18532 | -2.4757 | -0.28658 | 0.379144 | -0.72797 | 0.212525 | 0.32923 | 0.478114 | 0.4259 | 0.558685 |
| SL1344_RS00295_ | -3.81529 | -3.91191 | -3.66731 | 0.279977 | 0.736855 | 0.346279 | 0.354667 | 0.531541 | 0.666348 | 0.6452 | 0.747696 |
| SL1344_RS00300_ | -4.34295 | -4.45784 | -4.19402 | 0.173991 | 0.984283 | 0.158952 | 0.424384 | 0.636101 | 0.753958 | 0.7461 | 0.828051 |
| SL1344_RS00305_ | -3.0871 | -2.99758 | -3.20341 | -0.25935 | 0.790908 | -0.30784 | 0.372667 | 0.583987 | 0.716187 | 0.7154 | 0.805997 |
| SL1344_RS00310_ | -2.21804 | -2.38627 | -2.03374 | 0.356427 | 0.377309 | 0.905946 | 0.148568 | 0.2354 | 0.386011 | 0.3192 | 0.462018 |
| SL1344_RS00315_ | -2.85383 | -2.73004 | -3.058 | -0.28865 | 0.594229 | -0.44051 | 0.324 | 0.516248 | 0.657055 | 0.6321 | 0.742209 |
| SL1344_RS00320_ | -1.95383 | -1.97743 | -1.92938 | 0.067309 | 0.354119 | 0.184583 | 0.422 | 0.581001 | 0.704729 | 0.6909 | 0.782345 |
| SL1344_RS00325_ | 1.760728 | 1.697418 | 1.897276 | 0.164953 | 0.163528 | 0.953612 | 0.169333 | 0.239781 | 0.412678 | 0.3297 | 0.492579 |
| SL1344_RS00330_ | -3.00928 | -3.05413 | -2.94153 | 0.060937 | 0.650351 | 0.078822 | 0.46 | 0.672362 | 0.783251 | 0.775 | 0.851737 |
| SL1344_RS00335_ | -1.06224 | -1.20474 | -0.96224 | 0.210906 | 0.353322 | 0.593787 | 0.264 | 0.427616 | 0.583153 | 0.5389 | 0.668917 |
| SL1344_RS00340_ | 1.077504 | 0.931833 | 1.312803 | 0.366076 | 0.145586 | 2.441688 | 0.004665 | 0.031824 | 0.125246 | 0.1055 | 0.243588 |
| SL1344_RS00345_ | 1.888322 | 1.388877 | 2.402003 | 1.077655 | 0.299653 | 3.765743 | 4.68E-05 | 0.009794 | 0.075902 | 0.1 | 0.236592 |
| SL1344_RS00350_ | -2.86952 | -2.69755 | -3.02775 | -0.3374 | 0.427665 | -0.7184 | 0.208667 | 0.334753 | 0.480634 | 0.4255 | 0.557593 |
| SL1344_RS00355_ | -1.46981 | -1.35538 | -1.65028 | -0.2536 | 0.384748 | -0.62071 | 0.253165 | 0.390245 | 0.547331 | 0.5027 | 0.635948 |
| SL1344_RS00360_ | -2.00829 | -1.94275 | -2.08797 | -0.14773 | 0.334274 | -0.40573 | 0.331779 | 0.464589 | 0.60521 | 0.5784 | 0.691226 |
| SL1344_RS00365_ | -1.01623 | -1.04712 | -0.98373 | 0.062823 | 0.278033 | 0.21507 | 0.407062 | 0.590087 | 0.714605 | 0.6903 | 0.783118 |
| SL1344_RS00370_ | -0.00547 | -0.0191 | 0.005023 | 0.024148 | 0.192392 | 0.116281 | 0.435043 | 0.598344 | 0.720733 | 0.7056 | 0.794464 |
| SL1344_RS00375_ | -3.86129 | -3.58616 | -4.05983 | -0.5615 | 0.881163 | -0.61553 | 0.264667 | 0.3876 | 0.545323 | 0.511 | 0.637772 |
| SL1344_RS00380_ | -6.37416 | -6.06838 | -6.72014 | -0.81926 | 2.663487 | -0.25561 | 0.35 | 0.547799 | 0.689118 | 0.6878 | 0.782822 |
| SL1344_RS00385_ | -2.68959 | -3.11789 | -2.50279 | 0.594709 | 0.682014 | 0.789034 | 0.200667 | 0.288013 | 0.459911 | 0.394 | 0.548189 |
| SL1344_RS00390_ | -2.63218 | -2.86447 | -2.42087 | 0.437477 | 0.403563 | 1.024934 | 0.113333 | 0.19264 | 0.334801 | 0.2714 | 0.414694 |
| SL1344_RS00395_ | -2.62498 | -2.87583 | -2.16274 | 0.661174 | 0.523051 | 1.189628 | 0.109927 | 0.165578 | 0.316762 | 0.245 | 0.397432 |
| SL1344_RS00400_ | -3.28451 | -3.52158 | -2.36547 | 1.044575 | 0.779551 | 1.115754 | 0.166 | 0.207642 | 0.374304 | 0.3398 | 0.493736 |
| SL1344_RS00405_ | -1.47627 | -1.45087 | -1.50388 | 0.003412 | 0.414005 | 0.006002 | 0.495003 | 0.688139 | 0.79504 | 0.8096 | 0.873301 |
| SL1344_RS00410_ | -1.24559 | -1.19621 | -1.28029 | -0.08631 | 0.300509 | -0.25658 | 0.388 | 0.565326 | 0.693875 | 0.6886 | 0.780884 |
| SL1344_RS00415_ | 0.154967 | 0.029005 | 0.390878 | 0.347913 | 0.267887 | 1.22736 | 0.108594 | 0.137262 | 0.293432 | 0.2196 | 0.380454 |
| SL1344_RS00420_ | -1.09692 | -1.23672 | -0.95294 | 0.278609 | 0.238996 | 1.102782 | 0.111333 | 0.1773 | 0.320849 | 0.2569 | 0.402368 |
| SL1344_RS00425_ | -7.1322 | -7.62269 | -6.60241 | 0.813523 | 3.279979 | 0.202909 | 0.363091 | 0.593975 | 0.721149 | 0.691 | 0.784158 |
| SL1344_RS00430_ | -1.04846 | -0.92678 | -1.18913 | -0.26714 | 0.281552 | -0.92706 | 0.151899 | 0.231448 | 0.389732 | 0.3217 | 0.467808 |
| SL1344_RS00435_ | -2.48278 | -2.80281 | -2.24043 | 0.578054 | 0.513948 | 1.061507 | 0.098 | 0.169562 | 0.325092 | 0.2329 | 0.38368 |
| SL1344_RS00440_ | 0.001768 | -0.30512 | 0.381873 | 0.677679 | 0.227182 | 2.919738 | 4.68E-05 | 0.016676 | 0.092298 | 0.1004 | 0.237132 |
| SL1344_RS00450_ | 1.976011 | 1.814 | 2.063105 | 0.238613 | 0.123833 | 1.877623 | 0.026649 | 0.060017 | 0.180484 | 0.1237 | 0.266663 |
| SL1344_RS00455_ | 1.558214 | 1.454633 | 1.679754 | 0.225493 | 0.095739 | 2.322063 | 0.006001 | 0.033254 | 0.124141 | 0.1088 | 0.247668 |
| SL1344_RS00465_ | 2.379066 | 2.236855 | 2.61187 | 0.375954 | 0.104169 | 3.501393 | 4.68E-05 | 0.018756 | 0.098276 | 0.1 | 0.236592 |
| SL1344_RS00470_ | 3.563794 | 3.40448 | 3.723844 | 0.340481 | 0.08622 | 3.977591 | 4.68E-05 | 0.005953 | 0.059369 | 0.1 | 0.236592 |
| SL1344_RS00475_ | 3.558652 | 3.375511 | 3.753294 | 0.377175 | 0.05248 | 7.15033 | 4.68E-05 | 0.001632 | 0.035349 | 0.1 | 0.236592 |
| SL1344_RS00480_ | 1.181504 | 1.029356 | 1.515948 | 0.519925 | 0.363763 | 1.525476 | 0.017334 | 0.073948 | 0.209733 | 0.1139 | 0.254523 |
| SL1344_RS00485_ | 0.657441 | 0.593763 | 0.735041 | 0.13073 | 0.160019 | 0.777685 | 0.185876 | 0.298739 | 0.455547 | 0.3941 | 0.537137 |
| SL1344_RS00490_ | 1.731638 | 1.78318 | 1.625954 | -0.12495 | 0.189552 | -0.62159 | 0.273151 | 0.416499 | 0.585576 | 0.567 | 0.697962 |
| SL1344_RS00495_ | 0.771448 | 0.830403 | 0.718192 | -0.12569 | 0.214723 | -0.56659 | 0.247333 | 0.374847 | 0.540364 | 0.488 | 0.625743 |

| | | | | | | | | | | | |
|---|---|---|---|---|---|---|---|---|---|---|---|
| SL1344_RS00500_ | -2.10855 | -1.99759 | -2.25582 | -0.26032 | 0.387606 | -0.61967 | 0.247835 | 0.370464 | 0.51961 | 0.4702 | 0.599699 |
| SL1344_RS00505_ | -4.20216 | -4.35915 | -4.10321 | 0.246696 | 0.764472 | 0.31158 | 0.363758 | 0.524165 | 0.659825 | 0.6414 | 0.744431 |
| SL1344_RS00510_ | -2.28128 | -2.27723 | -2.28683 | 0.040104 | 0.565041 | 0.068495 | 0.469021 | 0.670889 | 0.783961 | 0.792 | 0.863631 |
| SL1344_RS00515_ | -2.11942 | -2.25873 | -1.83647 | 0.409865 | 0.55712 | 0.699344 | 0.225333 | 0.343032 | 0.508244 | 0.4464 | 0.594902 |
| SL1344_RS00520_ | -2.7603 | -2.88962 | -2.68329 | 0.197098 | 0.486024 | 0.358979 | 0.348434 | 0.512482 | 0.651094 | 0.6354 | 0.740928 |
| SL1344_RS00525_ | 1.259767 | 1.193086 | 1.378866 | 0.194702 | 0.139691 | 1.295861 | 0.068621 | 0.138626 | 0.290946 | 0.1952 | 0.342179 |
| SL1344_RS00530_ | 0.693142 | 0.435223 | 0.889921 | 0.4643 | 0.159882 | 2.836163 | 0.000673 | 0.025615 | 0.111625 | 0.1003 | 0.236996 |
| SL1344_RS00545_ | -0.22064 | -0.26262 | -0.15969 | 0.119942 | 0.220125 | 0.492309 | 0.288474 | 0.442404 | 0.588698 | 0.5674 | 0.678606 |
| SL1344_RS00550_ | -0.11827 | -0.11811 | -0.11837 | -0.00804 | 0.20959 | -0.04226 | 0.484344 | 0.622218 | 0.740682 | 0.7358 | 0.819405 |
| SL1344_RS00555_ | 4.40235 | 4.576052 | 4.273648 | -0.37519 | 0.193515 | -2.12924 | 4.68E-05 | 0.028183 | 0.126107 | 0.1 | 0.236592 |
| SL1344_RS00560_ | -1.09907 | -1.03775 | -1.15188 | -0.11116 | 0.249978 | -0.41993 | 0.317122 | 0.474731 | 0.612129 | 0.5764 | 0.686705 |
| SL1344_RS00565_ | -0.76039 | -0.71058 | -0.82112 | -0.11223 | 0.234261 | -0.46093 | 0.300667 | 0.441063 | 0.58359 | 0.5521 | 0.666726 |
| SL1344_RS00570_ | -0.87705 | -0.7106 | -1.08322 | -0.37263 | 0.261428 | -1.38427 | 0.054667 | 0.11348 | 0.251267 | 0.1748 | 0.320678 |
| SL1344_RS00575_ | -1.43598 | -1.4978 | -1.39588 | 0.096217 | 0.319356 | 0.289326 | 0.387333 | 0.555059 | 0.685216 | 0.6654 | 0.764595 |
| SL1344_RS00580_ | 0.081573 | 0.145256 | 0.012057 | -0.01469 | 0.508764 | -0.02987 | 0.487333 | 0.839768 | 0.904436 | 0.9409 | 0.964224 |
| SL1344_RS00590_ | 0.108979 | 0.066149 | 0.153307 | 0.090476 | 0.147839 | 0.574606 | 0.259333 | 0.377929 | 0.522347 | 0.4742 | 0.600608 |
| SL1344_RS00595_ | 0.277517 | 0.128085 | 0.454346 | 0.287754 | 0.224489 | 1.237247 | 0.095936 | 0.13462 | 0.285713 | 0.2041 | 0.361654 |
| SL1344_RS00600_ | 2.802961 | 2.730102 | 2.900817 | 0.175929 | 0.109146 | 1.569006 | 0.027982 | 0.093148 | 0.235157 | 0.1289 | 0.272622 |
| SL1344_RS00605_ | 3.373578 | 3.482503 | 3.212257 | -0.30314 | 0.219549 | -1.40654 | 0.017322 | 0.094048 | 0.242648 | 0.1181 | 0.260709 |
| SL1344_RS00610_ | 3.795848 | 3.784365 | 3.809942 | 0.012417 | 0.126108 | 0.097441 | 0.448 | 0.813196 | 0.887344 | 0.8449 | 0.905733 |
| SL1344_RS00615_ | 3.06839 | 3.045815 | 3.242661 | 0.187798 | 0.140241 | 0.866264 | 0.284667 | 0.330988 | 0.515632 | 0.5844 | 0.725106 |
| SL1344_RS00620_ | 5.542661 | 5.849036 | 5.454426 | -0.39136 | 0.05179 | -4.82283 | 4.68E-05 | 0.049633 | 0.170823 | 0.1 | 0.236592 |
| SL1344_RS00630_ | 3.16124 | 3.095815 | 3.198186 | 0.099233 | 0.07707 | 1.206004 | 0.094604 | 0.152388 | 0.307202 | 0.2244 | 0.377171 |
| SL1344_RS00635_ | 3.020818 | 2.948262 | 3.094582 | 0.147313 | 0.061478 | 2.344528 | 0.005997 | 0.031659 | 0.122715 | 0.1042 | 0.241725 |
| SL1344_RS00645_ | 3.087647 | 3.049293 | 3.157775 | 0.096582 | 0.105588 | 0.880502 | 0.176667 | 0.256564 | 0.430725 | 0.3536 | 0.51586 |
| SL1344_RS00650_ | 2.763158 | 2.778941 | 2.726683 | -0.04678 | 0.134968 | -0.27804 | 0.398 | 0.763962 | 0.853441 | 0.8236 | 0.893806 |
| SL1344_RS00655_ | 3.086714 | 3.075319 | 3.09362 | 0.014288 | 0.072253 | 0.176258 | 0.420667 | 0.627111 | 0.746737 | 0.7281 | 0.81664 |
| SL1344_RS00665_ | 2.515613 | 2.445854 | 2.669316 | 0.216496 | 0.089083 | 2.197771 | 0.008667 | 0.049593 | 0.160899 | 0.1098 | 0.248513 |
| SL1344_RS00670_ | 3.566955 | 3.522034 | 3.641168 | 0.083735 | 0.110414 | 0.773021 | 0.215856 | 0.320085 | 0.497266 | 0.4057 | 0.563122 |
| SL1344_RS00675_ | 4.360528 | 4.236996 | 4.383227 | 0.126819 | 0.09786 | 0.878667 | 0.247835 | 0.297112 | 0.482539 | 0.4883 | 0.647734 |
| SL1344_RS00680_ | 5.697438 | 5.6692 | 5.724299 | 0.024689 | 0.088945 | 0.278287 | 0.409727 | 0.69357 | 0.80776 | 0.877 | 0.923691 |
| SL1344_RS00685_ | 1.902594 | 2.007763 | 1.785109 | -0.22215 | 0.088972 | -2.43662 | 0.003332 | 0.028744 | 0.116142 | 0.1027 | 0.240034 |
| SL1344_RS00690_ | 4.465985 | 4.421922 | 4.506519 | 0.08027 | 0.061123 | 1.266956 | 0.074 | 0.120278 | 0.271237 | 0.186 | 0.341817 |
| SL1344_RS00695_ | -1.08161 | -1.19458 | -0.99704 | 0.241475 | 0.369053 | 0.61799 | 0.212525 | 0.331248 | 0.50243 | 0.4303 | 0.564318 |
| SL1344_RS00700_ | 0.385356 | 0.320359 | 0.426636 | 0.111896 | 0.217753 | 0.460377 | 0.295137 | 0.451857 | 0.607381 | 0.5931 | 0.709492 |
| SL1344_RS00705_ | 1.266319 | 1.232551 | 1.344421 | 0.103039 | 0.157699 | 0.592315 | 0.272485 | 0.436214 | 0.594062 | 0.5594 | 0.68891 |
| SL1344_RS00710_ | 1.203687 | 1.169208 | 1.240795 | 0.071959 | 0.109455 | 0.612331 | 0.249333 | 0.380716 | 0.523651 | 0.4813 | 0.605764 |
| SL1344_RS00715_ | 0.966783 | 1.046525 | 0.886162 | -0.18366 | 0.181681 | -0.96806 | 0.115257 | 0.196628 | 0.361972 | 0.268 | 0.417545 |
| SL1344_RS00720_ | 3.441331 | 3.195428 | 3.64082 | 0.444994 | 0.066059 | 6.574202 | 4.68E-05 | 0.00398 | 0.048603 | 0.1 | 0.236592 |
| SL1344_RS00725_ | -3.30735 | -3.30279 | -3.31229 | 0.003638 | 0.502407 | 0.010425 | 0.495003 | 0.548711 | 0.678817 | 0.6585 | 0.756839 |
| SL1344_RS00730_ | -4.55966 | -4.63191 | -4.50117 | 0.187823 | 0.97401 | 0.181224 | 0.421719 | 0.588637 | 0.715627 | 0.7113 | 0.798245 |
| SL1344_RS00735_ | -6.75463 | -6.06141 | -7.09884 | -0.84742 | 2.662997 | -0.24078 | 0.366 | 0.625695 | 0.743846 | 0.7116 | 0.8047 |
| SL1344_RS00740_ | 0.426201 | 0.482873 | 0.382803 | -0.10841 | 0.155249 | -0.65874 | 0.229333 | 0.352233 | 0.504281 | 0.457 | 0.585622 |
| SL1344_RS00745_ | 0.98179 | 0.797261 | 1.147317 | 0.351902 | 0.128803 | 2.703165 | 0.002001 | 0.025186 | 0.110046 | 0.1014 | 0.238344 |
| SL1344_RS00750_ | 1.615442 | 1.46416 | 1.772203 | 0.31467 | 0.12148 | 2.566866 | 0.002001 | 0.026866 | 0.114951 | 0.1016 | 0.238667 |

| | | | | | | | | | | | |
|---|---|---|---|---|---|---|---|---|---|---|---|
| SL1344_RS00755_ | -1.07297 | -1.05442 | -1.10281 | -0.06863 | 0.293352 | -0.21492 | 0.403333 | 0.579046 | 0.707989 | 0.7086 | 0.796608 |
| SL1344_RS00760_ | -2.48195 | -2.40893 | -2.65769 | -0.19567 | 0.729426 | -0.23591 | 0.409061 | 0.723363 | 0.824505 | 0.82 | 0.890543 |
| SL1344_RS00765_ | 0.476361 | 0.466976 | 0.494523 | 0.044623 | 0.181023 | 0.214392 | 0.408394 | 0.588365 | 0.71581 | 0.717 | 0.804075 |
| SL1344_RS00770_ | 2.694002 | 3.045248 | 2.243344 | -0.80111 | 0.107616 | -7.41996 | 4.68E-05 | 0.001086 | 0.0325 | 0.1 | 0.236592 |
| SL1344_RS00775_ | 5.468727 | 5.524774 | 5.385781 | -0.16732 | 0.091846 | -1.85196 | 0.001337 | 0.045365 | 0.160659 | 0.1008 | 0.237614 |
| SL1344_RS00785_ | 5.911362 | 6.005177 | 5.797009 | -0.20846 | 0.084664 | -2.39549 | 4.68E-05 | 0.029917 | 0.129525 | 0.1 | 0.236592 |
| SL1344_RS00790_ | -3.34151 | -3.00525 | -3.87421 | -0.71303 | 0.961452 | -0.71974 | 0.216522 | 0.33704 | 0.508717 | 0.443 | 0.591365 |
| SL1344_RS00795_ | -3.27111 | -3.36864 | -3.19271 | 0.233799 | 0.838824 | 0.254829 | 0.39507 | 0.612006 | 0.738101 | 0.7445 | 0.827656 |
| SL1344_RS00800_ | -1.4765 | -1.54501 | -1.38764 | 0.150447 | 0.285169 | 0.486715 | 0.290667 | 0.441575 | 0.582801 | 0.5462 | 0.66426 |
| SL1344_RS00805_ | 4.889835 | 4.394443 | 5.271637 | 0.876899 | 0.158914 | 5.639609 | 4.68E-05 | 0.012076 | 0.08455 | 0.1 | 0.236592 |
| SL1344_RS00810_ | 1.592411 | 1.769702 | 1.566037 | -0.1903 | 0.167944 | -0.69276 | 0.312667 | 0.454083 | 0.619435 | 0.6488 | 0.777654 |
| SL1344_RS00815_ | 2.58464 | 2.328182 | 2.74777 | 0.431421 | 0.160519 | 2.63051 | 4.68E-05 | 0.044406 | 0.158533 | 0.1 | 0.236592 |
| SL1344_RS00820_ | -0.85287 | -0.80476 | -0.89355 | -0.08137 | 0.236698 | -0.3341 | 0.357333 | 0.494848 | 0.632281 | 0.6098 | 0.717081 |
| SL1344_RS00825_ | 0.130188 | -0.01417 | 0.267333 | 0.286044 | 0.181529 | 1.505708 | 0.044667 | 0.095254 | 0.226584 | 0.1558 | 0.301263 |
| SL1344_RS00830_ | 0.920943 | 0.791372 | 1.072377 | 0.279239 | 0.144242 | 1.868578 | 0.016667 | 0.054691 | 0.166427 | 0.1221 | 0.263024 |
| SL1344_RS00835_ | 2.512333 | 2.491267 | 2.532532 | 0.071597 | 0.165106 | 0.430106 | 0.323333 | 0.478961 | 0.639802 | 0.6583 | 0.767322 |
| SL1344_RS00840_ | 1.085745 | 1.419627 | 0.577601 | -0.82479 | 0.177066 | -4.44817 | 4.68E-05 | 0.011007 | 0.076554 | 0.1 | 0.236592 |
| SL1344_RS00845_ | 0.860948 | 1.026983 | 0.421906 | -0.53633 | 0.280422 | -1.56666 | 0.054667 | 0.095029 | 0.241336 | 0.1516 | 0.303865 |
| SL1344_RS00850_ | -0.80327 | -0.7818 | -0.82936 | -0.05138 | 0.244534 | -0.17434 | 0.419054 | 0.580027 | 0.703753 | 0.6941 | 0.78415 |
| SL1344_RS00855_ | 1.620932 | 1.367942 | 1.948988 | 0.423874 | 0.404616 | 1.212262 | 0.110667 | 0.132193 | 0.2955 | 0.2052 | 0.374889 |
| SL1344_RS00860_ | 0.804403 | 0.639054 | 1.075799 | 0.42515 | 0.17468 | 2.337804 | 0.003998 | 0.034015 | 0.130767 | 0.1042 | 0.241893 |
| SL1344_RS00865_ | 1.409 | 1.576306 | 1.235847 | -0.34807 | 0.128674 | -2.63958 | 0.000673 | 0.022874 | 0.105477 | 0.1008 | 0.237659 |
| SL1344_RS00870_ | 2.990578 | 3.174804 | 2.747483 | -0.45058 | 0.107024 | -4.23989 | 4.68E-05 | 0.004642 | 0.053925 | 0.1 | 0.236592 |
| SL1344_RS00875_ | -0.14128 | -0.16231 | -0.12194 | 0.041519 | 0.172604 | 0.238004 | 0.401732 | 0.528057 | 0.660309 | 0.6431 | 0.743213 |
| SL1344_RS00880_ | 0.624986 | 0.350936 | 0.896952 | 0.498753 | 0.230517 | 2.146348 | 0.012 | 0.035853 | 0.137228 | 0.1095 | 0.249188 |
| SL1344_RS00885_ | -1.71346 | -1.67622 | -1.74832 | -0.07463 | 0.314694 | -0.21952 | 0.411059 | 0.541467 | 0.672684 | 0.648 | 0.749143 |
| SL1344_RS00890_ | -2.39635 | -2.02403 | -2.66182 | -0.61522 | 0.409497 | -1.43058 | 0.069333 | 0.108199 | 0.243416 | 0.1797 | 0.327827 |
| SL1344_RS00895_ | -5.11845 | -5.19365 | -5.04052 | 0.161708 | 0.928217 | 0.154268 | 0.429714 | 0.513667 | 0.645279 | 0.6222 | 0.724844 |
| SL1344_RS00900_ | -2.39176 | -2.20392 | -2.6759 | -0.48186 | 0.421621 | -1.08535 | 0.116667 | 0.194256 | 0.343872 | 0.2767 | 0.422785 |
| SL1344_RS00905_ | 0.135662 | -0.40229 | 0.436345 | 0.785679 | 0.348076 | 2.151853 | 0.004668 | 0.034944 | 0.13734 | 0.1039 | 0.241728 |
| SL1344_RS00910_ | 0.269624 | 0.221764 | 0.296887 | 0.078238 | 0.191713 | 0.373193 | 0.345769 | 0.526622 | 0.663992 | 0.6429 | 0.746946 |
| SL1344_RS00915_ | 1.779035 | 1.840416 | 1.725974 | -0.11653 | 0.103961 | -1.04489 | 0.113258 | 0.190938 | 0.341122 | 0.2651 | 0.410676 |
| SL1344_RS00920_ | 0.96697 | 0.987235 | 0.942097 | -0.04586 | 0.123454 | -0.35785 | 0.356 | 0.498028 | 0.634273 | 0.6126 | 0.716819 |
| SL1344_RS00925_ | 0.727827 | 0.768949 | 0.701817 | -0.06347 | 0.166873 | -0.36882 | 0.348 | 0.538234 | 0.673291 | 0.6526 | 0.755726 |
| SL1344_RS00930_ | 0.869649 | 0.918084 | 0.58193 | -0.32642 | 0.293723 | -0.81298 | 0.265157 | 0.317959 | 0.501546 | 0.5226 | 0.670405 |
| SL1344_RS00935_ | 2.887623 | 3.062307 | 2.68169 | -0.38017 | 0.066789 | -5.67155 | 4.68E-05 | 0.003491 | 0.045426 | 0.1 | 0.236592 |
| SL1344_RS00940_ | 0.256995 | 0.421822 | 0.110767 | -0.31518 | 0.181117 | -1.6592 | 0.027982 | 0.072058 | 0.194304 | 0.1347 | 0.278153 |
| SL1344_RS00945_ | 5.912427 | 5.883387 | 5.941298 | 0.020082 | 0.105549 | 0.170248 | 0.449034 | 0.882748 | 0.932319 | 0.9706 | 0.982561 |
| SL1344_RS00950_ | 1.236759 | 1.080288 | 1.344915 | 0.255282 | 0.12861 | 1.896448 | 0.020667 | 0.056734 | 0.171503 | 0.121 | 0.2629 |
| SL1344_RS00955_ | 0.391932 | 0.419645 | 0.360652 | -0.03615 | 0.206895 | -0.17163 | 0.431333 | 0.661991 | 0.775025 | 0.7723 | 0.847505 |
| SL1344_RS00960_ | 2.264577 | 2.275159 | 2.248871 | -0.02146 | 0.089307 | -0.21882 | 0.413333 | 0.60579 | 0.727916 | 0.7072 | 0.799298 |
| SL1344_RS00965_ | 3.349576 | 3.323977 | 3.410714 | 0.077534 | 0.085683 | 0.835489 | 0.189333 | 0.27108 | 0.441617 | 0.3776 | 0.535871 |
| SL1344_RS00970_ | -0.29895 | -0.4745 | -0.12693 | 0.348025 | 0.16247 | 2.066042 | 0.012667 | 0.043597 | 0.140561 | 0.1176 | 0.257119 |
| SL1344_RS00975_ | -2.62928 | -2.72527 | -2.57948 | 0.188023 | 0.609241 | 0.244693 | 0.386409 | 0.551268 | 0.691679 | 0.7035 | 0.795663 |
| SL1344_RS00980_ | -2.26982 | -2.15459 | -2.40579 | -0.25819 | 0.345125 | -0.68848 | 0.214 | 0.318623 | 0.464599 | 0.4183 | 0.549763 |

| | | | | | | | | | | | |
|---|---|---|---|---|---|---|---|---|---|---|---|
| SL1344_RS00985_ | -1.29283 | -1.23883 | -1.35148 | -0.1178 | 0.370956 | -0.32171 | 0.36 | 0.561487 | 0.695878 | 0.6706 | 0.772457 |
| SL1344_RS00990_ | -3.16674 | -3.20487 | -3.12591 | 0.08777 | 0.515708 | 0.155513 | 0.43 | 0.55179 | 0.680258 | 0.6641 | 0.760424 |
| SL1344_RS00995_ | -1.91467 | -1.97885 | -1.84579 | 0.136609 | 0.336856 | 0.385061 | 0.328 | 0.497173 | 0.632227 | 0.6063 | 0.712335 |
| SL1344_RS01000_ | -3.67589 | -3.73535 | -3.60748 | 0.144771 | 0.875212 | 0.153809 | 0.426667 | 0.644113 | 0.761155 | 0.7349 | 0.819949 |
| SL1344_RS01005_ | -3.58838 | -3.42516 | -3.78028 | -0.31032 | 0.699274 | -0.43194 | 0.321119 | 0.495554 | 0.637241 | 0.6091 | 0.720604 |
| SL1344_RS01010_ | -2.96083 | -2.76987 | -3.55797 | -0.75386 | 0.710003 | -0.90386 | 0.205863 | 0.269677 | 0.442334 | 0.3903 | 0.546879 |
| SL1344_RS01015_ | -3.5621 | -3.54396 | -3.58509 | -0.00823 | 0.663159 | -0.01002 | 0.494 | 0.604355 | 0.72651 | 0.7132 | 0.800327 |
| SL1344_RS01020_ | 0.361097 | 0.369022 | 0.346927 | -0.03859 | 0.245543 | -0.14335 | 0.435333 | 0.690936 | 0.798957 | 0.8236 | 0.885623 |
| SL1344_RS01025_ | 2.431241 | 2.37994 | 2.50827 | 0.131828 | 0.097785 | 1.260576 | 0.075949 | 0.135815 | 0.286187 | 0.2073 | 0.358944 |
| SL1344_RS01030_ | 1.007669 | 0.92412 | 1.083523 | 0.161834 | 0.110764 | 1.408668 | 0.064 | 0.116096 | 0.246456 | 0.1898 | 0.334473 |
| SL1344_RS01035_ | 1.276722 | 1.40566 | 1.138488 | -0.2713 | 0.140193 | -1.8794 | 0.011334 | 0.053516 | 0.166001 | 0.1155 | 0.255556 |
| SL1344_RS01040_ | -0.21431 | -0.01922 | -0.47748 | -0.46109 | 0.225815 | -1.93973 | 0.016 | 0.051996 | 0.163469 | 0.1137 | 0.253863 |
| SL1344_RS01045_ | 0.589357 | 0.676843 | 0.482217 | -0.19065 | 0.138747 | -1.30671 | 0.073951 | 0.135303 | 0.274756 | 0.2087 | 0.35559 |
| SL1344_RS01050_ | 2.081406 | 2.238193 | 2.011791 | -0.23352 | 0.215321 | -1.09979 | 0.110667 | 0.144038 | 0.307632 | 0.2188 | 0.38171 |
| SL1344_RS01055_ | 0.692034 | 0.515832 | 0.858192 | 0.389874 | 0.209452 | 1.860335 | 0.005997 | 0.049758 | 0.163168 | 0.1062 | 0.244524 |
| SL1344_RS01060_ | 3.373997 | 3.199396 | 3.516104 | 0.315126 | 0.066516 | 4.711612 | 4.68E-05 | 0.007162 | 0.061532 | 0.1 | 0.236592 |
| SL1344_RS01065_ | 2.320117 | 2.003438 | 2.468553 | 0.470577 | 0.195007 | 2.376725 | 4.68E-05 | 0.057495 | 0.182817 | 0.1 | 0.236592 |
| SL1344_RS01070_ | 5.005402 | 4.995208 | 5.142063 | 0.146503 | 0.14238 | 0.945463 | 0.231333 | 0.251618 | 0.438545 | 0.4796 | 0.628654 |
| SL1344_RS01075_ | 1.101681 | 1.10844 | 1.095457 | -0.00522 | 0.14614 | -0.03167 | 0.487009 | 0.661719 | 0.774047 | 0.77 | 0.846494 |
| SL1344_RS01080_ | 3.567787 | 3.536703 | 3.596738 | 0.06905 | 0.117602 | 0.579845 | 0.261159 | 0.378335 | 0.555115 | 0.5218 | 0.6607 |
| SL1344_RS01085_ | 1.218963 | 1.110822 | 1.322696 | 0.214024 | 0.124507 | 1.696019 | 0.025334 | 0.071435 | 0.191611 | 0.1331 | 0.2762 |
| SL1344_RS01090_ | 2.279178 | 2.304859 | 2.231594 | -0.06637 | 0.111929 | -0.55235 | 0.284667 | 0.456472 | 0.608735 | 0.5558 | 0.687505 |
| SL1344_RS01095_ | 5.869309 | 6.184825 | 5.613387 | -0.51996 | 0.159828 | -3.30477 | 4.68E-05 | 0.008848 | 0.073503 | 0.1 | 0.236592 |
| SL1344_RS01100_ | 6.333615 | 6.583289 | 6.226409 | -0.35592 | 0.11391 | -2.13948 | 4.68E-05 | 0.057825 | 0.185692 | 0.1 | 0.236592 |
| SL1344_RS01105_ | 2.72233 | 2.732046 | 2.714431 | -0.0074 | 0.133171 | -0.04943 | 0.480346 | 0.768417 | 0.856016 | 0.8523 | 0.905679 |
| SL1344_RS01110_ | 4.806522 | 4.89533 | 4.744941 | -0.14694 | 0.108563 | -1.37778 | 0.044 | 0.105021 | 0.257575 | 0.147 | 0.298603 |
| SL1344_RS01115_ | 1.701862 | 1.738623 | 1.657229 | -0.09135 | 0.124658 | -0.70086 | 0.205197 | 0.311451 | 0.478325 | 0.4211 | 0.557553 |
| SL1344_RS01120_ | 1.378335 | 1.491671 | 1.343476 | -0.14178 | 0.195174 | -0.62892 | 0.279813 | 0.408282 | 0.577392 | 0.5678 | 0.699199 |
| SL1344_RS01130_ | 4.101391 | 4.096008 | 4.110188 | 0.002863 | 0.086814 | 0.028571 | 0.489007 | 0.727833 | 0.829001 | 0.8705 | 0.920514 |
| SL1344_RS01135_ | 5.146551 | 5.042357 | 5.223575 | 0.161285 | 0.068688 | 2.293538 | 0.002002 | 0.027962 | 0.122655 | 0.1014 | 0.238468 |
| SL1344_RS01140_ | 5.264768 | 5.274539 | 5.253722 | 0.061062 | 0.157341 | 0.312299 | 0.426667 | 0.494003 | 0.662145 | 0.9463 | 0.967336 |
| SL1344_RS01145_ | 4.322902 | 4.333058 | 4.310199 | -0.04405 | 0.128936 | -0.37951 | 0.333333 | 0.504197 | 0.665317 | 0.6976 | 0.799121 |
| SL1344_RS01150_ | 2.178709 | 2.310162 | 1.992017 | -0.31026 | 0.101004 | -2.96403 | 0.000673 | 0.021293 | 0.103085 | 0.1003 | 0.236989 |
| SL1344_RS01155_ | 3.579906 | 3.695944 | 3.474187 | -0.24577 | 0.097644 | -2.53155 | 4.68E-05 | 0.022172 | 0.108276 | 0.1 | 0.236592 |
| SL1344_RS01160_ | 3.219505 | 3.344553 | 3.039656 | -0.32355 | 0.102079 | -3.23904 | 4.68E-05 | 0.021413 | 0.107019 | 0.1 | 0.236592 |
| SL1344_RS01165_ | 1.847099 | 1.917959 | 1.745418 | -0.17188 | 0.104571 | -1.54313 | 0.049967 | 0.090615 | 0.221111 | 0.1622 | 0.309386 |
| SL1344_RS01170_ | 3.348973 | 3.403822 | 3.27522 | -0.13615 | 0.095915 | -1.35681 | 0.061959 | 0.106504 | 0.252642 | 0.1713 | 0.324099 |
| SL1344_RS01175_ | 2.590868 | 2.677929 | 2.514944 | -0.1672 | 0.100277 | -1.63797 | 0.031979 | 0.073669 | 0.201877 | 0.134 | 0.278656 |
| SL1344_RS01180_ | 0.36367 | 0.422157 | 0.305763 | -0.11835 | 0.149969 | -0.73531 | 0.203864 | 0.322475 | 0.472411 | 0.42 | 0.553457 |
| SL1344_RS01185_ | 1.37491 | 1.220732 | 1.501619 | 0.305181 | 0.166638 | 1.821013 | 0.011326 | 0.052573 | 0.168344 | 0.1083 | 0.247382 |
| SL1344_RS01190_ | 0.168597 | -0.01293 | 0.257325 | 0.263636 | 0.273522 | 0.918613 | 0.156667 | 0.238205 | 0.413184 | 0.3198 | 0.477187 |
| SL1344_RS01195_ | 0.871563 | 0.781666 | 0.957463 | 0.177892 | 0.129956 | 1.299823 | 0.071952 | 0.130034 | 0.270044 | 0.1944 | 0.339721 |
| SL1344_RS01200_ | 3.181309 | 3.074349 | 3.418882 | 0.335666 | 0.088229 | 3.407689 | 4.68E-05 | 0.024656 | 0.114291 | 0.1 | 0.236592 |
| SL1344_RS01205_ | 2.499886 | 2.461372 | 2.58136 | 0.107995 | 0.205574 | 0.473221 | 0.314457 | 0.578314 | 0.722533 | 0.656 | 0.777869 |
| SL1344_RS01210_ | -0.41286 | -0.35933 | -0.44776 | -0.10447 | 0.295063 | -0.34685 | 0.358428 | 0.579673 | 0.710387 | 0.7069 | 0.797704 |

| | | | | | | | | | | |
|---|---|---|---|---|---|---|---|---|---|---|
| SL1344_RS01215_ | 0.374398 | 0.221301 | 0.564035 | 0.335277 | 0.19486 | 1.672088 | 0.039333 | 0.075524 | 0.202379 | 0.1448 | 0.292383 |
| SL1344_RS01220_ | 1.107283 | 1.020747 | 1.184037 | 0.166922 | 0.136818 | 1.164297 | 0.093937 | 0.156264 | 0.307024 | 0.2282 | 0.379079 |
| SL1344_RS01225_ | 3.60738 | 3.687562 | 3.569246 | -0.08123 | 0.113807 | -0.72028 | 0.226516 | 0.358409 | 0.5354 | 0.4633 | 0.617238 |
| SL1344_RS01230_ | 1.242095 | 1.323621 | 1.17533 | -0.16785 | 0.170886 | -0.96143 | 0.123917 | 0.216987 | 0.386869 | 0.2715 | 0.422911 |
| SL1344_RS01245_ | 0.623186 | 0.648634 | 0.599393 | -0.0625 | 0.174618 | -0.31803 | 0.357333 | 0.541274 | 0.676672 | 0.6696 | 0.767104 |
| SL1344_RS01250_ | 0.79214 | 0.921276 | 0.620061 | -0.30384 | 0.131776 | -2.25285 | 0.007995 | 0.041867 | 0.142044 | 0.1065 | 0.244789 |
| SL1344_RS01255_ | -0.12078 | -0.22536 | -0.04957 | 0.167105 | 0.264286 | 0.60072 | 0.25 | 0.402902 | 0.562402 | 0.5228 | 0.655042 |
| SL1344_RS01260_ | 0.81439 | 0.49098 | 0.866785 | 0.396633 | 0.770063 | 0.613708 | 0.226516 | 0.403224 | 0.585797 | 0.4033 | 0.58498 |
| SL1344_RS01265_ | 2.829323 | 3.004195 | 2.705128 | -0.30476 | 0.112125 | -2.71498 | 4.68E-05 | 0.022125 | 0.1076 | 0.1 | 0.236592 |
| SL1344_RS01270_ | 2.642821 | 2.990134 | 2.491301 | -0.46355 | 0.187997 | -2.28212 | 0.000673 | 0.034435 | 0.138264 | 0.1001 | 0.236731 |
| SL1344_RS01275_ | 1.927816 | 1.770592 | 2.071473 | 0.305167 | 0.575183 | 0.407986 | 0.4004 | 0.535209 | 0.69588 | 0.862 | 0.915581 |
| SL1344_RS01285_ | -1.59329 | -1.44014 | -1.90681 | -0.43865 | 0.369914 | -1.08582 | 0.121919 | 0.182549 | 0.340716 | 0.2728 | 0.424137 |
| SL1344_RS01290_ | 0.367927 | 0.297848 | 0.422071 | 0.145544 | 0.20831 | 0.692709 | 0.214524 | 0.318322 | 0.485257 | 0.4245 | 0.559483 |
| SL1344_RS01295_ | -0.78985 | -0.90712 | -0.59658 | 0.289909 | 0.28584 | 0.96746 | 0.150667 | 0.22727 | 0.387159 | 0.3184 | 0.471546 |
| SL1344_RS01300_ | -1.77242 | -1.86725 | -1.71544 | 0.137299 | 0.350036 | 0.349051 | 0.347333 | 0.544645 | 0.676024 | 0.6579 | 0.75926 |
| SL1344_RS01305_ | 1.494663 | 1.339373 | 1.621725 | 0.285018 | 0.11408 | 2.429522 | 0.002001 | 0.034723 | 0.130737 | 0.1008 | 0.237605 |
| SL1344_RS01310_ | -0.51224 | -0.77126 | -0.19243 | 0.539408 | 0.34247 | 1.541042 | 0.046 | 0.083372 | 0.218684 | 0.1459 | 0.295857 |
| SL1344_RS01315_ | 3.986435 | 4.28633 | 3.748112 | -0.53618 | 0.077987 | -6.80972 | 4.68E-05 | 0.00495 | 0.056045 | 0.1 | 0.236592 |
| SL1344_RS01320_ | -0.12892 | -0.14389 | -0.1168 | 0.018152 | 0.213273 | 0.08178 | 0.462667 | 0.624272 | 0.743256 | 0.7323 | 0.816139 |
| SL1344_RS01325_ | -0.13688 | -0.23226 | -0.00439 | 0.224429 | 0.178995 | 1.179232 | 0.099933 | 0.169119 | 0.313062 | 0.2491 | 0.395573 |
| SL1344_RS01330_ | -0.53663 | -0.48063 | -0.64718 | -0.15085 | 0.241274 | -0.56365 | 0.26982 | 0.437145 | 0.58539 | 0.5481 | 0.673061 |
| SL1344_RS01335_ | 1.514883 | 1.526037 | 1.486281 | -0.02143 | 0.203492 | -0.09651 | 0.467333 | 0.803311 | 0.880106 | 0.8903 | 0.933865 |
| SL1344_RS01340_ | 0.080753 | 0.096834 | 0.058745 | -0.00798 | 0.337658 | -0.02122 | 0.487009 | 0.770393 | 0.857405 | 0.8144 | 0.882836 |
| SL1344_RS01345_ | -2.01391 | -2.11876 | -1.92894 | 0.223014 | 0.489332 | 0.427351 | 0.313333 | 0.482034 | 0.6289 | 0.6137 | 0.723623 |
| SL1344_RS01350_ | -2.89298 | -2.96277 | -2.79373 | 0.154045 | 0.539232 | 0.268982 | 0.377333 | 0.5697 | 0.695539 | 0.6622 | 0.760825 |
| SL1344_RS01355_ | -1.61627 | -1.70046 | -1.53369 | 0.165709 | 0.271765 | 0.590279 | 0.254497 | 0.373505 | 0.517612 | 0.4763 | 0.599259 |
| SL1344_RS01360_ | -3.00047 | -2.97873 | -3.0265 | -0.08175 | 0.674737 | -0.1201 | 0.451699 | 0.665975 | 0.77854 | 0.7887 | 0.859134 |
| SL1344_RS01365_ | -1.71619 | -1.79757 | -1.61957 | 0.180426 | 0.30997 | 0.539989 | 0.278667 | 0.411537 | 0.555601 | 0.5178 | 0.638849 |
| SL1344_RS01370_ | -2.1464 | -2.13154 | -2.16408 | -0.04463 | 0.345877 | -0.11585 | 0.441706 | 0.541796 | 0.672595 | 0.6396 | 0.743111 |
| SL1344_RS01375_ | -6.35925 | -6.11 | -6.60128 | -0.48745 | 1.726989 | -0.26474 | 0.377082 | 0.528674 | 0.662511 | 0.6552 | 0.751008 |
| SL1344_RS01380_ | -0.54059 | -0.43981 | -0.6283 | -0.18704 | 0.196177 | -0.89406 | 0.152667 | 0.23592 | 0.38533 | 0.3192 | 0.463267 |
| SL1344_RS01385_ | -2.4832 | -2.58654 | -2.37856 | 0.207809 | 0.406857 | 0.49166 | 0.298667 | 0.453031 | 0.591385 | 0.5587 | 0.671119 |
| SL1344_RS01390_ | -1.46718 | -1.45205 | -1.49256 | -0.05153 | 0.322954 | -0.1449 | 0.429333 | 0.587319 | 0.711594 | 0.7029 | 0.791621 |
| SL1344_RS01395_ | -2.98922 | -2.91666 | -3.14233 | -0.23162 | 0.692795 | -0.29022 | 0.386409 | 0.578471 | 0.712067 | 0.7202 | 0.810227 |
| SL1344_RS01400_ | -3.96072 | -3.94723 | -3.97044 | 0.026118 | 0.775518 | 0.042324 | 0.487333 | 0.619542 | 0.740064 | 0.7362 | 0.819282 |
| SL1344_RS01405_ | -4.60096 | -4.4247 | -4.79725 | -0.37019 | 1.178741 | -0.27117 | 0.378667 | 0.582001 | 0.713128 | 0.7076 | 0.799433 |
| SL1344_RS01410_ | -5.19769 | -4.85953 | -5.43137 | -0.5037 | 1.220819 | -0.37277 | 0.325783 | 0.544438 | 0.67429 | 0.6332 | 0.739534 |
| SL1344_RS01415_ | -4.27937 | -4.35075 | -4.18361 | -0.03177 | 1.210519 | -0.02047 | 0.49 | 0.687962 | 0.79632 | 0.8268 | 0.886275 |
| SL1344_RS01420_ | -4.98425 | -4.60497 | -5.52133 | -0.39221 | 1.840906 | -0.20154 | 0.416 | 0.675597 | 0.7872 | 0.8015 | 0.868321 |
| SL1344_RS01425_ | -2.69189 | -2.68121 | -2.70276 | -0.04608 | 0.492506 | -0.08279 | 0.461026 | 0.610411 | 0.732078 | 0.726 | 0.811343 |
| SL1344_RS01430_ | -2.2746 | -2.20707 | -2.36342 | -0.15094 | 0.378572 | -0.39044 | 0.331779 | 0.4863 | 0.621885 | 0.5989 | 0.704691 |
| SL1344_RS01435_ | -3.90932 | -3.73037 | -4.12606 | -0.36921 | 0.5831 | -0.59887 | 0.24517 | 0.351972 | 0.496586 | 0.4557 | 0.581953 |
| SL1344_RS01440_ | -5.37228 | -4.93572 | -6.26029 | -1.32785 | 1.87753 | -0.66189 | 0.216667 | 0.316844 | 0.488024 | 0.4236 | 0.575933 |
| SL1344_RS01445_ | -0.68328 | -0.65165 | -0.74169 | -0.0774 | 0.243616 | -0.28564 | 0.377082 | 0.553159 | 0.684709 | 0.6632 | 0.763993 |
| SL1344_RS01450_ | -3.11578 | -3.14753 | -3.07835 | 0.062893 | 0.542676 | 0.103525 | 0.45503 | 0.5782 | 0.703795 | 0.6916 | 0.784731 |

| | | | | | | | | | | |
|---|---|---|---|---|---|---|---|---|---|---|
| SL1344_RS01455_ | -3.95393 | -4.17342 | -3.78146 | 0.319107 | 0.842363 | 0.356593 | 0.349333 | 0.535646 | 0.670485 | 0.6514 | 0.754105 |
| SL1344_RS01460_ | -2.58302 | -2.46834 | -2.79613 | -0.33558 | 0.526311 | -0.5685 | 0.266489 | 0.399636 | 0.556304 | 0.5243 | 0.648331 |
| SL1344_RS01465_ | -1.40588 | -1.42612 | -1.38639 | 0.025177 | 0.346696 | 0.0701 | 0.466667 | 0.641429 | 0.758946 | 0.7478 | 0.830453 |
| SL1344_RS01470_ | -4.46704 | -4.55801 | -4.36531 | 0.177057 | 0.927342 | 0.16622 | 0.422667 | 0.593072 | 0.716962 | 0.6955 | 0.787985 |
| SL1344_RS01475_ | 0.49595 | 0.543475 | 0.3542 | -0.15888 | 0.23824 | -0.54518 | 0.324 | 0.553537 | 0.694543 | 0.6722 | 0.785534 |
| SL1344_RS01480_ | 0.959335 | 1.138197 | 0.742676 | -0.34425 | 0.215532 | -1.56297 | 0.052 | 0.079865 | 0.213764 | 0.149 | 0.300478 |
| SL1344_RS01485_ | 0.332706 | 0.324163 | 0.349906 | 0.020367 | 0.197153 | 0.087461 | 0.46 | 0.673796 | 0.784881 | 0.7747 | 0.854232 |
| SL1344_RS01490_ | -0.99616 | -0.91766 | -1.09185 | -0.17799 | 0.235758 | -0.72267 | 0.213857 | 0.33621 | 0.485067 | 0.4357 | 0.568 |
| SL1344_RS01495_ | -3.33567 | -3.34292 | -3.32125 | 0.004998 | 0.79284 | 0.00966 | 0.495003 | 0.68556 | 0.793576 | 0.8075 | 0.873606 |
| SL1344_RS01500_ | -7.15085 | -6.83258 | -7.61041 | -0.8416 | 2.310447 | -0.28187 | 0.34044 | 0.48294 | 0.627948 | 0.6087 | 0.714065 |
| SL1344_RS01505_ | -3.47115 | -3.54892 | -3.41755 | 0.032618 | 0.771277 | 0.04227 | 0.481679 | 0.680633 | 0.789524 | 0.7839 | 0.857499 |
| SL1344_RS01515_ | -3.90656 | -3.62507 | -4.15979 | -0.56298 | 0.716938 | -0.75929 | 0.202532 | 0.311812 | 0.465446 | 0.4119 | 0.548758 |
| SL1344_RS01520_ | -6.94784 | -6.41151 | -7.56541 | -1.29785 | 2.536435 | -0.42735 | 0.283333 | 0.438689 | 0.591465 | 0.556 | 0.675248 |
| SL1344_RS01525_ | -1.01317 | -0.73049 | -1.46517 | -0.71231 | 0.432024 | -1.61115 | 0.042 | 0.075061 | 0.20649 | 0.138 | 0.285091 |
| SL1344_RS01530_ | -2.58292 | -2.48245 | -2.65584 | -0.19755 | 0.448725 | -0.39705 | 0.327781 | 0.483628 | 0.621555 | 0.6034 | 0.709636 |
| SL1344_RS01535_ | 0.081545 | -0.03649 | 0.209772 | 0.246159 | 0.254302 | 0.974526 | 0.139907 | 0.209813 | 0.375966 | 0.2844 | 0.441473 |
| SL1344_RS01540_ | -2.19293 | -2.27545 | -2.00974 | 0.221095 | 0.464869 | 0.396798 | 0.337775 | 0.57301 | 0.701156 | 0.656 | 0.765206 |
| SL1344_RS01545_ | -2.2425 | -2.04891 | -2.48566 | -0.45442 | 0.426119 | -1.02683 | 0.139333 | 0.217716 | 0.373056 | 0.3007 | 0.447716 |
| SL1344_RS01550_ | -3.56607 | -3.63864 | -3.50345 | 0.140449 | 0.686347 | 0.180092 | 0.423051 | 0.576194 | 0.703477 | 0.6833 | 0.777529 |
| SL1344_RS01565_ | -3.29902 | -3.35863 | -3.272 | 0.055084 | 0.590645 | 0.088458 | 0.462358 | 0.615002 | 0.737434 | 0.7243 | 0.813031 |
| SL1344_RS01570_ | 0.508323 | 0.412499 | 0.650263 | 0.263697 | 0.238526 | 1.074239 | 0.107333 | 0.18017 | 0.345452 | 0.2433 | 0.398205 |
| SL1344_RS01575_ | -1.06147 | -1.28263 | -0.66569 | 0.598286 | 0.350023 | 1.631166 | 0.039307 | 0.075594 | 0.204869 | 0.1416 | 0.288023 |
| SL1344_RS01580_ | 0.443366 | 0.261925 | 0.614906 | 0.333281 | 0.200851 | 1.620126 | 0.041306 | 0.079917 | 0.210546 | 0.1425 | 0.290731 |
| SL1344_RS01590_ | 2.252702 | 2.224173 | 2.301102 | 0.078749 | 0.094485 | 0.789655 | 0.17988 | 0.283791 | 0.442333 | 0.3908 | 0.528645 |
| SL1344_RS01595_ | 1.462793 | 1.386297 | 1.680331 | 0.270268 | 0.161154 | 1.474075 | 0.067955 | 0.109922 | 0.257004 | 0.1845 | 0.337443 |
| SL1344_RS01600_ | 1.032764 | 1.163782 | 0.939705 | -0.23431 | 0.159812 | -1.43821 | 0.05 | 0.109644 | 0.251882 | 0.1586 | 0.305578 |
| SL1344_RS01605_ | 1.194319 | 1.442776 | 0.896456 | -0.55427 | 0.145895 | -3.79186 | 4.68E-05 | 0.009049 | 0.068539 | 0.1 | 0.236592 |
| SL1344_RS01610_ | -1.19253 | -1.08345 | -1.34233 | -0.26322 | 0.261319 | -0.93717 | 0.136576 | 0.224119 | 0.37413 | 0.3147 | 0.453627 |
| SL1344_RS01615_ | -0.9158 | -0.93398 | -0.90402 | 0.026882 | 0.251701 | 0.091332 | 0.462667 | 0.57586 | 0.703652 | 0.693 | 0.785153 |
| SL1344_RS01620_ | 4.060263 | 3.773367 | 4.271761 | 0.498634 | 0.084161 | 5.860247 | 4.68E-05 | 0.005745 | 0.058653 | 0.1 | 0.236592 |
| SL1344_RS01625_ | 1.228019 | 1.504477 | 0.999873 | -0.50116 | 0.180388 | -2.74294 | 4.68E-05 | 0.017999 | 0.096834 | 0.1 | 0.236592 |
| SL1344_RS01635_ | 2.165513 | 2.203229 | 2.071055 | -0.11634 | 0.128727 | -0.79657 | 0.215856 | 0.311742 | 0.482297 | 0.4339 | 0.582891 |
| SL1344_RS01640_ | -1.41607 | -1.33385 | -1.52349 | -0.19354 | 0.291594 | -0.62942 | 0.237175 | 0.364931 | 0.514993 | 0.472 | 0.595762 |
| SL1344_RS01645_ | 1.44471 | 1.512329 | 1.379009 | -0.10336 | 0.171658 | -0.59841 | 0.269333 | 0.442134 | 0.599489 | 0.5458 | 0.67461 |
| SL1344_RS01650_ | 2.915704 | 2.880651 | 2.966806 | 0.059034 | 0.111499 | 0.509289 | 0.296 | 0.49694 | 0.647588 | 0.6082 | 0.726867 |
| SL1344_RS01655_ | -1.15841 | -1.15492 | -1.16499 | 0.047089 | 0.460031 | 0.093115 | 0.464667 | 0.71611 | 0.81737 | 0.822 | 0.885761 |
| SL1344_RS01660_ | -3.82956 | -3.77769 | -3.86097 | -0.08306 | 0.769719 | -0.09303 | 0.459694 | 0.606698 | 0.730495 | 0.7213 | 0.808594 |
| SL1344_RS01665_ | -5.75134 | -5.72751 | -5.7654 | 0.218537 | 2.575932 | 0.067263 | 0.453333 | 0.637358 | 0.763493 | 0.8463 | 0.900638 |
| SL1344_RS01670_ | -4.36443 | -4.45509 | -4.21618 | 0.173626 | 0.949474 | 0.153482 | 0.418667 | 0.633241 | 0.749541 | 0.734 | 0.818165 |
| SL1344_RS01675_ | 3.756959 | 3.562352 | 3.940948 | 0.413525 | 0.128987 | 3.254844 | 4.68E-05 | 0.008905 | 0.071894 | 0.1 | 0.236592 |
| SL1344_RS01680_ | -3.18573 | -3.31532 | -3.10742 | 0.176332 | 0.616929 | 0.271368 | 0.383744 | 0.577781 | 0.706652 | 0.6906 | 0.786241 |
| SL1344_RS01685_ | -2.53762 | -2.46777 | -2.60726 | -0.13693 | 0.498674 | -0.25759 | 0.391333 | 0.587392 | 0.713319 | 0.6968 | 0.788556 |
| SL1344_RS01690_ | -3.75198 | -3.8006 | -3.71988 | 0.08697 | 0.637217 | 0.121325 | 0.440667 | 0.568502 | 0.69687 | 0.6777 | 0.774413 |
| SL1344_RS01695_ | -2.18489 | -1.92138 | -2.46664 | -0.52458 | 0.483725 | -1.02821 | 0.123917 | 0.190834 | 0.351036 | 0.275 | 0.431112 |
| SL1344_RS01700_ | -0.81269 | -0.7402 | -0.88819 | -0.14672 | 0.258386 | -0.54236 | 0.265333 | 0.410936 | 0.558877 | 0.5252 | 0.645721 |

| | | | | | | | | | | | |
|---|---|---|---|---|---|---|---|---|---|---|---|
| SL1344_RS01705_ | -0.25454 | -0.30298 | -0.17794 | 0.096445 | 0.257073 | 0.348716 | 0.359333 | 0.596325 | 0.722261 | 0.694 | 0.79 |
| SL1344_RS01710_ | -2.45272 | -2.39338 | -2.49432 | -0.0717 | 0.481818 | -0.13384 | 0.450667 | 0.652651 | 0.766603 | 0.7506 | 0.833885 |
| SL1344_RS01715_ | -3.19934 | -2.97245 | -3.56517 | -0.57815 | 0.582046 | -0.90447 | 0.167222 | 0.249419 | 0.405958 | 0.3453 | 0.489765 |
| SL1344_RS01720_ | -2.07532 | -2.03326 | -2.13571 | -0.10934 | 0.372757 | -0.27399 | 0.384667 | 0.531881 | 0.666641 | 0.6429 | 0.745092 |
| SL1344_RS01725_ | -2.36624 | -2.34011 | -2.39265 | -0.02475 | 0.508641 | -0.04702 | 0.48 | 0.663623 | 0.77603 | 0.7816 | 0.854458 |
| SL1344_RS01730_ | -1.36429 | -1.78462 | -1.12605 | 0.652423 | 0.309082 | 1.972547 | 0.015334 | 0.056651 | 0.170059 | 0.1146 | 0.254624 |
| SL1344_RS01740_ | -5.9461 | -5.3101 | -6.49352 | -1.0858 | 1.807784 | -0.56757 | 0.243333 | 0.419453 | 0.572215 | 0.5172 | 0.646283 |
| SL1344_RS01745_ | -0.85303 | -0.8316 | -0.88343 | -0.08868 | 0.327666 | -0.24445 | 0.403333 | 0.588742 | 0.719476 | 0.7195 | 0.808365 |
| SL1344_RS01750_ | -2.06167 | -1.98397 | -2.11918 | -0.12787 | 0.357884 | -0.32774 | 0.369333 | 0.508207 | 0.643396 | 0.6143 | 0.72128 |
| SL1344_RS01755_ | -0.38368 | -0.48991 | -0.28182 | 0.207466 | 0.198758 | 1.00911 | 0.134577 | 0.21531 | 0.364671 | 0.3067 | 0.450567 |
| SL1344_RS01760_ | 0.601038 | 0.590125 | 0.622482 | 0.031043 | 0.189875 | 0.159553 | 0.427333 | 0.640943 | 0.757841 | 0.7598 | 0.837284 |
| SL1344_RS01765_ | -0.53316 | -0.44125 | -0.58493 | -0.08702 | 0.319227 | -0.25976 | 0.39507 | 0.688336 | 0.795186 | 0.7645 | 0.847898 |
| SL1344_RS01770_ | -4.77735 | -4.65583 | -4.96295 | -0.32689 | 1.536438 | -0.19962 | 0.403065 | 0.624675 | 0.748811 | 0.7631 | 0.841396 |
| SL1344_RS01775_ | -5.09391 | -4.53351 | -5.44274 | -0.81897 | 1.142435 | -0.65757 | 0.238667 | 0.379481 | 0.526794 | 0.468 | 0.602873 |
| SL1344_RS01780_ | -4.76376 | -4.19033 | -5.53356 | -1.35317 | 1.101369 | -1.15651 | 0.078 | 0.1558 | 0.305854 | 0.2062 | 0.352979 |
| SL1344_RS01785_ | -1.97315 | -1.97011 | -1.97634 | -0.02282 | 0.385929 | -0.05836 | 0.479333 | 0.6107 | 0.732626 | 0.7223 | 0.809055 |
| SL1344_RS01790_ | -3.15669 | -3.12838 | -3.19894 | 0.023468 | 0.784495 | 0.030136 | 0.490667 | 0.71807 | 0.817356 | 0.8113 | 0.878073 |
| SL1344_RS01795_ | -2.38198 | -2.3488 | -2.43544 | -0.0275 | 0.507978 | -0.04401 | 0.480667 | 0.675904 | 0.785205 | 0.7895 | 0.861151 |
| SL1344_RS01800_ | -4.60769 | -4.59339 | -4.61894 | -0.04682 | 1.074332 | -0.04472 | 0.479333 | 0.64688 | 0.763485 | 0.7579 | 0.838015 |
| SL1344_RS01805_ | -1.50136 | -1.74964 | -1.29179 | 0.44069 | 0.34344 | 1.219894 | 0.086667 | 0.140638 | 0.289012 | 0.2092 | 0.360674 |
| SL1344_RS01810_ | -2.52444 | -2.86159 | -2.28208 | 0.60087 | 0.494042 | 1.13999 | 0.077333 | 0.158838 | 0.313344 | 0.2034 | 0.349846 |
| SL1344_RS01815_ | -0.90253 | -1.04273 | -0.81488 | 0.255265 | 0.394672 | 0.636536 | 0.252 | 0.368007 | 0.535325 | 0.4956 | 0.636054 |
| SL1344_RS01820_ | -3.49442 | -3.4504 | -3.52003 | -0.05905 | 0.645333 | -0.07743 | 0.47 | 0.627733 | 0.746051 | 0.7269 | 0.814839 |
| SL1344_RS01825_ | -0.57285 | -0.44457 | -0.72176 | -0.28194 | 0.241131 | -1.12593 | 0.104667 | 0.175594 | 0.327616 | 0.2481 | 0.39781 |
| SL1344_RS01830_ | 1.174042 | 1.194925 | 1.153966 | -0.01892 | 0.164967 | -0.1167 | 0.455696 | 0.689109 | 0.796033 | 0.7949 | 0.863677 |
| SL1344_RS01835_ | 2.548667 | 2.588186 | 2.530717 | -0.04038 | 0.118756 | -0.28528 | 0.392 | 0.751138 | 0.842789 | 0.7841 | 0.867324 |
| SL1344_RS01840_ | -2.42249 | -2.80754 | -2.09361 | 0.716381 | 0.542204 | 1.301628 | 0.068621 | 0.12581 | 0.27262 | 0.1875 | 0.338783 |
| SL1344_RS01845_ | -1.08081 | -1.36143 | -0.82095 | 0.539908 | 0.224946 | 2.366085 | 0.005334 | 0.030978 | 0.119637 | 0.105 | 0.242949 |
| SL1344_RS01850_ | 0.341922 | 0.243784 | 0.620877 | 0.352357 | 0.248681 | 1.252716 | 0.097269 | 0.152199 | 0.313276 | 0.2295 | 0.384727 |
| SL1344_RS01855_ | -2.00572 | -2.36882 | -1.62841 | 0.744552 | 0.329588 | 2.156914 | 0.010667 | 0.037982 | 0.133811 | 0.1092 | 0.247938 |
| SL1344_RS01860_ | -0.8854 | -0.84799 | -0.90983 | -0.00823 | 0.393118 | -0.01913 | 0.491006 | 0.732169 | 0.830471 | 0.86 | 0.912 |
| SL1344_RS01865_ | -1.00603 | -1.01141 | -0.99994 | 0.012056 | 0.353987 | 0.030719 | 0.486 | 0.692053 | 0.797738 | 0.7961 | 0.864971 |
| SL1344_RS01870_ | 1.285083 | 1.105238 | 1.475769 | 0.430418 | 0.213723 | 1.990589 | 0.001336 | 0.038622 | 0.144619 | 0.1004 | 0.237118 |
| SL1344_RS01875_ | 3.369122 | 3.185627 | 3.770952 | 0.571723 | 0.105541 | 4.047201 | 4.68E-05 | 0.023368 | 0.114347 | 0.1 | 0.236592 |
| SL1344_RS01880_ | -0.13213 | -0.00762 | -0.26587 | -0.27064 | 0.207626 | -1.2475 | 0.074 | 0.132411 | 0.277369 | 0.197 | 0.344458 |
| SL1344_RS01885_ | -2.72378 | -2.72492 | -2.72306 | -0.01835 | 0.522834 | -0.03474 | 0.477682 | 0.638031 | 0.754235 | 0.746 | 0.827615 |
| SL1344_RS01890_ | -2.38162 | -2.46111 | -2.29053 | 0.159335 | 0.379578 | 0.386819 | 0.334444 | 0.487456 | 0.622926 | 0.5895 | 0.699743 |
| SL1344_RS01895_ | -1.75736 | -1.74759 | -1.76323 | -0.04939 | 0.456149 | -0.09067 | 0.456667 | 0.674888 | 0.785614 | 0.7945 | 0.863807 |
| SL1344_RS01900_ | -0.83254 | -0.86531 | -0.802 | 0.066727 | 0.232248 | 0.256718 | 0.382 | 0.542514 | 0.672733 | 0.6467 | 0.747323 |
| SL1344_RS01905_ | 3.01585 | 3.021652 | 3.013596 | -0.00698 | 0.094903 | -0.06575 | 0.475017 | 0.744272 | 0.838122 | 0.8407 | 0.90104 |
| SL1344_RS01910_ | 1.118222 | 1.12294 | 1.107306 | -0.01684 | 0.152253 | -0.10101 | 0.466 | 0.677893 | 0.787921 | 0.7823 | 0.857593 |
| SL1344_RS01915_ | -1.87107 | -1.94916 | -1.71688 | 0.218451 | 0.433415 | 0.471513 | 0.31046 | 0.516138 | 0.654678 | 0.619 | 0.731669 |
| SL1344_RS01930_ | -0.23423 | -0.46904 | 0.045402 | 0.508612 | 0.213014 | 2.311486 | 0.004668 | 0.032507 | 0.12618 | 0.1048 | 0.242954 |
| SL1344_RS01935_ | -1.61137 | -2.09977 | -1.27336 | 0.832327 | 0.326429 | 2.535667 | 0.001337 | 0.035757 | 0.130633 | 0.1013 | 0.238269 |
| SL1344_RS01940_ | 1.672644 | 1.558544 | 1.722713 | 0.160163 | 0.149241 | 1.0337 | 0.15523 | 0.223654 | 0.393128 | 0.3242 | 0.481852 |

| | | | | | | | | | | | |
|---|---|---|---|---|---|---|---|---|---|---|---|
| SL1344_RS01945_ | 1.87733 | 1.87342 | 1.881647 | -0.00037 | 0.119508 | -0.00207 | 0.498334 | 0.667537 | 0.779601 | 0.779 | 0.853985 |
| SL1344_RS01950_ | -1.69493 | -1.82676 | -1.31293 | 0.465128 | 0.530005 | 0.814433 | 0.210526 | 0.30492 | 0.476206 | 0.4161 | 0.575523 |
| SL1344_RS01955_ | -2.07924 | -2.17054 | -1.83433 | 0.312029 | 0.448064 | 0.573191 | 0.279813 | 0.466818 | 0.613338 | 0.5742 | 0.700304 |
| SL1344_RS01960_ | 4.069711 | 4.06919 | 4.07032 | -0.02673 | 0.180151 | -0.16741 | 0.42505 | 0.600057 | 0.743553 | 0.88 | 0.923737 |
| SL1344_RS01965_ | 0.823519 | 0.534468 | 1.090835 | 0.587865 | 0.191648 | 3.112574 | 4.68E-05 | 0.020685 | 0.104078 | 0.1 | 0.236592 |
| SL1344_RS01975_ | 2.557515 | 2.507567 | 2.636605 | 0.121287 | 0.121783 | 0.944144 | 0.156562 | 0.213699 | 0.383212 | 0.2998 | 0.462572 |
| SL1344_RS01980_ | -0.38068 | -0.24592 | -0.5035 | -0.25457 | 0.191497 | -1.24073 | 0.088667 | 0.146301 | 0.285275 | 0.2232 | 0.368612 |
| SL1344_RS01985_ | 0.705189 | 0.723118 | 0.677096 | -0.05772 | 0.170718 | -0.30631 | 0.367333 | 0.535593 | 0.672968 | 0.66 | 0.759014 |
| SL1344_RS01990_ | 1.260374 | 1.15123 | 1.403969 | 0.254485 | 0.145733 | 1.657101 | 0.039307 | 0.079242 | 0.208901 | 0.1371 | 0.28197 |
| SL1344_RS01995_ | 0.750081 | 0.780985 | 0.727988 | -0.04944 | 0.217528 | -0.207 | 0.412392 | 0.676335 | 0.787005 | 0.7954 | 0.865949 |
| SL1344_RS02000_ | 2.116929 | 2.110579 | 2.129489 | -0.0087 | 0.189129 | -0.04629 | 0.484344 | 0.782933 | 0.866649 | 0.9226 | 0.951782 |
| SL1344_RS02005_ | 1.168762 | 1.248911 | 1.04296 | -0.20191 | 0.127295 | -1.50838 | 0.053333 | 0.101466 | 0.234991 | 0.1682 | 0.314919 |
| SL1344_RS02010_ | 0.262506 | 0.073994 | 0.430499 | 0.365195 | 0.19053 | 1.892674 | 0.013991 | 0.049406 | 0.158974 | 0.1116 | 0.251075 |
| SL1344_RS02015_ | -0.50458 | -0.38038 | -0.63515 | -0.25399 | 0.200708 | -1.20811 | 0.086 | 0.150056 | 0.291316 | 0.2265 | 0.373734 |
| SL1344_RS02020_ | 1.650153 | 1.650416 | 1.650012 | -6.29E-05 | 0.092246 | -0.00104 | 0.499333 | 0.558688 | 0.686207 | 0.6674 | 0.765297 |
| SL1344_RS02025_ | -0.35692 | -0.25347 | -0.50083 | -0.2494 | 0.204985 | -1.14939 | 0.097935 | 0.165085 | 0.31413 | 0.2373 | 0.384455 |
| SL1344_RS02030_ | -7.24367 | -7.42457 | -7.14297 | 0.308332 | 2.557217 | 0.102018 | 0.452365 | 0.554614 | 0.692662 | 0.7001 | 0.792082 |
| SL1344_RS02045_ | 2.409459 | 2.533074 | 2.264372 | -0.2295 | 0.193202 | -1.21002 | 0.105333 | 0.133821 | 0.294142 | 0.2 | 0.364755 |
| SL1344_RS02050_ | 1.252414 | 1.323123 | 1.137272 | -0.17997 | 0.129283 | -1.30695 | 0.078614 | 0.129318 | 0.273881 | 0.2008 | 0.351434 |
| SL1344_RS02055_ | -0.85475 | -1.03826 | -0.68734 | 0.365186 | 0.275954 | 1.236511 | 0.065333 | 0.12893 | 0.273648 | 0.1885 | 0.337349 |
| SL1344_RS02060_ | 5.152474 | 5.08648 | 5.187071 | 0.146597 | 0.10753 | 1.421706 | 0.014667 | 0.081962 | 0.223077 | 0.1131 | 0.253341 |
| SL1344_RS02065_ | 2.097639 | 2.086953 | 2.112466 | -0.02121 | 0.185193 | -0.10665 | 0.463025 | 0.704383 | 0.811403 | 0.8916 | 0.931108 |
| SL1344_RS02070_ | 0.200278 | 0.476759 | -0.09781 | -0.59085 | 0.19269 | -3.05275 | 0.000673 | 0.015527 | 0.088036 | 0.1 | 0.236592 |
| SL1344_RS02075_ | 2.180916 | 2.296375 | 2.083065 | -0.2137 | 0.073293 | -2.84875 | 0.001336 | 0.018633 | 0.093754 | 0.1008 | 0.237657 |
| SL1344_RS02080_ | 2.984536 | 3.001817 | 2.949519 | -0.06041 | 0.110003 | -0.51342 | 0.294667 | 0.43955 | 0.602313 | 0.6101 | 0.726182 |
| SL1344_RS02085_ | 3.877442 | 3.873318 | 3.886495 | 0.011254 | 0.05635 | 0.187369 | 0.424384 | 0.621478 | 0.741687 | 0.7317 | 0.817939 |
| SL1344_RS02090_ | 2.795519 | 2.669739 | 2.891616 | 0.241687 | 0.116908 | 2.076758 | 0.002002 | 0.050378 | 0.166409 | 0.1007 | 0.237522 |
| SL1344_RS02095_ | -1.00393 | -0.92671 | -1.10787 | -0.19377 | 0.369483 | -0.51709 | 0.279147 | 0.423768 | 0.584477 | 0.558 | 0.680098 |
| SL1344_RS02100_ | -0.70605 | -0.69921 | -0.71586 | -0.03766 | 0.297144 | -0.11954 | 0.443038 | 0.634002 | 0.753063 | 0.7472 | 0.828923 |
| SL1344_RS02105_ | 1.366257 | 1.297923 | 1.42867 | 0.115036 | 0.148527 | 0.729587 | 0.212667 | 0.326047 | 0.491367 | 0.4311 | 0.575556 |
| SL1344_RS02110_ | 3.978364 | 3.962423 | 4.0177 | 0.052034 | 0.081911 | 0.589383 | 0.285333 | 0.460285 | 0.619252 | 0.5649 | 0.701858 |
| SL1344_RS02115_ | 3.892632 | 4.012056 | 3.862514 | -0.14338 | 0.245167 | -0.59582 | 0.334667 | 0.487964 | 0.657354 | 0.6988 | 0.820481 |
| SL1344_RS02120_ | -1.28863 | -1.58372 | -1.05257 | 0.526194 | 0.290359 | 1.739056 | 0.027333 | 0.066985 | 0.184275 | 0.1336 | 0.276813 |
| SL1344_RS02125_ | 1.534464 | 1.494091 | 1.624375 | 0.141278 | 0.219998 | 0.666621 | 0.220667 | 0.334297 | 0.509238 | 0.4181 | 0.579923 |
| SL1344_RS02130_ | 1.927148 | 1.814551 | 2.042845 | 0.232932 | 0.16267 | 1.406217 | 0.043305 | 0.105989 | 0.253116 | 0.1523 | 0.302205 |
| SL1344_RS02135_ | 1.952589 | 2.155622 | 1.899226 | -0.25345 | 0.310692 | -0.66832 | 0.319333 | 0.450155 | 0.624048 | 0.6575 | 0.78713 |
| SL1344_RS02145_ | 0.648895 | 0.603453 | 0.717252 | 0.110384 | 0.301907 | 0.352664 | 0.323118 | 0.600464 | 0.734216 | 0.6742 | 0.7832 |
| SL1344_RS02150_ | 0.523161 | 0.433725 | 0.605987 | 0.166574 | 0.18866 | 0.838725 | 0.177881 | 0.265767 | 0.428814 | 0.3551 | 0.504392 |
| SL1344_RS02155_ | 1.193463 | 0.759788 | 1.245199 | 0.455627 | 0.278603 | 0.894044 | 0.270486 | 0.280459 | 0.466827 | 0.5438 | 0.689296 |
| SL1344_RS02160_ | 2.694175 | 2.612102 | 2.769253 | 0.165873 | 0.131174 | 1.259379 | 0.068621 | 0.130572 | 0.287599 | 0.1746 | 0.328421 |
| SL1344_RS02165_ | 1.861488 | 1.956172 | 1.805962 | -0.16887 | 0.173592 | -0.93698 | 0.152667 | 0.217119 | 0.391419 | 0.3023 | 0.462867 |
| SL1344_RS02170_ | 1.083434 | 1.229377 | 1.014657 | -0.21915 | 0.183182 | -1.08059 | 0.124667 | 0.187704 | 0.354186 | 0.2622 | 0.41654 |
| SL1344_RS02175_ | 0.876084 | 0.837784 | 0.889768 | 0.03374 | 0.183594 | 0.179217 | 0.430667 | 0.686729 | 0.794962 | 0.7808 | 0.859599 |
| SL1344_RS02180_ | -1.69373 | -1.86629 | -1.57572 | 0.292611 | 0.308435 | 0.895802 | 0.166667 | 0.258427 | 0.411121 | 0.3532 | 0.493957 |
| SL1344_RS02185_ | -2.64154 | -2.36676 | -2.91672 | -0.55556 | 0.430465 | -1.21199 | 0.086 | 0.144228 | 0.284588 | 0.2144 | 0.360512 |

| | | | | | | | | | | | |
|---|---|---|---|---|---|---|---|---|---|---|---|
| SL1344_RS02190_ | -1.71044 | -1.65111 | -1.77195 | -0.13752 | 0.357786 | -0.34137 | 0.350667 | 0.540117 | 0.672888 | 0.6551 | 0.754848 |
| SL1344_RS02195_ | -1.66426 | -1.53354 | -1.75136 | -0.1869 | 0.423145 | -0.41063 | 0.322452 | 0.547867 | 0.682253 | 0.6441 | 0.753551 |
| SL1344_RS02200_ | 0.000672 | 0.046785 | -0.05246 | -0.09812 | 0.165582 | -0.55609 | 0.268667 | 0.404513 | 0.547297 | 0.5039 | 0.625353 |
| SL1344_RS02205_ | -3.31394 | -3.2569 | -3.38981 | -0.17018 | 0.709224 | -0.21595 | 0.410393 | 0.563699 | 0.697189 | 0.6909 | 0.78259 |
| SL1344_RS02210_ | -3.42194 | -3.22173 | -3.6528 | -0.43036 | 0.512341 | -0.76581 | 0.189207 | 0.291728 | 0.439 | 0.3807 | 0.514153 |
| SL1344_RS02215_ | 1.251056 | 1.109707 | 1.454877 | 0.336614 | 0.168888 | 1.950775 | 0.014657 | 0.045551 | 0.15548 | 0.1115 | 0.251495 |
| SL1344_RS02220_ | 0.546861 | 0.457043 | 0.822027 | 0.343703 | 0.209483 | 1.408578 | 0.082 | 0.123484 | 0.275086 | 0.2054 | 0.359049 |
| SL1344_RS02225_ | 2.892731 | 2.725284 | 3.023367 | 0.249673 | 0.25693 | 0.976798 | 0.131912 | 0.207669 | 0.388484 | 0.2355 | 0.406956 |
| SL1344_RS02230_ | 0.99428 | 0.929894 | 1.068703 | 0.142882 | 0.159769 | 0.846113 | 0.165223 | 0.258862 | 0.422822 | 0.3516 | 0.500575 |
| SL1344_RS02235_ | -1.58892 | -1.44671 | -1.71219 | -0.25188 | 0.383344 | -0.61742 | 0.257333 | 0.400714 | 0.555838 | 0.5138 | 0.642112 |
| SL1344_RS02240_ | -1.02346 | -0.67619 | -1.42554 | -0.77344 | 0.317378 | -2.41428 | 0.002667 | 0.030983 | 0.123804 | 0.1027 | 0.240015 |
| SL1344_RS02245_ | 4.2766 | 4.904529 | 3.519251 | -1.23526 | 0.238955 | -5.0526 | 4.68E-05 | 0.002012 | 0.041781 | 0.1 | 0.236592 |
| SL1344_RS02250_ | 2.579086 | 3.135087 | 1.795342 | -1.10105 | 0.45113 | -2.61888 | 4.68E-05 | 0.015395 | 0.094194 | 0.1 | 0.236592 |
| SL1344_RS02255_ | 3.036798 | 3.583584 | 2.216575 | -1.25356 | 0.346681 | -2.96111 | 4.68E-05 | 0.019505 | 0.105638 | 0.1 | 0.236592 |
| SL1344_RS02260_ | 4.646202 | 5.195332 | 3.686515 | -1.41975 | 0.282969 | -3.79879 | 4.68E-05 | 0.016982 | 0.099264 | 0.1 | 0.236592 |
| SL1344_RS02265_ | 4.339517 | 5.029508 | 3.311047 | -1.71933 | 0.249478 | -5.87714 | 4.68E-05 | 0.006863 | 0.066177 | 0.1 | 0.236592 |
| SL1344_RS02270_ | 1.549459 | 1.65462 | 1.415536 | -0.23914 | 0.102931 | -2.32013 | 0.003335 | 0.037622 | 0.134296 | 0.1049 | 0.242916 |
| SL1344_RS02275_ | 1.384209 | 1.571782 | 1.013293 | -0.54209 | 0.17839 | -2.75 | 4.68E-05 | 0.028137 | 0.121223 | 0.1001 | 0.236725 |
| SL1344_RS02280_ | 3.199685 | 2.969608 | 3.546789 | 0.482034 | 0.230453 | 2.085717 | 0.000673 | 0.034512 | 0.139649 | 0.1006 | 0.237396 |
| SL1344_RS02285_ | 5.34875 | 5.602526 | 5.257823 | -0.33613 | 0.11508 | -2.38771 | 4.68E-05 | 0.031116 | 0.132871 | 0.1 | 0.236592 |
| SL1344_RS02300_ | 5.240744 | 5.250087 | 5.230967 | -0.00481 | 0.123527 | -0.03276 | 0.488 | 0.895406 | 0.940054 | 0.9841 | 0.990529 |
| SL1344_RS02305_ | 7.609416 | 7.395318 | 7.803866 | 0.260761 | 0.224531 | 1.629631 | 0.108 | 0.072014 | 0.209994 | 0.1999 | 0.369894 |
| SL1344_RS02310_ | 3.109914 | 3.011502 | 3.19199 | 0.159211 | 0.095221 | 1.683676 | 0.041333 | 0.072651 | 0.202004 | 0.1396 | 0.288174 |
| SL1344_RS02315_ | -2.5335 | -2.79431 | -2.09329 | 0.695594 | 0.53194 | 1.265742 | 0.088608 | 0.146041 | 0.297553 | 0.2191 | 0.370555 |
| SL1344_RS02320_ | -0.97754 | -1.05811 | -0.89667 | 0.165234 | 0.235055 | 0.642302 | 0.232512 | 0.354151 | 0.502445 | 0.4545 | 0.583759 |
| SL1344_RS02325_ | 0.536793 | 0.580856 | 0.485491 | -0.11241 | 0.208302 | -0.51767 | 0.274484 | 0.436673 | 0.592532 | 0.5645 | 0.683652 |
| SL1344_RS02330_ | -0.71355 | -0.93675 | -0.49773 | 0.434443 | 0.219074 | 1.924563 | 0.016656 | 0.051357 | 0.158026 | 0.1177 | 0.258323 |
| SL1344_RS02335_ | -0.6809 | -0.53187 | -0.75697 | -0.1973 | 0.282134 | -0.62932 | 0.256667 | 0.411645 | 0.566791 | 0.5195 | 0.65122 |
| SL1344_RS02350_ | 1.10831 | 1.112674 | 1.101315 | -0.03743 | 0.183831 | -0.17357 | 0.414 | 0.585052 | 0.720377 | 0.7569 | 0.835112 |
| SL1344_RS02355_ | 1.14452 | 1.12121 | 1.177257 | 0.066072 | 0.152284 | 0.418383 | 0.306462 | 0.476964 | 0.624416 | 0.6104 | 0.716929 |
| SL1344_RS02360_ | -5.03749 | -4.90853 | -5.23368 | -0.33546 | 1.097628 | -0.2643 | 0.369087 | 0.536684 | 0.671364 | 0.6525 | 0.753961 |
| SL1344_RS02365_ | -1.17311 | -1.21809 | -1.09909 | 0.079161 | 0.343244 | 0.239385 | 0.399734 | 0.640351 | 0.757646 | 0.7464 | 0.830351 |
| SL1344_RS02370_ | 1.0162 | 1.002952 | 1.036433 | 0.030796 | 0.124631 | 0.236266 | 0.393738 | 0.567792 | 0.693208 | 0.6675 | 0.763837 |
| SL1344_RS02375_ | 2.580932 | 2.595599 | 2.571903 | -0.00583 | 0.114808 | -0.04619 | 0.474667 | 0.739386 | 0.834424 | 0.8442 | 0.902001 |
| SL1344_RS02380_ | -0.04635 | -0.11006 | 0.042029 | 0.151776 | 0.190173 | 0.750068 | 0.208 | 0.320594 | 0.47229 | 0.4154 | 0.552544 |
| SL1344_RS02385_ | 0.96463 | 0.939896 | 0.98422 | 0.048461 | 0.111763 | 0.386059 | 0.335776 | 0.466011 | 0.604882 | 0.5738 | 0.685025 |
| SL1344_RS02390_ | -1.61896 | -1.81901 | -1.4571 | 0.418827 | 0.448742 | 0.917093 | 0.125916 | 0.220856 | 0.390956 | 0.2852 | 0.434854 |
| SL1344_RS02395_ | 0.10482 | -0.07058 | 0.273022 | 0.421385 | 0.286901 | 1.421521 | 0.023334 | 0.096923 | 0.239739 | 0.1324 | 0.27515 |
| SL1344_RS02400_ | 2.01896 | 2.540073 | 1.566743 | -1.04434 | 0.303 | -3.34272 | 4.68E-05 | 0.006234 | 0.063192 | 0.1 | 0.236592 |
| SL1344_RS02405_ | 1.308308 | 1.219159 | 1.385045 | 0.172406 | 0.127695 | 1.304914 | 0.065956 | 0.129199 | 0.272989 | 0.1835 | 0.328498 |
| SL1344_RS02420_ | 4.369746 | 4.280912 | 4.455262 | 0.172349 | 0.052008 | 3.262798 | 4.68E-05 | 0.013261 | 0.082008 | 0.1001 | 0.23673 |
| SL1344_RS02425_ | 2.926046 | 2.940662 | 2.908854 | -0.03111 | 0.077583 | -0.37635 | 0.355097 | 0.548132 | 0.680145 | 0.6559 | 0.755948 |
| SL1344_RS02430_ | 1.493883 | 1.532354 | 1.450946 | -0.08549 | 0.114672 | -0.71069 | 0.216667 | 0.325903 | 0.479161 | 0.4321 | 0.563786 |
| SL1344_RS02435_ | 2.568196 | 2.573755 | 2.562381 | -0.02552 | 0.110749 | -0.23227 | 0.393738 | 0.631549 | 0.752809 | 0.7726 | 0.846696 |
| SL1344_RS02440_ | 1.549245 | 1.565239 | 1.519625 | -0.048 | 0.130355 | -0.34773 | 0.353333 | 0.549019 | 0.682905 | 0.6703 | 0.769213 |

| | | | | | | | | | | | |
|---|---|---|---|---|---|---|---|---|---|---|---|
| SL1344_RS02445_ | -1.0333 | -1.13726 | -0.92521 | 0.109838 | 0.454751 | 0.237182 | 0.405333 | 0.704445 | 0.809278 | 0.8082 | 0.874219 |
| SL1344_RS02450_ | -0.71137 | -0.52803 | -0.87294 | -0.34688 | 0.2922 | -1.15069 | 0.111333 | 0.16337 | 0.318253 | 0.2398 | 0.393671 |
| SL1344_RS02455_ | -1.6437 | -1.55048 | -1.75499 | -0.1954 | 0.398456 | -0.45356 | 0.314457 | 0.481007 | 0.624756 | 0.5931 | 0.706963 |
| SL1344_RS02460_ | 0.132256 | 0.469455 | -0.19193 | -0.68491 | 0.21866 | -3.198 | 4.68E-05 | 0.016605 | 0.092363 | 0.1 | 0.236592 |
| SL1344_RS02465_ | 2.635686 | 2.6391 | 2.630894 | -0.02431 | 0.115951 | -0.17255 | 0.411059 | 0.562851 | 0.704775 | 0.7273 | 0.815066 |
| SL1344_RS02470_ | 2.694052 | 2.741957 | 2.615542 | -0.13001 | 0.086939 | -1.40537 | 0.055297 | 0.115816 | 0.259963 | 0.1713 | 0.318224 |
| SL1344_RS02475_ | 2.891966 | 3.011642 | 2.76601 | -0.26126 | 0.117848 | -2.2041 | 0.000673 | 0.035603 | 0.13842 | 0.1014 | 0.238454 |
| SL1344_RS02480_ | 5.715215 | 5.788916 | 5.664279 | -0.11248 | 0.072553 | -1.56031 | 0.052 | 0.071479 | 0.205716 | 0.1457 | 0.297329 |
| SL1344_RS02485_ | 3.658176 | 3.789265 | 3.535724 | -0.24625 | 0.065478 | -3.72932 | 4.68E-05 | 0.009128 | 0.070033 | 0.1 | 0.236592 |
| SL1344_RS02490_ | 1.336131 | 1.302781 | 1.386072 | 0.101738 | 0.152723 | 0.59982 | 0.246667 | 0.359293 | 0.52567 | 0.4962 | 0.625044 |
| SL1344_RS02495_ | -1.47845 | -1.57152 | -1.36054 | 0.226766 | 0.362077 | 0.557441 | 0.274484 | 0.393498 | 0.54901 | 0.5168 | 0.641594 |
| SL1344_RS02500_ | -0.14815 | -0.16446 | -0.13218 | -0.005 | 0.260263 | -0.01694 | 0.492338 | 0.685368 | 0.792553 | 0.8165 | 0.877828 |
| SL1344_RS02505_ | 2.910353 | 3.044558 | 2.699143 | -0.35098 | 0.103305 | -3.36097 | 4.68E-05 | 0.014752 | 0.087976 | 0.1 | 0.236592 |
| SL1344_RS02510_ | 0.366565 | 0.388785 | 0.326095 | -0.05636 | 0.191671 | -0.26824 | 0.377082 | 0.59416 | 0.720644 | 0.7149 | 0.803973 |
| SL1344_RS02520_ | 0.074646 | 0.164854 | -0.04126 | -0.21367 | 0.174374 | -1.16862 | 0.092667 | 0.163936 | 0.311837 | 0.2372 | 0.382194 |
| SL1344_RS02525_ | -0.12104 | -0.01961 | -0.19483 | -0.16305 | 0.239695 | -0.63403 | 0.24517 | 0.385656 | 0.54244 | 0.491 | 0.625158 |
| SL1344_RS02530_ | -1.40893 | -1.33357 | -1.54779 | -0.19621 | 0.329435 | -0.53825 | 0.287808 | 0.455688 | 0.600086 | 0.568 | 0.687283 |
| SL1344_RS02540_ | 0.582635 | 0.463755 | 0.768214 | 0.2855 | 0.228683 | 1.19719 | 0.102667 | 0.151687 | 0.311359 | 0.2173 | 0.375606 |
| SL1344_RS02545_ | 1.0423 | 1.05721 | 1.016286 | -0.02152 | 0.174207 | -0.11324 | 0.459333 | 0.734147 | 0.830587 | 0.8052 | 0.878026 |
| SL1344_RS02550_ | 1.102871 | 1.092598 | 1.107455 | 0.009391 | 0.221912 | 0.033925 | 0.486342 | 0.782626 | 0.865829 | 0.8767 | 0.924363 |
| SL1344_RS02555_ | -0.73424 | -0.57057 | -0.89588 | -0.16201 | 0.470274 | -0.35615 | 0.355763 | 0.611401 | 0.740576 | 0.7285 | 0.815299 |
| SL1344_RS02560_ | -0.17139 | -0.27167 | -0.07283 | 0.206981 | 0.211765 | 0.91436 | 0.151899 | 0.232804 | 0.388421 | 0.3282 | 0.470989 |
| SL1344_RS02565_ | 2.851951 | 2.887706 | 2.791343 | -0.09275 | 0.0742 | -1.17207 | 0.109261 | 0.161517 | 0.311568 | 0.2443 | 0.394268 |
| SL1344_RS02570_ | 0.391124 | 0.474482 | 0.360647 | -0.096 | 0.20306 | -0.40974 | 0.345103 | 0.57377 | 0.706007 | 0.6863 | 0.789999 |
| SL1344_RS02575_ | -0.2958 | -0.27636 | -0.31782 | -0.0467 | 0.231274 | -0.18122 | 0.415057 | 0.598334 | 0.722401 | 0.7225 | 0.806587 |
| SL1344_RS02580_ | -0.64389 | -0.75819 | -0.41298 | 0.288074 | 0.354151 | 0.751216 | 0.23 | 0.327803 | 0.4982 | 0.4379 | 0.58706 |
| SL1344_RS02585_ | 1.525492 | 1.465457 | 1.682544 | 0.222155 | 0.158601 | 1.280416 | 0.083333 | 0.152081 | 0.314055 | 0.2077 | 0.359512 |
| SL1344_RS02590_ | -0.26186 | -0.2606 | -0.2647 | 0.024473 | 0.284579 | 0.081936 | 0.472352 | 0.692995 | 0.800651 | 0.8009 | 0.872965 |
| SL1344_RS02595_ | -0.88821 | -0.70625 | -1.03085 | -0.32352 | 0.255466 | -1.20446 | 0.09527 | 0.152668 | 0.297478 | 0.2317 | 0.380134 |
| SL1344_RS02600_ | -1.07033 | -0.94124 | -1.27287 | -0.31099 | 0.295156 | -0.98504 | 0.143238 | 0.221875 | 0.377147 | 0.3104 | 0.460022 |
| SL1344_RS02605_ | -0.59582 | -0.50793 | -0.70076 | -0.1871 | 0.265804 | -0.6523 | 0.233333 | 0.367288 | 0.524848 | 0.4724 | 0.607699 |
| SL1344_RS02610_ | 1.93823 | 1.949209 | 1.927158 | -0.02823 | 0.169953 | -0.18183 | 0.418388 | 0.69159 | 0.800467 | 0.8019 | 0.870521 |
| SL1344_RS02615_ | -1.79379 | -1.90141 | -1.68634 | 0.221383 | 0.289185 | 0.730768 | 0.201333 | 0.309568 | 0.454134 | 0.4067 | 0.538218 |
| SL1344_RS02620_ | -0.80215 | -0.91901 | -0.49792 | 0.391597 | 0.276393 | 1.244644 | 0.102598 | 0.156136 | 0.310814 | 0.2507 | 0.40352 |
| SL1344_RS02625_ | 1.998581 | 1.965582 | 2.017304 | 0.04925 | 0.104176 | 0.425649 | 0.316667 | 0.502238 | 0.641299 | 0.6124 | 0.722126 |
| SL1344_RS02630_ | -1.96452 | -2.29351 | -1.70289 | 0.596384 | 0.309315 | 1.835136 | 0.024667 | 0.062426 | 0.17208 | 0.1271 | 0.268263 |
| SL1344_RS02635_ | -2.80782 | -3.13893 | -2.49445 | 0.685623 | 0.536124 | 1.20053 | 0.091939 | 0.149754 | 0.298796 | 0.2245 | 0.373719 |
| SL1344_RS02640_ | -1.99617 | -2.17997 | -1.77421 | 0.433177 | 0.416663 | 1.010862 | 0.134577 | 0.203898 | 0.362689 | 0.2914 | 0.44049 |
| SL1344_RS02645_ | -2.67133 | -2.6143 | -2.76968 | -0.17343 | 0.541061 | -0.30944 | 0.363758 | 0.545088 | 0.68048 | 0.672 | 0.769183 |
| SL1344_RS02650_ | -3.74082 | -3.83764 | -3.6628 | 0.175575 | 0.757488 | 0.218386 | 0.398667 | 0.598054 | 0.723564 | 0.7058 | 0.7969 |
| SL1344_RS02655_ | -0.71729 | -0.65988 | -0.76764 | -0.11016 | 0.226594 | -0.45462 | 0.296 | 0.448262 | 0.588707 | 0.5491 | 0.666294 |
| SL1344_RS02660_ | -3.2647 | -3.21042 | -3.34724 | -0.14477 | 0.590298 | -0.22481 | 0.397735 | 0.589636 | 0.714713 | 0.7074 | 0.796138 |
| SL1344_RS02665_ | -2.31272 | -2.2723 | -2.35534 | -0.10448 | 0.531678 | -0.18746 | 0.418388 | 0.605698 | 0.729584 | 0.7147 | 0.802753 |
| SL1344_RS02670_ | 0.260724 | 0.17458 | 0.368822 | 0.206785 | 0.182144 | 1.082267 | 0.108667 | 0.181265 | 0.33599 | 0.2512 | 0.39956 |
| SL1344_RS02675_ | -1.84925 | -2.15369 | -1.66527 | 0.479849 | 0.405109 | 1.157754 | 0.120586 | 0.17329 | 0.330614 | 0.2571 | 0.410978 |

| | | | | | | | | | | | |
|---|---|---|---|---|---|---|---|---|---|---|---|
| SL1344_RS02680_ | -2.37375 | -2.53176 | -2.19958 | 0.32287 | 0.45497 | 0.669425 | 0.225333 | 0.361228 | 0.517101 | 0.4602 | 0.59687 |
| SL1344_RS02685_ | -2.12715 | -2.02114 | -2.25976 | -0.22091 | 0.388817 | -0.55014 | 0.279147 | 0.422576 | 0.570565 | 0.5276 | 0.65079 |
| SL1344_RS02690_ | -3.14604 | -3.19204 | -3.09849 | 0.091498 | 0.515916 | 0.176426 | 0.421053 | 0.55203 | 0.681327 | 0.6608 | 0.757094 |
| SL1344_RS02695_ | -4.03837 | -4.27288 | -3.7709 | 0.533851 | 0.86163 | 0.587037 | 0.253165 | 0.381774 | 0.536195 | 0.5046 | 0.630701 |
| SL1344_RS02700_ | -2.37164 | -2.27824 | -2.42574 | -0.17706 | 0.454958 | -0.34219 | 0.355763 | 0.538862 | 0.672416 | 0.653 | 0.753822 |
| SL1344_RS02705_ | 0.869216 | 1.026875 | 0.700221 | -0.32877 | 0.19803 | -1.66664 | 0.022652 | 0.075081 | 0.205378 | 0.127 | 0.271307 |
| SL1344_RS02710_ | -0.80655 | -0.5181 | -1.18255 | -0.66414 | 0.206032 | -3.19209 | 4.68E-05 | 0.017226 | 0.088419 | 0.1002 | 0.236857 |
| SL1344_RS02715_ | 1.614909 | 1.71642 | 1.543126 | -0.24094 | 0.237564 | -1.02838 | 0.107262 | 0.17221 | 0.342252 | 0.2325 | 0.390689 |
| SL1344_RS02720_ | 3.956435 | 4.03967 | 3.870565 | -0.18027 | 0.086947 | -2.11468 | 0.000673 | 0.05293 | 0.172848 | 0.1004 | 0.237122 |
| SL1344_RS02730_ | -1.08852 | -1.03485 | -1.13762 | -0.08043 | 0.339732 | -0.23545 | 0.395736 | 0.637394 | 0.755121 | 0.7402 | 0.823246 |
| SL1344_RS02735_ | -1.78215 | -1.6516 | -1.97006 | -0.35375 | 0.400529 | -0.85565 | 0.148667 | 0.259732 | 0.42466 | 0.3329 | 0.475727 |
| SL1344_RS02740_ | -2.6712 | -2.44723 | -2.98265 | -0.5392 | 0.46307 | -1.10902 | 0.104667 | 0.178629 | 0.32771 | 0.2571 | 0.400386 |
| SL1344_RS02745_ | 1.479334 | 1.665906 | 1.409786 | -0.23611 | 0.149616 | -1.35823 | 0.092605 | 0.133997 | 0.288817 | 0.2158 | 0.371495 |
| SL1344_RS02750_ | 2.05512 | 2.076102 | 2.032632 | -0.04956 | 0.099538 | -0.46593 | 0.2998 | 0.453361 | 0.598697 | 0.5705 | 0.683803 |
| SL1344_RS02760_ | -2.61597 | -2.99069 | -2.19503 | 0.827307 | 0.456542 | 1.768421 | 0.019987 | 0.063918 | 0.180677 | 0.1254 | 0.266788 |
| SL1344_RS02765_ | -3.35241 | -3.55483 | -3.18882 | 0.36696 | 0.533397 | 0.640802 | 0.238 | 0.345614 | 0.492067 | 0.4427 | 0.570252 |
| SL1344_RS02770_ | -2.02926 | -2.43435 | -1.71724 | 0.699834 | 0.391916 | 1.726713 | 0.028667 | 0.072184 | 0.194728 | 0.1327 | 0.276405 |
| SL1344_RS02775_ | -2.18868 | -2.13711 | -2.28008 | -0.14488 | 0.420679 | -0.29757 | 0.372 | 0.540415 | 0.673784 | 0.6597 | 0.758659 |
| SL1344_RS02790_ | -5.87188 | -6.09325 | -5.64195 | 0.444069 | 1.16328 | 0.353014 | 0.343771 | 0.46927 | 0.605824 | 0.572 | 0.683844 |
| SL1344_RS02795_ | -5.51056 | -4.90187 | -6.03363 | -1.30678 | 1.65832 | -0.74097 | 0.195333 | 0.297164 | 0.461182 | 0.4047 | 0.544309 |
| SL1344_RS02800_ | -3.21021 | -2.84593 | -3.37132 | -0.48823 | 0.902239 | -0.48535 | 0.318667 | 0.544384 | 0.685788 | 0.6387 | 0.755853 |
| SL1344_RS02805_ | -3.13281 | -3.47221 | -2.88362 | 0.258501 | 1.093433 | 0.235044 | 0.418 | 0.704509 | 0.810821 | 0.8521 | 0.905878 |
| SL1344_RS02815_ | -1.22817 | -0.86833 | -1.90385 | -1.03474 | 0.500134 | -2.03192 | 0.007329 | 0.044381 | 0.15438 | 0.1057 | 0.243975 |
| SL1344_RS02820_ | -2.18169 | -1.77348 | -3.32262 | -1.51834 | 0.676216 | -1.94359 | 0.008001 | 0.066996 | 0.195775 | 0.1081 | 0.247177 |
| SL1344_RS02825_ | -4.58651 | -3.59376 | -5.53358 | -1.89462 | 1.126567 | -1.62028 | 0.025983 | 0.081751 | 0.209979 | 0.1309 | 0.27449 |
| SL1344_RS02830_ | -5.43065 | -5.02581 | -6.28393 | -1.20395 | 1.67426 | -0.63177 | 0.229333 | 0.331687 | 0.500373 | 0.4489 | 0.591264 |
| SL1344_RS02835_ | -5.76844 | -5.53234 | -6.19792 | -0.51116 | 1.877252 | -0.24013 | 0.387333 | 0.592534 | 0.722758 | 0.7345 | 0.821044 |
| SL1344_RS02840_ | -0.96932 | -0.94761 | -1.01346 | -0.02775 | 0.347542 | -0.07331 | 0.470667 | 0.705279 | 0.80818 | 0.7818 | 0.861124 |
| SL1344_RS02845_ | -1.21646 | -1.36136 | -1.08719 | 0.269893 | 0.254099 | 1.019454 | 0.128667 | 0.208176 | 0.358391 | 0.2946 | 0.439637 |
| SL1344_RS02850_ | -1.26978 | -1.23702 | -1.31854 | -0.0805 | 0.286296 | -0.26367 | 0.384667 | 0.554981 | 0.683516 | 0.6497 | 0.751523 |
| SL1344_RS02855_ | -5.67897 | -6.04639 | -5.31149 | 0.638997 | 1.578916 | 0.375358 | 0.328 | 0.518266 | 0.65553 | 0.6227 | 0.731011 |
| SL1344_RS02860_ | -3.49029 | -3.32017 | -3.68469 | -0.38723 | 0.671364 | -0.55846 | 0.250667 | 0.391764 | 0.542717 | 0.5203 | 0.637717 |
| SL1344_RS02865_ | 2.372157 | 3.022815 | 2.156051 | -0.84603 | 0.136721 | -4.02637 | 4.68E-05 | 0.045163 | 0.161819 | 0.1 | 0.236592 |
| SL1344_RS02870_ | 0.824225 | 0.961837 | 0.733119 | -0.21394 | 0.16351 | -1.255 | 0.090667 | 0.140479 | 0.289848 | 0.224 | 0.37838 |
| SL1344_RS02875_ | 0.983872 | 0.919142 | 1.06331 | 0.143322 | 0.14813 | 0.952378 | 0.154 | 0.226419 | 0.385933 | 0.3191 | 0.466915 |
| SL1344_RS02880_ | 0.76295 | 0.625006 | 0.858953 | 0.229825 | 0.142041 | 1.553079 | 0.044667 | 0.08974 | 0.21866 | 0.1543 | 0.300033 |
| SL1344_RS02885_ | 1.368733 | 1.236908 | 1.520149 | 0.299359 | 0.136305 | 2.166656 | 0.002002 | 0.048172 | 0.157452 | 0.103 | 0.24021 |
| SL1344_RS02890_ | -2.91929 | -2.81683 | -3.09581 | -0.26059 | 0.594424 | -0.41052 | 0.332445 | 0.50677 | 0.649523 | 0.6302 | 0.739396 |
| SL1344_RS02895_ | -3.93347 | -3.81199 | -4.07406 | -0.27762 | 0.648763 | -0.397 | 0.330446 | 0.459536 | 0.599626 | 0.5663 | 0.679099 |
| SL1344_RS02900_ | -4.53132 | -4.29038 | -4.66163 | -0.31234 | 0.926813 | -0.31685 | 0.363758 | 0.555975 | 0.685809 | 0.666 | 0.765007 |
| SL1344_RS02905_ | -5.14488 | -5.39888 | -4.93658 | 0.458595 | 1.031601 | 0.413584 | 0.321119 | 0.470589 | 0.609885 | 0.575 | 0.686651 |
| SL1344_RS02915_ | -5.43468 | -5.40392 | -5.45553 | -0.13997 | 1.583047 | -0.07639 | 0.463333 | 0.63544 | 0.753512 | 0.7562 | 0.832696 |
| SL1344_RS02920_ | 1.849399 | 1.829554 | 1.874071 | 0.049453 | 0.115613 | 0.402791 | 0.333111 | 0.50375 | 0.645654 | 0.6312 | 0.736805 |
| SL1344_RS02925_ | -0.82288 | -0.83902 | -0.80752 | 0.031538 | 0.217483 | 0.13542 | 0.439041 | 0.539607 | 0.669038 | 0.6532 | 0.751398 |
| SL1344_RS02930_ | 0.210769 | 0.271328 | 0.016099 | -0.23148 | 0.218288 | -0.93422 | 0.186667 | 0.261866 | 0.429141 | 0.3913 | 0.541707 |

| | | | | | | | | | | | |
|---|---|---|---|---|---|---|---|---|---|---|---|
| SL1344_RS02935_ | -1.17314 | -1.19027 | -1.15615 | 0.012884 | 0.430281 | 0.03262 | 0.486667 | 0.732691 | 0.829567 | 0.8034 | 0.872794 |
| SL1344_RS02940_ | -1.79927 | -1.75883 | -1.85365 | -0.09805 | 0.328114 | -0.2688 | 0.371086 | 0.517923 | 0.652301 | 0.6367 | 0.73855 |
| SL1344_RS02945_ | -1.28157 | -1.47167 | -0.96601 | 0.489623 | 0.317975 | 1.403369 | 0.072618 | 0.111855 | 0.251604 | 0.1879 | 0.337115 |
| SL1344_RS02950_ | -4.41648 | -4.32065 | -4.51626 | -0.20286 | 0.706027 | -0.28029 | 0.373333 | 0.493193 | 0.626711 | 0.6029 | 0.707828 |
| SL1344_RS02955_ | -0.85037 | -1.01984 | -0.60066 | 0.411453 | 0.26576 | 1.465979 | 0.067289 | 0.108024 | 0.244393 | 0.1814 | 0.330175 |
| SL1344_RS02960_ | -2.76309 | -2.79277 | -2.74426 | -0.04468 | 0.68043 | -0.06551 | 0.471019 | 0.688661 | 0.795664 | 0.8008 | 0.870253 |
| SL1344_RS02965_ | -5.74301 | -5.37484 | -6.24609 | -0.70318 | 2.077158 | -0.32674 | 0.353333 | 0.570428 | 0.705705 | 0.6968 | 0.793119 |
| SL1344_RS02975_ | 1.13702 | 0.967139 | 1.256132 | 0.284458 | 0.125538 | 2.205414 | 0.007329 | 0.038673 | 0.137959 | 0.1059 | 0.244168 |
| SL1344_RS02980_ | -4.18861 | -4.18262 | -4.19324 | 0.029795 | 0.817476 | 0.0347 | 0.484344 | 0.590701 | 0.7149 | 0.7068 | 0.794484 |
| SL1344_RS02985_ | -2.50374 | -2.35858 | -2.75854 | -0.41218 | 0.552863 | -0.68425 | 0.203864 | 0.31995 | 0.485971 | 0.4255 | 0.562176 |
| SL1344_RS02990_ | -3.13186 | -3.0843 | -3.20737 | -0.14411 | 0.562212 | -0.22637 | 0.394667 | 0.537544 | 0.670588 | 0.6521 | 0.751732 |
| SL1344_RS02995_ | -2.21512 | -2.09513 | -2.37963 | -0.2874 | 0.40788 | -0.66737 | 0.228667 | 0.35345 | 0.505283 | 0.468 | 0.59707 |
| SL1344_RS03000_ | -2.16925 | -2.20748 | -2.13923 | 0.070033 | 0.386369 | 0.153279 | 0.428667 | 0.583255 | 0.708805 | 0.6997 | 0.790494 |
| SL1344_RS03005_ | -2.82516 | -2.9329 | -2.72491 | 0.200453 | 0.438984 | 0.429421 | 0.306 | 0.428984 | 0.570588 | 0.5323 | 0.651029 |
| SL1344_RS03020_ | -2.5946 | -2.59755 | -2.58925 | 0.009206 | 0.412194 | 0.018677 | 0.493333 | 0.544868 | 0.67659 | 0.654 | 0.754934 |
| SL1344_RS03025_ | -3.18644 | -3.19112 | -3.18281 | 0.022625 | 0.573554 | 0.036898 | 0.482345 | 0.604742 | 0.727636 | 0.7152 | 0.804065 |
| SL1344_RS03030_ | 4.030927 | 3.975575 | 4.130408 | 0.151314 | 0.107474 | 1.425876 | 0.06 | 0.08895 | 0.231198 | 0.1587 | 0.311845 |
| SL1344_RS03035_ | 0.953148 | 0.901776 | 1.032719 | 0.14069 | 0.152086 | 0.898836 | 0.151233 | 0.242343 | 0.40614 | 0.3271 | 0.470381 |
| SL1344_RS03040_ | -0.83187 | -0.69075 | -0.94108 | -0.27526 | 0.286334 | -0.89216 | 0.159893 | 0.24356 | 0.40216 | 0.3358 | 0.479284 |
| SL1344_RS03045_ | -0.19122 | 0.053898 | -0.40857 | -0.47211 | 0.224765 | -2.03773 | 0.008667 | 0.043593 | 0.148685 | 0.1086 | 0.24753 |
| SL1344_RS03050_ | -0.07299 | -0.07477 | -0.07047 | 0.009306 | 0.20173 | 0.047008 | 0.477333 | 0.617904 | 0.736942 | 0.7277 | 0.812361 |
| SL1344_RS03055_ | -0.48579 | -0.26434 | -0.73895 | -0.46358 | 0.21446 | -2.12007 | 0.009334 | 0.042247 | 0.143393 | 0.1125 | 0.251953 |
| SL1344_RS03060_ | 0.260362 | 0.238624 | 0.278209 | 0.027813 | 0.209558 | 0.118163 | 0.444 | 0.666222 | 0.778508 | 0.7601 | 0.840674 |
| SL1344_RS03065_ | 1.147548 | 1.230897 | 1.072518 | -0.15729 | 0.117065 | -1.27845 | 0.077333 | 0.131943 | 0.27006 | 0.2068 | 0.354126 |
| SL1344_RS03070_ | 4.679031 | 4.695191 | 4.658473 | -0.02018 | 0.144579 | -0.13634 | 0.449333 | 0.841392 | 0.906235 | 0.9679 | 0.980359 |
| SL1344_RS03075_ | 1.997071 | 2.016876 | 1.961621 | -0.06071 | 0.104105 | -0.54124 | 0.279813 | 0.418755 | 0.571766 | 0.5369 | 0.655331 |
| SL1344_RS03080_ | -5.53773 | -5.42323 | -5.64808 | -0.03373 | 1.829633 | -0.01832 | 0.491006 | 0.671764 | 0.784343 | 0.7993 | 0.866545 |
| SL1344_RS03085_ | -1.18036 | -1.22883 | -1.13471 | 0.115533 | 0.350917 | 0.318599 | 0.36 | 0.543771 | 0.680595 | 0.6716 | 0.77 |
| SL1344_RS03090_ | -2.81942 | -2.91612 | -2.60215 | 0.254213 | 0.617765 | 0.358316 | 0.342 | 0.603985 | 0.725382 | 0.6811 | 0.785776 |
| SL1344_RS03095_ | 0.839551 | 0.92889 | 0.612176 | -0.32393 | 0.264266 | -1.15521 | 0.11992 | 0.16152 | 0.329082 | 0.2403 | 0.403673 |
| SL1344_RS03100_ | 5.29175 | 5.270676 | 5.306841 | 0.078808 | 0.141906 | 0.586639 | 0.263824 | 0.332866 | 0.520293 | 0.5345 | 0.682975 |
| SL1344_RS03105_ | -1.32528 | -1.53783 | -1.09708 | 0.43893 | 0.271274 | 1.544004 | 0.046667 | 0.092416 | 0.220241 | 0.1617 | 0.307316 |
| SL1344_RS03110_ | 0.717342 | 0.84621 | 0.644645 | -0.18776 | 0.160455 | -1.07233 | 0.136667 | 0.198366 | 0.355014 | 0.2964 | 0.448413 |
| SL1344_RS03115_ | 2.932972 | 2.927103 | 2.93991 | 0.03197 | 0.119802 | 0.262902 | 0.393071 | 0.576317 | 0.7141 | 0.765 | 0.840917 |
| SL1344_RS03120_ | -0.82362 | 0.007799 | -1.22321 | -1.16337 | 0.422553 | -2.55677 | 4.68E-05 | 0.032248 | 0.131826 | 0.1004 | 0.237124 |
| SL1344_RS03125_ | -4.82997 | -4.8605 | -4.80812 | 0.187752 | 1.500951 | 0.084565 | 0.45 | 0.576241 | 0.715033 | 0.7519 | 0.831261 |
| SL1344_RS03130_ | -5.21043 | -5.37595 | -4.97344 | 0.287718 | 1.459407 | 0.174224 | 0.412667 | 0.654691 | 0.767268 | 0.7306 | 0.818189 |
| SL1344_RS03135_ | -4.3168 | -4.31809 | -4.31489 | -0.01702 | 0.831527 | -0.01909 | 0.491006 | 0.598269 | 0.722674 | 0.7116 | 0.800914 |
| SL1344_RS03140_ | -5.08943 | -4.87431 | -5.31442 | -0.52052 | 1.292882 | -0.39224 | 0.323333 | 0.497408 | 0.639087 | 0.6337 | 0.733859 |
| SL1344_RS03145_ | -6.93581 | -7.26709 | -6.65076 | 0.317208 | 3.686968 | 0.067811 | 0.462667 | 0.667932 | 0.781937 | 0.8144 | 0.877253 |
| SL1344_RS03150_ | -4.7731 | -4.40299 | -5.20883 | -0.7913 | 0.977056 | -0.75867 | 0.185876 | 0.294897 | 0.45061 | 0.3946 | 0.533642 |
| SL1344_RS03165_ | -0.2536 | -0.63969 | 0.136815 | 0.798856 | 0.226258 | 3.447284 | 4.68E-05 | 0.012092 | 0.078801 | 0.1 | 0.236592 |
| SL1344_RS03175_ | 5.592121 | 5.992465 | 5.401922 | -0.59213 | 0.145752 | -3.34502 | 4.68E-05 | 0.017566 | 0.100612 | 0.1 | 0.236592 |
| SL1344_RS03180_ | -0.50365 | -0.59132 | -0.33685 | 0.285802 | 0.383499 | 0.729917 | 0.232 | 0.327707 | 0.49971 | 0.4434 | 0.594952 |
| SL1344_RS03195_ | 2.723341 | 2.816254 | 2.508325 | -0.29802 | 0.098958 | -2.725 | 4.68E-05 | 0.034713 | 0.134774 | 0.1 | 0.236592 |

| | | | | | | | | | | | |
|---|---|---|---|---|---|---|---|---|---|---|---|
| SL1344_RS03200_ | -0.74698 | -1.29111 | -0.56677 | 0.682481 | 0.354647 | 1.61054 | 0.047333 | 0.10204 | 0.247699 | 0.158 | 0.30614 |
| SL1344_RS03205_ | 0.314645 | 0.583927 | -0.09153 | -0.67381 | 0.177998 | -3.76245 | 4.68E-05 | 0.012583 | 0.078013 | 0.1 | 0.236592 |
| SL1344_RS03210_ | 3.201093 | 3.359184 | 2.941181 | -0.44167 | 0.176776 | -2.64598 | 4.68E-05 | 0.014912 | 0.091841 | 0.1 | 0.236592 |
| SL1344_RS03215_ | 2.342541 | 2.368257 | 2.181761 | -0.175 | 0.129315 | -0.98224 | 0.216667 | 0.262502 | 0.441951 | 0.4539 | 0.603632 |
| SL1344_RS03225_ | 1.05657 | 0.968709 | 1.124072 | 0.132203 | 0.170593 | 0.736537 | 0.212525 | 0.33078 | 0.49597 | 0.4259 | 0.572526 |
| SL1344_RS03230_ | 1.612017 | 1.613768 | 1.61057 | -0.00296 | 0.096498 | -0.02564 | 0.486667 | 0.57252 | 0.697287 | 0.6796 | 0.771995 |
| SL1344_RS03235_ | 1.107124 | 1.263689 | 0.831067 | -0.41501 | 0.168658 | -2.29126 | 0.005331 | 0.036864 | 0.137802 | 0.1039 | 0.241735 |
| SL1344_RS03240_ | 0.881008 | 1.076665 | 0.708262 | -0.36714 | 0.130974 | -2.71982 | 0.002002 | 0.021036 | 0.099978 | 0.1017 | 0.238694 |
| SL1344_RS03245_ | -1.48952 | -1.15345 | -1.88136 | -0.72896 | 0.297223 | -2.42213 | 0.005334 | 0.030846 | 0.120394 | 0.1062 | 0.244286 |
| SL1344_RS03250_ | -0.38563 | -0.47117 | -0.28363 | 0.199965 | 0.258354 | 0.740745 | 0.20453 | 0.315142 | 0.475879 | 0.4149 | 0.555992 |
| SL1344_RS03255_ | 0.329411 | 0.412826 | 0.259919 | -0.17442 | 0.203905 | -0.84921 | 0.179333 | 0.269437 | 0.434956 | 0.3809 | 0.524485 |
| SL1344_RS03260_ | 0.275788 | 0.221622 | 0.379404 | 0.149657 | 0.265492 | 0.550339 | 0.268 | 0.443159 | 0.601319 | 0.5247 | 0.664889 |
| SL1344_RS03265_ | 2.023041 | 2.038482 | 1.986272 | -0.06113 | 0.157177 | -0.33152 | 0.373085 | 0.572885 | 0.712778 | 0.7345 | 0.824874 |
| SL1344_RS03270_ | 2.827402 | 2.813301 | 2.845206 | 0.022679 | 0.100447 | 0.202904 | 0.4 | 0.688686 | 0.797101 | 0.7648 | 0.846414 |
| SL1344_RS03275_ | 4.203745 | 4.202719 | 4.20507 | 0.005349 | 0.046546 | 0.112935 | 0.447702 | 0.598358 | 0.723039 | 0.7132 | 0.802625 |
| SL1344_RS03280_ | -5.04735 | -4.90923 | -5.25239 | -0.38068 | 1.225712 | -0.2731 | 0.358667 | 0.527035 | 0.664428 | 0.6454 | 0.745121 |
| SL1344_RS03285_ | -2.2509 | -2.30971 | -2.19711 | 0.115611 | 0.396694 | 0.263842 | 0.38 | 0.545503 | 0.677731 | 0.6574 | 0.756378 |
| SL1344_RS03290_ | -0.96626 | -1.06705 | -0.85275 | 0.204537 | 0.26548 | 0.716325 | 0.213333 | 0.329583 | 0.483949 | 0.4301 | 0.566688 |
| SL1344_RS03295_ | 1.842315 | 1.696793 | 1.968267 | 0.268105 | 0.107615 | 2.46625 | 0.004001 | 0.029181 | 0.119519 | 0.1038 | 0.241548 |
| SL1344_RS03300_ | 1.961526 | 1.325096 | 2.478632 | 1.16233 | 0.198349 | 5.90878 | 4.68E-05 | 0.002702 | 0.04383 | 0.1 | 0.236592 |
| SL1344_RS03305_ | -1.83059 | -1.8562 | -1.79232 | 0.088359 | 0.364522 | 0.221608 | 0.404667 | 0.569189 | 0.695779 | 0.6807 | 0.775304 |
| SL1344_RS03310_ | -3.52141 | -3.85927 | -3.31474 | 0.453068 | 0.895702 | 0.475749 | 0.308461 | 0.4873 | 0.638872 | 0.6018 | 0.721475 |
| SL1344_RS03315_ | -2.11431 | -2.14828 | -2.08139 | 0.084916 | 0.436459 | 0.179671 | 0.414 | 0.583967 | 0.710886 | 0.7069 | 0.794584 |
| SL1344_RS03320_ | -3.36105 | -3.38232 | -3.34342 | 0.037231 | 0.601647 | 0.058333 | 0.470353 | 0.605442 | 0.726735 | 0.7154 | 0.802914 |
| SL1344_RS03325_ | -1.4993 | -1.515 | -1.48441 | 0.025314 | 0.438261 | 0.049587 | 0.476667 | 0.709129 | 0.811629 | 0.7879 | 0.861912 |
| SL1344_RS03330_ | 0.851349 | 0.871968 | 0.836487 | -0.02937 | 0.167661 | -0.16391 | 0.429714 | 0.643435 | 0.760624 | 0.7523 | 0.833441 |
| SL1344_RS03335_ | 1.268196 | 1.353374 | 1.18121 | -0.18005 | 0.131747 | -1.3179 | 0.065956 | 0.128161 | 0.272918 | 0.1892 | 0.336893 |
| SL1344_RS03340_ | 1.301331 | 1.409111 | 1.196771 | -0.25729 | 0.202064 | -1.23739 | 0.062625 | 0.120706 | 0.272365 | 0.1747 | 0.324537 |
| SL1344_RS03350_ | 0.259826 | 0.569584 | -0.1838 | -0.7514 | 0.203467 | -3.52959 | 4.68E-05 | 0.015535 | 0.087742 | 0.1 | 0.236592 |
| SL1344_RS03355_ | 0.414581 | 0.907921 | -0.17154 | -1.06902 | 0.186598 | -5.71789 | 4.68E-05 | 0.00319 | 0.044612 | 0.1 | 0.236592 |
| SL1344_RS03360_ | 6.761382 | 7.032874 | 6.570262 | -0.50583 | 0.15706 | -2.7166 | 4.68E-05 | 0.060243 | 0.190341 | 0.1 | 0.236592 |
| SL1344_RS03365_ | 2.509636 | 2.540378 | 2.468107 | -0.07265 | 0.073245 | -0.96234 | 0.139241 | 0.220274 | 0.370228 | 0.3093 | 0.450664 |
| SL1344_RS03370_ | 2.986972 | 3.015202 | 2.955586 | -0.06862 | 0.078365 | -0.85346 | 0.178548 | 0.257452 | 0.422054 | 0.3619 | 0.506295 |
| SL1344_RS03375_ | 2.202099 | 2.160765 | 2.286736 | 0.123326 | 0.116355 | 1.005992 | 0.139333 | 0.218615 | 0.384428 | 0.3099 | 0.462801 |
| SL1344_RS03380_ | 2.760706 | 2.803359 | 2.715756 | -0.09846 | 0.093649 | -1.02735 | 0.110593 | 0.185407 | 0.345097 | 0.2649 | 0.412876 |
| SL1344_RS03385_ | 2.62151 | 2.750987 | 2.539323 | -0.20787 | 0.099664 | -1.99382 | 0.014667 | 0.045578 | 0.154759 | 0.1114 | 0.251264 |
| SL1344_RS03395_ | -2.49918 | -2.45227 | -2.55215 | -0.10416 | 0.441459 | -0.23897 | 0.402398 | 0.554445 | 0.685007 | 0.6603 | 0.760994 |
| SL1344_RS03400_ | 1.927098 | 1.999077 | 1.864376 | -0.19524 | 0.202468 | -1.00693 | 0.093937 | 0.171926 | 0.340702 | 0.2321 | 0.381855 |
| SL1344_RS03405_ | 1.669985 | 1.706085 | 1.444285 | -0.24939 | 0.31891 | -0.67709 | 0.260667 | 0.400932 | 0.580287 | 0.4922 | 0.655716 |
| SL1344_RS03410_ | 1.701672 | 1.830614 | 1.579678 | -0.26803 | 0.134465 | -1.97375 | 0.006663 | 0.046483 | 0.156067 | 0.1052 | 0.243357 |
| SL1344_RS03415_ | 2.228418 | 2.270508 | 2.057757 | -0.20606 | 0.253527 | -0.64956 | 0.313791 | 0.461319 | 0.633085 | 0.6295 | 0.764396 |
| SL1344_RS03420_ | 3.24815 | 3.275027 | 3.076731 | -0.19386 | 0.196552 | -0.8263 | 0.247835 | 0.333498 | 0.519602 | 0.4608 | 0.630467 |
| SL1344_RS03425_ | 3.214094 | 3.264459 | 3.020909 | -0.21091 | 0.263961 | -0.88591 | 0.196 | 0.216179 | 0.39813 | 0.3521 | 0.531231 |
| SL1344_RS03430_ | 2.373623 | 2.682053 | 2.16762 | -0.44097 | 0.273284 | -1.54858 | 0.068 | 0.080806 | 0.222236 | 0.1605 | 0.317257 |
| SL1344_RS03440_ | 1.443733 | 1.582249 | 1.304733 | -0.27899 | 0.094361 | -2.91079 | 0.000673 | 0.018375 | 0.091561 | 0.1008 | 0.237649 |

| | | | | | | | | | | | |
|---|---|---|---|---|---|---|---|---|---|---|---|
| SL1344_RS03445_ | 1.788814 | 1.73693 | 1.870273 | 0.131579 | 0.121884 | 1.061354 | 0.143904 | 0.207445 | 0.367945 | 0.291 | 0.446686 |
| SL1344_RS03450_ | 2.291848 | 2.288216 | 2.293382 | -0.00482 | 0.093818 | -0.05284 | 0.482 | 0.652406 | 0.766195 | 0.7629 | 0.840327 |
| SL1344_RS03455_ | 1.372009 | 1.240136 | 1.534195 | 0.291966 | 0.129387 | 2.225062 | 0.005997 | 0.032537 | 0.127283 | 0.1037 | 0.241359 |
| SL1344_RS03460_ | 3.806363 | 3.638031 | 4.049686 | 0.381256 | 0.113359 | 3.333709 | 4.68E-05 | 0.009479 | 0.073685 | 0.1 | 0.236592 |
| SL1344_RS03465_ | 3.574954 | 3.600515 | 3.449706 | -0.1365 | 0.09848 | -1.05003 | 0.186542 | 0.219597 | 0.397312 | 0.3609 | 0.52755 |
| SL1344_RS03475_ | 0.207342 | -0.16241 | 0.44556 | 0.629522 | 0.266516 | 2.33264 | 4.68E-05 | 0.057557 | 0.179978 | 0.1003 | 0.236983 |
| SL1344_RS03480_ | -1.17221 | -1.20189 | -1.13106 | 0.078526 | 0.280146 | 0.270995 | 0.378667 | 0.545424 | 0.675949 | 0.655 | 0.752816 |
| SL1344_RS03490_ | -1.86582 | -1.9255 | -1.82174 | 0.092818 | 0.355636 | 0.256982 | 0.39 | 0.553836 | 0.684606 | 0.6673 | 0.764877 |
| SL1344_RS03495_ | -0.22408 | -0.22316 | -0.2245 | 0.002886 | 0.22807 | 0.013062 | 0.494667 | 0.654487 | 0.765615 | 0.7657 | 0.840008 |
| SL1344_RS03505_ | 2.696714 | 3.098449 | 2.38409 | -0.6647 | 0.223924 | -2.9204 | 4.68E-05 | 0.01649 | 0.096349 | 0.1 | 0.236592 |
| SL1344_RS03510_ | -1.96686 | -1.8419 | -2.09832 | -0.35553 | 0.572676 | -0.57918 | 0.252498 | 0.355096 | 0.52557 | 0.5103 | 0.637938 |
| SL1344_RS03515_ | 0.661145 | 0.716775 | 0.590403 | -0.14228 | 0.194174 | -0.72388 | 0.201333 | 0.306239 | 0.474049 | 0.4187 | 0.555665 |
| SL1344_RS03520_ | 1.830922 | 1.950785 | 1.700157 | -0.24857 | 0.088387 | -2.71926 | 0.002001 | 0.021124 | 0.099562 | 0.1012 | 0.238077 |
| SL1344_RS03525_ | 4.371503 | 4.323046 | 4.425753 | 0.104055 | 0.047894 | 2.086969 | 0.009994 | 0.043306 | 0.148021 | 0.1104 | 0.249746 |
| SL1344_RS03530_ | 1.04859 | 1.083691 | 1.013395 | -0.14127 | 0.255811 | -0.5603 | 0.278 | 0.366332 | 0.538745 | 0.571 | 0.687616 |
| SL1344_RS03535_ | 2.458003 | 2.428382 | 2.579132 | 0.174171 | 0.305286 | 0.633659 | 0.234 | 0.350621 | 0.536048 | 0.4238 | 0.600277 |
| SL1344_RS03540_ | 2.944054 | 2.979601 | 2.893932 | 0.093105 | 0.336701 | 0.306717 | 0.442 | 0.727969 | 0.833787 | 0.9889 | 0.99327 |
| SL1344_RS03545_ | -6.18548 | -6.34773 | -6.04576 | 0.282246 | 1.744486 | 0.147456 | 0.430667 | 0.585602 | 0.716317 | 0.7184 | 0.808418 |
| SL1344_RS03550_ | -0.24531 | -0.42336 | -0.16415 | 0.248351 | 0.265325 | 0.864013 | 0.187333 | 0.27473 | 0.445042 | 0.3836 | 0.532123 |
| SL1344_RS03555_ | 0.750068 | 0.734921 | 0.775251 | 0.05562 | 0.160841 | 0.319188 | 0.371086 | 0.539801 | 0.674265 | 0.6658 | 0.760559 |
| SL1344_RS03560_ | -2.0289 | -1.99867 | -2.04456 | -0.03965 | 0.425889 | -0.08196 | 0.464667 | 0.63972 | 0.756605 | 0.7507 | 0.832728 |
| SL1344_RS03565_ | -2.58168 | -2.45567 | -2.68686 | -0.21779 | 0.504919 | -0.40497 | 0.336 | 0.518993 | 0.654976 | 0.6304 | 0.736644 |
| SL1344_RS03570_ | -3.76668 | -3.54367 | -3.90054 | -0.44136 | 0.84339 | -0.50174 | 0.299333 | 0.452594 | 0.602346 | 0.5812 | 0.695549 |
| SL1344_RS03575_ | -8.88287 | -8.85729 | -8.90879 | 0.02283 | 3.84365 | 0.006704 | 0.497333 | 0.555544 | 0.687699 | 0.686 | 0.779036 |
| SL1344_RS03580_ | 1.380156 | 1.356008 | 1.421939 | 0.063434 | 0.280767 | 0.253566 | 0.36509 | 0.71161 | 0.816734 | 0.7093 | 0.808366 |
| SL1344_RS03585_ | -0.1959 | -0.35614 | -0.07242 | 0.284089 | 0.193485 | 1.398698 | 0.06 | 0.109881 | 0.248402 | 0.1705 | 0.31791 |
| SL1344_RS03590_ | -1.13297 | -1.15376 | -1.10846 | 0.050934 | 0.23114 | 0.21683 | 0.408 | 0.516319 | 0.648208 | 0.6347 | 0.734104 |
| SL1344_RS03595_ | 1.327881 | 1.124693 | 1.452546 | 0.323791 | 0.131342 | 2.366795 | 0.003335 | 0.040001 | 0.142715 | 0.1033 | 0.240884 |
| SL1344_RS03600_ | 0.868761 | 0.699864 | 1.067779 | 0.320624 | 0.231303 | 1.367492 | 0.079333 | 0.110052 | 0.258328 | 0.1814 | 0.339369 |
| SL1344_RS03605_ | 1.288734 | 1.137026 | 1.44352 | 0.306554 | 0.099728 | 3.007394 | 0.000673 | 0.016288 | 0.087763 | 0.1002 | 0.23686 |
| SL1344_RS03610_ | 0.988479 | 0.888714 | 1.081357 | 0.192935 | 0.128275 | 1.451087 | 0.044 | 0.095987 | 0.22607 | 0.1594 | 0.304132 |
| SL1344_RS03615_ | -4.84494 | -5.04269 | -4.72164 | 0.346624 | 1.044416 | 0.280897 | 0.356429 | 0.505408 | 0.645446 | 0.6224 | 0.725222 |
| SL1344_RS03620_ | -1.52414 | -1.30815 | -2.03754 | -0.70271 | 0.459058 | -1.45084 | 0.071952 | 0.113353 | 0.261547 | 0.1904 | 0.344246 |
| SL1344_RS03625_ | -4.27061 | -4.04253 | -4.6777 | -0.60366 | 0.858041 | -0.64699 | 0.234 | 0.351527 | 0.508574 | 0.4732 | 0.606498 |
| SL1344_RS03630_ | -2.9755 | -2.91498 | -3.02714 | -0.13437 | 0.57407 | -0.22294 | 0.403333 | 0.58178 | 0.707335 | 0.7049 | 0.792961 |
| SL1344_RS03640_ | -4.87374 | -4.34252 | -5.28357 | -0.8897 | 1.042676 | -0.79173 | 0.193333 | 0.302219 | 0.454999 | 0.388 | 0.529742 |
| SL1344_RS03645_ | -3.5839 | -3.46692 | -3.74472 | -0.27197 | 0.59065 | -0.44062 | 0.303131 | 0.445447 | 0.587324 | 0.5592 | 0.672864 |
| SL1344_RS03650_ | -4.16362 | -4.1298 | -4.19823 | -0.2096 | 1.035669 | -0.20124 | 0.413333 | 0.619717 | 0.740875 | 0.7731 | 0.843748 |
| SL1344_RS03655_ | -4.18673 | -4.1789 | -4.19138 | -0.04332 | 0.817285 | -0.05253 | 0.475017 | 0.623127 | 0.741957 | 0.7279 | 0.812635 |
| SL1344_RS03660_ | -3.02439 | -2.91471 | -3.14765 | -0.21996 | 0.596465 | -0.35443 | 0.352667 | 0.534673 | 0.669637 | 0.6497 | 0.751202 |
| SL1344_RS03665_ | -4.26724 | -4.13374 | -4.41665 | -0.30303 | 0.806796 | -0.34542 | 0.347102 | 0.487933 | 0.629774 | 0.6117 | 0.717758 |
| SL1344_RS03670_ | -2.87543 | -2.66595 | -2.9864 | -0.35013 | 0.681461 | -0.45645 | 0.305796 | 0.489035 | 0.636549 | 0.6063 | 0.722976 |
| SL1344_RS03675_ | -1.37443 | -1.34733 | -1.42133 | -0.04434 | 0.375044 | -0.11508 | 0.456667 | 0.678598 | 0.787717 | 0.779 | 0.857294 |
| SL1344_RS03680_ | 0.797639 | 0.967684 | 0.651019 | -0.31514 | 0.12832 | -2.38519 | 0.003998 | 0.030095 | 0.118816 | 0.1043 | 0.241989 |
| SL1344_RS03685_ | 0.229102 | 0.226939 | 0.233019 | 0.002733 | 0.17252 | 0.017742 | 0.493005 | 0.588902 | 0.714544 | 0.7113 | 0.799956 |

| | | | | | | | | | | | |
|---|---|---|---|---|---|---|---|---|---|---|---|
| SL1344_RS03690_ | 5.547588 | 5.674588 | 5.326076 | -0.30279 | 0.132721 | -2.44308 | 4.68E-05 | 0.021754 | 0.11091 | 0.1 | 0.236592 |
| SL1344_RS03695_ | -3.72735 | -3.59026 | -3.87051 | -0.35575 | 0.896329 | -0.34372 | 0.347333 | 0.543477 | 0.682499 | 0.678 | 0.774999 |
| SL1344_RS03705_ | 3.125454 | 3.21267 | 2.900055 | -0.28581 | 0.219146 | -1.15848 | 0.128581 | 0.165828 | 0.338854 | 0.2341 | 0.404901 |
| SL1344_RS03710_ | 4.718096 | 4.729681 | 4.705882 | 0.078238 | 0.245043 | 0.323117 | 0.422385 | 0.641826 | 0.775299 | 0.9397 | 0.964242 |
| SL1344_RS03715_ | 5.193368 | 5.13286 | 5.254607 | 0.048691 | 0.14706 | 0.401293 | 0.358 | 0.559764 | 0.712441 | 0.7747 | 0.855086 |
| SL1344_RS03720_ | 6.301724 | 6.335987 | 6.241004 | -0.04946 | 0.165077 | -0.4091 | 0.288667 | 0.616907 | 0.757277 | 0.5872 | 0.730649 |
| SL1344_RS03725_ | 6.207988 | 6.214036 | 6.203769 | 0.000832 | 0.067713 | 0.012753 | 0.496 | 0.77347 | 0.861997 | 0.9565 | 0.97302 |
| SL1344_RS03730_ | 5.616677 | 5.62211 | 5.612444 | -0.01629 | 0.080842 | -0.17716 | 0.416667 | 0.582483 | 0.729038 | 0.8633 | 0.914007 |
| SL1344_RS03735_ | 5.984254 | 6.000753 | 5.95532 | -0.04097 | 0.101393 | -0.39356 | 0.357761 | 0.696351 | 0.812202 | 0.7549 | 0.850119 |
| SL1344_RS03740_ | 2.527068 | 2.544507 | 2.513146 | 0.057689 | 0.454906 | 0.157578 | 0.426 | 0.773621 | 0.864578 | 0.9091 | 0.942511 |
| SL1344_RS03745_ | 7.020599 | 6.931555 | 7.111956 | 0.183463 | 0.049168 | 3.813743 | 4.68E-05 | 0.011956 | 0.08235 | 0.1 | 0.236592 |
| SL1344_RS03750_ | 5.924376 | 5.762051 | 6.071141 | 0.312288 | 0.084934 | 3.501297 | 4.68E-05 | 0.069671 | 0.206052 | 0.1 | 0.236592 |
| SL1344_RS03755_ | 5.47978 | 5.316438 | 5.711723 | 0.324012 | 0.163122 | 2.144724 | 4.68E-05 | 0.030024 | 0.130827 | 0.1 | 0.236592 |
| SL1344_RS03760_ | 5.700132 | 5.558486 | 5.800327 | 0.237455 | 0.06647 | 3.454913 | 4.68E-05 | 0.014927 | 0.090971 | 0.1 | 0.236592 |
| SL1344_RS03765_ | 4.788335 | 4.906202 | 4.712938 | -0.19934 | 0.080798 | -2.51791 | 4.68E-05 | 0.037245 | 0.143585 | 0.1 | 0.236592 |
| SL1344_RS03770_ | 0.472269 | 0.59798 | 0.321544 | -0.27763 | 0.154992 | -1.7228 | 0.028 | 0.076199 | 0.198372 | 0.134 | 0.276307 |
| SL1344_RS03775_ | 1.783017 | 1.845089 | 1.559748 | -0.26991 | 0.141309 | -1.54589 | 0.057295 | 0.111772 | 0.260863 | 0.1691 | 0.315404 |
| SL1344_RS03780_ | 1.347329 | 1.314355 | 1.380308 | 0.059507 | 0.129345 | 0.448465 | 0.321785 | 0.518187 | 0.654305 | 0.6237 | 0.730461 |
| SL1344_RS03790_ | 4.637994 | 4.751708 | 4.620967 | -0.11716 | 0.109348 | -0.69625 | 0.309333 | 0.447003 | 0.621654 | 0.6433 | 0.776771 |
| SL1344_RS03795_ | 6.039465 | 6.1002 | 5.953568 | -0.12092 | 0.131553 | -0.91237 | 0.136667 | 0.220583 | 0.404669 | 0.2426 | 0.414783 |
| SL1344_RS03800_ | 4.805106 | 4.84475 | 4.653065 | -0.15135 | 0.122275 | -0.66566 | 0.333333 | 0.494192 | 0.662535 | 0.7 | 0.822108 |
| SL1344_RS03805_ | 1.408736 | 1.405812 | 1.41101 | -0.07596 | 0.246805 | -0.31029 | 0.388667 | 0.615388 | 0.746122 | 0.827 | 0.887139 |
| SL1344_RS03810_ | 1.888203 | 1.932163 | 1.767563 | -0.14788 | 0.147802 | -0.91105 | 0.184667 | 0.26846 | 0.443447 | 0.3766 | 0.532616 |
| SL1344_RS03815_ | 0.9024 | 0.97406 | 0.845837 | -0.13266 | 0.135557 | -0.91454 | 0.141905 | 0.236283 | 0.391736 | 0.3222 | 0.465113 |
| SL1344_RS03820_ | 1.020152 | 1.111064 | 0.932218 | -0.1854 | 0.155619 | -1.12896 | 0.093333 | 0.161827 | 0.313569 | 0.232 | 0.382784 |
| SL1344_RS03825_ | 0.372779 | 0.469812 | 0.268995 | -0.20136 | 0.136979 | -1.40808 | 0.062625 | 0.11324 | 0.244993 | 0.181 | 0.325308 |
| SL1344_RS03830_ | -1.74078 | -2.03059 | -1.60202 | 0.403471 | 0.387723 | 0.967479 | 0.159227 | 0.236524 | 0.400924 | 0.3378 | 0.488542 |
| SL1344_RS03835_ | -0.2786 | -0.37339 | -0.19831 | 0.170705 | 0.177822 | 0.913514 | 0.148 | 0.238466 | 0.385636 | 0.3252 | 0.464953 |
| SL1344_RS03840_ | 0.203804 | 0.103447 | 0.297857 | 0.200223 | 0.160936 | 1.15191 | 0.105263 | 0.161759 | 0.30795 | 0.2373 | 0.383924 |
| SL1344_RS03845_ | 0.823117 | 0.302372 | 1.136275 | 0.824906 | 0.221533 | 3.693238 | 4.68E-05 | 0.008396 | 0.067889 | 0.1 | 0.236592 |
| SL1344_RS03850_ | 2.39914 | 2.392366 | 2.406928 | -0.00957 | 0.129669 | -0.07856 | 0.467333 | 0.69798 | 0.804284 | 0.839 | 0.895785 |
| SL1344_RS03855_ | -0.55536 | -0.55329 | -0.56062 | -0.01448 | 0.255128 | -0.0483 | 0.478348 | 0.658914 | 0.771341 | 0.7647 | 0.843269 |
| SL1344_RS03860_ | -0.34697 | -0.26719 | -0.44842 | -0.17755 | 0.205206 | -0.81246 | 0.187209 | 0.27731 | 0.431847 | 0.3794 | 0.518261 |
| SL1344_RS03865_ | 0.079507 | 0.138236 | 0.038919 | -0.09572 | 0.200639 | -0.44298 | 0.320453 | 0.489164 | 0.628817 | 0.5989 | 0.709469 |
| SL1344_RS03870_ | -2.32932 | -2.26632 | -2.40921 | -0.21456 | 0.541384 | -0.37628 | 0.332445 | 0.517162 | 0.660623 | 0.6712 | 0.765414 |
| SL1344_RS03875_ | -4.25443 | -3.86122 | -4.61133 | -0.66978 | 1.063533 | -0.59415 | 0.242 | 0.406083 | 0.56066 | 0.4968 | 0.630646 |
| SL1344_RS03880_ | -5.36085 | -5.3765 | -5.3402 | 0.029469 | 1.37993 | 0.020157 | 0.49034 | 0.629386 | 0.748601 | 0.7495 | 0.829837 |
| SL1344_RS03885_ | -2.34309 | -2.34237 | -2.3436 | -0.02642 | 0.45368 | -0.05647 | 0.479014 | 0.628892 | 0.746902 | 0.7423 | 0.82416 |
| SL1344_RS03890_ | -2.62735 | -2.49763 | -2.75408 | -0.31544 | 0.588188 | -0.49598 | 0.293804 | 0.450513 | 0.600647 | 0.5863 | 0.697793 |
| SL1344_RS03895_ | -1.18104 | -1.02005 | -1.35925 | -0.33893 | 0.23746 | -1.36872 | 0.056629 | 0.114332 | 0.242523 | 0.1812 | 0.325069 |
| SL1344_RS03900_ | -2.36967 | -2.35214 | -2.40438 | -0.08498 | 0.474923 | -0.16221 | 0.429047 | 0.588865 | 0.716006 | 0.7134 | 0.799536 |
| SL1344_RS03905_ | -2.73821 | -2.89305 | -2.65649 | 0.165517 | 0.644207 | 0.247706 | 0.394 | 0.639148 | 0.759482 | 0.7606 | 0.841533 |
| SL1344_RS03915_ | 2.467009 | 2.432695 | 2.584926 | 0.142932 | 0.103303 | 1.168524 | 0.137908 | 0.187379 | 0.35291 | 0.2923 | 0.44683 |
| SL1344_RS03920_ | 1.824677 | 1.695019 | 1.939528 | 0.244262 | 0.100696 | 2.375799 | 0.004001 | 0.030814 | 0.121955 | 0.1024 | 0.2398 |
| SL1344_RS03925_ | 1.017553 | 0.748649 | 1.206236 | 0.462144 | 0.129696 | 3.520867 | 4.68E-05 | 0.014852 | 0.084231 | 0.1 | 0.236592 |

| | | | | | | | | | | | |
|---|---|---|---|---|---|---|---|---|---|---|---|
| SL1344_RS03930_ | 0.98912 | 0.698954 | 1.183251 | 0.496517 | 0.247366 | 2.022959 | 0.002668 | 0.036432 | 0.140135 | 0.1029 | 0.240414 |
| SL1344_RS03935_ | -3.38013 | -3.50625 | -3.21294 | 0.285911 | 0.569978 | 0.48777 | 0.296469 | 0.449395 | 0.590574 | 0.5422 | 0.662284 |
| SL1344_RS03940_ | 1.537743 | 1.536804 | 1.539213 | 0.002495 | 0.125045 | 0.018565 | 0.491672 | 0.640598 | 0.755359 | 0.7492 | 0.82808 |
| SL1344_RS03945_ | 0.882243 | 0.917111 | 0.837435 | -0.07726 | 0.137256 | -0.53772 | 0.275816 | 0.437196 | 0.581304 | 0.5475 | 0.665485 |
| SL1344_RS03950_ | 3.837252 | 4.051398 | 3.541803 | -0.73801 | 0.375363 | -2.08508 | 4.68E-05 | 0.022787 | 0.114206 | 0.1 | 0.236592 |
| SL1344_RS03970_ | 0.824542 | 0.687639 | 1.038591 | 0.335887 | 0.263898 | 1.271037 | 0.077282 | 0.121498 | 0.274882 | 0.1857 | 0.343247 |
| SL1344_RS03975_ | 1.603077 | 1.411151 | 1.802241 | 0.388726 | 0.102786 | 3.704498 | 4.68E-05 | 0.009263 | 0.068648 | 0.1 | 0.236592 |
| SL1344_RS03980_ | 2.969836 | 2.890371 | 3.03874 | 0.113843 | 0.16748 | 0.643978 | 0.231333 | 0.357397 | 0.537727 | 0.4599 | 0.610105 |
| SL1344_RS03985_ | 1.959491 | 1.9314 | 2.005598 | 0.067198 | 0.113088 | 0.565922 | 0.287333 | 0.447265 | 0.59823 | 0.5665 | 0.689864 |
| SL1344_RS03990_ | 1.401434 | 1.274973 | 1.508478 | 0.224605 | 0.123795 | 1.795616 | 0.027315 | 0.064457 | 0.182004 | 0.1326 | 0.277372 |
| SL1344_RS03995_ | 0.729917 | 0.623844 | 0.832874 | 0.233484 | 0.179273 | 1.268943 | 0.066667 | 0.129086 | 0.277133 | 0.1939 | 0.34217 |
| SL1344_RS04000_ | 0.78341 | 0.960821 | 0.531604 | -0.43067 | 0.148487 | -2.80962 | 4.68E-05 | 0.021699 | 0.102441 | 0.1003 | 0.236991 |
| SL1344_RS04005_ | 1.11241 | 1.241123 | 0.991226 | -0.23272 | 0.15759 | -1.42567 | 0.069287 | 0.10671 | 0.246577 | 0.1775 | 0.329793 |
| SL1344_RS04010_ | 0.698634 | 0.588434 | 0.778122 | 0.185655 | 0.138162 | 1.28148 | 0.079333 | 0.140193 | 0.280502 | 0.2178 | 0.363556 |
| SL1344_RS04015_ | -2.87237 | -2.86325 | -2.88209 | -0.02073 | 0.428591 | -0.04307 | 0.479333 | 0.553107 | 0.682723 | 0.6711 | 0.765909 |
| SL1344_RS04020_ | -2.81401 | -2.69024 | -3.10099 | -0.32886 | 0.650772 | -0.47012 | 0.326667 | 0.518395 | 0.663151 | 0.6401 | 0.75291 |
| SL1344_RS04025_ | -3.15742 | -2.84288 | -3.35964 | -0.48274 | 0.531734 | -0.83814 | 0.172552 | 0.273452 | 0.428363 | 0.3732 | 0.516573 |
| SL1344_RS04030_ | -3.67276 | -3.76864 | -3.56857 | 0.164398 | 0.686244 | 0.22541 | 0.391073 | 0.577837 | 0.703769 | 0.6795 | 0.776135 |
| SL1344_RS04035_ | -3.86332 | -4.14618 | -3.72222 | 0.380216 | 0.727594 | 0.464978 | 0.299134 | 0.454941 | 0.601162 | 0.5741 | 0.691646 |
| SL1344_RS04040_ | 2.28894 | 2.391582 | 2.241198 | -0.14634 | 0.109646 | -1.20067 | 0.109927 | 0.157836 | 0.314829 | 0.2337 | 0.387487 |
| SL1344_RS04045_ | -2.18933 | -1.92972 | -2.40568 | -0.46495 | 0.373085 | -1.18093 | 0.104 | 0.162753 | 0.307263 | 0.2423 | 0.390119 |
| SL1344_RS04050_ | 2.039865 | 2.025614 | 2.080151 | 0.045081 | 0.118404 | 0.323469 | 0.38 | 0.644381 | 0.762702 | 0.7358 | 0.829298 |
| SL1344_RS04055_ | 1.01397 | 0.885508 | 1.152385 | 0.267375 | 0.138647 | 1.896292 | 0.01599 | 0.055423 | 0.168066 | 0.1182 | 0.259542 |
| SL1344_RS04065_ | 2.84583 | 2.859467 | 2.829372 | -0.04826 | 0.104605 | -0.40826 | 0.30513 | 0.435948 | 0.59934 | 0.6072 | 0.714605 |
| SL1344_RS04070_ | 2.188167 | 2.25043 | 2.101868 | -0.11586 | 0.143758 | -0.80715 | 0.205863 | 0.304255 | 0.47839 | 0.3942 | 0.551068 |
| SL1344_RS04085_ | 4.21408 | 4.212758 | 4.215928 | 0.019985 | 0.081349 | 0.238504 | 0.4004 | 0.664285 | 0.778883 | 0.7942 | 0.863087 |
| SL1344_RS04090_ | -3.92011 | -3.61247 | -4.4644 | -0.93095 | 1.033813 | -0.81577 | 0.138 | 0.245466 | 0.418232 | 0.2947 | 0.44317 |
| SL1344_RS04095_ | -3.85205 | -3.78956 | -3.96573 | -0.15462 | 0.699582 | -0.2003 | 0.414 | 0.579324 | 0.706533 | 0.6879 | 0.784724 |
| SL1344_RS04100_ | -3.17542 | -3.16049 | -3.18961 | -0.02818 | 0.479665 | -0.0473 | 0.476349 | 0.561589 | 0.688834 | 0.6701 | 0.766909 |
| SL1344_RS04105_ | -0.82302 | -1.06595 | -0.66051 | 0.37861 | 0.298285 | 1.212236 | 0.109927 | 0.158168 | 0.310844 | 0.2392 | 0.392546 |
| SL1344_RS04110_ | -1.22728 | -1.42627 | -1.06126 | 0.361271 | 0.238848 | 1.458733 | 0.06 | 0.103324 | 0.230931 | 0.1743 | 0.318251 |
| SL1344_RS04115_ | -2.32835 | -2.44606 | -2.22561 | 0.22278 | 0.430049 | 0.491892 | 0.302465 | 0.454848 | 0.598576 | 0.5648 | 0.681935 |
| SL1344_RS04120_ | 3.223346 | 3.166615 | 3.363955 | 0.154989 | 0.224308 | 0.772013 | 0.195869 | 0.288137 | 0.473048 | 0.3652 | 0.543662 |
| SL1344_RS04125_ | 0.360328 | 0.203622 | 0.576033 | 0.341834 | 0.202035 | 1.623131 | 0.046636 | 0.080785 | 0.209926 | 0.1504 | 0.298557 |
| SL1344_RS04130_ | 0.527707 | 0.336294 | 0.68335 | 0.360126 | 0.194892 | 1.812942 | 0.013991 | 0.055774 | 0.171283 | 0.116 | 0.256881 |
| SL1344_RS04135_ | 1.369585 | 1.147265 | 1.578431 | 0.43813 | 0.130243 | 3.404929 | 4.68E-05 | 0.012997 | 0.081136 | 0.1 | 0.236592 |
| SL1344_RS04140_ | 0.793378 | 0.655594 | 0.945857 | 0.292138 | 0.150416 | 1.888063 | 0.018 | 0.05364 | 0.165282 | 0.1197 | 0.260932 |
| SL1344_RS04145_ | 0.33157 | 0.277238 | 0.410029 | 0.138728 | 0.242545 | 0.539172 | 0.288667 | 0.460082 | 0.61434 | 0.58 | 0.701559 |
| SL1344_RS04150_ | -1.51213 | -1.41881 | -1.62427 | -0.21182 | 0.345267 | -0.58141 | 0.251166 | 0.395364 | 0.547318 | 0.5202 | 0.641134 |
| SL1344_RS04160_ | 0.85379 | 0.90569 | 0.795363 | -0.10992 | 0.129294 | -0.81256 | 0.192 | 0.289648 | 0.442394 | 0.3912 | 0.529056 |
| SL1344_RS04165_ | -1.79351 | -1.72954 | -1.82173 | -0.05717 | 0.442734 | -0.11602 | 0.456362 | 0.702736 | 0.806726 | 0.7956 | 0.869177 |
| SL1344_RS04170_ | -3.89824 | -4.10603 | -3.75633 | 0.306539 | 0.708212 | 0.414185 | 0.324667 | 0.480931 | 0.621956 | 0.586 | 0.699427 |
| SL1344_RS04175_ | -0.49373 | -0.77845 | -0.22151 | 0.550372 | 0.228176 | 2.356653 | 0.005331 | 0.031935 | 0.124185 | 0.1056 | 0.243626 |
| SL1344_RS04180_ | -0.41947 | -0.35621 | -0.44989 | -0.09135 | 0.235101 | -0.34219 | 0.354667 | 0.55033 | 0.6806 | 0.6631 | 0.763138 |
| SL1344_RS04185_ | -0.42107 | -0.54493 | -0.29898 | 0.246902 | 0.187637 | 1.228067 | 0.09 | 0.144215 | 0.282904 | 0.2221 | 0.368262 |

| | | | | | | | | | | | |
|---|---|---|---|---|---|---|---|---|---|---|---|
| SL1344_RS04195_ | 0.683295 | 0.902131 | 0.352514 | -0.54469 | 0.188285 | -2.80004 | 0.000673 | 0.022028 | 0.10569 | 0.1003 | 0.236993 |
| SL1344_RS04200_ | 3.020982 | 3.521318 | 2.780622 | -0.70976 | 0.140831 | -3.78178 | 4.68E-05 | 0.025725 | 0.120285 | 0.1 | 0.236592 |
| SL1344_RS04205_ | 4.867254 | 4.700049 | 5.095536 | 0.395346 | 0.110397 | 3.691752 | 4.68E-05 | 0.021893 | 0.111184 | 0.1 | 0.236592 |
| SL1344_RS04210_ | -0.70146 | -0.64006 | -0.75819 | -0.13436 | 0.251054 | -0.49387 | 0.281812 | 0.442874 | 0.588966 | 0.5678 | 0.682245 |
| SL1344_RS04220_ | 0.044261 | 0.170787 | -0.07019 | -0.2378 | 0.1784 | -1.2701 | 0.084 | 0.138387 | 0.279691 | 0.2136 | 0.361944 |
| SL1344_RS04225_ | -0.63369 | -0.45522 | -0.83194 | -0.38845 | 0.233968 | -1.60602 | 0.036667 | 0.085839 | 0.213018 | 0.1441 | 0.287837 |
| SL1344_RS04230_ | 0.408414 | 0.486756 | 0.300275 | -0.18737 | 0.165465 | -1.04991 | 0.113258 | 0.185606 | 0.338287 | 0.265 | 0.413756 |
| SL1344_RS04235_ | 1.082544 | 1.265908 | 0.868563 | -0.39585 | 0.141792 | -2.73825 | 0.002668 | 0.021482 | 0.102753 | 0.1013 | 0.238321 |
| SL1344_RS04240_ | 1.244742 | 1.486184 | 1.068595 | -0.40975 | 0.139295 | -2.89837 | 0.000673 | 0.019 | 0.097808 | 0.1003 | 0.236996 |
| SL1344_RS04245_ | -3.20906 | -3.17261 | -3.23673 | -0.07033 | 0.516497 | -0.12647 | 0.441333 | 0.547663 | 0.677094 | 0.6511 | 0.751056 |
| SL1344_RS04250_ | -5.29316 | -5.30271 | -5.28153 | 0.000461 | 1.016966 | 0.00031 | 0.499667 | 0.566605 | 0.693613 | 0.6749 | 0.769562 |
| SL1344_RS04255_ | -1.94888 | -1.82631 | -2.11322 | -0.29499 | 0.357073 | -0.76565 | 0.183877 | 0.289058 | 0.445194 | 0.39 | 0.526611 |
| SL1344_RS04260_ | 0.564958 | 0.813798 | 0.426947 | -0.37554 | 0.179979 | -1.97505 | 0.017322 | 0.056219 | 0.171989 | 0.1202 | 0.260974 |
| SL1344_RS04265_ | -1.43715 | -1.65225 | -1.27391 | 0.39386 | 0.292407 | 1.26 | 0.073951 | 0.13544 | 0.275212 | 0.1999 | 0.34578 |
| SL1344_RS04270_ | -3.06593 | -3.32609 | -2.96582 | 0.287781 | 0.737077 | 0.337989 | 0.370667 | 0.660252 | 0.776655 | 0.7445 | 0.838711 |
| SL1344_RS04275_ | 0.852384 | 0.853456 | 0.851566 | -0.002 | 0.144522 | -0.01251 | 0.494667 | 0.620555 | 0.740441 | 0.7365 | 0.820651 |
| SL1344_RS04280_ | 0.669221 | 0.710834 | 0.634614 | -0.07422 | 0.14182 | -0.48171 | 0.298 | 0.446142 | 0.588807 | 0.5452 | 0.662902 |
| SL1344_RS04290_ | 2.673934 | 3.245822 | 2.303125 | -0.92389 | 0.138302 | -5.98353 | 4.68E-05 | 0.007409 | 0.066712 | 0.1 | 0.236592 |
| SL1344_RS04295_ | 2.337248 | 2.790016 | 2.040449 | -0.72108 | 0.154057 | -4.38329 | 4.68E-05 | 0.009747 | 0.074218 | 0.1 | 0.236592 |
| SL1344_RS04300_ | 1.749408 | 2.129945 | 1.298012 | -0.89367 | 0.227397 | -4.01841 | 4.68E-05 | 0.004132 | 0.052541 | 0.1 | 0.236592 |
| SL1344_RS04305_ | 1.284772 | 1.528273 | 0.908827 | -0.62696 | 0.195312 | -3.0883 | 4.68E-05 | 0.01913 | 0.099543 | 0.1 | 0.236592 |
| SL1344_RS04320_ | -3.89047 | -3.70316 | -4.14731 | -0.44193 | 0.715117 | -0.59417 | 0.262492 | 0.39384 | 0.545601 | 0.5114 | 0.636985 |
| SL1344_RS04325_ | -3.19728 | -2.9192 | -3.4075 | -0.54207 | 0.661244 | -0.75244 | 0.2 | 0.304367 | 0.466707 | 0.4159 | 0.553587 |
| SL1344_RS04330_ | -2.5662 | -2.34535 | -2.82828 | -0.55736 | 0.610421 | -0.89355 | 0.169887 | 0.240621 | 0.407495 | 0.325 | 0.476929 |
| SL1344_RS04335_ | -4.68786 | -4.63113 | -4.79456 | -0.17205 | 1.022685 | -0.14432 | 0.435333 | 0.593438 | 0.718456 | 0.7142 | 0.802925 |
| SL1344_RS04340_ | -1.17693 | -1.13146 | -1.22322 | -0.08903 | 0.236308 | -0.35114 | 0.339773 | 0.464633 | 0.60265 | 0.5697 | 0.6797 |
| SL1344_RS04345_ | -2.11737 | -2.03592 | -2.21798 | -0.18814 | 0.363194 | -0.47187 | 0.29447 | 0.433729 | 0.57931 | 0.55 | 0.665504 |
| SL1344_RS04350_ | -1.58999 | -1.64772 | -1.54279 | 0.056213 | 0.430059 | 0.121789 | 0.442 | 0.678075 | 0.788437 | 0.7775 | 0.853654 |
| SL1344_RS04355_ | -0.60267 | -0.52477 | -0.65872 | -0.1305 | 0.285817 | -0.41915 | 0.318 | 0.506131 | 0.647056 | 0.6103 | 0.724399 |
| SL1344_RS04360_ | -1.65495 | -1.86862 | -1.47369 | 0.394905 | 0.290325 | 1.303205 | 0.07 | 0.12672 | 0.264629 | 0.1959 | 0.341918 |
| SL1344_RS04365_ | 1.397053 | 1.418279 | 1.382299 | -0.07929 | 0.206868 | -0.36281 | 0.361333 | 0.579622 | 0.7147 | 0.735 | 0.820473 |
| SL1344_RS04370_ | 2.118862 | 1.92586 | 2.456181 | 0.517548 | 0.121667 | 3.85226 | 4.68E-05 | 0.017255 | 0.095645 | 0.1 | 0.236592 |
| SL1344_RS04375_ | 1.471291 | 1.634774 | 1.367643 | -0.25936 | 0.136057 | -1.81784 | 0.016667 | 0.057959 | 0.175352 | 0.1216 | 0.263672 |
| SL1344_RS04380_ | 0.632615 | 0.557449 | 0.727285 | 0.175829 | 0.163446 | 1.022665 | 0.12525 | 0.19716 | 0.353686 | 0.2765 | 0.42528 |
| SL1344_RS04385_ | 0.168205 | 0.085583 | 0.255089 | 0.16958 | 0.175204 | 0.931723 | 0.146569 | 0.237388 | 0.390243 | 0.3269 | 0.468558 |
| SL1344_RS04390_ | -0.14871 | -0.16933 | -0.10173 | 0.076694 | 0.24308 | 0.307777 | 0.378 | 0.5672 | 0.700891 | 0.7046 | 0.79681 |
| SL1344_RS04395_ | -0.09054 | 0.020207 | -0.18337 | -0.20787 | 0.179594 | -1.07917 | 0.107333 | 0.177849 | 0.325998 | 0.2517 | 0.399301 |
| SL1344_RS04400_ | -2.12961 | -2.05224 | -2.18845 | -0.13801 | 0.450036 | -0.27869 | 0.380667 | 0.582395 | 0.709229 | 0.6854 | 0.780182 |
| SL1344_RS04405_ | 1.083629 | 0.82797 | 1.235475 | 0.40124 | 0.138364 | 2.80936 | 0.000673 | 0.027456 | 0.115608 | 0.1006 | 0.237402 |
| SL1344_RS04410_ | -2.08725 | -2.68155 | -1.80439 | 0.868901 | 0.504968 | 1.61194 | 0.024 | 0.09433 | 0.233777 | 0.1293 | 0.271152 |
| SL1344_RS04415_ | 3.142506 | 3.280023 | 2.983325 | -0.30251 | 0.081354 | -3.72675 | 4.68E-05 | 0.008409 | 0.067625 | 0.1 | 0.236592 |
| SL1344_RS04420_ | -1.65828 | -1.71416 | -1.53534 | 0.196609 | 0.408625 | 0.437986 | 0.328667 | 0.505561 | 0.649319 | 0.6315 | 0.739642 |
| SL1344_RS04425_ | -0.02538 | -0.23782 | 0.229166 | 0.461705 | 0.195447 | 2.336712 | 0.006663 | 0.033836 | 0.127827 | 0.1061 | 0.244013 |
| SL1344_RS04430_ | 0.052911 | -0.01834 | 0.110964 | 0.141532 | 0.217284 | 0.619588 | 0.256667 | 0.382247 | 0.539566 | 0.5025 | 0.633356 |
| SL1344_RS04435_ | 0.529746 | 0.52087 | 0.539254 | 0.007157 | 0.171792 | 0.039127 | 0.481679 | 0.633484 | 0.75278 | 0.7536 | 0.833771 |

| | | | | | | | | | | |
|---|---|---|---|---|---|---|---|---|---|---|
| SL1344_RS04440_ | -1.31651 | -0.99913 | -1.51263 | -0.48102 | 0.327949 | -1.34739 | 0.086667 | 0.122935 | 0.266802 | 0.1994 | 0.352335 |
| SL1344_RS04450_ | -2.25217 | -2.05131 | -2.43798 | -0.36057 | 0.443156 | -0.77172 | 0.203864 | 0.32016 | 0.476014 | 0.4208 | 0.560965 |
| SL1344_RS04455_ | -1.08586 | -1.14335 | -1.01427 | 0.156301 | 0.324382 | 0.445727 | 0.318 | 0.483129 | 0.626569 | 0.6158 | 0.722575 |
| SL1344_RS04460_ | -0.98157 | -0.90515 | -1.02739 | -0.15609 | 0.307013 | -0.49165 | 0.293804 | 0.438297 | 0.590226 | 0.5774 | 0.687211 |
| SL1344_RS04465_ | -0.24627 | -0.09132 | -0.37265 | -0.28241 | 0.181647 | -1.49177 | 0.05 | 0.100401 | 0.229441 | 0.1703 | 0.315776 |
| SL1344_RS04470_ | -3.25756 | -3.11691 | -3.42646 | -0.29726 | 0.579117 | -0.49271 | 0.277149 | 0.430939 | 0.576706 | 0.5416 | 0.659616 |
| SL1344_RS04475_ | -3.72626 | -3.84344 | -3.58311 | 0.253953 | 0.782695 | 0.308937 | 0.375333 | 0.560744 | 0.692273 | 0.6741 | 0.772018 |
| SL1344_RS04480_ | -0.90517 | -1.01194 | -0.76259 | 0.242618 | 0.275639 | 0.838521 | 0.179333 | 0.26933 | 0.428461 | 0.3634 | 0.510166 |
| SL1344_RS04485_ | -1.50693 | -1.67098 | -1.3957 | 0.26359 | 0.368556 | 0.66457 | 0.231333 | 0.363158 | 0.521993 | 0.4711 | 0.608527 |
| SL1344_RS04495_ | 1.749898 | 1.74606 | 1.753129 | 0.063945 | 0.180935 | 0.317897 | 0.386667 | 0.555438 | 0.698009 | 0.7859 | 0.853121 |
| SL1344_RS04500_ | 2.907645 | 2.86915 | 3.122763 | 0.248721 | 0.165384 | 1.337025 | 0.101932 | 0.127431 | 0.287589 | 0.2283 | 0.390173 |
| SL1344_RS04505_ | 1.649419 | 1.800927 | 1.607987 | -0.17134 | 0.143146 | -1.04769 | 0.162667 | 0.220496 | 0.389818 | 0.3478 | 0.505189 |
| SL1344_RS04510_ | 2.137715 | 2.046558 | 2.22405 | 0.180852 | 0.092933 | 1.915138 | 0.012659 | 0.054292 | 0.165799 | 0.1146 | 0.254464 |
| SL1344_RS04515_ | 0.659904 | 0.561993 | 0.757525 | 0.17389 | 0.269675 | 0.634798 | 0.242667 | 0.370944 | 0.543169 | 0.4779 | 0.626543 |
| SL1344_RS04520_ | 1.517146 | 1.410659 | 1.661629 | 0.263251 | 0.149358 | 1.728687 | 0.02 | 0.063859 | 0.186016 | 0.1176 | 0.258649 |
| SL1344_RS04525_ | 0.6532 | 0.728806 | 0.566054 | -0.16502 | 0.142035 | -1.09797 | 0.104667 | 0.168841 | 0.315515 | 0.2461 | 0.392561 |
| SL1344_RS04530_ | 1.089212 | 0.977624 | 1.212784 | 0.241825 | 0.128201 | 1.853121 | 0.017988 | 0.062992 | 0.179158 | 0.1206 | 0.262155 |
| SL1344_RS04535_ | 1.366959 | 1.278176 | 1.456027 | 0.175616 | 0.133565 | 1.295683 | 0.076616 | 0.129207 | 0.276223 | 0.1941 | 0.345986 |
| SL1344_RS04540_ | 0.584316 | 0.462549 | 0.66677 | 0.199674 | 0.142008 | 1.327773 | 0.08 | 0.134392 | 0.274625 | 0.214 | 0.361849 |
| SL1344_RS04545_ | 0.194319 | 0.16243 | 0.224283 | 0.063287 | 0.147581 | 0.398138 | 0.325333 | 0.473408 | 0.607556 | 0.5827 | 0.68954 |
| SL1344_RS04550_ | 0.050931 | 0.034774 | 0.069919 | 0.040287 | 0.209127 | 0.195087 | 0.417722 | 0.616017 | 0.736197 | 0.7193 | 0.806313 |
| SL1344_RS04555_ | 1.092002 | 1.137117 | 1.051299 | -0.08996 | 0.135483 | -0.61632 | 0.248 | 0.382472 | 0.533102 | 0.5031 | 0.624182 |
| SL1344_RS04560_ | 1.653476 | 1.608112 | 1.711465 | 0.106105 | 0.104194 | 0.958255 | 0.136576 | 0.22029 | 0.371362 | 0.3074 | 0.449686 |
| SL1344_RS04565_ | 2.726019 | 2.889897 | 2.603309 | -0.28811 | 0.161196 | -1.86894 | 0.007329 | 0.057096 | 0.179914 | 0.1055 | 0.243829 |
| SL1344_RS04570_ | 1.265155 | 1.194396 | 1.34339 | 0.078526 | 0.337358 | 0.260665 | 0.371086 | 0.72047 | 0.824392 | 0.7659 | 0.84921 |
| SL1344_RS04575_ | 1.938452 | 1.790583 | 2.027907 | 0.256324 | 0.173694 | 1.43439 | 0.036667 | 0.094091 | 0.237502 | 0.1396 | 0.286948 |
| SL1344_RS04580_ | 4.495396 | 4.701958 | 4.377791 | -0.31893 | 0.19216 | -1.4548 | 0.061959 | 0.116561 | 0.276124 | 0.1547 | 0.309553 |
| SL1344_RS04585_ | 1.864176 | 1.840882 | 1.92811 | 0.091123 | 0.133636 | 0.640083 | 0.275333 | 0.414447 | 0.57682 | 0.5591 | 0.68345 |
| SL1344_RS04590_ | 4.252562 | 4.165367 | 4.29598 | 0.110203 | 0.094185 | 1.16108 | 0.136576 | 0.163953 | 0.3326 | 0.2407 | 0.410189 |
| SL1344_RS04600_ | -1.64339 | -1.73172 | -1.50424 | 0.205386 | 0.346426 | 0.542769 | 0.272 | 0.433714 | 0.581339 | 0.5426 | 0.665369 |
| SL1344_RS04605_ | -2.04467 | -2.26295 | -1.91412 | 0.339557 | 0.390435 | 0.781996 | 0.196667 | 0.293651 | 0.448561 | 0.3914 | 0.533462 |
| SL1344_RS04610_ | -1.38342 | -1.31929 | -1.45898 | -0.18837 | 0.387726 | -0.46817 | 0.300667 | 0.455011 | 0.607329 | 0.6077 | 0.715635 |
| SL1344_RS04615_ | -0.23572 | -0.18698 | -0.27932 | -0.08261 | 0.215715 | -0.37701 | 0.337775 | 0.525555 | 0.661483 | 0.6351 | 0.740014 |
| SL1344_RS04620_ | -2.07279 | -2.02329 | -2.15935 | -0.13295 | 0.433975 | -0.28245 | 0.367333 | 0.539539 | 0.67553 | 0.6696 | 0.765785 |
| SL1344_RS04625_ | -1.34297 | -1.6165 | -1.24641 | 0.349588 | 0.359273 | 0.854613 | 0.203333 | 0.291683 | 0.458275 | 0.4146 | 0.561974 |
| SL1344_RS04630_ | 2.519392 | 2.953368 | 1.981937 | -0.98568 | 0.150593 | -6.61277 | 4.68E-05 | 0.000965 | 0.032299 | 0.1 | 0.236592 |
| SL1344_RS04635_ | 2.460518 | 2.826226 | 2.078165 | -0.70672 | 0.168715 | -4.27125 | 4.68E-05 | 0.004448 | 0.053546 | 0.1 | 0.236592 |
| SL1344_RS04640_ | 1.183283 | 1.102911 | 1.263399 | 0.148486 | 0.141962 | 1.005963 | 0.141905 | 0.216001 | 0.375922 | 0.3007 | 0.454253 |
| SL1344_RS04645_ | 2.228284 | 2.143662 | 2.451999 | 0.299413 | 0.129613 | 1.967038 | 0.009994 | 0.070186 | 0.200535 | 0.112 | 0.251582 |
| SL1344_RS04650_ | 2.267362 | 2.206548 | 2.296697 | 0.073645 | 0.148285 | 0.469989 | 0.309793 | 0.568787 | 0.707453 | 0.6308 | 0.754563 |
| SL1344_RS04655_ | 3.785914 | 3.882365 | 3.763339 | -0.10591 | 0.0889 | -1.03293 | 0.176549 | 0.221863 | 0.396545 | 0.3538 | 0.517115 |
| SL1344_RS04665_ | 4.405107 | 4.359021 | 4.464447 | 0.111636 | 0.059885 | 1.886888 | 0.002667 | 0.078066 | 0.213193 | 0.1033 | 0.240766 |
| SL1344_RS04675_ | 2.690511 | 2.633538 | 2.757737 | 0.12356 | 0.073964 | 1.65123 | 0.035976 | 0.081307 | 0.207576 | 0.1423 | 0.28681 |
| SL1344_RS04680_ | 4.383481 | 4.325966 | 4.425412 | 0.107153 | 0.064141 | 1.578036 | 0.020653 | 0.075454 | 0.207485 | 0.121 | 0.262737 |
| SL1344_RS04700_ | -0.39452 | -0.78579 | 0.346144 | 1.085057 | 0.292143 | 3.388227 | 4.68E-05 | 0.020761 | 0.104928 | 0.1 | 0.236592 |

| | | | | | | | | | | | |
|---|---|---|---|---|---|---|---|---|---|---|---|
| SL1344_RS04705_ | 0.016542 | -0.05693 | 0.115789 | 0.162721 | 0.226424 | 0.670874 | 0.238 | 0.372713 | 0.529861 | 0.475 | 0.612499 |
| SL1344_RS04715_ | 3.057074 | 3.131309 | 2.923582 | -0.20993 | 0.113192 | -1.85156 | 0.014667 | 0.04846 | 0.162649 | 0.1131 | 0.253541 |
| SL1344_RS04720_ | -0.22646 | -0.04258 | -0.45663 | -0.41305 | 0.20239 | -1.95222 | 0.014657 | 0.050565 | 0.158587 | 0.1143 | 0.254144 |
| SL1344_RS04725_ | -0.02487 | 0.14618 | -0.24391 | -0.38469 | 0.16889 | -2.22094 | 0.005334 | 0.038911 | 0.136187 | 0.1075 | 0.246313 |
| SL1344_RS04730_ | 6.205218 | 6.0847 | 6.362173 | 0.270716 | 0.068217 | 3.77202 | 4.68E-05 | 0.01226 | 0.083813 | 0.1 | 0.236592 |
| SL1344_RS04740_ | 0.578822 | 0.709388 | 0.320639 | -0.38097 | 0.201757 | -1.76543 | 0.027982 | 0.066709 | 0.191044 | 0.1279 | 0.271537 |
| SL1344_RS04745_ | -0.96083 | -1.13821 | -0.80535 | 0.343865 | 0.280804 | 1.169239 | 0.093333 | 0.15803 | 0.310064 | 0.2304 | 0.380146 |
| SL1344_RS04750_ | 2.794213 | 2.753062 | 2.834523 | 0.072916 | 0.096507 | 0.7213 | 0.209194 | 0.334623 | 0.499567 | 0.4292 | 0.575259 |
| SL1344_RS04755_ | 1.52343 | 1.430871 | 1.661465 | 0.219068 | 0.135677 | 1.583771 | 0.046636 | 0.085037 | 0.216989 | 0.1536 | 0.302017 |
| SL1344_RS04760_ | -0.46323 | -0.48473 | -0.44247 | 0.043971 | 0.186592 | 0.216909 | 0.398667 | 0.520386 | 0.650856 | 0.6168 | 0.721209 |
| SL1344_RS04765_ | 1.657375 | 1.849322 | 1.454546 | -0.38823 | 0.123306 | -3.17443 | 4.68E-05 | 0.013617 | 0.082789 | 0.1 | 0.236592 |
| SL1344_RS04770_ | 6.618653 | 6.744181 | 6.435948 | -0.18276 | 0.207361 | -1.41514 | 0.111333 | 0.10113 | 0.254566 | 0.2 | 0.370027 |
| SL1344_RS04780_ | -1.13399 | -1.14939 | -1.1166 | 0.056217 | 0.366893 | 0.153223 | 0.437708 | 0.655063 | 0.770713 | 0.7664 | 0.844092 |
| SL1344_RS04785_ | 2.325802 | 2.418066 | 2.190117 | -0.18867 | 0.166917 | -1.11406 | 0.124 | 0.171741 | 0.340845 | 0.2378 | 0.405586 |
| SL1344_RS04790_ | 1.390107 | 1.501797 | 1.251455 | -0.26393 | 0.141335 | -1.84007 | 0.007995 | 0.070621 | 0.19642 | 0.1086 | 0.247185 |
| SL1344_RS04795_ | -0.44591 | -0.46161 | -0.42987 | 0.035058 | 0.211052 | 0.148969 | 0.432667 | 0.577616 | 0.702946 | 0.6838 | 0.777196 |
| SL1344_RS04800_ | -1.17495 | -1.39429 | -0.99384 | 0.421992 | 0.360433 | 1.108048 | 0.094604 | 0.160471 | 0.318159 | 0.2176 | 0.369711 |
| SL1344_RS04805_ | 1.312816 | 1.275547 | 1.35998 | 0.098001 | 0.139888 | 0.671316 | 0.229181 | 0.342884 | 0.504543 | 0.4605 | 0.592967 |
| SL1344_RS04810_ | 1.862825 | 1.73542 | 1.990438 | 0.261726 | 0.128044 | 1.961674 | 0.004665 | 0.046219 | 0.15646 | 0.1081 | 0.247118 |
| SL1344_RS04815_ | -0.87376 | -0.84253 | -0.92015 | -0.07998 | 0.294102 | -0.23621 | 0.4 | 0.636128 | 0.753312 | 0.7329 | 0.819467 |
| SL1344_RS04820_ | 1.217794 | 1.706178 | 1.034576 | -0.64162 | 0.290346 | -1.95084 | 0.007329 | 0.056254 | 0.179956 | 0.1051 | 0.243346 |
| SL1344_RS04825_ | 2.434209 | 2.686607 | 2.201745 | -0.49437 | 0.106944 | -4.50523 | 4.68E-05 | 0.007172 | 0.063146 | 0.1 | 0.236592 |
| SL1344_RS04830_ | 2.553627 | 2.863184 | 2.276779 | -0.58181 | 0.076294 | -7.52341 | 4.68E-05 | 0.001526 | 0.03454 | 0.1 | 0.236592 |
| SL1344_RS04840_ | 3.087777 | 3.053231 | 3.142577 | 0.095651 | 0.084201 | 1.056748 | 0.118 | 0.190174 | 0.35517 | 0.2594 | 0.411997 |
| SL1344_RS04845_ | 2.097008 | 2.020793 | 2.173629 | 0.160256 | 0.106999 | 1.463206 | 0.041972 | 0.101414 | 0.241668 | 0.1439 | 0.289901 |
| SL1344_RS04850_ | 2.81081 | 2.672047 | 2.917127 | 0.195837 | 0.145481 | 1.383394 | 0.083278 | 0.099628 | 0.246292 | 0.1748 | 0.334634 |
| SL1344_RS04855_ | 3.885111 | 3.79146 | 4.043107 | 0.241215 | 0.11967 | 2.080008 | 0.002002 | 0.032556 | 0.133554 | 0.1029 | 0.240443 |
| SL1344_RS04860_ | 5.334478 | 5.347727 | 5.323957 | 0.016238 | 0.148105 | 0.101237 | 0.452 | 0.692641 | 0.810112 | 0.9565 | 0.973571 |
| SL1344_RS04865_ | 4.456031 | 4.444876 | 4.471672 | 0.046 | 0.074588 | 0.584194 | 0.255829 | 0.354544 | 0.530213 | 0.5355 | 0.66176 |
| SL1344_RS04870_ | -0.38971 | -0.4235 | -0.31889 | 0.095865 | 0.227737 | 0.376458 | 0.341772 | 0.541257 | 0.675615 | 0.6534 | 0.758452 |
| SL1344_RS04875_ | -2.97142 | -3.09432 | -2.8704 | 0.228747 | 0.463297 | 0.456795 | 0.303333 | 0.443224 | 0.583463 | 0.5443 | 0.660096 |
| SL1344_RS04880_ | -1.86879 | -1.79164 | -1.93914 | -0.15015 | 0.356357 | -0.39046 | 0.325333 | 0.469448 | 0.612047 | 0.5824 | 0.69421 |
| SL1344_RS04885_ | -3.06092 | -2.78265 | -3.29487 | -0.50936 | 0.531848 | -0.89498 | 0.154 | 0.247874 | 0.400846 | 0.3394 | 0.480488 |
| SL1344_RS04890_ | 1.302004 | 1.344465 | 1.259879 | -0.0751 | 0.121898 | -0.59534 | 0.254667 | 0.398643 | 0.546352 | 0.5038 | 0.628406 |
| SL1344_RS04895_ | 2.093738 | 2.055257 | 2.181122 | 0.12273 | 0.101763 | 1.122978 | 0.124667 | 0.197296 | 0.357139 | 0.2866 | 0.435066 |
| SL1344_RS04900_ | -3.6459 | -3.96575 | -3.36011 | 0.611795 | 0.549092 | 1.031468 | 0.125916 | 0.199821 | 0.342342 | 0.2766 | 0.418877 |
| SL1344_RS04905_ | -4.97997 | -5.14501 | -4.89621 | 0.181635 | 1.137406 | 0.145879 | 0.433711 | 0.603273 | 0.726253 | 0.7247 | 0.809709 |
| SL1344_RS04910_ | -3.16408 | -3.49242 | -2.82587 | 0.662957 | 0.511606 | 1.232651 | 0.088608 | 0.145493 | 0.285371 | 0.2185 | 0.364713 |
| SL1344_RS04915_ | -2.28231 | -2.50014 | -1.96496 | 0.51303 | 0.453168 | 1.077319 | 0.120667 | 0.189445 | 0.342978 | 0.27 | 0.42086 |
| SL1344_RS04920_ | -2.59453 | -3.93637 | -1.93799 | 1.978523 | 0.641941 | 2.920665 | 4.68E-05 | 0.027476 | 0.117179 | 0.1 | 0.236592 |
| SL1344_RS04925_ | -3.55304 | -4.78166 | -2.73346 | 2.0518 | 0.606964 | 3.286006 | 0.000673 | 0.015465 | 0.08421 | 0.1 | 0.236592 |
| SL1344_RS04930_ | 2.385548 | 2.468878 | 2.193856 | -0.2618 | 0.136772 | -1.8334 | 0.021986 | 0.059481 | 0.181835 | 0.1215 | 0.263742 |
| SL1344_RS04935_ | -3.43799 | -4.02704 | -2.96814 | 1.058046 | 0.513233 | 1.975394 | 0.017322 | 0.051002 | 0.154605 | 0.1166 | 0.256155 |
| SL1344_RS04940_ | -3.01292 | -3.09369 | -2.92385 | 0.199123 | 0.524765 | 0.347314 | 0.352 | 0.501099 | 0.640326 | 0.6264 | 0.729583 |
| SL1344_RS04945_ | -2.25524 | -2.42143 | -2.11726 | 0.31973 | 0.51526 | 0.589346 | 0.25 | 0.389579 | 0.546744 | 0.5067 | 0.636891 |

| | | | | | | | | | | |
|---|---|---|---|---|---|---|---|---|---|---|
| SL1344_RS04950_ | -3.47946 | -3.87841 | -3.13857 | 0.703214 | 0.698449 | 0.944936 | 0.148568 | 0.225915 | 0.383659 | 0.3084 | 0.459471 |
| SL1344_RS04955_ | -4.41096 | -4.6493 | -4.08064 | 0.544292 | 1.087803 | 0.481135 | 0.298468 | 0.48055 | 0.626854 | 0.5897 | 0.707844 |
| SL1344_RS04960_ | -1.76834 | -1.73844 | -1.79756 | -0.07773 | 0.370007 | -0.20683 | 0.41439 | 0.583612 | 0.709098 | 0.7075 | 0.795162 |
| SL1344_RS04965_ | 2.527895 | 2.414737 | 2.649876 | 0.250007 | 0.102992 | 2.417009 | 4.68E-05 | 0.043532 | 0.152215 | 0.1002 | 0.236852 |
| SL1344_RS04970_ | -0.76684 | -0.63508 | -0.9392 | -0.32908 | 0.302438 | -0.99874 | 0.108 | 0.184028 | 0.342319 | 0.2632 | 0.412748 |
| SL1344_RS04975_ | -0.38039 | -0.30588 | -0.48631 | -0.18011 | 0.202922 | -0.85365 | 0.176 | 0.269435 | 0.423607 | 0.3645 | 0.502836 |
| SL1344_RS04980_ | -1.66046 | -1.64118 | -1.69364 | -0.03339 | 0.452197 | -0.06244 | 0.473684 | 0.709593 | 0.812137 | 0.8056 | 0.875775 |
| SL1344_RS04985_ | -6.78603 | -6.04716 | -8.35277 | -2.42412 | 3.218337 | -0.69708 | 0.187209 | 0.279366 | 0.452367 | 0.379 | 0.532426 |
| SL1344_RS04990_ | -4.683 | -4.80807 | -4.5479 | 0.24875 | 0.88484 | 0.27371 | 0.388 | 0.520706 | 0.653923 | 0.6222 | 0.727485 |
| SL1344_RS04995_ | -6.82552 | -8.53663 | -6.03605 | 2.440652 | 2.754744 | 0.790225 | 0.138574 | 0.238872 | 0.407885 | 0.2982 | 0.446808 |
| SL1344_RS05000_ | -2.96434 | -3.10776 | -2.76133 | 0.327343 | 0.714514 | 0.411904 | 0.335333 | 0.54067 | 0.677781 | 0.6541 | 0.758711 |
| SL1344_RS05005_ | 0.153615 | 0.162461 | 0.146634 | -0.02079 | 0.168633 | -0.12283 | 0.446369 | 0.571161 | 0.698787 | 0.6947 | 0.785178 |
| SL1344_RS05010_ | -6.69366 | -7.34244 | -6.38605 | 0.821384 | 2.783082 | 0.253663 | 0.364424 | 0.5337 | 0.677918 | 0.691 | 0.786135 |
| SL1344_RS05015_ | -0.45674 | -0.4568 | -0.45674 | 0.008478 | 0.230253 | 0.038219 | 0.483333 | 0.626984 | 0.746337 | 0.7373 | 0.820868 |
| SL1344_RS05020_ | -1.0432 | -0.82835 | -1.25967 | -0.47024 | 0.31612 | -1.44851 | 0.042667 | 0.098331 | 0.235704 | 0.1552 | 0.29967 |
| SL1344_RS05025_ | -6.4114 | -6.24547 | -6.59085 | -0.14779 | 3.703506 | -0.03435 | 0.474 | 0.681976 | 0.795379 | 0.8181 | 0.881471 |
| SL1344_RS05030_ | -2.96749 | -3.18471 | -2.80844 | 0.377233 | 0.543142 | 0.66303 | 0.233178 | 0.349627 | 0.503932 | 0.4574 | 0.588977 |
| SL1344_RS05035_ | -4.07367 | -4.01086 | -4.11792 | -0.09825 | 0.722829 | -0.12847 | 0.441333 | 0.577245 | 0.702163 | 0.6801 | 0.775112 |
| SL1344_RS05040_ | -2.14857 | -2.11789 | -2.18345 | -0.06937 | 0.359605 | -0.17165 | 0.428381 | 0.558175 | 0.684727 | 0.6653 | 0.760527 |
| SL1344_RS05045_ | -1.31494 | -1.4349 | -1.14561 | 0.265163 | 0.315965 | 0.811792 | 0.191206 | 0.302176 | 0.459192 | 0.4005 | 0.544294 |
| SL1344_RS05050_ | -4.652 | -5.16984 | -4.42154 | 0.670367 | 1.20451 | 0.483087 | 0.302 | 0.47102 | 0.62218 | 0.6112 | 0.725888 |
| SL1344_RS05055_ | -1.49849 | -1.62521 | -1.3946 | 0.417055 | 0.597994 | 0.701416 | 0.226667 | 0.304103 | 0.480274 | 0.4743 | 0.609675 |
| SL1344_RS05060_ | -0.39961 | -0.68136 | -0.25704 | 0.388118 | 0.299068 | 1.174973 | 0.112 | 0.154877 | 0.313984 | 0.2279 | 0.386899 |
| SL1344_RS05065_ | -3.6969 | -3.86929 | -3.57571 | 0.310394 | 0.687662 | 0.40522 | 0.316667 | 0.479207 | 0.62122 | 0.5989 | 0.707127 |
| SL1344_RS05070_ | -4.73879 | -5.45141 | -4.2542 | 1.197426 | 1.017469 | 1.106898 | 0.100667 | 0.174907 | 0.328752 | 0.2458 | 0.395337 |
| SL1344_RS05075_ | -2.98872 | -3.02438 | -2.94697 | 0.073807 | 0.548915 | 0.123061 | 0.434377 | 0.578684 | 0.70561 | 0.6984 | 0.789623 |
| SL1344_RS05080_ | -3.61179 | -3.83603 | -3.31414 | 0.590151 | 0.734585 | 0.772171 | 0.193333 | 0.299649 | 0.459511 | 0.4127 | 0.548263 |
| SL1344_RS05085_ | -1.96938 | -2.02614 | -1.88323 | 0.186811 | 0.422519 | 0.418376 | 0.329333 | 0.480404 | 0.624661 | 0.6071 | 0.715542 |
| SL1344_RS05090_ | -2.5032 | -2.70006 | -2.27128 | 0.423474 | 0.422787 | 0.942596 | 0.142 | 0.227367 | 0.378141 | 0.313 | 0.455881 |
| SL1344_RS05095_ | -3.35412 | -3.73312 | -3.09893 | 0.616063 | 0.702434 | 0.810094 | 0.19054 | 0.281224 | 0.445619 | 0.3873 | 0.534655 |
| SL1344_RS05100_ | -0.03194 | -0.13407 | 0.03721 | 0.172804 | 0.190818 | 0.857148 | 0.168667 | 0.26237 | 0.42074 | 0.3612 | 0.501304 |
| SL1344_RS05105_ | -3.11768 | -3.16028 | -3.06718 | 0.101084 | 0.498613 | 0.189537 | 0.419333 | 0.545267 | 0.675093 | 0.6513 | 0.751658 |
| SL1344_RS05110_ | 1.226168 | 1.368992 | 1.141906 | -0.23409 | 0.35433 | -0.67817 | 0.236509 | 0.334696 | 0.517219 | 0.4407 | 0.606946 |
| SL1344_RS05115_ | 2.719604 | 2.696323 | 2.748333 | 0.057337 | 0.089447 | 0.638709 | 0.246502 | 0.381589 | 0.539972 | 0.4979 | 0.63091 |
| SL1344_RS05120_ | -1.46682 | -1.58472 | -1.35919 | 0.220997 | 0.262623 | 0.774171 | 0.182545 | 0.287338 | 0.435232 | 0.3768 | 0.509997 |
| SL1344_RS05125_ | -2.22514 | -2.25531 | -2.16952 | 0.057237 | 0.410366 | 0.126619 | 0.445703 | 0.60837 | 0.727359 | 0.7082 | 0.797851 |
| SL1344_RS05130_ | -1.38588 | -1.4973 | -1.28071 | 0.208227 | 0.296735 | 0.674124 | 0.219853 | 0.332007 | 0.485112 | 0.4342 | 0.569516 |
| SL1344_RS05135_ | -0.33412 | -0.36018 | -0.30353 | 0.04714 | 0.223947 | 0.193426 | 0.413724 | 0.599563 | 0.723445 | 0.7121 | 0.799985 |
| SL1344_RS05140_ | -3.10156 | -2.71739 | -3.4017 | -0.68163 | 0.578402 | -1.11618 | 0.116 | 0.18086 | 0.333167 | 0.2633 | 0.411405 |
| SL1344_RS05165_ | -3.55173 | -3.85645 | -3.31917 | 0.494362 | 0.688725 | 0.698473 | 0.216 | 0.336645 | 0.492431 | 0.4367 | 0.572836 |
| SL1344_RS05175_ | -0.65014 | -0.59243 | -0.74505 | -0.14093 | 0.372047 | -0.37447 | 0.341106 | 0.577652 | 0.712028 | 0.668 | 0.774967 |
| SL1344_RS05190_ | 3.891208 | 3.70972 | 3.958996 | 0.237154 | 0.091879 | 2.134764 | 0.002002 | 0.054173 | 0.175537 | 0.1019 | 0.239052 |
| SL1344_RS05195_ | 0.472122 | 0.5251 | 0.327693 | -0.16774 | 0.213254 | -0.68694 | 0.253165 | 0.383587 | 0.547912 | 0.5166 | 0.656179 |
| SL1344_RS05200_ | 1.468375 | 1.636706 | 1.327599 | -0.30565 | 0.102329 | -2.91125 | 0.000673 | 0.018796 | 0.094711 | 0.1006 | 0.237388 |
| SL1344_RS05205_ | -0.35762 | -0.34408 | -0.37117 | -0.02415 | 0.209917 | -0.1146 | 0.452365 | 0.590395 | 0.713826 | 0.699 | 0.790928 |

| | | | | | | | | | | | |
|---|---|---|---|---|---|---|---|---|---|---|---|
| SL1344_RS05210_ | 1.715998 | 1.850364 | 1.596467 | -0.25013 | 0.133677 | -1.83572 | 0.019334 | 0.055393 | 0.17126 | 0.1188 | 0.260547 |
| SL1344_RS05215_ | 2.375837 | 2.573003 | 2.239285 | -0.31265 | 0.132308 | -2.30957 | 0.003998 | 0.032218 | 0.130801 | 0.1033 | 0.240982 |
| SL1344_RS05220_ | 1.262762 | 1.23726 | 1.319363 | 0.073834 | 0.174875 | 0.362532 | 0.341333 | 0.562283 | 0.699121 | 0.6739 | 0.779464 |
| SL1344_RS05225_ | 1.84243 | 1.743136 | 1.910462 | 0.169345 | 0.121099 | 1.329285 | 0.061293 | 0.121498 | 0.268865 | 0.1738 | 0.323036 |
| SL1344_RS05230_ | 1.469141 | 1.346702 | 1.523583 | 0.161066 | 0.182224 | 0.87343 | 0.182 | 0.265035 | 0.437785 | 0.3621 | 0.526122 |
| SL1344_RS05235_ | 4.369479 | 4.383591 | 4.342586 | -0.07879 | 0.130658 | -0.64718 | 0.242 | 0.311711 | 0.496748 | 0.4661 | 0.61806 |
| SL1344_RS05240_ | 2.525584 | 2.913337 | 2.149264 | -0.80613 | 0.177814 | -4.59913 | 4.68E-05 | 0.003194 | 0.047306 | 0.1 | 0.236592 |
| SL1344_RS05245_ | 1.952781 | 2.067167 | 1.721297 | -0.32636 | 0.159941 | -1.90422 | 0.013991 | 0.056447 | 0.17732 | 0.1136 | 0.254206 |
| SL1344_RS05250_ | 0.885984 | 0.910745 | 0.861228 | -0.05605 | 0.159036 | -0.33044 | 0.352 | 0.542552 | 0.677113 | 0.6755 | 0.770055 |
| SL1344_RS05255_ | 7.919781 | 7.797475 | 8.172485 | 0.37508 | 0.130597 | 3.065946 | 4.68E-05 | 0.00677 | 0.065872 | 0.1 | 0.236592 |
| SL1344_RS05260_ | 1.617216 | 1.944112 | 1.24294 | -0.67362 | 0.146761 | -4.53833 | 4.68E-05 | 0.004103 | 0.05074 | 0.1 | 0.236592 |
| SL1344_RS05265_ | -0.11641 | -0.16547 | -0.04935 | 0.113363 | 0.183308 | 0.57005 | 0.247169 | 0.392615 | 0.540859 | 0.5004 | 0.626187 |
| SL1344_RS05270_ | 1.181829 | 1.04992 | 1.369974 | 0.255339 | 0.217304 | 1.179241 | 0.115257 | 0.157165 | 0.318461 | 0.2296 | 0.394067 |
| SL1344_RS05275_ | -1.25463 | -1.18951 | -1.3264 | -0.14741 | 0.285903 | -0.47802 | 0.296469 | 0.460411 | 0.601772 | 0.5743 | 0.686691 |
| SL1344_RS05285_ | 2.574616 | 2.473981 | 2.780072 | 0.329436 | 0.174704 | 1.909655 | 0.002002 | 0.042529 | 0.153449 | 0.1027 | 0.240028 |
| SL1344_RS05290_ | -1.88125 | -2.2965 | -1.48905 | 0.812422 | 0.354341 | 2.224717 | 0.005997 | 0.035891 | 0.130624 | 0.1068 | 0.245342 |
| SL1344_RS05295_ | 2.110731 | 1.78478 | 2.457986 | 0.471982 | 0.332355 | 1.75317 | 0.041972 | 0.047817 | 0.165647 | 0.1334 | 0.281045 |
| SL1344_RS05300_ | 2.574304 | 2.573911 | 2.574574 | -0.00751 | 0.142707 | -0.0409 | 0.478 | 0.684724 | 0.796422 | 0.8323 | 0.892706 |
| SL1344_RS05305_ | 0.806955 | 0.804168 | 0.815057 | 0.007238 | 0.180045 | 0.034759 | 0.484344 | 0.705126 | 0.807202 | 0.8088 | 0.874858 |
| SL1344_RS05310_ | -1.13898 | -1.23034 | -1.01924 | 0.200452 | 0.259566 | 0.744378 | 0.214 | 0.320904 | 0.470252 | 0.4167 | 0.550243 |
| SL1344_RS05315_ | -0.21813 | -0.41916 | -0.05818 | 0.361739 | 0.169791 | 2.033234 | 0.014667 | 0.049656 | 0.153341 | 0.1192 | 0.259076 |
| SL1344_RS05320_ | -1.78488 | -2.15287 | -1.28293 | 0.850589 | 0.350133 | 2.332019 | 0.004668 | 0.032324 | 0.125606 | 0.106 | 0.244181 |
| SL1344_RS05325_ | 0.992037 | 1.26422 | 0.829834 | -0.42537 | 0.159939 | -2.5626 | 0.001337 | 0.031237 | 0.126015 | 0.1016 | 0.23867 |
| SL1344_RS05330_ | 4.000608 | 4.037555 | 3.812221 | -0.21984 | 0.074082 | -1.70852 | 0.018 | 0.099456 | 0.249065 | 0.1186 | 0.259199 |
| SL1344_RS05335_ | 1.448593 | 1.739816 | 1.404508 | -0.35037 | 0.379868 | -0.69548 | 0.299333 | 0.419609 | 0.598476 | 0.5972 | 0.739855 |
| SL1344_RS05340_ | -1.78234 | -2.09878 | -1.47617 | 0.635522 | 0.356214 | 1.736195 | 0.020667 | 0.067678 | 0.186004 | 0.132 | 0.274991 |
| SL1344_RS05345_ | -2.01659 | -1.32979 | -2.75206 | -1.45612 | 0.419172 | -3.41683 | 4.68E-05 | 0.010691 | 0.074315 | 0.1 | 0.236592 |
| SL1344_RS05350_ | -5.32184 | -4.70951 | -6.47956 | -1.37625 | 2.796558 | -0.43875 | 0.257162 | 0.414611 | 0.585599 | 0.5276 | 0.662982 |
| SL1344_RS05365_ | 1.852283 | 2.061837 | 1.353817 | -0.71417 | 0.320437 | -2.44058 | 4.68E-05 | 0.017237 | 0.098386 | 0.1 | 0.236592 |
| SL1344_RS05370_ | -3.14549 | -3.75569 | -2.81621 | 1.003194 | 0.861728 | 1.098134 | 0.098 | 0.171173 | 0.336661 | 0.2273 | 0.383236 |
| SL1344_RS05375_ | 1.209798 | 0.973948 | 1.408577 | 0.448517 | 0.15924 | 2.784517 | 4.68E-05 | 0.027161 | 0.117297 | 0.1 | 0.236592 |
| SL1344_RS05380_ | -0.62232 | -0.66882 | -0.5802 | 0.083143 | 0.194002 | 0.398475 | 0.316667 | 0.446145 | 0.585398 | 0.5473 | 0.663267 |
| SL1344_RS05385_ | -0.51035 | -0.59322 | -0.43373 | 0.152925 | 0.262819 | 0.559183 | 0.269154 | 0.427301 | 0.579268 | 0.5356 | 0.66126 |
| SL1344_RS05390_ | -0.51209 | -0.67447 | -0.40989 | 0.252104 | 0.235408 | 1.019998 | 0.141239 | 0.206662 | 0.362441 | 0.3022 | 0.450886 |
| SL1344_RS05395_ | -1.90301 | -1.87626 | -1.94417 | -0.08129 | 0.388328 | -0.19346 | 0.419054 | 0.581422 | 0.708355 | 0.6918 | 0.783622 |
| SL1344_RS05400_ | -2.54941 | -2.55673 | -2.53786 | 0.000953 | 0.455216 | 0.001229 | 0.499667 | 0.61512 | 0.735443 | 0.7183 | 0.807386 |
| SL1344_RS05405_ | -1.25619 | -1.23949 | -1.27985 | -0.03797 | 0.291994 | -0.11558 | 0.442 | 0.612862 | 0.733155 | 0.7228 | 0.808867 |
| SL1344_RS05410_ | -1.15086 | -1.05214 | -1.20933 | -0.1392 | 0.284414 | -0.44775 | 0.30513 | 0.502113 | 0.639 | 0.5978 | 0.712556 |
| SL1344_RS05415_ | -0.86169 | -0.88373 | -0.8311 | 0.050558 | 0.240194 | 0.201486 | 0.412 | 0.554765 | 0.683893 | 0.6628 | 0.761286 |
| SL1344_RS05420_ | -1.72933 | -1.67312 | -1.78666 | -0.11249 | 0.362488 | -0.28218 | 0.369087 | 0.558799 | 0.688743 | 0.6734 | 0.768707 |
| SL1344_RS05425_ | -1.64402 | -1.6509 | -1.63738 | -0.00052 | 0.331923 | -0.00129 | 0.499001 | 0.612397 | 0.732209 | 0.7211 | 0.807798 |
| SL1344_RS05430_ | -2.234 | -2.18349 | -2.28098 | -0.10416 | 0.360396 | -0.27335 | 0.383333 | 0.512121 | 0.645712 | 0.6245 | 0.727238 |
| SL1344_RS05435_ | -2.51171 | -2.39663 | -2.61849 | -0.21765 | 0.392527 | -0.52391 | 0.274484 | 0.423391 | 0.563953 | 0.5208 | 0.639914 |
| SL1344_RS05440_ | -1.70893 | -1.70329 | -1.71691 | -0.00661 | 0.319042 | -0.02196 | 0.489674 | 0.569669 | 0.697548 | 0.6799 | 0.774615 |
| SL1344_RS05445_ | -2.1265 | -2.14294 | -2.09643 | 0.025143 | 0.394124 | 0.054443 | 0.478667 | 0.617735 | 0.739782 | 0.7313 | 0.818315 |

| | | | | | | | | | | |
|---|---|---|---|---|---|---|---|---|---|---|
| SL1344_RS05450_ | -0.84542 | -0.79787 | -0.89618 | -0.086 | 0.266383 | -0.29617 | 0.369333 | 0.577554 | 0.702672 | 0.6833 | 0.777925 |
| SL1344_RS05455_ | -4.49483 | -4.03794 | -4.85503 | -0.52584 | 1.461933 | -0.33572 | 0.354667 | 0.637607 | 0.756391 | 0.7186 | 0.810058 |
| SL1344_RS05460_ | 1.790791 | 1.491503 | 2.076261 | 0.563939 | 0.214722 | 2.599866 | 4.68E-05 | 0.018527 | 0.100964 | 0.1 | 0.236592 |
| SL1344_RS05465_ | 2.138983 | 1.963392 | 2.299749 | 0.336124 | 0.07399 | 4.562072 | 4.68E-05 | 0.005516 | 0.05387 | 0.1 | 0.236592 |
| SL1344_RS05470_ | 0.203347 | 0.252932 | 0.153063 | -0.13248 | 0.227135 | -0.54906 | 0.282667 | 0.419851 | 0.575379 | 0.5679 | 0.684359 |
| SL1344_RS05475_ | -1.41321 | -1.35989 | -1.43855 | -0.05295 | 0.375652 | -0.12208 | 0.450366 | 0.695746 | 0.801537 | 0.791 | 0.865259 |
| SL1344_RS05480_ | -2.8154 | -2.93655 | -2.72615 | 0.279672 | 0.652428 | 0.378893 | 0.306667 | 0.451075 | 0.606637 | 0.6024 | 0.710787 |
| SL1344_RS05485_ | -2.08911 | -2.21339 | -1.99437 | 0.203513 | 0.376386 | 0.485906 | 0.296667 | 0.449575 | 0.590999 | 0.5541 | 0.671732 |
| SL1344_RS05490_ | 2.892785 | 2.801861 | 3.061176 | 0.251097 | 0.081966 | 2.868583 | 4.68E-05 | 0.025799 | 0.113869 | 0.1003 | 0.237 |
| SL1344_RS05495_ | 3.555873 | 3.295558 | 4.1763 | 0.845813 | 0.158122 | 3.980624 | 4.68E-05 | 0.030969 | 0.133047 | 0.1 | 0.236592 |
| SL1344_RS05500_ | 2.454695 | 2.059249 | 2.827676 | 0.791774 | 0.176379 | 4.612138 | 4.68E-05 | 0.010044 | 0.075419 | 0.1 | 0.236592 |
| SL1344_RS05505_ | 0.938598 | 1.152274 | 0.643559 | -0.50972 | 0.146697 | -3.40584 | 0.000673 | 0.014271 | 0.08275 | 0.1001 | 0.236724 |
| SL1344_RS05510_ | -0.83747 | -0.75995 | -0.94777 | -0.19349 | 0.231666 | -0.7742 | 0.198534 | 0.297344 | 0.450476 | 0.394 | 0.529859 |
| SL1344_RS05515_ | 3.102559 | 3.1628 | 3.063888 | -0.09106 | 0.090381 | -0.9357 | 0.151899 | 0.225606 | 0.394944 | 0.3122 | 0.470479 |
| SL1344_RS05525_ | 3.317854 | 3.264034 | 3.354151 | 0.109542 | 0.130496 | 0.857609 | 0.158561 | 0.238337 | 0.416931 | 0.3207 | 0.483559 |
| SL1344_RS05530_ | 1.963437 | 2.227151 | 1.701577 | -0.5235 | 0.092287 | -5.65166 | 4.68E-05 | 0.00294 | 0.043341 | 0.1 | 0.236592 |
| SL1344_RS05540_ | 1.58785 | 1.613678 | 1.564745 | -0.04884 | 0.095306 | -0.46949 | 0.292472 | 0.432889 | 0.574693 | 0.537 | 0.65473 |
| SL1344_RS05545_ | -0.6678 | -0.77191 | -0.56282 | 0.204362 | 0.205194 | 0.939501 | 0.144667 | 0.223956 | 0.370345 | 0.3104 | 0.451047 |
| SL1344_RS05550_ | -0.87255 | -0.81119 | -0.94922 | -0.13889 | 0.211778 | -0.59929 | 0.248 | 0.372751 | 0.515601 | 0.4778 | 0.601595 |
| SL1344_RS05555_ | -1.43965 | -1.76303 | -1.12764 | 0.624027 | 0.363337 | 1.660131 | 0.034644 | 0.074403 | 0.199854 | 0.1335 | 0.278202 |
| SL1344_RS05560_ | -1.45371 | -1.84912 | -1.08983 | 0.754799 | 0.295697 | 2.563647 | 0.003332 | 0.028632 | 0.116109 | 0.1017 | 0.238864 |
| SL1344_RS05565_ | -1.97497 | -2.44327 | -1.62666 | 0.797333 | 0.413773 | 1.871467 | 0.023334 | 0.063014 | 0.181279 | 0.1234 | 0.265953 |
| SL1344_RS05570_ | 0.668835 | 0.441902 | 0.841343 | 0.409457 | 0.185668 | 2.131013 | 0.007334 | 0.040071 | 0.143023 | 0.1057 | 0.243927 |
| SL1344_RS05580_ | -0.25005 | -0.10087 | -0.40853 | -0.32263 | 0.280072 | -1.10193 | 0.11459 | 0.175784 | 0.337326 | 0.2417 | 0.399113 |
| SL1344_RS05585_ | -5.79812 | -5.58893 | -6.03816 | -0.4851 | 1.135776 | -0.38148 | 0.33 | 0.441509 | 0.582271 | 0.5532 | 0.665967 |
| SL1344_RS05590_ | 1.7471 | 1.723879 | 1.760877 | 0.050091 | 0.121507 | 0.382102 | 0.334 | 0.499269 | 0.644832 | 0.6392 | 0.740756 |
| SL1344_RS05595_ | 1.553622 | 1.518103 | 1.599061 | 0.082654 | 0.101589 | 0.736797 | 0.198534 | 0.307169 | 0.461294 | 0.4122 | 0.546813 |
| SL1344_RS05600_ | 0.819787 | 0.702113 | 0.91466 | 0.21911 | 0.13154 | 1.612546 | 0.035333 | 0.089187 | 0.217604 | 0.1421 | 0.285463 |
| SL1344_RS05605_ | 0.522374 | 0.501748 | 0.532926 | 0.003568 | 0.234493 | 0.012307 | 0.495333 | 0.721583 | 0.821264 | 0.8172 | 0.883169 |
| SL1344_RS05610_ | -2.10449 | -2.09605 | -2.11367 | 0.022174 | 0.490861 | 0.035182 | 0.483678 | 0.675869 | 0.785754 | 0.7876 | 0.857778 |
| SL1344_RS05615_ | -4.81909 | -4.84351 | -4.78627 | 0.048324 | 0.930395 | 0.047973 | 0.474667 | 0.583385 | 0.707996 | 0.6982 | 0.788207 |
| SL1344_RS05620_ | -4.04824 | -4.31388 | -3.64344 | 0.686134 | 0.767773 | 0.807449 | 0.186667 | 0.281292 | 0.438957 | 0.382 | 0.524882 |
| SL1344_RS05625_ | -2.95024 | -2.80707 | -3.27428 | -0.34755 | 0.843638 | -0.36298 | 0.367755 | 0.595202 | 0.729501 | 0.7423 | 0.83086 |
| SL1344_RS05635_ | -2.67283 | -2.54508 | -2.80468 | -0.26494 | 0.40683 | -0.59978 | 0.260667 | 0.36595 | 0.511402 | 0.4771 | 0.599827 |
| SL1344_RS05650_ | 2.463081 | 2.787335 | 2.023316 | -0.78287 | 0.136722 | -5.75538 | 4.68E-05 | 0.00202 | 0.039944 | 0.1 | 0.236592 |
| SL1344_RS05660_ | -0.84397 | -0.98269 | -0.74324 | 0.245762 | 0.264469 | 0.869587 | 0.159333 | 0.244267 | 0.403371 | 0.3334 | 0.474993 |
| SL1344_RS05670_ | 4.159822 | 4.089379 | 4.215475 | 0.127965 | 0.072775 | 1.70124 | 0.022667 | 0.071075 | 0.201569 | 0.1209 | 0.263692 |
| SL1344_RS05675_ | -2.23608 | -2.12332 | -2.35455 | -0.2268 | 0.378217 | -0.58749 | 0.250667 | 0.395268 | 0.540188 | 0.4963 | 0.619701 |
| SL1344_RS05680_ | 1.456126 | 1.366935 | 1.829156 | 0.430004 | 0.205953 | 1.462114 | 0.059333 | 0.113589 | 0.268136 | 0.1706 | 0.323234 |
| SL1344_RS05685_ | -0.7721 | -0.92883 | -0.49689 | 0.394622 | 0.341904 | 1.108604 | 0.127249 | 0.17379 | 0.335225 | 0.259 | 0.419665 |
| SL1344_RS05690_ | 1.408406 | 1.57616 | 1.183467 | -0.38918 | 0.113616 | -3.29862 | 4.68E-05 | 0.01513 | 0.08529 | 0.1 | 0.236592 |
| SL1344_RS05695_ | 0.21327 | 0.409967 | 0.06808 | -0.33525 | 0.192137 | -1.71135 | 0.037333 | 0.07338 | 0.19733 | 0.139 | 0.284256 |
| SL1344_RS05700_ | -1.88645 | -1.8505 | -1.92385 | -0.06958 | 0.309996 | -0.20574 | 0.415333 | 0.522494 | 0.655643 | 0.6284 | 0.731042 |
| SL1344_RS05705_ | -3.66618 | -3.52663 | -3.79167 | -0.25944 | 0.597028 | -0.40732 | 0.313791 | 0.467376 | 0.607593 | 0.5811 | 0.69167 |
| SL1344_RS05710_ | -4.08252 | -4.25231 | -3.92178 | 0.269234 | 0.927831 | 0.266844 | 0.370667 | 0.594764 | 0.720507 | 0.7086 | 0.799816 |

| | | | | | | | | | | | |
|---|---|---|---|---|---|---|---|---|---|---|---|
| SL1344_RS05715_ | 1.638791 | 1.583666 | 1.711776 | 0.134672 | 0.107884 | 1.207607 | 0.086 | 0.152395 | 0.299277 | 0.214 | 0.359003 |
| SL1344_RS05730_ | 2.461995 | 2.569139 | 2.371884 | -0.20253 | 0.089558 | -2.24019 | 0.003332 | 0.038829 | 0.139607 | 0.1043 | 0.242232 |
| SL1344_RS05735_ | 2.420605 | 2.355628 | 2.497339 | 0.138841 | 0.092938 | 1.437625 | 0.050633 | 0.103685 | 0.241516 | 0.1735 | 0.32336 |
| SL1344_RS05740_ | 2.839504 | 2.694012 | 3.087405 | 0.387109 | 0.150957 | 2.596552 | 4.68E-05 | 0.018008 | 0.099016 | 0.1 | 0.236592 |
| SL1344_RS05745_ | 0.166838 | 0.183312 | 0.13943 | -0.04851 | 0.178492 | -0.25515 | 0.386 | 0.546904 | 0.677264 | 0.6538 | 0.751206 |
| SL1344_RS05750_ | 1.767771 | 1.48355 | 1.999625 | 0.515481 | 0.128819 | 3.910687 | 4.68E-05 | 0.012078 | 0.078677 | 0.1 | 0.236592 |
| SL1344_RS05755_ | 2.187417 | 1.835735 | 2.524923 | 0.692455 | 0.185642 | 3.70601 | 4.68E-05 | 0.006348 | 0.062133 | 0.1 | 0.236592 |
| SL1344_RS05760_ | 2.283695 | 1.981736 | 2.565881 | 0.594324 | 0.129314 | 4.751268 | 4.68E-05 | 0.008812 | 0.068581 | 0.1 | 0.236592 |
| SL1344_RS05765_ | 0.913265 | 0.852511 | 0.985193 | 0.135554 | 0.129628 | 0.996216 | 0.145903 | 0.219504 | 0.368507 | 0.3019 | 0.444997 |
| SL1344_RS05780_ | 4.616415 | 4.949518 | 4.139701 | -0.82396 | 0.181378 | -4.75683 | 4.68E-05 | 0.001417 | 0.037522 | 0.1 | 0.236592 |
| SL1344_RS05785_ | 4.224448 | 4.773971 | 3.670611 | -1.12277 | 0.109104 | -10.271 | 4.68E-05 | 0.000174 | 0.022554 | 0.1 | 0.236592 |
| SL1344_RS05790_ | 3.980501 | 4.524094 | 3.542424 | -0.96186 | 0.088178 | -10.2342 | 4.68E-05 | 0.00223 | 0.042169 | 0.1 | 0.236592 |
| SL1344_RS05800_ | 5.804192 | 6.196404 | 5.504705 | -0.68619 | 0.112651 | -6.19928 | 4.68E-05 | 0.000494 | 0.028464 | 0.1 | 0.236592 |
| SL1344_RS05810_ | 5.805971 | 6.326751 | 5.435402 | -0.89535 | 0.094333 | -8.544 | 4.68E-05 | 0.004744 | 0.056948 | 0.1 | 0.236592 |
| SL1344_RS05815_ | 3.898257 | 4.282258 | 3.456543 | -0.77394 | 0.14961 | -5.27216 | 4.68E-05 | 0.001114 | 0.034677 | 0.1 | 0.236592 |
| SL1344_RS05825_ | 3.641995 | 3.935472 | 3.320515 | -0.60172 | 0.100014 | -6.08581 | 4.68E-05 | 0.003535 | 0.048341 | 0.1 | 0.236592 |
| SL1344_RS05845_ | -6.09209 | -6.31884 | -5.92609 | 0.416096 | 1.481707 | 0.257412 | 0.381333 | 0.513623 | 0.65114 | 0.6323 | 0.733909 |
| SL1344_RS05850_ | 0.595993 | 0.722675 | 0.45949 | -0.26022 | 0.127074 | -1.99508 | 0.017322 | 0.048258 | 0.149333 | 0.1179 | 0.257868 |
| SL1344_RS05855_ | -0.03059 | -0.04153 | -0.02532 | 0.03217 | 0.238018 | 0.12115 | 0.447035 | 0.64581 | 0.763401 | 0.7652 | 0.842852 |
| SL1344_RS05860_ | -1.14456 | -1.39709 | -0.97924 | 0.422839 | 0.267056 | 1.491524 | 0.044637 | 0.096967 | 0.228013 | 0.1583 | 0.303503 |
| SL1344_RS05865_ | 1.162984 | 1.165316 | 1.161556 | -0.01391 | 0.148622 | -0.0866 | 0.46036 | 0.664596 | 0.776742 | 0.7795 | 0.852488 |
| SL1344_RS05870_ | 4.282489 | 4.450553 | 4.137409 | -0.27774 | 0.098806 | -2.81742 | 4.68E-05 | 0.01233 | 0.083256 | 0.1 | 0.236592 |
| SL1344_RS05875_ | 5.242247 | 5.27125 | 5.215447 | 0.001478 | 0.141855 | 0.010901 | 0.497668 | 0.89846 | 0.941979 | 0.9973 | 0.998398 |
| SL1344_RS05880_ | 3.571675 | 3.588836 | 3.39815 | -0.18916 | 0.13925 | -0.91775 | 0.249167 | 0.272763 | 0.459091 | 0.5187 | 0.665945 |
| SL1344_RS05885_ | 2.274553 | 2.340043 | 2.205181 | -0.12987 | 0.093518 | -1.30065 | 0.077282 | 0.124773 | 0.26772 | 0.1991 | 0.349887 |
| SL1344_RS05890_ | 3.098352 | 3.069994 | 3.120117 | 0.026394 | 0.105246 | 0.239131 | 0.403731 | 0.679907 | 0.790292 | 0.7901 | 0.860689 |
| SL1344_RS05900_ | 6.439814 | 6.496168 | 6.31586 | -0.17759 | 0.118942 | -1.57622 | 0.000673 | 0.092273 | 0.241254 | 0.1007 | 0.237534 |
| SL1344_RS05905_ | 5.634568 | 5.701189 | 5.620422 | -0.06924 | 0.078408 | -0.73312 | 0.277815 | 0.399179 | 0.577414 | 0.544 | 0.696477 |
| SL1344_RS05910_ | 0.662188 | 0.680196 | 0.640759 | -0.06862 | 0.214048 | -0.28967 | 0.357761 | 0.51568 | 0.662803 | 0.6846 | 0.775368 |
| SL1344_RS05915_ | 0.875689 | 0.938825 | 0.810603 | -0.12985 | 0.113575 | -1.068 | 0.118667 | 0.188479 | 0.332049 | 0.2689 | 0.411199 |
| SL1344_RS05920_ | 1.050427 | 1.126228 | 0.990155 | -0.14109 | 0.164926 | -0.82135 | 0.17988 | 0.273779 | 0.439114 | 0.3813 | 0.531159 |
| SL1344_RS05925_ | 1.260552 | 1.290724 | 1.233816 | -0.05861 | 0.110915 | -0.49312 | 0.288 | 0.430382 | 0.57527 | 0.5438 | 0.661752 |
| SL1344_RS05930_ | 1.210149 | 1.219468 | 1.193301 | -0.01795 | 0.134122 | -0.12063 | 0.453333 | 0.656599 | 0.771079 | 0.7527 | 0.837021 |
| SL1344_RS05935_ | 5.244984 | 4.948745 | 5.619319 | 0.670035 | 0.069721 | 9.529829 | 4.68E-05 | 0.004044 | 0.05333 | 0.1 | 0.236592 |
| SL1344_RS05940_ | -1.90816 | -1.85254 | -2.00411 | -0.10412 | 0.428241 | -0.2258 | 0.415723 | 0.640837 | 0.758775 | 0.7507 | 0.833944 |
| SL1344_RS05945_ | 1.784637 | 1.783334 | 1.788134 | 0.003307 | 0.143482 | 0.022574 | 0.494337 | 0.72747 | 0.825801 | 0.8211 | 0.888392 |
| SL1344_RS05950_ | 1.801334 | 1.743778 | 1.850202 | 0.115134 | 0.114724 | 0.996622 | 0.132 | 0.213814 | 0.375624 | 0.2844 | 0.431253 |
| SL1344_RS05955_ | 1.769074 | 1.644923 | 1.893725 | 0.264268 | 0.139177 | 1.865339 | 0.004668 | 0.062152 | 0.1847 | 0.1073 | 0.245782 |
| SL1344_RS05960_ | 1.246343 | 1.054701 | 1.370701 | 0.303285 | 0.173752 | 1.698274 | 0.031313 | 0.069889 | 0.196599 | 0.1304 | 0.274957 |
| SL1344_RS05965_ | 1.740201 | 1.652696 | 1.785736 | 0.11912 | 0.180022 | 0.648429 | 0.233333 | 0.368319 | 0.53987 | 0.4537 | 0.609361 |
| SL1344_RS05970_ | 3.532513 | 3.457808 | 3.601572 | 0.146327 | 0.07287 | 1.981714 | 0.012667 | 0.047966 | 0.159275 | 0.1104 | 0.250248 |
| SL1344_RS05975_ | 1.904052 | 1.795744 | 1.942394 | 0.138388 | 0.11416 | 1.06946 | 0.142667 | 0.201964 | 0.364868 | 0.309 | 0.460235 |
| SL1344_RS05980_ | -1.09691 | -1.27863 | -0.94864 | 0.330638 | 0.27984 | 1.138737 | 0.086667 | 0.154516 | 0.304004 | 0.2158 | 0.364264 |
| SL1344_RS05985_ | 2.672098 | 2.821731 | 2.594818 | -0.21881 | 0.085679 | -2.36238 | 0.004668 | 0.040726 | 0.144315 | 0.1049 | 0.242819 |
| SL1344_RS05990_ | 0.143453 | 0.522694 | 0.001185 | -0.54739 | 0.470166 | -1.26815 | 0.053333 | 0.118877 | 0.275844 | 0.1592 | 0.309865 |

| | | | | | | | | | | |
|---|---|---|---|---|---|---|---|---|---|---|
| SL1344_RS05995_ | 0.4977 | 0.513979 | 0.4818 | -0.03512 | 0.135877 | -0.23778 | 0.396667 | 0.527946 | 0.65875 | 0.632 | 0.733523 |
| SL1344_RS06000_ | 2.936323 | 2.822363 | 3.065691 | 0.242853 | 0.068536 | 3.458212 | 4.68E-05 | 0.011118 | 0.074868 | 0.1 | 0.236592 |
| SL1344_RS06005_ | 1.176511 | 1.309702 | 1.053073 | -0.25324 | 0.12618 | -1.96226 | 0.013334 | 0.048672 | 0.155794 | 0.1171 | 0.257648 |
| SL1344_RS06010_ | 1.237133 | 1.176494 | 1.316044 | 0.141866 | 0.116298 | 1.14202 | 0.112592 | 0.172569 | 0.321055 | 0.258 | 0.405081 |
| SL1344_RS06015_ | 1.545954 | 1.434523 | 1.67202 | 0.238692 | 0.103254 | 2.258279 | 0.004668 | 0.033588 | 0.126908 | 0.1048 | 0.242722 |
| SL1344_RS06020_ | 2.117178 | 2.039816 | 2.181363 | 0.153812 | 0.113785 | 1.298166 | 0.061959 | 0.122213 | 0.269332 | 0.1713 | 0.3191 |
| SL1344_RS06025_ | 2.439135 | 2.359441 | 2.497722 | 0.150246 | 0.135719 | 1.07846 | 0.110667 | 0.160506 | 0.32377 | 0.231 | 0.390875 |
| SL1344_RS06030_ | 2.458701 | 2.341633 | 2.561223 | 0.247266 | 0.131779 | 1.873048 | 0.002667 | 0.059669 | 0.182301 | 0.1029 | 0.24029 |
| SL1344_RS06035_ | 0.300014 | 0.400208 | 0.143501 | -0.24987 | 0.218453 | -1.08715 | 0.108 | 0.174744 | 0.335092 | 0.2478 | 0.402465 |
| SL1344_RS06045_ | 0.420531 | 0.550259 | 0.244891 | -0.31517 | 0.175864 | -1.71289 | 0.022652 | 0.073183 | 0.196057 | 0.1236 | 0.26503 |
| SL1344_RS06050_ | 0.872682 | 1.016464 | 0.668602 | -0.34728 | 0.128601 | -2.59072 | 0.002667 | 0.026624 | 0.112228 | 0.1034 | 0.240812 |
| SL1344_RS06055_ | 3.13735 | 3.061127 | 3.380821 | 0.325161 | 0.245842 | 1.438072 | 0.026649 | 0.072585 | 0.208767 | 0.128 | 0.272859 |
| SL1344_RS06070_ | 2.348159 | 2.162413 | 2.610912 | 0.443004 | 0.146103 | 2.853423 | 4.68E-05 | 0.025922 | 0.118129 | 0.1001 | 0.236725 |
| SL1344_RS06085_ | 2.115655 | 2.031166 | 2.205526 | 0.200658 | 0.186592 | 1.044745 | 0.128667 | 0.169095 | 0.337012 | 0.2439 | 0.406712 |
| SL1344_RS06090_ | 1.459181 | 1.408255 | 1.532002 | 0.126426 | 0.210489 | 0.581671 | 0.269333 | 0.399325 | 0.570395 | 0.5476 | 0.680318 |
| SL1344_RS06095_ | 2.380798 | 2.399826 | 2.364253 | -0.03298 | 0.103031 | -0.30385 | 0.375083 | 0.607124 | 0.732053 | 0.6993 | 0.795668 |
| SL1344_RS06100_ | -0.12097 | -0.22236 | -0.02015 | 0.199789 | 0.171553 | 1.095025 | 0.098601 | 0.169477 | 0.31352 | 0.2489 | 0.39359 |
| SL1344_RS06105_ | -0.49893 | -0.52583 | -0.45739 | 0.089743 | 0.283065 | 0.301549 | 0.369754 | 0.556476 | 0.688536 | 0.6799 | 0.774702 |
| SL1344_RS06110_ | -0.36778 | -0.44894 | -0.15154 | 0.284697 | 0.449237 | 0.641104 | 0.219853 | 0.375548 | 0.548124 | 0.4407 | 0.606314 |
| SL1344_RS06115_ | 6.065459 | 6.101671 | 6.038465 | -0.04266 | 0.100334 | -0.48597 | 0.261333 | 0.494149 | 0.659901 | 0.5054 | 0.664045 |
| SL1344_RS06120_ | 0.83664 | 1.280526 | 0.536656 | -0.69849 | 0.253676 | -2.73919 | 4.68E-05 | 0.017695 | 0.097336 | 0.1001 | 0.236732 |
| SL1344_RS06125_ | -5.28282 | -5.80518 | -3.99599 | 1.74733 | 2.332802 | 0.689129 | 0.223333 | 0.3677 | 0.534784 | 0.4441 | 0.601715 |
| SL1344_RS06130_ | -3.62032 | -3.20067 | -4.1577 | -0.97026 | 0.75131 | -1.25865 | 0.082667 | 0.132658 | 0.279866 | 0.2013 | 0.353039 |
| SL1344_RS06135_ | 1.1677 | 1.4377 | 0.865675 | -0.68866 | 0.367932 | -1.98539 | 4.68E-05 | 0.046783 | 0.162541 | 0.1 | 0.236592 |
| SL1344_RS06140_ | -0.04515 | -0.31082 | 0.097881 | 0.392751 | 0.244852 | 1.525029 | 0.044667 | 0.100035 | 0.241494 | 0.1539 | 0.300845 |
| SL1344_RS06150_ | -3.8406 | -4.16899 | -3.47049 | 0.765406 | 0.75834 | 0.976568 | 0.137333 | 0.214811 | 0.369861 | 0.3042 | 0.448103 |
| SL1344_RS06155_ | -7.53175 | -7.50248 | -7.57759 | -0.18244 | 2.50407 | -0.05828 | 0.46569 | 0.559058 | 0.691089 | 0.6928 | 0.783678 |
| SL1344_RS06160_ | -2.62352 | -3.13062 | -2.22204 | 0.925054 | 0.469881 | 1.912585 | 0.01599 | 0.055884 | 0.168005 | 0.1139 | 0.25405 |
| SL1344_RS06165_ | 1.990979 | 1.350232 | 2.412642 | 1.013843 | 0.260287 | 3.612496 | 4.68E-05 | 0.015344 | 0.092694 | 0.1 | 0.236592 |
| SL1344_RS06170_ | -2.68065 | -2.77362 | -2.58566 | 0.206652 | 0.488447 | 0.40393 | 0.328448 | 0.493438 | 0.633192 | 0.5994 | 0.709813 |
| SL1344_RS06180_ | 2.20391 | 1.955592 | 2.420034 | 0.476398 | 0.157382 | 3.067164 | 4.68E-05 | 0.024631 | 0.114201 | 0.1 | 0.236592 |
| SL1344_RS06185_ | -3.29224 | -3.39379 | -3.23681 | 0.195342 | 0.761832 | 0.235922 | 0.38974 | 0.57186 | 0.70634 | 0.713 | 0.80207 |
| SL1344_RS06190_ | -2.73002 | -2.71521 | -2.7525 | -0.0279 | 0.553534 | -0.05174 | 0.481013 | 0.664992 | 0.776622 | 0.7648 | 0.844277 |
| SL1344_RS06195_ | 1.909087 | 1.715562 | 2.389764 | 0.661483 | 0.279803 | 2.318726 | 4.68E-05 | 0.030995 | 0.1315 | 0.1 | 0.236592 |
| SL1344_RS06200_ | -1.65117 | -1.74421 | -1.512 | 0.288025 | 0.461243 | 0.575149 | 0.23984 | 0.371442 | 0.538129 | 0.5109 | 0.636514 |
| SL1344_RS06205_ | 3.629088 | 3.69851 | 3.500147 | -0.222 | 0.263201 | -0.95649 | 0.138574 | 0.19828 | 0.37906 | 0.2478 | 0.42009 |
| SL1344_RS06210_ | -4.224 | -3.66755 | -4.82151 | -1.23268 | 1.111253 | -1.04683 | 0.109261 | 0.178274 | 0.339393 | 0.2312 | 0.38516 |
| SL1344_RS06215_ | -0.52024 | -0.65794 | -0.37799 | 0.277249 | 0.209584 | 1.273326 | 0.08 | 0.137323 | 0.277382 | 0.2135 | 0.361462 |
| SL1344_RS06220_ | 1.893873 | 1.630938 | 2.110549 | 0.479508 | 0.098288 | 4.814851 | 4.68E-05 | 0.004901 | 0.052571 | 0.1 | 0.236592 |
| SL1344_RS06225_ | 0.545972 | -0.65089 | 1.551694 | 2.216844 | 0.234336 | 9.52708 | 4.68E-05 | 0.004702 | 0.053719 | 0.1 | 0.236592 |
| SL1344_RS06230_ | 2.461295 | 2.628177 | 2.292273 | -0.40495 | 0.215974 | -1.80231 | 4.68E-05 | 0.098772 | 0.250259 | 0.1 | 0.236592 |
| SL1344_RS06235_ | 0.899364 | 1.083622 | 0.721729 | -0.38994 | 0.216494 | -1.82391 | 4.68E-05 | 0.097974 | 0.246707 | 0.1009 | 0.2378 |
| SL1344_RS06240_ | -1.03877 | -0.93756 | -1.10532 | -0.16695 | 0.302358 | -0.51458 | 0.296469 | 0.45854 | 0.606389 | 0.5744 | 0.693968 |
| SL1344_RS06245_ | -0.5299 | -0.54249 | -0.51564 | 0.021833 | 0.211863 | 0.106903 | 0.452365 | 0.577916 | 0.703706 | 0.6908 | 0.782372 |
| SL1344_RS06250_ | -0.66253 | -0.63181 | -0.68164 | -0.05483 | 0.239529 | -0.22613 | 0.405333 | 0.570624 | 0.696699 | 0.685 | 0.777854 |

| | | | | | | | | | | | |
|---|---|---|---|---|---|---|---|---|---|---|---|
| SL1344_RS06255_ | -0.77246 | -0.90129 | -0.58301 | 0.310319 | 0.225503 | 1.315034 | 0.081279 | 0.134611 | 0.275106 | 0.2115 | 0.359097 |
| SL1344_RS06260_ | -0.76584 | -1.48662 | -0.19883 | 1.281605 | 0.244964 | 5.182648 | 4.68E-05 | 0.004546 | 0.050329 | 0.1 | 0.236592 |
| SL1344_RS06265_ | -1.90129 | -2.96352 | -1.23068 | 1.760807 | 0.515737 | 3.345164 | 4.68E-05 | 0.013203 | 0.081519 | 0.1 | 0.236592 |
| SL1344_RS06270_ | -1.0676 | -1.24998 | -0.90964 | 0.34037 | 0.241028 | 1.36403 | 0.071952 | 0.124503 | 0.258021 | 0.195 | 0.340402 |
| SL1344_RS06275_ | -0.1313 | -0.26441 | 0.06801 | 0.320267 | 0.208935 | 1.433324 | 0.06529 | 0.109161 | 0.248381 | 0.1816 | 0.33178 |
| SL1344_RS06280_ | -3.41333 | -3.57072 | -3.2207 | 0.329255 | 0.647531 | 0.465659 | 0.299134 | 0.483667 | 0.624371 | 0.5833 | 0.698447 |
| SL1344_RS06285_ | -0.15218 | -0.114 | -0.18225 | -0.05215 | 0.239181 | -0.21122 | 0.416389 | 0.649389 | 0.763346 | 0.7582 | 0.837244 |
| SL1344_RS06290_ | 1.489964 | 1.358787 | 1.796774 | 0.420416 | 0.147597 | 2.567668 | 0.001337 | 0.040746 | 0.147645 | 0.102 | 0.239039 |
| SL1344_RS06295_ | -4.30494 | -4.38487 | -4.23097 | 0.141809 | 0.70316 | 0.197423 | 0.412 | 0.526421 | 0.657561 | 0.6314 | 0.733443 |
| SL1344_RS06300_ | 0.777461 | 0.687413 | 0.870072 | 0.179673 | 0.169197 | 1.018914 | 0.134577 | 0.203456 | 0.361788 | 0.2852 | 0.436155 |
| SL1344_RS06305_ | 0.976507 | 0.827323 | 1.096122 | 0.282542 | 0.171929 | 1.646999 | 0.013325 | 0.083372 | 0.216972 | 0.1159 | 0.25592 |
| SL1344_RS06310_ | -1.21492 | -0.94913 | -1.38161 | -0.42047 | 0.329501 | -1.185 | 0.092 | 0.146042 | 0.297803 | 0.2196 | 0.372869 |
| SL1344_RS06315_ | 0.905744 | 0.89464 | 0.911644 | 0.02319 | 0.201214 | 0.109509 | 0.442667 | 0.677166 | 0.788107 | 0.7822 | 0.856356 |
| SL1344_RS06320_ | -2.27958 | -2.66654 | -2.05922 | 0.60685 | 0.506662 | 1.128545 | 0.099933 | 0.1775 | 0.335733 | 0.2445 | 0.39537 |
| SL1344_RS06325_ | -1.11203 | -1.45777 | -0.74507 | 0.781898 | 0.385713 | 2.047586 | 0.003335 | 0.04611 | 0.156944 | 0.1024 | 0.23968 |
| SL1344_RS06330_ | 2.193301 | 1.750725 | 2.912931 | 1.095709 | 0.212495 | 4.464287 | 4.68E-05 | 0.012828 | 0.085809 | 0.1 | 0.236592 |
| SL1344_RS06335_ | 0.64929 | 0.414886 | 0.914073 | 0.505245 | 0.167882 | 2.97362 | 4.68E-05 | 0.016483 | 0.090406 | 0.1002 | 0.236858 |
| SL1344_RS06340_ | 1.582332 | 1.427125 | 1.922155 | 0.471386 | 0.215876 | 2.156218 | 0.002001 | 0.030787 | 0.128569 | 0.1014 | 0.238446 |
| SL1344_RS06345_ | 0.509306 | 0.467501 | 0.652917 | 0.166106 | 0.236501 | 0.5649 | 0.316667 | 0.511714 | 0.661313 | 0.6369 | 0.76015 |
| SL1344_RS06350_ | -2.13152 | -1.99118 | -2.41219 | -0.38828 | 0.687156 | -0.49038 | 0.293138 | 0.436405 | 0.602548 | 0.5888 | 0.71164 |
| SL1344_RS06355_ | -0.75876 | -0.68487 | -0.81049 | -0.14334 | 0.247157 | -0.51187 | 0.283145 | 0.424914 | 0.572151 | 0.533 | 0.650098 |
| SL1344_RS06360_ | -1.4628 | -1.4747 | -1.44409 | 0.053021 | 0.394658 | 0.130234 | 0.445703 | 0.657752 | 0.770245 | 0.765 | 0.841362 |
| SL1344_RS06365_ | -4.86069 | -4.99641 | -4.73514 | 0.254714 | 0.869691 | 0.272089 | 0.381746 | 0.501335 | 0.637886 | 0.6106 | 0.718621 |
| SL1344_RS06370_ | 0.458475 | 0.124326 | 0.768312 | 0.639864 | 0.1528 | 4.115191 | 4.68E-05 | 0.006968 | 0.060667 | 0.1 | 0.236592 |
| SL1344_RS06375_ | 0.696201 | 0.36246 | 0.930833 | 0.567759 | 0.125671 | 4.422711 | 4.68E-05 | 0.006734 | 0.057915 | 0.1 | 0.236592 |
| SL1344_RS06380_ | 1.897613 | 1.436383 | 2.379346 | 0.926158 | 0.15928 | 5.841218 | 4.68E-05 | 0.00164 | 0.037376 | 0.1 | 0.236592 |
| SL1344_RS06385_ | 2.566776 | 2.036893 | 3.169497 | 1.142245 | 0.121189 | 9.589729 | 4.68E-05 | 0.00126 | 0.033534 | 0.1 | 0.236592 |
| SL1344_RS06390_ | 4.491358 | 4.409237 | 4.591352 | 0.192273 | 0.129302 | 1.462412 | 0.037309 | 0.078122 | 0.217587 | 0.1324 | 0.279777 |
| SL1344_RS06395_ | -1.99542 | -1.90027 | -2.13175 | -0.22428 | 0.361538 | -0.58025 | 0.256 | 0.397223 | 0.547533 | 0.5126 | 0.6363 |
| SL1344_RS06400_ | 0.646682 | 0.558989 | 1.004444 | 0.424889 | 0.239157 | 1.413697 | 0.074 | 0.129617 | 0.287536 | 0.1938 | 0.344872 |
| SL1344_RS06405_ | 1.492827 | 1.443643 | 1.597261 | 0.147346 | 0.178771 | 0.769143 | 0.231179 | 0.335963 | 0.509835 | 0.4599 | 0.606547 |
| SL1344_RS06410_ | 6.999165 | 6.922323 | 7.07666 | 0.157836 | 0.044134 | 3.390994 | 4.68E-05 | 0.082205 | 0.2259 | 0.1 | 0.236592 |
| SL1344_RS06415_ | 2.326515 | 2.37672 | 2.280091 | -0.11228 | 0.108342 | -0.97068 | 0.131333 | 0.201665 | 0.363926 | 0.2893 | 0.438681 |
| SL1344_RS06420_ | 2.905148 | 2.919453 | 2.892519 | -0.05793 | 0.1211 | -0.45009 | 0.319121 | 0.471345 | 0.63014 | 0.6701 | 0.771215 |
| SL1344_RS06425_ | 0.317627 | 0.107122 | 0.728484 | 0.5844 | 0.232072 | 2.351986 | 0.005334 | 0.035887 | 0.137118 | 0.1029 | 0.240278 |
| SL1344_RS06430_ | 0.273754 | 0.080772 | 0.424033 | 0.339405 | 0.187935 | 1.712521 | 0.021986 | 0.064687 | 0.183276 | 0.1276 | 0.26986 |
| SL1344_RS06435_ | 1.809757 | 1.712692 | 1.899988 | 0.196551 | 0.12721 | 1.510004 | 0.029333 | 0.095295 | 0.234518 | 0.1301 | 0.273547 |
| SL1344_RS06440_ | 1.313246 | 0.937447 | 1.379578 | 0.418635 | 0.21541 | 1.292955 | 0.099267 | 0.147027 | 0.313058 | 0.2335 | 0.391207 |
| SL1344_RS06445_ | 1.951049 | 1.779365 | 2.113556 | 0.332739 | 0.115146 | 2.794289 | 0.000673 | 0.019652 | 0.100132 | 0.1004 | 0.237122 |
| SL1344_RS06450_ | 2.758216 | 2.620487 | 2.903164 | 0.275544 | 0.08227 | 3.411036 | 4.68E-05 | 0.012431 | 0.079158 | 0.1003 | 0.236996 |
| SL1344_RS06455_ | 1.28487 | 1.208088 | 1.36591 | 0.156984 | 0.118519 | 1.285539 | 0.072618 | 0.129473 | 0.270547 | 0.2023 | 0.351062 |
| SL1344_RS06465_ | -1.64379 | -2.01632 | -1.2845 | 0.724781 | 0.318679 | 2.197195 | 0.008667 | 0.03871 | 0.136127 | 0.1103 | 0.248933 |
| SL1344_RS06470_ | 1.599941 | 1.356895 | 1.857115 | 0.508264 | 0.13825 | 3.63817 | 4.68E-05 | 0.008523 | 0.067951 | 0.1 | 0.236592 |
| SL1344_RS06475_ | -1.15439 | -1.30409 | -1.01177 | 0.289654 | 0.261036 | 1.039481 | 0.124584 | 0.196527 | 0.344181 | 0.2772 | 0.422307 |
| SL1344_RS06480_ | -1.66776 | -1.76429 | -1.55386 | 0.250344 | 0.420807 | 0.566455 | 0.273151 | 0.419577 | 0.575833 | 0.5529 | 0.674319 |

| | | | | | | | | | | | |
|---|---|---|---|---|---|---|---|---|---|---|---|
| SL1344_RS06485_ | -0.64944 | -0.90978 | -0.50678 | 0.416667 | 0.276834 | 1.397744 | 0.046 | 0.118578 | 0.264233 | 0.1598 | 0.304665 |
| SL1344_RS06490_ | -0.88654 | -1.52583 | -0.66519 | 0.823553 | 0.396645 | 1.797696 | 0.019987 | 0.081703 | 0.218557 | 0.1199 | 0.261429 |
| SL1344_RS06495_ | -1.11994 | -1.52167 | -0.77646 | 0.751542 | 0.23988 | 3.059722 | 0.000673 | 0.018425 | 0.092197 | 0.1006 | 0.237387 |
| SL1344_RS06500_ | 0.668832 | 0.518517 | 0.766829 | 0.245425 | 0.168971 | 1.369065 | 0.078614 | 0.123242 | 0.267296 | 0.1953 | 0.346772 |
| SL1344_RS06505_ | 0.406522 | 0.553052 | 0.250703 | -0.32611 | 0.195979 | -1.61587 | 0.027315 | 0.080012 | 0.210002 | 0.1314 | 0.274521 |
| SL1344_RS06510_ | 2.145242 | 1.916342 | 2.360805 | 0.44628 | 0.077148 | 5.709243 | 4.68E-05 | 0.003573 | 0.045615 | 0.1 | 0.236592 |
| SL1344_RS06515_ | 3.803987 | 3.341561 | 4.323143 | 0.973252 | 0.085434 | 11.26699 | 4.68E-05 | 0.000915 | 0.031039 | 0.1 | 0.236592 |
| SL1344_RS06520_ | 0.983014 | 0.672394 | 1.256522 | 0.608434 | 0.172424 | 3.511316 | 4.68E-05 | 0.009514 | 0.071305 | 0.1 | 0.236592 |
| SL1344_RS06525_ | 0.81374 | 0.420486 | 1.214878 | 0.826814 | 0.194557 | 4.284038 | 4.68E-05 | 0.011678 | 0.079761 | 0.1 | 0.236592 |
| SL1344_RS06530_ | -0.39748 | -0.65105 | 0.122141 | 0.742612 | 0.275809 | 2.464292 | 0.002001 | 0.034302 | 0.133659 | 0.1016 | 0.238729 |
| SL1344_RS06535_ | 0.390007 | -0.00878 | 0.779606 | 0.785098 | 0.169017 | 4.603996 | 4.68E-05 | 0.005254 | 0.05426 | 0.1 | 0.236592 |
| SL1344_RS06540_ | -1.07456 | -1.32246 | -0.57358 | 0.713898 | 0.31697 | 2.083301 | 0.015323 | 0.050089 | 0.161078 | 0.115 | 0.255011 |
| SL1344_RS06545_ | 0.722124 | 0.466037 | 1.097221 | 0.652823 | 0.198747 | 3.261747 | 4.68E-05 | 0.015772 | 0.090642 | 0.1 | 0.236592 |
| SL1344_RS06550_ | 2.707688 | 2.214746 | 3.192405 | 0.975767 | 0.093767 | 10.33781 | 4.68E-05 | 0.000518 | 0.026347 | 0.1 | 0.236592 |
| SL1344_RS06555_ | -1.23731 | -1.24717 | -1.22421 | 0.026997 | 0.287577 | 0.094557 | 0.457695 | 0.590241 | 0.713746 | 0.6868 | 0.78177 |
| SL1344_RS06565_ | -1.53137 | -1.27978 | -1.6705 | -0.36975 | 0.400824 | -0.85915 | 0.187333 | 0.272085 | 0.436409 | 0.3662 | 0.520032 |
| SL1344_RS06570_ | 1.930485 | 1.886201 | 2.0115 | 0.110111 | 0.183647 | 0.599983 | 0.248501 | 0.404693 | 0.573569 | 0.4772 | 0.631724 |
| SL1344_RS06575_ | 0.402187 | 0.218522 | 0.52368 | 0.318421 | 0.203303 | 1.469665 | 0.036 | 0.105084 | 0.246786 | 0.1392 | 0.282793 |
| SL1344_RS06580_ | 1.823642 | 1.430465 | 2.355017 | 0.904469 | 0.131196 | 6.596386 | 4.68E-05 | 0.004349 | 0.052238 | 0.1 | 0.236592 |
| SL1344_RS06605_ | -1.31747 | -1.1656 | -1.48633 | -0.35201 | 0.350063 | -0.95367 | 0.134667 | 0.221142 | 0.382526 | 0.2996 | 0.445947 |
| SL1344_RS06615_ | -3.41068 | -3.77665 | -3.06372 | 0.785674 | 0.863735 | 0.855631 | 0.147901 | 0.240042 | 0.409429 | 0.3094 | 0.461156 |
| SL1344_RS06630_ | 7.025072 | 7.073023 | 7.008397 | -0.09993 | 0.095965 | -1.09879 | 0.035333 | 0.160477 | 0.333986 | 0.1377 | 0.283349 |
| SL1344_RS06635_ | 6.883307 | 6.907224 | 6.847671 | -0.1815 | 0.203601 | -0.94311 | 0.147333 | 0.20908 | 0.392385 | 0.2744 | 0.446888 |
| SL1344_RS06640_ | 7.165469 | 7.201123 | 7.025193 | -0.17873 | 0.142909 | -1.4229 | 0.000673 | 0.07496 | 0.214374 | 0.1007 | 0.23752 |
| SL1344_RS06645_ | 2.079611 | 2.08879 | 2.072996 | -0.03103 | 0.127394 | -0.25007 | 0.406667 | 0.603915 | 0.730009 | 0.7421 | 0.823977 |
| SL1344_RS06650_ | 4.100225 | 3.901505 | 4.18316 | 0.274079 | 0.131782 | 1.752141 | 0.011992 | 0.081658 | 0.223383 | 0.1123 | 0.252912 |
| SL1344_RS06660_ | 0.245997 | 0.253912 | 0.234737 | -0.02635 | 0.177531 | -0.13642 | 0.438 | 0.592482 | 0.716344 | 0.7117 | 0.799039 |
| SL1344_RS06665_ | 0.12256 | -0.13685 | 0.401609 | 0.546926 | 0.186998 | 2.934934 | 0.000673 | 0.019379 | 0.097126 | 0.1008 | 0.237598 |
| SL1344_RS06670_ | 0.098827 | -0.21138 | 0.360288 | 0.575655 | 0.179263 | 3.199493 | 4.68E-05 | 0.018867 | 0.094368 | 0.1 | 0.236592 |
| SL1344_RS06675_ | 0.239662 | -0.04392 | 0.631605 | 0.659178 | 0.194735 | 3.266783 | 0.000673 | 0.014108 | 0.084029 | 0.1 | 0.236592 |
| SL1344_RS06685_ | 0.665672 | 0.39579 | 0.825566 | 0.418522 | 0.173654 | 2.205569 | 0.004668 | 0.039531 | 0.141868 | 0.1055 | 0.243716 |
| SL1344_RS06690_ | -2.04348 | -2.89353 | -1.83537 | 0.985231 | 1.146472 | 0.694447 | 0.2998 | 0.380491 | 0.562652 | 0.62 | 0.75501 |
| SL1344_RS06695_ | 1.113191 | 1.026686 | 1.187906 | 0.160771 | 0.111072 | 1.390189 | 0.062667 | 0.113303 | 0.245031 | 0.1809 | 0.326215 |
| SL1344_RS06700_ | 0.707882 | 0.382536 | 0.929205 | 0.558157 | 0.18808 | 2.945441 | 0.000673 | 0.031781 | 0.127885 | 0.1 | 0.236592 |
| SL1344_RS06705_ | 5.85774 | 5.691542 | 5.9347 | 0.243088 | 0.078776 | 2.882403 | 4.68E-05 | 0.045245 | 0.161917 | 0.1 | 0.236592 |
| SL1344_RS06710_ | -1.58895 | -1.63399 | -1.54583 | 0.074705 | 0.342131 | 0.205824 | 0.412 | 0.59737 | 0.721504 | 0.6938 | 0.787737 |
| SL1344_RS06715_ | -4.71755 | -5.10461 | -4.1864 | 0.848698 | 1.016813 | 0.764217 | 0.192 | 0.314363 | 0.468518 | 0.4012 | 0.543171 |
| SL1344_RS06720_ | -2.78381 | -2.992 | -2.67252 | 0.29402 | 0.516592 | 0.523655 | 0.291333 | 0.442654 | 0.591786 | 0.5579 | 0.67947 |
| SL1344_RS06730_ | -4.70779 | -4.4574 | -4.96511 | -0.59005 | 1.142459 | -0.46849 | 0.286 | 0.428884 | 0.582517 | 0.5568 | 0.672317 |
| SL1344_RS06735_ | -3.13932 | -3.24306 | -3.01546 | 0.228158 | 0.496464 | 0.424564 | 0.318454 | 0.470894 | 0.609273 | 0.5728 | 0.685604 |
| SL1344_RS06740_ | -0.6918 | -0.61004 | -0.79853 | -0.17664 | 0.298339 | -0.56551 | 0.266 | 0.409614 | 0.563292 | 0.5245 | 0.652443 |
| SL1344_RS06745_ | -2.78146 | -2.76027 | -2.82093 | -0.03381 | 0.507624 | -0.07012 | 0.472 | 0.614294 | 0.734474 | 0.7253 | 0.812178 |
| SL1344_RS06750_ | 0.661103 | 0.552602 | 0.801167 | 0.25368 | 0.192272 | 1.291538 | 0.073333 | 0.124988 | 0.272379 | 0.1933 | 0.345744 |
| SL1344_RS06755_ | -0.75364 | -0.81827 | -0.62175 | 0.186031 | 0.261501 | 0.627901 | 0.255829 | 0.413417 | 0.562711 | 0.5179 | 0.646935 |
| SL1344_RS06760_ | -2.30506 | -2.34131 | -2.26009 | 0.101359 | 0.482306 | 0.195086 | 0.42 | 0.622731 | 0.742616 | 0.739 | 0.821577 |

| | | | | | | | | | | |
|---|---|---|---|---|---|---|---|---|---|---|
| SL1344_RS06765_ | -3.17256 | -3.33649 | -2.9814 | 0.385385 | 0.596106 | 0.599868 | 0.258667 | 0.389624 | 0.542559 | 0.5035 | 0.628635 |
| SL1344_RS06770_ | -4.28451 | -4.51751 | -4.06575 | 0.457282 | 0.810972 | 0.50556 | 0.283811 | 0.430812 | 0.576083 | 0.5283 | 0.649419 |
| SL1344_RS06775_ | 1.056693 | 0.84413 | 1.304882 | 0.460941 | 0.124243 | 3.656266 | 0.000673 | 0.009779 | 0.069939 | 0.1 | 0.236592 |
| SL1344_RS06780_ | 2.562941 | 2.43678 | 2.658328 | 0.219696 | 0.076833 | 2.842381 | 0.000673 | 0.020626 | 0.099662 | 0.1009 | 0.237785 |
| SL1344_RS06785_ | 0.494161 | 0.211277 | 0.631489 | 0.39685 | 0.273109 | 1.414958 | 0.065956 | 0.104964 | 0.252041 | 0.1719 | 0.326372 |
| SL1344_RS06790_ | 2.608338 | 2.990959 | 2.325067 | -0.64905 | 0.196839 | -3.13311 | 4.68E-05 | 0.01751 | 0.098743 | 0.1 | 0.236592 |
| SL1344_RS06795_ | 0.061211 | -0.26992 | 0.419829 | 0.695658 | 0.192483 | 3.575812 | 4.68E-05 | 0.010126 | 0.072058 | 0.1 | 0.236592 |
| SL1344_RS06800_ | -2.98711 | -3.78925 | -2.45357 | 1.342401 | 0.488864 | 2.66505 | 0.002002 | 0.028093 | 0.113369 | 0.1018 | 0.238934 |
| SL1344_RS06805_ | -0.65244 | -1.16738 | -0.26071 | 0.910369 | 0.209013 | 4.245742 | 4.68E-05 | 0.007651 | 0.061142 | 0.1 | 0.236592 |
| SL1344_RS06810_ | -0.50017 | -0.72128 | -0.33531 | 0.381282 | 0.226966 | 1.566904 | 0.037333 | 0.080499 | 0.205962 | 0.1472 | 0.29289 |
| SL1344_RS06815_ | -0.32981 | -0.68148 | -0.06482 | 0.617773 | 0.206142 | 2.948792 | 4.68E-05 | 0.019402 | 0.095483 | 0.1002 | 0.236856 |
| SL1344_RS06820_ | -0.52846 | -0.8062 | -0.27475 | 0.528395 | 0.221032 | 2.317865 | 0.003335 | 0.029996 | 0.120218 | 0.1036 | 0.241164 |
| SL1344_RS06825_ | -1.725 | -1.92211 | -1.4114 | 0.500624 | 0.346326 | 1.377508 | 0.075333 | 0.123731 | 0.265143 | 0.1973 | 0.348133 |
| SL1344_RS06830_ | 0.894115 | 0.666867 | 1.141774 | 0.487668 | 0.165776 | 2.961891 | 4.68E-05 | 0.018718 | 0.097255 | 0.1001 | 0.236727 |
| SL1344_RS06835_ | 1.143741 | 1.104008 | 1.235669 | 0.106004 | 0.171691 | 0.556453 | 0.295137 | 0.483631 | 0.631789 | 0.5898 | 0.714759 |
| SL1344_RS06840_ | 10.60281 | 10.49668 | 10.60705 | 0.155992 | 0.169213 | 1.060207 | 0.224 | 0.15325 | 0.325791 | 0.3988 | 0.581059 |
| SL1344_RS06845_ | 2.899251 | 2.648685 | 3.223429 | 0.573214 | 0.093377 | 6.125847 | 4.68E-05 | 0.004238 | 0.051186 | 0.1 | 0.236592 |
| SL1344_RS06850_ | -0.08565 | -0.67912 | 0.480092 | 1.159866 | 0.16606 | 6.98535 | 4.68E-05 | 0.002298 | 0.03852 | 0.1 | 0.236592 |
| SL1344_RS06855_ | -1.81357 | -2.15826 | -1.33616 | 0.810289 | 0.428576 | 1.861784 | 0.016 | 0.059427 | 0.177573 | 0.1198 | 0.260997 |
| SL1344_RS06860_ | -0.13331 | -0.89403 | 0.354898 | 1.242138 | 0.246135 | 4.993476 | 4.68E-05 | 0.007585 | 0.062684 | 0.1 | 0.236592 |
| SL1344_RS06865_ | -0.50859 | -1.04945 | 0.037049 | 1.099199 | 0.254136 | 4.278358 | 4.68E-05 | 0.005863 | 0.057509 | 0.1 | 0.236592 |
| SL1344_RS06900_ | 2.632237 | 2.369333 | 2.929964 | 0.567776 | 0.079992 | 7.0129 | 4.68E-05 | 0.002213 | 0.039409 | 0.1 | 0.236592 |
| SL1344_RS06905_ | -0.28186 | -0.4118 | -0.1465 | 0.268973 | 0.170176 | 1.499433 | 0.053298 | 0.097986 | 0.22218 | 0.1663 | 0.311679 |
| SL1344_RS06925_ | -3.25982 | -3.09276 | -3.41103 | -0.30146 | 0.63317 | -0.44833 | 0.309333 | 0.489899 | 0.629307 | 0.583 | 0.698475 |
| SL1344_RS06930_ | -2.96268 | -2.9933 | -2.93572 | 0.029992 | 0.522098 | 0.051057 | 0.479014 | 0.613072 | 0.732519 | 0.7253 | 0.809751 |
| SL1344_RS06935_ | -3.26681 | -3.25032 | -3.28546 | -0.05762 | 0.730901 | -0.0656 | 0.47 | 0.660229 | 0.773987 | 0.7714 | 0.846206 |
| SL1344_RS06940_ | -2.66993 | -2.56685 | -2.72411 | -0.23803 | 0.558834 | -0.38788 | 0.342 | 0.499773 | 0.642765 | 0.6359 | 0.736372 |
| SL1344_RS06945_ | -1.92992 | -1.95829 | -1.889 | 0.046308 | 0.387413 | 0.108681 | 0.451033 | 0.637584 | 0.753142 | 0.7303 | 0.816797 |
| SL1344_RS06950_ | -2.47799 | -2.4742 | -2.48155 | -0.01138 | 0.442908 | -0.02387 | 0.487675 | 0.604281 | 0.726191 | 0.7089 | 0.798772 |
| SL1344_RS06955_ | -0.90632 | -1.12254 | -0.52863 | 0.563656 | 0.290194 | 1.84873 | 0.023318 | 0.060775 | 0.179343 | 0.1268 | 0.269849 |
| SL1344_RS06960_ | -2.63735 | -2.87856 | -2.388 | 0.492424 | 0.466248 | 0.996604 | 0.127915 | 0.204444 | 0.356267 | 0.2937 | 0.440112 |
| SL1344_RS06965_ | -2.58221 | -2.9355 | -2.28409 | 0.652964 | 0.486285 | 1.291096 | 0.076667 | 0.128537 | 0.272477 | 0.1896 | 0.338946 |
| SL1344_RS06970_ | -2.08692 | -2.11367 | -2.05803 | 0.066171 | 0.355582 | 0.168289 | 0.429047 | 0.558057 | 0.685835 | 0.6688 | 0.763646 |
| SL1344_RS06975_ | -2.11981 | -2.2253 | -2.00926 | 0.221191 | 0.408646 | 0.528406 | 0.278481 | 0.433155 | 0.577711 | 0.5406 | 0.659778 |
| SL1344_RS06980_ | -1.39194 | -1.60494 | -1.14931 | 0.442194 | 0.312539 | 1.375921 | 0.068 | 0.12183 | 0.26164 | 0.1929 | 0.342644 |
| SL1344_RS06990_ | -4.50146 | -3.93239 | -5.12366 | -1.18586 | 0.892554 | -1.26969 | 0.074667 | 0.131751 | 0.273142 | 0.1964 | 0.343504 |
| SL1344_RS06995_ | -5.21334 | -3.84886 | -6.6427 | -2.75121 | 1.363775 | -1.95128 | 0.012 | 0.056254 | 0.169141 | 0.1095 | 0.248737 |
| SL1344_RS07000_ | -3.05149 | -2.74892 | -3.76899 | -0.96103 | 0.740878 | -1.16203 | 0.111925 | 0.172053 | 0.335477 | 0.2481 | 0.404727 |
| SL1344_RS07005_ | -3.0551 | -2.47581 | -3.80339 | -1.32885 | 0.549317 | -2.36518 | 0.005997 | 0.036227 | 0.131114 | 0.1043 | 0.242087 |
| SL1344_RS07010_ | -3.61054 | -3.58881 | -3.6537 | 0.105238 | 1.049834 | 0.096056 | 0.462 | 0.714221 | 0.816908 | 0.8448 | 0.898943 |
| SL1344_RS07015_ | -1.70024 | -1.75796 | -1.63924 | 0.046465 | 0.539104 | 0.083023 | 0.464 | 0.734157 | 0.830542 | 0.8231 | 0.886102 |
| SL1344_RS07020_ | -4.39811 | -4.58418 | -4.24987 | 0.33504 | 0.804657 | 0.370648 | 0.333333 | 0.466007 | 0.606763 | 0.5722 | 0.683841 |
| SL1344_RS07025_ | -1.56971 | -1.61998 | -1.48551 | 0.141898 | 0.35098 | 0.379886 | 0.336 | 0.51317 | 0.650187 | 0.6309 | 0.735888 |
| SL1344_RS07030_ | -0.82094 | -0.95295 | -0.71827 | 0.283098 | 0.344467 | 0.757974 | 0.170553 | 0.261611 | 0.434032 | 0.3536 | 0.497386 |
| SL1344_RS07035_ | -2.22132 | -2.45271 | -1.96378 | 0.51734 | 0.398638 | 1.211646 | 0.076 | 0.14415 | 0.288936 | 0.2123 | 0.360167 |

| | | | | | | | | | | | |
|---|---|---|---|---|---|---|---|---|---|---|---|
| SL1344_RS07040_ | -2.20445 | -2.54486 | -1.88735 | 0.659063 | 0.401679 | 1.636038 | 0.031979 | 0.079036 | 0.203495 | 0.1377 | 0.280632 |
| SL1344_RS07045_ | -0.85555 | -1.09907 | -0.43668 | 0.62241 | 0.360198 | 1.627471 | 0.037975 | 0.079088 | 0.212039 | 0.1409 | 0.2882 |
| SL1344_RS07055_ | -4.73595 | -4.79629 | -4.68171 | 0.146904 | 0.907317 | 0.152093 | 0.442372 | 0.553703 | 0.684078 | 0.6683 | 0.765217 |
| SL1344_RS07060_ | -3.29869 | -3.21368 | -3.40106 | -0.18652 | 0.548612 | -0.31183 | 0.361333 | 0.527176 | 0.659553 | 0.6331 | 0.736424 |
| SL1344_RS07065_ | -2.60535 | -2.68404 | -2.49212 | 0.211263 | 0.503026 | 0.406142 | 0.339107 | 0.513876 | 0.650784 | 0.6294 | 0.734754 |
| SL1344_RS07070_ | -1.11949 | -1.16136 | -1.01912 | 0.155429 | 0.360269 | 0.401553 | 0.334667 | 0.515475 | 0.657263 | 0.6614 | 0.76042 |
| SL1344_RS07075_ | -0.88031 | -1.02338 | -0.76571 | 0.326527 | 0.466731 | 0.711677 | 0.201333 | 0.287616 | 0.465927 | 0.3848 | 0.534724 |
| SL1344_RS07080_ | 1.981575 | 1.955841 | 2.00099 | 0.058505 | 0.128103 | 0.420028 | 0.301333 | 0.46025 | 0.613771 | 0.6022 | 0.714079 |
| SL1344_RS07085_ | 1.545044 | 1.16162 | 1.708152 | 0.530684 | 0.213739 | 2.281904 | 0.000673 | 0.039358 | 0.146882 | 0.1006 | 0.237397 |
| SL1344_RS07090_ | 2.699003 | 2.483313 | 2.845165 | 0.327818 | 0.149066 | 2.087619 | 0.008661 | 0.044518 | 0.156619 | 0.1068 | 0.245589 |
| SL1344_RS07095_ | -0.49333 | -0.5294 | -0.46642 | 0.056473 | 0.22541 | 0.244613 | 0.391333 | 0.564729 | 0.692594 | 0.6788 | 0.772578 |
| SL1344_RS07100_ | -1.10388 | -1.33381 | -0.82847 | 0.487831 | 0.325246 | 1.454924 | 0.054 | 0.09832 | 0.234593 | 0.1645 | 0.314211 |
| SL1344_RS07115_ | 1.763549 | 1.636883 | 1.969524 | 0.329772 | 0.171934 | 1.929828 | 0.008001 | 0.043504 | 0.152997 | 0.1079 | 0.24677 |
| SL1344_RS07120_ | 4.078076 | 4.365618 | 3.965056 | -0.39721 | 0.087584 | -3.27338 | 4.68E-05 | 0.03828 | 0.147674 | 0.1 | 0.236592 |
| SL1344_RS07125_ | 0.498838 | 0.467759 | 0.535759 | 0.068417 | 0.135307 | 0.471188 | 0.295333 | 0.423227 | 0.564261 | 0.5265 | 0.64606 |
| SL1344_RS07130_ | 2.254784 | 2.109997 | 2.388321 | 0.288163 | 0.107908 | 2.596131 | 0.000673 | 0.023455 | 0.108649 | 0.1004 | 0.237122 |
| SL1344_RS07135_ | 1.446505 | 1.308187 | 1.557157 | 0.247077 | 0.110076 | 2.173657 | 0.009334 | 0.038222 | 0.136487 | 0.1079 | 0.246574 |
| SL1344_RS07140_ | -0.60546 | -0.49096 | -0.68178 | -0.18468 | 0.222068 | -0.7687 | 0.205333 | 0.32132 | 0.472592 | 0.4176 | 0.555755 |
| SL1344_RS07145_ | -1.61565 | -1.98846 | -1.20645 | 0.807539 | 0.336894 | 2.424344 | 0.001336 | 0.03331 | 0.128336 | 0.1021 | 0.239112 |
| SL1344_RS07150_ | 1.212527 | 0.956611 | 1.495319 | 0.549242 | 0.151355 | 3.651314 | 4.68E-05 | 0.009261 | 0.070011 | 0.1 | 0.236592 |
| SL1344_RS07155_ | 1.028372 | 0.77595 | 1.390385 | 0.609014 | 0.201828 | 2.947784 | 4.68E-05 | 0.023277 | 0.110456 | 0.1 | 0.236592 |
| SL1344_RS07160_ | -0.35228 | -0.54898 | 0.04361 | 0.558826 | 0.242446 | 2.184719 | 0.01066 | 0.047685 | 0.157422 | 0.1103 | 0.249615 |
| SL1344_RS07165_ | -0.80527 | -0.91639 | -0.70935 | 0.208179 | 0.208671 | 0.944656 | 0.15523 | 0.237158 | 0.38263 | 0.3232 | 0.462831 |
| SL1344_RS07170_ | -2.8141 | -2.95871 | -2.5839 | 0.307907 | 0.645779 | 0.437749 | 0.313333 | 0.54915 | 0.683524 | 0.6451 | 0.754764 |
| SL1344_RS07180_ | 3.132598 | 2.722023 | 3.811388 | 1.081866 | 0.121422 | 7.706829 | 4.68E-05 | 0.011567 | 0.082572 | 0.1 | 0.236592 |
| SL1344_RS07185_ | 0.345116 | 0.287944 | 0.442355 | 0.154532 | 0.169453 | 0.854191 | 0.179333 | 0.286111 | 0.442616 | 0.3876 | 0.530521 |
| SL1344_RS07190_ | 0.386098 | 0.021155 | 0.576942 | 0.5412 | 0.185813 | 2.811569 | 0.002001 | 0.03079 | 0.123846 | 0.1006 | 0.237387 |
| SL1344_RS07195_ | 1.256724 | 1.238354 | 1.274849 | 0.033837 | 0.133273 | 0.246093 | 0.4 | 0.589749 | 0.715621 | 0.6979 | 0.790355 |
| SL1344_RS07200_ | 3.508732 | 3.499121 | 3.531167 | 0.031251 | 0.07586 | 0.357701 | 0.361333 | 0.605015 | 0.731276 | 0.7237 | 0.815426 |
| SL1344_RS07205_ | 1.486609 | 1.523639 | 1.451982 | -0.07327 | 0.105032 | -0.66585 | 0.219187 | 0.348391 | 0.495791 | 0.4492 | 0.575956 |
| SL1344_RS07210_ | 2.745439 | 2.4603 | 3.019731 | 0.579341 | 0.164703 | 3.622299 | 4.68E-05 | 0.014328 | 0.089487 | 0.1 | 0.236592 |
| SL1344_RS07215_ | 2.71566 | 2.64009 | 2.798705 | 0.155696 | 0.080027 | 1.846515 | 0.022 | 0.056885 | 0.170958 | 0.1225 | 0.264805 |
| SL1344_RS07220_ | 0.900994 | 0.831852 | 1.00308 | 0.170814 | 0.130446 | 1.251538 | 0.102 | 0.156171 | 0.299487 | 0.2374 | 0.385631 |
| SL1344_RS07225_ | 0.931661 | 0.860607 | 0.992294 | 0.147314 | 0.17944 | 0.785226 | 0.187875 | 0.279478 | 0.444277 | 0.3841 | 0.525291 |
| SL1344_RS07230_ | 0.453502 | 0.401758 | 0.516061 | 0.116558 | 0.138031 | 0.795807 | 0.197868 | 0.299744 | 0.445615 | 0.3962 | 0.529297 |
| SL1344_RS07235_ | 0.580185 | 0.541074 | 0.630636 | 0.097692 | 0.148316 | 0.618572 | 0.235843 | 0.373541 | 0.522596 | 0.4878 | 0.610981 |
| SL1344_RS07240_ | 1.603502 | 1.62415 | 1.540445 | -0.08745 | 0.213103 | -0.34107 | 0.382412 | 0.663206 | 0.782807 | 0.7904 | 0.871156 |
| SL1344_RS07245_ | 0.127096 | 0.269395 | 0.066321 | -0.18719 | 0.204829 | -0.80822 | 0.218 | 0.332711 | 0.492387 | 0.4389 | 0.582195 |
| SL1344_RS07250_ | 0.774065 | 0.940118 | 0.552283 | -0.39535 | 0.183683 | -2.08292 | 0.006663 | 0.040605 | 0.144214 | 0.1077 | 0.24628 |
| SL1344_RS07255_ | 1.28265 | 1.389374 | 1.204275 | -0.18771 | 0.141648 | -1.28322 | 0.077333 | 0.127786 | 0.274417 | 0.1921 | 0.341811 |
| SL1344_RS07260_ | 0.361349 | 0.469438 | 0.125858 | -0.33358 | 0.18989 | -1.60204 | 0.052632 | 0.096384 | 0.233773 | 0.1587 | 0.305629 |
| SL1344_RS07265_ | -1.2283 | -0.82521 | -1.45503 | -0.58671 | 0.38797 | -1.48708 | 0.061293 | 0.098625 | 0.239868 | 0.1674 | 0.318902 |
| SL1344_RS07270_ | 2.111045 | 1.675074 | 2.418748 | 0.719499 | 0.162626 | 4.30997 | 4.68E-05 | 0.007781 | 0.066271 | 0.1 | 0.236592 |
| SL1344_RS07275_ | 1.816081 | 1.780588 | 1.845442 | 0.086492 | 0.164524 | 0.519651 | 0.291333 | 0.431367 | 0.59514 | 0.5944 | 0.711786 |
| SL1344_RS07285_ | 2.874096 | 2.588952 | 3.085165 | 0.495405 | 0.101754 | 4.700274 | 4.68E-05 | 0.010637 | 0.076007 | 0.1 | 0.236592 |

| | | | | | | | | | | |
|---|---|---|---|---|---|---|---|---|---|---|
| SL1344_RS07290_ | 1.982961 | 1.767732 | 2.189422 | 0.426074 | 0.102413 | 4.032313 | 4.68E-05 | 0.007772 | 0.064194 | 0.1 | 0.236592 |
| SL1344_RS07295_ | 4.595862 | 4.264471 | 4.901539 | 0.661913 | 0.092484 | 7.342089 | 4.68E-05 | 0.002999 | 0.046527 | 0.1 | 0.236592 |
| SL1344_RS07300_ | 3.120654 | 2.863653 | 3.357281 | 0.493218 | 0.079157 | 6.26728 | 4.68E-05 | 0.001875 | 0.038064 | 0.1 | 0.236592 |
| SL1344_RS07305_ | -0.67729 | -0.72867 | -0.61159 | 0.122886 | 0.229151 | 0.489889 | 0.292472 | 0.429325 | 0.57133 | 0.5412 | 0.657452 |
| SL1344_RS07310_ | -0.59272 | -0.7188 | -0.48589 | 0.24233 | 0.267351 | 0.882953 | 0.172667 | 0.257643 | 0.419955 | 0.3483 | 0.494657 |
| SL1344_RS07315_ | -2.86978 | -3.14218 | -2.62655 | 0.505995 | 0.405001 | 1.195259 | 0.094604 | 0.158579 | 0.294094 | 0.2382 | 0.380383 |
| SL1344_RS07320_ | 0.178245 | 0.280719 | 0.081981 | -0.20821 | 0.22075 | -0.92143 | 0.147333 | 0.23145 | 0.39284 | 0.3224 | 0.469733 |
| SL1344_RS07325_ | -5.31402 | -4.70495 | -5.70132 | -0.79239 | 1.676525 | -0.43706 | 0.30513 | 0.556561 | 0.687895 | 0.6101 | 0.730351 |
| SL1344_RS07330_ | 2.135892 | 2.23485 | 2.026692 | -0.23548 | 0.133575 | -1.7922 | 0.006001 | 0.05649 | 0.174854 | 0.1088 | 0.247683 |
| SL1344_RS07335_ | -1.83059 | -1.80708 | -1.84801 | -0.04066 | 0.35088 | -0.10765 | 0.454364 | 0.59726 | 0.720672 | 0.7105 | 0.797026 |
| SL1344_RS07340_ | -1.0429 | -1.0911 | -0.95943 | 0.105131 | 0.301104 | 0.323765 | 0.37575 | 0.609924 | 0.731752 | 0.6997 | 0.797221 |
| SL1344_RS07345_ | 3.081931 | 2.908986 | 3.240248 | 0.345643 | 0.119972 | 2.906812 | 4.68E-05 | 0.018995 | 0.101123 | 0.1 | 0.236592 |
| SL1344_RS07350_ | 3.174429 | 2.872119 | 3.518781 | 0.632358 | 0.126728 | 5.040277 | 4.68E-05 | 0.002789 | 0.044865 | 0.1 | 0.236592 |
| SL1344_RS07355_ | 3.417381 | 2.959254 | 3.767897 | 0.80522 | 0.090929 | 8.739724 | 4.68E-05 | 0.002762 | 0.043985 | 0.1 | 0.236592 |
| SL1344_RS07365_ | -0.26989 | -0.50894 | -0.08583 | 0.450736 | 0.309977 | 1.458966 | 0.014 | 0.104184 | 0.250392 | 0.1159 | 0.255854 |
| SL1344_RS07370_ | 0.432446 | -0.02031 | 0.828421 | 0.933568 | 0.296267 | 3.214335 | 4.68E-05 | 0.012551 | 0.081678 | 0.1 | 0.236592 |
| SL1344_RS07375_ | 0.881856 | 0.623801 | 1.080932 | 0.470479 | 0.153846 | 3.033781 | 4.68E-05 | 0.02694 | 0.115355 | 0.1 | 0.236592 |
| SL1344_RS07380_ | -0.97299 | -1.26113 | -0.72625 | 0.540378 | 0.22951 | 2.288694 | 0.007995 | 0.036494 | 0.132047 | 0.1065 | 0.244494 |
| SL1344_RS07385_ | 0.343745 | -0.28802 | 0.793295 | 1.0958 | 0.220564 | 4.982538 | 4.68E-05 | 0.004842 | 0.05293 | 0.1 | 0.236592 |
| SL1344_RS07390_ | -1.45152 | -1.39615 | -1.50198 | -0.11415 | 0.299914 | -0.37573 | 0.353098 | 0.501058 | 0.637161 | 0.6196 | 0.723425 |
| SL1344_RS07400_ | 1.995597 | 1.591597 | 2.374867 | 0.809622 | 0.15017 | 5.488017 | 4.68E-05 | 0.002688 | 0.043846 | 0.1 | 0.236592 |
| SL1344_RS07405_ | -1.03992 | -1.11567 | -0.98237 | 0.141076 | 0.273306 | 0.485944 | 0.295803 | 0.441717 | 0.587628 | 0.5537 | 0.669199 |
| SL1344_RS07410_ | -0.34651 | -0.7906 | -0.00219 | 0.778395 | 0.222203 | 3.498088 | 4.68E-05 | 0.012365 | 0.077699 | 0.1 | 0.236592 |
| SL1344_RS07415_ | -2.10622 | -2.38119 | -1.77927 | 0.572548 | 0.413854 | 1.335371 | 0.086 | 0.128747 | 0.272904 | 0.2002 | 0.351915 |
| SL1344_RS07420_ | -2.6052 | -2.99968 | -2.33242 | 0.671788 | 0.501962 | 1.273174 | 0.057333 | 0.129073 | 0.273132 | 0.1781 | 0.323695 |
| SL1344_RS07425_ | -2.83756 | -3.00162 | -2.67585 | 0.320274 | 0.44427 | 0.679715 | 0.219333 | 0.33665 | 0.48211 | 0.4321 | 0.562143 |
| SL1344_RS07430_ | 0.501418 | 0.487901 | 0.51716 | 0.025016 | 0.13446 | 0.174581 | 0.415723 | 0.519545 | 0.651407 | 0.6178 | 0.722075 |
| SL1344_RS07435_ | 0.456646 | 0.439058 | 0.479615 | 0.035337 | 0.155789 | 0.214565 | 0.401333 | 0.549312 | 0.679177 | 0.6604 | 0.759023 |
| SL1344_RS07440_ | -2.12051 | -2.41909 | -1.83126 | 0.591629 | 0.395091 | 1.432868 | 0.054667 | 0.108018 | 0.2429 | 0.1756 | 0.321128 |
| SL1344_RS07445_ | -0.13984 | -0.42638 | 0.272551 | 0.718012 | 0.272222 | 2.638879 | 4.68E-05 | 0.037926 | 0.142773 | 0.1 | 0.236592 |
| SL1344_RS07450_ | 1.068936 | 0.476786 | 1.567026 | 1.149784 | 0.275834 | 4.305022 | 4.68E-05 | 0.005674 | 0.059276 | 0.1 | 0.236592 |
| SL1344_RS07455_ | 1.520739 | 1.311958 | 1.793154 | 0.472366 | 0.144344 | 3.218728 | 4.68E-05 | 0.012413 | 0.080515 | 0.1 | 0.236592 |
| SL1344_RS07460_ | 0.063852 | -0.0374 | 0.259891 | 0.279255 | 0.224569 | 1.184178 | 0.114667 | 0.171038 | 0.328062 | 0.2559 | 0.410009 |
| SL1344_RS07465_ | 2.768233 | 2.74089 | 2.781731 | 0.038949 | 0.132376 | 0.273694 | 0.403731 | 0.672667 | 0.788026 | 0.824 | 0.888169 |
| SL1344_RS07470_ | 2.479946 | 2.490331 | 2.476244 | -0.00234 | 0.146198 | -0.01232 | 0.492 | 0.751614 | 0.845008 | 0.8621 | 0.916191 |
| SL1344_RS07475_ | -2.3755 | -2.40987 | -2.34973 | 0.057831 | 0.381705 | 0.149334 | 0.437708 | 0.557677 | 0.687399 | 0.6767 | 0.772113 |
| SL1344_RS07480_ | -0.64582 | -0.32661 | -0.88864 | -0.54032 | 0.299873 | -1.72766 | 0.03531 | 0.068509 | 0.193831 | 0.131 | 0.276213 |
| SL1344_RS07485_ | -0.8261 | -0.717 | -1.09207 | -0.33635 | 0.339741 | -0.85929 | 0.200667 | 0.283128 | 0.453869 | 0.398 | 0.55234 |
| SL1344_RS07490_ | -0.25602 | -0.18335 | -0.43663 | -0.19812 | 0.302753 | -0.58528 | 0.295803 | 0.459904 | 0.61751 | 0.5849 | 0.713117 |
| SL1344_RS07495_ | 0.251514 | 0.229227 | 0.289666 | 0.065824 | 0.252893 | 0.253728 | 0.394404 | 0.672146 | 0.784239 | 0.767 | 0.846817 |
| SL1344_RS07500_ | 3.691433 | 3.704758 | 3.669403 | -0.03565 | 0.065479 | -0.51407 | 0.291139 | 0.458313 | 0.607145 | 0.5769 | 0.693306 |
| SL1344_RS07505_ | 0.713399 | 0.617567 | 0.783691 | 0.171116 | 0.147247 | 1.075137 | 0.097935 | 0.175936 | 0.327507 | 0.2482 | 0.393244 |
| SL1344_RS07510_ | 2.802038 | 2.561483 | 3.039229 | 0.481998 | 0.076414 | 6.229534 | 4.68E-05 | 0.002959 | 0.043526 | 0.1 | 0.236592 |
| SL1344_RS07515_ | 1.378825 | 1.141477 | 1.550172 | 0.399824 | 0.13662 | 2.785772 | 4.68E-05 | 0.021509 | 0.103206 | 0.1005 | 0.237259 |
| SL1344_RS07520_ | -2.03456 | -2.16297 | -1.91316 | 0.249952 | 0.321318 | 0.728951 | 0.209333 | 0.321518 | 0.467031 | 0.4139 | 0.546424 |

| | | | | | | | | | | | |
|---|---|---|---|---|---|---|---|---|---|---|---|
| SL1344_RS07525_ | -2.22394 | -2.63424 | -1.88697 | 0.740942 | 0.351458 | 2.024746 | 0.016667 | 0.046355 | 0.148174 | 0.116 | 0.256375 |
| SL1344_RS07530_ | -0.52705 | -0.86499 | -0.25471 | 0.607488 | 0.207405 | 2.85006 | 0.001336 | 0.019782 | 0.0972 | 0.1007 | 0.237472 |
| SL1344_RS07535_ | 1.953416 | 1.96757 | 1.933187 | -0.10239 | 0.336949 | -0.31823 | 0.362667 | 0.635474 | 0.767517 | 0.7618 | 0.848328 |
| SL1344_RS07540_ | -0.75079 | -0.85052 | -0.58983 | 0.236717 | 0.268473 | 0.821136 | 0.193205 | 0.304584 | 0.463933 | 0.4004 | 0.546546 |
| SL1344_RS07550_ | -1.52735 | -1.77655 | -1.40543 | 0.377781 | 0.398741 | 0.851551 | 0.175883 | 0.264935 | 0.433281 | 0.3743 | 0.5181 |
| SL1344_RS07555_ | -1.30768 | -1.35866 | -1.24622 | 0.133303 | 0.322336 | 0.3921 | 0.326667 | 0.488871 | 0.629654 | 0.6149 | 0.720089 |
| SL1344_RS07560_ | -1.50577 | -1.38905 | -1.63221 | -0.2542 | 0.332982 | -0.7005 | 0.197333 | 0.319828 | 0.475672 | 0.4245 | 0.55977 |
| SL1344_RS07565_ | 0.412102 | 0.245403 | 0.553917 | 0.307314 | 0.137605 | 2.172245 | 0.009328 | 0.037992 | 0.133315 | 0.109 | 0.247737 |
| SL1344_RS07570_ | -0.80367 | -0.68707 | -1.02731 | -0.36232 | 0.317835 | -1.02443 | 0.117255 | 0.197883 | 0.363593 | 0.2705 | 0.418085 |
| SL1344_RS07575_ | -1.24532 | -1.07455 | -1.39861 | -0.31657 | 0.252558 | -1.21154 | 0.08994 | 0.155485 | 0.297192 | 0.228 | 0.375056 |
| SL1344_RS07580_ | -0.5574 | -0.85473 | -0.33522 | 0.547909 | 0.304339 | 1.755551 | 0.016667 | 0.062312 | 0.183836 | 0.1178 | 0.258749 |
| SL1344_RS07585_ | -1.02331 | -1.05556 | -0.99082 | 0.057952 | 0.226486 | 0.238706 | 0.393333 | 0.500406 | 0.635806 | 0.6135 | 0.717893 |
| SL1344_RS07590_ | -1.13545 | -1.15891 | -1.11371 | 0.048492 | 0.235618 | 0.185756 | 0.419054 | 0.522904 | 0.65389 | 0.6378 | 0.738887 |
| SL1344_RS07595_ | -0.89169 | -0.91154 | -0.87237 | 0.043071 | 0.25459 | 0.15427 | 0.435333 | 0.594641 | 0.717563 | 0.7076 | 0.794039 |
| SL1344_RS07600_ | -1.4556 | -0.64043 | -2.50779 | -1.88639 | 0.37091 | -5.07151 | 4.68E-05 | 0.003759 | 0.047575 | 0.1 | 0.236592 |
| SL1344_RS07605_ | 0.399397 | 0.630422 | 0.06058 | -0.5939 | 0.233779 | -2.57262 | 0.000673 | 0.023209 | 0.109979 | 0.1004 | 0.237134 |
| SL1344_RS07610_ | -1.76643 | -1.73802 | -1.80815 | -0.11406 | 0.445317 | -0.21997 | 0.392 | 0.575096 | 0.707748 | 0.7047 | 0.793858 |
| SL1344_RS07615_ | -0.39357 | -0.26945 | -0.48715 | -0.22089 | 0.218537 | -0.93762 | 0.14 | 0.236649 | 0.392166 | 0.3274 | 0.474091 |
| SL1344_RS07620_ | 0.44806 | 0.471617 | 0.419155 | -0.05794 | 0.189957 | -0.3035 | 0.373085 | 0.56291 | 0.693668 | 0.6783 | 0.773521 |
| SL1344_RS07625_ | -1.03118 | -0.83756 | -1.18999 | -0.34923 | 0.278085 | -1.17116 | 0.101266 | 0.153632 | 0.302063 | 0.2359 | 0.386831 |
| SL1344_RS07630_ | -0.81895 | -0.79167 | -0.84525 | -0.05292 | 0.215461 | -0.23199 | 0.4 | 0.52389 | 0.655663 | 0.628 | 0.731743 |
| SL1344_RS07635_ | 0.342328 | 0.434212 | 0.224435 | -0.21569 | 0.178371 | -1.14138 | 0.074 | 0.158972 | 0.309502 | 0.2215 | 0.368108 |
| SL1344_RS07640_ | 0.494681 | 0.534179 | 0.445757 | -0.10354 | 0.175501 | -0.54423 | 0.263824 | 0.398997 | 0.551078 | 0.5215 | 0.642536 |
| SL1344_RS07645_ | 0.94068 | 0.911503 | 0.974887 | 0.100565 | 0.206219 | 0.470614 | 0.31 | 0.450491 | 0.605921 | 0.6185 | 0.726393 |
| SL1344_RS07650_ | -0.04084 | -0.01353 | -0.0607 | -0.04485 | 0.175928 | -0.22355 | 0.4004 | 0.542841 | 0.672995 | 0.6608 | 0.757466 |
| SL1344_RS07655_ | -0.38327 | -0.52045 | -0.24383 | 0.279934 | 0.227234 | 1.202776 | 0.083278 | 0.140738 | 0.28628 | 0.2069 | 0.353999 |
| SL1344_RS07660_ | 1.687594 | 1.608725 | 1.740392 | 0.110808 | 0.145853 | 0.729862 | 0.21519 | 0.328749 | 0.495666 | 0.4234 | 0.571398 |
| SL1344_RS07665_ | 0.057392 | -0.19235 | 0.249946 | 0.448446 | 0.193096 | 2.323116 | 0.002667 | 0.041381 | 0.142206 | 0.1028 | 0.240171 |
| SL1344_RS07670_ | -1.01674 | -1.40532 | -0.74171 | 0.651827 | 0.272069 | 2.309418 | 0.008667 | 0.037814 | 0.137385 | 0.1063 | 0.244409 |
| SL1344_RS07675_ | -1.88746 | -1.95318 | -1.8407 | 0.119871 | 0.327183 | 0.351473 | 0.353333 | 0.499578 | 0.635426 | 0.6034 | 0.710446 |
| SL1344_RS07680_ | -2.55484 | -2.47963 | -2.62765 | -0.14849 | 0.408337 | -0.33428 | 0.355763 | 0.50189 | 0.637165 | 0.6102 | 0.718541 |
| SL1344_RS07685_ | -2.36712 | -2.47499 | -2.28622 | 0.192724 | 0.43364 | 0.404693 | 0.320453 | 0.463876 | 0.605598 | 0.5778 | 0.688126 |
| SL1344_RS07690_ | 0.530336 | 0.607384 | 0.332505 | -0.22238 | 0.320235 | -0.64733 | 0.267333 | 0.405994 | 0.578279 | 0.5233 | 0.670871 |
| SL1344_RS07695_ | -0.23879 | -0.18975 | -0.33675 | -0.15006 | 0.325925 | -0.44263 | 0.317788 | 0.527024 | 0.672613 | 0.6489 | 0.76042 |
| SL1344_RS07700_ | -0.06064 | 0.070429 | -0.12657 | -0.1911 | 0.211717 | -0.84159 | 0.202667 | 0.304781 | 0.464525 | 0.4101 | 0.554051 |
| SL1344_RS07705_ | 0.667471 | 0.761387 | 0.51502 | -0.25879 | 0.221799 | -1.14209 | 0.086 | 0.167622 | 0.332874 | 0.1964 | 0.347594 |
| SL1344_RS07710_ | 0.061024 | 0.130747 | -0.00839 | -0.14314 | 0.228936 | -0.6168 | 0.255829 | 0.396549 | 0.552541 | 0.5088 | 0.639065 |
| SL1344_RS07715_ | -2.01154 | -1.87282 | -2.18957 | -0.30916 | 0.339353 | -0.87209 | 0.168554 | 0.262433 | 0.4124 | 0.3522 | 0.493252 |
| SL1344_RS07720_ | -2.23951 | -2.2071 | -2.29631 | -0.08753 | 0.405514 | -0.1813 | 0.418388 | 0.594333 | 0.718249 | 0.7037 | 0.794833 |
| SL1344_RS07725_ | -1.05018 | -0.97507 | -1.11602 | -0.13994 | 0.230726 | -0.57988 | 0.263158 | 0.400087 | 0.54198 | 0.4971 | 0.619729 |
| SL1344_RS07735_ | -0.86719 | -0.57969 | -1.32982 | -0.74507 | 0.252504 | -2.90789 | 4.68E-05 | 0.022941 | 0.104302 | 0.1002 | 0.236855 |
| SL1344_RS07740_ | 1.793142 | 2.139658 | 1.261603 | -0.87787 | 0.123562 | -6.96463 | 4.68E-05 | 0.004724 | 0.052073 | 0.1 | 0.236592 |
| SL1344_RS07745_ | -0.43177 | -0.33996 | -0.53191 | -0.1818 | 0.209586 | -0.8313 | 0.187333 | 0.281388 | 0.434071 | 0.3732 | 0.514635 |
| SL1344_RS07750_ | -0.88078 | -0.91888 | -0.85609 | 0.062918 | 0.287986 | 0.191052 | 0.407333 | 0.62525 | 0.745704 | 0.7373 | 0.820744 |
| SL1344_RS07755_ | -0.01144 | 0.057448 | -0.07203 | -0.13925 | 0.260318 | -0.51451 | 0.281146 | 0.441409 | 0.595852 | 0.5584 | 0.683091 |

| | | | | | | | | | | | |
|---|---|---|---|---|---|---|---|---|---|---|---|
| SL1344_RS07760_ | 0.900343 | 0.758246 | 1.051082 | 0.315569 | 0.17262 | 1.843346 | 0.009334 | 0.063364 | 0.185545 | 0.1102 | 0.249448 |
| SL1344_RS07765_ | 0.693234 | 0.440189 | 1.006779 | 0.570462 | 0.185231 | 3.11327 | 4.68E-05 | 0.015725 | 0.089414 | 0.1 | 0.236592 |
| SL1344_RS07770_ | 0.925477 | 0.828952 | 1.429512 | 0.596171 | 0.932135 | 0.678784 | 0.224667 | 0.278447 | 0.466602 | 0.3992 | 0.581496 |
| SL1344_RS07775_ | -1.63907 | -1.80652 | -1.48436 | 0.323613 | 0.346594 | 0.901384 | 0.154667 | 0.237751 | 0.396963 | 0.3287 | 0.475977 |
| SL1344_RS07780_ | 2.2316 | 2.202324 | 2.258377 | 0.00952 | 0.306587 | 0.031695 | 0.47968 | 0.856531 | 0.915757 | 0.8881 | 0.931675 |
| SL1344_RS07785_ | 0.968203 | 0.693329 | 1.254707 | 0.57205 | 0.185915 | 3.010047 | 4.68E-05 | 0.013362 | 0.084057 | 0.1 | 0.236592 |
| SL1344_RS07790_ | -1.73456 | -1.86007 | -1.5639 | 0.317315 | 0.391387 | 0.753512 | 0.19054 | 0.297413 | 0.457874 | 0.3982 | 0.539102 |
| SL1344_RS07800_ | 3.836914 | 3.897638 | 3.816879 | -0.06401 | 0.094239 | -0.39196 | 0.347768 | 0.819857 | 0.891987 | 0.733 | 0.841446 |
| SL1344_RS07805_ | 1.815561 | 2.019853 | 1.532967 | -0.52549 | 0.174181 | -3.07953 | 4.68E-05 | 0.011376 | 0.079248 | 0.1 | 0.236592 |
| SL1344_RS07810_ | 4.056893 | 4.099398 | 3.905606 | -0.22021 | 0.20575 | -1.15776 | 0.057295 | 0.151295 | 0.322111 | 0.1607 | 0.312291 |
| SL1344_RS07815_ | 2.080837 | 1.884558 | 2.122498 | 0.223309 | 0.278582 | 0.637261 | 0.333777 | 0.486375 | 0.654665 | 0.7001 | 0.821083 |
| SL1344_RS07820_ | 3.934923 | 4.019417 | 3.919533 | -0.10062 | 0.140254 | -0.46984 | 0.33511 | 0.762124 | 0.856501 | 0.7057 | 0.825442 |
| SL1344_RS07825_ | 2.945664 | 3.22088 | 2.800395 | -0.3998 | 0.176953 | -2.32721 | 4.68E-05 | 0.024384 | 0.115516 | 0.1001 | 0.236726 |
| SL1344_RS07830_ | -1.40798 | -1.37658 | -1.43342 | -0.05166 | 0.353768 | -0.1467 | 0.431046 | 0.632425 | 0.749773 | 0.734 | 0.817026 |
| SL1344_RS07835_ | 9.523233 | 9.524582 | 9.521734 | -0.00218 | 0.094718 | -0.01513 | 0.488 | 0.816696 | 0.894062 | 0.997 | 0.998214 |
| SL1344_RS07840_ | 0.985727 | 1.166077 | 0.835422 | -0.32885 | 0.230684 | -1.37555 | 0.055963 | 0.111558 | 0.261518 | 0.1619 | 0.314501 |
| SL1344_RS07845_ | 2.607979 | 3.072817 | 2.114795 | -0.16792 | 1.154276 | -0.12949 | 0.443333 | 0.849879 | 0.914627 | 0.9994 | 0.999641 |
| SL1344_RS07850_ | 1.377043 | 1.606492 | 1.316978 | -0.23512 | 0.378251 | -0.3302 | 0.334 | 0.920496 | 0.955132 | 0.7036 | 0.824253 |
| SL1344_RS07855_ | -1.77376 | -1.6051 | -1.90705 | -0.31041 | 0.333042 | -0.87742 | 0.173333 | 0.251374 | 0.406581 | 0.3429 | 0.485815 |
| SL1344_RS07860_ | 0.272659 | 0.434248 | 0.071902 | -0.37929 | 0.194218 | -1.92408 | 0.007329 | 0.06249 | 0.181705 | 0.1077 | 0.245929 |
| SL1344_RS07865_ | -0.76241 | -0.56633 | -1.09214 | -0.51641 | 0.252779 | -1.96699 | 0.017334 | 0.060863 | 0.176408 | 0.12 | 0.26082 |
| SL1344_RS07870_ | -2.23826 | -2.11078 | -2.33881 | -0.20917 | 0.476832 | -0.40323 | 0.325783 | 0.526102 | 0.661984 | 0.6211 | 0.732512 |
| SL1344_RS07875_ | -1.396 | -1.29871 | -1.53196 | -0.11246 | 0.501098 | -0.22835 | 0.419054 | 0.693383 | 0.802008 | 0.8032 | 0.872865 |
| SL1344_RS07880_ | 0.284528 | 0.421144 | 0.144417 | -0.27706 | 0.168127 | -1.59434 | 0.039973 | 0.084599 | 0.211711 | 0.1484 | 0.29362 |
| SL1344_RS07885_ | -0.09629 | 0.096399 | -0.2865 | -0.36848 | 0.23057 | -1.53753 | 0.051333 | 0.093039 | 0.227813 | 0.1609 | 0.310583 |
| SL1344_RS07890_ | 3.030798 | 3.05616 | 2.966717 | -0.08627 | 0.093871 | -0.81321 | 0.213333 | 0.296524 | 0.469 | 0.4191 | 0.566577 |
| SL1344_RS07895_ | 0.369234 | 0.631211 | -0.0114 | -0.64936 | 0.203551 | -3.14272 | 4.68E-05 | 0.0239 | 0.109467 | 0.1 | 0.236592 |
| SL1344_RS07900_ | -3.7831 | -3.50881 | -4.11791 | -0.58493 | 0.789782 | -0.69333 | 0.221852 | 0.336273 | 0.494668 | 0.4344 | 0.574001 |
| SL1344_RS07905_ | -0.39141 | -0.19357 | -0.53931 | -0.33196 | 0.292719 | -1.11656 | 0.115257 | 0.16661 | 0.326574 | 0.2458 | 0.404481 |
| SL1344_RS07910_ | 0.307685 | 0.083454 | 0.422153 | 0.314961 | 0.211475 | 1.390797 | 0.071952 | 0.118174 | 0.26536 | 0.1878 | 0.340583 |
| SL1344_RS07915_ | -1.31346 | -1.38055 | -1.21296 | 0.155295 | 0.318974 | 0.427956 | 0.317788 | 0.504861 | 0.643632 | 0.6104 | 0.723129 |
| SL1344_RS07920_ | 1.736402 | 1.641216 | 1.901373 | 0.272589 | 0.152947 | 1.748543 | 0.018654 | 0.072076 | 0.201556 | 0.1209 | 0.262751 |
| SL1344_RS07925_ | 0.813425 | 0.795796 | 0.826484 | 0.012715 | 0.162356 | 0.079126 | 0.466667 | 0.656501 | 0.768819 | 0.7646 | 0.841197 |
| SL1344_RS07930_ | 0.84438 | 0.838172 | 0.857756 | 0.008068 | 0.174565 | 0.044237 | 0.486 | 0.693609 | 0.7991 | 0.7923 | 0.864307 |
| SL1344_RS07935_ | -0.71676 | -0.69939 | -0.74864 | -0.03583 | 0.269812 | -0.13801 | 0.447035 | 0.635228 | 0.752476 | 0.7497 | 0.830259 |
| SL1344_RS07955_ | -2.46884 | -2.24447 | -2.91356 | -0.64116 | 0.606749 | -1.02091 | 0.139907 | 0.201009 | 0.367618 | 0.2905 | 0.447643 |
| SL1344_RS07960_ | 1.757173 | 1.551129 | 1.949487 | 0.389903 | 0.128839 | 2.954405 | 0.000673 | 0.015802 | 0.089414 | 0.1002 | 0.236862 |
| SL1344_RS07965_ | 1.437141 | 1.513671 | 1.356575 | -0.15912 | 0.092853 | -1.6352 | 0.031979 | 0.075947 | 0.192414 | 0.1438 | 0.285938 |
| SL1344_RS07970_ | 1.139944 | 1.289908 | 1.001134 | -0.30584 | 0.144414 | -2.10598 | 0.004001 | 0.048938 | 0.159985 | 0.1044 | 0.242125 |
| SL1344_RS07975_ | -2.04582 | -2.08848 | -1.99873 | 0.097456 | 0.398236 | 0.210001 | 0.409727 | 0.56973 | 0.696447 | 0.6821 | 0.775351 |
| SL1344_RS07980_ | -0.8673 | -0.9277 | -0.81001 | 0.124959 | 0.275184 | 0.419174 | 0.317333 | 0.499022 | 0.638048 | 0.6088 | 0.715876 |
| SL1344_RS07990_ | -2.12774 | -1.92653 | -2.33849 | -0.41047 | 0.330187 | -1.18919 | 0.08994 | 0.153252 | 0.29207 | 0.2254 | 0.370122 |
| SL1344_RS07995_ | 1.622923 | 1.701396 | 1.545366 | -0.15203 | 0.124638 | -1.15625 | 0.104667 | 0.160974 | 0.313708 | 0.2375 | 0.390883 |
| SL1344_RS08000_ | 0.771573 | 0.816736 | 0.651027 | -0.14834 | 0.193622 | -0.69241 | 0.236667 | 0.360006 | 0.525909 | 0.4879 | 0.625774 |
| SL1344_RS08005_ | -0.00311 | 0.114715 | -0.09601 | -0.21107 | 0.174605 | -1.13788 | 0.105333 | 0.171555 | 0.318676 | 0.2522 | 0.399598 |

| | | | | | | | | | | |
|---|---|---|---|---|---|---|---|---|---|---|
| SL1344_RS08010_ | 3.402328 | 3.430802 | 3.383389 | -0.04745 | 0.056851 | -0.75742 | 0.207333 | 0.306085 | 0.46034 | 0.4054 | 0.542944 |
| SL1344_RS08015_ | 1.836465 | 1.616976 | 1.90447 | 0.215078 | 0.240544 | 0.78889 | 0.244667 | 0.335753 | 0.517477 | 0.4695 | 0.625151 |
| SL1344_RS08020_ | 0.929483 | 1.025438 | 0.790804 | -0.24767 | 0.181156 | -1.32709 | 0.061959 | 0.116241 | 0.263745 | 0.1748 | 0.325265 |
| SL1344_RS08025_ | -1.06065 | -1.16281 | -1.00373 | 0.159848 | 0.281634 | 0.494307 | 0.307129 | 0.436857 | 0.586301 | 0.5556 | 0.674798 |
| SL1344_RS08030_ | -0.39336 | -0.41172 | -0.36275 | 0.064443 | 0.251563 | 0.23957 | 0.398 | 0.593851 | 0.718594 | 0.7098 | 0.797499 |
| SL1344_RS08035_ | -0.79706 | -0.87531 | -0.69638 | 0.173663 | 0.237461 | 0.69349 | 0.227182 | 0.350077 | 0.501364 | 0.4515 | 0.584858 |
| SL1344_RS08040_ | -2.10479 | -2.28108 | -1.95108 | 0.329443 | 0.372247 | 0.864168 | 0.164557 | 0.252214 | 0.405149 | 0.338 | 0.478949 |
| SL1344_RS08045_ | -0.66619 | -0.73779 | -0.59716 | 0.146189 | 0.277166 | 0.519957 | 0.276482 | 0.432608 | 0.585302 | 0.5613 | 0.679942 |
| SL1344_RS08050_ | -1.2334 | -1.28235 | -1.17841 | 0.109267 | 0.275047 | 0.377274 | 0.344 | 0.489075 | 0.628532 | 0.5988 | 0.708587 |
| SL1344_RS08055_ | -2.35327 | -2.33009 | -2.37429 | -0.04017 | 0.354724 | -0.10701 | 0.456362 | 0.545559 | 0.674995 | 0.6601 | 0.756422 |
| SL1344_RS08060_ | -1.64304 | -1.76137 | -1.44027 | 0.300458 | 0.337385 | 0.84041 | 0.188541 | 0.296207 | 0.449912 | 0.3981 | 0.540007 |
| SL1344_RS08065_ | -0.78724 | -0.74557 | -0.81577 | -0.05784 | 0.284947 | -0.18641 | 0.415333 | 0.631859 | 0.74947 | 0.7411 | 0.823122 |
| SL1344_RS08070_ | -0.84588 | -0.90097 | -0.67988 | 0.198322 | 0.303848 | 0.53966 | 0.304464 | 0.514094 | 0.656897 | 0.6259 | 0.745476 |
| SL1344_RS08075_ | -1.3239 | -1.25315 | -1.43694 | -0.19322 | 0.30037 | -0.5896 | 0.259161 | 0.384366 | 0.536342 | 0.4986 | 0.622718 |
| SL1344_RS08080_ | -4.25156 | -4.21559 | -4.26912 | -0.02731 | 0.889914 | -0.03113 | 0.487675 | 0.639067 | 0.755019 | 0.7577 | 0.835879 |
| SL1344_RS08090_ | 1.39675 | 1.13622 | 1.733154 | 0.593669 | 0.159965 | 3.760076 | 4.68E-05 | 0.008655 | 0.068218 | 0.1 | 0.236592 |
| SL1344_RS08095_ | 2.136097 | 1.850763 | 2.412269 | 0.572445 | 0.110787 | 5.18692 | 4.68E-05 | 0.006889 | 0.0612 | 0.1 | 0.236592 |
| SL1344_RS08100_ | 0.680225 | 1.187738 | 0.138205 | -1.0968 | 0.243332 | -4.67242 | 4.68E-05 | 0.007311 | 0.065372 | 0.1 | 0.236592 |
| SL1344_RS08110_ | 0.17266 | -0.12526 | 0.486073 | 0.591982 | 0.241277 | 2.423014 | 0.001337 | 0.026864 | 0.117735 | 0.1016 | 0.238715 |
| SL1344_RS08115_ | -2.28971 | -2.28965 | -2.28984 | 0.007286 | 0.408 | 0.015817 | 0.493671 | 0.595648 | 0.719546 | 0.7085 | 0.797473 |
| SL1344_RS08120_ | -3.57709 | -3.82433 | -3.38892 | 0.207602 | 2.972423 | 0.080543 | 0.443333 | 0.895321 | 0.94007 | 0.8662 | 0.919597 |
| SL1344_RS08125_ | -4.25995 | -3.90244 | -4.60125 | -0.70301 | 1.26199 | -0.52643 | 0.259161 | 0.408815 | 0.571433 | 0.5082 | 0.642456 |
| SL1344_RS08130_ | -4.77644 | -4.13462 | -5.45464 | -1.36949 | 0.934555 | -1.4089 | 0.060626 | 0.110649 | 0.245462 | 0.1764 | 0.322099 |
| SL1344_RS08135_ | -1.19898 | -0.81571 | -1.63642 | -0.81467 | 0.243127 | -3.31237 | 4.68E-05 | 0.013823 | 0.080048 | 0.1002 | 0.236856 |
| SL1344_RS08140_ | -1.89375 | -1.53998 | -2.41135 | -0.8721 | 0.353258 | -2.38352 | 0.004668 | 0.034848 | 0.130089 | 0.1026 | 0.240004 |
| SL1344_RS08145_ | -3.23066 | -2.97527 | -3.54787 | -0.53563 | 0.679831 | -0.76555 | 0.198534 | 0.299793 | 0.462143 | 0.3997 | 0.544884 |
| SL1344_RS08150_ | -4.22179 | -3.89505 | -4.71247 | -0.82282 | 0.787491 | -0.97367 | 0.121253 | 0.214205 | 0.368293 | 0.2865 | 0.431115 |
| SL1344_RS08155_ | -1.92734 | -1.78592 | -2.05413 | -0.27194 | 0.349287 | -0.74259 | 0.20453 | 0.315238 | 0.464483 | 0.4055 | 0.539064 |
| SL1344_RS08160_ | 1.897855 | 2.163231 | 1.549116 | -0.61879 | 0.099956 | -6.19888 | 4.68E-05 | 0.004312 | 0.04927 | 0.1 | 0.236592 |
| SL1344_RS08165_ | 0.866076 | 0.928828 | 0.822556 | -0.11446 | 0.154742 | -0.70387 | 0.211859 | 0.328004 | 0.488588 | 0.439 | 0.576409 |
| SL1344_RS08170_ | 0.777435 | 0.560368 | 0.878879 | 0.282773 | 0.204674 | 1.283879 | 0.08994 | 0.135088 | 0.291026 | 0.2075 | 0.365664 |
| SL1344_RS08175_ | 2.734091 | 2.715554 | 2.759028 | 0.042436 | 0.132097 | 0.316167 | 0.364 | 0.597137 | 0.730136 | 0.7368 | 0.823738 |
| SL1344_RS08185_ | -0.91819 | -0.91381 | -0.92058 | -0.02921 | 0.280196 | -0.10111 | 0.461026 | 0.605794 | 0.727842 | 0.7091 | 0.799833 |
| SL1344_RS08190_ | -1.48061 | -1.12555 | -1.65604 | -0.53133 | 0.402436 | -1.2304 | 0.102667 | 0.155834 | 0.313548 | 0.2339 | 0.386615 |
| SL1344_RS08195_ | -1.38007 | -1.21795 | -1.49967 | -0.2815 | 0.280375 | -0.96487 | 0.151233 | 0.235141 | 0.385031 | 0.3272 | 0.469048 |
| SL1344_RS08200_ | 1.434178 | 1.493468 | 1.29367 | -0.17767 | 0.161987 | -0.97098 | 0.160667 | 0.223762 | 0.392288 | 0.3258 | 0.481832 |
| SL1344_RS08215_ | 1.654956 | 1.654216 | 1.657034 | -0.01076 | 0.145963 | -0.0656 | 0.472 | 0.688705 | 0.797119 | 0.805 | 0.873706 |
| SL1344_RS08220_ | -3.39647 | -3.69543 | -3.17749 | 0.408634 | 0.785071 | 0.490421 | 0.281333 | 0.468386 | 0.617673 | 0.5746 | 0.695987 |
| SL1344_RS08225_ | 1.276818 | 1.171469 | 1.422961 | 0.248603 | 0.116018 | 2.066224 | 0.015334 | 0.049642 | 0.157467 | 0.1163 | 0.256321 |
| SL1344_RS08230_ | -2.64784 | -2.49098 | -2.75945 | -0.30174 | 0.487591 | -0.56231 | 0.257162 | 0.400501 | 0.552729 | 0.5111 | 0.634977 |
| SL1344_RS08235_ | 4.959 | 5.023392 | 4.82914 | -0.19139 | 0.045266 | -4.05938 | 4.68E-05 | 0.008374 | 0.066948 | 0.1 | 0.236592 |
| SL1344_RS08240_ | 0.844762 | 0.763235 | 0.960996 | 0.189558 | 0.135543 | 1.294014 | 0.086 | 0.143316 | 0.285897 | 0.222 | 0.372054 |
| SL1344_RS08245_ | -0.8215 | -0.85124 | -0.78413 | 0.088383 | 0.319551 | 0.249767 | 0.39 | 0.602424 | 0.726951 | 0.7206 | 0.808198 |
| SL1344_RS08250_ | -2.04679 | -1.91207 | -2.24804 | -0.329 | 0.358221 | -0.83392 | 0.180546 | 0.269354 | 0.423118 | 0.3661 | 0.506616 |
| SL1344_RS08255_ | 1.41859 | 1.555851 | 1.238939 | -0.32237 | 0.11563 | -2.79184 | 0.001336 | 0.026809 | 0.112996 | 0.1012 | 0.237983 |

| | | | | | | | | | | | |
|---|---|---|---|---|---|---|---|---|---|---|---|
| SL1344_RS08260_ | 0.660739 | 0.663439 | 0.659889 | -0.00421 | 0.155622 | -0.02441 | 0.489333 | 0.641494 | 0.756076 | 0.7436 | 0.826385 |
| SL1344_RS08270_ | 0.489289 | 0.566512 | 0.367358 | -0.20224 | 0.213551 | -0.9064 | 0.148 | 0.223466 | 0.389126 | 0.3069 | 0.463735 |
| SL1344_RS08275_ | 1.864542 | 1.889036 | 1.847643 | -0.08707 | 0.273114 | -0.32277 | 0.33511 | 0.619278 | 0.752076 | 0.6984 | 0.800665 |
| SL1344_RS08280_ | 1.2491 | 1.227688 | 1.276252 | 0.027773 | 0.172865 | 0.162744 | 0.435043 | 0.713723 | 0.814811 | 0.7976 | 0.869379 |
| SL1344_RS08290_ | 3.96588 | 3.906911 | 4.099895 | 0.189004 | 0.090356 | 2.103469 | 0.007329 | 0.038151 | 0.143786 | 0.1044 | 0.242265 |
| SL1344_RS08295_ | 3.242134 | 3.61743 | 2.997828 | -0.59339 | 0.239844 | -2.59705 | 4.68E-05 | 0.013194 | 0.087118 | 0.1 | 0.236592 |
| SL1344_RS08300_ | 0.292314 | 0.24041 | 0.367295 | 0.107885 | 0.203212 | 0.496546 | 0.305333 | 0.493241 | 0.636878 | 0.5943 | 0.712629 |
| SL1344_RS08305_ | 0.038794 | 0.025586 | 0.047868 | -0.00622 | 0.245579 | -0.0256 | 0.492 | 0.700198 | 0.804418 | 0.8036 | 0.871586 |
| SL1344_RS08310_ | -0.75506 | -0.81621 | -0.69164 | 0.126425 | 0.2316 | 0.522879 | 0.270486 | 0.41372 | 0.557666 | 0.5246 | 0.642308 |
| SL1344_RS08315_ | -1.82199 | -1.81538 | -1.82593 | -0.01456 | 0.347473 | -0.03435 | 0.487333 | 0.601598 | 0.724592 | 0.7123 | 0.801647 |
| SL1344_RS08320_ | -3.55733 | -3.11373 | -3.94348 | -0.7992 | 0.6211 | -1.2146 | 0.098667 | 0.159583 | 0.305276 | 0.2342 | 0.384631 |
| SL1344_RS08325_ | -4.4197 | -4.55044 | -4.32589 | 0.181963 | 0.947533 | 0.158094 | 0.428 | 0.595517 | 0.721093 | 0.7138 | 0.801288 |
| SL1344_RS08330_ | -2.91977 | -2.51359 | -3.26874 | -0.79689 | 0.575697 | -1.36975 | 0.054667 | 0.113592 | 0.255201 | 0.1689 | 0.314544 |
| SL1344_RS08335_ | -3.91354 | -3.80423 | -4.10011 | -0.36542 | 1.022329 | -0.31957 | 0.348434 | 0.551903 | 0.69124 | 0.7032 | 0.79507 |
| SL1344_RS08340_ | -4.31109 | -4.11041 | -4.56799 | -0.48147 | 0.79402 | -0.57254 | 0.255829 | 0.390085 | 0.538408 | 0.4913 | 0.61606 |
| SL1344_RS08345_ | 3.333194 | 3.440638 | 3.171264 | -0.27135 | 0.075731 | -3.56006 | 4.68E-05 | 0.009998 | 0.072112 | 0.1 | 0.236592 |
| SL1344_RS08350_ | -0.42224 | -0.47398 | -0.35909 | 0.122021 | 0.250964 | 0.468815 | 0.297135 | 0.472983 | 0.618096 | 0.5823 | 0.698494 |
| SL1344_RS08355_ | -1.70294 | -1.72142 | -1.68994 | 0.043295 | 0.326586 | 0.129987 | 0.442372 | 0.573345 | 0.701436 | 0.7031 | 0.79294 |
| SL1344_RS08360_ | -0.4851 | -0.56388 | -0.36483 | 0.189719 | 0.219785 | 0.812151 | 0.196667 | 0.30588 | 0.457552 | 0.407 | 0.544538 |
| SL1344_RS08365_ | 0.313039 | 0.285379 | 0.337599 | 0.051446 | 0.167768 | 0.291224 | 0.369087 | 0.540355 | 0.670536 | 0.6433 | 0.744532 |
| SL1344_RS08370_ | -1.66071 | -1.66905 | -1.63815 | 0.057058 | 0.435449 | 0.130127 | 0.447333 | 0.672195 | 0.783573 | 0.7966 | 0.865836 |
| SL1344_RS08375_ | -2.84757 | -3.14001 | -2.6753 | 0.452992 | 0.535891 | 0.776707 | 0.195869 | 0.301126 | 0.461269 | 0.4097 | 0.5497 |
| SL1344_RS08380_ | 3.041258 | 3.071389 | 2.99808 | -0.06598 | 0.078174 | -0.78185 | 0.211333 | 0.327142 | 0.48737 | 0.425 | 0.568109 |
| SL1344_RS08385_ | -0.33092 | -0.19325 | -0.44813 | -0.29876 | 0.288029 | -1.04899 | 0.110667 | 0.197486 | 0.365731 | 0.2482 | 0.398366 |
| SL1344_RS08390_ | 0.70146 | 0.773229 | 0.5945 | -0.17883 | 0.16585 | -1.00886 | 0.131246 | 0.201116 | 0.359924 | 0.2798 | 0.430236 |
| SL1344_RS08395_ | 4.330632 | 4.471997 | 4.038359 | -0.43045 | 0.130963 | -2.84474 | 4.68E-05 | 0.03978 | 0.150894 | 0.1 | 0.236592 |
| SL1344_RS08400_ | 1.898069 | 1.779323 | 1.998871 | 0.231102 | 0.126122 | 1.778314 | 0.021986 | 0.06063 | 0.180571 | 0.118 | 0.259251 |
| SL1344_RS08405_ | 1.691055 | 1.67444 | 1.714081 | 0.034363 | 0.125819 | 0.253079 | 0.390667 | 0.60609 | 0.730952 | 0.7044 | 0.799344 |
| SL1344_RS08410_ | 1.231478 | 1.300497 | 1.200387 | -0.08936 | 0.132809 | -0.61365 | 0.255333 | 0.411229 | 0.564908 | 0.5274 | 0.658733 |
| SL1344_RS08415_ | 2.278886 | 2.290143 | 2.267527 | -0.02295 | 0.081334 | -0.26552 | 0.381746 | 0.537969 | 0.670271 | 0.6485 | 0.749741 |
| SL1344_RS08420_ | -0.76285 | -0.77687 | -0.74975 | 0.028697 | 0.208464 | 0.126898 | 0.446 | 0.540862 | 0.672618 | 0.6581 | 0.757 |
| SL1344_RS08425_ | -0.2836 | -0.22412 | -0.34536 | -0.12419 | 0.233439 | -0.51126 | 0.282478 | 0.447819 | 0.592782 | 0.5608 | 0.674672 |
| SL1344_RS08430_ | -1.26987 | -1.21972 | -1.33057 | -0.11333 | 0.268552 | -0.39652 | 0.339107 | 0.468221 | 0.607599 | 0.588 | 0.69725 |
| SL1344_RS08435_ | 2.330046 | 2.455677 | 2.244997 | -0.21923 | 0.138141 | -1.53977 | 0.018 | 0.10138 | 0.249185 | 0.1201 | 0.261715 |
| SL1344_RS08440_ | 0.088542 | 0.117211 | 0.065794 | -0.04646 | 0.173643 | -0.25391 | 0.385333 | 0.537645 | 0.668597 | 0.6463 | 0.746399 |
| SL1344_RS08445_ | 1.323202 | 1.388743 | 1.254582 | -0.13625 | 0.138978 | -0.93778 | 0.137242 | 0.214881 | 0.375268 | 0.3003 | 0.452211 |
| SL1344_RS08450_ | 0.682133 | 0.651875 | 0.723999 | 0.077979 | 0.164124 | 0.450764 | 0.329333 | 0.4993 | 0.639796 | 0.6115 | 0.721998 |
| SL1344_RS08455_ | 0.480421 | 0.489414 | 0.471315 | -0.05367 | 0.219251 | -0.23203 | 0.399734 | 0.601877 | 0.730026 | 0.7666 | 0.839459 |
| SL1344_RS08460_ | 0.783591 | 0.767634 | 0.803775 | 0.03818 | 0.17464 | 0.205037 | 0.399734 | 0.60869 | 0.731033 | 0.719 | 0.805772 |
| SL1344_RS08465_ | 0.579074 | 0.529459 | 0.635959 | 0.094838 | 0.215889 | 0.44516 | 0.306667 | 0.521497 | 0.663861 | 0.6199 | 0.735444 |
| SL1344_RS08470_ | -1.08745 | -1.25946 | -0.93874 | 0.355275 | 0.328459 | 1.018699 | 0.110667 | 0.197263 | 0.360919 | 0.2462 | 0.394571 |
| SL1344_RS08475_ | -2.17051 | -1.98225 | -2.40562 | -0.41568 | 0.370831 | -1.07888 | 0.117255 | 0.184436 | 0.330566 | 0.2646 | 0.410044 |
| SL1344_RS08480_ | -7.04131 | -6.93925 | -7.15264 | -0.19365 | 3.021221 | -0.05266 | 0.469021 | 0.629236 | 0.751363 | 0.7478 | 0.828238 |
| SL1344_RS08485_ | -1.35751 | -1.19162 | -1.48483 | -0.30619 | 0.280941 | -1.02183 | 0.124584 | 0.209465 | 0.360854 | 0.2985 | 0.442378 |
| SL1344_RS08490_ | 4.7384 | 4.951994 | 4.544105 | -0.40554 | 0.108501 | -3.66654 | 4.68E-05 | 0.007628 | 0.068009 | 0.1 | 0.236592 |

| | | | | | | | | | | | |
|---|---|---|---|---|---|---|---|---|---|---|---|
| SL1344_RS08500_ | 3.07515 | 3.540337 | 2.622511 | -0.81896 | 0.191643 | -4.43258 | 4.68E-05 | 0.002198 | 0.04256 | 0.1 | 0.236592 |
| SL1344_RS08505_ | 0.893132 | 0.84517 | 1.058167 | 0.197085 | 0.170358 | 0.955792 | 0.193871 | 0.258425 | 0.426451 | 0.3935 | 0.544529 |
| SL1344_RS08510_ | 1.306555 | 1.344518 | 1.274708 | -0.11207 | 0.195017 | -0.53703 | 0.271819 | 0.382486 | 0.553091 | 0.5549 | 0.674483 |
| SL1344_RS08515_ | 1.169828 | 1.300509 | 1.003936 | -0.30476 | 0.14854 | -1.97147 | 0.006001 | 0.055434 | 0.170649 | 0.107 | 0.24539 |
| SL1344_RS08520_ | 2.428945 | 2.44923 | 2.402008 | -0.11197 | 0.249012 | -0.474 | 0.327115 | 0.457489 | 0.627015 | 0.6732 | 0.78333 |
| SL1344_RS08525_ | -0.44836 | -0.10268 | -0.56028 | -0.4102 | 0.312539 | -1.12991 | 0.148568 | 0.190825 | 0.356399 | 0.301 | 0.457775 |
| SL1344_RS08530_ | -0.29376 | -0.17025 | -0.57405 | -0.40256 | 0.272197 | -1.36498 | 0.059294 | 0.131976 | 0.285975 | 0.1806 | 0.328576 |
| SL1344_RS08535_ | 1.987363 | 2.112348 | 1.811714 | -0.31301 | 0.142957 | -2.15315 | 0.002001 | 0.036008 | 0.138117 | 0.1017 | 0.238864 |
| SL1344_RS08540_ | 2.212126 | 2.403658 | 1.923794 | -0.47761 | 0.109474 | -4.28832 | 4.68E-05 | 0.013786 | 0.083655 | 0.1 | 0.236592 |
| SL1344_RS08545_ | -0.02506 | -0.14439 | 0.065622 | 0.202042 | 0.191922 | 1.002156 | 0.138574 | 0.216574 | 0.371659 | 0.2995 | 0.449111 |
| SL1344_RS08550_ | 2.010679 | 2.062837 | 1.939753 | -0.14123 | 0.137155 | -1.01568 | 0.1006 | 0.192407 | 0.358169 | 0.2531 | 0.403517 |
| SL1344_RS08555_ | 2.329729 | 2.556492 | 2.084224 | -0.47191 | 0.096971 | -4.8401 | 4.68E-05 | 0.005425 | 0.054669 | 0.1 | 0.236592 |
| SL1344_RS08560_ | 1.342022 | 1.097235 | 1.588873 | 0.507012 | 0.148126 | 3.439527 | 4.68E-05 | 0.012687 | 0.081284 | 0.1 | 0.236592 |
| SL1344_RS08570_ | 3.655225 | 3.652861 | 3.661886 | 0.004002 | 0.070413 | 0.048512 | 0.482345 | 0.71449 | 0.815756 | 0.7985 | 0.872734 |
| SL1344_RS08575_ | 2.974321 | 3.227524 | 2.811661 | -0.42824 | 0.150298 | -2.83918 | 4.68E-05 | 0.0124 | 0.083454 | 0.1 | 0.236592 |
| SL1344_RS08580_ | 1.483655 | 1.364356 | 1.609054 | 0.20258 | 0.202762 | 0.993916 | 0.143904 | 0.207701 | 0.380588 | 0.2762 | 0.442527 |
| SL1344_RS08590_ | 1.758 | 1.691385 | 1.843991 | 0.155529 | 0.09948 | 1.491488 | 0.048 | 0.100774 | 0.232672 | 0.1635 | 0.307717 |
| SL1344_RS08595_ | 1.419063 | 1.566538 | 1.296301 | -0.27754 | 0.155163 | -1.75573 | 0.022667 | 0.060577 | 0.181078 | 0.1187 | 0.260442 |
| SL1344_RS08600_ | 1.785559 | 1.897152 | 1.660159 | -0.23675 | 0.114624 | -2.00841 | 0.01066 | 0.047311 | 0.155607 | 0.1129 | 0.253039 |
| SL1344_RS08605_ | 1.576767 | 1.655475 | 1.513043 | -0.15184 | 0.119883 | -1.2344 | 0.07 | 0.141624 | 0.290929 | 0.1921 | 0.337743 |
| SL1344_RS08610_ | -2.19215 | -2.18878 | -2.19717 | -0.01732 | 0.419924 | -0.03869 | 0.482667 | 0.615754 | 0.736849 | 0.7228 | 0.809486 |
| SL1344_RS08615_ | -1.28622 | -1.04889 | -1.52503 | -0.47114 | 0.266508 | -1.69823 | 0.032 | 0.070471 | 0.188352 | 0.1361 | 0.279479 |
| SL1344_RS08620_ | -0.26694 | -0.16539 | -0.41823 | -0.26658 | 0.270464 | -0.96363 | 0.147235 | 0.212329 | 0.376795 | 0.3004 | 0.458407 |
| SL1344_RS08625_ | -0.0722 | 0.057316 | -0.22022 | -0.28045 | 0.167725 | -1.61302 | 0.036667 | 0.082464 | 0.204349 | 0.146 | 0.289105 |
| SL1344_RS08630_ | 0.177294 | 0.275662 | 0.071902 | -0.20066 | 0.142961 | -1.35578 | 0.069953 | 0.117111 | 0.247573 | 0.1921 | 0.336736 |
| SL1344_RS08635_ | -0.99348 | -1.19271 | -0.75275 | 0.448261 | 0.247209 | 1.748315 | 0.028 | 0.068183 | 0.185298 | 0.1278 | 0.269483 |
| SL1344_RS08640_ | -3.2024 | -3.44806 | -3.02857 | 0.428694 | 0.528534 | 0.761497 | 0.189207 | 0.288355 | 0.441929 | 0.3783 | 0.515951 |
| SL1344_RS08645_ | -3.54803 | -3.83554 | -3.2447 | 0.611295 | 0.598215 | 0.979278 | 0.142667 | 0.226198 | 0.376107 | 0.311 | 0.452414 |
| SL1344_RS08650_ | -1.27703 | -1.42108 | -1.11303 | 0.330498 | 0.336863 | 0.964292 | 0.146667 | 0.218373 | 0.378143 | 0.3045 | 0.453047 |
| SL1344_RS08655_ | 6.245429 | 5.930489 | 6.508548 | 0.577169 | 0.277968 | 2.308682 | 4.68E-05 | 0.030099 | 0.131587 | 0.1 | 0.236592 |
| SL1344_RS08660_ | 0.656315 | 0.667409 | 0.647735 | -0.0206 | 0.138489 | -0.13176 | 0.440373 | 0.568593 | 0.695807 | 0.6816 | 0.775371 |
| SL1344_RS08665_ | 2.415314 | 2.715871 | 2.310961 | -0.39805 | 0.126039 | -2.49904 | 4.68E-05 | 0.052983 | 0.17338 | 0.1009 | 0.237742 |
| SL1344_RS08670_ | 1.920785 | 2.09322 | 1.836666 | -0.24677 | 0.107549 | -2.13501 | 0.015334 | 0.051305 | 0.162834 | 0.1126 | 0.252126 |
| SL1344_RS08675_ | 0.27524 | 0.596796 | -0.17859 | -0.7709 | 0.181186 | -4.23558 | 4.68E-05 | 0.009119 | 0.067857 | 0.1 | 0.236592 |
| SL1344_RS08680_ | -0.69994 | -0.70471 | -0.69268 | -0.00985 | 0.285878 | -0.03665 | 0.486 | 0.669735 | 0.780649 | 0.7731 | 0.850683 |
| SL1344_RS08685_ | 1.110531 | 0.972807 | 1.356831 | 0.398391 | 0.218426 | 1.902225 | 0.006663 | 0.050006 | 0.165648 | 0.1087 | 0.247568 |
| SL1344_RS08690_ | 0.873948 | 1.008753 | 0.704754 | -0.30517 | 0.138165 | -2.16461 | 0.006663 | 0.041782 | 0.142111 | 0.1076 | 0.246071 |
| SL1344_RS08695_ | 0.987264 | 0.943354 | 1.115886 | 0.15723 | 0.195533 | 0.696668 | 0.251832 | 0.362556 | 0.531545 | 0.4895 | 0.634591 |
| SL1344_RS08700_ | 0.013297 | 0.032299 | -0.01211 | -0.05125 | 0.196104 | -0.24384 | 0.398 | 0.553187 | 0.682905 | 0.6717 | 0.76555 |
| SL1344_RS08705_ | 2.391032 | 2.345751 | 2.427013 | 0.095201 | 0.161313 | 0.570513 | 0.254667 | 0.360667 | 0.540481 | 0.5056 | 0.647025 |
| SL1344_RS08710_ | 2.302078 | 2.191383 | 2.40419 | 0.216058 | 0.10846 | 1.969986 | 0.007995 | 0.053103 | 0.168391 | 0.1077 | 0.246645 |
| SL1344_RS08715_ | 1.673001 | 1.514869 | 1.831679 | 0.293523 | 0.14544 | 1.984967 | 0.019334 | 0.04535 | 0.154613 | 0.1168 | 0.258757 |
| SL1344_RS08725_ | 3.895651 | 3.897927 | 3.894556 | -0.00234 | 0.056362 | -0.04099 | 0.482 | 0.645991 | 0.759927 | 0.7435 | 0.826927 |
| SL1344_RS08730_ | -0.54542 | -0.39767 | -0.69123 | -0.2976 | 0.202754 | -1.42585 | 0.059294 | 0.113153 | 0.242632 | 0.1816 | 0.325602 |
| SL1344_RS08735_ | 4.721016 | 4.704245 | 4.730917 | 0.042701 | 0.083138 | 0.522531 | 0.272485 | 0.402367 | 0.575997 | 0.5563 | 0.689828 |

| | | | | | | | | | | | |
|---|---|---|---|---|---|---|---|---|---|---|---|
| SL1344_RS08740_ | 1.014239 | 1.147707 | 0.968025 | -0.15015 | 0.177993 | -0.71703 | 0.258494 | 0.383073 | 0.545583 | 0.507 | 0.647605 |
| SL1344_RS08750_ | 3.824418 | 3.89131 | 3.74706 | -0.09426 | 0.163157 | -0.60589 | 0.258667 | 0.406337 | 0.584224 | 0.5035 | 0.646991 |
| SL1344_RS08755_ | 1.45053 | 1.462354 | 1.441593 | -0.01752 | 0.112831 | -0.13917 | 0.439041 | 0.584177 | 0.709291 | 0.6828 | 0.778464 |
| SL1344_RS08760_ | 1.729515 | 1.753839 | 1.692401 | -0.06433 | 0.105307 | -0.58444 | 0.248 | 0.375544 | 0.528258 | 0.4989 | 0.619358 |
| SL1344_RS08765_ | 0.653523 | 0.911042 | 0.290121 | -0.61094 | 0.166776 | -3.58876 | 4.68E-05 | 0.013742 | 0.081836 | 0.1001 | 0.236728 |
| SL1344_RS08770_ | 2.460066 | 2.877505 | 2.049948 | -0.83847 | 0.118925 | -7.08606 | 4.68E-05 | 0.002448 | 0.042109 | 0.1 | 0.236592 |
| SL1344_RS08775_ | 0.205451 | 0.386939 | -0.19493 | -0.57365 | 0.22362 | -2.33351 | 0.003335 | 0.045711 | 0.155177 | 0.1029 | 0.240416 |
| SL1344_RS08780_ | -0.94603 | -0.72593 | -1.22609 | -0.48373 | 0.288478 | -1.59077 | 0.044 | 0.081693 | 0.208998 | 0.1472 | 0.293308 |
| SL1344_RS08785_ | 0.236474 | 0.129036 | 0.376087 | 0.253116 | 0.170377 | 1.405144 | 0.046667 | 0.106515 | 0.242033 | 0.1701 | 0.316077 |
| SL1344_RS08815_ | 1.135351 | 1.602791 | 0.674164 | -0.93989 | 0.163329 | -5.6606 | 4.68E-05 | 0.002695 | 0.043597 | 0.1 | 0.236592 |
| SL1344_RS08825_ | 0.241823 | 0.278046 | 0.21658 | -0.07126 | 0.194321 | -0.34034 | 0.348667 | 0.521403 | 0.658604 | 0.6498 | 0.747711 |
| SL1344_RS08830_ | 2.099504 | 2.129302 | 2.036844 | -0.08394 | 0.099884 | -0.75653 | 0.218521 | 0.332235 | 0.491745 | 0.4429 | 0.584174 |
| SL1344_RS08840_ | -0.25325 | -0.21166 | -0.29074 | -0.05484 | 0.241165 | -0.2306 | 0.396 | 0.617425 | 0.739027 | 0.7259 | 0.814266 |
| SL1344_RS08845_ | 3.705113 | 3.75433 | 3.625551 | -0.12646 | 0.095015 | -1.24213 | 0.085943 | 0.139554 | 0.299939 | 0.1868 | 0.343807 |
| SL1344_RS08850_ | 1.309568 | 1.427963 | 1.204505 | -0.22508 | 0.108312 | -2.02089 | 0.011334 | 0.046262 | 0.148635 | 0.1141 | 0.253877 |
| SL1344_RS08855_ | -0.35792 | -0.18076 | -0.67832 | -0.483 | 0.244747 | -1.8417 | 0.022667 | 0.065733 | 0.186495 | 0.1253 | 0.267697 |
| SL1344_RS08860_ | 1.236226 | 1.392499 | 1.051285 | -0.357 | 0.163899 | -2.15197 | 0.002667 | 0.038535 | 0.140824 | 0.1017 | 0.238811 |
| SL1344_RS08865_ | 1.697249 | 1.886563 | 1.362336 | -0.5117 | 0.143693 | -3.41063 | 4.68E-05 | 0.020756 | 0.103517 | 0.1 | 0.236592 |
| SL1344_RS08870_ | 3.217545 | 3.387442 | 3.043142 | -0.34574 | 0.060953 | -5.57673 | 4.68E-05 | 0.003376 | 0.045468 | 0.1 | 0.236592 |
| SL1344_RS08880_ | 3.763195 | 3.974195 | 3.54415 | -0.53659 | 0.208154 | -2.74408 | 4.68E-05 | 0.013816 | 0.089271 | 0.1 | 0.236592 |
| SL1344_RS08885_ | 5.005103 | 5.140865 | 4.737258 | -0.40897 | 0.117038 | -3.50758 | 4.68E-05 | 0.013617 | 0.088237 | 0.1 | 0.236592 |
| SL1344_RS08890_ | 1.492435 | 1.732578 | 1.213061 | -0.53026 | 0.141805 | -3.76886 | 4.68E-05 | 0.013713 | 0.083167 | 0.1 | 0.236592 |
| SL1344_RS08895_ | 2.055822 | 1.61193 | 2.180739 | 0.586135 | 0.303543 | 1.845204 | 0.002001 | 0.058778 | 0.185554 | 0.102 | 0.239159 |
| SL1344_RS08900_ | 0.151147 | 0.373594 | -0.08813 | -0.47952 | 0.214973 | -2.17101 | 0.002002 | 0.036393 | 0.135618 | 0.1046 | 0.242457 |
| SL1344_RS08905_ | 3.219508 | 3.327219 | 3.092643 | -0.2349 | 0.058703 | -3.97422 | 4.68E-05 | 0.008443 | 0.064803 | 0.1 | 0.236592 |
| SL1344_RS08910_ | 0.755546 | 0.854517 | 0.524908 | -0.31374 | 0.168074 | -1.69383 | 0.038 | 0.088651 | 0.223398 | 0.1494 | 0.295076 |
| SL1344_RS08915_ | -3.63952 | -3.62911 | -3.65497 | -0.02611 | 0.855674 | -0.02535 | 0.488341 | 0.68184 | 0.790589 | 0.7931 | 0.863957 |
| SL1344_RS08920_ | -1.84121 | -1.82075 | -1.86095 | -0.04791 | 0.317534 | -0.13428 | 0.443704 | 0.542025 | 0.671814 | 0.6398 | 0.741817 |
| SL1344_RS08925_ | -4.05007 | -3.96966 | -4.18111 | -0.19918 | 0.770977 | -0.23366 | 0.392405 | 0.555897 | 0.687666 | 0.6746 | 0.772085 |
| SL1344_RS08930_ | -3.95672 | -3.5693 | -4.81952 | -1.2394 | 0.902757 | -1.26725 | 0.064 | 0.137786 | 0.290106 | 0.1763 | 0.322953 |
| SL1344_RS08935_ | -4.5544 | -4.53952 | -4.57306 | -0.05396 | 0.934868 | -0.05048 | 0.476 | 0.596581 | 0.719932 | 0.7164 | 0.802363 |
| SL1344_RS08940_ | -3.64454 | -3.51014 | -3.79796 | -0.26376 | 0.679526 | -0.36706 | 0.333777 | 0.4944 | 0.63413 | 0.6164 | 0.722381 |
| SL1344_RS08945_ | -2.54311 | -2.57408 | -2.51598 | 0.059412 | 0.409407 | 0.140799 | 0.439707 | 0.552633 | 0.681021 | 0.6627 | 0.760911 |
| SL1344_RS08950_ | -2.36385 | -2.14827 | -2.65702 | -0.5043 | 0.40365 | -1.1984 | 0.099933 | 0.164181 | 0.310881 | 0.2351 | 0.380813 |
| SL1344_RS08955_ | -2.50071 | -2.24925 | -2.82379 | -0.5709 | 0.393541 | -1.37753 | 0.066 | 0.121233 | 0.253917 | 0.1911 | 0.33608 |
| SL1344_RS08960_ | 4.90482 | 5.161642 | 4.596371 | -0.49831 | 0.21275 | -2.41794 | 4.68E-05 | 0.016647 | 0.098141 | 0.1 | 0.236592 |
| SL1344_RS08965_ | 2.334948 | 2.188596 | 2.575449 | 0.378508 | 0.092858 | 3.944677 | 4.68E-05 | 0.009468 | 0.070073 | 0.1 | 0.236592 |
| SL1344_RS08970_ | 0.228302 | -0.09584 | 0.722264 | 0.780119 | 0.350252 | 2.235219 | 0.002001 | 0.0288 | 0.124811 | 0.1019 | 0.239136 |
| SL1344_RS08985_ | -0.13999 | -0.07938 | -0.21925 | -0.13938 | 0.194172 | -0.67603 | 0.233844 | 0.362235 | 0.514372 | 0.4648 | 0.595512 |
| SL1344_RS08995_ | 1.68852 | 2.236048 | 0.947555 | -1.27966 | 0.152276 | -8.31365 | 4.68E-05 | 0.00314 | 0.045372 | 0.1 | 0.236592 |
| SL1344_RS09000_ | 1.747058 | 2.382675 | 1.022279 | -1.32955 | 0.17682 | -7.49496 | 4.68E-05 | 0.001449 | 0.034906 | 0.1 | 0.236592 |
| SL1344_RS09005_ | 2.086918 | 1.699383 | 2.391678 | 0.704603 | 0.144958 | 4.820878 | 4.68E-05 | 0.003364 | 0.047683 | 0.1 | 0.236592 |
| SL1344_RS09010_ | 2.552491 | 2.513554 | 2.648112 | 0.142407 | 0.147098 | 0.904879 | 0.171219 | 0.243772 | 0.42182 | 0.3368 | 0.497006 |
| SL1344_RS09015_ | 1.657734 | 1.679411 | 1.620685 | -0.0446 | 0.126487 | -0.33184 | 0.369754 | 0.584863 | 0.712982 | 0.699 | 0.791977 |
| SL1344_RS09020_ | 1.909494 | 2.13857 | 1.697113 | -0.44932 | 0.1443 | -3.10319 | 4.68E-05 | 0.020824 | 0.104472 | 0.1 | 0.236592 |

| | | | | | | | | | | | |
|---|---|---|---|---|---|---|---|---|---|---|---|
| SL1344_RS09025_ | -2.11561 | -2.02448 | -2.18691 | -0.2634 | 0.521036 | -0.471 | 0.313125 | 0.450071 | 0.604457 | 0.6144 | 0.720203 |
| SL1344_RS09030_ | 2.831395 | 2.538949 | 3.1231 | 0.446025 | 0.226117 | 2.309287 | 0.000673 | 0.02086 | 0.107925 | 0.1003 | 0.236999 |
| SL1344_RS09035_ | -0.19374 | -0.26991 | -0.09423 | 0.179466 | 0.192742 | 0.871951 | 0.172 | 0.27651 | 0.429279 | 0.3801 | 0.520859 |
| SL1344_RS09040_ | 0.549425 | 0.577669 | 0.520029 | -0.05397 | 0.126359 | -0.39661 | 0.325117 | 0.445423 | 0.585156 | 0.5477 | 0.662563 |
| SL1344_RS09045_ | 1.717686 | 1.737355 | 1.692411 | -0.04655 | 0.093225 | -0.44637 | 0.303797 | 0.4634 | 0.603268 | 0.5674 | 0.681522 |
| SL1344_RS09050_ | 0.653696 | 0.647795 | 0.662365 | 0.036205 | 0.187836 | 0.192972 | 0.428 | 0.639327 | 0.757211 | 0.7703 | 0.844416 |
| SL1344_RS09055_ | 2.817968 | 3.018649 | 2.607969 | -0.43687 | 0.106976 | -4.07866 | 4.68E-05 | 0.005228 | 0.056484 | 0.1 | 0.236592 |
| SL1344_RS09060_ | 4.334283 | 4.373938 | 4.290025 | -0.02015 | 0.12878 | -0.19032 | 0.428381 | 0.728552 | 0.83089 | 0.8766 | 0.920875 |
| SL1344_RS09065_ | 3.987654 | 4.001088 | 3.959283 | -0.03354 | 0.078796 | -0.37183 | 0.361333 | 0.656173 | 0.774335 | 0.7273 | 0.825053 |
| SL1344_RS09070_ | -0.67484 | -0.20474 | -1.28979 | -1.07434 | 0.222667 | -4.83993 | 4.68E-05 | 0.006849 | 0.058253 | 0.1 | 0.236592 |
| SL1344_RS09085_ | 1.156821 | 1.505364 | 0.722972 | -0.78006 | 0.122193 | -6.20619 | 4.68E-05 | 0.003538 | 0.045434 | 0.1 | 0.236592 |
| SL1344_RS09090_ | 0.937915 | 1.425989 | 0.460755 | -0.98512 | 0.184162 | -5.41963 | 4.68E-05 | 0.006072 | 0.059787 | 0.1 | 0.236592 |
| SL1344_RS09095_ | 1.496645 | 1.881075 | 1.068007 | -0.82272 | 0.117514 | -6.94713 | 4.68E-05 | 0.002431 | 0.040081 | 0.1 | 0.236592 |
| SL1344_RS09100_ | 2.682647 | 3.094313 | 2.26542 | -0.8584 | 0.133379 | -6.41032 | 4.68E-05 | 0.001179 | 0.034129 | 0.1 | 0.236592 |
| SL1344_RS09105_ | 0.780692 | 1.080794 | 0.435233 | -0.66714 | 0.27045 | -2.53789 | 4.68E-05 | 0.029493 | 0.126458 | 0.1001 | 0.236729 |
| SL1344_RS09115_ | 0.858606 | 1.103122 | 0.601872 | -0.49947 | 0.116513 | -4.26535 | 4.68E-05 | 0.006916 | 0.059311 | 0.1 | 0.236592 |
| SL1344_RS09125_ | -1.30338 | -1.22032 | -1.4048 | -0.17353 | 0.304184 | -0.55704 | 0.273817 | 0.424347 | 0.57141 | 0.5141 | 0.638835 |
| SL1344_RS09130_ | 0.775515 | 1.01435 | 0.597923 | -0.4135 | 0.188453 | -2.10029 | 0.006667 | 0.039226 | 0.141895 | 0.1078 | 0.246577 |
| SL1344_RS09135_ | -0.48137 | -0.22968 | -0.78793 | -0.56137 | 0.205931 | -2.72345 | 0.000673 | 0.024038 | 0.105682 | 0.1015 | 0.238538 |
| SL1344_RS09140_ | 4.459041 | 4.475513 | 4.447582 | -0.05028 | 0.110035 | -0.44421 | 0.319121 | 0.463947 | 0.632778 | 0.6755 | 0.785015 |
| SL1344_RS09145_ | 3.898389 | 3.81088 | 3.958821 | 0.080737 | 0.140008 | 0.699363 | 0.231333 | 0.379479 | 0.557996 | 0.4443 | 0.595511 |
| SL1344_RS09150_ | 5.120014 | 5.088899 | 5.195551 | 0.085524 | 0.115245 | 0.819018 | 0.189333 | 0.26254 | 0.447185 | 0.3502 | 0.528061 |
| SL1344_RS09155_ | -0.47606 | -0.18867 | -0.68835 | -0.49274 | 0.236455 | -2.03663 | 0.014 | 0.049272 | 0.158565 | 0.1142 | 0.254671 |
| SL1344_RS09160_ | 0.181441 | 0.710449 | -0.18534 | -0.90957 | 0.25802 | -3.51416 | 4.68E-05 | 0.009348 | 0.071222 | 0.1 | 0.236592 |
| SL1344_RS09165_ | -0.54334 | -0.16465 | -1.15991 | -1.00831 | 0.357581 | -2.75871 | 4.68E-05 | 0.039537 | 0.144926 | 0.1001 | 0.236726 |
| SL1344_RS09170_ | -0.01907 | 0.423425 | -0.5514 | -0.98296 | 0.205718 | -4.73619 | 4.68E-05 | 0.006101 | 0.056746 | 0.1 | 0.236592 |
| SL1344_RS09175_ | 5.614129 | 5.912942 | 5.210743 | -0.65407 | 0.174731 | -3.60982 | 4.68E-05 | 0.008867 | 0.073571 | 0.1 | 0.236592 |
| SL1344_RS09180_ | 4.94647 | 5.254204 | 4.377556 | -0.88397 | 0.225192 | -3.92317 | 4.68E-05 | 0.004669 | 0.056632 | 0.1 | 0.236592 |
| SL1344_RS09185_ | 1.55935 | 1.713488 | 1.501733 | -0.19307 | 0.143999 | -1.21328 | 0.127333 | 0.163214 | 0.322406 | 0.2594 | 0.41722 |
| SL1344_RS09190_ | 1.533711 | 1.269173 | 1.582472 | 0.300988 | 0.221642 | 1.084246 | 0.167222 | 0.202886 | 0.380002 | 0.3337 | 0.498961 |
| SL1344_RS09200_ | 3.208668 | 3.414685 | 3.099701 | -0.30164 | 0.118797 | -2.51754 | 0.000673 | 0.021285 | 0.106802 | 0.1 | 0.236592 |
| SL1344_RS09205_ | -1.17611 | -0.90643 | -1.46142 | -0.61462 | 0.399277 | -1.54963 | 0.020653 | 0.084217 | 0.218648 | 0.126 | 0.268742 |
| SL1344_RS09210_ | 3.475633 | 3.800998 | 3.189355 | -0.60911 | 0.092066 | -6.68762 | 4.68E-05 | 0.001283 | 0.034577 | 0.1 | 0.236592 |
| SL1344_RS09235_ | 2.68318 | 2.836855 | 2.491417 | -0.34446 | 0.059574 | -5.75644 | 4.68E-05 | 0.002918 | 0.042503 | 0.1 | 0.236592 |
| SL1344_RS09245_ | 1.246445 | 1.468836 | 0.845321 | -0.62653 | 0.209161 | -2.92569 | 4.68E-05 | 0.024352 | 0.112721 | 0.1 | 0.236592 |
| SL1344_RS09250_ | -1.17374 | -1.07294 | -1.30552 | -0.23446 | 0.249129 | -0.89726 | 0.157229 | 0.244991 | 0.395547 | 0.3378 | 0.476888 |
| SL1344_RS09260_ | 0.725959 | 1.100457 | 0.312508 | -0.83189 | 0.320699 | -2.58792 | 4.68E-05 | 0.03189 | 0.131448 | 0.1 | 0.236592 |
| SL1344_RS09270_ | -3.32145 | -3.14995 | -3.4588 | -0.32185 | 0.610531 | -0.49119 | 0.29447 | 0.445614 | 0.590153 | 0.5577 | 0.673558 |
| SL1344_RS09275_ | 0.757782 | 1.095236 | 0.207104 | -0.86035 | 0.255025 | -3.31904 | 4.68E-05 | 0.012422 | 0.081404 | 0.1 | 0.236592 |
| SL1344_RS09285_ | -0.99598 | -0.83389 | -1.57457 | -0.70859 | 0.370979 | -1.54216 | 0.052 | 0.108803 | 0.25749 | 0.1646 | 0.312747 |
| SL1344_RS09290_ | -0.27998 | -0.04842 | -0.45868 | -0.40462 | 0.205607 | -1.87466 | 0.023984 | 0.056529 | 0.167877 | 0.1265 | 0.268941 |
| SL1344_RS09295_ | -2.68938 | -2.64287 | -2.74081 | -0.10835 | 0.517932 | -0.20101 | 0.409061 | 0.567632 | 0.698025 | 0.6885 | 0.781999 |
| SL1344_RS09300_ | -3.96355 | -4.05445 | -3.8705 | 0.168463 | 0.685795 | 0.214689 | 0.398667 | 0.553978 | 0.682734 | 0.6639 | 0.760928 |
| SL1344_RS09305_ | -1.17646 | -0.85858 | -1.59247 | -0.73992 | 0.263678 | -2.74266 | 0.001336 | 0.023904 | 0.105462 | 0.1007 | 0.237527 |
| SL1344_RS09310_ | -4.77682 | -4.51699 | -5.23215 | -0.72454 | 1.068105 | -0.63211 | 0.232512 | 0.350451 | 0.509453 | 0.4705 | 0.600141 |

| | | | | | | | | | | | |
|---|---|---|---|---|---|---|---|---|---|---|---|
| SL1344_RS09315_ | -2.2715 | -2.19089 | -2.40442 | -0.22962 | 0.440325 | -0.47686 | 0.276667 | 0.406391 | 0.559269 | 0.5372 | 0.652991 |
| SL1344_RS09320_ | 1.798786 | 1.863344 | 1.732208 | -0.23674 | 0.235599 | -1.00785 | 0.111259 | 0.17095 | 0.339757 | 0.2413 | 0.391352 |
| SL1344_RS09325_ | -3.25658 | -2.96007 | -3.53689 | -0.62915 | 0.632746 | -0.95125 | 0.138667 | 0.214429 | 0.370751 | 0.2995 | 0.445319 |
| SL1344_RS09335_ | -4.59911 | -4.14994 | -5.24958 | -1.10554 | 0.833336 | -1.23072 | 0.089274 | 0.143628 | 0.28414 | 0.21 | 0.355478 |
| SL1344_RS09340_ | -1.63433 | -1.05988 | -2.16564 | -1.10112 | 0.389714 | -2.78838 | 0.000673 | 0.019957 | 0.099905 | 0.1002 | 0.236855 |
| SL1344_RS09345_ | -5.03494 | -4.20385 | -5.9543 | -1.75596 | 1.195154 | -1.36779 | 0.052 | 0.107739 | 0.246226 | 0.166 | 0.312923 |
| SL1344_RS09350_ | -5.00803 | -4.41837 | -5.8065 | -1.38578 | 0.981279 | -1.32375 | 0.064624 | 0.120728 | 0.257495 | 0.1848 | 0.330134 |
| SL1344_RS09355_ | -3.70261 | -3.49886 | -4.0118 | -0.55306 | 0.712613 | -0.714 | 0.19 | 0.300669 | 0.457808 | 0.3914 | 0.52632 |
| SL1344_RS09360_ | -1.82814 | -1.54863 | -1.97444 | -0.38801 | 0.409602 | -0.85987 | 0.173218 | 0.269452 | 0.43317 | 0.3679 | 0.517902 |
| SL1344_RS09365_ | 1.03192 | 0.74269 | 1.30903 | 0.564719 | 0.476057 | 1.149092 | 0.096667 | 0.16776 | 0.341816 | 0.1896 | 0.355286 |
| SL1344_RS09370_ | 1.70898 | 2.126198 | 1.172892 | -0.9583 | 0.123368 | -7.72104 | 4.68E-05 | 0.003839 | 0.04827 | 0.1 | 0.236592 |
| SL1344_RS09380_ | 2.375489 | 2.731836 | 2.011588 | -0.72897 | 0.100665 | -7.28056 | 4.68E-05 | 0.002102 | 0.039144 | 0.1 | 0.236592 |
| SL1344_RS09385_ | -0.50618 | -0.03704 | -0.81927 | -0.78062 | 0.418268 | -1.90686 | 0.005997 | 0.046593 | 0.158929 | 0.1069 | 0.245653 |
| SL1344_RS09390_ | 0.046905 | 0.184536 | -0.01725 | -0.19951 | 0.246917 | -0.72718 | 0.229847 | 0.340686 | 0.510101 | 0.4677 | 0.609178 |
| SL1344_RS09395_ | 1.179579 | 1.44677 | 0.86039 | -0.5837 | 0.12731 | -4.62046 | 4.68E-05 | 0.00615 | 0.057078 | 0.1 | 0.236592 |
| SL1344_RS09400_ | 2.029195 | 2.116342 | 1.890464 | -0.22773 | 0.111476 | -1.99156 | 0.006663 | 0.062243 | 0.182936 | 0.1097 | 0.248348 |
| SL1344_RS09405_ | 0.796502 | 1.093 | 0.535707 | -0.56366 | 0.180554 | -3.16598 | 0.000673 | 0.014915 | 0.086934 | 0.1 | 0.236592 |
| SL1344_RS09410_ | 1.988957 | 2.186965 | 1.723031 | -0.45484 | 0.113228 | -3.96346 | 4.68E-05 | 0.009924 | 0.071587 | 0.1 | 0.236592 |
| SL1344_RS09420_ | 0.304357 | 0.720332 | -0.25432 | -0.98245 | 0.218865 | -4.49517 | 4.68E-05 | 0.010487 | 0.073113 | 0.1 | 0.236592 |
| SL1344_RS09425_ | 2.581969 | 2.693459 | 2.377743 | -0.34086 | 0.16054 | -2.15738 | 0.000673 | 0.032252 | 0.13199 | 0.1008 | 0.237594 |
| SL1344_RS09430_ | 1.363733 | 1.548075 | 1.183978 | -0.36674 | 0.158105 | -2.25894 | 0.005997 | 0.031924 | 0.128407 | 0.1035 | 0.241035 |
| SL1344_RS09435_ | 3.327183 | 3.550026 | 3.047017 | -0.55106 | 0.188891 | -3.1535 | 4.68E-05 | 0.013352 | 0.087377 | 0.1 | 0.236592 |
| SL1344_RS09440_ | 1.805046 | 1.831403 | 1.767313 | -0.06195 | 0.092 | -0.62086 | 0.256496 | 0.382187 | 0.530929 | 0.4938 | 0.620014 |
| SL1344_RS09455_ | 2.350631 | 2.722116 | 2.111438 | -0.60235 | 0.113919 | -5.03613 | 4.68E-05 | 0.008083 | 0.066907 | 0.1 | 0.236592 |
| SL1344_RS09460_ | 1.460712 | 1.662312 | 1.150803 | -0.50704 | 0.123117 | -3.95602 | 4.68E-05 | 0.012246 | 0.078277 | 0.1 | 0.236592 |
| SL1344_RS09465_ | 0.512256 | 0.810955 | 0.097399 | -0.71405 | 0.163104 | -4.30376 | 4.68E-05 | 0.010519 | 0.071681 | 0.1 | 0.236592 |
| SL1344_RS09470_ | 0.644774 | 0.881226 | 0.253311 | -0.63265 | 0.206341 | -2.97985 | 4.68E-05 | 0.014913 | 0.087665 | 0.1 | 0.236592 |
| SL1344_RS09485_ | -3.76192 | -2.66819 | -4.18352 | -1.39003 | 1.01131 | -1.24237 | 0.113258 | 0.153307 | 0.313363 | 0.2442 | 0.401629 |
| SL1344_RS09490_ | -1.80313 | -1.47703 | -2.37471 | -0.86788 | 0.421915 | -1.97497 | 0.012667 | 0.057841 | 0.173831 | 0.1147 | 0.25511 |
| SL1344_RS09495_ | 2.642905 | 2.963641 | 2.309747 | -0.64409 | 0.126924 | -5.06475 | 4.68E-05 | 0.003295 | 0.047377 | 0.1 | 0.236592 |
| SL1344_RS09500_ | 2.61597 | 2.930345 | 2.409955 | -0.51938 | 0.193278 | -2.78198 | 4.68E-05 | 0.040626 | 0.152277 | 0.1 | 0.236592 |
| SL1344_RS09510_ | 3.802571 | 4.184879 | 3.428552 | -0.7915 | 0.116047 | -6.92126 | 4.68E-05 | 0.000613 | 0.029444 | 0.1 | 0.236592 |
| SL1344_RS09515_ | 0.554504 | 0.772862 | 0.291102 | -0.51772 | 0.221025 | -2.38515 | 4.68E-05 | 0.034423 | 0.133171 | 0.1002 | 0.236859 |
| SL1344_RS09520_ | -0.12926 | 0.240036 | -0.35942 | -0.57472 | 0.222792 | -2.46193 | 0.002001 | 0.031825 | 0.127136 | 0.1029 | 0.240403 |
| SL1344_RS09525_ | 1.92393 | 2.27059 | 1.699243 | -0.5677 | 0.156455 | -3.43858 | 4.68E-05 | 0.022702 | 0.110967 | 0.1 | 0.236592 |
| SL1344_RS09535_ | 1.888789 | 2.020291 | 1.721714 | -0.30898 | 0.118521 | -2.61187 | 0.000673 | 0.03904 | 0.142717 | 0.1 | 0.236592 |
| SL1344_RS09540_ | 0.850903 | 1.158804 | 0.474488 | -0.69039 | 0.195763 | -3.55621 | 4.68E-05 | 0.008627 | 0.068675 | 0.1 | 0.236592 |
| SL1344_RS09545_ | -0.72065 | -0.17334 | -1.18094 | -1.00601 | 0.235304 | -4.27333 | 4.68E-05 | 0.006568 | 0.059223 | 0.1 | 0.236592 |
| SL1344_RS09550_ | 2.6789 | 3.187623 | 2.190966 | -0.96013 | 0.153553 | -6.30982 | 4.68E-05 | 0.001654 | 0.037347 | 0.1 | 0.236592 |
| SL1344_RS09555_ | -1.76071 | -1.05369 | -1.99268 | -0.88526 | 0.449823 | -1.63999 | 0.050633 | 0.095046 | 0.236177 | 0.1586 | 0.305328 |
| SL1344_RS09635_ | -0.57104 | -0.48008 | -0.72194 | -0.23982 | 0.225046 | -0.99416 | 0.136667 | 0.222411 | 0.376597 | 0.3183 | 0.462688 |
| SL1344_RS09640_ | 1.920582 | 2.208915 | 1.87462 | -0.29524 | 0.198598 | -0.95339 | 0.226 | 0.258098 | 0.441112 | 0.4458 | 0.606935 |
| SL1344_RS09645_ | 1.109965 | 1.165406 | 1.064332 | -0.11094 | 0.177402 | -0.59181 | 0.273151 | 0.404963 | 0.567862 | 0.5392 | 0.666331 |
| SL1344_RS09650_ | -0.02389 | 0.025445 | -0.07709 | -0.10435 | 0.290997 | -0.3451 | 0.352 | 0.5712 | 0.705776 | 0.673 | 0.775796 |
| SL1344_RS09655_ | -7.19747 | -6.71573 | -7.58806 | -1.23486 | 2.771534 | -0.36595 | 0.314457 | 0.49248 | 0.636301 | 0.6265 | 0.731147 |

| | | | | | | | | | | | |
|---|---|---|---|---|---|---|---|---|---|---|---|
| SL1344_RS09660_ | -0.91875 | -0.58163 | -1.17768 | -0.58951 | 0.28213 | -2.0295 | 0.012667 | 0.047667 | 0.15595 | 0.1155 | 0.255966 |
| SL1344_RS09670_ | -0.62416 | -0.43951 | -0.75838 | -0.32102 | 0.213548 | -1.39395 | 0.063291 | 0.112342 | 0.247952 | 0.1882 | 0.334019 |
| SL1344_RS09680_ | 5.070861 | 5.167671 | 4.697092 | -0.46434 | 0.130673 | -2.21659 | 4.68E-05 | 0.043066 | 0.157972 | 0.1 | 0.236592 |
| SL1344_RS09685_ | -1.09673 | -0.64798 | -1.54666 | -0.87683 | 0.302497 | -2.77548 | 0.001336 | 0.019326 | 0.097884 | 0.1011 | 0.238062 |
| SL1344_RS09690_ | 0.124885 | 0.377552 | -0.25305 | -0.62741 | 0.164262 | -3.7465 | 4.68E-05 | 0.011436 | 0.073552 | 0.1 | 0.236592 |
| SL1344_RS09695_ | 2.481678 | 2.577991 | 2.35929 | -0.23428 | 0.121112 | -1.8779 | 0.003998 | 0.062908 | 0.187759 | 0.1036 | 0.241177 |
| SL1344_RS09700_ | 1.614746 | 1.83866 | 1.340568 | -0.49457 | 0.195482 | -2.46063 | 0.000673 | 0.02782 | 0.121489 | 0.1004 | 0.237129 |
| SL1344_RS09705_ | 1.105954 | 1.239558 | 0.963289 | -0.28007 | 0.101725 | -2.68887 | 0.002002 | 0.02011 | 0.096877 | 0.1013 | 0.238216 |
| SL1344_RS09710_ | -0.56213 | -0.20578 | -1.05606 | -0.85067 | 0.215488 | -3.98669 | 4.68E-05 | 0.010554 | 0.070834 | 0.1 | 0.236592 |
| SL1344_RS09715_ | 1.567916 | 1.659764 | 1.098027 | -0.57082 | 0.459579 | -1.35229 | 0.068667 | 0.090954 | 0.237978 | 0.1683 | 0.321574 |
| SL1344_RS09725_ | 2.071345 | 2.352089 | 1.79474 | -0.51519 | 0.183533 | -2.7834 | 4.68E-05 | 0.014756 | 0.089946 | 0.1 | 0.236592 |
| SL1344_RS09730_ | 2.425171 | 2.699538 | 2.140915 | -0.56679 | 0.08973 | -6.33274 | 4.68E-05 | 0.002588 | 0.041928 | 0.1 | 0.236592 |
| SL1344_RS09805_ | 6.770758 | 7.182301 | 6.39048 | -0.80872 | 0.127017 | -6.42909 | 4.68E-05 | 0.000881 | 0.032969 | 0.1 | 0.236592 |
| SL1344_RS09810_ | 5.330368 | 5.712693 | 4.946434 | -0.79468 | 0.082863 | -9.74472 | 4.68E-05 | 0.000436 | 0.026709 | 0.1 | 0.236592 |
| SL1344_RS09820_ | 0.811481 | 0.934856 | 0.690279 | -0.27869 | 0.217869 | -1.3027 | 0.049333 | 0.127828 | 0.282274 | 0.1578 | 0.304937 |
| SL1344_RS09850_ | 5.407089 | 5.697718 | 5.037234 | -0.67055 | 0.097142 | -6.95548 | 4.68E-05 | 0.002021 | 0.041084 | 0.1 | 0.236592 |
| SL1344_RS09855_ | 4.577881 | 4.811416 | 4.276502 | -0.53522 | 0.076626 | -6.97248 | 4.68E-05 | 0.001043 | 0.03268 | 0.1 | 0.236592 |
| SL1344_RS09860_ | 4.112017 | 4.350843 | 3.776254 | -0.56876 | 0.153154 | -3.77285 | 4.68E-05 | 0.004224 | 0.053878 | 0.1 | 0.236592 |
| SL1344_RS09875_ | 4.387314 | 4.557155 | 4.110576 | -0.45023 | 0.100381 | -4.44618 | 4.68E-05 | 0.010588 | 0.077709 | 0.1 | 0.236592 |
| SL1344_RS09885_ | 4.524754 | 4.876398 | 4.148032 | -0.7273 | 0.071695 | -10.0975 | 4.68E-05 | 0.000531 | 0.026796 | 0.1 | 0.236592 |
| SL1344_RS09890_ | 3.88135 | 4.096994 | 3.556833 | -0.54857 | 0.114718 | -4.89301 | 4.68E-05 | 0.001927 | 0.040384 | 0.1 | 0.236592 |
| SL1344_RS09895_ | 3.07273 | 3.37583 | 2.693663 | -0.6937 | 0.1001 | -7.01446 | 4.68E-05 | 0.001256 | 0.033957 | 0.1 | 0.236592 |
| SL1344_RS09900_ | 3.20699 | 3.608518 | 2.803233 | -0.81886 | 0.099136 | -8.28163 | 4.68E-05 | 0.001543 | 0.036277 | 0.1 | 0.236592 |
| SL1344_RS09910_ | 1.055954 | 1.372134 | 0.620497 | -0.76069 | 0.2467 | -3.05212 | 4.68E-05 | 0.012431 | 0.082276 | 0.1 | 0.236592 |
| SL1344_RS09920_ | 1.299641 | 1.211202 | 1.706246 | 0.470425 | 0.238197 | 1.369583 | 0.083944 | 0.129332 | 0.289889 | 0.192 | 0.346625 |
| SL1344_RS09925_ | -0.8704 | -1.46853 | -0.5068 | 0.946172 | 0.349589 | 2.658425 | 4.68E-05 | 0.022802 | 0.108016 | 0.1005 | 0.237251 |
| SL1344_RS09930_ | -1.37792 | -1.63366 | -1.11518 | 0.506543 | 0.308491 | 1.592805 | 0.041306 | 0.084984 | 0.210271 | 0.1511 | 0.297367 |
| SL1344_RS09935_ | 0.374952 | 0.09408 | 0.590641 | 0.49556 | 0.140739 | 3.438455 | 4.68E-05 | 0.012887 | 0.077218 | 0.1002 | 0.236863 |
| SL1344_RS09940_ | 3.175224 | 2.811888 | 3.764712 | 0.876921 | 0.185963 | 3.89607 | 4.68E-05 | 0.015552 | 0.094256 | 0.1 | 0.236592 |
| SL1344_RS09945_ | 0.165172 | 0.149374 | 0.204314 | 0.04326 | 0.236149 | 0.167527 | 0.427049 | 0.684489 | 0.79282 | 0.7816 | 0.858643 |
| SL1344_RS09950_ | -1.59638 | -1.71125 | -1.43248 | 0.293714 | 0.420888 | 0.654212 | 0.239333 | 0.365403 | 0.528707 | 0.4861 | 0.623645 |
| SL1344_RS09955_ | -0.13901 | -0.42678 | 0.084404 | 0.50564 | 0.206842 | 2.358757 | 0.003998 | 0.032086 | 0.125495 | 0.1031 | 0.240626 |
| SL1344_RS09960_ | 1.050247 | 0.950796 | 1.192273 | 0.241426 | 0.114781 | 1.98756 | 0.020667 | 0.052028 | 0.158857 | 0.1221 | 0.262986 |
| SL1344_RS09965_ | -1.44667 | -1.50017 | -1.37753 | 0.096961 | 0.356275 | 0.243176 | 0.394404 | 0.608197 | 0.730768 | 0.7193 | 0.807162 |
| SL1344_RS09970_ | -3.94793 | -3.78061 | -4.18129 | -0.38778 | 0.808869 | -0.44909 | 0.312 | 0.481711 | 0.626356 | 0.5954 | 0.710684 |
| SL1344_RS09975_ | 0.152353 | 0.358652 | -0.12767 | -0.44548 | 0.252515 | -1.72135 | 0.037975 | 0.06766 | 0.193019 | 0.1341 | 0.280722 |
| SL1344_RS09985_ | -0.34023 | -0.38233 | -0.30857 | 0.077312 | 0.219216 | 0.343395 | 0.345103 | 0.506159 | 0.64419 | 0.6278 | 0.731275 |
| SL1344_RS09990_ | -2.15849 | -2.15093 | -2.16544 | -0.00137 | 0.398288 | -0.0032 | 0.496667 | 0.604394 | 0.72702 | 0.7101 | 0.799807 |
| SL1344_RS09995_ | -5.19521 | -5.12763 | -5.30175 | -0.1589 | 1.098487 | -0.14224 | 0.439707 | 0.573936 | 0.702637 | 0.6922 | 0.7851 |
| SL1344_RS10000_ | -2.62923 | -2.33461 | -3.0242 | -0.67778 | 0.486396 | -1.30164 | 0.082 | 0.13086 | 0.275924 | 0.2009 | 0.351479 |
| SL1344_RS10005_ | -1.42025 | -1.30821 | -1.54847 | -0.23684 | 0.282002 | -0.78624 | 0.191333 | 0.297051 | 0.447146 | 0.3906 | 0.528165 |
| SL1344_RS10010_ | -1.61975 | -1.6224 | -1.61695 | 0.005123 | 0.270268 | 0.020228 | 0.489674 | 0.52715 | 0.660241 | 0.6403 | 0.743395 |
| SL1344_RS10015_ | -3.4615 | -3.34404 | -3.66055 | -0.33841 | 0.641271 | -0.4711 | 0.296469 | 0.432202 | 0.582077 | 0.5578 | 0.671468 |
| SL1344_RS10020_ | -3.14238 | -3.15832 | -3.1279 | 1.64E-05 | 0.652409 | 0.000193 | 0.5 | 0.659076 | 0.773161 | 0.7675 | 0.845842 |
| SL1344_RS10025_ | -2.63408 | -2.4887 | -2.76468 | -0.27645 | 0.404169 | -0.62158 | 0.241839 | 0.361973 | 0.507913 | 0.4609 | 0.589193 |

| | | | | | | | | | | | |
|---|---|---|---|---|---|---|---|---|---|---|---|
| SL1344_RS10030_ | -3.56704 | -3.39191 | -3.7949 | -0.43423 | 0.678811 | -0.60874 | 0.247169 | 0.368555 | 0.525524 | 0.4892 | 0.613869 |
| SL1344_RS10035_ | -2.66523 | -2.48904 | -2.78165 | -0.27465 | 0.461677 | -0.54158 | 0.268 | 0.437744 | 0.578554 | 0.5391 | 0.658926 |
| SL1344_RS10040_ | -1.47249 | -1.38738 | -1.54097 | -0.15319 | 0.289965 | -0.48798 | 0.293804 | 0.429247 | 0.572187 | 0.536 | 0.652402 |
| SL1344_RS10045_ | -1.68685 | -1.55801 | -1.76113 | -0.17608 | 0.357579 | -0.45641 | 0.304464 | 0.494667 | 0.635371 | 0.5978 | 0.713397 |
| SL1344_RS10050_ | 0.129198 | 0.106017 | 0.163381 | 0.057195 | 0.192267 | 0.28109 | 0.381079 | 0.570189 | 0.699465 | 0.6753 | 0.773913 |
| SL1344_RS10055_ | -1.93433 | -2.03434 | -1.88352 | 0.125236 | 0.450993 | 0.262725 | 0.392 | 0.605937 | 0.733099 | 0.7323 | 0.820524 |
| SL1344_RS10060_ | -1.46179 | -1.39707 | -1.54002 | -0.15157 | 0.312336 | -0.45115 | 0.304 | 0.453242 | 0.598005 | 0.5738 | 0.685882 |
| SL1344_RS10065_ | -0.69429 | -0.63215 | -0.78447 | -0.17189 | 0.272814 | -0.59015 | 0.253831 | 0.382307 | 0.539016 | 0.5117 | 0.632792 |
| SL1344_RS10070_ | -2.59027 | -2.36521 | -2.7862 | -0.38063 | 0.563445 | -0.63607 | 0.245836 | 0.389463 | 0.547246 | 0.4892 | 0.625161 |
| SL1344_RS10075_ | -2.81867 | -2.53015 | -3.17571 | -0.62645 | 0.477113 | -1.26666 | 0.076 | 0.132864 | 0.274362 | 0.2041 | 0.35228 |
| SL1344_RS10080_ | -3.52375 | -3.33605 | -3.73364 | -0.40601 | 0.555227 | -0.67975 | 0.222518 | 0.335016 | 0.481241 | 0.4351 | 0.564606 |
| SL1344_RS10085_ | -3.17016 | -2.92713 | -3.40364 | -0.45269 | 0.574167 | -0.73095 | 0.200533 | 0.318828 | 0.475718 | 0.4146 | 0.553726 |
| SL1344_RS10090_ | -3.01815 | -2.74363 | -3.27182 | -0.59387 | 0.88839 | -0.63193 | 0.187209 | 0.309791 | 0.486395 | 0.386 | 0.537341 |
| SL1344_RS10095_ | -0.98436 | -0.8035 | -1.19499 | -0.4025 | 0.289475 | -1.2928 | 0.057333 | 0.121718 | 0.265039 | 0.1748 | 0.321588 |
| SL1344_RS10100_ | -2.50567 | -2.16965 | -2.68153 | -0.4823 | 0.468693 | -0.93689 | 0.171219 | 0.254662 | 0.41127 | 0.3553 | 0.503399 |
| SL1344_RS10105_ | -1.88001 | -1.50004 | -2.14367 | -0.61841 | 0.380008 | -1.52616 | 0.045303 | 0.091075 | 0.221674 | 0.1614 | 0.30953 |
| SL1344_RS10110_ | -4.0132 | -3.68083 | -4.33742 | -0.63866 | 0.734118 | -0.84157 | 0.172 | 0.275942 | 0.430346 | 0.3658 | 0.50745 |
| SL1344_RS10115_ | -0.92348 | -0.69042 | -1.16837 | -0.47884 | 0.247119 | -1.89079 | 0.015334 | 0.054217 | 0.163608 | 0.1203 | 0.260859 |
| SL1344_RS10120_ | -2.62286 | -2.39787 | -2.85883 | -0.48245 | 0.482457 | -0.94637 | 0.150566 | 0.225437 | 0.380586 | 0.3146 | 0.457801 |
| SL1344_RS10130_ | -1.0811 | -0.91462 | -1.23158 | -0.23519 | 0.396673 | -0.56974 | 0.272667 | 0.439128 | 0.596269 | 0.5435 | 0.672902 |
| SL1344_RS10135_ | -2.50115 | -2.27118 | -2.74895 | -0.46619 | 0.376553 | -1.21717 | 0.089333 | 0.15287 | 0.292824 | 0.2303 | 0.376159 |
| SL1344_RS10140_ | -2.026 | -1.88916 | -2.22716 | -0.38775 | 0.488923 | -0.73405 | 0.207861 | 0.307042 | 0.47438 | 0.4197 | 0.557952 |
| SL1344_RS10145_ | -2.37879 | -2.07721 | -2.60176 | -0.56444 | 0.487698 | -1.10321 | 0.099333 | 0.17631 | 0.331053 | 0.2496 | 0.398371 |
| SL1344_RS10150_ | -3.15756 | -2.72901 | -3.56303 | -0.84996 | 0.570067 | -1.44393 | 0.042638 | 0.098871 | 0.231832 | 0.1619 | 0.307843 |
| SL1344_RS10155_ | -1.74878 | -1.62474 | -1.87943 | -0.24967 | 0.337332 | -0.6819 | 0.223185 | 0.344069 | 0.494036 | 0.4508 | 0.580574 |
| SL1344_RS10160_ | -1.09279 | -0.88437 | -1.42168 | -0.54948 | 0.338643 | -1.59729 | 0.026 | 0.09883 | 0.239315 | 0.129 | 0.271293 |
| SL1344_RS10165_ | -1.56047 | -1.44007 | -1.70274 | -0.2621 | 0.282033 | -0.86791 | 0.159227 | 0.263967 | 0.412367 | 0.3627 | 0.499402 |
| SL1344_RS10170_ | -1.43782 | -1.37487 | -1.50664 | -0.16065 | 0.325852 | -0.46645 | 0.305333 | 0.464541 | 0.608284 | 0.5824 | 0.692694 |
| SL1344_RS10175_ | -2.72705 | -2.55548 | -2.8213 | -0.28956 | 0.625357 | -0.4556 | 0.311333 | 0.501843 | 0.646045 | 0.6207 | 0.733265 |
| SL1344_RS10180_ | -1.86643 | -2.03513 | -1.7975 | 0.206847 | 0.411331 | 0.469191 | 0.319121 | 0.503631 | 0.64768 | 0.6141 | 0.73203 |
| SL1344_RS10185_ | -2.58636 | -2.64885 | -2.49097 | 0.13644 | 0.521952 | 0.230734 | 0.396667 | 0.611526 | 0.733756 | 0.7075 | 0.799131 |
| SL1344_RS10190_ | -1.51134 | -1.43422 | -1.61842 | -0.21419 | 0.418181 | -0.49312 | 0.293333 | 0.446165 | 0.600447 | 0.5881 | 0.70336 |
| SL1344_RS10195_ | -3.34677 | -3.3257 | -3.37343 | -0.03708 | 0.499309 | -0.06131 | 0.469687 | 0.549361 | 0.678261 | 0.6551 | 0.754264 |
| SL1344_RS10200_ | 1.343936 | 1.634437 | 1.184199 | -0.43647 | 0.255793 | -1.52913 | 0.048667 | 0.101122 | 0.250929 | 0.1431 | 0.293155 |
| SL1344_RS10205_ | -0.86933 | -0.82624 | -0.97552 | -0.15857 | 0.313384 | -0.45506 | 0.308667 | 0.477926 | 0.626225 | 0.6114 | 0.722855 |
| SL1344_RS10210_ | -1.78691 | -1.7227 | -1.82894 | -0.05768 | 0.422241 | -0.12309 | 0.452365 | 0.671233 | 0.781256 | 0.767 | 0.844744 |
| SL1344_RS10215_ | 3.510494 | 3.530308 | 3.487905 | -0.05315 | 0.08718 | -0.59128 | 0.254 | 0.367337 | 0.535417 | 0.4908 | 0.627849 |
| SL1344_RS10225_ | -0.27173 | -0.27675 | -0.26642 | 0.004204 | 0.204364 | 0.022881 | 0.490667 | 0.612784 | 0.733069 | 0.7376 | 0.818552 |
| SL1344_RS10230_ | 1.153532 | 1.165058 | 1.133773 | -0.01507 | 0.144303 | -0.10383 | 0.459694 | 0.655634 | 0.770197 | 0.7652 | 0.843464 |
| SL1344_RS10235_ | -2.68876 | -2.84393 | -2.52133 | 0.327832 | 0.387343 | 0.784949 | 0.197202 | 0.295033 | 0.440441 | 0.384 | 0.517637 |
| SL1344_RS10240_ | -4.04384 | -4.18007 | -3.9005 | 0.299904 | 0.658486 | 0.403132 | 0.32 | 0.464201 | 0.604683 | 0.5722 | 0.684611 |
| SL1344_RS10245_ | -2.6109 | -2.65196 | -2.59433 | 0.051166 | 0.558288 | 0.079351 | 0.466356 | 0.661132 | 0.775667 | 0.7774 | 0.855211 |
| SL1344_RS10250_ | -1.95038 | -2.12232 | -1.78279 | 0.331704 | 0.301654 | 1.058273 | 0.123917 | 0.199136 | 0.3404 | 0.2733 | 0.415912 |
| SL1344_RS10255_ | -1.16996 | -1.29709 | -1.05802 | 0.244902 | 0.240596 | 0.965565 | 0.141333 | 0.226606 | 0.375141 | 0.3121 | 0.454328 |
| SL1344_RS10260_ | -1.51251 | -1.63576 | -1.40557 | 0.224229 | 0.28031 | 0.736003 | 0.198534 | 0.323408 | 0.471777 | 0.4183 | 0.552275 |

| | | | | | | | | | | | |
|---|---|---|---|---|---|---|---|---|---|---|---|
| SL1344_RS10265_ | -2.30101 | -2.27867 | -2.31842 | 0.016313 | 0.568188 | 0.030772 | 0.486667 | 0.6972 | 0.803999 | 0.8134 | 0.878917 |
| SL1344_RS10270_ | -1.44019 | -1.51626 | -1.34453 | 0.208633 | 0.377479 | 0.535274 | 0.283333 | 0.433368 | 0.586192 | 0.5734 | 0.687387 |
| SL1344_RS10275_ | 0.896993 | 0.801103 | 1.255985 | 0.441468 | 0.210856 | 1.597837 | 0.041306 | 0.102735 | 0.251128 | 0.145 | 0.290527 |
| SL1344_RS10280_ | 3.118603 | 2.862422 | 3.317876 | 0.453925 | 0.085606 | 5.325437 | 4.68E-05 | 0.007074 | 0.062085 | 0.1 | 0.236592 |
| SL1344_RS10285_ | -1.11147 | -1.14199 | -1.05983 | 0.031163 | 0.60552 | 0.047557 | 0.482345 | 0.811837 | 0.886636 | 0.9124 | 0.945475 |
| SL1344_RS10290_ | -5.03284 | -5.75801 | -4.75156 | 0.995957 | 1.34701 | 0.614564 | 0.246502 | 0.357397 | 0.522613 | 0.4921 | 0.626803 |
| SL1344_RS10295_ | -5.67299 | -5.40055 | -6.64277 | -1.17291 | 1.801112 | -0.53113 | 0.282 | 0.407687 | 0.570398 | 0.5771 | 0.700396 |
| SL1344_RS10300_ | -3.96561 | -3.78001 | -4.31682 | -0.41504 | 1.125853 | -0.36173 | 0.365333 | 0.603857 | 0.732801 | 0.7115 | 0.808574 |
| SL1344_RS10305_ | -2.83977 | -2.73262 | -2.96386 | -0.23004 | 0.42018 | -0.50591 | 0.289333 | 0.417693 | 0.558148 | 0.5212 | 0.638978 |
| SL1344_RS10310_ | -4.9647 | -4.90311 | -5.03957 | -0.22849 | 1.622662 | -0.12196 | 0.426 | 0.619744 | 0.747965 | 0.7908 | 0.860458 |
| SL1344_RS10315_ | -2.05329 | -2.09674 | -2.0269 | 0.074913 | 0.361359 | 0.178526 | 0.423333 | 0.571675 | 0.699034 | 0.6805 | 0.776272 |
| SL1344_RS10320_ | 1.944289 | 1.971015 | 1.930366 | -0.03696 | 0.104202 | -0.33341 | 0.364667 | 0.564455 | 0.69587 | 0.6756 | 0.774746 |
| SL1344_RS10325_ | -3.13804 | -3.09296 | -3.17886 | -0.09462 | 0.488512 | -0.18161 | 0.427049 | 0.531205 | 0.662572 | 0.6388 | 0.741576 |
| SL1344_RS10330_ | 2.169408 | 2.05799 | 2.333176 | 0.281445 | 0.137912 | 2.044265 | 0.004668 | 0.039659 | 0.144894 | 0.1034 | 0.241011 |
| SL1344_RS10335_ | -2.55976 | -2.59273 | -2.51939 | 0.095959 | 0.501065 | 0.184249 | 0.41972 | 0.603333 | 0.72743 | 0.717 | 0.805323 |
| SL1344_RS10340_ | -0.42113 | -0.75418 | -0.29115 | 0.440883 | 0.262463 | 1.475239 | 0.053298 | 0.109138 | 0.253251 | 0.1704 | 0.317704 |
| SL1344_RS10345_ | 0.354811 | 0.003559 | 0.628153 | 0.626652 | 0.147095 | 4.293109 | 4.68E-05 | 0.007822 | 0.062325 | 0.1 | 0.236592 |
| SL1344_RS10350_ | 1.220291 | 1.049163 | 1.332026 | 0.26216 | 0.265194 | 0.948973 | 0.146569 | 0.2208 | 0.399057 | 0.2823 | 0.448732 |
| SL1344_RS10355_ | -0.71038 | -0.76483 | -0.62427 | 0.130499 | 0.536249 | 0.23761 | 0.368421 | 0.711572 | 0.815945 | 0.7356 | 0.827249 |
| SL1344_RS10360_ | -0.83695 | -1.12317 | -0.51503 | 0.608336 | 0.221576 | 2.645217 | 0.002668 | 0.022603 | 0.10363 | 0.1023 | 0.239481 |
| SL1344_RS10365_ | -1.54199 | -1.57603 | -1.48288 | 0.04704 | 0.46545 | 0.094976 | 0.456667 | 0.724135 | 0.823283 | 0.8254 | 0.888623 |
| SL1344_RS10370_ | -1.95481 | -2.43492 | -1.54686 | 0.888196 | 0.298409 | 2.920821 | 0.001336 | 0.01937 | 0.093901 | 0.1006 | 0.237394 |
| SL1344_RS10375_ | 1.076611 | 0.683327 | 1.484241 | 0.815768 | 0.143537 | 5.635121 | 4.68E-05 | 0.002927 | 0.043933 | 0.1 | 0.236592 |
| SL1344_RS10380_ | 1.555034 | 1.170658 | 1.88505 | 0.717945 | 0.154533 | 4.677448 | 4.68E-05 | 0.005946 | 0.057845 | 0.1 | 0.236592 |
| SL1344_RS10390_ | 0.168967 | -0.23347 | 0.579043 | 0.906818 | 0.335513 | 2.720544 | 0.000673 | 0.017608 | 0.097795 | 0.1 | 0.236592 |
| SL1344_RS10395_ | 1.516459 | 1.112827 | 1.92532 | 0.810285 | 0.147686 | 5.396666 | 4.68E-05 | 0.003172 | 0.04576 | 0.1 | 0.236592 |
| SL1344_RS10400_ | 0.977941 | 0.450022 | 1.394062 | 0.914651 | 0.191905 | 4.621299 | 4.68E-05 | 0.005466 | 0.055899 | 0.1 | 0.236592 |
| SL1344_RS10405_ | 0.992935 | 0.263967 | 1.460114 | 1.19564 | 0.230676 | 4.997481 | 4.68E-05 | 0.005166 | 0.055293 | 0.1 | 0.236592 |
| SL1344_RS10410_ | -0.15015 | -0.92974 | 0.191287 | 1.11425 | 0.391418 | 2.597059 | 4.68E-05 | 0.041553 | 0.151837 | 0.1 | 0.236592 |
| SL1344_RS10415_ | 1.153187 | 0.586355 | 1.652895 | 1.085609 | 0.178855 | 6.190949 | 4.68E-05 | 0.007823 | 0.065686 | 0.1 | 0.236592 |
| SL1344_RS10420_ | 1.248774 | 0.64646 | 1.7282 | 1.091934 | 0.215308 | 5.079328 | 4.68E-05 | 0.002499 | 0.042861 | 0.1 | 0.236592 |
| SL1344_RS10430_ | 1.741716 | 1.260189 | 2.217532 | 0.997237 | 0.190535 | 5.260846 | 4.68E-05 | 0.002085 | 0.041042 | 0.1 | 0.236592 |
| SL1344_RS10435_ | 1.796858 | 1.213205 | 2.315717 | 1.106243 | 0.219127 | 5.083821 | 4.68E-05 | 0.002424 | 0.04343 | 0.1 | 0.236592 |
| SL1344_RS10440_ | 1.564519 | 0.993758 | 2.168902 | 1.176257 | 0.196386 | 6.108585 | 4.68E-05 | 0.002612 | 0.042781 | 0.1 | 0.236592 |
| SL1344_RS10445_ | 1.167357 | 0.324644 | 1.65485 | 1.363596 | 0.341696 | 4.082549 | 4.68E-05 | 0.00319 | 0.048151 | 0.1 | 0.236592 |
| SL1344_RS10450_ | 0.754707 | 0.234299 | 1.197958 | 0.995259 | 0.202414 | 4.916174 | 4.68E-05 | 0.006241 | 0.05854 | 0.1 | 0.236592 |
| SL1344_RS10455_ | 0.595296 | 0.227997 | 0.973999 | 0.879638 | 0.345722 | 2.574672 | 4.68E-05 | 0.01943 | 0.102156 | 0.1 | 0.236592 |
| SL1344_RS10470_ | 1.569368 | 1.060561 | 1.954543 | 0.806752 | 0.241337 | 3.340746 | 4.68E-05 | 0.009131 | 0.07189 | 0.1 | 0.236592 |
| SL1344_RS10475_ | 1.616432 | 1.18719 | 2.045796 | 0.878202 | 0.229418 | 3.83918 | 4.68E-05 | 0.006857 | 0.063526 | 0.1 | 0.236592 |
| SL1344_RS10480_ | 1.09496 | 0.741211 | 1.607679 | 0.847775 | 0.275792 | 3.089806 | 4.68E-05 | 0.011266 | 0.079757 | 0.1 | 0.236592 |
| SL1344_RS10485_ | -1.62312 | -2.06272 | -1.4101 | 0.637356 | 0.366853 | 1.601226 | 0.041306 | 0.092517 | 0.225625 | 0.1501 | 0.295788 |
| SL1344_RS10490_ | -3.45712 | -5.47342 | -2.28926 | 3.303077 | 0.933507 | 3.534518 | 4.68E-05 | 0.021645 | 0.09895 | 0.1 | 0.236592 |
| SL1344_RS10495_ | -2.75264 | -4.18811 | -1.71225 | 2.518458 | 0.573888 | 4.462929 | 4.68E-05 | 0.005796 | 0.05614 | 0.1 | 0.236592 |
| SL1344_RS10515_ | -2.18228 | -2.90419 | -1.48283 | 1.413841 | 0.468359 | 2.945707 | 0.000673 | 0.017419 | 0.092414 | 0.1003 | 0.23699 |
| SL1344_RS10520_ | -5.03777 | -6.51009 | -4.26689 | 2.25897 | 1.295448 | 1.613611 | 0.02 | 0.093617 | 0.227845 | 0.129 | 0.270682 |

| | | | | | | | | | | | |
|---|---|---|---|---|---|---|---|---|---|---|---|
| SL1344_RS10530_ | -3.25988 | -3.59463 | -2.9647 | 0.632643 | 0.501949 | 1.196379 | 0.087275 | 0.149044 | 0.285547 | 0.2268 | 0.369981 |
| SL1344_RS10535_ | -3.23381 | -3.33419 | -3.08839 | 0.271371 | 0.76873 | 0.331889 | 0.35976 | 0.575485 | 0.705191 | 0.6806 | 0.7778 |
| SL1344_RS10540_ | -3.64874 | -3.49707 | -3.79038 | -0.28878 | 0.581281 | -0.47604 | 0.3 | 0.447805 | 0.586741 | 0.5446 | 0.659262 |
| SL1344_RS10545_ | -4.9444 | -4.79561 | -5.10613 | -0.31976 | 0.934669 | -0.32332 | 0.357095 | 0.489568 | 0.626152 | 0.5916 | 0.701906 |
| SL1344_RS10550_ | -3.43289 | -3.35083 | -3.54913 | -0.19268 | 0.547131 | -0.33866 | 0.359094 | 0.498673 | 0.635543 | 0.6109 | 0.718256 |
| SL1344_RS10555_ | -2.71208 | -2.79755 | -2.62586 | 0.141897 | 0.519569 | 0.266846 | 0.383744 | 0.570648 | 0.69939 | 0.6804 | 0.776776 |
| SL1344_RS10560_ | -2.8733 | -2.71551 | -3.10034 | -0.36137 | 0.562275 | -0.60381 | 0.237841 | 0.384241 | 0.538551 | 0.4935 | 0.623926 |
| SL1344_RS10565_ | -2.55466 | -2.35485 | -2.8488 | -0.48157 | 0.471896 | -0.95664 | 0.14457 | 0.227229 | 0.381909 | 0.3073 | 0.454429 |
| SL1344_RS10570_ | -3.66567 | -3.48683 | -3.78457 | -0.32536 | 0.734525 | -0.38879 | 0.336 | 0.531764 | 0.667075 | 0.6392 | 0.745021 |
| SL1344_RS10575_ | -4.98128 | -4.20752 | -5.53608 | -1.30867 | 1.179194 | -1.05817 | 0.125333 | 0.198147 | 0.355061 | 0.2809 | 0.432056 |
| SL1344_RS10580_ | -4.2228 | -3.70637 | -4.82238 | -1.11948 | 0.774597 | -1.38964 | 0.062667 | 0.119315 | 0.255485 | 0.1818 | 0.328271 |
| SL1344_RS10585_ | -1.5604 | -1.51623 | -1.60325 | -0.08581 | 0.289845 | -0.27908 | 0.38 | 0.506735 | 0.640828 | 0.6147 | 0.718405 |
| SL1344_RS10590_ | -0.75252 | -0.77026 | -0.73616 | 0.050146 | 0.416215 | 0.120832 | 0.434 | 0.694333 | 0.801584 | 0.7815 | 0.856272 |
| SL1344_RS10595_ | 3.667249 | 3.59388 | 3.701861 | 0.104676 | 0.071483 | 1.383759 | 0.071286 | 0.117663 | 0.263661 | 0.1915 | 0.344485 |
| SL1344_RS10600_ | 1.689688 | 1.729684 | 1.556686 | -0.15879 | 0.146032 | -0.93695 | 0.193871 | 0.256392 | 0.42839 | 0.3854 | 0.540906 |
| SL1344_RS10640_ | 2.455789 | 2.569471 | 2.269169 | -0.21601 | 0.308498 | -0.65529 | 0.272 | 0.405653 | 0.586094 | 0.5408 | 0.690609 |
| SL1344_RS10645_ | 0.58978 | 0.244587 | 0.914105 | 0.685726 | 0.17863 | 3.842525 | 4.68E-05 | 0.008633 | 0.067658 | 0.1 | 0.236592 |
| SL1344_RS10650_ | 1.967012 | 1.78735 | 2.418169 | 0.608512 | 0.156719 | 2.79642 | 4.68E-05 | 0.038969 | 0.147736 | 0.1 | 0.236592 |
| SL1344_RS10655_ | -0.31021 | -0.72521 | 0.031693 | 0.748738 | 0.205325 | 3.666946 | 4.68E-05 | 0.011442 | 0.074707 | 0.1 | 0.236592 |
| SL1344_RS10660_ | -0.0645 | -0.55919 | 0.290851 | 0.858992 | 0.251872 | 3.316033 | 4.68E-05 | 0.018101 | 0.094896 | 0.1 | 0.236592 |
| SL1344_RS10665_ | -0.0521 | -0.47826 | 0.279161 | 0.752841 | 0.195457 | 3.814294 | 4.68E-05 | 0.008571 | 0.066365 | 0.1 | 0.236592 |
| SL1344_RS10670_ | 0.948738 | 0.692163 | 1.257798 | 0.571554 | 0.157669 | 3.528777 | 4.68E-05 | 0.012477 | 0.080066 | 0.1 | 0.236592 |
| SL1344_RS10675_ | -0.14717 | -0.54 | 0.310984 | 0.860967 | 0.224475 | 3.848105 | 4.68E-05 | 0.010792 | 0.075052 | 0.1 | 0.236592 |
| SL1344_RS10680_ | 0.402935 | 0.094164 | 0.63077 | 0.539624 | 0.148408 | 3.576996 | 4.68E-05 | 0.013379 | 0.079443 | 0.1001 | 0.236726 |
| SL1344_RS10685_ | 0.842811 | 0.545889 | 1.13097 | 0.5885 | 0.140108 | 4.15322 | 4.68E-05 | 0.008351 | 0.064663 | 0.1 | 0.236592 |
| SL1344_RS10690_ | 0.421829 | 0.071757 | 0.723621 | 0.649437 | 0.161196 | 3.967435 | 4.68E-05 | 0.008627 | 0.066153 | 0.1 | 0.236592 |
| SL1344_RS10700_ | 1.984184 | 1.757995 | 2.225166 | 0.465822 | 0.099239 | 4.632758 | 4.68E-05 | 0.004798 | 0.052482 | 0.1 | 0.236592 |
| SL1344_RS10705_ | 3.040133 | 2.576523 | 3.462356 | 0.888201 | 0.127524 | 6.940622 | 4.68E-05 | 0.000835 | 0.03138 | 0.1 | 0.236592 |
| SL1344_RS10710_ | 2.499081 | 2.170811 | 2.748531 | 0.583278 | 0.128488 | 4.561478 | 4.68E-05 | 0.004044 | 0.050909 | 0.1 | 0.236592 |
| SL1344_RS10715_ | 2.456362 | 2.09153 | 2.74433 | 0.667303 | 0.150284 | 4.444606 | 4.68E-05 | 0.005932 | 0.059222 | 0.1 | 0.236592 |
| SL1344_RS10720_ | 2.761928 | 2.502887 | 3.006495 | 0.504956 | 0.08184 | 6.203317 | 4.68E-05 | 0.002423 | 0.040879 | 0.1 | 0.236592 |
| SL1344_RS10725_ | 2.30536 | 2.04096 | 2.477334 | 0.442057 | 0.149008 | 2.891468 | 4.68E-05 | 0.031454 | 0.130561 | 0.1 | 0.236592 |
| SL1344_RS10730_ | 2.187437 | 2.030377 | 2.320851 | 0.302486 | 0.14459 | 2.145997 | 4.68E-05 | 0.079082 | 0.217372 | 0.1001 | 0.236722 |
| SL1344_RS10735_ | 2.555756 | 2.380674 | 2.692165 | 0.321059 | 0.158627 | 2.050211 | 0.001336 | 0.052704 | 0.172202 | 0.1009 | 0.237778 |
| SL1344_RS10740_ | 2.454214 | 2.215819 | 2.649935 | 0.439848 | 0.093189 | 4.784836 | 4.68E-05 | 0.010531 | 0.074015 | 0.1 | 0.236592 |
| SL1344_RS10745_ | 3.14303 | 2.899216 | 3.397453 | 0.518221 | 0.094194 | 5.513748 | 4.68E-05 | 0.00263 | 0.04348 | 0.1 | 0.236592 |
| SL1344_RS10750_ | 4.49783 | 4.218897 | 4.737734 | 0.520699 | 0.136493 | 3.91199 | 4.68E-05 | 0.011075 | 0.080317 | 0.1 | 0.236592 |
| SL1344_RS10755_ | 2.980947 | 2.644947 | 3.246621 | 0.626372 | 0.195356 | 3.24542 | 4.68E-05 | 0.019613 | 0.104568 | 0.1 | 0.236592 |
| SL1344_RS10760_ | 2.328791 | 1.85773 | 2.659854 | 0.790952 | 0.235399 | 3.295175 | 4.68E-05 | 0.02434 | 0.116529 | 0.1 | 0.236592 |
| SL1344_RS10765_ | 2.498301 | 2.165781 | 2.700654 | 0.520964 | 0.132304 | 3.490005 | 4.68E-05 | 0.020023 | 0.103663 | 0.1 | 0.236592 |
| SL1344_RS10770_ | 2.065595 | 1.782537 | 2.228918 | 0.45001 | 0.165777 | 2.599251 | 4.68E-05 | 0.035521 | 0.138765 | 0.1 | 0.236592 |
| SL1344_RS10775_ | 1.660957 | 1.34591 | 1.809542 | 0.444865 | 0.172223 | 2.34373 | 0.001337 | 0.034414 | 0.135651 | 0.1015 | 0.238537 |
| SL1344_RS10780_ | 2.126174 | 1.869599 | 2.350668 | 0.482331 | 0.103588 | 4.656172 | 4.68E-05 | 0.008882 | 0.067331 | 0.1 | 0.236592 |
| SL1344_RS10785_ | 2.169273 | 2.054536 | 2.297607 | 0.24271 | 0.089671 | 2.639236 | 0.001337 | 0.022753 | 0.105164 | 0.1016 | 0.238677 |
| SL1344_RS10790_ | 3.122677 | 3.154145 | 3.078361 | -0.06094 | 0.157127 | -0.39532 | 0.316456 | 0.590767 | 0.730221 | 0.636 | 0.758311 |

| | | | | | | | | | | | |
|---|---|---|---|---|---|---|---|---|---|---|---|
| SL1344_RS10795_ | -2.2766 | -2.02518 | -2.43305 | -0.41934 | 0.405475 | -0.95756 | 0.157333 | 0.239461 | 0.392838 | 0.3308 | 0.473576 |
| SL1344_RS10800_ | -2.66007 | -2.4902 | -3.00272 | -0.48094 | 0.575244 | -0.79243 | 0.203198 | 0.302392 | 0.469489 | 0.4179 | 0.562178 |
| SL1344_RS10805_ | -2.69195 | -2.96046 | -2.59297 | 0.313724 | 0.568824 | 0.479635 | 0.322667 | 0.515174 | 0.658398 | 0.648 | 0.761104 |
| SL1344_RS10810_ | -2.75624 | -2.82515 | -2.70076 | 0.124005 | 0.457704 | 0.268342 | 0.387333 | 0.529074 | 0.662436 | 0.6402 | 0.743857 |
| SL1344_RS10815_ | -2.93426 | -2.68826 | -3.12823 | -0.44246 | 0.62359 | -0.67028 | 0.222518 | 0.355586 | 0.515851 | 0.4534 | 0.592856 |
| SL1344_RS10820_ | -3.28753 | -3.33645 | -3.25274 | 0.096096 | 0.512261 | 0.171075 | 0.421333 | 0.52867 | 0.660886 | 0.6427 | 0.743333 |
| SL1344_RS10825_ | -2.39209 | -2.45583 | -2.32224 | 0.129378 | 0.345091 | 0.360143 | 0.345103 | 0.463338 | 0.600361 | 0.5657 | 0.67736 |
| SL1344_RS10830_ | -2.45961 | -2.35332 | -2.57068 | -0.20339 | 0.381508 | -0.51108 | 0.271819 | 0.392441 | 0.536086 | 0.4932 | 0.615851 |
| SL1344_RS10835_ | -3.63911 | -3.77697 | -3.49803 | 0.291037 | 0.554819 | 0.480755 | 0.285333 | 0.42874 | 0.569318 | 0.5377 | 0.652863 |
| SL1344_RS10840_ | -2.58919 | -2.3991 | -2.71004 | -0.28648 | 0.44464 | -0.57514 | 0.263158 | 0.419705 | 0.563767 | 0.5265 | 0.649658 |
| SL1344_RS10845_ | -2.7343 | -2.68282 | -2.79671 | -0.06004 | 0.606995 | -0.10141 | 0.463691 | 0.679559 | 0.787072 | 0.7926 | 0.859674 |
| SL1344_RS10850_ | -5.2845 | -5.20327 | -5.35849 | -0.14014 | 0.964429 | -0.12702 | 0.444 | 0.532151 | 0.66431 | 0.6433 | 0.744371 |
| SL1344_RS10855_ | -3.97745 | -3.71747 | -4.28926 | -0.56863 | 0.786682 | -0.68591 | 0.22052 | 0.337311 | 0.494058 | 0.4457 | 0.581617 |
| SL1344_RS10860_ | -2.92538 | -3.12385 | -2.77536 | 0.331415 | 0.587696 | 0.525718 | 0.293138 | 0.430727 | 0.580191 | 0.5431 | 0.66385 |
| SL1344_RS10865_ | -2.55485 | -2.46968 | -2.59828 | -0.15084 | 0.502347 | -0.27932 | 0.391333 | 0.58911 | 0.714787 | 0.7004 | 0.791955 |
| SL1344_RS10870_ | -3.9111 | -3.86341 | -3.95094 | 0.007039 | 0.871258 | 0.006653 | 0.497002 | 0.648492 | 0.76427 | 0.7627 | 0.839424 |
| SL1344_RS10875_ | -2.97295 | -3.11835 | -2.72894 | 0.365852 | 0.591816 | 0.55365 | 0.274 | 0.454019 | 0.599982 | 0.5533 | 0.676803 |
| SL1344_RS10880_ | -2.06743 | -2.07623 | -2.06105 | -0.03192 | 0.442735 | -0.06483 | 0.470353 | 0.655335 | 0.769136 | 0.7765 | 0.850861 |
| SL1344_RS10885_ | -5.02638 | -5.41521 | -4.60227 | 0.81245 | 1.039253 | 0.727845 | 0.202 | 0.303357 | 0.456403 | 0.3972 | 0.534838 |
| SL1344_RS10890_ | -3.69437 | -4.06732 | -3.20113 | 0.857001 | 0.661024 | 1.257101 | 0.085943 | 0.145574 | 0.290539 | 0.2239 | 0.372252 |
| SL1344_RS10895_ | 0.713494 | 0.484818 | 0.854103 | 0.383724 | 0.207353 | 1.949497 | 0.003998 | 0.092725 | 0.235831 | 0.1054 | 0.243371 |
| SL1344_RS10900_ | 3.261823 | 3.094339 | 3.46898 | 0.364193 | 0.10931 | 3.270043 | 4.68E-05 | 0.009334 | 0.072579 | 0.1 | 0.236592 |
| SL1344_RS10905_ | 1.691907 | 1.558863 | 1.850968 | 0.266013 | 0.147187 | 1.772782 | 0.026649 | 0.056691 | 0.175508 | 0.1255 | 0.269845 |
| SL1344_RS10910_ | 0.672946 | 0.775879 | 0.445979 | -0.31538 | 0.224858 | -1.31022 | 0.083944 | 0.123923 | 0.275562 | 0.2047 | 0.361765 |
| SL1344_RS10915_ | 1.230678 | 1.015144 | 1.505288 | 0.483791 | 0.141506 | 3.285954 | 4.68E-05 | 0.013086 | 0.081439 | 0.1002 | 0.23686 |
| SL1344_RS10920_ | -2.47433 | -2.65166 | -2.32229 | 0.330621 | 0.368987 | 0.835248 | 0.170553 | 0.267674 | 0.413926 | 0.3563 | 0.492407 |
| SL1344_RS10930_ | -1.28844 | -1.31669 | -1.24439 | 0.086079 | 0.283676 | 0.2678 | 0.377082 | 0.533104 | 0.666605 | 0.6399 | 0.741307 |
| SL1344_RS10935_ | -0.58396 | -0.89152 | -0.36456 | 0.533645 | 0.297099 | 1.782758 | 0.019334 | 0.058941 | 0.177505 | 0.1214 | 0.263193 |
| SL1344_RS10940_ | -0.63488 | -0.78215 | -0.52107 | 0.271539 | 0.272542 | 0.951866 | 0.133333 | 0.216408 | 0.375747 | 0.3018 | 0.449063 |
| SL1344_RS10945_ | -0.73437 | -0.79195 | -0.67135 | 0.068445 | 0.330436 | 0.210702 | 0.419054 | 0.669686 | 0.780028 | 0.7735 | 0.848157 |
| SL1344_RS10950_ | -0.02313 | -0.01471 | -0.03367 | -0.01898 | 0.161667 | -0.10872 | 0.452667 | 0.544059 | 0.674923 | 0.6628 | 0.758777 |
| SL1344_RS10955_ | 0.48502 | 0.445041 | 0.551091 | 0.097953 | 0.159098 | 0.556973 | 0.283145 | 0.435059 | 0.583157 | 0.5384 | 0.66446 |
| SL1344_RS10960_ | -3.92479 | -4.02055 | -3.83345 | 0.17729 | 0.617794 | 0.268607 | 0.382667 | 0.509913 | 0.64252 | 0.6167 | 0.721877 |
| SL1344_RS10965_ | -3.54888 | -3.72341 | -3.21753 | 0.52924 | 0.819908 | 0.61035 | 0.263158 | 0.399593 | 0.556425 | 0.5236 | 0.650841 |
| SL1344_RS10970_ | -0.88442 | -1.04817 | -0.81462 | 0.185524 | 0.365958 | 0.463291 | 0.356667 | 0.614959 | 0.741497 | 0.7198 | 0.818932 |
| SL1344_RS10975_ | -0.45217 | -0.44316 | -0.46208 | -0.00965 | 0.250364 | -0.03427 | 0.487333 | 0.654995 | 0.768527 | 0.7631 | 0.840282 |
| SL1344_RS10980_ | -3.81076 | -3.98778 | -3.69533 | 0.321925 | 0.708096 | 0.408513 | 0.302667 | 0.455901 | 0.600895 | 0.5761 | 0.689264 |
| SL1344_RS10985_ | -3.2066 | -2.96784 | -3.46163 | -0.48304 | 0.558207 | -0.83538 | 0.178667 | 0.274561 | 0.42623 | 0.359 | 0.500407 |
| SL1344_RS10990_ | -2.35453 | -2.11268 | -2.50335 | -0.40142 | 0.51707 | -0.70027 | 0.221186 | 0.33608 | 0.498623 | 0.4587 | 0.594749 |
| SL1344_RS10995_ | -5.92605 | -5.89971 | -5.93838 | -0.05926 | 1.69143 | -0.0284 | 0.489674 | 0.634761 | 0.753925 | 0.7589 | 0.840088 |
| SL1344_RS11000_ | 0.012898 | -0.2585 | 0.195041 | 0.451991 | 0.186312 | 2.360705 | 0.006001 | 0.037655 | 0.135237 | 0.1058 | 0.243976 |
| SL1344_RS11005_ | 2.598022 | 1.900367 | 2.969727 | 1.043601 | 0.151574 | 5.733066 | 4.68E-05 | 0.015393 | 0.093582 | 0.1 | 0.236592 |
| SL1344_RS11010_ | 0.112398 | 0.420988 | -0.22109 | -0.65176 | 0.188344 | -3.48223 | 4.68E-05 | 0.010809 | 0.073676 | 0.1 | 0.236592 |
| SL1344_RS11015_ | 0.187056 | 0.381369 | -0.05642 | -0.4642 | 0.226565 | -2.03702 | 0.004001 | 0.061735 | 0.183676 | 0.1022 | 0.239457 |
| SL1344_RS11020_ | -0.16017 | 0.073692 | -0.46006 | -0.54262 | 0.231826 | -2.27938 | 0.002667 | 0.042798 | 0.147082 | 0.1016 | 0.238722 |

| | | | | | | | | | | | |
|---|---|---|---|---|---|---|---|---|---|---|---|
| SL1344_RS11025_ | 0.700455 | 0.604726 | 0.815723 | 0.211029 | 0.142257 | 1.430524 | 0.051299 | 0.109011 | 0.24437 | 0.1708 | 0.317017 |
| SL1344_RS11030_ | -1.10494 | -1.45736 | -0.73632 | 0.719881 | 0.302686 | 2.380082 | 0.002668 | 0.031456 | 0.124991 | 0.1039 | 0.241738 |
| SL1344_RS11035_ | -1.48735 | -1.75058 | -0.8603 | 0.830469 | 0.404307 | 1.843235 | 0.028667 | 0.065433 | 0.189313 | 0.1292 | 0.272128 |
| SL1344_RS11040_ | -0.01033 | -0.03049 | 0.002426 | 0.041454 | 0.191595 | 0.19079 | 0.416389 | 0.587667 | 0.712155 | 0.7044 | 0.792872 |
| SL1344_RS11045_ | -2.4453 | -2.4982 | -2.41078 | 0.106803 | 0.44726 | 0.219305 | 0.397735 | 0.552528 | 0.685274 | 0.6783 | 0.773467 |
| SL1344_RS11050_ | -1.47271 | -1.52441 | -1.42941 | 0.118222 | 0.361792 | 0.296093 | 0.363333 | 0.545544 | 0.677628 | 0.671 | 0.764437 |
| SL1344_RS11055_ | -3.39982 | -3.3092 | -3.68286 | -0.34619 | 0.834918 | -0.37335 | 0.364 | 0.586076 | 0.720151 | 0.7232 | 0.820063 |
| SL1344_RS11060_ | 0.559483 | 0.499167 | 0.660924 | 0.168692 | 0.237752 | 0.686789 | 0.240506 | 0.35087 | 0.521696 | 0.4701 | 0.611 |
| SL1344_RS11065_ | -1.39634 | -1.44822 | -1.29771 | 0.118298 | 0.39633 | 0.264542 | 0.384 | 0.661356 | 0.774241 | 0.7462 | 0.833027 |
| SL1344_RS11070_ | 2.952727 | 2.892102 | 2.991014 | 0.096192 | 0.088263 | 1.049612 | 0.121333 | 0.184959 | 0.345526 | 0.2725 | 0.431391 |
| SL1344_RS11080_ | -2.09166 | -1.9673 | -2.20129 | -0.23947 | 0.476007 | -0.49376 | 0.304667 | 0.475979 | 0.621498 | 0.5859 | 0.702146 |
| SL1344_RS11085_ | -3.20818 | -2.96882 | -3.48271 | -0.52464 | 0.612613 | -0.81781 | 0.171219 | 0.262152 | 0.421526 | 0.3573 | 0.502784 |
| SL1344_RS11090_ | 0.462142 | 0.616602 | 0.364187 | -0.23896 | 0.183295 | -1.24092 | 0.096667 | 0.150503 | 0.301752 | 0.234 | 0.388742 |
| SL1344_RS11095_ | 0.977811 | 0.959008 | 0.991097 | 0.034712 | 0.120271 | 0.267521 | 0.391073 | 0.525318 | 0.65509 | 0.6304 | 0.731322 |
| SL1344_RS11100_ | 1.605207 | 1.64334 | 1.543747 | -0.09404 | 0.118811 | -0.74744 | 0.196 | 0.309706 | 0.467001 | 0.4165 | 0.554885 |
| SL1344_RS11105_ | -0.63006 | -1.16802 | -0.18945 | 0.981498 | 0.263258 | 3.677673 | 4.68E-05 | 0.009777 | 0.070442 | 0.1 | 0.236592 |
| SL1344_RS11110_ | 0.183235 | 0.221522 | 0.126297 | -0.08091 | 0.186628 | -0.41122 | 0.314667 | 0.509692 | 0.648418 | 0.6206 | 0.730048 |
| SL1344_RS11115_ | -0.60167 | -0.90473 | -0.33819 | 0.569165 | 0.246897 | 2.257134 | 0.004665 | 0.036699 | 0.133861 | 0.1063 | 0.244191 |
| SL1344_RS11120_ | 0.069959 | -0.34771 | 0.414768 | 0.75761 | 0.184981 | 4.000971 | 4.68E-05 | 0.008582 | 0.066162 | 0.1 | 0.236592 |
| SL1344_RS11125_ | 0.844923 | 0.386377 | 1.310524 | 0.919131 | 0.153566 | 6.022495 | 4.68E-05 | 0.002344 | 0.04055 | 0.1 | 0.236592 |
| SL1344_RS11130_ | 1.054418 | 0.661995 | 1.452399 | 0.750592 | 0.198195 | 3.765712 | 4.68E-05 | 0.007506 | 0.065369 | 0.1 | 0.236592 |
| SL1344_RS11135_ | 1.658708 | 1.414055 | 1.920115 | 0.521145 | 0.121519 | 4.32145 | 4.68E-05 | 0.008762 | 0.068265 | 0.1 | 0.236592 |
| SL1344_RS11140_ | 2.405166 | 2.115702 | 2.705211 | 0.591376 | 0.096965 | 6.137966 | 4.68E-05 | 0.002482 | 0.041153 | 0.1 | 0.236592 |
| SL1344_RS11145_ | 1.195711 | 1.123576 | 1.272499 | 0.147046 | 0.108917 | 1.278302 | 0.080613 | 0.138587 | 0.276795 | 0.2116 | 0.357561 |
| SL1344_RS11150_ | -2.03675 | -2.16959 | -1.84598 | 0.344099 | 0.385058 | 0.862245 | 0.172552 | 0.274635 | 0.429609 | 0.3747 | 0.511928 |
| SL1344_RS11155_ | -2.07045 | -2.20722 | -1.95882 | 0.271671 | 0.433608 | 0.596036 | 0.255163 | 0.372741 | 0.530612 | 0.4883 | 0.615102 |
| SL1344_RS11160_ | -0.83246 | -1.21986 | -0.44027 | 0.780135 | 0.217572 | 3.552356 | 4.68E-05 | 0.010019 | 0.070237 | 0.1001 | 0.236722 |
| SL1344_RS11165_ | -2.30009 | -2.39879 | -2.17947 | 0.214671 | 0.359284 | 0.559808 | 0.254667 | 0.391351 | 0.536054 | 0.4932 | 0.616602 |
| SL1344_RS11170_ | -1.08065 | -1.17834 | -0.98616 | 0.189188 | 0.239723 | 0.743583 | 0.200533 | 0.298535 | 0.447906 | 0.3987 | 0.533795 |
| SL1344_RS11175_ | -0.40676 | -0.43404 | -0.386 | 0.049722 | 0.288632 | 0.158861 | 0.429714 | 0.657836 | 0.771106 | 0.7537 | 0.834548 |
| SL1344_RS11180_ | -1.95948 | -1.90156 | -2.0055 | -0.1011 | 0.344403 | -0.27927 | 0.389074 | 0.52692 | 0.658115 | 0.6383 | 0.739265 |
| SL1344_RS11185_ | -2.26353 | -2.3918 | -2.11477 | 0.255442 | 0.443288 | 0.522349 | 0.269154 | 0.437761 | 0.584906 | 0.5444 | 0.664859 |
| SL1344_RS11190_ | -2.60324 | -2.79159 | -2.40003 | 0.430892 | 0.55528 | 0.73447 | 0.207333 | 0.312009 | 0.475097 | 0.4191 | 0.560187 |
| SL1344_RS11195_ | -2.86293 | -2.4412 | -3.23938 | -0.85618 | 0.57729 | -1.44859 | 0.038667 | 0.092656 | 0.227651 | 0.1418 | 0.286085 |
| SL1344_RS11200_ | -1.01302 | -0.66974 | -1.33959 | -0.68704 | 0.290208 | -2.28901 | 0.004001 | 0.032193 | 0.12655 | 0.1035 | 0.241106 |
| SL1344_RS11205_ | -0.73086 | -0.64743 | -0.88898 | -0.22513 | 0.258124 | -0.80034 | 0.207333 | 0.313664 | 0.47157 | 0.4206 | 0.561467 |
| SL1344_RS11210_ | -0.10726 | -0.19485 | -0.05315 | 0.123617 | 0.217507 | 0.544605 | 0.285809 | 0.437757 | 0.587427 | 0.5512 | 0.674714 |
| SL1344_RS11215_ | 1.305296 | 1.383038 | 1.19771 | -0.18704 | 0.11628 | -1.58651 | 0.042638 | 0.09137 | 0.219919 | 0.1525 | 0.296583 |
| SL1344_RS11220_ | 1.663444 | 1.607768 | 1.751882 | 0.145285 | 0.140352 | 0.981023 | 0.145333 | 0.204114 | 0.368207 | 0.2896 | 0.446037 |
| SL1344_RS11225_ | 0.243672 | 0.054513 | 0.421355 | 0.359987 | 0.185098 | 1.86918 | 0.023984 | 0.058217 | 0.17342 | 0.1259 | 0.26862 |
| SL1344_RS11230_ | 0.651319 | 0.35115 | 0.920178 | 0.566465 | 0.1329 | 4.257723 | 4.68E-05 | 0.007083 | 0.059995 | 0.1 | 0.236592 |
| SL1344_RS11235_ | 1.867529 | 1.564114 | 2.186205 | 0.607874 | 0.115711 | 5.13849 | 4.68E-05 | 0.003734 | 0.048422 | 0.1 | 0.236592 |
| SL1344_RS11240_ | 0.380448 | 0.234185 | 0.530136 | 0.288024 | 0.179989 | 1.51307 | 0.058 | 0.093755 | 0.226927 | 0.1611 | 0.310141 |
| SL1344_RS11245_ | 1.139189 | 0.996568 | 1.193166 | 0.166991 | 0.202973 | 0.782518 | 0.211193 | 0.31323 | 0.486058 | 0.4196 | 0.578244 |
| SL1344_RS11250_ | 3.242374 | 3.202053 | 3.274821 | 0.080395 | 0.137915 | 0.551014 | 0.285333 | 0.422962 | 0.593483 | 0.5707 | 0.705184 |

| | | | | | | | | | | | |
|---|---|---|---|---|---|---|---|---|---|---|---|
| SL1344_RS11255_ | -0.71658 | -0.79217 | -0.59936 | 0.143733 | 0.359143 | 0.381193 | 0.337775 | 0.597539 | 0.726313 | 0.6844 | 0.786185 |
| SL1344_RS11260_ | 0.565681 | 0.568753 | 0.558951 | 0.01231 | 0.193531 | 0.06127 | 0.473333 | 0.664778 | 0.777732 | 0.7731 | 0.850077 |
| SL1344_RS11265_ | 0.928767 | 0.824482 | 0.980432 | 0.148646 | 0.14001 | 0.956058 | 0.156562 | 0.231417 | 0.390583 | 0.3241 | 0.472446 |
| SL1344_RS11270_ | 3.003724 | 2.991467 | 3.01933 | 0.040093 | 0.099525 | 0.386323 | 0.339107 | 0.521213 | 0.667491 | 0.6722 | 0.770524 |
| SL1344_RS11275_ | -1.93336 | -2.04279 | -1.8577 | 0.186525 | 0.404475 | 0.446132 | 0.304 | 0.466986 | 0.611225 | 0.589 | 0.702161 |
| SL1344_RS11280_ | -0.82343 | -0.85992 | -0.79171 | 0.06177 | 0.223918 | 0.269002 | 0.37575 | 0.511602 | 0.645765 | 0.6212 | 0.72753 |
| SL1344_RS11285_ | -4.13774 | -4.20142 | -4.02626 | 0.272533 | 1.011268 | 0.25743 | 0.402398 | 0.615153 | 0.737322 | 0.734 | 0.817845 |
| SL1344_RS11290_ | -2.53299 | -3.2548 | -2.06853 | 1.187166 | 0.471515 | 2.445841 | 0.002002 | 0.034159 | 0.128648 | 0.1035 | 0.240946 |
| SL1344_RS11295_ | -1.82996 | -1.71417 | -1.98823 | -0.27703 | 0.332537 | -0.79028 | 0.196536 | 0.294179 | 0.448862 | 0.3852 | 0.521572 |
| SL1344_RS11300_ | 1.799549 | 1.789262 | 1.803805 | 0.018036 | 0.161502 | 0.098889 | 0.461026 | 0.742253 | 0.836829 | 0.8592 | 0.911621 |
| SL1344_RS11305_ | 1.358726 | 1.44162 | 1.300854 | -0.15216 | 0.196019 | -0.7606 | 0.191872 | 0.282212 | 0.455439 | 0.3761 | 0.533352 |
| SL1344_RS11310_ | 1.269482 | 0.583976 | 1.478166 | 0.87948 | 0.193343 | 2.906491 | 4.68E-05 | 0.051376 | 0.172459 | 0.1 | 0.236592 |
| SL1344_RS11315_ | 1.05899 | 0.96946 | 1.202733 | 0.225322 | 0.13725 | 1.594719 | 0.043305 | 0.086054 | 0.214379 | 0.1533 | 0.299077 |
| SL1344_RS11320_ | -0.94965 | -1.02243 | -0.83091 | 0.150802 | 0.331709 | 0.419636 | 0.327115 | 0.5457 | 0.681326 | 0.6488 | 0.755939 |
| SL1344_RS11325_ | -1.9785 | -2.20692 | -1.84333 | 0.332325 | 0.457702 | 0.700414 | 0.235843 | 0.357714 | 0.517929 | 0.4618 | 0.602236 |
| SL1344_RS11330_ | -2.3735 | -2.3154 | -2.44279 | -0.1182 | 0.405164 | -0.26831 | 0.385743 | 0.529983 | 0.663078 | 0.6337 | 0.737937 |
| SL1344_RS11335_ | -2.13134 | -2.05484 | -2.21996 | -0.17416 | 0.378724 | -0.40642 | 0.318 | 0.477837 | 0.61951 | 0.5958 | 0.706181 |
| SL1344_RS11340_ | -3.04418 | -3.0283 | -3.08013 | -0.0833 | 0.669318 | -0.11234 | 0.456 | 0.626621 | 0.74853 | 0.7527 | 0.832262 |
| SL1344_RS11345_ | -6.17363 | -5.92126 | -6.36765 | -0.47609 | 1.510014 | -0.28805 | 0.371333 | 0.528546 | 0.659222 | 0.6435 | 0.742217 |
| SL1344_RS11350_ | -3.18415 | -3.63397 | -2.96756 | 0.599679 | 0.742145 | 0.72388 | 0.230667 | 0.341796 | 0.508026 | 0.455 | 0.597838 |
| SL1344_RS11355_ | 1.981993 | 2.055102 | 1.900693 | -0.15681 | 0.097001 | -1.56922 | 0.038641 | 0.086881 | 0.21442 | 0.1473 | 0.291678 |
| SL1344_RS11360_ | -0.19398 | -0.38784 | 0.103248 | 0.47189 | 0.235091 | 1.91405 | 0.022 | 0.054329 | 0.168744 | 0.1213 | 0.263187 |
| SL1344_RS11365_ | 0.995833 | 0.987776 | 1.004679 | -0.02032 | 0.23109 | -0.08904 | 0.464357 | 0.724736 | 0.823737 | 0.8457 | 0.900235 |
| SL1344_RS11375_ | 1.488973 | 1.357524 | 1.667851 | 0.304397 | 0.132861 | 2.210241 | 0.007995 | 0.034578 | 0.132247 | 0.1057 | 0.244139 |
| SL1344_RS11380_ | 1.358519 | 1.1395 | 1.557906 | 0.42033 | 0.104301 | 4.031191 | 4.68E-05 | 0.008872 | 0.066903 | 0.1 | 0.236592 |
| SL1344_RS11385_ | 0.92113 | 0.710174 | 1.165023 | 0.46055 | 0.147791 | 3.071976 | 4.68E-05 | 0.018022 | 0.094577 | 0.1 | 0.236592 |
| SL1344_RS11390_ | 0.521593 | 0.169531 | 0.769608 | 0.600886 | 0.141333 | 4.237843 | 4.68E-05 | 0.007878 | 0.062576 | 0.1 | 0.236592 |
| SL1344_RS11395_ | 1.396214 | 1.106388 | 1.647312 | 0.54374 | 0.154049 | 3.553014 | 4.68E-05 | 0.009432 | 0.071559 | 0.1 | 0.236592 |
| SL1344_RS11400_ | 1.887842 | 2.005613 | 1.783394 | -0.23142 | 0.113061 | -2.01213 | 0.007334 | 0.050513 | 0.162405 | 0.1048 | 0.242727 |
| SL1344_RS11405_ | -2.6189 | -2.74928 | -2.41277 | 0.348063 | 0.50437 | 0.643805 | 0.248 | 0.385612 | 0.539273 | 0.502 | 0.628628 |
| SL1344_RS11410_ | 0.792644 | 0.7154 | 0.849578 | 0.145964 | 0.176437 | 0.793069 | 0.156 | 0.258114 | 0.42502 | 0.3418 | 0.483396 |
| SL1344_RS11415_ | 0.46946 | 0.550431 | 0.411962 | -0.14083 | 0.187499 | -0.7013 | 0.21519 | 0.334138 | 0.491778 | 0.4356 | 0.574721 |
| SL1344_RS11420_ | 0.420902 | 0.296317 | 0.485295 | 0.187423 | 0.199748 | 0.871379 | 0.185876 | 0.267468 | 0.434025 | 0.382 | 0.529758 |
| SL1344_RS11425_ | 3.55086 | 3.791367 | 3.31459 | -0.49587 | 0.134009 | -3.70709 | 4.68E-05 | 0.006204 | 0.06238 | 0.1 | 0.236592 |
| SL1344_RS11430_ | 0.45009 | 0.146122 | 0.716874 | 0.570572 | 0.194752 | 2.879599 | 0.000673 | 0.02181 | 0.104022 | 0.1002 | 0.236862 |
| SL1344_RS11435_ | 0.755045 | 0.735133 | 0.768097 | 0.03984 | 0.154472 | 0.231929 | 0.405333 | 0.564192 | 0.692504 | 0.6762 | 0.772234 |
| SL1344_RS11440_ | 0.498798 | 0.49991 | 0.496482 | -0.00337 | 0.168726 | -0.02354 | 0.490667 | 0.636479 | 0.753351 | 0.7328 | 0.819302 |
| SL1344_RS11445_ | 2.331172 | 2.267566 | 2.387557 | 0.14959 | 0.143448 | 1.022642 | 0.101266 | 0.171318 | 0.337595 | 0.239 | 0.393415 |
| SL1344_RS11450_ | -0.17135 | -0.17083 | -0.17164 | 0.093991 | 0.510912 | 0.192435 | 0.424384 | 0.659701 | 0.781775 | 0.848 | 0.902166 |
| SL1344_RS11455_ | -3.96606 | -4.23105 | -3.76607 | 0.461927 | 0.792349 | 0.535429 | 0.281146 | 0.418595 | 0.569689 | 0.5392 | 0.660785 |
| SL1344_RS11460_ | -6.05363 | -5.73742 | -6.42153 | -0.72371 | 1.539051 | -0.42969 | 0.297801 | 0.439648 | 0.584467 | 0.5557 | 0.668135 |
| SL1344_RS11465_ | 1.904021 | 1.780689 | 2.007124 | 0.235241 | 0.140807 | 1.694725 | 0.03 | 0.070572 | 0.198405 | 0.1312 | 0.275504 |
| SL1344_RS11470_ | -4.74741 | -4.96736 | -4.65063 | 0.249892 | 1.292498 | 0.182786 | 0.42505 | 0.635786 | 0.757806 | 0.7781 | 0.854767 |
| SL1344_RS11475_ | -3.15244 | -3.22027 | -3.01341 | 0.239593 | 0.744738 | 0.297454 | 0.378667 | 0.627338 | 0.747211 | 0.738 | 0.821799 |
| SL1344_RS11480_ | -2.93247 | -3.15509 | -2.72533 | 0.422628 | 0.437674 | 0.927753 | 0.150566 | 0.232464 | 0.382116 | 0.3181 | 0.459513 |

| | | | | | | | | | | | |
|---|---|---|---|---|---|---|---|---|---|---|---|
| SL1344_RS11485_ | -4.50761 | -4.61621 | -4.39056 | 0.247625 | 0.832274 | 0.285128 | 0.380667 | 0.513324 | 0.648631 | 0.6265 | 0.730599 |
| SL1344_RS11490_ | -5.48337 | -6.27854 | -4.9579 | 1.327092 | 1.193813 | 1.053002 | 0.116589 | 0.194265 | 0.342873 | 0.2747 | 0.417843 |
| SL1344_RS11495_ | 1.638441 | 1.442097 | 1.932222 | 0.486266 | 0.118394 | 3.945463 | 4.68E-05 | 0.01007 | 0.071937 | 0.1 | 0.236592 |
| SL1344_RS11500_ | -2.88033 | -2.83021 | -2.93495 | -0.11517 | 0.443688 | -0.22963 | 0.392667 | 0.503972 | 0.637617 | 0.6103 | 0.715056 |
| SL1344_RS11505_ | -2.9166 | -2.92937 | -2.90146 | 0.052269 | 0.553878 | 0.082586 | 0.461692 | 0.631343 | 0.749392 | 0.7486 | 0.828143 |
| SL1344_RS11510_ | -3.79627 | -3.86694 | -3.74737 | 0.120426 | 0.800996 | 0.155634 | 0.423333 | 0.621439 | 0.742396 | 0.7395 | 0.822089 |
| SL1344_RS11515_ | -0.04454 | -0.07713 | 0.00385 | 0.089455 | 0.171434 | 0.497307 | 0.3 | 0.44474 | 0.584784 | 0.549 | 0.663698 |
| SL1344_RS11520_ | -5.7119 | -6.02235 | -5.44242 | 0.530586 | 1.299718 | 0.377749 | 0.327115 | 0.484531 | 0.623855 | 0.589 | 0.698788 |
| SL1344_RS11525_ | -3.12055 | -3.3181 | -2.93524 | 0.368375 | 0.478775 | 0.738966 | 0.198667 | 0.313444 | 0.461366 | 0.4017 | 0.535994 |
| SL1344_RS11535_ | -3.98307 | -4.58057 | -3.77321 | 0.752072 | 0.908596 | 0.716796 | 0.217189 | 0.324253 | 0.492307 | 0.4581 | 0.600202 |
| SL1344_RS11540_ | 1.490942 | 1.476743 | 1.500597 | 0.048257 | 0.156488 | 0.284548 | 0.376 | 0.56222 | 0.697398 | 0.7198 | 0.803247 |
| SL1344_RS11545_ | 3.16769 | 2.80438 | 3.439458 | 0.638824 | 0.14505 | 4.417298 | 4.68E-05 | 0.00354 | 0.049461 | 0.1 | 0.236592 |
| SL1344_RS11550_ | -0.33791 | -0.62593 | -0.0002 | 0.653077 | 0.333605 | 1.888945 | 0.008661 | 0.052031 | 0.169142 | 0.1077 | 0.246697 |
| SL1344_RS11555_ | 1.739245 | 1.458235 | 1.930847 | 0.442696 | 0.325913 | 1.342381 | 0.088667 | 0.106109 | 0.259346 | 0.1831 | 0.346449 |
| SL1344_RS11560_ | 1.01972 | 0.823175 | 1.130179 | 0.318521 | 0.245146 | 1.304471 | 0.072618 | 0.113184 | 0.263985 | 0.1788 | 0.333578 |
| SL1344_RS11565_ | -1.76771 | -1.92476 | -1.67529 | 0.23295 | 0.452915 | 0.495174 | 0.290667 | 0.451308 | 0.602856 | 0.573 | 0.694039 |
| SL1344_RS11570_ | 0.101528 | 0.109566 | 0.090755 | 0.012489 | 0.28167 | 0.046672 | 0.48 | 0.728619 | 0.826856 | 0.8405 | 0.898235 |
| SL1344_RS11585_ | 3.692412 | 3.307958 | 4.048228 | 0.743024 | 0.14109 | 5.238526 | 4.68E-05 | 0.008955 | 0.072706 | 0.1 | 0.236592 |
| SL1344_RS11595_ | 5.142399 | 4.899397 | 5.512119 | 0.587353 | 0.10394 | 5.06579 | 4.68E-05 | 0.006128 | 0.062333 | 0.1 | 0.236592 |
| SL1344_RS11600_ | 4.267135 | 3.969518 | 4.550752 | 0.588974 | 0.065364 | 9.232829 | 4.68E-05 | 0.003847 | 0.050476 | 0.1 | 0.236592 |
| SL1344_RS11605_ | 6.583519 | 6.314447 | 6.833636 | 0.526983 | 0.03997 | 13.59519 | 4.68E-05 | 0.002016 | 0.041075 | 0.1 | 0.236592 |
| SL1344_RS11610_ | 3.064941 | 2.927171 | 3.235126 | 0.29866 | 0.088051 | 3.345476 | 0.000673 | 0.012043 | 0.079371 | 0.1 | 0.236592 |
| SL1344_RS11615_ | 4.125187 | 3.835831 | 4.37899 | 0.531206 | 0.070582 | 7.450573 | 4.68E-05 | 0.001254 | 0.03382 | 0.1 | 0.236592 |
| SL1344_RS11620_ | 1.006757 | 0.189927 | 1.6317 | 1.446755 | 0.181549 | 8.088623 | 4.68E-05 | 0.004308 | 0.050941 | 0.1 | 0.236592 |
| SL1344_RS11625_ | -0.64578 | -0.56663 | -0.75165 | -0.1719 | 0.249357 | -0.65119 | 0.232512 | 0.366757 | 0.521638 | 0.4667 | 0.602233 |
| SL1344_RS11630_ | 0.095118 | 0.151719 | 0.030674 | -0.15158 | 0.243148 | -0.60726 | 0.239174 | 0.371272 | 0.53684 | 0.4844 | 0.615188 |
| SL1344_RS11635_ | 1.00262 | 1.112377 | 0.901143 | -0.2102 | 0.127812 | -1.49425 | 0.051299 | 0.098913 | 0.229812 | 0.1692 | 0.315543 |
| SL1344_RS11640_ | 0.049559 | 0.314675 | -0.21667 | -0.57367 | 0.30312 | -1.87731 | 0.005334 | 0.054038 | 0.171681 | 0.1052 | 0.243371 |
| SL1344_RS11670_ | -1.38248 | -1.34313 | -1.41921 | -0.18312 | 0.551763 | -0.28383 | 0.361333 | 0.481303 | 0.644656 | 0.7031 | 0.794263 |
| SL1344_RS11675_ | -3.61024 | -3.43843 | -3.72998 | -0.26516 | 1.031979 | -0.23909 | 0.404 | 0.692957 | 0.79961 | 0.8066 | 0.872377 |
| SL1344_RS11680_ | -2.76641 | -2.39901 | -3.09558 | -0.7255 | 0.547451 | -1.2504 | 0.075283 | 0.131677 | 0.277674 | 0.1992 | 0.348456 |
| SL1344_RS11685_ | 1.866712 | 1.988706 | 1.735454 | -0.25608 | 0.100337 | -2.47787 | 0.005331 | 0.029352 | 0.119379 | 0.1027 | 0.240123 |
| SL1344_RS11690_ | 4.106662 | 4.104248 | 4.109779 | 0.020491 | 0.133373 | 0.147072 | 0.425716 | 0.695136 | 0.808044 | 0.8653 | 0.915112 |
| SL1344_RS11695_ | 2.987592 | 2.94804 | 3.119507 | 0.152842 | 0.119851 | 1.182037 | 0.133245 | 0.162017 | 0.327638 | 0.27 | 0.435328 |
| SL1344_RS11700_ | 2.422595 | 2.402277 | 2.450183 | 0.048704 | 0.110762 | 0.403224 | 0.332445 | 0.533575 | 0.67237 | 0.6523 | 0.756708 |
| SL1344_RS11705_ | -0.43014 | -0.42092 | -0.44188 | 0.013941 | 0.280844 | 0.049612 | 0.480346 | 0.677662 | 0.786426 | 0.7925 | 0.861219 |
| SL1344_RS11710_ | -0.37037 | -0.35544 | -0.3789 | -0.01088 | 0.256437 | -0.0366 | 0.482 | 0.650874 | 0.765323 | 0.7691 | 0.844278 |
| SL1344_RS11715_ | 7.83496 | 7.589923 | 8.009377 | 0.418193 | 0.109857 | 3.906286 | 4.68E-05 | 0.014758 | 0.092905 | 0.1 | 0.236592 |
| SL1344_RS11720_ | 7.083967 | 6.913402 | 7.21698 | 0.306617 | 0.05286 | 5.789392 | 4.68E-05 | 0.017712 | 0.100955 | 0.1 | 0.236592 |
| SL1344_RS11725_ | 5.434397 | 5.10909 | 5.80192 | 0.63579 | 0.127166 | 5.265181 | 4.68E-05 | 0.000912 | 0.033112 | 0.1 | 0.236592 |
| SL1344_RS11730_ | 4.915939 | 4.58364 | 5.31841 | 0.731561 | 0.06299 | 11.36818 | 4.68E-05 | 0.002044 | 0.041222 | 0.1 | 0.236592 |
| SL1344_RS11735_ | 5.109058 | 4.793267 | 5.428482 | 0.702948 | 0.116494 | 6.092699 | 4.68E-05 | 0.000703 | 0.030722 | 0.1 | 0.236592 |
| SL1344_RS11740_ | -4.92798 | -4.81293 | -5.14103 | -0.42341 | 1.51455 | -0.23308 | 0.371086 | 0.52436 | 0.672927 | 0.6948 | 0.787268 |
| SL1344_RS11745_ | -1.91339 | -1.88318 | -1.94981 | -0.06342 | 0.371097 | -0.14691 | 0.427333 | 0.596157 | 0.719146 | 0.7013 | 0.791337 |
| SL1344_RS11750_ | -0.25715 | -0.46396 | -0.11022 | 0.383384 | 0.279087 | 1.319396 | 0.027315 | 0.136389 | 0.293886 | 0.1321 | 0.274545 |

| | | | | | | | | | | | |
|---|---|---|---|---|---|---|---|---|---|---|---|
| SL1344_RS11755_ | -1.52531 | -1.82402 | -1.31144 | 0.536634 | 0.412329 | 1.2334 | 0.063291 | 0.13836 | 0.293225 | 0.1662 | 0.313737 |
| SL1344_RS11760_ | -0.57597 | -0.91468 | -0.22714 | 0.700826 | 0.361451 | 1.969412 | 0.001337 | 0.055321 | 0.174237 | 0.1034 | 0.241123 |
| SL1344_RS11765_ | -1.63324 | -1.71872 | -1.50364 | 0.194196 | 0.353662 | 0.500682 | 0.295803 | 0.473499 | 0.615688 | 0.579 | 0.694587 |
| SL1344_RS11770_ | 1.196219 | 1.057979 | 1.310836 | 0.259676 | 0.126188 | 2.014099 | 0.006663 | 0.051224 | 0.160852 | 0.1078 | 0.246164 |
| SL1344_RS11775_ | 0.516369 | 0.547784 | 0.435448 | -0.09235 | 0.244712 | -0.23008 | 0.387333 | 0.808357 | 0.883704 | 0.7889 | 0.874139 |
| SL1344_RS11780_ | 0.322299 | 0.175243 | 0.458467 | 0.284117 | 0.148938 | 1.871128 | 0.017988 | 0.058921 | 0.169997 | 0.1215 | 0.262354 |
| SL1344_RS11785_ | 1.846415 | 1.687149 | 2.045679 | 0.346559 | 0.144414 | 2.368747 | 0.004665 | 0.028941 | 0.122558 | 0.1032 | 0.240862 |
| SL1344_RS11790_ | 2.620771 | 2.585227 | 2.690963 | 0.107215 | 0.128898 | 0.824878 | 0.190667 | 0.26903 | 0.443578 | 0.3632 | 0.526682 |
| SL1344_RS11795_ | 3.163967 | 3.020357 | 3.397443 | 0.370848 | 0.136832 | 2.776039 | 4.68E-05 | 0.01479 | 0.089961 | 0.1 | 0.236592 |
| SL1344_RS11800_ | 4.15305 | 3.969276 | 4.50554 | 0.530182 | 0.114934 | 3.787307 | 4.68E-05 | 0.018885 | 0.103277 | 0.1 | 0.236592 |
| SL1344_RS11805_ | 2.760829 | 2.599988 | 2.983636 | 0.356631 | 0.121964 | 2.859762 | 4.68E-05 | 0.016051 | 0.092592 | 0.1001 | 0.236719 |
| SL1344_RS11810_ | 2.764037 | 2.5542 | 3.044817 | 0.483009 | 0.092083 | 5.180729 | 4.68E-05 | 0.006993 | 0.062616 | 0.1 | 0.236592 |
| SL1344_RS11815_ | 1.21237 | 0.954437 | 1.504631 | 0.546797 | 0.129945 | 4.147859 | 4.68E-05 | 0.006836 | 0.060884 | 0.1 | 0.236592 |
| SL1344_RS11820_ | -0.19338 | -0.2612 | -0.1248 | 0.136611 | 0.199404 | 0.647034 | 0.236509 | 0.369116 | 0.520386 | 0.475 | 0.605264 |
| SL1344_RS11825_ | 1.826045 | 1.758409 | 1.911933 | 0.170722 | 0.154512 | 1.057546 | 0.116 | 0.175187 | 0.338518 | 0.2574 | 0.413569 |
| SL1344_RS11830_ | 2.485068 | 2.332705 | 2.62175 | 0.286167 | 0.130485 | 2.191861 | 0.004001 | 0.032063 | 0.130476 | 0.1021 | 0.239385 |
| SL1344_RS11840_ | 2.260715 | 2.012544 | 2.518252 | 0.507548 | 0.0937 | 5.381989 | 4.68E-05 | 0.003264 | 0.045351 | 0.1 | 0.236592 |
| SL1344_RS11845_ | 1.045445 | 0.945435 | 1.165329 | 0.225322 | 0.12912 | 1.702463 | 0.023984 | 0.075616 | 0.198227 | 0.1332 | 0.274564 |
| SL1344_RS11850_ | 2.52185 | 2.441033 | 2.562187 | 0.120458 | 0.11519 | 1.016626 | 0.143333 | 0.195348 | 0.361104 | 0.2911 | 0.452004 |
| SL1344_RS11855_ | 1.237261 | 1.270601 | 1.207161 | -0.09148 | 0.211463 | -0.42567 | 0.30513 | 0.454658 | 0.616661 | 0.6059 | 0.722362 |
| SL1344_RS11860_ | 3.708035 | 3.431578 | 4.089108 | 0.648256 | 0.139137 | 4.599747 | 4.68E-05 | 0.004666 | 0.055117 | 0.1 | 0.236592 |
| SL1344_RS11865_ | -0.91544 | -1.25961 | -0.69554 | 0.460447 | 0.49963 | 0.908301 | 0.150566 | 0.226486 | 0.401159 | 0.3046 | 0.472271 |
| SL1344_RS11870_ | 0.366734 | 0.451981 | 0.300814 | -0.15331 | 0.20949 | -0.69584 | 0.228667 | 0.349056 | 0.511455 | 0.4562 | 0.597054 |
| SL1344_RS11880_ | -1.00284 | -1.04758 | -0.95264 | 0.087447 | 0.268882 | 0.309059 | 0.363091 | 0.549393 | 0.678413 | 0.6506 | 0.751079 |
| SL1344_RS11885_ | 4.402395 | 4.073994 | 4.703299 | 0.634384 | 0.147896 | 4.43792 | 4.68E-05 | 0.00725 | 0.066719 | 0.1 | 0.236592 |
| SL1344_RS11890_ | 3.747893 | 3.41714 | 3.989843 | 0.596962 | 0.157648 | 3.520394 | 4.68E-05 | 0.03234 | 0.135772 | 0.1 | 0.236592 |
| SL1344_RS11895_ | 4.886263 | 4.647439 | 5.02786 | 0.378857 | 0.146412 | 2.775163 | 4.68E-05 | 0.00946 | 0.075267 | 0.1 | 0.236592 |
| SL1344_RS11900_ | 2.429052 | 2.287986 | 2.66756 | 0.399467 | 0.188115 | 2.04217 | 0.002002 | 0.044291 | 0.157695 | 0.101 | 0.237945 |
| SL1344_RS11905_ | 3.714246 | 3.53469 | 3.865712 | 0.298821 | 0.118603 | 2.460711 | 4.68E-05 | 0.022764 | 0.111506 | 0.1 | 0.236592 |
| SL1344_RS11910_ | 3.431016 | 3.227798 | 3.571639 | 0.357828 | 0.139376 | 2.700145 | 4.68E-05 | 0.016317 | 0.094921 | 0.1 | 0.236592 |
| SL1344_RS11915_ | 4.138157 | 3.900189 | 4.342539 | 0.409009 | 0.134399 | 3.135999 | 4.68E-05 | 0.007862 | 0.069177 | 0.1 | 0.236592 |
| SL1344_RS11920_ | 5.704013 | 5.623764 | 5.867858 | 0.243107 | 0.101991 | 2.256672 | 4.68E-05 | 0.04501 | 0.16111 | 0.1001 | 0.236722 |
| SL1344_RS11925_ | 4.733635 | 4.61998 | 4.861795 | 0.23921 | 0.139376 | 1.69296 | 0.007334 | 0.066802 | 0.199878 | 0.107 | 0.245932 |
| SL1344_RS11930_ | 3.579798 | 3.439353 | 3.665464 | 0.258736 | 0.15381 | 1.754319 | 0.002001 | 0.047864 | 0.164833 | 0.1029 | 0.240467 |
| SL1344_RS11935_ | 5.044108 | 5.052214 | 5.033796 | -0.00399 | 0.107678 | -0.0255 | 0.484 | 0.620064 | 0.759341 | 0.9694 | 0.98133 |
| SL1344_RS11940_ | 3.95674 | 3.969745 | 3.773275 | -0.18929 | 0.075439 | -0.89408 | 0.293333 | 0.279077 | 0.465539 | 0.5948 | 0.732824 |
| SL1344_RS11945_ | 3.645515 | 3.725774 | 3.47099 | -0.24603 | 0.093335 | -2.39914 | 0.002001 | 0.037114 | 0.141982 | 0.1011 | 0.237996 |
| SL1344_RS11950_ | -3.79517 | -3.98917 | -3.64039 | 0.351043 | 0.723144 | 0.44483 | 0.307129 | 0.453475 | 0.597238 | 0.5637 | 0.679069 |
| SL1344_RS11960_ | 2.134357 | 2.097858 | 2.182151 | 0.089807 | 0.113788 | 0.762266 | 0.201865 | 0.312796 | 0.476479 | 0.4182 | 0.559679 |
| SL1344_RS11965_ | -0.17498 | -0.35312 | -0.09072 | 0.232881 | 0.236924 | 0.869169 | 0.175333 | 0.256841 | 0.422066 | 0.364 | 0.515333 |
| SL1344_RS11970_ | 1.4581 | 1.157726 | 1.681883 | 0.517457 | 0.125383 | 4.072789 | 4.68E-05 | 0.00848 | 0.066614 | 0.1 | 0.236592 |
| SL1344_RS11975_ | 2.673576 | 2.420542 | 2.934392 | 0.505512 | 0.08594 | 5.773302 | 4.68E-05 | 0.004244 | 0.050259 | 0.1 | 0.236592 |
| SL1344_RS11980_ | 1.520995 | 1.358287 | 1.651299 | 0.292559 | 0.108552 | 2.686324 | 0.000673 | 0.026804 | 0.113142 | 0.1012 | 0.238178 |
| SL1344_RS11985_ | 1.262934 | 1.236188 | 1.300292 | 0.080986 | 0.163659 | 0.452219 | 0.309793 | 0.449345 | 0.603308 | 0.5926 | 0.706555 |
| SL1344_RS12000_ | 0.056074 | 0.247002 | -0.09669 | -0.34607 | 0.222618 | -1.55756 | 0.042638 | 0.088414 | 0.221569 | 0.1459 | 0.292194 |

| | | | | | | | | | | | |
|---|---|---|---|---|---|---|---|---|---|---|---|
| SL1344_RS12005_ | 1.822708 | 1.67564 | 1.923251 | 0.247723 | 0.123559 | 1.956592 | 0.011334 | 0.058513 | 0.17635 | 0.1086 | 0.247643 |
| SL1344_RS12010_ | 0.385541 | 0.14301 | 0.532297 | 0.377161 | 0.188855 | 1.941108 | 0.013991 | 0.055541 | 0.169854 | 0.1188 | 0.259982 |
| SL1344_RS12015_ | 1.044409 | 0.837897 | 1.27429 | 0.433829 | 0.156792 | 2.679468 | 4.68E-05 | 0.021646 | 0.104962 | 0.101 | 0.237915 |
| SL1344_RS12020_ | -1.04409 | -1.26244 | -0.82092 | 0.441678 | 0.23654 | 1.770813 | 0.026649 | 0.064534 | 0.17957 | 0.1308 | 0.273104 |
| SL1344_RS12025_ | -1.02303 | -1.15887 | -0.90064 | 0.260229 | 0.267832 | 0.92704 | 0.141239 | 0.220853 | 0.376809 | 0.3088 | 0.454596 |
| SL1344_RS12030_ | -6.36312 | -6.22363 | -6.6543 | -0.4769 | 2.302062 | -0.15685 | 0.403065 | 0.547444 | 0.690294 | 0.7243 | 0.811987 |
| SL1344_RS12035_ | 0.757878 | 0.516013 | 0.998561 | 0.484058 | 0.177039 | 2.714938 | 4.68E-05 | 0.020417 | 0.101876 | 0.1004 | 0.237117 |
| SL1344_RS12040_ | 2.391144 | 2.413556 | 2.365857 | -0.03767 | 0.114314 | -0.31941 | 0.371333 | 0.606224 | 0.731536 | 0.7195 | 0.808534 |
| SL1344_RS12045_ | 1.174634 | 1.130455 | 1.221059 | 0.091571 | 0.130106 | 0.671531 | 0.231845 | 0.352251 | 0.507623 | 0.464 | 0.596473 |
| SL1344_RS12050_ | 0.723363 | 0.522394 | 0.843251 | 0.319614 | 0.177727 | 1.693913 | 0.026649 | 0.075124 | 0.202614 | 0.13 | 0.273454 |
| SL1344_RS12055_ | -2.3561 | -2.75297 | -2.0021 | 0.807982 | 0.545332 | 1.399968 | 0.038641 | 0.102799 | 0.243059 | 0.1493 | 0.294492 |
| SL1344_RS12060_ | 0.652244 | 0.30524 | 0.934943 | 0.644156 | 0.170157 | 3.723054 | 4.68E-05 | 0.011209 | 0.074911 | 0.1 | 0.236592 |
| SL1344_RS12065_ | -0.81151 | -1.03634 | -0.54399 | 0.482752 | 0.23758 | 2.032051 | 0.018 | 0.048505 | 0.154573 | 0.1162 | 0.256941 |
| SL1344_RS12070_ | -1.05926 | -1.19914 | -0.89875 | 0.312054 | 0.274214 | 1.054963 | 0.114667 | 0.182681 | 0.334445 | 0.2603 | 0.406692 |
| SL1344_RS12075_ | -0.84131 | -0.89914 | -0.79488 | 0.117823 | 0.273576 | 0.3835 | 0.336442 | 0.510939 | 0.649376 | 0.6335 | 0.736223 |
| SL1344_RS12080_ | 2.564136 | 2.347307 | 2.731838 | 0.396676 | 0.12114 | 3.146373 | 4.68E-05 | 0.052547 | 0.174173 | 0.1 | 0.236592 |
| SL1344_RS12085_ | -1.25378 | -1.2432 | -1.27099 | -0.04584 | 0.307315 | -0.13848 | 0.438 | 0.61042 | 0.73166 | 0.7218 | 0.808369 |
| SL1344_RS12090_ | 1.024373 | 0.992489 | 1.040173 | 0.053784 | 0.168059 | 0.299524 | 0.382 | 0.564646 | 0.69894 | 0.7007 | 0.793089 |
| SL1344_RS12095_ | 0.888612 | 0.899426 | 0.883748 | -0.04069 | 0.207052 | -0.18085 | 0.42 | 0.648146 | 0.766481 | 0.7929 | 0.862592 |
| SL1344_RS12100_ | -0.40463 | -0.48487 | -0.24607 | 0.230096 | 0.364021 | 0.620172 | 0.238 | 0.386108 | 0.553487 | 0.461 | 0.616004 |
| SL1344_RS12105_ | -1.59133 | -1.53143 | -1.66288 | -0.13187 | 0.283463 | -0.42102 | 0.313333 | 0.447424 | 0.587142 | 0.5449 | 0.66276 |
| SL1344_RS12110_ | -2.32267 | -2.26622 | -2.4266 | -0.13334 | 0.451918 | -0.28111 | 0.382412 | 0.574966 | 0.703828 | 0.6841 | 0.782184 |
| SL1344_RS12115_ | 1.499022 | 1.544682 | 1.473167 | -0.10285 | 0.214189 | -0.45585 | 0.304667 | 0.498459 | 0.654571 | 0.6306 | 0.74996 |
| SL1344_RS12120_ | 2.27075 | 2.303969 | 2.204337 | -0.07196 | 0.159974 | -0.41078 | 0.359333 | 0.628696 | 0.755061 | 0.7377 | 0.832097 |
| SL1344_RS12125_ | 0.980382 | 1.057614 | 0.927507 | -0.1306 | 0.129819 | -0.9383 | 0.148667 | 0.235471 | 0.389357 | 0.3238 | 0.467558 |
| SL1344_RS12130_ | 2.520662 | 2.496856 | 2.545446 | 0.068613 | 0.113845 | 0.598742 | 0.265333 | 0.38224 | 0.545669 | 0.5222 | 0.648229 |
| SL1344_RS12135_ | 2.376696 | 2.351018 | 2.414692 | 0.075789 | 0.136541 | 0.527574 | 0.279333 | 0.434689 | 0.596408 | 0.5718 | 0.699304 |
| SL1344_RS12140_ | 3.703923 | 3.732051 | 3.690473 | -0.03236 | 0.076398 | -0.36228 | 0.358667 | 0.645798 | 0.764511 | 0.7312 | 0.826348 |
| SL1344_RS12145_ | 0.506922 | 0.408859 | 0.583496 | 0.18475 | 0.150449 | 1.13501 | 0.093937 | 0.164001 | 0.310993 | 0.2305 | 0.374917 |
| SL1344_RS12155_ | 1.627379 | 1.428366 | 1.68795 | 0.24304 | 0.144281 | 1.42842 | 0.065333 | 0.118436 | 0.268778 | 0.189 | 0.340483 |
| SL1344_RS12160_ | 2.357379 | 2.368273 | 2.334005 | -0.02103 | 0.107072 | -0.16872 | 0.431712 | 0.702664 | 0.807896 | 0.7829 | 0.864356 |
| SL1344_RS12170_ | 0.960501 | 0.900848 | 1.047678 | 0.142504 | 0.135485 | 1.020948 | 0.139241 | 0.212935 | 0.365336 | 0.2964 | 0.44385 |
| SL1344_RS12175_ | -4.56673 | -4.77895 | -4.36566 | 0.396863 | 0.743117 | 0.484525 | 0.284667 | 0.408396 | 0.547966 | 0.5115 | 0.628198 |
| SL1344_RS12180_ | -5.31852 | -5.36037 | -5.25912 | 0.116376 | 1.263545 | 0.078785 | 0.461026 | 0.635014 | 0.752699 | 0.7409 | 0.824099 |
| SL1344_RS12185_ | -5.95849 | -5.3863 | -6.40343 | -0.93311 | 1.540051 | -0.55418 | 0.243333 | 0.413603 | 0.557146 | 0.5014 | 0.628452 |
| SL1344_RS12190_ | -4.6197 | -4.899 | -4.40947 | 0.476142 | 0.838645 | 0.511031 | 0.279333 | 0.39324 | 0.539478 | 0.4993 | 0.61934 |
| SL1344_RS12195_ | -1.46956 | -1.32062 | -1.62137 | -0.31842 | 0.320951 | -0.95281 | 0.14 | 0.226152 | 0.381259 | 0.3201 | 0.462441 |
| SL1344_RS12200_ | 4.762805 | 4.608416 | 4.798842 | 0.180476 | 0.156301 | 0.972055 | 0.193205 | 0.230561 | 0.415332 | 0.362 | 0.540235 |
| SL1344_RS12205_ | 1.470171 | 1.46389 | 1.482717 | 0.030912 | 0.126327 | 0.212241 | 0.414667 | 0.601342 | 0.726216 | 0.7167 | 0.805811 |
| SL1344_RS12210_ | 0.978856 | 0.948995 | 1.021517 | 0.095502 | 0.158006 | 0.541262 | 0.260493 | 0.400824 | 0.559292 | 0.5237 | 0.647619 |
| SL1344_RS12215_ | -0.24801 | -0.36959 | -0.05292 | 0.299489 | 0.284941 | 1.012386 | 0.140573 | 0.205038 | 0.368623 | 0.2873 | 0.44729 |
| SL1344_RS12220_ | -0.13536 | -0.15508 | -0.11961 | -0.04074 | 0.34698 | -0.1156 | 0.453333 | 0.731187 | 0.829015 | 0.8711 | 0.917358 |
| SL1344_RS12225_ | 0.624745 | 0.511198 | 0.710522 | 0.195993 | 0.152698 | 1.199709 | 0.088608 | 0.148633 | 0.293458 | 0.22 | 0.369426 |
| SL1344_RS12230_ | 1.56773 | 1.367393 | 1.771749 | 0.404225 | 0.094037 | 4.283506 | 4.68E-05 | 0.006148 | 0.056824 | 0.1 | 0.236592 |
| SL1344_RS12235_ | 1.260757 | 1.1177 | 1.330497 | 0.203658 | 0.1457 | 1.346491 | 0.085333 | 0.140362 | 0.292916 | 0.2142 | 0.367136 |

| | | | | | | | | | | | |
|---|---|---|---|---|---|---|---|---|---|---|---|
| SL1344_RS12240_ | -0.55904 | -0.6804 | -0.31196 | 0.34577 | 0.240099 | 1.358545 | 0.083333 | 0.135454 | 0.280912 | 0.2137 | 0.362425 |
| SL1344_RS12245_ | 2.53823 | 2.461714 | 2.586803 | 0.122673 | 0.079721 | 1.488173 | 0.059294 | 0.099543 | 0.231881 | 0.1678 | 0.315108 |
| SL1344_RS12250_ | 1.603175 | 1.473572 | 1.736462 | 0.282254 | 0.146925 | 1.892774 | 0.007334 | 0.047459 | 0.157742 | 0.1073 | 0.245719 |
| SL1344_RS12260_ | 3.505068 | 3.425904 | 3.595579 | 0.17751 | 0.075954 | 2.413075 | 4.68E-05 | 0.047304 | 0.160522 | 0.1002 | 0.236859 |
| SL1344_RS12265_ | 2.553058 | 2.58952 | 2.341186 | -0.22683 | 0.179049 | -0.60029 | 0.334 | 0.572128 | 0.722312 | 0.7 | 0.822042 |
| SL1344_RS12270_ | 3.178657 | 3.240029 | 3.15496 | -0.07034 | 0.101574 | -0.57265 | 0.301133 | 0.50171 | 0.65477 | 0.6273 | 0.750156 |
| SL1344_RS12275_ | -0.31889 | -0.63912 | -0.02755 | 0.6199 | 0.174942 | 3.492124 | 4.68E-05 | 0.010835 | 0.073252 | 0.1 | 0.236592 |
| SL1344_RS12280_ | -2.89779 | -2.98405 | -2.77827 | 0.210271 | 0.903934 | 0.215701 | 0.402 | 0.712065 | 0.813915 | 0.8052 | 0.872798 |
| SL1344_RS12285_ | -1.10451 | -1.20356 | -0.87367 | 0.293957 | 0.322938 | 0.824175 | 0.194 | 0.295999 | 0.45814 | 0.4032 | 0.549368 |
| SL1344_RS12290_ | 0.782296 | 0.534431 | 1.082395 | 0.543146 | 0.140061 | 3.757752 | 4.68E-05 | 0.009904 | 0.070568 | 0.1 | 0.236592 |
| SL1344_RS12295_ | 0.895891 | 0.611098 | 1.157608 | 0.549462 | 0.140428 | 3.921178 | 4.68E-05 | 0.008283 | 0.066077 | 0.1 | 0.236592 |
| SL1344_RS12300_ | -0.37315 | -0.61216 | -0.19423 | 0.415871 | 0.197938 | 2.052343 | 0.009328 | 0.047473 | 0.151981 | 0.1108 | 0.249453 |
| SL1344_RS12305_ | -2.55069 | -2.85217 | -2.32155 | 0.555417 | 0.533361 | 0.983984 | 0.134 | 0.20536 | 0.36307 | 0.2997 | 0.448419 |
| SL1344_RS12310_ | -2.16924 | -2.28083 | -2.03892 | 0.2401 | 0.444046 | 0.492917 | 0.292667 | 0.452836 | 0.596184 | 0.5582 | 0.675672 |
| SL1344_RS12315_ | 0.261753 | -0.06684 | 0.430511 | 0.513254 | 0.353611 | 1.496054 | 0.043305 | 0.079538 | 0.216274 | 0.1436 | 0.291827 |
| SL1344_RS12320_ | -6.33544 | -6.43017 | -6.23661 | 0.190992 | 1.389613 | 0.11604 | 0.441039 | 0.511891 | 0.646329 | 0.6195 | 0.723182 |
| SL1344_RS12325_ | -1.98728 | -1.93301 | -2.07087 | -0.16571 | 0.407763 | -0.36357 | 0.342438 | 0.504916 | 0.645651 | 0.6305 | 0.733907 |
| SL1344_RS12330_ | 2.233393 | 2.094258 | 2.31076 | 0.225103 | 0.132656 | 1.633834 | 0.015323 | 0.090358 | 0.231184 | 0.1147 | 0.254889 |
| SL1344_RS12335_ | 2.35641 | 2.224088 | 2.542807 | 0.309633 | 0.090809 | 3.272185 | 4.68E-05 | 0.013497 | 0.082238 | 0.1 | 0.236592 |
| SL1344_RS12340_ | -0.22269 | -0.51855 | -0.08677 | 0.417217 | 0.246694 | 1.544429 | 0.050633 | 0.096465 | 0.234817 | 0.1603 | 0.308523 |
| SL1344_RS12345_ | 0.847298 | 0.708532 | 0.997609 | 0.293727 | 0.158304 | 1.844415 | 0.022652 | 0.056858 | 0.170258 | 0.1231 | 0.265104 |
| SL1344_RS12350_ | 0.692951 | 0.550406 | 0.818922 | 0.257505 | 0.163175 | 1.502155 | 0.052632 | 0.090988 | 0.224078 | 0.1557 | 0.304609 |
| SL1344_RS12355_ | 0.837686 | 0.778873 | 0.908252 | 0.152979 | 0.18217 | 0.798122 | 0.187209 | 0.274541 | 0.441247 | 0.3792 | 0.526057 |
| SL1344_RS12360_ | -0.48341 | -0.61007 | -0.31959 | 0.330225 | 0.280288 | 1.169183 | 0.069287 | 0.164972 | 0.326927 | 0.1846 | 0.330206 |
| SL1344_RS12365_ | 2.039823 | 2.079906 | 1.975628 | -0.05915 | 0.166589 | -0.34144 | 0.387075 | 0.680007 | 0.791271 | 0.7851 | 0.861957 |
| SL1344_RS12370_ | 2.06441 | 1.945721 | 2.122823 | 0.16068 | 0.144871 | 1.023673 | 0.143238 | 0.203259 | 0.371979 | 0.2927 | 0.451753 |
| SL1344_RS12375_ | 0.556546 | 0.605434 | 0.300995 | -0.27219 | 0.285627 | -0.637 | 0.322667 | 0.516588 | 0.673364 | 0.6708 | 0.79754 |
| SL1344_RS12380_ | 2.482883 | 2.633807 | 2.348709 | -0.30694 | 0.131664 | -2.43666 | 4.68E-05 | 0.039113 | 0.146076 | 0.1 | 0.236592 |
| SL1344_RS12385_ | 2.004126 | 1.652483 | 2.304727 | 0.636964 | 0.145283 | 4.308143 | 4.68E-05 | 0.008024 | 0.066144 | 0.1 | 0.236592 |
| SL1344_RS12390_ | 2.10734 | 1.938442 | 2.523429 | 0.535483 | 0.258129 | 1.959265 | 0.002668 | 0.045187 | 0.160238 | 0.1026 | 0.240068 |
| SL1344_RS12395_ | 3.400454 | 3.376354 | 3.452293 | 0.06554 | 0.111892 | 0.602436 | 0.257162 | 0.437384 | 0.601431 | 0.5015 | 0.653966 |
| SL1344_RS12400_ | 2.028681 | 2.170283 | 1.887017 | -0.27156 | 0.147767 | -1.84825 | 0.02465 | 0.05668 | 0.176254 | 0.1208 | 0.264049 |
| SL1344_RS12405_ | 1.676579 | 1.673439 | 1.682295 | -0.02029 | 0.229585 | -0.08248 | 0.467022 | 0.756267 | 0.848848 | 0.9093 | 0.943737 |
| SL1344_RS12410_ | 2.593295 | 2.618126 | 2.553968 | -0.01807 | 0.168237 | -0.08586 | 0.465333 | 0.813648 | 0.88735 | 0.9202 | 0.951946 |
| SL1344_RS12415_ | 1.537266 | 1.534237 | 1.540781 | 0.020656 | 0.143533 | 0.135628 | 0.441706 | 0.643494 | 0.760672 | 0.7718 | 0.845241 |
| SL1344_RS12420_ | -2.01136 | -2.00859 | -2.01252 | -0.01893 | 0.4416 | -0.04309 | 0.483011 | 0.651841 | 0.76695 | 0.7711 | 0.84581 |
| SL1344_RS12425_ | -6.98737 | -6.81838 | -7.13157 | -0.35813 | 1.937655 | -0.17688 | 0.422385 | 0.54028 | 0.672551 | 0.6663 | 0.76388 |
| SL1344_RS12430_ | -1.63105 | -1.60559 | -1.68247 | -0.03932 | 0.380504 | -0.08623 | 0.462358 | 0.679643 | 0.788152 | 0.7733 | 0.852104 |
| SL1344_RS12435_ | -4.71294 | -4.47017 | -5.05949 | -0.64138 | 1.183348 | -0.48051 | 0.265333 | 0.399781 | 0.559768 | 0.5282 | 0.650306 |
| SL1344_RS12440_ | -2.88594 | -2.98991 | -2.71273 | 0.308643 | 0.53903 | 0.538909 | 0.286476 | 0.431369 | 0.578079 | 0.5525 | 0.670459 |
| SL1344_RS12445_ | -0.96502 | -0.85154 | -1.07272 | -0.22654 | 0.281848 | -0.75149 | 0.197333 | 0.302757 | 0.459527 | 0.4027 | 0.542161 |
| SL1344_RS12450_ | 0.601073 | 0.530955 | 0.706056 | 0.174122 | 0.162561 | 0.995543 | 0.134 | 0.206038 | 0.363801 | 0.2883 | 0.441185 |
| SL1344_RS12455_ | 0.507274 | 0.229989 | 0.613439 | 0.365025 | 0.197515 | 1.635627 | 0.048634 | 0.088368 | 0.224557 | 0.1457 | 0.291899 |
| SL1344_RS12465_ | 3.590884 | 3.54952 | 3.652802 | 0.107087 | 0.082339 | 1.279277 | 0.068 | 0.117798 | 0.265742 | 0.1825 | 0.334428 |
| SL1344_RS12480_ | -5.21209 | -4.86237 | -5.7517 | -1.04068 | 2.01373 | -0.43136 | 0.231179 | 0.366692 | 0.542619 | 0.4681 | 0.600824 |

| | | | | | | | | | | | |
|---|---|---|---|---|---|---|---|---|---|---|---|
| SL1344_RS12485_ | 3.567076 | 3.451996 | 3.722831 | 0.273899 | 0.080669 | 3.343213 | 4.68E-05 | 0.016574 | 0.093152 | 0.1 | 0.236592 |
| SL1344_RS12490_ | 6.346372 | 6.229952 | 6.569528 | 0.343364 | 0.142487 | 2.548711 | 4.68E-05 | 0.018584 | 0.103498 | 0.1 | 0.236592 |
| SL1344_RS12495_ | 5.062184 | 4.784618 | 5.297583 | 0.505449 | 0.069418 | 7.201772 | 4.68E-05 | 0.00302 | 0.046202 | 0.1 | 0.236592 |
| SL1344_RS12500_ | -2.07988 | -2.38406 | -1.88613 | 0.495909 | 0.409553 | 1.143624 | 0.101333 | 0.163642 | 0.315775 | 0.2309 | 0.379528 |
| SL1344_RS12510_ | -1.06711 | -1.08093 | -1.05035 | 0.034373 | 0.255769 | 0.139822 | 0.446 | 0.550916 | 0.6798 | 0.6614 | 0.757677 |
| SL1344_RS12515_ | -1.22378 | -1.35884 | -1.08566 | 0.284933 | 0.311409 | 0.897559 | 0.165333 | 0.254543 | 0.411426 | 0.3531 | 0.49572 |
| SL1344_RS12520_ | -2.2755 | -2.52589 | -2.1888 | 0.285053 | 0.545046 | 0.422378 | 0.362425 | 0.633087 | 0.756417 | 0.7288 | 0.829734 |
| SL1344_RS12525_ | -2.07665 | -2.2842 | -1.84574 | 0.426201 | 0.355818 | 1.128585 | 0.111259 | 0.175377 | 0.32364 | 0.2589 | 0.40672 |
| SL1344_RS12555_ | 3.825725 | 3.417762 | 4.043273 | 0.614037 | 0.13468 | 3.903114 | 4.68E-05 | 0.017498 | 0.099363 | 0.1 | 0.236592 |
| SL1344_RS12560_ | 2.742627 | 2.681421 | 2.793125 | 0.101057 | 0.093654 | 1.015692 | 0.125916 | 0.194193 | 0.356081 | 0.2678 | 0.426176 |
| SL1344_RS12565_ | 0.765706 | 0.800725 | 0.722322 | -0.08062 | 0.130082 | -0.57756 | 0.255163 | 0.39717 | 0.542352 | 0.5008 | 0.622326 |
| SL1344_RS12570_ | 0.906491 | 0.925556 | 0.880374 | -0.06366 | 0.215712 | -0.27039 | 0.389333 | 0.610992 | 0.739718 | 0.7607 | 0.840186 |
| SL1344_RS12575_ | 1.387713 | 1.40478 | 1.343628 | -0.04781 | 0.169089 | -0.2549 | 0.406667 | 0.699147 | 0.804732 | 0.7825 | 0.863516 |
| SL1344_RS12580_ | -0.60261 | -0.58172 | -0.62183 | -0.04041 | 0.190243 | -0.19521 | 0.412667 | 0.519108 | 0.64949 | 0.6259 | 0.727409 |
| SL1344_RS12585_ | -0.71819 | -0.79287 | -0.63859 | 0.159755 | 0.283787 | 0.522145 | 0.265823 | 0.418565 | 0.572481 | 0.5404 | 0.662691 |
| SL1344_RS12590_ | -2.45249 | -2.69749 | -2.20051 | 0.501866 | 0.561207 | 0.885469 | 0.163333 | 0.243985 | 0.409085 | 0.3199 | 0.470684 |
| SL1344_RS12595_ | 0.179335 | 0.128932 | 0.230173 | 0.099953 | 0.145316 | 0.653244 | 0.229847 | 0.352417 | 0.497666 | 0.449 | 0.576916 |
| SL1344_RS12600_ | -1.51339 | -1.79467 | -1.26004 | 0.529888 | 0.324703 | 1.594962 | 0.038641 | 0.085611 | 0.212227 | 0.1478 | 0.291959 |
| SL1344_RS12605_ | -1.31901 | -1.52251 | -0.92193 | 0.585298 | 0.379613 | 1.438614 | 0.052632 | 0.099767 | 0.239514 | 0.1745 | 0.325061 |
| SL1344_RS12610_ | -1.27885 | -1.55389 | -1.03771 | 0.513187 | 0.34669 | 1.385282 | 0.067289 | 0.110546 | 0.251972 | 0.1771 | 0.328639 |
| SL1344_RS12615_ | -2.59796 | -2.75932 | -2.49155 | 0.264376 | 0.435083 | 0.573635 | 0.265333 | 0.3883 | 0.537273 | 0.5092 | 0.630741 |
| SL1344_RS12620_ | -1.79139 | -1.91937 | -1.64136 | 0.279119 | 0.359099 | 0.725065 | 0.213857 | 0.331081 | 0.485485 | 0.4279 | 0.564572 |
| SL1344_RS12630_ | -1.79139 | -1.999 | -1.63664 | 0.367928 | 0.362087 | 0.963206 | 0.129913 | 0.21101 | 0.366691 | 0.2836 | 0.428358 |
| SL1344_RS12635_ | -3.45276 | -3.57025 | -3.25791 | 0.318872 | 0.656781 | 0.427854 | 0.321119 | 0.516466 | 0.653078 | 0.6191 | 0.728587 |
| SL1344_RS12640_ | -2.21171 | -2.39458 | -2.0809 | 0.298473 | 0.378994 | 0.760113 | 0.193205 | 0.307805 | 0.461629 | 0.4053 | 0.544451 |
| SL1344_RS12645_ | -1.80189 | -1.92034 | -1.73316 | 0.168608 | 0.336211 | 0.469319 | 0.318454 | 0.46772 | 0.609546 | 0.5815 | 0.696435 |
| SL1344_RS12650_ | -3.92557 | -3.86089 | -4.01795 | -0.16493 | 0.760391 | -0.20236 | 0.408667 | 0.56457 | 0.695677 | 0.6845 | 0.779216 |
| SL1344_RS12655_ | -2.26359 | -2.47041 | -2.09318 | 0.391032 | 0.514731 | 0.742099 | 0.199201 | 0.299414 | 0.462459 | 0.4165 | 0.559235 |
| SL1344_RS12660_ | -2.81377 | -2.87206 | -2.77259 | 0.090967 | 0.621063 | 0.153724 | 0.432 | 0.641652 | 0.758787 | 0.7586 | 0.836369 |
| SL1344_RS12665_ | -4.18619 | -4.25817 | -4.10797 | 0.097777 | 0.971459 | 0.089327 | 0.462 | 0.666464 | 0.777218 | 0.7779 | 0.850665 |
| SL1344_RS12670_ | -3.24575 | -3.2464 | -3.2455 | 0.011416 | 0.610374 | 0.016358 | 0.492667 | 0.634656 | 0.752841 | 0.7496 | 0.831167 |
| SL1344_RS12675_ | -3.85513 | -4.33816 | -3.50815 | 0.868285 | 0.752265 | 1.038971 | 0.1006 | 0.187342 | 0.343102 | 0.2454 | 0.391194 |
| SL1344_RS12690_ | 5.058259 | 4.943827 | 5.171754 | 0.212576 | 0.114292 | 1.784307 | 0.008661 | 0.053374 | 0.176152 | 0.1077 | 0.246882 |
| SL1344_RS12695_ | 0.447963 | 0.148574 | 0.742161 | 0.595693 | 0.155316 | 3.788102 | 4.68E-05 | 0.008976 | 0.067607 | 0.1 | 0.236592 |
| SL1344_RS12700_ | 1.485714 | 1.129453 | 1.955108 | 0.810912 | 0.145241 | 5.565518 | 4.68E-05 | 0.002826 | 0.043896 | 0.1 | 0.236592 |
| SL1344_RS12705_ | 1.419312 | 1.345938 | 1.67947 | 0.338848 | 0.214794 | 1.370255 | 0.084 | 0.139369 | 0.301981 | 0.1989 | 0.354169 |
| SL1344_RS12710_ | -1.44352 | -1.51089 | -1.39423 | 0.117593 | 0.295729 | 0.37037 | 0.346 | 0.505982 | 0.640569 | 0.6264 | 0.728468 |
| SL1344_RS12715_ | -0.11853 | -0.06109 | -0.20732 | -0.16715 | 0.223262 | -0.70253 | 0.194667 | 0.305842 | 0.471024 | 0.3945 | 0.531284 |
| SL1344_RS12720_ | -0.70265 | -0.88496 | -0.49404 | 0.396709 | 0.247661 | 1.583755 | 0.033311 | 0.085294 | 0.212591 | 0.1428 | 0.285528 |
| SL1344_RS12725_ | 0.050925 | -0.36945 | 0.366709 | 0.693193 | 0.287454 | 2.331734 | 0.002668 | 0.02926 | 0.123977 | 0.1026 | 0.240066 |
| SL1344_RS12730_ | 0.411876 | 0.093789 | 0.706649 | 0.626933 | 0.184463 | 3.379784 | 4.68E-05 | 0.014235 | 0.084245 | 0.1 | 0.236592 |
| SL1344_RS12735_ | 0.241472 | 0.275445 | 0.196652 | -0.07959 | 0.162118 | -0.47113 | 0.29447 | 0.444697 | 0.587992 | 0.5637 | 0.676195 |
| SL1344_RS12740_ | -0.24381 | -0.23258 | -0.25519 | -0.02655 | 0.209623 | -0.11345 | 0.453333 | 0.587574 | 0.711271 | 0.6985 | 0.787847 |
| SL1344_RS12745_ | 1.755978 | 1.682811 | 1.80728 | 0.137416 | 0.118326 | 1.123814 | 0.081333 | 0.174893 | 0.334318 | 0.2202 | 0.367181 |
| SL1344_RS12750_ | 1.083522 | 0.927623 | 1.442825 | 0.495721 | 0.188784 | 2.270453 | 0.003998 | 0.044031 | 0.154728 | 0.1029 | 0.240389 |

| | | | | | | | | | | |
|---|---|---|---|---|---|---|---|---|---|---|
| SL1344_RS12755_ | 1.414215 | 1.333726 | 1.555956 | 0.21571 | 0.114382 | 1.742815 | 0.032667 | 0.072068 | 0.193537 | 0.1428 | 0.286606 |
| SL1344_RS12760_ | 1.185451 | 1.008848 | 1.296062 | 0.275143 | 0.197829 | 1.353371 | 0.068 | 0.10489 | 0.251064 | 0.1769 | 0.332528 |
| SL1344_RS12765_ | 1.681664 | 1.530521 | 1.806097 | 0.27036 | 0.156854 | 1.667917 | 0.031979 | 0.065903 | 0.191442 | 0.1294 | 0.274249 |
| SL1344_RS12770_ | 4.425054 | 4.416336 | 4.434593 | 0.014384 | 0.048847 | 0.278611 | 0.382667 | 0.601068 | 0.725345 | 0.708 | 0.799194 |
| SL1344_RS12775_ | 3.598282 | 3.586582 | 3.612047 | 0.024683 | 0.063966 | 0.367528 | 0.346436 | 0.541613 | 0.676564 | 0.6573 | 0.758172 |
| SL1344_RS12780_ | 1.832963 | 1.840966 | 1.823494 | -0.03129 | 0.128695 | -0.21651 | 0.4004 | 0.598795 | 0.724028 | 0.7281 | 0.812 |
| SL1344_RS12785_ | 3.133382 | 3.162521 | 3.096624 | -0.04874 | 0.120431 | -0.38855 | 0.332 | 0.574535 | 0.713148 | 0.6731 | 0.781151 |
| SL1344_RS12790_ | -2.18673 | -2.27289 | -2.10811 | 0.164154 | 0.359051 | 0.421496 | 0.311333 | 0.474479 | 0.613314 | 0.5859 | 0.696268 |
| SL1344_RS12795_ | 0.573915 | 0.548392 | 0.616978 | 0.060474 | 0.183536 | 0.296355 | 0.382 | 0.620293 | 0.741566 | 0.7161 | 0.810233 |
| SL1344_RS12800_ | 1.844616 | 1.800135 | 1.896224 | 0.100125 | 0.103226 | 0.90802 | 0.152 | 0.248219 | 0.403118 | 0.3414 | 0.484277 |
| SL1344_RS12805_ | 1.994158 | 1.955663 | 2.102962 | 0.143674 | 0.118072 | 1.039771 | 0.146667 | 0.211543 | 0.37954 | 0.3158 | 0.467274 |
| SL1344_RS12810_ | 1.280953 | 1.374475 | 1.171477 | -0.21612 | 0.14524 | -1.45683 | 0.038667 | 0.096803 | 0.233852 | 0.1411 | 0.28541 |
| SL1344_RS12815_ | 1.818662 | 1.860511 | 1.791455 | -0.03957 | 0.180402 | -0.19325 | 0.425716 | 0.763453 | 0.852138 | 0.8717 | 0.920191 |
| SL1344_RS12820_ | 3.491242 | 3.458614 | 3.522817 | 0.009973 | 0.159127 | 0.067441 | 0.478667 | 0.829024 | 0.897535 | 0.9382 | 0.962952 |
| SL1344_RS12825_ | 1.334857 | 1.615331 | 1.209601 | -0.38602 | 0.178152 | -1.94924 | 0.016 | 0.057456 | 0.177804 | 0.1134 | 0.253457 |
| SL1344_RS12830_ | 1.093432 | 1.171156 | 1.009651 | -0.16304 | 0.105857 | -1.46539 | 0.051299 | 0.102113 | 0.229421 | 0.1714 | 0.314776 |
| SL1344_RS12835_ | 2.600119 | 2.33275 | 2.814748 | 0.482487 | 0.102335 | 4.717162 | 4.68E-05 | 0.010045 | 0.072852 | 0.1 | 0.236592 |
| SL1344_RS12840_ | 1.896046 | 1.680091 | 2.210868 | 0.541649 | 0.137882 | 3.987552 | 4.68E-05 | 0.00802 | 0.066706 | 0.1 | 0.236592 |
| SL1344_RS12845_ | 0.002953 | -0.5121 | 0.274413 | 0.778574 | 0.231173 | 3.176617 | 4.68E-05 | 0.024924 | 0.11211 | 0.1003 | 0.236932 |
| SL1344_RS12850_ | -1.99884 | -2.26914 | -1.85366 | 0.397101 | 0.529623 | 0.698205 | 0.227182 | 0.332327 | 0.500116 | 0.4448 | 0.592339 |
| SL1344_RS12855_ | 1.324678 | 1.504724 | 1.112334 | -0.40294 | 0.235432 | -1.72753 | 0.013991 | 0.064463 | 0.192498 | 0.1145 | 0.255582 |
| SL1344_RS12860_ | -3.17065 | -3.13286 | -3.21746 | -0.09508 | 0.515688 | -0.17406 | 0.426667 | 0.554357 | 0.685143 | 0.6661 | 0.762565 |
| SL1344_RS12865_ | -4.73893 | -4.85895 | -4.65026 | 0.209913 | 0.825498 | 0.246539 | 0.393071 | 0.497219 | 0.632918 | 0.6055 | 0.710495 |
| SL1344_RS12870_ | 2.914296 | 2.910258 | 2.917191 | 0.027492 | 0.1397 | 0.195626 | 0.431046 | 0.680052 | 0.794574 | 0.852 | 0.905041 |
| SL1344_RS12875_ | 1.794813 | 1.896677 | 1.698824 | -0.20542 | 0.269533 | -0.73813 | 0.207861 | 0.294018 | 0.477314 | 0.3988 | 0.564425 |
| SL1344_RS12880_ | 0.501328 | 0.672793 | 0.361022 | -0.31092 | 0.20453 | -1.51158 | 0.051965 | 0.094543 | 0.230744 | 0.1574 | 0.305215 |
| SL1344_RS12885_ | 0.532698 | 0.628637 | 0.417404 | -0.21337 | 0.172565 | -1.20197 | 0.084 | 0.146253 | 0.297046 | 0.2172 | 0.368998 |
| SL1344_RS12890_ | 1.789513 | 1.743443 | 1.840133 | 0.098296 | 0.094884 | 0.972025 | 0.1499 | 0.226061 | 0.379978 | 0.3154 | 0.459663 |
| SL1344_RS12900_ | -1.66884 | -1.55126 | -1.8001 | -0.25678 | 0.306566 | -0.79314 | 0.188667 | 0.28479 | 0.437576 | 0.3824 | 0.519721 |
| SL1344_RS12905_ | -3.06765 | -2.69219 | -3.31069 | -0.60553 | 0.597158 | -0.94777 | 0.142572 | 0.227692 | 0.384976 | 0.3237 | 0.471723 |
| SL1344_RS12910_ | -0.87473 | -0.66535 | -1.1248 | -0.45625 | 0.210949 | -2.11069 | 0.01066 | 0.039948 | 0.136628 | 0.1095 | 0.248295 |
| SL1344_RS12915_ | 1.87711 | 1.948579 | 1.812541 | -0.14114 | 0.170404 | -0.79907 | 0.186542 | 0.281861 | 0.459203 | 0.3769 | 0.533292 |
| SL1344_RS12920_ | 3.012548 | 2.965233 | 3.084184 | 0.12294 | 0.078092 | 1.509136 | 0.038641 | 0.100572 | 0.23929 | 0.1489 | 0.294595 |
| SL1344_RS12925_ | 2.608854 | 2.582883 | 2.649677 | 0.070052 | 0.182577 | 0.399066 | 0.302667 | 0.552218 | 0.699916 | 0.6192 | 0.743918 |
| SL1344_RS12930_ | 2.76026 | 2.800286 | 2.735723 | -0.06902 | 0.093295 | -0.70512 | 0.227182 | 0.332183 | 0.495739 | 0.4454 | 0.583955 |
| SL1344_RS12935_ | 3.414076 | 3.214278 | 3.535079 | 0.32876 | 0.163381 | 1.992637 | 0.001336 | 0.035668 | 0.141211 | 0.1006 | 0.23739 |
| SL1344_RS12940_ | 2.445561 | 2.406636 | 2.481642 | 0.081519 | 0.101695 | 0.779122 | 0.191333 | 0.289976 | 0.455705 | 0.3818 | 0.52753 |
| SL1344_RS12945_ | 1.878594 | 2.027125 | 1.658625 | -0.36249 | 0.111948 | -3.16448 | 4.68E-05 | 0.017851 | 0.094565 | 0.1001 | 0.236729 |
| SL1344_RS12950_ | 3.762815 | 3.950132 | 3.345505 | -0.56324 | 0.226739 | -1.96336 | 0.000673 | 0.051768 | 0.174159 | 0.1003 | 0.236998 |
| SL1344_RS12955_ | -0.07223 | -0.08606 | -0.05475 | 0.031615 | 0.230798 | 0.129493 | 0.446667 | 0.648769 | 0.763954 | 0.751 | 0.832508 |
| SL1344_RS12960_ | 0.770562 | 0.809209 | 0.733478 | -0.07592 | 0.139847 | -0.52532 | 0.279147 | 0.421751 | 0.568346 | 0.5348 | 0.653749 |
| SL1344_RS12965_ | 0.065672 | 0.058519 | 0.072958 | 0.014173 | 0.165373 | 0.077242 | 0.464357 | 0.575647 | 0.701415 | 0.6764 | 0.772011 |
| SL1344_RS12970_ | 1.892588 | 1.893818 | 1.891307 | -0.00256 | 0.09194 | -0.02641 | 0.486342 | 0.589474 | 0.713522 | 0.7085 | 0.797008 |
| SL1344_RS12975_ | 0.963447 | 0.853798 | 1.087642 | 0.237198 | 0.154913 | 1.480102 | 0.042638 | 0.095685 | 0.230688 | 0.152 | 0.297464 |
| SL1344_RS12980_ | 3.550892 | 3.362532 | 3.741996 | 0.386279 | 0.064312 | 6.017638 | 4.68E-05 | 0.003412 | 0.046227 | 0.1 | 0.236592 |

| | | | | | | | | | | |
|---|---|---|---|---|---|---|---|---|---|---|
| SL1344_RS12985_ | 2.770966 | 2.605476 | 3.040486 | 0.446132 | 0.176933 | 2.694146 | 4.68E-05 | 0.019471 | 0.103066 | 0.1 | 0.236592 |
| SL1344_RS12990_ | -1.61043 | -1.63338 | -1.58909 | 0.053092 | 0.340011 | 0.159484 | 0.437042 | 0.602642 | 0.72501 | 0.7157 | 0.801699 |
| SL1344_RS12995_ | 0.870932 | 1.065521 | 0.795543 | -0.23953 | 0.217944 | -1.02586 | 0.160667 | 0.210334 | 0.379732 | 0.3078 | 0.471193 |
| SL1344_RS13000_ | 1.912412 | 1.980768 | 1.840858 | -0.15134 | 0.117853 | -1.21614 | 0.06529 | 0.129833 | 0.276349 | 0.1888 | 0.338489 |
| SL1344_RS13005_ | 1.352766 | 1.358307 | 1.345264 | 0.031161 | 0.186415 | 0.152459 | 0.437708 | 0.69429 | 0.801343 | 0.8516 | 0.90268 |
| SL1344_RS13010_ | 1.396654 | 1.354503 | 1.436935 | 0.087115 | 0.119585 | 0.670386 | 0.231179 | 0.343009 | 0.498736 | 0.4618 | 0.591809 |
| SL1344_RS13015_ | 2.112462 | 2.104033 | 2.124561 | 0.015442 | 0.088599 | 0.16858 | 0.421333 | 0.583813 | 0.707425 | 0.6825 | 0.777602 |
| SL1344_RS13025_ | 2.324064 | 2.333835 | 2.317492 | -0.00847 | 0.239436 | -0.04279 | 0.478 | 0.84645 | 0.909179 | 0.8752 | 0.922239 |
| SL1344_RS13030_ | 1.733543 | 1.745898 | 1.722383 | -0.03782 | 0.137555 | -0.2518 | 0.38974 | 0.607466 | 0.733022 | 0.7484 | 0.828645 |
| SL1344_RS13035_ | 2.52525 | 2.563888 | 2.460447 | -0.10782 | 0.105979 | -0.96342 | 0.149234 | 0.226527 | 0.393416 | 0.3143 | 0.467985 |
| SL1344_RS13040_ | 0.6907 | 0.858529 | 0.516386 | -0.33797 | 0.154361 | -2.13983 | 0.008667 | 0.038159 | 0.136735 | 0.1081 | 0.246924 |
| SL1344_RS13045_ | 1.753621 | 1.830005 | 1.652644 | -0.18165 | 0.122847 | -1.40077 | 0.055963 | 0.10346 | 0.2428 | 0.1697 | 0.319328 |
| SL1344_RS13050_ | 1.215426 | 1.447703 | 0.857389 | -0.59976 | 0.189321 | -3.26234 | 4.68E-05 | 0.02819 | 0.122025 | 0.1 | 0.236592 |
| SL1344_RS13075_ | -0.27073 | -0.31392 | -0.17378 | 0.135151 | 0.288433 | 0.420838 | 0.344 | 0.558243 | 0.694907 | 0.6782 | 0.779633 |
| SL1344_RS13080_ | -0.75567 | -0.80387 | -0.65352 | 0.113012 | 0.306148 | 0.328773 | 0.372 | 0.620111 | 0.741201 | 0.7008 | 0.798707 |
| SL1344_RS13085_ | 0.944739 | 0.242904 | 1.641213 | 1.385747 | 0.167332 | 8.191883 | 4.68E-05 | 0.001116 | 0.032346 | 0.1 | 0.236592 |
| SL1344_RS13090_ | 0.592374 | 0.81871 | 0.431207 | -0.38654 | 0.155636 | -2.38159 | 0.003332 | 0.033045 | 0.127437 | 0.1044 | 0.242251 |
| SL1344_RS13095_ | 5.242487 | 4.955852 | 5.442776 | 0.49248 | 0.09593 | 5.177825 | 4.68E-05 | 0.001189 | 0.035246 | 0.1 | 0.236592 |
| SL1344_RS13100_ | -0.03975 | 0.035017 | -0.10914 | -0.15193 | 0.184679 | -0.77106 | 0.191333 | 0.297324 | 0.454126 | 0.3992 | 0.538059 |
| SL1344_RS13105_ | 1.042916 | 1.022801 | 1.075589 | 0.062657 | 0.146647 | 0.412445 | 0.335776 | 0.493436 | 0.635659 | 0.6232 | 0.726634 |
| SL1344_RS13115_ | 1.904301 | 1.865995 | 2.190865 | 0.296815 | 0.174963 | 0.983288 | 0.228 | 0.246363 | 0.427965 | 0.4453 | 0.602821 |
| SL1344_RS13120_ | -0.93825 | -1.07138 | -0.82012 | 0.236671 | 0.308373 | 0.706782 | 0.229333 | 0.342824 | 0.500788 | 0.4455 | 0.585001 |
| SL1344_RS13125_ | 1.722671 | 1.707182 | 1.736723 | 0.046485 | 0.148567 | 0.302757 | 0.377748 | 0.587794 | 0.717701 | 0.7306 | 0.814722 |
| SL1344_RS13130_ | 2.156371 | 2.123374 | 2.188493 | 0.063727 | 0.082852 | 0.707431 | 0.200667 | 0.318738 | 0.468907 | 0.4198 | 0.552322 |
| SL1344_RS13135_ | 0.829368 | 0.783929 | 0.879932 | 0.129409 | 0.191823 | 0.624954 | 0.237333 | 0.352381 | 0.516779 | 0.4976 | 0.622684 |
| SL1344_RS13140_ | 1.081724 | 1.062669 | 1.100954 | -0.00692 | 0.210582 | -0.02727 | 0.484667 | 0.724369 | 0.824273 | 0.8565 | 0.907725 |
| SL1344_RS13145_ | 2.249488 | 2.23692 | 2.274901 | 0.024021 | 0.147571 | 0.141118 | 0.449333 | 0.786871 | 0.868766 | 0.8589 | 0.914604 |
| SL1344_RS13150_ | -2.53956 | -2.55237 | -2.52388 | 0.053717 | 0.570252 | 0.092976 | 0.458361 | 0.656551 | 0.770279 | 0.7711 | 0.845273 |
| SL1344_RS13155_ | 0.565687 | 0.640458 | 0.516085 | -0.12721 | 0.169447 | -0.71437 | 0.222518 | 0.344704 | 0.501641 | 0.4507 | 0.584451 |
| SL1344_RS13160_ | -0.09817 | -0.20741 | 0.080079 | 0.282791 | 0.188716 | 1.443851 | 0.058 | 0.109032 | 0.243144 | 0.18 | 0.327659 |
| SL1344_RS13165_ | 1.301635 | 1.142013 | 1.468608 | 0.359691 | 0.189198 | 1.836671 | 0.005334 | 0.050002 | 0.164772 | 0.1054 | 0.243761 |
| SL1344_RS13170_ | 0.694711 | 0.541246 | 1.158024 | 0.61029 | 0.396043 | 1.678889 | 0.014 | 0.056659 | 0.180015 | 0.1144 | 0.255036 |
| SL1344_RS13175_ | -0.30502 | -0.71426 | -0.04089 | 0.676171 | 0.278712 | 2.329454 | 0.002002 | 0.029641 | 0.122844 | 0.1026 | 0.239955 |
| SL1344_RS13180_ | -0.94998 | -0.91427 | -1.0328 | -0.07704 | 0.32685 | -0.23203 | 0.416 | 0.671344 | 0.784364 | 0.771 | 0.853283 |
| SL1344_RS13185_ | -0.67565 | -0.51838 | -0.83761 | -0.3254 | 0.284844 | -1.09542 | 0.104667 | 0.165415 | 0.32179 | 0.2419 | 0.396117 |
| SL1344_RS13190_ | -4.28843 | -4.01058 | -4.58493 | -0.57305 | 0.680489 | -0.78003 | 0.182545 | 0.291446 | 0.437977 | 0.3794 | 0.514904 |
| SL1344_RS13195_ | -2.19837 | -2.13948 | -2.26818 | -0.11603 | 0.563692 | -0.20899 | 0.410667 | 0.644731 | 0.763848 | 0.7645 | 0.841997 |
| SL1344_RS13200_ | -3.94268 | -3.58642 | -4.20794 | -0.5964 | 0.845803 | -0.62071 | 0.23984 | 0.382105 | 0.535439 | 0.4748 | 0.609105 |
| SL1344_RS13210_ | 1.003686 | 0.884015 | 1.139362 | 0.25093 | 0.12791 | 1.95126 | 0.017334 | 0.052034 | 0.159592 | 0.1217 | 0.262146 |
| SL1344_RS13215_ | 1.200404 | 1.113626 | 1.295914 | 0.180507 | 0.116937 | 1.512192 | 0.046667 | 0.095842 | 0.22402 | 0.1632 | 0.307651 |
| SL1344_RS13220_ | 2.428411 | 2.357963 | 2.471392 | 0.127055 | 0.117609 | 0.997713 | 0.117255 | 0.192723 | 0.360039 | 0.2537 | 0.406849 |
| SL1344_RS13225_ | 1.774587 | 1.70587 | 1.832975 | 0.10336 | 0.162208 | 0.630998 | 0.240506 | 0.383931 | 0.551692 | 0.4912 | 0.633261 |
| SL1344_RS13230_ | 3.2821 | 3.29538 | 3.274067 | -0.01149 | 0.092867 | -0.09879 | 0.45503 | 0.731961 | 0.830888 | 0.8147 | 0.886155 |
| SL1344_RS13235_ | 3.408003 | 3.408364 | 3.407327 | 0.004856 | 0.098782 | 0.053249 | 0.472352 | 0.747427 | 0.840287 | 0.8159 | 0.881355 |
| SL1344_RS13240_ | 0.617279 | 0.517199 | 0.684581 | 0.169293 | 0.141241 | 1.12532 | 0.107928 | 0.176854 | 0.326229 | 0.2592 | 0.40719 |

| | | | | | | | | | | |
|---|---|---|---|---|---|---|---|---|---|---|
| SL1344_RS13245_ | 1.000574 | 0.773687 | 1.097906 | 0.317251 | 0.151595 | 1.964498 | 0.015323 | 0.062955 | 0.183995 | 0.1172 | 0.257486 |
| SL1344_RS13250_ | -4.29581 | -4.13564 | -4.45372 | -0.39858 | 0.956993 | -0.38327 | 0.332 | 0.504003 | 0.645 | 0.6199 | 0.728576 |
| SL1344_RS13255_ | -4.30973 | -4.01401 | -4.7485 | -0.75101 | 0.81985 | -0.82711 | 0.157333 | 0.249414 | 0.40478 | 0.344 | 0.481957 |
| SL1344_RS13260_ | -3.49607 | -3.21796 | -3.67201 | -0.39941 | 0.778148 | -0.46373 | 0.297801 | 0.515527 | 0.654165 | 0.5982 | 0.718138 |
| SL1344_RS13265_ | -1.7279 | -1.76883 | -1.69301 | 0.071177 | 0.370141 | 0.170395 | 0.423333 | 0.595694 | 0.719187 | 0.7026 | 0.794059 |
| SL1344_RS13270_ | -3.63375 | -3.44561 | -3.95763 | -0.57141 | 0.882652 | -0.5869 | 0.247333 | 0.380814 | 0.543798 | 0.5146 | 0.644439 |
| SL1344_RS13275_ | -1.38134 | -1.57286 | -1.227 | 0.351086 | 0.343944 | 0.965988 | 0.133333 | 0.209699 | 0.368925 | 0.2962 | 0.446081 |
| SL1344_RS13280_ | -2.38315 | -2.62162 | -2.16919 | 0.451216 | 0.367428 | 1.155207 | 0.097935 | 0.169934 | 0.313983 | 0.2467 | 0.392336 |
| SL1344_RS13285_ | -2.87105 | -2.85325 | -2.90255 | -0.00863 | 0.55979 | -0.01232 | 0.495333 | 0.644675 | 0.760068 | 0.7658 | 0.841107 |
| SL1344_RS13290_ | -2.35798 | -2.65338 | -1.83818 | 0.77229 | 0.653078 | 1.127161 | 0.120586 | 0.17234 | 0.336818 | 0.2511 | 0.409833 |
| SL1344_RS13295_ | -3.71837 | -3.82871 | -3.59486 | 0.204388 | 0.672992 | 0.284138 | 0.369333 | 0.53486 | 0.667618 | 0.6357 | 0.739334 |
| SL1344_RS13300_ | -0.23013 | -0.5396 | -0.12483 | 0.39531 | 0.392426 | 0.932331 | 0.175217 | 0.226128 | 0.402517 | 0.3324 | 0.499482 |
| SL1344_RS13305_ | -4.45015 | -4.72992 | -4.25473 | 0.502542 | 0.870505 | 0.524643 | 0.285143 | 0.422129 | 0.568912 | 0.53 | 0.652155 |
| SL1344_RS13310_ | -4.21036 | -4.32709 | -4.12479 | 0.299808 | 1.08644 | 0.238729 | 0.371752 | 0.552819 | 0.691432 | 0.6908 | 0.78381 |
| SL1344_RS13315_ | -0.34697 | -0.30038 | -0.44244 | -0.13581 | 0.228104 | -0.54285 | 0.291805 | 0.447029 | 0.593492 | 0.5618 | 0.683397 |
| SL1344_RS13320_ | -2.41879 | -2.38148 | -2.45424 | -0.10301 | 0.534411 | -0.17522 | 0.419054 | 0.622347 | 0.743984 | 0.7396 | 0.821564 |
| SL1344_RS13325_ | 3.970862 | 4.02991 | 3.922529 | -0.05253 | 0.157758 | -0.35901 | 0.393071 | 0.705914 | 0.816577 | 0.8387 | 0.901155 |
| SL1344_RS13330_ | -2.33654 | -2.41523 | -2.24632 | 0.174073 | 0.387796 | 0.418522 | 0.329333 | 0.463983 | 0.604219 | 0.5658 | 0.679757 |
| SL1344_RS13335_ | -2.53235 | -2.72574 | -2.29136 | 0.417465 | 0.42839 | 0.932548 | 0.151899 | 0.247 | 0.399504 | 0.3304 | 0.476019 |
| SL1344_RS13340_ | -3.00386 | -3.0561 | -2.95197 | 0.087465 | 0.608469 | 0.137452 | 0.438374 | 0.636662 | 0.754612 | 0.7441 | 0.825924 |
| SL1344_RS13345_ | -1.34336 | -1.49933 | -1.21031 | 0.289601 | 0.254202 | 1.041899 | 0.124667 | 0.199781 | 0.344232 | 0.2811 | 0.42265 |
| SL1344_RS13350_ | -2.13982 | -2.39403 | -1.9458 | 0.446738 | 0.36568 | 1.167079 | 0.098601 | 0.162084 | 0.306141 | 0.2361 | 0.380184 |
| SL1344_RS13355_ | -1.42727 | -1.72248 | -1.11389 | 0.603409 | 0.291849 | 1.99415 | 0.011992 | 0.047769 | 0.153035 | 0.1159 | 0.256193 |
| SL1344_RS13360_ | -0.85933 | -1.10803 | -0.59308 | 0.520657 | 0.241862 | 2.11531 | 0.007334 | 0.039899 | 0.138881 | 0.1082 | 0.24678 |
| SL1344_RS13365_ | -3.16485 | -3.4456 | -3.01027 | 0.45241 | 0.725704 | 0.590993 | 0.248 | 0.380983 | 0.54279 | 0.5149 | 0.64419 |
| SL1344_RS13370_ | -1.58787 | -1.66954 | -1.49009 | 0.182789 | 0.383635 | 0.452913 | 0.314667 | 0.492988 | 0.635806 | 0.6071 | 0.71866 |
| SL1344_RS13375_ | -3.7284 | -4.30348 | -3.31901 | 1.003295 | 0.715316 | 1.360278 | 0.060626 | 0.121634 | 0.263641 | 0.1697 | 0.315834 |
| SL1344_RS13380_ | -0.78898 | -0.94377 | -0.67006 | 0.28517 | 0.246704 | 1.073066 | 0.12 | 0.187542 | 0.339658 | 0.2641 | 0.410465 |
| SL1344_RS13385_ | -3.34182 | -3.69636 | -2.79651 | 0.873679 | 0.590431 | 1.424824 | 0.065956 | 0.114447 | 0.252003 | 0.1881 | 0.337355 |
| SL1344_RS13390_ | -2.43868 | -2.70301 | -2.31591 | 0.377364 | 0.496908 | 0.691533 | 0.247169 | 0.365664 | 0.523619 | 0.4841 | 0.617809 |
| SL1344_RS13395_ | -7.12874 | -6.95137 | -7.2977 | -0.27541 | 3.377556 | -0.06053 | 0.457029 | 0.638126 | 0.759254 | 0.7753 | 0.849586 |
| SL1344_RS13400_ | 0.749091 | 0.842289 | 0.499319 | -0.34601 | 0.25897 | -1.21695 | 0.086 | 0.152005 | 0.3158 | 0.2066 | 0.362486 |
| SL1344_RS13405_ | -2.58446 | -2.92745 | -2.27612 | 0.646994 | 0.407687 | 1.536546 | 0.043333 | 0.091548 | 0.216607 | 0.1526 | 0.296881 |
| SL1344_RS13410_ | -2.49504 | -2.87326 | -2.10612 | 0.545779 | 0.801412 | 0.692766 | 0.217189 | 0.364855 | 0.53434 | 0.4467 | 0.595843 |
| SL1344_RS13415_ | -1.8174 | -1.64322 | -2.01753 | -0.37683 | 0.366629 | -1.0029 | 0.13 | 0.205729 | 0.360486 | 0.2955 | 0.442802 |
| SL1344_RS13420_ | -3.1296 | -3.44044 | -2.91421 | 0.478422 | 0.668123 | 0.659699 | 0.235177 | 0.360687 | 0.518615 | 0.4578 | 0.597466 |
| SL1344_RS13425_ | -5.7752 | -5.76798 | -5.77731 | 0.004642 | 1.65152 | 0.002817 | 0.498334 | 0.652632 | 0.769113 | 0.7791 | 0.855025 |
| SL1344_RS13430_ | -4.50466 | -5.04306 | -4.19911 | 0.833375 | 0.999051 | 0.751062 | 0.201199 | 0.296285 | 0.456837 | 0.399 | 0.539576 |
| SL1344_RS13435_ | -5.28721 | -5.46446 | -5.03887 | 0.450983 | 1.242654 | 0.331418 | 0.353333 | 0.541522 | 0.673787 | 0.6489 | 0.75149 |
| SL1344_RS13440_ | 0.661854 | 0.621284 | 0.940895 | 0.312977 | 0.249534 | 0.865871 | 0.268 | 0.326341 | 0.505772 | 0.5343 | 0.67686 |
| SL1344_RS13450_ | -0.28362 | 0.040379 | -0.57939 | -0.61953 | 0.209291 | -2.92816 | 4.68E-05 | 0.016603 | 0.090757 | 0.1004 | 0.237128 |
| SL1344_RS13455_ | 0.975619 | 0.848705 | 1.064135 | 0.209606 | 0.17243 | 1.107369 | 0.125333 | 0.178551 | 0.341207 | 0.2554 | 0.412919 |
| SL1344_RS13460_ | -2.56212 | -2.72465 | -2.44702 | 0.271418 | 0.563494 | 0.463494 | 0.308 | 0.492911 | 0.637361 | 0.5967 | 0.714216 |
| SL1344_RS13465_ | -1.75145 | -1.83745 | -1.61022 | 0.218928 | 0.444945 | 0.458894 | 0.315333 | 0.503227 | 0.646577 | 0.6105 | 0.724844 |
| SL1344_RS13470_ | -2.89543 | -3.08969 | -2.65547 | 0.464748 | 0.527643 | 0.816544 | 0.171333 | 0.26879 | 0.425969 | 0.3661 | 0.507273 |

| | | | | | | | | | | | |
|---|---|---|---|---|---|---|---|---|---|---|---|
| SL1344_RS13475_ | -2.13403 | -2.55475 | -1.96742 | 0.569319 | 0.449878 | 1.118145 | 0.12 | 0.175963 | 0.335105 | 0.2623 | 0.412931 |
| SL1344_RS13480_ | -2.97286 | -3.30677 | -2.70328 | 0.596072 | 0.603238 | 0.901481 | 0.168554 | 0.244757 | 0.405572 | 0.3381 | 0.486739 |
| SL1344_RS13485_ | -1.60491 | -1.8047 | -1.13947 | 0.559575 | 0.573 | 0.959127 | 0.168667 | 0.230487 | 0.40308 | 0.3208 | 0.486929 |
| SL1344_RS13490_ | -1.80042 | -2.1359 | -1.59139 | 0.532739 | 0.370965 | 1.393186 | 0.060626 | 0.114461 | 0.2532 | 0.1759 | 0.323041 |
| SL1344_RS13495_ | -2.45748 | -2.7966 | -2.25742 | 0.523596 | 0.489121 | 1.030625 | 0.126 | 0.200191 | 0.360184 | 0.2893 | 0.440381 |
| SL1344_RS13500_ | 2.388199 | 2.443444 | 2.360047 | -0.0852 | 0.147917 | -0.5925 | 0.246502 | 0.402788 | 0.570686 | 0.4811 | 0.633989 |
| SL1344_RS13505_ | 4.947763 | 4.956815 | 4.939345 | -0.00452 | 0.09843 | -0.04611 | 0.482 | 0.855076 | 0.914846 | 0.9661 | 0.979551 |
| SL1344_RS13510_ | -3.03958 | -3.09158 | -2.99592 | 0.116776 | 0.856223 | 0.126764 | 0.439333 | 0.687121 | 0.795622 | 0.7782 | 0.85317 |
| SL1344_RS13515_ | -1.90521 | -2.17738 | -1.58731 | 0.613969 | 0.357498 | 1.645564 | 0.030667 | 0.077294 | 0.201096 | 0.1382 | 0.280755 |
| SL1344_RS13520_ | -0.49999 | -0.8292 | -0.32241 | 0.499506 | 0.226962 | 2.038183 | 0.01 | 0.055499 | 0.169348 | 0.114 | 0.254302 |
| SL1344_RS13525_ | -2.79734 | -3.06195 | -2.40959 | 0.615574 | 0.583351 | 0.990616 | 0.150667 | 0.219623 | 0.379539 | 0.3139 | 0.467164 |
| SL1344_RS13530_ | -4.64275 | -5.10298 | -4.21779 | 0.91159 | 0.877876 | 1.030336 | 0.137242 | 0.212169 | 0.361098 | 0.2946 | 0.437128 |
| SL1344_RS13535_ | -1.99052 | -2.15033 | -1.79445 | 0.372095 | 0.55587 | 0.653714 | 0.220667 | 0.342753 | 0.510921 | 0.453 | 0.59635 |
| SL1344_RS13540_ | -0.87864 | -0.8137 | -0.91931 | -0.08759 | 0.25956 | -0.3108 | 0.360426 | 0.570274 | 0.698483 | 0.6715 | 0.771122 |
| SL1344_RS13545_ | 0.623596 | 0.489644 | 0.790493 | 0.31013 | 0.157136 | 1.913215 | 0.014657 | 0.056485 | 0.170333 | 0.1149 | 0.25525 |
| SL1344_RS13550_ | 2.318958 | 2.044193 | 2.499232 | 0.463399 | 0.138841 | 3.348367 | 4.68E-05 | 0.02885 | 0.124748 | 0.1 | 0.236592 |
| SL1344_RS13555_ | 2.885888 | 2.697119 | 3.111871 | 0.436825 | 0.153532 | 2.706616 | 4.68E-05 | 0.049178 | 0.168344 | 0.1 | 0.236592 |
| SL1344_RS13560_ | 2.857657 | 2.794688 | 2.984983 | 0.1754 | 0.115829 | 1.450735 | 0.056629 | 0.094137 | 0.235519 | 0.1652 | 0.317602 |
| SL1344_RS13565_ | -1.47944 | -1.51319 | -1.43554 | 0.078407 | 0.270832 | 0.270203 | 0.383744 | 0.5199 | 0.652421 | 0.6322 | 0.73349 |
| SL1344_RS13570_ | 0.072454 | 0.189014 | -0.03685 | -0.24503 | 0.197792 | -1.1717 | 0.083944 | 0.153456 | 0.303771 | 0.2222 | 0.369771 |
| SL1344_RS13575_ | 2.780483 | 2.769961 | 2.788601 | 0.154592 | 0.288409 | 0.281657 | 0.328 | 0.385422 | 0.570632 | 0.7045 | 0.798798 |
| SL1344_RS13580_ | -0.3955 | -0.46453 | -0.32725 | 0.10631 | 0.264064 | 0.390905 | 0.344437 | 0.558523 | 0.691404 | 0.6762 | 0.774421 |
| SL1344_RS13590_ | 6.537751 | 6.66485 | 6.517693 | -0.12925 | 0.145653 | -0.98368 | 0.190667 | 0.166581 | 0.341372 | 0.3511 | 0.530456 |
| SL1344_RS13595_ | 2.109185 | 1.967287 | 2.23952 | 0.276178 | 0.095539 | 2.891388 | 4.68E-05 | 0.017762 | 0.093578 | 0.1004 | 0.237123 |
| SL1344_RS13605_ | 1.140199 | 1.302957 | 0.965352 | -0.34914 | 0.148967 | -2.3034 | 0.001337 | 0.029928 | 0.12269 | 0.1019 | 0.239069 |
| SL1344_RS13610_ | -0.00957 | 0.042467 | -0.05813 | -0.10349 | 0.158991 | -0.61431 | 0.251333 | 0.369216 | 0.513574 | 0.473 | 0.597639 |
| SL1344_RS13615_ | 2.996367 | 2.795829 | 3.23711 | 0.436215 | 0.095942 | 4.483979 | 4.68E-05 | 0.004153 | 0.051003 | 0.1 | 0.236592 |
| SL1344_RS13625_ | -0.43206 | -0.65934 | -0.289 | 0.353761 | 0.24778 | 1.37262 | 0.080667 | 0.122809 | 0.267966 | 0.1981 | 0.350743 |
| SL1344_RS13630_ | -0.25584 | -0.32404 | -0.20775 | 0.11981 | 0.195827 | 0.570808 | 0.259161 | 0.385913 | 0.535179 | 0.5039 | 0.626469 |
| SL1344_RS13635_ | -3.24567 | -3.94963 | -2.77999 | 1.196685 | 0.642036 | 1.800018 | 0.008667 | 0.072937 | 0.19697 | 0.1106 | 0.250108 |
| SL1344_RS13645_ | 4.092932 | 3.996136 | 4.179989 | 0.187715 | 0.475822 | 0.269978 | 0.438374 | 0.578818 | 0.730357 | 0.9943 | 0.996614 |
| SL1344_RS13650_ | -0.85669 | -0.7167 | -1.04144 | -0.33948 | 0.285689 | -1.11815 | 0.082 | 0.168073 | 0.323929 | 0.2219 | 0.370367 |
| SL1344_RS13655_ | 1.208853 | 1.05185 | 1.339416 | 0.318199 | 0.788599 | 0.428734 | 0.286 | 0.523147 | 0.686258 | 0.5686 | 0.716956 |
| SL1344_RS13660_ | 5.195106 | 5.151097 | 5.323859 | 0.153262 | 0.108566 | 1.495583 | 0.055963 | 0.078625 | 0.217975 | 0.1536 | 0.306676 |
| SL1344_RS13665_ | 2.279266 | 2.1991 | 2.360185 | 0.131384 | 0.158168 | 0.811968 | 0.184544 | 0.26634 | 0.442999 | 0.3489 | 0.513421 |
| SL1344_RS13670_ | 2.71179 | 2.597431 | 2.75388 | 0.120098 | 0.149635 | 0.707481 | 0.258 | 0.37333 | 0.549618 | 0.5197 | 0.665486 |
| SL1344_RS13675_ | 2.187619 | 2.284227 | 2.136565 | -0.1563 | 0.138585 | -1.09373 | 0.106596 | 0.18417 | 0.352615 | 0.2483 | 0.403976 |
| SL1344_RS13680_ | 6.599002 | 6.459601 | 6.856264 | 0.399022 | 0.149655 | 2.841112 | 4.68E-05 | 0.007496 | 0.068721 | 0.1 | 0.236592 |
| SL1344_RS13685_ | 1.818551 | 1.972069 | 1.79099 | -0.16637 | 0.138484 | -0.85842 | 0.256496 | 0.339027 | 0.512406 | 0.5223 | 0.665311 |
| SL1344_RS13705_ | -2.28067 | -2.64712 | -1.84645 | 0.89851 | 0.612469 | 1.412467 | 0.038667 | 0.09595 | 0.236108 | 0.1456 | 0.29214 |
| SL1344_RS13710_ | -0.49106 | -0.84177 | -0.23219 | 0.592167 | 0.274841 | 2.087986 | 0.008661 | 0.045209 | 0.153253 | 0.1068 | 0.245552 |
| SL1344_RS13715_ | 2.409527 | 2.434459 | 2.356778 | -0.07049 | 0.102509 | -0.62455 | 0.275816 | 0.43141 | 0.585319 | 0.5443 | 0.673717 |
| SL1344_RS13720_ | 2.561725 | 2.57954 | 2.546321 | -0.05035 | 0.155184 | -0.30644 | 0.345333 | 0.557269 | 0.700852 | 0.6789 | 0.781876 |
| SL1344_RS13725_ | 1.685676 | 1.746546 | 1.584069 | -0.13307 | 0.255991 | -0.48287 | 0.326449 | 0.523195 | 0.679085 | 0.6862 | 0.79373 |
| SL1344_RS13730_ | -5.78579 | -5.66285 | -6.02026 | -0.32284 | 1.484329 | -0.19958 | 0.403333 | 0.55594 | 0.690972 | 0.6884 | 0.782937 |

| | | | | | | | | | | |
|---|---|---|---|---|---|---|---|---|---|---|
| SL1344_RS13735_ | -5.01562 | -5.26426 | -4.70834 | 0.590809 | 1.052805 | 0.516636 | 0.289807 | 0.432409 | 0.575934 | 0.5323 | 0.653591 |
| SL1344_RS13745_ | -5.01326 | -5.8061 | -3.34897 | 2.236198 | 1.313793 | 1.635571 | 0.048667 | 0.082108 | 0.21246 | 0.1573 | 0.304792 |
| SL1344_RS13750_ | -6.06346 | -5.28374 | -7.15278 | -1.84345 | 1.491708 | -1.17848 | 0.080613 | 0.151753 | 0.293832 | 0.2148 | 0.36092 |
| SL1344_RS13755_ | -7.03923 | -6.8743 | -7.15215 | -0.0859 | 2.448297 | -0.02512 | 0.486667 | 0.601478 | 0.728406 | 0.7279 | 0.81474 |
| SL1344_RS13760_ | -5.25933 | -5.15673 | -5.37368 | -0.16423 | 1.181857 | -0.13343 | 0.439041 | 0.570421 | 0.699565 | 0.6838 | 0.779076 |
| SL1344_RS13765_ | -3.61605 | -3.55323 | -3.77307 | -0.13965 | 0.815525 | -0.15404 | 0.429047 | 0.664787 | 0.777638 | 0.7658 | 0.849724 |
| SL1344_RS13770_ | -3.33603 | -3.39515 | -3.28138 | 0.108181 | 0.571206 | 0.176213 | 0.421333 | 0.567618 | 0.693542 | 0.6747 | 0.769451 |
| SL1344_RS13775_ | -1.36226 | -1.39283 | -1.32837 | -0.03932 | 0.45821 | -0.08563 | 0.469687 | 0.702182 | 0.80618 | 0.8502 | 0.901295 |
| SL1344_RS13780_ | 2.879371 | 2.842159 | 3.078953 | 0.219543 | 0.123405 | 1.19277 | 0.146 | 0.170115 | 0.340871 | 0.2914 | 0.454287 |
| SL1344_RS13785_ | 5.113315 | 5.303719 | 4.918389 | -0.23249 | 0.209055 | -1.69723 | 0.091273 | 0.065836 | 0.198987 | 0.1853 | 0.350317 |
| SL1344_RS13790_ | 5.080819 | 5.299438 | 4.851072 | -0.3203 | 0.196373 | -2.09974 | 4.68E-05 | 0.028503 | 0.127406 | 0.1002 | 0.236858 |
| SL1344_RS13800_ | 3.814188 | 4.083397 | 3.500537 | -0.38728 | 0.277866 | -1.85573 | 0.025317 | 0.045031 | 0.161443 | 0.1228 | 0.267028 |
| SL1344_RS13805_ | 2.894096 | 2.769222 | 3.122801 | 0.292088 | 0.174156 | 1.642141 | 0.047333 | 0.068589 | 0.200443 | 0.1413 | 0.291353 |
| SL1344_RS13810_ | 0.577125 | 0.797438 | 0.388793 | -0.40038 | 0.15296 | -2.56461 | 0.002002 | 0.025718 | 0.111423 | 0.1018 | 0.238986 |
| SL1344_RS13815_ | 0.792761 | 0.777982 | 0.809724 | 0.037002 | 0.14262 | 0.247756 | 0.393738 | 0.575225 | 0.701724 | 0.685 | 0.779891 |
| SL1344_RS13820_ | -1.2546 | -1.28732 | -1.22785 | 0.054941 | 0.2476 | 0.20937 | 0.40573 | 0.505371 | 0.63928 | 0.6182 | 0.722005 |
| SL1344_RS13825_ | 4.646626 | 4.659135 | 4.512585 | -0.14805 | 0.184185 | -0.81322 | 0.249167 | 0.284664 | 0.47346 | 0.4885 | 0.640098 |
| SL1344_RS13830_ | 1.289166 | 1.33187 | 1.258181 | -0.06792 | 0.215998 | -0.29935 | 0.35443 | 0.658314 | 0.775961 | 0.7134 | 0.810317 |
| SL1344_RS13840_ | 2.283709 | 2.299584 | 2.269902 | -0.02692 | 0.109535 | -0.23498 | 0.397735 | 0.634705 | 0.754029 | 0.734 | 0.819995 |
| SL1344_RS13845_ | 1.69936 | 1.696142 | 1.706617 | 0.00271 | 0.152572 | 0.016353 | 0.494337 | 0.729526 | 0.826654 | 0.8399 | 0.897458 |
| SL1344_RS13850_ | 2.600111 | 2.653547 | 2.523406 | -0.11697 | 0.111267 | -0.99896 | 0.156 | 0.212832 | 0.380742 | 0.2963 | 0.45714 |
| SL1344_RS13855_ | 2.473192 | 2.742956 | 2.227217 | -0.43061 | 0.187994 | -2.4576 | 4.68E-05 | 0.019067 | 0.10204 | 0.1006 | 0.237388 |
| SL1344_RS13860_ | 0.082501 | 0.0752 | 0.0997 | 0.037709 | 0.231272 | 0.142617 | 0.44437 | 0.679461 | 0.789674 | 0.7843 | 0.859981 |
| SL1344_RS13865_ | -3.10784 | -3.04547 | -3.17343 | -0.16537 | 0.606889 | -0.24749 | 0.38441 | 0.567605 | 0.696727 | 0.6878 | 0.778768 |
| SL1344_RS13870_ | -2.22908 | -2.16745 | -2.29072 | -0.12656 | 0.379237 | -0.3136 | 0.36509 | 0.499305 | 0.632915 | 0.6051 | 0.710824 |
| SL1344_RS13875_ | -2.43983 | -2.58608 | -2.35663 | 0.225844 | 0.46405 | 0.443432 | 0.303131 | 0.463511 | 0.610303 | 0.5797 | 0.69669 |
| SL1344_RS13880_ | -3.25831 | -3.22666 | -3.27424 | 0.03665 | 0.800382 | 0.037991 | 0.481679 | 0.668933 | 0.783286 | 0.812 | 0.877675 |
| SL1344_RS13885_ | -2.96549 | -2.63423 | -3.30444 | -0.65636 | 0.570245 | -1.12845 | 0.109333 | 0.168867 | 0.320748 | 0.2451 | 0.394728 |
| SL1344_RS13890_ | -4.00765 | -4.17666 | -3.86059 | 0.308061 | 0.701951 | 0.426477 | 0.328448 | 0.460638 | 0.60237 | 0.5652 | 0.680339 |
| SL1344_RS13895_ | -1.62356 | -1.63833 | -1.61412 | 0.03179 | 0.545402 | 0.052211 | 0.476 | 0.756435 | 0.846977 | 0.8633 | 0.913525 |
| SL1344_RS13900_ | -4.42547 | -4.39399 | -4.44674 | -0.04845 | 0.762434 | -0.05555 | 0.470667 | 0.557161 | 0.684332 | 0.6578 | 0.755154 |
| SL1344_RS13905_ | -3.64363 | -3.6512 | -3.6327 | -0.00507 | 0.683577 | -0.00948 | 0.496336 | 0.611403 | 0.734183 | 0.7271 | 0.814282 |
| SL1344_RS13910_ | -4.45857 | -4.39037 | -4.50869 | -0.05232 | 1.033339 | -0.04736 | 0.480346 | 0.663871 | 0.775864 | 0.7646 | 0.843562 |
| SL1344_RS13915_ | -3.44203 | -3.4689 | -3.4137 | 0.042906 | 0.656459 | 0.05927 | 0.473018 | 0.608499 | 0.730336 | 0.7166 | 0.804493 |
| SL1344_RS13920_ | -5.62348 | -4.77405 | -6.0976 | -1.05032 | 2.263896 | -0.41124 | 0.299333 | 0.583291 | 0.711753 | 0.6182 | 0.74089 |
| SL1344_RS13930_ | -1.45288 | -1.54474 | -1.39521 | 0.068119 | 0.472364 | 0.130354 | 0.446369 | 0.737656 | 0.833128 | 0.8218 | 0.887871 |
| SL1344_RS13935_ | -2.20397 | -2.08232 | -2.28282 | -0.15398 | 0.525618 | -0.26599 | 0.385743 | 0.646138 | 0.76131 | 0.7193 | 0.811685 |
| SL1344_RS13940_ | -4.29383 | -4.14897 | -4.44135 | -0.29183 | 0.794607 | -0.33427 | 0.357095 | 0.508376 | 0.643466 | 0.624 | 0.727109 |
| SL1344_RS13950_ | -5.78809 | -6.45178 | -5.27029 | 1.182346 | 1.246674 | 0.898569 | 0.157895 | 0.251185 | 0.398057 | 0.3309 | 0.469771 |
| SL1344_RS13955_ | -4.91953 | -5.18182 | -4.7469 | 0.453754 | 1.126708 | 0.372336 | 0.333333 | 0.491148 | 0.633689 | 0.606 | 0.716149 |
| SL1344_RS13965_ | -5.31254 | -4.88039 | -5.82782 | -0.91153 | 1.131363 | -0.72918 | 0.178 | 0.28833 | 0.442565 | 0.3733 | 0.511384 |
| SL1344_RS13970_ | -4.58846 | -4.7463 | -4.46742 | 0.28915 | 0.830302 | 0.318233 | 0.358428 | 0.504088 | 0.641847 | 0.6241 | 0.72767 |
| SL1344_RS13975_ | -5.16687 | -4.98838 | -5.34464 | -0.35158 | 0.950786 | -0.32019 | 0.350667 | 0.471519 | 0.610559 | 0.5881 | 0.696956 |
| SL1344_RS13980_ | -4.55688 | -4.74661 | -4.34491 | 0.137305 | 1.390398 | 0.086605 | 0.453698 | 0.702548 | 0.807809 | 0.8323 | 0.891046 |
| SL1344_RS13985_ | -6.14894 | -6.06766 | -6.27462 | -0.10371 | 1.799636 | -0.05197 | 0.475017 | 0.614525 | 0.738748 | 0.7437 | 0.828911 |

| | | | | | | | | | | | |
|---|---|---|---|---|---|---|---|---|---|---|---|
| SL1344_RS14000_ | -4.17097 | -4.00341 | -4.26167 | -0.2649 | 1.005182 | -0.23959 | 0.403333 | 0.638767 | 0.75525 | 0.7464 | 0.829038 |
| SL1344_RS14005_ | -3.62981 | -3.56784 | -3.74373 | -0.14769 | 0.694414 | -0.19849 | 0.409061 | 0.589115 | 0.715549 | 0.7118 | 0.801546 |
| SL1344_RS14010_ | -4.37694 | -4.57939 | -4.21791 | 0.406346 | 1.135665 | 0.329774 | 0.354667 | 0.531894 | 0.671673 | 0.6544 | 0.75547 |
| SL1344_RS14015_ | -3.40915 | -3.38557 | -3.43068 | -0.17151 | 0.918515 | -0.17918 | 0.415057 | 0.586319 | 0.721353 | 0.7628 | 0.837411 |
| SL1344_RS14020_ | -4.10446 | -4.19262 | -3.97892 | 0.283016 | 0.86402 | 0.307389 | 0.366422 | 0.555348 | 0.686857 | 0.6831 | 0.775849 |
| SL1344_RS14025_ | 2.315935 | 2.174912 | 2.43837 | 0.328949 | 0.197632 | 1.814488 | 0.001337 | 0.05202 | 0.171252 | 0.1008 | 0.237657 |
| SL1344_RS14030_ | 2.357585 | 2.196668 | 2.641611 | 0.437205 | 0.164119 | 2.461943 | 4.68E-05 | 0.035429 | 0.13962 | 0.1 | 0.236592 |
| SL1344_RS14035_ | 0.786488 | 0.945188 | 0.409619 | -0.45421 | 0.328299 | -1.32592 | 0.098 | 0.120718 | 0.277972 | 0.1999 | 0.363324 |
| SL1344_RS14040_ | 3.18697 | 3.385516 | 2.772825 | -0.58079 | 0.248855 | -1.87005 | 0.001337 | 0.063891 | 0.195389 | 0.1013 | 0.238319 |
| SL1344_RS14045_ | 2.970934 | 3.109146 | 2.732439 | -0.36483 | 0.210194 | -1.55789 | 0.02998 | 0.098728 | 0.249318 | 0.1294 | 0.275844 |
| SL1344_RS14050_ | -0.28409 | -0.09083 | -0.51187 | -0.45655 | 0.300295 | -1.53317 | 0.019334 | 0.096101 | 0.237522 | 0.1203 | 0.26112 |
| SL1344_RS14055_ | -0.65728 | -0.25924 | -1.11293 | -0.87332 | 0.275956 | -3.05962 | 4.68E-05 | 0.015601 | 0.088124 | 0.1001 | 0.236723 |
| SL1344_RS14060_ | -1.55941 | -1.52819 | -1.60306 | -0.09112 | 0.330125 | -0.26506 | 0.388667 | 0.557166 | 0.688219 | 0.6755 | 0.77008 |
| SL1344_RS14065_ | -3.24101 | -3.22217 | -3.25133 | -0.0103 | 0.59823 | -0.01141 | 0.494667 | 0.614613 | 0.735288 | 0.7256 | 0.810967 |
| SL1344_RS14070_ | -6.06768 | -5.69433 | -7.18385 | -1.32254 | 2.544386 | -0.43814 | 0.300667 | 0.417728 | 0.585872 | 0.5902 | 0.713511 |
| SL1344_RS14075_ | -5.65381 | -5.3318 | -6.10175 | -0.73397 | 1.40161 | -0.48841 | 0.286 | 0.440543 | 0.586675 | 0.5639 | 0.678738 |
| SL1344_RS14080_ | -4.96793 | -4.74053 | -5.19938 | -0.35506 | 1.82716 | -0.17518 | 0.409333 | 0.694331 | 0.800795 | 0.7756 | 0.85101 |
| SL1344_RS14085_ | 1.470186 | 1.492183 | 1.454613 | -0.03701 | 0.102893 | -0.33567 | 0.36 | 0.514079 | 0.645755 | 0.6249 | 0.726751 |
| SL1344_RS14090_ | -4.269 | -4.45862 | -4.02312 | 0.409159 | 0.854068 | 0.441021 | 0.309793 | 0.470663 | 0.612583 | 0.5752 | 0.692349 |
| SL1344_RS14095_ | -4.50098 | -3.99376 | -5.24531 | -1.27934 | 0.929948 | -1.30647 | 0.066667 | 0.132612 | 0.278849 | 0.1818 | 0.327876 |
| SL1344_RS14100_ | 0.104831 | 0.06679 | 0.144202 | 0.056957 | 0.219952 | 0.248689 | 0.397735 | 0.614393 | 0.736477 | 0.7261 | 0.812313 |
| SL1344_RS14105_ | 0.920378 | 0.824138 | 1.055937 | 0.22638 | 0.147003 | 1.500383 | 0.055333 | 0.097789 | 0.232272 | 0.1682 | 0.315689 |
| SL1344_RS14110_ | 0.41995 | 0.45092 | 0.370515 | -0.0533 | 0.212094 | -0.22588 | 0.413724 | 0.672578 | 0.783369 | 0.7722 | 0.851112 |
| SL1344_RS14115_ | -0.66349 | -0.6488 | -0.70071 | -0.0394 | 0.284704 | -0.12385 | 0.456362 | 0.681211 | 0.790199 | 0.7869 | 0.862293 |
| SL1344_RS14120_ | 2.807904 | 2.605126 | 2.946129 | 0.300938 | 0.137723 | 2.187972 | 0.005331 | 0.033259 | 0.134274 | 0.1049 | 0.243113 |
| SL1344_RS14125_ | -0.28097 | -0.66265 | -0.11538 | 0.550522 | 0.383759 | 1.449927 | 0.050633 | 0.088219 | 0.228377 | 0.1581 | 0.308458 |
| SL1344_RS14130_ | 0.660172 | 0.575365 | 0.810602 | 0.272297 | 0.269498 | 0.980257 | 0.145903 | 0.206071 | 0.379103 | 0.2883 | 0.449236 |
| SL1344_RS14135_ | 1.48082 | 1.515583 | 1.464767 | -0.03055 | 0.151399 | -0.16957 | 0.422385 | 0.723071 | 0.822735 | 0.7865 | 0.865493 |
| SL1344_RS14140_ | -3.93089 | -4.00751 | -3.82655 | 0.160552 | 0.733325 | 0.191902 | 0.416389 | 0.58348 | 0.707376 | 0.6956 | 0.786108 |
| SL1344_RS14150_ | -3.74715 | -3.92494 | -3.55257 | 0.356154 | 0.609137 | 0.570635 | 0.264 | 0.385337 | 0.529196 | 0.4867 | 0.611228 |
| SL1344_RS14155_ | -3.9686 | -4.08789 | -3.83421 | 0.247798 | 0.692741 | 0.343886 | 0.355097 | 0.50449 | 0.637269 | 0.6102 | 0.713936 |
| SL1344_RS14165_ | -4.07963 | -4.12834 | -4.04114 | 0.100717 | 0.745581 | 0.129091 | 0.439333 | 0.565551 | 0.692076 | 0.6696 | 0.766384 |
| SL1344_RS14170_ | -3.62989 | -3.92461 | -3.26081 | 0.682408 | 0.881292 | 0.738788 | 0.19054 | 0.301056 | 0.467012 | 0.4103 | 0.555054 |
| SL1344_RS14175_ | 0.188808 | 0.176154 | 0.202899 | 0.029653 | 0.146556 | 0.183496 | 0.424384 | 0.525332 | 0.656613 | 0.6383 | 0.737697 |
| SL1344_RS14180_ | 0.620886 | 0.279712 | 0.960392 | 0.608391 | 0.25589 | 2.352171 | 0.005331 | 0.025131 | 0.114715 | 0.1031 | 0.240731 |
| SL1344_RS14190_ | -1.53489 | -1.65584 | -0.89346 | 0.688716 | 0.503016 | 1.041428 | 0.196 | 0.234406 | 0.410223 | 0.3948 | 0.550903 |
| SL1344_RS14195_ | 1.055674 | 1.301248 | 0.837648 | -0.46261 | 0.141953 | -3.19721 | 4.68E-05 | 0.014501 | 0.084929 | 0.1002 | 0.236856 |
| SL1344_RS14200_ | 0.531293 | 0.497401 | 0.621512 | 0.119513 | 0.207557 | 0.525091 | 0.314457 | 0.496009 | 0.644581 | 0.6235 | 0.740454 |
| SL1344_RS14205_ | -4.05198 | -4.20229 | -3.76555 | 0.358694 | 1.065985 | 0.290785 | 0.377082 | 0.656764 | 0.770497 | 0.7282 | 0.820634 |
| SL1344_RS14210_ | -2.73027 | -2.71771 | -2.73315 | -0.0027 | 0.658071 | -0.00397 | 0.498 | 0.703566 | 0.807579 | 0.802 | 0.874398 |
| SL1344_RS14215_ | 2.208902 | 2.075253 | 2.250313 | 0.163358 | 0.103991 | 1.33604 | 0.096 | 0.143776 | 0.298533 | 0.2382 | 0.389179 |
| SL1344_RS14220_ | 0.736997 | 0.618919 | 0.942111 | 0.312372 | 0.152187 | 2.002866 | 0.016656 | 0.054714 | 0.168431 | 0.1172 | 0.258168 |
| SL1344_RS14225_ | 1.134429 | 1.223975 | 1.081971 | -0.13223 | 0.127103 | -0.99269 | 0.153231 | 0.233436 | 0.388261 | 0.3319 | 0.480447 |
| SL1344_RS14230_ | -3.91481 | -3.97179 | -3.86731 | 0.121772 | 0.877088 | 0.12359 | 0.436667 | 0.625993 | 0.744948 | 0.732 | 0.816681 |
| SL1344_RS14235_ | -4.71512 | -4.67255 | -4.76219 | -0.11103 | 0.926977 | -0.10756 | 0.453698 | 0.577967 | 0.703359 | 0.6981 | 0.787338 |

| | | | | | | | | | | | |
|---|---|---|---|---|---|---|---|---|---|---|---|
| SL1344_RS14240_ | -5.32896 | -5.2169 | -5.4315 | -0.24328 | 1.378653 | -0.14686 | 0.426 | 0.609283 | 0.733465 | 0.7216 | 0.808372 |
| SL1344_RS14245_ | -3.77598 | -4.04574 | -3.50908 | 0.492714 | 0.786544 | 0.584028 | 0.24517 | 0.403485 | 0.555912 | 0.5022 | 0.632854 |
| SL1344_RS14250_ | -3.1273 | -2.99645 | -3.27976 | -0.28882 | 0.527481 | -0.489 | 0.297333 | 0.423918 | 0.570738 | 0.5374 | 0.655656 |
| SL1344_RS14255_ | 0.792427 | 0.939958 | 0.646588 | -0.2931 | 0.150171 | -1.95968 | 0.013334 | 0.050596 | 0.158557 | 0.1152 | 0.255274 |
| SL1344_RS14260_ | -2.52481 | -2.9624 | -2.32278 | 0.582677 | 0.539842 | 0.988501 | 0.136576 | 0.210465 | 0.37539 | 0.3024 | 0.458509 |
| SL1344_RS14265_ | -4.55088 | -4.54563 | -4.55669 | -0.03275 | 0.913163 | -0.02797 | 0.483333 | 0.592751 | 0.717134 | 0.6981 | 0.790164 |
| SL1344_RS14270_ | -3.93906 | -3.84469 | -4.04259 | -0.18018 | 0.653401 | -0.24106 | 0.386 | 0.518318 | 0.651325 | 0.6302 | 0.731484 |
| SL1344_RS14275_ | -3.40274 | -3.26252 | -3.53861 | -0.36211 | 0.843566 | -0.43154 | 0.318667 | 0.491824 | 0.639588 | 0.6357 | 0.740214 |
| SL1344_RS14280_ | -1.10042 | -1.61703 | -0.86938 | 0.726426 | 0.37434 | 1.82145 | 0.009994 | 0.075074 | 0.205502 | 0.1147 | 0.254741 |
| SL1344_RS14285_ | -4.14822 | -4.50322 | -3.90379 | 0.599162 | 0.845258 | 0.666806 | 0.212 | 0.332192 | 0.490491 | 0.4375 | 0.572607 |
| SL1344_RS14290_ | -3.14945 | -3.17479 | -3.12882 | 0.048267 | 0.484619 | 0.090824 | 0.457029 | 0.545772 | 0.67482 | 0.6542 | 0.753946 |
| SL1344_RS14295_ | -1.26236 | -1.20887 | -1.31938 | -0.13701 | 0.336421 | -0.38785 | 0.339773 | 0.50479 | 0.64659 | 0.6441 | 0.744122 |
| SL1344_RS14300_ | -0.34553 | -0.32681 | -0.35746 | -0.01367 | 0.233383 | -0.05533 | 0.474667 | 0.656839 | 0.771218 | 0.7617 | 0.841788 |
| SL1344_RS14305_ | 0.887696 | 0.726517 | 0.972718 | 0.247557 | 0.168013 | 1.369118 | 0.058 | 0.119356 | 0.266969 | 0.1697 | 0.317634 |
| SL1344_RS14310_ | -3.65426 | -3.73631 | -3.58155 | 0.164711 | 0.977179 | 0.151985 | 0.438667 | 0.688782 | 0.79703 | 0.8082 | 0.87416 |
| SL1344_RS14315_ | -3.06291 | -2.98902 | -3.14461 | -0.14951 | 0.512216 | -0.28237 | 0.373333 | 0.518719 | 0.650105 | 0.6206 | 0.723356 |
| SL1344_RS14320_ | 1.082688 | 1.13121 | 1.012134 | -0.12306 | 0.133008 | -0.86648 | 0.175883 | 0.268704 | 0.427558 | 0.3656 | 0.507997 |
| SL1344_RS14325_ | 2.962365 | 2.989888 | 2.868609 | -0.10135 | 0.115261 | -0.76053 | 0.252 | 0.364799 | 0.536594 | 0.5096 | 0.654117 |
| SL1344_RS14330_ | -1.53581 | -1.47037 | -1.60671 | -0.13439 | 0.339892 | -0.37983 | 0.336442 | 0.515886 | 0.653 | 0.627 | 0.733489 |
| SL1344_RS14335_ | -2.3098 | -2.33238 | -2.29293 | 0.044249 | 0.482356 | 0.086135 | 0.458361 | 0.643977 | 0.759494 | 0.7439 | 0.826827 |
| SL1344_RS14345_ | 9.735423 | 5.78499 | 9.778457 | 3.994463 | 0.235067 | 0.976232 | 0.333333 | 0.19698 | 0.379046 | 0.7 | 0.822108 |
| SL1344_RS14350_ | -1.98446 | -2.43962 | -1.62981 | 0.8161 | 0.385175 | 2.009672 | 0.007334 | 0.051193 | 0.159638 | 0.1103 | 0.249267 |
| SL1344_RS14360_ | -0.3409 | -0.18086 | -0.71059 | -0.50346 | 0.270355 | -1.68667 | 0.031979 | 0.082927 | 0.216434 | 0.1364 | 0.280535 |
| SL1344_RS14365_ | -2.47172 | -2.70575 | -2.21931 | 0.519097 | 0.502197 | 1.022515 | 0.127249 | 0.199536 | 0.357975 | 0.2789 | 0.428144 |
| SL1344_RS14370_ | -2.55337 | -2.4849 | -2.62432 | -0.15452 | 0.491021 | -0.2961 | 0.367089 | 0.532342 | 0.667862 | 0.6507 | 0.750968 |
| SL1344_RS14375_ | -0.50542 | -0.36985 | -0.65516 | -0.29595 | 0.243836 | -1.16401 | 0.093937 | 0.155985 | 0.306705 | 0.2314 | 0.381779 |
| SL1344_RS14380_ | -2.5849 | -2.34818 | -2.96605 | -0.62211 | 0.467701 | -1.24439 | 0.088608 | 0.141803 | 0.288573 | 0.2133 | 0.361828 |
| SL1344_RS14385_ | -0.04974 | -0.21441 | 0.010519 | 0.189794 | 0.29024 | 0.570355 | 0.291805 | 0.447268 | 0.609227 | 0.6018 | 0.725931 |
| SL1344_RS14390_ | 3.068634 | 3.145139 | 2.924208 | -0.2147 | 0.080842 | -2.49183 | 0.002002 | 0.037607 | 0.140255 | 0.1018 | 0.238996 |
| SL1344_RS14395_ | 3.3203 | 3.339054 | 3.194755 | -0.13836 | 0.126093 | -0.80628 | 0.276667 | 0.337884 | 0.521191 | 0.5715 | 0.71348 |
| SL1344_RS14400_ | -0.65123 | -0.62292 | -0.67349 | -0.02292 | 0.275571 | -0.07754 | 0.470353 | 0.671377 | 0.782292 | 0.7777 | 0.853135 |
| SL1344_RS14405_ | -0.33765 | -0.24269 | -0.44149 | -0.19999 | 0.20127 | -0.94403 | 0.14 | 0.226202 | 0.379777 | 0.3115 | 0.45621 |
| SL1344_RS14410_ | 0.889956 | 1.018349 | 0.705811 | -0.30846 | 0.137098 | -2.16767 | 0.009328 | 0.042185 | 0.144211 | 0.11 | 0.249263 |
| SL1344_RS14415_ | 0.448759 | 0.746943 | 0.149695 | -0.59385 | 0.138134 | -4.26766 | 4.68E-05 | 0.006619 | 0.058989 | 0.1 | 0.236592 |
| SL1344_RS14420_ | -2.9304 | -3.06281 | -2.84548 | 0.225918 | 0.536005 | 0.386852 | 0.337109 | 0.498323 | 0.638294 | 0.6131 | 0.719561 |
| SL1344_RS14425_ | -0.32895 | -0.33502 | -0.32151 | 0.025972 | 0.225379 | 0.107407 | 0.454364 | 0.624563 | 0.742804 | 0.7355 | 0.818469 |
| SL1344_RS14430_ | -1.99724 | -2.0173 | -1.97807 | 0.020646 | 0.380219 | 0.053122 | 0.474 | 0.606275 | 0.726635 | 0.7208 | 0.804445 |
| SL1344_RS14435_ | 0.338791 | 0.338727 | 0.338967 | -0.00579 | 0.172127 | -0.02912 | 0.486342 | 0.622231 | 0.742474 | 0.7353 | 0.819652 |
| SL1344_RS14440_ | -1.92344 | -1.7566 | -2.03877 | -0.27004 | 0.358181 | -0.70535 | 0.224517 | 0.352076 | 0.503564 | 0.4537 | 0.587055 |
| SL1344_RS14445_ | -1.98871 | -1.97893 | -2.00443 | 0.005229 | 0.425343 | 0.008992 | 0.496 | 0.671918 | 0.781943 | 0.7806 | 0.855784 |
| SL1344_RS14450_ | -1.33246 | -1.19272 | -1.49474 | -0.31878 | 0.328046 | -0.9278 | 0.135333 | 0.22935 | 0.387916 | 0.3225 | 0.469356 |
| SL1344_RS14455_ | -0.84739 | -0.63195 | -1.08186 | -0.44969 | 0.21925 | -1.97441 | 0.019987 | 0.053309 | 0.159863 | 0.1195 | 0.259937 |
| SL1344_RS14460_ | -1.31538 | -1.1936 | -1.3748 | -0.13706 | 0.378516 | -0.32667 | 0.362425 | 0.633418 | 0.752027 | 0.7146 | 0.811226 |
| SL1344_RS14465_ | -2.08801 | -1.95683 | -2.20632 | -0.26091 | 0.349481 | -0.69994 | 0.223333 | 0.326488 | 0.477908 | 0.4298 | 0.561983 |
| SL1344_RS14470_ | 0.286668 | -0.35052 | 0.703039 | 1.040779 | 0.21088 | 4.877469 | 4.68E-05 | 0.006618 | 0.059325 | 0.1 | 0.236592 |

| | | | | | | | | | | | |
|---|---|---|---|---|---|---|---|---|---|---|---|
| SL1344_RS14475_ | -0.90551 | -1.31441 | -0.5622 | 0.756472 | 0.208348 | 3.513694 | 4.68E-05 | 0.010803 | 0.071832 | 0.1001 | 0.236727 |
| SL1344_RS14480_ | -1.90264 | -1.8639 | -1.97927 | -0.11289 | 0.39211 | -0.26068 | 0.38974 | 0.578854 | 0.708538 | 0.7026 | 0.795214 |
| SL1344_RS14485_ | -0.52033 | -0.50647 | -0.53608 | -0.0312 | 0.205838 | -0.15937 | 0.439707 | 0.555312 | 0.684385 | 0.6714 | 0.766859 |
| SL1344_RS14490_ | 2.076885 | 2.429771 | 1.939838 | -0.46515 | 0.146927 | -2.50276 | 0.000673 | 0.045007 | 0.158631 | 0.1005 | 0.237254 |
| SL1344_RS14495_ | 2.767305 | 2.749725 | 2.865556 | 0.113387 | 0.169649 | 0.556585 | 0.334444 | 0.555353 | 0.705393 | 0.7075 | 0.824763 |
| SL1344_RS14500_ | 2.957297 | 2.445047 | 3.364342 | 0.918541 | 0.113172 | 8.014235 | 4.68E-05 | 0.00353 | 0.04851 | 0.1 | 0.236592 |
| SL1344_RS14505_ | -2.17885 | -2.01712 | -2.30246 | -0.28635 | 0.426391 | -0.63707 | 0.23984 | 0.375824 | 0.529077 | 0.4808 | 0.609465 |
| SL1344_RS14510_ | 1.644712 | 1.214988 | 2.124352 | 0.901125 | 0.294105 | 3.037262 | 4.68E-05 | 0.009138 | 0.073631 | 0.1 | 0.236592 |
| SL1344_RS14515_ | 0.35466 | 0.398541 | 0.328431 | -0.04775 | 0.262689 | -0.1736 | 0.433333 | 0.751921 | 0.843954 | 0.8245 | 0.88935 |
| SL1344_RS14520_ | -2.45169 | -2.40943 | -2.50639 | -0.08756 | 0.428346 | -0.17078 | 0.422 | 0.579651 | 0.705721 | 0.7011 | 0.791389 |
| SL1344_RS14525_ | -5.22196 | -5.3847 | -4.96707 | 0.378877 | 1.571007 | 0.209823 | 0.382667 | 0.631184 | 0.751424 | 0.7217 | 0.810497 |
| SL1344_RS14530_ | -5.59889 | -5.41394 | -5.77301 | -0.39251 | 1.263319 | -0.2835 | 0.364424 | 0.536793 | 0.668479 | 0.6501 | 0.749064 |
| SL1344_RS14540_ | -2.33503 | -2.19593 | -2.44436 | -0.24627 | 0.369931 | -0.64029 | 0.243171 | 0.367752 | 0.513647 | 0.477 | 0.600754 |
| SL1344_RS14545_ | -2.15158 | -2.1726 | -2.11111 | 0.038992 | 0.417738 | 0.090943 | 0.462667 | 0.635639 | 0.750778 | 0.7343 | 0.81955 |
| SL1344_RS14550_ | -0.58559 | -0.47836 | -0.71369 | -0.23644 | 0.243497 | -0.91961 | 0.164 | 0.244222 | 0.401402 | 0.3306 | 0.479323 |
| SL1344_RS14555_ | -0.95992 | -1.25573 | -0.73093 | 0.513538 | 0.298326 | 1.663537 | 0.032 | 0.072914 | 0.196915 | 0.1357 | 0.279758 |
| SL1344_RS14560_ | 0.094546 | 0.11431 | 0.07775 | -0.03319 | 0.154514 | -0.21483 | 0.411059 | 0.521018 | 0.65317 | 0.6278 | 0.731518 |
| SL1344_RS14565_ | -1.12304 | -0.95963 | -1.24715 | -0.30784 | 0.348479 | -0.83124 | 0.180546 | 0.273688 | 0.437654 | 0.3767 | 0.521869 |
| SL1344_RS14570_ | 1.744777 | 1.739382 | 1.749983 | -0.01462 | 0.245139 | -0.05929 | 0.473333 | 0.797989 | 0.87711 | 0.9089 | 0.944369 |
| SL1344_RS14575_ | 1.562535 | 1.556524 | 1.567095 | 0.008778 | 0.105024 | 0.071281 | 0.466667 | 0.587564 | 0.711819 | 0.6945 | 0.787368 |
| SL1344_RS14585_ | -1.91696 | -1.97122 | -1.85523 | 0.141918 | 0.551744 | 0.242304 | 0.390667 | 0.631626 | 0.75109 | 0.751 | 0.831065 |
| SL1344_RS14600_ | 1.58261 | 1.558614 | 1.690212 | 0.128024 | 0.173929 | 0.580178 | 0.328448 | 0.534868 | 0.68369 | 0.6756 | 0.794514 |
| SL1344_RS14605_ | 1.979052 | 1.959456 | 2.03599 | 0.068951 | 0.175623 | 0.302199 | 0.390406 | 0.788171 | 0.870224 | 0.8032 | 0.882796 |
| SL1344_RS14610_ | -1.12183 | -1.2402 | -1.01517 | 0.233091 | 0.39714 | 0.567755 | 0.262492 | 0.422979 | 0.581458 | 0.5428 | 0.672366 |
| SL1344_RS14615_ | 0.130817 | 0.166941 | 0.078359 | -0.12609 | 0.379495 | -0.3405 | 0.357761 | 0.579451 | 0.718591 | 0.727 | 0.819565 |
| SL1344_RS14620_ | 1.241762 | 1.360846 | 1.137358 | -0.16538 | 0.319827 | -0.48019 | 0.339773 | 0.561009 | 0.708533 | 0.7161 | 0.813668 |
| SL1344_RS14625_ | 2.342964 | 2.396268 | 2.294773 | -0.25872 | 0.278385 | -0.92618 | 0.135243 | 0.195081 | 0.372068 | 0.2911 | 0.444683 |
| SL1344_RS14630_ | 4.540367 | 4.590753 | 4.42242 | -0.20558 | 0.169085 | -1.41701 | 0.005331 | 0.087392 | 0.23254 | 0.1061 | 0.244504 |
| SL1344_RS14640_ | 4.141857 | 3.90749 | 4.288488 | 0.362443 | 0.1712 | 2.101383 | 4.68E-05 | 0.033066 | 0.136617 | 0.1 | 0.236592 |
| SL1344_RS14645_ | -1.23074 | -1.20891 | -1.25841 | -0.04958 | 0.39226 | -0.11383 | 0.453698 | 0.691811 | 0.798807 | 0.8055 | 0.871642 |
| SL1344_RS14655_ | 0.864592 | 0.832528 | 0.904778 | 0.087275 | 0.222782 | 0.381754 | 0.348667 | 0.545312 | 0.686809 | 0.69 | 0.786643 |
| SL1344_RS14660_ | 1.045603 | 1.001487 | 1.088357 | 0.09159 | 0.11318 | 0.747207 | 0.201865 | 0.30701 | 0.451896 | 0.4042 | 0.534771 |
| SL1344_RS14675_ | -0.79875 | -0.34271 | -1.03649 | -0.69705 | 0.338017 | -1.87902 | 0.009328 | 0.069471 | 0.198735 | 0.1095 | 0.248458 |
| SL1344_RS14680_ | 1.443362 | 1.927338 | 1.098864 | -0.84084 | 0.245275 | -3.46682 | 4.68E-05 | 0.017044 | 0.097142 | 0.1 | 0.236592 |
| SL1344_RS14685_ | -0.02752 | -0.08897 | 0.004271 | 0.092455 | 0.209466 | 0.390801 | 0.336442 | 0.500815 | 0.642184 | 0.6254 | 0.73288 |
| SL1344_RS14690_ | -0.43762 | -0.49425 | -0.3831 | 0.108165 | 0.185338 | 0.562705 | 0.26449 | 0.386326 | 0.528574 | 0.4988 | 0.619078 |
| SL1344_RS14700_ | 1.795293 | 1.628436 | 1.963024 | 0.330557 | 0.104129 | 3.180512 | 0.000673 | 0.014767 | 0.085301 | 0.1003 | 0.236989 |
| SL1344_RS14705_ | 0.207421 | 0.244548 | 0.105204 | -0.11882 | 0.218999 | -0.47776 | 0.316 | 0.491486 | 0.640041 | 0.6304 | 0.743429 |
| SL1344_RS14710_ | -2.55718 | -2.61679 | -2.49843 | 0.127305 | 0.475915 | 0.254721 | 0.392 | 0.560084 | 0.689462 | 0.6708 | 0.765931 |
| SL1344_RS14715_ | -1.97822 | -2.07752 | -1.9152 | 0.120335 | 0.484194 | 0.234962 | 0.407333 | 0.637972 | 0.755596 | 0.7485 | 0.831541 |
| SL1344_RS14720_ | 0.971252 | 0.948239 | 0.988229 | 0.043499 | 0.134708 | 0.304249 | 0.369754 | 0.540871 | 0.673795 | 0.6561 | 0.75639 |
| SL1344_RS14725_ | -3.75799 | -3.69104 | -3.83427 | -0.1719 | 0.665891 | -0.22102 | 0.406 | 0.558778 | 0.686341 | 0.6825 | 0.77364 |
| SL1344_RS14730_ | -1.2828 | -1.17935 | -1.41436 | -0.23515 | 0.256746 | -0.86042 | 0.163891 | 0.258062 | 0.407392 | 0.3507 | 0.487691 |
| SL1344_RS14735_ | -1.98537 | -1.9323 | -2.03347 | -0.09388 | 0.351058 | -0.25843 | 0.379333 | 0.531382 | 0.66492 | 0.6407 | 0.742926 |
| SL1344_RS14740_ | -2.21436 | -2.11501 | -2.39438 | -0.25886 | 0.431933 | -0.56301 | 0.259827 | 0.405506 | 0.55743 | 0.5351 | 0.656975 |

| | | | | | | | | | | | |
|---|---|---|---|---|---|---|---|---|---|---|---|
| SL1344_RS14745_ | -1.9577 | -1.92102 | -2.02052 | -0.08048 | 0.367955 | -0.20566 | 0.406 | 0.568814 | 0.697013 | 0.6756 | 0.773832 |
| SL1344_RS14750_ | -2.90802 | -3.00057 | -2.80876 | 0.219809 | 0.559091 | 0.364597 | 0.340667 | 0.500241 | 0.638556 | 0.617 | 0.722347 |
| SL1344_RS14765_ | -1.87474 | -1.64873 | -2.07251 | -0.41705 | 0.31863 | -1.23612 | 0.093271 | 0.152202 | 0.290399 | 0.2271 | 0.373666 |
| SL1344_RS14770_ | -4.39451 | -4.40147 | -4.38293 | 0.007548 | 1.053664 | 0.005717 | 0.496 | 0.667439 | 0.779296 | 0.787 | 0.858314 |
| SL1344_RS14775_ | -4.44157 | -4.91699 | -4.01025 | 0.872778 | 0.950216 | 0.869604 | 0.15 | 0.243982 | 0.400851 | 0.3261 | 0.472913 |
| SL1344_RS14780_ | 2.595632 | 2.601384 | 2.585223 | -0.06823 | 0.241996 | -0.30208 | 0.373333 | 0.543827 | 0.698744 | 0.7885 | 0.862104 |
| SL1344_RS14785_ | 2.234804 | 2.194287 | 2.273266 | 0.084191 | 0.09008 | 0.865321 | 0.162 | 0.258251 | 0.416452 | 0.3489 | 0.492312 |
| SL1344_RS14790_ | 0.930304 | 0.888317 | 0.961688 | 0.074748 | 0.154653 | 0.458382 | 0.316 | 0.489313 | 0.633325 | 0.6062 | 0.71679 |
| SL1344_RS14795_ | 2.279061 | 2.235281 | 2.333357 | 0.103568 | 0.104838 | 0.944949 | 0.146569 | 0.217185 | 0.379728 | 0.3067 | 0.457651 |
| SL1344_RS14800_ | 2.379292 | 2.343654 | 2.417003 | 0.079395 | 0.097418 | 0.794213 | 0.189333 | 0.288023 | 0.452719 | 0.3891 | 0.531775 |
| SL1344_RS14805_ | 1.377579 | 1.370563 | 1.383318 | 0.011636 | 0.095628 | 0.121887 | 0.448 | 0.534106 | 0.663442 | 0.6424 | 0.740201 |
| SL1344_RS14810_ | -2.23174 | -2.19485 | -2.29925 | -0.06507 | 0.474758 | -0.13231 | 0.439707 | 0.651284 | 0.766453 | 0.7536 | 0.83656 |
| SL1344_RS14815_ | -2.38376 | -2.1177 | -2.70879 | -0.5892 | 0.395991 | -1.44357 | 0.048 | 0.106577 | 0.238189 | 0.1694 | 0.313647 |
| SL1344_RS14820_ | -2.52386 | -2.54575 | -2.48918 | 0.076688 | 0.452635 | 0.164334 | 0.430667 | 0.584402 | 0.707585 | 0.6962 | 0.786409 |
| SL1344_RS14825_ | -2.73538 | -2.60002 | -2.93353 | -0.28373 | 0.572649 | -0.48511 | 0.305796 | 0.484226 | 0.62894 | 0.6059 | 0.719459 |
| SL1344_RS14830_ | -1.33225 | -1.1716 | -1.45749 | -0.30993 | 0.317204 | -0.92211 | 0.144 | 0.240502 | 0.397468 | 0.3257 | 0.468283 |
| SL1344_RS14850_ | 3.261324 | 3.60296 | 2.959908 | -0.64516 | 0.166672 | -3.75728 | 4.68E-05 | 0.007998 | 0.069534 | 0.1 | 0.236592 |
| SL1344_RS14855_ | 3.119011 | 3.620494 | 2.869726 | -0.74018 | 0.103404 | -5.98963 | 4.68E-05 | 0.017357 | 0.099354 | 0.1 | 0.236592 |
| SL1344_RS14900_ | 3.624619 | 3.8365 | 3.583482 | -0.27758 | 0.260786 | -1.14351 | 0.075333 | 0.148745 | 0.318988 | 0.1848 | 0.340588 |
| SL1344_RS14905_ | 2.196692 | 2.507979 | 1.892431 | -0.61437 | 0.193007 | -3.19442 | 4.68E-05 | 0.01865 | 0.101269 | 0.1 | 0.236592 |
| SL1344_RS14910_ | 2.424241 | 2.806958 | 2.121713 | -0.68071 | 0.210967 | -3.12527 | 4.68E-05 | 0.013987 | 0.088987 | 0.1 | 0.236592 |
| SL1344_RS15010_ | 2.304265 | 2.537295 | 2.071088 | -0.50797 | 0.140333 | -3.68316 | 4.68E-05 | 0.007052 | 0.064111 | 0.1 | 0.236592 |
| SL1344_RS15015_ | 1.811225 | 1.915533 | 1.61124 | -0.30641 | 0.212588 | -1.44862 | 0.022652 | 0.130699 | 0.292634 | 0.1184 | 0.260665 |
| SL1344_RS15020_ | -0.44995 | -0.51803 | -0.40468 | 0.124518 | 0.442403 | 0.278188 | 0.364424 | 0.671416 | 0.78767 | 0.7643 | 0.84706 |
| SL1344_RS15025_ | -2.50292 | -2.20679 | -2.95828 | -0.76249 | 0.436591 | -1.65827 | 0.030646 | 0.079304 | 0.203071 | 0.1415 | 0.284283 |
| SL1344_RS15030_ | 1.494088 | 1.308116 | 1.567543 | 0.255272 | 0.152972 | 1.508731 | 0.058 | 0.114522 | 0.263211 | 0.1603 | 0.30788 |
| SL1344_RS15035_ | 1.727405 | 1.619262 | 1.813824 | 0.200304 | 0.131432 | 1.488165 | 0.04064 | 0.091773 | 0.227331 | 0.151 | 0.298141 |
| SL1344_RS15040_ | -5.21881 | -5.20419 | -5.24412 | -0.10326 | 1.530776 | -0.05462 | 0.469687 | 0.612764 | 0.741808 | 0.769 | 0.847816 |
| SL1344_RS15045_ | -2.34122 | -2.26143 | -2.44493 | -0.16541 | 0.448705 | -0.33556 | 0.356 | 0.538583 | 0.673686 | 0.643 | 0.74832 |
| SL1344_RS15050_ | -4.83637 | -4.98219 | -4.69833 | 0.280439 | 0.858725 | 0.289665 | 0.357095 | 0.482256 | 0.619609 | 0.587 | 0.696779 |
| SL1344_RS15055_ | 2.655282 | 2.545256 | 2.763387 | 0.22444 | 0.083705 | 2.654339 | 4.68E-05 | 0.023098 | 0.106851 | 0.1004 | 0.237133 |
| SL1344_RS15060_ | -2.36362 | -2.37356 | -2.35431 | -0.01287 | 0.475172 | -0.02442 | 0.489007 | 0.635722 | 0.753904 | 0.7586 | 0.83666 |
| SL1344_RS15065_ | -2.38074 | -2.31486 | -2.43886 | -0.15734 | 0.498222 | -0.3025 | 0.361093 | 0.561625 | 0.694136 | 0.6815 | 0.776943 |
| SL1344_RS15070_ | -1.68366 | -1.63628 | -1.73259 | -0.10111 | 0.291465 | -0.32699 | 0.355333 | 0.479422 | 0.617504 | 0.5943 | 0.703111 |
| SL1344_RS15075_ | 0.331848 | 0.120504 | 0.512844 | 0.404957 | 0.232949 | 1.682491 | 0.018 | 0.075521 | 0.206199 | 0.118 | 0.259744 |
| SL1344_RS15080_ | -1.21155 | -1.50727 | -0.96341 | 0.574443 | 0.365655 | 1.531259 | 0.038667 | 0.094538 | 0.232648 | 0.1422 | 0.287598 |
| SL1344_RS15085_ | -1.3081 | -1.45907 | -1.1747 | 0.299437 | 0.458496 | 0.663976 | 0.234 | 0.359849 | 0.529385 | 0.4708 | 0.613203 |
| SL1344_RS15090_ | -1.27796 | -1.51057 | -1.12039 | 0.39436 | 0.417436 | 0.922909 | 0.151233 | 0.230738 | 0.396964 | 0.3185 | 0.476049 |
| SL1344_RS15095_ | -0.29769 | -0.4034 | -0.12858 | 0.275353 | 0.225367 | 1.133217 | 0.116589 | 0.175265 | 0.328619 | 0.2534 | 0.404291 |
| SL1344_RS15100_ | -0.59627 | -0.63572 | -0.5543 | 0.121288 | 0.294255 | 0.399103 | 0.336 | 0.51442 | 0.655797 | 0.6633 | 0.759866 |
| SL1344_RS15105_ | 0.520277 | 0.472953 | 0.611572 | 0.102062 | 0.228166 | 0.406985 | 0.346436 | 0.586347 | 0.717034 | 0.6952 | 0.794865 |
| SL1344_RS15110_ | 0.288671 | 0.151162 | 0.400628 | 0.285796 | 0.244626 | 1.142733 | 0.060626 | 0.154 | 0.314201 | 0.1891 | 0.336181 |
| SL1344_RS15115_ | -4.63319 | -4.78833 | -4.52436 | 0.237006 | 0.975344 | 0.228527 | 0.390406 | 0.561664 | 0.691469 | 0.6759 | 0.77098 |
| SL1344_RS15120_ | -2.06591 | -2.00298 | -2.11801 | -0.15861 | 0.460733 | -0.30114 | 0.361093 | 0.567318 | 0.69887 | 0.7046 | 0.792918 |
| SL1344_RS15125_ | -3.39452 | -3.13635 | -3.63576 | -0.47703 | 0.571049 | -0.77335 | 0.194 | 0.311497 | 0.462943 | 0.4072 | 0.5448 |

| | | | | | | | | | | | |
|---|---|---|---|---|---|---|---|---|---|---|---|
| SL1344_RS15140_ | 1.929665 | 1.889029 | 1.972582 | 0.145569 | 0.209436 | 0.689141 | 0.203198 | 0.281668 | 0.462045 | 0.4148 | 0.555424 |
| SL1344_RS15145_ | 1.754026 | 1.621231 | 1.856162 | 0.239298 | 0.11121 | 2.140501 | 0.004665 | 0.048621 | 0.156729 | 0.107 | 0.245357 |
| SL1344_RS15150_ | 2.017072 | 1.926276 | 2.124317 | 0.162292 | 0.145781 | 1.092117 | 0.127249 | 0.173222 | 0.337331 | 0.2479 | 0.411658 |
| SL1344_RS15155_ | 0.420263 | 0.314801 | 0.488954 | 0.17132 | 0.179834 | 0.884968 | 0.161333 | 0.251436 | 0.411021 | 0.3509 | 0.498153 |
| SL1344_RS15160_ | 0.3292 | 0.457681 | 0.23875 | -0.23596 | 0.242834 | -0.91276 | 0.164 | 0.237717 | 0.406047 | 0.3328 | 0.485836 |
| SL1344_RS15165_ | 0.78429 | 0.814768 | 0.741922 | -0.05861 | 0.184216 | -0.29881 | 0.376416 | 0.629005 | 0.748884 | 0.7279 | 0.815541 |
| SL1344_RS15170_ | 0.837004 | 2.237923 | -0.86921 | -3.12267 | 0.235545 | -13.4474 | 4.68E-05 | 0.001509 | 0.034983 | 0.1 | 0.236592 |
| SL1344_RS15190_ | 0.769681 | 0.900136 | 0.581091 | -0.31858 | 0.155883 | -1.98059 | 0.01599 | 0.055689 | 0.168335 | 0.1159 | 0.256135 |
| SL1344_RS15195_ | -2.08749 | -2.13665 | -2.04602 | 0.087669 | 0.33841 | 0.244437 | 0.389074 | 0.511762 | 0.645648 | 0.6212 | 0.725495 |
| SL1344_RS15200_ | -0.19912 | -0.2137 | -0.18067 | 0.028052 | 0.202466 | 0.126472 | 0.444 | 0.585275 | 0.710379 | 0.6941 | 0.786251 |
| SL1344_RS15205_ | -0.64806 | -0.6141 | -0.69223 | -0.08379 | 0.216708 | -0.34611 | 0.339773 | 0.497158 | 0.63232 | 0.6056 | 0.712438 |
| SL1344_RS15210_ | -0.90018 | -0.90979 | -0.8909 | 0.02052 | 0.284984 | 0.067549 | 0.471686 | 0.638939 | 0.754683 | 0.7374 | 0.820857 |
| SL1344_RS15215_ | -0.16395 | -0.11711 | -0.2753 | -0.12252 | 0.268151 | -0.40876 | 0.331779 | 0.544873 | 0.684644 | 0.6593 | 0.769066 |
| SL1344_RS15220_ | -1.56486 | -1.55187 | -1.57829 | -0.01897 | 0.307193 | -0.05274 | 0.477015 | 0.580894 | 0.706417 | 0.6938 | 0.786356 |
| SL1344_RS15225_ | -0.42269 | -0.32298 | -0.54788 | -0.22819 | 0.215795 | -0.99324 | 0.118667 | 0.202748 | 0.354286 | 0.2847 | 0.426585 |
| SL1344_RS15230_ | -0.56131 | -0.27577 | -0.75135 | -0.45573 | 0.234378 | -1.87036 | 0.02465 | 0.060787 | 0.17677 | 0.1299 | 0.273041 |
| SL1344_RS15260_ | -0.02038 | -0.22593 | 0.105389 | 0.334062 | 0.203647 | 1.558088 | 0.035976 | 0.092969 | 0.225345 | 0.1467 | 0.290817 |
| SL1344_RS15265_ | 1.485957 | 1.563484 | 1.408763 | -0.16527 | 0.126557 | -1.25266 | 0.077948 | 0.136419 | 0.283396 | 0.2017 | 0.349512 |
| SL1344_RS15270_ | 5.795973 | 5.708066 | 5.848543 | 0.160131 | 0.092329 | 1.920769 | 4.68E-05 | 0.039815 | 0.150237 | 0.1001 | 0.236726 |
| SL1344_RS15275_ | 3.864037 | 3.895138 | 3.84299 | -0.05412 | 0.054322 | -0.93762 | 0.149234 | 0.230245 | 0.389378 | 0.3203 | 0.463679 |
| SL1344_RS15280_ | 1.103071 | 1.119123 | 1.092421 | -0.01309 | 0.146194 | -0.08134 | 0.462358 | 0.653183 | 0.767377 | 0.7667 | 0.842977 |
| SL1344_RS15285_ | 0.734098 | 0.711401 | 0.811902 | 0.085481 | 0.252223 | 0.281435 | 0.391739 | 0.773275 | 0.85884 | 0.7976 | 0.878531 |
| SL1344_RS15290_ | 1.618191 | 1.632263 | 1.595677 | -0.05874 | 0.154811 | -0.34142 | 0.334444 | 0.473032 | 0.630621 | 0.6376 | 0.739862 |
| SL1344_RS15295_ | 3.905826 | 3.797566 | 4.01175 | 0.231079 | 0.08041 | 2.974654 | 4.68E-05 | 0.024779 | 0.11486 | 0.1 | 0.236592 |
| SL1344_RS15310_ | 1.407786 | 1.10155 | 1.888446 | 0.799117 | 0.247304 | 3.187596 | 4.68E-05 | 0.011035 | 0.078988 | 0.1 | 0.236592 |
| SL1344_RS15315_ | 3.592431 | 3.250977 | 4.145246 | 0.795586 | 0.321751 | 2.851485 | 4.68E-05 | 0.00866 | 0.072899 | 0.1 | 0.236592 |
| SL1344_RS15320_ | 2.784513 | 2.452685 | 3.164018 | 0.718105 | 0.107082 | 6.951309 | 4.68E-05 | 0.004293 | 0.052538 | 0.1 | 0.236592 |
| SL1344_RS15325_ | 3.840238 | 3.540105 | 4.120189 | 0.586795 | 0.082868 | 7.326511 | 4.68E-05 | 0.014477 | 0.091016 | 0.1 | 0.236592 |
| SL1344_RS15330_ | 1.526802 | 1.329951 | 1.660264 | 0.320597 | 0.147381 | 2.018923 | 0.009994 | 0.049654 | 0.162297 | 0.1115 | 0.251443 |
| SL1344_RS15355_ | 1.079189 | 0.978325 | 1.181679 | 0.209451 | 0.144013 | 1.42984 | 0.052632 | 0.101773 | 0.23834 | 0.163 | 0.310406 |
| SL1344_RS15360_ | 3.948785 | 3.720571 | 4.139295 | 0.425159 | 0.080446 | 5.32955 | 4.68E-05 | 0.008292 | 0.06848 | 0.1 | 0.236592 |
| SL1344_RS15365_ | 5.949798 | 5.956597 | 5.94045 | -0.01315 | 0.074024 | -0.1103 | 0.448 | 0.828467 | 0.898488 | 0.9268 | 0.956588 |
| SL1344_RS15370_ | 5.406873 | 5.372608 | 5.581122 | 0.205642 | 0.063278 | 1.597728 | 0.005334 | 0.105699 | 0.260325 | 0.1074 | 0.246257 |
| SL1344_RS15375_ | 0.028857 | 0.003982 | 0.051992 | 0.048142 | 0.167615 | 0.270076 | 0.385333 | 0.50799 | 0.64434 | 0.612 | 0.717405 |
| SL1344_RS15380_ | -0.93707 | -1.31071 | -0.73464 | 0.532122 | 0.349391 | 1.416978 | 0.069953 | 0.105843 | 0.249528 | 0.1721 | 0.323635 |
| SL1344_RS15385_ | -2.04494 | -2.12791 | -1.9729 | 0.150919 | 0.325772 | 0.424134 | 0.308 | 0.440878 | 0.582091 | 0.5515 | 0.666762 |
| SL1344_RS15390_ | -2.16214 | -2.57466 | -1.76325 | 0.812905 | 0.350499 | 2.242955 | 0.008661 | 0.036379 | 0.131133 | 0.1083 | 0.246592 |
| SL1344_RS15395_ | -0.1875 | -0.7534 | 0.166343 | 0.915261 | 0.227539 | 3.931345 | 4.68E-05 | 0.014377 | 0.084359 | 0.1 | 0.236592 |
| SL1344_RS15400_ | -1.02555 | -1.38981 | -0.60164 | 0.788761 | 0.236933 | 3.239906 | 0.000673 | 0.013039 | 0.078746 | 0.1003 | 0.236991 |
| SL1344_RS15405_ | -0.8601 | -1.12825 | -0.56335 | 0.581269 | 0.328891 | 1.762118 | 0.018 | 0.062213 | 0.182928 | 0.1162 | 0.25661 |
| SL1344_RS15410_ | 0.810707 | 0.662769 | 0.964215 | 0.298753 | 0.124755 | 2.289293 | 0.009994 | 0.035276 | 0.128035 | 0.1091 | 0.247737 |
| SL1344_RS15415_ | 2.087985 | 2.146269 | 2.00278 | -0.1458 | 0.10161 | -1.38263 | 0.068667 | 0.110049 | 0.250506 | 0.175 | 0.32435 |
| SL1344_RS15420_ | -0.10061 | 0.071474 | -0.31517 | -0.4033 | 0.223008 | -1.75494 | 0.015334 | 0.074581 | 0.20114 | 0.1141 | 0.253814 |
| SL1344_RS15425_ | 0.683068 | 0.796761 | 0.574838 | -0.2226 | 0.123736 | -1.74174 | 0.028667 | 0.070658 | 0.185544 | 0.1417 | 0.283439 |
| SL1344_RS15430_ | 3.045693 | 3.009774 | 3.176006 | 0.170694 | 0.138836 | 1.124255 | 0.116 | 0.179969 | 0.353477 | 0.2474 | 0.406835 |

| | | | | | | | | | | | |
|---|---|---|---|---|---|---|---|---|---|---|---|
| SL1344_RS15435_ | 0.610155 | 0.597387 | 0.63038 | 0.035632 | 0.136808 | 0.250493 | 0.389333 | 0.514588 | 0.647936 | 0.6233 | 0.727973 |
| SL1344_RS15440_ | -0.23523 | -0.37458 | -0.06925 | 0.301033 | 0.189633 | 1.528762 | 0.044637 | 0.088303 | 0.21523 | 0.1547 | 0.300171 |
| SL1344_RS15445_ | 0.361085 | 0.395985 | 0.332497 | -0.08664 | 0.199393 | -0.4192 | 0.33511 | 0.48438 | 0.630446 | 0.6356 | 0.736825 |
| SL1344_RS15450_ | 0.54214 | 0.531743 | 0.564748 | 0.023227 | 0.205444 | 0.10371 | 0.462667 | 0.716558 | 0.816489 | 0.8012 | 0.873846 |
| SL1344_RS15455_ | 2.250694 | 2.336741 | 2.099335 | -0.23716 | 0.139939 | -1.60949 | 0.035976 | 0.082899 | 0.219131 | 0.1361 | 0.282131 |
| SL1344_RS15460_ | -0.52576 | -0.29551 | -0.85284 | -0.55483 | 0.226164 | -2.39268 | 0.003332 | 0.034744 | 0.129755 | 0.1056 | 0.243584 |
| SL1344_RS15465_ | 0.79596 | 0.967651 | 0.582302 | -0.3856 | 0.130063 | -2.91493 | 0.000673 | 0.019614 | 0.095732 | 0.1005 | 0.237203 |
| SL1344_RS15475_ | 0.327665 | 0.238035 | 0.526423 | 0.260507 | 0.219961 | 1.100786 | 0.135909 | 0.192361 | 0.353975 | 0.2867 | 0.445804 |
| SL1344_RS15480_ | 1.032702 | 0.924953 | 1.151196 | 0.222565 | 0.126911 | 1.688535 | 0.030646 | 0.071514 | 0.192054 | 0.1376 | 0.281866 |
| SL1344_RS15485_ | 2.257204 | 2.162771 | 2.368742 | 0.200639 | 0.111113 | 1.766406 | 0.02465 | 0.063746 | 0.185885 | 0.1252 | 0.268792 |
| SL1344_RS15490_ | 1.871849 | 1.807314 | 1.925153 | 0.1182 | 0.104433 | 1.0817 | 0.132 | 0.188709 | 0.341491 | 0.2708 | 0.42154 |
| SL1344_RS15495_ | 1.672056 | 1.560363 | 1.817141 | 0.253315 | 0.099938 | 2.450391 | 0.003332 | 0.029959 | 0.119964 | 0.1041 | 0.241887 |
| SL1344_RS15500_ | -3.0865 | -3.12588 | -3.06608 | 0.10215 | 0.630342 | 0.146338 | 0.435043 | 0.616587 | 0.740892 | 0.7437 | 0.82801 |
| SL1344_RS15505_ | -3.19369 | -3.26067 | -3.12562 | 0.130345 | 0.457535 | 0.261445 | 0.375083 | 0.489641 | 0.623047 | 0.5978 | 0.704893 |
| SL1344_RS15510_ | -2.94023 | -2.90191 | -2.99018 | -0.11751 | 0.686669 | -0.16619 | 0.423051 | 0.634705 | 0.754604 | 0.7537 | 0.833777 |
| SL1344_RS15515_ | -3.97016 | -3.78301 | -4.11313 | -0.25083 | 0.850576 | -0.26372 | 0.383744 | 0.62359 | 0.742848 | 0.7181 | 0.807872 |
| SL1344_RS15520_ | -4.5956 | -4.26197 | -4.75079 | -0.37322 | 1.191549 | -0.24219 | 0.385743 | 0.689328 | 0.794752 | 0.7348 | 0.829892 |
| SL1344_RS15525_ | 1.71875 | 1.727445 | 1.696314 | -0.03393 | 0.152421 | -0.20252 | 0.422385 | 0.677982 | 0.790135 | 0.8071 | 0.876382 |
| SL1344_RS15530_ | 1.218045 | 1.247473 | 1.202781 | -0.05674 | 0.15585 | -0.32877 | 0.357761 | 0.545408 | 0.682791 | 0.6877 | 0.78322 |
| SL1344_RS15535_ | 3.547127 | 3.472598 | 3.667041 | 0.19867 | 0.082283 | 2.331979 | 0.002002 | 0.033904 | 0.133499 | 0.1014 | 0.238455 |
| SL1344_RS15540_ | 3.418159 | 3.381989 | 3.438841 | 0.090751 | 0.154014 | 0.618618 | 0.270667 | 0.375858 | 0.556145 | 0.5345 | 0.68039 |
| SL1344_RS15545_ | -2.26411 | -2.3265 | -2.21823 | 0.094444 | 0.405912 | 0.233537 | 0.407062 | 0.559021 | 0.688247 | 0.6703 | 0.767325 |
| SL1344_RS15550_ | 1.614028 | 1.877844 | 1.490273 | -0.36949 | 0.170802 | -2.05813 | 0.005997 | 0.044594 | 0.155284 | 0.1089 | 0.248015 |
| SL1344_RS15555_ | 1.512093 | 1.702497 | 1.260818 | -0.44477 | 0.12347 | -3.58215 | 4.68E-05 | 0.017564 | 0.092926 | 0.1 | 0.236592 |
| SL1344_RS15560_ | 0.411311 | -0.21213 | 0.839829 | 1.057867 | 0.303363 | 3.514322 | 4.68E-05 | 0.007057 | 0.064483 | 0.1 | 0.236592 |
| SL1344_RS15565_ | 0.731877 | 0.599424 | 0.930275 | 0.334197 | 0.160799 | 2.017531 | 0.011334 | 0.053488 | 0.166516 | 0.1101 | 0.249422 |
| SL1344_RS15570_ | 1.069041 | 0.898074 | 1.247595 | 0.323287 | 0.169153 | 1.843651 | 0.023334 | 0.0531 | 0.167768 | 0.1235 | 0.267056 |
| SL1344_RS15575_ | 2.093253 | 2.024856 | 2.168814 | 0.141878 | 0.099346 | 1.375762 | 0.063333 | 0.10867 | 0.246892 | 0.1723 | 0.322175 |
| SL1344_RS15580_ | 0.889381 | 0.865828 | 0.908842 | 0.084354 | 0.206825 | 0.376272 | 0.349767 | 0.520366 | 0.6653 | 0.6962 | 0.787162 |
| SL1344_RS15585_ | 0.756093 | 0.668268 | 1.046457 | 0.347266 | 0.201191 | 1.399543 | 0.095333 | 0.129 | 0.284095 | 0.2131 | 0.366667 |
| SL1344_RS15590_ | -1.68299 | -1.2194 | -2.09901 | -0.84378 | 0.413004 | -2.02159 | 0.018654 | 0.045499 | 0.153118 | 0.1139 | 0.254745 |
| SL1344_RS15595_ | -0.69614 | -0.8334 | -0.5587 | 0.275379 | 0.223835 | 1.176311 | 0.097333 | 0.157342 | 0.300805 | 0.2302 | 0.377763 |
| SL1344_RS15600_ | 0.709789 | 0.616968 | 0.874336 | 0.227104 | 0.217291 | 0.954452 | 0.157229 | 0.22006 | 0.387762 | 0.3064 | 0.468773 |
| SL1344_RS15605_ | -0.89537 | -1.0515 | -0.73425 | 0.312645 | 0.228029 | 1.318886 | 0.074667 | 0.129771 | 0.264401 | 0.1967 | 0.341057 |
| SL1344_RS15610_ | -1.17542 | -1.47574 | -0.80483 | 0.690612 | 0.341925 | 1.95656 | 0.008661 | 0.046757 | 0.155353 | 0.1104 | 0.249569 |
| SL1344_RS15615_ | -2.08046 | -2.49918 | -1.77239 | 0.717467 | 0.388708 | 1.804964 | 0.023334 | 0.067559 | 0.186507 | 0.1278 | 0.269679 |
| SL1344_RS15620_ | 1.568279 | 1.696202 | 1.411922 | -0.28631 | 0.097943 | -2.8368 | 0.001336 | 0.022497 | 0.101952 | 0.1011 | 0.237856 |
| SL1344_RS15625_ | 0.500496 | 0.691707 | 0.267236 | -0.42921 | 0.164983 | -2.55231 | 0.000673 | 0.030381 | 0.121797 | 0.1009 | 0.237793 |
| SL1344_RS15630_ | -1.86855 | -1.74801 | -1.95754 | -0.23778 | 0.411136 | -0.56658 | 0.261333 | 0.396966 | 0.551291 | 0.5219 | 0.643455 |
| SL1344_RS15635_ | -0.72794 | -0.72702 | -0.72835 | -0.0094 | 0.236083 | -0.03961 | 0.481679 | 0.585821 | 0.711797 | 0.6987 | 0.789726 |
| SL1344_RS15640_ | 0.739912 | 0.805526 | 0.651108 | -0.10981 | 0.303073 | -0.33723 | 0.382412 | 0.65957 | 0.779725 | 0.8297 | 0.890998 |
| SL1344_RS15645_ | -1.91499 | -1.94824 | -1.87756 | 0.054267 | 0.477465 | 0.112659 | 0.452365 | 0.695976 | 0.801615 | 0.801 | 0.869169 |
| SL1344_RS15650_ | -1.76719 | -1.99905 | -1.69133 | 0.3134 | 0.52921 | 0.504261 | 0.291333 | 0.434309 | 0.599663 | 0.605 | 0.724767 |
| SL1344_RS15655_ | -3.93205 | -4.12817 | -3.72746 | 0.408751 | 0.815201 | 0.457184 | 0.303797 | 0.480877 | 0.623251 | 0.5969 | 0.707884 |
| SL1344_RS15660_ | -3.03877 | -3.04491 | -3.03509 | 0.008161 | 0.459823 | 0.016619 | 0.491333 | 0.553398 | 0.682311 | 0.6738 | 0.767019 |

| | | | | | | | | | | | |
|---|---|---|---|---|---|---|---|---|---|---|---|
| SL1344_RS15665_ | -3.54062 | -3.42026 | -3.74537 | -0.29617 | 0.688542 | -0.38805 | 0.340667 | 0.511016 | 0.650936 | 0.6324 | 0.740554 |
| SL1344_RS15670_ | -2.63559 | -2.61106 | -2.67081 | -0.05673 | 0.459596 | -0.11672 | 0.440373 | 0.577887 | 0.703361 | 0.6928 | 0.78319 |
| SL1344_RS15675_ | -4.40099 | -4.01253 | -4.73495 | -0.62724 | 0.971821 | -0.59628 | 0.241333 | 0.398346 | 0.546025 | 0.495 | 0.623028 |
| SL1344_RS15680_ | -0.26617 | -0.14565 | -0.39793 | -0.26074 | 0.217906 | -1.18792 | 0.094667 | 0.157565 | 0.305051 | 0.2294 | 0.375717 |
| SL1344_RS15685_ | -3.46853 | -3.4268 | -3.49965 | -0.12141 | 0.808451 | -0.13248 | 0.442667 | 0.669525 | 0.780376 | 0.78 | 0.852906 |
| SL1344_RS15690_ | 2.371167 | 2.214852 | 2.436221 | 0.217665 | 0.127496 | 1.580483 | 0.027333 | 0.098715 | 0.24373 | 0.131 | 0.273961 |
| SL1344_RS15695_ | 2.238305 | 2.238272 | 2.238451 | 0.024218 | 0.152928 | 0.12034 | 0.424 | 0.538609 | 0.69248 | 0.7903 | 0.858509 |
| SL1344_RS15700_ | -6.80084 | -6.26566 | -7.43124 | -1.16514 | 1.686432 | -0.66066 | 0.212667 | 0.322225 | 0.46793 | 0.4176 | 0.547413 |
| SL1344_RS15705_ | -2.62843 | -2.84764 | -2.39901 | 0.45993 | 0.49669 | 0.848428 | 0.157333 | 0.253527 | 0.411489 | 0.3431 | 0.488814 |
| SL1344_RS15710_ | 0.399152 | 0.451623 | 0.352703 | -0.09864 | 0.258829 | -0.35044 | 0.35 | 0.578815 | 0.712693 | 0.7019 | 0.795776 |
| SL1344_RS15715_ | 1.285341 | 1.323657 | 1.19415 | -0.12718 | 0.143089 | -0.81191 | 0.205333 | 0.30044 | 0.467761 | 0.4161 | 0.560618 |
| SL1344_RS15720_ | 1.518677 | 1.546226 | 1.49831 | -0.14066 | 0.259409 | -0.54907 | 0.296 | 0.36995 | 0.5468 | 0.6196 | 0.729166 |
| SL1344_RS15725_ | 4.356536 | 4.086658 | 4.466376 | 0.378727 | 0.151169 | 2.506358 | 4.68E-05 | 0.056305 | 0.182336 | 0.1 | 0.236592 |
| SL1344_RS15730_ | 3.202681 | 3.239147 | 3.181339 | -0.06717 | 0.119354 | -0.57105 | 0.267333 | 0.424656 | 0.591621 | 0.5472 | 0.686175 |
| SL1344_RS15735_ | 2.822687 | 2.784886 | 2.949499 | 0.153001 | 0.108094 | 1.196 | 0.118588 | 0.1652 | 0.330021 | 0.2556 | 0.412209 |
| SL1344_RS15740_ | 2.014266 | 1.983282 | 2.13698 | 0.139055 | 0.182107 | 0.55403 | 0.325333 | 0.550834 | 0.699265 | 0.6855 | 0.804477 |
| SL1344_RS15745_ | 1.168113 | 1.198216 | 1.119573 | -0.08854 | 0.144281 | -0.59286 | 0.251832 | 0.378777 | 0.534231 | 0.492 | 0.613772 |
| SL1344_RS15750_ | 1.139251 | 1.001616 | 1.257276 | 0.242068 | 0.15371 | 1.552747 | 0.044667 | 0.08707 | 0.219893 | 0.1544 | 0.303134 |
| SL1344_RS15755_ | 2.150758 | 2.058582 | 2.337976 | 0.267229 | 0.117933 | 2.152901 | 0.006667 | 0.052113 | 0.167337 | 0.1098 | 0.248822 |
| SL1344_RS15760_ | 1.18285 | 1.101642 | 1.261411 | 0.17249 | 0.139857 | 1.18545 | 0.095333 | 0.153537 | 0.306088 | 0.2267 | 0.376102 |
| SL1344_RS15765_ | -1.28247 | -1.37171 | -1.19121 | 0.171682 | 0.385051 | 0.435121 | 0.314 | 0.505675 | 0.649286 | 0.6201 | 0.731272 |
| SL1344_RS15770_ | 2.559994 | 2.394882 | 2.726099 | 0.341607 | 0.10606 | 3.255851 | 4.68E-05 | 0.023025 | 0.110058 | 0.1 | 0.236592 |
| SL1344_RS15775_ | 1.512322 | 1.414419 | 1.556672 | 0.136555 | 0.123317 | 1.020904 | 0.156 | 0.230133 | 0.391889 | 0.3295 | 0.48155 |
| SL1344_RS15780_ | 1.068503 | 0.863454 | 1.23849 | 0.377799 | 0.179895 | 2.03917 | 0.005331 | 0.048603 | 0.160995 | 0.1045 | 0.242408 |
| SL1344_RS15785_ | 0.808479 | 0.61164 | 0.97565 | 0.377021 | 0.159775 | 2.311863 | 0.004001 | 0.034106 | 0.130932 | 0.1034 | 0.240955 |
| SL1344_RS15790_ | -2.19342 | -2.29803 | -2.10078 | 0.203237 | 0.339719 | 0.564022 | 0.265333 | 0.382929 | 0.526401 | 0.4822 | 0.604609 |
| SL1344_RS15795_ | 6.118153 | 5.901443 | 6.287015 | 0.381604 | 0.059274 | 6.379538 | 4.68E-05 | 0.000996 | 0.032907 | 0.1 | 0.236592 |
| SL1344_RS15800_ | 4.968981 | 4.740474 | 5.113585 | 0.371585 | 0.088295 | 4.309031 | 4.68E-05 | 0.003433 | 0.049028 | 0.1 | 0.236592 |
| SL1344_RS15810_ | 1.805684 | 1.695803 | 1.888528 | 0.193714 | 0.10653 | 1.783369 | 0.024 | 0.068697 | 0.190702 | 0.1266 | 0.26835 |
| SL1344_RS15815_ | 2.353174 | 2.187812 | 2.453866 | 0.264191 | 0.094024 | 2.742591 | 0.000673 | 0.024371 | 0.109346 | 0.1011 | 0.237998 |
| SL1344_RS15820_ | 2.829754 | 2.784018 | 2.880578 | 0.094101 | 0.075388 | 1.210089 | 0.102 | 0.153632 | 0.30314 | 0.2306 | 0.381195 |
| SL1344_RS15825_ | 2.238091 | 2.219915 | 2.26269 | 0.049414 | 0.090955 | 0.50878 | 0.283811 | 0.439038 | 0.584974 | 0.5572 | 0.670786 |
| SL1344_RS15830_ | 3.155902 | 3.124987 | 3.254417 | 0.122176 | 0.158244 | 0.811784 | 0.202 | 0.265408 | 0.445409 | 0.3598 | 0.536288 |
| SL1344_RS15835_ | 0.975922 | 0.896042 | 1.151377 | 0.260893 | 0.187917 | 1.29358 | 0.058 | 0.13786 | 0.296581 | 0.1692 | 0.317415 |
| SL1344_RS15840_ | 1.740357 | 1.555504 | 1.888362 | 0.323334 | 0.173417 | 1.852365 | 0.015323 | 0.055963 | 0.175726 | 0.1157 | 0.25722 |
| SL1344_RS15845_ | 3.265918 | 2.937871 | 3.537503 | 0.570232 | 0.116144 | 4.846743 | 4.68E-05 | 0.004013 | 0.050776 | 0.1 | 0.236592 |
| SL1344_RS15850_ | -0.28595 | -0.38294 | -0.21278 | 0.159118 | 0.228279 | 0.662899 | 0.23984 | 0.362863 | 0.518651 | 0.4682 | 0.602928 |
| SL1344_RS15855_ | -3.45106 | -3.42888 | -3.47014 | 0.007248 | 0.75794 | 0.013198 | 0.494337 | 0.661641 | 0.773252 | 0.7681 | 0.843692 |
| SL1344_RS15860_ | 1.704851 | 1.501102 | 1.862162 | 0.357085 | 0.136169 | 2.548826 | 0.001337 | 0.027833 | 0.118648 | 0.1005 | 0.237247 |
| SL1344_RS15865_ | -1.35701 | -1.13544 | -1.59136 | -0.4654 | 0.277287 | -1.6317 | 0.03531 | 0.079029 | 0.200354 | 0.1426 | 0.285228 |
| SL1344_RS15870_ | 1.775928 | 1.54214 | 1.912904 | 0.363233 | 0.11851 | 2.961657 | 4.68E-05 | 0.024316 | 0.11 | 0.1007 | 0.237464 |
| SL1344_RS15875_ | 5.826249 | 5.552243 | 6.142337 | 0.591498 | 0.073691 | 8.145893 | 4.68E-05 | 0.000317 | 0.025208 | 0.1 | 0.236592 |
| SL1344_RS15880_ | 4.253066 | 3.833903 | 4.480523 | 0.641255 | 0.108936 | 5.191809 | 4.68E-05 | 0.017318 | 0.09939 | 0.1 | 0.236592 |
| SL1344_RS15890_ | -0.71682 | -0.59529 | -0.81235 | -0.22009 | 0.297431 | -0.68642 | 0.219187 | 0.343918 | 0.505289 | 0.4396 | 0.582335 |
| SL1344_RS15895_ | 0.399117 | 0.19455 | 0.586427 | 0.393712 | 0.155593 | 2.485197 | 0.002667 | 0.028188 | 0.115937 | 0.1022 | 0.239528 |

| | | | | | | | | | | | |
|---|---|---|---|---|---|---|---|---|---|---|---|
| SL1344_RS15900_ | 0.292974 | 0.132482 | 0.573727 | 0.381714 | 0.284863 | 1.299 | 0.093271 | 0.130279 | 0.285609 | 0.2044 | 0.364382 |
| SL1344_RS15905_ | 0.651885 | 0.566065 | 0.791216 | 0.207676 | 0.17884 | 1.087477 | 0.135333 | 0.193844 | 0.35137 | 0.2781 | 0.433463 |
| SL1344_RS15910_ | -0.43672 | -0.76336 | -0.26106 | 0.508763 | 0.311652 | 1.548752 | 0.029333 | 0.103752 | 0.249471 | 0.1318 | 0.275408 |
| SL1344_RS15915_ | 5.422409 | 5.362322 | 5.526662 | 0.19854 | 0.102341 | 2.042284 | 4.68E-05 | 0.036346 | 0.143436 | 0.1 | 0.236592 |
| SL1344_RS15920_ | 1.336083 | 1.244763 | 1.39373 | 0.140368 | 0.13866 | 0.988558 | 0.151333 | 0.228637 | 0.390153 | 0.315 | 0.468493 |
| SL1344_RS15925_ | 3.409814 | 3.286496 | 3.42576 | 0.139913 | 0.142886 | 0.820849 | 0.280667 | 0.330609 | 0.514966 | 0.5624 | 0.704547 |
| SL1344_RS15930_ | -1.94861 | -1.98934 | -1.89516 | 0.098958 | 0.347499 | 0.26414 | 0.38441 | 0.546033 | 0.676104 | 0.6524 | 0.753863 |
| SL1344_RS15935_ | -2.58035 | -2.32541 | -2.93752 | -0.59795 | 0.484681 | -1.19326 | 0.09527 | 0.156232 | 0.304888 | 0.2322 | 0.383153 |
| SL1344_RS15940_ | -1.24565 | -1.34158 | -1.14768 | 0.193351 | 0.263593 | 0.708383 | 0.217333 | 0.339166 | 0.486759 | 0.4349 | 0.566926 |
| SL1344_RS15945_ | -0.87814 | -0.62077 | -1.01804 | -0.41136 | 0.299658 | -1.29522 | 0.081279 | 0.147598 | 0.302639 | 0.2021 | 0.350976 |
| SL1344_RS15950_ | -2.30083 | -2.30772 | -2.29601 | -0.06999 | 0.593584 | -0.12528 | 0.447333 | 0.683843 | 0.792762 | 0.8004 | 0.868356 |
| SL1344_RS15955_ | -2.1707 | -2.17903 | -2.15874 | 0.028121 | 0.385193 | 0.064823 | 0.470353 | 0.593833 | 0.716995 | 0.7086 | 0.796582 |
| SL1344_RS15960_ | 2.174314 | 2.069963 | 2.389275 | 0.30583 | 0.125197 | 2.264845 | 0.004668 | 0.042046 | 0.149427 | 0.1053 | 0.243521 |
| SL1344_RS15965_ | -3.2487 | -3.30913 | -3.16498 | 0.110865 | 0.683172 | 0.146575 | 0.435043 | 0.652261 | 0.765375 | 0.763 | 0.83797 |
| SL1344_RS15970_ | 2.97312 | 2.921708 | 3.002316 | 0.091965 | 0.130144 | 0.742603 | 0.2 | 0.28767 | 0.464878 | 0.3812 | 0.541644 |
| SL1344_RS15975_ | -0.48726 | -0.30055 | -0.60106 | -0.28869 | 0.248793 | -1.10713 | 0.115257 | 0.180797 | 0.336231 | 0.2635 | 0.415155 |
| SL1344_RS15985_ | 3.500616 | 3.816455 | 3.386515 | -0.44138 | 0.189238 | -2.39234 | 4.68E-05 | 0.033071 | 0.136611 | 0.1 | 0.236592 |
| SL1344_RS15990_ | 1.230023 | 1.00294 | 1.529806 | 0.527529 | 0.165582 | 3.1885 | 4.68E-05 | 0.012352 | 0.08039 | 0.1 | 0.236592 |
| SL1344_RS15995_ | -1.88919 | -1.77892 | -2.00238 | -0.27155 | 0.481379 | -0.53291 | 0.274484 | 0.412528 | 0.569873 | 0.5334 | 0.658782 |
| SL1344_RS16000_ | -1.13617 | -0.8079 | -1.46753 | -0.63529 | 0.323564 | -1.91468 | 0.019321 | 0.051379 | 0.163335 | 0.1177 | 0.259301 |
| SL1344_RS16005_ | 0.921836 | 0.94727 | 0.883067 | -0.0731 | 0.168006 | -0.41041 | 0.323784 | 0.504078 | 0.646204 | 0.6291 | 0.735342 |
| SL1344_RS16015_ | 0.506486 | 0.609104 | 0.426952 | -0.18175 | 0.176807 | -0.97561 | 0.137908 | 0.217712 | 0.37562 | 0.3053 | 0.453678 |
| SL1344_RS16020_ | 0.098764 | 0.361301 | 0.034954 | -0.30312 | 0.25837 | -0.93449 | 0.203198 | 0.260341 | 0.435485 | 0.404 | 0.558523 |
| SL1344_RS16025_ | -1.87682 | -2.03984 | -1.80233 | 0.184588 | 0.445847 | 0.354416 | 0.358667 | 0.588864 | 0.7186 | 0.6976 | 0.798015 |
| SL1344_RS16030_ | -1.97031 | -1.78512 | -2.10712 | -0.30913 | 0.339017 | -0.85496 | 0.168 | 0.267874 | 0.415396 | 0.3577 | 0.496839 |
| SL1344_RS16035_ | 0.615382 | 0.676005 | 0.525547 | -0.13158 | 0.206572 | -0.62579 | 0.260667 | 0.413853 | 0.5705 | 0.5096 | 0.646914 |
| SL1344_RS16040_ | 1.113352 | 1.096584 | 1.132362 | 0.03103 | 0.177039 | 0.167581 | 0.424 | 0.65528 | 0.77115 | 0.7437 | 0.829603 |
| SL1344_RS16045_ | 0.651132 | 0.55342 | 0.740328 | 0.202464 | 0.163883 | 1.168657 | 0.094667 | 0.162405 | 0.314738 | 0.2281 | 0.376499 |
| SL1344_RS16050_ | 0.902124 | 0.946296 | 0.866043 | -0.07848 | 0.126312 | -0.57134 | 0.253333 | 0.395323 | 0.5404 | 0.4971 | 0.620751 |
| SL1344_RS16055_ | 1.475133 | 1.542336 | 1.413361 | -0.1306 | 0.123131 | -1.00927 | 0.123917 | 0.198055 | 0.354611 | 0.2868 | 0.435335 |
| SL1344_RS16060_ | -0.14325 | -0.31212 | 0.09538 | 0.370352 | 0.269728 | 1.311284 | 0.095333 | 0.130342 | 0.281215 | 0.2033 | 0.360343 |
| SL1344_RS16065_ | 5.877615 | 5.53379 | 6.136933 | 0.59909 | 0.066583 | 8.984345 | 4.68E-05 | 0.000652 | 0.029026 | 0.1 | 0.236592 |
| SL1344_RS16070_ | 1.712066 | 1.628981 | 1.956284 | 0.32276 | 0.197424 | 1.506594 | 0.042 | 0.090852 | 0.233551 | 0.146 | 0.293194 |
| SL1344_RS16075_ | 2.42474 | 2.124724 | 2.709374 | 0.521321 | 0.235634 | 2.210742 | 0.000673 | 0.028885 | 0.126566 | 0.1004 | 0.237128 |
| SL1344_RS16080_ | 1.220213 | 1.148887 | 1.284865 | 0.120526 | 0.151756 | 0.731875 | 0.208 | 0.320018 | 0.48316 | 0.4125 | 0.559927 |
| SL1344_RS16085_ | 0.105006 | 0.143113 | 0.060938 | -0.09325 | 0.199438 | -0.42518 | 0.312458 | 0.4795 | 0.622022 | 0.6032 | 0.711227 |
| SL1344_RS16090_ | 1.175171 | 1.061886 | 1.486424 | 0.397155 | 0.19735 | 1.644946 | 0.042638 | 0.089805 | 0.231423 | 0.1431 | 0.290108 |
| SL1344_RS16095_ | 1.252972 | 1.115878 | 1.378858 | 0.258564 | 0.131445 | 1.869415 | 0.021319 | 0.052792 | 0.164748 | 0.1174 | 0.258951 |
| SL1344_RS16100_ | 2.884385 | 2.693466 | 3.021965 | 0.336262 | 0.109491 | 3.053848 | 4.68E-05 | 0.025791 | 0.116135 | 0.1 | 0.236592 |
| SL1344_RS16105_ | 0.665486 | 0.473781 | 0.796117 | 0.28968 | 0.222633 | 1.21989 | 0.105333 | 0.144608 | 0.301614 | 0.2103 | 0.370006 |
| SL1344_RS16110_ | -5.18679 | -4.98271 | -5.48975 | -0.49838 | 1.095816 | -0.40132 | 0.31046 | 0.453205 | 0.59548 | 0.5624 | 0.675811 |
| SL1344_RS16115_ | -2.46603 | -2.50067 | -2.41367 | 0.120709 | 0.542215 | 0.210662 | 0.414667 | 0.614124 | 0.735105 | 0.7274 | 0.812611 |
| SL1344_RS16120_ | -1.36171 | -1.0067 | -1.55842 | -0.48458 | 0.455463 | -1.04483 | 0.141905 | 0.195541 | 0.361805 | 0.2741 | 0.43679 |
| SL1344_RS16125_ | -2.38095 | -2.11148 | -2.67508 | -0.57188 | 0.387533 | -1.39641 | 0.061333 | 0.114023 | 0.247288 | 0.1818 | 0.327436 |
| SL1344_RS16130_ | -0.37448 | -0.56781 | -0.30397 | 0.242524 | 0.241064 | 0.913038 | 0.179333 | 0.26582 | 0.429182 | 0.3803 | 0.526428 |

| | | | | | | | | | | | |
|---|---|---|---|---|---|---|---|---|---|---|---|
| SL1344_RS16135_ | -1.67673 | -1.82479 | -1.42157 | 0.352785 | 0.46021 | 0.714887 | 0.227333 | 0.342718 | 0.507735 | 0.4309 | 0.582096 |
| SL1344_RS16140_ | -0.54919 | -0.64658 | -0.46769 | 0.164806 | 0.279471 | 0.559704 | 0.267155 | 0.442804 | 0.592452 | 0.543 | 0.66997 |
| SL1344_RS16145_ | -0.08418 | -0.08278 | -0.08689 | -0.00253 | 0.193971 | -0.01354 | 0.492667 | 0.627254 | 0.744286 | 0.7407 | 0.822972 |
| SL1344_RS16150_ | 0.247044 | 0.122294 | 0.452138 | 0.316741 | 0.172879 | 1.750106 | 0.037333 | 0.07496 | 0.19738 | 0.1425 | 0.286847 |
| SL1344_RS16155_ | -2.91636 | -3.03901 | -2.83085 | 0.169238 | 0.534772 | 0.297928 | 0.371752 | 0.56557 | 0.694318 | 0.6804 | 0.77715 |
| SL1344_RS16160_ | -1.5338 | -1.49895 | -1.5594 | -0.07992 | 0.366095 | -0.19781 | 0.419333 | 0.611419 | 0.733663 | 0.7204 | 0.80783 |
| SL1344_RS16165_ | -1.39412 | -1.47931 | -1.33689 | 0.148751 | 0.30334 | 0.457553 | 0.31 | 0.450682 | 0.595182 | 0.5654 | 0.680872 |
| SL1344_RS16170_ | -0.71811 | -0.60092 | -0.80202 | -0.22752 | 0.266376 | -0.80581 | 0.177881 | 0.268096 | 0.429741 | 0.3594 | 0.498067 |
| SL1344_RS16175_ | -1.40991 | -1.26675 | -1.6238 | -0.36623 | 0.413805 | -0.85035 | 0.186542 | 0.267465 | 0.435112 | 0.375 | 0.527438 |
| SL1344_RS16180_ | -0.74681 | -0.74369 | -0.74997 | -0.01015 | 0.215532 | -0.0483 | 0.478 | 0.554077 | 0.684015 | 0.669 | 0.766288 |
| SL1344_RS16185_ | -0.84884 | -1.04351 | -0.66649 | 0.377501 | 0.228689 | 1.586115 | 0.044637 | 0.084338 | 0.209528 | 0.1461 | 0.2907 |
| SL1344_RS16190_ | -0.55217 | -0.58401 | -0.50026 | 0.0863 | 0.34669 | 0.238972 | 0.404 | 0.668876 | 0.780737 | 0.7797 | 0.854917 |
| SL1344_RS16195_ | -0.39826 | -0.4074 | -0.38235 | 0.002083 | 0.386576 | 0.004474 | 0.496667 | 0.727476 | 0.828763 | 0.867 | 0.918245 |
| SL1344_RS16200_ | -1.63388 | -1.82442 | -1.4675 | 0.355278 | 0.29411 | 1.156975 | 0.098 | 0.165719 | 0.307597 | 0.2455 | 0.390584 |
| SL1344_RS16205_ | -2.92314 | -3.21533 | -2.6163 | 0.578843 | 0.499641 | 1.089238 | 0.113258 | 0.184843 | 0.334561 | 0.2626 | 0.410971 |
| SL1344_RS16210_ | 1.588132 | 1.130473 | 1.807612 | 0.665671 | 0.14158 | 4.396868 | 4.68E-05 | 0.013941 | 0.085425 | 0.1 | 0.236592 |
| SL1344_RS16215_ | 2.210471 | 1.792631 | 2.450914 | 0.644939 | 0.176591 | 3.364005 | 4.68E-05 | 0.023643 | 0.113463 | 0.1 | 0.236592 |
| SL1344_RS16220_ | 1.350726 | 0.857403 | 1.738495 | 0.880098 | 0.126489 | 6.823551 | 4.68E-05 | 0.003375 | 0.044572 | 0.1 | 0.236592 |
| SL1344_RS16225_ | 1.817912 | 1.29519 | 2.096166 | 0.784025 | 0.154014 | 4.564285 | 4.68E-05 | 0.013921 | 0.086575 | 0.1 | 0.236592 |
| SL1344_RS16235_ | 1.872231 | 1.596476 | 2.240697 | 0.635247 | 0.143954 | 4.217329 | 4.68E-05 | 0.012923 | 0.08361 | 0.1 | 0.236592 |
| SL1344_RS16240_ | 0.977285 | 0.747302 | 1.184702 | 0.437524 | 0.119322 | 3.54475 | 0.000673 | 0.009997 | 0.06951 | 0.1 | 0.236592 |
| SL1344_RS16245_ | -1.67981 | -1.91985 | -1.44674 | 0.470808 | 0.284883 | 1.580499 | 0.037975 | 0.083283 | 0.205374 | 0.1453 | 0.289115 |
| SL1344_RS16250_ | -1.65923 | -1.67203 | -1.64162 | 0.007548 | 0.401021 | 0.018169 | 0.492338 | 0.670653 | 0.781009 | 0.7807 | 0.854149 |
| SL1344_RS16255_ | 1.827021 | 1.785905 | 1.893469 | 0.112546 | 0.246283 | 0.469102 | 0.286 | 0.492092 | 0.652223 | 0.5702 | 0.70939 |
| SL1344_RS16260_ | 1.360898 | 1.279731 | 1.423051 | 0.141238 | 0.128877 | 1.037141 | 0.115257 | 0.189783 | 0.343938 | 0.272 | 0.421897 |
| SL1344_RS16265_ | 1.691673 | 1.65289 | 1.742321 | 0.095929 | 0.116494 | 0.784103 | 0.206529 | 0.29606 | 0.458379 | 0.4013 | 0.544187 |
| SL1344_RS16270_ | 3.017637 | 2.821646 | 3.13347 | 0.313623 | 0.099729 | 3.088872 | 4.68E-05 | 0.027006 | 0.119295 | 0.1 | 0.236592 |
| SL1344_RS16275_ | 4.281762 | 4.14765 | 4.435363 | 0.318371 | 0.144174 | 2.274893 | 4.68E-05 | 0.044776 | 0.160495 | 0.1 | 0.236592 |
| SL1344_RS16280_ | 3.122807 | 2.990822 | 3.319562 | 0.342065 | 0.180623 | 1.820013 | 4.68E-05 | 0.102531 | 0.255843 | 0.1 | 0.236592 |
| SL1344_RS16285_ | 2.721967 | 2.538965 | 2.980752 | 0.489937 | 0.244825 | 2.188762 | 4.68E-05 | 0.033517 | 0.136876 | 0.1 | 0.236592 |
| SL1344_RS16290_ | 3.11404 | 2.94153 | 3.25259 | 0.300678 | 0.116388 | 2.454191 | 0.000673 | 0.025949 | 0.117446 | 0.1006 | 0.237399 |
| SL1344_RS16305_ | 2.384092 | 2.618271 | 2.173823 | -0.46434 | 0.106184 | -4.34832 | 4.68E-05 | 0.005712 | 0.056929 | 0.1 | 0.236592 |
| SL1344_RS16310_ | 3.466836 | 3.841674 | 2.922573 | -0.9118 | 0.098059 | -8.9142 | 4.68E-05 | 0.003232 | 0.047256 | 0.1 | 0.236592 |
| SL1344_RS16315_ | 1.453288 | 1.898658 | 0.845573 | -1.07209 | 0.178746 | -6.08978 | 4.68E-05 | 0.001641 | 0.037444 | 0.1 | 0.236592 |
| SL1344_RS16320_ | 1.875703 | 2.380692 | 1.293636 | -1.10587 | 0.136337 | -8.06881 | 4.68E-05 | 0.002074 | 0.038434 | 0.1 | 0.236592 |
| SL1344_RS16325_ | 2.778434 | 2.598963 | 2.945613 | 0.358114 | 0.105402 | 3.258673 | 4.68E-05 | 0.055721 | 0.180173 | 0.1 | 0.236592 |
| SL1344_RS16330_ | -0.85406 | -0.96112 | -0.81436 | 0.1144 | 0.357332 | 0.250209 | 0.405063 | 0.72757 | 0.826447 | 0.7789 | 0.864965 |
| SL1344_RS16335_ | 0.096298 | -0.38272 | 0.378099 | 0.689518 | 0.495661 | 1.444294 | 0.073285 | 0.088191 | 0.2318 | 0.1638 | 0.320894 |
| SL1344_RS16340_ | -0.18534 | -0.85685 | 0.430412 | 1.178644 | 0.733814 | 1.527698 | 0.05996 | 0.080813 | 0.222377 | 0.157 | 0.312658 |
| SL1344_RS16350_ | 1.62714 | 1.542539 | 1.962619 | 0.423793 | 0.232437 | 1.511849 | 0.052 | 0.118411 | 0.27606 | 0.1548 | 0.303327 |
| SL1344_RS16355_ | 0.435189 | 0.397287 | 0.552826 | 0.130564 | 0.217701 | 0.453145 | 0.339107 | 0.657383 | 0.772996 | 0.7063 | 0.816943 |
| SL1344_RS16360_ | 0.60688 | 0.555301 | 0.66644 | 0.108496 | 0.127169 | 0.824331 | 0.170667 | 0.273866 | 0.421639 | 0.3696 | 0.505338 |
| SL1344_RS16365_ | 1.027897 | 0.701103 | 1.328913 | 0.62779 | 0.103702 | 5.926263 | 4.68E-05 | 0.003079 | 0.04275 | 0.1 | 0.236592 |
| SL1344_RS16370_ | -1.84101 | -1.83719 | -1.8456 | -0.02003 | 0.3621 | -0.05079 | 0.476667 | 0.621249 | 0.741331 | 0.73 | 0.816213 |
| SL1344_RS16375_ | 0.723393 | 0.767647 | 0.665598 | -0.10037 | 0.15281 | -0.61061 | 0.261333 | 0.39035 | 0.542092 | 0.4897 | 0.620688 |

| | | | | | | | | | | | |
|---|---|---|---|---|---|---|---|---|---|---|---|
| SL1344_RS16380_ | -3.28991 | -3.3527 | -3.25283 | 0.093151 | 0.573217 | 0.14375 | 0.442667 | 0.603386 | 0.72643 | 0.7099 | 0.800385 |
| SL1344_RS16385_ | 1.45547 | 1.533263 | 1.227444 | -0.27206 | 0.229607 | -1.06933 | 0.149234 | 0.199227 | 0.37493 | 0.2923 | 0.459348 |
| SL1344_RS16390_ | -0.80156 | -0.81114 | -0.78852 | 0.038046 | 0.289106 | 0.122493 | 0.445037 | 0.642443 | 0.758263 | 0.761 | 0.839059 |
| SL1344_RS16395_ | -3.26011 | -3.46224 | -3.11125 | 0.352037 | 0.589725 | 0.541035 | 0.254497 | 0.384689 | 0.534749 | 0.4994 | 0.621824 |
| SL1344_RS16400_ | -0.96146 | -0.97524 | -0.94353 | 0.041621 | 0.27775 | 0.142966 | 0.435333 | 0.612142 | 0.73351 | 0.7229 | 0.8092 |
| SL1344_RS16405_ | 1.972942 | 1.799172 | 2.131165 | 0.331538 | 0.085427 | 3.908713 | 4.68E-05 | 0.008521 | 0.064982 | 0.1 | 0.236592 |
| SL1344_RS16410_ | 0.321528 | 0.383044 | 0.24909 | -0.13589 | 0.164415 | -0.77834 | 0.203864 | 0.301558 | 0.455794 | 0.3843 | 0.526734 |
| SL1344_RS16415_ | 3.024164 | 3.001562 | 3.043426 | 0.04464 | 0.066552 | 0.609481 | 0.244 | 0.377418 | 0.52955 | 0.4872 | 0.614793 |
| SL1344_RS16420_ | 2.149206 | 2.120583 | 2.161789 | 0.053764 | 0.126675 | 0.372599 | 0.338667 | 0.491901 | 0.644318 | 0.645 | 0.747402 |
| SL1344_RS16425_ | 2.903526 | 2.755827 | 2.981987 | 0.196553 | 0.285786 | 0.719164 | 0.221186 | 0.309486 | 0.496493 | 0.4021 | 0.577717 |
| SL1344_RS16440_ | -0.15211 | -0.12239 | -0.18342 | 0.01956 | 0.313601 | 0.064469 | 0.473333 | 0.73234 | 0.829328 | 0.8767 | 0.920001 |
| SL1344_RS16445_ | 2.769022 | 2.776033 | 2.761579 | -0.01605 | 0.094941 | -0.16681 | 0.426 | 0.662911 | 0.776134 | 0.7816 | 0.853912 |
| SL1344_RS16450_ | 3.380776 | 3.343741 | 3.441007 | 0.111627 | 0.113814 | 1.016825 | 0.11 | 0.180434 | 0.350752 | 0.2303 | 0.389736 |
| SL1344_RS16455_ | 2.044577 | 2.055445 | 2.026845 | -0.02727 | 0.100657 | -0.2601 | 0.388408 | 0.555745 | 0.687712 | 0.6761 | 0.77333 |
| SL1344_RS16460_ | 3.188275 | 3.106204 | 3.278374 | 0.184551 | 0.090618 | 1.998987 | 0.001337 | 0.051478 | 0.167305 | 0.1015 | 0.238589 |
| SL1344_RS16465_ | 1.993115 | 1.730196 | 2.23477 | 0.533642 | 0.139018 | 3.951127 | 4.68E-05 | 0.008304 | 0.068254 | 0.1 | 0.236592 |
| SL1344_RS16470_ | 0.334236 | 0.370966 | 0.236796 | -0.10119 | 0.240384 | -0.34259 | 0.38 | 0.663211 | 0.780358 | 0.7652 | 0.854211 |
| SL1344_RS16480_ | 3.126213 | 3.029625 | 3.22982 | 0.225314 | 0.151213 | 1.590578 | 0.005334 | 0.088404 | 0.232308 | 0.1055 | 0.243741 |
| SL1344_RS16485_ | 3.476011 | 3.220673 | 3.679175 | 0.448271 | 0.080964 | 5.484794 | 4.68E-05 | 0.002661 | 0.043206 | 0.1 | 0.236592 |
| SL1344_RS16490_ | 0.364153 | 0.511305 | 0.173406 | -0.32682 | 0.235245 | -1.35919 | 0.062625 | 0.11797 | 0.267604 | 0.1738 | 0.326767 |
| SL1344_RS16495_ | 0.964969 | 0.921415 | 1.014449 | 0.090998 | 0.123685 | 0.718788 | 0.222 | 0.329806 | 0.480005 | 0.4318 | 0.565429 |
| SL1344_RS16500_ | 0.43963 | -0.07997 | 1.177943 | 1.157989 | 0.343132 | 3.381871 | 4.68E-05 | 0.00828 | 0.069526 | 0.1 | 0.236592 |
| SL1344_RS16505_ | -0.98803 | -1.15975 | -0.84579 | 0.321172 | 0.272913 | 1.101097 | 0.11 | 0.173977 | 0.325165 | 0.2509 | 0.399842 |
| SL1344_RS16510_ | -2.30048 | -2.50564 | -2.09372 | 0.413769 | 0.342789 | 1.144424 | 0.100667 | 0.169044 | 0.30873 | 0.2422 | 0.386032 |
| SL1344_RS16515_ | -0.21925 | -0.17295 | -0.27427 | -0.12252 | 0.284714 | -0.42125 | 0.330667 | 0.516048 | 0.659761 | 0.6318 | 0.741327 |
| SL1344_RS16520_ | 2.570985 | 2.558821 | 2.580931 | 0.096433 | 0.220421 | 0.39921 | 0.334667 | 0.40281 | 0.582867 | 0.6962 | 0.789673 |
| SL1344_RS16525_ | 2.306912 | 2.284909 | 2.406162 | 0.10722 | 0.115985 | 0.764184 | 0.252498 | 0.366024 | 0.534895 | 0.5171 | 0.658741 |
| SL1344_RS16530_ | 1.948164 | 1.363462 | 2.482085 | 1.187765 | 0.193576 | 6.141436 | 4.68E-05 | 0.001116 | 0.033745 | 0.1 | 0.236592 |
| SL1344_RS16535_ | -2.82607 | -2.74607 | -2.95387 | -0.15706 | 0.572669 | -0.26596 | 0.392 | 0.610873 | 0.734698 | 0.723 | 0.813478 |
| SL1344_RS16540_ | -0.07568 | -0.09331 | -0.04014 | 0.034139 | 0.297685 | 0.097382 | 0.460667 | 0.770049 | 0.856426 | 0.8513 | 0.908077 |
| SL1344_RS16545_ | 1.930608 | 1.771574 | 2.11582 | 0.349498 | 0.117296 | 3.018032 | 4.68E-05 | 0.022276 | 0.106709 | 0.1001 | 0.236729 |
| SL1344_RS16550_ | 2.350896 | 2.248743 | 2.550167 | 0.261628 | 0.157365 | 1.529646 | 0.071286 | 0.094723 | 0.238698 | 0.1648 | 0.320184 |
| SL1344_RS16555_ | 1.367817 | 1.218532 | 1.47852 | 0.265361 | 0.131686 | 1.943895 | 0.012667 | 0.056054 | 0.169981 | 0.1125 | 0.252454 |
| SL1344_RS16560_ | 0.560363 | 0.597046 | 0.514357 | -0.08496 | 0.165755 | -0.48103 | 0.291333 | 0.443648 | 0.590045 | 0.5672 | 0.681418 |
| SL1344_RS16565_ | 1.582765 | 1.525077 | 1.694157 | 0.172325 | 0.152494 | 1.056021 | 0.133245 | 0.200967 | 0.368621 | 0.2757 | 0.432755 |
| SL1344_RS16570_ | 1.068416 | 1.028253 | 1.091975 | 0.049427 | 0.172838 | 0.276031 | 0.393071 | 0.640228 | 0.758683 | 0.753 | 0.83459 |
| SL1344_RS16575_ | -1.34929 | -1.49389 | -1.16186 | 0.325885 | 0.280941 | 1.062397 | 0.122585 | 0.19018 | 0.336942 | 0.276 | 0.421114 |
| SL1344_RS16580_ | 2.21428 | 2.093306 | 2.311613 | 0.218024 | 0.074387 | 2.855093 | 0.001336 | 0.020113 | 0.096436 | 0.1009 | 0.237739 |
| SL1344_RS16585_ | 1.215945 | 1.176708 | 1.270271 | 0.101082 | 0.147918 | 0.643505 | 0.231845 | 0.355326 | 0.516833 | 0.4541 | 0.591386 |
| SL1344_RS16590_ | 4.650034 | 5.252866 | 4.149328 | -1.102 | 0.198613 | -5.65643 | 4.68E-05 | 0.005927 | 0.062238 | 0.1 | 0.236592 |
| SL1344_RS16595_ | 3.481008 | 3.621717 | 3.278605 | -0.34041 | 0.080262 | -4.10501 | 4.68E-05 | 0.012286 | 0.080415 | 0.1 | 0.236592 |
| SL1344_RS16600_ | 4.663412 | 4.666541 | 4.661558 | -0.00592 | 0.063842 | -0.09654 | 0.459027 | 0.761807 | 0.851583 | 0.8771 | 0.924352 |
| SL1344_RS16605_ | -0.37326 | -0.84078 | 0.136406 | 0.85881 | 0.391804 | 2.17822 | 0.006663 | 0.02893 | 0.124463 | 0.1043 | 0.242306 |
| SL1344_RS16610_ | -2.71294 | -2.62492 | -2.93476 | -0.26705 | 0.593952 | -0.38974 | 0.361093 | 0.584119 | 0.714049 | 0.6953 | 0.796171 |
| SL1344_RS16615_ | -1.88109 | -1.94499 | -1.84648 | 0.0663 | 0.401433 | 0.14584 | 0.443704 | 0.656538 | 0.771694 | 0.7571 | 0.841024 |

| | | | | | | | | | | |
|---|---|---|---|---|---|---|---|---|---|---|
| SL1344_RS16620_ | 0.122349 | -0.26532 | 0.58259 | 0.825835 | 0.230516 | 3.563417 | 4.68E-05 | 0.011274 | 0.077109 | 0.1 | 0.236592 |
| SL1344_RS16635_ | -0.7604 | -0.98302 | -0.56205 | 0.471396 | 0.363104 | 1.312629 | 0.055963 | 0.124323 | 0.277159 | 0.1634 | 0.311814 |
| SL1344_RS16640_ | 0.33975 | 0.329067 | 0.353973 | 0.030778 | 0.159711 | 0.181112 | 0.427333 | 0.558569 | 0.686348 | 0.6649 | 0.761337 |
| SL1344_RS16645_ | 0.398433 | 0.419252 | 0.366016 | -0.06122 | 0.180205 | -0.30544 | 0.36 | 0.537164 | 0.671786 | 0.6616 | 0.758631 |
| SL1344_RS16650_ | 0.478553 | 0.339187 | 0.55759 | 0.220481 | 0.196225 | 1.053122 | 0.124667 | 0.189322 | 0.349499 | 0.2707 | 0.423946 |
| SL1344_RS16655_ | -2.70589 | -2.47196 | -2.82211 | -0.30909 | 0.578403 | -0.46891 | 0.307333 | 0.515691 | 0.655578 | 0.6055 | 0.72459 |
| SL1344_RS16660_ | 2.067456 | 1.8589 | 2.499816 | 0.622945 | 0.144288 | 3.988194 | 4.68E-05 | 0.01348 | 0.08548 | 0.1 | 0.236592 |
| SL1344_RS16665_ | -0.04407 | -0.10491 | 0.017072 | 0.120713 | 0.172424 | 0.655202 | 0.236667 | 0.351669 | 0.502153 | 0.4616 | 0.592844 |
| SL1344_RS16675_ | 2.105194 | 2.056976 | 2.154257 | 0.11512 | 0.119421 | 0.910063 | 0.145333 | 0.220338 | 0.384858 | 0.3203 | 0.467781 |
| SL1344_RS16680_ | 1.205074 | 1.088389 | 1.29514 | 0.202095 | 0.133061 | 1.433385 | 0.063333 | 0.107952 | 0.246876 | 0.1725 | 0.321731 |
| SL1344_RS16685_ | 2.965417 | 2.848248 | 3.339883 | 0.475607 | 0.262573 | 1.764332 | 0.002667 | 0.056629 | 0.182418 | 0.1039 | 0.241783 |
| SL1344_RS16690_ | 2.953237 | 2.828991 | 3.182018 | 0.332211 | 0.125218 | 2.358795 | 0.001337 | 0.037182 | 0.142799 | 0.1012 | 0.238188 |
| SL1344_RS16695_ | 3.47289 | 3.19783 | 3.730657 | 0.516006 | 0.08366 | 6.110333 | 4.68E-05 | 0.001636 | 0.037121 | 0.1 | 0.236592 |
| SL1344_RS16700_ | 3.544781 | 3.401817 | 3.696473 | 0.255478 | 0.131043 | 1.97611 | 0.009994 | 0.036919 | 0.142776 | 0.1084 | 0.247789 |
| SL1344_RS16705_ | -4.10408 | -4.26013 | -3.98557 | 0.244711 | 0.809673 | 0.283107 | 0.373751 | 0.543448 | 0.677953 | 0.6721 | 0.768151 |
| SL1344_RS16710_ | 0.432447 | 0.064956 | 0.818207 | 0.734896 | 0.190021 | 3.812964 | 4.68E-05 | 0.008013 | 0.066288 | 0.1 | 0.236592 |
| SL1344_RS16715_ | 2.057722 | 1.910422 | 2.215424 | 0.306475 | 0.105014 | 2.86519 | 0.000673 | 0.019225 | 0.097414 | 0.1006 | 0.237384 |
| SL1344_RS16720_ | -0.07243 | -0.27808 | 0.166225 | 0.459222 | 0.218468 | 2.068344 | 0.006667 | 0.04289 | 0.147121 | 0.1083 | 0.246933 |
| SL1344_RS16725_ | -0.74566 | -1.02824 | -0.47819 | 0.557596 | 0.269309 | 2.00704 | 0.008667 | 0.046882 | 0.153818 | 0.1121 | 0.251832 |
| SL1344_RS16730_ | 0.865901 | 0.624354 | 1.144058 | 0.525461 | 0.196633 | 2.610415 | 0.000673 | 0.024619 | 0.113414 | 0.1005 | 0.237259 |
| SL1344_RS16745_ | 0.593887 | 0.841945 | 0.54223 | -0.28056 | 0.279706 | -0.76891 | 0.268667 | 0.366027 | 0.542682 | 0.5309 | 0.679673 |
| SL1344_RS16750_ | 1.375408 | 1.646414 | 1.171838 | -0.3294 | 0.304414 | -1.28513 | 0.110667 | 0.136542 | 0.298227 | 0.2075 | 0.3756 |
| SL1344_RS16755_ | 0.374193 | 0.641273 | 0.24885 | -0.38003 | 0.343844 | -1.015 | 0.146667 | 0.208267 | 0.385291 | 0.2716 | 0.434727 |
| SL1344_RS16775_ | 2.392811 | 2.136609 | 2.713706 | 0.59963 | 0.211384 | 2.811949 | 4.68E-05 | 0.012712 | 0.085084 | 0.1 | 0.236592 |
| SL1344_RS16780_ | 4.118862 | 3.79171 | 4.580526 | 0.742598 | 0.156233 | 4.613791 | 4.68E-05 | 0.004548 | 0.055245 | 0.1 | 0.236592 |
| SL1344_RS16785_ | 3.366751 | 3.138128 | 3.745696 | 0.595928 | 0.115075 | 4.489091 | 4.68E-05 | 0.014402 | 0.089925 | 0.1 | 0.236592 |
| SL1344_RS16790_ | 2.780223 | 2.500908 | 3.213797 | 0.682173 | 0.134678 | 4.76838 | 4.68E-05 | 0.007884 | 0.067603 | 0.1 | 0.236592 |
| SL1344_RS16795_ | 1.430931 | 1.222728 | 1.678289 | 0.450129 | 0.141353 | 3.117794 | 4.68E-05 | 0.015723 | 0.089904 | 0.1001 | 0.23672 |
| SL1344_RS16800_ | 1.833957 | 1.800369 | 1.978876 | 0.16908 | 0.182203 | 0.722119 | 0.282478 | 0.384285 | 0.559543 | 0.5583 | 0.698915 |
| SL1344_RS16805_ | -1.77286 | -1.89497 | -1.60974 | 0.320057 | 0.508171 | 0.592847 | 0.242505 | 0.376488 | 0.542413 | 0.468 | 0.61098 |
| SL1344_RS16810_ | -2.42357 | -2.70562 | -2.14906 | 0.555714 | 0.379068 | 1.433472 | 0.069333 | 0.113625 | 0.246421 | 0.1874 | 0.332945 |
| SL1344_RS16815_ | -2.06095 | -2.44915 | -1.81376 | 0.62264 | 0.399035 | 1.527787 | 0.052 | 0.100073 | 0.23463 | 0.1674 | 0.314244 |
| SL1344_RS16820_ | -2.88215 | -3.22366 | -2.62314 | 0.592989 | 0.515885 | 1.079898 | 0.113333 | 0.181628 | 0.333122 | 0.261 | 0.410756 |
| SL1344_RS16825_ | -2.85028 | -2.72738 | -3.05813 | -0.26892 | 0.630476 | -0.40867 | 0.347102 | 0.557026 | 0.692111 | 0.6749 | 0.779561 |
| SL1344_RS16830_ | -4.94383 | -4.61019 | -5.33647 | -0.70889 | 0.991053 | -0.65047 | 0.212667 | 0.340915 | 0.493621 | 0.4377 | 0.568813 |
| SL1344_RS16835_ | -3.6417 | -4.08282 | -3.26961 | 0.830134 | 0.594671 | 1.321164 | 0.075949 | 0.128115 | 0.26585 | 0.1995 | 0.346684 |
| SL1344_RS16840_ | -1.28495 | -1.35648 | -1.22383 | 0.215991 | 0.484016 | 0.433279 | 0.317788 | 0.475401 | 0.631937 | 0.6378 | 0.744851 |
| SL1344_RS16845_ | -3.64104 | -3.53877 | -3.69616 | -0.09808 | 0.869218 | -0.1032 | 0.454 | 0.68796 | 0.795312 | 0.7857 | 0.860539 |
| SL1344_RS16850_ | 1.692463 | 1.573575 | 1.816187 | 0.245437 | 0.105149 | 2.31806 | 0.002002 | 0.032468 | 0.12561 | 0.1027 | 0.240102 |
| SL1344_RS16855_ | 0.592439 | 0.62631 | 0.558917 | -0.06811 | 0.13528 | -0.45225 | 0.302 | 0.429796 | 0.571963 | 0.5359 | 0.653639 |
| SL1344_RS16860_ | 3.029362 | 3.020936 | 3.038652 | 0.008633 | 0.085199 | 0.088895 | 0.463691 | 0.68324 | 0.792405 | 0.7954 | 0.864816 |
| SL1344_RS16865_ | 1.942878 | 1.862943 | 2.019373 | 0.156007 | 0.092926 | 1.642654 | 0.041333 | 0.077491 | 0.200621 | 0.1429 | 0.286909 |
| SL1344_RS16870_ | 1.257656 | 1.250855 | 1.263777 | 0.023906 | 0.149366 | 0.155481 | 0.433711 | 0.640984 | 0.757241 | 0.7609 | 0.837215 |
| SL1344_RS16875_ | 2.713013 | 2.707957 | 2.720827 | 0.012132 | 0.086009 | 0.132908 | 0.447035 | 0.650806 | 0.766813 | 0.761 | 0.839737 |
| SL1344_RS16880_ | -1.14956 | -1.23202 | -1.04376 | 0.211388 | 0.27617 | 0.703625 | 0.217189 | 0.328275 | 0.481947 | 0.4437 | 0.574485 |

| | | | | | | | | | | | |
|---|---|---|---|---|---|---|---|---|---|---|---|
| SL1344_RS16885_ | -0.38482 | -0.71678 | -0.0495 | 0.671723 | 0.199094 | 3.351682 | 0.000673 | 0.013029 | 0.079199 | 0.1002 | 0.23686 |
| SL1344_RS16890_ | -1.13551 | -0.91538 | -1.21575 | -0.25823 | 0.348604 | -0.64995 | 0.267821 | 0.437968 | 0.591744 | 0.5512 | 0.684477 |
| SL1344_RS16895_ | -1.06611 | -1.00198 | -1.12269 | -0.0457 | 0.497651 | -0.08499 | 0.46036 | 0.779384 | 0.863102 | 0.842 | 0.901664 |
| SL1344_RS16900_ | 2.868213 | 2.611177 | 3.184919 | 0.521791 | 0.176654 | 2.86694 | 4.68E-05 | 0.016567 | 0.09577 | 0.1 | 0.236592 |
| SL1344_RS16905_ | 2.93426 | 2.597293 | 3.053908 | 0.439308 | 0.17661 | 2.123888 | 0.000673 | 0.043086 | 0.156199 | 0.1003 | 0.236991 |
| SL1344_RS16910_ | 1.149472 | 1.269627 | 1.038218 | -0.25298 | 0.181026 | -1.37347 | 0.039307 | 0.111559 | 0.259299 | 0.1443 | 0.290778 |
| SL1344_RS16915_ | 1.649327 | 1.601353 | 1.751594 | 0.137811 | 0.163951 | 0.804851 | 0.204 | 0.29873 | 0.470947 | 0.4073 | 0.562096 |
| SL1344_RS16920_ | 0.397555 | 0.347281 | 0.525841 | 0.167093 | 0.182949 | 0.789261 | 0.214667 | 0.337762 | 0.496324 | 0.452 | 0.594565 |
| SL1344_RS16925_ | -3.52293 | -3.51059 | -3.54182 | -0.14531 | 0.804656 | -0.15853 | 0.430667 | 0.610299 | 0.735817 | 0.7413 | 0.82245 |
| SL1344_RS16930_ | -4.54213 | -4.49259 | -4.6453 | -0.12945 | 0.961831 | -0.12751 | 0.445703 | 0.604543 | 0.728422 | 0.733 | 0.818832 |
| SL1344_RS16935_ | -1.03127 | -0.99545 | -1.06162 | -0.0663 | 0.241753 | -0.26133 | 0.4004 | 0.529097 | 0.66225 | 0.6482 | 0.747034 |
| SL1344_RS16940_ | 1.550059 | 1.825 | 1.080158 | -0.73978 | 0.241239 | -2.98048 | 4.68E-05 | 0.035439 | 0.140485 | 0.1 | 0.236592 |
| SL1344_RS16945_ | 5.769176 | 5.777793 | 5.763633 | -0.01331 | 0.068019 | -0.15697 | 0.436667 | 0.851291 | 0.912476 | 0.8929 | 0.936187 |
| SL1344_RS16950_ | 4.851107 | 4.854115 | 4.846292 | -0.04955 | 0.115121 | -0.42326 | 0.341333 | 0.447563 | 0.62003 | 0.735 | 0.823787 |
| SL1344_RS16955_ | 5.178599 | 5.479933 | 4.953875 | -0.52127 | 0.144122 | -3.66799 | 4.68E-05 | 0.007275 | 0.067437 | 0.1 | 0.236592 |
| SL1344_RS16960_ | 2.462722 | 2.654576 | 2.375556 | -0.26913 | 0.105831 | -2.30376 | 0.003335 | 0.047798 | 0.160518 | 0.1037 | 0.241344 |
| SL1344_RS16965_ | 1.610244 | 1.729269 | 1.569098 | -0.1334 | 0.15083 | -0.8176 | 0.221852 | 0.323489 | 0.492808 | 0.4505 | 0.599054 |
| SL1344_RS16970_ | 5.277505 | 5.364809 | 5.269922 | -0.09566 | 0.093567 | -0.80417 | 0.306 | 0.332556 | 0.518644 | 0.6336 | 0.765778 |
| SL1344_RS16980_ | 1.588484 | 1.826984 | 1.410687 | -0.4149 | 0.103718 | -3.98145 | 4.68E-05 | 0.009462 | 0.068822 | 0.1 | 0.236592 |
| SL1344_RS16985_ | 1.727413 | 1.857939 | 1.608842 | -0.26533 | 0.247447 | -1.0868 | 0.083333 | 0.183135 | 0.35887 | 0.1837 | 0.341969 |
| SL1344_RS16990_ | -0.25665 | -0.28012 | -0.23089 | 0.047196 | 0.207677 | 0.203825 | 0.41 | 0.586657 | 0.709884 | 0.6921 | 0.784119 |
| SL1344_RS16995_ | 0.47974 | 0.769243 | 0.162201 | -0.60779 | 0.140867 | -4.27564 | 4.68E-05 | 0.006601 | 0.058435 | 0.1 | 0.236592 |
| SL1344_RS17000_ | 1.817811 | 1.854015 | 1.736289 | -0.15306 | 0.331904 | -0.49977 | 0.262492 | 0.467356 | 0.63659 | 0.5026 | 0.66125 |
| SL1344_RS17005_ | 3.944752 | 4.027828 | 3.893218 | -0.14732 | 0.09053 | -1.56913 | 0.025334 | 0.078396 | 0.213191 | 0.1288 | 0.272657 |
| SL1344_RS17010_ | 2.829619 | 2.691467 | 2.960293 | 0.271691 | 0.127519 | 2.097063 | 0.004001 | 0.036944 | 0.141303 | 0.1036 | 0.241351 |
| SL1344_RS17015_ | 0.475209 | 0.504572 | 0.439092 | -0.09601 | 0.225432 | -0.42047 | 0.312667 | 0.491765 | 0.639121 | 0.6492 | 0.748441 |
| SL1344_RS17030_ | 3.104131 | 3.074265 | 3.145744 | 0.062315 | 0.16636 | 0.318335 | 0.402398 | 0.677809 | 0.79604 | 0.8778 | 0.924743 |
| SL1344_RS17035_ | 1.546588 | 2.28707 | 1.229254 | -1.05863 | 0.266979 | -3.38739 | 4.68E-05 | 0.031257 | 0.133285 | 0.1 | 0.236592 |
| SL1344_RS17040_ | 2.228319 | 2.435285 | 1.974864 | -0.4551 | 0.098389 | -4.54937 | 4.68E-05 | 0.005727 | 0.056431 | 0.1 | 0.236592 |
| SL1344_RS17045_ | 0.758837 | 1.16913 | 0.583577 | -0.56024 | 0.263264 | -2.00172 | 0.007995 | 0.046295 | 0.159497 | 0.1079 | 0.246873 |
| SL1344_RS17050_ | 1.547686 | 1.640799 | 1.49713 | -0.11391 | 0.197795 | -0.52304 | 0.312 | 0.481014 | 0.639421 | 0.6263 | 0.744437 |
| SL1344_RS17060_ | 0.539998 | 0.643687 | 0.33322 | -0.30487 | 0.174147 | -1.66386 | 0.04064 | 0.083552 | 0.213984 | 0.1441 | 0.288437 |
| SL1344_RS17065_ | 3.313238 | 3.495732 | 3.226277 | -0.26045 | 0.114095 | -2.24477 | 0.002002 | 0.029439 | 0.125189 | 0.1012 | 0.2382 |
| SL1344_RS17070_ | 3.677624 | 4.022115 | 3.497478 | -0.51506 | 0.085486 | -5.26808 | 4.68E-05 | 0.013358 | 0.08644 | 0.1 | 0.236592 |
| SL1344_RS17075_ | 3.222451 | 3.232894 | 3.199196 | -0.03062 | 0.079868 | -0.34384 | 0.363333 | 0.600094 | 0.728585 | 0.7273 | 0.818415 |
| SL1344_RS17080_ | -3.14819 | -2.94468 | -3.32912 | -0.40332 | 0.601264 | -0.59933 | 0.233178 | 0.368561 | 0.524687 | 0.4714 | 0.605496 |
| SL1344_RS17085_ | 2.558347 | 2.592847 | 2.522738 | -0.06814 | 0.073722 | -0.86969 | 0.177333 | 0.268697 | 0.422894 | 0.3575 | 0.501093 |
| SL1344_RS17090_ | -0.92405 | -0.81777 | -1.11544 | -0.28045 | 0.301454 | -0.85856 | 0.176549 | 0.272634 | 0.435607 | 0.3722 | 0.521896 |
| SL1344_RS17095_ | 2.359848 | 2.306611 | 2.45053 | 0.143392 | 0.093086 | 1.48499 | 0.057961 | 0.10652 | 0.246167 | 0.1653 | 0.31188 |
| SL1344_RS17100_ | 2.164177 | 2.068027 | 2.274822 | 0.219148 | 0.113499 | 1.88401 | 0.007329 | 0.050991 | 0.162747 | 0.1089 | 0.247827 |
| SL1344_RS17105_ | 0.728251 | 0.582595 | 0.825797 | 0.249991 | 0.167074 | 1.466354 | 0.036642 | 0.109704 | 0.250801 | 0.1505 | 0.294816 |
| SL1344_RS17110_ | 0.74768 | 0.57055 | 0.78284 | 0.190555 | 0.210037 | 0.677836 | 0.300667 | 0.447285 | 0.610181 | 0.618 | 0.748733 |
| SL1344_RS17115_ | 1.185528 | 1.263971 | 1.075085 | -0.1546 | 0.184459 | -0.80898 | 0.193871 | 0.295555 | 0.466227 | 0.393 | 0.547414 |
| SL1344_RS17120_ | 1.69299 | 1.749632 | 1.668304 | -0.08835 | 0.177331 | -0.45901 | 0.33511 | 0.516031 | 0.666304 | 0.662 | 0.771551 |
| SL1344_RS17125_ | 2.186736 | 2.114079 | 2.22477 | 0.113279 | 0.117238 | 0.899367 | 0.175333 | 0.248356 | 0.417802 | 0.3408 | 0.491513 |

| | | | | | | | | | | | |
|---|---|---|---|---|---|---|---|---|---|---|---|
| SL1344_RS17130_ | 2.292636 | 2.246393 | 2.368114 | 0.08789 | 0.160178 | 0.524268 | 0.291333 | 0.487533 | 0.643872 | 0.5807 | 0.711366 |
| SL1344_RS17135_ | 2.023188 | 2.033533 | 2.018385 | -0.01202 | 0.091408 | -0.12436 | 0.445037 | 0.608396 | 0.728855 | 0.7086 | 0.800011 |
| SL1344_RS17140_ | 2.434348 | 2.408966 | 2.460665 | 0.047591 | 0.116185 | 0.397323 | 0.336667 | 0.547425 | 0.6868 | 0.6554 | 0.762017 |
| SL1344_RS17145_ | 2.298956 | 2.247136 | 2.372042 | 0.121568 | 0.077169 | 1.502119 | 0.051299 | 0.10075 | 0.229866 | 0.1748 | 0.319399 |
| SL1344_RS17150_ | 3.621406 | 3.595747 | 3.683684 | 0.089396 | 0.077449 | 1.067521 | 0.116667 | 0.193723 | 0.36203 | 0.2666 | 0.419958 |
| SL1344_RS17155_ | 4.291699 | 4.11779 | 4.473066 | 0.366289 | 0.104986 | 3.530123 | 4.68E-05 | 0.020728 | 0.10756 | 0.1 | 0.236592 |
| SL1344_RS17160_ | 2.621668 | 2.535627 | 2.71732 | 0.186396 | 0.081003 | 2.26305 | 0.002001 | 0.037853 | 0.136884 | 0.1025 | 0.239765 |
| SL1344_RS17165_ | 3.251069 | 3.131507 | 3.322015 | 0.185449 | 0.06715 | 2.655565 | 0.002667 | 0.029131 | 0.119572 | 0.1025 | 0.239812 |
| SL1344_RS17170_ | 0.982938 | 0.938545 | 1.047061 | 0.104149 | 0.120663 | 0.794767 | 0.182545 | 0.298237 | 0.448286 | 0.3943 | 0.531546 |
| SL1344_RS17175_ | -1.63169 | -2.12364 | -1.24846 | 0.871678 | 0.290732 | 3.029292 | 0.000673 | 0.01759 | 0.09053 | 0.1006 | 0.237396 |
| SL1344_RS17180_ | 2.238197 | 2.592118 | 2.202289 | -0.36308 | 0.185823 | -1.00268 | 0.202667 | 0.213818 | 0.394447 | 0.4059 | 0.569669 |
| SL1344_RS17185_ | 2.591416 | 2.719088 | 2.424232 | -0.30101 | 0.245382 | -1.2185 | 0.087333 | 0.129741 | 0.29192 | 0.1872 | 0.350351 |
| SL1344_RS17190_ | 2.857385 | 2.819736 | 2.899068 | 0.047888 | 0.18416 | 0.237214 | 0.415057 | 0.823642 | 0.894345 | 0.8923 | 0.935564 |
| SL1344_RS17195_ | -0.64734 | -0.41445 | -0.92241 | -0.51073 | 0.197633 | -2.50448 | 0.002668 | 0.026301 | 0.110089 | 0.1033 | 0.240817 |
| SL1344_RS17200_ | -4.97539 | -4.64533 | -5.50117 | -0.85298 | 1.103215 | -0.70793 | 0.188541 | 0.310501 | 0.471253 | 0.4048 | 0.5411 |
| SL1344_RS17205_ | 1.28139 | 1.172658 | 1.396351 | 0.221988 | 0.10018 | 2.187447 | 0.008001 | 0.038136 | 0.133802 | 0.1085 | 0.24702 |
| SL1344_RS17210_ | 0.134864 | 0.020835 | 0.229887 | 0.220504 | 0.247317 | 0.890936 | 0.165889 | 0.246251 | 0.414366 | 0.3374 | 0.491555 |
| SL1344_RS17215_ | -2.8719 | -2.96712 | -2.72904 | 0.190313 | 0.663596 | 0.255362 | 0.388 | 0.657541 | 0.771202 | 0.7274 | 0.819652 |
| SL1344_RS17220_ | -1.29244 | -1.23239 | -1.4511 | -0.17452 | 0.394125 | -0.40304 | 0.347102 | 0.590365 | 0.721699 | 0.6934 | 0.798362 |
| SL1344_RS17225_ | 0.800344 | 0.771932 | 0.816281 | 0.041632 | 0.1442 | 0.265911 | 0.38974 | 0.573149 | 0.7026 | 0.6776 | 0.777095 |
| SL1344_RS17230_ | 0.993955 | 0.763483 | 1.231493 | 0.412089 | 0.209288 | 1.919676 | 0.022667 | 0.047647 | 0.160136 | 0.1213 | 0.26471 |
| SL1344_RS17235_ | 0.287492 | 0.141649 | 0.545289 | 0.391384 | 0.169237 | 2.188989 | 0.011992 | 0.043622 | 0.147498 | 0.1111 | 0.250371 |
| SL1344_RS17240_ | 0.239016 | 0.034957 | 0.460839 | 0.419594 | 0.199992 | 2.070593 | 0.009994 | 0.042513 | 0.147343 | 0.1109 | 0.250643 |
| SL1344_RS17245_ | 1.291354 | 1.160302 | 1.402408 | 0.263694 | 0.158345 | 1.613727 | 0.016667 | 0.082154 | 0.214748 | 0.1209 | 0.261959 |
| SL1344_RS17250_ | 1.126385 | 0.85654 | 1.317075 | 0.449594 | 0.191194 | 2.286166 | 0.001337 | 0.027441 | 0.119673 | 0.1021 | 0.239393 |
| SL1344_RS17255_ | 2.333603 | 2.329941 | 2.339631 | 0.025467 | 0.115824 | 0.209507 | 0.406 | 0.609043 | 0.735426 | 0.758 | 0.835854 |
| SL1344_RS17260_ | 3.927002 | 3.986679 | 3.845058 | -0.1501 | 0.073921 | -1.87132 | 0.000673 | 0.11624 | 0.273552 | 0.1003 | 0.236989 |
| SL1344_RS17270_ | -0.13994 | -0.39966 | 0.076852 | 0.477021 | 0.169185 | 2.756324 | 0.001337 | 0.022108 | 0.101452 | 0.1013 | 0.238334 |
| SL1344_RS17275_ | 5.339691 | 5.626609 | 5.030355 | -0.59664 | 0.067461 | -8.79019 | 4.68E-05 | 0.002542 | 0.04386 | 0.1 | 0.236592 |
| SL1344_RS17280_ | 4.255321 | 4.539122 | 3.814204 | -0.70496 | 0.095854 | -6.6219 | 4.68E-05 | 0.006589 | 0.063761 | 0.1 | 0.236592 |
| SL1344_RS17285_ | 1.224386 | 1.193074 | 1.252802 | 0.059474 | 0.112942 | 0.52943 | 0.284667 | 0.421862 | 0.564782 | 0.5373 | 0.652875 |
| SL1344_RS17290_ | 3.305156 | 3.234184 | 3.464908 | 0.212456 | 0.102617 | 1.88758 | 0.011334 | 0.054892 | 0.174776 | 0.1148 | 0.255735 |
| SL1344_RS17295_ | 3.19794 | 2.889357 | 3.427159 | 0.525106 | 0.13289 | 3.966958 | 4.68E-05 | 0.005879 | 0.059862 | 0.1 | 0.236592 |
| SL1344_RS17300_ | 1.641905 | 1.594916 | 1.702082 | 0.125667 | 0.15434 | 0.799467 | 0.18521 | 0.277122 | 0.44848 | 0.3856 | 0.530301 |
| SL1344_RS17305_ | -2.66637 | -2.69342 | -2.64944 | 0.034 | 0.538186 | 0.054506 | 0.477015 | 0.654401 | 0.76932 | 0.7567 | 0.841093 |
| SL1344_RS17310_ | -1.80771 | -1.86152 | -1.72289 | 0.124739 | 0.346757 | 0.33925 | 0.35443 | 0.548603 | 0.678005 | 0.6456 | 0.750365 |
| SL1344_RS17315_ | -0.98445 | -0.79677 | -1.2937 | -0.49442 | 0.30867 | -1.53096 | 0.041972 | 0.094245 | 0.230018 | 0.1523 | 0.297768 |
| SL1344_RS17320_ | -4.67985 | -4.65942 | -4.71193 | -0.05948 | 1.052631 | -0.05315 | 0.472 | 0.622139 | 0.74311 | 0.7364 | 0.82073 |
| SL1344_RS17325_ | -1.98875 | -2.17921 | -1.87797 | 0.29084 | 0.423742 | 0.62672 | 0.247835 | 0.376657 | 0.535039 | 0.4928 | 0.624296 |
| SL1344_RS17330_ | -2.17792 | -2.162 | -2.19618 | -0.02782 | 0.348151 | -0.07568 | 0.467688 | 0.566364 | 0.692094 | 0.6752 | 0.769071 |
| SL1344_RS17335_ | -3.03053 | -3.12848 | -2.88486 | 0.221033 | 0.584994 | 0.32273 | 0.351765 | 0.566565 | 0.694531 | 0.6637 | 0.765234 |
| SL1344_RS17340_ | 0.360805 | 0.309283 | 0.489276 | 0.159243 | 0.238021 | 0.579681 | 0.289807 | 0.459713 | 0.616951 | 0.5949 | 0.718607 |
| SL1344_RS17345_ | 0.605224 | 0.652986 | 0.475104 | -0.16979 | 0.189766 | -0.79959 | 0.219853 | 0.31472 | 0.481794 | 0.4365 | 0.580562 |
| SL1344_RS17350_ | 5.053947 | 5.059218 | 5.044154 | -0.02343 | 0.142952 | -0.18983 | 0.396 | 0.631621 | 0.767148 | 0.8104 | 0.879594 |
| SL1344_RS17355_ | 2.058984 | 2.067955 | 2.047956 | -0.02749 | 0.120661 | -0.231 | 0.399734 | 0.631266 | 0.75154 | 0.7359 | 0.820616 |

| | | | | | | | | | | |
|---|---|---|---|---|---|---|---|---|---|---|
| SL1344_RS17360_ | -2.99359 | -3.05688 | -2.91415 | 0.298827 | 0.903688 | 0.321745 | 0.367755 | 0.586904 | 0.719893 | 0.7344 | 0.8203 |
| SL1344_RS17365_ | -0.8286 | -0.65607 | -1.06233 | -0.40445 | 0.242534 | -1.65479 | 0.037309 | 0.082786 | 0.208143 | 0.1482 | 0.292467 |
| SL1344_RS17370_ | -0.69283 | -1.23408 | -0.24122 | 1.031348 | 0.313394 | 3.362236 | 4.68E-05 | 0.010849 | 0.075797 | 0.1 | 0.236592 |
| SL1344_RS17375_ | -0.72642 | -0.74267 | -0.71238 | 0.020643 | 0.250285 | 0.079168 | 0.46569 | 0.624051 | 0.743285 | 0.7335 | 0.817614 |
| SL1344_RS17380_ | -1.58758 | -1.58629 | -1.59067 | -0.0045 | 0.265587 | -0.01183 | 0.496336 | 0.529107 | 0.659361 | 0.6342 | 0.734984 |
| SL1344_RS17385_ | -5.11683 | -4.86496 | -5.46726 | -0.73229 | 1.584422 | -0.40653 | 0.303333 | 0.456828 | 0.61411 | 0.608 | 0.71734 |
| SL1344_RS17390_ | -0.0014 | -0.04785 | 0.062766 | 0.121468 | 0.206517 | 0.543064 | 0.271153 | 0.418035 | 0.570082 | 0.544 | 0.663851 |
| SL1344_RS17395_ | 1.896675 | 1.67907 | 2.102558 | 0.425537 | 0.097059 | 4.344472 | 4.68E-05 | 0.006304 | 0.058089 | 0.1 | 0.236592 |
| SL1344_RS17400_ | 2.974121 | 2.850003 | 3.075975 | 0.230905 | 0.078701 | 2.943563 | 4.68E-05 | 0.023594 | 0.108145 | 0.1004 | 0.237127 |
| SL1344_RS17405_ | 2.849554 | 2.801562 | 2.874321 | 0.093216 | 0.141165 | 0.665546 | 0.227182 | 0.321963 | 0.505072 | 0.4464 | 0.592092 |
| SL1344_RS17410_ | 1.770225 | 1.788574 | 1.738892 | -0.06352 | 0.133077 | -0.44824 | 0.305796 | 0.434387 | 0.593664 | 0.5841 | 0.694839 |
| SL1344_RS17415_ | 1.364959 | 1.338994 | 1.39325 | 0.083704 | 0.155803 | 0.512616 | 0.290473 | 0.429019 | 0.584581 | 0.5907 | 0.7004 |
| SL1344_RS17420_ | 2.235844 | 2.279739 | 2.169576 | -0.10399 | 0.129915 | -0.79545 | 0.192 | 0.290105 | 0.459837 | 0.3844 | 0.541714 |
| SL1344_RS17425_ | 3.879261 | 4.058686 | 3.670498 | -0.40069 | 0.079291 | -5.14758 | 4.68E-05 | 0.00401 | 0.050642 | 0.1 | 0.236592 |
| SL1344_RS17430_ | -6.41304 | -6.26623 | -6.59291 | -0.29339 | 1.395167 | -0.18995 | 0.403065 | 0.496198 | 0.629421 | 0.5968 | 0.703586 |
| SL1344_RS17440_ | 2.502016 | 2.359815 | 2.640873 | 0.266423 | 0.136455 | 1.931852 | 0.013991 | 0.046955 | 0.159671 | 0.1122 | 0.252738 |
| SL1344_RS17445_ | 0.706642 | 0.617306 | 0.783596 | 0.167545 | 0.126921 | 1.285176 | 0.081279 | 0.145331 | 0.28286 | 0.2231 | 0.367408 |
| SL1344_RS17450_ | -0.25186 | -0.29195 | -0.18895 | 0.129395 | 0.246371 | 0.493694 | 0.302465 | 0.435668 | 0.586546 | 0.5717 | 0.684664 |
| SL1344_RS17455_ | 2.890778 | 2.933493 | 2.696061 | -0.23516 | 0.133007 | -1.37599 | 0.079281 | 0.133255 | 0.294314 | 0.1898 | 0.341798 |
| SL1344_RS17460_ | 4.137742 | 4.129634 | 4.151577 | 0.030801 | 0.068393 | 0.412518 | 0.32445 | 0.449406 | 0.611813 | 0.6293 | 0.734341 |
| SL1344_RS17465_ | -3.65405 | -3.15044 | -4.24113 | -1.04547 | 0.690196 | -1.44469 | 0.055297 | 0.104082 | 0.240143 | 0.1624 | 0.311126 |
| SL1344_RS17470_ | 0.169933 | 0.275784 | 0.04318 | -0.24036 | 0.183857 | -1.25792 | 0.078667 | 0.136918 | 0.283662 | 0.1999 | 0.350034 |
| SL1344_RS17475_ | 0.873345 | 0.914875 | 0.799613 | -0.11863 | 0.170503 | -0.64206 | 0.253831 | 0.393483 | 0.5525 | 0.5165 | 0.648064 |
| SL1344_RS17480_ | 2.387739 | 2.661294 | 1.937717 | -0.72171 | 0.215052 | -3.42917 | 4.68E-05 | 0.006092 | 0.061509 | 0.1 | 0.236592 |
| SL1344_RS17490_ | -0.83631 | -0.76641 | -1.00479 | -0.20968 | 0.302545 | -0.65526 | 0.250667 | 0.398558 | 0.555421 | 0.5151 | 0.649926 |
| SL1344_RS17495_ | -0.10545 | 0.141203 | -0.24678 | -0.37449 | 0.252826 | -1.41127 | 0.062667 | 0.105154 | 0.246966 | 0.1791 | 0.331655 |
| SL1344_RS17500_ | 0.237707 | 0.177338 | 0.270423 | 0.071508 | 0.198928 | 0.335337 | 0.371333 | 0.595023 | 0.719414 | 0.6993 | 0.795105 |
| SL1344_RS17505_ | -3.40294 | -3.22064 | -3.59808 | -0.36702 | 0.532319 | -0.68908 | 0.228667 | 0.337347 | 0.48443 | 0.4382 | 0.567007 |
| SL1344_RS17510_ | -3.70231 | -3.76055 | -3.62445 | 0.115981 | 0.817041 | 0.131493 | 0.440373 | 0.647639 | 0.76273 | 0.7472 | 0.829434 |
| SL1344_RS17515_ | -0.49898 | -0.48781 | -0.51049 | -0.02488 | 0.227074 | -0.10985 | 0.450366 | 0.607758 | 0.728697 | 0.7128 | 0.801395 |
| SL1344_RS17520_ | -1.07102 | -1.21158 | -0.93547 | 0.283297 | 0.266389 | 1.033372 | 0.11992 | 0.201307 | 0.351967 | 0.2847 | 0.427674 |
| SL1344_RS17530_ | -0.86456 | -1.02218 | -0.77854 | 0.272597 | 0.535514 | 0.511708 | 0.279813 | 0.448288 | 0.612777 | 0.5416 | 0.684055 |
| SL1344_RS17540_ | 3.061925 | 2.616389 | 3.179974 | 0.562431 | 0.66099 | 0.741345 | 0.218521 | 0.335919 | 0.525193 | 0.4027 | 0.58451 |
| SL1344_RS17545_ | -2.54837 | -2.27086 | -2.77344 | -0.48803 | 0.568008 | -0.81981 | 0.167888 | 0.268228 | 0.432433 | 0.3529 | 0.504493 |
| SL1344_RS17550_ | 0.69488 | 0.476856 | 0.842867 | 0.366004 | 0.969117 | 0.476353 | 0.261159 | 0.478039 | 0.65022 | 0.5008 | 0.662752 |
| SL1344_RS17555_ | 1.248361 | 1.252503 | 1.244405 | -0.00907 | 0.160341 | -0.05546 | 0.469687 | 0.694014 | 0.799111 | 0.7841 | 0.856692 |
| SL1344_RS17560_ | 0.682438 | 0.378757 | 0.816297 | 0.385412 | 0.241872 | 1.480261 | 0.065956 | 0.101581 | 0.24718 | 0.1688 | 0.322375 |
| SL1344_RS17565_ | 1.53173 | 1.437516 | 1.640531 | 0.206712 | 0.108371 | 1.814806 | 0.022667 | 0.063646 | 0.179565 | 0.1284 | 0.270759 |
| SL1344_RS17570_ | 0.438731 | 0.234724 | 0.488371 | 0.213804 | 0.262876 | 0.646697 | 0.294667 | 0.437594 | 0.60303 | 0.595 | 0.727948 |
| SL1344_RS17575_ | 3.42447 | 3.29236 | 3.652003 | 0.366375 | 0.126322 | 2.884967 | 4.68E-05 | 0.023956 | 0.114634 | 0.1 | 0.236592 |
| SL1344_RS17580_ | 4.009847 | 3.937414 | 4.304709 | 0.340576 | 0.25041 | 1.380198 | 0.095333 | 0.100433 | 0.252734 | 0.1824 | 0.344295 |
| SL1344_RS17585_ | 1.822146 | 1.523445 | 2.004091 | 0.471419 | 0.122003 | 3.659555 | 4.68E-05 | 0.01474 | 0.086479 | 0.1 | 0.236592 |
| SL1344_RS17590_ | 2.656009 | 2.562193 | 2.80052 | 0.232741 | 0.087404 | 2.583313 | 0.001337 | 0.027075 | 0.116432 | 0.1022 | 0.239407 |
| SL1344_RS17595_ | 2.623513 | 2.636871 | 2.608597 | -0.03158 | 0.077834 | -0.37414 | 0.346436 | 0.514323 | 0.649104 | 0.6252 | 0.728271 |
| SL1344_RS17600_ | -0.08098 | -0.18244 | 0.022606 | 0.205507 | 0.176859 | 1.122586 | 0.111259 | 0.178708 | 0.324642 | 0.2593 | 0.405763 |

| | | | | | | | | | | |
|---|---|---|---|---|---|---|---|---|---|---|
| SL1344_RS17605_ | 0.892009 | 0.729861 | 1.037415 | 0.302009 | 0.162097 | 1.765857 | 0.022667 | 0.062159 | 0.181243 | 0.1229 | 0.265495 |
| SL1344_RS17610_ | 3.066872 | 2.825767 | 3.216786 | 0.386317 | 0.096147 | 3.93441 | 4.68E-05 | 0.009095 | 0.070302 | 0.1 | 0.236592 |
| SL1344_RS17615_ | 0.436305 | 0.424523 | 0.451819 | 0.021713 | 0.382605 | 0.058013 | 0.462667 | 0.809675 | 0.884475 | 0.8251 | 0.888606 |
| SL1344_RS17620_ | 1.143197 | 1.100984 | 1.266283 | 0.152501 | 0.177373 | 0.759584 | 0.244504 | 0.350945 | 0.520897 | 0.4863 | 0.628694 |
| SL1344_RS17625_ | 0.123306 | 0.101604 | 0.158259 | 0.044143 | 0.232376 | 0.181786 | 0.43038 | 0.673531 | 0.782736 | 0.7745 | 0.850411 |
| SL1344_RS17630_ | 5.519471 | 5.531418 | 5.318049 | -0.21189 | 0.13483 | -0.95577 | 0.285143 | 0.249189 | 0.436656 | 0.5716 | 0.715812 |
| SL1344_RS17635_ | 6.751217 | 6.827318 | 6.635524 | -0.15142 | 0.175322 | -0.86869 | 0.133245 | 0.237953 | 0.42466 | 0.236 | 0.408244 |
| SL1344_RS17640_ | 6.163917 | 6.221502 | 6.108799 | -0.11323 | 0.114179 | -0.92421 | 0.124667 | 0.222223 | 0.406268 | 0.2311 | 0.402383 |
| SL1344_RS17645_ | 3.949009 | 3.941639 | 3.958594 | 0.131401 | 0.277364 | 0.401416 | 0.348 | 0.396945 | 0.580429 | 0.7337 | 0.828774 |
| SL1344_RS17650_ | 4.756035 | 4.835962 | 4.681649 | -0.0943 | 0.152193 | -0.76676 | 0.188541 | 0.312975 | 0.499531 | 0.3357 | 0.505606 |
| SL1344_RS17655_ | 5.131841 | 5.201537 | 5.111559 | -0.08084 | 0.15373 | -0.57293 | 0.242667 | 0.456559 | 0.630182 | 0.4453 | 0.61856 |
| SL1344_RS17660_ | 6.797248 | 6.844554 | 6.734433 | -0.03089 | 0.192857 | -0.22701 | 0.372418 | 0.669725 | 0.796148 | 0.8149 | 0.888049 |
| SL1344_RS17665_ | 4.996046 | 4.987783 | 5.001504 | 0.016181 | 0.197808 | 0.083143 | 0.450366 | 0.780387 | 0.869767 | 0.9214 | 0.95335 |
| SL1344_RS17670_ | 3.919598 | 3.953418 | 3.881603 | -0.03789 | 0.233629 | -0.16015 | 0.398667 | 0.80848 | 0.886901 | 0.8542 | 0.913237 |
| SL1344_RS17675_ | 4.929118 | 5.020477 | 4.847841 | -0.12869 | 0.220741 | -0.51874 | 0.288667 | 0.415726 | 0.596806 | 0.57 | 0.716984 |
| SL1344_RS17680_ | 4.849127 | 4.936716 | 4.729396 | -0.11856 | 0.209136 | -0.74457 | 0.187209 | 0.32476 | 0.512862 | 0.3514 | 0.524277 |
| SL1344_RS17685_ | 5.490687 | 5.701816 | 5.368132 | -0.20775 | 0.235315 | -1.2873 | 0.108594 | 0.133446 | 0.29926 | 0.2002 | 0.370238 |
| SL1344_RS17690_ | 5.291244 | 5.484882 | 5.090709 | -0.22825 | 0.223139 | -1.60232 | 0.112 | 0.079522 | 0.221483 | 0.1987 | 0.368294 |
| SL1344_RS17700_ | 5.989478 | 6.111596 | 5.682347 | -0.32519 | 0.201066 | -1.48719 | 0.093937 | 0.088135 | 0.235178 | 0.1836 | 0.348157 |
| SL1344_RS17705_ | 5.458564 | 5.641988 | 5.224155 | -0.32329 | 0.171631 | -2.08766 | 4.68E-05 | 0.030586 | 0.132011 | 0.1 | 0.236592 |
| SL1344_RS17710_ | 5.616369 | 5.844816 | 5.297623 | -0.36353 | 0.232779 | -1.85089 | 0.006663 | 0.044527 | 0.161108 | 0.1045 | 0.242604 |
| SL1344_RS17715_ | 6.131885 | 6.334773 | 6.021506 | -0.32146 | 0.174904 | -1.83965 | 4.68E-05 | 0.040905 | 0.153924 | 0.1 | 0.236592 |
| SL1344_RS17720_ | 5.089393 | 5.377017 | 5.012583 | -0.36664 | 0.216007 | -1.68858 | 0.003335 | 0.082098 | 0.225749 | 0.102 | 0.239257 |
| SL1344_RS17725_ | 6.190695 | 6.486049 | 6.092261 | -0.39476 | 0.152625 | -2.50364 | 4.68E-05 | 0.063123 | 0.195094 | 0.1 | 0.236592 |
| SL1344_RS17730_ | 6.04427 | 6.284776 | 5.907505 | -0.25357 | 0.263074 | -1.3101 | 0.109261 | 0.126184 | 0.289933 | 0.2 | 0.370027 |
| SL1344_RS17735_ | 5.182789 | 5.498385 | 5.065834 | -0.33121 | 0.311923 | -1.36688 | 0.108594 | 0.107739 | 0.26402 | 0.2002 | 0.370241 |
| SL1344_RS17740_ | 4.990246 | 5.198615 | 4.836314 | -0.21733 | 0.233301 | -1.4023 | 0.115923 | 0.10652 | 0.261813 | 0.1999 | 0.369894 |
| SL1344_RS17745_ | 5.620863 | 5.819047 | 5.378478 | -0.20382 | 0.318995 | -1.29074 | 0.107333 | 0.177452 | 0.355558 | 0.2 | 0.370027 |
| SL1344_RS17750_ | 5.61028 | 5.821614 | 5.495456 | -0.25416 | 0.180067 | -1.6908 | 0.042667 | 0.060087 | 0.18915 | 0.1376 | 0.286748 |
| SL1344_RS17755_ | 5.480797 | 5.806154 | 5.317362 | -0.3253 | 0.302775 | -1.47267 | 0.113924 | 0.086887 | 0.233337 | 0.1997 | 0.36963 |
| SL1344_RS17760_ | 5.461635 | 5.84775 | 5.331601 | -0.5047 | 0.255795 | -1.92864 | 4.68E-05 | 0.04397 | 0.160121 | 0.1 | 0.236592 |
| SL1344_RS17765_ | 4.261734 | 4.465915 | 3.86586 | -0.49957 | 0.21801 | -1.98275 | 4.68E-05 | 0.044953 | 0.161578 | 0.1 | 0.236592 |
| SL1344_RS17770_ | -4.74932 | -4.91009 | -4.58052 | 0.340884 | 0.799052 | 0.395255 | 0.327115 | 0.438879 | 0.578714 | 0.5416 | 0.656335 |
| SL1344_RS17775_ | 3.333793 | 2.956475 | 3.870414 | 0.871344 | 0.176816 | 4.803525 | 4.68E-05 | 0.003905 | 0.051699 | 0.1 | 0.236592 |
| SL1344_RS17780_ | -1.59562 | -1.6026 | -1.59007 | -0.13402 | 0.56157 | -0.21619 | 0.41972 | 0.657142 | 0.775014 | 0.8263 | 0.884972 |
| SL1344_RS17785_ | 6.855915 | 6.868887 | 6.839332 | -0.03004 | 0.131915 | -0.26328 | 0.387742 | 0.778659 | 0.868882 | 0.8632 | 0.91834 |
| SL1344_RS17790_ | 7.365627 | 7.460814 | 7.157666 | -0.17785 | 0.214921 | -1.17661 | 0.119333 | 0.179742 | 0.358454 | 0.2004 | 0.370453 |
| SL1344_RS17795_ | 5.109023 | 5.298314 | 4.744322 | -0.39836 | 0.215357 | -1.59903 | 0.054 | 0.075335 | 0.215106 | 0.1541 | 0.30871 |
| SL1344_RS17800_ | 4.580869 | 4.699526 | 4.113254 | -0.53189 | 0.18777 | -1.5179 | 0.005997 | 0.091279 | 0.239928 | 0.1054 | 0.243782 |
| SL1344_RS17805_ | 0.691031 | 0.972179 | 0.628752 | -0.31784 | 0.203078 | -1.18276 | 0.143238 | 0.183231 | 0.350826 | 0.3021 | 0.456482 |
| SL1344_RS17810_ | 0.60613 | 0.455921 | 0.694544 | 0.233511 | 0.166463 | 1.318156 | 0.079947 | 0.133285 | 0.278887 | 0.203 | 0.354395 |
| SL1344_RS17815_ | 1.121088 | 0.988728 | 1.203989 | 0.22301 | 0.203017 | 1.083435 | 0.114667 | 0.162381 | 0.325267 | 0.2415 | 0.401493 |
| SL1344_RS17820_ | 1.564753 | 1.65585 | 1.493045 | -0.15446 | 0.133367 | -1.08261 | 0.125916 | 0.181607 | 0.339751 | 0.2614 | 0.416677 |
| SL1344_RS17825_ | 4.452614 | 4.445492 | 4.458426 | 0.019579 | 0.073906 | 0.250526 | 0.392 | 0.649238 | 0.768419 | 0.7641 | 0.843373 |
| SL1344_RS17830_ | 0.683762 | 0.764468 | 0.447948 | -0.31525 | 0.29132 | -0.96214 | 0.150566 | 0.221562 | 0.399175 | 0.3011 | 0.462742 |

| | | | | | | | | | | | |
|---|---|---|---|---|---|---|---|---|---|---|---|
| SL1344_RS17835_ | 3.679222 | 3.63156 | 3.7107 | 0.082161 | 0.074468 | 1.063104 | 0.118588 | 0.178706 | 0.340986 | 0.2557 | 0.414403 |
| SL1344_RS17840_ | -0.52807 | -0.52632 | -0.52883 | -0.01555 | 0.234012 | -0.06161 | 0.477682 | 0.621037 | 0.740459 | 0.729 | 0.814426 |
| SL1344_RS17845_ | 1.126572 | 1.166695 | 1.058703 | -0.11501 | 0.163391 | -0.63763 | 0.253165 | 0.367165 | 0.530251 | 0.4883 | 0.622084 |
| SL1344_RS17850_ | -1.2391 | -1.02239 | -1.51421 | -0.49639 | 0.24684 | -1.92954 | 0.015334 | 0.053126 | 0.159452 | 0.116 | 0.255504 |
| SL1344_RS17855_ | 2.064983 | 2.046803 | 2.081866 | -0.02626 | 0.178909 | -0.1383 | 0.451033 | 0.700898 | 0.807721 | 0.9063 | 0.939763 |
| SL1344_RS17860_ | -3.48143 | -3.35906 | -3.59274 | -0.2315 | 0.60615 | -0.33724 | 0.353098 | 0.520063 | 0.652435 | 0.6288 | 0.731411 |
| SL1344_RS17865_ | 1.261103 | 0.943209 | 1.58768 | 0.558496 | 0.237354 | 2.425995 | 0.001337 | 0.022025 | 0.108703 | 0.102 | 0.23927 |
| SL1344_RS17870_ | 1.434503 | 1.380859 | 1.496738 | 0.114922 | 0.100599 | 1.065404 | 0.111925 | 0.185617 | 0.330756 | 0.2689 | 0.414587 |
| SL1344_RS17875_ | -0.07988 | -0.33961 | 0.182383 | 0.523789 | 0.163926 | 3.159658 | 4.68E-05 | 0.013823 | 0.081008 | 0.1002 | 0.236856 |
| SL1344_RS17880_ | 1.534965 | 1.387637 | 1.682803 | 0.301991 | 0.119405 | 2.522948 | 0.003332 | 0.030383 | 0.121797 | 0.1018 | 0.238937 |
| SL1344_RS17885_ | 1.866609 | 1.863297 | 1.869684 | -0.02381 | 0.157239 | -0.14303 | 0.441706 | 0.703839 | 0.808493 | 0.8378 | 0.894675 |
| SL1344_RS17895_ | 2.946197 | 2.946303 | 2.946149 | 0.010638 | 0.092323 | 0.100828 | 0.458361 | 0.686597 | 0.795043 | 0.8025 | 0.870055 |
| SL1344_RS17900_ | 0.504287 | 0.469889 | 0.53853 | 0.071947 | 0.130482 | 0.501511 | 0.295333 | 0.415482 | 0.556194 | 0.5136 | 0.632227 |
| SL1344_RS17905_ | -0.0624 | -0.13722 | 0.000528 | 0.135298 | 0.182719 | 0.678683 | 0.208 | 0.334146 | 0.485129 | 0.4405 | 0.570946 |
| SL1344_RS17910_ | -0.66862 | -1.01362 | -0.38377 | 0.646979 | 0.271902 | 2.32953 | 0.003998 | 0.033731 | 0.129882 | 0.1023 | 0.239551 |
| SL1344_RS17915_ | -1.31191 | -1.65907 | -1.08837 | 0.571383 | 0.299243 | 1.824342 | 0.015323 | 0.068249 | 0.187339 | 0.1213 | 0.26162 |
| SL1344_RS17920_ | 1.891756 | 2.009001 | 1.704057 | -0.29584 | 0.11306 | -2.50186 | 0.003998 | 0.031873 | 0.12631 | 0.1039 | 0.241684 |
| SL1344_RS17925_ | -1.79421 | -1.66908 | -1.96105 | -0.28059 | 0.366535 | -0.78457 | 0.199867 | 0.319857 | 0.47698 | 0.4208 | 0.559285 |
| SL1344_RS17930_ | 2.614344 | 2.77851 | 2.36287 | -0.4099 | 0.088808 | -4.46484 | 4.68E-05 | 0.009725 | 0.070765 | 0.1 | 0.236592 |
| SL1344_RS17940_ | 0.122072 | 0.133135 | 0.112074 | -0.04322 | 0.228519 | -0.17803 | 0.434 | 0.617259 | 0.740024 | 0.7413 | 0.823092 |
| SL1344_RS17965_ | 3.452243 | 3.218302 | 3.515679 | 0.285651 | 0.109497 | 1.723291 | 0.01066 | 0.082612 | 0.223994 | 0.1123 | 0.252309 |
| SL1344_RS17970_ | 2.246778 | 2.101601 | 2.390477 | 0.288113 | 0.082143 | 3.484685 | 4.68E-05 | 0.010724 | 0.073163 | 0.1001 | 0.236728 |
| SL1344_RS17975_ | 1.836833 | 1.662531 | 2.003051 | 0.356803 | 0.139982 | 2.565472 | 4.68E-05 | 0.031613 | 0.127942 | 0.1001 | 0.236727 |
| SL1344_RS17985_ | 3.959973 | 3.915149 | 4.035084 | 0.130781 | 0.110319 | 1.27012 | 0.051299 | 0.132965 | 0.293806 | 0.1591 | 0.309809 |
| SL1344_RS17990_ | 3.260702 | 3.211846 | 3.282374 | 0.06262 | 0.1285 | 0.396953 | 0.372 | 0.690622 | 0.80348 | 0.784 | 0.86885 |
| SL1344_RS17995_ | 2.598939 | 2.666423 | 2.490474 | -0.15913 | 0.1061 | -1.38945 | 0.072667 | 0.109642 | 0.254015 | 0.1829 | 0.337776 |
| SL1344_RS18000_ | -0.91037 | -1.06504 | -0.71697 | 0.348993 | 0.315046 | 1.03642 | 0.124 | 0.185165 | 0.34257 | 0.2664 | 0.419729 |
| SL1344_RS18005_ | -2.92843 | -3.13591 | -2.76299 | 0.376711 | 0.510493 | 0.675805 | 0.230667 | 0.34302 | 0.499276 | 0.4491 | 0.583559 |
| SL1344_RS18010_ | -2.94261 | -2.77943 | -3.31028 | -0.51794 | 0.616192 | -0.77002 | 0.207861 | 0.296534 | 0.460782 | 0.4082 | 0.551545 |
| SL1344_RS18015_ | -2.90201 | -2.85753 | -2.95428 | -0.09547 | 0.430448 | -0.19189 | 0.41439 | 0.526584 | 0.657139 | 0.634 | 0.73446 |
| SL1344_RS18020_ | -2.65233 | -2.5642 | -2.73897 | -0.17555 | 0.431968 | -0.3954 | 0.330446 | 0.482298 | 0.617532 | 0.581 | 0.690955 |
| SL1344_RS18025_ | 1.901383 | 1.917712 | 1.874476 | -0.0543 | 0.108502 | -0.45268 | 0.308 | 0.450334 | 0.598808 | 0.5666 | 0.681603 |
| SL1344_RS18030_ | 0.544428 | 0.574394 | 0.51346 | -0.06118 | 0.166038 | -0.33795 | 0.351099 | 0.528268 | 0.662994 | 0.649 | 0.74973 |
| SL1344_RS18035_ | 3.163113 | 3.233014 | 3.114683 | -0.12025 | 0.074147 | -1.60121 | 0.037309 | 0.089481 | 0.223666 | 0.1447 | 0.289854 |
| SL1344_RS18040_ | 0.297314 | 0.457629 | 0.083484 | -0.32384 | 0.257491 | -1.22869 | 0.103333 | 0.146066 | 0.30342 | 0.2143 | 0.376028 |
| SL1344_RS18045_ | -0.05041 | 0.009631 | -0.10941 | -0.11703 | 0.193764 | -0.56969 | 0.263333 | 0.411472 | 0.559711 | 0.5199 | 0.643605 |
| SL1344_RS18050_ | 1.369943 | 1.38398 | 1.362413 | -0.01083 | 0.127701 | -0.07761 | 0.468 | 0.660927 | 0.772365 | 0.769 | 0.846878 |
| SL1344_RS18055_ | 0.533868 | 0.502862 | 0.547743 | 0.029485 | 0.19973 | 0.137682 | 0.448667 | 0.697984 | 0.804138 | 0.7951 | 0.869022 |
| SL1344_RS18060_ | 5.85766 | 5.8877 | 5.803213 | -0.05628 | 0.127106 | -0.43002 | 0.337333 | 0.624951 | 0.762403 | 0.694 | 0.812774 |
| SL1344_RS18075_ | 0.142133 | 0.26202 | -0.16217 | -0.39314 | 0.249834 | -1.38249 | 0.069333 | 0.121765 | 0.273459 | 0.1933 | 0.347785 |
| SL1344_RS18080_ | 2.434265 | 2.483236 | 2.375388 | -0.11334 | 0.090111 | -1.19938 | 0.08994 | 0.149106 | 0.297258 | 0.2218 | 0.37114 |
| SL1344_RS18090_ | 0.630509 | 0.433801 | 0.795175 | 0.36461 | 0.127712 | 2.775738 | 0.000673 | 0.020873 | 0.097872 | 0.101 | 0.237873 |
| SL1344_RS18095_ | -0.54901 | -0.66447 | -0.41873 | 0.262749 | 0.23729 | 1.05245 | 0.104597 | 0.183145 | 0.336536 | 0.2552 | 0.401622 |
| SL1344_RS18100_ | -0.12914 | -0.15475 | -0.10548 | 0.045198 | 0.196763 | 0.213611 | 0.404 | 0.562203 | 0.689076 | 0.6724 | 0.767165 |
| SL1344_RS18105_ | -0.14977 | -0.36231 | 0.066368 | 0.424185 | 0.233332 | 1.76449 | 0.023984 | 0.062288 | 0.180877 | 0.1264 | 0.269774 |

| | | | | | | | | | | |
|---|---|---|---|---|---|---|---|---|---|---|
| SL1344_RS18110_ | 0.104814 | 0.052227 | 0.179097 | 0.08449 | 0.43014 | 0.168089 | 0.445333 | 0.838672 | 0.904183 | 0.9115 | 0.947366 |
| SL1344_RS18115_ | 2.369284 | 2.324255 | 2.428774 | 0.076515 | 0.218306 | 0.353575 | 0.329333 | 0.627696 | 0.758654 | 0.708 | 0.810836 |
| SL1344_RS18120_ | -0.14296 | -0.22156 | -0.08229 | 0.143008 | 0.199538 | 0.665671 | 0.23451 | 0.367895 | 0.519689 | 0.471 | 0.603099 |
| SL1344_RS18125_ | 1.32362 | 1.048858 | 1.780632 | 0.723018 | 0.230055 | 3.203845 | 4.68E-05 | 0.00942 | 0.07289 | 0.1 | 0.236592 |
| SL1344_RS18130_ | 1.691875 | 1.390285 | 2.141855 | 0.746038 | 0.145063 | 5.011264 | 4.68E-05 | 0.009347 | 0.072363 | 0.1 | 0.236592 |
| SL1344_RS18140_ | 1.221217 | 1.178524 | 1.251997 | 0.06408 | 0.13963 | 0.438608 | 0.327115 | 0.518827 | 0.657511 | 0.6268 | 0.738998 |
| SL1344_RS18145_ | 2.051734 | 2.059239 | 2.041628 | -0.0011 | 0.208696 | -0.0057 | 0.497668 | 0.815179 | 0.888302 | 0.9125 | 0.945819 |
| SL1344_RS18150_ | 0.201402 | 0.313403 | 0.071465 | -0.24027 | 0.153798 | -1.46195 | 0.052 | 0.097124 | 0.225077 | 0.1665 | 0.311585 |
| SL1344_RS18155_ | 0.234004 | 0.240429 | 0.226416 | -0.02082 | 0.205473 | -0.08703 | 0.457029 | 0.643875 | 0.759853 | 0.7654 | 0.840538 |
| SL1344_RS18160_ | -0.61825 | -0.46676 | -0.70116 | -0.22608 | 0.259664 | -0.81652 | 0.198534 | 0.301271 | 0.46345 | 0.4137 | 0.556285 |
| SL1344_RS18165_ | 0.215016 | 0.162339 | 0.274499 | 0.115436 | 0.164992 | 0.670831 | 0.225849 | 0.353099 | 0.50444 | 0.4533 | 0.583882 |
| SL1344_RS18170_ | 1.745553 | 1.616322 | 1.854608 | 0.256713 | 0.15774 | 1.672579 | 0.005331 | 0.10351 | 0.252207 | 0.1059 | 0.244236 |
| SL1344_RS18175_ | 1.910573 | 2.027818 | 1.758526 | -0.27728 | 0.120465 | -2.27364 | 0.003332 | 0.048639 | 0.160008 | 0.102 | 0.239204 |
| SL1344_RS18180_ | 0.711897 | 0.794897 | 0.622854 | -0.16752 | 0.175948 | -0.88592 | 0.166 | 0.250496 | 0.414536 | 0.3354 | 0.488 |
| SL1344_RS18185_ | 7.369273 | 7.371097 | 7.365958 | 0.044726 | 0.109135 | 0.327749 | 0.39507 | 0.480999 | 0.652261 | 0.8812 | 0.92439 |
| SL1344_RS18190_ | -1.53034 | -1.62802 | -1.41198 | 0.211706 | 0.300512 | 0.684832 | 0.232667 | 0.35105 | 0.499454 | 0.4471 | 0.576445 |
| SL1344_RS18195_ | 0.600864 | 0.451773 | 0.8258 | 0.356212 | 0.173879 | 1.949587 | 0.017322 | 0.051744 | 0.163334 | 0.1206 | 0.262843 |
| SL1344_RS18205_ | -2.73824 | -2.81505 | -2.64933 | 0.159174 | 0.407092 | 0.371169 | 0.333333 | 0.473467 | 0.610575 | 0.5831 | 0.692247 |
| SL1344_RS18210_ | -1.70669 | -1.95355 | -1.53602 | 0.385768 | 0.542529 | 0.691231 | 0.224 | 0.342011 | 0.512745 | 0.4513 | 0.597378 |
| SL1344_RS18215_ | -1.5656 | -1.71742 | -1.39217 | 0.326975 | 0.275477 | 1.13596 | 0.110593 | 0.175198 | 0.318126 | 0.2493 | 0.39356 |
| SL1344_RS18220_ | 2.695688 | 2.64986 | 2.732444 | 0.106664 | 0.137183 | 0.788578 | 0.202 | 0.286764 | 0.462753 | 0.4257 | 0.574567 |
| SL1344_RS18225_ | 3.516043 | 3.193909 | 3.82465 | 0.639822 | 0.07115 | 9.044453 | 4.68E-05 | 0.001498 | 0.035219 | 0.1 | 0.236592 |
| SL1344_RS18230_ | 2.199064 | 1.82856 | 2.670496 | 0.825344 | 0.207469 | 4.048006 | 4.68E-05 | 0.003831 | 0.05133 | 0.1 | 0.236592 |
| SL1344_RS18235_ | 2.336049 | 1.989755 | 2.691066 | 0.673912 | 0.164155 | 4.088272 | 4.68E-05 | 0.004866 | 0.055758 | 0.1 | 0.236592 |
| SL1344_RS18240_ | 2.805317 | 2.482231 | 3.069345 | 0.588881 | 0.081809 | 7.094142 | 4.68E-05 | 0.002968 | 0.04329 | 0.1 | 0.236592 |
| SL1344_RS18245_ | 3.011542 | 2.673561 | 3.236787 | 0.5481 | 0.102603 | 5.223442 | 4.68E-05 | 0.003956 | 0.049707 | 0.1 | 0.236592 |
| SL1344_RS18250_ | 2.863242 | 2.706027 | 3.024419 | 0.341265 | 0.109591 | 3.157616 | 4.68E-05 | 0.019453 | 0.100491 | 0.1 | 0.236592 |
| SL1344_RS18255_ | -0.47982 | -0.52458 | -0.4299 | 0.09572 | 0.234048 | 0.386635 | 0.333111 | 0.509211 | 0.647375 | 0.6188 | 0.728202 |
| SL1344_RS18270_ | -0.57829 | -0.68775 | -0.47268 | 0.216919 | 0.178878 | 1.149135 | 0.105929 | 0.176072 | 0.315598 | 0.2561 | 0.397864 |
| SL1344_RS18275_ | 0.514418 | 0.361396 | 0.712091 | 0.351705 | 0.154945 | 2.191194 | 0.004668 | 0.03813 | 0.136807 | 0.1057 | 0.243812 |
| SL1344_RS18280_ | 0.027269 | 0.086847 | -0.07091 | -0.14501 | 0.201653 | -0.69009 | 0.244 | 0.369345 | 0.523497 | 0.4729 | 0.607629 |
| SL1344_RS18285_ | -2.32659 | -2.84475 | -2.14833 | 0.679063 | 0.62964 | 0.959894 | 0.161892 | 0.232554 | 0.406753 | 0.3293 | 0.483279 |
| SL1344_RS18290_ | -3.08651 | -3.2999 | -2.91716 | 0.486461 | 0.753668 | 0.616071 | 0.251333 | 0.370739 | 0.533692 | 0.4976 | 0.630605 |
| SL1344_RS18295_ | -3.47879 | -3.67702 | -3.2772 | 0.403965 | 0.647363 | 0.60796 | 0.257162 | 0.381888 | 0.535793 | 0.4975 | 0.624698 |
| SL1344_RS18300_ | -1.21403 | -1.42855 | -0.93932 | 0.47864 | 0.297406 | 1.55403 | 0.045969 | 0.088044 | 0.217186 | 0.1542 | 0.30155 |
| SL1344_RS18305_ | 0.52535 | 0.417699 | 0.688482 | 0.258136 | 0.188862 | 1.319805 | 0.082 | 0.127485 | 0.274954 | 0.2033 | 0.357297 |
| SL1344_RS18310_ | 1.538752 | 1.233927 | 1.773387 | 0.558708 | 0.151014 | 3.67024 | 4.68E-05 | 0.009203 | 0.070913 | 0.1 | 0.236592 |
| SL1344_RS18315_ | 0.357037 | 0.011938 | 0.784909 | 0.772947 | 0.251811 | 3.062949 | 4.68E-05 | 0.013448 | 0.084442 | 0.1 | 0.236592 |
| SL1344_RS18325_ | -0.36379 | -0.5009 | -0.23564 | 0.286228 | 0.283347 | 0.976463 | 0.129333 | 0.202061 | 0.365856 | 0.2891 | 0.442765 |
| SL1344_RS18330_ | -0.3042 | -0.35619 | -0.26682 | 0.09416 | 0.26577 | 0.331025 | 0.358428 | 0.56753 | 0.698474 | 0.6857 | 0.779642 |
| SL1344_RS18335_ | 2.891531 | 2.597723 | 3.137894 | 0.533341 | 0.118539 | 4.464645 | 4.68E-05 | 0.003764 | 0.050237 | 0.1 | 0.236592 |
| SL1344_RS18340_ | -0.93256 | -1.09207 | -0.71814 | 0.378594 | 0.289131 | 1.25034 | 0.092667 | 0.148914 | 0.299421 | 0.2207 | 0.37162 |
| SL1344_RS18345_ | -0.80692 | -1.02246 | -0.60817 | 0.396868 | 0.355454 | 1.073775 | 0.105929 | 0.170642 | 0.332426 | 0.2411 | 0.399469 |
| SL1344_RS18355_ | -1.80835 | -1.84222 | -1.77882 | 0.070952 | 0.329853 | 0.201332 | 0.410667 | 0.548704 | 0.680036 | 0.6683 | 0.764752 |
| SL1344_RS18360_ | -1.37009 | -1.28777 | -1.44167 | -0.15313 | 0.320376 | -0.45921 | 0.311792 | 0.486293 | 0.627174 | 0.6007 | 0.711176 |

| | | | | | | | | | | | |
|---|---|---|---|---|---|---|---|---|---|---|---|
| SL1344_RS18365_ | -1.90205 | -1.93469 | -1.86428 | 0.034694 | 0.460784 | 0.077305 | 0.464357 | 0.67707 | 0.786373 | 0.7788 | 0.852146 |
| SL1344_RS18370_ | -1.42544 | -1.29224 | -1.59558 | -0.30036 | 0.263286 | -1.1039 | 0.112592 | 0.180173 | 0.325368 | 0.261 | 0.40673 |
| SL1344_RS18375_ | 0.410788 | 0.441423 | 0.381793 | -0.06923 | 0.241868 | -0.27653 | 0.366422 | 0.602828 | 0.730869 | 0.704 | 0.799944 |
| SL1344_RS18380_ | 0.468705 | 0.500489 | 0.435329 | -0.06815 | 0.155667 | -0.41303 | 0.32 | 0.469348 | 0.609238 | 0.5761 | 0.686801 |
| SL1344_RS18385_ | 0.081645 | 0.217514 | -0.09345 | -0.32384 | 0.218664 | -1.40532 | 0.060667 | 0.107642 | 0.248509 | 0.1711 | 0.319723 |
| SL1344_RS18390_ | 4.866029 | 4.892136 | 4.774438 | -0.11004 | 0.074275 | -1.46252 | 0.075949 | 0.096886 | 0.242519 | 0.1835 | 0.338181 |
| SL1344_RS18395_ | 2.955638 | 2.742055 | 3.14268 | 0.400107 | 0.085246 | 4.651558 | 4.68E-05 | 0.004469 | 0.051879 | 0.1 | 0.236592 |
| SL1344_RS18400_ | 1.928753 | 1.819079 | 2.091824 | 0.258976 | 0.142714 | 1.770463 | 0.031333 | 0.063672 | 0.187442 | 0.1308 | 0.276104 |
| SL1344_RS18405_ | 2.439575 | 2.358887 | 2.71041 | 0.32374 | 0.155081 | 1.639248 | 0.049967 | 0.08958 | 0.233434 | 0.1456 | 0.294581 |
| SL1344_RS18415_ | 0.309587 | 0.286608 | 0.325285 | 0.020199 | 0.22954 | 0.086817 | 0.462 | 0.71085 | 0.812571 | 0.8265 | 0.887484 |
| SL1344_RS18420_ | -0.58214 | -0.41461 | -0.79266 | -0.38134 | 0.281817 | -1.32082 | 0.071286 | 0.122017 | 0.267428 | 0.186 | 0.336004 |
| SL1344_RS18425_ | 1.230185 | 1.101392 | 1.301917 | 0.189527 | 0.159421 | 1.130974 | 0.117921 | 0.172763 | 0.331751 | 0.2581 | 0.416509 |
| SL1344_RS18440_ | -0.70437 | -0.70586 | -0.70394 | -0.02874 | 0.308162 | -0.08966 | 0.456 | 0.676073 | 0.785334 | 0.7889 | 0.859336 |
| SL1344_RS18445_ | -1.09874 | -0.714 | -1.30125 | -0.51636 | 0.388345 | -1.26322 | 0.103265 | 0.142821 | 0.299131 | 0.2228 | 0.38194 |
| SL1344_RS18450_ | 3.198288 | 3.213481 | 3.185627 | -0.01867 | 0.093417 | -0.19216 | 0.421719 | 0.690852 | 0.797131 | 0.8024 | 0.869597 |
| SL1344_RS18455_ | 0.39557 | 0.23797 | 0.699398 | 0.447837 | 0.194428 | 2.147037 | 0.010667 | 0.04892 | 0.159859 | 0.1108 | 0.249723 |
| SL1344_RS18460_ | -0.4434 | -0.67306 | -0.2606 | 0.411426 | 0.204953 | 1.976388 | 0.010667 | 0.056293 | 0.166814 | 0.1161 | 0.256384 |
| SL1344_RS18465_ | 0.399132 | 0.382516 | 0.407437 | 0.011686 | 0.217069 | 0.052185 | 0.482 | 0.720419 | 0.821413 | 0.8175 | 0.885889 |
| SL1344_RS18470_ | -0.08874 | -0.07838 | -0.09372 | -0.02603 | 0.223138 | -0.10582 | 0.458667 | 0.651018 | 0.764596 | 0.7534 | 0.834489 |
| SL1344_RS18475_ | 0.602672 | 0.390197 | 0.788426 | 0.398669 | 0.133495 | 2.928612 | 0.000673 | 0.018397 | 0.092939 | 0.1006 | 0.237395 |
| SL1344_RS18480_ | 2.25063 | 2.141899 | 2.436205 | 0.294536 | 0.108417 | 2.519996 | 0.001337 | 0.035564 | 0.13614 | 0.1008 | 0.237652 |
| SL1344_RS18485_ | 1.413992 | 1.269081 | 1.780489 | 0.490149 | 0.178498 | 2.325872 | 0.002002 | 0.051168 | 0.169175 | 0.1026 | 0.240062 |
| SL1344_RS18490_ | 0.751848 | 0.68656 | 0.840986 | 0.150892 | 0.152607 | 0.909891 | 0.145237 | 0.237448 | 0.394763 | 0.329 | 0.475608 |
| SL1344_RS18495_ | 3.231497 | 3.313647 | 3.081763 | -0.23191 | 0.098722 | -2.30746 | 0.000673 | 0.054492 | 0.175026 | 0.1006 | 0.237337 |
| SL1344_RS18500_ | 1.387762 | 1.079316 | 1.820135 | 0.719621 | 0.161302 | 4.273816 | 4.68E-05 | 0.008472 | 0.067916 | 0.1 | 0.236592 |
| SL1344_RS18505_ | 6.012795 | 5.973993 | 6.090501 | 0.177753 | 0.151239 | 1.23705 | 0.020653 | 0.102229 | 0.255366 | 0.1204 | 0.263642 |
| SL1344_RS18510_ | 3.117217 | 2.768159 | 3.399338 | 0.65621 | 0.11368 | 5.814509 | 4.68E-05 | 0.003 | 0.045093 | 0.1 | 0.236592 |
| SL1344_RS18515_ | 0.459676 | 0.216037 | 0.651876 | 0.434913 | 0.142279 | 3.016241 | 0.000673 | 0.018305 | 0.092082 | 0.1008 | 0.237565 |
| SL1344_RS18520_ | 2.621063 | 2.318481 | 2.915924 | 0.603291 | 0.102798 | 5.860192 | 4.68E-05 | 0.002409 | 0.041404 | 0.1 | 0.236592 |
| SL1344_RS18525_ | 1.378934 | 1.449634 | 1.307783 | -0.14376 | 0.105763 | -1.30599 | 0.08461 | 0.136471 | 0.276316 | 0.2141 | 0.359894 |
| SL1344_RS18530_ | 0.810802 | 0.847193 | 0.779206 | -0.07906 | 0.151331 | -0.49065 | 0.290667 | 0.433582 | 0.583433 | 0.5567 | 0.673861 |
| SL1344_RS18535_ | 2.855294 | 2.758722 | 2.933759 | 0.184396 | 0.085885 | 2.067583 | 0.003335 | 0.052753 | 0.166845 | 0.104 | 0.241858 |
| SL1344_RS18540_ | 0.288111 | 0.208208 | 0.607108 | 0.480087 | 0.662292 | 0.78044 | 0.212525 | 0.268436 | 0.454252 | 0.3855 | 0.565847 |
| SL1344_RS18545_ | 1.140428 | 0.93383 | 1.48196 | 0.588326 | 0.46444 | 1.2197 | 0.097333 | 0.130568 | 0.293805 | 0.1934 | 0.357855 |
| SL1344_RS18550_ | 0.732629 | 0.435415 | 1.192816 | 0.796225 | 0.587137 | 1.307982 | 0.079333 | 0.107145 | 0.262108 | 0.1688 | 0.328073 |
| SL1344_RS18555_ | 2.155134 | 2.124002 | 2.618597 | 0.643269 | 0.752408 | 0.939102 | 0.224 | 0.186093 | 0.365506 | 0.3983 | 0.580518 |
| SL1344_RS18560_ | 0.233722 | 0.276766 | 0.166652 | -0.11402 | 0.176378 | -0.61513 | 0.256667 | 0.37275 | 0.526097 | 0.484 | 0.610014 |
| SL1344_RS18565_ | 0.786644 | 0.635677 | 0.921613 | 0.301125 | 0.182769 | 1.601913 | 0.026 | 0.075409 | 0.203306 | 0.1308 | 0.273839 |
| SL1344_RS18570_ | 4.25662 | 4.332468 | 4.208889 | -0.0664 | 0.125874 | -0.59967 | 0.26449 | 0.429503 | 0.602983 | 0.5306 | 0.6713 |
| SL1344_RS18575_ | -5.03188 | -4.72732 | -5.45056 | -0.72842 | 0.944183 | -0.71778 | 0.20986 | 0.305795 | 0.457181 | 0.4035 | 0.536688 |
| SL1344_RS18580_ | -2.93512 | -2.80909 | -3.04452 | -0.24233 | 0.494714 | -0.45644 | 0.303131 | 0.456037 | 0.59695 | 0.5637 | 0.677627 |
| SL1344_RS18585_ | -0.74897 | -0.75076 | -0.74733 | -0.04685 | 0.345643 | -0.13342 | 0.447702 | 0.677195 | 0.787074 | 0.7998 | 0.866856 |
| SL1344_RS18590_ | 0.374628 | 0.226217 | 0.496875 | 0.285062 | 0.172153 | 1.578834 | 0.030646 | 0.083598 | 0.211255 | 0.1433 | 0.287134 |
| SL1344_RS18595_ | -0.55502 | -0.57573 | -0.53755 | 0.039495 | 0.220122 | 0.173789 | 0.42 | 0.574539 | 0.701507 | 0.6877 | 0.779847 |
| SL1344_RS18600_ | 2.294892 | 2.64133 | 2.17245 | -0.44934 | 0.16289 | -2.22675 | 0.002667 | 0.049432 | 0.166914 | 0.1015 | 0.238542 |

| | | | | | | | | | | | |
|---|---|---|---|---|---|---|---|---|---|---|---|
| SL1344_RS18610_ | 0.960992 | 0.679107 | 1.232545 | 0.576724 | 0.166176 | 3.424689 | 4.68E-05 | 0.010127 | 0.072953 | 0.1 | 0.236592 |
| SL1344_RS18615_ | 1.890566 | 1.840806 | 1.915963 | 0.081754 | 0.159047 | 0.525069 | 0.279147 | 0.470064 | 0.625468 | 0.5745 | 0.703309 |
| SL1344_RS18620_ | 5.150182 | 5.005117 | 5.439771 | 0.451454 | 0.161787 | 2.856954 | 4.68E-05 | 0.011547 | 0.082708 | 0.1 | 0.236592 |
| SL1344_RS18625_ | 1.67454 | 1.664712 | 1.680582 | 0.02278 | 0.108733 | 0.207667 | 0.417055 | 0.587477 | 0.712984 | 0.7056 | 0.793849 |
| SL1344_RS18630_ | 2.818331 | 2.545185 | 3.147606 | 0.570146 | 0.124637 | 4.638506 | 4.68E-05 | 0.003989 | 0.050995 | 0.1 | 0.236592 |
| SL1344_RS18635_ | 1.363625 | 1.01683 | 1.772695 | 0.75653 | 0.140449 | 5.226645 | 4.68E-05 | 0.005392 | 0.055683 | 0.1 | 0.236592 |
| SL1344_RS18640_ | 1.791866 | 1.3975 | 2.206605 | 0.816663 | 0.131947 | 6.271118 | 4.68E-05 | 0.001934 | 0.038637 | 0.1 | 0.236592 |
| SL1344_RS18655_ | -0.87323 | -1.27131 | -0.492 | 0.774348 | 0.236266 | 3.216664 | 4.68E-05 | 0.013339 | 0.080607 | 0.1 | 0.236592 |
| SL1344_RS18660_ | 1.128825 | 0.812005 | 1.477613 | 0.666772 | 0.119002 | 5.500219 | 4.68E-05 | 0.004207 | 0.048391 | 0.1 | 0.236592 |
| SL1344_RS18665_ | -0.48101 | -0.93414 | -0.11936 | 0.816259 | 0.17914 | 4.455363 | 4.68E-05 | 0.005789 | 0.054078 | 0.1 | 0.236592 |
| SL1344_RS18670_ | 1.553862 | 1.193457 | 1.994302 | 0.777819 | 0.159765 | 4.832753 | 4.68E-05 | 0.004116 | 0.050987 | 0.1 | 0.236592 |
| SL1344_RS18675_ | 3.694361 | 3.711919 | 3.665953 | -0.03308 | 0.158272 | -0.12002 | 0.400667 | 0.701382 | 0.817001 | 0.8191 | 0.892234 |
| SL1344_RS18680_ | -4.92592 | -4.9248 | -4.93094 | -0.06191 | 1.157682 | -0.04502 | 0.482 | 0.629652 | 0.749192 | 0.7468 | 0.827739 |
| SL1344_RS18685_ | -0.94929 | -0.80174 | -1.18747 | -0.3987 | 0.29326 | -1.28109 | 0.068667 | 0.141057 | 0.292762 | 0.1861 | 0.33428 |
| SL1344_RS18690_ | 0.697632 | 0.555849 | 0.831282 | 0.278195 | 0.167896 | 1.646159 | 0.027982 | 0.073704 | 0.199717 | 0.1312 | 0.274673 |
| SL1344_RS18695_ | 0.709972 | 0.621986 | 0.79203 | 0.169363 | 0.117584 | 1.353145 | 0.069953 | 0.124595 | 0.257481 | 0.1948 | 0.340142 |
| SL1344_RS18700_ | 0.948019 | 0.812581 | 1.081579 | 0.279397 | 0.153394 | 1.800544 | 0.018 | 0.061811 | 0.179594 | 0.1192 | 0.260547 |
| SL1344_RS18705_ | 0.082401 | -0.11034 | 0.276407 | 0.393981 | 0.17564 | 2.159746 | 0.005334 | 0.03767 | 0.135179 | 0.1068 | 0.245291 |
| SL1344_RS18710_ | 2.798602 | 2.584726 | 3.051017 | 0.472727 | 0.095967 | 4.935184 | 4.68E-05 | 0.003775 | 0.048604 | 0.1 | 0.236592 |
| SL1344_RS18715_ | -2.33876 | -2.26731 | -2.42883 | -0.18124 | 0.414429 | -0.41065 | 0.320453 | 0.473327 | 0.612917 | 0.5888 | 0.697811 |
| SL1344_RS18720_ | -1.63591 | -1.66009 | -1.59956 | 0.075602 | 0.329985 | 0.212933 | 0.409061 | 0.573401 | 0.701143 | 0.6817 | 0.776943 |
| SL1344_RS18725_ | -1.07502 | -0.94893 | -1.16125 | -0.21598 | 0.270412 | -0.75841 | 0.20986 | 0.32705 | 0.482323 | 0.4309 | 0.567186 |
| SL1344_RS18735_ | 0.209189 | 0.036853 | 0.386948 | 0.147889 | 0.599084 | 0.258661 | 0.421719 | 0.712707 | 0.821772 | 0.9215 | 0.951985 |
| SL1344_RS18740_ | 1.275843 | 1.17443 | 1.449426 | 0.2753 | 0.176502 | 1.462004 | 0.058628 | 0.09779 | 0.239882 | 0.1581 | 0.307939 |
| SL1344_RS18745_ | -2.04903 | -2.00597 | -2.15584 | -0.11428 | 0.442817 | -0.22461 | 0.40573 | 0.626653 | 0.747215 | 0.7274 | 0.818275 |
| SL1344_RS18750_ | -2.30656 | -2.35916 | -2.22743 | 0.092491 | 0.459802 | 0.19284 | 0.415057 | 0.61706 | 0.736891 | 0.7181 | 0.807708 |
| SL1344_RS18755_ | -1.61911 | -1.54356 | -1.69468 | -0.13098 | 0.339863 | -0.35642 | 0.35 | 0.513852 | 0.651805 | 0.618 | 0.727964 |
| SL1344_RS18760_ | -4.87899 | -4.30147 | -5.46583 | -1.16391 | 0.903358 | -1.21658 | 0.096 | 0.150138 | 0.289125 | 0.2218 | 0.36794 |
| SL1344_RS18765_ | -3.72289 | -3.6688 | -3.79879 | -0.19915 | 0.902515 | -0.20103 | 0.404667 | 0.596775 | 0.725087 | 0.7284 | 0.813236 |
| SL1344_RS18770_ | -0.93398 | -1.29445 | -0.70838 | 0.575072 | 0.284562 | 1.930144 | 0.016656 | 0.054722 | 0.168976 | 0.1131 | 0.253023 |
| SL1344_RS18775_ | 1.143883 | 1.213281 | 1.021244 | -0.20204 | 0.164735 | -1.15539 | 0.085277 | 0.15541 | 0.313639 | 0.2069 | 0.357051 |
| SL1344_RS18780_ | 0.985008 | 0.994633 | 0.97907 | -0.05154 | 0.218457 | -0.24364 | 0.406396 | 0.649191 | 0.766849 | 0.8042 | 0.869464 |
| SL1344_RS18785_ | 1.953474 | 1.954199 | 1.952807 | -0.00213 | 0.082205 | -0.02522 | 0.486 | 0.546416 | 0.674475 | 0.6506 | 0.749583 |
| SL1344_RS18790_ | 0.787291 | 0.625009 | 0.903098 | 0.265766 | 0.197204 | 1.310572 | 0.075949 | 0.128219 | 0.280759 | 0.1856 | 0.340455 |
| SL1344_RS18795_ | 1.615363 | 1.483228 | 1.930723 | 0.416974 | 0.172964 | 2.052909 | 0.011334 | 0.057407 | 0.179093 | 0.1113 | 0.250985 |
| SL1344_RS18800_ | 2.992665 | 2.851629 | 3.096639 | 0.239968 | 0.083916 | 2.780375 | 4.68E-05 | 0.022938 | 0.107131 | 0.1002 | 0.23686 |
| SL1344_RS18805_ | 2.180688 | 1.822095 | 2.510193 | 0.689491 | 0.108479 | 6.334556 | 4.68E-05 | 0.001849 | 0.037738 | 0.1 | 0.236592 |
| SL1344_RS18815_ | 0.996022 | 0.612024 | 1.553823 | 0.7847 | 0.374673 | 2.253411 | 0.001336 | 0.025745 | 0.11921 | 0.1008 | 0.237666 |
| SL1344_RS18820_ | 0.291179 | 0.372749 | 0.241524 | -0.12742 | 0.183911 | -0.64298 | 0.243333 | 0.392151 | 0.544826 | 0.5042 | 0.6333 |
| SL1344_RS18825_ | 0.437705 | 0.647102 | 0.303407 | -0.24738 | 0.317408 | -0.8075 | 0.197202 | 0.3041 | 0.48 | 0.3836 | 0.541798 |
| SL1344_RS18830_ | -1.89063 | -1.89282 | -1.88946 | -0.00284 | 0.333399 | -0.00984 | 0.496336 | 0.58753 | 0.71166 | 0.6973 | 0.78757 |
| SL1344_RS18835_ | 3.631467 | 3.554868 | 3.673536 | 0.13039 | 0.098626 | 1.306178 | 0.05996 | 0.11166 | 0.261699 | 0.1737 | 0.325011 |
| SL1344_RS18840_ | 1.664473 | 1.762773 | 1.628634 | -0.1089 | 0.146795 | -0.64738 | 0.267155 | 0.425823 | 0.584982 | 0.5419 | 0.680035 |
| SL1344_RS18845_ | -1.33791 | -1.41799 | -1.24262 | 0.213045 | 0.3658 | 0.583011 | 0.273817 | 0.420144 | 0.575272 | 0.5562 | 0.675352 |
| SL1344_RS18850_ | -1.18521 | -1.05516 | -1.37228 | -0.31306 | 0.28427 | -1.01793 | 0.118667 | 0.194974 | 0.348343 | 0.2771 | 0.42358 |

| | | | | | | | | | | |
|---|---|---|---|---|---|---|---|---|---|---|
| SL1344_RS18855_ | -2.40652 | -2.64853 | -2.06068 | 0.566165 | 0.420388 | 1.298166 | 0.085943 | 0.141592 | 0.28543 | 0.2155 | 0.363257 |
| SL1344_RS18860_ | 0.083931 | 0.038925 | 0.145643 | 0.105361 | 0.211211 | 0.477479 | 0.285143 | 0.462744 | 0.610327 | 0.5807 | 0.697609 |
| SL1344_RS18865_ | 0.045114 | -0.00688 | 0.079573 | 0.073218 | 0.214198 | 0.326031 | 0.362425 | 0.571832 | 0.702236 | 0.6742 | 0.774448 |
| SL1344_RS18870_ | -0.02523 | -0.25498 | 0.299261 | 0.550415 | 0.198988 | 2.673527 | 0.001336 | 0.026601 | 0.113983 | 0.1014 | 0.238349 |
| SL1344_RS18875_ | 3.129214 | 3.322348 | 3.032422 | -0.27647 | 0.108933 | -2.39726 | 0.000673 | 0.027688 | 0.121099 | 0.1004 | 0.237123 |
| SL1344_RS18880_ | -2.80154 | -3.01339 | -2.39898 | 0.590376 | 0.503147 | 1.055449 | 0.137242 | 0.201906 | 0.355434 | 0.2962 | 0.447443 |
| SL1344_RS18885_ | -1.44997 | -1.56441 | -1.30611 | 0.262129 | 0.292571 | 0.824921 | 0.177333 | 0.28244 | 0.434226 | 0.3779 | 0.515099 |
| SL1344_RS18890_ | -0.03003 | 0.171984 | -0.26515 | -0.4347 | 0.221655 | -1.84582 | 0.018654 | 0.056853 | 0.17199 | 0.1193 | 0.260826 |
| SL1344_RS18895_ | 1.32336 | 1.270866 | 1.441117 | 0.169262 | 0.14377 | 1.090996 | 0.12 | 0.191909 | 0.353923 | 0.2724 | 0.422404 |
| SL1344_RS18900_ | 0.94982 | 0.972059 | 0.916631 | -0.04446 | 0.154273 | -0.27261 | 0.385077 | 0.606357 | 0.730131 | 0.7224 | 0.809924 |
| SL1344_RS18905_ | -2.48271 | -2.42676 | -2.53328 | -0.11173 | 0.448568 | -0.23841 | 0.397333 | 0.56807 | 0.69592 | 0.6814 | 0.775969 |
| SL1344_RS18910_ | -4.18941 | -3.8864 | -4.36974 | -0.43336 | 0.99677 | -0.3995 | 0.324 | 0.532414 | 0.668941 | 0.6281 | 0.739102 |
| SL1344_RS18915_ | -3.28998 | -3.32729 | -3.24053 | 0.072456 | 0.497872 | 0.132378 | 0.44 | 0.529996 | 0.6632 | 0.6354 | 0.73905 |
| SL1344_RS18920_ | -5.73799 | -5.52439 | -5.94431 | -0.42587 | 1.345085 | -0.29933 | 0.360667 | 0.516126 | 0.650737 | 0.627 | 0.730573 |
| SL1344_RS18925_ | -4.33146 | -4.26879 | -4.38427 | -0.17368 | 1.177337 | -0.1331 | 0.429333 | 0.661002 | 0.776574 | 0.7699 | 0.846439 |
| SL1344_RS18930_ | -3.78961 | -3.49846 | -4.11361 | -0.62325 | 0.590952 | -0.96509 | 0.143238 | 0.218385 | 0.363371 | 0.3006 | 0.441429 |
| SL1344_RS18935_ | -2.97763 | -2.69869 | -3.20171 | -0.48626 | 0.491927 | -0.94533 | 0.149333 | 0.235617 | 0.389624 | 0.3229 | 0.47011 |
| SL1344_RS18945_ | -3.82312 | -3.44403 | -4.20831 | -0.75209 | 0.63134 | -1.12981 | 0.103333 | 0.172804 | 0.316519 | 0.2515 | 0.39669 |
| SL1344_RS18950_ | -3.35227 | -3.19279 | -3.53989 | -0.35327 | 0.491877 | -0.68522 | 0.234667 | 0.336974 | 0.482706 | 0.441 | 0.568931 |
| SL1344_RS18955_ | 0.044077 | -0.05599 | 0.150753 | 0.241743 | 0.238889 | 0.977651 | 0.120667 | 0.192861 | 0.35525 | 0.2776 | 0.427532 |
| SL1344_RS18960_ | -0.23091 | -0.14641 | -0.33587 | -0.18651 | 0.183924 | -0.94232 | 0.130667 | 0.214958 | 0.364993 | 0.3055 | 0.448432 |
| SL1344_RS18965_ | 1.466671 | 1.065347 | 1.860946 | 0.814763 | 0.186432 | 4.413467 | 4.68E-05 | 0.009848 | 0.072769 | 0.1 | 0.236592 |
| SL1344_RS18970_ | -0.27415 | -0.52895 | 0.016087 | 0.539433 | 0.234183 | 2.231185 | 0.006663 | 0.035133 | 0.132553 | 0.1055 | 0.24373 |
| SL1344_RS18975_ | 2.562745 | 2.421029 | 2.643209 | 0.218379 | 0.087358 | 2.392835 | 0.004668 | 0.037305 | 0.136795 | 0.1046 | 0.242514 |
| SL1344_RS18980_ | 2.706901 | 2.642638 | 2.72582 | 0.074541 | 0.137898 | 0.335933 | 0.360667 | 0.83873 | 0.904386 | 0.7597 | 0.857452 |
| SL1344_RS18985_ | 2.202776 | 2.019004 | 2.38719 | 0.36805 | 0.092412 | 3.915805 | 4.68E-05 | 0.007775 | 0.063785 | 0.1 | 0.236592 |
| SL1344_RS18990_ | 3.495488 | 3.135998 | 3.711511 | 0.572629 | 0.149131 | 3.873732 | 4.68E-05 | 0.003909 | 0.051761 | 0.1 | 0.236592 |
| SL1344_RS18995_ | 2.14307 | 2.000948 | 2.249847 | 0.248831 | 0.092387 | 2.614539 | 0.002002 | 0.026469 | 0.113035 | 0.1023 | 0.239565 |
| SL1344_RS19005_ | -1.73871 | -3.27436 | -0.87081 | 2.382566 | 0.504618 | 4.600855 | 4.68E-05 | 0.012826 | 0.080411 | 0.1 | 0.236592 |
| SL1344_RS19010_ | 1.368581 | 1.368912 | 1.36785 | -0.01052 | 0.18667 | -0.04796 | 0.480667 | 0.736041 | 0.833217 | 0.8351 | 0.898824 |
| SL1344_RS19015_ | -5.94445 | -5.50078 | -6.39176 | -1.00075 | 1.52712 | -0.58847 | 0.24517 | 0.378711 | 0.530013 | 0.4824 | 0.609035 |
| SL1344_RS19020_ | -0.28109 | -0.32474 | -0.24117 | 0.078789 | 0.189587 | 0.378207 | 0.349101 | 0.495696 | 0.629669 | 0.606 | 0.711638 |
| SL1344_RS19025_ | -0.61256 | -0.44879 | -0.72722 | -0.28123 | 0.296701 | -0.89957 | 0.171885 | 0.251566 | 0.418433 | 0.349 | 0.500872 |
| SL1344_RS19030_ | -2.33951 | -2.24606 | -2.42784 | -0.17644 | 0.42458 | -0.38222 | 0.327781 | 0.504037 | 0.638333 | 0.6145 | 0.719174 |
| SL1344_RS19040_ | -1.19372 | -1.0437 | -1.30395 | -0.30539 | 0.381575 | -0.80486 | 0.172 | 0.273193 | 0.444233 | 0.3678 | 0.510456 |
| SL1344_RS19045_ | 0.372855 | 0.388424 | 0.35811 | -0.04478 | 0.185484 | -0.23713 | 0.401066 | 0.583901 | 0.710651 | 0.7094 | 0.796767 |
| SL1344_RS19050_ | 1.121604 | 0.704149 | 1.490679 | 0.791136 | 0.152696 | 5.254286 | 4.68E-05 | 0.00974 | 0.071071 | 0.1 | 0.236592 |
| SL1344_RS19055_ | -0.08015 | -0.50517 | 0.298891 | 0.815544 | 0.186656 | 4.322204 | 4.68E-05 | 0.008289 | 0.064321 | 0.1 | 0.236592 |
| SL1344_RS19065_ | 3.933556 | 3.818989 | 4.020166 | 0.195188 | 0.073747 | 2.616323 | 0.000673 | 0.019693 | 0.101113 | 0.1006 | 0.237393 |
| SL1344_RS19070_ | 4.475979 | 4.359175 | 4.488733 | 0.124108 | 0.083475 | 0.760303 | 0.313791 | 0.378144 | 0.559928 | 0.6498 | 0.781073 |
| SL1344_RS19075_ | 2.48816 | 2.482897 | 2.493591 | 0.010499 | 0.091229 | 0.119021 | 0.444667 | 0.63366 | 0.751393 | 0.7376 | 0.822553 |
| SL1344_RS19080_ | 2.201467 | 2.213344 | 2.177195 | -0.02399 | 0.114327 | -0.18257 | 0.428381 | 0.713367 | 0.815007 | 0.7852 | 0.86476 |
| SL1344_RS19085_ | 2.511982 | 2.504317 | 2.519187 | 0.002724 | 0.111631 | 0.023547 | 0.49 | 0.730819 | 0.826552 | 0.8249 | 0.887122 |
| SL1344_RS19090_ | 2.39559 | 2.509965 | 2.26708 | -0.24226 | 0.085996 | -2.7497 | 0.001337 | 0.021329 | 0.1023 | 0.101 | 0.237924 |
| SL1344_RS19095_ | 1.082058 | 0.955804 | 1.156748 | 0.199712 | 0.146444 | 1.29368 | 0.078667 | 0.137693 | 0.286606 | 0.2019 | 0.351238 |

| | | | | | | | | | | |
|---|---|---|---|---|---|---|---|---|---|---|---|
| SL1344_RS19100_ | -0.93334 | -0.96606 | -0.91118 | 0.083108 | 0.291245 | 0.262725 | 0.383078 | 0.562429 | 0.693416 | 0.6857 | 0.779124 |
| SL1344_RS19105_ | 4.676685 | 4.433201 | 4.973087 | 0.545394 | 0.102269 | 5.496325 | 4.68E-05 | 0.002437 | 0.043277 | 0.1 | 0.236592 |
| SL1344_RS19110_ | 3.846955 | 3.558097 | 4.101895 | 0.541758 | 0.058687 | 9.245241 | 4.68E-05 | 0.000982 | 0.030349 | 0.1 | 0.236592 |
| SL1344_RS19115_ | 2.951832 | 2.900757 | 3.015506 | 0.122563 | 0.116349 | 1.046829 | 0.118588 | 0.176863 | 0.343028 | 0.2578 | 0.418783 |
| SL1344_RS19120_ | 2.680115 | 2.616427 | 2.749308 | 0.132264 | 0.060221 | 2.122515 | 0.009334 | 0.041413 | 0.138391 | 0.1133 | 0.25269 |
| SL1344_RS19125_ | 1.927122 | 1.853403 | 1.962528 | 0.086808 | 0.147703 | 0.509882 | 0.318 | 0.527767 | 0.673737 | 0.6348 | 0.754274 |
| SL1344_RS19130_ | 2.285553 | 2.311718 | 2.265943 | -0.04915 | 0.148701 | -0.34619 | 0.343105 | 0.593346 | 0.726101 | 0.6944 | 0.794033 |
| SL1344_RS19135_ | 1.736936 | 1.845884 | 1.675784 | -0.16471 | 0.113369 | -1.33182 | 0.080667 | 0.132734 | 0.279734 | 0.2088 | 0.359934 |
| SL1344_RS19140_ | 1.060295 | 0.942699 | 1.19164 | 0.264582 | 0.157659 | 1.636192 | 0.022 | 0.077882 | 0.20715 | 0.1235 | 0.264969 |
| SL1344_RS19145_ | 0.467924 | 0.497233 | 0.433429 | -0.06496 | 0.140501 | -0.43751 | 0.303797 | 0.441698 | 0.583742 | 0.5554 | 0.670583 |
| SL1344_RS19150_ | 1.582746 | 1.608731 | 1.548176 | -0.06512 | 0.113097 | -0.54114 | 0.278 | 0.429024 | 0.576328 | 0.544 | 0.66209 |
| SL1344_RS19155_ | 1.835016 | 1.872556 | 1.80589 | -0.07891 | 0.121138 | -0.58142 | 0.253165 | 0.369574 | 0.528639 | 0.4866 | 0.615248 |
| SL1344_RS19160_ | 1.867497 | 1.929729 | 1.835663 | -0.09515 | 0.11944 | -0.72866 | 0.215333 | 0.317551 | 0.483162 | 0.4365 | 0.576388 |
| SL1344_RS19165_ | 0.230291 | 0.238902 | 0.219882 | -0.0271 | 0.195657 | -0.12387 | 0.442667 | 0.618871 | 0.739731 | 0.732 | 0.816511 |
| SL1344_RS19170_ | 1.819281 | 1.809259 | 1.827466 | 0.024839 | 0.14804 | 0.151524 | 0.432378 | 0.685121 | 0.792967 | 0.7821 | 0.856434 |
| SL1344_RS19175_ | 2.233933 | 2.253721 | 2.219873 | -0.02748 | 0.09079 | -0.28385 | 0.379747 | 0.57868 | 0.70459 | 0.6799 | 0.777493 |
| SL1344_RS19180_ | 1.869523 | 1.982479 | 1.764515 | -0.22072 | 0.090923 | -2.40281 | 0.003998 | 0.029297 | 0.118614 | 0.1034 | 0.241019 |
| SL1344_RS19185_ | 2.208626 | 2.244658 | 2.172079 | -0.07366 | 0.109303 | -0.65504 | 0.236509 | 0.360402 | 0.521631 | 0.4784 | 0.614055 |
| SL1344_RS19190_ | 1.805762 | 1.795331 | 1.824242 | 0.034479 | 0.133607 | 0.237617 | 0.409061 | 0.632034 | 0.752612 | 0.7644 | 0.841923 |
| SL1344_RS19195_ | 0.206395 | 0.247102 | 0.159516 | -0.08791 | 0.179365 | -0.46787 | 0.304667 | 0.467737 | 0.609284 | 0.5812 | 0.694185 |
| SL1344_RS19200_ | 5.422428 | 5.742637 | 5.076044 | -0.74951 | 0.181116 | -3.939 | 4.68E-05 | 0.007434 | 0.068195 | 0.1 | 0.236592 |
| SL1344_RS19205_ | 4.791394 | 5.11672 | 4.360995 | -0.80306 | 0.211434 | -4.0539 | 4.68E-05 | 0.002488 | 0.045231 | 0.1 | 0.236592 |
| SL1344_RS19210_ | -0.1601 | -0.18904 | -0.12953 | 0.06039 | 0.1706 | 0.323142 | 0.353333 | 0.49284 | 0.627809 | 0.5874 | 0.697723 |
| SL1344_RS19215_ | 1.958258 | 2.023525 | 1.885703 | -0.14792 | 0.12048 | -1.21481 | 0.086 | 0.143169 | 0.294539 | 0.209 | 0.360885 |
| SL1344_RS19220_ | 1.870901 | 1.805909 | 1.90468 | 0.091036 | 0.123921 | 0.67873 | 0.231333 | 0.363026 | 0.523032 | 0.466 | 0.607508 |
| SL1344_RS19225_ | 1.529295 | 1.723061 | 1.425303 | -0.28663 | 0.125284 | -2.12314 | 0.007334 | 0.045258 | 0.152167 | 0.1126 | 0.252062 |
| SL1344_RS19230_ | 0.186394 | -0.06554 | 0.509585 | 0.595747 | 0.218466 | 2.69736 | 4.68E-05 | 0.025249 | 0.11358 | 0.1005 | 0.237204 |
| SL1344_RS19235_ | -0.29529 | -0.33954 | -0.24954 | 0.10455 | 0.222333 | 0.443604 | 0.319787 | 0.470694 | 0.616978 | 0.5953 | 0.707986 |
| SL1344_RS19240_ | 2.542445 | 2.549417 | 2.535781 | 0.021716 | 0.132922 | 0.159645 | 0.441039 | 0.675321 | 0.787936 | 0.8681 | 0.913057 |
| SL1344_RS19245_ | -0.57555 | -0.61135 | -0.52903 | 0.080233 | 0.196322 | 0.379255 | 0.323333 | 0.474852 | 0.611861 | 0.5784 | 0.688741 |
| SL1344_RS19250_ | -2.229 | -2.31646 | -2.16349 | 0.189756 | 0.498342 | 0.359832 | 0.349767 | 0.520825 | 0.661289 | 0.6542 | 0.75341 |
| SL1344_RS19255_ | 0.210482 | 0.252008 | 0.094621 | -0.16543 | 0.267113 | -0.58151 | 0.302667 | 0.435788 | 0.600705 | 0.5983 | 0.720564 |
| SL1344_RS19260_ | -0.90474 | -0.89154 | -0.92845 | -0.03413 | 0.245982 | -0.11522 | 0.45 | 0.594658 | 0.718183 | 0.7052 | 0.796335 |
| SL1344_RS19265_ | 1.642263 | 1.636807 | 1.645912 | 0.009151 | 0.107695 | 0.083873 | 0.463025 | 0.614962 | 0.735532 | 0.7285 | 0.813778 |
| SL1344_RS19270_ | 2.131409 | 2.133257 | 2.130242 | -0.00331 | 0.135554 | -0.02746 | 0.488 | 0.722132 | 0.821847 | 0.8037 | 0.872425 |
| SL1344_RS19275_ | 3.644064 | 3.672223 | 3.607611 | -0.04343 | 0.099276 | -0.43898 | 0.331333 | 0.53983 | 0.68413 | 0.658 | 0.764447 |
| SL1344_RS19280_ | 1.559923 | 1.602773 | 1.520403 | -0.08324 | 0.104167 | -0.77321 | 0.182667 | 0.285763 | 0.440363 | 0.3816 | 0.519338 |
| SL1344_RS19285_ | 1.870233 | 1.862171 | 1.879845 | -0.00015 | 0.128049 | -0.00076 | 0.499333 | 0.692917 | 0.799143 | 0.8094 | 0.873582 |
| SL1344_RS19290_ | 1.817007 | 2.422528 | 1.488286 | -0.91144 | 0.15895 | -4.77078 | 4.68E-05 | 0.015898 | 0.094374 | 0.1 | 0.236592 |
| SL1344_RS19295_ | 2.015968 | 2.659585 | 1.725652 | -0.91853 | 0.147833 | -4.51009 | 4.68E-05 | 0.036218 | 0.143774 | 0.1 | 0.236592 |
| SL1344_RS19300_ | -0.84717 | -0.79733 | -0.88426 | -0.07973 | 0.228322 | -0.32632 | 0.36509 | 0.531283 | 0.662049 | 0.6275 | 0.732703 |
| SL1344_RS19305_ | 2.597569 | 2.512191 | 2.774494 | 0.242493 | 0.113502 | 1.958386 | 0.014657 | 0.05541 | 0.174438 | 0.1138 | 0.25433 |
| SL1344_RS19310_ | -1.10995 | -1.2692 | -0.97865 | 0.291268 | 0.258949 | 1.053007 | 0.112 | 0.184194 | 0.333597 | 0.2594 | 0.405954 |
| SL1344_RS19315_ | -2.53353 | -2.42954 | -2.58847 | -0.18004 | 0.546566 | -0.31412 | 0.366422 | 0.581589 | 0.70911 | 0.7004 | 0.793271 |
| SL1344_RS19320_ | -1.07416 | -1.13884 | -1.00885 | 0.03372 | 0.451423 | 0.072652 | 0.472667 | 0.746208 | 0.840139 | 0.8674 | 0.914605 |

| | | | | | | | | | | | |
|---|---|---|---|---|---|---|---|---|---|---|---|
| SL1344_RS19325_ | -2.77248 | -2.90841 | -2.71051 | 0.155509 | 0.551415 | 0.271888 | 0.389074 | 0.605802 | 0.728667 | 0.7096 | 0.802417 |
| SL1344_RS19330_ | 2.03891 | 2.228036 | 1.992062 | -0.23704 | 0.180964 | -1.20989 | 0.094667 | 0.178294 | 0.351755 | 0.2198 | 0.371845 |
| SL1344_RS19335_ | 0.451651 | 0.323311 | 0.622307 | 0.297539 | 0.156382 | 1.841606 | 0.018 | 0.061318 | 0.176187 | 0.1207 | 0.261049 |
| SL1344_RS19340_ | 1.46228 | 1.504274 | 1.400755 | -0.09507 | 0.12389 | -0.74396 | 0.199201 | 0.316605 | 0.475218 | 0.412 | 0.555229 |
| SL1344_RS19345_ | -4.86877 | -4.80165 | -4.93651 | -0.1422 | 0.969657 | -0.13125 | 0.439707 | 0.576923 | 0.701644 | 0.6848 | 0.777408 |
| SL1344_RS19350_ | -0.92532 | -0.89013 | -0.97499 | -0.02335 | 0.438465 | -0.04947 | 0.477333 | 0.758803 | 0.849165 | 0.8707 | 0.918874 |
| SL1344_RS19355_ | -2.0562 | -2.29344 | -1.81034 | 0.490767 | 0.358087 | 1.310625 | 0.067289 | 0.123808 | 0.261113 | 0.1933 | 0.34018 |
| SL1344_RS19360_ | -1.99739 | -2.06592 | -1.92136 | 0.14767 | 0.303057 | 0.466519 | 0.309127 | 0.432675 | 0.573113 | 0.537 | 0.653055 |
| SL1344_RS19365_ | -1.39946 | -1.1939 | -1.67818 | -0.47911 | 0.279898 | -1.62714 | 0.038 | 0.078415 | 0.200253 | 0.1423 | 0.285748 |
| SL1344_RS19370_ | 0.763257 | 0.727355 | 0.96067 | 0.21197 | 0.211148 | 0.749792 | 0.271333 | 0.394579 | 0.564062 | 0.5623 | 0.700958 |
| SL1344_RS19375_ | 0.185643 | 0.29309 | 0.097627 | -0.18993 | 0.163735 | -1.11271 | 0.116589 | 0.180426 | 0.328617 | 0.2596 | 0.407133 |
| SL1344_RS19380_ | -2.25786 | -1.81116 | -2.73666 | -0.92529 | 0.442239 | -2.02926 | 0.010667 | 0.044069 | 0.148177 | 0.1119 | 0.251932 |
| SL1344_RS19385_ | 0.121444 | 0.244861 | 0.021545 | -0.25543 | 0.247176 | -1.01994 | 0.102667 | 0.187591 | 0.351655 | 0.2442 | 0.391233 |
| SL1344_RS19390_ | -1.03141 | -1.22896 | -0.95661 | 0.254259 | 0.327051 | 0.690728 | 0.244 | 0.364373 | 0.526983 | 0.4983 | 0.632619 |
| SL1344_RS19395_ | -0.61821 | -0.7239 | -0.56859 | 0.124861 | 0.293294 | 0.383397 | 0.341106 | 0.568239 | 0.701419 | 0.6832 | 0.78529 |
| SL1344_RS19400_ | -1.07838 | -1.20454 | -0.95448 | 0.242388 | 0.249831 | 0.929363 | 0.144667 | 0.226573 | 0.374991 | 0.3112 | 0.455271 |
| SL1344_RS19405_ | -1.93621 | -2.18714 | -1.66346 | 0.523448 | 0.345426 | 1.454091 | 0.064 | 0.104773 | 0.238332 | 0.173 | 0.320489 |
| SL1344_RS19410_ | -2.35643 | -2.70161 | -2.06244 | 0.653574 | 0.565853 | 1.110143 | 0.090606 | 0.160471 | 0.31774 | 0.2335 | 0.388 |
| SL1344_RS19415_ | -3.10453 | -3.25707 | -2.94785 | 0.287189 | 0.556813 | 0.454843 | 0.309333 | 0.466504 | 0.609914 | 0.5656 | 0.684286 |
| SL1344_RS19420_ | -4.17891 | -4.07703 | -4.27556 | -0.18988 | 0.802233 | -0.22483 | 0.398401 | 0.568677 | 0.695715 | 0.6754 | 0.77084 |
| SL1344_RS19425_ | 2.323733 | 2.239376 | 2.390878 | 0.161169 | 0.107686 | 1.507196 | 0.035976 | 0.106236 | 0.249513 | 0.1412 | 0.284236 |
| SL1344_RS19430_ | -2.16719 | -2.68555 | -1.66969 | 1.151066 | 0.626185 | 1.829613 | 0.002668 | 0.049559 | 0.164385 | 0.1043 | 0.242143 |
| SL1344_RS19435_ | 0.814355 | 0.645465 | 0.948205 | 0.319397 | 0.163876 | 1.857394 | 0.008001 | 0.064557 | 0.185936 | 0.1083 | 0.247191 |
| SL1344_RS19445_ | -0.43064 | -0.49078 | -0.39391 | 0.092228 | 0.339628 | 0.242172 | 0.387742 | 0.635367 | 0.758031 | 0.7748 | 0.850391 |
| SL1344_RS19450_ | 0.259375 | 0.12891 | 0.348238 | 0.236094 | 0.237597 | 0.974878 | 0.112 | 0.201009 | 0.369227 | 0.2535 | 0.405248 |
| SL1344_RS19455_ | -2.02494 | -2.10559 | -1.94502 | 0.16181 | 0.37579 | 0.413964 | 0.325333 | 0.487682 | 0.624851 | 0.5966 | 0.704091 |
| SL1344_RS19460_ | -2.54026 | -3.30319 | -2.37553 | 0.89119 | 0.684297 | 1.023479 | 0.164557 | 0.212102 | 0.3858 | 0.3318 | 0.485881 |
| SL1344_RS19465_ | 0.197211 | -0.02297 | 0.423228 | 0.447021 | 0.168748 | 2.616476 | 0.003332 | 0.025666 | 0.110764 | 0.1025 | 0.23978 |
| SL1344_RS19470_ | -0.09982 | -0.32644 | 0.095533 | 0.429988 | 0.187664 | 2.210194 | 0.004665 | 0.040014 | 0.139959 | 0.104 | 0.241763 |
| SL1344_RS19475_ | -0.2964 | -0.34407 | -0.25693 | 0.085673 | 0.19156 | 0.427866 | 0.32445 | 0.462704 | 0.60223 | 0.5693 | 0.681518 |
| SL1344_RS19480_ | 0.172568 | 0.081449 | 0.22444 | 0.129703 | 0.195501 | 0.612851 | 0.253165 | 0.400838 | 0.553719 | 0.507 | 0.638228 |
| SL1344_RS19485_ | 1.913198 | 1.778834 | 2.013362 | 0.24606 | 0.17259 | 1.414603 | 0.03 | 0.11495 | 0.268057 | 0.1295 | 0.273638 |
| SL1344_RS19490_ | -0.28205 | -0.39642 | -0.20623 | 0.135216 | 0.291582 | 0.447731 | 0.321785 | 0.535542 | 0.675382 | 0.6402 | 0.748366 |
| SL1344_RS19495_ | -1.85677 | -2.03473 | -1.58172 | 0.432746 | 0.382631 | 1.07554 | 0.126582 | 0.203854 | 0.360877 | 0.291 | 0.442559 |
| SL1344_RS19500_ | -0.77883 | -0.79206 | -0.76793 | 0.039191 | 0.279019 | 0.125782 | 0.440373 | 0.617515 | 0.738845 | 0.7312 | 0.814605 |
| SL1344_RS19505_ | -0.47778 | -0.33906 | -0.64302 | -0.30811 | 0.204815 | -1.41302 | 0.052 | 0.106803 | 0.240511 | 0.1687 | 0.314985 |
| SL1344_RS19510_ | -1.20386 | -0.93586 | -1.4777 | -0.54052 | 0.235932 | -2.22591 | 0.008661 | 0.039376 | 0.135429 | 0.1116 | 0.250682 |
| SL1344_RS19515_ | 0.333128 | 0.242312 | 0.492464 | 0.238589 | 0.203612 | 1.122681 | 0.124 | 0.178265 | 0.336502 | 0.2651 | 0.421052 |
| SL1344_RS19520_ | -0.26731 | -0.53046 | 0.008469 | 0.539357 | 0.184186 | 2.909253 | 0.000673 | 0.017891 | 0.092007 | 0.1006 | 0.237389 |
| SL1344_RS19525_ | -0.01336 | -0.19834 | 0.180809 | 0.398066 | 0.210948 | 1.931608 | 0.011992 | 0.054105 | 0.166492 | 0.1119 | 0.251293 |
| SL1344_RS19530_ | -0.48676 | -0.48316 | -0.48998 | -0.00605 | 0.200588 | -0.03249 | 0.487675 | 0.567488 | 0.693378 | 0.6825 | 0.774228 |
| SL1344_RS19535_ | -1.10518 | -0.82751 | -1.25107 | -0.41303 | 0.310646 | -1.23889 | 0.091333 | 0.156337 | 0.309667 | 0.2241 | 0.374857 |
| SL1344_RS19540_ | 0.773628 | 1.017481 | 0.48574 | -0.53148 | 0.13068 | -3.98255 | 4.68E-05 | 0.007892 | 0.063173 | 0.1 | 0.236592 |
| SL1344_RS19545_ | 2.693925 | 2.965679 | 2.657438 | -0.31665 | 0.32447 | -0.5865 | 0.333111 | 0.605565 | 0.749385 | 0.7 | 0.822108 |
| SL1344_RS19550_ | 1.419732 | 1.619624 | 1.188159 | -0.44959 | 0.257116 | -1.77442 | 0.003335 | 0.082778 | 0.223825 | 0.1024 | 0.239776 |

| | | | | | | | | | | | |
|---|---|---|---|---|---|---|---|---|---|---|---|
| SL1344_RS19555_ | -1.6069 | -1.58703 | -1.62287 | -0.03674 | 0.346828 | -0.09019 | 0.459027 | 0.6137 | 0.733973 | 0.7201 | 0.806175 |
| SL1344_RS19565_ | 0.558231 | 0.42244 | 0.717425 | 0.293802 | 0.157385 | 1.776795 | 0.020667 | 0.063082 | 0.180724 | 0.1245 | 0.266657 |
| SL1344_RS19570_ | -1.20074 | -1.30715 | -1.07457 | 0.2393 | 0.288793 | 0.789039 | 0.198534 | 0.3022 | 0.456406 | 0.4014 | 0.539122 |
| SL1344_RS19575_ | -2.1071 | -2.19302 | -2.04023 | 0.142541 | 0.36183 | 0.367621 | 0.33511 | 0.478054 | 0.618895 | 0.6046 | 0.713284 |
| SL1344_RS19580_ | 0.13209 | -0.1084 | 0.331147 | 0.440644 | 0.163802 | 2.616803 | 0.002667 | 0.026696 | 0.112169 | 0.1035 | 0.240782 |
| SL1344_RS19585_ | 1.896502 | 1.818841 | 1.969405 | 0.15209 | 0.087886 | 1.674565 | 0.030646 | 0.075706 | 0.195579 | 0.1394 | 0.282879 |
| SL1344_RS19590_ | 0.720653 | 0.647668 | 0.786532 | 0.139738 | 0.121905 | 1.05502 | 0.125333 | 0.193614 | 0.337044 | 0.2809 | 0.424017 |
| SL1344_RS19595_ | 0.428896 | 0.348488 | 0.532556 | 0.14124 | 0.233626 | 0.591904 | 0.267155 | 0.430662 | 0.588799 | 0.5346 | 0.666034 |
| SL1344_RS19600_ | 1.850029 | 1.540382 | 2.105693 | 0.55848 | 0.14496 | 3.845744 | 4.68E-05 | 0.007877 | 0.065859 | 0.1 | 0.236592 |
| SL1344_RS19605_ | 2.238832 | 2.92623 | 1.703368 | -1.20913 | 0.268842 | -4.34021 | 4.68E-05 | 0.008032 | 0.06969 | 0.1 | 0.236592 |
| SL1344_RS19610_ | 2.808452 | 2.910067 | 2.721964 | -0.21753 | 0.168509 | -1.36433 | 0.02998 | 0.12217 | 0.281183 | 0.1231 | 0.266607 |
| SL1344_RS19615_ | 1.019632 | 0.954494 | 1.308935 | 0.319717 | 0.22748 | 1.116037 | 0.165333 | 0.204127 | 0.37896 | 0.3313 | 0.489833 |
| SL1344_RS19620_ | 0.881065 | 0.902187 | 0.857517 | -0.04356 | 0.133884 | -0.3035 | 0.352667 | 0.522317 | 0.653406 | 0.6253 | 0.729241 |
| SL1344_RS19625_ | 0.87503 | 0.95005 | 0.803696 | -0.16508 | 0.175155 | -0.89485 | 0.157895 | 0.241718 | 0.407463 | 0.3413 | 0.491914 |
| SL1344_RS19630_ | -1.58301 | -1.656 | -1.50056 | 0.229878 | 0.458586 | 0.480121 | 0.300667 | 0.458415 | 0.613375 | 0.6214 | 0.73003 |
| SL1344_RS19635_ | 0.741772 | 0.57064 | 0.962638 | 0.381529 | 0.20554 | 1.779167 | 0.030646 | 0.063389 | 0.185696 | 0.1243 | 0.267802 |
| SL1344_RS19640_ | -0.34466 | -0.3935 | -0.27067 | 0.125688 | 0.295339 | 0.401519 | 0.342438 | 0.550633 | 0.687841 | 0.6714 | 0.77238 |
| SL1344_RS19645_ | -2.01957 | -2.24408 | -1.88503 | 0.358368 | 0.377388 | 0.88815 | 0.165223 | 0.250661 | 0.406728 | 0.3477 | 0.489602 |
| SL1344_RS19650_ | 0.433738 | 0.245577 | 0.540436 | 0.267717 | 0.398559 | 0.5905 | 0.276667 | 0.401744 | 0.578877 | 0.5424 | 0.686354 |
| SL1344_RS19655_ | -0.36552 | -0.56839 | -0.08602 | 0.487159 | 0.361047 | 1.315854 | 0.072 | 0.112857 | 0.26284 | 0.1779 | 0.333701 |
| SL1344_RS19660_ | -0.32813 | -0.68094 | -0.0128 | 0.61528 | 0.351349 | 1.701765 | 0.028 | 0.067398 | 0.194936 | 0.1276 | 0.2731 |
| SL1344_RS19665_ | 3.075291 | 3.258094 | 2.888678 | -0.36374 | 0.116094 | -3.12237 | 4.68E-05 | 0.013322 | 0.085167 | 0.1 | 0.236592 |
| SL1344_RS19670_ | -2.99092 | -3.13888 | -2.84042 | 0.309035 | 0.448539 | 0.636087 | 0.245836 | 0.355224 | 0.502592 | 0.4646 | 0.589272 |
| SL1344_RS19675_ | -0.85939 | -0.84133 | -0.8789 | -0.05466 | 0.281989 | -0.17252 | 0.426382 | 0.608304 | 0.730614 | 0.7301 | 0.812001 |
| SL1344_RS19680_ | -2.5461 | -2.67203 | -2.42053 | 0.262944 | 0.58898 | 0.423208 | 0.32445 | 0.512288 | 0.653291 | 0.6293 | 0.736696 |
| SL1344_RS19685_ | 2.219255 | 2.067186 | 2.278754 | 0.214008 | 0.145105 | 1.344015 | 0.034 | 0.140133 | 0.30242 | 0.1435 | 0.288204 |
| SL1344_RS19690_ | -0.11626 | -0.2079 | 0.066881 | 0.246593 | 0.33665 | 0.703368 | 0.223851 | 0.338248 | 0.510353 | 0.4243 | 0.583431 |
| SL1344_RS19695_ | 0.216445 | 0.160767 | 0.251449 | 0.094711 | 0.171959 | 0.51489 | 0.283145 | 0.432057 | 0.577548 | 0.5396 | 0.656904 |
| SL1344_RS19700_ | -1.9526 | -2.08752 | -1.89621 | 0.194296 | 0.43844 | 0.413265 | 0.336442 | 0.499261 | 0.644578 | 0.635 | 0.742224 |
| SL1344_RS19705_ | -1.32666 | -1.37995 | -1.26025 | 0.085755 | 0.437315 | 0.185936 | 0.411059 | 0.687614 | 0.796173 | 0.759 | 0.841967 |
| SL1344_RS19710_ | -2.62939 | -2.92097 | -2.30707 | 0.611134 | 0.454745 | 1.27066 | 0.085943 | 0.139394 | 0.279731 | 0.214 | 0.36211 |
| SL1344_RS19715_ | -0.16277 | -0.26164 | -0.06558 | 0.2162 | 0.260585 | 0.81947 | 0.173333 | 0.268012 | 0.435145 | 0.3504 | 0.497986 |
| SL1344_RS19720_ | -0.75693 | -0.85082 | -0.5462 | 0.298596 | 0.298225 | 0.890719 | 0.185333 | 0.271095 | 0.436839 | 0.3846 | 0.533123 |
| SL1344_RS19725_ | 0.934177 | 0.745903 | 1.034088 | 0.240359 | 0.24854 | 0.927854 | 0.159227 | 0.23215 | 0.407668 | 0.3142 | 0.477375 |
| SL1344_RS19730_ | -2.36106 | -2.50033 | -2.22072 | 0.284489 | 0.5156 | 0.532494 | 0.281333 | 0.428937 | 0.583367 | 0.5378 | 0.664436 |
| SL1344_RS19735_ | -0.93342 | -1.04402 | -0.80132 | 0.24379 | 0.251445 | 0.896208 | 0.154667 | 0.26339 | 0.417366 | 0.3563 | 0.497867 |
| SL1344_RS19740_ | 1.623215 | 1.379903 | 2.004856 | 0.602541 | 0.1642 | 3.476806 | 4.68E-05 | 0.017948 | 0.097479 | 0.1 | 0.236592 |
| SL1344_RS19745_ | 4.239708 | 4.239133 | 4.240015 | 0.008941 | 0.068988 | 0.127022 | 0.442372 | 0.689372 | 0.796998 | 0.8035 | 0.870071 |
| SL1344_RS19750_ | 1.620013 | 1.612664 | 1.627051 | 0.015095 | 0.102293 | 0.151044 | 0.440373 | 0.583991 | 0.70932 | 0.6897 | 0.782472 |
| SL1344_RS19755_ | 3.368584 | 3.360583 | 3.382331 | 0.028321 | 0.094723 | 0.28285 | 0.383744 | 0.633495 | 0.75527 | 0.7569 | 0.837588 |
| SL1344_RS19765_ | 3.95807 | 4.405141 | 3.560993 | -0.80305 | 0.239639 | -3.29687 | 4.68E-05 | 0.006421 | 0.064066 | 0.1 | 0.236592 |
| SL1344_RS19770_ | 1.895545 | 2.177898 | 1.455163 | -0.66099 | 0.401438 | -1.64285 | 0.022667 | 0.055186 | 0.179918 | 0.12 | 0.263339 |
| SL1344_RS19775_ | -0.0665 | 0.440479 | -0.27739 | -0.69523 | 0.252835 | -2.32129 | 0.003332 | 0.053241 | 0.172454 | 0.1025 | 0.239901 |
| SL1344_RS19780_ | 4.057021 | 4.239931 | 3.799076 | -0.42438 | 0.141186 | -2.98782 | 4.68E-05 | 0.00977 | 0.076123 | 0.1 | 0.236592 |
| SL1344_RS19785_ | 0.286802 | 0.267727 | 0.30035 | 0.050134 | 0.231356 | 0.190696 | 0.417055 | 0.624874 | 0.748044 | 0.7508 | 0.83255 |

| | | | | | | | | | | | |
|---|---|---|---|---|---|---|---|---|---|---|---|
| SL1344_RS19790_ | -1.02959 | -0.80272 | -1.56084 | -0.75158 | 0.366807 | -1.86028 | 0.016 | 0.078229 | 0.211004 | 0.1163 | 0.256875 |
| SL1344_RS19795_ | -1.63827 | -1.42359 | -1.89793 | -0.6338 | 0.589756 | -1.06338 | 0.099933 | 0.16341 | 0.327625 | 0.2378 | 0.39141 |
| SL1344_RS19800_ | 1.446304 | 1.447246 | 1.445851 | 0.017888 | 0.159819 | 0.105754 | 0.443038 | 0.633317 | 0.754631 | 0.7679 | 0.843559 |
| SL1344_RS19805_ | 1.48034 | 1.372904 | 1.573088 | 0.201931 | 0.10139 | 1.976476 | 0.013334 | 0.052589 | 0.159875 | 0.117 | 0.257125 |
| SL1344_RS19810_ | -2.59143 | -2.64132 | -2.53917 | 0.054257 | 0.585451 | 0.07935 | 0.466356 | 0.672048 | 0.781495 | 0.7785 | 0.851506 |
| SL1344_RS19815_ | -0.81667 | -0.84971 | -0.77994 | 0.069212 | 0.215742 | 0.289191 | 0.367089 | 0.499758 | 0.634575 | 0.6036 | 0.710304 |
| SL1344_RS19820_ | -0.74918 | -1.16585 | -0.31036 | 0.854033 | 0.194864 | 4.329166 | 4.68E-05 | 0.006388 | 0.057199 | 0.1 | 0.236592 |
| SL1344_RS19825_ | 2.130177 | 1.865222 | 2.36023 | 0.488391 | 0.107402 | 4.442338 | 4.68E-05 | 0.005625 | 0.056263 | 0.1 | 0.236592 |
| SL1344_RS19830_ | 1.124253 | 1.276409 | 1.038989 | -0.22448 | 0.155495 | -1.34911 | 0.076 | 0.119903 | 0.266532 | 0.1917 | 0.345299 |
| SL1344_RS19835_ | 0.950209 | 1.125927 | 0.734674 | -0.37869 | 0.160168 | -2.30939 | 0.006001 | 0.031668 | 0.126211 | 0.1064 | 0.244997 |
| SL1344_RS19840_ | 1.640273 | 1.521203 | 1.794536 | 0.269492 | 0.107866 | 2.418599 | 0.003998 | 0.02971 | 0.120445 | 0.104 | 0.241773 |
| SL1344_RS19845_ | 0.678876 | 0.600436 | 0.712323 | 0.098583 | 0.181185 | 0.484203 | 0.31046 | 0.483894 | 0.631143 | 0.6039 | 0.721495 |
| SL1344_RS19850_ | 0.193346 | 0.183641 | 0.198141 | 0.015037 | 0.235057 | 0.06621 | 0.472667 | 0.687262 | 0.793695 | 0.7912 | 0.860945 |
| SL1344_RS19855_ | 0.23152 | 0.284898 | 0.170966 | -0.07174 | 0.25811 | -0.24921 | 0.395333 | 0.662718 | 0.776875 | 0.7656 | 0.842484 |
| SL1344_RS19860_ | 1.108209 | 1.194328 | 1.051183 | -0.14111 | 0.115097 | -1.13867 | 0.106667 | 0.176102 | 0.323099 | 0.2609 | 0.406789 |
| SL1344_RS19865_ | -1.34532 | -1.4669 | -1.24886 | 0.21698 | 0.263955 | 0.768162 | 0.191872 | 0.293033 | 0.441392 | 0.3859 | 0.522076 |
| SL1344_RS19870_ | -4.14396 | -4.22991 | -3.93728 | 0.352105 | 0.964661 | 0.327437 | 0.368421 | 0.591499 | 0.716061 | 0.7001 | 0.791186 |
| SL1344_RS19875_ | 0.32502 | 0.173067 | 0.467838 | 0.295261 | 0.139063 | 2.029793 | 0.014657 | 0.045246 | 0.145639 | 0.1169 | 0.256723 |
| SL1344_RS19880_ | 4.817943 | 4.752255 | 4.857914 | 0.109097 | 0.057127 | 1.877107 | 0.009328 | 0.065505 | 0.191818 | 0.1068 | 0.245384 |
| SL1344_RS19885_ | 4.184812 | 4.236946 | 4.155299 | -0.09328 | 0.08772 | -1.09561 | 0.086609 | 0.159927 | 0.326299 | 0.2045 | 0.361204 |
| SL1344_RS19890_ | -1.12866 | -1.15249 | -1.11268 | 0.033374 | 0.312114 | 0.103824 | 0.455696 | 0.640655 | 0.757528 | 0.7452 | 0.828352 |
| SL1344_RS19900_ | 6.221698 | 5.971224 | 6.455412 | 0.465622 | 0.092318 | 4.982251 | 4.68E-05 | 0.001793 | 0.040235 | 0.1 | 0.236592 |
| SL1344_RS19905_ | 5.782416 | 5.631011 | 5.967717 | 0.414175 | 0.14876 | 2.871487 | 4.68E-05 | 0.010512 | 0.079252 | 0.1 | 0.236592 |
| SL1344_RS19910_ | 7.063898 | 6.909745 | 7.270794 | 0.383082 | 0.103185 | 3.902815 | 4.68E-05 | 0.007447 | 0.06828 | 0.1 | 0.236592 |
| SL1344_RS19915_ | 5.419916 | 5.394172 | 5.461993 | 0.069448 | 0.111043 | 0.582959 | 0.283811 | 0.423067 | 0.601594 | 0.5514 | 0.6986 |
| SL1344_RS19920_ | 5.80827 | 5.702752 | 5.845197 | 0.140272 | 0.090255 | 1.371458 | 0.099267 | 0.132902 | 0.297126 | 0.1936 | 0.356785 |
| SL1344_RS19925_ | 4.875397 | 4.771821 | 4.893658 | 0.123505 | 0.149107 | 0.881794 | 0.18521 | 0.185786 | 0.362759 | 0.3398 | 0.51748 |
| SL1344_RS19930_ | 4.724663 | 4.713057 | 4.745486 | 0.027446 | 0.092767 | 0.284534 | 0.378 | 0.746597 | 0.842269 | 0.8011 | 0.876607 |
| SL1344_RS19935_ | 3.006026 | 3.0444 | 2.83427 | -0.17541 | 0.156392 | -0.84975 | 0.244 | 0.307187 | 0.491385 | 0.4765 | 0.633311 |
| SL1344_RS19945_ | 1.951882 | 1.873086 | 2.09945 | 0.221463 | 0.135757 | 1.511 | 0.045333 | 0.089352 | 0.226222 | 0.1588 | 0.308082 |
| SL1344_RS19950_ | 1.718056 | 1.945291 | 1.388869 | -0.53847 | 0.212725 | -2.44546 | 0.001336 | 0.0373 | 0.1433 | 0.1002 | 0.23686 |
| SL1344_RS19955_ | -0.24333 | -0.31354 | -0.18891 | 0.125139 | 0.22303 | 0.539394 | 0.274667 | 0.423647 | 0.576337 | 0.5407 | 0.661947 |
| SL1344_RS19960_ | 4.233069 | 4.040881 | 6.201523 | 2.065995 | 0.257828 | 1.235868 | 0.002667 | 0.144004 | 0.314098 | 0.1014 | 0.238458 |
| SL1344_RS19965_ | 0.747594 | 0.643574 | 0.88573 | 0.240757 | 0.156822 | 1.475381 | 0.055333 | 0.100059 | 0.234881 | 0.1692 | 0.318234 |
| SL1344_RS19970_ | 1.859133 | 1.784289 | 1.936522 | 0.153901 | 0.084491 | 1.700568 | 0.036 | 0.071787 | 0.189053 | 0.1378 | 0.281794 |
| SL1344_RS19980_ | 0.027336 | 0.401014 | -0.19081 | -0.5735 | 0.216579 | -2.52201 | 0.003998 | 0.033103 | 0.130153 | 0.1021 | 0.239331 |
| SL1344_RS19985_ | -0.43704 | -0.41757 | -0.46471 | -0.05452 | 0.210035 | -0.23298 | 0.394 | 0.539889 | 0.672058 | 0.6557 | 0.753455 |
| SL1344_RS19990_ | 0.534138 | 0.525744 | 0.540685 | 0.014653 | 0.134793 | 0.093361 | 0.459694 | 0.5521 | 0.681216 | 0.6683 | 0.764718 |
| SL1344_RS19995_ | 4.608335 | 4.476644 | 4.750151 | 0.258222 | 0.080248 | 3.258511 | 4.68E-05 | 0.009024 | 0.072427 | 0.1 | 0.236592 |
| SL1344_RS20000_ | 1.938722 | 1.740586 | 2.12035 | 0.378845 | 0.105741 | 3.595412 | 4.68E-05 | 0.010004 | 0.071449 | 0.1 | 0.236592 |
| SL1344_RS20005_ | 1.620074 | 1.368204 | 1.886365 | 0.499017 | 0.148228 | 3.303658 | 4.68E-05 | 0.010814 | 0.075965 | 0.1 | 0.236592 |
| SL1344_RS20020_ | 1.119537 | 1.009624 | 1.303032 | 0.248803 | 0.24531 | 0.941513 | 0.152667 | 0.22903 | 0.405181 | 0.308 | 0.473635 |
| SL1344_RS20025_ | -0.91872 | -1.12101 | -0.70469 | 0.409112 | 0.811317 | 0.509627 | 0.270667 | 0.429378 | 0.604375 | 0.5435 | 0.691529 |
| SL1344_RS20030_ | -1.97135 | -1.78723 | -2.35724 | -0.57104 | 0.470321 | -1.14776 | 0.096602 | 0.171111 | 0.331773 | 0.2254 | 0.37542 |
| SL1344_RS20035_ | 2.696937 | 2.05824 | 2.917908 | 0.865249 | 0.668353 | 0.959449 | 0.185876 | 0.257708 | 0.446194 | 0.3179 | 0.494525 |

| | | | | | | | | | | | |
|---|---|---|---|---|---|---|---|---|---|---|---|
| SL1344_RS20045_ | 2.376795 | 2.397373 | 2.331321 | -0.06923 | 0.301492 | -0.20988 | 0.402398 | 0.775572 | 0.865027 | 0.8512 | 0.911804 |
| SL1344_RS20050_ | 2.978892 | 3.073283 | 2.706305 | -0.25104 | 0.415667 | -0.72946 | 0.228667 | 0.361374 | 0.547672 | 0.4105 | 0.590723 |
| SL1344_RS20055_ | 0.879706 | 0.863394 | 0.907028 | 0.006885 | 0.230042 | 0.025965 | 0.489007 | 0.75515 | 0.846094 | 0.8703 | 0.918652 |
| SL1344_RS20060_ | 4.026085 | 3.796138 | 4.09683 | 0.301395 | 0.140224 | 1.797187 | 0.005997 | 0.093552 | 0.242253 | 0.1042 | 0.242191 |
| SL1344_RS20065_ | -1.65011 | -1.46225 | -1.95507 | -0.49121 | 0.311482 | -1.45931 | 0.051299 | 0.105964 | 0.239399 | 0.1699 | 0.314228 |
| SL1344_RS20070_ | 2.94789 | 2.949707 | 2.94605 | 0.002378 | 0.214034 | 0.01112 | 0.497002 | 0.8069 | 0.884213 | 0.9706 | 0.982516 |
| SL1344_RS20075_ | 0.041416 | 0.181732 | -0.0535 | -0.23288 | 0.183152 | -1.23211 | 0.105929 | 0.157116 | 0.303132 | 0.2338 | 0.381799 |
| SL1344_RS20085_ | 1.348666 | 1.319202 | 1.379298 | 0.071477 | 0.187275 | 0.363204 | 0.339773 | 0.541752 | 0.684584 | 0.6647 | 0.768468 |
| SL1344_RS20090_ | 1.718853 | 1.685758 | 1.746614 | 0.065633 | 0.167463 | 0.362199 | 0.339333 | 0.543085 | 0.685928 | 0.6796 | 0.779312 |
| SL1344_RS20095_ | 2.182145 | 2.266686 | 2.092663 | -0.17513 | 0.083491 | -2.02615 | 0.016 | 0.044549 | 0.146623 | 0.1137 | 0.253778 |
| SL1344_RS20100_ | -0.63004 | -0.50756 | -0.75127 | -0.24295 | 0.192409 | -1.23661 | 0.093937 | 0.156699 | 0.294054 | 0.2337 | 0.376766 |
| SL1344_RS20105_ | -0.72904 | -0.71305 | -0.74441 | -0.05655 | 0.306168 | -0.17433 | 0.424384 | 0.63243 | 0.752769 | 0.7647 | 0.841254 |
| SL1344_RS20110_ | 0.334926 | 0.246062 | 0.413949 | 0.166357 | 0.157621 | 0.983716 | 0.14 | 0.218692 | 0.371677 | 0.3051 | 0.45144 |
| SL1344_RS20115_ | 3.082941 | 3.467026 | 2.84222 | -0.62492 | 0.22789 | -2.87816 | 4.68E-05 | 0.038894 | 0.149375 | 0.1 | 0.236592 |
| SL1344_RS20120_ | 1.626469 | 1.528361 | 1.709912 | 0.182207 | 0.114562 | 1.535948 | 0.050633 | 0.093932 | 0.227402 | 0.1567 | 0.303805 |
| SL1344_RS20125_ | -1.19722 | -1.23262 | -1.16216 | 0.069452 | 0.235453 | 0.26176 | 0.389074 | 0.495965 | 0.629946 | 0.6059 | 0.711993 |
| SL1344_RS20130_ | 1.30324 | 1.266055 | 1.343971 | 0.080196 | 0.121317 | 0.625766 | 0.246667 | 0.367341 | 0.519546 | 0.475 | 0.603618 |
| SL1344_RS20135_ | 3.033566 | 3.048522 | 3.022197 | -0.03194 | 0.081562 | -0.38178 | 0.345103 | 0.537611 | 0.673758 | 0.6736 | 0.769066 |
| SL1344_RS20140_ | 2.955719 | 3.045437 | 2.910706 | -0.13374 | 0.079184 | -1.61165 | 0.042638 | 0.098085 | 0.236167 | 0.1525 | 0.297256 |
| SL1344_RS20145_ | 5.00165 | 5.008506 | 4.990239 | -0.02432 | 0.10042 | -0.2363 | 0.390406 | 0.732244 | 0.83378 | 0.7936 | 0.867979 |
| SL1344_RS20150_ | 4.940381 | 5.044073 | 4.818126 | -0.21681 | 0.055047 | -3.87443 | 4.68E-05 | 0.007347 | 0.06486 | 0.1 | 0.236592 |
| SL1344_RS20155_ | 2.592878 | 2.532258 | 2.655259 | 0.129429 | 0.094951 | 1.299434 | 0.069333 | 0.123734 | 0.271752 | 0.1845 | 0.33574 |
| SL1344_RS20160_ | 3.021612 | 2.977185 | 3.078361 | 0.099495 | 0.059122 | 1.649626 | 0.032645 | 0.080801 | 0.203299 | 0.1463 | 0.290394 |
| SL1344_RS20165_ | 2.212911 | 2.200812 | 2.223187 | 0.023286 | 0.087302 | 0.248217 | 0.387075 | 0.582039 | 0.706672 | 0.6881 | 0.780757 |
| SL1344_RS20175_ | 1.948423 | 1.842738 | 2.058911 | 0.231453 | 0.122832 | 1.850208 | 0.015323 | 0.057151 | 0.175497 | 0.1121 | 0.251947 |
| SL1344_RS20180_ | 0.862992 | 0.700601 | 1.001946 | 0.315128 | 0.163342 | 1.90398 | 0.009328 | 0.061543 | 0.18094 | 0.1084 | 0.247368 |
| SL1344_RS20185_ | 1.03474 | 0.932463 | 1.198761 | 0.27892 | 0.184863 | 1.478848 | 0.049301 | 0.095012 | 0.23453 | 0.1563 | 0.305271 |
| SL1344_RS20190_ | 1.737605 | 1.640212 | 1.837002 | 0.201606 | 0.119648 | 1.696559 | 0.023334 | 0.072023 | 0.196744 | 0.1243 | 0.266368 |
| SL1344_RS20195_ | 1.246816 | 1.193306 | 1.284297 | 0.092824 | 0.122214 | 0.713551 | 0.211193 | 0.326908 | 0.481446 | 0.4354 | 0.5666 |
| SL1344_RS20200_ | 0.834849 | 0.733015 | 0.901163 | 0.171775 | 0.144726 | 1.105867 | 0.102598 | 0.177722 | 0.332507 | 0.2488 | 0.395412 |
| SL1344_RS20205_ | 1.706506 | 1.647331 | 1.801421 | 0.152377 | 0.110421 | 1.293706 | 0.085333 | 0.135275 | 0.281057 | 0.204 | 0.352718 |
| SL1344_RS20210_ | 0.991068 | 0.9199 | 1.070272 | 0.151477 | 0.125684 | 1.160939 | 0.099267 | 0.168616 | 0.312046 | 0.2423 | 0.387679 |
| SL1344_RS20215_ | 1.801386 | 1.649262 | 1.872211 | 0.205863 | 0.169771 | 1.19558 | 0.096602 | 0.140614 | 0.298196 | 0.2255 | 0.386849 |
| SL1344_RS20220_ | 1.182023 | 1.164598 | 1.201898 | -0.01661 | 0.247851 | -0.06889 | 0.472 | 0.784694 | 0.867381 | 0.9032 | 0.939837 |
| SL1344_RS20225_ | 1.456431 | 1.485304 | 1.418106 | -0.0313 | 0.235477 | -0.12165 | 0.456667 | 0.802449 | 0.879811 | 0.8768 | 0.924989 |
| SL1344_RS20230_ | 2.53072 | 2.579979 | 2.351314 | -0.20052 | 0.182223 | -0.88397 | 0.228 | 0.292608 | 0.477049 | 0.436 | 0.599219 |
| SL1344_RS20235_ | 2.963155 | 3.13279 | 2.617426 | -0.25467 | 0.573586 | -0.77597 | 0.214 | 0.31848 | 0.507535 | 0.3934 | 0.575308 |
| SL1344_RS20240_ | 2.941097 | 2.792681 | 3.064233 | 0.275678 | 0.075245 | 3.726977 | 4.68E-05 | 0.009331 | 0.069657 | 0.1 | 0.236592 |
| SL1344_RS20245_ | 2.518174 | 2.456602 | 2.613631 | 0.147176 | 0.100198 | 1.391987 | 0.067955 | 0.102558 | 0.243085 | 0.1746 | 0.32724 |
| SL1344_RS20250_ | 2.378928 | 2.25995 | 2.478483 | 0.219255 | 0.089329 | 2.39134 | 0.002668 | 0.030718 | 0.122908 | 0.1031 | 0.240713 |
| SL1344_RS20255_ | 2.811001 | 2.695547 | 2.88795 | 0.192865 | 0.0735 | 2.502177 | 0.003335 | 0.031437 | 0.12409 | 0.1019 | 0.23912 |
| SL1344_RS20260_ | 3.585474 | 3.51912 | 3.639494 | 0.123593 | 0.060826 | 1.993165 | 0.009994 | 0.047128 | 0.15513 | 0.1107 | 0.250073 |
| SL1344_RS20265_ | -4.15923 | -4.30855 | -3.94452 | 0.289411 | 0.980155 | 0.297209 | 0.356667 | 0.597883 | 0.721591 | 0.6917 | 0.786833 |
| SL1344_RS20270_ | -4.79033 | -4.79365 | -4.78839 | 0.147824 | 1.200274 | 0.115416 | 0.453698 | 0.66553 | 0.776285 | 0.7775 | 0.853605 |
| SL1344_RS20275_ | -4.05293 | -4.03818 | -4.07214 | -0.09038 | 0.855876 | -0.09328 | 0.461026 | 0.624961 | 0.744216 | 0.7328 | 0.815991 |

| | | | | | | | | | | |
|---|---|---|---|---|---|---|---|---|---|---|
| SL1344_RS20280_ | 1.130165 | 1.075266 | 1.364004 | 0.25838 | 0.189193 | 1.079043 | 0.178667 | 0.210848 | 0.383634 | 0.3536 | 0.513125 |
| SL1344_RS20285_ | -4.20134 | -4.27429 | -4.09802 | 0.318803 | 1.000437 | 0.283549 | 0.384667 | 0.565724 | 0.697403 | 0.6999 | 0.78988 |
| SL1344_RS20290_ | 1.382702 | 1.454458 | 1.34569 | -0.09067 | 0.228472 | -0.3423 | 0.37 | 0.70315 | 0.810727 | 0.7903 | 0.870722 |
| SL1344_RS20295_ | 2.288032 | 2.25197 | 2.401974 | 0.137326 | 0.121419 | 0.99592 | 0.185876 | 0.243983 | 0.413633 | 0.3615 | 0.518947 |
| SL1344_RS20300_ | 1.906866 | 1.85213 | 2.01573 | 0.15048 | 0.121028 | 1.13042 | 0.113924 | 0.16339 | 0.32109 | 0.251 | 0.408035 |
| SL1344_RS20305_ | 1.893347 | 1.80982 | 1.979635 | 0.170974 | 0.080607 | 2.032481 | 0.014667 | 0.046981 | 0.148563 | 0.1157 | 0.255947 |
| SL1344_RS20310_ | 0.612411 | 0.596074 | 0.629914 | 0.034648 | 0.12449 | 0.255999 | 0.386409 | 0.494522 | 0.629771 | 0.6014 | 0.707185 |
| SL1344_RS20315_ | 2.905618 | 3.052231 | 2.820775 | -0.22074 | 0.132325 | -1.66938 | 0.036 | 0.062884 | 0.188043 | 0.1311 | 0.276683 |
| SL1344_RS20325_ | -3.7981 | -3.81831 | -3.76967 | 0.044128 | 0.646401 | 0.061602 | 0.472352 | 0.575168 | 0.701763 | 0.6884 | 0.781224 |
| SL1344_RS20330_ | -5.23146 | -5.3229 | -5.15378 | -0.14876 | 1.948225 | -0.06808 | 0.461026 | 0.670289 | 0.784861 | 0.8366 | 0.891863 |
| SL1344_RS20335_ | -0.92729 | -0.91195 | -0.94284 | -0.0328 | 0.232794 | -0.13452 | 0.44 | 0.538593 | 0.669874 | 0.6525 | 0.752507 |
| SL1344_RS20340_ | -3.13482 | -2.82856 | -3.63981 | -0.82764 | 0.691482 | -1.09358 | 0.111925 | 0.175242 | 0.334788 | 0.246 | 0.399118 |
| SL1344_RS20345_ | 2.777846 | 2.706718 | 2.820958 | 0.10913 | 0.103936 | 1.02348 | 0.140573 | 0.206952 | 0.372007 | 0.2935 | 0.451373 |
| SL1344_RS20350_ | 1.931186 | 1.927568 | 1.942599 | 0.012592 | 0.116591 | 0.105457 | 0.464 | 0.686817 | 0.795148 | 0.7863 | 0.863282 |
| SL1344_RS20355_ | 0.079993 | 0.210659 | -0.04754 | -0.2597 | 0.154381 | -1.62032 | 0.044667 | 0.08607 | 0.207026 | 0.1573 | 0.301329 |
| SL1344_RS20365_ | 1.317279 | 1.331803 | 1.291946 | -0.0256 | 0.151711 | -0.16369 | 0.435333 | 0.664942 | 0.778565 | 0.7776 | 0.852596 |
| SL1344_RS20370_ | 1.075417 | 1.012713 | 1.150971 | 0.132371 | 0.148529 | 0.860196 | 0.175333 | 0.270359 | 0.432903 | 0.3744 | 0.522191 |
| SL1344_RS20375_ | -1.46019 | -1.51758 | -1.43142 | 0.056318 | 0.36707 | 0.150353 | 0.441333 | 0.659391 | 0.772658 | 0.7528 | 0.837096 |
| SL1344_RS20390_ | -0.95918 | -0.94101 | -0.98711 | -0.05034 | 0.438211 | -0.11385 | 0.455696 | 0.718402 | 0.819741 | 0.8426 | 0.898762 |
| SL1344_RS20395_ | 1.407012 | 1.320899 | 1.449222 | 0.118624 | 0.126405 | 0.901677 | 0.19054 | 0.278524 | 0.443167 | 0.3841 | 0.534217 |
| SL1344_RS20400_ | 3.494689 | 3.521274 | 3.468797 | -0.0639 | 0.078902 | -0.77662 | 0.19054 | 0.28755 | 0.455433 | 0.4062 | 0.549416 |
| SL1344_RS20405_ | 1.037939 | 1.046128 | 1.024164 | -0.05161 | 0.210377 | -0.22389 | 0.391739 | 0.660252 | 0.77616 | 0.7864 | 0.858186 |
| SL1344_RS20410_ | 2.260053 | 2.267407 | 2.254324 | 0.00342 | 0.121321 | 0.025113 | 0.488341 | 0.71247 | 0.814711 | 0.8348 | 0.893964 |
| SL1344_RS20415_ | 1.871743 | 1.707892 | 2.025335 | 0.322143 | 0.120842 | 2.645713 | 0.002001 | 0.023622 | 0.109089 | 0.1004 | 0.237126 |
| SL1344_RS20425_ | 3.685149 | 3.779825 | 3.666287 | -0.11287 | 0.121794 | -0.76382 | 0.264 | 0.353853 | 0.535775 | 0.5365 | 0.686642 |
| SL1344_RS20430_ | 3.571612 | 3.544522 | 3.607514 | 0.038699 | 0.128809 | 0.270462 | 0.4 | 0.694837 | 0.805908 | 0.8644 | 0.913485 |
| SL1344_RS20435_ | 2.331406 | 2.411761 | 2.267735 | -0.15161 | 0.151425 | -1.04698 | 0.099933 | 0.195527 | 0.369629 | 0.2286 | 0.386873 |
| SL1344_RS20440_ | 0.009899 | -0.03009 | 0.047469 | 0.085167 | 0.198668 | 0.443621 | 0.318667 | 0.46957 | 0.611923 | 0.5943 | 0.700892 |
| SL1344_RS20445_ | -0.0627 | 0.124633 | -0.11818 | -0.21708 | 0.252714 | -0.7435 | 0.25 | 0.364171 | 0.530513 | 0.4981 | 0.640158 |
| SL1344_RS20450_ | 3.571509 | 3.693971 | 3.464347 | -0.23157 | 0.056566 | -4.09534 | 4.68E-05 | 0.007163 | 0.06248 | 0.1 | 0.236592 |
| SL1344_RS20455_ | 2.658181 | 2.647586 | 2.667665 | 0.006374 | 0.094399 | 0.063806 | 0.465023 | 0.676081 | 0.785388 | 0.785 | 0.857088 |
| SL1344_RS20460_ | 0.064863 | 0.106676 | 0.04133 | -0.06593 | 0.19863 | -0.30534 | 0.364424 | 0.548651 | 0.680601 | 0.6622 | 0.76091 |
| SL1344_RS20470_ | 0.988917 | 1.224524 | 0.67731 | -0.57066 | 0.206415 | -2.7648 | 4.68E-05 | 0.019013 | 0.099883 | 0.1 | 0.236592 |
| SL1344_RS20475_ | 3.076993 | 3.026516 | 3.133776 | 0.11841 | 0.088482 | 1.335316 | 0.039333 | 0.120772 | 0.270287 | 0.1459 | 0.290706 |
| SL1344_RS20480_ | 2.613219 | 2.533396 | 2.689451 | 0.177628 | 0.15093 | 1.177121 | 0.085277 | 0.138393 | 0.298775 | 0.1966 | 0.353941 |
| SL1344_RS20485_ | 1.33786 | 1.25637 | 1.406072 | 0.149466 | 0.129275 | 1.119626 | 0.111925 | 0.172651 | 0.326315 | 0.2517 | 0.402101 |
| SL1344_RS20495_ | -1.72875 | -1.73335 | -1.72602 | 0.069394 | 0.558257 | 0.129442 | 0.441333 | 0.660188 | 0.775934 | 0.7953 | 0.863776 |
| SL1344_RS20500_ | -1.05649 | -0.82242 | -1.41485 | -0.69959 | 0.638978 | -1.08119 | 0.081945 | 0.168819 | 0.340184 | 0.1942 | 0.350785 |
| SL1344_RS20505_ | -1.37644 | -1.22505 | -1.59391 | -0.32478 | 0.547739 | -0.55481 | 0.267821 | 0.401482 | 0.571997 | 0.5306 | 0.6647 |
| SL1344_RS20510_ | 0.008258 | -0.24865 | 0.082136 | 0.344416 | 0.628941 | 0.490446 | 0.333333 | 0.487036 | 0.655248 | 0.694 | 0.812346 |
| SL1344_RS20520_ | 0.194911 | 0.120707 | 0.444463 | 0.297955 | 0.275921 | 0.979703 | 0.176549 | 0.227878 | 0.398654 | 0.3436 | 0.50572 |
| SL1344_RS20530_ | 1.842131 | 1.814391 | 1.866986 | 0.080838 | 0.16325 | 0.481373 | 0.291139 | 0.418015 | 0.583765 | 0.5944 | 0.707233 |
| SL1344_RS20535_ | 2.634146 | 2.526049 | 2.717917 | 0.189294 | 0.106414 | 1.797446 | 0.02465 | 0.062504 | 0.184317 | 0.125 | 0.268417 |
| SL1344_RS20540_ | 2.598475 | 2.425591 | 2.666036 | 0.225803 | 0.123284 | 1.686615 | 0.034 | 0.078295 | 0.212728 | 0.1359 | 0.281776 |
| SL1344_RS20545_ | 1.35427 | 1.470432 | 1.260026 | -0.33443 | 0.288506 | -1.14302 | 0.070667 | 0.134639 | 0.296235 | 0.1814 | 0.334012 |

| | | | | | | | | | | | |
|---|---|---|---|---|---|---|---|---|---|---|---|
| SL1344_RS20550_ | -2.16167 | -1.95177 | -2.40264 | -0.44532 | 0.368495 | -1.16702 | 0.099267 | 0.158242 | 0.299887 | 0.2297 | 0.375821 |
| SL1344_RS20555_ | 3.128635 | 2.896939 | 3.313621 | 0.426138 | 0.11164 | 3.821498 | 4.68E-05 | 0.021467 | 0.108052 | 0.1 | 0.236592 |
| SL1344_RS20560_ | 2.034786 | 2.068881 | 1.993347 | -0.09317 | 0.136107 | -0.66526 | 0.224517 | 0.336133 | 0.508333 | 0.4634 | 0.601005 |
| SL1344_RS20565_ | 8.443884 | 8.511785 | 8.37501 | 0.014446 | 0.218799 | 0.07624 | 0.4497 | 0.876927 | 0.930504 | 0.9997 | 0.999827 |
| SL1344_RS20570_ | 2.189063 | 2.162576 | 2.336545 | 0.156688 | 0.134637 | 0.863443 | 0.253165 | 0.329268 | 0.506319 | 0.509 | 0.658048 |
| SL1344_RS20575_ | 2.771435 | 2.784283 | 2.761957 | -0.02243 | 0.067662 | -0.30216 | 0.357761 | 0.509334 | 0.644907 | 0.6139 | 0.721826 |
| SL1344_RS20580_ | 0.606389 | 0.591921 | 0.637416 | 0.0331 | 0.159812 | 0.193774 | 0.420386 | 0.624425 | 0.743789 | 0.7209 | 0.81272 |
| SL1344_RS20585_ | -0.31275 | -0.31926 | -0.30569 | 0.016032 | 0.226833 | 0.069453 | 0.468354 | 0.628895 | 0.747912 | 0.7378 | 0.821802 |
| SL1344_RS20590_ | 2.957956 | 3.062581 | 2.923686 | -0.1406 | 0.168575 | -0.87286 | 0.185333 | 0.240018 | 0.419864 | 0.3489 | 0.52442 |
| SL1344_RS20595_ | 3.808473 | 4.059756 | 3.569784 | -0.4469 | 0.112567 | -4.00181 | 4.68E-05 | 0.003857 | 0.051249 | 0.1 | 0.236592 |
| SL1344_RS20600_ | -1.36608 | -1.20691 | -1.55393 | -0.35061 | 0.28212 | -1.11803 | 0.105263 | 0.16608 | 0.314731 | 0.2423 | 0.391327 |
| SL1344_RS20605_ | -3.20317 | -3.04202 | -3.31184 | -0.257 | 0.665529 | -0.33826 | 0.349101 | 0.569967 | 0.699951 | 0.6693 | 0.770771 |
| SL1344_RS20610_ | -3.30053 | -3.49899 | -3.11249 | 0.293593 | 0.774403 | 0.365984 | 0.349333 | 0.569956 | 0.700935 | 0.6773 | 0.774315 |
| SL1344_RS20615_ | -2.49635 | -2.49745 | -2.49476 | 0.004472 | 0.397029 | 0.011995 | 0.494337 | 0.576433 | 0.7003 | 0.6799 | 0.773137 |
| SL1344_RS20620_ | -2.94812 | -2.89978 | -3.04789 | -0.17544 | 0.612575 | -0.27017 | 0.382667 | 0.545996 | 0.681455 | 0.6745 | 0.76955 |
| SL1344_RS20625_ | -1.94168 | -2.15542 | -1.76704 | 0.399935 | 0.344565 | 1.080862 | 0.104597 | 0.183235 | 0.332914 | 0.2503 | 0.396885 |
| SL1344_RS20635_ | -1.40067 | -1.28159 | -1.5749 | -0.30581 | 0.313814 | -0.94745 | 0.141333 | 0.225791 | 0.381344 | 0.2991 | 0.441908 |
| SL1344_RS20640_ | -0.116 | -0.26793 | -0.01371 | 0.255019 | 0.217742 | 1.136833 | 0.09 | 0.169381 | 0.323274 | 0.2264 | 0.373623 |
| SL1344_RS20645_ | -0.50698 | -0.5269 | -0.4763 | 0.044774 | 0.240962 | 0.170875 | 0.426667 | 0.613775 | 0.734516 | 0.7179 | 0.805664 |
| SL1344_RS20650_ | -1.73497 | -1.68093 | -1.79659 | -0.10775 | 0.318706 | -0.31715 | 0.356429 | 0.514291 | 0.649538 | 0.6201 | 0.725783 |
| SL1344_RS20655_ | -1.38558 | -1.22911 | -1.61675 | -0.38689 | 0.31964 | -1.1037 | 0.107262 | 0.171192 | 0.323608 | 0.2539 | 0.403935 |
| SL1344_RS20660_ | -2.45955 | -2.55457 | -2.37584 | 0.18398 | 0.460067 | 0.372724 | 0.338667 | 0.515794 | 0.65124 | 0.6358 | 0.737168 |
| SL1344_RS20665_ | -3.032 | -2.81312 | -3.34107 | -0.54285 | 0.519911 | -0.98303 | 0.124667 | 0.2116 | 0.362845 | 0.2846 | 0.42784 |
| SL1344_RS20670_ | -1.80079 | -1.85775 | -1.74072 | 0.122468 | 0.300738 | 0.380347 | 0.333111 | 0.465911 | 0.604946 | 0.5683 | 0.681022 |
| SL1344_RS20675_ | 1.677138 | 1.571477 | 1.764695 | 0.196622 | 0.146914 | 1.29066 | 0.062667 | 0.120561 | 0.269234 | 0.1713 | 0.321502 |
| SL1344_RS20680_ | 1.778519 | 1.865925 | 1.675311 | -0.18577 | 0.112767 | -1.5921 | 0.048634 | 0.088308 | 0.218674 | 0.1566 | 0.304882 |
| SL1344_RS20685_ | 1.91506 | 1.892028 | 1.937302 | 0.042587 | 0.092371 | 0.42392 | 0.316667 | 0.466352 | 0.605864 | 0.5741 | 0.684938 |
| SL1344_RS20690_ | 1.028136 | 1.01125 | 1.077193 | 0.059941 | 0.200483 | 0.26661 | 0.403333 | 0.700953 | 0.806502 | 0.7902 | 0.86919 |
| SL1344_RS20695_ | 2.189519 | 2.191325 | 2.188738 | -0.04754 | 0.195229 | -0.25416 | 0.401333 | 0.608583 | 0.741427 | 0.8082 | 0.87236 |
| SL1344_RS20700_ | 0.911371 | 0.95907 | 0.861241 | -0.15409 | 0.257886 | -0.55438 | 0.283145 | 0.417237 | 0.58576 | 0.5874 | 0.713467 |
| SL1344_RS20705_ | 0.268734 | 0.369816 | 0.067327 | -0.32242 | 0.308673 | -1.0198 | 0.141905 | 0.192709 | 0.363232 | 0.2785 | 0.442153 |
| SL1344_RS20710_ | -0.09471 | 0.033179 | -0.20763 | -0.2423 | 0.19563 | -1.19541 | 0.091333 | 0.163696 | 0.311787 | 0.2354 | 0.381813 |
| SL1344_RS20715_ | -0.69894 | -0.83201 | -0.58912 | 0.239463 | 0.24179 | 0.940891 | 0.145903 | 0.230603 | 0.386053 | 0.3257 | 0.46998 |
| SL1344_RS20720_ | -0.37346 | -0.42887 | -0.27001 | 0.14879 | 0.326511 | 0.421083 | 0.334667 | 0.555349 | 0.694235 | 0.6563 | 0.767013 |
| SL1344_RS20725_ | 3.321389 | 3.477818 | 3.214934 | -0.25516 | 0.103952 | -2.41273 | 0.001336 | 0.026085 | 0.117452 | 0.101 | 0.237928 |
| SL1344_RS20730_ | 3.741442 | 3.958218 | 3.658549 | -0.28955 | 0.149249 | -1.52003 | 0.045303 | 0.110488 | 0.266426 | 0.144 | 0.295053 |
| SL1344_RS20735_ | 3.560995 | 3.809986 | 3.509298 | -0.29072 | 0.096422 | -1.64291 | 0.015334 | 0.097295 | 0.246857 | 0.1154 | 0.256416 |
| SL1344_RS20740_ | 4.833753 | 4.989692 | 4.660257 | -0.292 | 0.125595 | -2.35321 | 4.68E-05 | 0.019312 | 0.104317 | 0.1 | 0.236592 |
| SL1344_RS20745_ | 4.07828 | 4.286312 | 3.887385 | -0.3988 | 0.094308 | -4.21521 | 4.68E-05 | 0.003918 | 0.05151 | 0.1 | 0.236592 |
| SL1344_RS20750_ | 0.737095 | 0.730114 | 0.747416 | 0.004047 | 0.188311 | 0.020686 | 0.491333 | 0.695672 | 0.801035 | 0.7958 | 0.86656 |
| SL1344_RS20755_ | -1.94691 | -1.88991 | -2.00306 | -0.1143 | 0.314765 | -0.32904 | 0.356429 | 0.494312 | 0.630923 | 0.6094 | 0.715411 |
| SL1344_RS20760_ | -3.01586 | -2.95291 | -3.06019 | -0.12531 | 0.724224 | -0.15989 | 0.427333 | 0.660003 | 0.77284 | 0.7602 | 0.838225 |
| SL1344_RS20765_ | -1.57839 | -1.44406 | -1.6933 | -0.25464 | 0.29344 | -0.81131 | 0.188 | 0.285946 | 0.434887 | 0.3865 | 0.521367 |
| SL1344_RS20770_ | -0.74576 | -0.52201 | -1.04414 | -0.52289 | 0.218161 | -2.36685 | 0.004665 | 0.03387 | 0.12542 | 0.1054 | 0.243237 |
| SL1344_RS20775_ | 0.730428 | 0.769098 | 0.703663 | -0.08334 | 0.229308 | -0.35214 | 0.365756 | 0.551811 | 0.692655 | 0.701 | 0.795507 |

| | | | | | | | | | | | |
|---|---|---|---|---|---|---|---|---|---|---|---|
| SL1344_RS20780_ | 0.098893 | 0.04965 | 0.186943 | 0.138953 | 0.212833 | 0.600922 | 0.273333 | 0.405673 | 0.56076 | 0.53 | 0.6542 |
| SL1344_RS20785_ | 0.096316 | -0.06999 | 0.31165 | 0.377425 | 0.199926 | 1.867244 | 0.027315 | 0.060319 | 0.177077 | 0.1242 | 0.266591 |
| SL1344_RS20790_ | -1.62816 | -1.82481 | -1.4552 | 0.366076 | 0.319907 | 1.083618 | 0.115333 | 0.180312 | 0.328458 | 0.2587 | 0.405491 |
| SL1344_RS20795_ | -2.82364 | -2.82307 | -2.82504 | -0.01778 | 0.499323 | -0.02832 | 0.487675 | 0.599437 | 0.721626 | 0.7049 | 0.794297 |
| SL1344_RS20800_ | -3.24001 | -3.09796 | -3.38536 | -0.30299 | 0.55777 | -0.49088 | 0.291333 | 0.431792 | 0.576816 | 0.5444 | 0.662558 |
| SL1344_RS20805_ | -0.54564 | -0.66564 | -0.45218 | 0.212289 | 0.206716 | 0.983565 | 0.145237 | 0.220608 | 0.371696 | 0.3031 | 0.449334 |
| SL1344_RS20810_ | -0.74685 | -1.06817 | -0.43899 | 0.628259 | 0.220226 | 2.773372 | 0.001337 | 0.019557 | 0.096945 | 0.1014 | 0.238395 |
| SL1344_RS20815_ | -2.72509 | -2.83608 | -2.64856 | 0.181291 | 0.448696 | 0.378667 | 0.332 | 0.485011 | 0.622826 | 0.5901 | 0.702412 |
| SL1344_RS20820_ | -6.85704 | -6.61763 | -7.12244 | -0.5201 | 1.699959 | -0.28447 | 0.369333 | 0.469574 | 0.609793 | 0.5841 | 0.692941 |
| SL1344_RS20825_ | -3.91573 | -3.96032 | -3.89365 | -0.02277 | 0.855277 | -0.02702 | 0.489007 | 0.658767 | 0.771055 | 0.7766 | 0.85039 |
| SL1344_RS20830_ | -2.37938 | -2.18344 | -2.49227 | -0.29786 | 0.437952 | -0.62921 | 0.252498 | 0.389359 | 0.541165 | 0.4972 | 0.627281 |
| SL1344_RS20835_ | -4.41231 | -4.6233 | -4.24302 | 0.386569 | 0.833129 | 0.429198 | 0.326449 | 0.462271 | 0.604491 | 0.5726 | 0.687052 |
| SL1344_RS20840_ | -3.58449 | -3.88111 | -3.34545 | 0.613225 | 0.768932 | 0.733923 | 0.193205 | 0.302494 | 0.467205 | 0.4028 | 0.539264 |
| SL1344_RS20845_ | 1.53375 | 1.509533 | 1.549653 | 0.038056 | 0.28576 | 0.128511 | 0.448 | 0.738447 | 0.838209 | 0.8953 | 0.936956 |
| SL1344_RS20850_ | 0.406714 | 0.432453 | 0.37105 | -0.05928 | 0.17304 | -0.32484 | 0.360426 | 0.530602 | 0.665623 | 0.6449 | 0.748529 |
| SL1344_RS20855_ | 1.161344 | 1.195701 | 1.119547 | -0.07731 | 0.124667 | -0.58683 | 0.253831 | 0.400631 | 0.548264 | 0.5106 | 0.633135 |
| SL1344_RS20875_ | 1.314589 | 1.247305 | 1.375374 | 0.127231 | 0.095338 | 1.267715 | 0.083278 | 0.145273 | 0.278226 | 0.2215 | 0.365072 |
| SL1344_RS20880_ | 2.198258 | 1.917453 | 2.299635 | 0.357437 | 0.225548 | 1.587751 | 0.04064 | 0.075048 | 0.211218 | 0.1397 | 0.288187 |
| SL1344_RS20890_ | 2.747101 | 2.6874 | 2.811263 | 0.095493 | 0.132271 | 0.730089 | 0.214524 | 0.341239 | 0.514202 | 0.4312 | 0.58282 |
| SL1344_RS20895_ | -0.59133 | -0.62956 | -0.55442 | 0.055266 | 0.273691 | 0.190551 | 0.413333 | 0.622767 | 0.742951 | 0.7357 | 0.819888 |
| SL1344_RS20900_ | -0.90027 | -1.15178 | -0.71406 | 0.442656 | 0.247332 | 1.734627 | 0.02465 | 0.067884 | 0.185542 | 0.1267 | 0.268555 |
| SL1344_RS20905_ | 1.5019 | 1.444668 | 1.580685 | 0.157427 | 0.160809 | 0.952229 | 0.131333 | 0.223823 | 0.391484 | 0.2938 | 0.442569 |
| SL1344_RS20910_ | 1.297651 | 1.301699 | 1.293911 | -0.00296 | 0.133829 | -0.02199 | 0.493005 | 0.644542 | 0.760533 | 0.7612 | 0.838422 |
| SL1344_RS20915_ | -3.6125 | -3.38825 | -3.84183 | -0.43771 | 0.615532 | -0.67096 | 0.231179 | 0.356018 | 0.50462 | 0.4482 | 0.58079 |
| SL1344_RS20920_ | 2.121347 | 2.331308 | 1.89011 | -0.40874 | 0.139424 | -2.88474 | 4.68E-05 | 0.014899 | 0.089122 | 0.1001 | 0.236725 |
| SL1344_RS20925_ | 3.247343 | 3.409178 | 3.102249 | -0.36951 | 0.162757 | -2.41114 | 4.68E-05 | 0.019504 | 0.103876 | 0.1 | 0.236592 |
| SL1344_RS20935_ | 3.277 | 3.432523 | 3.177456 | -0.2512 | 0.085492 | -2.80493 | 4.68E-05 | 0.021974 | 0.10643 | 0.1 | 0.236592 |
| SL1344_RS20940_ | 2.756661 | 2.874807 | 2.694444 | -0.1952 | 0.157289 | -1.2669 | 0.057333 | 0.115531 | 0.269264 | 0.1649 | 0.317199 |
| SL1344_RS20945_ | 2.732989 | 2.919899 | 2.471287 | -0.44997 | 0.081695 | -5.50534 | 4.68E-05 | 0.003451 | 0.045929 | 0.1 | 0.236592 |
| SL1344_RS20950_ | 3.889724 | 4.155197 | 3.5473 | -0.60293 | 0.06501 | -9.14886 | 4.68E-05 | 0.002072 | 0.038964 | 0.1 | 0.236592 |
| SL1344_RS20955_ | 4.210132 | 4.52609 | 3.950366 | -0.56467 | 0.189544 | -3.12806 | 4.68E-05 | 0.007498 | 0.068145 | 0.1 | 0.236592 |
| SL1344_RS20960_ | 1.749636 | 1.941732 | 1.533911 | -0.40601 | 0.096147 | -4.13738 | 4.68E-05 | 0.007092 | 0.060903 | 0.1 | 0.236592 |
| SL1344_RS20965_ | 3.445938 | 3.724767 | 3.081081 | -0.65944 | 0.127224 | -5.17236 | 4.68E-05 | 0.002754 | 0.04468 | 0.1 | 0.236592 |
| SL1344_RS20970_ | 4.313024 | 4.201834 | 4.456767 | 0.257093 | 0.066868 | 3.749861 | 4.68E-05 | 0.016133 | 0.092833 | 0.1 | 0.236592 |
| SL1344_RS20975_ | -0.57837 | -0.56215 | -0.60863 | -0.02176 | 0.278904 | -0.06969 | 0.471333 | 0.684699 | 0.793074 | 0.781 | 0.857516 |
| SL1344_RS20980_ | -0.00146 | -0.10375 | 0.135344 | 0.231752 | 0.205525 | 1.087414 | 0.116667 | 0.18517 | 0.341397 | 0.2613 | 0.414821 |
| SL1344_RS20985_ | 1.84873 | 1.809644 | 1.931474 | 0.121676 | 0.133302 | 0.834873 | 0.192667 | 0.278796 | 0.449848 | 0.3915 | 0.538716 |
| SL1344_RS20990_ | 2.945766 | 2.97121 | 2.907088 | -0.07413 | 0.082561 | -0.85963 | 0.149234 | 0.245509 | 0.413305 | 0.321 | 0.463199 |
| SL1344_RS20995_ | 8.423789 | 8.296805 | 8.584424 | 0.286672 | 0.062197 | 4.552962 | 4.68E-05 | 0.016521 | 0.097841 | 0.1 | 0.236592 |
| SL1344_RS21000_ | 6.451764 | 6.407764 | 6.490781 | 0.076207 | 0.139854 | 0.41137 | 0.379333 | 0.562626 | 0.716936 | 0.8281 | 0.893795 |
| SL1344_RS21005_ | 3.978503 | 3.731564 | 4.153473 | 0.390194 | 0.160536 | 2.410254 | 4.68E-05 | 0.02284 | 0.113079 | 0.1 | 0.236592 |
| SL1344_RS21010_ | 3.636125 | 3.782367 | 3.613085 | -0.14928 | 0.094965 | -0.99334 | 0.209194 | 0.245337 | 0.426396 | 0.4301 | 0.586434 |
| SL1344_RS21015_ | -0.09236 | -0.05176 | -0.18806 | -0.11578 | 0.239325 | -0.41847 | 0.333777 | 0.542065 | 0.680468 | 0.6541 | 0.764398 |
| SL1344_RS21020_ | 4.040975 | 4.034251 | 4.045312 | -6.55E-05 | 0.076066 | -0.00071 | 0.499333 | 0.718591 | 0.820248 | 0.8334 | 0.895204 |
| SL1344_RS21025_ | 1.695732 | 1.747036 | 1.562499 | -0.18312 | 0.141794 | -1.18347 | 0.094 | 0.172812 | 0.336419 | 0.241 | 0.392576 |

| | | | | | | | | | | |
|---|---|---|---|---|---|---|---|---|---|---|
| SL1344_RS21030_ | 2.214962 | 2.149287 | 2.35271 | 0.195862 | 0.114216 | 1.598145 | 0.042 | 0.095238 | 0.235501 | 0.1557 | 0.303576 |
| SL1344_RS21035_ | 2.087414 | 2.081786 | 2.09598 | 0.004171 | 0.107619 | 0.033679 | 0.483678 | 0.671776 | 0.782802 | 0.7724 | 0.851673 |
| SL1344_RS21040_ | 1.664873 | 1.701143 | 1.64952 | -0.03598 | 0.151706 | -0.18988 | 0.423051 | 0.762564 | 0.851531 | 0.8025 | 0.879745 |
| SL1344_RS21045_ | 4.15504 | 4.183861 | 3.808315 | -0.35818 | 0.265935 | -1.26432 | 0.157229 | 0.124836 | 0.287548 | 0.289 | 0.462479 |
| SL1344_RS21050_ | -1.8183 | -1.73841 | -1.86301 | -0.16 | 0.424617 | -0.36677 | 0.349333 | 0.539777 | 0.676607 | 0.6689 | 0.767415 |
| SL1344_RS21055_ | -1.00099 | -0.97422 | -1.03124 | -0.05999 | 0.215689 | -0.25552 | 0.39507 | 0.503858 | 0.638036 | 0.6207 | 0.724221 |
| SL1344_RS21075_ | -3.88895 | -3.8494 | -3.92618 | -0.07854 | 0.707186 | -0.10106 | 0.464357 | 0.593893 | 0.717575 | 0.6985 | 0.790644 |
| SL1344_RS21080_ | -3.57414 | -3.36535 | -3.82616 | -0.48741 | 0.689543 | -0.66616 | 0.214524 | 0.337072 | 0.495724 | 0.4362 | 0.569 |
| SL1344_RS21085_ | 0.382569 | 0.377817 | 0.385732 | 0.040255 | 0.229777 | 0.154712 | 0.425716 | 0.619876 | 0.744158 | 0.7754 | 0.847659 |
| SL1344_RS21090_ | -0.32353 | -0.22387 | -0.4503 | -0.22114 | 0.217787 | -0.95676 | 0.152565 | 0.225767 | 0.382216 | 0.316 | 0.465029 |
| SL1344_RS21100_ | 3.513658 | 3.286163 | 3.694784 | 0.408902 | 0.08405 | 4.837068 | 4.68E-05 | 0.003535 | 0.048258 | 0.1 | 0.236592 |
| SL1344_RS21105_ | -2.21206 | -2.05711 | -2.3311 | -0.26627 | 0.377774 | -0.67293 | 0.236667 | 0.365577 | 0.517019 | 0.4735 | 0.602514 |
| SL1344_RS21115_ | 1.388995 | 1.322098 | 1.540323 | 0.200397 | 0.163521 | 1.183879 | 0.118 | 0.161132 | 0.319969 | 0.248 | 0.407262 |
| SL1344_RS21120_ | 0.910255 | 0.613415 | 1.120957 | 0.519815 | 0.163314 | 3.216981 | 4.68E-05 | 0.022649 | 0.106176 | 0.1 | 0.236592 |
| SL1344_RS21125_ | -1.54669 | -1.79291 | -1.22812 | 0.47699 | 0.510552 | 0.927267 | 0.161226 | 0.238557 | 0.40975 | 0.314 | 0.476564 |
| SL1344_RS21130_ | -0.71787 | -0.94536 | -0.12492 | 0.746644 | 0.386307 | 1.724497 | 0.038641 | 0.079908 | 0.215264 | 0.1377 | 0.284156 |
| SL1344_RS21135_ | 1.098671 | 0.899041 | 1.291401 | 0.393426 | 0.153271 | 2.496363 | 0.003332 | 0.025669 | 0.11346 | 0.1019 | 0.239069 |
| SL1344_RS21140_ | 0.966092 | 0.898788 | 1.139256 | 0.218243 | 0.176196 | 1.104172 | 0.138667 | 0.191353 | 0.354478 | 0.2898 | 0.446468 |
| SL1344_RS21145_ | -0.64709 | -0.96852 | -0.44315 | 0.512808 | 0.290122 | 1.722793 | 0.025334 | 0.074256 | 0.202584 | 0.1276 | 0.271433 |
| SL1344_RS21150_ | -0.4287 | -0.52594 | -0.34788 | 0.166657 | 0.242426 | 0.646362 | 0.241839 | 0.379403 | 0.53197 | 0.4782 | 0.610755 |
| SL1344_RS21155_ | 2.563325 | 2.514968 | 2.624335 | 0.110665 | 0.120885 | 0.922948 | 0.149234 | 0.228085 | 0.399512 | 0.3035 | 0.460797 |
| SL1344_RS21160_ | 3.169475 | 3.102755 | 3.29325 | 0.183512 | 0.074702 | 2.29353 | 0.005334 | 0.038896 | 0.141087 | 0.1051 | 0.24285 |
| SL1344_RS21165_ | 1.235077 | 0.974556 | 1.489055 | 0.514839 | 0.111061 | 4.563022 | 4.68E-05 | 0.005799 | 0.055702 | 0.1 | 0.236592 |
| SL1344_RS21170_ | -1.73566 | -1.75202 | -1.70971 | 0.048082 | 0.314155 | 0.140475 | 0.438374 | 0.559635 | 0.685908 | 0.6631 | 0.760904 |
| SL1344_RS21175_ | -0.77319 | -0.70444 | -0.93394 | -0.19309 | 0.317527 | -0.53206 | 0.306462 | 0.504674 | 0.651374 | 0.6215 | 0.738711 |
| SL1344_RS21180_ | 0.811252 | 0.816494 | 0.807235 | -0.02292 | 0.205946 | -0.10334 | 0.457695 | 0.690755 | 0.798289 | 0.8123 | 0.877498 |
| SL1344_RS21185_ | 3.228398 | 3.128897 | 3.31637 | 0.206568 | 0.106349 | 2.008741 | 0.000673 | 0.05418 | 0.173991 | 0.1011 | 0.237994 |
| SL1344_RS21195_ | 2.50975 | 2.492741 | 2.538969 | 0.065747 | 0.183803 | 0.379711 | 0.31046 | 0.537185 | 0.687718 | 0.6323 | 0.751135 |
| SL1344_RS21200_ | 1.949833 | 1.815245 | 2.012434 | 0.188572 | 0.147193 | 1.185207 | 0.115333 | 0.153599 | 0.314369 | 0.2418 | 0.402544 |
| SL1344_RS21205_ | -0.29744 | -0.21788 | -0.39454 | -0.17132 | 0.203487 | -0.78531 | 0.182667 | 0.291436 | 0.445848 | 0.3959 | 0.536196 |
| SL1344_RS21210_ | 2.889895 | 2.710769 | 3.016589 | 0.317158 | 0.158352 | 1.966739 | 0.003332 | 0.043979 | 0.156335 | 0.1042 | 0.24219 |
| SL1344_RS21215_ | 2.197425 | 2.12458 | 2.257559 | 0.128288 | 0.159715 | 0.799652 | 0.186542 | 0.280747 | 0.455881 | 0.3753 | 0.535094 |
| SL1344_RS21220_ | -1.75017 | -2.15662 | -1.58917 | 0.467756 | 0.717667 | 0.602142 | 0.272485 | 0.447353 | 0.615865 | 0.533 | 0.682679 |
| SL1344_RS21225_ | -0.52674 | -0.30893 | -0.8144 | -0.54737 | 0.355301 | -1.53123 | 0.037333 | 0.082558 | 0.21771 | 0.1385 | 0.284447 |
| SL1344_RS21230_ | -2.4723 | -2.15341 | -2.92018 | -0.84032 | 0.779672 | -1.0628 | 0.122667 | 0.172848 | 0.339076 | 0.2488 | 0.411472 |
| SL1344_RS21235_ | -0.19353 | -0.57244 | -0.10555 | 0.497068 | 0.753377 | 0.593495 | 0.328667 | 0.490043 | 0.658124 | 0.6835 | 0.806281 |
| SL1344_RS21245_ | 0.978032 | 0.980791 | 0.975884 | 0.002001 | 0.169114 | 0.009969 | 0.494667 | 0.680779 | 0.789697 | 0.7827 | 0.857444 |
| SL1344_RS21250_ | 1.479164 | 1.346937 | 1.581423 | 0.237046 | 0.120737 | 1.914868 | 0.012667 | 0.057768 | 0.171942 | 0.1156 | 0.255595 |
| SL1344_RS21255_ | 0.920314 | 0.885887 | 0.970411 | 0.103252 | 0.156774 | 0.612307 | 0.248501 | 0.374428 | 0.532786 | 0.5025 | 0.623256 |
| SL1344_RS21260_ | -5.9533 | -5.93314 | -5.96315 | 0.02405 | 1.595735 | 0.01365 | 0.492 | 0.616613 | 0.739252 | 0.7305 | 0.815842 |
| SL1344_RS21265_ | 1.084388 | 1.12115 | 1.009315 | -0.08809 | 0.277936 | -0.30026 | 0.387075 | 0.79401 | 0.874406 | 0.8047 | 0.882693 |
| SL1344_RS21270_ | 1.401889 | 1.420107 | 1.37373 | -0.04617 | 0.282195 | -0.17212 | 0.4 | 0.763601 | 0.852747 | 0.779 | 0.85588 |
| SL1344_RS21275_ | 3.809672 | 3.9806 | 3.724458 | -0.32652 | 0.204892 | -1.7756 | 4.68E-05 | 0.046702 | 0.163929 | 0.1001 | 0.236729 |
| SL1344_RS21280_ | 2.96414 | 3.081089 | 2.883238 | -0.27572 | 0.259276 | -1.05651 | 0.125916 | 0.159612 | 0.331454 | 0.2244 | 0.39263 |
| SL1344_RS21285_ | 7.423008 | 7.415236 | 7.429558 | 0.034292 | 0.117786 | 0.3466 | 0.328 | 0.560734 | 0.715099 | 0.6832 | 0.800399 |

| | | | | | | | | | | | |
|---|---|---|---|---|---|---|---|---|---|---|---|
| SL1344_RS21290_ | 3.188558 | 3.320148 | 2.984669 | -0.33739 | 0.102415 | -3.27979 | 4.68E-05 | 0.02744 | 0.121657 | 0.1 | 0.236592 |
| SL1344_RS21300_ | 4.496517 | 4.781372 | 4.259934 | -0.30213 | 0.340884 | -1.38635 | 0.104 | 0.12105 | 0.282563 | 0.2 | 0.370027 |
| SL1344_RS21305_ | 5.853052 | 6.168676 | 5.664266 | -0.26922 | 0.283659 | -1.29488 | 0.108594 | 0.145494 | 0.315701 | 0.2 | 0.370027 |
| SL1344_RS21310_ | 6.338978 | 6.473324 | 6.092431 | -0.36819 | 0.174698 | -2.27072 | 4.68E-05 | 0.017825 | 0.101495 | 0.1 | 0.236592 |
| SL1344_RS21315_ | 7.501059 | 7.736656 | 7.359582 | -0.38789 | 0.13494 | -3.1007 | 4.68E-05 | 0.016311 | 0.097404 | 0.1 | 0.236592 |
| SL1344_RS21320_ | 6.372678 | 6.384153 | 6.312256 | -0.07353 | 0.14185 | -0.59384 | 0.224 | 0.426124 | 0.60556 | 0.4033 | 0.584994 |
| SL1344_RS21325_ | 6.769626 | 6.779257 | 6.761352 | 0.000412 | 0.1641 | 0.002643 | 0.497333 | 0.919903 | 0.955064 | 0.9859 | 0.99147 |
| SL1344_RS21330_ | -2.0501 | -1.98485 | -2.14756 | -0.09242 | 0.822312 | -0.10834 | 0.450366 | 0.766618 | 0.854705 | 0.8932 | 0.932525 |
| SL1344_RS21335_ | -5.14759 | -5.00569 | -5.25857 | -0.27108 | 1.356916 | -0.18381 | 0.414 | 0.611273 | 0.734366 | 0.7245 | 0.811381 |
| SL1344_RS21340_ | -0.25924 | -0.2905 | -0.2254 | 0.066236 | 0.228009 | 0.263158 | 0.372 | 0.572857 | 0.700271 | 0.6868 | 0.77857 |
| SL1344_RS21345_ | -3.13274 | -2.99156 | -3.27985 | -0.28552 | 0.44837 | -0.58593 | 0.252498 | 0.373756 | 0.517095 | 0.4769 | 0.60083 |
| SL1344_RS21350_ | -0.22351 | -0.21796 | -0.22831 | 0.005399 | 0.300567 | 0.014923 | 0.49034 | 0.710671 | 0.813572 | 0.7829 | 0.858039 |
| SL1344_RS21355_ | -1.63705 | -1.71536 | -1.58391 | 0.143803 | 0.353549 | 0.36904 | 0.334444 | 0.491246 | 0.632009 | 0.6098 | 0.716274 |
| SL1344_RS21360_ | -3.23524 | -3.21429 | -3.25242 | -0.03665 | 0.513061 | -0.06244 | 0.470353 | 0.548962 | 0.677518 | 0.6564 | 0.754542 |
| SL1344_RS21365_ | -1.91952 | -1.93704 | -1.90866 | 0.043041 | 0.415624 | 0.082346 | 0.45503 | 0.636361 | 0.755086 | 0.7578 | 0.836027 |
| SL1344_RS21370_ | -2.36684 | -2.49203 | -2.12099 | 0.368541 | 0.60012 | 0.548563 | 0.288474 | 0.462499 | 0.613546 | 0.5783 | 0.700097 |
| SL1344_RS21375_ | -1.05424 | -1.20842 | -0.93252 | 0.258685 | 0.302998 | 0.812718 | 0.196 | 0.292922 | 0.45392 | 0.3918 | 0.536589 |
| SL1344_RS21380_ | 0.307034 | 0.183181 | 0.414112 | 0.235236 | 0.158317 | 1.413717 | 0.063957 | 0.113575 | 0.248401 | 0.1767 | 0.323393 |
| SL1344_RS21390_ | 2.156393 | 2.051384 | 2.224748 | 0.115536 | 0.169501 | 0.703354 | 0.228 | 0.356048 | 0.528209 | 0.44 | 0.588499 |
| SL1344_RS21395_ | 1.391695 | 1.341393 | 1.498462 | 0.157644 | 0.1303 | 1.140808 | 0.124667 | 0.190751 | 0.347891 | 0.2781 | 0.427855 |
| SL1344_RS21400_ | 1.543954 | 1.501695 | 1.617042 | 0.118201 | 0.162464 | 0.710208 | 0.216522 | 0.335635 | 0.503866 | 0.4386 | 0.589768 |
| SL1344_RS21420_ | 1.080267 | 0.811485 | 1.24877 | 0.410113 | 0.180671 | 2.17052 | 0.007329 | 0.046052 | 0.156841 | 0.1073 | 0.246161 |
| SL1344_RS21425_ | 1.407296 | 1.279326 | 1.660543 | 0.375561 | 0.204069 | 1.782514 | 0.017322 | 0.057509 | 0.179307 | 0.1209 | 0.263102 |
| SL1344_RS21430_ | 0.726859 | 0.695951 | 0.805806 | 0.080101 | 0.196835 | 0.346685 | 0.366422 | 0.696319 | 0.802074 | 0.7438 | 0.83972 |
| SL1344_RS21435_ | 0.632927 | 0.740555 | 0.4477 | -0.2958 | 0.179137 | -1.59319 | 0.037309 | 0.089401 | 0.222736 | 0.1423 | 0.287327 |
| SL1344_RS21440_ | -1.24521 | -1.67463 | -1.01776 | 0.445128 | 0.947806 | 0.528334 | 0.260493 | 0.470403 | 0.639597 | 0.5208 | 0.675321 |
| SL1344_RS21445_ | -5.33197 | -5.79344 | -5.06276 | 0.701005 | 1.358813 | 0.488943 | 0.289141 | 0.439661 | 0.590483 | 0.5607 | 0.682343 |
| SL1344_RS21450_ | 0.804399 | 0.321551 | 0.87715 | 0.552092 | 0.498392 | 0.801647 | 0.279147 | 0.360147 | 0.546163 | 0.562 | 0.711812 |
| SL1344_RS21460_ | -4.53807 | -4.3475 | -4.74474 | -0.39976 | 0.758768 | -0.47985 | 0.287142 | 0.420057 | 0.561439 | 0.5202 | 0.638639 |
| SL1344_RS21465_ | 0.884563 | 0.618699 | 1.141279 | 0.467253 | 0.267685 | 1.747662 | 0.027982 | 0.055529 | 0.176211 | 0.1254 | 0.270318 |
| SL1344_RS21475_ | -0.7084 | -1.00009 | -0.55844 | 0.428578 | 0.299634 | 1.360107 | 0.065956 | 0.124344 | 0.273976 | 0.1785 | 0.329071 |
| SL1344_RS21480_ | -0.4159 | -0.43628 | -0.37988 | 0.061697 | 0.225488 | 0.256445 | 0.388667 | 0.560656 | 0.690505 | 0.6766 | 0.772037 |
| SL1344_RS21485_ | -0.29697 | -0.24951 | -0.34056 | -0.08969 | 0.22911 | -0.38216 | 0.337775 | 0.521482 | 0.657749 | 0.6269 | 0.733241 |
| SL1344_RS21490_ | 0.340985 | 0.170303 | 0.536531 | 0.321047 | 0.254002 | 1.231681 | 0.098667 | 0.143447 | 0.300078 | 0.212 | 0.372622 |
| SL1344_RS21495_ | 1.144189 | 1.111582 | 1.167082 | -0.00397 | 0.310095 | -0.01318 | 0.496 | 0.81851 | 0.891059 | 0.9004 | 0.938333 |
| SL1344_RS21505_ | 1.239977 | 1.119432 | 1.34955 | 0.230843 | 0.121003 | 1.885442 | 0.021986 | 0.054875 | 0.165503 | 0.1193 | 0.259973 |
| SL1344_RS21510_ | 2.724306 | 2.672466 | 2.937859 | 0.207024 | 0.176154 | 0.628073 | 0.332445 | 0.555028 | 0.708994 | 0.697 | 0.819698 |
| SL1344_RS21515_ | -4.29397 | -4.43888 | -4.18418 | 0.239548 | 0.780897 | 0.273555 | 0.377748 | 0.528972 | 0.662397 | 0.6477 | 0.747754 |
| SL1344_RS21520_ | 0.668444 | 0.74757 | 0.609815 | -0.12862 | 0.153663 | -0.78579 | 0.191872 | 0.306918 | 0.462563 | 0.4029 | 0.544767 |
| SL1344_RS21525_ | 1.819565 | 1.71168 | 1.923002 | 0.210587 | 0.095524 | 2.135463 | 0.009334 | 0.04308 | 0.144346 | 0.1133 | 0.25334 |
| SL1344_RS21530_ | 0.977638 | 0.817081 | 1.127201 | 0.298573 | 0.283062 | 1.044669 | 0.126667 | 0.175608 | 0.345813 | 0.2434 | 0.41055 |
| SL1344_RS21535_ | -0.76953 | -0.65701 | -0.9039 | -0.25593 | 0.283491 | -0.8638 | 0.164557 | 0.251381 | 0.412633 | 0.3393 | 0.484829 |
| SL1344_RS21540_ | -2.03037 | -2.10671 | -1.93646 | 0.205945 | 0.514153 | 0.387013 | 0.33 | 0.514769 | 0.657726 | 0.6315 | 0.740828 |
| SL1344_RS21545_ | -4.91846 | -5.28079 | -4.66717 | 0.525117 | 1.270811 | 0.392115 | 0.335333 | 0.517818 | 0.65867 | 0.6318 | 0.740154 |
| SL1344_RS21550_ | -1.30018 | -1.3486 | -1.2136 | 0.103371 | 0.338273 | 0.280088 | 0.385743 | 0.608362 | 0.731528 | 0.7066 | 0.801406 |

| | | | | | | | | | | | |
|---|---|---|---|---|---|---|---|---|---|---|---|
| SL1344_RS21555_ | -2.49691 | -2.64345 | -2.35889 | 0.31605 | 0.502073 | 0.604062 | 0.253333 | 0.382104 | 0.53518 | 0.5063 | 0.629121 |
| SL1344_RS21560_ | -2.07697 | -2.28936 | -1.86205 | 0.444579 | 0.40407 | 1.065683 | 0.122585 | 0.191754 | 0.344921 | 0.2705 | 0.419968 |
| SL1344_RS21565_ | -4.22139 | -4.16162 | -4.3035 | -0.11172 | 0.723203 | -0.12571 | 0.437042 | 0.564321 | 0.688489 | 0.6667 | 0.76269 |
| SL1344_RS21570_ | -3.37015 | -3.18443 | -3.547 | -0.29363 | 0.731807 | -0.38544 | 0.338441 | 0.553303 | 0.685466 | 0.6689 | 0.767716 |
| SL1344_RS21575_ | -5.08851 | -5.48847 | -4.64638 | 0.847872 | 1.117295 | 0.728259 | 0.204667 | 0.330873 | 0.482848 | 0.4327 | 0.565038 |
| SL1344_RS21580_ | -0.0631 | 0.058052 | -0.24797 | -0.30318 | 0.181004 | -1.58575 | 0.046636 | 0.090341 | 0.219436 | 0.1548 | 0.300355 |
| SL1344_RS21585_ | -1.77024 | -1.49027 | -2.11284 | -0.62521 | 0.318644 | -1.91699 | 0.017334 | 0.059483 | 0.170984 | 0.1191 | 0.260019 |
| SL1344_RS21590_ | -2.27016 | -2.14547 | -2.41262 | -0.31622 | 0.540287 | -0.53729 | 0.273817 | 0.42015 | 0.57625 | 0.5592 | 0.678692 |
| SL1344_RS21595_ | -1.27443 | -1.3713 | -1.20675 | 0.192065 | 0.337691 | 0.526028 | 0.273151 | 0.401385 | 0.558447 | 0.538 | 0.654989 |
| SL1344_RS21600_ | -2.20422 | -2.20132 | -2.2072 | -0.00074 | 0.473588 | -0.0013 | 0.498667 | 0.650533 | 0.765841 | 0.7663 | 0.842305 |
| SL1344_RS21605_ | -3.88397 | -3.69235 | -4.41154 | -0.68623 | 0.865853 | -0.69869 | 0.241173 | 0.352003 | 0.516068 | 0.4837 | 0.620987 |
| SL1344_RS21610_ | -2.22635 | -2.24064 | -2.21094 | 0.021125 | 0.426585 | 0.04972 | 0.474667 | 0.602543 | 0.727302 | 0.7123 | 0.803289 |
| SL1344_RS21615_ | -1.8702 | -1.83482 | -1.92346 | -0.04462 | 0.45282 | -0.08754 | 0.469021 | 0.673671 | 0.783766 | 0.7937 | 0.862615 |
| SL1344_RS21620_ | -1.58423 | -1.44831 | -1.69493 | -0.24705 | 0.290746 | -0.79208 | 0.192 | 0.288522 | 0.437734 | 0.3844 | 0.519794 |
| SL1344_RS21625_ | -3.08418 | -3.07763 | -3.08851 | -0.0726 | 0.682784 | -0.10983 | 0.454 | 0.652007 | 0.767488 | 0.7777 | 0.852599 |
| SL1344_RS21630_ | -1.67462 | -1.55876 | -1.80218 | -0.20318 | 0.39656 | -0.49099 | 0.293138 | 0.477618 | 0.621973 | 0.5897 | 0.70527 |
| SL1344_RS21635_ | -4.91772 | -5.30015 | -4.50609 | 0.944093 | 1.226445 | 0.722789 | 0.195333 | 0.301871 | 0.464771 | 0.415 | 0.553028 |
| SL1344_RS21640_ | -5.11819 | -5.12455 | -5.11059 | -0.03134 | 1.207929 | -0.02245 | 0.489333 | 0.606301 | 0.730406 | 0.724 | 0.808686 |
| SL1344_RS21645_ | -4.52722 | -4.69446 | -4.42517 | 0.111448 | 1.25342 | 0.074647 | 0.471019 | 0.711061 | 0.813701 | 0.8137 | 0.881969 |
| SL1344_RS21650_ | -4.48264 | -4.64076 | -4.38479 | 0.129691 | 1.077337 | 0.110303 | 0.448 | 0.660769 | 0.773332 | 0.7652 | 0.844562 |
| SL1344_RS21655_ | -4.13852 | -4.00044 | -4.24396 | -0.33091 | 0.889957 | -0.37403 | 0.352 | 0.532456 | 0.667165 | 0.6608 | 0.756515 |
| SL1344_RS21660_ | -0.56976 | -0.63743 | -0.45469 | 0.184449 | 0.237525 | 0.718715 | 0.211193 | 0.339037 | 0.495281 | 0.454 | 0.584998 |
| SL1344_RS21665_ | 1.36109 | 1.17062 | 1.534216 | 0.384487 | 0.154782 | 2.485897 | 0.000673 | 0.033053 | 0.129459 | 0.1002 | 0.23686 |
| SL1344_RS21670_ | 3.728321 | 3.587674 | 3.857628 | 0.273117 | 0.056831 | 4.7508 | 4.68E-05 | 0.006363 | 0.05839 | 0.1 | 0.236592 |
| SL1344_RS21675_ | -0.03454 | 0.200544 | -0.35073 | -0.56789 | 0.253472 | -2.28954 | 4.68E-05 | 0.055664 | 0.174275 | 0.1001 | 0.236727 |
| SL1344_RS21680_ | -3.21499 | -3.21523 | -3.21496 | -0.00911 | 0.607503 | -0.01503 | 0.493671 | 0.613405 | 0.734504 | 0.718 | 0.805398 |
| SL1344_RS21685_ | -3.54928 | -3.42015 | -3.70403 | -0.28088 | 0.563383 | -0.49886 | 0.294 | 0.422025 | 0.565951 | 0.5339 | 0.65092 |
| SL1344_RS21690_ | 0.070973 | 0.043992 | 0.123884 | 0.089113 | 0.20342 | 0.382279 | 0.353333 | 0.532161 | 0.668709 | 0.6461 | 0.750062 |
| SL1344_RS21695_ | -1.57646 | -1.4829 | -1.64252 | -0.15662 | 0.3202 | -0.46836 | 0.296667 | 0.452531 | 0.596711 | 0.5625 | 0.678412 |
| SL1344_RS21710_ | 3.530902 | 3.22233 | 3.827223 | 0.610788 | 0.06804 | 8.996962 | 4.68E-05 | 0.001513 | 0.034721 | 0.1 | 0.236592 |
| SL1344_RS21715_ | 1.189495 | 0.758921 | 1.486118 | 0.725718 | 0.116829 | 6.100824 | 4.68E-05 | 0.00236 | 0.039921 | 0.1 | 0.236592 |
| SL1344_RS21725_ | 1.473238 | 1.088223 | 1.812921 | 0.736847 | 0.133892 | 5.610973 | 4.68E-05 | 0.010395 | 0.074094 | 0.1 | 0.236592 |
| SL1344_RS21730_ | 0.534266 | 0.424195 | 0.71045 | 0.25378 | 0.22513 | 1.09546 | 0.121333 | 0.174212 | 0.33643 | 0.2588 | 0.420331 |
| SL1344_RS21735_ | 0.37895 | 0.158357 | 0.59842 | 0.431002 | 0.183362 | 2.305441 | 0.002668 | 0.030424 | 0.123258 | 0.1041 | 0.242006 |
| SL1344_RS21740_ | 4.069486 | 4.03136 | 4.129578 | 0.080173 | 0.105942 | 0.751265 | 0.223333 | 0.327206 | 0.507758 | 0.4288 | 0.586642 |
| SL1344_RS21745_ | -0.11705 | -0.13181 | -0.10766 | 0.027346 | 0.198151 | 0.130527 | 0.441333 | 0.606186 | 0.728955 | 0.7191 | 0.807771 |
| SL1344_RS21755_ | 0.611089 | 0.695752 | 0.512258 | -0.20091 | 0.18841 | -1.02886 | 0.106596 | 0.190672 | 0.352922 | 0.25 | 0.400124 |
| SL1344_RS21760_ | 1.145657 | 1.074409 | 1.326949 | 0.285708 | 0.26451 | 1.054661 | 0.136667 | 0.172245 | 0.341321 | 0.2692 | 0.436586 |
| SL1344_RS21765_ | 3.91676 | 3.827676 | 4.263076 | 0.435283 | 0.118306 | 2.129351 | 4.68E-05 | 0.066676 | 0.200292 | 0.1 | 0.236592 |
| SL1344_RS21770_ | 0.385327 | 0.462372 | 0.30866 | -0.14949 | 0.147687 | -0.9432 | 0.137333 | 0.219135 | 0.368685 | 0.3086 | 0.450402 |
| SL1344_RS21775_ | 0.527794 | 0.507974 | 0.571063 | 0.058599 | 0.205964 | 0.267198 | 0.402 | 0.637185 | 0.756903 | 0.749 | 0.834559 |
| SL1344_RS21780_ | 2.573815 | 2.702282 | 2.492115 | -0.20634 | 0.082835 | -2.43303 | 0.005997 | 0.034873 | 0.132092 | 0.1036 | 0.241287 |
| SL1344_RS21785_ | -1.69326 | -1.6862 | -1.70899 | -0.02739 | 0.420815 | -0.07085 | 0.472667 | 0.665845 | 0.779061 | 0.7808 | 0.857092 |
| SL1344_RS21790_ | 1.608793 | 1.340657 | 1.925705 | 0.570864 | 0.140327 | 3.985957 | 4.68E-05 | 0.008207 | 0.067203 | 0.1 | 0.236592 |
| SL1344_RS21795_ | 1.875442 | 1.918134 | 1.826134 | -0.08557 | 0.139425 | -0.6149 | 0.26 | 0.407634 | 0.568342 | 0.5178 | 0.653903 |

| | | | | | | | | | | | |
|---|---|---|---|---|---|---|---|---|---|---|---|
| SL1344_RS21800_ | 1.084063 | 1.015524 | 1.138986 | 0.144053 | 0.173739 | 0.808879 | 0.162 | 0.261155 | 0.434336 | 0.3403 | 0.487616 |
| SL1344_RS21805_ | 1.210284 | 1.095042 | 1.381158 | 0.299785 | 0.15549 | 1.919165 | 0.005331 | 0.066634 | 0.192398 | 0.1071 | 0.245498 |
| SL1344_RS21810_ | 3.588354 | 3.647364 | 3.563412 | -0.09234 | 0.107473 | -0.86828 | 0.172552 | 0.232615 | 0.407714 | 0.3351 | 0.501516 |
| SL1344_RS21815_ | 1.85564 | 1.697639 | 2.048805 | 0.318546 | 0.149114 | 2.082039 | 0.012659 | 0.039188 | 0.143709 | 0.1132 | 0.254063 |
| SL1344_RS21820_ | 1.059089 | 0.878709 | 1.334582 | 0.442128 | 0.149706 | 2.804993 | 0.000673 | 0.022935 | 0.106902 | 0.1003 | 0.236993 |
| SL1344_RS21825_ | -4.23931 | -4.2025 | -4.27157 | -0.1086 | 0.856922 | -0.12339 | 0.450366 | 0.601312 | 0.725094 | 0.7229 | 0.808285 |
| SL1344_RS21830_ | -5.71559 | -5.39078 | -6.12569 | -0.66151 | 1.703236 | -0.36426 | 0.326449 | 0.501595 | 0.643549 | 0.6106 | 0.719079 |
| SL1344_RS21835_ | 2.828379 | 2.908081 | 2.678047 | -0.22455 | 0.096895 | -2.22905 | 0.005331 | 0.050344 | 0.164903 | 0.1042 | 0.242016 |
| SL1344_RS21840_ | -3.66698 | -3.67065 | -3.6605 | -0.00386 | 0.686605 | -0.00568 | 0.498 | 0.623957 | 0.741656 | 0.7276 | 0.812565 |
| SL1344_RS21845_ | 3.241567 | 3.312189 | 3.212982 | -0.09172 | 0.074269 | -1.09624 | 0.149234 | 0.209095 | 0.372468 | 0.3133 | 0.468193 |
| SL1344_RS21880_ | -2.61089 | -2.54235 | -2.6805 | -0.13966 | 0.55715 | -0.22759 | 0.392 | 0.604627 | 0.728367 | 0.7118 | 0.801349 |
| SL1344_RS21885_ | -0.78463 | -0.67878 | -0.91277 | -0.2344 | 0.258054 | -0.87881 | 0.181213 | 0.276508 | 0.432082 | 0.3729 | 0.516266 |
| SL1344_RS21890_ | -1.1361 | -0.97311 | -1.28063 | -0.30472 | 0.259154 | -1.12516 | 0.113924 | 0.176038 | 0.322018 | 0.2602 | 0.406619 |
| SL1344_RS21895_ | -1.19308 | -1.11343 | -1.27969 | -0.17546 | 0.256116 | -0.65289 | 0.234667 | 0.352341 | 0.499479 | 0.46 | 0.586615 |
| SL1344_RS21900_ | -0.05611 | -0.27353 | 0.154282 | 0.431848 | 0.196721 | 2.141051 | 0.004665 | 0.040316 | 0.141353 | 0.1063 | 0.244739 |
| SL1344_RS21905_ | 0.702175 | 0.857948 | 0.486652 | -0.37138 | 0.169769 | -2.0885 | 0.007995 | 0.044405 | 0.150171 | 0.1068 | 0.245173 |
| SL1344_RS21910_ | 1.866393 | 1.769565 | 1.970605 | 0.206955 | 0.10307 | 1.968681 | 0.01 | 0.051305 | 0.16117 | 0.1134 | 0.252726 |
| SL1344_RS21915_ | 1.234561 | 1.178284 | 1.286334 | 0.108427 | 0.152356 | 0.68307 | 0.224 | 0.346023 | 0.507692 | 0.456 | 0.594889 |
| SL1344_RS21920_ | -3.73232 | -3.67946 | -3.79963 | -0.06183 | 0.884169 | -0.0585 | 0.474667 | 0.688919 | 0.795843 | 0.7889 | 0.860667 |
| SL1344_RS21925_ | -2.22651 | -2.03809 | -2.39967 | -0.36451 | 0.336563 | -1.01641 | 0.135243 | 0.197684 | 0.342077 | 0.275 | 0.417419 |
| SL1344_RS21930_ | 0.155755 | 0.100814 | 0.236065 | 0.148335 | 0.187974 | 0.749552 | 0.193871 | 0.311328 | 0.467427 | 0.4077 | 0.540686 |
| SL1344_RS21935_ | -2.8213 | -2.77879 | -2.83749 | -0.10804 | 0.691385 | -0.14078 | 0.448368 | 0.69456 | 0.800578 | 0.803 | 0.873108 |
| SL1344_RS21940_ | 1.812874 | 1.91068 | 1.616466 | -0.29407 | 0.171898 | -1.63109 | 0.023318 | 0.092437 | 0.235221 | 0.1231 | 0.265983 |
| SL1344_RS21945_ | -0.97336 | -1.30169 | -0.75443 | 0.543004 | 0.263356 | 1.960886 | 0.020653 | 0.052101 | 0.161277 | 0.116 | 0.256506 |
| SL1344_RS21950_ | 4.076622 | 3.824776 | 4.28976 | 0.458439 | 0.122942 | 3.702377 | 4.68E-05 | 0.013943 | 0.088464 | 0.1 | 0.236592 |
| SL1344_RS21965_ | 2.365749 | 1.958645 | 2.629605 | 0.665148 | 0.144596 | 4.514593 | 4.68E-05 | 0.006639 | 0.061594 | 0.1 | 0.236592 |
| SL1344_RS21970_ | 1.831497 | 1.436515 | 2.171491 | 0.71564 | 0.228235 | 3.095679 | 4.68E-05 | 0.014557 | 0.089287 | 0.1 | 0.236592 |
| SL1344_RS21980_ | 2.894607 | 2.931201 | 2.863411 | -0.10603 | 0.159095 | -0.66287 | 0.225333 | 0.308871 | 0.489871 | 0.4574 | 0.605913 |
| SL1344_RS21985_ | 0.080499 | -0.33906 | 0.309352 | 0.631116 | 0.227653 | 2.669664 | 0.001337 | 0.034367 | 0.131922 | 0.1016 | 0.238688 |
| SL1344_RS21990_ | 2.273947 | 2.305734 | 2.233897 | -0.09484 | 0.132748 | -0.70894 | 0.213857 | 0.308603 | 0.481702 | 0.4505 | 0.587136 |
| SL1344_RS21995_ | -0.06612 | -0.11921 | 0.011391 | 0.12873 | 0.188547 | 0.615627 | 0.253831 | 0.386439 | 0.538534 | 0.4987 | 0.626999 |
| SL1344_RS22000_ | 0.571704 | 0.651455 | 0.376125 | -0.25746 | 0.202734 | -1.15638 | 0.11992 | 0.173727 | 0.335864 | 0.2631 | 0.421032 |
| SL1344_RS22005_ | -0.98896 | -1.08151 | -0.92065 | 0.166808 | 0.343565 | 0.4604 | 0.296 | 0.489081 | 0.634177 | 0.5908 | 0.708781 |
| SL1344_RS22010_ | -1.02249 | -1.30502 | -0.83898 | 0.487172 | 0.32917 | 1.392837 | 0.041306 | 0.11867 | 0.266635 | 0.1506 | 0.296691 |
| SL1344_RS22015_ | 2.133456 | 2.24172 | 2.108195 | -0.12159 | 0.134186 | -0.72567 | 0.26982 | 0.395473 | 0.564038 | 0.5642 | 0.699961 |
| SL1344_RS22020_ | 1.775566 | 1.397789 | 2.200003 | 0.791025 | 0.169624 | 4.665846 | 4.68E-05 | 0.004072 | 0.050819 | 0.1 | 0.236592 |
| SL1344_RS22030_ | 2.306388 | 2.018123 | 2.558171 | 0.558136 | 0.126452 | 4.531627 | 4.68E-05 | 0.005423 | 0.056792 | 0.1 | 0.236592 |
| SL1344_RS22035_ | 2.219654 | 1.968757 | 2.440093 | 0.483346 | 0.13327 | 3.613105 | 4.68E-05 | 0.011844 | 0.079055 | 0.1 | 0.236592 |
| SL1344_RS22040_ | 2.482929 | 2.346907 | 2.68616 | 0.350125 | 0.135333 | 2.489511 | 4.68E-05 | 0.031535 | 0.130493 | 0.1004 | 0.237116 |
| SL1344_RS22045_ | 0.478335 | 0.153818 | 0.703024 | 0.549473 | 0.17614 | 3.033061 | 4.68E-05 | 0.022939 | 0.105991 | 0.1005 | 0.237252 |
| SL1344_RS22050_ | 2.208306 | 1.976653 | 2.492117 | 0.517898 | 0.154192 | 3.385722 | 4.68E-05 | 0.008924 | 0.071083 | 0.1 | 0.236592 |
| SL1344_RS22055_ | 0.841046 | 0.703689 | 1.039544 | 0.347398 | 0.167699 | 2.045686 | 0.005331 | 0.057016 | 0.174415 | 0.1077 | 0.246313 |
| SL1344_RS22060_ | -0.87463 | -1.08239 | -0.65044 | 0.431472 | 0.231775 | 1.80196 | 0.019987 | 0.058247 | 0.169389 | 0.123 | 0.264401 |
| SL1344_RS22065_ | 1.197265 | 0.946397 | 1.492983 | 0.47147 | 0.228191 | 2.126168 | 0.008001 | 0.033713 | 0.134566 | 0.1084 | 0.247791 |
| SL1344_RS22070_ | 0.089709 | -0.85333 | 0.643754 | 1.484817 | 0.242473 | 6.00284 | 4.68E-05 | 0.008068 | 0.065787 | 0.1 | 0.236592 |

| | | | | | | | | | | |
|---|---|---|---|---|---|---|---|---|---|---|
| SL1344_RS22075_ | -1.24932 | -1.87487 | -0.85735 | 1.031057 | 0.35221 | 2.834516 | 0.000673 | 0.018169 | 0.09517 | 0.1001 | 0.236727 |
| SL1344_RS22080_ | 3.798596 | 3.662108 | 3.83513 | 0.193601 | 0.150981 | 1.30102 | 0.067955 | 0.099331 | 0.248585 | 0.1752 | 0.328396 |
| SL1344_RS22085_ | 3.303435 | 3.124536 | 3.393639 | 0.273857 | 0.144957 | 1.813065 | 0.007334 | 0.063586 | 0.192659 | 0.1077 | 0.246742 |
| SL1344_RS22090_ | 2.378257 | 2.209251 | 2.687265 | 0.478225 | 0.174503 | 2.648576 | 0.000673 | 0.04368 | 0.157309 | 0.1 | 0.236592 |
| SL1344_RS22095_ | 1.131495 | 1.080953 | 1.238361 | 0.151428 | 0.201441 | 0.717078 | 0.227848 | 0.345576 | 0.515189 | 0.4426 | 0.596163 |
| SL1344_RS22100_ | 1.58823 | 1.575503 | 1.600488 | 0.046983 | 0.186094 | 0.235489 | 0.397069 | 0.648674 | 0.769099 | 0.8036 | 0.871112 |
| SL1344_RS22105_ | 2.419341 | 1.884895 | 2.893252 | 1.025529 | 0.130631 | 7.947289 | 4.68E-05 | 0.000798 | 0.030276 | 0.1 | 0.236592 |
| SL1344_RS22110_ | 0.514881 | -0.08709 | 1.000402 | 1.06935 | 0.216439 | 4.846983 | 4.68E-05 | 0.003986 | 0.049728 | 0.1 | 0.236592 |
| SL1344_RS22115_ | 0.628755 | -0.00539 | 1.106277 | 1.135514 | 0.250499 | 4.529835 | 4.68E-05 | 0.004818 | 0.053942 | 0.1 | 0.236592 |
| SL1344_RS22120_ | 0.791522 | 0.3022 | 1.183764 | 0.873882 | 0.175051 | 4.943909 | 4.68E-05 | 0.00442 | 0.050525 | 0.1 | 0.236592 |
| SL1344_RS22125_ | -2.54769 | -2.39414 | -2.73399 | -0.3469 | 0.394492 | -0.81955 | 0.171333 | 0.271875 | 0.419275 | 0.3587 | 0.493943 |
| SL1344_RS22130_ | 0.244819 | 0.510117 | -0.07756 | -0.61586 | 0.261693 | -2.39723 | 0.000673 | 0.027724 | 0.120803 | 0.1009 | 0.237745 |
| SL1344_RS22155_ | -6.31222 | -6.42231 | -6.17594 | 0.19314 | 1.588963 | 0.114848 | 0.439333 | 0.548663 | 0.67773 | 0.6617 | 0.758596 |
| SL1344_RS22160_ | 0.154131 | 0.224393 | 0.08751 | -0.14738 | 0.225444 | -0.62118 | 0.235177 | 0.365745 | 0.52312 | 0.4781 | 0.610054 |
| SL1344_RS22165_ | 0.955373 | 1.021438 | 0.878599 | -0.19227 | 0.214332 | -0.89123 | 0.151899 | 0.227942 | 0.399136 | 0.3364 | 0.485221 |
| SL1344_RS22170_ | 1.82105 | 1.761511 | 1.842361 | 0.079569 | 0.133941 | 0.535334 | 0.298667 | 0.450778 | 0.607931 | 0.5924 | 0.713704 |
| SL1344_RS22175_ | -3.6811 | -3.34069 | -4.18392 | -0.82988 | 0.646694 | -1.21419 | 0.088667 | 0.157011 | 0.303037 | 0.2273 | 0.374123 |
| SL1344_RS22180_ | -1.47144 | -1.38938 | -1.65737 | -0.23948 | 0.366211 | -0.60368 | 0.267333 | 0.416925 | 0.5716 | 0.538 | 0.666697 |
| SL1344_RS22185_ | -2.36753 | -2.44455 | -2.31614 | 0.116676 | 0.53359 | 0.209851 | 0.407062 | 0.643017 | 0.761066 | 0.7412 | 0.827264 |
| SL1344_RS22190_ | 0.819776 | 0.745827 | 0.916735 | 0.18671 | 0.184933 | 0.985589 | 0.125333 | 0.205355 | 0.3714 | 0.2855 | 0.438232 |
| SL1344_RS22195_ | 0.69912 | 0.650261 | 0.75189 | 0.101808 | 0.129618 | 0.752451 | 0.205197 | 0.312661 | 0.461645 | 0.407 | 0.542747 |
| SL1344_RS22200_ | -0.16471 | -0.20112 | -0.11548 | 0.08765 | 0.188327 | 0.426171 | 0.302 | 0.457461 | 0.597328 | 0.5679 | 0.678947 |
| SL1344_RS22205_ | 4.166338 | 3.664803 | 4.660752 | 0.967783 | 0.151996 | 6.39177 | 4.68E-05 | 0.001301 | 0.035887 | 0.1 | 0.236592 |
| SL1344_RS22210_ | 7.712663 | 7.523997 | 7.879235 | 0.355003 | 0.037541 | 9.48776 | 4.68E-05 | 0.005815 | 0.061683 | 0.1 | 0.236592 |
| SL1344_RS22215_ | 5.506409 | 5.475141 | 5.539935 | 0.045779 | 0.227693 | 0.173775 | 0.450667 | 0.939846 | 0.966475 | 1 | 1 |
| SL1344_RS22220_ | 2.270081 | 2.430508 | 2.005268 | -0.41937 | 0.135556 | -3.01935 | 4.68E-05 | 0.013371 | 0.084319 | 0.1 | 0.236592 |
| SL1344_RS22225_ | 0.429577 | 0.214381 | 0.586165 | 0.369239 | 0.141403 | 2.53712 | 0.002668 | 0.027656 | 0.113969 | 0.1032 | 0.240519 |
| SL1344_RS22235_ | 8.22624 | 8.255381 | 8.194946 | -0.0095 | 0.127981 | -0.11296 | 0.418 | 0.754375 | 0.853265 | 0.8719 | 0.920738 |
| SL1344_RS22240_ | 4.252682 | 4.269484 | 4.230158 | 0.020654 | 0.139333 | 0.157451 | 0.454667 | 0.817026 | 0.889986 | 0.9697 | 0.981352 |
| SL1344_RS22245_ | -1.18501 | -1.29856 | -1.09966 | 0.119459 | 0.477214 | 0.242205 | 0.397735 | 0.672452 | 0.784183 | 0.7897 | 0.859186 |
| SL1344_RS22250_ | 0.310399 | 0.332082 | 0.293161 | -0.04153 | 0.160621 | -0.23203 | 0.39 | 0.556292 | 0.685339 | 0.6575 | 0.756938 |
| SL1344_RS22255_ | 2.882074 | 2.939886 | 2.741146 | -0.17633 | 0.158328 | -1.02124 | 0.139241 | 0.193212 | 0.367804 | 0.2729 | 0.442238 |
| SL1344_RS22260_ | -2.675 | -2.72347 | -2.63333 | 0.01714 | 0.651262 | 0.02498 | 0.489007 | 0.691625 | 0.799353 | 0.805 | 0.872933 |
| SL1344_RS22265_ | 4.249115 | 4.26924 | 4.227448 | -0.02546 | 0.166302 | -0.14655 | 0.427049 | 0.814871 | 0.890044 | 0.8932 | 0.936026 |
| SL1344_RS22275_ | -1.35891 | -1.16709 | -1.49798 | -0.32357 | 0.335636 | -0.88703 | 0.175217 | 0.257749 | 0.419493 | 0.3468 | 0.496375 |
| SL1344_RS22280_ | 0.207363 | 0.014222 | 0.571392 | 0.539956 | 0.220307 | 2.327675 | 0.004001 | 0.043505 | 0.151088 | 0.1042 | 0.241879 |
| SL1344_RS22285_ | 4.825018 | 4.448982 | 5.051568 | 0.607254 | 0.127756 | 4.636297 | 4.68E-05 | 0.010378 | 0.078243 | 0.1 | 0.236592 |
| SL1344_RS22290_ | 3.835021 | 3.561687 | 4.094841 | 0.53813 | 0.058129 | 9.266081 | 4.68E-05 | 0.00149 | 0.034131 | 0.1 | 0.236592 |
| SL1344_RS22295_ | 5.296941 | 4.994559 | 5.523796 | 0.517071 | 0.087898 | 5.854421 | 4.68E-05 | 0.001834 | 0.039499 | 0.1 | 0.236592 |
| SL1344_RS22300_ | 6.364276 | 6.154922 | 6.604636 | 0.458686 | 0.103562 | 4.638413 | 4.68E-05 | 0.00276 | 0.046386 | 0.1 | 0.236592 |
| SL1344_RS22305_ | 0.851116 | 0.840029 | 0.867457 | 0.035202 | 0.164461 | 0.20891 | 0.413724 | 0.603854 | 0.726632 | 0.7167 | 0.80356 |
| SL1344_RS22310_ | -1.40801 | -1.29781 | -1.5228 | -0.23056 | 0.255989 | -0.8339 | 0.166556 | 0.279075 | 0.424539 | 0.3724 | 0.50764 |
| SL1344_RS22315_ | 0.228793 | 0.128714 | 0.333366 | 0.198448 | 0.165288 | 1.130102 | 0.098667 | 0.163227 | 0.309154 | 0.2381 | 0.387728 |
| SL1344_RS22320_ | 2.710804 | 2.556177 | 2.877089 | 0.324206 | 0.090364 | 3.59215 | 4.68E-05 | 0.009025 | 0.069563 | 0.1 | 0.236592 |
| SL1344_RS22325_ | 1.744557 | 1.556673 | 1.908829 | 0.351879 | 0.089715 | 3.849622 | 4.68E-05 | 0.009143 | 0.066436 | 0.1 | 0.236592 |

| | | | | | | | | | | | |
|---|---|---|---|---|---|---|---|---|---|---|---|
| SL1344_RS22330_ | 1.818697 | 1.722864 | 1.897402 | 0.164095 | 0.116258 | 1.344128 | 0.073951 | 0.123979 | 0.270727 | 0.1925 | 0.345683 |
| SL1344_RS22335_ | 1.462115 | 1.234022 | 1.736064 | 0.480823 | 0.165325 | 2.908318 | 0.000673 | 0.014846 | 0.088353 | 0.1001 | 0.236723 |
| SL1344_RS22340_ | 1.06853 | 0.630308 | 1.201521 | 0.34637 | 0.472918 | 0.735455 | 0.278 | 0.373622 | 0.557004 | 0.5584 | 0.702274 |
| SL1344_RS22345_ | 2.004448 | 1.959646 | 2.053708 | 0.063044 | 0.145493 | 0.430543 | 0.324667 | 0.548462 | 0.68728 | 0.6479 | 0.755463 |
| SL1344_RS22350_ | 0.790644 | 0.808822 | 0.779515 | -0.05111 | 0.192906 | -0.25856 | 0.381333 | 0.58774 | 0.719014 | 0.7403 | 0.82129 |
| SL1344_RS22355_ | -1.65776 | -1.82102 | -1.47549 | 0.347995 | 0.300509 | 1.091805 | 0.118588 | 0.190045 | 0.335477 | 0.2659 | 0.411647 |
| SL1344_RS22360_ | -0.228 | -0.16344 | -0.3023 | -0.14379 | 0.183222 | -0.73348 | 0.218521 | 0.317944 | 0.466181 | 0.4125 | 0.544978 |
| SL1344_RS22365_ | 1.087586 | 0.954324 | 1.271684 | 0.320592 | 0.140345 | 2.202636 | 0.004001 | 0.042159 | 0.146505 | 0.1042 | 0.241992 |
| SL1344_RS22370_ | -0.29921 | -0.55019 | -0.03716 | 0.511496 | 0.170703 | 2.950119 | 0.001337 | 0.018461 | 0.093236 | 0.101 | 0.237939 |
| SL1344_RS22375_ | 2.012213 | 1.934562 | 2.103171 | 0.170768 | 0.104381 | 1.551932 | 0.043305 | 0.090079 | 0.222564 | 0.1528 | 0.299592 |
| SL1344_RS22380_ | 2.326415 | 2.230033 | 2.463442 | 0.229273 | 0.088517 | 2.507777 | 0.003335 | 0.03035 | 0.122394 | 0.1033 | 0.240893 |
| SL1344_RS22385_ | 4.166837 | 3.989748 | 4.377311 | 0.363673 | 0.131319 | 2.639109 | 0.000673 | 0.025684 | 0.119841 | 0.1001 | 0.236726 |
| SL1344_RS22395_ | 4.128556 | 4.057715 | 4.349122 | 0.280351 | 0.12779 | 2.028761 | 0.002001 | 0.045693 | 0.161035 | 0.1007 | 0.23753 |
| SL1344_RS22400_ | 3.780883 | 3.614021 | 3.96928 | 0.346381 | 0.087793 | 3.933451 | 4.68E-05 | 0.005167 | 0.056748 | 0.1 | 0.236592 |
| SL1344_RS22405_ | 4.588974 | 4.485726 | 4.700082 | 0.244714 | 0.078819 | 3.250885 | 4.68E-05 | 0.008888 | 0.071675 | 0.1 | 0.236592 |
| SL1344_RS22410_ | -2.357 | -2.30148 | -2.41289 | -0.10528 | 0.357928 | -0.28299 | 0.371752 | 0.495944 | 0.62959 | 0.6046 | 0.7099 |
| SL1344_RS22415_ | 5.094711 | 5.132612 | 5.058317 | -0.08764 | 0.050993 | -1.69584 | 0.008667 | 0.066438 | 0.192226 | 0.1091 | 0.247957 |
| SL1344_RS22420_ | 1.503753 | 1.379044 | 1.64914 | 0.269506 | 0.102853 | 2.556184 | 0.003335 | 0.026406 | 0.111224 | 0.1036 | 0.241024 |
| SL1344_RS22425_ | 4.031229 | 3.930957 | 4.257306 | 0.32806 | 0.150163 | 2.167783 | 4.68E-05 | 0.032281 | 0.1347 | 0.1 | 0.236592 |
| SL1344_RS22430_ | 2.326748 | 2.280795 | 2.454188 | 0.130875 | 0.181541 | 0.586548 | 0.326 | 0.58515 | 0.72666 | 0.6769 | 0.799816 |
| SL1344_RS22435_ | -4.16824 | -4.32726 | -4.02864 | 0.294197 | 0.667093 | 0.421079 | 0.314 | 0.438086 | 0.577942 | 0.539 | 0.655385 |
| SL1344_RS22440_ | -2.25662 | -2.31544 | -2.17137 | 0.165855 | 0.434304 | 0.359797 | 0.346 | 0.53285 | 0.666822 | 0.6568 | 0.755071 |
| SL1344_RS22445_ | -2.37029 | -2.25582 | -2.48366 | -0.21888 | 0.390633 | -0.55382 | 0.273151 | 0.408485 | 0.552078 | 0.5055 | 0.62709 |
| SL1344_RS22450_ | -3.26245 | -3.45067 | -3.10455 | 0.327577 | 0.525713 | 0.585033 | 0.247835 | 0.372667 | 0.519495 | 0.4741 | 0.601756 |
| SL1344_RS22455_ | -3.80165 | -4.01755 | -3.57122 | 0.462065 | 0.682097 | 0.648229 | 0.241333 | 0.356057 | 0.507432 | 0.4553 | 0.585546 |
| SL1344_RS22460_ | -2.79691 | -2.68934 | -3.05432 | -0.26541 | 0.732934 | -0.31634 | 0.386409 | 0.689319 | 0.799417 | 0.7775 | 0.862404 |
| SL1344_RS22465_ | -0.6094 | -0.63637 | -0.57488 | 0.151234 | 0.368318 | 0.403126 | 0.342667 | 0.521571 | 0.668242 | 0.7083 | 0.797884 |
| SL1344_RS22470_ | 0.698644 | 0.444984 | 1.214219 | 0.745999 | 0.198447 | 3.411644 | 4.68E-05 | 0.019125 | 0.100505 | 0.1 | 0.236592 |
| SL1344_RS22475_ | 1.123156 | 0.229234 | 1.839338 | 1.630722 | 0.197098 | 8.363356 | 4.68E-05 | 0.000696 | 0.028755 | 0.1 | 0.236592 |
| SL1344_RS22480_ | 2.759593 | 1.883425 | 3.708078 | 1.803231 | 0.292896 | 6.138376 | 4.68E-05 | 0.001295 | 0.035856 | 0.1 | 0.236592 |
| SL1344_RS22485_ | 0.175344 | -0.08394 | 0.522372 | 0.563581 | 0.260939 | 2.12351 | 0.008001 | 0.037436 | 0.139909 | 0.1105 | 0.250398 |
| SL1344_RS22490_ | 0.960389 | 0.950088 | 0.967777 | 0.016772 | 0.178582 | 0.089778 | 0.462358 | 0.682688 | 0.790536 | 0.7634 | 0.84228 |
| SL1344_RS22495_ | -1.28924 | -1.31651 | -1.2542 | 0.058938 | 0.292482 | 0.186805 | 0.415057 | 0.589327 | 0.712937 | 0.6951 | 0.786915 |
| SL1344_RS22500_ | -1.62031 | -1.7962 | -1.45718 | 0.339967 | 0.329346 | 0.979502 | 0.138574 | 0.216457 | 0.371694 | 0.3009 | 0.449156 |
| SL1344_RS22505_ | -3.06291 | -3.4697 | -2.69161 | 0.781281 | 0.471938 | 1.593353 | 0.041306 | 0.082578 | 0.203952 | 0.1479 | 0.29129 |
| SL1344_RS22510_ | -2.20831 | -2.59668 | -1.91754 | 0.708065 | 0.497968 | 1.38114 | 0.037333 | 0.114818 | 0.259501 | 0.1514 | 0.296026 |
| SL1344_RS22520_ | -1.3654 | -1.53924 | -1.20288 | 0.326758 | 0.29823 | 1.029178 | 0.132667 | 0.202112 | 0.35526 | 0.2823 | 0.431851 |
| SL1344_RS22525_ | -0.79241 | -0.89097 | -0.68747 | 0.202259 | 0.270808 | 0.696276 | 0.225183 | 0.349621 | 0.505403 | 0.4502 | 0.586595 |
| SL1344_RS22530_ | -1.69337 | -1.87364 | -1.49227 | 0.383127 | 0.323052 | 1.136461 | 0.111333 | 0.167616 | 0.314292 | 0.2461 | 0.39335 |
| SL1344_RS22535_ | 2.076033 | 2.16228 | 1.702224 | -0.44609 | 0.228506 | -1.39157 | 0.074667 | 0.130967 | 0.294242 | 0.1806 | 0.337393 |
| SL1344_RS22540_ | 3.911796 | 4.026103 | 3.527927 | -0.45679 | 0.240519 | -1.34889 | 0.109261 | 0.125939 | 0.289218 | 0.1972 | 0.365618 |
| SL1344_RS22545_ | 4.233685 | 4.37773 | 3.886186 | -0.47285 | 0.172665 | -1.73197 | 0.006663 | 0.085485 | 0.230818 | 0.1044 | 0.24245 |
| SL1344_RS22550_ | 3.40848 | 3.465816 | 3.032978 | -0.41026 | 0.140799 | -1.16313 | 0.121253 | 0.167293 | 0.342295 | 0.23 | 0.396501 |
| SL1344_RS22555_ | 4.786935 | 4.953098 | 4.412444 | -0.41719 | 0.202381 | -1.6263 | 0.025334 | 0.073912 | 0.212649 | 0.1249 | 0.269835 |
| SL1344_RS22560_ | -1.16448 | -1.11488 | -1.21604 | -0.11474 | 0.275607 | -0.38115 | 0.332445 | 0.493748 | 0.628531 | 0.5989 | 0.704487 |

| | | | | | | | | | | | |
|---|---|---|---|---|---|---|---|---|---|---|---|
| SL1344_RS22565_ | 1.883232 | 1.584002 | 2.120566 | 0.538981 | 0.086826 | 6.202609 | 4.68E-05 | 0.002333 | 0.03928 | 0.1 | 0.236592 |
| SL1344_RS22570_ | 1.260253 | 0.994165 | 1.35604 | 0.377506 | 0.336477 | 1.113154 | 0.067333 | 0.210209 | 0.392555 | 0.1722 | 0.3258 |
| SL1344_RS22575_ | 1.966628 | 2.014147 | 1.90564 | -0.12557 | 0.153463 | -0.79381 | 0.194667 | 0.27615 | 0.450229 | 0.3781 | 0.529108 |
| SL1344_RS22580_ | -1.23691 | -1.24716 | -1.2209 | 0.03498 | 0.304803 | 0.111699 | 0.450667 | 0.608972 | 0.731707 | 0.7255 | 0.812391 |
| SL1344_RS22585_ | -1.09213 | -0.87276 | -1.42774 | -0.54554 | 0.317067 | -1.64142 | 0.025983 | 0.079554 | 0.208367 | 0.1365 | 0.281057 |
| SL1344_RS22590_ | -2.31847 | -2.18517 | -2.4722 | -0.29239 | 0.3399 | -0.81152 | 0.187875 | 0.283276 | 0.432027 | 0.3835 | 0.518389 |
| SL1344_RS22595_ | -1.06491 | -1.07739 | -1.05066 | 0.03221 | 0.290475 | 0.112908 | 0.450667 | 0.606393 | 0.72826 | 0.7175 | 0.803828 |
| SL1344_RS22600_ | 3.286325 | 3.259425 | 3.319711 | 0.067973 | 0.07285 | 0.874063 | 0.158561 | 0.235111 | 0.399747 | 0.3331 | 0.480629 |
| SL1344_RS22610_ | 0.400304 | 0.384667 | 0.421788 | -0.04175 | 0.282915 | -0.14025 | 0.437708 | 0.714045 | 0.816821 | 0.8809 | 0.922987 |
| SL1344_RS22615_ | 4.839465 | 5.183643 | 4.408366 | -0.72764 | 0.162768 | -4.67971 | 4.68E-05 | 0.00171 | 0.039678 | 0.1 | 0.236592 |
| SL1344_RS22620_ | 2.18515 | 2.200457 | 2.171188 | -0.02918 | 0.096503 | -0.2908 | 0.37 | 0.582341 | 0.70876 | 0.6816 | 0.776876 |
| SL1344_RS22625_ | 1.890688 | 2.00092 | 1.760547 | -0.25633 | 0.125959 | -2.02957 | 0.006667 | 0.045996 | 0.154336 | 0.1044 | 0.242187 |
| SL1344_RS22630_ | 1.818176 | 1.430638 | 2.175061 | 0.770434 | 0.144131 | 5.349549 | 4.68E-05 | 0.002975 | 0.045258 | 0.1 | 0.236592 |
| SL1344_RS22635_ | 3.28856 | 3.05292 | 3.567571 | 0.512437 | 0.110358 | 4.698394 | 4.68E-05 | 0.011374 | 0.080024 | 0.1 | 0.236592 |
| SL1344_RS22640_ | 2.485769 | 2.568737 | 2.38207 | -0.17598 | 0.228583 | -0.76308 | 0.205863 | 0.294468 | 0.478938 | 0.3916 | 0.558419 |
| SL1344_RS22645_ | -2.43593 | -2.40082 | -2.4717 | -0.07306 | 0.450897 | -0.16767 | 0.428667 | 0.575703 | 0.702909 | 0.6963 | 0.786345 |
| SL1344_RS22650_ | -2.91228 | -2.68426 | -3.11161 | -0.4465 | 0.519157 | -0.8102 | 0.19054 | 0.285064 | 0.439496 | 0.3881 | 0.526881 |
| SL1344_RS22655_ | 3.809129 | 3.870862 | 3.70223 | -0.17644 | 0.087385 | -2.048 | 0.002002 | 0.066823 | 0.196574 | 0.101 | 0.237863 |
| SL1344_RS22660_ | 4.202785 | 4.031369 | 4.356544 | 0.369802 | 0.173874 | 2.336157 | 4.68E-05 | 0.025677 | 0.120202 | 0.1 | 0.236592 |
| SL1344_RS22665_ | 2.364581 | 2.183361 | 2.430869 | 0.232172 | 0.158233 | 1.412466 | 0.074 | 0.098547 | 0.244754 | 0.1808 | 0.33671 |
| SL1344_RS22670_ | -3.04407 | -3.17065 | -2.93908 | 0.238588 | 0.720623 | 0.316781 | 0.368667 | 0.560526 | 0.696103 | 0.6791 | 0.776945 |
| SL1344_RS22675_ | 2.007537 | 2.068715 | 1.942248 | -0.13085 | 0.163446 | -0.78072 | 0.197202 | 0.279945 | 0.455391 | 0.3756 | 0.532431 |
| SL1344_RS22680_ | -0.96085 | -1.15263 | -0.80099 | 0.363213 | 0.292684 | 1.204895 | 0.096602 | 0.150541 | 0.30182 | 0.2183 | 0.370931 |
| SL1344_RS22685_ | -7.07248 | -6.94308 | -7.17611 | -0.25558 | 3.191933 | -0.05486 | 0.461026 | 0.635335 | 0.757273 | 0.756 | 0.835367 |
| SL1344_RS22690_ | -0.20683 | -0.44693 | -0.04802 | 0.388255 | 0.189387 | 1.989026 | 0.016667 | 0.055876 | 0.16683 | 0.1212 | 0.262192 |
| SL1344_RS22695_ | -0.44759 | -0.46645 | -0.42215 | 0.02683 | 0.261362 | 0.092729 | 0.456667 | 0.668042 | 0.778959 | 0.7684 | 0.845488 |
| SL1344_RS22700_ | -0.87299 | -1.01755 | -0.75685 | 0.319551 | 0.356791 | 0.833082 | 0.159893 | 0.249179 | 0.419634 | 0.3487 | 0.495048 |
| SL1344_RS22705_ | -2.72432 | -3.02489 | -2.22262 | 0.800454 | 0.550807 | 1.369204 | 0.071286 | 0.122387 | 0.267504 | 0.1944 | 0.343995 |
| SL1344_RS22710_ | -4.38769 | -4.36519 | -4.40463 | -0.09013 | 1.367206 | -0.05518 | 0.479333 | 0.730221 | 0.82763 | 0.8368 | 0.896931 |
| SL1344_RS22715_ | -1.62489 | -1.69504 | -1.54698 | 0.16932 | 0.389081 | 0.414941 | 0.33 | 0.516798 | 0.656021 | 0.6455 | 0.747068 |
| SL1344_RS22720_ | -1.52857 | -1.37587 | -1.63106 | -0.25145 | 0.313868 | -0.74246 | 0.212667 | 0.339805 | 0.490099 | 0.4386 | 0.57041 |
| SL1344_RS22725_ | -0.63794 | -0.95995 | -0.44644 | 0.52194 | 0.332555 | 1.533724 | 0.034 | 0.077069 | 0.208467 | 0.141 | 0.287306 |
| SL1344_RS22730_ | -2.12291 | -2.18206 | -2.07493 | 0.076395 | 0.473872 | 0.166569 | 0.431046 | 0.660784 | 0.774296 | 0.7637 | 0.841516 |
| SL1344_RS22735_ | -0.20317 | -0.31101 | -0.10397 | 0.228839 | 0.242821 | 0.90184 | 0.159227 | 0.230812 | 0.395462 | 0.3211 | 0.46922 |
| SL1344_RS22740_ | -1.52234 | -1.56539 | -1.47811 | 0.084658 | 0.305944 | 0.268042 | 0.382667 | 0.540097 | 0.671421 | 0.6489 | 0.750363 |
| SL1344_RS22745_ | -2.27313 | -2.19576 | -2.35978 | -0.1694 | 0.395235 | -0.38429 | 0.330667 | 0.487836 | 0.625399 | 0.5969 | 0.706716 |
| SL1344_RS22750_ | -1.29486 | -1.07067 | -1.5015 | -0.42733 | 0.254372 | -1.63387 | 0.036 | 0.079435 | 0.200772 | 0.1483 | 0.291799 |
| SL1344_RS22755_ | -2.5564 | -2.73063 | -2.40901 | 0.402514 | 0.61685 | 0.610934 | 0.241839 | 0.357424 | 0.521465 | 0.4897 | 0.619214 |
| SL1344_RS22760_ | -1.9507 | -2.08101 | -1.84069 | 0.255811 | 0.375825 | 0.658488 | 0.230667 | 0.342917 | 0.498755 | 0.4544 | 0.582929 |
| SL1344_RS22765_ | 0.574715 | 0.44631 | 0.683323 | 0.240741 | 0.135752 | 1.734974 | 0.026 | 0.071256 | 0.189718 | 0.1304 | 0.272416 |
| SL1344_RS22770_ | 2.710534 | 2.641603 | 2.778989 | 0.141492 | 0.086523 | 1.562161 | 0.030667 | 0.076497 | 0.202324 | 0.1382 | 0.282196 |
| SL1344_RS22775_ | 2.901284 | 2.959736 | 2.823545 | -0.17702 | 0.138811 | -1.2782 | 0.053298 | 0.111265 | 0.261722 | 0.1615 | 0.310663 |
| SL1344_RS22780_ | -3.9033 | -4.03773 | -3.76472 | 0.280274 | 0.668148 | 0.374254 | 0.334444 | 0.482675 | 0.618451 | 0.5891 | 0.698864 |
| SL1344_RS22785_ | -2.54546 | -2.72802 | -2.20246 | 0.49305 | 0.567946 | 0.785847 | 0.205197 | 0.314834 | 0.475177 | 0.4237 | 0.568257 |
| SL1344_RS22790_ | -3.03258 | -3.16403 | -2.91848 | 0.22739 | 0.538498 | 0.401415 | 0.316667 | 0.460853 | 0.602089 | 0.5628 | 0.678346 |

| | | | | | | | | | | | |
|---|---|---|---|---|---|---|---|---|---|---|---|
| SL1344_RS22795_ | -2.31762 | -2.43343 | -2.1653 | 0.323596 | 0.46422 | 0.675452 | 0.23984 | 0.35266 | 0.511111 | 0.4712 | 0.59741 |
| SL1344_RS22800_ | -2.42086 | -2.53922 | -2.28434 | 0.25848 | 0.380136 | 0.635076 | 0.227182 | 0.357824 | 0.501624 | 0.45 | 0.577241 |
| SL1344_RS22805_ | -1.89564 | -1.89444 | -1.89721 | -0.00729 | 0.305658 | -0.02266 | 0.49034 | 0.541753 | 0.671583 | 0.6488 | 0.748063 |
| SL1344_RS22810_ | -1.88667 | -2.23293 | -1.55022 | 0.687441 | 0.327397 | 2.024037 | 0.011992 | 0.043344 | 0.144983 | 0.1109 | 0.250276 |
| SL1344_RS22815_ | -1.70282 | -1.98788 | -1.54384 | 0.467083 | 0.423736 | 1.042866 | 0.096667 | 0.18916 | 0.352915 | 0.2396 | 0.387708 |
| SL1344_RS22820_ | 0.698131 | 0.558387 | 0.81888 | 0.252451 | 0.184175 | 1.352293 | 0.063333 | 0.115292 | 0.2597 | 0.1759 | 0.327715 |
| SL1344_RS22825_ | 0.171026 | 0.294076 | 0.057977 | -0.23399 | 0.188296 | -1.20485 | 0.093271 | 0.154024 | 0.301824 | 0.2231 | 0.374048 |
| SL1344_RS22830_ | 0.163691 | 0.182409 | 0.145507 | -0.0353 | 0.15525 | -0.21892 | 0.397069 | 0.521714 | 0.653829 | 0.6218 | 0.72647 |
| SL1344_RS22835_ | -0.20768 | -0.30589 | 0.082961 | 0.359958 | 0.226618 | 1.354473 | 0.099933 | 0.137021 | 0.288689 | 0.2334 | 0.384144 |
| SL1344_RS22840_ | 3.061572 | 2.816956 | 3.282784 | 0.438804 | 0.11793 | 3.697175 | 4.68E-05 | 0.008785 | 0.070464 | 0.1 | 0.236592 |
| SL1344_RS22845_ | -5.3491 | -5.55351 | -5.17078 | 0.430801 | 1.380941 | 0.287685 | 0.356 | 0.544153 | 0.678406 | 0.6586 | 0.757556 |
| SL1344_RS22850_ | -0.49284 | -0.5297 | -0.4492 | 0.141224 | 0.327479 | 0.400961 | 0.323118 | 0.466136 | 0.622099 | 0.6542 | 0.753264 |
| SL1344_RS22855_ | 0.803411 | 0.758945 | 0.839943 | 0.083891 | 0.141735 | 0.563168 | 0.257828 | 0.41002 | 0.554894 | 0.5183 | 0.637656 |
| SL1344_RS22860_ | -1.50794 | -1.49103 | -1.51972 | -0.02681 | 0.302842 | -0.08414 | 0.465023 | 0.59298 | 0.715714 | 0.7029 | 0.792609 |
| SL1344_RS22865_ | -0.15364 | -0.24252 | -0.07373 | 0.170641 | 0.17438 | 0.901216 | 0.153231 | 0.242626 | 0.390471 | 0.3302 | 0.470022 |
| SL1344_RS22870_ | 1.647206 | 1.566959 | 1.717622 | 0.163459 | 0.157164 | 1.019667 | 0.111333 | 0.192105 | 0.358615 | 0.2537 | 0.408993 |
| SL1344_RS22875_ | 5.64163 | 5.46809 | 5.802017 | 0.355617 | 0.081487 | 4.494974 | 4.68E-05 | 0.006518 | 0.063473 | 0.1 | 0.236592 |
| SL1344_RS22880_ | -0.72273 | -0.95902 | -0.50355 | 0.459088 | 0.235229 | 1.91043 | 0.018654 | 0.056098 | 0.167024 | 0.1211 | 0.261701 |
| SL1344_RS22885_ | -0.18895 | -0.46973 | -0.01216 | 0.453921 | 0.231011 | 1.846848 | 0.022652 | 0.063788 | 0.184919 | 0.1206 | 0.262476 |
| SL1344_RS22890_ | -3.93508 | -3.89244 | -3.99208 | -0.14173 | 0.982038 | -0.13649 | 0.437708 | 0.662259 | 0.775956 | 0.7839 | 0.854923 |
| SL1344_RS22895_ | -1.97942 | -2.24773 | -1.77567 | 0.467031 | 0.559176 | 0.780707 | 0.203198 | 0.286411 | 0.457943 | 0.3816 | 0.537913 |
| SL1344_RS22900_ | -2.14239 | -2.25223 | -2.00654 | 0.269459 | 0.431837 | 0.577474 | 0.255333 | 0.39866 | 0.549344 | 0.5163 | 0.638814 |
| SL1344_RS22905_ | -3.19483 | -3.79885 | -2.65513 | 1.135869 | 0.774937 | 1.381826 | 0.074617 | 0.109927 | 0.254952 | 0.1682 | 0.319787 |
| SL1344_RS22910_ | -4.51535 | -5.03218 | -4.03119 | 0.981226 | 1.041847 | 0.873242 | 0.161892 | 0.247856 | 0.408864 | 0.3301 | 0.480048 |
| SL1344_RS22915_ | -3.87809 | -4.51664 | -3.37608 | 1.223419 | 0.941183 | 1.25499 | 0.078 | 0.130418 | 0.279597 | 0.2009 | 0.353254 |
| SL1344_RS22920_ | 0.233636 | -0.05852 | 0.351953 | 0.364997 | 0.296981 | 1.156955 | 0.121919 | 0.162614 | 0.327171 | 0.2509 | 0.414945 |
| SL1344_RS22925_ | -1.29759 | -1.05299 | -1.54196 | -0.48643 | 0.284263 | -1.68696 | 0.031333 | 0.077328 | 0.199863 | 0.1442 | 0.287712 |
| SL1344_RS22930_ | 2.016133 | 1.679537 | 2.167279 | 0.440517 | 0.232343 | 1.580489 | 0.048 | 0.092747 | 0.239751 | 0.1437 | 0.294788 |
| SL1344_RS22935_ | 0.244851 | 0.060779 | 0.402802 | 0.340657 | 0.154927 | 2.187349 | 0.008001 | 0.037309 | 0.133814 | 0.1073 | 0.245496 |
| SL1344_RS22940_ | -1.27483 | -1.35035 | -1.21694 | 0.19609 | 0.422601 | 0.462123 | 0.318 | 0.499303 | 0.647259 | 0.6376 | 0.745392 |
| SL1344_RS22945_ | 0.253795 | 0.346609 | 0.018442 | -0.31925 | 0.25284 | -1.12584 | 0.113924 | 0.16788 | 0.331715 | 0.2528 | 0.411122 |
| SL1344_RS22950_ | -1.46462 | -1.34137 | -1.55143 | -0.22792 | 0.366985 | -0.57189 | 0.279147 | 0.411745 | 0.56722 | 0.5273 | 0.654635 |
| SL1344_RS22955_ | 4.45546 | 4.476743 | 4.434116 | -0.03527 | 0.06976 | -0.48585 | 0.295803 | 0.480615 | 0.636325 | 0.5922 | 0.715907 |
| SL1344_RS22960_ | 1.756509 | 1.737809 | 1.769642 | 0.032144 | 0.113012 | 0.286416 | 0.378414 | 0.597396 | 0.721244 | 0.7134 | 0.800562 |
| SL1344_RS22965_ | 3.981535 | 3.994518 | 3.965094 | -0.0243 | 0.086587 | -0.26152 | 0.385077 | 0.64967 | 0.770168 | 0.7776 | 0.854533 |
| SL1344_RS22970_ | 1.941965 | 1.972188 | 1.918589 | -0.06067 | 0.13197 | -0.43167 | 0.319333 | 0.473947 | 0.627349 | 0.6151 | 0.727703 |
| SL1344_RS22975_ | 2.301646 | 2.187755 | 2.452249 | 0.276035 | 0.151056 | 1.880605 | 0.004001 | 0.062143 | 0.186528 | 0.105 | 0.24282 |
| SL1344_RS22980_ | 1.933373 | 1.7736 | 2.091381 | 0.332118 | 0.134223 | 2.542419 | 0.000673 | 0.039791 | 0.14513 | 0.1001 | 0.236727 |
| SL1344_RS22985_ | -1.00307 | -1.17745 | -0.84195 | 0.334185 | 0.230489 | 1.402789 | 0.059333 | 0.110308 | 0.241681 | 0.1791 | 0.324166 |
| SL1344_RS22995_ | -1.99464 | -2.27225 | -1.78089 | 0.508734 | 0.393691 | 1.224062 | 0.074 | 0.141973 | 0.289678 | 0.1993 | 0.344548 |
| SL1344_RS23000_ | -2.11753 | -2.11244 | -2.12433 | -0.01001 | 0.392041 | -0.02012 | 0.491006 | 0.598623 | 0.72117 | 0.7089 | 0.797567 |
| SL1344_RS23010_ | 1.104208 | 0.938599 | 1.303064 | 0.363433 | 0.125483 | 2.91871 | 0.001336 | 0.019017 | 0.095215 | 0.1007 | 0.237523 |
| SL1344_RS23015_ | -1.44879 | -1.3308 | -1.62251 | -0.18352 | 0.861596 | -0.21607 | 0.401333 | 0.844824 | 0.908473 | 0.8182 | 0.891615 |
| SL1344_RS23020_ | -5.61976 | -5.52905 | -5.68672 | 0.07768 | 2.261957 | 0.025213 | 0.480667 | 0.614626 | 0.746772 | 0.7985 | 0.867964 |
| SL1344_RS23025_ | 2.340467 | 2.187963 | 2.510155 | 0.295411 | 0.189599 | 1.533111 | 0.054 | 0.080294 | 0.218278 | 0.1475 | 0.299283 |

| | | | | | | | | | | |
|---|---|---|---|---|---|---|---|---|---|---|
| SL1344_RS23030_ | 2.343575 | 2.210878 | 2.539787 | 0.2739 | 0.165924 | 1.656192 | 0.040667 | 0.065062 | 0.192773 | 0.1407 | 0.290223 |
| SL1344_RS23035_ | 2.898099 | 2.871042 | 2.924484 | 0.057445 | 0.088666 | 0.613461 | 0.240667 | 0.370493 | 0.5307 | 0.4923 | 0.623287 |
| SL1344_RS23040_ | 3.117573 | 3.158777 | 3.099898 | -0.05306 | 0.076644 | -0.62895 | 0.266 | 0.408148 | 0.563867 | 0.5218 | 0.655156 |
| SL1344_RS23045_ | 1.824707 | 1.889692 | 1.739894 | -0.16179 | 0.137933 | -1.15001 | 0.087941 | 0.147698 | 0.302533 | 0.2142 | 0.366693 |
| SL1344_RS23050_ | 2.113872 | 2.139894 | 2.08095 | -0.05946 | 0.10078 | -0.54431 | 0.278667 | 0.428754 | 0.579966 | 0.5539 | 0.675374 |
| SL1344_RS23055_ | 2.96577 | 2.874888 | 3.063579 | 0.19253 | 0.082907 | 2.302584 | 0.002002 | 0.033496 | 0.13085 | 0.1031 | 0.24067 |
| SL1344_RS23060_ | 3.694929 | 3.562591 | 3.777002 | 0.211255 | 0.06207 | 3.199062 | 4.68E-05 | 0.017117 | 0.092363 | 0.1 | 0.236592 |
| SL1344_RS23065_ | 0.973903 | 0.900057 | 1.11261 | 0.214432 | 0.183006 | 1.122708 | 0.121919 | 0.176299 | 0.339102 | 0.2586 | 0.416075 |
| SL1344_RS23070_ | 0.66844 | 0.71399 | 0.546245 | -0.14774 | 0.185612 | -0.71427 | 0.236 | 0.365329 | 0.52509 | 0.4795 | 0.618691 |
| SL1344_RS23075_ | -5.66049 | -6.08152 | -5.24077 | 0.763366 | 1.687299 | 0.412435 | 0.287142 | 0.475107 | 0.620906 | 0.5589 | 0.68085 |
| SL1344_RS23080_ | 1.71779 | 1.845477 | 1.574506 | -0.28985 | 0.135552 | -2.15427 | 0.002002 | 0.03951 | 0.142279 | 0.1016 | 0.238677 |
| SL1344_RS23085_ | 1.214991 | 1.045915 | 1.367138 | 0.354334 | 0.168653 | 2.138954 | 0.002001 | 0.038618 | 0.14171 | 0.1016 | 0.238727 |
| SL1344_RS23090_ | -0.72323 | -0.97412 | -0.54241 | 0.441762 | 0.327985 | 1.344761 | 0.066622 | 0.111932 | 0.256913 | 0.177 | 0.329028 |
| SL1344_RS23095_ | -2.59602 | -2.48359 | -2.71663 | -0.25869 | 0.473066 | -0.52725 | 0.276482 | 0.436779 | 0.580907 | 0.5498 | 0.665214 |
| SL1344_RS23100_ | 2.718028 | 2.493916 | 2.981011 | 0.467587 | 0.166373 | 2.775338 | 4.68E-05 | 0.014976 | 0.090672 | 0.1 | 0.236592 |
| SL1344_RS23105_ | -2.43868 | -2.5426 | -2.30004 | 0.203546 | 0.480727 | 0.403377 | 0.317333 | 0.524151 | 0.659553 | 0.6198 | 0.72998 |
| SL1344_RS23110_ | -1.30464 | -1.35682 | -1.15612 | 0.161615 | 0.413165 | 0.308054 | 0.378667 | 0.718575 | 0.818681 | 0.767 | 0.856349 |
| SL1344_RS23115_ | -1.03872 | -1.02138 | -1.05449 | -0.04025 | 0.247261 | -0.14019 | 0.433333 | 0.548252 | 0.680213 | 0.669 | 0.765061 |
| SL1344_RS23120_ | 2.614598 | 2.683853 | 2.577401 | -0.10086 | 0.09494 | -0.99133 | 0.158667 | 0.223232 | 0.387833 | 0.323 | 0.478772 |
| SL1344_RS23125_ | -2.14923 | -2.04408 | -2.27047 | -0.23482 | 0.415904 | -0.53381 | 0.29 | 0.433104 | 0.57882 | 0.5436 | 0.661656 |
| SL1344_RS23130_ | -4.29397 | -4.47161 | -4.10369 | 0.360347 | 0.736666 | 0.456066 | 0.307129 | 0.457371 | 0.595859 | 0.5556 | 0.671183 |
| SL1344_RS23140_ | -0.93633 | -1.05638 | -0.62355 | 0.404147 | 0.63388 | 0.67101 | 0.216667 | 0.328446 | 0.508403 | 0.3976 | 0.570221 |
| SL1344_RS23145_ | -1.03237 | -1.06921 | -0.9857 | 0.066454 | 0.296391 | 0.206909 | 0.418667 | 0.60196 | 0.725563 | 0.7111 | 0.799797 |
| SL1344_RS23150_ | 0.593089 | 0.360887 | 0.815543 | 0.453791 | 0.145898 | 3.077848 | 0.000673 | 0.016277 | 0.08851 | 0.1004 | 0.237132 |
| SL1344_RS23155_ | -2.96767 | -3.36518 | -2.5994 | 0.77219 | 0.688185 | 1.090362 | 0.107262 | 0.174843 | 0.335685 | 0.2414 | 0.398238 |
| SL1344_RS23160_ | -3.45156 | -3.84155 | -2.90183 | 0.709152 | 1.080642 | 0.648237 | 0.246 | 0.399448 | 0.563883 | 0.4773 | 0.622834 |
| SL1344_RS23165_ | -4.67259 | -4.72438 | -4.59665 | 0.137129 | 0.883537 | 0.152317 | 0.439707 | 0.562428 | 0.689764 | 0.6699 | 0.763981 |
| SL1344_RS23170_ | -1.9334 | -1.66655 | -2.04243 | -0.34427 | 0.430023 | -0.72217 | 0.244 | 0.378068 | 0.536519 | 0.4951 | 0.633452 |
| SL1344_RS23175_ | 0.167666 | 0.098463 | 0.247049 | 0.172263 | 0.233255 | 0.694685 | 0.204 | 0.318906 | 0.483811 | 0.4326 | 0.573526 |
| SL1344_RS23180_ | 1.019038 | 1.076031 | 0.936696 | -0.15909 | 0.179593 | -0.87668 | 0.125333 | 0.234217 | 0.405939 | 0.2903 | 0.435916 |
| SL1344_RS23185_ | -1.30571 | -1.49665 | -1.05952 | 0.448069 | 0.308712 | 1.370584 | 0.052632 | 0.114039 | 0.253797 | 0.1789 | 0.325874 |
| SL1344_RS23190_ | 1.937132 | 1.321704 | 2.552657 | 1.235546 | 0.159566 | 7.743277 | 4.68E-05 | 0.000817 | 0.0306 | 0.1 | 0.236592 |
| SL1344_RS23195_ | -4.26266 | -4.57124 | -4.04031 | 0.531791 | 0.726017 | 0.681593 | 0.220667 | 0.340699 | 0.486838 | 0.4292 | 0.560241 |
| SL1344_RS23200_ | -5.22896 | -5.71509 | -4.52907 | 1.020982 | 2.104723 | 0.443479 | 0.262667 | 0.494103 | 0.642901 | 0.5246 | 0.66539 |
| SL1344_RS23205_ | -0.44494 | -0.0253 | -0.77898 | -0.78735 | 0.362628 | -2.1992 | 0.001337 | 0.053866 | 0.174477 | 0.1013 | 0.238261 |
| SL1344_RS23210_ | -2.59529 | -2.53457 | -2.65908 | -0.10727 | 0.503405 | -0.20129 | 0.413058 | 0.610955 | 0.732545 | 0.7227 | 0.810075 |
| SL1344_RS23215_ | 0.829334 | 0.823315 | 0.834981 | 0.012012 | 0.131706 | 0.08944 | 0.458 | 0.56048 | 0.688698 | 0.6723 | 0.768135 |
| SL1344_RS23220_ | 1.200899 | 1.165916 | 1.24775 | 0.088928 | 0.151332 | 0.561863 | 0.267821 | 0.418002 | 0.573308 | 0.5466 | 0.66864 |
| SL1344_RS23225_ | 3.296009 | 3.203516 | 3.393003 | 0.18953 | 0.077011 | 2.399963 | 0.001336 | 0.028312 | 0.119983 | 0.1015 | 0.238606 |
| SL1344_RS23230_ | 0.772038 | 0.52445 | 1.026234 | 0.504246 | 0.199452 | 2.495224 | 4.68E-05 | 0.026485 | 0.116509 | 0.1006 | 0.237389 |
| SL1344_RS23235_ | 0.335196 | 0.180497 | 0.442911 | 0.264113 | 0.205526 | 1.229412 | 0.081279 | 0.148804 | 0.303498 | 0.2066 | 0.356473 |
| SL1344_RS23240_ | 0.306379 | 0.167045 | 0.417838 | 0.218813 | 0.235236 | 0.905192 | 0.169887 | 0.250567 | 0.41978 | 0.3354 | 0.492921 |
| SL1344_RS23245_ | 1.907663 | 1.939069 | 1.809862 | -0.11453 | 0.132579 | -0.76377 | 0.228514 | 0.335966 | 0.503622 | 0.472 | 0.615001 |
| SL1344_RS23250_ | -1.07433 | -1.0553 | -1.08515 | -0.05068 | 0.458668 | -0.10255 | 0.461692 | 0.748591 | 0.841734 | 0.8672 | 0.917472 |
| SL1344_RS23265_ | 2.082262 | 2.04694 | 2.117934 | 0.059767 | 0.14014 | 0.405139 | 0.331333 | 0.541374 | 0.683237 | 0.6485 | 0.757942 |

| | | | | | | | | | | |
|---|---|---|---|---|---|---|---|---|---|---|
| SL1344_RS23270_ | 0.810869 | -0.00185 | 1.409335 | 1.38593 | 0.207503 | 6.587491 | 4.68E-05 | 0.002699 | 0.042131 | 0.1 | 0.236592 |
| SL1344_RS23275_ | 0.491962 | -0.32092 | 1.191368 | 1.545081 | 0.224152 | 6.974802 | 4.68E-05 | 0.00279 | 0.042115 | 0.1 | 0.236592 |
| SL1344_RS23280_ | 1.017225 | 0.115854 | 1.67442 | 1.556863 | 0.167295 | 9.313517 | 4.68E-05 | 0.002616 | 0.041649 | 0.1 | 0.236592 |
| SL1344_RS23285_ | 1.31218 | 0.566729 | 2.06968 | 1.499837 | 0.134246 | 11.17836 | 4.68E-05 | 0.000524 | 0.025654 | 0.1 | 0.236592 |
| SL1344_RS23290_ | 1.631299 | 0.827118 | 2.250107 | 1.432653 | 0.169909 | 8.482092 | 4.68E-05 | 0.000689 | 0.02835 | 0.1 | 0.236592 |
| SL1344_RS23295_ | 3.572027 | 2.810281 | 3.978869 | 1.157879 | 0.102831 | 9.636101 | 4.68E-05 | 0.014933 | 0.09311 | 0.1 | 0.236592 |
| SL1344_RS23300_ | 3.832044 | 3.701865 | 3.998687 | 0.300344 | 0.052668 | 5.715323 | 4.68E-05 | 0.003241 | 0.045113 | 0.1 | 0.236592 |
| SL1344_RS23305_ | 0.549795 | 0.475409 | 0.697379 | 0.21507 | 0.160993 | 1.199037 | 0.113258 | 0.162579 | 0.312753 | 0.2493 | 0.400336 |
| SL1344_RS23310_ | 1.82978 | 1.811908 | 1.845883 | 0.009744 | 0.144699 | 0.058485 | 0.470667 | 0.70311 | 0.806918 | 0.8187 | 0.879998 |
| SL1344_RS23315_ | 0.932507 | 0.920187 | 0.940057 | 0.001216 | 0.168054 | 0.007048 | 0.497668 | 0.681094 | 0.791041 | 0.7832 | 0.861498 |
| SL1344_RS23320_ | 0.460405 | 0.377781 | 0.580193 | 0.211698 | 0.230198 | 0.871084 | 0.156667 | 0.237904 | 0.407478 | 0.3209 | 0.481208 |
| SL1344_RS23325_ | 0.129312 | 0.134903 | 0.124804 | -0.00122 | 0.18942 | -0.00667 | 0.497668 | 0.621139 | 0.739662 | 0.7318 | 0.815211 |
| SL1344_RS23330_ | -7.15667 | -6.5964 | -8.0285 | -1.41241 | 2.08188 | -0.59338 | 0.222 | 0.339918 | 0.49611 | 0.438 | 0.569317 |
| SL1344_RS23335_ | -5.55373 | -5.49433 | -5.64897 | -0.16157 | 1.33748 | -0.10747 | 0.445333 | 0.569369 | 0.700458 | 0.694 | 0.787562 |
| SL1344_RS23340_ | 1.11085 | 0.892397 | 1.381347 | 0.478432 | 0.172135 | 2.750018 | 0.000673 | 0.018376 | 0.097409 | 0.1005 | 0.237259 |
| SL1344_RS23345_ | -2.43693 | -2.4624 | -2.41394 | 0.105691 | 0.816894 | 0.122912 | 0.428667 | 0.563856 | 0.709027 | 0.755 | 0.834872 |
| SL1344_RS23350_ | -0.56358 | -0.39827 | -0.65413 | -0.24135 | 0.226054 | -0.9952 | 0.154667 | 0.237556 | 0.392326 | 0.3399 | 0.486694 |
| SL1344_RS23355_ | -4.29721 | -4.13361 | -4.46823 | -0.34713 | 0.952224 | -0.3394 | 0.348434 | 0.529739 | 0.666838 | 0.6483 | 0.750064 |
| SL1344_RS23360_ | 2.771094 | 2.86119 | 2.619936 | -0.2245 | 0.1041 | -2.07364 | 0.010667 | 0.044797 | 0.153776 | 0.1134 | 0.2538 |
| SL1344_RS23365_ | 4.671973 | 4.706299 | 4.494983 | -0.21267 | 0.176789 | -1.24481 | 0.017334 | 0.173601 | 0.350455 | 0.1155 | 0.255782 |
| SL1344_RS23370_ | 1.153175 | 1.383761 | 1.055151 | -0.31464 | 0.146754 | -1.94898 | 0.018 | 0.062378 | 0.183335 | 0.1222 | 0.26317 |
| SL1344_RS23375_ | 0.114328 | 0.143349 | 0.081163 | -0.06459 | 0.179713 | -0.3396 | 0.357333 | 0.506012 | 0.641776 | 0.6159 | 0.720774 |
| SL1344_RS23380_ | -0.22682 | -0.12207 | -0.32775 | -0.20076 | 0.188013 | -1.02914 | 0.124 | 0.196732 | 0.345342 | 0.2755 | 0.420107 |
| SL1344_RS23385_ | -0.27388 | -0.09767 | -0.39455 | -0.27186 | 0.270561 | -0.97166 | 0.162558 | 0.232532 | 0.398975 | 0.3266 | 0.481752 |
| SL1344_RS23390_ | 1.230091 | 1.261591 | 1.180631 | -0.07743 | 0.133261 | -0.54695 | 0.286667 | 0.453628 | 0.601069 | 0.5667 | 0.687216 |
| SL1344_RS23395_ | 2.245921 | 2.320225 | 2.192111 | -0.13887 | 0.118858 | -1.12958 | 0.105333 | 0.160556 | 0.32075 | 0.2341 | 0.389736 |
| SL1344_RS23400_ | 3.547655 | 3.334593 | 3.761168 | 0.357996 | 0.138727 | 2.765222 | 4.68E-05 | 0.012632 | 0.084484 | 0.1 | 0.236592 |
| SL1344_RS23410_ | 0.745734 | 0.652469 | 0.962507 | 0.30869 | 0.178242 | 1.573979 | 0.049967 | 0.105263 | 0.248397 | 0.1593 | 0.305209 |
| SL1344_RS23415_ | 0.045532 | -0.02752 | 0.135152 | 0.165443 | 0.190543 | 0.828235 | 0.172667 | 0.276678 | 0.432903 | 0.3707 | 0.51108 |
| SL1344_RS23420_ | 1.057072 | 0.971064 | 1.153825 | 0.179302 | 0.121852 | 1.429959 | 0.052 | 0.103971 | 0.2361 | 0.1697 | 0.316227 |
| SL1344_RS23425_ | 1.93019 | 1.852291 | 2.017488 | 0.166419 | 0.105087 | 1.533463 | 0.044 | 0.09046 | 0.221149 | 0.158 | 0.303606 |
| SL1344_RS23430_ | 1.035078 | 0.989164 | 1.095667 | 0.099402 | 0.152181 | 0.608804 | 0.247169 | 0.390796 | 0.544373 | 0.485 | 0.618776 |
| SL1344_RS23435_ | 1.156261 | 1.094898 | 1.207462 | 0.114196 | 0.142117 | 0.77339 | 0.203864 | 0.306077 | 0.464896 | 0.4055 | 0.547842 |
| SL1344_RS23440_ | 2.860762 | 2.783038 | 2.880893 | 0.08894 | 0.137083 | 0.478334 | 0.340667 | 0.635809 | 0.764179 | 0.7156 | 0.826971 |
| SL1344_RS23445_ | 3.401807 | 3.366671 | 3.464895 | 0.143022 | 0.165036 | 0.863257 | 0.191872 | 0.236271 | 0.417801 | 0.3489 | 0.517688 |
| SL1344_RS23450_ | -1.30091 | -1.27487 | -1.31968 | -0.02438 | 0.317814 | -0.07242 | 0.468354 | 0.629671 | 0.748653 | 0.7326 | 0.818526 |
| SL1344_RS23455_ | -1.22621 | -1.28479 | -1.18428 | 0.104721 | 0.270905 | 0.347281 | 0.353764 | 0.505095 | 0.640963 | 0.6122 | 0.720517 |
| SL1344_RS23460_ | -4.88994 | -5.5678 | -4.34472 | 1.189684 | 1.157115 | 0.97278 | 0.138 | 0.211979 | 0.370595 | 0.2936 | 0.443007 |
| SL1344_RS23465_ | -2.88232 | -2.87309 | -2.89505 | 0.000319 | 0.540343 | 0.000902 | 0.5 | 0.617614 | 0.73829 | 0.7369 | 0.819623 |
| SL1344_RS23470_ | -3.32633 | -3.40013 | -3.27283 | 0.127903 | 0.5588 | 0.209877 | 0.413058 | 0.545776 | 0.676595 | 0.665 | 0.761484 |
| SL1344_RS23480_ | -1.03159 | -0.88163 | -1.19049 | -0.3106 | 0.23819 | -1.23628 | 0.082667 | 0.137903 | 0.277078 | 0.2102 | 0.356296 |
| SL1344_RS23485_ | 0.831839 | 0.826041 | 0.837836 | 0.014614 | 0.132332 | 0.108002 | 0.452365 | 0.571033 | 0.695285 | 0.6744 | 0.77044 |
| SL1344_RS23495_ | 2.303379 | 2.192034 | 2.427072 | 0.248844 | 0.111962 | 2.176501 | 0.001337 | 0.037902 | 0.139938 | 0.1021 | 0.239323 |
| SL1344_RS23500_ | 4.624439 | 4.5713 | 4.644315 | 0.073101 | 0.076998 | 0.902134 | 0.142667 | 0.23769 | 0.41894 | 0.293 | 0.450359 |
| SL1344_RS23505_ | 2.13244 | 1.928075 | 2.288954 | 0.359037 | 0.099145 | 3.614603 | 4.68E-05 | 0.01047 | 0.072815 | 0.1 | 0.236592 |

| | | | | | | | | | | | |
|---|---|---|---|---|---|---|---|---|---|---|---|
| SL1344_RS23510_ | 0.63451 | 0.527382 | 0.757409 | 0.22358 | 0.151146 | 1.431335 | 0.058628 | 0.106689 | 0.242931 | 0.172 | 0.320162 |
| SL1344_RS23515_ | -0.46836 | -0.29303 | -0.75124 | -0.41992 | 0.334114 | -1.19125 | 0.095936 | 0.151343 | 0.311194 | 0.2184 | 0.375729 |
| SL1344_RS23520_ | 1.548513 | 1.662367 | 1.456518 | -0.19802 | 0.13638 | -1.45156 | 0.058628 | 0.107698 | 0.248013 | 0.1772 | 0.329141 |
| SL1344_RS23525_ | 3.420579 | 3.30591 | 3.524522 | 0.235253 | 0.092832 | 2.504575 | 4.68E-05 | 0.021461 | 0.10666 | 0.1002 | 0.236854 |
| SL1344_RS23530_ | 1.589879 | 1.532999 | 1.651585 | 0.099603 | 0.21539 | 0.468766 | 0.309127 | 0.512969 | 0.666489 | 0.6237 | 0.745168 |
| SL1344_RS23535_ | 2.181327 | 2.146508 | 2.205785 | 0.039547 | 0.119886 | 0.299704 | 0.373333 | 0.621433 | 0.743878 | 0.7357 | 0.820713 |
| SL1344_RS23540_ | 0.901473 | 0.914098 | 0.88377 | -0.02813 | 0.134356 | -0.19011 | 0.427333 | 0.57751 | 0.703211 | 0.6794 | 0.775792 |
| SL1344_RS23545_ | -0.65615 | -0.63447 | -0.6789 | -0.03451 | 0.220816 | -0.15254 | 0.436667 | 0.565316 | 0.69215 | 0.6734 | 0.768695 |
| SL1344_RS23550_ | -2.48361 | -2.58157 | -2.34441 | 0.248892 | 0.444753 | 0.532988 | 0.277815 | 0.423794 | 0.571237 | 0.5453 | 0.663874 |
| SL1344_RS23555_ | -4.36522 | -4.45429 | -4.27237 | 0.172721 | 0.763667 | 0.211691 | 0.403731 | 0.535575 | 0.664144 | 0.6397 | 0.740044 |
| SL1344_RS23560_ | -2.14302 | -2.26596 | -2.0277 | 0.244451 | 0.37313 | 0.623209 | 0.243171 | 0.36412 | 0.512887 | 0.4696 | 0.597817 |
| SL1344_RS23565_ | -3.96991 | -4.00677 | -3.94597 | 0.024753 | 1.23303 | 0.019104 | 0.490667 | 0.748814 | 0.841687 | 0.852 | 0.906237 |
| SL1344_RS23570_ | -3.01486 | -2.94612 | -3.10062 | -0.17143 | 0.565241 | -0.27785 | 0.363333 | 0.539313 | 0.673172 | 0.656 | 0.754809 |
| SL1344_RS23575_ | 0.339201 | 0.55546 | 0.163902 | -0.36 | 0.221812 | -1.5475 | 0.058667 | 0.086903 | 0.221752 | 0.1576 | 0.309512 |
| SL1344_RS23580_ | -1.43307 | -1.50756 | -1.36642 | 0.173572 | 0.428864 | 0.392009 | 0.33511 | 0.506141 | 0.651596 | 0.6506 | 0.753024 |
| SL1344_RS23590_ | -5.86965 | -5.56692 | -6.15002 | -0.62267 | 2.737468 | -0.2069 | 0.358667 | 0.559306 | 0.700197 | 0.674 | 0.773261 |
| SL1344_RS23595_ | -0.6875 | -0.82495 | -0.57932 | 0.244617 | 0.266159 | 0.875339 | 0.169887 | 0.251612 | 0.410528 | 0.3425 | 0.489928 |
| SL1344_RS23640_ | 0.828847 | 0.696516 | 0.942682 | 0.140753 | 0.32422 | 0.453798 | 0.297333 | 0.503512 | 0.660796 | 0.5982 | 0.723126 |
| SL1344_RS23645_ | -0.92527 | -0.81756 | -1.08226 | -0.26355 | 0.279738 | -0.88568 | 0.166 | 0.254465 | 0.414279 | 0.3542 | 0.499292 |
| SL1344_RS23650_ | 1.580112 | 1.555067 | 1.59863 | 0.021738 | 0.168664 | 0.11859 | 0.455696 | 0.739085 | 0.83421 | 0.8481 | 0.902837 |
| SL1344_RS23655_ | 0.609429 | 0.593088 | 0.633694 | 0.052912 | 0.167647 | 0.287918 | 0.366 | 0.543314 | 0.675521 | 0.6703 | 0.764612 |
| SL1344_RS23660_ | 1.554834 | 1.504715 | 1.625018 | 0.11363 | 0.121011 | 0.862398 | 0.172552 | 0.270939 | 0.429702 | 0.3749 | 0.52006 |
| SL1344_RS23665_ | -0.74271 | -0.49234 | -1.13078 | -0.62517 | 0.290519 | -2.10872 | 0.01066 | 0.045412 | 0.151566 | 0.1122 | 0.251675 |
| SL1344_RS23670_ | 5.023451 | 4.692531 | 5.306732 | 0.52012 | 0.157817 | 3.5211 | 4.68E-05 | 0.004401 | 0.055115 | 0.1 | 0.236592 |
| SL1344_RS23675_ | 2.87539 | 2.715052 | 3.276283 | 0.544836 | 0.222877 | 2.338709 | 4.68E-05 | 0.030265 | 0.130403 | 0.1 | 0.236592 |
| SL1344_RS23680_ | 2.092642 | 1.839604 | 2.283516 | 0.421248 | 0.182719 | 2.277781 | 0.001336 | 0.025889 | 0.117965 | 0.101 | 0.237939 |
| SL1344_RS23685_ | -1.02741 | -1.12148 | -0.93446 | 0.216006 | 0.383404 | 0.553765 | 0.254 | 0.36718 | 0.534119 | 0.4771 | 0.614483 |
| SL1344_RS23690_ | -2.457 | -2.46628 | -2.44082 | -0.0044 | 0.513757 | -0.00805 | 0.498 | 0.66202 | 0.774423 | 0.77 | 0.847414 |
| SL1344_RS23695_ | -2.6754 | -2.56043 | -2.77162 | -0.19628 | 0.499362 | -0.3743 | 0.330446 | 0.505193 | 0.642286 | 0.6142 | 0.722341 |
| SL1344_RS23700_ | -1.92473 | -1.95341 | -1.89695 | 0.037656 | 0.388072 | 0.08725 | 0.459027 | 0.632801 | 0.750598 | 0.7331 | 0.818953 |
| SL1344_RS23705_ | -3.17266 | -3.40975 | -3.0415 | 0.366307 | 0.737239 | 0.476219 | 0.297801 | 0.454923 | 0.611605 | 0.5941 | 0.712644 |
| SL1344_RS23710_ | -0.64699 | -0.8942 | -0.45985 | 0.441562 | 0.280149 | 1.471081 | 0.042638 | 0.09387 | 0.231195 | 0.1474 | 0.293782 |
| SL1344_RS23715_ | -2.55456 | -2.61267 | -2.48579 | 0.126705 | 0.391056 | 0.320468 | 0.366422 | 0.499411 | 0.63418 | 0.6141 | 0.718026 |
| SL1344_RS23720_ | -2.22702 | -2.0642 | -2.33807 | -0.22301 | 0.489315 | -0.42158 | 0.333111 | 0.560119 | 0.693243 | 0.6579 | 0.765707 |
| SL1344_RS23725_ | -2.99836 | -2.69343 | -3.36638 | -0.67288 | 0.467706 | -1.38692 | 0.065956 | 0.116842 | 0.249947 | 0.1869 | 0.333041 |
| SL1344_RS23730_ | -2.22353 | -2.18136 | -2.26402 | -0.07305 | 0.3568 | -0.19739 | 0.409727 | 0.520135 | 0.65351 | 0.6277 | 0.732398 |
| SL1344_RS23735_ | -3.20762 | -3.02037 | -3.40786 | -0.37369 | 0.597353 | -0.57654 | 0.265333 | 0.40462 | 0.555108 | 0.5136 | 0.638922 |
| SL1344_RS23740_ | -3.74158 | -3.82193 | -3.67242 | 0.127805 | 0.997113 | 0.114925 | 0.43571 | 0.672034 | 0.783913 | 0.7616 | 0.841044 |
| SL1344_RS23745_ | -1.17225 | -1.10751 | -1.24475 | -0.14369 | 0.270311 | -0.49245 | 0.283811 | 0.439201 | 0.580292 | 0.5407 | 0.657521 |
| SL1344_RS23750_ | -4.04198 | -3.97077 | -4.16398 | -0.25031 | 0.967538 | -0.23692 | 0.395736 | 0.596823 | 0.725603 | 0.7417 | 0.824004 |
| SL1344_RS23755_ | -3.70513 | -3.40276 | -3.93689 | -0.56547 | 0.757971 | -0.71548 | 0.222518 | 0.328611 | 0.485155 | 0.4308 | 0.566758 |
| SL1344_RS23760_ | -4.3918 | -4.32464 | -4.46331 | -0.16715 | 1.016476 | -0.14222 | 0.423051 | 0.630252 | 0.748849 | 0.7312 | 0.816803 |
| SL1344_RS23765_ | -1.46929 | -1.37794 | -1.57642 | -0.18995 | 0.339747 | -0.49789 | 0.293138 | 0.455017 | 0.601086 | 0.5615 | 0.681646 |
| SL1344_RS23770_ | -2.64134 | -2.56814 | -2.73184 | -0.19064 | 0.470795 | -0.3816 | 0.337109 | 0.492005 | 0.631098 | 0.6001 | 0.709125 |
| SL1344_RS23775_ | -1.65577 | -1.17918 | -2.50915 | -1.33364 | 0.485677 | -2.65562 | 4.68E-05 | 0.034895 | 0.134413 | 0.1001 | 0.236727 |

| | | | | | | | | | | |
|---|---|---|---|---|---|---|---|---|---|---|
| SL1344_RS23780_ | -3.07154 | -3.22448 | -2.92886 | 0.305198 | 0.479039 | 0.598852 | 0.255163 | 0.367736 | 0.513545 | 0.4624 | 0.589243 |
| SL1344_RS23785_ | -2.39949 | -2.61576 | -1.997 | 0.537562 | 0.640319 | 0.800833 | 0.202 | 0.309098 | 0.475023 | 0.4013 | 0.554578 |
| SL1344_RS23790_ | -4.10167 | -4.19039 | -4.05518 | 0.018341 | 0.962346 | 0.017951 | 0.492 | 0.682102 | 0.790118 | 0.7797 | 0.856627 |
| SL1344_RS23795_ | -3.47684 | -3.80176 | -3.08795 | 0.698715 | 0.55589 | 1.175912 | 0.102 | 0.160716 | 0.303424 | 0.2417 | 0.386867 |
| SL1344_RS23800_ | -3.51443 | -3.79416 | -3.15837 | 0.664042 | 0.7135 | 0.904152 | 0.162667 | 0.248173 | 0.407701 | 0.3423 | 0.488731 |
| SL1344_RS23805_ | -3.81118 | -3.69695 | -3.94696 | -0.34646 | 0.964155 | -0.34742 | 0.348434 | 0.542534 | 0.681616 | 0.6889 | 0.781486 |
| SL1344_RS23810_ | -1.69576 | -1.17316 | -2.47182 | -1.27945 | 0.421576 | -2.96135 | 4.68E-05 | 0.022984 | 0.105564 | 0.1001 | 0.236726 |
| SL1344_RS23815_ | -1.24985 | -0.95614 | -1.502 | -0.5824 | 0.326976 | -1.7193 | 0.018654 | 0.071317 | 0.195627 | 0.1192 | 0.259722 |
| SL1344_RS23820_ | -4.71189 | -4.63637 | -4.7541 | -0.16499 | 0.995555 | -0.15231 | 0.431046 | 0.600852 | 0.724313 | 0.7075 | 0.799298 |
| SL1344_RS23825_ | -3.60959 | -3.61982 | -3.59946 | 0.026748 | 0.650208 | 0.038425 | 0.484344 | 0.595678 | 0.718822 | 0.7037 | 0.792434 |
| SL1344_RS23830_ | -2.80998 | -2.85089 | -2.77209 | 0.076299 | 0.418261 | 0.166917 | 0.428 | 0.501783 | 0.634999 | 0.6094 | 0.714287 |
| SL1344_RS23840_ | -3.01791 | -2.97548 | -3.06312 | -0.08025 | 0.466286 | -0.16637 | 0.417722 | 0.534551 | 0.666203 | 0.647 | 0.746435 |
| SL1344_RS23845_ | -4.5723 | -3.96564 | -5.0781 | -1.1355 | 1.000439 | -1.05366 | 0.133245 | 0.188815 | 0.344078 | 0.2676 | 0.420139 |
| SL1344_RS23850_ | -4.73614 | -4.54868 | -4.893 | -0.40062 | 0.98281 | -0.3708 | 0.334667 | 0.499721 | 0.6345 | 0.6118 | 0.715692 |
| SL1344_RS23855_ | -6.80176 | -7.62026 | -6.09179 | 1.46656 | 2.524481 | 0.481654 | 0.245836 | 0.391969 | 0.548635 | 0.4841 | 0.612948 |
| SL1344_RS23860_ | -1.80478 | -1.72371 | -1.98639 | -0.27599 | 0.429052 | -0.57067 | 0.265823 | 0.385528 | 0.546236 | 0.5214 | 0.648617 |
| SL1344_RS23865_ | -3.83477 | -3.4482 | -4.09069 | -0.63157 | 0.795199 | -0.74251 | 0.212 | 0.324128 | 0.482205 | 0.4274 | 0.566432 |
| SL1344_RS23870_ | -4.43761 | -4.52838 | -4.3626 | 0.150547 | 0.703577 | 0.19767 | 0.411726 | 0.514322 | 0.644733 | 0.6262 | 0.727374 |
| SL1344_RS23875_ | -2.5026 | -2.45304 | -2.54876 | -0.09356 | 0.446476 | -0.19588 | 0.413058 | 0.578971 | 0.706145 | 0.6958 | 0.788024 |
| SL1344_RS23880_ | -5.94108 | -5.95133 | -5.93554 | 0.042847 | 1.323122 | 0.026382 | 0.489007 | 0.549884 | 0.678848 | 0.6547 | 0.753555 |
| SL1344_RS23885_ | -6.50044 | -6.43463 | -6.60629 | -0.2609 | 2.151022 | -0.08478 | 0.437333 | 0.566841 | 0.703821 | 0.7224 | 0.807865 |
| SL1344_RS23890_ | -6.83717 | -7.41453 | -6.45523 | 0.968915 | 2.608048 | 0.33012 | 0.330667 | 0.497348 | 0.644359 | 0.6294 | 0.736638 |
| SL1344_RS23895_ | -6.39828 | -5.95207 | -6.92525 | -0.96629 | 1.715449 | -0.4884 | 0.270486 | 0.424735 | 0.572661 | 0.5331 | 0.651443 |
| SL1344_RS23900_ | -5.8771 | -6.42016 | -4.9992 | 1.218152 | 1.584796 | 0.721717 | 0.211859 | 0.355532 | 0.504851 | 0.4396 | 0.580745 |
| SL1344_RS23905_ | -7.87937 | -7.40265 | -8.42017 | -0.95281 | 3.923191 | -0.20643 | 0.37042 | 0.56157 | 0.697311 | 0.6991 | 0.791663 |
| SL1344_RS23910_ | -0.66783 | -0.72831 | -0.63333 | 0.094708 | 0.347111 | 0.246909 | 0.39507 | 0.63235 | 0.754549 | 0.7726 | 0.848331 |
| SL1344_RS23915_ | -0.55117 | -0.58335 | -0.4971 | 0.114195 | 0.262094 | 0.408386 | 0.336442 | 0.491723 | 0.635871 | 0.6303 | 0.73433 |
| SL1344_RS23920_ | 3.250823 | 3.211203 | 3.275818 | 0.074315 | 0.10629 | 0.738035 | 0.173884 | 0.299661 | 0.482031 | 0.3552 | 0.499623 |
| SL1344_RS23925_ | 2.239398 | 2.242839 | 2.2376 | -0.02175 | 0.162429 | -0.13021 | 0.447333 | 0.745528 | 0.839641 | 0.8615 | 0.912075 |
| SL1344_RS23930_ | 3.500702 | 3.337672 | 3.653817 | 0.342447 | 0.115944 | 2.92546 | 4.68E-05 | 0.01242 | 0.08325 | 0.1 | 0.236592 |
| SL1344_RS23935_ | 1.025001 | 1.02532 | 1.024965 | -0.00111 | 0.145118 | -0.00704 | 0.496667 | 0.639282 | 0.756018 | 0.752 | 0.832856 |
| SL1344_RS23940_ | 0.640023 | 0.628695 | 0.648921 | 0.072729 | 0.288714 | 0.233971 | 0.393738 | 0.537541 | 0.688095 | 0.7753 | 0.847849 |
| SL1344_RS23945_ | 0.575578 | 0.484144 | 0.733821 | 0.257523 | 0.191786 | 1.267731 | 0.08 | 0.1451 | 0.299703 | 0.1993 | 0.349527 |
| SL1344_RS23950_ | 2.544341 | 2.510106 | 2.589338 | 0.055272 | 0.194321 | 0.301515 | 0.338441 | 0.687766 | 0.800731 | 0.7042 | 0.806538 |
| SL1344_RS23955_ | -0.05859 | -0.09913 | -0.03848 | 0.046783 | 0.218942 | 0.188426 | 0.418 | 0.632166 | 0.750985 | 0.739 | 0.824709 |
| SL1344_RS23960_ | 2.915742 | 2.825687 | 2.992577 | 0.166901 | 0.060919 | 2.707121 | 0.001337 | 0.022329 | 0.102633 | 0.1025 | 0.239645 |
| SL1344_RS23965_ | 2.523748 | 2.496295 | 2.556231 | 0.063768 | 0.084242 | 0.719445 | 0.222 | 0.313923 | 0.474994 | 0.4288 | 0.565233 |
| SL1344_RS23970_ | 3.129664 | 3.0551 | 3.228366 | 0.179163 | 0.109499 | 1.622551 | 0.02465 | 0.085087 | 0.22383 | 0.1247 | 0.268389 |
| SL1344_RS23975_ | 2.259307 | 2.061681 | 2.4678 | 0.334429 | 0.182786 | 1.924637 | 0.02 | 0.041056 | 0.15013 | 0.1177 | 0.260166 |
| SL1344_RS23980_ | -2.45982 | -2.47675 | -2.44855 | 0.047054 | 0.413326 | 0.113279 | 0.453698 | 0.585506 | 0.711327 | 0.6965 | 0.788779 |
| SL1344_RS23985_ | -4.78875 | -4.71088 | -4.89552 | -0.1884 | 0.887568 | -0.19765 | 0.408 | 0.529524 | 0.662482 | 0.6412 | 0.741693 |
| SL1344_RS23990_ | -6.06255 | -5.96983 | -6.15738 | -0.14382 | 3.772975 | -0.03422 | 0.47 | 0.69979 | 0.808497 | 0.8016 | 0.870091 |
| SL1344_RS23995_ | -2.81458 | -3.31086 | -2.37452 | 0.939157 | 0.428934 | 2.075738 | 0.010667 | 0.042284 | 0.140874 | 0.1137 | 0.253215 |
| SL1344_RS24000_ | -1.18933 | -1.35341 | -0.94821 | 0.392534 | 0.271675 | 1.352837 | 0.080613 | 0.129924 | 0.26835 | 0.2074 | 0.356606 |
| SL1344_RS24005_ | -5.57525 | -5.45767 | -5.73581 | -0.37868 | 1.662069 | -0.18451 | 0.409061 | 0.61086 | 0.737079 | 0.7545 | 0.833335 |

| | | | | | | | | | | |
|---|---|---|---|---|---|---|---|---|---|---|
| SL1344_RS24010_ | -4.86622 | -4.81312 | -4.91514 | -0.33156 | 1.452857 | -0.19388 | 0.406667 | 0.595122 | 0.725908 | 0.7551 | 0.832763 |
| SL1344_RS24015_ | -2.0116 | -2.17953 | -1.917 | 0.239554 | 0.453707 | 0.513994 | 0.288 | 0.455939 | 0.608071 | 0.5855 | 0.705967 |
| SL1344_RS24020_ | 4.07647 | 4.045103 | 4.134644 | 0.11197 | 0.129238 | 0.899565 | 0.167333 | 0.220317 | 0.400279 | 0.3129 | 0.482749 |
| SL1344_RS24025_ | -5.59684 | -6.13752 | -4.95359 | 1.262908 | 1.332418 | 0.902688 | 0.156562 | 0.250074 | 0.404306 | 0.3397 | 0.482528 |
| SL1344_RS24030_ | -4.71336 | -4.30384 | -5.07679 | -0.76693 | 1.196365 | -0.62032 | 0.223185 | 0.385479 | 0.540462 | 0.4677 | 0.605644 |
| SL1344_RS24035_ | -1.66223 | -1.84519 | -1.47164 | 0.368044 | 0.303742 | 1.164176 | 0.096667 | 0.161425 | 0.304293 | 0.2329 | 0.378064 |
| SL1344_RS24040_ | -0.9086 | -0.94819 | -0.83995 | 0.147521 | 0.329771 | 0.421598 | 0.331113 | 0.471728 | 0.6227 | 0.6217 | 0.725975 |
| SL1344_RS24045_ | 0.823213 | 0.586049 | 0.940738 | 0.342293 | 0.167072 | 1.835819 | 0.022652 | 0.067836 | 0.191614 | 0.1263 | 0.268691 |
| SL1344_RS24050_ | 1.482672 | 1.423017 | 1.567669 | 0.146682 | 0.115266 | 1.216333 | 0.087275 | 0.146305 | 0.293016 | 0.2166 | 0.362925 |
| SL1344_RS24055_ | 1.418338 | 1.326136 | 1.476273 | 0.143761 | 0.109817 | 1.238731 | 0.081945 | 0.146807 | 0.290595 | 0.2193 | 0.366665 |
| SL1344_RS24060_ | 1.320483 | 1.286916 | 1.360175 | 0.072763 | 0.10102 | 0.67487 | 0.223333 | 0.349617 | 0.495983 | 0.4385 | 0.56817 |
| SL1344_RS24065_ | 1.613396 | 1.539864 | 1.729839 | 0.230431 | 0.195292 | 1.172304 | 0.08 | 0.14383 | 0.305951 | 0.1945 | 0.348215 |
| SL1344_RS24070_ | 1.409923 | 1.445633 | 1.36767 | -0.06482 | 0.236184 | -0.26108 | 0.391333 | 0.649161 | 0.771713 | 0.8004 | 0.87014 |
| SL1344_RS24075_ | 1.311792 | 1.488723 | 1.188679 | -0.3002 | 0.140644 | -2.08908 | 0.008667 | 0.049011 | 0.159552 | 0.1093 | 0.24832 |
| SL1344_RS24080_ | -2.38031 | -2.67913 | -2.19564 | 0.485222 | 0.435332 | 1.091191 | 0.115257 | 0.184362 | 0.335369 | 0.2632 | 0.408751 |
| SL1344_RS24085_ | 3.831404 | 3.794815 | 3.857414 | 0.064459 | 0.052029 | 1.168387 | 0.085943 | 0.156827 | 0.305971 | 0.2268 | 0.375885 |
| SL1344_RS24090_ | -3.84084 | -3.57155 | -4.09209 | -0.52042 | 0.675903 | -0.73456 | 0.203864 | 0.329291 | 0.480509 | 0.4239 | 0.560402 |
| SL1344_RS24095_ | -2.26627 | -2.3965 | -2.07501 | 0.306697 | 0.656021 | 0.448271 | 0.293138 | 0.504917 | 0.652802 | 0.5923 | 0.717297 |
| SL1344_RS24100_ | -5.35418 | -4.83464 | -6.08123 | -1.24339 | 1.052372 | -1.06546 | 0.105929 | 0.180932 | 0.328648 | 0.2507 | 0.394968 |
| SL1344_RS24105_ | -0.45234 | -0.55043 | -0.36979 | 0.179319 | 0.19276 | 0.879909 | 0.157333 | 0.245278 | 0.39403 | 0.3414 | 0.481404 |
| SL1344_RS24110_ | -2.03566 | -2.17681 | -1.923 | 0.264735 | 0.354402 | 0.712135 | 0.201865 | 0.320189 | 0.472319 | 0.4308 | 0.561081 |
| SL1344_RS24115_ | -2.38439 | -2.49964 | -2.29482 | 0.2573 | 0.579095 | 0.420705 | 0.322452 | 0.505109 | 0.650261 | 0.6444 | 0.747731 |
| SL1344_RS24120_ | -1.70783 | -1.59131 | -1.78639 | -0.17339 | 0.626102 | -0.26906 | 0.369087 | 0.671075 | 0.784598 | 0.7299 | 0.821836 |
| SL1344_RS24125_ | 2.495873 | 2.473748 | 2.515753 | 0.009291 | 0.157802 | 0.059116 | 0.476667 | 0.785635 | 0.867741 | 0.8953 | 0.933809 |
| SL1344_RS24130_ | -2.08978 | -2.06677 | -2.10382 | -0.02795 | 0.418002 | -0.06506 | 0.476349 | 0.639927 | 0.75675 | 0.7513 | 0.834871 |
| SL1344_RS24135_ | -1.72703 | -1.81533 | -1.68163 | 0.162285 | 0.467425 | 0.286123 | 0.370667 | 0.555633 | 0.694095 | 0.7102 | 0.801479 |
| SL1344_RS24140_ | -2.93942 | -2.74121 | -3.12593 | -0.37161 | 0.554084 | -0.62455 | 0.235843 | 0.366213 | 0.519967 | 0.472 | 0.604008 |
| SL1344_RS24145_ | -3.24957 | -3.34 | -3.13646 | 0.278491 | 0.85327 | 0.301703 | 0.373751 | 0.586927 | 0.717491 | 0.719 | 0.808435 |
| SL1344_RS24150_ | -0.50084 | -0.93261 | -0.26744 | 0.683607 | 0.334289 | 1.946398 | 0.008661 | 0.064331 | 0.189327 | 0.111 | 0.250542 |
| SL1344_RS24155_ | -0.14546 | -0.27286 | 0.08771 | 0.363186 | 0.274697 | 1.235082 | 0.082612 | 0.13294 | 0.285253 | 0.2042 | 0.357291 |
| SL1344_RS24160_ | -5.36024 | -5.37329 | -5.33977 | 0.004897 | 1.092162 | 0.004918 | 0.498334 | 0.569471 | 0.697005 | 0.6933 | 0.783175 |
| SL1344_RS24165_ | 1.823645 | 1.954107 | 1.70869 | -0.26513 | 0.209576 | -1.37443 | 0.025334 | 0.135785 | 0.298853 | 0.1219 | 0.264424 |
| SL1344_RS24170_ | 1.841284 | 2.076555 | 1.709276 | -0.43298 | 0.294044 | -1.58752 | 0.005334 | 0.092286 | 0.239492 | 0.1036 | 0.241131 |
| SL1344_RS24175_ | 3.861537 | 4.142291 | 3.830776 | -0.30499 | 0.278161 | -1.14527 | 0.101266 | 0.100859 | 0.253231 | 0.22 | 0.381365 |
| SL1344_RS24180_ | 2.715781 | 2.764345 | 2.693648 | -0.05653 | 0.151192 | -0.26311 | 0.388667 | 0.842223 | 0.906609 | 0.8227 | 0.894637 |
| SL1344_RS24185_ | -0.06109 | -0.25225 | 0.089828 | 0.336695 | 0.206329 | 1.574818 | 0.041306 | 0.083617 | 0.212013 | 0.1465 | 0.292247 |
| SL1344_RS24190_ | 0.034693 | -0.03215 | 0.233544 | 0.210614 | 0.298024 | 0.622871 | 0.273151 | 0.439132 | 0.601692 | 0.5574 | 0.695075 |
| SL1344_RS24195_ | -0.41745 | -0.60048 | -0.21833 | 0.398461 | 0.232494 | 1.680128 | 0.021334 | 0.070804 | 0.193049 | 0.1261 | 0.268419 |
| SL1344_RS24200_ | -5.20876 | -5.01423 | -5.37267 | -0.35729 | 1.311007 | -0.22341 | 0.387333 | 0.589861 | 0.714563 | 0.6943 | 0.787184 |
| SL1344_RS24205_ | 2.014336 | 2.20604 | 1.851089 | -0.41789 | 0.252264 | -1.75686 | 4.68E-05 | 0.081678 | 0.223133 | 0.1006 | 0.237392 |
| SL1344_RS24210_ | 0.766645 | 0.775027 | 0.762221 | 0.019272 | 0.234179 | 0.087886 | 0.468 | 0.707953 | 0.812236 | 0.8354 | 0.893119 |
| SL1344_RS24215_ | 0.862316 | 0.793672 | 0.993361 | 0.178146 | 0.170742 | 0.978412 | 0.156 | 0.221922 | 0.386435 | 0.313 | 0.470821 |
| SL1344_RS24220_ | 2.206834 | 2.146714 | 2.310779 | 0.167795 | 0.11823 | 1.347874 | 0.064 | 0.131985 | 0.28483 | 0.1847 | 0.333933 |
| SL1344_RS24225_ | 1.097665 | 1.040675 | 1.160211 | 0.118032 | 0.113651 | 0.986413 | 0.132667 | 0.215426 | 0.363622 | 0.3052 | 0.448914 |
| SL1344_RS24230_ | 0.018705 | -0.08693 | 0.169431 | 0.269734 | 0.286386 | 0.906728 | 0.164667 | 0.239232 | 0.410318 | 0.3348 | 0.493157 |

| | | | | | | | | | | | |
|---|---|---|---|---|---|---|---|---|---|---|---|
| SL1344_RS24235_ | 0.804103 | 0.701163 | 0.914404 | 0.252125 | 0.2333 | 1.100128 | 0.1 | 0.165857 | 0.33039 | 0.2337 | 0.389293 |
| SL1344_RS24240_ | 1.763629 | 1.671279 | 1.847744 | 0.165662 | 0.155049 | 1.032042 | 0.134577 | 0.195694 | 0.36239 | 0.269 | 0.431143 |
| SL1344_RS24245_ | 1.860241 | 1.750731 | 1.974082 | 0.226852 | 0.091287 | 2.414871 | 0.003332 | 0.028661 | 0.116435 | 0.1033 | 0.240512 |
| SL1344_RS24250_ | 2.556369 | 2.830861 | 2.424021 | -0.4054 | 0.158962 | -2.42064 | 4.68E-05 | 0.02732 | 0.121809 | 0.1001 | 0.236729 |
| SL1344_RS24255_ | -1.66685 | -1.48585 | -1.86582 | -0.3811 | 0.302817 | -1.1826 | 0.100667 | 0.164895 | 0.307062 | 0.2422 | 0.387525 |
| SL1344_RS24260_ | -2.2052 | -2.38848 | -1.97192 | 0.398473 | 0.389885 | 0.94152 | 0.153231 | 0.238887 | 0.387768 | 0.3291 | 0.471304 |
| SL1344_RS24265_ | 0.306756 | 0.280795 | 0.333498 | 0.043236 | 0.276038 | 0.159249 | 0.424 | 0.700218 | 0.806338 | 0.7698 | 0.848139 |
| SL1344_RS24270_ | -0.85148 | -0.91055 | -0.78361 | 0.128861 | 0.230219 | 0.495921 | 0.284 | 0.431173 | 0.57137 | 0.5349 | 0.65288 |
| SL1344_RS24275_ | -0.71112 | -0.66357 | -0.75658 | -0.09139 | 0.206023 | -0.4177 | 0.309127 | 0.449402 | 0.58697 | 0.5551 | 0.668082 |
| SL1344_RS24280_ | 0.010096 | 0.078794 | -0.10834 | -0.17801 | 0.199895 | -0.85384 | 0.18 | 0.275343 | 0.433613 | 0.3735 | 0.516822 |
| SL1344_RS24285_ | 2.07628 | 2.097962 | 2.033254 | -0.05038 | 0.155652 | -0.30346 | 0.39 | 0.707299 | 0.81249 | 0.7771 | 0.859343 |
| SL1344_RS24290_ | 1.880763 | 1.904085 | 1.855968 | -0.00674 | 0.161429 | -0.04257 | 0.485333 | 0.760827 | 0.849295 | 0.8766 | 0.920126 |
| SL1344_RS24295_ | 2.389335 | 2.483968 | 2.285711 | -0.21207 | 0.104431 | -1.98323 | 0.009328 | 0.048383 | 0.158746 | 0.1067 | 0.244996 |
| SL1344_RS24300_ | 1.663477 | 1.671362 | 1.659258 | -0.03116 | 0.156041 | -0.18798 | 0.427049 | 0.645682 | 0.764772 | 0.7842 | 0.856697 |
| SL1344_RS24305_ | -0.15326 | 0.28596 | -0.64515 | -0.94906 | 0.295129 | -3.22346 | 4.68E-05 | 0.014449 | 0.086097 | 0.1 | 0.236592 |
| SL1344_RS24310_ | -2.29421 | -1.99949 | -2.50404 | -0.49067 | 0.406312 | -1.11277 | 0.117921 | 0.184089 | 0.332615 | 0.2679 | 0.41602 |
| SL1344_RS24315_ | -6.34346 | -5.94018 | -6.82071 | -0.90418 | 1.411596 | -0.61349 | 0.233333 | 0.366923 | 0.513682 | 0.4601 | 0.58633 |
| SL1344_RS24320_ | -6.54867 | -6.64245 | -6.41515 | 0.194196 | 1.804616 | 0.106666 | 0.446 | 0.571201 | 0.702065 | 0.6877 | 0.78161 |
| SL1344_RS24325_ | -3.85478 | -3.7201 | -4.0781 | -0.37378 | 0.732712 | -0.47839 | 0.296 | 0.460224 | 0.605374 | 0.5797 | 0.694989 |
| SL1344_RS24330_ | -2.82159 | -2.62279 | -3.00851 | -0.39812 | 0.45214 | -0.84229 | 0.173333 | 0.270963 | 0.421273 | 0.3577 | 0.497393 |
| SL1344_RS24335_ | -4.31777 | -4.75375 | -3.97609 | 0.778286 | 0.776161 | 0.927657 | 0.145333 | 0.236765 | 0.390233 | 0.3209 | 0.463705 |
| SL1344_RS24340_ | -4.476 | -4.12861 | -4.89551 | -0.70341 | 1.11596 | -0.62221 | 0.251832 | 0.388073 | 0.543962 | 0.4913 | 0.623673 |
| SL1344_RS24345_ | -3.74713 | -3.69783 | -3.78803 | -0.10364 | 0.74368 | -0.13478 | 0.444667 | 0.610634 | 0.731071 | 0.7286 | 0.811481 |
| SL1344_RS24350_ | -4.68681 | -5.0515 | -4.33481 | 0.736666 | 0.859884 | 0.813312 | 0.183211 | 0.287032 | 0.437872 | 0.3793 | 0.515488 |
| SL1344_RS24355_ | -5.60768 | -5.96232 | -5.22069 | 0.815309 | 1.410034 | 0.554866 | 0.261333 | 0.392208 | 0.544177 | 0.5124 | 0.635908 |
| SL1344_RS24360_ | -4.38077 | -4.39103 | -4.37306 | 0.021518 | 1.029874 | 0.017447 | 0.492338 | 0.667629 | 0.781117 | 0.7923 | 0.864597 |
| SL1344_RS24365_ | -4.72819 | -4.50255 | -5.01686 | -0.49166 | 0.869691 | -0.52023 | 0.263333 | 0.396647 | 0.542248 | 0.489 | 0.613267 |
| SL1344_RS24370_ | -1.40863 | -1.42814 | -1.38925 | 0.043916 | 0.300768 | 0.124534 | 0.44437 | 0.602354 | 0.725255 | 0.7105 | 0.800601 |
| SL1344_RS24380_ | -1.56189 | -1.64659 | -1.4823 | 0.160065 | 0.280682 | 0.539078 | 0.271333 | 0.39055 | 0.535812 | 0.4961 | 0.618295 |
| SL1344_RS24385_ | -0.86939 | -0.87013 | -0.86862 | 0.007569 | 0.257166 | 0.023865 | 0.488341 | 0.611924 | 0.732971 | 0.7141 | 0.802754 |
| SL1344_RS24390_ | -5.79985 | -5.93733 | -5.59692 | 0.35025 | 1.36511 | 0.231803 | 0.393738 | 0.56163 | 0.688841 | 0.6557 | 0.755645 |
| SL1344_RS24395_ | -2.38295 | -2.23735 | -2.85203 | -0.59111 | 0.649052 | -0.80177 | 0.215333 | 0.305311 | 0.48054 | 0.446 | 0.592499 |
| SL1344_RS24400_ | -4.65402 | -4.37002 | -4.96551 | -0.66737 | 1.16452 | -0.51815 | 0.267333 | 0.424952 | 0.576077 | 0.517 | 0.642623 |
| SL1344_RS24405_ | -5.78874 | -6.17713 | -5.36314 | 0.925711 | 1.540777 | 0.56439 | 0.270486 | 0.411719 | 0.560565 | 0.5217 | 0.644631 |
| SL1344_RS24410_ | 1.395121 | 1.368388 | 1.424258 | 0.084467 | 0.180129 | 0.468794 | 0.3 | 0.443177 | 0.603549 | 0.6167 | 0.726549 |
| SL1344_RS24415_ | -1.81105 | -2.17741 | -1.44773 | 0.756586 | 0.39965 | 1.840259 | 0.015334 | 0.05765 | 0.173633 | 0.1151 | 0.254953 |
| SL1344_RS24420_ | -0.57898 | -0.61938 | -0.52078 | 0.060593 | 0.336792 | 0.161326 | 0.430667 | 0.722347 | 0.82111 | 0.8161 | 0.882586 |
| SL1344_RS24425_ | -4.03016 | -3.86175 | -4.21473 | -0.4227 | 0.912825 | -0.44054 | 0.296667 | 0.459078 | 0.607893 | 0.5872 | 0.697563 |
| SL1344_RS24430_ | 2.6143 | 2.9555 | 2.379105 | -0.5503 | 0.193385 | -2.81137 | 4.68E-05 | 0.015987 | 0.094246 | 0.1 | 0.236592 |
| SL1344_RS24435_ | 0.689678 | 0.744693 | 0.574147 | -0.15296 | 0.187841 | -0.73144 | 0.227333 | 0.346277 | 0.50915 | 0.4552 | 0.597861 |
| SL1344_RS24440_ | 2.863808 | 2.937663 | 2.791896 | -0.13568 | 0.095921 | -1.32653 | 0.08 | 0.115745 | 0.262454 | 0.1834 | 0.338456 |
| SL1344_RS24445_ | 0.632961 | 0.48151 | 0.741193 | 0.252838 | 0.153206 | 1.550256 | 0.049967 | 0.09065 | 0.221747 | 0.1567 | 0.303783 |
| SL1344_RS24450_ | -0.91867 | -1.22524 | -0.71784 | 0.511111 | 0.342463 | 1.462467 | 0.042667 | 0.107629 | 0.251901 | 0.1488 | 0.296514 |
| SL1344_RS24455_ | -2.20397 | -2.49517 | -1.90902 | 0.587337 | 0.347746 | 1.567318 | 0.046 | 0.083706 | 0.205619 | 0.155 | 0.298733 |
| SL1344_RS24460_ | 8.353536 | 8.55269 | 8.08447 | -0.45124 | 0.078594 | -5.32181 | 4.68E-05 | 0.004221 | 0.054634 | 0.1 | 0.236592 |

| | | | | | | | | | | | |
|---|---|---|---|---|---|---|---|---|---|---|---|
| SL1344_RS24465_ | -0.00902 | -0.10511 | 0.065952 | 0.039211 | 0.464983 | 0.080812 | 0.472 | 0.815285 | 0.888667 | 0.9447 | 0.966193 |
| SL1344_RS24470_ | 6.931225 | 6.872707 | 7.023326 | 0.156282 | 0.16369 | 1.142316 | 0.110593 | 0.144169 | 0.313661 | 0.1994 | 0.369062 |
| SL1344_RS24475_ | 1.598152 | 1.716265 | 1.319732 | -0.38296 | 0.140968 | -2.47866 | 0.003335 | 0.044222 | 0.153133 | 0.1018 | 0.238941 |
| SL1344_RS24480_ | 3.662542 | 3.614789 | 3.704202 | 0.098637 | 0.067442 | 1.419295 | 0.035976 | 0.10792 | 0.250865 | 0.1386 | 0.281436 |
| SL1344_RS24485_ | 0.088961 | -0.03953 | 0.439531 | 0.448754 | 0.232996 | 1.668711 | 0.046636 | 0.088337 | 0.225566 | 0.1474 | 0.293709 |
| SL1344_RS24490_ | -1.54648 | -1.84633 | -1.43215 | 0.394939 | 0.364876 | 0.937134 | 0.173218 | 0.247222 | 0.412985 | 0.3652 | 0.513585 |
| SL1344_RS24495_ | -1.05726 | -1.2128 | -0.89692 | 0.328231 | 0.330023 | 0.934218 | 0.141333 | 0.223061 | 0.386375 | 0.2983 | 0.449081 |
| SL1344_RS24500_ | 0.430433 | 0.132846 | 0.677986 | 0.544685 | 0.213897 | 2.503145 | 0.002002 | 0.028635 | 0.120474 | 0.1018 | 0.238982 |
| SL1344_RS24505_ | -1.16207 | -1.34264 | -0.9335 | 0.394389 | 0.342965 | 1.118436 | 0.11459 | 0.176822 | 0.335449 | 0.2533 | 0.410125 |
| SL1344_RS24510_ | -3.93895 | -4.45622 | -3.53023 | 0.936648 | 0.647065 | 1.37908 | 0.056 | 0.110326 | 0.241879 | 0.1709 | 0.314459 |
| SL1344_RS24515_ | -0.90847 | -1.45867 | -0.54493 | 0.944203 | 0.416454 | 2.18001 | 0.001336 | 0.043302 | 0.152036 | 0.1008 | 0.237659 |
| SL1344_RS24520_ | -1.86712 | -2.10181 | -1.63718 | 0.46809 | 0.346527 | 1.305744 | 0.073333 | 0.128591 | 0.268114 | 0.1978 | 0.345036 |
| SL1344_RS24525_ | -0.41177 | -0.53316 | -0.28737 | 0.300918 | 0.336055 | 0.881998 | 0.134 | 0.233384 | 0.409703 | 0.2821 | 0.436169 |
| SL1344_RS24530_ | -0.49149 | -1.06425 | -0.126 | 0.95188 | 0.294271 | 3.250613 | 4.68E-05 | 0.026832 | 0.116352 | 0.1 | 0.236592 |
| SL1344_RS24535_ | -1.33646 | -1.72821 | -0.97841 | 0.759187 | 0.251159 | 2.917219 | 0.001336 | 0.017281 | 0.088738 | 0.1006 | 0.237341 |
| SL1344_RS24540_ | -1.32972 | -1.66116 | -1.0631 | 0.603378 | 0.27544 | 2.094043 | 0.010667 | 0.042801 | 0.143662 | 0.1094 | 0.24827 |
| SL1344_RS24545_ | -2.01963 | -2.36942 | -1.71337 | 0.653162 | 0.438801 | 1.429333 | 0.06 | 0.109323 | 0.249088 | 0.1678 | 0.316727 |
| SL1344_RS24550_ | 0.886789 | 0.581743 | 1.178891 | 0.599358 | 0.163157 | 3.714686 | 4.68E-05 | 0.011436 | 0.075928 | 0.1 | 0.236592 |
| SL1344_RS24555_ | -1.16777 | -1.46457 | -0.91304 | 0.546253 | 0.257495 | 2.063613 | 0.009328 | 0.041627 | 0.140634 | 0.112 | 0.25069 |
| SL1344_RS24560_ | -0.83906 | -0.95968 | -0.75341 | 0.234393 | 0.290448 | 0.777425 | 0.173218 | 0.275645 | 0.439842 | 0.3678 | 0.505824 |
| SL1344_RS24565_ | -1.33339 | -1.56313 | -1.16881 | 0.398269 | 0.307748 | 1.231011 | 0.077333 | 0.142465 | 0.290054 | 0.2011 | 0.34891 |
| SL1344_RS24570_ | -2.60119 | -2.71955 | -2.46534 | 0.304536 | 0.638579 | 0.447286 | 0.292 | 0.464786 | 0.616738 | 0.5968 | 0.712188 |
| SL1344_RS24575_ | -1.5955 | -1.01375 | -2.21655 | -1.23454 | 0.432099 | -2.8291 | 0.000673 | 0.019951 | 0.099944 | 0.1004 | 0.237119 |
| SL1344_RS24580_ | 3.039727 | 3.689234 | 2.423385 | -1.25342 | 0.112677 | -11.0891 | 4.68E-05 | 0.000632 | 0.028289 | 0.1 | 0.236592 |
| SL1344_RS24585_ | 1.044176 | 1.597345 | 0.563754 | -1.02753 | 0.188672 | -5.39412 | 4.68E-05 | 0.002631 | 0.043091 | 0.1 | 0.236592 |
| SL1344_RS24590_ | -0.64773 | -0.52415 | -0.77952 | -0.25433 | 0.22806 | -1.07609 | 0.109927 | 0.189475 | 0.339991 | 0.2639 | 0.411182 |
| SL1344_RS24595_ | -4.56315 | -4.77664 | -4.37333 | 0.397804 | 0.91366 | 0.395036 | 0.321785 | 0.478682 | 0.618692 | 0.5865 | 0.69789 |
| SL1344_RS24600_ | -1.60982 | -1.89068 | -1.39404 | 0.474351 | 0.369114 | 1.206269 | 0.088 | 0.142003 | 0.291174 | 0.2107 | 0.364311 |
| SL1344_RS24605_ | -0.05206 | -0.3932 | 0.188204 | 0.582592 | 0.190828 | 3.010478 | 4.68E-05 | 0.02255 | 0.103156 | 0.1004 | 0.23713 |
| SL1344_RS24610_ | -0.0969 | -0.21514 | 0.0393 | 0.249183 | 0.181349 | 1.358038 | 0.070667 | 0.124786 | 0.263132 | 0.1982 | 0.346434 |
| SL1344_RS24615_ | -0.27185 | -0.36163 | -0.15068 | 0.227637 | 0.288415 | 0.750269 | 0.207333 | 0.308161 | 0.476471 | 0.4119 | 0.56051 |
| SL1344_RS24620_ | -0.05481 | -0.35246 | 0.172812 | 0.528808 | 0.212108 | 2.485948 | 0.002002 | 0.030457 | 0.122023 | 0.1019 | 0.239126 |
| SL1344_RS24625_ | -1.45356 | -1.80576 | -1.27135 | 0.528729 | 0.413519 | 1.194139 | 0.081945 | 0.156932 | 0.316707 | 0.2184 | 0.370955 |
| SL1344_RS24630_ | -0.14391 | -0.29276 | -0.0049 | 0.291027 | 0.167882 | 1.665333 | 0.036667 | 0.073905 | 0.192729 | 0.1395 | 0.282183 |
| SL1344_RS24635_ | 0.142137 | -0.14575 | 0.3496 | 0.51732 | 0.237657 | 2.118855 | 0.004665 | 0.048144 | 0.159197 | 0.103 | 0.24044 |
| SL1344_RS24640_ | -2.29426 | -2.69241 | -1.97396 | 0.734058 | 0.458567 | 1.568496 | 0.033311 | 0.086473 | 0.21707 | 0.1323 | 0.274702 |
| SL1344_RS24645_ | -0.90314 | -1.11412 | -0.75842 | 0.374497 | 0.306501 | 1.168847 | 0.078 | 0.158736 | 0.315361 | 0.2005 | 0.349573 |
| SL1344_RS24650_ | -3.02908 | -3.1612 | -2.84016 | 0.30109 | 0.507277 | 0.554786 | 0.262 | 0.421244 | 0.565945 | 0.5285 | 0.650203 |
| SL1344_RS24655_ | -3.08435 | -3.34512 | -2.85009 | 0.495428 | 0.503217 | 0.938944 | 0.150566 | 0.240515 | 0.391532 | 0.3277 | 0.470398 |
| SL1344_RS24660_ | 0.405831 | 0.537151 | 0.238129 | -0.31484 | 0.183271 | -1.68919 | 0.019321 | 0.081754 | 0.210213 | 0.1261 | 0.267544 |
| SL1344_RS24665_ | 0.036189 | 0.054022 | 0.010238 | -0.03815 | 0.256792 | -0.14282 | 0.44 | 0.679996 | 0.789405 | 0.7848 | 0.857051 |
| SL1344_RS24670_ | 0.338645 | 0.51446 | 0.158579 | -0.35481 | 0.137856 | -2.53614 | 0.003332 | 0.026272 | 0.11025 | 0.1026 | 0.239935 |
| SL1344_RS24675_ | -1.841 | -1.57847 | -2.10169 | -0.51579 | 0.395965 | -1.24739 | 0.089333 | 0.144067 | 0.293538 | 0.2162 | 0.368274 |
| SL1344_RS24690_ | 5.750886 | 6.233804 | 5.391074 | -0.82905 | 0.128338 | -6.3043 | 4.68E-05 | 0.002217 | 0.043162 | 0.1 | 0.236592 |
| SL1344_RS24695_ | 2.746355 | 3.213026 | 2.227789 | -1.00208 | 0.115961 | -8.68189 | 4.68E-05 | 0.001467 | 0.034727 | 0.1 | 0.236592 |

| | | | | | | | | | | | |
|---|---|---|---|---|---|---|---|---|---|---|---|
| SL1344_RS24700_ | 2.953681 | 3.453709 | 2.421062 | -1.0531 | 0.10361 | -10.2073 | 4.68E-05 | 0.00043 | 0.02533 | 0.1 | 0.236592 |
| SL1344_RS24705_ | 2.764556 | 3.243019 | 2.426149 | -0.81695 | 0.155603 | -5.20726 | 4.68E-05 | 0.008432 | 0.070523 | 0.1 | 0.236592 |
| SL1344_RS24710_ | 2.490704 | 3.080124 | 2.009017 | -1.04916 | 0.178288 | -5.84031 | 4.68E-05 | 0.002551 | 0.0436 | 0.1 | 0.236592 |
| SL1344_RS24730_ | 4.577645 | 4.898662 | 4.305207 | -0.58212 | 0.090115 | -6.54024 | 4.68E-05 | 0.000733 | 0.030584 | 0.1 | 0.236592 |
| SL1344_RS24735_ | 5.943852 | 6.121905 | 5.754173 | -0.37833 | 0.055667 | -7.10934 | 4.68E-05 | 0.001973 | 0.039911 | 0.1 | 0.236592 |
| SL1344_RS24740_ | -4.26306 | -4.58241 | -4.04776 | 0.411533 | 1.25187 | 0.328549 | 0.357333 | 0.58593 | 0.718661 | 0.7022 | 0.796952 |
| SL1344_RS24745_ | -6.25854 | -5.56308 | -8.24883 | -2.65796 | 2.76325 | -0.86941 | 0.131246 | 0.215934 | 0.387041 | 0.2759 | 0.428962 |
| SL1344_RS24750_ | -1.86578 | -1.8158 | -1.96402 | -0.14474 | 0.435569 | -0.33189 | 0.373751 | 0.570318 | 0.701451 | 0.6914 | 0.786757 |
| SL1344_RS24755_ | 0.245577 | 0.274032 | 0.195202 | -0.07619 | 0.193352 | -0.35287 | 0.355097 | 0.547947 | 0.68194 | 0.6668 | 0.767129 |
| SL1344_RS24760_ | -3.5266 | -3.0867 | -4.59385 | -1.43491 | 1.01292 | -1.29199 | 0.082667 | 0.128949 | 0.283999 | 0.1908 | 0.34564 |
| SL1344_RS24780_ | -2.08892 | -1.85526 | -2.31854 | -0.49148 | 0.432338 | -1.1053 | 0.103265 | 0.17302 | 0.324961 | 0.2503 | 0.398477 |
| SL1344_RS24785_ | 2.469004 | 2.533558 | 2.409453 | -0.14409 | 0.154225 | -0.88434 | 0.148568 | 0.224422 | 0.401688 | 0.2959 | 0.457323 |
| SL1344_RS24790_ | 2.386513 | 2.532051 | 2.305416 | -0.22781 | 0.12419 | -1.78021 | 0.019987 | 0.05877 | 0.178815 | 0.124 | 0.266697 |
| SL1344_RS24795_ | -4.04145 | -3.72621 | -4.23408 | -0.45929 | 0.843462 | -0.46265 | 0.29447 | 0.505291 | 0.641899 | 0.5867 | 0.705056 |
| SL1344_RS24800_ | -7.63632 | -8.05835 | -7.15145 | 0.794116 | 3.000411 | 0.213242 | 0.370667 | 0.562722 | 0.692859 | 0.6615 | 0.761795 |
| SL1344_RS24805_ | -4.33922 | -4.10717 | -4.52208 | -0.41027 | 0.908247 | -0.40519 | 0.319333 | 0.504848 | 0.6435 | 0.6006 | 0.714261 |
| SL1344_RS24810_ | -3.91753 | -3.74015 | -4.78762 | -1.00139 | 1.054587 | -0.77343 | 0.240506 | 0.318757 | 0.494618 | 0.4926 | 0.636171 |
| SL1344_RS24820_ | 4.111582 | 4.345342 | 3.846203 | -0.53527 | 0.188332 | -2.9454 | 4.68E-05 | 0.022422 | 0.112611 | 0.1 | 0.236592 |
| SL1344_RS24825_ | 1.102773 | 1.298683 | 0.825767 | -0.49044 | 0.187528 | -2.62779 | 0.000673 | 0.034004 | 0.133007 | 0.1002 | 0.236855 |
| SL1344_RS24830_ | -2.67752 | -2.46172 | -3.1075 | -0.62599 | 0.583355 | -0.98053 | 0.148568 | 0.219526 | 0.383329 | 0.3144 | 0.467071 |
| SL1344_RS24835_ | -5.31817 | -4.82712 | -5.90477 | -1.05751 | 0.990014 | -1.01042 | 0.127915 | 0.202294 | 0.346735 | 0.2844 | 0.425606 |
| SL1344_RS24840_ | -6.18712 | -5.60334 | -7.17273 | -1.60208 | 1.941379 | -0.73974 | 0.139907 | 0.256946 | 0.424778 | 0.2975 | 0.441961 |
| SL1344_RS24845_ | -6.58623 | -6.27806 | -6.87098 | -0.56945 | 1.722846 | -0.29571 | 0.358 | 0.51786 | 0.652763 | 0.6245 | 0.730934 |
| SL1344_RS24850_ | -3.8079 | -3.77377 | -3.85567 | -0.10225 | 0.682704 | -0.1345 | 0.435333 | 0.570344 | 0.699054 | 0.6897 | 0.77978 |
| SL1344_RS24860_ | 0.754105 | 0.705475 | 0.819722 | 0.116314 | 0.321719 | 0.345303 | 0.351765 | 0.521955 | 0.676551 | 0.7211 | 0.811311 |
| SL1344_RS24865_ | -2.01842 | -2.22529 | -1.73477 | 0.530652 | 0.458264 | 1.096324 | 0.109333 | 0.174269 | 0.331255 | 0.2518 | 0.403762 |
| SL1344_RS24875_ | -6.14849 | -6.26863 | -5.99419 | 0.488106 | 1.9311 | 0.234636 | 0.395333 | 0.579233 | 0.70893 | 0.7165 | 0.802692 |
| SL1344_RS24885_ | -4.48222 | -4.52744 | -4.43956 | 0.145778 | 0.925871 | 0.144627 | 0.443333 | 0.607109 | 0.727657 | 0.7149 | 0.801909 |
| SLnc0002_ | -2.95983 | -3.19899 | -2.67393 | 0.525724 | 0.440021 | 1.097861 | 0.110667 | 0.17734 | 0.321294 | 0.255 | 0.399989 |
| SLnc0010_ | -2.99447 | -2.77861 | -3.18778 | -0.42677 | 0.555272 | -0.72183 | 0.205333 | 0.312801 | 0.469657 | 0.4101 | 0.54908 |
| SLnc0010a_ | -0.45334 | -0.34001 | -0.6536 | -0.2904 | 0.315408 | -0.88275 | 0.163225 | 0.245599 | 0.41449 | 0.343 | 0.49717 |
| SLnc0014_ | 3.121676 | 3.332942 | 2.870066 | -0.5007 | 0.142074 | -3.55172 | 4.68E-05 | 0.007628 | 0.067333 | 0.1 | 0.236592 |
| SLnc0022_ | -0.1935 | 0.111205 | -0.30845 | -0.36217 | 0.298516 | -1.09666 | 0.138574 | 0.185446 | 0.351466 | 0.2782 | 0.440013 |
| SLnc0030_ | 1.907543 | 1.879533 | 1.925442 | 0.030352 | 0.164648 | 0.172916 | 0.425716 | 0.708312 | 0.811714 | 0.8303 | 0.890227 |
| SLnc0030a_ | -1.36236 | -1.40496 | -1.31245 | 0.107794 | 0.380987 | 0.269395 | 0.378 | 0.593724 | 0.720449 | 0.7171 | 0.805387 |
| SLnc0031_ | -3.26207 | -3.44695 | -3.04835 | 0.488927 | 0.687506 | 0.675061 | 0.222667 | 0.334533 | 0.495979 | 0.4551 | 0.587131 |
| SLnc0035_ | 4.10308 | 3.788994 | 4.332648 | 0.544304 | 0.105514 | 5.188442 | 4.68E-05 | 0.014749 | 0.091842 | 0.1 | 0.236592 |
| SLnc0039_ | 3.781544 | 3.763161 | 3.849544 | 0.185325 | 0.253912 | 0.765944 | 0.226 | 0.289212 | 0.476727 | 0.4185 | 0.596305 |
| SLnc0048_ | -0.74677 | -0.92965 | -0.47173 | 0.450661 | 0.239224 | 1.774239 | 0.03 | 0.066511 | 0.18564 | 0.1336 | 0.276865 |
| SLnc1001_ | -4.5225 | -4.39725 | -4.629 | -0.21155 | 0.84929 | -0.22192 | 0.401333 | 0.551595 | 0.680619 | 0.6564 | 0.755915 |
| SLnc1002_ | -2.25136 | -1.63325 | -2.4966 | -0.84183 | 0.773554 | -1.13211 | 0.117255 | 0.158481 | 0.322251 | 0.2466 | 0.410614 |
| SLnc1006_ | 0.237846 | 0.551275 | -0.06407 | -0.56888 | 0.278261 | -2.04994 | 0.01 | 0.042119 | 0.149735 | 0.1094 | 0.249128 |
| SLnc1008_ | -3.6644 | -3.52556 | -3.73573 | -0.12973 | 0.962618 | -0.10941 | 0.456362 | 0.717985 | 0.820102 | 0.8168 | 0.885536 |
| SLnc1013_ | -2.31605 | -2.48747 | -2.13757 | 0.346829 | 0.518994 | 0.630741 | 0.2505 | 0.387852 | 0.544437 | 0.4894 | 0.622185 |
| SLnc1014_ | -2.01705 | -2.25038 | -1.78889 | 0.462021 | 0.363164 | 1.197542 | 0.092667 | 0.153333 | 0.29782 | 0.2311 | 0.379301 |

| | | | | | | | | | | | |
|---|---|---|---|---|---|---|---|---|---|---|---|
| SLnc1015_ | -2.11664 | -2.53227 | -1.83365 | 0.719547 | 0.421824 | 1.687417 | 0.024667 | 0.086532 | 0.218664 | 0.1242 | 0.264995 |
| SLnc1016_ | -2.45893 | -2.51807 | -2.4108 | 0.100513 | 0.356106 | 0.262317 | 0.386409 | 0.492894 | 0.6301 | 0.6036 | 0.710778 |
| SLnc1017_ | -1.58694 | -1.7628 | -1.52093 | 0.209489 | 0.384014 | 0.470739 | 0.328448 | 0.526665 | 0.668152 | 0.6454 | 0.75913 |
| SLnc1018_ | -3.52078 | -3.92071 | -3.03289 | 0.884967 | 0.649639 | 1.248602 | 0.08461 | 0.139868 | 0.283725 | 0.2092 | 0.358207 |
| SLnc1019_ | -3.55431 | -3.67778 | -3.33547 | 0.255572 | 0.852955 | 0.271684 | 0.387742 | 0.652972 | 0.765741 | 0.7308 | 0.820581 |
| SLnc1021_ | -1.82693 | -2.02701 | -1.47449 | 0.578081 | 0.505426 | 1.104492 | 0.121919 | 0.182573 | 0.347259 | 0.2608 | 0.416019 |
| SLnc1022_ | -2.68677 | -2.72997 | -2.59548 | 0.079811 | 0.567996 | 0.126496 | 0.446369 | 0.688142 | 0.793536 | 0.7765 | 0.854414 |
| SLnc1024_ | -2.52392 | -2.64443 | -2.35794 | 0.302443 | 0.464869 | 0.614358 | 0.243171 | 0.379578 | 0.530728 | 0.488 | 0.614789 |
| SLnc1025_ | -2.55219 | -2.84323 | -2.39557 | 0.437165 | 0.664843 | 0.628281 | 0.249167 | 0.378067 | 0.543986 | 0.4971 | 0.637798 |
| SLnc1026_ | -2.64207 | -2.77221 | -2.55938 | 0.189709 | 0.51755 | 0.335109 | 0.358667 | 0.550484 | 0.68434 | 0.6561 | 0.761344 |
| SLnc1027_ | -2.89192 | -2.57624 | -3.20228 | -0.71965 | 0.68767 | -1.02056 | 0.116589 | 0.185749 | 0.347779 | 0.2671 | 0.41763 |
| SLnc1028_ | -1.87745 | -2.08398 | -1.48328 | 0.580605 | 0.40729 | 1.292162 | 0.090667 | 0.138496 | 0.285774 | 0.2201 | 0.371181 |
| SLnc1029_ | -4.66117 | -4.14936 | -5.5142 | -1.33917 | 1.278439 | -0.98466 | 0.127249 | 0.199221 | 0.363689 | 0.2638 | 0.416757 |
| sRNA1_ | 2.418951 | 2.485977 | 2.357565 | -0.13917 | 0.126984 | -1.11635 | 0.063291 | 0.166837 | 0.333116 | 0.185 | 0.335839 |
| sRNA10_ | 1.315905 | 1.337388 | 1.284071 | -0.04938 | 0.12968 | -0.35517 | 0.346436 | 0.549735 | 0.683873 | 0.6505 | 0.755467 |
| sRNA11_ | -3.26836 | -3.38269 | -3.04883 | 0.289794 | 0.699886 | 0.370198 | 0.341106 | 0.602311 | 0.724346 | 0.6913 | 0.790724 |
| sRNA12_ | -5.07321 | -5.05023 | -5.13082 | -0.04204 | 1.066107 | -0.03873 | 0.481333 | 0.596342 | 0.719996 | 0.7196 | 0.806225 |
| sRNA15_ | -6.66635 | -7.2401 | -6.22742 | 1.07738 | 1.993824 | 0.453653 | 0.263824 | 0.398717 | 0.556941 | 0.5147 | 0.634707 |
| sRNA16_ | -0.99241 | -1.05605 | -0.87509 | 0.324088 | 0.640906 | 0.482789 | 0.2998 | 0.476512 | 0.63922 | 0.6139 | 0.739288 |
| sRNA18_ | -5.96602 | -6.56197 | -5.55454 | 1.107539 | 2.042257 | 0.466382 | 0.235333 | 0.369573 | 0.538117 | 0.4718 | 0.599632 |
| sRNA19_ | -9.41222 | -9.90044 | -8.91874 | 0.873845 | 4.148449 | 0.188127 | 0.39507 | 0.505581 | 0.645599 | 0.6285 | 0.733078 |
| sRNA2_ | 2.270477 | 2.580995 | 1.939622 | -0.53597 | 0.284824 | -2.02687 | 0.004001 | 0.035928 | 0.142523 | 0.1048 | 0.243 |
| sRNA21_ | -0.07843 | -0.00982 | -0.15982 | -0.16893 | 0.263252 | -0.60367 | 0.252 | 0.387936 | 0.552343 | 0.5308 | 0.658737 |
| sRNA22_ | -2.40052 | -2.38593 | -2.43062 | -0.05203 | 0.522615 | -0.08927 | 0.465333 | 0.677396 | 0.786473 | 0.7817 | 0.856395 |
| sRNA23_ | -5.93133 | -5.25153 | -6.41961 | -1.10886 | 1.639949 | -0.60115 | 0.236509 | 0.39207 | 0.543428 | 0.4727 | 0.606904 |
| sRNA24_ | -5.61107 | -6.14344 | -5.33952 | 0.770262 | 1.416523 | 0.491784 | 0.294667 | 0.439686 | 0.58933 | 0.5658 | 0.68475 |
| sRNA25_ | -7.21426 | -7.32193 | -7.12854 | 0.439283 | 3.80387 | 0.095505 | 0.444 | 0.65994 | 0.777599 | 0.8192 | 0.882309 |
| sRNA26_ | -6.38884 | -5.86125 | -6.9923 | -1.06112 | 1.720932 | -0.55883 | 0.253165 | 0.390879 | 0.541136 | 0.5021 | 0.625952 |
| sRNA28_ | -5.94598 | -5.09039 | -6.65896 | -1.46018 | 1.685318 | -0.79741 | 0.183211 | 0.303565 | 0.457544 | 0.3714 | 0.51853 |
| sRNA29_ | -5.35515 | -5.5927 | -5.20962 | 0.38007 | 1.431924 | 0.231132 | 0.396 | 0.578722 | 0.70897 | 0.701 | 0.793894 |
| sRNA31_ | -4.48278 | -4.40218 | -4.58972 | -0.2063 | 0.945158 | -0.19564 | 0.412392 | 0.576799 | 0.706312 | 0.7018 | 0.792063 |
| sRNA32_ | -5.19597 | -5.16021 | -5.236 | -0.10927 | 1.134847 | -0.08887 | 0.454364 | 0.580799 | 0.707441 | 0.704 | 0.792707 |
| sRNA33_ | -3.1977 | -3.24994 | -3.13784 | 0.112992 | 0.545879 | 0.194028 | 0.419333 | 0.565167 | 0.692492 | 0.6732 | 0.769327 |
| sRNA5_ | -5.35236 | -5.57232 | -5.06875 | 0.520242 | 1.286953 | 0.347404 | 0.344 | 0.525905 | 0.660455 | 0.6309 | 0.735283 |
| sRNA7_ | -2.59709 | -2.68956 | -2.49895 | -0.00752 | 0.815872 | -0.00942 | 0.494 | 0.75297 | 0.845521 | 0.896 | 0.933882 |
| STnc10_ | -8.43074 | -6.88278 | -9.78761 | -2.62236 | 3.72957 | -0.62325 | 0.21 | 0.370146 | 0.514831 | 0.4184 | 0.555307 |
| STnc1010_ | -2.35266 | -2.35441 | -2.35083 | 0.033637 | 0.568557 | 0.052355 | 0.479014 | 0.697746 | 0.802533 | 0.7939 | 0.864668 |
| STnc1030_ | -2.97071 | -2.83219 | -3.18676 | -0.30375 | 0.621211 | -0.47511 | 0.321333 | 0.511498 | 0.652626 | 0.629 | 0.737526 |
| STnc1040_ | -5.83753 | -6.10002 | -5.52472 | 0.531373 | 1.395974 | 0.334279 | 0.353098 | 0.520424 | 0.652315 | 0.622 | 0.726499 |
| STnc1065_ | 1.899878 | 2.725705 | 1.105795 | -1.62925 | 0.265721 | -6.14795 | 4.68E-05 | 0.001499 | 0.036784 | 0.1 | 0.236592 |
| STnc1070_ | 0.202613 | 0.196836 | 0.205301 | -0.00181 | 0.199071 | -0.00924 | 0.496 | 0.657047 | 0.768786 | 0.7586 | 0.836398 |
| STnc1080_ | -4.77643 | -4.84065 | -4.7457 | 0.227402 | 1.459489 | 0.121933 | 0.425333 | 0.553132 | 0.699065 | 0.7345 | 0.818621 |
| STnc1090_ | -1.06014 | -1.14544 | -0.96886 | 0.175201 | 0.280485 | 0.589876 | 0.247169 | 0.377991 | 0.527866 | 0.4818 | 0.610993 |
| STnc1100_ | -3.72418 | -3.73446 | -3.71273 | -0.10231 | 1.027612 | -0.09754 | 0.46036 | 0.711645 | 0.814589 | 0.8096 | 0.876482 |
| STnc1130_ | -0.34925 | -0.0383 | -0.76017 | -0.7319 | 0.2626 | -2.71561 | 0.000673 | 0.022765 | 0.106842 | 0.1003 | 0.237001 |

| | | | | | | | | | | | |
|---|---|---|---|---|---|---|---|---|---|---|---|
| STnc1150_ | 1.295816 | 1.309095 | 1.284067 | -0.07338 | 0.222951 | -0.32422 | 0.364667 | 0.502761 | 0.657987 | 0.7261 | 0.810506 |
| STnc1160_ | 1.649166 | 1.731106 | 1.6132 | -0.20158 | 0.334168 | -0.63943 | 0.249833 | 0.359916 | 0.541461 | 0.4862 | 0.645624 |
| STnc1170_ | -1.53237 | -1.84509 | -1.34281 | 0.490698 | 0.34092 | 1.352015 | 0.067955 | 0.123852 | 0.266959 | 0.1895 | 0.337155 |
| STnc1180_ | -2.10197 | -2.32388 | -1.89851 | 0.427203 | 0.314266 | 1.2975 | 0.074 | 0.135997 | 0.269021 | 0.2082 | 0.352128 |
| STnc1190_ | 4.329156 | 4.33688 | 4.32298 | -0.00386 | 0.154824 | -0.02184 | 0.484344 | 0.891697 | 0.937678 | 0.9508 | 0.970788 |
| STnc1200_ | -2.10245 | -2.11924 | -2.07403 | 0.022687 | 0.417043 | 0.044852 | 0.481013 | 0.634801 | 0.752673 | 0.7379 | 0.821774 |
| STnc1210_ | -3.04024 | -3.20162 | -2.87735 | 0.334843 | 0.567064 | 0.559475 | 0.256667 | 0.405787 | 0.557455 | 0.5111 | 0.637492 |
| STnc1220_ | -6.81273 | -6.94457 | -6.61234 | 0.696704 | 2.661758 | 0.222178 | 0.388 | 0.573362 | 0.706779 | 0.7295 | 0.812149 |
| STnc1275_ | -1.0527 | -0.68873 | -1.53364 | -0.84336 | 0.240165 | -3.44883 | 4.68E-05 | 0.014927 | 0.082858 | 0.1 | 0.236592 |
| STnc1280_ | -2.00186 | -2.18873 | -1.85753 | 0.329813 | 0.325796 | 0.939923 | 0.143238 | 0.229796 | 0.377502 | 0.3184 | 0.459862 |
| STnc1285_ | 2.913075 | 3.623666 | 2.31723 | -1.31997 | 0.349442 | -3.73249 | 4.68E-05 | 0.003618 | 0.051664 | 0.1 | 0.236592 |
| STnc1290_ | 0.274928 | 0.59498 | 0.013288 | -0.45839 | 0.3819 | -1.24041 | 0.102598 | 0.137941 | 0.299216 | 0.2014 | 0.366887 |
| STnc130_ | -4.54415 | -4.3439 | -4.69231 | -0.30383 | 1.170142 | -0.22621 | 0.403065 | 0.631945 | 0.750685 | 0.7229 | 0.812422 |
| STnc1320_ | -4.19126 | -4.46857 | -3.91552 | 0.561545 | 0.74638 | 0.679724 | 0.217189 | 0.345665 | 0.495771 | 0.4358 | 0.568723 |
| STnc1340_ | 0.491362 | 0.666186 | 0.339787 | -0.32571 | 0.151528 | -2.07066 | 0.012 | 0.045456 | 0.149044 | 0.1125 | 0.251877 |
| STnc1380_ | -4.00361 | -4.04706 | -3.9746 | 0.094689 | 0.820078 | 0.103252 | 0.454364 | 0.609111 | 0.731309 | 0.7233 | 0.810871 |
| STnc1385_ | 3.410584 | 3.467357 | 3.361359 | -0.05591 | 0.212501 | -0.24552 | 0.41972 | 0.831298 | 0.899922 | 0.9076 | 0.94478 |
| STnc1390_ | -4.79625 | -4.55601 | -5.01913 | -0.44233 | 0.846544 | -0.4722 | 0.283811 | 0.419069 | 0.561884 | 0.5228 | 0.639504 |
| STnc140_ | -4.34662 | -4.89874 | -3.94292 | 0.963775 | 0.885106 | 1.017065 | 0.111925 | 0.189667 | 0.341255 | 0.2537 | 0.39862 |
| STnc1400_ | -0.1851 | 0.156194 | -0.44276 | -0.59777 | 0.280467 | -2.13247 | 0.005997 | 0.040696 | 0.145945 | 0.1045 | 0.242409 |
| STnc1405_ | 3.071824 | 3.429881 | 2.894266 | -0.52553 | 0.190946 | -2.47934 | 4.68E-05 | 0.033074 | 0.136402 | 0.1 | 0.236592 |
| STnc1410_ | -2.92665 | -3.41509 | -2.58554 | 0.835472 | 0.485098 | 1.665034 | 0.025334 | 0.078368 | 0.199903 | 0.1368 | 0.278459 |
| STnc1460_ | 5.804275 | 6.042278 | 5.753528 | -0.29047 | 0.206841 | -1.52764 | 4.68E-05 | 0.135069 | 0.302042 | 0.1 | 0.236592 |
| STnc1490_ | -0.6212 | -0.5181 | -0.71799 | -0.20208 | 0.254752 | -0.7609 | 0.200667 | 0.299283 | 0.455757 | 0.4027 | 0.541029 |
| STnc150_ | -0.49185 | -0.56823 | -0.22949 | 0.305905 | 0.34895 | 0.762982 | 0.240667 | 0.350474 | 0.522733 | 0.4864 | 0.634689 |
| STnc1500_ | -2.82746 | -2.81187 | -2.84617 | -0.03761 | 0.432999 | -0.08893 | 0.465023 | 0.535167 | 0.666741 | 0.6489 | 0.746942 |
| STnc1510_ | -0.81115 | -0.65382 | -0.93097 | -0.33685 | 0.405837 | -0.78493 | 0.189333 | 0.269711 | 0.441521 | 0.3824 | 0.535649 |
| STnc1530_ | -5.61908 | -5.73368 | -5.54171 | 0.227174 | 1.214628 | 0.154008 | 0.434667 | 0.536847 | 0.669308 | 0.6522 | 0.751362 |
| STnc1540_ | -1.877 | -1.828 | -1.9116 | -0.08029 | 0.362408 | -0.19946 | 0.415333 | 0.569834 | 0.697124 | 0.6834 | 0.776603 |
| STnc1545_ | 0.952762 | 0.963368 | 0.934107 | -0.0332 | 0.206902 | -0.14221 | 0.432667 | 0.703792 | 0.808485 | 0.8263 | 0.888157 |
| STnc1550_ | -0.16576 | 0.741042 | -0.7593 | -1.31659 | 0.503829 | -2.6175 | 4.68E-05 | 0.022286 | 0.110883 | 0.1 | 0.236592 |
| STnc1560_ | 3.350538 | 3.258562 | 3.43286 | 0.175717 | 0.163594 | 1.082529 | 0.105263 | 0.186042 | 0.362917 | 0.1936 | 0.356508 |
| STnc1565_ | 0.115735 | 0.20554 | -0.13965 | -0.31592 | 0.272995 | -1.05065 | 0.159227 | 0.213788 | 0.385496 | 0.3132 | 0.471533 |
| STnc1580_ | -2.24769 | -2.31696 | -2.18223 | 0.134271 | 0.359427 | 0.342259 | 0.350667 | 0.49162 | 0.627656 | 0.5958 | 0.705933 |
| STnc1590_ | 2.238767 | 2.108737 | 2.385015 | 0.185048 | 0.229587 | 0.872557 | 0.145333 | 0.245414 | 0.426007 | 0.2822 | 0.451764 |
| STnc1600_ | -1.10266 | -1.09519 | -1.11437 | -0.03011 | 0.298221 | -0.09543 | 0.46 | 0.637059 | 0.753673 | 0.7358 | 0.82014 |
| STnc1640_ | -1.1429 | -1.31334 | -1.03472 | 0.308475 | 0.306593 | 0.977444 | 0.125333 | 0.224214 | 0.386261 | 0.2924 | 0.436557 |
| STnc1650_ | -5.96586 | -6.32378 | -5.60132 | 0.715389 | 1.526853 | 0.455337 | 0.303797 | 0.452478 | 0.595347 | 0.5718 | 0.683594 |
| STnc1660_ | -0.08263 | -0.12778 | -0.00459 | 0.14124 | 0.238405 | 0.565127 | 0.262 | 0.400818 | 0.558761 | 0.5269 | 0.649406 |
| STnc1670_ | -0.20657 | -0.35616 | -0.07277 | 0.283659 | 0.203031 | 1.299779 | 0.075283 | 0.131417 | 0.275685 | 0.1955 | 0.344237 |
| STnc1675_ | 5.723388 | 5.80402 | 5.581038 | -0.13212 | 0.218292 | -0.88678 | 0.158561 | 0.254005 | 0.441702 | 0.308 | 0.484734 |
| STnc1680_ | -0.24766 | -0.25096 | -0.24466 | 0.018706 | 0.451433 | 0.051667 | 0.475333 | 0.787462 | 0.869081 | 0.84 | 0.89866 |
| STnc1685_ | 6.675895 | 4.060236 | 6.680604 | 2.619687 | 0.058181 | 0.939352 | 0.333333 | 0.21154 | 0.395698 | 0.7 | 0.822108 |
| STnc1690_ | -3.31978 | -3.23856 | -3.42433 | -0.25883 | 0.732274 | -0.32251 | 0.343771 | 0.517114 | 0.659707 | 0.6533 | 0.75178 |
| STnc170_ | -4.51012 | -3.52796 | -6.83098 | -3.29153 | 1.42009 | -2.23035 | 0.001336 | 0.052501 | 0.163901 | 0.1006 | 0.237389 |

| | | | | | | | | | | | |
|---|---|---|---|---|---|---|---|---|---|---|---|
| STnc1700_ | -4.45293 | -4.37739 | -4.52236 | -0.12614 | 0.77401 | -0.14926 | 0.426 | 0.534558 | 0.665273 | 0.6422 | 0.742487 |
| STnc1710_ | 0.018189 | 0.200104 | -0.03595 | -0.19873 | 0.324518 | -0.46504 | 0.330667 | 0.623442 | 0.751584 | 0.6959 | 0.809714 |
| STnc1760_ | -3.10004 | -3.05259 | -3.13882 | -0.08532 | 0.498465 | -0.16834 | 0.426 | 0.559879 | 0.687606 | 0.6748 | 0.769704 |
| STnc1820_ | -1.95644 | -1.90731 | -2.01472 | -0.11244 | 0.47023 | -0.22037 | 0.396667 | 0.605768 | 0.730584 | 0.7132 | 0.803043 |
| STnc1830_ | -2.76341 | -2.77563 | -2.74211 | 0.04386 | 0.455475 | 0.096471 | 0.455333 | 0.572071 | 0.697859 | 0.6914 | 0.781937 |
| STnc1870_ | -7.25013 | -8.35853 | -6.79014 | 1.324982 | 3.705134 | 0.277748 | 0.373333 | 0.543605 | 0.689288 | 0.7306 | 0.82678 |
| STnc1880_ | -3.22936 | -3.09501 | -3.34545 | -0.27218 | 0.569921 | -0.45565 | 0.306667 | 0.475119 | 0.614959 | 0.5884 | 0.697175 |
| STnc1890_ | -5.14352 | -5.31396 | -4.97735 | 0.301523 | 1.3205 | 0.197492 | 0.403065 | 0.606508 | 0.730117 | 0.7169 | 0.805447 |
| STnc190_ | -0.89303 | -0.70771 | -1.11509 | -0.49673 | 0.475815 | -1.12018 | 0.064 | 0.172634 | 0.343005 | 0.1796 | 0.328096 |
| STnc1920_ | -0.06099 | 0.025022 | -0.1162 | -0.09554 | 0.264035 | -0.34809 | 0.356667 | 0.595312 | 0.723703 | 0.6973 | 0.79345 |
| STnc1990_ | -1.31903 | -1.27349 | -1.35453 | -0.08915 | 0.298007 | -0.27314 | 0.383333 | 0.548964 | 0.679612 | 0.6673 | 0.761635 |
| STnc20_ | -2.5761 | -2.82955 | -2.20907 | 0.587015 | 0.458939 | 1.182422 | 0.106 | 0.165295 | 0.313427 | 0.2519 | 0.40365 |
| STnc200_ | -1.54393 | -1.36215 | -1.80594 | -0.467 | 0.469933 | -0.94739 | 0.139907 | 0.213186 | 0.382624 | 0.2943 | 0.453007 |
| STnc2000_ | 0.078418 | 0.454927 | -0.36452 | -0.82945 | 0.210719 | -3.92959 | 4.68E-05 | 0.007649 | 0.0647 | 0.1 | 0.236592 |
| STnc2010_ | 0.013476 | -0.37322 | 0.26864 | 0.667033 | 0.301974 | 2.200283 | 0.003332 | 0.031799 | 0.129195 | 0.1022 | 0.239414 |
| STnc2030_ | -2.24672 | -2.48003 | -1.95849 | 0.423557 | 0.613009 | 0.661656 | 0.244667 | 0.386751 | 0.549613 | 0.4704 | 0.614183 |
| STnc2040_ | 0.27852 | 0.364689 | -1.27145 | -1.61977 | 0.836465 | -1.26881 | 0.134667 | 0.12897 | 0.293343 | 0.2706 | 0.438781 |
| STnc205_ | 5.312566 | 6.134658 | 4.459859 | -1.77286 | 0.185704 | -9.45467 | 4.68E-05 | 0.00098 | 0.033793 | 0.1 | 0.236592 |
| STnc2070_ | -0.01017 | 0.301629 | -0.38471 | -0.4905 | 0.661245 | -0.76282 | 0.18521 | 0.30621 | 0.491765 | 0.3463 | 0.512632 |
| STnc2080_ | 0.415136 | 0.317303 | 0.470505 | 0.24417 | 0.393897 | 0.618733 | 0.267333 | 0.360821 | 0.539836 | 0.5272 | 0.671836 |
| STnc2090_ | 5.567711 | 5.405948 | 5.972484 | 0.555778 | 0.087725 | 3.590021 | 4.68E-05 | 0.046379 | 0.164831 | 0.1 | 0.236592 |
| STnc210_ | -7.45851 | -6.50579 | -8.83576 | -2.33845 | 2.833203 | -0.78422 | 0.155333 | 0.262773 | 0.424152 | 0.3339 | 0.47903 |
| STnc2100_ | -3.12672 | -3.14799 | -3.08935 | 0.062467 | 0.740839 | 0.078438 | 0.464667 | 0.703331 | 0.806665 | 0.8095 | 0.874402 |
| STnc2110_ | -1.83306 | -1.49833 | -2.15156 | -0.73909 | 0.508777 | -1.54226 | 0.01599 | 0.107066 | 0.255386 | 0.1195 | 0.260635 |
| STnc2130_ | -1.21607 | -1.25444 | -1.13753 | 0.1063 | 0.315219 | 0.313493 | 0.377333 | 0.583473 | 0.70981 | 0.6799 | 0.779422 |
| STnc2160_ | -4.65394 | -4.47827 | -4.78264 | -0.25412 | 1.113056 | -0.19867 | 0.410667 | 0.616729 | 0.737108 | 0.72 | 0.806722 |
| STnc2170_ | -3.63482 | -3.75793 | -3.56536 | 0.251988 | 0.768908 | 0.284826 | 0.370667 | 0.522357 | 0.666339 | 0.6617 | 0.762127 |
| STnc2175_ | -1.87592 | -1.82455 | -1.92057 | -0.10311 | 0.334767 | -0.29212 | 0.373085 | 0.514584 | 0.649495 | 0.6172 | 0.72325 |
| STnc2180_ | -0.08587 | -0.00358 | -0.38294 | -0.46391 | 0.597013 | -0.81198 | 0.194 | 0.236082 | 0.417672 | 0.3636 | 0.536902 |
| STnc220_ | -4.86805 | -4.58266 | -5.17694 | -0.5834 | 0.957782 | -0.57671 | 0.242 | 0.390334 | 0.538227 | 0.4918 | 0.615066 |
| STnc230_ | -6.93025 | -6.30012 | -7.58008 | -1.33147 | 2.5755 | -0.45445 | 0.267155 | 0.416613 | 0.571816 | 0.5217 | 0.644981 |
| STnc240_ | -2.2608 | -2.24422 | -2.28471 | -0.03927 | 0.392042 | -0.09347 | 0.461333 | 0.580813 | 0.705842 | 0.691 | 0.784438 |
| STnc260_ | -6.94721 | -6.68532 | -7.3674 | -0.69652 | 2.480052 | -0.2349 | 0.374 | 0.513592 | 0.657886 | 0.6578 | 0.759158 |
| STnc280_ | -5.5432 | -5.94639 | -5.20067 | 0.75614 | 1.252077 | 0.540172 | 0.262492 | 0.384202 | 0.53318 | 0.487 | 0.610264 |
| STnc290_ | -0.30373 | -0.23705 | -0.39828 | -0.16885 | 0.238122 | -0.65519 | 0.227182 | 0.335451 | 0.496255 | 0.446 | 0.581932 |
| STnc30_ | -0.42572 | -0.47126 | -0.37666 | 0.12329 | 0.246055 | 0.473075 | 0.312667 | 0.465902 | 0.613694 | 0.6061 | 0.714834 |
| STnc300_ | -7.61371 | -7.62834 | -7.6002 | -0.00579 | 3.217137 | -0.00104 | 0.499667 | 0.606568 | 0.733172 | 0.7348 | 0.81841 |
| STnc3000_ | -4.63911 | -4.7991 | -4.448 | 0.352399 | 0.801031 | 0.409704 | 0.319121 | 0.457904 | 0.597521 | 0.5587 | 0.67268 |
| STnc3020_ | -5.58144 | -5.22395 | -6.06172 | -0.8183 | 1.343297 | -0.56395 | 0.260493 | 0.397729 | 0.551017 | 0.5098 | 0.63674 |
| STnc3030_ | -3.53378 | -3.04359 | -3.85131 | -0.78905 | 0.668227 | -1.10274 | 0.112667 | 0.178792 | 0.333136 | 0.2626 | 0.413836 |
| STnc3040_ | -5.32645 | -5.27606 | -5.37948 | -0.06739 | 1.317349 | -0.04141 | 0.481679 | 0.628478 | 0.747829 | 0.7456 | 0.827304 |
| STnc3060_ | -4.10932 | -4.13841 | -4.07056 | 0.071917 | 0.752142 | 0.093335 | 0.458361 | 0.601657 | 0.722246 | 0.6999 | 0.790975 |
| STnc3070_ | -1.55621 | -1.78899 | -1.19307 | 0.56373 | 0.417805 | 1.296707 | 0.083333 | 0.122908 | 0.271916 | 0.197 | 0.352418 |
| STnc3080_ | -6.94125 | -7.16852 | -6.82384 | 0.434992 | 2.51133 | 0.143856 | 0.427333 | 0.572591 | 0.707897 | 0.7221 | 0.811153 |
| STnc3090_ | -5.03639 | -4.5582 | -5.6605 | -1.31984 | 1.964608 | -0.61544 | 0.150667 | 0.294463 | 0.474971 | 0.3182 | 0.464119 |

| | | | | | | | | | | | |
|---|---|---|---|---|---|---|---|---|---|---|---|
| STnc310_ | -4.97723 | -4.4235 | -5.55708 | -1.15329 | 0.959151 | -1.11549 | 0.1006 | 0.167566 | 0.312248 | 0.2375 | 0.384627 |
| STnc3100_ | 0.923234 | 0.833546 | 1.057894 | 0.247125 | 0.200915 | 1.225936 | 0.064667 | 0.148732 | 0.30992 | 0.1729 | 0.320996 |
| STnc3110_ | -0.22912 | 0.021368 | -0.29906 | -0.29851 | 0.286422 | -0.8714 | 0.20986 | 0.289243 | 0.463789 | 0.4182 | 0.571241 |
| STnc3120_ | -1.51461 | -1.72759 | -1.30541 | 0.479669 | 0.427401 | 1.07852 | 0.106596 | 0.167531 | 0.326558 | 0.2402 | 0.390008 |
| STnc3150_ | -2.33309 | -1.98827 | -2.69385 | -0.77774 | 0.540935 | -1.43815 | 0.046 | 0.100329 | 0.240136 | 0.1495 | 0.294604 |
| STnc3160_ | -3.99371 | -4.05298 | -3.91 | 0.175123 | 0.821813 | 0.201395 | 0.414 | 0.593598 | 0.717687 | 0.695 | 0.786423 |
| STnc3170_ | -0.92749 | -0.98891 | -0.84937 | -0.0217 | 0.518735 | -0.04875 | 0.483011 | 0.779389 | 0.863859 | 0.9199 | 0.949493 |
| STnc3180_ | -4.01969 | -4.00519 | -4.03615 | 0.01197 | 0.84461 | 0.015269 | 0.49567 | 0.626595 | 0.746727 | 0.7474 | 0.828046 |
| STnc320_ | -2.37253 | -2.26887 | -2.54022 | -0.26864 | 0.520681 | -0.49508 | 0.300466 | 0.474339 | 0.621546 | 0.5957 | 0.711952 |
| STnc3210_ | -6.52366 | -6.66284 | -6.38812 | 0.237131 | 1.712568 | 0.126377 | 0.440373 | 0.556984 | 0.687523 | 0.6705 | 0.765655 |
| STnc3220_ | -0.82266 | -0.81124 | -0.83572 | -0.0458 | 0.361101 | -0.11778 | 0.441706 | 0.671518 | 0.783412 | 0.7683 | 0.844212 |
| STnc3230_ | -2.98213 | -3.03142 | -2.8952 | 0.093865 | 0.624631 | 0.129163 | 0.4497 | 0.660372 | 0.773734 | 0.7474 | 0.831975 |
| STnc3270_ | -3.30001 | -3.23922 | -3.37963 | -0.13717 | 0.548415 | -0.22582 | 0.387742 | 0.529762 | 0.66429 | 0.6422 | 0.743785 |
| STnc3280_ | -3.38003 | -3.36092 | -3.40455 | 0.013012 | 0.734882 | 0.016044 | 0.493005 | 0.671761 | 0.781101 | 0.7813 | 0.854307 |
| STnc3290_ | -2.15132 | -2.14211 | -2.16196 | -0.07086 | 0.61146 | -0.11503 | 0.439707 | 0.639401 | 0.761877 | 0.7806 | 0.85425 |
| STnc3300_ | -3.29758 | -3.27546 | -3.32883 | -0.04059 | 0.597199 | -0.06496 | 0.471019 | 0.615669 | 0.736328 | 0.7278 | 0.81449 |
| STnc3310_ | 2.423152 | 2.532261 | 2.243239 | -0.30107 | 0.172561 | -1.72211 | 0.014 | 0.053648 | 0.173807 | 0.1129 | 0.253385 |
| STnc3330_ | -4.92586 | -4.97252 | -4.88771 | 0.018096 | 1.394799 | 0.012445 | 0.494 | 0.684961 | 0.793356 | 0.807 | 0.873726 |
| STnc3340_ | -4.24642 | -4.33782 | -4.16757 | 0.172926 | 0.949794 | 0.174185 | 0.423718 | 0.612528 | 0.734923 | 0.722 | 0.80921 |
| STnc3350_ | -2.97142 | -2.92264 | -2.99887 | -0.10785 | 0.499735 | -0.18559 | 0.429333 | 0.584218 | 0.709612 | 0.7024 | 0.792029 |
| STnc340_ | -5.04315 | -3.8584 | -5.93578 | -1.97628 | 1.364993 | -1.40411 | 0.060667 | 0.105614 | 0.24458 | 0.1735 | 0.324628 |
| STnc3400_ | -5.21625 | -5.38157 | -5.04961 | 0.36525 | 1.121857 | 0.298097 | 0.356667 | 0.515881 | 0.650368 | 0.6355 | 0.736003 |
| STnc3410_ | -0.39938 | -0.59184 | -0.21313 | 0.380765 | 0.273637 | 1.334173 | 0.064624 | 0.120057 | 0.267122 | 0.1811 | 0.333511 |
| STnc3420_ | -2.722 | -2.81988 | -2.62468 | 0.203908 | 0.556525 | 0.348907 | 0.354 | 0.54893 | 0.683083 | 0.6703 | 0.76839 |
| STnc3430_ | -0.54188 | -1.86557 | 0.350682 | 2.236645 | 0.369741 | 6.349916 | 4.68E-05 | 0.011802 | 0.079917 | 0.1 | 0.236592 |
| STnc3440_ | -3.32781 | -3.77646 | -3.0395 | 0.772059 | 0.725359 | 1.011723 | 0.098 | 0.191954 | 0.351581 | 0.2469 | 0.392731 |
| STnc3460_ | -0.16526 | -0.28279 | -0.07945 | 0.200005 | 0.239994 | 0.804506 | 0.18521 | 0.291113 | 0.454268 | 0.3887 | 0.533853 |
| STnc3480_ | -3.80058 | -3.05865 | -4.07383 | -0.96041 | 0.834887 | -1.01293 | 0.172667 | 0.238571 | 0.40055 | 0.3513 | 0.502649 |
| STnc3490_ | -5.14918 | -4.86968 | -5.65576 | -0.82936 | 1.54887 | -0.49496 | 0.267155 | 0.398931 | 0.560756 | 0.5295 | 0.652928 |
| STnc350_ | -1.95507 | -2.08445 | -1.85387 | 0.220444 | 0.401719 | 0.522494 | 0.280667 | 0.422222 | 0.571514 | 0.5213 | 0.647403 |
| STnc3500_ | -3.42096 | -3.42594 | -3.4137 | -0.01038 | 0.642751 | -0.01525 | 0.492667 | 0.620928 | 0.740713 | 0.7259 | 0.811965 |
| STnc3510_ | -0.69781 | -0.53461 | -0.80071 | -0.29427 | 0.378409 | -0.7313 | 0.221852 | 0.323181 | 0.494238 | 0.4419 | 0.587502 |
| STnc3540_ | 3.79617 | 3.846341 | 3.585186 | -0.26609 | 0.22311 | -1.26435 | 0.028648 | 0.152707 | 0.324245 | 0.1236 | 0.266407 |
| STnc3550_ | -2.09751 | -2.50927 | -1.48468 | 0.994394 | 0.389279 | 2.467125 | 0.003998 | 0.028299 | 0.116723 | 0.1039 | 0.241617 |
| STnc3580_ | -3.23595 | -3.6283 | -2.9743 | 0.633704 | 0.66625 | 0.892948 | 0.153897 | 0.240708 | 0.401936 | 0.3227 | 0.469079 |
| STnc360_ | -6.29159 | -6.24455 | -6.38067 | -0.15276 | 1.664556 | -0.08627 | 0.464357 | 0.59418 | 0.718949 | 0.7056 | 0.795592 |
| STnc3600_ | -1.57813 | -0.62103 | -3.04566 | -2.48207 | 0.57818 | -4.32028 | 4.68E-05 | 0.005997 | 0.057501 | 0.1 | 0.236592 |
| STnc3610_ | -0.88774 | -0.84067 | -0.93569 | -0.09255 | 0.211925 | -0.40217 | 0.316667 | 0.436003 | 0.572932 | 0.54 | 0.652775 |
| STnc3620_ | -3.28147 | -3.30331 | -3.26816 | 0.017923 | 0.560613 | 0.030787 | 0.486 | 0.580746 | 0.70638 | 0.6998 | 0.789859 |
| STnc3630_ | -1.03566 | -1.12384 | -0.94782 | 0.20707 | 0.343039 | 0.566179 | 0.26 | 0.398881 | 0.556913 | 0.5324 | 0.656595 |
| STnc3640_ | -1.85121 | -1.4968 | -2.46485 | -0.93561 | 0.5185 | -1.70105 | 0.020667 | 0.078098 | 0.20969 | 0.1232 | 0.265701 |
| STnc3680_ | 1.815294 | 1.853781 | 1.787103 | -0.06334 | 0.118964 | -0.5036 | 0.29447 | 0.452112 | 0.601305 | 0.5692 | 0.68836 |
| STnc370_ | -6.44624 | -5.94218 | -8.33291 | -2.27868 | 3.199265 | -0.59201 | 0.24 | 0.321818 | 0.498304 | 0.475 | 0.616899 |
| STnc3700_ | -3.69602 | -3.66259 | -3.72523 | -0.06366 | 0.678763 | -0.08978 | 0.463025 | 0.60191 | 0.723563 | 0.7046 | 0.794715 |
| STnc3720_ | -3.13714 | -3.48153 | -2.86378 | 0.627385 | 0.478723 | 1.239838 | 0.090667 | 0.145089 | 0.282235 | 0.2201 | 0.363211 |

| | | | | | | | | | | | |
|---|---|---|---|---|---|---|---|---|---|---|---|
| STnc3730_ | -1.21828 | -1.18466 | -1.24971 | -0.06195 | 0.236065 | -0.24796 | 0.390667 | 0.497139 | 0.63297 | 0.6152 | 0.719292 |
| STnc3750_ | -3.27404 | -3.39831 | -3.16224 | 0.250373 | 0.538902 | 0.440551 | 0.309127 | 0.441312 | 0.585413 | 0.5538 | 0.66672 |
| STnc3770_ | -2.60814 | -2.73092 | -2.39293 | 0.311887 | 0.522794 | 0.560918 | 0.276482 | 0.453611 | 0.601144 | 0.5743 | 0.693105 |
| STnc3790_ | -4.84509 | -4.46193 | -5.33785 | -0.87315 | 0.855208 | -0.96003 | 0.130667 | 0.214058 | 0.362494 | 0.301 | 0.443725 |
| STnc380_ | -5.10006 | -5.24492 | -4.88975 | 0.374541 | 1.048646 | 0.334735 | 0.351765 | 0.505271 | 0.642513 | 0.6223 | 0.726119 |
| STnc3800_ | -1.23339 | -1.28109 | -1.17976 | 0.097936 | 0.23113 | 0.389194 | 0.324 | 0.440968 | 0.581218 | 0.5472 | 0.662145 |
| STnc3830_ | -2.4222 | -2.8632 | -2.10424 | 0.761044 | 0.366791 | 2.019123 | 0.016656 | 0.052235 | 0.157917 | 0.1154 | 0.255239 |
| STnc3850_ | -3.55799 | -3.79094 | -3.29515 | 0.489606 | 0.571837 | 0.787014 | 0.184667 | 0.286822 | 0.43663 | 0.3789 | 0.516722 |
| STnc3860_ | -2.74483 | -2.76437 | -2.72555 | 0.071401 | 0.57291 | 0.115878 | 0.45503 | 0.647523 | 0.762606 | 0.7555 | 0.833906 |
| STnc3870_ | -0.58339 | -0.33911 | -0.8188 | -0.41545 | 0.420573 | -0.96077 | 0.137333 | 0.205669 | 0.378106 | 0.279 | 0.446366 |
| STnc3880_ | -3.92427 | -3.98414 | -3.86607 | 0.139522 | 0.640693 | 0.196771 | 0.411726 | 0.526512 | 0.657483 | 0.6315 | 0.734255 |
| STnc3890_ | -3.54386 | -3.32112 | -3.70536 | -0.45213 | 0.732398 | -0.5915 | 0.258494 | 0.386191 | 0.53967 | 0.5078 | 0.632188 |
| STnc390_ | 1.660674 | 1.379578 | 2.040232 | 0.62723 | 0.220292 | 2.799071 | 4.68E-05 | 0.018605 | 0.100554 | 0.1 | 0.236592 |
| STnc3920_ | -6.85685 | -6.46079 | -8.30692 | -1.8172 | 3.117102 | -0.47029 | 0.292472 | 0.391618 | 0.560599 | 0.57 | 0.697729 |
| STnc3950_ | -1.73704 | -1.44397 | -1.99781 | -0.55772 | 0.330967 | -1.57971 | 0.042638 | 0.08443 | 0.211105 | 0.1476 | 0.292591 |
| STnc3960_ | -1.84442 | -1.99837 | -1.78396 | 0.166404 | 0.465596 | 0.280284 | 0.397333 | 0.697791 | 0.804192 | 0.7618 | 0.852317 |
| STnc3970_ | -4.78736 | -4.88845 | -4.70164 | 0.198493 | 1.029193 | 0.176767 | 0.417722 | 0.576458 | 0.70549 | 0.6964 | 0.78736 |
| STnc3990_ | -1.33571 | -1.39895 | -1.22408 | 0.169303 | 0.421065 | 0.371115 | 0.341772 | 0.582787 | 0.713308 | 0.681 | 0.782583 |
| STnc40_ | 0.569355 | 0.594376 | 0.545747 | -0.03548 | 0.257819 | -0.13097 | 0.434667 | 0.72498 | 0.823875 | 0.82 | 0.882796 |
| STnc4000_ | -4.09425 | -4.33785 | -3.90212 | 0.453713 | 0.709836 | 0.594948 | 0.253165 | 0.383049 | 0.528079 | 0.4892 | 0.611218 |
| STnc4010_ | -4.2602 | -5.04371 | -3.79924 | 1.239442 | 0.799818 | 1.441098 | 0.049301 | 0.103779 | 0.237527 | 0.1629 | 0.307718 |
| STnc4020_ | -7.75433 | -7.54768 | -7.99353 | -0.47682 | 2.352525 | -0.15478 | 0.407062 | 0.515055 | 0.653437 | 0.6349 | 0.737841 |
| STnc4040_ | -1.28041 | -1.52207 | -1.04485 | 0.501876 | 0.3341 | 1.472875 | 0.049967 | 0.099602 | 0.237018 | 0.1551 | 0.300867 |
| STnc4070_ | -4.04794 | -3.9321 | -4.11761 | -0.15621 | 0.968382 | -0.15195 | 0.439333 | 0.664445 | 0.775561 | 0.7674 | 0.844314 |
| STnc4080_ | -4.26703 | -4.29407 | -4.24551 | -0.03279 | 0.964373 | -0.03247 | 0.485333 | 0.651926 | 0.766221 | 0.7673 | 0.842604 |
| STnc410_ | -5.98292 | -6.29317 | -5.77628 | 0.479775 | 1.323944 | 0.351015 | 0.342 | 0.472629 | 0.614508 | 0.5865 | 0.696663 |
| STnc4100_ | -3.53851 | -2.90488 | -4.34909 | -1.44546 | 0.707474 | -1.95026 | 0.014657 | 0.05379 | 0.165309 | 0.1124 | 0.252048 |
| STnc4110_ | -3.95039 | -3.75063 | -4.16606 | -0.37633 | 0.921759 | -0.39344 | 0.341106 | 0.546821 | 0.68214 | 0.6565 | 0.760072 |
| STnc4120_ | -2.4212 | -2.54247 | -2.2865 | 0.260596 | 0.409791 | 0.584436 | 0.251333 | 0.378585 | 0.525549 | 0.4784 | 0.604838 |
| STnc4130_ | -2.33713 | -1.95781 | -2.73262 | -0.79577 | 0.416114 | -1.84259 | 0.021986 | 0.05552 | 0.167632 | 0.1192 | 0.25984 |
| STnc4140_ | -2.51313 | -2.39084 | -2.57497 | -0.13483 | 0.491956 | -0.2421 | 0.38974 | 0.623025 | 0.740589 | 0.7121 | 0.804746 |
| STnc4150_ | -6.47489 | -6.42913 | -6.52042 | -0.01245 | 1.834322 | -0.00946 | 0.496 | 0.589818 | 0.714367 | 0.7015 | 0.792044 |
| STnc4160_ | -1.75232 | -1.65183 | -1.84794 | -0.18975 | 0.281328 | -0.62334 | 0.246 | 0.358299 | 0.503526 | 0.4661 | 0.591069 |
| STnc4170_ | -5.97591 | -5.72111 | -6.34327 | -0.58401 | 1.718927 | -0.32181 | 0.345769 | 0.518508 | 0.658108 | 0.6424 | 0.74636 |
| STnc4190_ | -7.24634 | -6.65856 | -8.38947 | -1.67581 | 2.809665 | -0.53566 | 0.245333 | 0.362366 | 0.526343 | 0.4805 | 0.611721 |
| STnc420_ | -5.9667 | -5.11874 | -6.83456 | -1.83048 | 1.567414 | -1.13122 | 0.088608 | 0.160887 | 0.307218 | 0.2286 | 0.374438 |
| STnc4200_ | -3.45884 | -3.50906 | -3.41712 | 0.17356 | 0.983397 | 0.177985 | 0.422667 | 0.676626 | 0.788104 | 0.7875 | 0.859453 |
| STnc4210_ | 0.450166 | 0.470273 | 0.436893 | -0.01657 | 0.193385 | -0.08446 | 0.456 | 0.676318 | 0.785671 | 0.7829 | 0.857262 |
| STnc4220_ | -0.58613 | -0.61074 | -0.51085 | 0.095858 | 0.361763 | 0.233555 | 0.412667 | 0.730167 | 0.827633 | 0.794 | 0.873031 |
| STnc4230_ | -2.04498 | -2.13621 | -1.97431 | 0.18402 | 0.50852 | 0.334245 | 0.35443 | 0.587656 | 0.716725 | 0.7053 | 0.798287 |
| STnc4240_ | -1.24361 | -1.19156 | -1.31434 | -0.05569 | 0.397448 | -0.13685 | 0.449034 | 0.689778 | 0.797243 | 0.8116 | 0.87593 |
| STnc4250_ | 2.542833 | 2.675669 | 2.394383 | -0.32194 | 0.149466 | -2.30442 | 4.68E-05 | 0.035041 | 0.137674 | 0.1 | 0.236592 |
| STnc4260_ | -5.73987 | -5.37387 | -6.36537 | -1.05872 | 1.935214 | -0.44374 | 0.221186 | 0.355031 | 0.527432 | 0.447 | 0.576578 |
| STnc430_ | -5.62674 | -5.47659 | -5.86429 | -0.56327 | 2.098984 | -0.20693 | 0.361093 | 0.494533 | 0.652025 | 0.6826 | 0.777769 |
| STnc440_ | -1.20826 | -1.16676 | -1.26935 | -0.15177 | 0.523909 | -0.3066 | 0.369333 | 0.63049 | 0.754918 | 0.7327 | 0.821801 |

| | | | | | | | | | | | |
|---|---|---|---|---|---|---|---|---|---|---|---|
| STnc450_ | -0.45402 | -0.40513 | -0.52858 | -0.16968 | 0.385239 | -0.41307 | 0.308667 | 0.497645 | 0.650383 | 0.6268 | 0.743318 |
| STnc460_ | -8.09839 | -6.96301 | -9.82961 | -2.87497 | 3.054694 | -0.83382 | 0.143904 | 0.233733 | 0.385491 | 0.2974 | 0.439475 |
| STnc470_ | -1.70466 | -1.7965 | -1.59722 | 0.198991 | 0.310205 | 0.617144 | 0.251832 | 0.391163 | 0.53851 | 0.4974 | 0.62023 |
| STnc475_ | 5.240464 | 5.625262 | 5.112041 | -0.5069 | 0.092261 | -3.93978 | 4.68E-05 | 0.037788 | 0.147667 | 0.1 | 0.236592 |
| STnc480_ | -3.0222 | -3.29276 | -2.83547 | 0.461693 | 0.602505 | 0.695624 | 0.212667 | 0.32393 | 0.48438 | 0.4273 | 0.566239 |
| STnc50_ | -6.35005 | -6.08132 | -6.60009 | -0.41651 | 1.724515 | -0.21448 | 0.378667 | 0.552095 | 0.681753 | 0.6551 | 0.753705 |
| STnc500_ | -1.84433 | -1.87607 | -1.80626 | 0.08197 | 0.322089 | 0.229794 | 0.397333 | 0.539386 | 0.672874 | 0.6504 | 0.751393 |
| STnc510_ | -1.18651 | -1.5466 | -0.85485 | 0.685627 | 0.273133 | 2.428605 | 0.005331 | 0.028898 | 0.117953 | 0.1042 | 0.242076 |
| STnc520_ | -0.9287 | -0.9463 | -0.9161 | 0.140447 | 0.488931 | 0.288739 | 0.389333 | 0.608978 | 0.739156 | 0.7887 | 0.859317 |
| STnc530_ | -4.3602 | -4.63588 | -3.92767 | 0.702607 | 0.938242 | 0.693666 | 0.244667 | 0.37539 | 0.531608 | 0.4898 | 0.620808 |
| STnc540_ | 3.824795 | 3.837434 | 3.798729 | -0.0357 | 0.145558 | -0.2234 | 0.407728 | 0.855152 | 0.914709 | 0.8623 | 0.918273 |
| STnc590_ | 0.384754 | 0.582648 | 0.330517 | -0.23094 | 0.294798 | -0.59686 | 0.318 | 0.539624 | 0.688816 | 0.6622 | 0.784823 |
| STnc60_ | -9.33656 | -8.90118 | -9.79139 | -0.75715 | 3.994991 | -0.17464 | 0.408 | 0.506859 | 0.643616 | 0.6231 | 0.726296 |
| STnc600_ | 3.759545 | 4.021383 | 3.516527 | -0.54482 | 0.181415 | -2.97912 | 4.68E-05 | 0.032695 | 0.136607 | 0.1 | 0.236592 |
| STnc610_ | 0.501995 | 0.637584 | 0.30785 | -0.38368 | 0.350427 | -1.18973 | 0.063333 | 0.142934 | 0.306725 | 0.1681 | 0.319389 |
| STnc620_ | 1.41405 | 1.441717 | 1.267089 | -0.16084 | 0.197698 | -0.58488 | 0.332667 | 0.557719 | 0.703945 | 0.6983 | 0.816464 |
| STnc640_ | -2.25362 | -1.52578 | -2.62967 | -1.04263 | 0.475696 | -2.04648 | 0.017334 | 0.051432 | 0.163863 | 0.1169 | 0.257346 |
| STnc650_ | 3.244174 | 3.360133 | 3.188731 | -0.13242 | 0.152396 | -0.88574 | 0.175333 | 0.246617 | 0.427114 | 0.3237 | 0.49565 |
| STnc660_ | 1.939528 | 1.619167 | 2.083242 | 0.43467 | 0.182494 | 2.173425 | 0.001337 | 0.043314 | 0.154265 | 0.1031 | 0.240718 |
| STnc670_ | 2.667089 | 2.988183 | 2.34119 | -0.64734 | 0.060716 | -10.5359 | 4.68E-05 | 0.000645 | 0.026449 | 0.1 | 0.236592 |
| STnc680_ | 2.975108 | 2.733581 | 3.253857 | 0.501872 | 0.112981 | 4.404893 | 4.68E-05 | 0.004242 | 0.052542 | 0.1 | 0.236592 |
| STnc690_ | -3.2182 | -3.16482 | -3.28489 | -0.11845 | 0.706222 | -0.15693 | 0.429714 | 0.632958 | 0.752193 | 0.7456 | 0.827984 |
| STnc695_ | -1.31922 | -1.63406 | -0.75053 | 0.85462 | 0.350106 | 2.313708 | 0.005331 | 0.038879 | 0.141392 | 0.1054 | 0.243516 |
| STnc70_ | 2.67868 | 2.749068 | 2.560576 | -0.17355 | 0.251549 | -0.59365 | 0.291333 | 0.390591 | 0.572666 | 0.5887 | 0.717406 |
| STnc720_ | -1.53519 | -2.31106 | -0.6784 | 1.538266 | 0.48038 | 3.218243 | 4.68E-05 | 0.011337 | 0.078333 | 0.1001 | 0.236722 |
| STnc730_ | 1.145857 | 1.308495 | 0.980781 | -0.16521 | 0.3792 | -0.52259 | 0.283811 | 0.468441 | 0.635686 | 0.5767 | 0.706946 |
| STnc740_ | -5.96195 | -6.08657 | -5.77758 | 0.427911 | 1.58802 | 0.249477 | 0.393333 | 0.575778 | 0.70239 | 0.6927 | 0.784198 |
| STnc750_ | -1.05913 | -0.82822 | -1.17748 | -0.28802 | 0.432681 | -0.65743 | 0.243333 | 0.38148 | 0.547137 | 0.4768 | 0.624926 |
| STnc760_ | 0.362935 | 0.29283 | 0.420818 | 0.12649 | 0.321811 | 0.381556 | 0.325333 | 0.55017 | 0.694893 | 0.6495 | 0.761864 |
| STnc770_ | 2.974448 | 2.990867 | 2.961349 | 0.019774 | 0.231767 | 0.093552 | 0.458667 | 0.666658 | 0.790168 | 0.9232 | 0.952725 |
| STnc780_ | -0.09806 | -0.03106 | -0.15516 | -0.15172 | 0.248794 | -0.57982 | 0.266667 | 0.405429 | 0.561547 | 0.5437 | 0.661977 |
| STnc790_ | 0.219188 | 0.532289 | 0.005914 | -0.52638 | 0.23359 | -2.18996 | 0.003335 | 0.041472 | 0.14727 | 0.1042 | 0.242118 |
| STnc800_ | 2.810827 | 2.868708 | 2.633443 | -0.2291 | 0.181464 | -1.24276 | 0.063957 | 0.169901 | 0.343496 | 0.1725 | 0.326128 |
| STnc810_ | -1.38582 | -1.71893 | -1.01743 | 0.671712 | 0.381875 | 1.709825 | 0.035976 | 0.066802 | 0.190278 | 0.1324 | 0.277405 |
| STnc835_ | 3.603751 | 3.684544 | 3.467398 | -0.14195 | 0.281683 | -0.42794 | 0.343333 | 0.739931 | 0.843224 | 0.7192 | 0.833462 |
| STnc840_ | 2.462414 | 2.921651 | 1.905254 | -0.98115 | 0.258449 | -3.72453 | 4.68E-05 | 0.005233 | 0.058801 | 0.1 | 0.236592 |
| STnc850_ | -3.10224 | -3.13463 | -3.0732 | 0.051821 | 0.524979 | 0.092054 | 0.451033 | 0.571523 | 0.697206 | 0.681 | 0.773562 |
| STnc870_ | 1.031663 | 0.744883 | 1.171582 | 0.282875 | 0.38141 | 0.815288 | 0.207861 | 0.314681 | 0.498594 | 0.3806 | 0.546547 |
| STnc880_ | -1.19259 | -1.11134 | -1.27671 | -0.02931 | 0.512967 | -0.058 | 0.47435 | 0.768533 | 0.855451 | 0.889 | 0.928986 |
| STnc890_ | -0.36978 | -0.27363 | -0.43317 | -0.15998 | 0.377437 | -0.42017 | 0.317333 | 0.53639 | 0.682268 | 0.6345 | 0.751471 |
| STnc920_ | -0.0048 | 0.019172 | -0.01814 | -0.00921 | 0.231314 | -0.03887 | 0.485676 | 0.695143 | 0.800132 | 0.7912 | 0.864517 |
| STnc925_ | -0.43324 | -0.47176 | -0.37111 | 0.090908 | 0.306742 | 0.27174 | 0.385333 | 0.631688 | 0.753181 | 0.738 | 0.825197 |
| STnc930_ | -4.83437 | -4.5891 | -5.20427 | -0.52181 | 1.393915 | -0.35749 | 0.353764 | 0.556642 | 0.694314 | 0.6916 | 0.788148 |
| STnc940_ | -2.96701 | -3.05607 | -2.89441 | 0.164946 | 0.646806 | 0.231288 | 0.402667 | 0.618089 | 0.738855 | 0.7187 | 0.807057 |
| STnc950_ | -4.5879 | -4.59476 | -4.57977 | -0.14057 | 1.35861 | -0.09757 | 0.457695 | 0.708034 | 0.811228 | 0.8123 | 0.877231 |

| | | | | | | | | | | | |
|---|---|---|---|---|---|---|---|---|---|---|---|
| STnc970_ | -1.09617 | -1.1015 | -1.09102 | 0.015945 | 0.240114 | 0.055347 | 0.472 | 0.550717 | 0.681061 | 0.6618 | 0.759868 |
| StyR.103a_ | -8.70462 | -8.88114 | -8.49391 | 0.499172 | 4.038834 | 0.103991 | 0.433045 | 0.559854 | 0.693535 | 0.6913 | 0.784484 |
| StyR.103b_ | -8.91618 | -8.06654 | -9.96855 | -1.69557 | 3.568566 | -0.41002 | 0.287808 | 0.436329 | 0.579092 | 0.5287 | 0.647045 |
| StyR.103c_ | -8.37786 | -6.95283 | -9.72001 | -2.50578 | 3.799768 | -0.63094 | 0.204 | 0.378043 | 0.523398 | 0.4336 | 0.568191 |
| StyR.103g_ | -8.90093 | -8.90709 | -8.89244 | 0.009219 | 3.959143 | 0.002083 | 0.499001 | 0.562643 | 0.693114 | 0.6833 | 0.777712 |
| StyR.122_ | -4.64264 | -4.93121 | -4.34113 | 0.548818 | 0.996922 | 0.503467 | 0.276482 | 0.458265 | 0.600832 | 0.5557 | 0.674299 |
| StyR.126_ | -1.71734 | -1.72092 | -1.71365 | 0.006521 | 0.300131 | 0.020967 | 0.488341 | 0.547107 | 0.675151 | 0.6544 | 0.753062 |
| StyR.128_ | -5.03905 | -4.92181 | -5.16841 | -0.2714 | 0.920433 | -0.27211 | 0.373751 | 0.485245 | 0.623124 | 0.5992 | 0.70708 |
| StyR.134_ | -0.53175 | -0.53943 | -0.51981 | 0.017616 | 0.250816 | 0.062532 | 0.468354 | 0.620975 | 0.742039 | 0.7355 | 0.820547 |
| StyR.137_ | -8.84777 | -8.90188 | -8.78601 | 0.07689 | 3.916257 | 0.014554 | 0.492667 | 0.544398 | 0.680306 | 0.6871 | 0.778811 |
| StyR.150_ | -4.79339 | -5.13464 | -4.58638 | 0.515751 | 1.023014 | 0.478891 | 0.293333 | 0.440202 | 0.588486 | 0.5548 | 0.675506 |
| StyR.169_ | -1.13144 | -1.36343 | -0.95413 | 0.403472 | 0.247218 | 1.607068 | 0.039333 | 0.088724 | 0.212926 | 0.1552 | 0.299074 |
| StyR.172_ | -8.22056 | -8.92541 | -7.61178 | 1.186252 | 3.523376 | 0.294401 | 0.336667 | 0.524311 | 0.659936 | 0.6335 | 0.73588 |
| StyR.178_ | -8.82972 | -8.8306 | -8.82814 | -0.06805 | 3.950795 | -0.01254 | 0.493005 | 0.552 | 0.686333 | 0.6872 | 0.780427 |
| StyR.195_ | -3.48958 | -3.74426 | -3.25507 | 0.488483 | 0.519249 | 0.884675 | 0.157895 | 0.243584 | 0.391449 | 0.332 | 0.470913 |
| StyR.196_ | -6.52427 | -7.27925 | -6.11625 | 1.119692 | 2.451034 | 0.387825 | 0.289141 | 0.415171 | 0.577957 | 0.5509 | 0.676107 |
| StyR.199_ | -3.58971 | -3.63767 | -3.51641 | 0.130714 | 0.724052 | 0.176869 | 0.424667 | 0.611916 | 0.733769 | 0.7251 | 0.809811 |
| StyR.215a_ | 1.221695 | 1.27179 | 0.896104 | -0.36657 | 0.323814 | -0.79444 | 0.275816 | 0.333289 | 0.518781 | 0.5395 | 0.692516 |
| StyR.215b_ | 0.709754 | 0.804255 | 0.654282 | -0.14524 | 0.232387 | -0.61008 | 0.259827 | 0.403819 | 0.56977 | 0.5277 | 0.665163 |
| StyR.215c_ | -3.36171 | -3.28082 | -3.43574 | -0.20186 | 0.673739 | -0.26393 | 0.378414 | 0.565505 | 0.694355 | 0.6924 | 0.782605 |
| StyR.215d_ | -1.30132 | -1.11674 | -1.76688 | -0.59215 | 0.440692 | -1.22412 | 0.104597 | 0.148196 | 0.307835 | 0.2269 | 0.385415 |
| StyR.215e_ | -3.0376 | -2.75932 | -3.29648 | -0.45303 | 0.855157 | -0.52833 | 0.294667 | 0.48029 | 0.634126 | 0.6005 | 0.721921 |
| StyR.215f_ | -0.7759 | -0.86554 | -0.67497 | 0.249905 | 0.384392 | 0.610161 | 0.248 | 0.365548 | 0.532983 | 0.5185 | 0.648426 |
| StyR.215g_ | -3.67402 | -3.41169 | -3.83598 | -0.36983 | 0.694565 | -0.47707 | 0.299134 | 0.500317 | 0.638539 | 0.5878 | 0.705563 |
| StyR.219_ | -5.02917 | -4.73042 | -5.37499 | -0.65595 | 0.9008 | -0.68978 | 0.22052 | 0.329054 | 0.472964 | 0.4231 | 0.550624 |
| StyR.222_ | -6.20399 | -7.30952 | -5.79076 | 1.481916 | 2.28805 | 0.537492 | 0.224667 | 0.339292 | 0.509208 | 0.4476 | 0.5832 |
| StyR.234_ | -5.89857 | -5.84885 | -5.92919 | -0.02775 | 1.574585 | -0.01431 | 0.492 | 0.617862 | 0.73912 | 0.7228 | 0.809897 |
| StyR.243_ | -0.42209 | -0.29614 | -0.814 | -0.50828 | 0.381264 | -1.15009 | 0.100667 | 0.175348 | 0.345654 | 0.2312 | 0.387475 |
| StyR.248_ | -1.69654 | -1.95564 | -1.40544 | 0.565225 | 0.339101 | 1.621171 | 0.031979 | 0.079366 | 0.203913 | 0.1376 | 0.280805 |
| StyR.24a_ | -4.12452 | -4.08659 | -4.15386 | -0.01697 | 1.061525 | -0.01386 | 0.492338 | 0.69337 | 0.799222 | 0.8082 | 0.873088 |
| StyR.24b_ | -2.06559 | -1.18617 | -3.36834 | -2.19777 | 0.457331 | -4.84259 | 4.68E-05 | 0.012486 | 0.076059 | 0.1 | 0.236592 |
| StyR.24c_ | -3.35374 | -2.56609 | -4.60351 | -2.05061 | 0.702972 | -2.91174 | 0.000673 | 0.031621 | 0.122295 | 0.1 | 0.236592 |
| StyR.24d_ | -4.66012 | -5.01224 | -4.3807 | 0.627017 | 0.813174 | 0.72001 | 0.20453 | 0.321045 | 0.466003 | 0.4166 | 0.54859 |
| StyR.24e_ | -4.67517 | -3.60162 | -5.38699 | -1.63118 | 1.127384 | -1.41509 | 0.064624 | 0.109874 | 0.24902 | 0.1822 | 0.332982 |
| StyR.24f_ | -2.97186 | -2.70107 | -3.42347 | -0.73398 | 0.6533 | -1.02515 | 0.116667 | 0.190918 | 0.353986 | 0.2586 | 0.410335 |
| StyR.24g_ | -4.19493 | -3.41662 | -5.69069 | -2.27712 | 0.891928 | -2.46214 | 0.000673 | 0.03729 | 0.13252 | 0.1013 | 0.238116 |
| StyR.250_ | -4.66151 | -4.37023 | -5.0076 | -0.63827 | 0.828524 | -0.73349 | 0.209194 | 0.321119 | 0.468384 | 0.4225 | 0.553685 |
| StyR.252_ | -1.70462 | -1.73051 | -1.65878 | 0.094541 | 0.387028 | 0.241307 | 0.406396 | 0.600501 | 0.72556 | 0.7156 | 0.805561 |
| StyR.254_ | -5.02459 | -5.45834 | -4.70046 | 0.833614 | 1.334101 | 0.527758 | 0.223333 | 0.351894 | 0.519008 | 0.4643 | 0.592927 |
| StyR.262_ | -0.71582 | -0.77115 | -0.63405 | 0.138775 | 0.235399 | 0.541981 | 0.267821 | 0.423383 | 0.569087 | 0.5301 | 0.652523 |
| StyR.264_ | -3.46116 | -3.63636 | -3.25037 | 0.378669 | 0.556837 | 0.63561 | 0.238667 | 0.357013 | 0.505646 | 0.4621 | 0.591526 |
| StyR.27_ | -2.32469 | -3.10288 | -1.97218 | 1.079289 | 0.581472 | 1.66923 | 0.030667 | 0.079605 | 0.213316 | 0.1323 | 0.277095 |
| StyR.277_ | 0.598443 | 0.454765 | 0.70749 | 0.253468 | 0.22641 | 1.066833 | 0.114667 | 0.172771 | 0.335699 | 0.2485 | 0.407996 |
| StyR.281_ | -4.4023 | -4.40059 | -4.40352 | -0.07771 | 1.0058 | -0.07048 | 0.468 | 0.623417 | 0.743785 | 0.7368 | 0.821212 |
| StyR.285_ | -3.75852 | -3.87873 | -3.63511 | 0.234985 | 0.677558 | 0.330114 | 0.352667 | 0.53541 | 0.668524 | 0.6503 | 0.751226 |

| | | | | | | | | | | | |
|---|---|---|---|---|---|---|---|---|---|---|---|
| StyR.287_ | -0.42785 | -0.49821 | -0.03346 | 0.440052 | 0.454674 | 0.681097 | 0.298667 | 0.442944 | 0.613245 | 0.6138 | 0.751404 |
| StyR.29_ | 2.010673 | 2.601428 | 1.590238 | -0.95701 | 0.237907 | -3.83576 | 4.68E-05 | 0.00957 | 0.07473 | 0.1 | 0.236592 |
| StyR.291_ | -1.79046 | -1.82687 | -1.77017 | 0.02136 | 0.376664 | 0.051132 | 0.480346 | 0.62901 | 0.746309 | 0.7443 | 0.826487 |
| StyR.292_ | -6.80843 | -6.64458 | -7.0852 | -0.53604 | 2.141291 | -0.20222 | 0.38 | 0.495905 | 0.643829 | 0.6341 | 0.736802 |
| StyR.293_ | -2.76469 | -2.85806 | -2.61718 | 0.229223 | 0.50295 | 0.415521 | 0.326 | 0.509207 | 0.644611 | 0.608 | 0.7178 |
| StyR.295_ | -4.36309 | -4.35355 | -4.37553 | -0.00868 | 0.7588 | -0.00935 | 0.497002 | 0.568313 | 0.695052 | 0.6809 | 0.774628 |
| StyR.296_ | 3.284895 | 3.343044 | 3.16893 | -0.16043 | 0.269463 | -0.61922 | 0.232 | 0.350481 | 0.536041 | 0.4249 | 0.599883 |
| StyR.3_ | -1.78514 | -1.7019 | -1.84094 | -0.09541 | 0.433458 | -0.20572 | 0.413058 | 0.65823 | 0.77115 | 0.7487 | 0.832504 |
| StyR.302_ | -8.3809 | -8.90269 | -7.9176 | 0.882496 | 3.58288 | 0.2059 | 0.385333 | 0.51393 | 0.653808 | 0.635 | 0.739881 |
| StyR.305_ | -1.27468 | -0.84982 | -1.81033 | -0.907 | 0.498029 | -1.82107 | 0.022 | 0.052752 | 0.169951 | 0.1208 | 0.264065 |
| StyR.306_ | 1.048654 | 0.96305 | 1.236919 | 0.262279 | 0.161675 | 1.498472 | 0.062667 | 0.103382 | 0.245017 | 0.1734 | 0.323801 |
| StyR.315_ | -6.62012 | -6.65922 | -6.5945 | 0.006611 | 3.244521 | 0.002067 | 0.498667 | 0.667236 | 0.782745 | 0.7877 | 0.859793 |
| StyR.322_ | -5.63402 | -5.39597 | -5.95961 | -0.55249 | 1.280926 | -0.41276 | 0.319121 | 0.458726 | 0.602473 | 0.5831 | 0.693322 |
| StyR.327_ | -4.41118 | -4.38169 | -4.44712 | 0.108451 | 1.150775 | 0.104691 | 0.464667 | 0.659624 | 0.774109 | 0.7951 | 0.863758 |
| StyR.328_ | -6.02941 | -5.82035 | -6.36066 | -0.63449 | 2.139633 | -0.23464 | 0.343771 | 0.493822 | 0.64861 | 0.6489 | 0.751892 |
| StyR.329a_ | -1.15091 | -1.43671 | -0.86425 | 0.508405 | 0.378695 | 1.270574 | 0.093271 | 0.129983 | 0.279206 | 0.2055 | 0.362391 |
| StyR.329b_ | -3.66437 | -3.75181 | -3.53687 | 0.164449 | 0.787992 | 0.181106 | 0.422385 | 0.659527 | 0.772198 | 0.7433 | 0.829979 |
| StyR.336_ | -3.66398 | -4.07839 | -3.21097 | 0.880097 | 0.648226 | 1.318062 | 0.06529 | 0.12568 | 0.264771 | 0.1937 | 0.341349 |
| StyR.341_ | -7.4707 | -7.19758 | -7.62873 | -0.6971 | 3.559158 | -0.17029 | 0.412 | 0.634203 | 0.753707 | 0.766 | 0.842149 |
| StyR.354_ | -7.37524 | -7.21419 | -7.58799 | -0.49498 | 2.465869 | -0.15446 | 0.40573 | 0.532833 | 0.671503 | 0.6728 | 0.76798 |
| StyR.358_ | -4.76383 | -4.73662 | -4.79805 | -0.07272 | 0.861823 | -0.07424 | 0.461333 | 0.548427 | 0.678763 | 0.6688 | 0.764876 |
| StyR.362_ | -5.45673 | -5.84142 | -5.20766 | 0.63173 | 1.558508 | 0.357847 | 0.311126 | 0.450467 | 0.603488 | 0.5853 | 0.698621 |
| StyR.369_ | -6.90163 | -5.6361 | -7.97548 | -2.1623 | 2.458329 | -0.81773 | 0.150667 | 0.259814 | 0.419832 | 0.3145 | 0.466567 |
| StyR.377_ | 1.536656 | 1.486525 | 1.590729 | 0.067708 | 0.244825 | 0.269074 | 0.393333 | 0.683074 | 0.796567 | 0.8245 | 0.887756 |
| StyR.378_ | -8.00842 | -8.06339 | -7.96355 | 0.033279 | 2.901568 | 0.009882 | 0.493005 | 0.565134 | 0.696486 | 0.6849 | 0.776918 |
| StyR.44a_ | -1.71333 | -1.54271 | -1.95007 | -0.27756 | 0.613295 | -0.41524 | 0.337109 | 0.560667 | 0.701403 | 0.6951 | 0.792378 |
| StyR.44b_ | 0.109735 | 0.164341 | -0.04605 | -0.17431 | 0.265708 | -0.56185 | 0.307333 | 0.487399 | 0.641525 | 0.6362 | 0.755012 |
| StyR.44c_ | -2.8253 | -2.84969 | -2.7848 | 0.006209 | 0.726935 | 0.007592 | 0.496667 | 0.705855 | 0.810457 | 0.8283 | 0.889182 |
| StyR.44d_ | -2.66105 | -2.4742 | -2.88101 | -0.42593 | 0.572689 | -0.68473 | 0.207195 | 0.323075 | 0.487083 | 0.4412 | 0.579309 |
| StyR.44e_ | -4.63992 | -4.60389 | -4.67836 | -0.10423 | 0.958908 | -0.10159 | 0.4497 | 0.584413 | 0.708416 | 0.6893 | 0.781127 |
| StyR.44f_ | -4.71516 | -4.651 | -4.79923 | 0.215865 | 1.989216 | 0.103868 | 0.453333 | 0.730211 | 0.830034 | 0.886 | 0.928503 |
| StyR.44g_ | -4.4345 | -3.97578 | -4.76141 | -0.63769 | 1.20794 | -0.477 | 0.299333 | 0.509237 | 0.650226 | 0.5825 | 0.706421 |
| StyR.51_ | 0.927169 | 1.068701 | 0.750655 | -0.46422 | 0.339676 | -1.3954 | 0.014667 | 0.09126 | 0.235223 | 0.1136 | 0.253674 |
| StyR.55_ | 3.928284 | 4.056405 | 3.542918 | -0.44677 | 0.197573 | -1.37959 | 0.101333 | 0.126358 | 0.289798 | 0.1888 | 0.355048 |
| StyR.8_ | 2.125533 | 2.266751 | 1.720507 | -0.51722 | 0.193156 | -2.07713 | 0.002002 | 0.059681 | 0.186333 | 0.1023 | 0.239604 |
| StyR.88_ | -0.47371 | -0.31615 | -0.73659 | -0.43947 | 0.306507 | -1.39957 | 0.056 | 0.112649 | 0.259551 | 0.1622 | 0.311474 |
| StyR.90_ | -5.58723 | -5.27439 | -5.89147 | -0.6583 | 1.245618 | -0.47685 | 0.292667 | 0.434147 | 0.579718 | 0.5442 | 0.661219 |
| StyR.99_ | -1.31159 | -1.31032 | -1.31289 | -0.0274 | 0.453209 | -0.05648 | 0.47435 | 0.700931 | 0.807614 | 0.8271 | 0.891426 |
| tp2_ | -6.1837 | -6.05725 | -6.27031 | -0.22346 | 1.760059 | -0.10937 | 0.447702 | 0.626895 | 0.745357 | 0.7355 | 0.818123 |
| tp28_ | -0.86684 | -0.49218 | -1.16911 | -0.668 | 0.279214 | -2.32778 | 0.004668 | 0.029518 | 0.121312 | 0.1027 | 0.240082 |
| tpk3_ | -0.55625 | -0.402 | -0.77135 | -0.43704 | 0.373752 | -1.18146 | 0.078 | 0.137249 | 0.293735 | 0.193 | 0.345476 |
| tpke11_ | 3.698762 | 3.714894 | 3.682606 | -0.03634 | 0.156203 | -0.22895 | 0.382412 | 0.721906 | 0.82659 | 0.8008 | 0.874482 |
| tpke70_ | -1.75101 | -2.21852 | -1.63151 | 0.563263 | 0.504373 | 0.947067 | 0.177215 | 0.242177 | 0.416816 | 0.3619 | 0.513177 |
| zipA.leader_ | 1.630389 | 1.770236 | 1.555698 | -0.21238 | 0.159257 | -1.23933 | 0.08 | 0.144852 | 0.304948 | 0.202 | 0.356881 |

### Table S3

## EE

| Gene | Manual_Name | Mean_exp | wt_exp | tnpA_exp | diff.btw | diff.win | effect | BH.q |
|---|---|---|---|---|---|---|---|---|
| SL1344_RS18810_cspA | cspA | 6.419062741 | 7.097225009 | 5.642287591 | -1.488111047 | 0.208209484 | -7.06767878 | 0.014046512 |
| SL1344_RS15245_cysI | cysI | 4.056164126 | 4.686254003 | 3.347055128 | -1.395832738 | 0.204092842 | -6.906043779 | 0.046492757 |
| SL1344_RS18655_ | yhjR | -0.175635645 | 0.420255151 | -0.936960747 | -1.371876014 | 0.284105542 | -4.805558158 | 0.044874286 |
| SL1344_RS06145_cspH | cspH | 0.453623017 | 0.974281324 | -0.326425646 | -1.31680604 | 0.31948141 | -4.138720827 | 0.048052754 |
| SL1344_RS10925_yegD | yegD | 1.105670054 | 1.732708717 | 0.448425334 | -1.286800292 | 0.160963891 | -7.967381483 | 0.021527603 |
| SL1344_RS15250_cysJ | cysJ | 3.961420778 | 4.518189381 | 3.292039642 | -1.276365694 | 0.1891843 | -6.836986975 | 0.034873732 |
| SL1344_RS22040_ | eptA | 3.174909785 | 3.722117569 | 2.449211881 | -1.267158258 | 0.119887168 | -10.43306078 | 0.019521133 |
| SL1344_RS15255_ptpS | ptpS | 1.914132541 | 2.522176035 | 1.267214467 | -1.26413147 | 0.116482635 | -10.82576655 | 0.014528149 |
| SL1344_RS12540_cysW | cysW | 2.395311822 | 2.967392289 | 1.74293836 | -1.231334168 | 0.124414141 | -9.766974013 | 0.02611489 |
| SL1344_RS21985_ | yjcO | 0.733936926 | 1.302722343 | 0.071026303 | -1.227867633 | 0.16945707 | -7.180261678 | 0.026149853 |
| SL1344_RS10700_ | ugd | 1.00308898 | 1.537990928 | 0.380413972 | -1.15610963 | 0.14200896 | -8.162686998 | 0.024494651 |
| SL1344_RS18650_yhjQ | yhjQ | 1.410054478 | 1.923803022 | 0.787398733 | -1.137689652 | 0.119280818 | -9.489672369 | 0.019448682 |
| SL1344_RS18135_malT | malT | 2.670235292 | 3.201441739 | 2.074507441 | -1.135073597 | 0.101569693 | -11.26541326 | 0.026045236 |
| SL1344_RS12535_cysA | cysA | 2.374848878 | 2.896759145 | 1.854442429 | -1.130127519 | 0.233502511 | -4.926656611 | 0.038562095 |
| SL1344_RS23255_yjiY | yjiY | 2.067857536 | 2.595633757 | 1.522075098 | -1.118925448 | 0.198855483 | -5.605948099 | 0.027689682 |
| SL1344_RS10695_wzzB | wzzB | 2.656564197 | 3.189828041 | 2.123876797 | -1.075360559 | 0.156566045 | -6.930816896 | 0.028953611 |
| bcsA_bcsA | bcsA | 2.004876475 | 2.491151912 | 1.462831503 | -1.042287125 | 0.122861587 | -8.608564075 | 0.023263476 |
| IsrE_isrE | isrE | -0.173774023 | 0.285604687 | -0.751933646 | -1.039982266 | 0.196107162 | -5.201415676 | 0.038049167 |
| SL1344_RS07340_ | ydgI | 2.452740525 | 2.907362981 | 1.894330745 | -1.024775617 | 0.121449548 | -8.453049834 | 0.020437285 |
| SL1344_RS09100_ | yoaB | 3.144017303 | 2.639373618 | 3.66649123 | 1.006263192 | 0.132748776 | 7.564152963 | 0.023325851 |
| SL1344_RS05780_ | flgA | 4.029801777 | 3.576962075 | 4.583639094 | 1.008607651 | 0.11579271 | 8.494163025 | 0.041110794 |
| SL1344_RS05785_ | flgB | 3.973523968 | 3.496875889 | 4.523586055 | 1.016581265 | 0.109591592 | 9.207515444 | 0.019511564 |
| SL1344_RS05815_ | flgH | 3.445989793 | 2.874819218 | 3.906542298 | 1.023076772 | 0.136631169 | 7.460242717 | 0.020330782 |
| SL1344_RS09225_yebR | yebR | 3.718558062 | 3.199604481 | 4.234652987 | 1.04112789 | 0.069281914 | 15.02926838 | 0.010918994 |
| SL1344_RS09095_ | yoaA | 2.462091382 | 1.933616787 | 2.978931712 | 1.041181092 | 0.094314296 | 11.05197723 | 0.014451685 |
| SL1344_RS05790_ | flgC | 3.537710028 | 3.001742434 | 4.05199206 | 1.041910424 | 0.074261154 | 13.98464132 | 0.010865385 |
| SL1344_RS03690_ | gltA | 3.728976152 | 3.147925467 | 4.190727357 | 1.045723005 | 0.103720696 | 10.01729337 | 0.019069151 |
| flgF_flgF | flgF | 4.59508316 | 4.105600206 | 5.169695436 | 1.053106889 | 0.067903461 | 14.99793395 | 0.022081965 |
| SL1344_RS05800_ | flgE | 4.885416065 | 4.408407516 | 5.470073318 | 1.053790504 | 0.064668177 | 15.79503915 | 0.027562432 |
| SL1344_RS07740_ | STM1554 | 1.475745489 | 0.832106621 | 1.897774495 | 1.056958012 | 0.149692772 | 6.850728617 | 0.041195799 |
| SL1344_RS09370_ | STM1873 | 2.746360326 | 2.136551719 | 3.193945875 | 1.064875792 | 0.130324294 | 8.211022743 | 0.022955514 |
| SL1344_RS15800_ | gcvH | 1.941878303 | 1.327121114 | 2.43282326 | 1.069604813 | 0.180625722 | 5.903380874 | 0.028144936 |
| SL1344_RS23265_ | STM4534 | 1.338709519 | 0.778542431 | 1.86143451 | 1.089521975 | 0.150337644 | 7.179003978 | 0.025271759 |
| SL1344_RS09150_ | STM1832 | 2.807353952 | 2.265611805 | 3.368920386 | 1.096496043 | 0.16147347 | 6.822095241 | 0.022676963 |
| SL1344_RS07805_ | adhP | 1.503038193 | 0.90073481 | 1.987865449 | 1.101896684 | 0.16912114 | 6.46362015 | 0.029102611 |
| SL1344_RS18600_ | yhjG | 2.515971863 | 1.953137607 | 3.080778592 | 1.117755114 | 0.109643032 | 10.22339074 | 0.015306698 |
| flgD_flgD | flgD | 4.188347548 | 3.627937578 | 4.731515601 | 1.119249887 | 0.121873544 | 9.311773336 | 0.026210129 |
| SL1344_RS24700_ | SL1344_RS24700 | 3.391146397 | 2.848051222 | 3.970362108 | 1.121484362 | 0.105939593 | 10.62738386 | 0.016859288 |
| SL1344_RS09420_ | purT | 0.483272819 | -0.036455482 | 1.102424262 | 1.122801263 | 0.222942407 | 4.9093638 | 0.045013561 |
| SL1344_RS05810_ | flgG | 5.322672406 | 4.795727729 | 5.926962459 | 1.130926504 | 0.077553843 | 14.29196198 | 0.026137317 |
| SL1344_RS09405_ | yebE | 0.969512581 | 0.366512837 | 1.491535214 | 1.131027984 | 0.176294543 | 6.541322189 | 0.039388961 |
| SL1344_RS09400_ | ptrB | 1.428854248 | 0.877220891 | 2.016499122 | 1.131281496 | 0.131544266 | 8.591019714 | 0.019198097 |
| SL1344_RS24695_ | SL1344_RS24695 | 3.154313982 | 2.572006456 | 3.718378658 | 1.14507897 | 0.127504459 | 8.963596752 | 0.015782944 |
| rpsV_rpsV | rpsV | 1.805865304 | 1.278198915 | 2.458976333 | 1.145222536 | 0.197587593 | 5.724977832 | 0.034093618 |

| | | | | | | | | |
|---|---|---|---|---|---|---|---|---|
| SL1344_RS02865_ | STM0567 | 2.191308179 | 1.56526536 | 2.728549631 | 1.154670559 | 0.175387203 | 6.533385724 | 0.023184017 |
| SL1344_RS09375_membrane.protein | STM1874 | 2.59139072 | 1.968837202 | 3.123232828 | 1.161655302 | 0.104941061 | 10.95922902 | 0.0140794 |
| SL1344_RS22270_rflM | rflM | 2.869984061 | 2.319415566 | 3.445287596 | 1.164674856 | 0.179126237 | 6.557579275 | 0.027991014 |
| SL1344_RS08100_ | ydcI | 0.966232427 | 0.28584774 | 1.452500709 | 1.165156381 | 0.157404971 | 7.326805296 | 0.030017308 |
| SL1344_RS24820_ | yncL | 3.510508891 | 2.909135295 | 4.160676767 | 1.20724756 | 0.174522804 | 6.817833521 | 0.028618604 |
| SL1344_RS09545_ | yecM | 0.905197678 | 0.240927699 | 1.444045942 | 1.207413411 | 0.169032331 | 7.150695522 | 0.032247258 |
| SL1344_RS16310_ | STM3154 | 3.323443559 | 2.654324924 | 3.861409786 | 1.209591146 | 0.122264025 | 10.05082747 | 0.038820858 |
| SLnc0014_ | SLnc0014 | 0.789820856 | 0.084169824 | 1.30706173 | 1.220894307 | 0.194569553 | 6.259811614 | 0.046883662 |
| SL1344_RS14885_hilD | hilD | 2.377187541 | 1.765622162 | 2.917304772 | 1.221617672 | 0.210001924 | 5.902644703 | 0.023782471 |
| SL1344_RS16325_ | yghA | 1.882296643 | 1.263606555 | 2.48915854 | 1.230675199 | 0.160152667 | 7.705366021 | 0.01994067 |
| SL1344_RS09290_ | STM1859 | -1.004119291 | -1.665272554 | -0.426129966 | 1.23456574 | 0.280478715 | 4.304154955 | 0.049350378 |
| SL1344_RS09540_ | cutC | 1.242112218 | 0.631870796 | 1.886644314 | 1.243544764 | 0.139455819 | 8.841092661 | 0.018737903 |
| SL1344_RS09795_lysine.N.methylase | fliB | 2.939225739 | 2.245828739 | 3.509861644 | 1.255046319 | 0.184938424 | 6.792064962 | 0.023670409 |
| SL1344_RS08975_ycgR | ycgR | 3.077798169 | 2.419497872 | 3.666727476 | 1.256569859 | 0.095413197 | 13.16561674 | 0.01071182 |
| SL1344_RS16750_ | tdcE | 1.128497179 | 0.43723571 | 1.716721458 | 1.274162927 | 0.145058153 | 8.757947792 | 0.020263742 |
| SL1344_RS09760_yecC | yecC | 1.481362513 | 0.798515317 | 2.078202304 | 1.274841766 | 0.180394433 | 7.082569435 | 0.023568646 |
| STnc840_ | STnc840 | 1.959239357 | 1.272219215 | 2.558733575 | 1.277608432 | 0.169340343 | 7.611889864 | 0.019827995 |
| SL1344_RS09770_yedO | yedO | 1.528714396 | 0.873812395 | 2.132566892 | 1.279905843 | 0.211983096 | 6.145200452 | 0.033918072 |
| SL1344_RS09410_ | yebF | 1.312456198 | 0.619513186 | 1.91404377 | 1.29144454 | 0.145487336 | 8.950364423 | 0.019047414 |
| SL1344_RS16320_ | STM3156 | 1.99112977 | 1.304017801 | 2.622152166 | 1.326039822 | 0.113655994 | 11.71801523 | 0.012745591 |
| SL1344_RS05645_ymdA | ymdA | 1.040439866 | 0.249280718 | 1.66012414 | 1.340417466 | 0.267435362 | 4.971683887 | 0.036502328 |
| SL1344_RS05770_flgN | flgN | 4.588268723 | 3.910110795 | 5.260829108 | 1.341469121 | 0.138971468 | 9.61821181 | 0.012752666 |
| SL1344_RS16315_ | STM3155 | 1.991665067 | 1.214730231 | 2.572824279 | 1.346353041 | 0.149194096 | 8.877880664 | 0.034752948 |
| SL1344_RS16295_DUF2623.domain.containing.protein | yghW | 1.760898319 | 1.132006326 | 2.544147984 | 1.362099174 | 0.215723239 | 6.205765485 | 0.028346021 |
| SL1344_RS05775_flgM | flgM | 4.062044313 | 3.37721498 | 4.764763807 | 1.396879636 | 0.140937201 | 10.02597513 | 0.026335032 |
| flgL_flgL | flgL | 5.222227736 | 4.523908198 | 5.926007174 | 1.400794137 | 0.053833249 | 26.19795106 | 0.007909151 |
| SL1344_RS03965_modC | modC | 3.162052666 | 2.45883454 | 3.87228866 | 1.404521791 | 0.110874414 | 12.68339556 | 0.014375108 |
| SL1344_RS07945_srfB | srfB | 3.941243697 | 3.222339444 | 4.645787207 | 1.412616277 | 0.149043599 | 9.380226472 | 0.016223079 |
| SL1344_RS06460_DUF1496.domain.containing.protein | STM1300 | 2.338065514 | 1.607537276 | 3.029339879 | 1.422534346 | 0.192821339 | 7.493069104 | 0.029723294 |
| SL1344_RS08990_ycgO | ycgO | 3.185836132 | 2.491779411 | 3.935023308 | 1.424764616 | 0.124239944 | 11.36350759 | 0.012360327 |
| SL1344_RS16300_methyl.accepting.chemotaxis.protein | STM3152 | 4.491344177 | 3.800061179 | 5.243814121 | 1.425787671 | 0.113734546 | 12.49423159 | 0.009288144 |
| SL1344_RS24580_ | SL1344_RS24580 | 1.524403395 | 0.69579905 | 2.12818127 | 1.428800318 | 0.137905266 | 10.281803 | 0.021893522 |
| SL1344_RS07940_hypothetical.protein | srfA | 3.551981359 | 2.844849036 | 4.268887735 | 1.432641682 | 0.146668364 | 9.811859936 | 0.030125021 |
| RyeF_ryeF | RyeF | 0.453566074 | -0.307320228 | 1.133138641 | 1.439201608 | 0.171474545 | 8.381320097 | 0.02085438 |
| SL1344_RS07950_srfC | srfC | 3.547016733 | 2.842379854 | 4.27518789 | 1.43997097 | 0.12664869 | 11.39233849 | 0.023149596 |
| SL1344_RS23260_tsr | tsr | 6.495358272 | 5.784158144 | 7.236024179 | 1.44105851 | 0.058443698 | 24.56445079 | 0.004366308 |
| SL1344_RS09560_glycoside.hydrolase.family.105.protein | STM1911 | 1.706991799 | 0.908972887 | 2.397269187 | 1.491281532 | 0.143101581 | 10.47049142 | 0.014373563 |
| modB_modB | modB | 2.661557259 | 1.939182771 | 3.458710901 | 1.49350576 | 0.135045782 | 10.9038722 | 0.017374703 |
| SL1344_RS11875_cheV | cheV | 4.125388988 | 3.362715758 | 4.837653773 | 1.506334705 | 0.109813924 | 13.90262856 | 0.013115267 |
| SL1344_RS05640_csgC | csgC | 0.976996543 | 0.178561915 | 1.608132002 | 1.511012943 | 0.300181865 | 5.149707983 | 0.039620996 |
| SL1344_RS18605_yhjH | yhjH | 4.006014591 | 3.200422236 | 4.795318718 | 1.517188192 | 0.189557008 | 8.08753487 | 0.01215761 |
| SraC_sraC | SraC | 3.412784354 | 2.511625464 | 4.017645085 | 1.517567166 | 0.219345377 | 6.832444608 | 0.025623652 |
| SL1344_RS05830_flgK | flgK | 5.48249124 | 4.696100979 | 6.237171908 | 1.520414367 | 0.069572651 | 21.95421887 | 0.004533891 |
| SL1344_RS14845_hilC | hilC | 2.001293974 | 1.22961975 | 2.758533573 | 1.534440364 | 0.213840994 | 7.266082718 | 0.038548633 |
| SL1344_RS16630_aer | aer | 2.165888621 | 1.338086331 | 2.900965209 | 1.538151745 | 0.200264204 | 7.644209677 | 0.027769122 |

| | | | | | | | | |
|---|---|---|---|---|---|---|---|---|
| SL1344_RS21960_nrfC | nrfC | 0.174866838 | -0.753517624 | 0.824665373 | 1.576726583 | 0.248267655 | 6.269139336 | 0.029574253 |
| SL1344_RS09565_flhE | flhE | 2.301557338 | 1.453344377 | 3.103185093 | 1.58235673 | 0.196109509 | 8.013426434 | 0.018292334 |
| SL1344_RS03955_modA | modA | 2.852457744 | 2.048243761 | 3.636485974 | 1.587469869 | 0.104869295 | 15.04353678 | 0.010146909 |
| StyR.305_ | StyR.305 | -1.037292468 | -1.887203065 | -0.292261718 | 1.594256964 | 0.284190631 | 5.578246205 | 0.037661378 |
| SL1344_RS16760_tdcC | tdcC | 1.130860337 | 0.273382915 | 1.886877934 | 1.616080292 | 0.161161287 | 10.01936348 | 0.019257218 |
| SL1344_RS09625_flhC | flhC | 3.984253677 | 3.110802381 | 4.711947848 | 1.627266893 | 0.132592009 | 12.20765094 | 0.011773641 |
| SL1344_RS06065_hypothetical.protein | STM3032 | 0.844358877 | -0.048497354 | 1.625823591 | 1.672955444 | 0.238843919 | 7.081164764 | 0.033251317 |
| SL1344_RS21955_nrfB | nrfB | -0.006362656 | -0.897808298 | 0.770519689 | 1.689144703 | 0.229714663 | 7.299945164 | 0.022467006 |
| InvR_invR | InvR | 2.491244036 | 1.557807708 | 3.254172862 | 1.707186946 | 0.220992542 | 7.970731659 | 0.048147822 |
| SL1344_RS16625_methyl.accepting.chemotaxis.protein | STM3216 | 5.202906586 | 4.383048865 | 6.133739964 | 1.745146533 | 0.072053801 | 23.23335138 | 0.019931416 |
| STnc3480_ | STnc3480 | -1.248781577 | -2.198794665 | -0.480545065 | 1.748002585 | 0.385357013 | 4.480641052 | 0.043859897 |
| SL1344_RS09570_flhA | flhA | 4.396241638 | 3.523909141 | 5.298935349 | 1.750764731 | 0.148069545 | 11.65176313 | 0.015431941 |
| SL1344_RS14355_iroB | iroB | -0.683541091 | -1.548743051 | 0.325388138 | 1.807573259 | 0.389868399 | 4.567343227 | 0.045446087 |
| SL1344_RS09775_fliY | fliY | 3.519804505 | 2.605492461 | 4.423625937 | 1.8120605 | 0.091714775 | 19.84384309 | 0.007299907 |
| STnc600_ | STnc600 | 0.995777 | -0.151716553 | 1.651475766 | 1.816017867 | 0.296942055 | 5.934040542 | 0.043598535 |
| SL1344_RS09280_N.acetyltransferase | STM1857 | -0.103957665 | -1.091435017 | 0.744164262 | 1.825437165 | 0.305126502 | 6.010539767 | 0.042203885 |
| SL1344_RS16230_methyl.accepting.chemotaxis.protein | STM3138 | 2.16548682 | 1.100448581 | 3.000576917 | 1.901004377 | 0.154751023 | 12.36421511 | 0.031494751 |
| SL1344_RS09785_fliA | fliA | 5.545836614 | 4.631671583 | 6.482516976 | 1.910807628 | 0.170556176 | 11.34425348 | 0.014360643 |
| SL1344_RS09575_flhB | flhB | 4.011615192 | 3.074764659 | 5.013690734 | 1.917285917 | 0.115611269 | 16.39679935 | 0.010977844 |
| SL1344_RS09195_hypothetical.protein | STM1841 | -1.094272672 | -2.119651457 | -0.143684507 | 1.959326192 | 0.428207162 | 4.589435772 | 0.042394955 |
| SL1344_RS21865_HlyD.family.type.I.secretion.periplasmic.adaptor.subunit | siiD | -0.983469416 | -2.054224364 | -0.056688377 | 1.9778188 | 0.358244068 | 5.521630883 | 0.033906772 |
| STnc1405_ | STnc1405 | 0.117124101 | -0.918413438 | 1.05995444 | 2.017530243 | 0.264068823 | 7.752706736 | 0.021622798 |
| SL1344_RS09580_cheZ | cheZ | 5.344095707 | 4.319599453 | 6.288628513 | 2.027674712 | 0.173074143 | 11.90077242 | 0.012860661 |
| SL1344_RS08105_methyl.accepting.chemotaxis.protein | trg | 1.724570599 | 0.646934955 | 2.694411608 | 2.035191477 | 0.142868669 | 14.25616253 | 0.010960007 |
| SL1344_RS14860_orgA | orgA | 2.052963109 | 1.004755978 | 3.020835543 | 2.073637942 | 0.221278451 | 9.457268339 | 0.013502444 |
| SL1344_RS21855_siiB | siiB | -1.006959861 | -2.273669372 | -0.17372108 | 2.099873268 | 0.352077018 | 5.987093527 | 0.044391213 |
| SL1344_RS09780_fliZ | fliZ | 4.374024066 | 3.275929827 | 5.365909006 | 2.10218249 | 0.082471561 | 25.50883399 | 0.004634667 |
| SL1344_RS24725_IS91.family.transposase | SL1344_RS24725 | 0.791772113 | -0.418879083 | 1.694577913 | 2.142373223 | 0.239066419 | 9.026182196 | 0.016423523 |
| SL1344_RS09585_chemotaxis.regulator | cheY | 3.875979251 | 2.777772463 | 4.897820896 | 2.142883283 | 0.123922316 | 17.20640767 | 0.006634798 |
| SL1344_RS14920_sipA | sipA | 1.1345437 | -0.002838095 | 2.117925767 | 2.146530203 | 0.201971667 | 10.84802867 | 0.013636878 |
| SL1344_RS09590_chemotaxis.response.regulator.protein.glutamate.methylesterase | cheB | 3.785640167 | 2.734251455 | 4.890560207 | 2.155477294 | 0.194043159 | 11.13886326 | 0.013810541 |
| SL1344_RS18435_methyl.accepting.chemotaxis.citrate.transducer | tcp | 3.215709554 | 2.080109521 | 4.252169129 | 2.171417527 | 0.082611949 | 26.50373454 | 0.007834257 |
| STnc1285_ | STnc1285 | 2.531325255 | 1.414031333 | 3.610690248 | 2.186225905 | 0.2884833 | 7.563495233 | 0.046021047 |
| SL1344_RS09755_sdiA | sdiA | 2.316461944 | 1.145221925 | 3.388031482 | 2.217953869 | 0.17763552 | 12.44066321 | 0.011619053 |
| SL1344_RS14895_iagB | iagB | 0.490593634 | -0.795690406 | 1.358890311 | 2.218400274 | 0.332700585 | 6.737920779 | 0.024330525 |
| SL1344_RS09595_chemotaxis.protein.methyltransferase | cheR | 4.276969012 | 3.102008224 | 5.29559874 | 2.218814439 | 0.141643529 | 15.70235333 | 0.005571355 |
| RUF_125_ | cheM-cheR_IGR | 2.27289672 | 1.014857074 | 3.314760255 | 2.303885846 | 0.161421238 | 14.35085656 | 0.023903588 |
| SL1344_RS21850_siiA | siiA | -2.030717933 | -3.417798385 | -1.072829764 | 2.352665201 | 0.461687872 | 5.099693169 | 0.039649776 |
| SL1344_RS14840_spi4.transcriptional.regulator | sprB | -0.212758664 | -1.546407687 | 0.828219433 | 2.387181168 | 0.244206649 | 9.693109221 | 0.019394548 |
| RUF_219c_ | hilC-sprB_IGR | -0.437915969 | -1.737375986 | 0.645632625 | 2.395092641 | 0.373030012 | 6.346402666 | 0.026116961 |
| STnc205_ | STnc205 | 4.775777624 | 3.535626054 | 5.957002726 | 2.420647036 | 0.061165884 | 39.59002533 | 0.003438448 |
| motB_motB | motB | 5.1606932 | 3.915279527 | 6.386341578 | 2.434202409 | 0.133152282 | 18.32463247 | 0.004453934 |
| SL1344_RS09610_cheA | cheA | 6.251374771 | 5.087450782 | 7.521963039 | 2.436357858 | 0.15604642 | 15.71489294 | 0.004262471 |
| SL1344_RS14980_invC | invC | -0.126215919 | -1.436247212 | 1.025649803 | 2.442809027 | 0.249646491 | 9.791595798 | 0.017076322 |
| SL1344_RS14865_prgK | prgK | 2.009988149 | 0.695641671 | 3.171092611 | 2.484969702 | 0.179259219 | 13.95448227 | 0.011865632 |

| | | | | | | | | |
|---|---|---|---|---|---|---|---|---|
| SL1344_RS09605_CheW | CheW | 6.108829296 | 4.886034766 | 7.383245277 | 2.49046073 | 0.080004277 | 31.17887468 | 0.005890127 |
| SL1344_RS24715_mobA | mobA | 4.095642886 | 2.803286786 | 5.270092658 | 2.492733262 | 0.119804114 | 20.84744921 | 0.005594591 |
| spaN_spaN | spaN | -0.220006965 | -1.618936189 | 0.844193922 | 2.525143153 | 0.318283055 | 7.911793137 | 0.019945195 |
| SL1344_RS09675_hypothetical.protein | STM1934 | 2.22939568 | 0.914725207 | 3.414896258 | 2.535839001 | 0.205338606 | 12.48046964 | 0.013495271 |
| SL1344_RS10630_sopA | sopA | -0.71800026 | -2.191397152 | 0.308877974 | 2.537772168 | 0.405457535 | 6.278963215 | 0.035709128 |
| SL1344_RS09600_cheM | cheM | 5.064598122 | 3.7916604 | 6.324974701 | 2.547998652 | 0.118317071 | 21.58098578 | 0.012004357 |
| SL1344_RS24720_mobC | mobC | 1.866079893 | 0.318497624 | 2.930481108 | 2.602745094 | 0.237277291 | 10.60012456 | 0.037093481 |
| SL1344_RS14950_spaR | spaR | -1.318185988 | -2.652955992 | 0.027909043 | 2.626061566 | 0.42807728 | 6.115346478 | 0.030399477 |
| SL1344_RS06600_lpxR | lpxR | -0.384177069 | -1.984439309 | 0.685957349 | 2.680526135 | 0.300368482 | 8.940706809 | 0.030479233 |
| SL1344_RS09620_motA | motA | 5.007755774 | 3.634214683 | 6.393933904 | 2.700483576 | 0.19828057 | 13.68946538 | 0.005279752 |
| SL1344_RS14965_spaO | spaO | -0.47875027 | -2.157408938 | 0.642431312 | 2.799956436 | 0.33273923 | 8.456398132 | 0.029330153 |
| SL1344_RS14955_spaQ | spaQ | -1.133876649 | -2.771170661 | 0.068431665 | 2.810845869 | 0.375017287 | 7.508039557 | 0.027067618 |
| SL1344_RS14925_sipD | sipD | -0.085266264 | -1.769775083 | 1.066234888 | 2.842701841 | 0.430816665 | 6.442256975 | 0.030177049 |
| SL1344_RS14960_spaP | spaP | -0.63790011 | -2.419606163 | 0.507624045 | 2.928857998 | 0.386126113 | 7.61693687 | 0.04304549 |
| SL1344_RS22150_rtsA | rtsA | 0.452902666 | -1.062321105 | 1.792610775 | 2.941557787 | 0.437518983 | 6.899931821 | 0.031997126 |
| SL1344_RS14890_hilA | hilA | 0.46216544 | -1.068878221 | 1.844773165 | 2.94633598 | 0.320253329 | 9.255629251 | 0.020626093 |
| SL1344_RS14990_invA | invA | 0.682471625 | -0.936556786 | 2.080826505 | 3.018579431 | 0.204182441 | 14.85897979 | 0.010021995 |
| SL1344_RS14870_prgJ | prgJ | 0.822571265 | -1.005174376 | 2.295626704 | 3.299913064 | 0.213358023 | 15.38494333 | 0.02107073 |
| SL1344_RS05355_sigE | sigE | 0.800489393 | -1.01830937 | 2.337659121 | 3.366385307 | 0.186194022 | 18.14525321 | 0.011740449 |
| SL1344_RS14930_sipC | sipC | 2.327392381 | 0.538687681 | 3.927122235 | 3.384909767 | 0.123010798 | 27.62603278 | 0.005578486 |
| SL1344_RS14995_invE | invE | 0.132476992 | -1.92046863 | 1.56917593 | 3.502716262 | 0.304574765 | 11.45221763 | 0.021052468 |
| SL1344_RS22145_rtsB | rtsB | -1.2442888 | -3.49905731 | 0.089942354 | 3.589880254 | 0.489967752 | 7.179369027 | 0.03777489 |
| SL1344_RS14935_sipB | sipB | 1.488832336 | -0.557600678 | 3.145242582 | 3.712649722 | 0.252036627 | 14.72291981 | 0.018904594 |
| SL1344_RS14880_prgH | prgH | 1.097127461 | -1.013791909 | 2.692927343 | 3.712893743 | 0.208255375 | 17.90356227 | 0.017272341 |
| SL1344_RS14875_prgI | prgI | 0.06611795 | -2.181916865 | 1.556241612 | 3.75119875 | 0.432206482 | 8.557957982 | 0.043340865 |
| sopB_sopB | sopB | 0.751940032 | -1.34041714 | 2.419716275 | 3.797656393 | 0.280755375 | 13.55305934 | 0.010570109 |
| SL1344_RS14940_sicA | sicA | 0.104307709 | -2.088841248 | 1.710310908 | 3.800894935 | 0.299777447 | 12.76025132 | 0.025408169 |
| SL1344_RS15000_invG | invG | 0.481355901 | -1.625204761 | 2.254505709 | 3.874918355 | 0.234489853 | 16.4774541 | 0.012041437 |
| SL1344_RS15005_invF | invF | 0.526054695 | -1.758653107 | 2.230655994 | 4.028525175 | 0.394680239 | 10.26070057 | 0.041715541 |
| SL1344_RS13925_sopE | sopE | 0.434435894 | -2.428513668 | 2.474587091 | 4.933570915 | 0.361997062 | 13.68190033 | 0.023873407 |
| SL1344_RS09265_sopE2 | sopE2 | -0.715698333 | -3.649321815 | 1.299861847 | 4.959373015 | 0.432247897 | 11.43210645 | 0.025318374 |

# ME

| Gene | Mean_exp | wt_exp | tnpA_exp | diff.btw | diff.win | effect | BH.q |
|---|---|---|---|---|---|---|---|
| SL1344_RS09600_cheM | 5.313096221 | 3.945069949 | 6.327482473 | 2.293831679 | 0.282546149 | 8.058211331 | 0.044105446 |
| SL1344_RS09595_chemotaxis.protein.methyltransferase | 4.276339117 | 3.223302266 | 5.24220378 | 1.911027479 | 0.216183871 | 8.968186724 | 0.021658227 |
| SL1344_RS09590_chemotaxis.response.regulator.protein.glutamate.methylesterase | 3.81034308 | 2.943074892 | 4.757886144 | 1.817268639 | 0.135259953 | 13.47054511 | 0.04948947 |
| SL1344_RS09605_CheW | 6.259060363 | 5.362645786 | 7.397403443 | 2.029843943 | 0.136003083 | 14.49714706 | 0.043913517 |
| SL1344_RS09580_cheZ | 5.267001926 | 4.312117338 | 6.214357267 | 1.915775088 | 0.089085338 | 21.54586406 | 0.016101994 |
| flgD_flgD | 4.029160576 | 3.543110345 | 4.541567768 | 1.069428666 | 0.155134549 | 6.937712476 | 0.033432083 |
| SL1344_RS05830_flgK | 5.499988746 | 4.748293183 | 6.130492834 | 1.381092731 | 0.266639678 | 5.3729771 | 0.047211767 |
| SL1344_RS09575_flhB | 3.40364242 | 2.400015285 | 4.335952996 | 2.002389768 | 0.322813581 | 6.548941197 | 0.043415737 |
| SL1344_RS09785_fliA | 5.375667337 | 4.465469404 | 6.38016751 | 1.970516075 | 0.239510393 | 8.227138263 | 0.037943293 |
| SL1344_RS09780_fliZ | 4.662276312 | 3.665553667 | 5.54323743 | 1.953904365 | 0.210500175 | 9.405472143 | 0.020573912 |
| SL1344_RS09560_glycoside.hydrolase.family.105.protein | 1.050008137 | 0.239598326 | 1.645332862 | 1.427258739 | 0.189861285 | 7.644935239 | 0.041549685 |
| SL1344_RS14890_hilA | 1.262688343 | 0.193604829 | 2.359693773 | 2.108941992 | 0.237274655 | 8.911363504 | 0.03864478 |
| SL1344_RS06065_hypothetical.protein | 0.332317142 | -0.609470806 | 1.160606621 | 1.780848326 | 0.171435463 | 10.46275179 | 0.031454413 |
| SL1344_RS07940_hypothetical.protein | 3.225944222 | 2.474355735 | 4.108031717 | 1.565700497 | 0.206909881 | 7.389563634 | 0.044176783 |
| SL1344_RS09675_hypothetical.protein | 1.711741766 | 0.397263311 | 2.906395905 | 2.50262221 | 0.13858671 | 18.03988309 | 0.022712892 |
| SL1344_RS14895_iagB | 0.963783237 | -0.129471078 | 1.926038301 | 2.005583187 | 0.264220148 | 7.554433999 | 0.037514827 |
| SL1344_RS14990_invA | 1.202701029 | -0.230979315 | 2.347211476 | 2.601637527 | 0.160703753 | 16.14451736 | 0.020379878 |
| SL1344_RS14980_invC | 0.169792706 | -0.955937495 | 1.188272753 | 2.155684713 | 0.209955296 | 10.22857523 | 0.030325846 |
| SL1344_RS14995_invE | 0.985765192 | -0.528145204 | 2.067016902 | 2.597086472 | 0.182858837 | 14.04864359 | 0.042809879 |
| SL1344_RS15005_invF | 1.552501907 | 0.111887234 | 2.683617203 | 2.572675138 | 0.153054626 | 16.62891534 | 0.035732696 |
| SL1344_RS15000_invG | 1.282412599 | -0.128884332 | 2.498852504 | 2.627483174 | 0.216574572 | 12.12511595 | 0.024683533 |
| SL1344_RS16300_methyl.accepting.chemotaxis.protein | 3.866711648 | 3.159928376 | 4.699414803 | 1.388319244 | 0.309365566 | 4.905674559 | 0.048569076 |
| SL1344_RS16625_methyl.accepting.chemotaxis.protein | 5.546734684 | 4.741203605 | 6.114300215 | 1.425790743 | 0.326122421 | 4.678756646 | 0.04676688 |
| SL1344_RS24715_mobA | 4.149796836 | 2.807088799 | 5.138473469 | 2.282819124 | 0.279875515 | 8.184517919 | 0.038751025 |
| SL1344_RS09620_motA | 5.067574502 | 3.610429536 | 6.148330881 | 2.529978836 | 0.353937917 | 7.19106547 | 0.033232465 |
| motB_motB | 5.175884004 | 4.008267648 | 6.225983125 | 2.113708605 | 0.234824888 | 9.234705873 | 0.020176911 |
| SL1344_RS21955_nrfB | -0.046488915 | -1.073007671 | 0.851345065 | 1.915449109 | 0.236353621 | 8.066405143 | 0.040769193 |
| SL1344_RS14860_orgA | 2.082579366 | 0.979788748 | 3.096607515 | 2.122266421 | 0.172422874 | 12.21744399 | 0.022920521 |
| SL1344_RS14880_prgH | 1.702464212 | 0.255852773 | 2.942434555 | 2.759712699 | 0.302378095 | 9.215940202 | 0.02566909 |
| SL1344_RS14875_prgI | 0.385362699 | -1.067738114 | 1.876523752 | 2.912282041 | 0.392086361 | 7.378251797 | 0.037708502 |
| SL1344_RS14870_prgJ | 1.310280897 | -0.113750943 | 2.675676501 | 2.770031767 | 0.249815646 | 11.16039759 | 0.026857306 |
| SL1344_RS14865_prgK | 2.23497102 | 1.076568662 | 3.394485835 | 2.297131968 | 0.187406972 | 12.23103803 | 0.025807149 |
| SL1344_RS14940_sicA | 0.318794582 | -1.544216347 | 1.928106023 | 3.521955923 | 0.330409238 | 10.72445589 | 0.028904491 |
| SL1344_RS05355_sigE | 1.030820806 | -0.810625713 | 2.848520627 | 3.505004703 | 0.434269269 | 8.188222177 | 0.028026183 |
| SL1344_RS14920_sipA | 1.18534644 | -0.157104952 | 2.334105594 | 2.543344006 | 0.28757154 | 8.961169503 | 0.034807389 |
| SL1344_RS14935_sipB | 1.925299485 | 0.230556095 | 3.471767331 | 3.266159246 | 0.221811896 | 14.84577921 | 0.025874124 |
| SL1344_RS14930_sipC | 2.528367853 | 0.584523648 | 4.363357329 | 3.751749189 | 0.187015584 | 19.903222 | 0.015517636 |
| SL1344_RS14925_sipD | -0.060121071 | -1.753138958 | 1.251552029 | 3.093917003 | 0.400110577 | 7.942234095 | 0.037385896 |
| SL1344_RS10630_sopA | -0.935780218 | -2.11292451 | 0.053497257 | 2.199911481 | 0.307200939 | 7.098623599 | 0.046630313 |
| sopB_sopB | 1.136603764 | -0.76841136 | 2.883886376 | 3.656526249 | 0.313439593 | 11.66799894 | 0.025019864 |

| | | | | | | | |
|---|---|---|---|---|---|---|---|
| SL1344_RS13925_sopE | 0.925993217 | -1.2051904 | 2.770729463 | 3.976163808 | 0.417099785 | 9.393196846 | 0.032021139 |
| spaM_spaM | -0.597855633 | -2.364291728 | 0.739884071 | 3.013190629 | 0.412308182 | 7.270615877 | 0.049991225 |
| spaN_spaN | 0.114803244 | -1.363757898 | 1.169074271 | 2.550457082 | 0.255255827 | 9.909438287 | 0.038263414 |
| SL1344_RS14965_spaO | -0.21400462 | -1.640519892 | 1.019100782 | 2.623494503 | 0.300219854 | 8.715630905 | 0.0405136 |
| SL1344_RS07945_srfB | 3.65595235 | 2.893827174 | 4.391199128 | 1.469222317 | 0.193212322 | 7.481062172 | 0.036645274 |
| SL1344_RS07950_srfC | 3.256693227 | 2.565054643 | 4.022870507 | 1.427674662 | 0.12106283 | 11.72660473 | 0.031488517 |
| SL1344_RS10695_wzzB | 2.505508582 | 3.276316189 | 1.814397904 | -1.452513268 | 0.132800096 | -10.88080659 | 0.033861131 |
| SL1344_RS08975_ycgR | 2.67757148 | 2.091519381 | 3.345178453 | 1.155171306 | 0.225749143 | 5.30189738 | 0.047839266 |
| SL1344_RS09760_yecC | 1.188718045 | 0.53425333 | 1.806004693 | 1.288178453 | 0.149530375 | 8.599505296 | 0.038884828 |
| RUF_125_ | 2.388038256 | 1.352205132 | 3.339584482 | 1.973399189 | 0.198449943 | 9.905683186 | 0.026904216 |
| SL1344_RS03540_ | 0.577001654 | 1.386343822 | -0.276323217 | -1.655232862 | 0.171122298 | -9.701576847 | 0.035170415 |
| SL1344_RS05785_ | 3.870752099 | 3.316239735 | 4.376561284 | 1.072605181 | 0.085218576 | 12.66204721 | 0.027262654 |
| SL1344_RS09085_ | 1.551432232 | 1.007897498 | 2.119096564 | 1.098668021 | 0.122796331 | 8.907509162 | 0.040189327 |
| SL1344_RS09090_ | 2.034847314 | 1.439331725 | 2.518276502 | 1.080844425 | 0.094689274 | 11.27655577 | 0.031072277 |
| SL1344_RS09095_ | 2.006444168 | 1.477093564 | 2.49769367 | 1.031286926 | 0.104453763 | 9.782567894 | 0.03769026 |
| SL1344_RS09100_ | 3.353336304 | 2.887743084 | 3.922846788 | 1.029059057 | 0.208427832 | 5.091116143 | 0.049255693 |
| SL1344_RS09410_ | 1.289524313 | 0.682646126 | 1.872132599 | 1.188610595 | 0.118382805 | 10.07590544 | 0.03386129 |
| SL1344_RS11620_ | 0.658182927 | 1.423376305 | -0.146711531 | -1.59577532 | 0.224861097 | -7.159589847 | 0.044983876 |
| SL1344_RS21710_ | 4.935007906 | 5.412539993 | 4.369051658 | -1.005947189 | 0.142497633 | -7.142363386 | 0.037715134 |
| SL1344_RS24695_ | 3.025713482 | 2.35205251 | 3.567425787 | 1.219010362 | 0.123879184 | 9.82383718 | 0.027121731 |
| STnc1405_ | 0.55831788 | -0.602044345 | 1.478684853 | 2.094301851 | 0.201427769 | 10.41841754 | 0.03477911 |
| STnc205_ | 4.98474965 | 3.950127717 | 5.999965898 | 2.048914648 | 0.117739832 | 17.31534077 | 0.015515064 |
| STnc600_ | 1.661594428 | 0.839949837 | 2.49506933 | 1.656738803 | 0.18276322 | 9.029311886 | 0.04533566 |

# LE

| Gene | Manual_Name | Mean_exp | wt_exp | tnpA_exp | diff.btw | diff.win | effect | BH.q |
|---|---|---|---|---|---|---|---|---|
| SL1344_RS10625_phsA | phsA | 1.290927914 | 3.693277907 | -0.790674718 | -4.409864556 | 0.322869246 | -13.31190435 | 0.037090442 |
| SL1344_RS10620_phsB | phsB | -0.430655497 | 1.596570602 | -2.330699823 | -3.794306937 | 0.52652843 | -7.220972135 | 0.035289319 |
| SL1344_RS10615_phsC | phsC | 0.585564164 | 2.538986753 | -1.220250223 | -3.690230455 | 0.416198649 | -8.957771871 | 0.032475203 |
| SL1344_RS22480_ | yjfO | 2.759592948 | 3.708077879 | 1.883424721 | -1.803231057 | 0.292895553 | -6.138375627 | 0.035856236 |
| SL1344_RS22475_ | yjfN | 1.123156449 | 1.839337984 | 0.229234297 | -1.630722108 | 0.197098402 | -8.363355526 | 0.028755184 |
| SL1344_RS23280_ | STM4537 | 1.017224714 | 1.674419581 | 0.115854124 | -1.556863436 | 0.167295186 | -9.313516812 | 0.041648734 |
| SL1344_RS23275_ | STM4536 | 0.491961954 | 1.191367842 | -0.320924254 | -1.545081387 | 0.224151662 | -6.974801525 | 0.042114782 |
| SL1344_RS23285_ | STM4538 | 1.312180248 | 2.069679913 | 0.566729225 | -1.499836985 | 0.134246252 | -11.17835897 | 0.025654383 |
| SL1344_RS23290_ | STM4539 | 1.631299129 | 2.250107369 | 0.827117658 | -1.432652623 | 0.169909413 | -8.482092233 | 0.028349642 |
| SL1344_RS23270_ | STM4535 | 0.810868751 | 1.409335441 | -0.001853448 | -1.385930195 | 0.20750296 | -6.587491435 | 0.042131025 |
| SL1344_RS13085_ | csiE | 0.944739211 | 1.641213008 | 0.242903595 | -1.385747111 | 0.167332387 | -8.19188283 | 0.032346417 |
| SL1344_RS10445_ | cbiF | 1.167357118 | 1.654850384 | 0.324643757 | -1.363595899 | 0.341695565 | -4.08254862 | 0.048151477 |
| SL1344_RS23190_ | STM4519 | 1.937131713 | 2.552657094 | 1.321704006 | -1.235546034 | 0.159566226 | -7.743276659 | 0.030600322 |
| SL1344_RS16770_tdcA | tdcA | 0.352110649 | 0.907879897 | -0.287584993 | -1.19037552 | 0.167298173 | -7.106900206 | 0.036067292 |
| SL1344_RS16530_ | glgS | 1.948164456 | 2.482084677 | 1.36346203 | -1.187765233 | 0.193576444 | -6.141435596 | 0.033745 |
| SL1344_RS10440_ | cbiG | 1.564519163 | 2.168902308 | 0.993757993 | -1.176256651 | 0.196386179 | -6.108585007 | 0.042780604 |
| SL1344_RS03300_ | ybeL | 1.961526477 | 2.478631877 | 1.325095713 | -1.162330062 | 0.198349356 | -5.908780131 | 0.043829821 |
| SL1344_RS06850_ | orf48 | -0.085651912 | 0.480091827 | -0.679116973 | -1.159865622 | 0.166059511 | -6.985349692 | 0.038520359 |
| SL1344_RS06385_ | yeaG | 2.566775924 | 3.169496892 | 2.036892558 | -1.142244545 | 0.121188994 | -9.589728651 | 0.033533746 |
| SL1344_RS10435_ | cbiH | 1.796857889 | 2.315716592 | 1.213204619 | -1.106242576 | 0.2191265 | -5.083821277 | 0.043430205 |
| SL1344_RS10420_ | cbiL | 1.248774275 | 1.728199831 | 0.646460058 | -1.091933502 | 0.215308345 | -5.07932759 | 0.042860861 |
| cbiC_cbiC | cbiC | 0.928616058 | 1.476348443 | 0.402992042 | -1.086375356 | 0.14108781 | -7.714548646 | 0.034310769 |
| SL1344_RS06590_pfkB | pfkB | 2.887780474 | 3.481245669 | 2.316837777 | -1.086103375 | 0.194319383 | -5.649673732 | 0.035089123 |
| SL1344_RS22110_ | dmsB | 0.514880633 | 1.000401601 | -0.087086063 | -1.069350043 | 0.216439113 | -4.846982516 | 0.049728115 |
| SL1344_RS10425_sirohydrochlorin.cobaltochelatase | cbiK | 1.214825407 | 1.726034336 | 0.670379389 | -1.055144539 | 0.146313217 | -7.226216362 | 0.036441125 |
| SL1344_RS10695_wzzB | wzzB | 2.462760485 | 2.936530734 | 1.886404828 | -1.050663505 | 0.086409295 | -12.12890653 | 0.028743245 |
| SL1344_RS22105_ | STM4305 | 2.419340593 | 2.893252461 | 1.884894868 | -1.025529267 | 0.130630568 | -7.947289312 | 0.030276115 |
| SL1344_RS21865_HlyD.family.type.I.secretion.periplasmic.adaptor.subunit | siiD | 1.917744369 | 2.36309073 | 1.33509801 | -1.005485316 | 0.221357894 | -4.48302217 | 0.045179379 |
| SL1344_RS24695_ | SL1344_RS24695 | 2.746355029 | 2.227789288 | 3.213026061 | 1.002081799 | 0.115960746 | 8.68188871 | 0.034727074 |
| InvR_invR | InvR | 5.404928328 | 4.805160527 | 5.945874534 | 1.019319985 | 0.268534187 | 3.855327159 | 0.046065955 |
| SL1344_RS05355_sigE | sigE | 4.323092984 | 3.74827914 | 4.769050438 | 1.023187034 | 0.222417626 | 4.828671217 | 0.041222957 |
| SL1344_RS24585_ | SL1344_RS24585 | 1.044176305 | 0.563754447 | 1.597344812 | 1.027525179 | 0.188671759 | 5.394119373 | 0.043091167 |
| SL1344_RS24710_ | SL1344_RS24710 | 2.490704145 | 2.009016705 | 3.080123526 | 1.049157152 | 0.17828828 | 5.840312449 | 0.043600342 |
| SL1344_RS06460_DUF1496.domain.containing.protein | STM1300 | 3.390192966 | 2.786387131 | 3.841275441 | 1.051917535 | 0.185375215 | 5.678240534 | 0.040286489 |
| SL1344_RS24700_ | SL1344_RS24700 | 2.953680888 | 2.421062195 | 3.453709149 | 1.053098038 | 0.10361033 | 10.20727551 | 0.025329818 |
| SL1344_RS03355_ | gltJ | 0.414580862 | -0.171536572 | 0.907920961 | 1.069019847 | 0.186597597 | 5.71788682 | 0.044611576 |
| SL1344_RS16315_ | STM3155 | 1.45328832 | 0.845573115 | 1.898657716 | 1.072091051 | 0.178745906 | 6.089776862 | 0.037444089 |
| SL1344_RS09080_fadD | fadD | 1.399017352 | 0.854205751 | 1.929453237 | 1.076222737 | 0.141858544 | 7.685715728 | 0.031710652 |
| SL1344_RS16320_ | STM3156 | 1.875703166 | 1.293635524 | 2.380692359 | 1.105865458 | 0.136336645 | 8.068810059 | 0.038433947 |
| SL1344_RS05785_ | flgB | 4.224448339 | 3.670610701 | 4.773971062 | 1.122774568 | 0.109103879 | 10.27096706 | 0.022554233 |
| SL1344_RS05645_ymdA | ymdA | 2.420349002 | 1.825576894 | 2.926079152 | 1.136119117 | 0.145467704 | 7.938423512 | 0.038884303 |
| SL1344_RS09770_yedO | yedO | 1.885322799 | 1.312977413 | 2.459284375 | 1.158977357 | 0.126796767 | 9.166468473 | 0.031201053 |
| SL1344_RS09480_ruvA | ruvA | 1.015054916 | 0.382429949 | 1.573814237 | 1.185612152 | 0.161991946 | 7.172271196 | 0.033541733 |
| SL1344_RS11875_cheV | cheV | 5.196217878 | 4.590090999 | 5.863350763 | 1.198259288 | 0.139365281 | 8.564099926 | 0.027639008 |

| | | | | | | | | |
|---|---|---|---|---|---|---|---|---|
| modB_modB | modB | 3.518414083 | 2.97099389 | 4.203977494 | 1.209374645 | 0.134170345 | 8.751911436 | 0.03328311 |
| SL1344_RS03965_modC | modC | 3.566850508 | 2.9610365 | 4.166220372 | 1.215675768 | 0.127287345 | 9.693167659 | 0.032631438 |
| SL1344_RS09765_yecS | yecS | 0.873191283 | 0.24874491 | 1.47285961 | 1.233019599 | 0.190524351 | 6.508030486 | 0.038506067 |
| SL1344_RS02245_ | cyoE | 4.276600419 | 3.519250964 | 4.904528508 | 1.235256587 | 0.23895501 | 5.052596904 | 0.041781385 |
| SL1344_RS24580_ | SL1344_RS24580 | 3.039726962 | 2.423385179 | 3.689233705 | 1.253418116 | 0.112676976 | 11.08912706 | 0.028288612 |
| SL1344_RS08975_ycgR | ycgR | 2.419402747 | 1.818051357 | 3.00982625 | 1.266647095 | 0.202474787 | 6.446846431 | 0.038754307 |
| SL1344_RS09760_yecC | yecC | 1.853845917 | 1.164173691 | 2.432362368 | 1.267564203 | 0.097996518 | 12.94740036 | 0.024991158 |
| SL1344_RS08995_ | dadX | 1.688519591 | 0.947554594 | 2.236047684 | 1.279661527 | 0.152275682 | 8.313653039 | 0.045372178 |
| SL1344_RS07940_hypothetical.protein | srfA | 3.645148982 | 2.976912658 | 4.243485026 | 1.281720484 | 0.102237349 | 12.62880293 | 0.024378467 |
| SL1344_RS09585_chemotaxis.regulator | cheY | 4.351922531 | 3.715895142 | 4.935886686 | 1.299337853 | 0.175683914 | 7.402682525 | 0.04820805 |
| SL1344_RS09750_yecF | yecF | 1.135943398 | 0.522387497 | 1.839185048 | 1.308137334 | 0.173309983 | 7.607132269 | 0.037755277 |
| SL1344_RS09000_ | dadA | 1.747058484 | 1.022279461 | 2.382674705 | 1.329549601 | 0.176820272 | 7.494961633 | 0.034906097 |
| SL1344_RS09745_hypothetical.protein | SL1344_RS09745 | -0.464752284 | -1.213602211 | 0.133483548 | 1.339945211 | 0.226194157 | 5.868579537 | 0.045897343 |
| SL1344_RS08105_methyl.accepting.chemotaxis.protein | | 5.156913303 | 4.517491317 | 5.845325412 | 1.358894667 | 0.205447894 | 6.822032701 | 0.031998831 |
| SL1344_RS09565_flhE | trg | 2.53543265 | 1.829092897 | 3.211487552 | 1.381043162 | 0.110627953 | 12.49250734 | 0.022373592 |
| SL1344_RS09580_cheZ | flhE | 5.328427255 | 4.650421125 | 6.04442897 | 1.395645689 | 0.157961086 | 8.670680927 | 0.044167407 |
| SL1344_RS09775_fliY | cheZ | 4.712990801 | 4.00870389 | 5.372420151 | 1.409437984 | 0.158481843 | 9.042002814 | 0.040708355 |
| SraC_sraC | fliY | 2.330926463 | 1.652613081 | 3.071847462 | 1.41510178 | 0.174296059 | 7.928786327 | 0.037524426 |
| SL1344_RS09785_fliA | SraC | 5.690989698 | 5.053939332 | 6.497668929 | 1.435604839 | 0.206682295 | 6.991151905 | 0.025557471 |
| SL1344_RS09590_chemotaxis.response.regulator.protein.glutamate.methylesterase | fliA | 3.996498621 | 3.251592219 | 4.676904225 | 1.442872472 | 0.168936792 | 8.655221264 | 0.026482396 |
| SL1344_RS22270_rflM | cheB | 0.74122764 | 0.006201957 | 1.477707793 | 1.462650897 | 0.202270693 | 7.24809101 | 0.035343242 |
| SL1344_RS09575_flhB | rflM | 3.752462534 | 2.966941264 | 4.479707939 | 1.510889968 | 0.085285049 | 17.69542005 | 0.016940183 |
| SL1344_RS09570_flhA | flhB | 4.388665527 | 3.575598965 | 5.12361301 | 1.549796954 | 0.091359576 | 16.93363152 | 0.015992042 |
| SL1344_RS09600_cheM | flhA | 5.891596997 | 5.17307005 | 6.725627433 | 1.553500319 | 0.128287242 | 11.80359547 | 0.03167266 |
| SL1344_RS09780_fliZ | cheM | 5.406820262 | 4.652751267 | 6.172187178 | 1.583131424 | 0.135996088 | 11.69399399 | 0.018876115 |
| RUF_125_ | cheM-cheR_IGR | 2.953623987 | 2.157361169 | 3.707795138 | 1.584738886 | 0.148371827 | 10.859854 | 0.02443091 |
| STnc1065_ | STnc1065 | 1.899877983 | 1.105794586 | 2.725704695 | 1.629249828 | 0.265720543 | 6.147946012 | 0.036784369 |
| SL1344_RS09595_chemotaxis.protein.methyltransferase | STM3216 | 4.343066509 | 3.523798452 | 5.188383857 | 1.664142971 | 0.111487817 | 14.92597369 | 0.018203639 |
| SL1344_RS09610_cheA | cheR | 6.52031626 | 5.769991651 | 7.431432961 | 1.666513821 | 0.181007079 | 9.389917567 | 0.029987376 |
| SL1344_RS16295_DUF2623.domain.containing.protein | cheA | 0.764719741 | -0.027515467 | 1.714579823 | 1.68375929 | 0.269992044 | 6.172968615 | 0.042030236 |
| SL1344_RS09265_sopE2 | yghW | 2.40607184 | 1.610598533 | 3.363772513 | 1.696475256 | 0.211223324 | 7.68408027 | 0.042295914 |
| SL1344_RS06065_hypothetical.protein | sopE2 | 0.643969998 | -0.280004731 | 1.422543061 | 1.705315804 | 0.131325353 | 12.86168401 | 0.024222867 |
| SL1344_RS09605_CheW | STM3032 | 6.623055132 | 5.801331055 | 7.442738484 | 1.71336886 | 0.160344074 | 10.68251592 | 0.018779572 |
| SL1344_RS09755_sdiA | CheW | 3.076884069 | 2.199990002 | 3.914655748 | 1.722203327 | 0.171625212 | 10.09504406 | 0.030788317 |
| SL1344_RS12530_cysM | sdiA | 1.274891249 | 0.369167813 | 2.171785171 | 1.747947063 | 0.215373985 | 8.110409604 | 0.028279235 |
| STnc205_ | cysM | 5.312566217 | 4.459858842 | 6.134658024 | 1.77285852 | 0.185704375 | 9.454673244 | 0.033792724 |
| motB_motB | motB | 5.233258489 | 4.35067775 | 6.18577104 | 1.81491788 | 0.155292937 | 11.8898515 | 0.017204722 |
| SL1344_RS09620_motA | motA | 5.144830796 | 4.323410774 | 6.158313381 | 1.839751593 | 0.234807597 | 7.995530697 | 0.025048265 |
| SL1344_RS09675_hypothetical.protein | STM1934 | 2.431989215 | 1.465341892 | 3.347786768 | 1.874875207 | 0.103025247 | 18.13038772 | 0.017415182 |
| SL1344_RS07600_ | STM1528 | -1.455604734 | -2.507786628 | -0.640426747 | 1.886387458 | 0.370909806 | 5.071514064 | 0.047575339 |
| SL1344_RS24720_mobC | mobC | 2.242567363 | 0.999972364 | 3.216878283 | 2.193524038 | 0.28812943 | 7.544898194 | 0.049724669 |
| SL1344_RS24715_mobA | mobA | 3.691919699 | 2.559730435 | 4.756351548 | 2.237970089 | 0.138081079 | 16.43199719 | 0.022829744 |
| SL1344_RS15170_ | ygbE | 0.837004374 | -0.869208621 | 2.23792286 | 3.122671985 | 0.235545046 | 13.4474373 | 0.03498313 |
| SL1344_RS08495_yciW | yciW | 0.912870063 | -0.815250544 | 2.672466231 | 3.386107474 | 0.467180801 | 7.26139573 | 0.029716246 |
| SL1344_RS12475_cysK | cysK | 5.327924957 | 3.226004485 | 7.158138172 | 3.569932956 | 0.491525535 | 6.95373487 | 0.028858037 |

| | | | | | | | | |
|---|---|---|---|---|---|---|---|---|
| SL1344_RS15175_cysC | cysC | 2.649839189 | 0.052627943 | 4.491644056 | 4.339759337 | 0.42429819 | 9.794084095 | 0.049253092 |
| SL1344_RS12540_cysW | cysW | 2.659288361 | 0.177617695 | 4.663416832 | 4.357972459 | 0.49009167 | 8.242932623 | 0.040642441 |
| SL1344_RS15180_cysN | cysN | 3.439915028 | 0.735726338 | 5.631472264 | 4.693766652 | 0.571900688 | 7.239451659 | 0.040062142 |
| SL1344_RS12545_cysU | cysU | 1.772297127 | -1.141026054 | 4.010328545 | 4.864584154 | 0.746319136 | 6.647100101 | 0.036951149 |
| SL1344_RS15185_cysD | cysD | 2.11015451 | -1.059073116 | 4.79519241 | 5.466643955 | 0.742856263 | 7.770180233 | 0.027522797 |
| SL1344_RS12550_cysP | cysP | 2.315584549 | -0.935938925 | 5.397265852 | 5.640318538 | 0.919480178 | 6.703985561 | 0.025723293 |

IS91 family
transposase
RS09540
RS05800
cysA
csgC
ptpS
flgF
siiA
RS07340
ryeF
RS07740
RS09195
RS14840
SLnc0014
RS05810
iroB
yhjH
hisF
RS09370
RS16325
isrE
spaQ
rtsA
flhC
RS03690
RS02865
nrfC
RS09290
cysJ
lpxR
RS05780
spaR
spaP
tdcC
ycgO
rtsB
RS15800
cspA
RS18435
STnc840
STnc1285

asmA
RS09150
yebR
flgN
siiB
aer
malT
tsr
RS09405
RS22040
RS07805
bcsA
RS16230
RS21985
yjiY
hilC
bcsR
rpsV
RS09280
flgL
STnc3480
flgM
RS05815
cysI
RS09400
styR
RS05790
RS24820
hilD
yegD
RS09420
RS23265
modA
RS09795
RS09375
RS16310
RUF_219c
RS08100
cspH
RS09545
yhjQ

RS16750

ME

RS09085
RS09090
spaM
malE
RS11620
RS03540

LE

RS24710
phsA
RS24585
RS09000
RS23190
RS23270
cysK
cbiC
RS23285
RS10445
RS10420
tdcA
yecS
yecF
RS10425
RS22110
RS22475
RS03355
RS22480
ruvA
RS15170
cysM
cysN
phsC
STnc1065
RS06850
RS02245
fadD
RS07600
RS23280
RS13085
cysD
yciW
phsB
RS22105
cysU
cysC
cysP
pfkB
RS06385
RS03300

RS10435
RS08995
RS10440
RS16530
RS23275
RS23290
RS09745

invG
sopB
orgA
STnc600
sopA
iagB
RS09100
sicA
invC
RS09560
prgJ
RS16625
spaN
srfC
hilA
invF
prgI
RS16300
sopE
prgK
sipA
flgK
nrfB
sipB
STnc1405
prgH
invE
invA
RS09095
sipD
RS09410
sipC
spaO
flgD
srfB

EE + LE

RS16315
mobC
RS16295
yedO
cysW
invR
RS21865
modB
fliY
RS24580
RS09585
ymdA
RS24700
sraC
rflM
sopE2
cheA
RS08105
modC
RS16320
cheV
sdiA
flhE
flhA
RS06460

EE + ME + LE

fliA
cheW
cheM
RS09590
RS06065
yecC
motA
ycgR
motB
flhB
mobA
RS24695
STnc205
RS05785
cheZ
sigE
RUF_125
fliZ
wzzB
RS09595
RS09675
RS07940

### Table S5

| Rank | Energy [kJ/mol] | z-Score | Interaction [dot-bracket] | mRNA [Start] | mRNA [End] | sRNA | Gene Annotation | Locus Tag | Strand | Genomic Coordinates |
|---|---|---|---|---|---|---|---|---|---|---|
| 1. | -18.41 | -4.47 | ((((((((((((((((.((((((&)))))))))))))))))))))) | 42 | 66 | 3-26 | L idonate transport protein | STM4482 idnT | - | 4727312-4725793 |
| 2. | -17.84 | -4.21 | ((((((((.((((((((&)))))))).)))))))) | -71 | -53 | 2-20 | soluble pyridine nucleotide transhydrogenase | STM4126 udhA | - | 4345580-4343980 |
| 3. | -16.94 | -3.81 | (((.((((((((((((((((((&)))))))...))))))))))).))) | 492 | 514 | 1-26 | putative regulatory protein | STM1704 yciT | + | 1798712-1799664 |
| 4. | -16.85 | -3.77 | ((((((((((((((((&)))).)))))))))))) | 735 | 750 | 1-17 | putative inner membrane protein | STM4443 | + | 4684778-4685754 |
| 5. | -16.83 | -3.76 | ((((((((...((((..((((((((&)))))..))))..))))....))))))))) | 73 | 99 | 4-33 | putative sugar kinase | STM4024.S yihV | + | 4234619-4235721 |
| 6. | -16.80 | -3.75 | ((((((((.(((((((((((((&)))..)))))))..)))...))))))))) | 1262 | 1284 | 4-32 | tRNA(Ile) lysidine synthetase | STM0236 mesJ | + | 278471-279963 |
| 7. | -16.75 | -3.73 | ((.((((((((((((&)))))))))))).)) | -22 | -7 | 25-40 | transposase | STM0946 Art200 | + | 1024966-1025624 |
| 8. | -16.75 | -3.73 | ((.((((((((((((&)))))))))))).)) | -22 | -7 | 25-40 | transposase for IS200 | STM4311 | + | 4559216-4559874 |
| 9. | -16.74 | -3.72 | ((.((((((((((((&)))))))))))).)) | -21 | -6 | 25-40 | transposase for IS200 | STM3479 | - | 3635818-3635160 |
| 10. | -16.74 | -3.72 | ((.((((((((((((&)))))))))))).)) | -21 | -6 | 25-40 | transposase | STM3032 | - | 3194976-3194318 |
| 11. | -16.74 | -3.72 | ((.((((((((((((&)))))))))))).)) | -21 | -6 | 25-40 | transposase | STM2471 | - | 2579989-2579331 |
| 12. | -16.65 | -3.68 | (((((((.((((((((&)))))))).))))))) | 123 | 138 | 25-40 | phage tail component L like protein | STM2592 | - | 2743043-2742145 |
| 13. | -16.60 | -3.66 | ((.((((((((((((&)))))))))))).)) | 30 | 45 | 25-40 | transposase for IS200 | STM1957 | - | 2046444-2045735 |
| 14. | -16.40 | -3.57 | (((((((.(((((((((((...((((((&))))))...))))))))))).))))))) | -21 | 6 | 14-41 | putative glutathione S transferase | STM0862 yliJ | - | 935453-934627 |
| 15. | -16.40 | -3.57 | ((((((((((((((&)))))))))))))) | 1419 | 1432 | 4-17 | potassium efflux protein KefA | STM0478 aefA | + | 534326-537888 |
| 16. | -16.34 | -3.54 | (((((((...((((((((.....(((((&)))))......))).))))))..))))))) | 2783 | 2809 | 1-28 | putative minor tail protein | STM0918 | + | 988425-991786 |
| 17. | -16.20 | -3.48 | ((((((((.((((((((&)))))))).)))))))) | 13 | 29 | 1-17 | tail assembly protein K like | STM2591 | - | 2742200-2741401 |
| 18. | -16.03 | -3.4 | ((((((((((((((.(((((.((((((&))))...)).))))))))))))))))))) | -96 | -70 | 4-32 | putative cytoplasmic protein | STM2329 | - | 2440245-2439893 |
| 19. | -16.02 | -3.4 | (((((((..(((((((..(((((((((((&))))))..))))))))))))..))))))) | -132 | -104 | 2-30 | 3 octaprenyl 4 hydroxybenzoate carboxy lyase | STM2356 ubiX | - | 2467351-2466582 |
| 20. | -16.01 | -3.39 | (((((((((((.(((((((.....(((((&)))))...))))))))))))))))))) | 71 | 99 | 2-27 | hypothetical protein STM4562 | STM4562 | + | 4816407-4816786 |
| 21. | -15.98 | -3.38 | (((((((..(((((((((...((((&))))...))))))))))))))))) | 1218 | 1242 | 4-26 | putative dehydrogenase | STM0858 | + | 930132-932017 |
| 22. | -15.90 | -3.35 | ((((((.........((((((((((&))))).))))))......)))))) | 355 | 380 | 4-28 | 30S ribosomal protein S5 | STM3423 rpsE | - | 3588690-3587987 |
| 23. | -15.90 | -3.35 | ((((....(((.((.((((((((((&)))))))))).)).)))...)))) | 416 | 441 | 24-48 | putative dithiobiotin synthetase | STM1489 ynfK | + | 1566247-1567142 |
| 24. | -15.82 | -3.31 | ((((((((((((((((&)))).)))))))))))) | -53 | -38 | 1-17 | putative inner membrane protein | STM4444 | + | 4685566-4686410 |
| 25. | -15.78 | -3.29 | (((((((..(((((((((&))))))))).))))))) | 338 | 355 | 1-17 | 50S ribosomal protein L7/L12 | STM4152 rplL | + | 4365025-4365590 |
| 26. | -15.77 | -3.29 | ((((((.((((((((((((((((&))))...))))))).)))))))))) | 69 | 91 | 4-29 | putative regulatory protein | STM3084.S | - | 3248047-3247098 |
| 27. | -15.75 | -3.28 | (((((..((.((((.((((((((((((&)))))))))))))).))...))))) | 78 | 103 | 2-27 | glycerol diffusion | STM4087 glpF | - | 4296916-4295871 |
| 28. | -15.71 | -3.26 | (((((.(((((((((((.(((...((((((&))))))...))).))))))))))))))) | 444 | 473 | 4-32 | putative amidohydrolase | STM3133 | - | 3294223-3293145 |
| 29. | -15.47 | -3.15 | (((((((.....(((.(((((((((&))))))))).)))......))))))) | 163 | 187 | 4-30 | putative phage antiterminator | STM2239 | - | 2339752-2339184 |
| 30. | -15.36 | -3.1 | ((((((((((((((((((.(((((&))))).))....))))))))))))))) | 640 | 663 | 23-50 | virulence protein | STM1395 ssaD | + | 1481676-1483087 |
| 31. | -15.31 | -3.08 | ((((((..(((..((.((((((((&))))))))..))..)))..))).))) | 73 | 96 | 25-50 | hypothetical protein STM2728 | STM2728 | - | 2870315-2869927 |
| 32. | -15.22 | -3.04 | ((((((((((((((((&))))).))))))))))) | 323 | 338 | 11-27 | F0F1 ATP synthase subunit I | STM3872 atpI | - | 4080756-4080176 |
| 33. | -15.21 | -3.04 | (((((.((((((((((.((((((&))))))......))))))))).))))) | 1656 | 1677 | 12-37 | ferrioxamine receptor | STM0364 foxA | + | 412463-414771 |
| 34. | -15.18 | -3.02 | (((((((..(((((((((&))))))))).))))))) | 802 | 819 | 1-17 | putative phosphoribulokinase | STM3464 prkB | + | 3613768-3614837 |
| 35. | -15.16 | -3.01 | (((((((.(((.(((((((((((&)))))))..)))))......))).))))))) | 1338 | 1360 | 1-29 | hypothetical protein STM2013 | STM2013 yeeO | - | 2096034-2094377 |
| 36. | -15.09 | -2.98 | ((..(((((((((((&)))))))))))..)) | 2846 | 2860 | 26-40 | aminoglycoside/multidrug efflux system | STM2481 acrD | + | 2593447-2596760 |
| 37. | -15.06 | -2.97 | (((((..((((((((...(((((&)))))..)))))))).....))))) | 600 | 622 | 4-28 | hypothetical protein STM1989 | STM1989 yedI | - | 2072993-2071882 |
| 38. | -15.03 | -2.96 | ((((((((...((((((.((((..(((((&))))).)))).))))))....)))))))) | -189 | -161 | 14-42 | 4 hydroxyphenylacetate catabolism | STM1101 hpaG | + | 1188021-1189510 |
| 39. | -15.01 | -2.95 | (((((((((((((...((((((&)))....))).))))))))))))) | 722 | 743 | 4-27 | putative glutamic dehyrogenase like protein | STM1795 | + | 1893331-1894856 |
